# Analysis of solid-state heparin samples by ATR-FTIR spectroscopy

**DOI:** 10.1101/538074

**Authors:** Anthony Devlin, Courtney J. Mycroft-West, Jeremy E. Turnbull, Marco Guerrini, Edwin A. Yates, Mark A. Skidmore

## Abstract

The widely used anticoagulant pharmaceutical, heparin, is a polydisperse, heterogeneous polysaccharide. Heparin is one of the essential medicines defined by the World Health Organisation but, during 2007-2008, was the subject of adulteration. The intrinsic heterogeneity and variability of heparin makes it a challenge to monitor its purity by conventional means. This has led to the adoption of alternative approaches for its analysis and quality control, some of which are based on multivariate analysis of ^1^H NMR spectra, or exploit correlation techniques. Such NMR spectroscopy-based analyses, however, require costly and technically demanding NMR instrumentation. Here, an alternative approach based on the use of attenuated total reflectance Fourier transform infrared spectroscopy (FTIR-ATR) combined with multivariate analysis is proposed. FTIR-ATR employs more affordable and easy-to-use technology and, when combined with multivariate analysis of the resultant spectra, readily differentiates between glycosaminoglycans of different types, between heparin samples of distinct animal origins and enables the detection of both known heparin contaminants, such as over-sulphated chondroitin sulfate (OSCS), as well as other alien sulphated polysaccharides in heparin samples to a degree of sensitivity comparable to that achievable by NMR. The approach will permit the rapid and cost-effective monitoring of pharmaceutical heparin at any stage of the production process and indeed, in principle, the quality control of any heterogeneous or variable material.

## Introduction

Unlike the majority of pharmaceutical agents, the widely used clinical anticoagulant heparin lacks a unique, defined structure. As a polydisperse, polysaccharide natural product derived from animal tissue, primarily the intestines of pigs at present, heparin is inherently varied in structure. Although variable, the typical composition of heparin can, nevertheless, be defined within bounds, in terms of the substitution of its underlying repeating disaccharide unit (comprising 1-4 linked uronate - glucosamine disaccharides), its substitution with a variety of O-, N-sulfates and N-acetyl groups in the α-D-glucosamine residues, and its uronic acid content (α-L-iduronate or β-D-glucoronate).^1,2^ The structural diversity of pharmaceutical heparin is further compounded by variations between individual animals, as well as the practice of amalgamating the material obtained from many individual animals before processing. The processing stage can also introduce structural modifications, depending on which method of preparation is being employed and whether unfractionated or low molecular weight heparin (LMWH) is being prepared. These consist of modifications to its constituent saccharides that include base-catalysed epoxide formation,^3^ de-O- and or N-sulfation,^4^ β-elimination and oxidation^5,6^, amongst others especially if the process is not carefully controlled. The need for sensitive methods with which to analyse heparin was highlighted by the global contamination of pharmaceutical heparin in 2007-8,^7,8,9^ caused by the introduction of an unnatural chemically modified GAG (over-sulfated chondroitin sulfate; OSCS) into the supply chain, that resulted in approximately 150 deaths and 350 other adverse events in the USA alone.^10^

The first challenge facing those who wish to analyse pharmaceutical heparin therefore involves being able to identify alien (i.e. non-heparin) agents within heterogeneous heparin samples. The second challenge is to be able to detect heparins from different animal or distinct tissue sources. This is particularly relevant given the re-introduction of bovine heparin into the US market.

In terms of the approaches that need to be adopted, the challenges of detecting a known alien substance and that of product quality control, which may include detection of previously unknown alien substances, are distinct, even if they are based on the application of similar principles. Since the adulteration of commercial pharmaceutical heparin in 2007-8, there have been many attempts to characterise^11^ and provide methods for, the detection of the major contaminating substance that was identified on that occasion,^9,12^ over-sulfated chondroitin sulfate (OSCS), but far fewer approaches have been reported that address the more demanding challenge of detecting *any* potential contaminant. The first attempts of which we are aware to address this depended first on changing the framework in which the purity of heparin was discussed, to recognise the fundamentally variable nature of the product. Then, it was necessary to design an approach which was sufficiently flexible and, at the same time generally applicable, to be able to detect *any* alien substance.

The first step was to view heparin, not as an individual product in a similar way to a single molecule pharmaceutical agent, paracetamol say, but rather as a group of subtly distinct substances that can be grouped in terms of their similarity using non-parametric approaches, for example, principal component analysis (PCA). This allowed extant, accepted pharmaceutical heparin samples to be grouped, depending on the degree to which their ^1^H NMR spectra resembled one another to form a library of accepted *bona fide* heparins.^13^ The second stage required the correlation of ^1^H NMR spectral features within a group of these structurally varied, *bona fide* heparin library samples to form a matrix, which then allowed comparison with the test sample of interest, and revealed any differences between the test sample and the library, from which a decision could be made concerning the provenance of the test sample.^14^ It is worth emphasising that it was never the aim of either of these approaches to detect contamination with OSCS in particular, rather to define an approach for the detection, in principle, of *any* alien species, among samples of variable composition. While this approach has been developed further in a series of papers by the same authors to include two dimensional (^1^H-^13^C) NMR,^15,16^ these approaches, nevertheless, still require access to a high-field NMR spectrometer and skilled technical assistance to meet the standards required to ensure reproducibility.

Other spectroscopic methods, including FTIR and Raman spectroscopy have been applied to the question of heparin purity in the past (reviewed in^12^) but, the present approach utilises attenuated total reflectance Fourier Transform infrared spectroscopy (ATR-FTIR), which enables rapid, facile analysis of samples in either their dry, powdered state or in solution, and is readily applicable to quality control of both the final pharmaceutical product and intermediates arising during the production process. Here, advantage is taken of the ability of ATR-FTIR to examine dry samples in order to simplify considerably the sample preparation.

The approach adopted in this work is based on the detection of chemical structures by Fourier transform infrared spectroscopy (FTIR). For larger molecules, the total number of vibrational modes is high and the number of infrared spectral bands and their complex superposition are further increased by overtones which correspond, in the classical picture, to harmonics of the fundamental vibrational modes. An additional nuance is provided by internal hydrogen bonds between constituent groups of the molecule which, since they change the dipoles of interacting groups, can also alter subtly the frequency of their vibrational modes. The combined effect of these factors is that FTIR spectra of biological macromolecules comprise complex, but characteristic spectra, which are very sensitive to changes in their detailed chemical composition. It is this feature of FTIR spectra that is exploited in the present approach.

A particular advantage of ATR-FTIR over other forms is that, following the simple adaptation of a standard FTIR instrument, the spectra of samples in the solid state can be recorded, obviating the need for expensive instrumentation, such as that associated with NMR spectroscopy. Additionally, the method avoids the need for samples to be prepared in deuterated water, thereby eliminating another potential source of variation. Sample preparation and data collection are simple and, furthermore, are suitable for adaptation to a portable format, which will be convenient for producers and regulators alike.

## Results

To begin exploring the viability of FTIR-ATR as a means of differentiating heparin samples, a large library of different GAG polysaccharides was first created. It was important to include a range of GAG polysaccharides, since they are structurally heterogeneous, both between species and between batches from the same species and this variability must be captured within the library if useful comparisons are to be made.^13^ The heparin samples that form the library, even though they are diverse, nevertheless possess similar structural characteristics which can provide useful metrics either for the grouping of samples, or for the purposes of classification and separating samples from analytical motives. An outline of the procedure is as follows (more details are provided in Methods): ATR-FTIR spectra of a series of GAG polysaccharide samples were recorded using a standard laboratory ATR-FTIR instrument. The GAG series consisted of: 176 heparins (Hps) (comprising 69 porcine mucosal heparins (PMHs), 55 bovine mucosal heparins (BMHs), 33 ovine mucosal heparins (OMHs) and 19 bovine lung heparins (BLHs), 31 heparan sulfates (HS)), 29 chondroitin sulfates (13 CS-As and 16 CS-Cs), 21 dermatan sulfates (DS), 10 hyaluronic acids (HAs) and 8 semi-synthetically modified polysaccharides including 6 over-sulfated chondroitin sulfates (OSCSs), 1 over-sulfated agarose sulfate (OSAS) and 1 dextran sulfate (DeS). These spectra were subjected to a preliminary smoothing procedure, followed by a baseline correction involving application of a custom fitted 7^th^-order polynomial baseline. The absorbance values of the resultant spectra were normalised and the variable regions located at >3600cm^-1^, below 700 cm^-1^ and between 2500 and 2000 cm^-1^ were removed to reduce the effects of environmental variation. The first derivatives of these spectra were then calculated and these were then subjected to principal component analysis (PCA) using singular value decomposition (SVD). The first few principal components, PC1 to PC4 covering ∼85% of variation (**Fig. 2.1**), were plotted allowing the different GAG groups to be distinguished.

**Figure 1.1.**
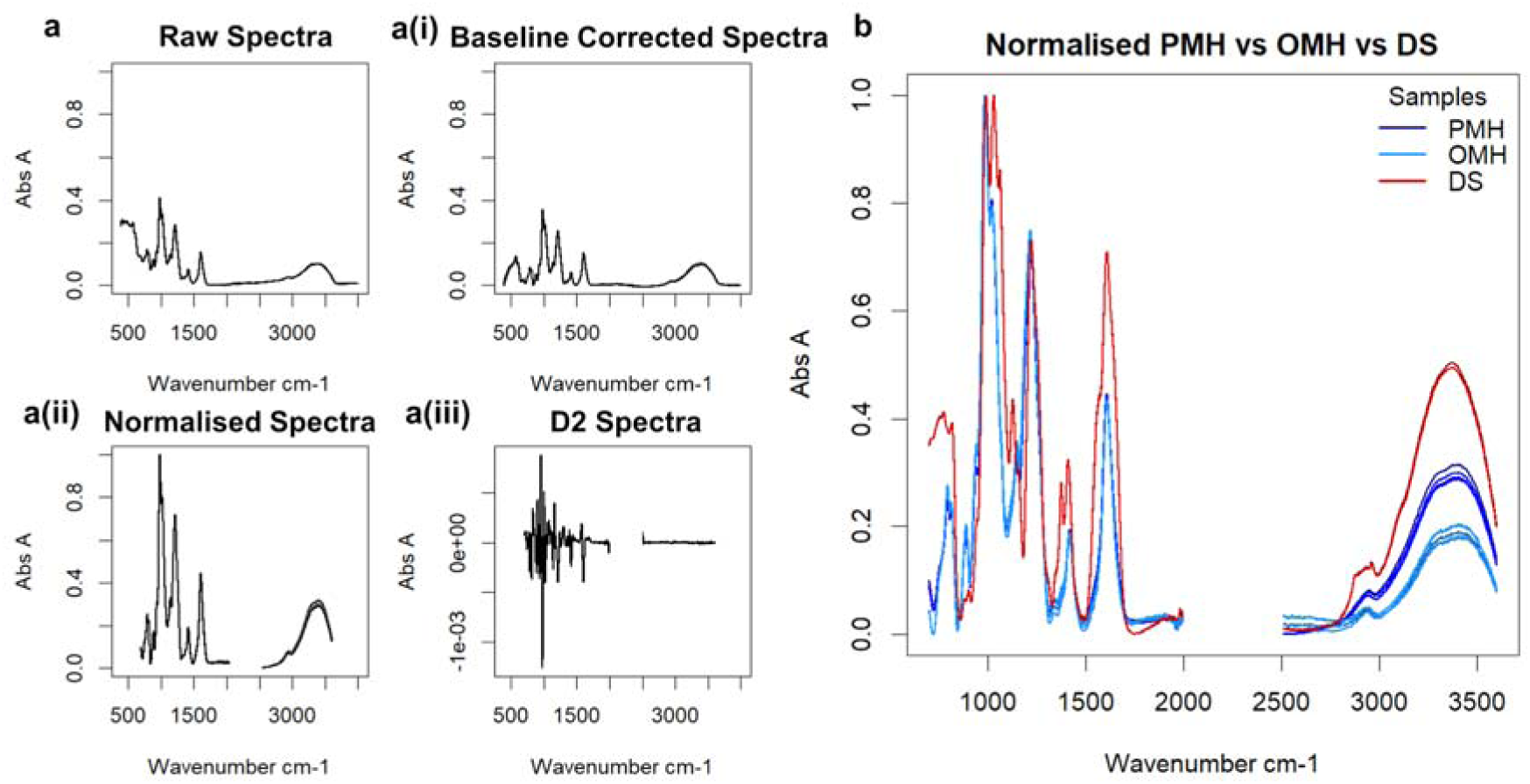
**a**. Steps in the ATR-FTIR spectral preparation following recording on a dry sample of pharmaceutical heparin. **a** The raw spectrum of a random heparin **ai** Baseline corrected (7^th^ order polynomial) spectrum **aii** Spectrum following normalisation and removal of variable regions <700 cm^-1^, > 3600 cm^-1^ and between 2000 and 2500 cm^-1^. **aiii** The second derivative of the resultant spectrum. **b**. Normalised ATR-FTIR spectra of a sample of PMH, an OMH and a DS (700-3600 cm^-1^) showing similarity between PMH and OMH in the regions 700-2000 cm^-1^, only differing significantly in the highly variable OH region 3000-3500 cm^-1^. In contrast, DS is distinct from either PMH or OMH in the region 700-2000 cm^-1^, corresponding to structural variations.

**Figure 2.1.**
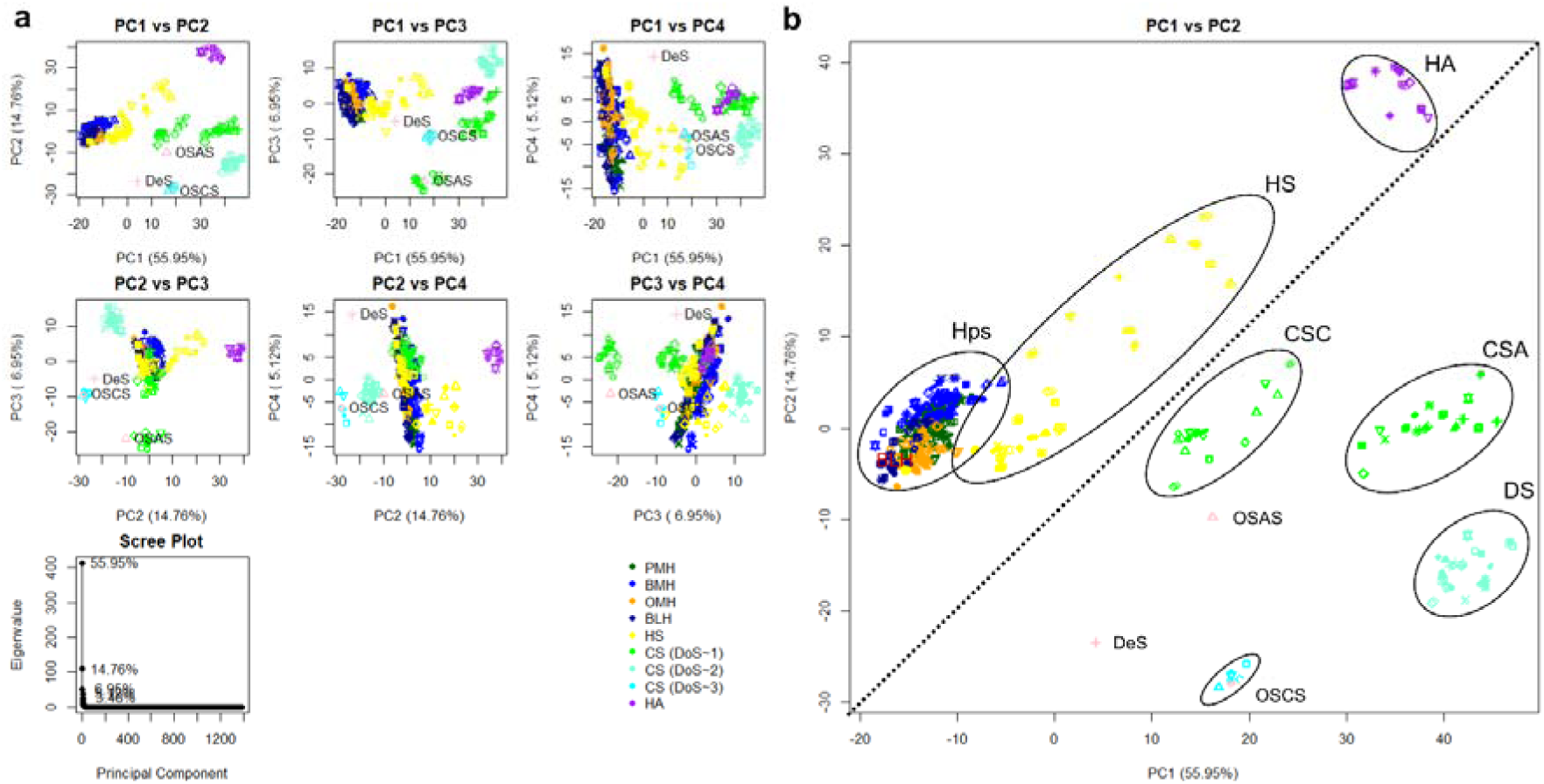
Principal component analysis of ATR-FTIR spectra of glycosaminoglycans. **a.** PCA scores and scree plot(s) for different GAG families. At least two components are needed to separate into family groups, meaning that earlier components appear to separate by gross structure, while later components separate by individual constituents shared by all the carbohydrate species. **b.** Score plot for PC1 vs PC2. GAG families are highlighted to show groupings and the plot is split in half diagonally to distinguish different glycosidic bond types (β 1-4 upper left vs β1-3/4 lower right). In all plots, 176 heparins, comprising: 69 PMH (dark green) 57 BMH (blue), 33 OMH (orange), 19 BLH (dark blue)), 31 HS (orange), 40 chondroitin (including 13 CS-A samples, 16 CS-C samples and 21 DS (green for monosulphated chondroitin samples and teal for disulphated chondroitin samples), 11 HA (purple) and 6 OSCS (including the OSCS selected for contamination study later (aquamarine)) samples, as well as an OSAS and DeS sample, also used in the contamination study (Fig. 2.3) are compared. All 5 repeats of each sample are shown and the polysaccharides selected for the contamination study are highlighted in lilac. The ellipses are for illustrative purposes only.

**Figure 2.2.**
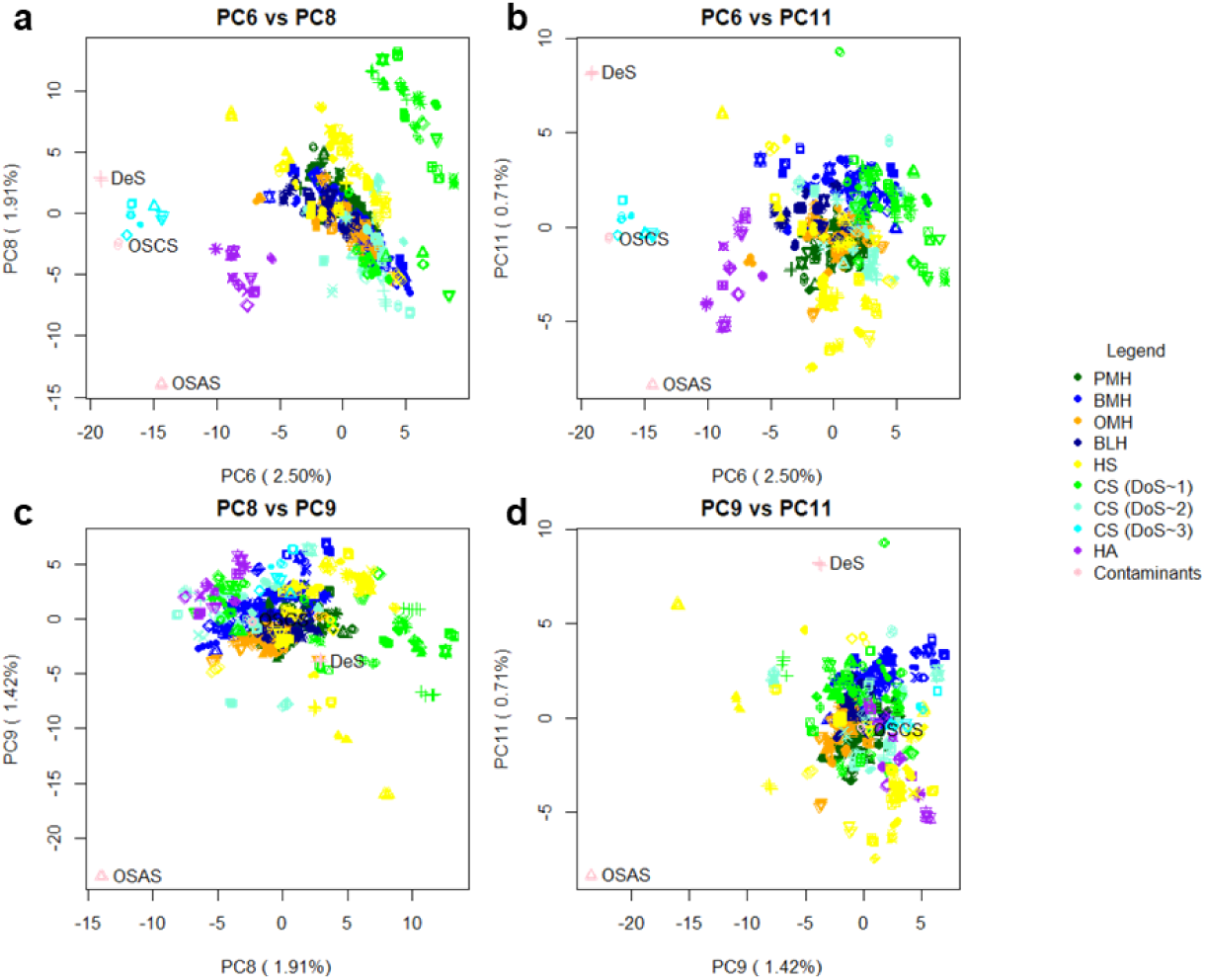
The lower principal components detect increasingly subtle differences, and can differentiate synthetically modified polysaccharides. Score plots for lower PCs following analysis of FTIR-ATR spectra of different GAG species and other polysaccharides. As the PCs cover progressively less variance, subtler changes become more apparent, amplifying the distinctness of relative outliers, such as OSCS, OSAS and DeS. All 5 repeats of each spectrum are shown. **a.** score plot of PC6 vs PC8 **b.** score plot of PC6 vs PC11 **c.** score plot of PC8 vs PC9 **d.** score plot of PC9 vs PC11.

Clear separations between samples comprising □-D-glucosamine and □-D-galactosamine residue-containing GAGs can be seen with PC1 and PC2 (**Figs.1, 2**) where HA, HS and Hp (GlcN containing) reside in the top left half of the plot, while the various CS samples (GalN-containing) remain in the bottom right. Within these two groups, across PC1, the type(s) of sulphation are separated; in the glucosamine region, three general regions can be seen - the purple HA (entirely unsulphated) group, the yellow variable HS group (HS consists of a combination of 6-O-sulphated, 2-O-sulphated and N-sulphated residues, the levels of which determine the location of a sample within this group) and the green-blue-orange Hp group, which contains the same sulphation types (albeit more frequently throughout the chain) and a final, rarer, 3-O-sulphation. The discrimination of sulphation type is clearer in the galactosamine group. Here, 6-O-Sulphated DS and CS-A are on the far right, while 4-O-sulphated CS-C is to the left. The OSCS lies closer to the 4-O-sulphated group, while DS (containing 2-O-sulphated IdoA) moves slightly further right and is better differentiated from CS-A. OSCS, also possessing 6-O- and 2-O-sulphation is positioned slightly to the left of the densest region of the CS-C data-point cluster. Across PC1, the ratio of N-sulfate (NS) to N-acetylation (N-Ac) is also distinguished, with samples not containing NS (i.e. HA and the CSs) being separated to the right, while NS-containing species (heparin and some HS samples) separate to the left.

PC2 appears to separate by degree of sulphation (DoS), orthogonal to the GlcN/ GalN groups. Again, the entirely unsulphated HA resides at the top, samples descending down to Hp with an average DoS of 2.3. Meanwhile, CS-A and CS-C (a DoS of ∼ 1) start at the top of the GalN region, moving down to DS in the middle with a DoS of ∼2 and finishing at OSCS with a DoS in excess of 3. PC3 may separate samples in terms of the detailed geometries of their sulfate and carboxylate groups, which both present relatively large signals in FTIR spectra, but are known to vary according to their disposition around the sugar pyranose rings.^17^ Through its association with a signal at ∼1000 cm^-1^ of the FTIR spectra, PC4 is likely to represent primarily the extent of solvation of free hydroxyl groups along the glycan backbone, which is highly variable and a function of environmental factors.

The early components separate the GAGs very well by gross structural features, however contained within this library are two additional carbohydrates - over-sulphated agarose sulphate (OSAS) and dextran sulphate (DeS). Agarose, a D-galactose-containing polysaccharide, in which every other D-galactose residue possesses an oxygen bridge between C1 and C3, is found in the GalN region (as expected), while DeS, a sulphated glucose polymer, consists primarily of β(1-6) glycosidic bonds (not found in the other GAGs) and, on average, each sugar residue has two sulphate groups. This di-sulphated character enables it to fit well within the group containing ∼2 sulphates of the GalN-containing groups, (around the DS-containing region). It may be expected that DeS would fall within the GlcN region, confirmed by PC2, however, it also possesses β(1-3) linkages, consistent with its appearance in the GalN (β(1-3), β(1-4))-containing group, but is separated from the rest. The relative contribution of the OSAS and DeS samples towards the entire library is very low, at 0.7% of the total number of spectra, so they are relatively dwarfed, however, later components can still separate them strongly from the GAG library. Their unique structural features (a β(1-6) bond and a C1-C3 oxygen bridge) are differentiated by PCs 6, 8, 9 and 11 (2.40%, 1.83%, 1.37% and 0.68% of the variance, respectively). PCs 8 and 9 separate OSAS entirely from the remainder of the polysaccharides, presumably on the basis of their C1-C3 oxygen bridges, while PC 11 separates DeS clearly from the rest of the polysaccharides, presumably on the basis of their β(1-6) bonds. Interestingly, PC 6 separates all artificially sulphated (OSCS, OSAS and DeS) polysaccharides from the rest of the GAG polysaccharides.

After establishing ATR-FTIR as a valid tool for distinguishing polysaccharides on the basis of their structural features, some of its potential practical uses were also investigated. First, the detection of heparin contaminated by the addition of either a chemically modified GAG or synthetically derivatised non-GAG polysaccharide species was investigated. One each of the OSCS polysaccharides and the PMH samples was selected at random, and contaminated with each other on a defined weight for weight basis. The ATR-FTIR spectra of the resulting contaminated mixtures were recorded, followed by PCA analysis as previously outlined, the results of which are shown in **Fig. 2.3a**. In general, as the percentage OSCS content increases, the location on the PC score plot of the sample moves further from the heparin library in a linear manner. At the lower range of contamination (≤ 2.5% w/w) this discriminatory power is lost and the contaminated samples spread out across the heparin library (**Fig. 2.3a(i-iv)**). Lower components are required for samples to be distinguished. PC 5 was needed to distinguish 1% OSCS contamination comfortably. Below 1% contamination, the contaminated sample appears at the edge of the library, with separation arguable at 0.5%, but not definitive. At 0.25% contamination the contaminated heparin is indistinguishable from other outlying heparin samples.

**Figure 2.3.**
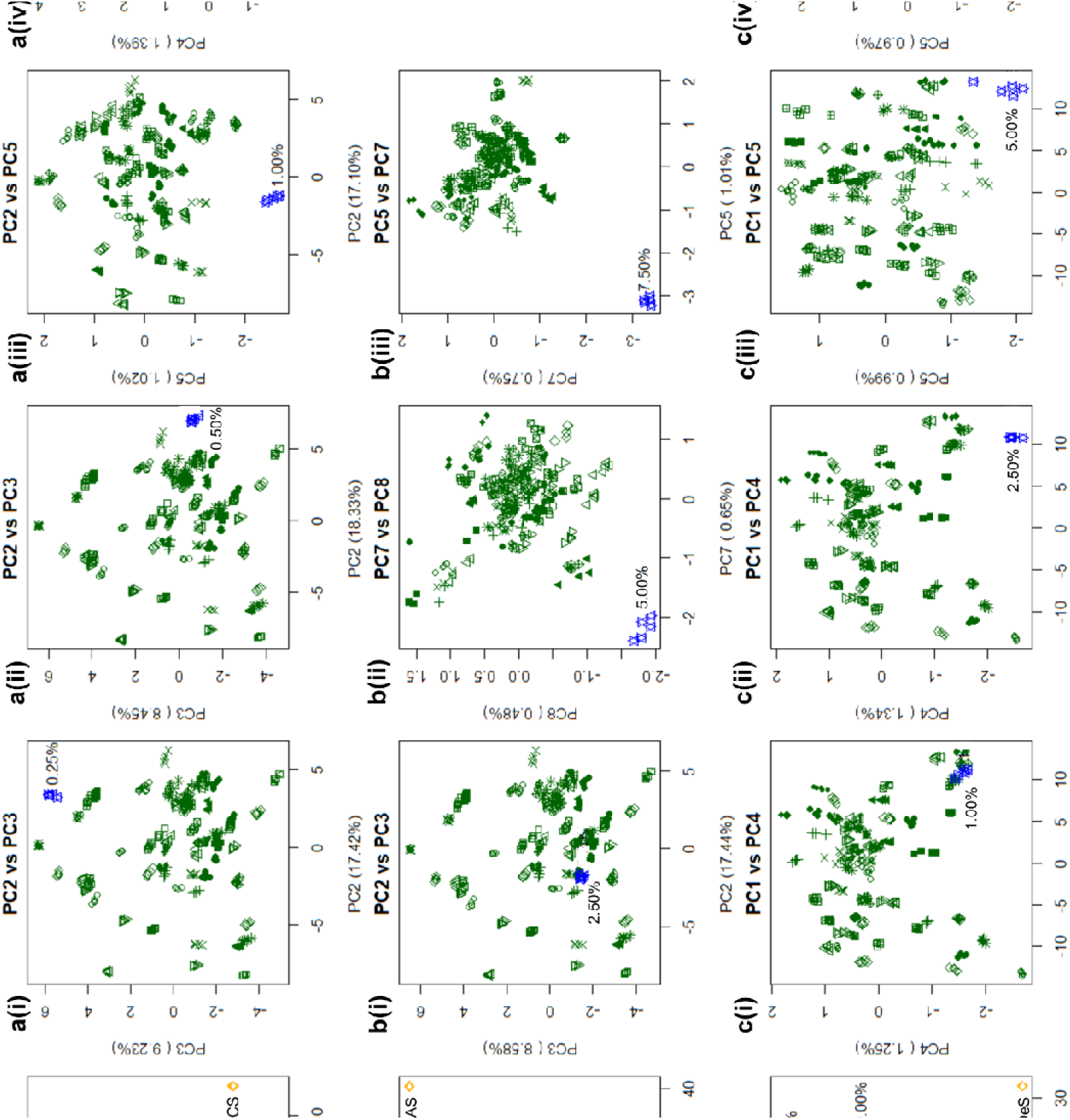
Detection of known contaminants in porcine mucosal heparin. PCA Score plots of FTIR-ATR spectra of the library of 69 bona fide PMH samples analysed in the presence of a randomly selected heparin contaminated deliberately with varying levels of OSCS (**a**), OSAS (**b**) and DeS (**c**) (confirmed to contain no other contaminants and mixed w/w% with 40%, 20%, 15%, 10%, 7.5%, 5%, 2.5%, 1%, 0.5%, 0.25%, 0.13% and 0.06% contaminant). Differentiation down to 0.5%, 5.0% and 2.5% contamination are achievable respectively. All 5 repeats for each sample are shown. **a.** score plot PC2 vs PC3 of heparin vs. the entire contamination series. **ai.** score plot of PC2 vs PC3 of heparin vs. 0.25% OSCS. **aii.** score plot PC2 vs PC3 of heparin vs 0.5% OSCS contamination. **aiii.** score plot PC2 vs PC5 of heparin vs 1.0% OSCS contaminated heparin. **aiv.** PC2 vs PC4 of heparin vs 2.5% OSCS contaminated heparin. **b.** score plot PC1 vs PC2 of heparin vs. the entire contamination series. **bi.** score plot of PC2 vs PC3 of heparin vs. 2.5% OSCS. **bii.** score plot PC7 vs PC8 of heparin vs 5.0% OSCS contamination. **biii.** Score plot PC5 vs PC7 of heparin vs 7.5% OSAS contamination. **c.** score plot PC1 vs PC2 of heparin vs. the entire contamination series. **ci.** score plot of PC1 vs PC4 of heparin vs. 1.0% DeS contamination. **cii.** score plot PC1 vs PC4 of heparin vs 2.5% DeS contamination. **ciii.** score plot PC1 vs PC5 of heparin vs 5.0% DeS contamination. **civ.** score plot PC4 vs PC5 of heparin vs 7.5% DeS contamination.

In previous work employing ^1^H NMR spectroscopy,^14^ PCA was able to distinguish 2.5% OSCS contamination. OSCS exhibits a well-defined difference in ^1^H NMR chemical shift values of signals from the N-acetyl group, making detection by multivariate analysis relatively straightforward. To present a stronger challenge, and to cover the possibility of any future contamination by non-GAG polysaccharides whose features may reside entirely under the mass of carbohydrate NMR signals, an N-acetyl free, chemically modified polysaccharide, OSAS, was employed. The same heparin sample that had been utilised previously was contaminated with OSAS in a comparable manner (**Fig. 2.3b**). The results were broadly similar to those for OSCS contamination, the entire contamination series extending progressively from the cluster representing the heparin samples and, as before, the lower percentage contaminants merging with heparin. Using PCs 5 and 7 and, 7 and 8, 7.5% and 5% contamination were separated strongly from the cluster of heparin samples. At 2.5% contamination, no separation was evident. Another carbohydrate DeS, made of a glucopyranose polysaccharide backbone, possessing a variable sulfation pattern and level, like heparin, was utilised in a manner akin to OSCS and OSAS (**Fig. 2.3c**). The results were similar to these contaminations, with the entire contamination series extending progressively from the heparin cluster. Using PCs 1 and 4, separation at 5% and 2.5% levels were achieved, while 1% contamination sits on the edge of the cluster.

More recently, heparin from different tissue types or animal species has entered the market, potentially, having been introduced deliberately. It will be important to be able to distinguish heparin from different animal sources. In fact, while bovine heparin was, until the 1990’s, widely used medicinally, it fell into disuse throughout much of the western world following the bovine spongiform encephalopathy (BSE) crisis. Nevertheless, in some countries where the use of pig products is sensitive for religious or cultural reasons, and throughout much of South America, bovine heparin continues to be used. Heparin from bovine sources is now due to be reintroduced into the North American market. It has been established that Hps from distinct animal sources exhibit different structural features^18,1^ and, in light of this, the same 176 Hps as used above_from different species and tissues were subjected to PCA following ATR-FTIR spectral acquisition.

Bovine mucosal heparin separates strongly from the rest of the heparins with PCs 2 and 3, however, OMH, PMH and BLH are separated less well (**Fig. 2.4**). Through PC 5, all heparin species form one large cluster, however, 4 nodes within this cluster are evident - each populated with a single Hp species, with BLH at the top, BMH to the right, PMH at the bottom left and OMH at the top left. PMH, OMH and BLH do not separate entirely, but do form distinct nodules. To further examine these samples, in **Figs. 2.4C** and **D**, OMH with PMH and OMH with BLH were compared respectively. Again, the two species formed nodes at opposing ends of their combined data clusters, with some cross-over between the two.

**Figure 2.4.**
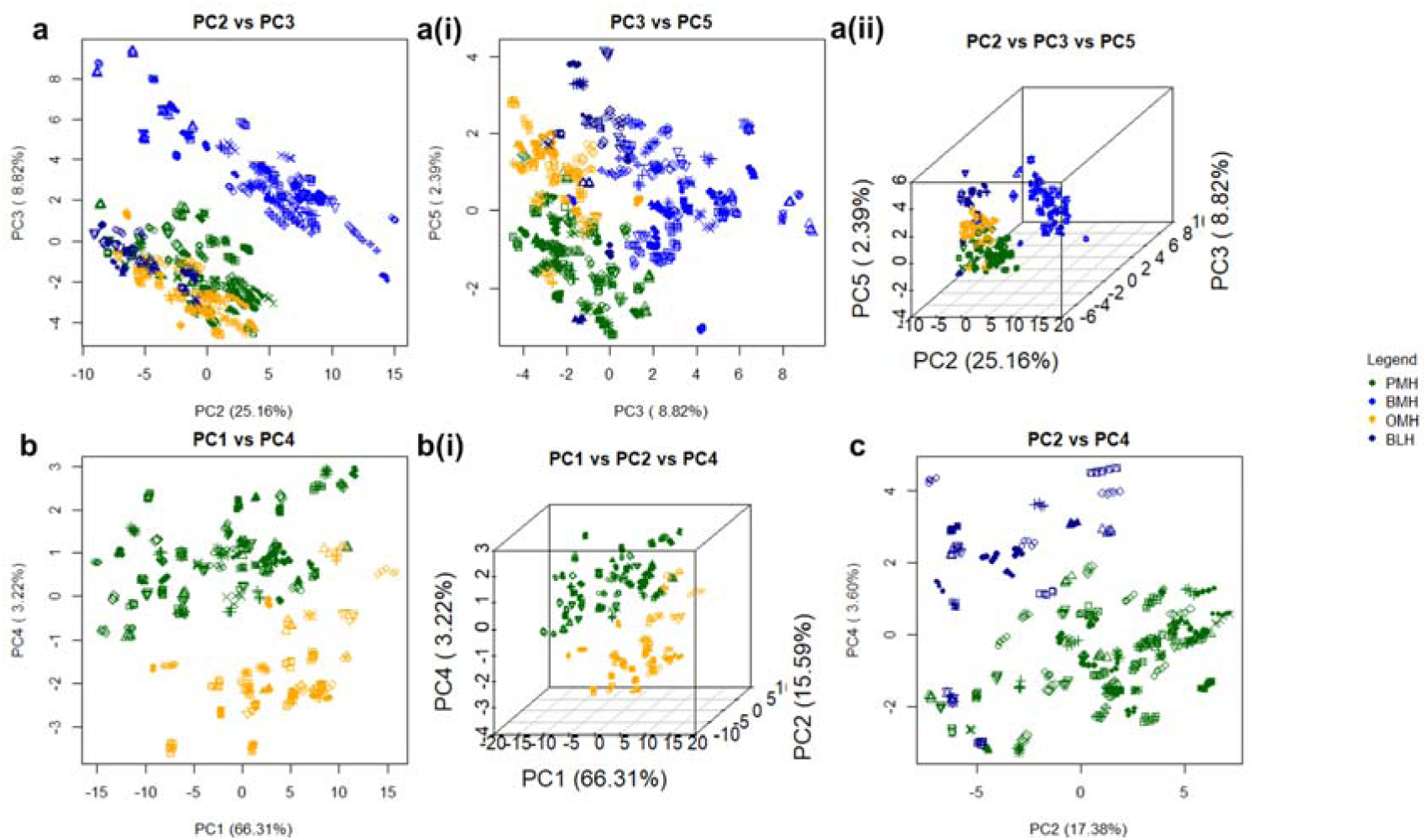
Separation of heparin in terms of species. PCA Score plots following analysis of ATR-FTIR spectra of heparin samples from different animal species. Samples of heparin from each animal species can be separated into their respective clusters. All 5 repeats of 69 PMH (dark green), 57 BMH (blue), 33 OMH (orange) and 19 BLH (dark blue) spectra were compared using PCA. **a.** All four species compared against each other. **a** score plot of PC2 vs PC3 **ai** score plot of PC3 vs PC5 **aii** 3D score plot of PC2 vs PC3 vs PC5. **b.** Spectra of samples of OMH and PMH are compared. **b** score plot of PC1 vs PC4 **bi** 3D score plot of PC1 vs PC2 vs PC4 respectively and for BLH vs PMH in **c.** Score plot of PC2 vs PC4 of PCA of spectra of BLH and PMH samples.

Finally, a series with PMH and BMH contamination was created; the PMH selected for the previous contamination series was employed, along with another randomly selected PMH and two randomly selected BMH samples from the library. These were contaminated with each other, to form two PMH:BMH contamination series containing 1%, 2.5%, 5%, 7.5%, 10%, 15%, 20% and 40% BMH. Given that the structure of heparins from different sources are, in broad terms, similar, comparison of any single inter-species contamination series will be highly dictated by the level of structural similarity between the parental materials. For example, if a bovine-like porcine heparin is contaminated with a bovine heparin, the separation between the two will be less, hence four contamination series were employed to counter this. When all four of the entire contamination series were plotted against the PMH library, the contamination series emerged from the library in a linear fashion according to the level of contamination or clustered tightly towards one edge of the library; the 40% contaminated sample being clearly removed from the main heparin cluster in all cases. When singular contaminated samples were plotted against the heparin library, the 40% contaminated sample could be detected in all samples while the 20% contaminated sample was separated in all series apart from series b (Fig.2.5b). And as low as 10% could be seen in series a (Fig.2.5a). Very low components were used in some cases to distinguish the species; in series **a**, these were PCs 10 and 5 covering 0.37% and 0.99% of the variance respectively and allowing separation of 20% and 40% contamination respectively; in series **b**, they were PCs 6 and 8 covering 0.43% and 0.95% respectively, allowing for separation of 40% contamination; in series **c**, they were PCs 4, 11, and 14 at 1.31%, 0.26%, and 0.16% variance allowing separation of 10%, 20% and 40% contamination respectively; and in series **d**, they were PCs 11 and 8 covering 0.26% and 0.44% of the variance, allowing for separation of 20% and 40% contamination respectively. Such small variations are expected, and arise from the high structural similarity of these molecules (**Fig. 2.5**).

**Fig. 2.5.**
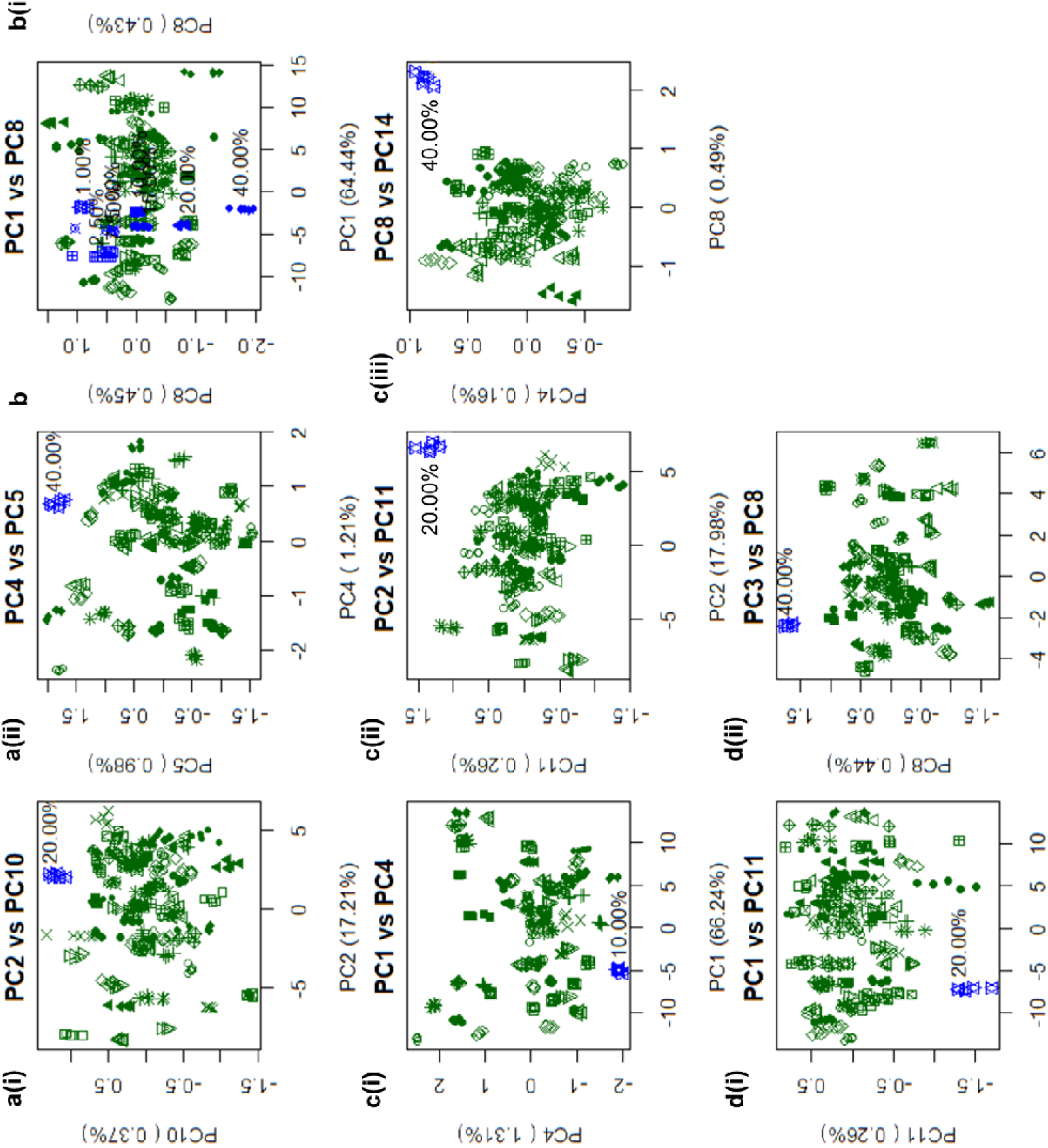
Detection of porcine heparin contaminated by bovine mucosal heparin. PCA score plots of FTIR-ATR spectra of the library of 69 bona fide heparin samples analysed in the presence of a randomly selected PMH contaminated deliberately with varying levels of a randomly selected BMH (confirmed to contain no other contaminants and mixed (w/w) with 40%, 20%, 15%, 10%, 7.5%, 5%, 2.5%, 1% BMH). Four contamination series (a:d) were created using the same contaminated PMH as before, among 3 other randomly selected PMH samples with 4 randomly selected BMH samples. Series a:d correspond to **Fig2**.**a-d**. For each, the first score plot shown represents the entire contamination series against the entire PMH library. 40% contamination is detected in all cases while 20% is detectable in series **a, b** and **d** and 10% is detectable in series **a**. **ai** score plot of PC2 vs PC10 of PMH(a) vs. 20.0% BMH(a) contamination. **aii** score plot of PC4 vs PC5 of PMH(a) vs. 40.0% BMH(a) contamination. **bi** score plot of PC6 vs PC8 of PMH(b) vs. 40.0% BMH(b) contamination. **ci** score plot of PC1 vs PC4 of PMH(c) vs. 10.0% BMH(c) contamination. **cii** score plot of PC2 vs PC11 of PMH(c) vs. 20.0% BMH(c) contamination. **ciii** score plot of PC8 vs PC14 of PMH(c) vs. 40.0% BMH(c) contamination. **di** score plot of PC1 vs PC11 of PMH(d) vs. 20.0% BMH(d) contamination. **dii** score plot of PC3 vs PC8 of PMH(d) vs. 40.0% BMH(d) contamination

## Discussion

Any attempt at the full assignment of the FTIR spectrum of a sample of heparin has yet to be reported. While FTIR and NMR spectra yield broadly complementary information, that available from FTIR currently lags someway behind that available from NMR. In part, this disparity is due to the availability of highly informative multi-dimensional experiments (e.g. COSY, TOCSY and HSQC) and high resolution instruments, which enable manipulation of pulse sequences and signal assignment to be achieved so successfully. It is also a consequence, however, of the superior signal dispersion of the NMR spectrum, especially in the ^13^C dimension, it being possible in some cases, to distinguish between identical chemical features (e.g. mono-sulphated glucosamine residues) that are embedded in distinct saccharide sequences.^19^ In contrast, FTIR signals from particular structural features are typically broad, super-imposed and more difficult to distinguish, although the changes observed in Amide I bands, for example, in different protein secondary structures, do demonstrate that FTIR is sufficiently sensitive to distinguish three-dimensional structures. There are also some features evident in FTIR spectra that are difficult to control and these arise, in particular, from variations in the solvation level, evident through changes in broad OH bands. These regions were eliminated, as far as possible, by excluding the relevant spectral regions (see Methods). A similar process is also conducted during the processing of NMR spectra to account for the deuterated water peak.^13,14^ In the present work, variation in hydration between samples has been minimised through standardised sample drying procedures prior to recording the spectra.

While some correlation experiments have been performed in the FTIR field^20^, true 2-dimensional FTIR spectroscopy remains a specialised topic. Nevertheless, some basic IR band assignments in heparin have been made, including vibrational modes associated with carboxyl group stretches, N-acetyl group modes, including C=O and N-H stretches that are analogous to Amide I and II bands in peptides, and N- and O-sulfate stretches, as well as some more general carbohydrate associated modes involving C-O-C deformations around the glycosidic linkages: in relation to the FTIR spectrum of heparin and its derivatives, the 800-950 cm^-1^ region contains complex C-O-S stretching modes; that at 890 cm^-1^ being attributed to C-O-S,^21^ glycosidic bond and C-O-S stretching at 937 cm^-1^ and the 6-O-S in glucosamine at 827 cm^-1^ and 820 cm^-1. 22,23^ Features at 800 cm^-1^ have been assigned to axial O-sulfates in L-Ido A residues^17^. Bands at 1230 cm^-1^ have been proposed to arise from S=O stretches,^24^ while those at 1430 cm^-1^ have been attributed to symmetric carbonyl stretching,^25^ and those at 1635 cm^-1^, to asymmetric stretches.^26^

The results presented in Figs 2.3 demonstrate that the differentiation of samples containing known contaminants using the present ATR-FTIR approach is achievable; 0.5%, 5.0% and 2.5% (w/w) of OSCS, OSAS and DeS respectively were readily distinguishable by visual inspection from a heparin library. These results are comparable to those achievable using NMR spectroscopy, which is normally considered to be a considerably higher resolution technique but is, undeniably, more technically demanding and expensive than ATR-FTIR, especially since the advent of competitive, portable laboratory FTIR instruments. It is also clear that there is a significant level of structural complexity within ATR-FTIR spectra that remains to be fully exploited, and for which simple band assignments cannot account; this will be the topic of future investigations. For example, the differentiation of N-acetyl GlcN and N-acetyl GalN residues, as well as information pertaining to glycosidic linkage geometry that is both subtle and complex, will all require detailed complementary studies to decipher. Despite these caveats, the original aim of this work, to ascertain whether it is possible to employ ATR-FTIR to distinguish GAG polysaccharides from distinct origins and to detect contaminants has been shown to be readily achievable.

To conclude, sample preparation for ATR-FTIR is straightforward and, since it can employ solid material, rather than require dissolution, this removes the need for very expensive NMR spectrometers, accurate solution preparation, or skilled NMR technical assistance, making the current approach both economical and accessible. The method is highly sensitive and can detect structural variation both between different GAG types, but also between GAGs and other polysaccharides. Notably, it is sufficiently sensitive to distinguish between samples of heparin derived from distinct species (e.g. PMH vs. BMH) and from the same species, (e.g. BLH vs. BMH). This is particularly relevant in the context of the re-introduction of bovine-derived heparin into the US heparin market and also of the need to remain vigilant regarding the addition of heparin from other species into existing heparin, for example, ovine sources, which would be difficult to detect by current FDA methods. The approach offers a means of readily controlling purity, origin and process that is beyond the requirements of current FDA regulations. It should appeal to manufacturers as a means of guaranteeing provenance and of providing confidence in their manufacturing processes at a fraction of the cost of NMR.

## Methods

### Polysaccharide samples

A range of suppliers and manufacturers were used, full details of which can be found in the supplementary data, tables 1-7.

### FTIR sample preparation

Prior to spectral acquisition a small amount of sample preparation was performed; 1-10 mg of dry sample was taken and 1 ml of deionised water added. The solution was frozen at −80°C and lyophilised overnight. It is important that the same drying technique is used for all samples to minimise variation from this source. Care was also taken to use the same type of tubes, to minimise variation arising from different drying rates.

### ATR-FTIR

Samples were recorded using a Bruker Alpha I spectrometer in the region of 4000 to 400 cm^-1^, for 32 scans at a resolution of 2 cm^-1^ (approx 70 seconds acquisition time), 5 times. A background spectrum was taken prior to each sample, using the same settings as for sample acquisition but with a completely clean stage. 1-10 mg of dried sample was placed upon the crystal stage of the instrument, ensuring that the entirety of the crystal was covered. Sufficient sample was employed to ensure that at least 5 *µ*m thickness was obtained, as this is the extent to which the effervescent ATR wave penetrates. The instrument stage was cleaned with acetone and dried between samples. Spectra were recorded using OPUS software (Bruker) and exported in CSV format.

### FTIR processing

All processing and subsequent analysis was performed on an Asus Vivobook Pro (M580VD-EB76), using an intel core i7-7700HQ. Spectra were imported into R studio v1.1.463 prior to a preliminary smooth, employing a Savitzky-Golay smoothing algorithm (*signal* package, ***sgolayfilter***), with a 21 neighbour, 2^nd^ degree polynomial smooth, facilitating the improved accuracy of later corrections.

### Baseline correction

While a background spectrum was taken prior to each new sample, perturbations in the air can still occur during spectral acquisition. To further reduce the effects of these perturbations, each individual smoothed spectra underwent a custom baseline correction using a 7^th^ order polynomial. First, the spectra were separated into 6 equally spaced regions (buckets), with the minimum absorbance value for each of these buckets and their relevant wavenumber (x axis) values taken. The start and end values for the spectrum were added to these values and, from the resultant 8 x-y pairs, the coefficients for a 7^th^ order polynomial were generated using the *base* R ***lm*** function. The baseline was calculated utilising the generated coefficients and the original x-axis and then subtracted from the smoothed spectrum.

### Pre-PCA preparation

To remove the effects of the variable quantity of sample used to record a spectrum (which cannot be regulated easily, because there is no control over the amount of sample placed into contact with the ATR crystal), the corrected spectra were normalised (0-1) using the equation

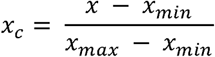

Where *x* is the value to be corrected, *x*_*c*_ is the resultant corrected value, *x*_*max*_ the maximum *x* value for the spectrum and *x*_*mln*_ the minimum *x* value for the spectrum.The normalised spectra had variable regions removed; these occur due to variable CO_2_ and H_2_O levels in the environment, (< 700 cm^-1^, between 2000 and 2500 cm^-1^ and >3600 cm^-1^). The second derivative was taken, again using the Savitzky-Golay algorithm but, this time, with 41 neighbours and a 2^nd^ order polynomial. The preliminary smooth is not always needed, but if omitted, then more neighbours are required for optimum output during this step. It was observed that a less aggressive smoothing procedure at the beginning of the process, removed anomalous baseline corrections entirely.

### PCA

The normalised and corrected matrix of intensities underwent PCA using singular value decomposition with the *base* ***prcomp*** function in R.

During this process, the matrix was mean centered, but not scaled in any other way. Through comparison of the scree and loading plots, suitable PC scores were chosen to plot against each other as x-y scatter graphs. Low PCs were employed, assuming that the 5 individual repeats across this component form compact groups.

### Defining optimum smoothing and correction parameters

To define the optimum smoothing parameters, different degrees of smoothing both in terms of neighbours and polynomial were applied to the spectra and the resultant PC scores compared. If the spectra were not smoothed sufficiently, their individual 5 repeats spread across the score plots i.e they are dissimilar, while if the samples are over-smoothed, all samples from diverse polysaccharide types intermingle, yielding no meaningful separation. The plots with the least smoothing and the most repeated sample grouping were taken forward, and these comprised 21 neighbours using a 2^nd^ order polynomial for the preliminary smooth, and 41 neighbours and a 2^nd^ order polynomial for the pre-differentiation smooth. The optimum baseline polynomial was also defined in a similar manner, using distinct polynomials in the range of 2^nd^ order to 9^th^ order. For an n^th^ order polynomial, the spectra were divided into n - 1 buckets, and the same script run, as above. Second and 3^rd^ order polynomials provide relatively poor baselines, often resulting in early or late baseline anomalies, in which alien peaks are introduced as a consequence of their effective rigidity. For 4^th^ order polynomials and higher, however, the baselines are adequate. A 7^th^ order polynomial was chosen because it produced the fewest unusable corrections, i.e samples whose baseline becomes more curved

### Preparation of spectral libraries

Prior to sample comparison through PCA, individual sample libraries were created for each polysaccharide species (PMH, OMH, BMH, BLH, CS-A, CS-C, DS, HA, HS and OSCS). Each polysaccharide library was compared to itself with PCA, and through use of the first 10 principal components, any outstanding samples (i.e any that appeared distinctly removed from the main data-cluster of the library) were removed from the spectral library. The aforementioned anomalies were due to the particularly unusual nature of the samples, with abnormal bands and/or correlations between bands intensities. Unique samples (OSAS and DeS) were also introduced, however, these contained relatively little heterogeneity, hence a library was unnecessary.

### Polysaccharide sample comparison

The various libraries were subsequently compared against each other in a series of PCA score plots, starting with all the libraries compared against each other, including two additional molecules (DeS and OSAS). To test the practical application of this method, three contamination series were created through the selection of a random PMH sample from the PMH library, and subsequent weight by weight contamination to create a series of contaminated samples at the levels of 0.0625%, 0.125%, 0.25%, 0.50%, 1.0%, 2.5%, 5.0%, 7.5%, 10.0%, 15.0%, 20.0% and 40.0%. Each series contained the same PMH sample, but a different contaminant, namely OSCS, OSAS or DeS. During this study, all contaminated samples were compared against the PMH library using PCA, followed by successive PCA for each individual contaminated species against the PMH library, where a sample contaminated at the 0.06% w/w/ level was first compared against all PMHs, then one at 0.125% w/w contamination was compared against all PMH *etc*.

All of the Hp species libraries, PMH, OMH, BMH and BLH, were compared to each other using PCA and following the analysis, owing to their clustering as opposed to separation, PMH with OMH, and PMH with BLH, underwent PCA. A series of four PMH:BMH contamination series were then created. The comparison of PMH and BMH was selected, as these two have the best separation and represent the most likely case of heparin contamination. The heparin used for previous contamination studies was utilised, along with three other randomly selected PMH samples. Four BMH samples were selected at random, and randomly matched with the four PMH samples. The PMH samples were contaminated (w/w) with their BMH partner, at the levels of 1.0%, 2.5%, 5.0%, 7.5%, 10.0%, 15.0%, 20.0% and 40.0%. All contaminated samples were compared against the PMH library using PCA, followed by successive PCAs for each individual contaminated species against the PMH library, where 1.0% was compared against all PMH samples, then 2.5% was compared against all PMH samples *etc*.

## Abbreviations

LMWH: low molecular weight heparin.
GAG: glycosaminoglycan.
OSCS: over-sulphated chondroitin sulphate.
PCA: principal component analysis.
NMR: nuclear magnetic resonance.
FTIR: Fourier transform infrared.
ATR: attenuated total reflectance.
PMH: porcine mucosal heparin.
BMH: bovine mucosal heparin.
OMH: ovine musocal heparin.
CS-A: chondroitin sulphate-A.
CS-C: chondroitin sulphate-C.
DS: dermatan sulphate.
DeS: dextran sulphate.
OSAS: over-sulphated agarose sulphate.
HS: heparan sulphate.
HA: hyaluronic acid.
Hp: heparin.
PC: principal component.
SVD: singular value decomposition.
DoS: degree of sulphation.
BSE: bovine spongiform encephalopathy.

