## Supplementary Data for "Analysis of solid-state heparin samples by ATR-FTIR spectroscopy"

**Graph Title Key -**
“Raw”: Raw data post 21neighbour, $2^{nd}$ polynomial Savitsky Golay smooth.

“Normalised (0-1)”: Raw spectra, following a custom $7^{th}$ order polynomial baseline correction and subsequent y-axis scaling between 0 and 1.

“D2”: second derivative of Normalised spectra, following a 41neighbour, $2^{nd}$ order polynomial Savitsky Golay smooth.

All PCA analyses contain a matrix plot, scree plot, loading plots for PCs 1:4, and score plots for PCs 1:5 as well as a page for any relevant preparation information.

**Contents:**

Page 2-6: Tables of used samples

Page 7-23: Raw spectra of PMH, OMH, BMH, BLH and HS.

Page 24-40: Normalised spectra of PMH, OMH, BMH, BLH and HS.

Page 41-57: D2 spectra of PMH, OMH, BMH, BLH and HS.

Page 58-62: Raw spectra of CS, DS, OSCS and HA.

Page 63-67: Normalised spectra of CS, DS, OSCS and HA.

Page 68-72: D2 spectra of CS, DS, OSCS and HA.

Page 73-75: Raw spectra of OSCS, OSAS and Des Selected for contamination, and subsequent contaminated samples.

Page 76-78: Normalised spectra of OSCS, OSAS and Des Selected for contamination, and subsequent contaminated samples.

Page 79-81: D2 spectra of OSCS, OSAS and Des Selected for contamination, and subsequent contaminated samples.

Page 82-84: Raw spectra of PMH contaminated with BMH, series a:d

Page 85-87: Normalised spectra of PMH contaminated with BMH, series a:d

Page 88-90: D2 spectra of PMH contaminated with BMH, series a:d

Page 91-109: PCA of Heps vs GAGs (“Everything”)

Page 110-349: PCA of OSCS contamination series, All vs PMH and each contaminated sample.

Page 350-589: PCA of OSAS contamination series, All vs PMH and each contaminated sample.

Page 590-879: PCA of DeS contamination series, All vs PMH and each contaminated sample.

Page 880-935: PCA of PMH vs BMH vs OMG vs BLH, PMH vs OMH and PMH vs BLH

Page 936-1584: PCA of PMH contaminated with BMH, All vs PMH and each contaminated sample, series a:d

*Table 1: Table of PMH's used and their source(s)*

| Sample Name | Supplier | Batch No. |
|---|---|---|
| Hep 3 | Acros | A0286020 |
| Hep 4 | Applichem | 1X007824 |
| Hep A | Applichem | 30009511 |
| Hep C | Applichem | 40011435 |
| Hep D | Calbiochem | 375093 |
| Hep E | Celsius | PH-89915 |
| Hep F | Bioberica | G3060 |
| Hep H | Leo Pharmaceuticals | DD6985 |
| Hep J | Sigma | H3393 |
| Hep K | Sigma | SLBQ1361V |
| Hep L | Sigma | SLBP2731V |
| Hep M | Sigma | SLBN9145V |
| Hep N | Sigma | SLBP6873V |
| Hep O | Alfa Aesar | 10198820 |
| Hep P | Acros | A0382416 |
| Hep Q | Sigma | SLBQ1351V |
| Hep R | Sigma | SLBP2731V |
| Hep S | Applichem | 6C015265 |
| Carsyn Hep | Carbosynth | YH093541501 |
| Hstd | EU Pharma Ref Std | Y001282 |
| RStd | Ronzoni Institute | G9850 |
| P1 | Ronzoni Institute | G12010 |
| P2 | Ronzoni Institute | G12007 |
| P3 | Ronzoni Institute | G12006 |
| P4 | Ronzoni Institute | A5363 |
| P5 | Ronzoni Institute | |
| P6 | Ronzoni Institute | A5675 |
| P7 | Ronzoni Institute | G7768 |
| P8 | Ronzoni Institute | G12009 |
| P9 | Ronzoni Institute | G12012 |
| P10 | Ronzoni Institute | G12013 |
| P11 | Ronzoni Institute | A353PSA |
| P12 | Ronzoni Institute | A3201QA |
| P13 | Ronzoni Institute | A3288QA |
| P14 | Ronzoni Institute | A5229 |
| P15 | Ronzoni Institute | A5595 |
| P16 | Ronzoni Institute | A5605 |
| P17 | Ronzoni Institute | G7765 |
| P19 | Ronzoni Institute | G10921 |
| P20 | Ronzoni Institute | G10922 |
| P21 | Ronzoni Institute | G11653 |
| P23 | Ronzoni Institute | G11657 |

| Sample Name | Supplier | Batch No. |
|---|---|---|
| P24 | Ronzoni Institute | G12011 |
| P25 | Ronzoni Institute | A3100A |
| P26 | Ronzoni Institute | A5668 |
| P27 | Ronzoni Institute | G13456 |
| P28 | Ronzoni Institute | G11429 |
| P29 | Ronzoni Institute | G11444 |
| P30 | Ronzoni Institute | G1143 |
| P31 | Ronzoni Institute | A3787 |
| P32 | Ronzoni Institute | A3934 |
| P33 | Ronzoni Institute | A4077 |
| P34 | Ronzoni Institute | A4427 |
| P35 | Ronzoni Institute | A4464 |
| P36 | Ronzoni Institute | A5302 |
| P37 | Ronzoni Institute | A5306 |
| P38 | Ronzoni Institute | A5319 |
| P39 | Ronzoni Institute | A5444 |
| P40 | Ronzoni Institute | A5468 |
| P41 | Ronzoni Institute | A5569 |
| P42 | Ronzoni Institute | A5613 |
| P43 | Ronzoni Institute | A5615 |
| P44 | Ronzoni Institute | A01106023 |
| P45 | Ronzoni Institute | G11627 |
| P47 | Ronzoni Institute | G11629 |
| P48 | Ronzoni Institute | G11661 |
| P49 | Ronzoni Institute | G13100 |
| P50 | Ronzoni Institute | G13425 |
| P51 | Ronzoni Institute | G13426 |
| P52 | Ronzoni Institute | G13427 |
| P53 | Ronzoni Institute | G6202 |
| P54 | Ronzoni Institute | G6203 |

*Table 2: Table of BMH's used and their source(s)*

| **Sample Name** | **Supplier** | **Batch No.** |
|---|---|---|
| B1 | Ronzoni Institute | G12019 |
| B2 | Ronzoni Institute | G12018 |
| B3 | Ronzoni Institute | G12015 |
| B4 | Ronzoni Institute | G12028 |
| B5 | Ronzoni Institute | G12022 |
| B6 | Ronzoni Institute | G7850 |
| B7 | Ronzoni Institute | G7847 |
| B9 | Ronzoni Institute | G7773 |
| B10 | Ronzoni Institute | G7848 |
| B11 | Ronzoni Institute | G7844 |
| B12 | Ronzoni Institute | G7841 |
| B13 | Ronzoni Institute | G7846 |
| B14 | Ronzoni Institute | G10803 |
| B15 | Ronzoni Institute | G7845 |
| B16 | Ronzoni Institute | G12014 |
| B18 | Ronzoni Institute | G12017 |
| B19 | Ronzoni Institute | G12020 |
| B20 | Ronzoni Institute | G12021 |
| B21 | Ronzoni Institute | G12662 |
| B22 | Ronzoni Institute | G12667 |
| B23 | Ronzoni Institute | G12668 |
| B24 | Ronzoni Institute | G12669 |
| B26 | Ronzoni Institute | G12804 |
| B27 | Ronzoni Institute | G13420 |
| B28 | Ronzoni Institute | G13421 |
| B29 | Ronzoni Institute | G13422 |
| B30 | Ronzoni Institute | G13423 |
| B31 | Ronzoni Institute | G12657 |
| B32 | Ronzoni Institute | G12658 |
| B33 | Ronzoni Institute | G12659 |
| B34 | Ronzoni Institute | G12661 |
| B35 | Ronzoni Institute | G12664 |
| B36 | Ronzoni Institute | G12665 |
| B37 | Ronzoni Institute | G12666 |
| B38 | Ronzoni Institute | G13420 |
| B39 | Ronzoni Institute | G13421 |
| B40 | Ronzoni Institute | G13422 |
| B41 | Ronzoni Institute | G13423 |
| B42 | Ronzoni Institute | G12660 |
| B43 | Ronzoni Institute | G12663 |
| B44 | Ronzoni Institute | G1467 |
| B45 | Ronzoni Institute | G1468 |
| B46 | Ronzoni Institute | G1469 |

| **Sample Name** | **Supplier** | **Batch No.** |
|---|---|---|
| B47 | Ronzoni Institute | G1470 |
| B48 | Ronzoni Institute | G1471 |
| B49 | Ronzoni Institute | G1472 |
| B50 | Ronzoni Institute | G1473 |
| B51 | Ronzoni Institute | G1474 |
| B52 | Ronzoni Institute | G1475 |
| B53 | Ronzoni Institute | G1476 |
| B60 | Ronzoni Institute | G12802 |
| B61 | Ronzoni Institute | G12803 |
| B62 | Ronzoni Institute | G13421 |
| B63 | Ronzoni Institute | G13422 |
| B64 | Ronzoni Institute | G13504 |
| B65 | Ronzoni Institute | G13505 |
| B66 | Ronzoni Institute | G13506 |

*Table 3: Table of OMH's used and their source(s)*

| Sample Name | Supplier | Batch No. |
|---|---|---|
| O1 | Ronzoni Institute | G12028 |
| O2 | Ronzoni Institute | G12026 |
| O3 | Ronzoni Institute | G12025 |
| O4 | Ronzoni Institute | G12024 |
| O5 | Ronzoni Institute | G12029 |
| O6 | Ronzoni Institute | G8117 |
| O7 | Ronzoni Institute | G11448 |
| O8 | Ronzoni Institute | G12077 |
| O10 | Ronzoni Institute | G1371 |
| O11 | Ronzoni Institute | G1372 |
| O12 | Ronzoni Institute | G1373 |
| O13 | Ronzoni Institute | G12205 |
| O14 | Ronzoni Institute | G12206 |
| O15 | Ronzoni Institute | G12207 |
| O16 | Ronzoni Institute | G12288 |
| O17 | Ronzoni Institute | G12301 |
| O18 | Ronzoni Institute | G12302 |
| O19 | Ronzoni Institute | G13096 |
| O20 | Ronzoni Institute | G13097 |
| O21 | Ronzoni Institute | G13098 |
| O24 | Ronzoni Institute | G13689 |
| O25 | Ronzoni Institute | G13690 |
| O26 | Ronzoni Institute | G13691 |
| O27 | Gifted by Ed Yates | NA |
| O28 | Ronzoni Institute | G11447 |
| O29 | Ronzoni Institute | G11449 |
| O30 | Ronzoni Institute | G11480 |
| O31 | Ronzoni Institute | G11482 |
| O32 | Ronzoni Institute | G11489 |
| O33 | Ronzoni Institute | G11936 |
| O34 | Ronzoni Institute | G11937 |
| O36 | Ronzoni Institute | G10786 |
| O38 | Ronzoni Institute | GUK |

*Table 4: Table of BLH's used and their source(s)*

| Sample Name | Supplier | Batch No. |
|---|---|---|
| BLH1 | Ronzoni Institute | G4141 |
| BLH2 | Ronzoni Institute | G46 |
| BLH3 | Ronzoni Institute | G65 |
| BLH4 | Ronzoni Institute | G121 |
| BLH5 | Ronzoni Institute | G11099 |
| BLH6 | Ronzoni Institute | G11100 |
| BLH7 | Ronzoni Institute | G11101 |
| BLH8 | Ronzoni Institute | G11388 |
| BLH9 | Ronzoni Institute | G11389 |
| BLH10 | Ronzoni Institute | G11463 |
| BLH11 | Ronzoni Institute | G12271 |
| BLH12 | Ronzoni Institute | G12272 |
| BLH13 | Ronzoni Institute | G12275 |
| BLH14 | Ronzoni Institute | G17273 |
| BLH15 | Ronzoni Institute | G17274 |
| BLH18 | Ronzoni Institute | G11757 |
| BLH19 | Ronzoni Institute | G11758 |
| BLH20 | Ronzoni Institute | G11759 |
| BLH21 | Ronzoni Institute | G10787 |

*Table 5: Table of HAs used and their source(s)*

| Sample Name | Supplier | Batch No. |
|---|---|---|
| HAstd | Ronzoni Institute | G9980 |
| HA1 | Ronzoni Institute | G13998 |
| HA2 | Ronzoni Institute | G1039 |
| HA3 | Ronzoni Institute | G1093 |
| HA4 | Ronzoni Institute | G1824 |
| HA5 | Ronzoni Institute | G5733 |
| HA6 | Ronzoni Institute | G28 |
| HA11 | Ronzoni Institute | G5673 |
| HA12 | Ronzoni Institute | G5674 |
| HA14 | Ronzoni Institute | G45 |

*Table 6: Table of HSs used and their source(s)*

| Sample Name | Supplier | Batch No. |
|---|---|---|
| PHS | Ronzoni Institute | G116 |
| OHS | Ronzoni Institute | G13014 |
| OHS | Ronzoni Institute | G13015 |
| HS1 | Ronzoni Institute | G108 |
| HS2 | Ronzoni Institute | G120 |
| HS3 | Ronzoni Institute | G1568 |
| HS4 | Ronzoni Institute | G41 |
| HS5 | Ronzoni Institute | G42 |
| HS8 | Ronzoni Institute | G101 |
| HS9 | Ronzoni Institute | G107 |
| HS12 | Ronzoni Institute | G1807 |
| HS14 | Ronzoni Institute | HHS-S |
| HS16 | Ronzoni Institute | G34 |
| HS17 | Ronzoni Institute | G67 |
| HS18 | Ronzoni Institute | G104 |
| HS21 | Ronzoni Institute | G5021 |
| HS22 | Ronzoni Institute | LOTUKa |
| HS23 | Ronzoni Institute | G1567 |
| HS24 | Ronzoni Institute | G7861 |
| HS25 | Ronzoni Institute | O258 |
| HS26 | Ronzoni Institute | OBGR |
| HS27 | Ronzoni Institute | G5547 |
| HS28 | Ronzoni Institute | HS141 |
| HS29 | Ronzoni Institute | HS9335 |
| HS30 | Ronzoni Institute | HS137 |
| HS31 | Ronzoni Institute | HS138 |
| HS32 | Ronzoni Institute | HS139 - 140 |
| HS33 | Ronzoni Institute | HS149 |
| HS34 | Ronzoni Institute | HS934 |
| HS35 | Ronzoni Institute | HS20594 |
| HS36 | Ronzoni Institute | HS271192 |

*Table 7: Table of Chondroitins used and their source(s)*

| Sample Name | Supplier | Batch No. |
|---|---|---|
| CSA1 | Carbosynth | YC152881701 |
| CSA2 | Ronzoni Institute | G920 |
| CSA3 | Ronzoni Institute | G1125B |
| CSA5 | Ronzoni Institute | G2426 |
| CSA7 | Ronzoni Institute | G167 |
| CSA8 | Ronzoni Institute | G3432 |
| CSA9 | Ronzoni Institute | G4090 |
| DSstd | Celsus | 10596 |
| DS3 | Ronzoni Institute | G1021 |
| DS6 | Ronzoni Institute | G33 |
| DS8 | Ronzoni Institute | G593 |
| DS9 | Ronzoni Institute | G5026 |
| DS10 | Ronzoni Institute | G6987 |
| DS11 | Ronzoni Institute | G904D51 |
| DS12 | Ronzoni Institute | DSB2 |
| DS13 | Ronzoni Institute | G904 |
| DS14 | Ronzoni Institute | G904/1 |
| DS16 | Ronzoni Institute | G904D44 |
| DS17 | Ronzoni Institute | G904D47 |
| DS18 | Ronzoni Institute | G904D54 |
| DS20 | Ronzoni Institute | G1084 |
| DS21 | Ronzoni Institute | G6322 |
| DS22 | Ronzoni Institute | GC180 |
| DS23 | Ronzoni Institute | GC185 |
| DS24 | Ronzoni Institute | GC181 |
| DS25 | Ronzoni Institute | GC182 |
| DS26 | Ronzoni Institute | A1908 |
| DS27 | Ronzoni Institute | DS52 |
| CSC2 | Ronzoni Institute | G1223 |
| CSC3 | Ronzoni Institute | G56 |
| CSC4 | Ronzoni Institute | G113 |
| CSC6 | Ronzoni Institute | MPS2 |
| CSC7 | Ronzoni Institute | MPS3 |
| CS1 | Ronzoni Institute | DFSTDO |
| CS2 | Ronzoni Institute | MPSTDN |
| CS4 | Ronzoni Institute | G267 |
| CS5 | Ronzoni Institute | G1381 |
| CS6 | Ronzoni Institute | G1673A |
| CS7 | Ronzoni Institute | G7222 |
| CS8 | Ronzoni Institute | G11251I |
| CS9 | Ronzoni Institute | P6098 |
| CS11 | Ronzoni Institute | G9175 |
| CS12 | Ronzoni Institute | G9176 |

| ***Sample Name*** | ***Supplier*** | ***Batch No.*** |
|---|---|---|
| CS13 | Ronzoni Institute | G9178 |
| CS15 | Ronzoni Institute | G8501 |
| CS16 | Ronzoni Institute | G8585 |
| CS17 | Ronzoni Institute | G8783 |
| CS18 | Ronzoni Institute | G9131 |
| CS19 | Ronzoni Institute | G9132 |
| CS20 | Ronzoni Institute | G9133 |
| CS21 | Ronzoni Institute | G91039 |
| CS22 | Ronzoni Institute | G6866 |
| OSCS | FDA | H0M191 |
| OSCS1a | Ronzoni Institute | G62439 (1) |
| OSCS1b | Ronzoni Institute | G62439 (2) |
| OSCS1c | Ronzoni Institute | G62439 (3) |
| OSCS2 | Ronzoni Institute | G6243b1 |
| OSCS3 | Ronzoni Institute | G6296 |
| OSCS4 | Ronzoni Institute | G6469 |
| OSAS | *Made in house* | NA |
| DeS | Fluka | 1053244-42507314 |

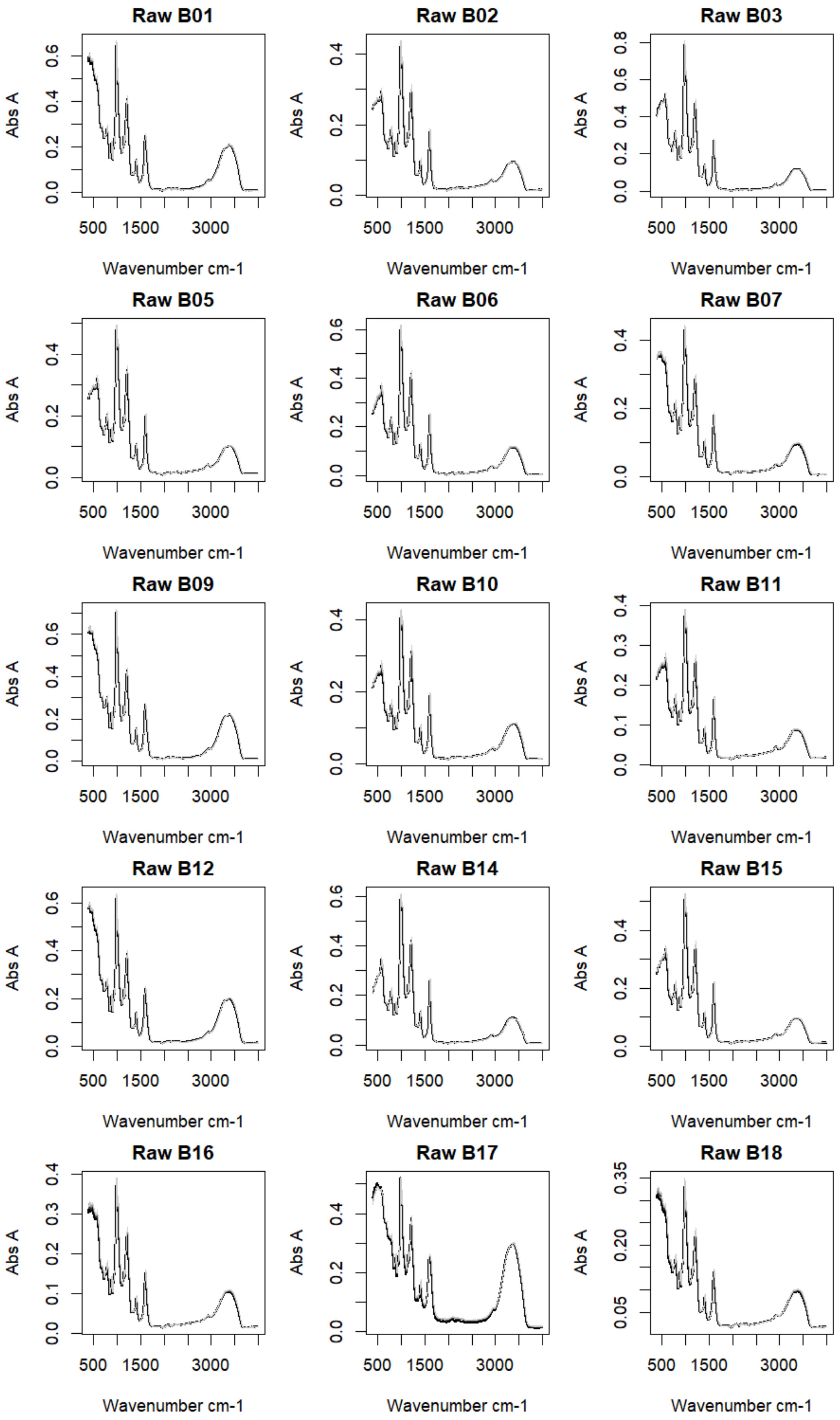
Raw B01
Raw B02
Raw B03
Raw B05
Raw B06
Raw B07
Raw B09
Raw B10
Raw B11
Raw B12
Raw B14
Raw B15
Raw B16
Raw B17
Raw B18
Abs A
Wavenumber cm-1
500
1500
3000

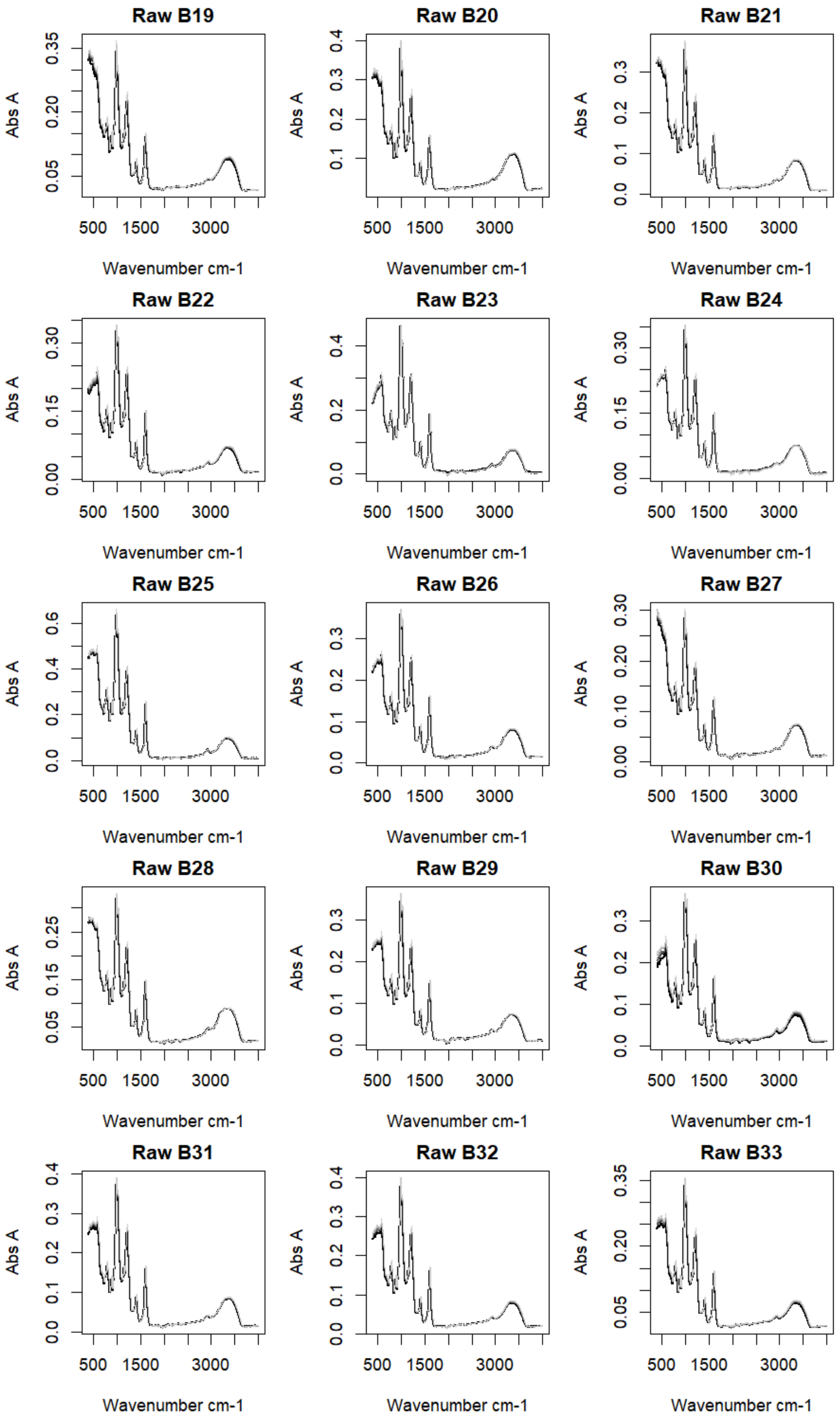
Raw B19
Abs A
0.05 0.20 0.35
500 1500 3000
Wavenumber cm-1
Raw B20
Abs A
0.1 0.2 0.3 0.4
500 1500 3000
Wavenumber cm-1
Raw B21
Abs A
0.0 0.1 0.2 0.3
500 1500 3000
Wavenumber cm-1
Raw B22
Abs A
0.00 0.15 0.30
500 1500 3000
Wavenumber cm-1
Raw B23
Abs A
0.0 0.2 0.4
500 1500 3000
Wavenumber cm-1
Raw B24
Abs A
0.00 0.15 0.30
500 1500 3000
Wavenumber cm-1
Raw B25
Abs A
0.0 0.2 0.4 0.6
500 1500 3000
Wavenumber cm-1
Raw B26
Abs A
0.0 0.1 0.2 0.3
500 1500 3000
Wavenumber cm-1
Raw B27
Abs A
0.00 0.10 0.20 0.30
500 1500 3000
Wavenumber cm-1
Raw B28
Abs A
0.05 0.15 0.25
500 1500 3000
Wavenumber cm-1
Raw B29
Abs A
0.0 0.1 0.2 0.3
500 1500 3000
Wavenumber cm-1
Raw B30
Abs A
0.0 0.1 0.2 0.3
500 1500 3000
Wavenumber cm-1
Raw B31
Abs A
0.0 0.1 0.2 0.3 0.4
500 1500 3000
Wavenumber cm-1
Raw B32
Abs A
0.0 0.1 0.2 0.3 0.4
500 1500 3000
Wavenumber cm-1
Raw B33
Abs A
0.05 0.20 0.35
500 1500 3000
Wavenumber cm-1

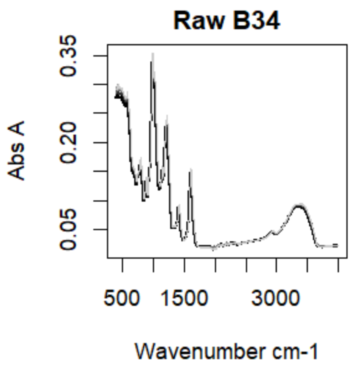
Raw B34
Abs A
0.05
0.20
0.35
500
1500
3000
Wavenumber cm-1

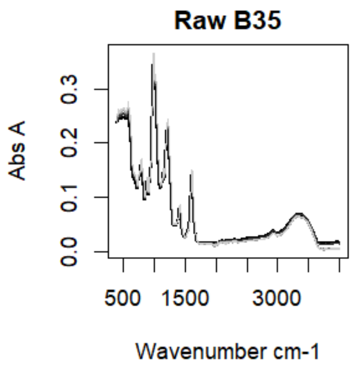
Raw B35
Abs A
0.0
0.1
0.2
0.3
500
1500
3000
Wavenumber cm-1

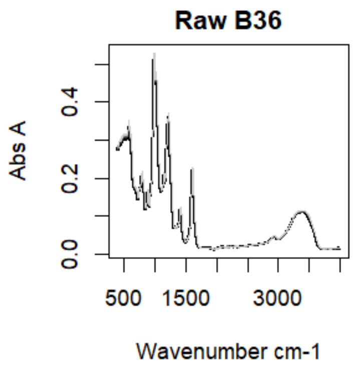
Raw B36
Abs A
0.0
0.2
0.4
500
1500
3000
Wavenumber cm-1

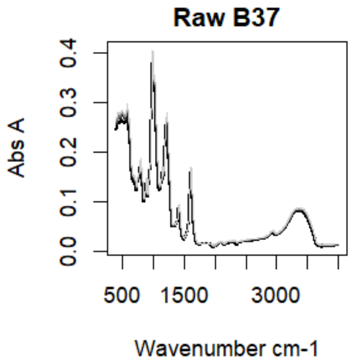
Raw B37
Abs A
0.0
0.1
0.2
0.3
0.4
500
1500
3000
Wavenumber cm-1

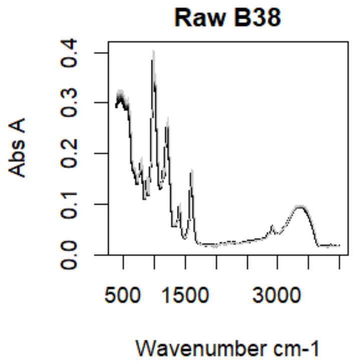
Raw B38
Abs A
0.0
0.1
0.2
0.3
0.4
500
1500
3000
Wavenumber cm-1

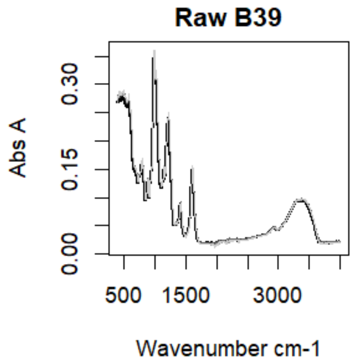
Raw B39
Abs A
0.00
0.15
0.30
500
1500
3000
Wavenumber cm-1

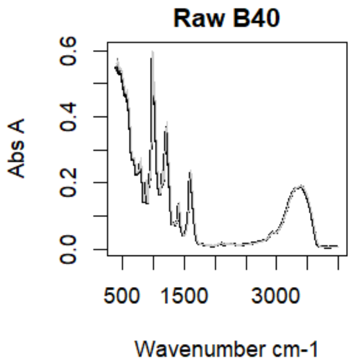
Raw B40
Abs A
0.0
0.2
0.4
0.6
500
1500
3000
Wavenumber cm-1

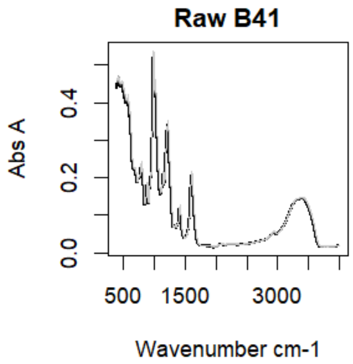
Raw B41
Abs A
0.0
0.2
0.4
500
1500
3000
Wavenumber cm-1

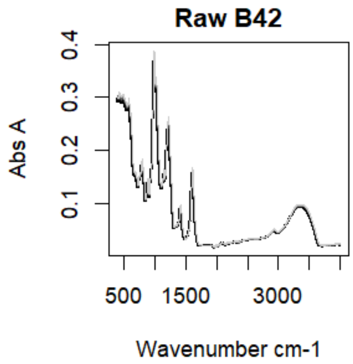
Raw B42
Abs A
0.1
0.2
0.3
0.4
500
1500
3000
Wavenumber cm-1

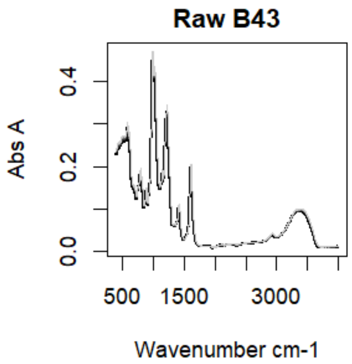
Raw B43
Abs A
0.0
0.2
0.4
500
1500
3000
Wavenumber cm-1

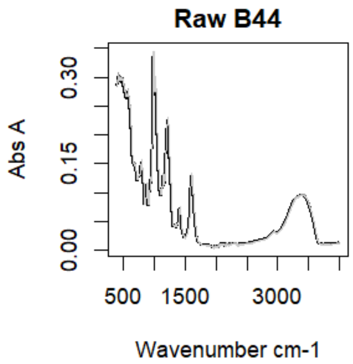
Raw B44
Abs A
0.00
0.15
0.30
500
1500
3000
Wavenumber cm-1

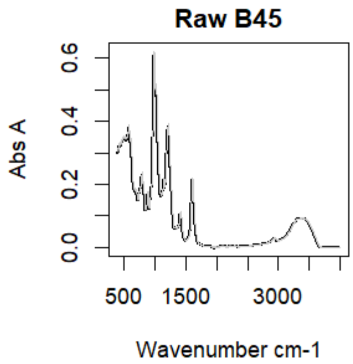
Raw B45
Abs A
0.0
0.2
0.4
0.6
500
1500
3000
Wavenumber cm-1

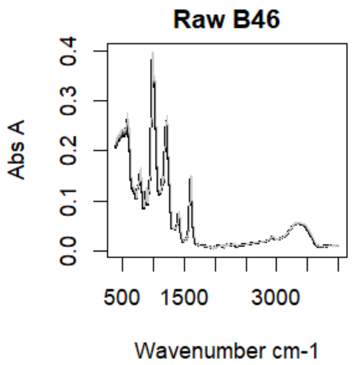
Raw B46
Abs A
0.0
0.1
0.2
0.3
0.4
500
1500
3000
Wavenumber cm-1

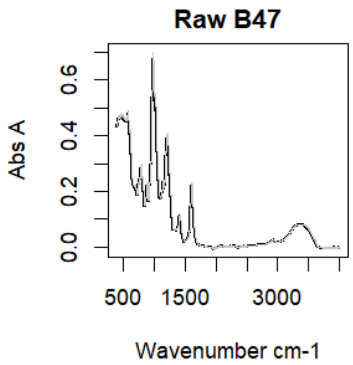
Raw B47
Abs A
0.0
0.2
0.4
0.6
500
1500
3000
Wavenumber cm-1

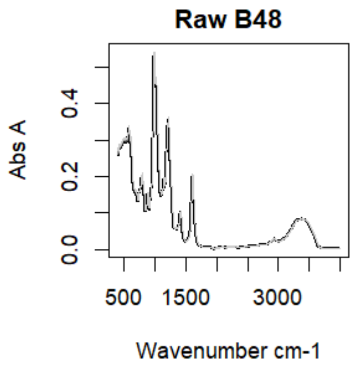
Raw B48
Abs A
0.0
0.2
0.4
500
1500
3000
Wavenumber cm-1

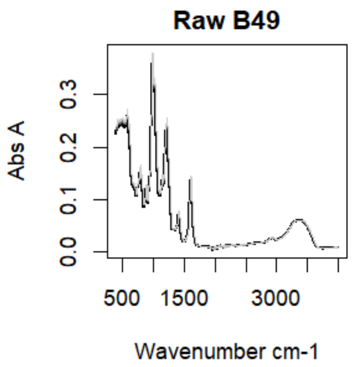
Raw B49
Abs A
0.0 0.1 0.2 0.3
500 1500 3000
Wavenumber cm-1

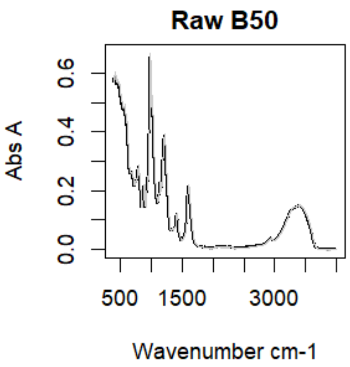
Raw B50
Abs A
0.0 0.2 0.4 0.6
500 1500 3000
Wavenumber cm-1

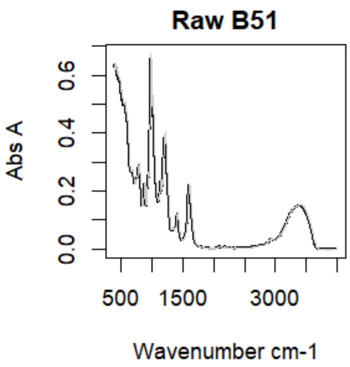
Raw B51
Abs A
0.0 0.2 0.4 0.6
500 1500 3000
Wavenumber cm-1

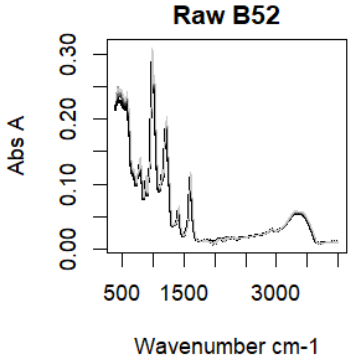
Raw B52
Abs A
0.00 0.10 0.20 0.30
500 1500 3000
Wavenumber cm-1

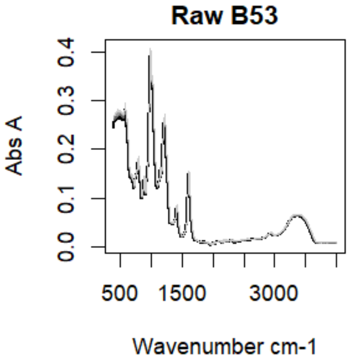
Raw B53
Abs A
0.0 0.1 0.2 0.3 0.4
500 1500 3000
Wavenumber cm-1

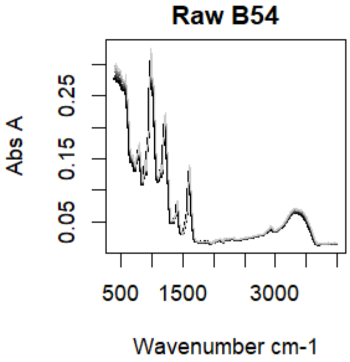
Raw B54
Abs A
0.05 0.15 0.25
500 1500 3000
Wavenumber cm-1

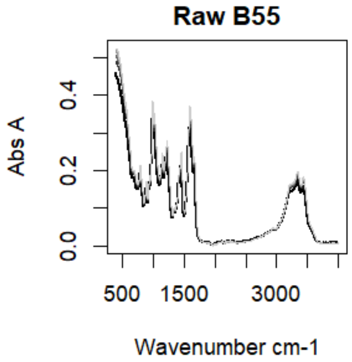
Raw B55
Abs A
0.0 0.2 0.4
500 1500 3000
Wavenumber cm-1

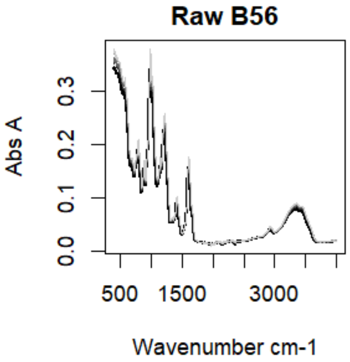
Raw B56
Abs A
0.0 0.1 0.2 0.3
500 1500 3000
Wavenumber cm-1

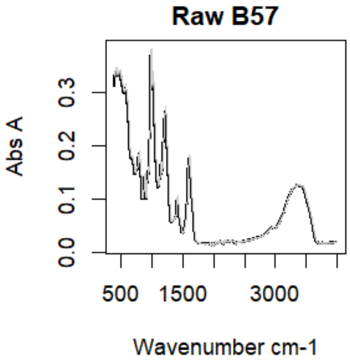
Raw B57
Abs A
0.0 0.1 0.2 0.3
500 1500 3000
Wavenumber cm-1

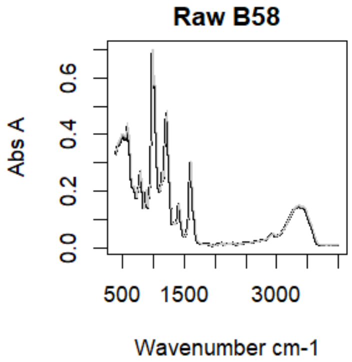
Raw B58
Abs A
0.0 0.2 0.4 0.6
500 1500 3000
Wavenumber cm-1

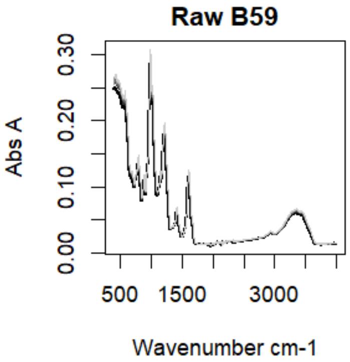
Raw B59
Abs A
0.00 0.10 0.20 0.30
500 1500 3000
Wavenumber cm-1

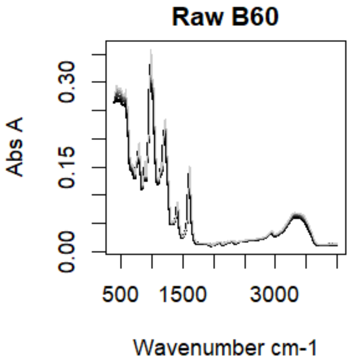
Raw B60
Abs A
0.00 0.15 0.30
500 1500 3000
Wavenumber cm-1

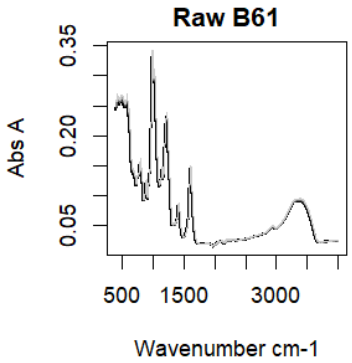
Raw B61
Abs A
0.05 0.20 0.35
500 1500 3000
Wavenumber cm-1

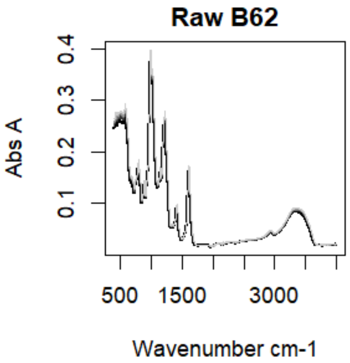
Raw B62
Abs A
0.1 0.2 0.3 0.4
500 1500 3000
Wavenumber cm-1

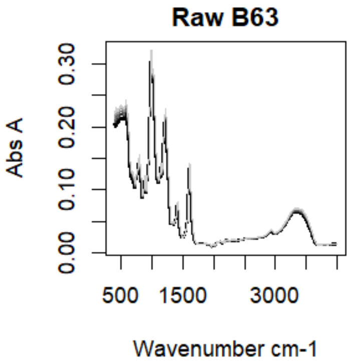
Raw B63
Abs A
0.00 0.10 0.20 0.30
500 1500 3000
Wavenumber cm-1

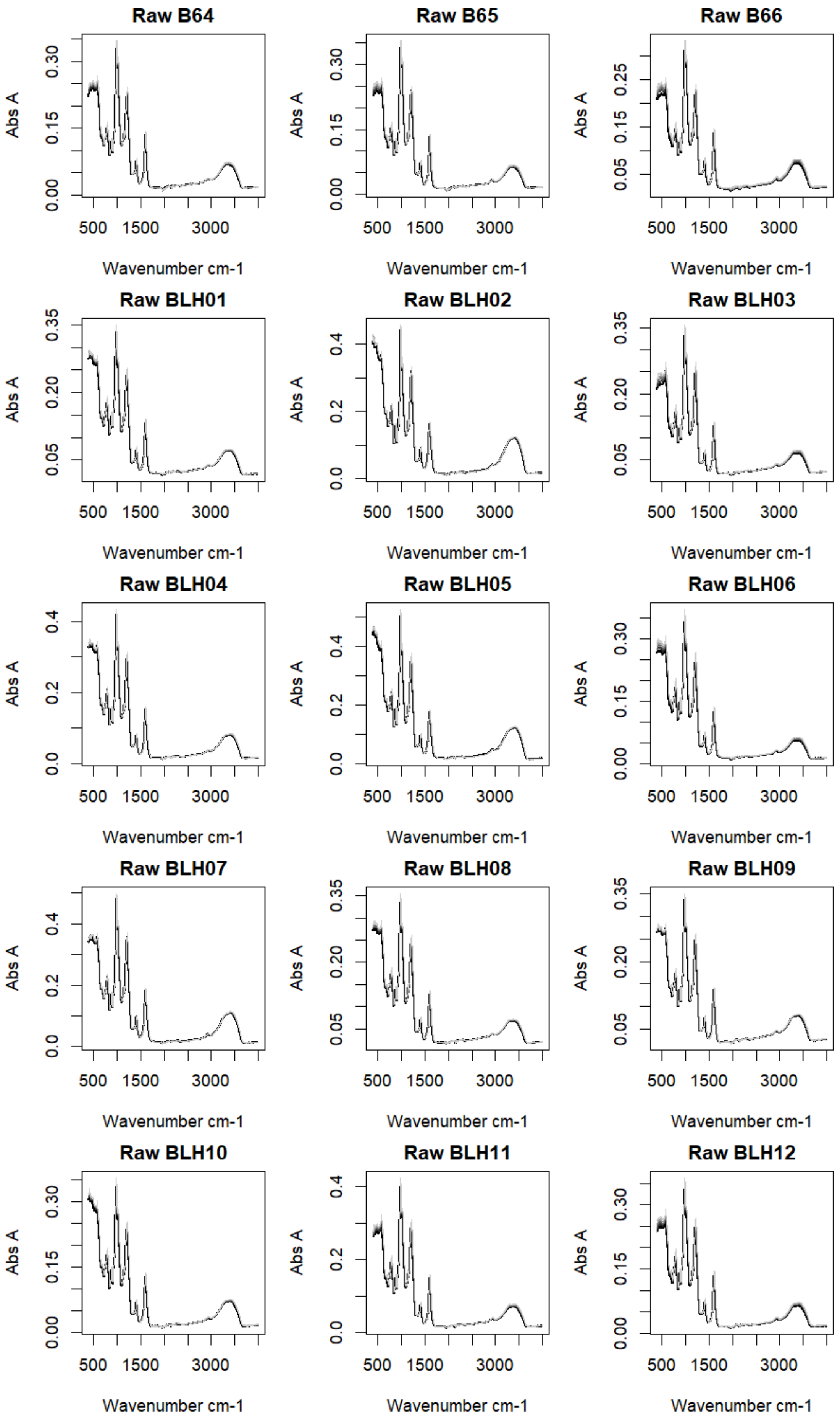
Raw B64
Raw B65
Raw B66
Raw BLH01
Raw BLH02
Raw BLH03
Raw BLH04
Raw BLH05
Raw BLH06
Raw BLH07
Raw BLH08
Raw BLH09
Raw BLH10
Raw BLH11
Raw BLH12
Abs A
Wavenumber cm-1
500 1500 3000

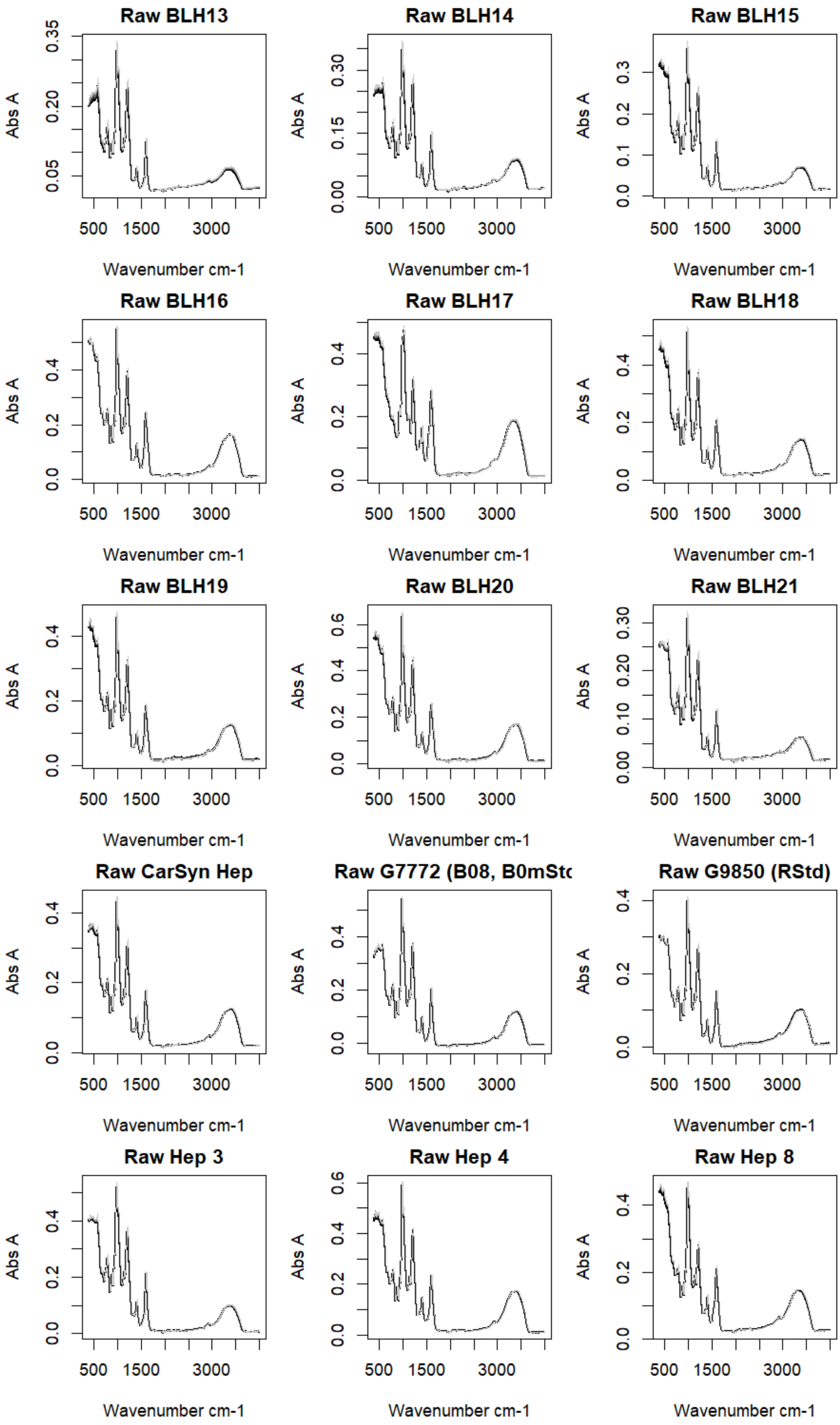
Raw BLH13
Raw BLH14
Raw BLH15
Raw BLH16
Raw BLH17
Raw BLH18
Raw BLH19
Raw BLH20
Raw BLH21
Raw CarSyn Hep
Raw G7772 (B08, B0mStc
Raw G9850 (RStd)
Raw Hep 3
Raw Hep 4
Raw Hep 8
Abs A
Wavenumber cm-1
500 1500 3000

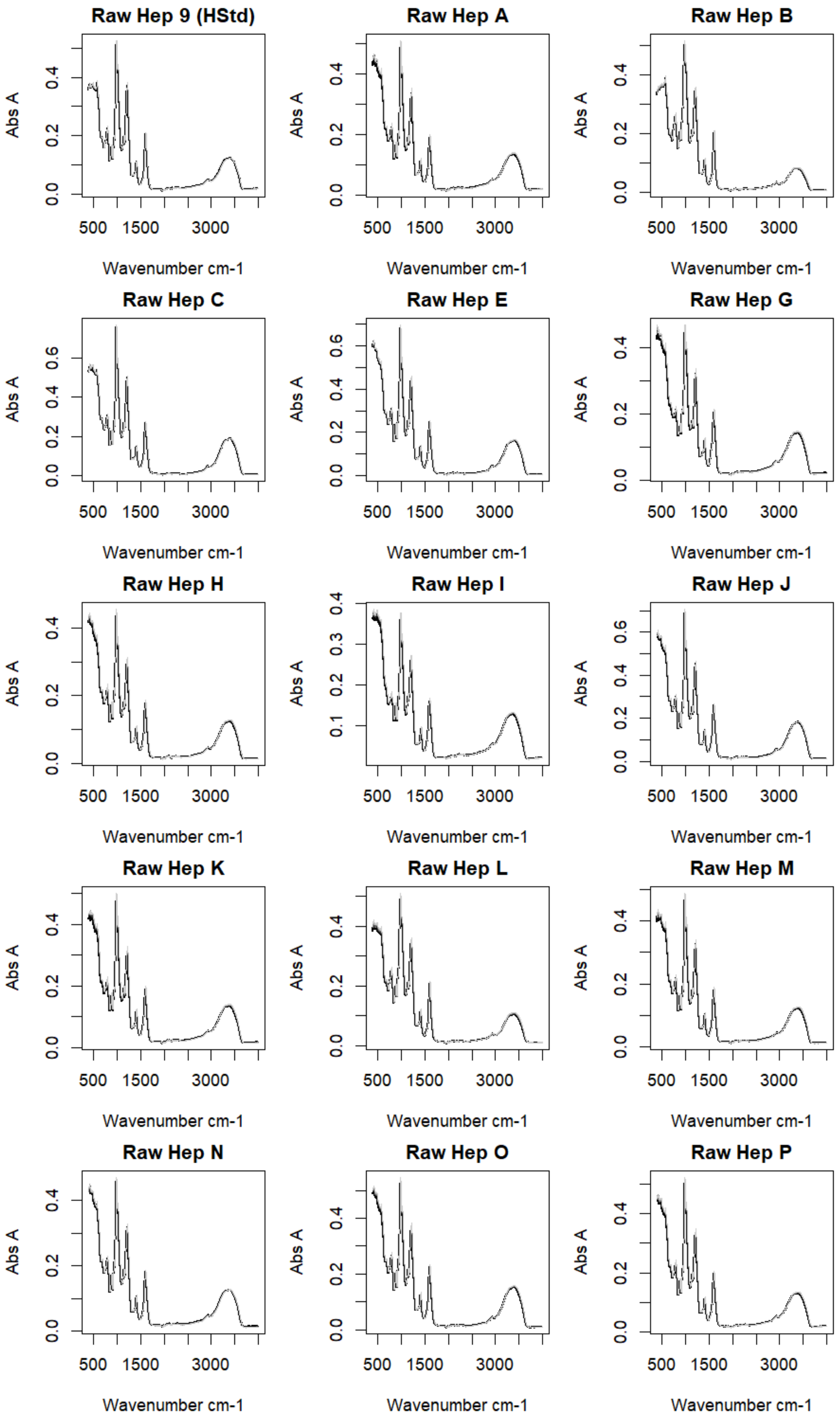
Raw Hep 9 (HStd)
Raw Hep A
Raw Hep B
Raw Hep C
Raw Hep E
Raw Hep G
Raw Hep H
Raw Hep I
Raw Hep J
Raw Hep K
Raw Hep L
Raw Hep M
Raw Hep N
Raw Hep O
Raw Hep P
Abs A
Wavenumber cm-1
500
1500
3000

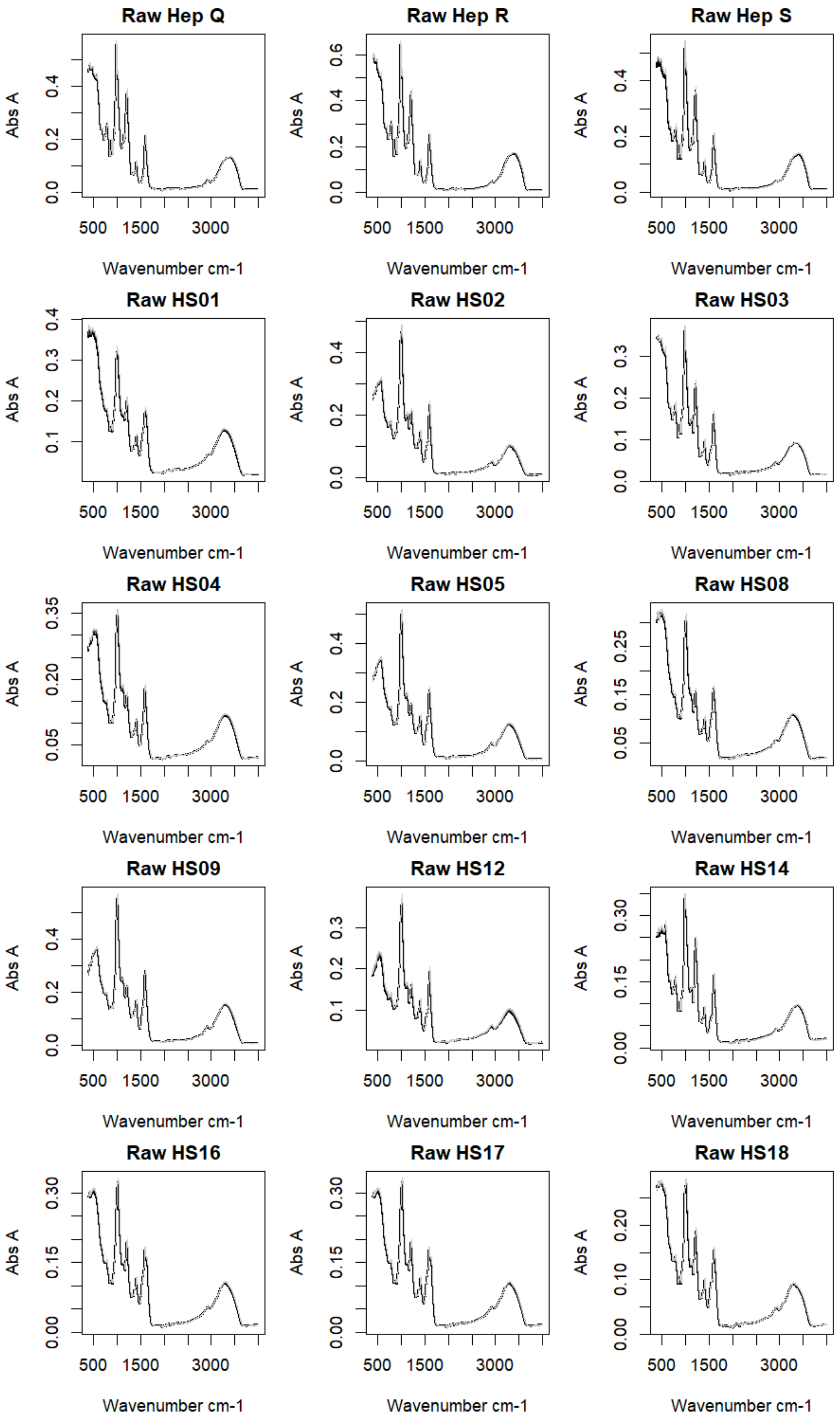
Raw Hep Q
Raw Hep R
Raw Hep S
Raw HS01
Raw HS02
Raw HS03
Raw HS04
Raw HS05
Raw HS08
Raw HS09
Raw HS12
Raw HS14
Raw HS16
Raw HS17
Raw HS18
Abs A
Wavenumber cm-1
500
1500
3000

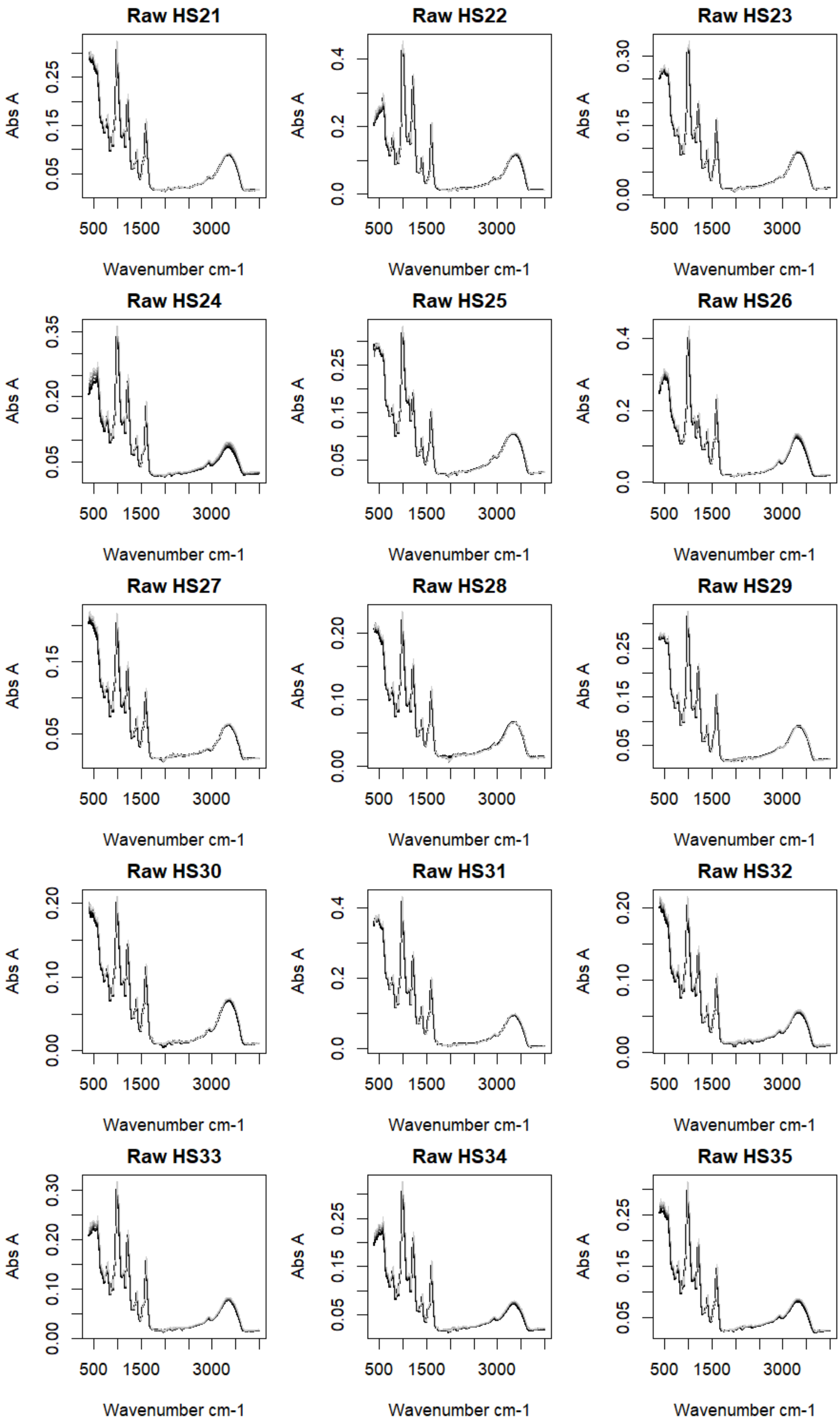
Raw HS21
Raw HS22
Raw HS23
Raw HS24
Raw HS25
Raw HS26
Raw HS27
Raw HS28
Raw HS29
Raw HS30
Raw HS31
Raw HS32
Raw HS33
Raw HS34
Raw HS35
Abs A
Wavenumber cm-1
500 1500 3000

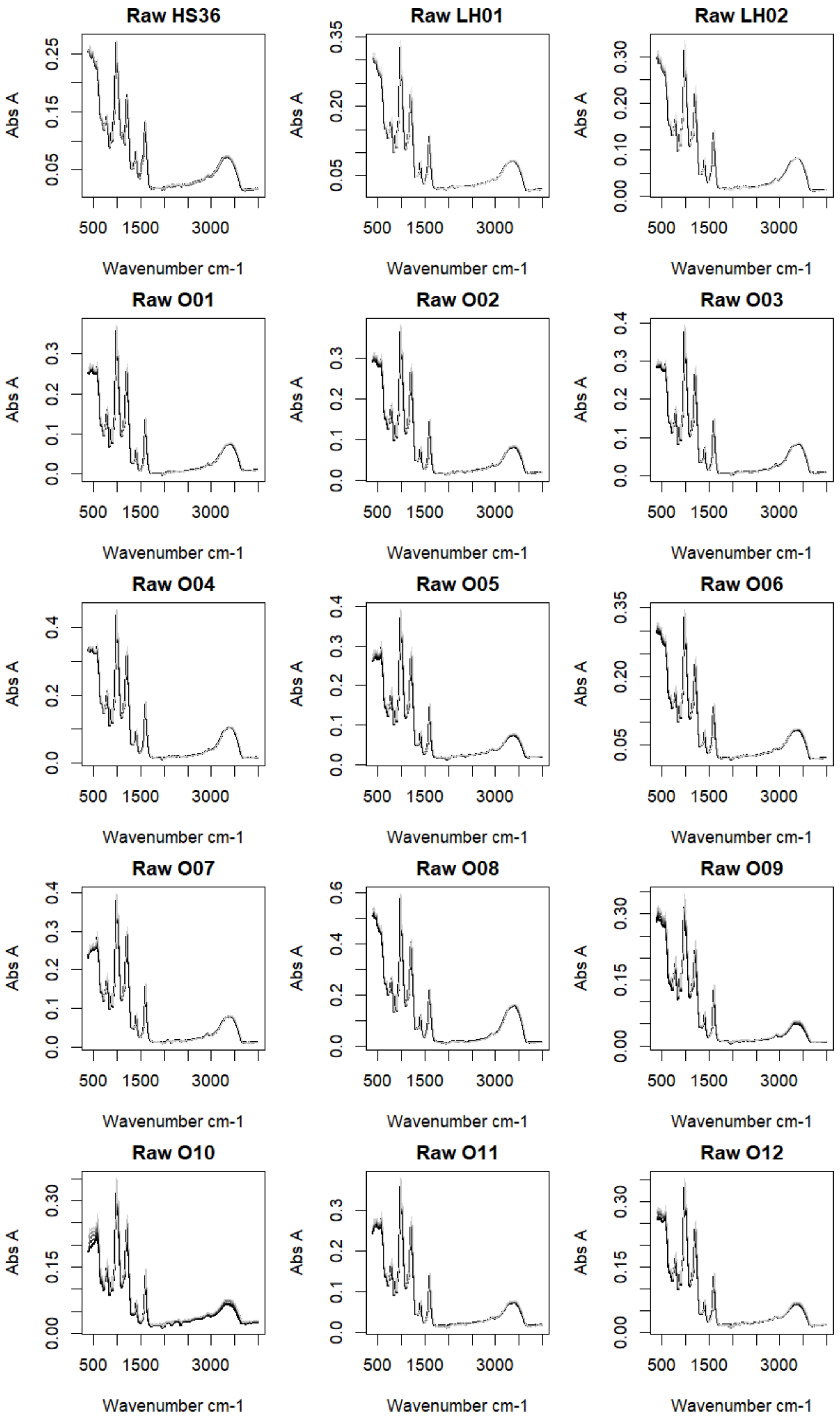
Raw HS36
Raw LH01
Raw LH02
Raw O01
Raw O02
Raw O03
Raw O04
Raw O05
Raw O06
Raw O07
Raw O08
Raw O09
Raw O10
Raw O11
Raw O12
Abs A
Wavenumber cm-1
500
1500
3000

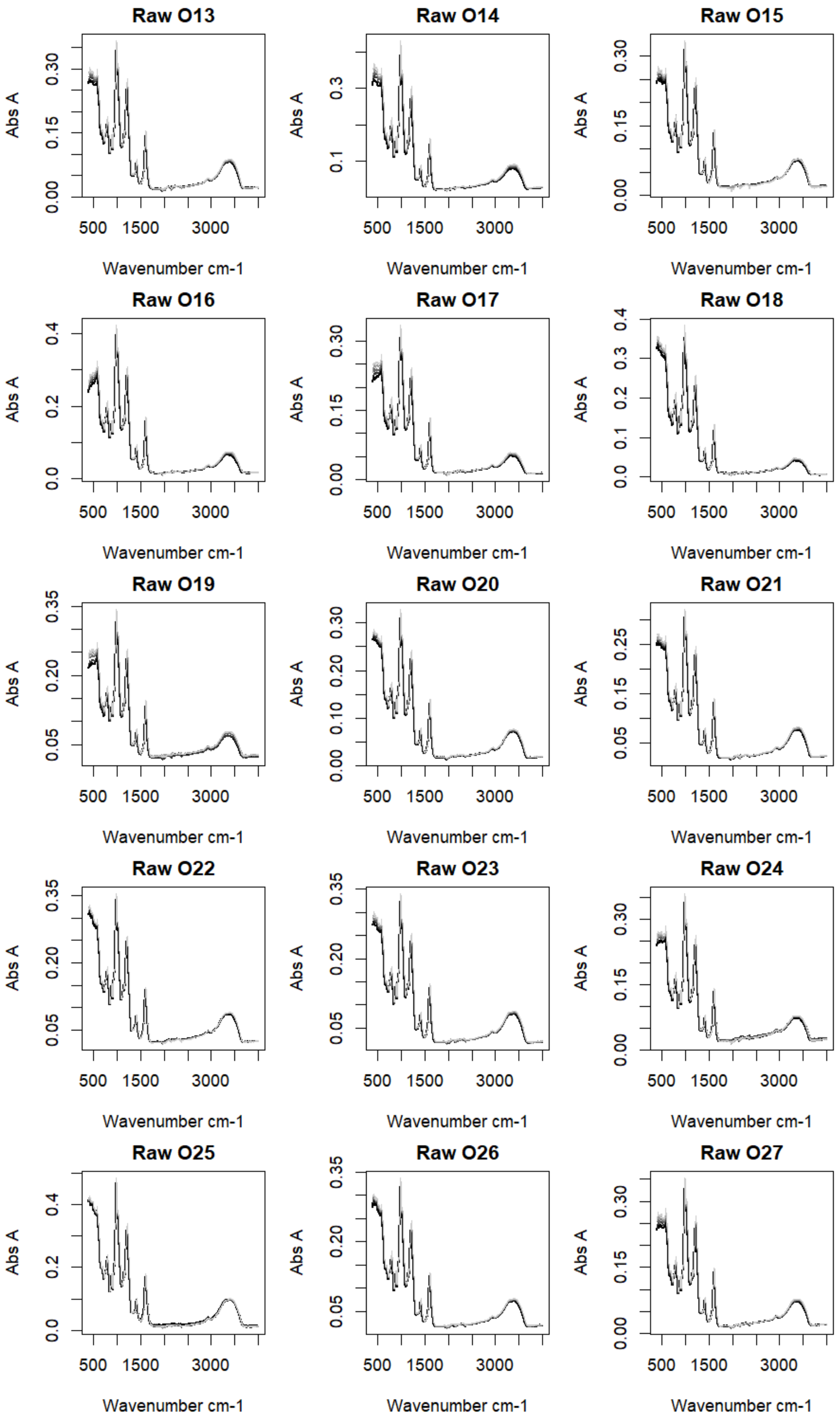
Raw O13
Raw O14
Raw O15
Raw O16
Raw O17
Raw O18
Raw O19
Raw O20
Raw O21
Raw O22
Raw O23
Raw O24
Raw O25
Raw O26
Raw O27
Abs A
Wavenumber cm-1
500
1500
3000

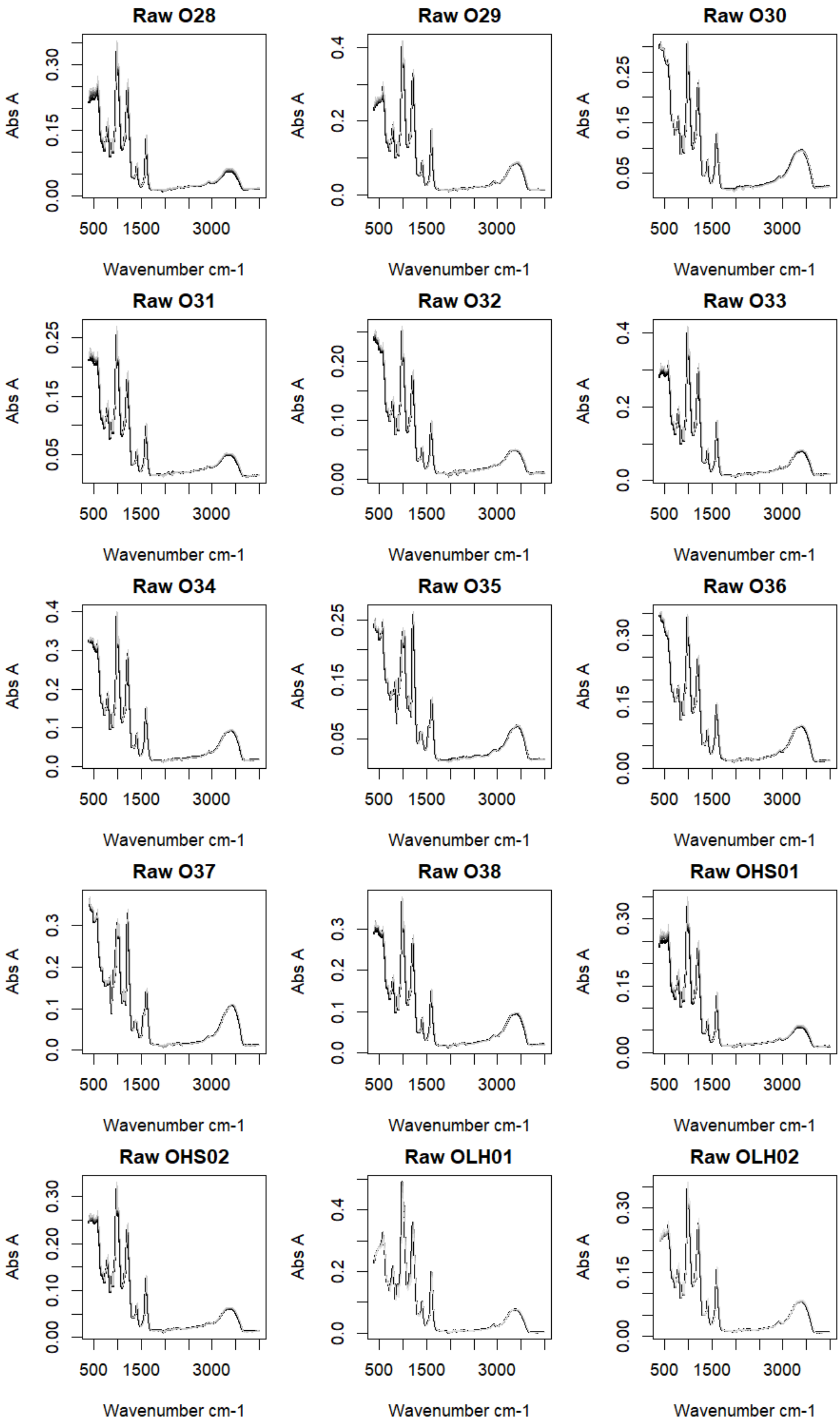
Raw O28
Raw O29
Raw O30
Raw O31
Raw O32
Raw O33
Raw O34
Raw O35
Raw O36
Raw O37
Raw O38
Raw OHS01
Raw OHS02
Raw OLH01
Raw OLH02
Abs A
Wavenumber cm-1
500 1500 3000

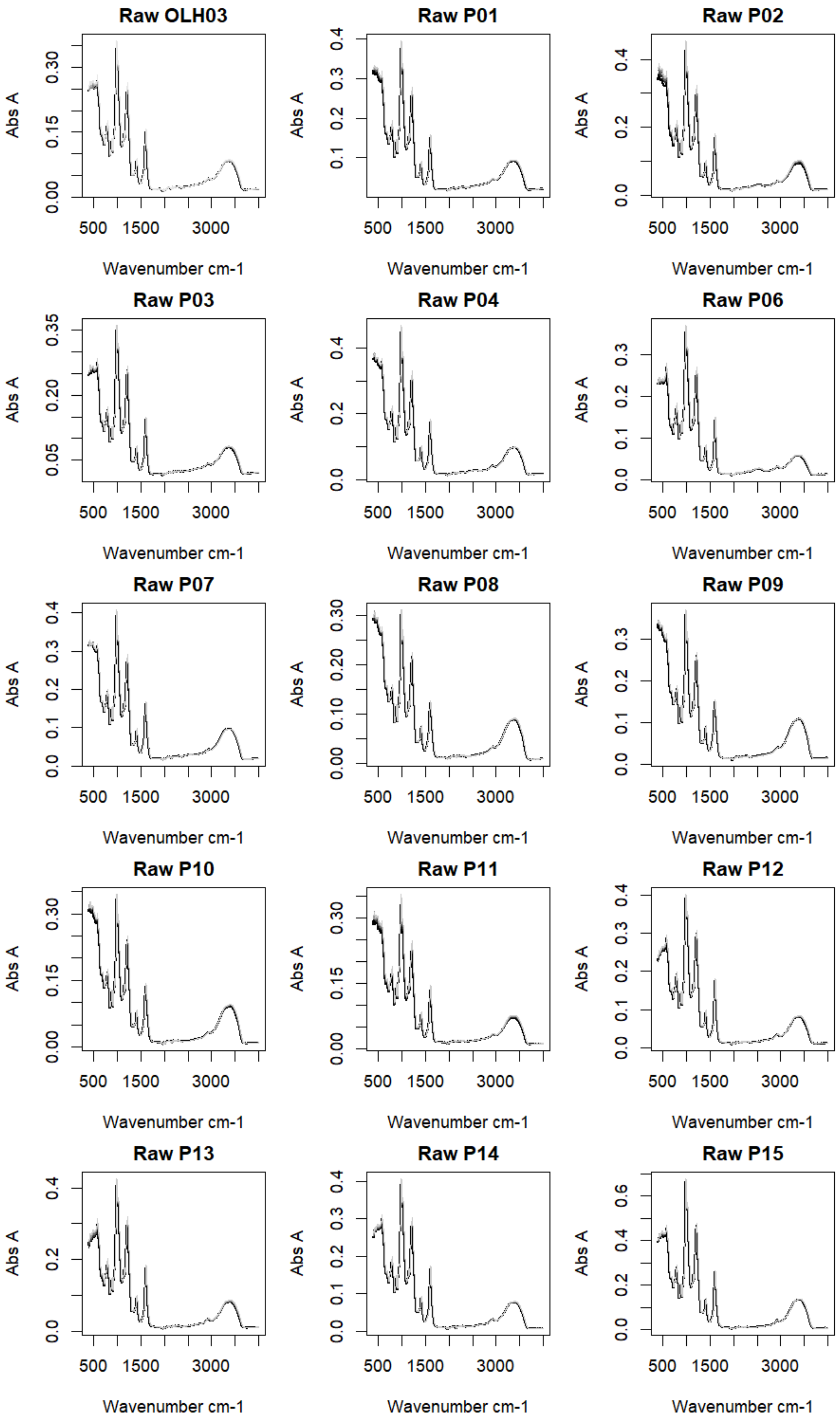
Raw OLH03
Abs A
0.00 0.15 0.30
500 1500 3000
Wavenumber cm-1
Raw P01
Abs A
0.1 0.2 0.3 0.4
500 1500 3000
Wavenumber cm-1
Raw P02
Abs A
0.0 0.2 0.4
500 1500 3000
Wavenumber cm-1
Raw P03
Abs A
0.05 0.20 0.35
500 1500 3000
Wavenumber cm-1
Raw P04
Abs A
0.0 0.2 0.4
500 1500 3000
Wavenumber cm-1
Raw P06
Abs A
0.0 0.1 0.2 0.3
500 1500 3000
Wavenumber cm-1
Raw P07
Abs A
0.0 0.1 0.2 0.3 0.4
500 1500 3000
Wavenumber cm-1
Raw P08
Abs A
0.00 0.10 0.20 0.30
500 1500 3000
Wavenumber cm-1
Raw P09
Abs A
0.0 0.1 0.2 0.3
500 1500 3000
Wavenumber cm-1
Raw P10
Abs A
0.00 0.15 0.30
500 1500 3000
Wavenumber cm-1
Raw P11
Abs A
0.00 0.15 0.30
500 1500 3000
Wavenumber cm-1
Raw P12
Abs A
0.0 0.1 0.2 0.3 0.4
500 1500 3000
Wavenumber cm-1
Raw P13
Abs A
0.0 0.2 0.4
500 1500 3000
Wavenumber cm-1
Raw P14
Abs A
0.0 0.1 0.2 0.3 0.4
500 1500 3000
Wavenumber cm-1
Raw P15
Abs A
0.0 0.2 0.4 0.6
500 1500 3000
Wavenumber cm-1

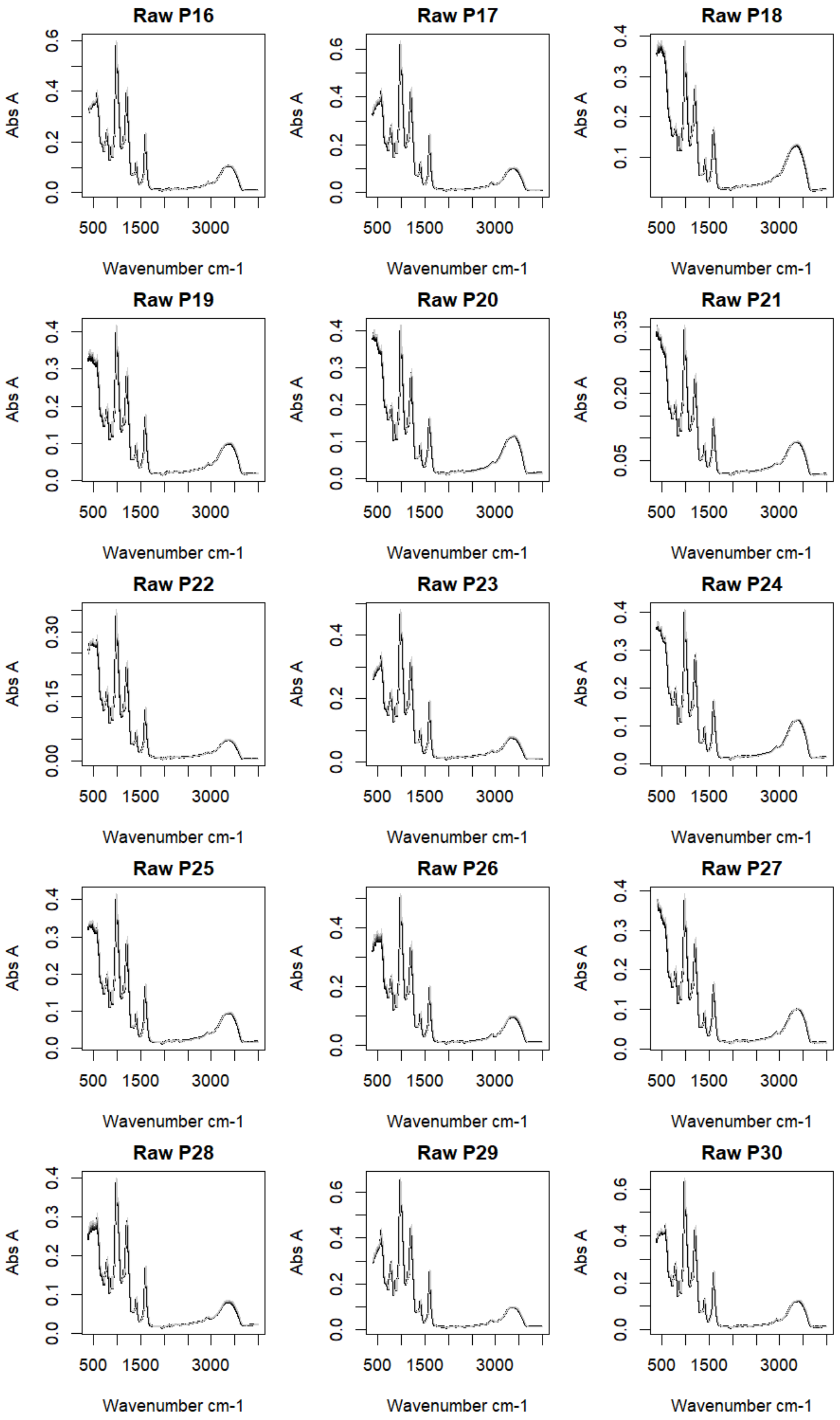
Raw P16
Raw P17
Raw P18
Raw P19
Raw P20
Raw P21
Raw P22
Raw P23
Raw P24
Raw P25
Raw P26
Raw P27
Raw P28
Raw P29
Raw P30
Abs A
Wavenumber cm-1
500 1500 3000

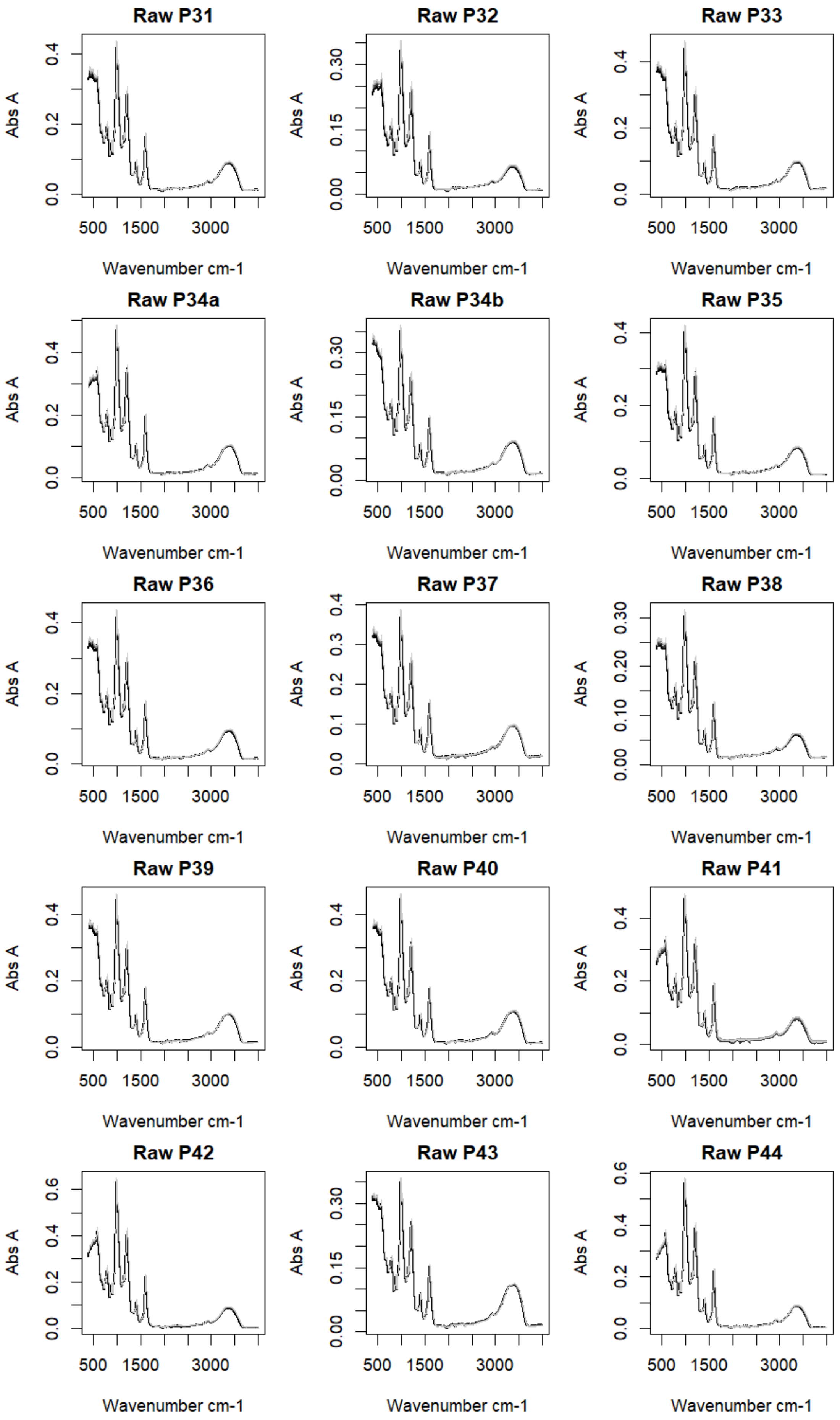
Raw P31
Raw P32
Raw P33
Raw P34a
Raw P34b
Raw P35
Raw P36
Raw P37
Raw P38
Raw P39
Raw P40
Raw P41
Raw P42
Raw P43
Raw P44
Abs A
Wavenumber cm-1
500 1500 3000

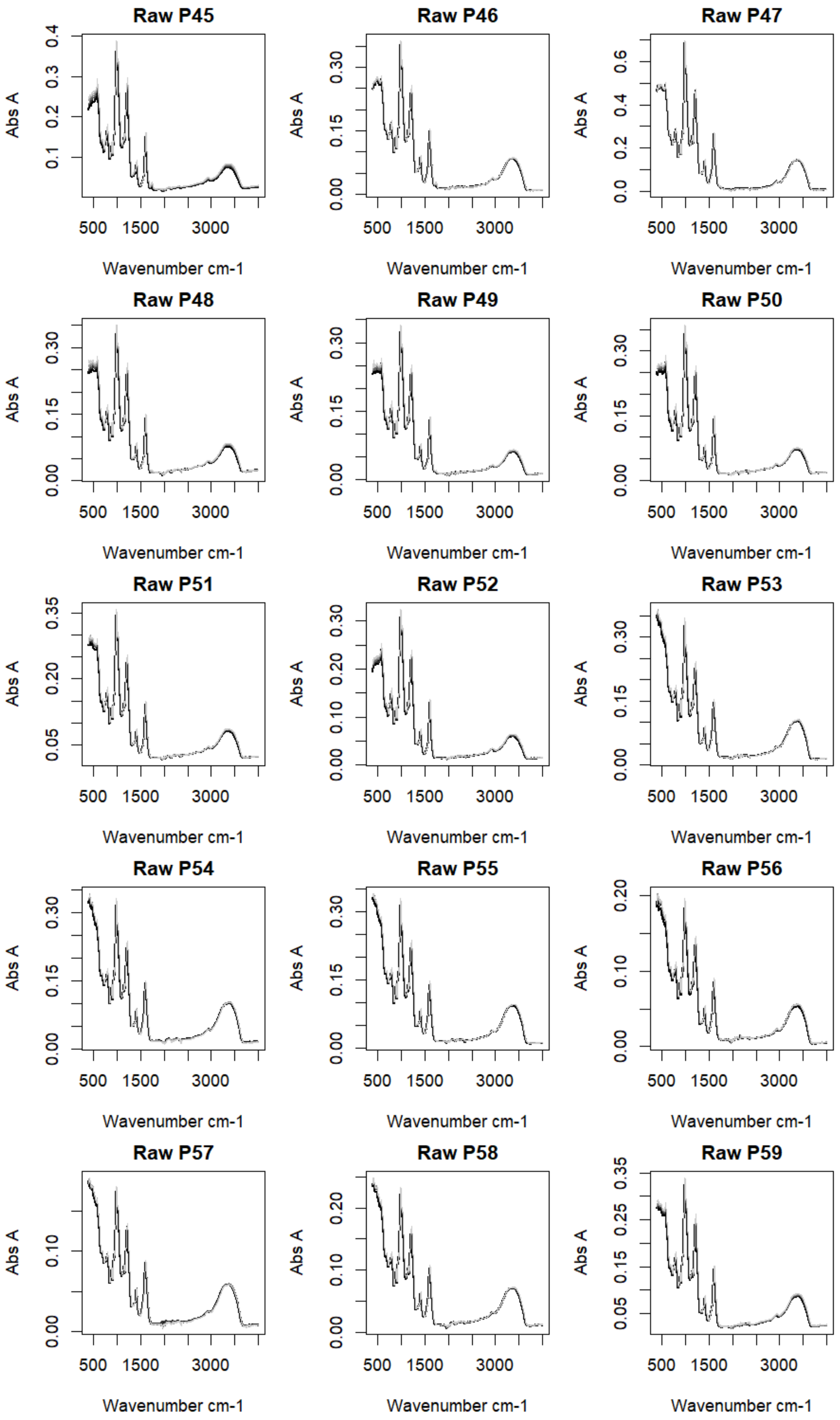

Raw P45
Raw P46
Raw P47
Raw P48
Raw P49
Raw P50
Raw P51
Raw P52
Raw P53
Raw P54
Raw P55
Raw P56
Raw P57
Raw P58
Raw P59
Abs A
500 1500 3000
Wavenumber cm-1

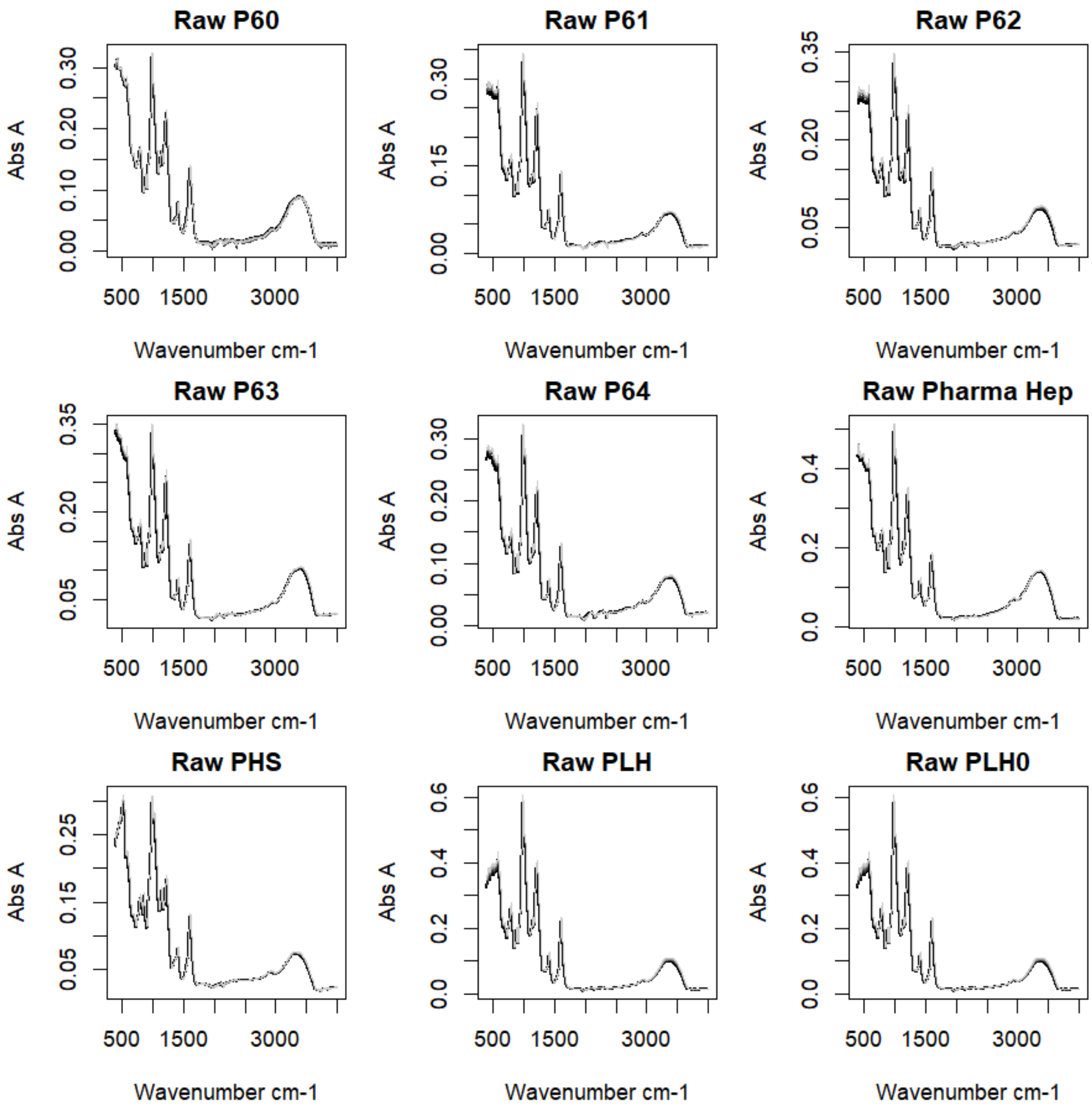
Raw P60
Raw P61
Raw P62
Raw P63
Raw P64
Raw Pharma Hep
Raw PHS
Raw PLH
Raw PLH0
Abs A
Wavenumber cm-1
500
1500
3000

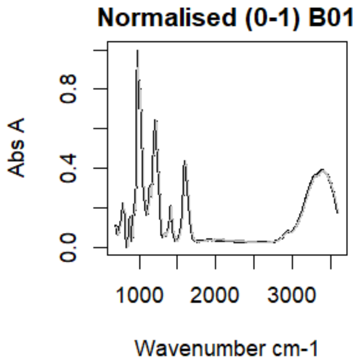
Normalised (0-1) B01
Abs A
0.0
0.4
0.8
1000
2000
3000
Wavenumber cm-1

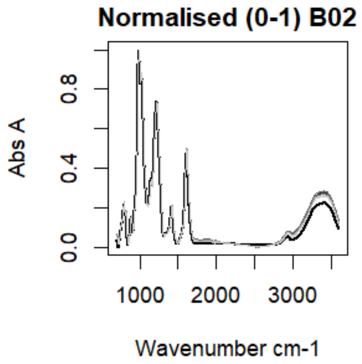
Normalised (0-1) B02
Abs A
0.0
0.4
0.8
1000
2000
3000
Wavenumber cm-1

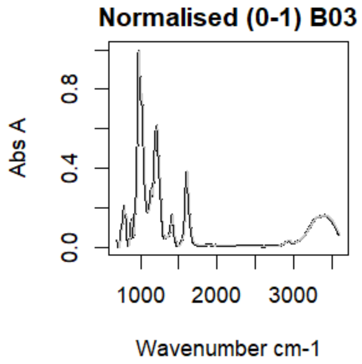
Normalised (0-1) B03
Abs A
0.0
0.4
0.8
1000
2000
3000
Wavenumber cm-1

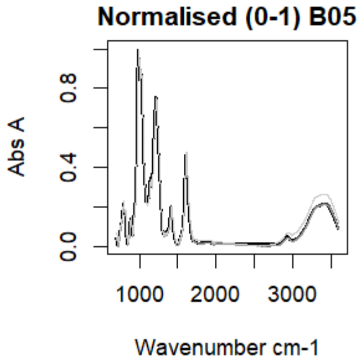
Normalised (0-1) B05
Abs A
0.0
0.4
0.8
1000
2000
3000
Wavenumber cm-1

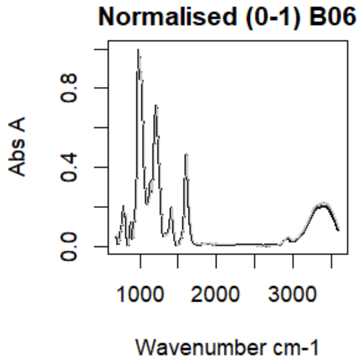
Normalised (0-1) B06
Abs A
0.0
0.4
0.8
1000
2000
3000
Wavenumber cm-1

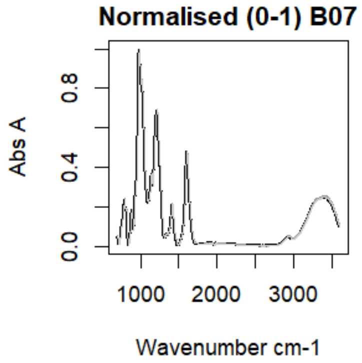
Normalised (0-1) B07
Abs A
0.0
0.4
0.8
1000
2000
3000
Wavenumber cm-1

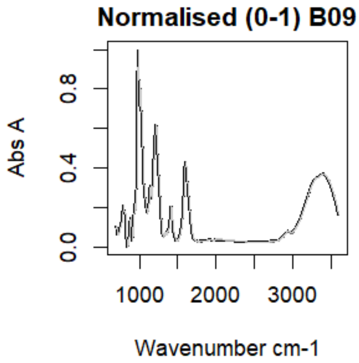
Normalised (0-1) B09
Abs A
0.0
0.4
0.8
1000
2000
3000
Wavenumber cm-1

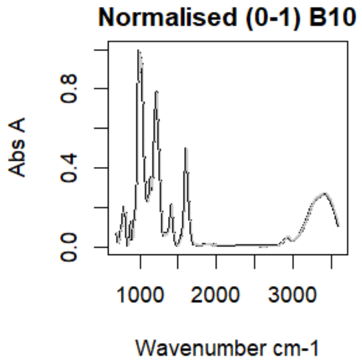
Normalised (0-1) B10
Abs A
0.0
0.4
0.8
1000
2000
3000
Wavenumber cm-1

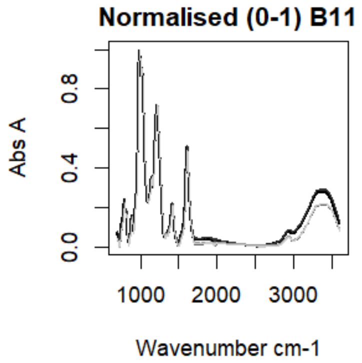
Normalised (0-1) B11
Abs A
0.0
0.4
0.8
1000
2000
3000
Wavenumber cm-1

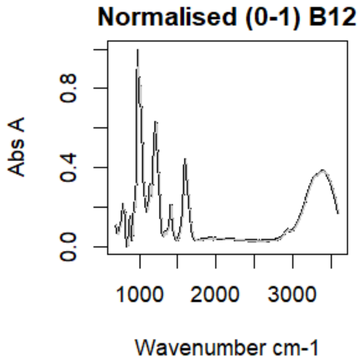
Normalised (0-1) B12
Abs A
0.0
0.4
0.8
1000
2000
3000
Wavenumber cm-1

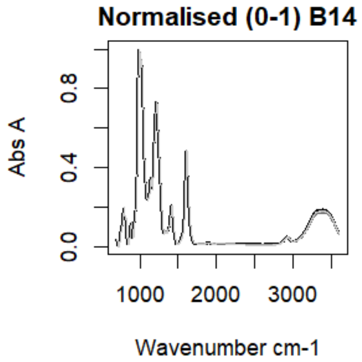
Normalised (0-1) B14
Abs A
0.0
0.4
0.8
1000
2000
3000
Wavenumber cm-1

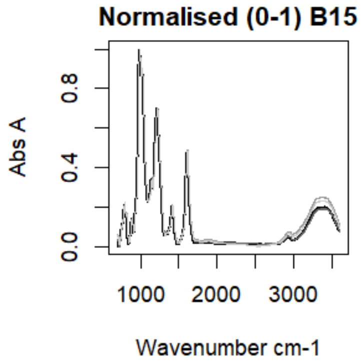
Normalised (0-1) B15
Abs A
0.0
0.4
0.8
1000
2000
3000
Wavenumber cm-1

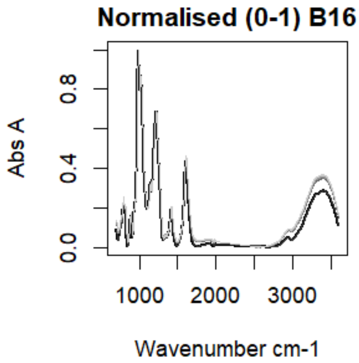
Normalised (0-1) B16
Abs A
0.0
0.4
0.8
1000
2000
3000
Wavenumber cm-1

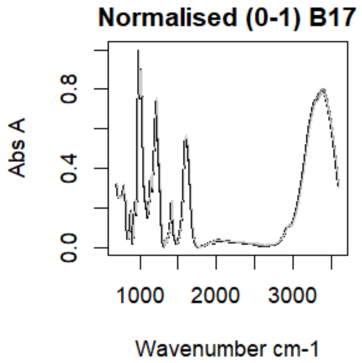
Normalised (0-1) B17
Abs A
0.0
0.4
0.8
1000
2000
3000
Wavenumber cm-1

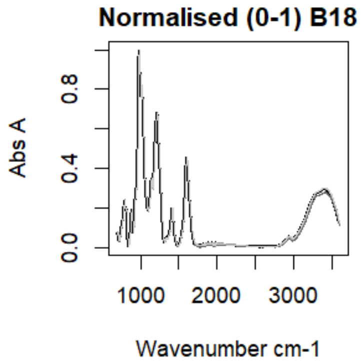
Normalised (0-1) B18
Abs A
0.0
0.4
0.8
1000
2000
3000
Wavenumber cm-1

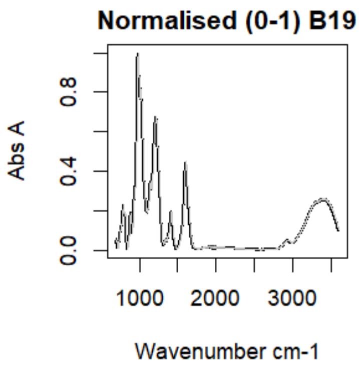
Normalised (0-1) B19
Abs A
0.0
0.4
0.8
1000
2000
3000
Wavenumber cm-1

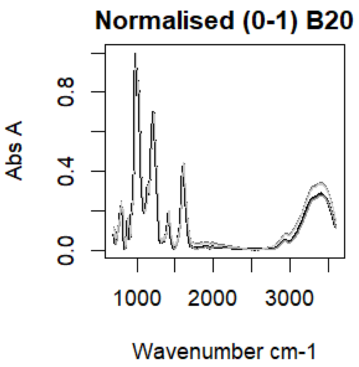
Normalised (0-1) B20
Abs A
0.0
0.4
0.8
1000
2000
3000
Wavenumber cm-1

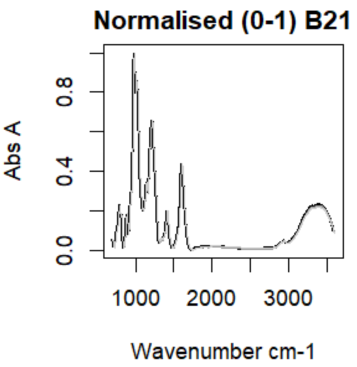
Normalised (0-1) B21
Abs A
0.0
0.4
0.8
1000
2000
3000
Wavenumber cm-1

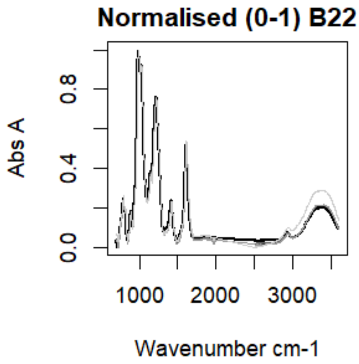
Normalised (0-1) B22
Abs A
0.0
0.4
0.8
1000
2000
3000
Wavenumber cm-1

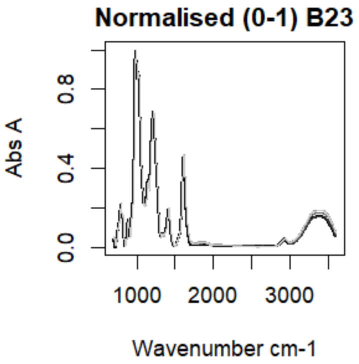
Normalised (0-1) B23
Abs A
0.0
0.4
0.8
1000
2000
3000
Wavenumber cm-1

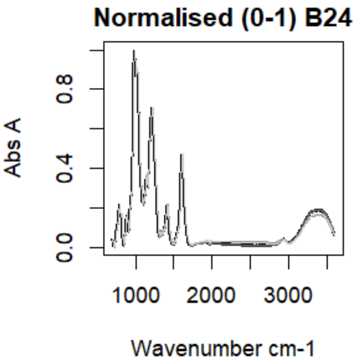
Normalised (0-1) B24
Abs A
0.0
0.4
0.8
1000
2000
3000
Wavenumber cm-1

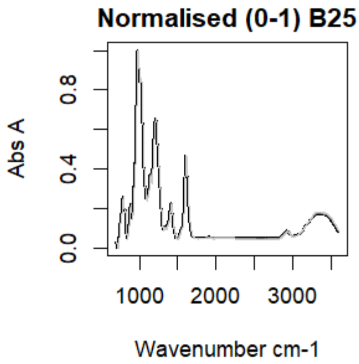
Normalised (0-1) B25
Abs A
0.0
0.4
0.8
1000
2000
3000
Wavenumber cm-1

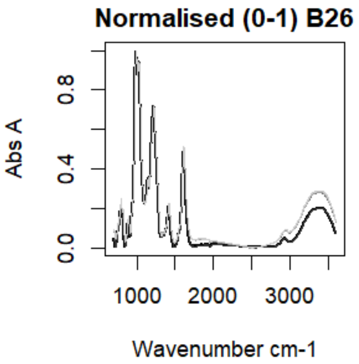
Normalised (0-1) B26
Abs A
0.0
0.4
0.8
1000
2000
3000
Wavenumber cm-1

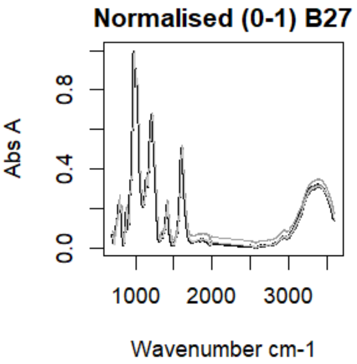
Normalised (0-1) B27
Abs A
0.0
0.4
0.8
1000
2000
3000
Wavenumber cm-1

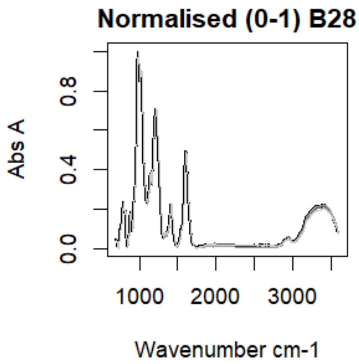
Normalised (0-1) B28
Abs A
0.0
0.4
0.8
1000
2000
3000
Wavenumber cm-1

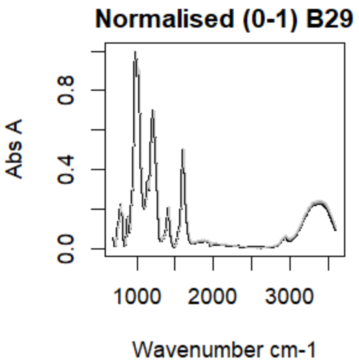
Normalised (0-1) B29
Abs A
0.0
0.4
0.8
1000
2000
3000
Wavenumber cm-1

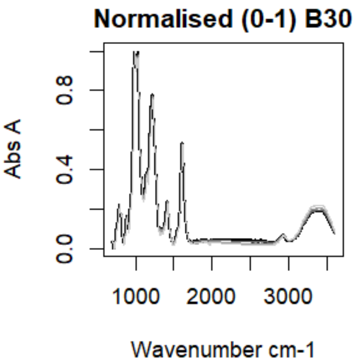
Normalised (0-1) B30
Abs A
0.0
0.4
0.8
1000
2000
3000
Wavenumber cm-1

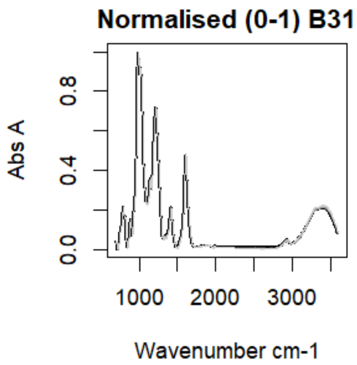
Normalised (0-1) B31
Abs A
0.0
0.4
0.8
1000
2000
3000
Wavenumber cm-1

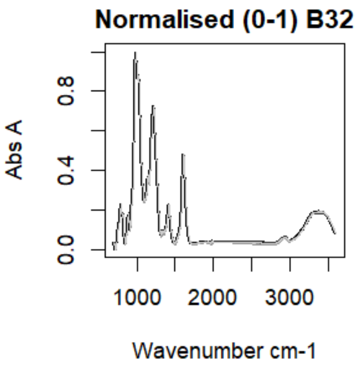
Normalised (0-1) B32
Abs A
0.0
0.4
0.8
1000
2000
3000
Wavenumber cm-1

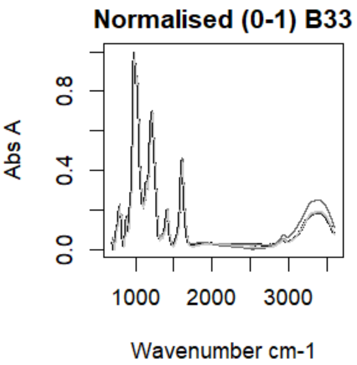
Normalised (0-1) B33
Abs A
0.0
0.4
0.8
1000
2000
3000
Wavenumber cm-1

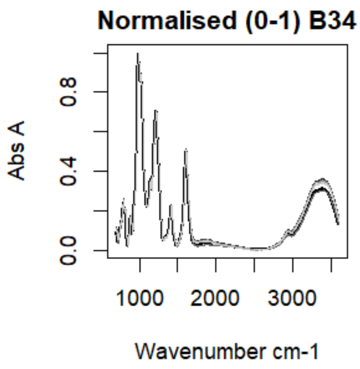
Normalised (0-1) B34
Abs A
0.0
0.4
0.8
1000
2000
3000
Wavenumber cm-1

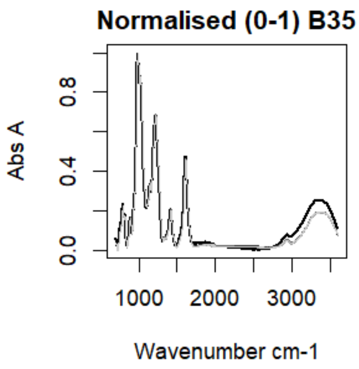
Normalised (0-1) B35
Abs A
0.0
0.4
0.8
1000
2000
3000
Wavenumber cm-1

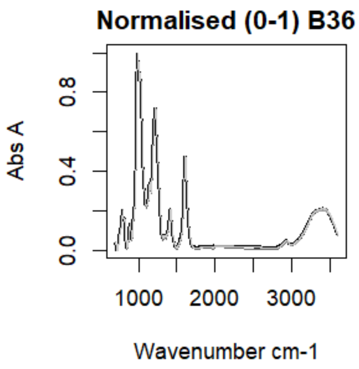
Normalised (0-1) B36
Abs A
0.0
0.4
0.8
1000
2000
3000
Wavenumber cm-1

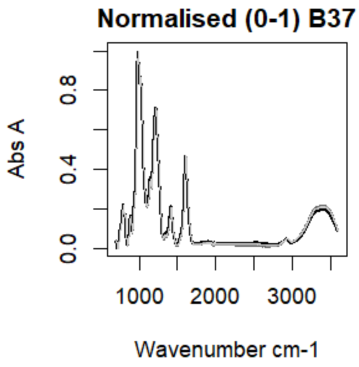
Normalised (0-1) B37
Abs A
0.0
0.4
0.8
1000
2000
3000
Wavenumber cm-1

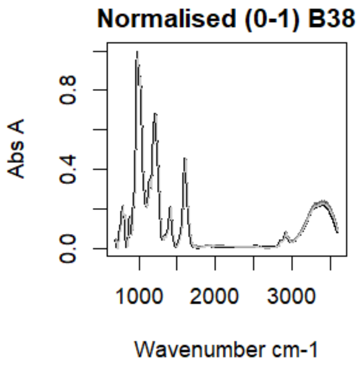
Normalised (0-1) B38
Abs A
0.0
0.4
0.8
1000
2000
3000
Wavenumber cm-1

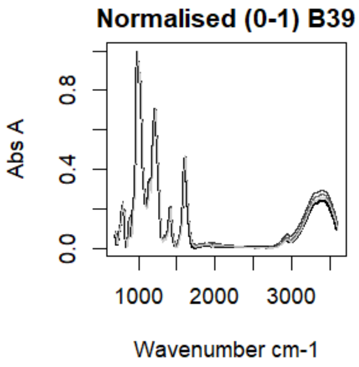
Normalised (0-1) B39
Abs A
0.0
0.4
0.8
1000
2000
3000
Wavenumber cm-1

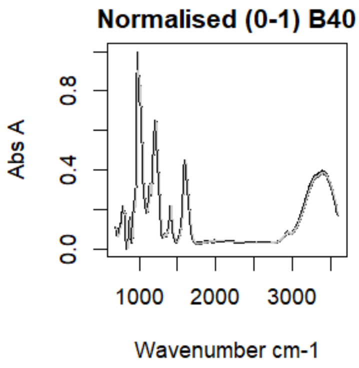
Normalised (0-1) B40
Abs A
0.0
0.4
0.8
1000
2000
3000
Wavenumber cm-1

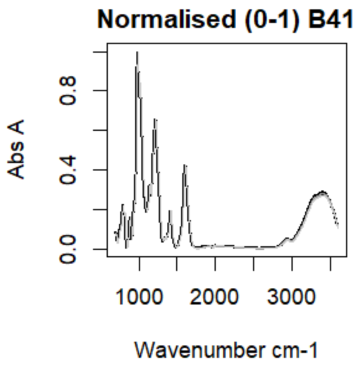
Normalised (0-1) B41
Abs A
0.0
0.4
0.8
1000
2000
3000
Wavenumber cm-1

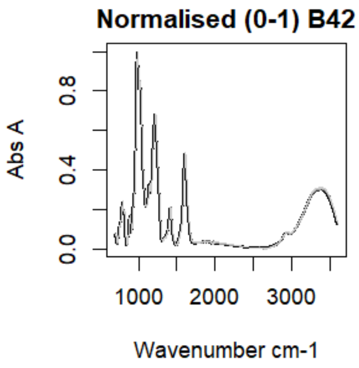
Normalised (0-1) B42
Abs A
0.0
0.4
0.8
1000
2000
3000
Wavenumber cm-1

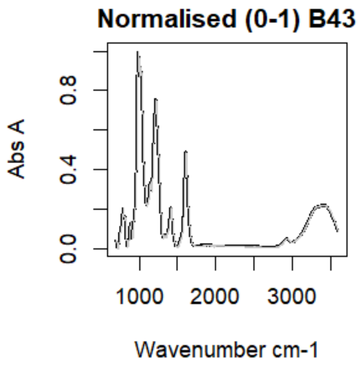
Normalised (0-1) B43
Abs A
0.0
0.4
0.8
1000
2000
3000
Wavenumber cm-1

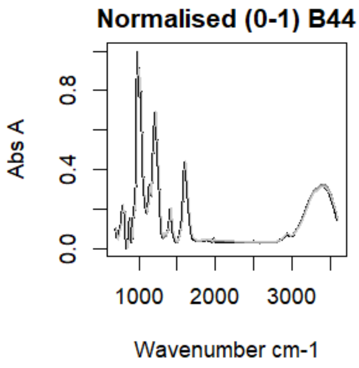
Normalised (0-1) B44
Abs A
0.0
0.4
0.8
1000
2000
3000
Wavenumber cm-1

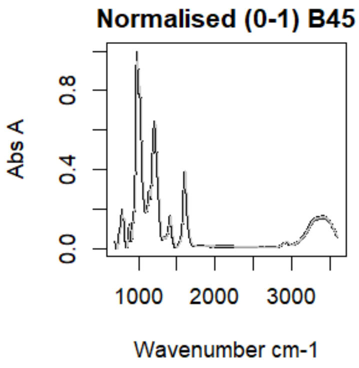
Normalised (0-1) B45
Abs A
0.0
0.4
0.8
1000
2000
3000
Wavenumber cm-1

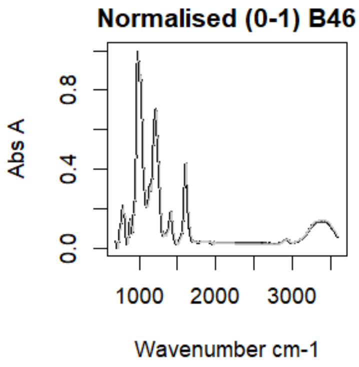
Normalised (0-1) B46
Abs A
0.0
0.4
0.8
1000
2000
3000
Wavenumber cm-1

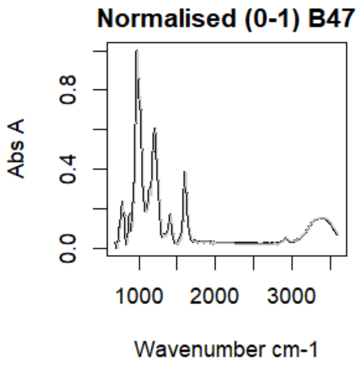
Normalised (0-1) B47
Abs A
0.0
0.4
0.8
1000
2000
3000
Wavenumber cm-1

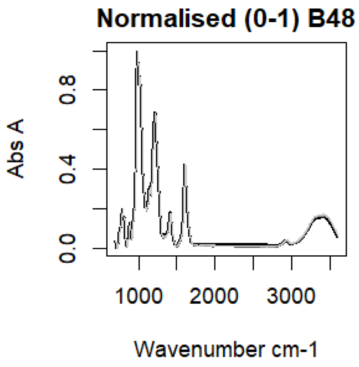
Normalised (0-1) B48
Abs A
0.0
0.4
0.8
1000
2000
3000
Wavenumber cm-1

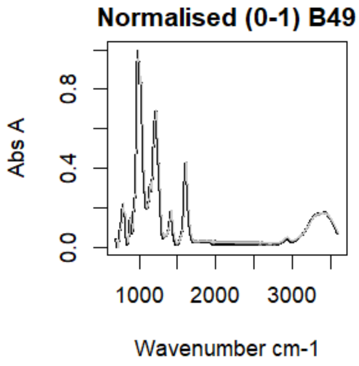
Normalised (0-1) B49
Abs A
0.0
0.4
0.8
1000
2000
3000
Wavenumber cm-1

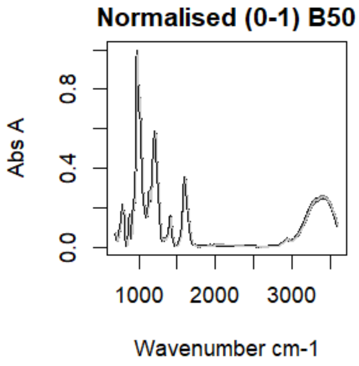
Normalised (0-1) B50
Abs A
0.0
0.4
0.8
1000
2000
3000
Wavenumber cm-1

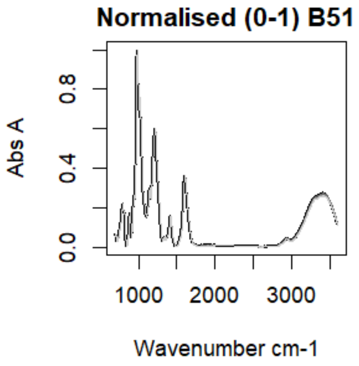
Normalised (0-1) B51
Abs A
0.0
0.4
0.8
1000
2000
3000
Wavenumber cm-1

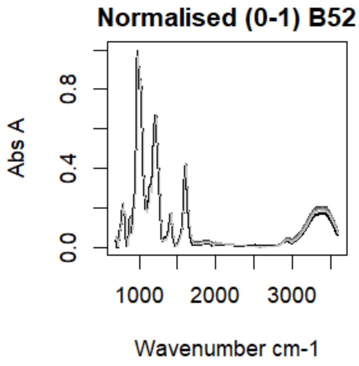
Normalised (0-1) B52
Abs A
0.0
0.4
0.8
1000
2000
3000
Wavenumber cm-1

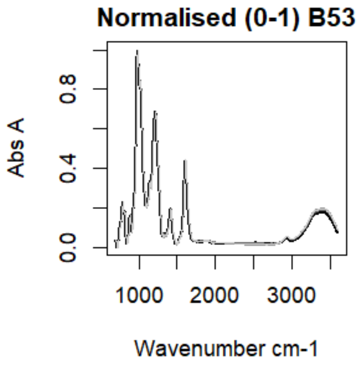
Normalised (0-1) B53
Abs A
0.0
0.4
0.8
1000
2000
3000
Wavenumber cm-1

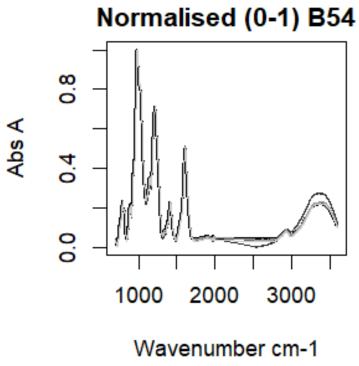
Normalised (0-1) B54
Abs A
0.0
0.4
0.8
1000
2000
3000
Wavenumber cm-1

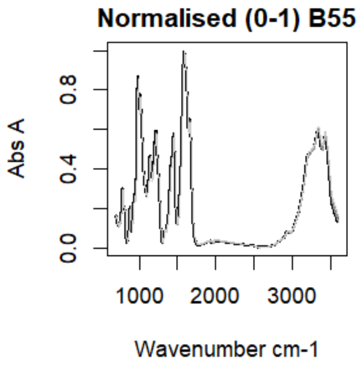
Normalised (0-1) B55
Abs A
0.0
0.4
0.8
1000
2000
3000
Wavenumber cm-1

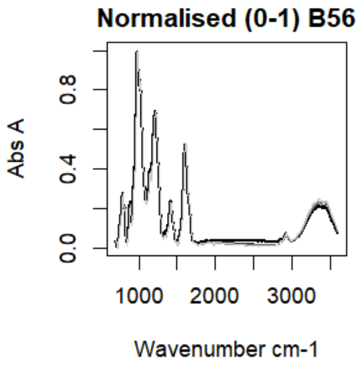
Normalised (0-1) B56
Abs A
0.0
0.4
0.8
1000
2000
3000
Wavenumber cm-1

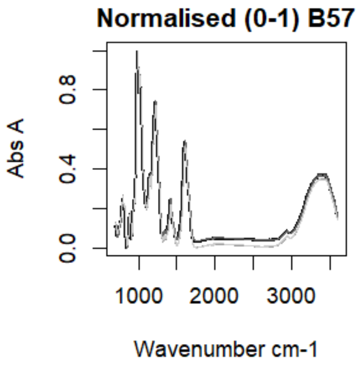
Normalised (0-1) B57
Abs A
0.0
0.4
0.8
1000
2000
3000
Wavenumber cm-1

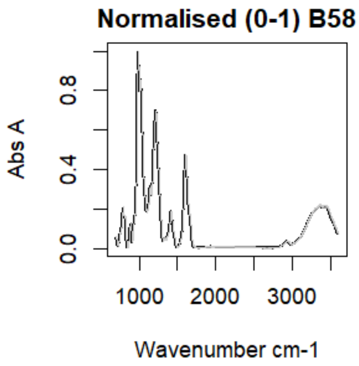
Normalised (0-1) B58
Abs A
0.0
0.4
0.8
1000
2000
3000
Wavenumber cm-1

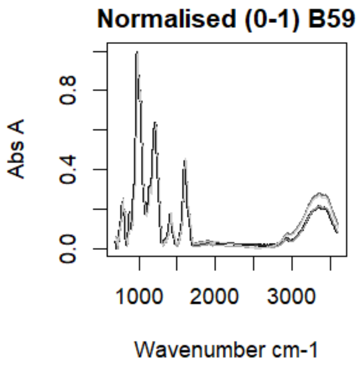
Normalised (0-1) B59
Abs A
0.0
0.4
0.8
1000
2000
3000
Wavenumber cm-1

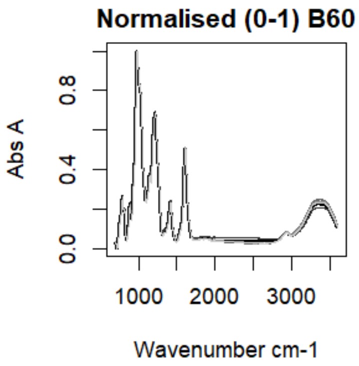
Normalised (0-1) B60
Abs A
0.0
0.4
0.8
1000
2000
3000
Wavenumber cm-1

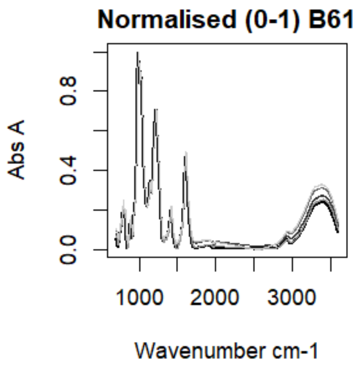
Normalised (0-1) B61
Abs A
0.0
0.4
0.8
1000
2000
3000
Wavenumber cm-1

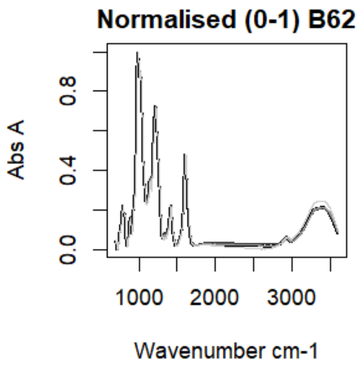
Normalised (0-1) B62
Abs A
0.0
0.4
0.8
1000
2000
3000
Wavenumber cm-1

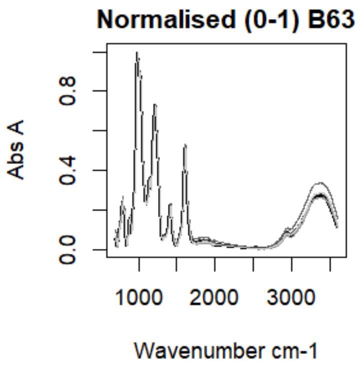
Normalised (0-1) B63
Abs A
0.0
0.4
0.8
1000
2000
3000
Wavenumber cm-1

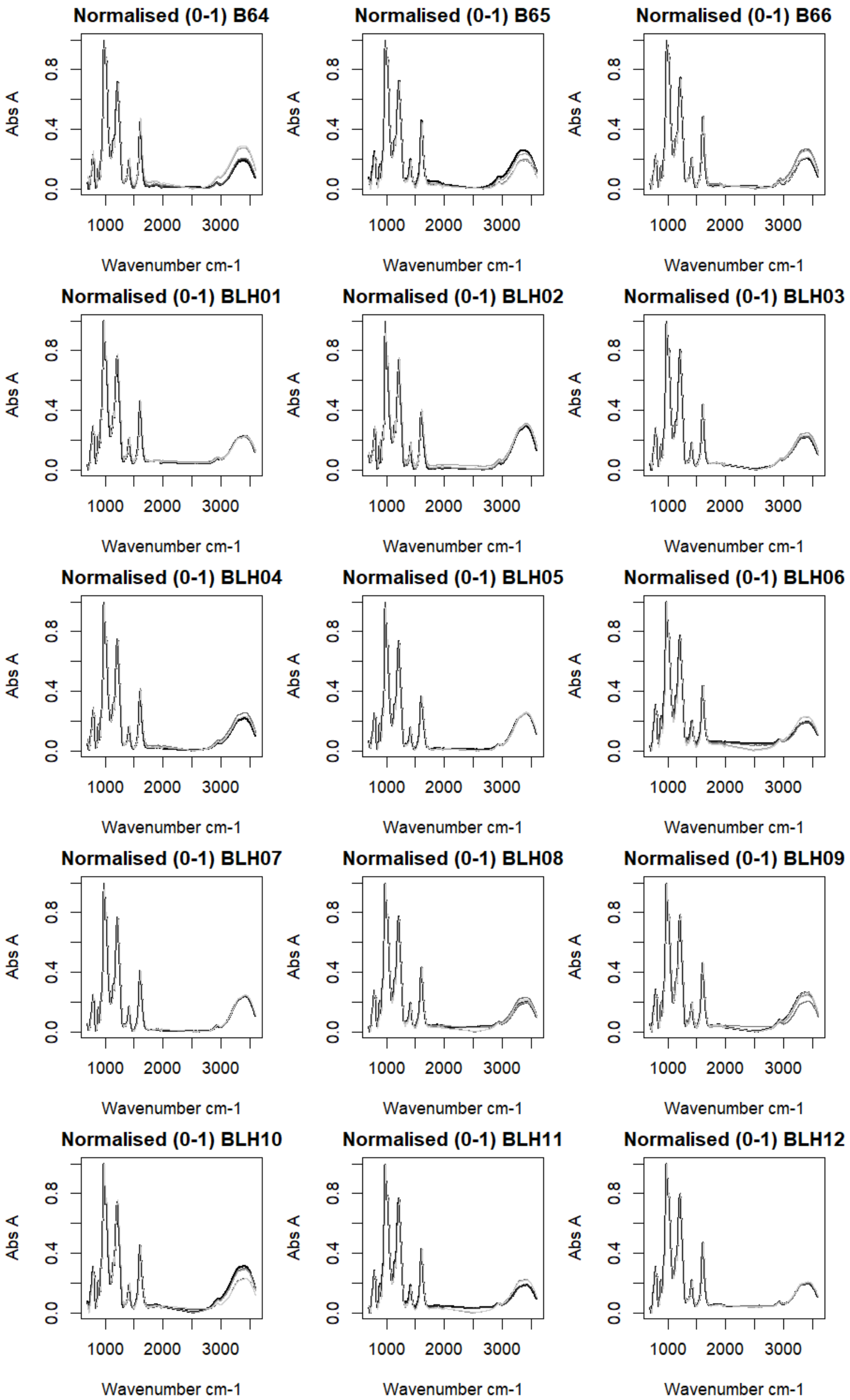
Normalised (0-1) B64
Normalised (0-1) B65
Normalised (0-1) B66
Normalised (0-1) BLH01
Normalised (0-1) BLH02
Normalised (0-1) BLH03
Normalised (0-1) BLH04
Normalised (0-1) BLH05
Normalised (0-1) BLH06
Normalised (0-1) BLH07
Normalised (0-1) BLH08
Normalised (0-1) BLH09
Normalised (0-1) BLH10
Normalised (0-1) BLH11
Normalised (0-1) BLH12
Abs A
0.0
0.4
0.8
1000
2000
3000
Wavenumber cm-1

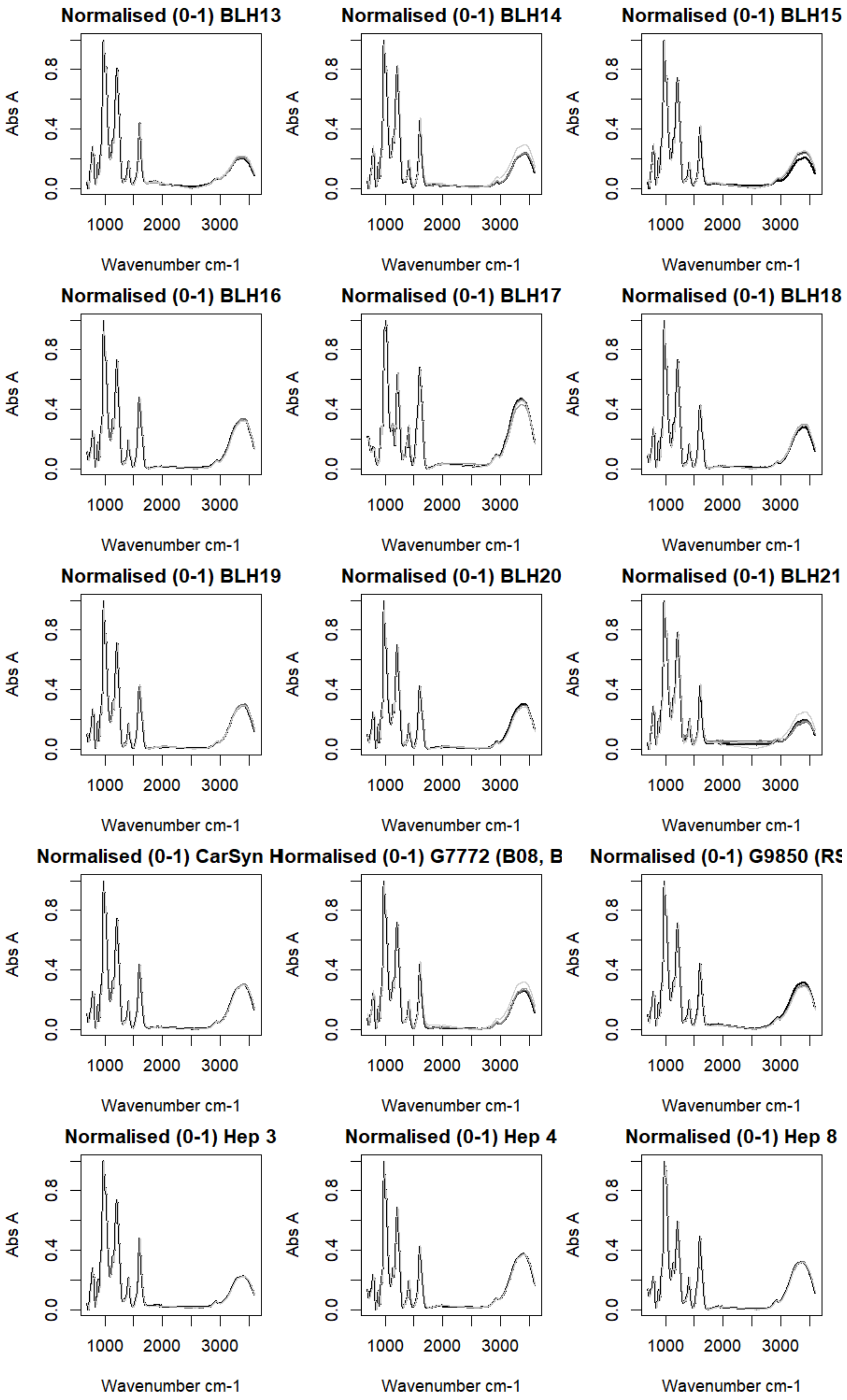

Normalised (0-1) BLH13
Normalised (0-1) BLH14
Normalised (0-1) BLH15
Normalised (0-1) BLH16
Normalised (0-1) BLH17
Normalised (0-1) BLH18
Normalised (0-1) BLH19
Normalised (0-1) BLH20
Normalised (0-1) BLH21
Normalised (0-1) CarSyn H
Normalised (0-1) G7772 (B08, B
Normalised (0-1) G9850 (RS
Normalised (0-1) Hep 3
Normalised (0-1) Hep 4
Normalised (0-1) Hep 8
Abs A
0.0
0.4
0.8
1000
2000
3000
Wavenumber cm-1

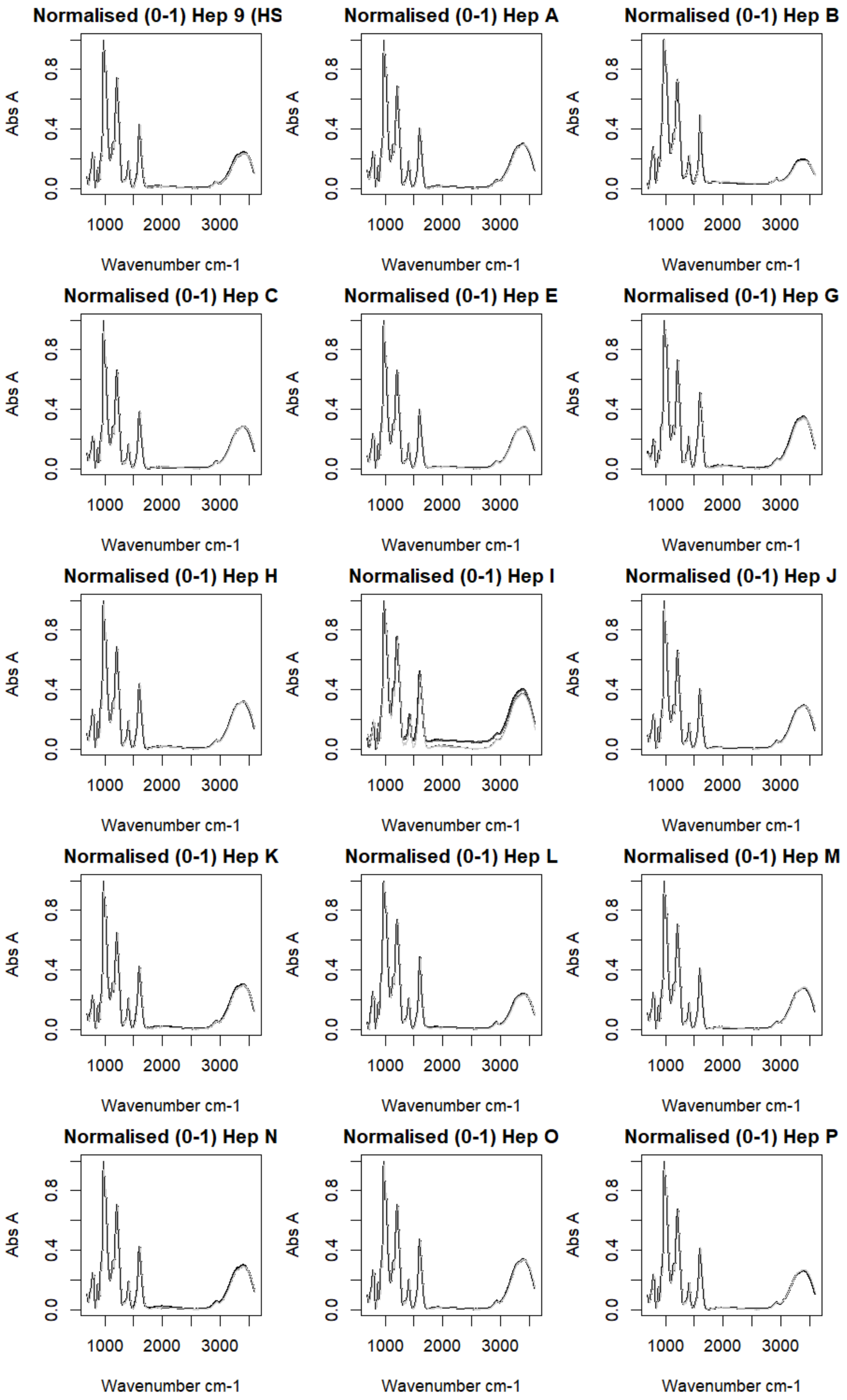
Normalised (0-1) Hep 9 (HS
Normalised (0-1) Hep A
Normalised (0-1) Hep B
Normalised (0-1) Hep C
Normalised (0-1) Hep E
Normalised (0-1) Hep G
Normalised (0-1) Hep H
Normalised (0-1) Hep I
Normalised (0-1) Hep J
Normalised (0-1) Hep K
Normalised (0-1) Hep L
Normalised (0-1) Hep M
Normalised (0-1) Hep N
Normalised (0-1) Hep O
Normalised (0-1) Hep P
Abs A
0.0
0.4
0.8
1000
2000
3000
Wavenumber cm-1

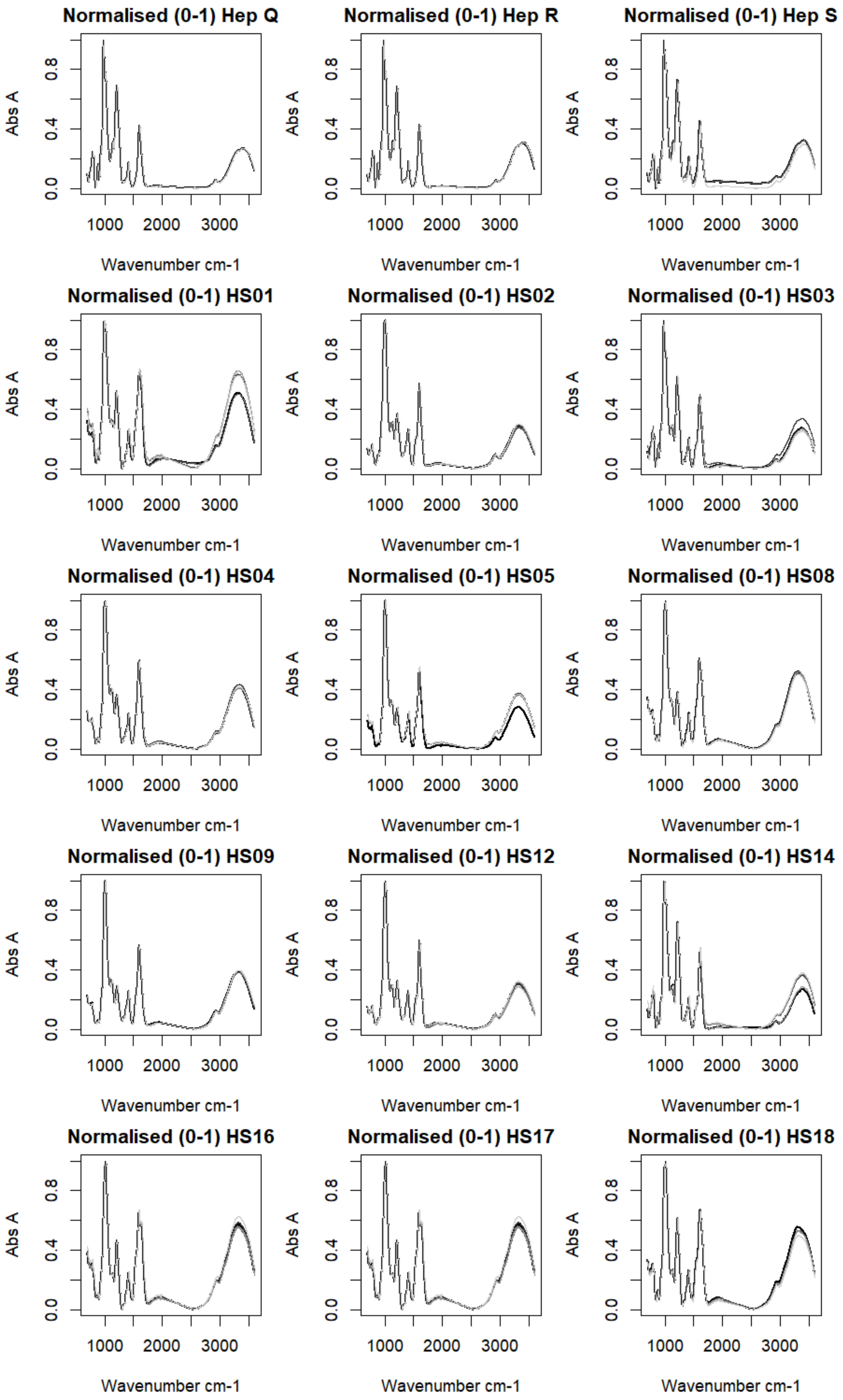
Normalised (0-1) Hep Q
Normalised (0-1) Hep R
Normalised (0-1) Hep S
Normalised (0-1) HS01
Normalised (0-1) HS02
Normalised (0-1) HS03
Normalised (0-1) HS04
Normalised (0-1) HS05
Normalised (0-1) HS08
Normalised (0-1) HS09
Normalised (0-1) HS12
Normalised (0-1) HS14
Normalised (0-1) HS16
Normalised (0-1) HS17
Normalised (0-1) HS18
Abs A
0.0
0.4
0.8
1000
2000
3000
Wavenumber cm-1

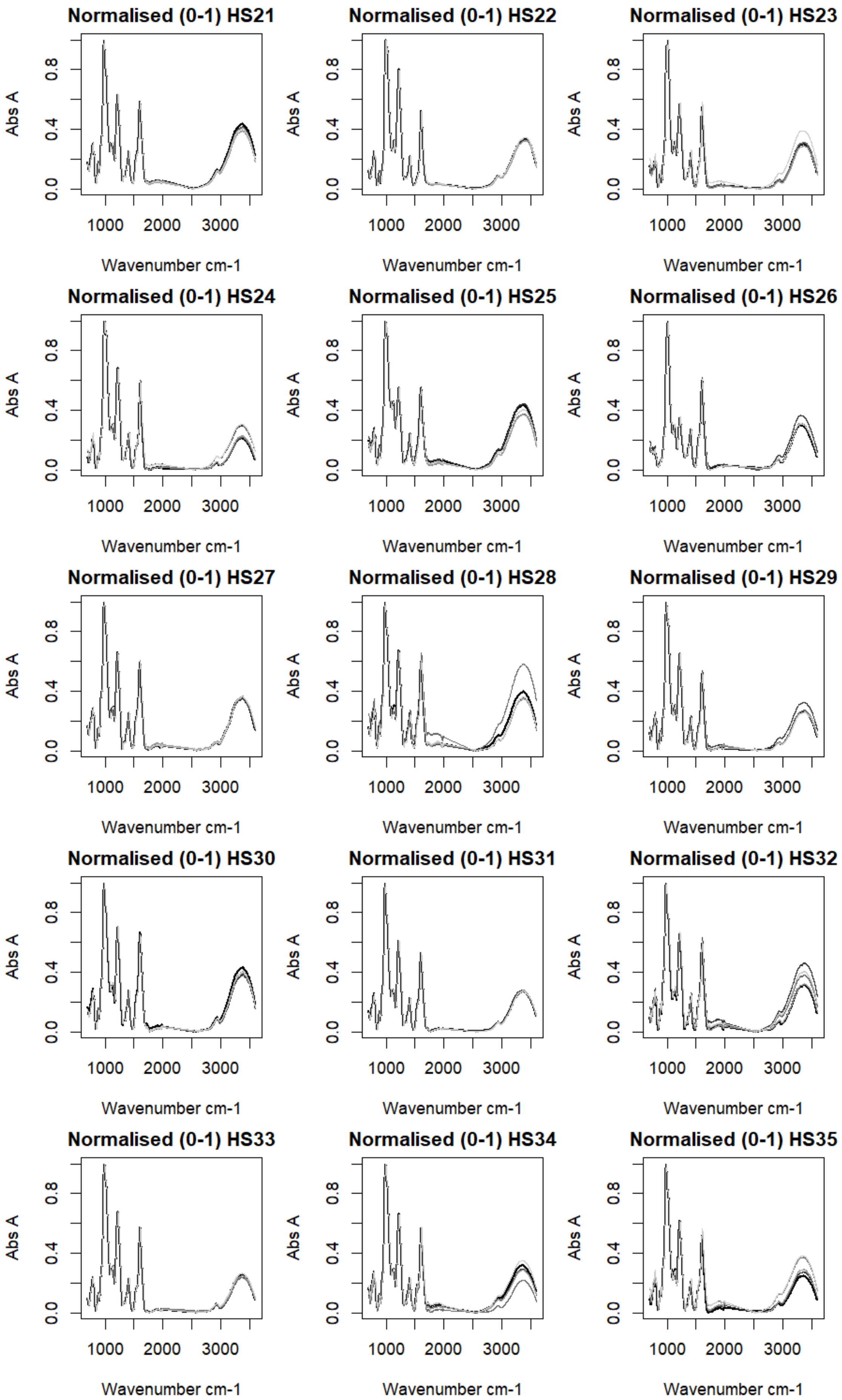
Normalised (0-1) HS21
Normalised (0-1) HS22
Normalised (0-1) HS23
Normalised (0-1) HS24
Normalised (0-1) HS25
Normalised (0-1) HS26
Normalised (0-1) HS27
Normalised (0-1) HS28
Normalised (0-1) HS29
Normalised (0-1) HS30
Normalised (0-1) HS31
Normalised (0-1) HS32
Normalised (0-1) HS33
Normalised (0-1) HS34
Normalised (0-1) HS35
Abs A
0.0
0.4
0.8
1000
2000
3000
Wavenumber cm-1

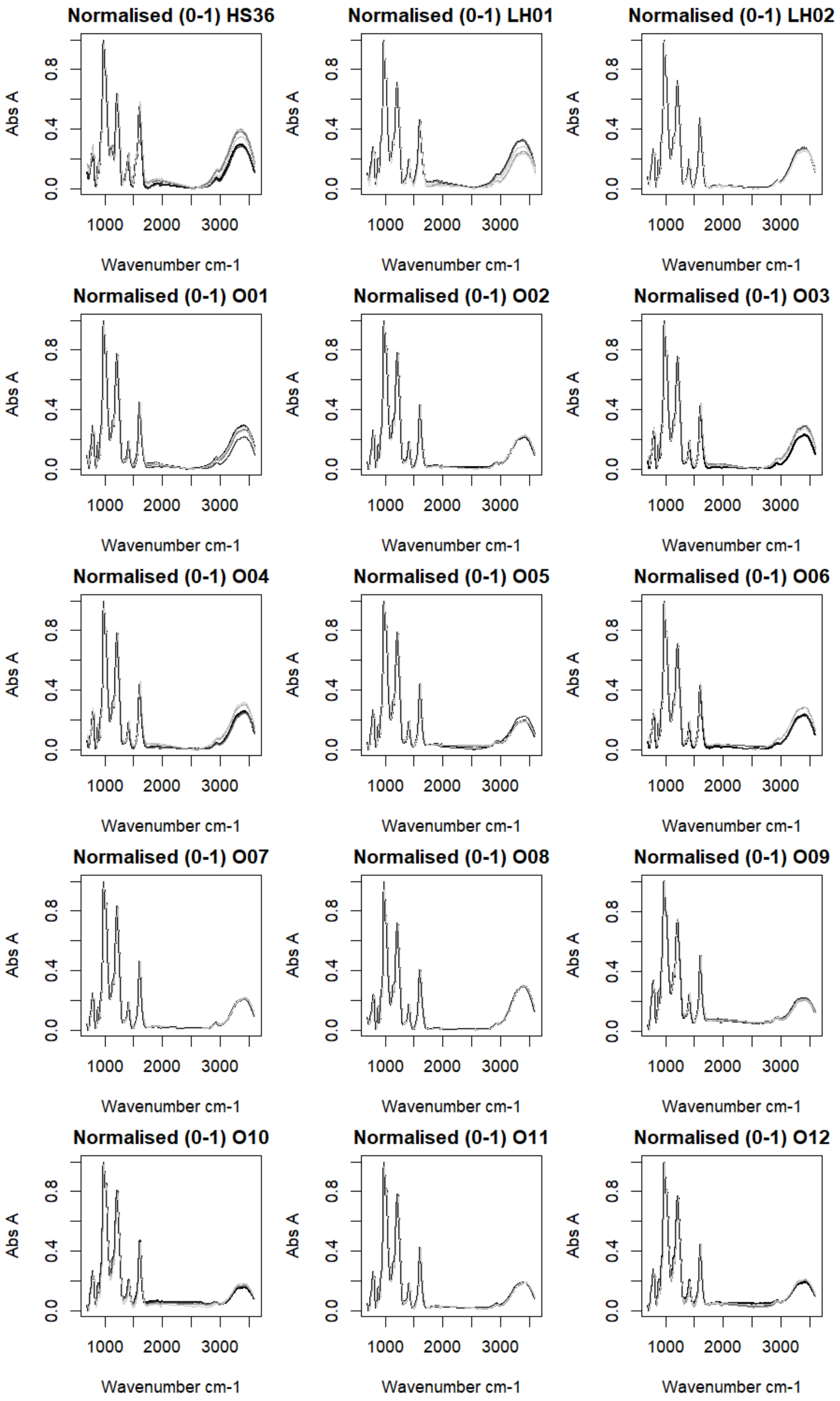
Normalised (0-1) HS36
Normalised (0-1) LH01
Normalised (0-1) LH02
Normalised (0-1) O01
Normalised (0-1) O02
Normalised (0-1) O03
Normalised (0-1) O04
Normalised (0-1) O05
Normalised (0-1) O06
Normalised (0-1) O07
Normalised (0-1) O08
Normalised (0-1) O09
Normalised (0-1) O10
Normalised (0-1) O11
Normalised (0-1) O12
Abs A
0.0
0.4
0.8
1000
2000
3000
Wavenumber cm-1

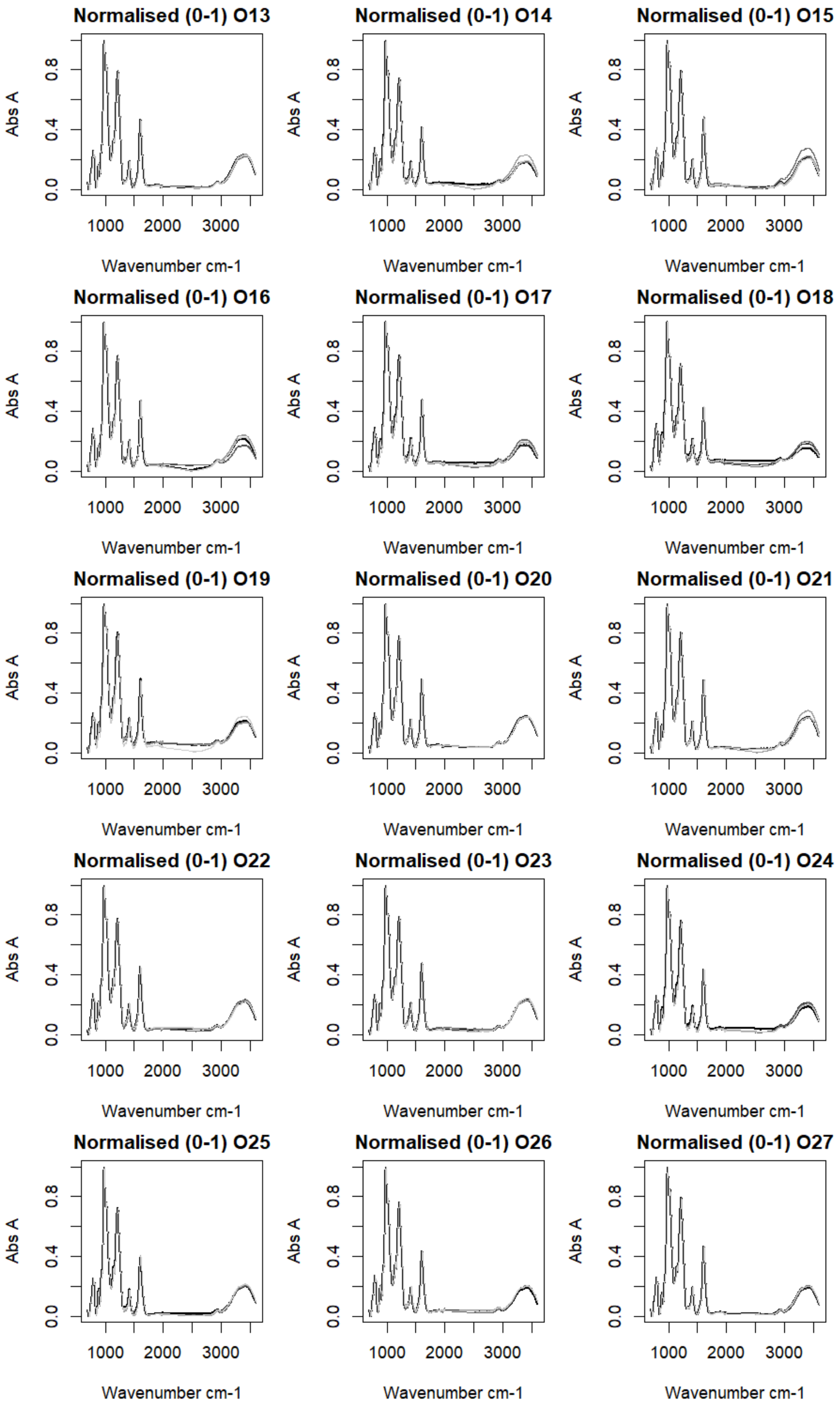

Normalised (0-1) O13
Normalised (0-1) O14
Normalised (0-1) O15
Normalised (0-1) O16
Normalised (0-1) O17
Normalised (0-1) O18
Normalised (0-1) O19
Normalised (0-1) O20
Normalised (0-1) O21
Normalised (0-1) O22
Normalised (0-1) O23
Normalised (0-1) O24
Normalised (0-1) O25
Normalised (0-1) O26
Normalised (0-1) O27
Abs A
0.0
0.4
0.8
1000
2000
3000
Wavenumber cm-1

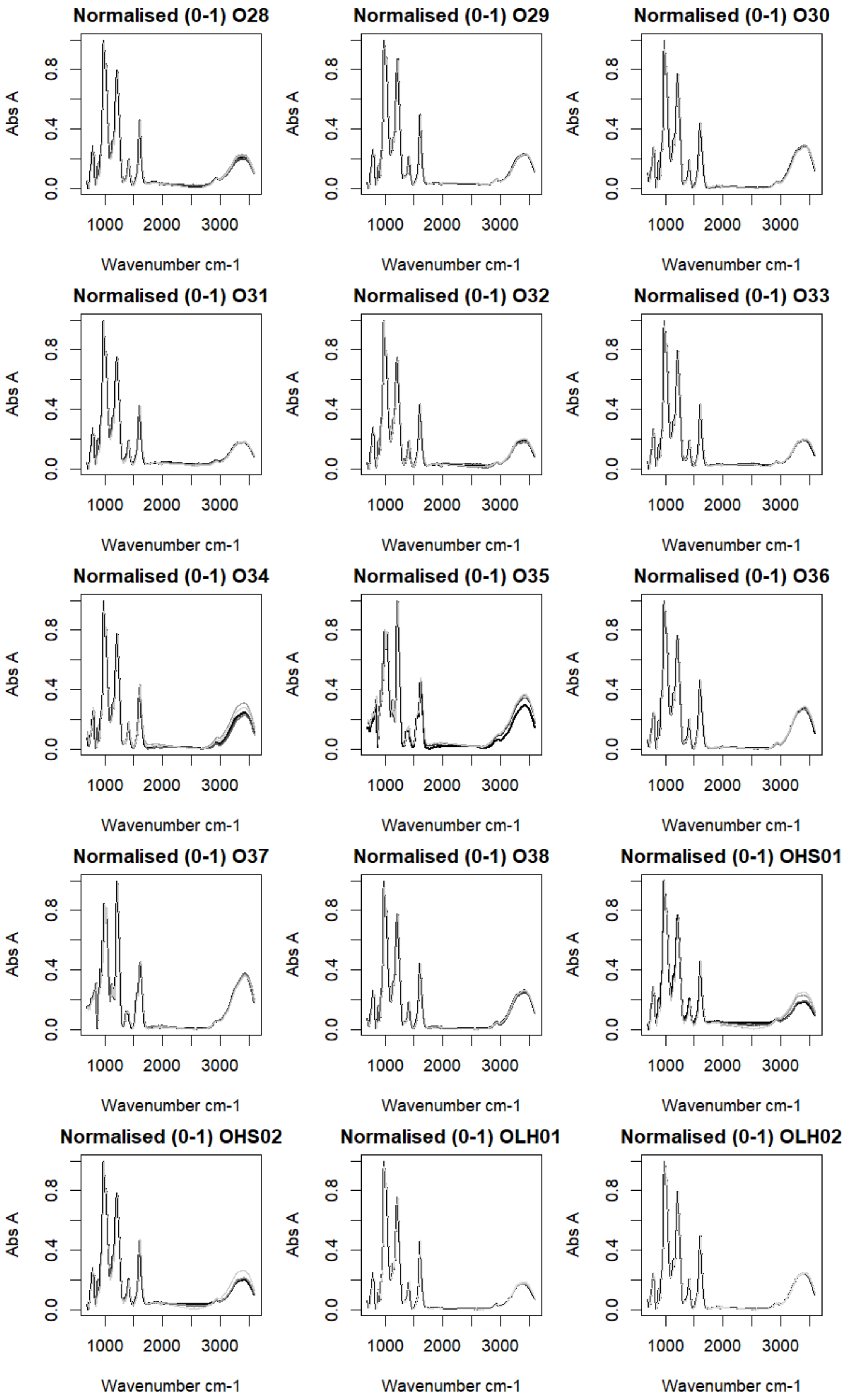
Normalised (0-1) O28
Normalised (0-1) O29
Normalised (0-1) O30
Normalised (0-1) O31
Normalised (0-1) O32
Normalised (0-1) O33
Normalised (0-1) O34
Normalised (0-1) O35
Normalised (0-1) O36
Normalised (0-1) O37
Normalised (0-1) O38
Normalised (0-1) OHS01
Normalised (0-1) OHS02
Normalised (0-1) OLH01
Normalised (0-1) OLH02
Abs A
0.0
0.4
0.8
1000
2000
3000
Wavenumber cm-1

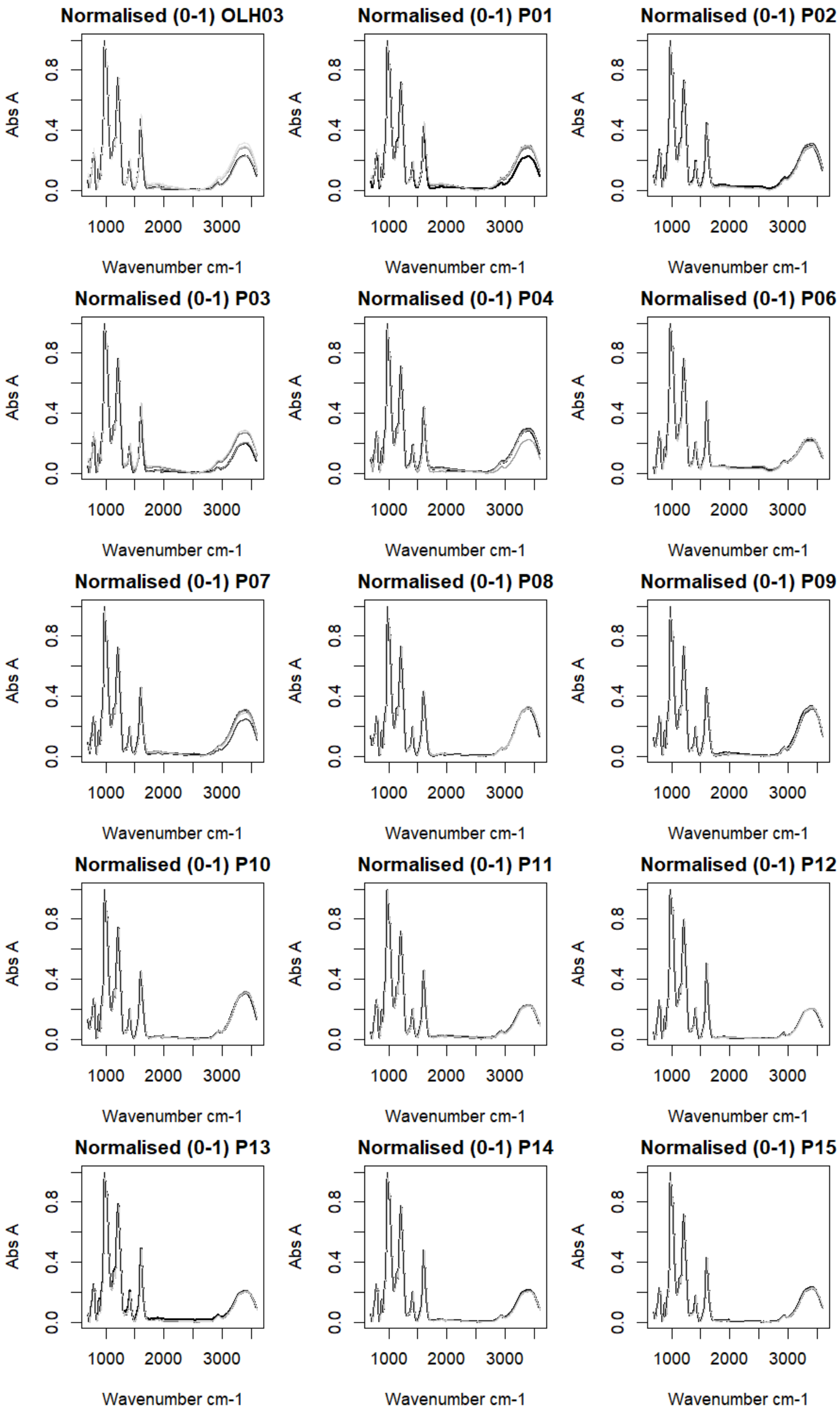
Normalised (0-1) OLH03
Normalised (0-1) P01
Normalised (0-1) P02
Normalised (0-1) P03
Normalised (0-1) P04
Normalised (0-1) P06
Normalised (0-1) P07
Normalised (0-1) P08
Normalised (0-1) P09
Normalised (0-1) P10
Normalised (0-1) P11
Normalised (0-1) P12
Normalised (0-1) P13
Normalised (0-1) P14
Normalised (0-1) P15
Abs A
0.0
0.4
0.8
1000
2000
3000
Wavenumber cm-1

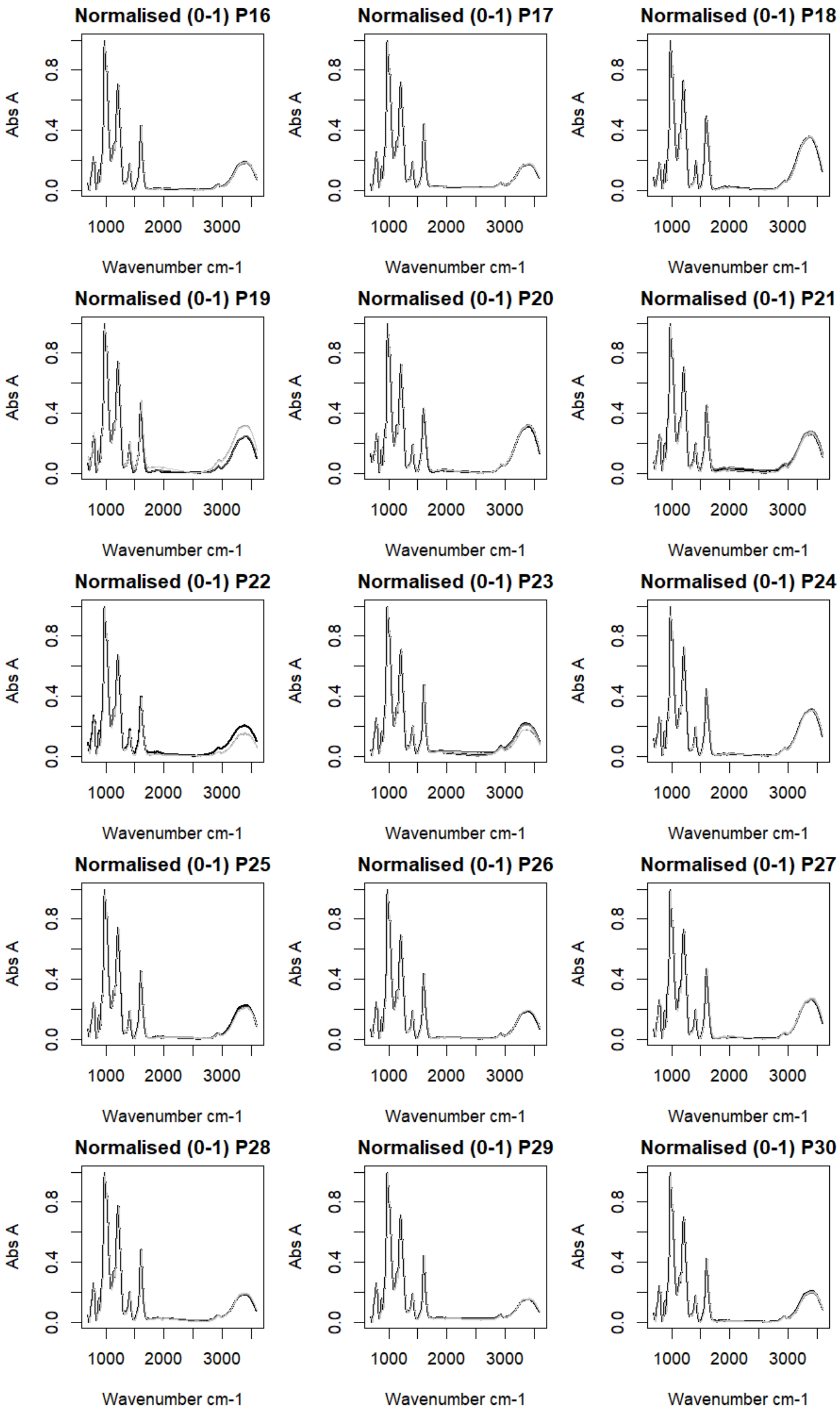
Normalised (0-1) P16
Normalised (0-1) P17
Normalised (0-1) P18
Normalised (0-1) P19
Normalised (0-1) P20
Normalised (0-1) P21
Normalised (0-1) P22
Normalised (0-1) P23
Normalised (0-1) P24
Normalised (0-1) P25
Normalised (0-1) P26
Normalised (0-1) P27
Normalised (0-1) P28
Normalised (0-1) P29
Normalised (0-1) P30
Abs A
0.0
0.4
0.8
1000
2000
3000
Wavenumber cm-1

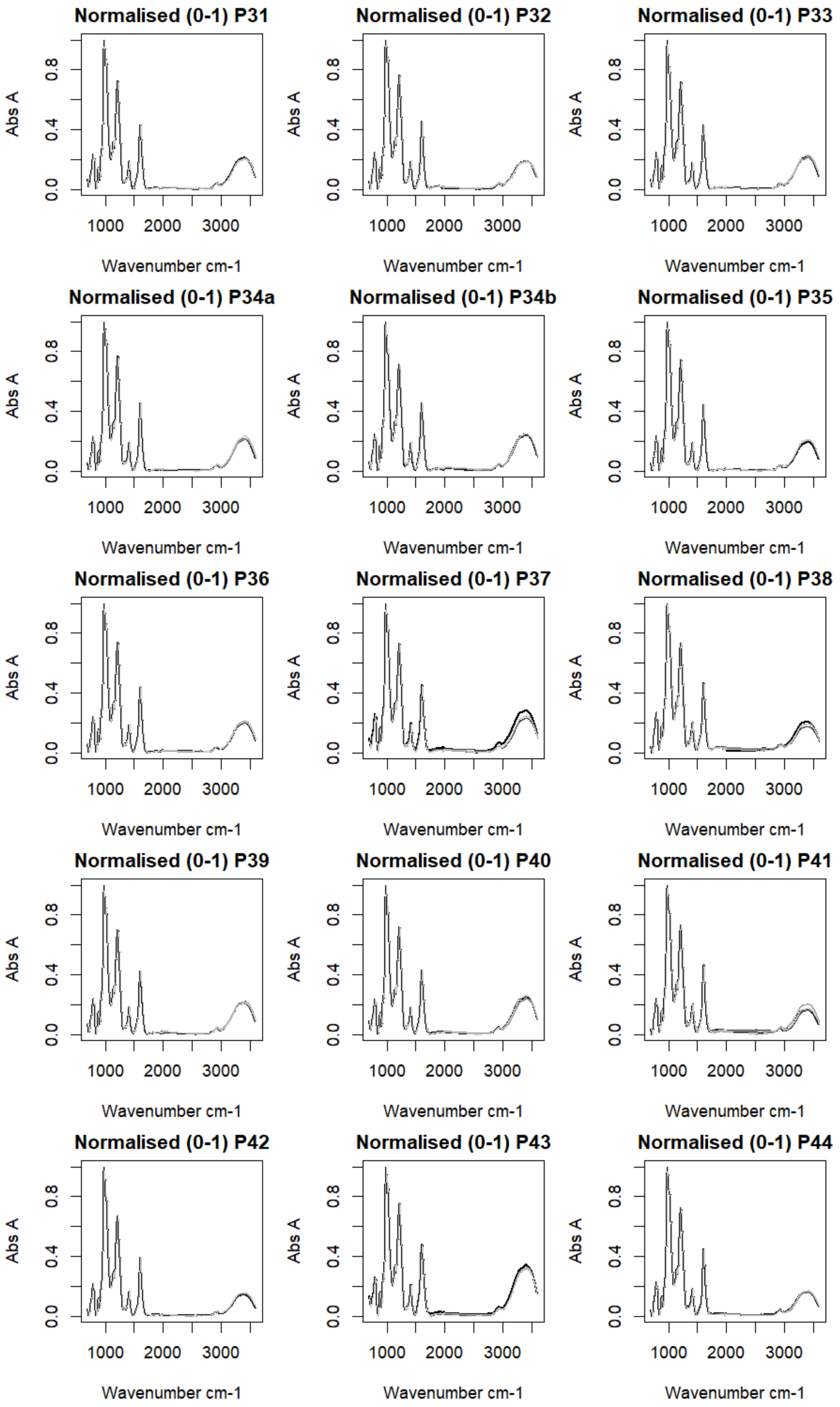
Normalised (0-1) P31
Normalised (0-1) P32
Normalised (0-1) P33
Normalised (0-1) P34a
Normalised (0-1) P34b
Normalised (0-1) P35
Normalised (0-1) P36
Normalised (0-1) P37
Normalised (0-1) P38
Normalised (0-1) P39
Normalised (0-1) P40
Normalised (0-1) P41
Normalised (0-1) P42
Normalised (0-1) P43
Normalised (0-1) P44
Abs A
0.0
0.4
0.8
1000
2000
3000
Wavenumber cm-1

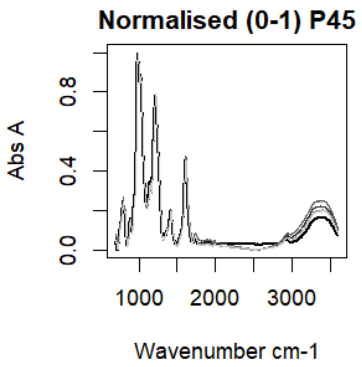
Normalised (0-1) P45
Abs A
0.0 0.4 0.8
1000 2000 3000
Wavenumber cm-1

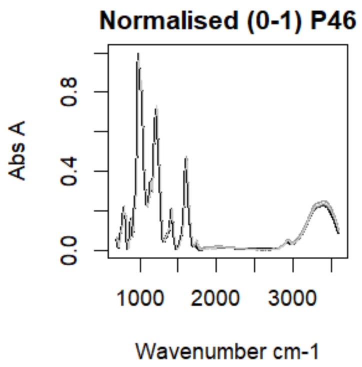
Normalised (0-1) P46
Abs A
0.0 0.4 0.8
1000 2000 3000
Wavenumber cm-1

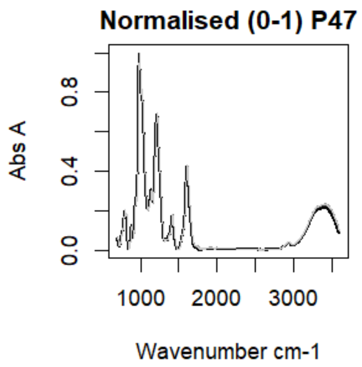
Normalised (0-1) P47
Abs A
0.0 0.4 0.8
1000 2000 3000
Wavenumber cm-1

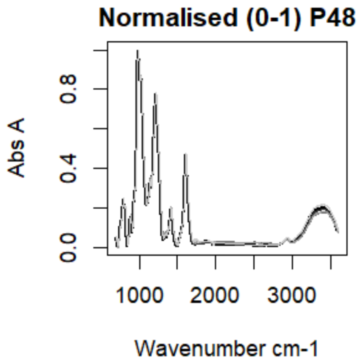
Normalised (0-1) P48
Abs A
0.0 0.4 0.8
1000 2000 3000
Wavenumber cm-1

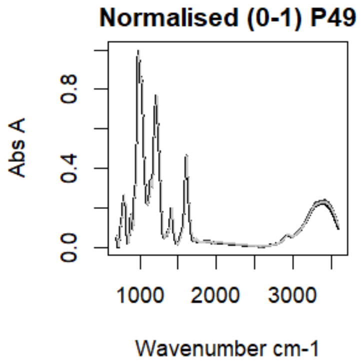
Normalised (0-1) P49
Abs A
0.0 0.4 0.8
1000 2000 3000
Wavenumber cm-1

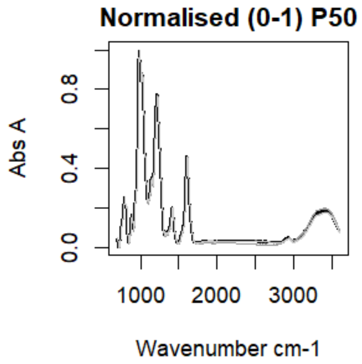
Normalised (0-1) P50
Abs A
0.0 0.4 0.8
1000 2000 3000
Wavenumber cm-1

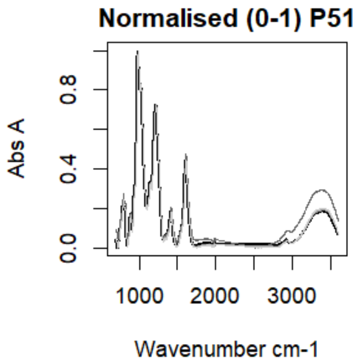
Normalised (0-1) P51
Abs A
0.0 0.4 0.8
1000 2000 3000
Wavenumber cm-1

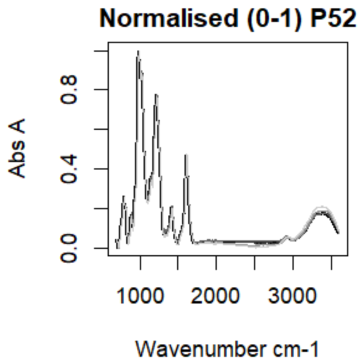
Normalised (0-1) P52
Abs A
0.0 0.4 0.8
1000 2000 3000
Wavenumber cm-1

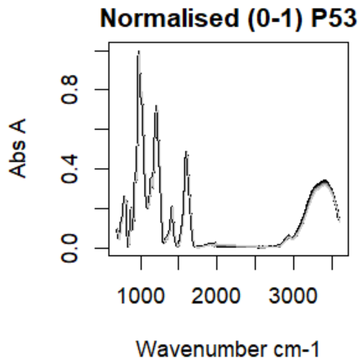
Normalised (0-1) P53
Abs A
0.0 0.4 0.8
1000 2000 3000
Wavenumber cm-1

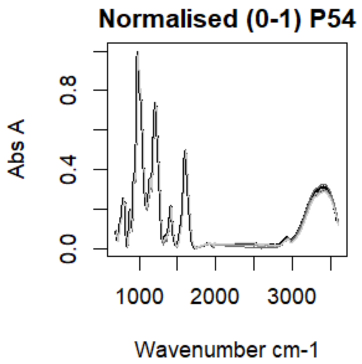
Normalised (0-1) P54
Abs A
0.0 0.4 0.8
1000 2000 3000
Wavenumber cm-1

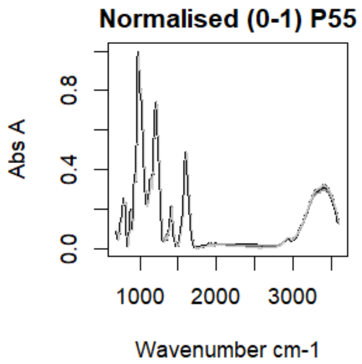
Normalised (0-1) P55
Abs A
0.0 0.4 0.8
1000 2000 3000
Wavenumber cm-1

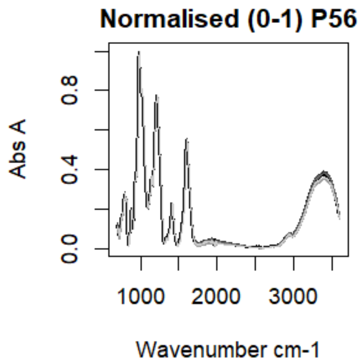
Normalised (0-1) P56
Abs A
0.0 0.4 0.8
1000 2000 3000
Wavenumber cm-1

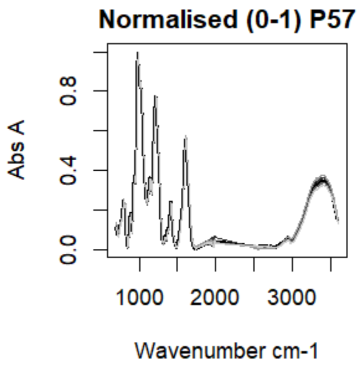
Normalised (0-1) P57
Abs A
0.0 0.4 0.8
1000 2000 3000
Wavenumber cm-1

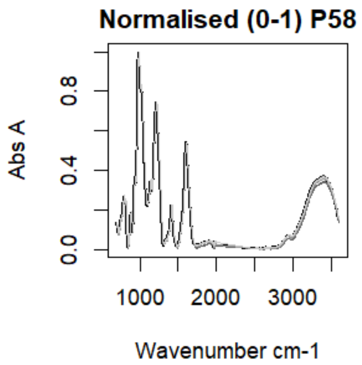
Normalised (0-1) P58
Abs A
0.0 0.4 0.8
1000 2000 3000
Wavenumber cm-1

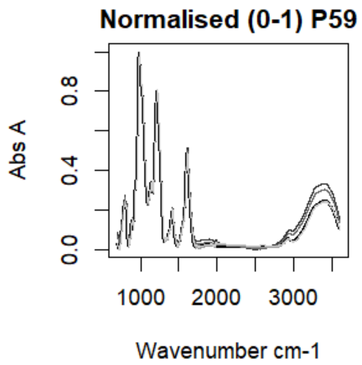
Normalised (0-1) P59
Abs A
0.0 0.4 0.8
1000 2000 3000
Wavenumber cm-1

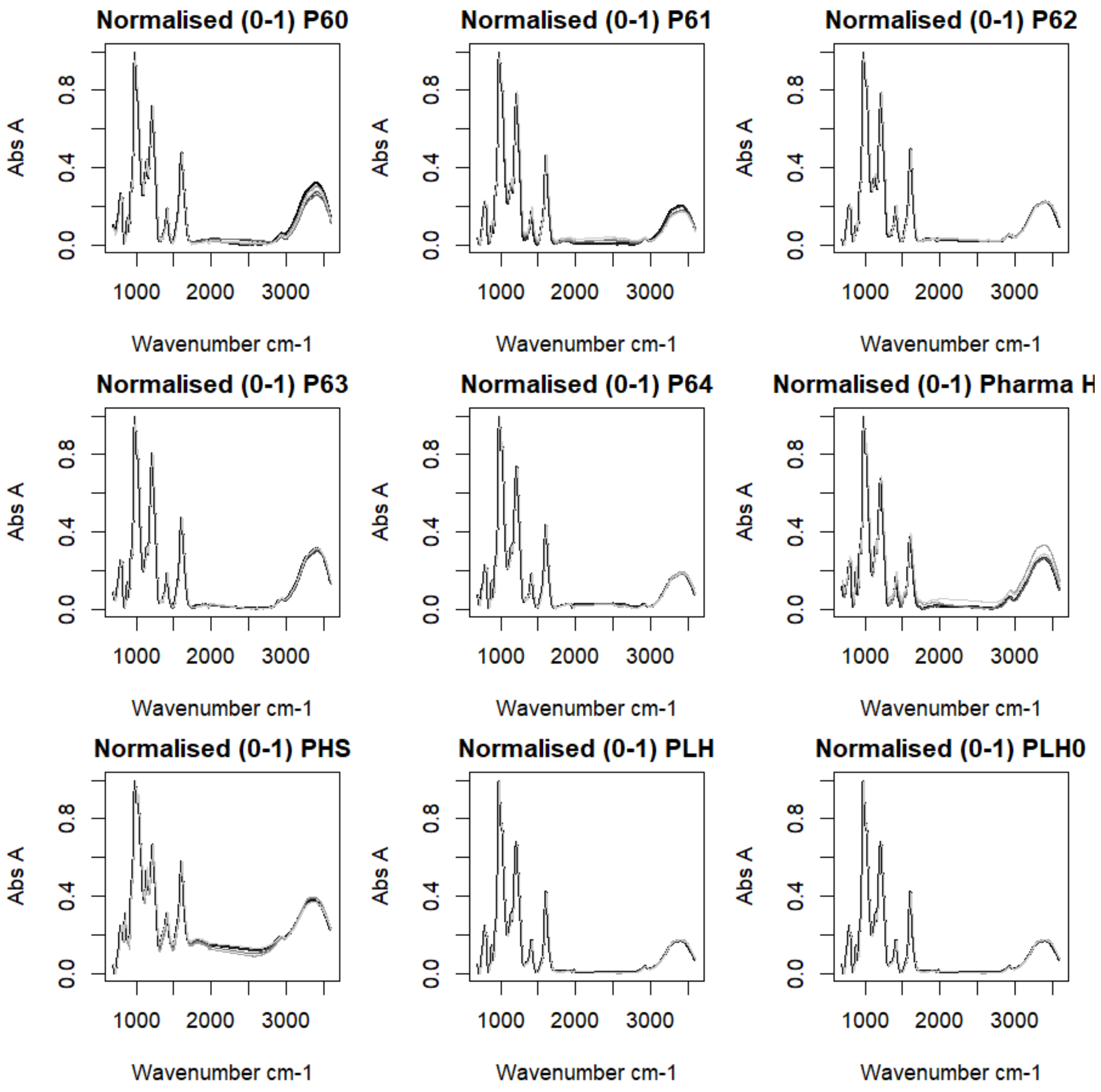
Normalised (0-1) P60
Normalised (0-1) P61
Normalised (0-1) P62
Normalised (0-1) P63
Normalised (0-1) P64
Normalised (0-1) Pharma H
Normalised (0-1) PHS
Normalised (0-1) PLH
Normalised (0-1) PLH0
Abs A
0.0
0.4
0.8
1000
2000
3000
Wavenumber cm-1

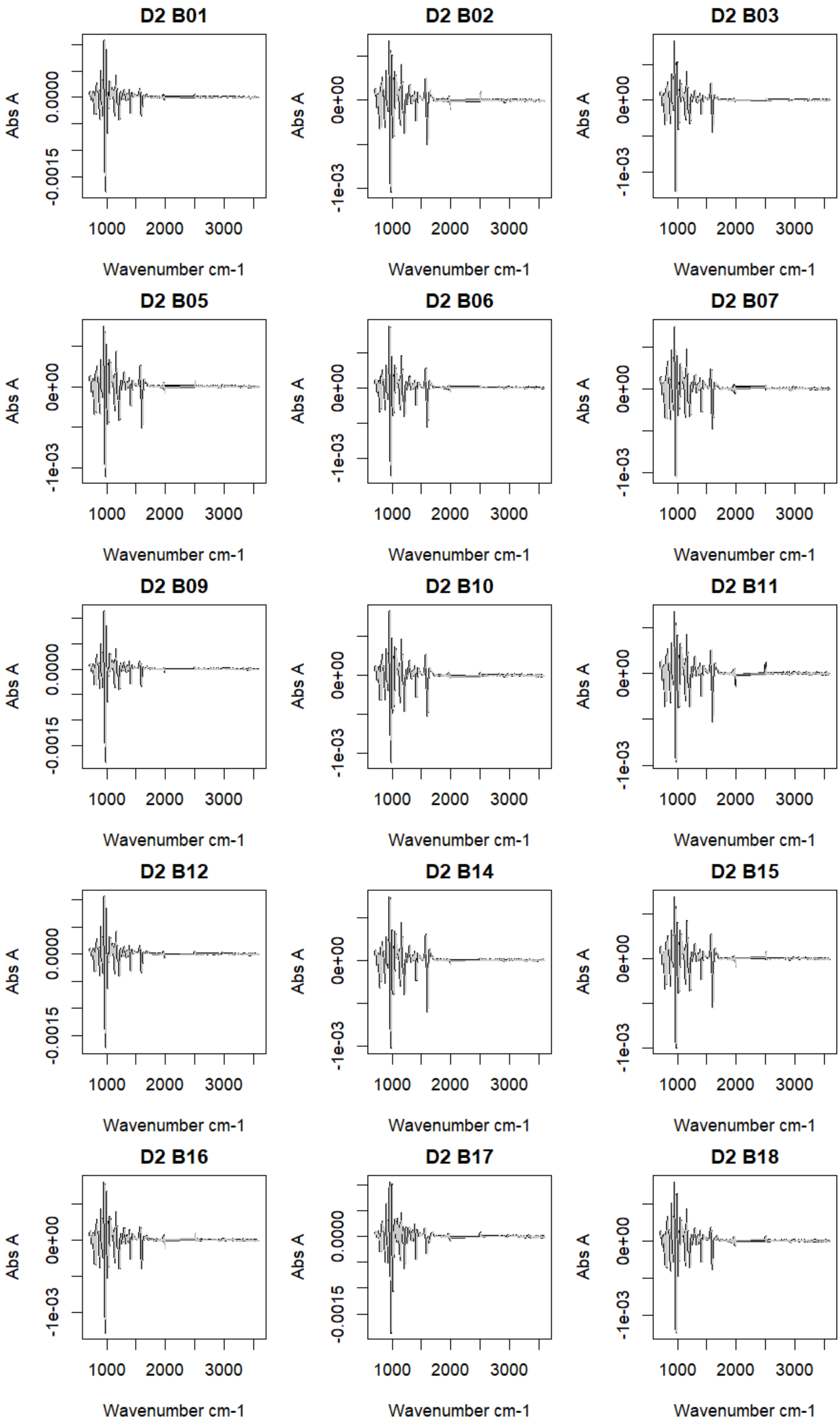
D2 B01
Abs A
0.0000
-0.0015
1000 2000 3000
Wavenumber cm-1
D2 B02
Abs A
0e+00
-1e-03
1000 2000 3000
Wavenumber cm-1
D2 B03
Abs A
0e+00
-1e-03
1000 2000 3000
Wavenumber cm-1
D2 B05
Abs A
0e+00
-1e-03
1000 2000 3000
Wavenumber cm-1
D2 B06
Abs A
0e+00
-1e-03
1000 2000 3000
Wavenumber cm-1
D2 B07
Abs A
0e+00
-1e-03
1000 2000 3000
Wavenumber cm-1
D2 B09
Abs A
0.0000
-0.0015
1000 2000 3000
Wavenumber cm-1
D2 B10
Abs A
0e+00
-1e-03
1000 2000 3000
Wavenumber cm-1
D2 B11
Abs A
0e+00
-1e-03
1000 2000 3000
Wavenumber cm-1
D2 B12
Abs A
0.0000
-0.0015
1000 2000 3000
Wavenumber cm-1
D2 B14
Abs A
0e+00
-1e-03
1000 2000 3000
Wavenumber cm-1
D2 B15
Abs A
0e+00
-1e-03
1000 2000 3000
Wavenumber cm-1
D2 B16
Abs A
0e+00
-1e-03
1000 2000 3000
Wavenumber cm-1
D2 B17
Abs A
0.0000
-0.0015
1000 2000 3000
Wavenumber cm-1
D2 B18
Abs A
0e+00
-1e-03
1000 2000 3000
Wavenumber cm-1

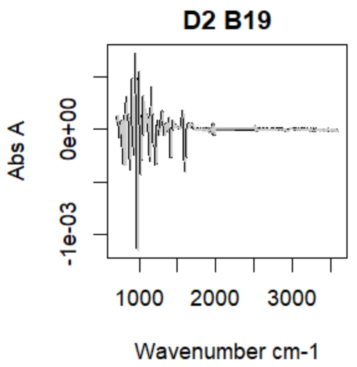
D2 B19
Abs A
0e+00
-1e-03
1000 2000 3000
Wavenumber cm-1

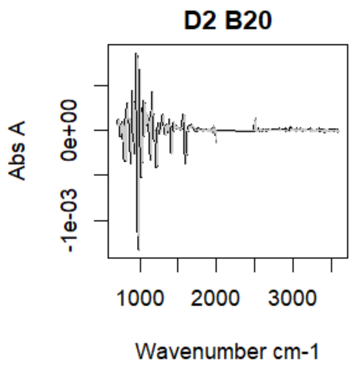
D2 B20
Abs A
0e+00
-1e-03
1000 2000 3000
Wavenumber cm-1

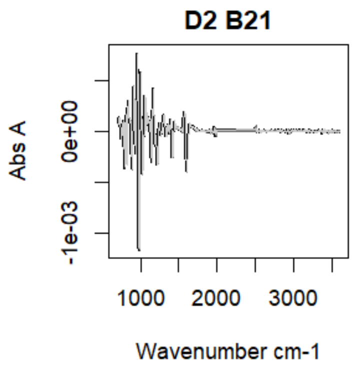
D2 B21
Abs A
0e+00
-1e-03
1000 2000 3000
Wavenumber cm-1

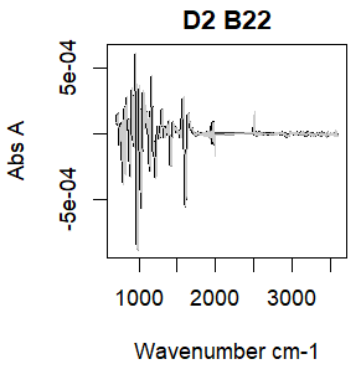
D2 B22
Abs A
5e-04
-5e-04
1000 2000 3000
Wavenumber cm-1

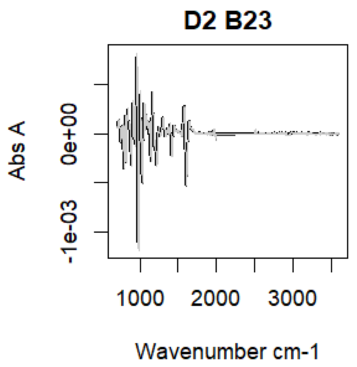
D2 B23
Abs A
0e+00
-1e-03
1000 2000 3000
Wavenumber cm-1

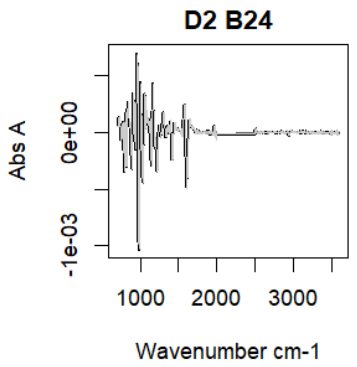
D2 B24
Abs A
0e+00
-1e-03
1000 2000 3000
Wavenumber cm-1

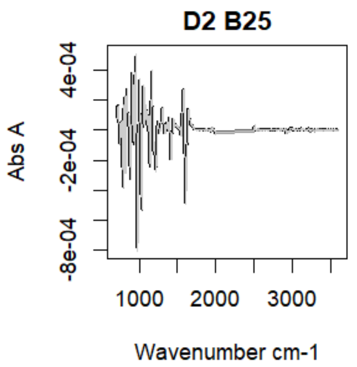
D2 B25
Abs A
4e-04
-2e-04
-8e-04
1000 2000 3000
Wavenumber cm-1

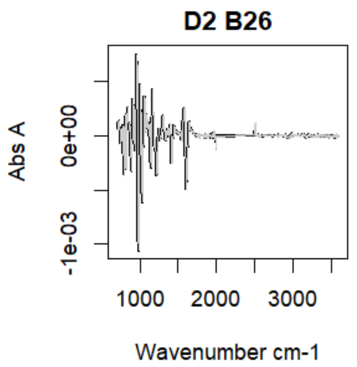
D2 B26
Abs A
0e+00
-1e-03
1000 2000 3000
Wavenumber cm-1

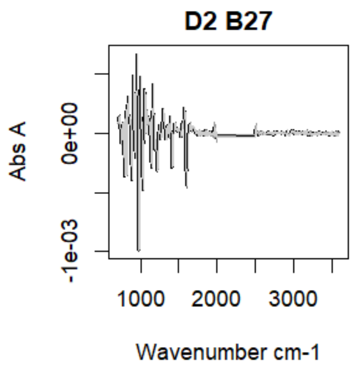
D2 B27
Abs A
0e+00
-1e-03
1000 2000 3000
Wavenumber cm-1

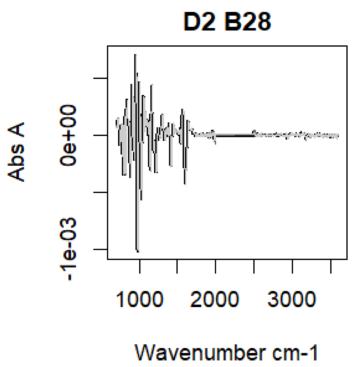
D2 B28
Abs A
0e+00
-1e-03
1000 2000 3000
Wavenumber cm-1

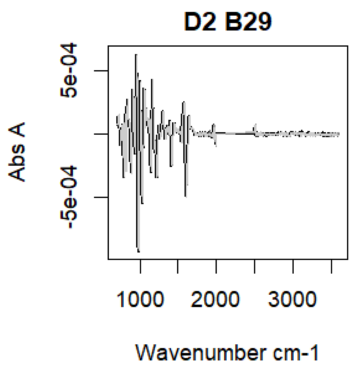
D2 B29
Abs A
5e-04
-5e-04
1000 2000 3000
Wavenumber cm-1

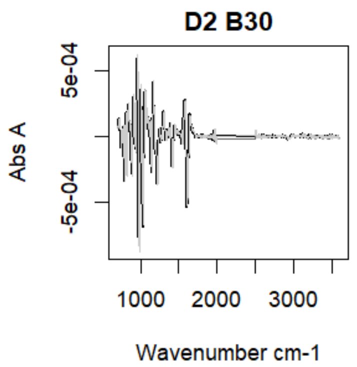
D2 B30
Abs A
5e-04
-5e-04
1000 2000 3000
Wavenumber cm-1

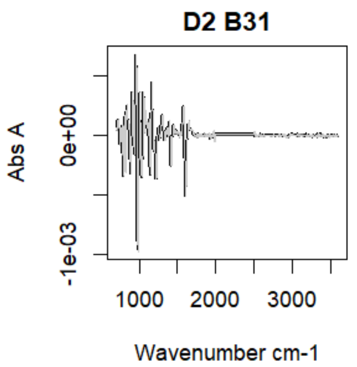
D2 B31
Abs A
0e+00
-1e-03
1000 2000 3000
Wavenumber cm-1

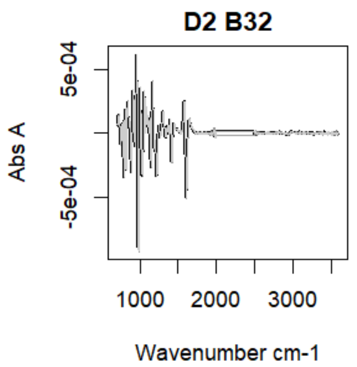
D2 B32
Abs A
5e-04
-5e-04
1000 2000 3000
Wavenumber cm-1

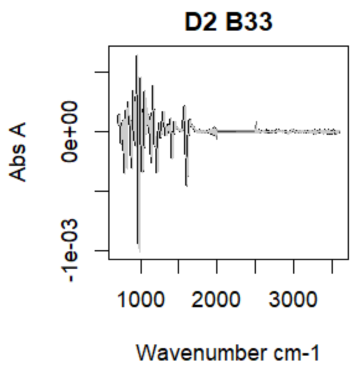
D2 B33
Abs A
0e+00
-1e-03
1000 2000 3000
Wavenumber cm-1

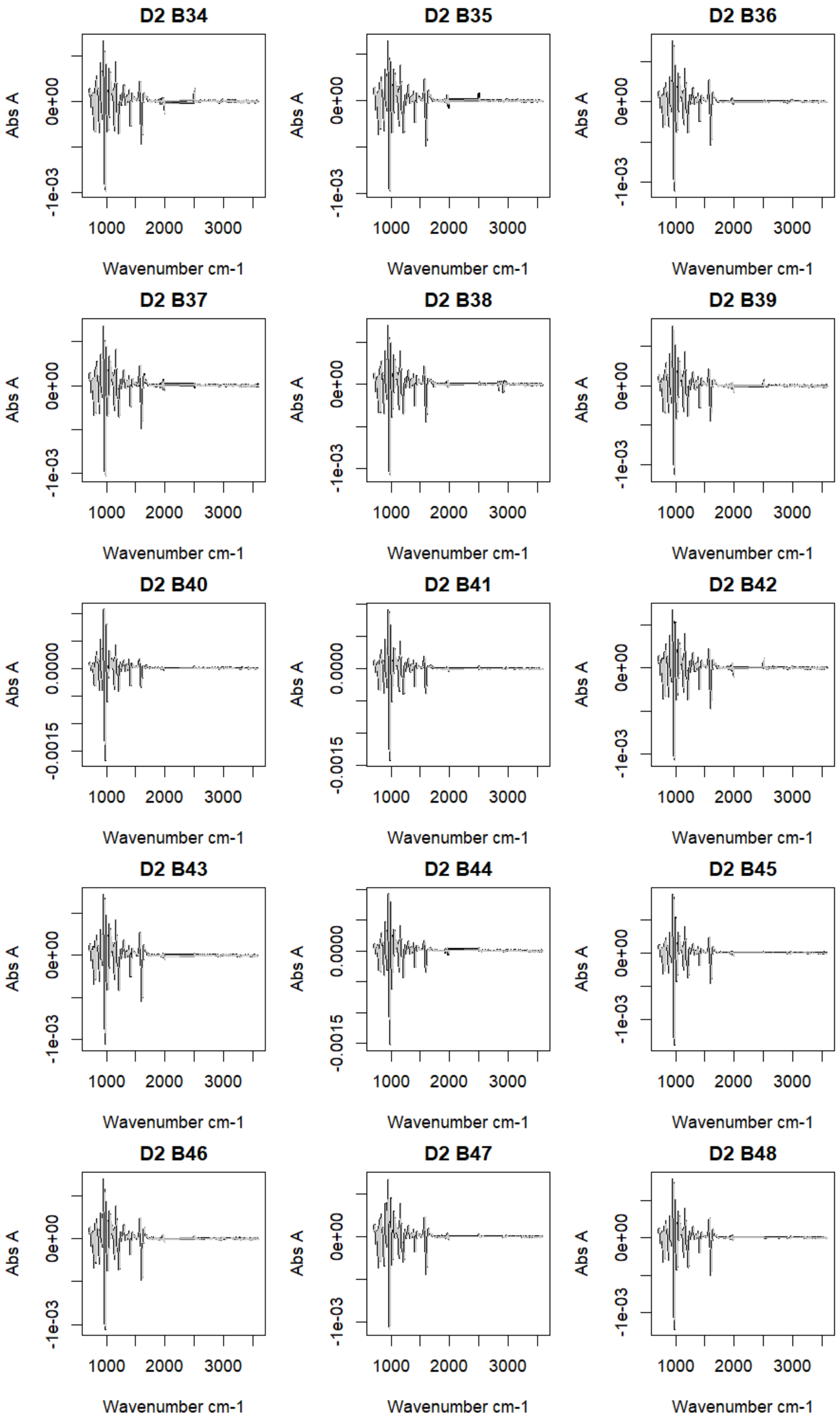
D2 B34
D2 B35
D2 B36
D2 B37
D2 B38
D2 B39
D2 B40
D2 B41
D2 B42
D2 B43
D2 B44
D2 B45
D2 B46
D2 B47
D2 B48
Abs A
0e+00
-1e-03
0.0000
-0.0015
1000
2000
3000
Wavenumber cm-1

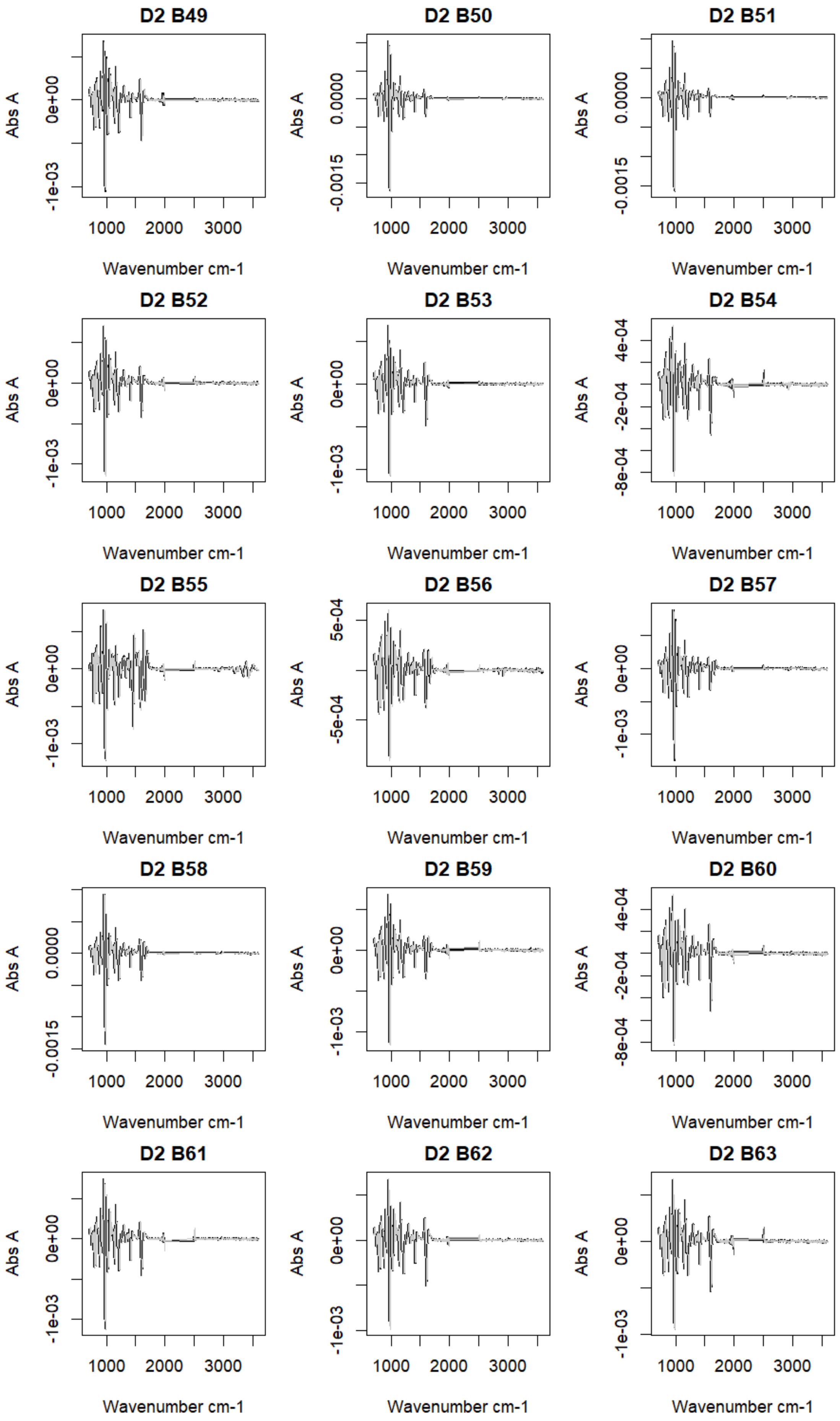
D2 B49
Abs A
0e+00
-1e-03
1000 2000 3000
Wavenumber cm-1
D2 B50
Abs A
0.0000
-0.0015
1000 2000 3000
Wavenumber cm-1
D2 B51
Abs A
0.0000
-0.0015
1000 2000 3000
Wavenumber cm-1
D2 B52
Abs A
0e+00
-1e-03
1000 2000 3000
Wavenumber cm-1
D2 B53
Abs A
0e+00
-1e-03
1000 2000 3000
Wavenumber cm-1
D2 B54
Abs A
4e-04
-2e-04
-8e-04
1000 2000 3000
Wavenumber cm-1
D2 B55
Abs A
0e+00
-1e-03
1000 2000 3000
Wavenumber cm-1
D2 B56
Abs A
5e-04
-5e-04
1000 2000 3000
Wavenumber cm-1
D2 B57
Abs A
0e+00
-1e-03
1000 2000 3000
Wavenumber cm-1
D2 B58
Abs A
0.0000
-0.0015
1000 2000 3000
Wavenumber cm-1
D2 B59
Abs A
0e+00
-1e-03
1000 2000 3000
Wavenumber cm-1
D2 B60
Abs A
4e-04
-2e-04
-8e-04
1000 2000 3000
Wavenumber cm-1
D2 B61
Abs A
0e+00
-1e-03
1000 2000 3000
Wavenumber cm-1
D2 B62
Abs A
0e+00
-1e-03
1000 2000 3000
Wavenumber cm-1
D2 B63
Abs A
0e+00
-1e-03
1000 2000 3000
Wavenumber cm-1

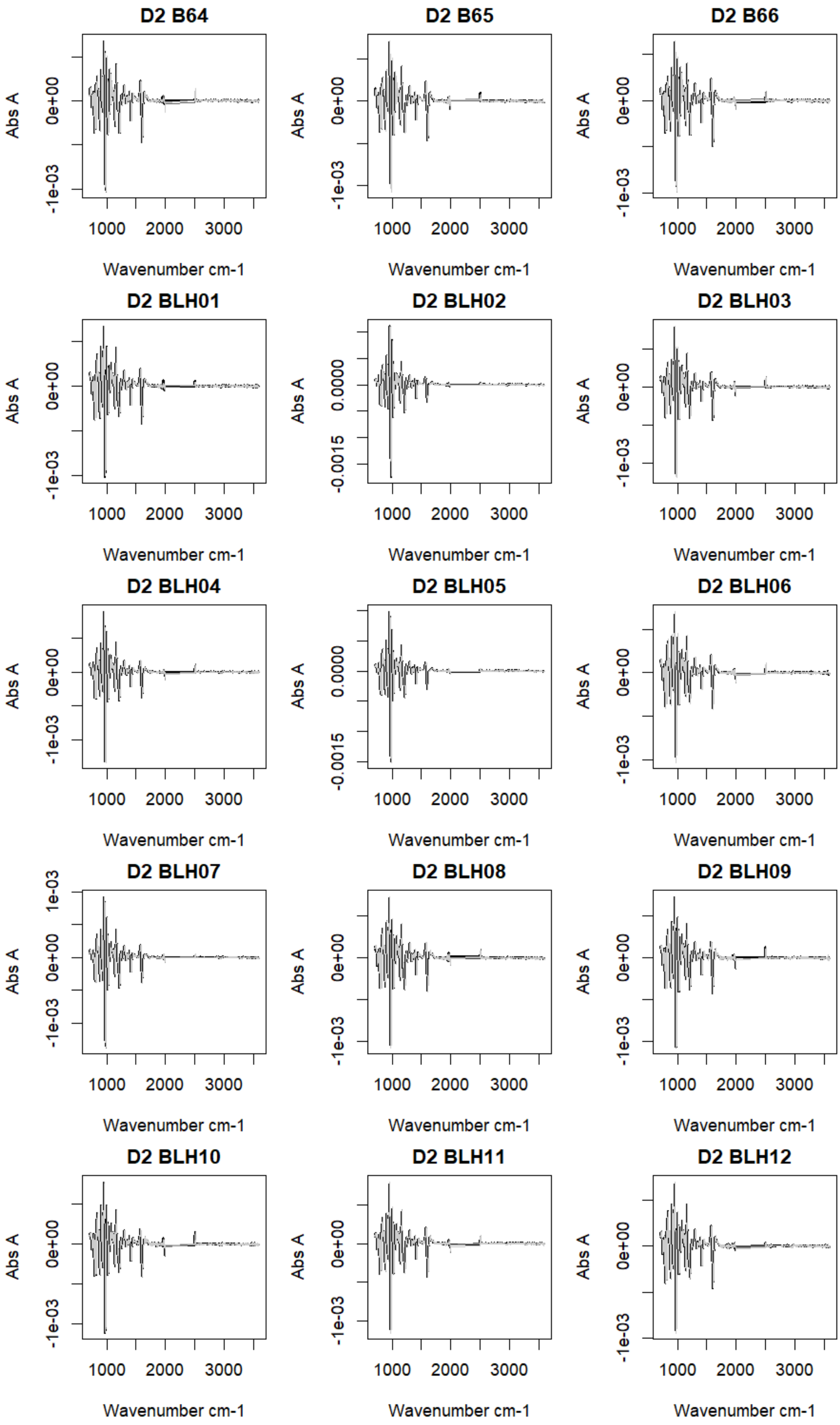
D2 B64
Abs A
0e+00
-1e-03
1000 2000 3000
Wavenumber cm-1
D2 B65
Abs A
0e+00
-1e-03
1000 2000 3000
Wavenumber cm-1
D2 B66
Abs A
0e+00
-1e-03
1000 2000 3000
Wavenumber cm-1
D2 BLH01
Abs A
0e+00
-1e-03
1000 2000 3000
Wavenumber cm-1
D2 BLH02
Abs A
0.0000
-0.0015
1000 2000 3000
Wavenumber cm-1
D2 BLH03
Abs A
0e+00
-1e-03
1000 2000 3000
Wavenumber cm-1
D2 BLH04
Abs A
0e+00
-1e-03
1000 2000 3000
Wavenumber cm-1
D2 BLH05
Abs A
0.0000
-0.0015
1000 2000 3000
Wavenumber cm-1
D2 BLH06
Abs A
0e+00
-1e-03
1000 2000 3000
Wavenumber cm-1
D2 BLH07
Abs A
1e-03
0e+00
-1e-03
1000 2000 3000
Wavenumber cm-1
D2 BLH08
Abs A
0e+00
-1e-03
1000 2000 3000
Wavenumber cm-1
D2 BLH09
Abs A
0e+00
-1e-03
1000 2000 3000
Wavenumber cm-1
D2 BLH10
Abs A
0e+00
-1e-03
1000 2000 3000
Wavenumber cm-1
D2 BLH11
Abs A
0e+00
-1e-03
1000 2000 3000
Wavenumber cm-1
D2 BLH12
Abs A
0e+00
-1e-03
1000 2000 3000
Wavenumber cm-1

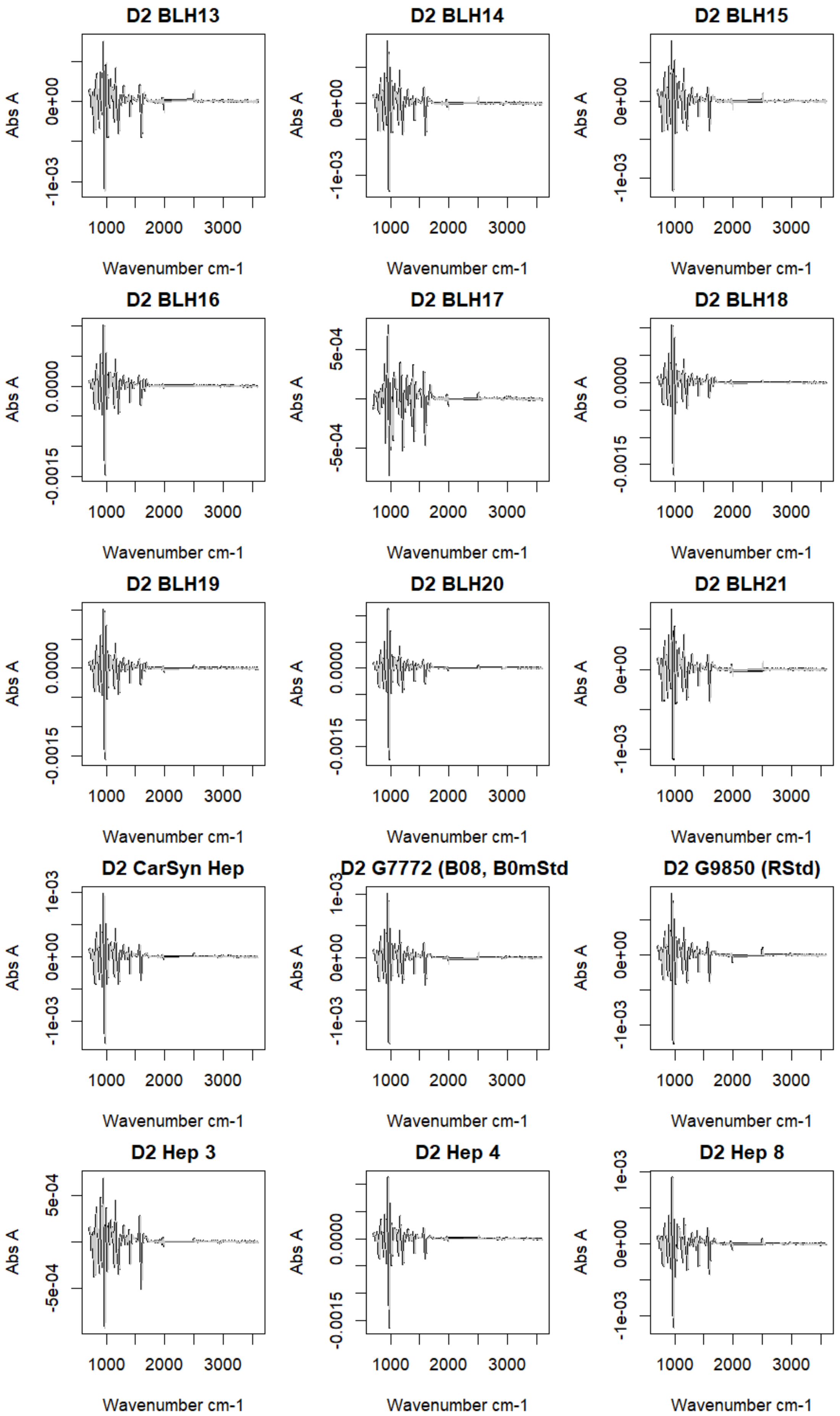
D2 BLH13
D2 BLH14
D2 BLH15
D2 BLH16
D2 BLH17
D2 BLH18
D2 BLH19
D2 BLH20
D2 BLH21
D2 CarSyn Hep
D2 G7772 (B08, B0mStd
D2 G9850 (RStd)
D2 Hep 3
D2 Hep 4
D2 Hep 8
Abs A
Wavenumber cm-1
1000
2000
3000
0e+00
-1e-03
1e-03
0.0000
-0.0015
5e-04
-5e-04

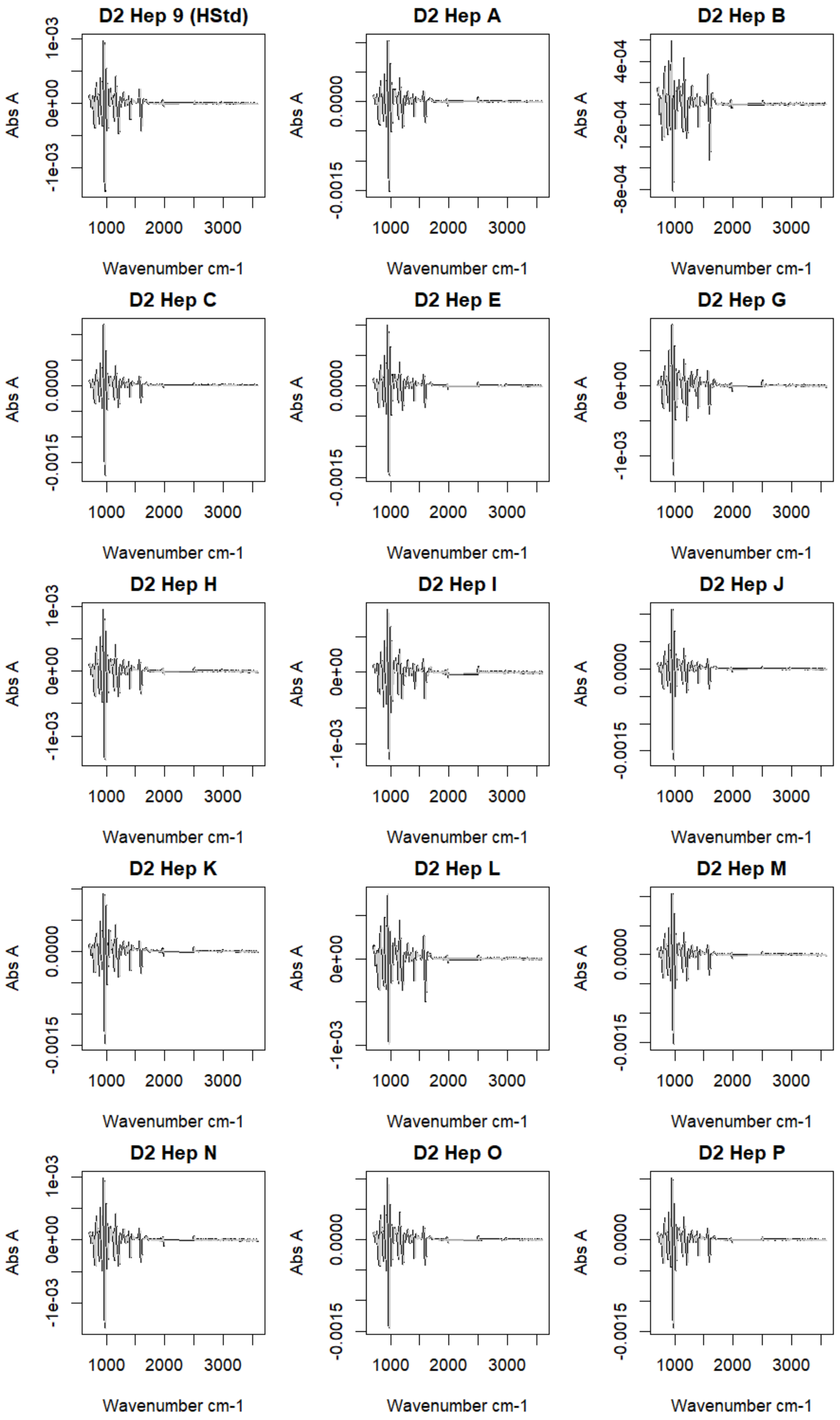

D2 Hep 9 (HStd)
D2 Hep A
D2 Hep B
D2 Hep C
D2 Hep E
D2 Hep G
D2 Hep H
D2 Hep I
D2 Hep J
D2 Hep K
D2 Hep L
D2 Hep M
D2 Hep N
D2 Hep O
D2 Hep P
Abs A
Wavenumber cm-1
1000
2000
3000
1e-03
0e+00
-1e-03
0.0000
-0.0015
4e-04
-2e-04
-8e-04

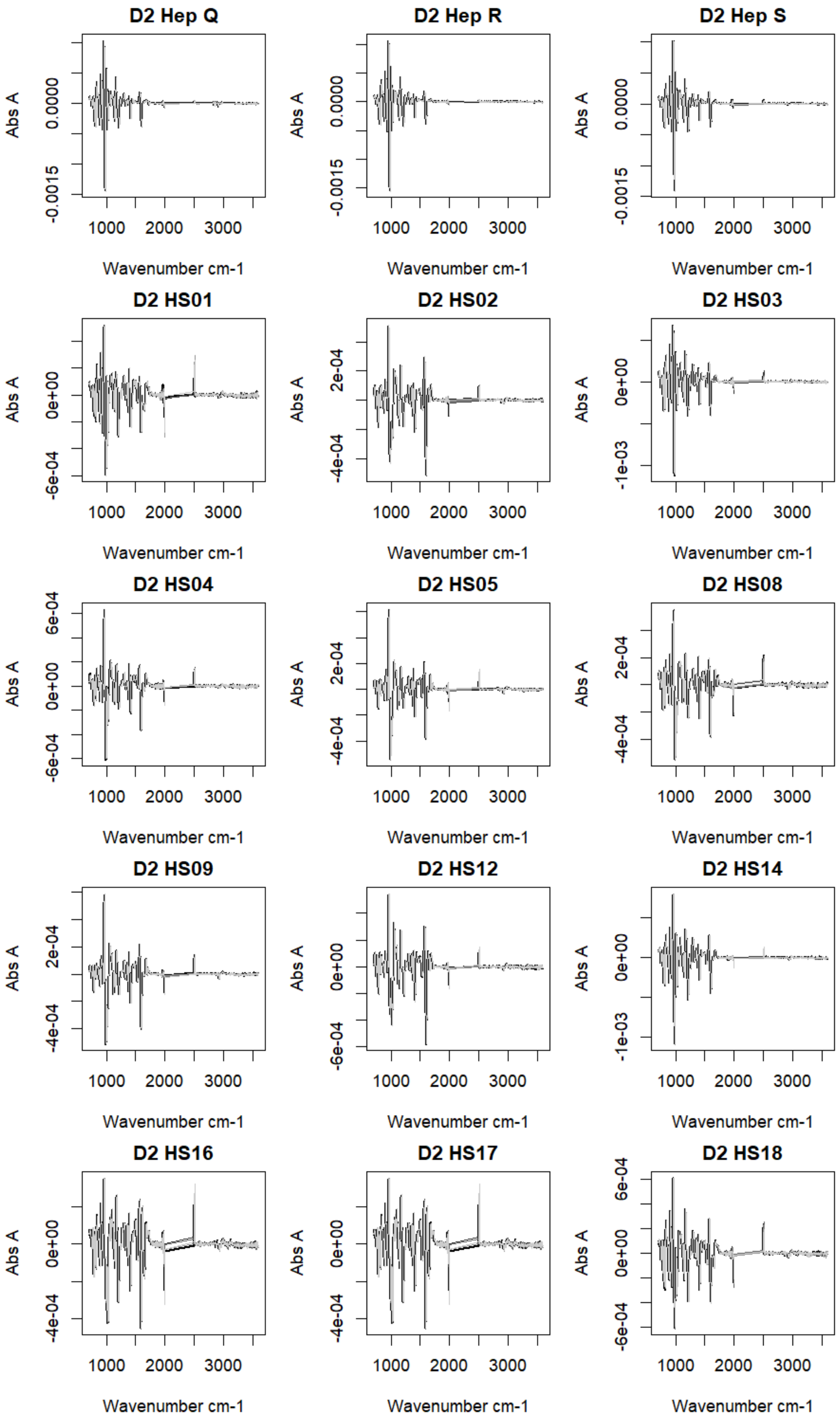
D2 Hep Q
D2 Hep R
D2 Hep S
D2 HS01
D2 HS02
D2 HS03
D2 HS04
D2 HS05
D2 HS08
D2 HS09
D2 HS12
D2 HS14
D2 HS16
D2 HS17
D2 HS18
Abs A
Wavenumber cm-1
1000
2000
3000
0.0000
-0.0015
0e+00
2e-04
-4e-04
6e-04
-6e-04
-1e-03

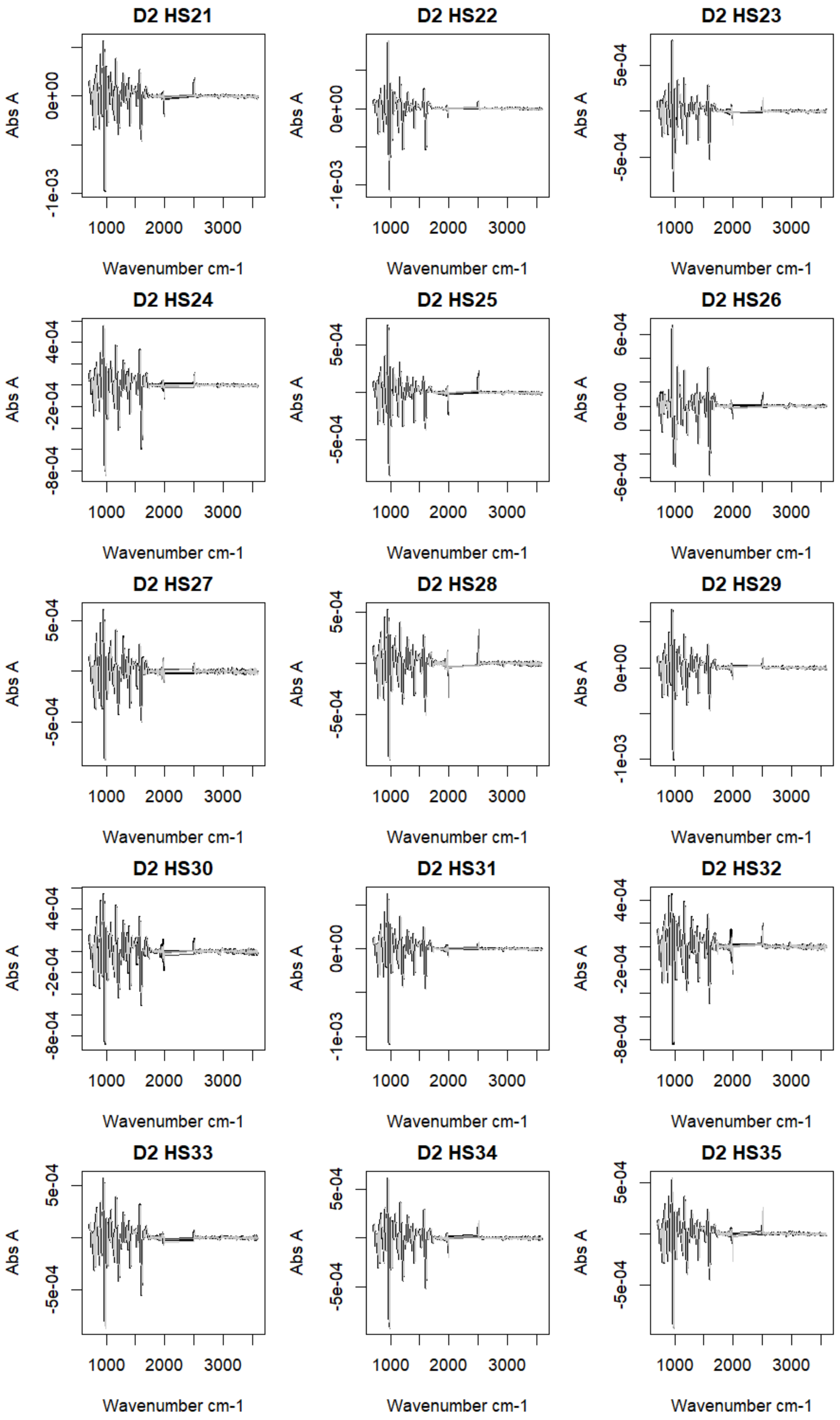
D2 HS21
Abs A
0e+00
-1e-03
1000 2000 3000
Wavenumber cm-1
D2 HS22
Abs A
0e+00
-1e-03
1000 2000 3000
Wavenumber cm-1
D2 HS23
Abs A
5e-04
-5e-04
1000 2000 3000
Wavenumber cm-1
D2 HS24
Abs A
4e-04
-2e-04
-8e-04
1000 2000 3000
Wavenumber cm-1
D2 HS25
Abs A
5e-04
-5e-04
1000 2000 3000
Wavenumber cm-1
D2 HS26
Abs A
6e-04
0e+00
-6e-04
1000 2000 3000
Wavenumber cm-1
D2 HS27
Abs A
5e-04
-5e-04
1000 2000 3000
Wavenumber cm-1
D2 HS28
Abs A
5e-04
-5e-04
1000 2000 3000
Wavenumber cm-1
D2 HS29
Abs A
0e+00
-1e-03
1000 2000 3000
Wavenumber cm-1
D2 HS30
Abs A
4e-04
-2e-04
-8e-04
1000 2000 3000
Wavenumber cm-1
D2 HS31
Abs A
0e+00
-1e-03
1000 2000 3000
Wavenumber cm-1
D2 HS32
Abs A
4e-04
-2e-04
-8e-04
1000 2000 3000
Wavenumber cm-1
D2 HS33
Abs A
5e-04
-5e-04
1000 2000 3000
Wavenumber cm-1
D2 HS34
Abs A
5e-04
-5e-04
1000 2000 3000
Wavenumber cm-1
D2 HS35
Abs A
5e-04
-5e-04
1000 2000 3000
Wavenumber cm-1

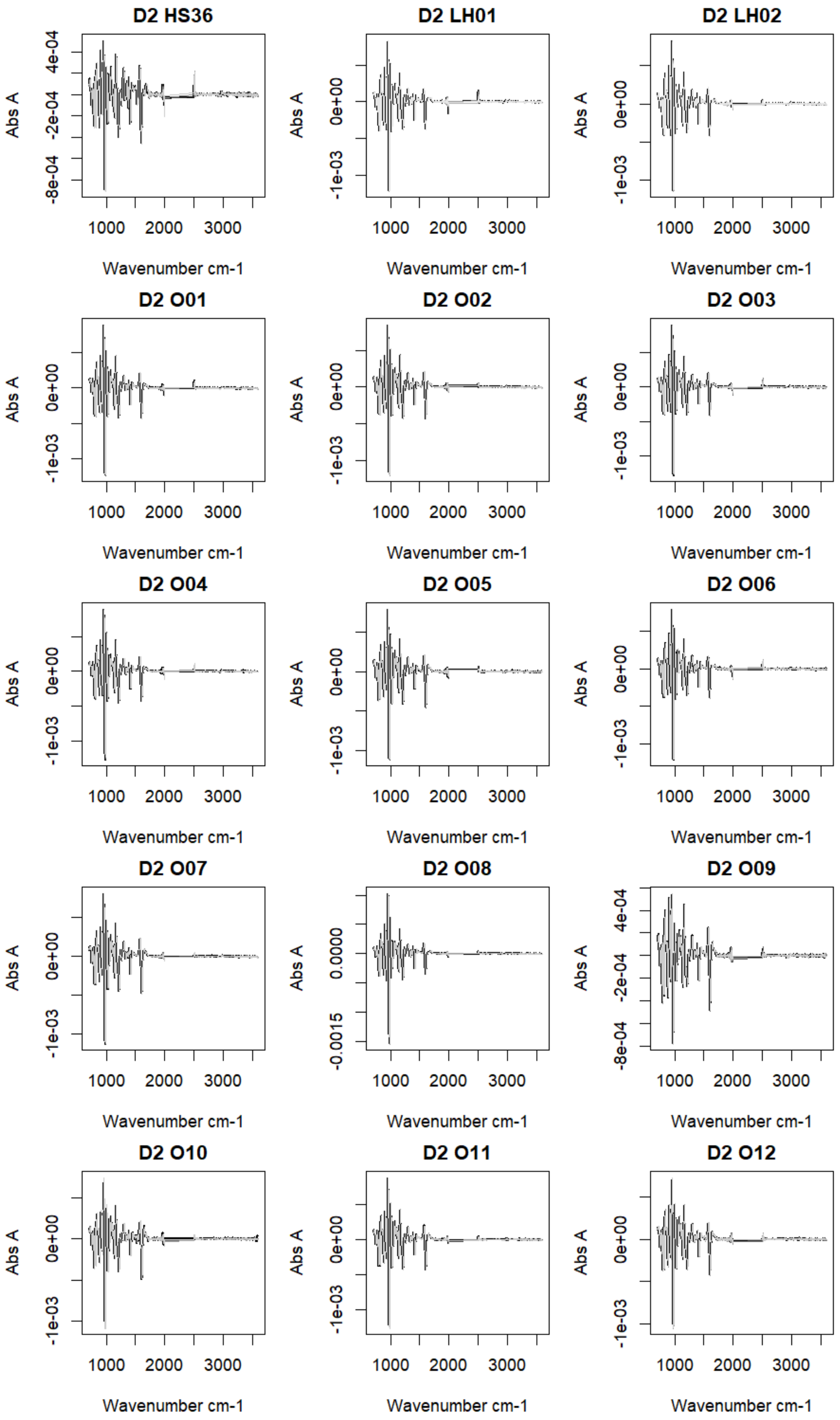
D2 HS36
Abs A
4e-04
-2e-04
-8e-04
1000
2000
3000
Wavenumber cm-1
D2 LH01
Abs A
0e+00
-1e-03
1000
2000
3000
Wavenumber cm-1
D2 LH02
Abs A
0e+00
-1e-03
1000
2000
3000
Wavenumber cm-1
D2 O01
Abs A
0e+00
-1e-03
1000
2000
3000
Wavenumber cm-1
D2 O02
Abs A
0e+00
-1e-03
1000
2000
3000
Wavenumber cm-1
D2 O03
Abs A
0e+00
-1e-03
1000
2000
3000
Wavenumber cm-1
D2 O04
Abs A
0e+00
-1e-03
1000
2000
3000
Wavenumber cm-1
D2 O05
Abs A
0e+00
-1e-03
1000
2000
3000
Wavenumber cm-1
D2 O06
Abs A
0e+00
-1e-03
1000
2000
3000
Wavenumber cm-1
D2 O07
Abs A
0e+00
-1e-03
1000
2000
3000
Wavenumber cm-1
D2 O08
Abs A
0.0000
-0.0015
1000
2000
3000
Wavenumber cm-1
D2 O09
Abs A
4e-04
-2e-04
-8e-04
1000
2000
3000
Wavenumber cm-1
D2 O10
Abs A
0e+00
-1e-03
1000
2000
3000
Wavenumber cm-1
D2 O11
Abs A
0e+00
-1e-03
1000
2000
3000
Wavenumber cm-1
D2 O12
Abs A
0e+00
-1e-03
1000
2000
3000
Wavenumber cm-1

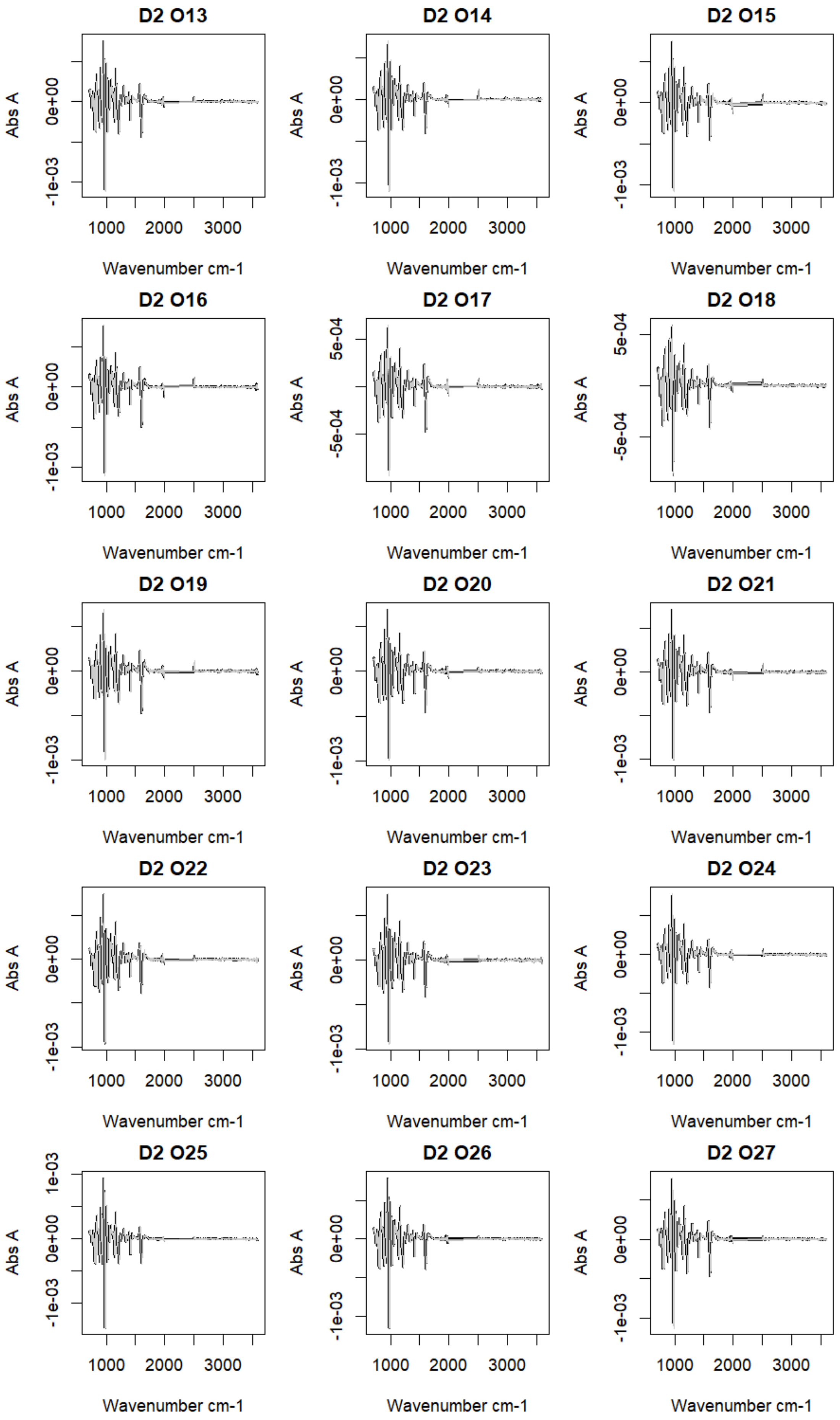
D2 O13
D2 O14
D2 O15
D2 O16
D2 O17
D2 O18
D2 O19
D2 O20
D2 O21
D2 O22
D2 O23
D2 O24
D2 O25
D2 O26
D2 O27
Abs A
Wavenumber cm-1
1000 2000 3000
0e+00
-1e-03
1e-03
5e-04
-5e-04

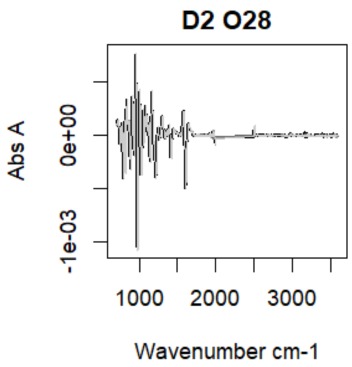
D2 O28
Abs A
0e+00
-1e-03
1000 2000 3000
Wavenumber cm-1

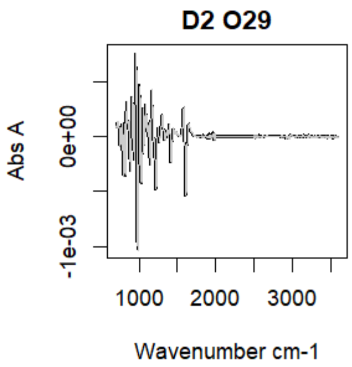
D2 O29
Abs A
0e+00
-1e-03
1000 2000 3000
Wavenumber cm-1

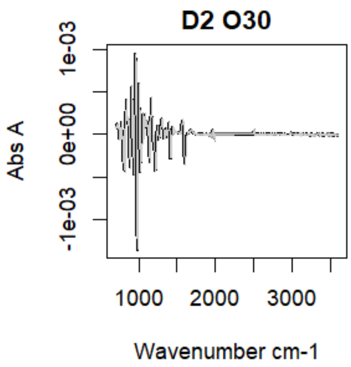
D2 O30
Abs A
1e-03
0e+00
-1e-03
1000 2000 3000
Wavenumber cm-1

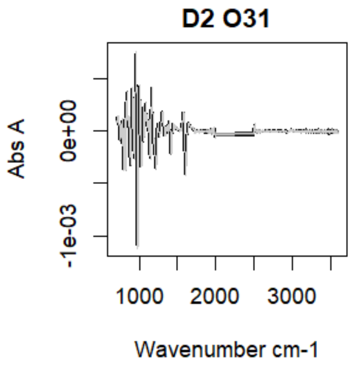
D2 O31
Abs A
0e+00
-1e-03
1000 2000 3000
Wavenumber cm-1

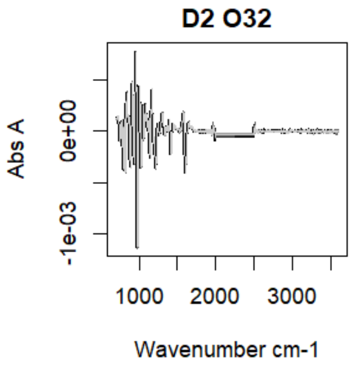
D2 O32
Abs A
0e+00
-1e-03
1000 2000 3000
Wavenumber cm-1

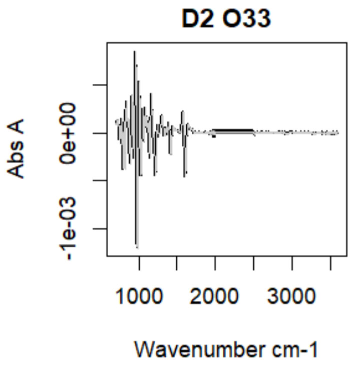
D2 O33
Abs A
0e+00
-1e-03
1000 2000 3000
Wavenumber cm-1

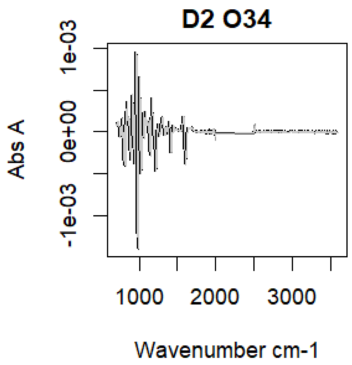
D2 O34
Abs A
1e-03
0e+00
-1e-03
1000 2000 3000
Wavenumber cm-1

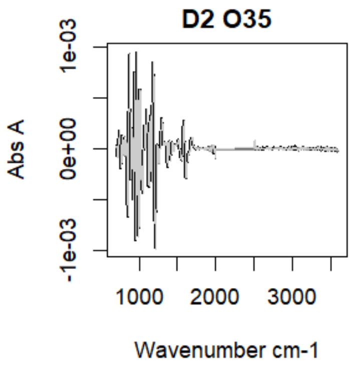
D2 O35
Abs A
1e-03
0e+00
-1e-03
1000 2000 3000
Wavenumber cm-1

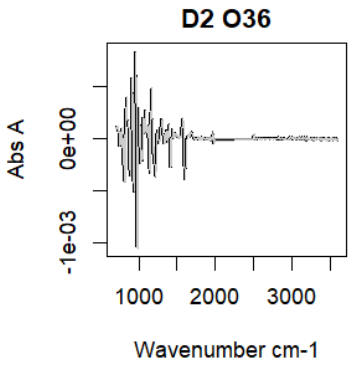
D2 O36
Abs A
0e+00
-1e-03
1000 2000 3000
Wavenumber cm-1

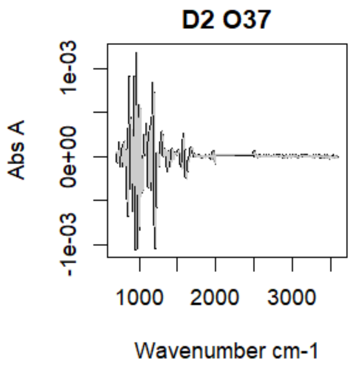
D2 O37
Abs A
1e-03
0e+00
-1e-03
1000 2000 3000
Wavenumber cm-1

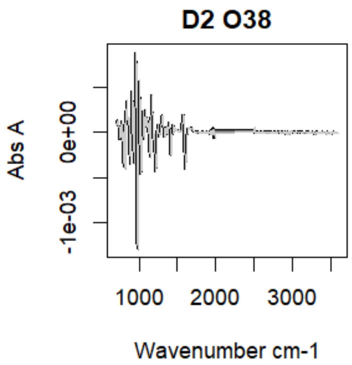
D2 O38
Abs A
0e+00
-1e-03
1000 2000 3000
Wavenumber cm-1

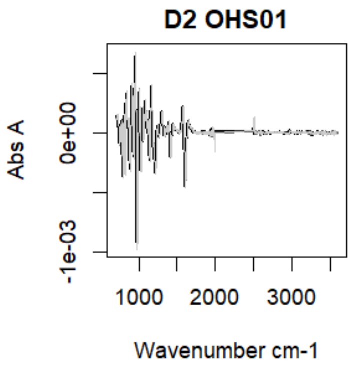
D2 OHS01
Abs A
0e+00
-1e-03
1000 2000 3000
Wavenumber cm-1

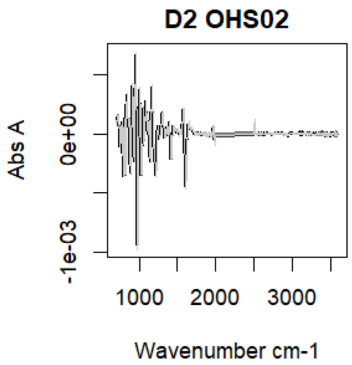
D2 OHS02
Abs A
0e+00
-1e-03
1000 2000 3000
Wavenumber cm-1

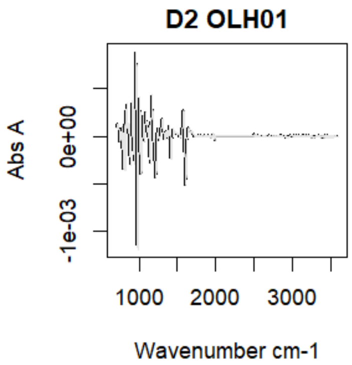
D2 OLH01
Abs A
0e+00
-1e-03
1000 2000 3000
Wavenumber cm-1

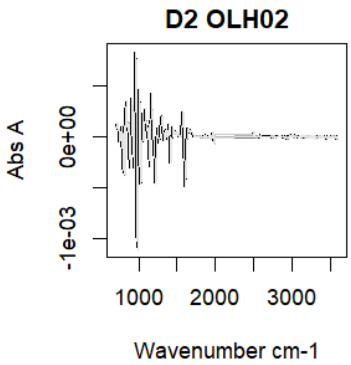
D2 OLH02
Abs A
0e+00
-1e-03
1000 2000 3000
Wavenumber cm-1

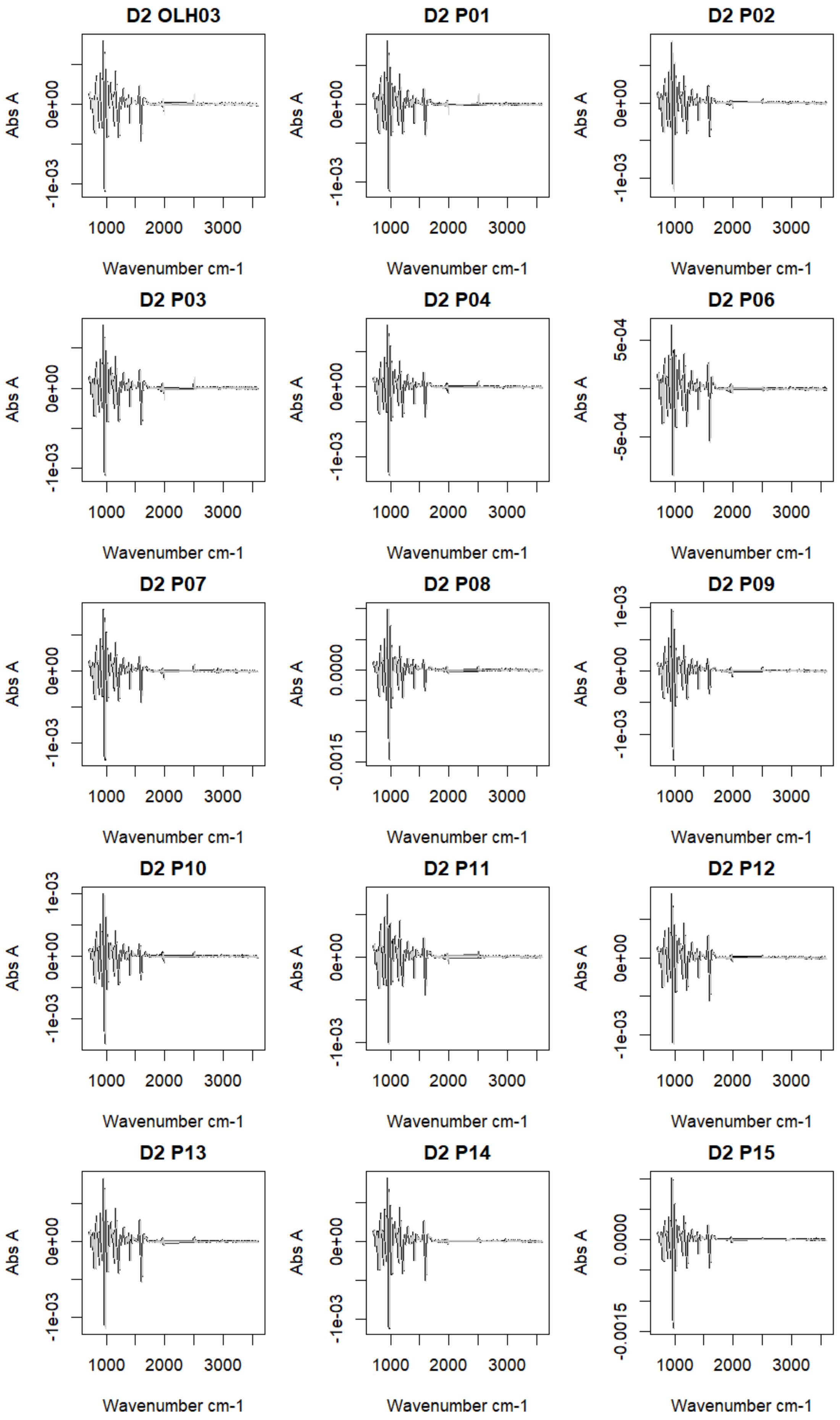
D2 OLH03
D2 P01
D2 P02
D2 P03
D2 P04
D2 P06
D2 P07
D2 P08
D2 P09
D2 P10
D2 P11
D2 P12
D2 P13
D2 P14
D2 P15
Abs A
Wavenumber cm-1
1000 2000 3000
-1e-03 0e+00 1e-03
-5e-04 5e-04
-0.0015 0.0000

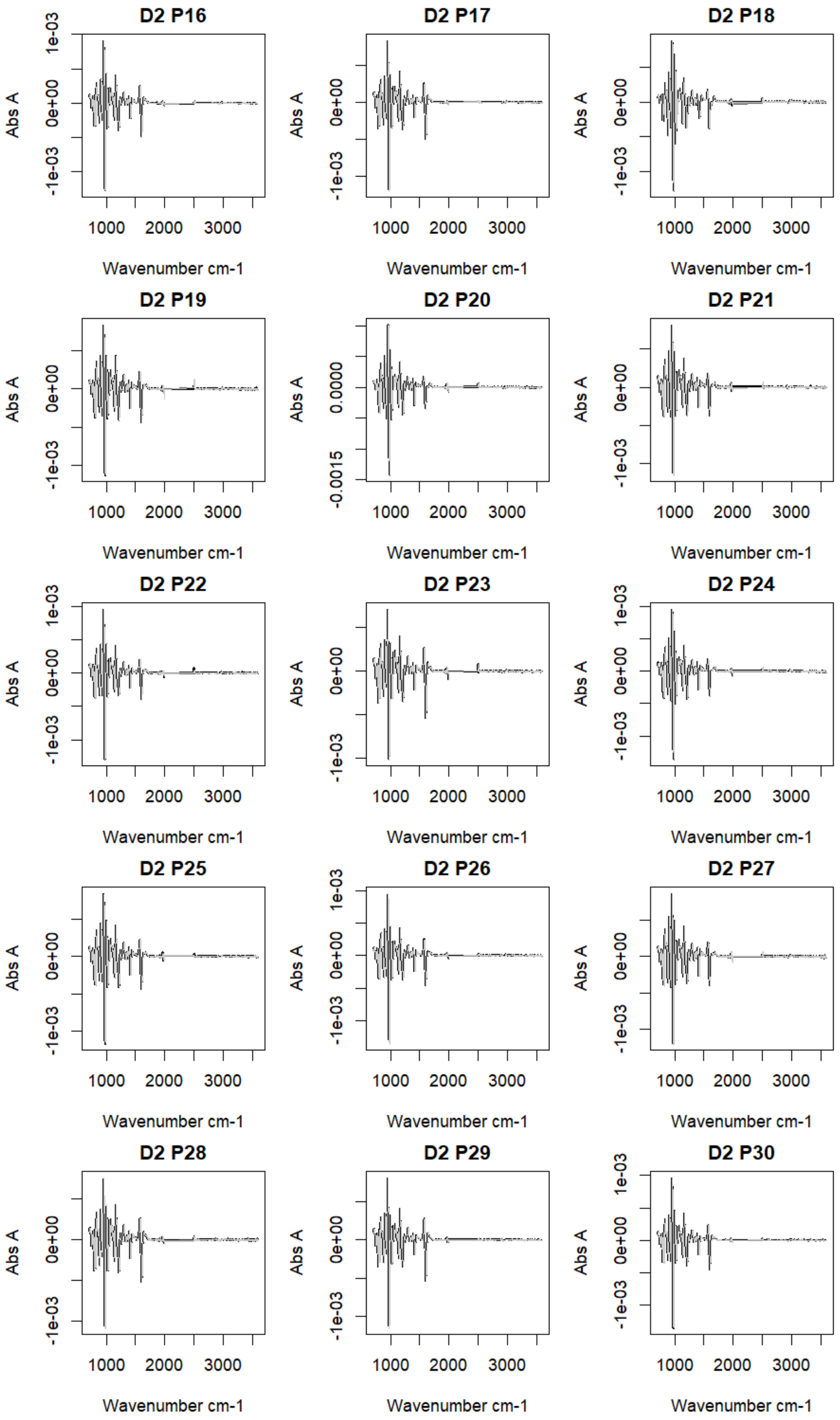
D2 P16
D2 P17
D2 P18
D2 P19
D2 P20
D2 P21
D2 P22
D2 P23
D2 P24
D2 P25
D2 P26
D2 P27
D2 P28
D2 P29
D2 P30
Abs A
Wavenumber cm-1
1000
2000
3000
1e-03
0e+00
-1e-03
0.0000
-0.0015

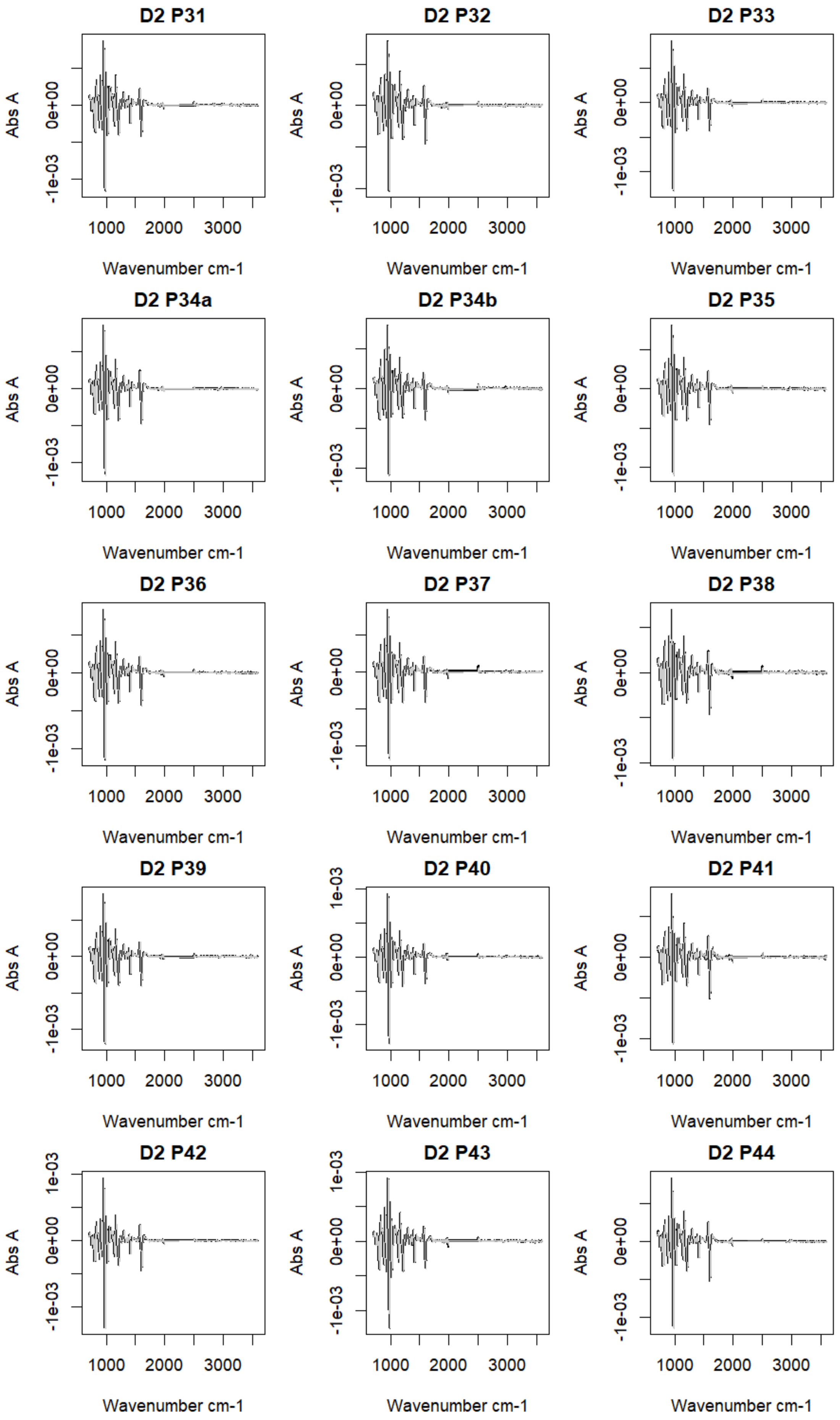
D2 P31
D2 P32
D2 P33
D2 P34a
D2 P34b
D2 P35
D2 P36
D2 P37
D2 P38
D2 P39
D2 P40
D2 P41
D2 P42
D2 P43
D2 P44
Abs A
1e-03
0e+00
-1e-03
1000
2000
3000
Wavenumber cm-1

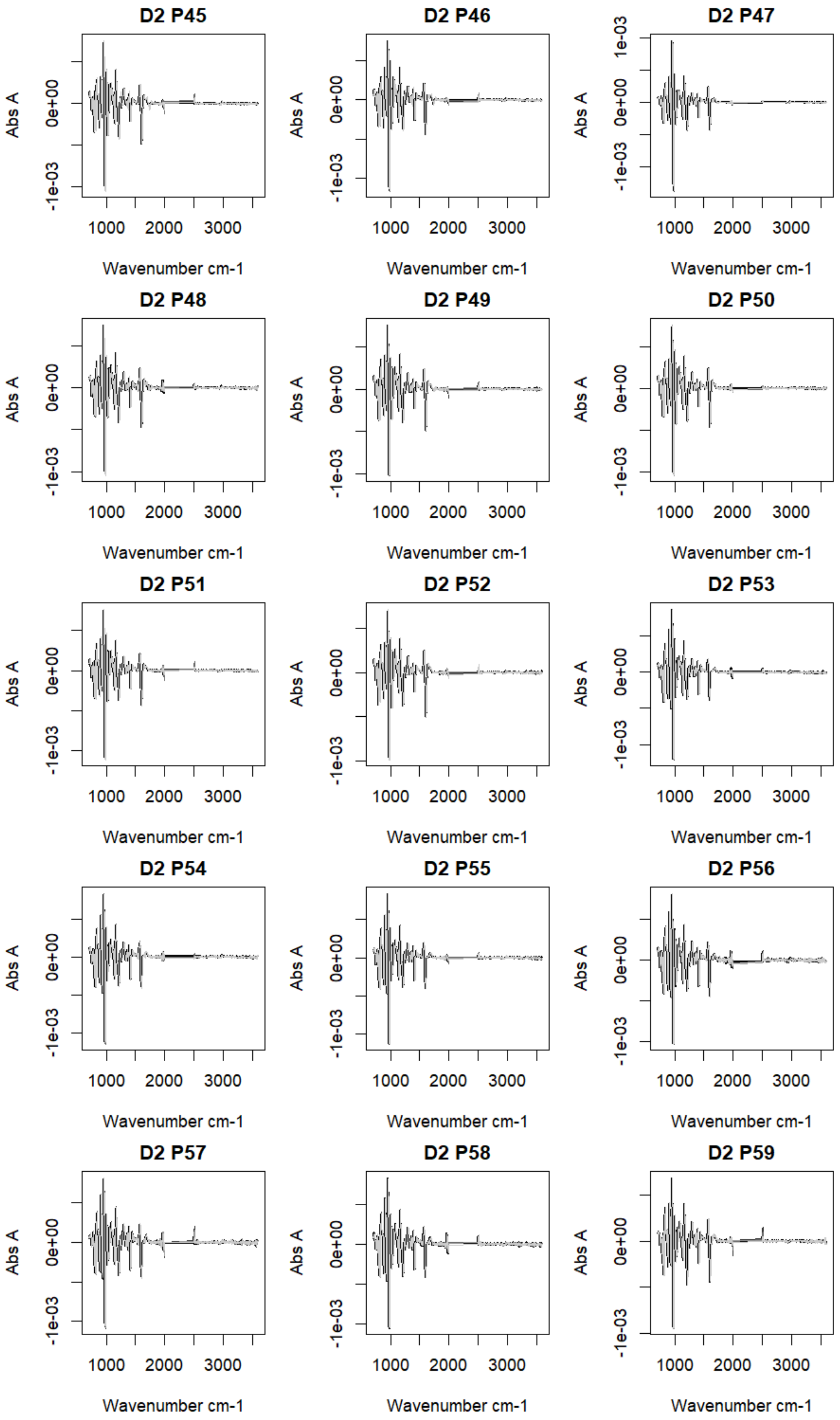
D2 P45
D2 P46
D2 P47
D2 P48
D2 P49
D2 P50
D2 P51
D2 P52
D2 P53
D2 P54
D2 P55
D2 P56
D2 P57
D2 P58
D2 P59
Abs A
1e-03
0e+00
-1e-03
1000
2000
3000
Wavenumber cm-1

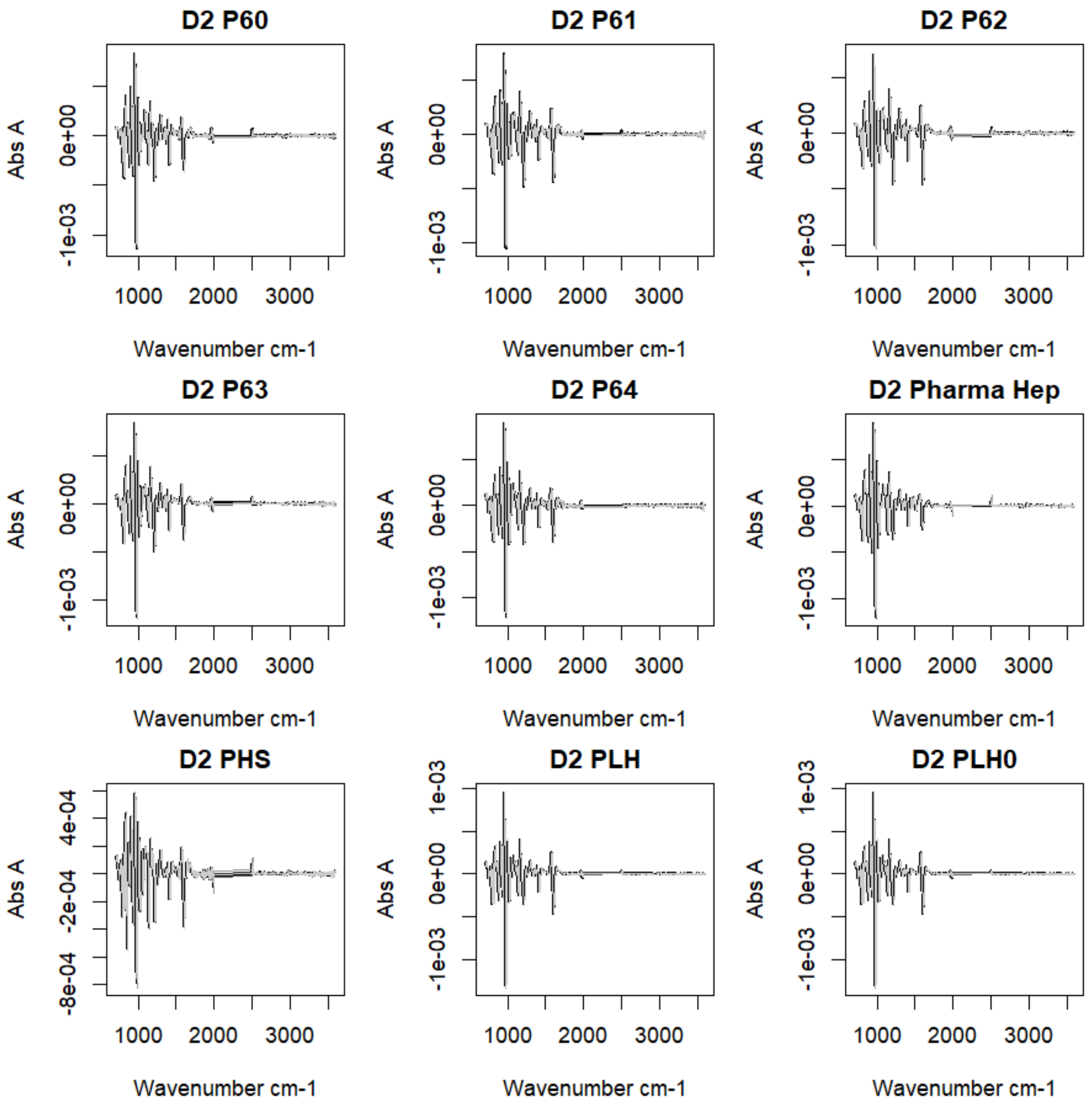
D2 P60
D2 P61
D2 P62
D2 P63
D2 P64
D2 Pharma Hep
D2 PHS
D2 PLH
D2 PLH0
Abs A
Wavenumber cm-1
1000
2000
3000
-1e-03
0e+00
1e-03
-8e-04
-2e-04
4e-04

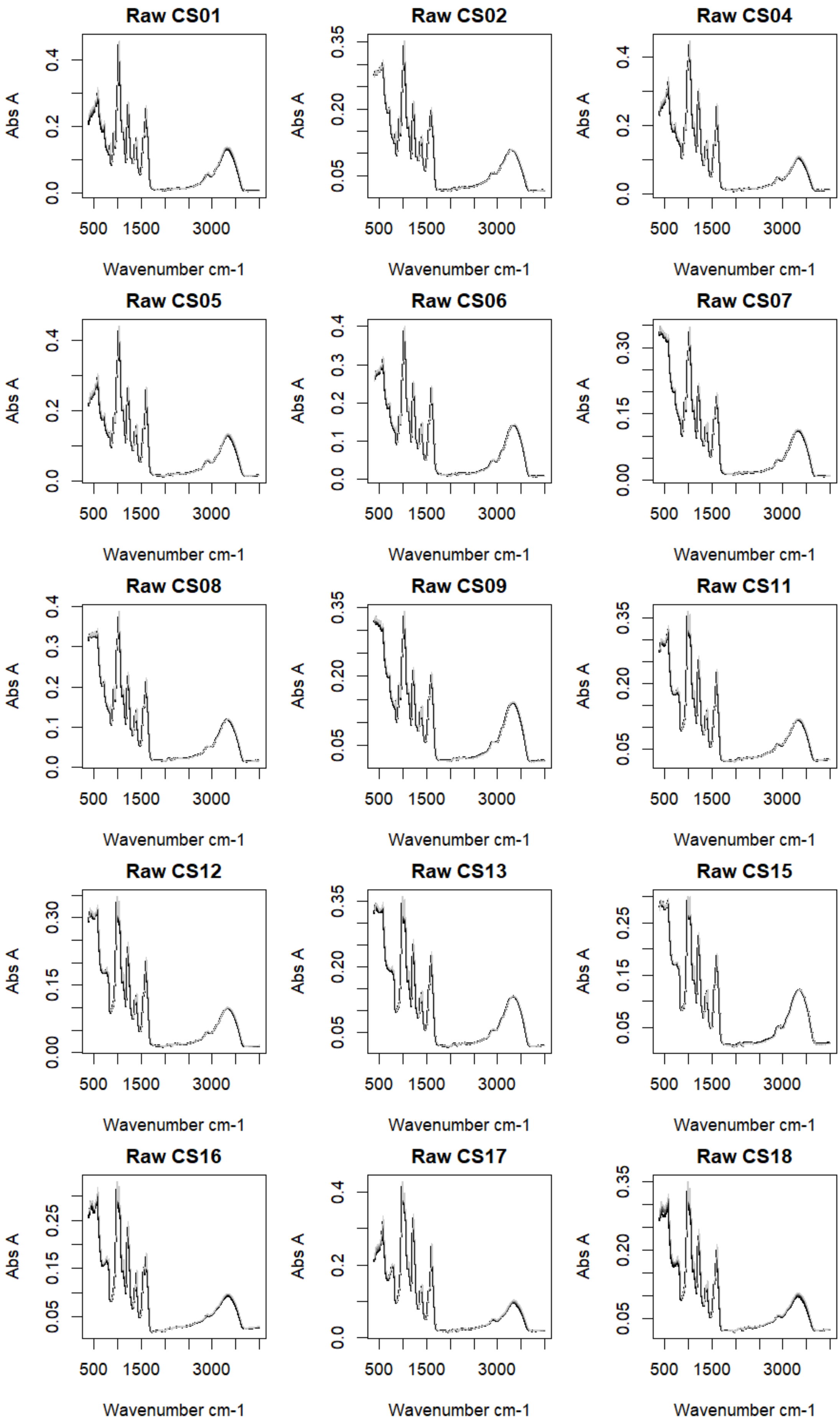

Raw CS01
Raw CS02
Raw CS04
Raw CS05
Raw CS06
Raw CS07
Raw CS08
Raw CS09
Raw CS11
Raw CS12
Raw CS13
Raw CS15
Raw CS16
Raw CS17
Raw CS18
Abs A
Wavenumber cm-1
500
1500
3000

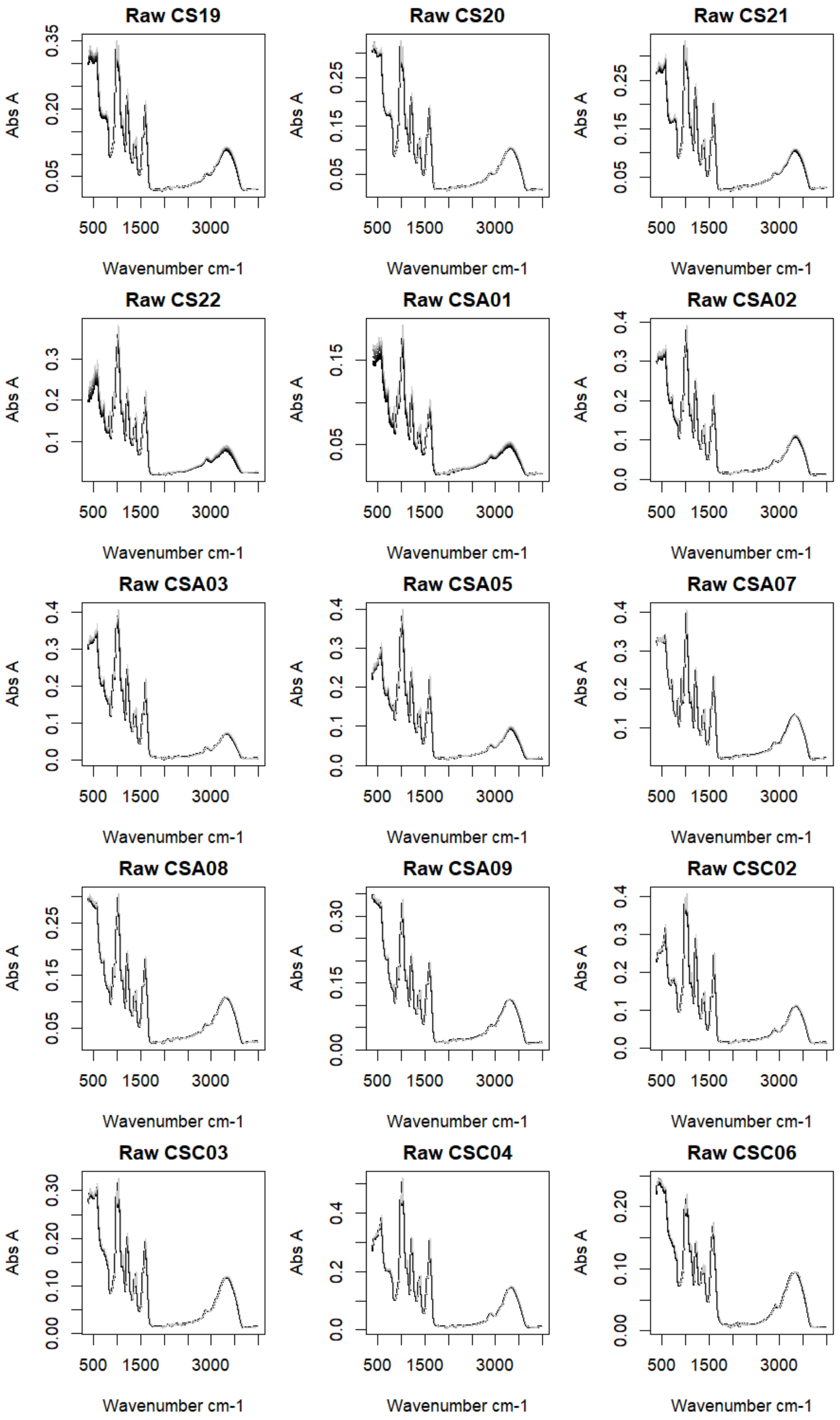

Raw CS19
Raw CS20
Raw CS21
Raw CS22
Raw CSA01
Raw CSA02
Raw CSA03
Raw CSA05
Raw CSA07
Raw CSA08
Raw CSA09
Raw CSC02
Raw CSC03
Raw CSC04
Raw CSC06
Abs A
Wavenumber cm-1
500 1500 3000

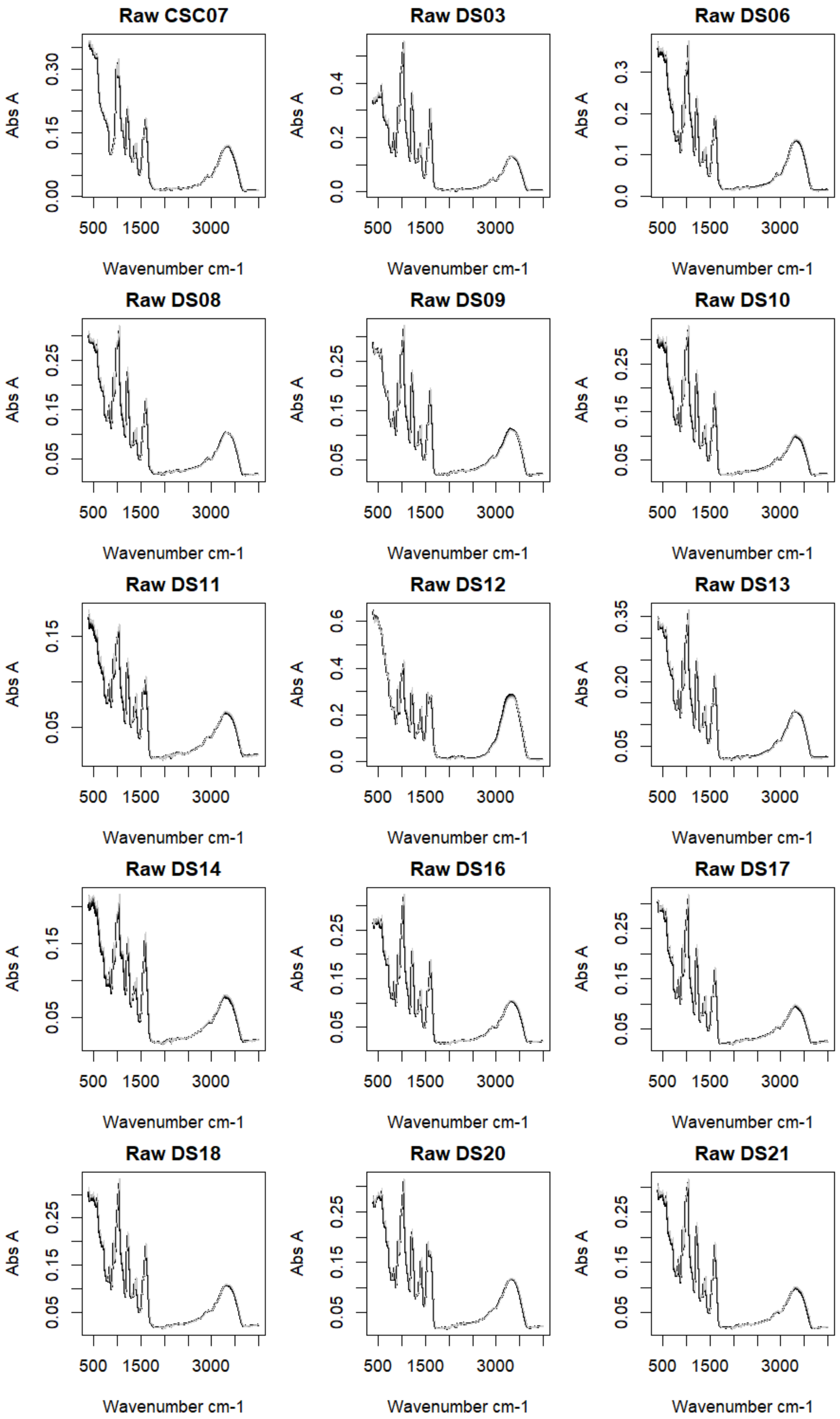

Raw CSC07
Raw DS03
Raw DS06
Raw DS08
Raw DS09
Raw DS10
Raw DS11
Raw DS12
Raw DS13
Raw DS14
Raw DS16
Raw DS17
Raw DS18
Raw DS20
Raw DS21
Abs A
Wavenumber cm-1
500
1500
3000

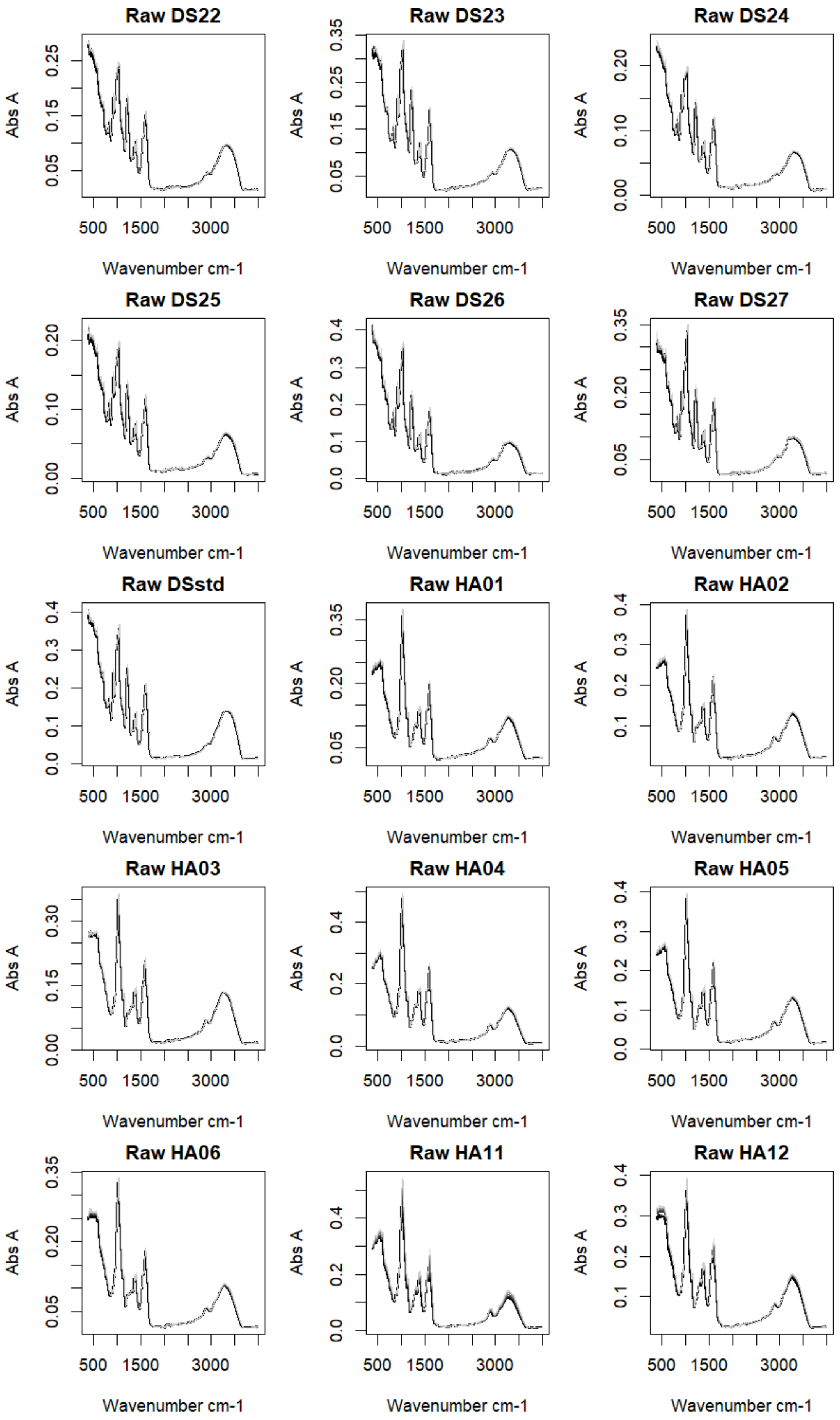
Raw DS22
Raw DS23
Raw DS24
Raw DS25
Raw DS26
Raw DS27
Raw DSstd
Raw HA01
Raw HA02
Raw HA03
Raw HA04
Raw HA05
Raw HA06
Raw HA11
Raw HA12
Abs A
Wavenumber cm-1
500 1500 3000

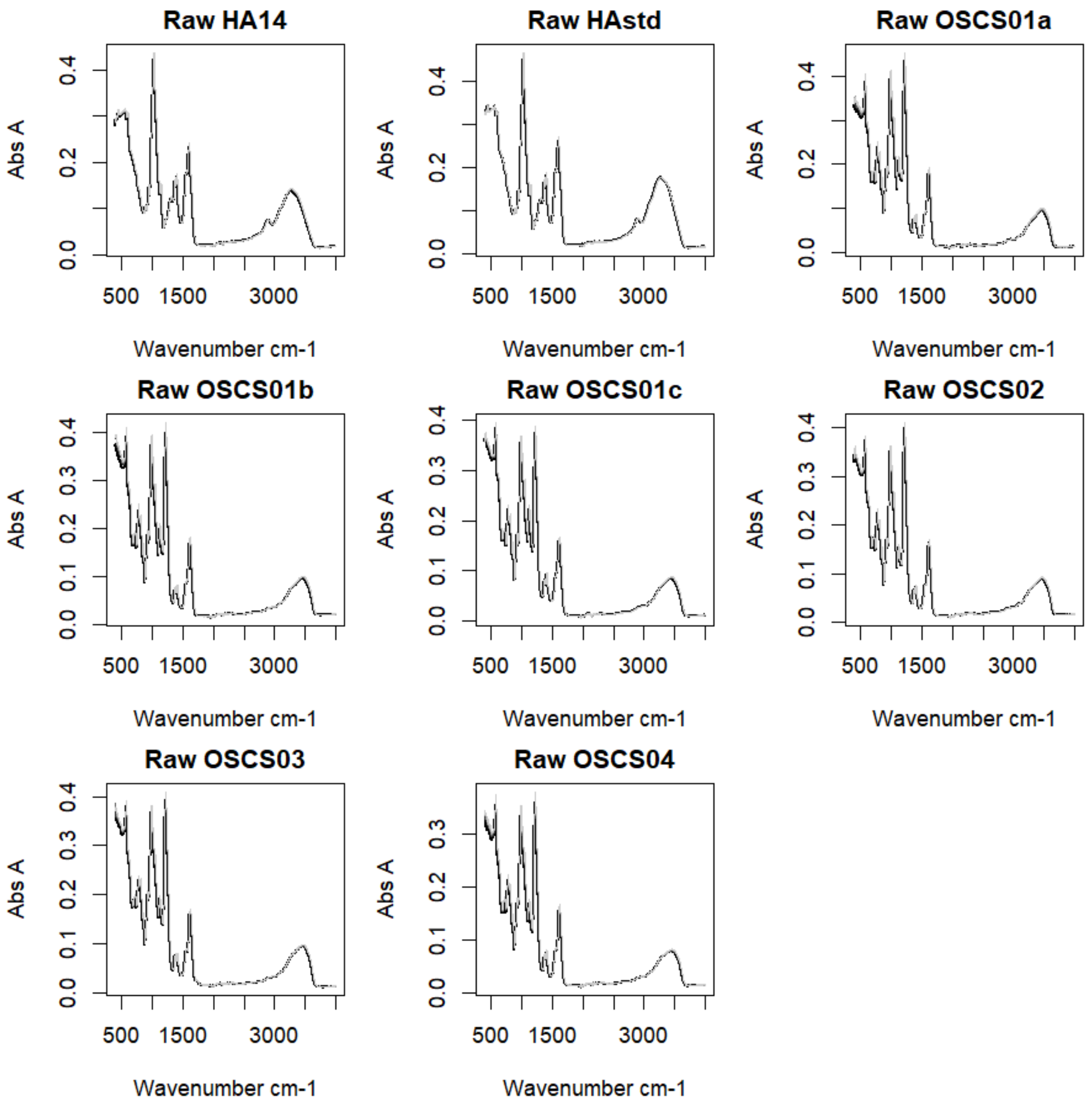
Raw HA14
Raw HAstd
Raw OSCS01a
Raw OSCS01b
Raw OSCS01c
Raw OSCS02
Raw OSCS03
Raw OSCS04
Abs A
Wavenumber cm-1
500
1500
3000
0.0
0.1
0.2
0.3
0.4

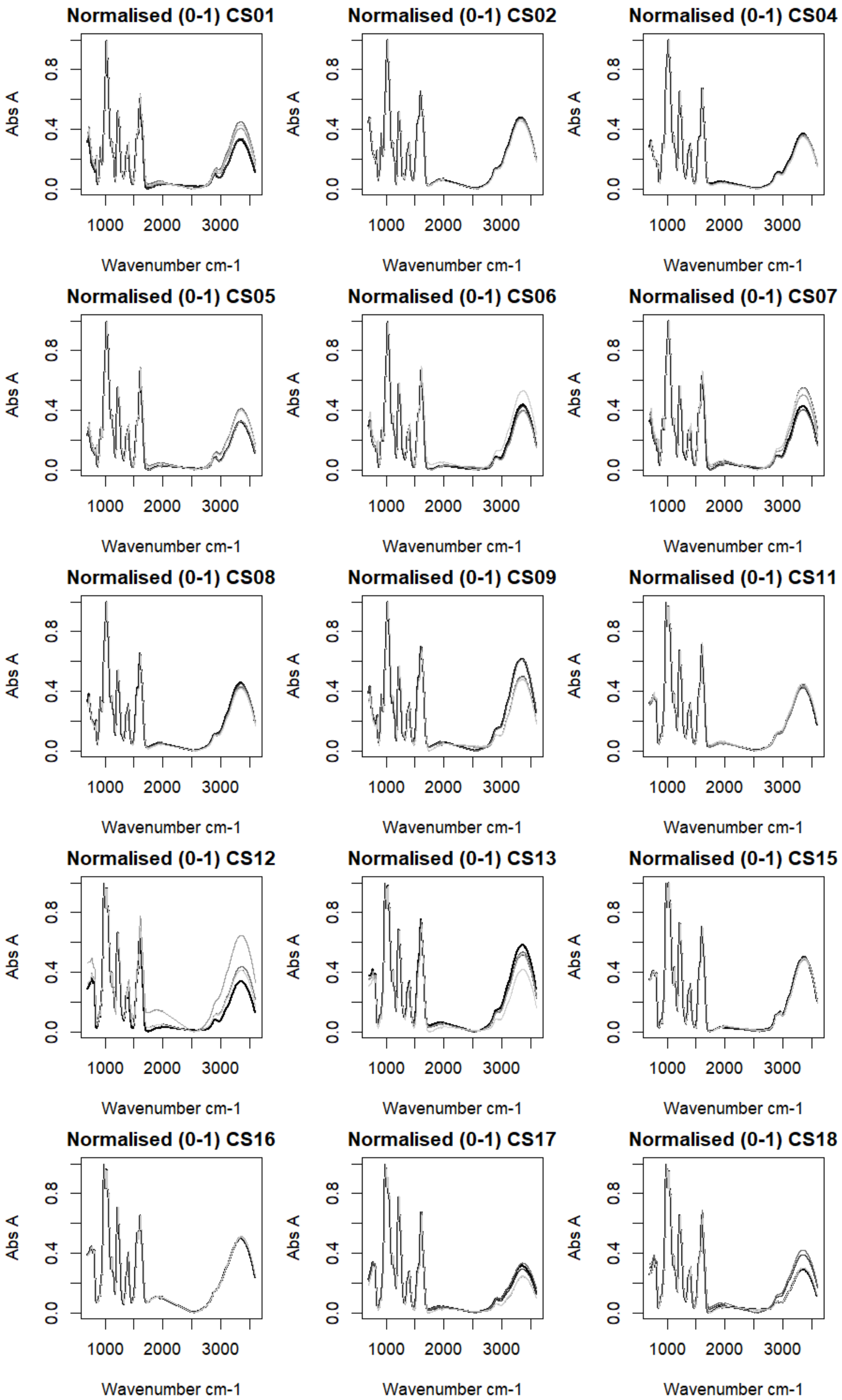
Normalised (0-1) CS01
Normalised (0-1) CS02
Normalised (0-1) CS04
Normalised (0-1) CS05
Normalised (0-1) CS06
Normalised (0-1) CS07
Normalised (0-1) CS08
Normalised (0-1) CS09
Normalised (0-1) CS11
Normalised (0-1) CS12
Normalised (0-1) CS13
Normalised (0-1) CS15
Normalised (0-1) CS16
Normalised (0-1) CS17
Normalised (0-1) CS18
Abs A
0.0
0.4
0.8
1000
2000
3000
Wavenumber cm-1

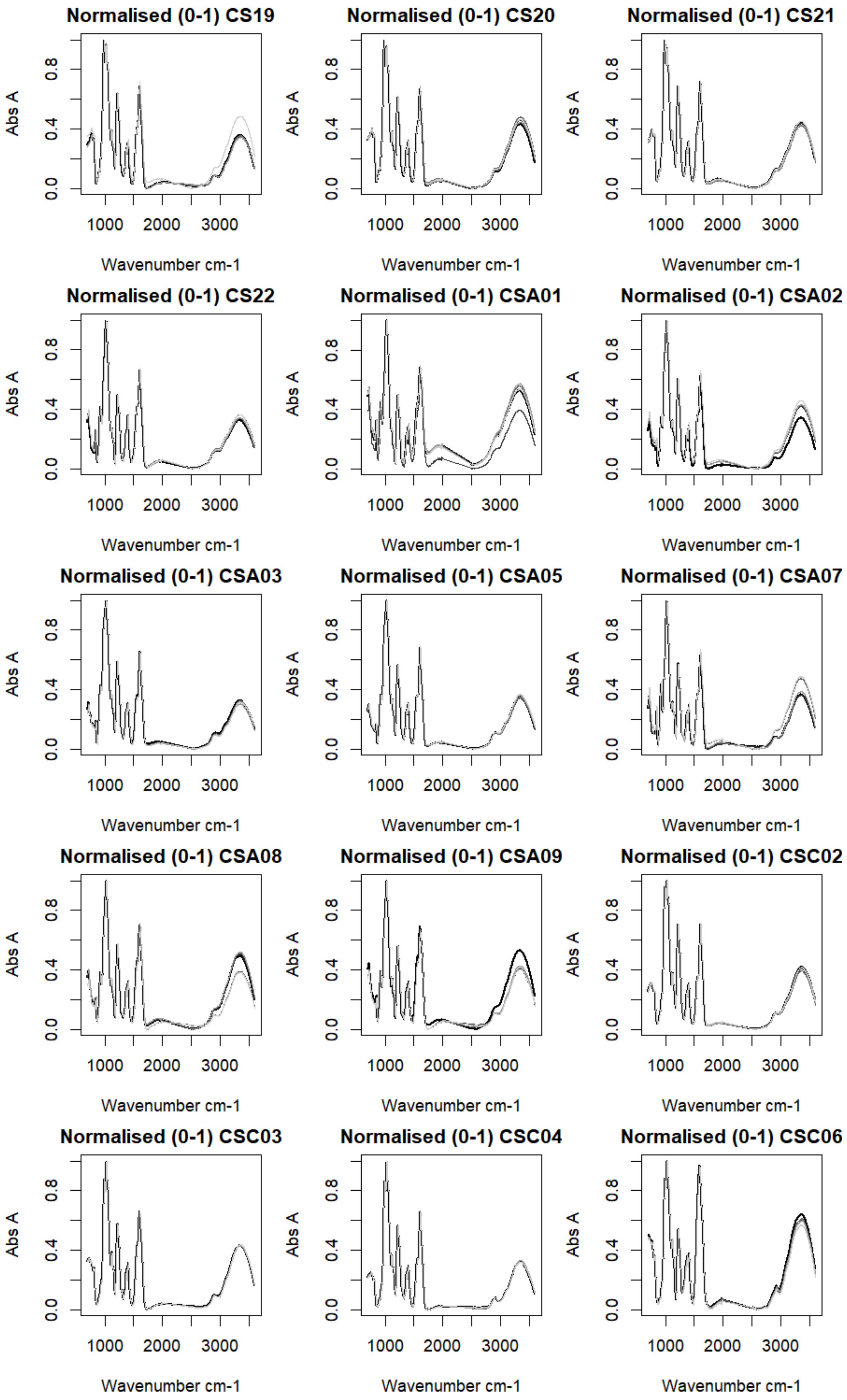
Normalised (0-1) CS19
Normalised (0-1) CS20
Normalised (0-1) CS21
Normalised (0-1) CS22
Normalised (0-1) CSA01
Normalised (0-1) CSA02
Normalised (0-1) CSA03
Normalised (0-1) CSA05
Normalised (0-1) CSA07
Normalised (0-1) CSA08
Normalised (0-1) CSA09
Normalised (0-1) CSC02
Normalised (0-1) CSC03
Normalised (0-1) CSC04
Normalised (0-1) CSC06
Abs A
0.0
0.4
0.8
1000
2000
3000
Wavenumber cm-1

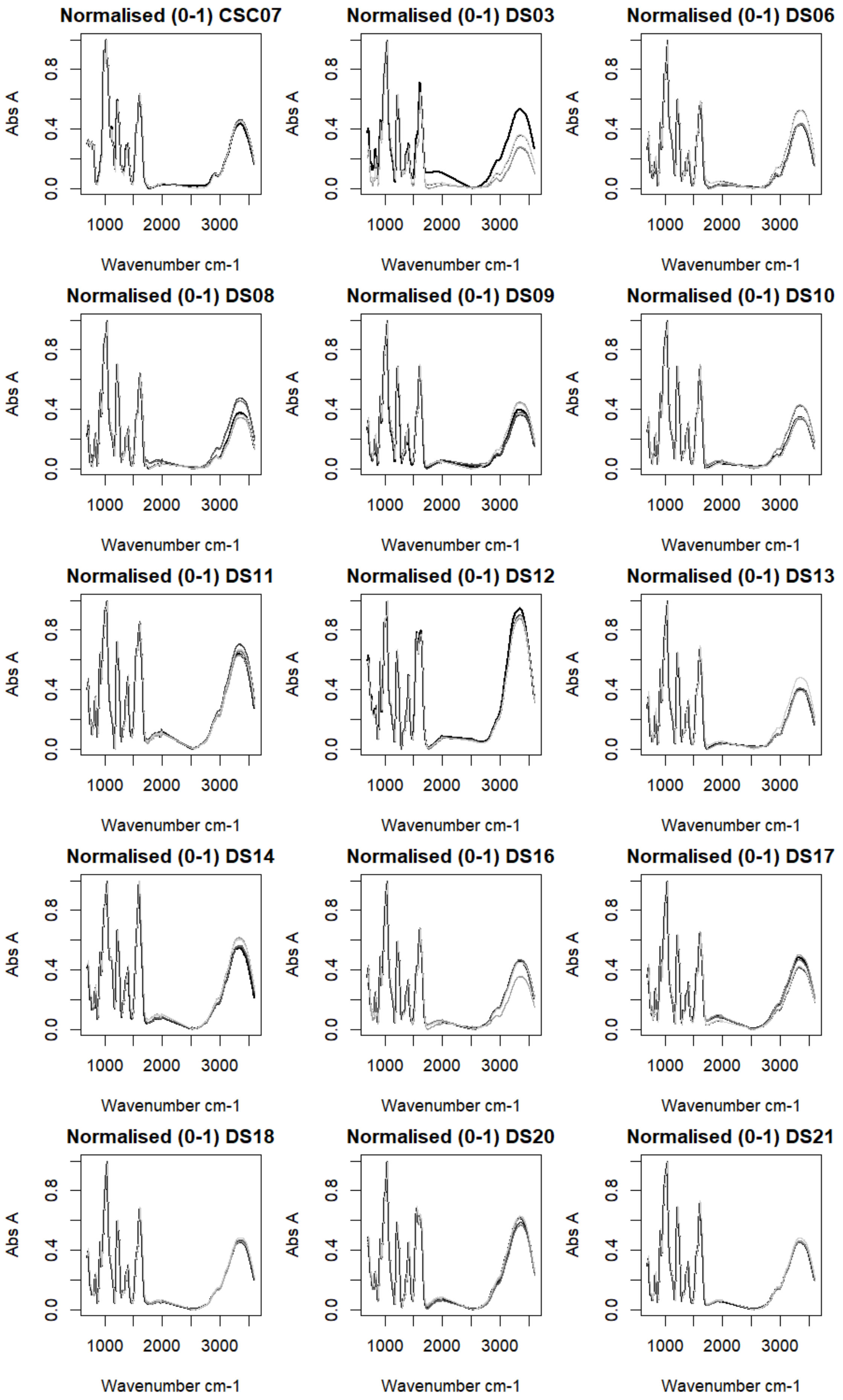
Normalised (0-1) CSC07
Normalised (0-1) DS03
Normalised (0-1) DS06
Normalised (0-1) DS08
Normalised (0-1) DS09
Normalised (0-1) DS10
Normalised (0-1) DS11
Normalised (0-1) DS12
Normalised (0-1) DS13
Normalised (0-1) DS14
Normalised (0-1) DS16
Normalised (0-1) DS17
Normalised (0-1) DS18
Normalised (0-1) DS20
Normalised (0-1) DS21
Abs A
0.0
0.4
0.8
1000
2000
3000
Wavenumber cm-1

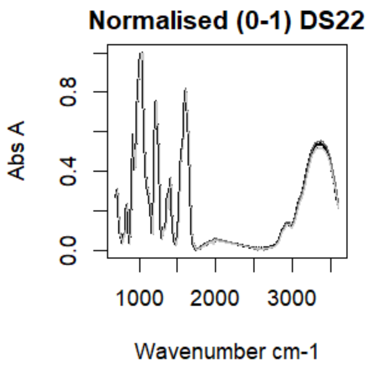
Normalised (0-1) DS22
Abs A
0.0
0.4
0.8
1000
2000
3000
Wavenumber cm-1

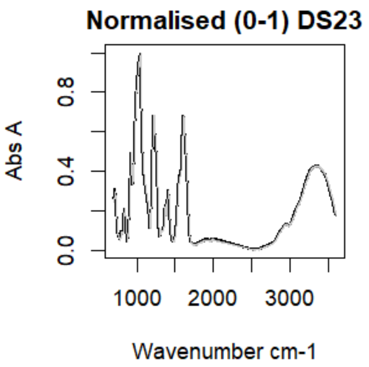
Normalised (0-1) DS23
Abs A
0.0
0.4
0.8
1000
2000
3000
Wavenumber cm-1

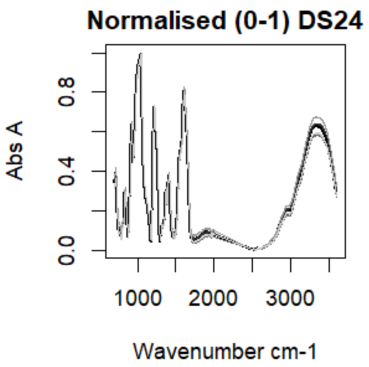
Normalised (0-1) DS24
Abs A
0.0
0.4
0.8
1000
2000
3000
Wavenumber cm-1

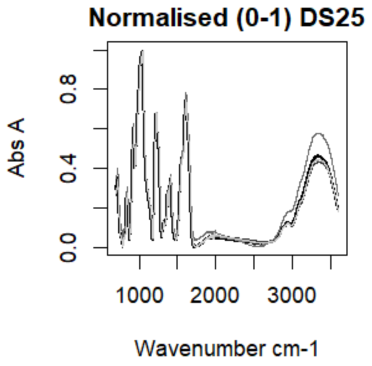
Normalised (0-1) DS25
Abs A
0.0
0.4
0.8
1000
2000
3000
Wavenumber cm-1

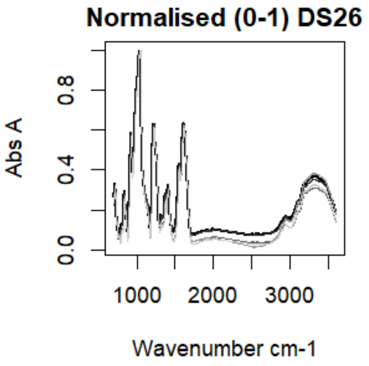
Normalised (0-1) DS26
Abs A
0.0
0.4
0.8
1000
2000
3000
Wavenumber cm-1

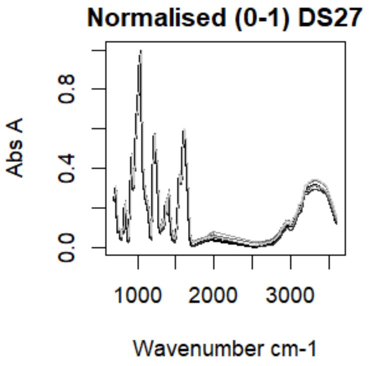
Normalised (0-1) DS27
Abs A
0.0
0.4
0.8
1000
2000
3000
Wavenumber cm-1

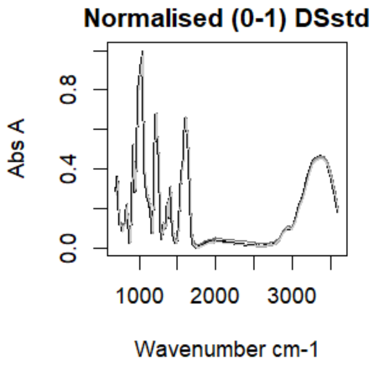
Normalised (0-1) DSstd
Abs A
0.0
0.4
0.8
1000
2000
3000
Wavenumber cm-1

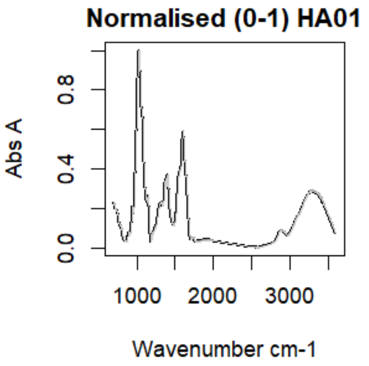
Normalised (0-1) HA01
Abs A
0.0
0.4
0.8
1000
2000
3000
Wavenumber cm-1

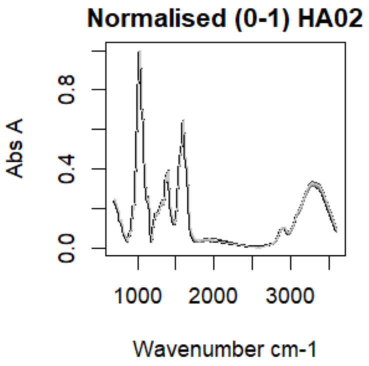
Normalised (0-1) HA02
Abs A
0.0
0.4
0.8
1000
2000
3000
Wavenumber cm-1

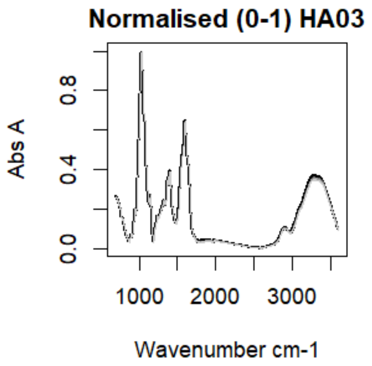
Normalised (0-1) HA03
Abs A
0.0
0.4
0.8
1000
2000
3000
Wavenumber cm-1

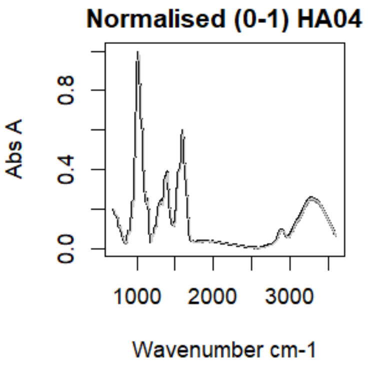
Normalised (0-1) HA04
Abs A
0.0
0.4
0.8
1000
2000
3000
Wavenumber cm-1

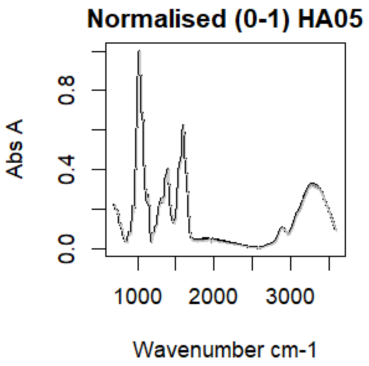
Normalised (0-1) HA05
Abs A
0.0
0.4
0.8
1000
2000
3000
Wavenumber cm-1

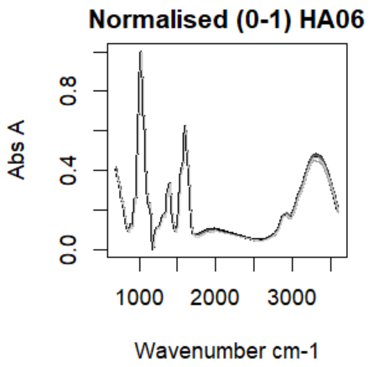
Normalised (0-1) HA06
Abs A
0.0
0.4
0.8
1000
2000
3000
Wavenumber cm-1

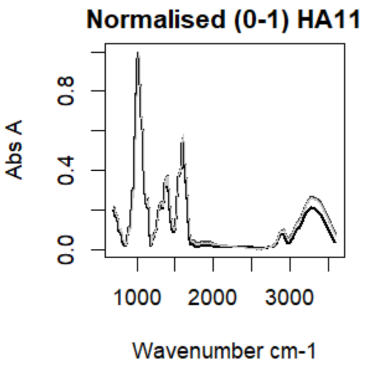
Normalised (0-1) HA11
Abs A
0.0
0.4
0.8
1000
2000
3000
Wavenumber cm-1

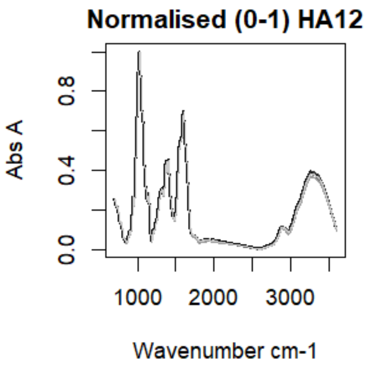
Normalised (0-1) HA12
Abs A
0.0
0.4
0.8
1000
2000
3000
Wavenumber cm-1

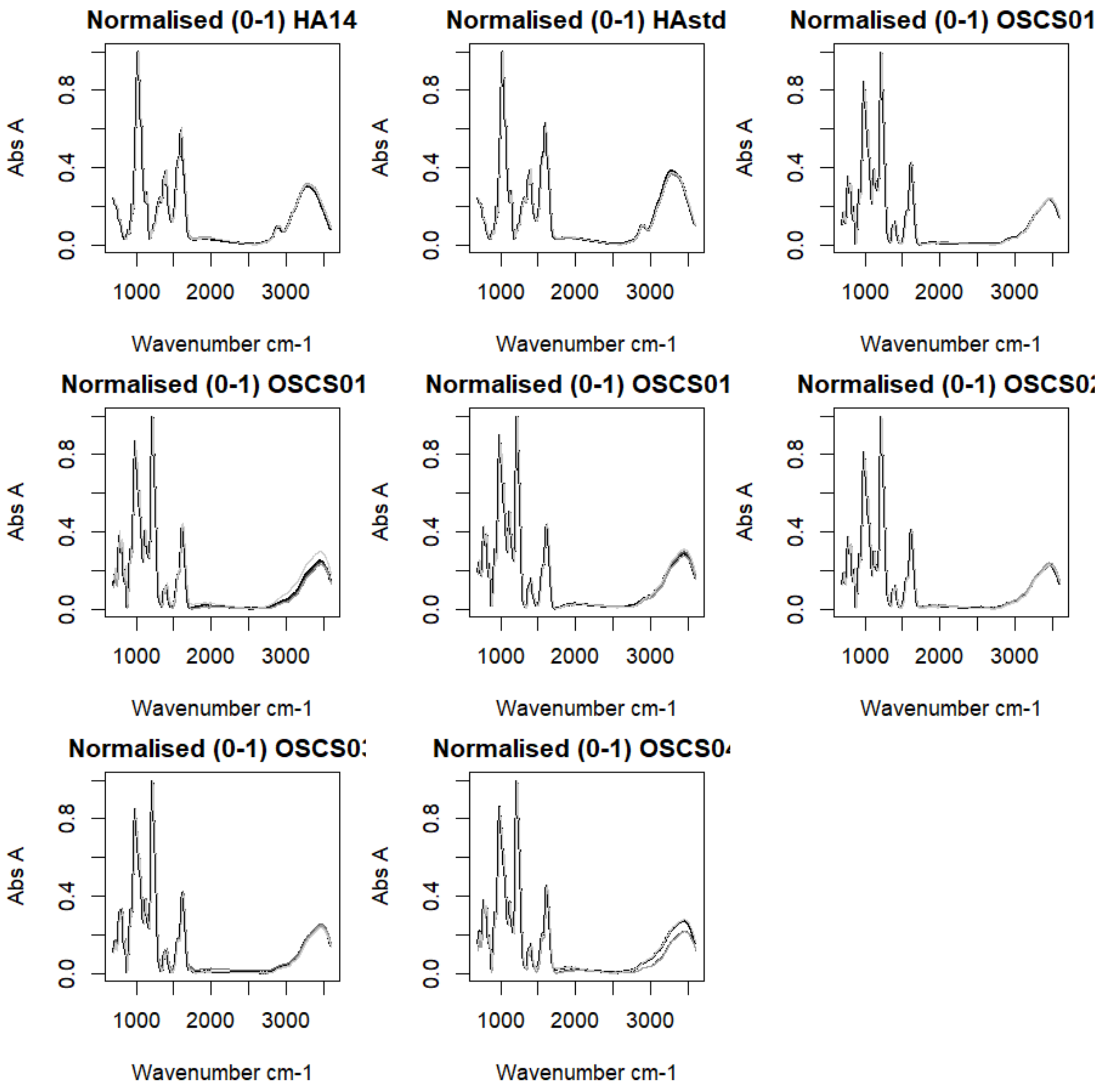
Normalised (0-1) HA14
Normalised (0-1) HAstd
Normalised (0-1) OSCS01
Normalised (0-1) OSCS01
Normalised (0-1) OSCS01
Normalised (0-1) OSCS0
Normalised (0-1) OSCS0
Normalised (0-1) OSCS0
Abs A
0.0
0.4
0.8
1000
2000
3000
Wavenumber cm-1

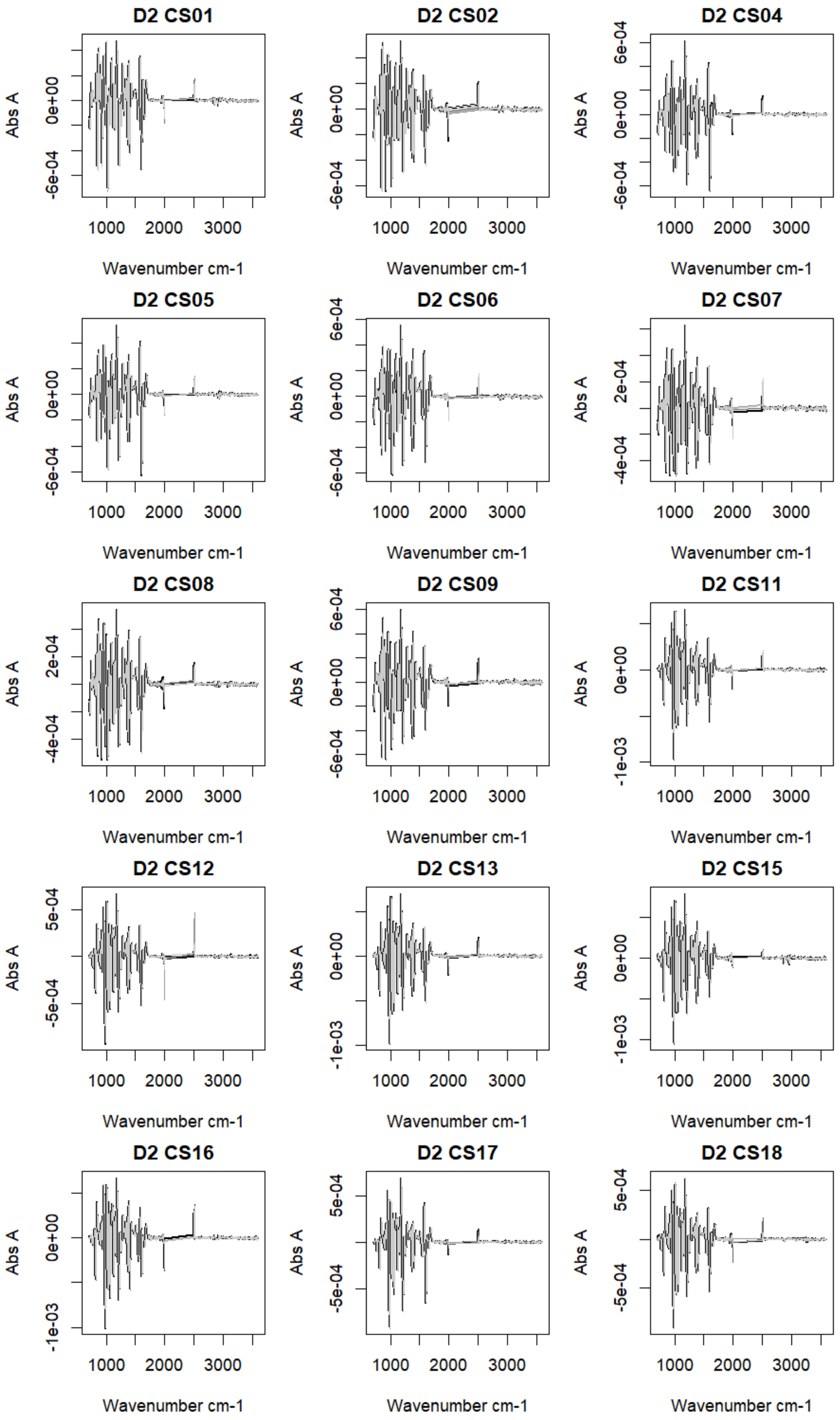
D2 CS01
D2 CS02
D2 CS04
D2 CS05
D2 CS06
D2 CS07
D2 CS08
D2 CS09
D2 CS11
D2 CS12
D2 CS13
D2 CS15
D2 CS16
D2 CS17
D2 CS18
Abs A
Wavenumber cm-1
1000
2000
3000
6e-04
0e+00
-6e-04
2e-04
-4e-04
-1e-03
5e-04
-5e-04

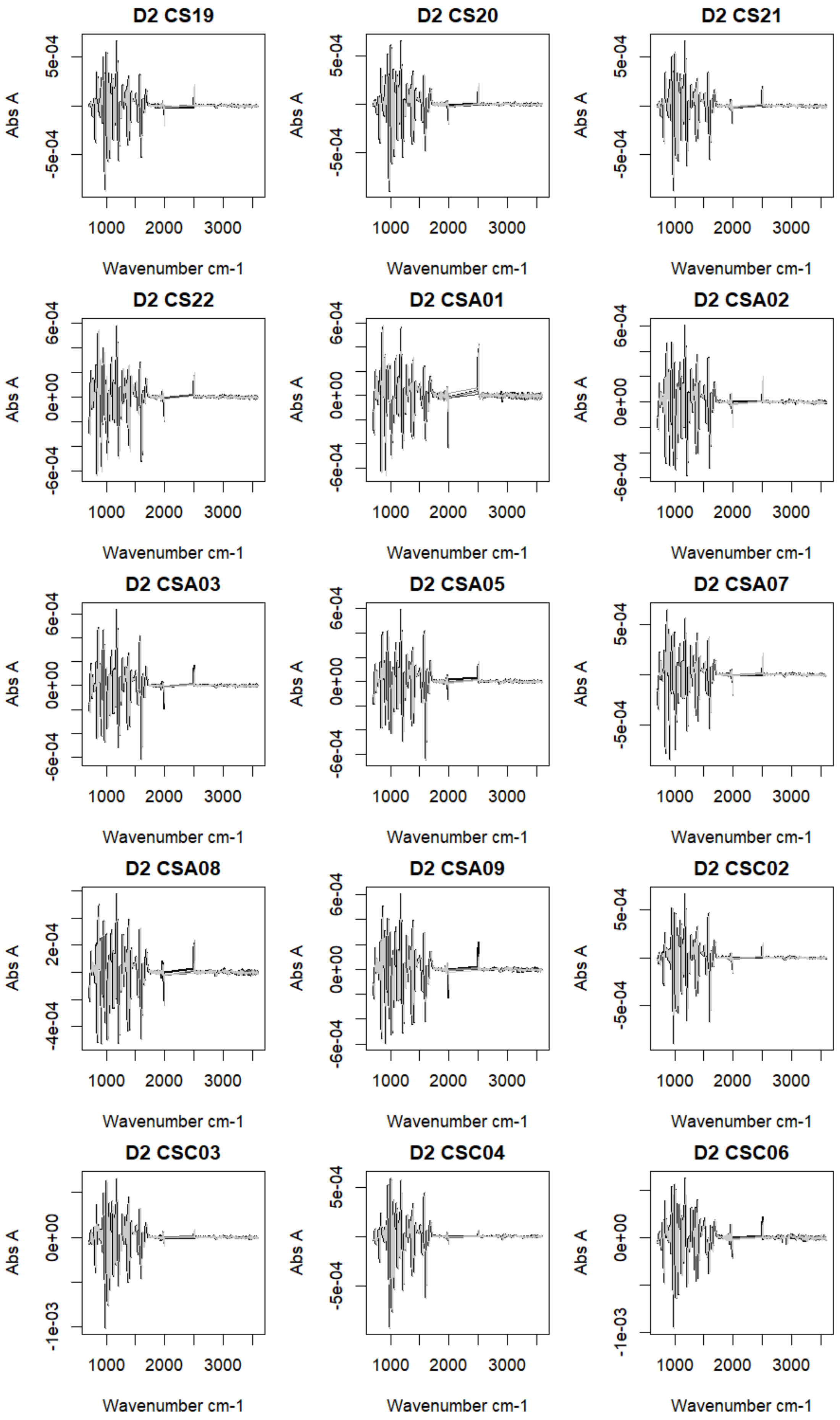
D2 CS19
D2 CS20
D2 CS21
D2 CS22
D2 CSA01
D2 CSA02
D2 CSA03
D2 CSA05
D2 CSA07
D2 CSA08
D2 CSA09
D2 CSC02
D2 CSC03
D2 CSC04
D2 CSC06
Abs A
Wavenumber cm-1
1000 2000 3000
5e-04
-5e-04
6e-04
0e+00
-6e-04
2e-04
-4e-04
-1e-03

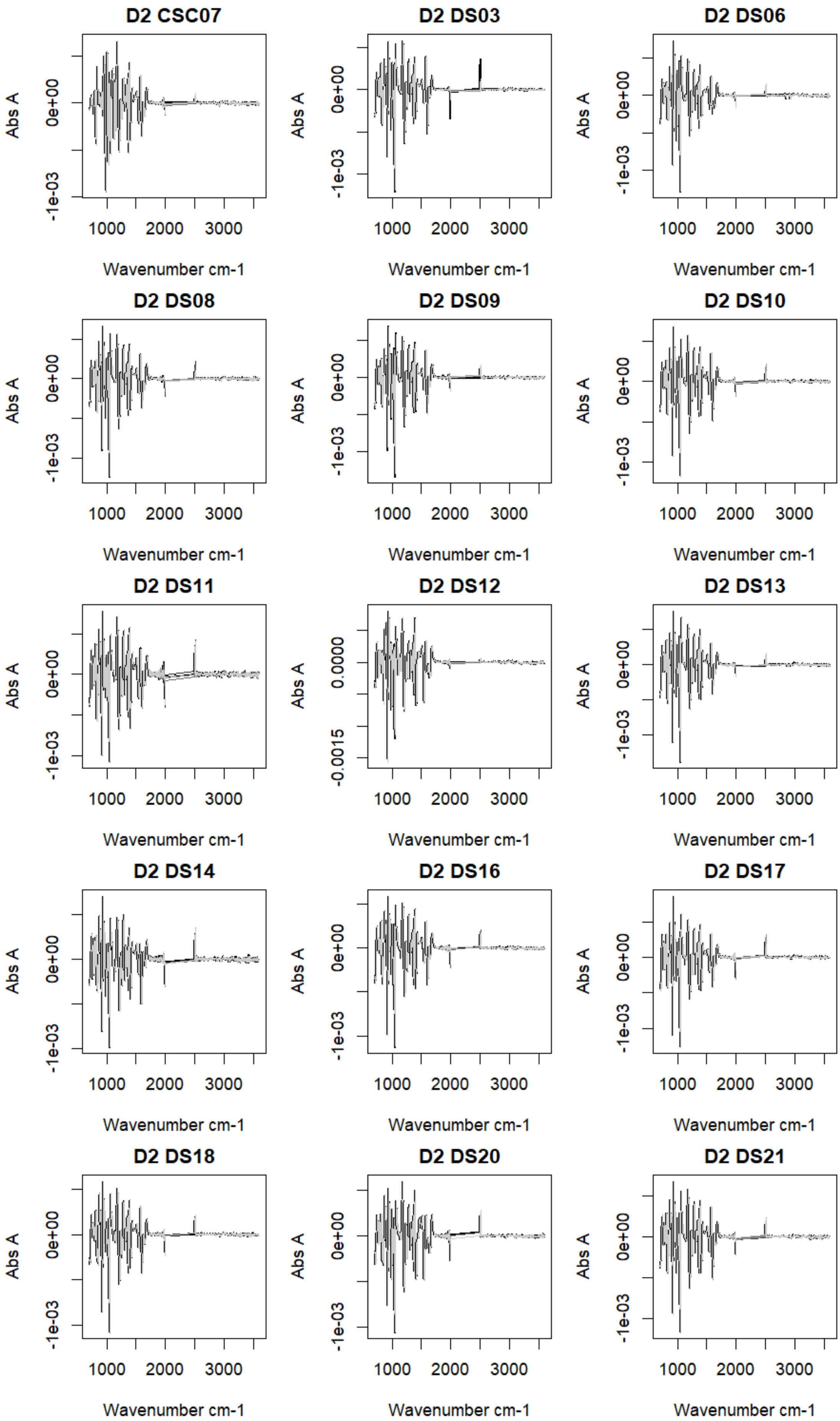

D2 CSC07
D2 DS03
D2 DS06
D2 DS08
D2 DS09
D2 DS10
D2 DS11
D2 DS12
D2 DS13
D2 DS14
D2 DS16
D2 DS17
D2 DS18
D2 DS20
D2 DS21
Abs A
0e+00
-1e-03
0.0000
-0.0015
1000
2000
3000
Wavenumber cm-1

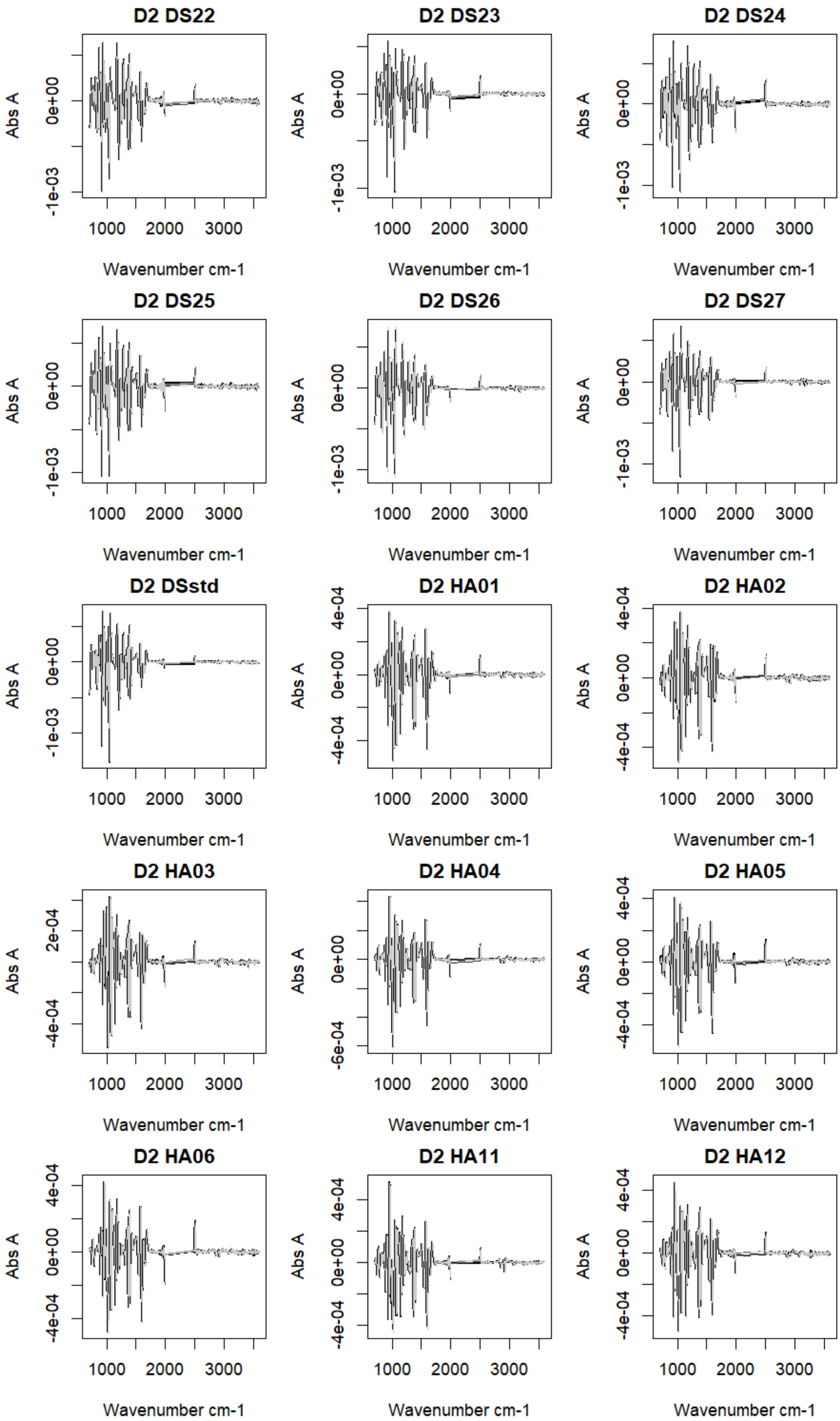
D2 DS22
D2 DS23
D2 DS24
D2 DS25
D2 DS26
D2 DS27
D2 DSstd
D2 HA01
D2 HA02
D2 HA03
D2 HA04
D2 HA05
D2 HA06
D2 HA11
D2 HA12
Abs A
Wavenumber cm-1
1000
2000
3000

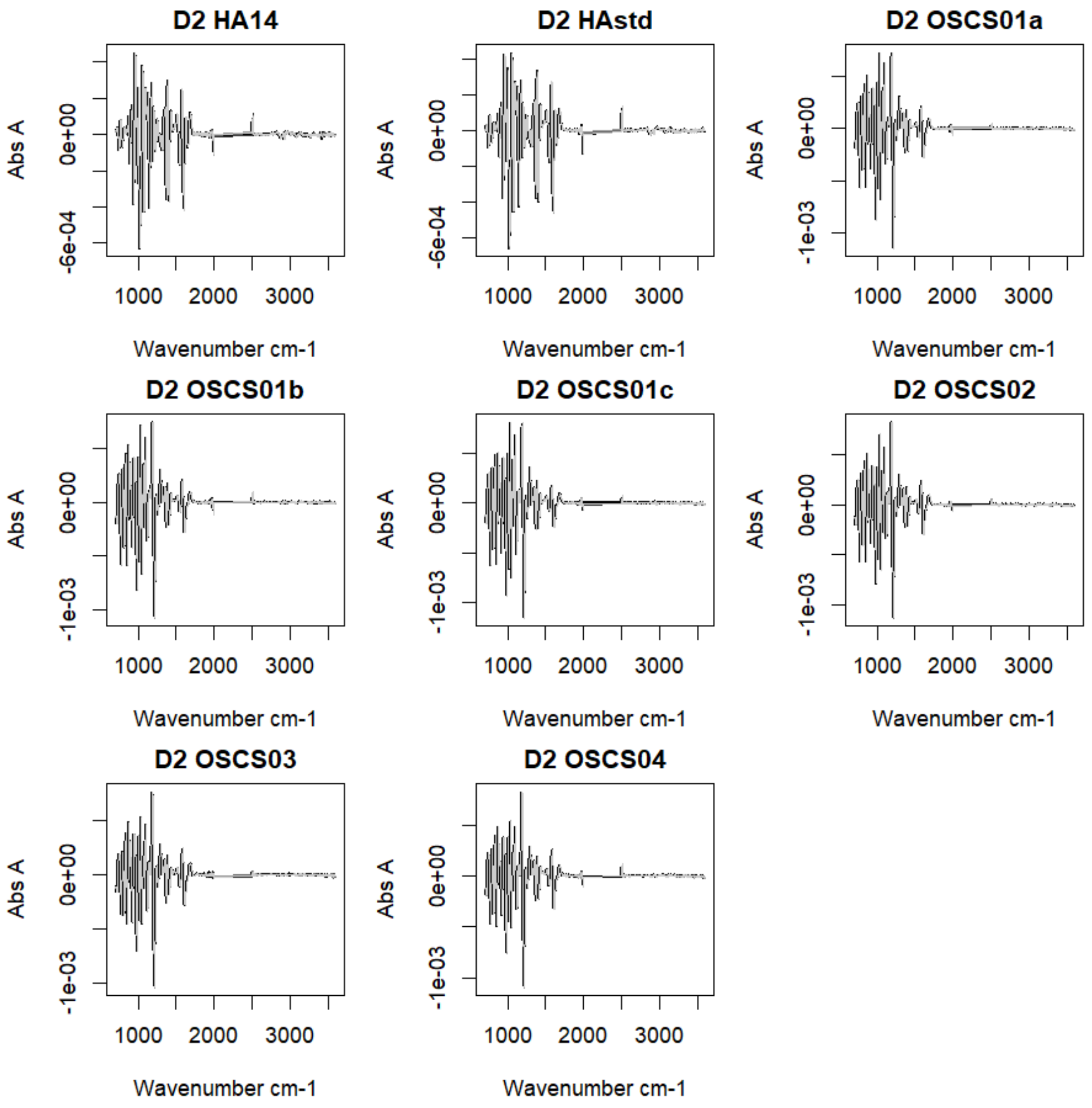
D2 HA14
D2 HAstd
D2 OSCS01a
D2 OSCS01b
D2 OSCS01c
D2 OSCS02
D2 OSCS03
D2 OSCS04
Abs A
Wavenumber cm-1
1000
2000
3000
0e+00
-6e-04
-1e-03

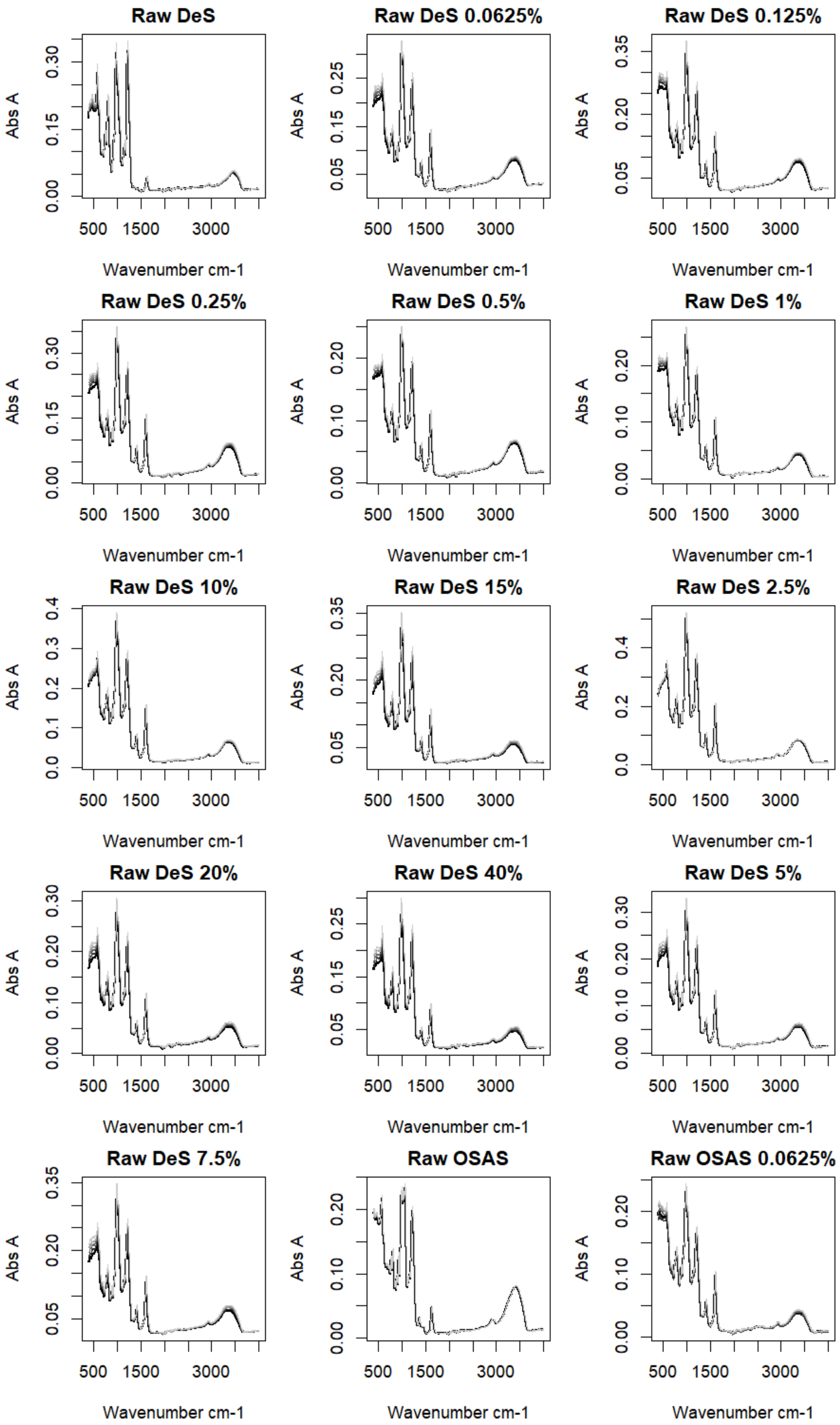
Raw DeS
Abs A
0.00 0.15 0.30
500 1500 3000
Wavenumber cm-1
Raw DeS 0.0625%
Abs A
0.05 0.15 0.25
500 1500 3000
Wavenumber cm-1
Raw DeS 0.125%
Abs A
0.05 0.20 0.35
500 1500 3000
Wavenumber cm-1
Raw DeS 0.25%
Abs A
0.00 0.15 0.30
500 1500 3000
Wavenumber cm-1
Raw DeS 0.5%
Abs A
0.00 0.10 0.20
500 1500 3000
Wavenumber cm-1
Raw DeS 1%
Abs A
0.00 0.10 0.20
500 1500 3000
Wavenumber cm-1
Raw DeS 10%
Abs A
0.0 0.1 0.2 0.3 0.4
500 1500 3000
Wavenumber cm-1
Raw DeS 15%
Abs A
0.05 0.20 0.35
500 1500 3000
Wavenumber cm-1
Raw DeS 2.5%
Abs A
0.0 0.2 0.4
500 1500 3000
Wavenumber cm-1
Raw DeS 20%
Abs A
0.00 0.10 0.20 0.30
500 1500 3000
Wavenumber cm-1
Raw DeS 40%
Abs A
0.05 0.15 0.25
500 1500 3000
Wavenumber cm-1
Raw DeS 5%
Abs A
0.00 0.10 0.20 0.30
500 1500 3000
Wavenumber cm-1
Raw DeS 7.5%
Abs A
0.05 0.20 0.35
500 1500 3000
Wavenumber cm-1
Raw OSAS
Abs A
0.00 0.10 0.20
500 1500 3000
Wavenumber cm-1
Raw OSAS 0.0625%
Abs A
0.00 0.10 0.20
500 1500 3000
Wavenumber cm-1

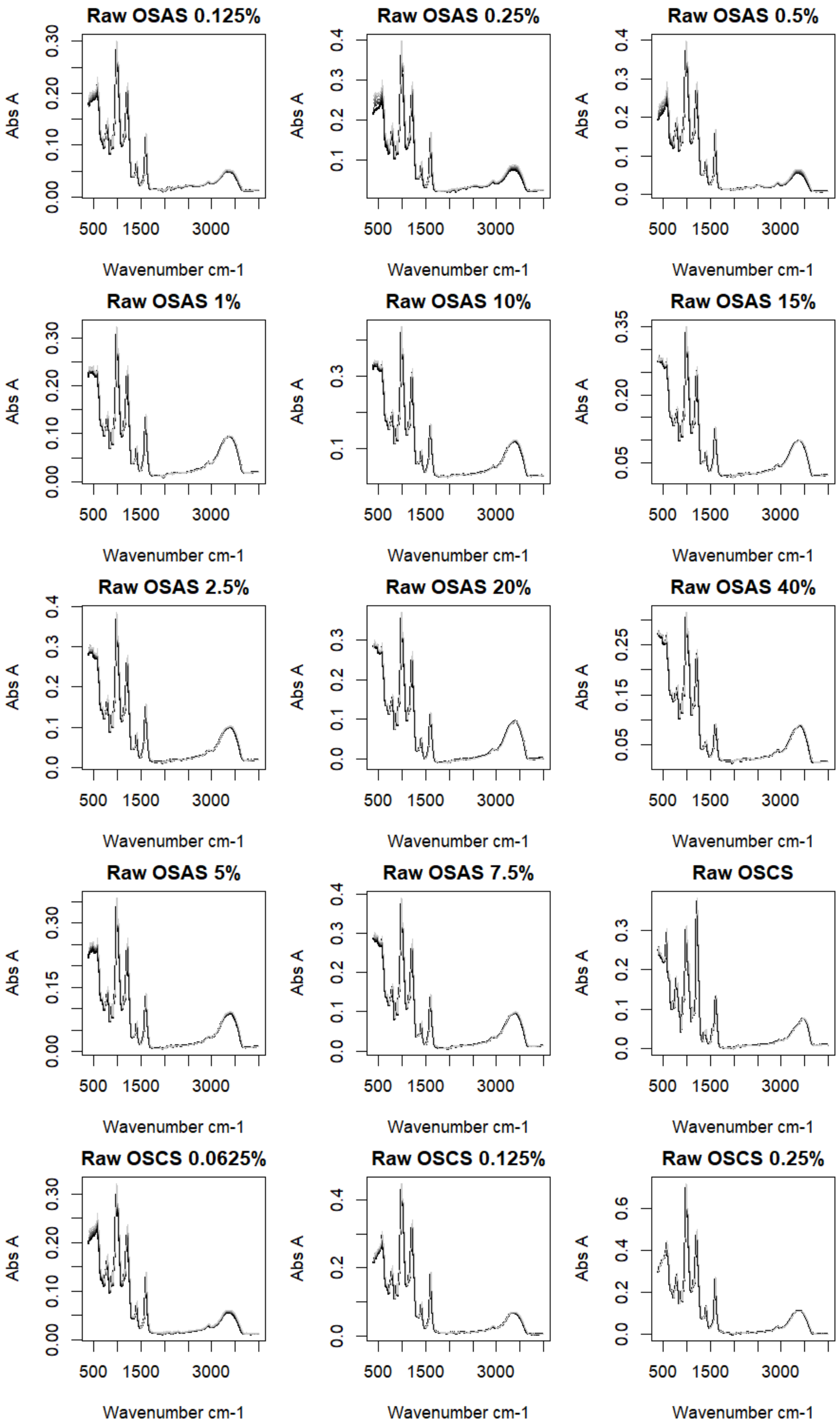
Raw OSAS 0.125%
Raw OSAS 0.25%
Raw OSAS 0.5%
Raw OSAS 1%
Raw OSAS 10%
Raw OSAS 15%
Raw OSAS 2.5%
Raw OSAS 20%
Raw OSAS 40%
Raw OSAS 5%
Raw OSAS 7.5%
Raw OSCS
Raw OSCS 0.0625%
Raw OSCS 0.125%
Raw OSCS 0.25%
Abs A
Wavenumber cm-1
500 1500 3000

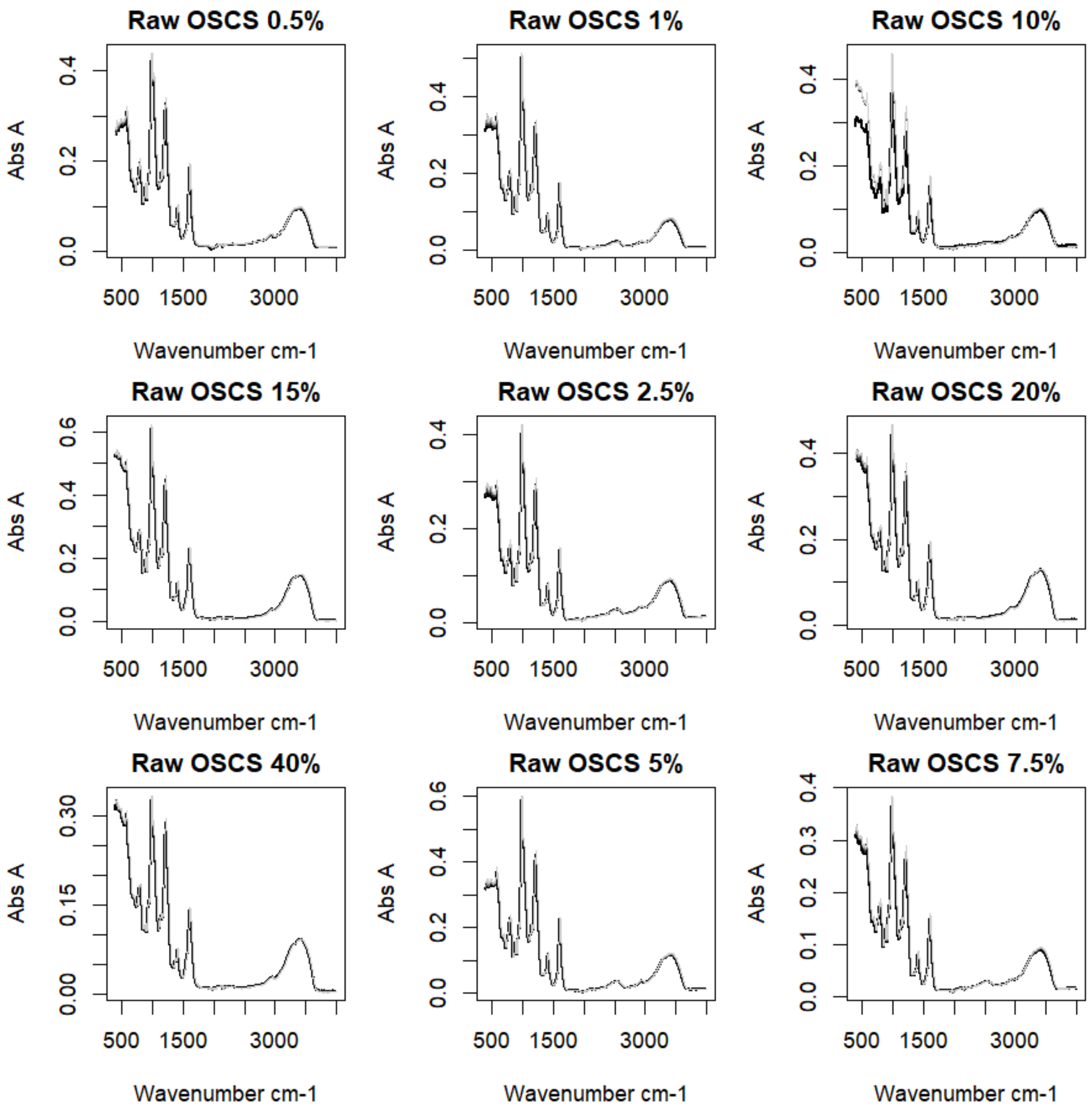
Raw OSCS 0.5%
Raw OSCS 1%
Raw OSCS 10%
Raw OSCS 15%
Raw OSCS 2.5%
Raw OSCS 20%
Raw OSCS 40%
Raw OSCS 5%
Raw OSCS 7.5%
Abs A
Wavenumber cm-1
500
1500
3000
0.0
0.1
0.2
0.3
0.4
0.6
0.00
0.15
0.30

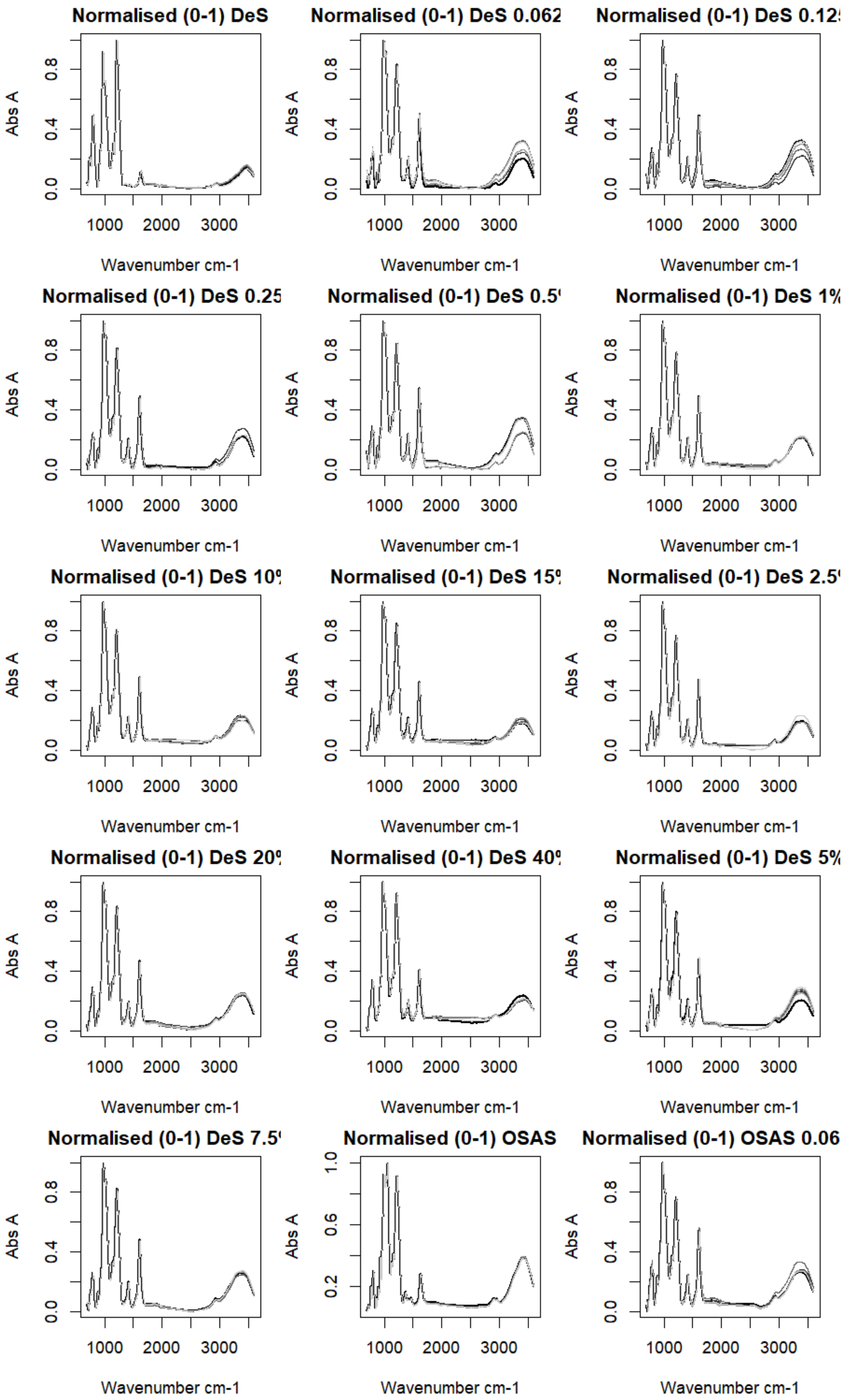

Normalised (0-1) DeS
Normalised (0-1) DeS 0.062
Normalised (0-1) DeS 0.12
Normalised (0-1) DeS 0.25
Normalised (0-1) DeS 0.5
Normalised (0-1) DeS 1%
Normalised (0-1) DeS 10
Normalised (0-1) DeS 15
Normalised (0-1) DeS 2.5
Normalised (0-1) DeS 20
Normalised (0-1) DeS 40
Normalised (0-1) DeS 5%
Normalised (0-1) DeS 7.5
Normalised (0-1) OSAS
Normalised (0-1) OSAS 0.06
Abs A
Wavenumber cm-1
1000
2000
3000
0.0
0.4
0.8
0.2
0.6
1.0

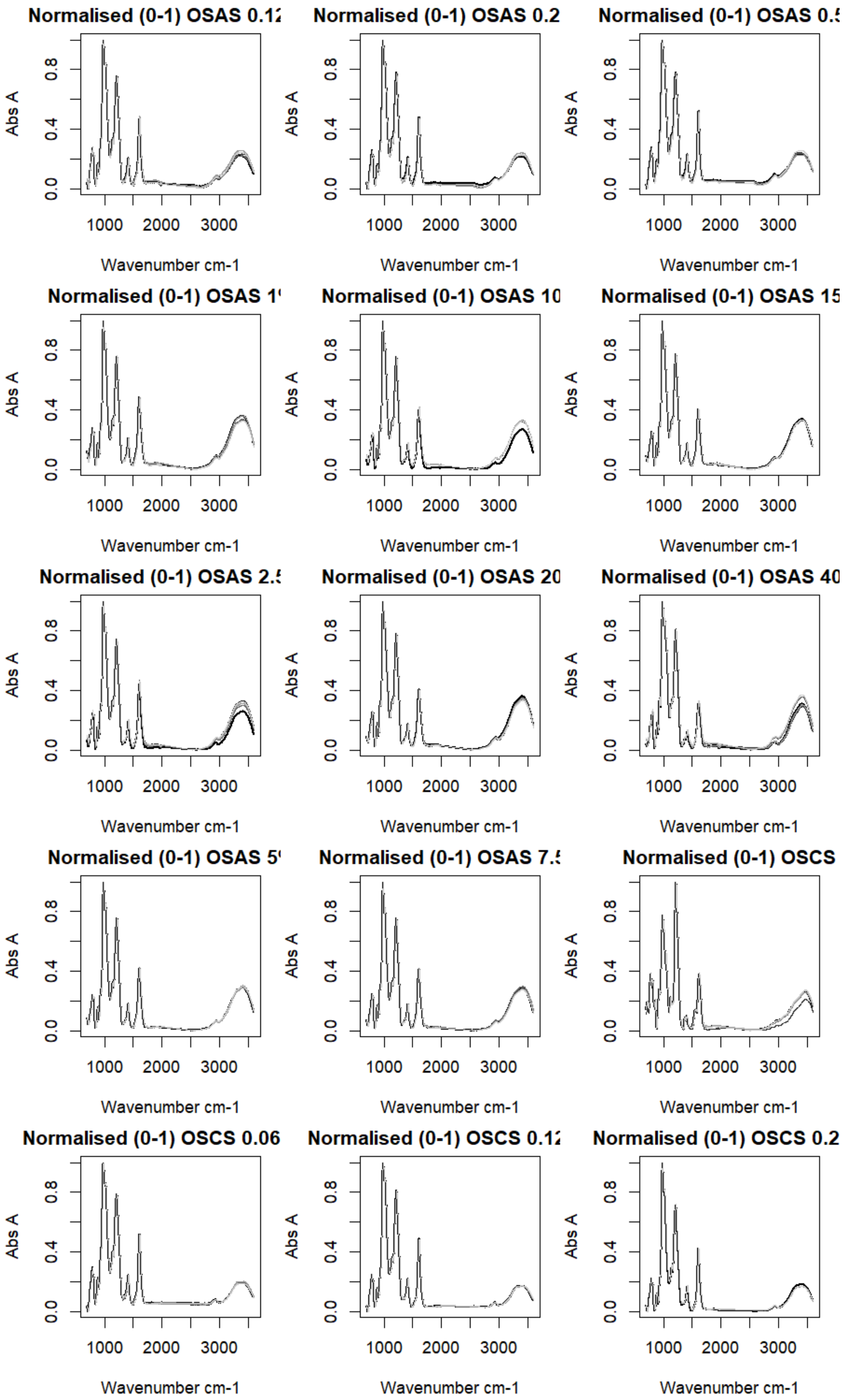
Normalised (0-1) OSAS 0.1
Normalised (0-1) OSAS 0.2
Normalised (0-1) OSAS 0.
Normalised (0-1) OSAS 1
Normalised (0-1) OSAS 10
Normalised (0-1) OSAS 15
Normalised (0-1) OSAS 2.
Normalised (0-1) OSAS 20
Normalised (0-1) OSAS 40
Normalised (0-1) OSAS 5
Normalised (0-1) OSAS 7.
Normalised (0-1) OSCS
Normalised (0-1) OSCS 0.06
Normalised (0-1) OSCS 0.1
Normalised (0-1) OSCS 0.2
Abs A
0.0
0.4
0.8
1000
2000
3000
Wavenumber cm-1

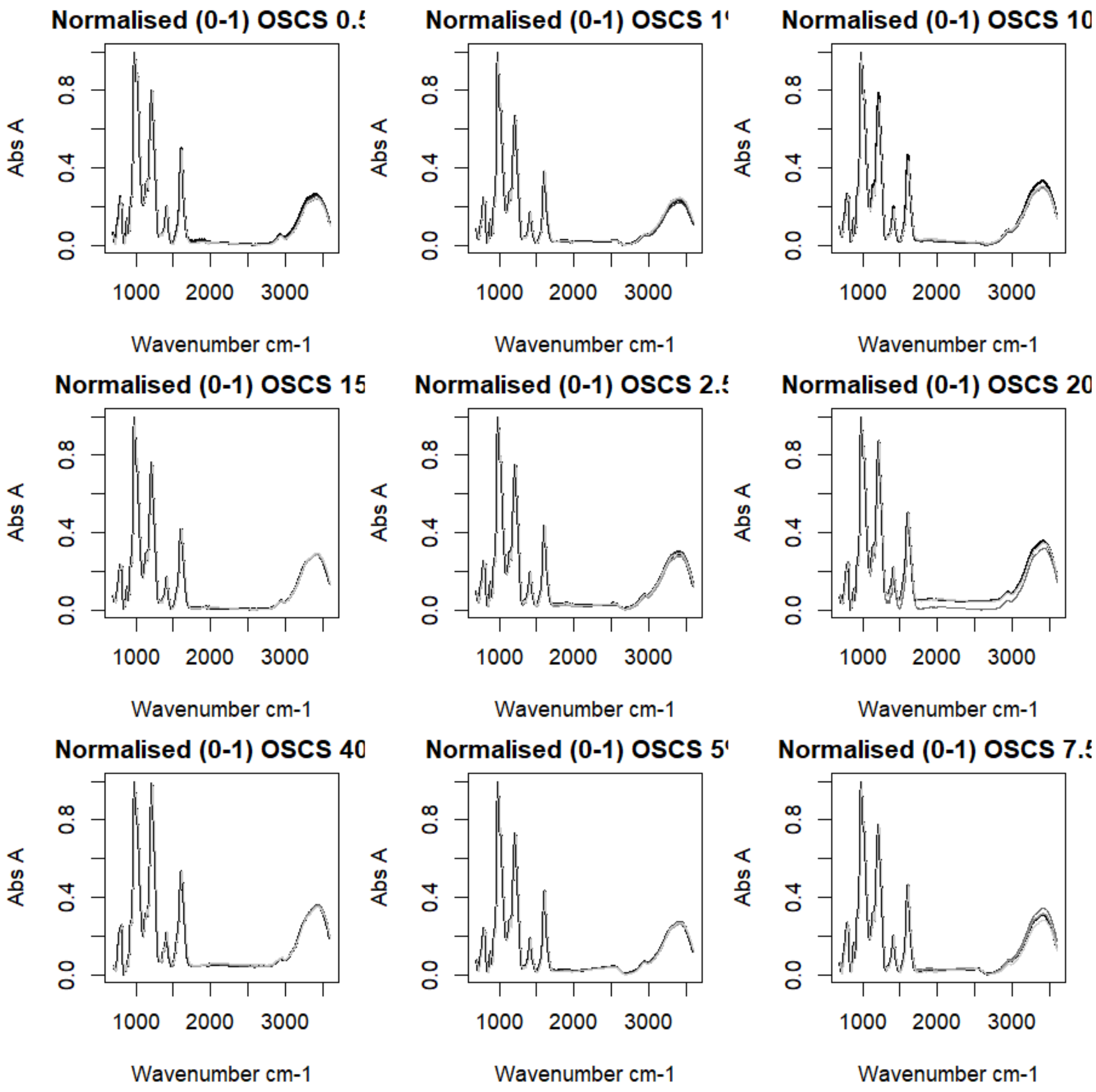
Normalised (0-1) OSCS 0.
Normalised (0-1) OSCS 1
Normalised (0-1) OSCS 10
Normalised (0-1) OSCS 15
Normalised (0-1) OSCS 2.
Normalised (0-1) OSCS 20
Normalised (0-1) OSCS 40
Normalised (0-1) OSCS 5
Normalised (0-1) OSCS 7.
Abs A
0.0
0.4
0.8
1000
2000
3000
Wavenumber cm-1

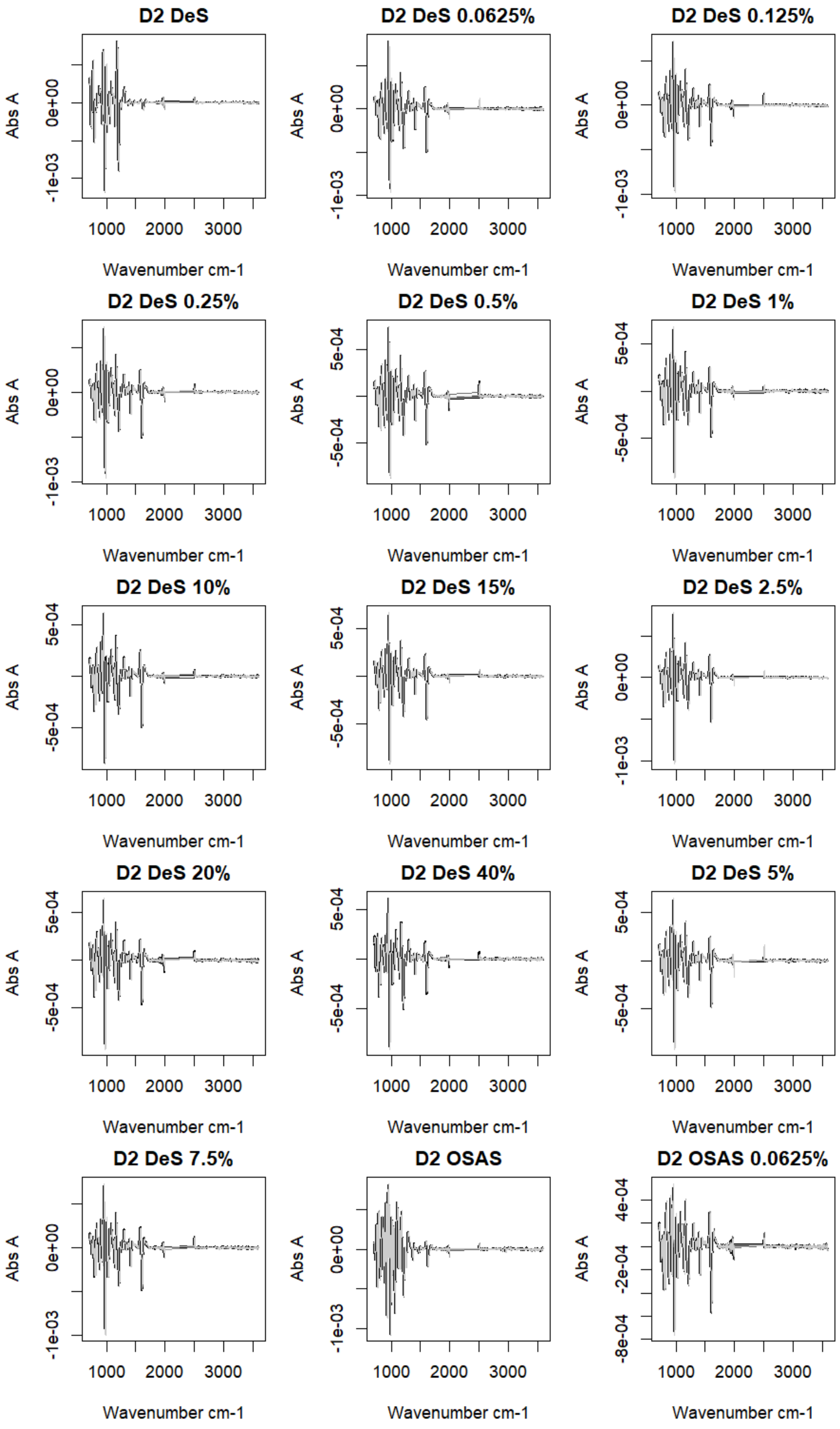
D2 DeS
D2 DeS 0.0625%
D2 DeS 0.125%
D2 DeS 0.25%
D2 DeS 0.5%
D2 DeS 1%
D2 DeS 10%
D2 DeS 15%
D2 DeS 2.5%
D2 DeS 20%
D2 DeS 40%
D2 DeS 5%
D2 DeS 7.5%
D2 OSAS
D2 OSAS 0.0625%
Abs A
Wavenumber cm-1
1000 2000 3000
-1e-03 0e+00
-5e-04 5e-04
-8e-04 -2e-04 4e-04

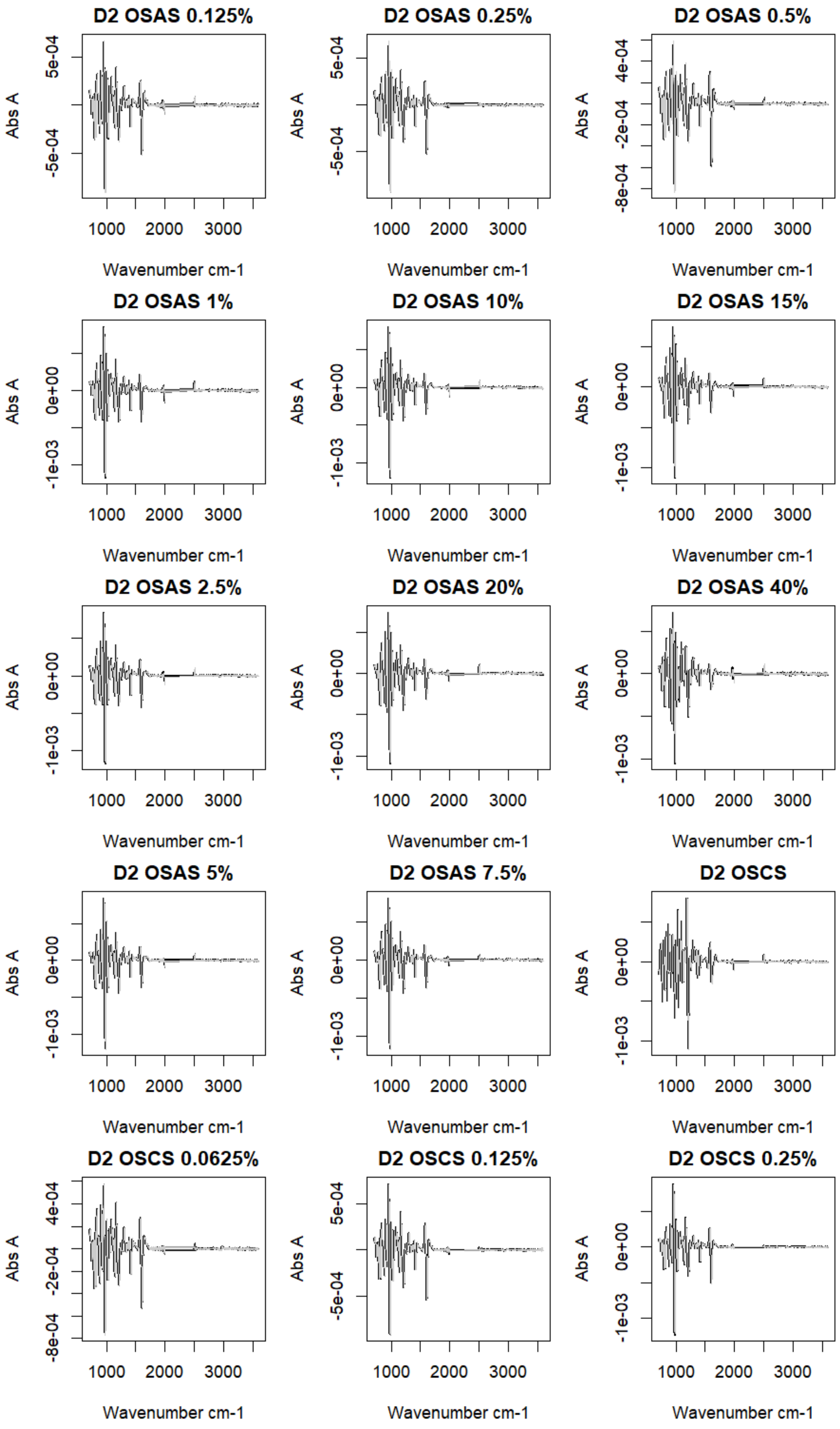
D2 OSAS 0.125%
D2 OSAS 0.25%
D2 OSAS 0.5%
D2 OSAS 1%
D2 OSAS 10%
D2 OSAS 15%
D2 OSAS 2.5%
D2 OSAS 20%
D2 OSAS 40%
D2 OSAS 5%
D2 OSAS 7.5%
D2 OSCS
D2 OSCS 0.0625%
D2 OSCS 0.125%
D2 OSCS 0.25%
Abs A
Wavenumber cm-1
1000 2000 3000
5e-04
-5e-04
4e-04
-2e-04
-8e-04
0e+00
-1e-03

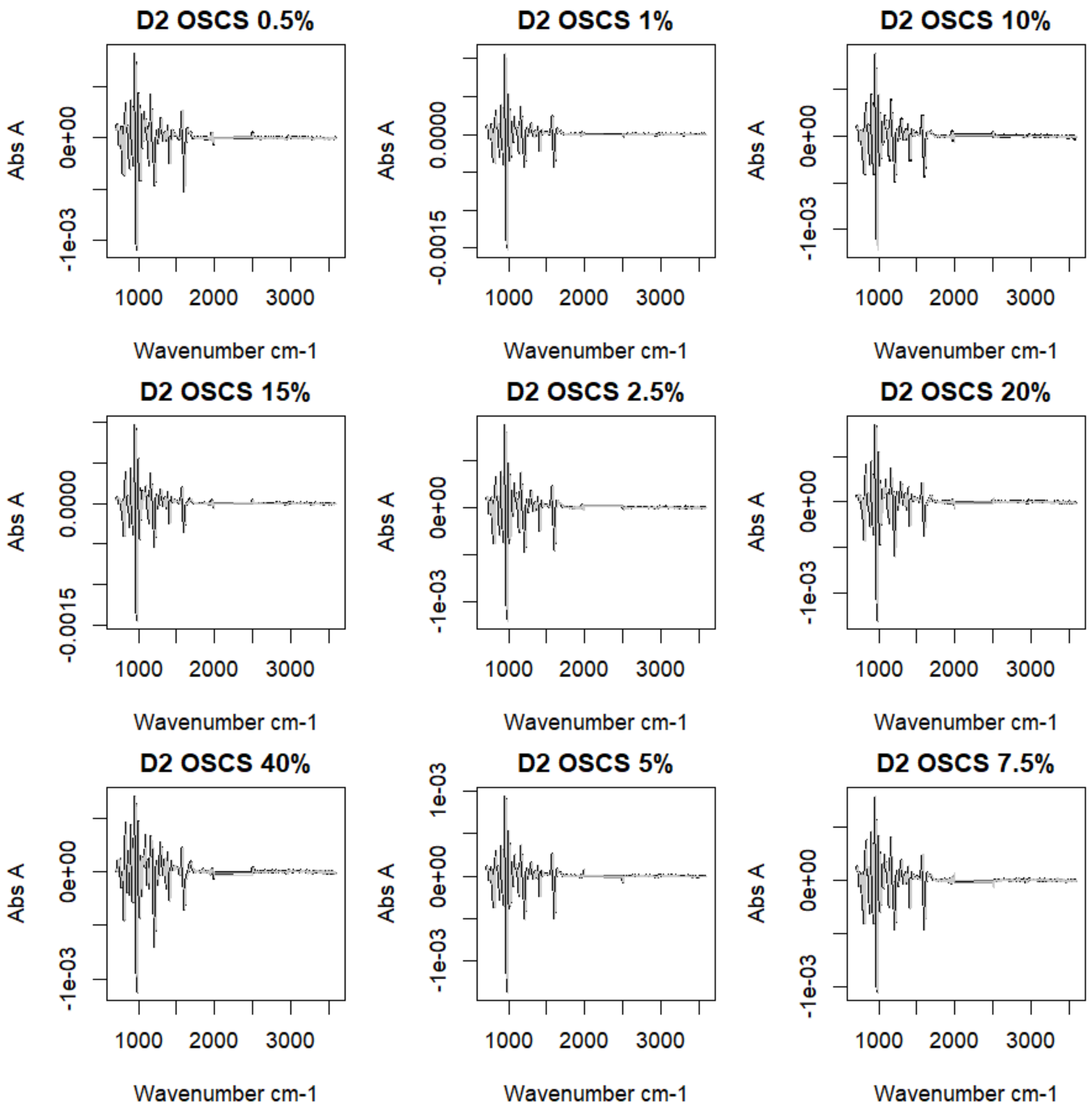
D2 OSCS 0.5%
D2 OSCS 1%
D2 OSCS 10%
D2 OSCS 15%
D2 OSCS 2.5%
D2 OSCS 20%
D2 OSCS 40%
D2 OSCS 5%
D2 OSCS 7.5%
Abs A
Wavenumber cm-1
1000
2000
3000
-1e-03
0e+00
1e-03
0.0000
-0.0015

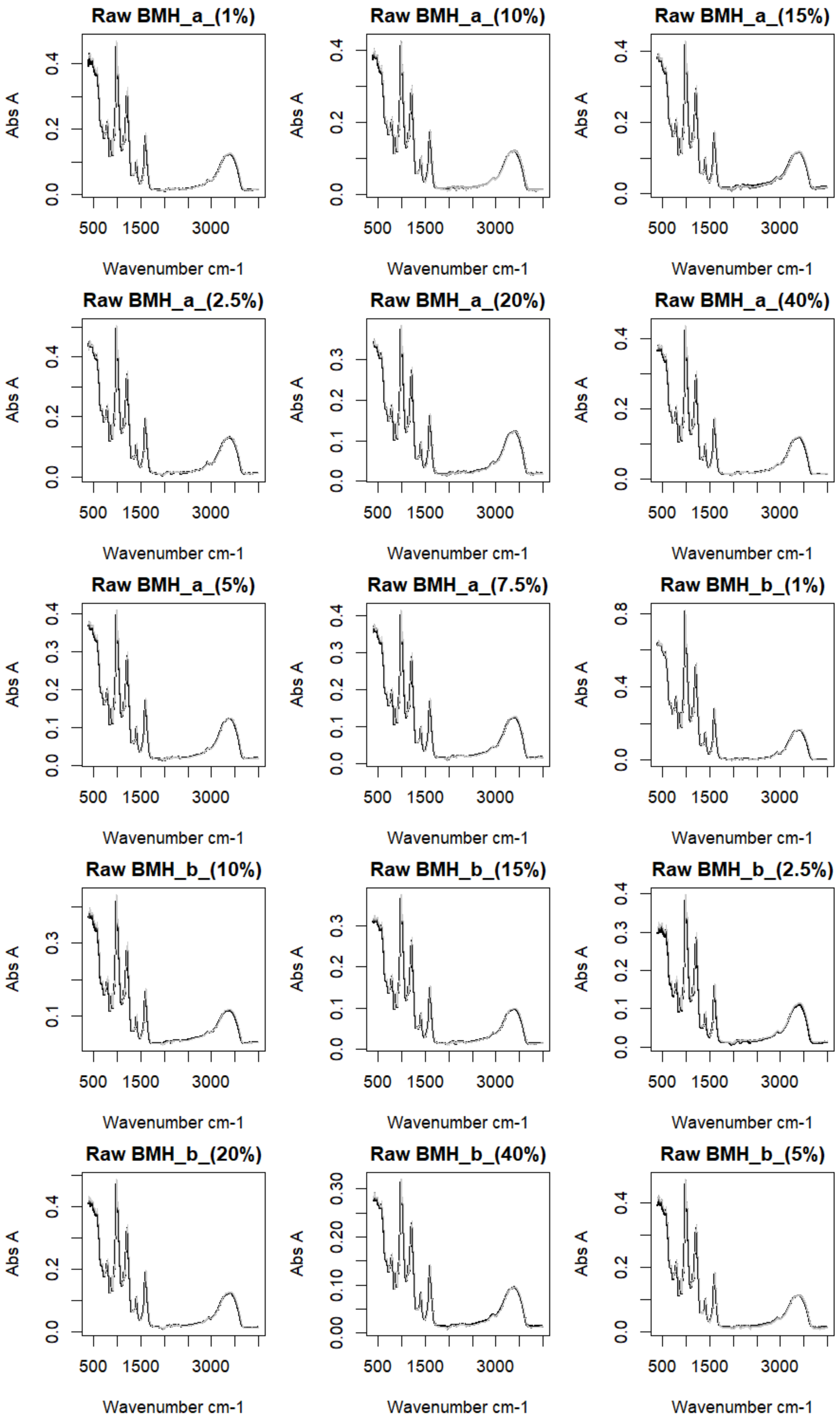
Raw BMH_a_(1%)
Raw BMH_a_(10%)
Raw BMH_a_(15%)
Raw BMH_a_(2.5%)
Raw BMH_a_(20%)
Raw BMH_a_(40%)
Raw BMH_a_(5%)
Raw BMH_a_(7.5%)
Raw BMH_b_(1%)
Raw BMH_b_(10%)
Raw BMH_b_(15%)
Raw BMH_b_(2.5%)
Raw BMH_b_(20%)
Raw BMH_b_(40%)
Raw BMH_b_(5%)
Abs A
Wavenumber cm-1

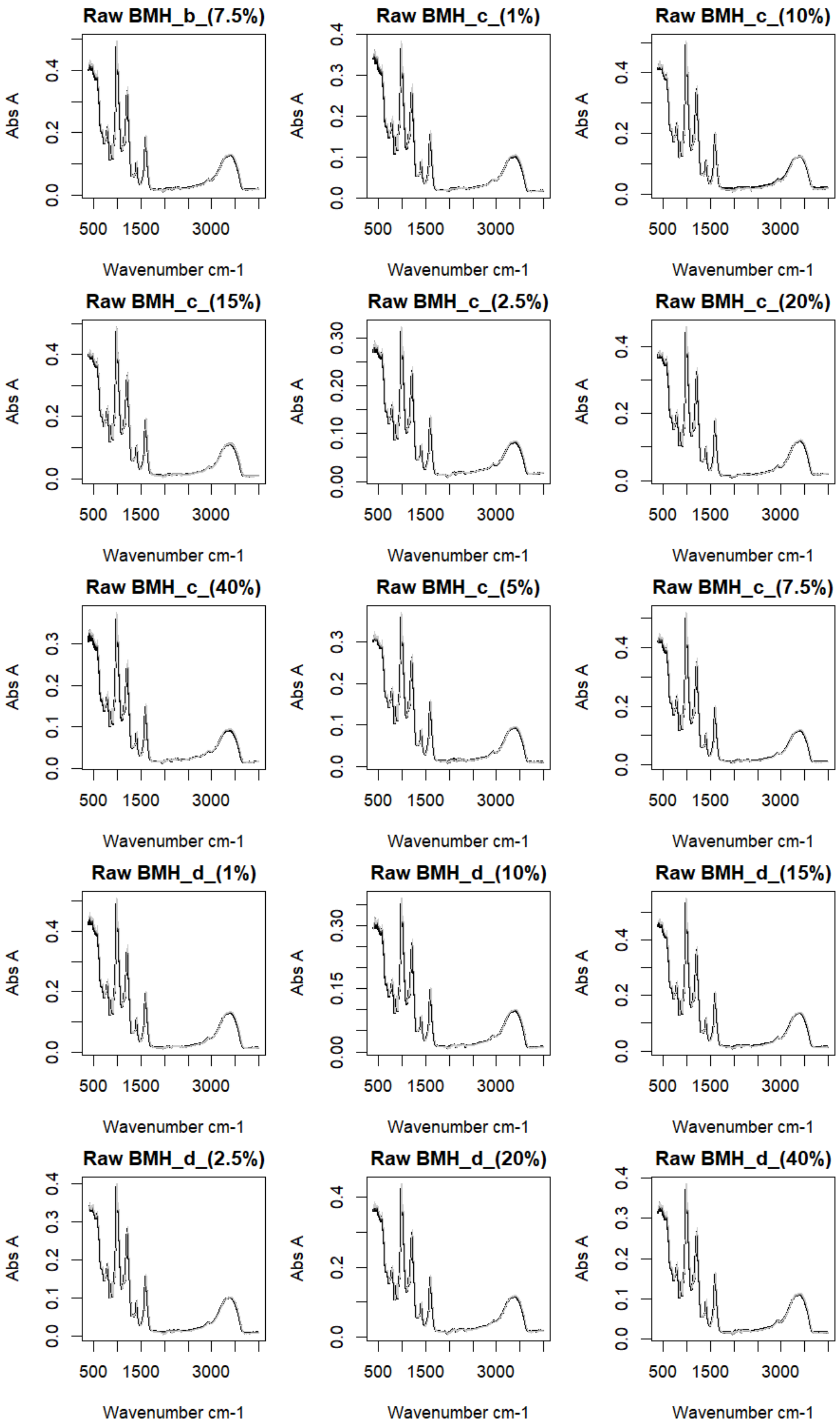

Raw BMH_b_(7.5%)
Raw BMH_c_(1%)
Raw BMH_c_(10%)
Raw BMH_c_(15%)
Raw BMH_c_(2.5%)
Raw BMH_c_(20%)
Raw BMH_c_(40%)
Raw BMH_c_(5%)
Raw BMH_c_(7.5%)
Raw BMH_d_(1%)
Raw BMH_d_(10%)
Raw BMH_d_(15%)
Raw BMH_d_(2.5%)
Raw BMH_d_(20%)
Raw BMH_d_(40%)
Abs A
Wavenumber cm-1
500
1500
3000

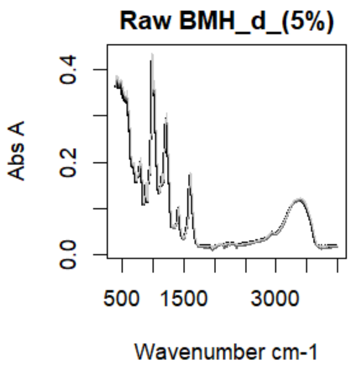

Raw BMH_d_(5%)
Abs A
0.0
0.2
0.4
500
1500
3000
Wavenumber cm-1


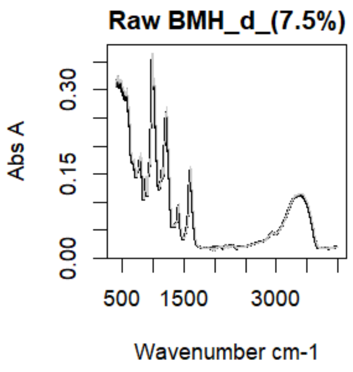

Raw BMH_d_(7.5%)
Abs A
0.00
0.15
0.30
500
1500
3000
Wavenumber cm-1

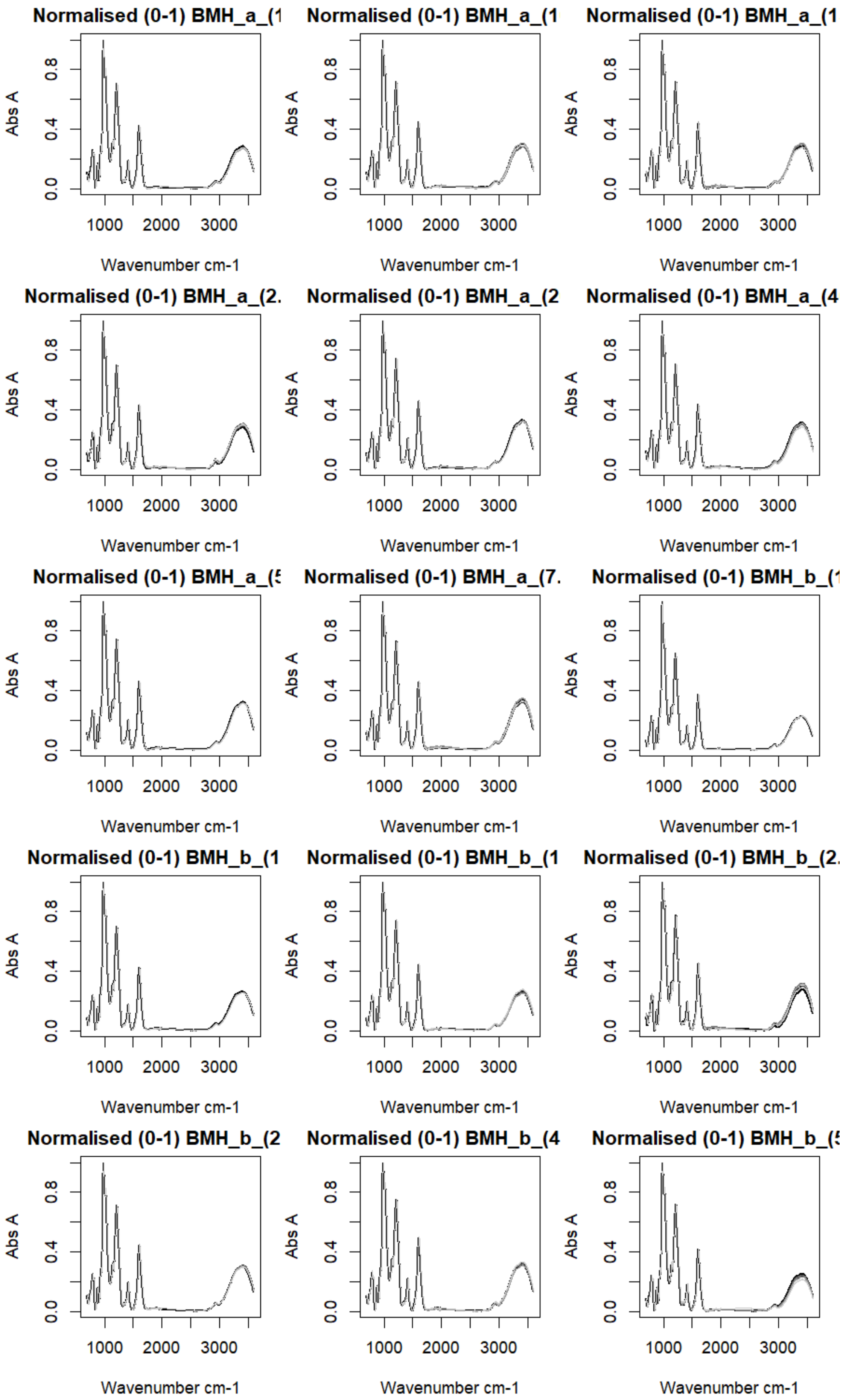
Normalised (0-1) BMH_a_(1
Normalised (0-1) BMH_a_(1
Normalised (0-1) BMH_a_(1
Normalised (0-1) BMH_a_(2.
Normalised (0-1) BMH_a_(2
Normalised (0-1) BMH_a_(4
Normalised (0-1) BMH_a_(5
Normalised (0-1) BMH_a_(7.
Normalised (0-1) BMH_b_(1
Normalised (0-1) BMH_b_(1
Normalised (0-1) BMH_b_(1
Normalised (0-1) BMH_b_(2.
Normalised (0-1) BMH_b_(2
Normalised (0-1) BMH_b_(4
Normalised (0-1) BMH_b_(5
Abs A
0.0
0.4
0.8
1000
2000
3000
Wavenumber cm-1

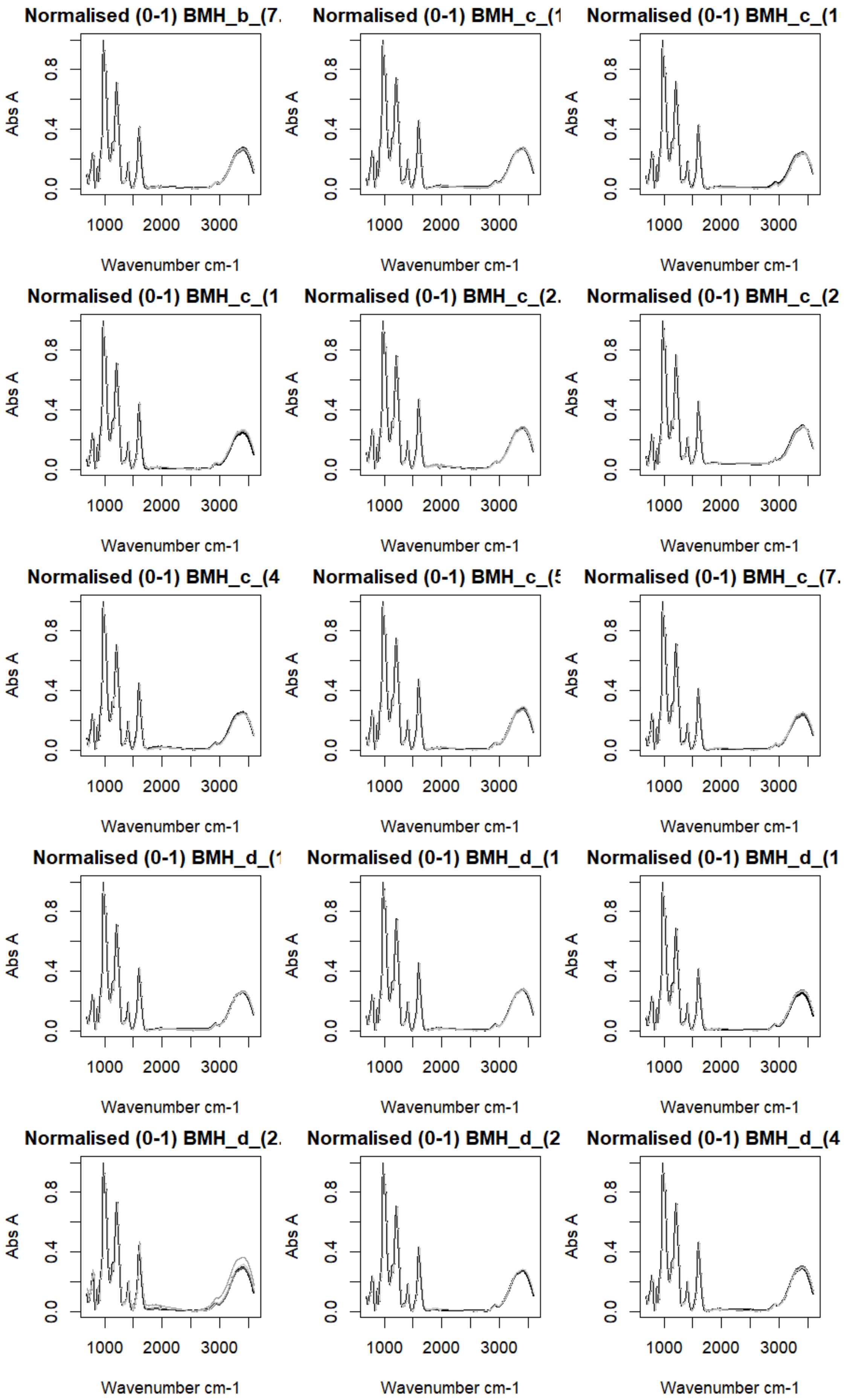
Normalised (0-1) BMH_b_(7.
Normalised (0-1) BMH_c_(1
Normalised (0-1) BMH_c_(1
Normalised (0-1) BMH_c_(1
Normalised (0-1) BMH_c_(2.
Normalised (0-1) BMH_c_(2
Normalised (0-1) BMH_c_(4
Normalised (0-1) BMH_c_(5
Normalised (0-1) BMH_c_(7.
Normalised (0-1) BMH_d_(1
Normalised (0-1) BMH_d_(1
Normalised (0-1) BMH_d_(1
Normalised (0-1) BMH_d_(2.
Normalised (0-1) BMH_d_(2
Normalised (0-1) BMH_d_(4
Abs A
0.0
0.4
0.8
1000
2000
3000
Wavenumber cm-1

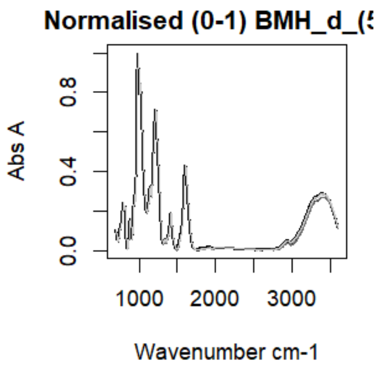
Normalised (0-1) BMH_d_(5
Abs A
0.0
0.4
0.8
1000
2000
3000
Wavenumber cm-1

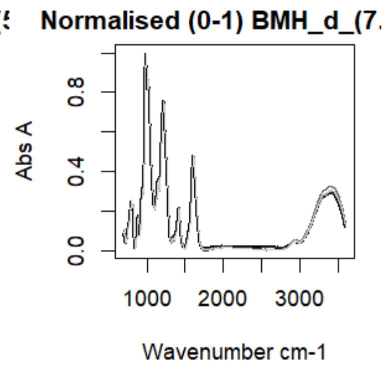
Normalised (0-1) BMH_d_(7.
Abs A
0.0
0.4
0.8
1000
2000
3000
Wavenumber cm-1

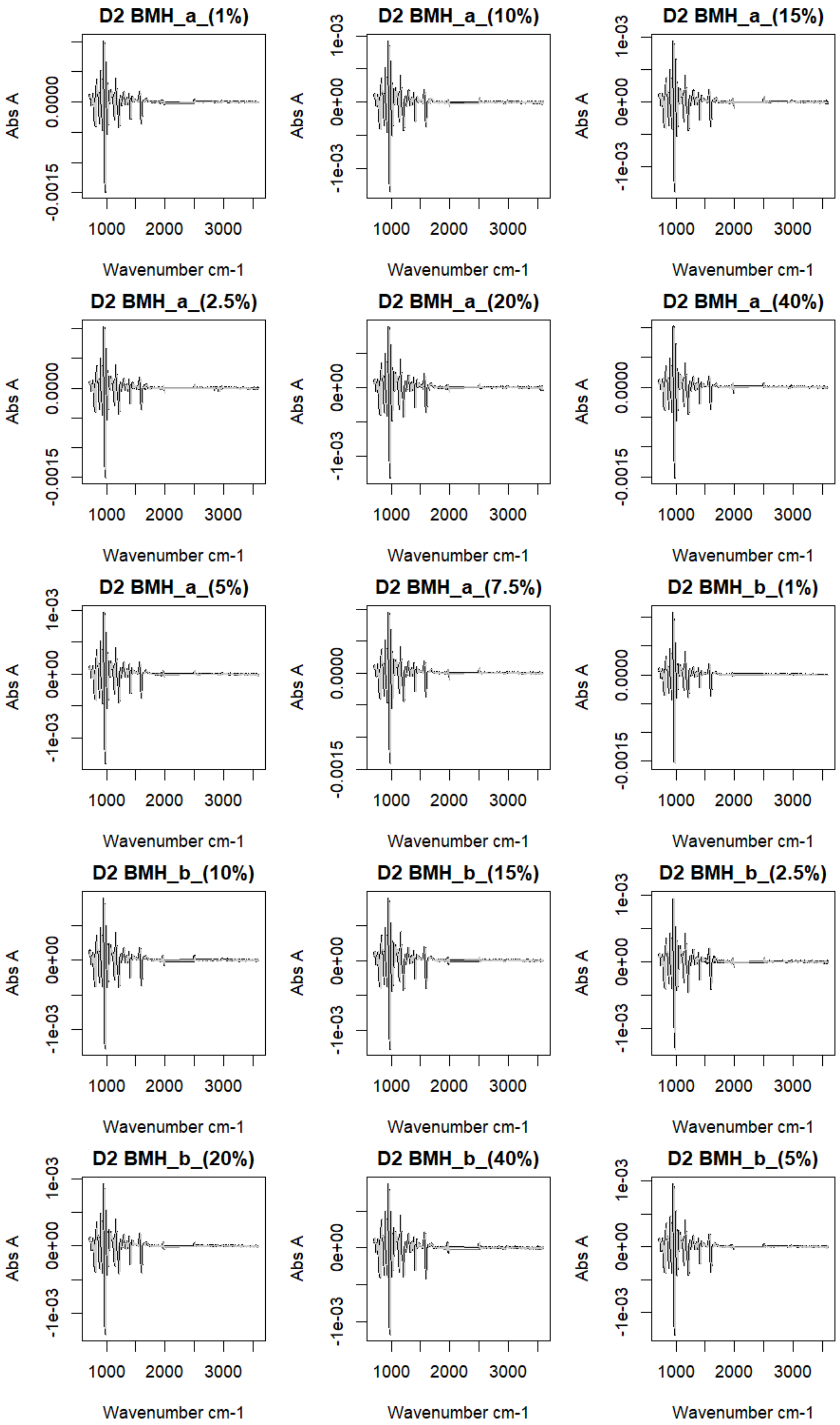
D2 BMH_a_(1%)
D2 BMH_a_(10%)
D2 BMH_a_(15%)
D2 BMH_a_(2.5%)
D2 BMH_a_(20%)
D2 BMH_a_(40%)
D2 BMH_a_(5%)
D2 BMH_a_(7.5%)
D2 BMH_b_(1%)
D2 BMH_b_(10%)
D2 BMH_b_(15%)
D2 BMH_b_(2.5%)
D2 BMH_b_(20%)
D2 BMH_b_(40%)
D2 BMH_b_(5%)
Abs A
Wavenumber cm-1
1000
2000
3000
0.0000
-0.0015
1e-03
0e+00
-1e-03

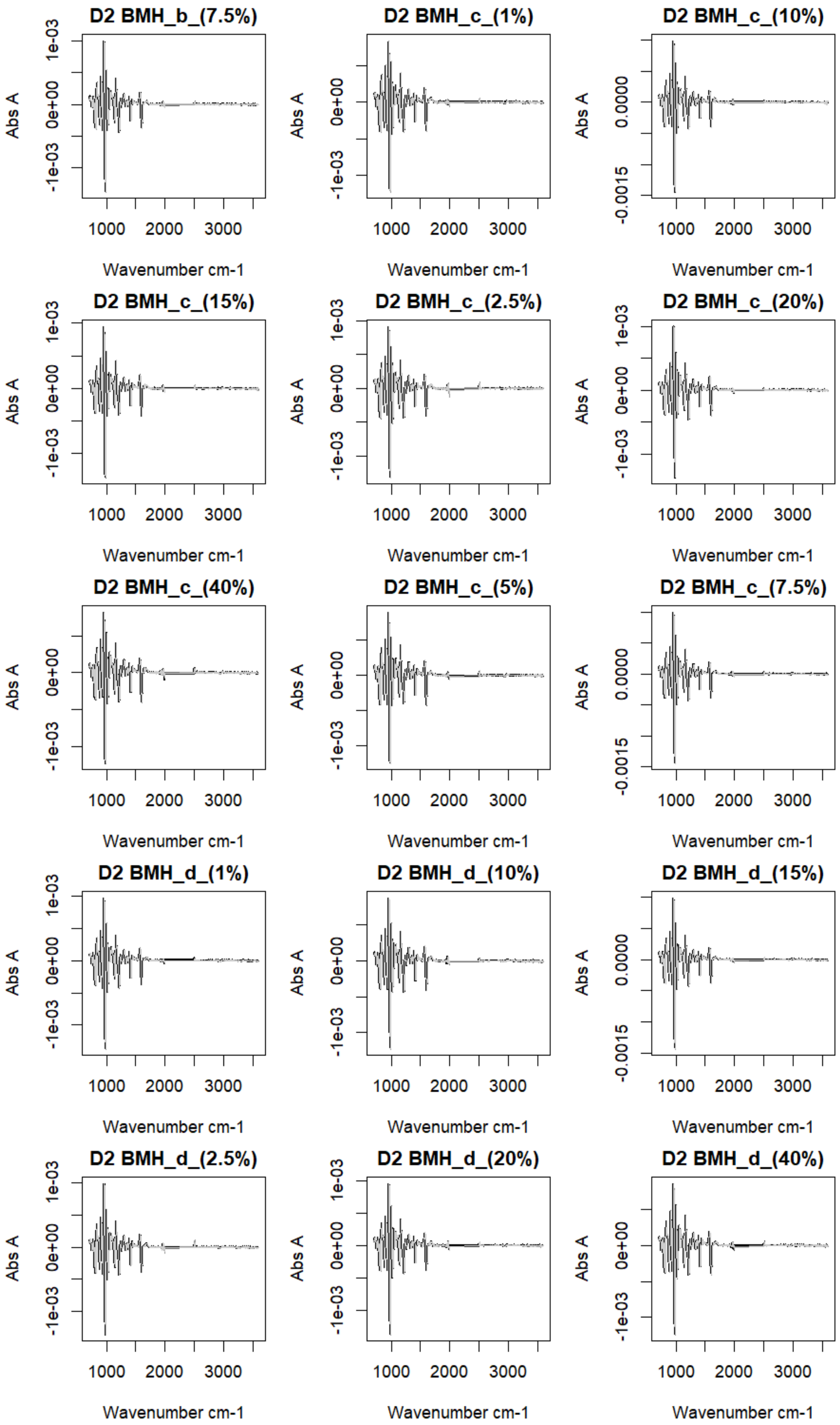
D2 BMH_b_(7.5%)
D2 BMH_c_(1%)
D2 BMH_c_(10%)
D2 BMH_c_(15%)
D2 BMH_c_(2.5%)
D2 BMH_c_(20%)
D2 BMH_c_(40%)
D2 BMH_c_(5%)
D2 BMH_c_(7.5%)
D2 BMH_d_(1%)
D2 BMH_d_(10%)
D2 BMH_d_(15%)
D2 BMH_d_(2.5%)
D2 BMH_d_(20%)
D2 BMH_d_(40%)
Abs A
Wavenumber cm-1
1000
2000
3000
-1e-03
0e+00
1e-03
-0.0015
0.0000

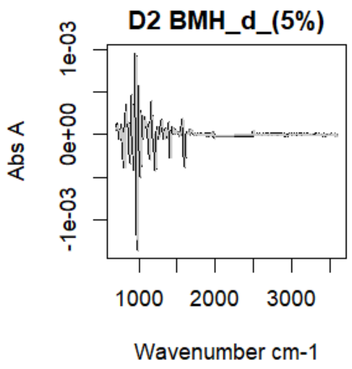
D2 BMH_d_(5%)
Abs A
1e-03
0e+00
-1e-03
1000
2000
3000
Wavenumber cm-1

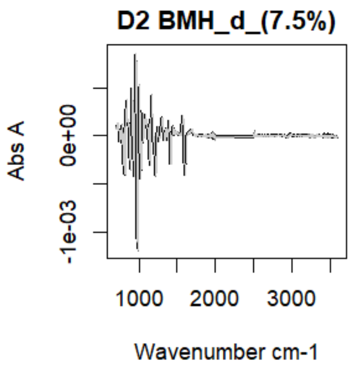
D2 BMH_d_(7.5%)
Abs A
0e+00
-1e-03
1000
2000
3000
Wavenumber cm-1

### EVerything New Code D2 7th

Anthony Devlin, Keele University

### PCA of PCA of Everything

Scaled Matrix PCAed

7th order polynomial baseline correction performed

Raw spectra smoothed to 21 Neighbours, 2nd order PN, Savtisky Golay

Normalisation between 0 and 100

D2 spectra smoothed to 41 Neighbours, 2nd order PN, Savtisky Golay

Regions Analysed: 3601−2501 and 2001−700cm−1

Samples Analysed: 24 BLH's 55 BMH's 31 C1's 21 C2's 6 C3's 3 Contam's
10 GAG's 31 HS's 33 OMH's 1 PLH's 69 PMH's

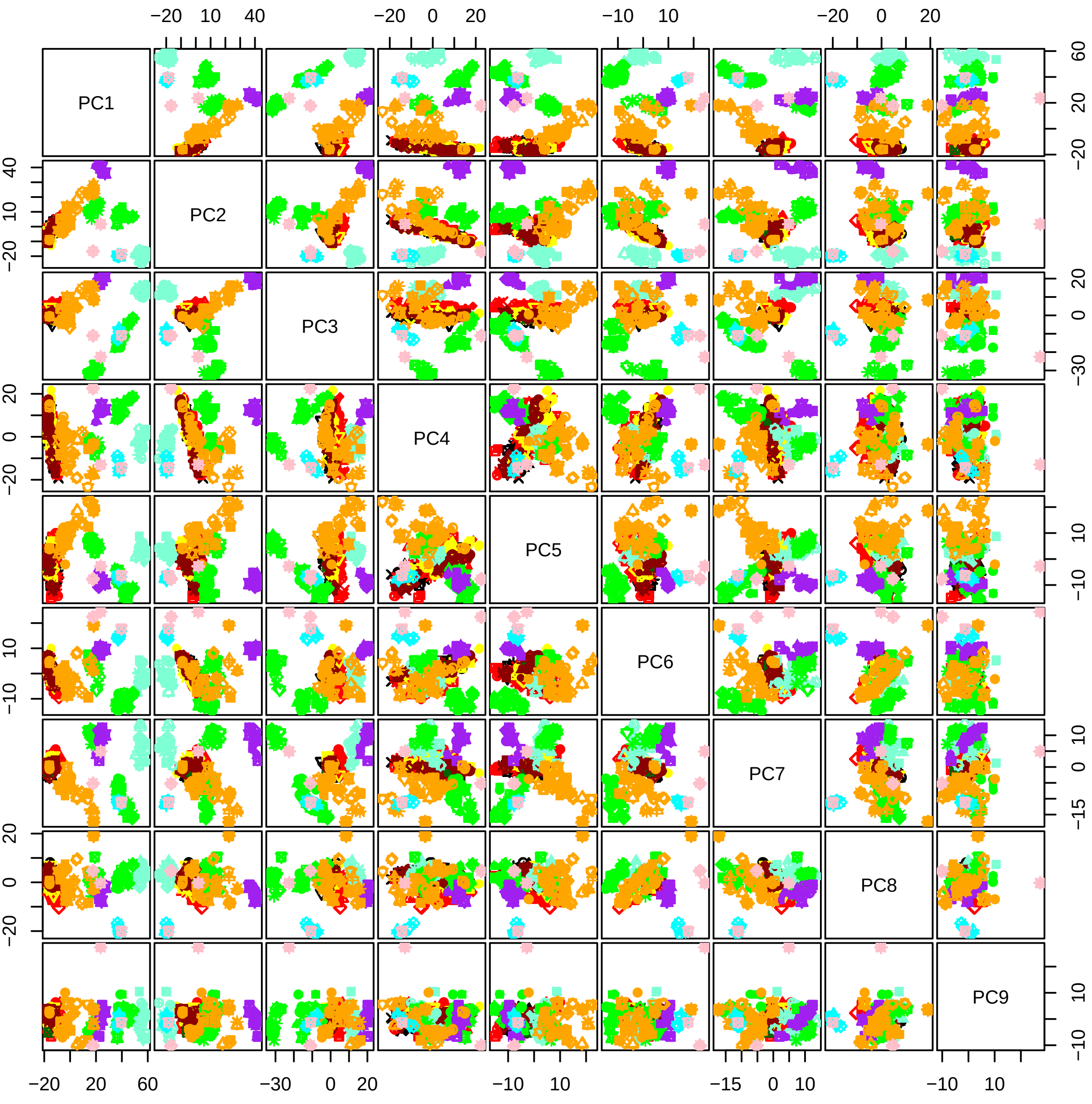
PC1
PC2
PC3
PC4
PC5
PC6
PC7
PC8
PC9
−20
10
40
−20
0
20
−10
10
−20
0
20
60
20
−20
20
0
−30
20
10
−10
10
0
−15
10
−10
−20
20
60
−30
0
20
−10
10
−15
0
10
−10
10

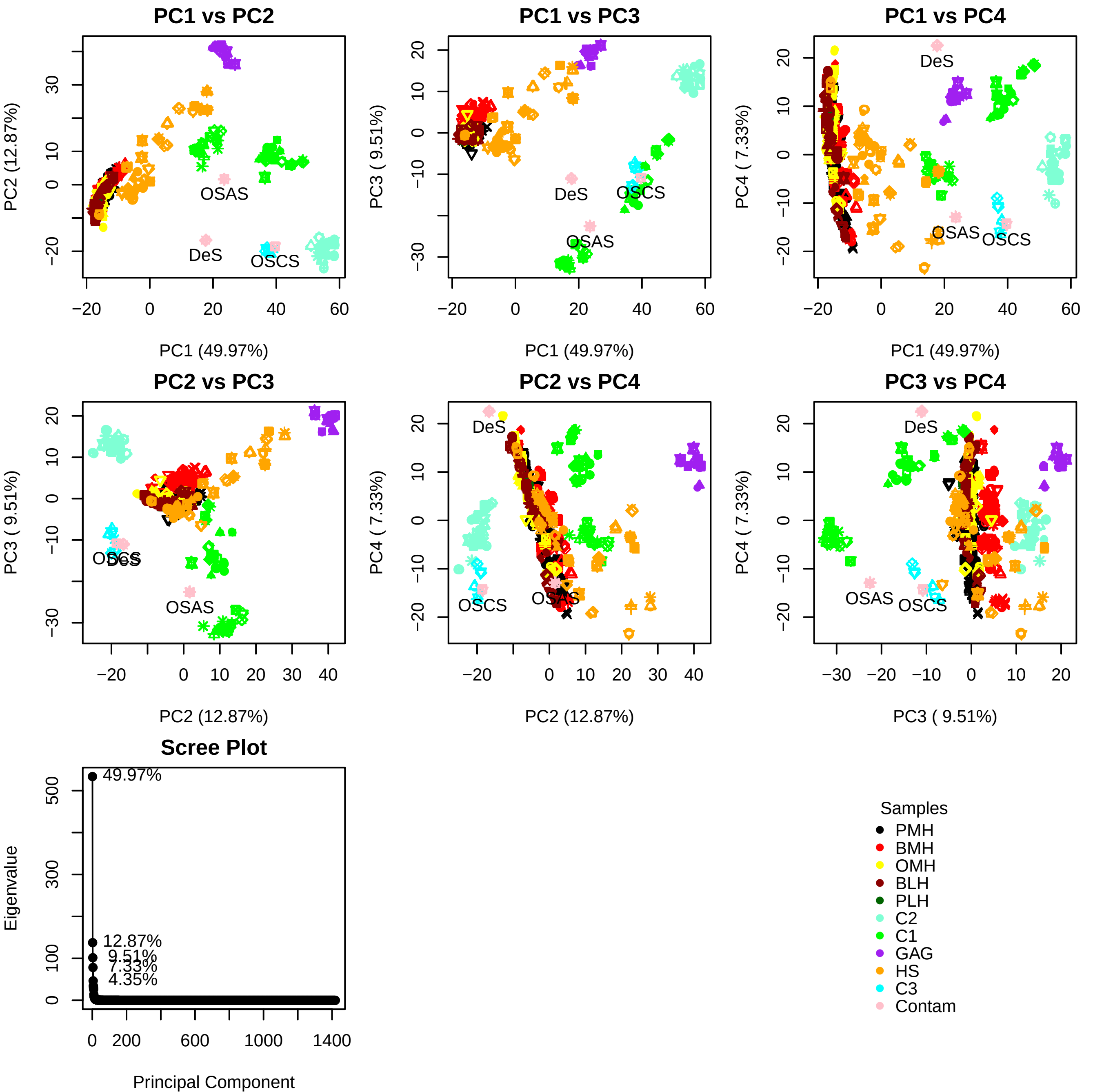

PC1 vs PC2
PC1 vs PC3
PC1 vs PC4
PC2 vs PC3
PC2 vs PC4
PC3 vs PC4
Scree Plot
PC1 (49.97%)
PC2 (12.87%)
PC3 ( 9.51%)
PC4 ( 7.33%)
OSAS
DeS
OSCS
Eigenvalue
Principal Component
49.97%
12.87%
9.51%
7.33%
4.35%
Samples
PMH
BMH
OMH
BLH
PLH
C2
C1
GAG
HS
C3
Contam

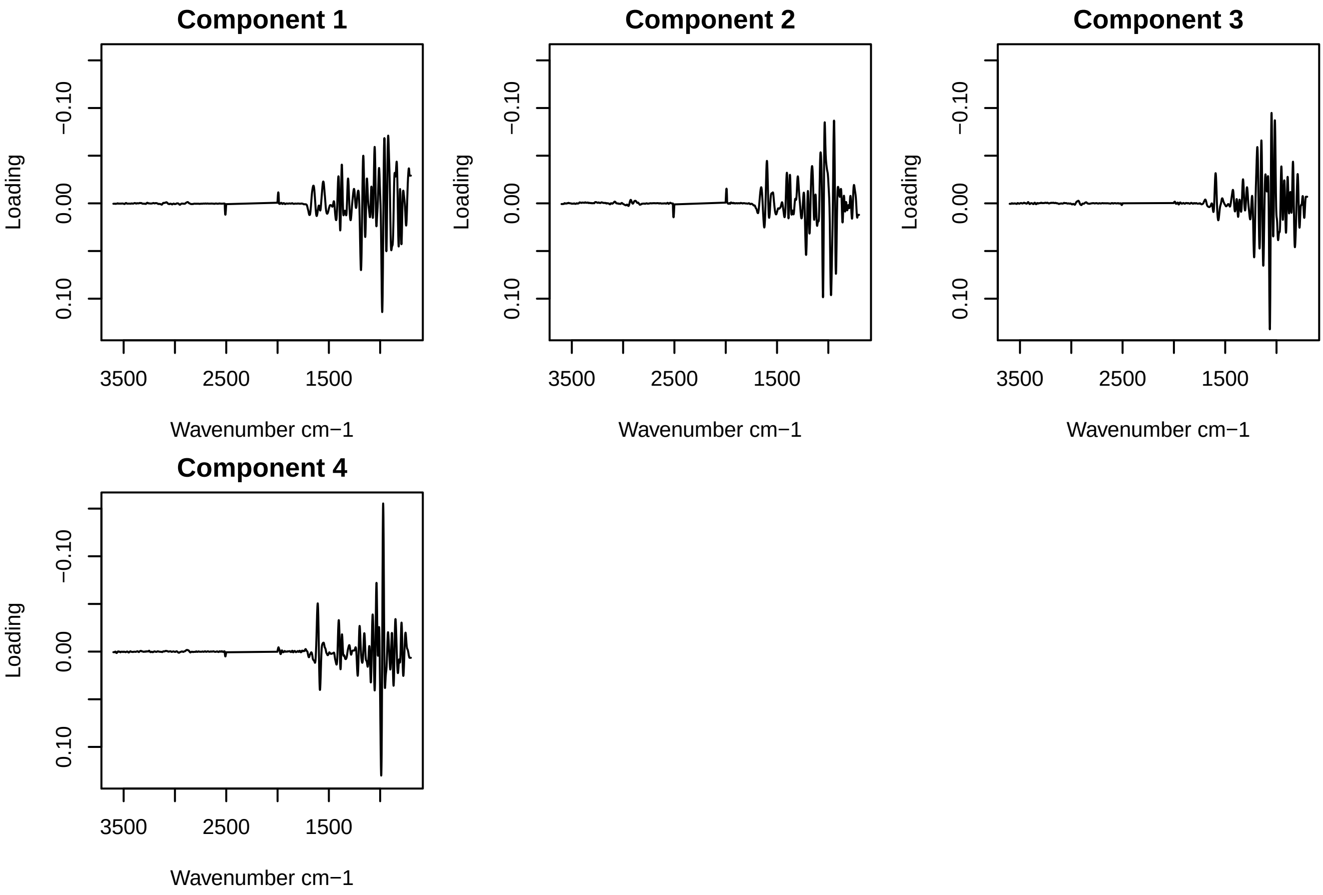
Component 1
Component 2
Component 3
Component 4
Loading
−0.10
0.00
0.10
3500
2500
1500
Wavenumber cm−1

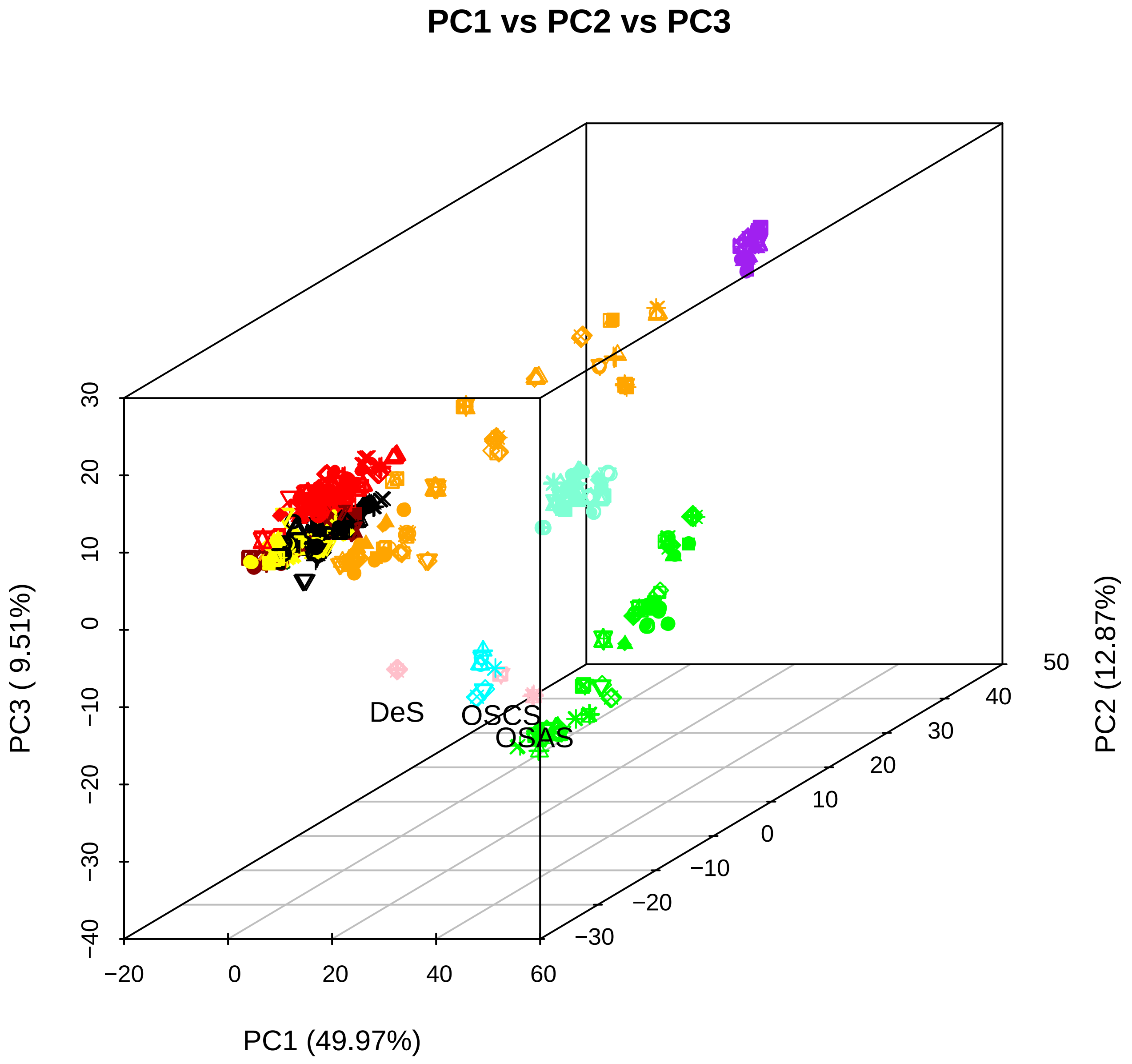
PC1 vs PC2 vs PC3
PC1 (49.97%)
PC2 (12.87%)
PC3 ( 9.51%)
−20
0
20
40
60
−30
−20
−10
0
10
20
30
40
50
−40
−30
−20
−10
0
10
20
30
DeS
OSCS
OSAS

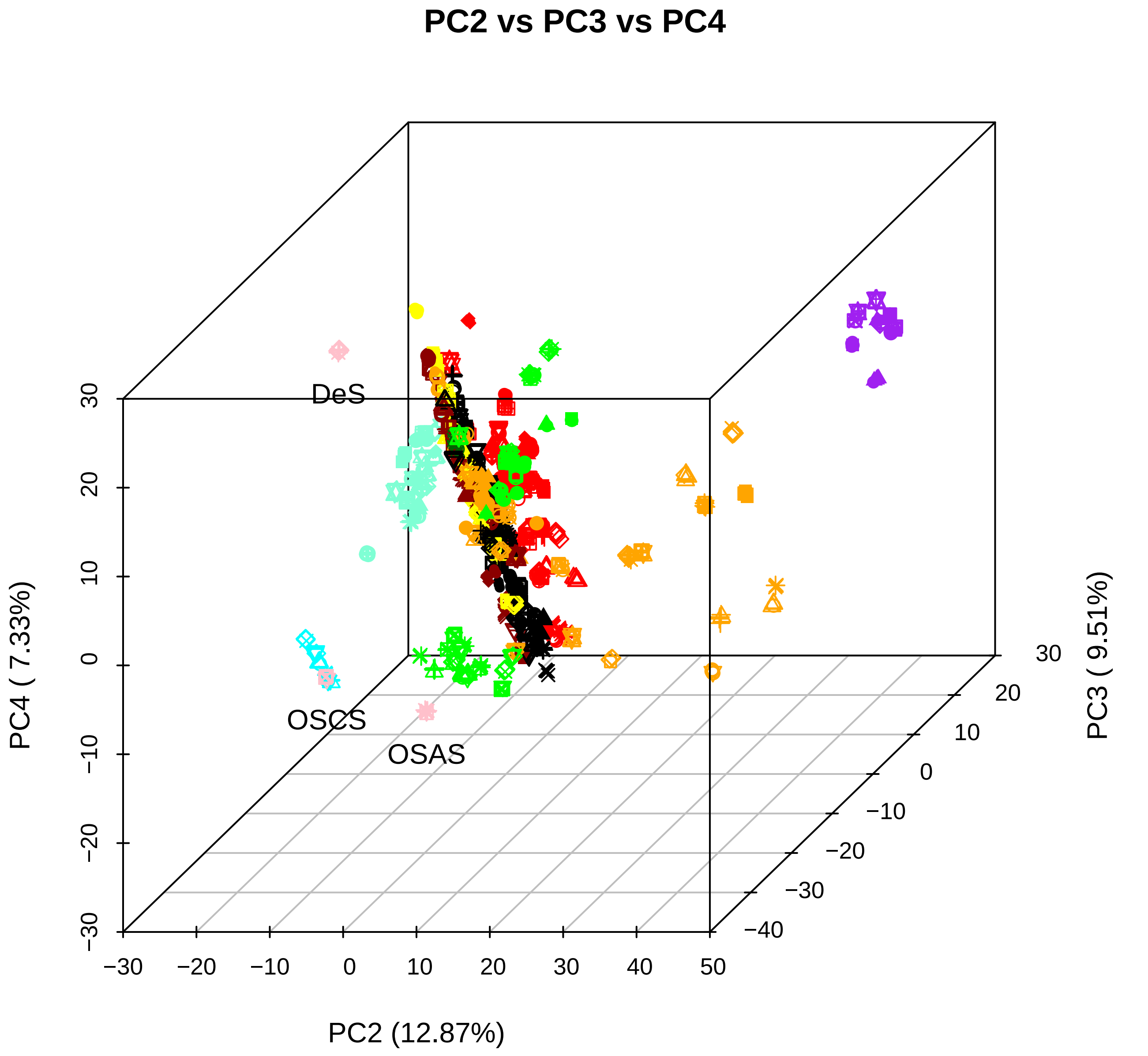
PC2 vs PC3 vs PC4
DeS
OSCS
OSAS
PC2 (12.87%)
PC3 ( 9.51%)
PC4 ( 7.33%)
−30
−20
−10
0
10
20
30
40
50
−40
−30
−20
−10
0
10
20
30

## PC1 vs PC2 vs PC4

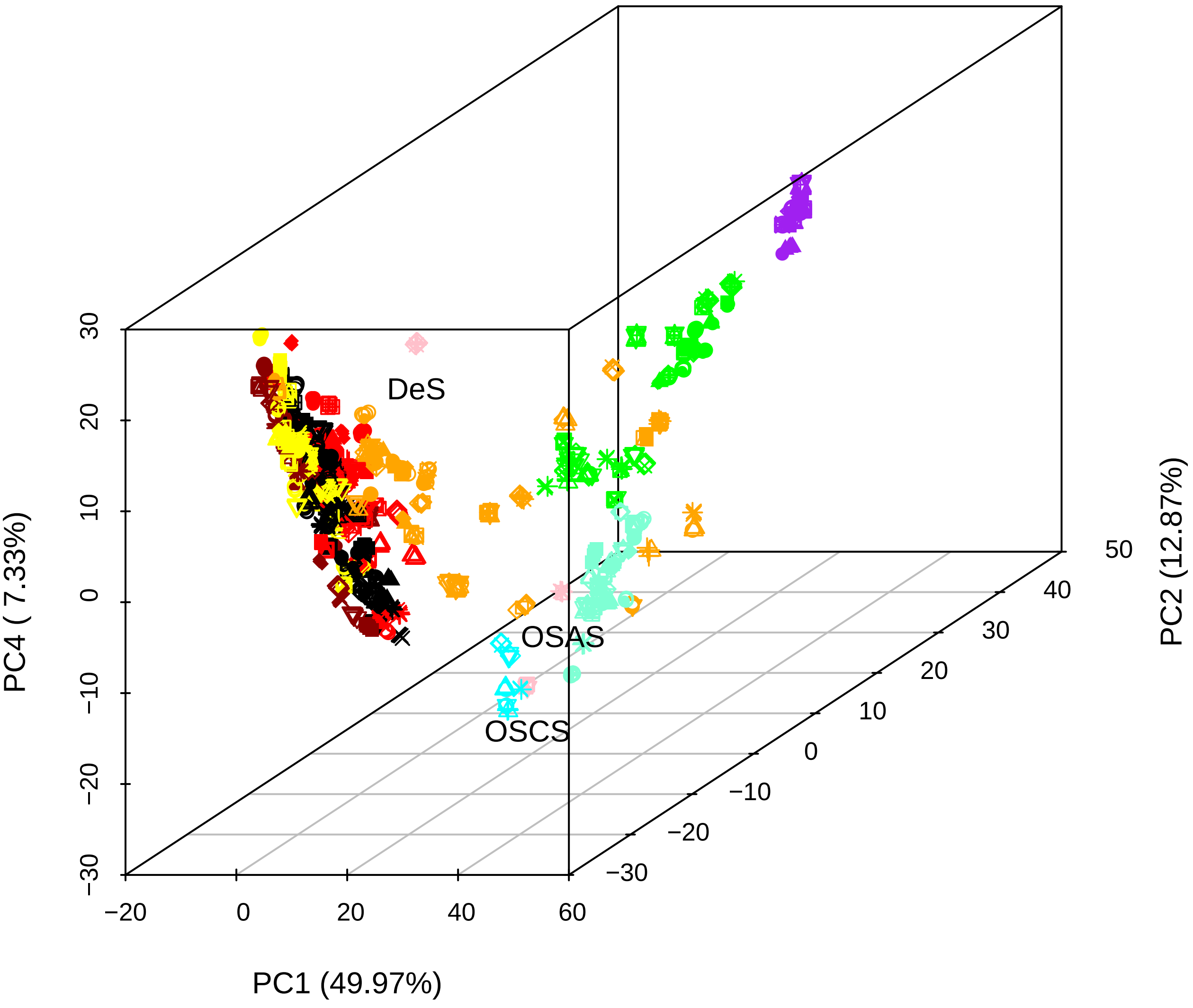

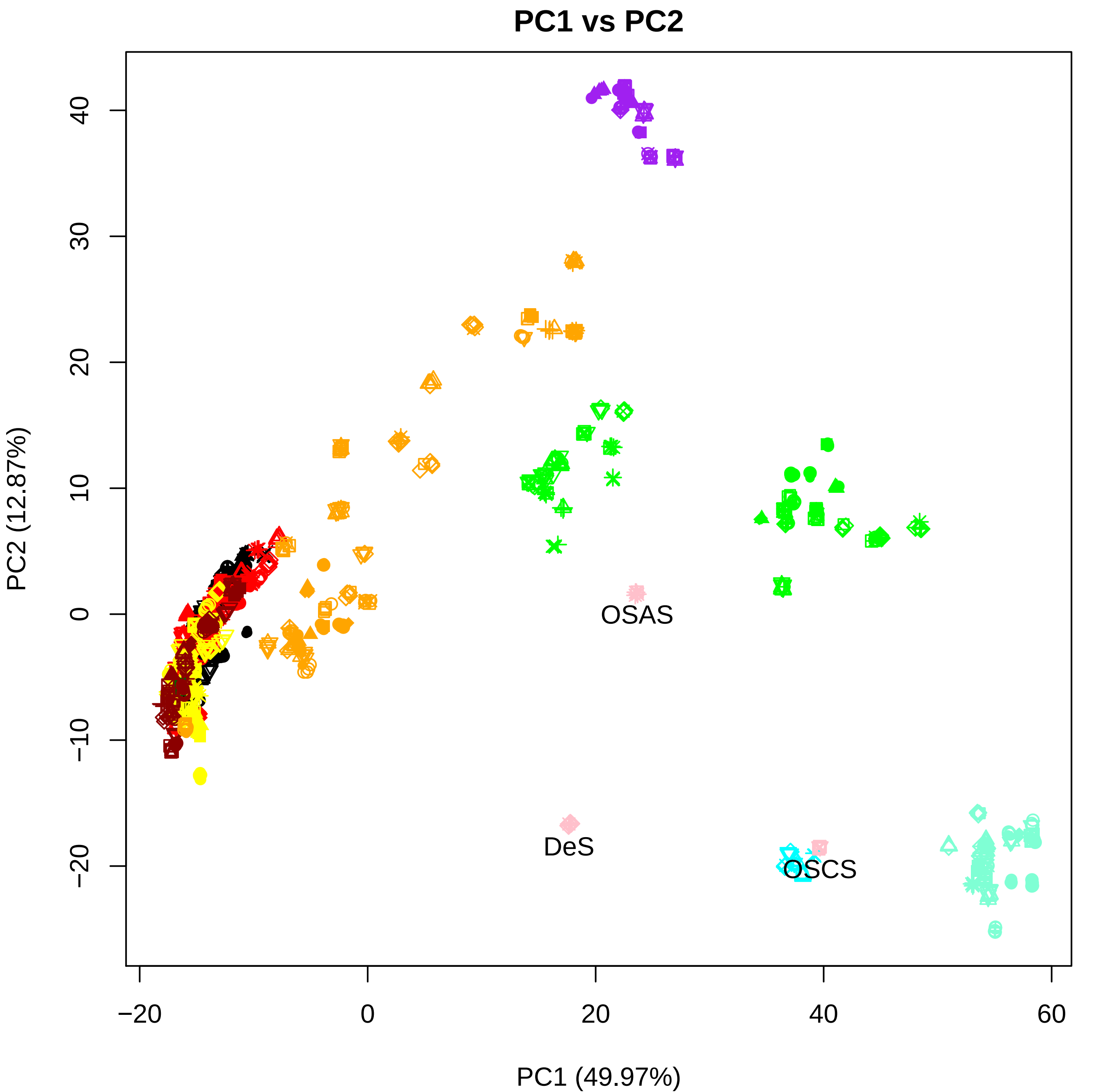
PC1 vs PC2
PC2 (12.87%)
PC1 (49.97%)
−20
0
20
40
60
−20
−10
0
10
20
30
40
OSAS
DeS
OSCS

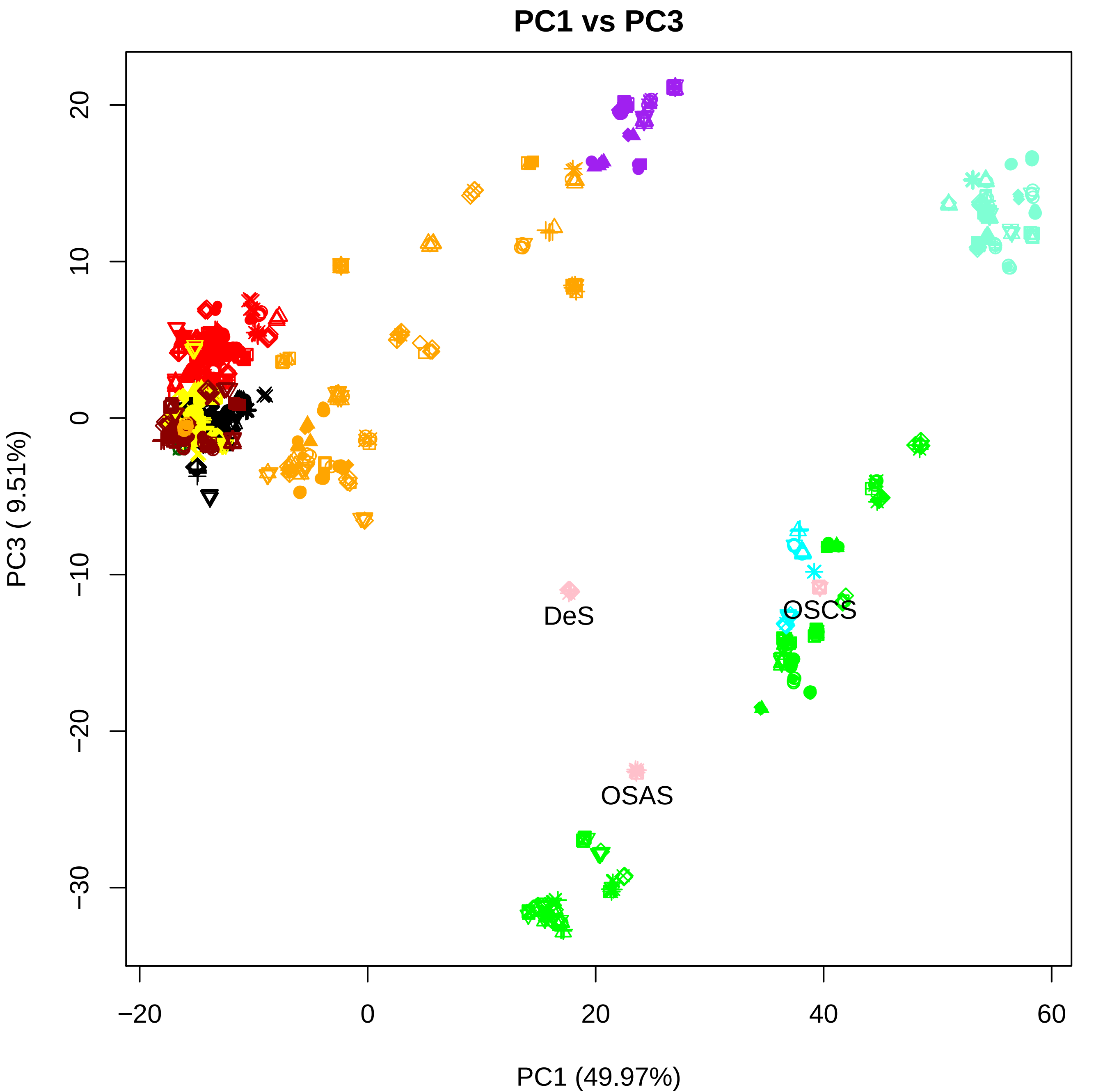
PC1 vs PC3
PC3 ( 9.51%)
PC1 (49.97%)
20
10
0
−10
−20
−30
−20
0
20
40
60
DeS
OSCS
OSAS

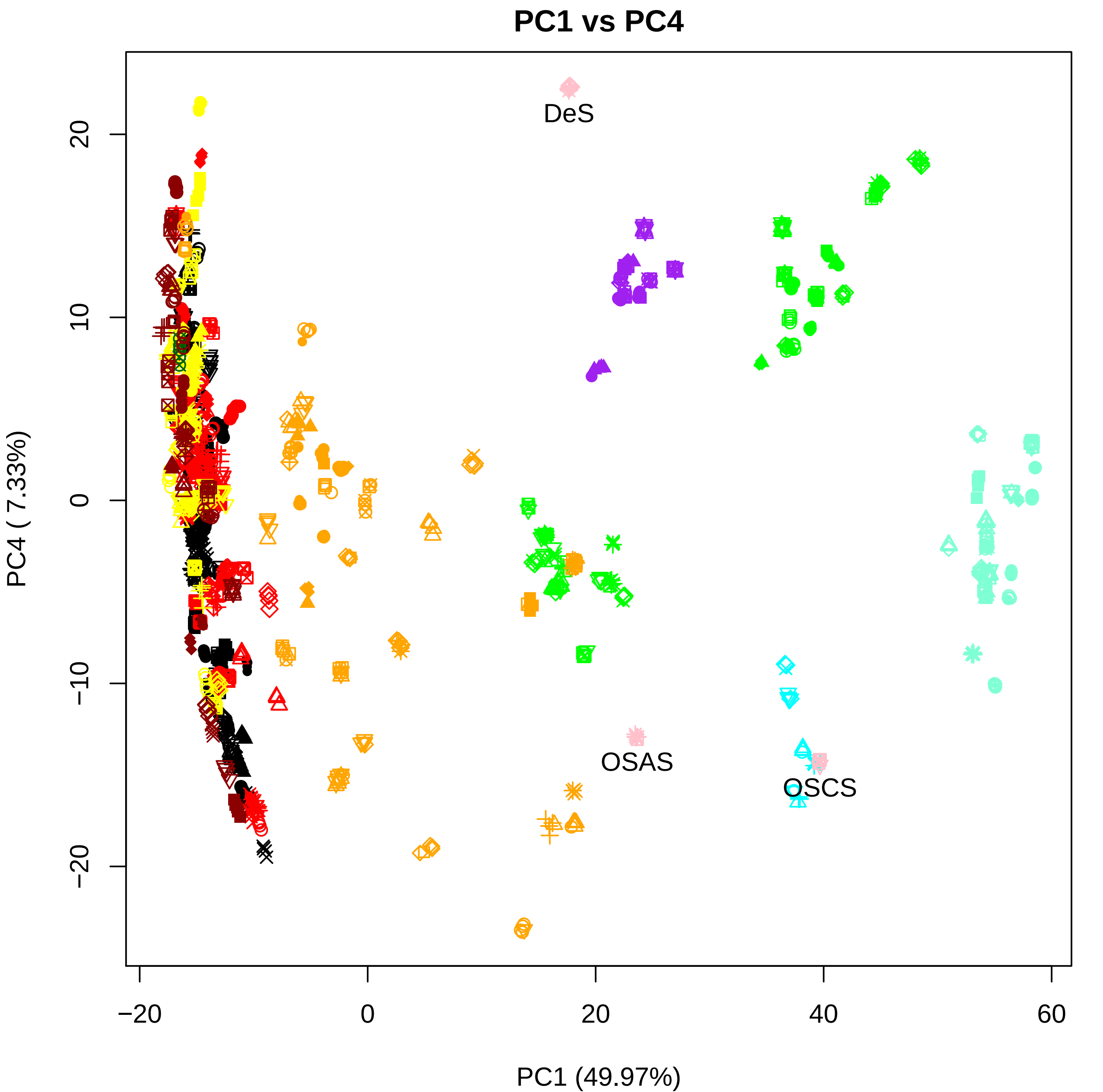

PC1 vs PC4
DeS
OSAS
OSCS
PC4 ( 7.33%)
PC1 (49.97%)
20
10
0
−10
−20
−20
0
20
40
60

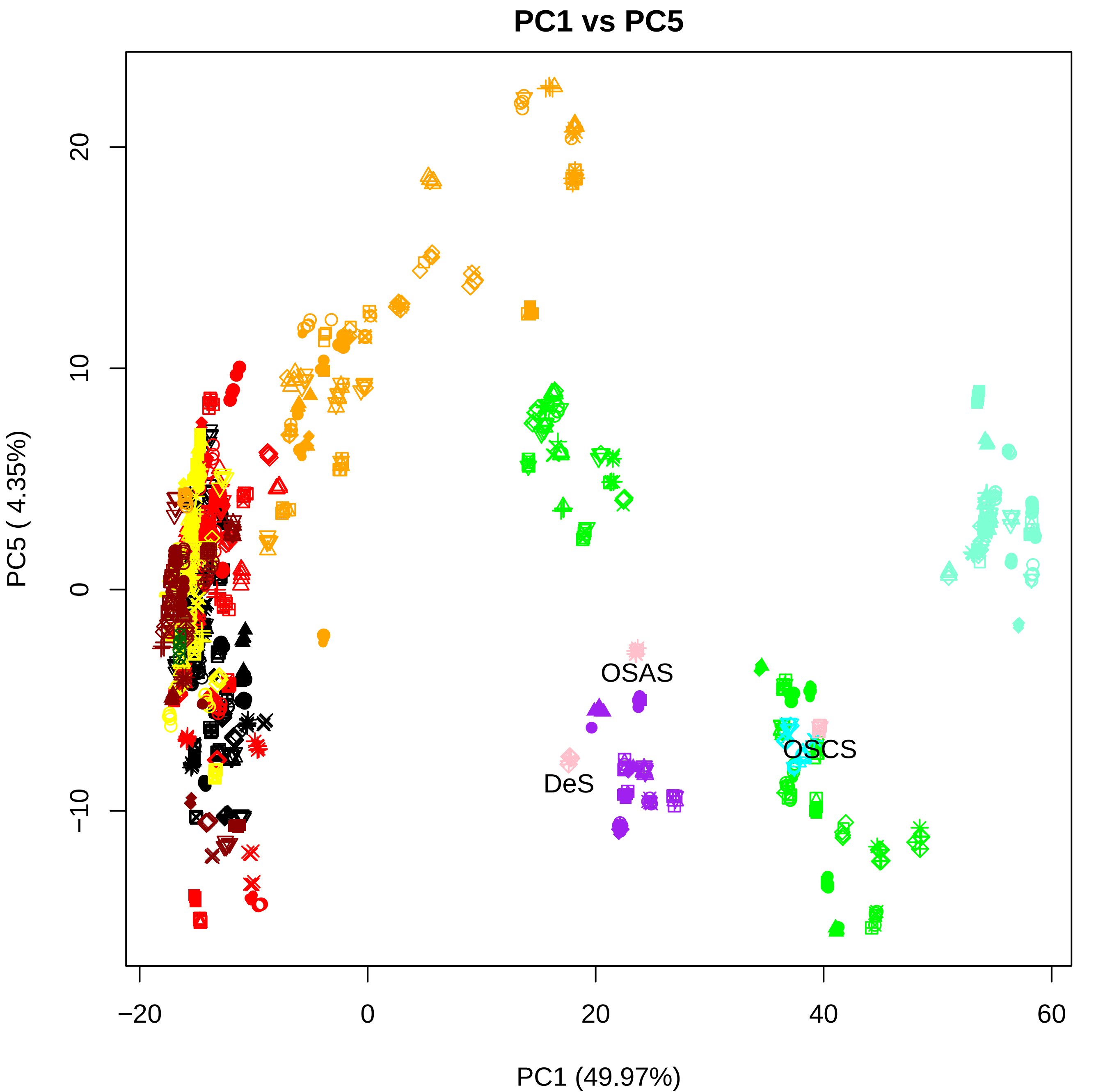
PC1 vs PC5
PC5 ( 4.35%)
PC1 (49.97%)
20
10
0
−10
−20
0
20
40
60
OSAS
DeS
OSCS

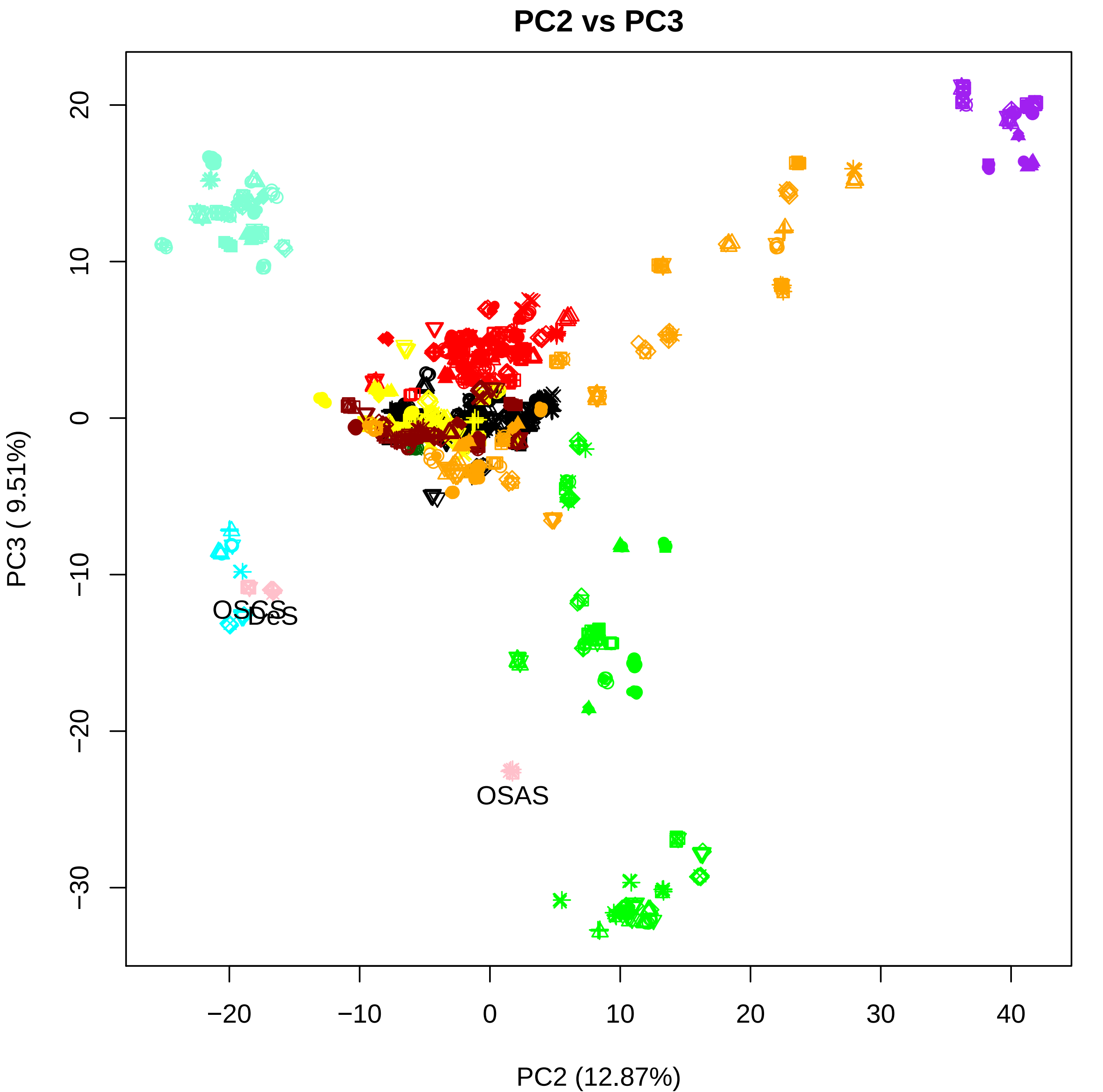
PC2 vs PC3
PC3 ( 9.51%)
PC2 (12.87%)
20
10
0
−10
−20
−30
−20
−10
0
10
20
30
40
OSCS
DeS
OSAS

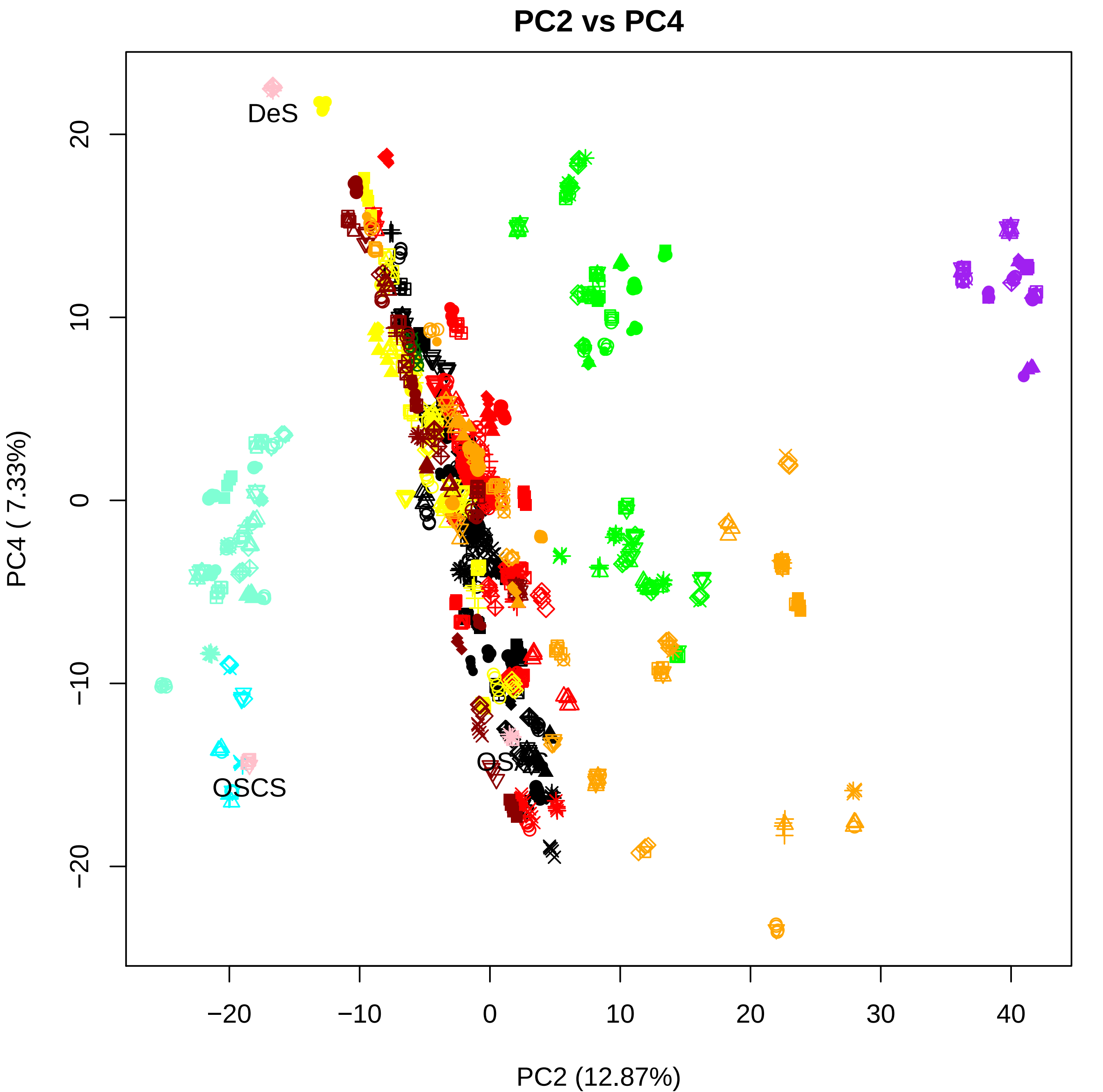
PC2 vs PC4
DeS
OSCS
PC4 ( 7.33%)
20
10
0
−10
−20
PC2 (12.87%)
−20
−10
0
10
20
30
40

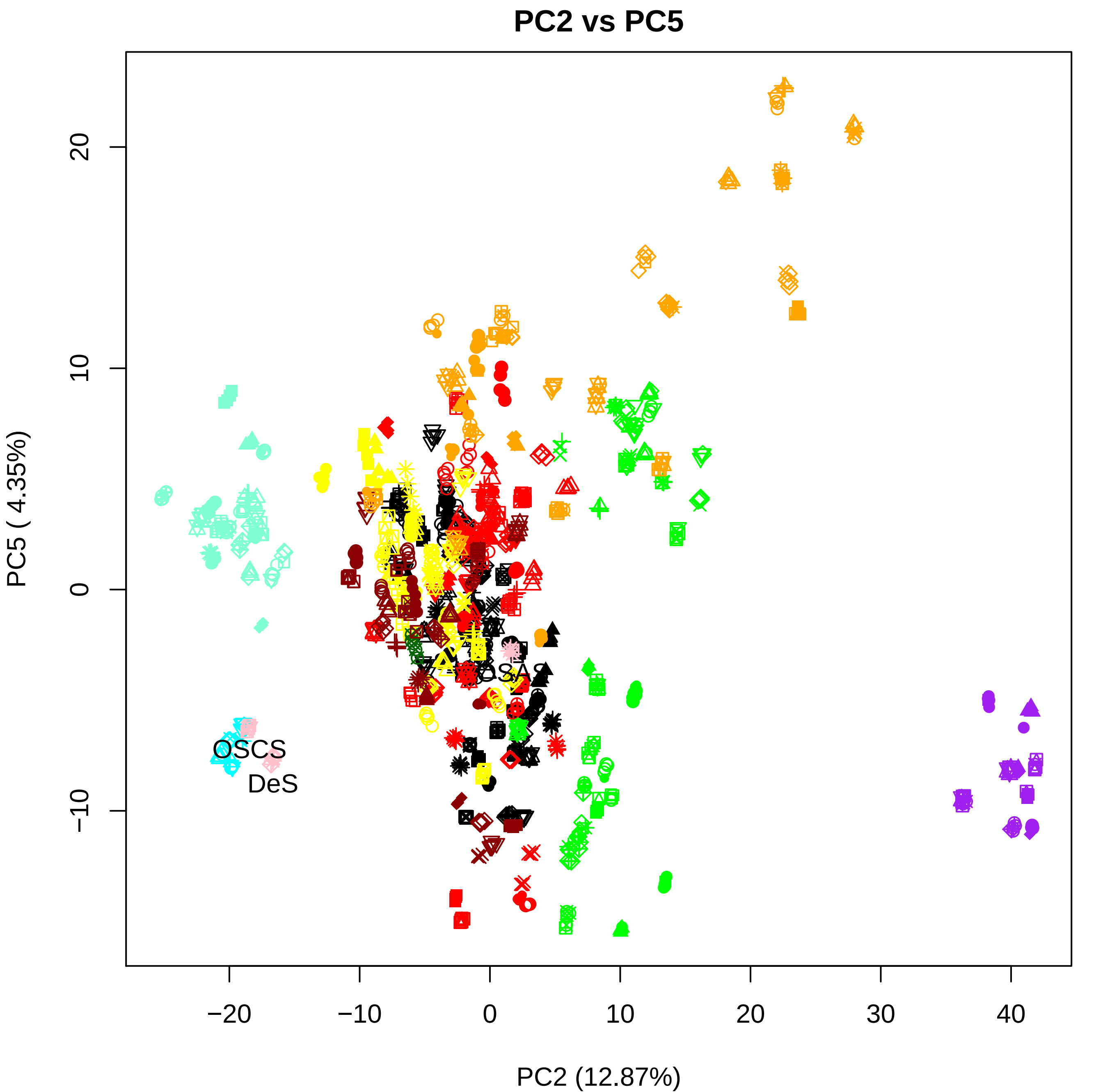
PC2 vs PC5
PC5 ( 4.35%)
PC2 (12.87%)
20
10
0
−10
−20
−10
0
10
20
30
40
OSCS
DeS
SAS

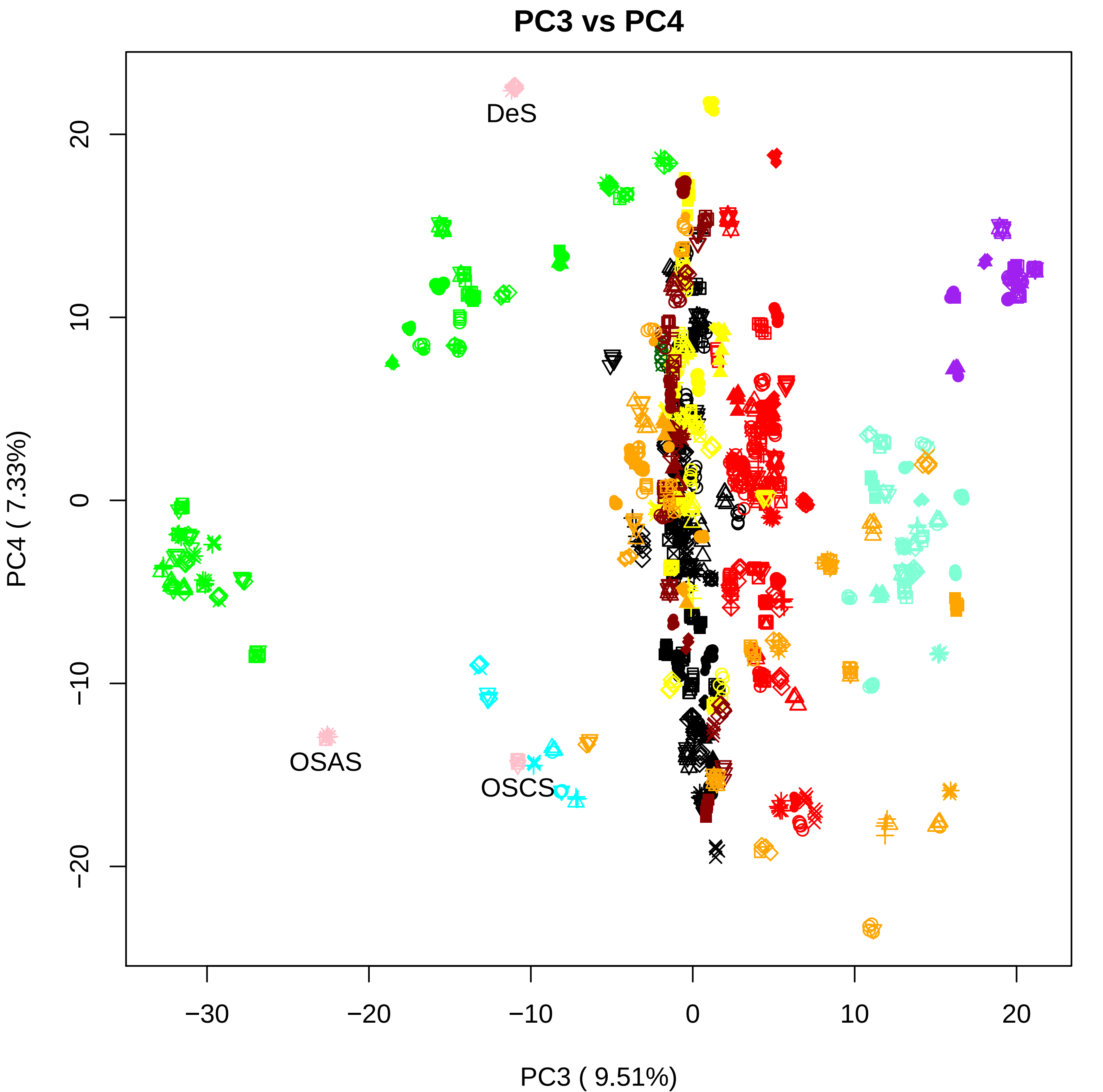
PC3 vs PC4
DeS
OSAS
OSCS
PC4 ( 7.33%)
PC3 ( 9.51%)
20
10
0
−10
−20
−30
−20
−10
0
10
20

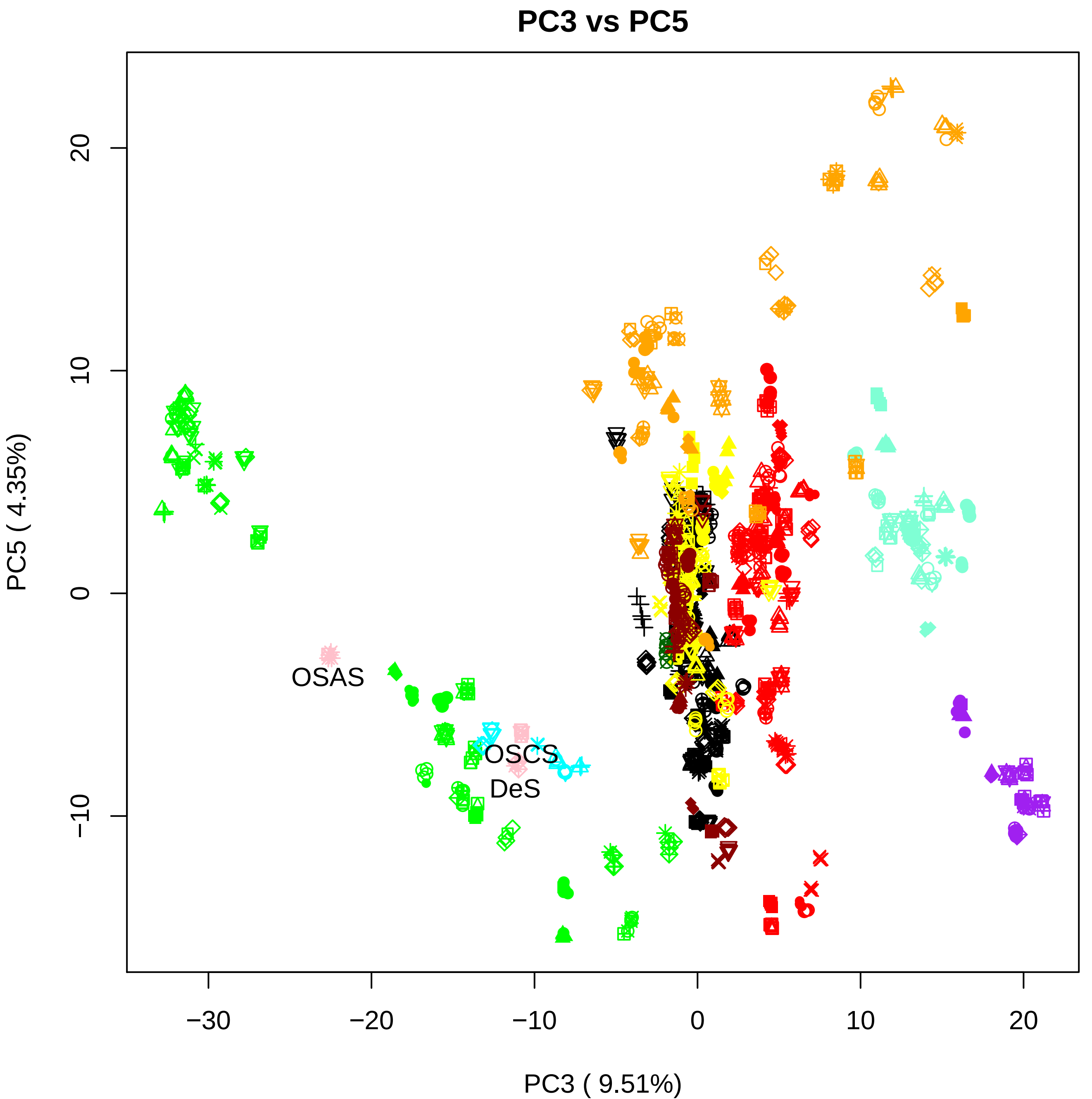
PC3 vs PC5
PC5 ( 4.35%)
PC3 ( 9.51%)
20
10
0
−10
−30
−20
−10
0
10
20
OSAS
OSCS
DeS

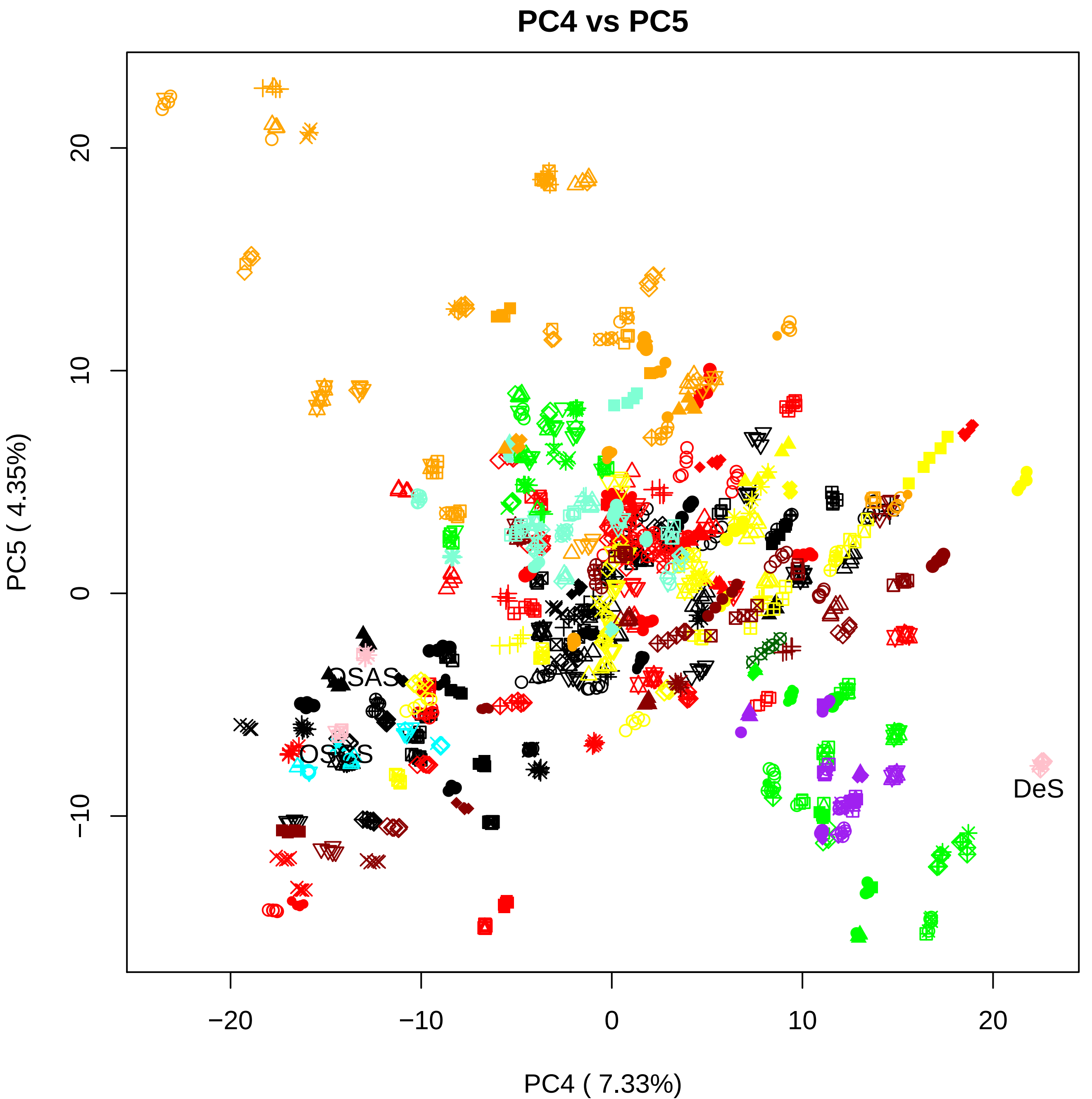
PC4 vs PC5
PC5 ( 4.35%)
PC4 ( 7.33%)
20
10
0
−10
−20
−10
0
10
20
OSAS
OSAS
DeS

### OSCS Contam 7th

Anthony Devlin, Keele University

### PCA of All  OSCS vs PMH

Scaled Matrix PCAed

7th order polynomial baseline correction performed

Raw spectra smoothed to 21 Neighbours, 2nd order PN, Savtisky Golay

Normalisation between 0 and 100

D2 spectra smoothed to 41 Neighbours, 2nd order PN, Savtisky Golay

Regions Analysed: 3601−2501 and 2001−700cm−1

Samples Analysed: 12 Contam's 1 OSCS's 45 PMH's

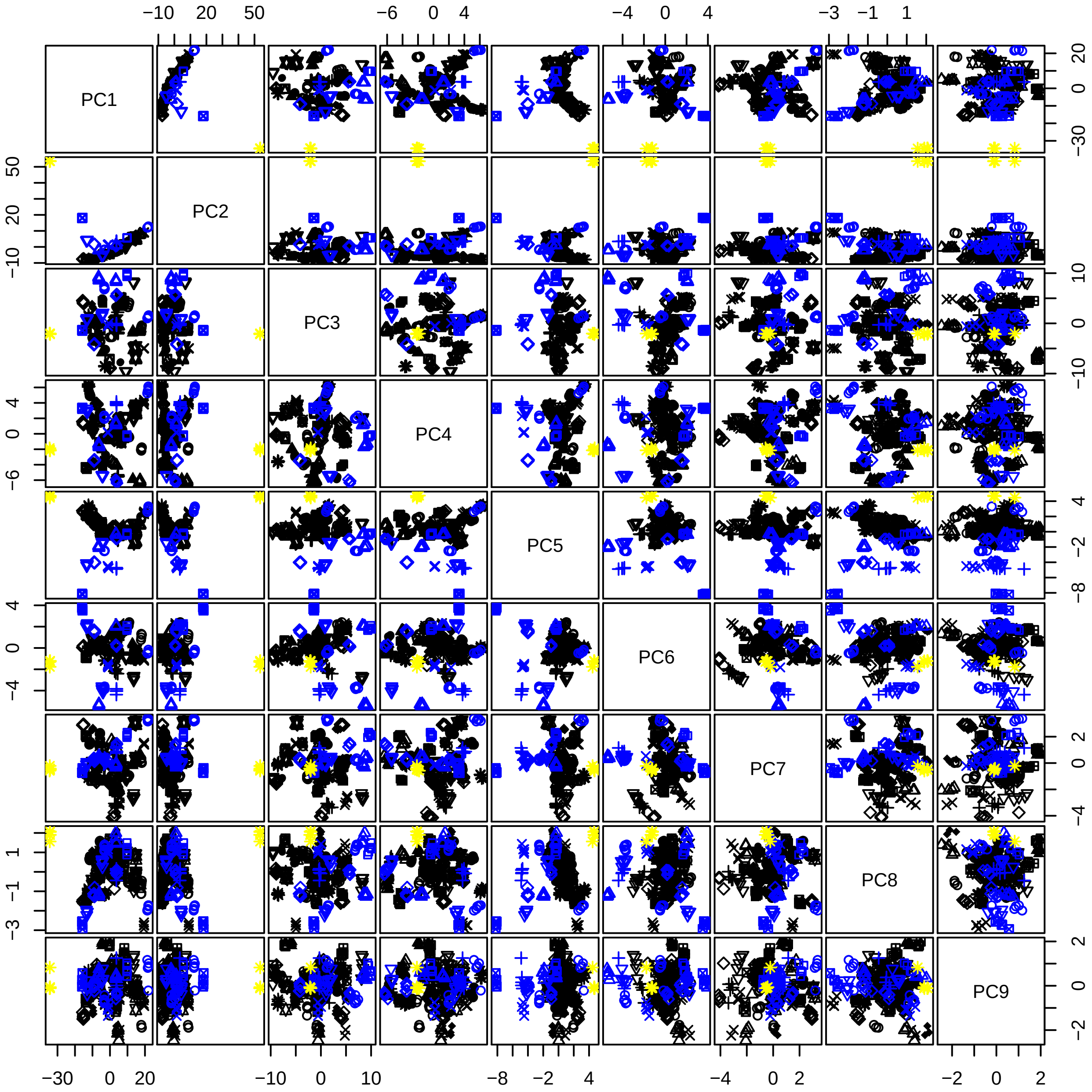
PC1
PC2
PC3
PC4
PC5
PC6
PC7
PC8
PC9
−10 20 50
−6 0 4
−4 0 4
−3 −1 1
20 0 −30
50 20 −10
10 0 −10
4 0 −6
4 −2 −8
4 0 −4
2 0 −4
1 −1 −3
2 0 −2
−30 0 20
−10 0 10
−8 −2 4
−4 0 2
−2 0 2

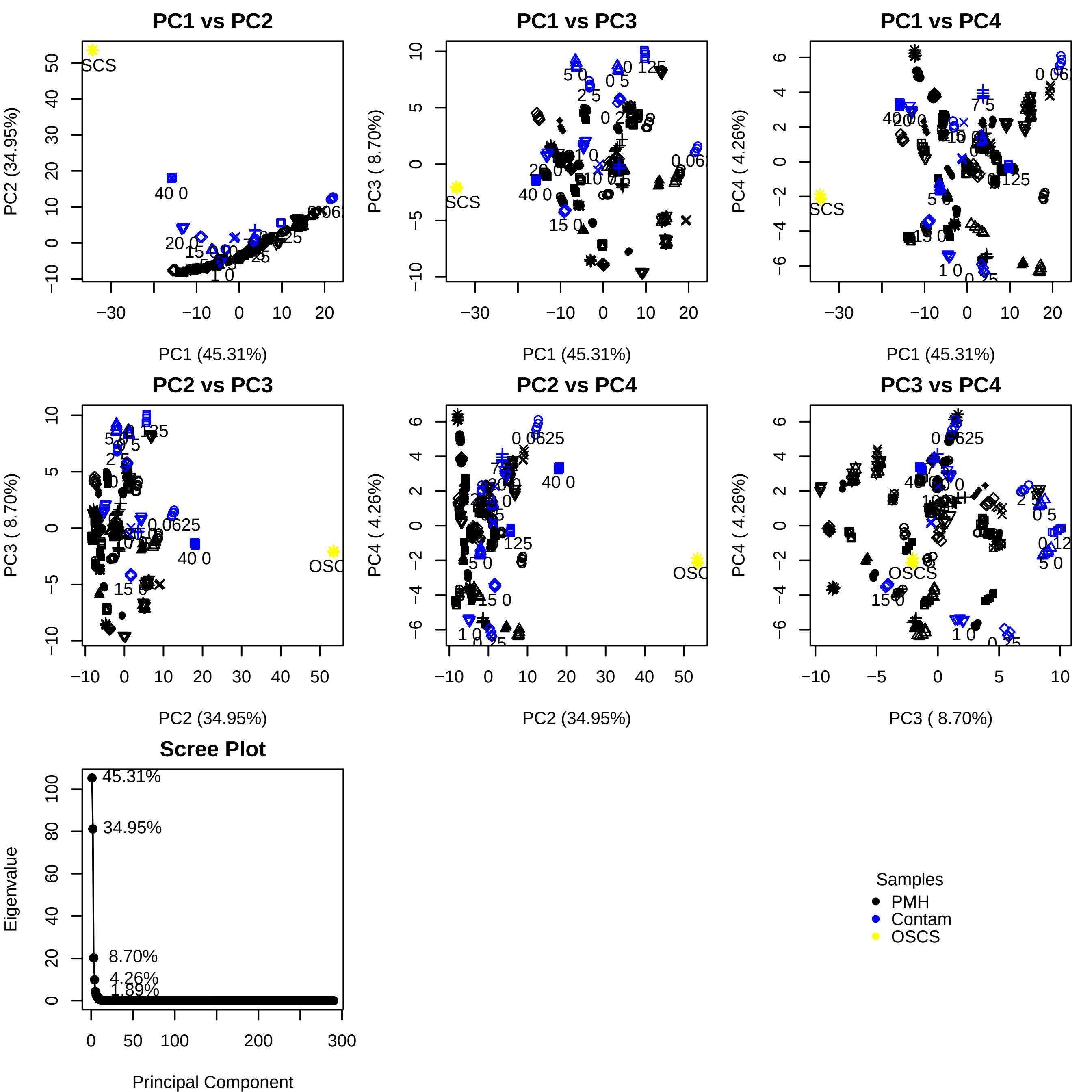

PC1 vs PC2
PC1 vs PC3
PC1 vs PC4
PC2 vs PC3
PC2 vs PC4
PC3 vs PC4
Scree Plot
PC1 (45.31%)
PC2 (34.95%)
PC3 ( 8.70%)
PC4 ( 4.26%)
Eigenvalue
Principal Component
45.31%
34.95%
8.70%
4.26%
1.89%
Samples
PMH
Contam
OSCS

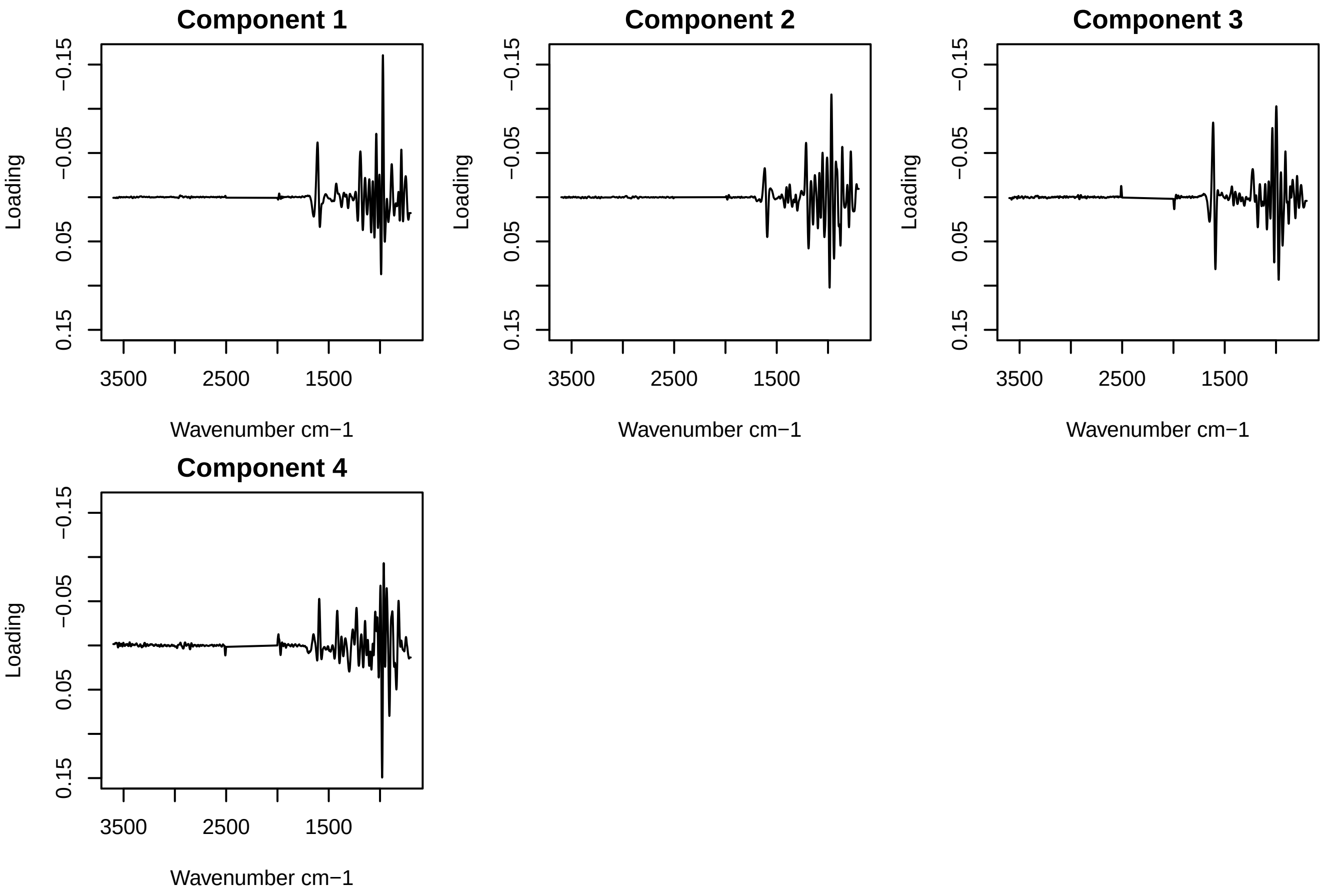
Component 1
Component 2
Component 3
Component 4
Loading
−0.15
−0.05
0.05
0.15
3500
2500
1500
Wavenumber cm−1

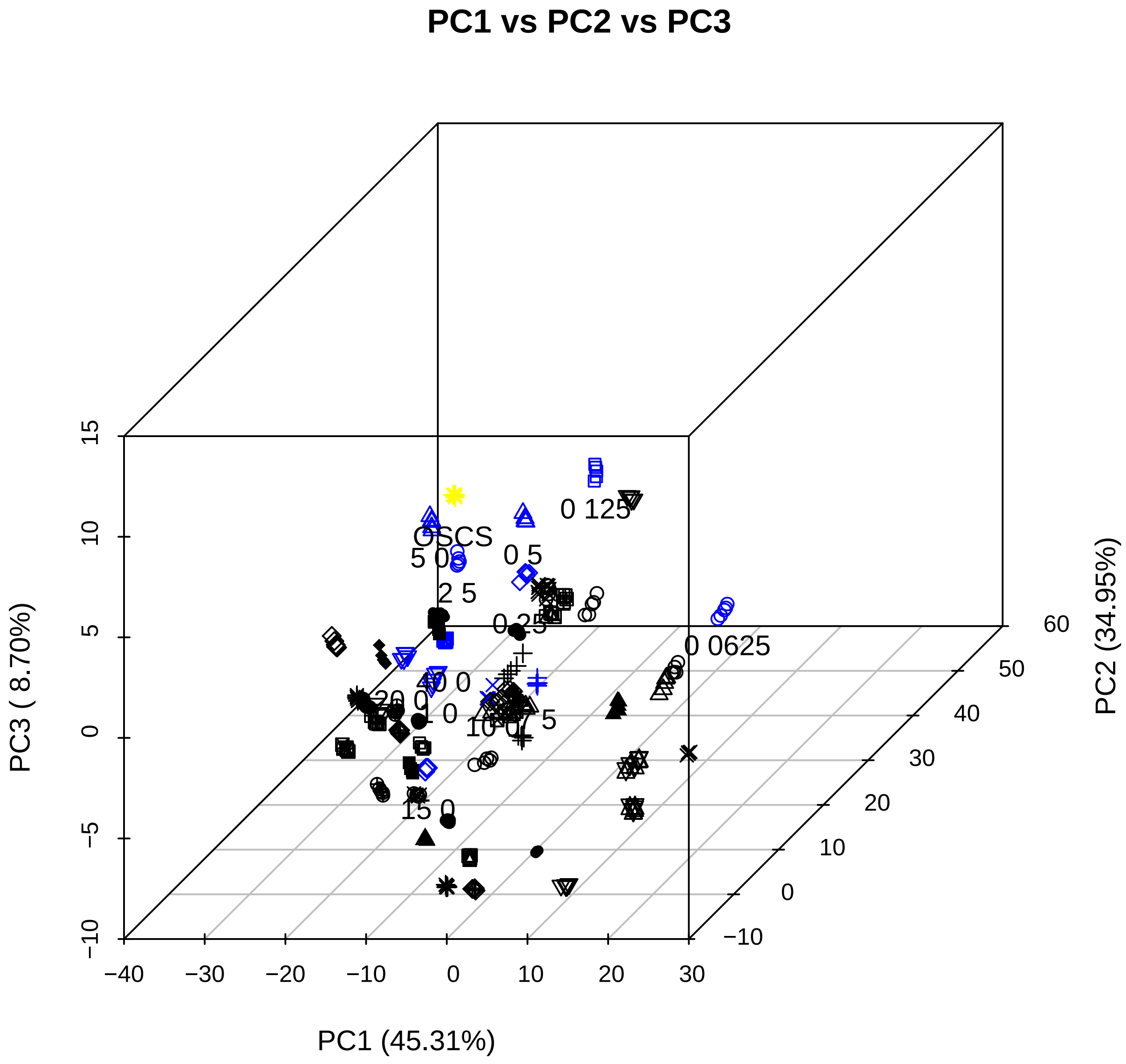

PC1 vs PC2 vs PC3
PC1 (45.31%)
PC2 (34.95%)
PC3 ( 8.70%)
−40
−30
−20
−10
0
10
20
30
−10
0
10
20
30
40
50
60
−10
−5
0
5
10
15
OSCS
0 125
5 0
0 5
2 5
0 25
0 0625
40 0
20 0
1 0
10 0
7 5
15 0

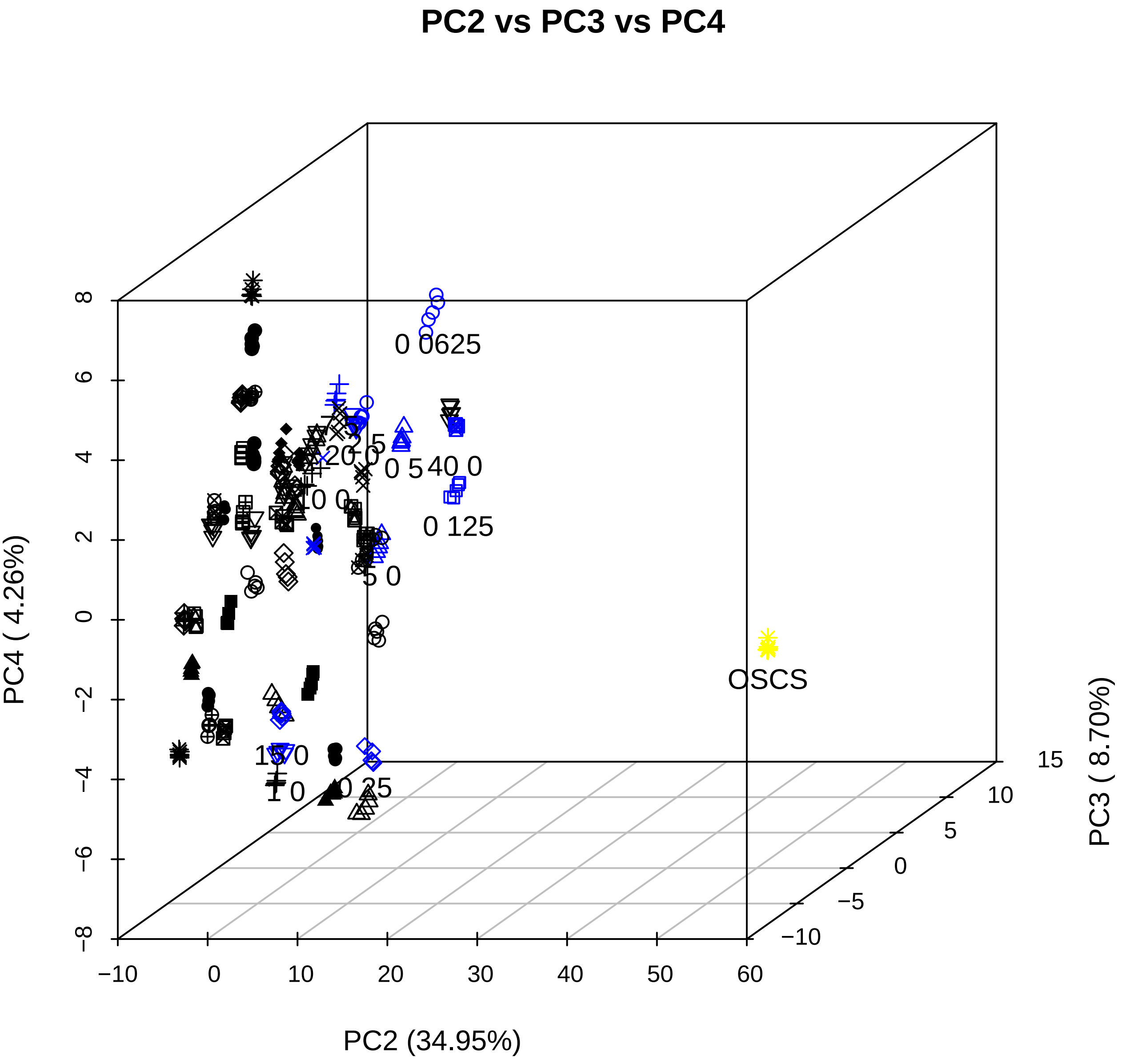

PC2 vs PC3 vs PC4
PC2 (34.95%)
PC3 ( 8.70%)
PC4 ( 4.26%)
0 0625
2 5
20 0
0 5
40 0
10 0
0 125
5 0
OSCS
15 0
1 0
0 25

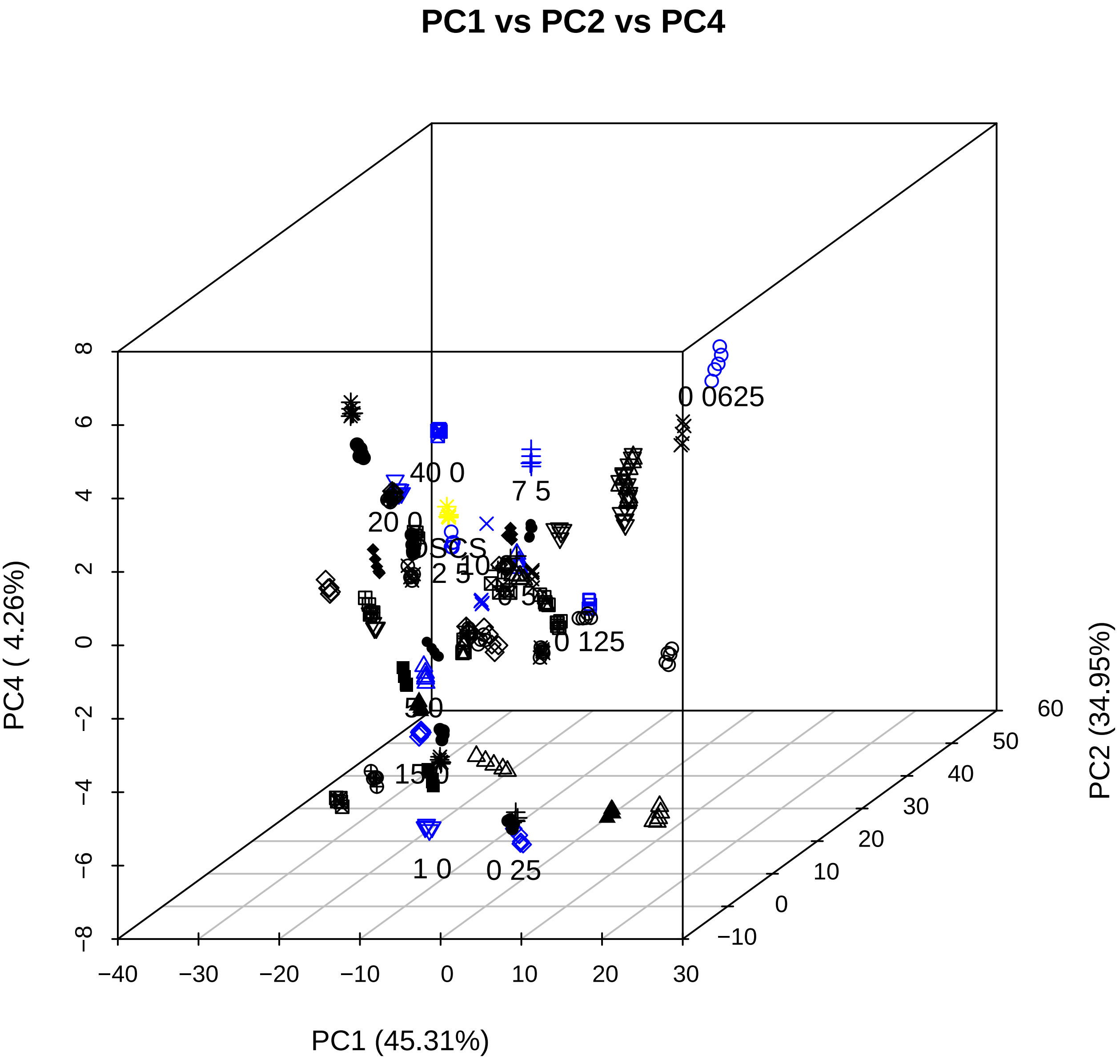
PC1 vs PC2 vs PC4
0 0625
40 0
7 5
20 0
DSCS
10
2 5
0 5
0 125
5 0
15 0
1 0
0 25
PC1 (45.31%)
PC2 (34.95%)
PC4 ( 4.26%)
−40
−30
−20
−10
0
10
20
30
−10
0
10
20
30
40
50
60
−8
−6
−4
−2
0
2
4
6
8

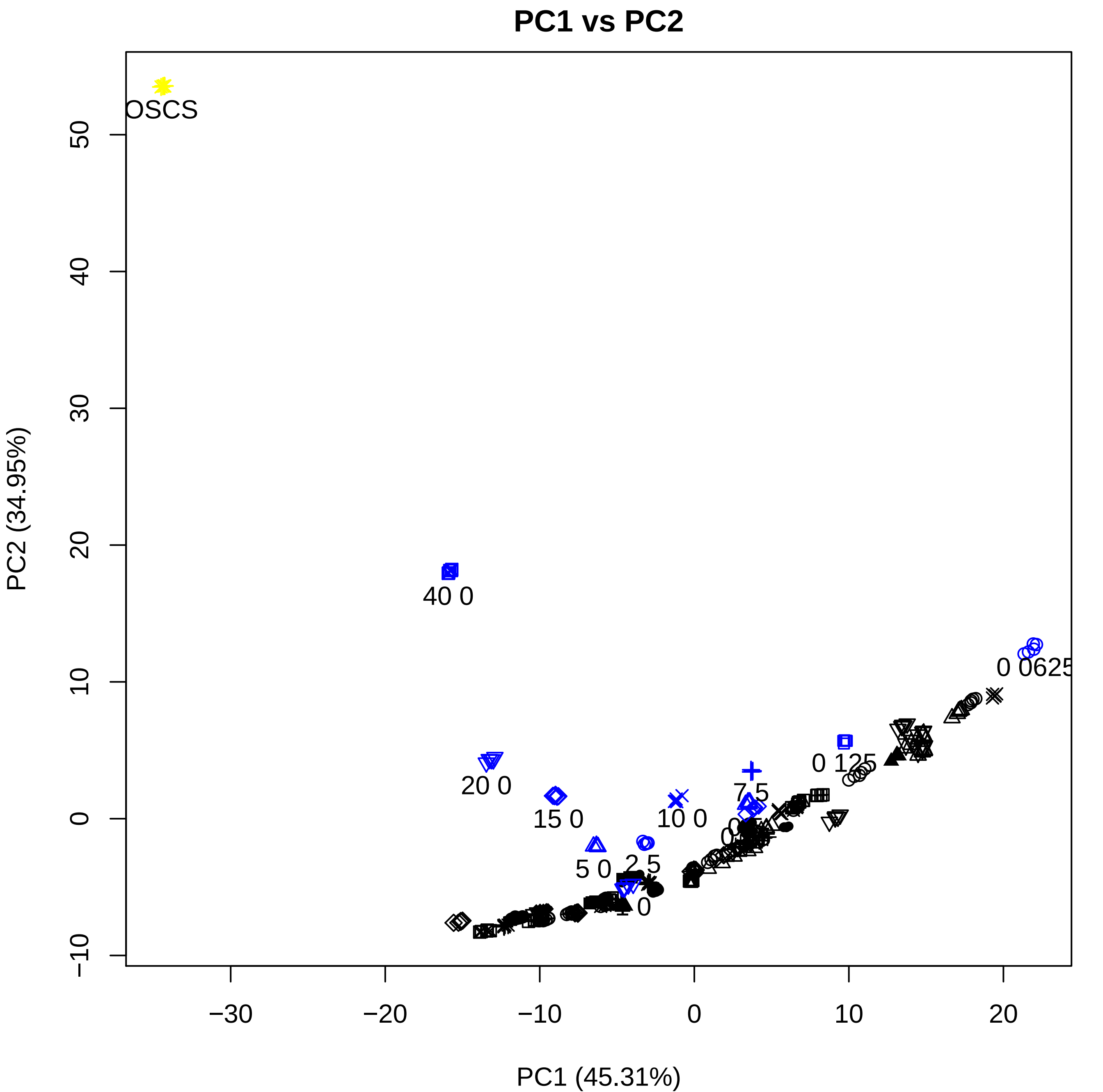
PC1 vs PC2
OSCS
40 0
20 0
15 0
10 0
7 5
5 0
2 5
0 0625
0 125
PC2 (34.95%)
PC1 (45.31%)
−30
−20
−10
0
10
20
30
40
50

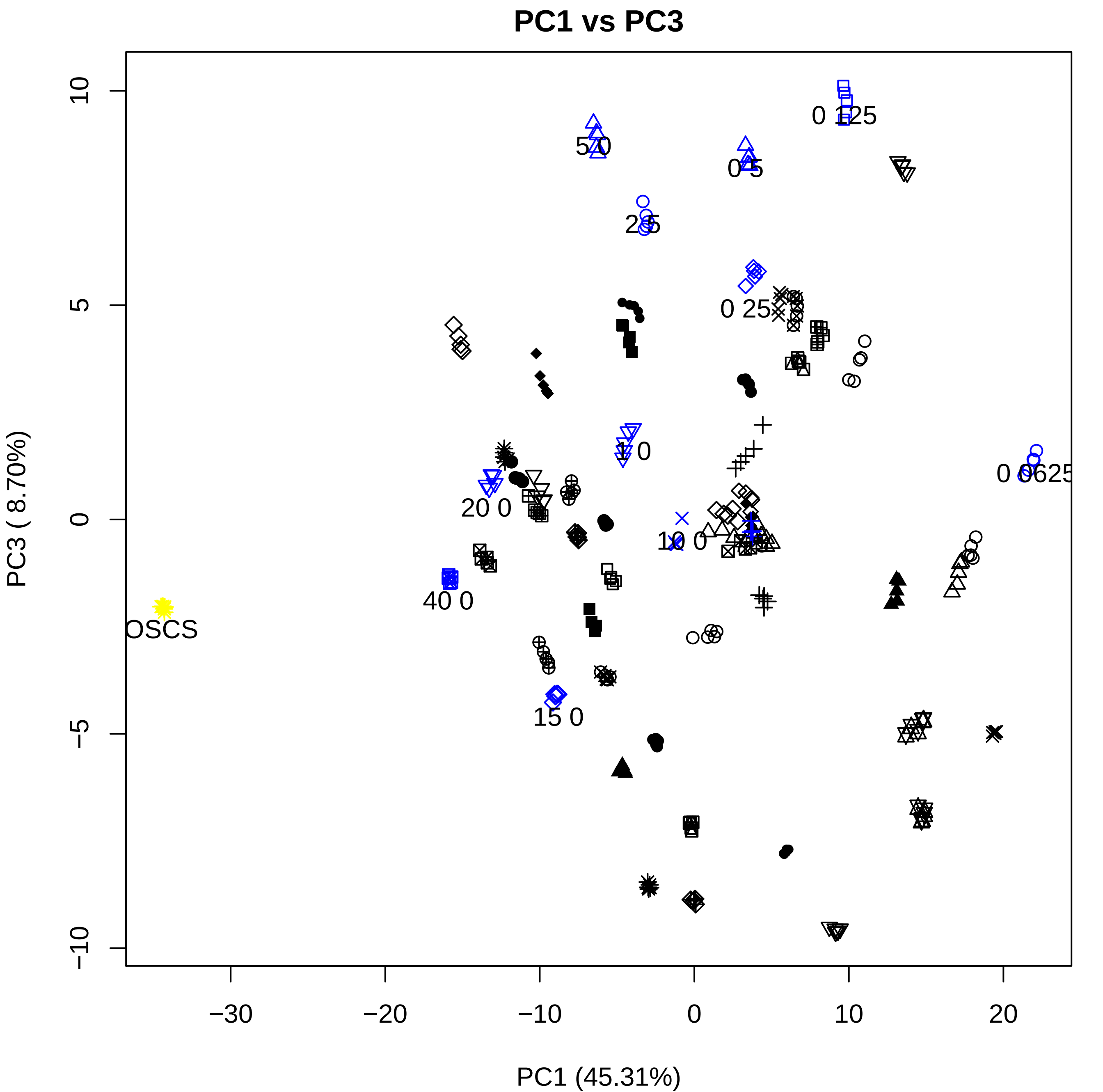
PC1 vs PC3
PC3 ( 8.70%)
PC1 (45.31%)
10
5
0
−5
−10
−30
−20
−10
0
10
20
0 125
5 0
0 5
2 5
0 25
1 0
0 0625
20 0
10 0
40 0
OSCS
15 0

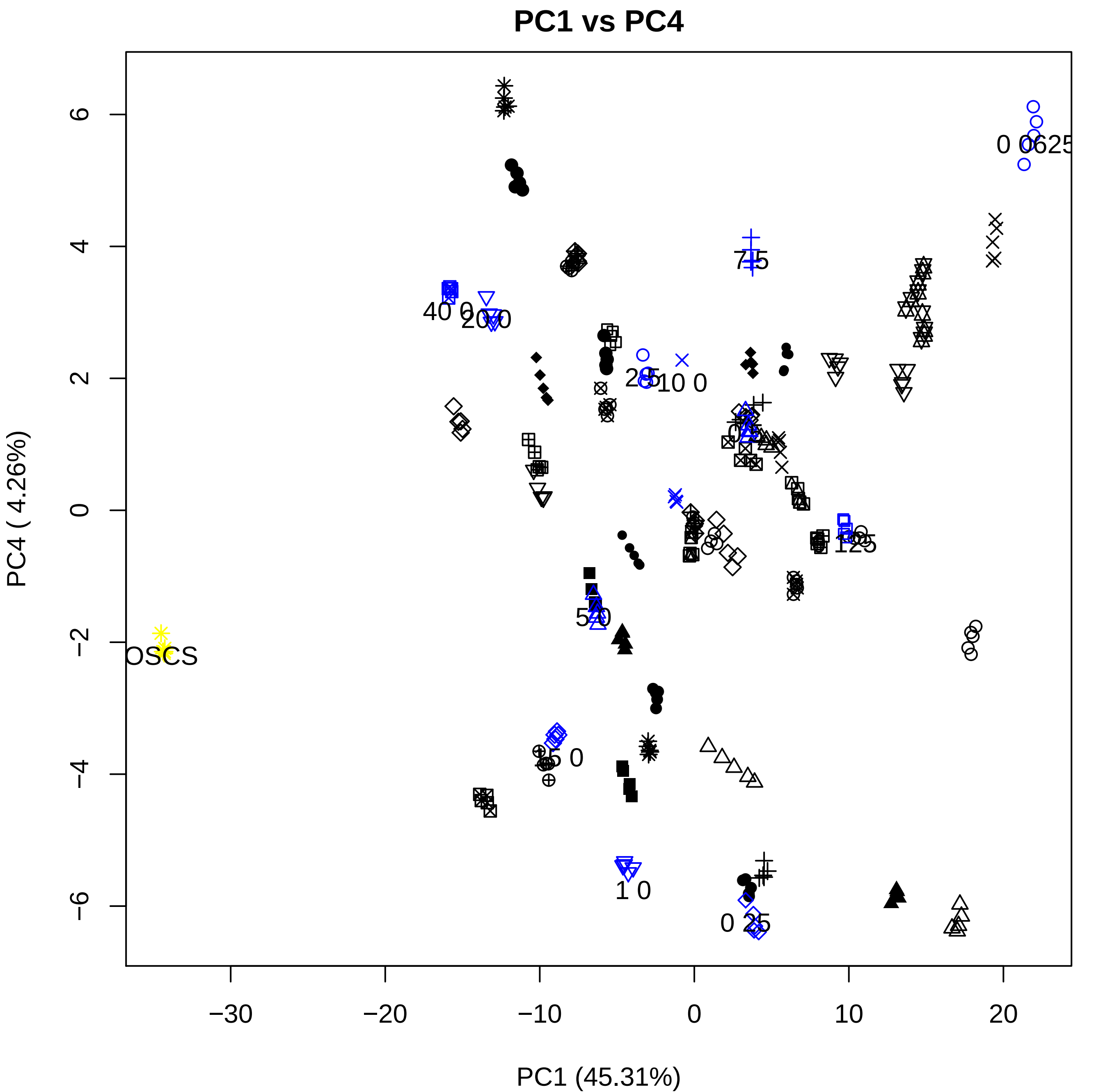

PC1 vs PC4
PC4 ( 4.26%)
PC1 (45.31%)
6
4
2
0
−2
−4
−6
−30
−20
−10
0
10
20
0 0625
7 5
40 0
20 0
2 5
10 0
1 25
5 0
OSCS
15 0
1 0
0 25

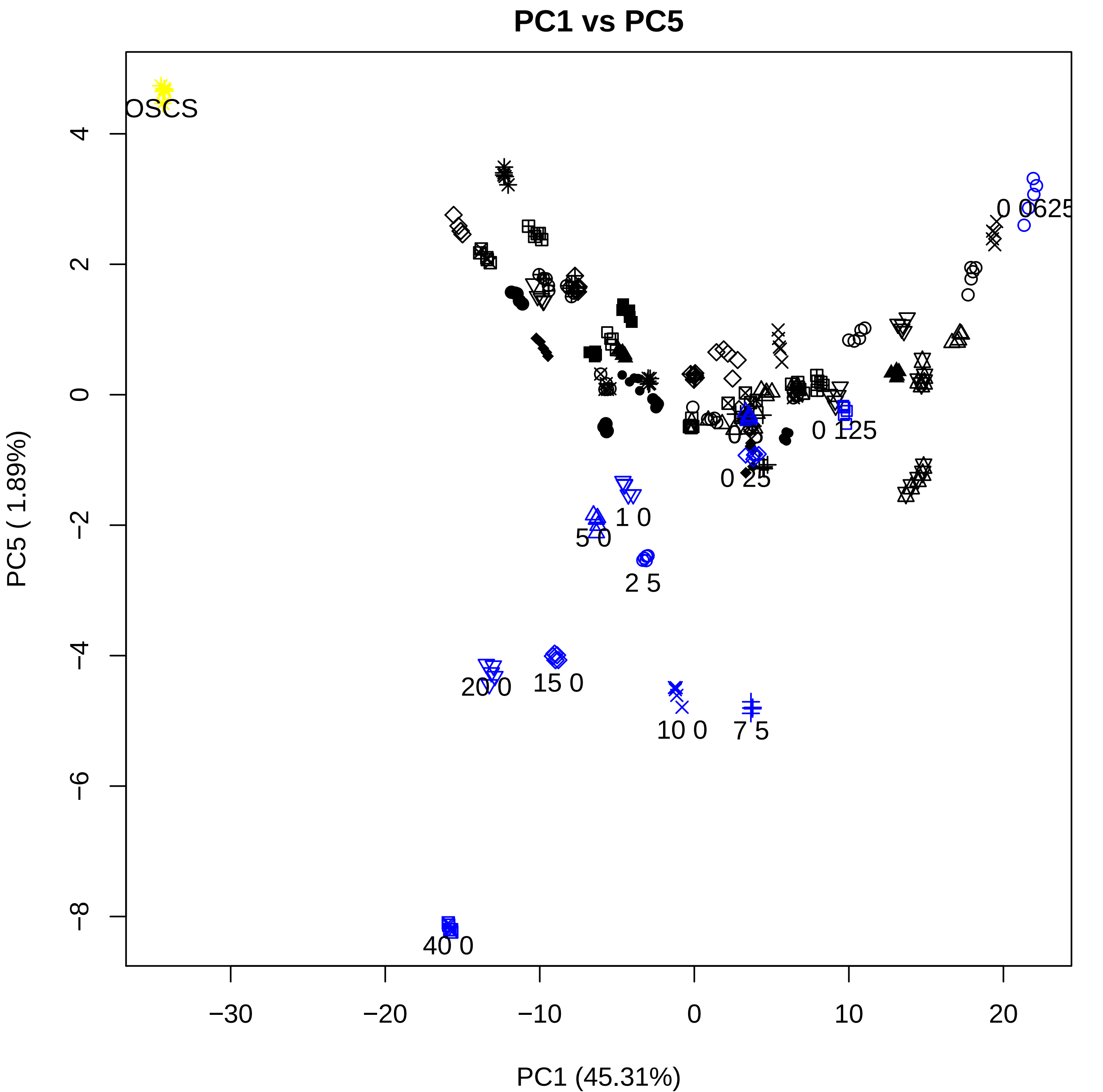
PC1 vs PC5
OSCS
0 0625
0 125
0 5
0 25
1 0
5 0
2 5
20 0
15 0
10 0
7 5
40 0
4
2
0
−2
−4
−6
−8
−30
−20
−10
0
10
20
PC1 (45.31%)
PC5 ( 1.89%)

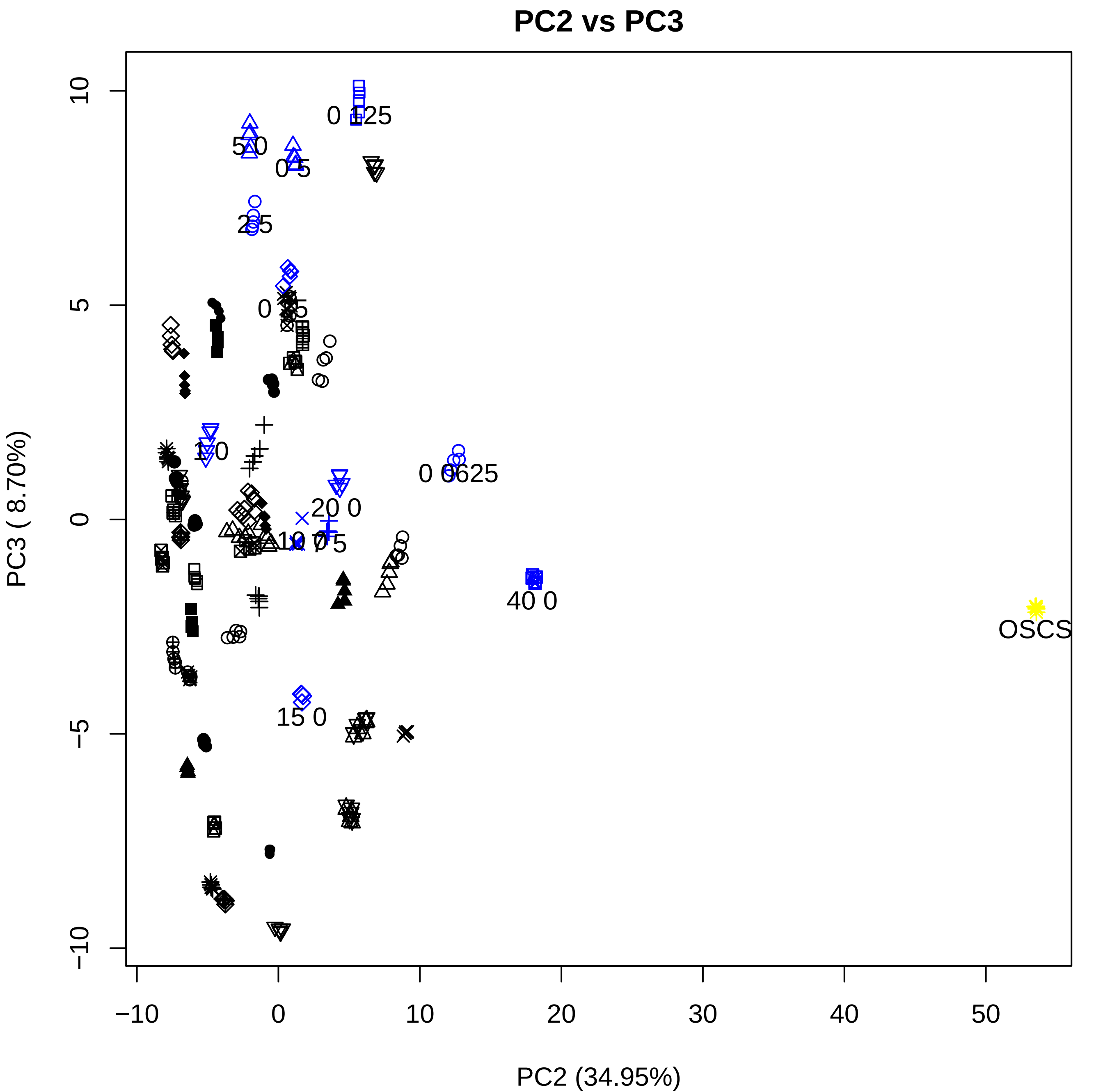

PC2 vs PC3
PC3 ( 8.70%)
PC2 (34.95%)
10
5
0
−5
−10
−10
0
10
20
30
40
50
0 125
5 0
0 5
2 5
1 0
0 0625
20 0
7 5
40 0
OSCS
15 0

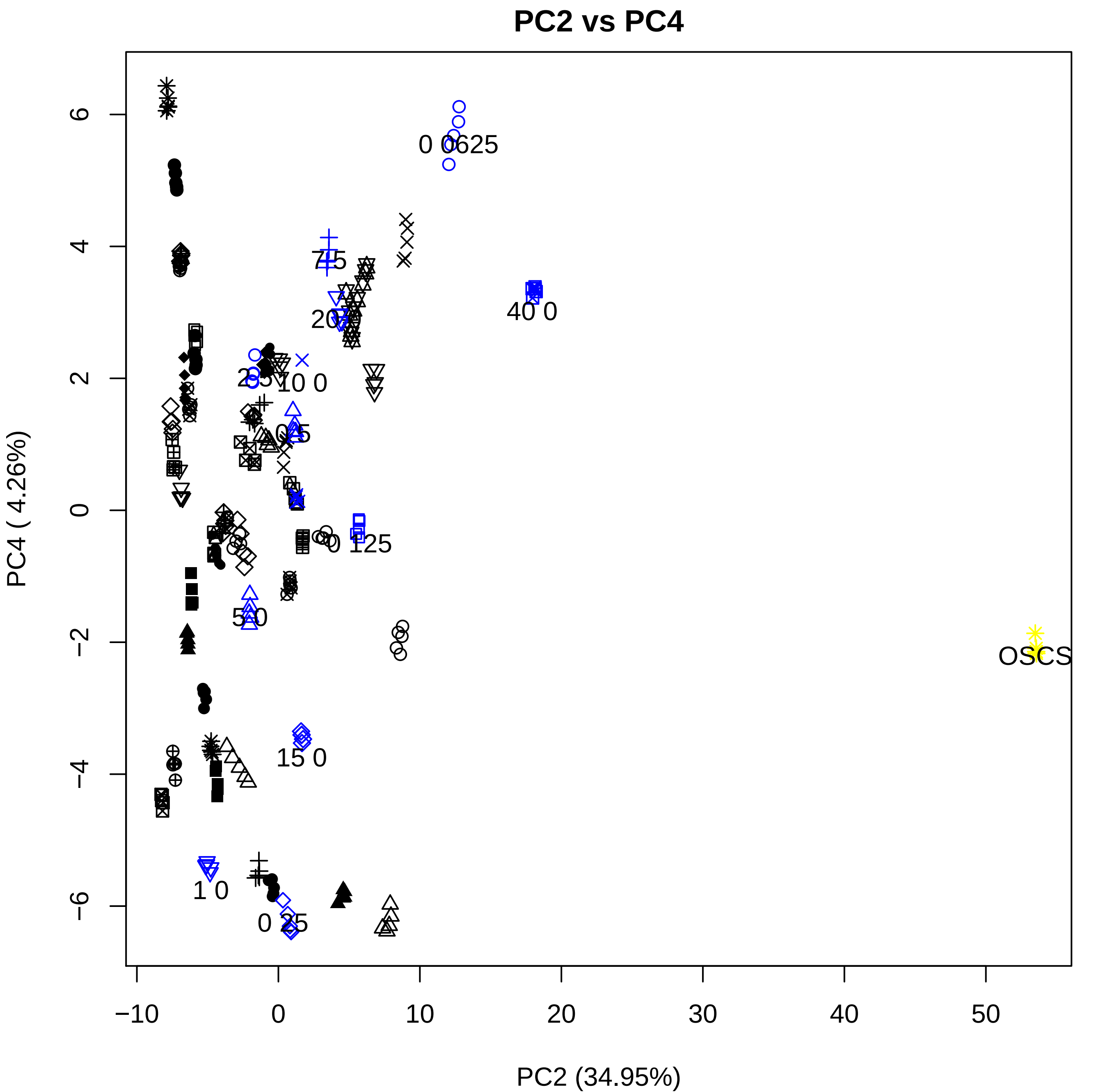
PC2 vs PC4
0 0625
7 5
40 0
10 0
0 125
5 0
15 0
1 0
0 25
OSCS
PC2 (34.95%)
PC4 ( 4.26%)
−10
0
10
20
30
40
50
6
4
2
0
−2
−4
−6

# PC2 vs PC5

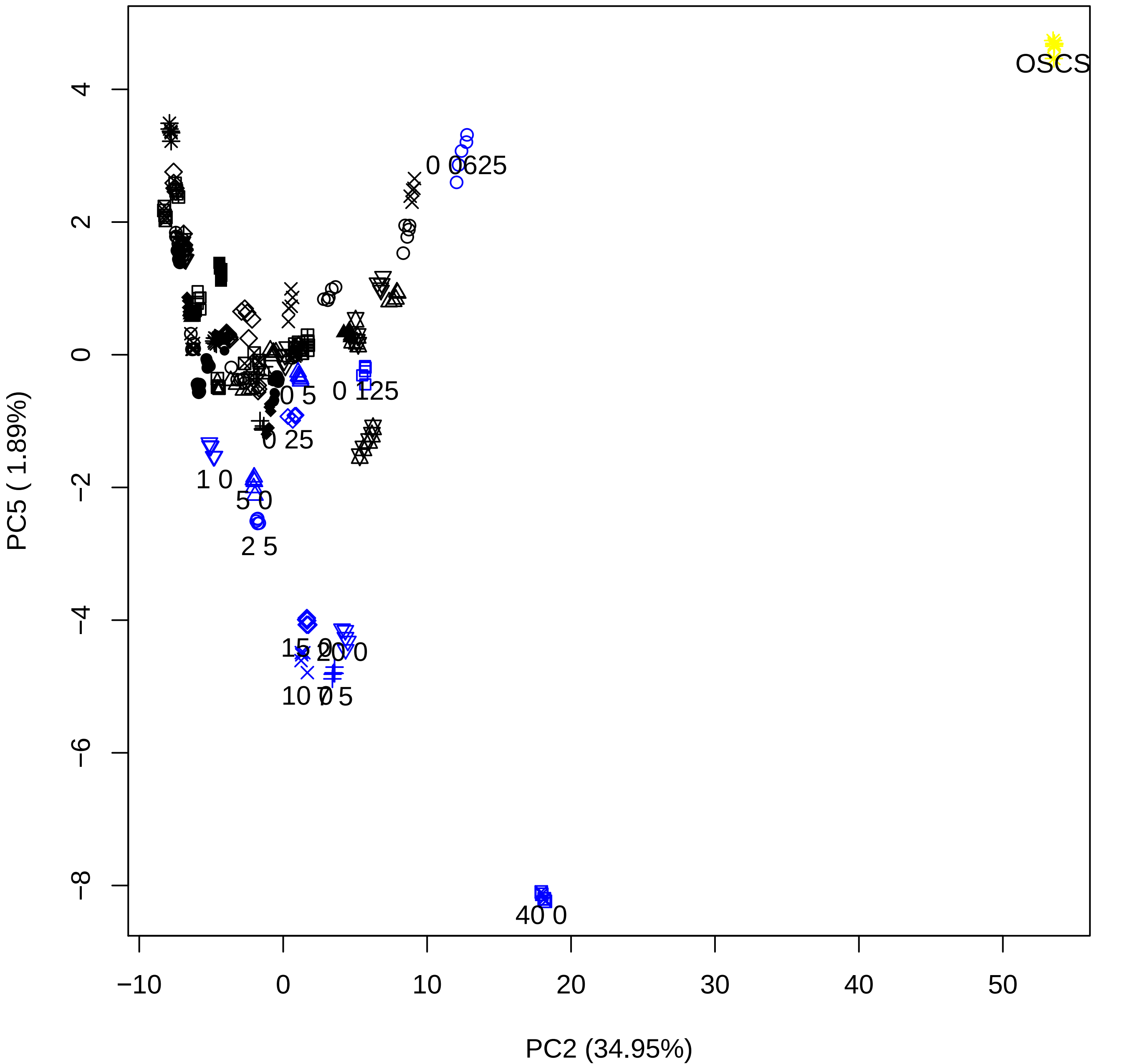

OSCS
0 0625
0 5
0 125
0 25
1 0
5 0
2 5
15 0
20 0
10 0
7 5
40 0
PC5 ( 1.89%)
PC2 (34.95%)

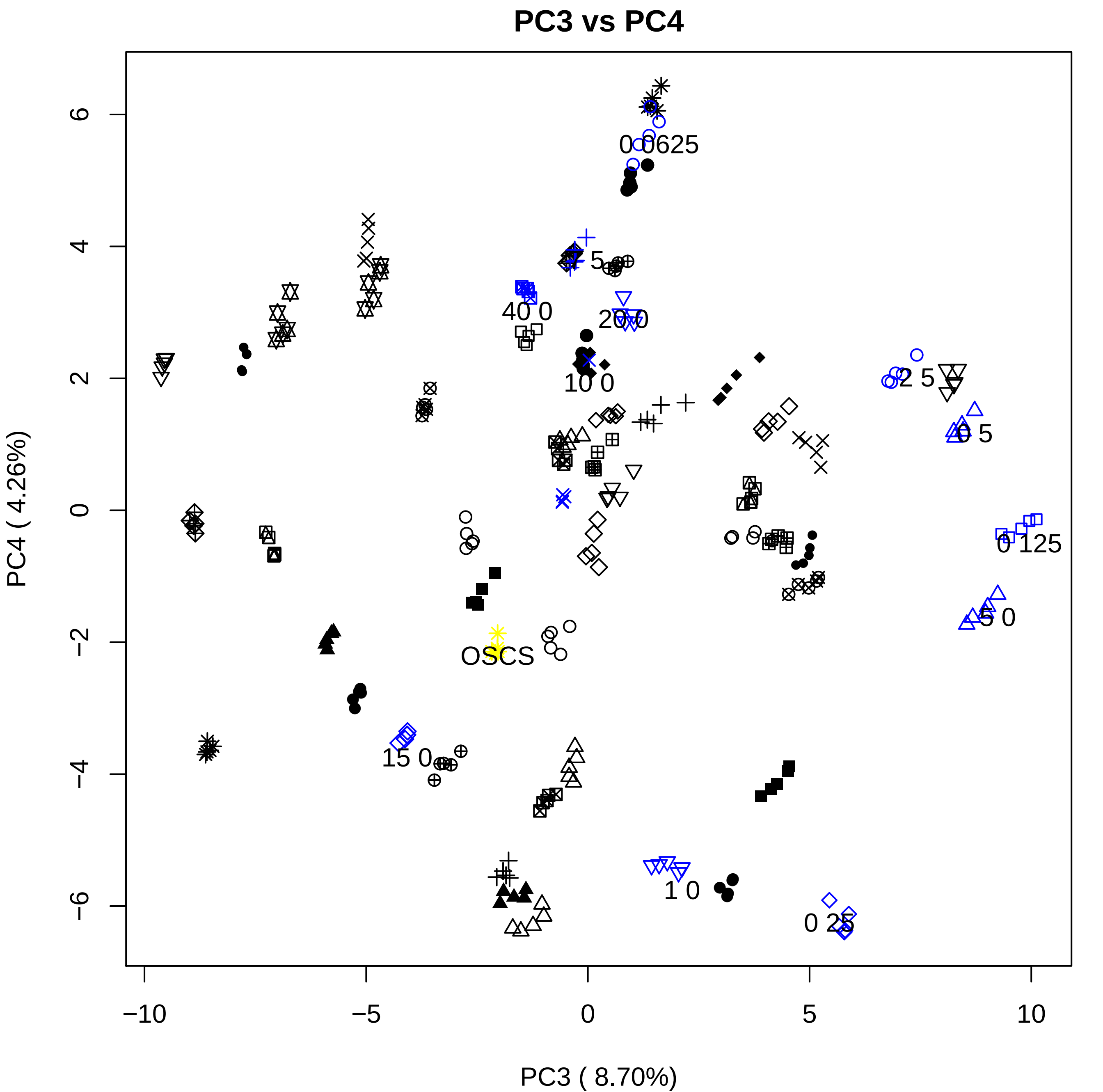
PC3 vs PC4
0 0625
5
40 0
20 0
10 0
2 5
0 5
0 125
5 0
OSCS
15 0
1 0
0 25
PC4 ( 4.26%)
PC3 ( 8.70%)
6
4
2
0
−2
−4
−6
−10
−5
0
5
10

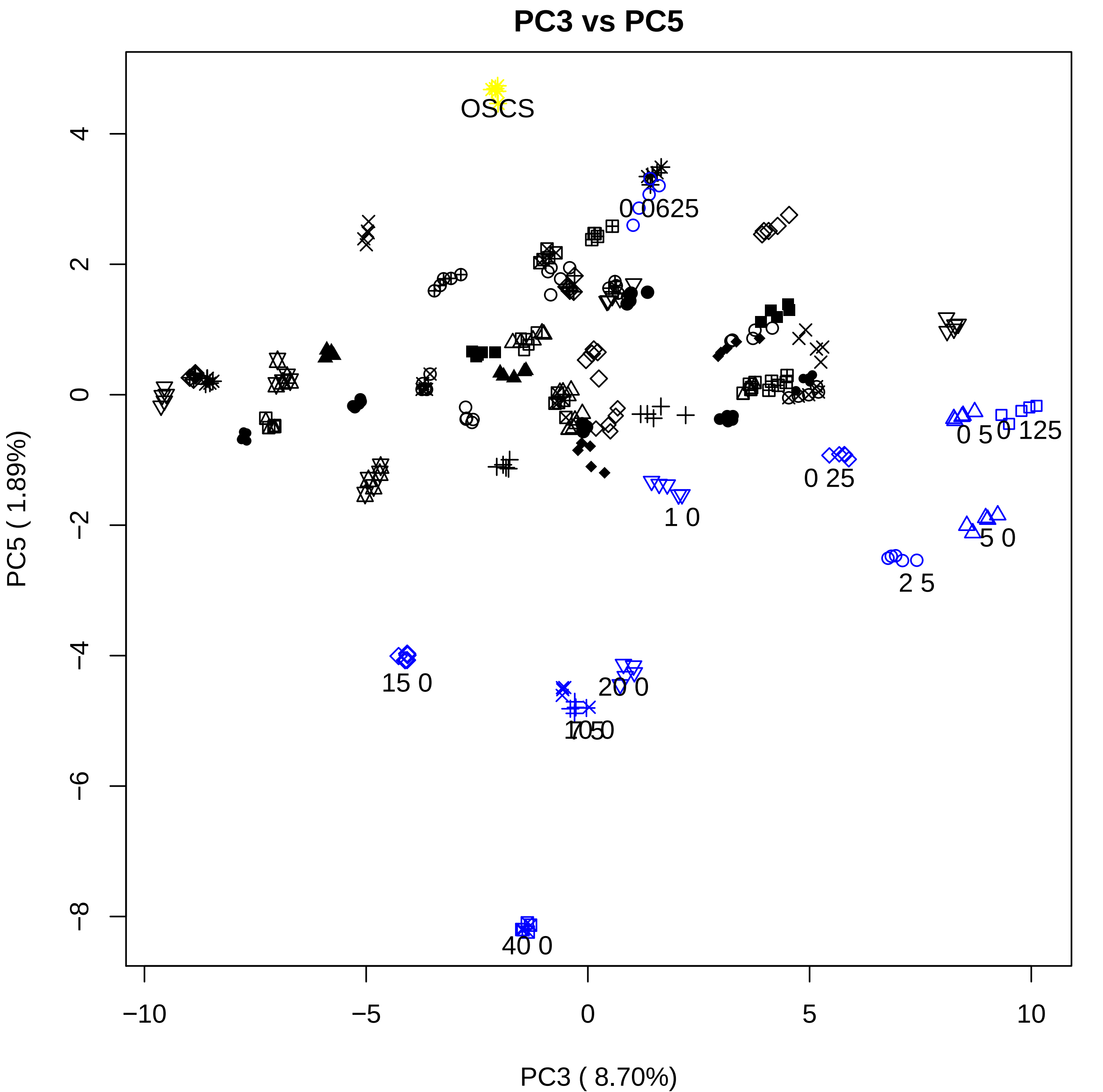
PC3 vs PC5
OSCS
0 0625
0 5
0 125
0 25
1 0
5 0
2 5
15 0
20 0
10 0
7 5
40 0
PC5 ( 1.89%)
PC3 ( 8.70%)
4
2
0
−2
−4
−6
−8
−10
−5
0
5
10

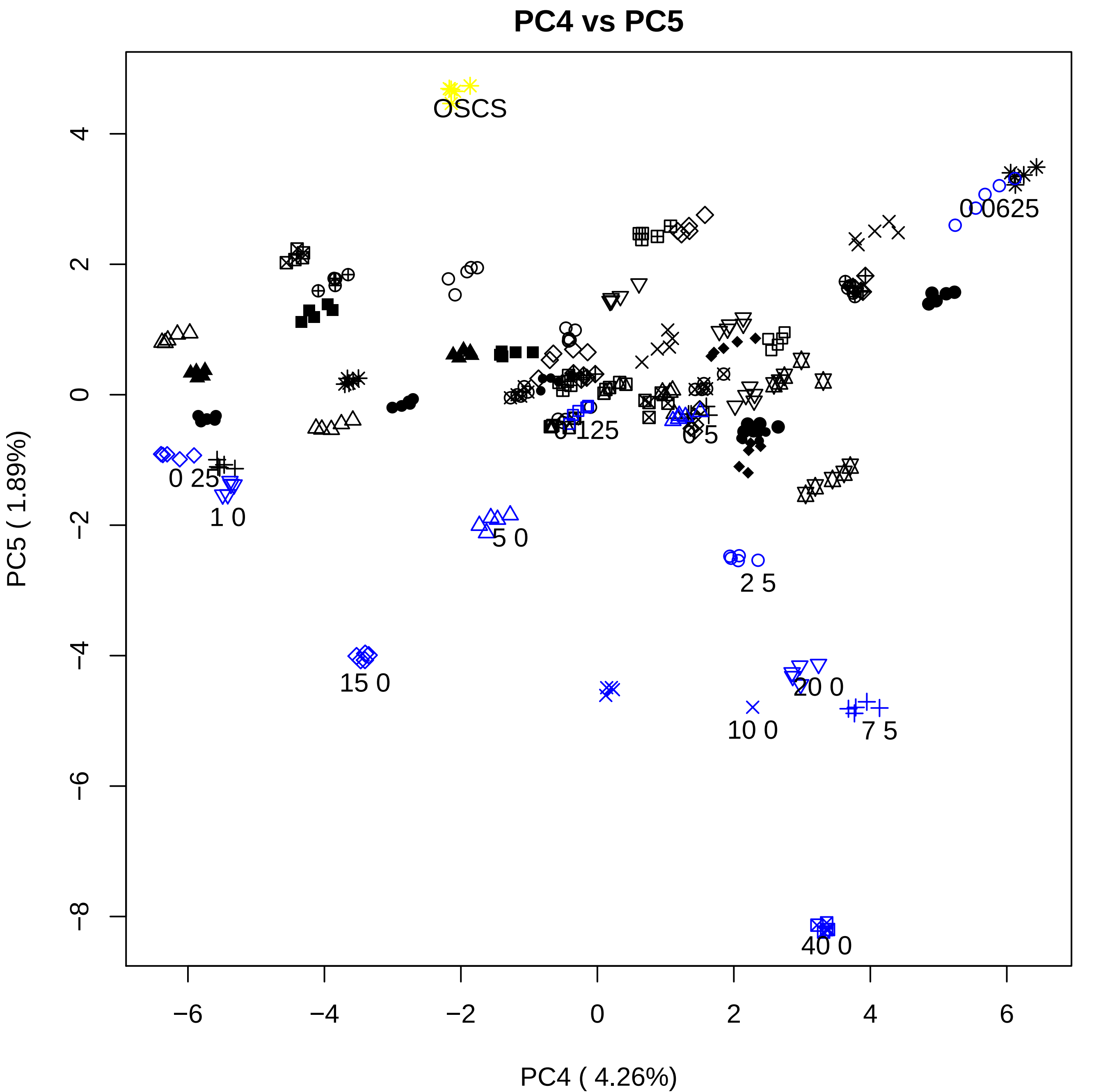

PC4 vs PC5
OSCS
0.0625
0.125
0.5
0.25
1.0
5.0
2.5
15.0
20.0
10.0
7.5
40.0
PC4 ( 4.26%)
PC5 ( 1.89%)
−6
−4
−2
0
2
4
6
−8

### PCA of 0.06%  OSCS vs PMH

Scaled Matrix PCAed

7th order polynomial baseline correction performed

Raw spectra smoothed to 21 Neighbours, 2nd order PN, Savtisky Golay

Normalisation between 0 and 100

D2 spectra smoothed to 41 Neighbours, 2nd order PN, Savtisky Golay

Regions Analysed: 3601−2501 and 2001−700cm−1

Samples Analysed: 1 Contam's 0 OSCS's 45 PMH's

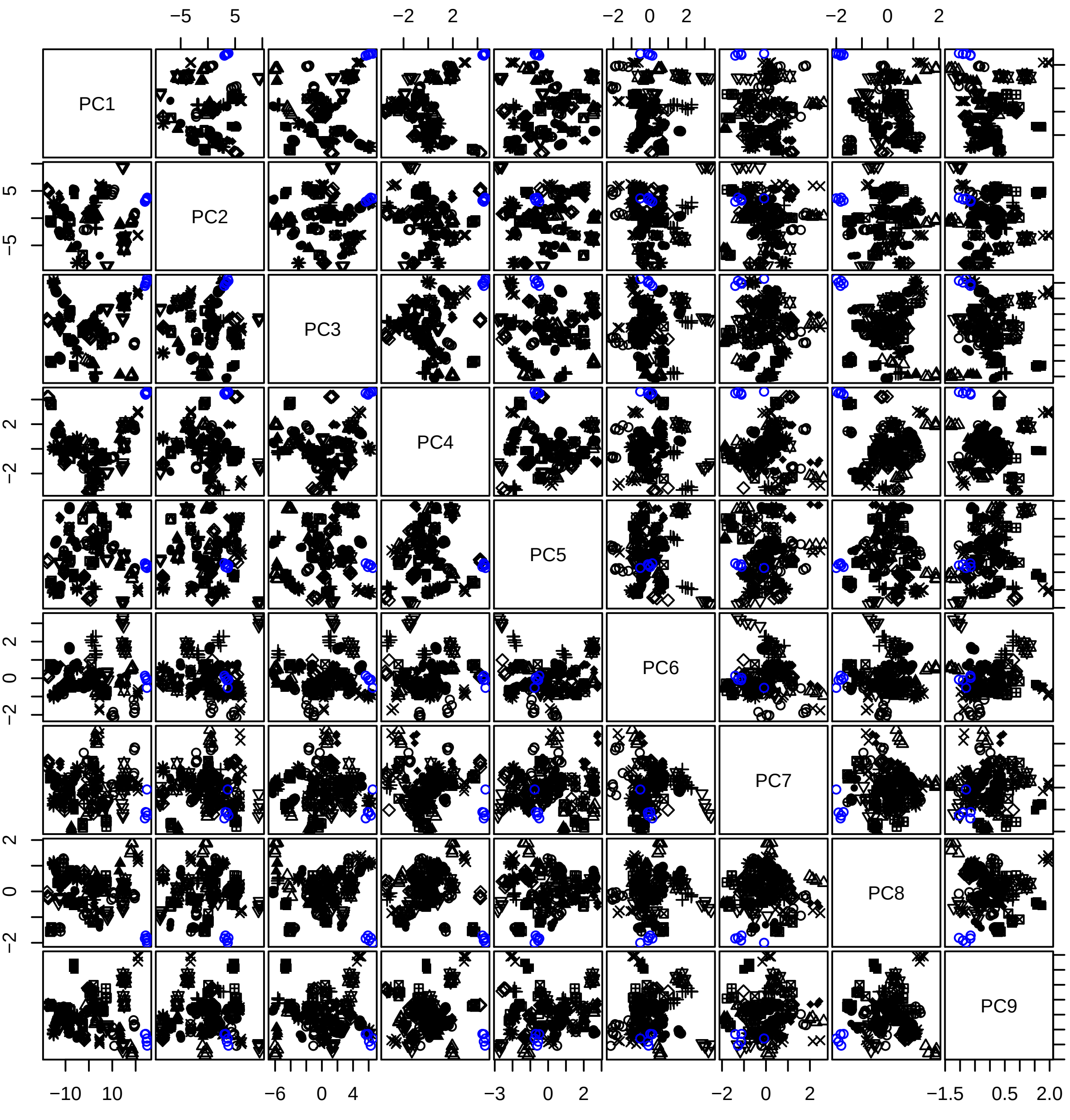
PC1
PC2
PC3
PC4
PC5
PC6
PC7
PC8
PC9

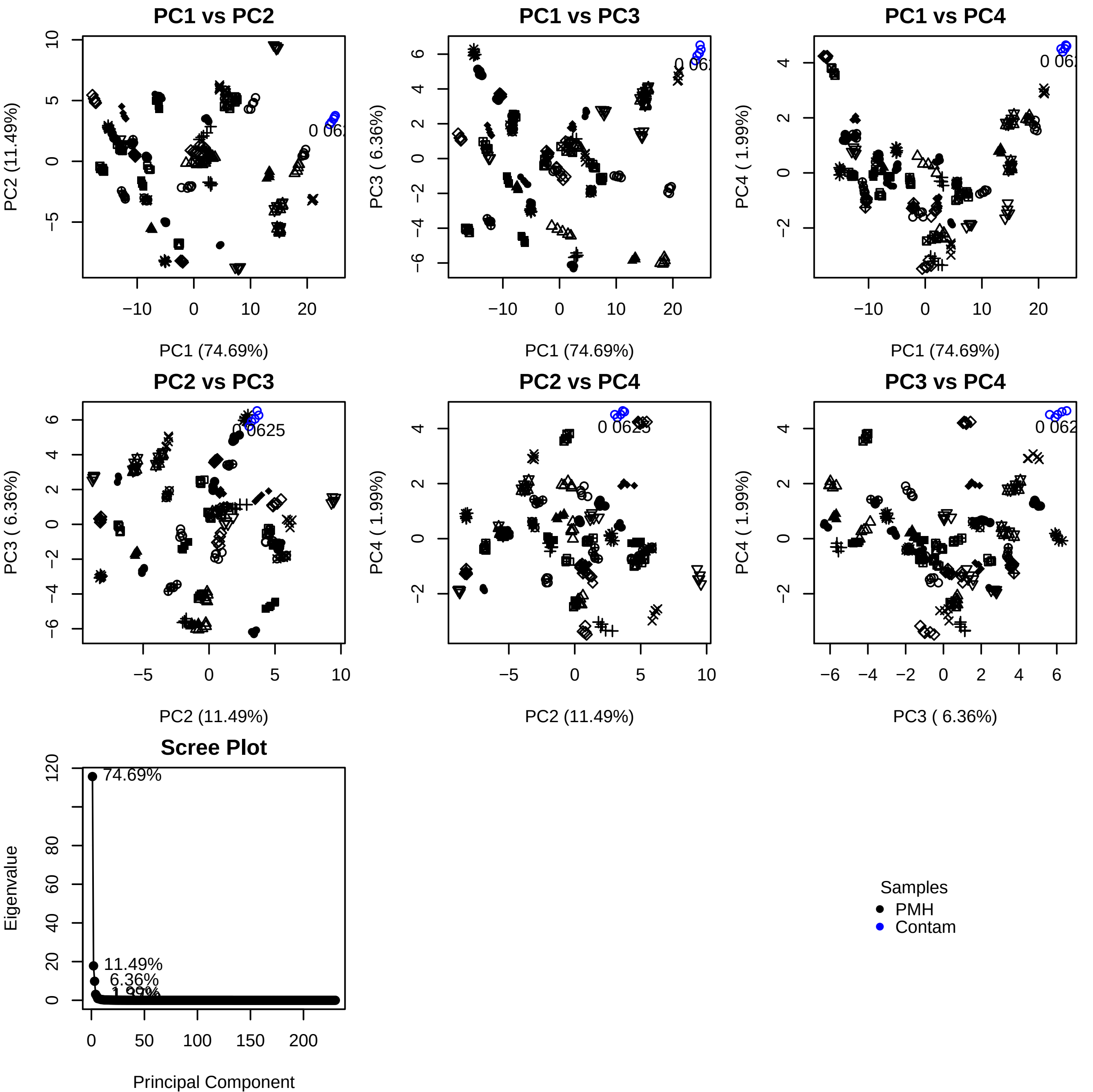
PC1 vs PC2
PC1 vs PC3
PC1 vs PC4
PC2 vs PC3
PC2 vs PC4
PC3 vs PC4
Scree Plot
PC1 (74.69%)
PC2 (11.49%)
PC3 ( 6.36%)
PC4 ( 1.99%)
0 0625
74.69%
11.49%
6.36%
Eigenvalue
Principal Component
Samples
PMH
Contam

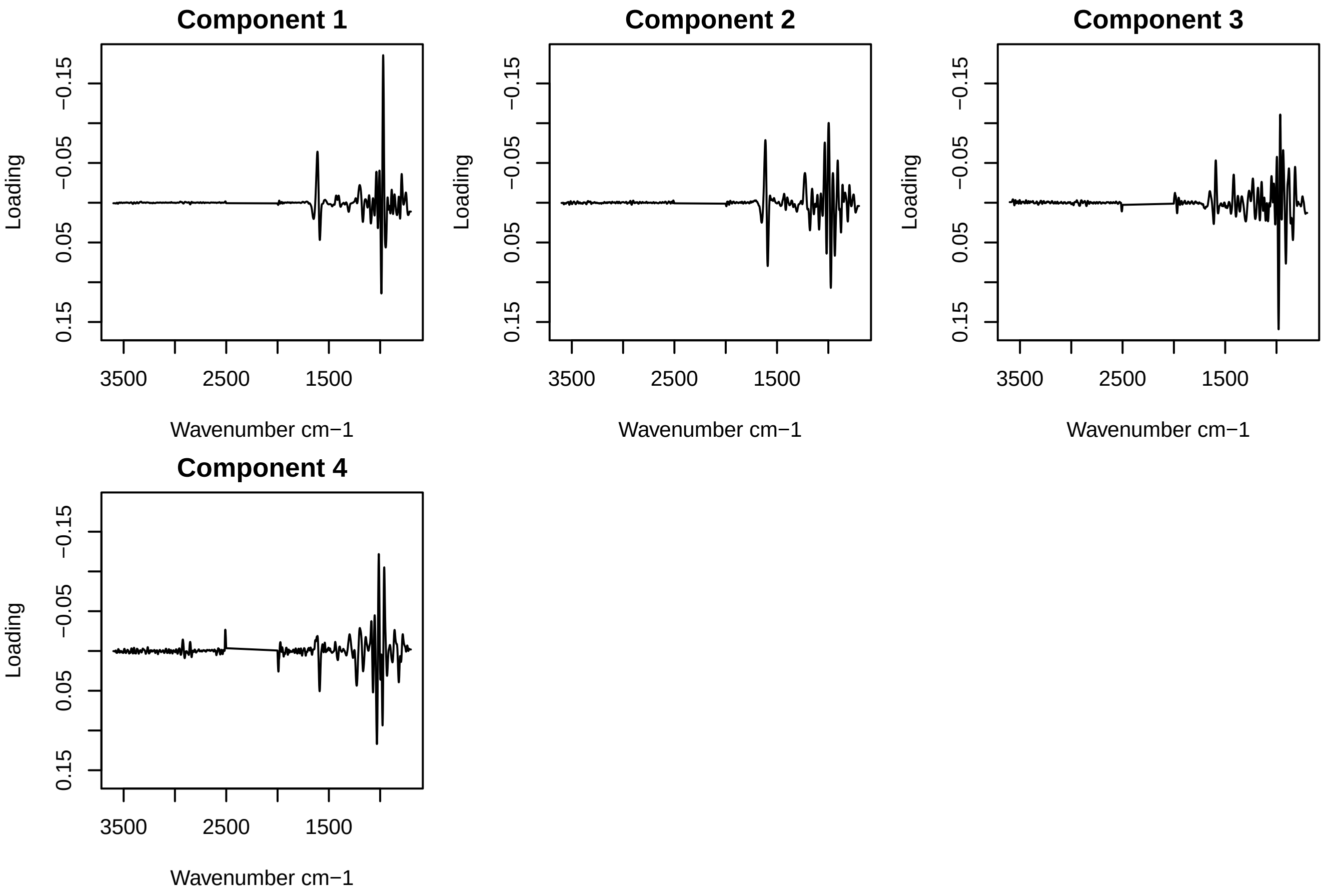
Component 1
Component 2
Component 3
Component 4
Loading
−0.15
−0.05
0.05
0.15
3500
2500
1500
Wavenumber cm−1

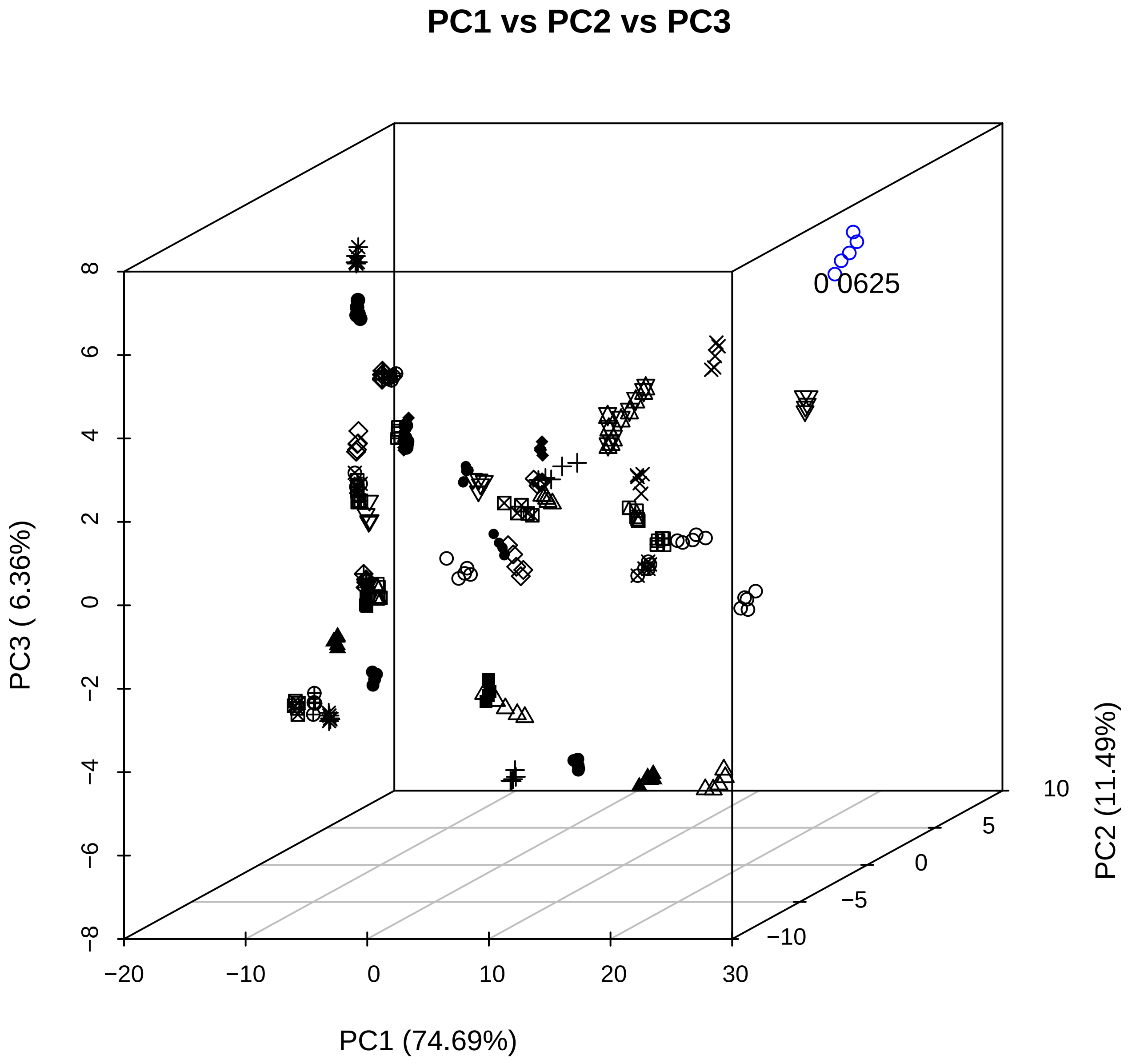
PC1 vs PC2 vs PC3
PC1 (74.69%)
PC2 (11.49%)
PC3 ( 6.36%)
0 0625
−20
−10
0
10
20
30
−10
−5
0
5
10
−8
−6
−4
−2
0
2
4
6
8

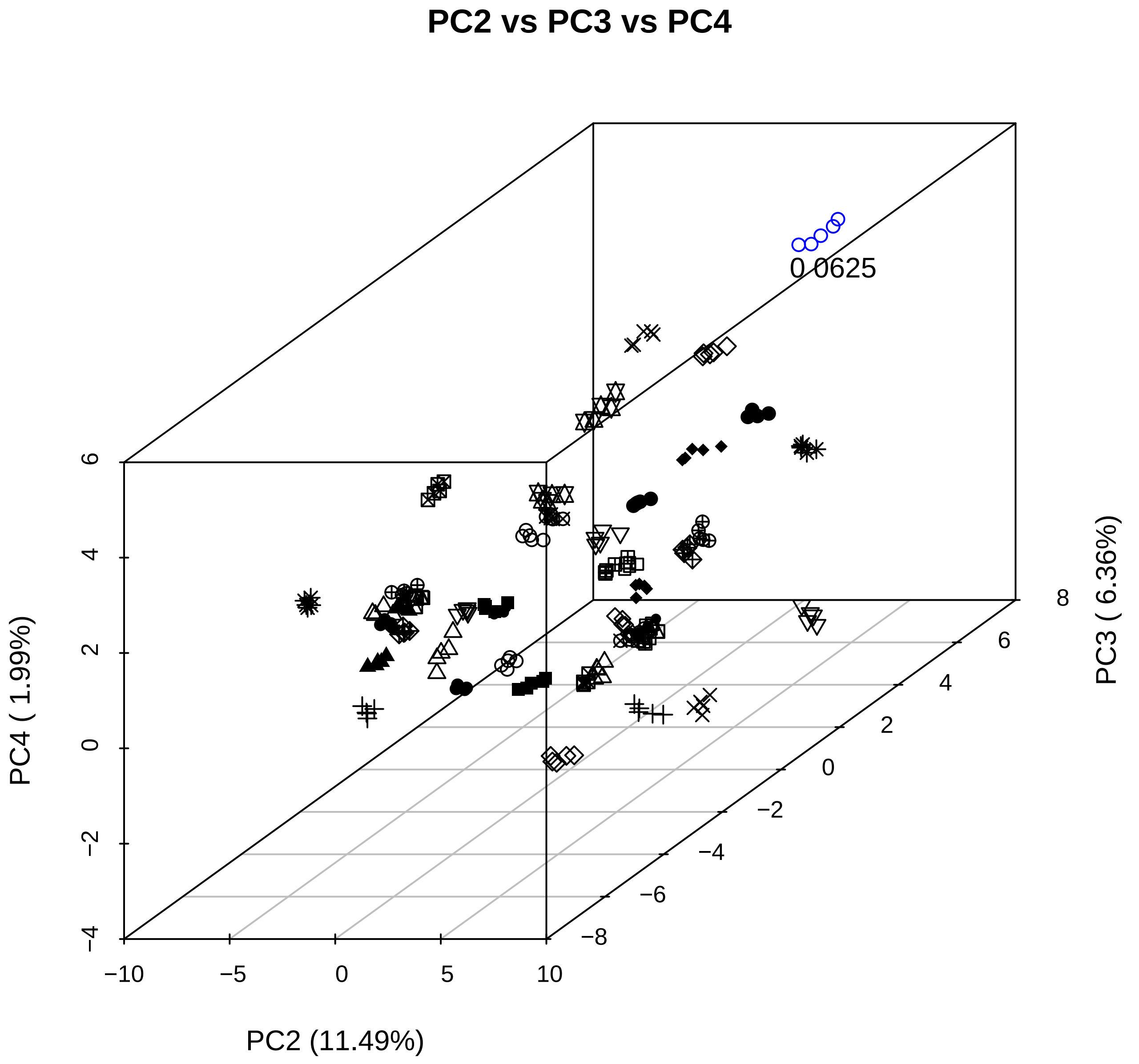
PC2 vs PC3 vs PC4
0.0625
PC2 (11.49%)
PC4 ( 1.99%)
PC3 ( 6.36%)
−10
−5
0
5
10
−4
−2
0
2
4
6
−8
−6
−4
−2
0
2
4
6
8

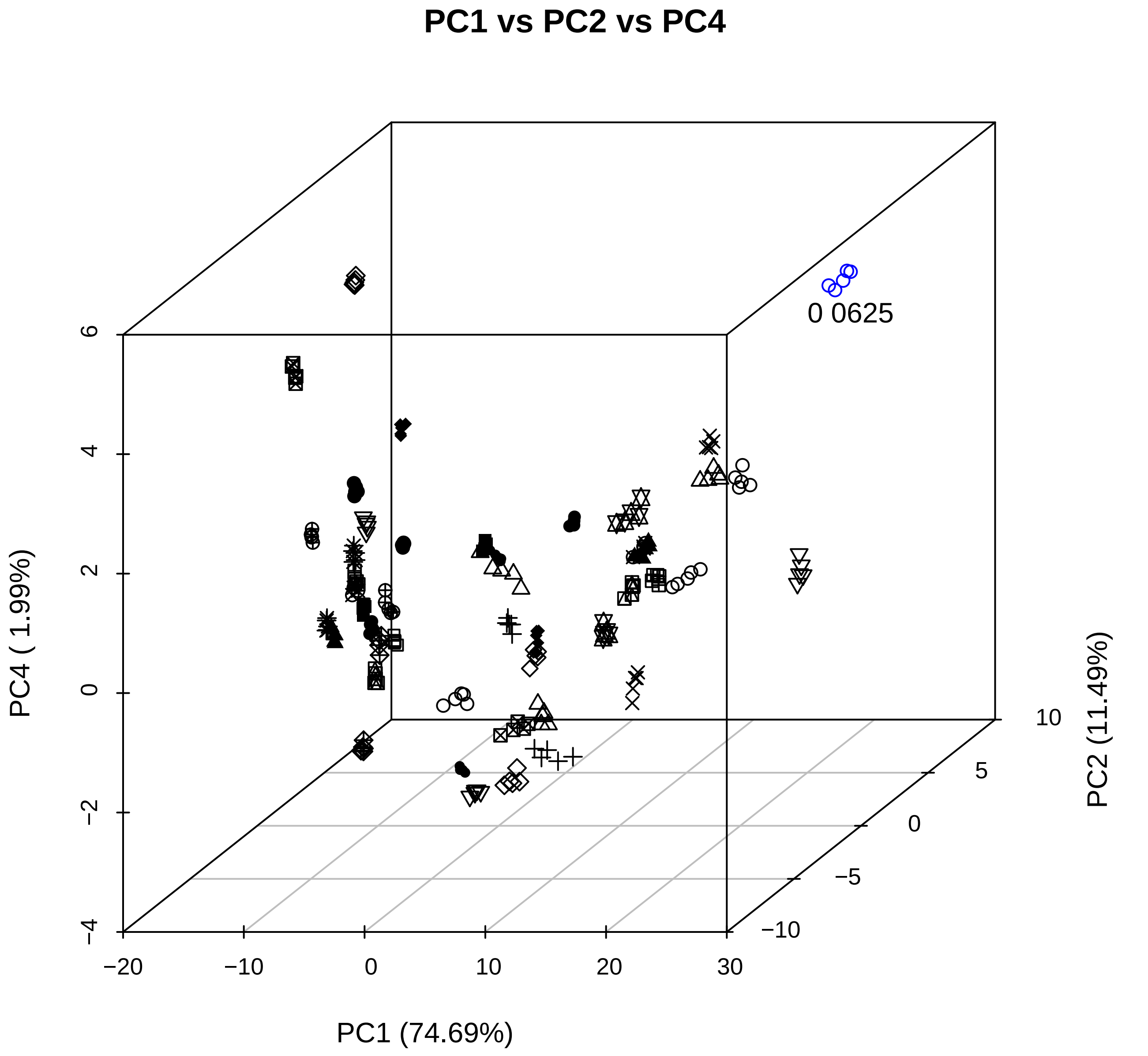
PC1 vs PC2 vs PC4
0 0625
PC4 ( 1.99%)
PC1 (74.69%)
PC2 (11.49%)
6
4
2
0
−2
−4
−20
−10
0
10
20
30
10
5
0
−5
−10

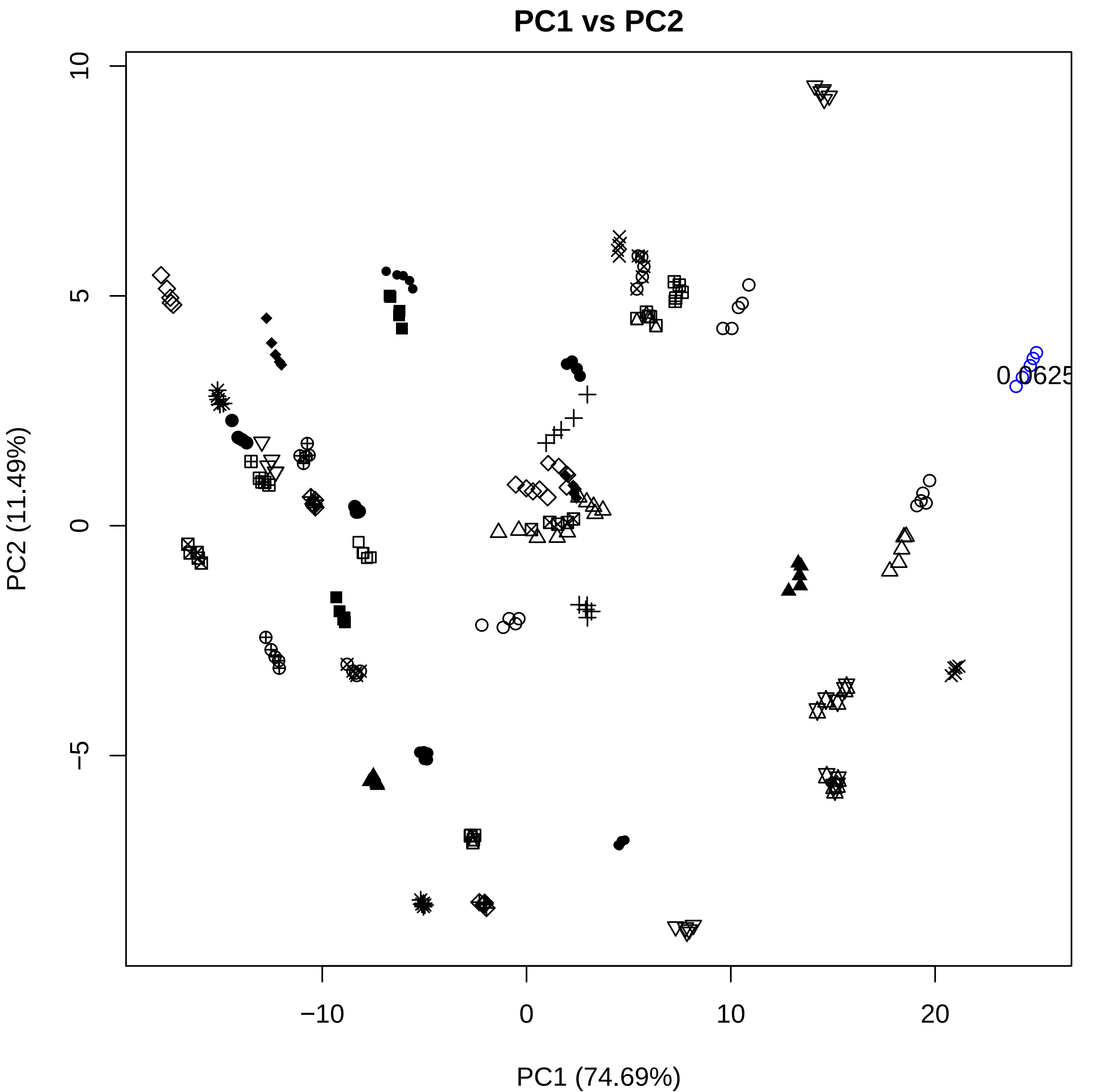
PC1 vs PC2
PC2 (11.49%)
PC1 (74.69%)
10
5
0
−5
−10
0
10
20
0.0625

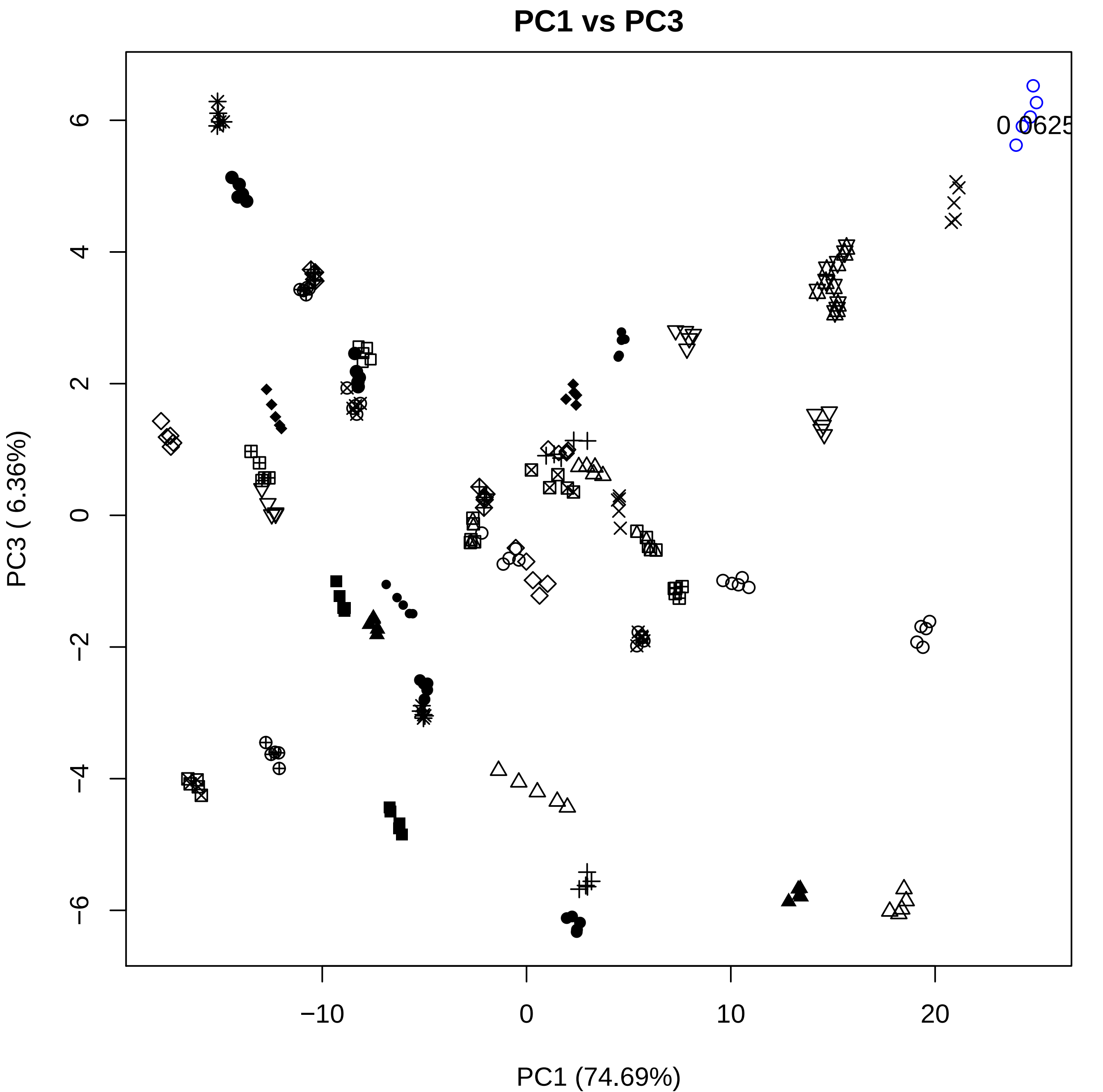
PC1 vs PC3
PC3 ( 6.36%)
PC1 (74.69%)
6
4
2
0
−2
−4
−6
−10
0
10
20
0.0625

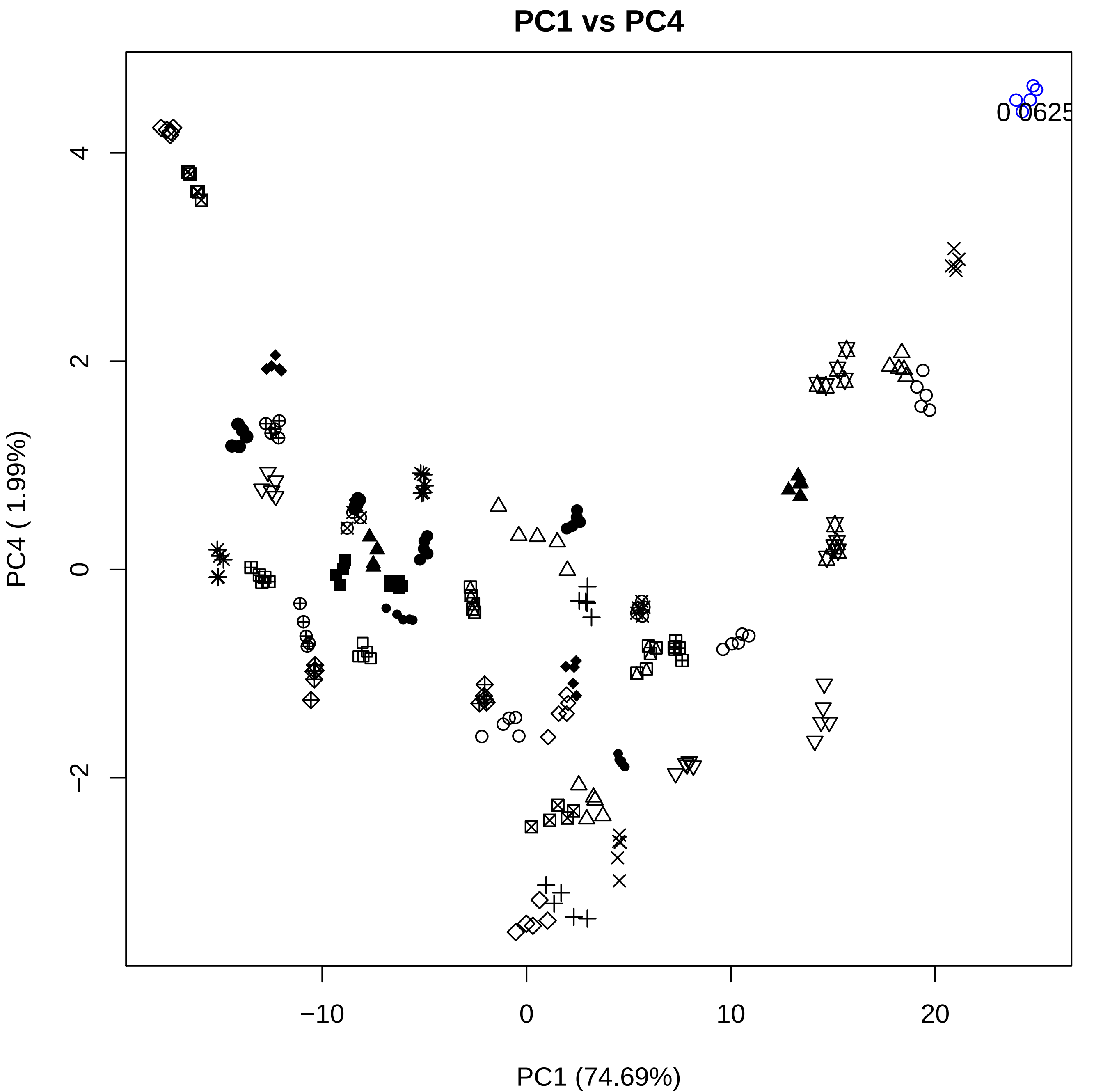
PC1 vs PC4
0 0625
4
2
0
−2
−10
0
10
20
PC4 ( 1.99%)
PC1 (74.69%)

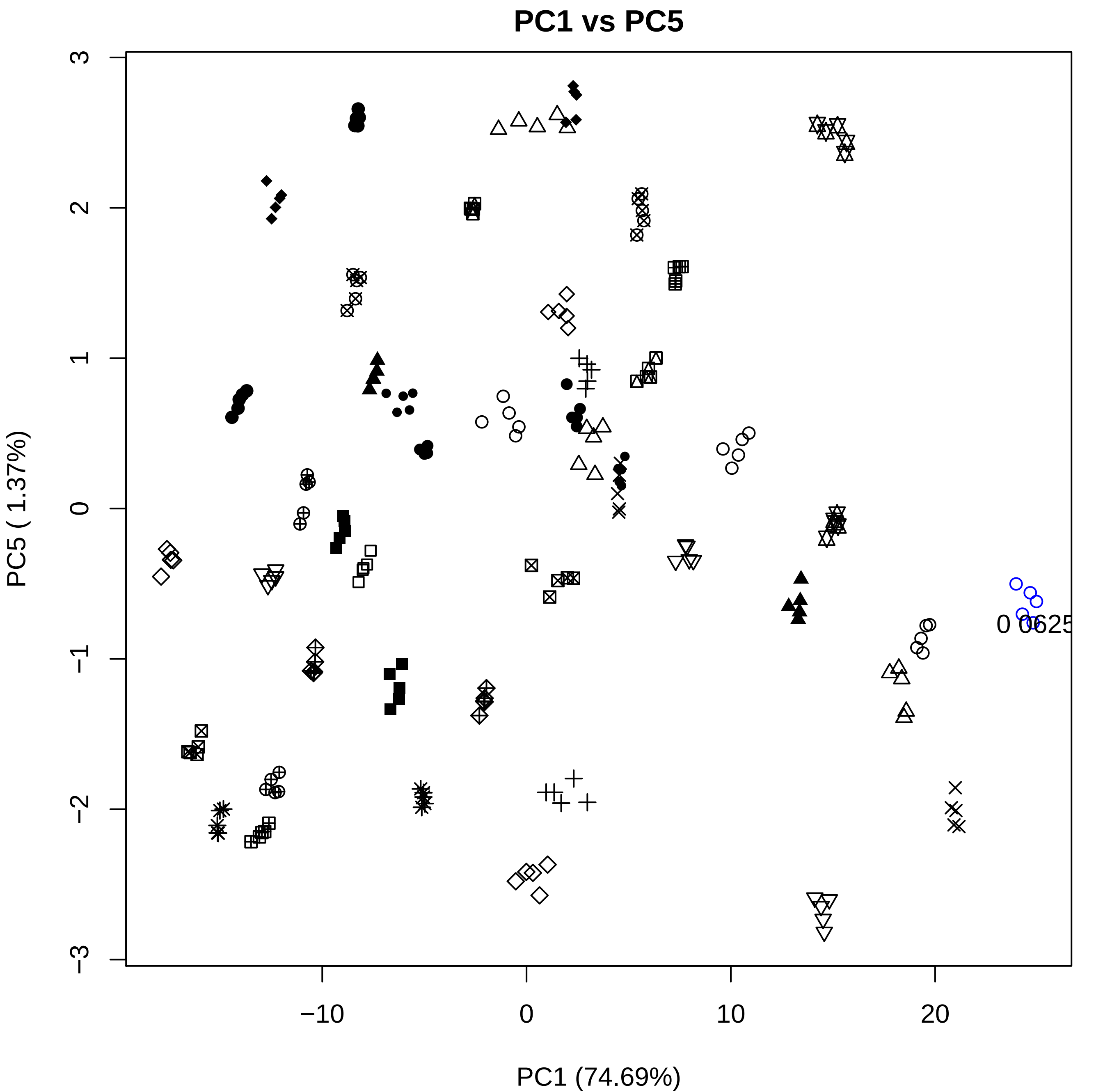

PC1 vs PC5
PC5 ( 1.37%)
PC1 (74.69%)
3
2
1
0
−1
−2
−3
−10
0
10
20
0 0625

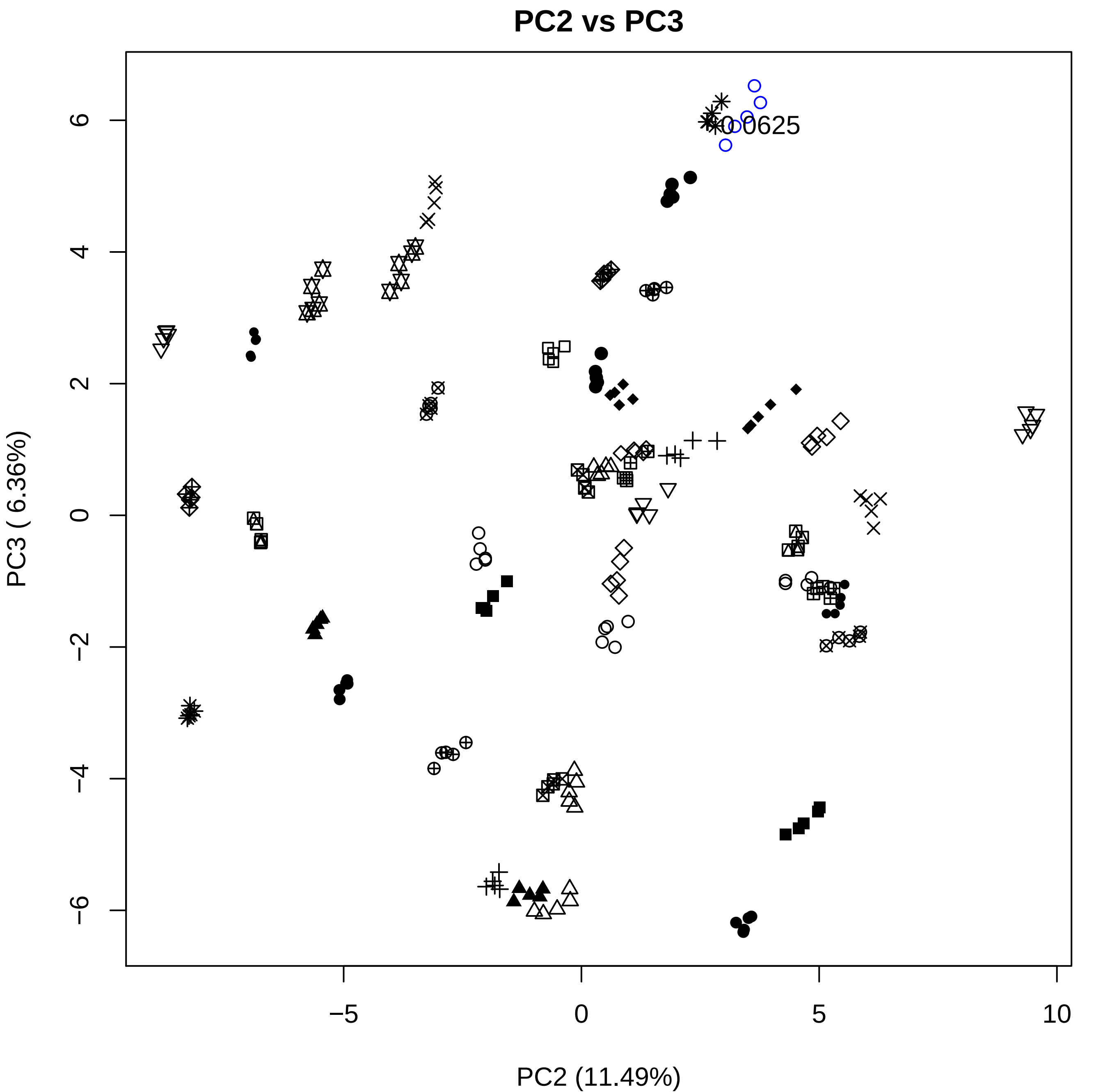

PC2 vs PC3
00625
PC3 ( 6.36%)
PC2 (11.49%)
−6
−4
−2
0
2
4
6
−5
0
5
10

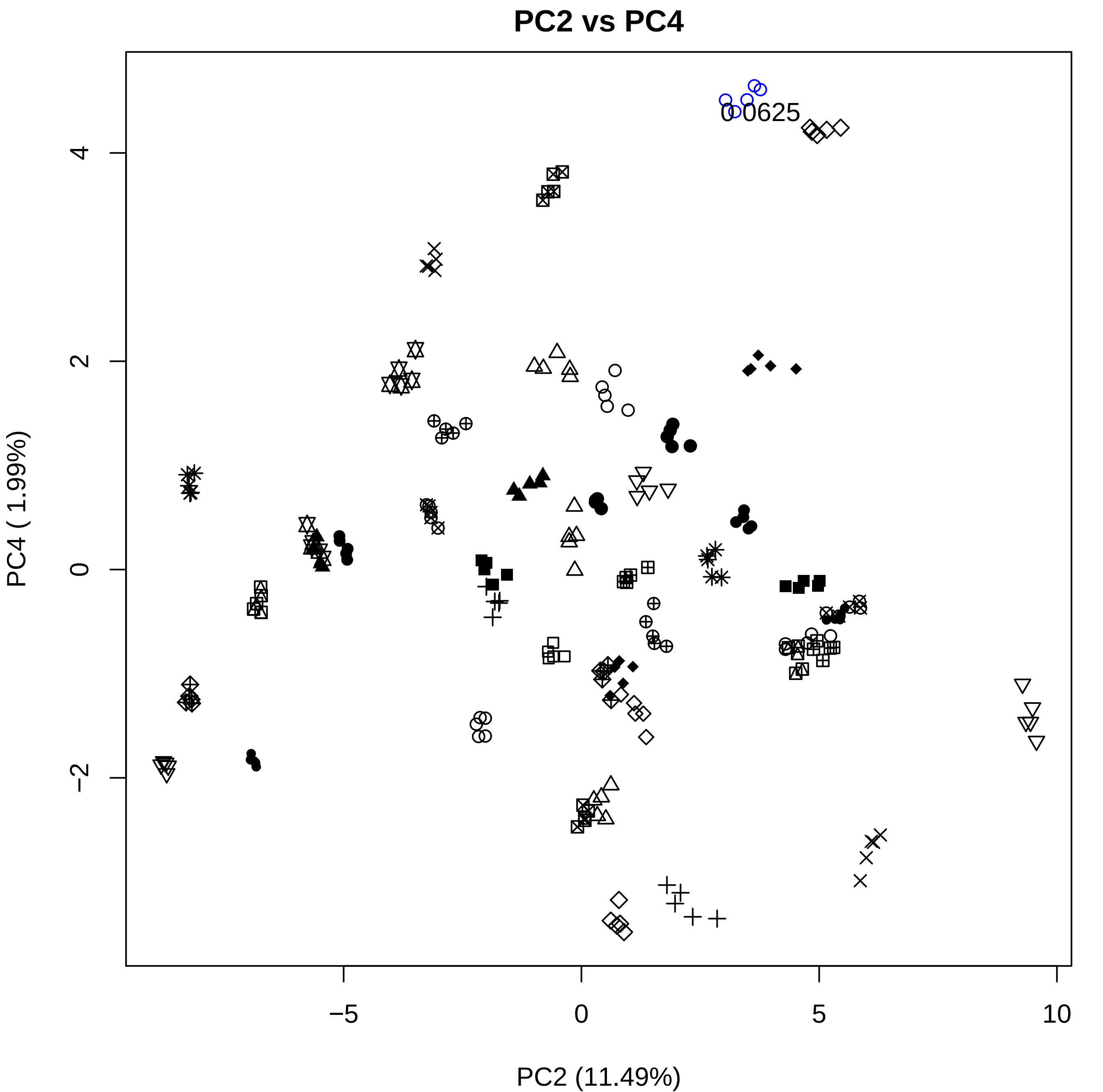
PC2 vs PC4
00625
PC4 ( 1.99%)
PC2 (11.49%)
−5
0
5
10
−2
0
2
4

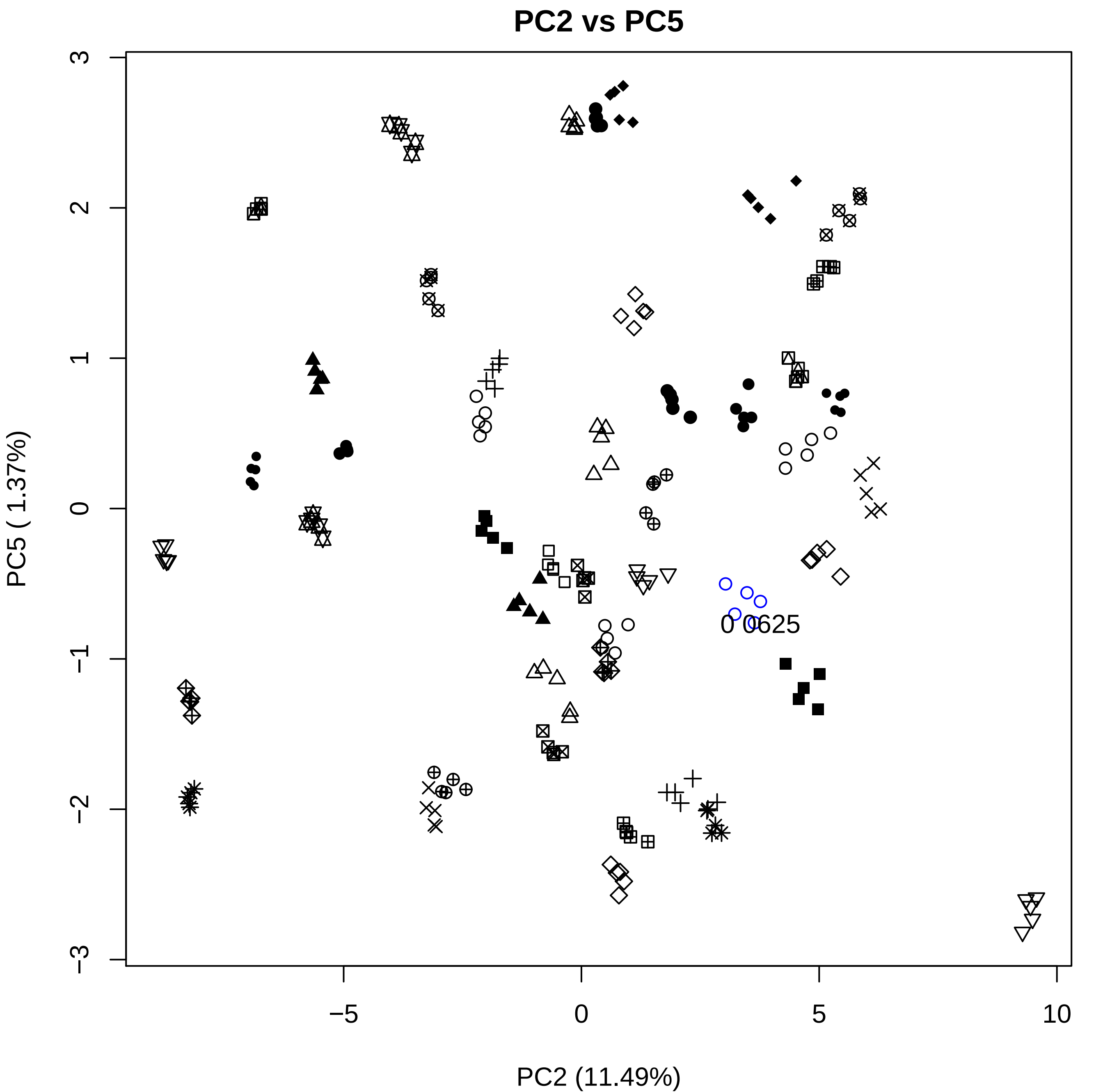

PC2 vs PC5
0.0625
PC2 (11.49%)
PC5 ( 1.37%)
−5
0
5
10
−3
−2
−1
0
1
2
3

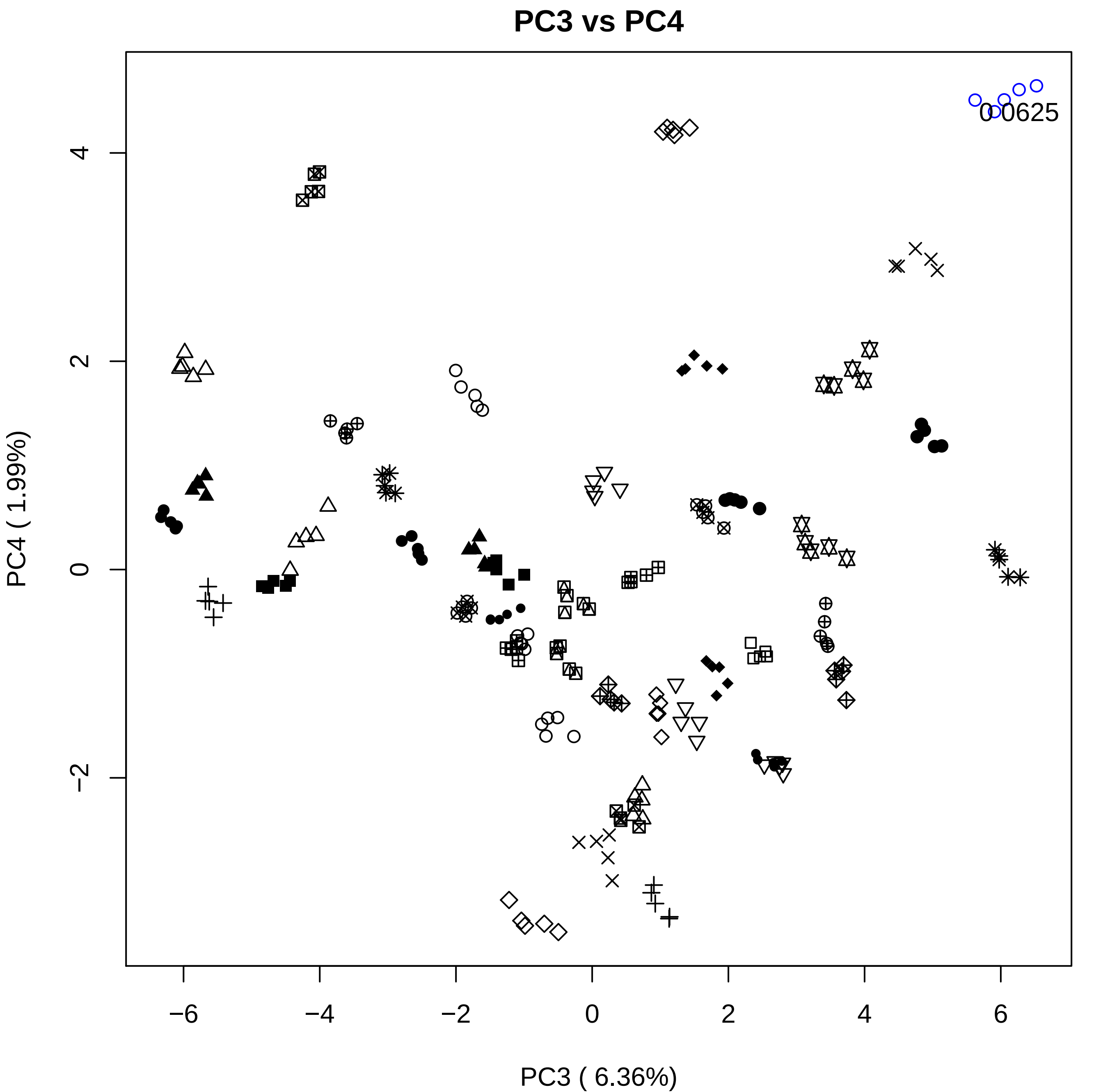
PC3 vs PC4
0.0625
PC4 ( 1.99%)
PC3 ( 6.36%)
−6
−4
−2
0
2
4
6
−2
0
2
4

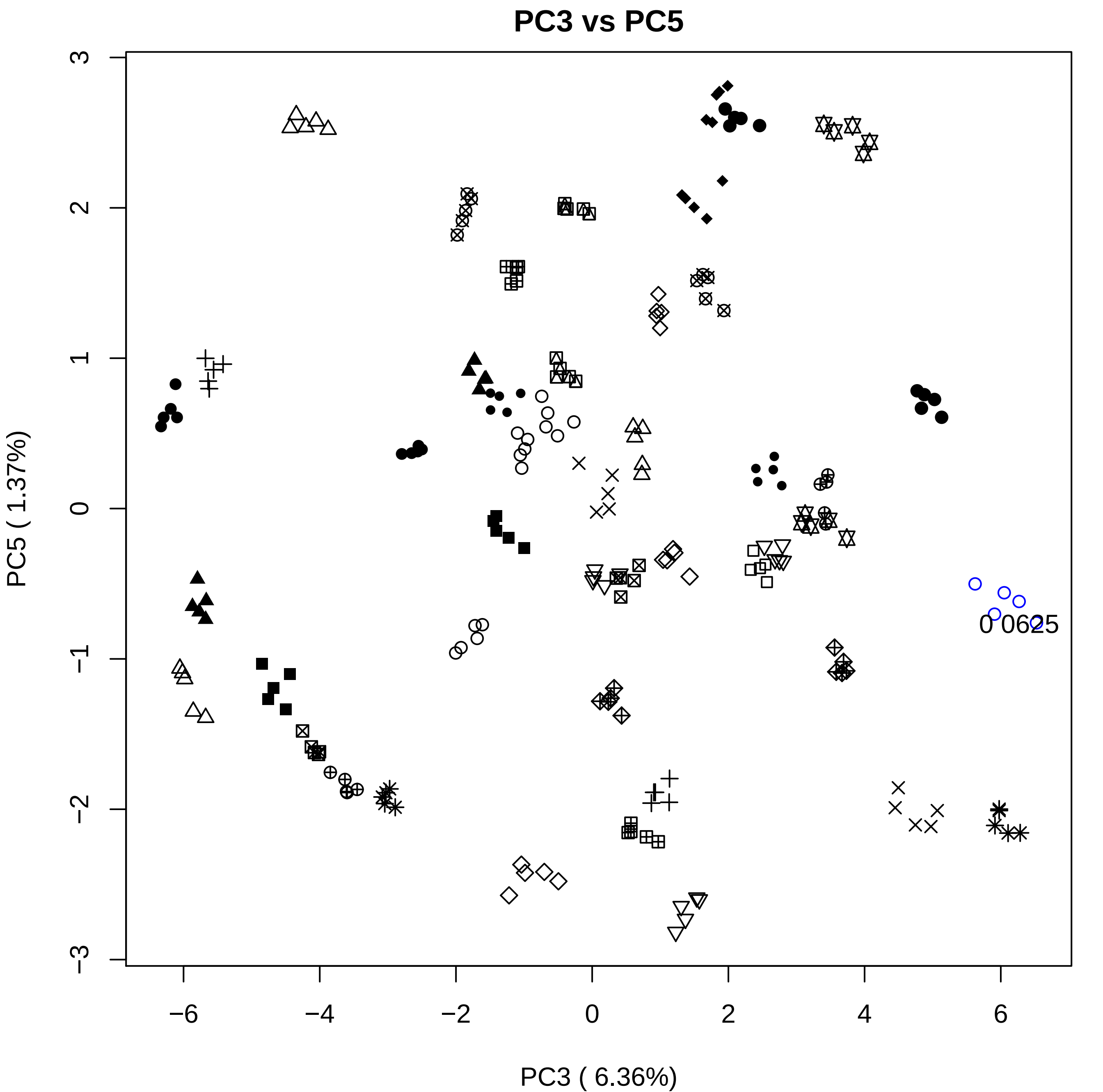
PC3 vs PC5
PC5 ( 1.37%)
PC3 ( 6.36%)
3
2
1
0
−1
−2
−3
−6
−4
−2
0
2
4
6
0 0625

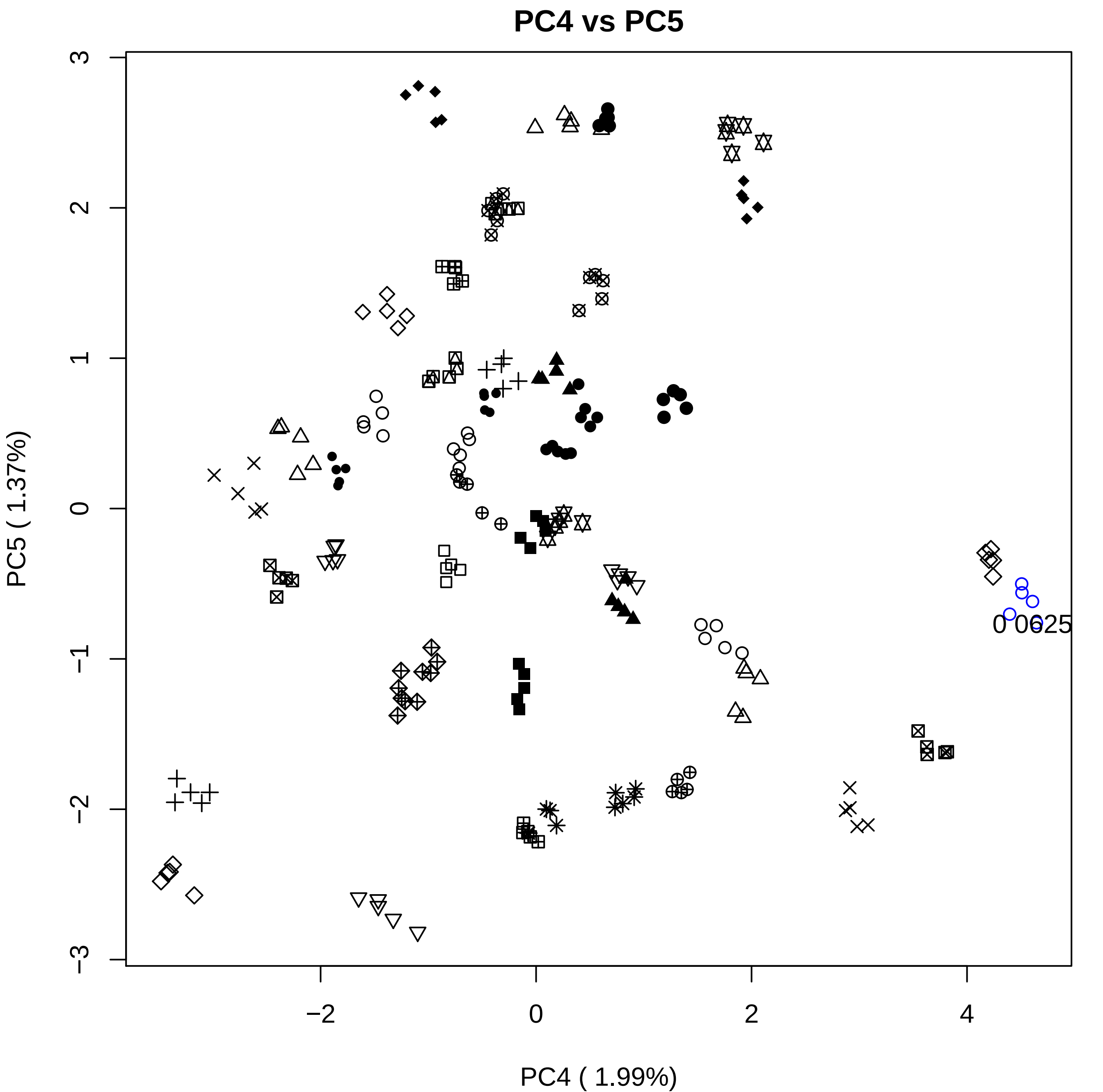
PC4 vs PC5
PC5 ( 1.37%)
PC4 ( 1.99%)
−3
−2
−1
0
1
2
3
−2
0
2
4
0.0625

### PCA of 0.12%  OSCS vs PMH

Scaled Matrix PCAed

7th order polynomial baseline correction performed

Raw spectra smoothed to 21 Neighbours, 2nd order PN, Savtisky Golay

Normalisation between 0 and 100

D2 spectra smoothed to 41 Neighbours, 2nd order PN, Savtisky Golay

Regions Analysed: 3601−2501 and 2001−700cm−1

Samples Analysed: 1 Contam's 0 OSCS's 45 PMH's

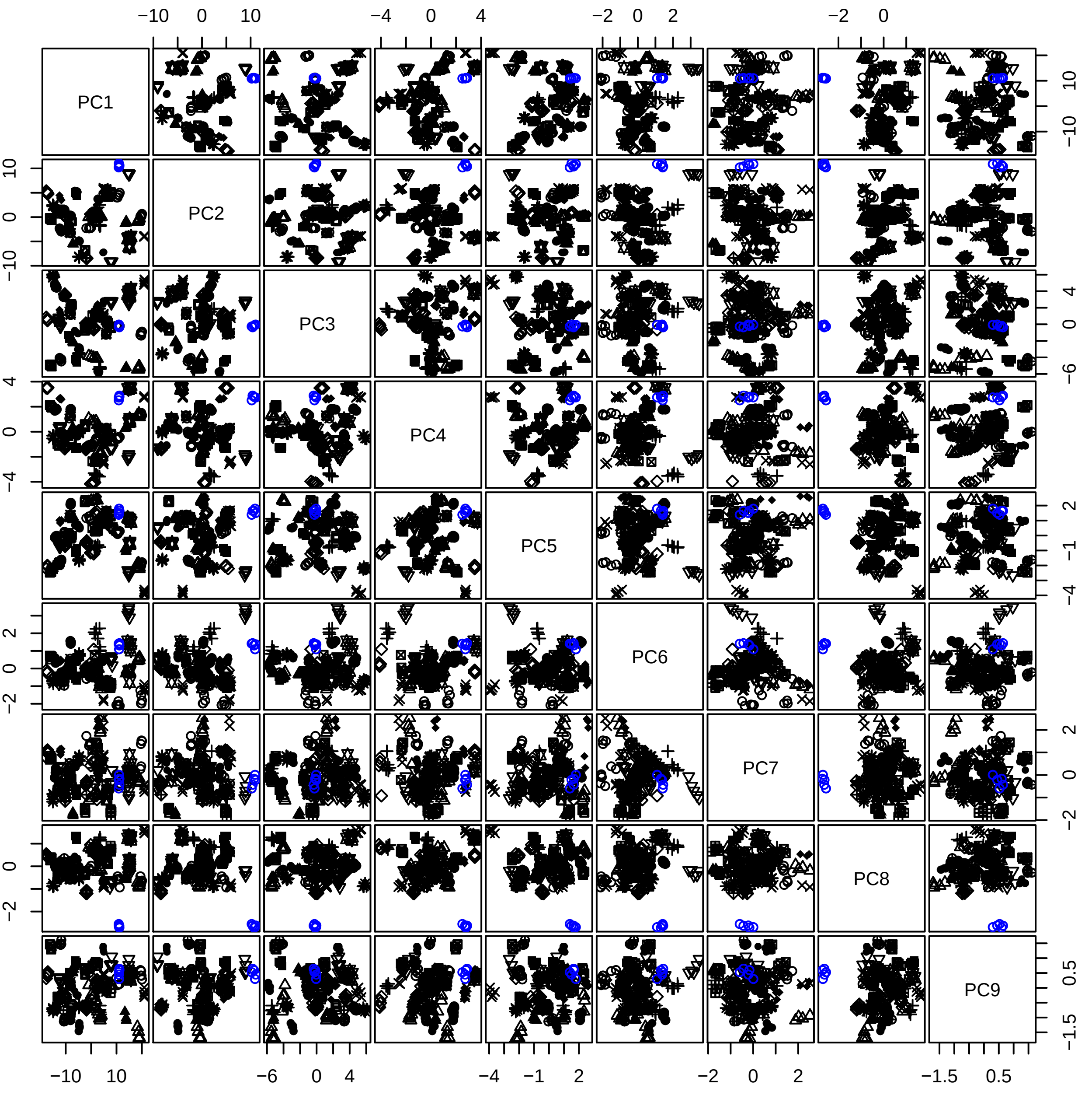
PC1
PC2
PC3
PC4
PC5
PC6
PC7
PC8
PC9
−10 0 10
−4 0 4
−2 0 2
−2 0
10 −10
10 0 −10
4 0 −6
2 −1 −4
2 0 −2
0.5 −1.5
−4 0 4
0 −2
−10 10
−6 0 4
−4 −1 2
−2 0 2
−1.5 0.5

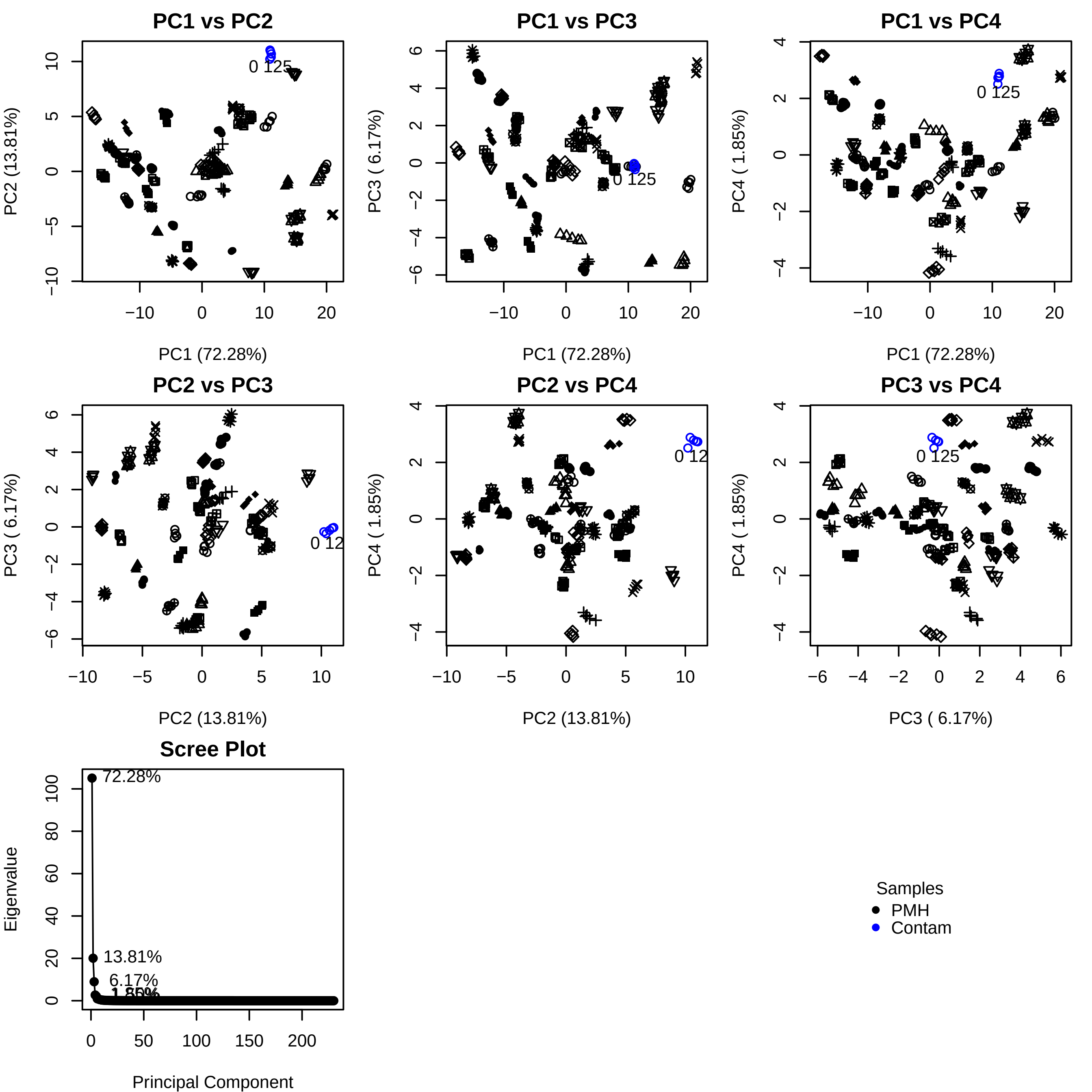
PC1 vs PC2
PC1 vs PC3
PC1 vs PC4
PC2 vs PC3
PC2 vs PC4
PC3 vs PC4
Scree Plot
PC1 (72.28%)
PC2 (13.81%)
PC3 ( 6.17%)
PC4 ( 1.85%)
0 125
72.28%
13.81%
6.17%
Eigenvalue
Principal Component
Samples
PMH
Contam

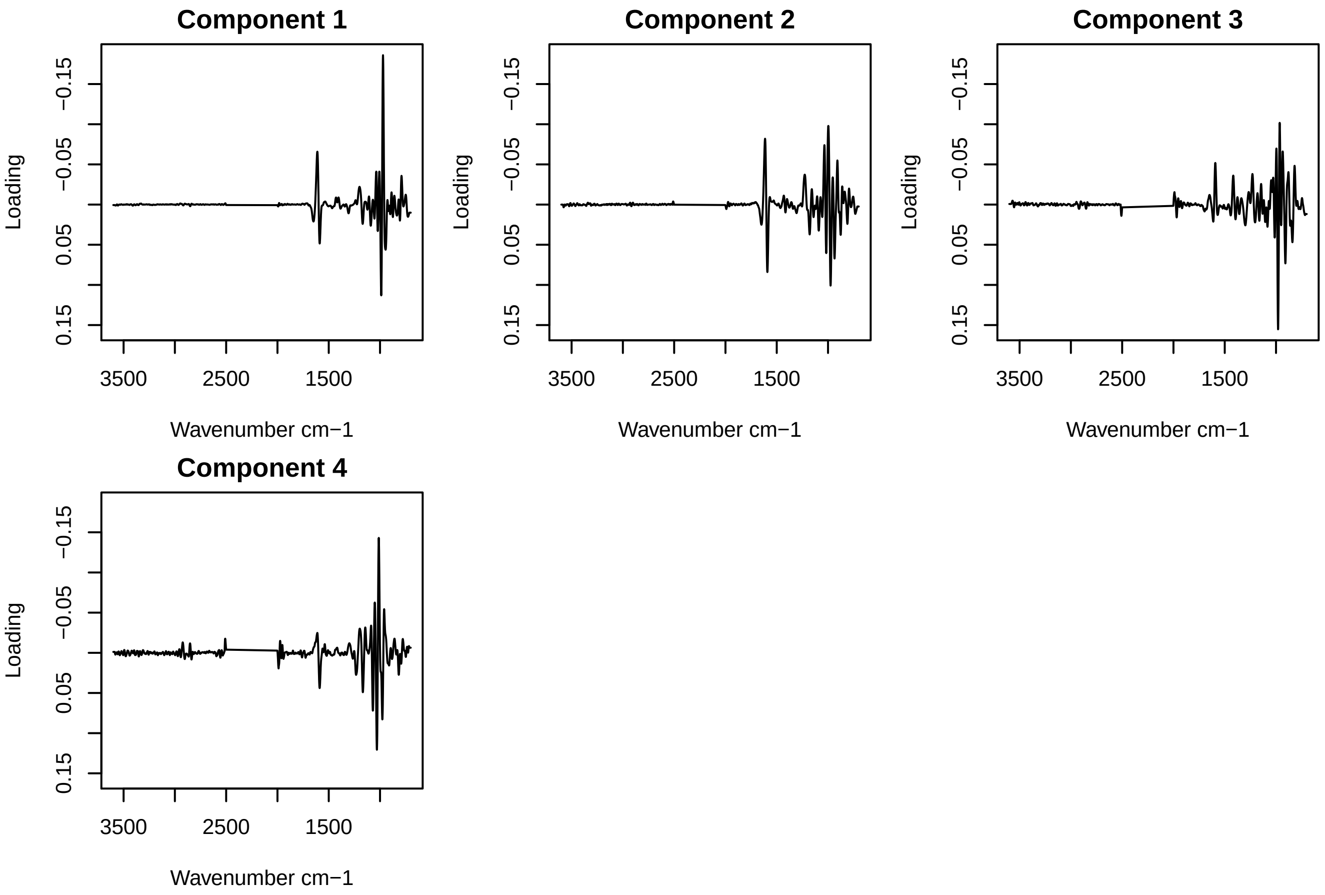
Component 1
Component 2
Component 3
Component 4
Loading
−0.15
−0.05
0.05
0.15
3500
2500
1500
Wavenumber cm−1

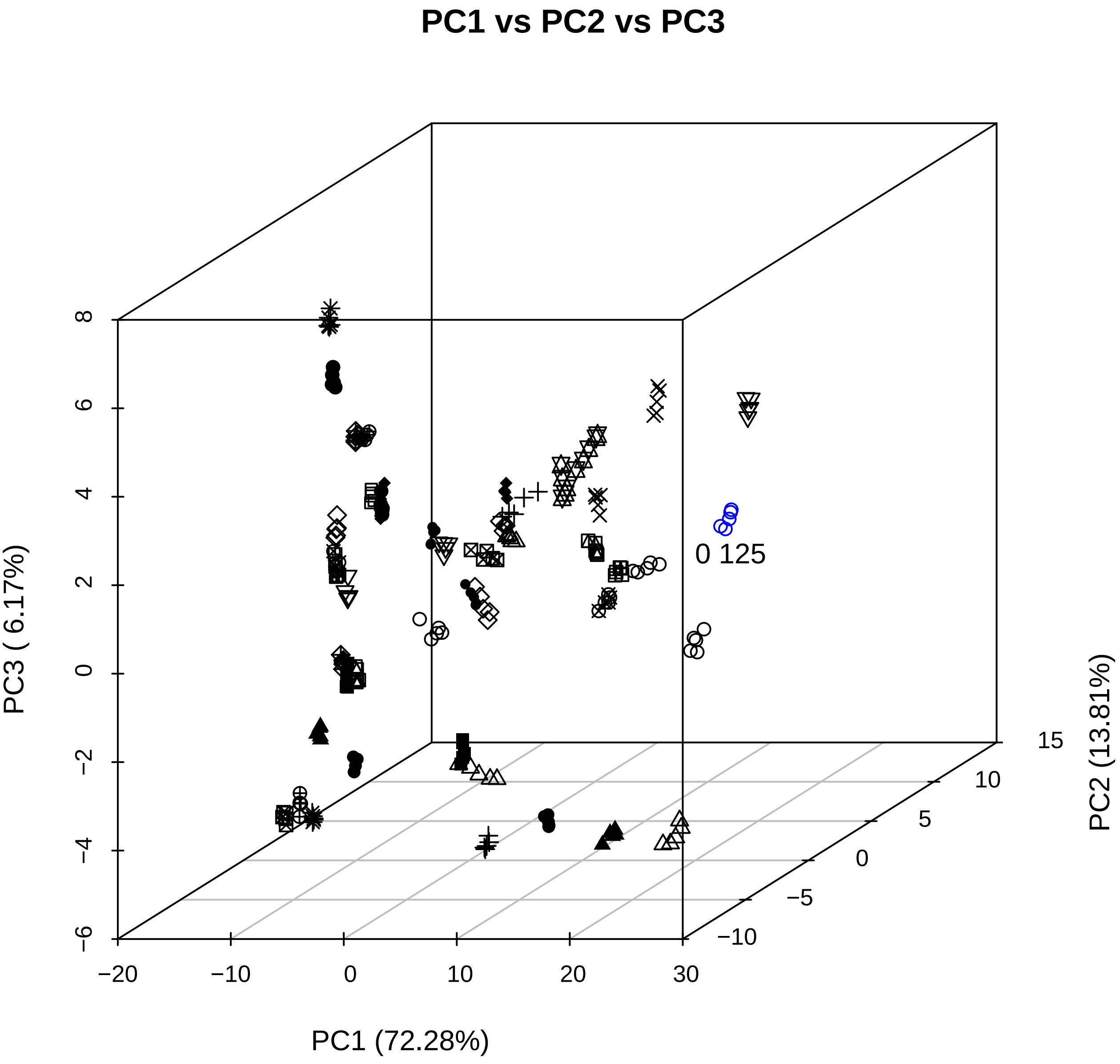
PC1 vs PC2 vs PC3
PC1 (72.28%)
PC2 (13.81%)
PC3 ( 6.17%)
0 125
−20
−10
0
10
20
30
−10
−5
0
5
10
15
−6
−4
−2
0
2
4
6
8

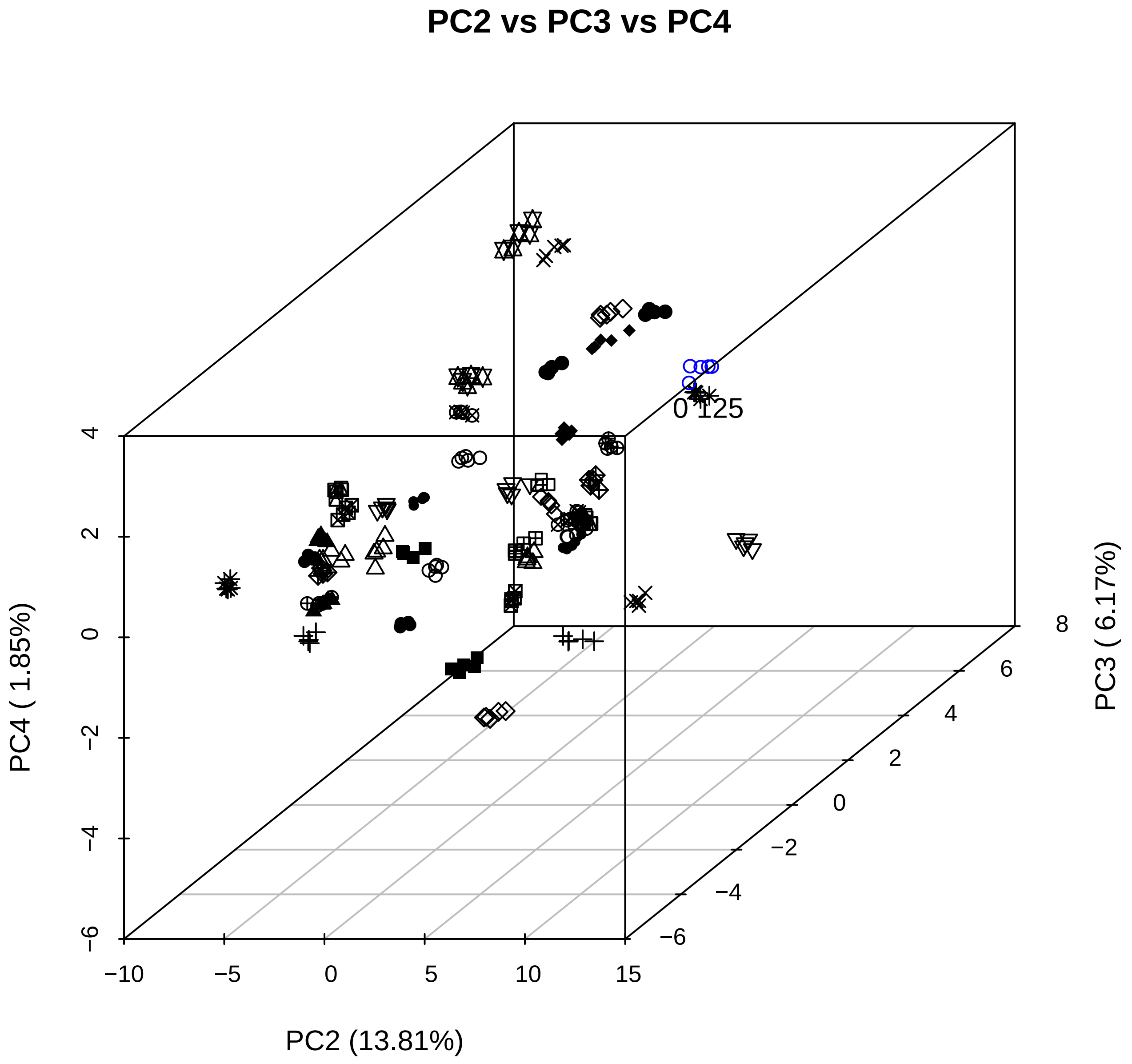
PC2 vs PC3 vs PC4
0 125
PC2 (13.81%)
PC3 ( 6.17%)
PC4 ( 1.85%)
−10
−5
0
5
10
15
−6
−4
−2
0
2
4
6
8
−6
−4
−2
0
2
4

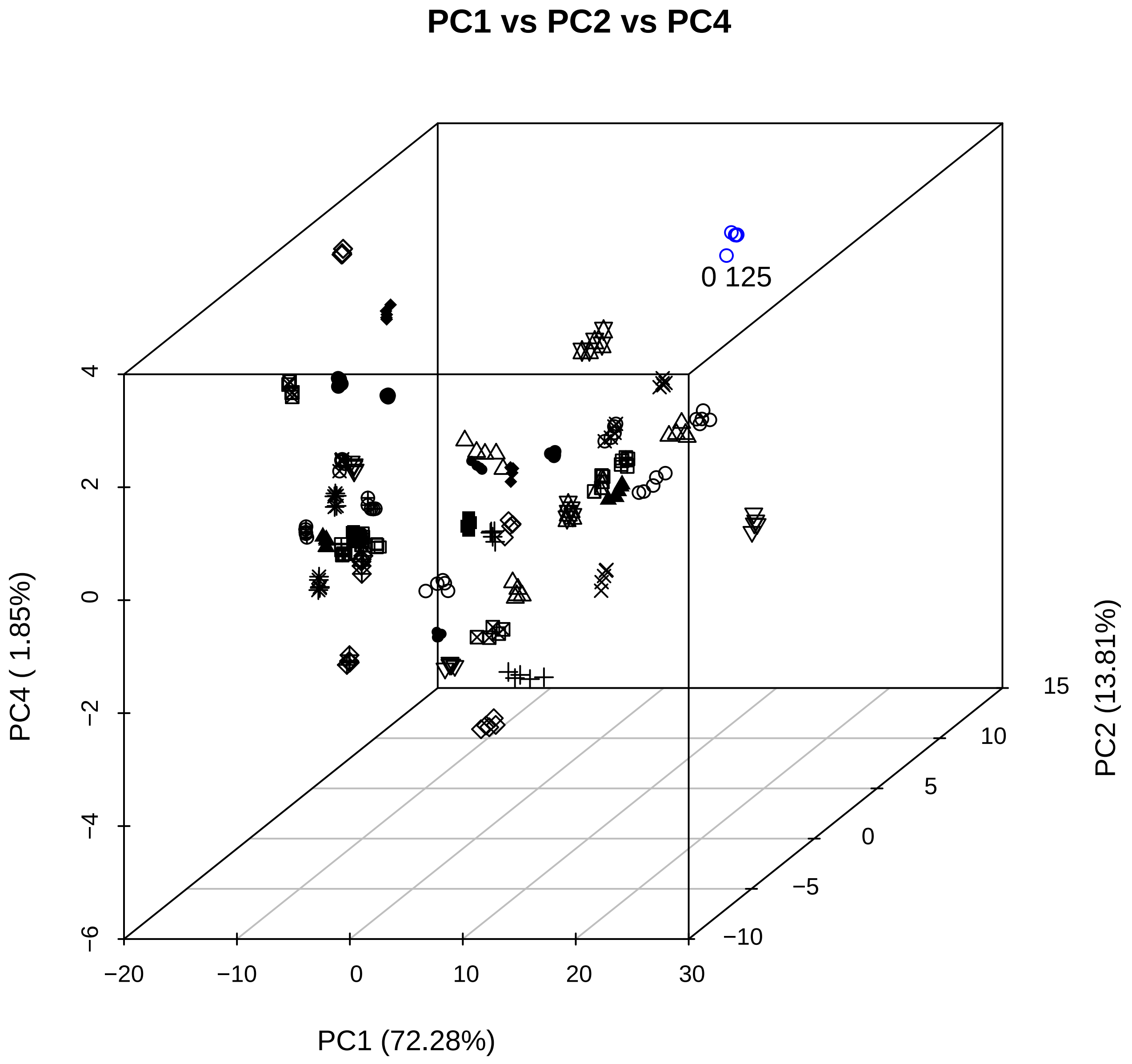

PC1 vs PC2 vs PC4
0 125
PC1 (72.28%)
−20
−10
0
10
20
30
PC2 (13.81%)
−10
−5
0
5
10
15
PC4 ( 1.85%)
−6
−4
−2
0
2
4

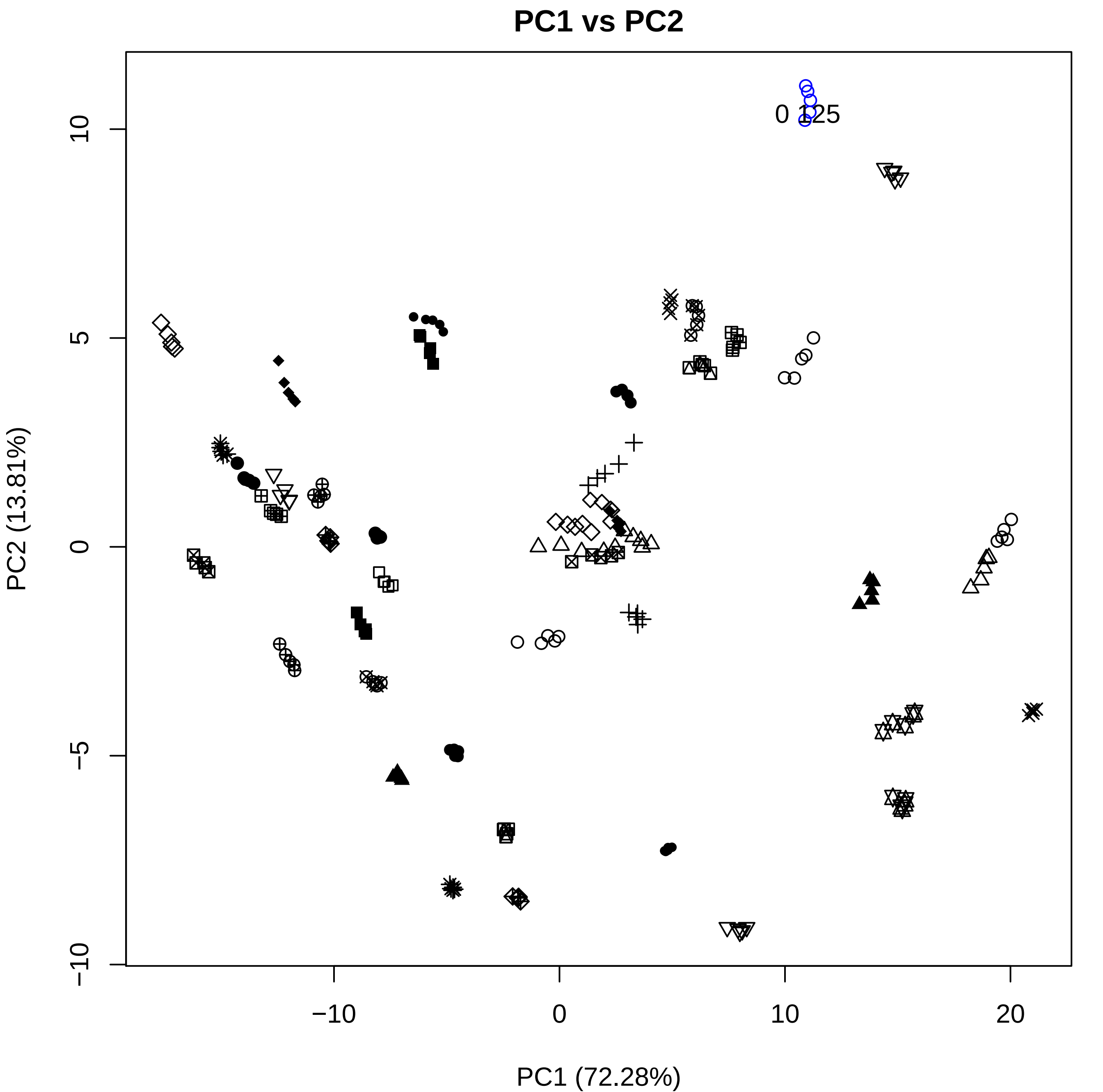

PC1 vs PC2
0 125
PC2 (13.81%)
PC1 (72.28%)
10
5
0
−5
−10
−10
0
10
20

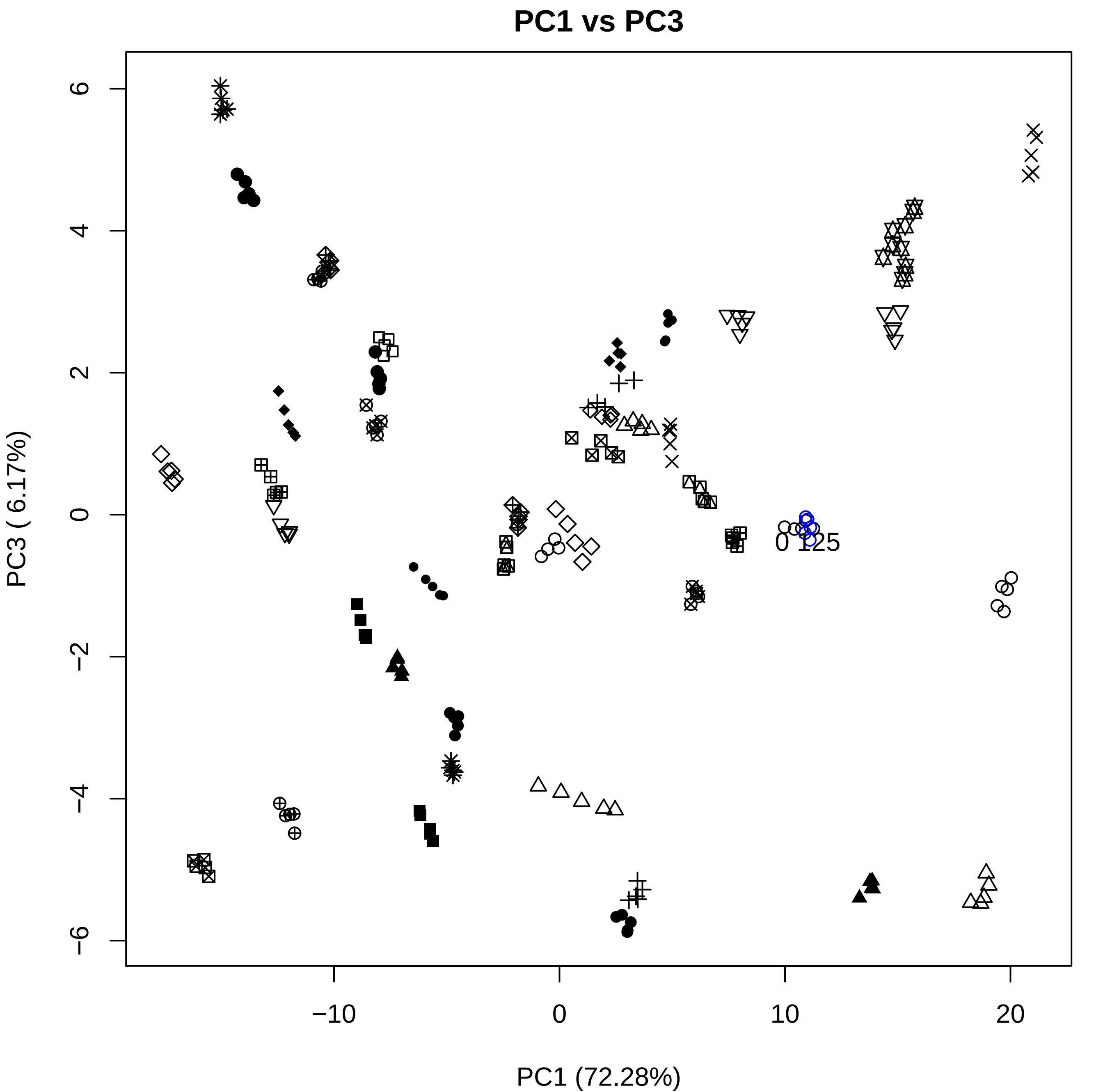
PC1 vs PC3
PC3 ( 6.17%)
PC1 (72.28%)
6
4
2
0
−2
−4
−6
−10
0
10
20
0 125

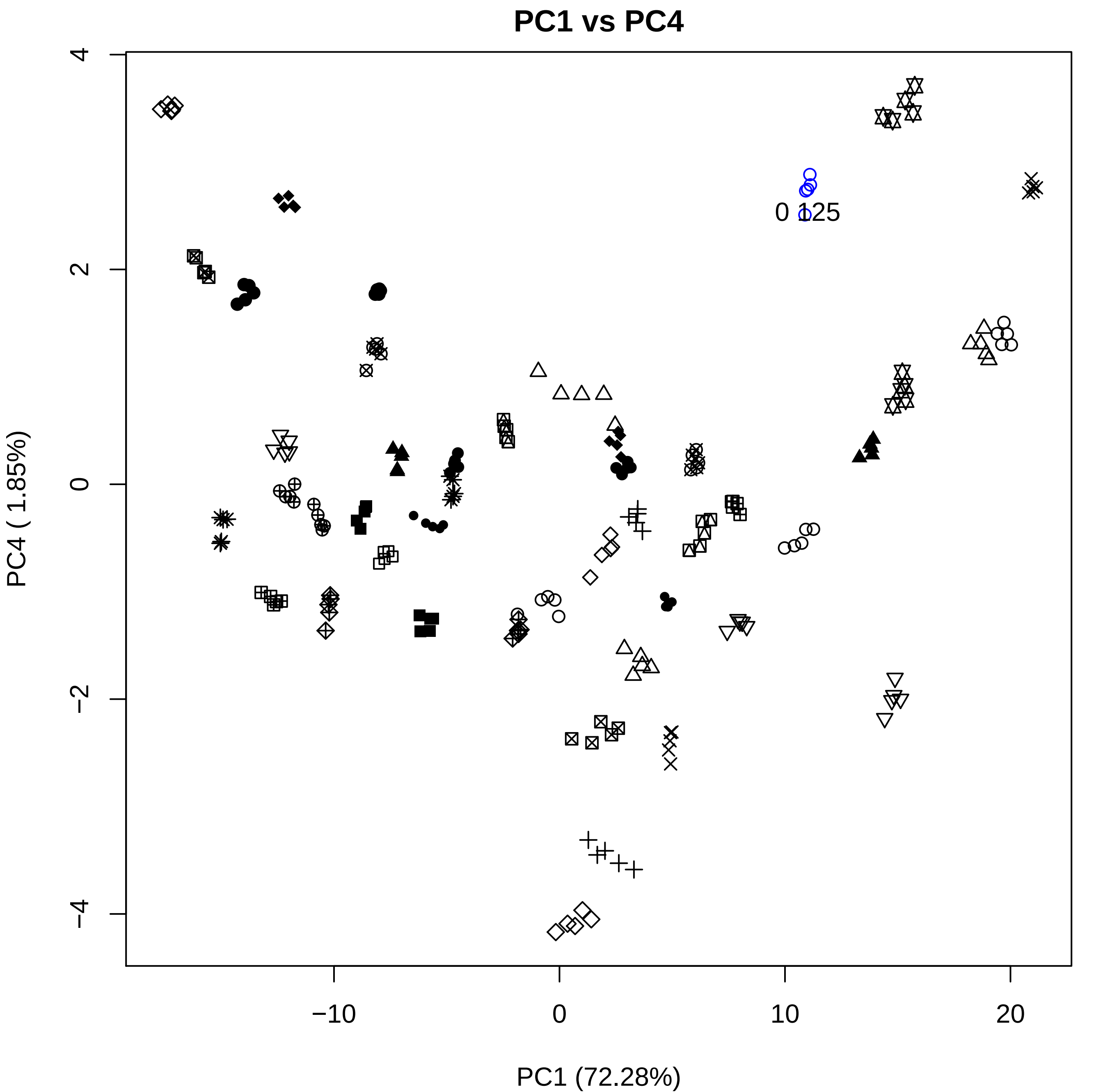

PC1 vs PC4
PC4 ( 1.85%)
PC1 (72.28%)
4
2
0
−2
−4
−10
0
10
20
0.125

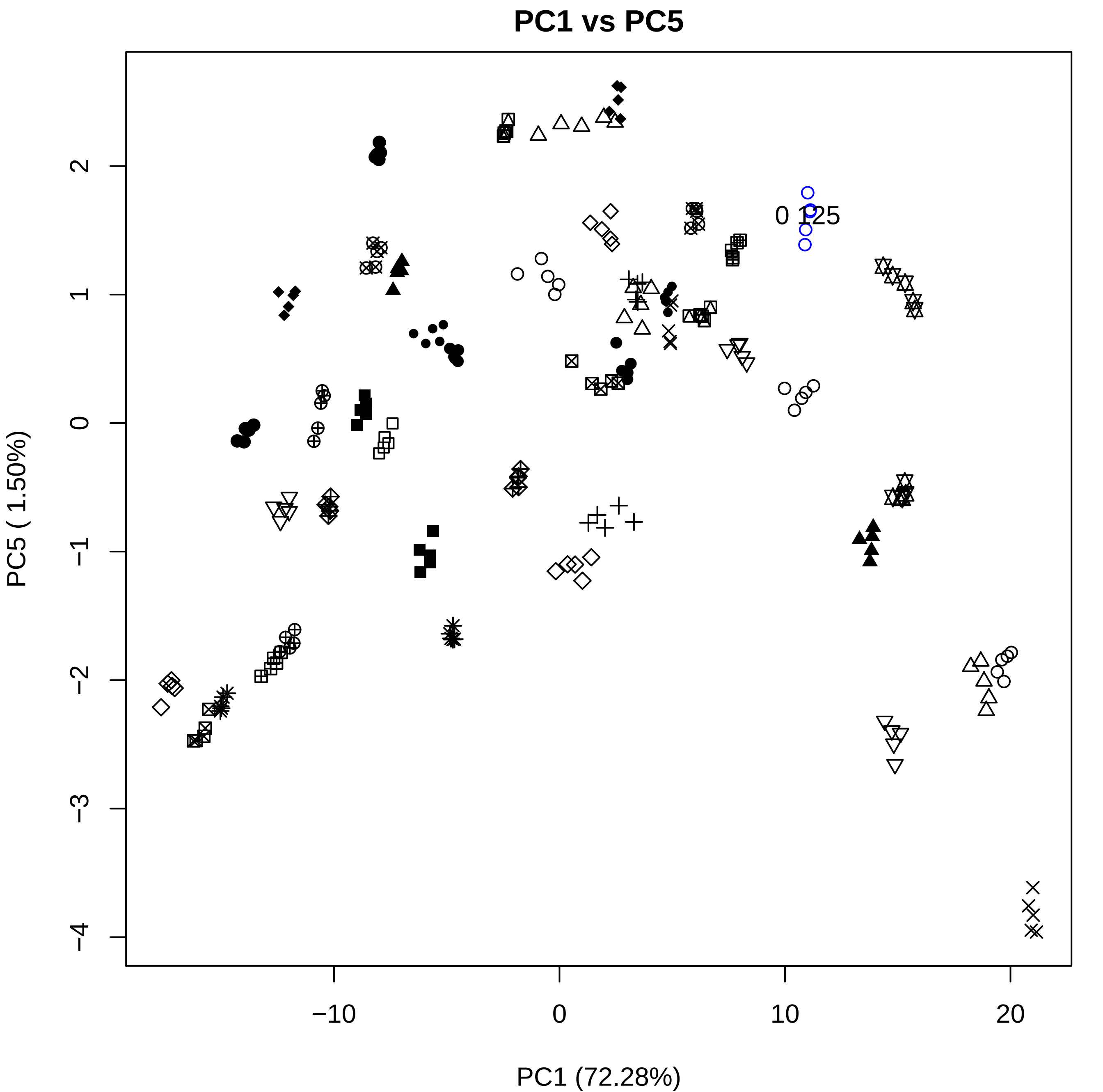
PC1 vs PC5
0 125
PC5 ( 1.50%)
PC1 (72.28%)
2
1
0
−1
−2
−3
−4
−10
0
10
20

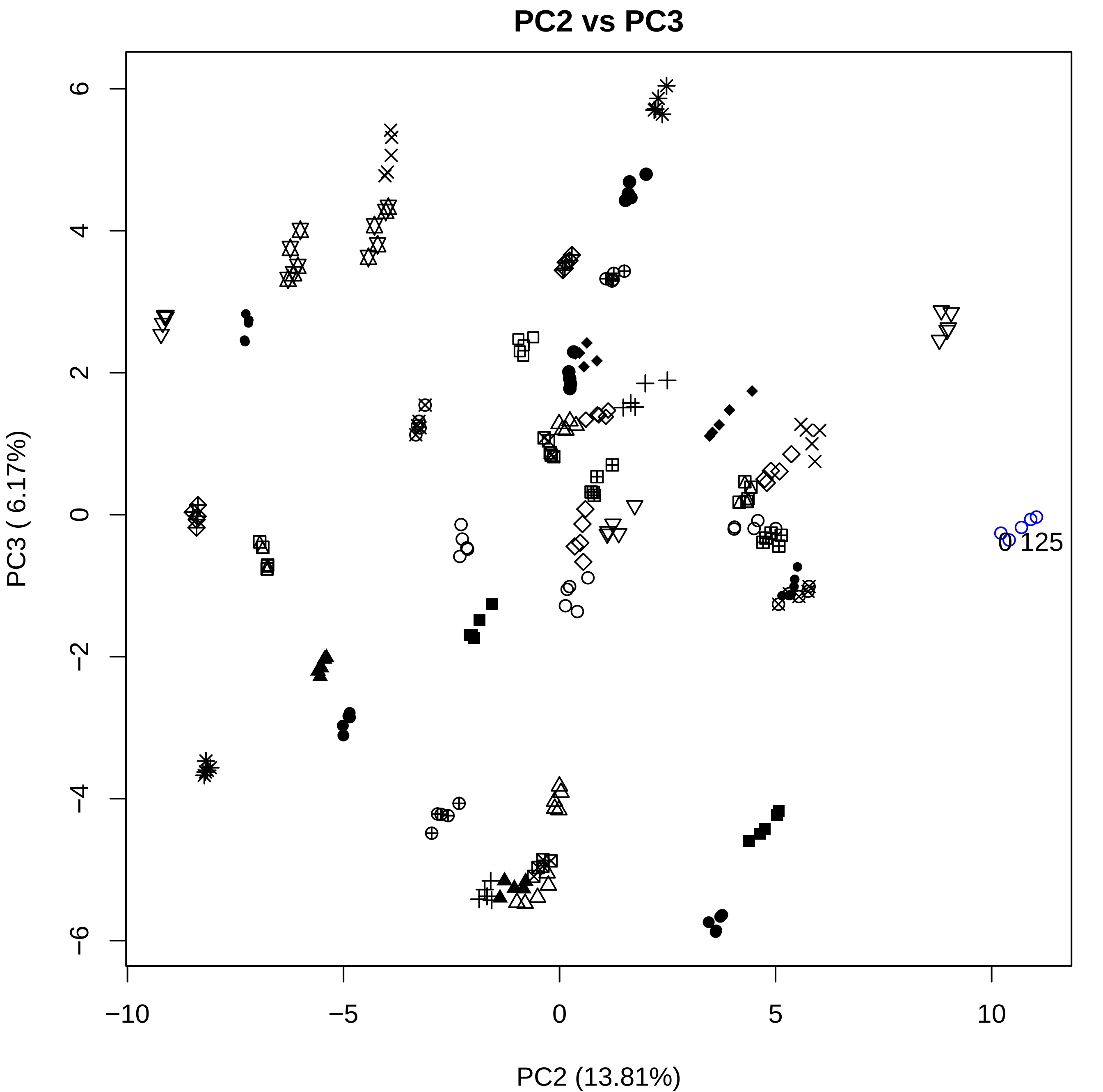

PC2 vs PC3
PC3 ( 6.17%)
PC2 (13.81%)
6
4
2
0
−2
−4
−6
−10
−5
0
5
10
0 125

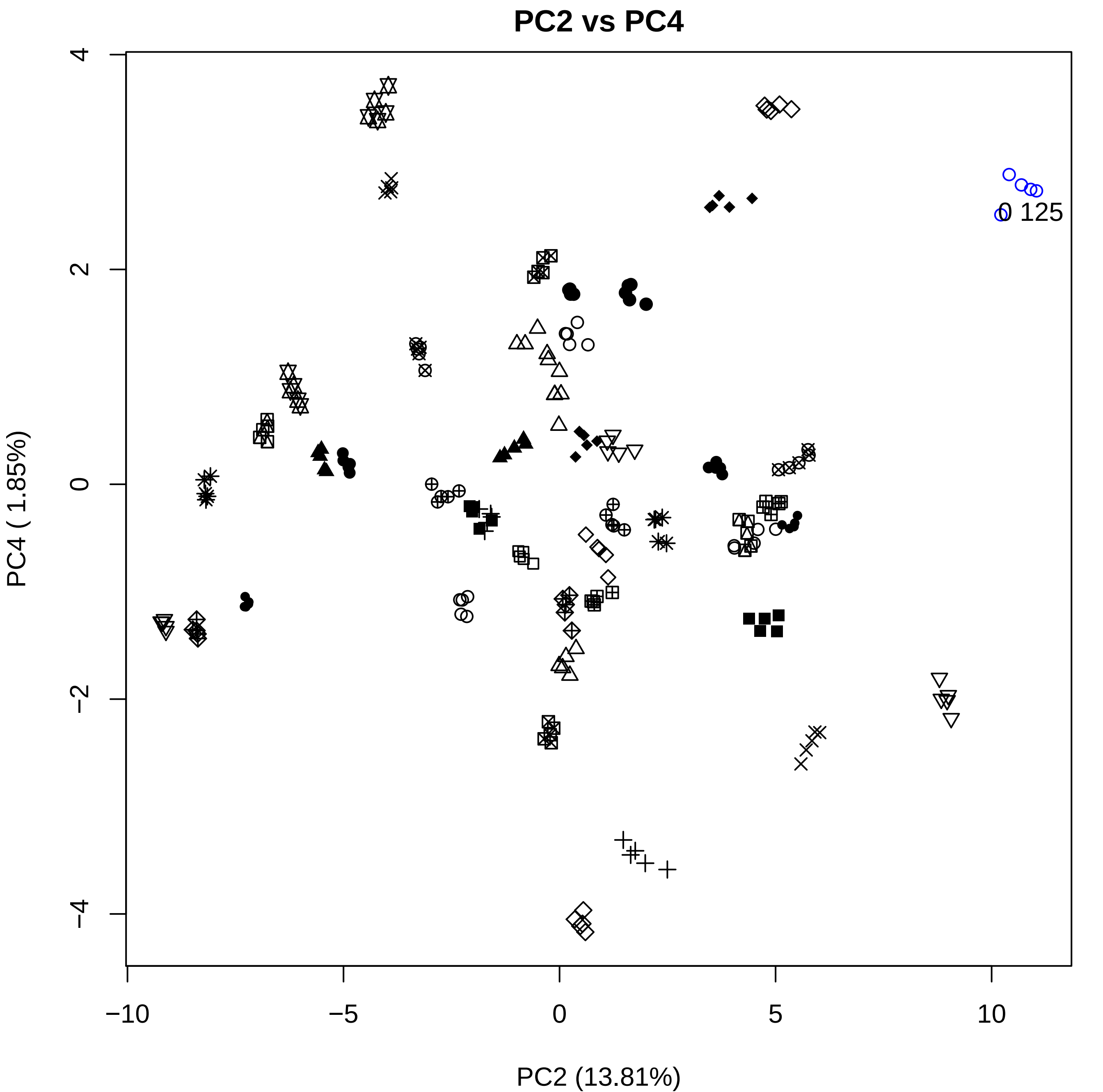
PC2 vs PC4
PC4 ( 1.85%)
PC2 (13.81%)
4
2
0
−2
−4
−10
−5
0
5
10
0 125

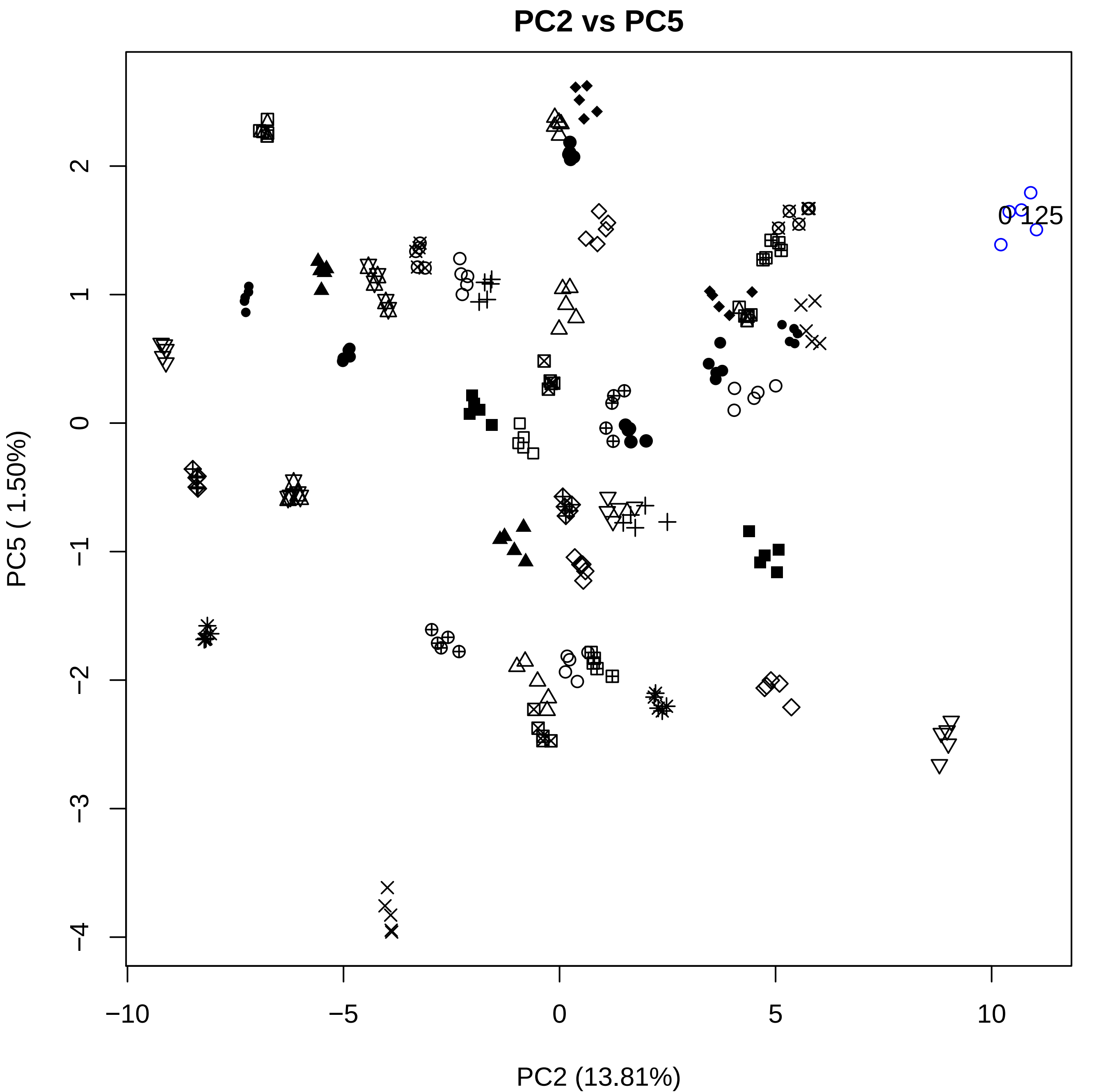

PC2 vs PC5
0 125
PC5 ( 1.50%)
PC2 (13.81%)
−10
−5
0
5
10
−4
−3
−2
−1
0
1
2

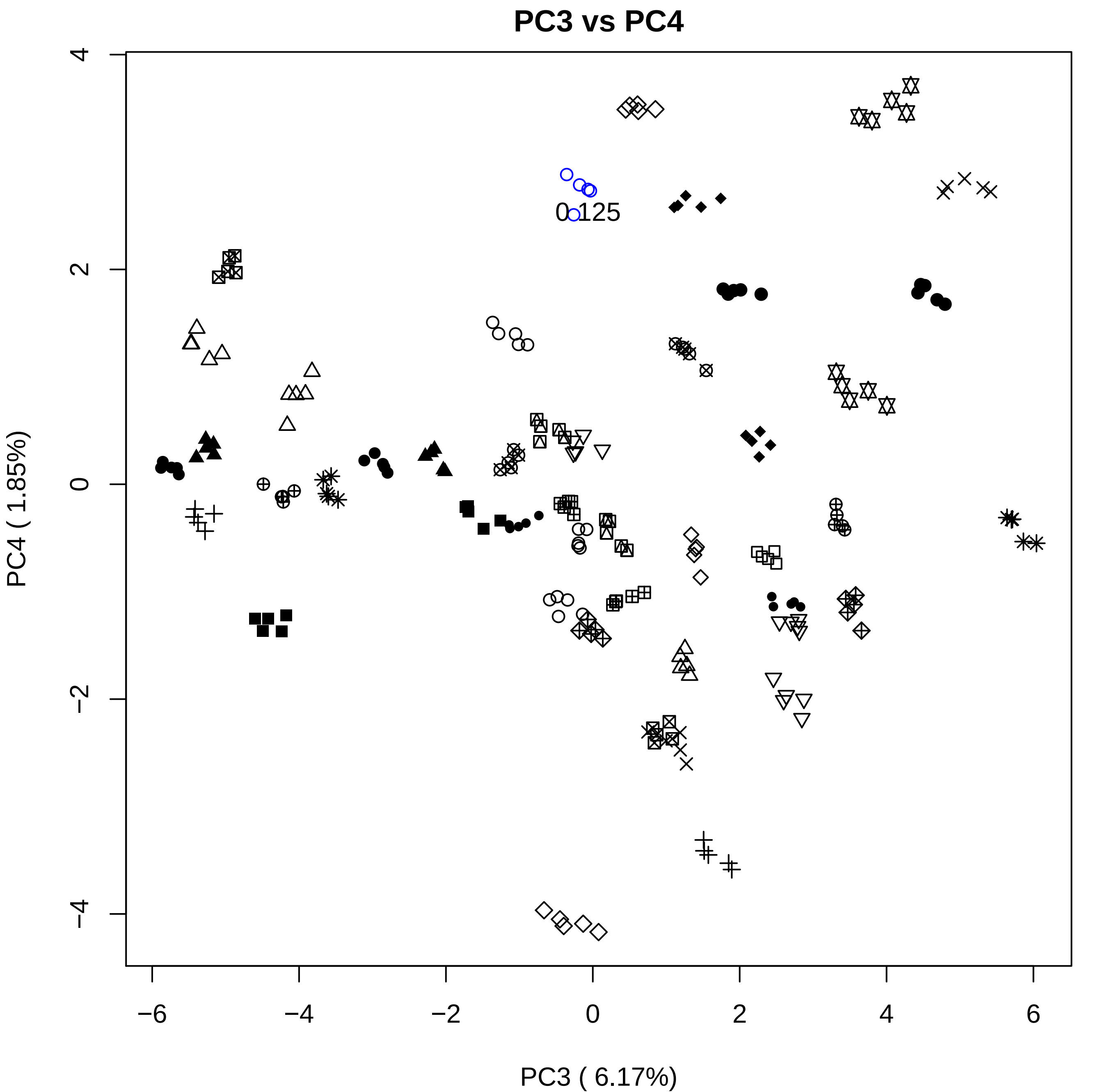
PC3 vs PC4
PC4 ( 1.85%)
PC3 ( 6.17%)
0.125
4
2
0
−2
−4
−6
−4
−2
0
2
4
6

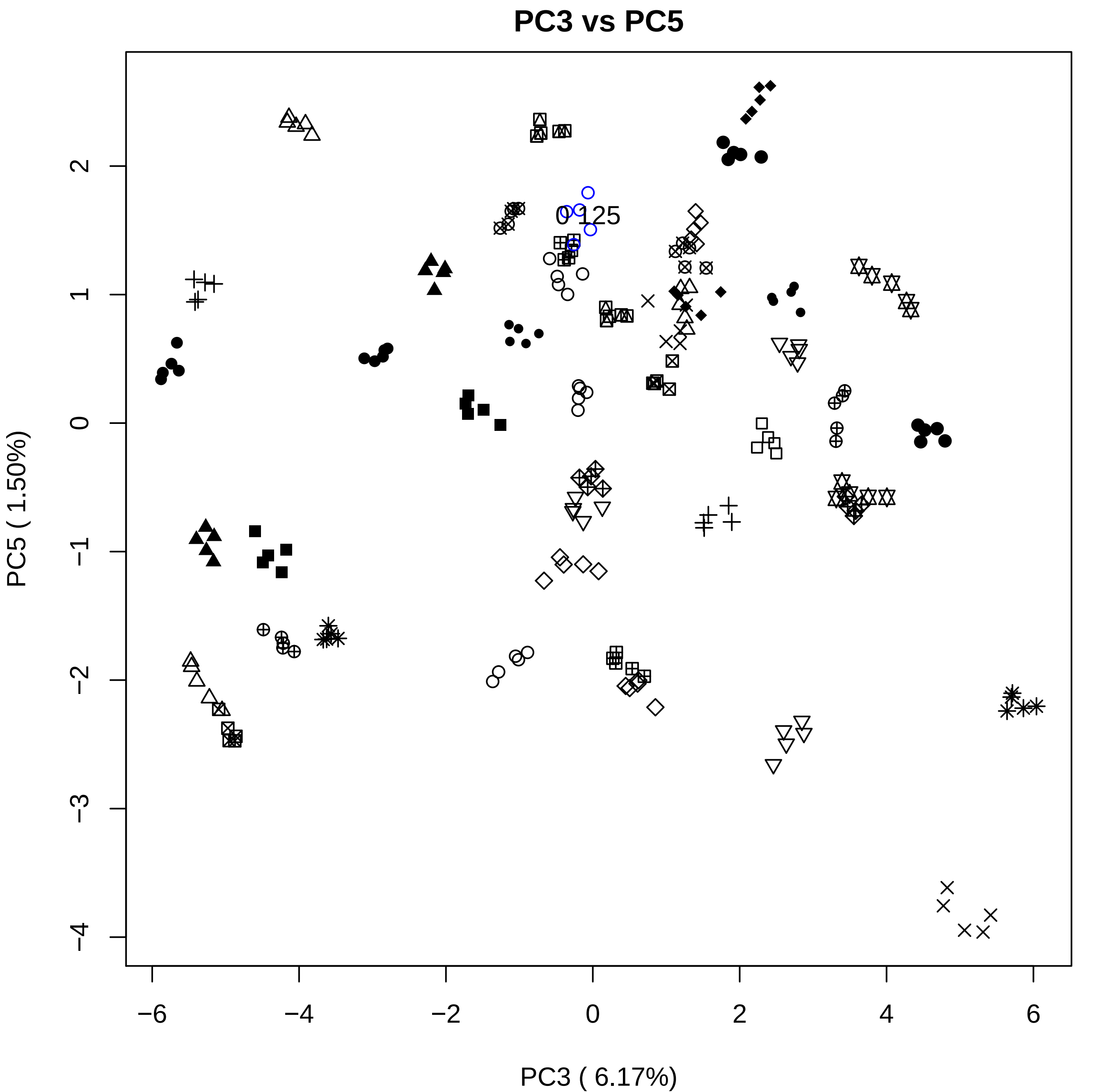
PC3 vs PC5
PC5 ( 1.50%)
PC3 ( 6.17%)
0 125
−6
−4
−2
0
2
4
6
2
1
0
−1
−2
−3
−4

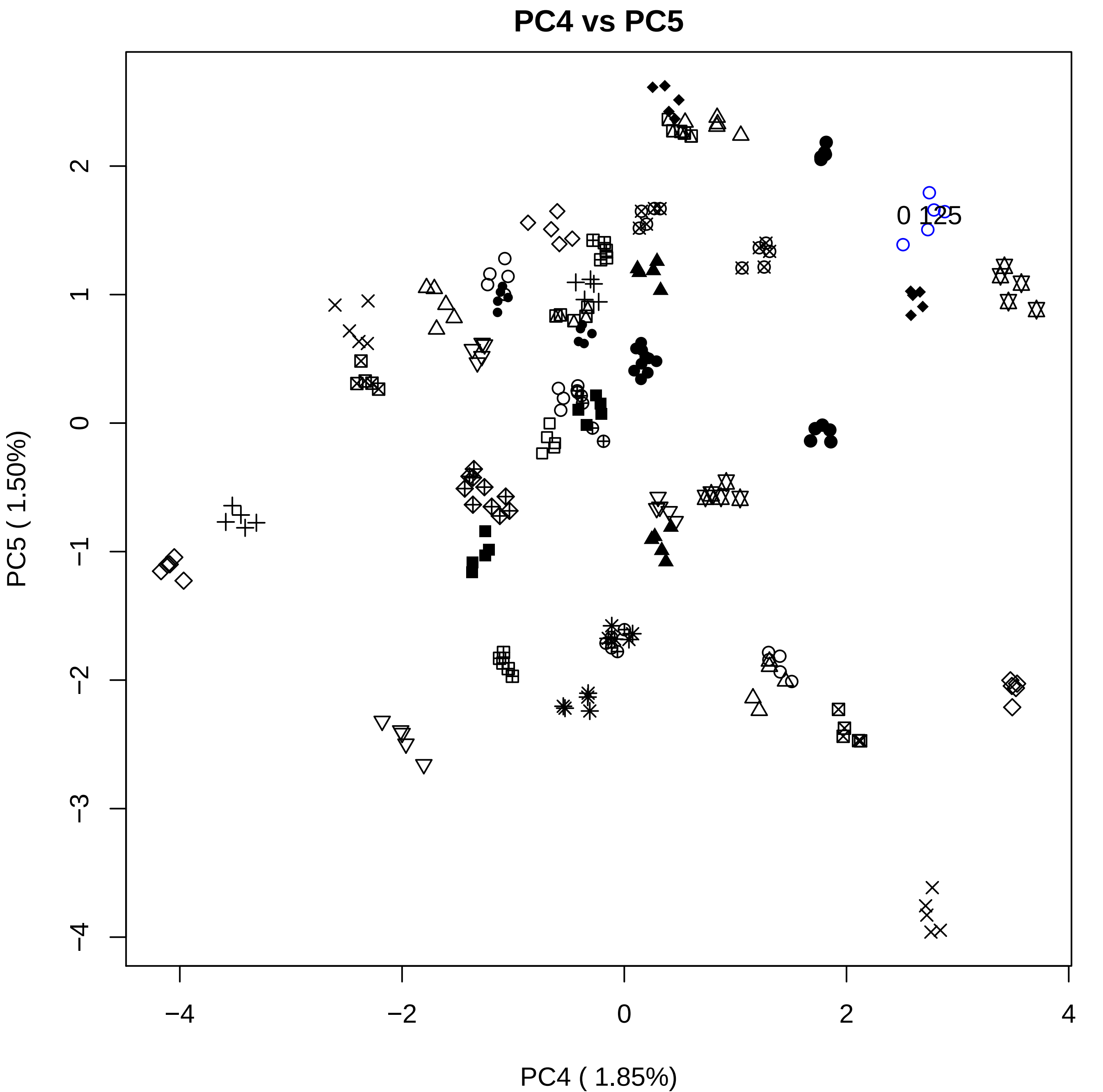
PC4 vs PC5
PC5 ( 1.50%)
PC4 ( 1.85%)
0 125
2
1
0
−1
−2
−3
−4
−4
−2
0
2
4

### PCA of 0.25%  OSCS vs PMH

Scaled Matrix PCAed

7th order polynomial baseline correction performed

Raw spectra smoothed to 21 Neighbours, 2nd order PN, Savtisky Golay

Normalisation between 0 and 100

D2 spectra smoothed to 41 Neighbours, 2nd order PN, Savtisky Golay

Regions Analysed: 3601−2501 and 2001−700cm−1

Samples Analysed: 1 Contam's 0 OSCS's 45 PMH's

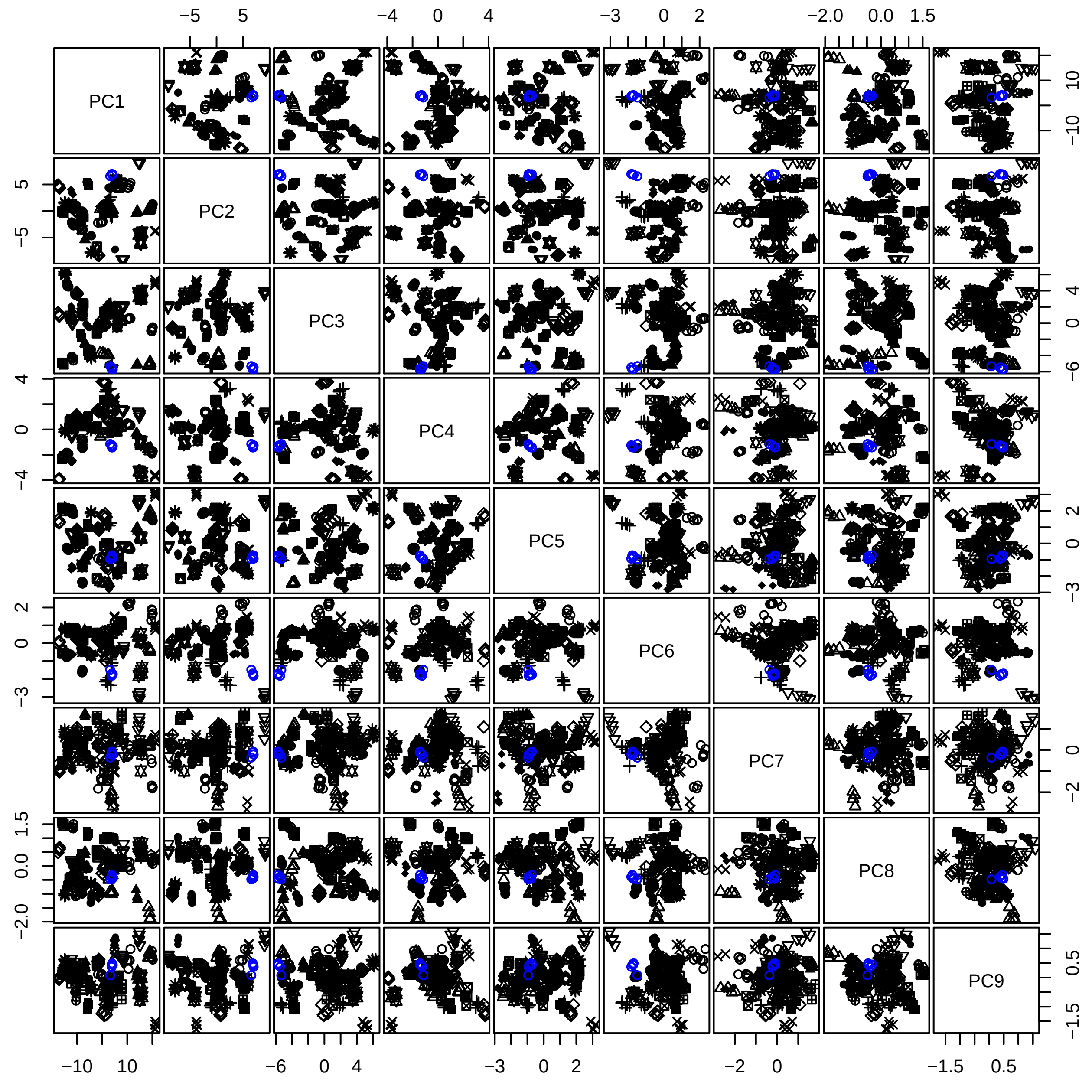
PC1
PC2
PC3
PC4
PC5
PC6
PC7
PC8
PC9

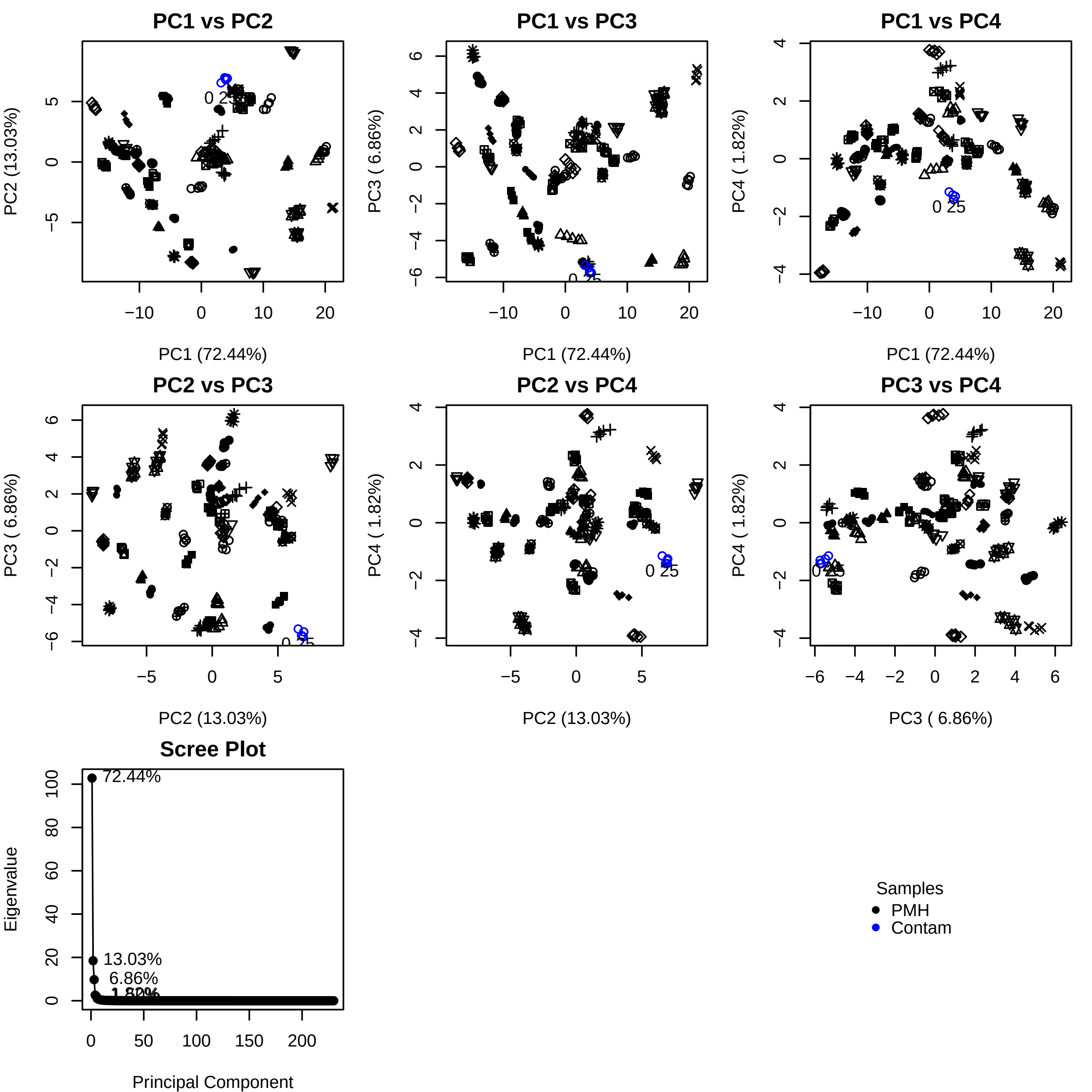
PC1 vs PC2
PC1 vs PC3
PC1 vs PC4
PC2 vs PC3
PC2 vs PC4
PC3 vs PC4
Scree Plot
PC1 (72.44%)
PC2 (13.03%)
PC3 ( 6.86%)
PC4 ( 1.82%)
Eigenvalue
Principal Component
72.44%
13.03%
6.86%
Samples
PMH
Contam

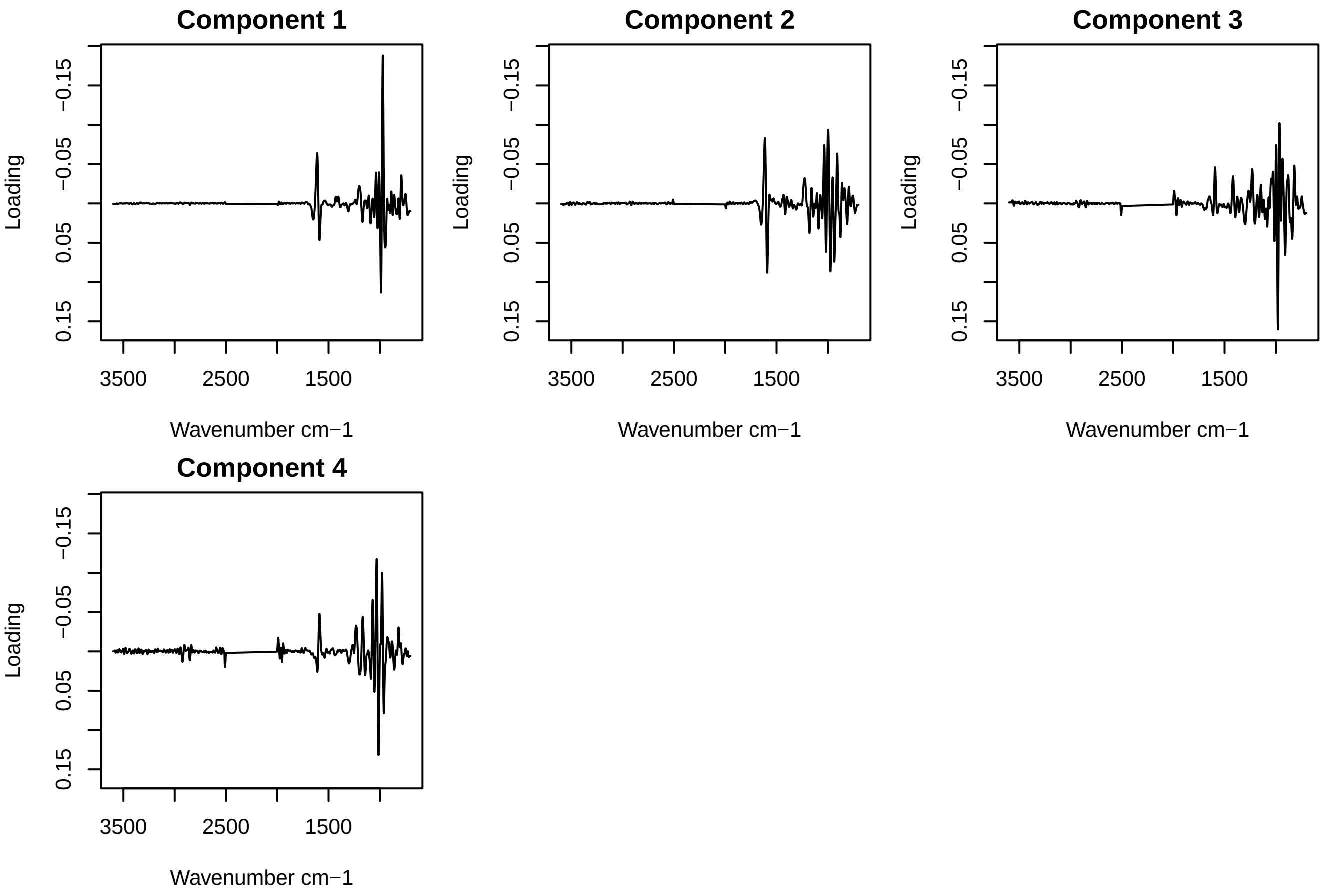
Component 1
Component 2
Component 3
Component 4
Loading
−0.15
−0.05
0.05
0.15
3500
2500
1500
Wavenumber cm−1

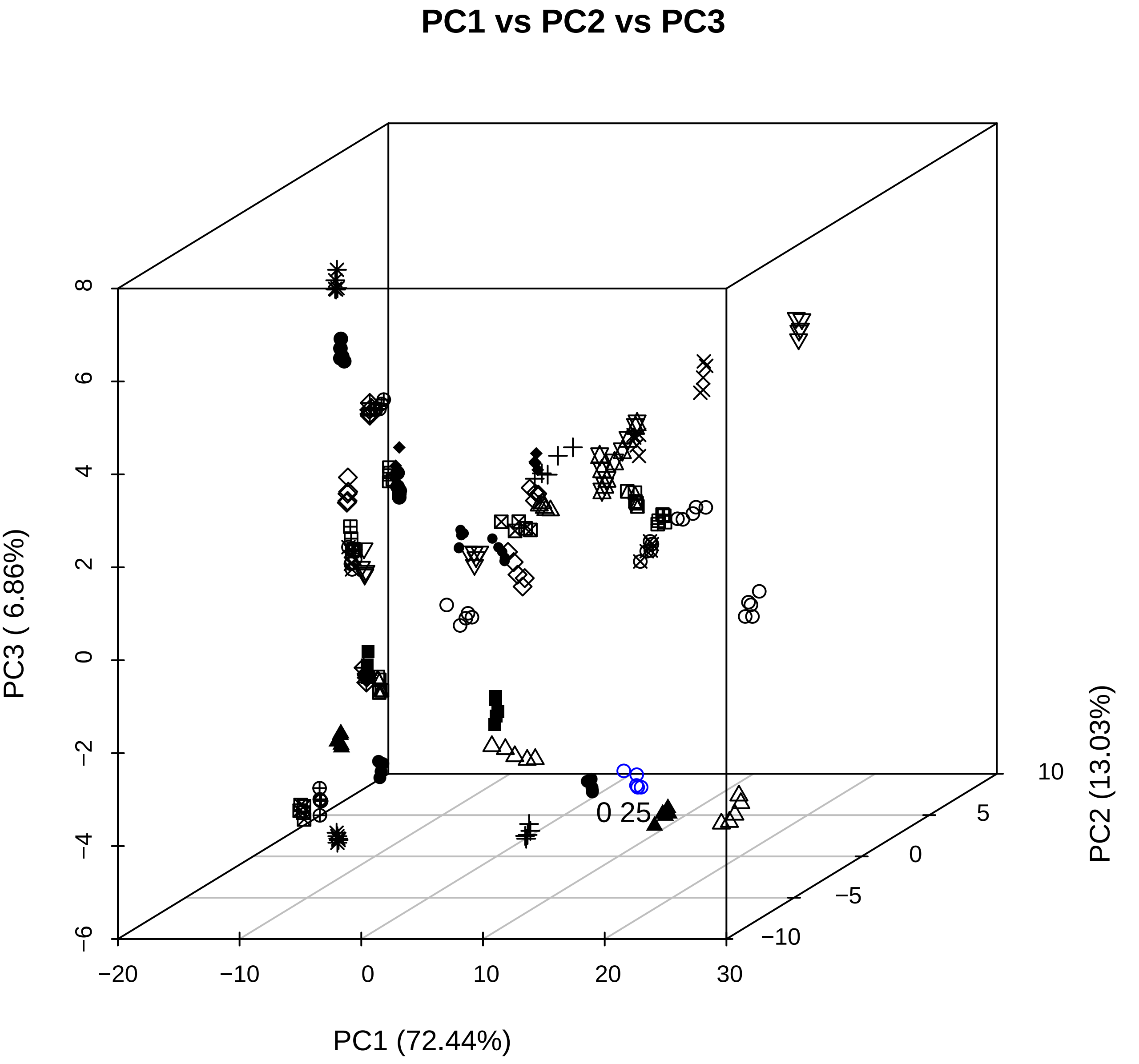
PC1 vs PC2 vs PC3
PC1 (72.44%)
PC2 (13.03%)
PC3 ( 6.86%)
−20
−10
0
10
20
30
−10
−5
0
5
10
−6
−4
−2
0
2
4
6
8
0 25

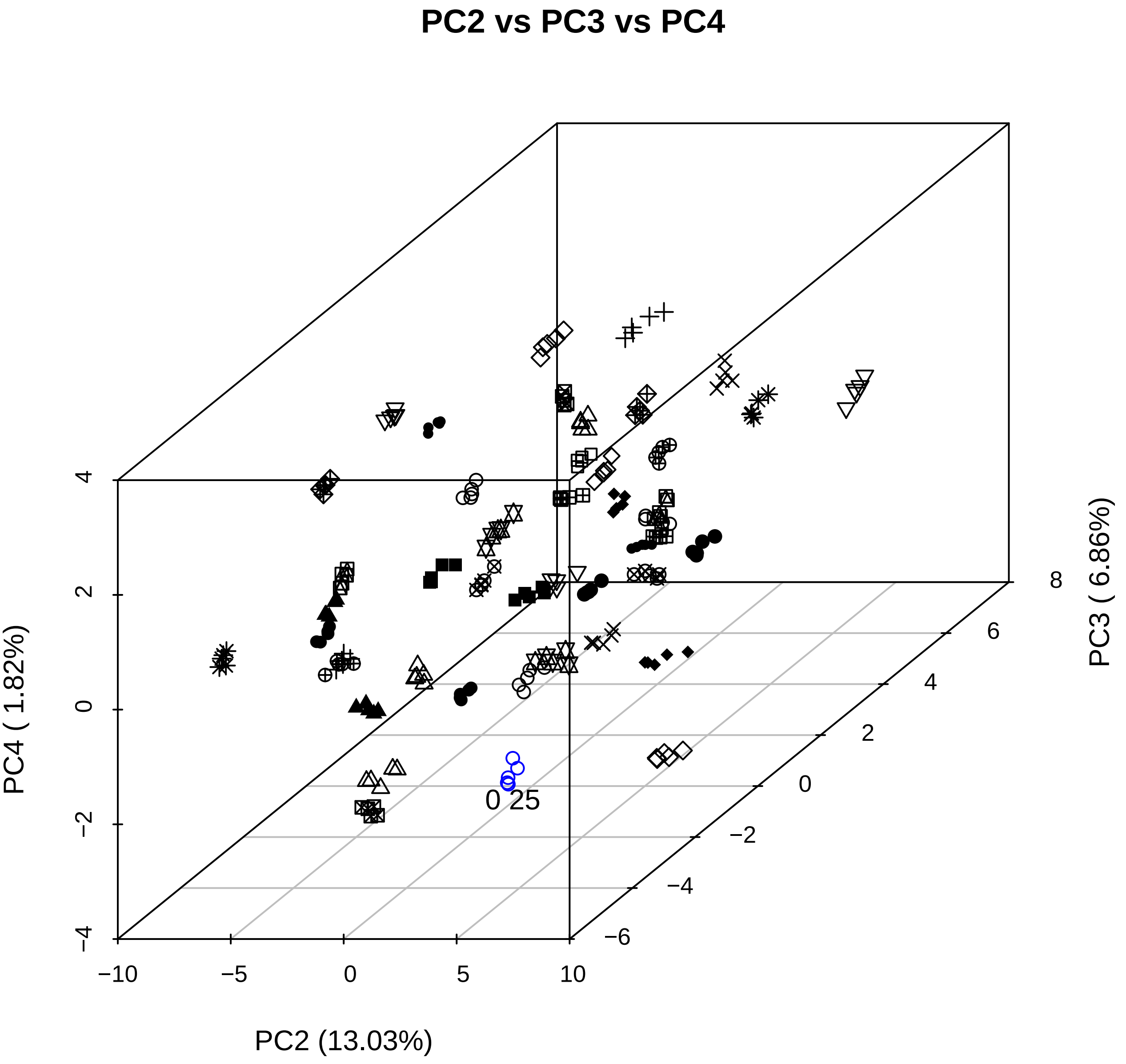

PC2 vs PC3 vs PC4
PC4 ( 1.82%)
PC3 ( 6.86%)
PC2 (13.03%)
0 25
−4
−2
0
2
4
−10
−5
0
5
10
−6
−4
−2
0
2
4
6
8

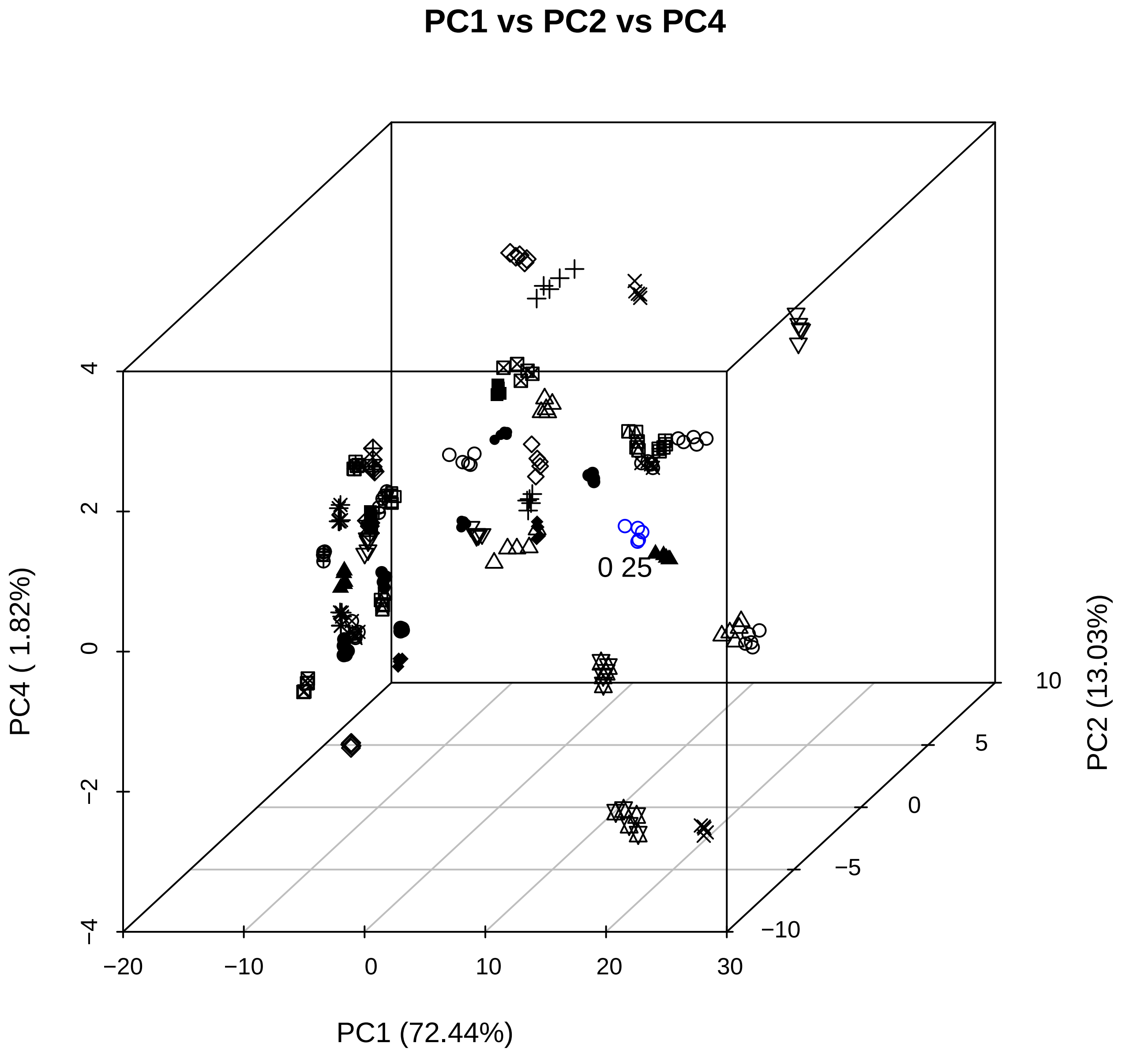
PC1 vs PC2 vs PC4
PC1 (72.44%)
PC2 (13.03%)
PC4 ( 1.82%)
−20
−10
0
10
20
30
−10
−5
0
5
10
−4
−2
0
2
4
0 25

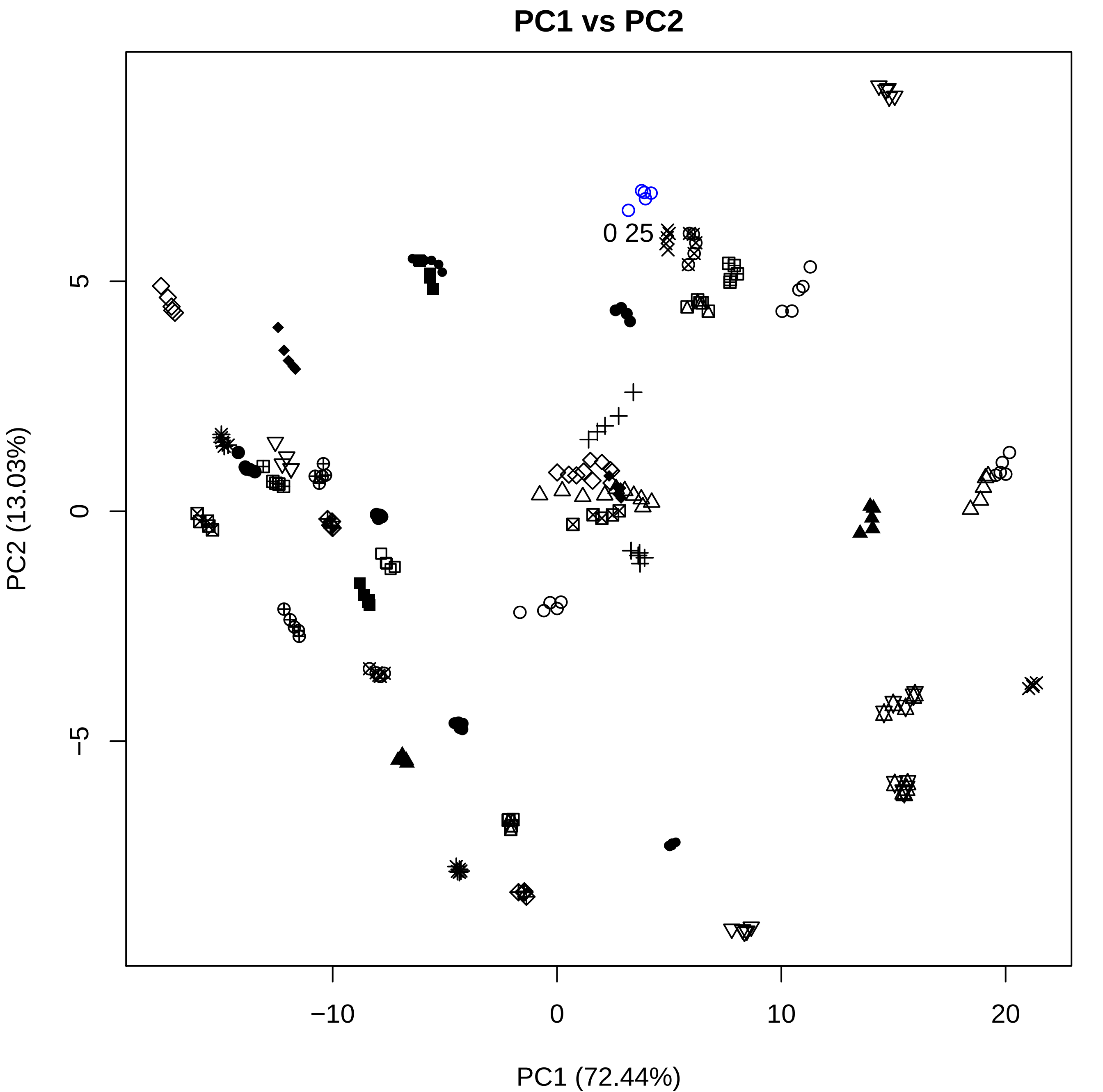

PC1 vs PC2
0 25
PC2 (13.03%)
5
0
−5
−10
0
10
20
PC1 (72.44%)

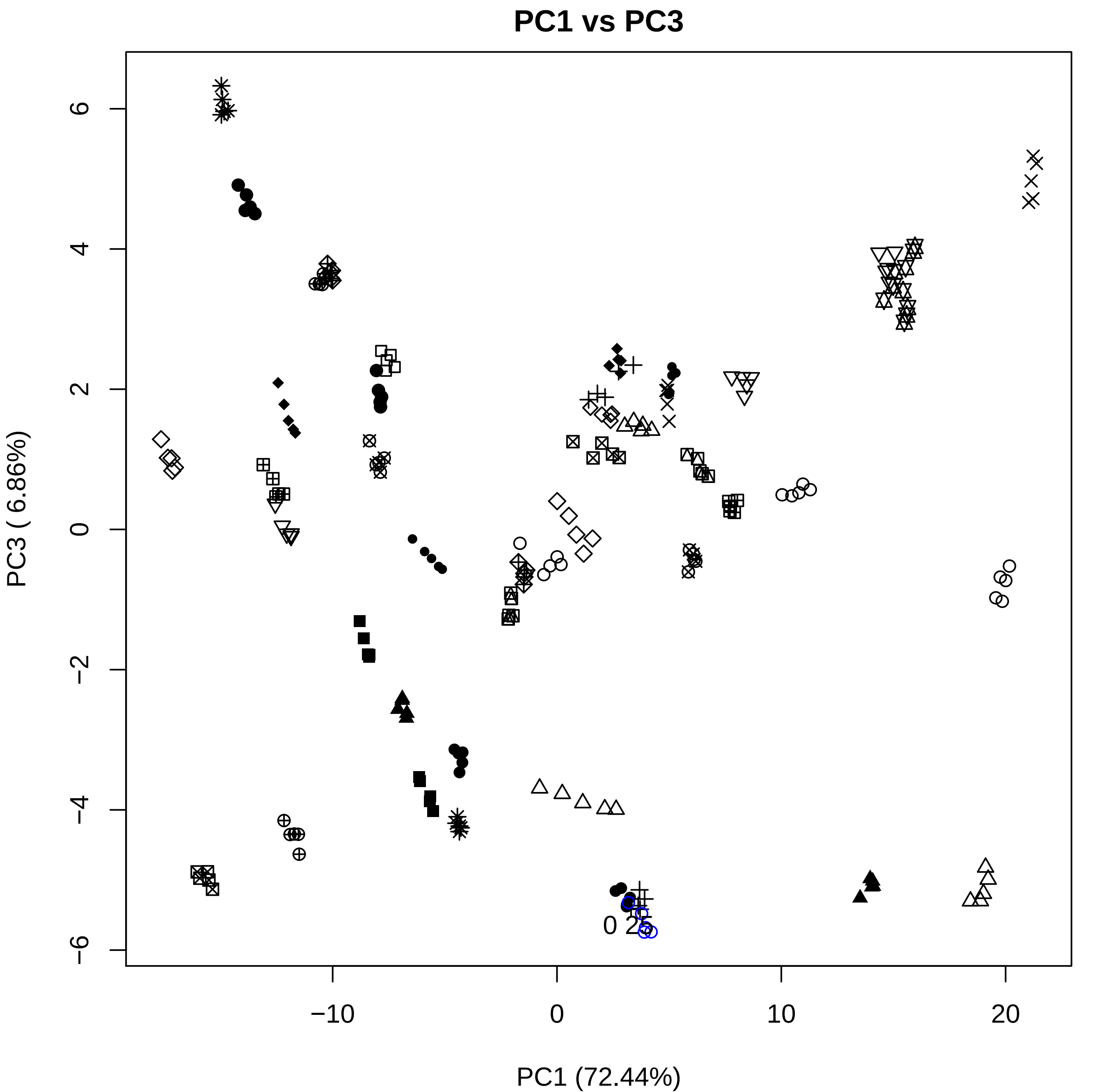

PC1 vs PC3
PC3 ( 6.86%)
PC1 (72.44%)
6
4
2
0
−2
−4
−6
−10
0
10
20
0 25

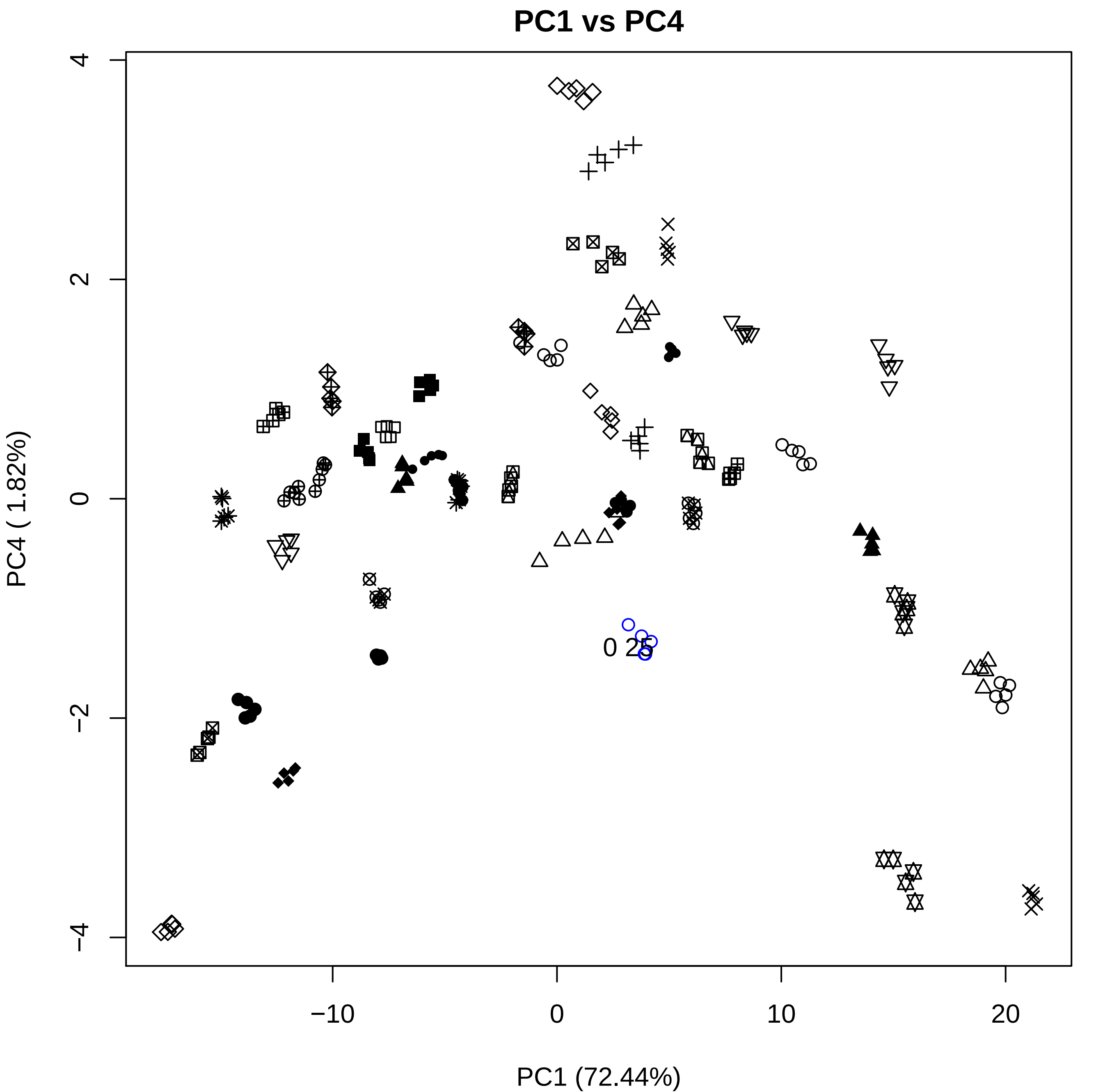
PC1 vs PC4
PC4 ( 1.82%)
PC1 (72.44%)
4
2
0
−2
−4
−10
0
10
20
0 25

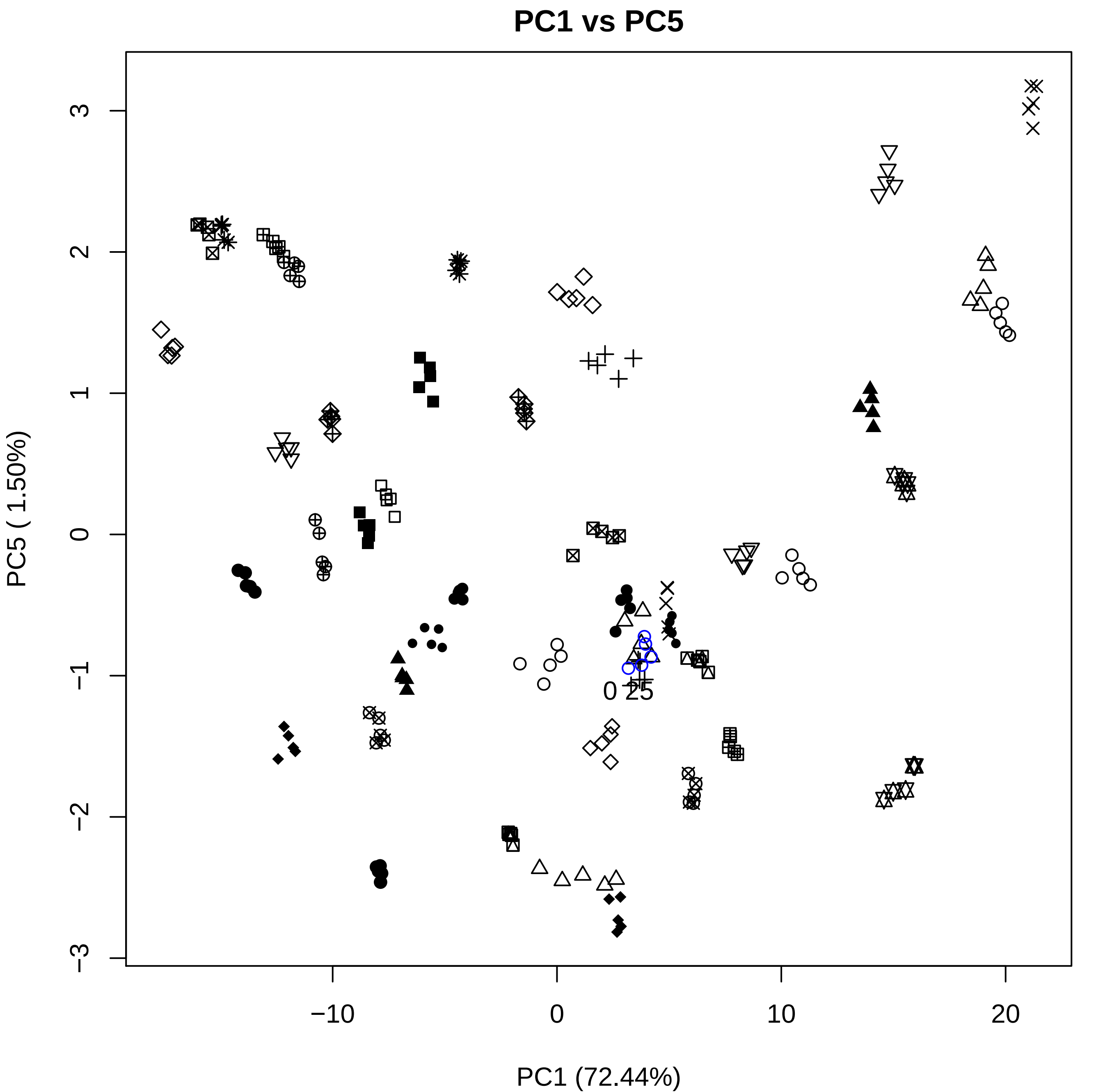
PC1 vs PC5
PC5 ( 1.50%)
PC1 (72.44%)
3
2
1
0
−1
−2
−3
−10
0
10
20
0 25

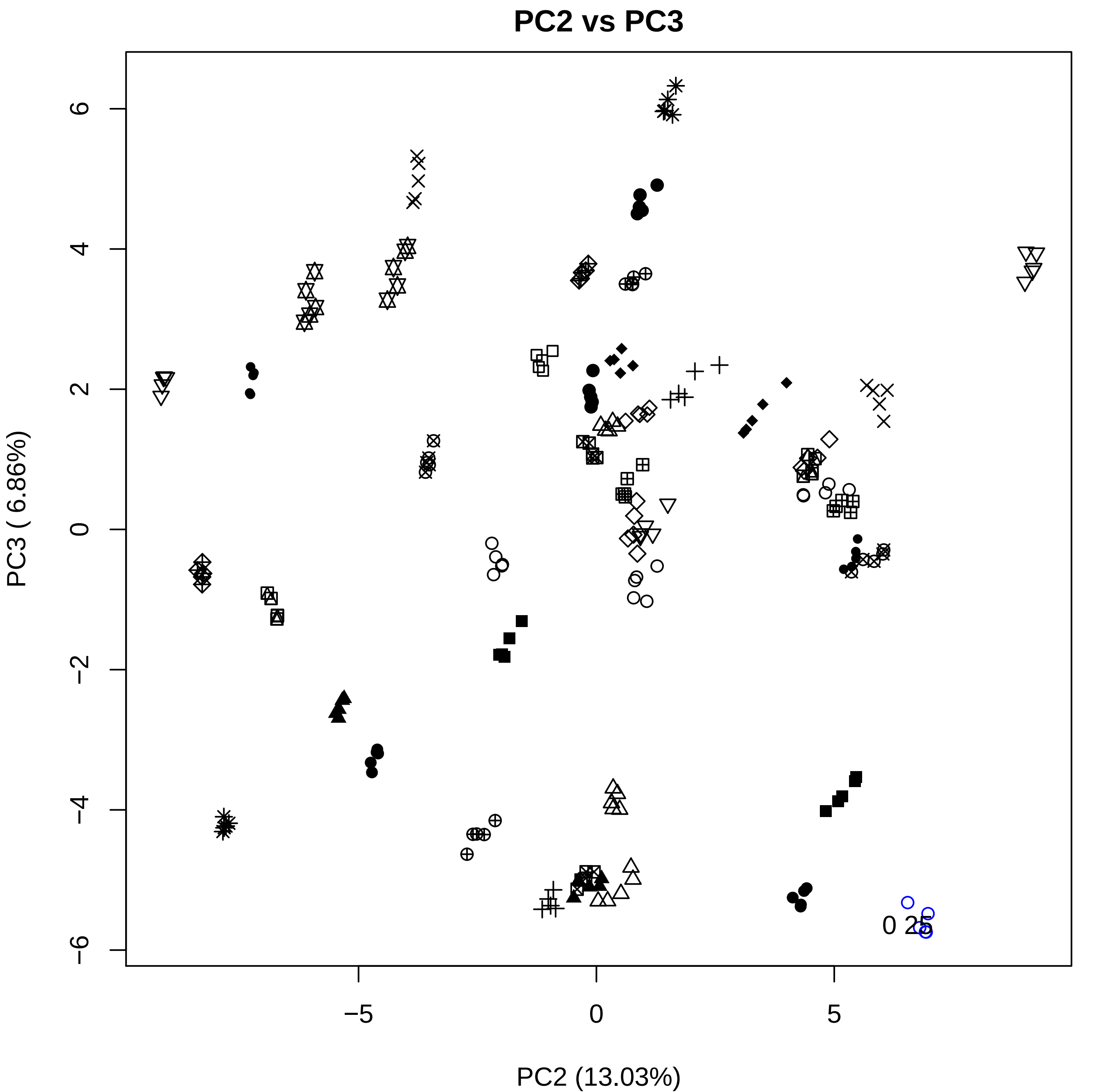
PC2 vs PC3
PC3 ( 6.86%)
PC2 (13.03%)
6
4
2
0
−2
−4
−6
−5
0
5
0 25

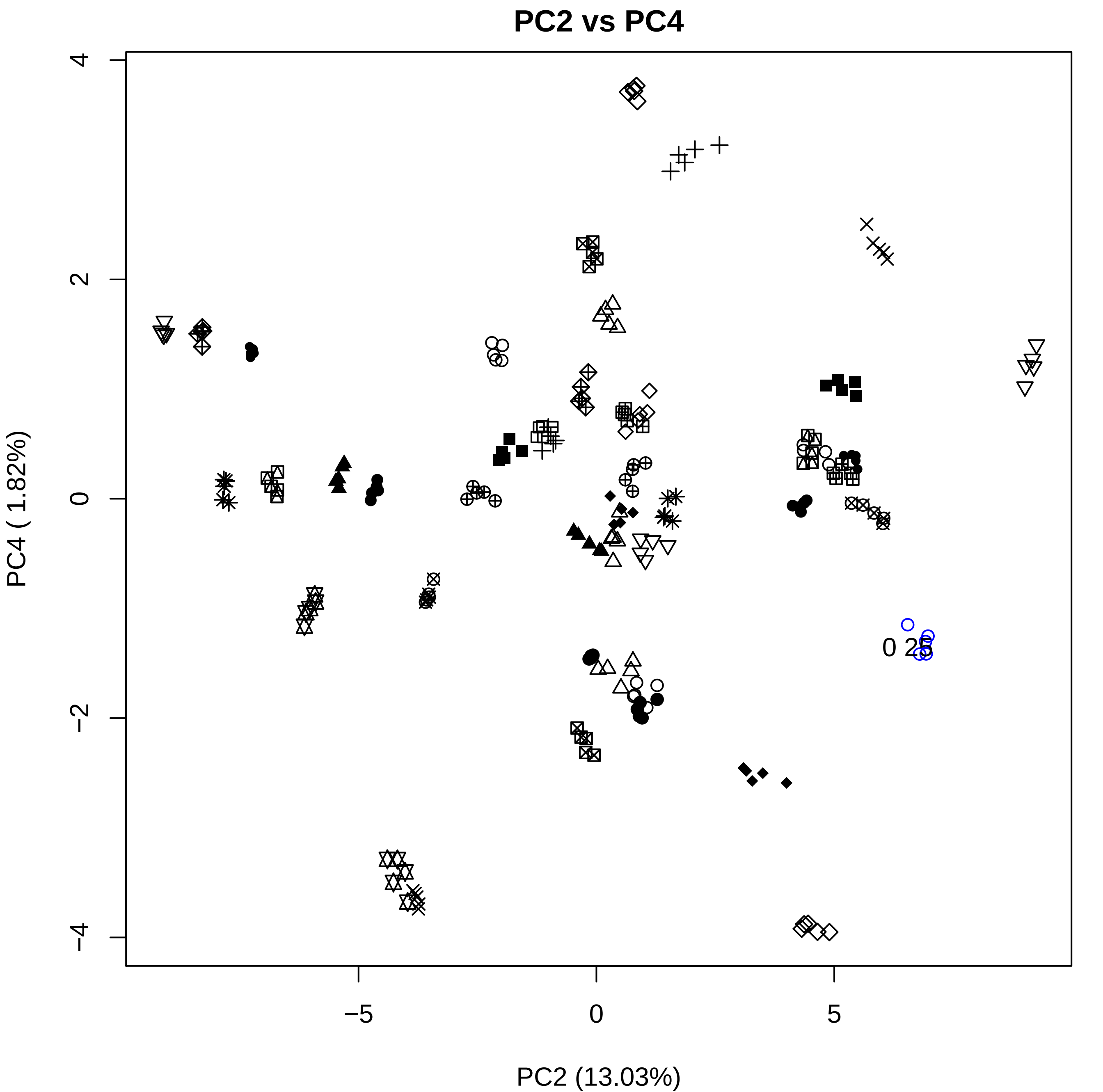
PC2 vs PC4
PC4 ( 1.82%)
PC2 (13.03%)
4
2
0
−2
−4
−5
0
5
0 25

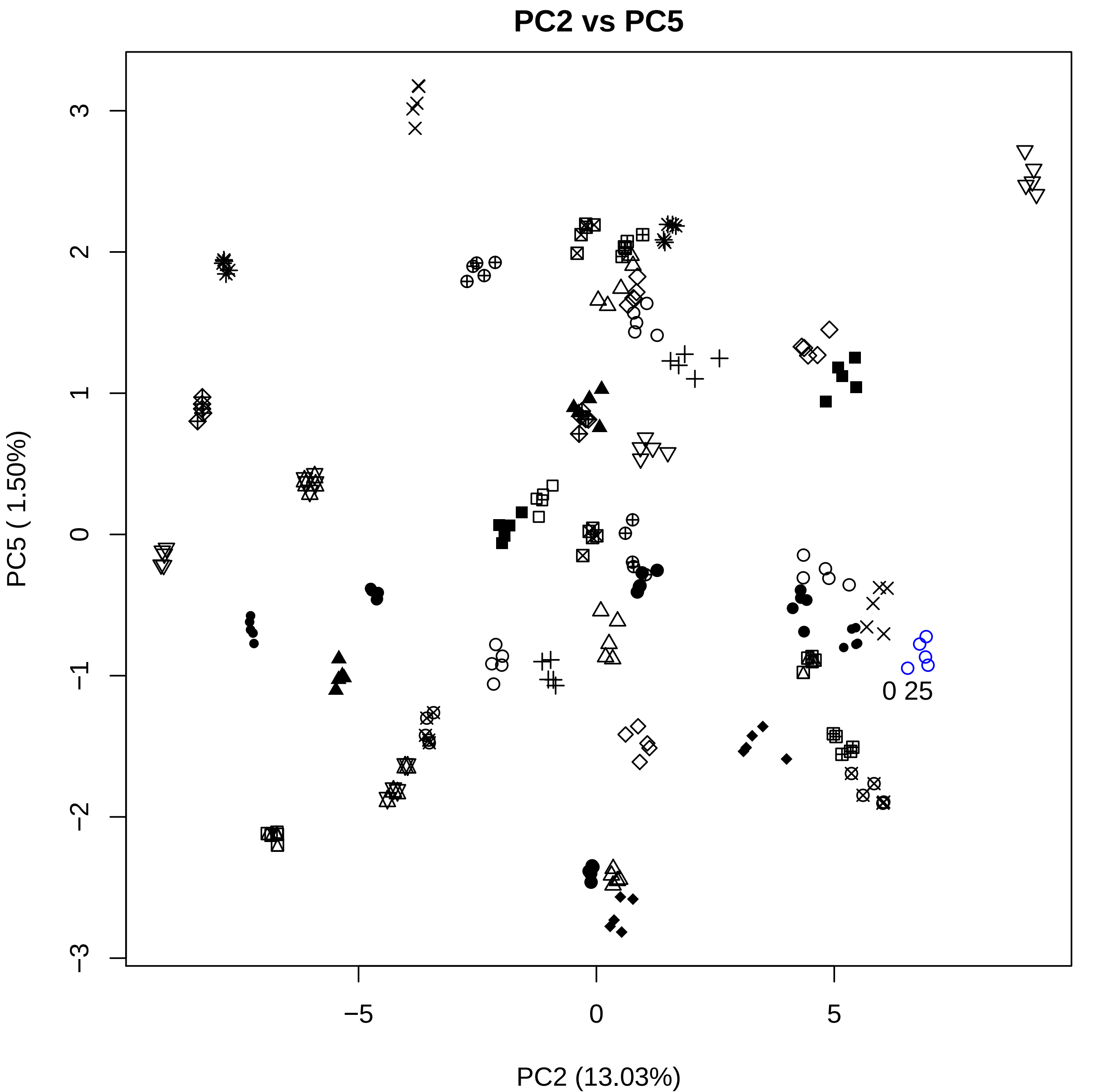

PC2 vs PC5
PC5 ( 1.50%)
PC2 (13.03%)
3
2
1
0
−1
−2
−3
−5
0
5
0 25

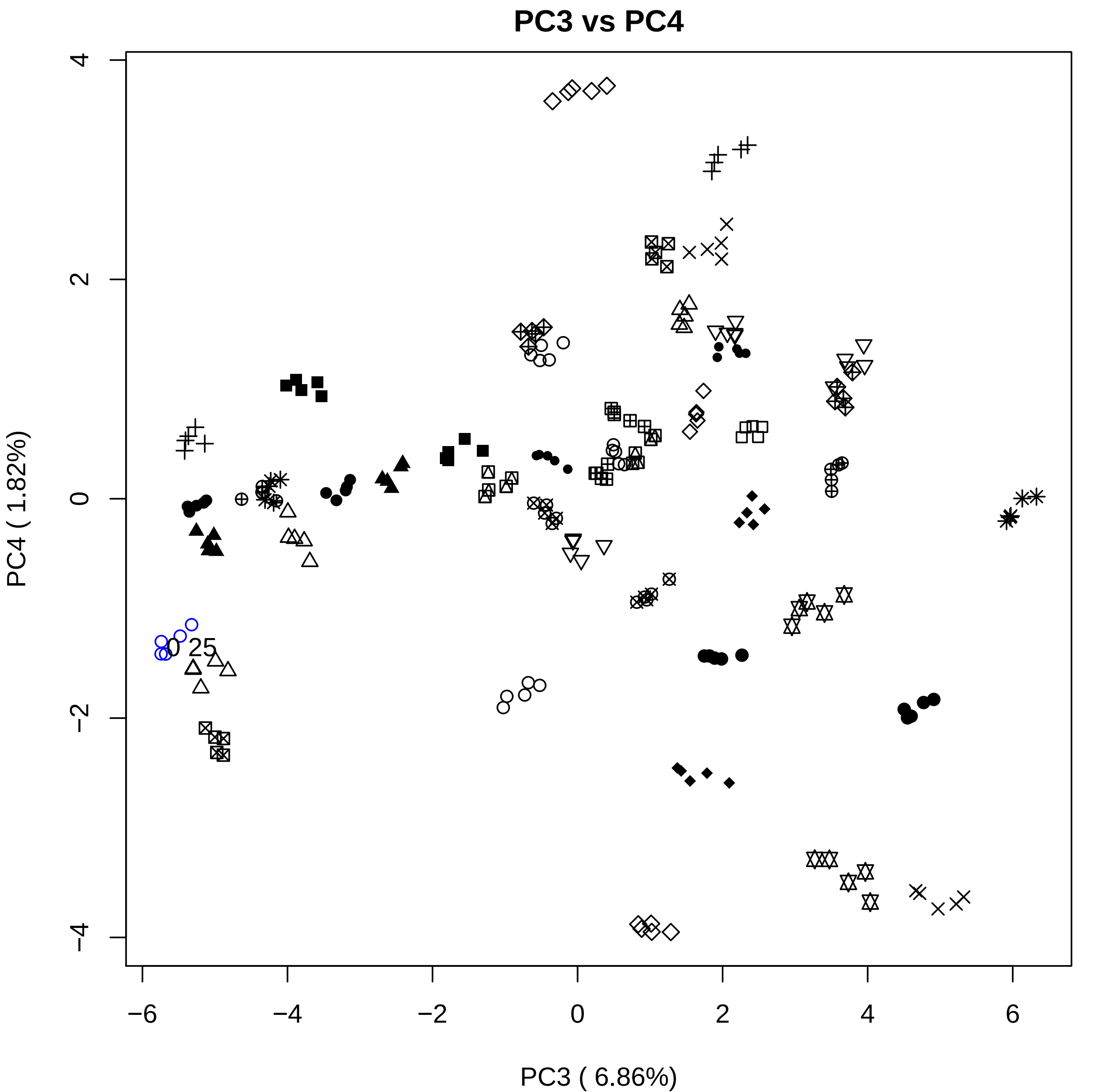

PC3 vs PC4
PC4 ( 1.82%)
PC3 ( 6.86%)
4
2
0
−2
−4
−6
−4
−2
0
2
4
6
0 25

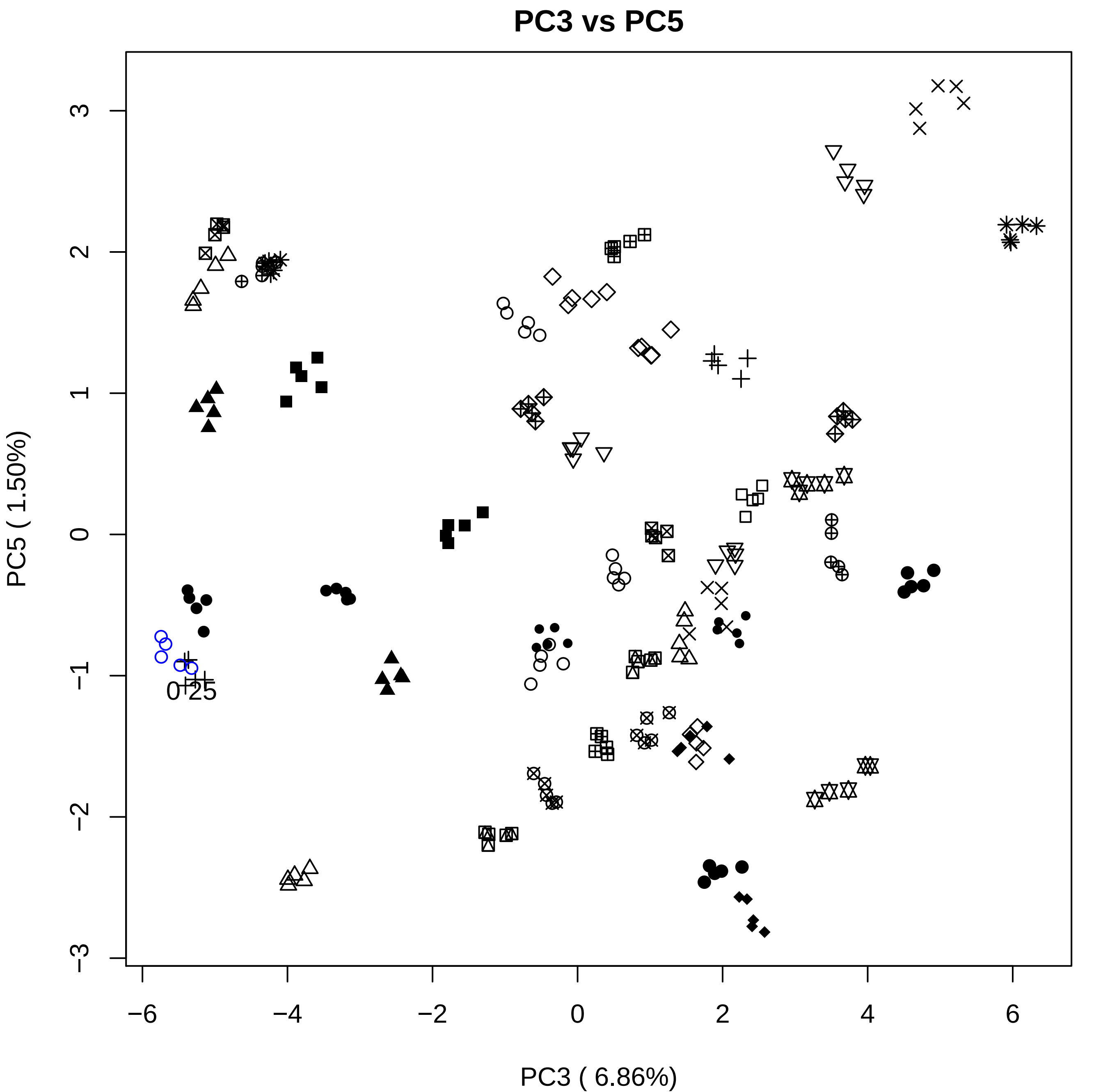
PC3 vs PC5
PC5 ( 1.50%)
PC3 ( 6.86%)
0·25
3
2
1
0
−1
−2
−3
−6
−4
−2
0
2
4
6

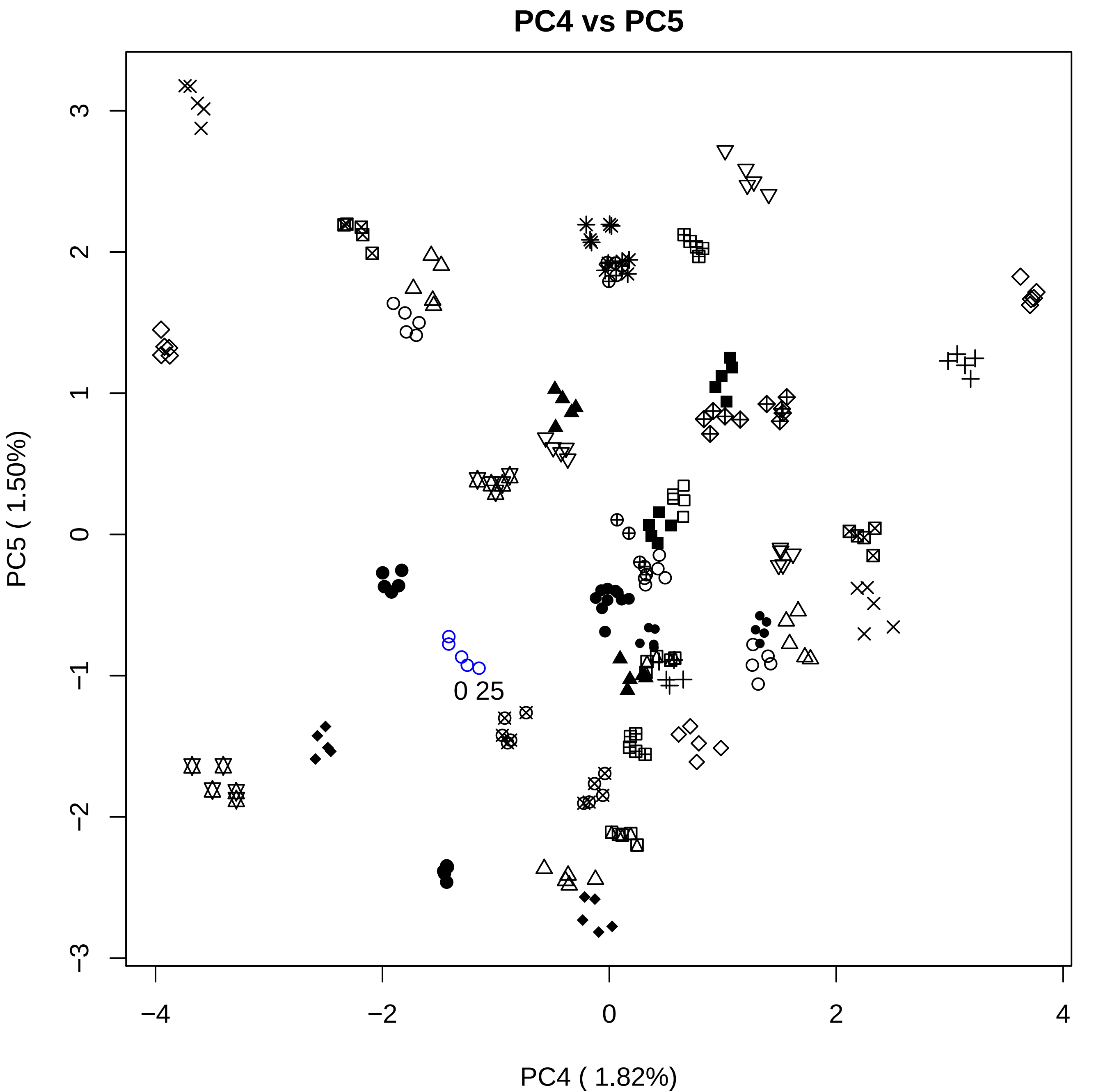

PC4 vs PC5
PC5 ( 1.50%)
PC4 ( 1.82%)
0 25
−4
−2
0
2
4
3
2
1
0
−1
−2
−3

### PCA of 0.5%  OSCS vs PMH

Scaled Matrix PCAed

7th order polynomial baseline correction performed

Raw spectra smoothed to 21 Neighbours, 2nd order PN, Savtisky Golay

Normalisation between 0 and 100

D2 spectra smoothed to 41 Neighbours, 2nd order PN, Savtisky Golay

Regions Analysed: 3601−2501 and 2001−700cm−1

Samples Analysed: 1 Contam's 0 OSCS's 45 PMH's

Keele University AD                                        Sun Dec 09 16:50:48 2018

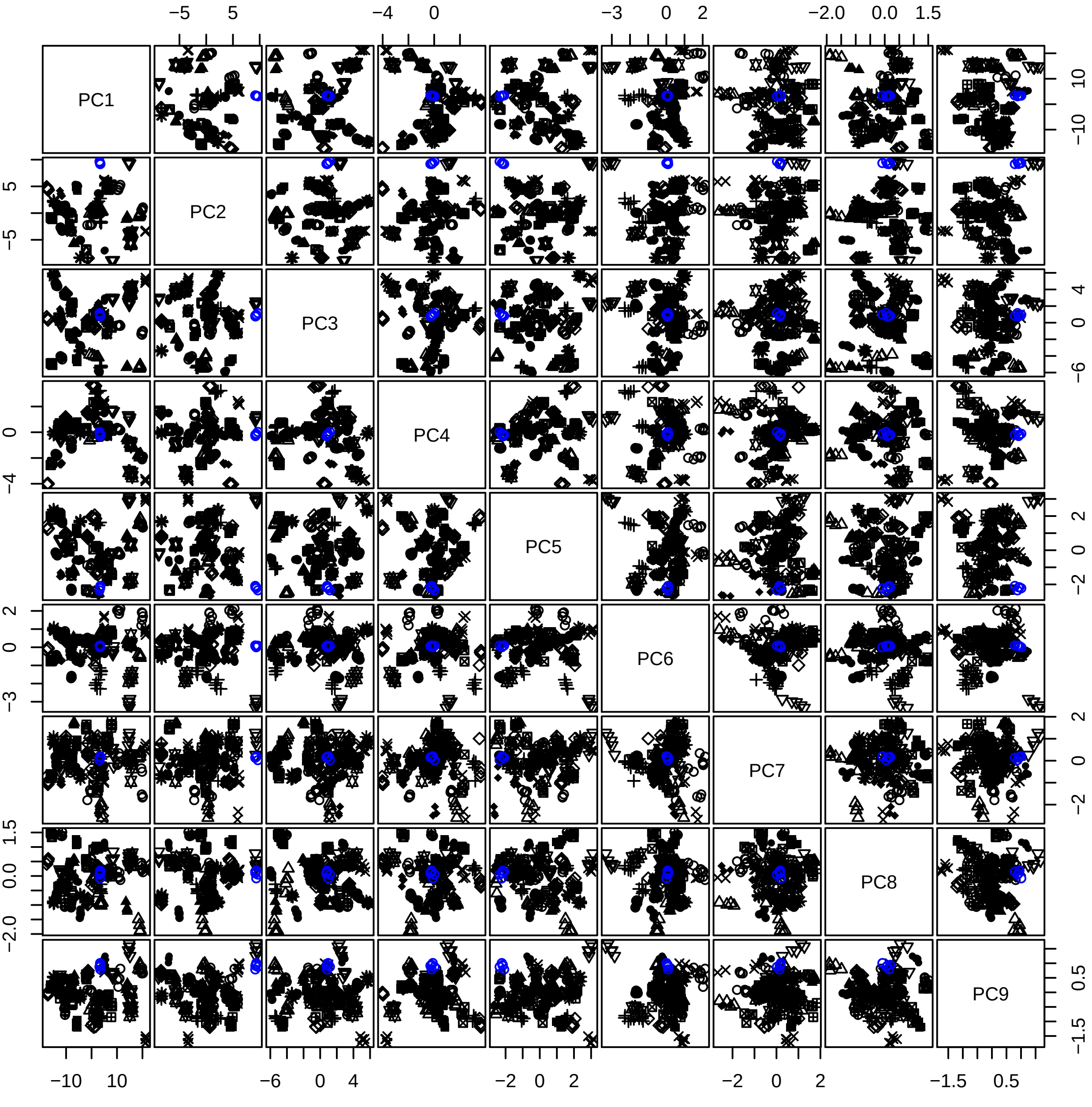
PC1
PC2
PC3
PC4
PC5
PC6
PC7
PC8
PC9

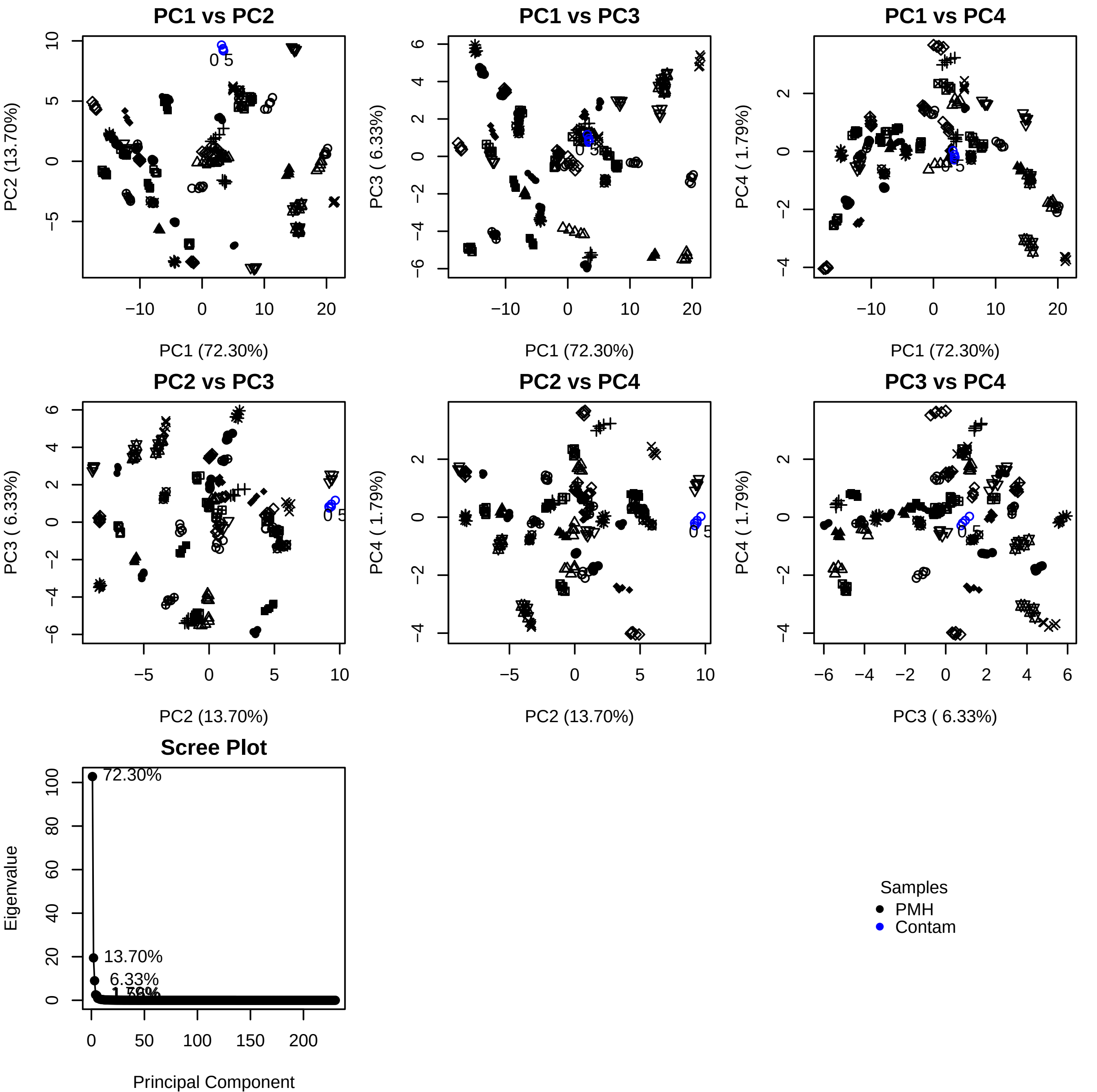
PC1 vs PC2
PC1 (72.30%)
PC2 (13.70%)
0 5
PC1 vs PC3
PC1 (72.30%)
PC3 ( 6.33%)
0 5
PC1 vs PC4
PC1 (72.30%)
PC4 ( 1.79%)
0 5
PC2 vs PC3
PC2 (13.70%)
PC3 ( 6.33%)
0 5
PC2 vs PC4
PC2 (13.70%)
PC4 ( 1.79%)
0 5
PC3 vs PC4
PC3 ( 6.33%)
PC4 ( 1.79%)
0 5
Scree Plot
Eigenvalue
Principal Component
72.30%
13.70%
6.33%
Samples
PMH
Contam

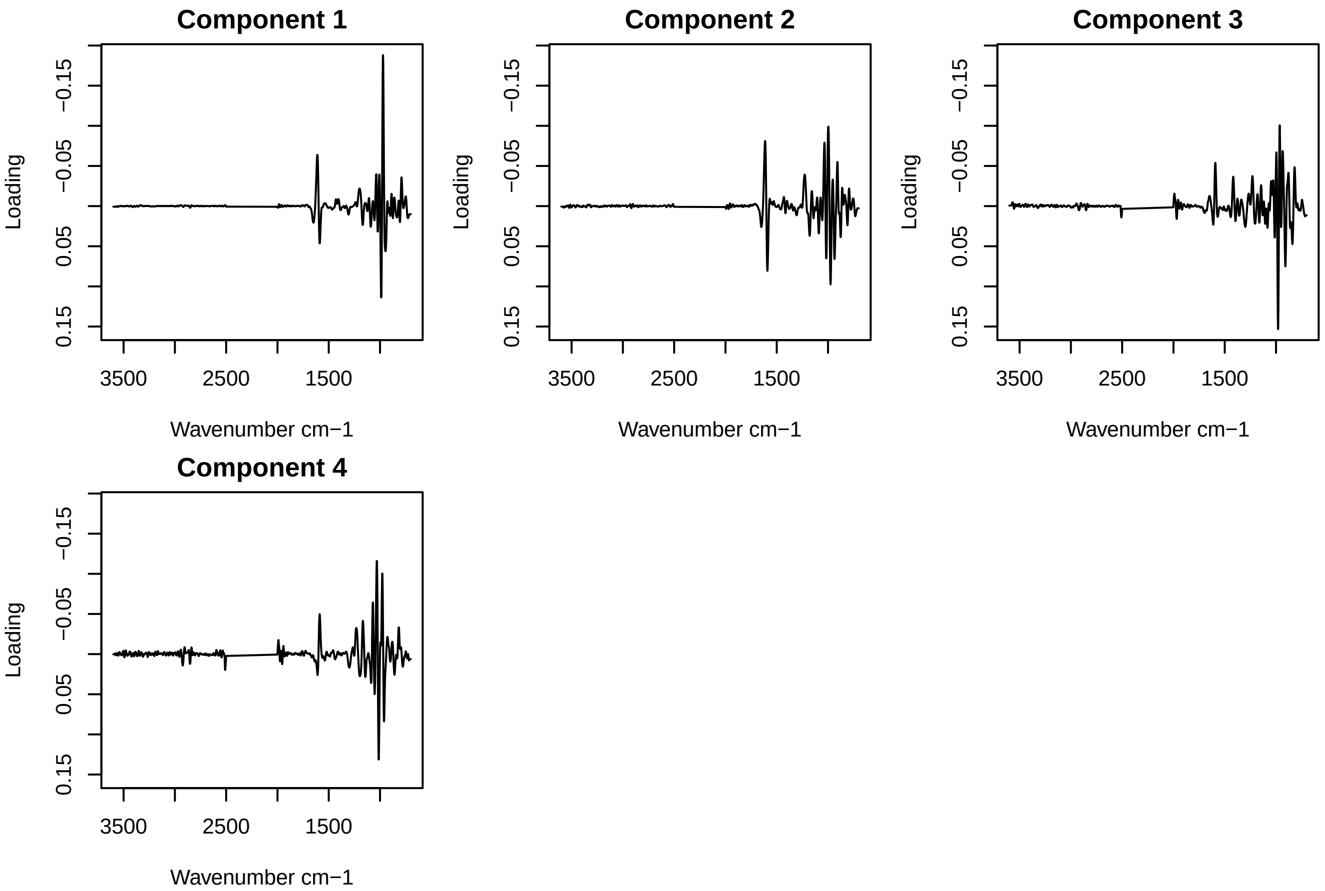
Component 1
Component 2
Component 3
Component 4
Loading
−0.15
−0.05
0.05
0.15
3500
2500
1500
Wavenumber cm−1

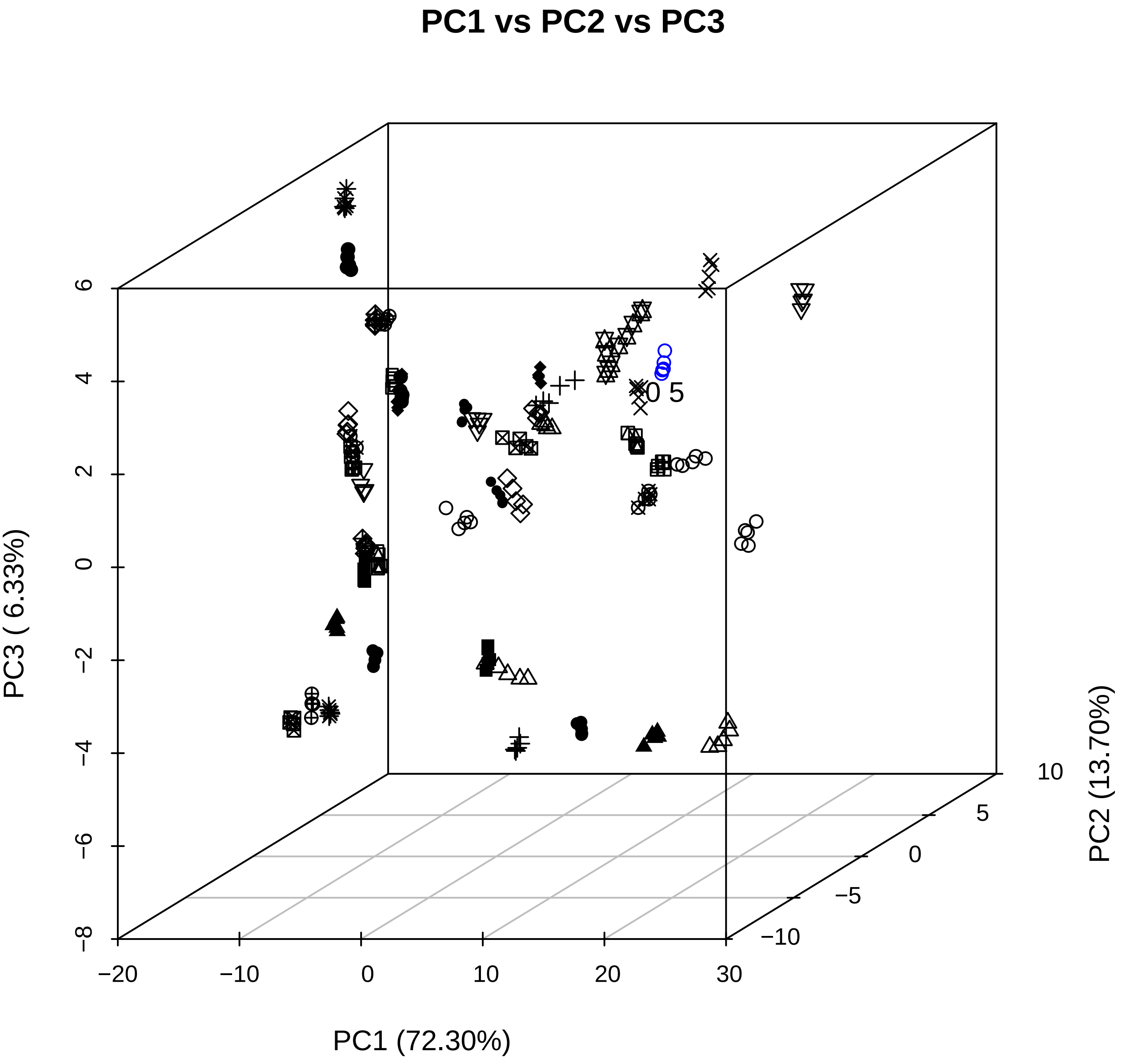
PC1 vs PC2 vs PC3
PC1 (72.30%)
PC2 (13.70%)
PC3 ( 6.33%)
−20
−10
0
10
20
30
−10
−5
0
5
10
−8
−6
−4
−2
0
2
4
6
0 5

# PC2 vs PC3 vs PC4

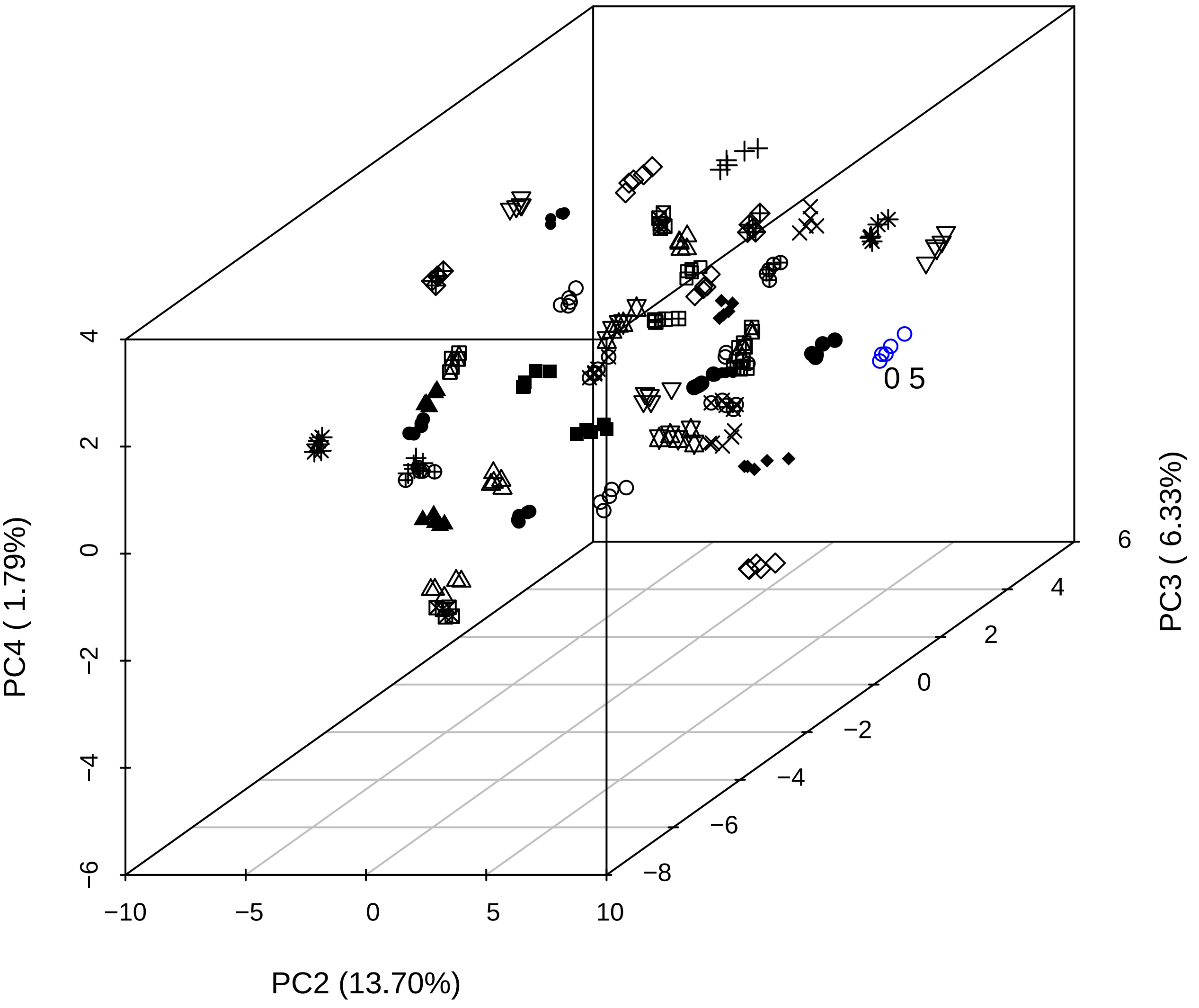

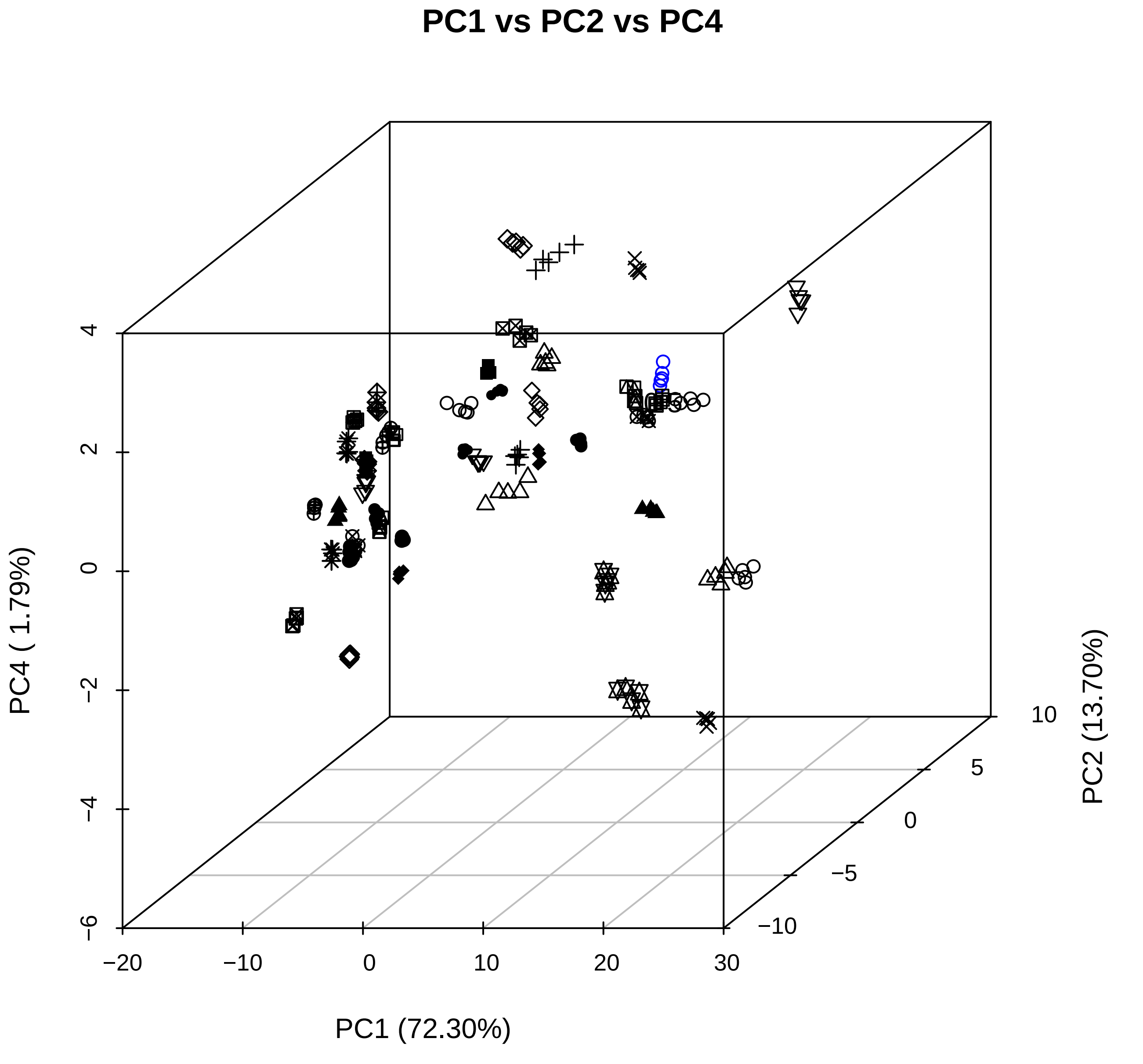
PC1 vs PC2 vs PC4
PC1 (72.30%)
PC2 (13.70%)
PC4 ( 1.79%)
−20
−10
0
10
20
30
−10
−5
0
5
10
−6
−4
−2
0
2
4

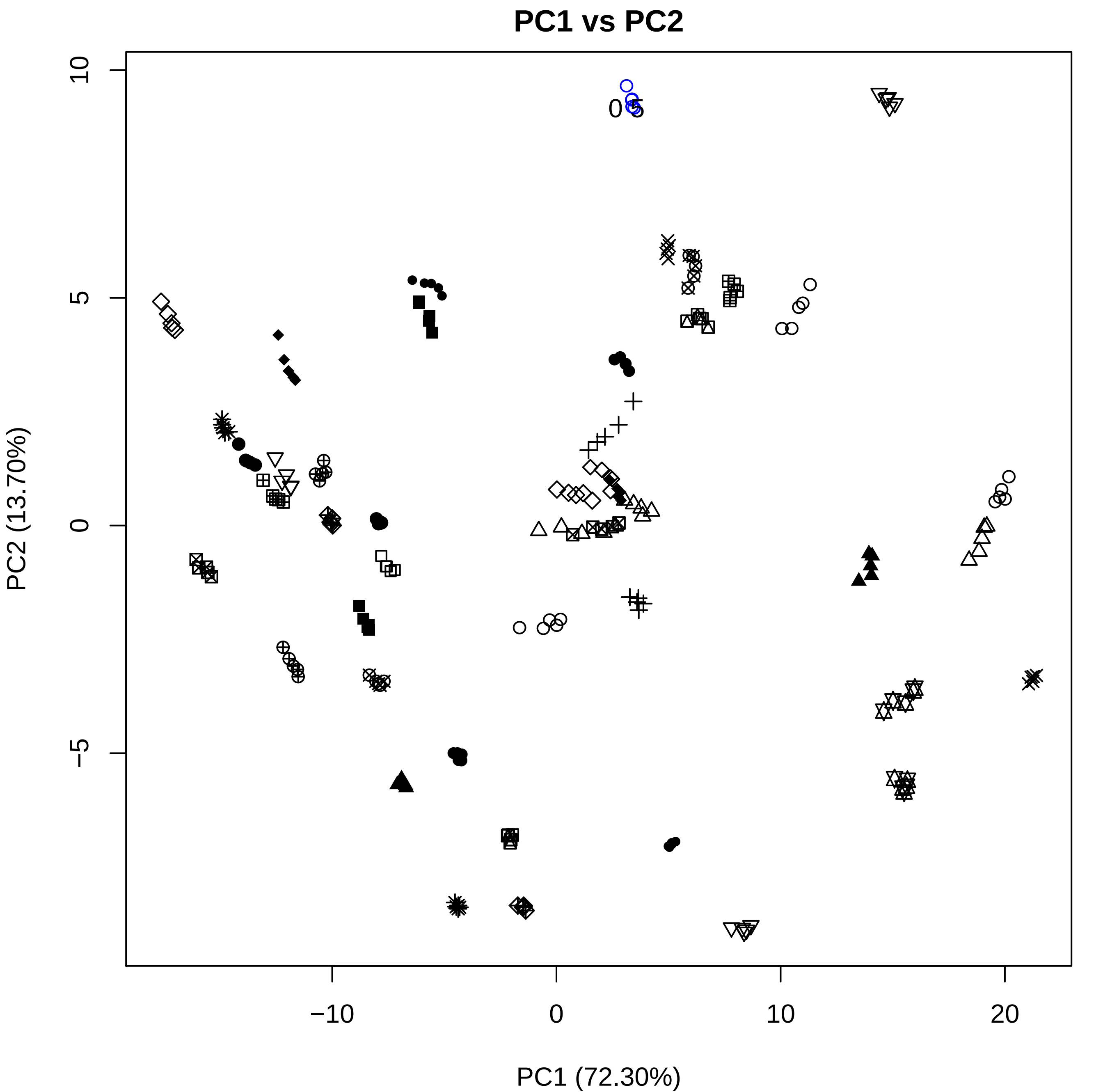

PC1 vs PC2
PC2 (13.70%)
PC1 (72.30%)
10
5
0
−5
−10
0
10
20
0 5

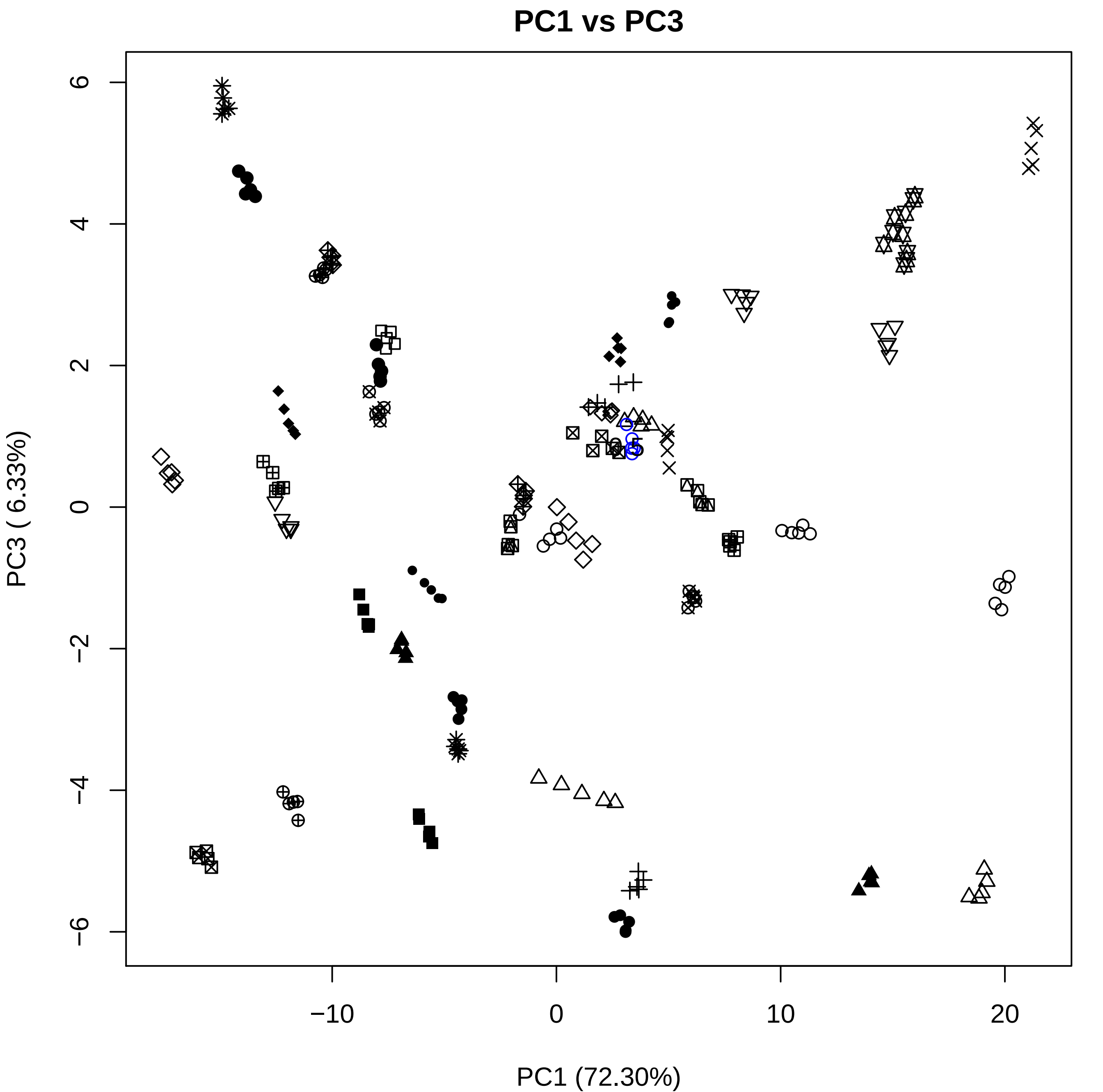
PC1 vs PC3
PC3 ( 6.33%)
PC1 (72.30%)
6
4
2
0
−2
−4
−6
−10
0
10
20

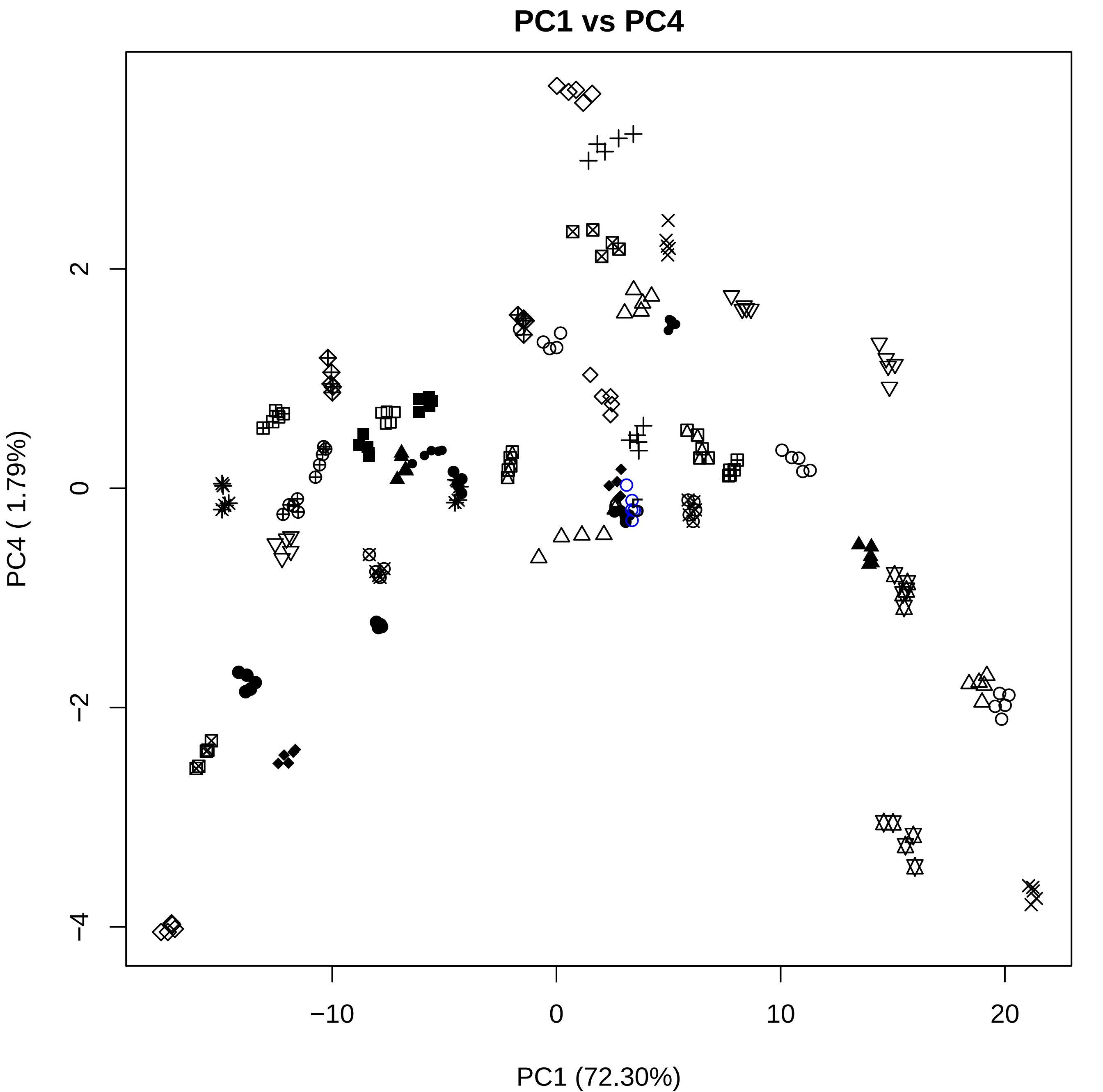
PC1 vs PC4
PC4 ( 1.79%)
PC1 (72.30%)
2
0
−2
−4
−10
0
10
20

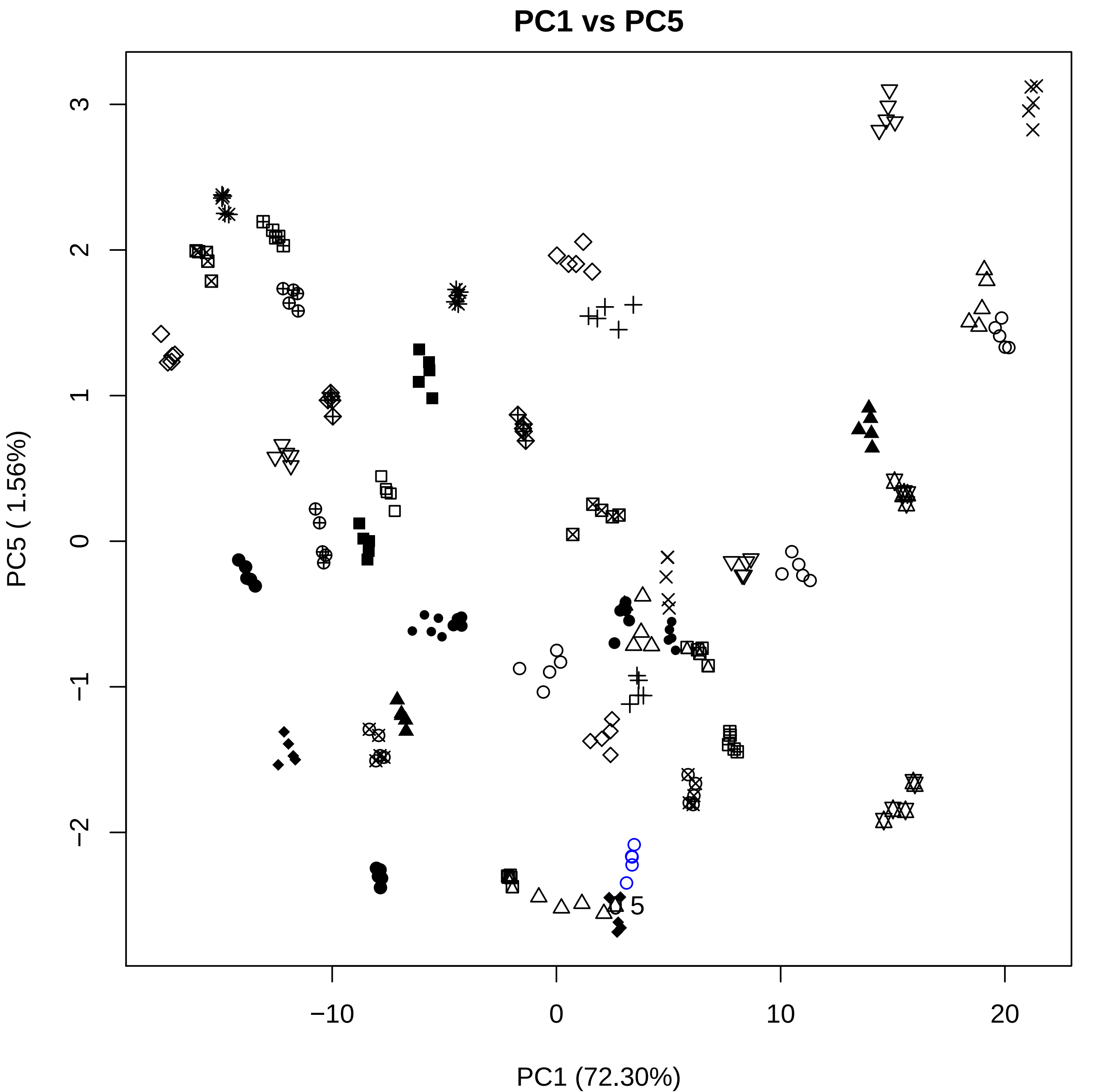

PC1 vs PC5
PC5 ( 1.56%)
PC1 (72.30%)
−10
0
10
20
−2
−1
0
1
2
3
5

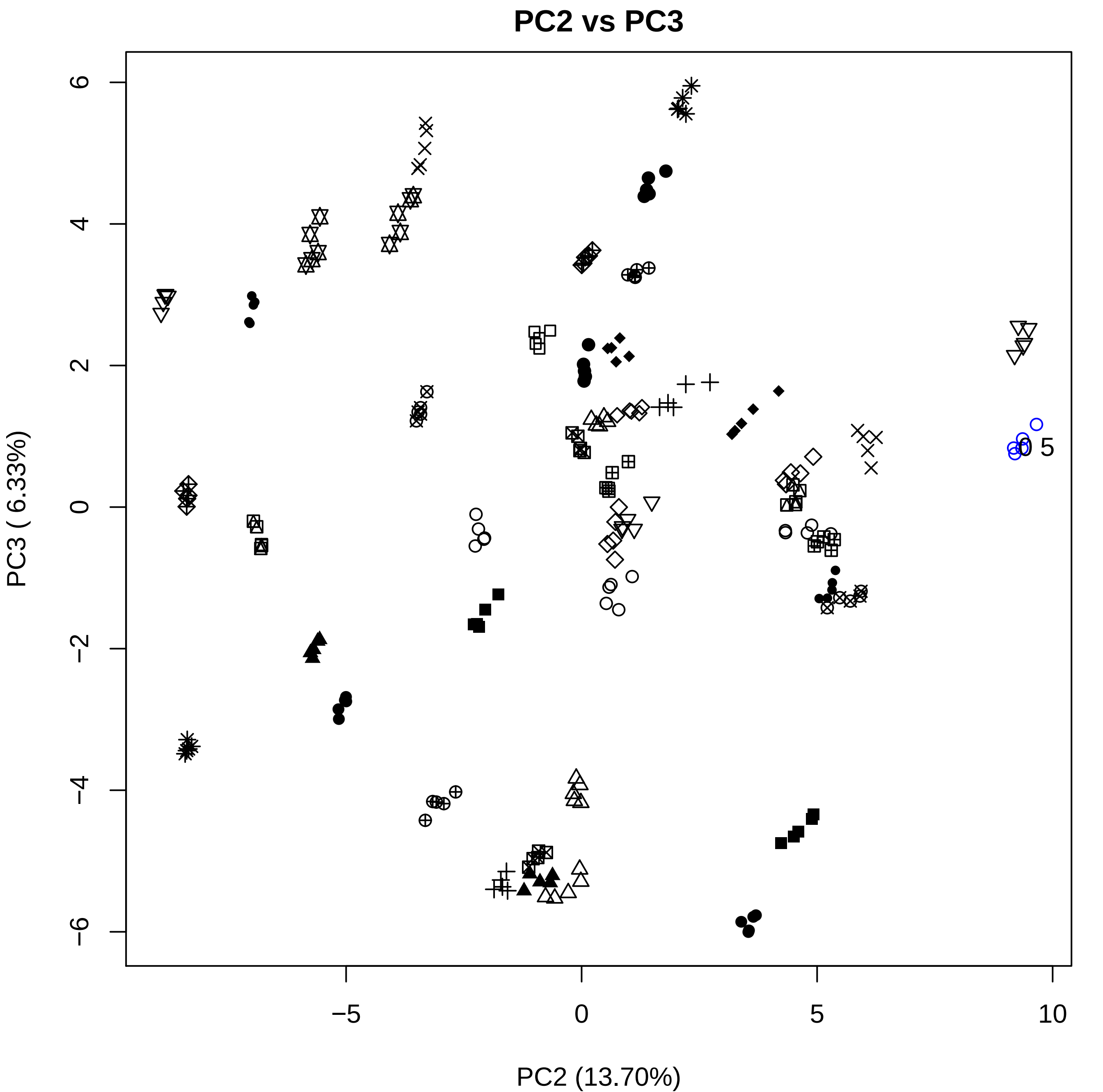
PC2 vs PC3
PC3 ( 6.33%)
PC2 (13.70%)
6
4
2
0
−2
−4
−6
−5
0
5
10
0 5

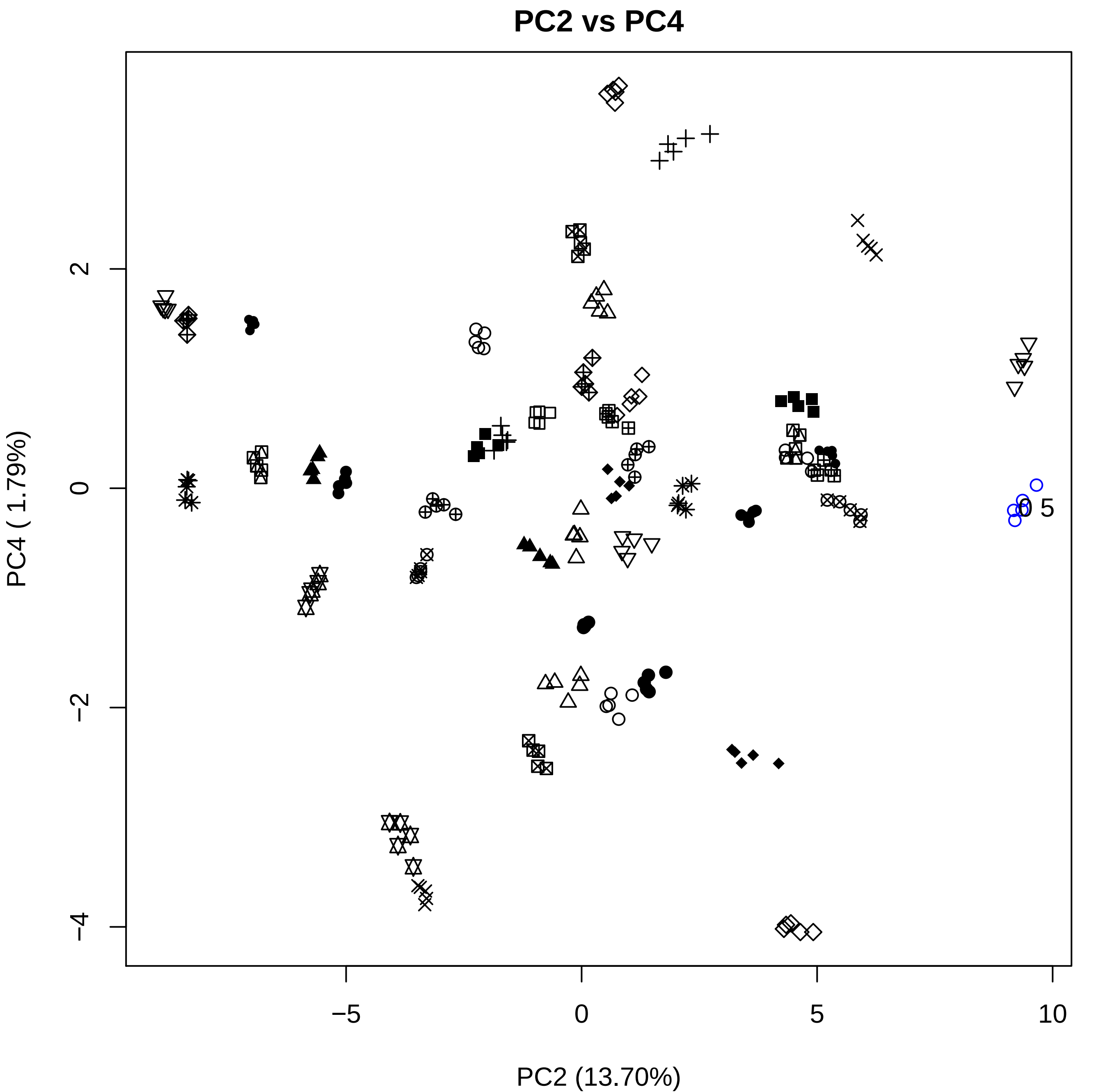

PC2 vs PC4
PC4 ( 1.79%)
PC2 (13.70%)
2
0
−2
−4
−5
0
5
10
0 5

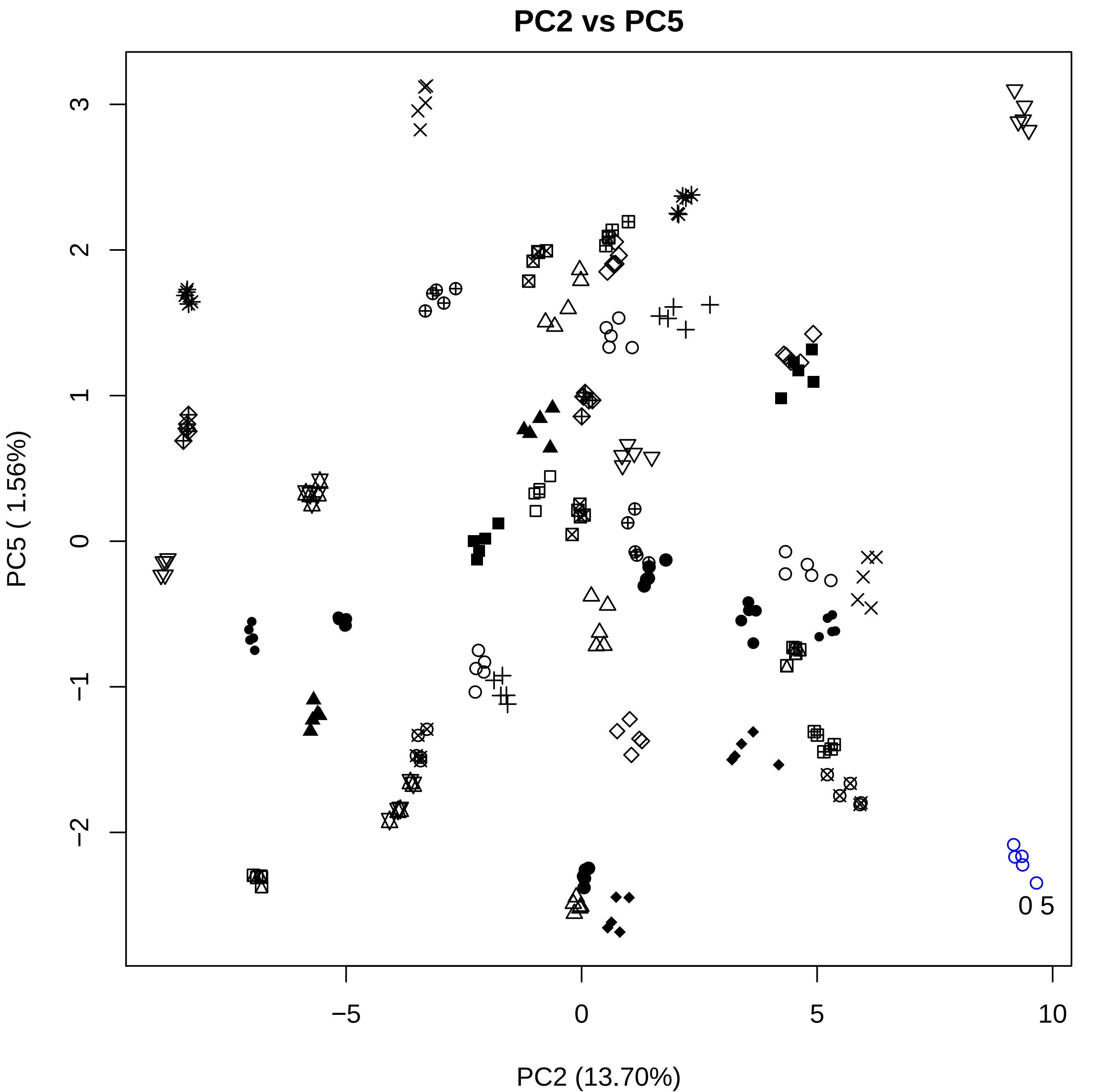
PC2 vs PC5
PC5 ( 1.56%)
PC2 (13.70%)
3
2
1
0
−1
−2
−5
0
5
10
0 5

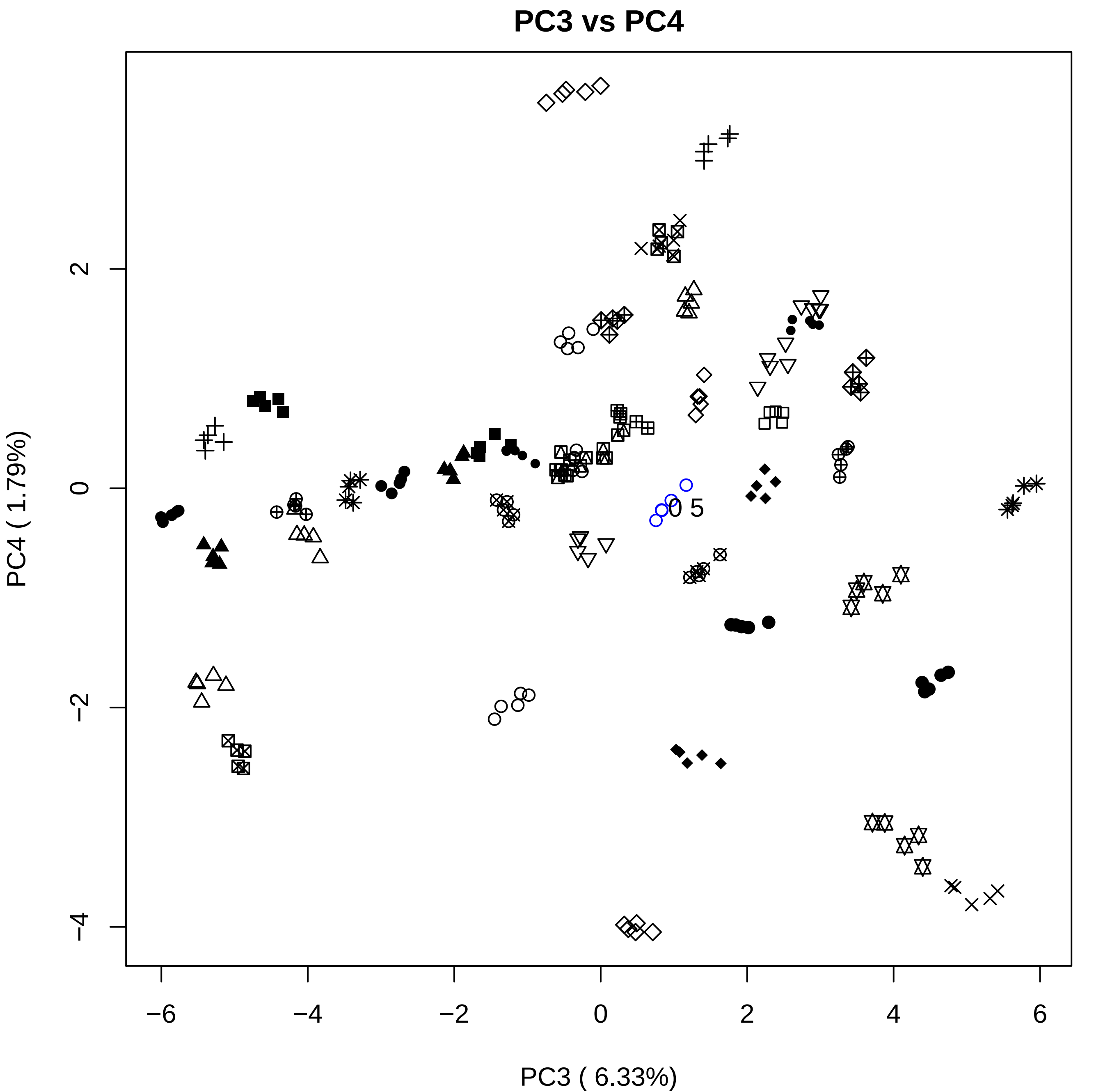
PC3 vs PC4
PC4 ( 1.79%)
PC3 ( 6.33%)
2
0
−2
−4
−6
−4
−2
0
2
4
6
0 5

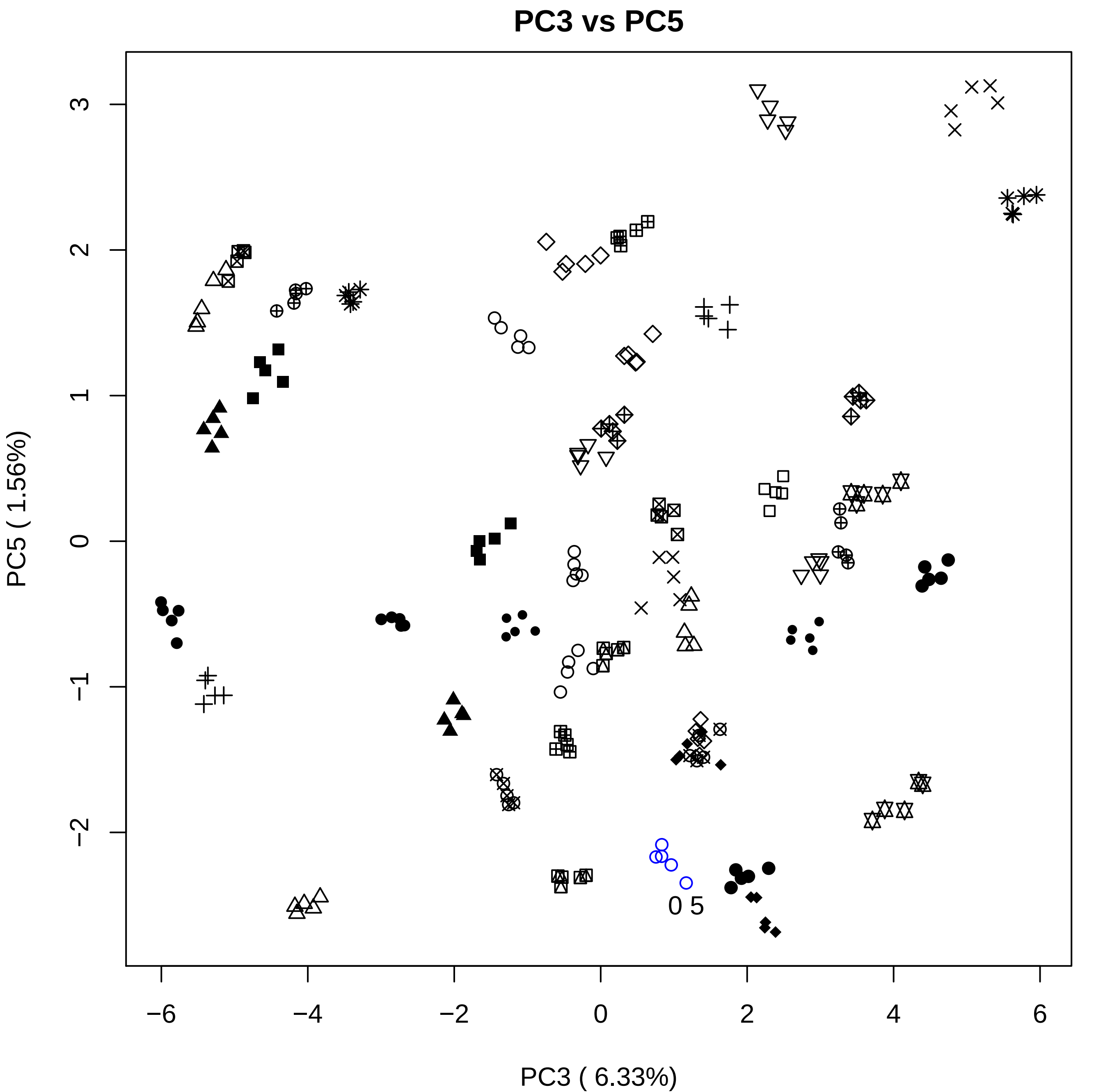
PC3 vs PC5
PC5 ( 1.56%)
PC3 ( 6.33%)
3
2
1
0
−1
−2
−6
−4
−2
0
2
4
6
0 5

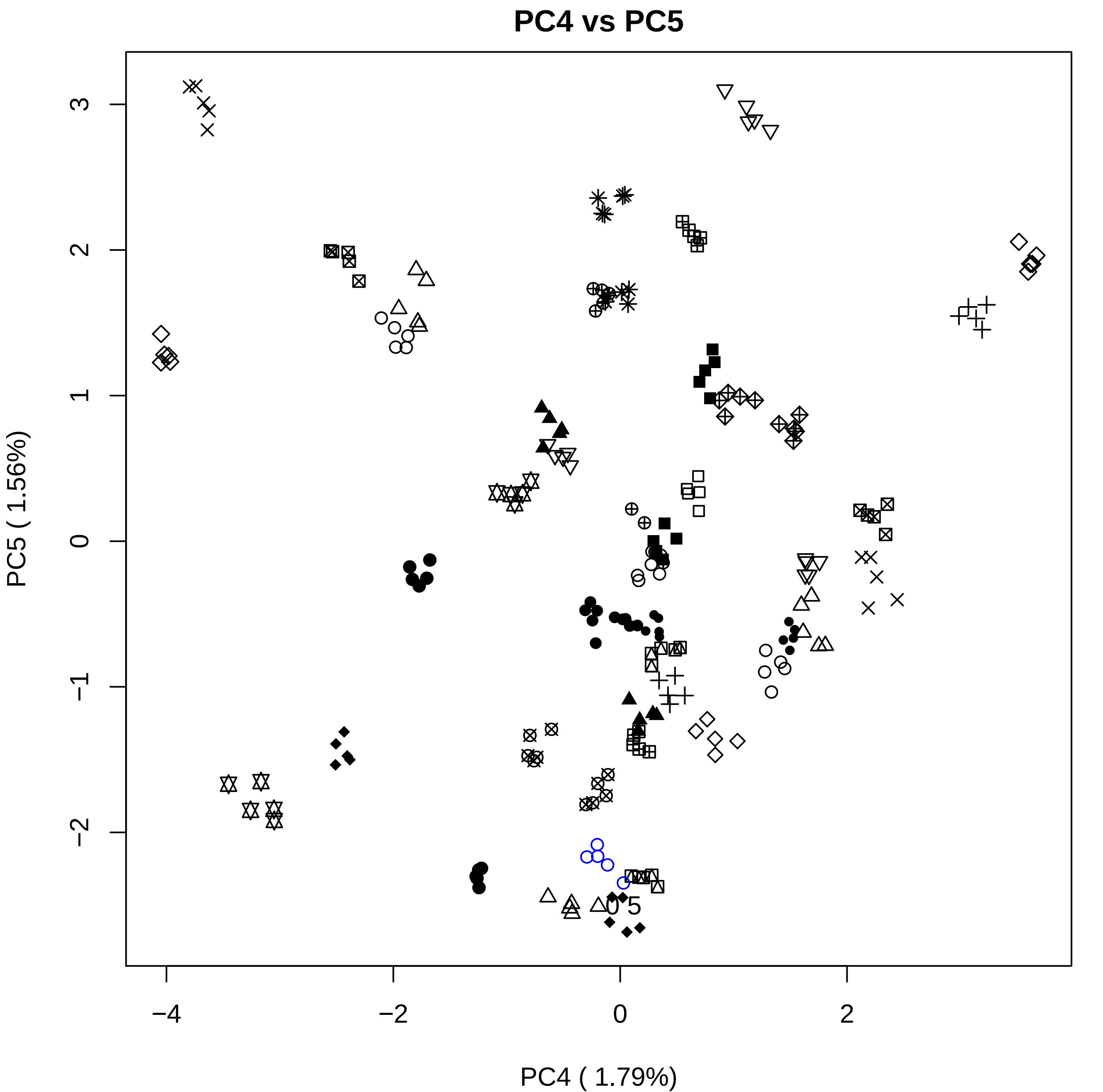
PC4 vs PC5
PC5 ( 1.56%)
PC4 ( 1.79%)
−4
−2
0
2
3
1
−1
0 5

### PCA of 1%  OSCS vs PMH

Scaled Matrix PCAed

7th order polynomial baseline correction performed

Raw spectra smoothed to 21 Neighbours, 2nd order PN, Savtisky Golay

Normalisation between 0 and 100

D2 spectra smoothed to 41 Neighbours, 2nd order PN, Savtisky Golay

Regions Analysed: 3601−2501 and 2001−700cm−1

Samples Analysed: 1 Contam's 0 OSCS's 45 PMH's

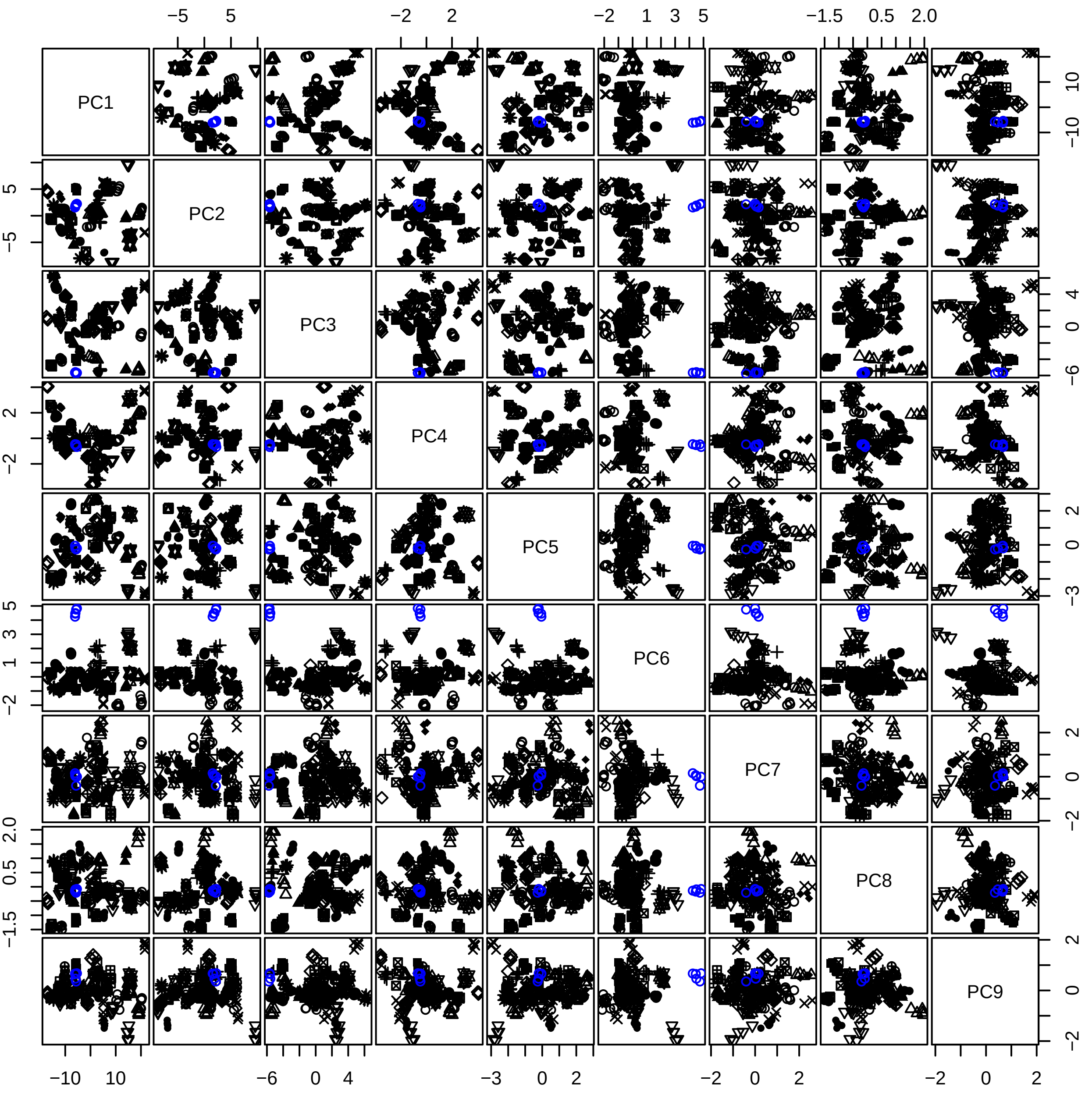
PC1
PC2
PC3
PC4
PC5
PC6
PC7
PC8
PC9
−5
5
−2
2
−2
1
3
5
−1.5
0.5
2.0
10
−10
5
−5
4
0
−6
2
−2
2
0
−3
5
3
1
−2
2
0
−2
2.0
0.5
−1.5
2
0
−2
−10
10
−6
0
4
−3
0
2
−2
0
2
−2
0
2

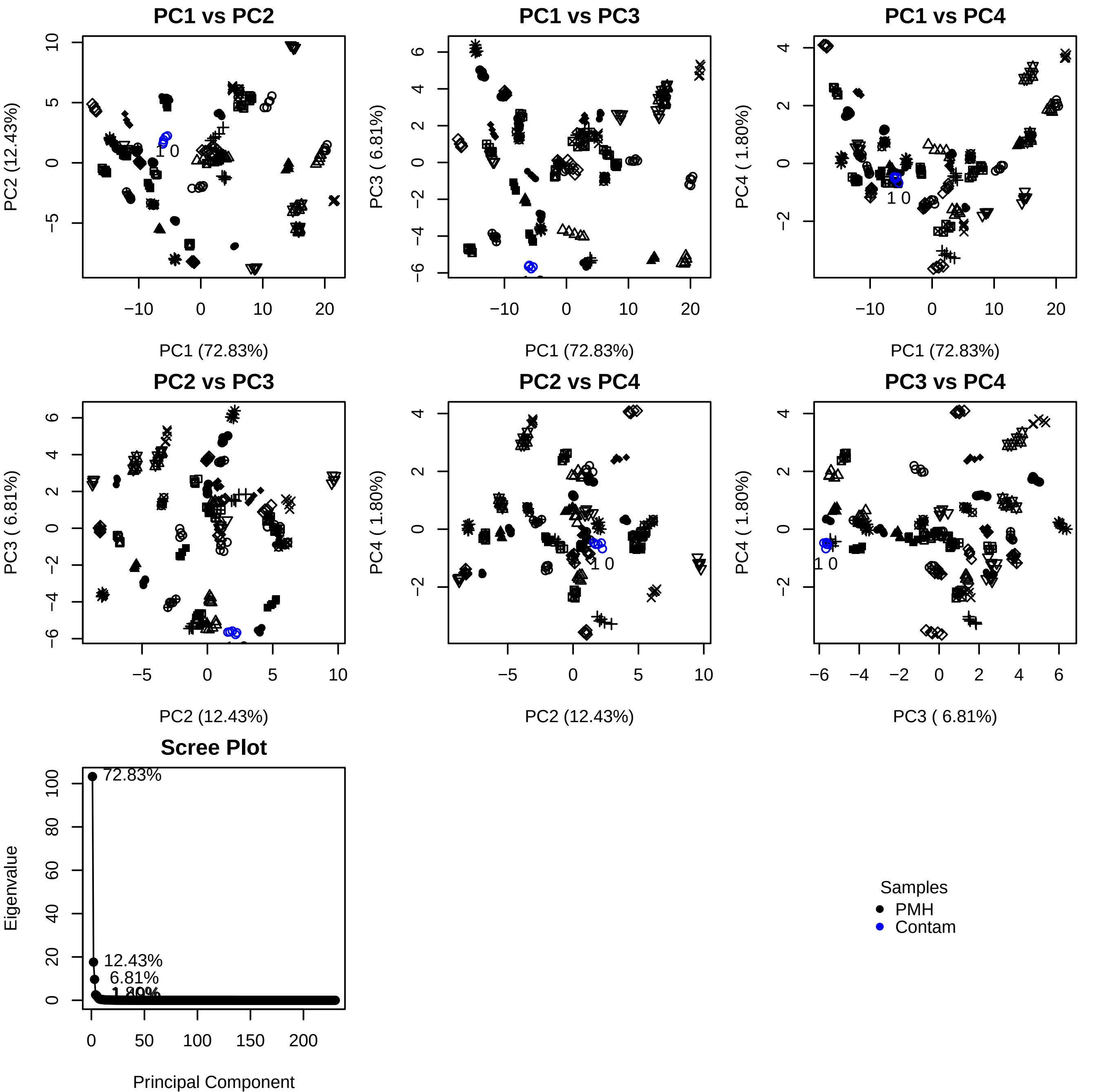
PC1 vs PC2
PC1 vs PC3
PC1 vs PC4
PC2 vs PC3
PC2 vs PC4
PC3 vs PC4
Scree Plot
PC1 (72.83%)
PC2 (12.43%)
PC3 ( 6.81%)
PC4 ( 1.80%)
72.83%
12.43%
6.81%
Eigenvalue
Principal Component
Samples
PMH
Contam

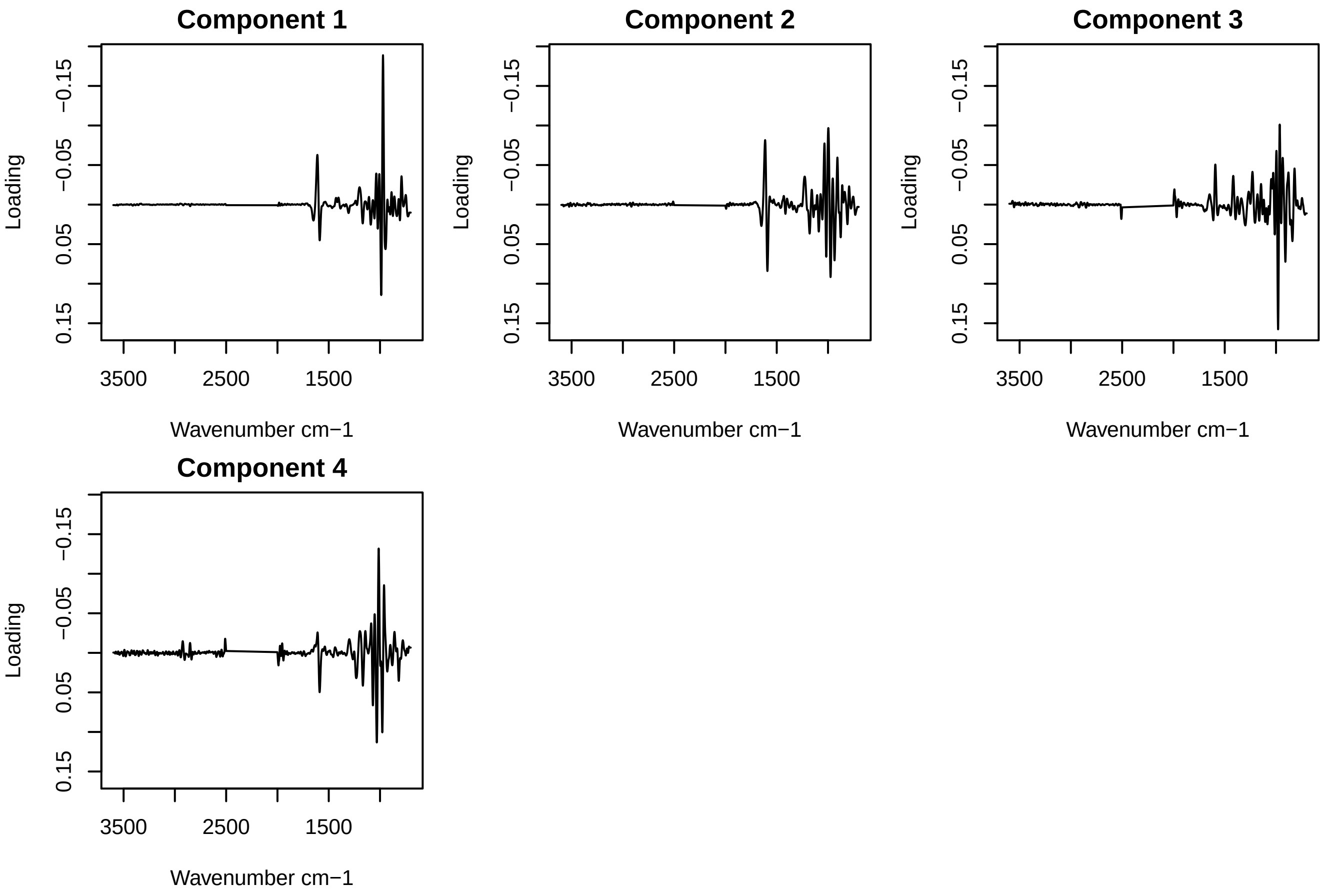
Component 1
Component 2
Component 3
Component 4
Loading
−0.15
−0.05
0.05
0.15
3500
2500
1500
Wavenumber cm−1

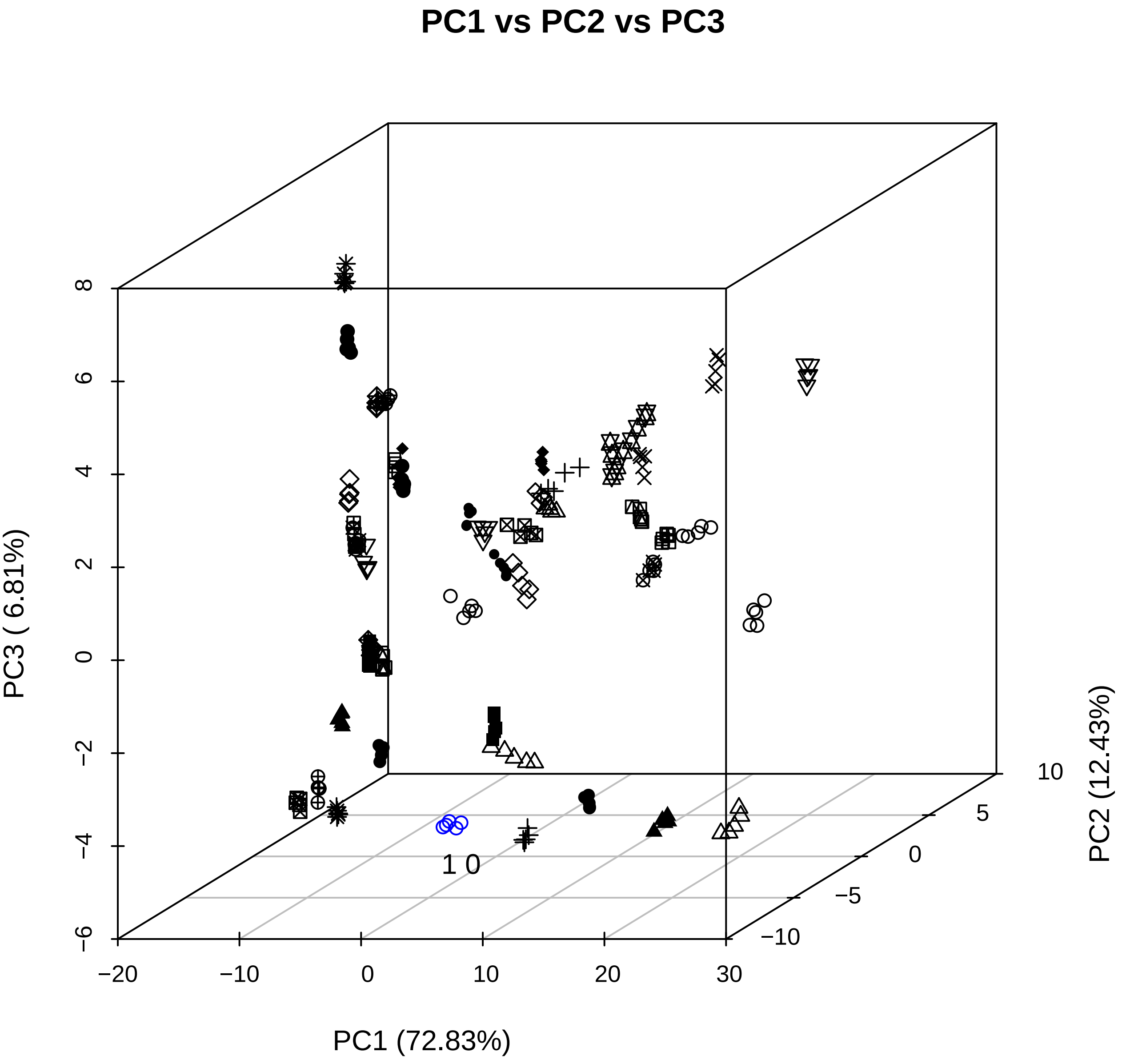

PC1 vs PC2 vs PC3
PC3 ( 6.81%)
PC1 (72.83%)
PC2 (12.43%)
8
6
4
2
0
−2
−4
−6
−20
−10
0
10
20
30
10
5
0
−5
−10
1 0

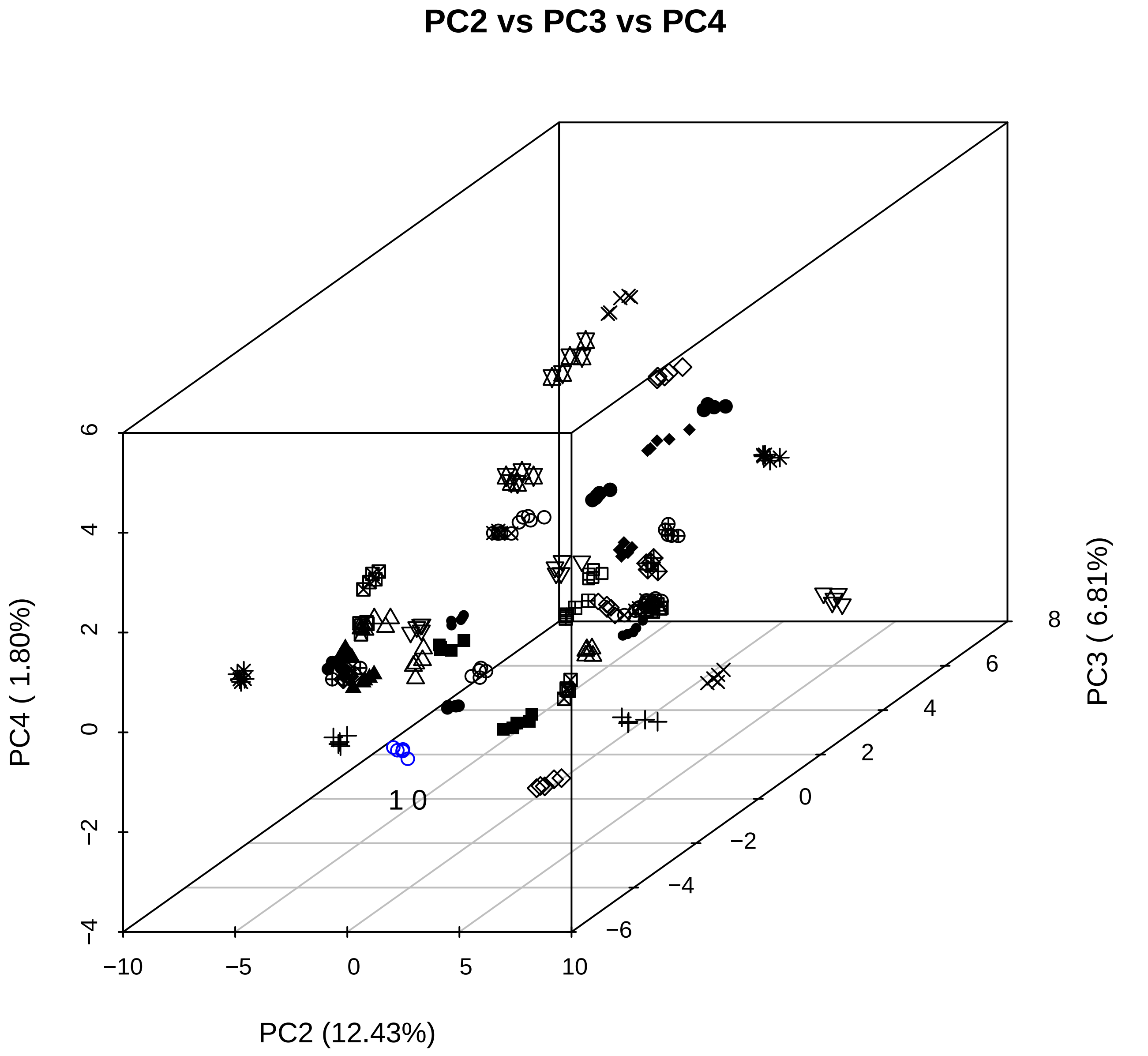
PC2 vs PC3 vs PC4
PC4 ( 1.80%)
PC2 (12.43%)
PC3 ( 6.81%)
−4
−2
0
2
4
6
−10
−5
0
5
10
−6
−4
−2
0
2
4
6
8
10

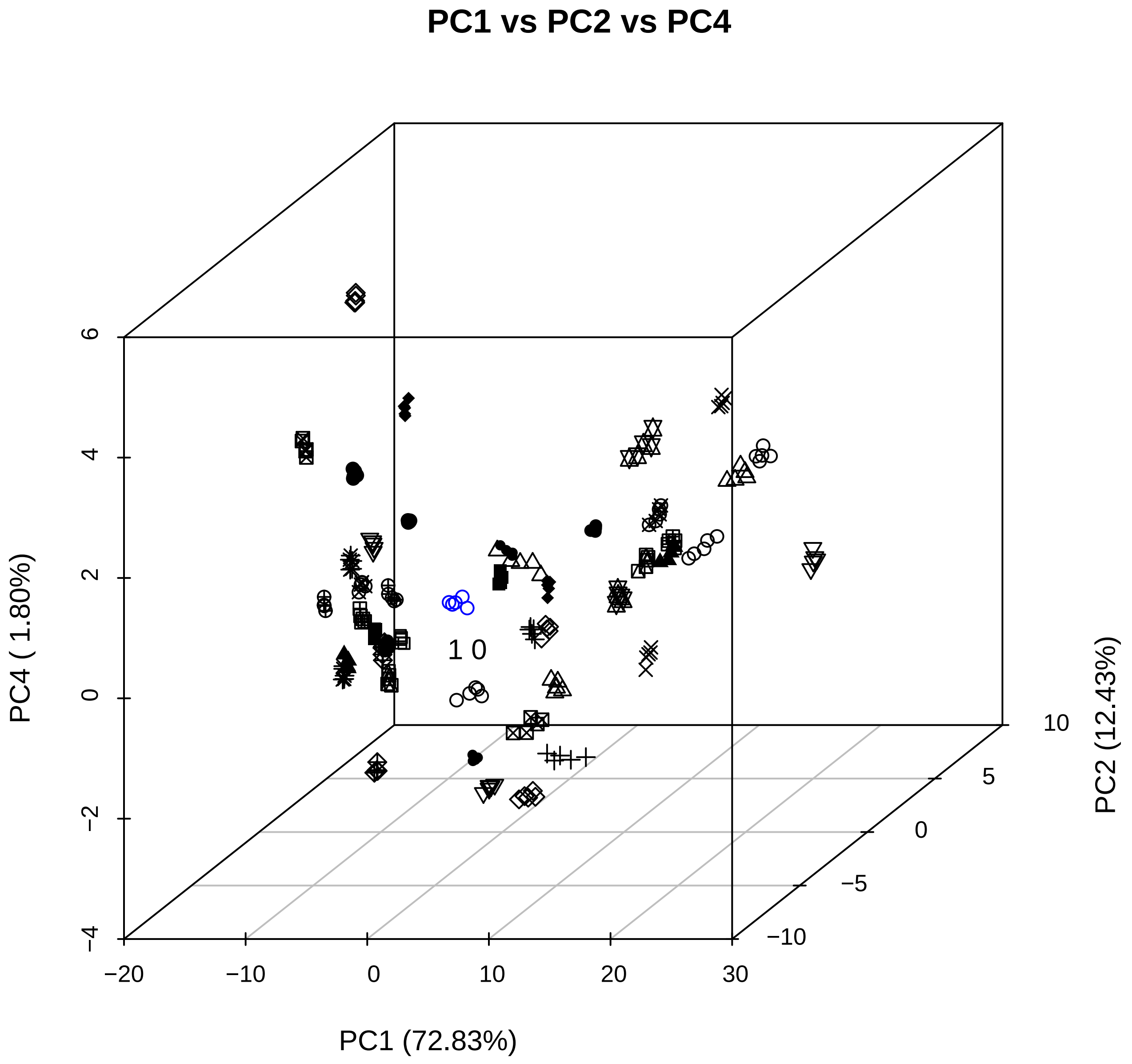

PC1 vs PC2 vs PC4
PC4 ( 1.80%)
PC1 (72.83%)
PC2 (12.43%)
−4
−2
0
2
4
6
−20
−10
0
10
20
30
−10
−5
0
5
10
1 0

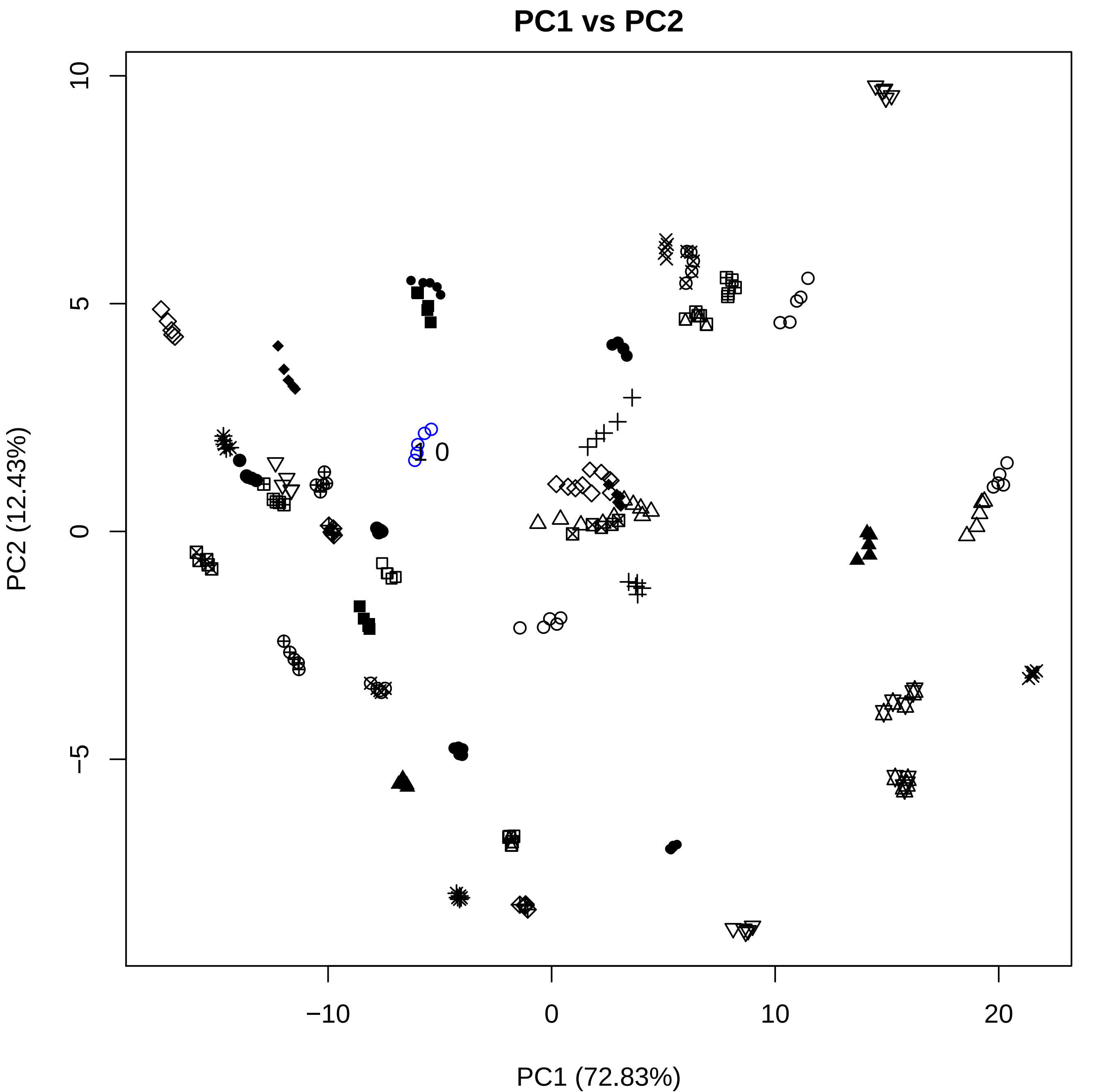
PC1 vs PC2
PC1 (72.83%)
PC2 (12.43%)
−10
0
10
20
−5
0
5
10
1 0

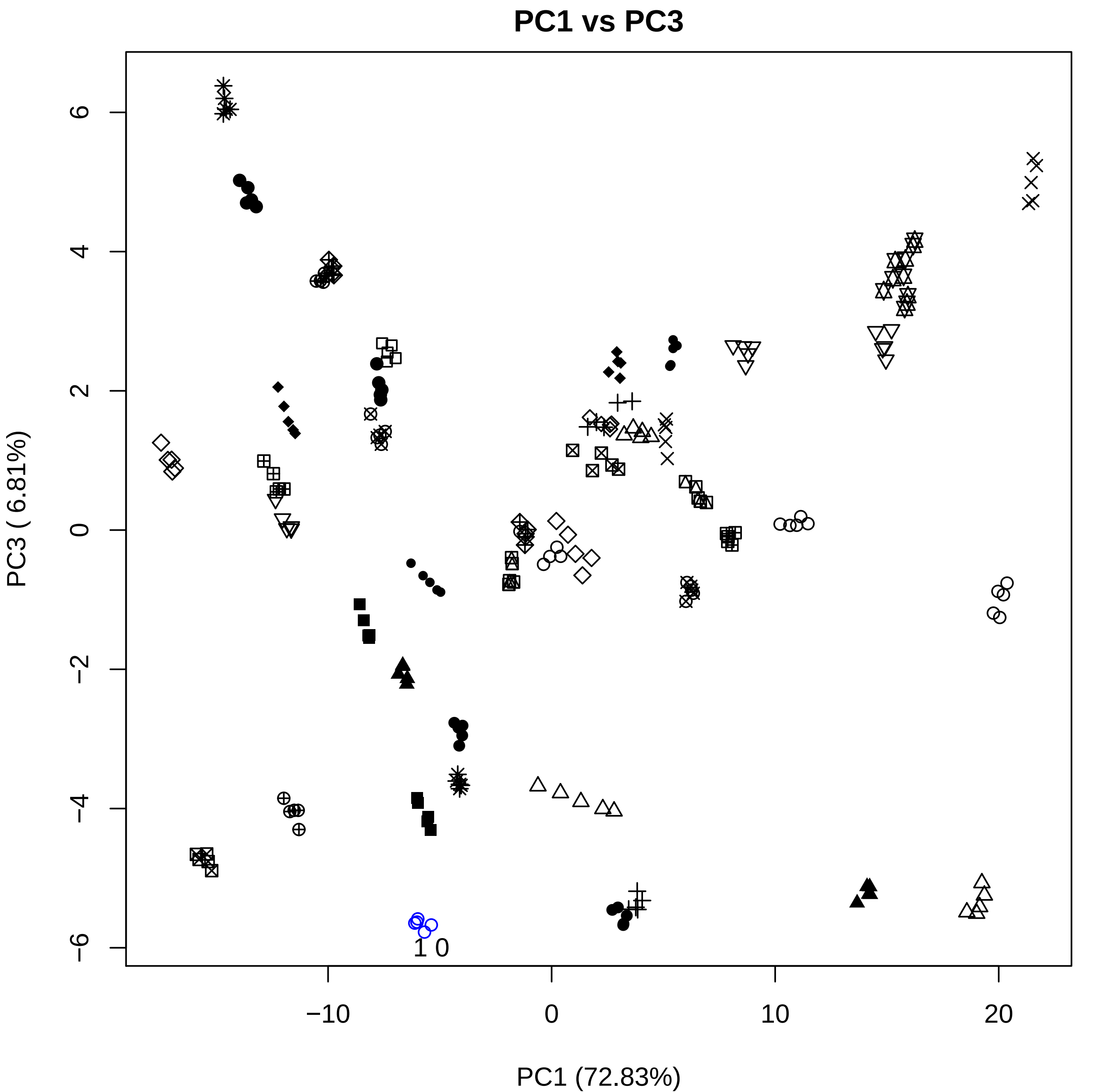
PC1 vs PC3
PC3 ( 6.81%)
PC1 (72.83%)
6
4
2
0
−2
−4
−6
−10
0
10
20
1 0

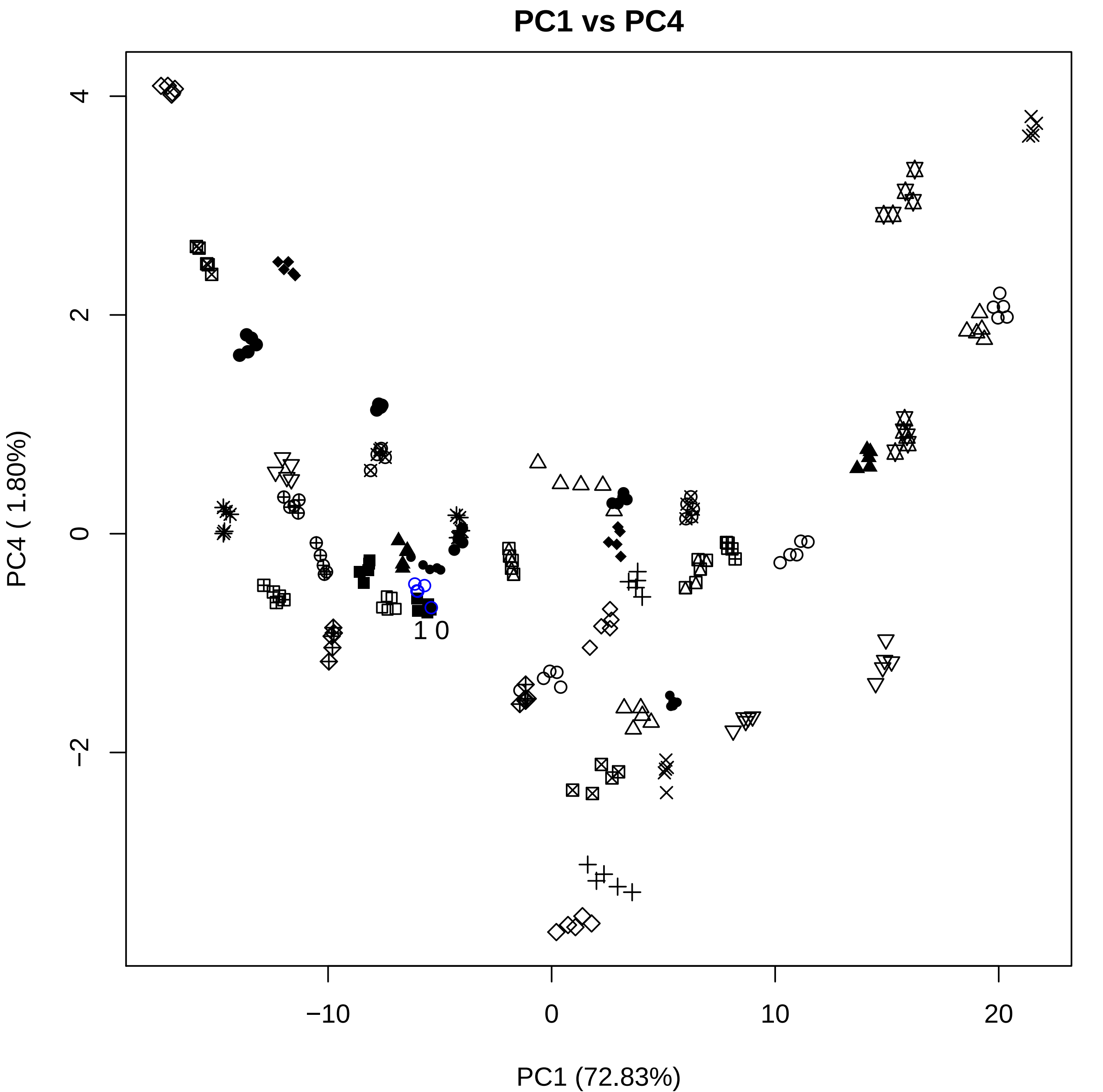

PC1 vs PC4
PC4 ( 1.80%)
PC1 (72.83%)
4
2
0
−2
−10
0
10
20
1 0

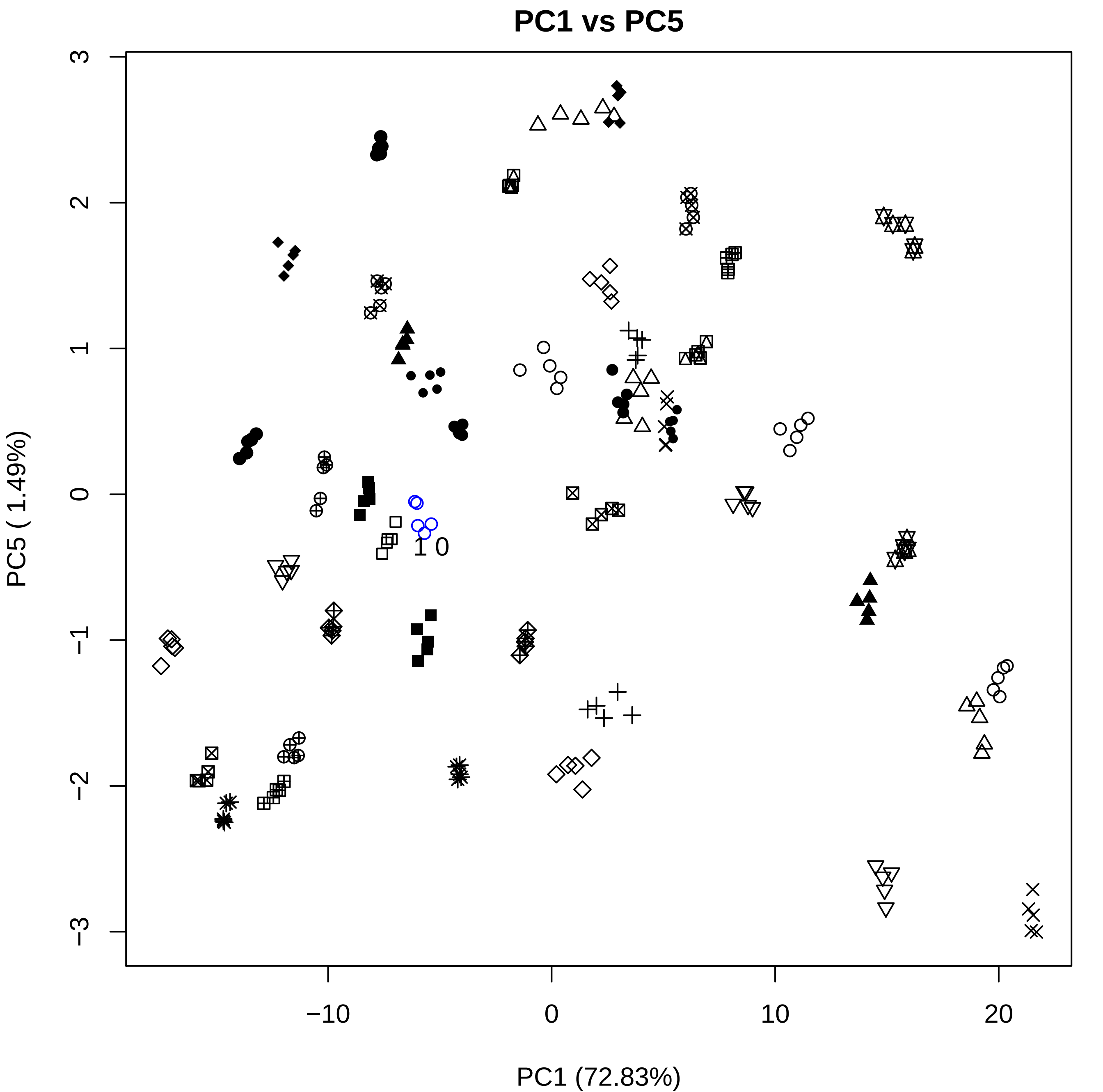
PC1 vs PC5
PC5 ( 1.49%)
PC1 (72.83%)
3
2
1
0
−1
−2
−3
−10
0
10
20
1 0

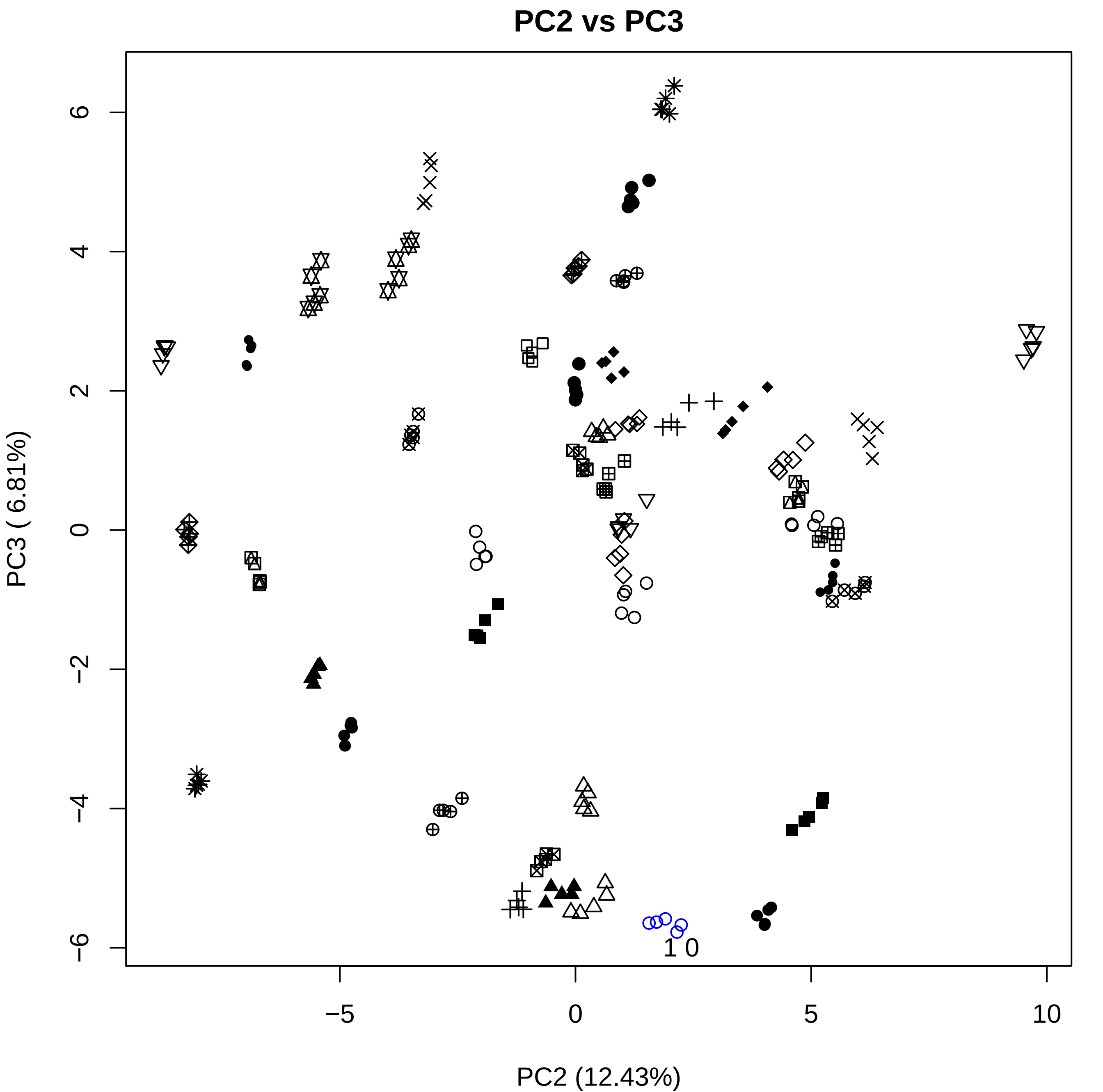
PC2 vs PC3
PC3 ( 6.81%)
PC2 (12.43%)
6
4
2
0
−2
−4
−6
−5
0
5
10
1 0

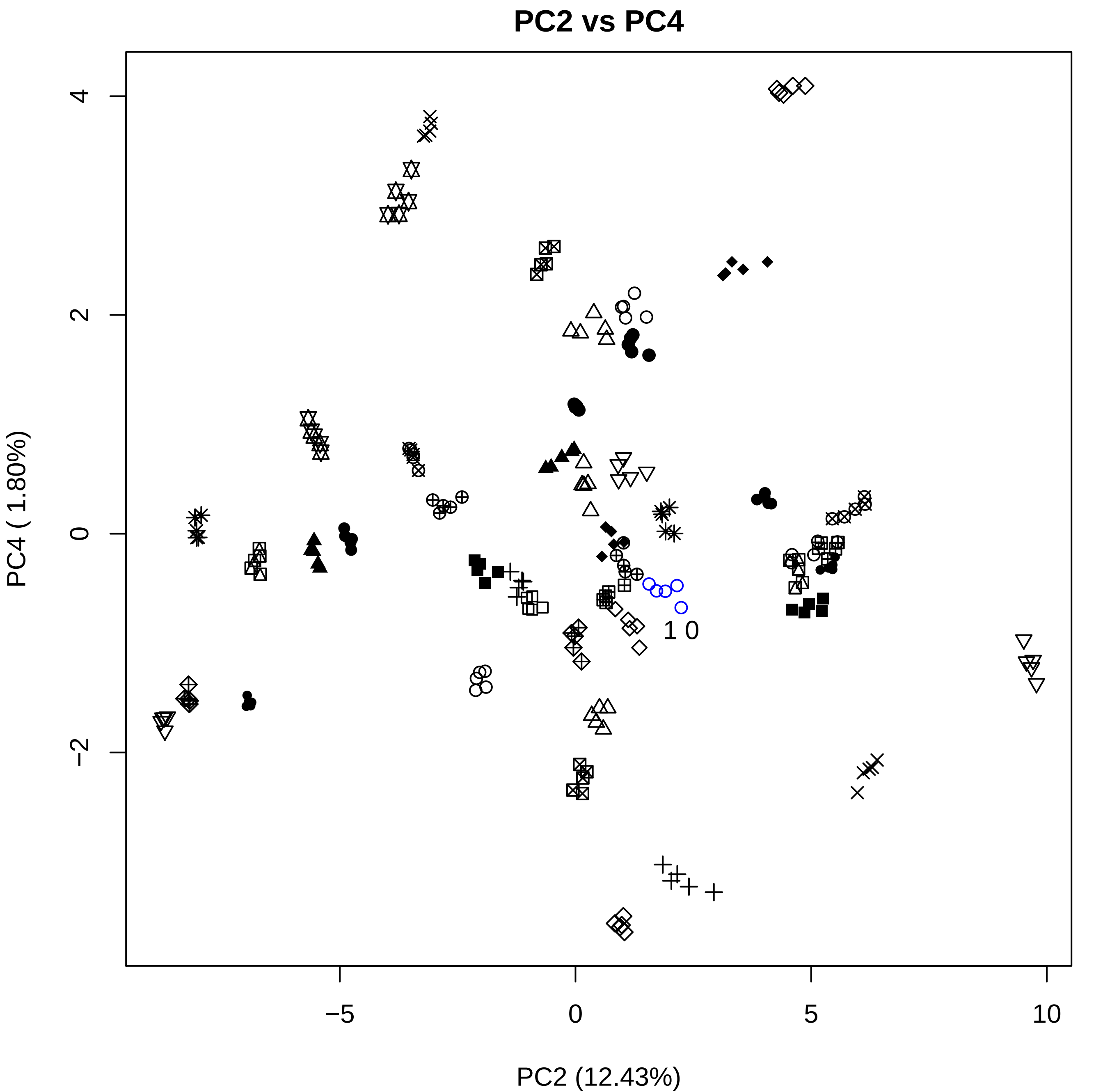
PC2 vs PC4
PC4 ( 1.80%)
PC2 (12.43%)
4
2
0
−2
−5
0
5
10
1 0

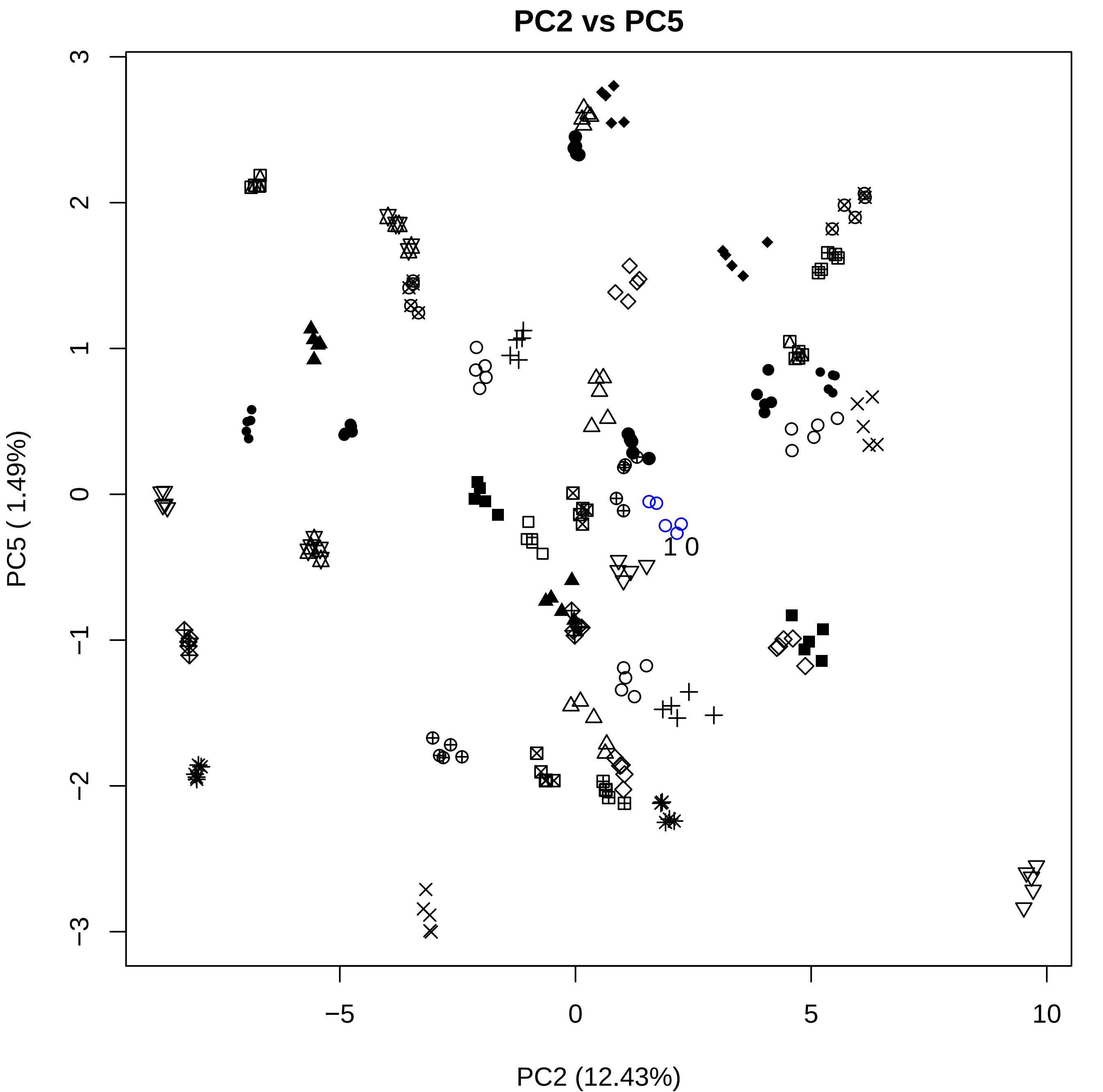
PC2 vs PC5
PC5 ( 1.49%)
PC2 (12.43%)
3
2
1
0
−1
−2
−3
−5
0
5
10
1 0

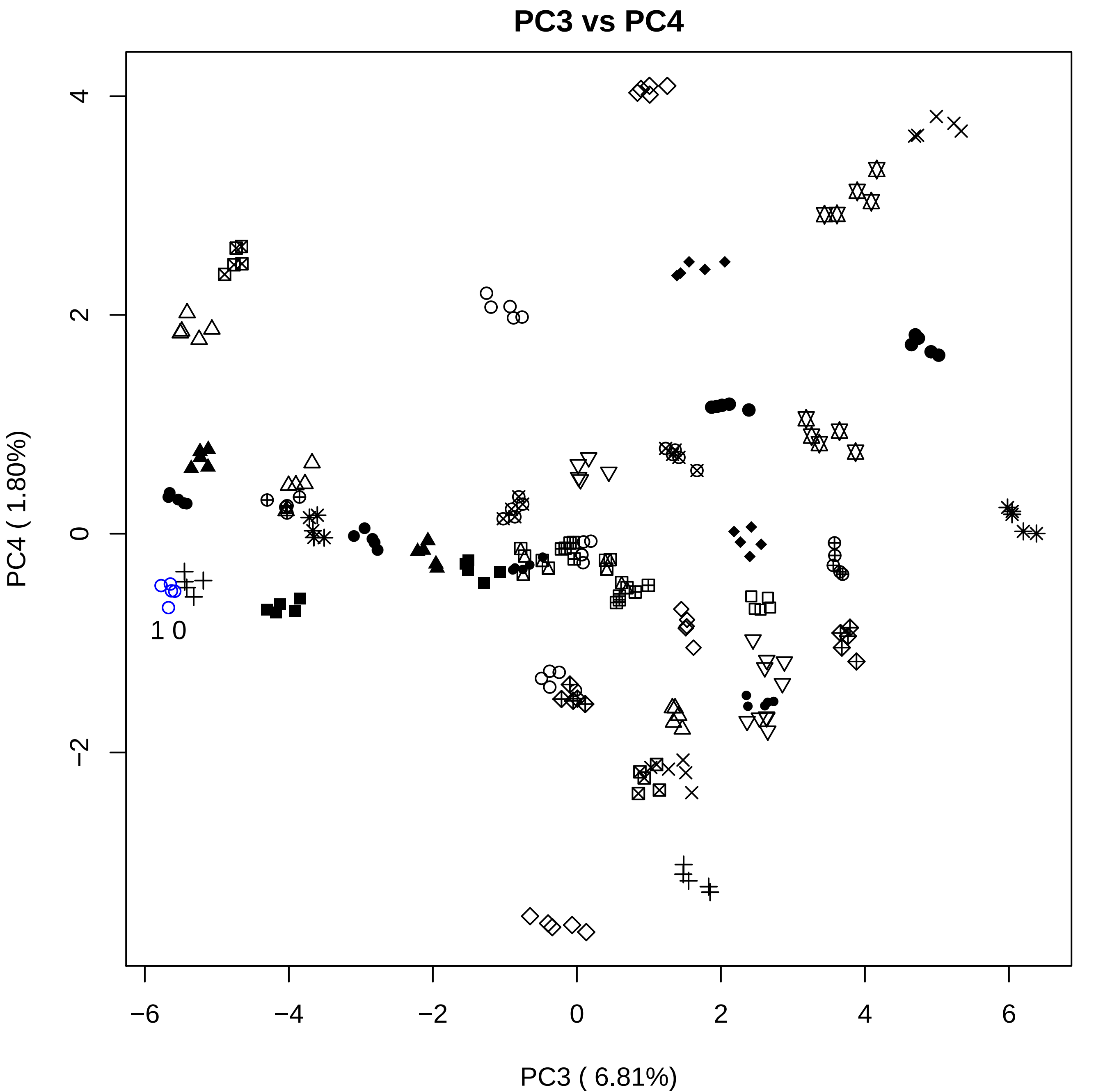
PC3 vs PC4
4
2
0
−2
PC4 ( 1.80%)
1 0
−6
−4
−2
0
2
4
6
PC3 ( 6.81%)

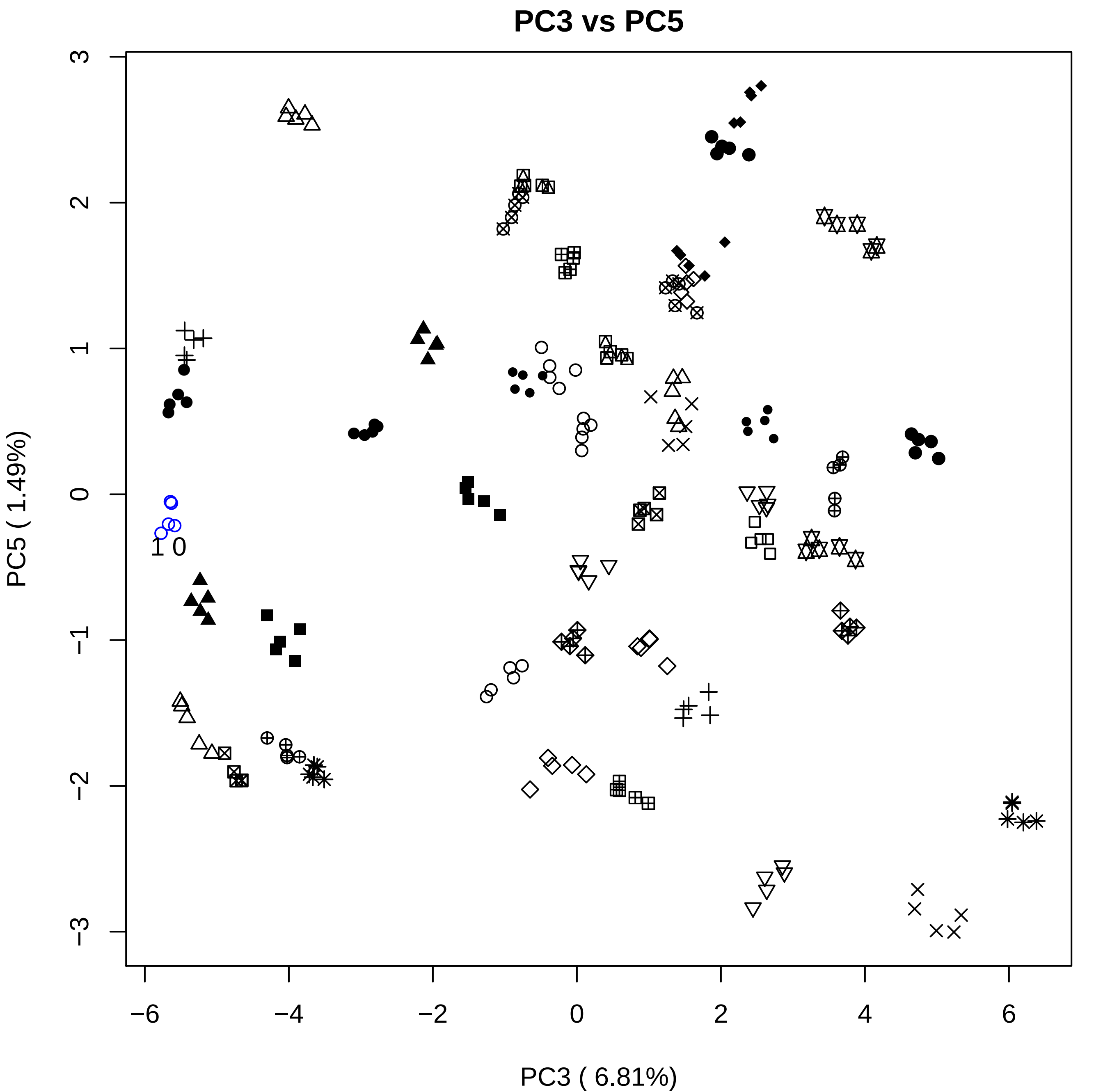
PC3 vs PC5
PC5 ( 1.49%)
PC3 ( 6.81%)
−6
−4
−2
0
2
4
6
−3
−2
−1
0
1
2
3
1 0

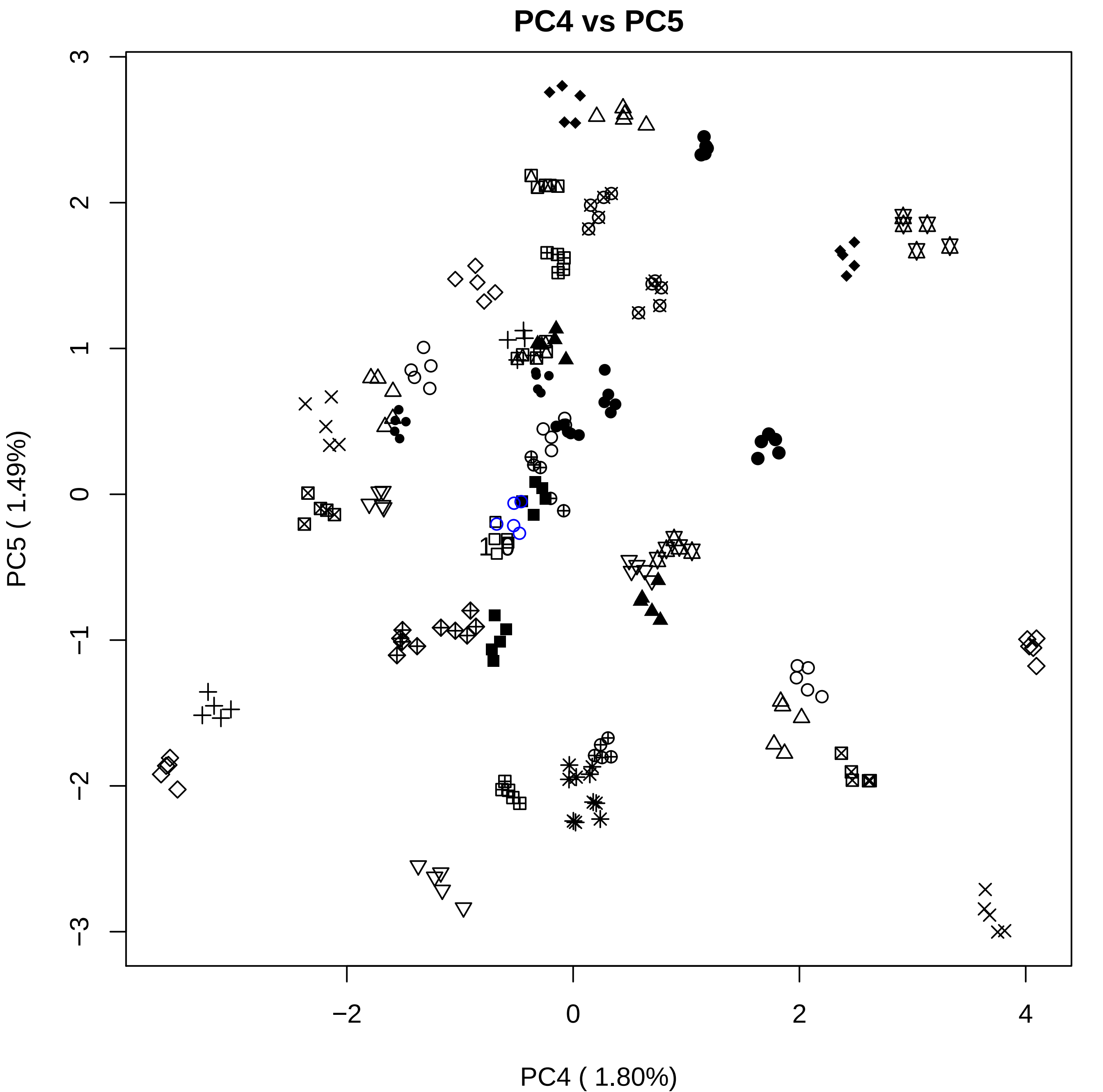
PC4 vs PC5
PC5 ( 1.49%)
PC4 ( 1.80%)
3
2
1
0
−1
−2
−3
−2
0
2
4
10

### PCA of 2.5%  OSCS vs PMH

Scaled Matrix PCAed

7th order polynomial baseline correction performed

Raw spectra smoothed to 21 Neighbours, 2nd order PN, Savtisky Golay

Normalisation between 0 and 100

D2 spectra smoothed to 41 Neighbours, 2nd order PN, Savtisky Golay

Regions Analysed: 3601−2501 and 2001−700cm−1

Samples Analysed: 1 Contam's 0 OSCS's 45 PMH's

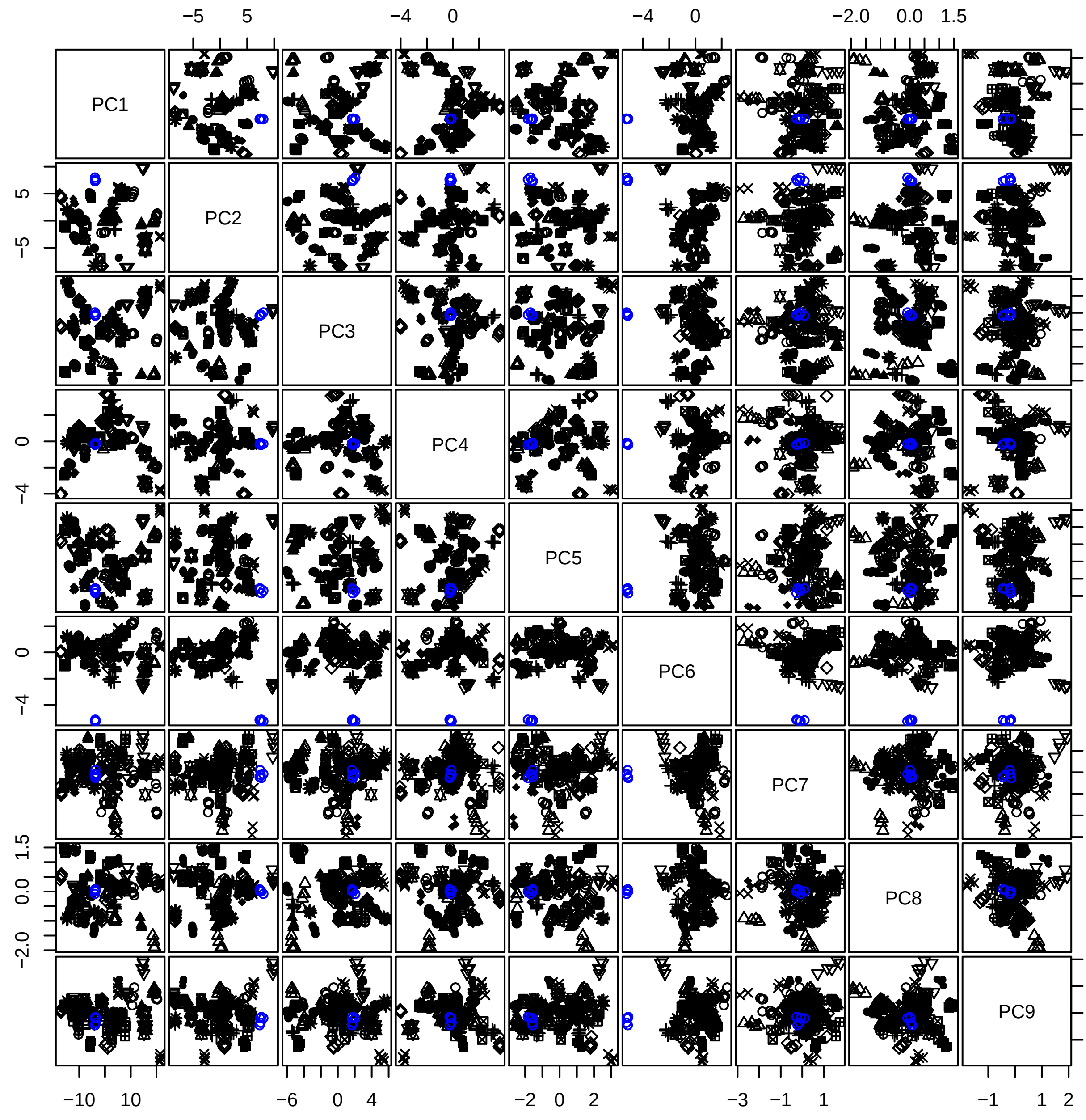

PC1
PC2
PC3
PC4
PC5
PC6
PC7
PC8
PC9
−5 5
−4 0
−4 0
−2.0 0.0 1.5
10 −10
5 −5
4 0 −6
0 −4
2 0 −2
0 −4
1 −1 −3
1.5 0.0 −2.0
2 1 −1
−10 10
−6 0 4
−2 0 2
−3 −1 1
−1 1 2

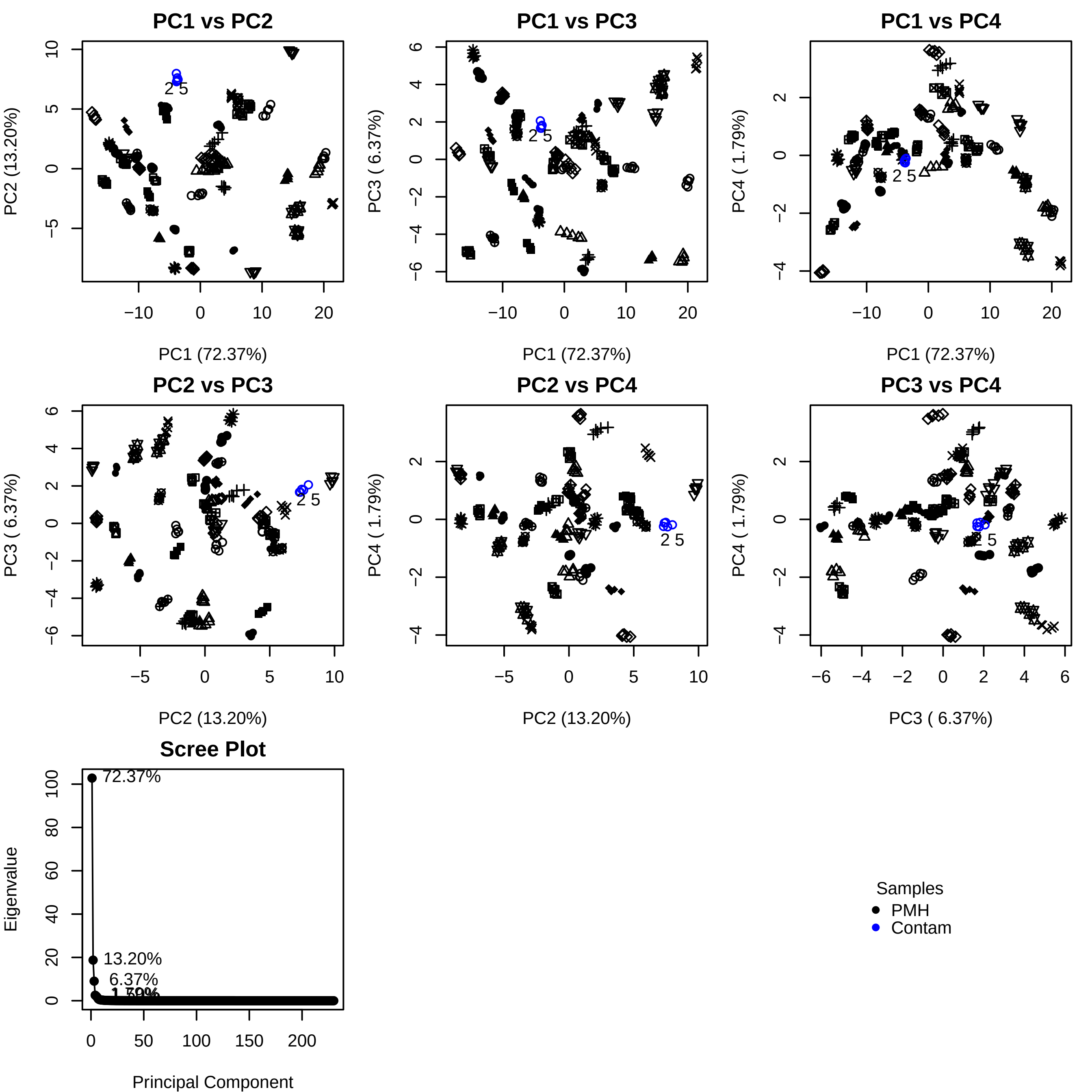

PC1 vs PC2
PC1 vs PC3
PC1 vs PC4
PC2 vs PC3
PC2 vs PC4
PC3 vs PC4
Scree Plot
PC1 (72.37%)
PC2 (13.20%)
PC3 ( 6.37%)
PC4 ( 1.79%)
Eigenvalue
Principal Component
72.37%
13.20%
6.37%
Samples
PMH
Contam

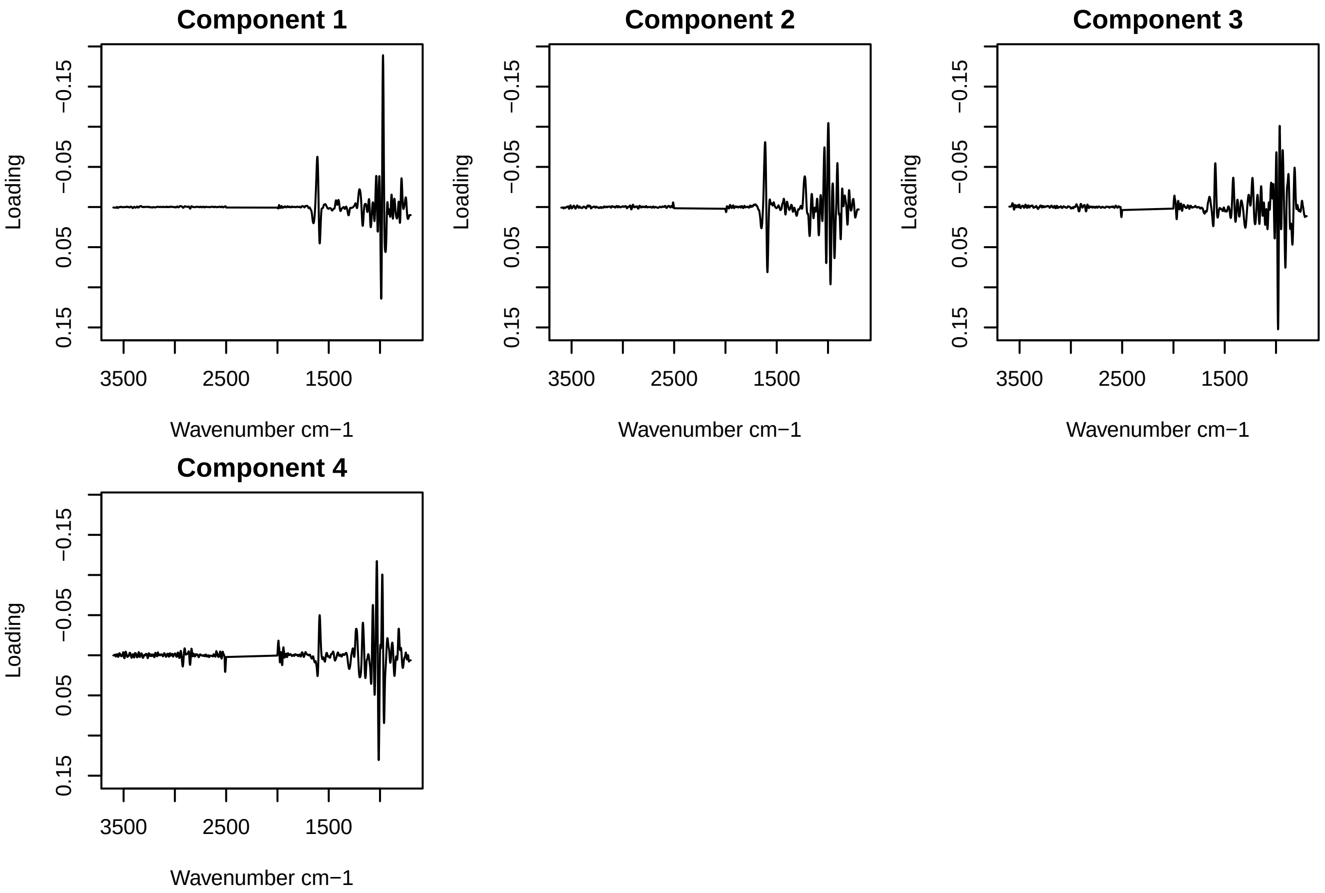
Component 1
Component 2
Component 3
Component 4
Loading
Wavenumber cm−1
−0.15
−0.05
0.05
0.15
3500
2500
1500

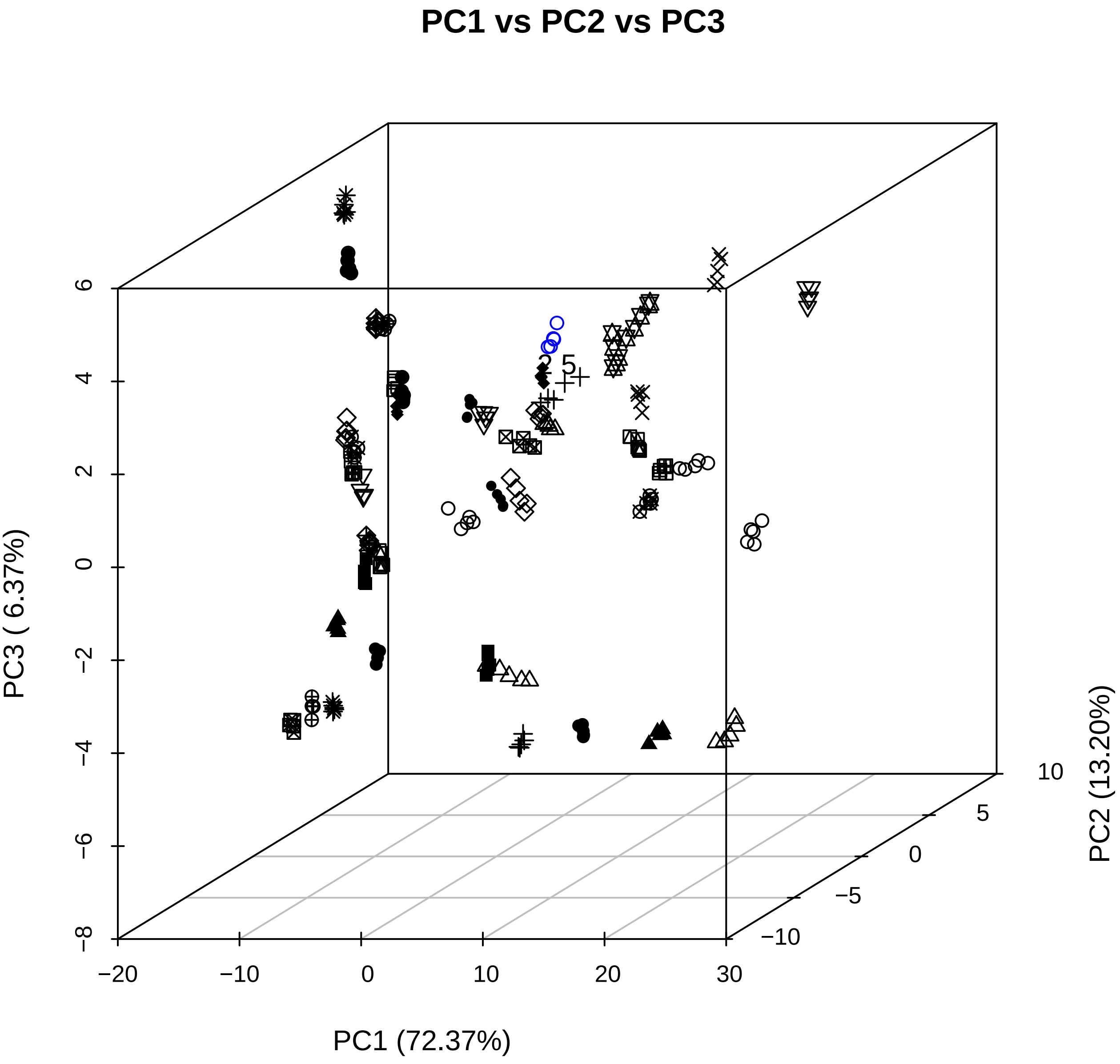
PC1 vs PC2 vs PC3
PC3 ( 6.37%)
PC1 (72.37%)
PC2 (13.20%)
6
4
2
0
−2
−4
−6
−8
−20
−10
0
10
20
30
10
5
0
−5
−10
2
5

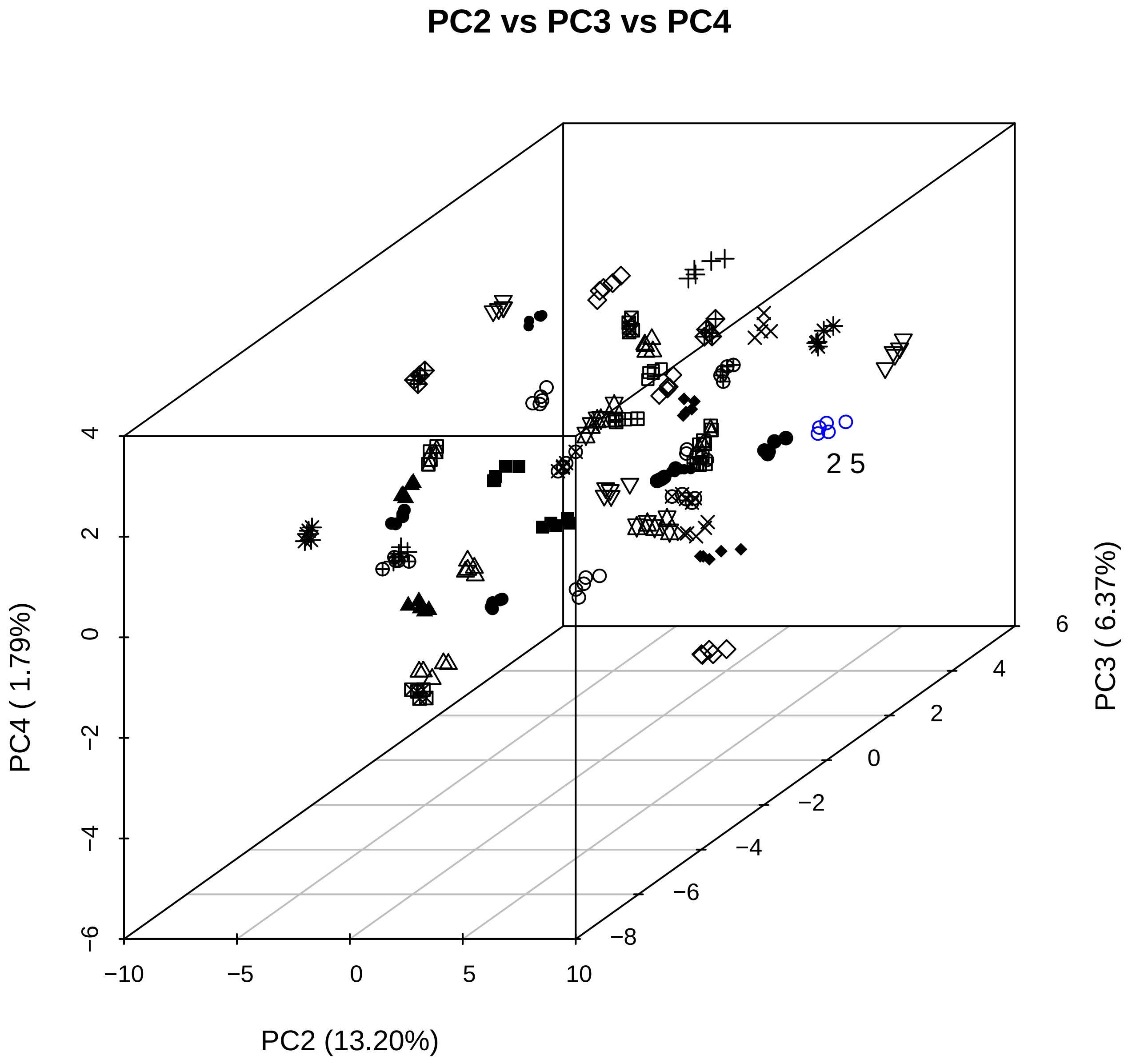
PC2 vs PC3 vs PC4
25
PC2 (13.20%)
PC3 ( 6.37%)
PC4 ( 1.79%)
−10
−5
0
5
10
−8
−6
−4
−2
0
2
4
6
−6
−4
−2
0
2
4

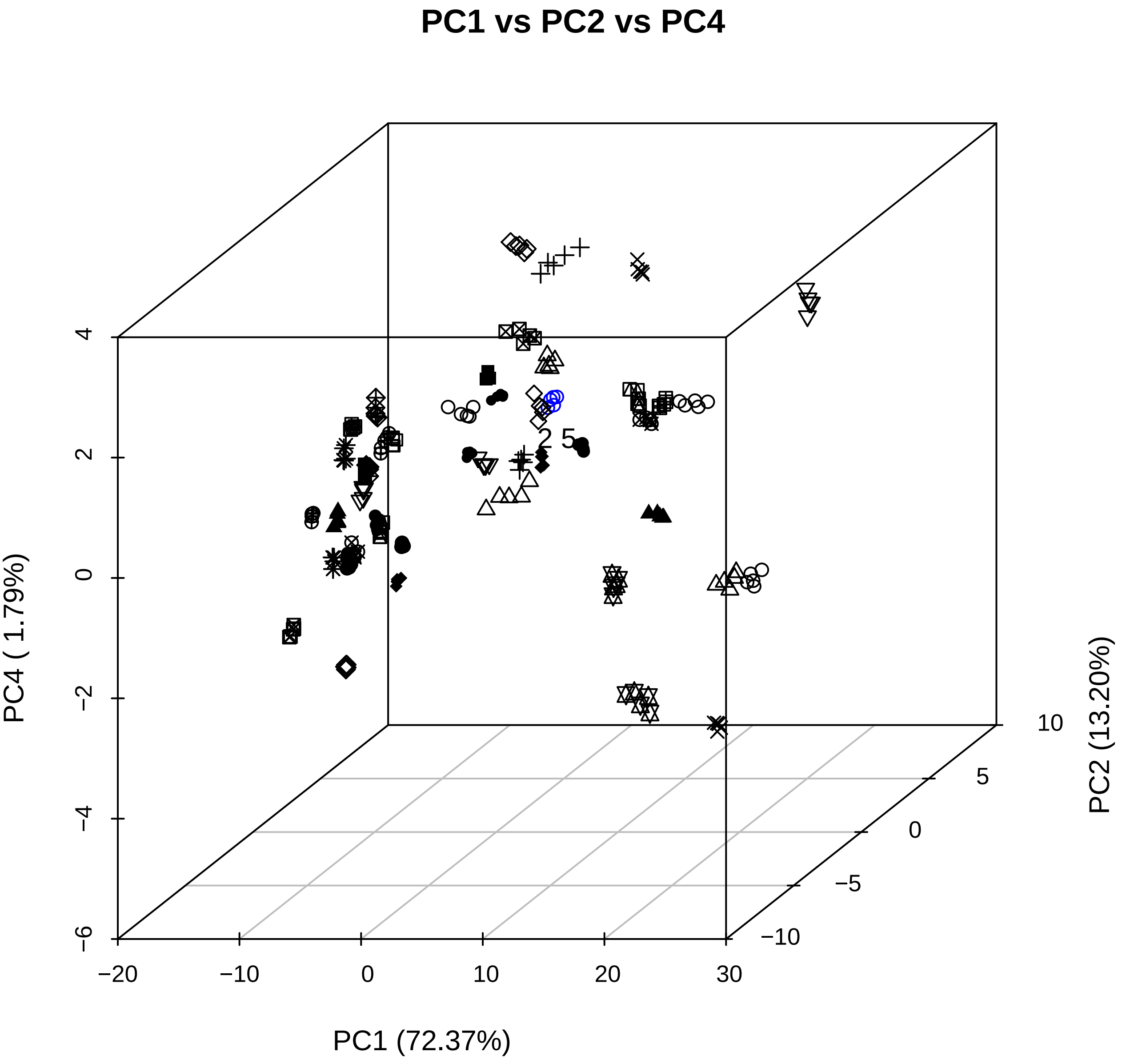
PC1 vs PC2 vs PC4
PC1 (72.37%)
PC2 (13.20%)
PC4 ( 1.79%)
−20
−10
0
10
20
30
−10
−5
0
5
10
−6
−4
−2
0
2
4
2
5

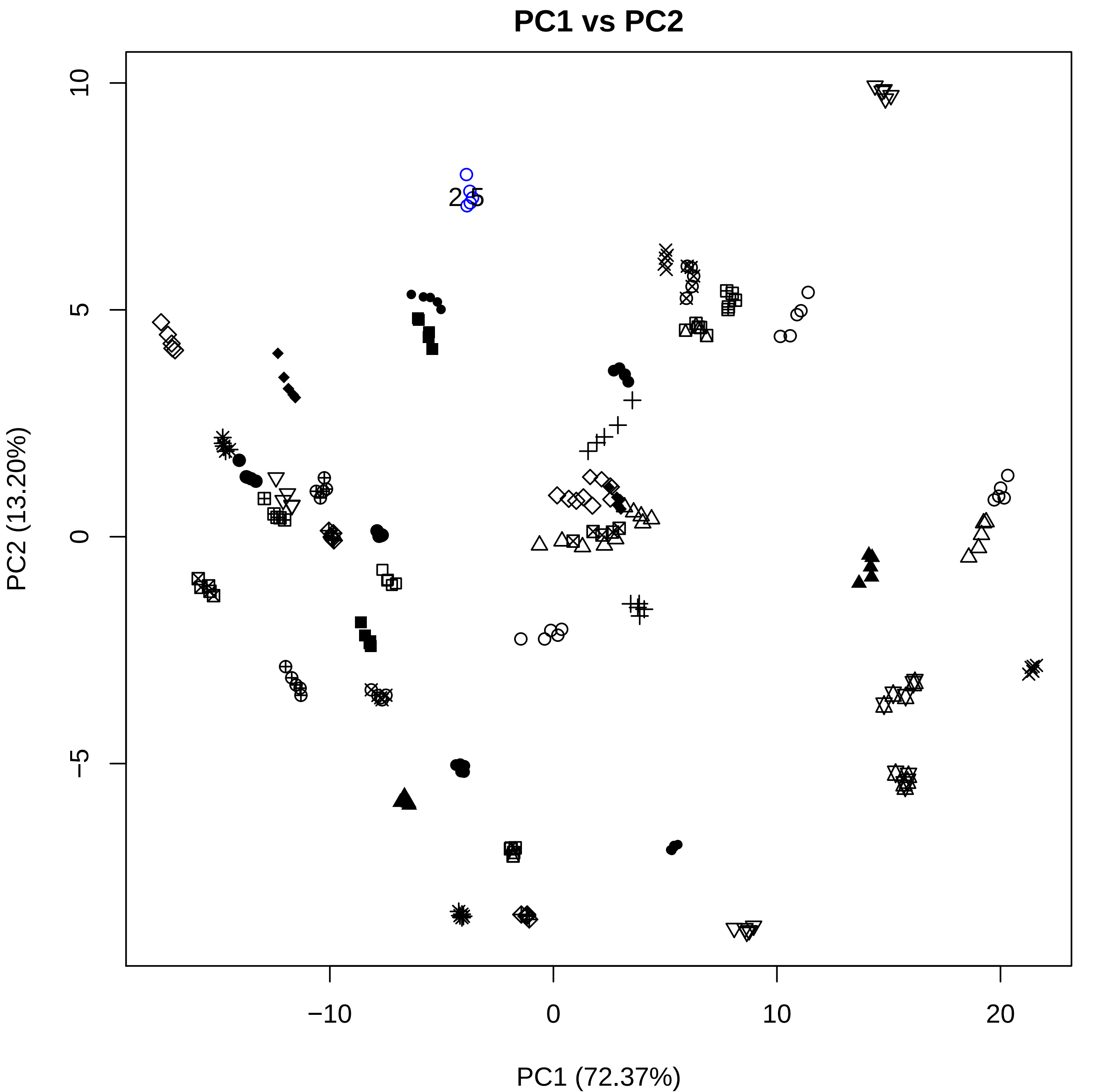

PC1 vs PC2
PC2 (13.20%)
PC1 (72.37%)
10
5
0
−5
−10
0
10
20
25

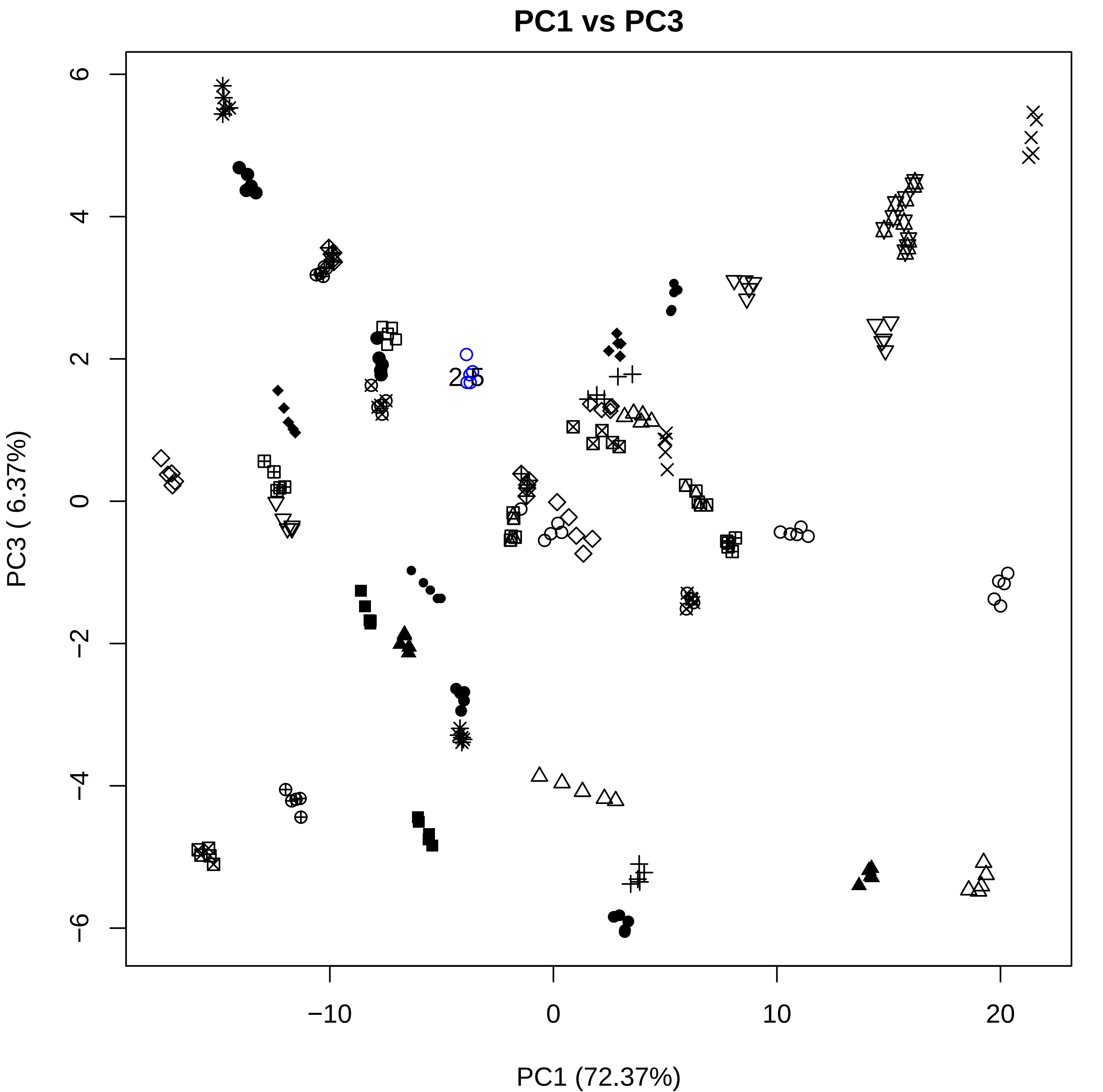
PC1 vs PC3
PC3 ( 6.37%)
6
4
2
0
−2
−4
−6
25
−10
0
10
20
PC1 (72.37%)

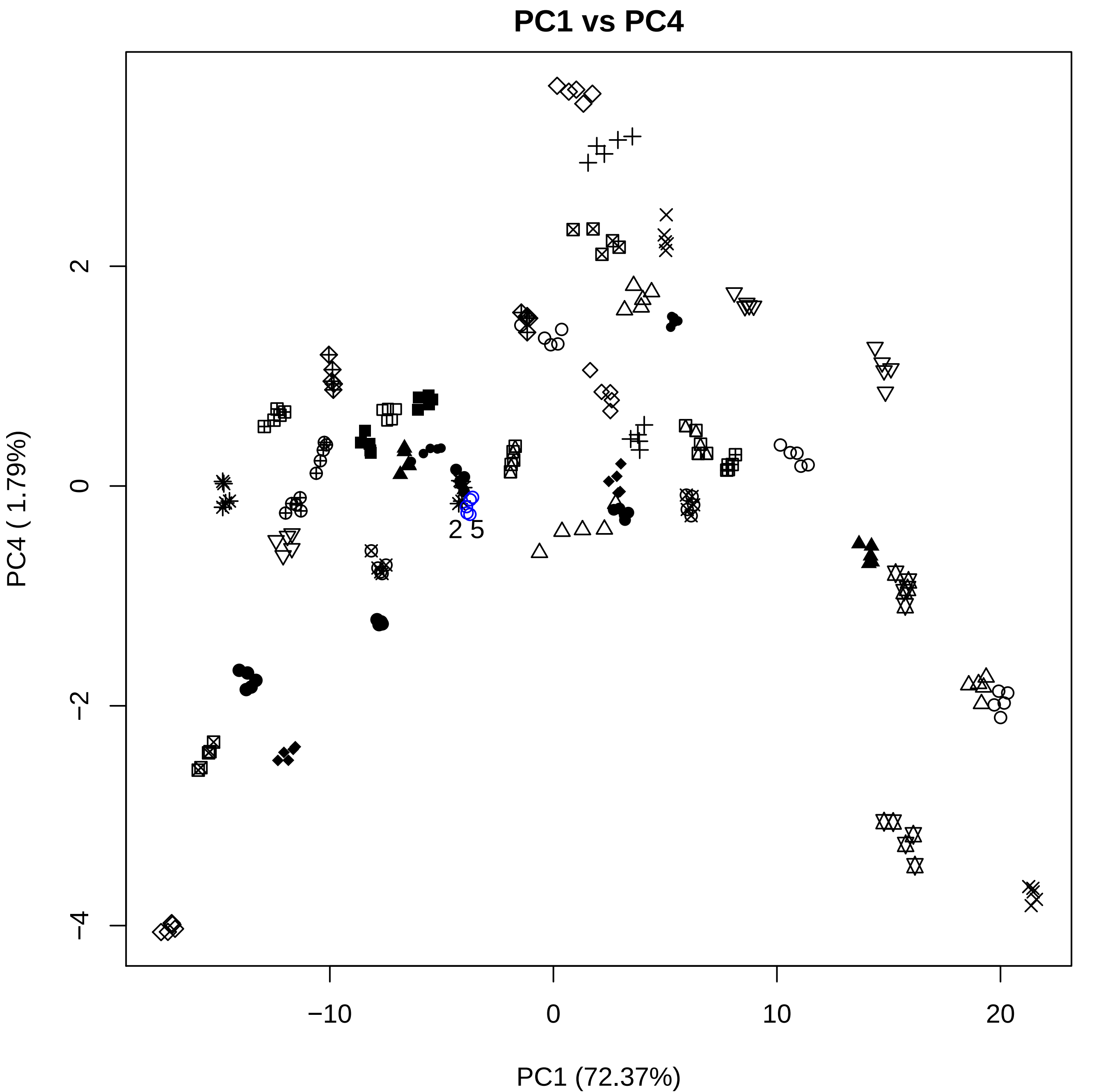
PC1 vs PC4
PC4 ( 1.79%)
PC1 (72.37%)
2
0
−2
−4
−10
0
10
20
2 5

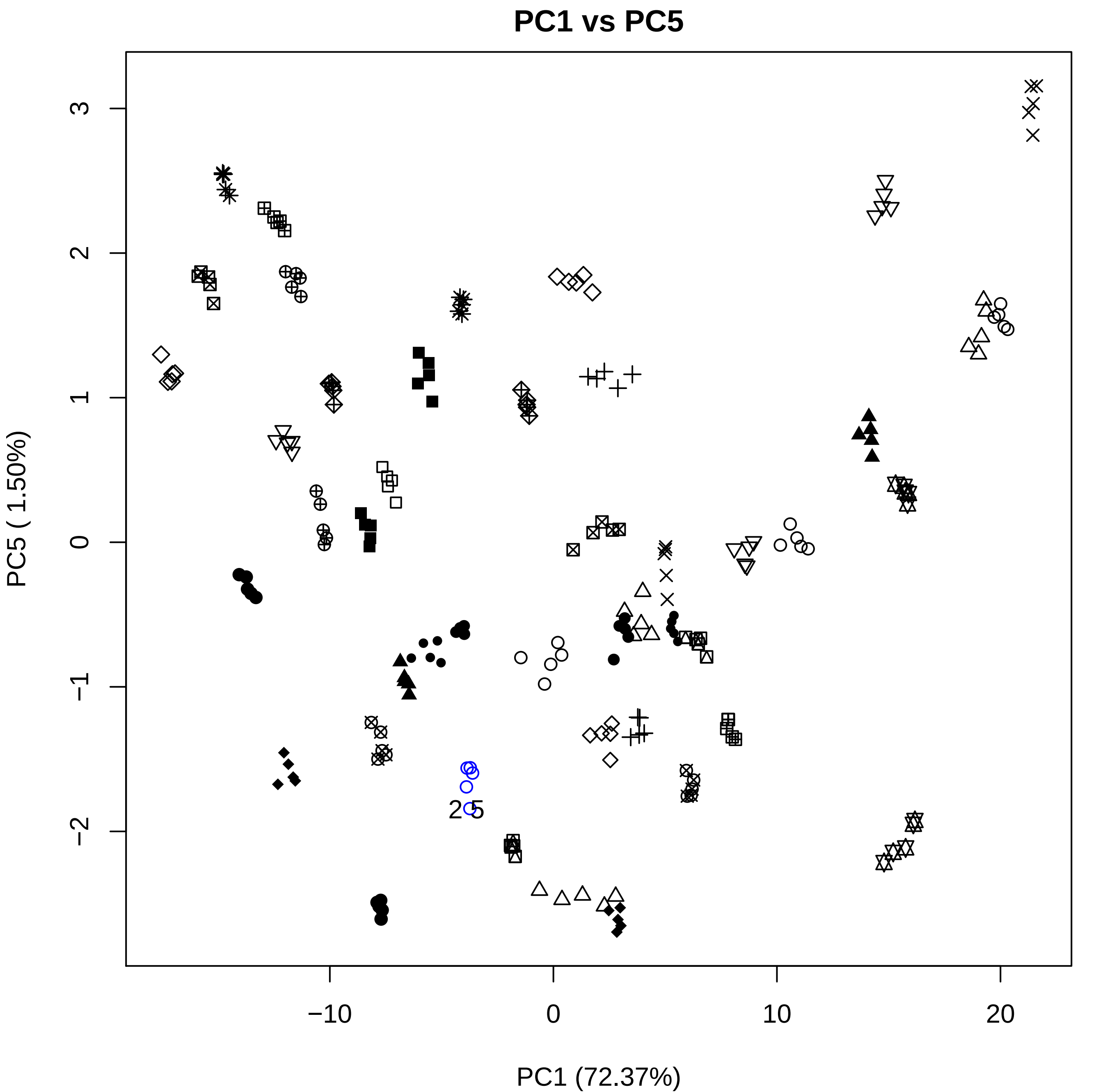
PC1 vs PC5
PC5 ( 1.50%)
PC1 (72.37%)
3
2
1
0
−1
−2
−10
0
10
20
25

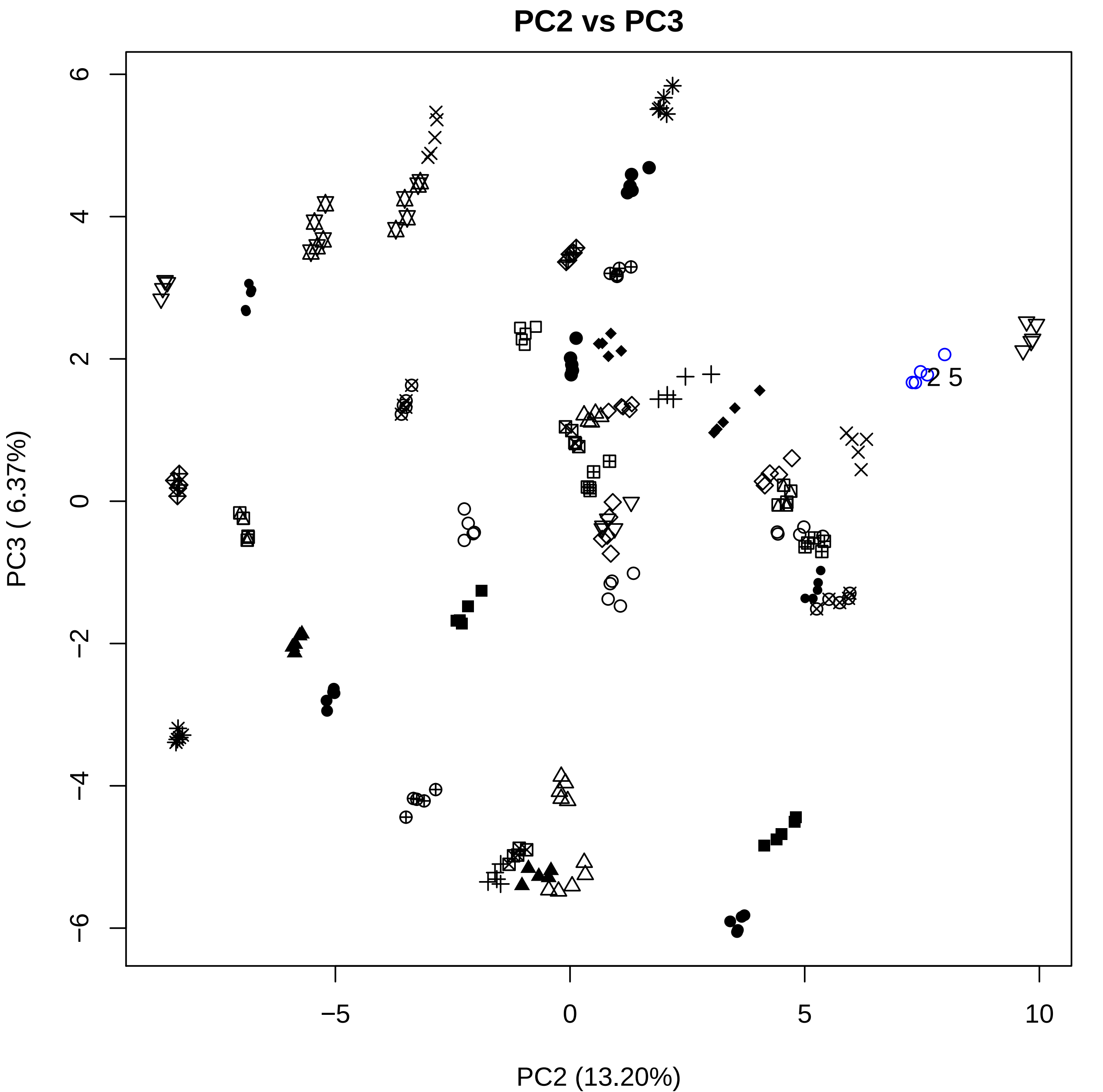

PC2 vs PC3
PC3 ( 6.37%)
PC2 (13.20%)
6
4
2
0
−2
−4
−6
−5
0
5
10
2 5

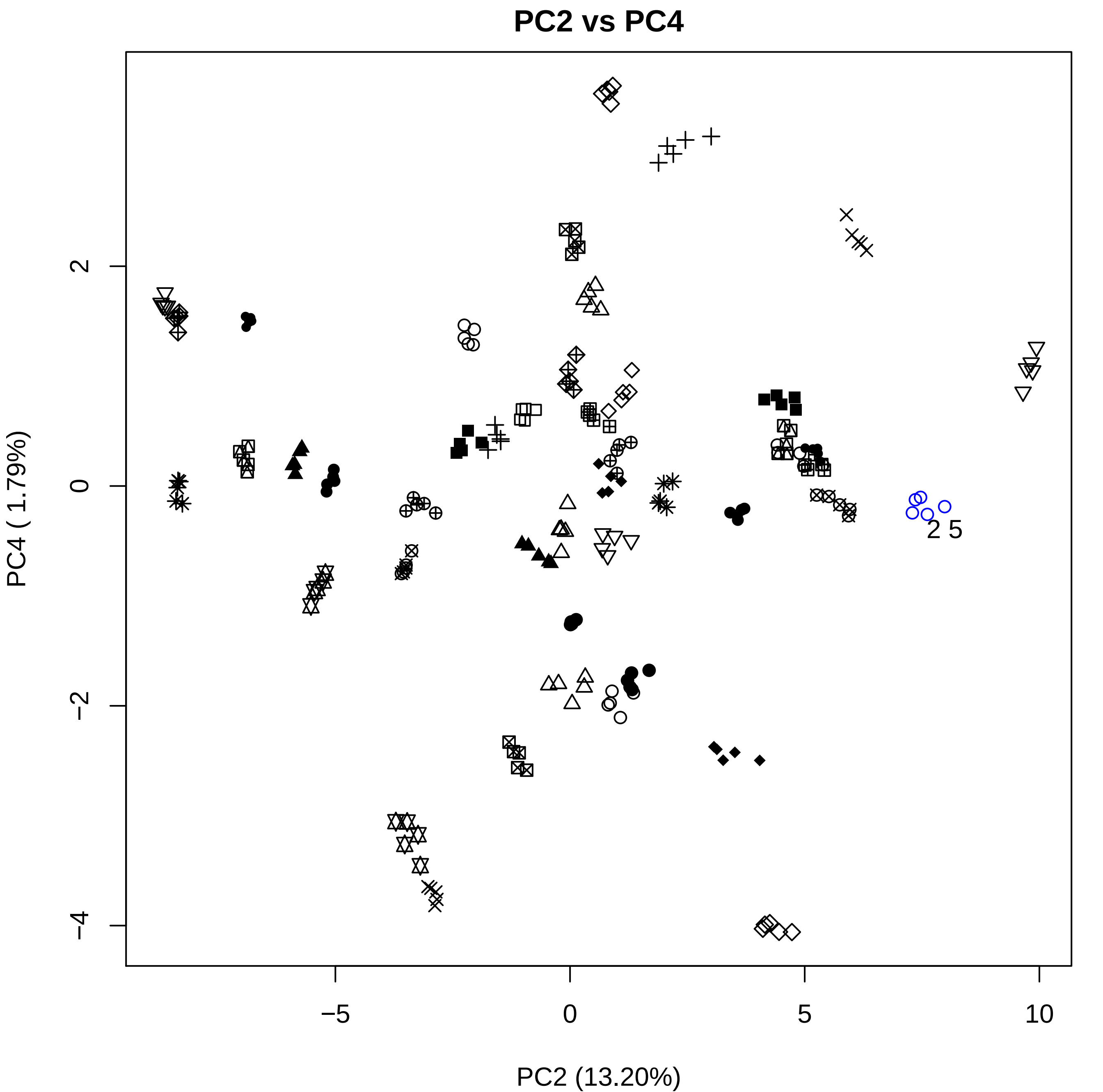
PC2 vs PC4
PC4 ( 1.79%)
PC2 (13.20%)
2
0
−2
−4
−5
0
5
10
2 5

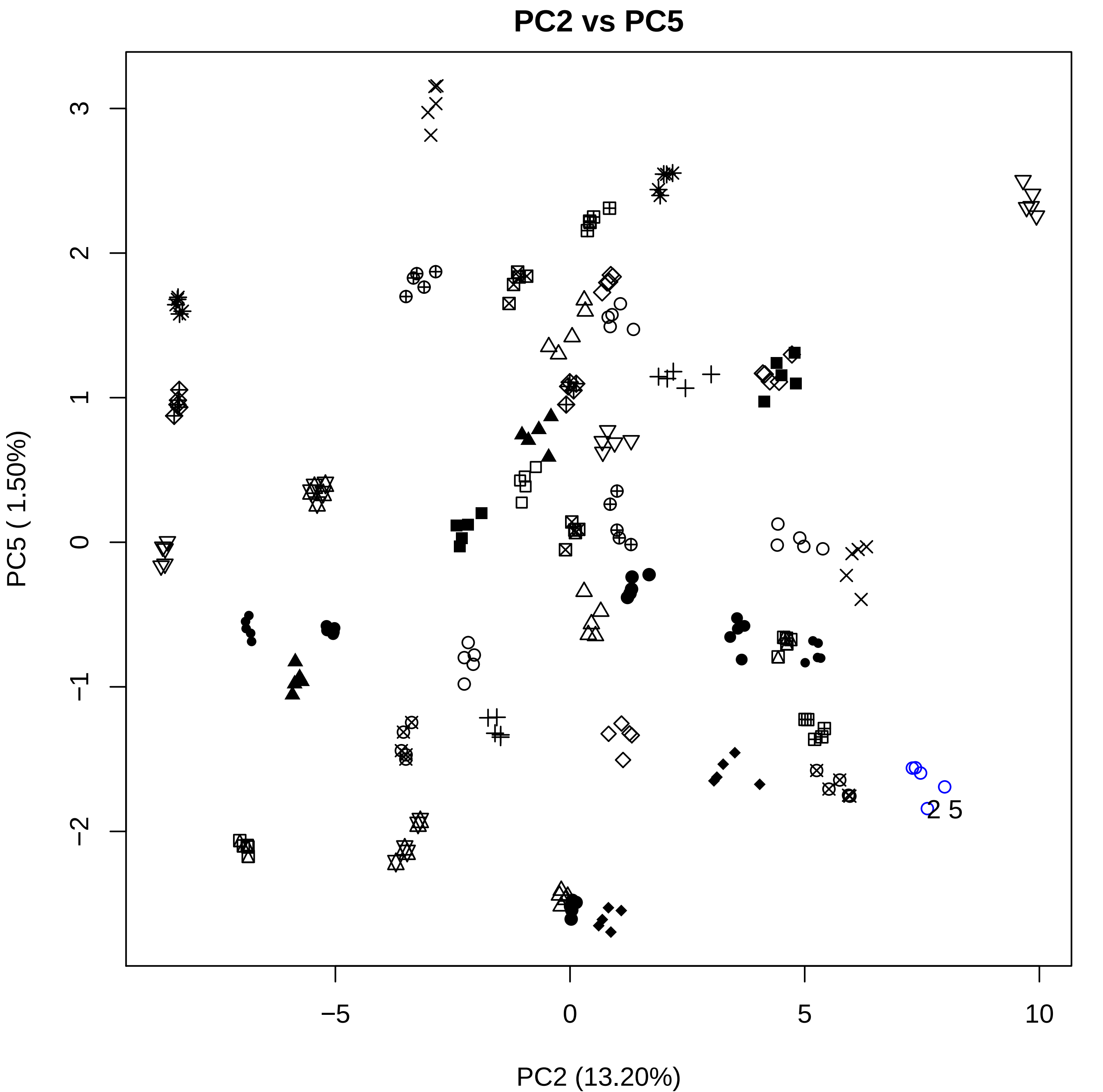
PC2 vs PC5
PC5 ( 1.50%)
PC2 (13.20%)
3
2
1
0
−1
−2
−5
0
5
10
2 5

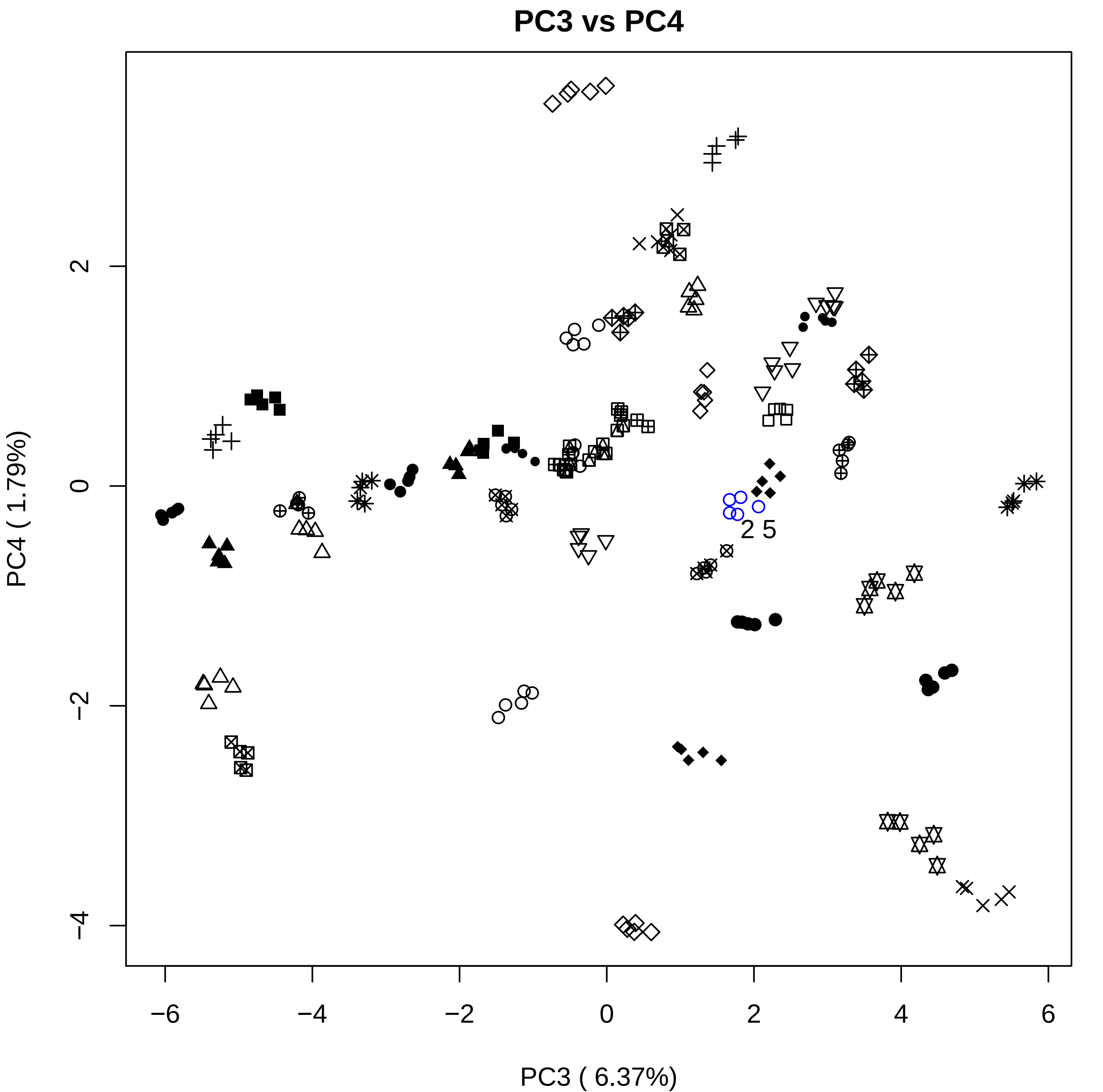
PC3 vs PC4
PC4 ( 1.79%)
PC3 ( 6.37%)
2
0
−2
−4
−6
−4
−2
0
2
4
6
2 5

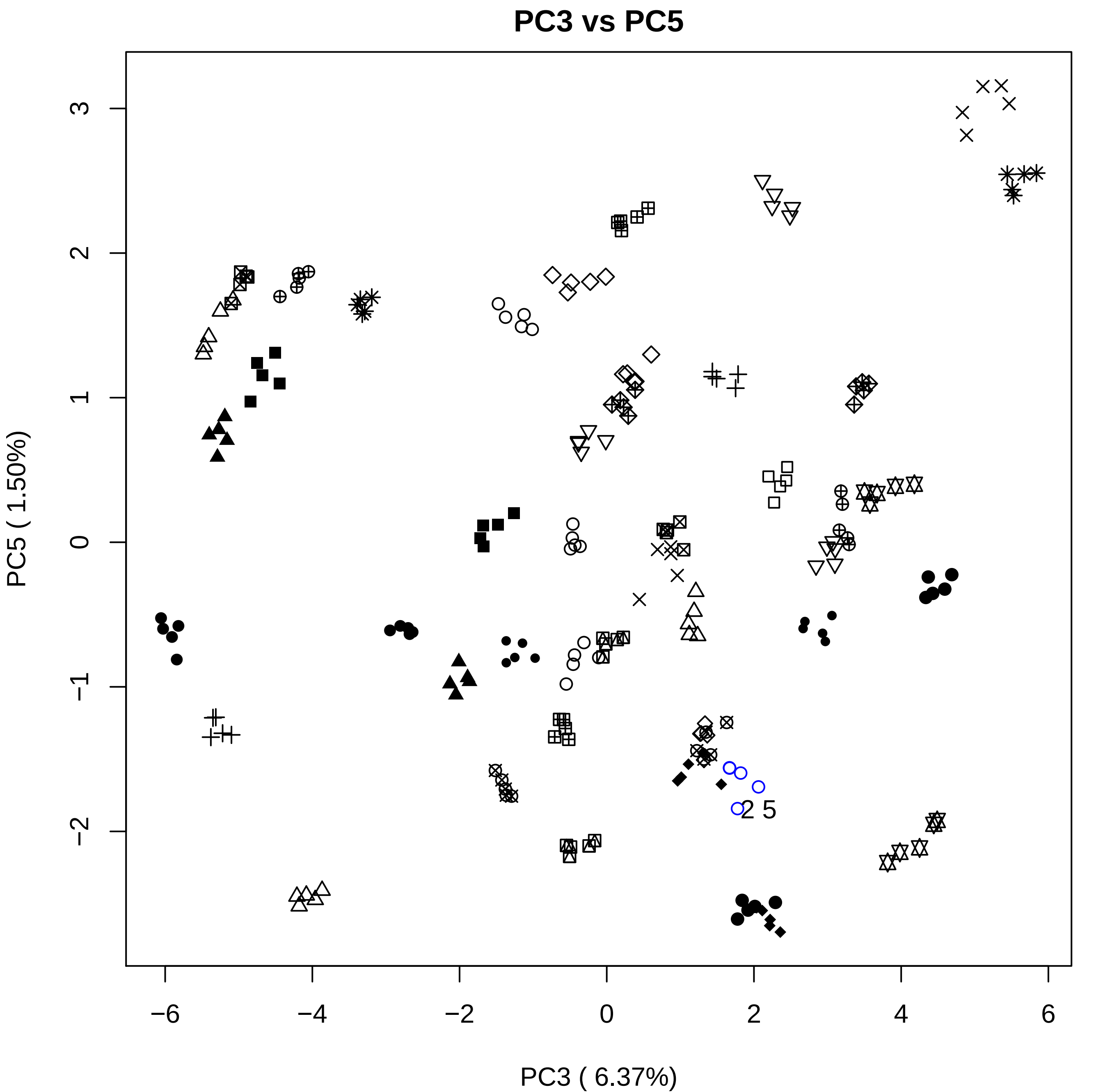
PC3 vs PC5
PC5 ( 1.50%)
PC3 ( 6.37%)
−6
−4
−2
0
2
4
6
3
2
1
0
−1
−2
2 5

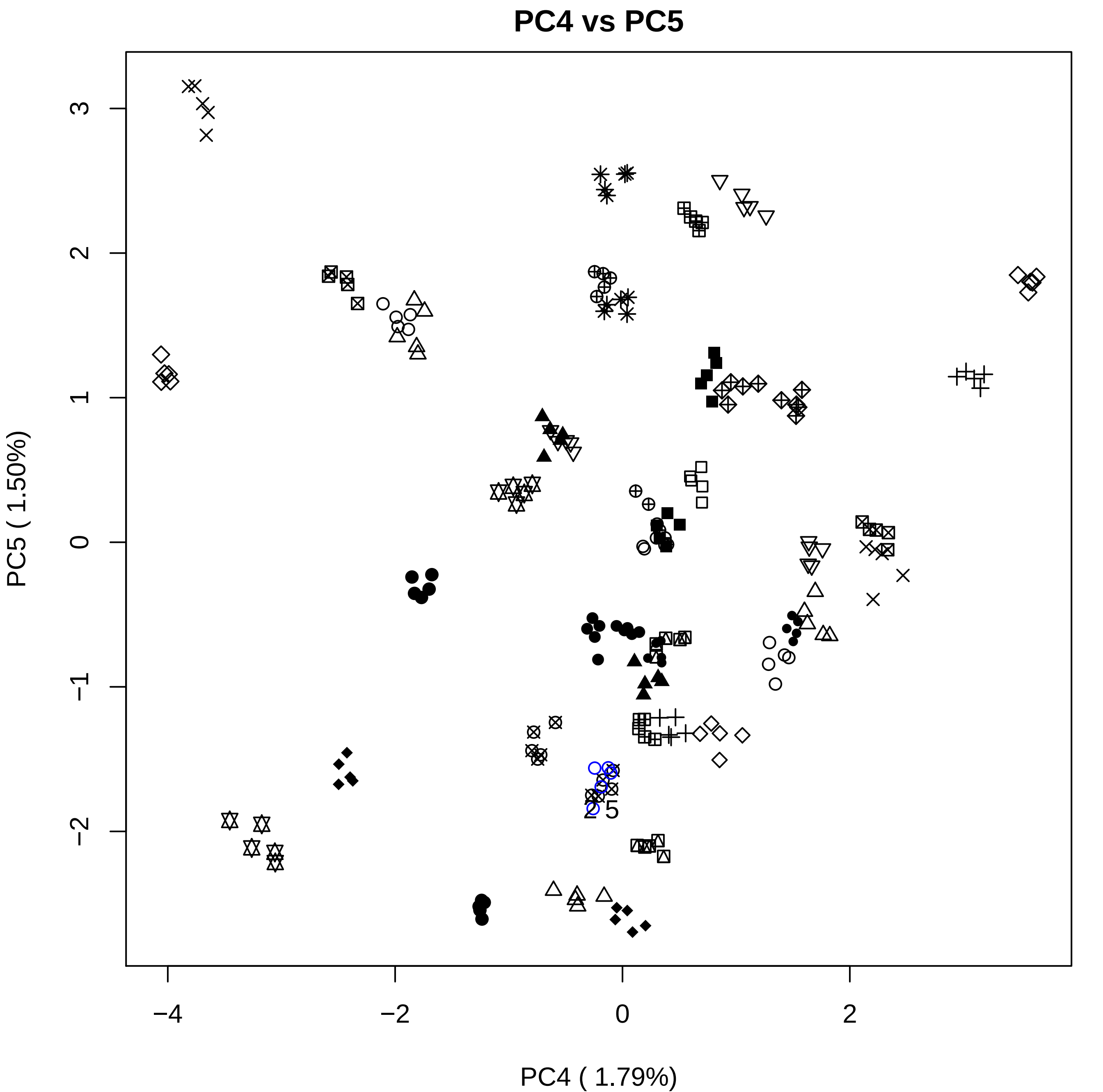
PC4 vs PC5
PC5 ( 1.50%)
PC4 ( 1.79%)
3
2
1
0
−1
−2
−4
−2
0
2
2
5

### PCA of 5%  OSCS vs PMH

Scaled Matrix PCAed

7th order polynomial baseline correction performed

Raw spectra smoothed to 21 Neighbours, 2nd order PN, Savtisky Golay

Normalisation between 0 and 100

D2 spectra smoothed to 41 Neighbours, 2nd order PN, Savtisky Golay

Regions Analysed: 3601−2501 and 2001−700cm−1

Samples Analysed: 1 Contam's 0 OSCS's 45 PMH's

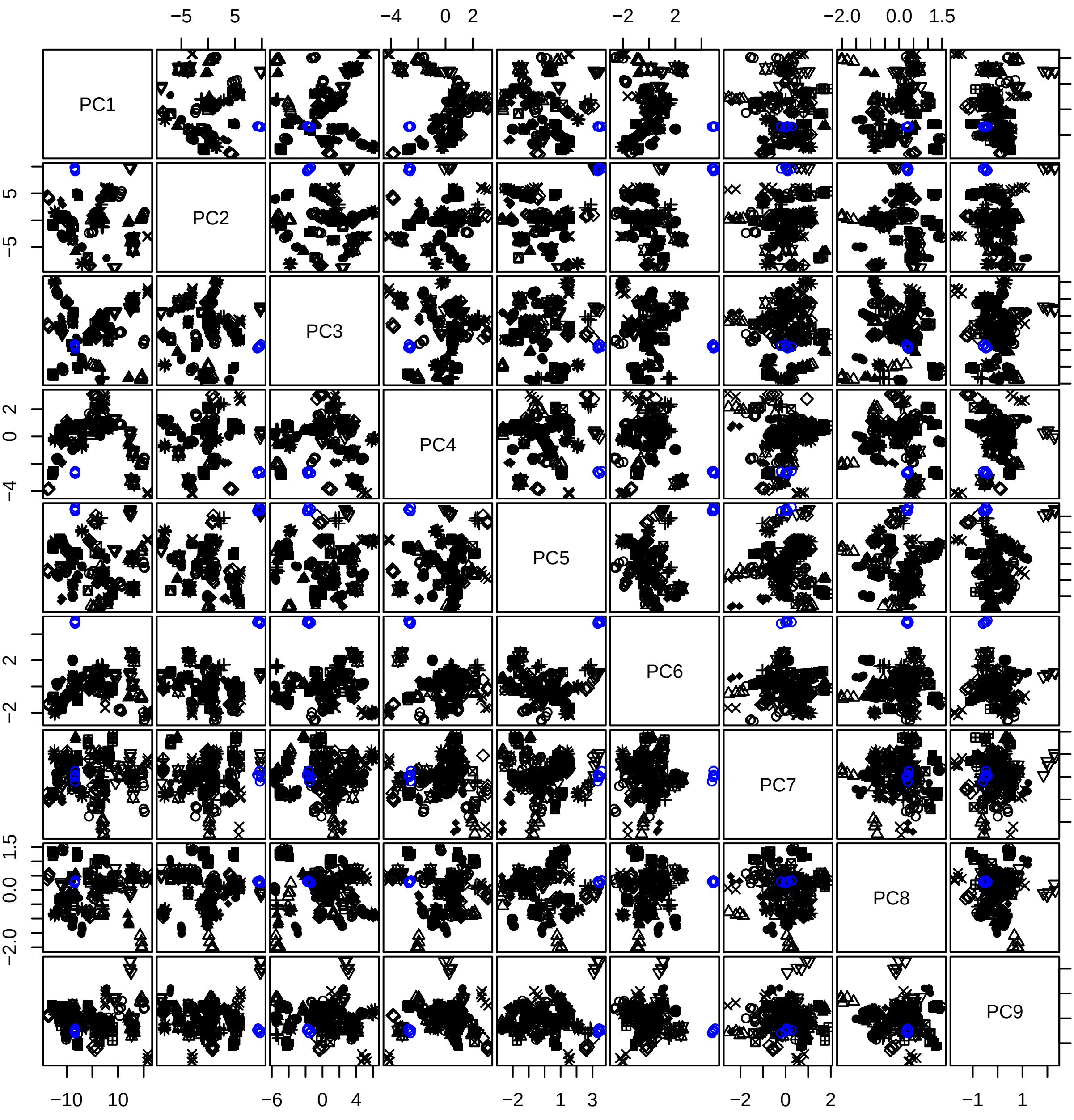
PC1
PC2
PC3
PC4
PC5
PC6
PC7
PC8
PC9
−5 5
−4 0 2
−2 2
−2.0 0.0 1.5
10 −10
5 −5
4 0 −6
2 0 −4
3 1 −2
2 −2
2 0 −2
1.5 0.0 −2.0
1 −1
−10 10
−6 0 4
−2 1 3
−2 0 2
−1 1

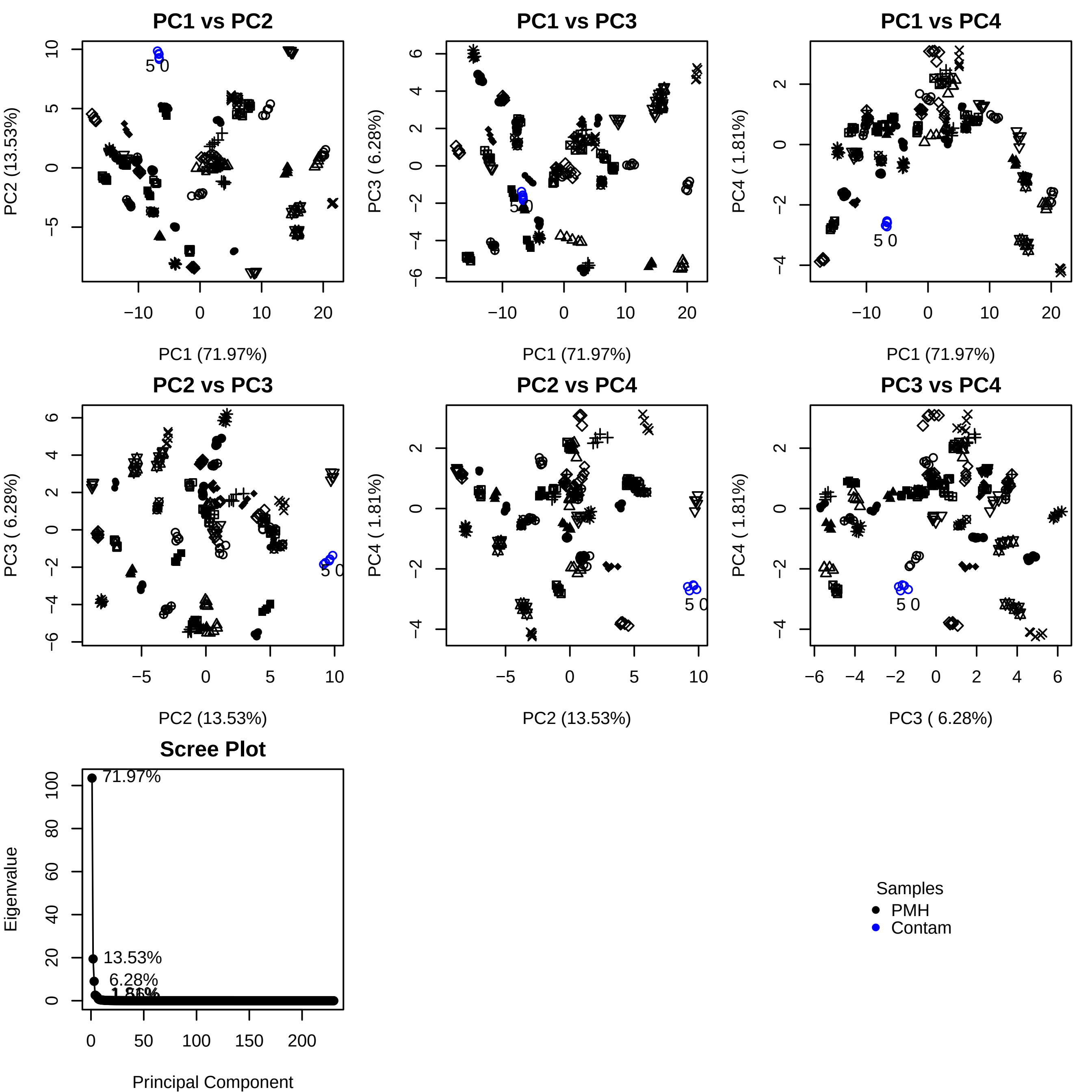

PC1 vs PC2
PC1 vs PC3
PC1 vs PC4
PC2 vs PC3
PC2 vs PC4
PC3 vs PC4
Scree Plot
PC1 (71.97%)
PC2 (13.53%)
PC3 ( 6.28%)
PC4 ( 1.81%)
Eigenvalue
Principal Component
71.97%
13.53%
6.28%
50
Samples
PMH
Contam

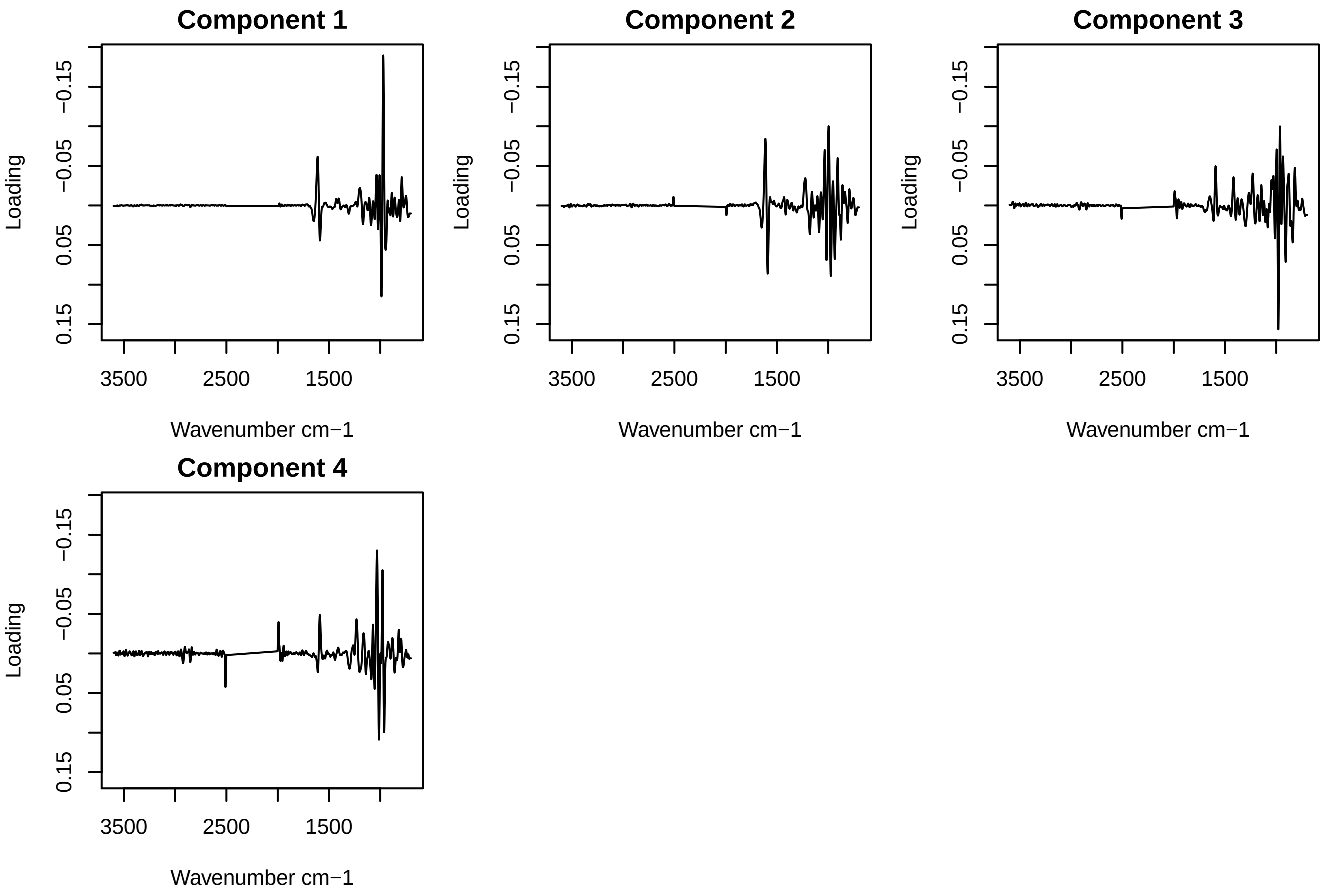
Component 1
Component 2
Component 3
Component 4
Loading
−0.15
−0.05
0.05
0.15
3500
2500
1500
Wavenumber cm−1

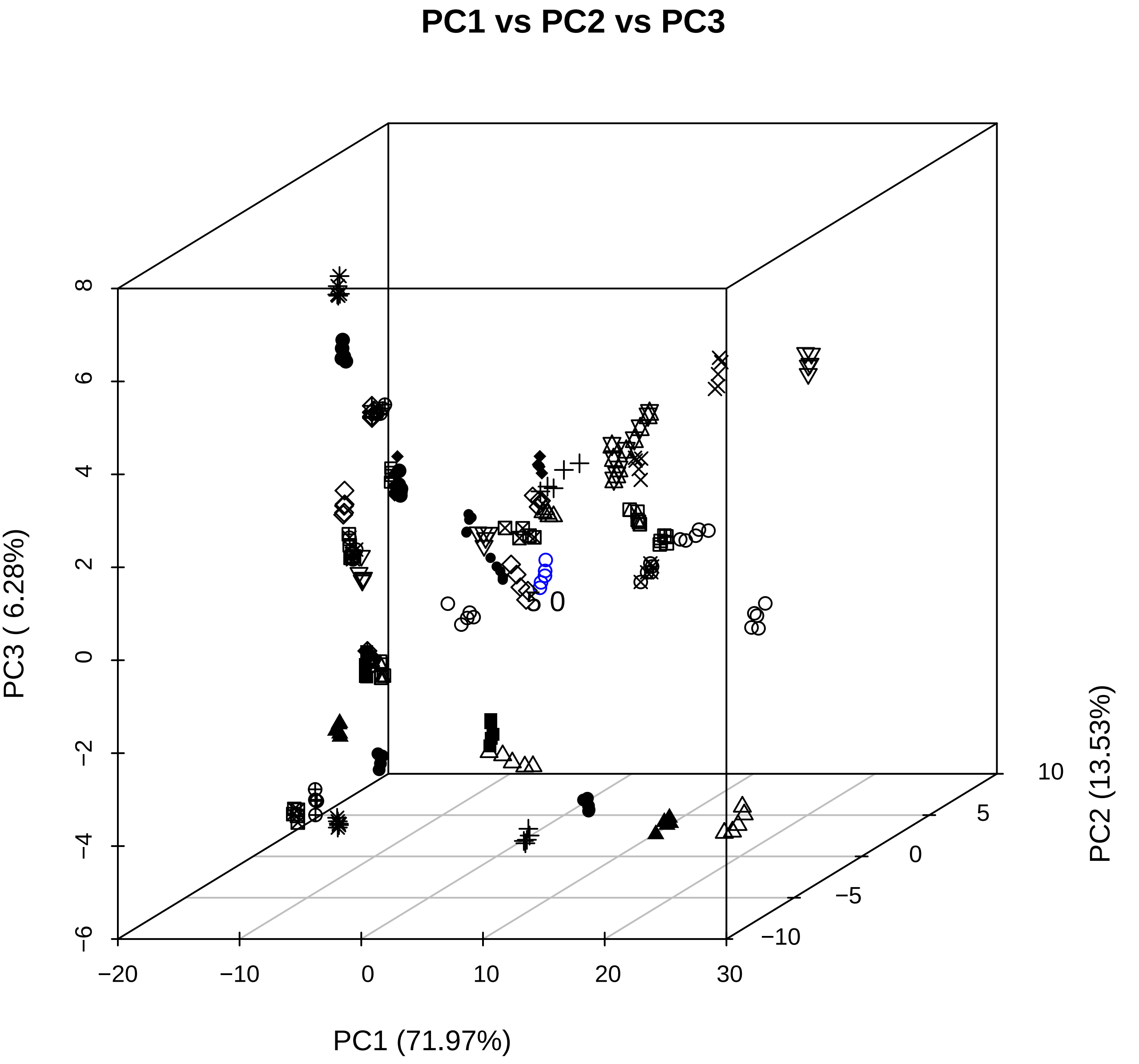
PC1 vs PC2 vs PC3
PC1 (71.97%)
PC2 (13.53%)
PC3 ( 6.28%)
−20
−10
0
10
20
30
−10
−5
0
5
10
−6
−4
−2
0
2
4
6
8

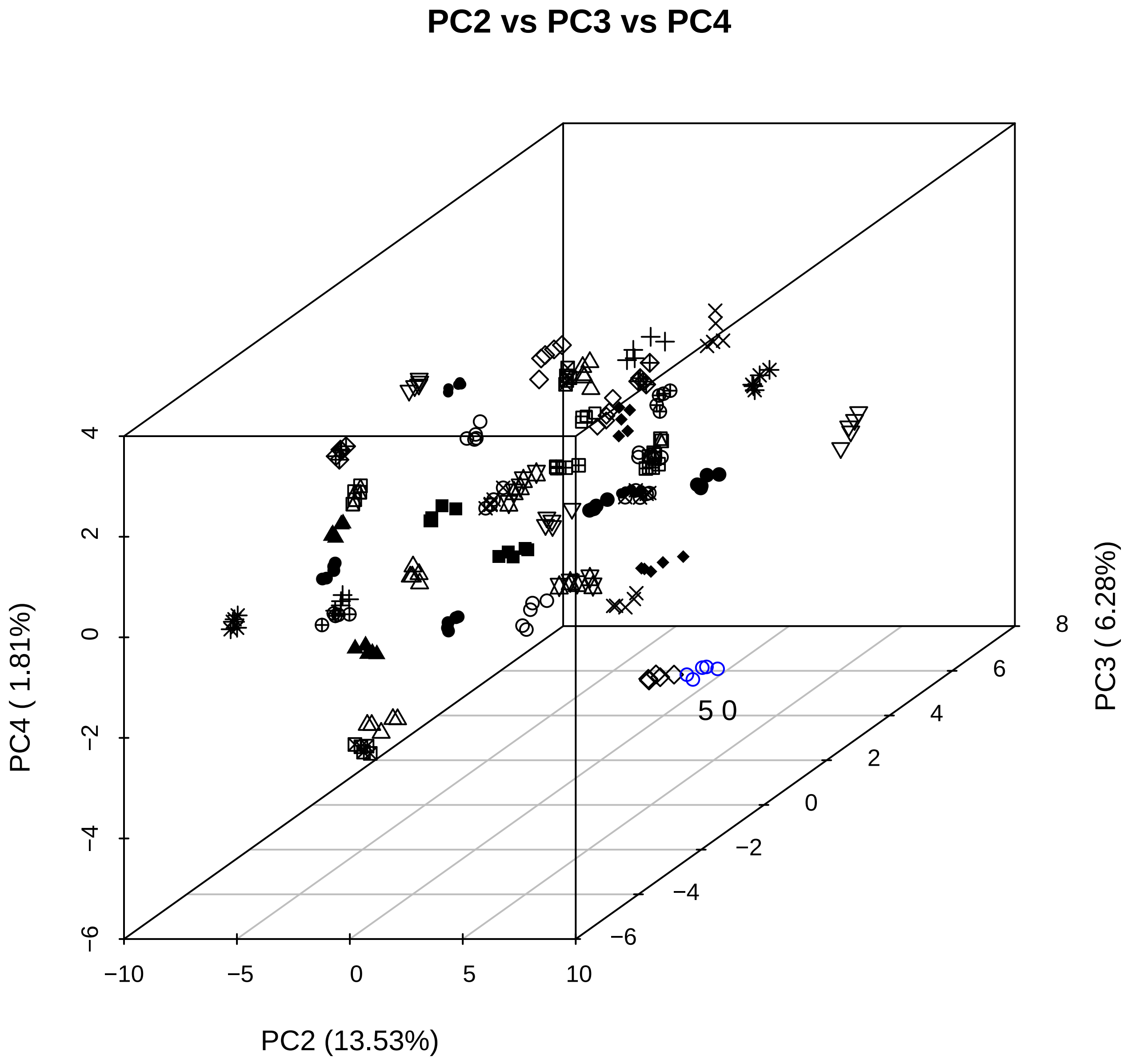

PC2 vs PC3 vs PC4
PC2 (13.53%)
PC3 ( 6.28%)
PC4 ( 1.81%)
−10
−5
0
5
10
−6
−4
−2
0
2
4
6
8
50

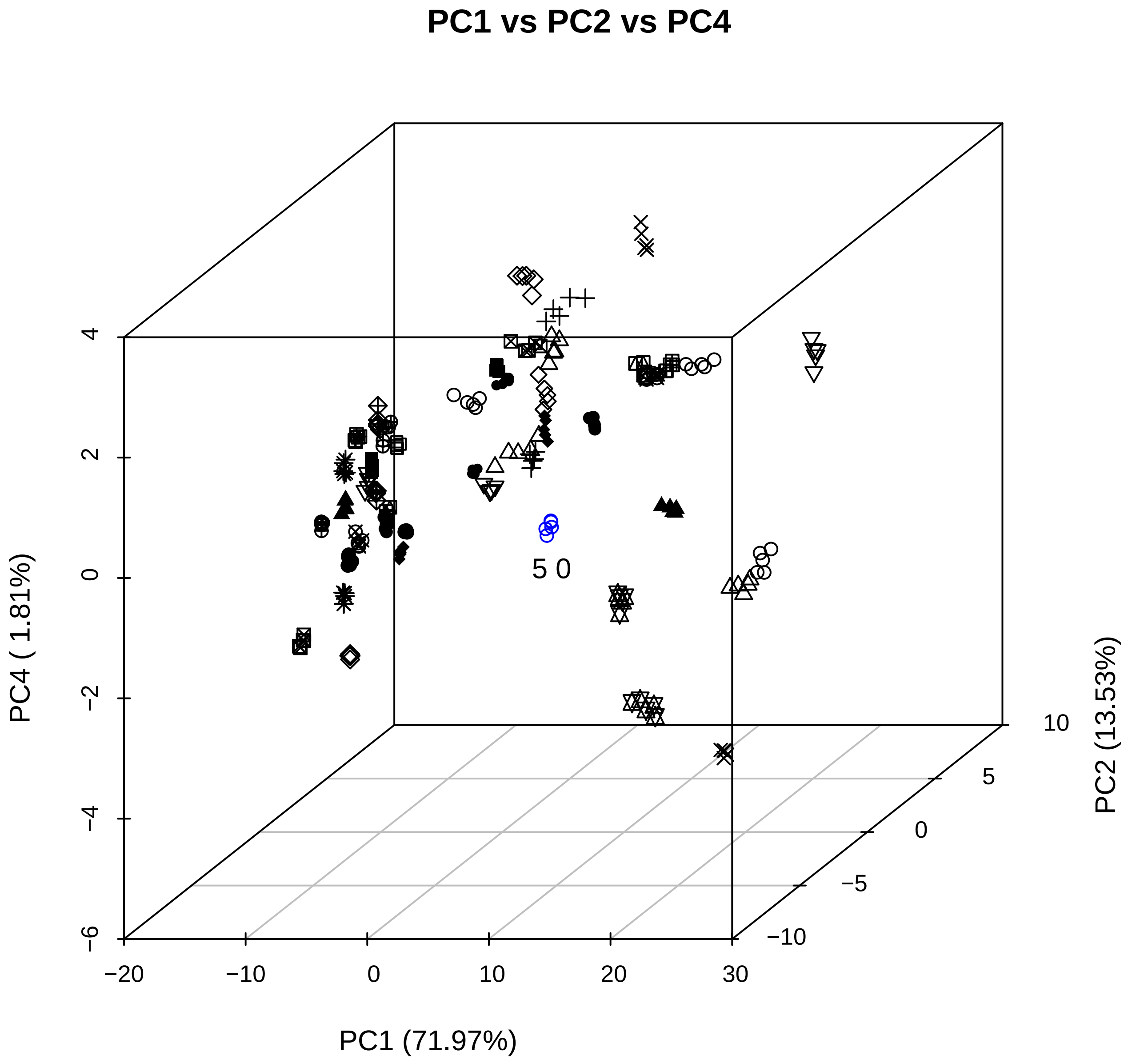

PC1 vs PC2 vs PC4
5 0
PC4 ( 1.81%)
4
2
0
−2
−4
−6
−20
−10
0
10
20
30
PC1 (71.97%)
PC2 (13.53%)
10
5
0
−5
−10

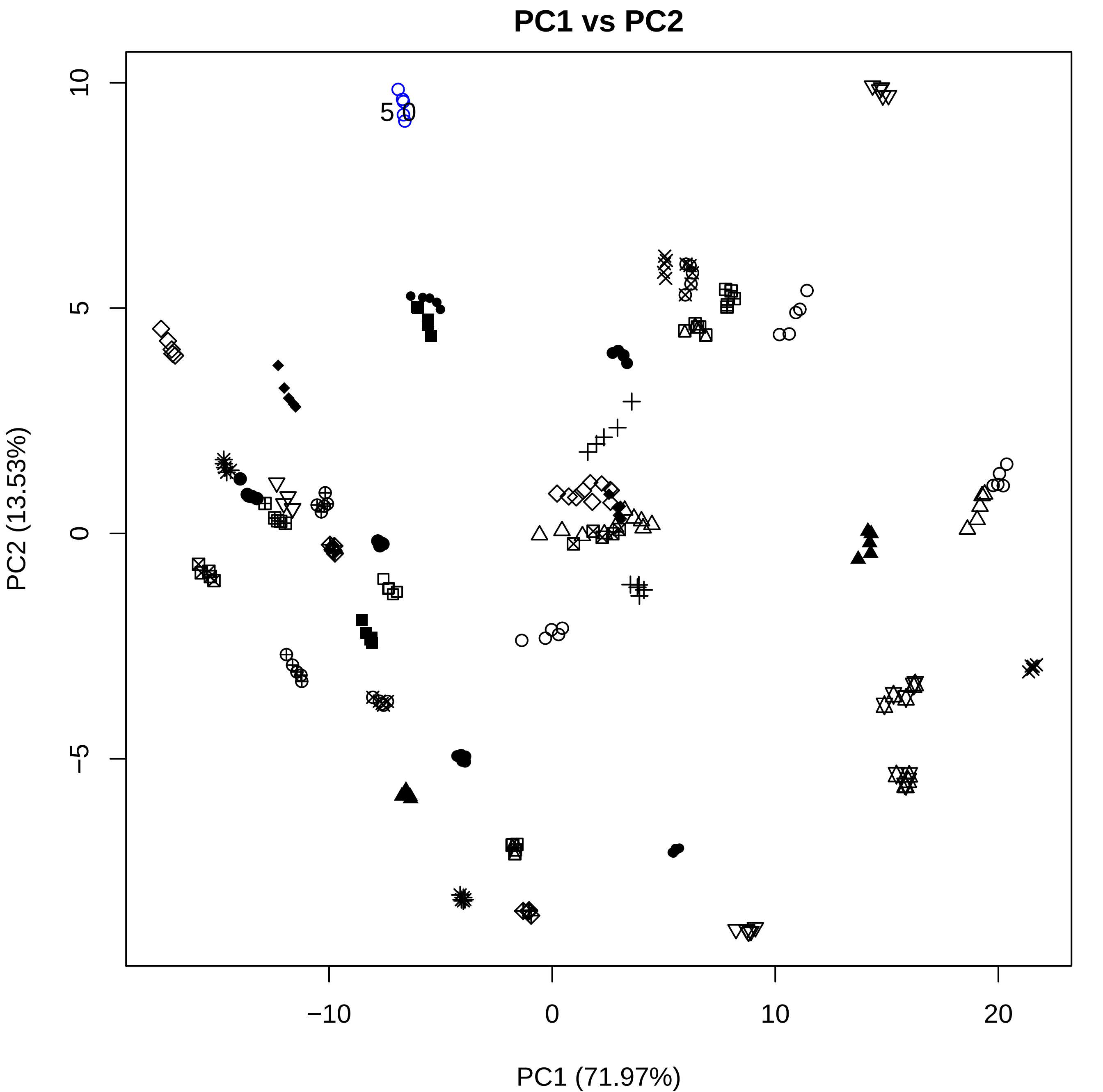

PC1 vs PC2
50
PC1 (71.97%)
PC2 (13.53%)
−10
0
10
20
10
5
0
−5

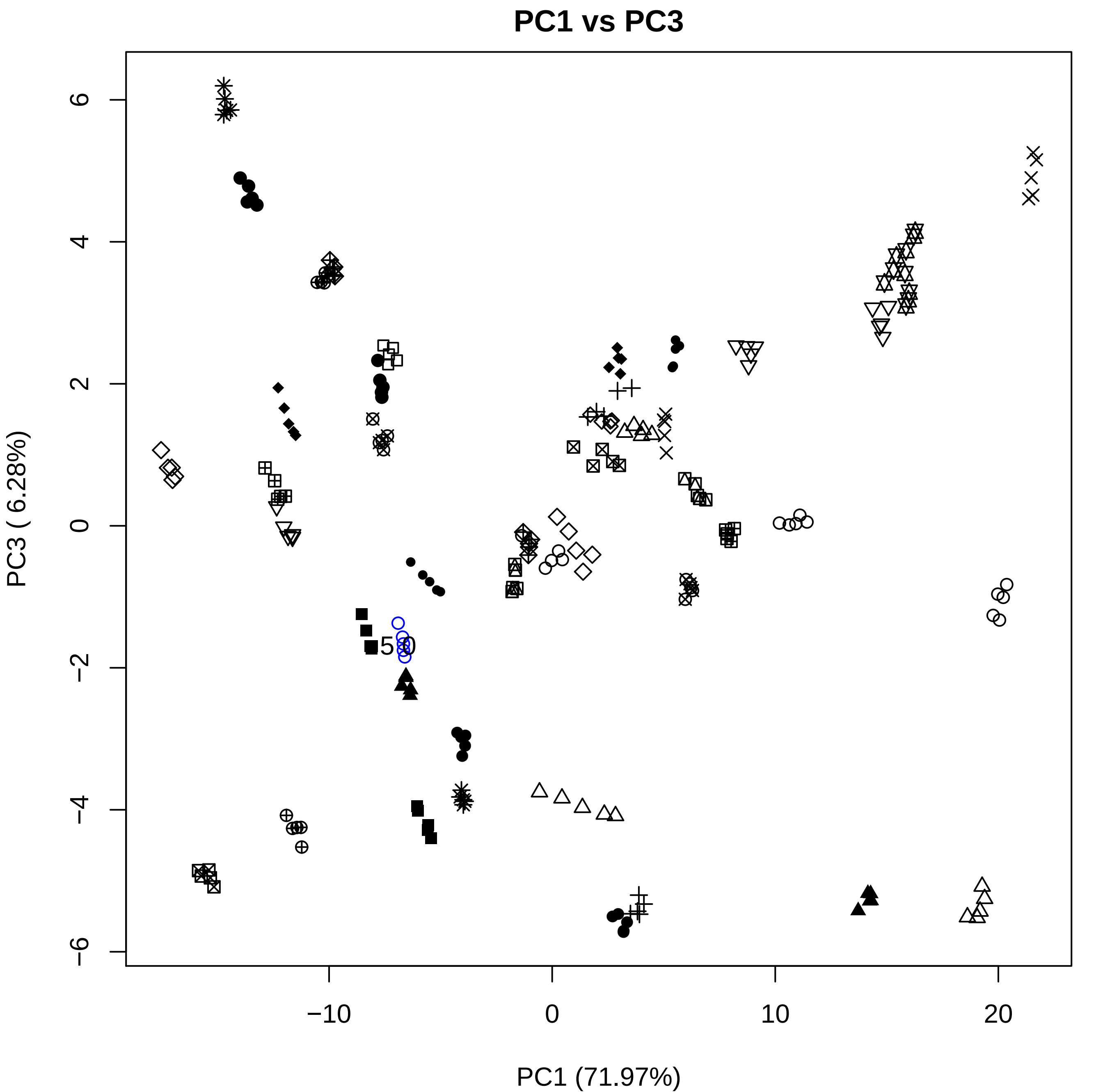
PC1 vs PC3
PC3 ( 6.28%)
PC1 (71.97%)
6
4
2
0
−2
−4
−6
−10
0
10
20
50

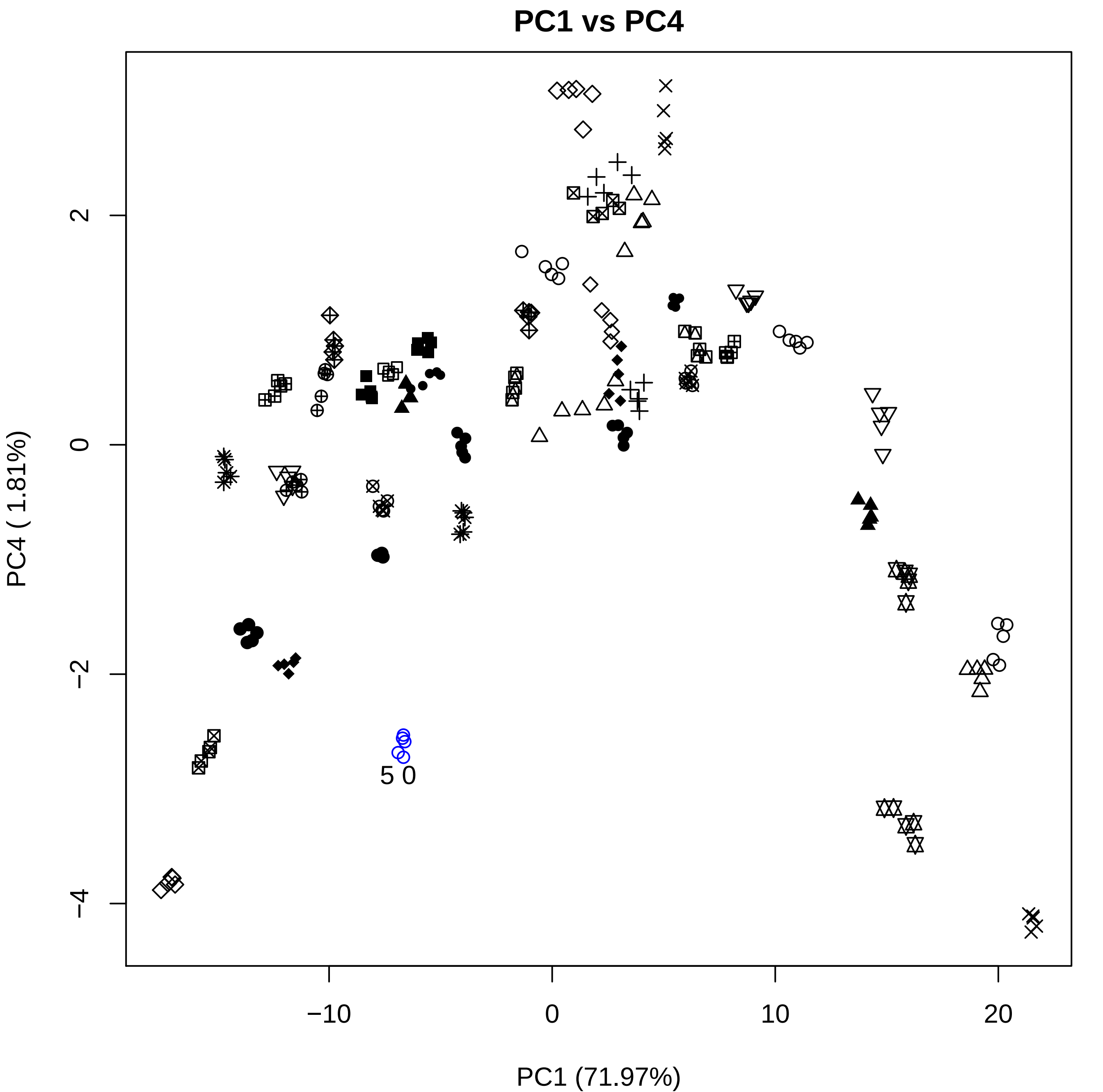
PC1 vs PC4
PC4 ( 1.81%)
PC1 (71.97%)
2
0
−2
−4
−10
0
10
20
5 0

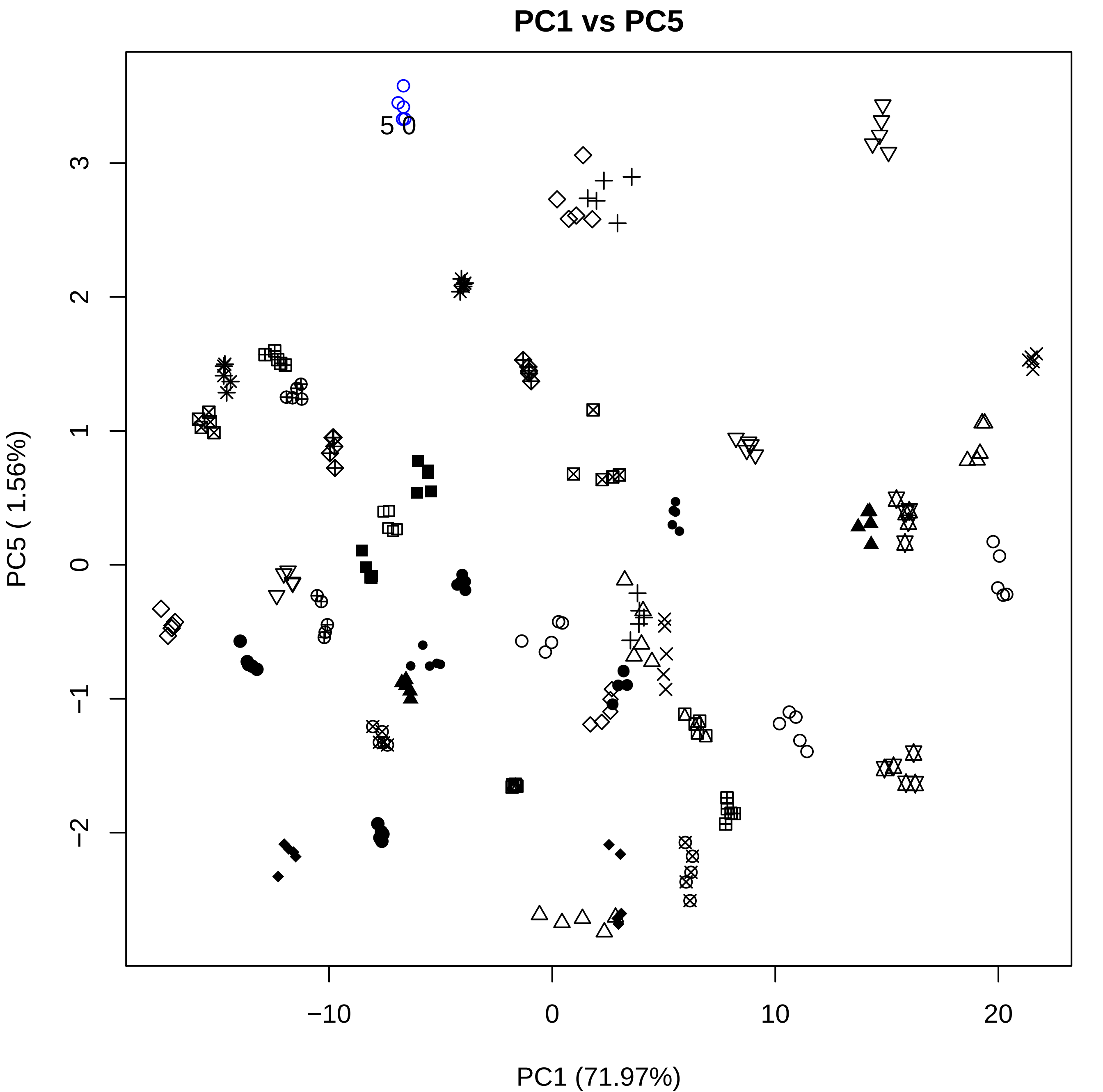

PC1 vs PC5
50
PC5 ( 1.56%)
PC1 (71.97%)
3
2
1
0
−1
−2
−10
0
10
20

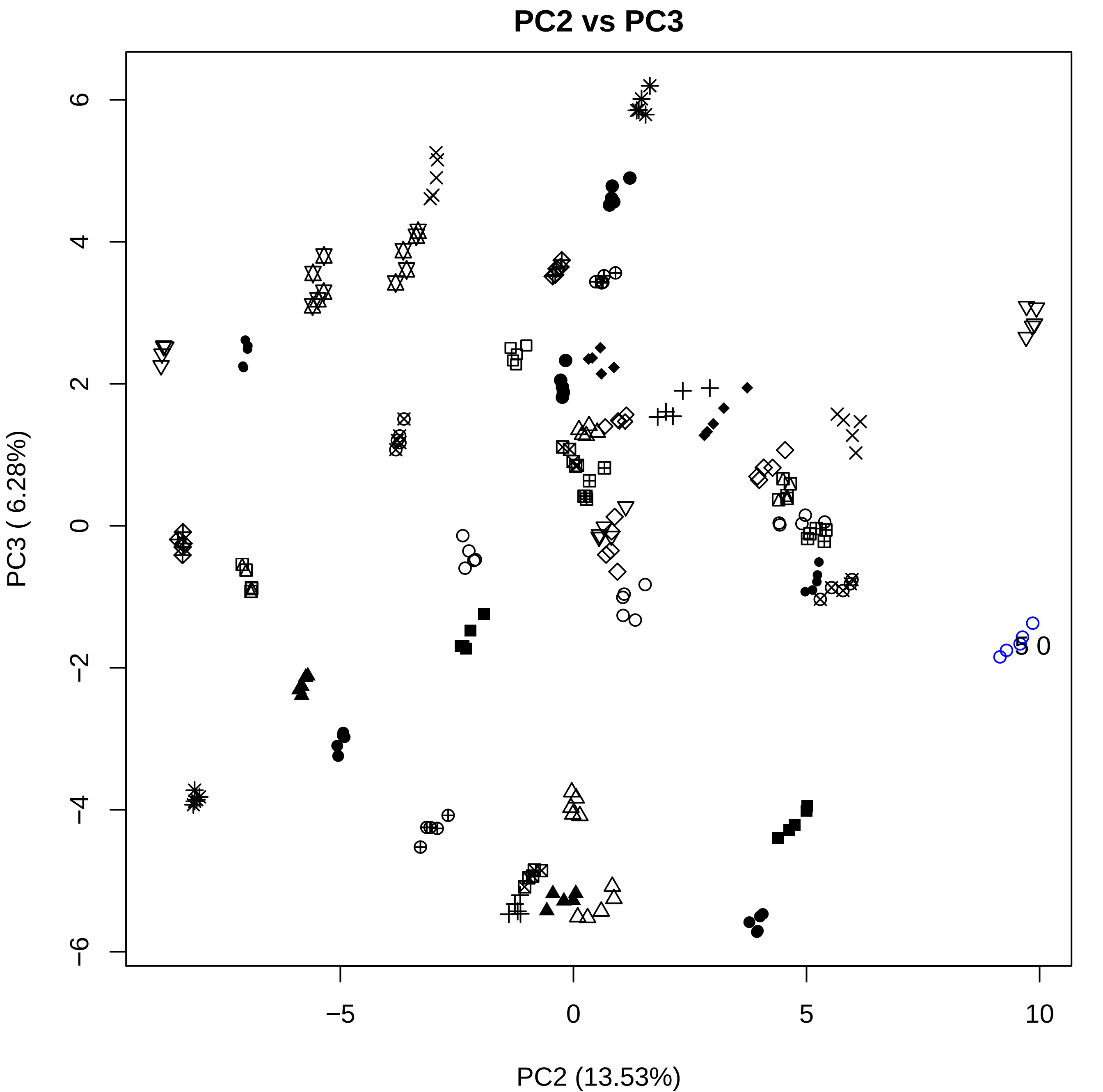

PC2 vs PC3
PC3 ( 6.28%)
PC2 (13.53%)
6
4
2
0
−2
−4
−6
−5
0
5
10
5 0

## PC2 vs PC4

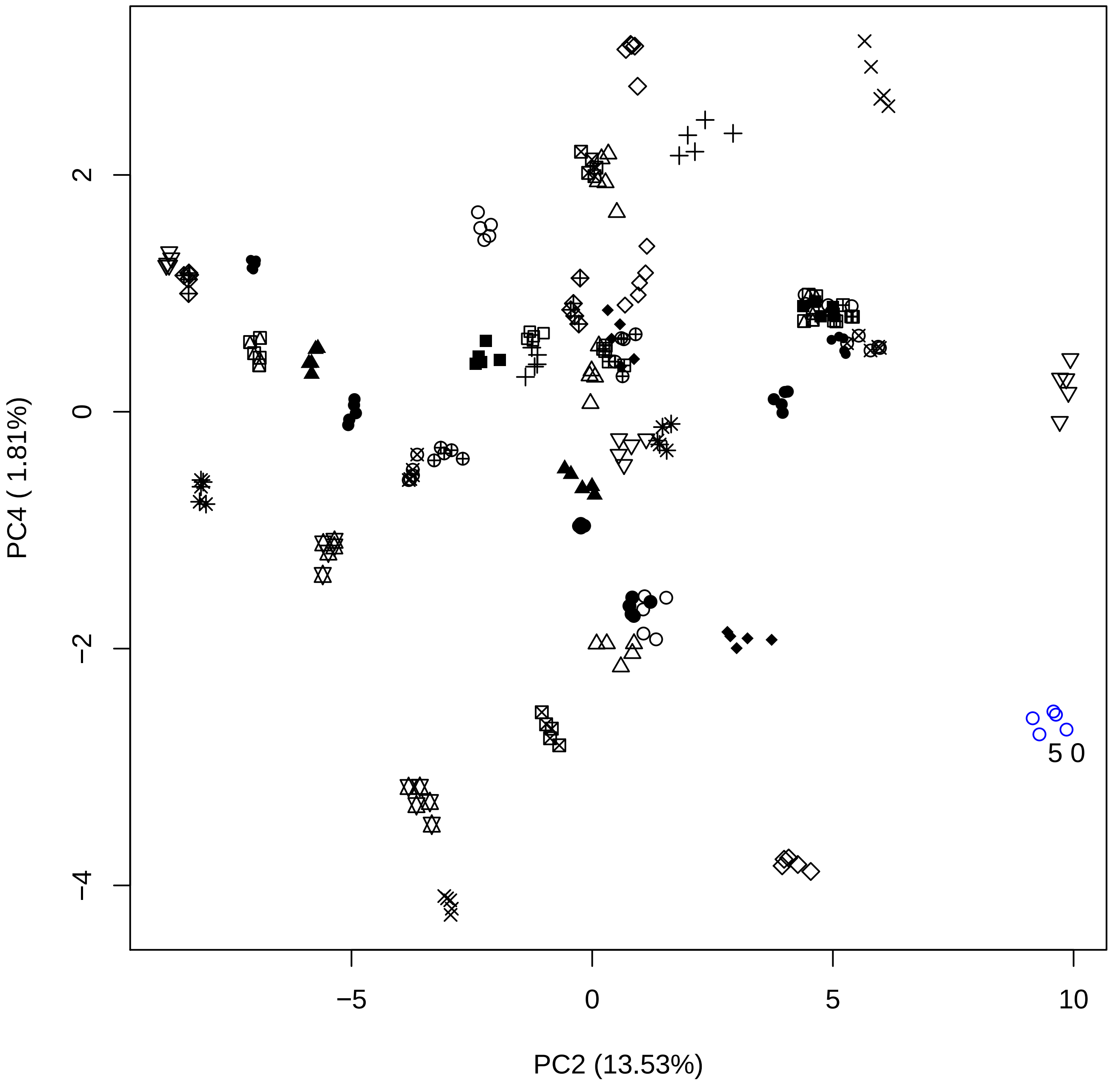

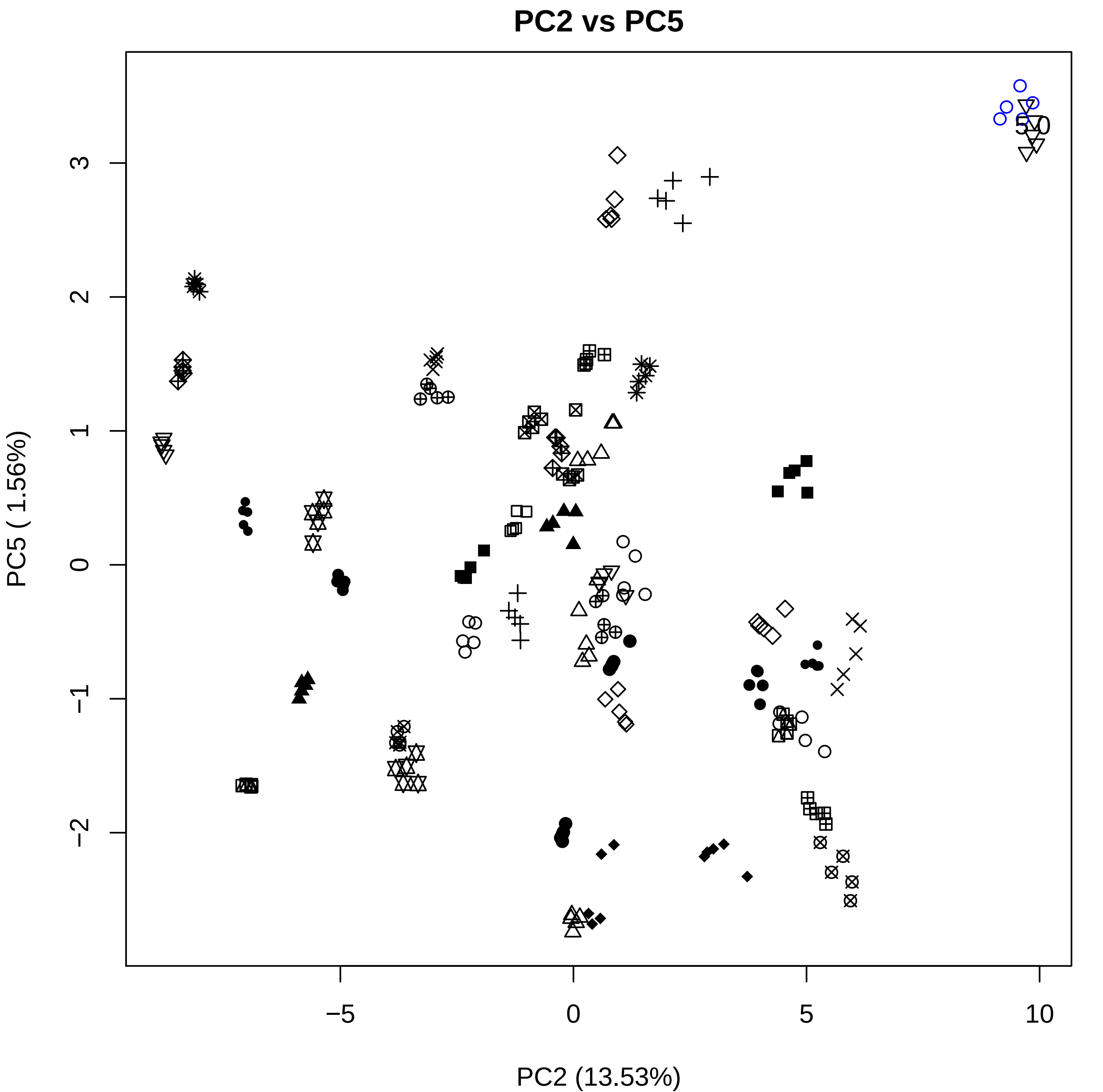

PC2 vs PC5
PC5 ( 1.56%)
PC2 (13.53%)
−5
0
5
10
−2
−1
0
1
2
3
50

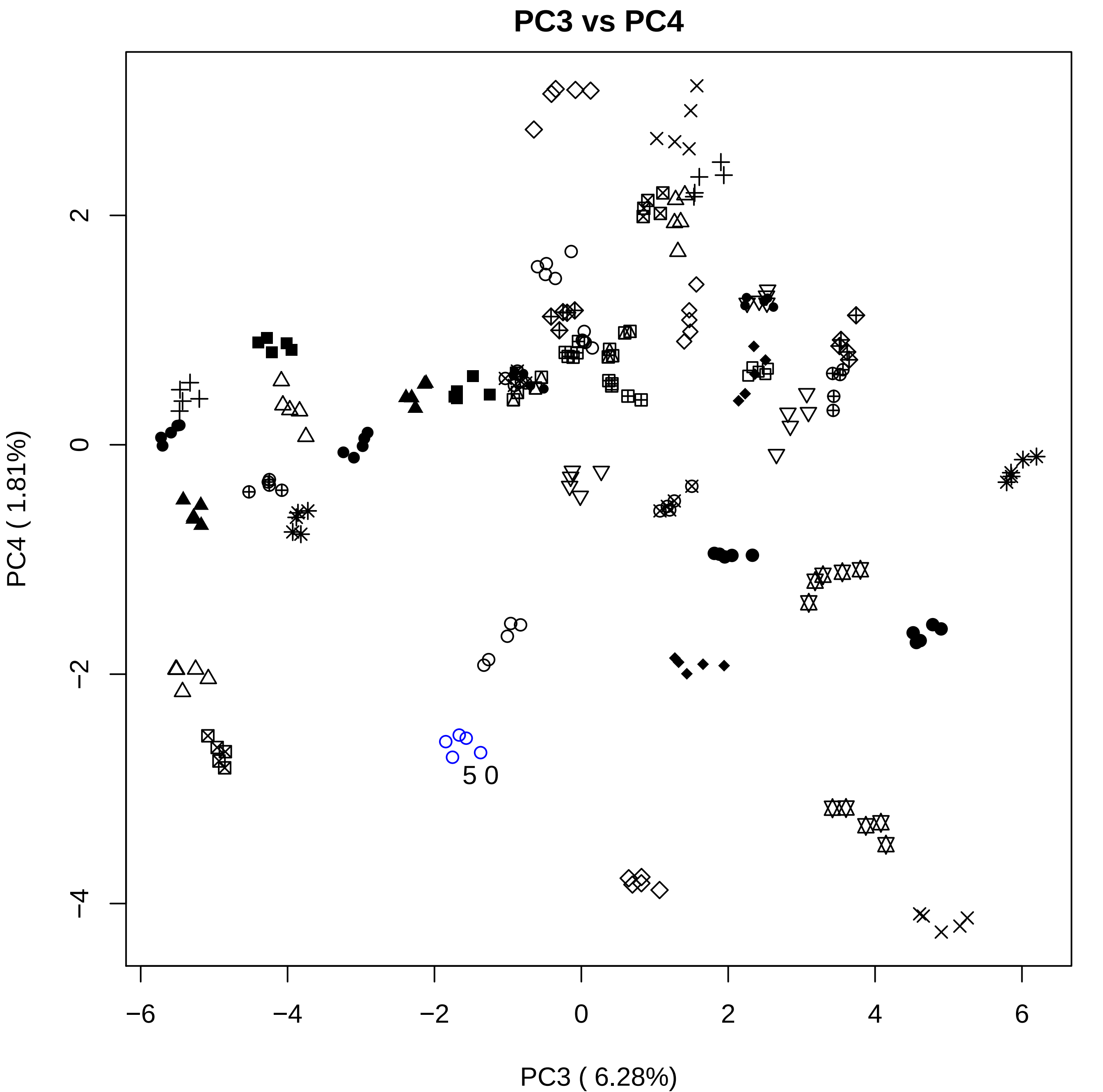
PC3 vs PC4
PC4 ( 1.81%)
PC3 ( 6.28%)
−6
−4
−2
0
2
4
6
50

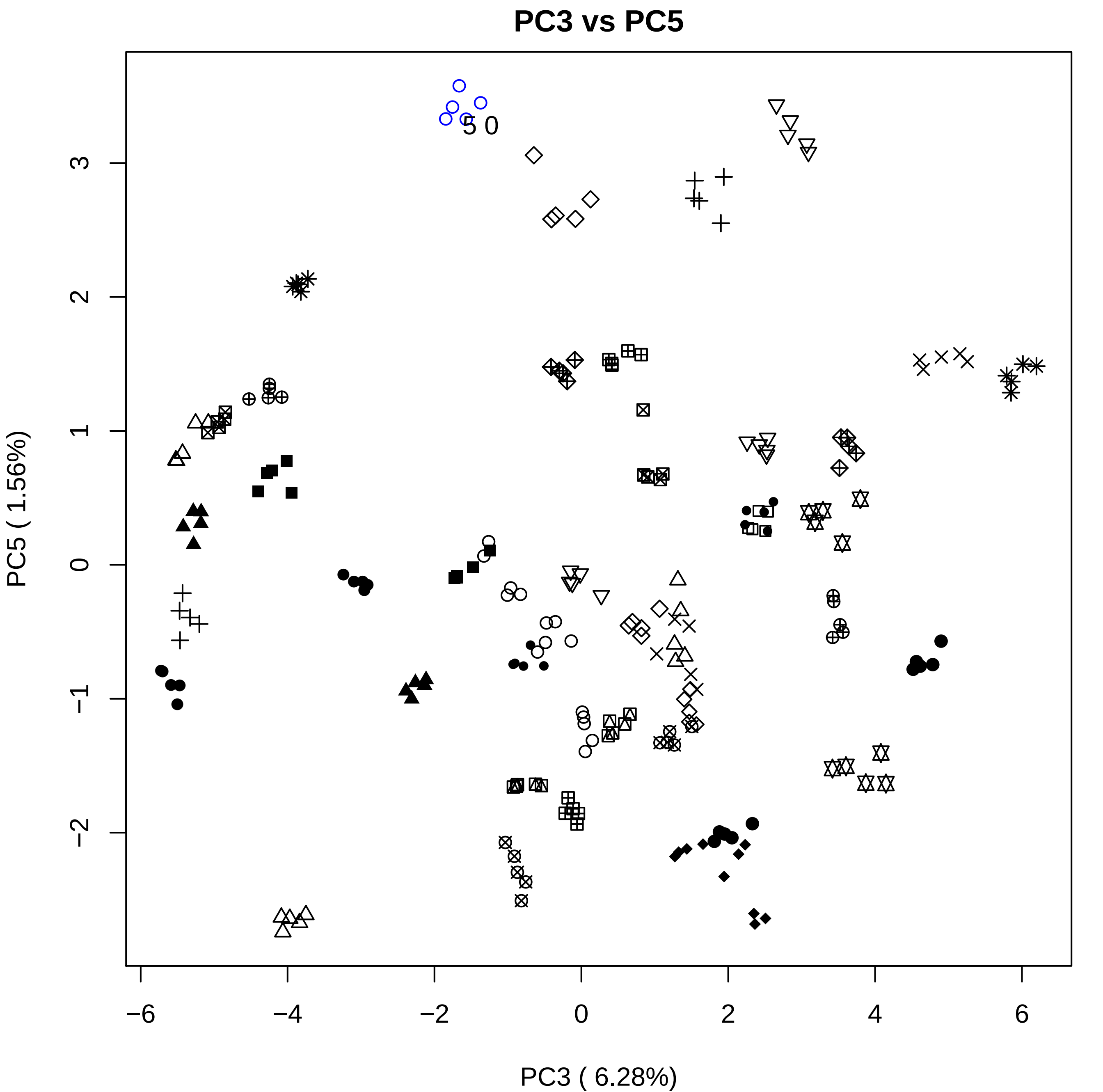

PC3 vs PC5
5 0
PC5 ( 1.56%)
PC3 ( 6.28%)
−6
−4
−2
0
2
4
6
−2
−1
0
1
2
3

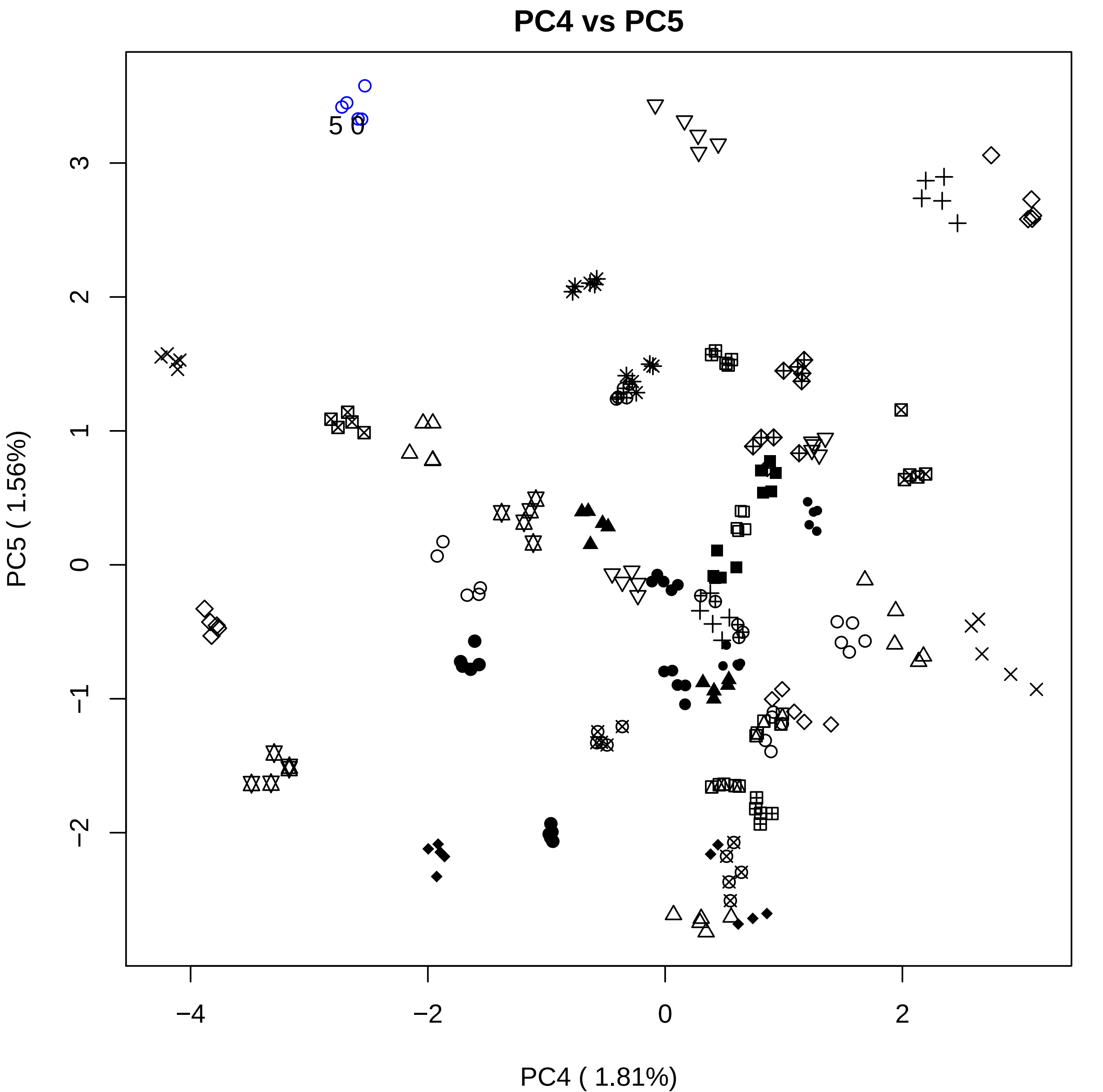

PC4 vs PC5
50
PC5 ( 1.56%)
PC4 ( 1.81%)
−4
−2
0
2
−2
−1
0
1
2
3

### PCA of 7.5%  OSCS vs PMH

Scaled Matrix PCAed

7th order polynomial baseline correction performed

Raw spectra smoothed to 21 Neighbours, 2nd order PN, Savtisky Golay

Normalisation between 0 and 100

D2 spectra smoothed to 41 Neighbours, 2nd order PN, Savtisky Golay

Regions Analysed: 3601−2501 and 2001−700cm−1

Samples Analysed: 1 Contam's 0 OSCS's 45 PMH's

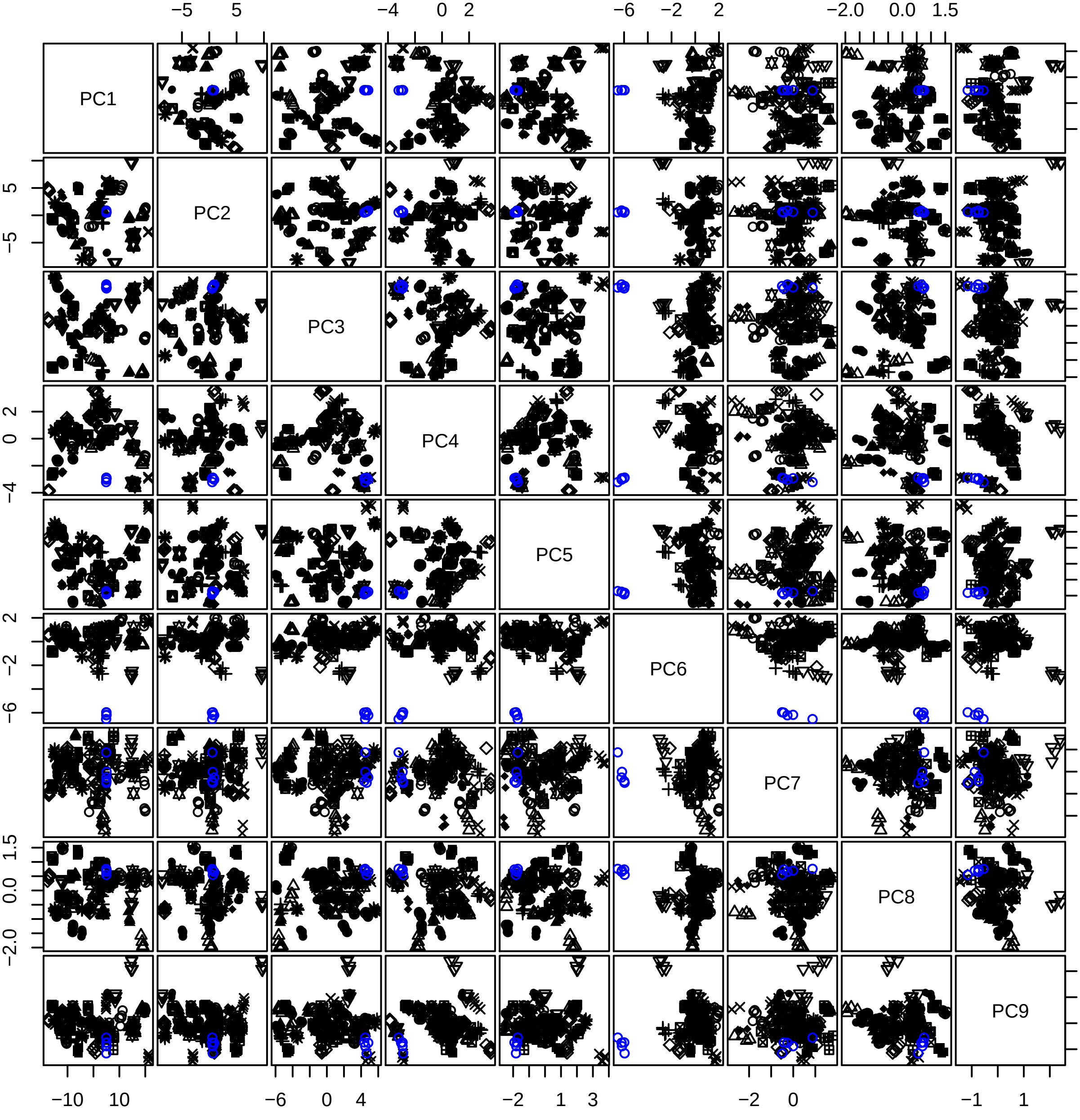
PC1
PC2
PC3
PC4
PC5
PC6
PC7
PC8
PC9

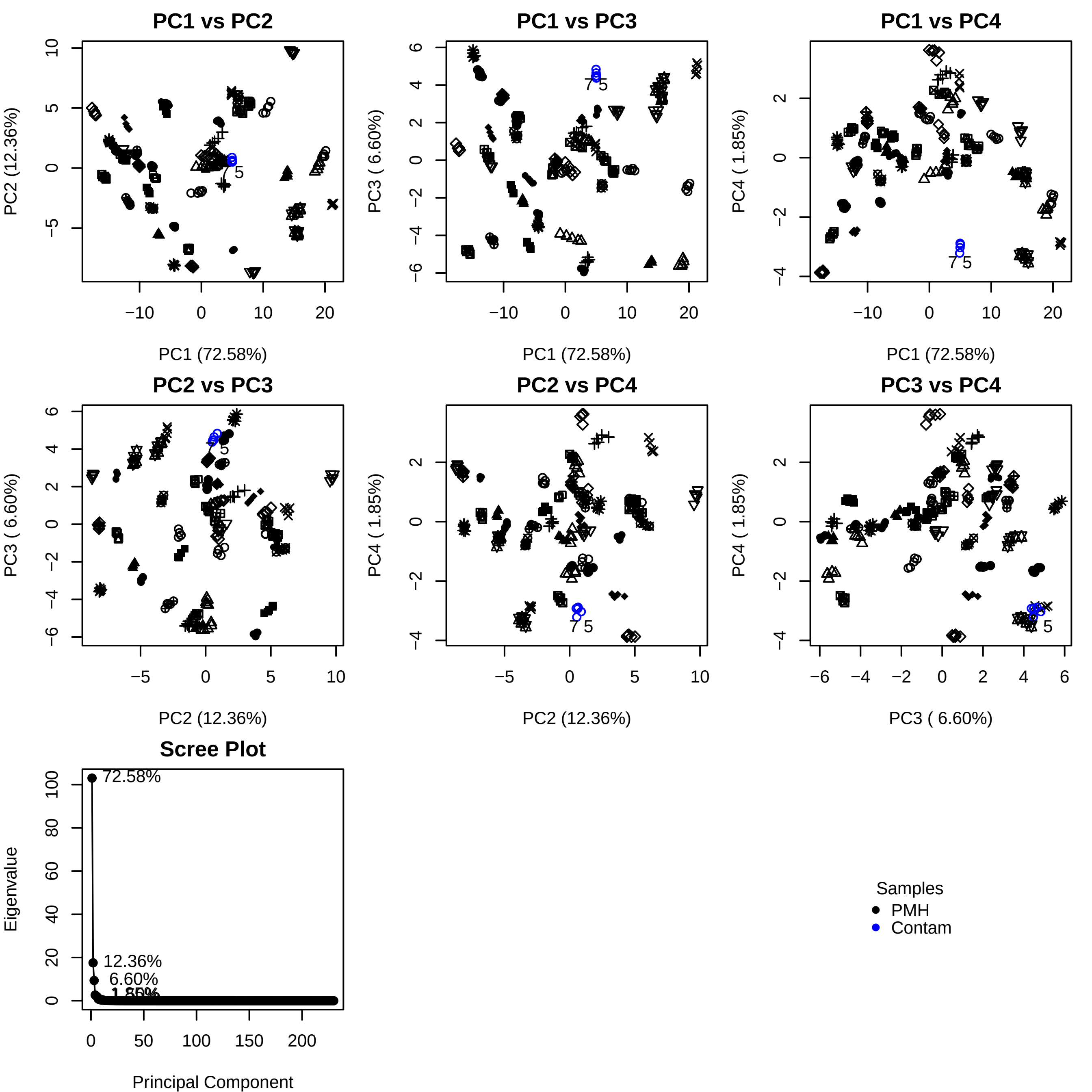

PC1 vs PC2
PC1 vs PC3
PC1 vs PC4
PC2 vs PC3
PC2 vs PC4
PC3 vs PC4
PC1 (72.58%)
PC2 (12.36%)
PC3 ( 6.60%)
PC4 ( 1.85%)
Scree Plot
Eigenvalue
Principal Component
72.58%
12.36%
6.60%
Samples
PMH
Contam

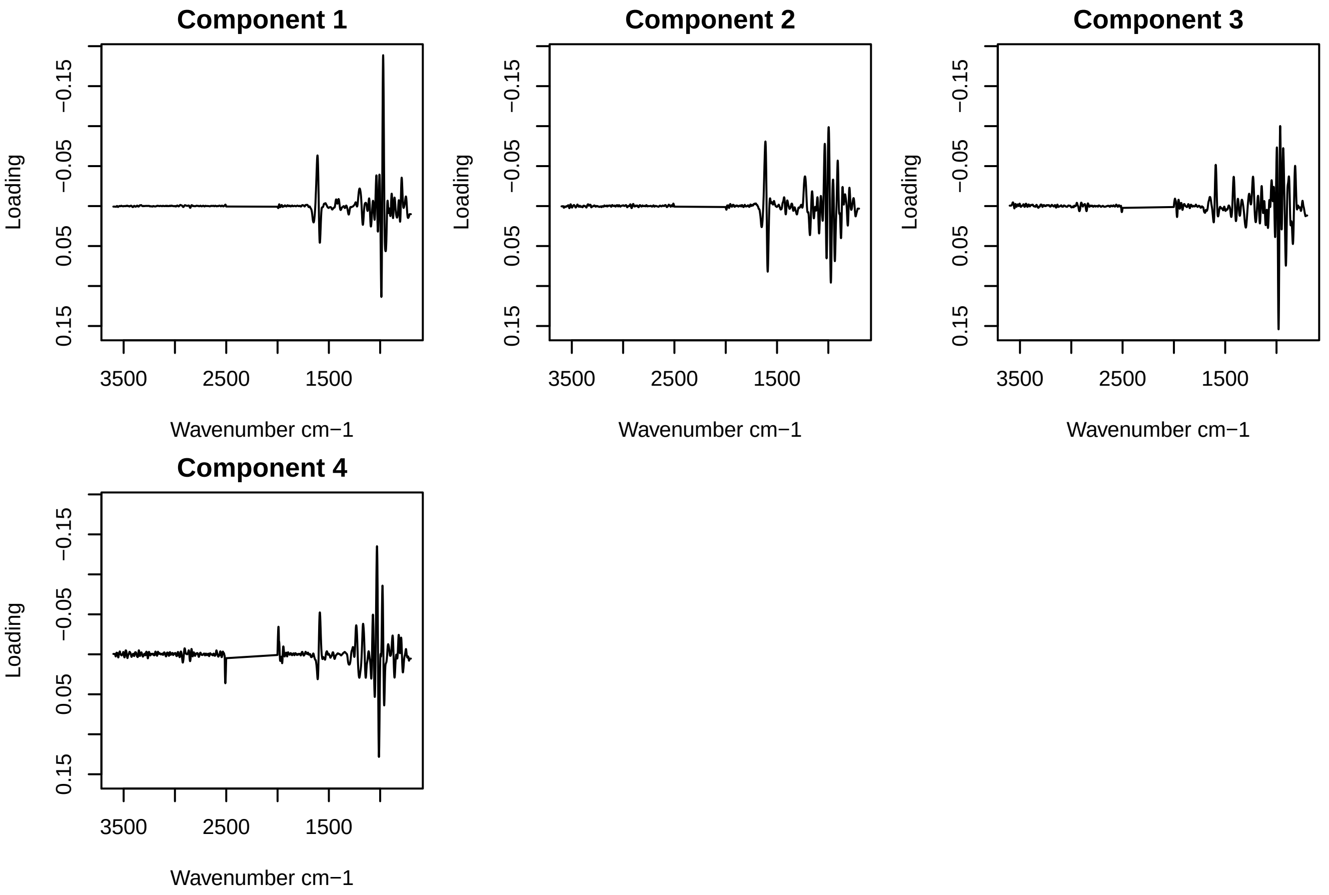
Component 1
Component 2
Component 3
Component 4
Loading
−0.15
−0.05
0.05
0.15
3500
2500
1500
Wavenumber cm−1

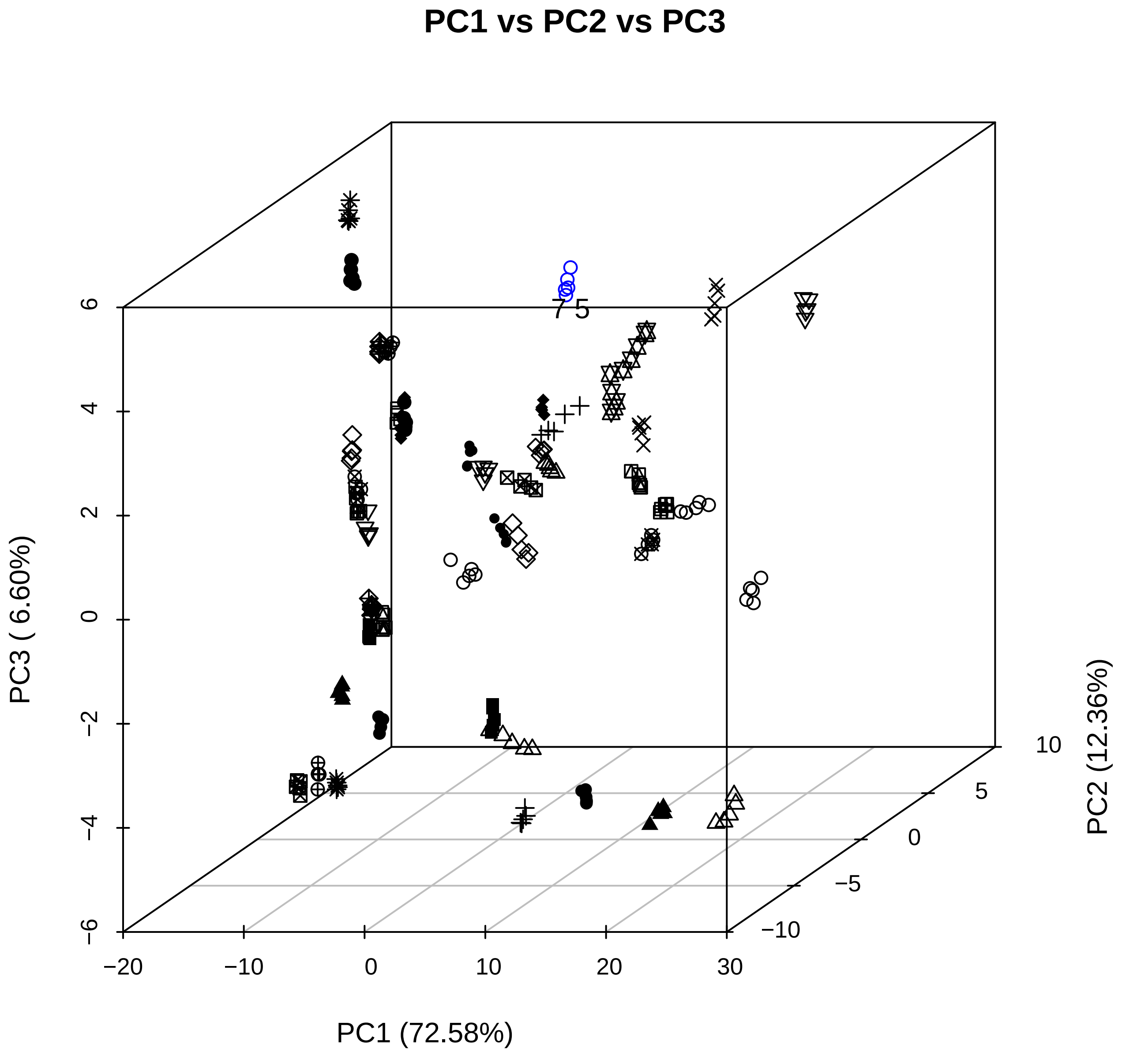
PC1 vs PC2 vs PC3
75
PC3 ( 6.60%)
6
4
2
0
−2
−4
−6
−20
−10
0
10
20
30
PC1 (72.58%)
−10
−5
0
5
10
PC2 (12.36%)

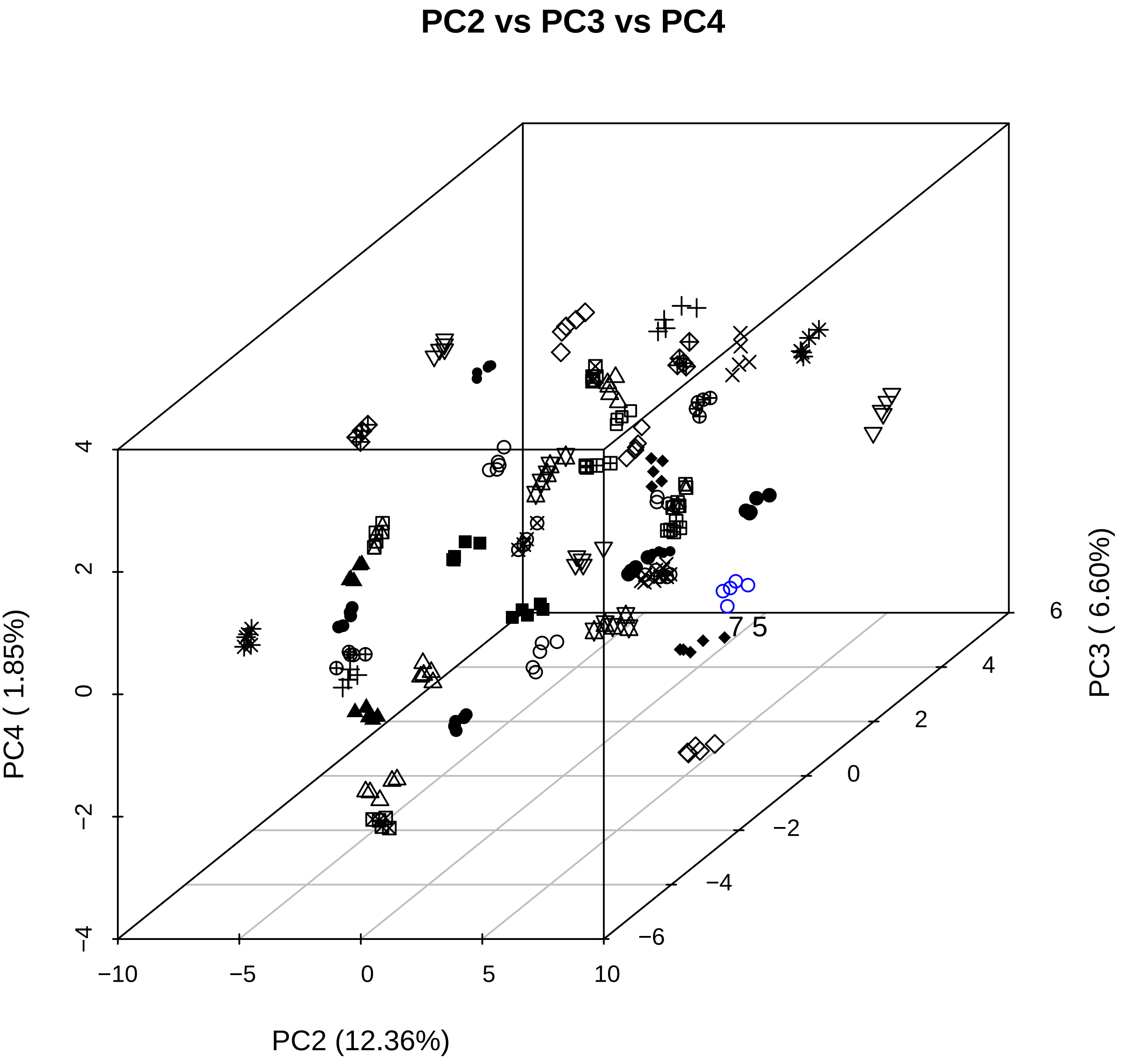

PC2 vs PC3 vs PC4
PC2 (12.36%)
PC4 ( 1.85%)
PC3 ( 6.60%)
−10
−5
0
5
10
−4
−2
0
2
4
−6
−4
−2
0
2
4
6
7 5

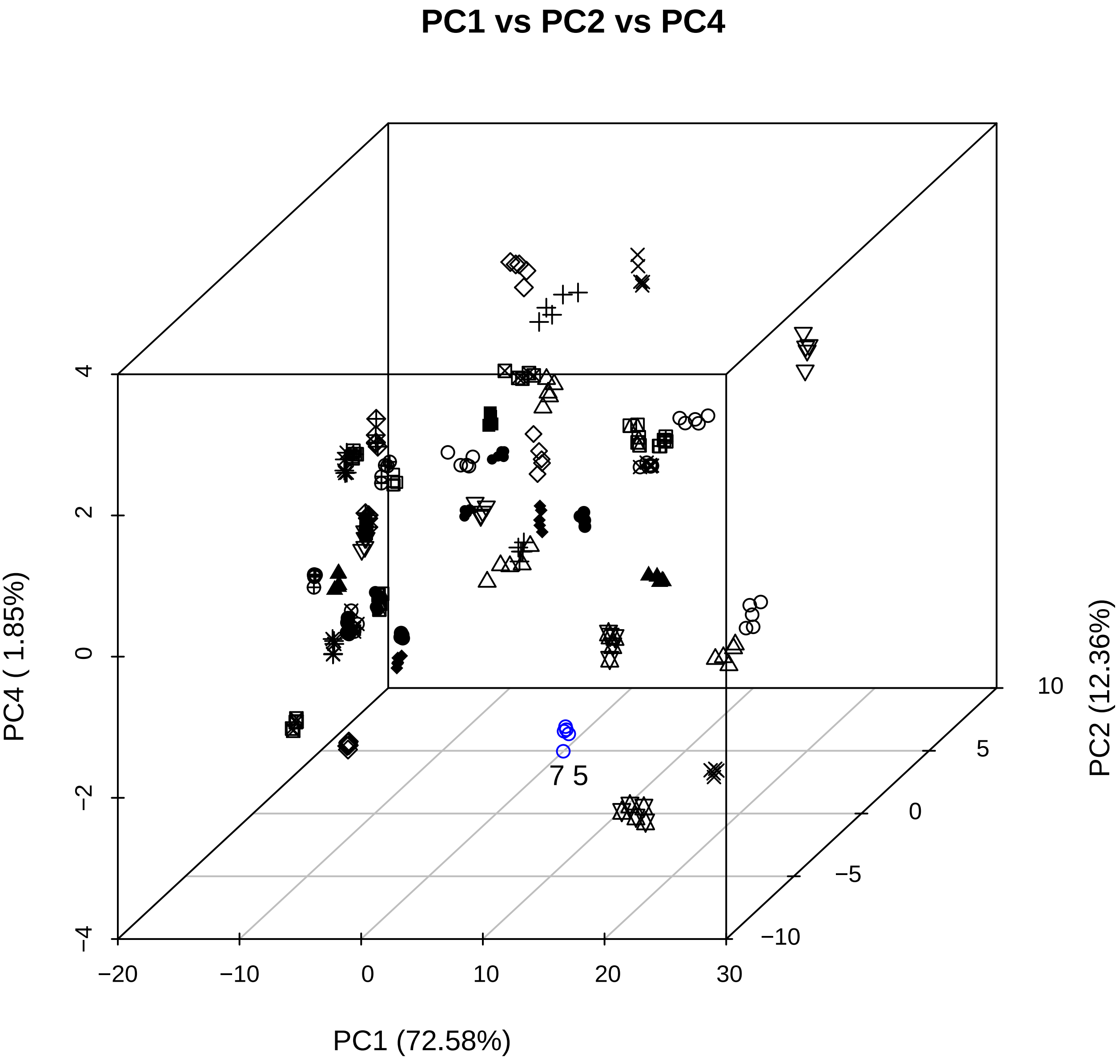
PC1 vs PC2 vs PC4
PC4 ( 1.85%)
PC1 (72.58%)
PC2 (12.36%)
4
2
0
−2
−4
−20
−10
0
10
20
30
10
5
0
−5
−10
7 5

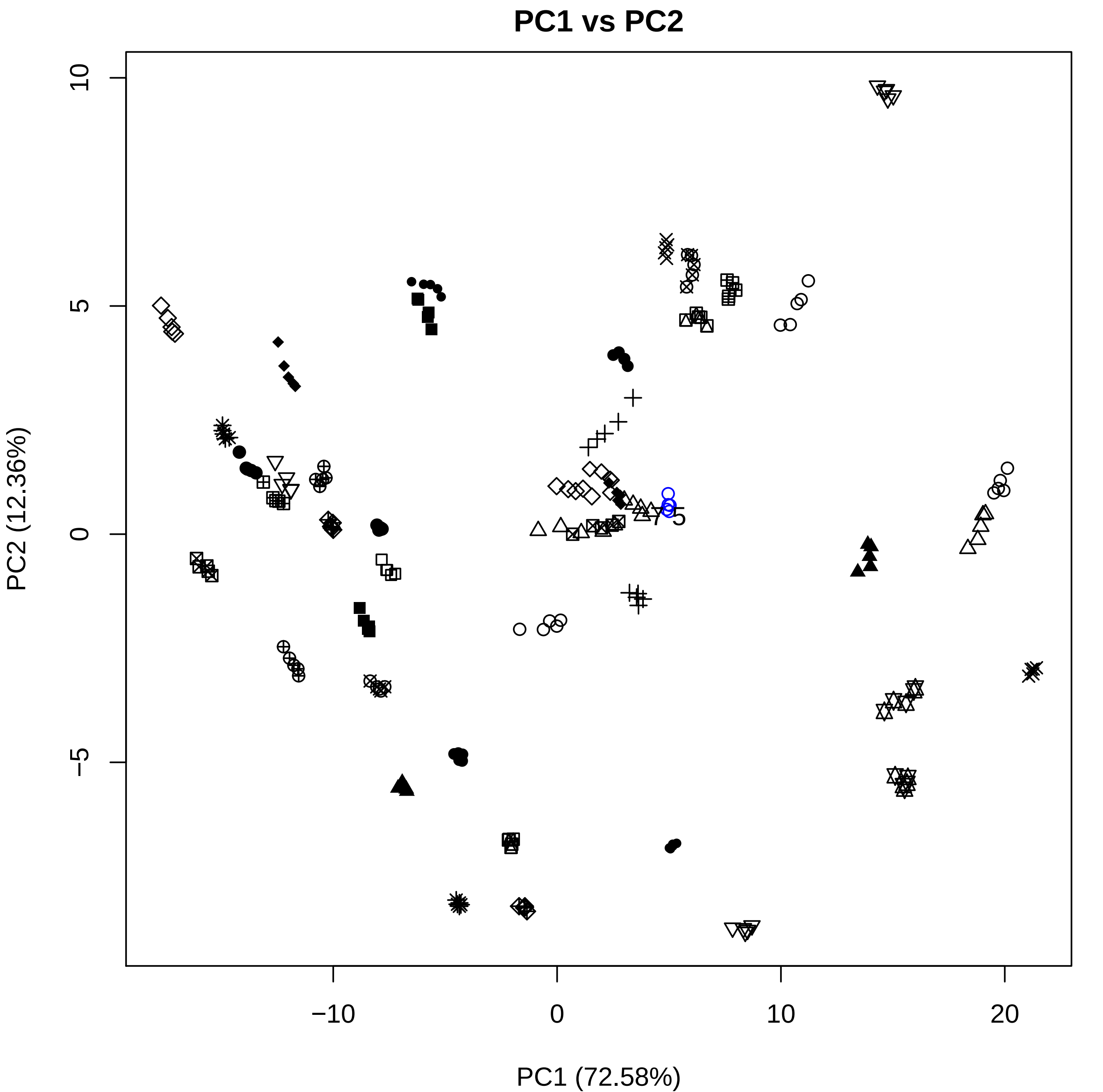
PC1 vs PC2
PC2 (12.36%)
PC1 (72.58%)
10
5
0
−5
−10
0
10
20
7
5

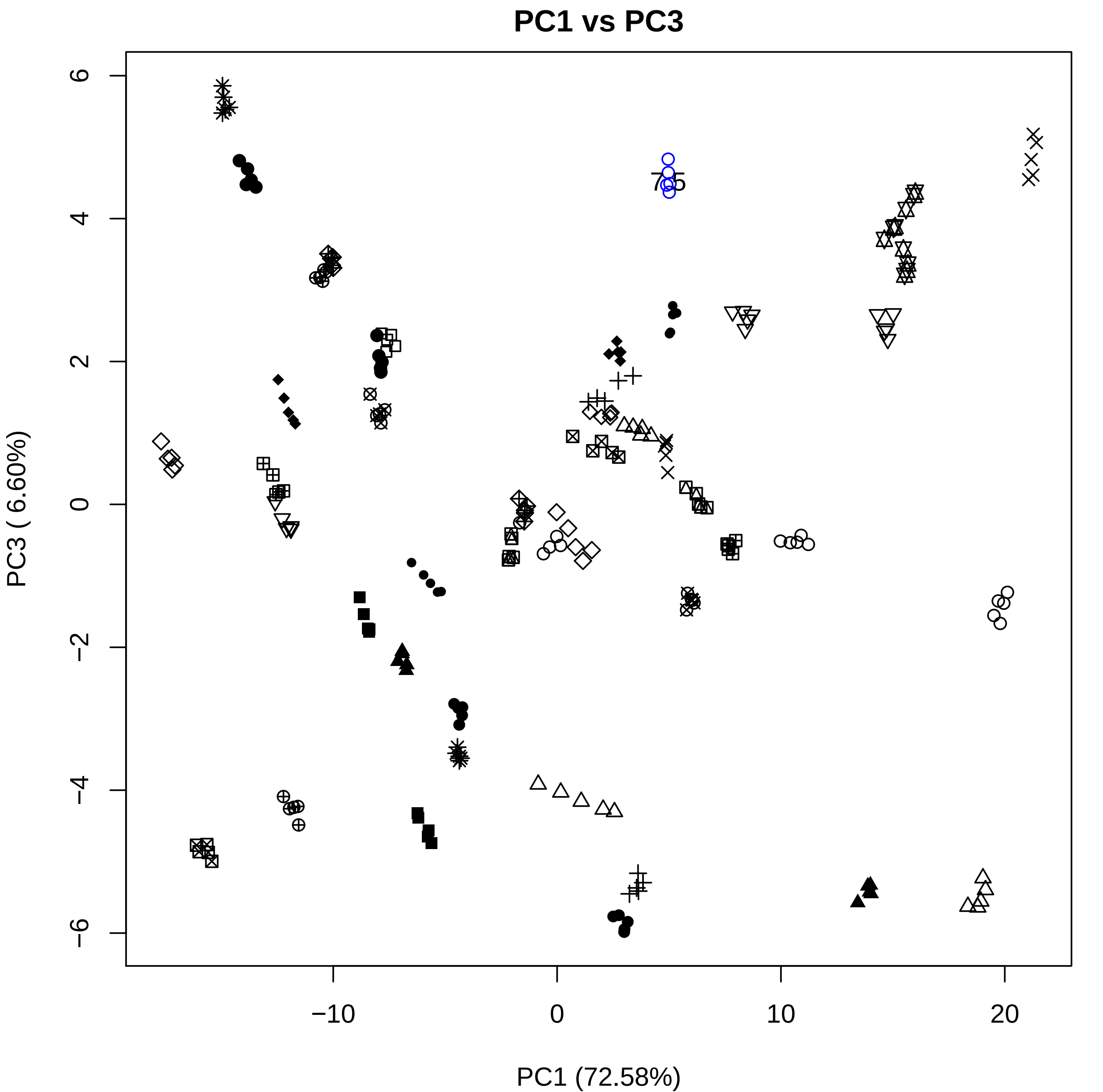
PC1 vs PC3
PC3 ( 6.60%)
PC1 (72.58%)
6
4
2
0
−2
−4
−6
−10
0
10
20
75

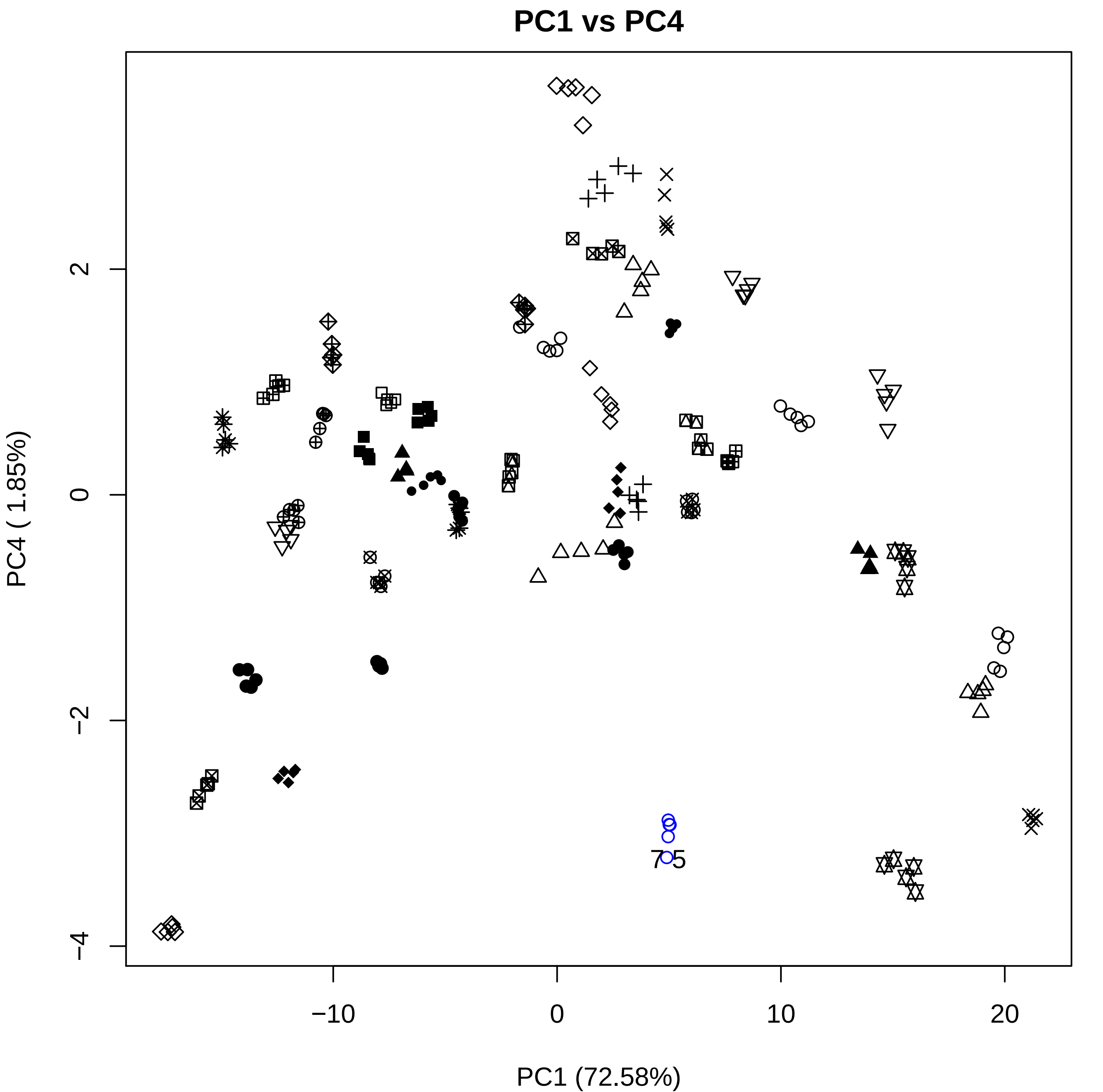
PC1 vs PC4
PC4 ( 1.85%)
PC1 (72.58%)
2
0
−2
−4
−10
0
10
20
75

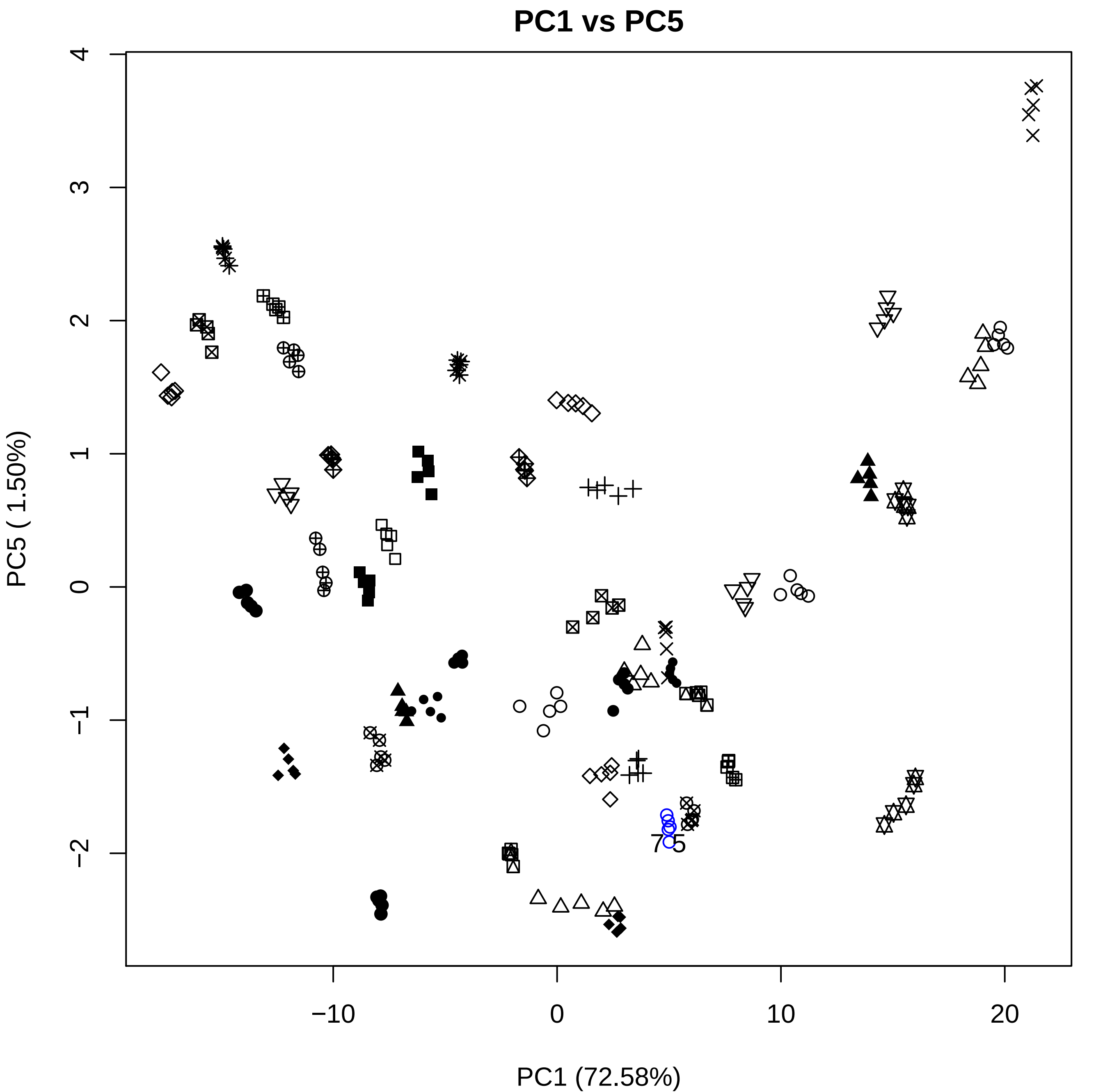
PC1 vs PC5
PC5 ( 1.50%)
PC1 (72.58%)
4
3
2
1
0
−1
−2
−10
0
10
20
75

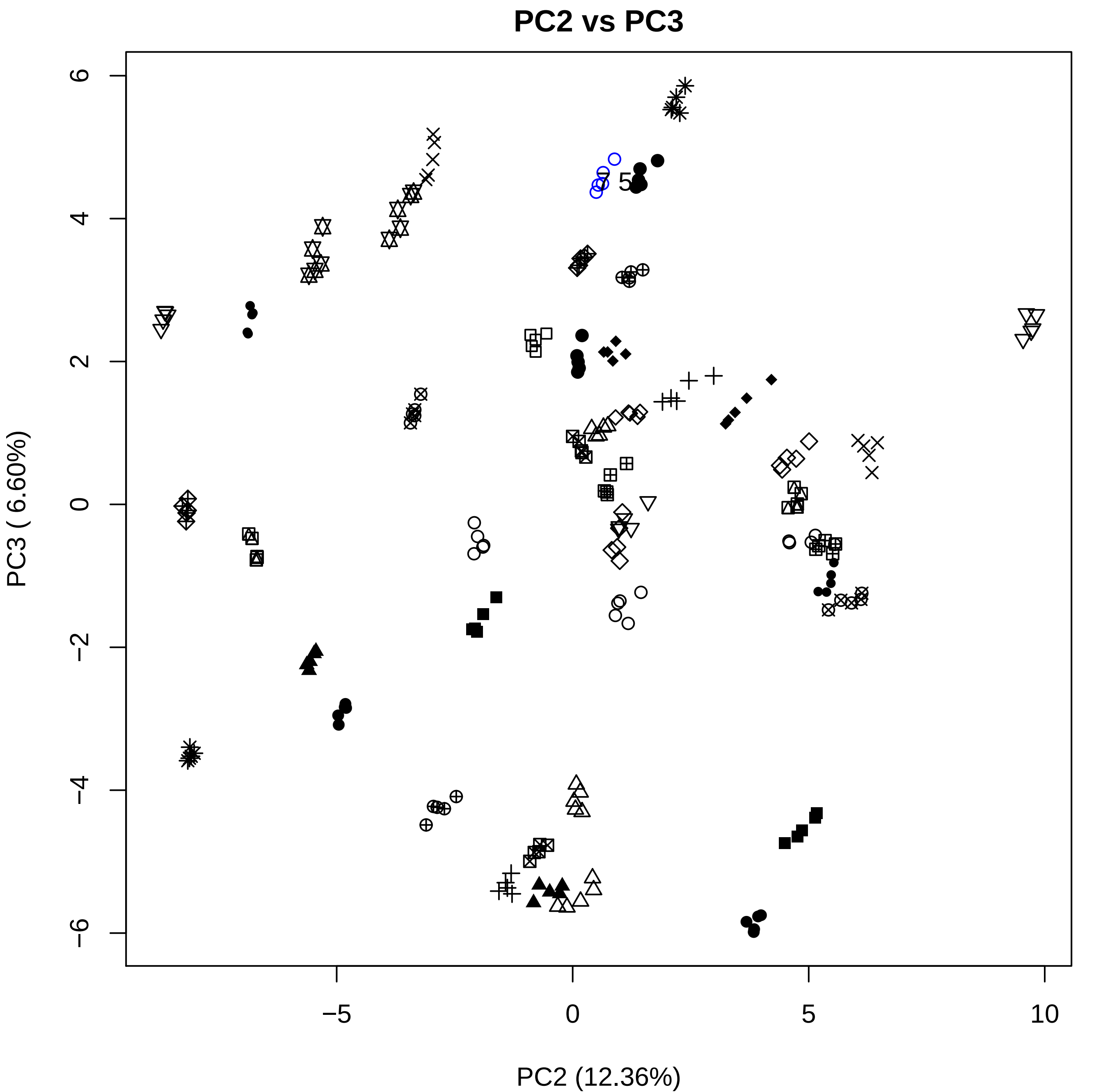

PC2 vs PC3
PC3 ( 6.60%)
PC2 (12.36%)
6
4
2
0
−2
−4
−6
−5
0
5
10
7
5

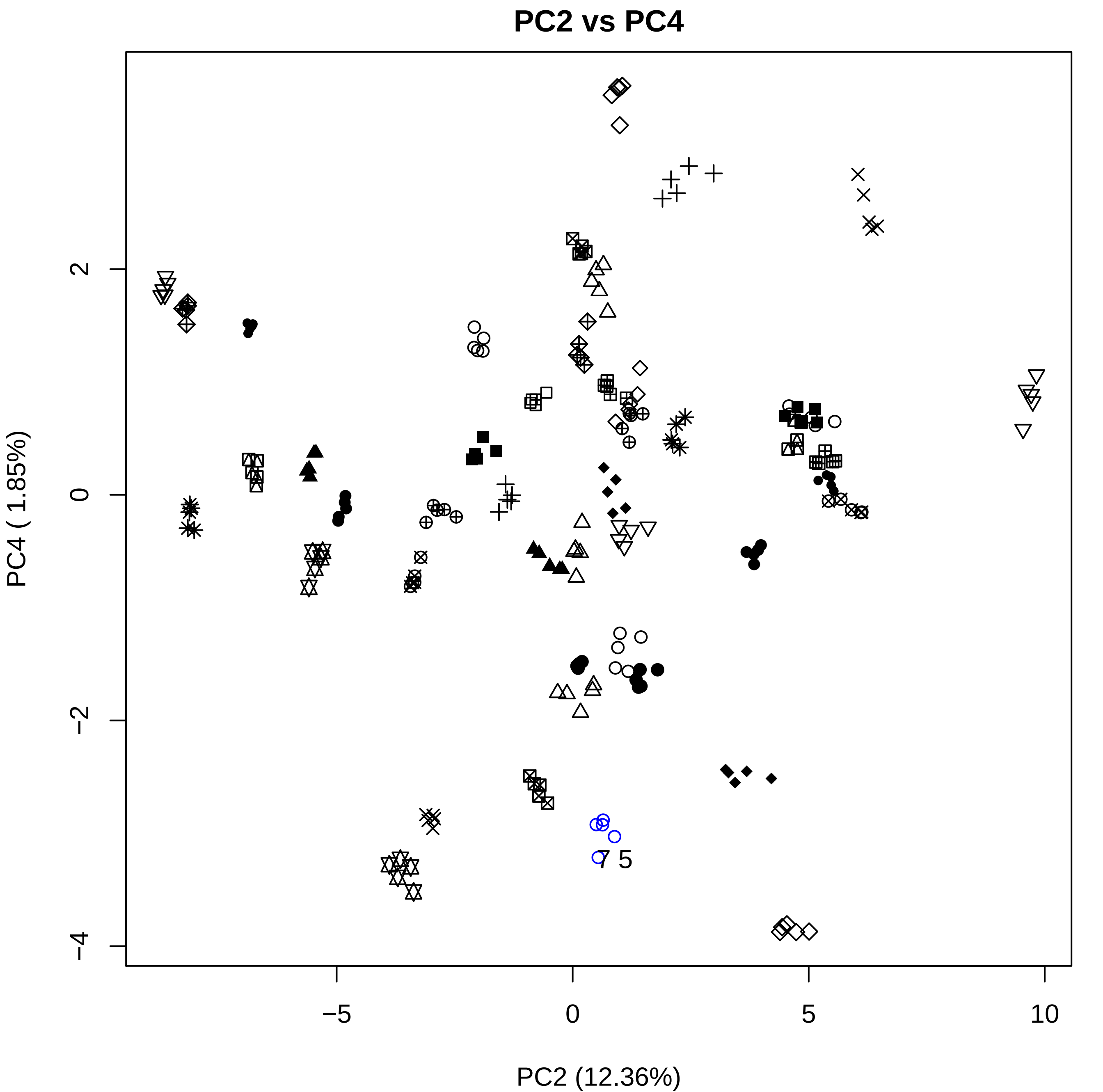
PC2 vs PC4
PC4 ( 1.85%)
PC2 (12.36%)
2
0
−2
−4
−5
0
5
10
7 5

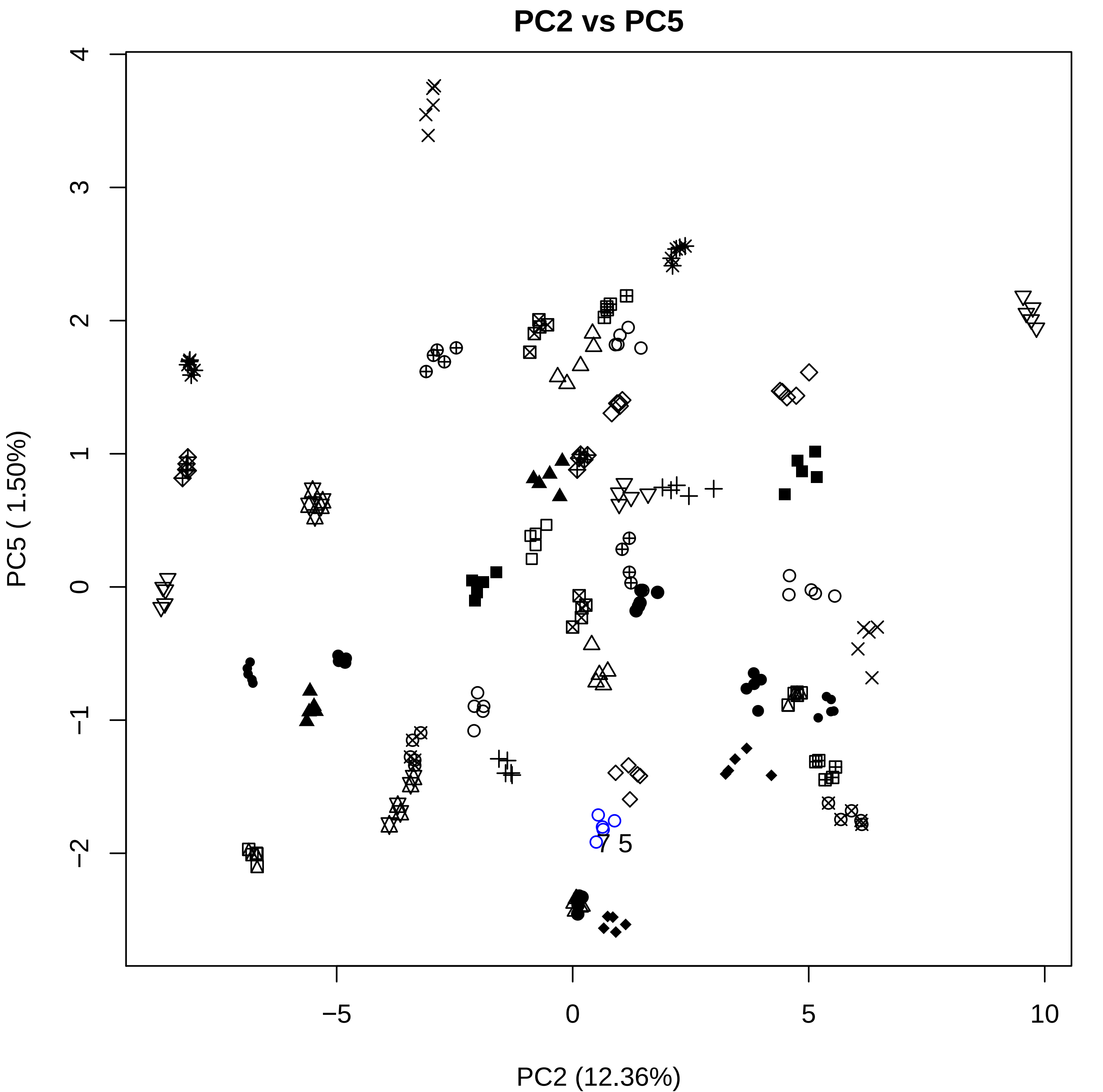
PC2 vs PC5
PC5 ( 1.50%)
PC2 (12.36%)
4
3
2
1
0
−1
−2
−5
0
5
10
7 5

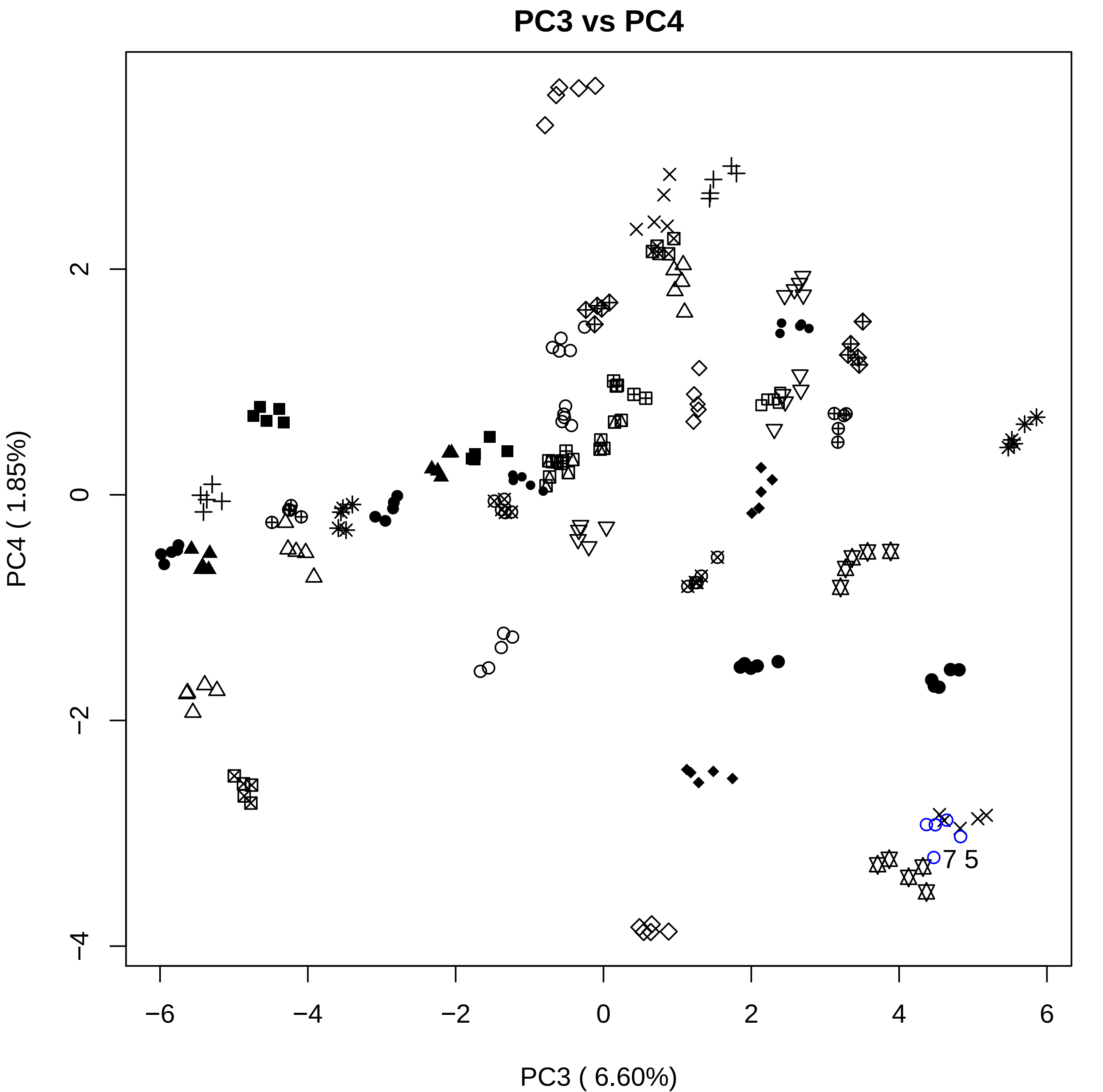

PC3 vs PC4
PC4 ( 1.85%)
PC3 ( 6.60%)
2
0
−2
−4
−6
−4
−2
0
2
4
6
7 5

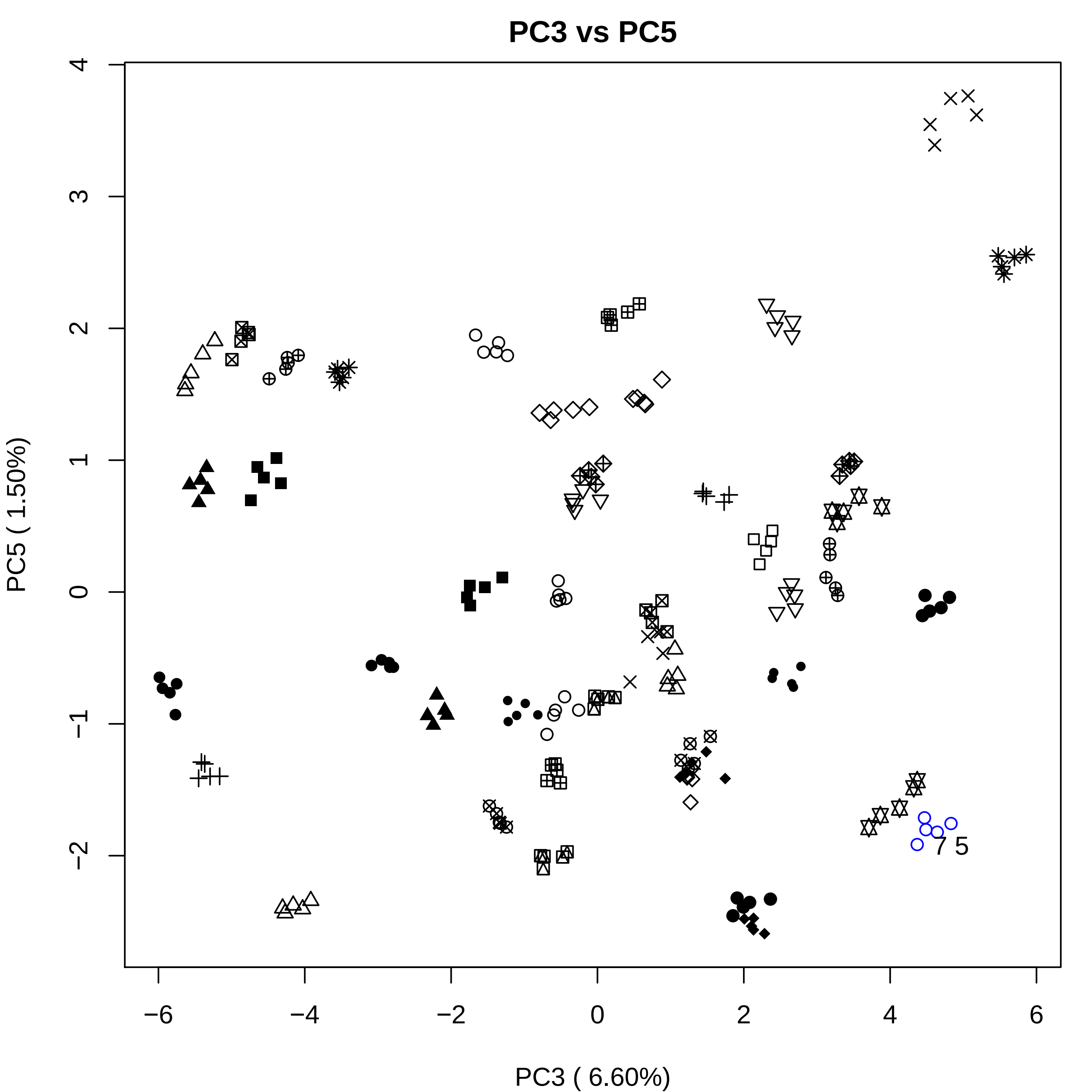
PC3 vs PC5
PC5 ( 1.50%)
PC3 ( 6.60%)
4
3
2
1
0
−1
−2
−6
−4
−2
0
2
4
6
7 5

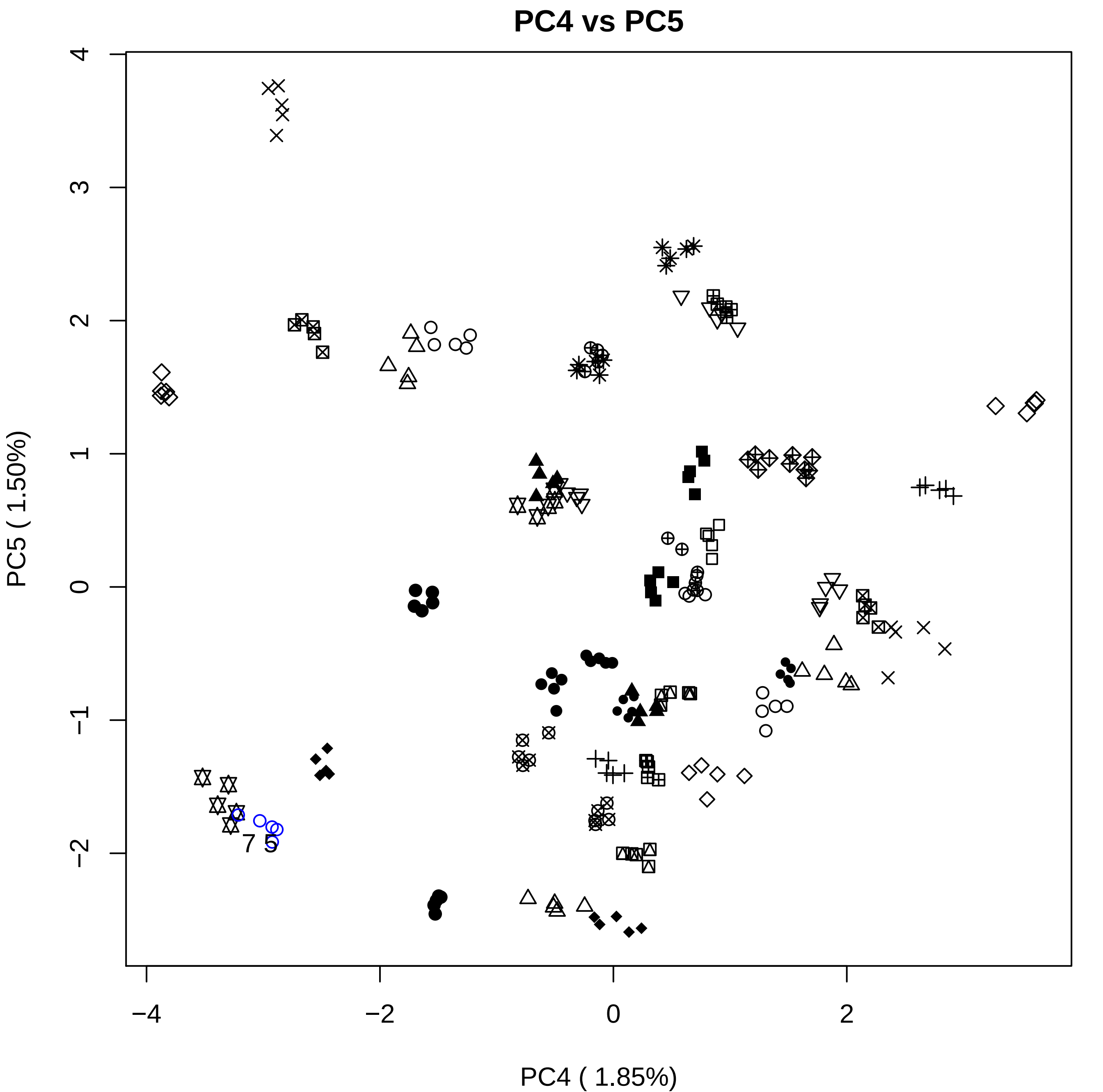

PC4 vs PC5
PC5 ( 1.50%)
PC4 ( 1.85%)
4
3
2
1
0
−1
−2
−4
−2
0
2
7
5

### PCA of 10%  OSCS vs PMH

Scaled Matrix PCAed

7th order polynomial baseline correction performed

Raw spectra smoothed to 21 Neighbours, 2nd order PN, Savtisky Golay

Normalisation between 0 and 100

D2 spectra smoothed to 41 Neighbours, 2nd order PN, Savtisky Golay

Regions Analysed: 3601−2501 and 2001−700cm−1

Samples Analysed: 1 Contam's 0 OSCS's 45 PMH's

Keele University AD    Sun Dec 09 16:51:14 2018

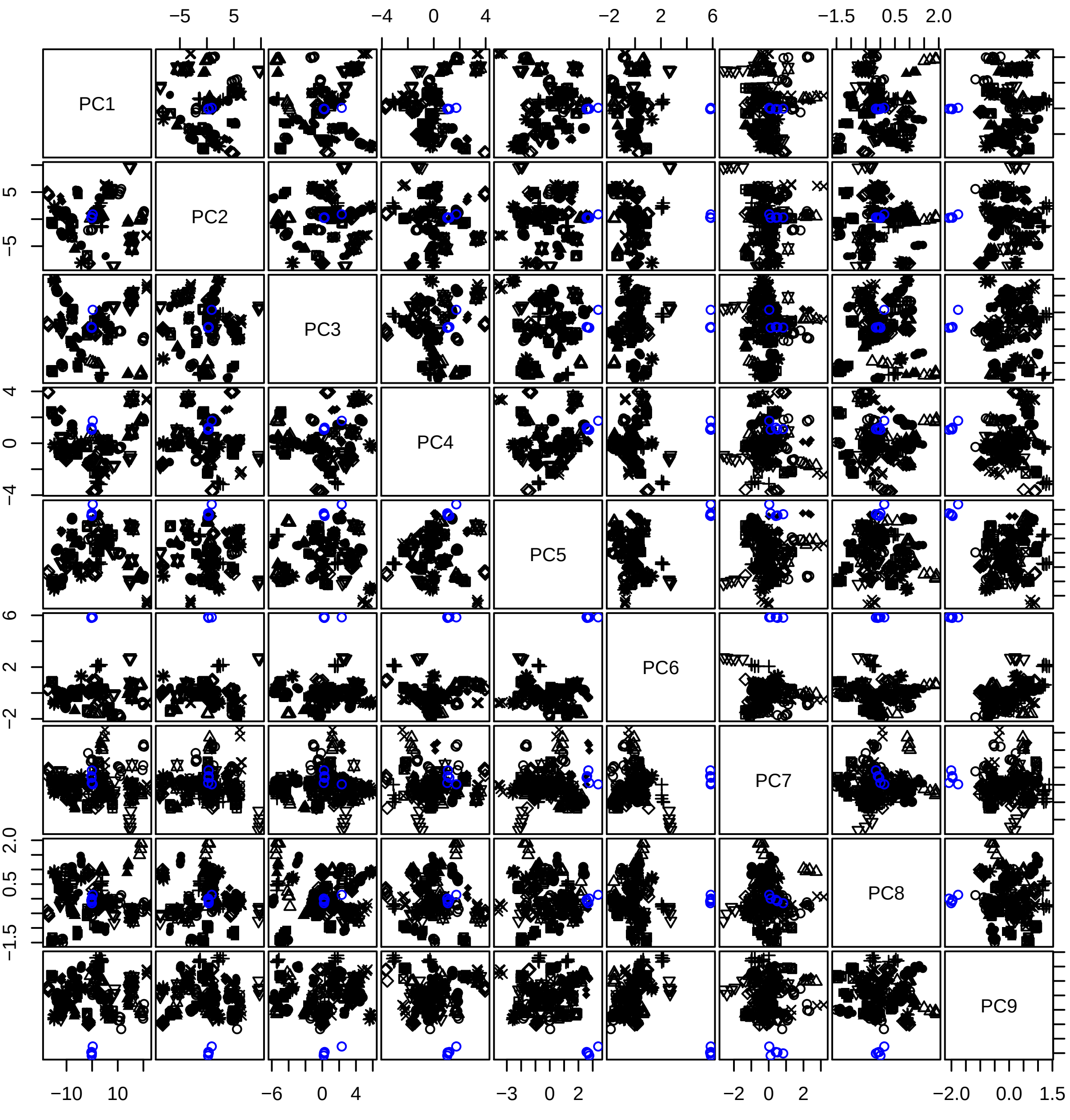
PC1
PC2
PC3
PC4
PC5
PC6
PC7
PC8
PC9

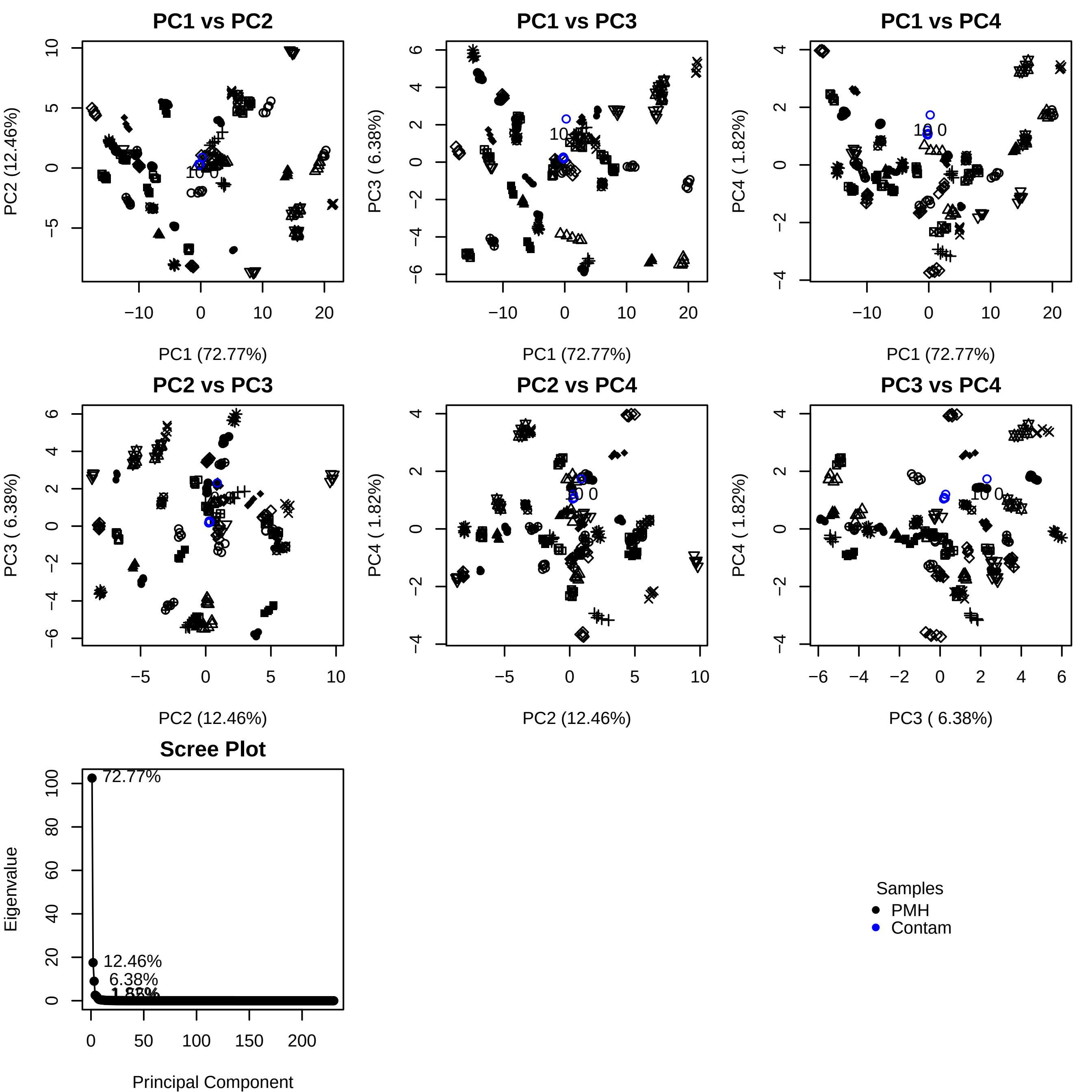
PC1 vs PC2
PC1 vs PC3
PC1 vs PC4
PC1 (72.77%)
PC2 (12.46%)
PC3 ( 6.38%)
PC4 ( 1.82%)
PC2 vs PC3
PC2 vs PC4
PC3 vs PC4
Scree Plot
Eigenvalue
Principal Component
72.77%
12.46%
6.38%
Samples
PMH
Contam

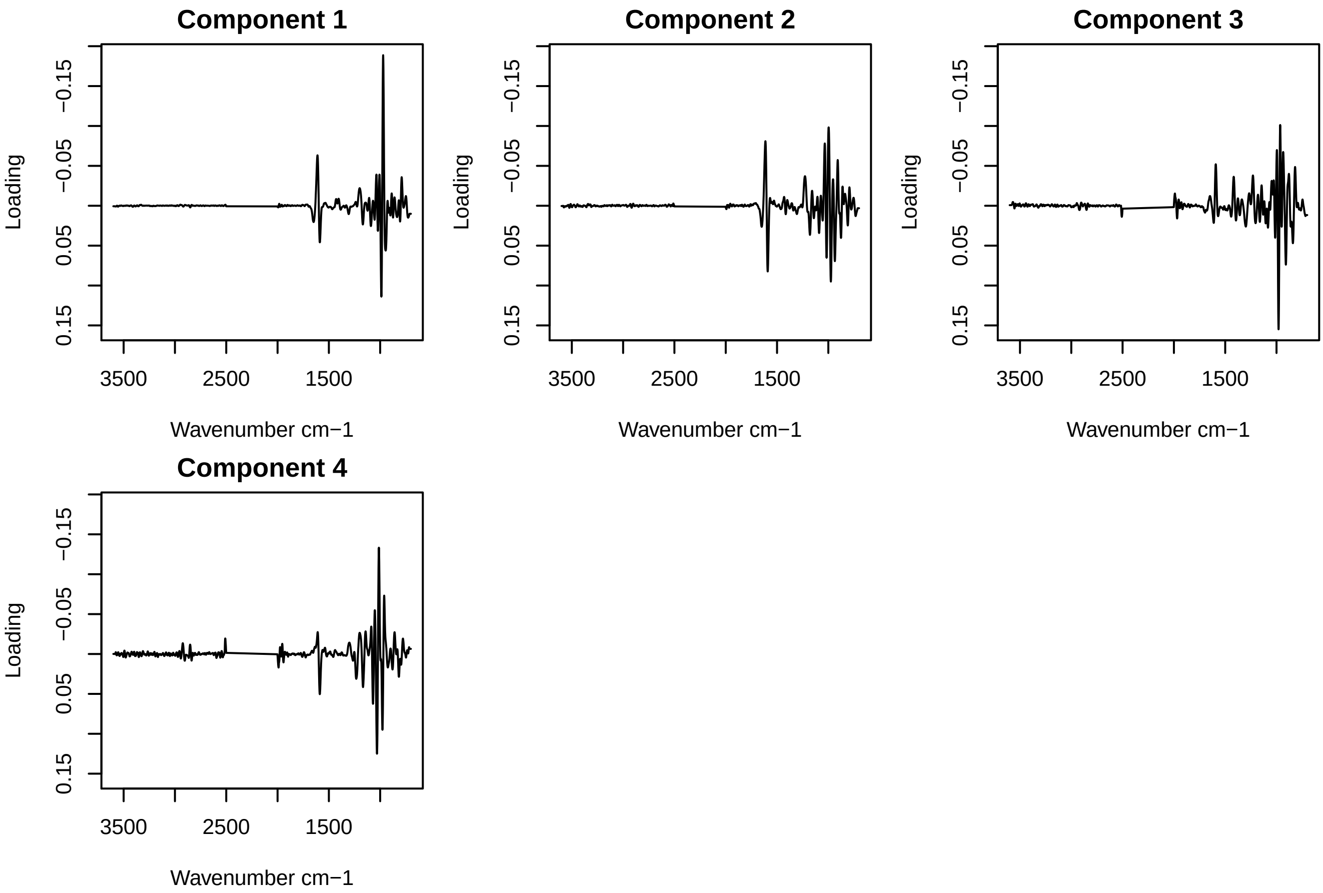
Component 1
Component 2
Component 3
Component 4
Loading
−0.15
−0.05
0.05
0.15
3500
2500
1500
Wavenumber cm−1

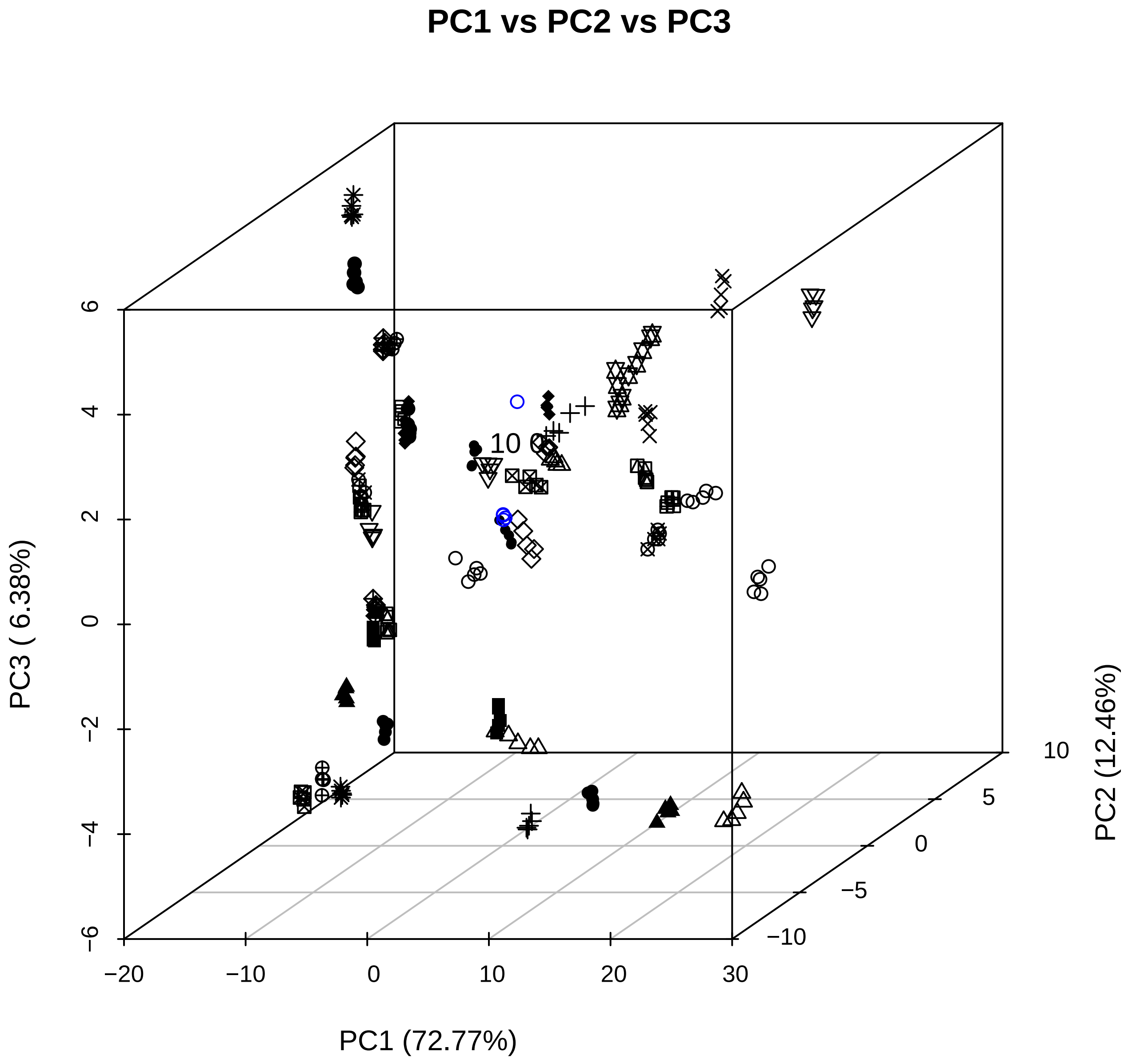
PC1 vs PC2 vs PC3
PC1 (72.77%)
PC2 (12.46%)
PC3 ( 6.38%)
−20
−10
0
10
20
30
−10
−5
0
5
10
−6
−4
−2
0
2
4
6
10 0

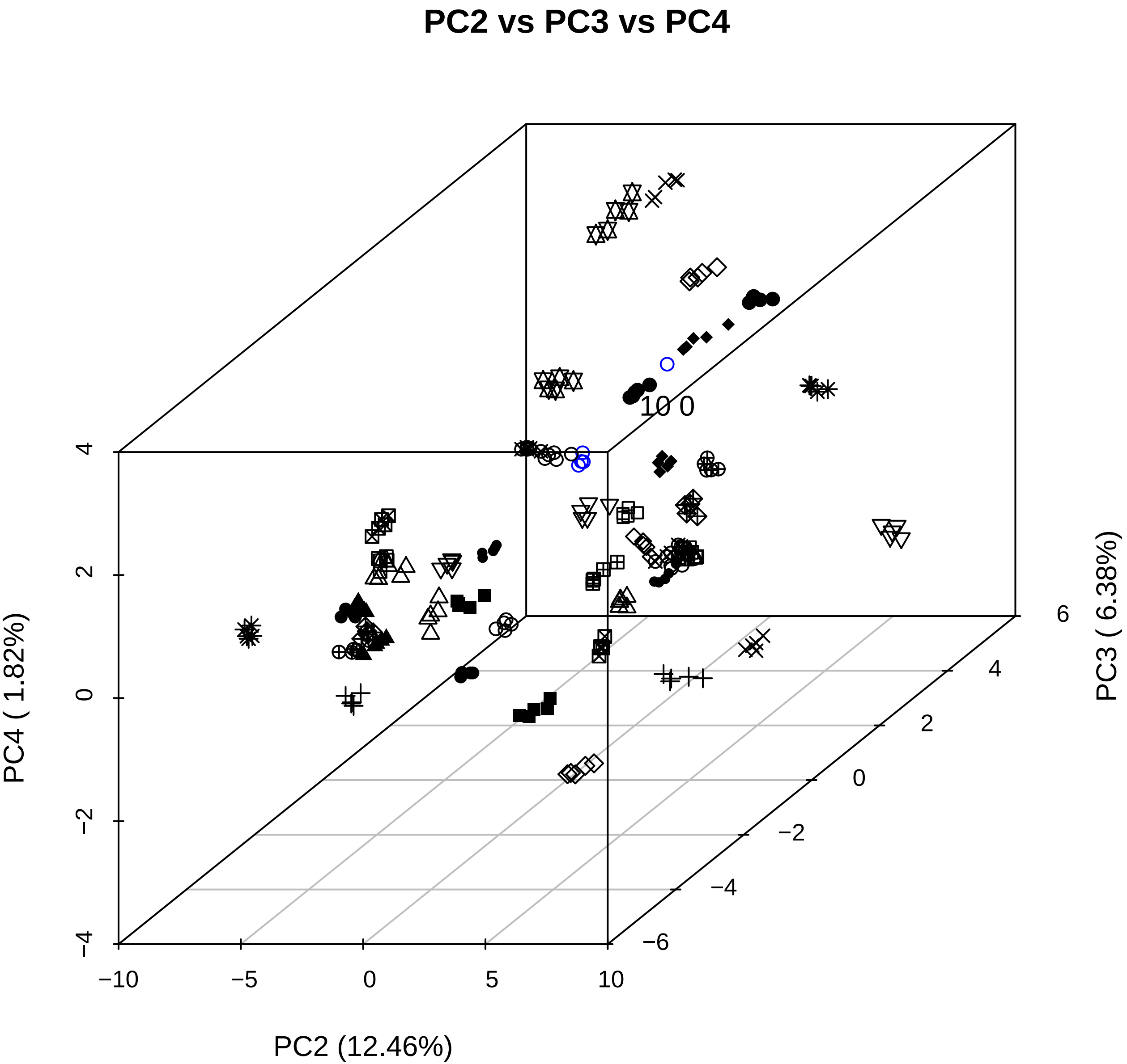
PC2 vs PC3 vs PC4
PC2 (12.46%)
PC4 ( 1.82%)
PC3 ( 6.38%)
−10
−5
0
5
10
−4
−2
0
2
4
−6
−4
−2
0
2
4
6
10 0

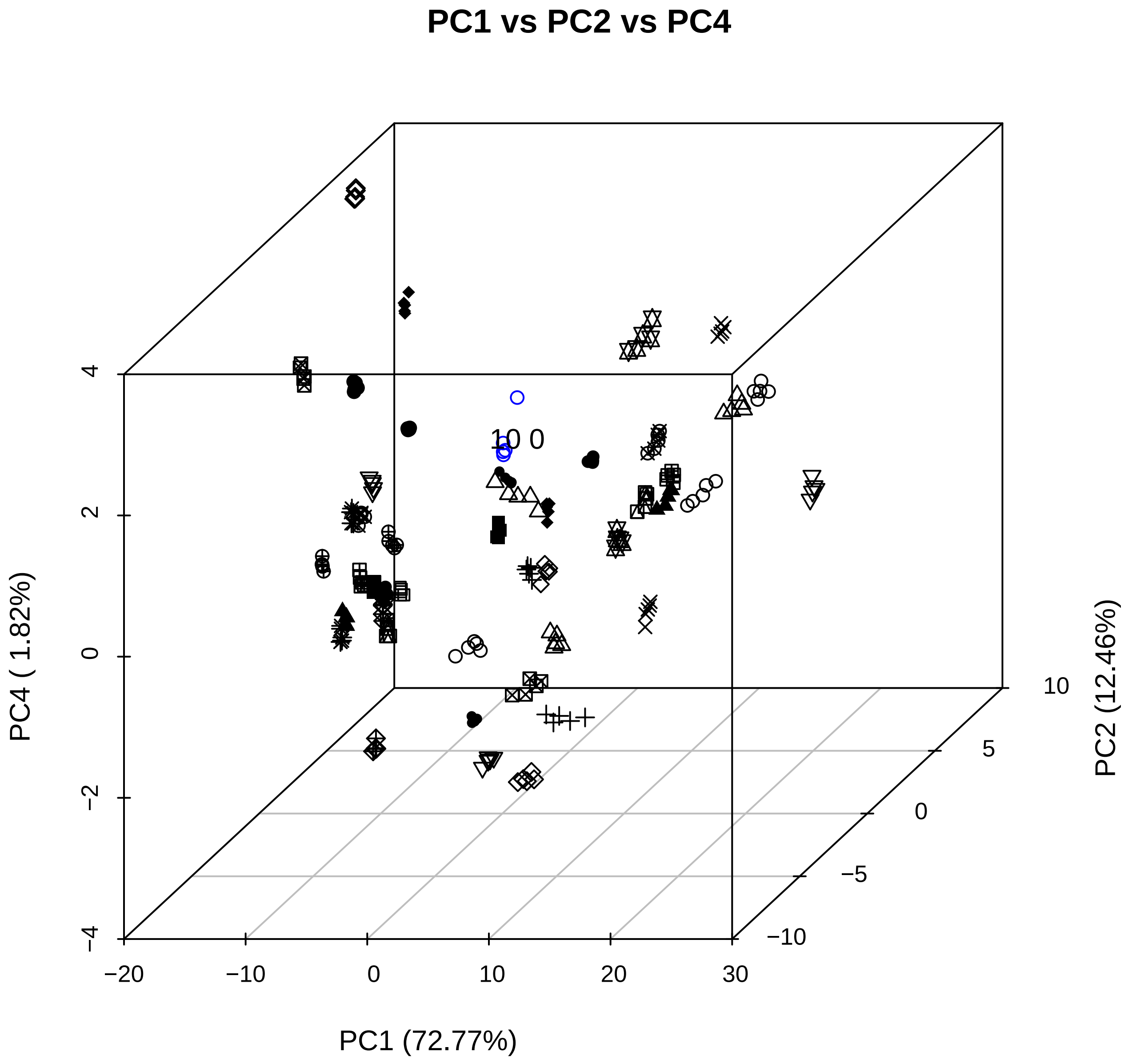
PC1 vs PC2 vs PC4
PC1 (72.77%)
PC2 (12.46%)
PC4 ( 1.82%)
−20
−10
0
10
20
30
−10
−5
0
5
10
−4
−2
0
2
4
10 0

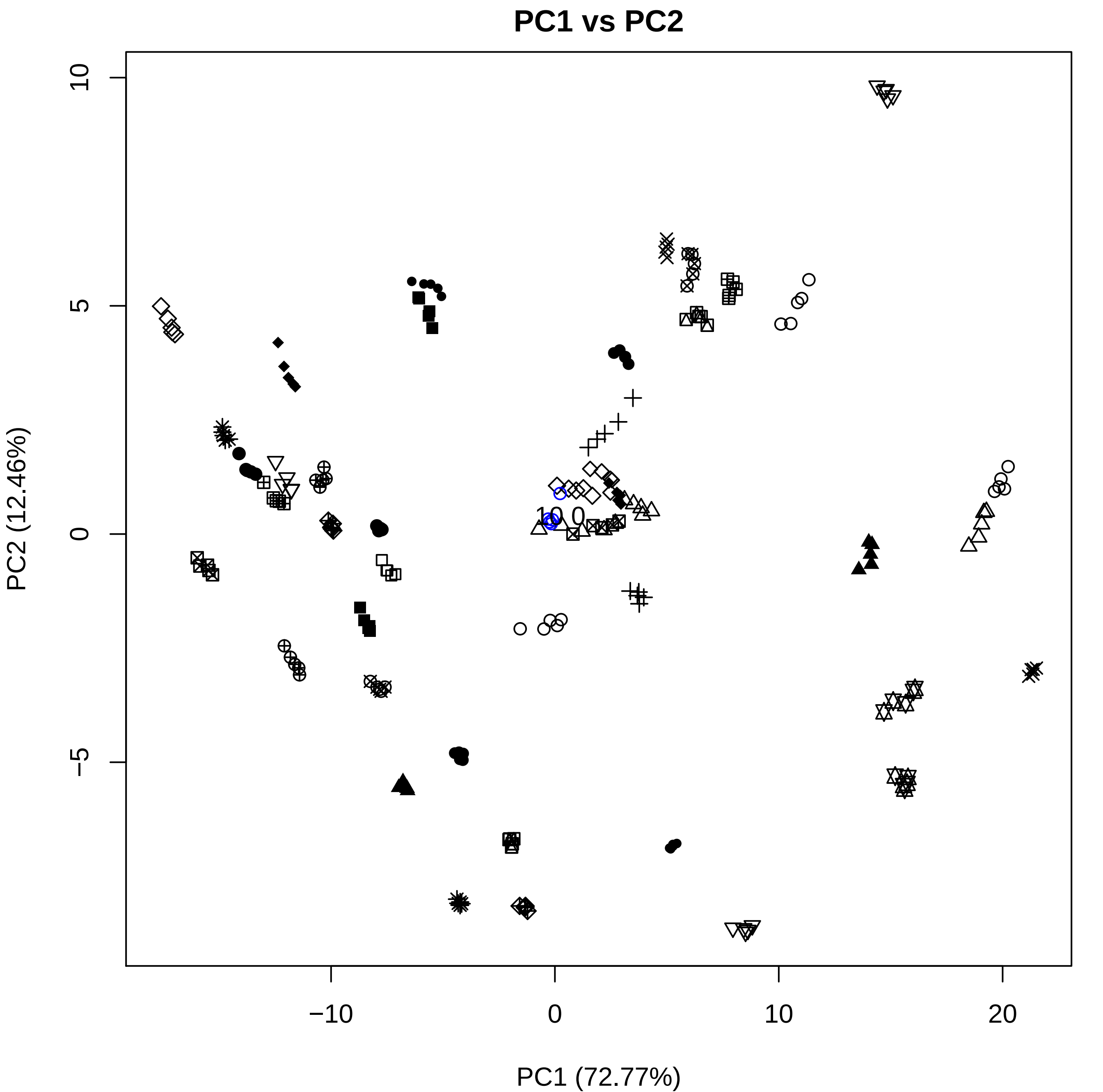
PC1 vs PC2
PC2 (12.46%)
PC1 (72.77%)
10
5
0
−5
−10
0
10
20
10 0

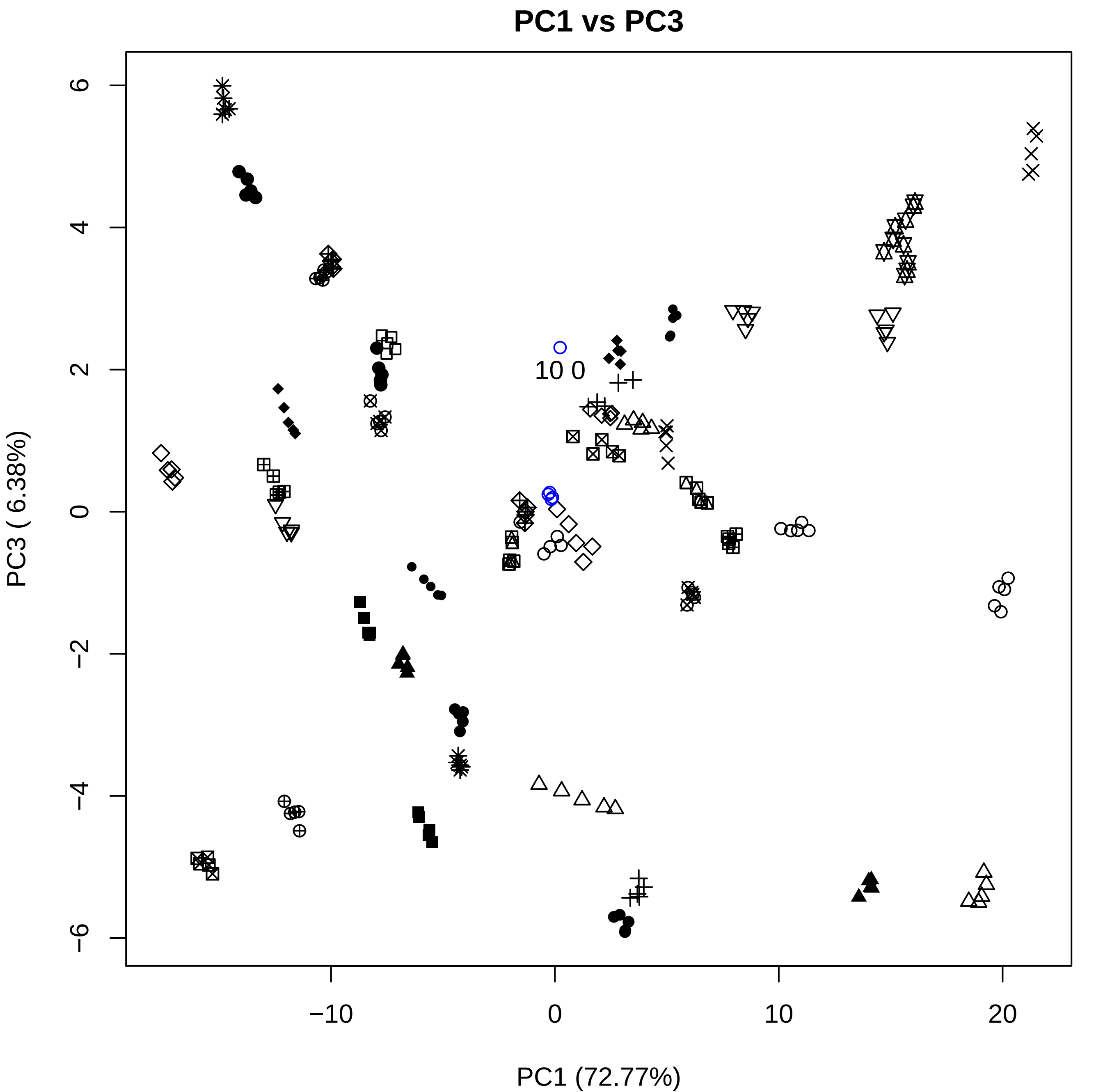
PC1 vs PC3
10 0
PC3 ( 6.38%)
PC1 (72.77%)
−10
0
10
20
−6
−4
−2
0
2
4
6

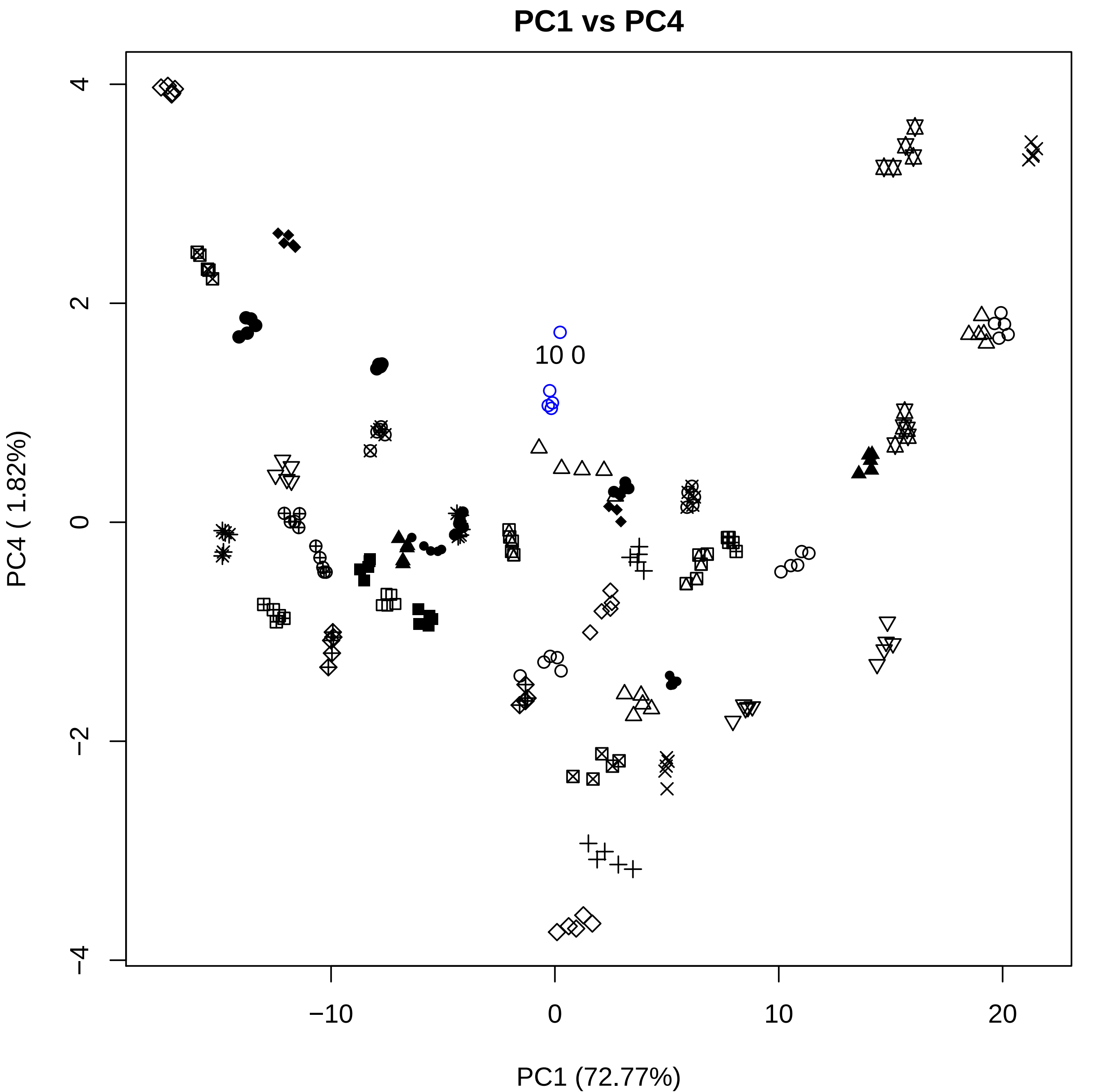
PC1 vs PC4
PC4 ( 1.82%)
PC1 (72.77%)
10 0
−4
−2
0
2
4
−10
0
10
20

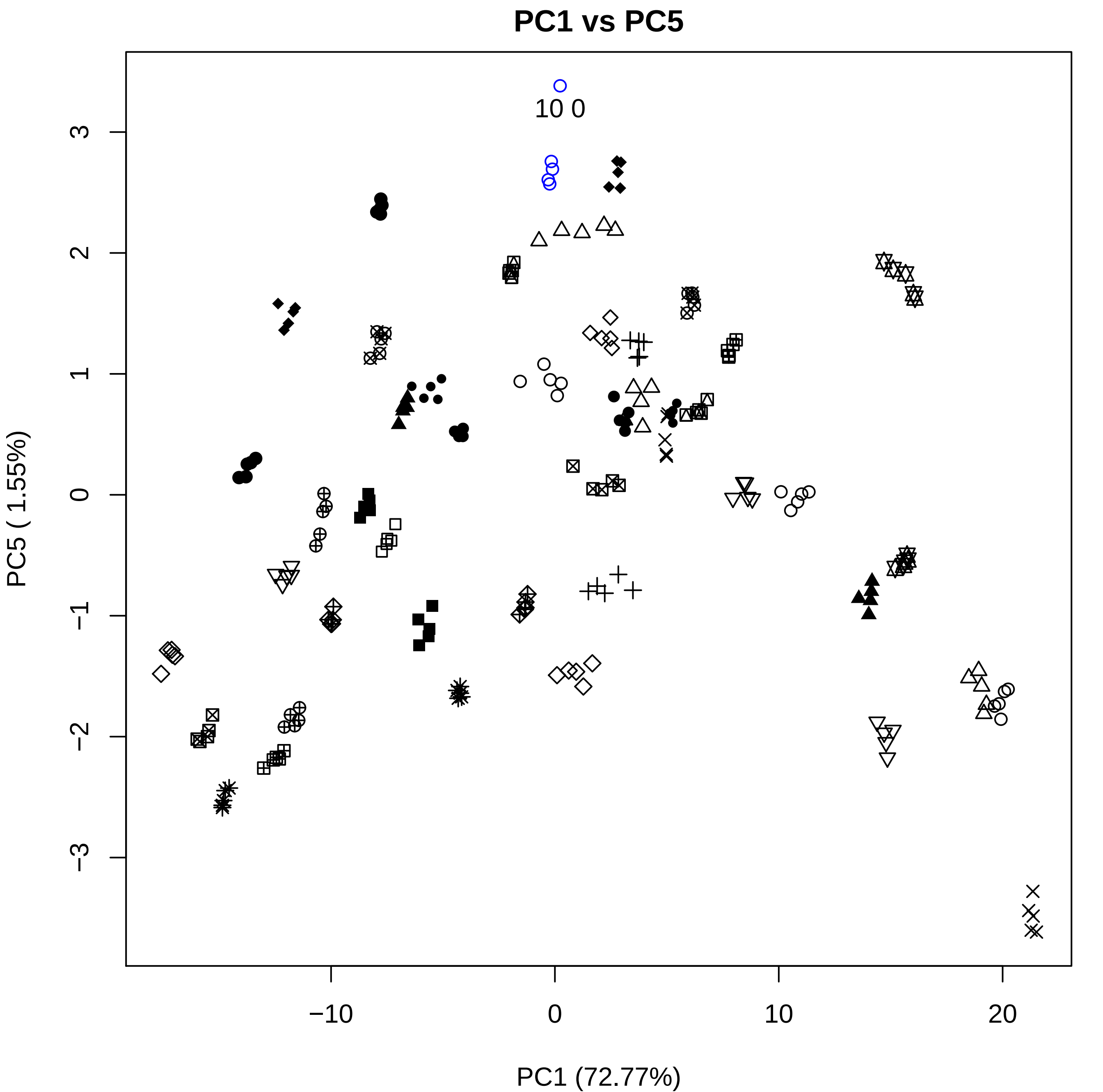
PC1 vs PC5
10 0
PC5 ( 1.55%)
PC1 (72.77%)
−10
0
10
20
−3
−2
−1
0
1
2
3

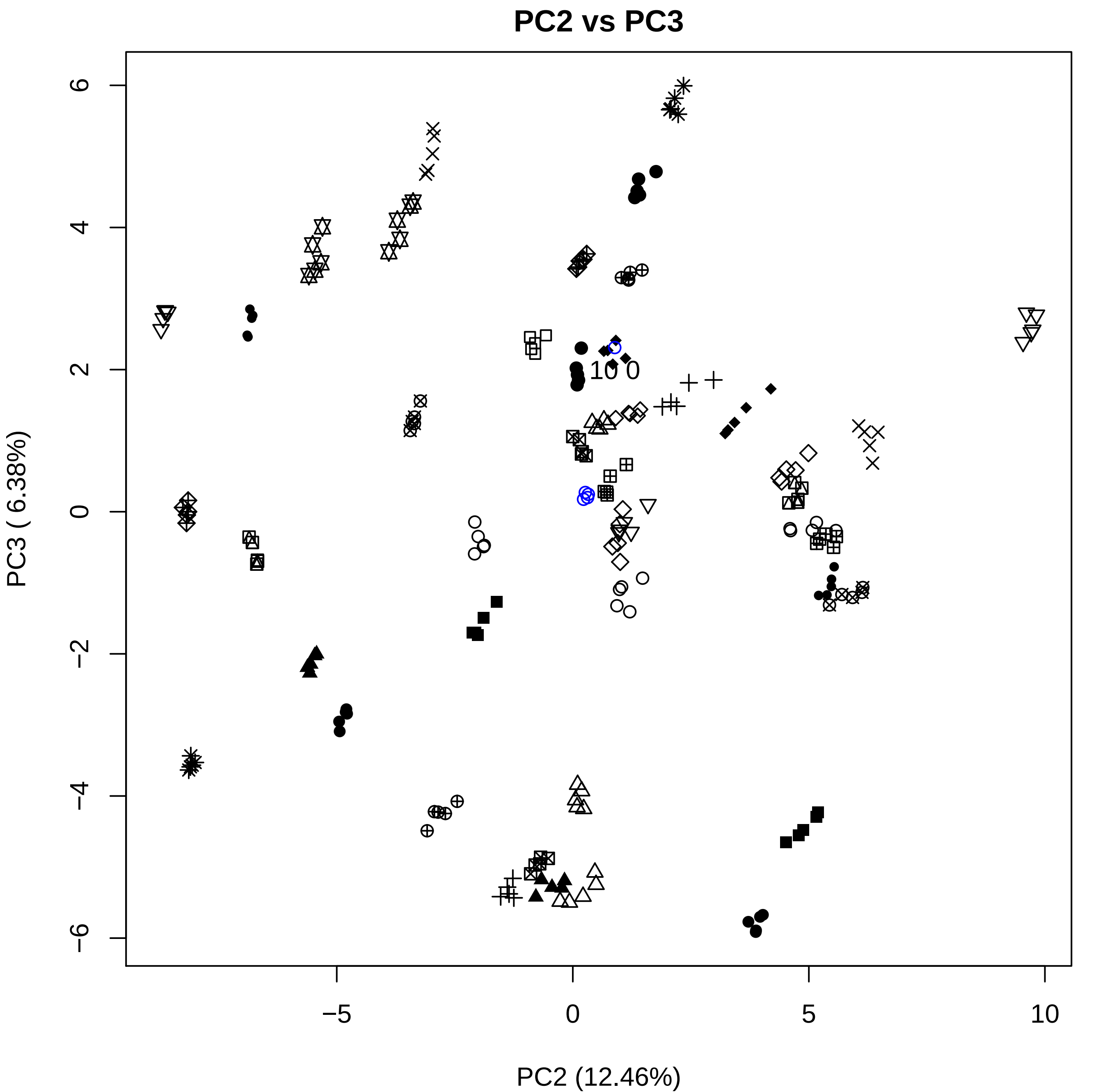

PC2 vs PC3
PC3 ( 6.38%)
PC2 (12.46%)
6
4
2
0
−2
−4
−6
−5
0
5
10
10 0

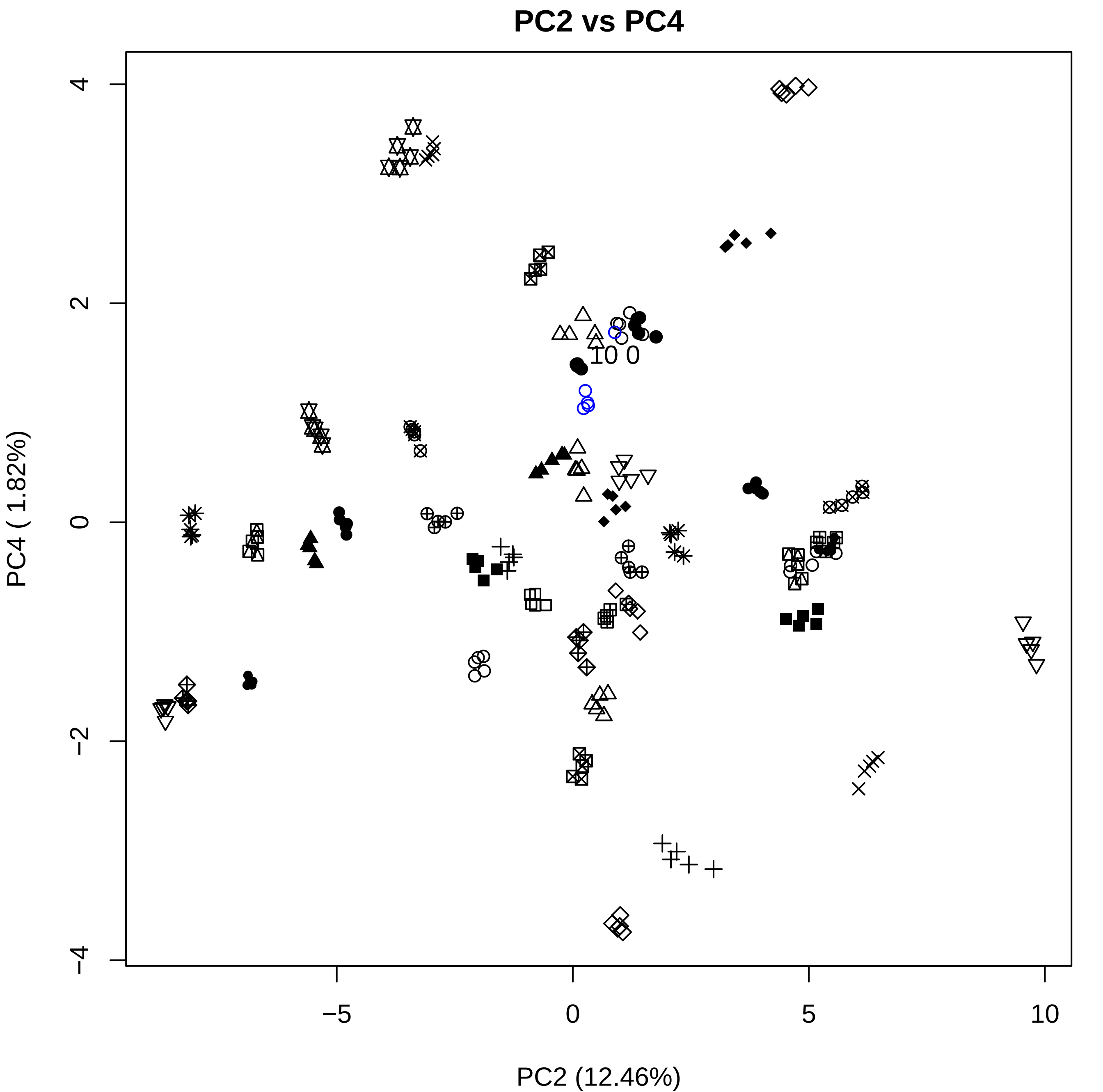
PC2 vs PC4
PC4 ( 1.82%)
PC2 (12.46%)
4
2
0
−2
−4
−5
0
5
10
10 0

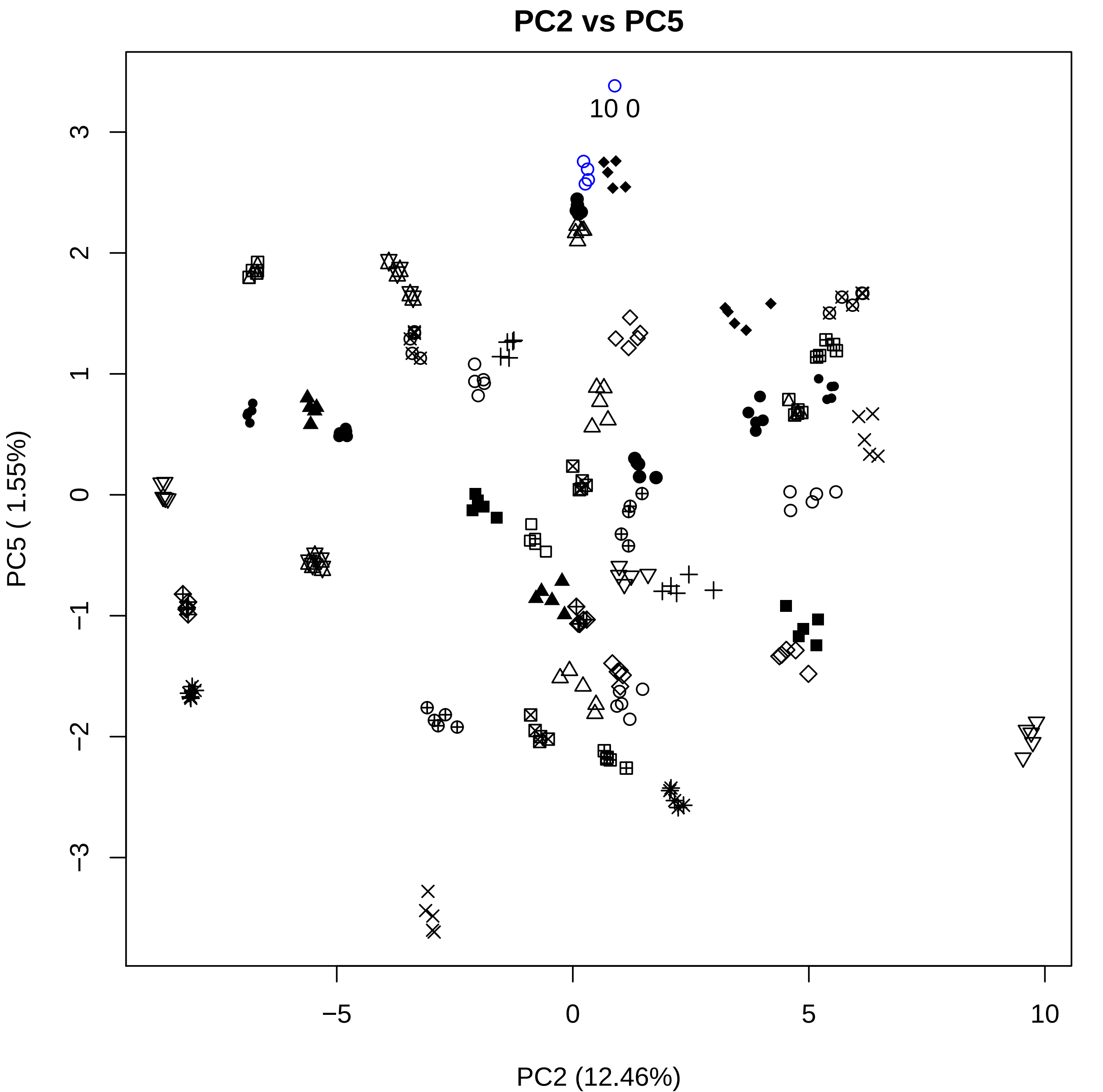
PC2 vs PC5
10 0
PC5 ( 1.55%)
PC2 (12.46%)
3
2
1
0
−1
−2
−3
−5
0
5
10

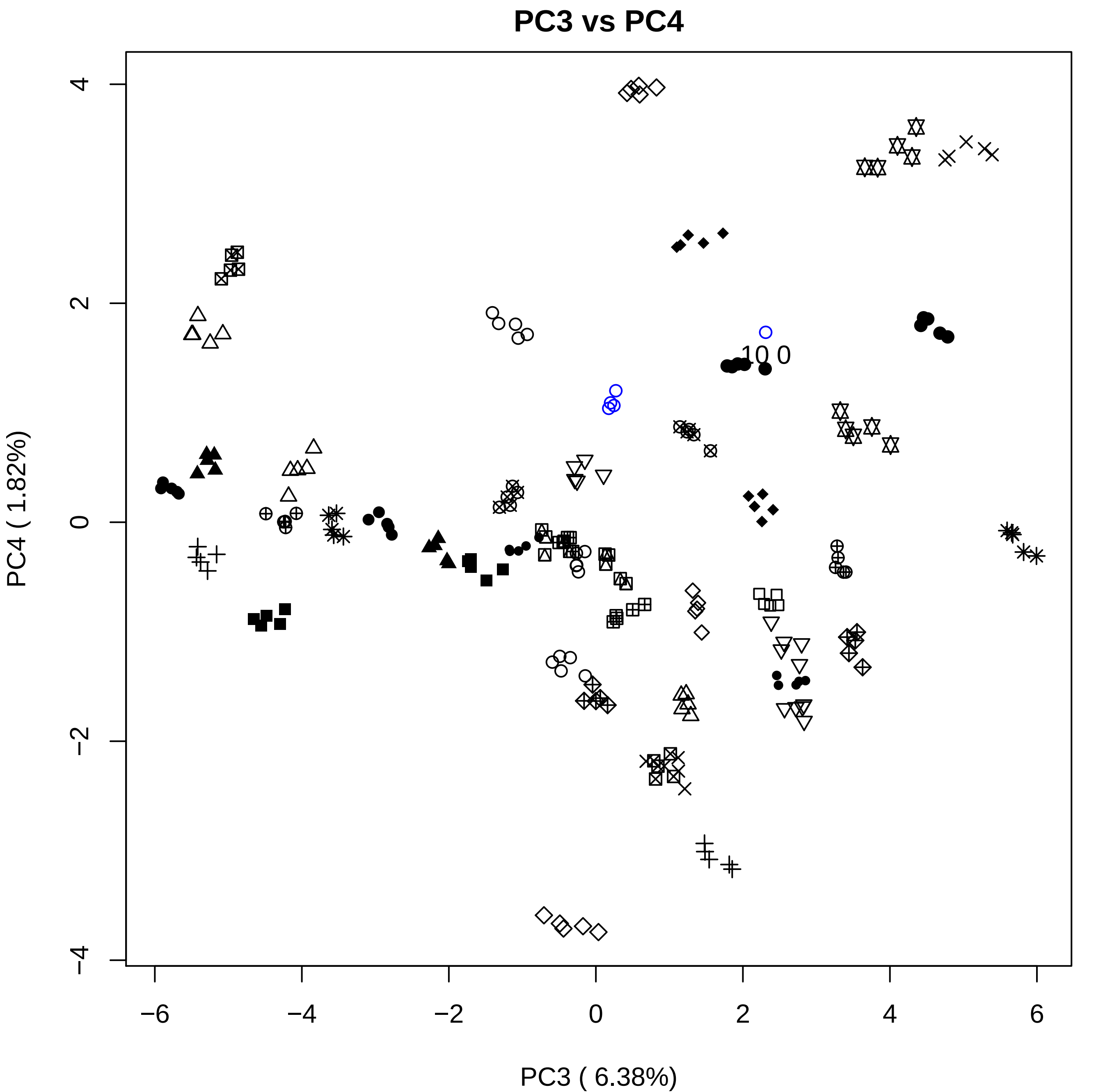
PC3 vs PC4
PC4 ( 1.82%)
PC3 ( 6.38%)
−6
−4
−2
0
2
4
6
10 0

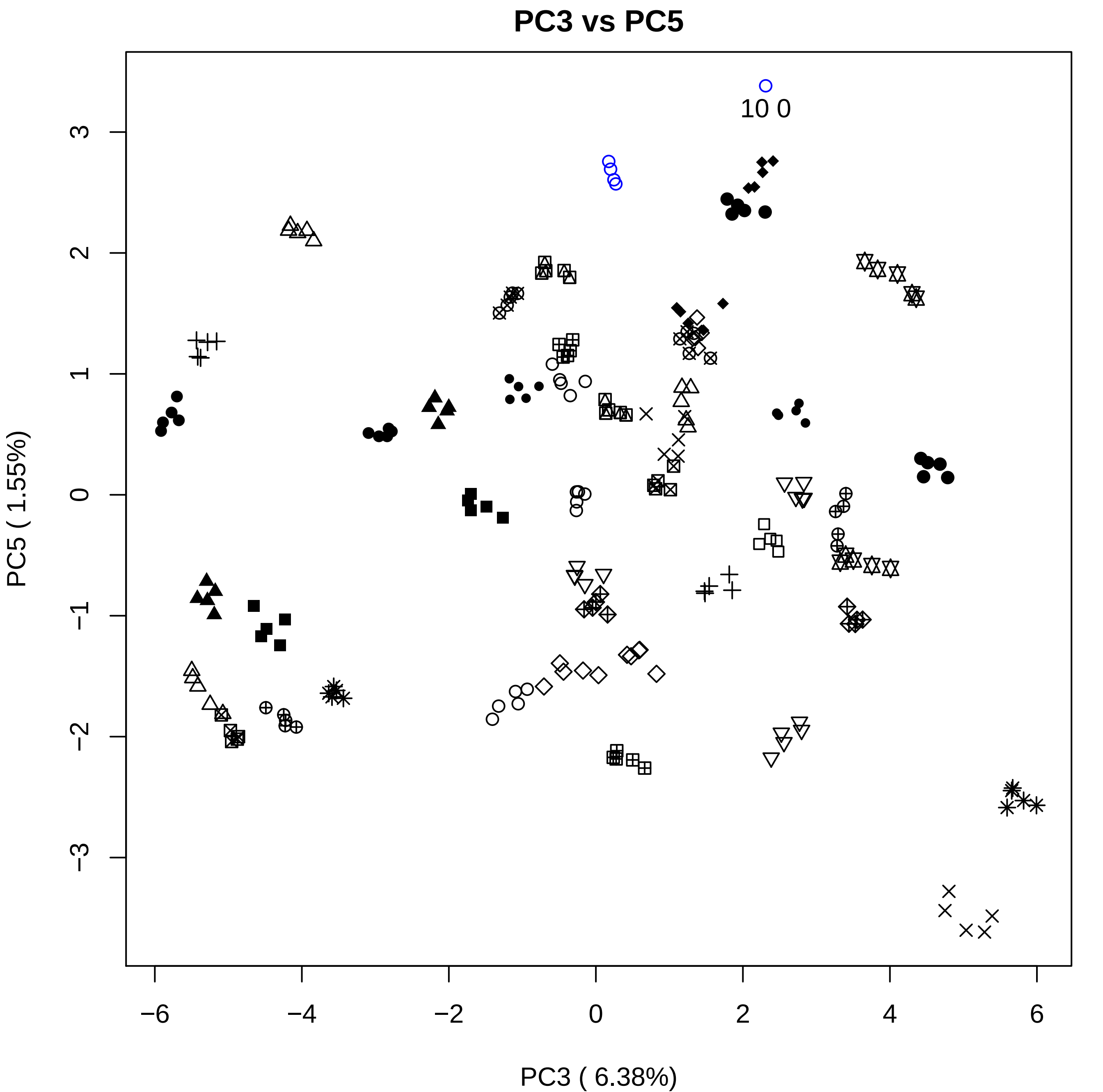

PC3 vs PC5
10 0
PC5 ( 1.55%)
PC3 ( 6.38%)
−6
−4
−2
0
2
4
6
−3
−2
−1
0
1
2
3

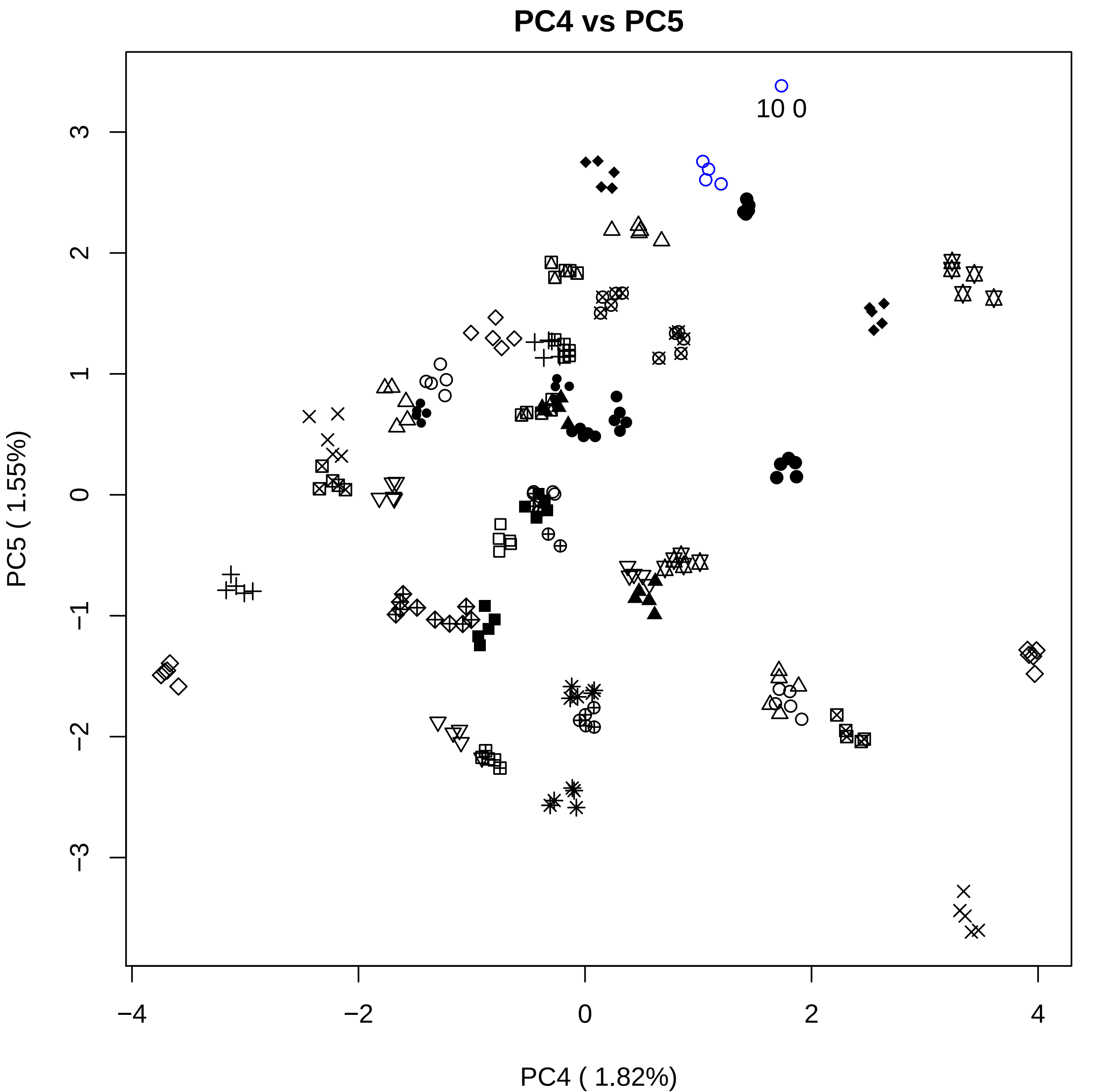
PC4 vs PC5
10 0
PC5 ( 1.55%)
PC4 ( 1.82%)
−4
−2
0
2
4
−3
−2
−1
0
1
2
3

### PCA of 15%  OSCS vs PMH

Scaled Matrix PCAed

7th order polynomial baseline correction performed

Raw spectra smoothed to 21 Neighbours, 2nd order PN, Savtisky Golay

Normalisation between 0 and 100

D2 spectra smoothed to 41 Neighbours, 2nd order PN, Savtisky Golay

Regions Analysed: 3601−2501 and 2001−700cm−1

Samples Analysed: 1 Contam's 0 OSCS's 45 PMH's

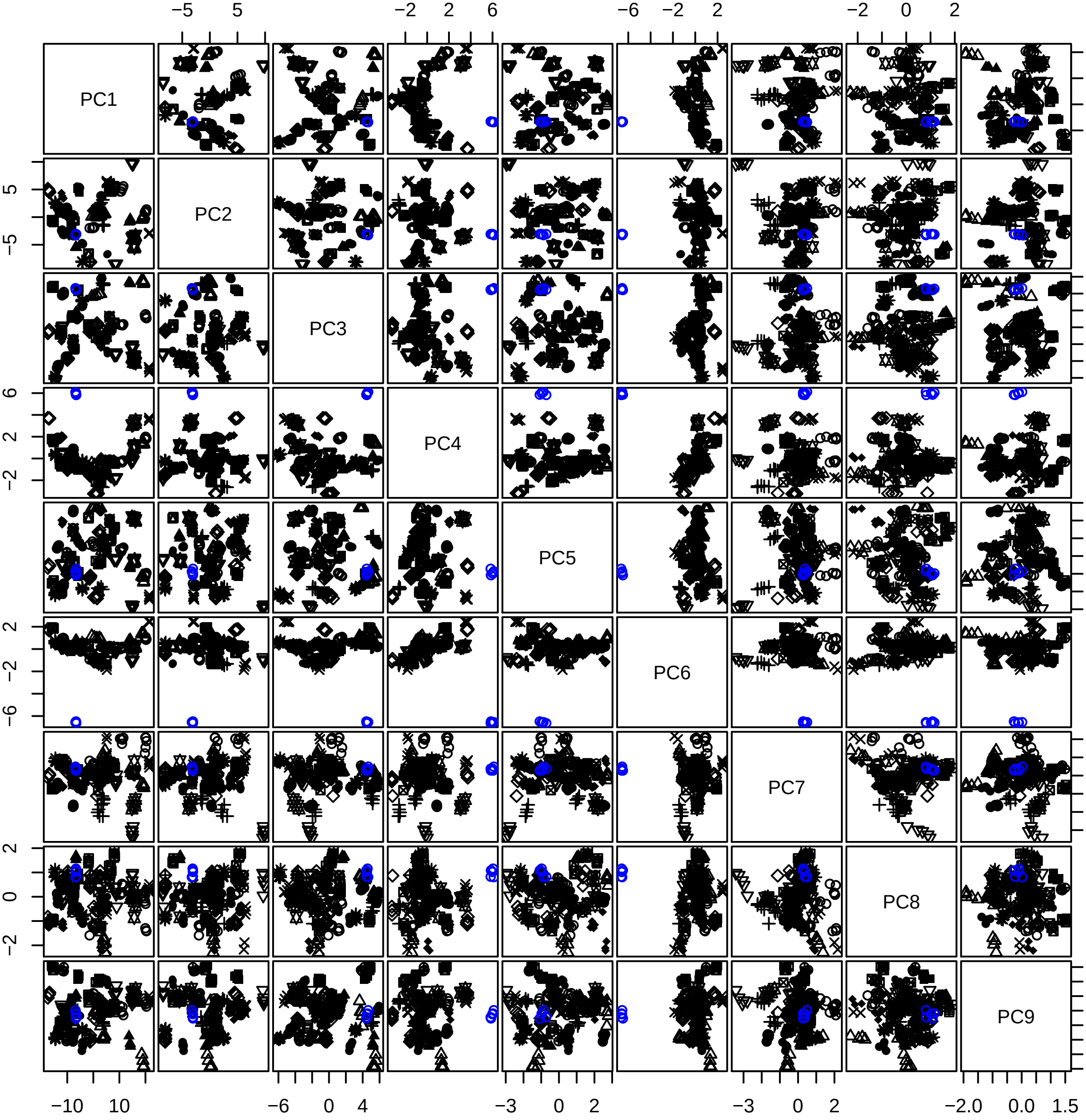
PC1
PC2
PC3
PC4
PC5
PC6
PC7
PC8
PC9
−5 5
−2 2 6
−6 −2 2
−2 0 2
10 −10
5 −5
4 0 −6
6 2 −2
2 0 −3
2 −2 −6
2 0 −3
2 0 −2
1.5 0.0 −2.0
−10 10
−6 0 4
−3 0 2
−3 0 2
−2.0 0.0 1.5

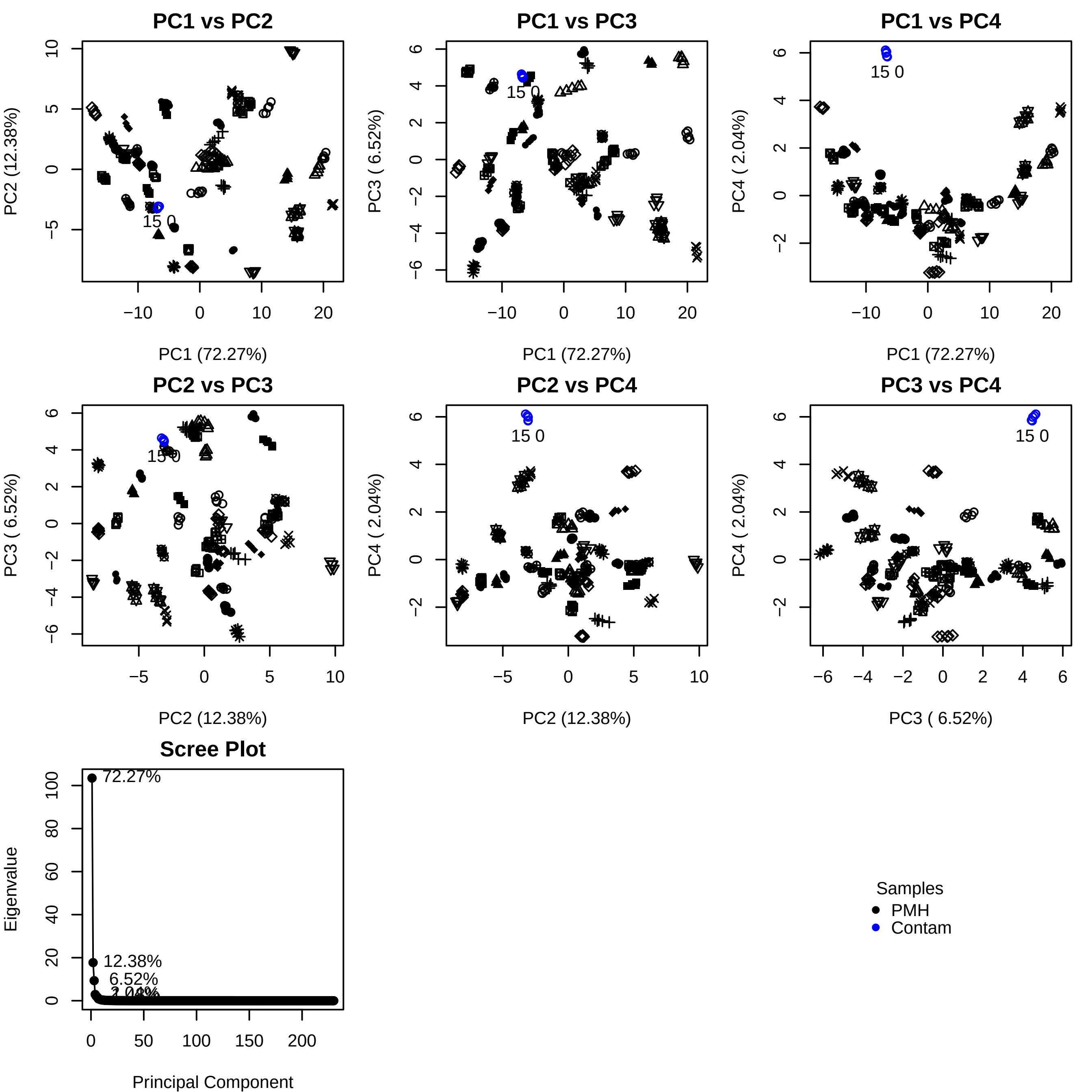
PC1 vs PC2
PC2 (12.38%)
PC1 (72.27%)
15 0
PC1 vs PC3
PC3 ( 6.52%)
PC1 (72.27%)
15 0
PC1 vs PC4
PC4 ( 2.04%)
PC1 (72.27%)
15 0
PC2 vs PC3
PC3 ( 6.52%)
PC2 (12.38%)
15 0
PC2 vs PC4
PC4 ( 2.04%)
PC2 (12.38%)
15 0
PC3 vs PC4
PC4 ( 2.04%)
PC3 ( 6.52%)
15 0
Scree Plot
Eigenvalue
Principal Component
72.27%
12.38%
6.52%
Samples
PMH
Contam

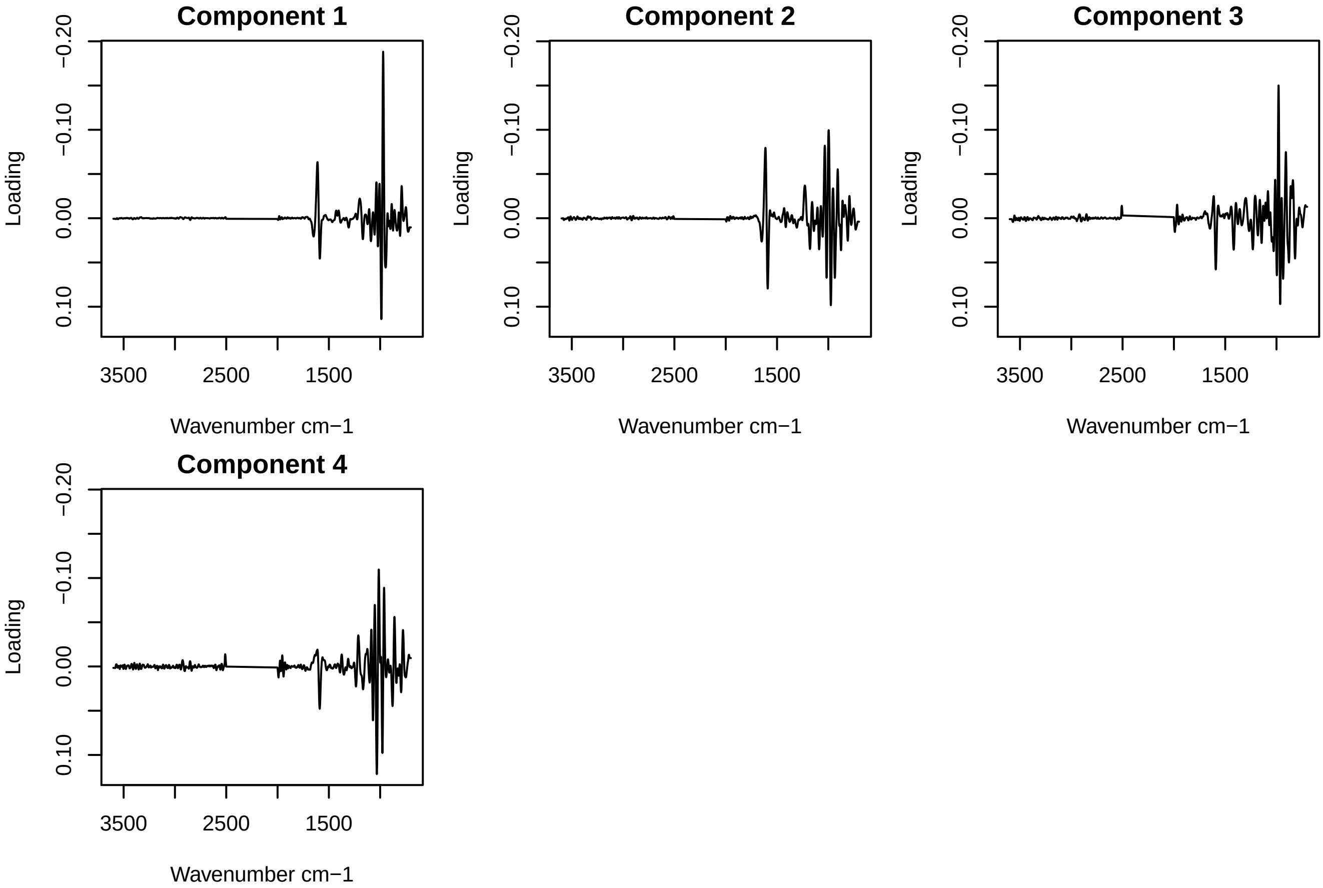
Component 1
Component 2
Component 3
Component 4
Loading
−0.20
−0.10
0.00
0.10
3500
2500
1500
Wavenumber cm−1

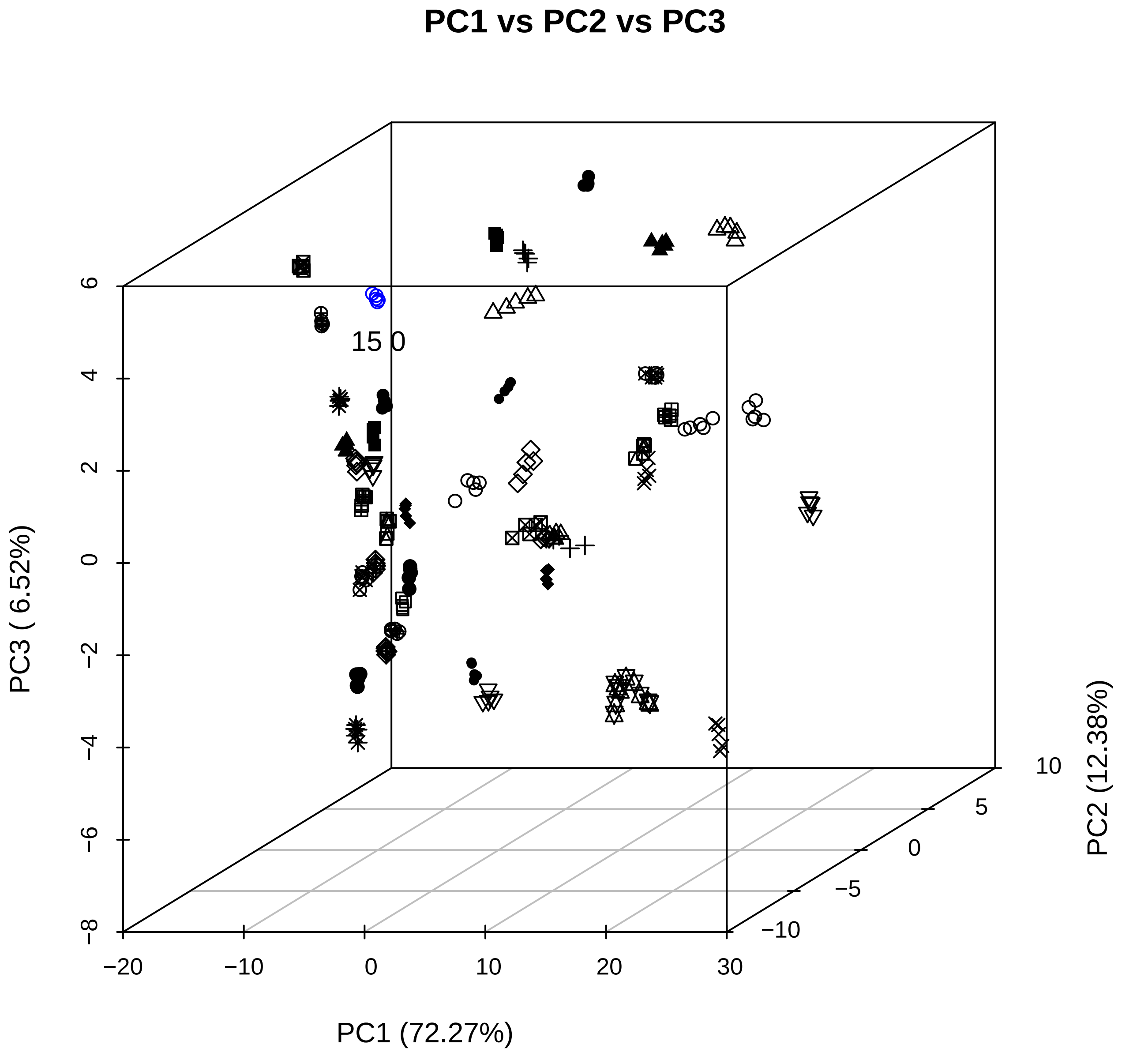
PC1 vs PC2 vs PC3
15 0
PC1 (72.27%)
PC2 (12.38%)
PC3 ( 6.52%)
−20
−10
0
10
20
30
−10
−5
0
5
10
−8
−6
−4
−2
0
2
4
6

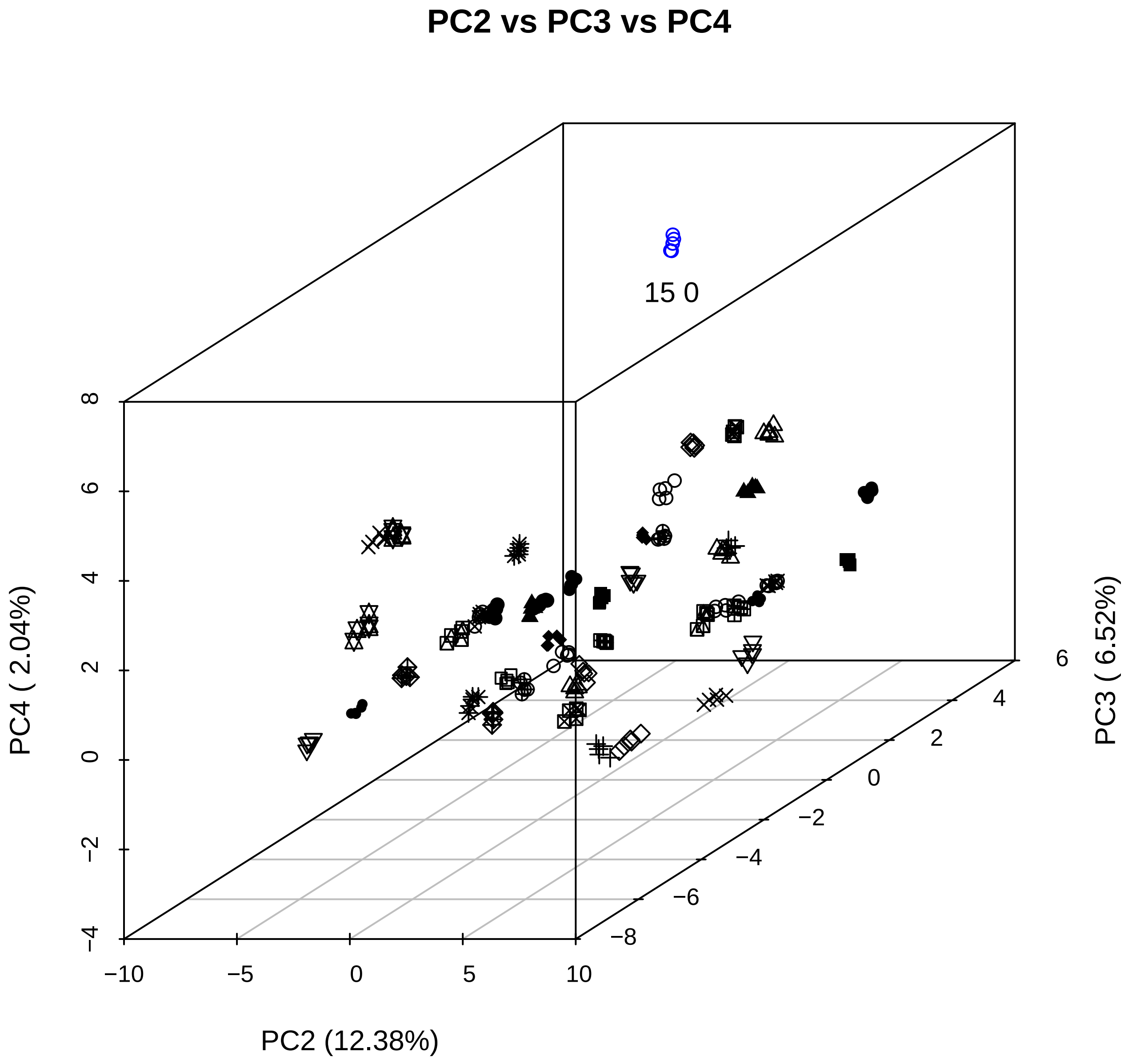

PC2 vs PC3 vs PC4
15 0
PC2 (12.38%)
PC3 ( 6.52%)
PC4 ( 2.04%)
−10
−5
0
5
10
−8
−6
−4
−2
0
2
4
6
−4
−2
0
2
4
6
8

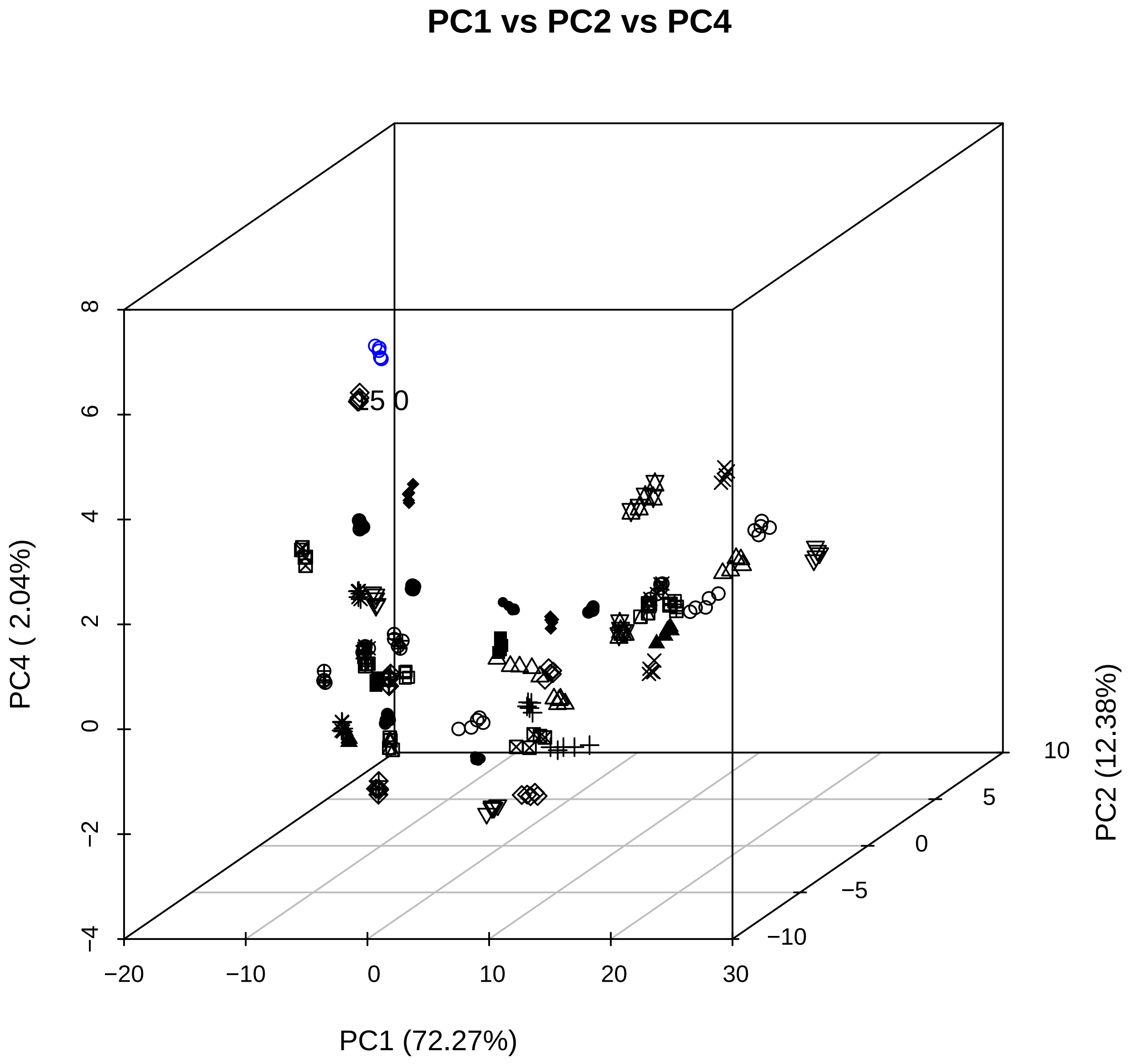
PC1 vs PC2 vs PC4
25 0
PC1 (72.27%)
PC2 (12.38%)
PC4 ( 2.04%)
−20
−10
0
10
20
30
−10
−5
0
5
10
−4
−2
0
2
4
6
8

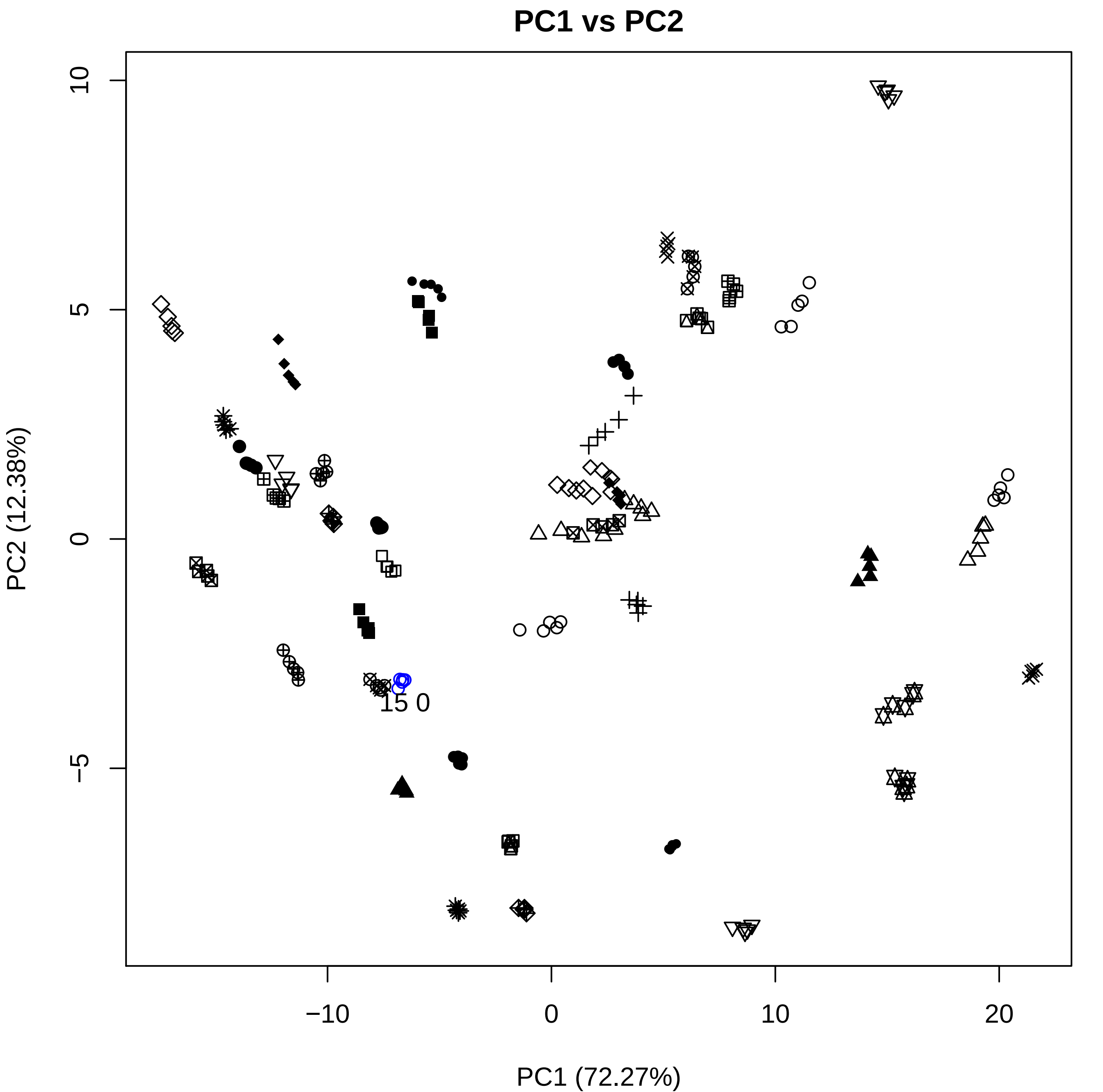
PC1 vs PC2
15 0
PC2 (12.38%)
PC1 (72.27%)
10
5
0
−5
−10
0
10
20

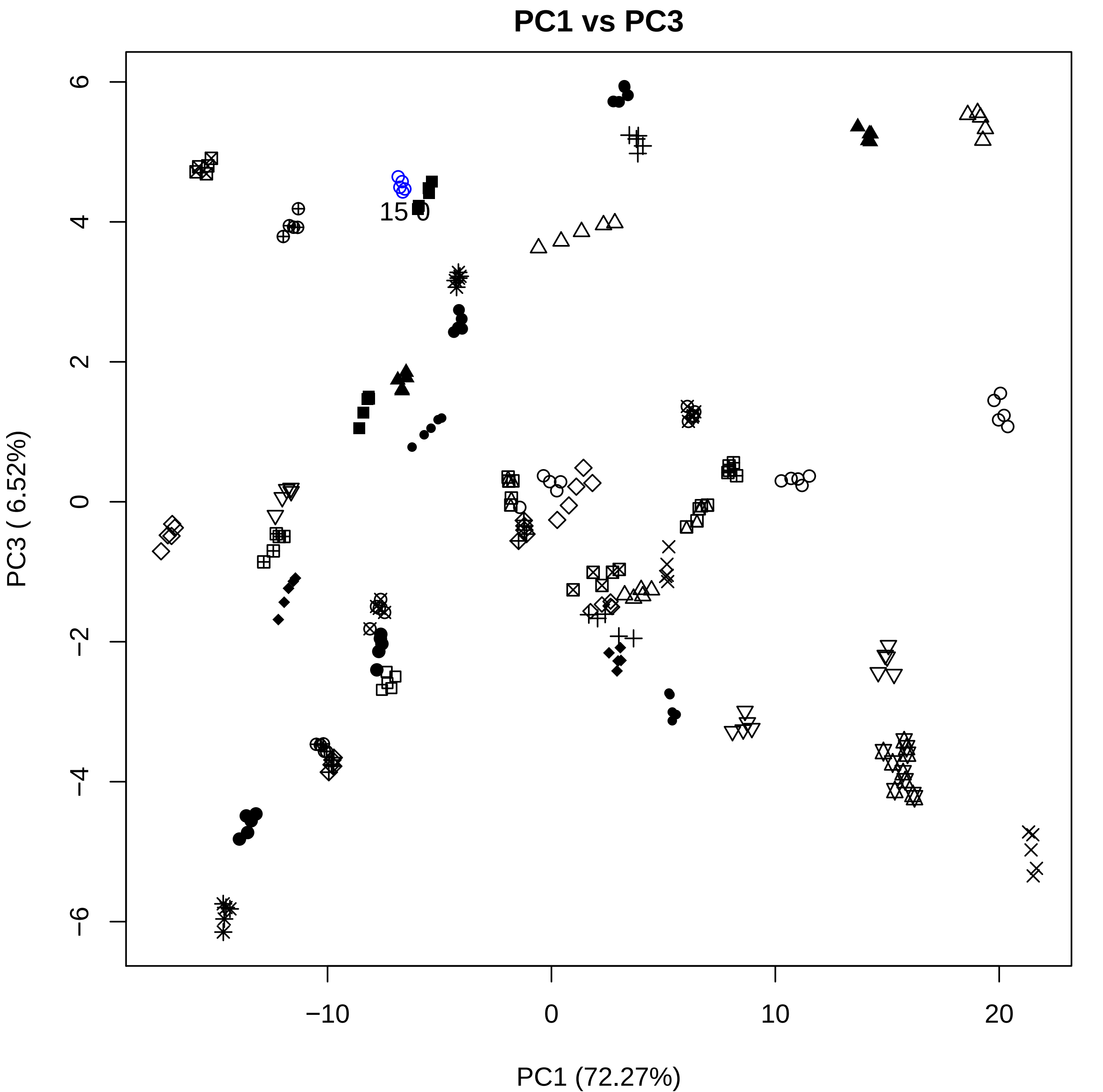
PC1 vs PC3
PC3 ( 6.52%)
PC1 (72.27%)
6
4
2
0
−2
−4
−6
−10
0
10
20
150

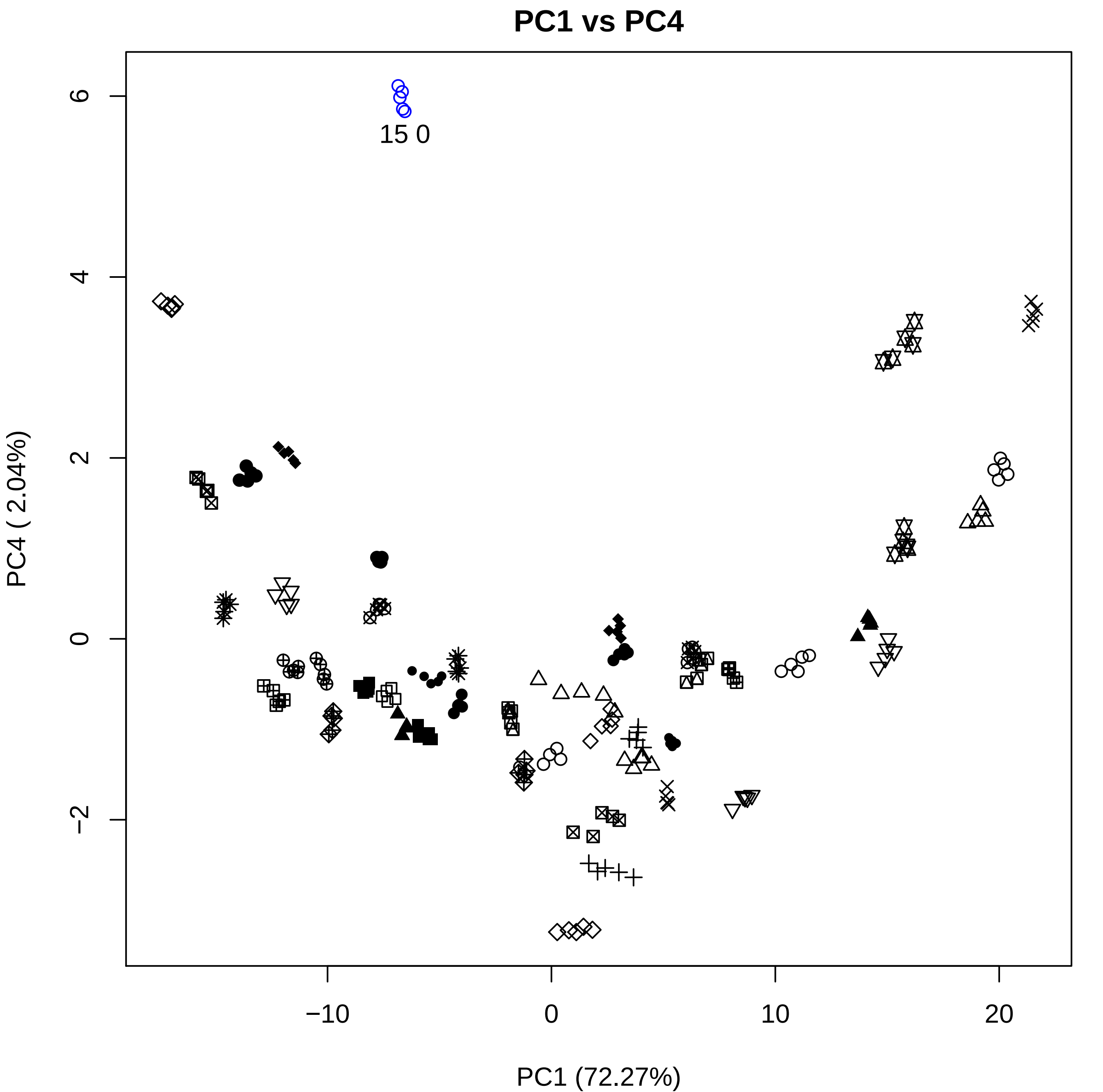

PC1 vs PC4
15 0
PC4 ( 2.04%)
PC1 (72.27%)
6
4
2
0
−2
−10
0
10
20

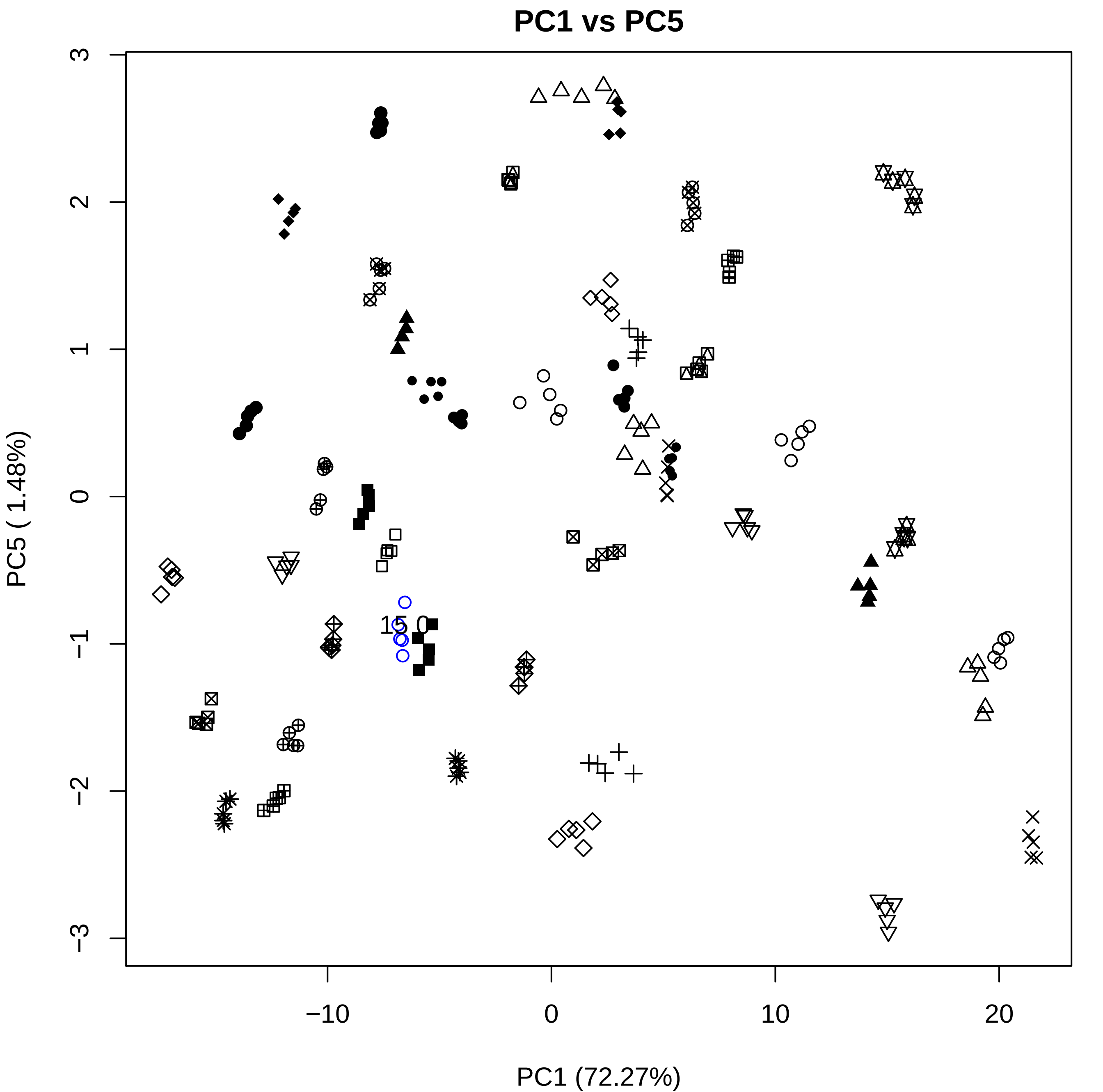
PC1 vs PC5
PC5 ( 1.48%)
PC1 (72.27%)
3
2
1
0
−1
−2
−3
−10
0
10
20
15.0

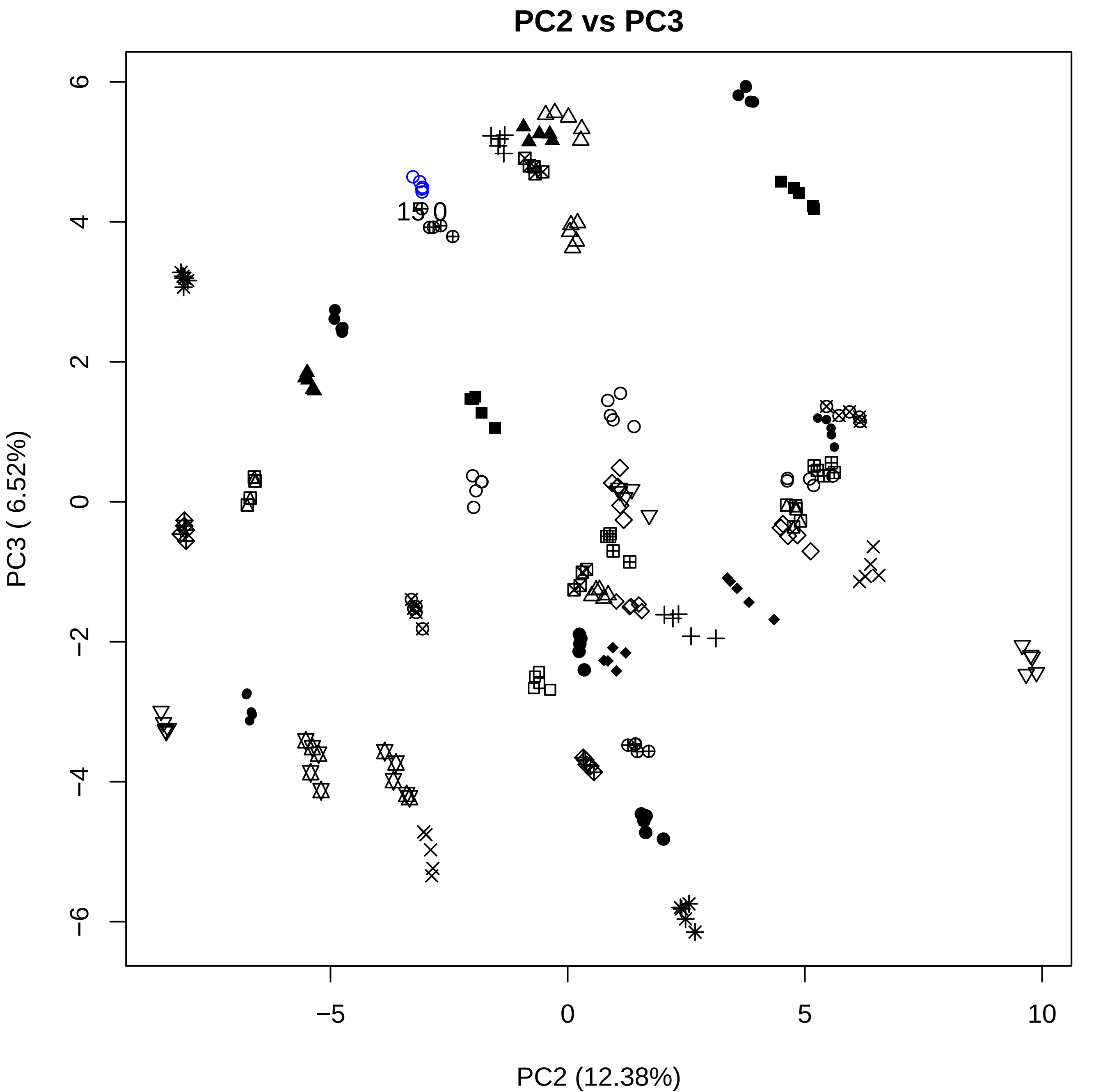

PC2 vs PC3
PC3 ( 6.52%)
PC2 (12.38%)
6
4
2
0
−2
−4
−6
−5
0
5
10
15
0

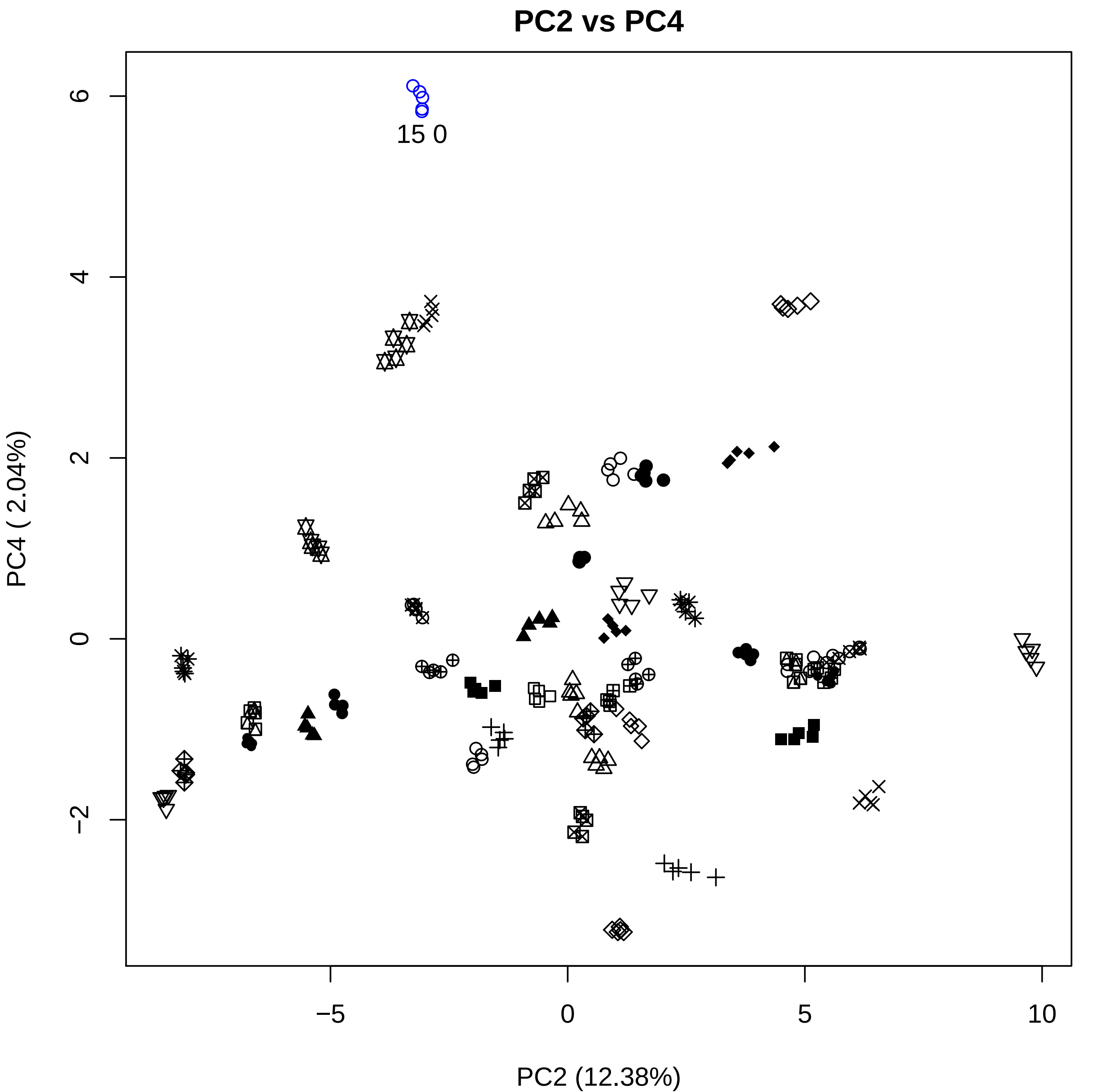

PC2 vs PC4
15 0
6
4
2
0
−2
PC4 ( 2.04%)
−5
0
5
10
PC2 (12.38%)

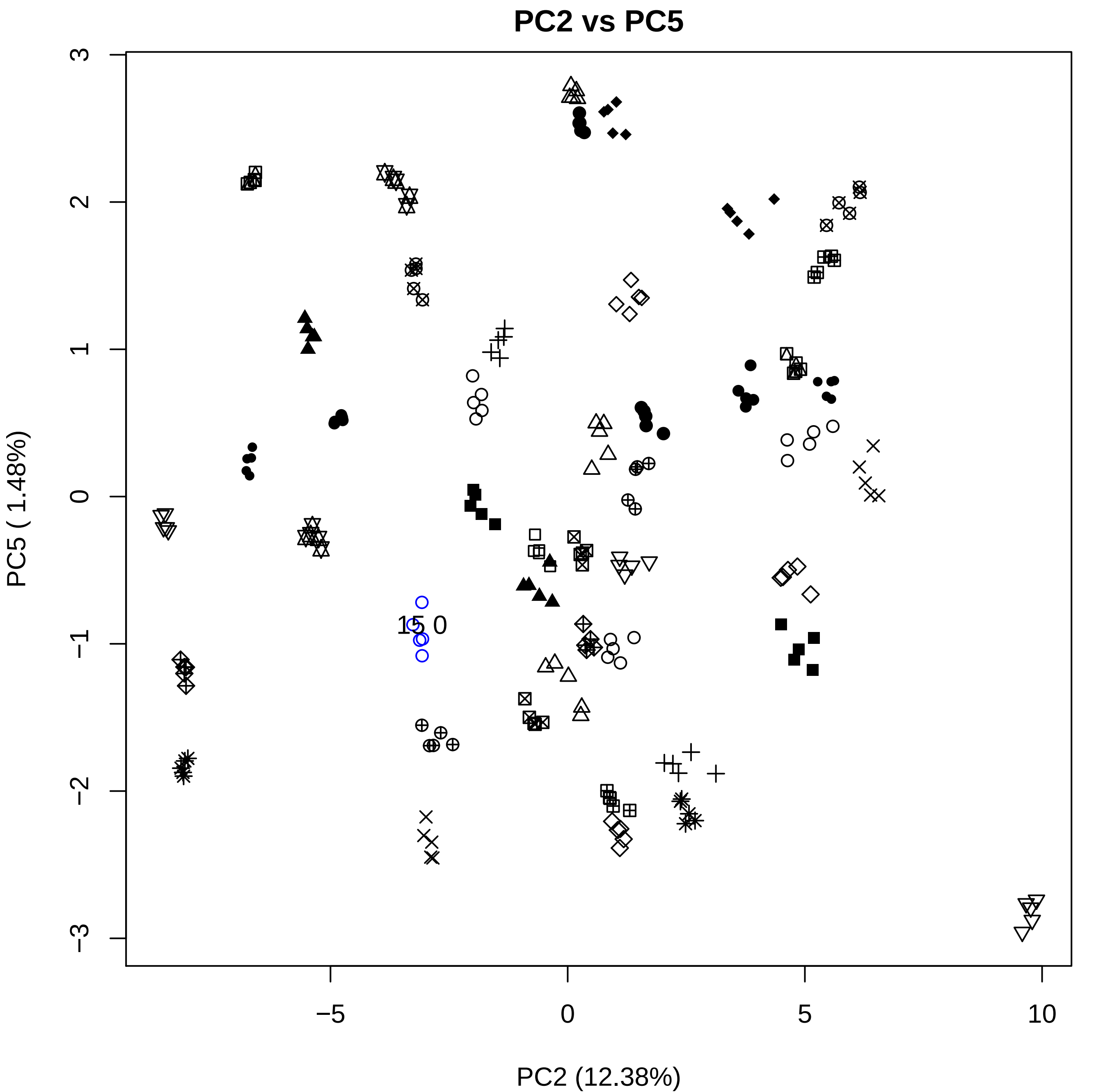

PC2 vs PC5
PC5 ( 1.48%)
PC2 (12.38%)
3
2
1
0
−1
−2
−3
−5
0
5
10
15 0

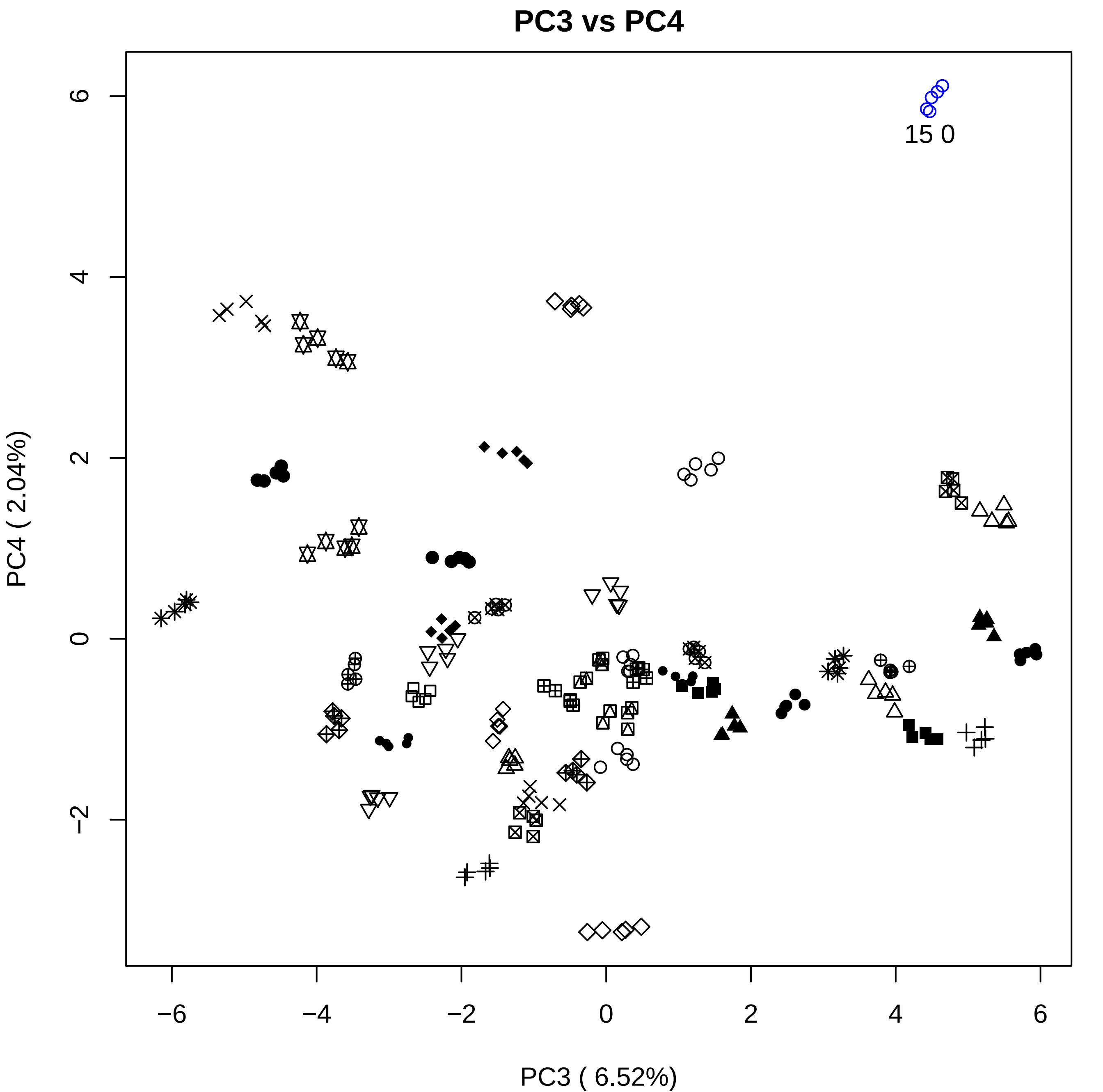

PC3 vs PC4
15 0
PC4 ( 2.04%)
PC3 ( 6.52%)
−6
−4
−2
0
2
4
6
−2
0
2
4
6

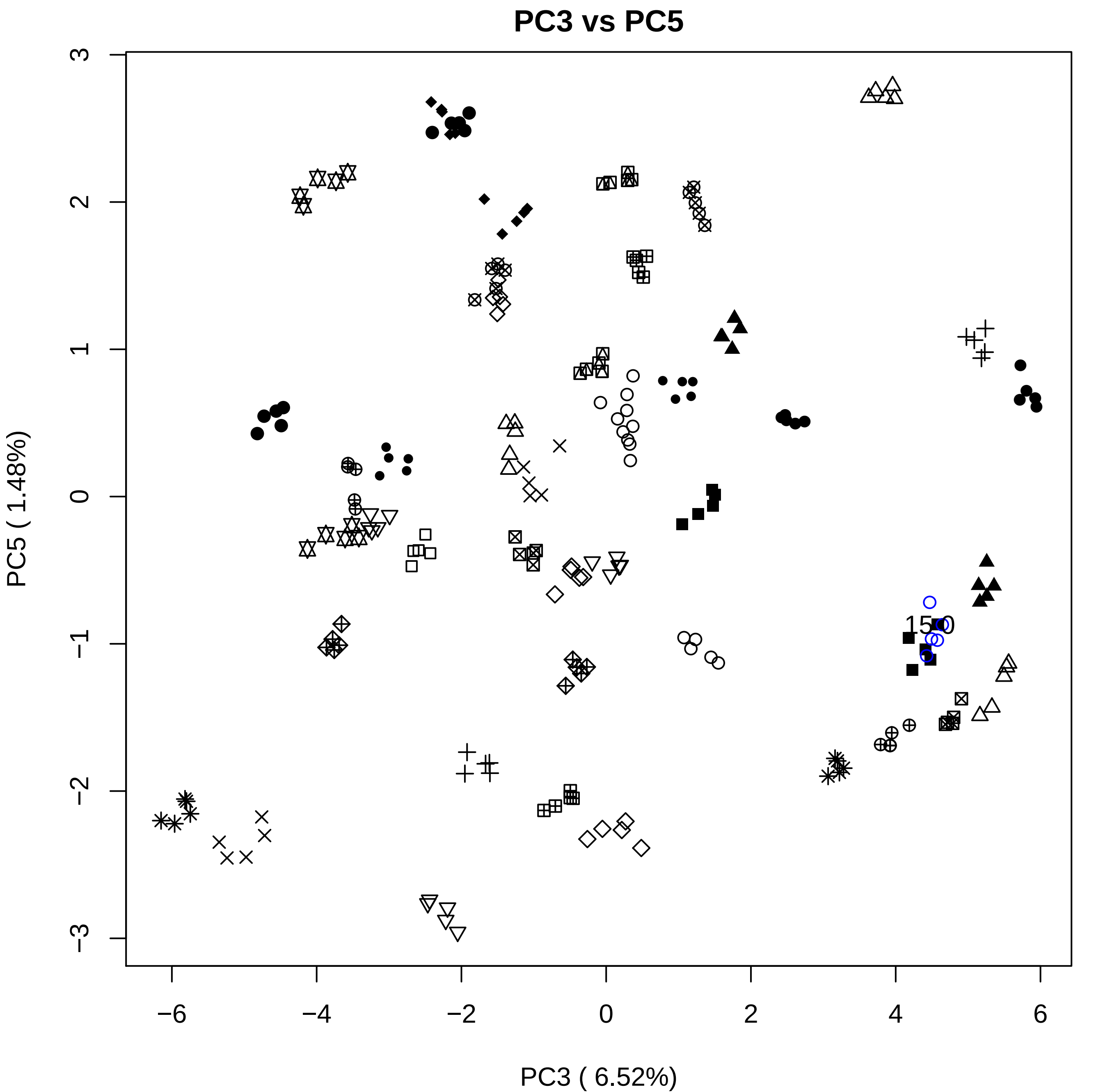
PC3 vs PC5
PC5 ( 1.48%)
PC3 ( 6.52%)
3
2
1
0
−1
−2
−3
−6
−4
−2
0
2
4
6
150

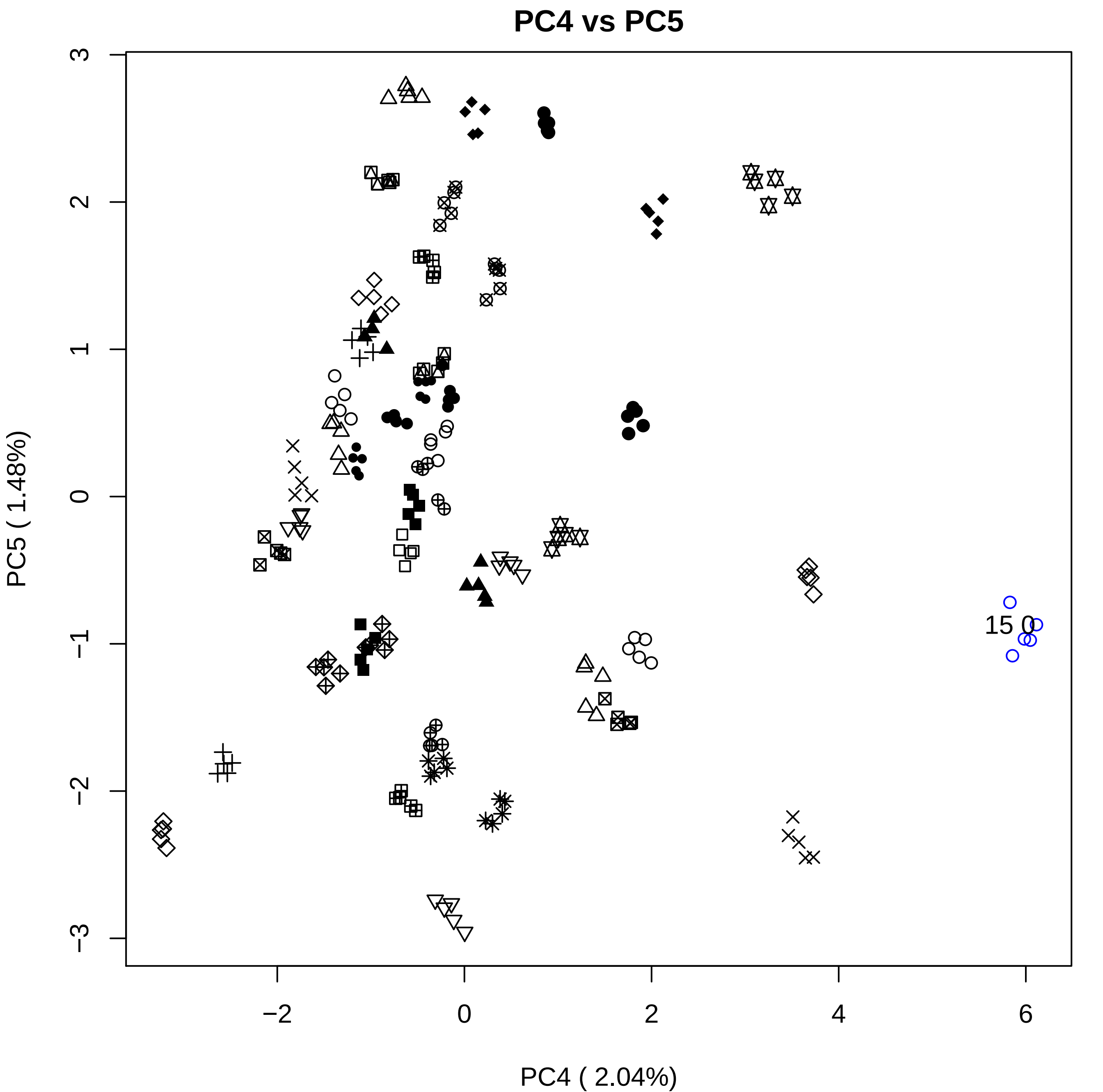
PC4 vs PC5
PC5 ( 1.48%)
PC4 ( 2.04%)
3
2
1
0
−1
−2
−3
−2
0
2
4
6
15 0

### PCA of 20%  OSCS vs PMH

Scaled Matrix PCAed

7th order polynomial baseline correction performed

Raw spectra smoothed to 21 Neighbours, 2nd order PN, Savtisky Golay

Normalisation between 0 and 100

D2 spectra smoothed to 41 Neighbours, 2nd order PN, Savtisky Golay

Regions Analysed: 3601−2501 and 2001−700cm−1

Samples Analysed: 1 Contam's 0 OSCS's 45 PMH's

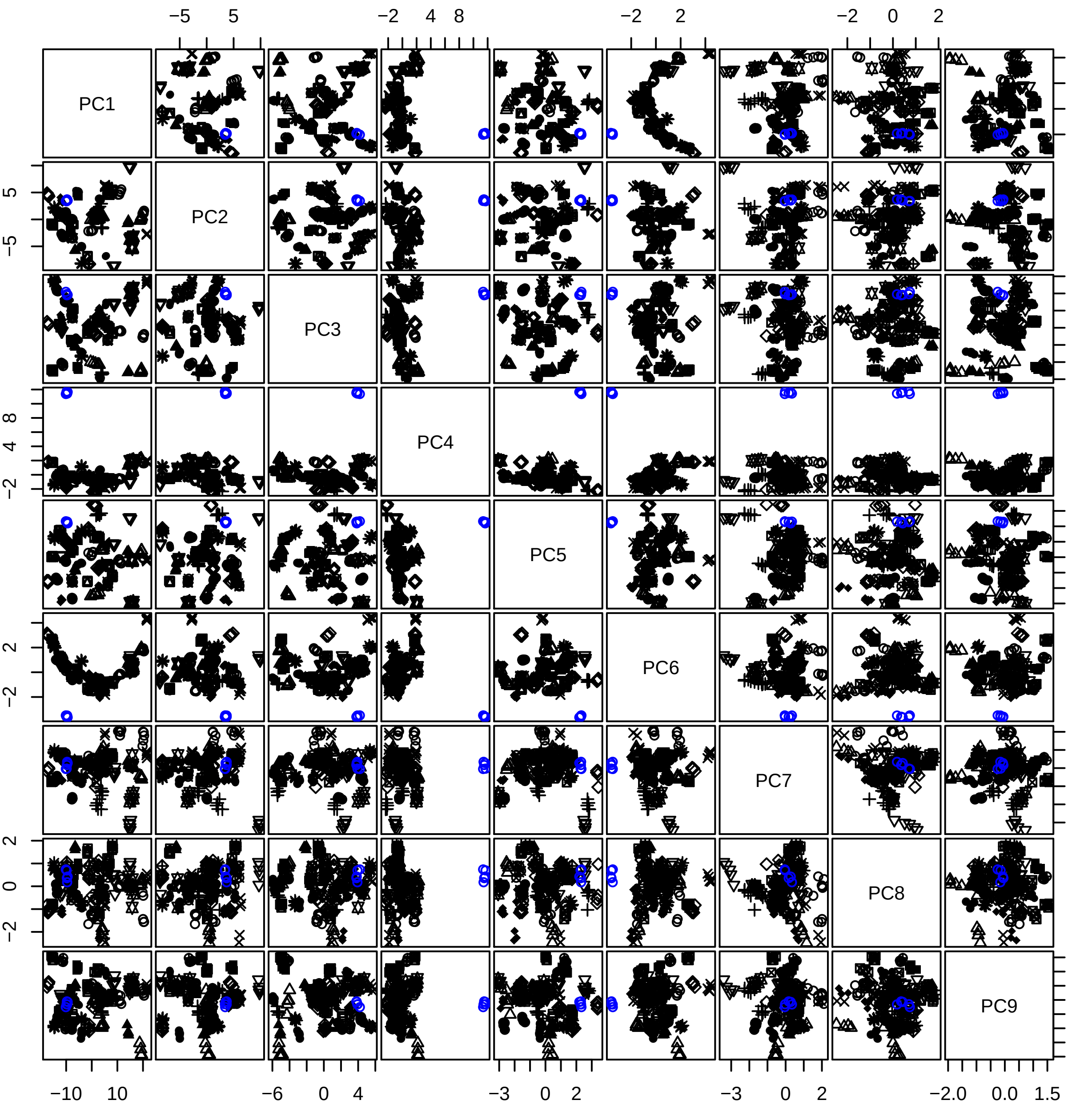
PC1
PC2
PC3
PC4
PC5
PC6
PC7
PC8
PC9
−5
5
−2
4
8
−2
2
−2
0
2
10
−10
5
−5
4
0
−6
8
4
−2
2
0
−3
2
−2
2
0
−3
2
0
−2
1.5
0.0
−2.0
−10
10
−6
0
4
−3
0
2
−3
0
2
−2.0
0.0
1.5

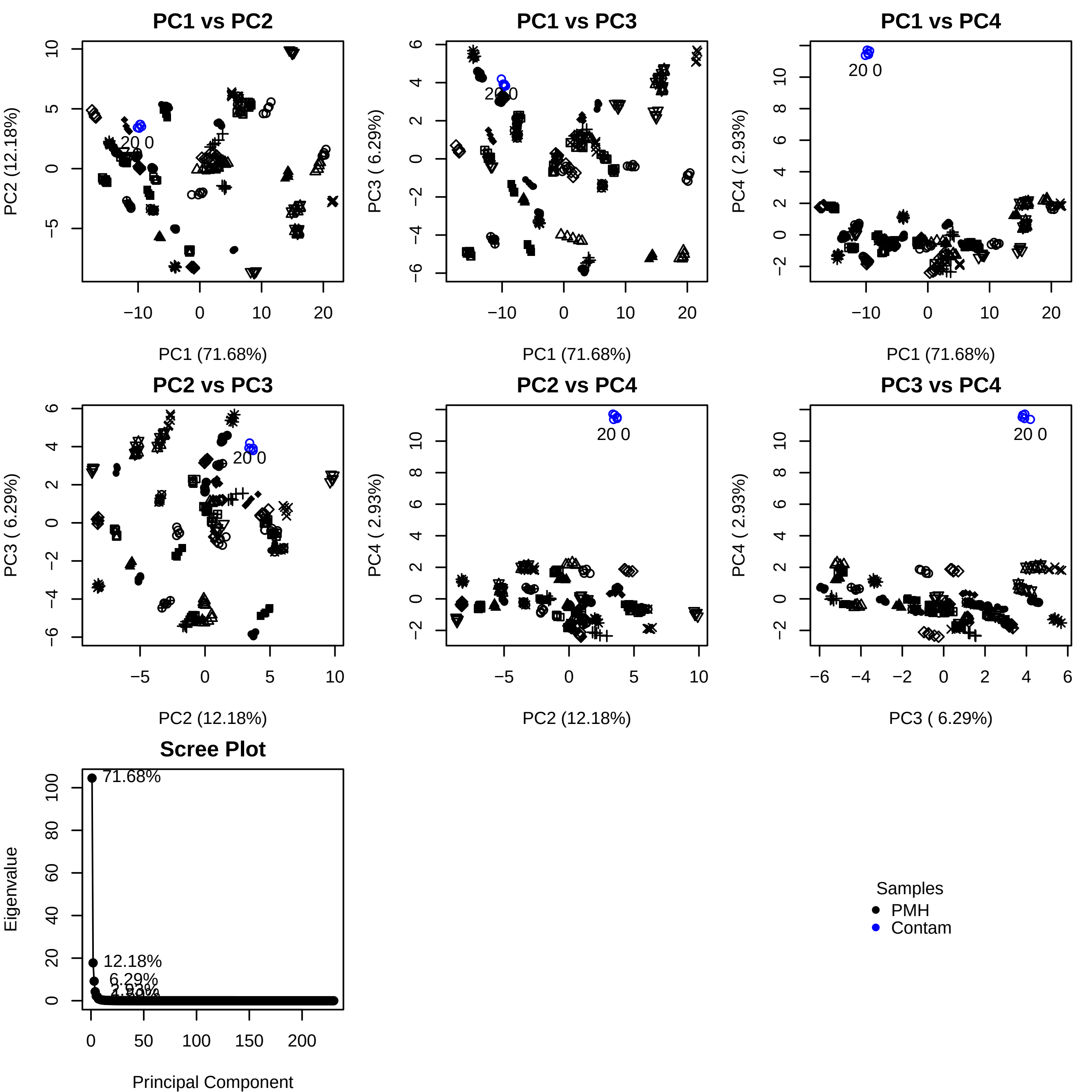

PC1 vs PC2
PC1 vs PC3
PC1 vs PC4
PC2 vs PC3
PC2 vs PC4
PC3 vs PC4
Scree Plot
PC1 (71.68%)
PC2 (12.18%)
PC3 ( 6.29%)
PC4 ( 2.93%)
20 0
71.68%
12.18%
6.29%
Eigenvalue
Principal Component
Samples
PMH
Contam

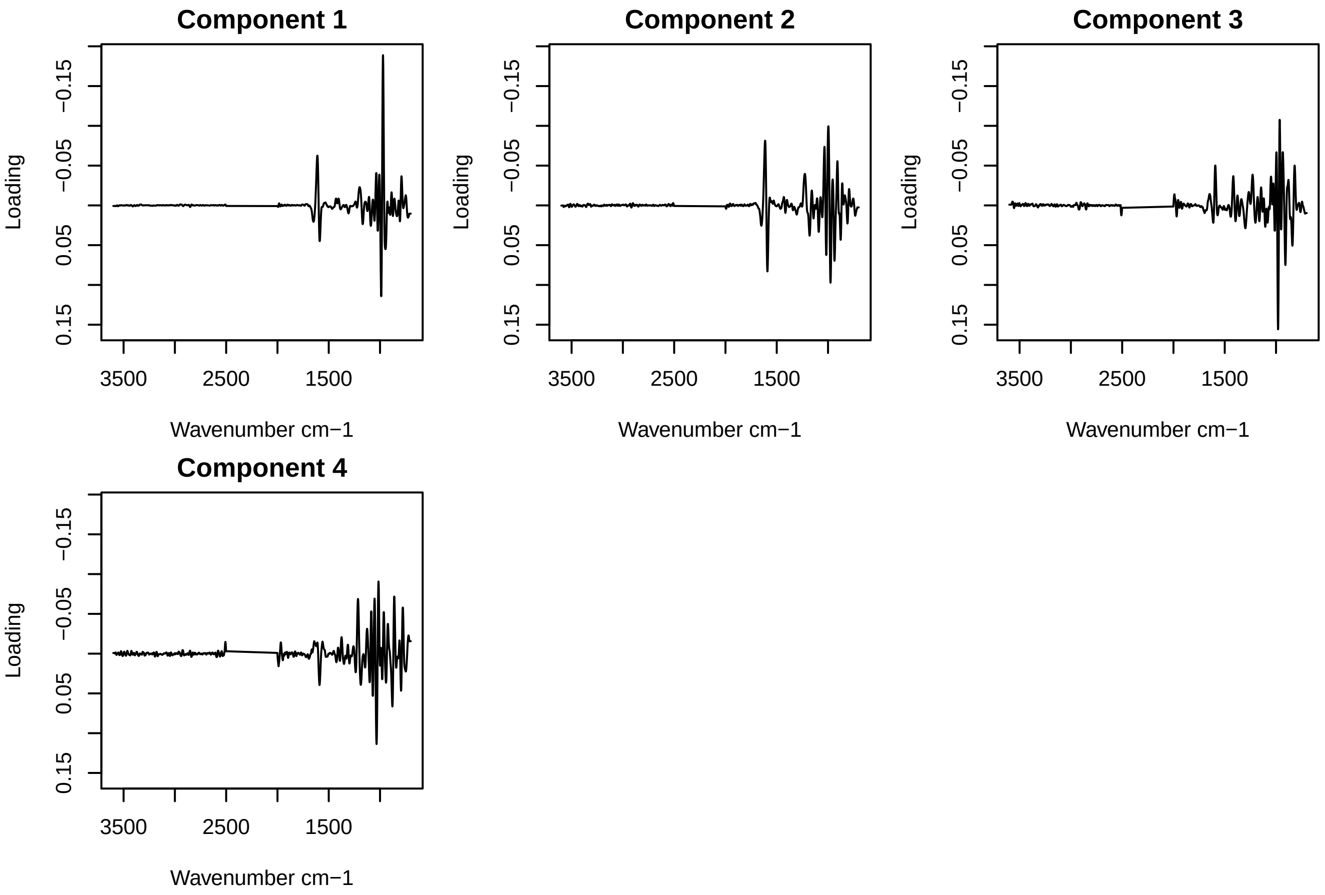
Component 1
Loading
−0.15
−0.05
0.05
0.15
3500
2500
1500
Wavenumber cm−1
Component 2
Loading
−0.15
−0.05
0.05
0.15
3500
2500
1500
Wavenumber cm−1
Component 3
Loading
−0.15
−0.05
0.05
0.15
3500
2500
1500
Wavenumber cm−1
Component 4
Loading
−0.15
−0.05
0.05
0.15
3500
2500
1500
Wavenumber cm−1

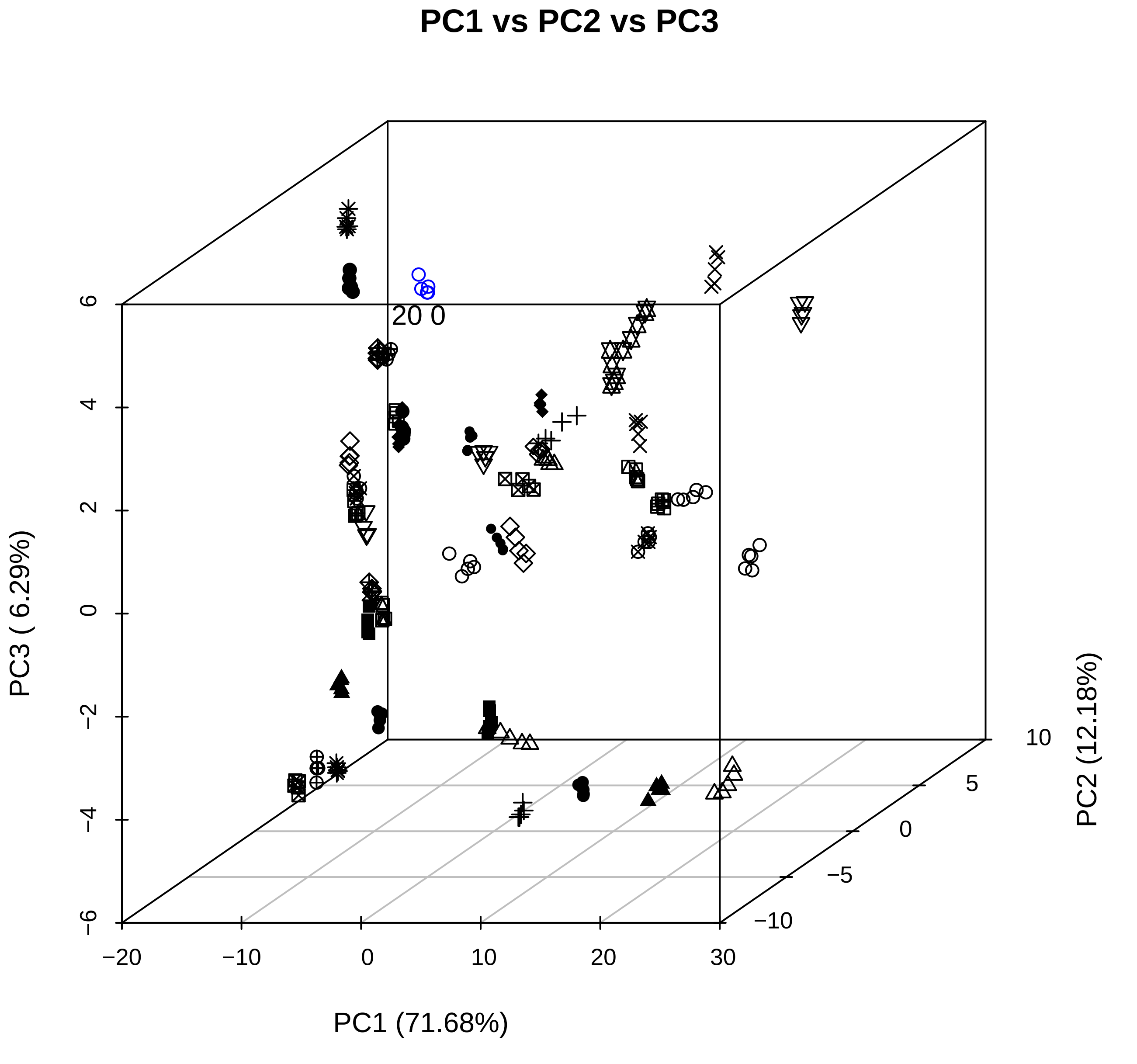
PC1 vs PC2 vs PC3
20 0
PC1 (71.68%)
PC2 (12.18%)
PC3 ( 6.29%)
−20
−10
0
10
20
30
−10
−5
0
5
10
−6
−4
−2
0
2
4
6

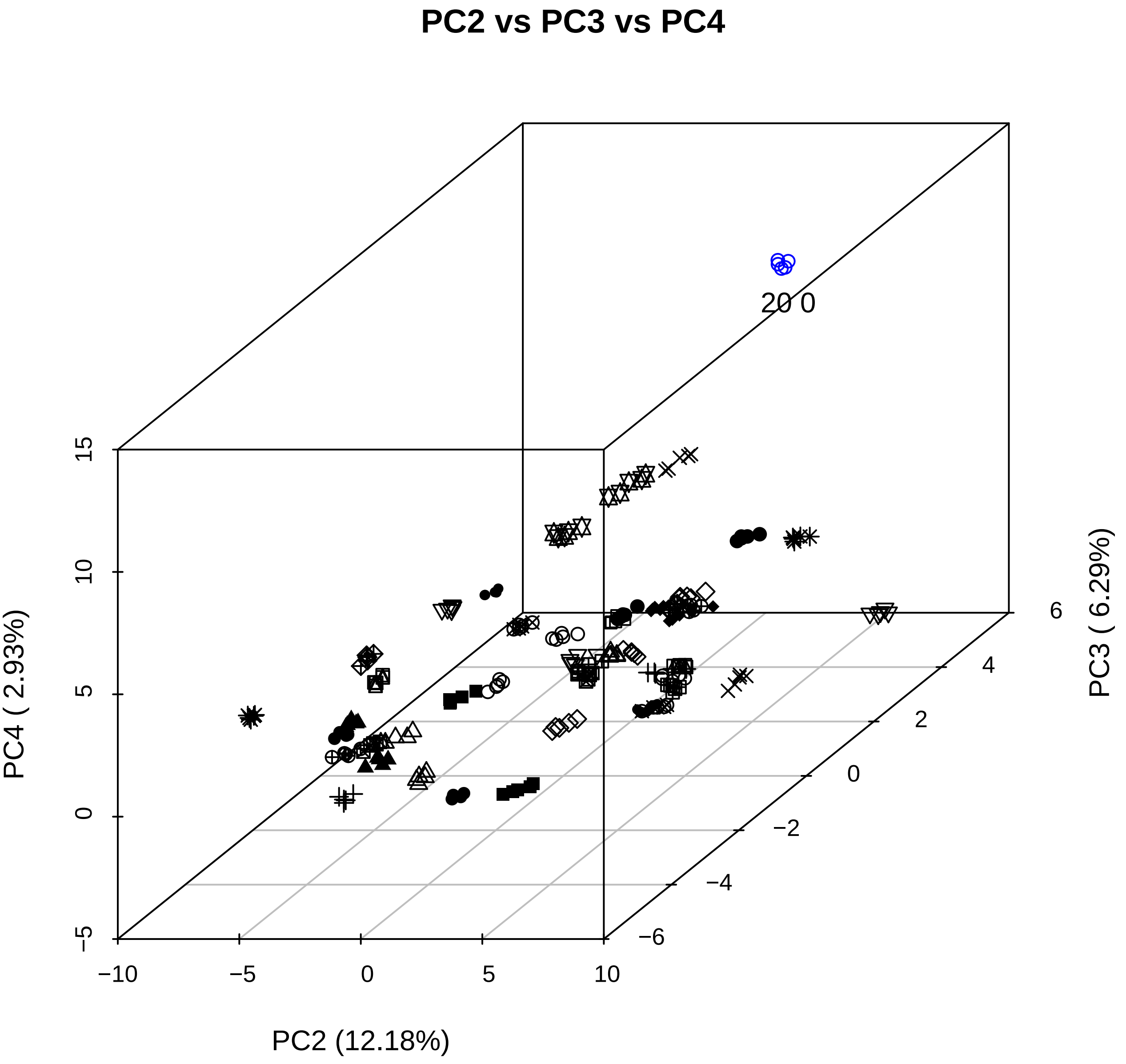
PC2 vs PC3 vs PC4
20 0
PC4 ( 2.93%)
PC3 ( 6.29%)
PC2 (12.18%)
15
10
5
0
−5
−10
−5
0
5
10
6
4
2
0
−2
−4
−6

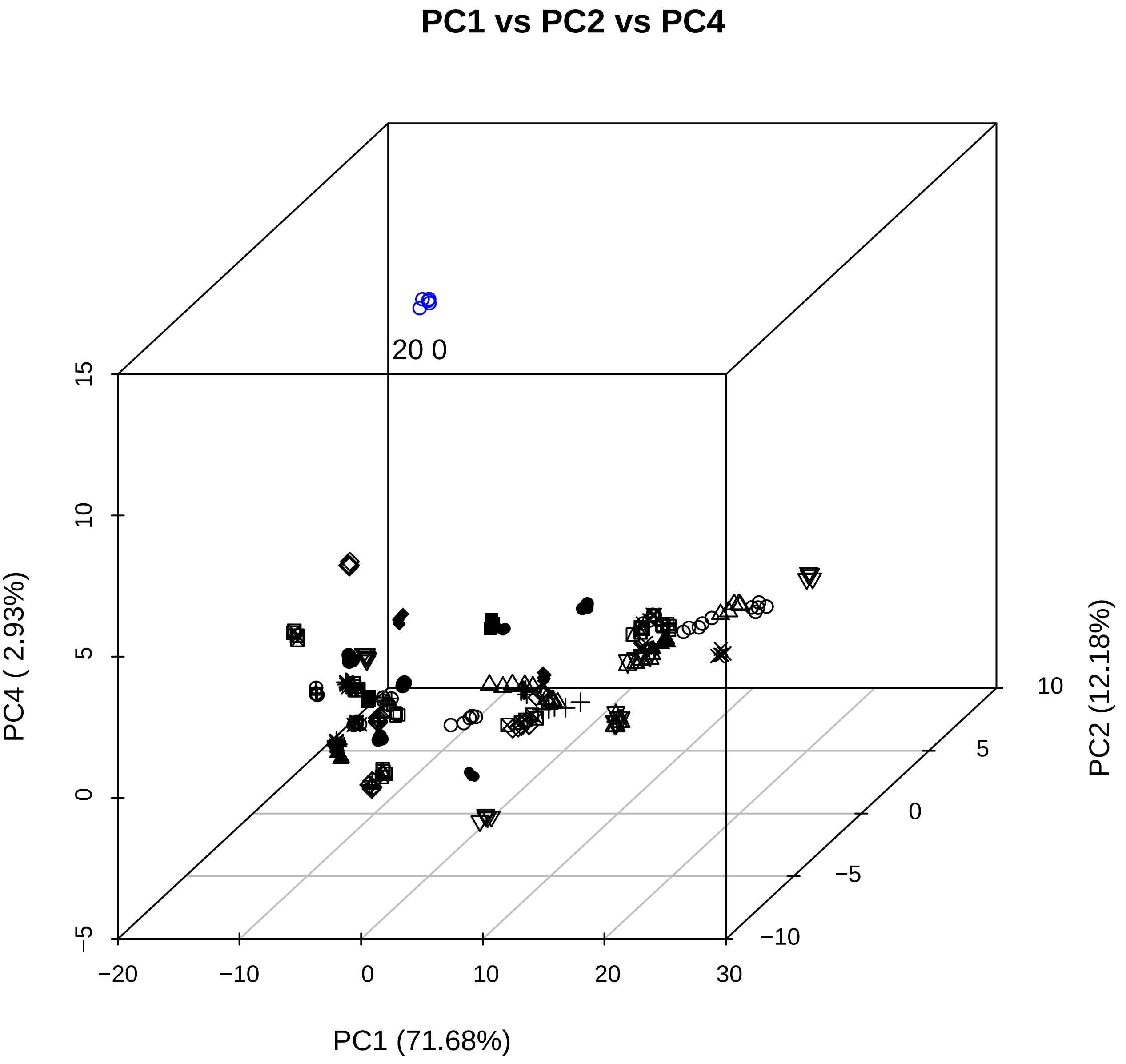
PC1 vs PC2 vs PC4
20 0
PC4 ( 2.93%)
15
10
5
0
−5
−20
−10
0
10
20
30
PC1 (71.68%)
PC2 (12.18%)
10
5
0
−5
−10

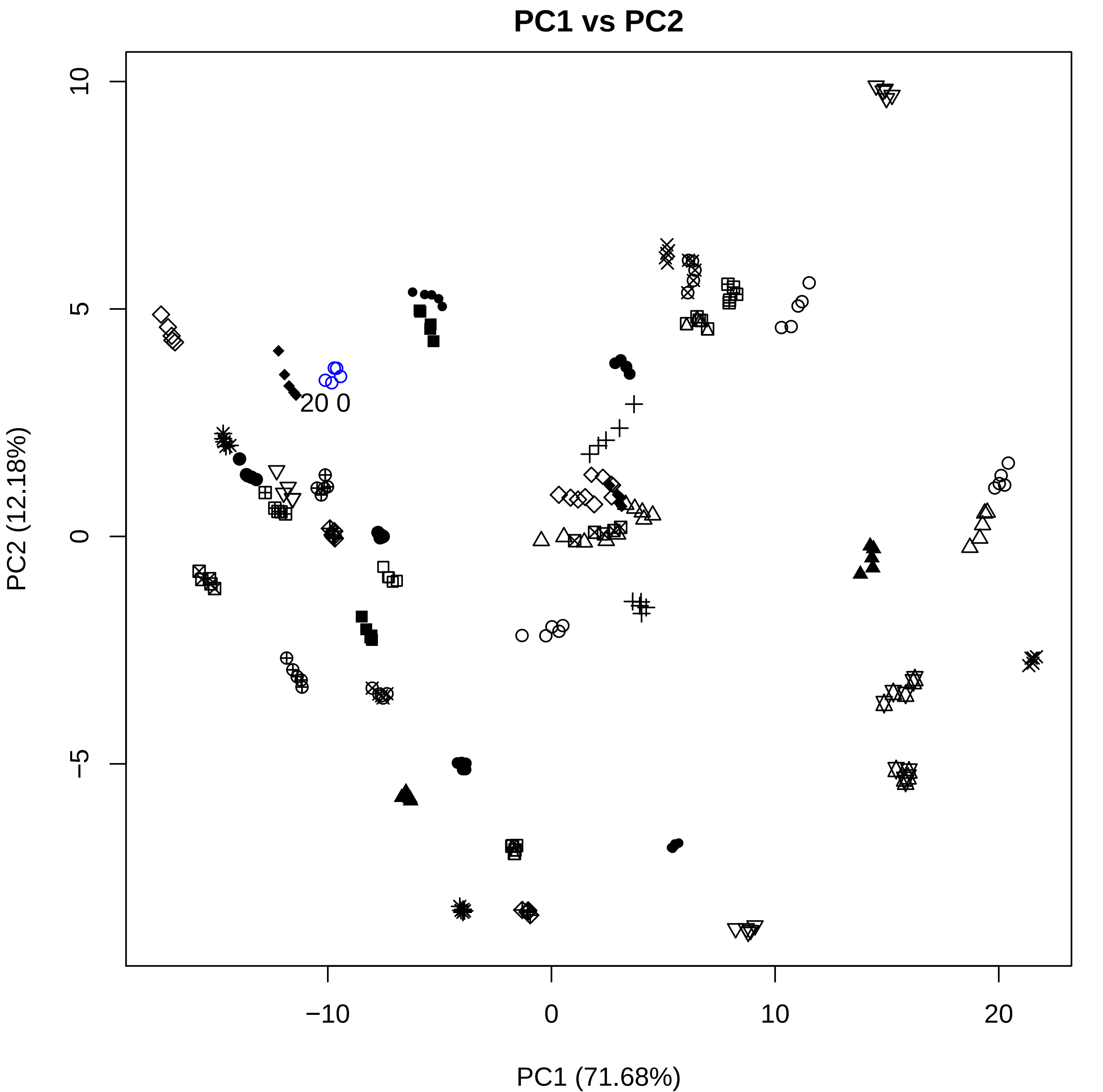

PC1 vs PC2
10
5
0
−5
PC2 (12.18%)
−10
0
10
20
PC1 (71.68%)
20 0

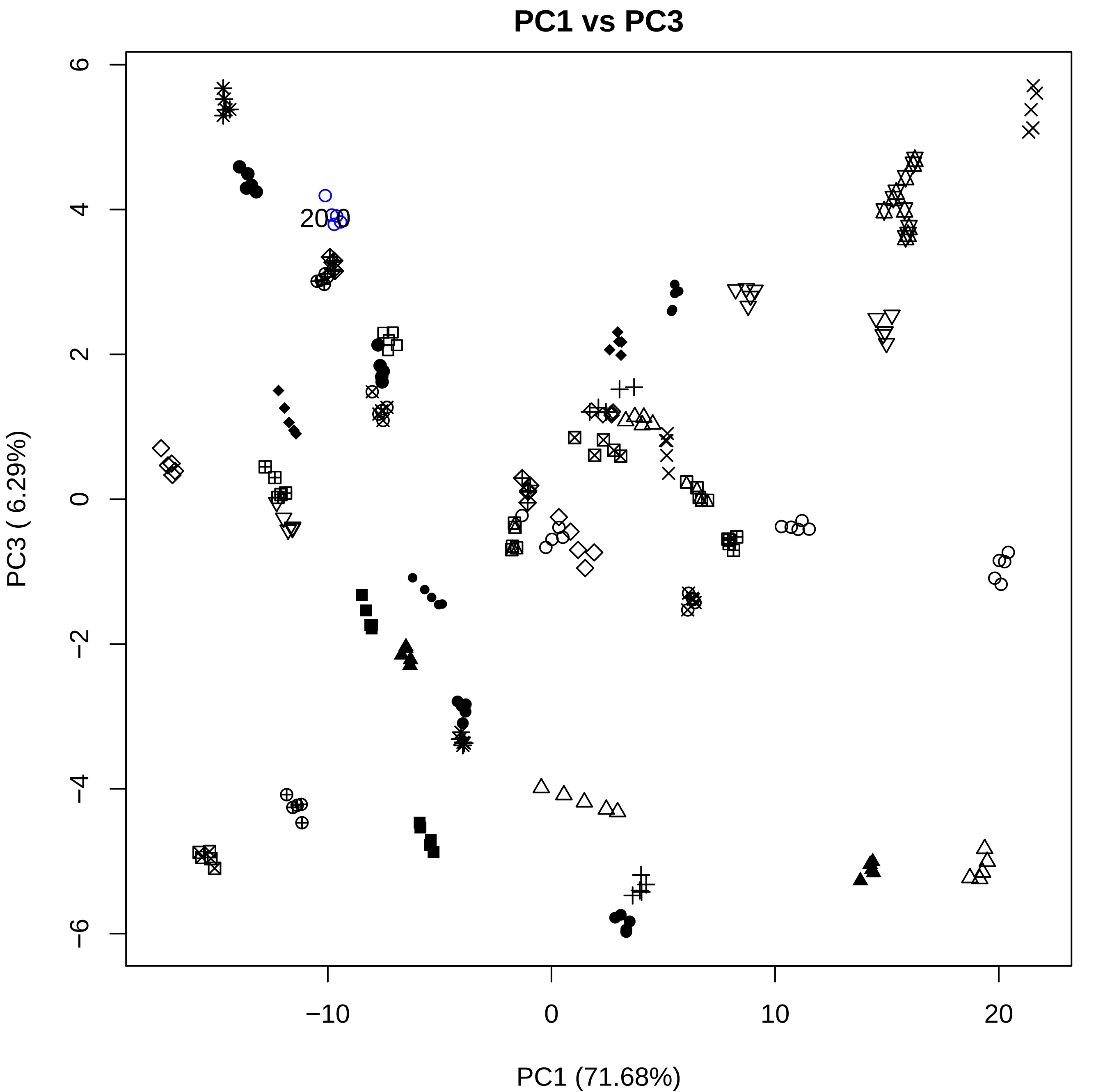
PC1 vs PC3
PC3 ( 6.29%)
PC1 (71.68%)
6
4
2
0
−2
−4
−6
−10
0
10
20

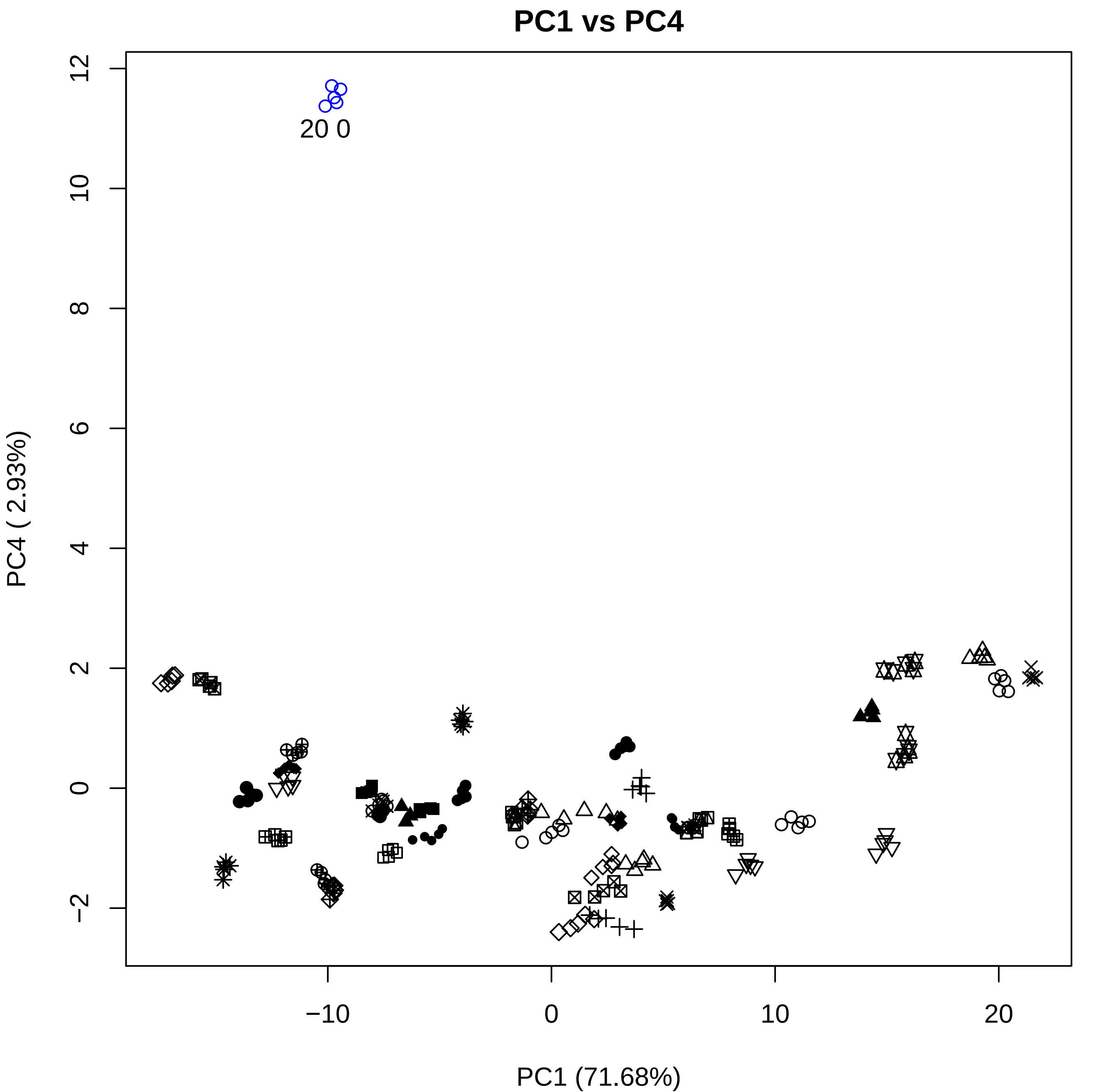

PC1 vs PC4
20 0
PC4 ( 2.93%)
PC1 (71.68%)
12
10
8
6
4
2
0
−2
−10
0
10
20

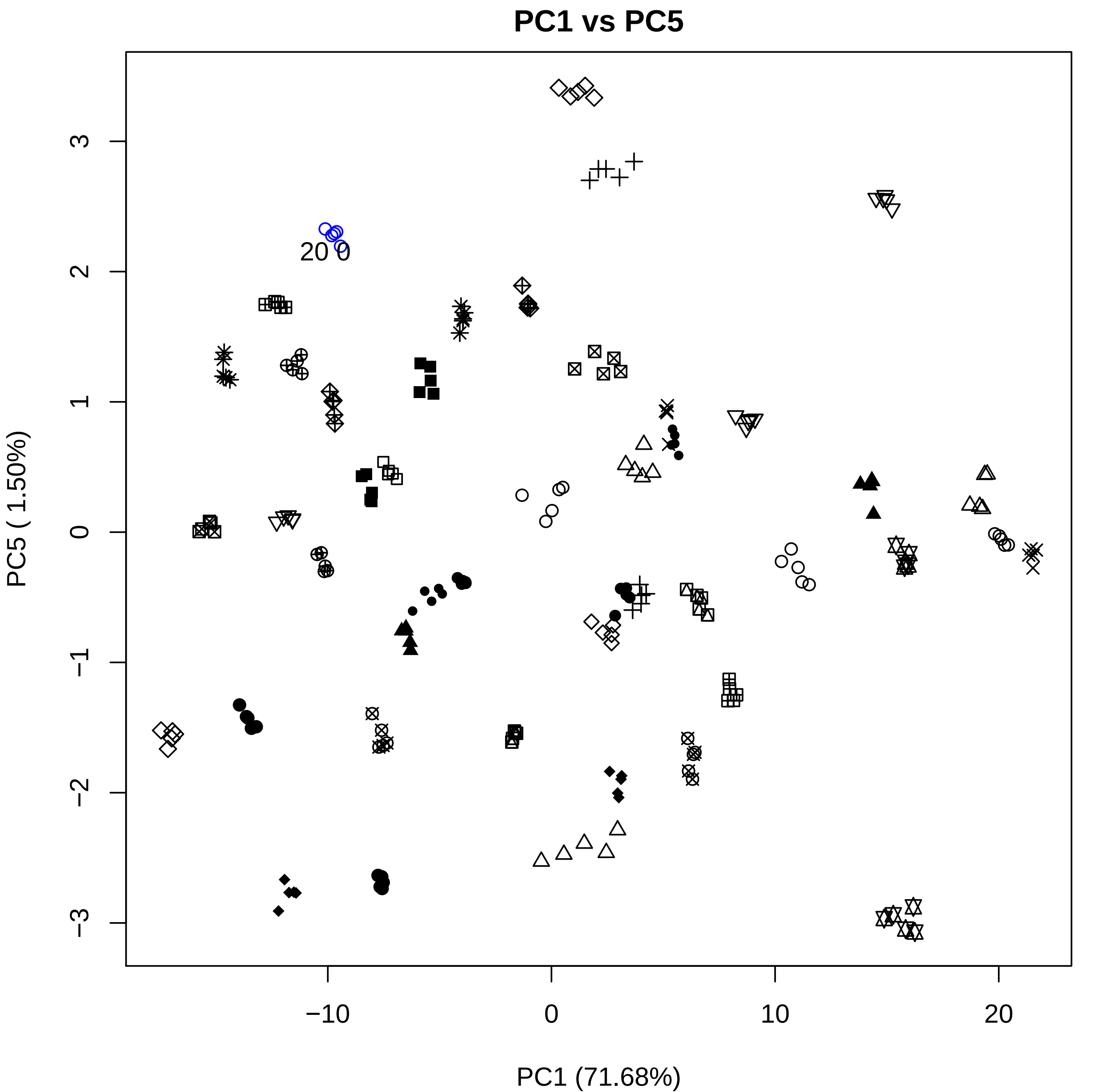

PC1 vs PC5
20 0
3
2
1
0
−1
−2
−3
−10
0
10
20
PC5 ( 1.50%)
PC1 (71.68%)

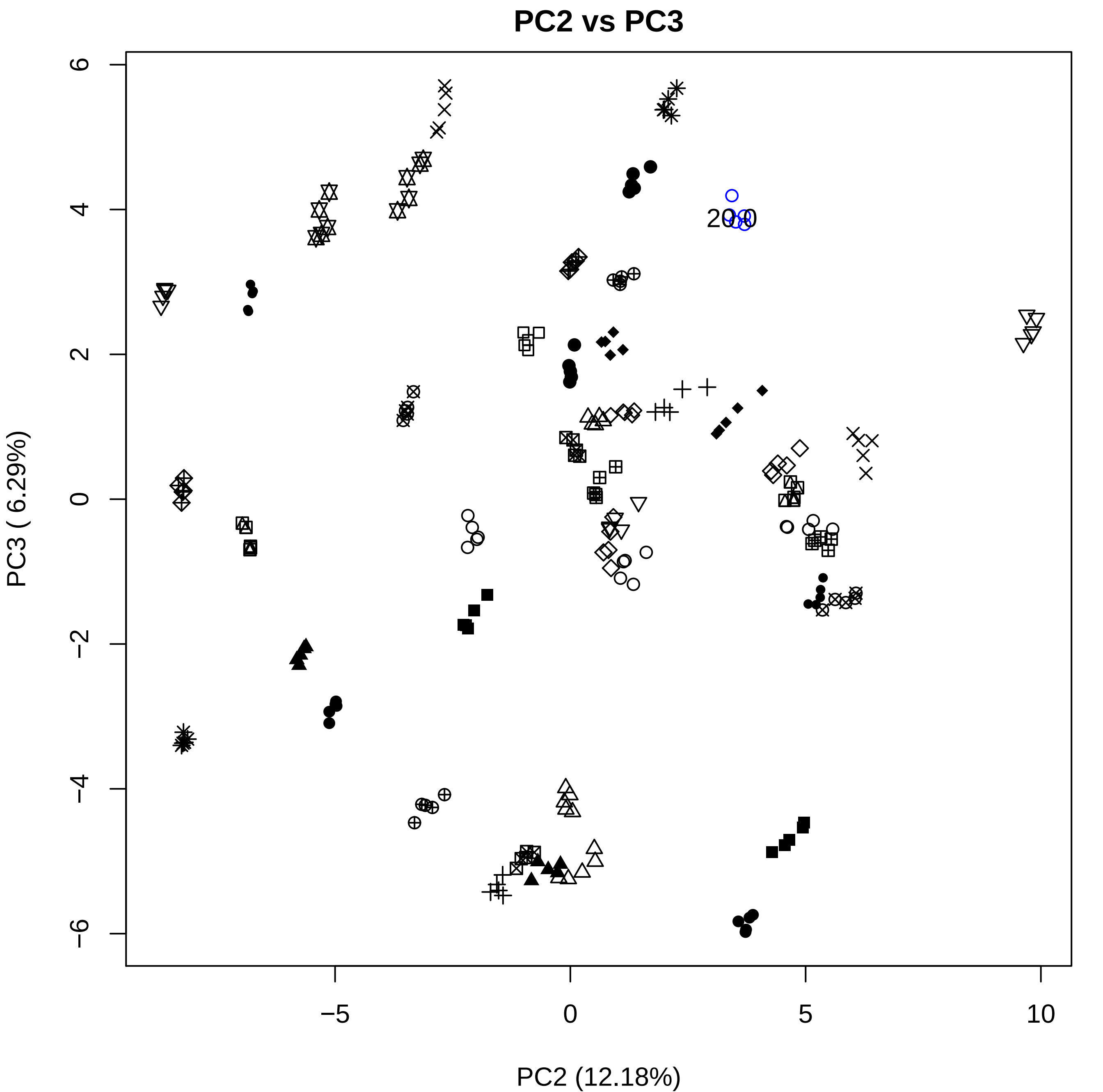

PC2 vs PC3
PC3 ( 6.29%)
PC2 (12.18%)
6
4
2
0
−2
−4
−6
−5
0
5
10
2010

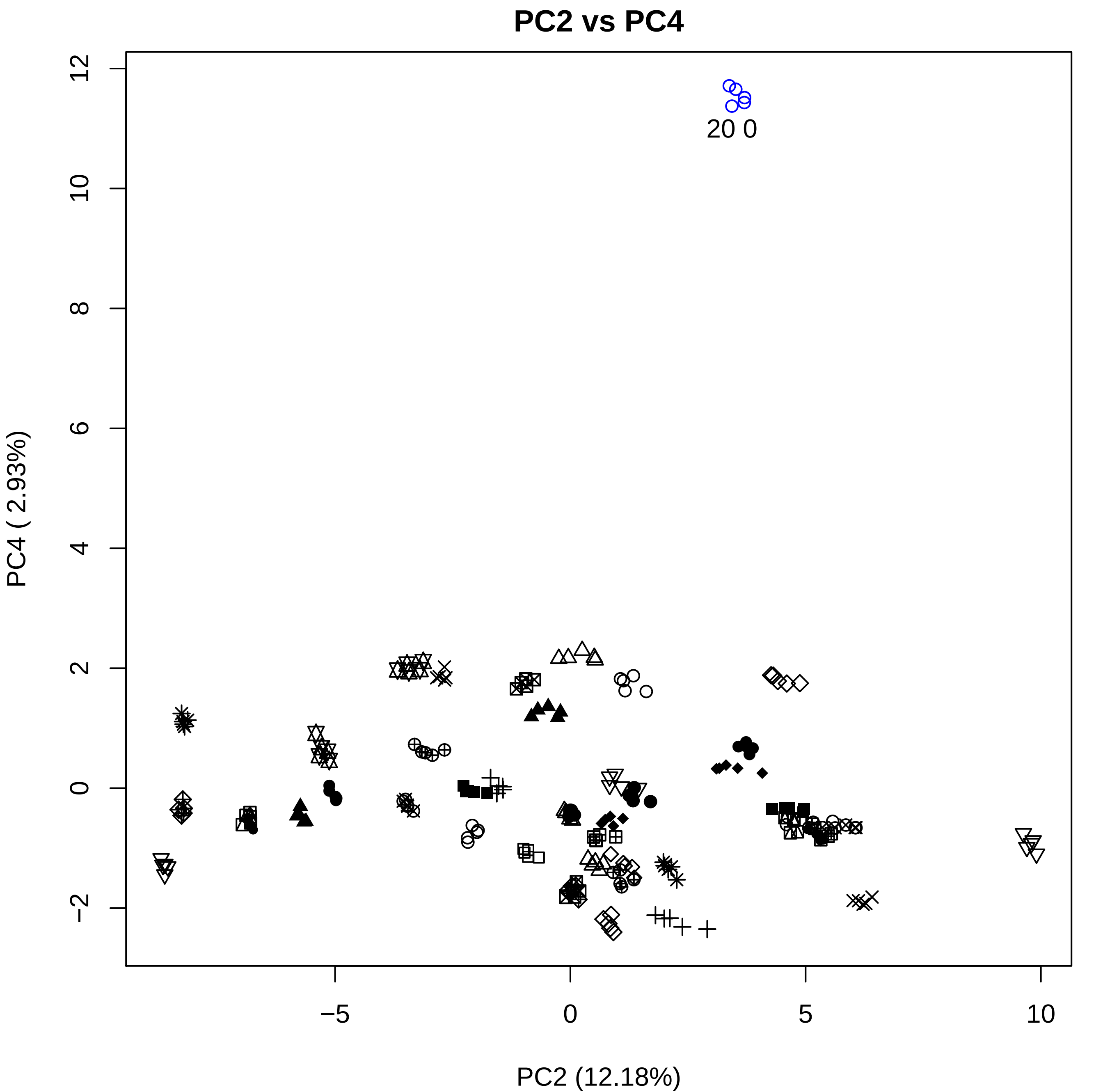

PC2 vs PC4
20 0
PC4 ( 2.93%)
PC2 (12.18%)
−2
0
2
4
6
8
10
12
−5
0
5
10

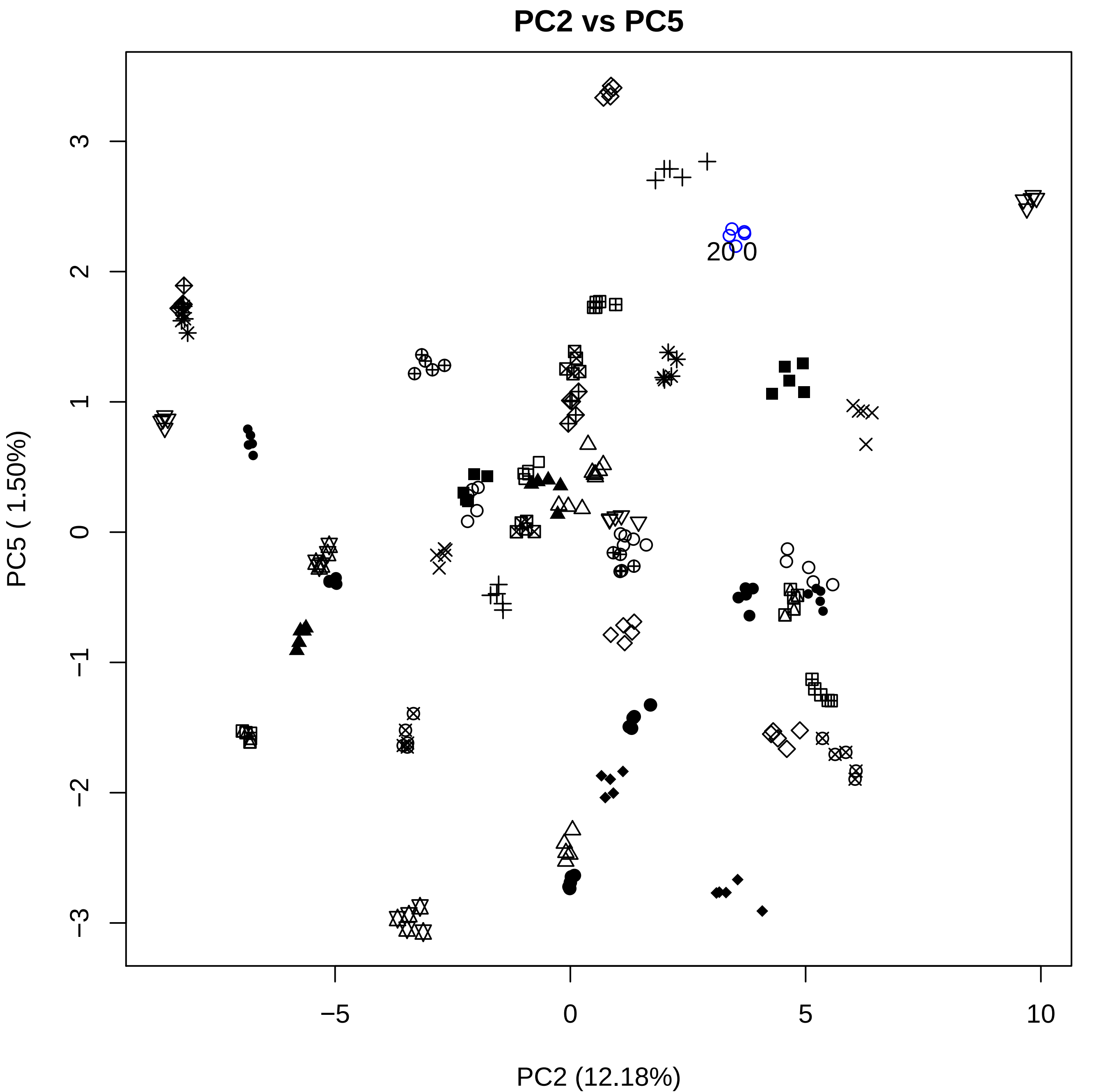
PC2 vs PC5
PC5 ( 1.50%)
PC2 (12.18%)
3
2
1
0
−1
−2
−3
−5
0
5
10
20 0

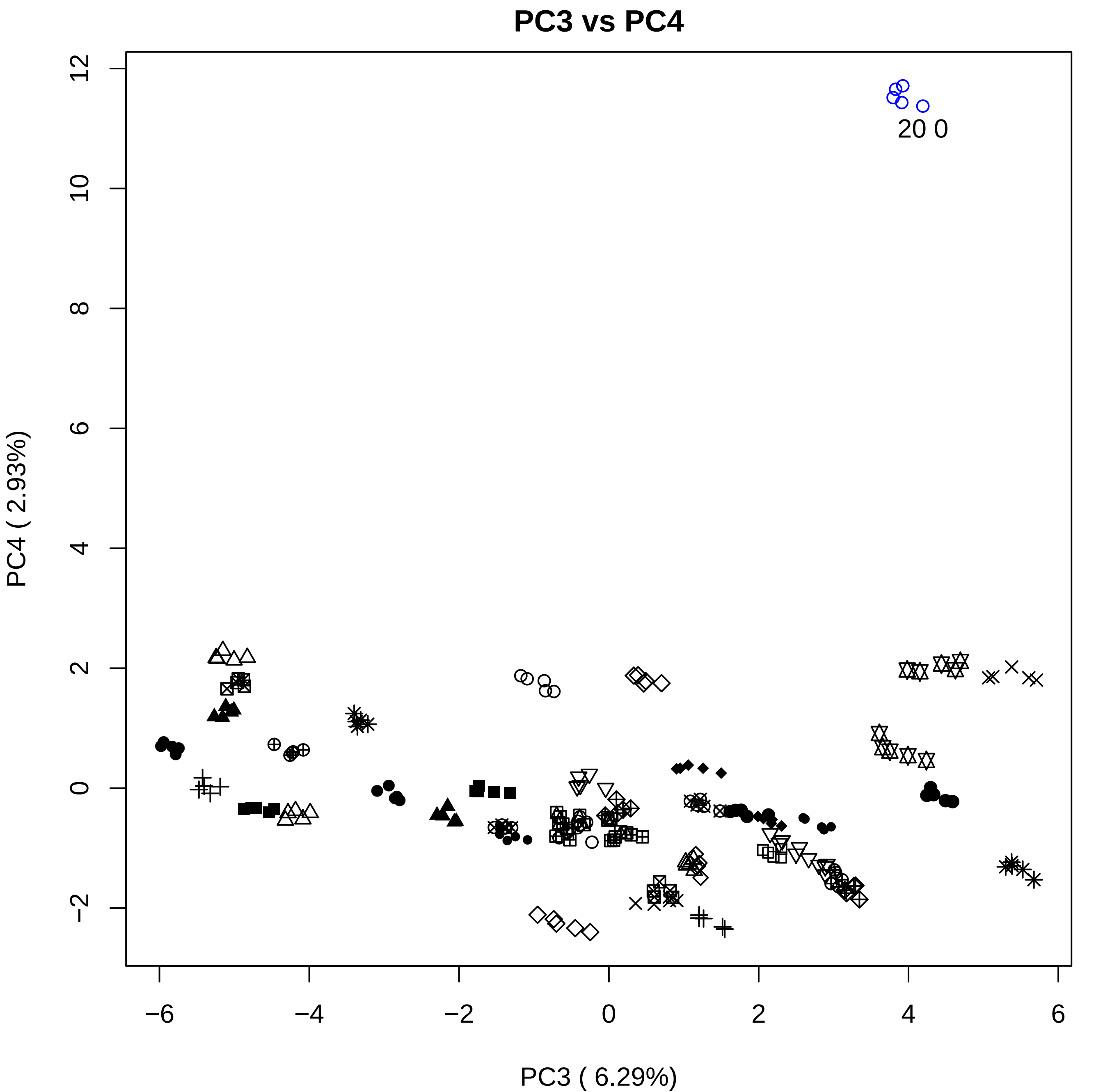
PC3 vs PC4
20 0
PC4 ( 2.93%)
PC3 ( 6.29%)
−2
0
2
4
6
8
10
12
−6
−4
−2
0
2
4
6

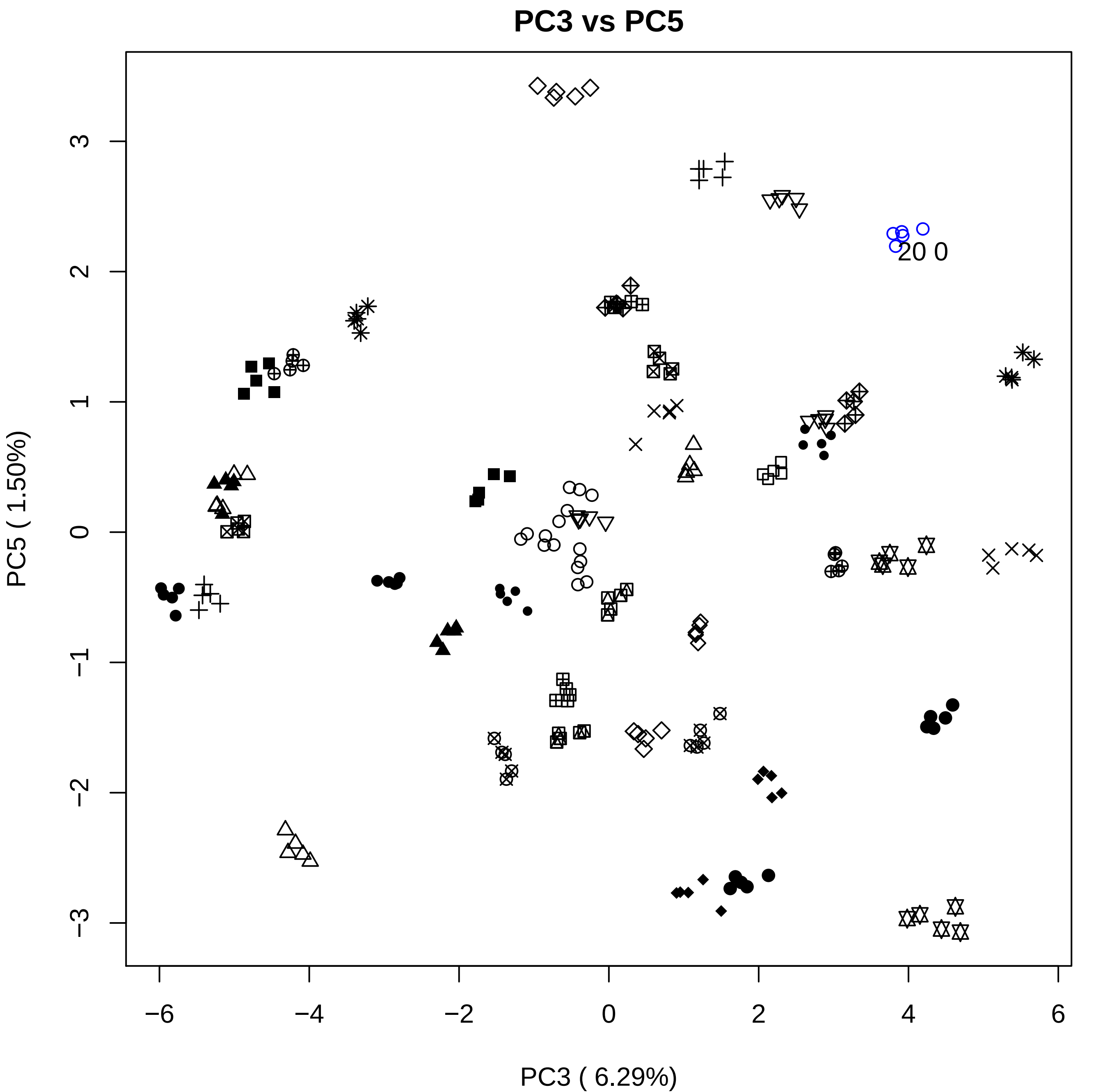
PC3 vs PC5
PC5 ( 1.50%)
PC3 ( 6.29%)
3
2
1
0
−1
−2
−3
−6
−4
−2
0
2
4
6
20 0

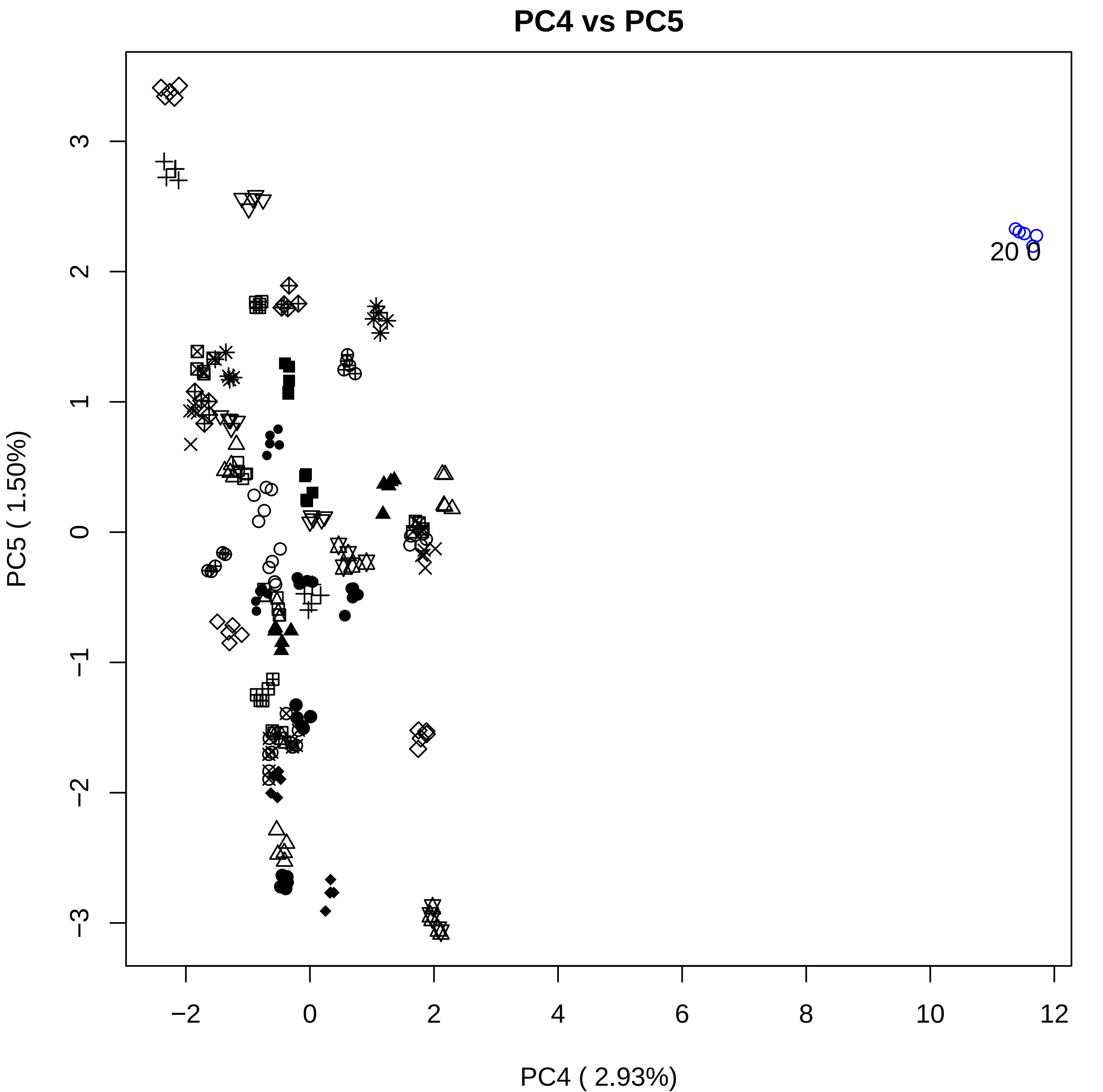
PC4 vs PC5
PC5 ( 1.50%)
PC4 ( 2.93%)
3
2
1
0
−1
−2
−3
−2
0
2
4
6
8
10
12
20 0

### PCA of 40%  OSCS vs PMH

Scaled Matrix PCAed

7th order polynomial baseline correction performed

Raw spectra smoothed to 21 Neighbours, 2nd order PN, Savtisky Golay

Normalisation between 0 and 100

D2 spectra smoothed to 41 Neighbours, 2nd order PN, Savtisky Golay

Regions Analysed: 3601−2501 and 2001−700cm−1

Samples Analysed: 1 Contam's 0 OSCS's 45 PMH's

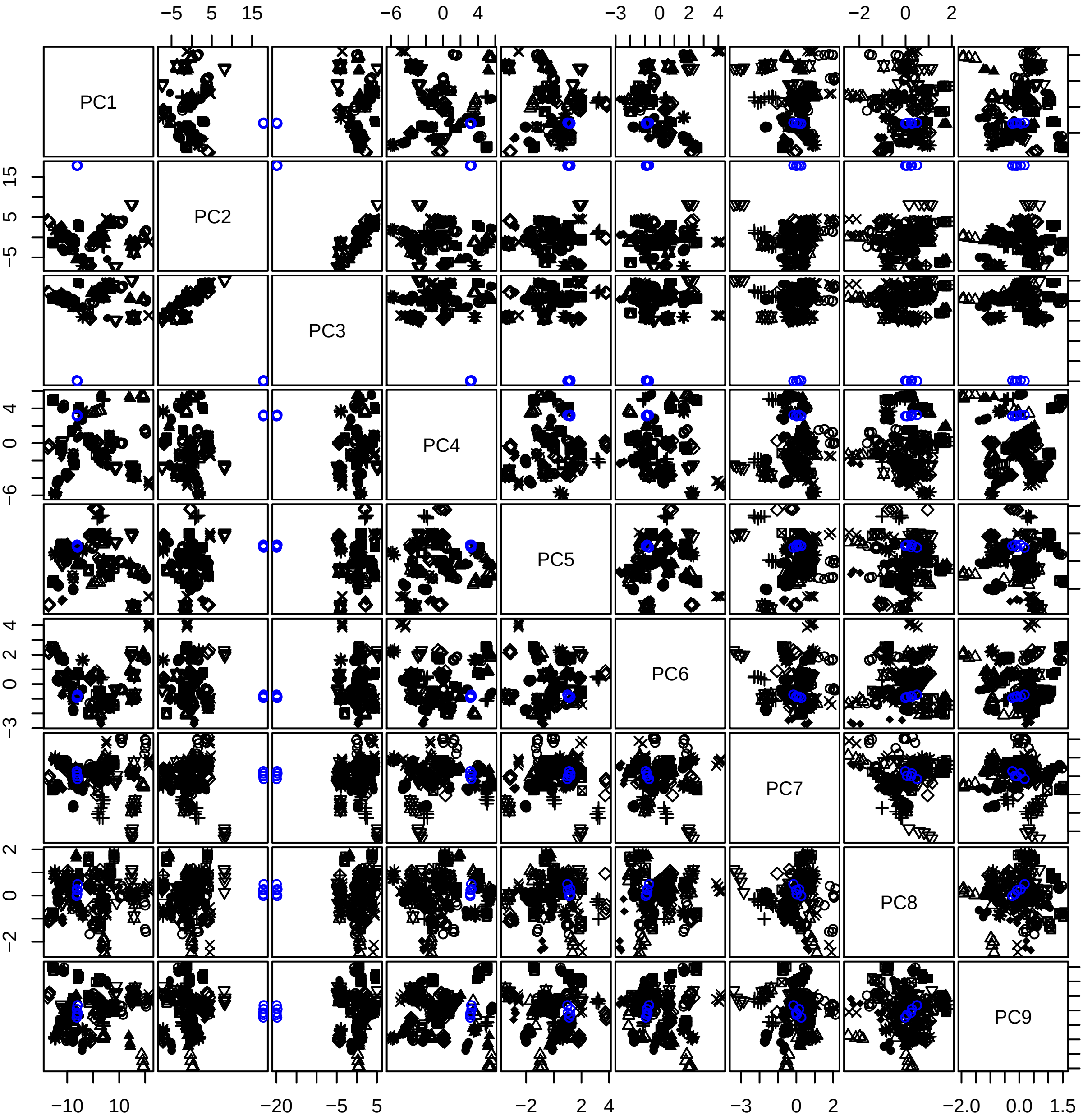
PC1
PC2
PC3
PC4
PC5
PC6
PC7
PC8
PC9
−5 5 15
−6 0 4
−3 0 2 4
−2 0 2
10
−10
15
5
−5
5
−5
−20
4
0
−6
4
2
−2
4
2
0
−3
2
0
−3
2
0
−2
1.5
0.0
−2.0
−10 10
−20 −5 5
−2 2 4
−3 0 2
−2.0 0.0 1.5

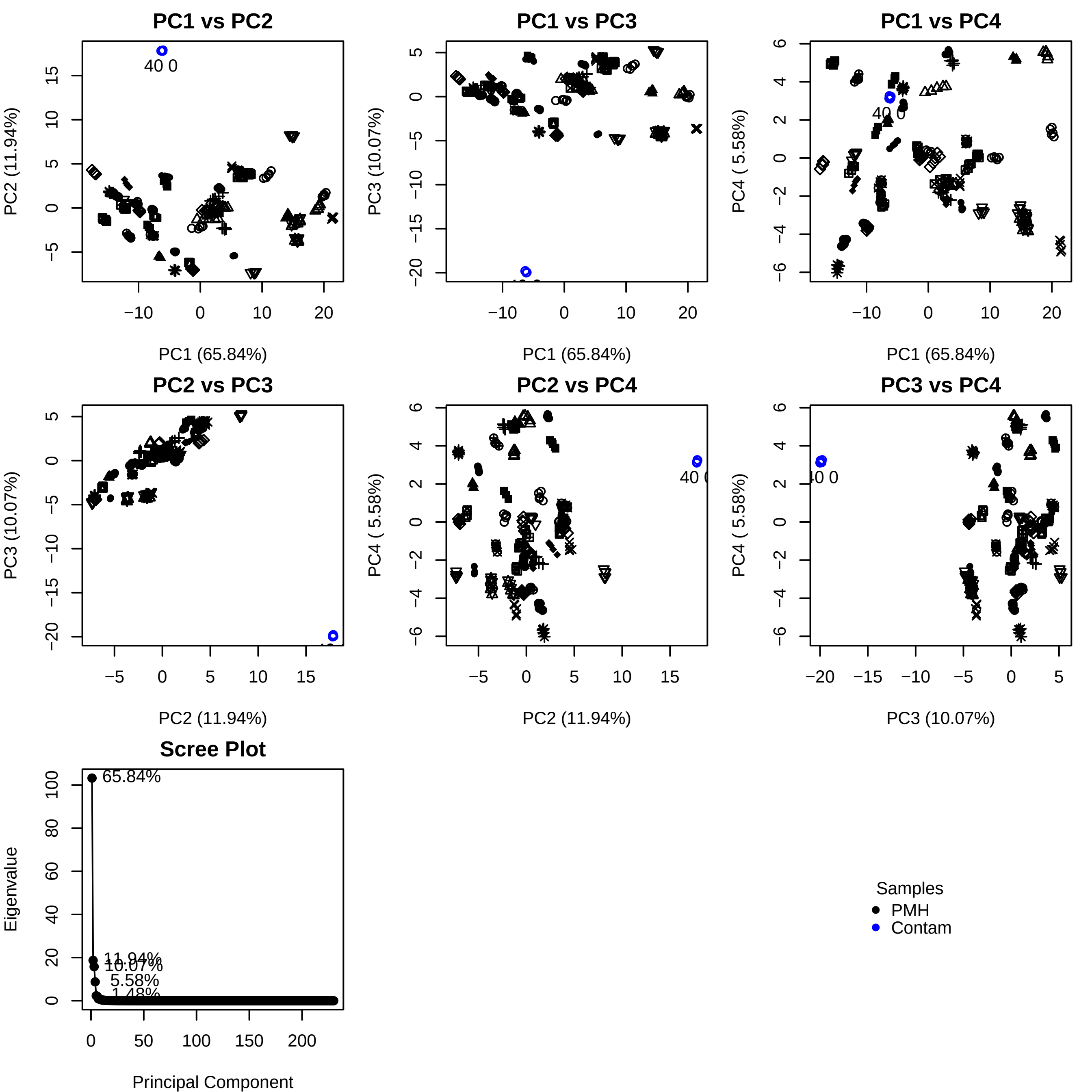

PC1 vs PC2
PC1 vs PC3
PC1 vs PC4
PC2 vs PC3
PC2 vs PC4
PC3 vs PC4
Scree Plot
PC1 (65.84%)
PC2 (11.94%)
PC3 (10.07%)
PC4 ( 5.58%)
Eigenvalue
Principal Component
65.84%
11.94%
10.07%
5.58%
1.48%
40 0
Samples
PMH
Contam

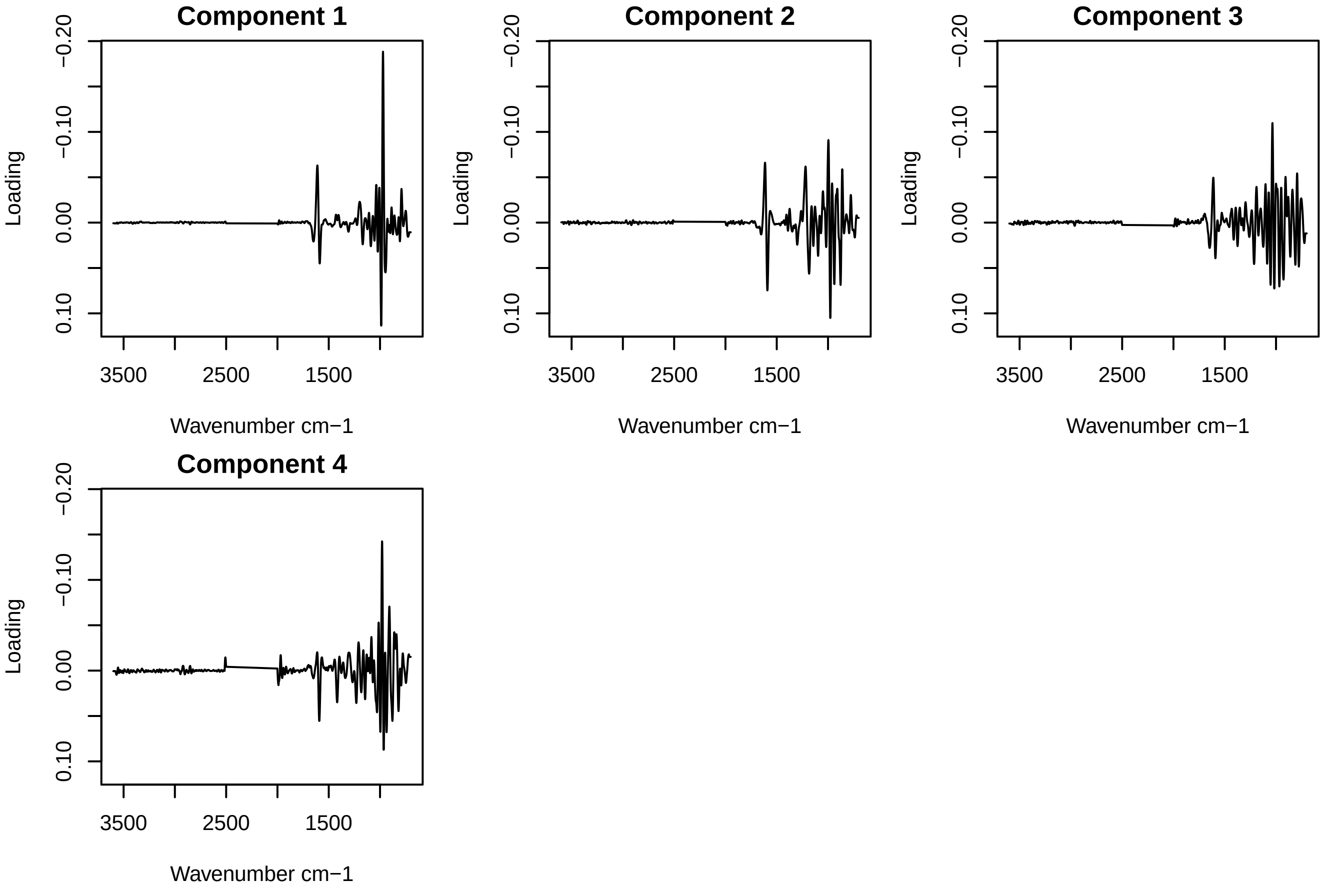
Component 1
Component 2
Component 3
Component 4
Loading
−0.20
−0.10
0.00
0.10
3500
2500
1500
Wavenumber cm−1

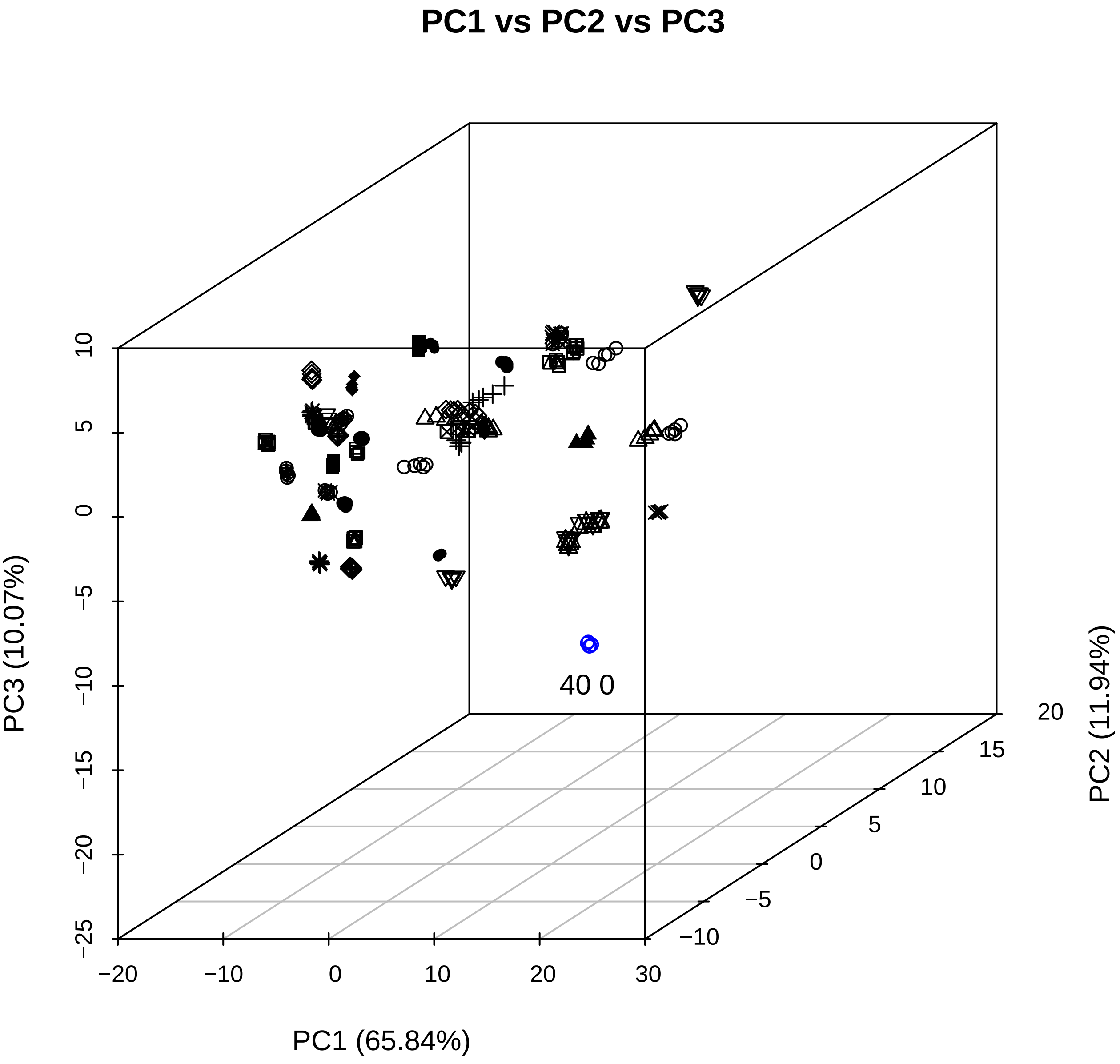

PC1 vs PC2 vs PC3
PC1 (65.84%)
PC2 (11.94%)
PC3 (10.07%)
−20
−10
0
10
20
30
−10
−5
0
5
10
15
20
−25
−20
−15
−10
−5
0
5
10
40 0

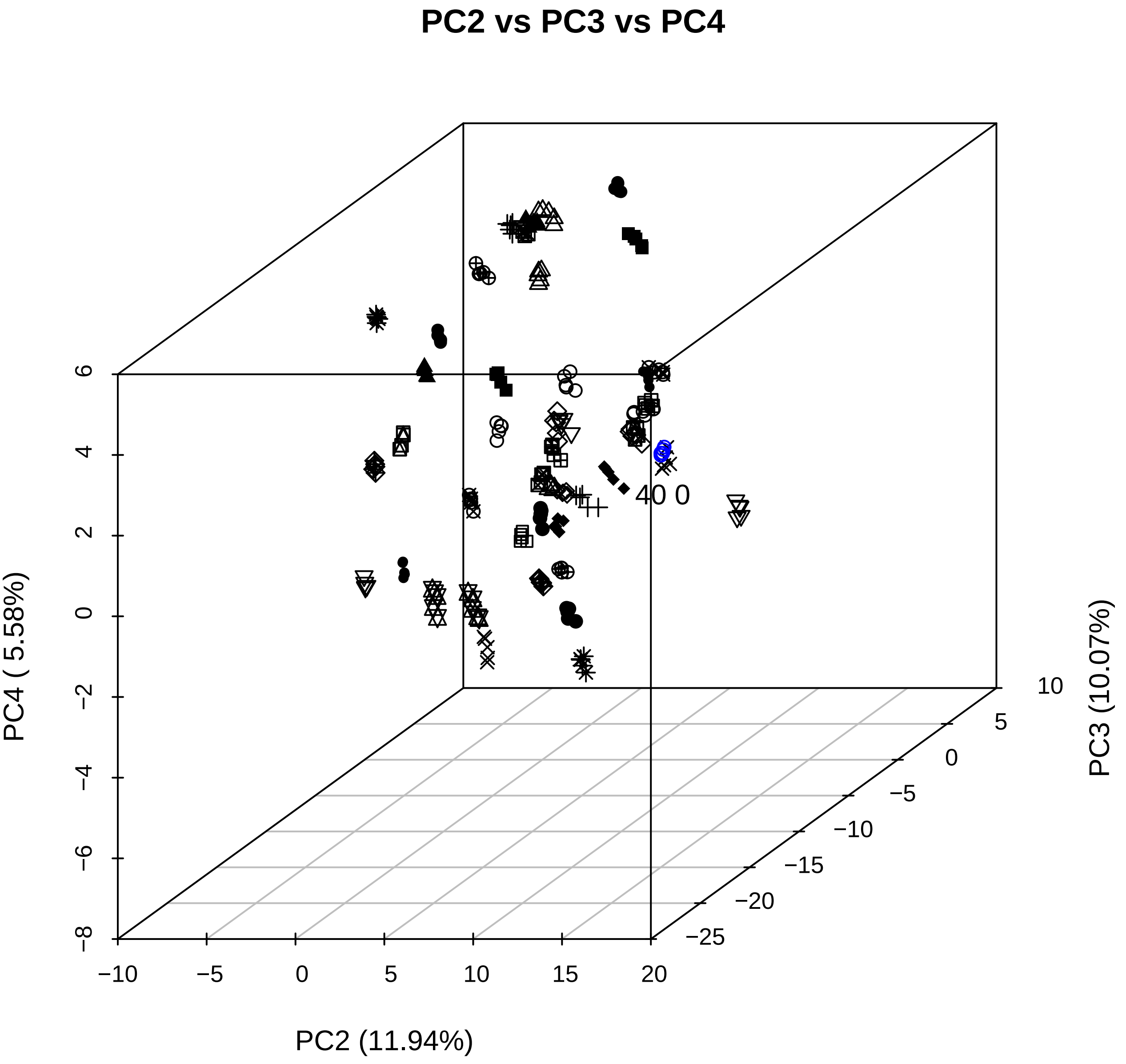
PC2 vs PC3 vs PC4
PC2 (11.94%)
PC3 (10.07%)
PC4 ( 5.58%)
−10
−5
0
5
10
15
20
−25
−20
−15
−10
−5
0
5
10
−8
−6
−4
−2
0
2
4
6
40 0

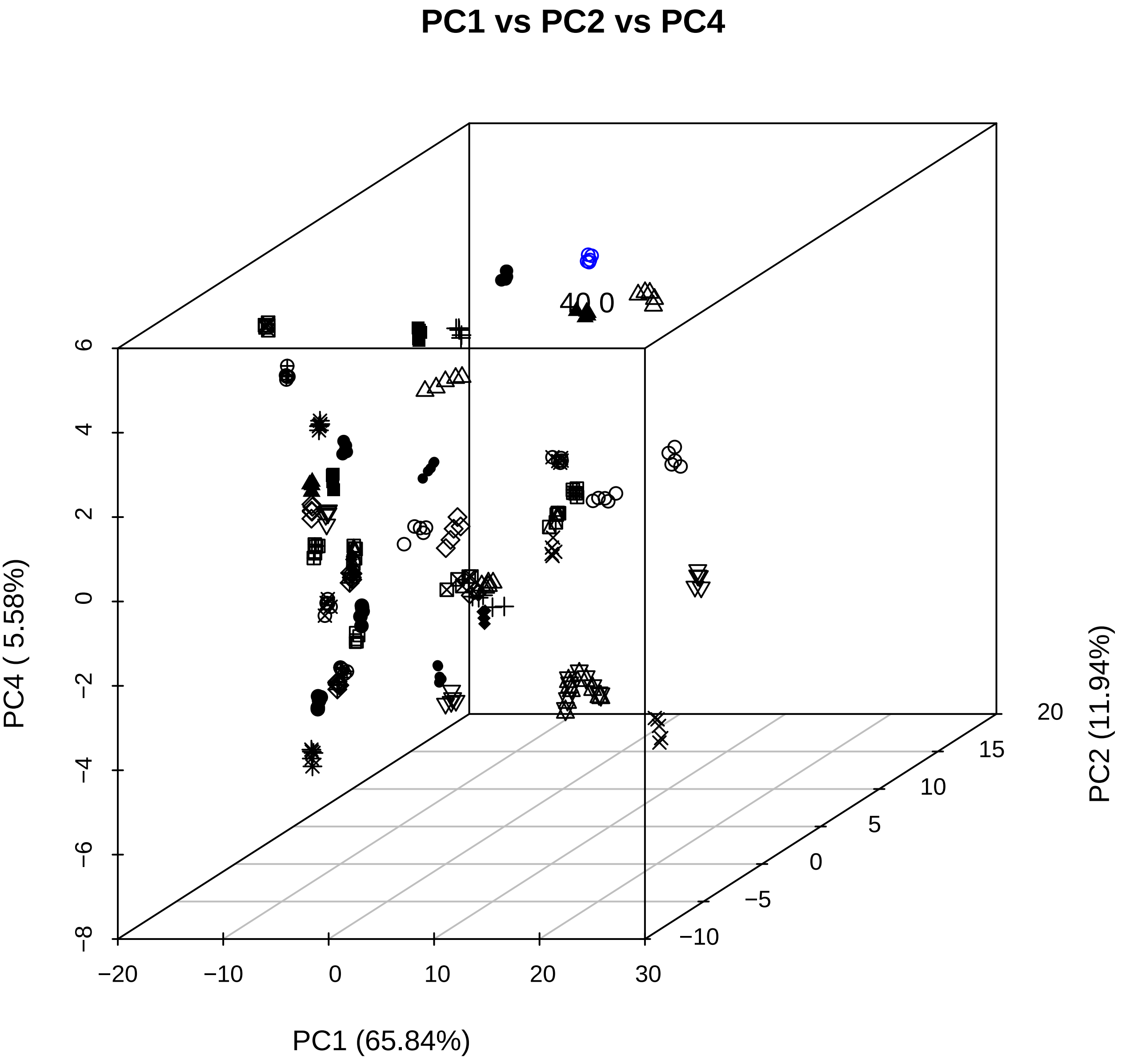
PC1 vs PC2 vs PC4
PC1 (65.84%)
PC2 (11.94%)
PC4 ( 5.58%)
−20
−10
0
10
20
30
−10
−5
0
5
10
15
20
−8
−6
−4
−2
0
2
4
6
40
0

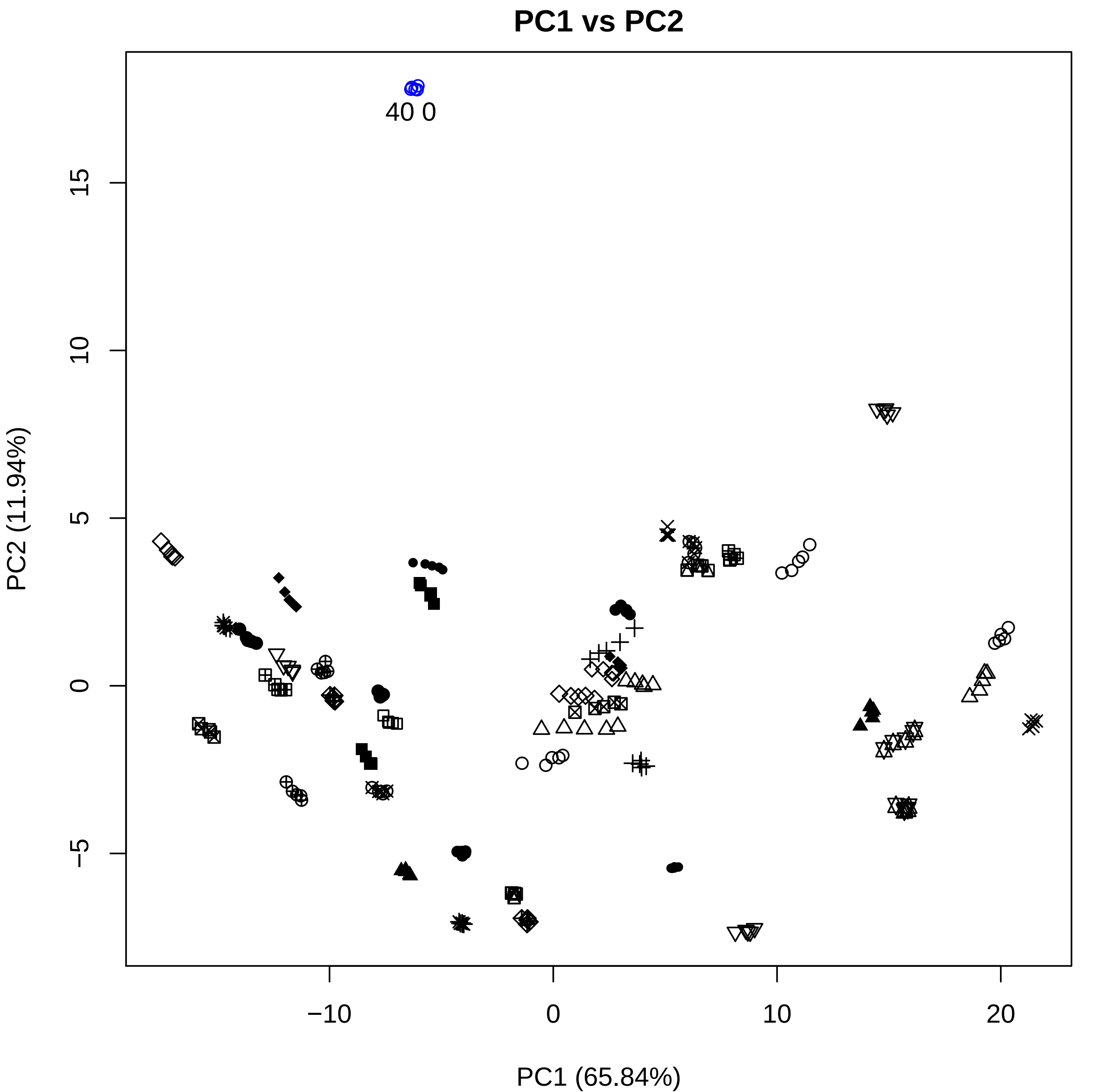

PC1 vs PC2
40 0
PC2 (11.94%)
PC1 (65.84%)
−10
0
10
20
−5
0
5
10
15

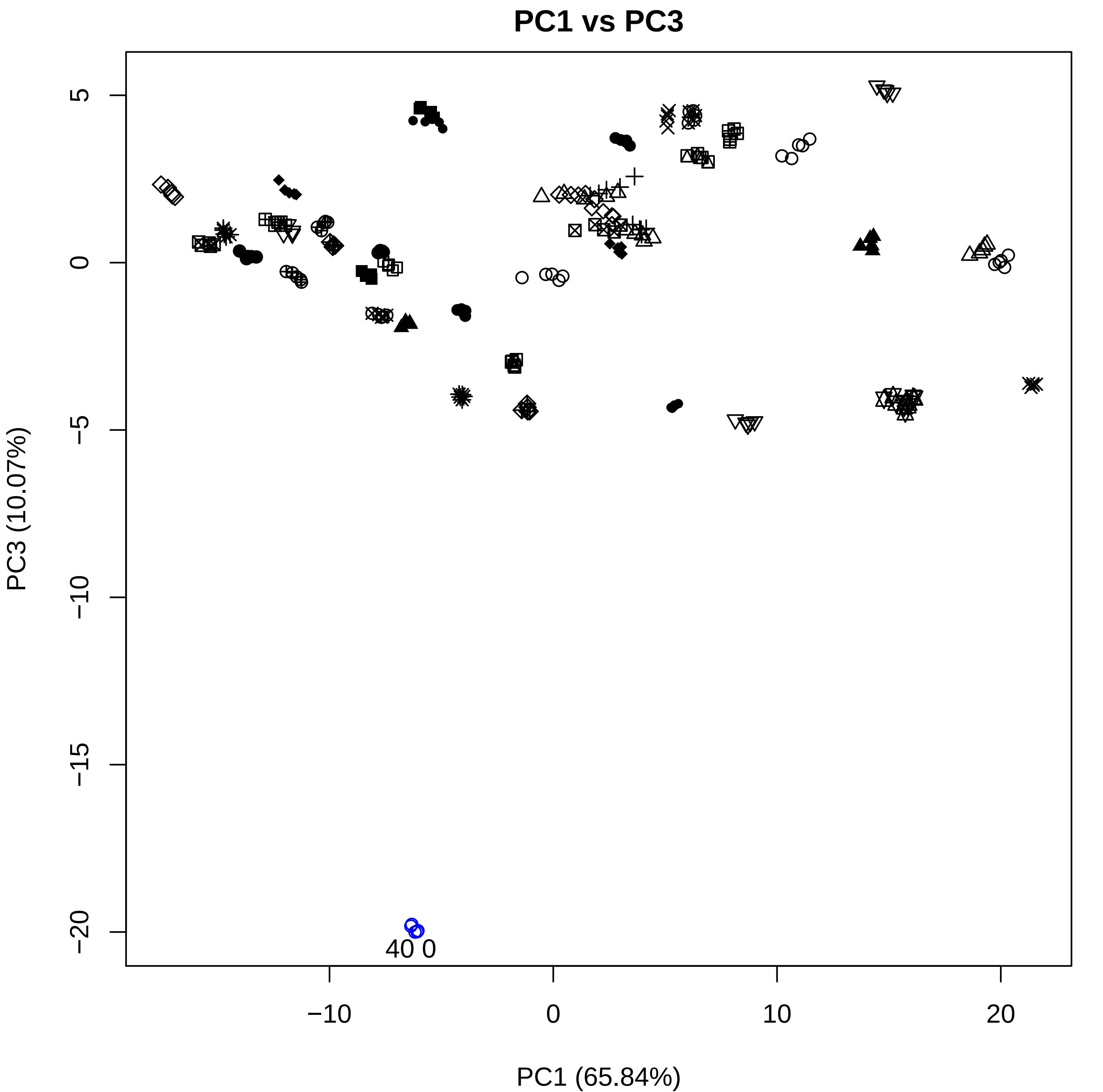
PC1 vs PC3
PC3 (10.07%)
PC1 (65.84%)
5
0
−5
−10
−15
−20
−10
0
10
20
40 0

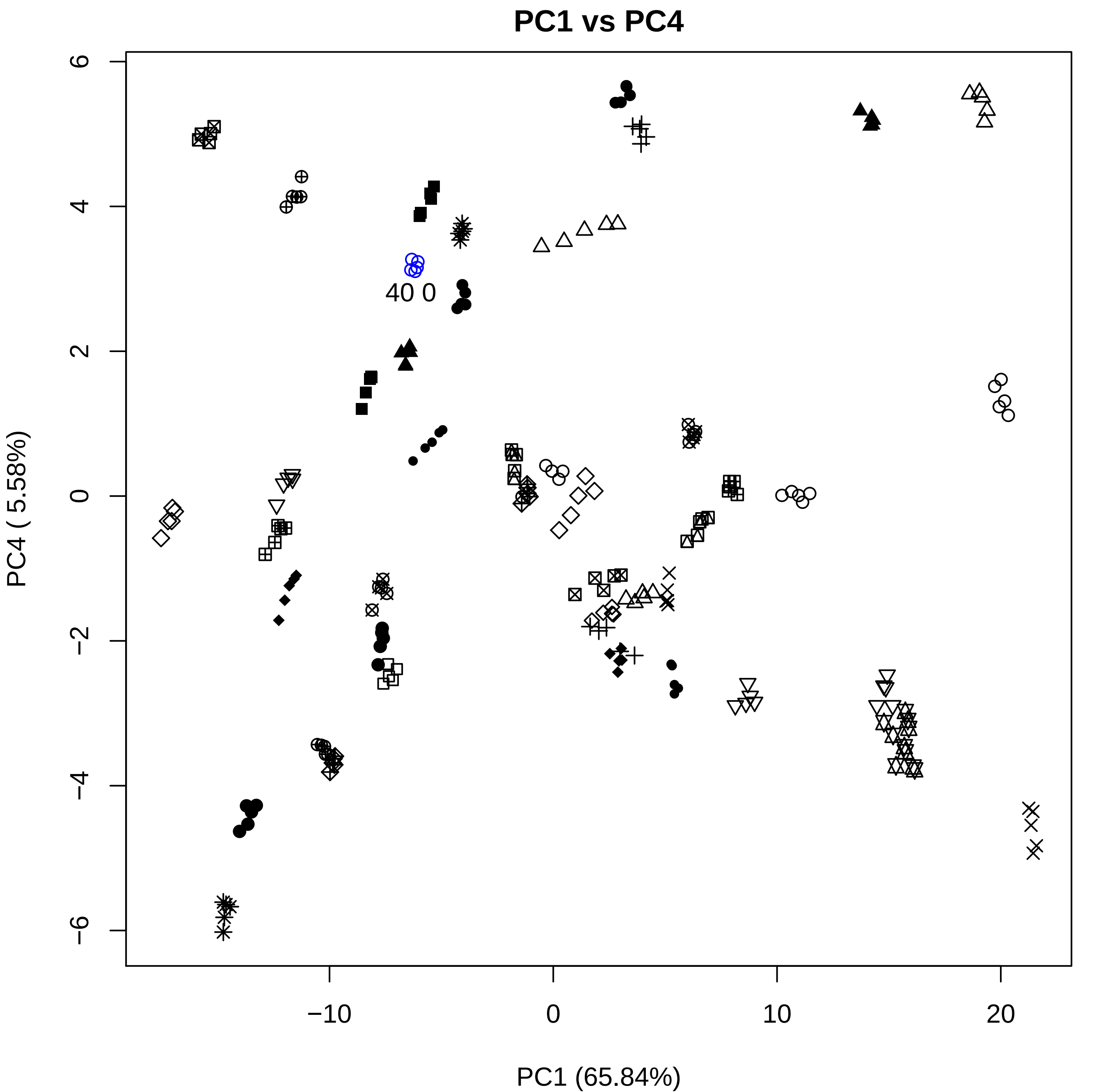
PC1 vs PC4
PC4 ( 5.58%)
PC1 (65.84%)
6
4
2
0
−2
−4
−6
−10
0
10
20
40 0

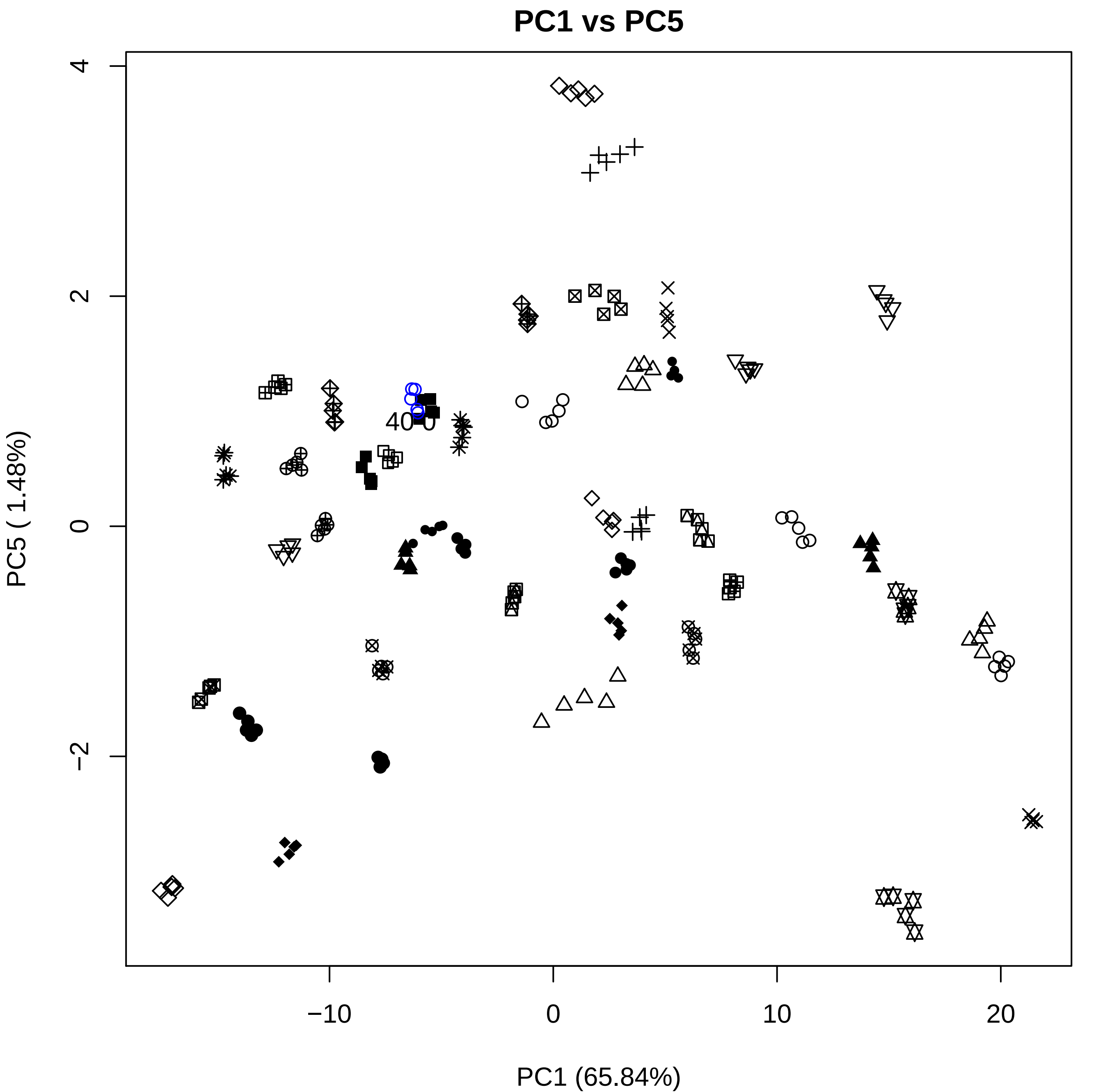
PC1 vs PC5
PC5 ( 1.48%)
PC1 (65.84%)
4
2
0
−2
−10
0
10
20
40

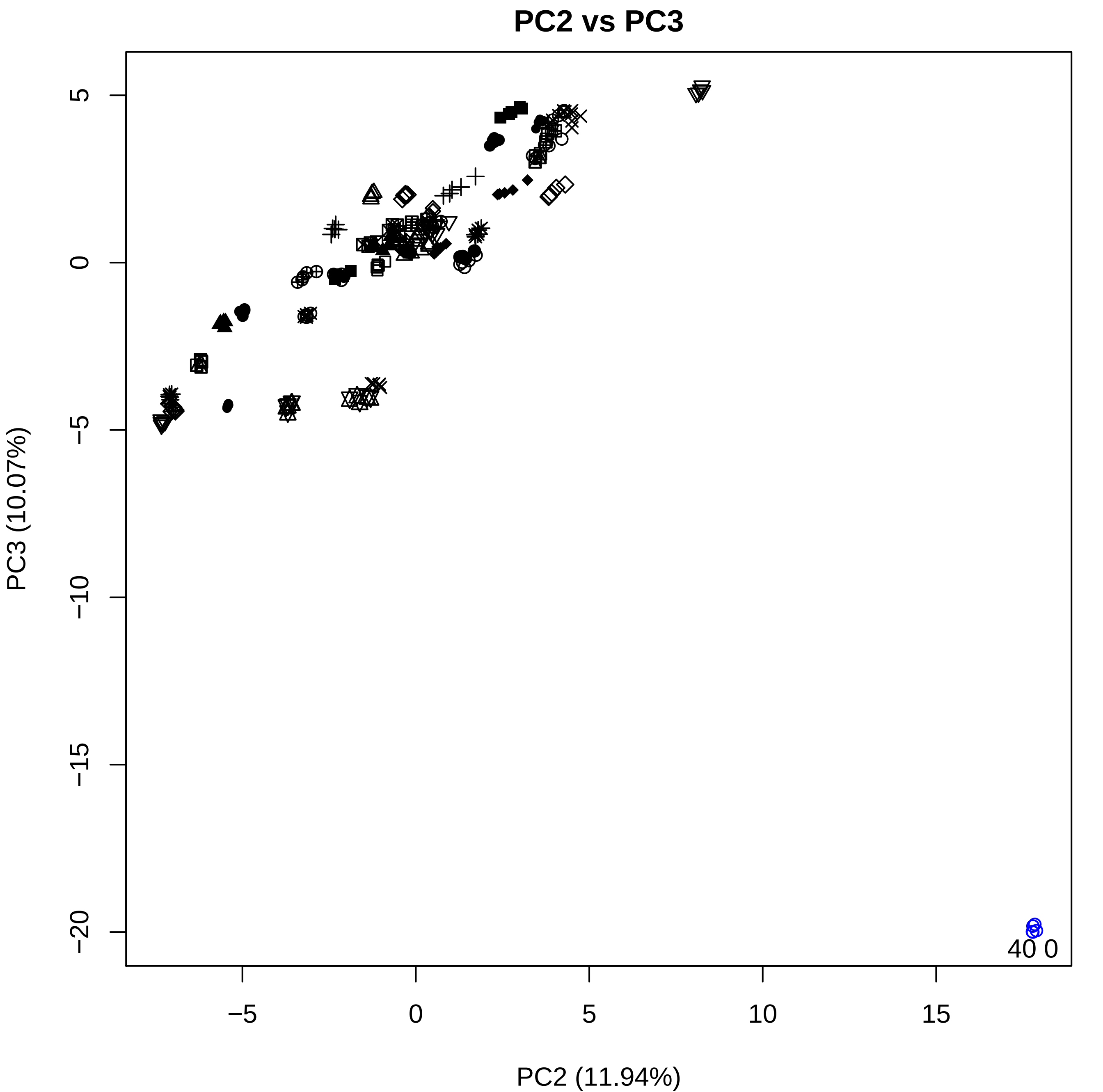
PC2 vs PC3
PC3 (10.07%)
PC2 (11.94%)
5
0
−5
−10
−15
−20
−5
0
5
10
15
40 0

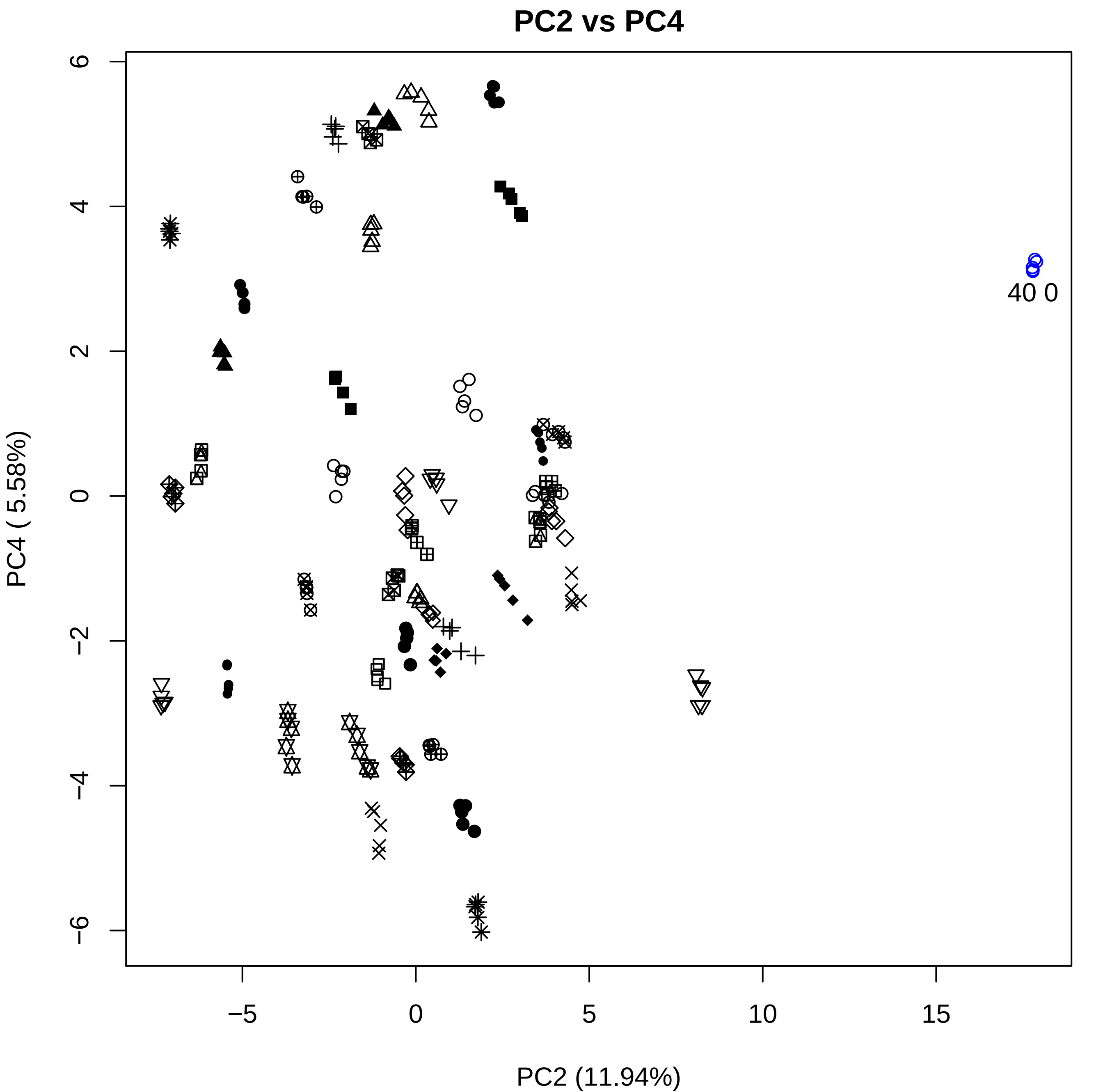
PC2 vs PC4
6
4
2
0
−2
−4
−6
PC4 ( 5.58%)
−5
0
5
10
15
PC2 (11.94%)
40 0

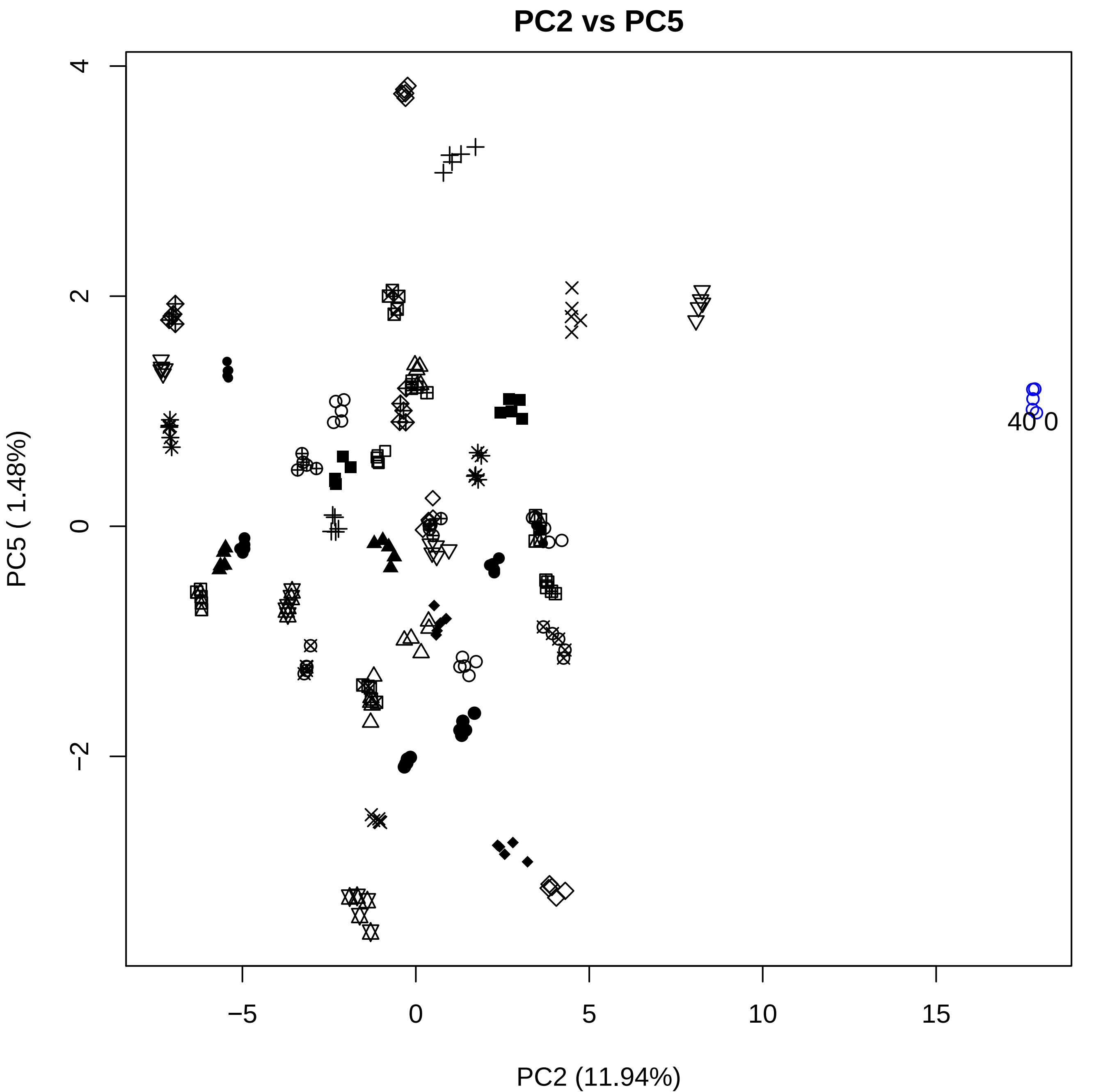
PC2 vs PC5
PC5 ( 1.48%)
PC2 (11.94%)
4
2
0
−2
−5
0
5
10
15
40 0

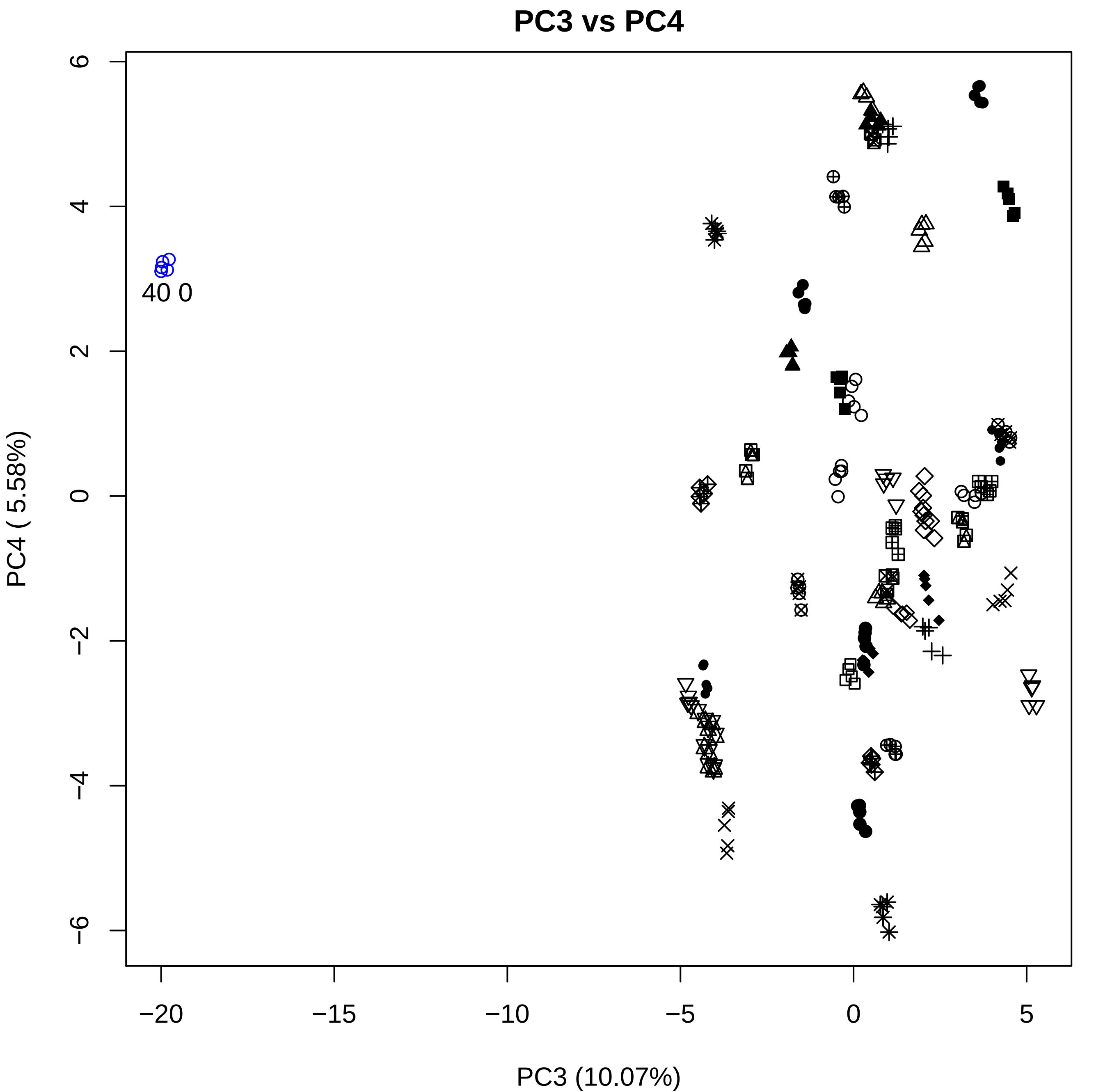

PC3 vs PC4
PC4 ( 5.58%)
PC3 (10.07%)
40 0
6
4
2
0
−2
−4
−6
−20
−15
−10
−5
0
5

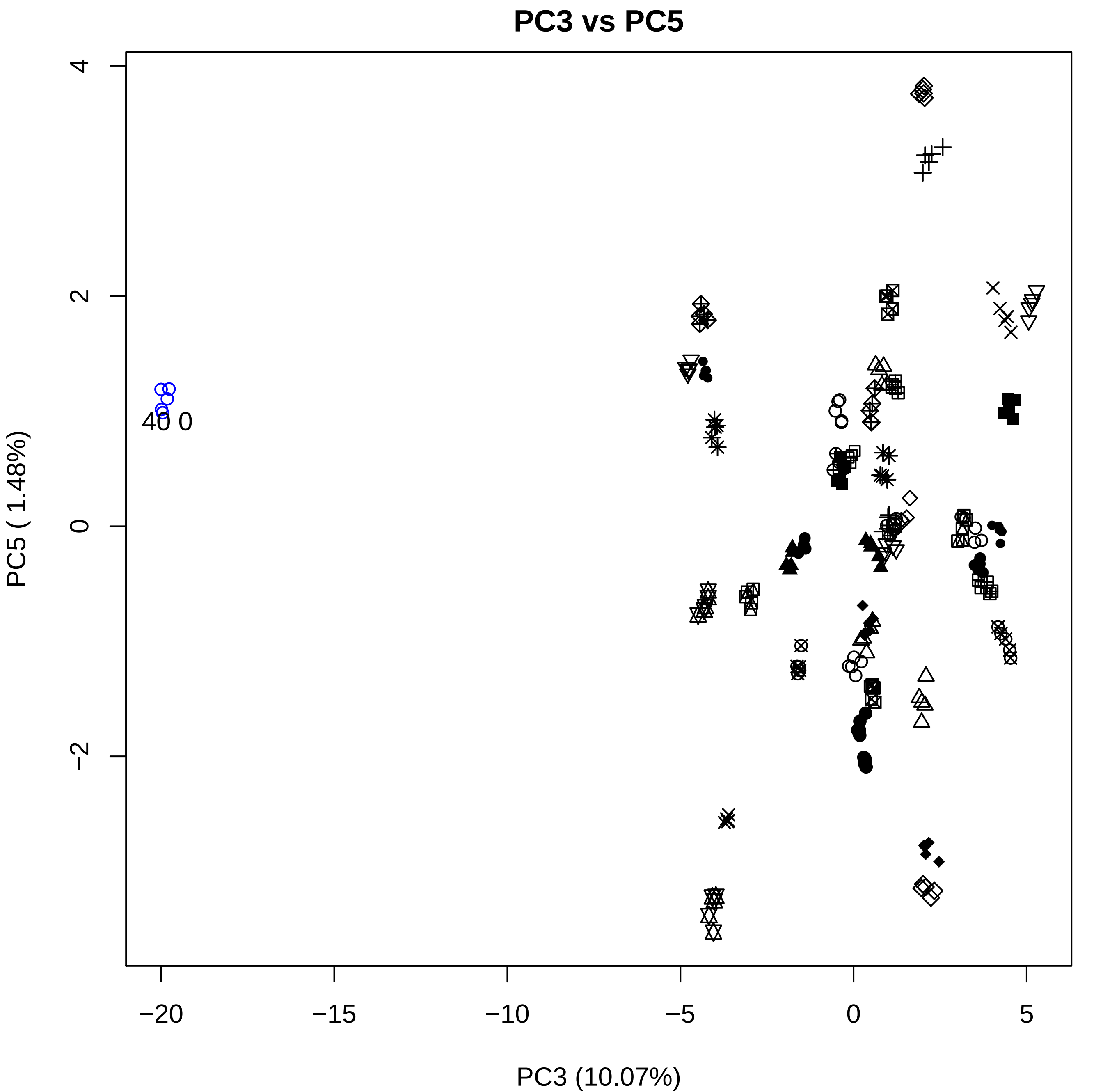

PC3 vs PC5
PC5 ( 1.48%)
PC3 (10.07%)
40 0
4
2
0
−2
−20
−15
−10
−5
0
5

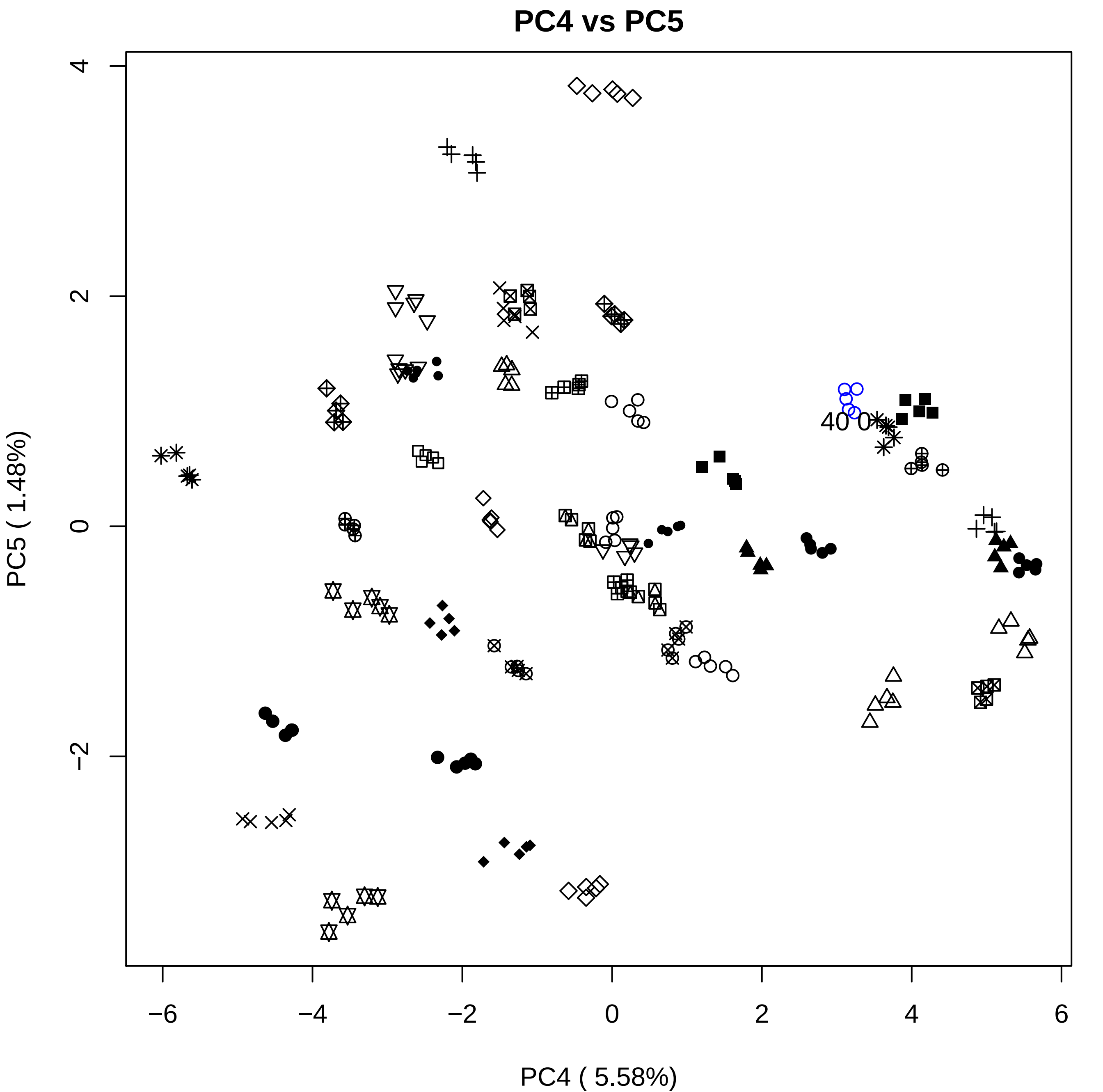
PC4 vs PC5
PC5 ( 1.48%)
PC4 ( 5.58%)
4
2
0
−2
−6
−4
−2
0
2
4
6
40 0

### PCA of 100%  OSCS vs PMH

Scaled Matrix PCAed

7th order polynomial baseline correction performed

Raw spectra smoothed to 21 Neighbours, 2nd order PN, Savtisky Golay

Normalisation between 0 and 100

D2 spectra smoothed to 41 Neighbours, 2nd order PN, Savtisky Golay

Regions Analysed: 3601−2501 and 2001−700cm−1

Samples Analysed: 0 Contam's 1 OSCS's 45 PMH's

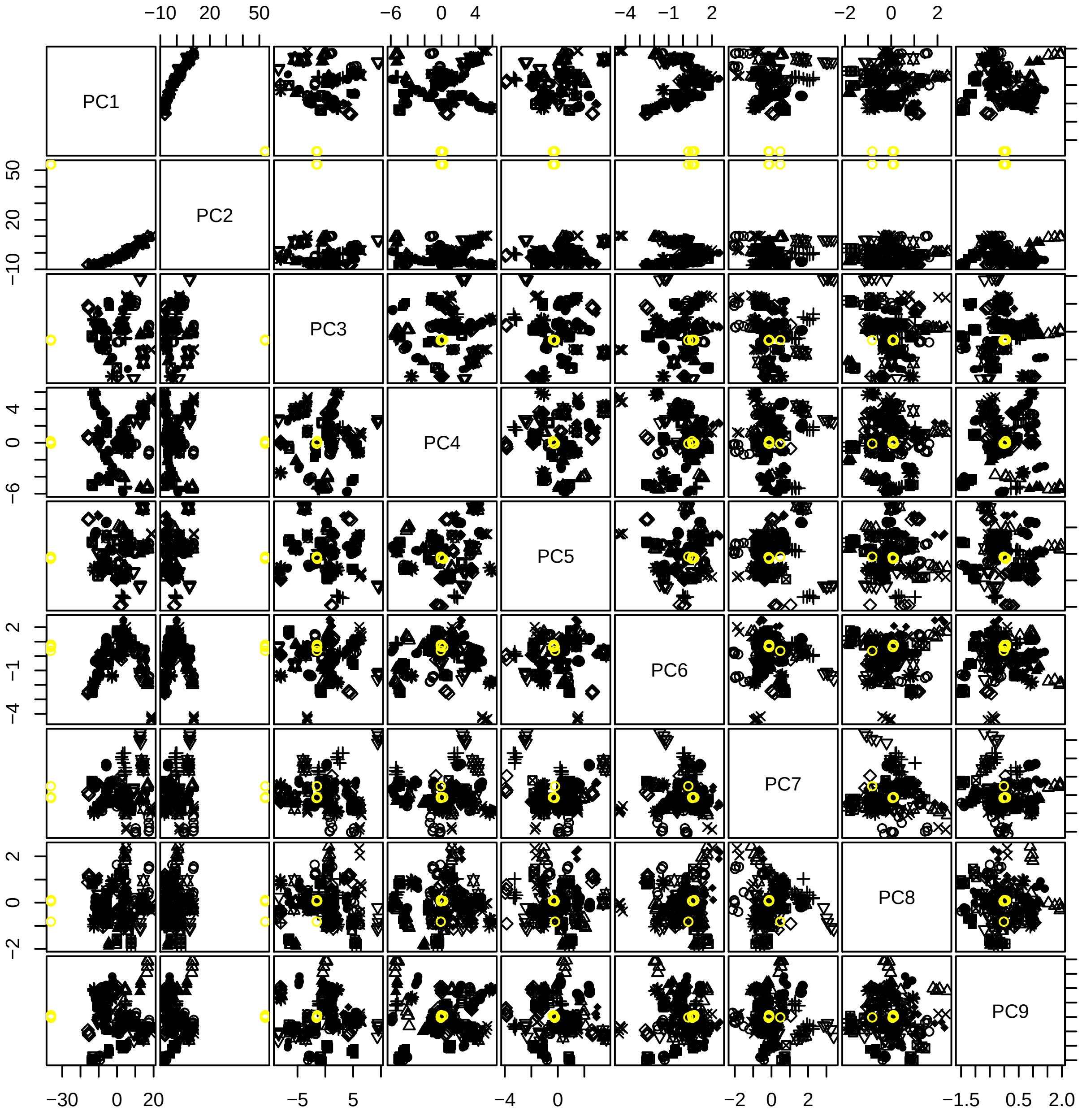
PC1
PC2
PC3
PC4
PC5
PC6
PC7
PC8
PC9

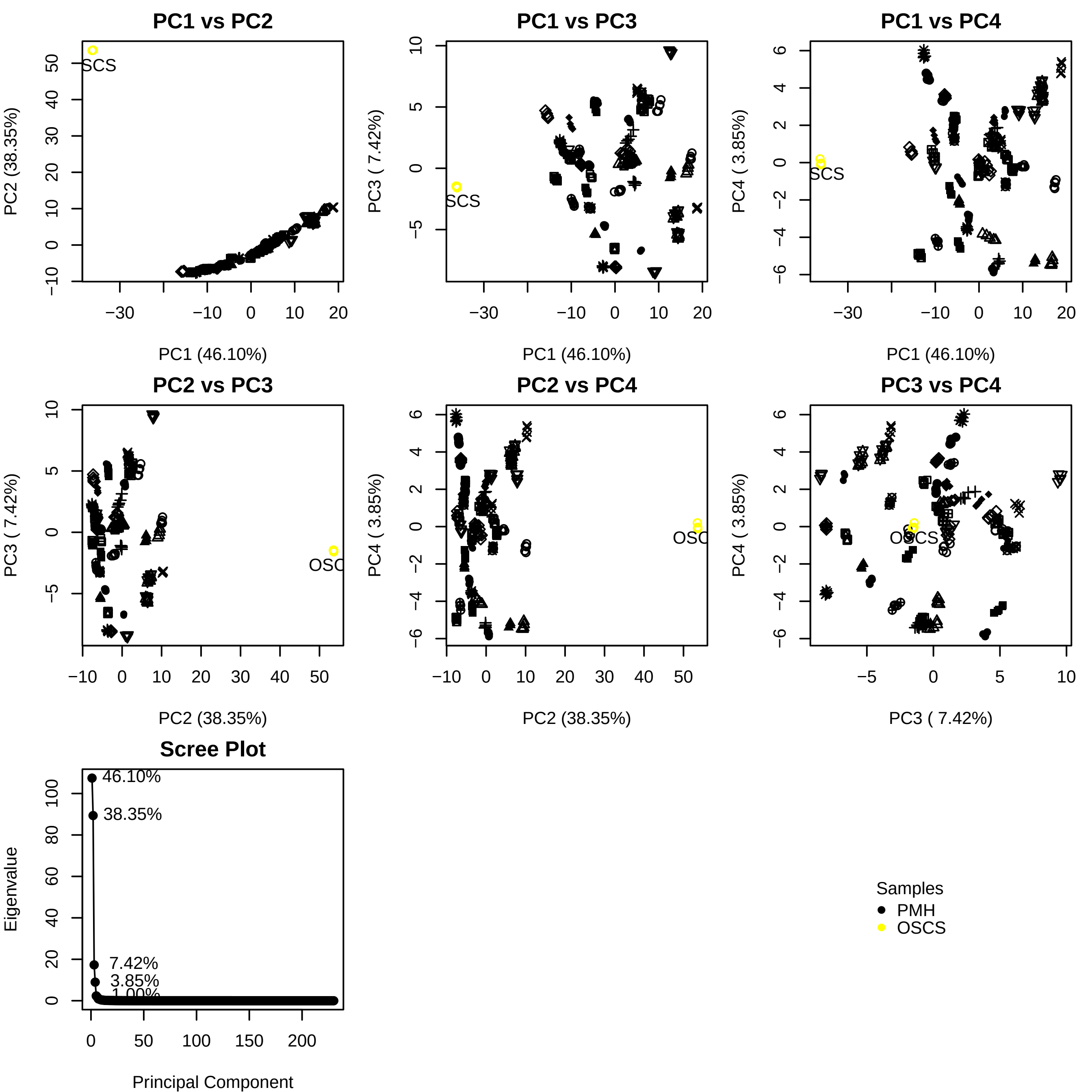

PC1 vs PC2
PC1 vs PC3
PC1 vs PC4
PC2 vs PC3
PC2 vs PC4
PC3 vs PC4
Scree Plot
PC1 (46.10%)
PC2 (38.35%)
PC3 ( 7.42%)
PC4 ( 3.85%)
SCS
OSC
OSCS
Eigenvalue
Principal Component
46.10%
38.35%
7.42%
3.85%
1.00%
Samples
PMH
OSCS

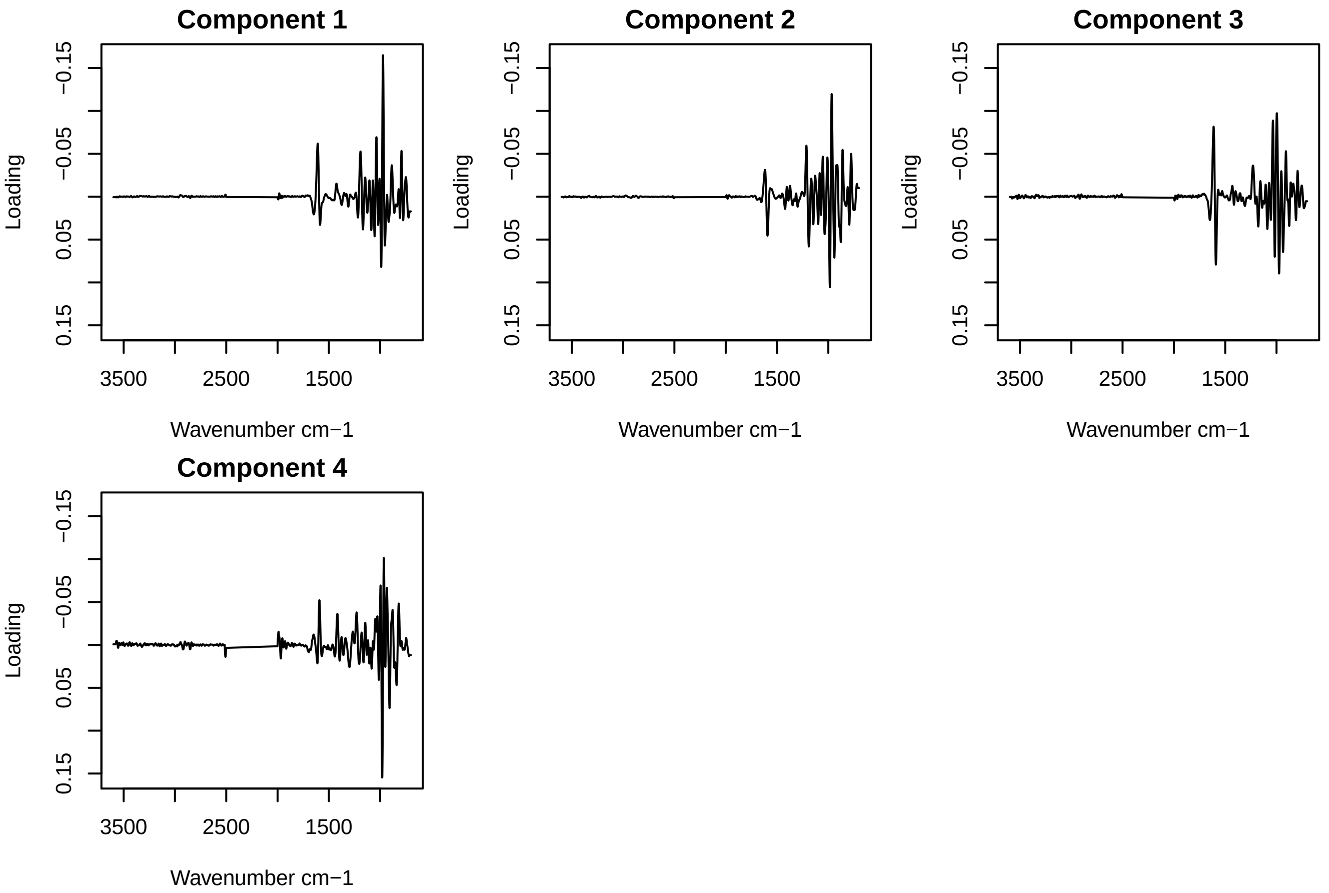
Component 1
Component 2
Component 3
Component 4
Loading
−0.15
−0.05
0.05
0.15
3500
2500
1500
Wavenumber cm−1

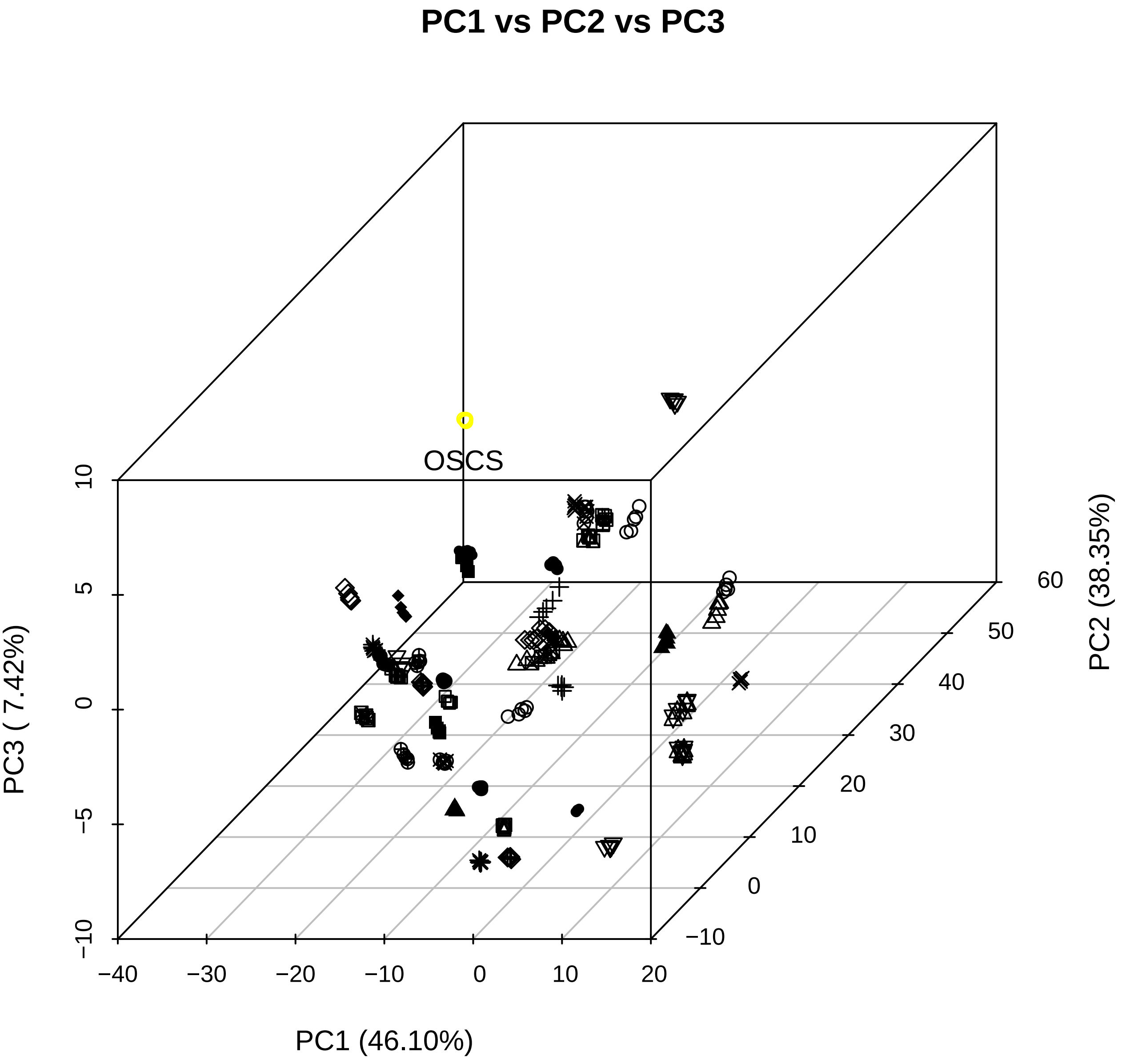
PC1 vs PC2 vs PC3
OSCS
PC1 (46.10%)
PC2 (38.35%)
PC3 ( 7.42%)
−40 −30 −20 −10 0 10 20
−10 0 10 20 30 40 50 60
−10 −5 0 5 10

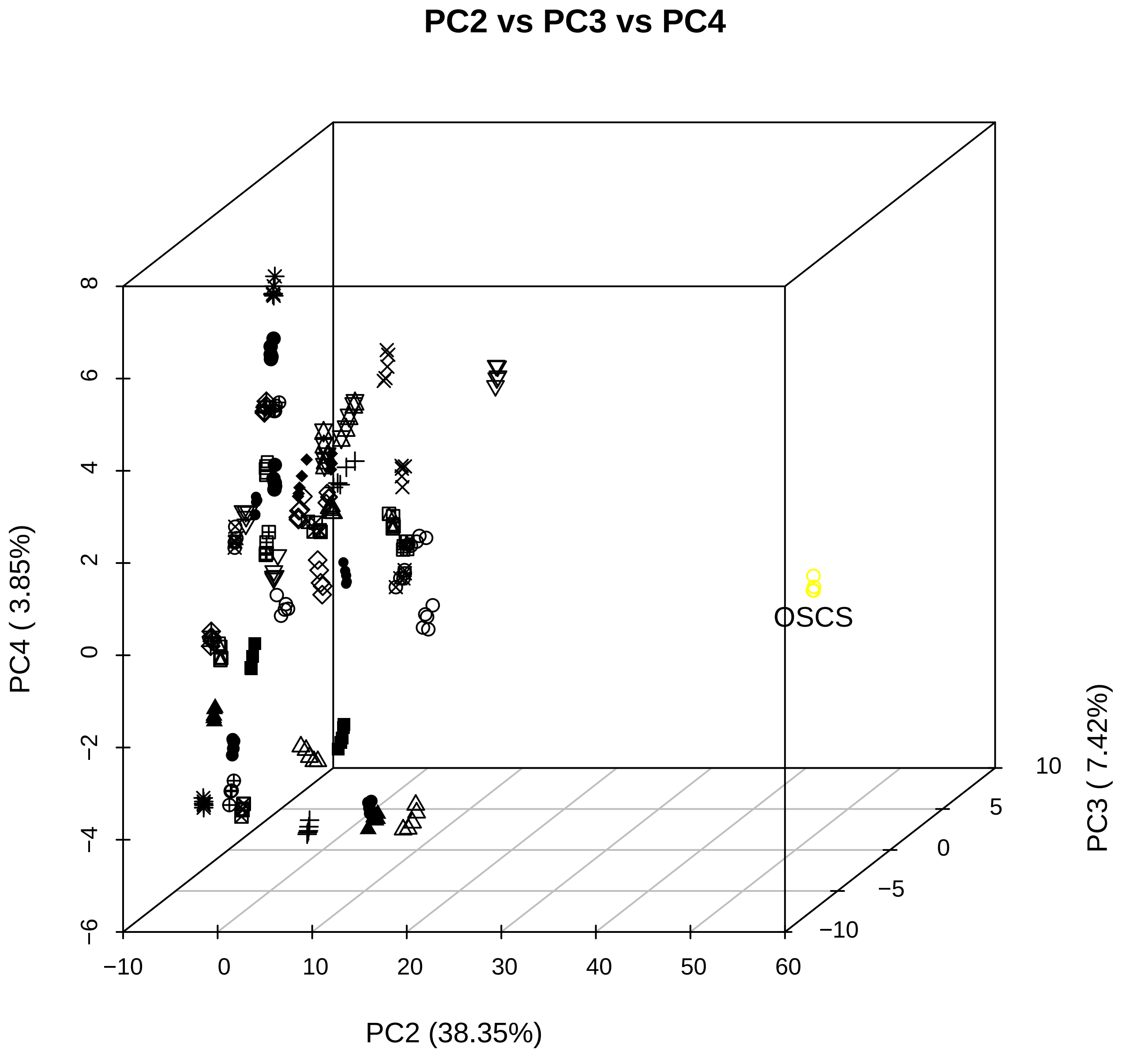
PC2 vs PC3 vs PC4
PC4 ( 3.85%)
PC2 (38.35%)
PC3 ( 7.42%)
OSCS
8
6
4
2
0
−2
−4
−6
−10
0
10
20
30
40
50
60
10
5
0
−5
−10

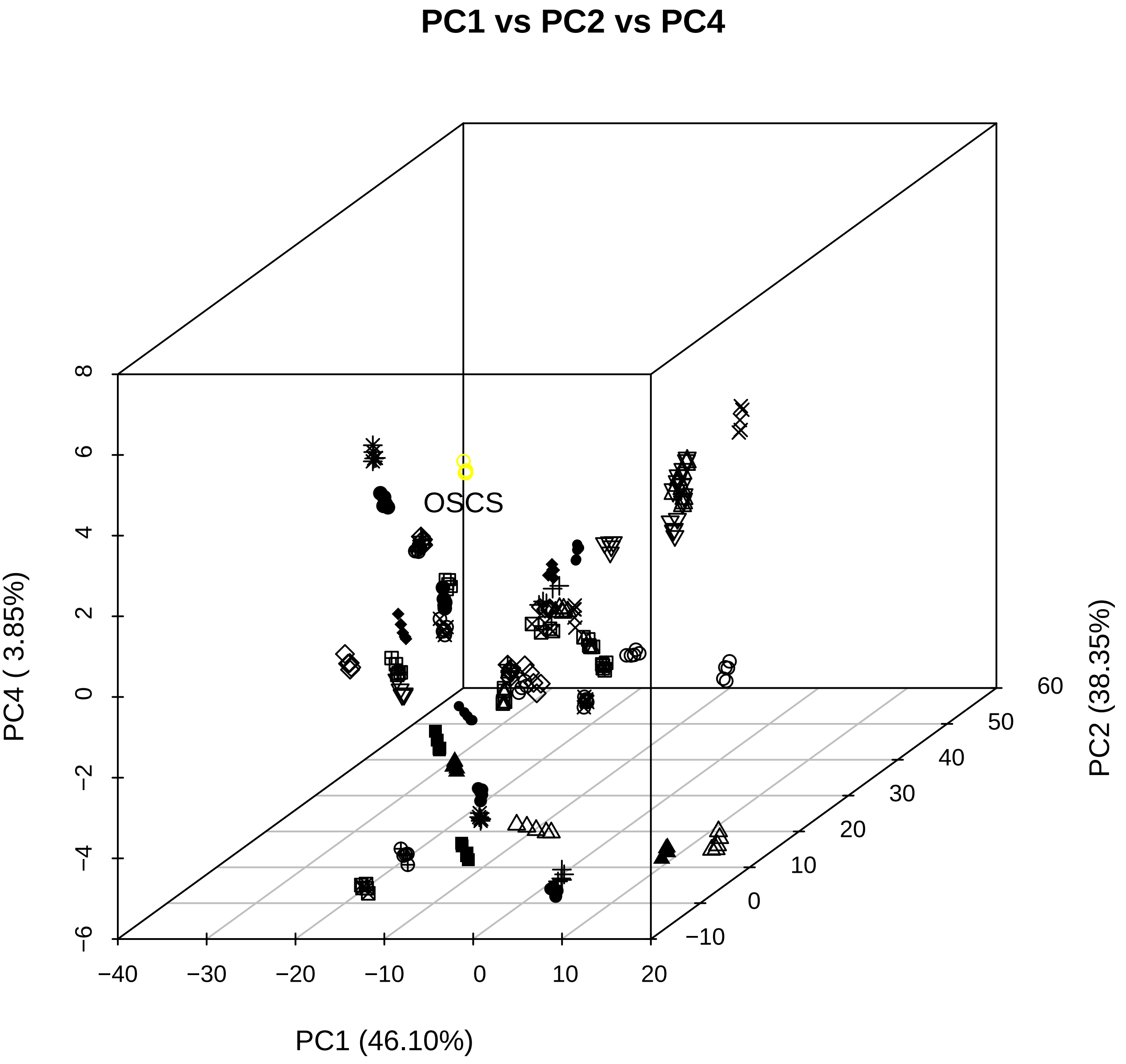

PC1 vs PC2 vs PC4
OSCS
PC4 ( 3.85%)
PC1 (46.10%)
PC2 (38.35%)
8
6
4
2
0
−2
−4
−6
−40
−30
−20
−10
0
10
20
60
50
40
30
20
10
0
−10

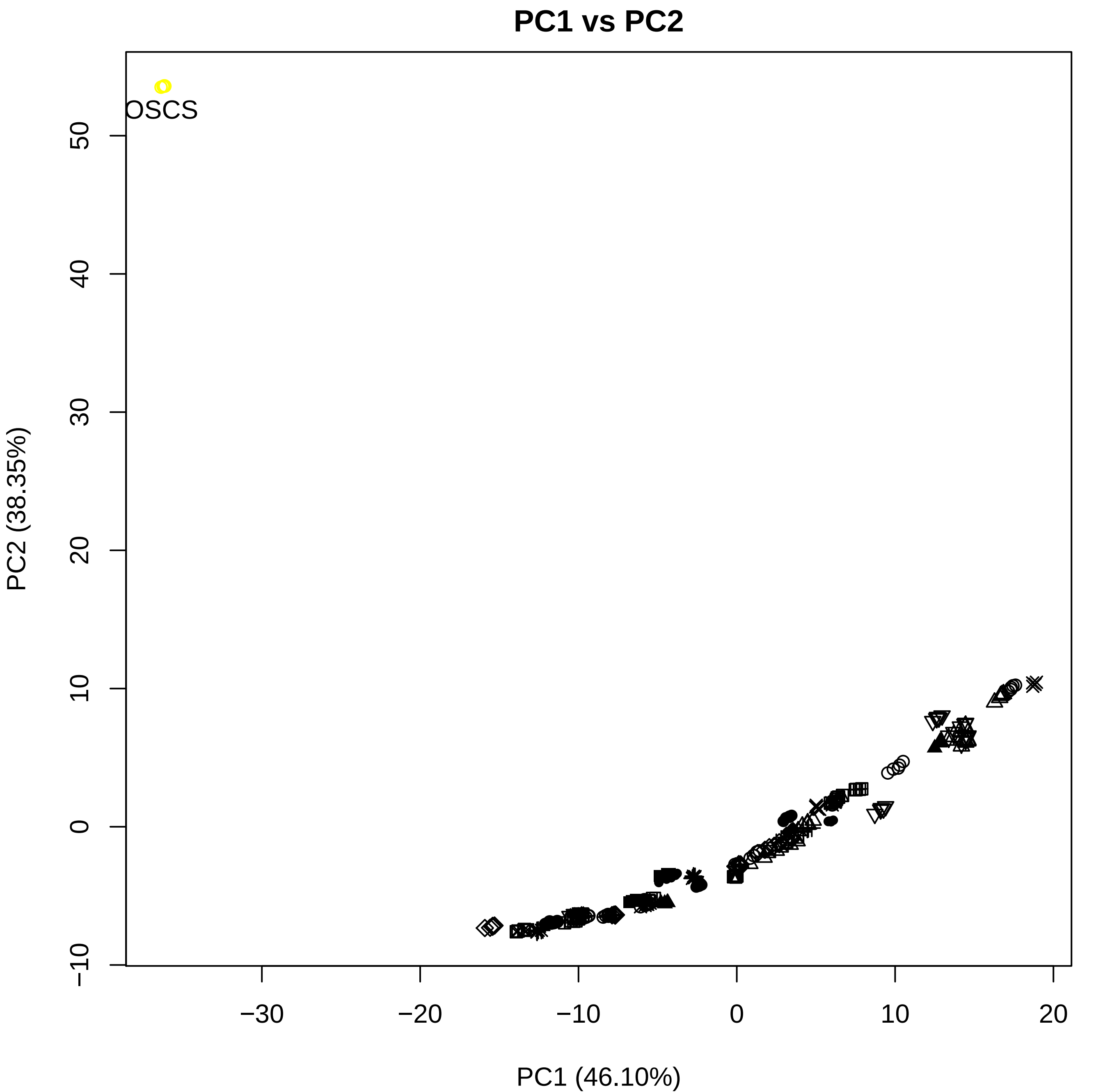
PC1 vs PC2
OSCS
PC2 (38.35%)
PC1 (46.10%)
−30
−20
−10
0
10
20
50
40
30
20
10
0
−10

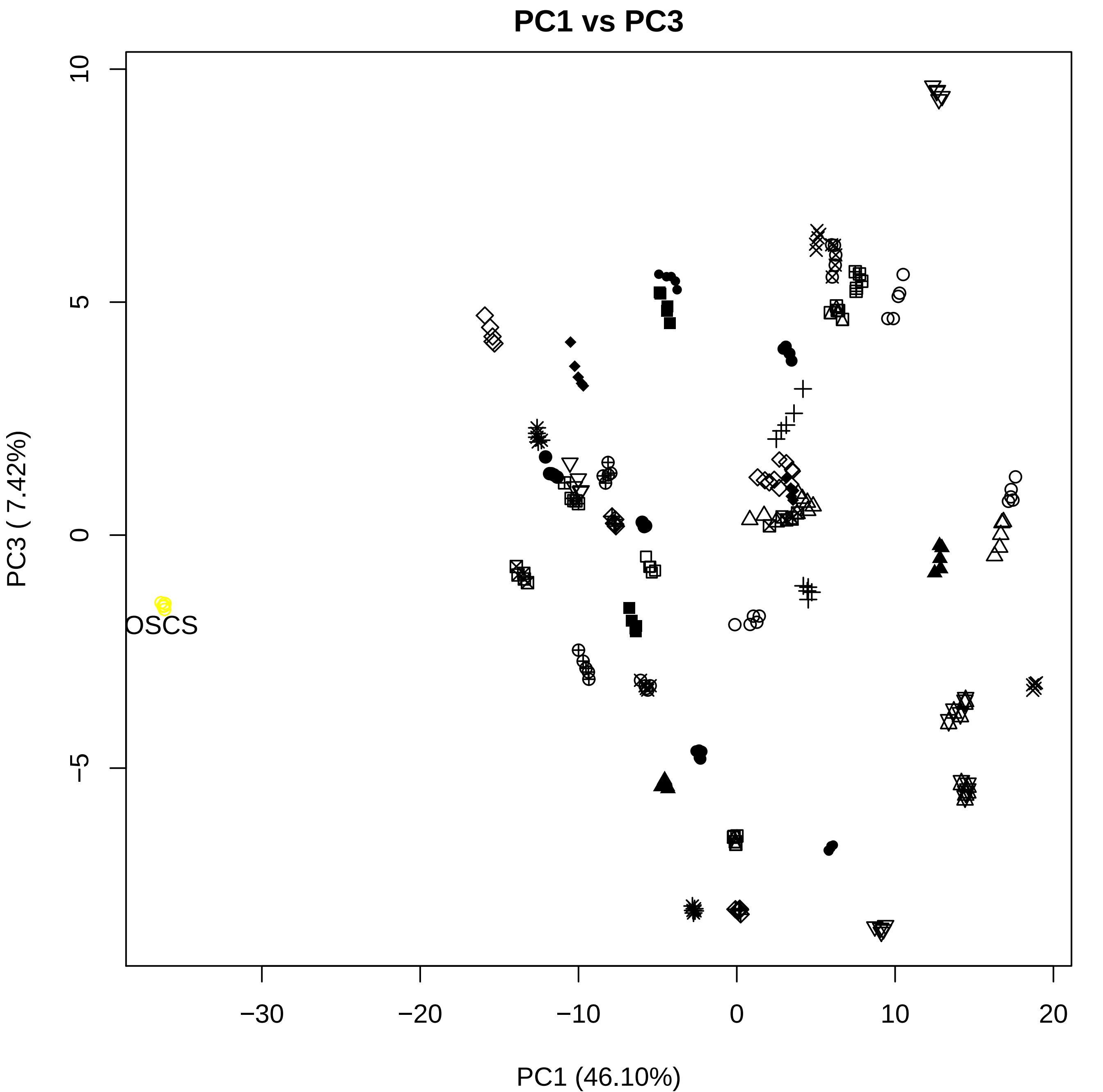

PC1 vs PC3
10
5
0
−5
PC3 ( 7.42%)
OSCS
−30
−20
−10
0
10
20
PC1 (46.10%)

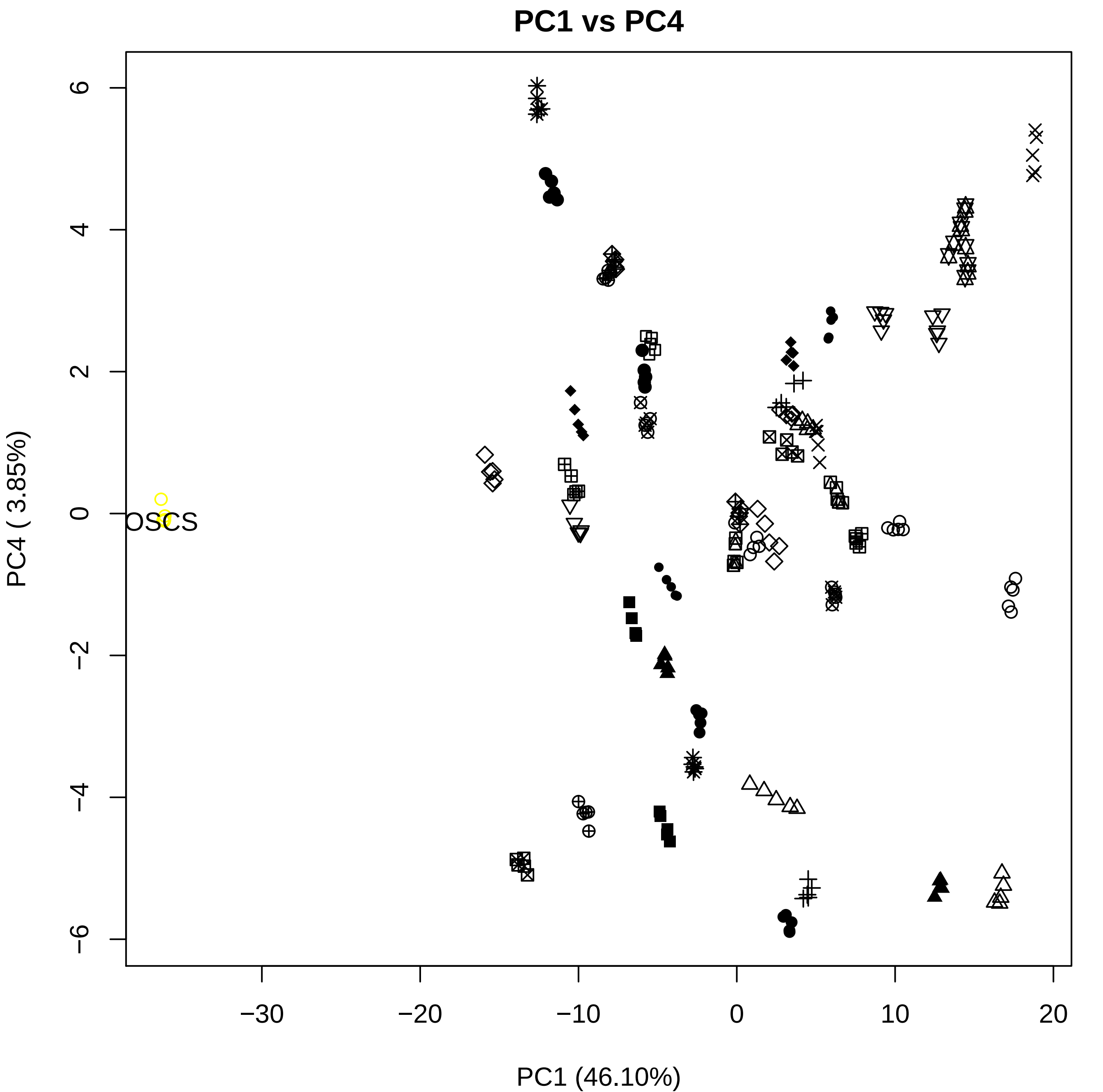
PC1 vs PC4
PC4 ( 3.85%)
PC1 (46.10%)
OSCS
6
4
2
0
−2
−4
−6
−30
−20
−10
0
10
20

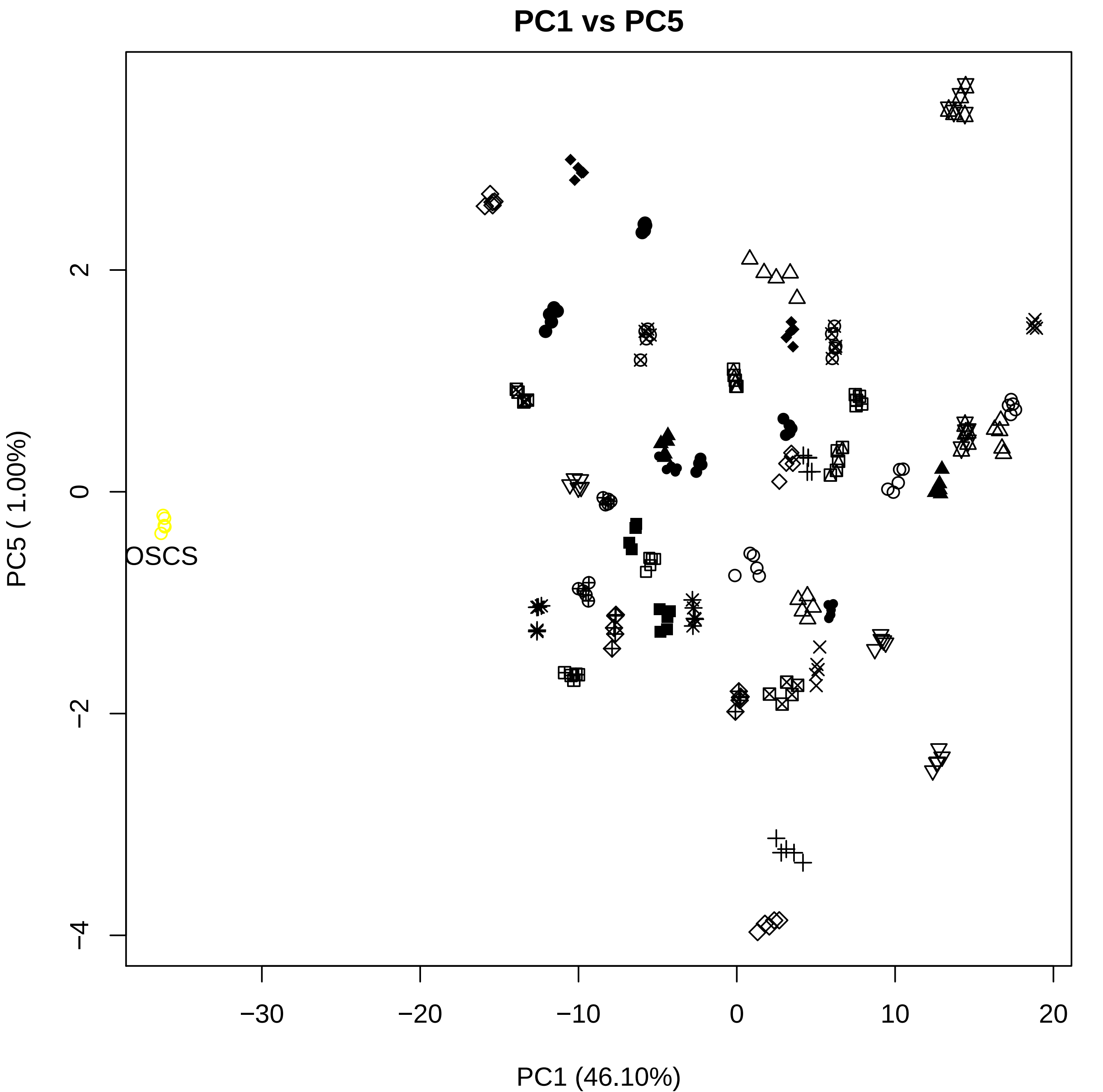
PC1 vs PC5
PC5 ( 1.00%)
PC1 (46.10%)
OSCS
2
0
−2
−4
−30
−20
−10
0
10
20

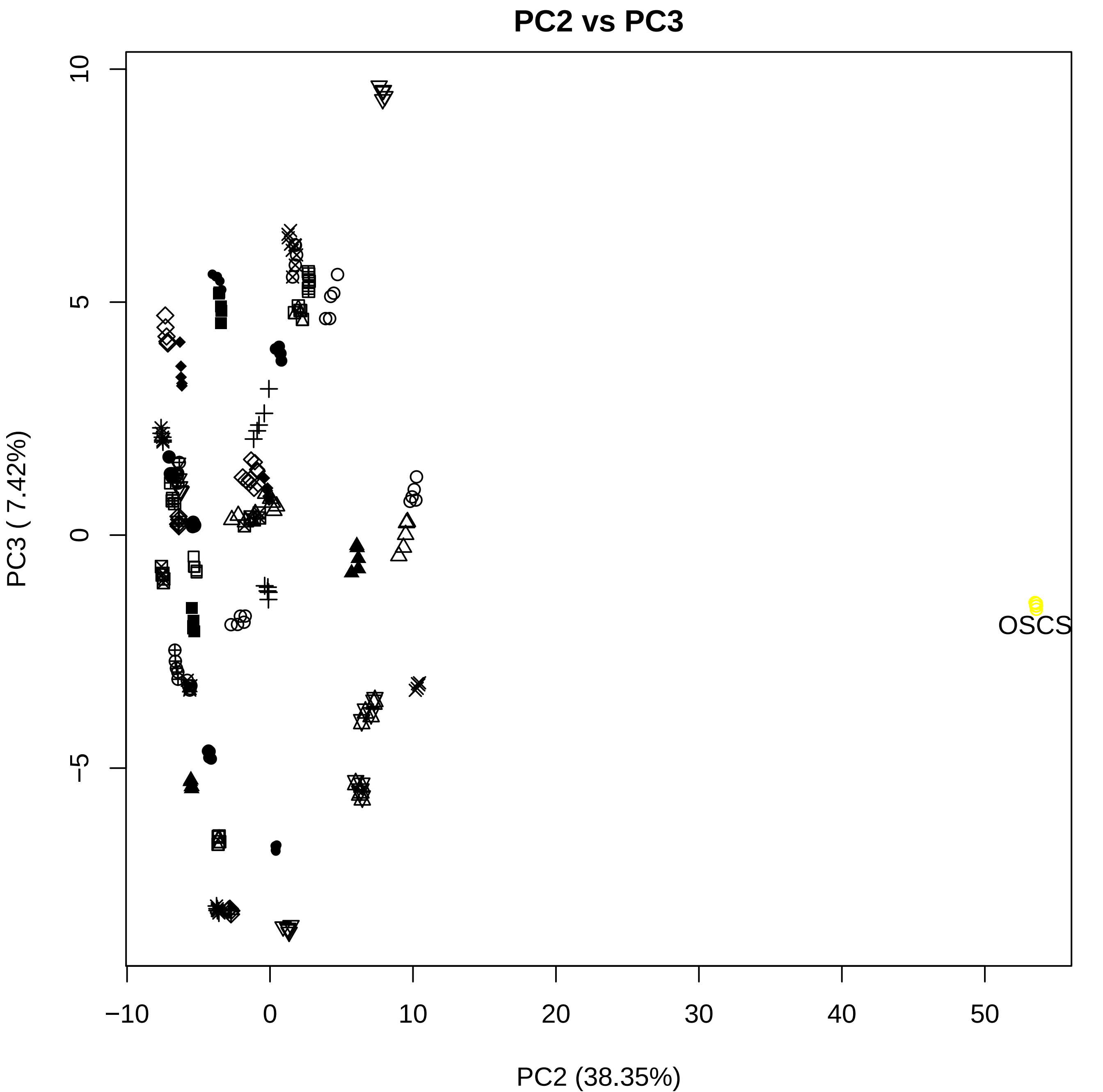
PC2 vs PC3
PC3 ( 7.42%)
10
5
0
−5
−10
0
10
20
30
40
50
PC2 (38.35%)
OSCS

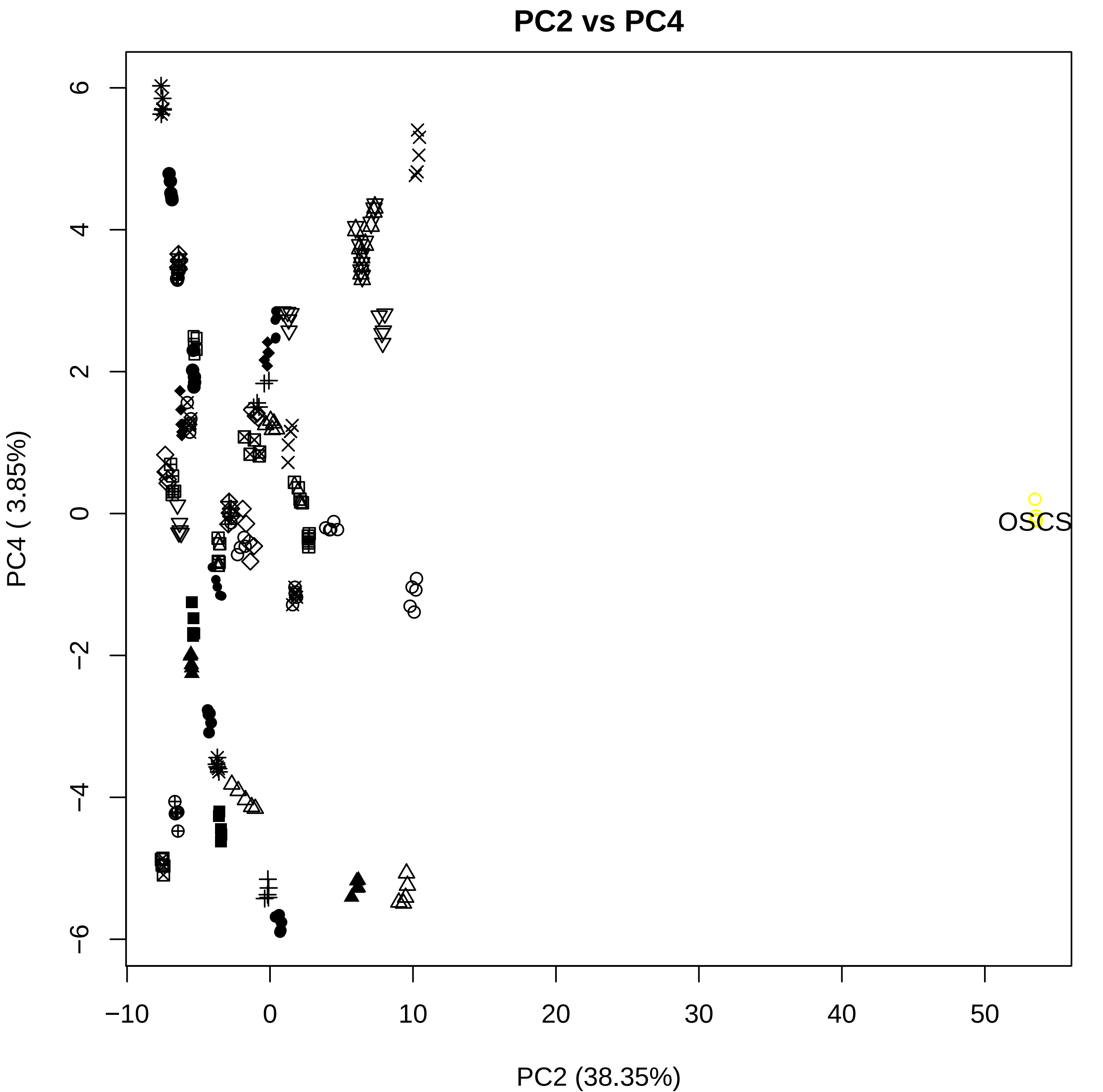

PC2 vs PC4
PC4 ( 3.85%)
PC2 (38.35%)
6
4
2
0
−2
−4
−6
−10
0
10
20
30
40
50
OSCS

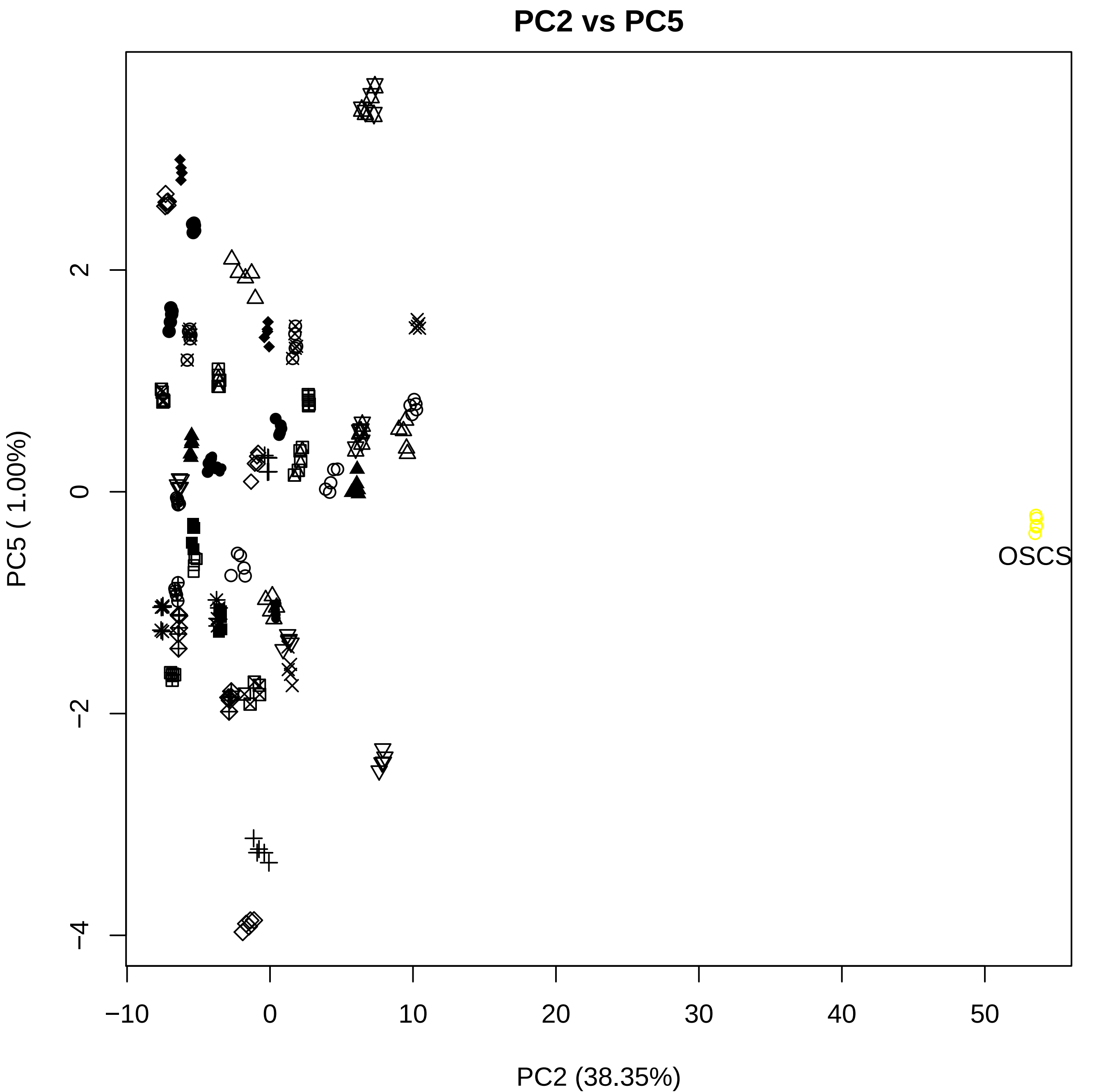

PC2 vs PC5
PC5 ( 1.00%)
PC2 (38.35%)
2
0
−2
−4
−10
0
10
20
30
40
50
OSCS

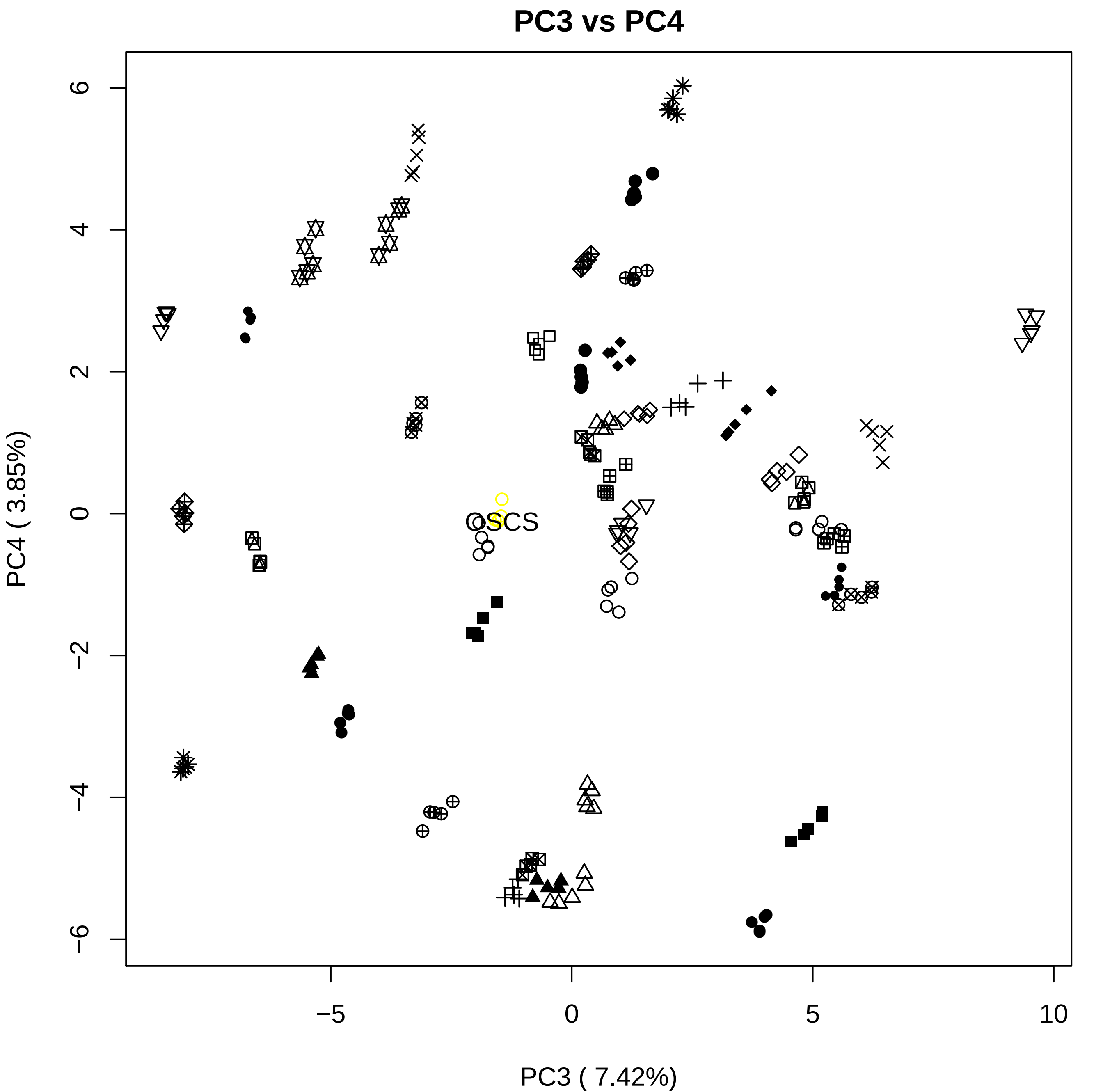

PC3 vs PC4
PC4 ( 3.85%)
PC3 ( 7.42%)
OSCS
−6
−4
−2
0
2
4
6
−5
0
5
10

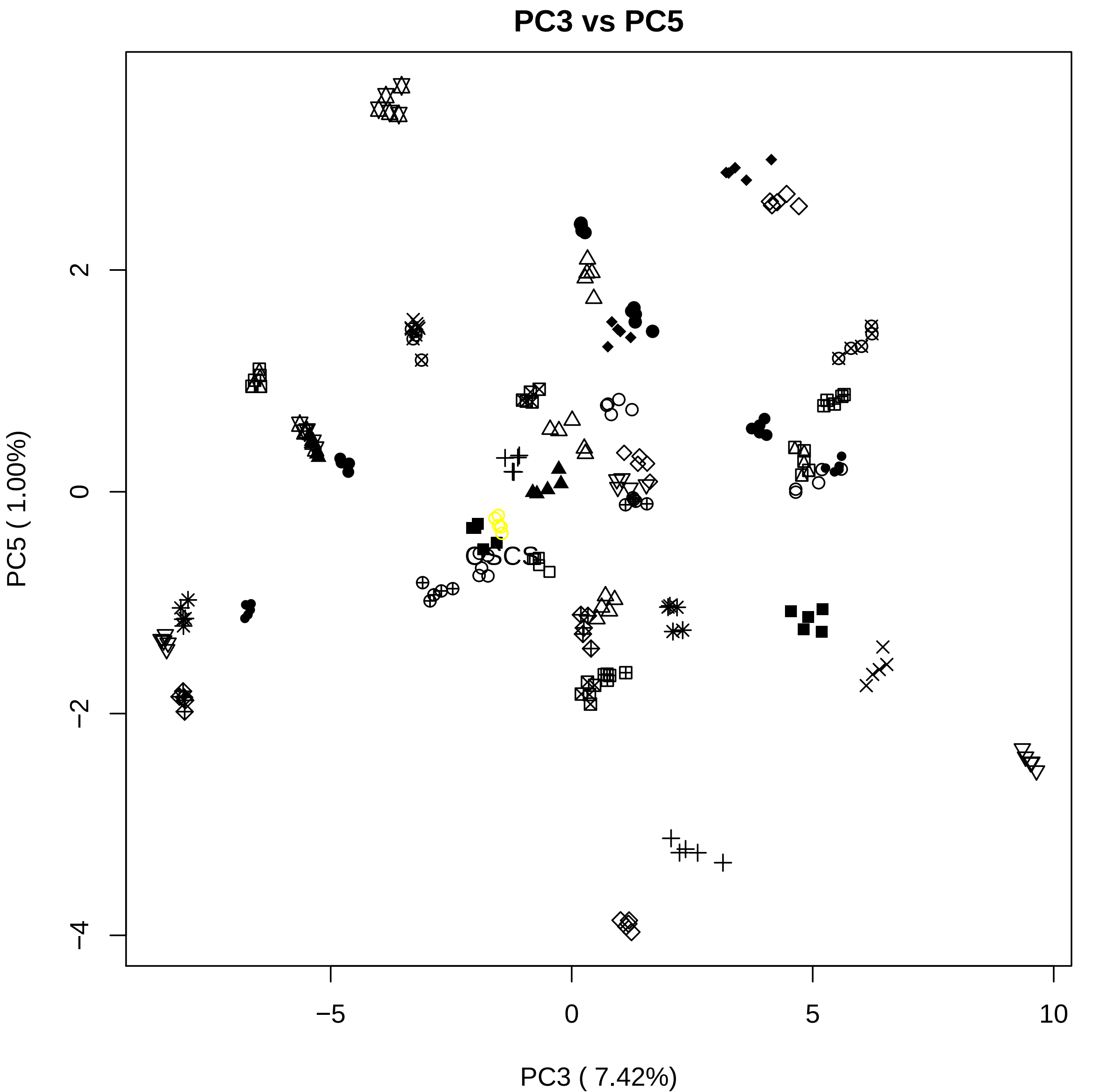
PC3 vs PC5
PC5 ( 1.00%)
PC3 ( 7.42%)
2
0
−2
−4
−5
0
5
10

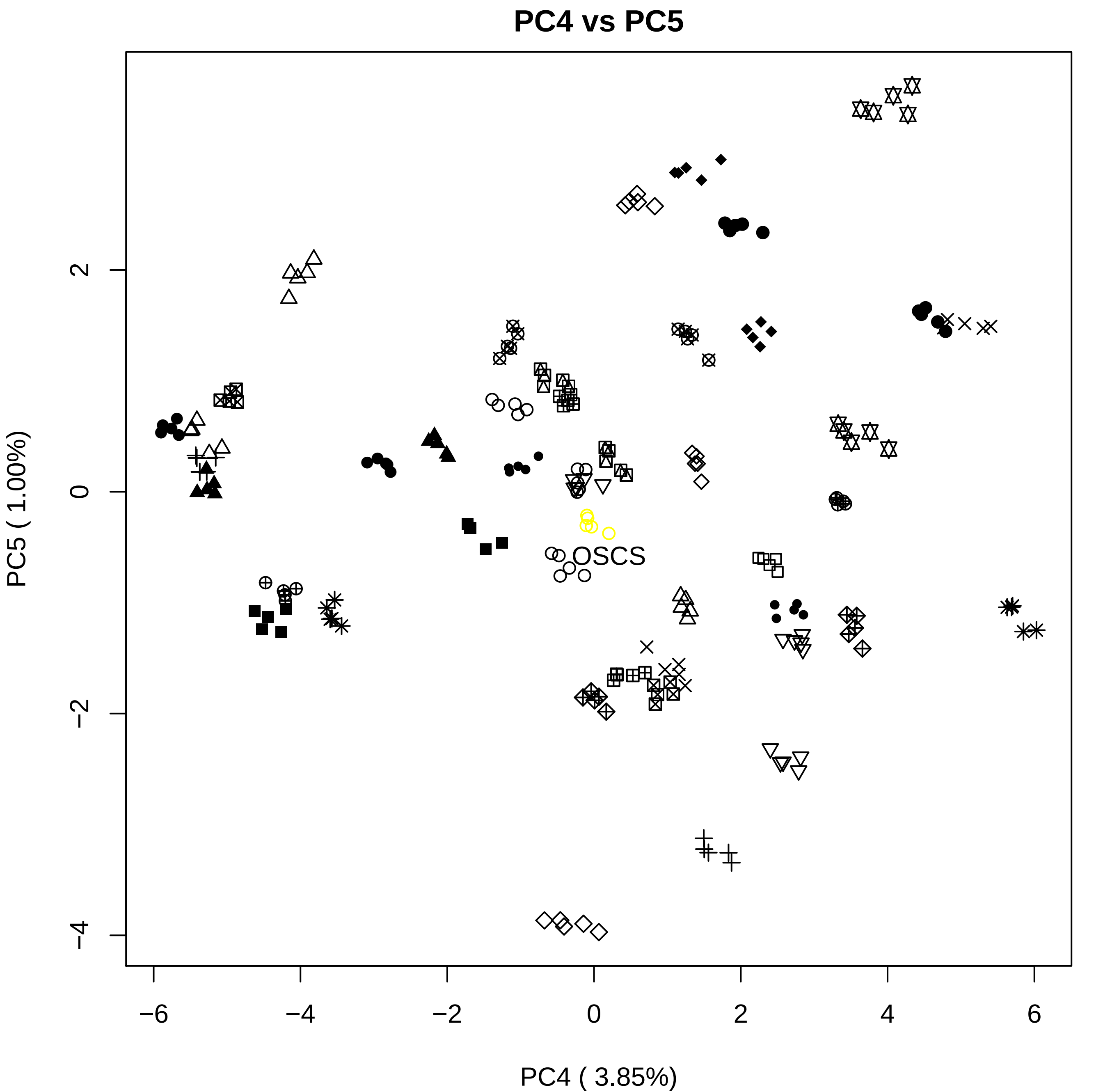
PC4 vs PC5
PC5 ( 1.00%)
PC4 ( 3.85%)
OSCS
−6
−4
−2
0
2
4
6
2
0
−2
−4

### OSAS Contam 7th

Anthony Devlin, Keele University

### PCA of All  OSAS vs PMH

Scaled Matrix PCAed

7th order polynomial baseline correction performed

Raw spectra smoothed to 21 Neighbours, 2nd order PN, Savtisky Golay

Normalisation between 0 and 100

D2 spectra smoothed to 41 Neighbours, 2nd order PN, Savtisky Golay

Regions Analysed: 3601−2501 and 2001−700cm−1

Samples Analysed: 12 Contam's 1 OSAS's 45 PMH's

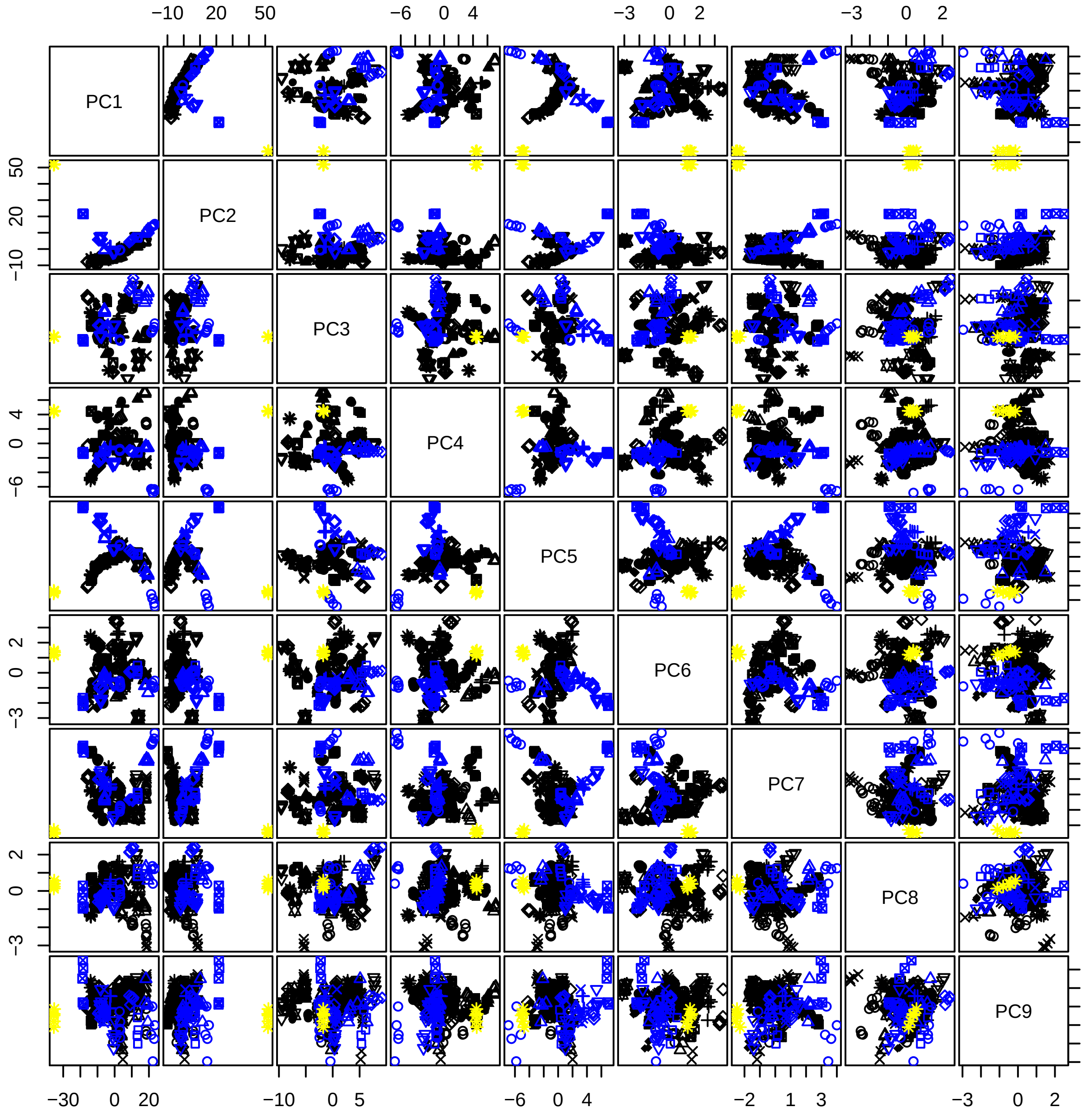
PC1
PC2
PC3
PC4
PC5
PC6
PC7
PC8
PC9
−10 20 50
−6 0 4
−3 0 2
−3 0 2
20 0 −30
50 20 −10
5 0 −10
4 0 −6
4 0 −6
2 0 −3
3 1 −2
2 0 −3
2 0 −3
−30 0 20
−10 0 5
−6 0 4
−2 1 3
−3 0 2

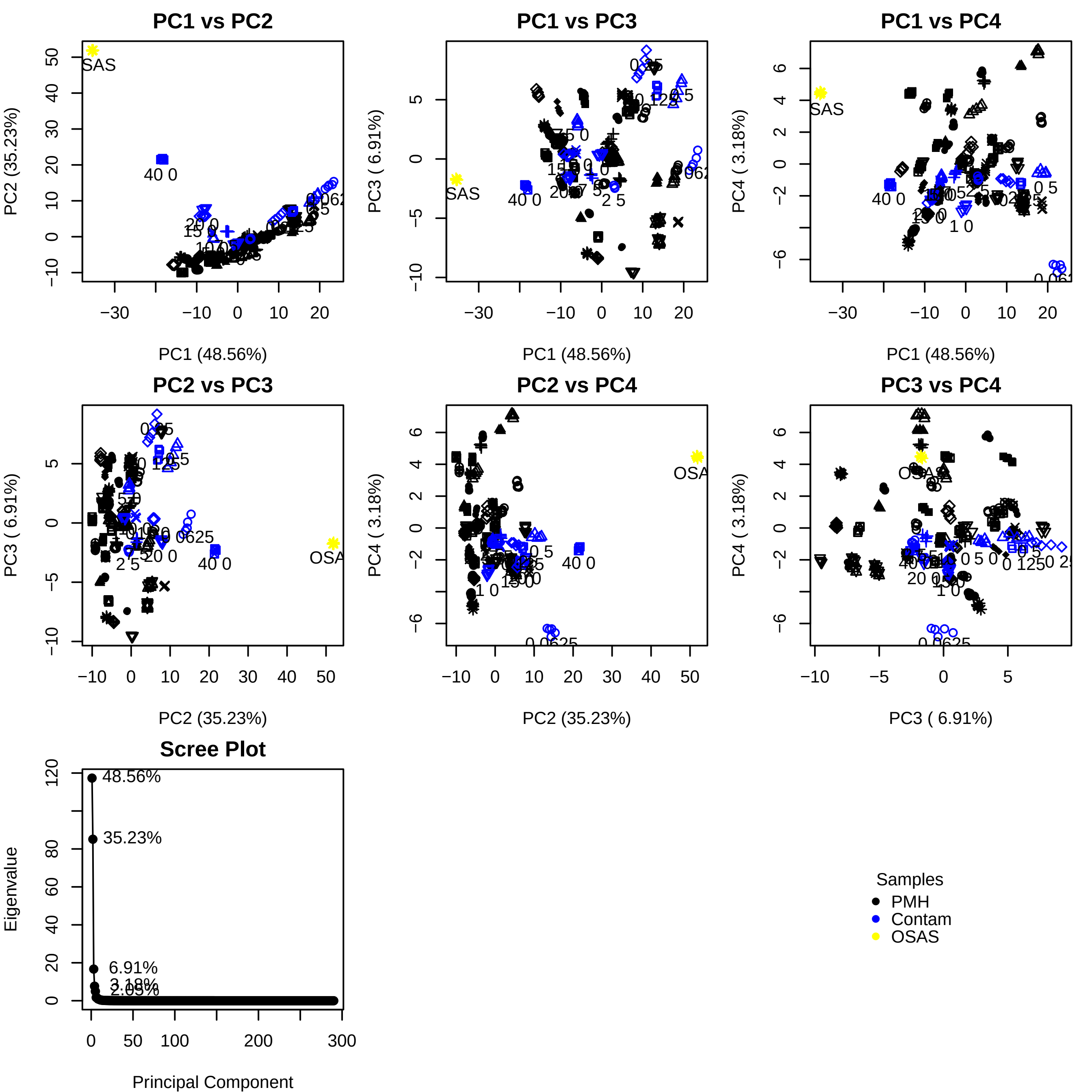
PC1 vs PC2
PC1 vs PC3
PC1 vs PC4
PC2 vs PC3
PC2 vs PC4
PC3 vs PC4
PC1 (48.56%)
PC2 (35.23%)
PC3 ( 6.91%)
PC4 ( 3.18%)
Scree Plot
48.56%
35.23%
6.91%
Eigenvalue
Principal Component
Samples
PMH
Contam
OSAS

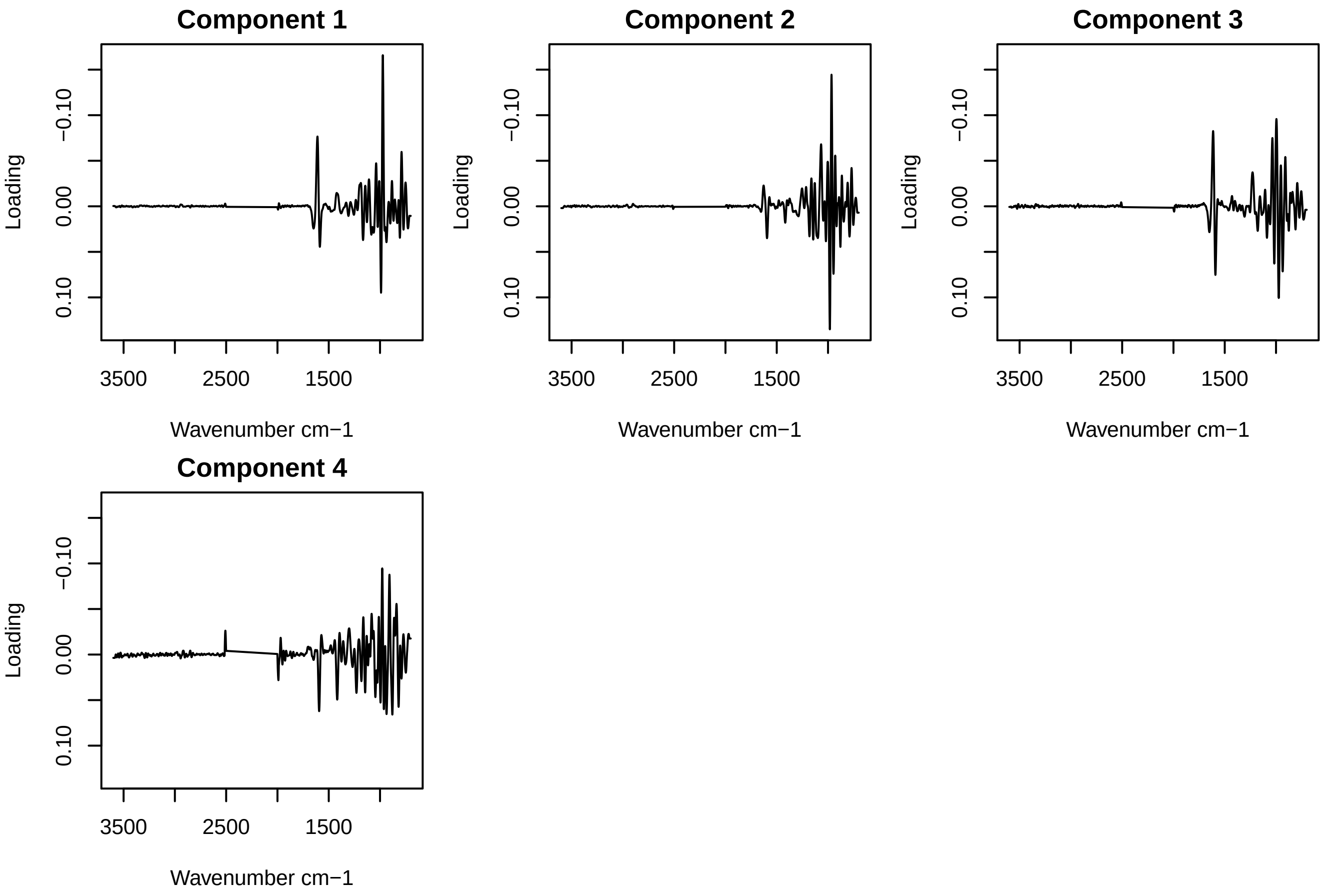

Component 1
Component 2
Component 3
Component 4
Loading
−0.10
0.00
0.10
3500
2500
1500
Wavenumber cm−1

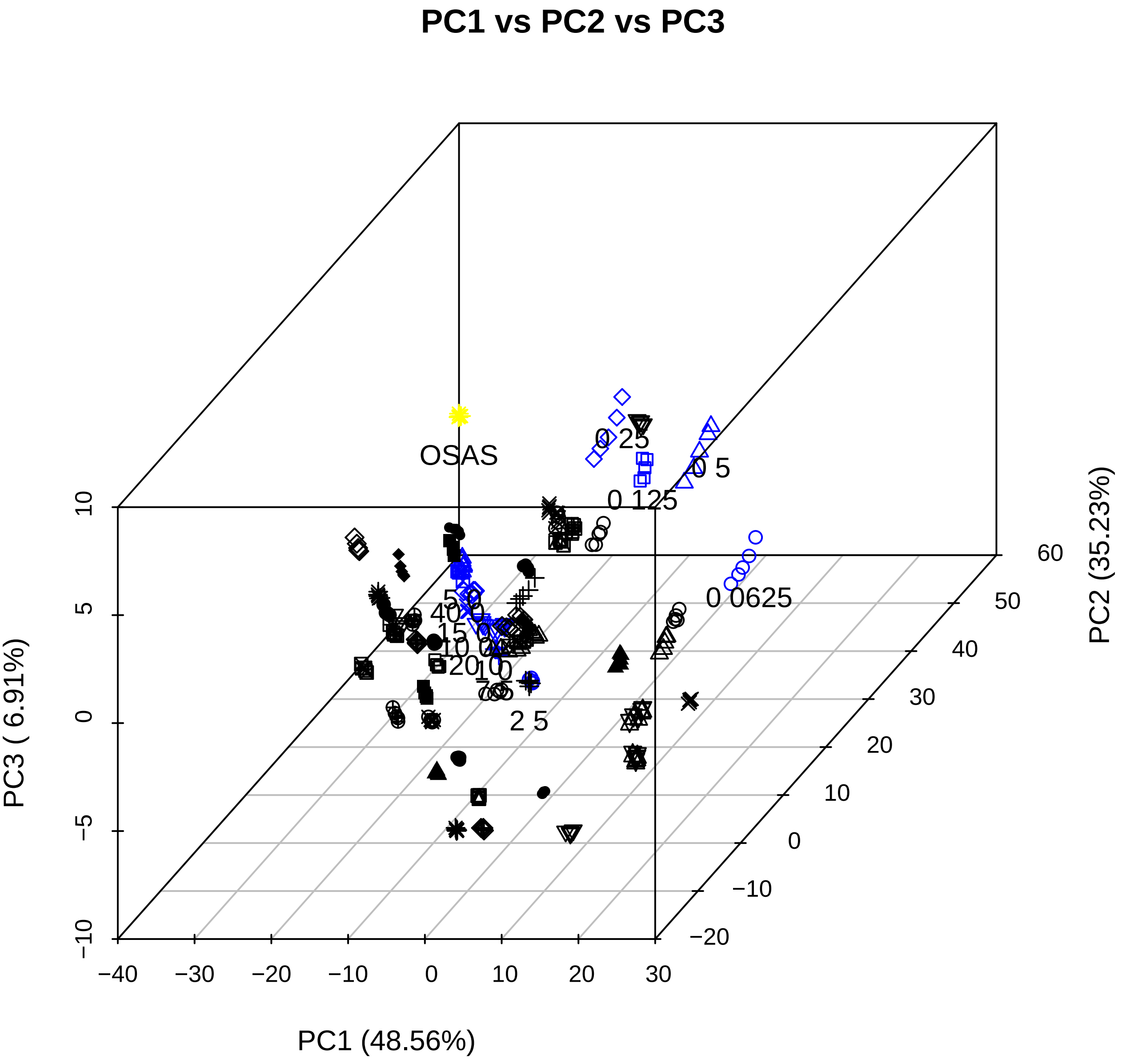

PC1 vs PC2 vs PC3
OSAS
0 25
0 5
0 125
0 0625
5 0
40
15
10
20
10
2 5
PC1 (48.56%)
PC2 (35.23%)
PC3 ( 6.91%)
−40
−30
−20
−10
0
10
20
30
−20
−10
0
10
20
30
40
50
60
−10
−5
0
5
10

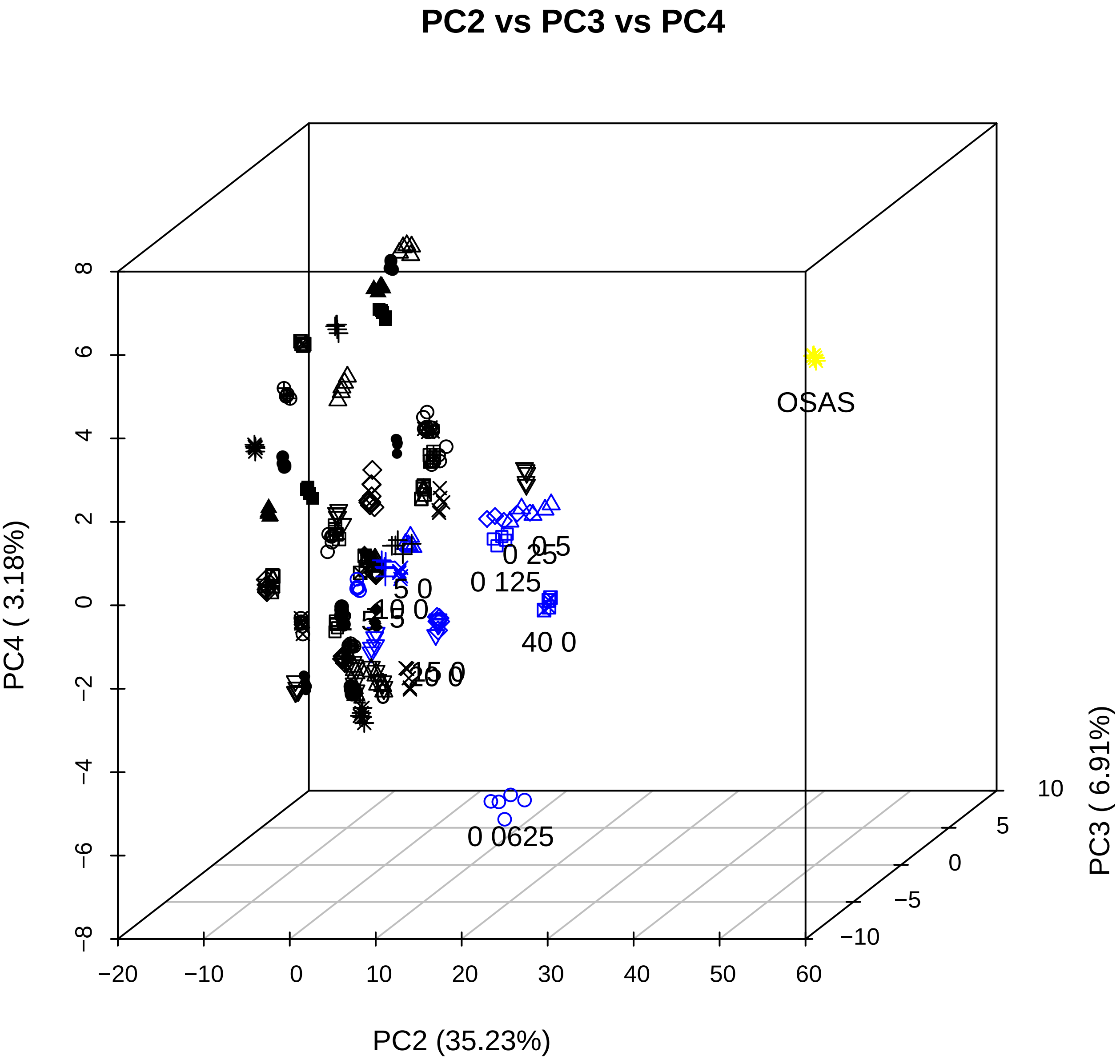
PC2 vs PC3 vs PC4
PC4 ( 3.18%)
PC2 (35.23%)
PC3 ( 6.91%)
−8
−6
−4
−2
0
2
4
6
8
−20
−10
0
10
20
30
40
50
60
−10
−5
0
5
10
OSAS
0 5
0 25
0 125
5 0
10 0
40 0
15 0
20 0
0 0625

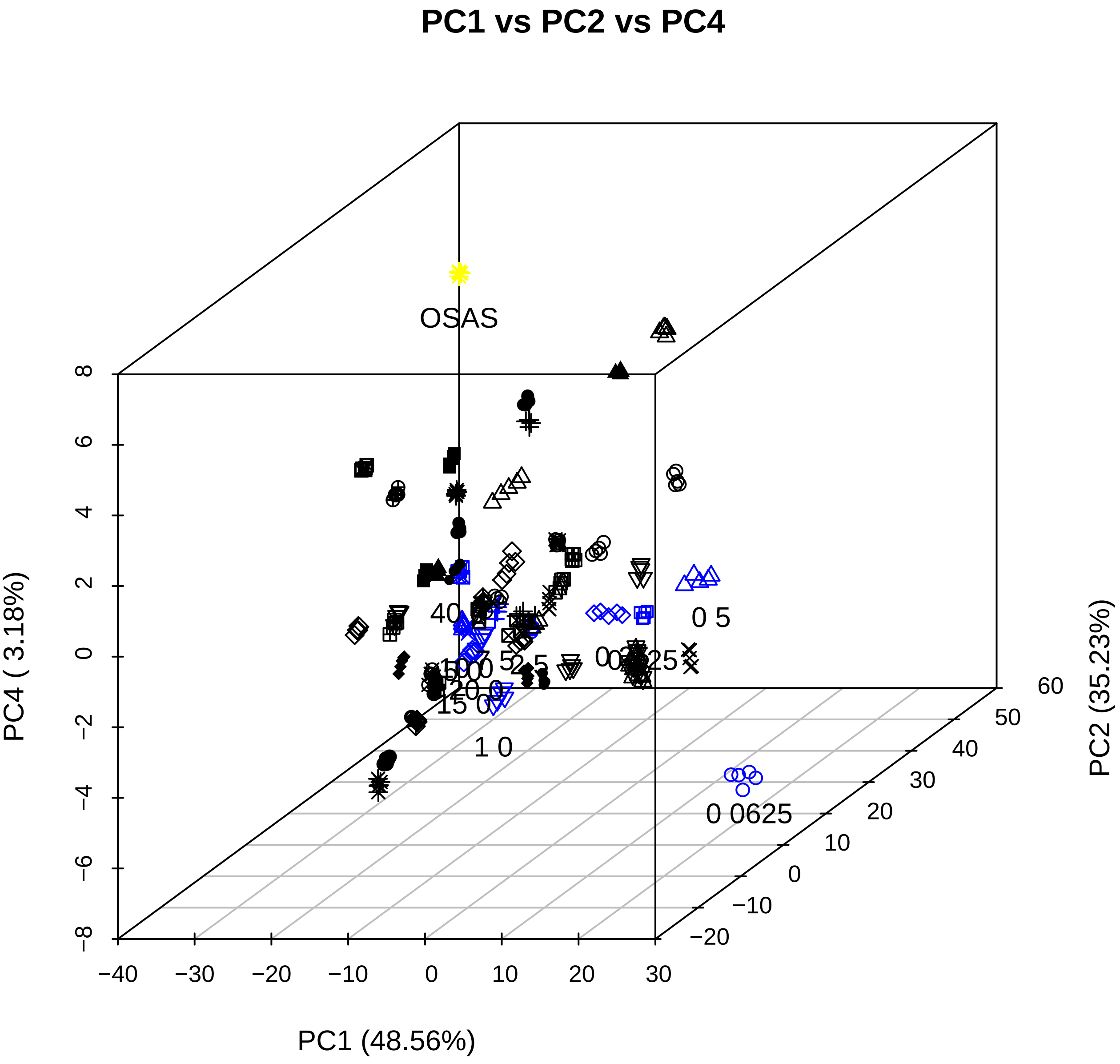
PC1 vs PC2 vs PC4
OSAS
0 5
1 0
0 0625
15 0
PC1 (48.56%)
PC2 (35.23%)
PC4 ( 3.18%)
−40
−30
−20
−10
0
10
20
30
−20
−10
0
10
20
30
40
50
60
−8
−6
−4
−2
0
2
4
6
8

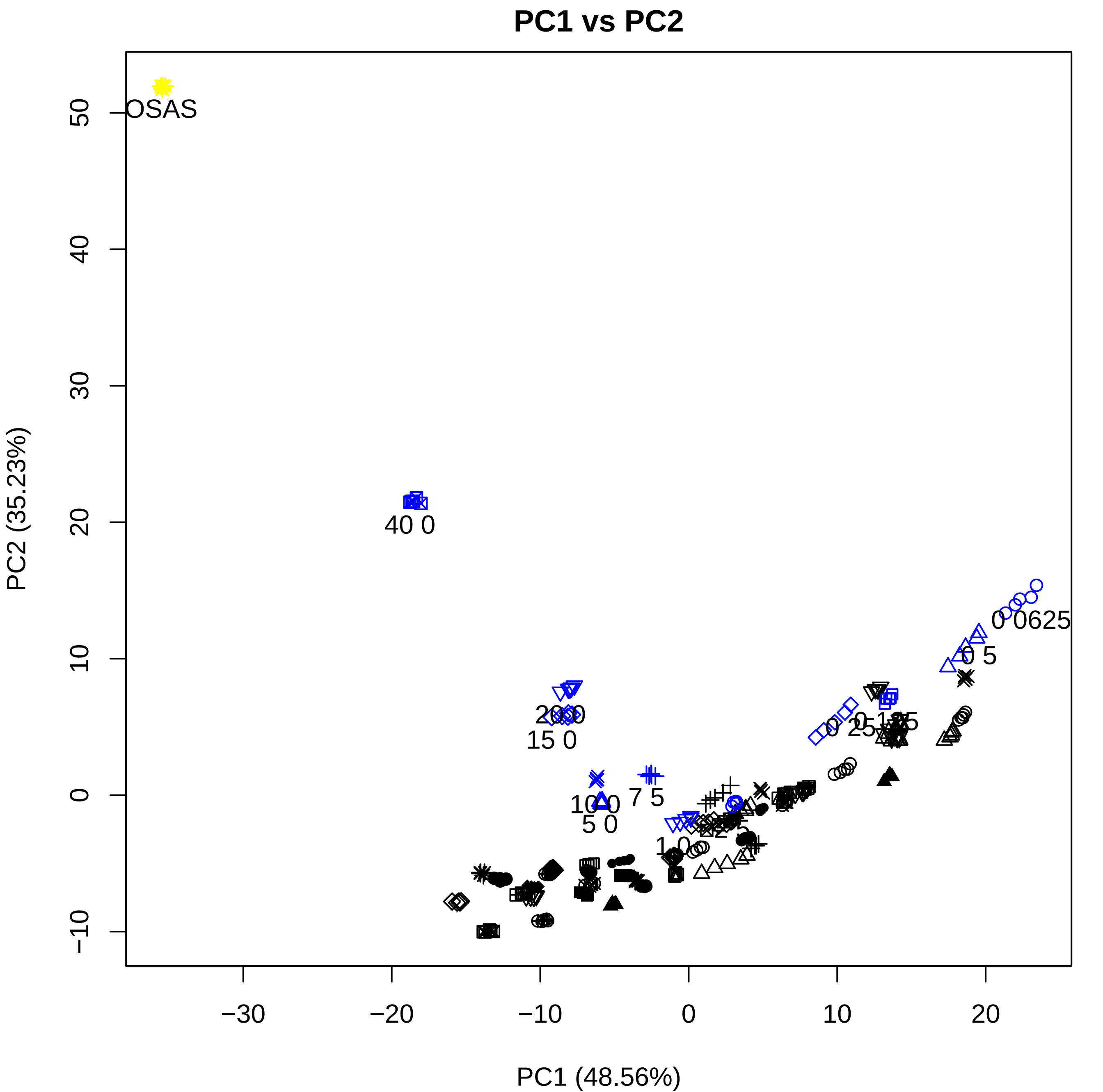

PC1 vs PC2
OSAS
40 0
20 0
15 0
10 0
7 5
5 0
0 0625
0 5
0 25
50
40
30
20
10
0
−10
−30
−20
−10
0
10
20
PC2 (35.23%)
PC1 (48.56%)

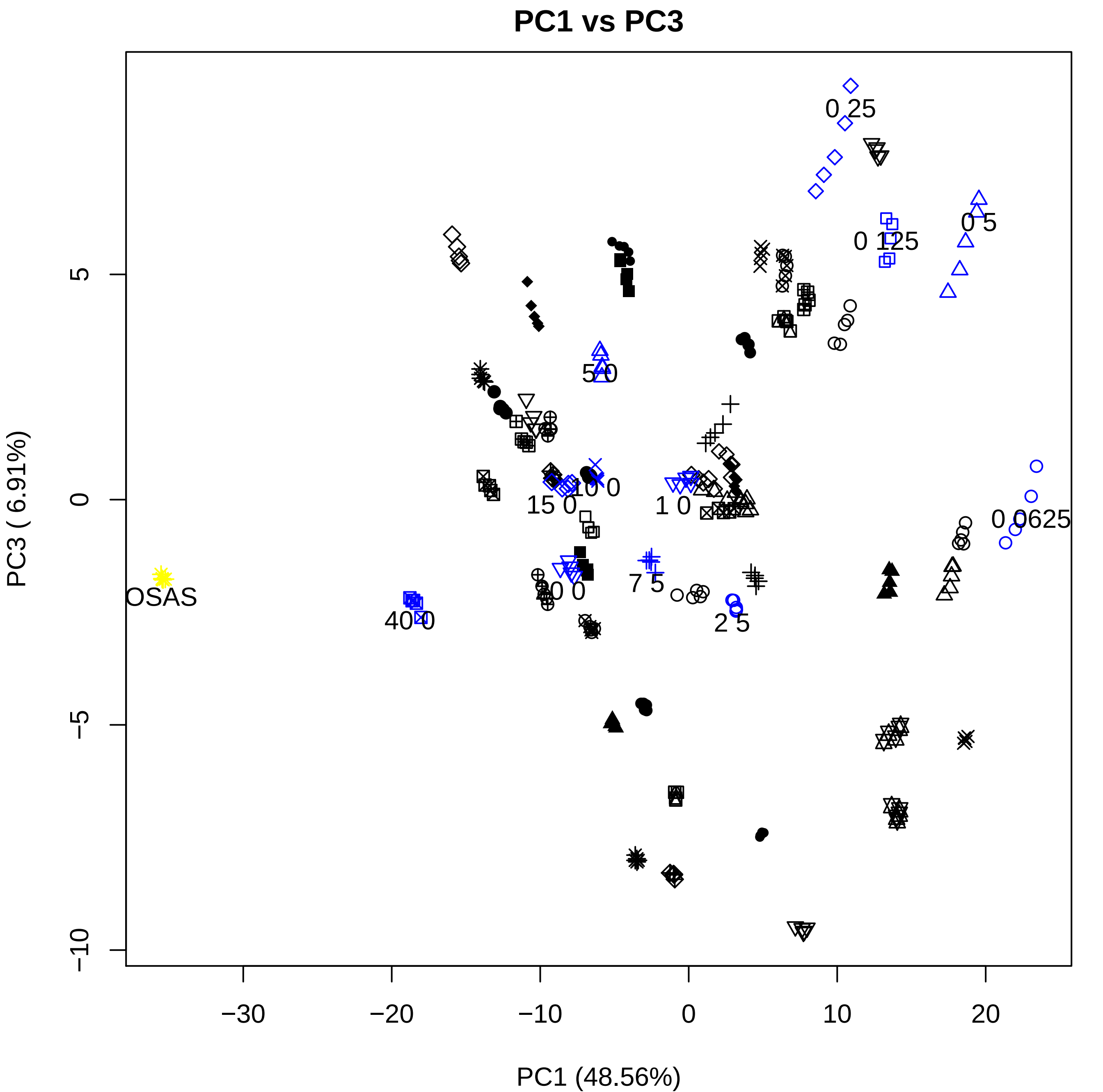
PC1 vs PC3
PC3 ( 6.91%)
PC1 (48.56%)
5
0
−5
−10
−30
−20
−10
0
10
20
0 25
0 5
0 125
0 0625
5 0
10 0
15 0
1 0
7 5
2 5
20 0
40 0
OSAS

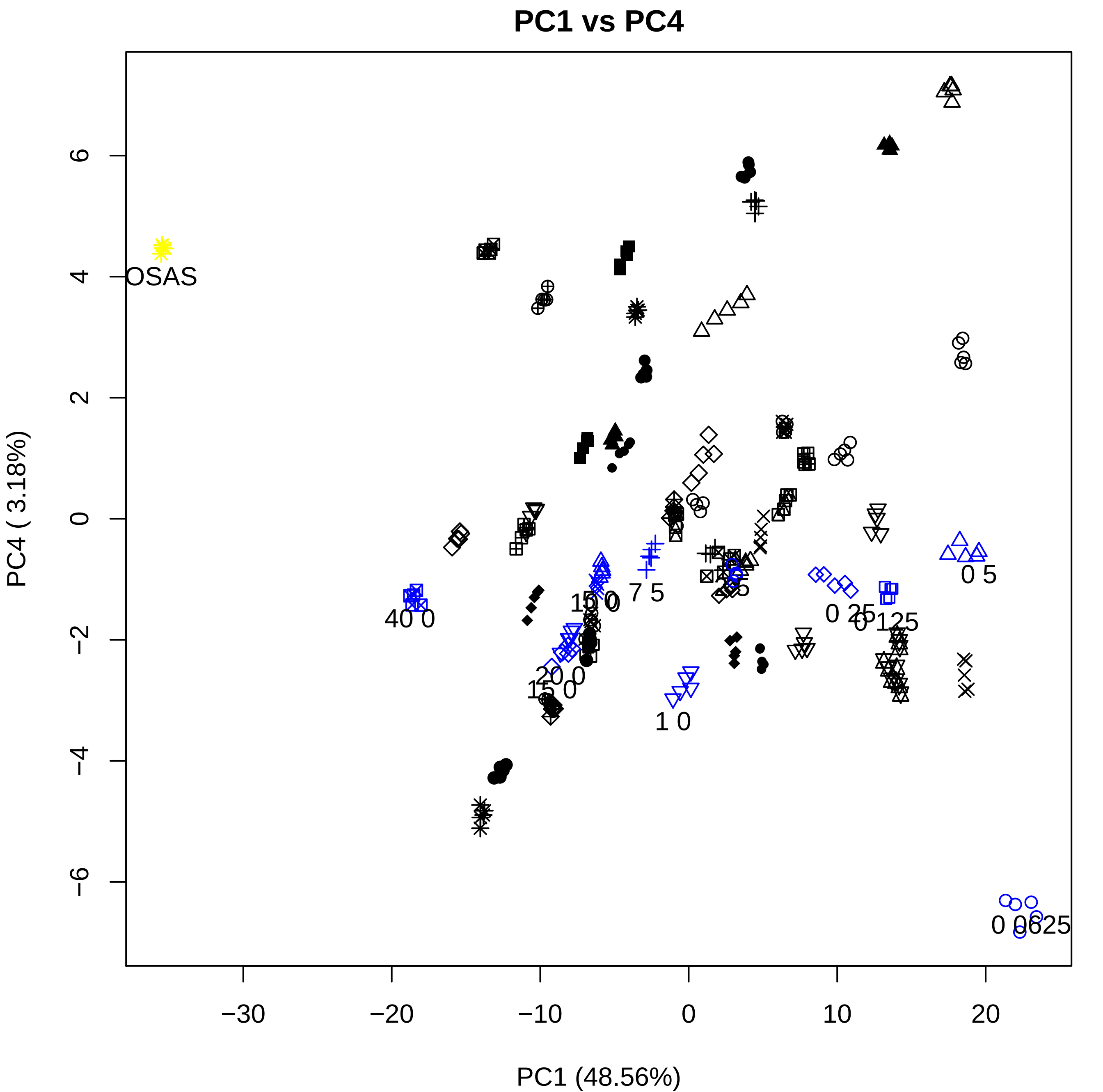
PC1 vs PC4
PC4 ( 3.18%)
PC1 (48.56%)
6
4
2
0
−2
−4
−6
−30
−20
−10
0
10
20
OSAS
40 0
7 5
0 5
0 25
0 125
20 0
15 0
1 0
0 0625

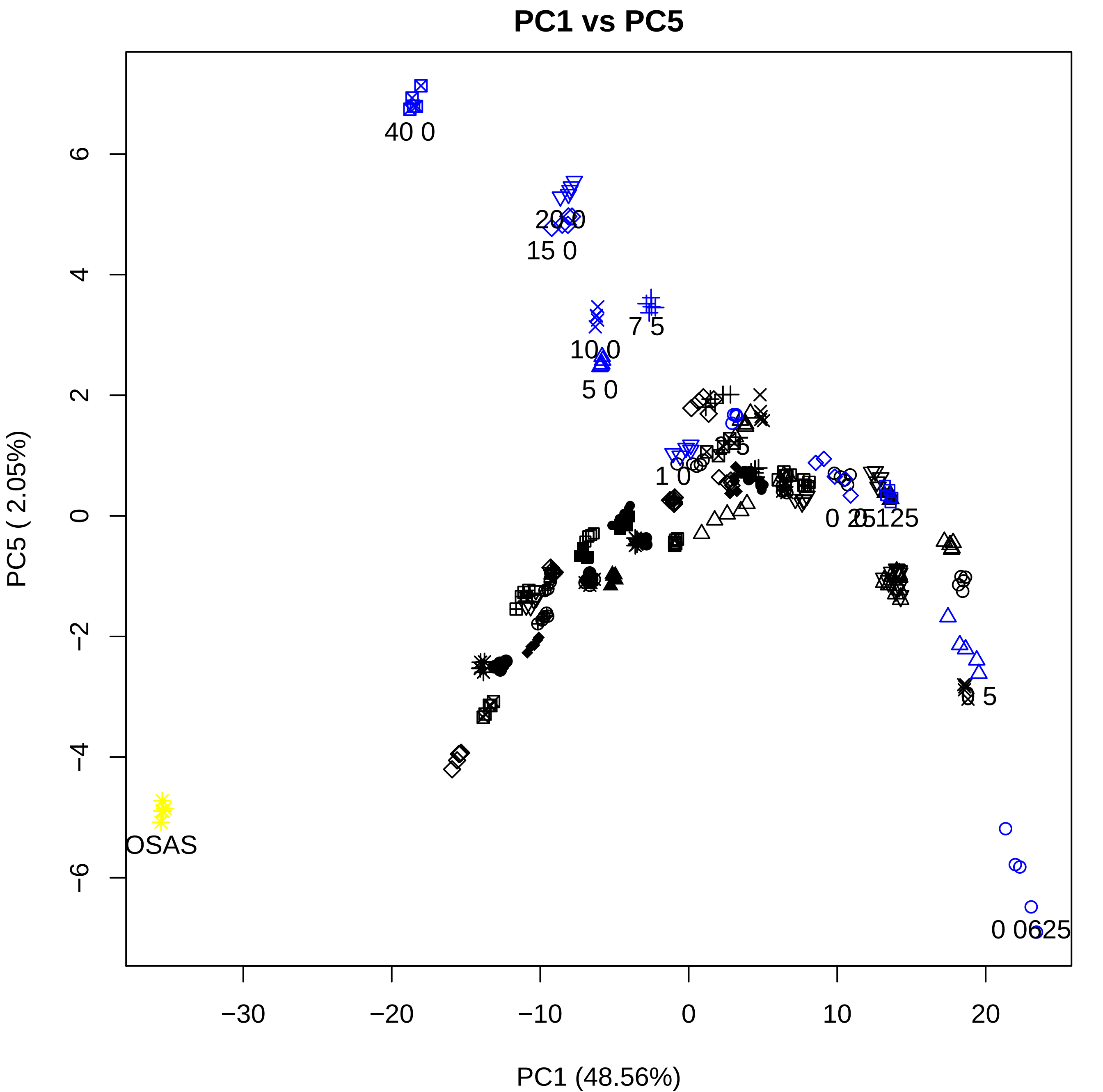

PC1 vs PC5
40 0
20 0
15 0
7 5
10 0
5 0
1 0
0 25
0 125
0 5
OSAS
0 0625
PC5 ( 2.05%)
PC1 (48.56%)
6
4
2
0
−2
−4
−6
−30
−20
−10
0
10
20

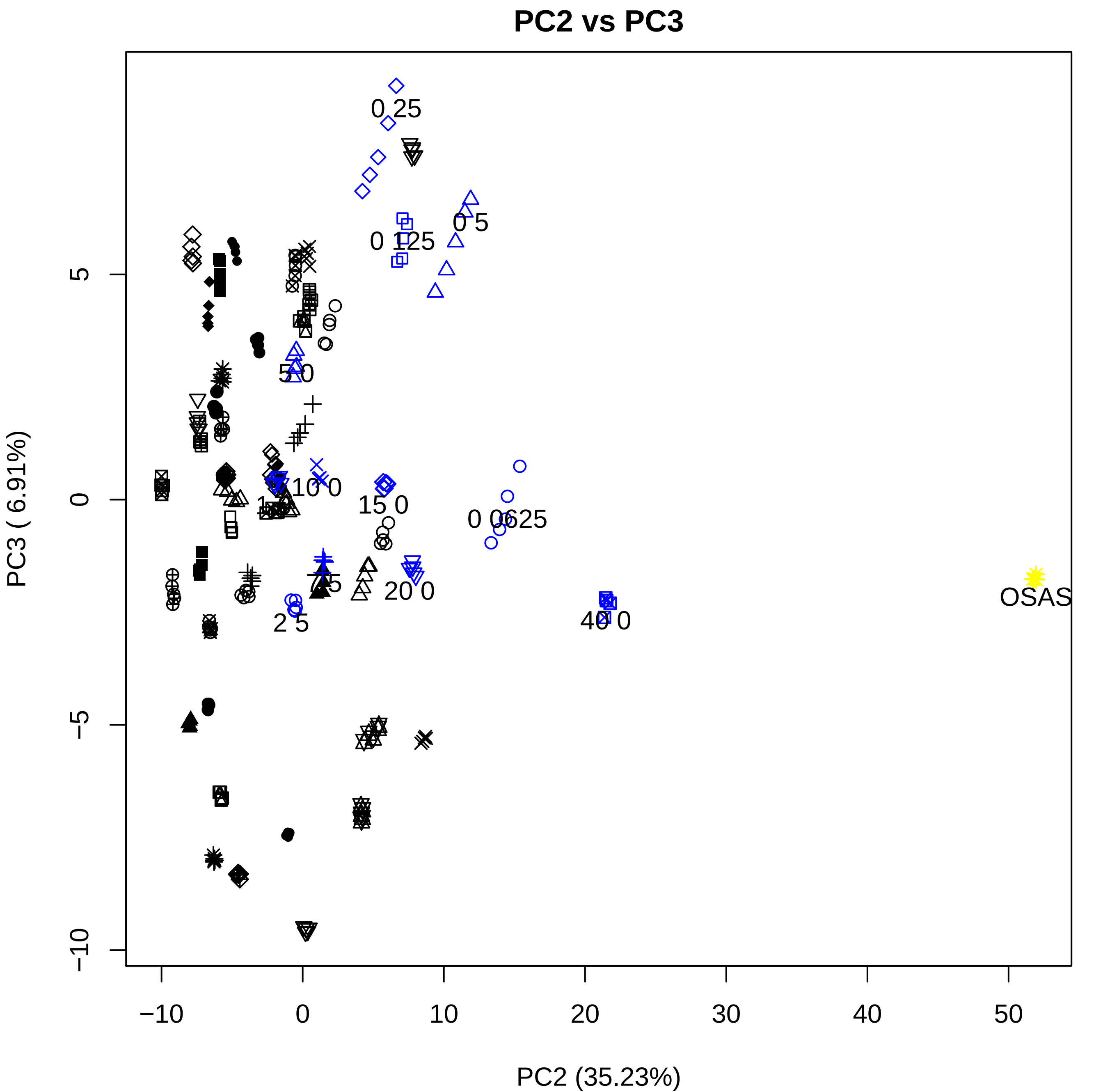

PC2 vs PC3
0 25
0 5
0 125
5 0
10 0
15 0
0 0625
20 0
2 5
40 0
OSAS
PC3 ( 6.91%)
PC2 (35.23%)
5
0
−5
−10
−10
0
10
20
30
40
50

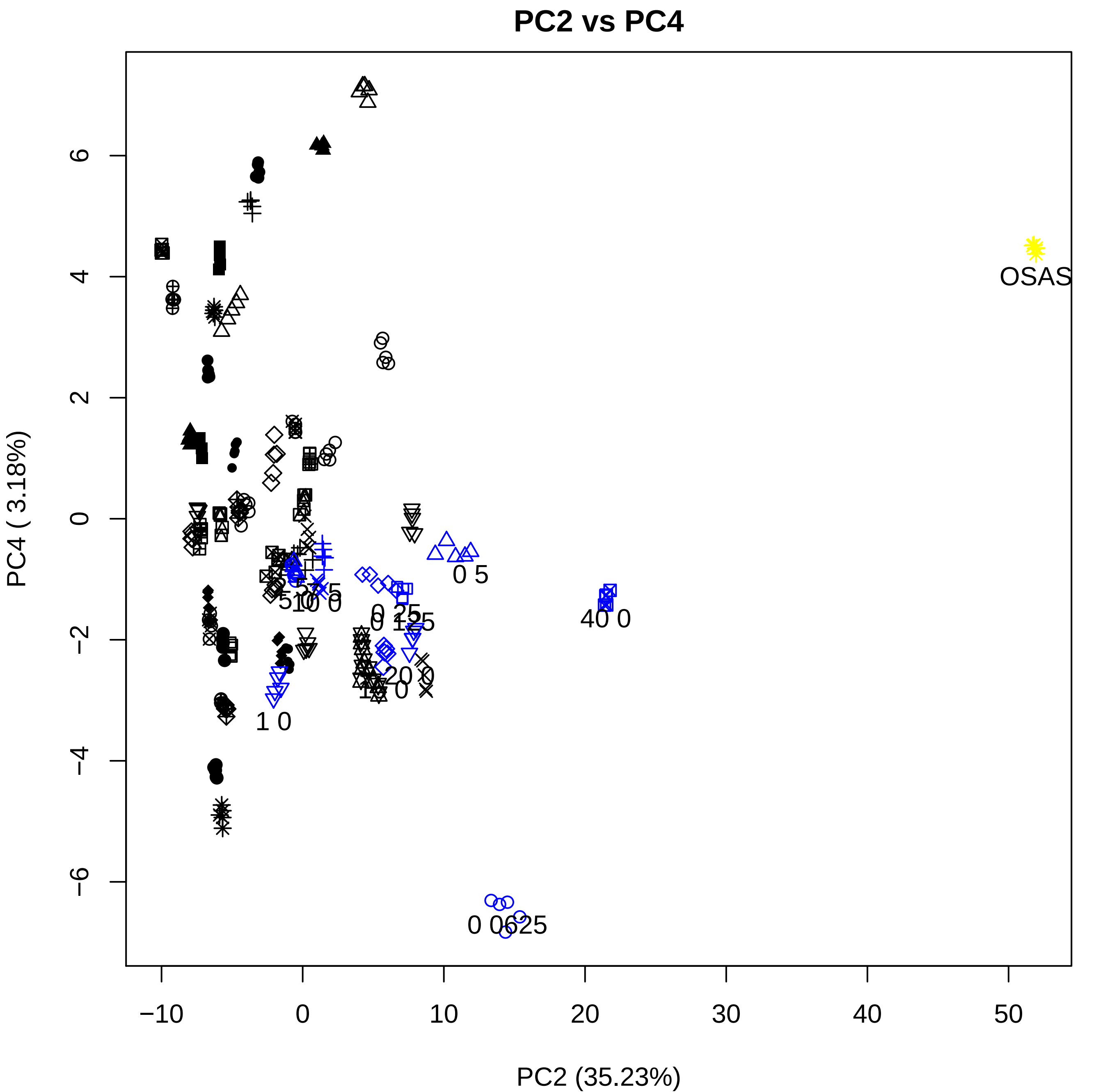
PC2 vs PC4
PC4 ( 3.18%)
PC2 (35.23%)
6
4
2
0
−2
−4
−6
−10
0
10
20
30
40
50
OSAS
0 5
40 0
1 0
0 0625

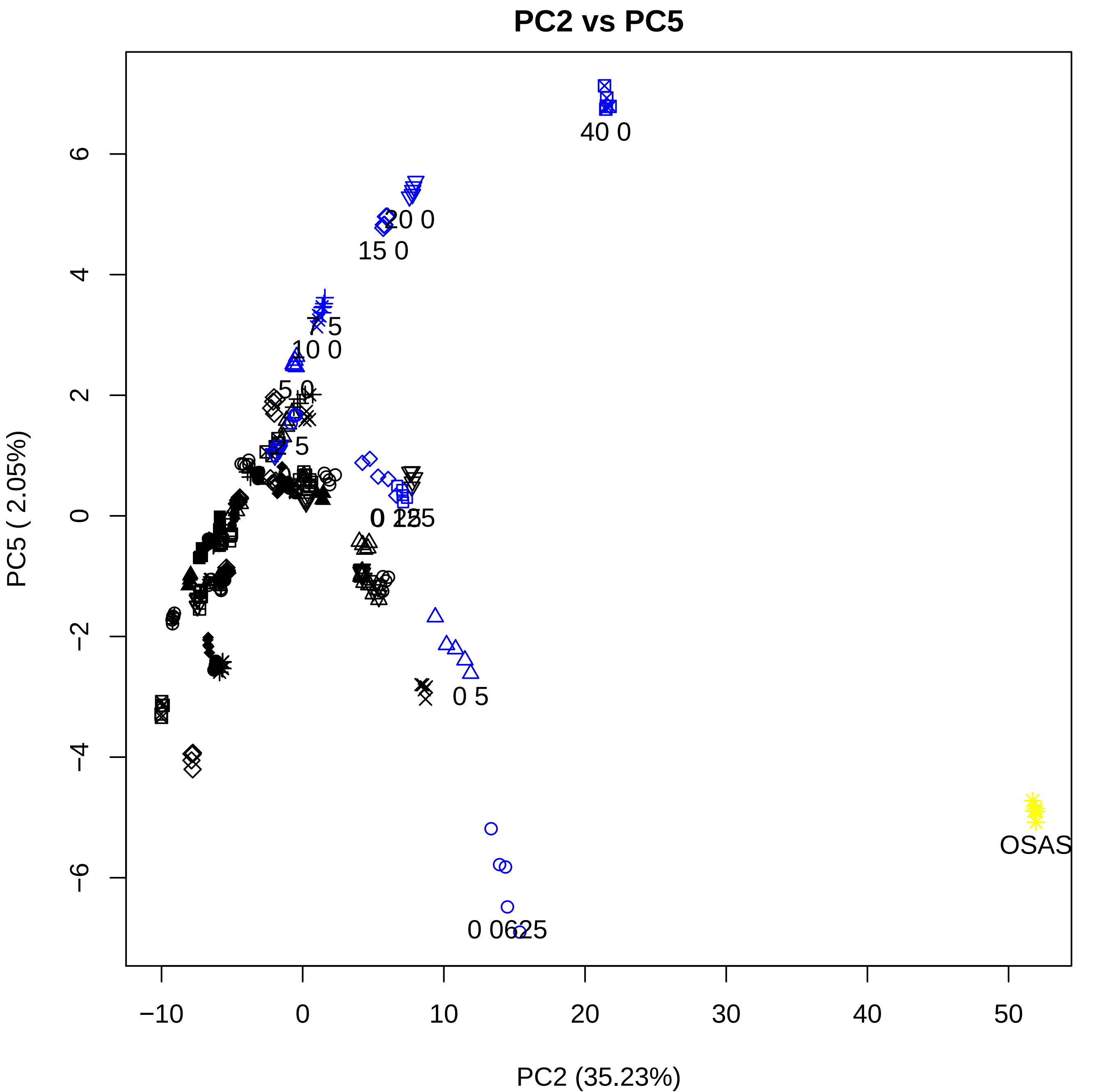

PC2 vs PC5
40 0
20 0
15 0
7 5
10 0
5 0
5
0 125
0 25
0 5
0 0625
OSAS
PC5 ( 2.05%)
PC2 (35.23%)
6
4
2
0
−2
−4
−6
−10
0
10
20
30
40
50

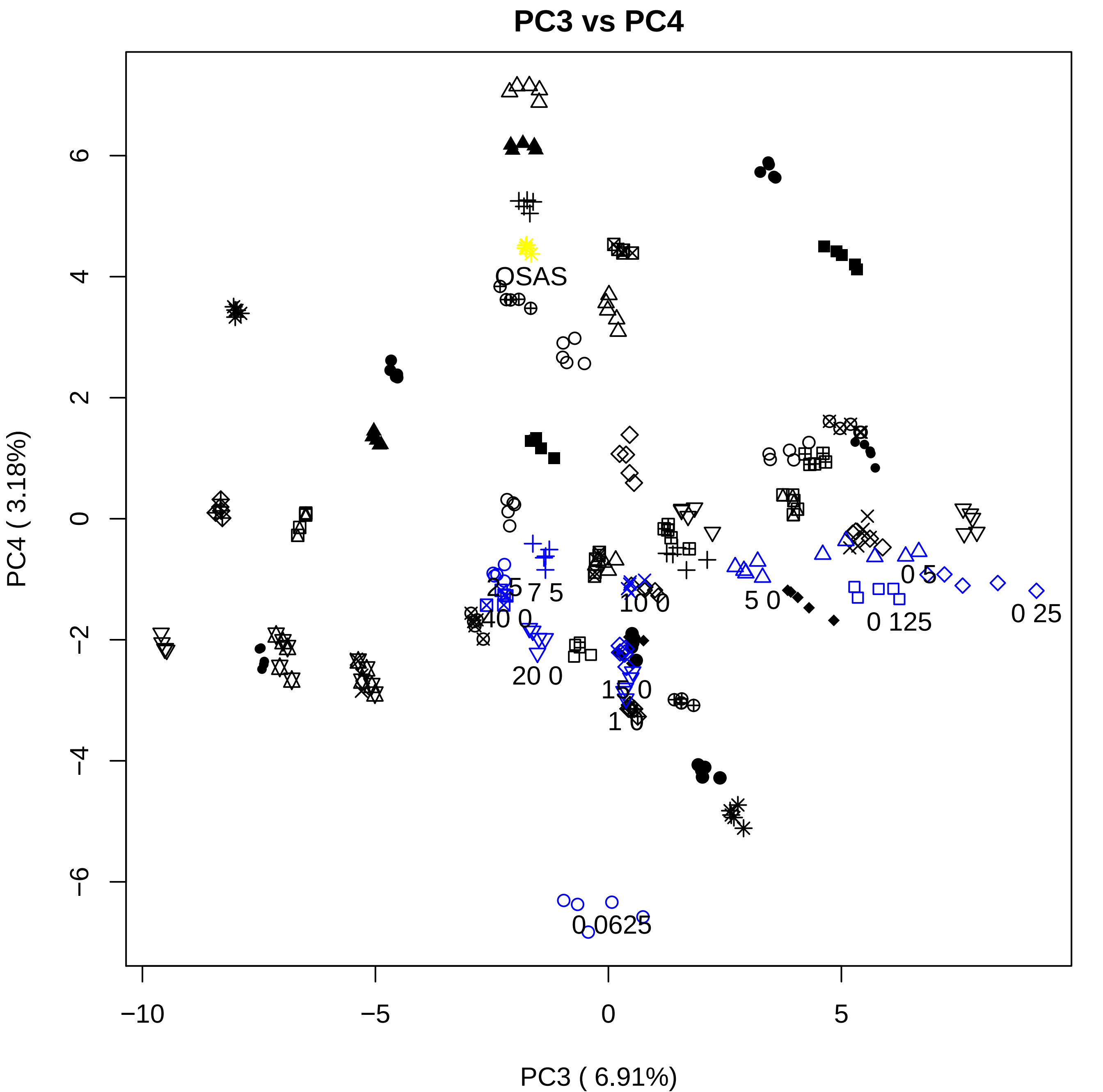

PC3 vs PC4
PC4 ( 3.18%)
PC3 ( 6.91%)
6
4
2
0
−2
−4
−6
−10
−5
0
5
OSAS
2.5
7.5
40.0
10.0
5.0
0.5
0.125
0.25
20.0
15.0
1.0
0.0625

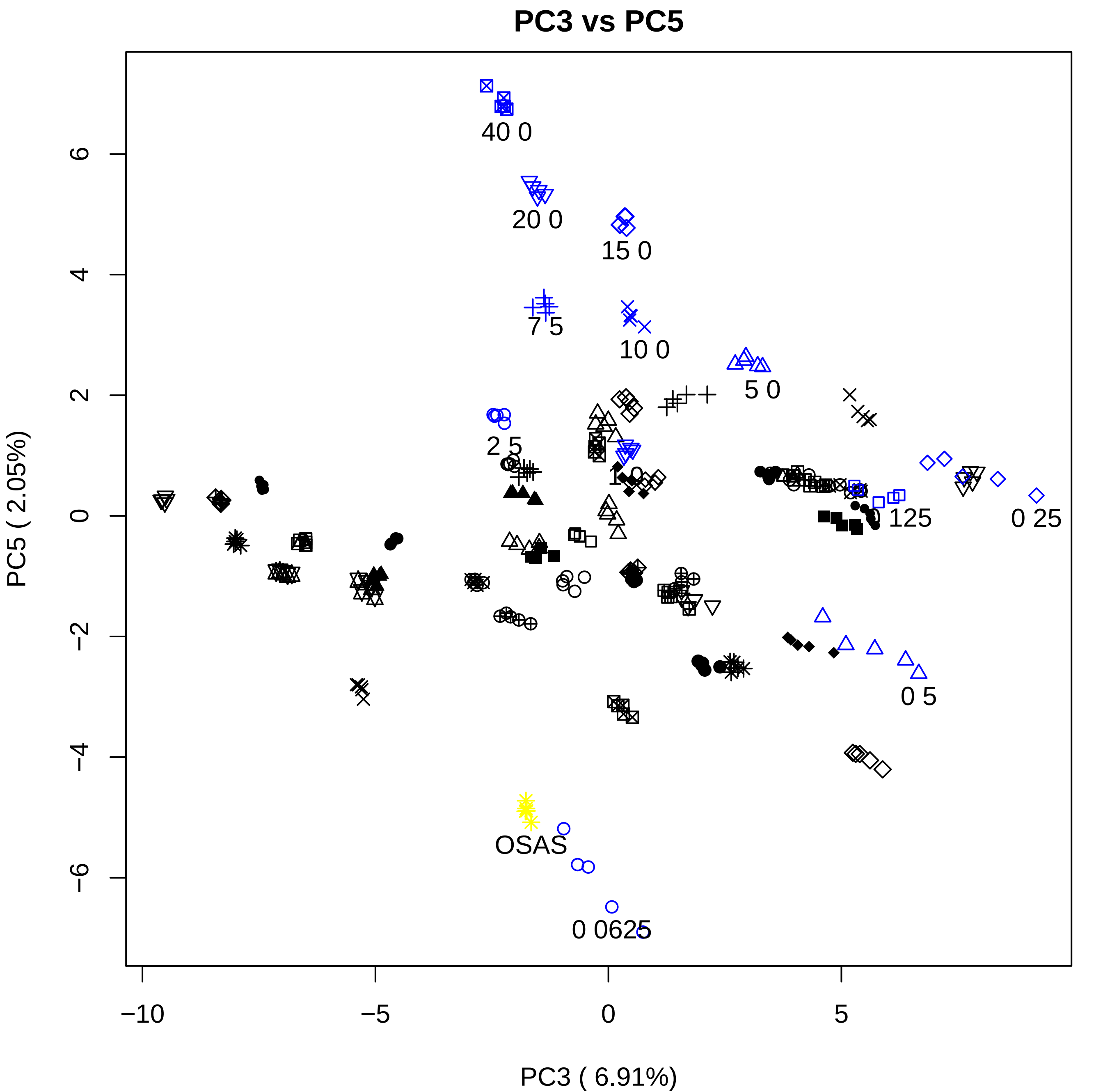
PC3 vs PC5
40 0
20 0
15 0
7 5
10 0
5 0
2 5
1 0
0 125
0 25
0 5
OSAS
0 0625
PC5 ( 2.05%)
6
4
2
0
−2
−4
−6
PC3 ( 6.91%)
−10
−5
0
5

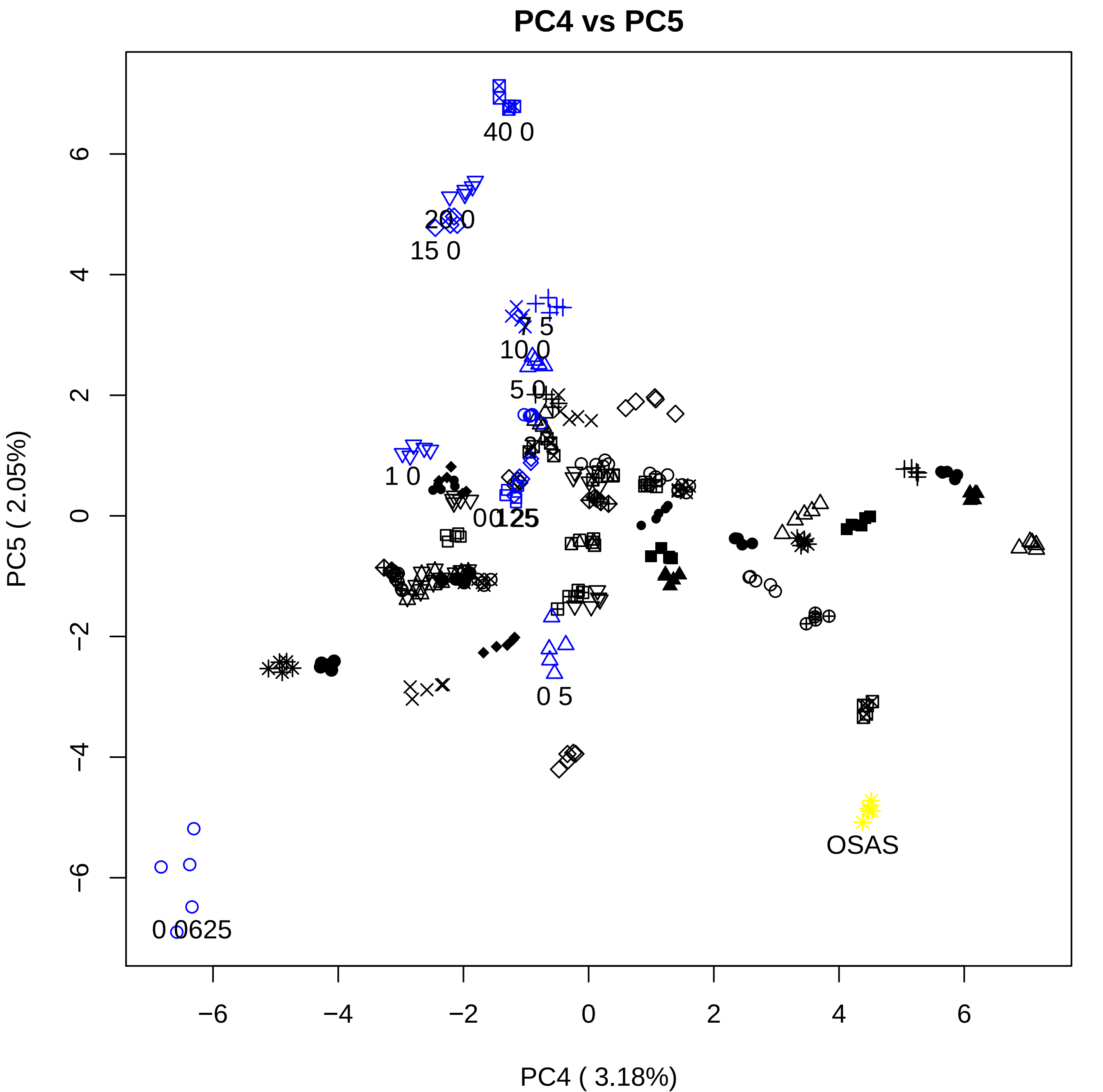

PC4 vs PC5
40 0
20 0
15 0
7 5
10 0
5 0
1 0
0 125
0 5
OSAS
0 0625
PC5 ( 2.05%)
PC4 ( 3.18%)
−6
−4
−2
0
2
4
6

### PCA of 0.06%  OSAS vs PMH

Scaled Matrix PCAed

7th order polynomial baseline correction performed

Raw spectra smoothed to 21 Neighbours, 2nd order PN, Savtisky Golay

Normalisation between 0 and 100

D2 spectra smoothed to 41 Neighbours, 2nd order PN, Savtisky Golay

Regions Analysed: 3601−2501 and 2001−700cm−1

Samples Analysed: 1 Contam's 0 OSAS's 45 PMH's

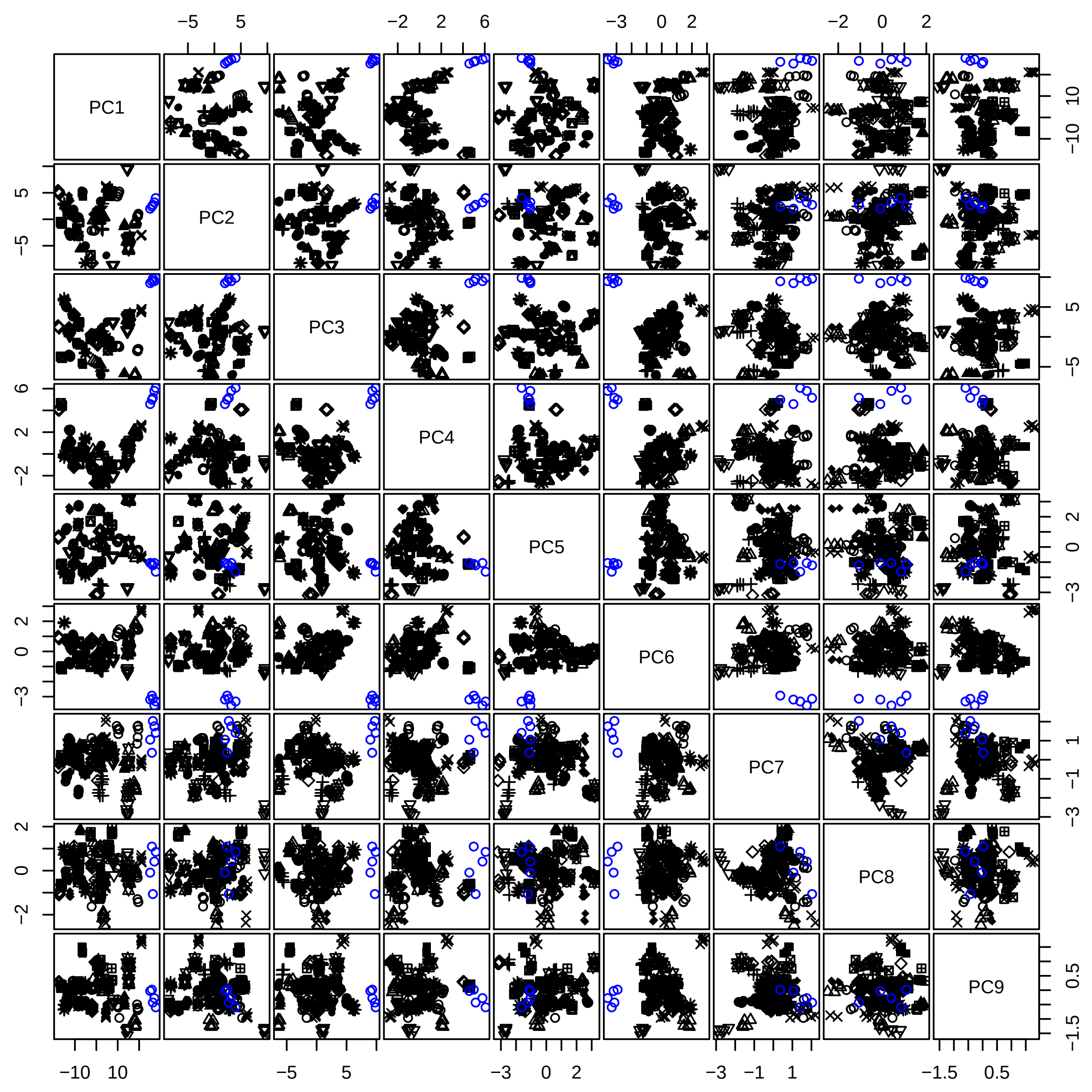
PC1
PC2
PC3
PC4
PC5
PC6
PC7
PC8
PC9
−5 5
−2 2 6
−3 0 2
−2 0 2
10 −10
5 −5
6 2 −2
2 0 −3
2 0 −2
1 −1 −3
0.5 −1.5
−10 10
−5 5
−3 0 2
−3 −1 1
−1.5 0.5

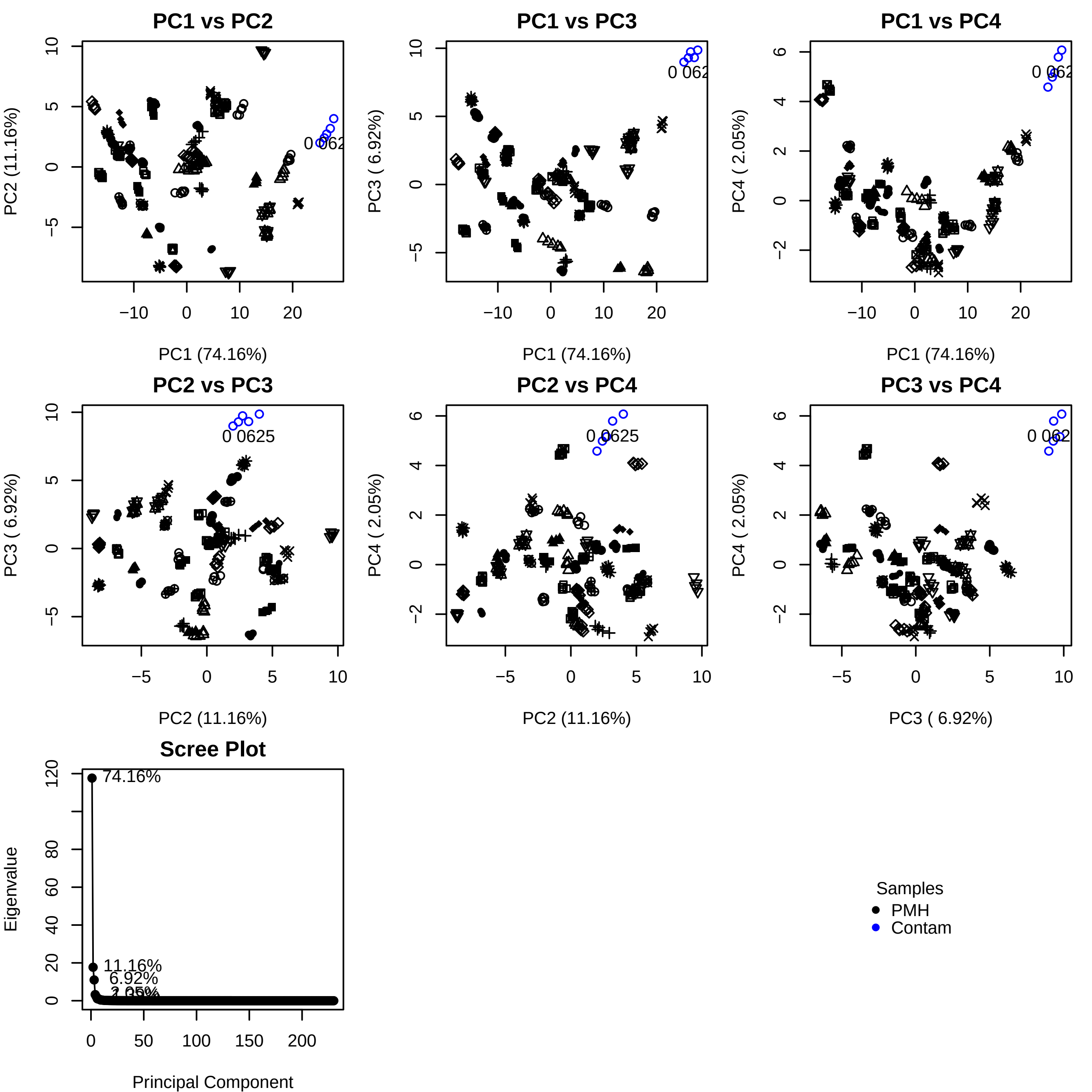

PC1 vs PC2
PC1 vs PC3
PC1 vs PC4
PC2 vs PC3
PC2 vs PC4
PC3 vs PC4
Scree Plot
PC1 (74.16%)
PC2 (11.16%)
PC3 ( 6.92%)
PC4 ( 2.05%)
0.0625
74.16%
11.16%
6.92%
Eigenvalue
Principal Component
Samples
PMH
Contam

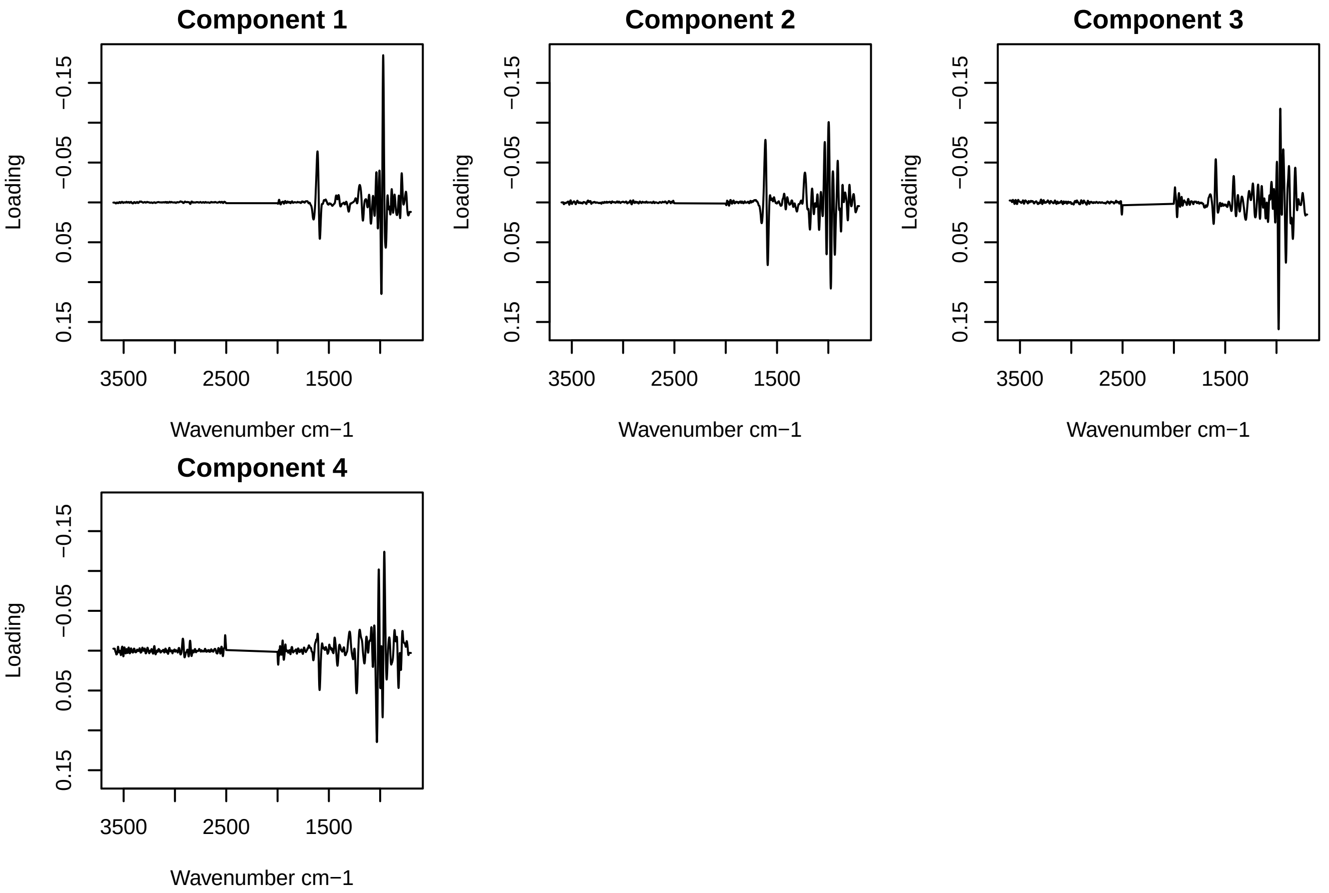
Component 1
Component 2
Component 3
Component 4
Loading
−0.15
−0.05
0.05
0.15
3500
2500
1500
Wavenumber cm−1

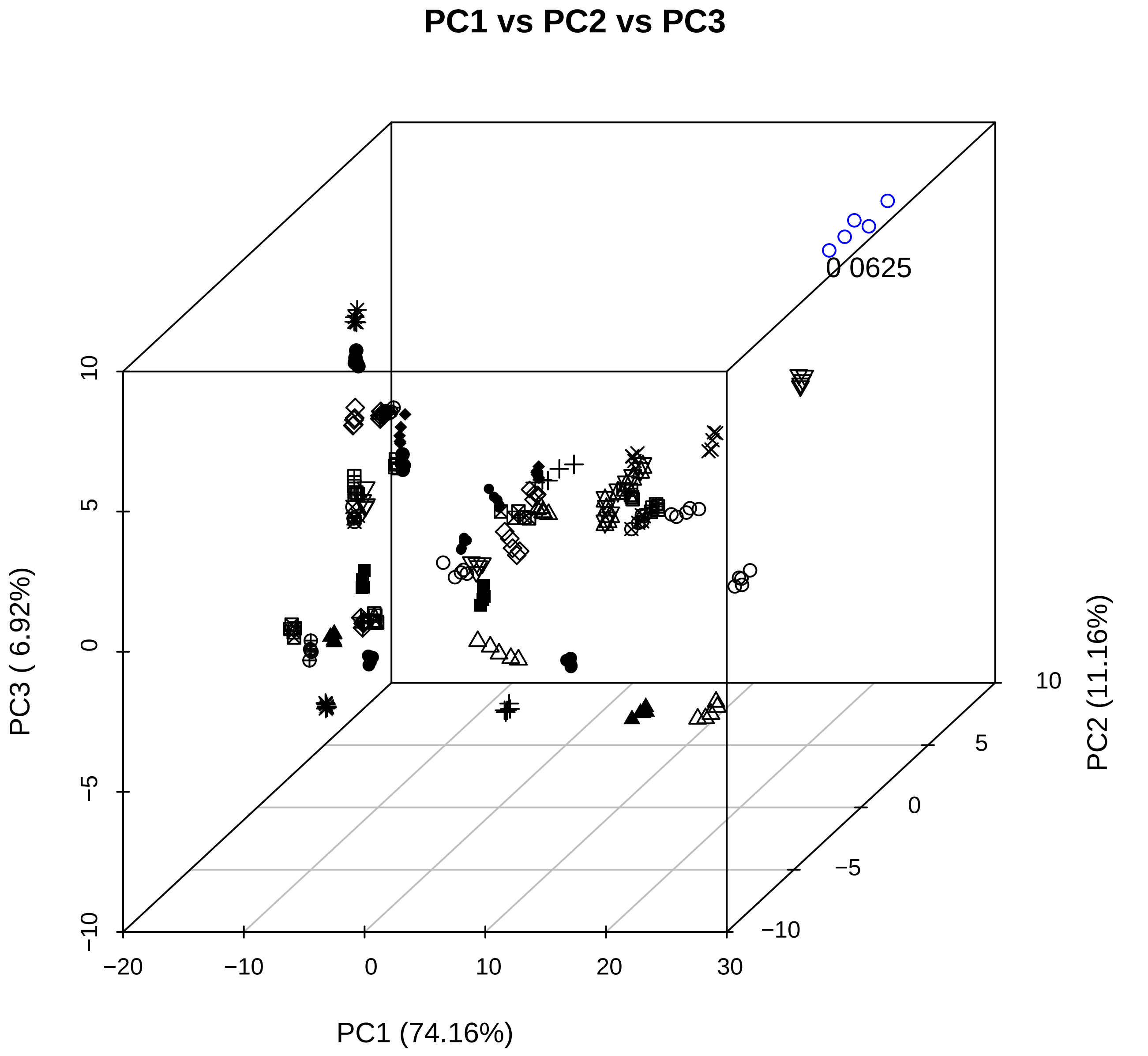
PC1 vs PC2 vs PC3
0 0625
PC1 (74.16%)
PC2 (11.16%)
PC3 ( 6.92%)
−20
−10
0
10
20
30
−10
−5
0
5
10

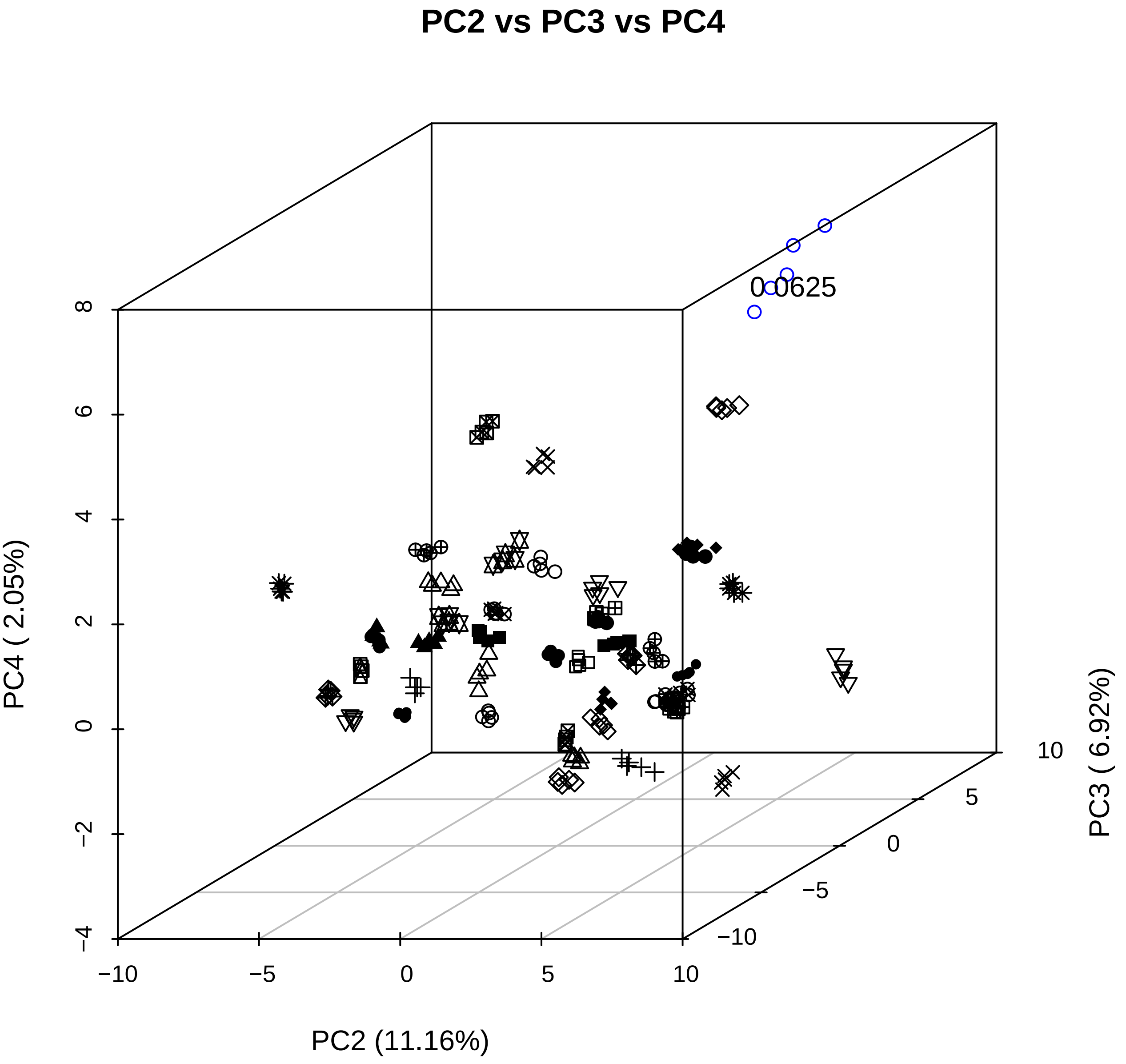

PC2 vs PC3 vs PC4
0.0625
PC2 (11.16%)
PC3 ( 6.92%)
PC4 ( 2.05%)
−10
−5
0
5
10
−4
−2
0
2
4
6
8

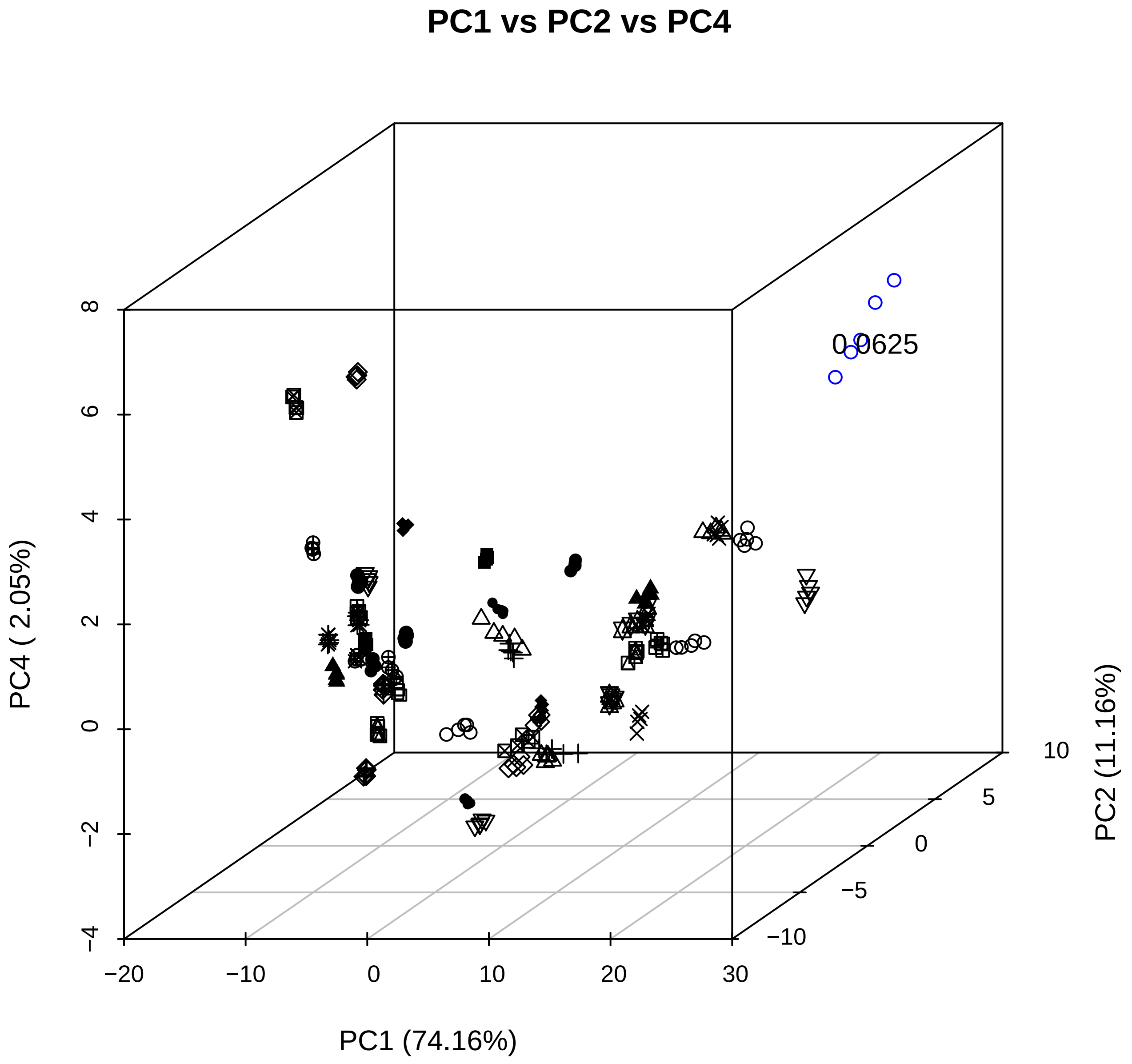
PC1 vs PC2 vs PC4
0.0625
PC1 (74.16%)
PC2 (11.16%)
PC4 ( 2.05%)
−20
−10
0
10
20
30
−10
−5
0
5
10
−4
−2
0
2
4
6
8

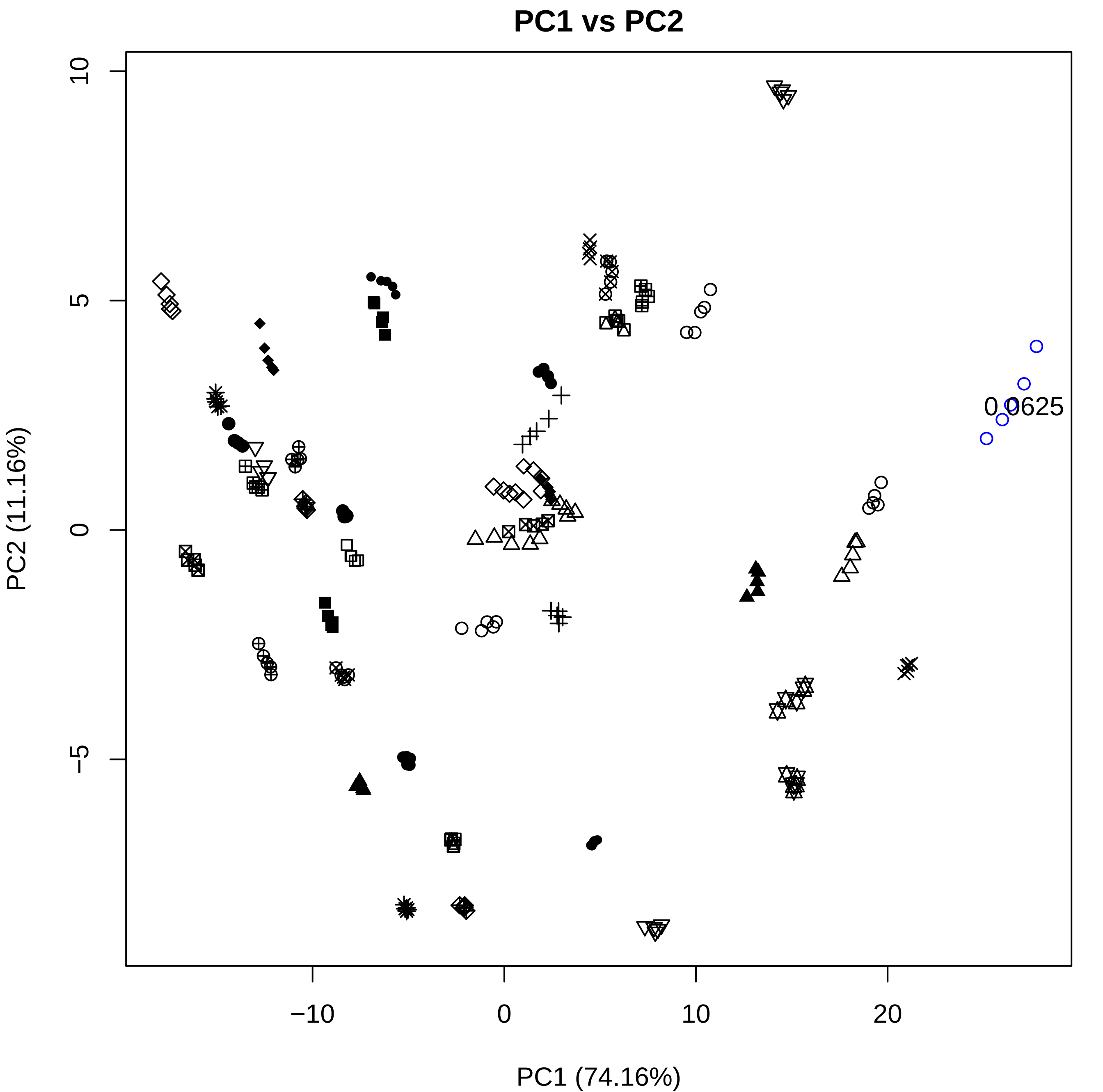
PC1 vs PC2
PC2 (11.16%)
PC1 (74.16%)
10
5
0
−5
−10
0
10
20
0.0625

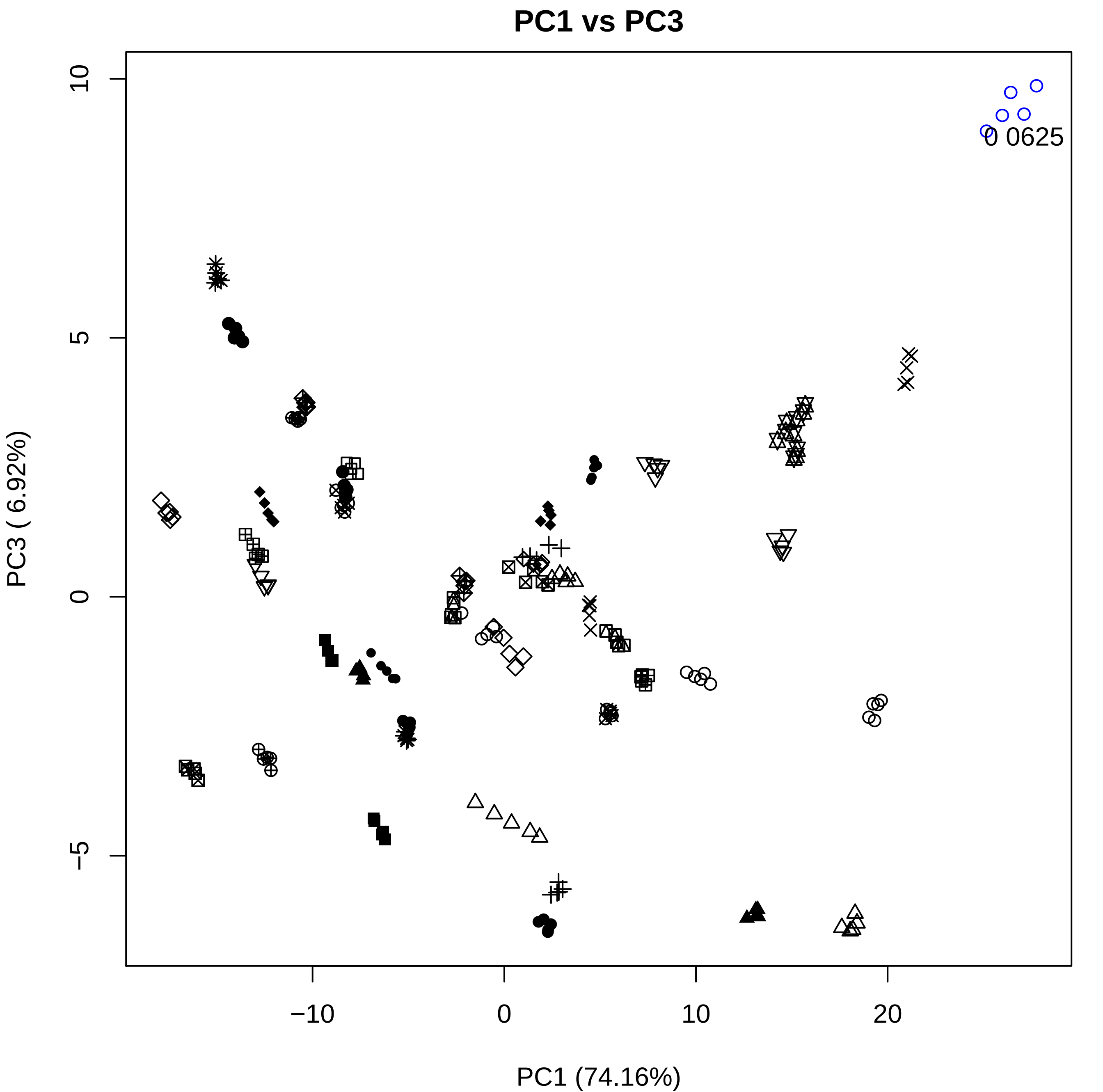
PC1 vs PC3
0 0625
10
5
0
−5
PC3 ( 6.92%)
−10
0
10
20
PC1 (74.16%)

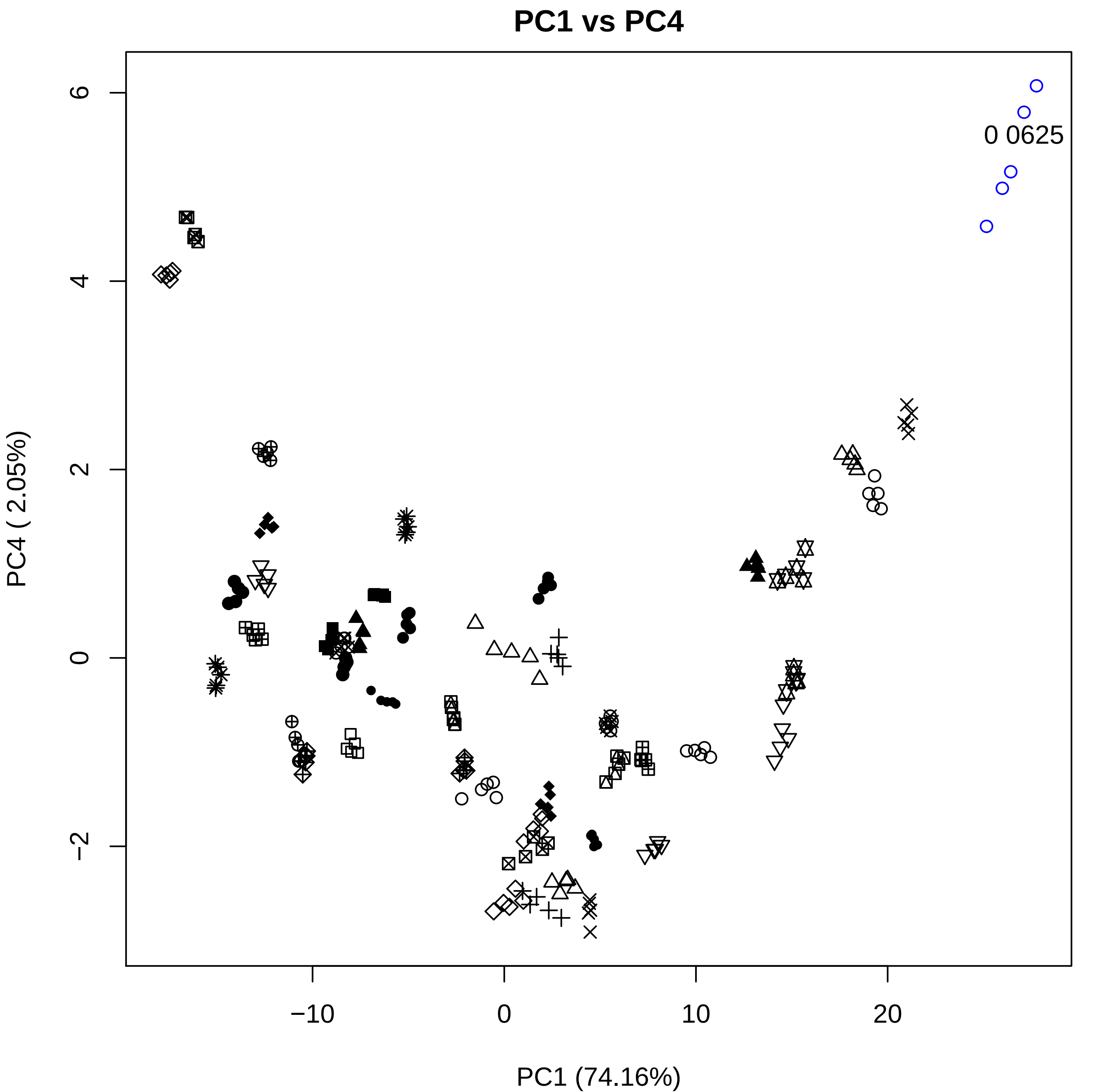

PC1 vs PC4
0 0625
PC4 ( 2.05%)
PC1 (74.16%)
−10
0
10
20
−2
0
2
4
6

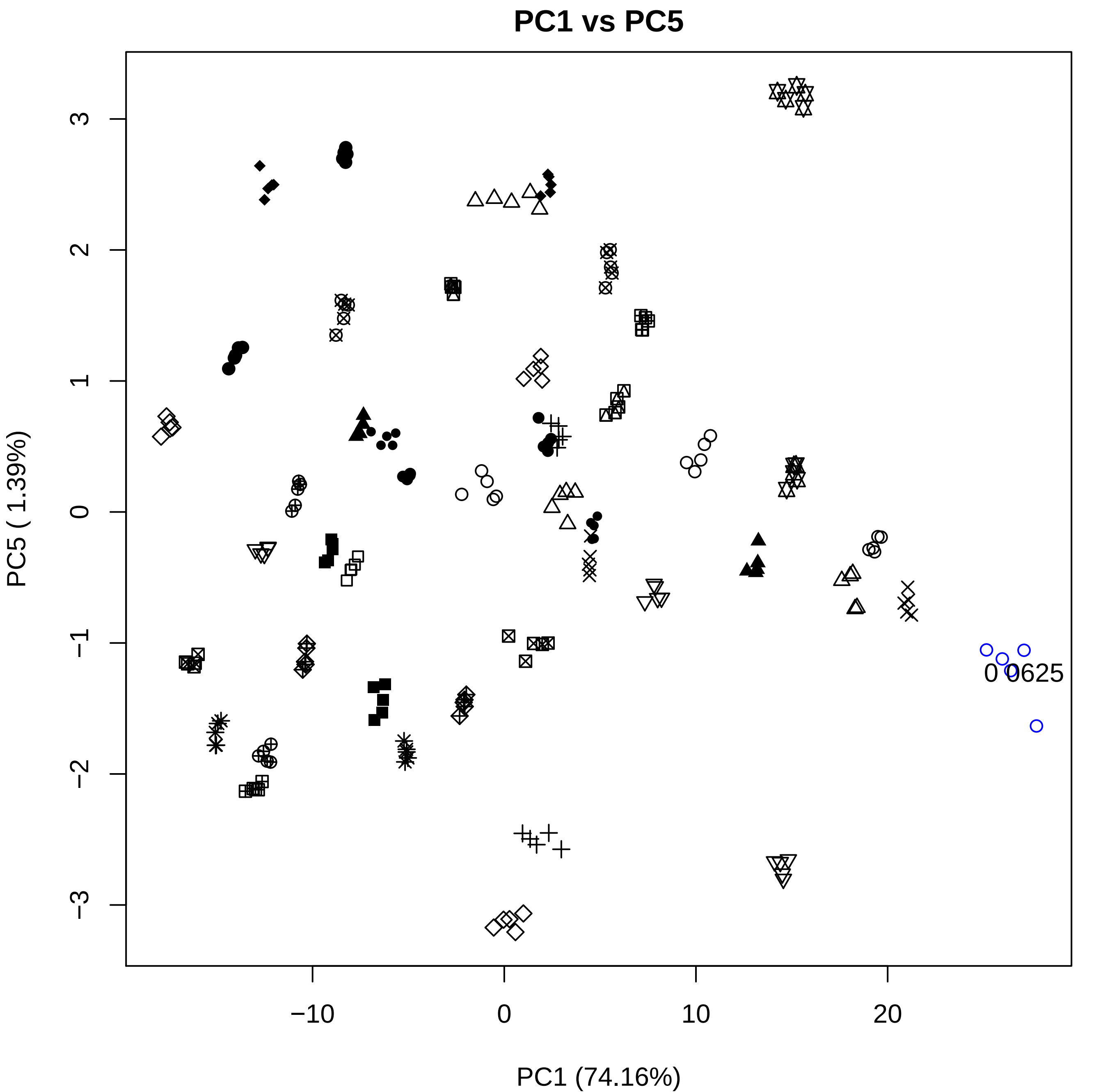

PC1 vs PC5
PC5 ( 1.39%)
PC1 (74.16%)
3
2
1
0
−1
−2
−3
−10
0
10
20
0 0625

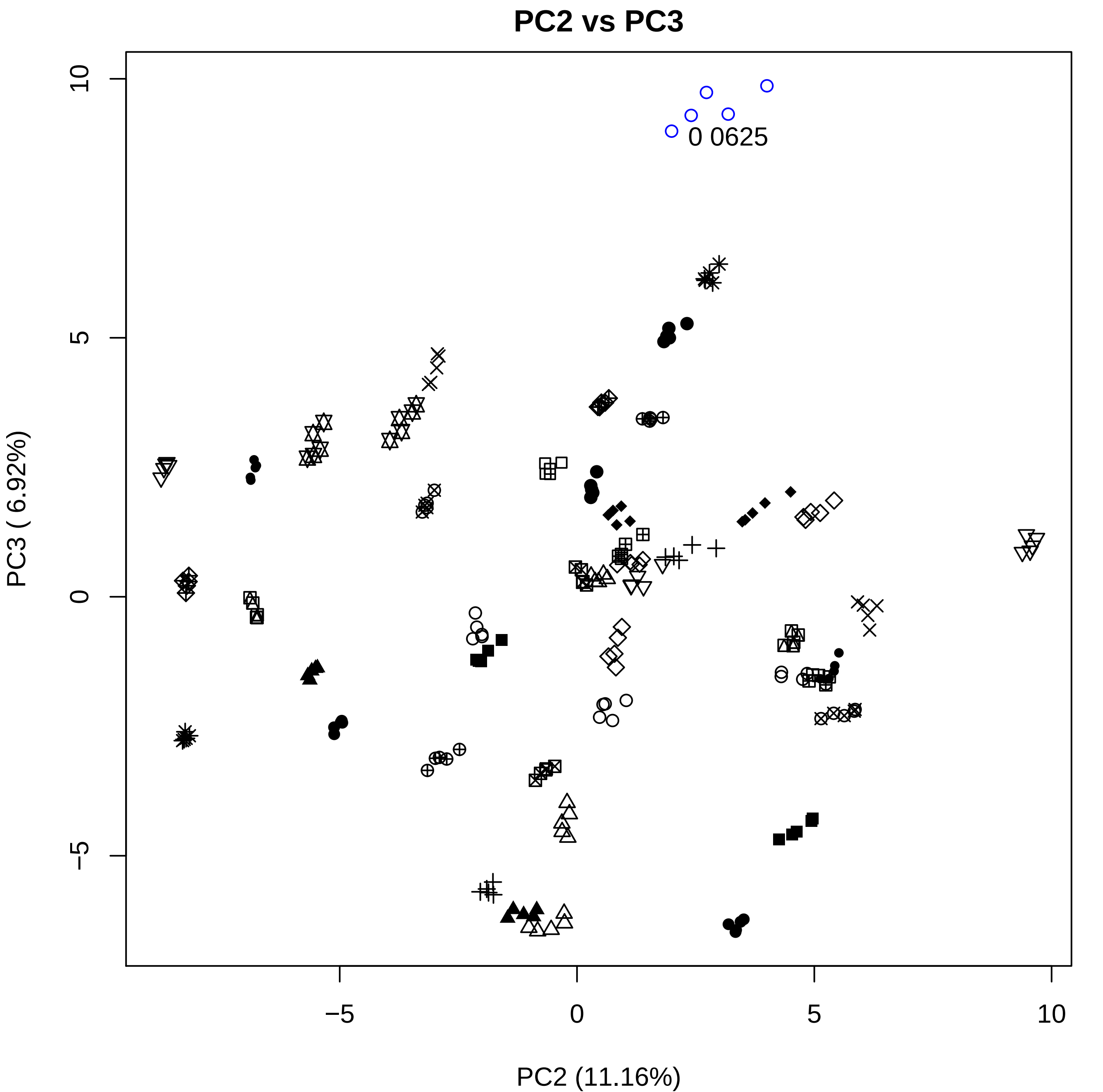
PC2 vs PC3
0 0625
PC3 ( 6.92%)
PC2 (11.16%)
10
5
0
−5
−5
0
5
10

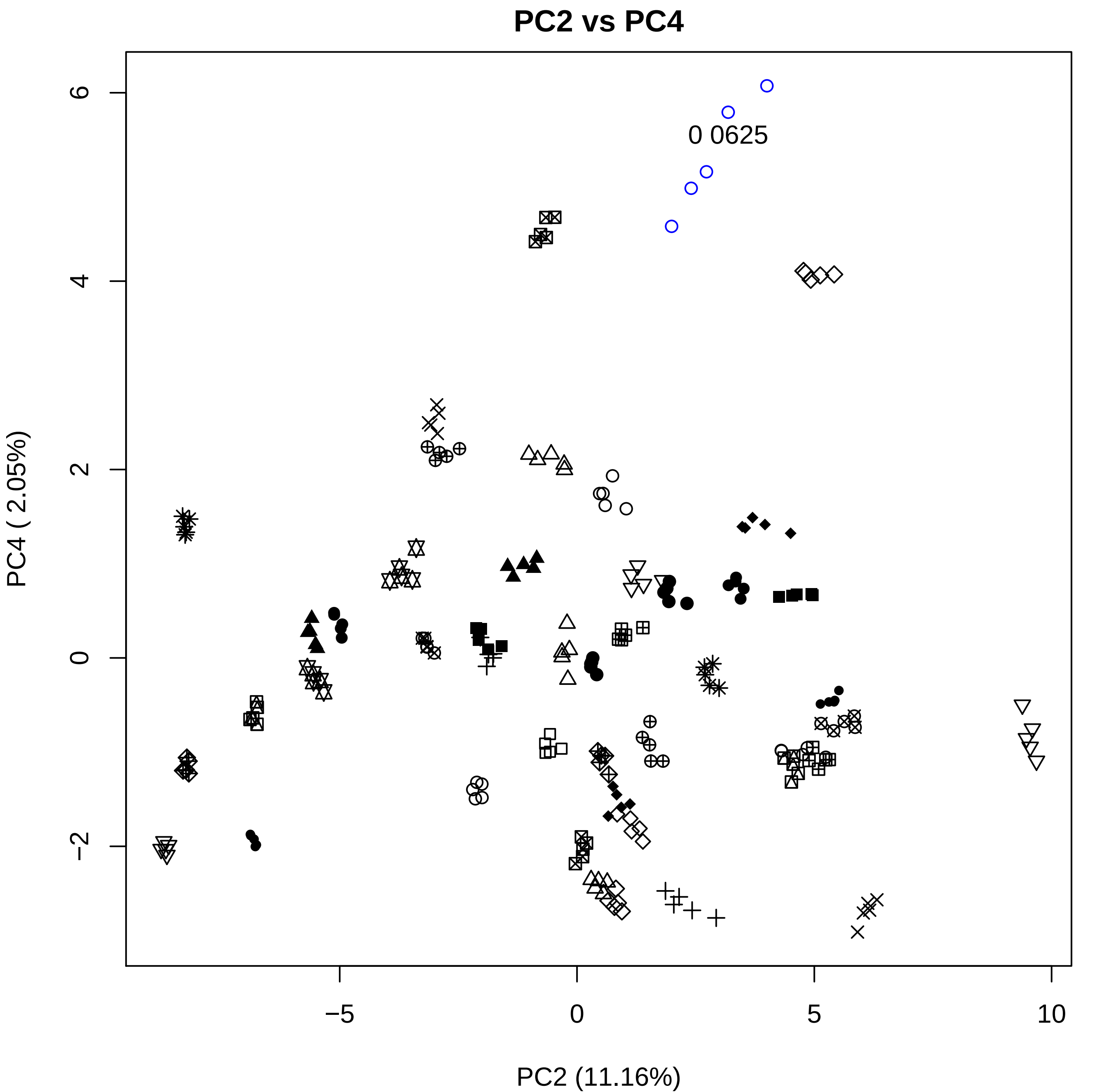
PC2 vs PC4
0 0625
PC4 ( 2.05%)
PC2 (11.16%)
−2
0
2
4
6
−5
0
5
10

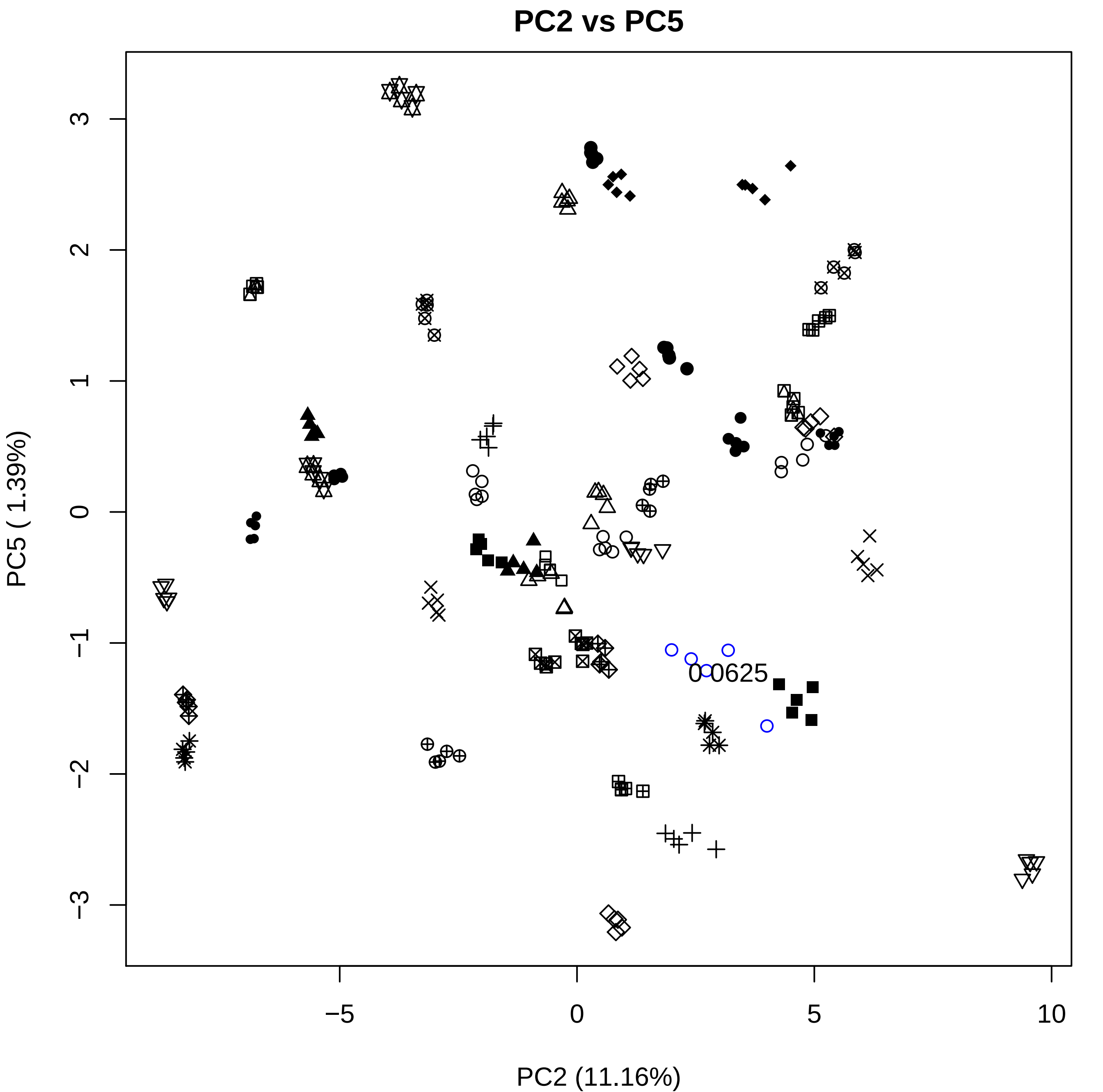
PC2 vs PC5
PC5 ( 1.39%)
PC2 (11.16%)
3
2
1
0
−1
−2
−3
−5
0
5
10
0.0625

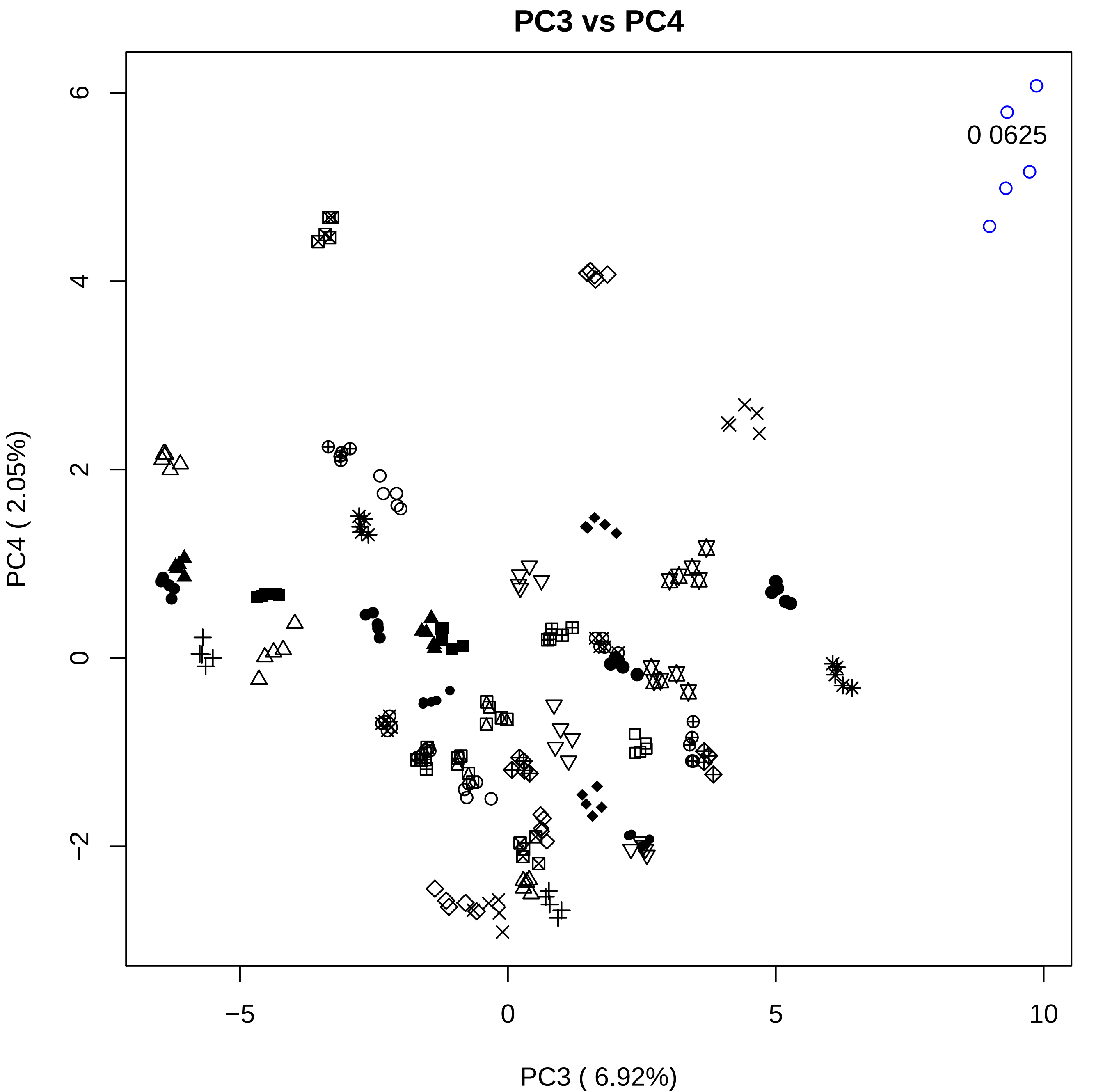
PC3 vs PC4
0 0625
PC4 ( 2.05%)
6
4
2
0
−2
−5
0
5
10
PC3 ( 6.92%)

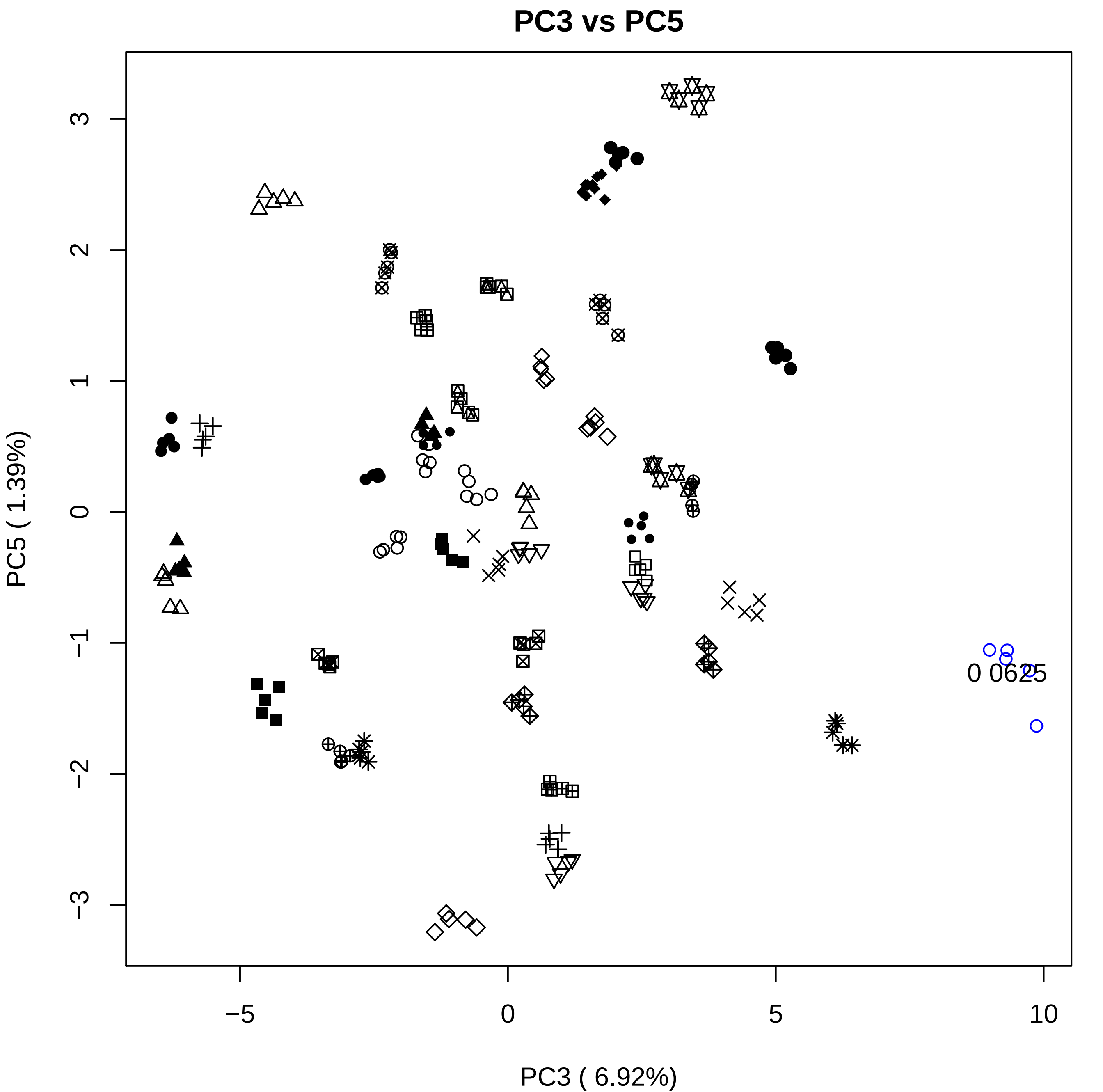

PC3 vs PC5
0 0625
PC5 ( 1.39%)
PC3 ( 6.92%)
−5
0
5
10
−3
−2
−1
0
1
2
3

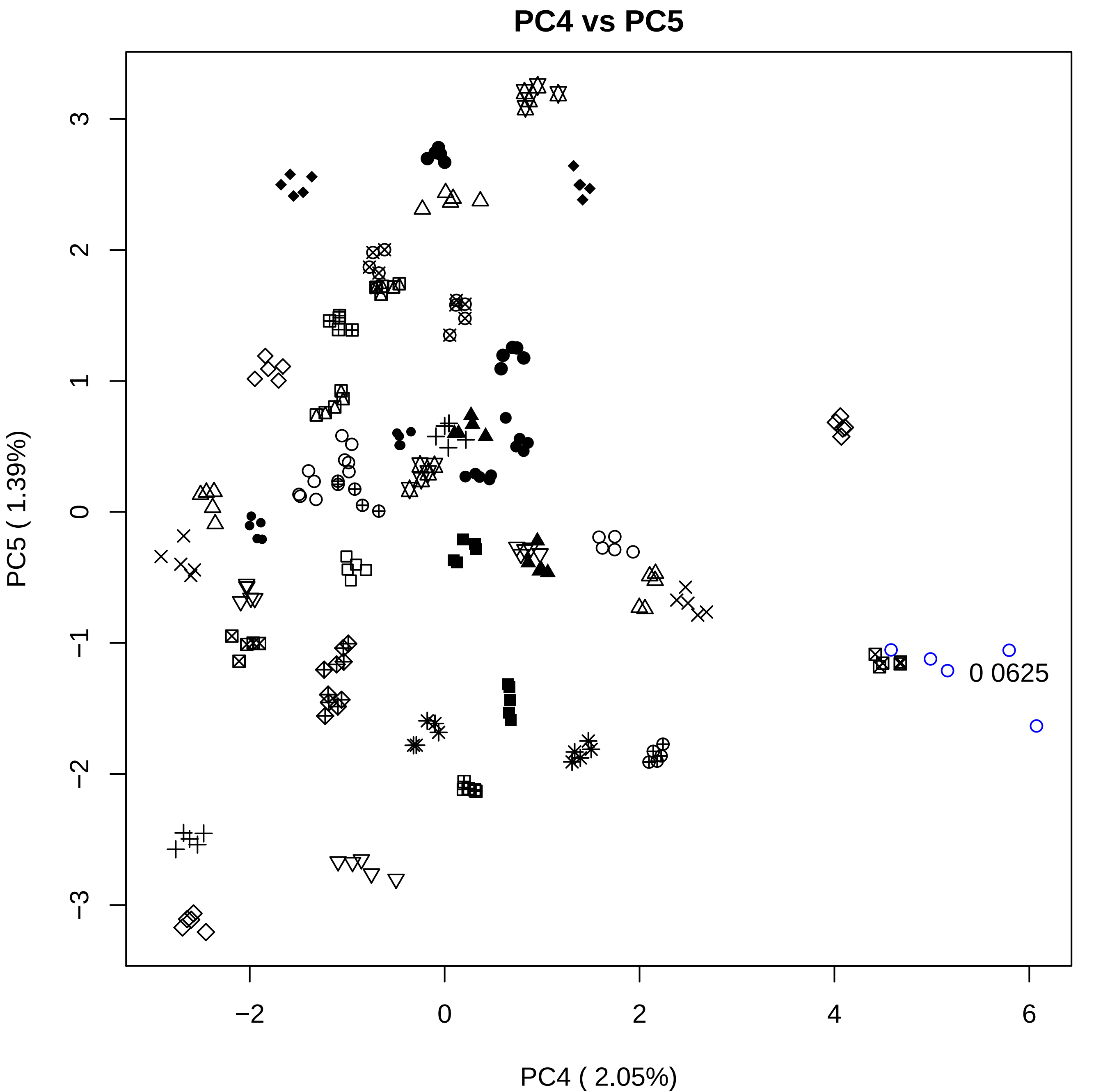
PC4 vs PC5
PC5 ( 1.39%)
PC4 ( 2.05%)
0 0625
3
2
1
0
−1
−2
−3
−2
0
2
4
6

### PCA of 0.12%  OSAS vs PMH

Scaled Matrix PCAed

7th order polynomial baseline correction performed

Raw spectra smoothed to 21 Neighbours, 2nd order PN, Savtisky Golay

Normalisation between 0 and 100

D2 spectra smoothed to 41 Neighbours, 2nd order PN, Savtisky Golay

Regions Analysed: 3601−2501 and 2001−700cm−1

Samples Analysed: 1 Contam's 0 OSAS's 45 PMH's

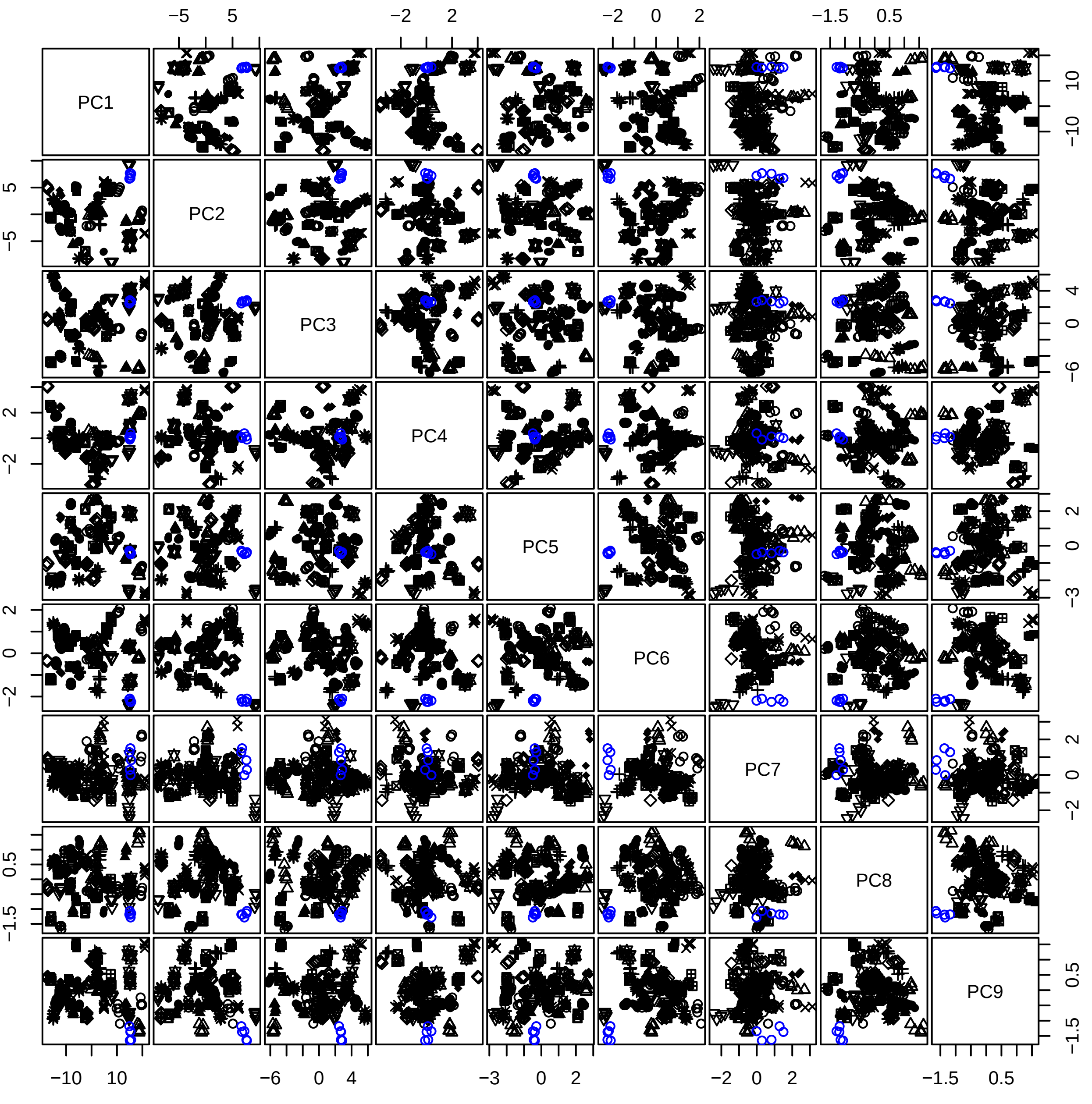
PC1
PC2
PC3
PC4
PC5
PC6
PC7
PC8
PC9

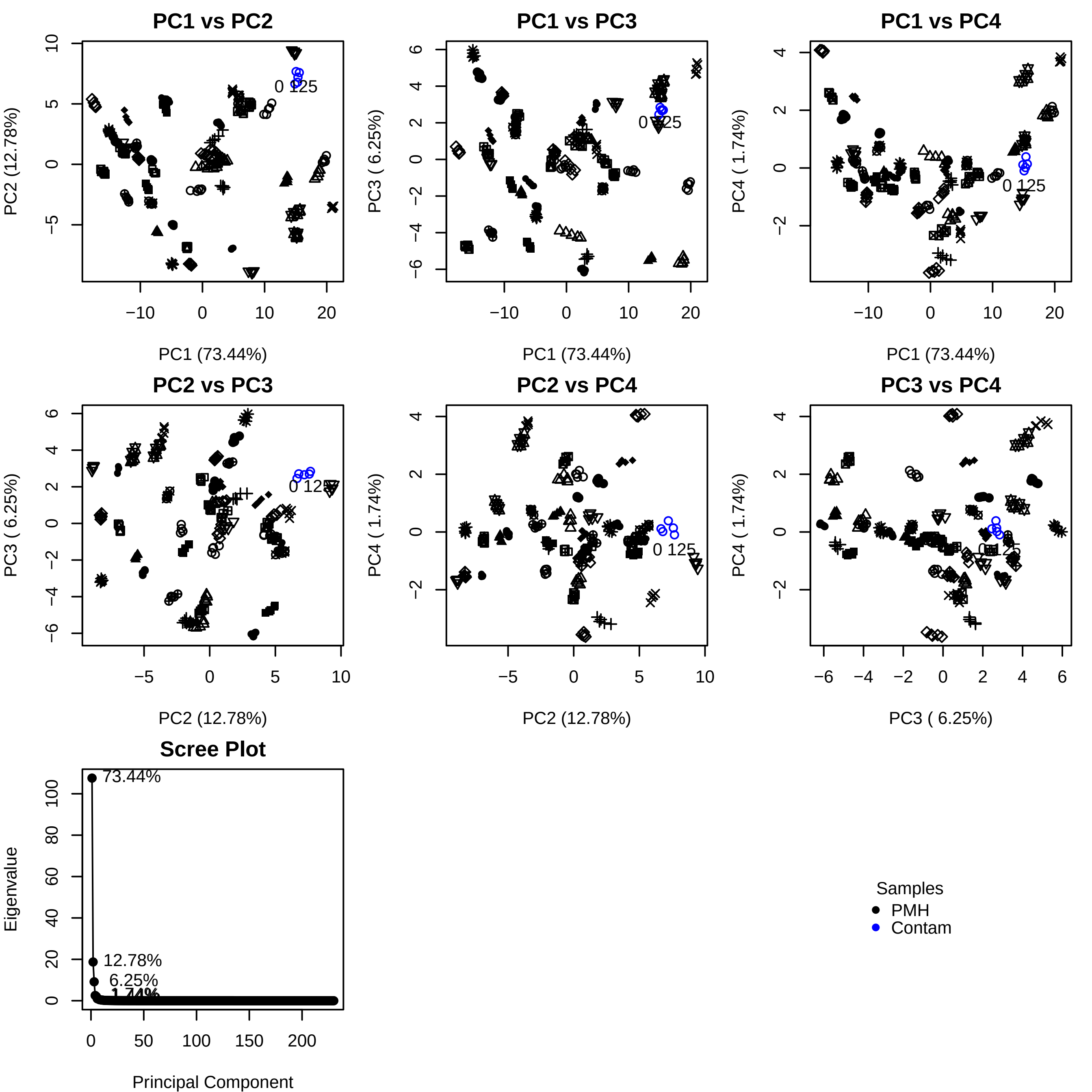

PC1 vs PC2
PC2 (12.78%)
PC1 (73.44%)
0 125
PC1 vs PC3
PC3 ( 6.25%)
PC1 (73.44%)
0 125
PC1 vs PC4
PC4 ( 1.74%)
PC1 (73.44%)
0 125
PC2 vs PC3
PC3 ( 6.25%)
PC2 (12.78%)
0 125
PC2 vs PC4
PC4 ( 1.74%)
PC2 (12.78%)
0 125
PC3 vs PC4
PC4 ( 1.74%)
PC3 ( 6.25%)
Scree Plot
Eigenvalue
Principal Component
73.44%
12.78%
6.25%
Samples
PMH
Contam

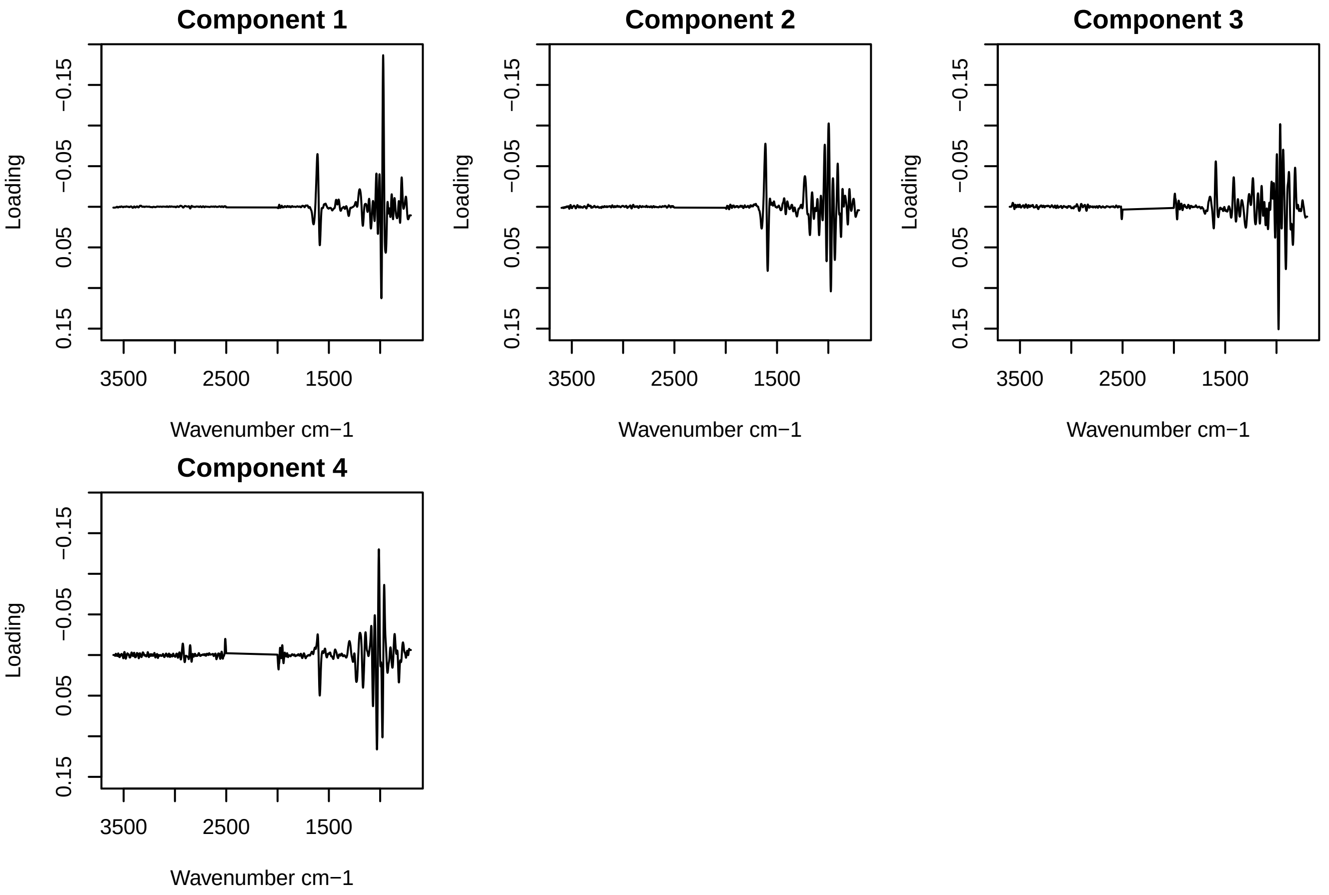
Component 1
Component 2
Component 3
Component 4
Loading
−0.15
−0.05
0.05
0.15
3500
2500
1500
Wavenumber cm−1

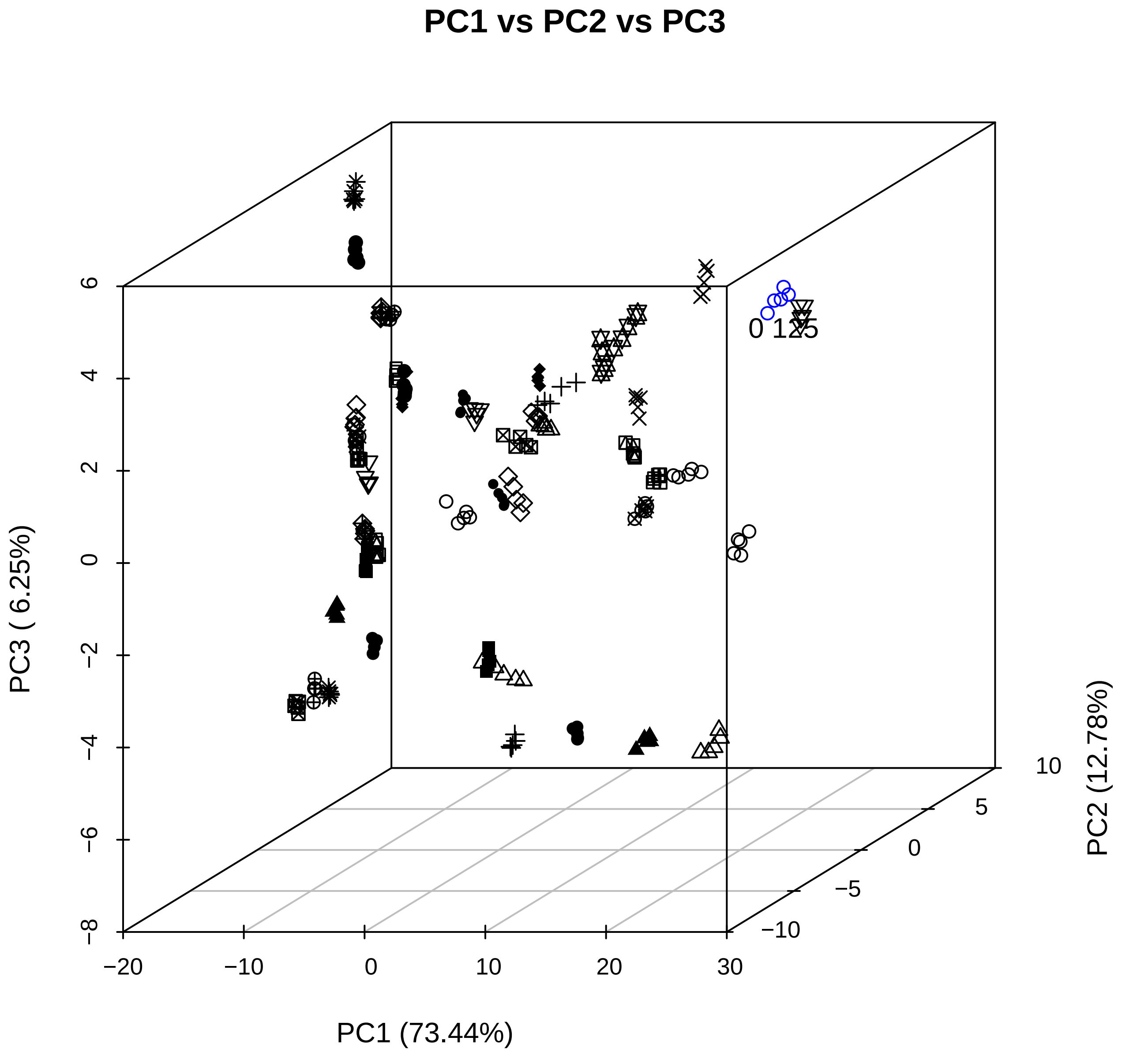

PC1 vs PC2 vs PC3
PC1 (73.44%)
PC2 (12.78%)
PC3 ( 6.25%)
−20
−10
0
10
20
30
−10
−5
0
5
10
−8
−6
−4
−2
0
2
4
6
0 125

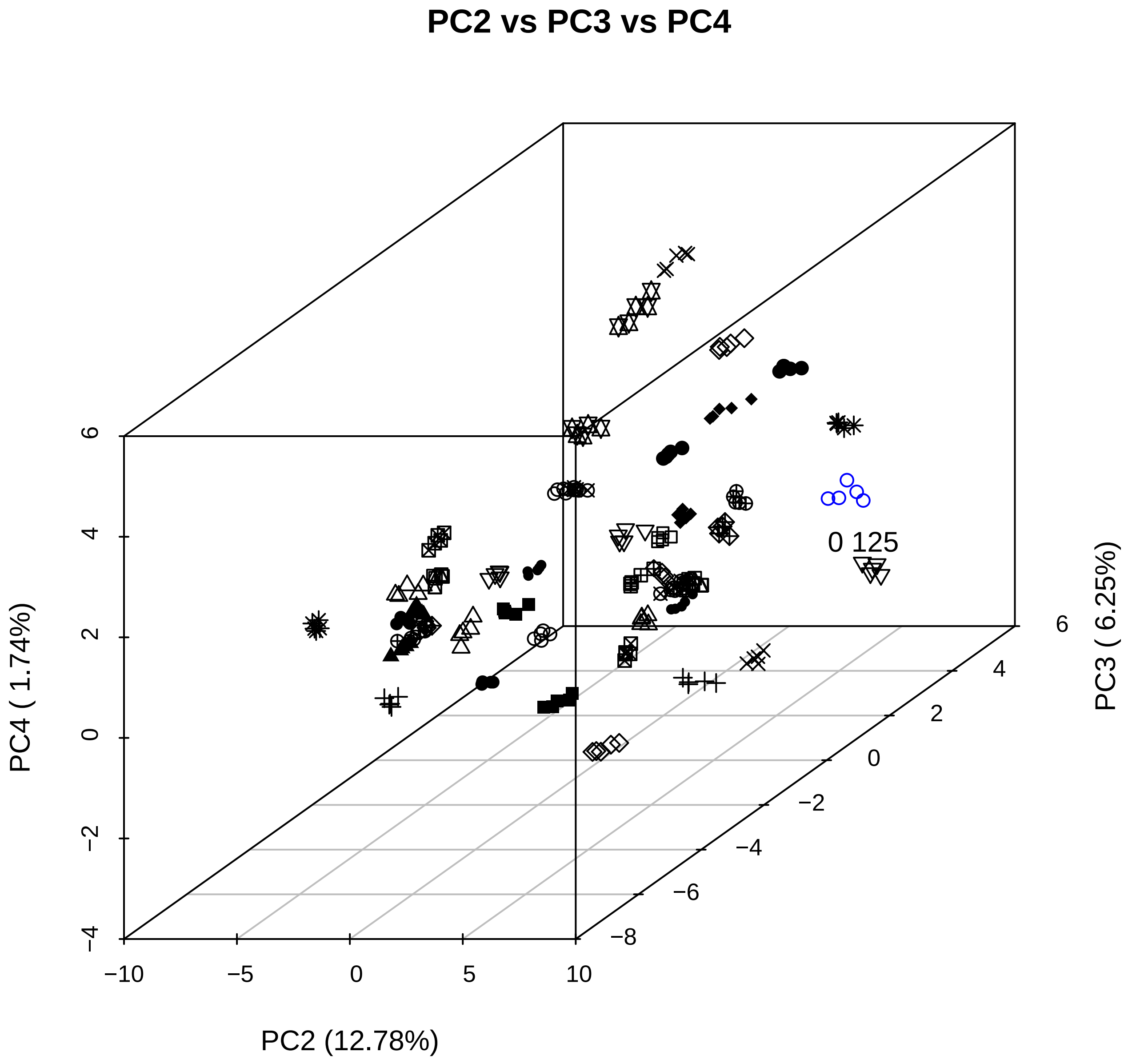
PC2 vs PC3 vs PC4
PC2 (12.78%)
PC4 ( 1.74%)
PC3 ( 6.25%)
0 125
−10
−5
0
5
10
−4
−2
0
2
4
6
−8
−6
−4
−2
0
2
4
6

## PC1 vs PC2 vs PC4

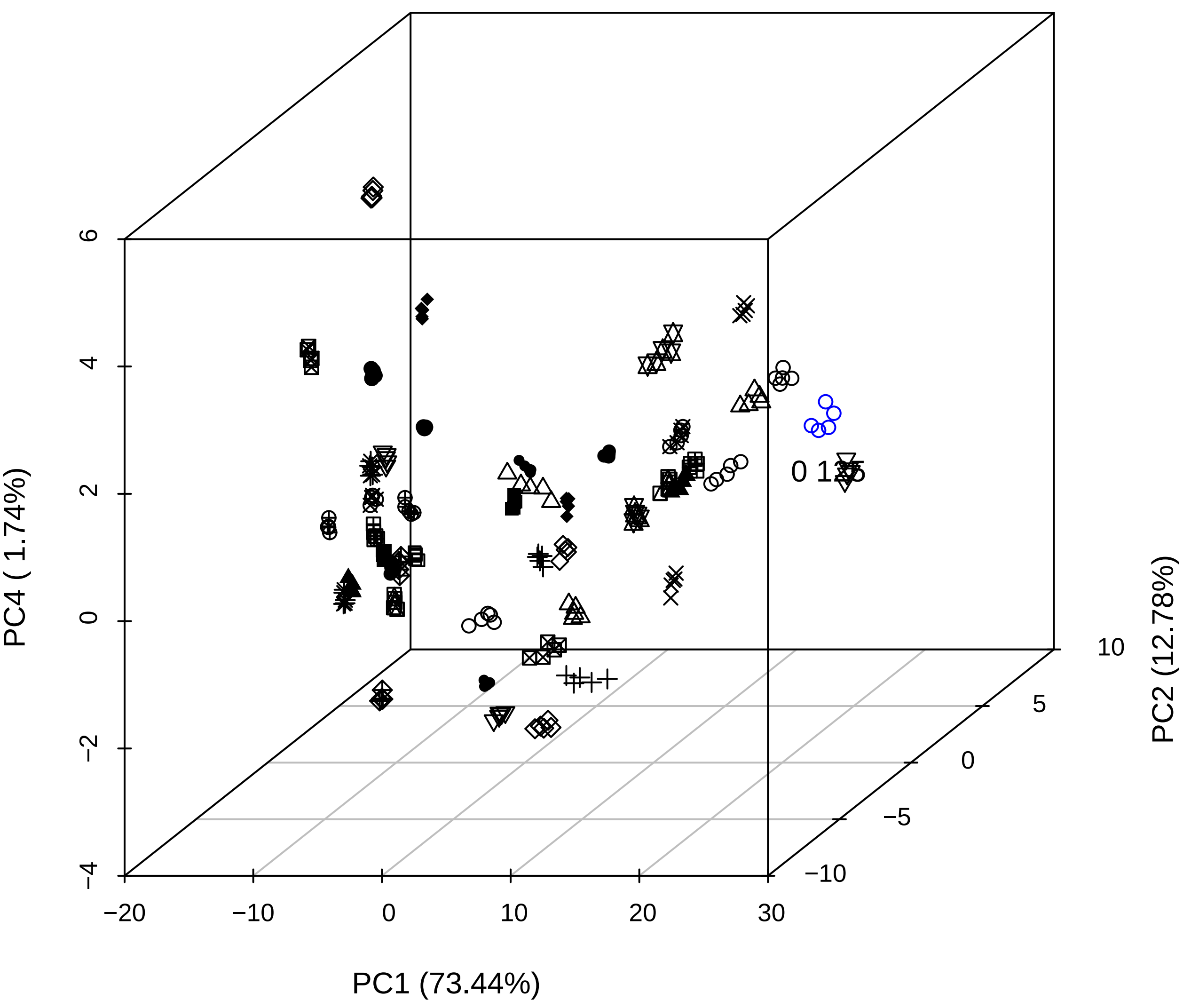

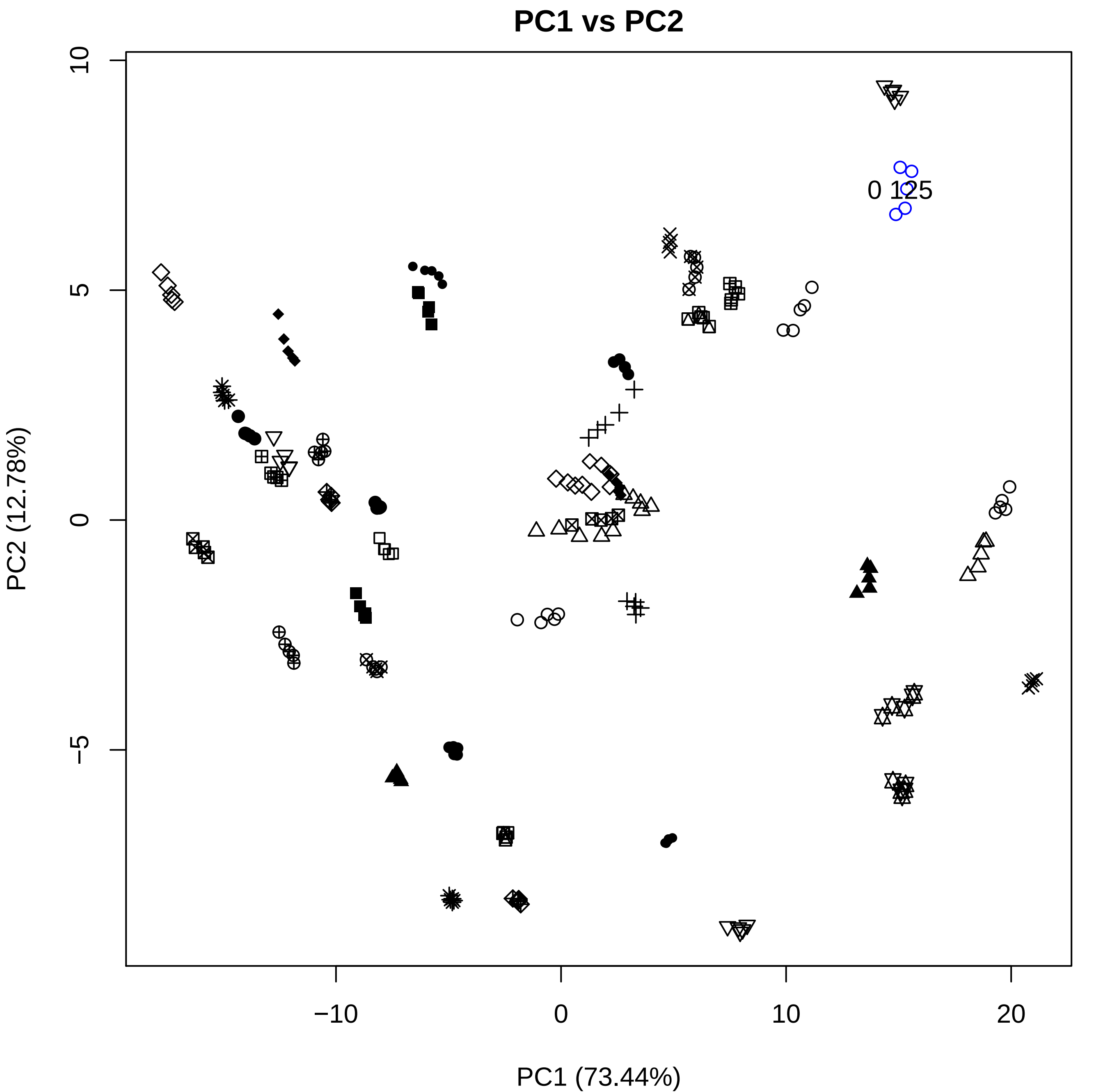
PC1 vs PC2
0 125
10
5
0
−5
−10
0
10
20
PC2 (12.78%)
PC1 (73.44%)

# PC1 vs PC3

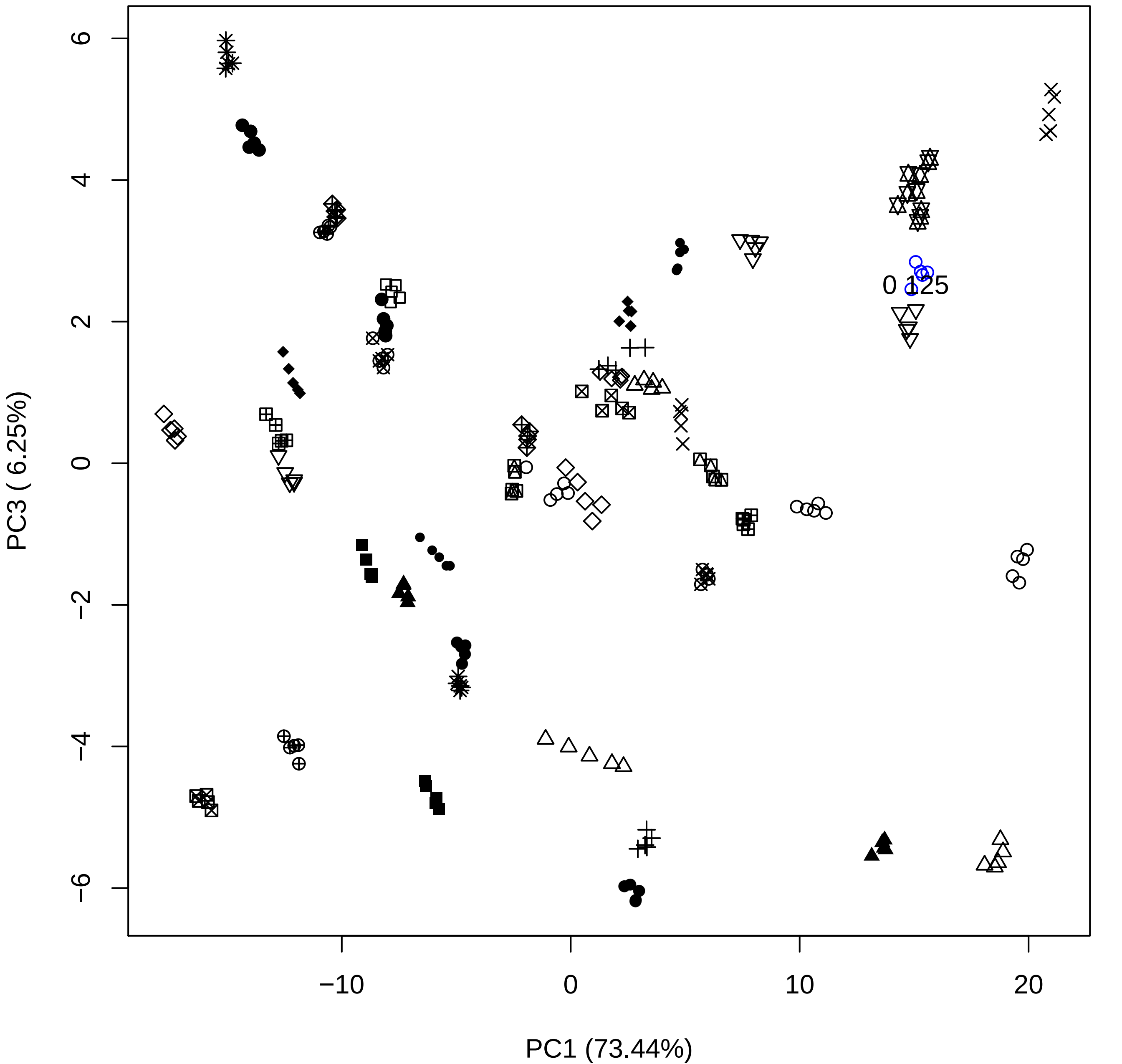

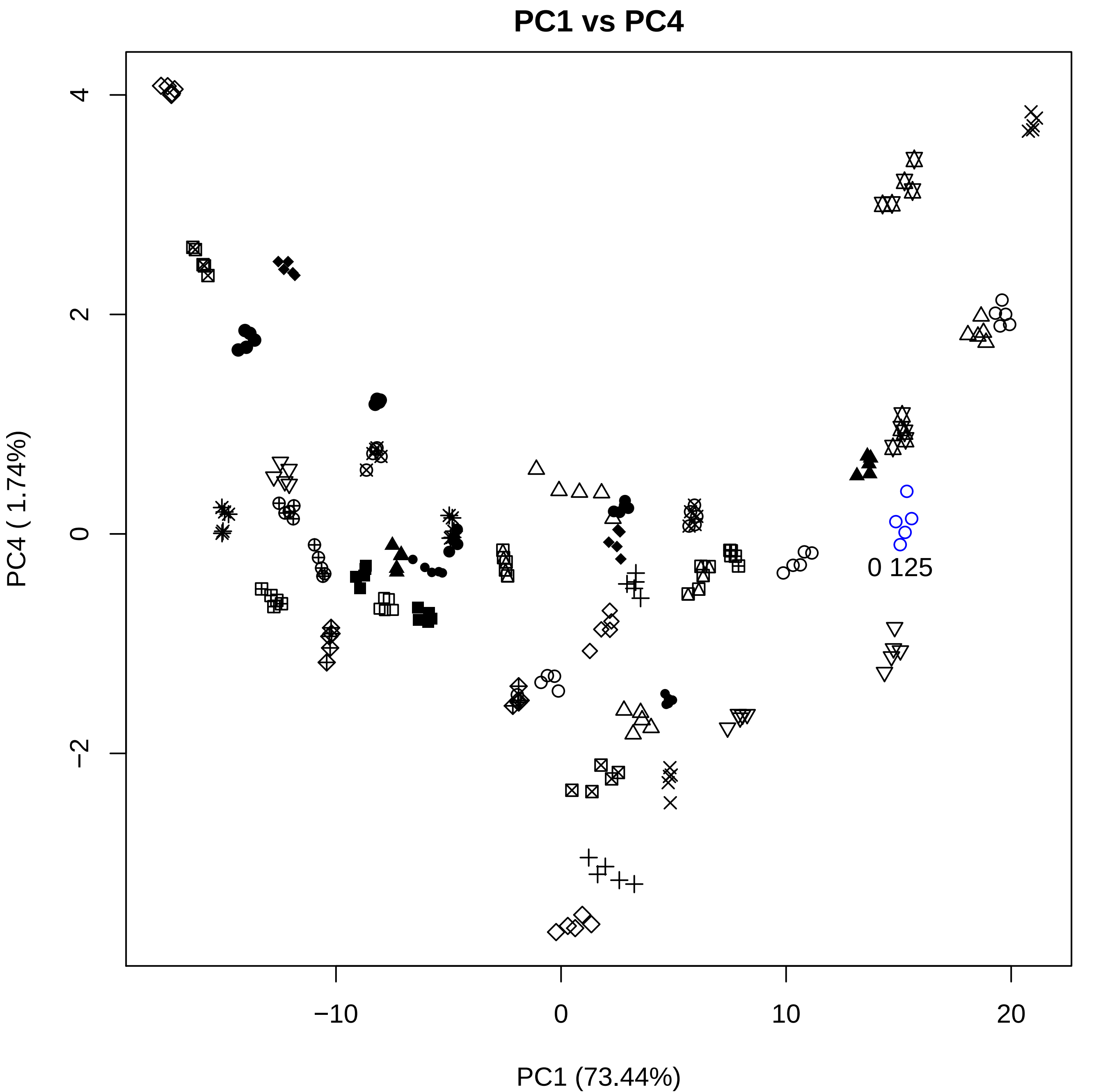
PC1 vs PC4
4
2
0
−2
PC4 ( 1.74%)
−10
0
10
20
PC1 (73.44%)
0 125

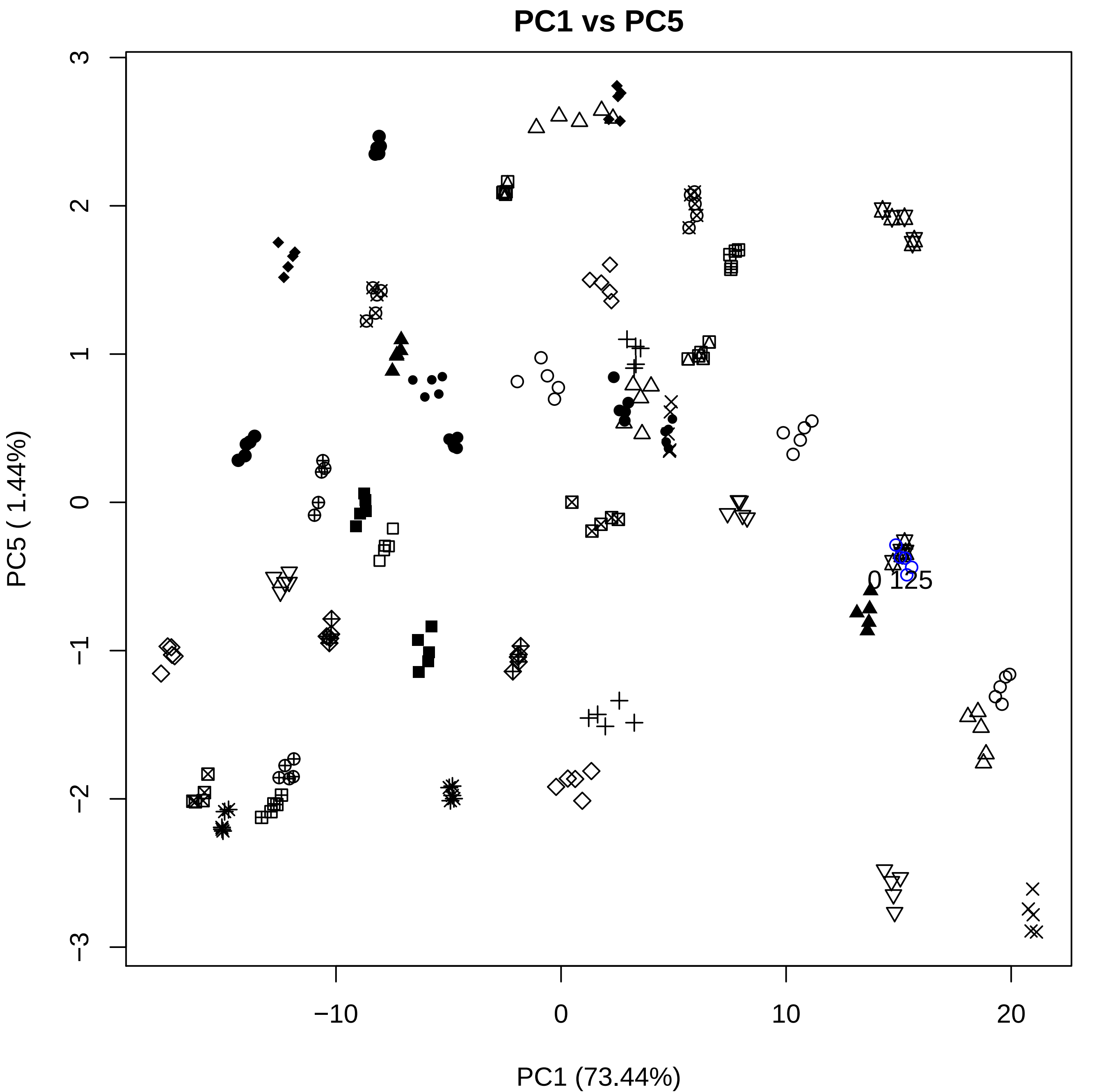
PC1 vs PC5
PC5 ( 1.44%)
PC1 (73.44%)
3
2
1
0
−1
−2
−3
−10
0
10
20
0 125

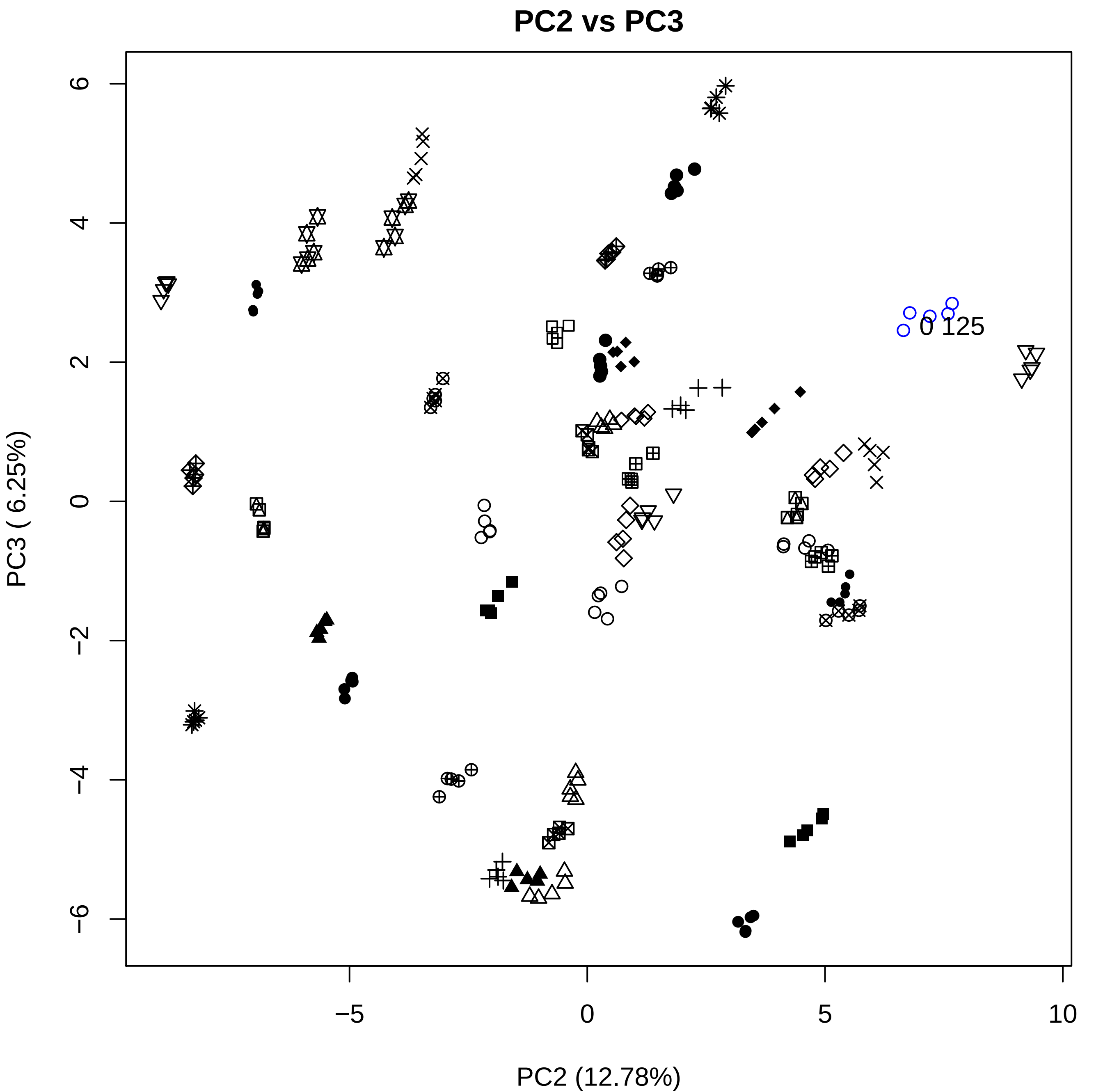
PC2 vs PC3
0 125
PC3 ( 6.25%)
PC2 (12.78%)
6
4
2
0
−2
−4
−6
−5
0
5
10

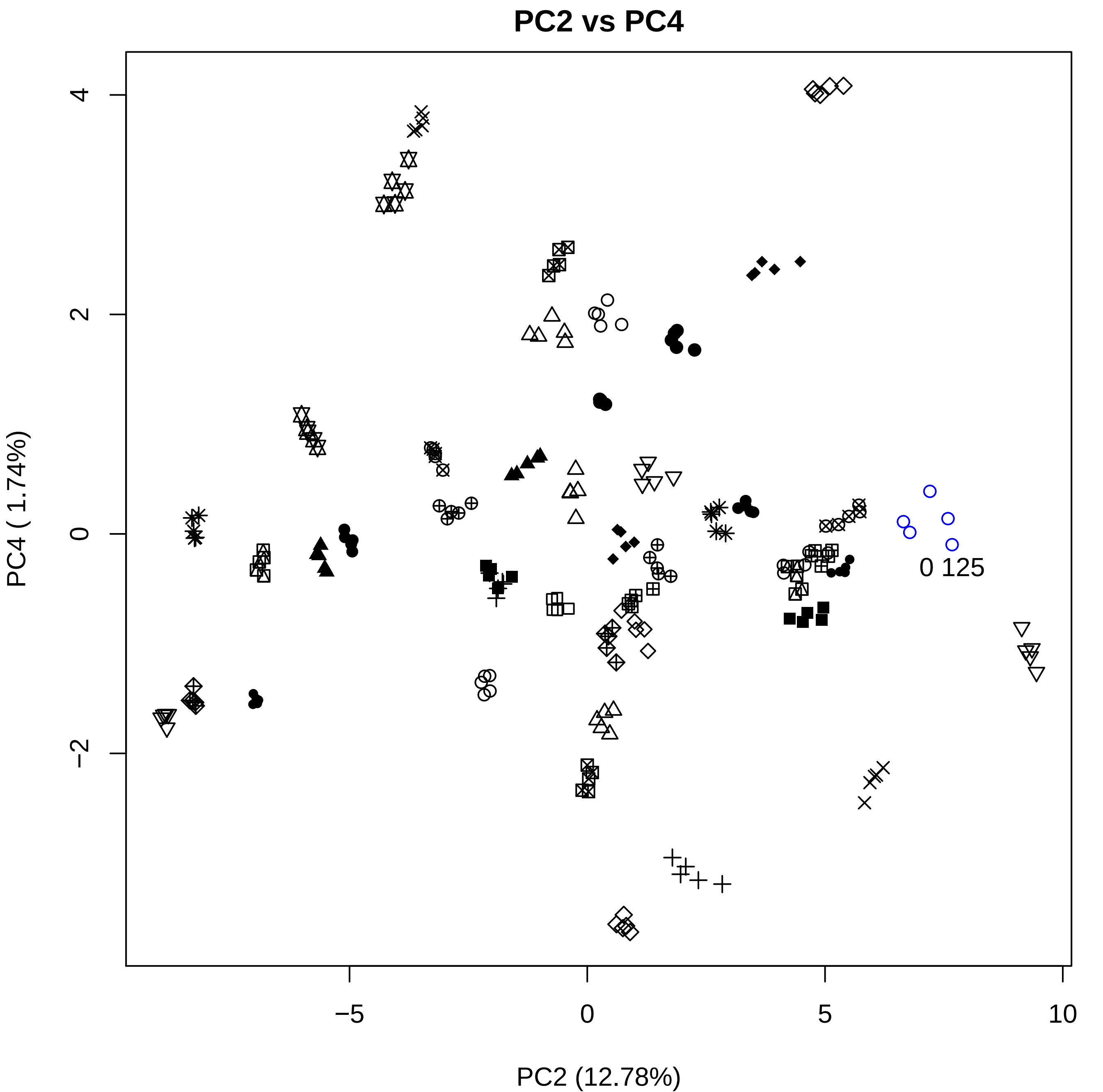
PC2 vs PC4
PC4 ( 1.74%)
PC2 (12.78%)
4
2
0
−2
−5
0
5
10
0 125

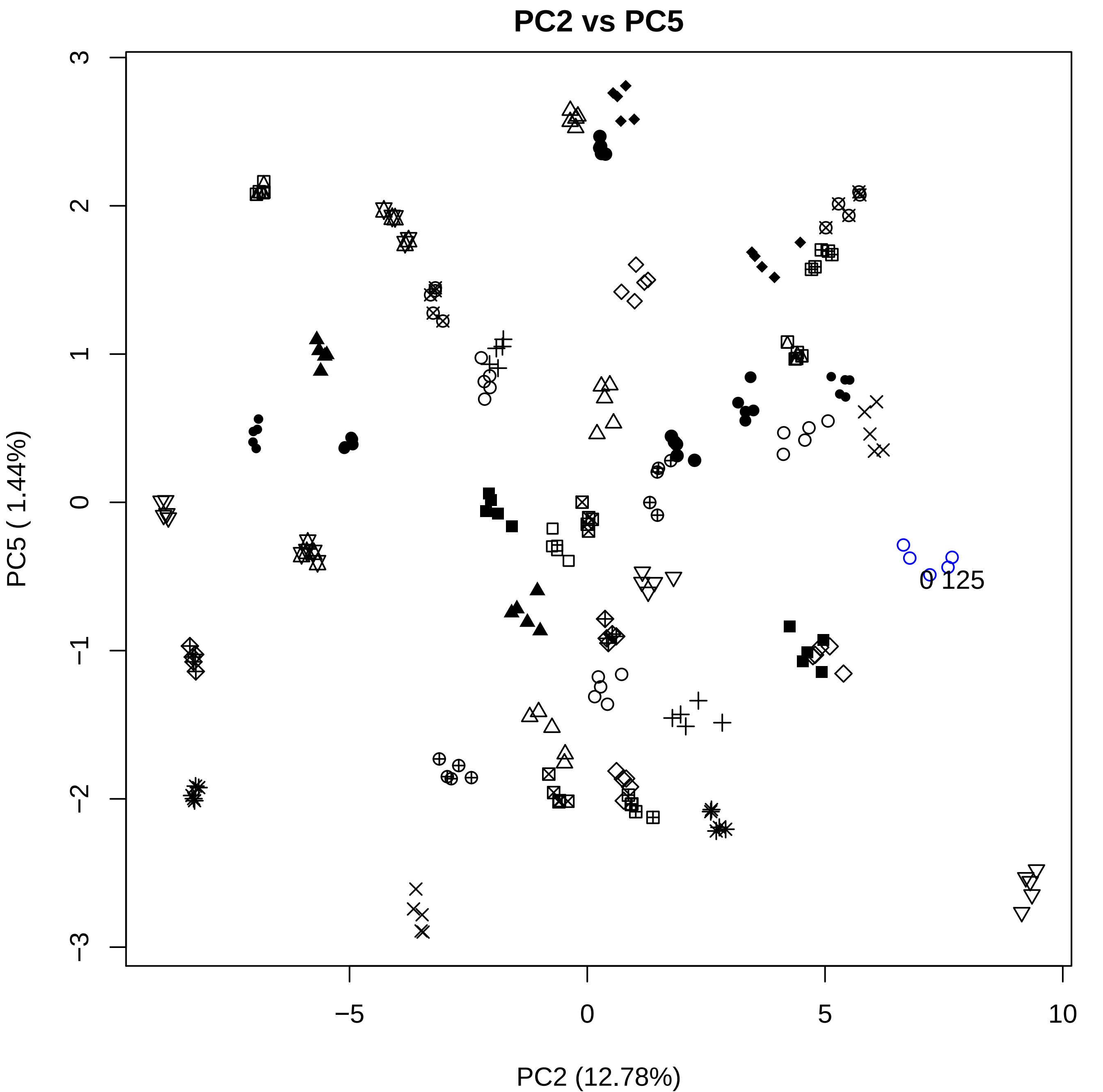
PC2 vs PC5
PC5 ( 1.44%)
PC2 (12.78%)
0 125
3
2
1
0
−1
−2
−3
−5
0
5
10

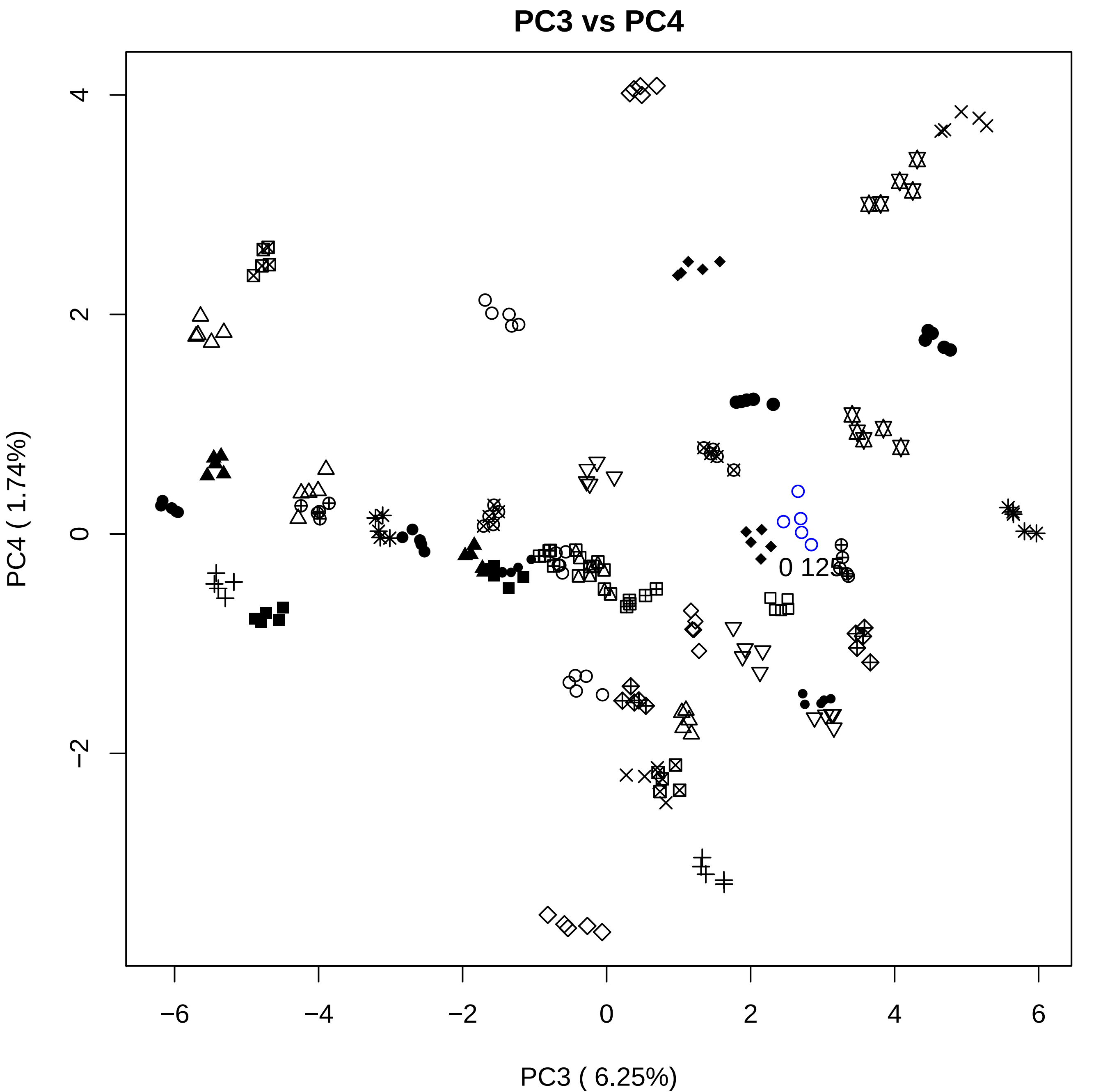
PC3 vs PC4
PC4 ( 1.74%)
PC3 ( 6.25%)
4
2
0
−2
−6
−4
−2
0
2
4
6
0 125

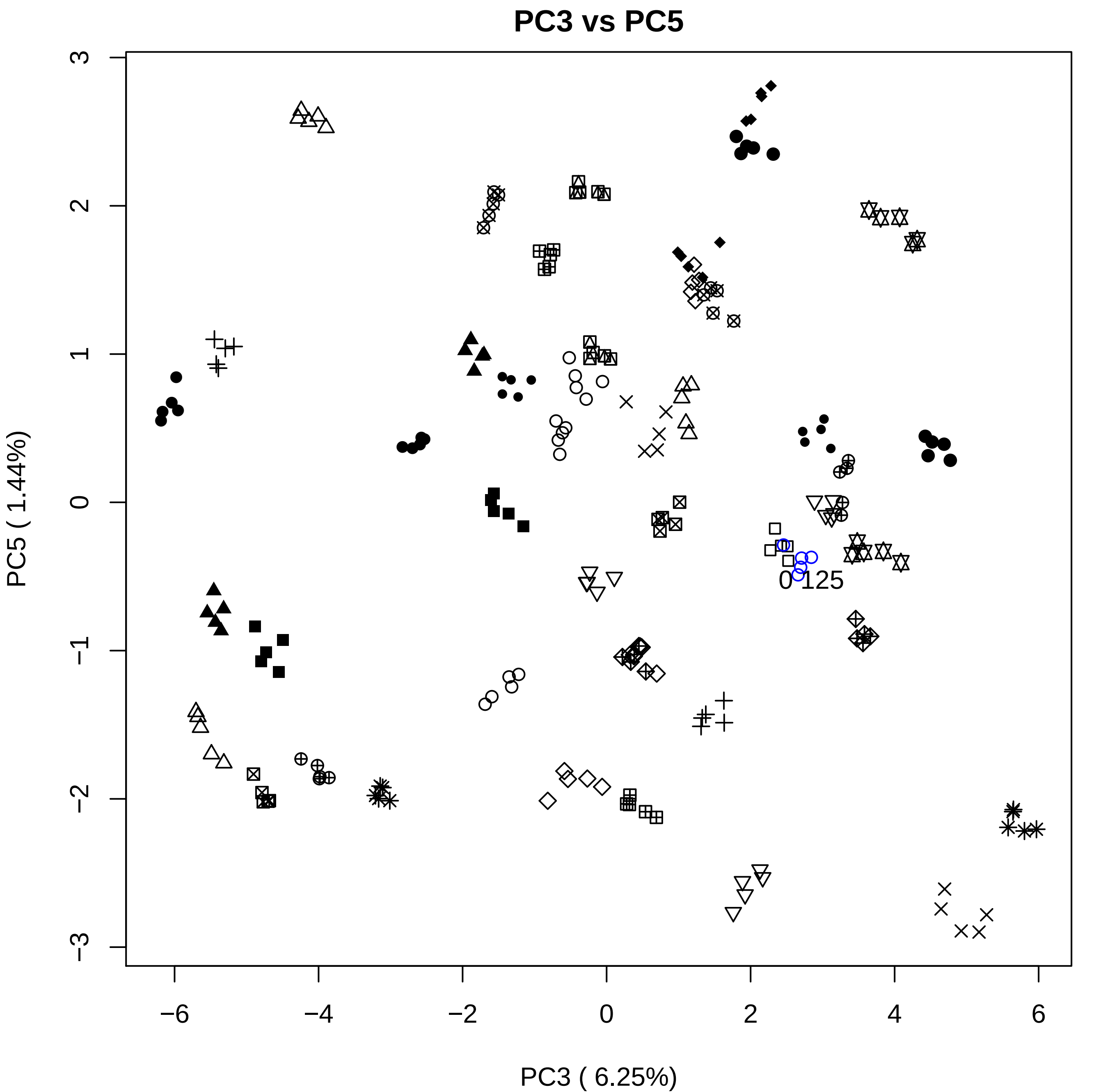
PC3 vs PC5
PC5 ( 1.44%)
PC3 ( 6.25%)
3
2
1
0
−1
−2
−3
−6
−4
−2
0
2
4
6
0 125

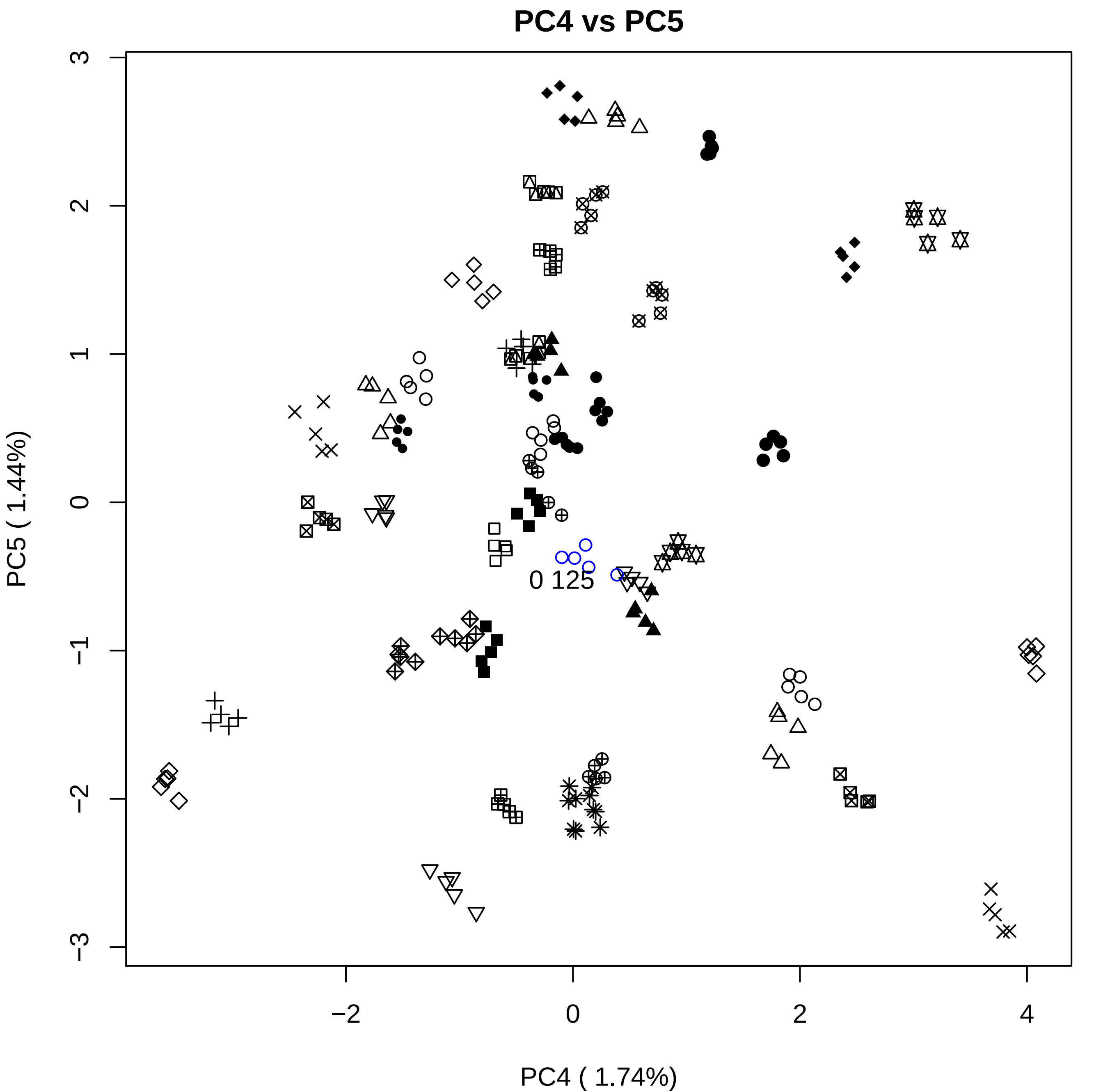
PC4 vs PC5
3
2
1
0
−1
−2
−3
PC5 ( 1.44%)
0 125
−2
0
2
4
PC4 ( 1.74%)

### PCA of 0.25%  OSAS vs PMH

Scaled Matrix PCAed

7th order polynomial baseline correction performed

Raw spectra smoothed to 21 Neighbours, 2nd order PN, Savtisky Golay

Normalisation between 0 and 100

D2 spectra smoothed to 41 Neighbours, 2nd order PN, Savtisky Golay

Regions Analysed: 3601−2501 and 2001−700cm−1

Samples Analysed: 1 Contam's 0 OSAS's 45 PMH's

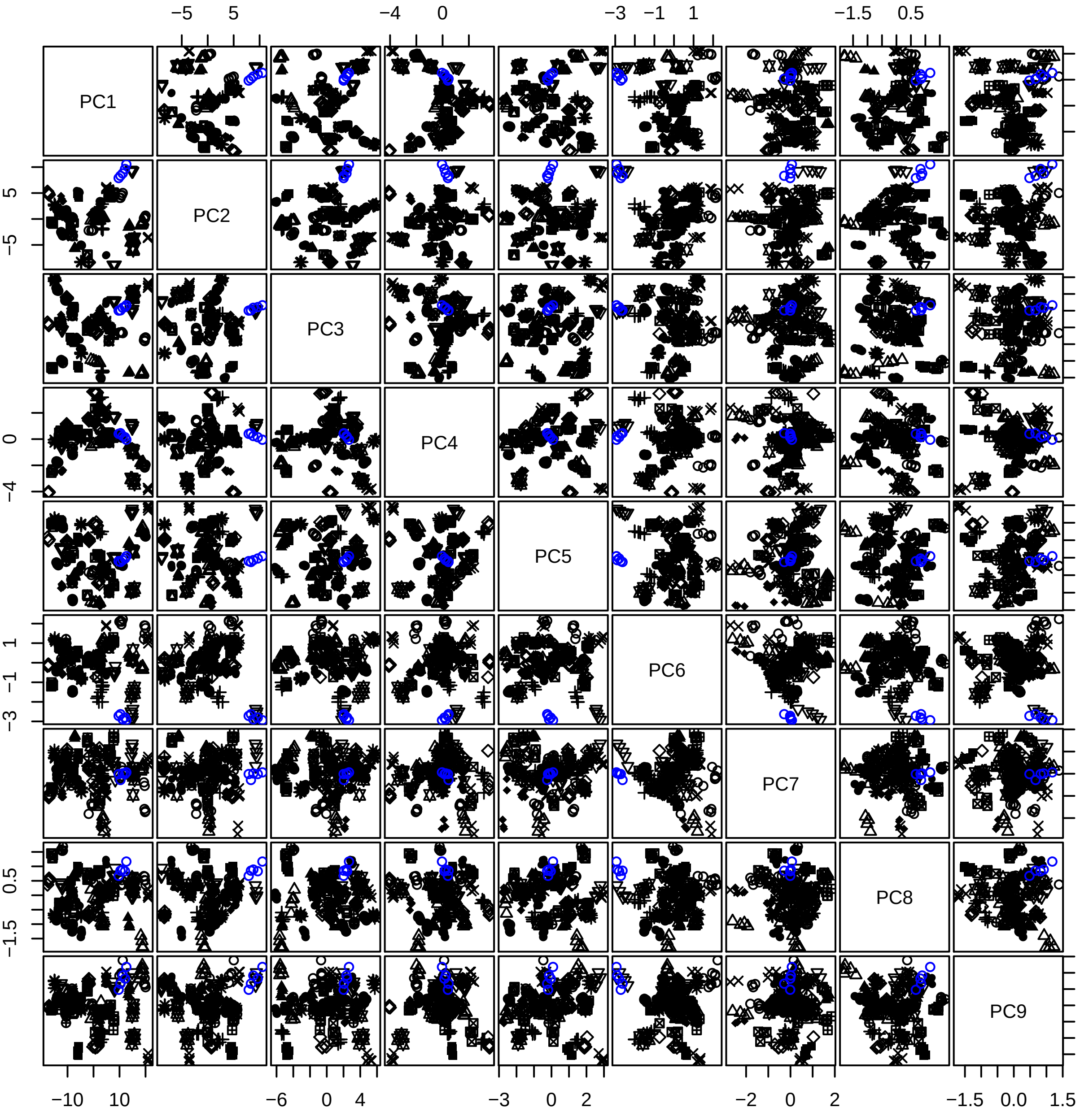
PC1
PC2
PC3
PC4
PC5
PC6
PC7
PC8
PC9

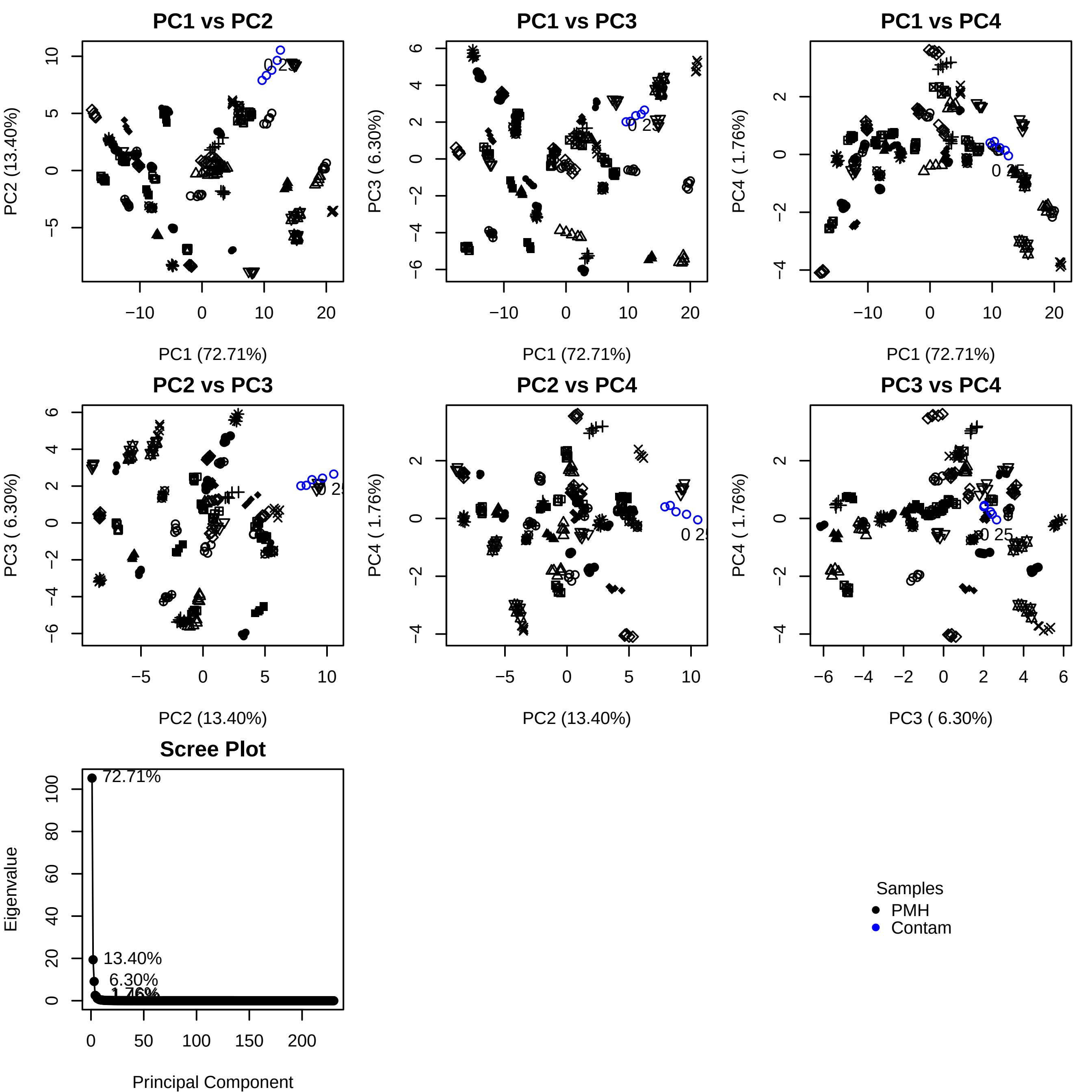
PC1 vs PC2
PC1 vs PC3
PC1 vs PC4
PC2 vs PC3
PC2 vs PC4
PC3 vs PC4
Scree Plot
PC1 (72.71%)
PC2 (13.40%)
PC3 ( 6.30%)
PC4 ( 1.76%)
72.71%
13.40%
6.30%
Eigenvalue
Principal Component
Samples
PMH
Contam

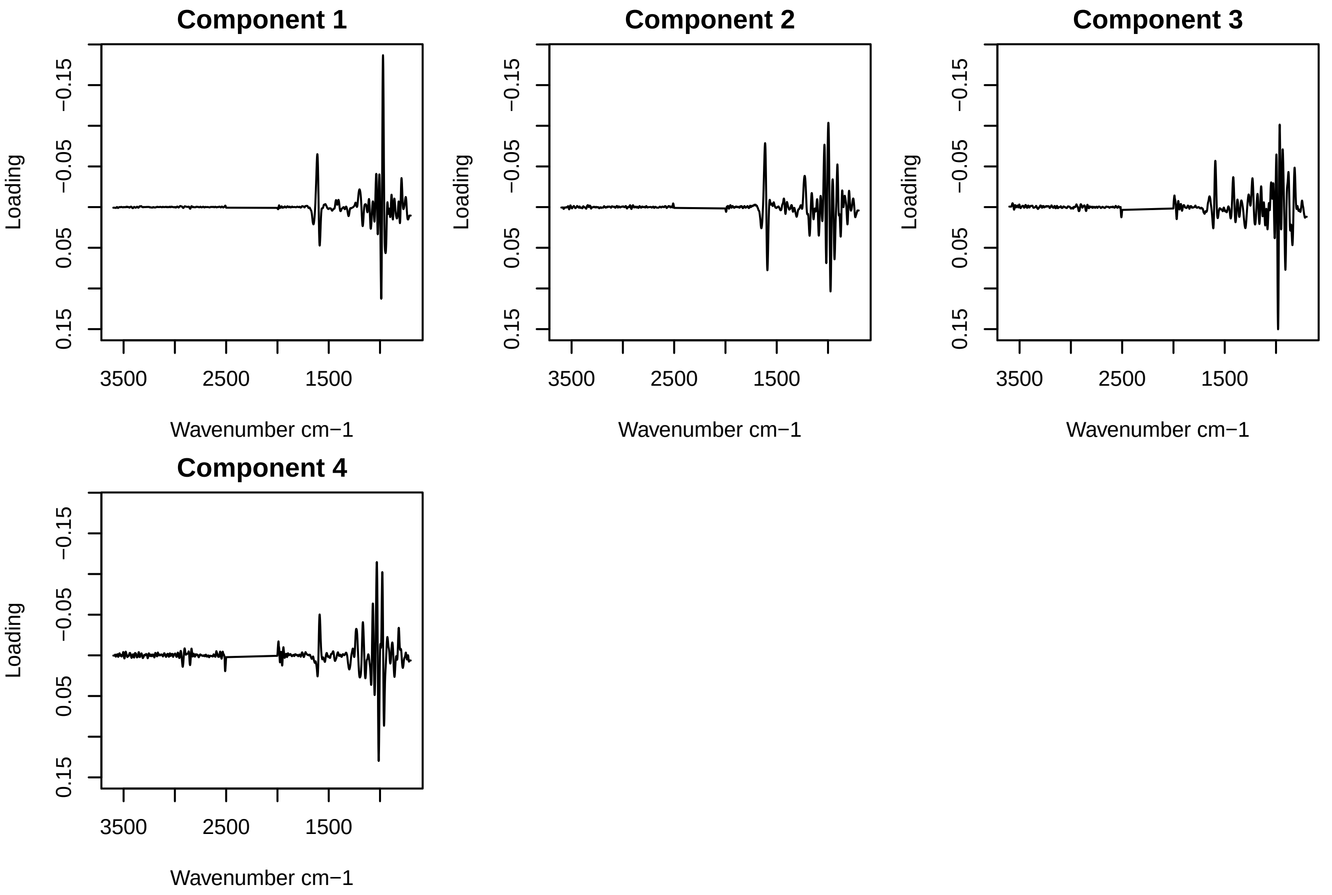
Component 1
Component 2
Component 3
Component 4
Loading
−0.15
−0.05
0.05
0.15
3500
2500
1500
Wavenumber cm−1

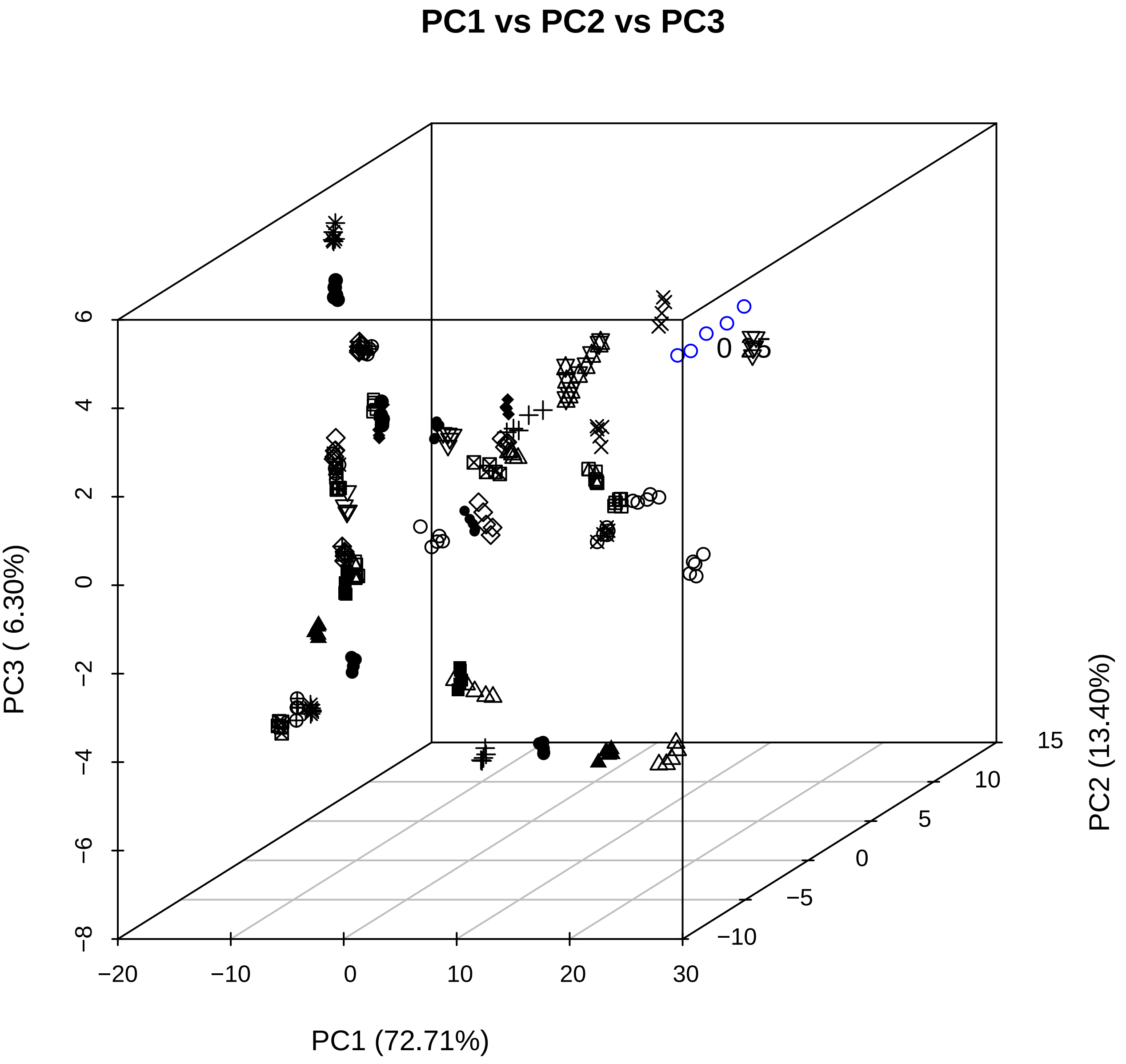
PC1 vs PC2 vs PC3
PC1 (72.71%)
PC2 (13.40%)
PC3 ( 6.30%)
−20
−10
0
10
20
30
−10
−5
0
5
10
15
−8
−6
−4
−2
0
2
4
6

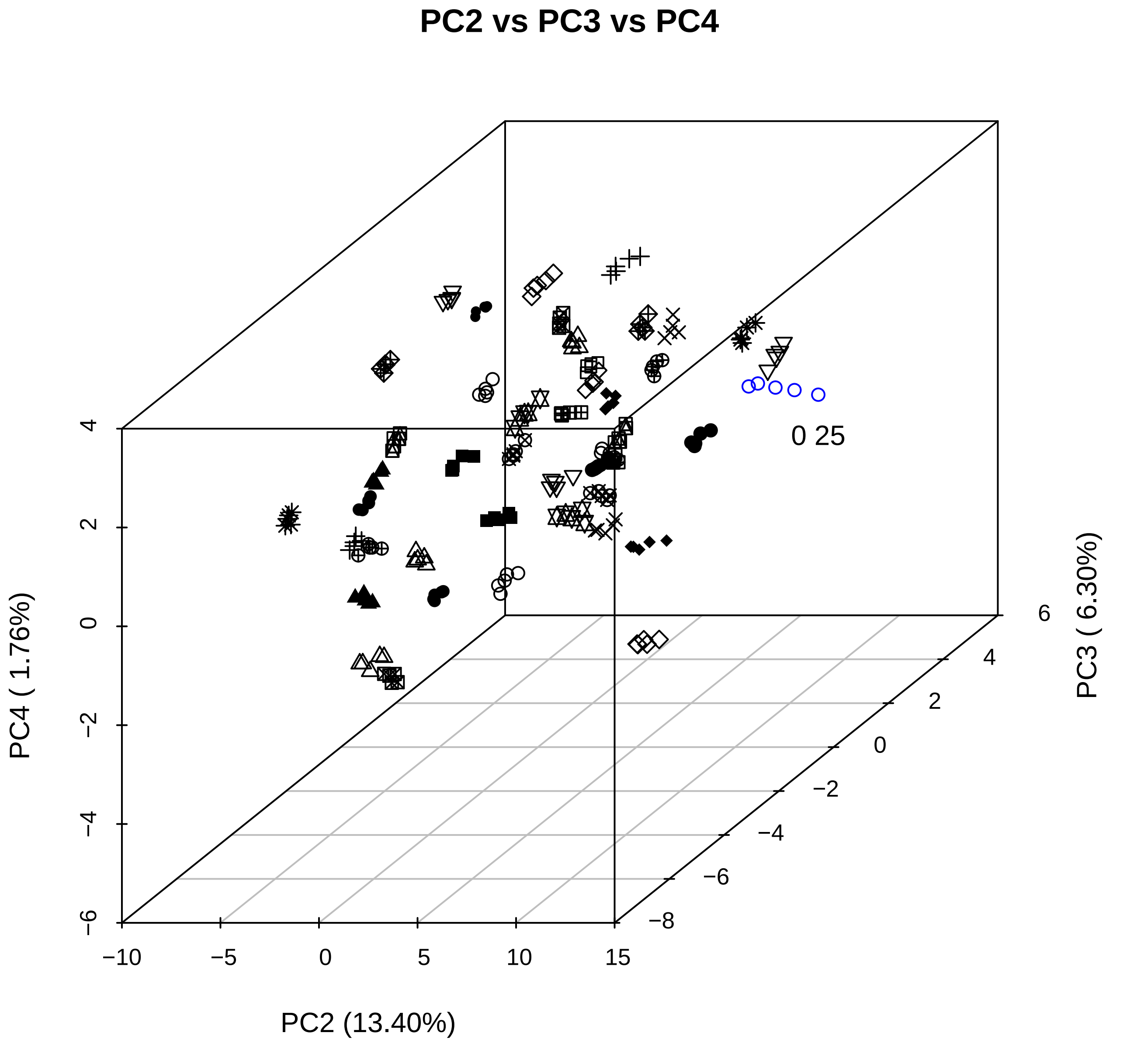

PC2 vs PC3 vs PC4
0 25
PC4 ( 1.76%)
4
2
0
−2
−4
−6
−10
−5
0
5
10
15
PC2 (13.40%)
PC3 ( 6.30%)
6
4
2
0
−2
−4
−6
−8

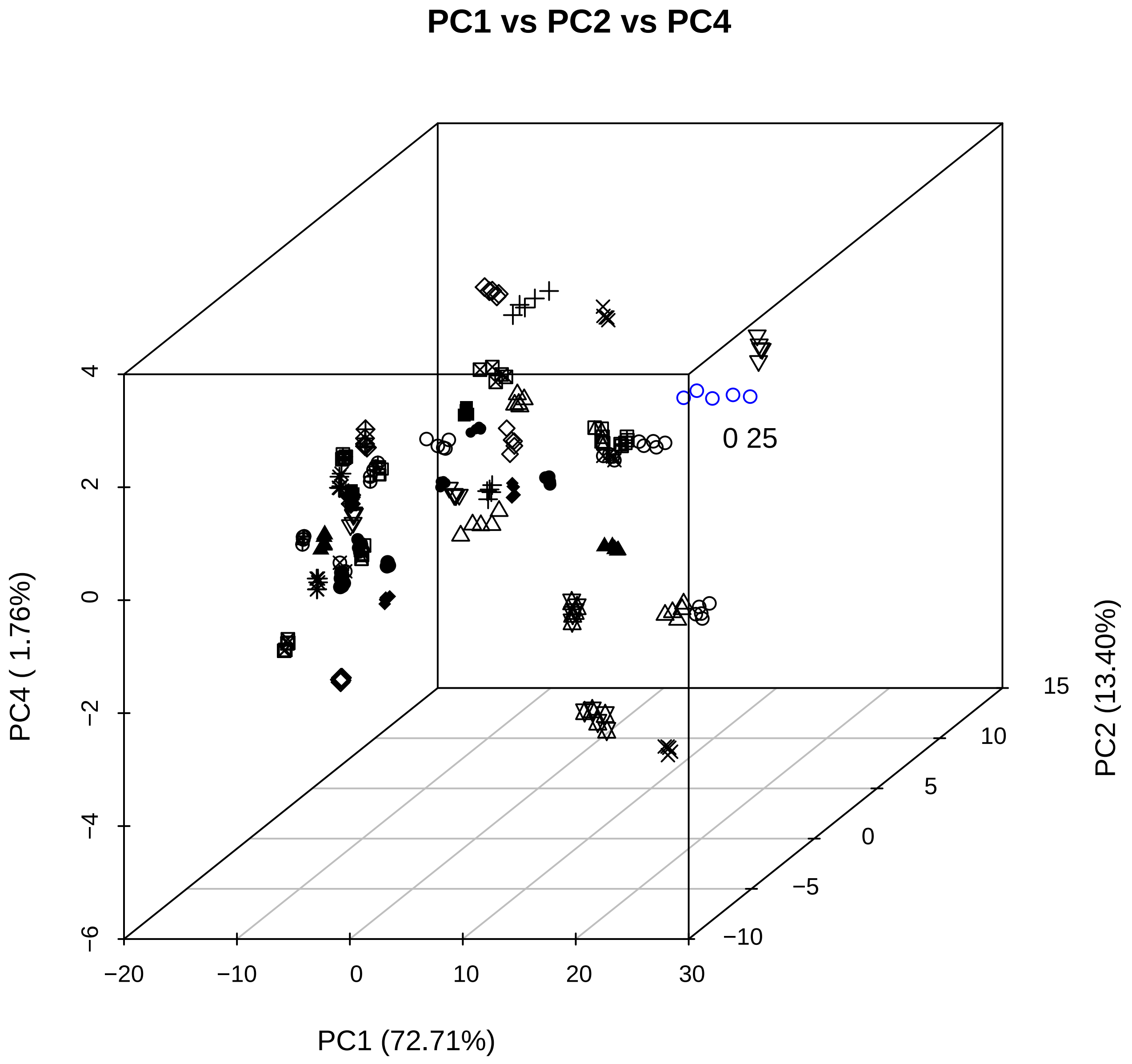
PC1 vs PC2 vs PC4
0 25
PC1 (72.71%)
PC2 (13.40%)
PC4 ( 1.76%)
−20
−10
0
10
20
30
−10
−5
0
5
10
15
−6
−4
−2
0
2
4

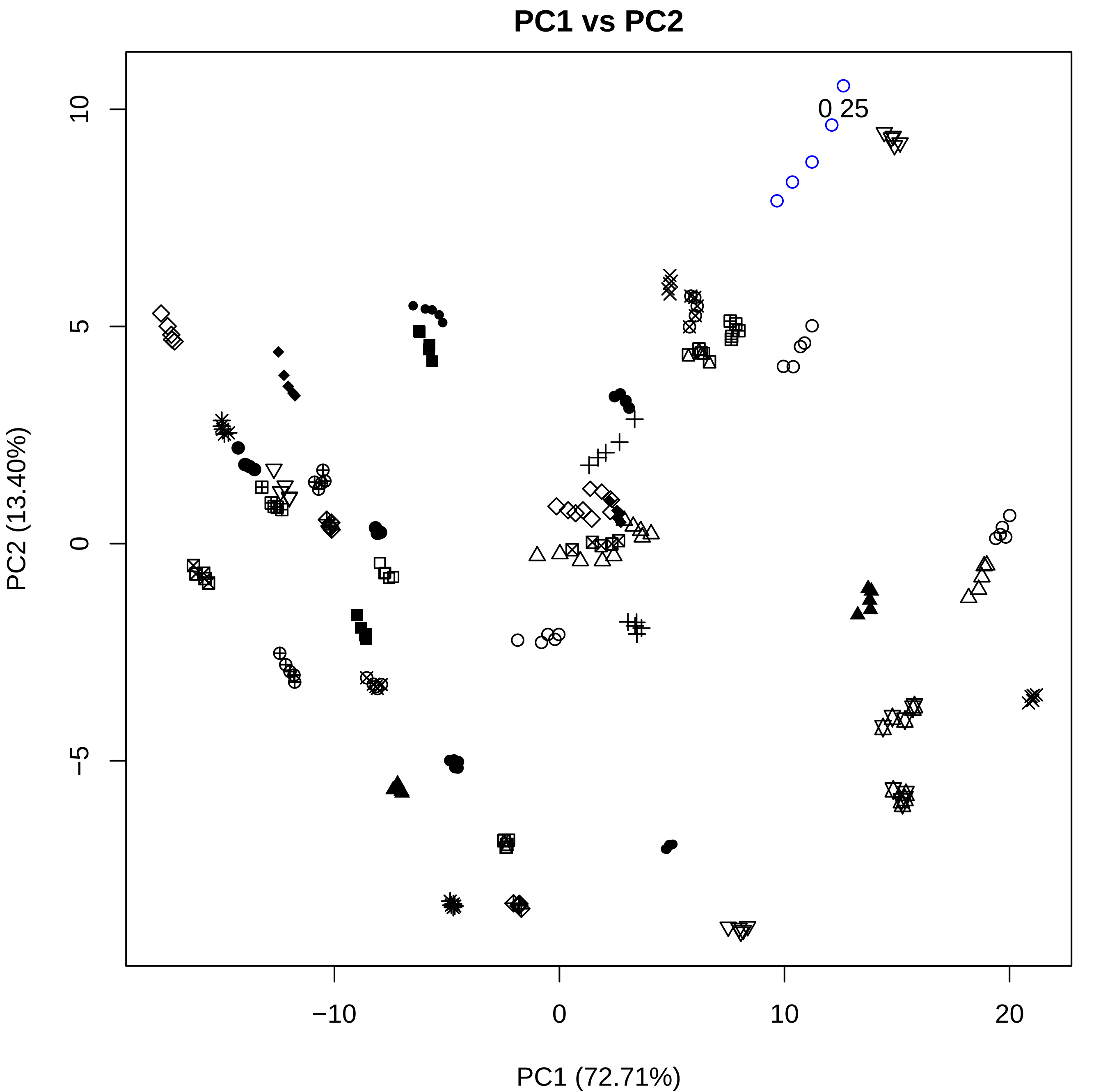
PC1 vs PC2
0 25
PC2 (13.40%)
PC1 (72.71%)
10
5
0
−5
−10
0
10
20

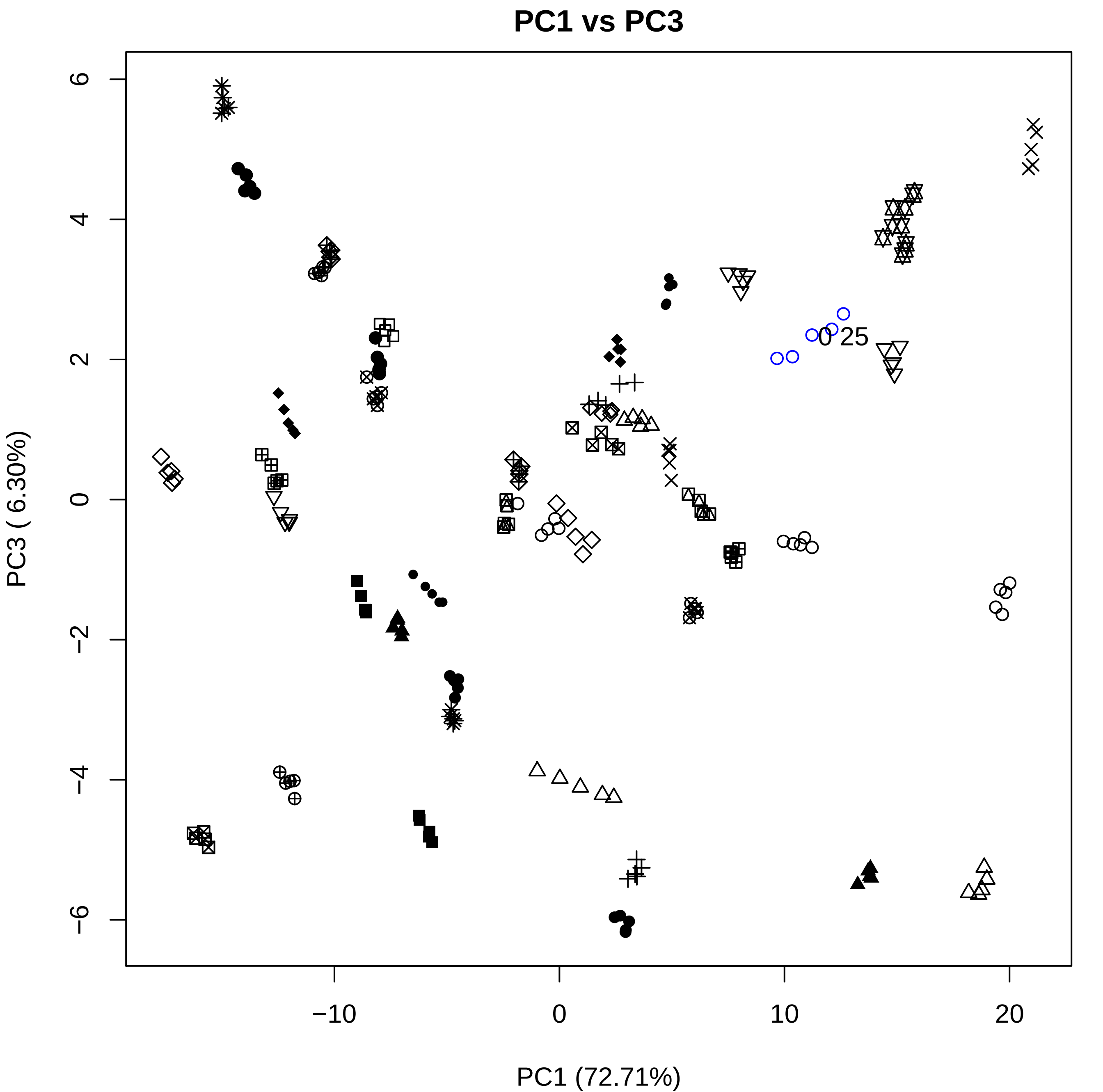
PC1 vs PC3
PC3 ( 6.30%)
PC1 (72.71%)
6
4
2
0
−2
−4
−6
−10
0
10
20
0 25

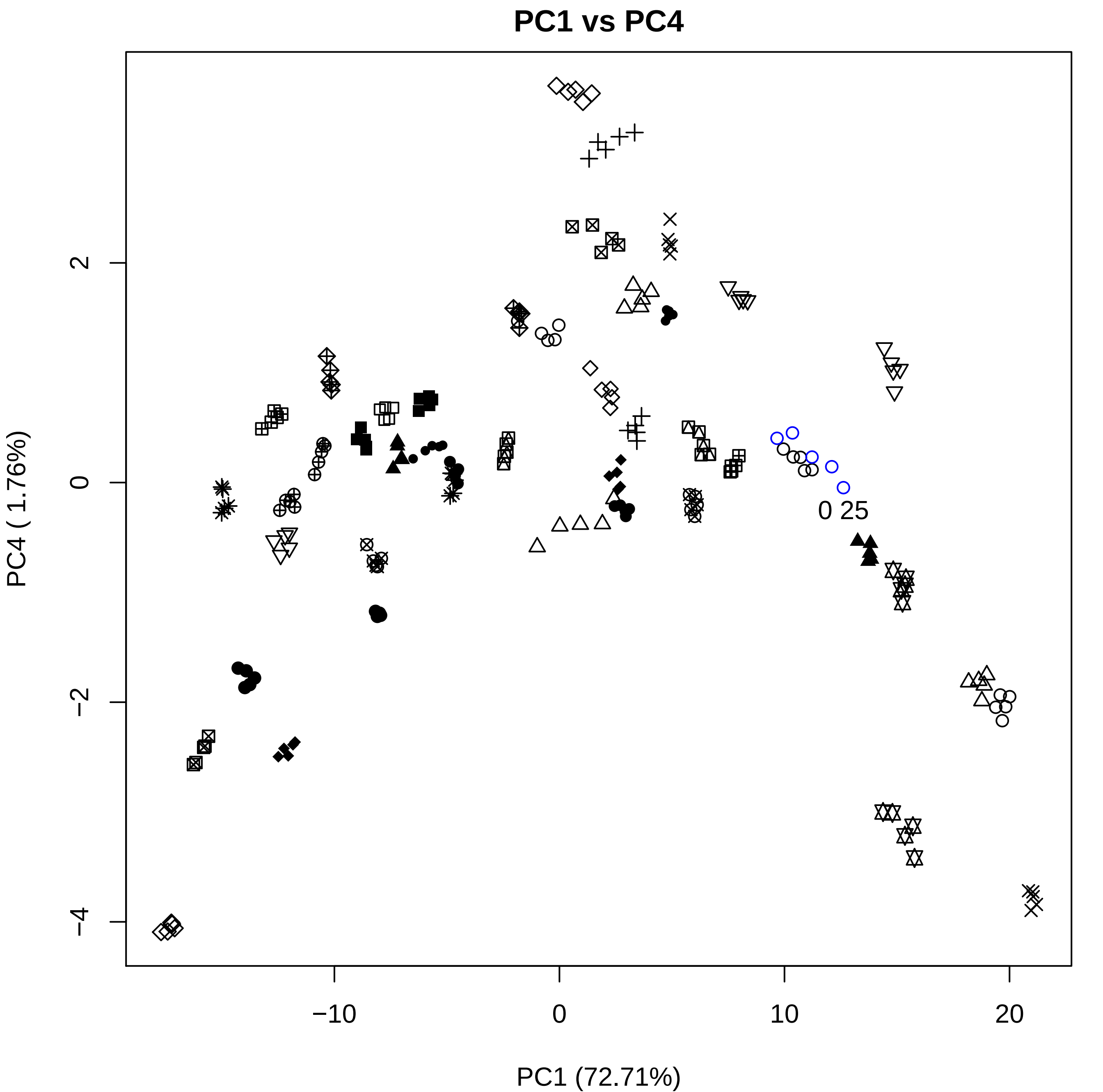
PC1 vs PC4
PC4 ( 1.76%)
PC1 (72.71%)
2
0
−2
−4
−10
0
10
20
0 25

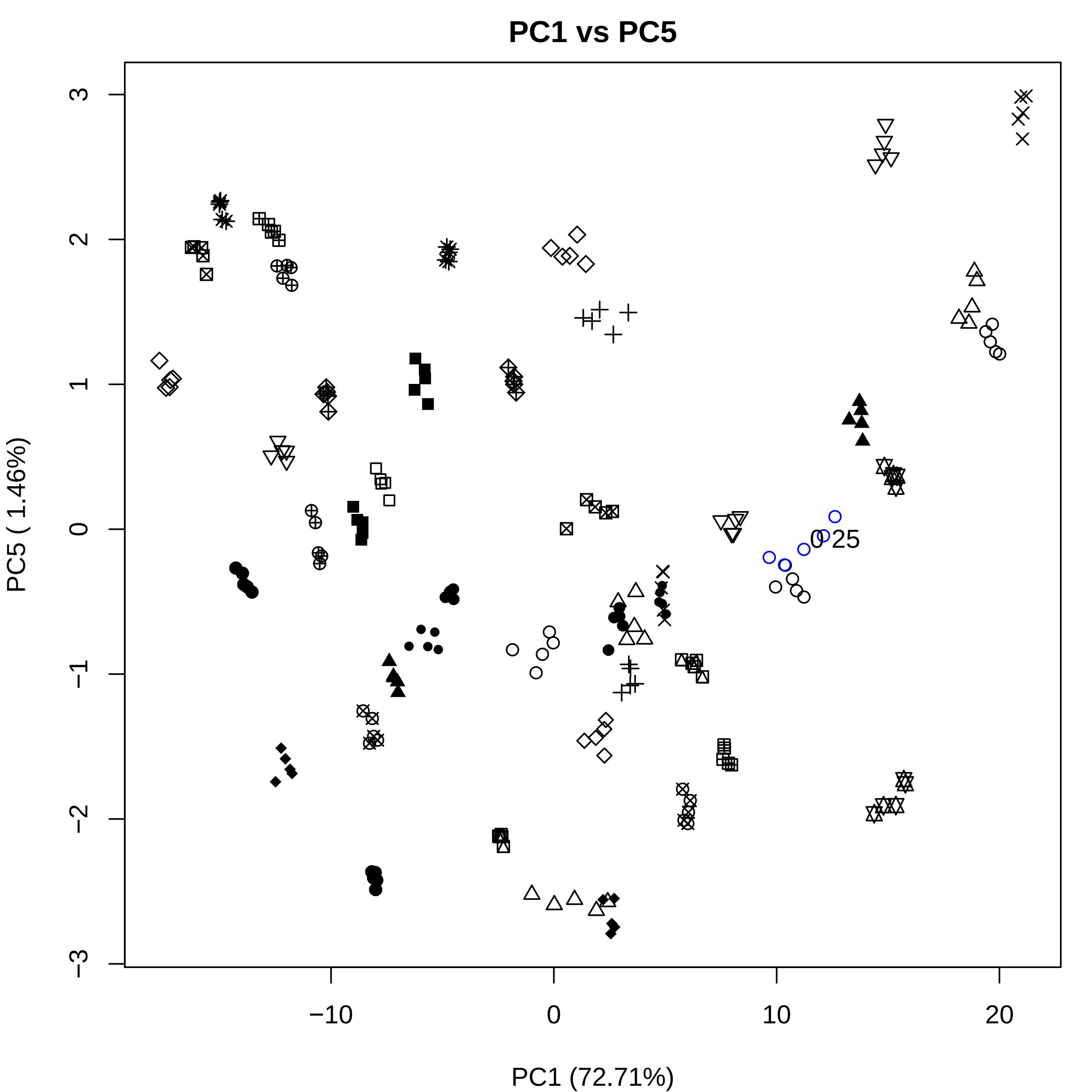

PC1 vs PC5
PC5 ( 1.46%)
PC1 (72.71%)
3
2
1
0
−1
−2
−3
−10
0
10
20
025

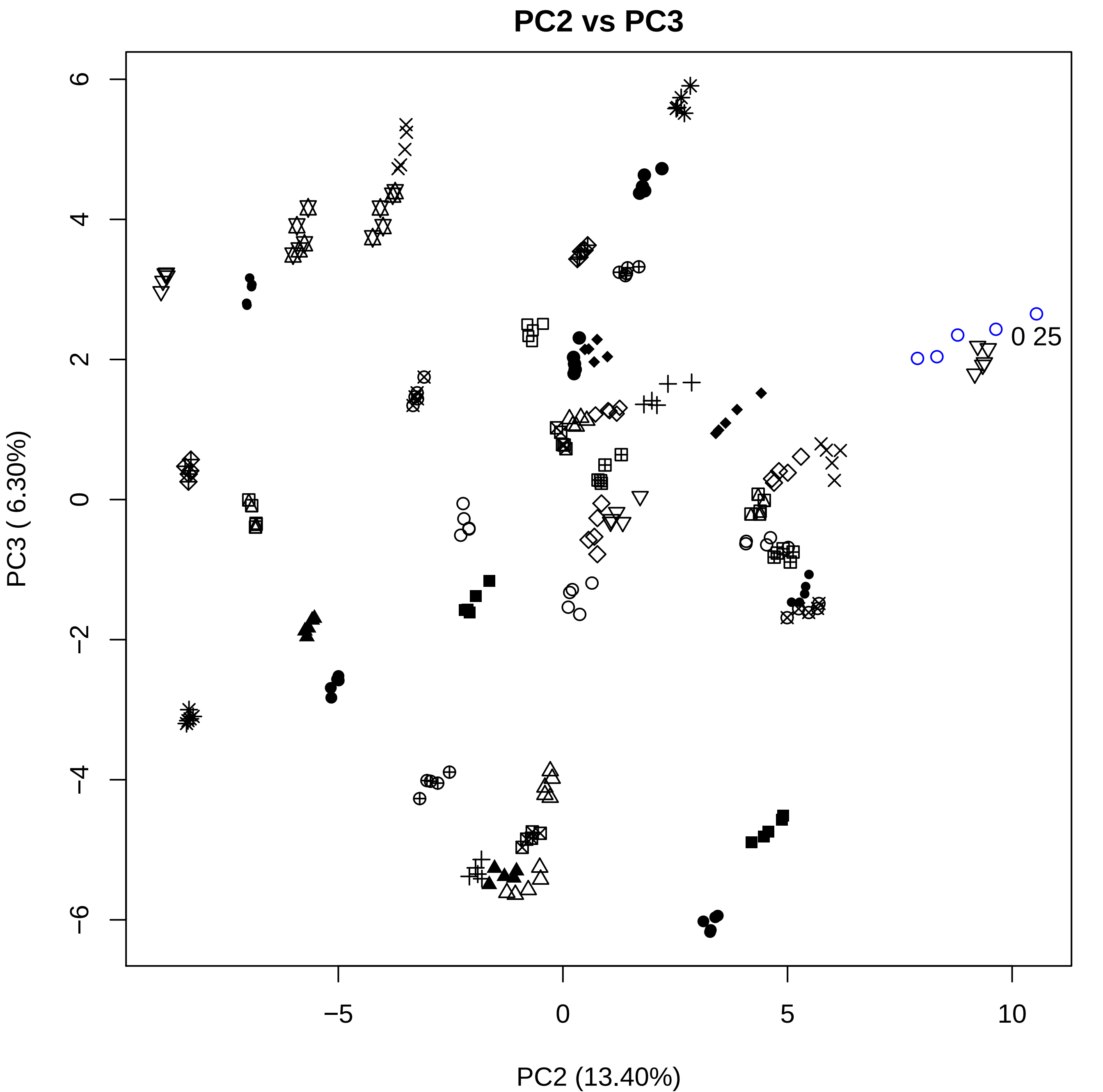
PC2 vs PC3
PC3 ( 6.30%)
PC2 (13.40%)
6
4
2
0
−2
−4
−6
−5
0
5
10
0 25

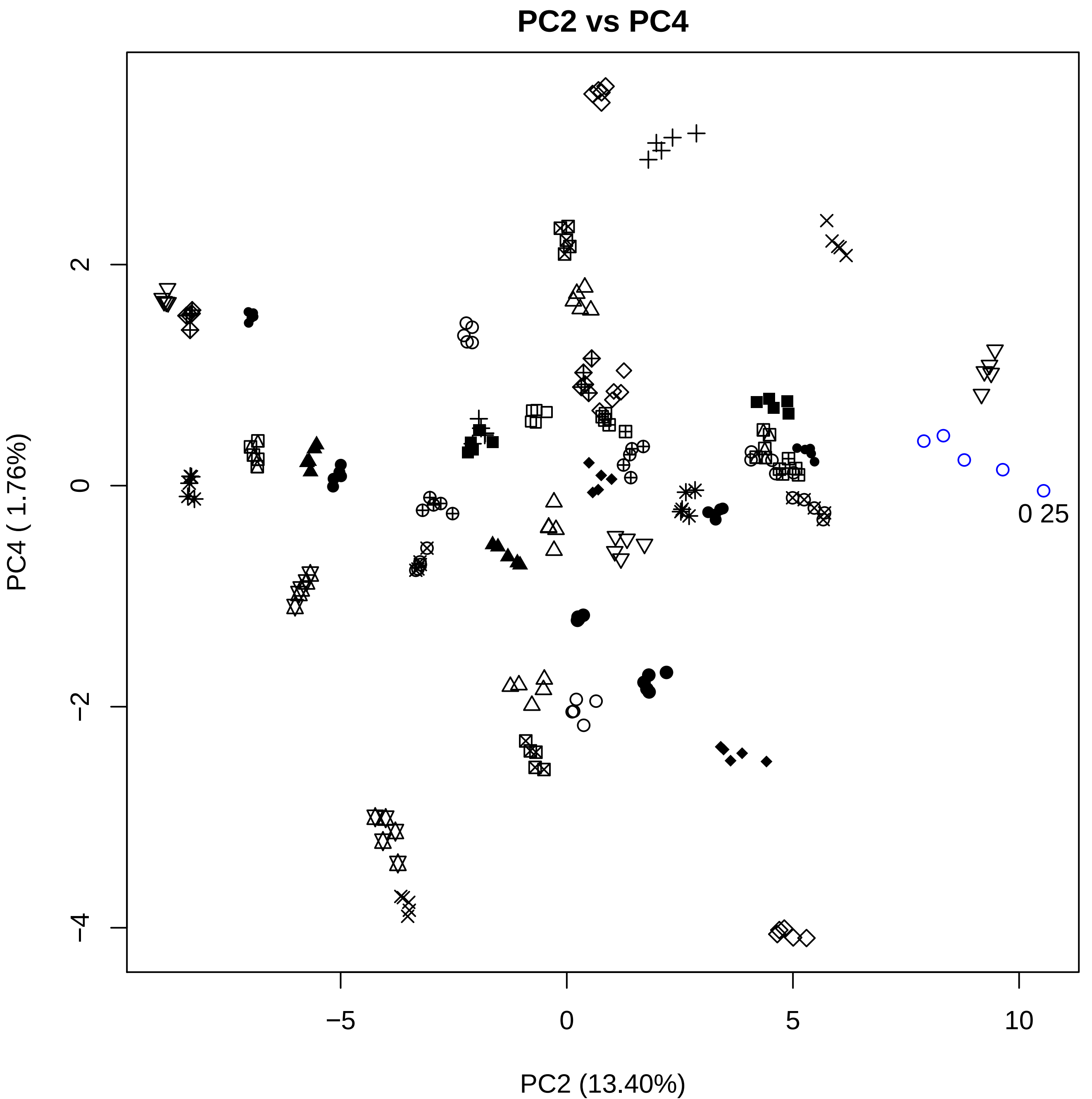
PC2 vs PC4
PC4 ( 1.76%)
PC2 (13.40%)
2
0
−2
−4
−5
0
5
10
0 25

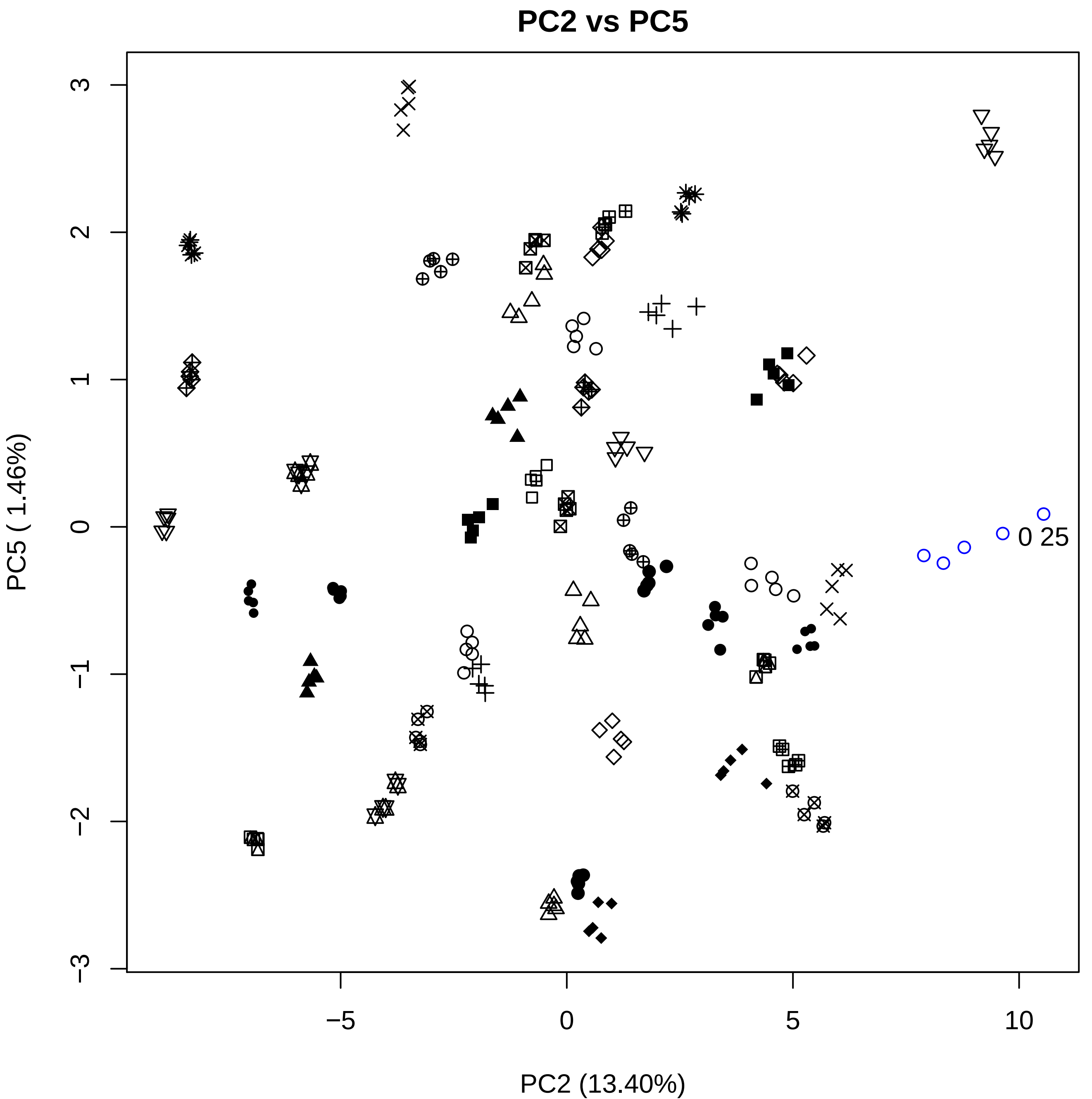
PC2 vs PC5
PC5 ( 1.46%)
PC2 (13.40%)
0 25
3
2
1
0
−1
−2
−3
−5
0
5
10

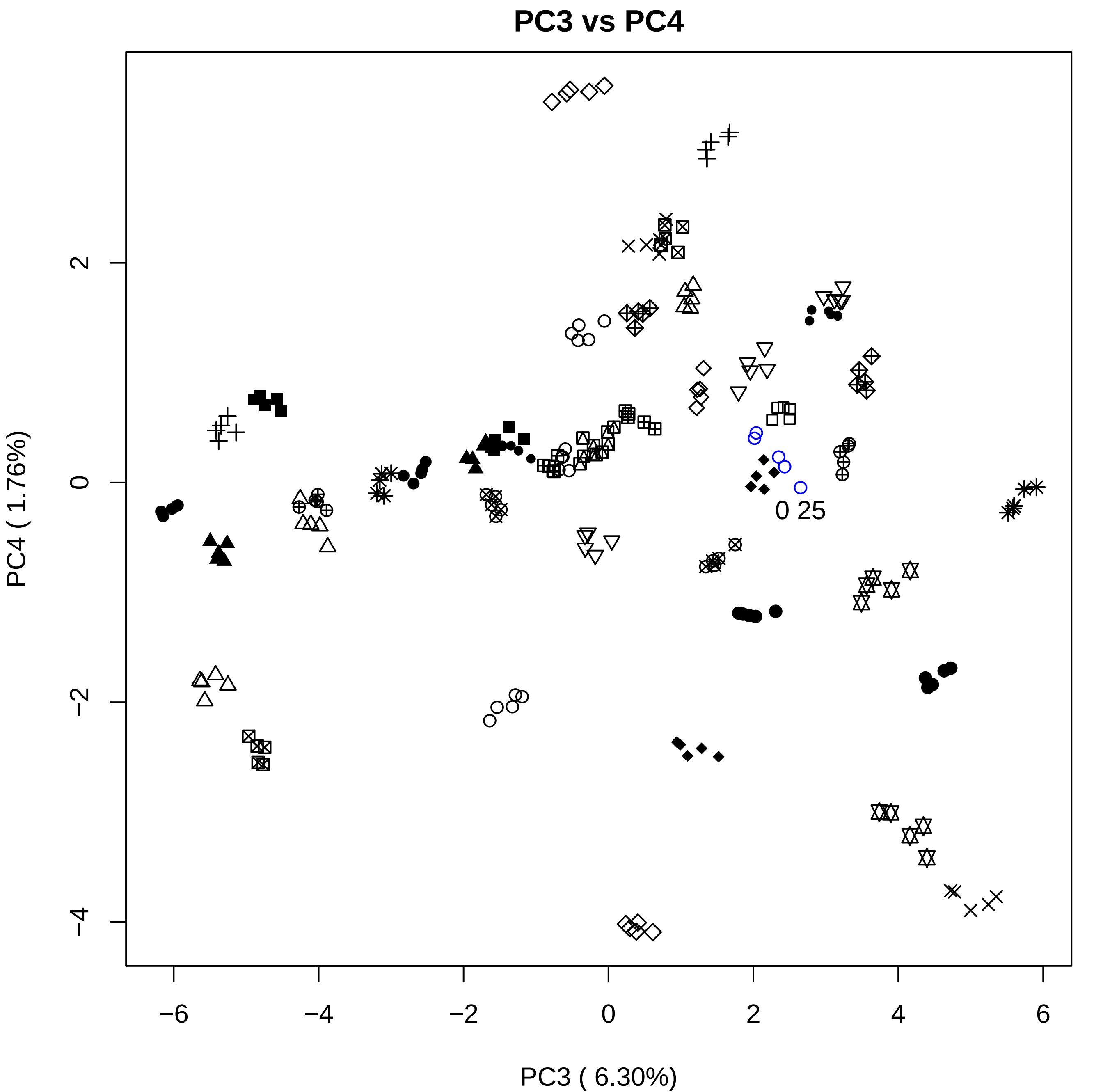
PC3 vs PC4
PC4 ( 1.76%)
PC3 ( 6.30%)
2
0
−2
−4
−6
−4
−2
0
2
4
6
0 25

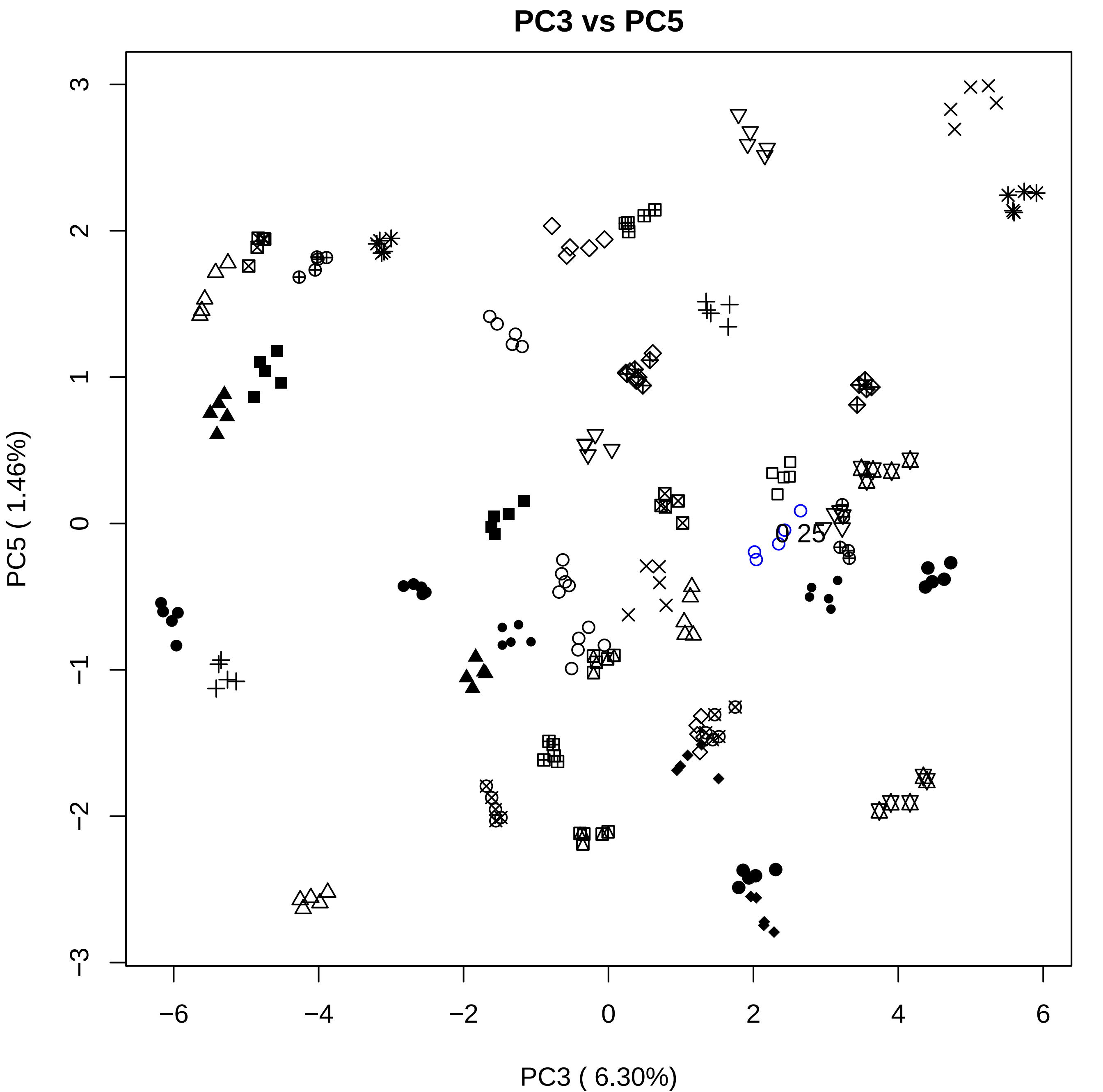

PC3 vs PC5
PC5 ( 1.46%)
PC3 ( 6.30%)
−6
−4
−2
0
2
4
6
−3
−2
−1
0
1
2
3
25

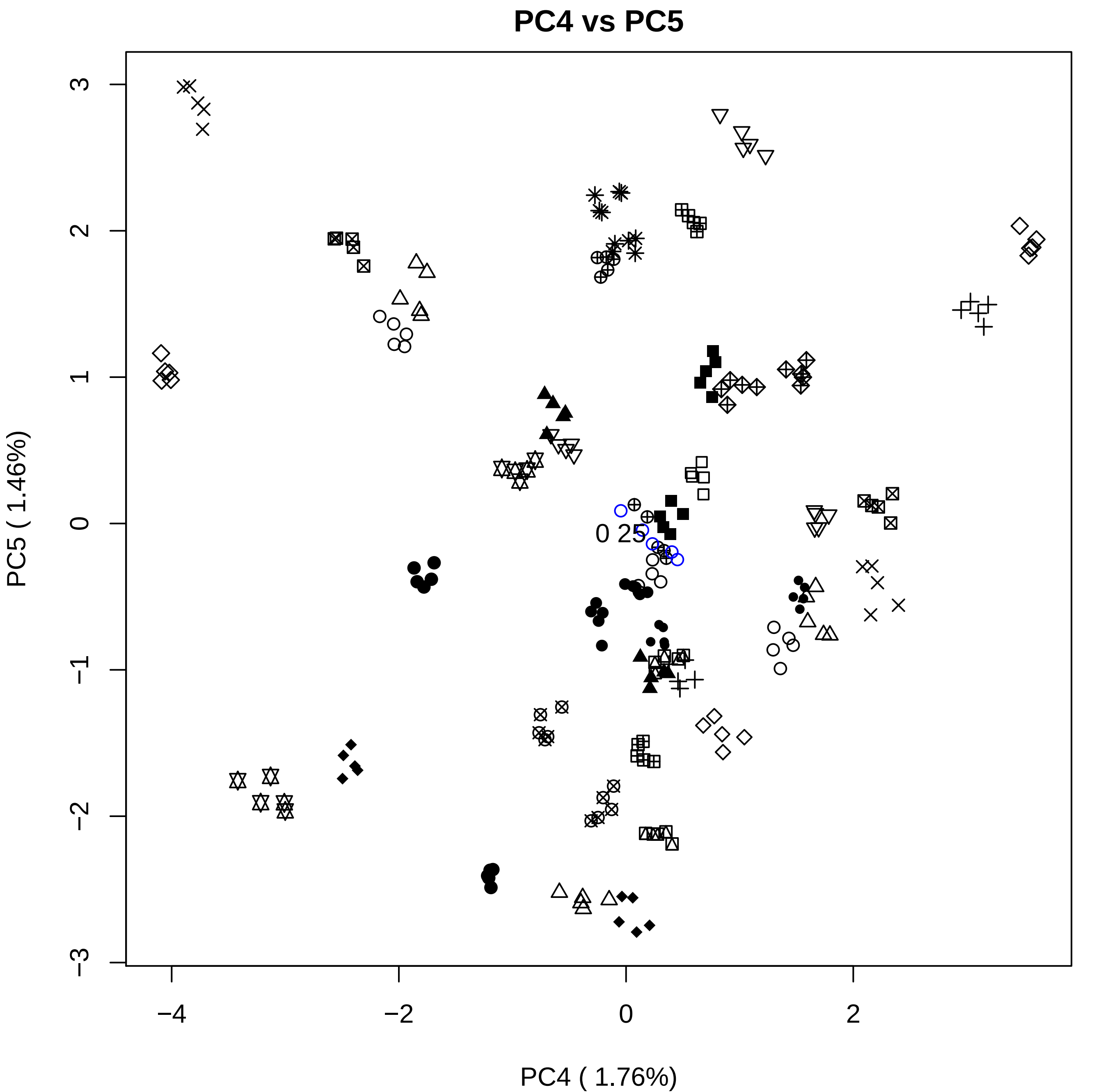
PC4 vs PC5
PC5 ( 1.46%)
PC4 ( 1.76%)
3
2
1
0
−1
−2
−3
−4
−2
0
2
0 25

### PCA of 0.5%  OSAS vs PMH

Scaled Matrix PCAed

7th order polynomial baseline correction performed

Raw spectra smoothed to 21 Neighbours, 2nd order PN, Savtisky Golay

Normalisation between 0 and 100

D2 spectra smoothed to 41 Neighbours, 2nd order PN, Savtisky Golay

Regions Analysed: 3601−2501 and 2001−700cm−1

Samples Analysed: 1 Contam's 0 OSAS's 45 PMH's

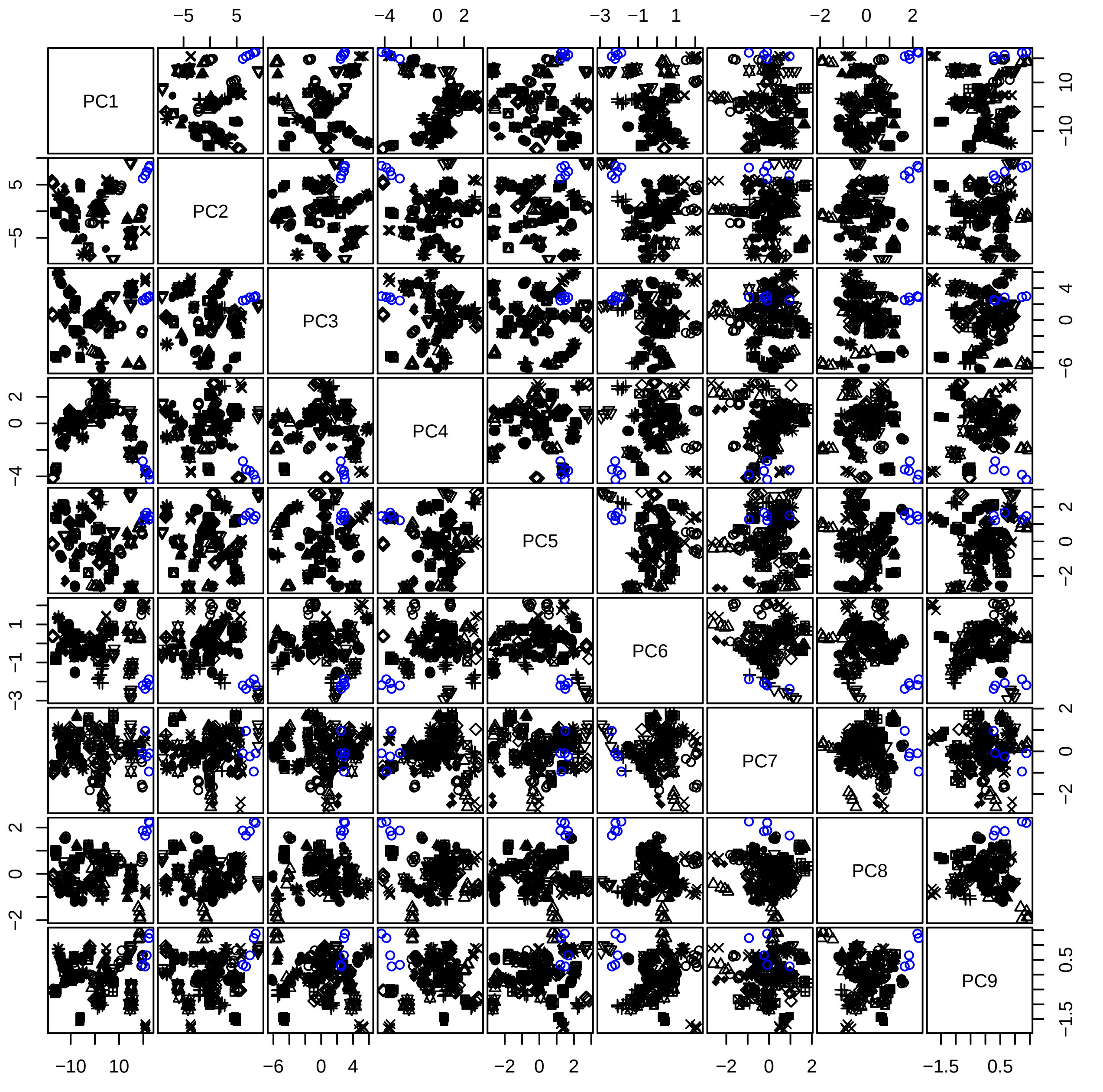
PC1
PC2
PC3
PC4
PC5
PC6
PC7
PC8
PC9

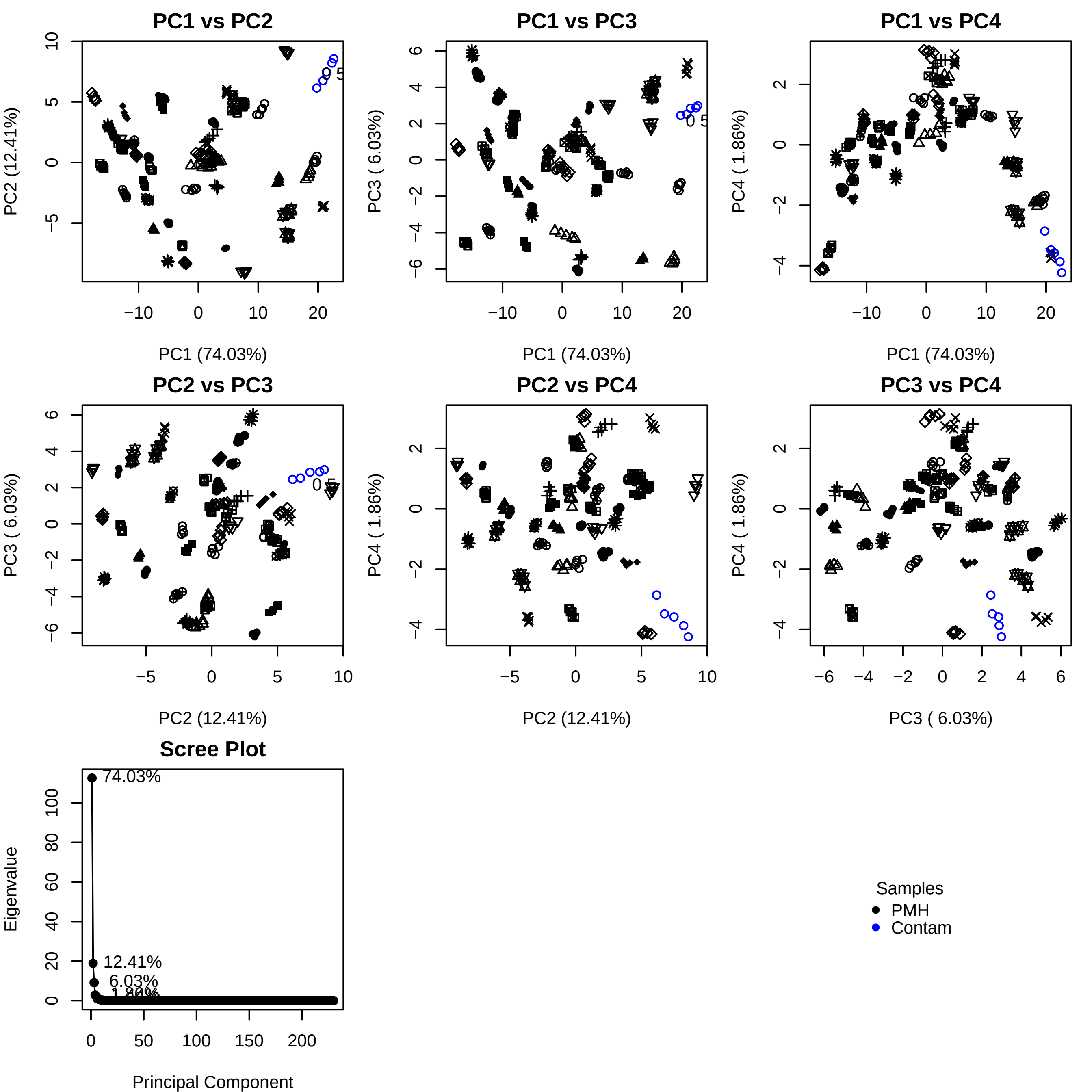
PC1 vs PC2
PC1 vs PC3
PC1 vs PC4
PC2 vs PC3
PC2 vs PC4
PC3 vs PC4
Scree Plot
PC1 (74.03%)
PC2 (12.41%)
PC3 ( 6.03%)
PC4 ( 1.86%)
Eigenvalue
Principal Component
74.03%
12.41%
6.03%
Samples
PMH
Contam

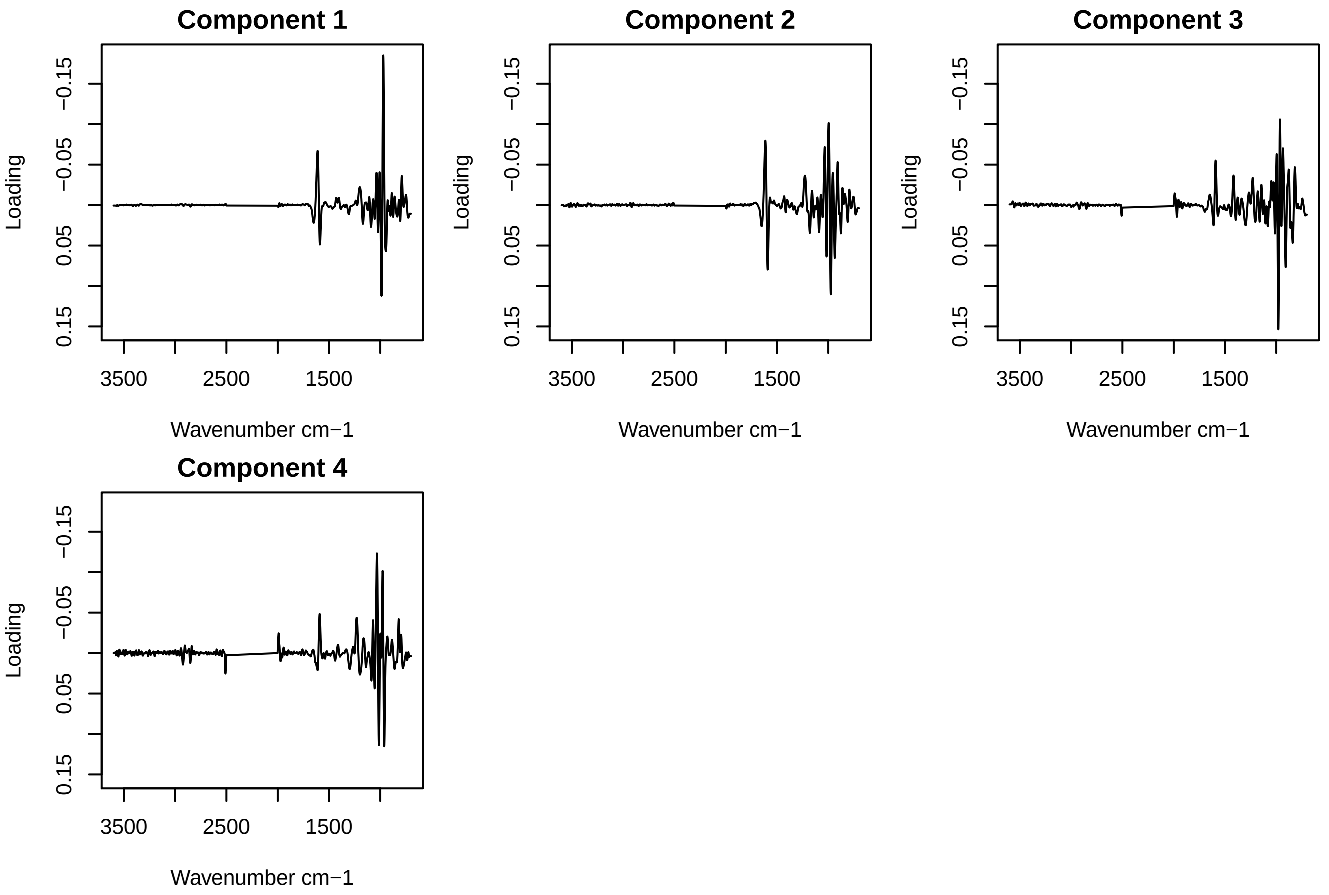

Component 1
Component 2
Component 3
Component 4
Loading
−0.15
−0.05
0.05
0.15
3500
2500
1500
Wavenumber cm−1

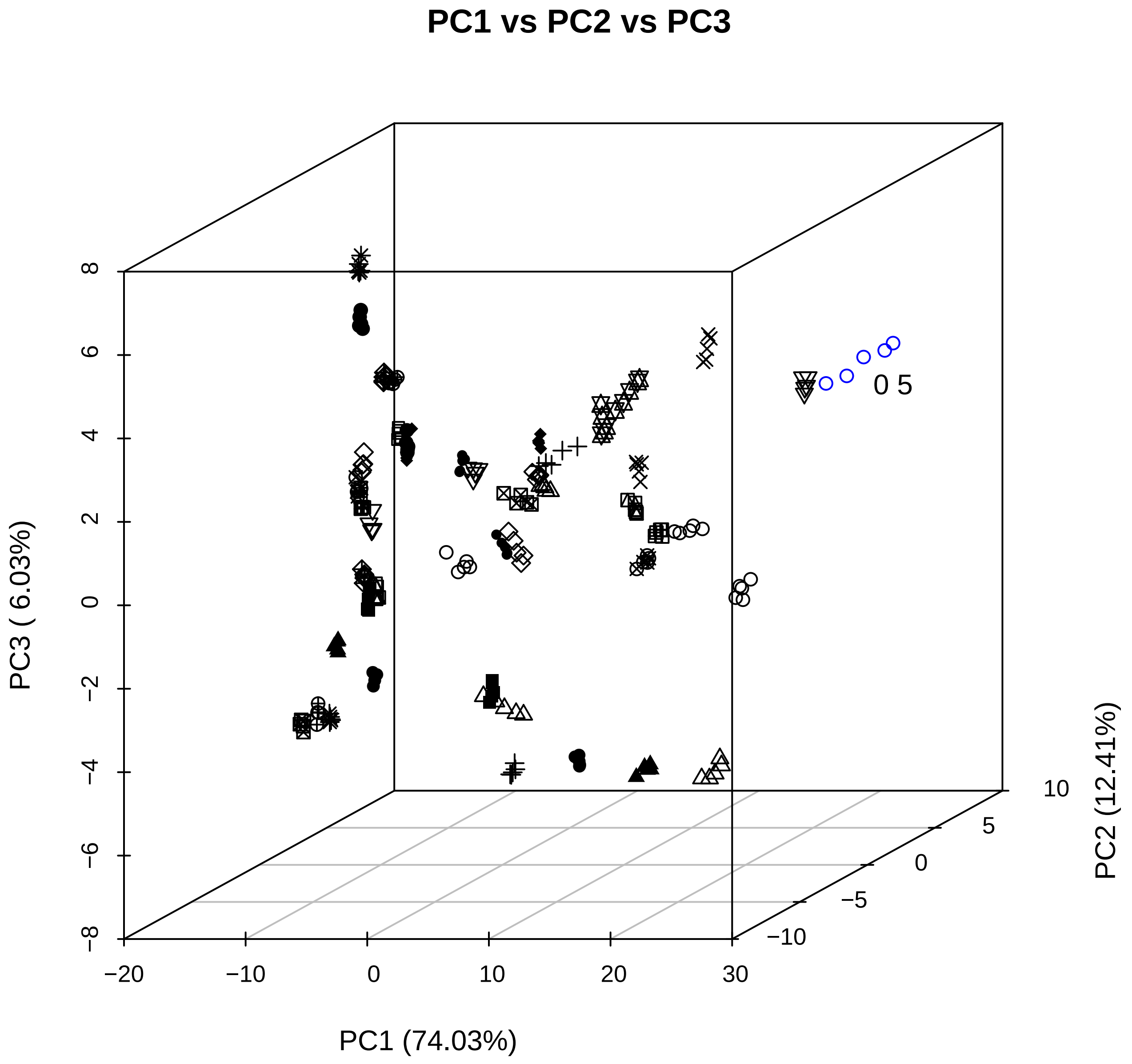
PC1 vs PC2 vs PC3
PC3 ( 6.03%)
PC1 (74.03%)
PC2 (12.41%)
0 5
8
6
4
2
0
−2
−4
−6
−8
−20
−10
0
10
20
30
10
5
0
−5
−10

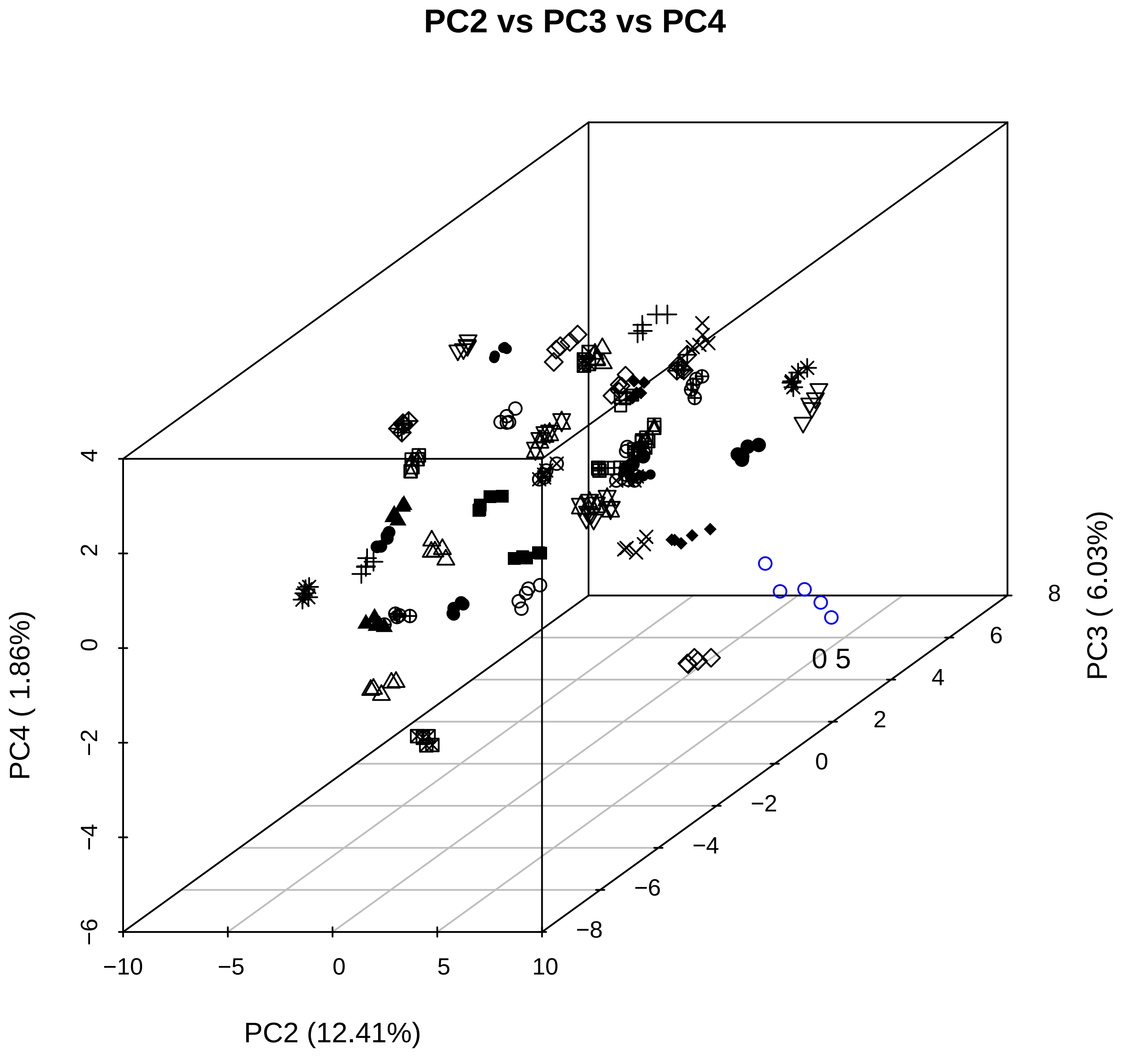
PC2 vs PC3 vs PC4
PC4 ( 1.86%)
4
2
0
−2
−4
−6
−10
−5
0
5
10
PC2 (12.41%)
PC3 ( 6.03%)
8
6
4
2
0
−2
−4
−6
−8
0 5

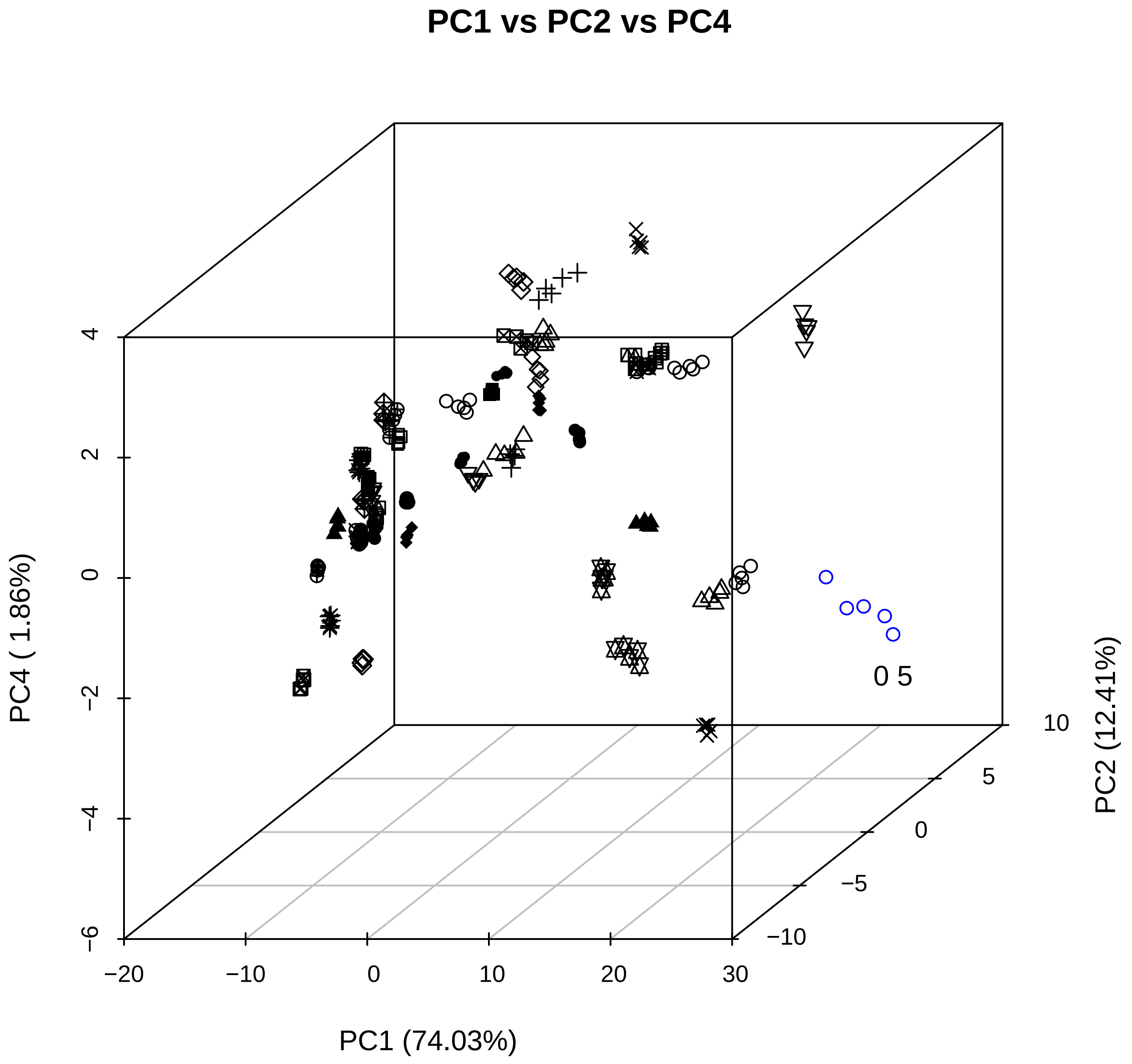

PC1 vs PC2 vs PC4
PC1 (74.03%)
PC2 (12.41%)
PC4 ( 1.86%)
−20
−10
0
10
20
30
−10
−5
0
5
10
−6
−4
−2
0
2
4
0 5

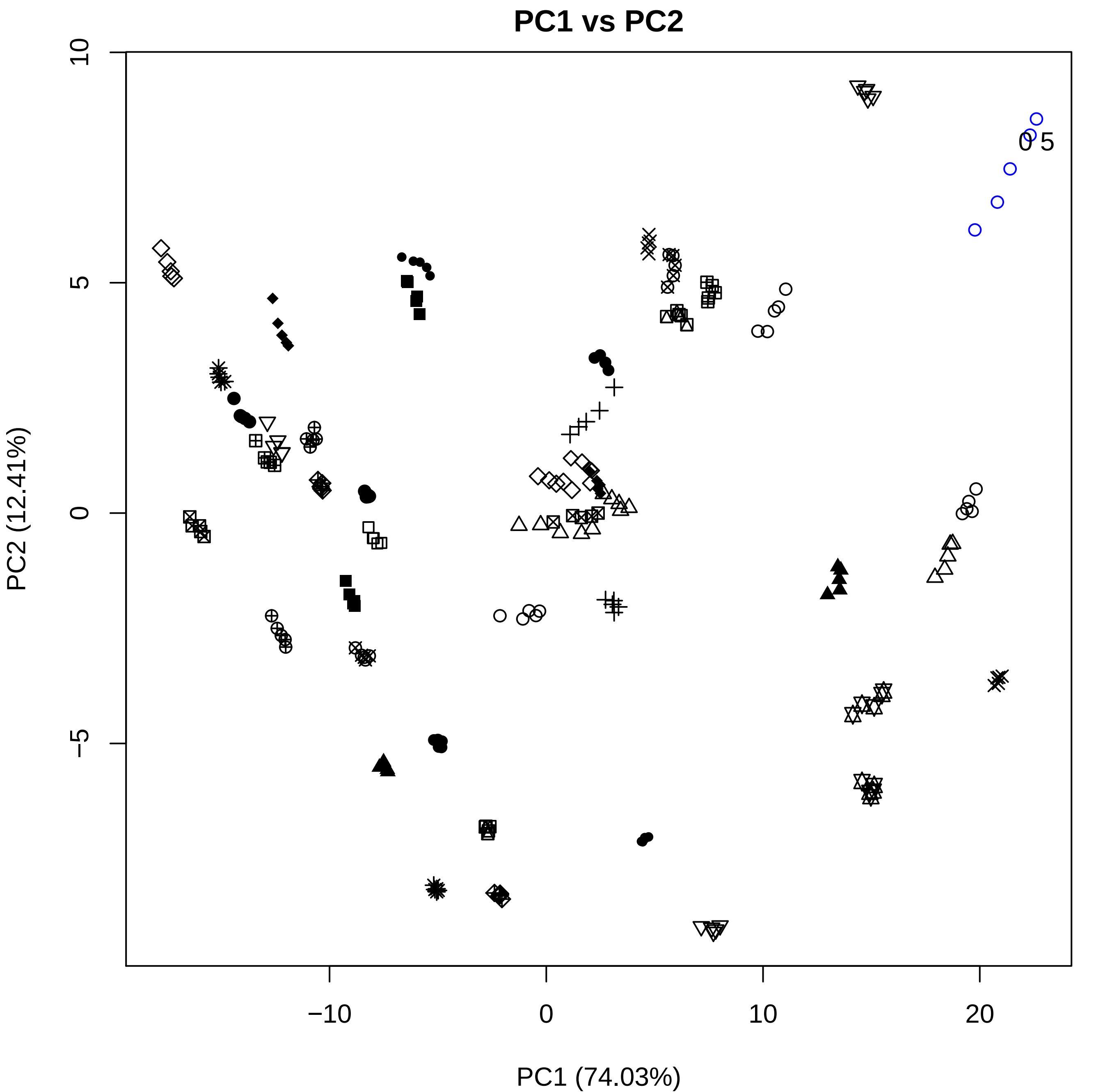
PC1 vs PC2
PC2 (12.41%)
PC1 (74.03%)
10
5
0
−5
−10
0
10
20
0 5

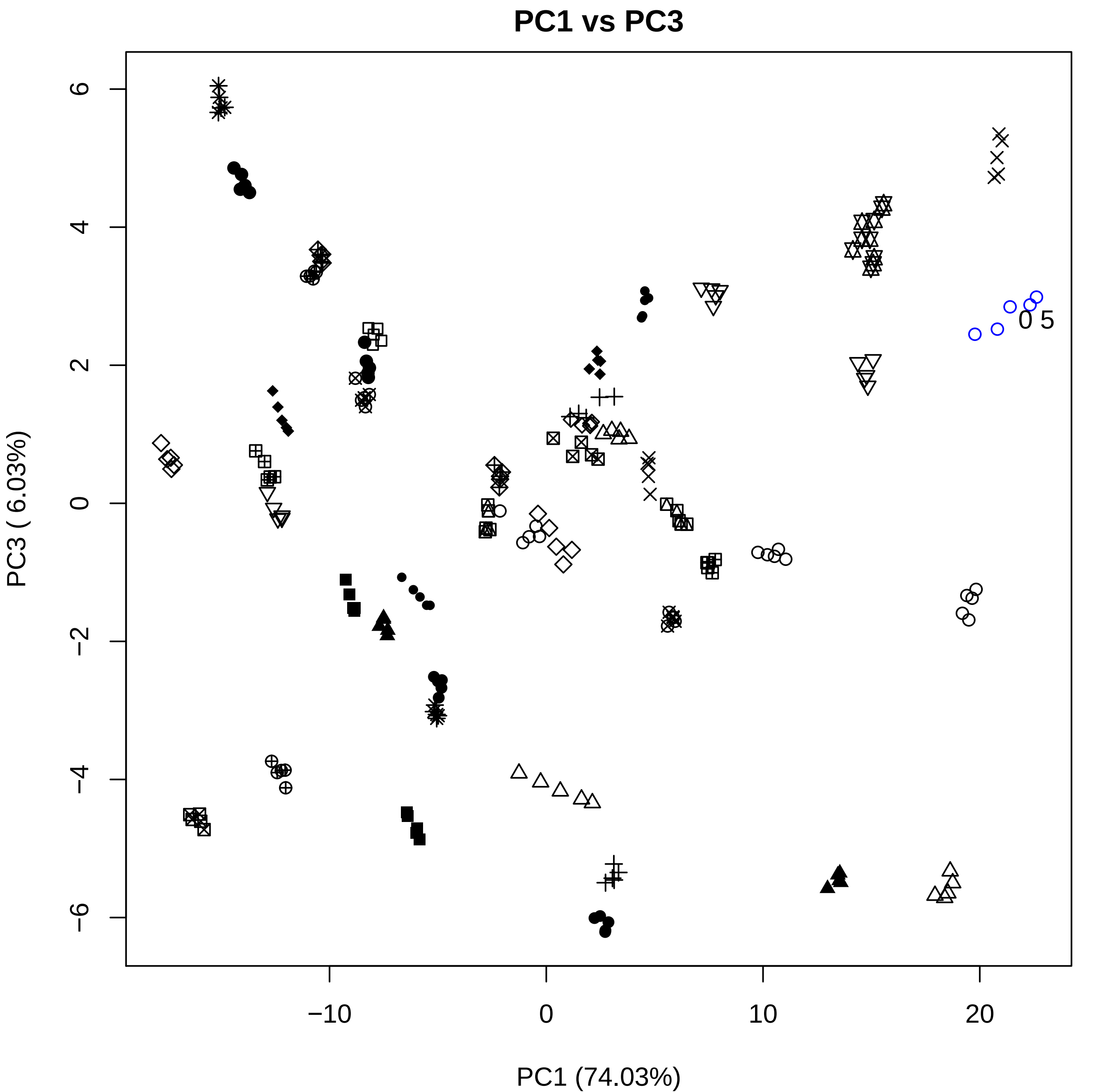
PC1 vs PC3
PC3 ( 6.03%)
PC1 (74.03%)
6
4
2
0
−2
−4
−6
−10
0
10
20
0 5

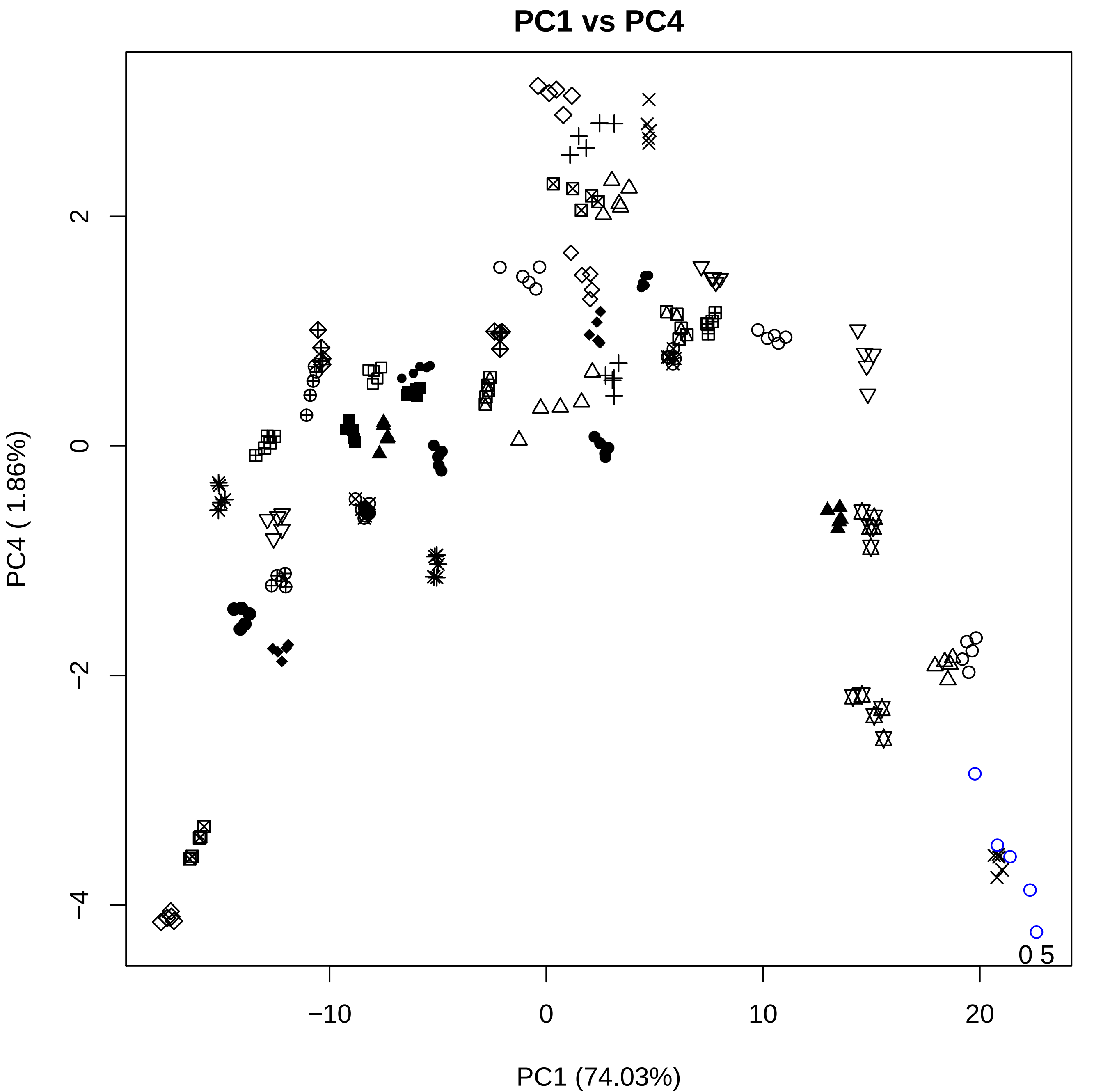
PC1 vs PC4
PC4 ( 1.86%)
PC1 (74.03%)
2
0
−2
−4
−10
0
10
20
0 5

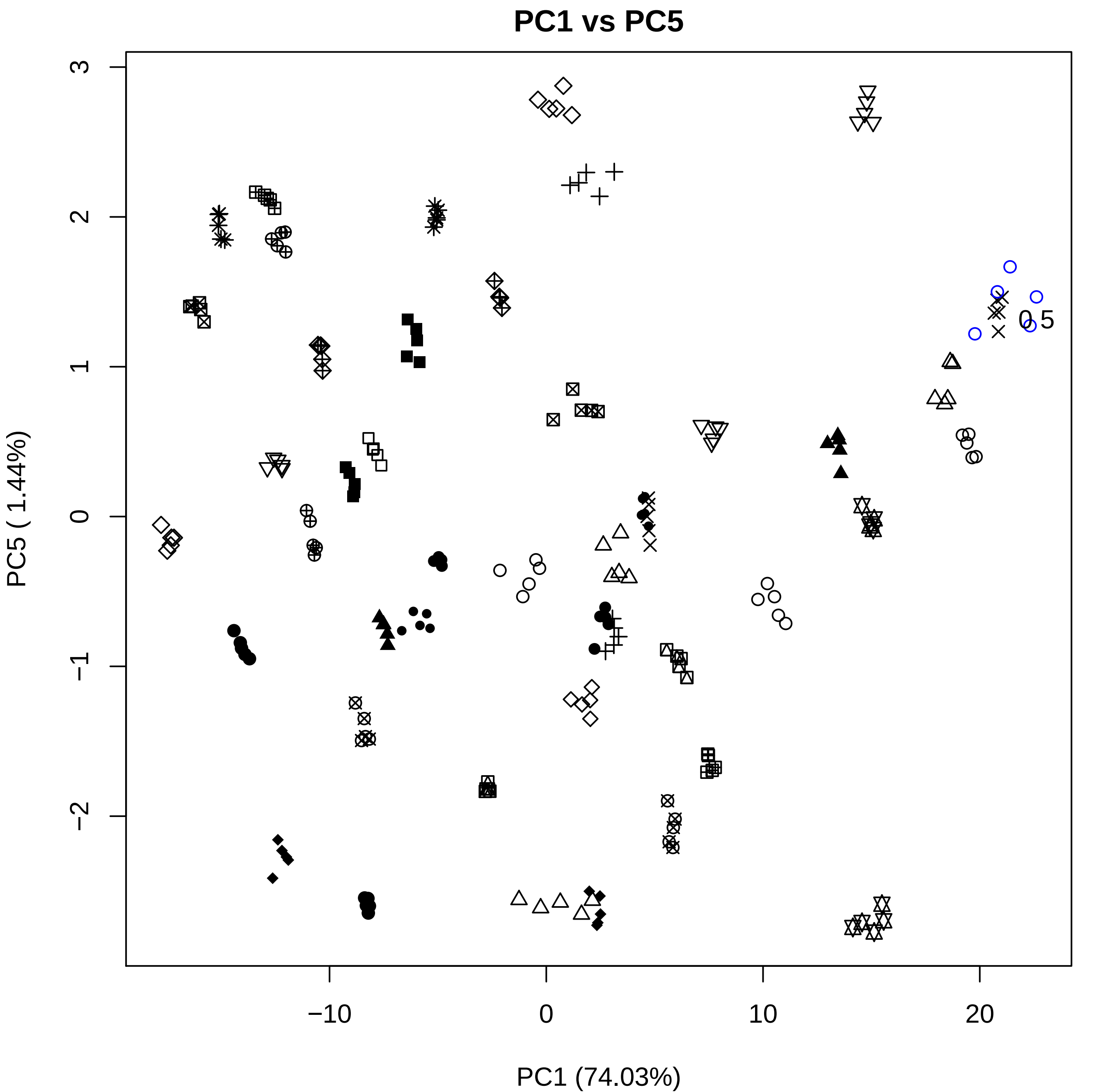
PC1 vs PC5
PC5 ( 1.44%)
PC1 (74.03%)
3
2
1
0
−1
−2
−10
0
10
20
0.5

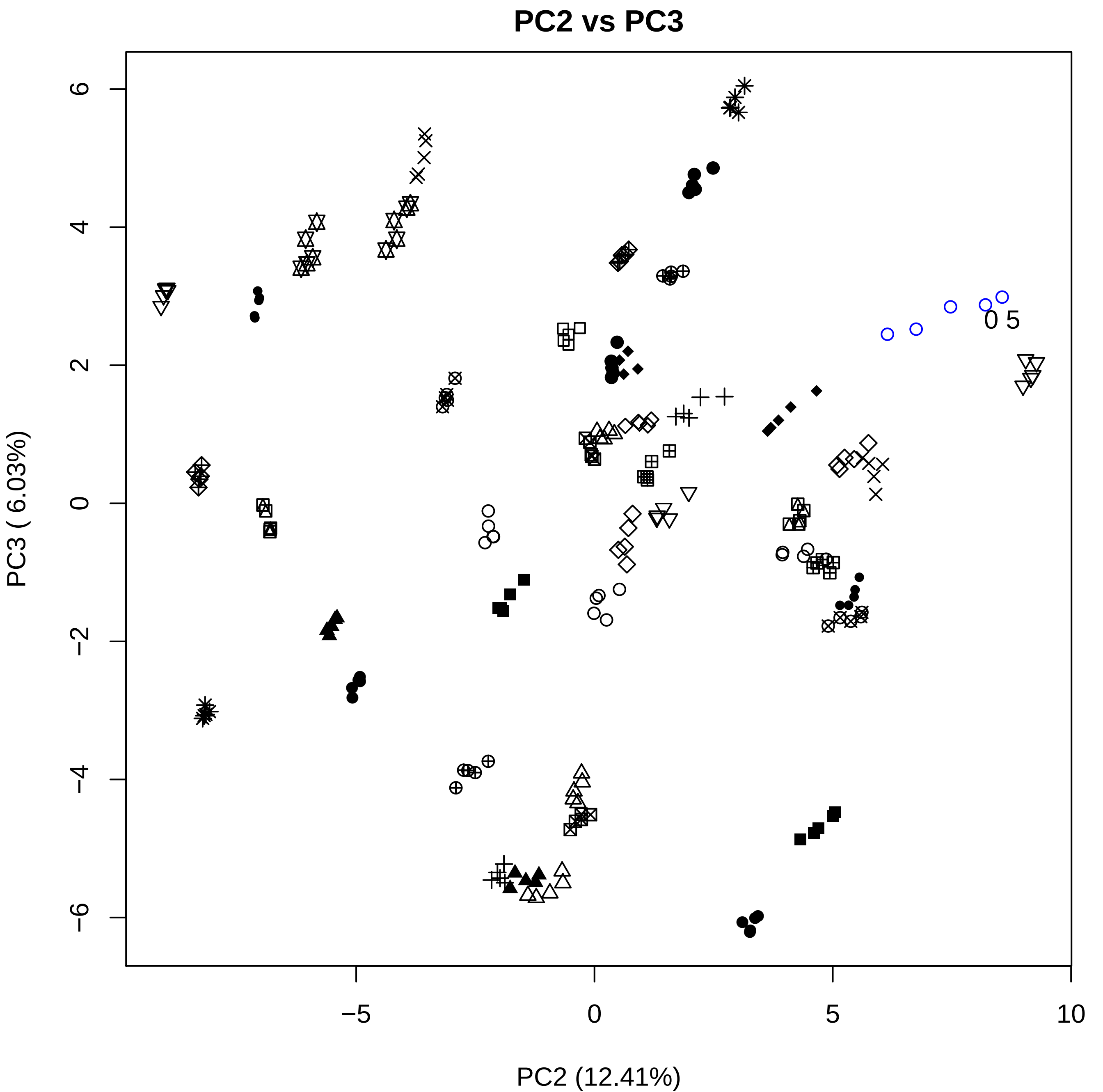
PC2 vs PC3
PC3 ( 6.03%)
PC2 (12.41%)
6
4
2
0
−2
−4
−6
−5
0
5
10
0 5

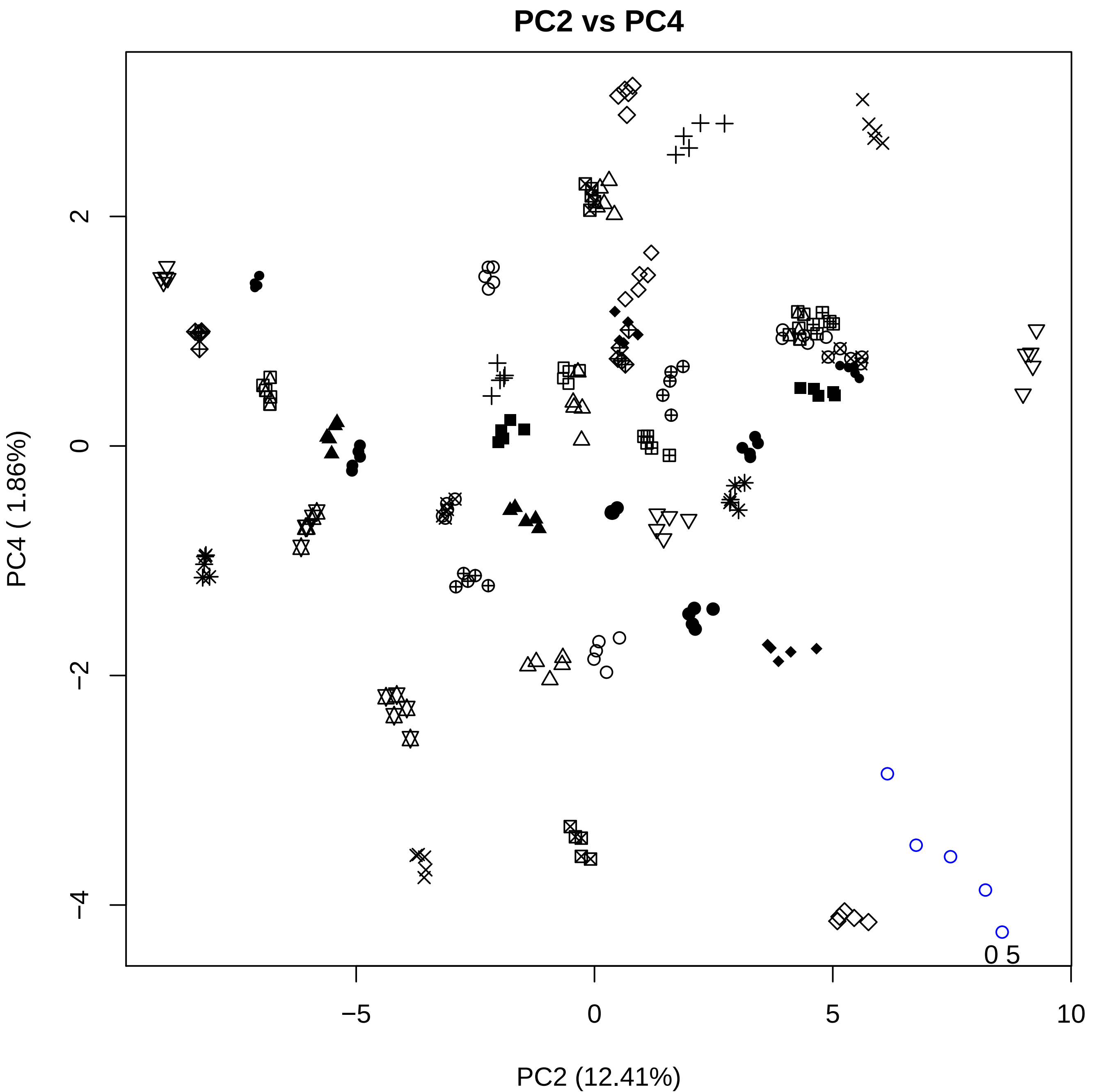
PC2 vs PC4
PC4 ( 1.86%)
PC2 (12.41%)
2
0
−2
−4
−5
0
5
10
0 5

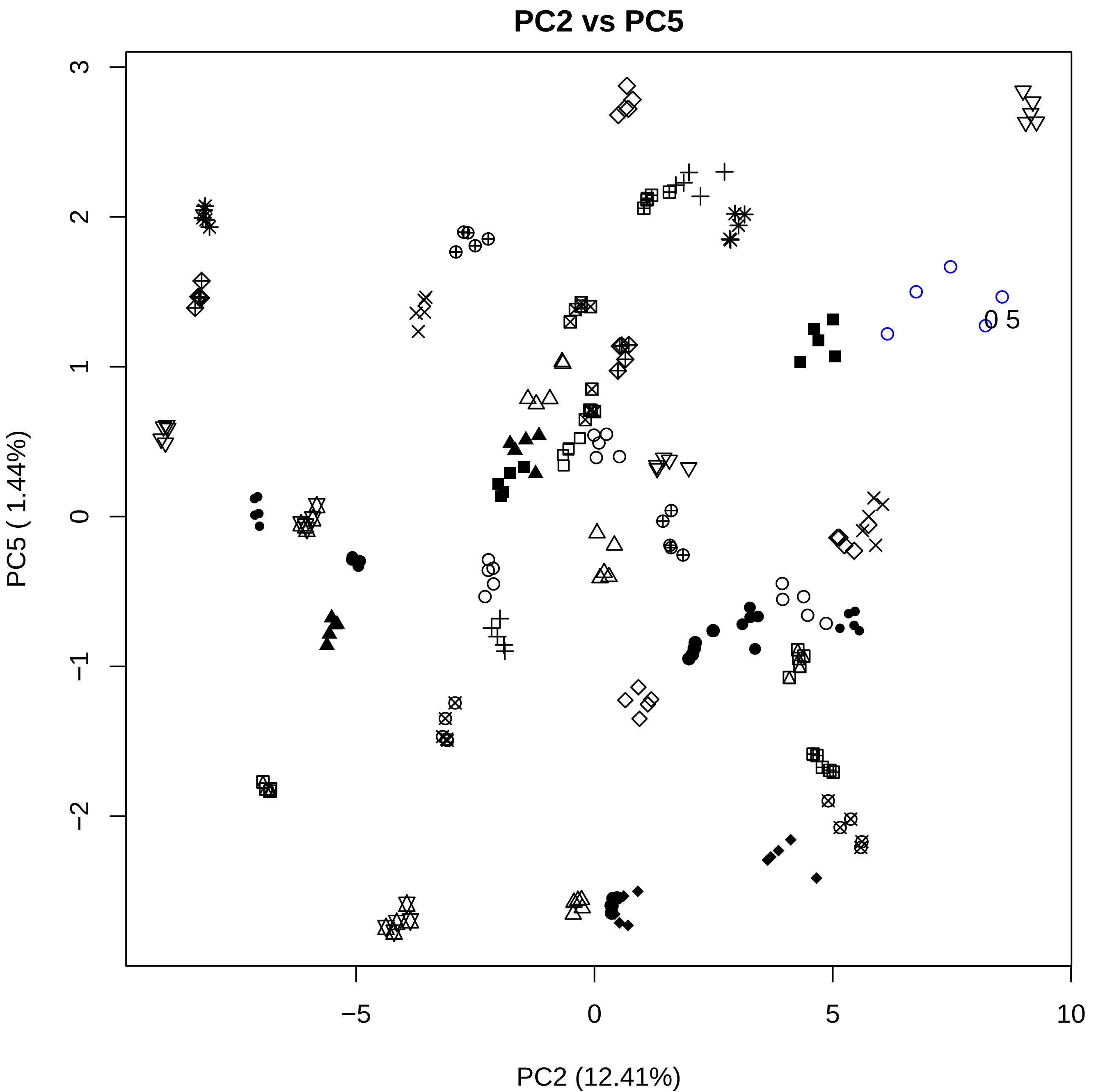
PC2 vs PC5
PC5 ( 1.44%)
PC2 (12.41%)
3
2
1
0
−1
−2
−5
0
5
10
0 5

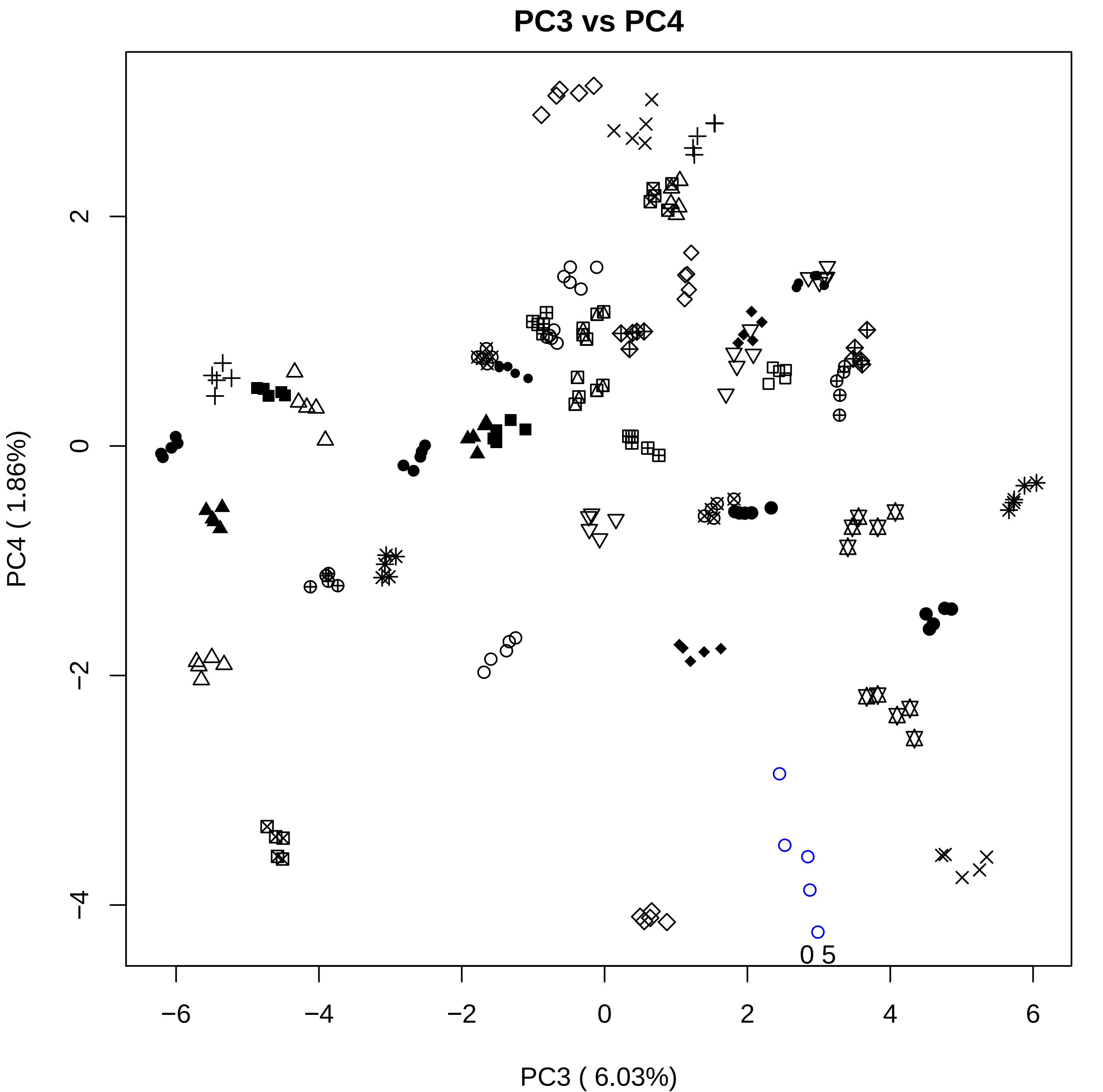
PC3 vs PC4
PC4 ( 1.86%)
PC3 ( 6.03%)
−6
−4
−2
0
2
4
6
2
0
−2
−4
0 5

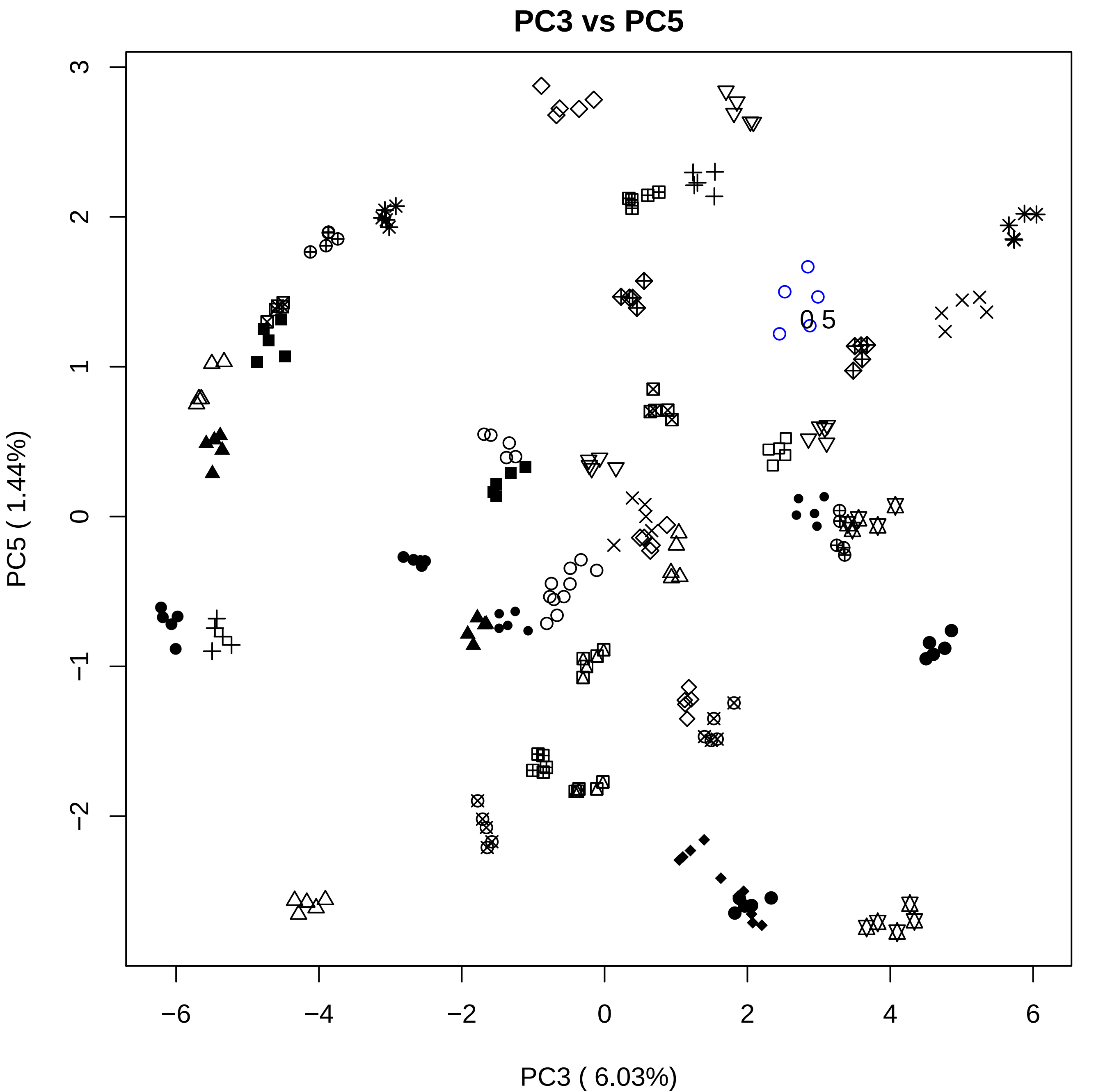
PC3 vs PC5
PC5 ( 1.44%)
PC3 ( 6.03%)
3
2
1
0
−1
−2
−6
−4
−2
0
2
4
6
0 5

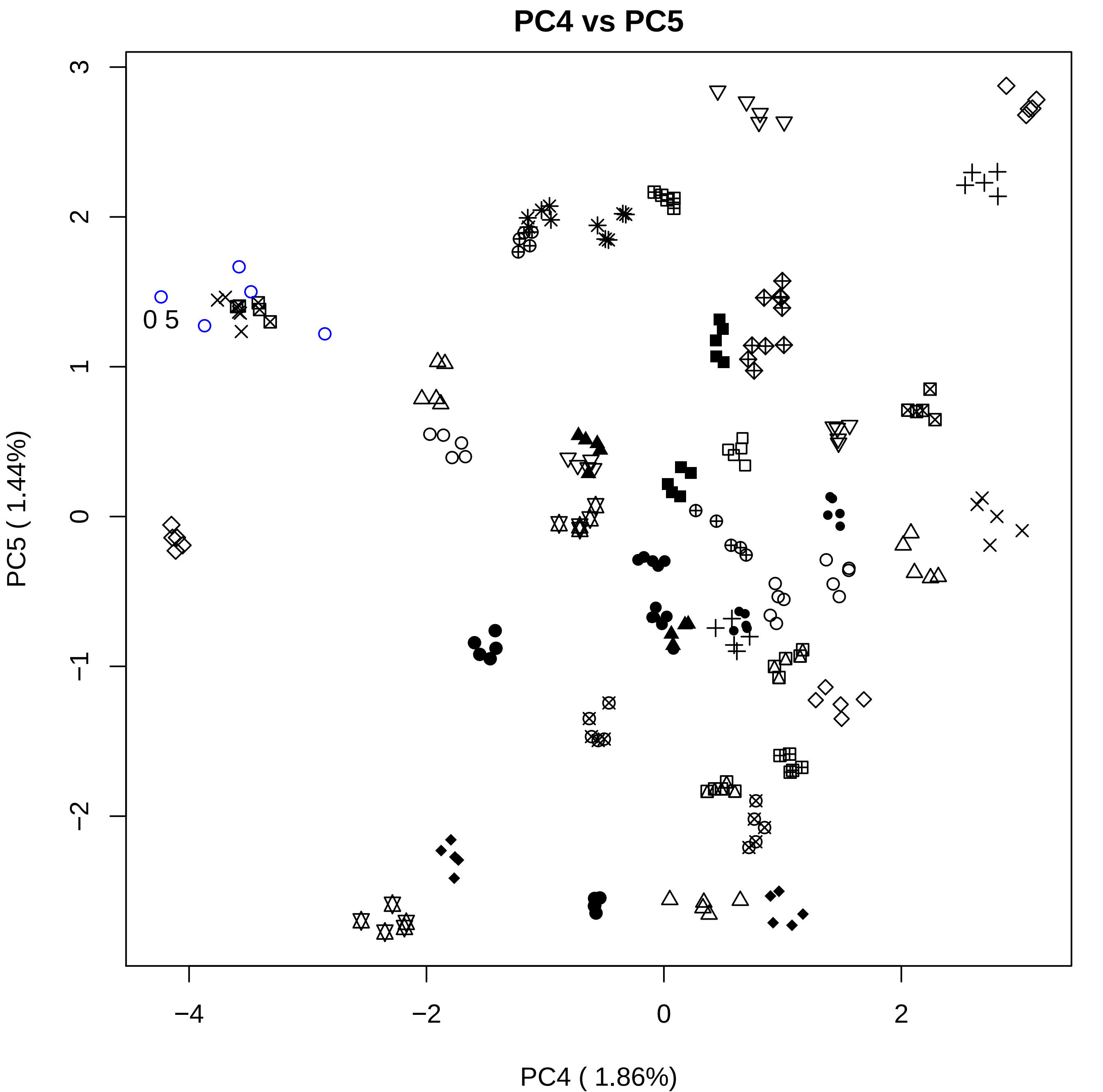
PC4 vs PC5
PC5 ( 1.44%)
PC4 ( 1.86%)
3
2
1
0
−1
−2
−4
−2
0
2
0 5

### PCA of 1%  OSAS vs PMH

Scaled Matrix PCAed

7th order polynomial baseline correction performed

Raw spectra smoothed to 21 Neighbours, 2nd order PN, Savtisky Golay

Normalisation between 0 and 100

D2 spectra smoothed to 41 Neighbours, 2nd order PN, Savtisky Golay

Regions Analysed: 3601−2501 and 2001−700cm−1

Samples Analysed: 1 Contam's 0 OSAS's 45 PMH's

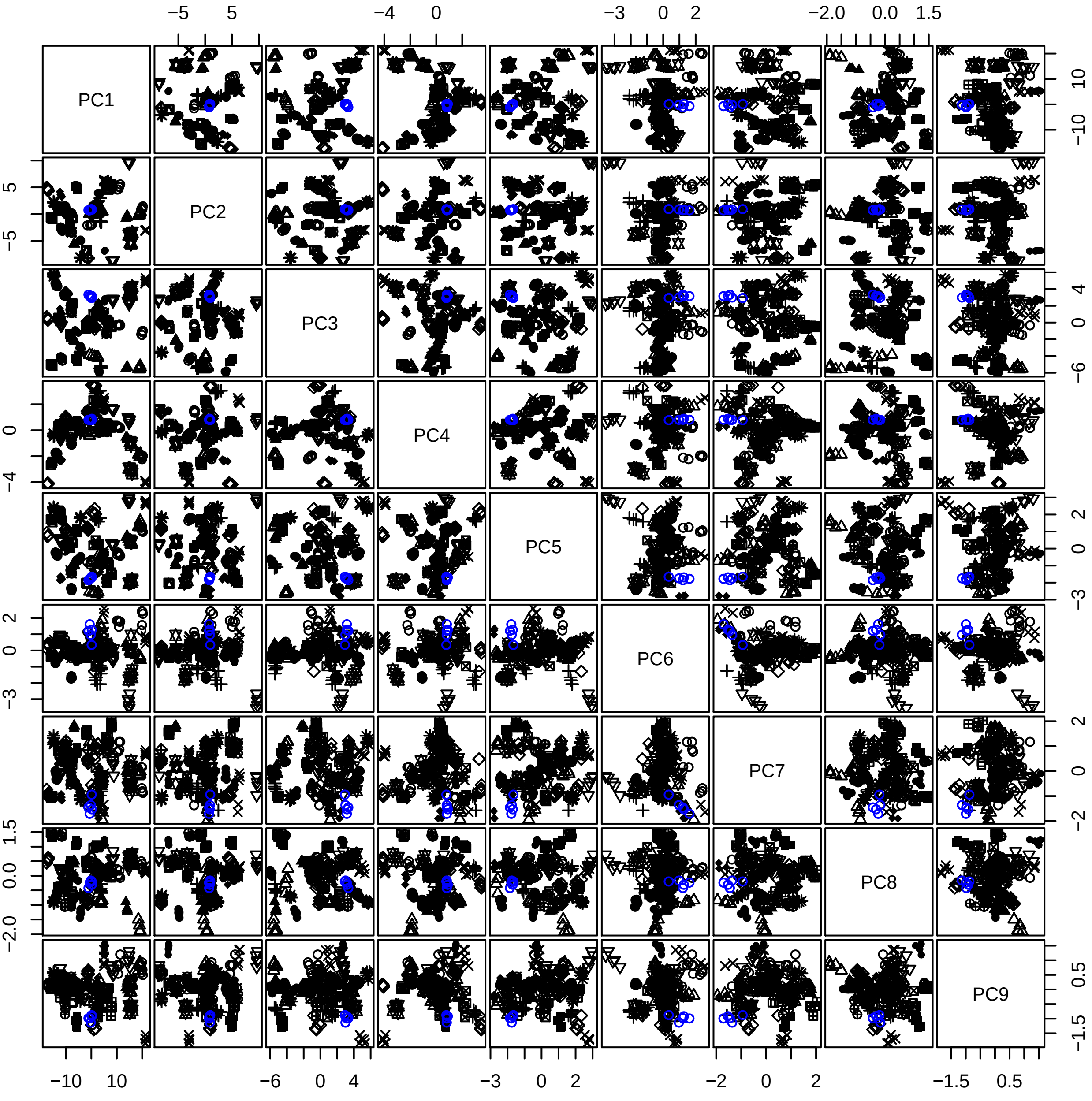
PC1
PC2
PC3
PC4
PC5
PC6
PC7
PC8
PC9

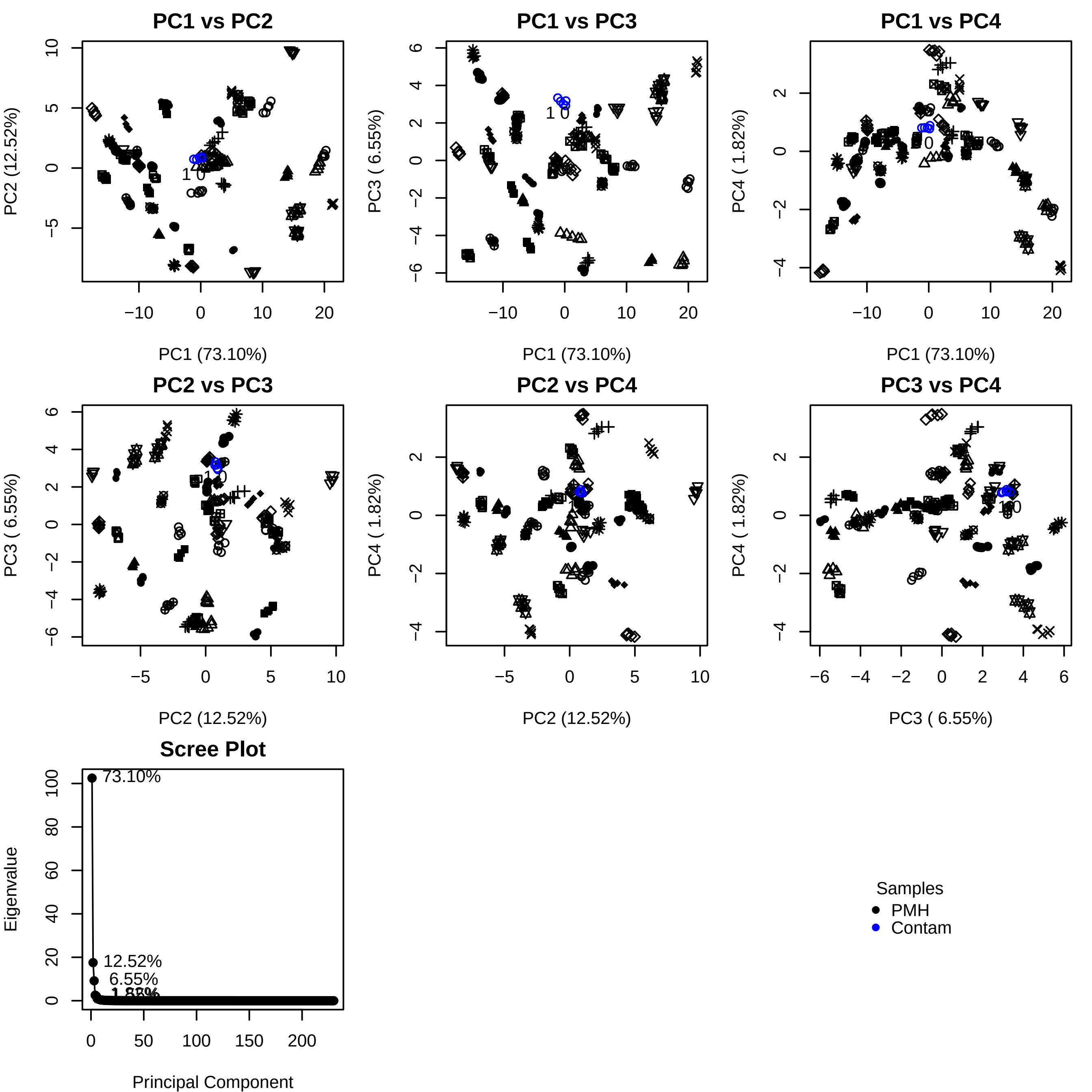
PC1 vs PC2
PC1 vs PC3
PC1 vs PC4
PC1 (73.10%)
PC2 (12.52%)
PC3 ( 6.55%)
PC4 ( 1.82%)
PC2 vs PC3
PC2 vs PC4
PC3 vs PC4
Scree Plot
Eigenvalue
Principal Component
73.10%
12.52%
6.55%
Samples
PMH
Contam

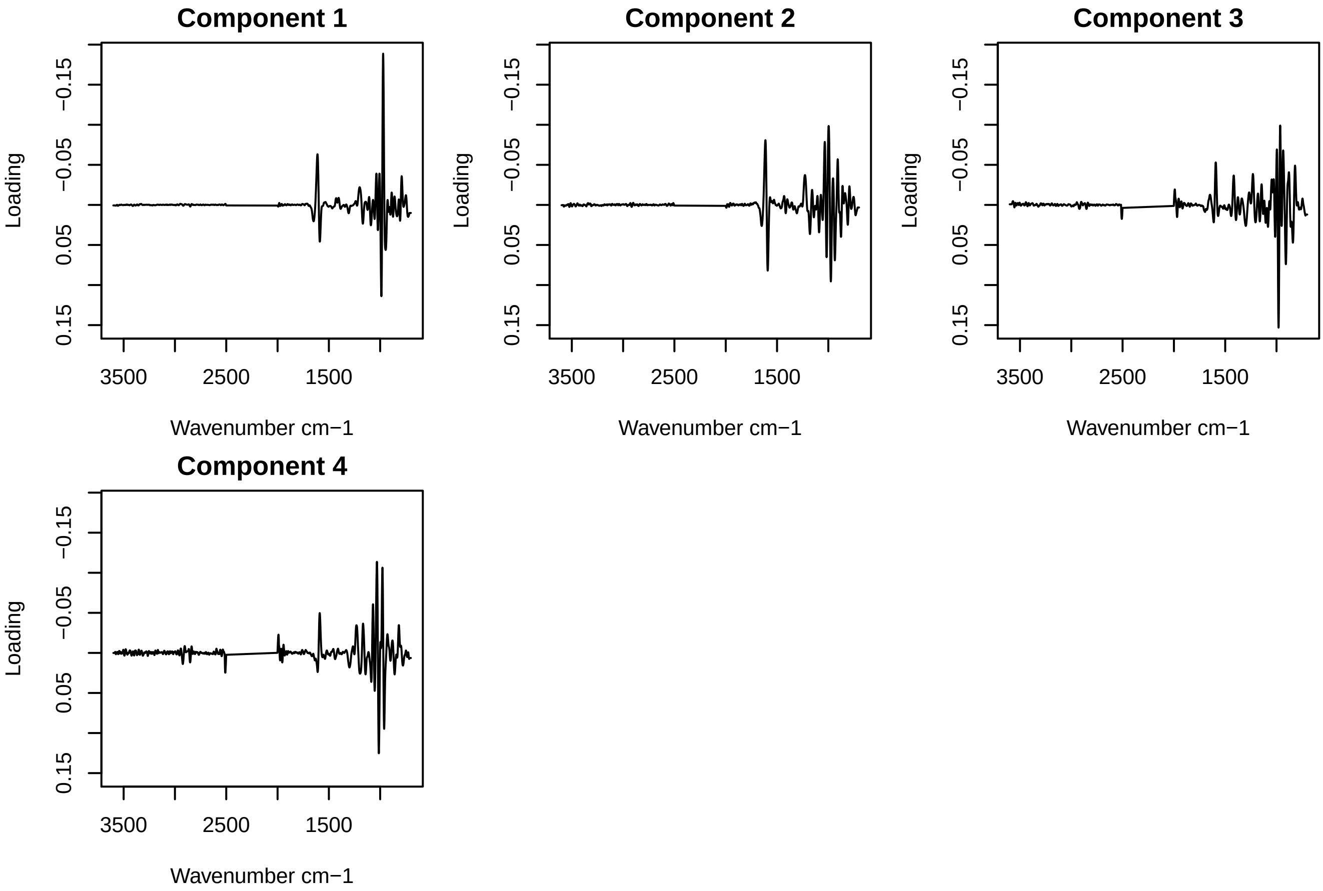
Component 1
Component 2
Component 3
Component 4
Loading
−0.15
−0.05
0.05
0.15
3500
2500
1500
Wavenumber cm−1

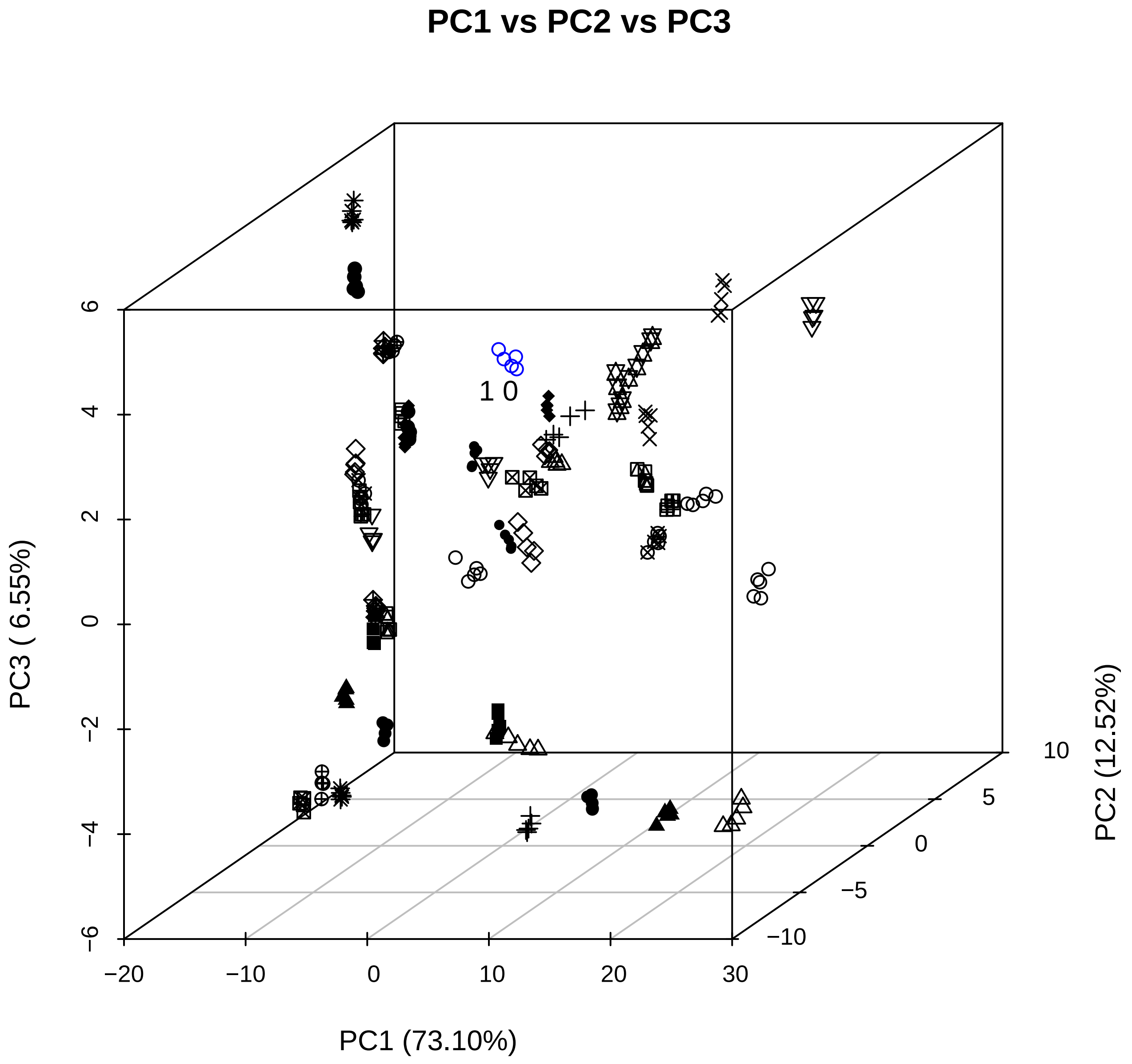

PC1 vs PC2 vs PC3
PC1 (73.10%)
PC2 (12.52%)
PC3 ( 6.55%)
−20
−10
0
10
20
30
−10
−5
0
5
10
−6
−4
−2
0
2
4
6
1 0

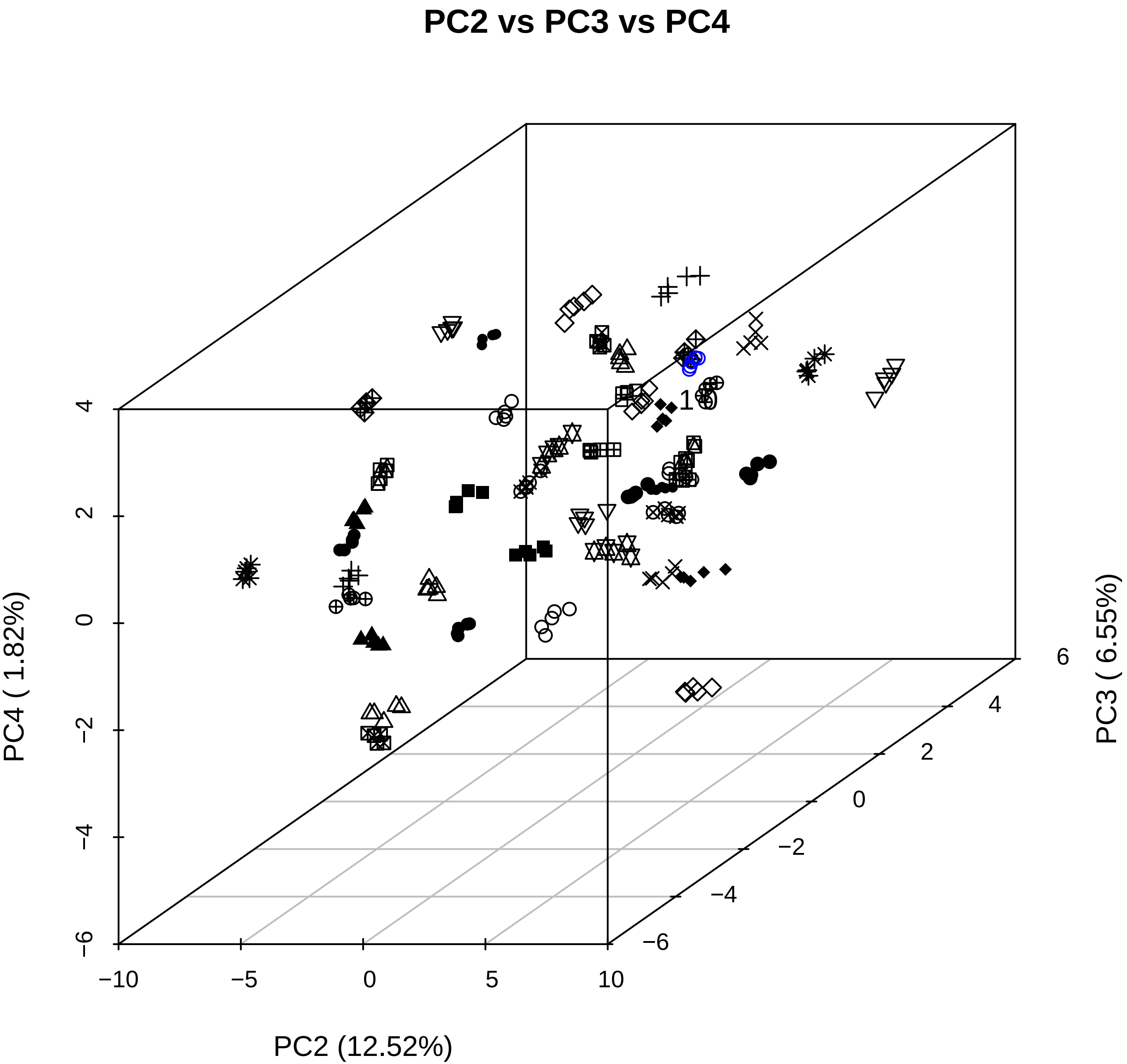
PC2 vs PC3 vs PC4
PC2 (12.52%)
PC3 ( 6.55%)
PC4 ( 1.82%)
−10
−5
0
5
10
−6
−4
−2
0
2
4
6
16

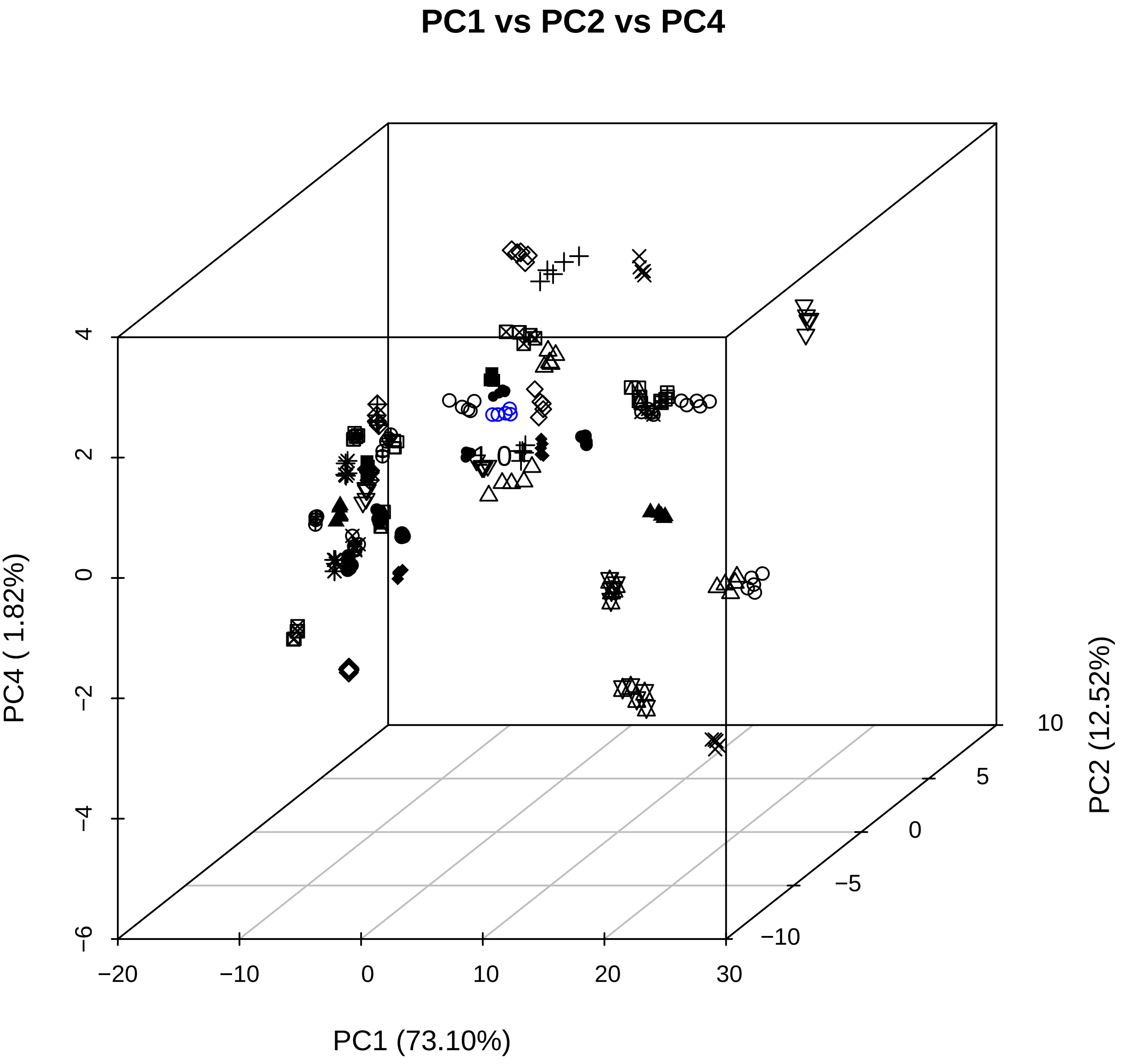

PC1 vs PC2 vs PC4
PC4 ( 1.82%)
PC1 (73.10%)
PC2 (12.52%)
4
2
0
−2
−4
−6
−20
−10
0
10
20
30
10
5
0
−5
−10

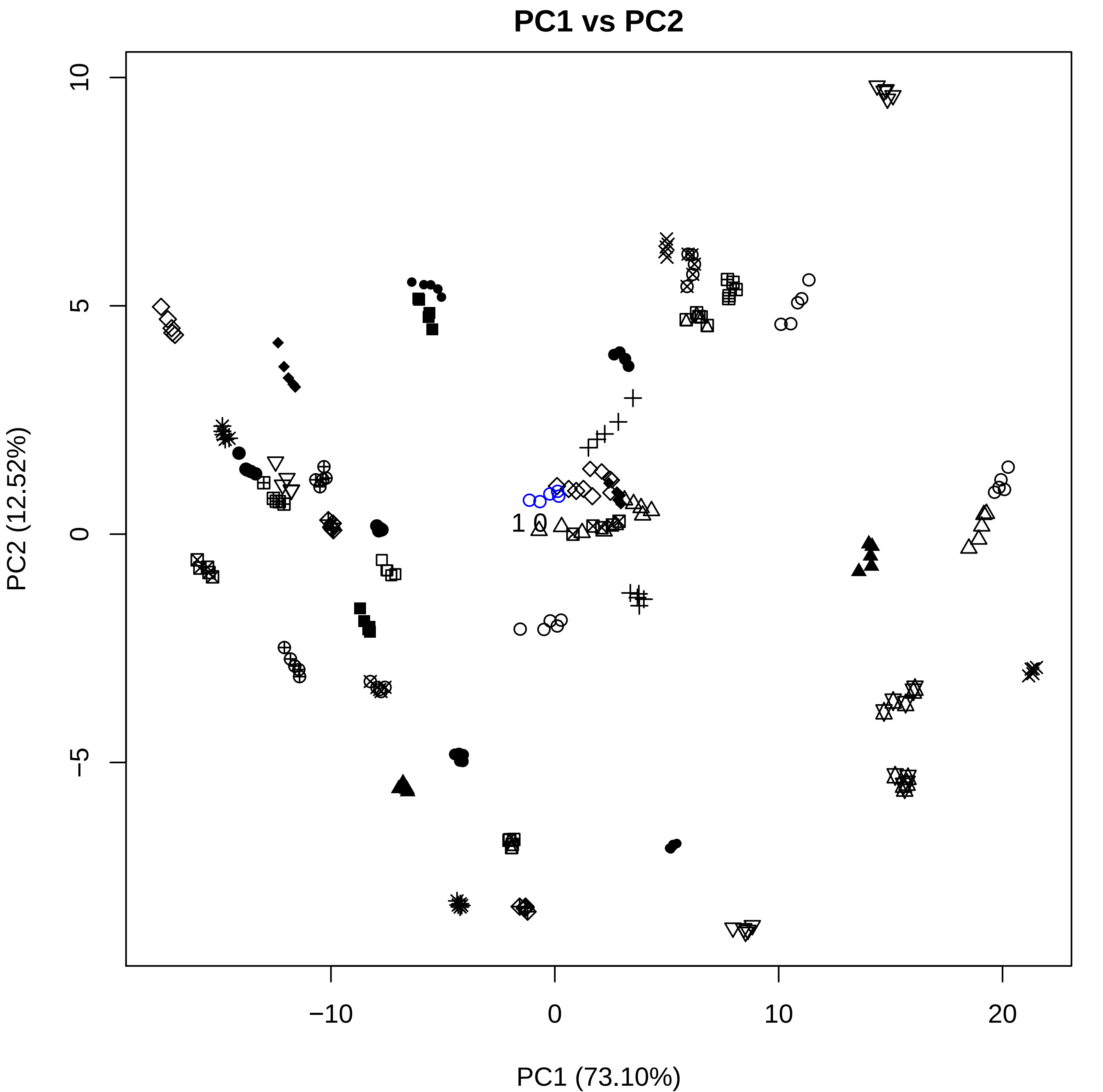
PC1 vs PC2
PC1 (73.10%)
PC2 (12.52%)
−10
0
10
20
−5
0
5
10
1 0

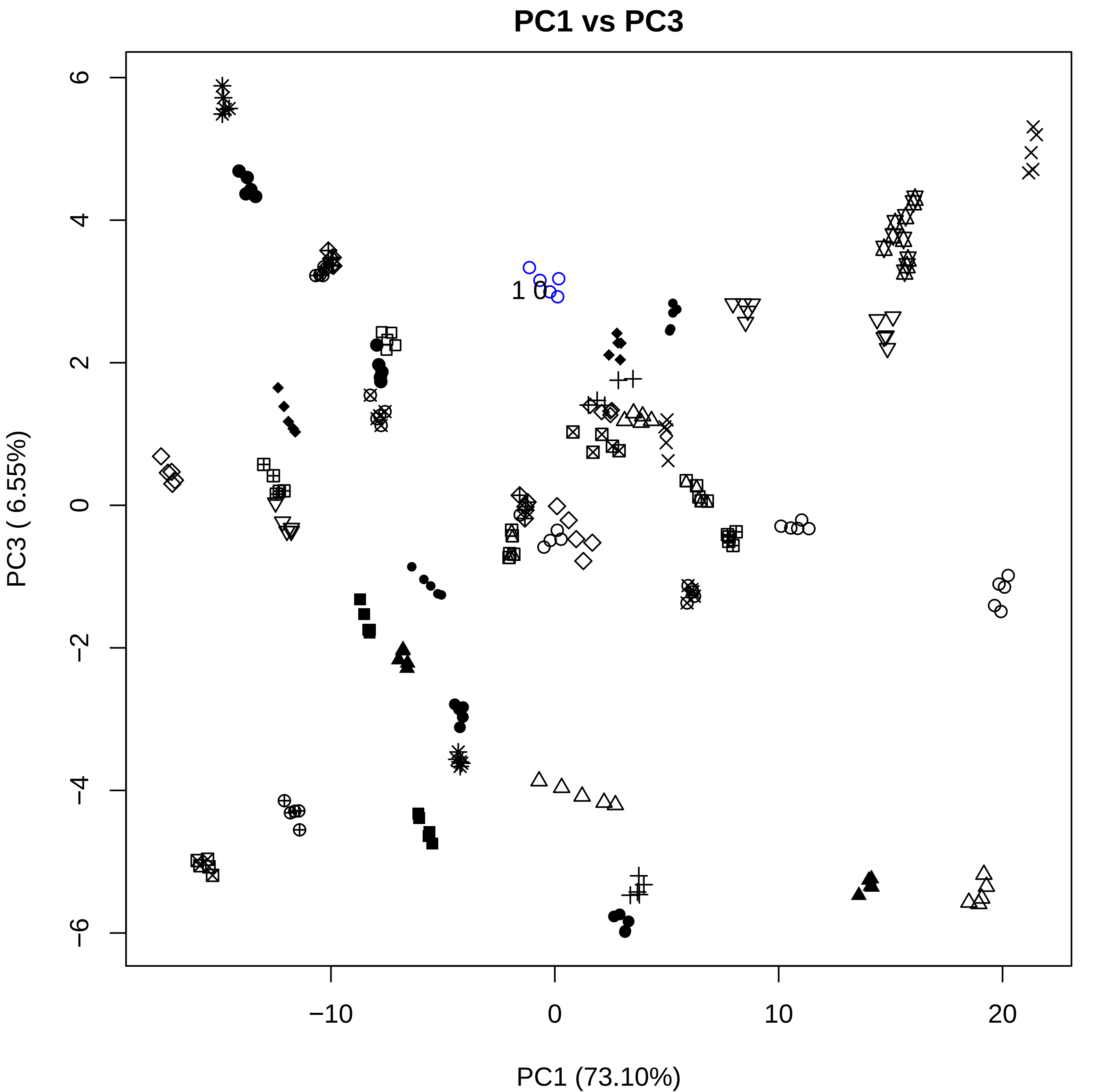

PC1 vs PC3
PC3 ( 6.55%)
PC1 (73.10%)
6
4
2
0
−2
−4
−6
−10
0
10
20
1 0

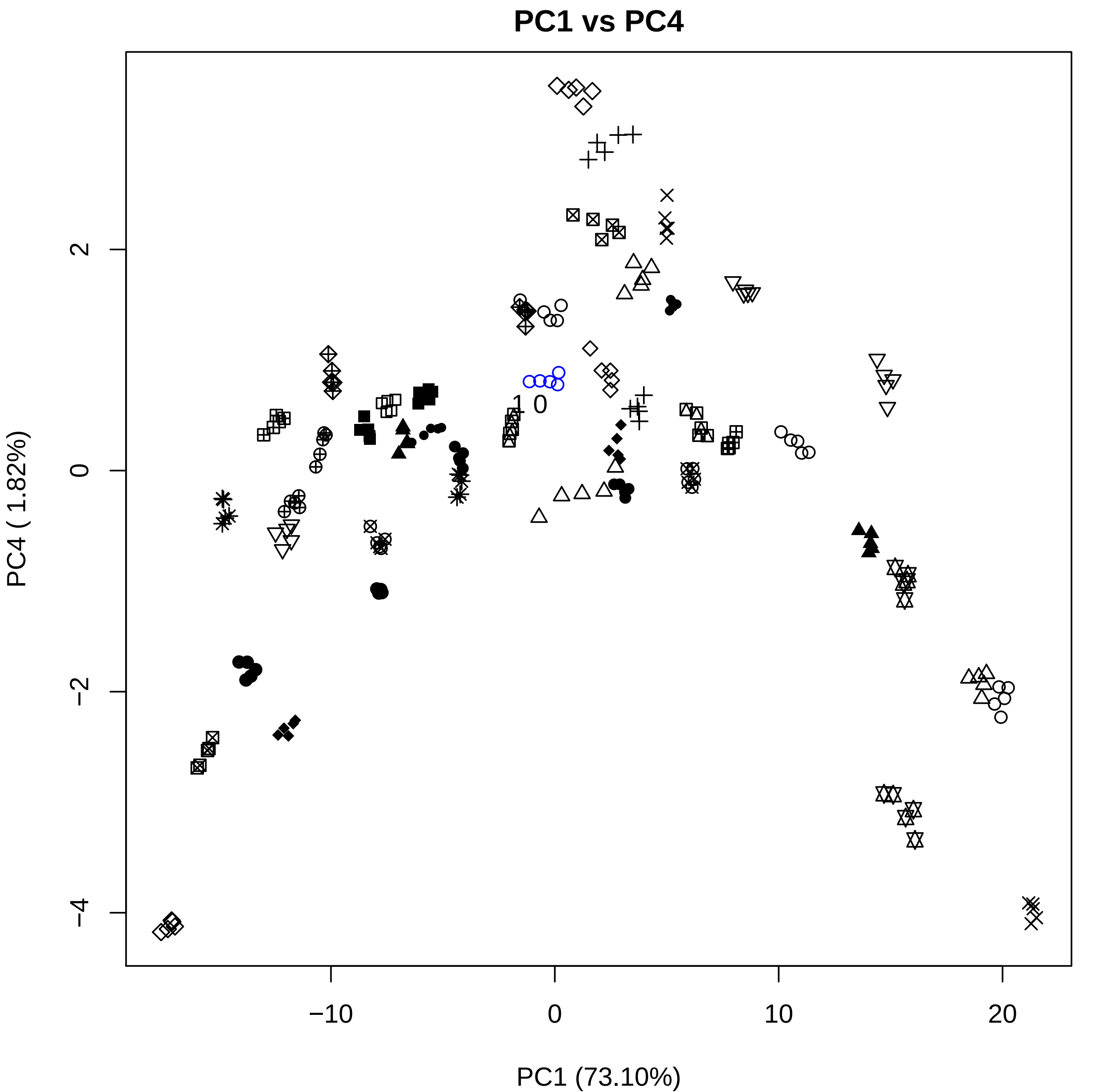
PC1 vs PC4
PC4 ( 1.82%)
PC1 (73.10%)
2
0
−2
−4
−10
0
10
20
1 0

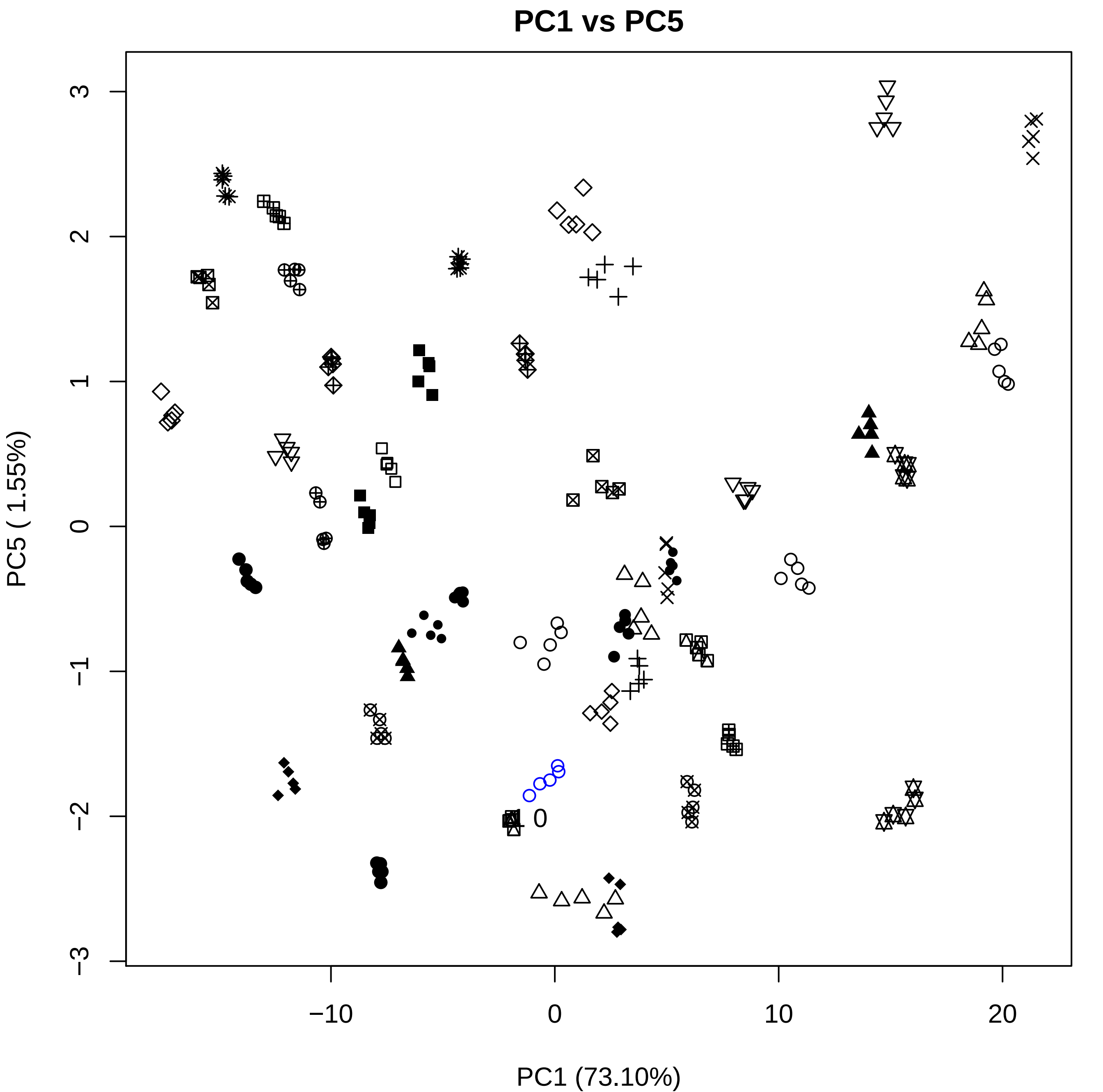
PC1 vs PC5
PC5 ( 1.55%)
PC1 (73.10%)
3
2
1
0
−1
−2
−3
−10
0
10
20
1 0

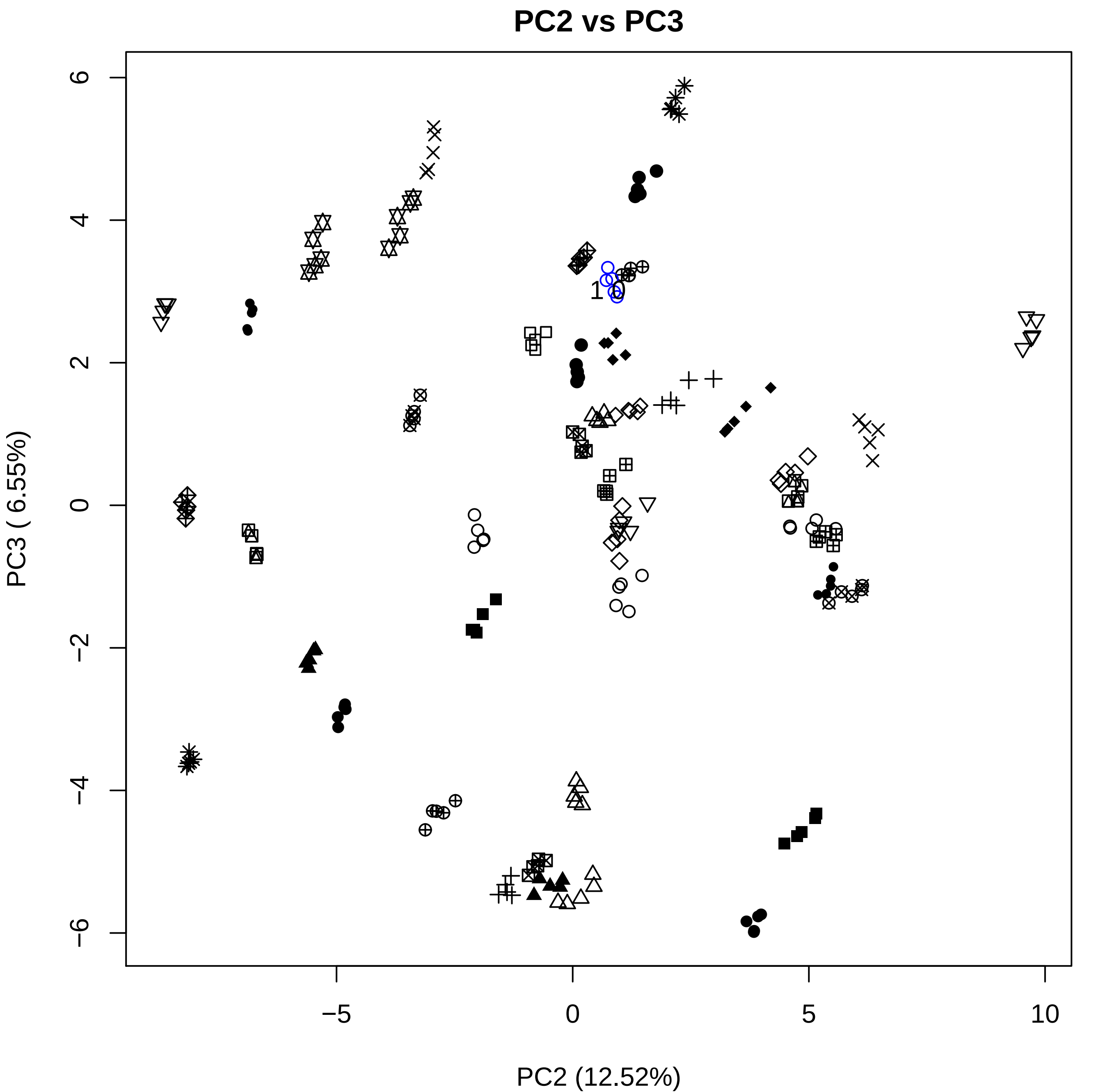
PC2 vs PC3
PC3 ( 6.55%)
PC2 (12.52%)
6
4
2
0
−2
−4
−6
−5
0
5
10

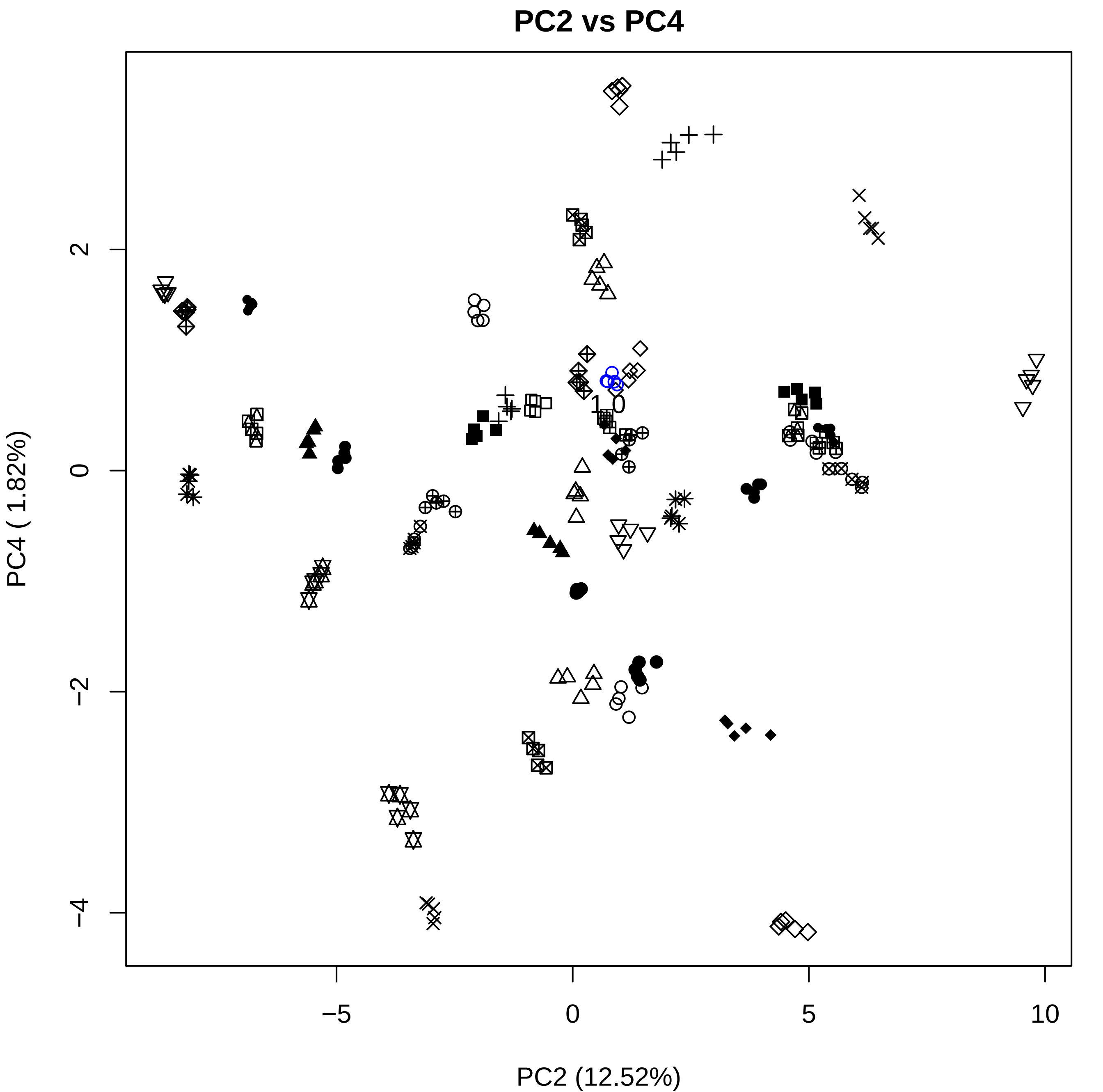
PC2 vs PC4
PC4 ( 1.82%)
PC2 (12.52%)
2
0
−2
−4
−5
0
5
10
10

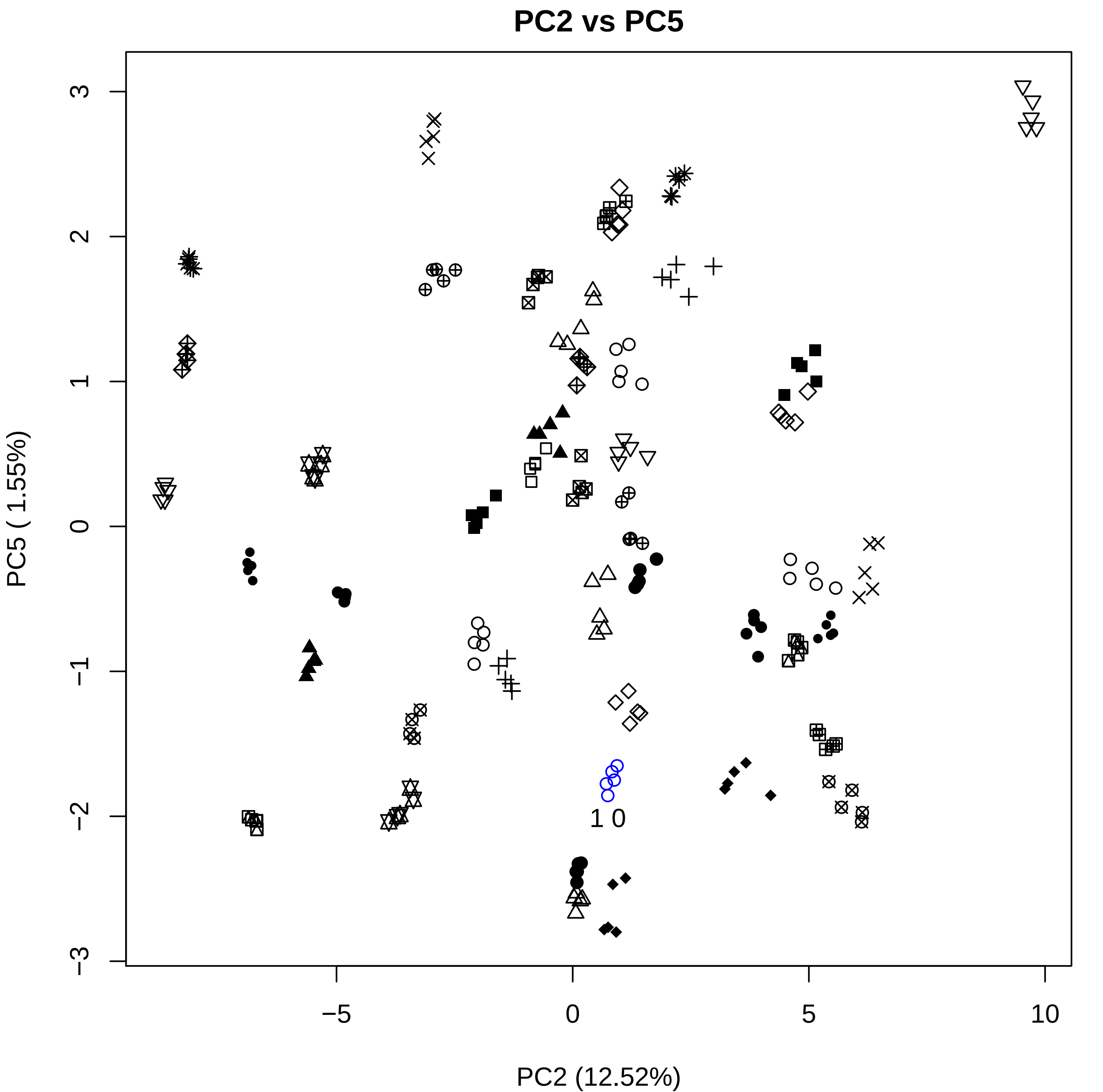
PC2 vs PC5
PC5 ( 1.55%)
PC2 (12.52%)
3
2
1
0
−1
−2
−3
−5
0
5
10
1 0

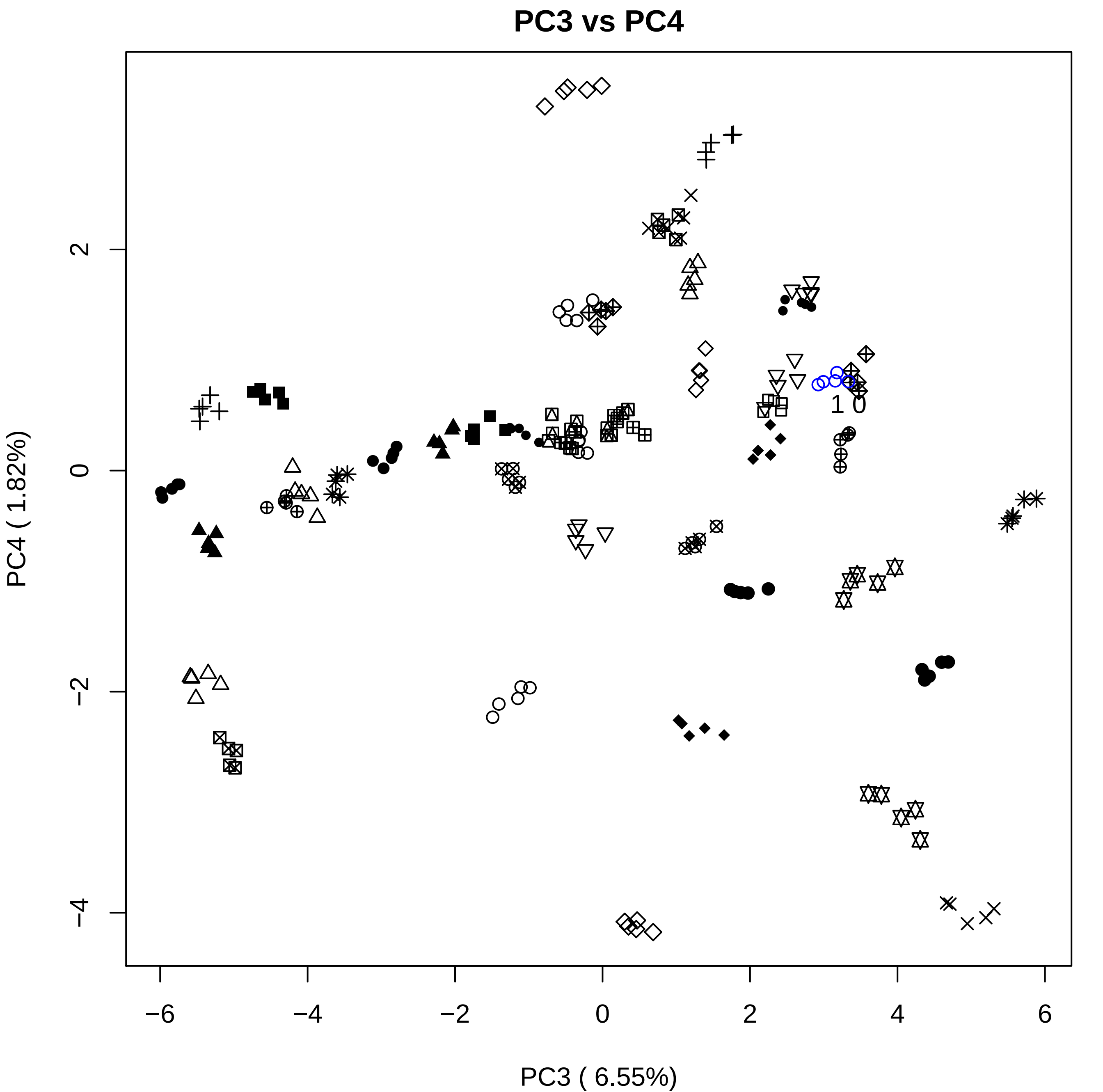
PC3 vs PC4
PC4 ( 1.82%)
PC3 ( 6.55%)
−6
−4
−2
0
2
4
6
2
0
−2
−4
1 0

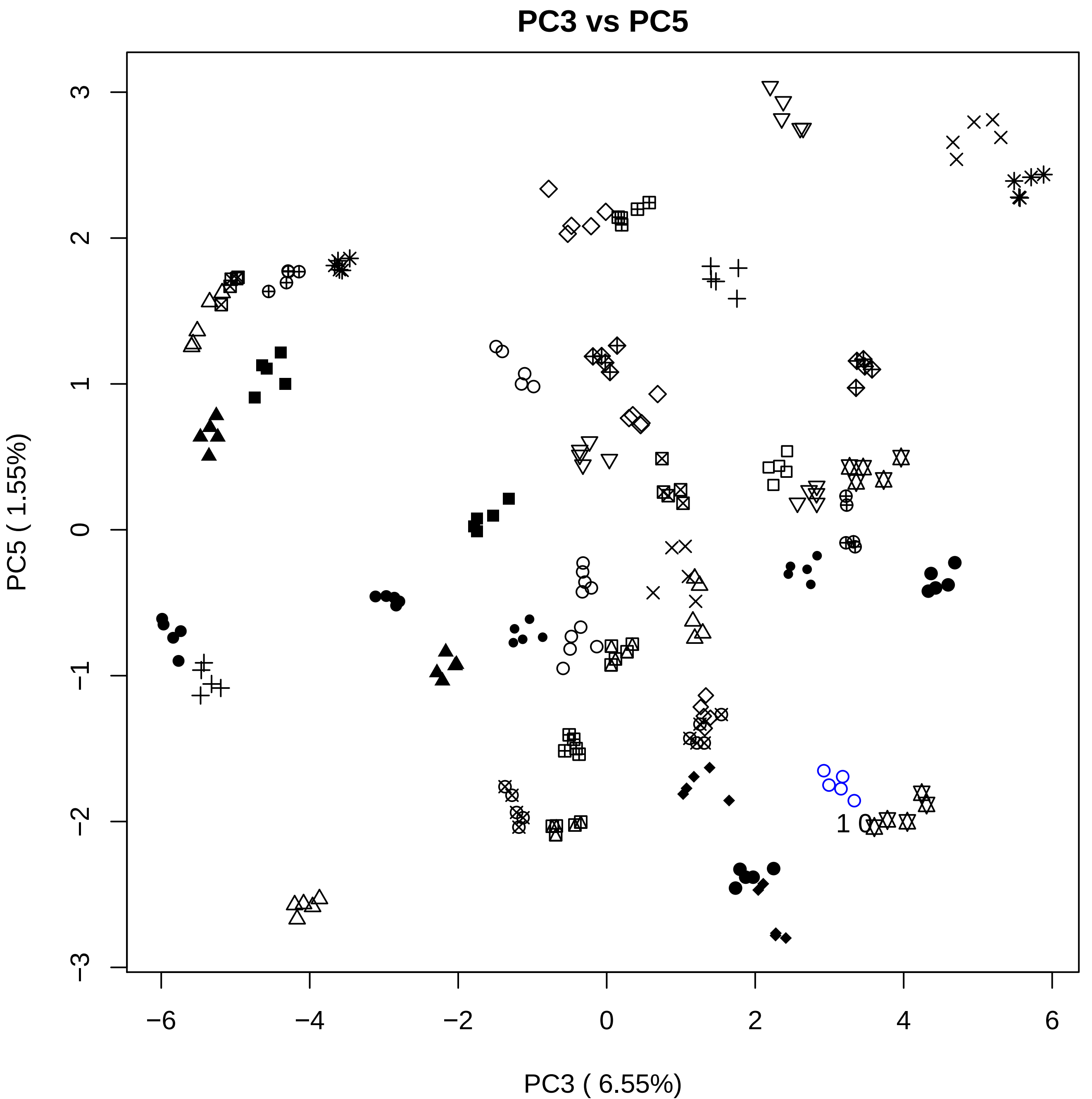
PC3 vs PC5
PC5 ( 1.55%)
PC3 ( 6.55%)
3
2
1
0
−1
−2
−3
−6
−4
−2
0
2
4
6
10

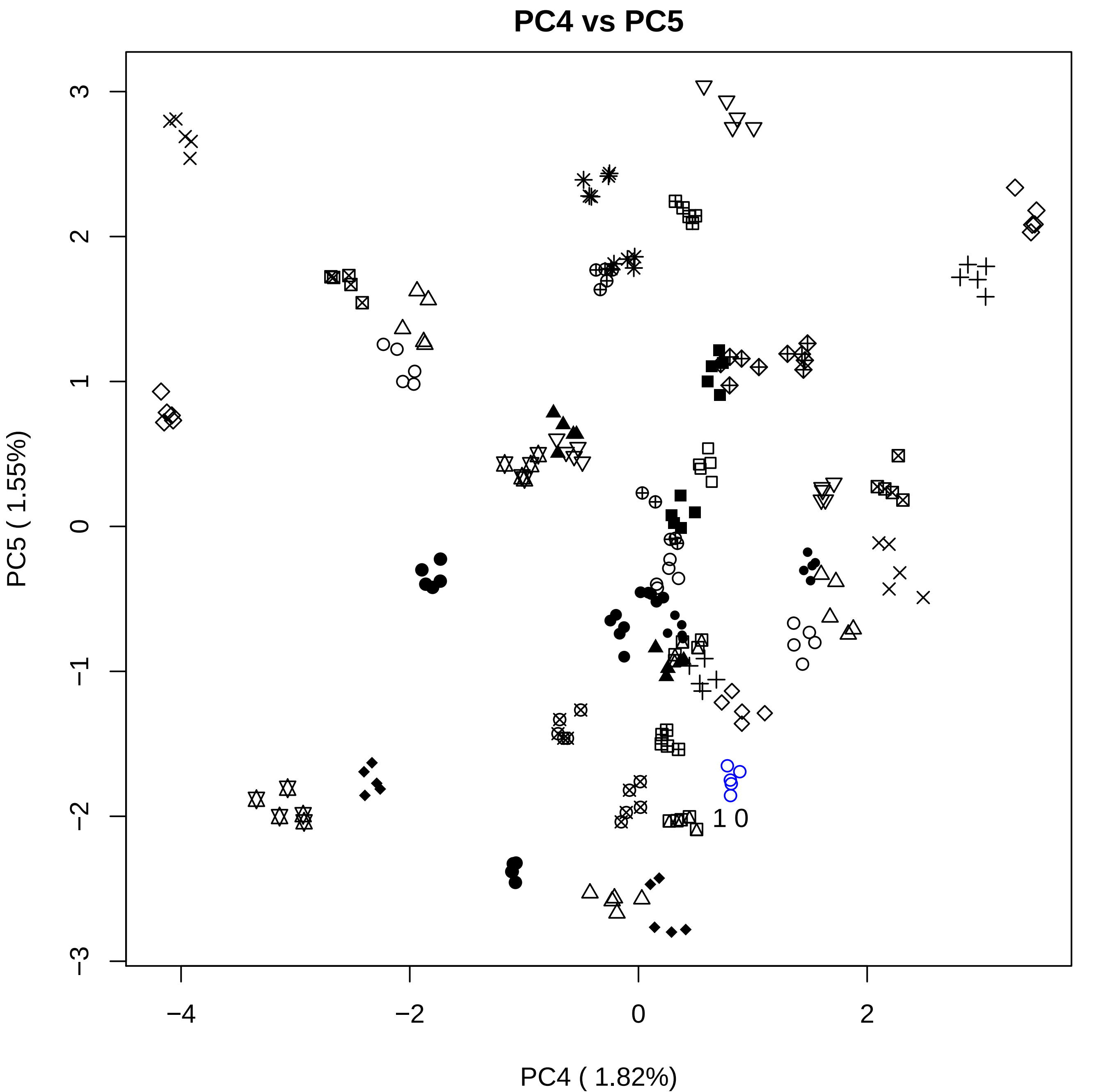
PC4 vs PC5
PC5 ( 1.55%)
PC4 ( 1.82%)
3
2
1
0
−1
−2
−3
−4
−2
0
2
1 0

### PCA of 2.5%  OSAS vs PMH

Scaled Matrix PCAed

7th order polynomial baseline correction performed

Raw spectra smoothed to 21 Neighbours, 2nd order PN, Savtisky Golay

Normalisation between 0 and 100

D2 spectra smoothed to 41 Neighbours, 2nd order PN, Savtisky Golay

Regions Analysed: 3601−2501 and 2001−700cm−1

Samples Analysed: 1 Contam's 0 OSAS's 45 PMH's

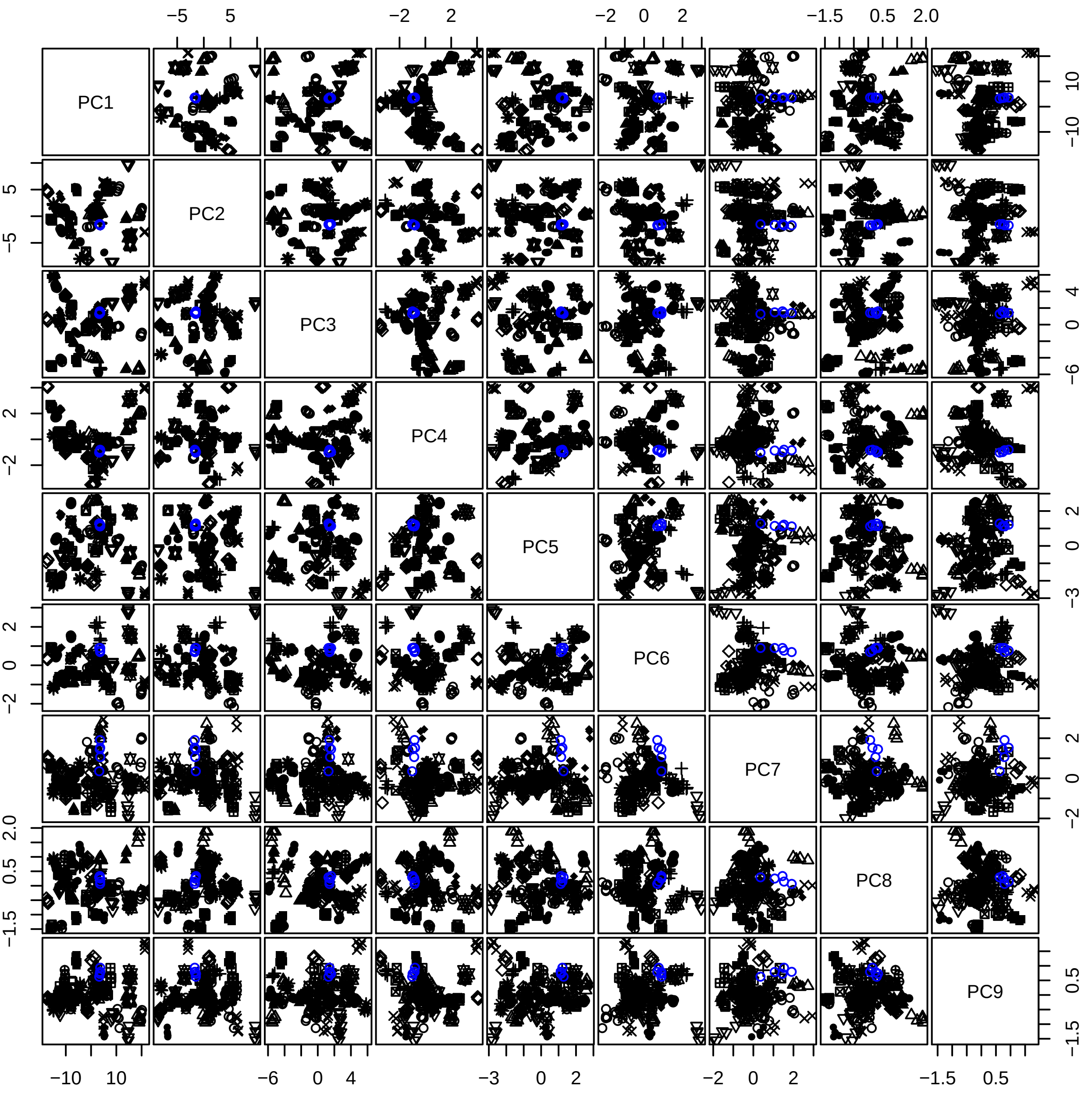
PC1
PC2
PC3
PC4
PC5
PC6
PC7
PC8
PC9

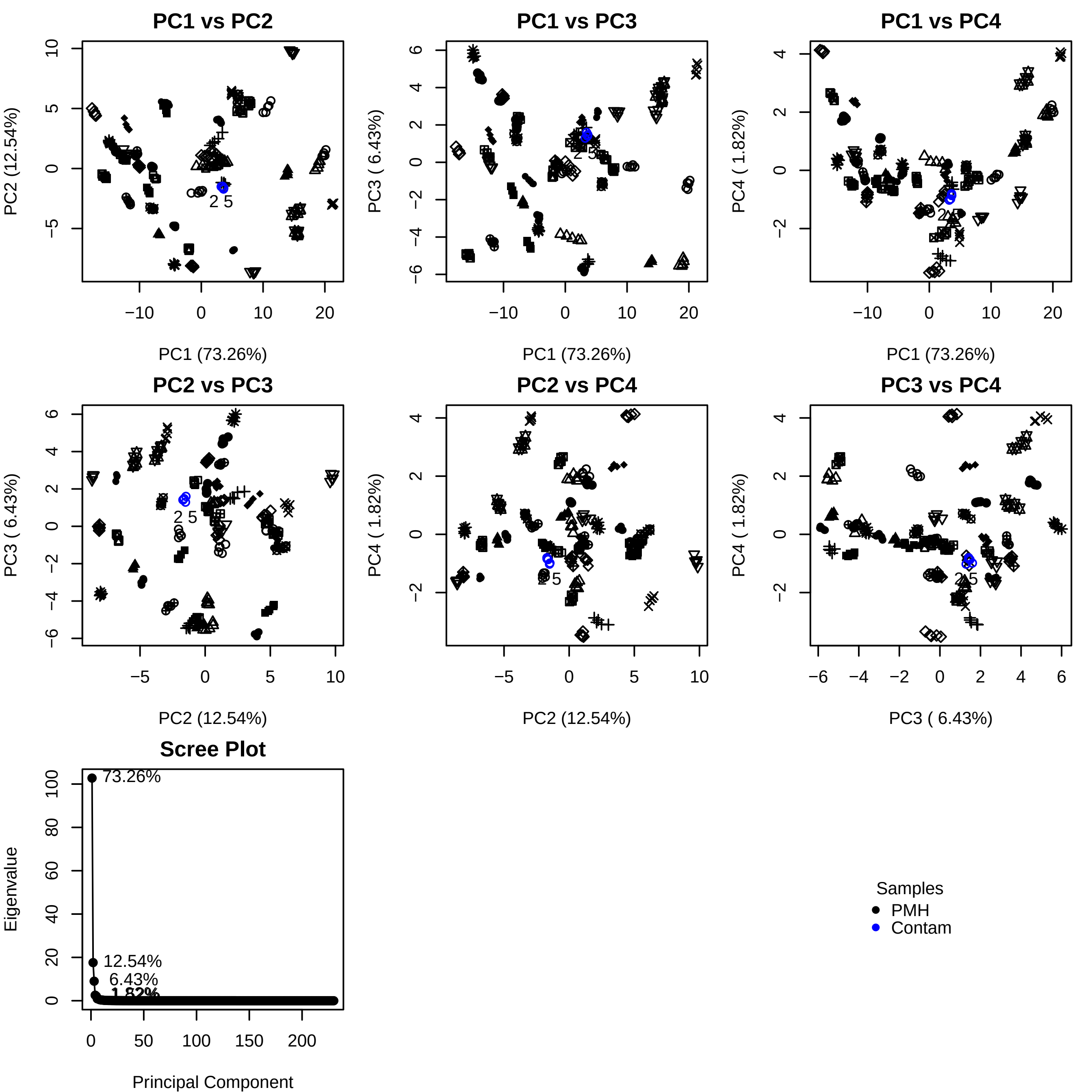
PC1 vs PC2
PC2 (12.54%)
PC1 (73.26%)
PC1 vs PC3
PC3 ( 6.43%)
PC1 (73.26%)
PC1 vs PC4
PC4 ( 1.82%)
PC1 (73.26%)
PC2 vs PC3
PC3 ( 6.43%)
PC2 (12.54%)
PC2 vs PC4
PC4 ( 1.82%)
PC2 (12.54%)
PC3 vs PC4
PC4 ( 1.82%)
PC3 ( 6.43%)
Scree Plot
Eigenvalue
Principal Component
73.26%
12.54%
6.43%
1.82%
Samples
PMH
Contam

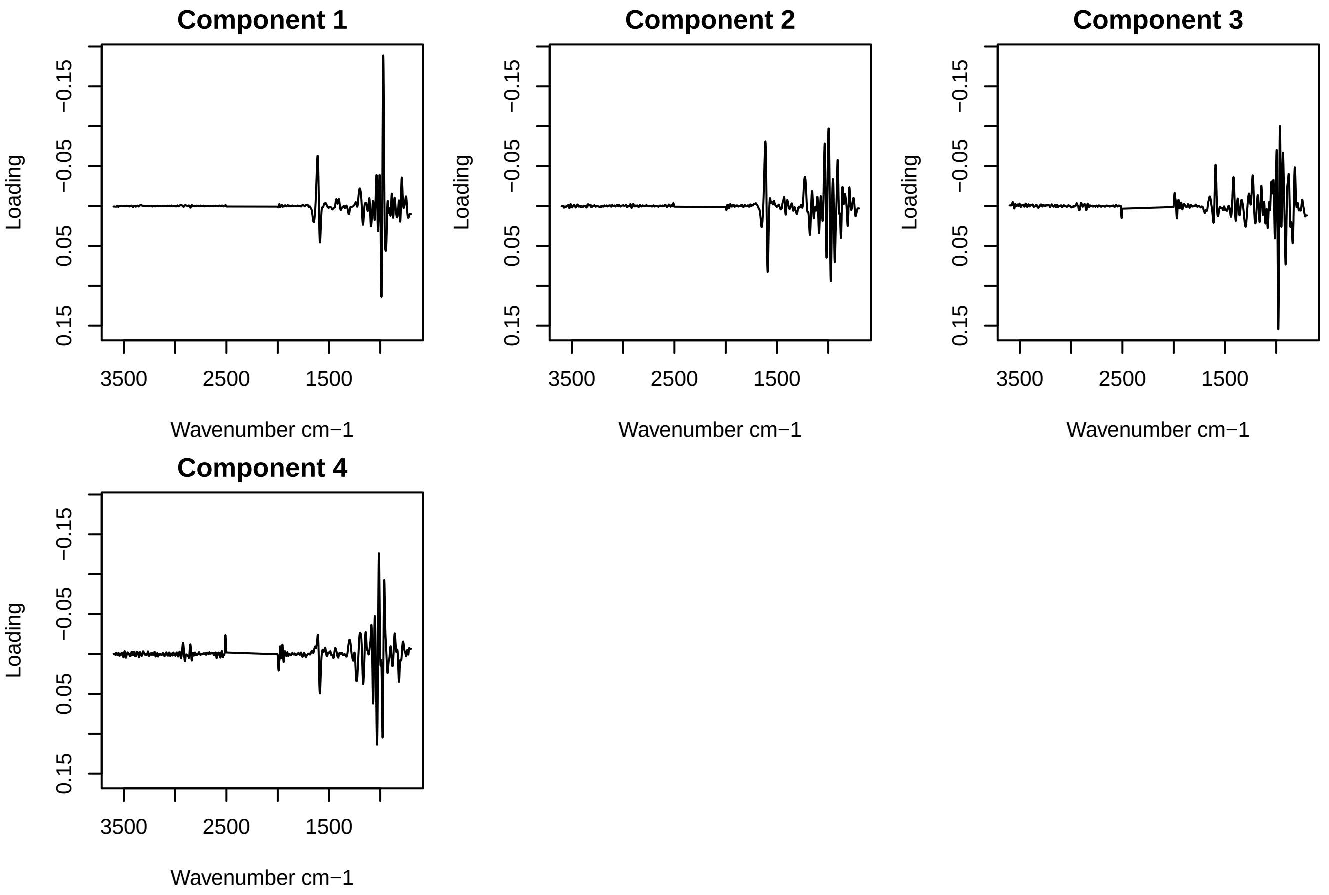
Component 1
Component 2
Component 3
Component 4
Loading
−0.15
−0.05
0.05
0.15
3500
2500
1500
Wavenumber cm−1

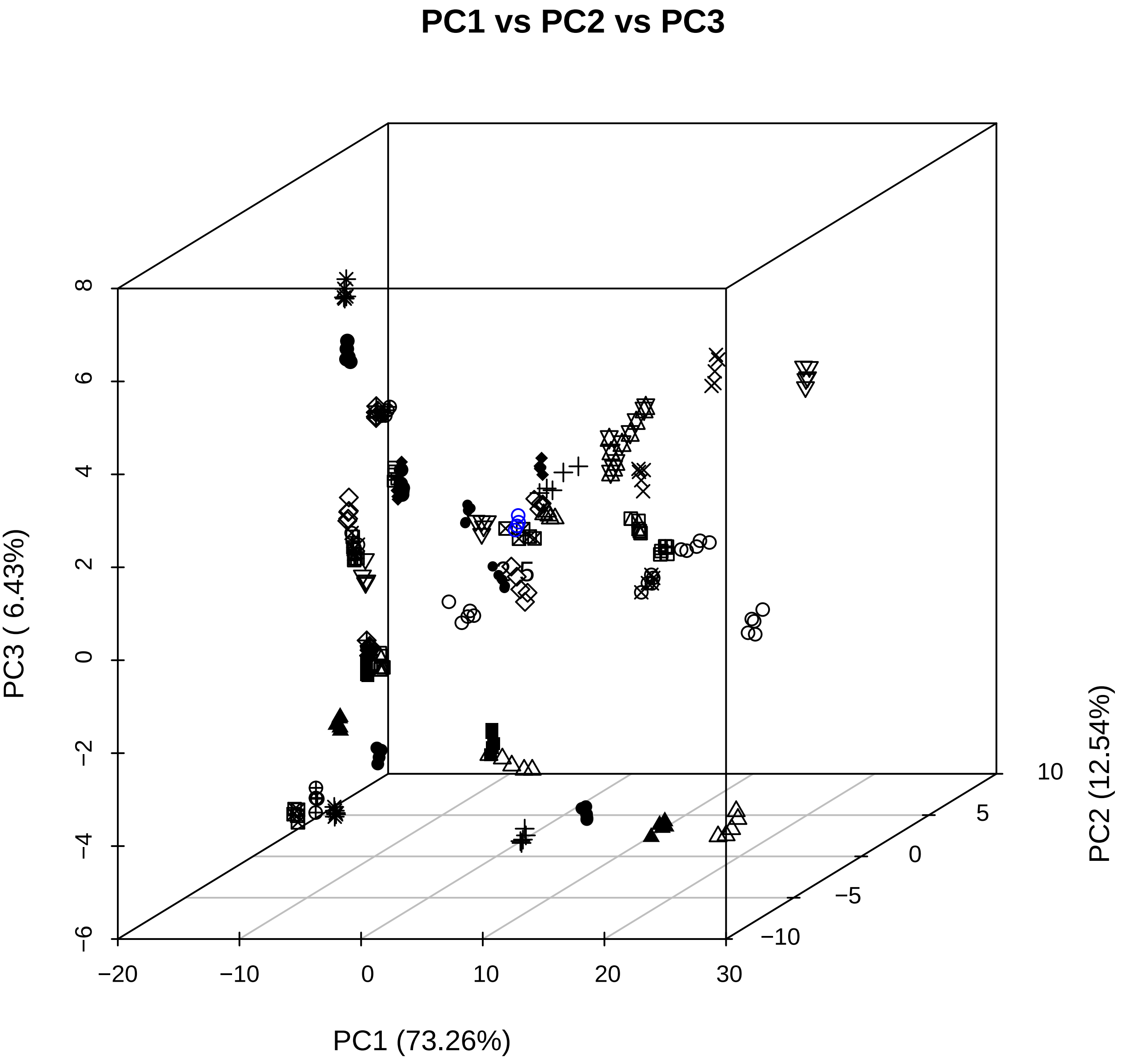
PC1 vs PC2 vs PC3
PC1 (73.26%)
PC2 (12.54%)
PC3 ( 6.43%)
−20
−10
0
10
20
30
−10
−5
0
5
10
−6
−4
−2
0
2
4
6
8

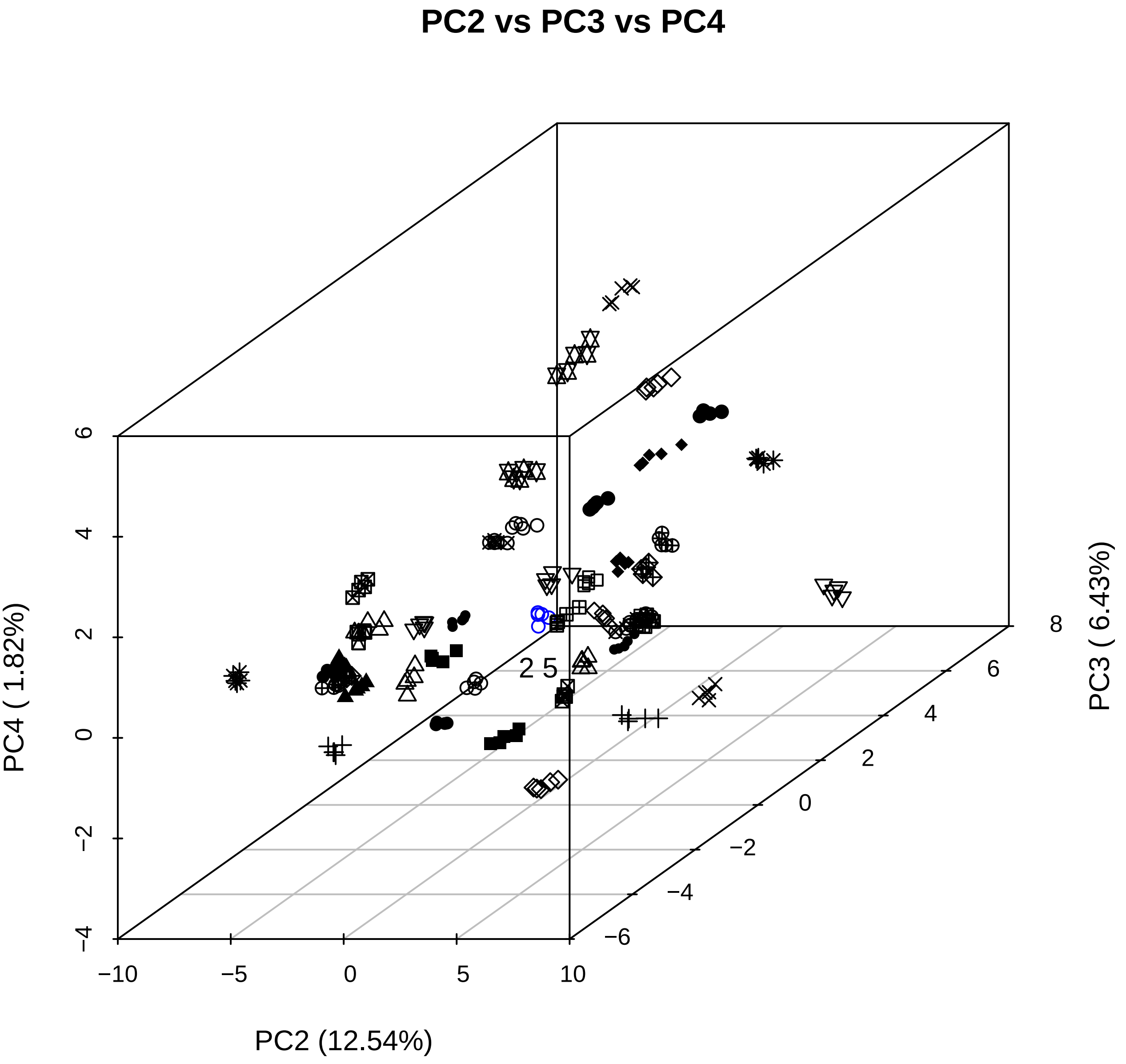
PC2 vs PC3 vs PC4
PC2 (12.54%)
PC3 ( 6.43%)
PC4 ( 1.82%)
−10
−5
0
5
10
−6
−4
−2
0
2
4
6
8
−4
−2
0
2
4
6
2 5

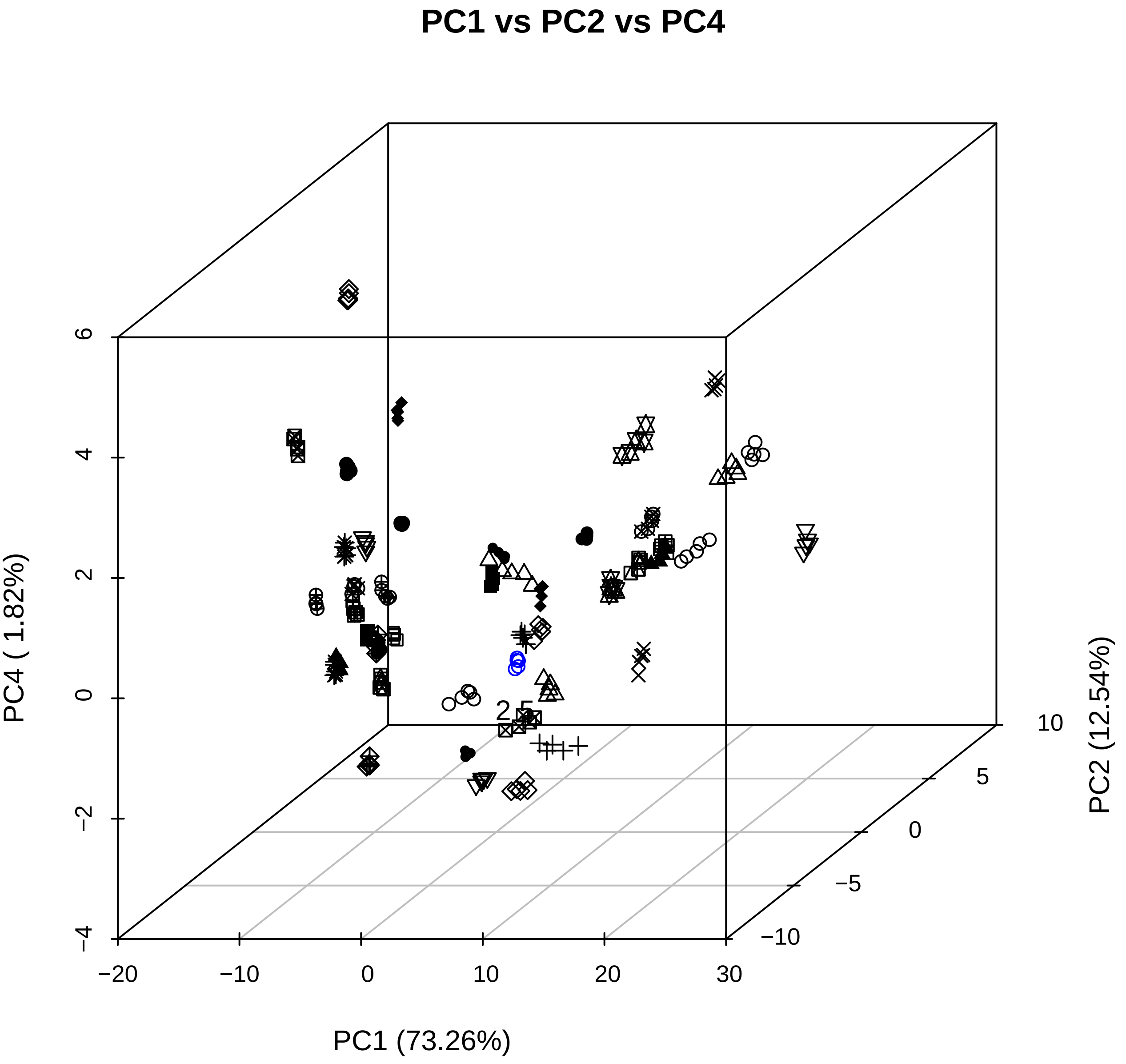

PC1 vs PC2 vs PC4
PC1 (73.26%)
PC2 (12.54%)
PC4 ( 1.82%)
−20
−10
0
10
20
30
−10
−5
0
5
10
−4
−2
0
2
4
6
2
5

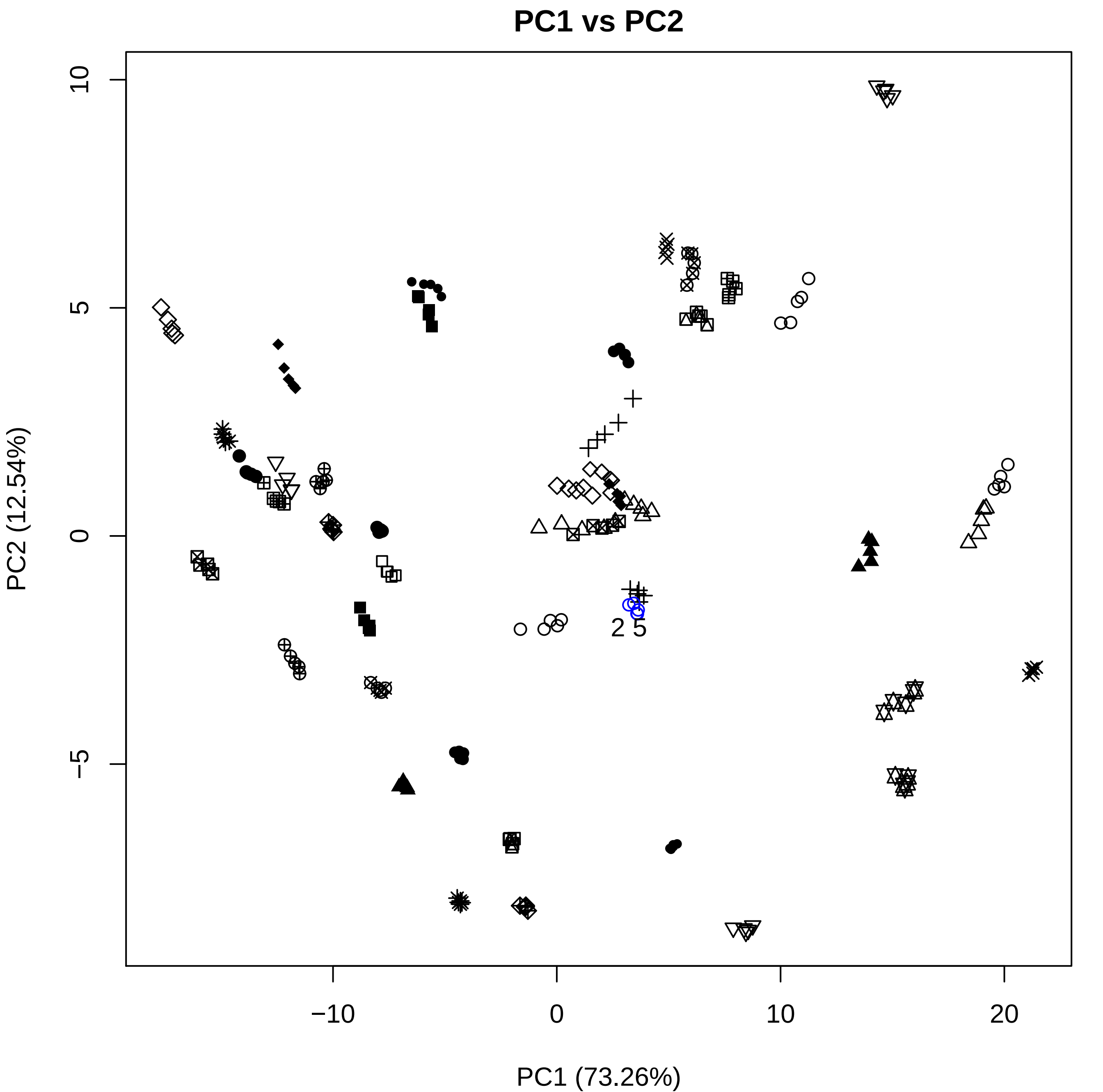
PC1 vs PC2
PC2 (12.54%)
10
5
0
−5
−10
0
10
20
2 5
PC1 (73.26%)

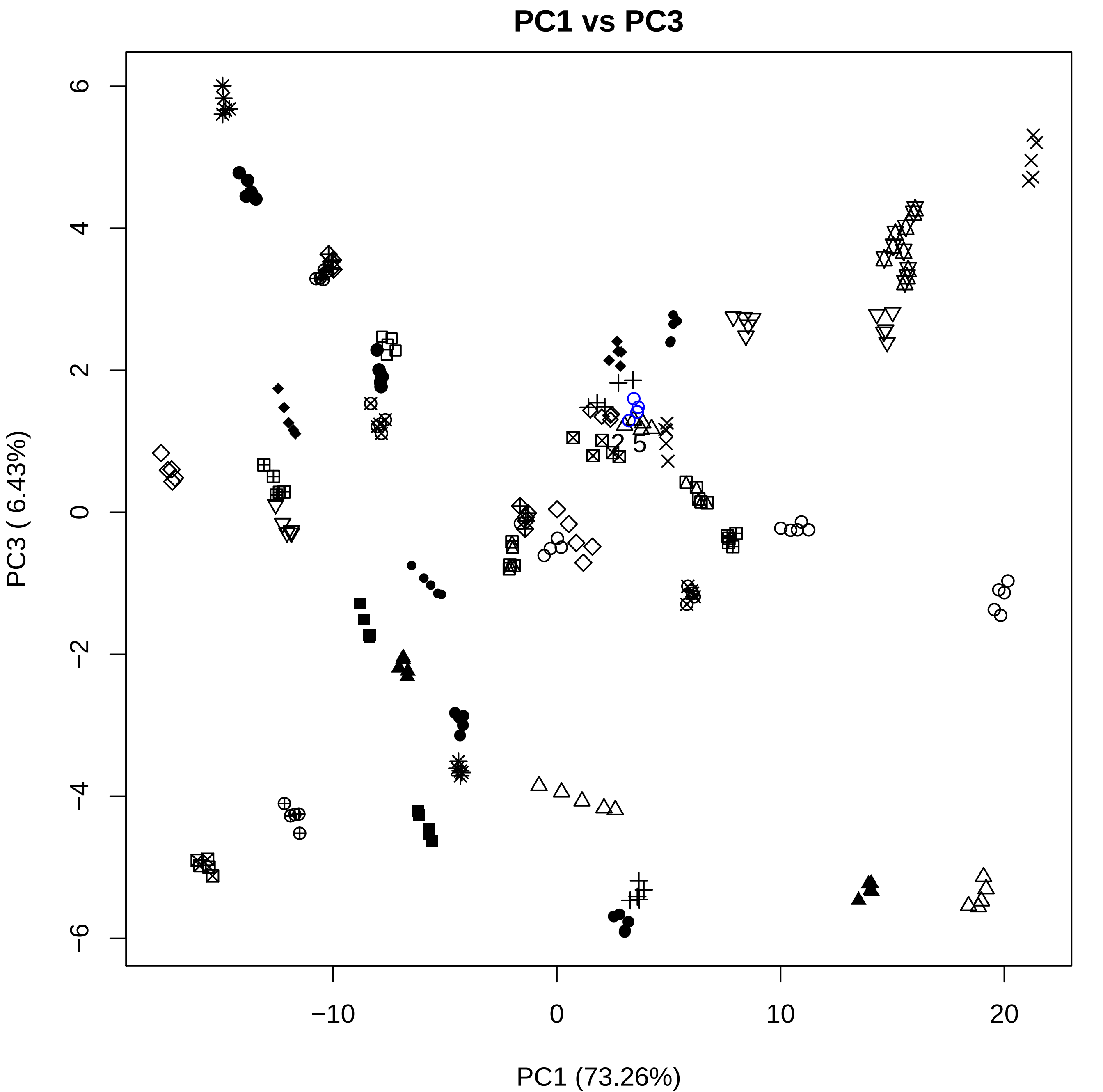
PC1 vs PC3
PC3 ( 6.43%)
6
4
2
0
−2
−4
−6
−10
0
10
20
PC1 (73.26%)
2
5

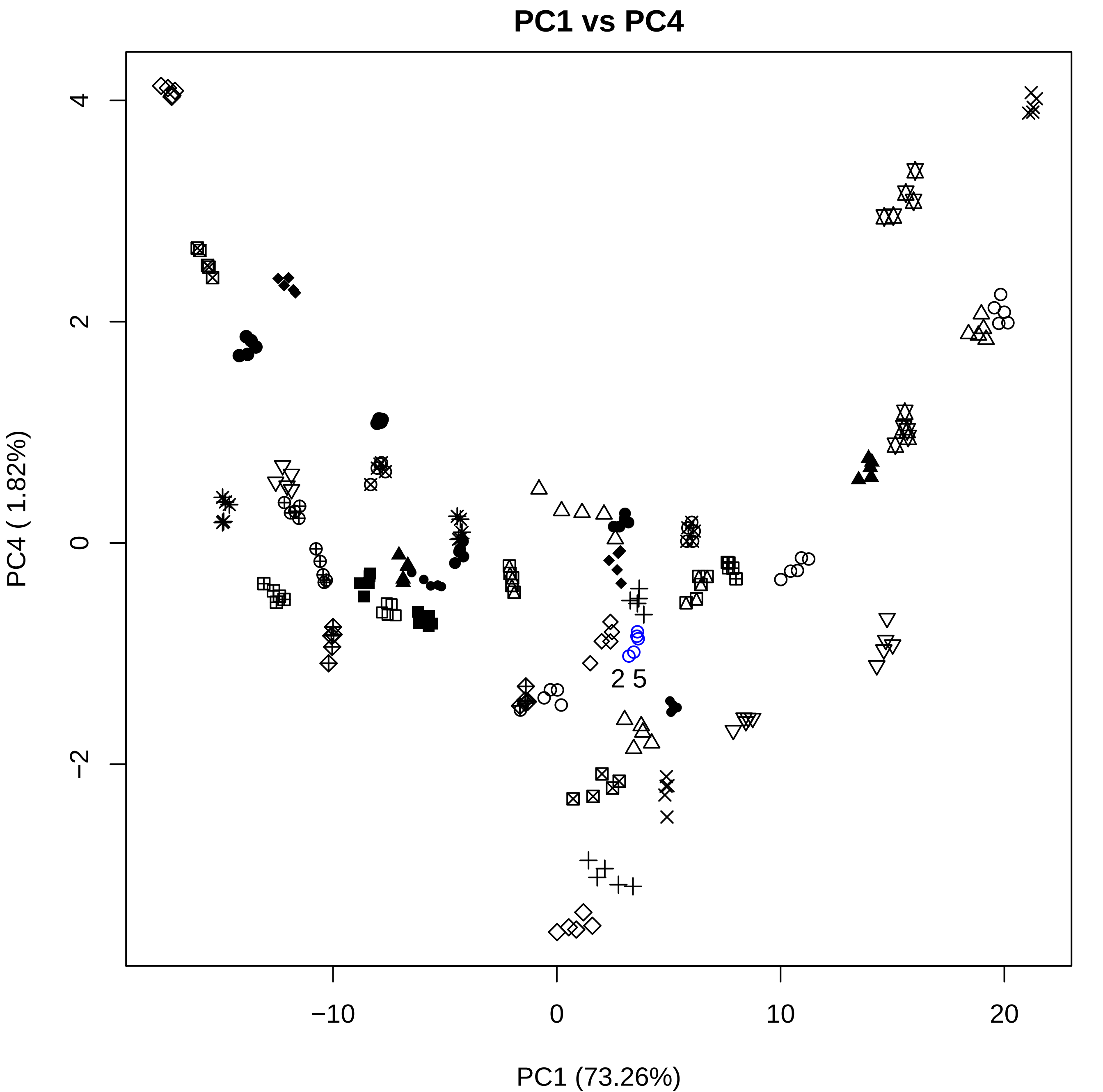
PC1 vs PC4
PC4 ( 1.82%)
PC1 (73.26%)
4
2
0
−2
−10
0
10
20
2 5

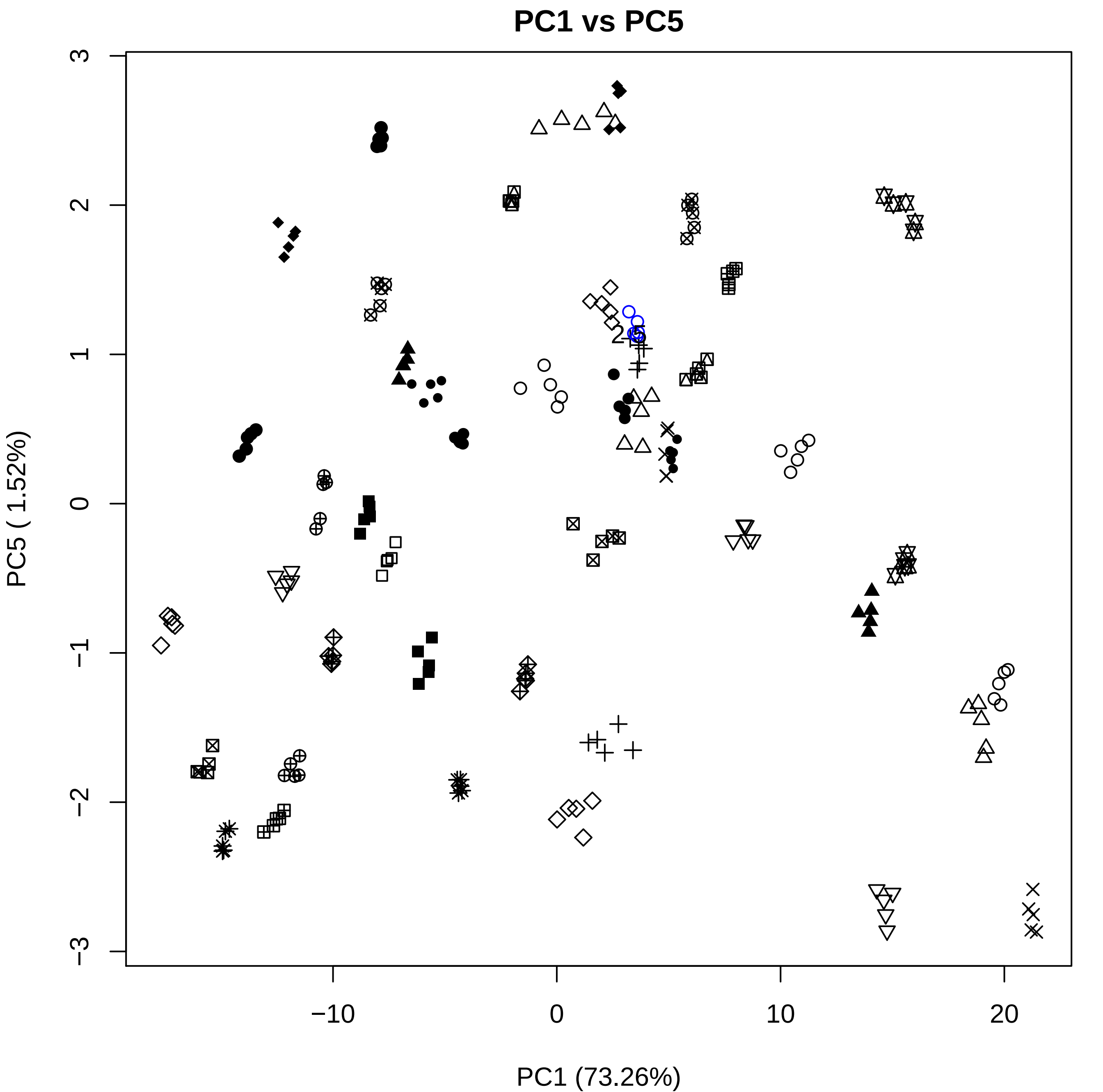
PC1 vs PC5
PC5 ( 1.52%)
PC1 (73.26%)
3
2
1
0
−1
−2
−3
−10
0
10
20

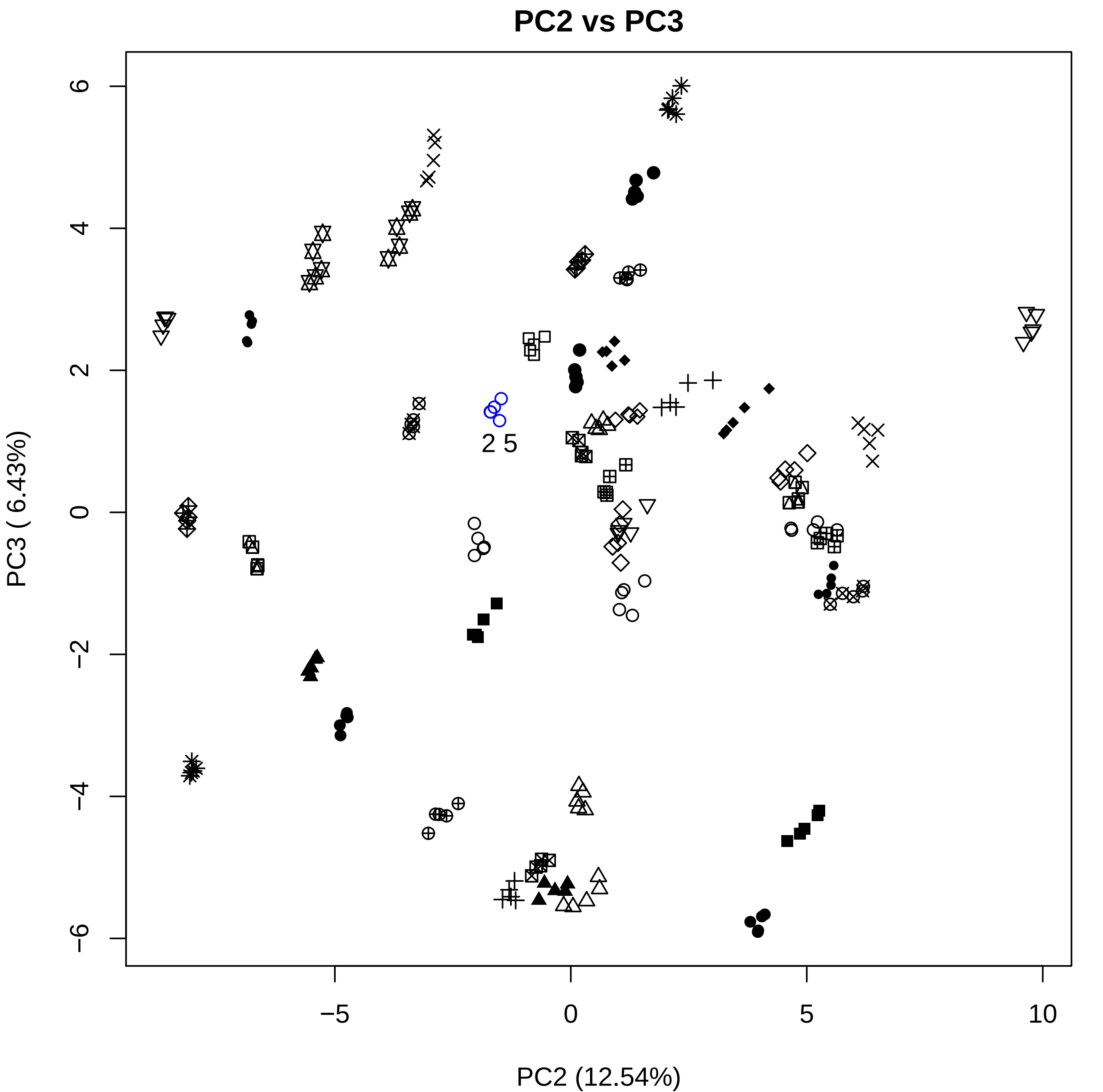
PC2 vs PC3
PC3 ( 6.43%)
PC2 (12.54%)
6
4
2
0
−2
−4
−6
−5
0
5
10
2 5

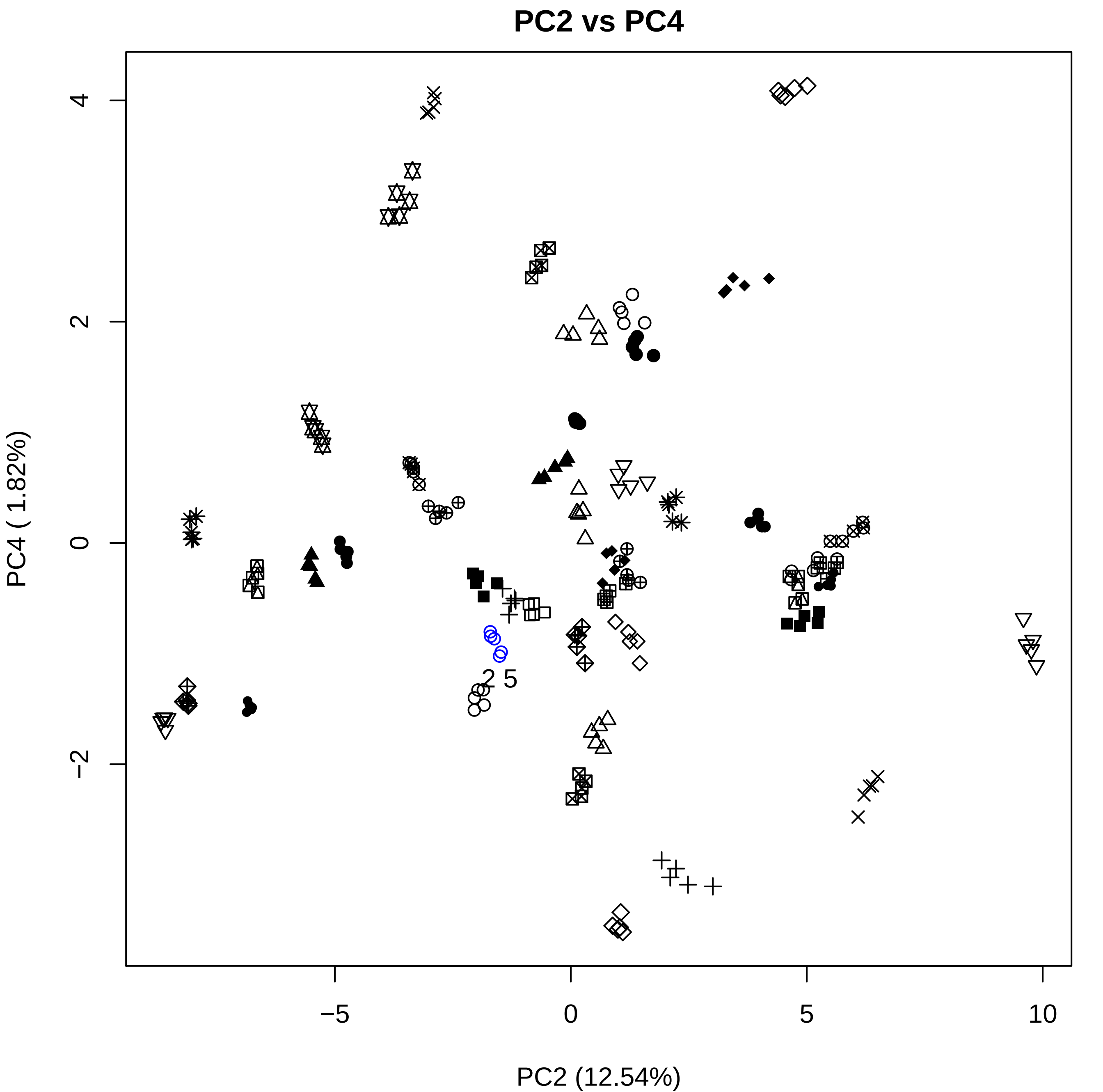
PC2 vs PC4
PC4 ( 1.82%)
PC2 (12.54%)
4
2
0
−2
−5
0
5
10
2 5

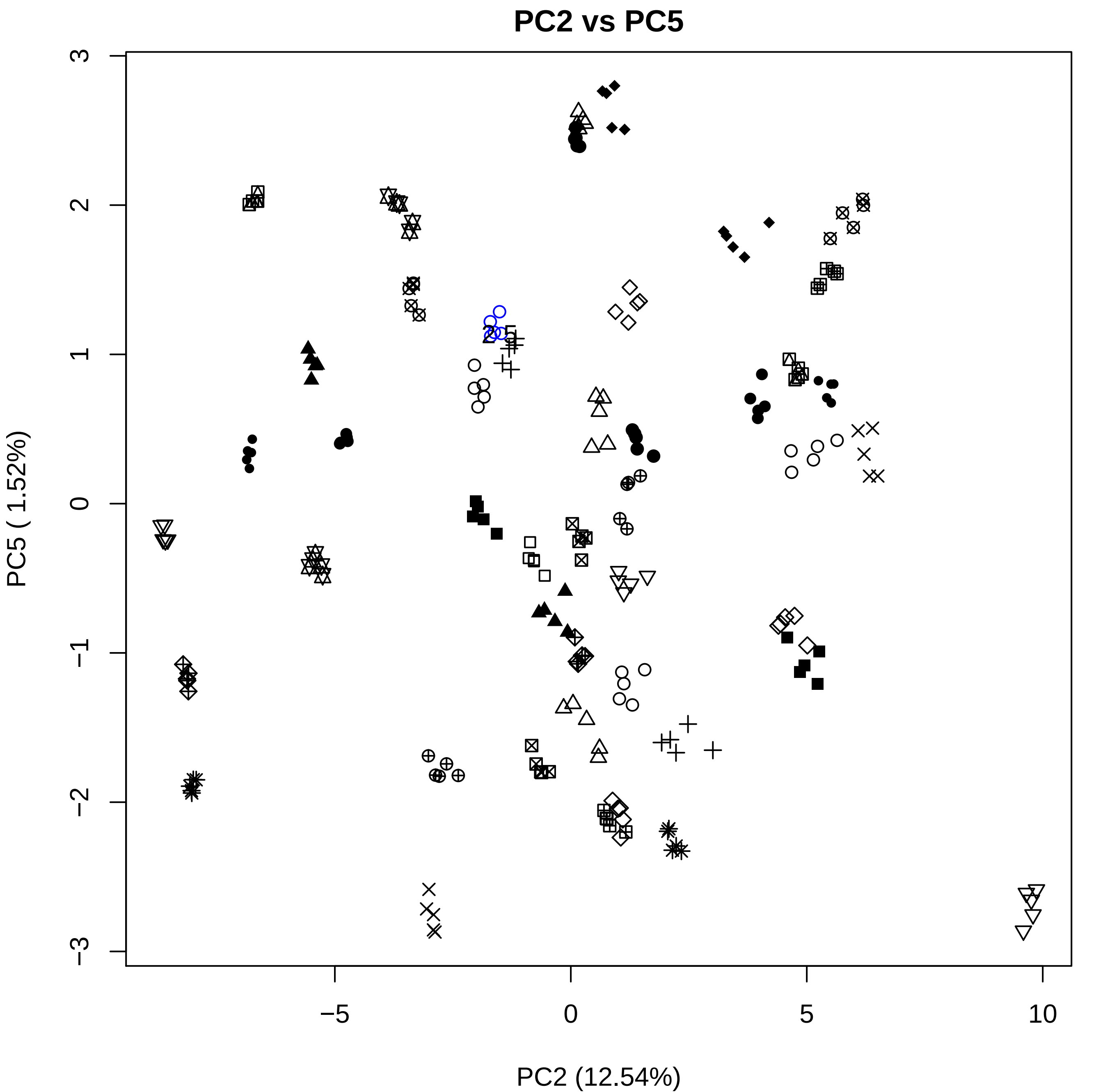
PC2 vs PC5
PC2 (12.54%)
PC5 ( 1.52%)
−5
0
5
10
−3
−2
−1
0
1
2
3

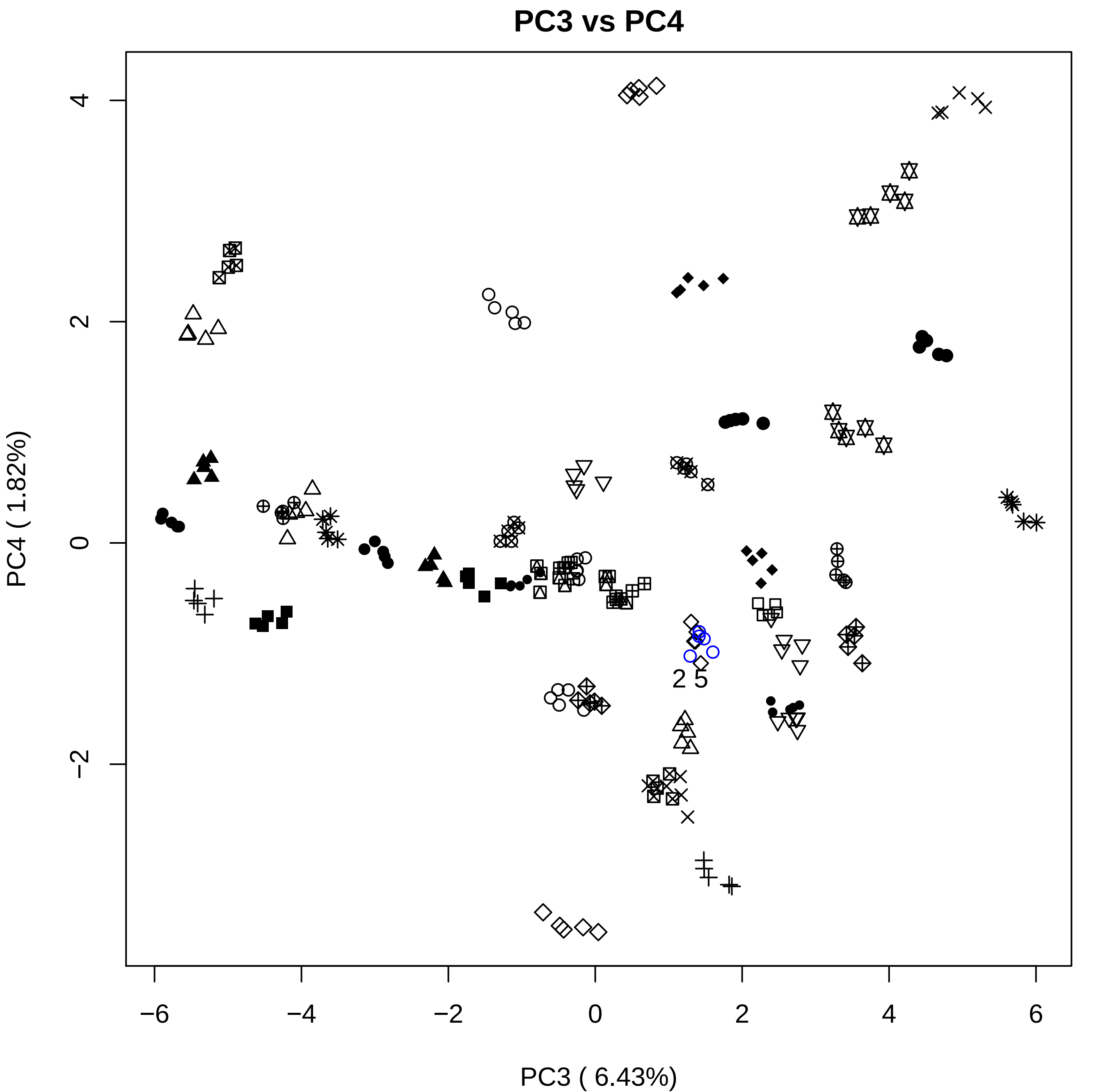
PC3 vs PC4
PC4 ( 1.82%)
PC3 ( 6.43%)
−6
−4
−2
0
2
4
6
4
2
0
−2
2 5

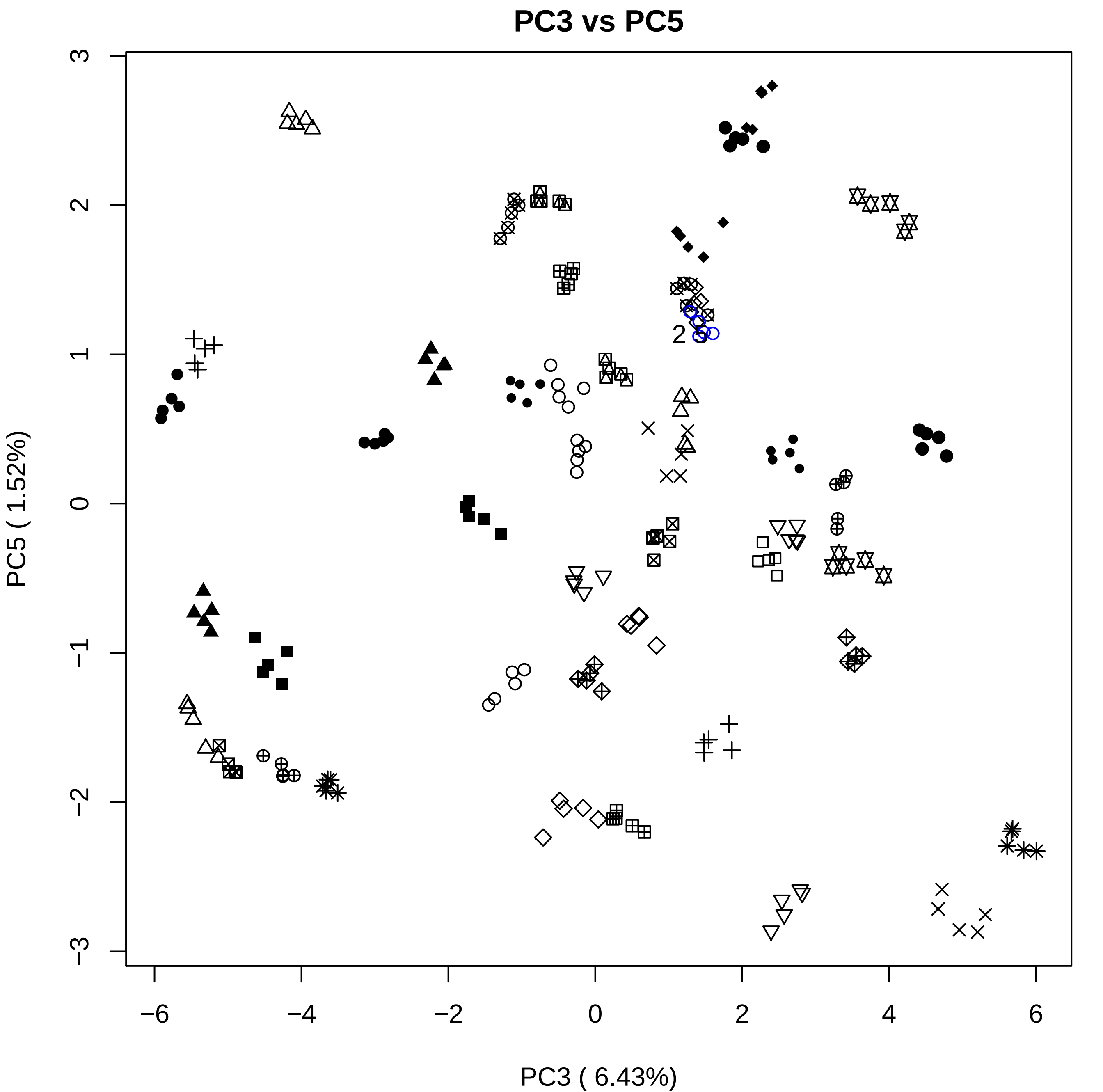
PC3 vs PC5
PC3 ( 6.43%)
PC5 ( 1.52%)
−6
−4
−2
0
2
4
6
−3
−2
−1
0
1
2
3
2 5

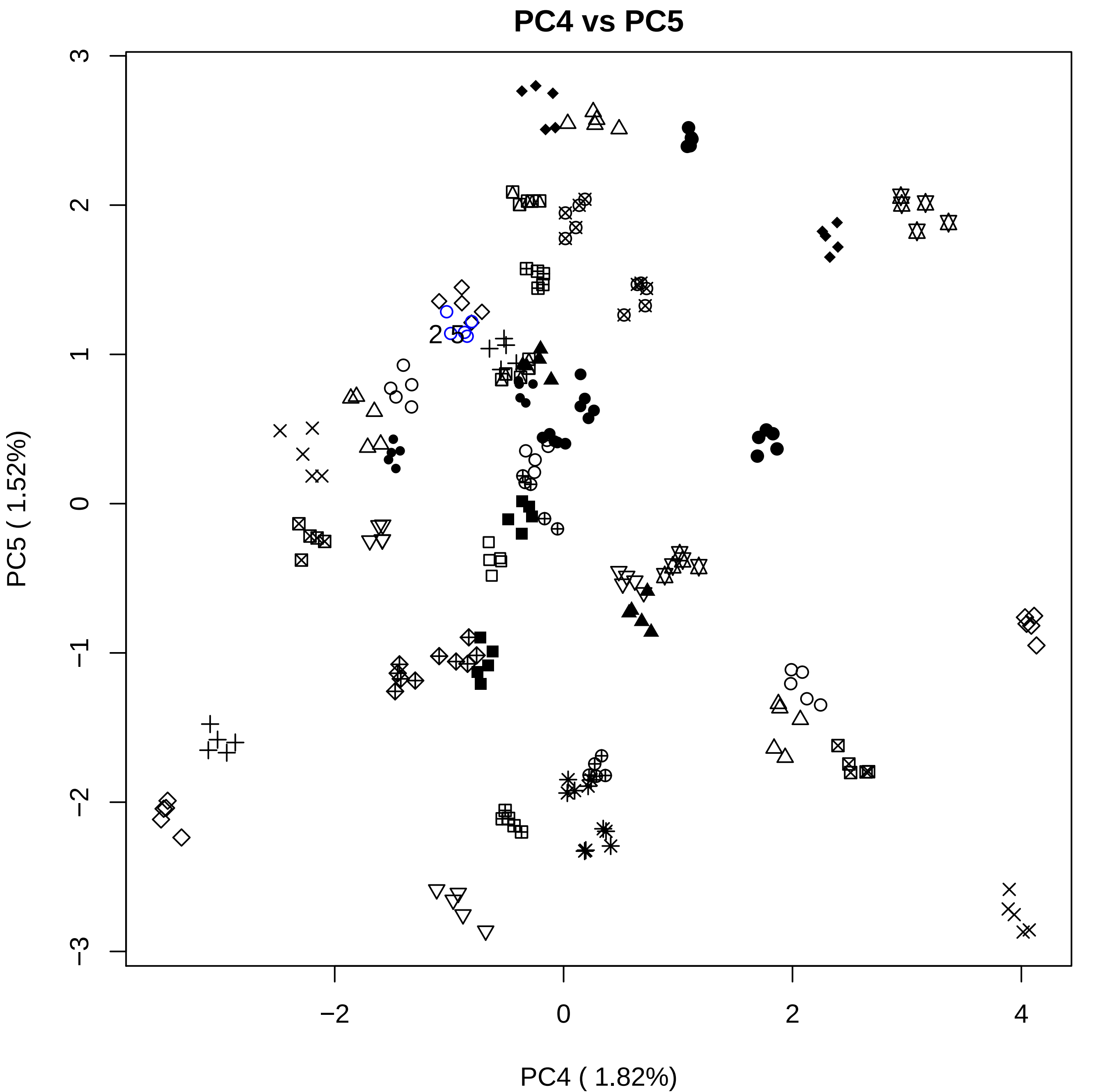
PC4 vs PC5
PC5 ( 1.52%)
PC4 ( 1.82%)
3
2
1
0
−1
−2
−3
−2
0
2
4
25

### PCA of 5%  OSAS vs PMH

Scaled Matrix PCAed

7th order polynomial baseline correction performed

Raw spectra smoothed to 21 Neighbours, 2nd order PN, Savtisky Golay

Normalisation between 0 and 100

D2 spectra smoothed to 41 Neighbours, 2nd order PN, Savtisky Golay

Regions Analysed: 3601−2501 and 2001−700cm−1

Samples Analysed: 1 Contam's 0 OSAS's 45 PMH's

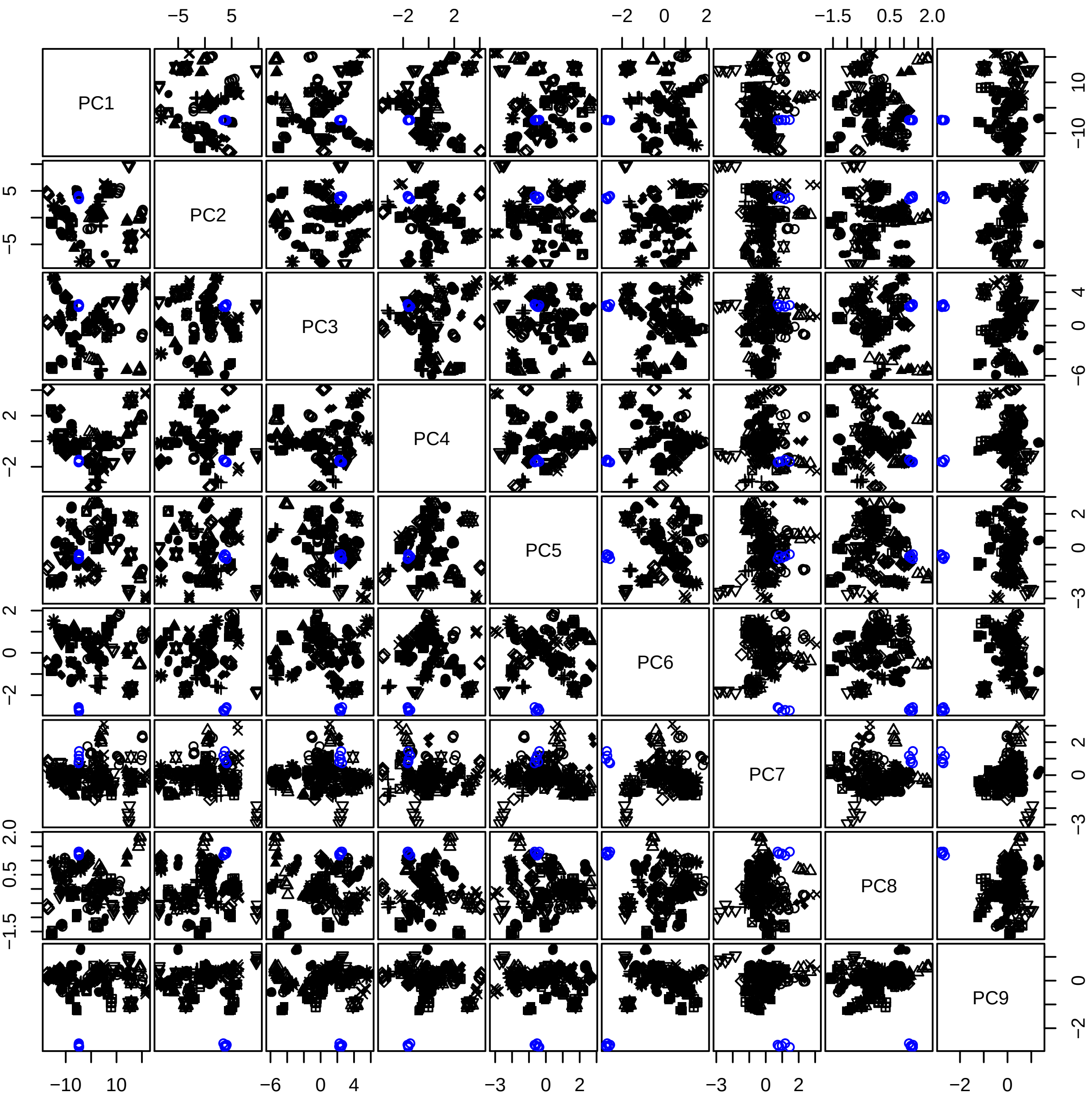
PC1
PC2
PC3
PC4
PC5
PC6
PC7
PC8
PC9

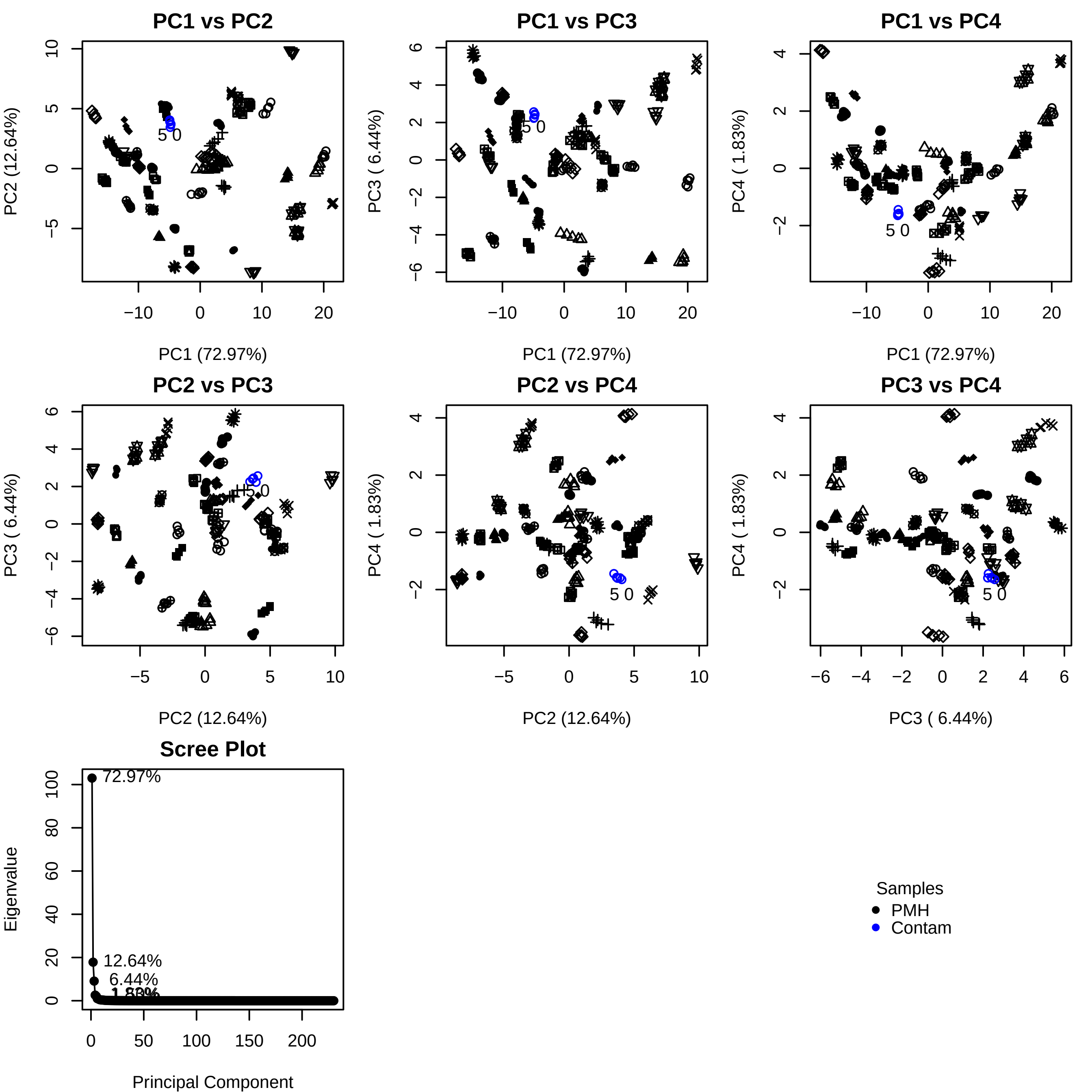

PC1 vs PC2
PC1 vs PC3
PC1 vs PC4
PC2 vs PC3
PC2 vs PC4
PC3 vs PC4
Scree Plot
PC1 (72.97%)
PC2 (12.64%)
PC3 ( 6.44%)
PC4 ( 1.83%)
5 0
Eigenvalue
Principal Component
72.97%
12.64%
6.44%
Samples
PMH
Contam

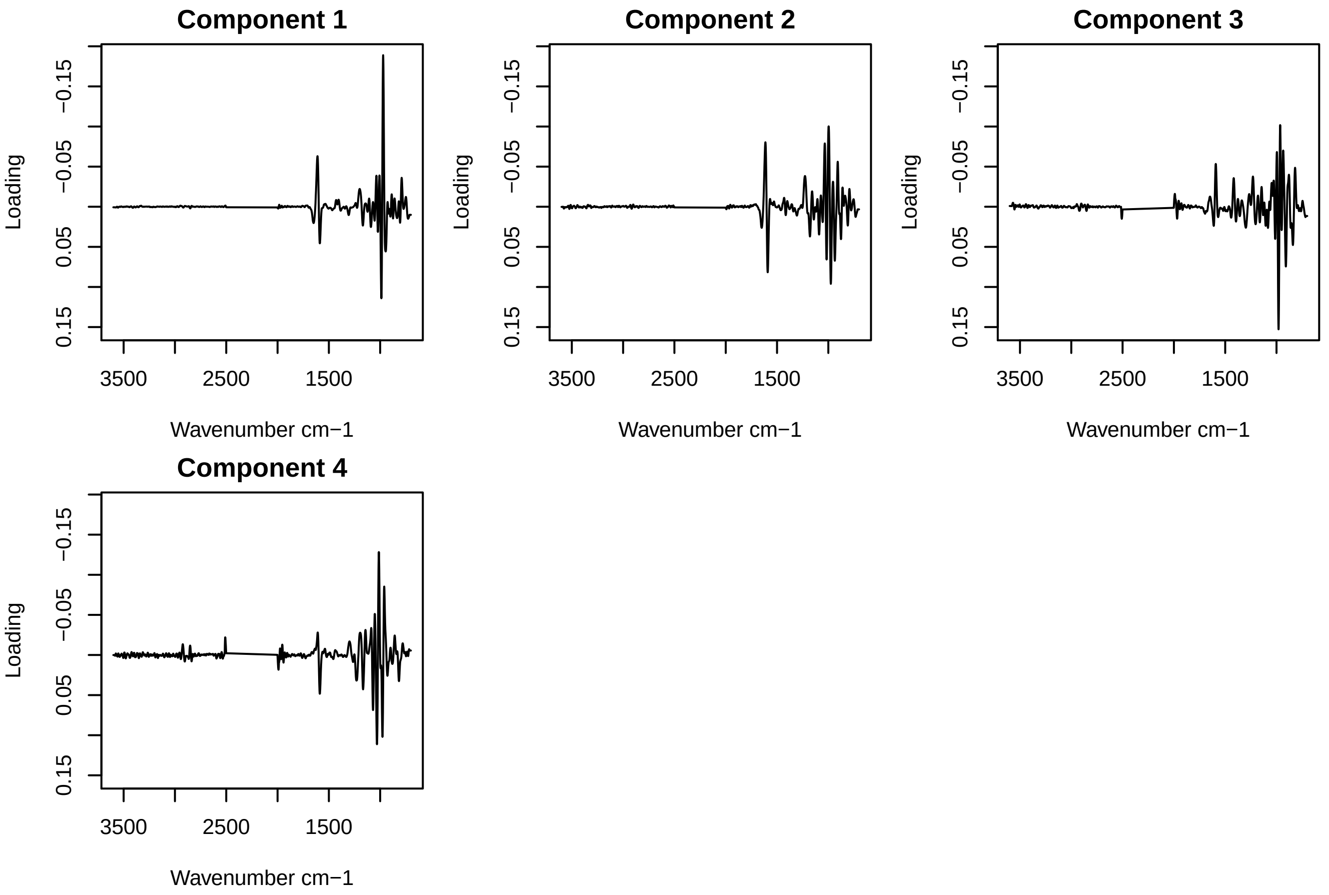
Component 1
Component 2
Component 3
Component 4
Loading
−0.15
−0.05
0.05
0.15
3500
2500
1500
Wavenumber cm−1

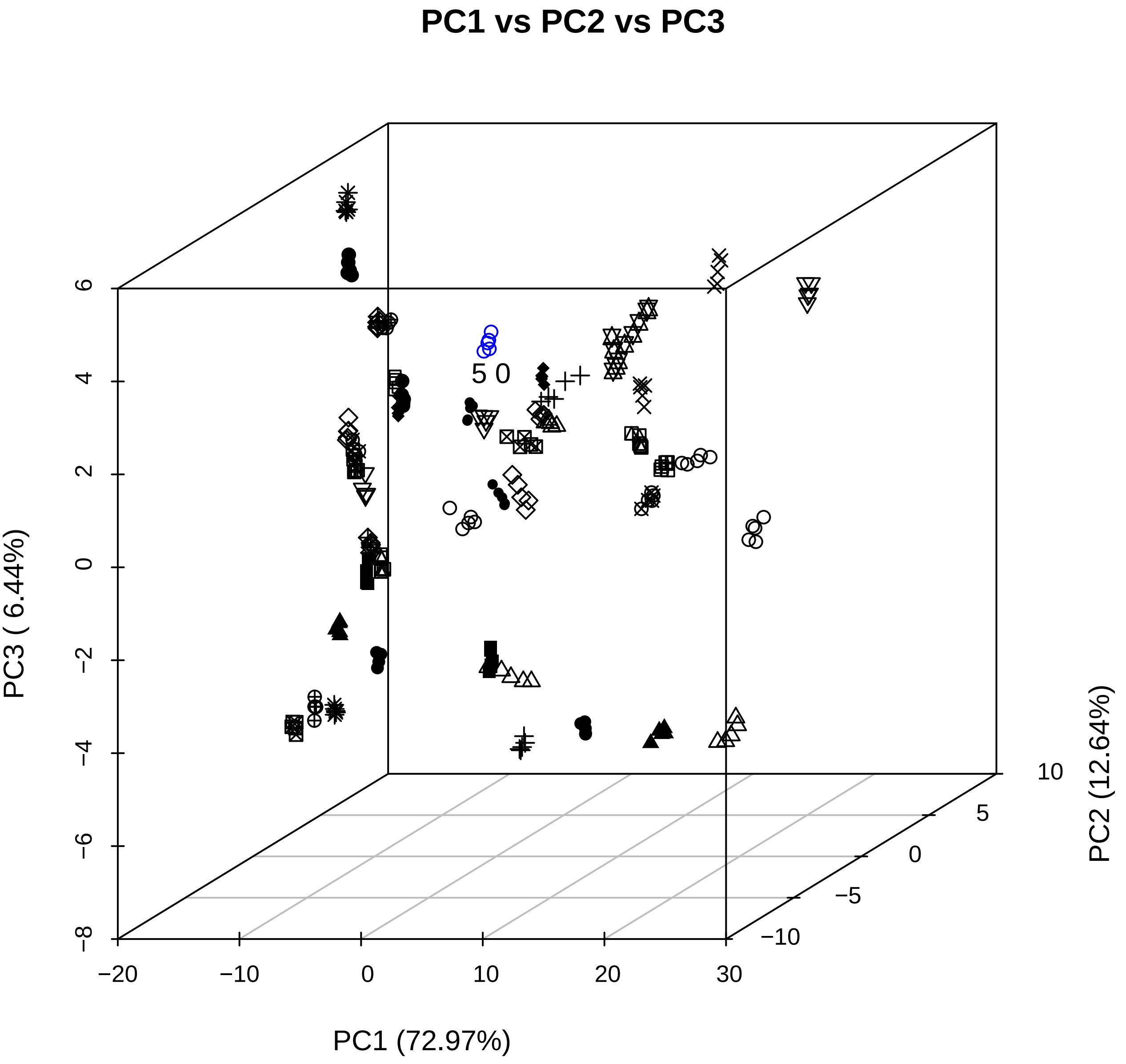
PC1 vs PC2 vs PC3
PC1 (72.97%)
PC2 (12.64%)
PC3 ( 6.44%)
−20
−10
0
10
20
30
−10
−5
0
5
10
−8
−6
−4
−2
0
2
4
6
5 0

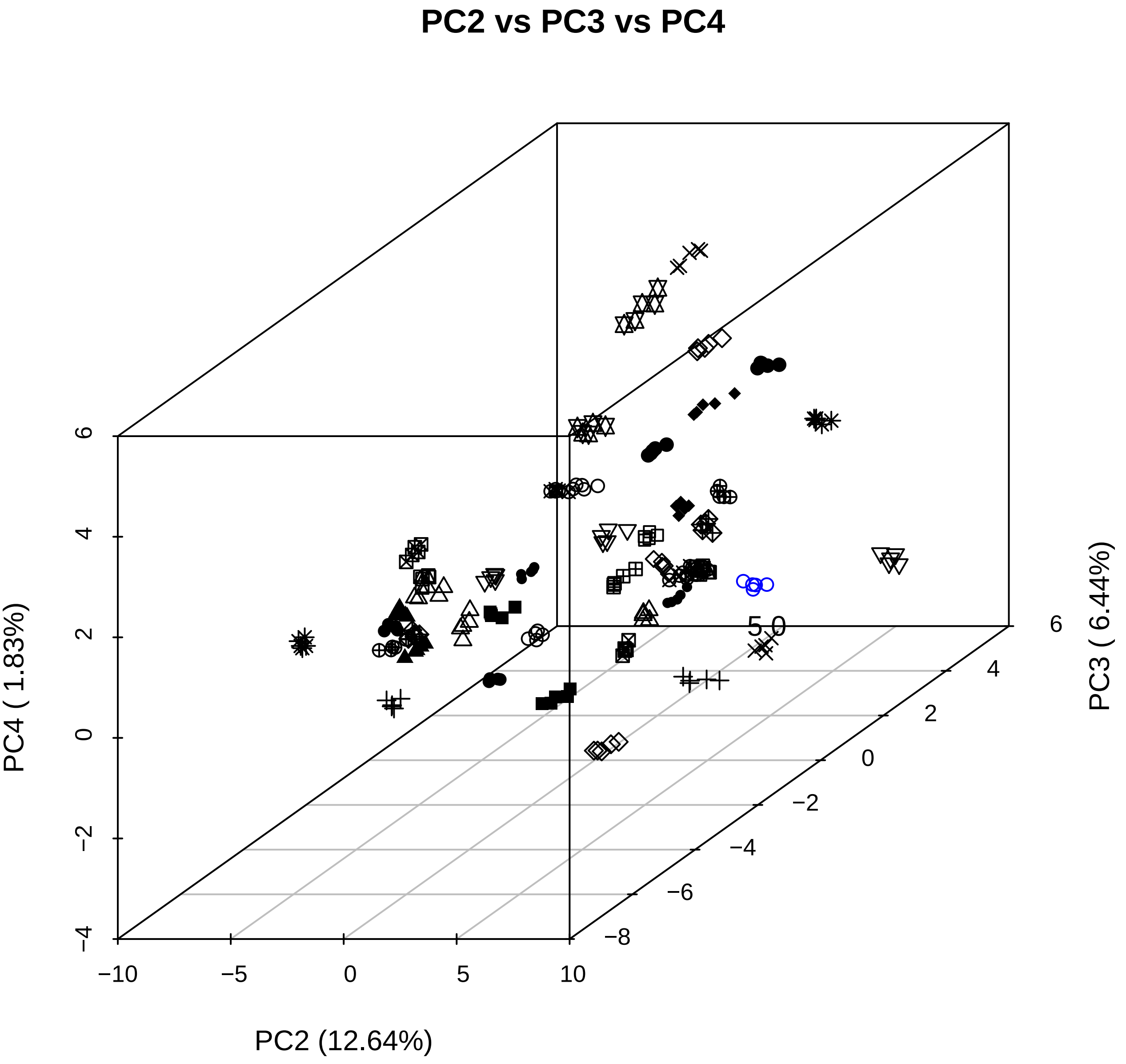
PC2 vs PC3 vs PC4
PC2 (12.64%)
PC3 ( 6.44%)
PC4 ( 1.83%)
−10
−5
0
5
10
−8
−6
−4
−2
0
2
4
6
−4
−2
0
2
4
6
5.0

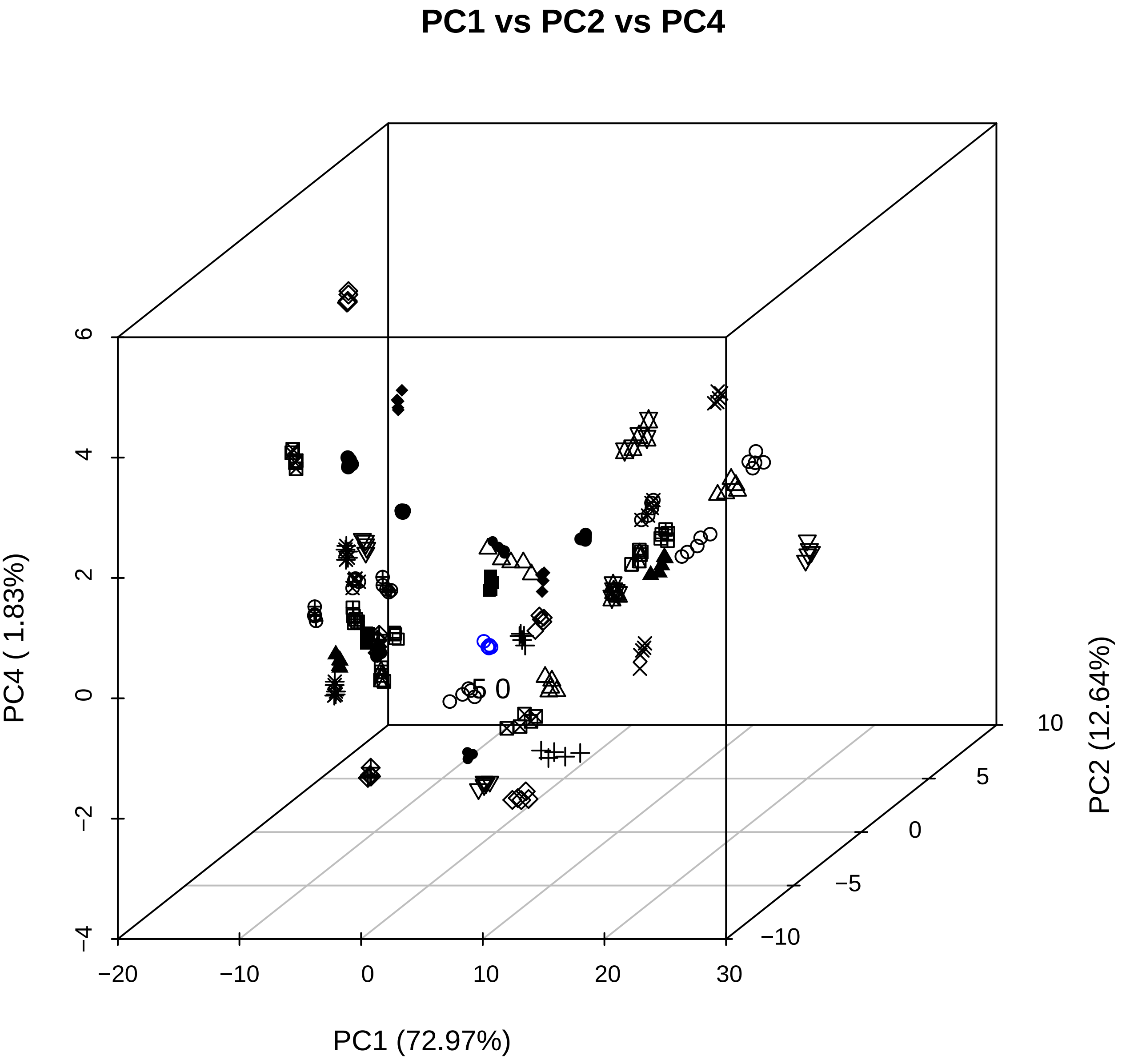
PC1 vs PC2 vs PC4
PC4 ( 1.83%)
PC1 (72.97%)
PC2 (12.64%)
−20
−10
0
10
20
30
−4
−2
0
2
4
6
−10
−5
0
5
10
5 0

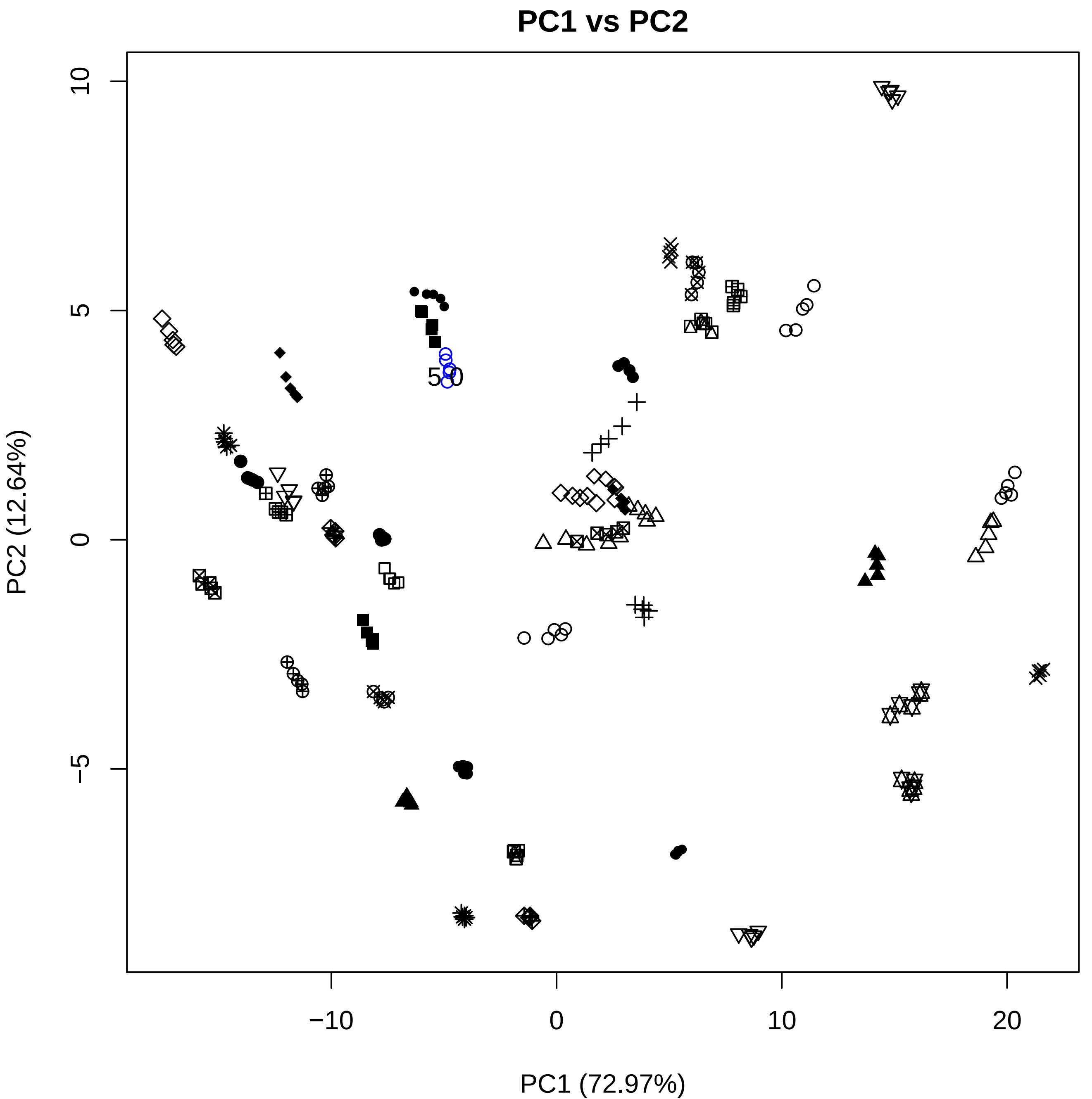
PC1 vs PC2
PC2 (12.64%)
PC1 (72.97%)
10
5
0
−5
−10
0
10
20
50

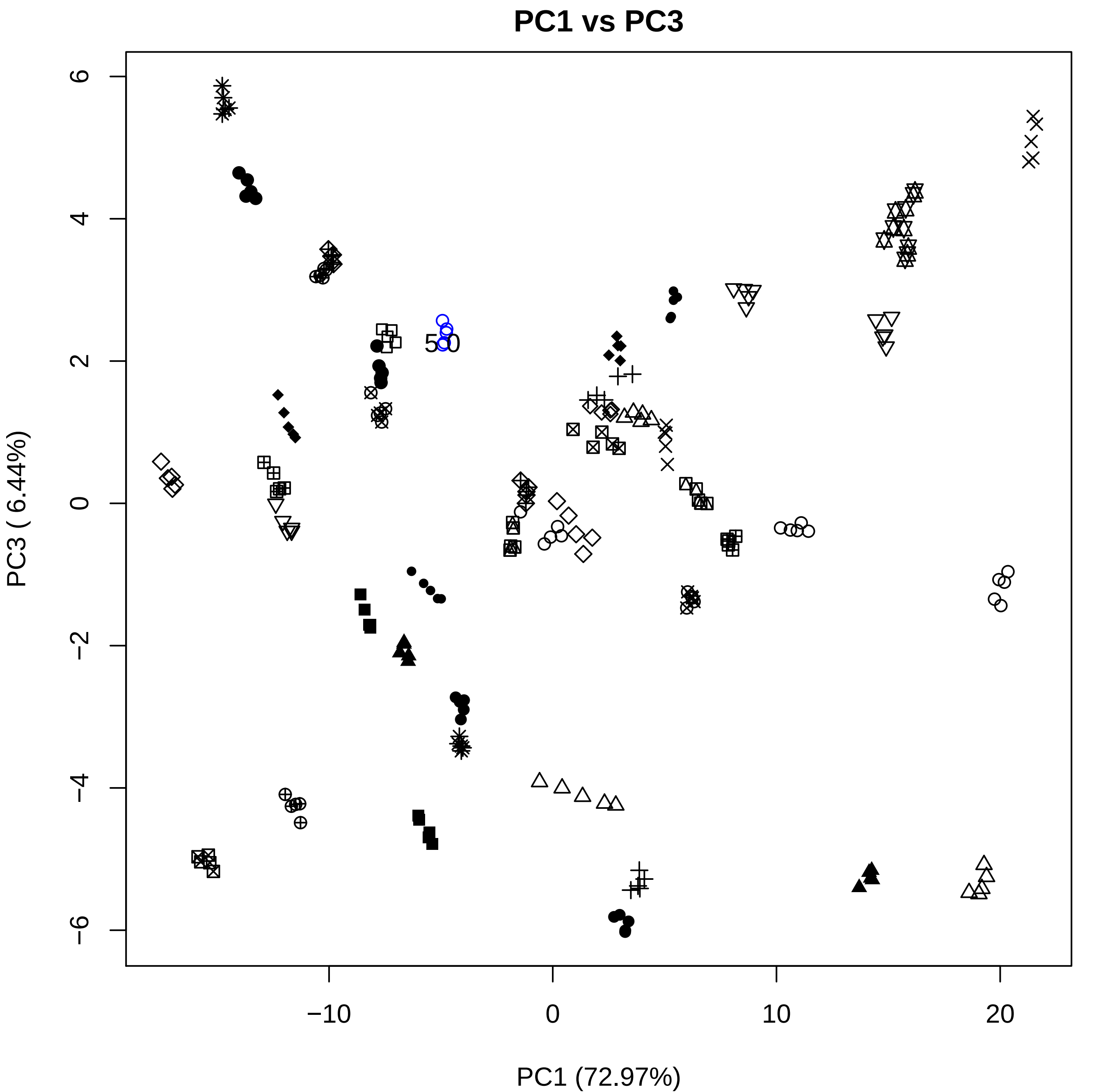
PC1 vs PC3
PC3 ( 6.44%)
PC1 (72.97%)
6
4
2
0
−2
−4
−6
−10
0
10
20
50

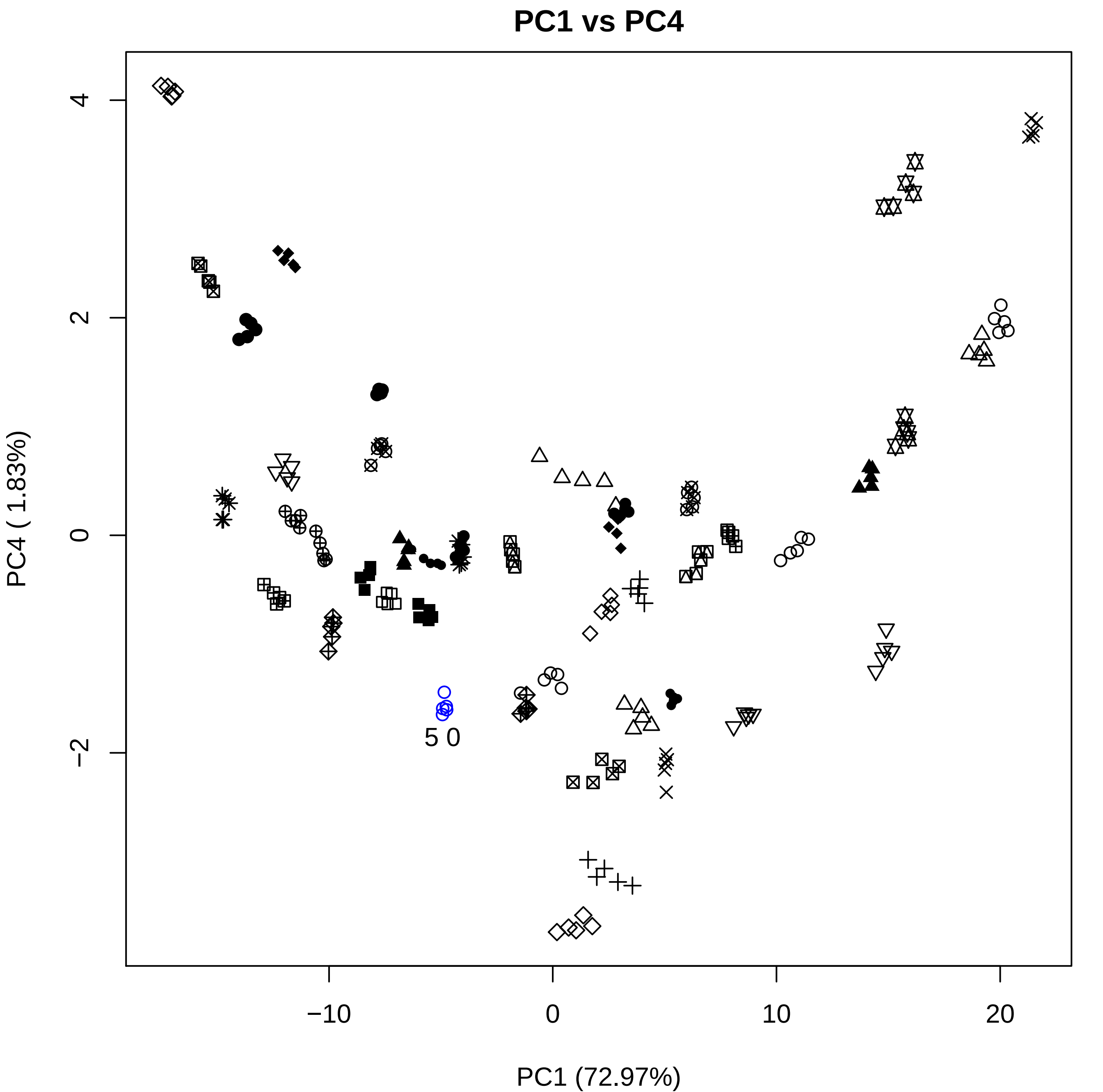
PC1 vs PC4
PC4 ( 1.83%)
PC1 (72.97%)
4
2
0
−2
−10
0
10
20
5 0

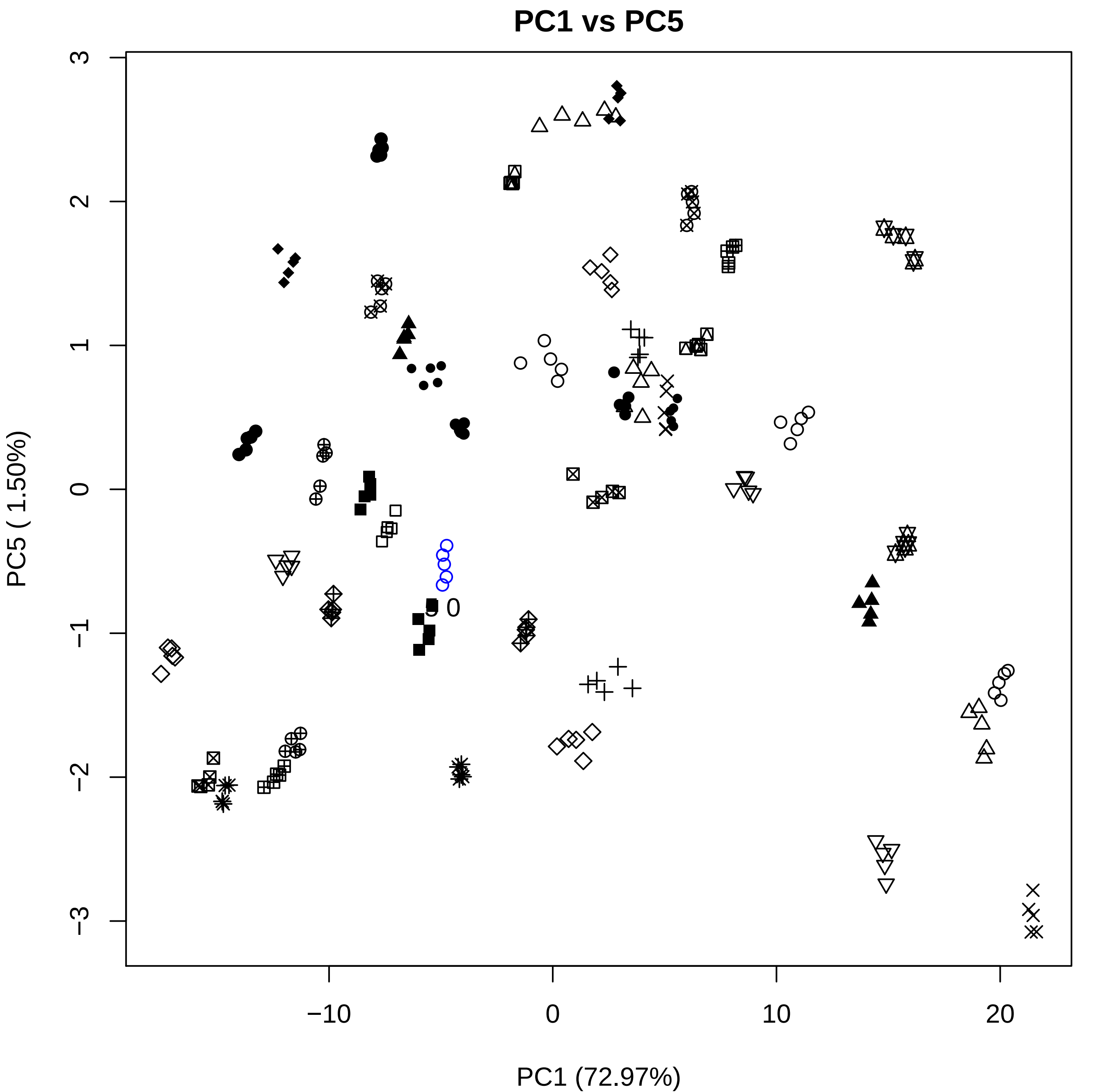

PC1 vs PC5
PC5 ( 1.50%)
PC1 (72.97%)
3
2
1
0
−1
−2
−3
−10
0
10
20

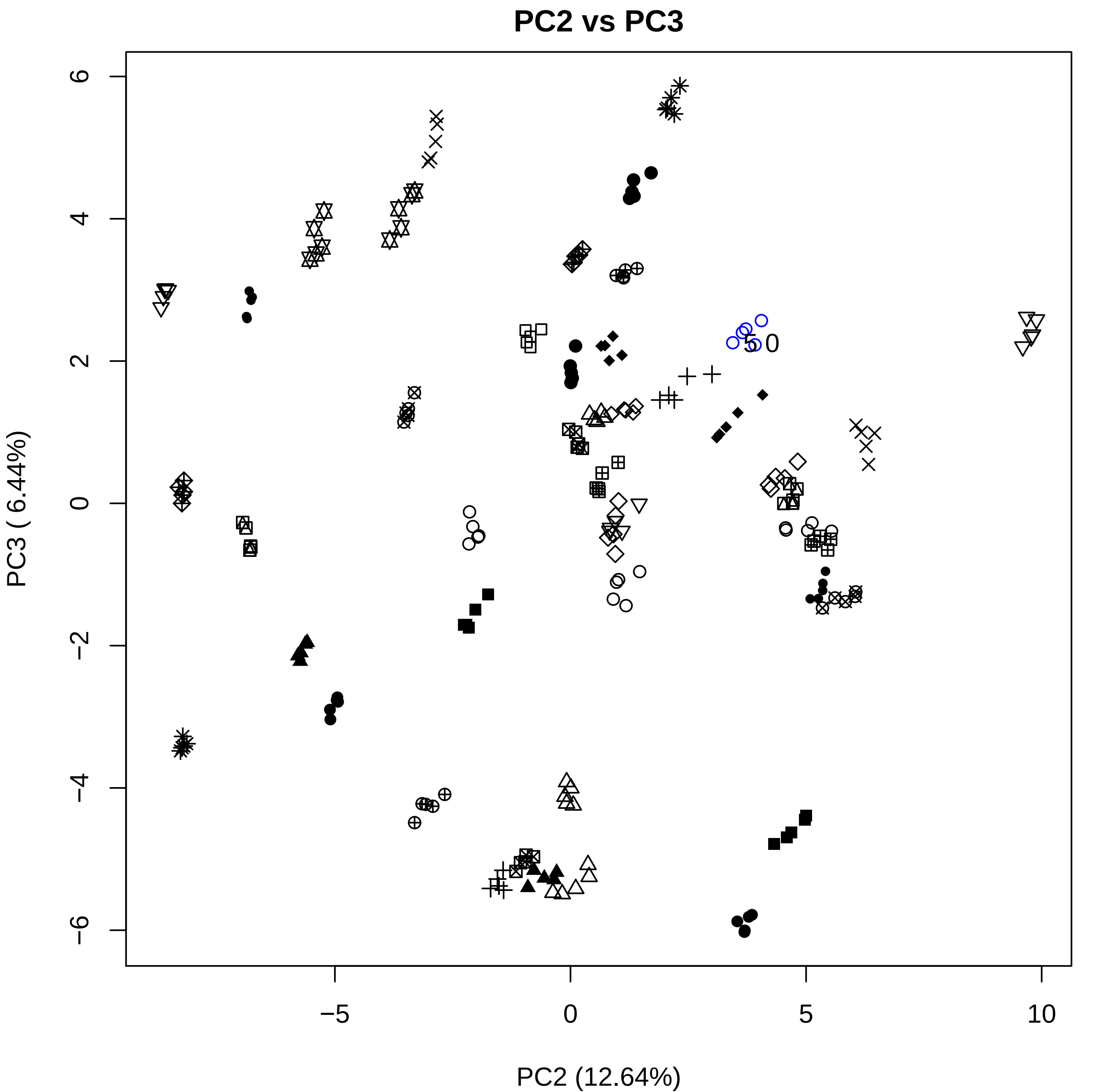
PC2 vs PC3
PC3 ( 6.44%)
PC2 (12.64%)
−6
−4
−2
0
2
4
6
−5
0
5
10
50

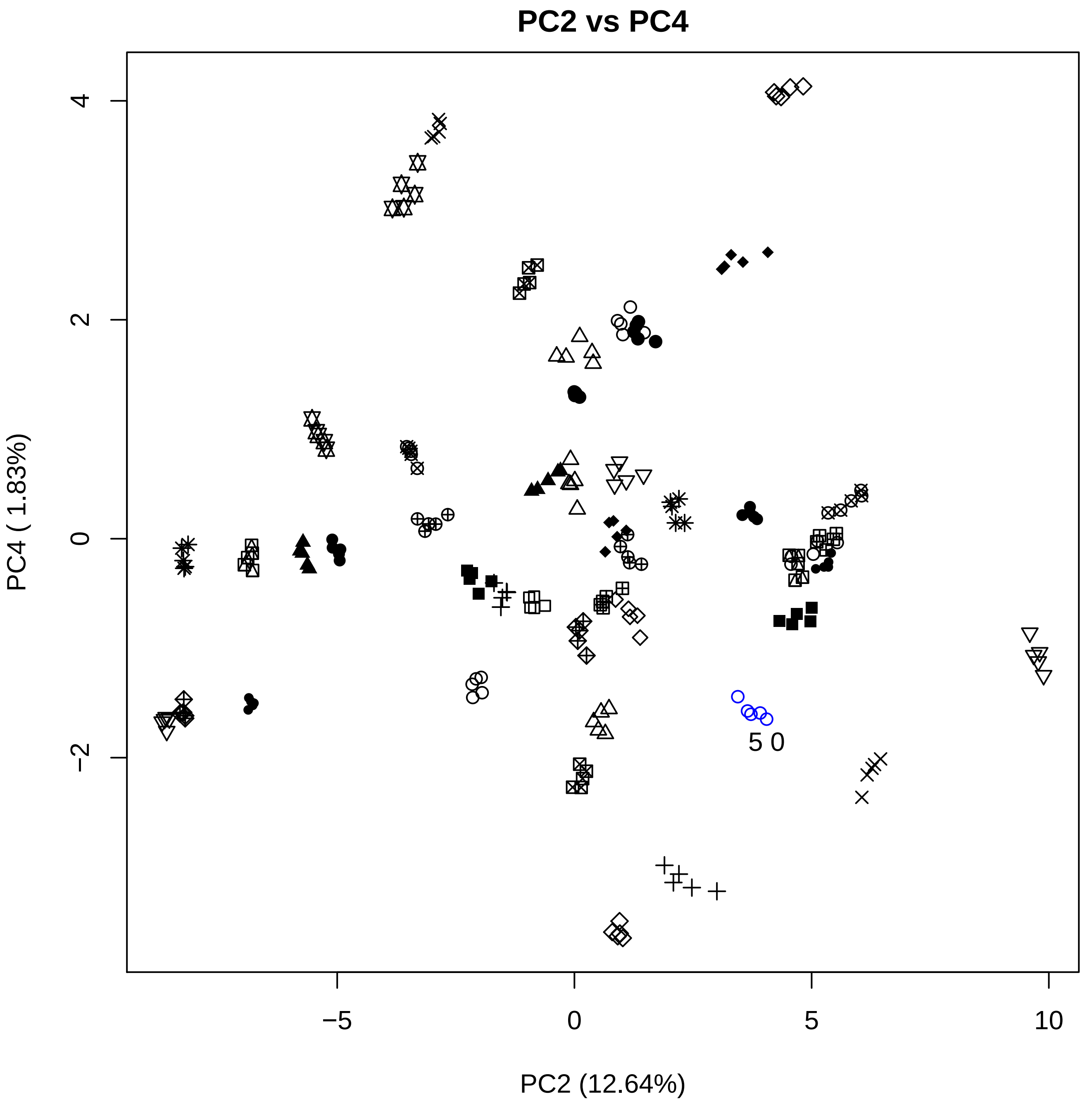

PC2 vs PC4
PC4 ( 1.83%)
PC2 (12.64%)
4
2
0
−2
−5
0
5
10
5 0

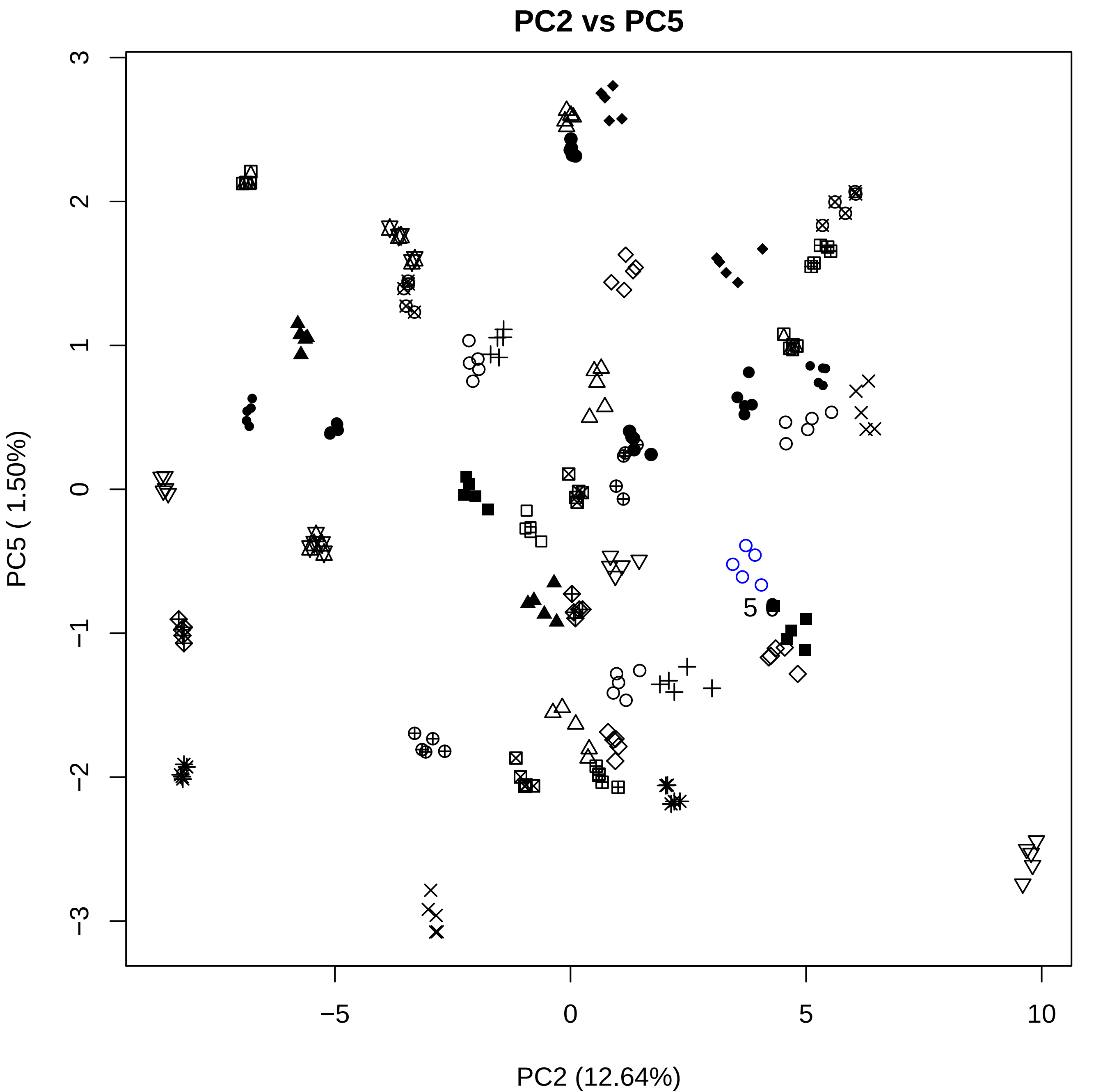
PC2 vs PC5
PC5 ( 1.50%)
PC2 (12.64%)
3
2
1
0
−1
−2
−3
−5
0
5
10
5

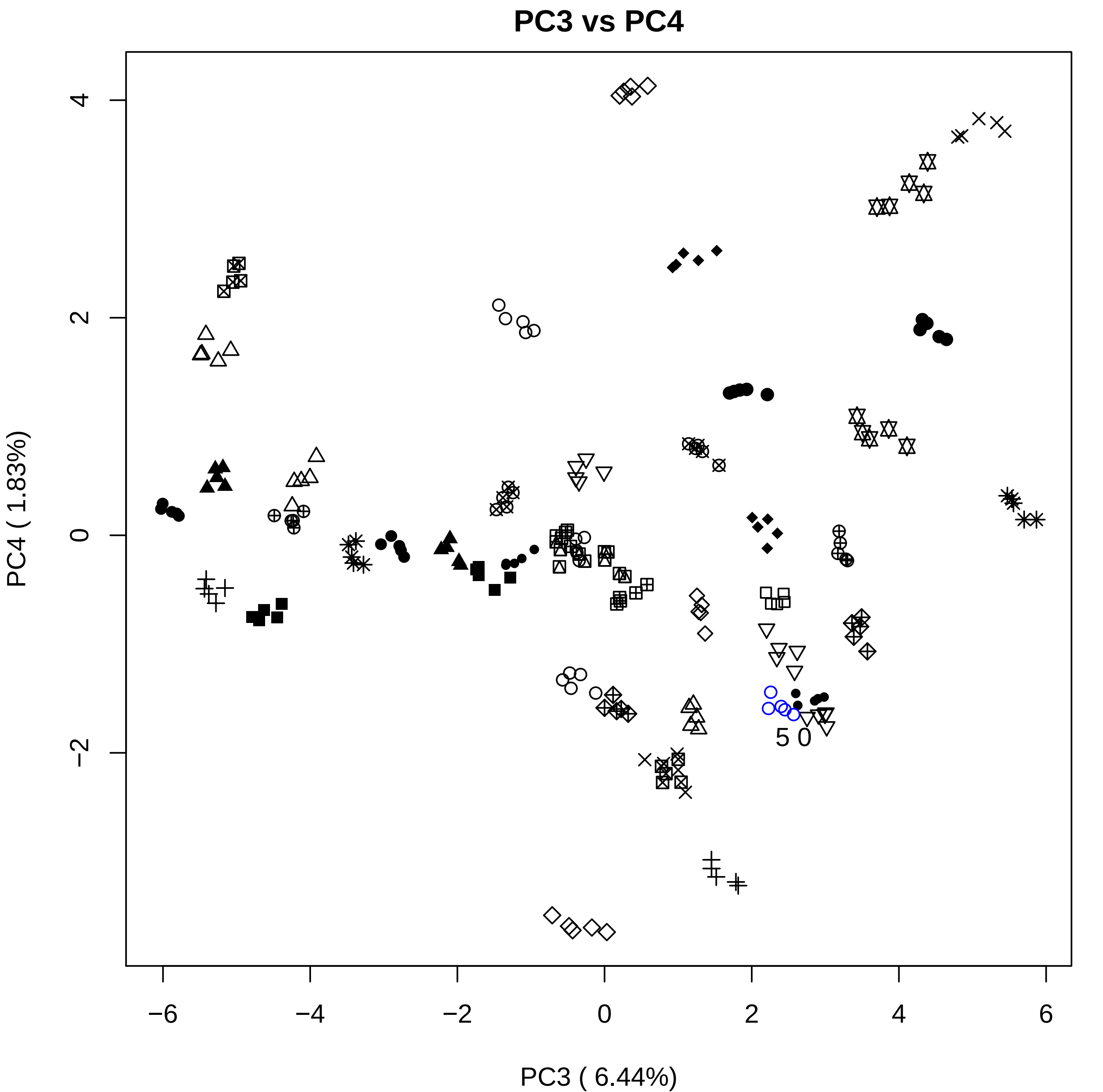
PC3 vs PC4
PC4 ( 1.83%)
PC3 ( 6.44%)
4
2
0
−2
−6
−4
−2
0
2
4
6
5 0

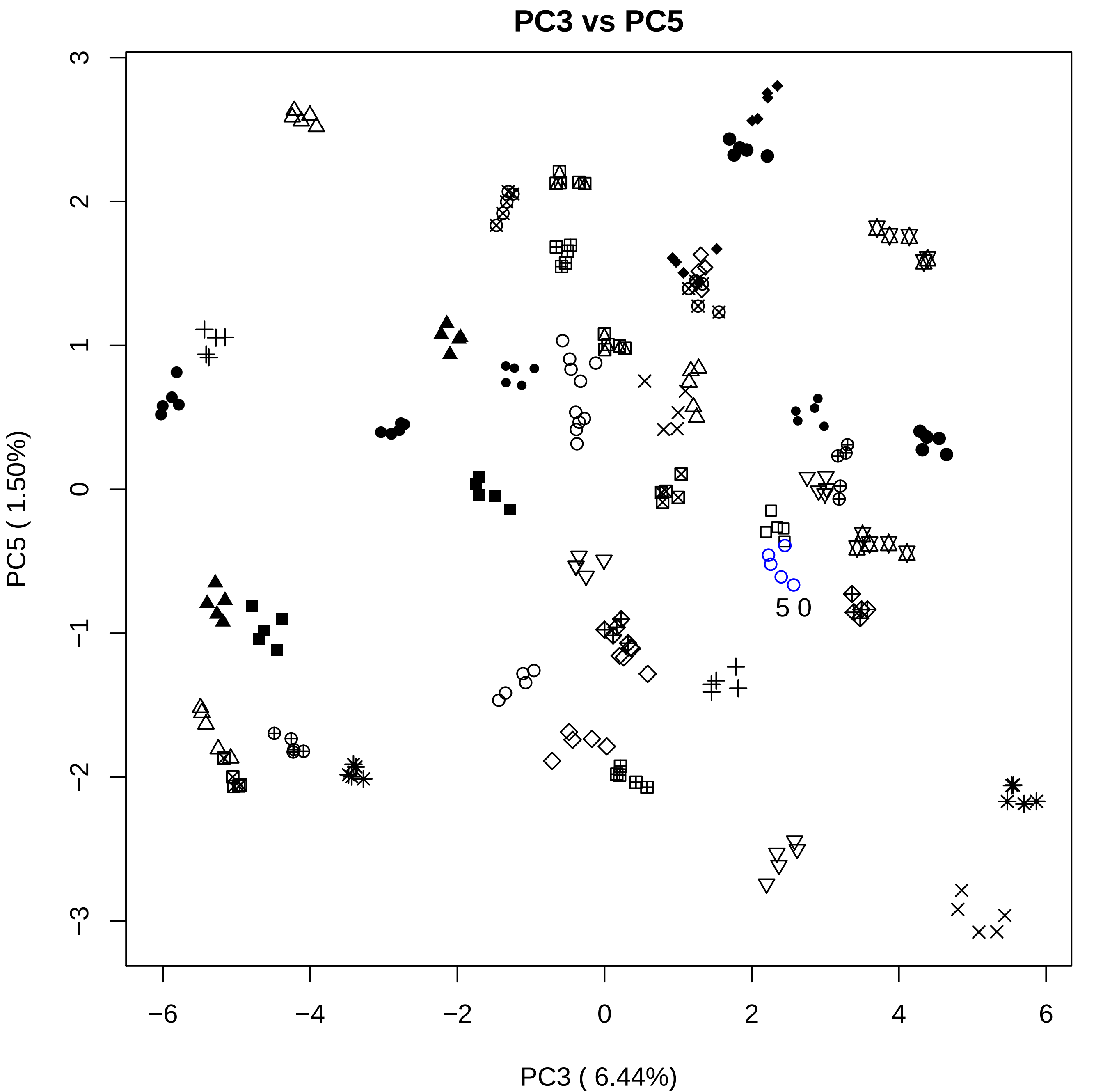
PC3 vs PC5
PC5 ( 1.50%)
PC3 ( 6.44%)
−6
−4
−2
0
2
4
6
−3
−2
−1
0
1
2
3
5 0

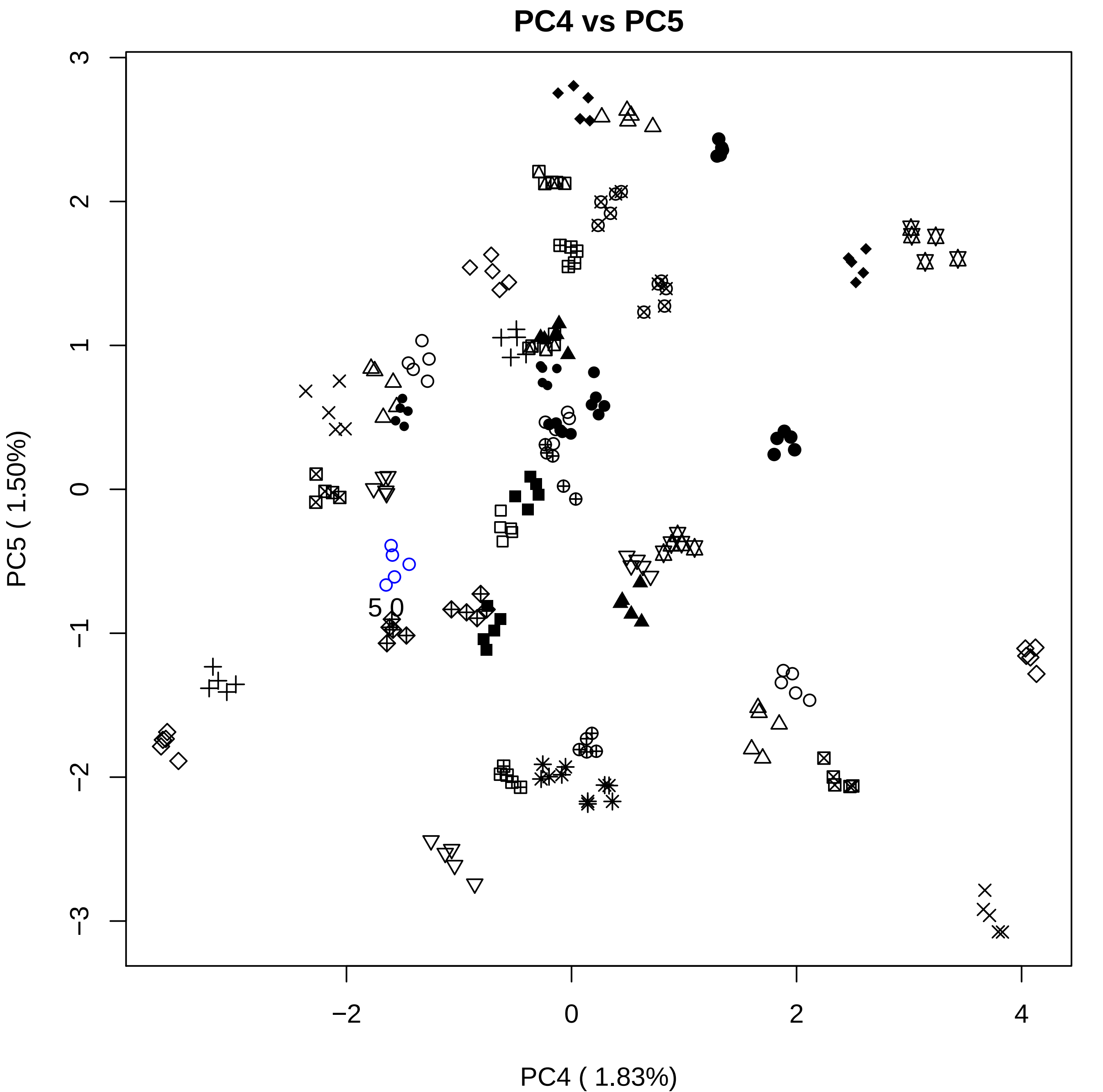
PC4 vs PC5
PC5 ( 1.50%)
PC4 ( 1.83%)
3
2
1
0
−1
−2
−3
−2
0
2
4
50

### PCA of 7.5%  OSAS vs PMH

Scaled Matrix PCAed

7th order polynomial baseline correction performed

Raw spectra smoothed to 21 Neighbours, 2nd order PN, Savtisky Golay

Normalisation between 0 and 100

D2 spectra smoothed to 41 Neighbours, 2nd order PN, Savtisky Golay

Regions Analysed: 3601−2501 and 2001−700cm−1

Samples Analysed: 1 Contam's 0 OSAS's 45 PMH's

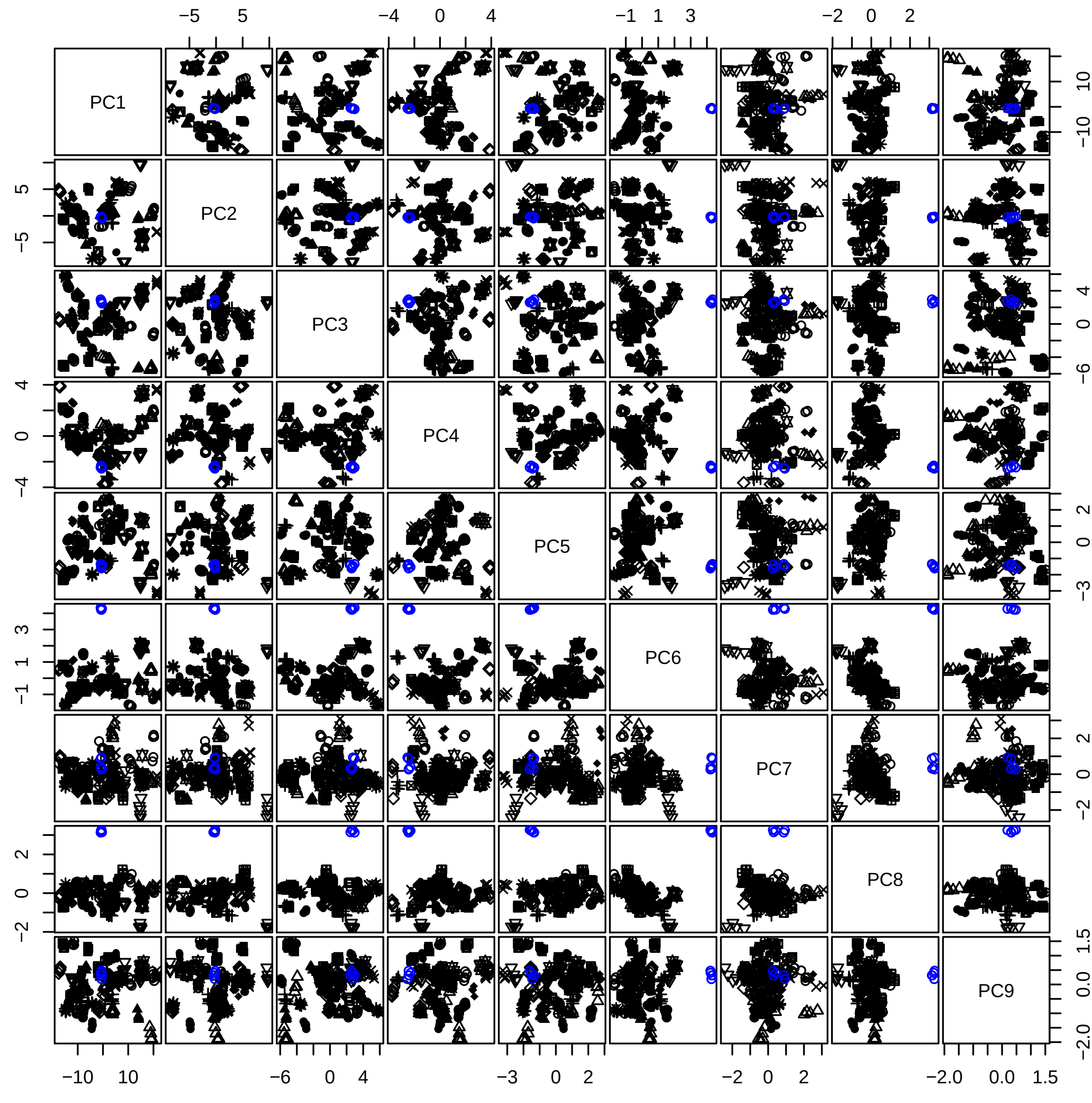

PC1
PC2
PC3
PC4
PC5
PC6
PC7
PC8
PC9

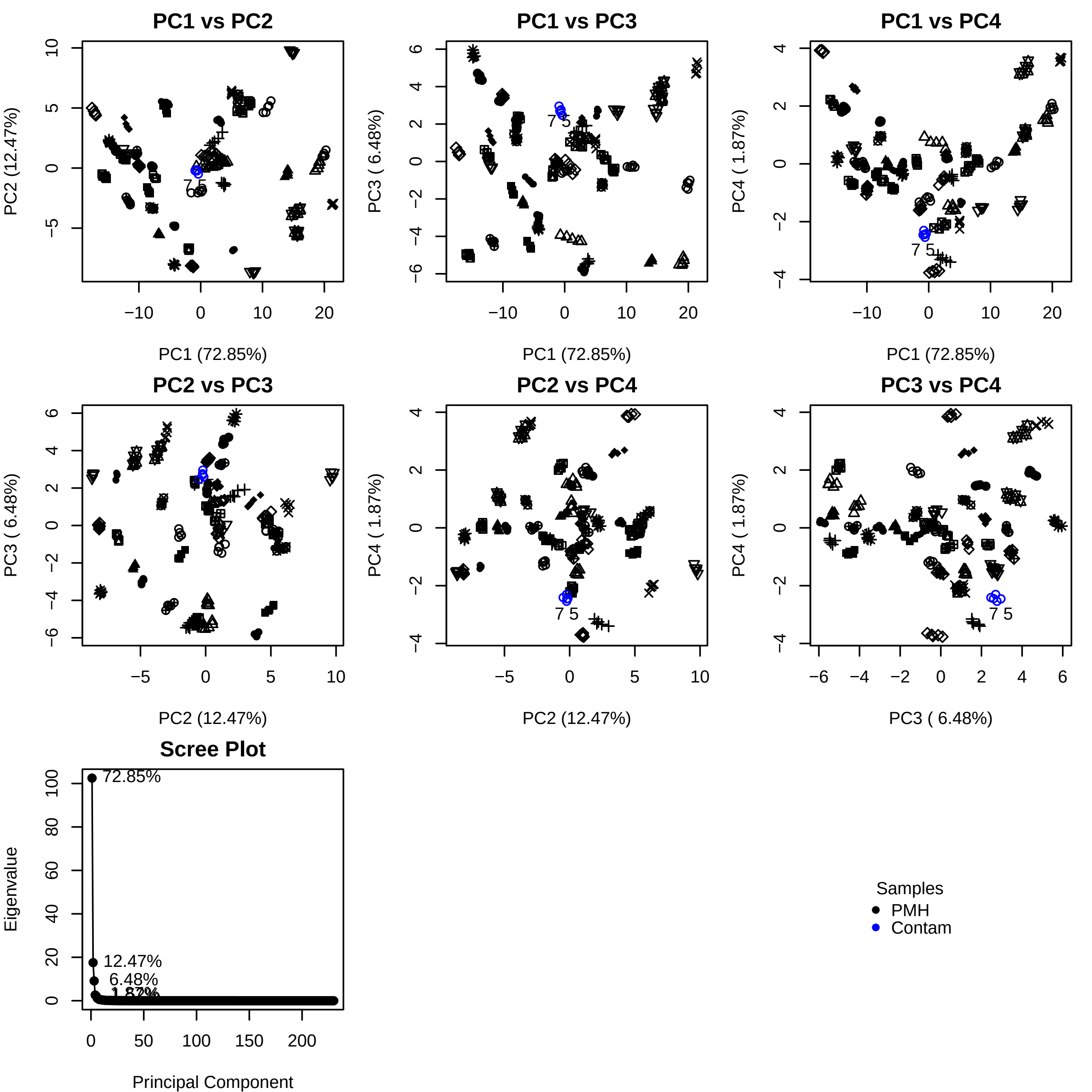

PC1 vs PC2
PC1 vs PC3
PC1 vs PC4
PC2 vs PC3
PC2 vs PC4
PC3 vs PC4
Scree Plot
PC1 (72.85%)
PC2 (12.47%)
PC3 ( 6.48%)
PC4 ( 1.87%)
7 5
72.85%
12.47%
6.48%
Eigenvalue
Principal Component
Samples
PMH
Contam

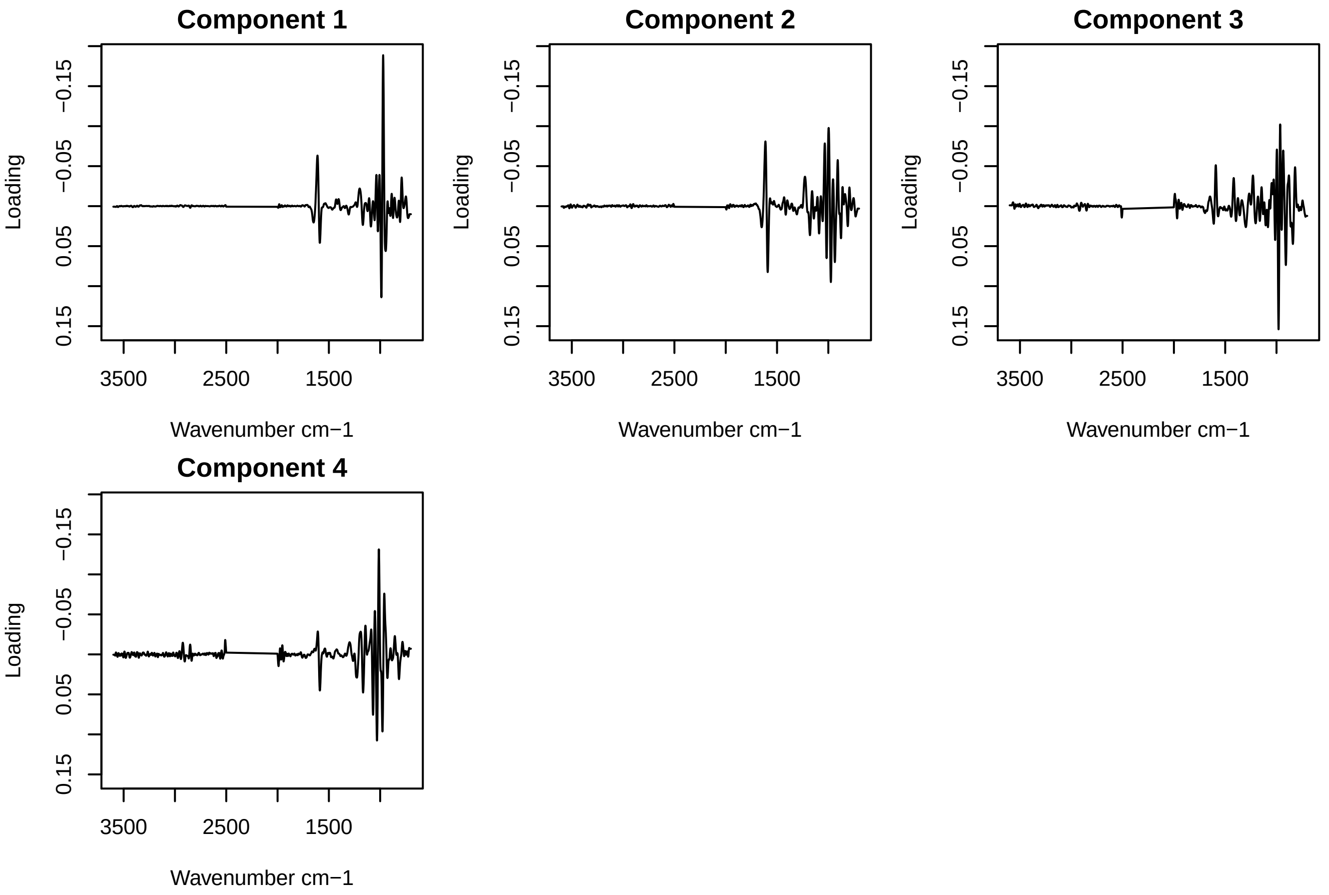
Component 1
Component 2
Component 3
Component 4
Loading
−0.15
−0.05
0.05
0.15
3500
2500
1500
Wavenumber cm−1

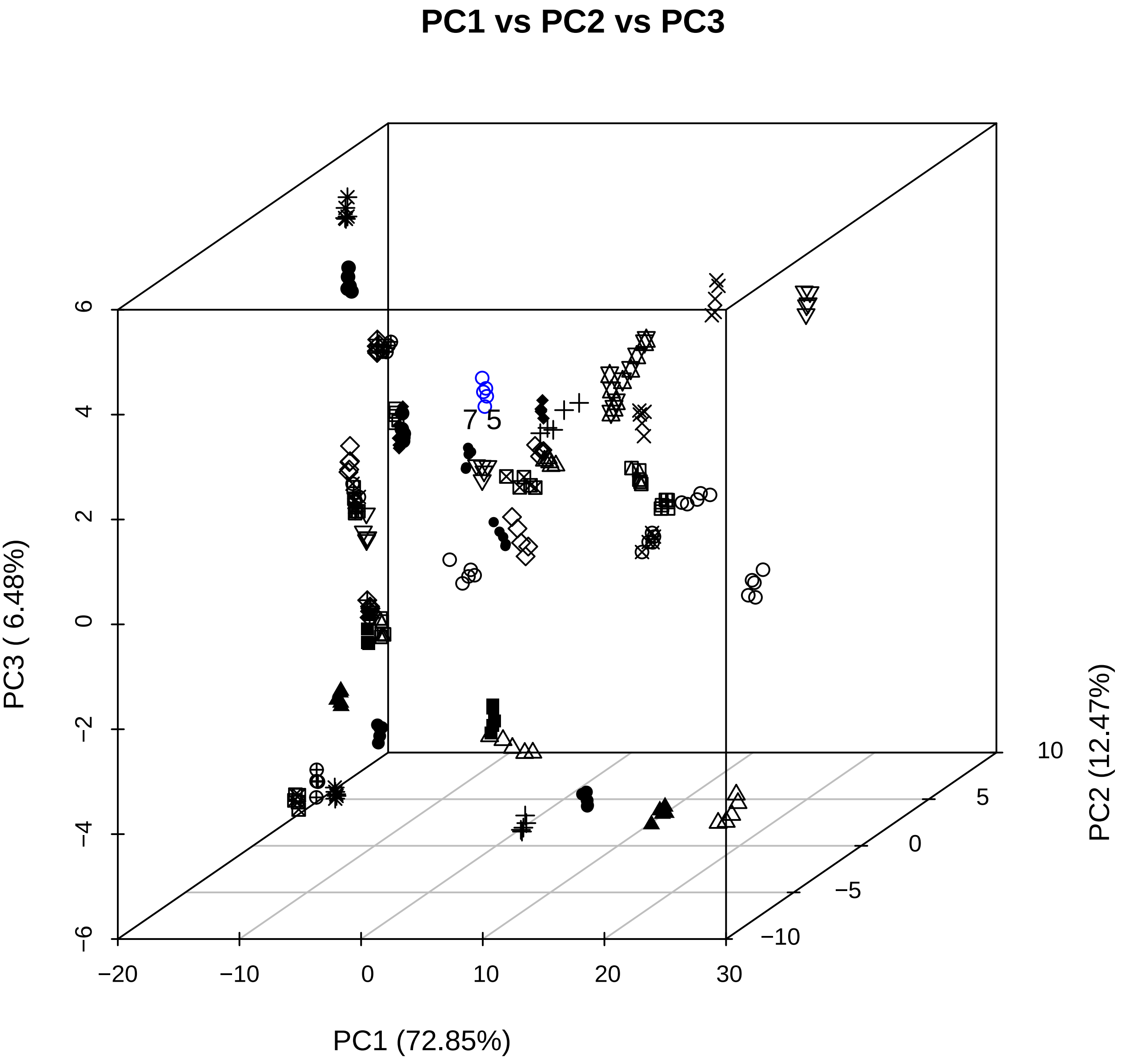
PC1 vs PC2 vs PC3
PC1 (72.85%)
PC2 (12.47%)
PC3 ( 6.48%)
−20
−10
0
10
20
30
−10
−5
0
5
10
−6
−4
−2
0
2
4
6
7 5

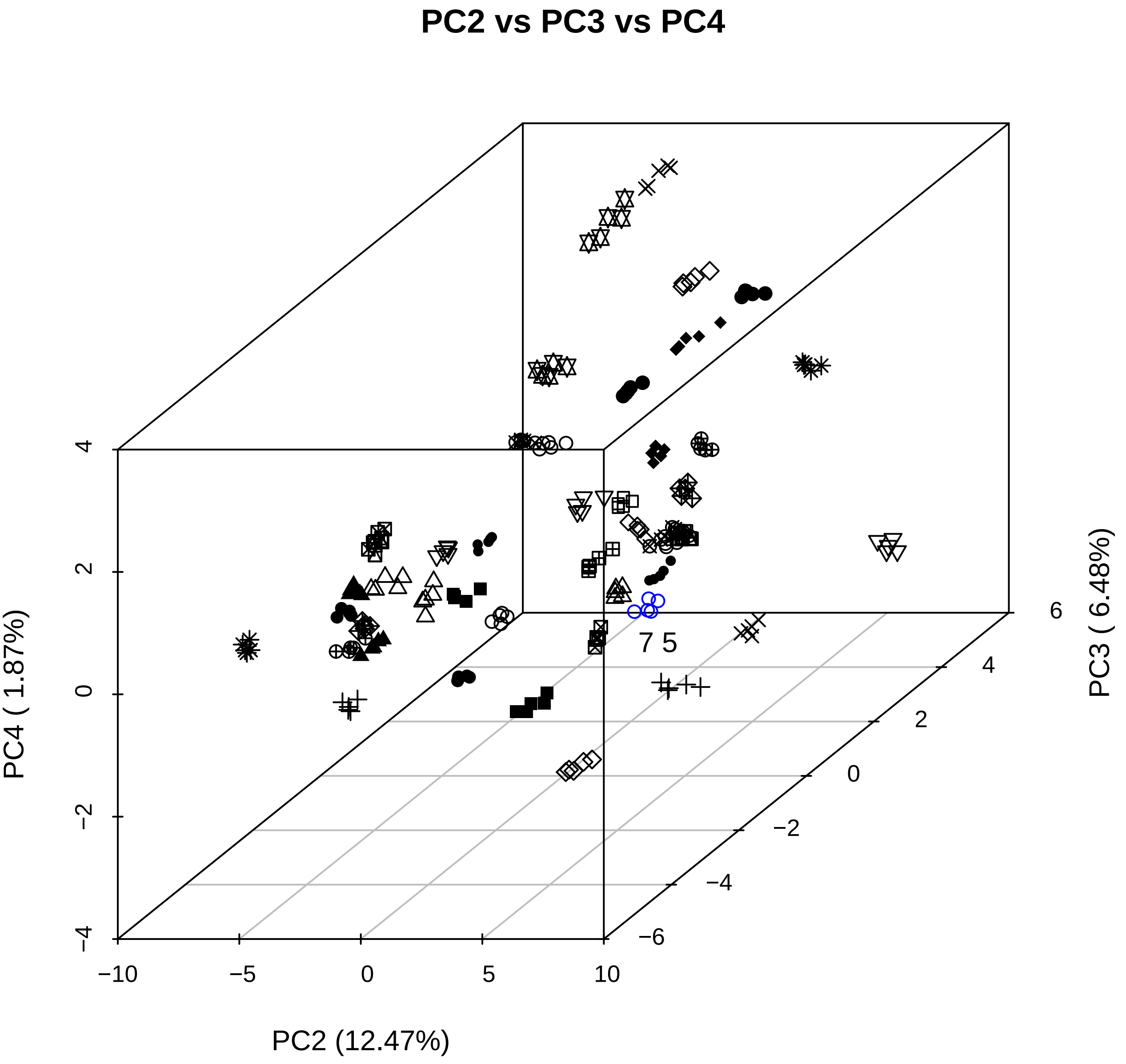

PC2 vs PC3 vs PC4
PC4 ( 1.87%)
PC3 ( 6.48%)
PC2 (12.47%)
−10
−5
0
5
10
−4
−2
0
2
4
−6
−4
−2
0
2
4
6
7 5

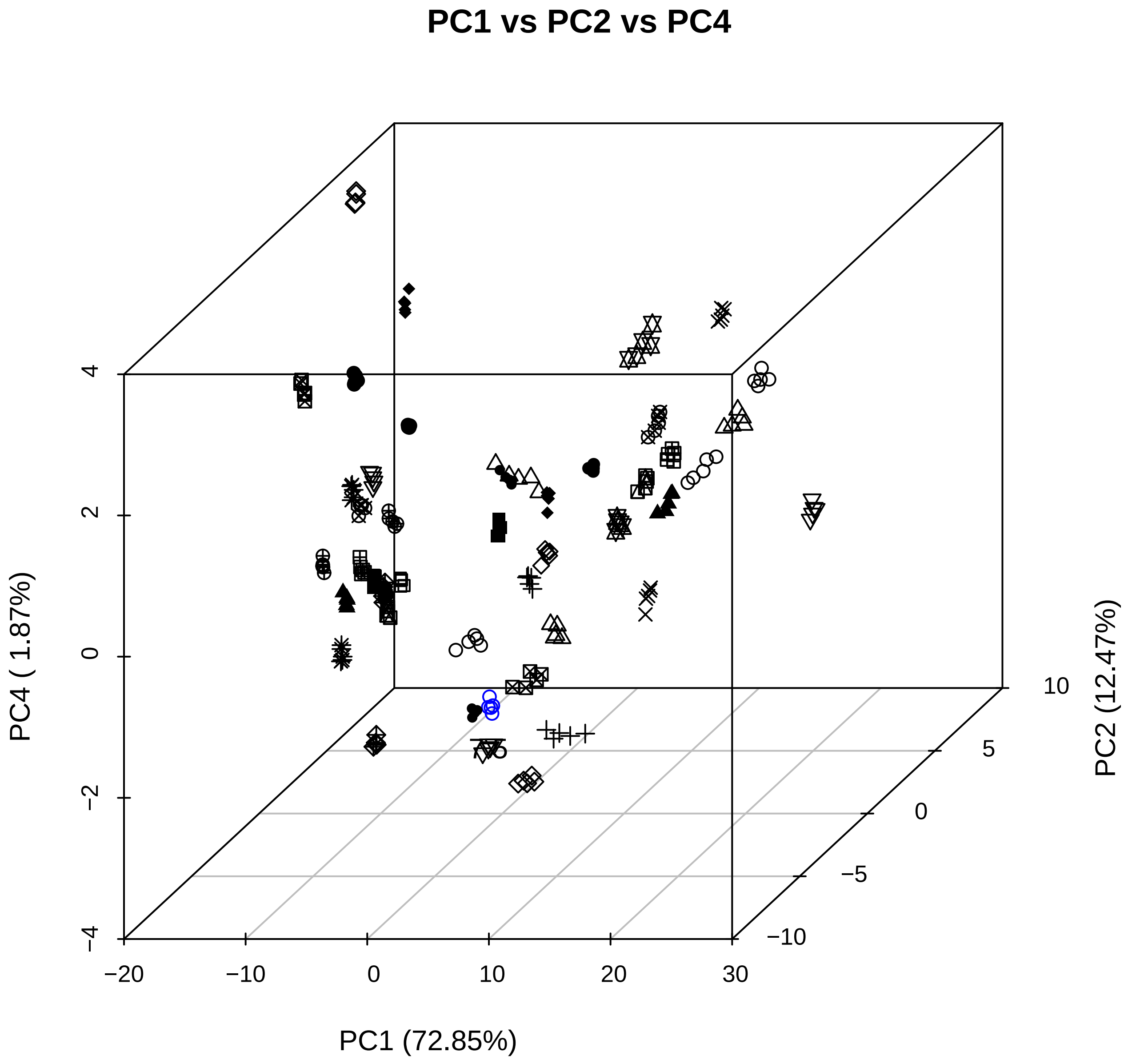
PC1 vs PC2 vs PC4
PC1 (72.85%)
PC2 (12.47%)
PC4 ( 1.87%)
−20
−10
0
10
20
30
−10
−5
0
5
10
−4
−2
0
2
4

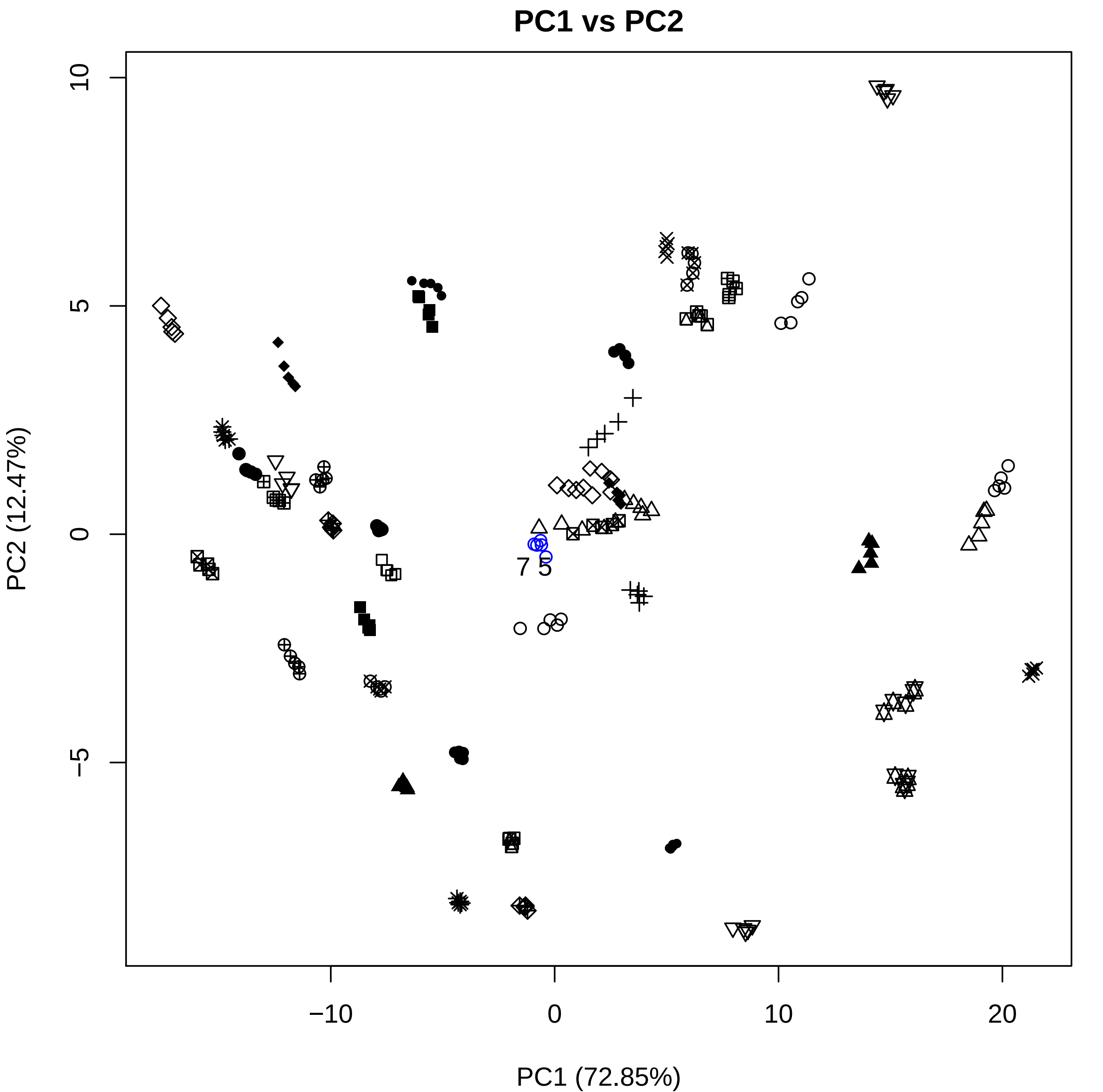
PC1 vs PC2
PC2 (12.47%)
PC1 (72.85%)
10
5
0
−5
−10
0
10
20
7 5

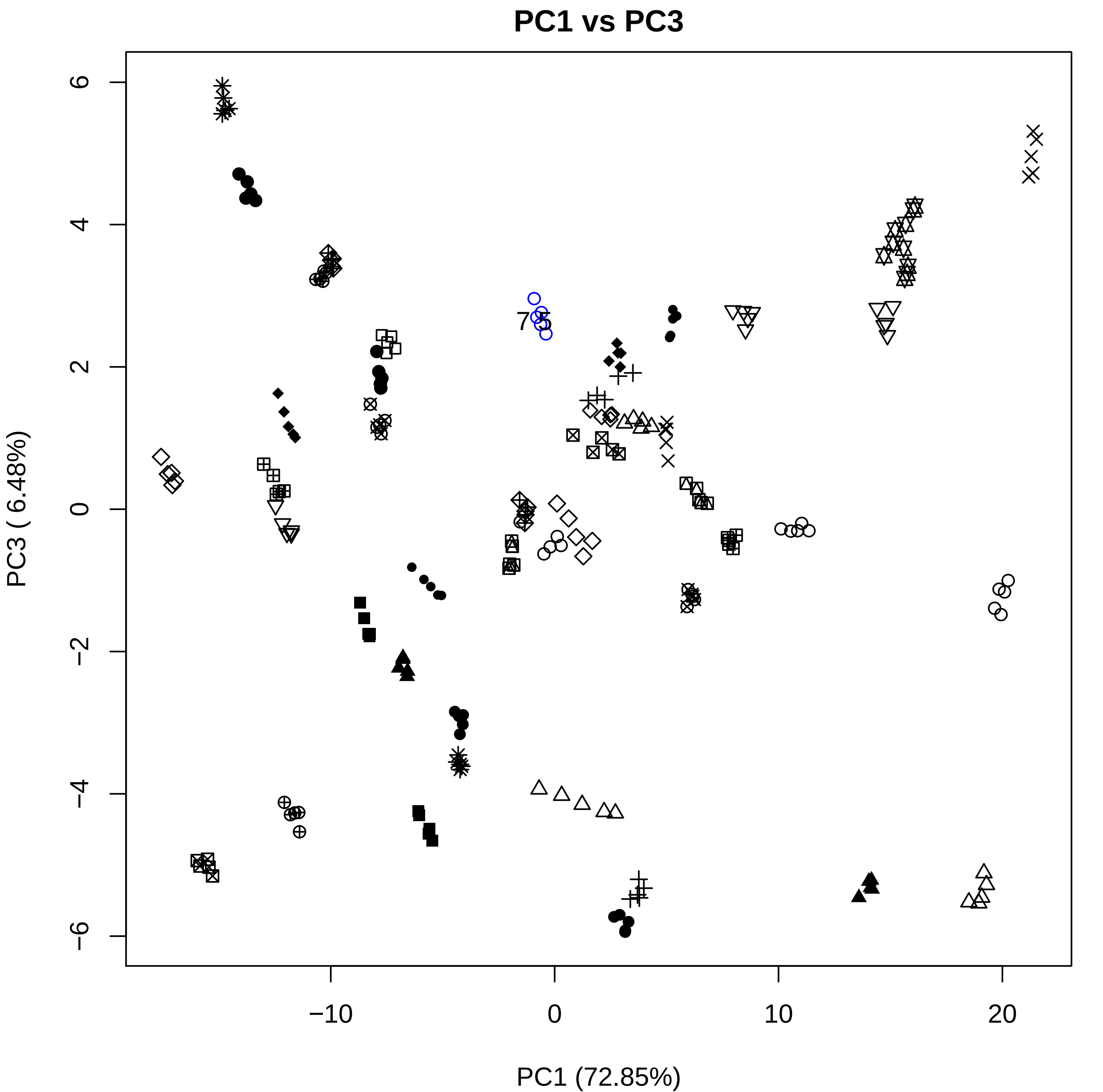
PC1 vs PC3
PC3 ( 6.48%)
PC1 (72.85%)
6
4
2
0
−2
−4
−6
−10
0
10
20
75

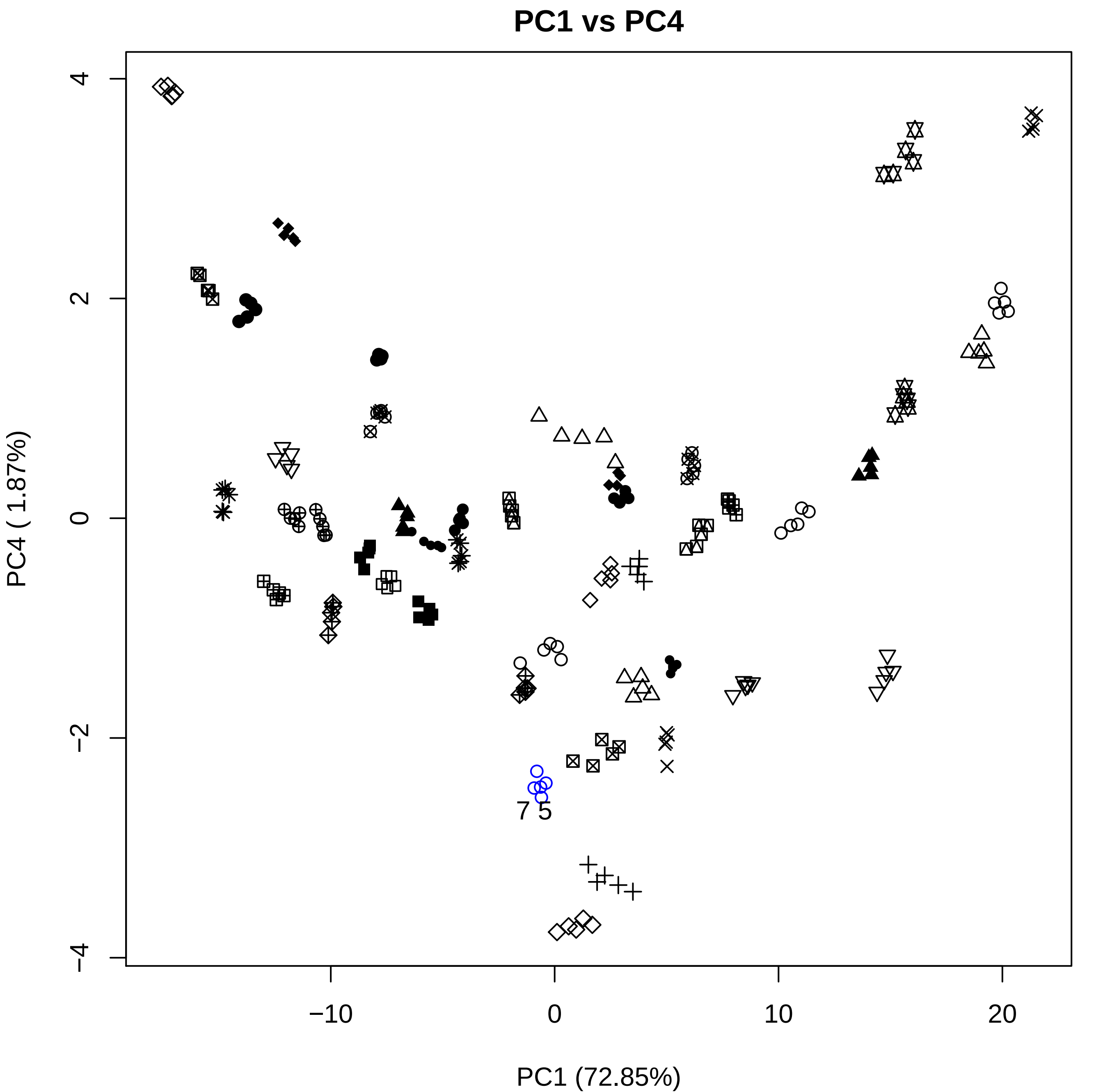
PC1 vs PC4
PC4 ( 1.87%)
PC1 (72.85%)
4
2
0
−2
−4
−10
0
10
20
7 5

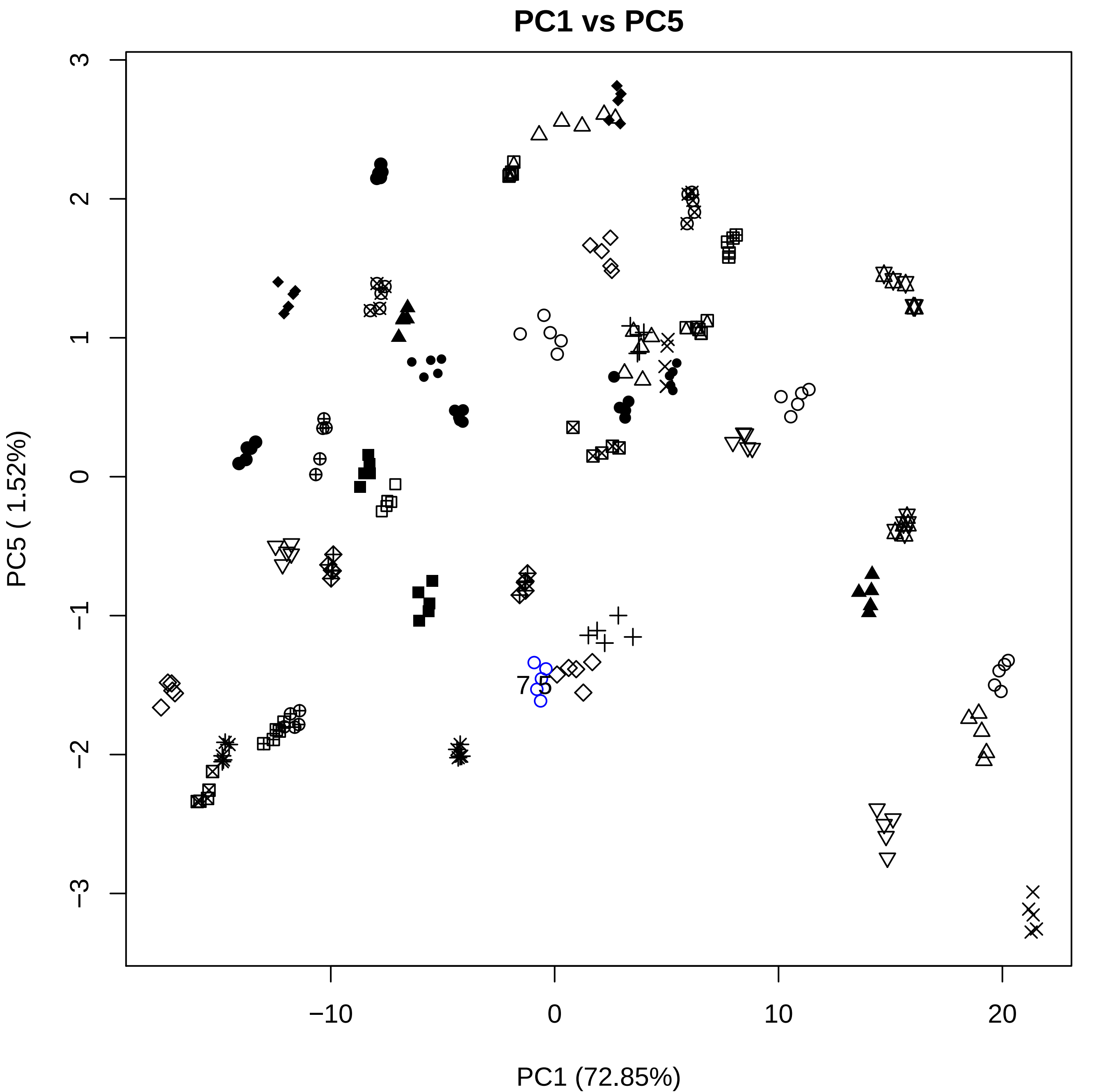
PC1 vs PC5
PC5 ( 1.52%)
PC1 (72.85%)
3
2
1
0
−1
−2
−3
−10
0
10
20
75

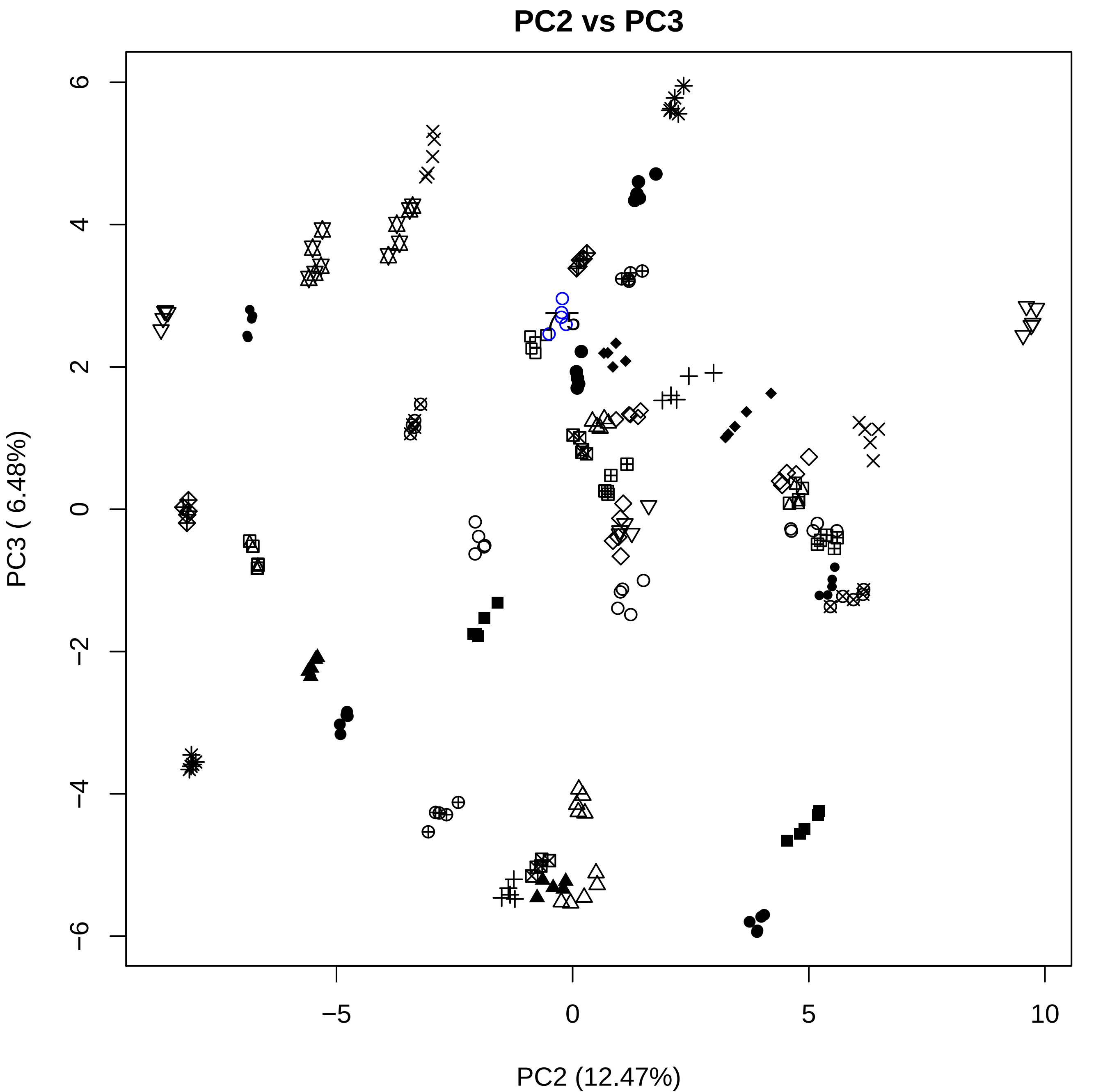

PC2 vs PC3
6
4
2
0
−2
−4
−6
PC3 ( 6.48%)
−5
0
5
10
PC2 (12.47%)

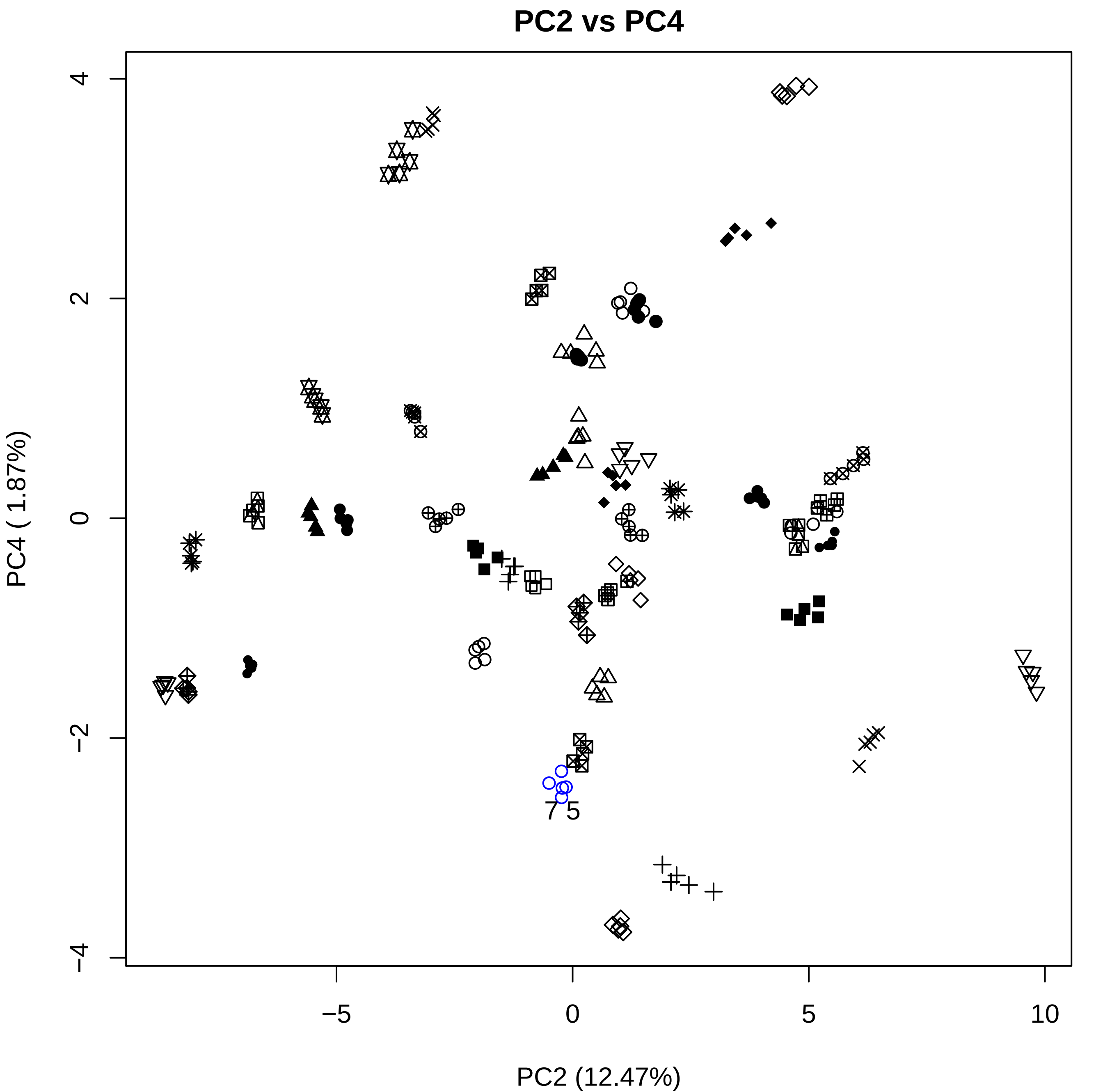
PC2 vs PC4
PC4 ( 1.87%)
PC2 (12.47%)
4
2
0
−2
−4
−5
0
5
10
7 5

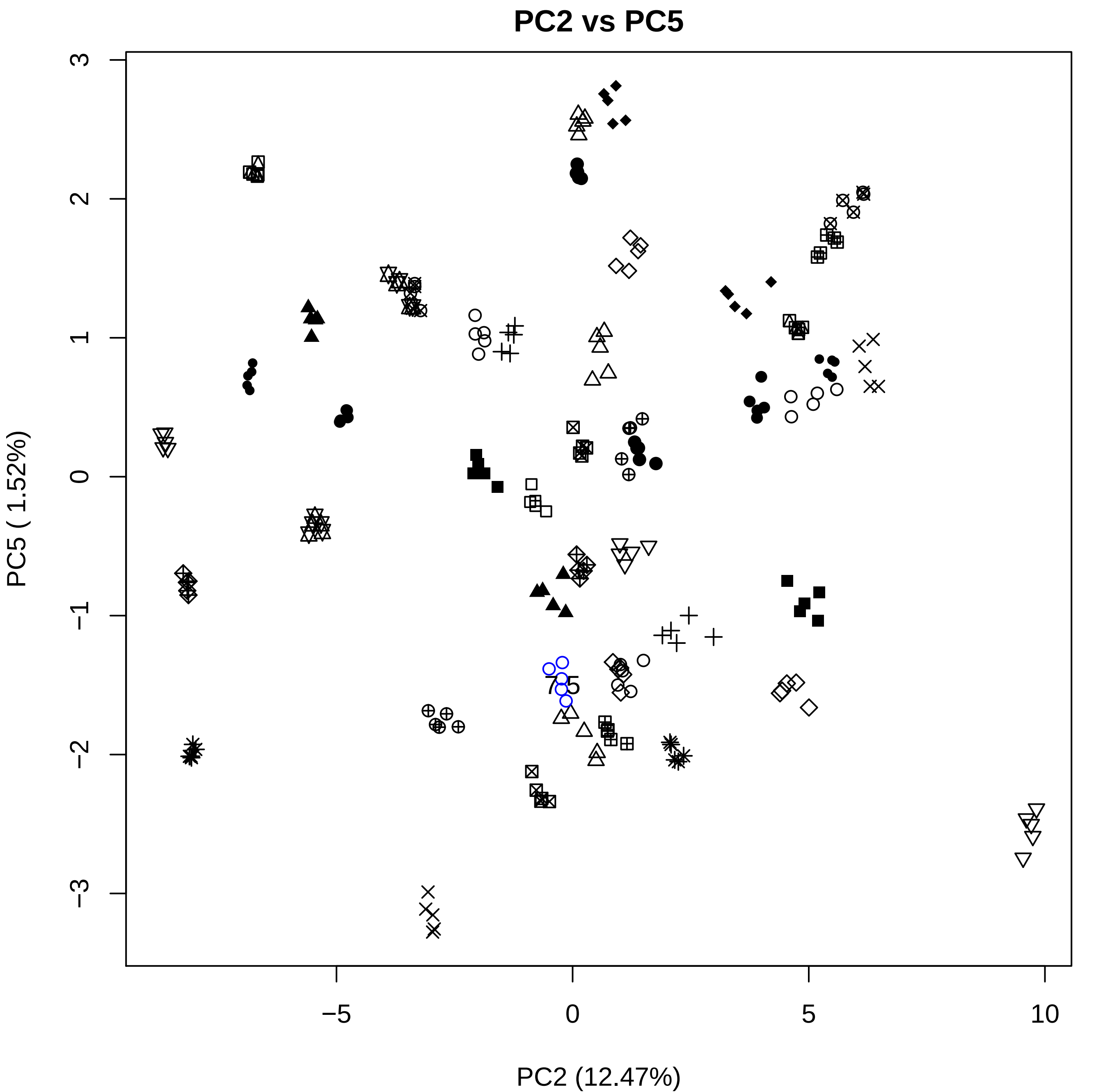
PC2 vs PC5
PC5 ( 1.52%)
PC2 (12.47%)
3
2
1
0
−1
−2
−3
−5
0
5
10
75

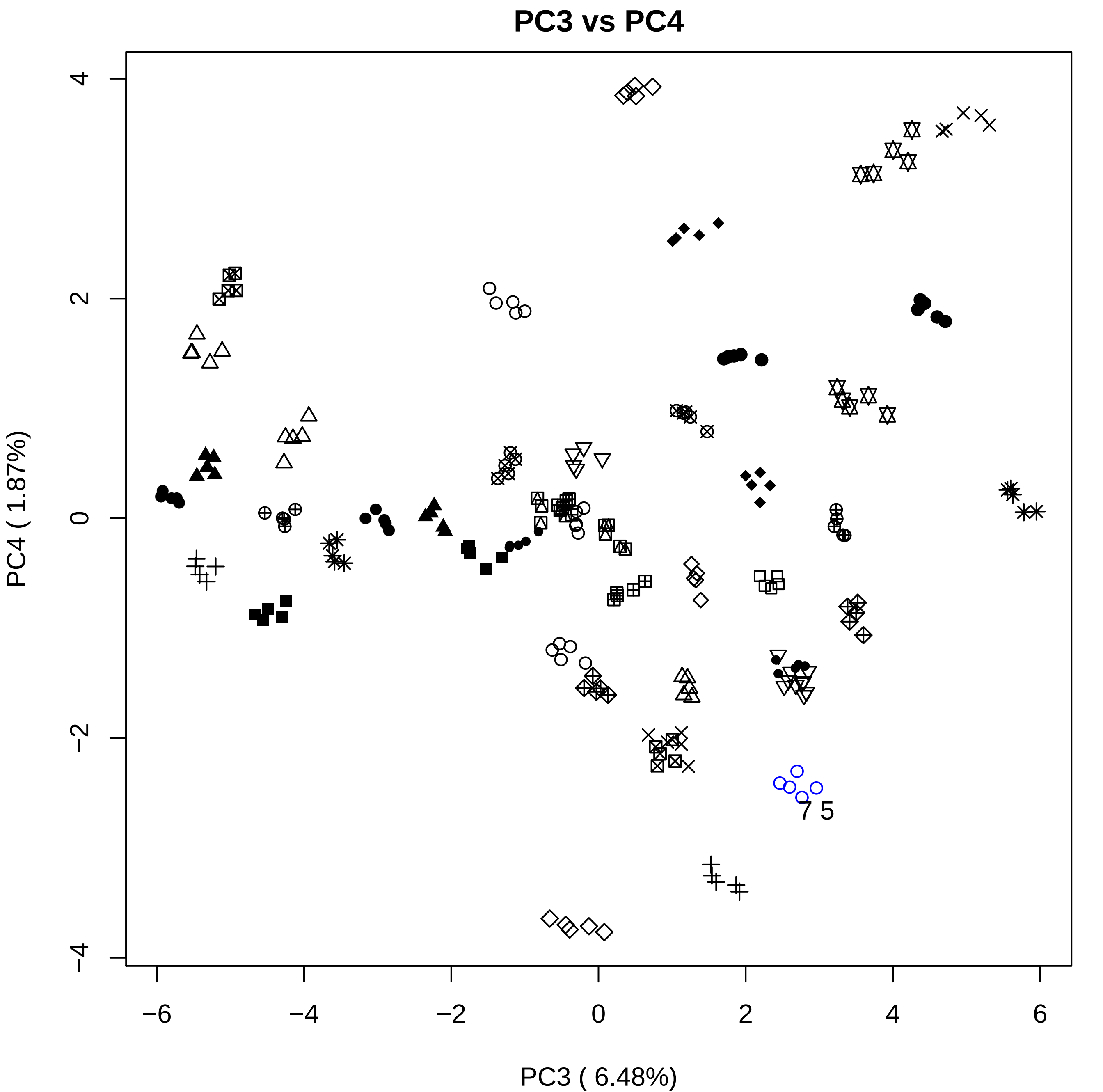
PC3 vs PC4
PC4 ( 1.87%)
PC3 ( 6.48%)
4
2
0
−2
−4
−6
−4
−2
0
2
4
6
7 5

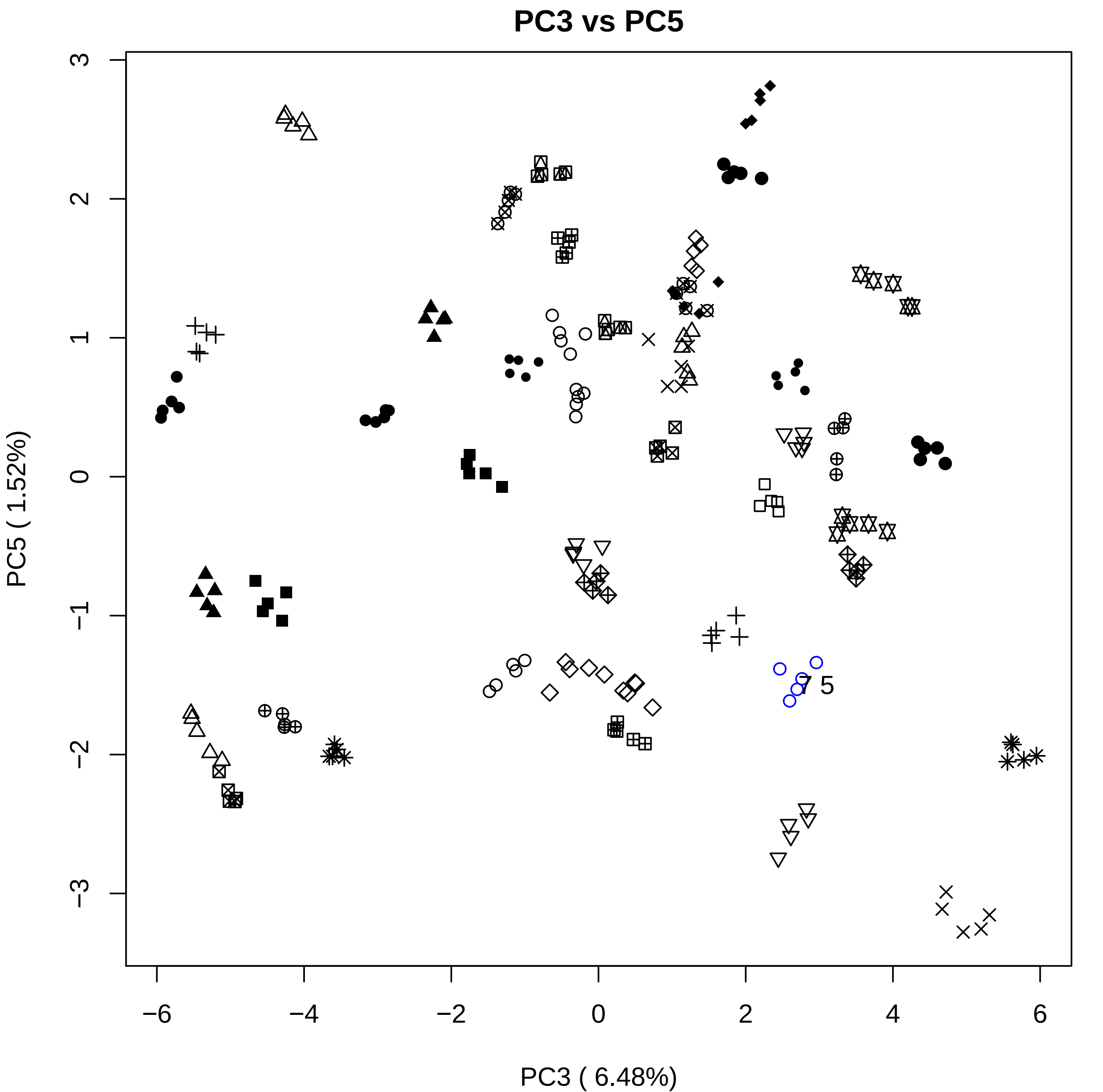
PC3 vs PC5
PC5 ( 1.52%)
PC3 ( 6.48%)
3
2
1
0
−1
−2
−3
−6
−4
−2
0
2
4
6
7 5

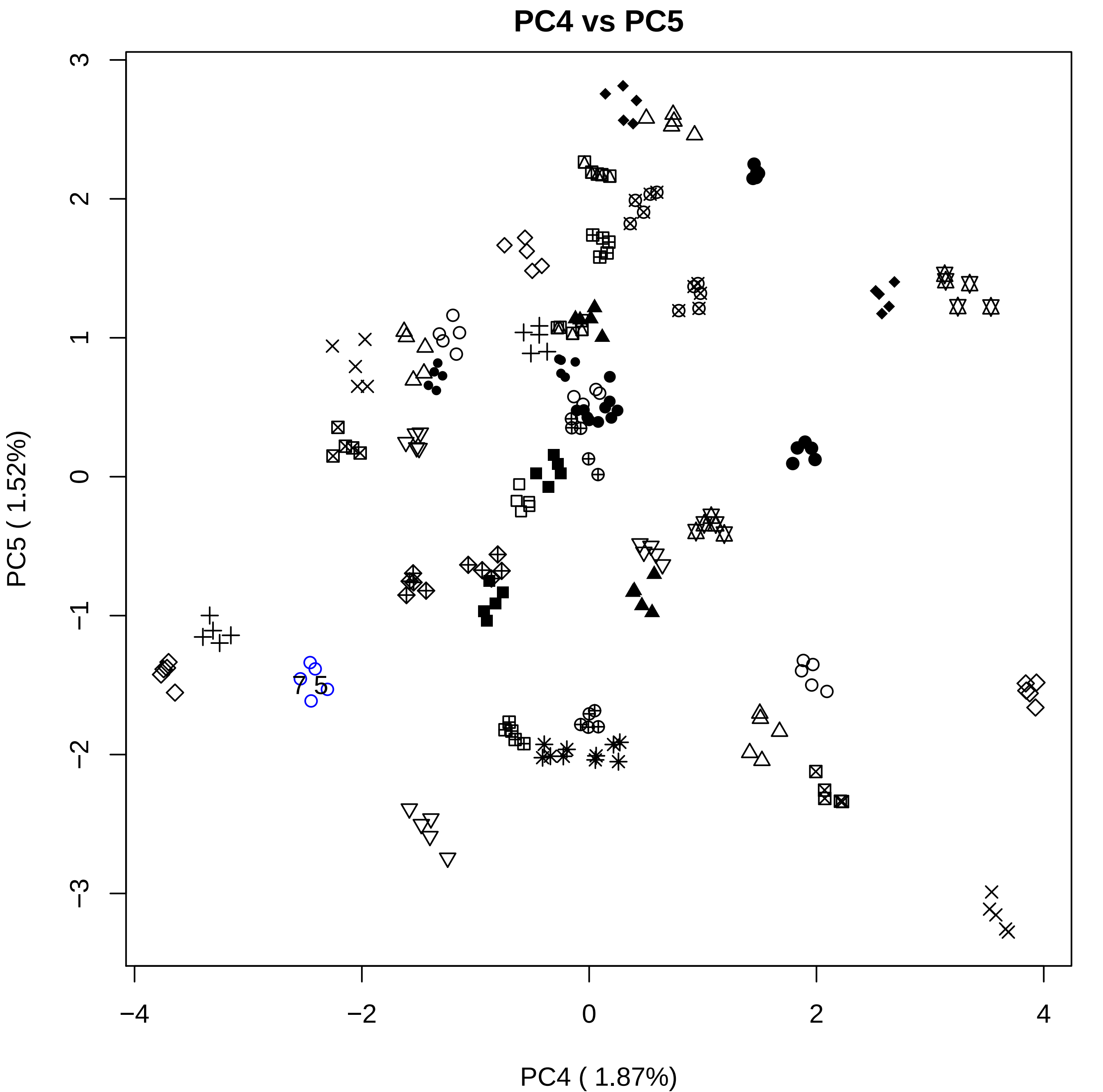

PC4 vs PC5
PC5 ( 1.52%)
PC4 ( 1.87%)
3
2
1
0
−1
−2
−3
−4
−2
0
2
4
7 5

### PCA of 10%  OSAS vs PMH

Scaled Matrix PCAed

7th order polynomial baseline correction performed

Raw spectra smoothed to 21 Neighbours, 2nd order PN, Savtisky Golay

Normalisation between 0 and 100

D2 spectra smoothed to 41 Neighbours, 2nd order PN, Savtisky Golay

Regions Analysed: 3601−2501 and 2001−700cm−1

Samples Analysed: 1 Contam's 0 OSAS's 45 PMH's

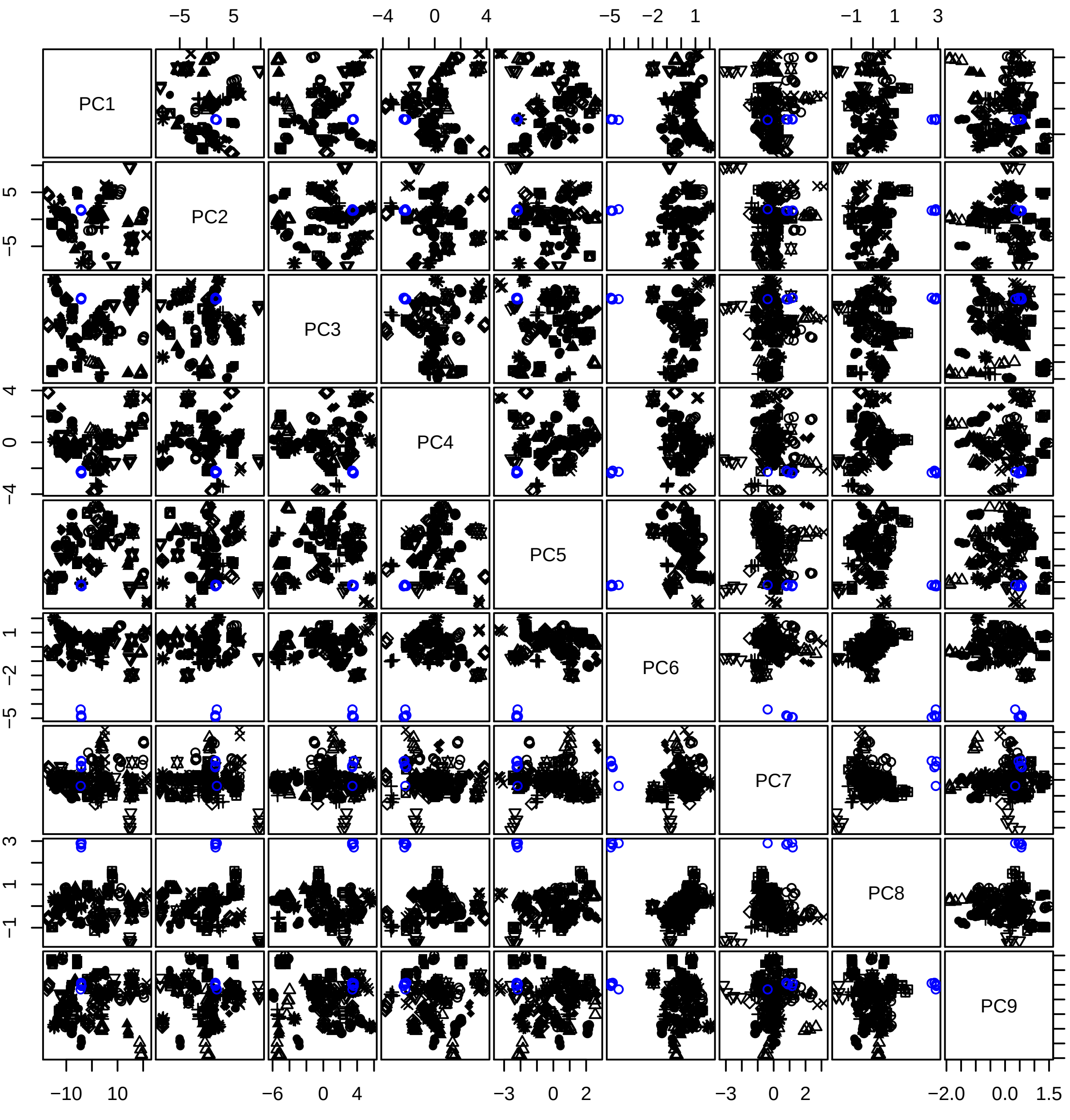
PC1
PC2
PC3
PC4
PC5
PC6
PC7
PC8
PC9

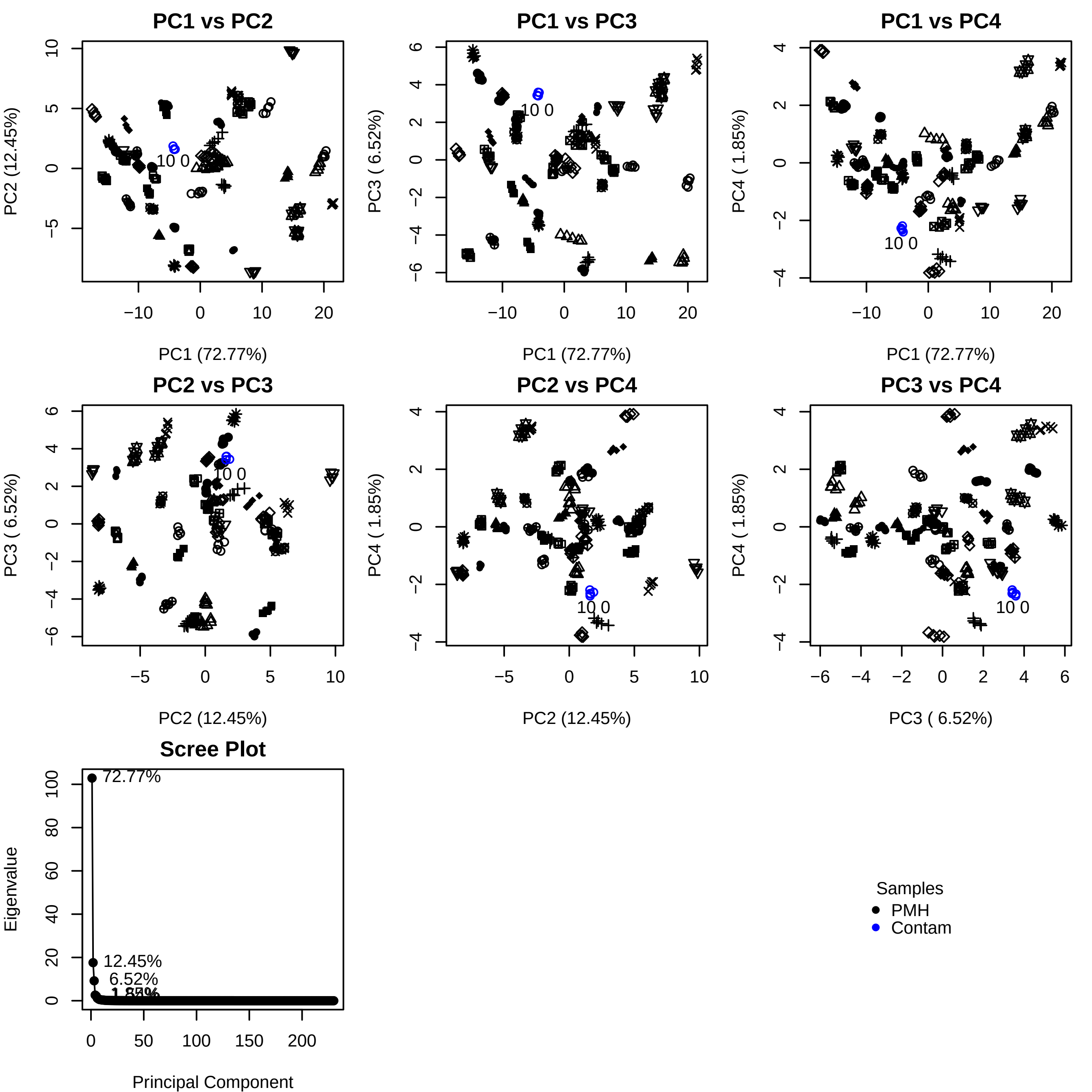

PC1 vs PC2
PC1 vs PC3
PC1 vs PC4
PC2 vs PC3
PC2 vs PC4
PC3 vs PC4
Scree Plot
PC1 (72.77%)
PC2 (12.45%)
PC3 ( 6.52%)
PC4 ( 1.85%)
10 0
Eigenvalue
Principal Component
72.77%
12.45%
6.52%
Samples
PMH
Contam

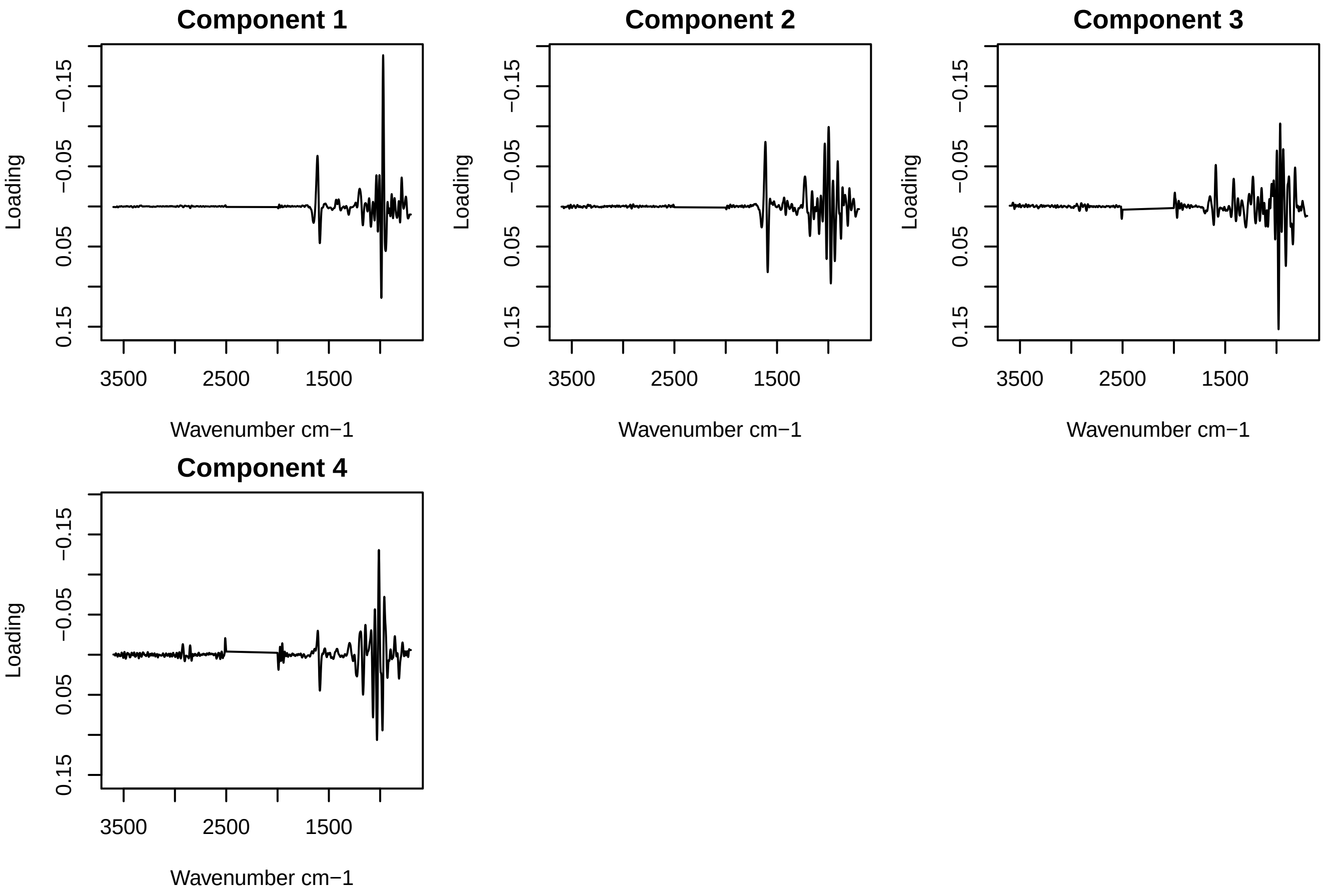
Component 1
Component 2
Component 3
Component 4
Loading
−0.15
−0.05
0.05
0.15
3500
2500
1500
Wavenumber cm−1

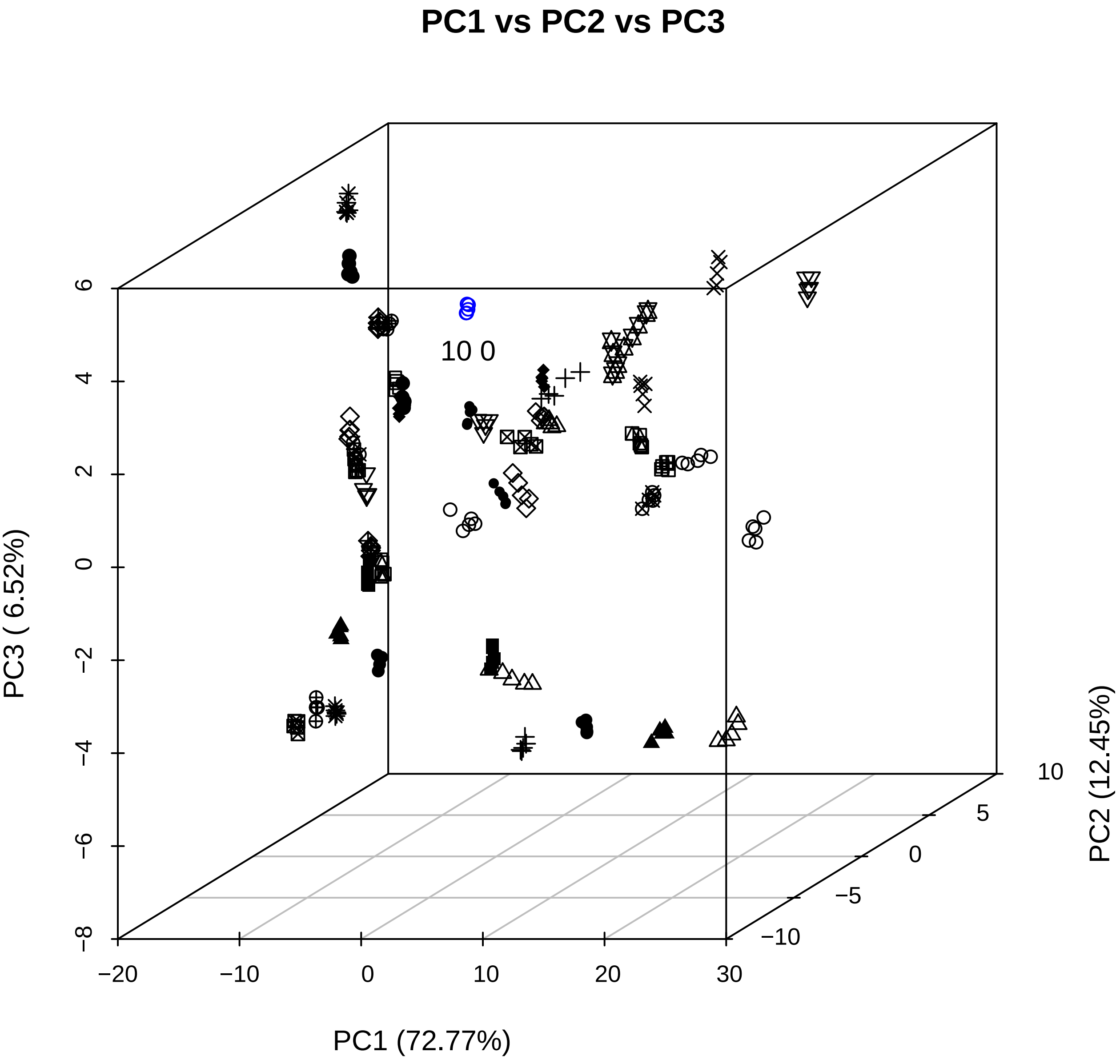
PC1 vs PC2 vs PC3
10 0
PC3 ( 6.52%)
−8
−6
−4
−2
0
2
4
6
PC1 (72.77%)
−20
−10
0
10
20
30
PC2 (12.45%)
−10
−5
0
5
10

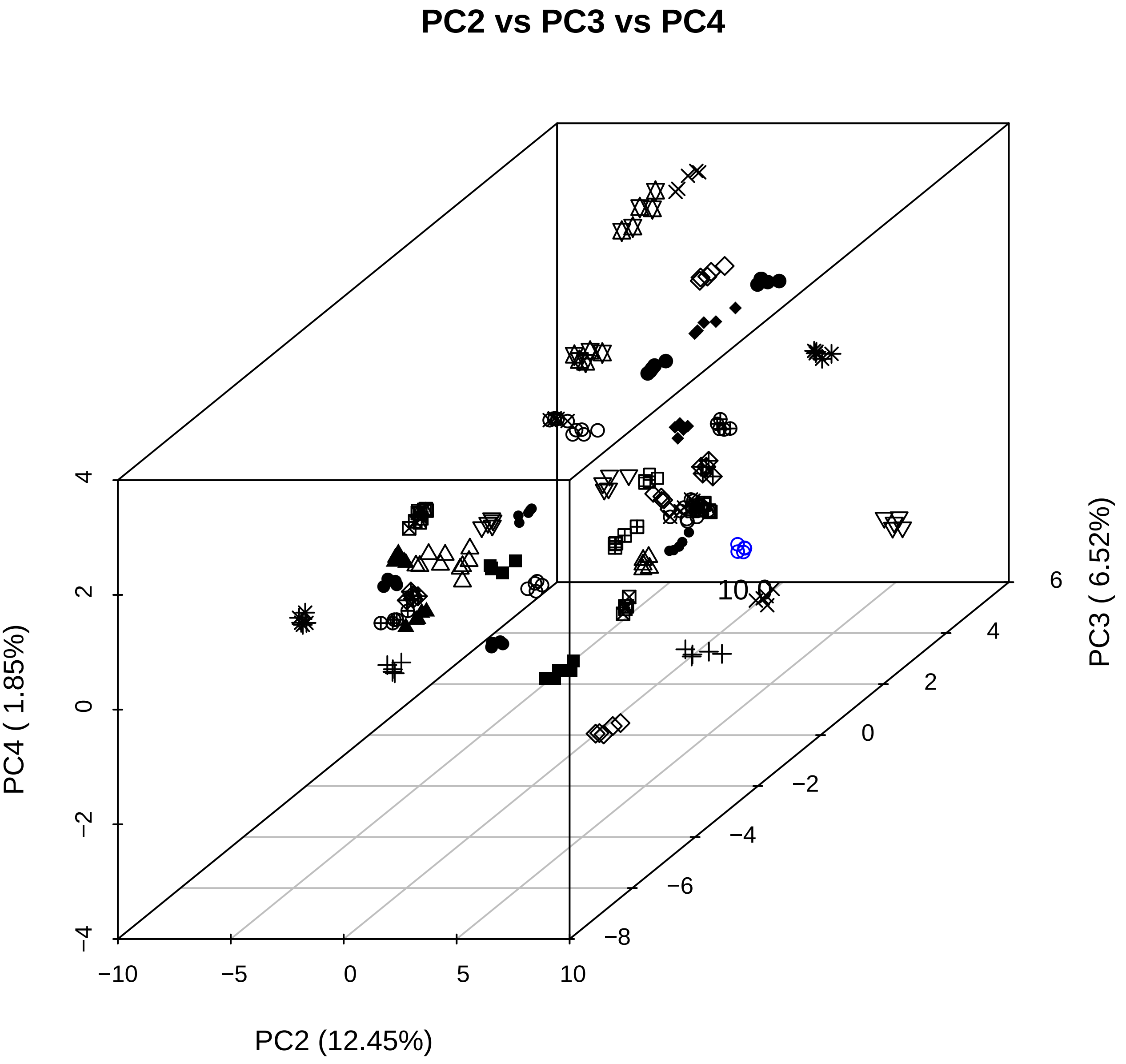

PC2 vs PC3 vs PC4
PC4 ( 1.85%)
PC2 (12.45%)
PC3 ( 6.52%)
10.0
−4
−2
0
2
4
−10
−5
0
5
10
−8
−6
−4
−2
0
2
4
6

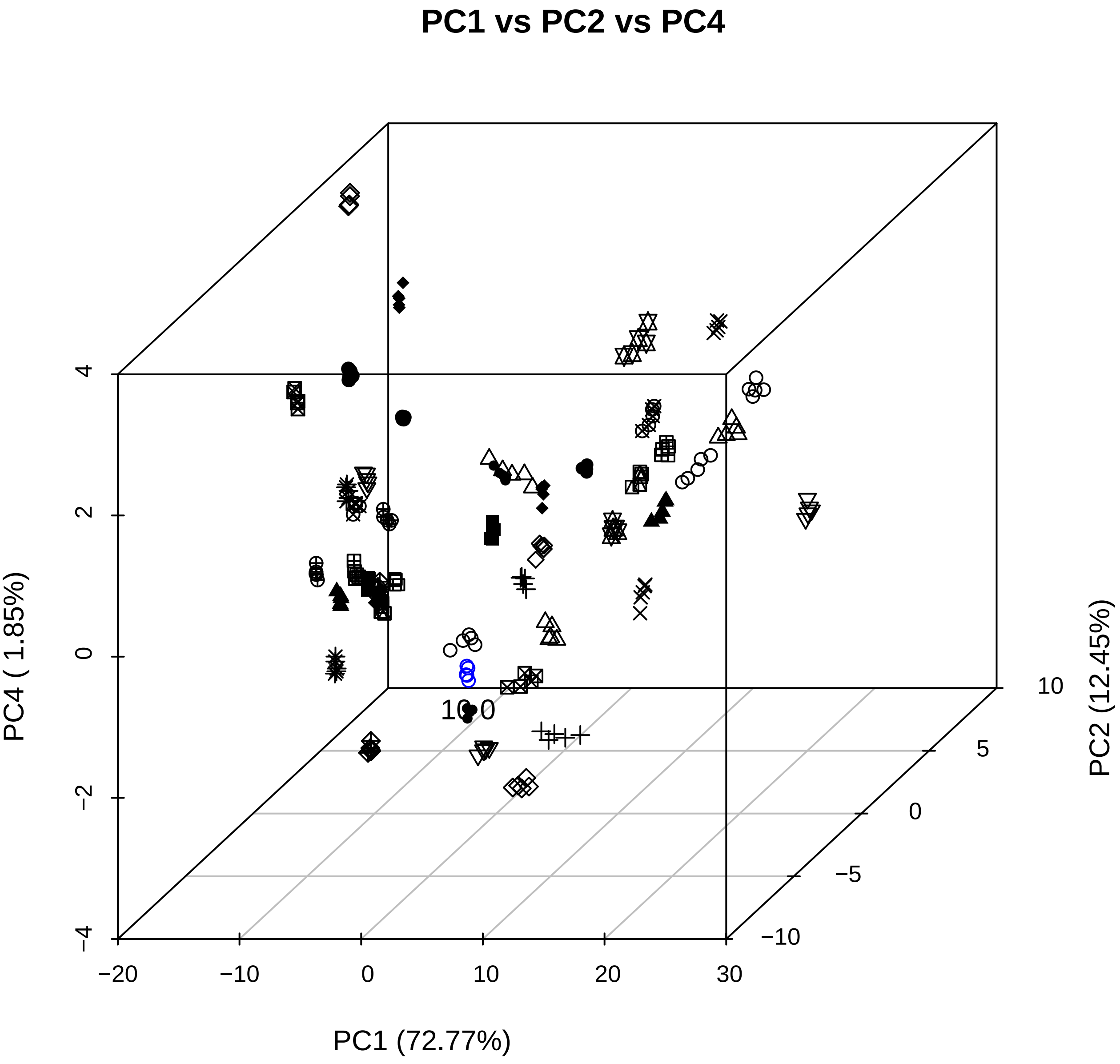
PC1 vs PC2 vs PC4
PC1 (72.77%)
PC2 (12.45%)
PC4 ( 1.85%)
−20
−10
0
10
20
30
−10
−5
0
5
10
−4
−2
0
2
4
100

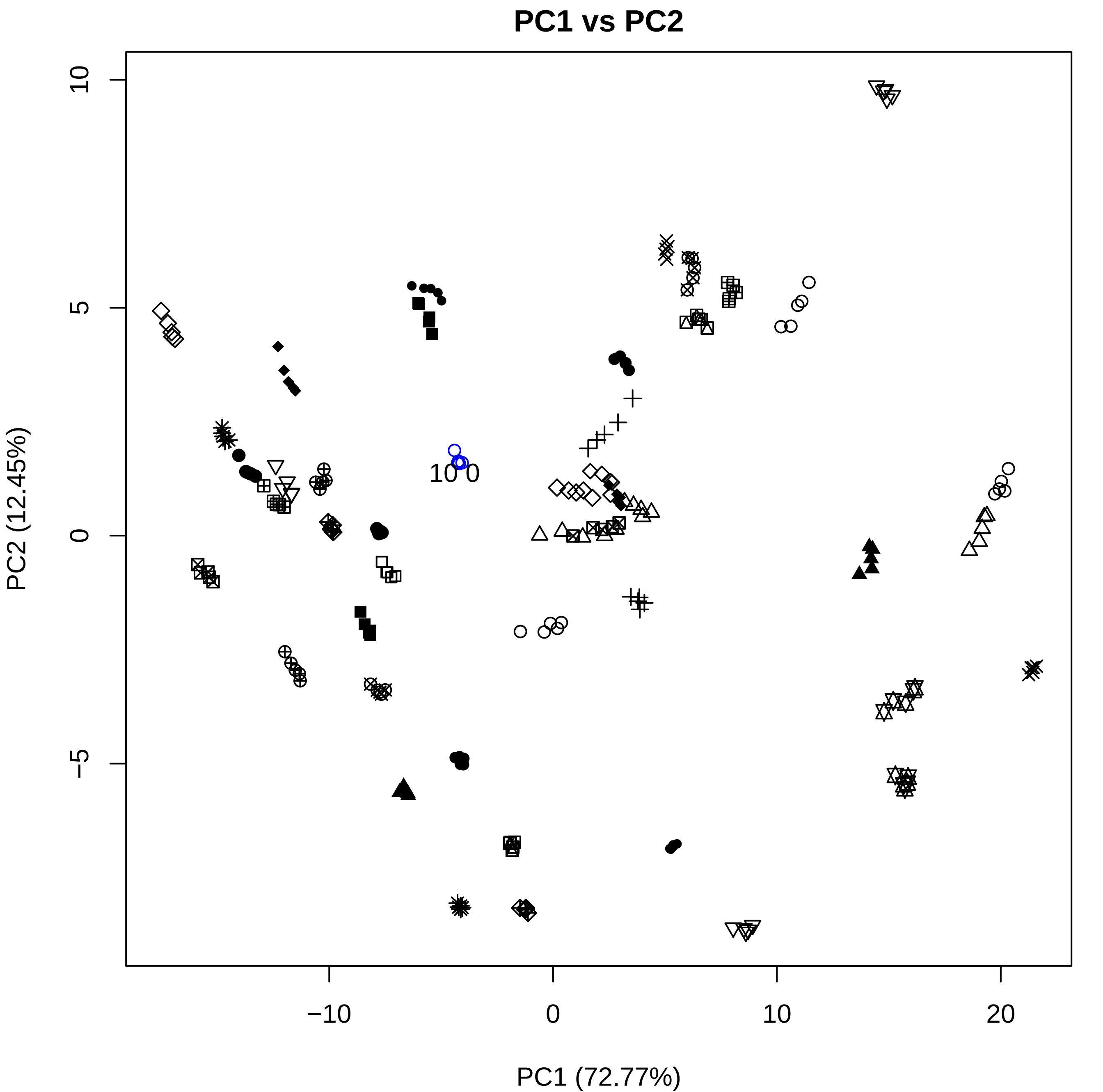
PC1 vs PC2
PC1 (72.77%)
PC2 (12.45%)
10 0
−10
0
10
20
−5
0
5
10

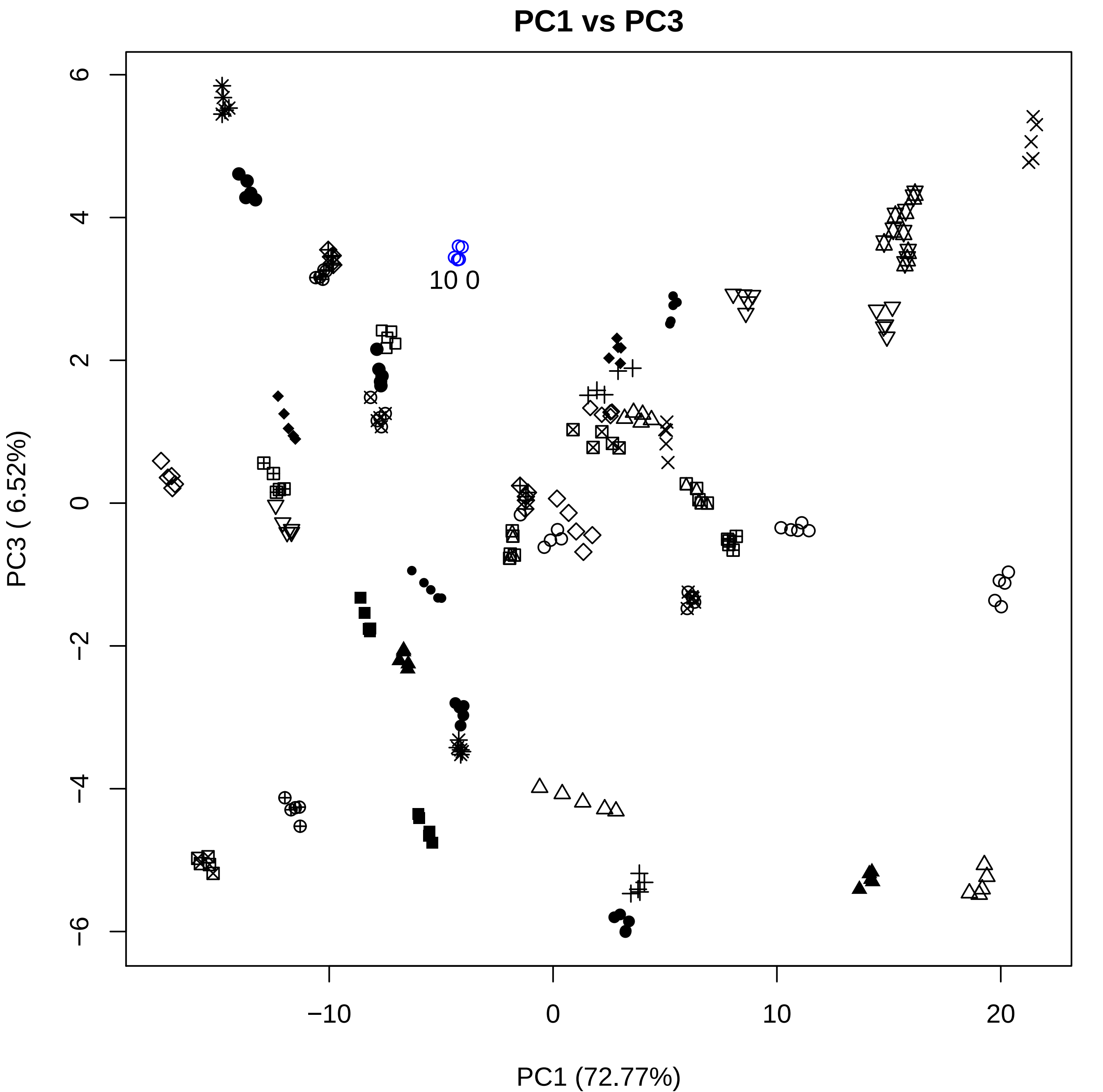

PC1 vs PC3
PC3 ( 6.52%)
PC1 (72.77%)
6
4
2
0
−2
−4
−6
−10
0
10
20
10 0

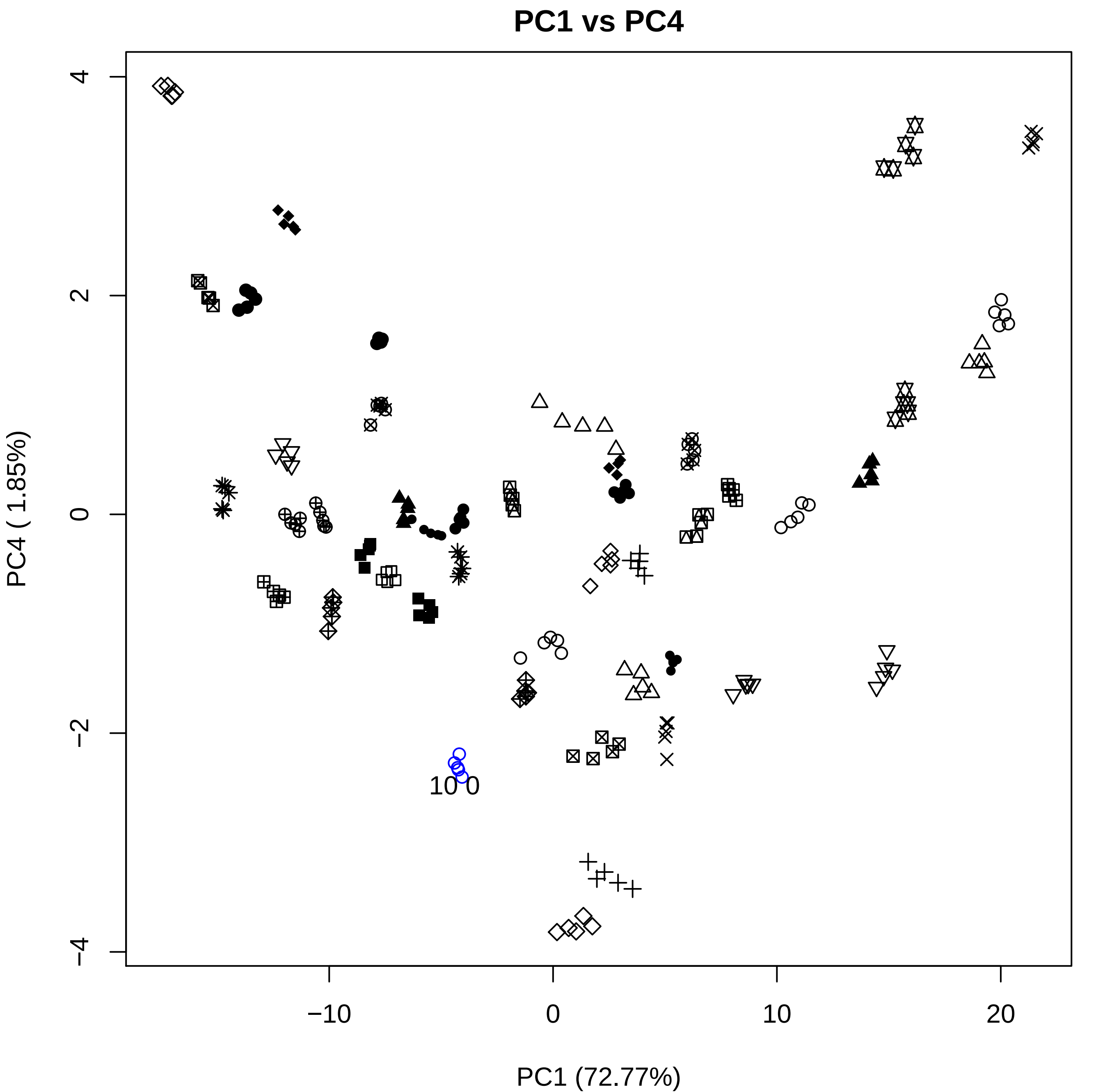
PC1 vs PC4
PC4 ( 1.85%)
PC1 (72.77%)
4
2
0
−2
−4
−10
0
10
20
10 0

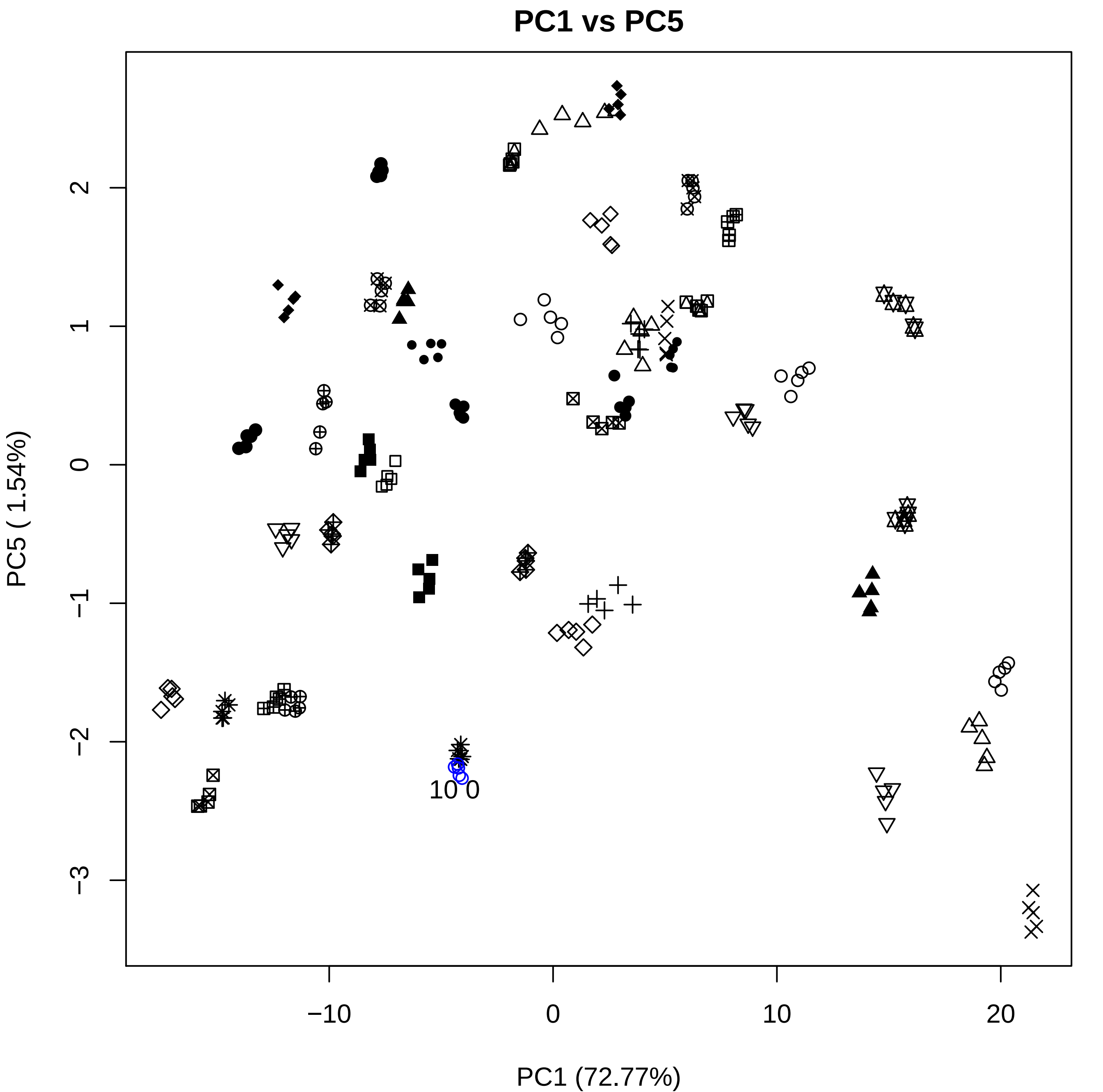
PC1 vs PC5
PC5 ( 1.54%)
PC1 (72.77%)
2
1
0
−1
−2
−3
−10
0
10
20
10 0

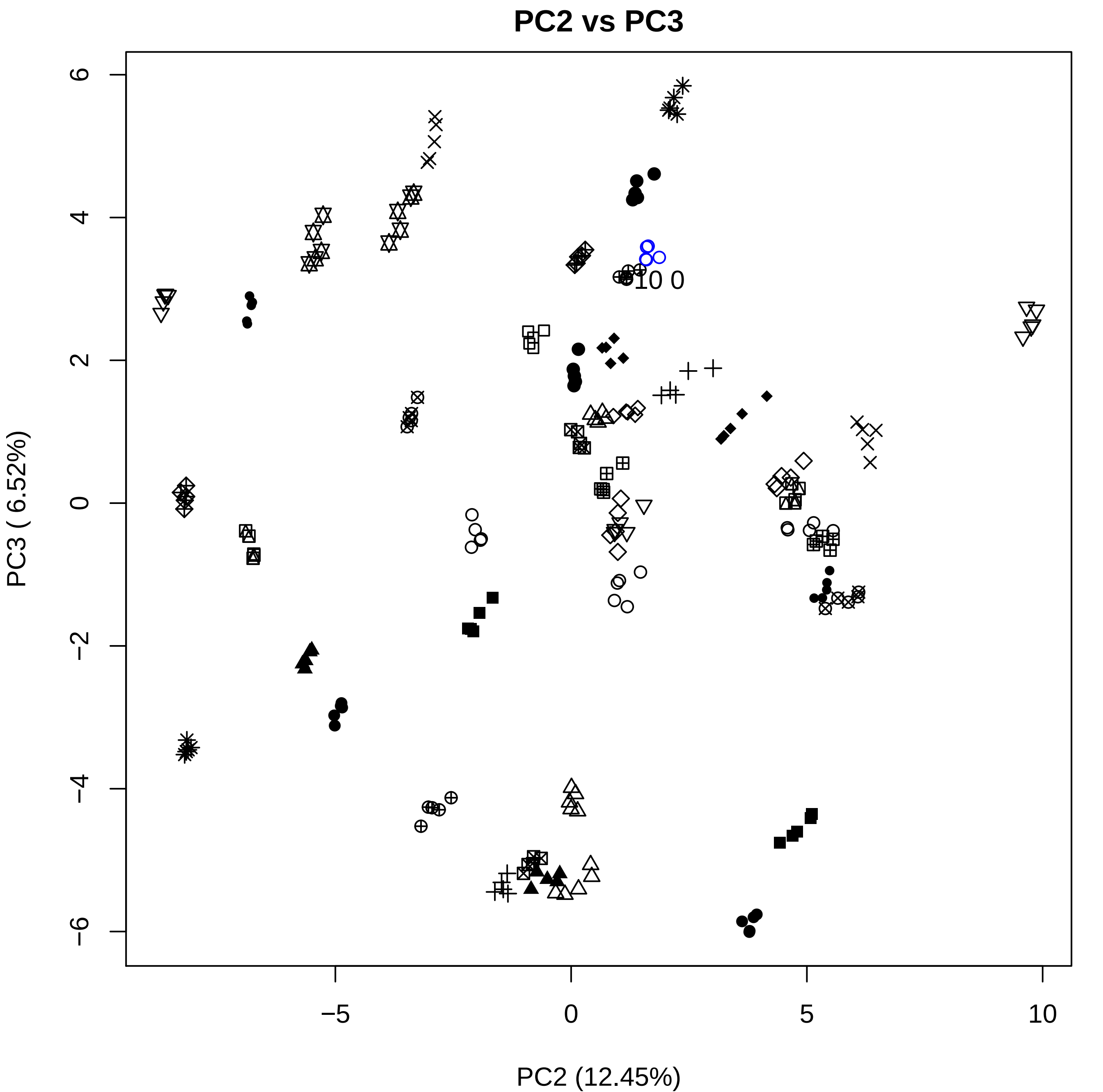
PC2 vs PC3
PC3 ( 6.52%)
PC2 (12.45%)
−5
0
5
10
−6
−4
−2
0
2
4
6
10 0

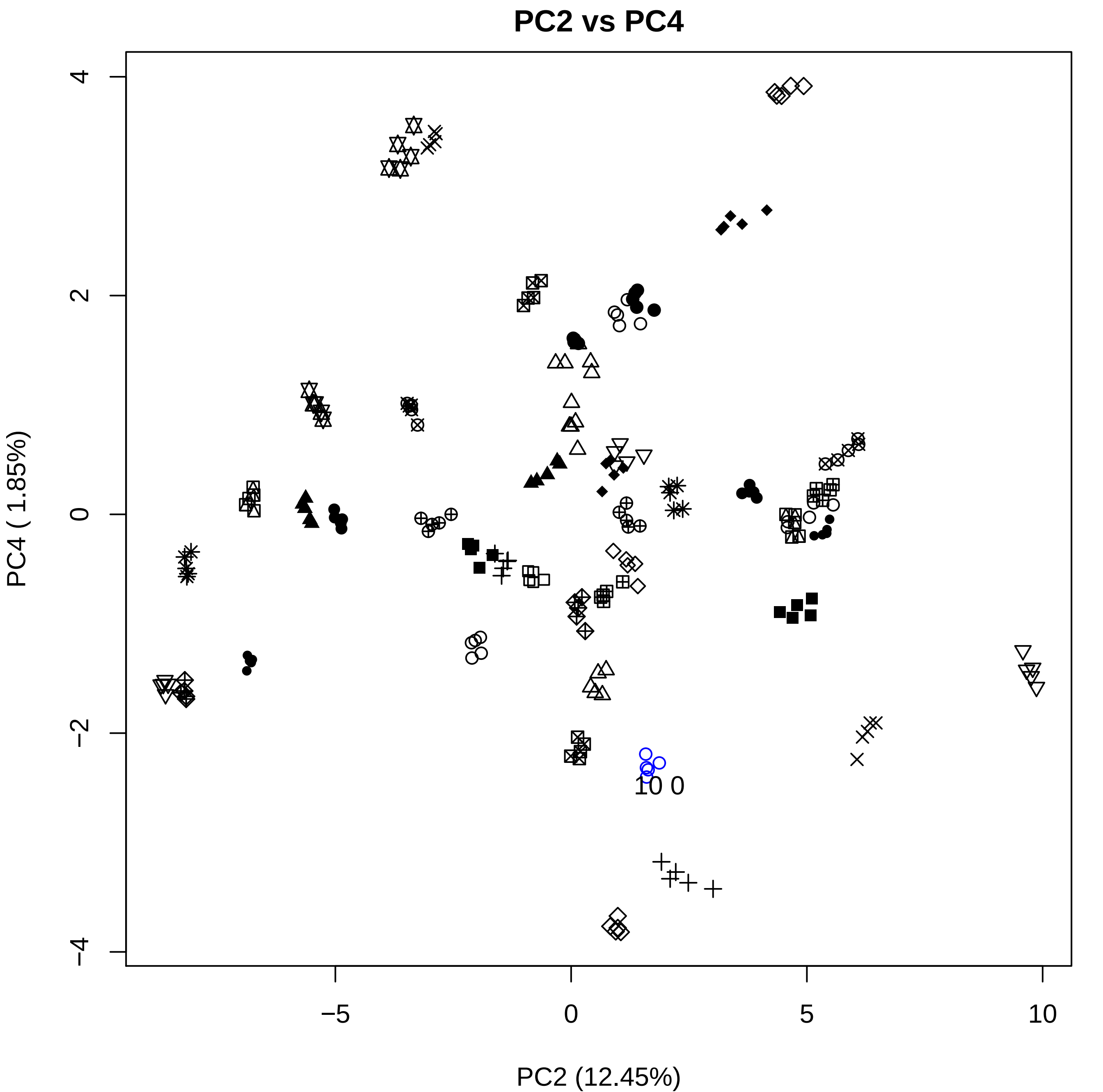
PC2 vs PC4
PC4 ( 1.85%)
PC2 (12.45%)
4
2
0
−2
−4
−5
0
5
10
10 0

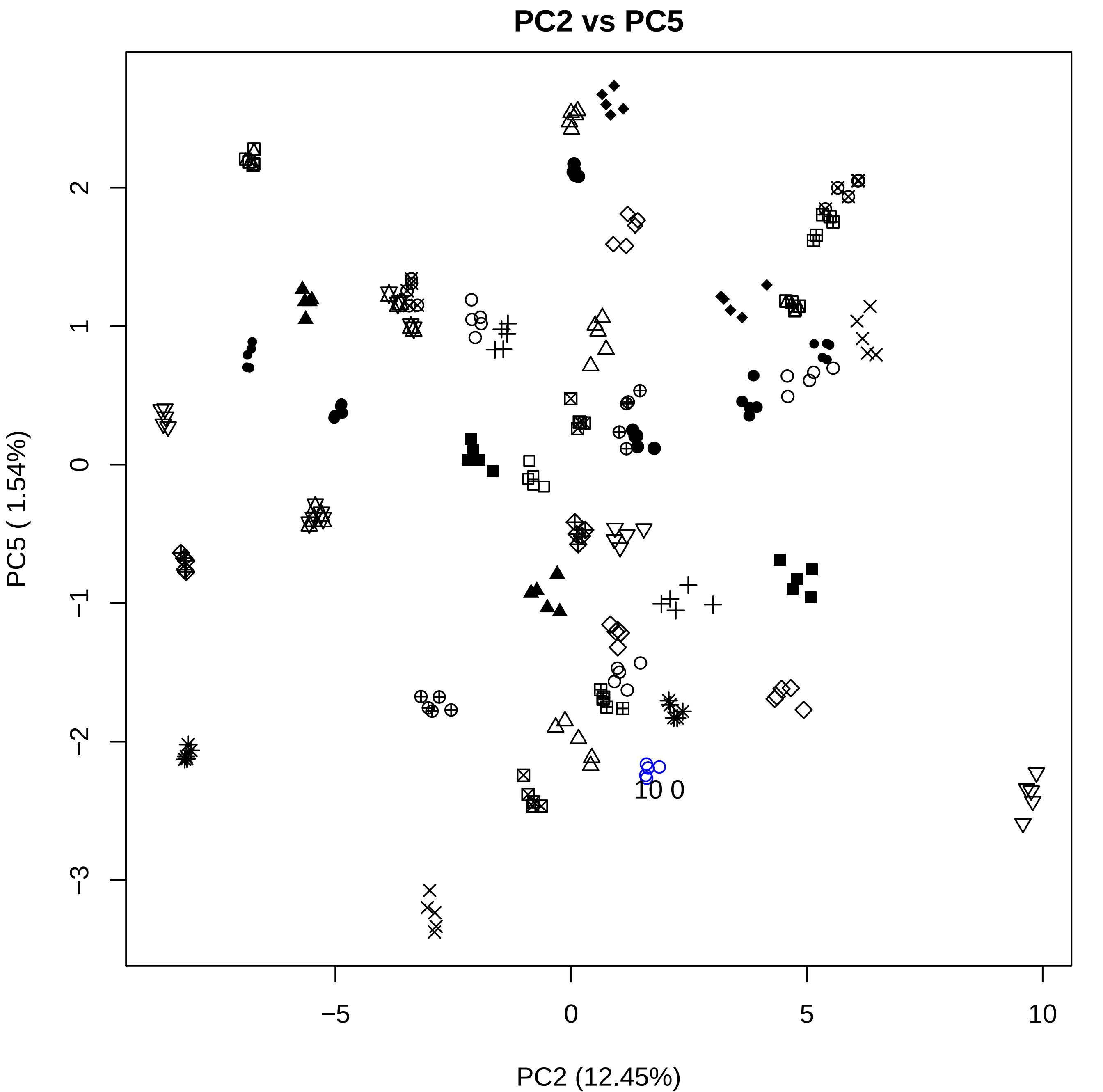

PC2 vs PC5
PC5 ( 1.54%)
PC2 (12.45%)
2
1
0
−1
−2
−3
−5
0
5
10
10 0

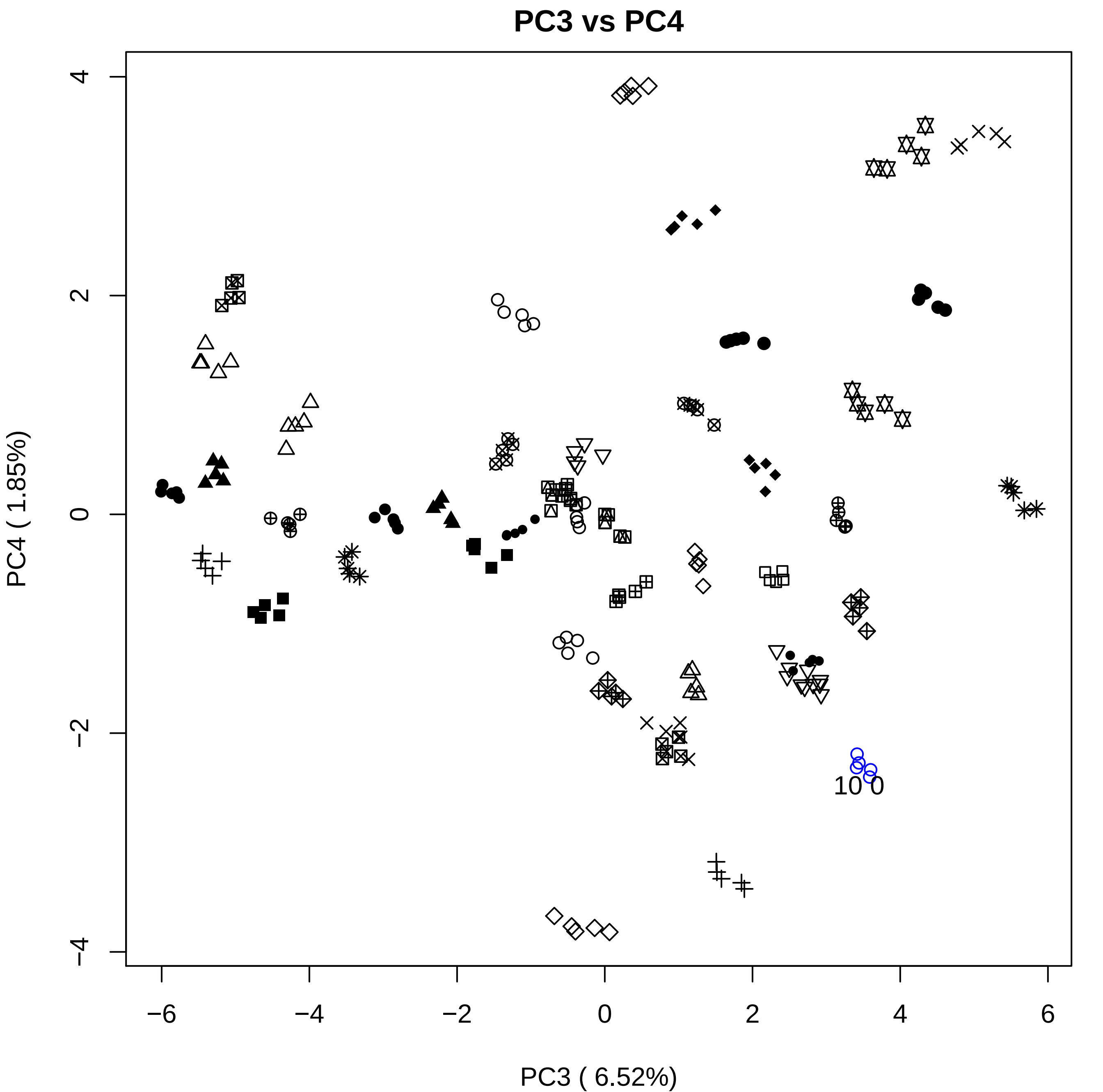

PC3 vs PC4
PC4 ( 1.85%)
PC3 ( 6.52%)
4
2
0
−2
−4
−6
−4
−2
0
2
4
6
10 0

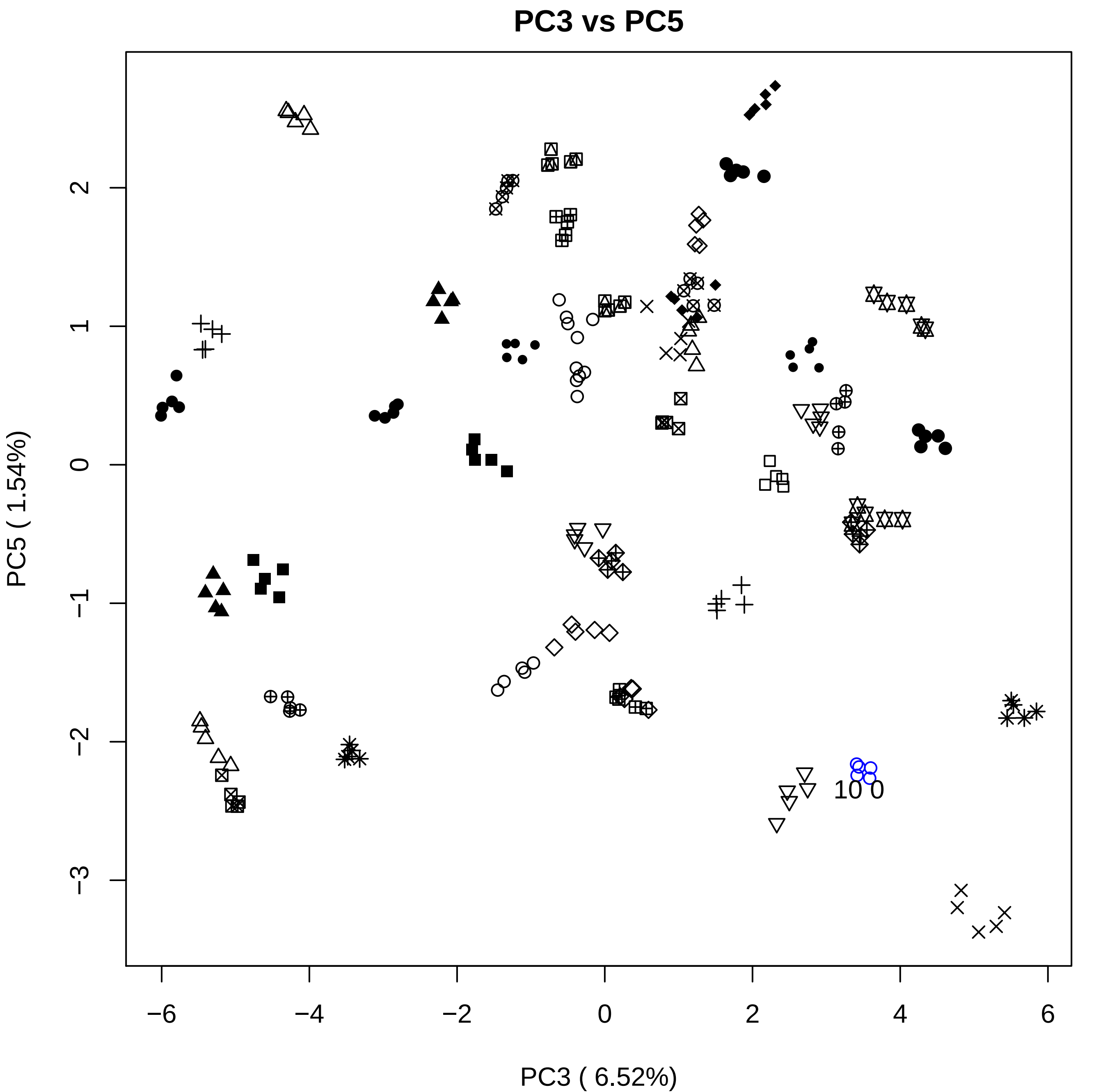
PC3 vs PC5
PC5 ( 1.54%)
PC3 ( 6.52%)
−6
−4
−2
0
2
4
6
−3
−2
−1
0
1
2
10 0

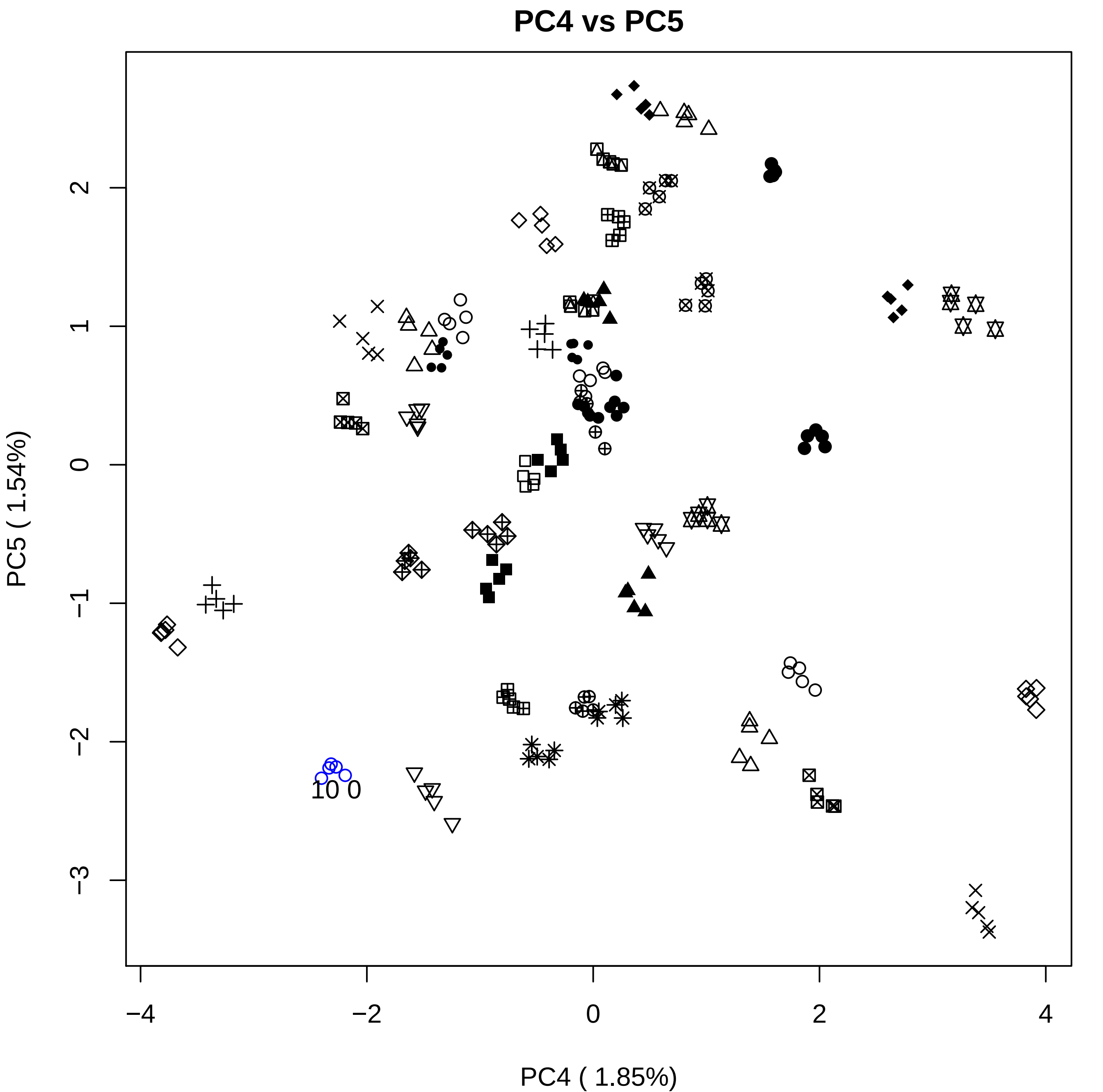

PC4 vs PC5
PC5 ( 1.54%)
PC4 ( 1.85%)
−4
−2
0
2
4
−3
−2
−1
0
1
2
10 0

### PCA of 15%  OSAS vs PMH

Scaled Matrix PCAed

7th order polynomial baseline correction performed

Raw spectra smoothed to 21 Neighbours, 2nd order PN, Savtisky Golay

Normalisation between 0 and 100

D2 spectra smoothed to 41 Neighbours, 2nd order PN, Savtisky Golay

Regions Analysed: 3601−2501 and 2001−700cm−1

Samples Analysed: 1 Contam's 0 OSAS's 45 PMH's

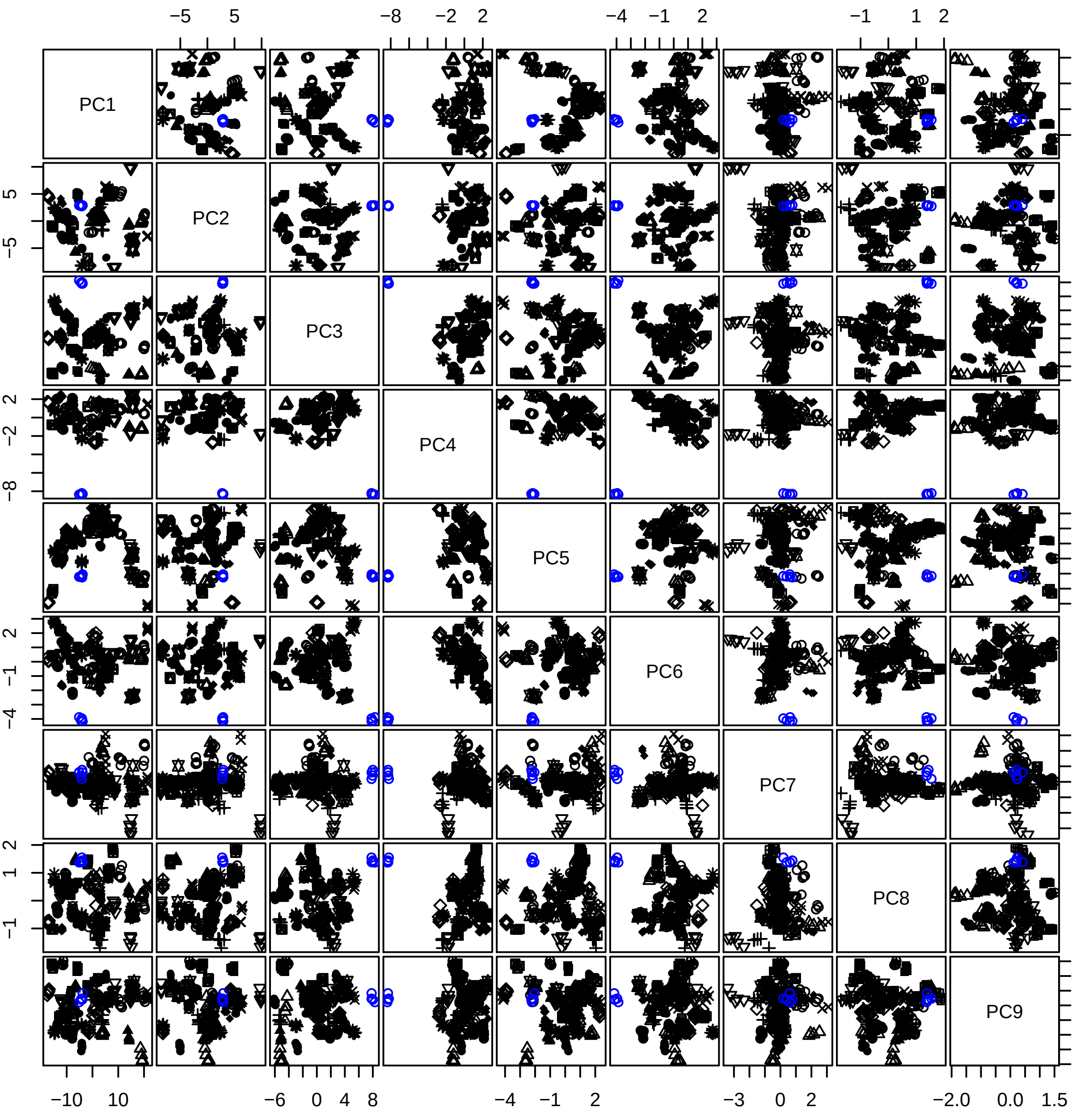
PC1
PC2
PC3
PC4
PC5
PC6
PC7
PC8
PC9
−5
5
−8
−2
2
−4
−1
2
−1
1
2
10
−10
5
−5
8
4
0
−6
2
−2
−8
2
−1
−4
2
−1
−4
2
0
−3
2
1
−1
1.5
0.0
−2.0
−10
10
−6
0
4
8
−4
−1
2
−3
0
2
−2.0
0.0
1.5

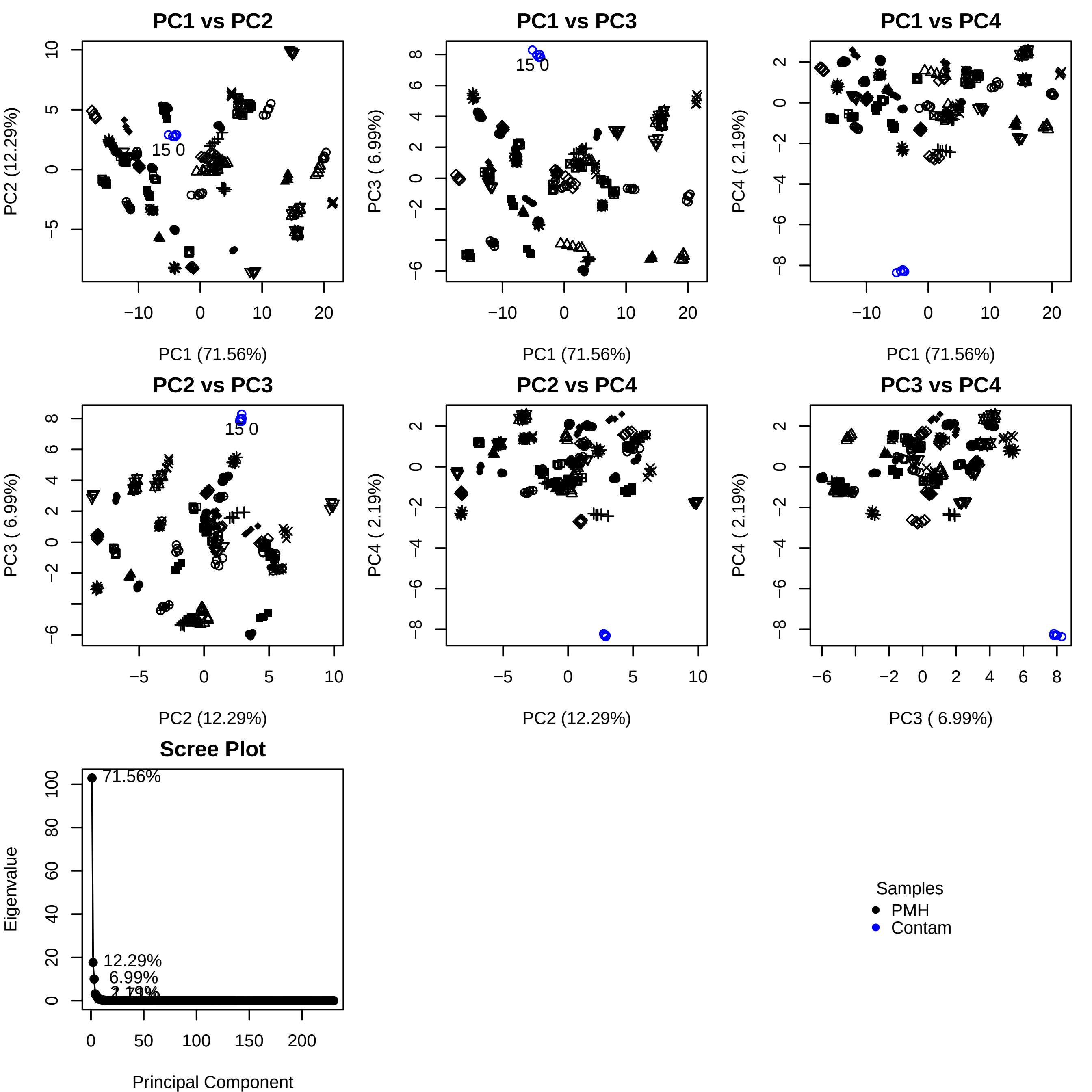
PC1 vs PC2
PC1 vs PC3
PC1 vs PC4
PC2 vs PC3
PC2 vs PC4
PC3 vs PC4
Scree Plot
PC1 (71.56%)
PC2 (12.29%)
PC3 ( 6.99%)
PC4 ( 2.19%)
15 0
Eigenvalue
Principal Component
71.56%
12.29%
6.99%
Samples
PMH
Contam

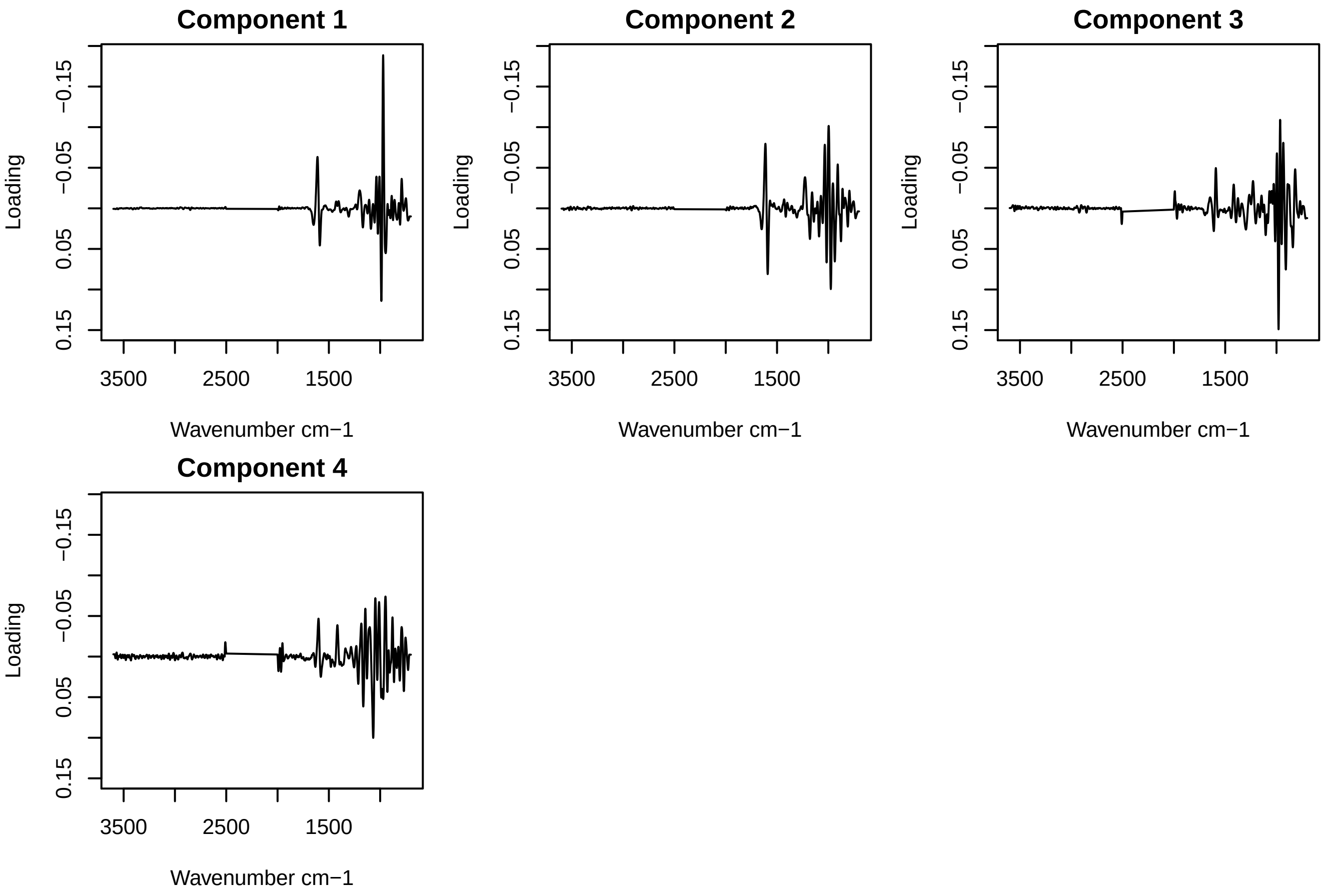

Component 1
Component 2
Component 3
Component 4
Loading
−0.15
−0.05
0.05
0.15
3500
2500
1500
Wavenumber cm−1

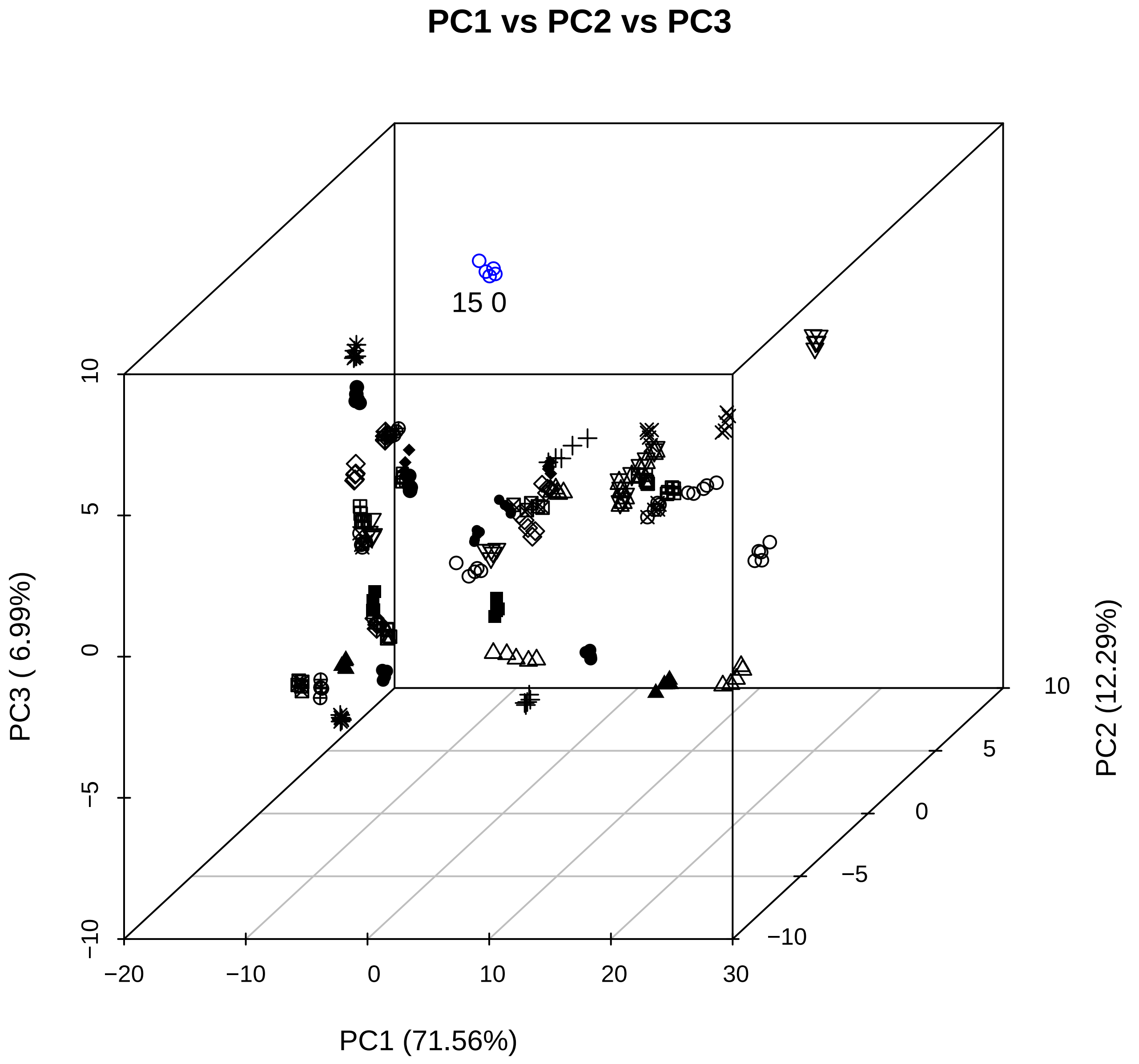
PC1 vs PC2 vs PC3
15 0
PC1 (71.56%)
PC2 (12.29%)
PC3 ( 6.99%)
−20
−10
0
10
20
30
−10
−5
0
5
10

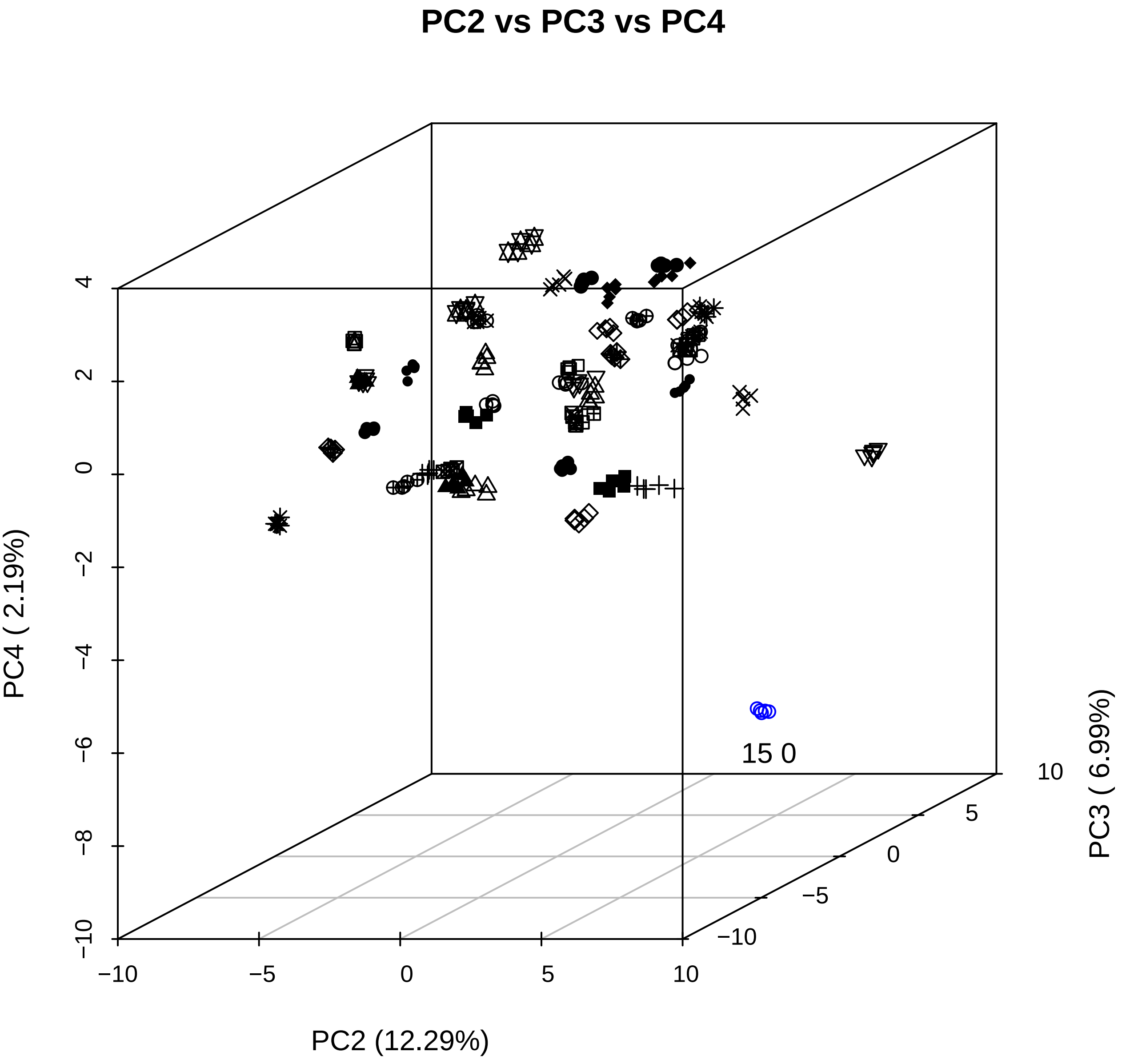
PC2 vs PC3 vs PC4
PC2 (12.29%)
PC3 ( 6.99%)
PC4 ( 2.19%)
−10
−5
0
5
10
−10
−8
−6
−4
−2
0
2
4
15 0

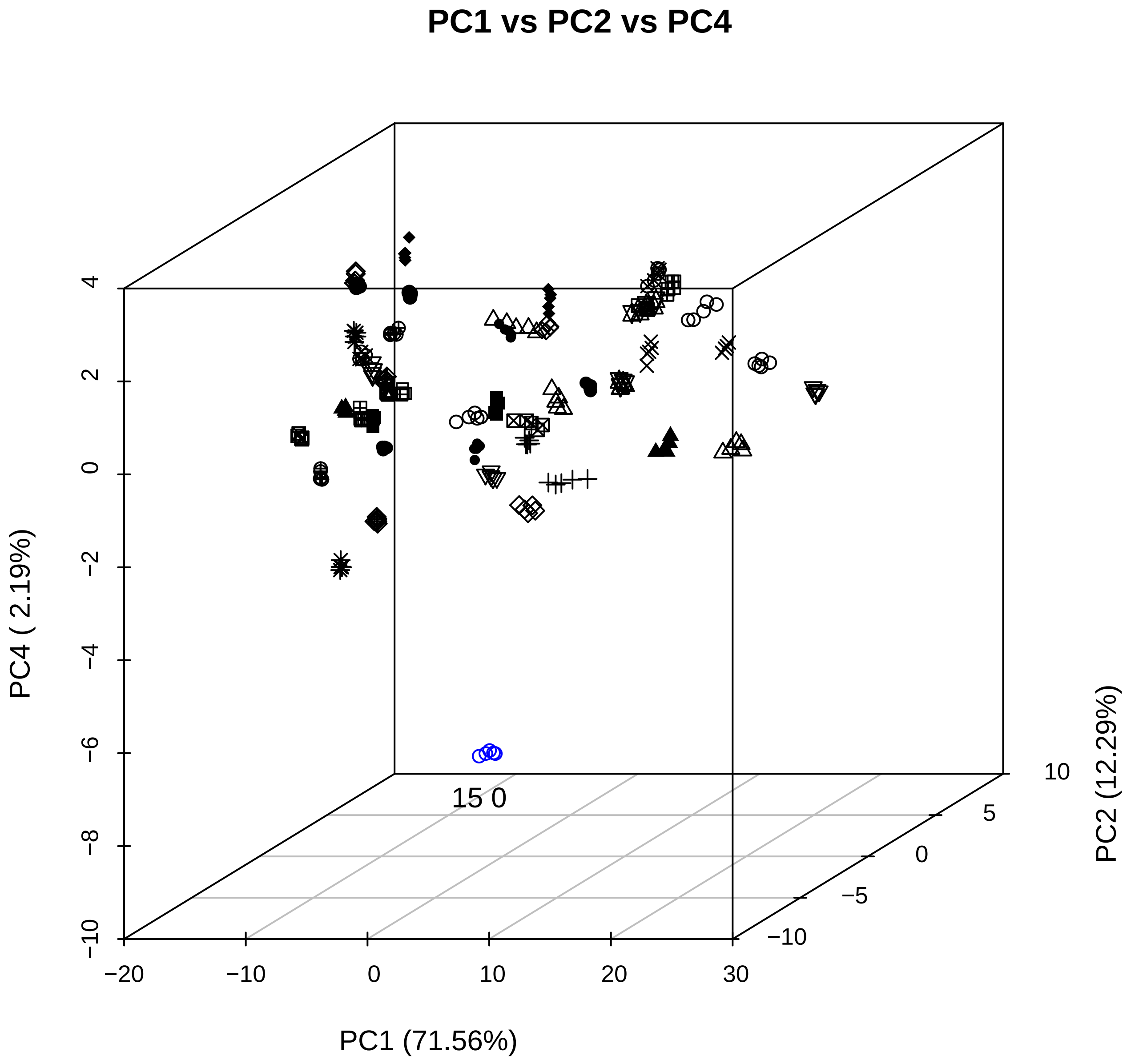

PC1 vs PC2 vs PC4
PC4 ( 2.19%)
4
2
0
−2
−4
−6
−8
−10
15 0
−20
−10
0
10
20
30
PC1 (71.56%)
10
5
0
−5
−10
PC2 (12.29%)

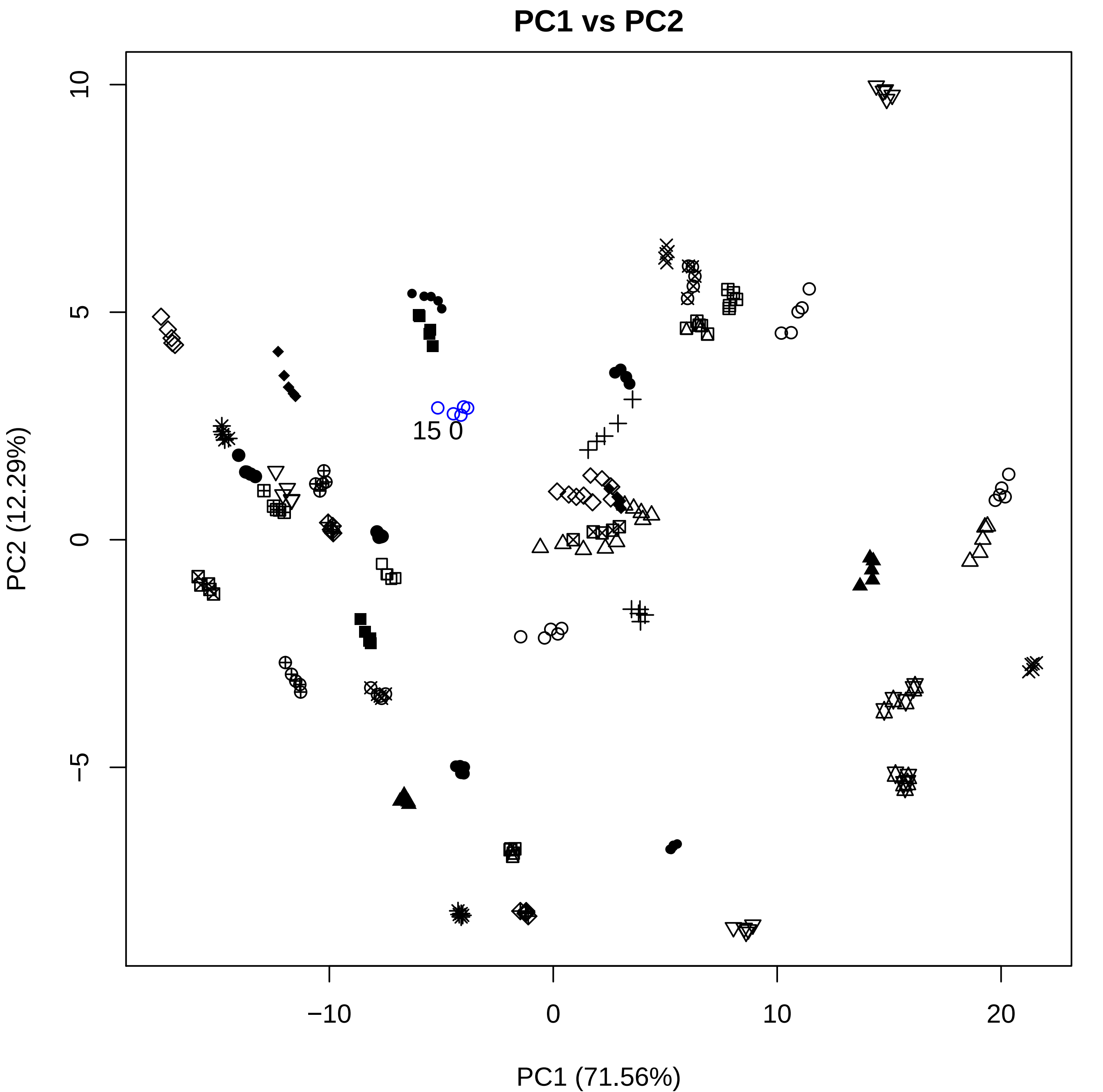

PC1 vs PC2
15 0
PC1 (71.56%)
PC2 (12.29%)
−10
0
10
20
−5
0
5
10

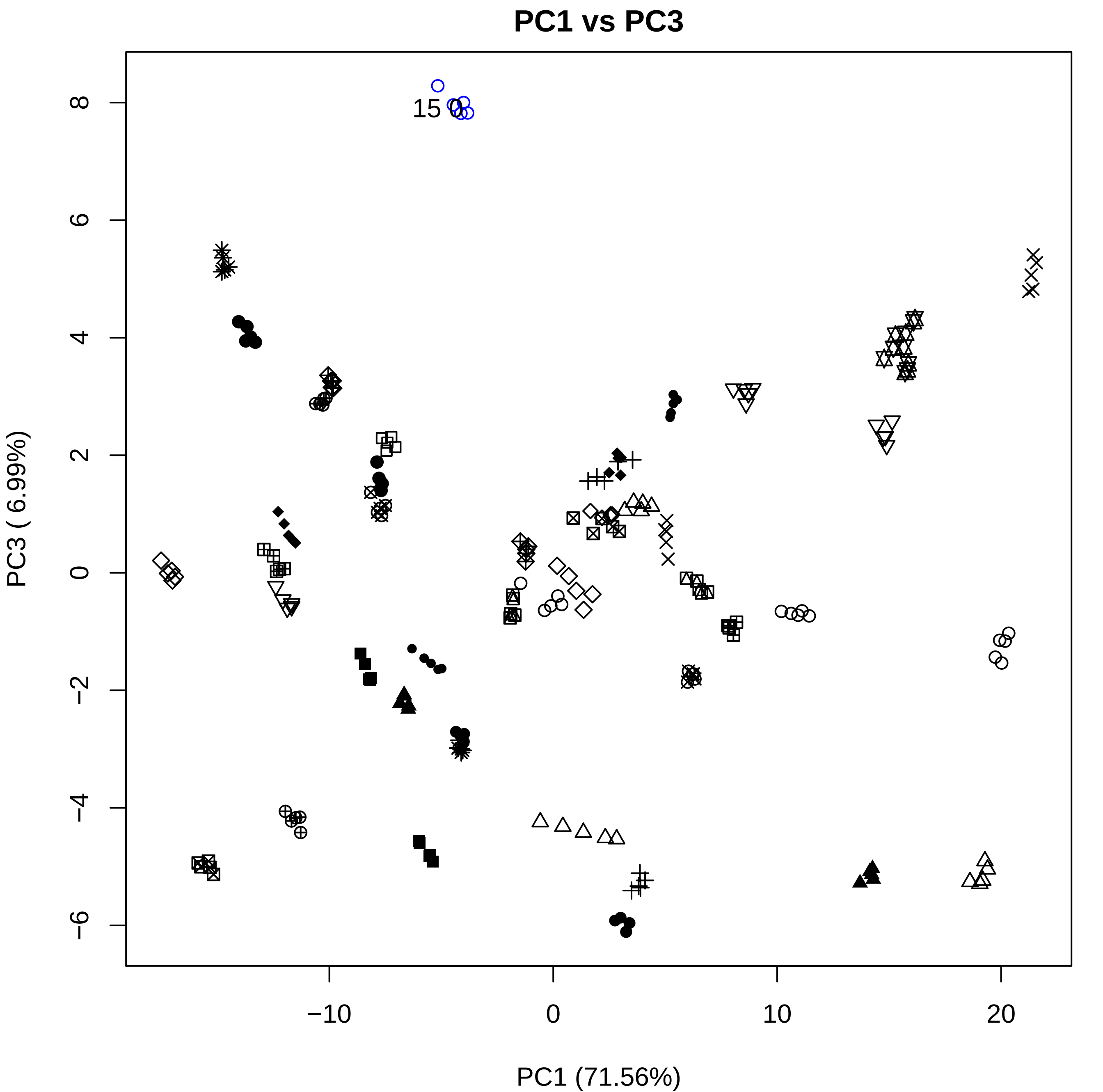
PC1 vs PC3
PC3 ( 6.99%)
PC1 (71.56%)
15 0
8
6
4
2
0
−2
−4
−6
−10
0
10
20

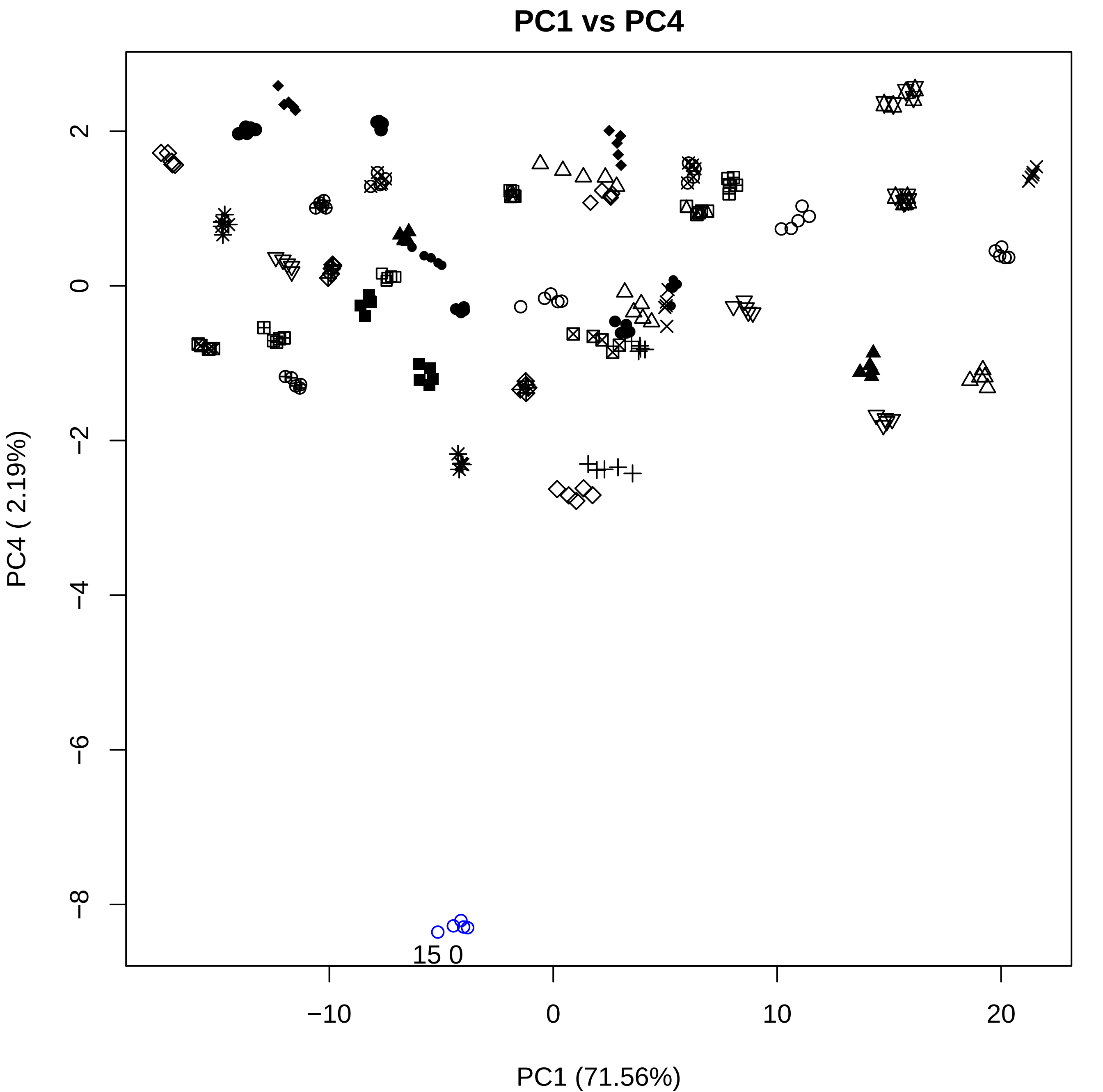
PC1 vs PC4
PC4 ( 2.19%)
PC1 (71.56%)
2
0
−2
−4
−6
−8
−10
0
10
20
15 0

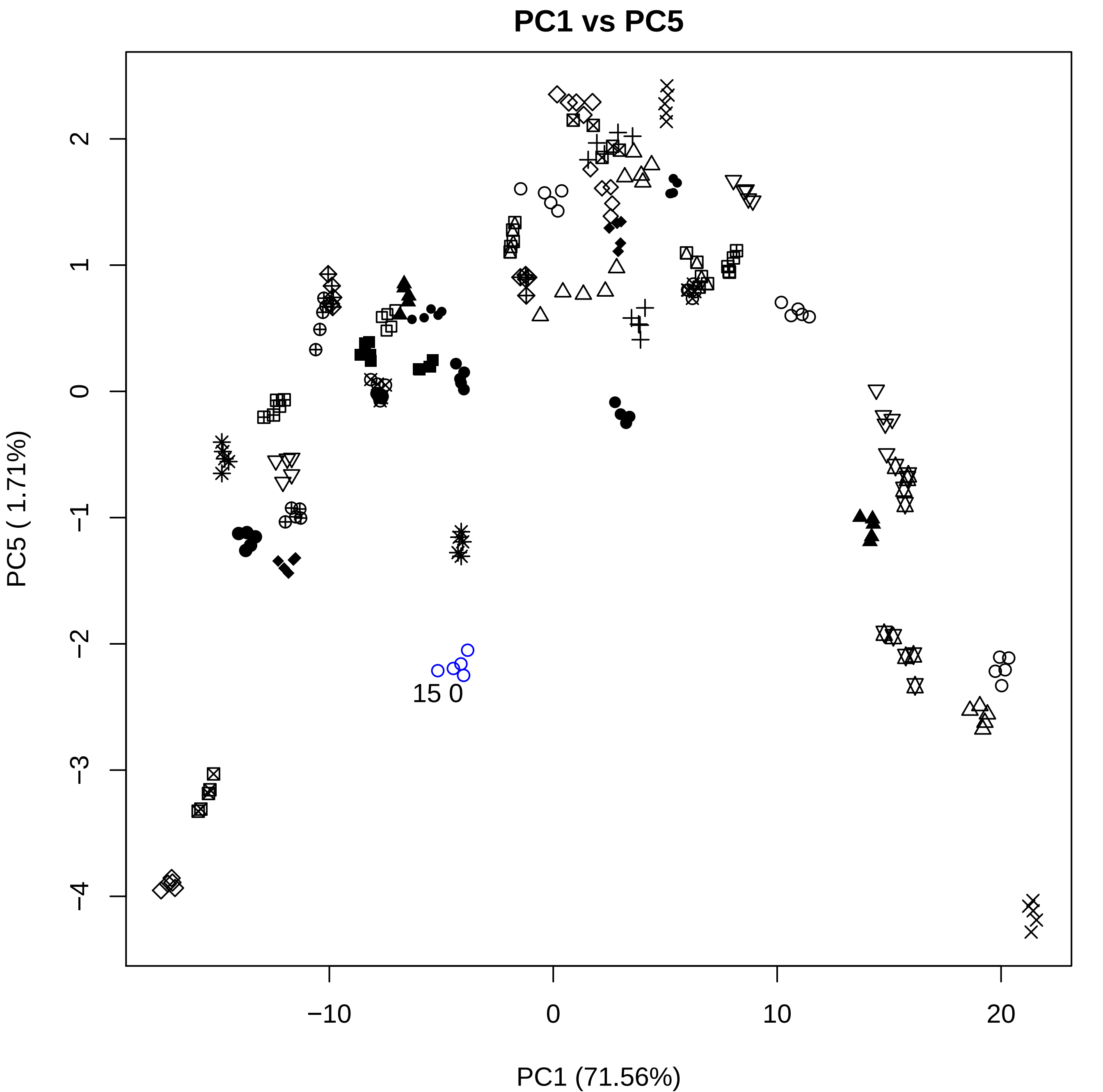
PC1 vs PC5
PC5 ( 1.71%)
PC1 (71.56%)
2
1
0
−1
−2
−3
−4
−10
0
10
20
15 0

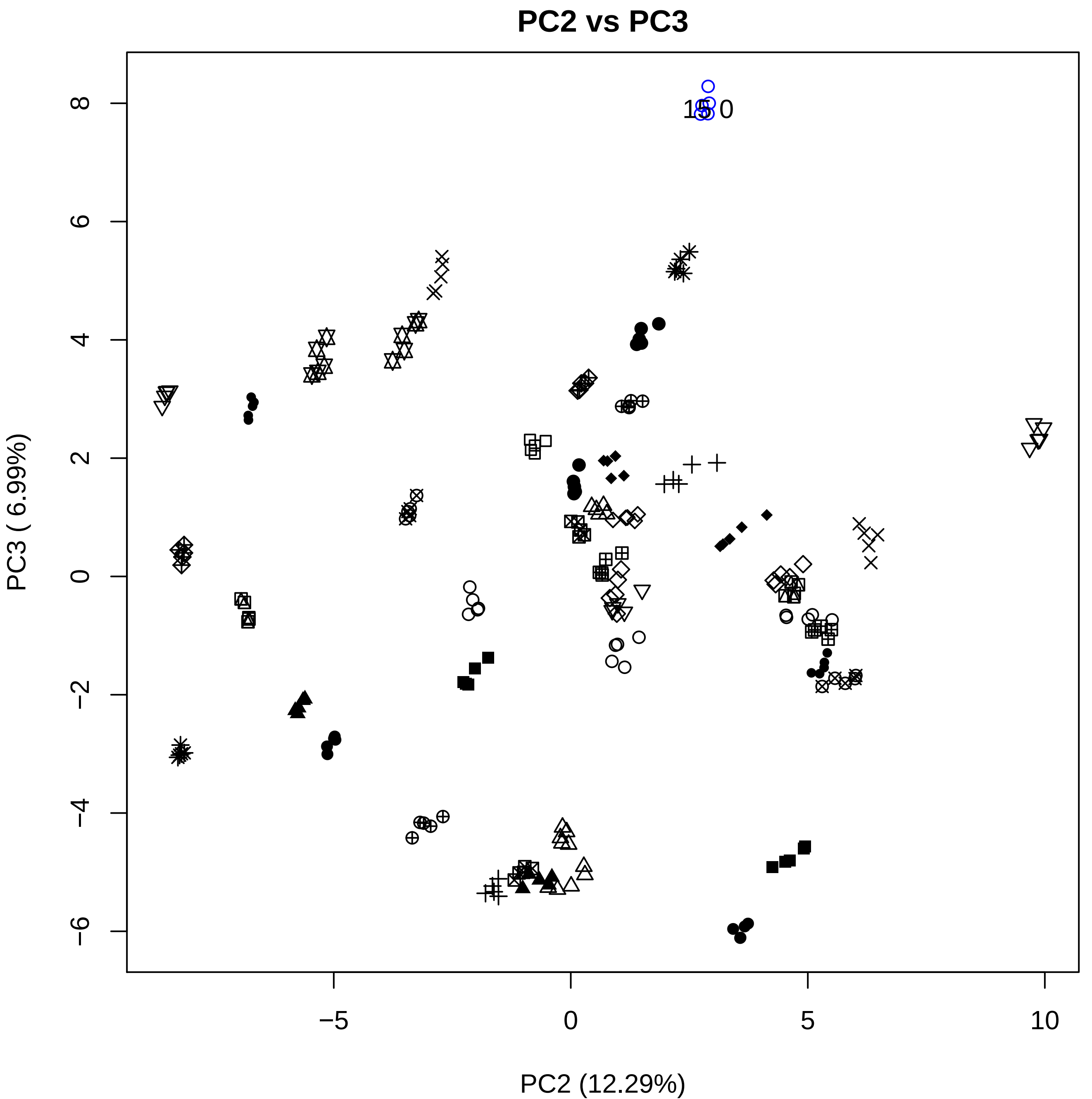
PC2 vs PC3
PC3 ( 6.99%)
PC2 (12.29%)
8
6
4
2
0
−2
−4
−6
−5
0
5
10
150

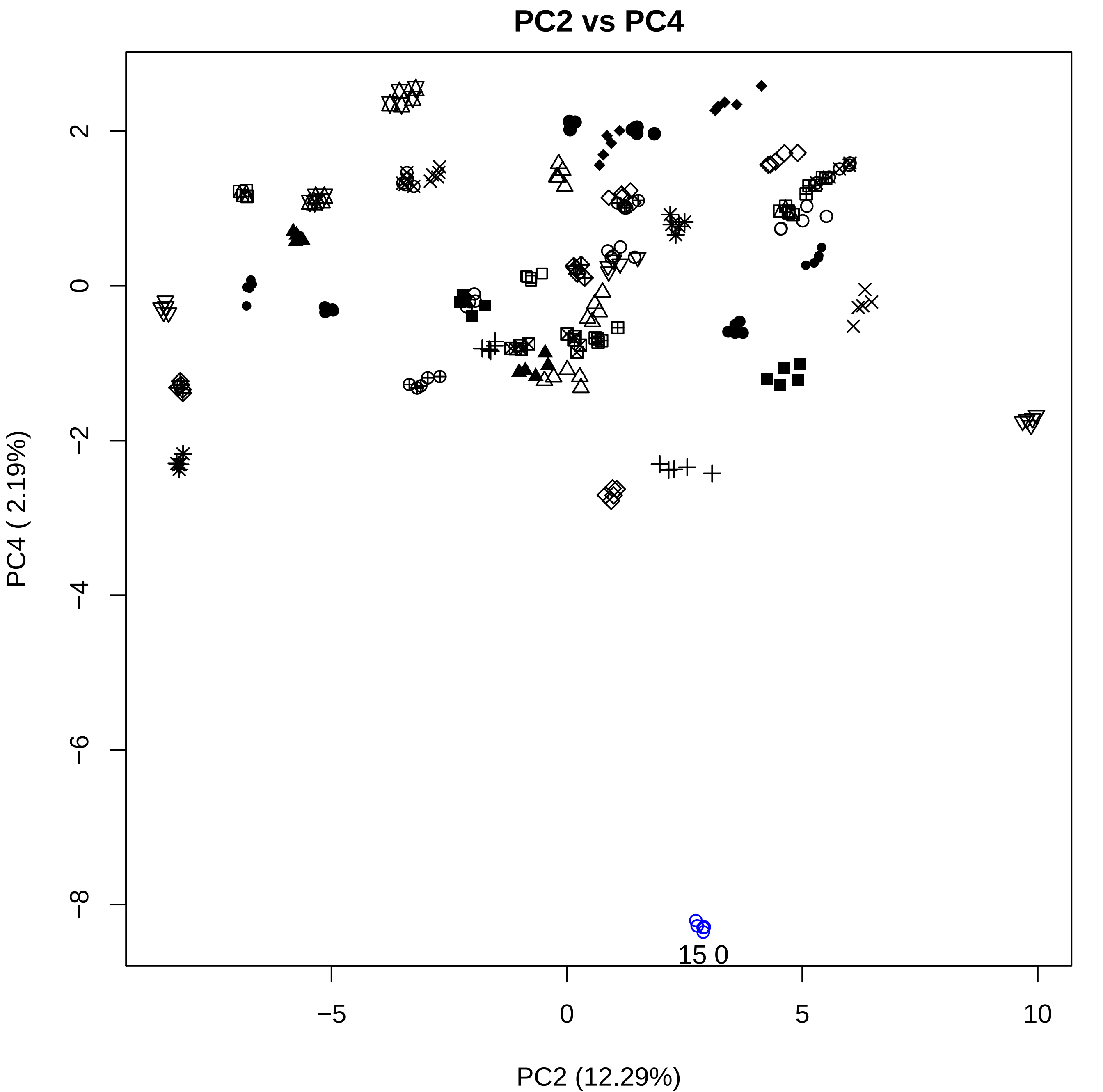
PC2 vs PC4
PC4 ( 2.19%)
PC2 (12.29%)
2
0
−2
−4
−6
−8
−5
0
5
10
15 0

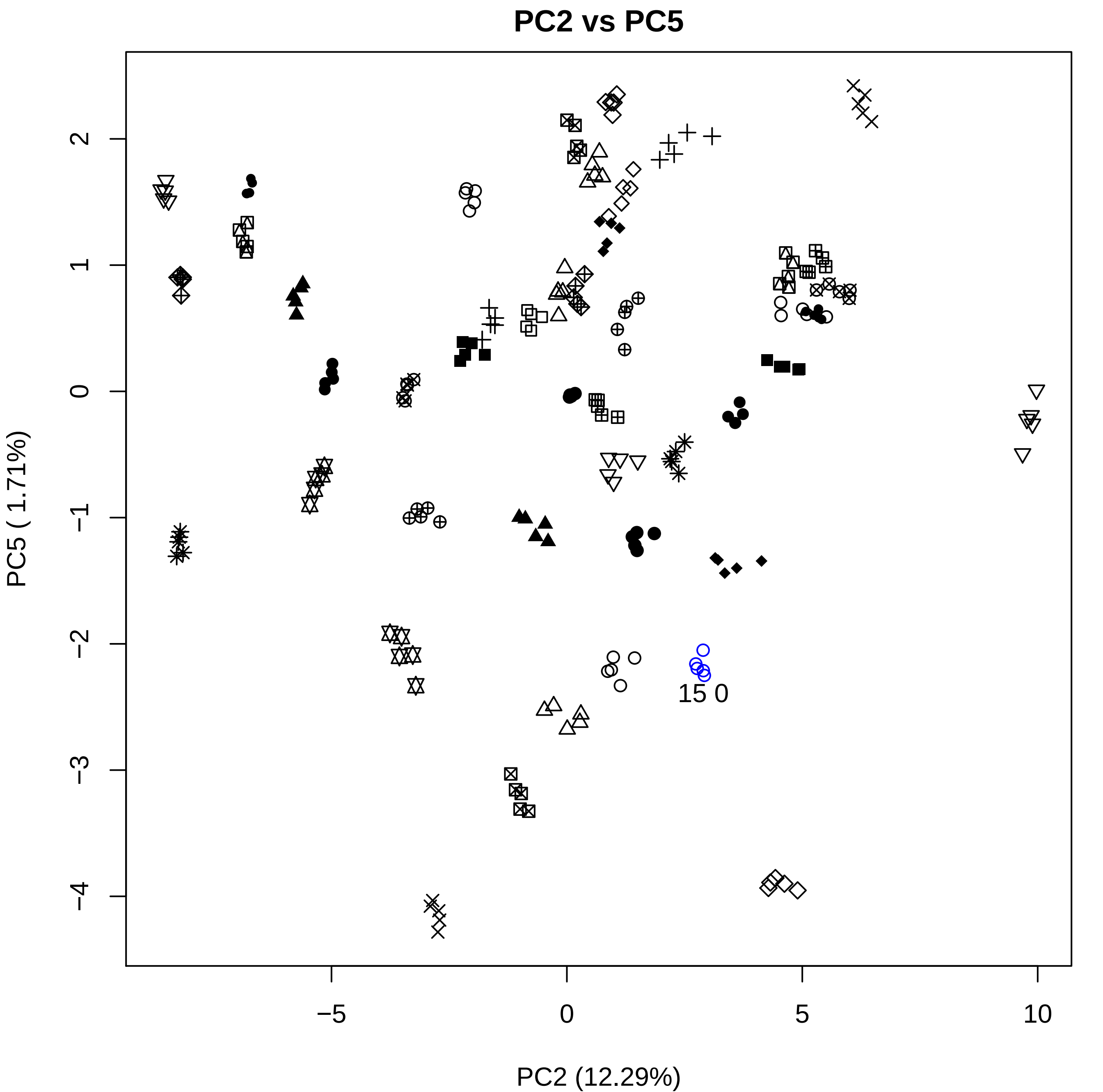
PC2 vs PC5
PC5 ( 1.71%)
PC2 (12.29%)
2
1
0
−1
−2
−3
−4
−5
0
5
10
15 0

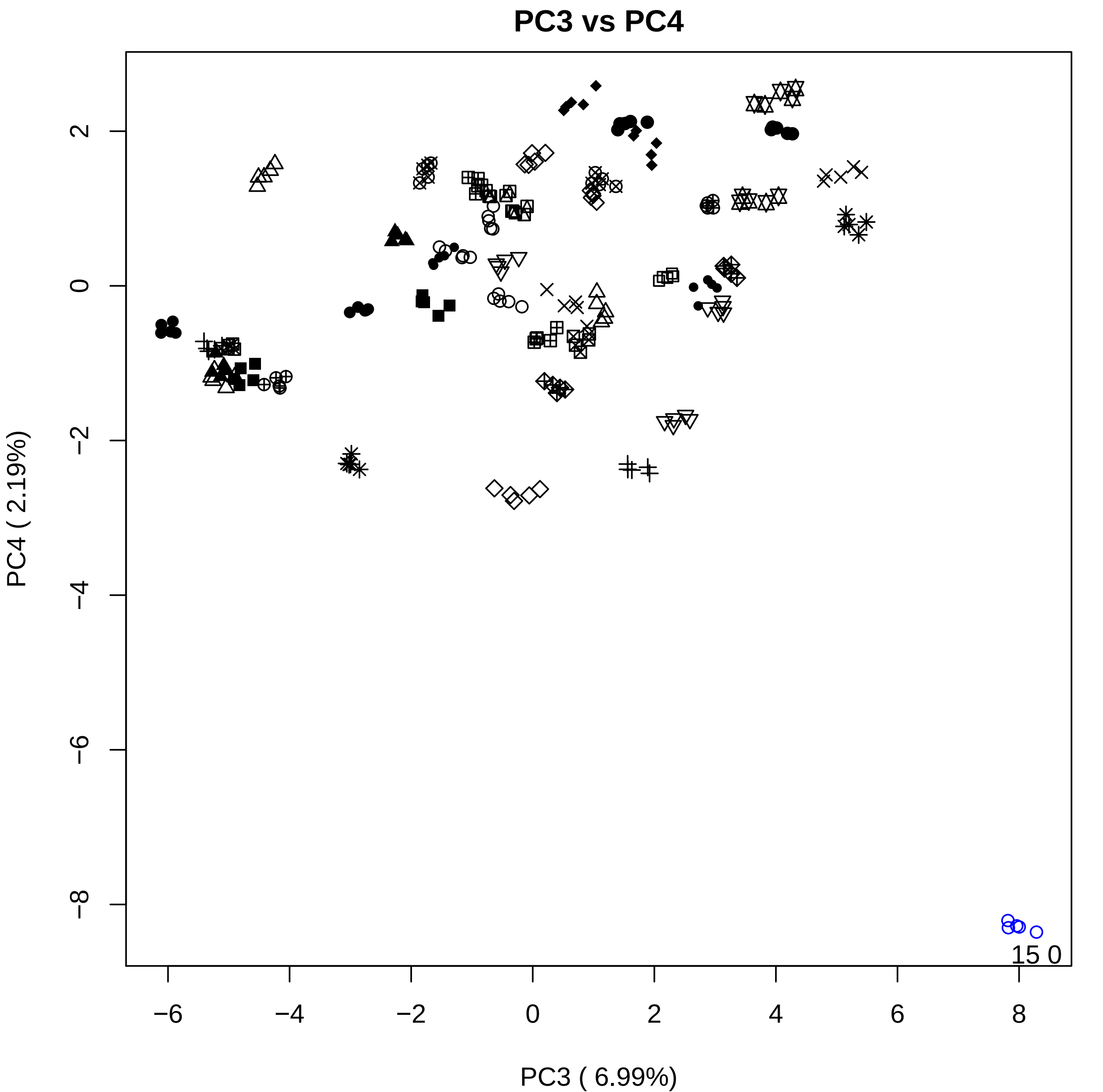
PC3 vs PC4
PC4 ( 2.19%)
PC3 ( 6.99%)
15 0
−6
−4
−2
0
2
4
6
8
−8
−6
−4
−2
0
2

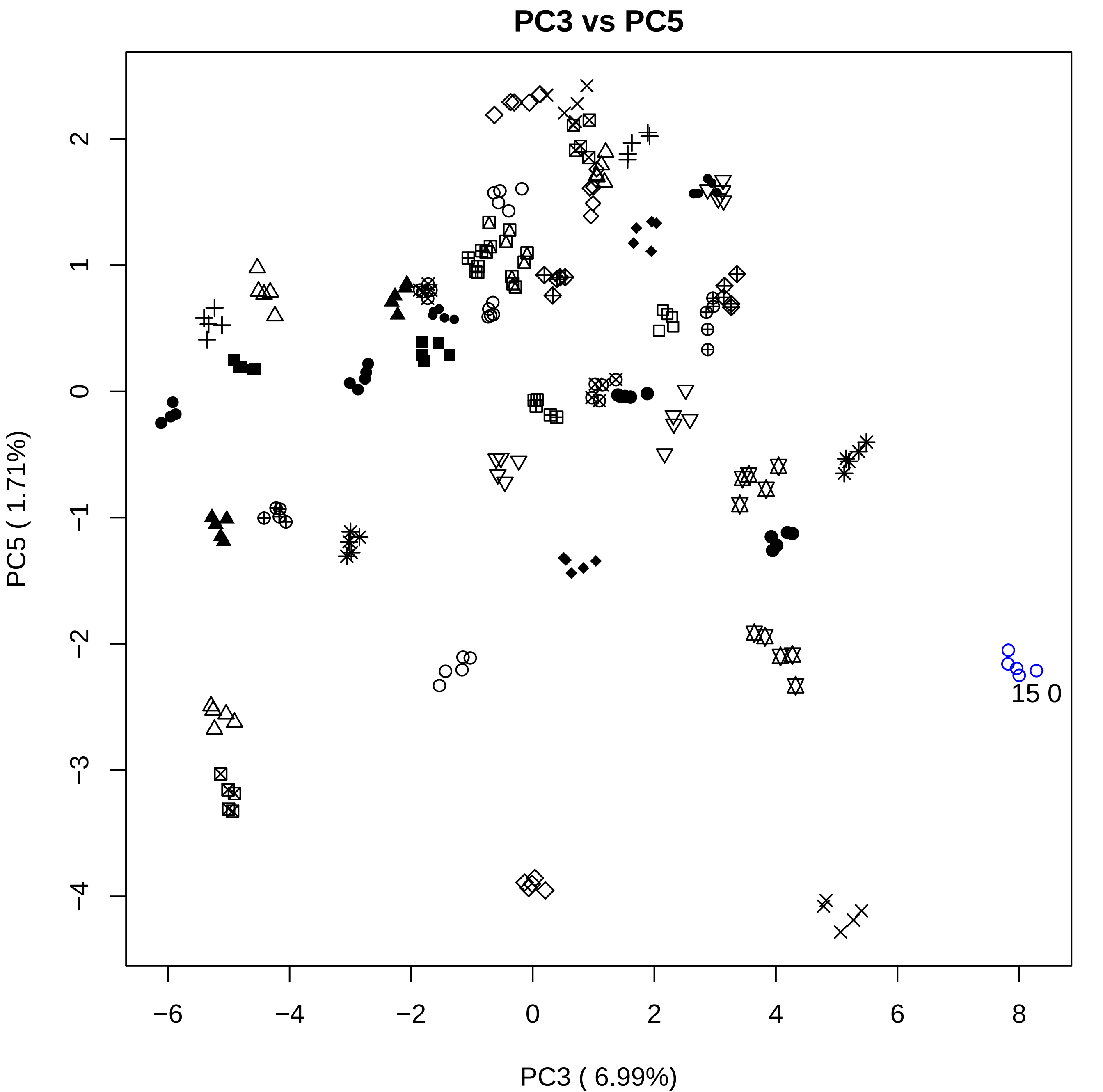

PC3 vs PC5
PC5 ( 1.71%)
PC3 ( 6.99%)
−6
−4
−2
0
2
4
6
8
−4
−3
−2
−1
0
1
2
15 0

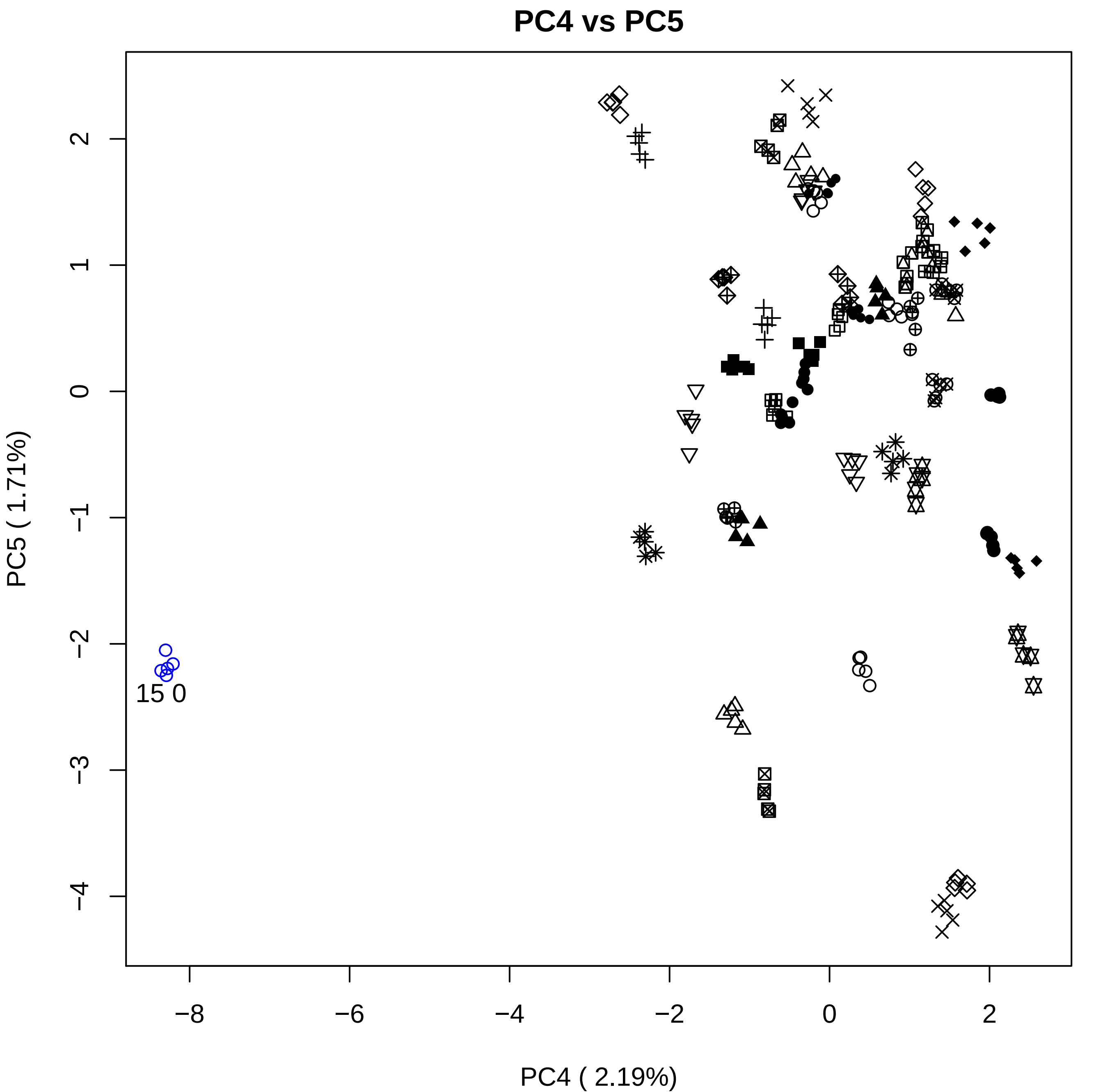

PC4 vs PC5
PC4 ( 2.19%)
PC5 ( 1.71%)
−8
−6
−4
−2
0
2
−4
−3
−2
−1
0
1
2
15 0

### PCA of 20%  OSAS vs PMH

Scaled Matrix PCAed

7th order polynomial baseline correction performed

Raw spectra smoothed to 21 Neighbours, 2nd order PN, Savtisky Golay

Normalisation between 0 and 100

D2 spectra smoothed to 41 Neighbours, 2nd order PN, Savtisky Golay

Regions Analysed: 3601−2501 and 2001−700cm−1

Samples Analysed: 1 Contam's 0 OSAS's 45 PMH's

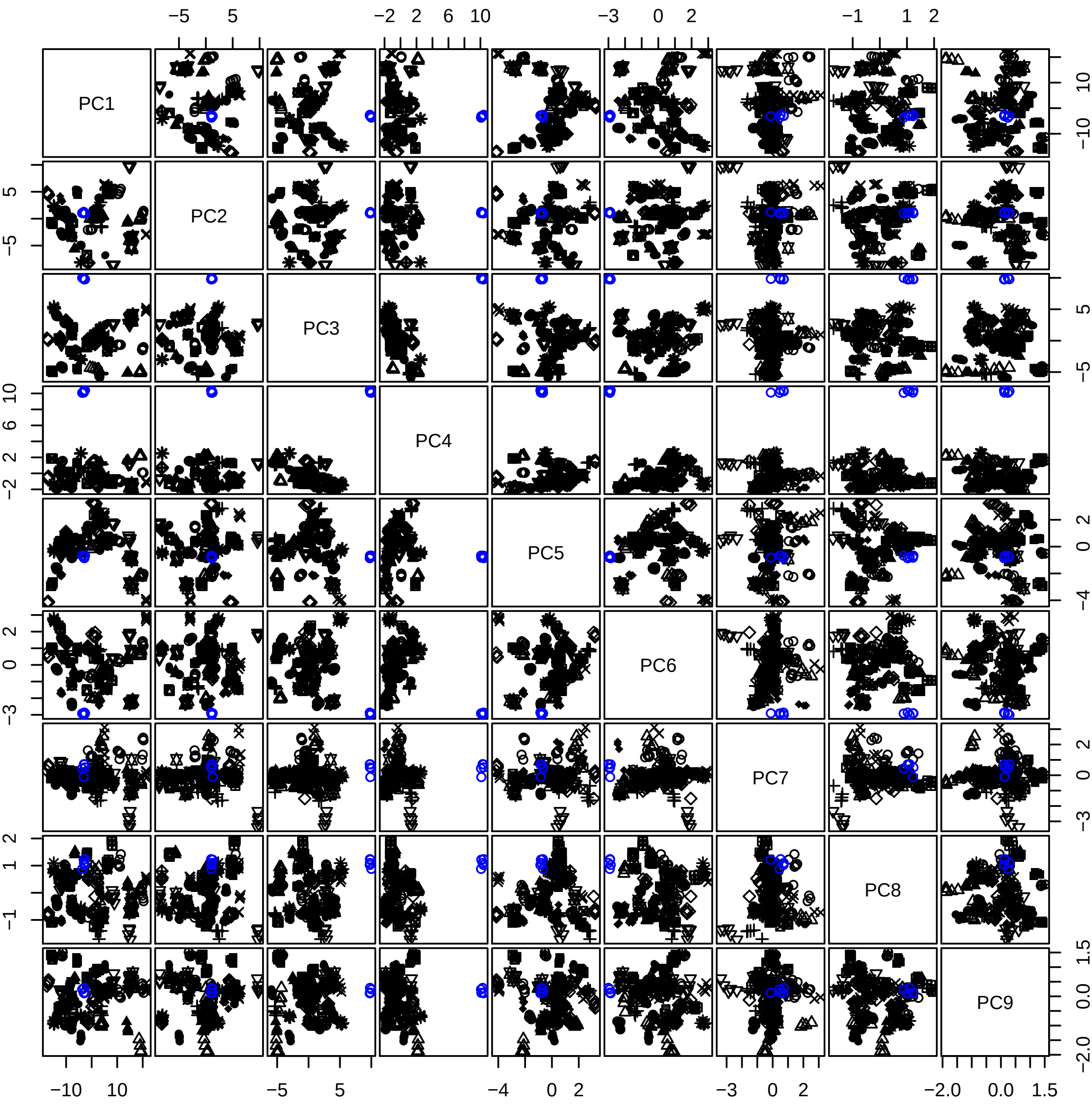
PC1
PC2
PC3
PC4
PC5
PC6
PC7
PC8
PC9

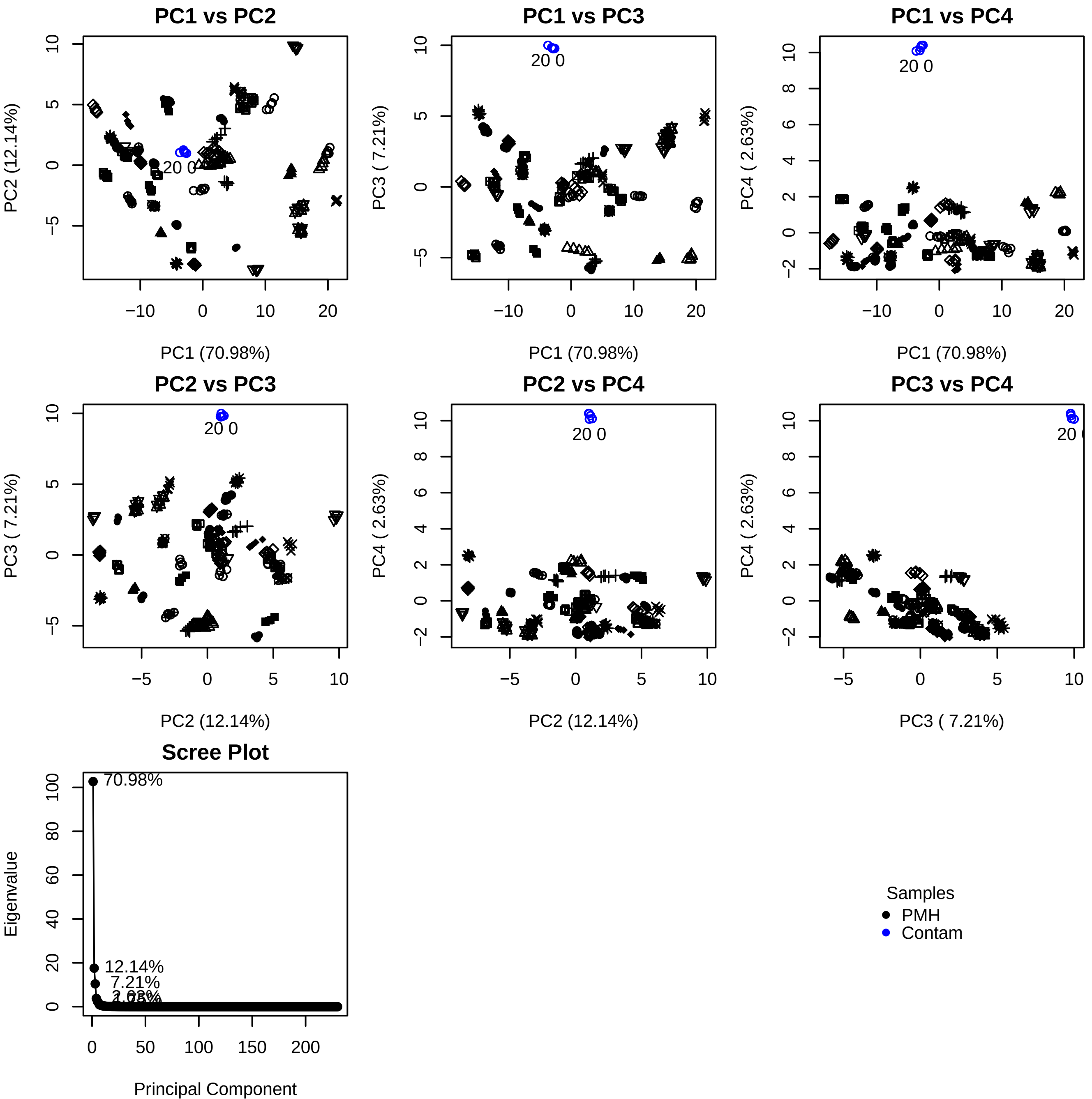
PC1 vs PC2
PC1 vs PC3
PC1 vs PC4
PC2 vs PC3
PC2 vs PC4
PC3 vs PC4
Scree Plot
PC1 (70.98%)
PC2 (12.14%)
PC3 ( 7.21%)
PC4 ( 2.63%)
20 0
70.98%
12.14%
7.21%
Eigenvalue
Principal Component
Samples
PMH
Contam

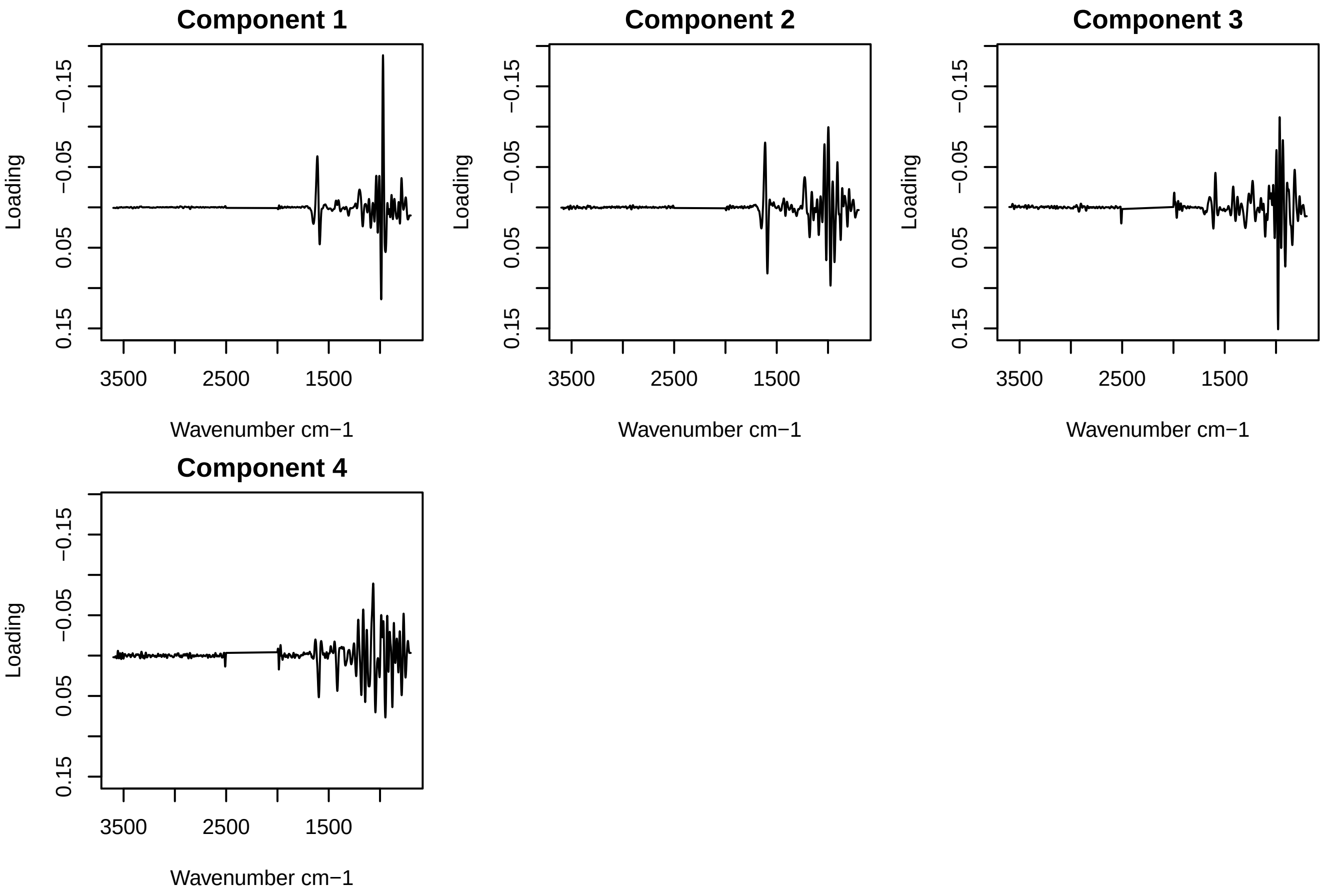
Component 1
Component 2
Component 3
Component 4
Loading
−0.15
−0.05
0.05
0.15
3500
2500
1500
Wavenumber cm−1

PC1 vs PC2 vs PC3

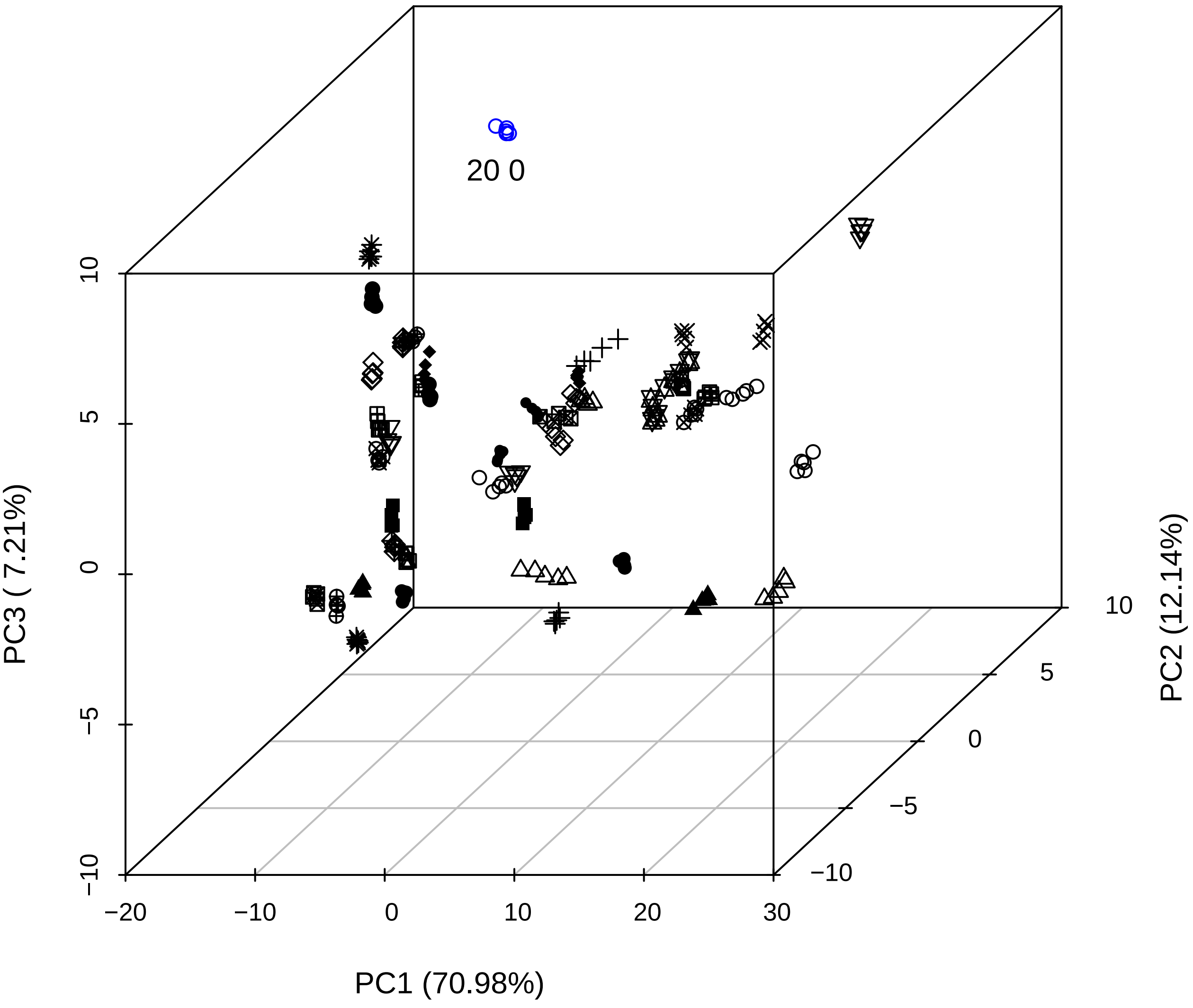

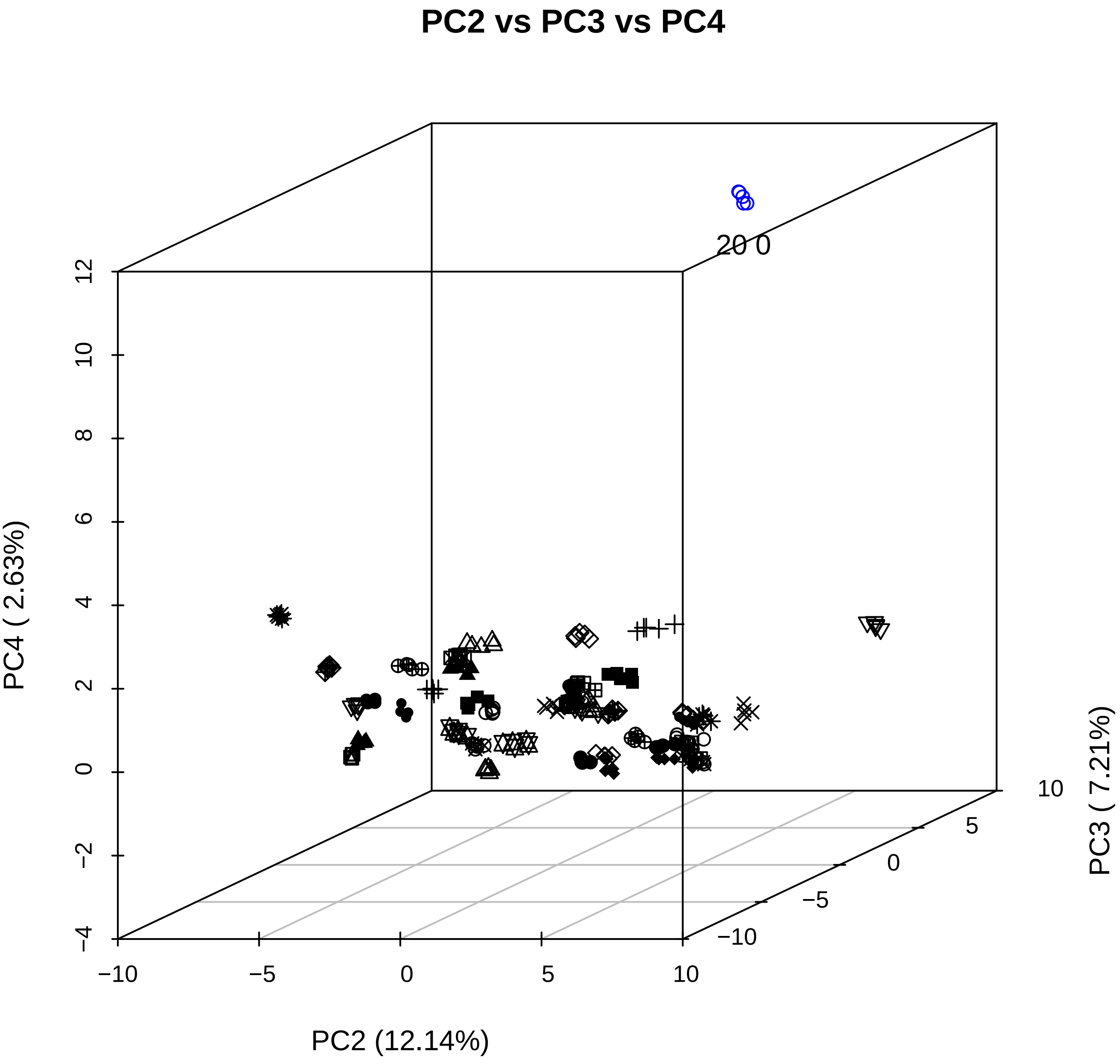

PC2 vs PC3 vs PC4
PC4 ( 2.63%)
PC2 (12.14%)
PC3 ( 7.21%)
−4
−2
0
2
4
6
8
10
12
−10
−5
0
5
10
−10
−5
0
5
10
20 0

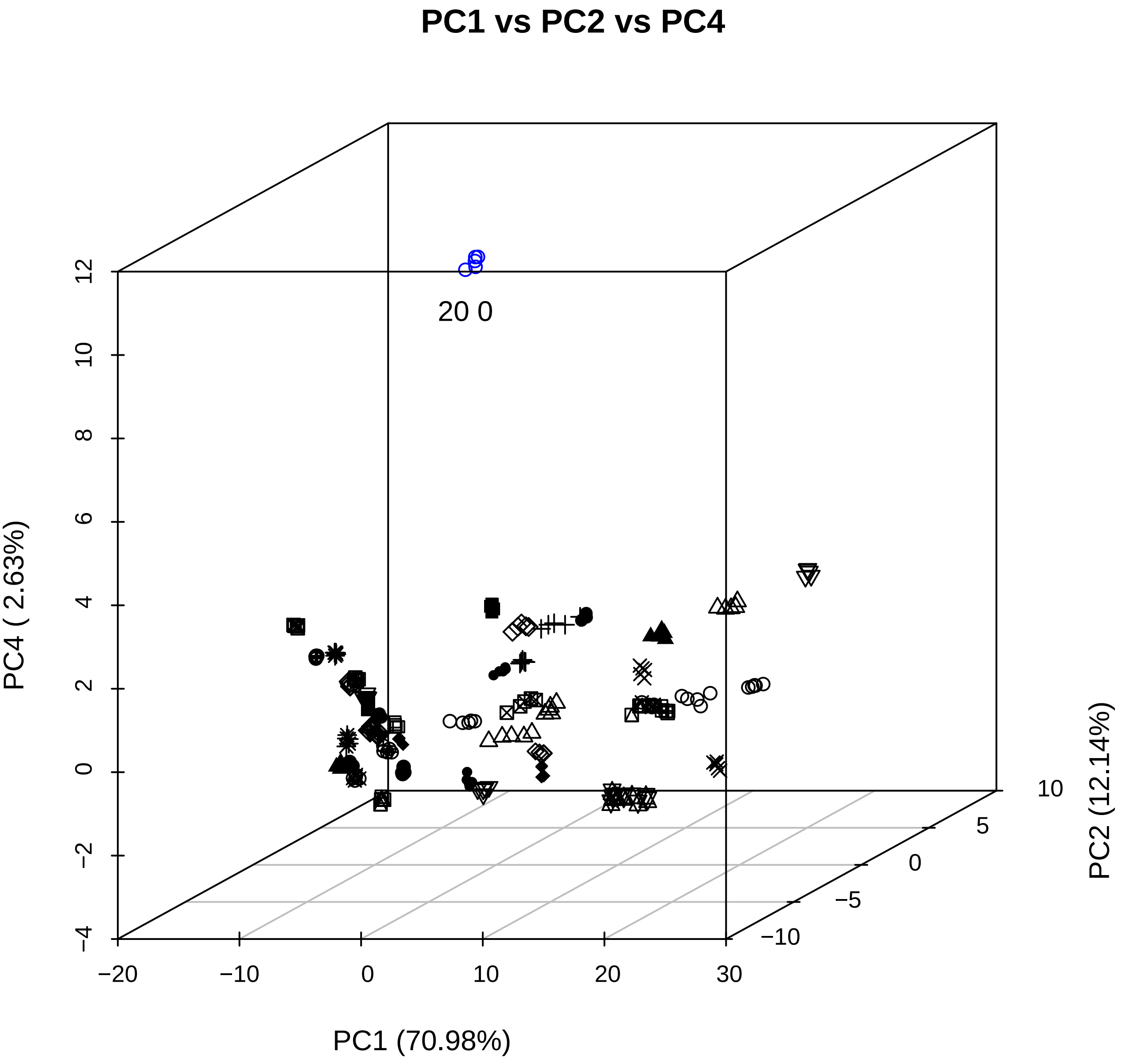

PC1 vs PC2 vs PC4
20 0
PC4 ( 2.63%)
12
10
8
6
4
2
0
−2
−4
−20
−10
0
10
20
30
PC1 (70.98%)
PC2 (12.14%)
10
5
0
−5
−10

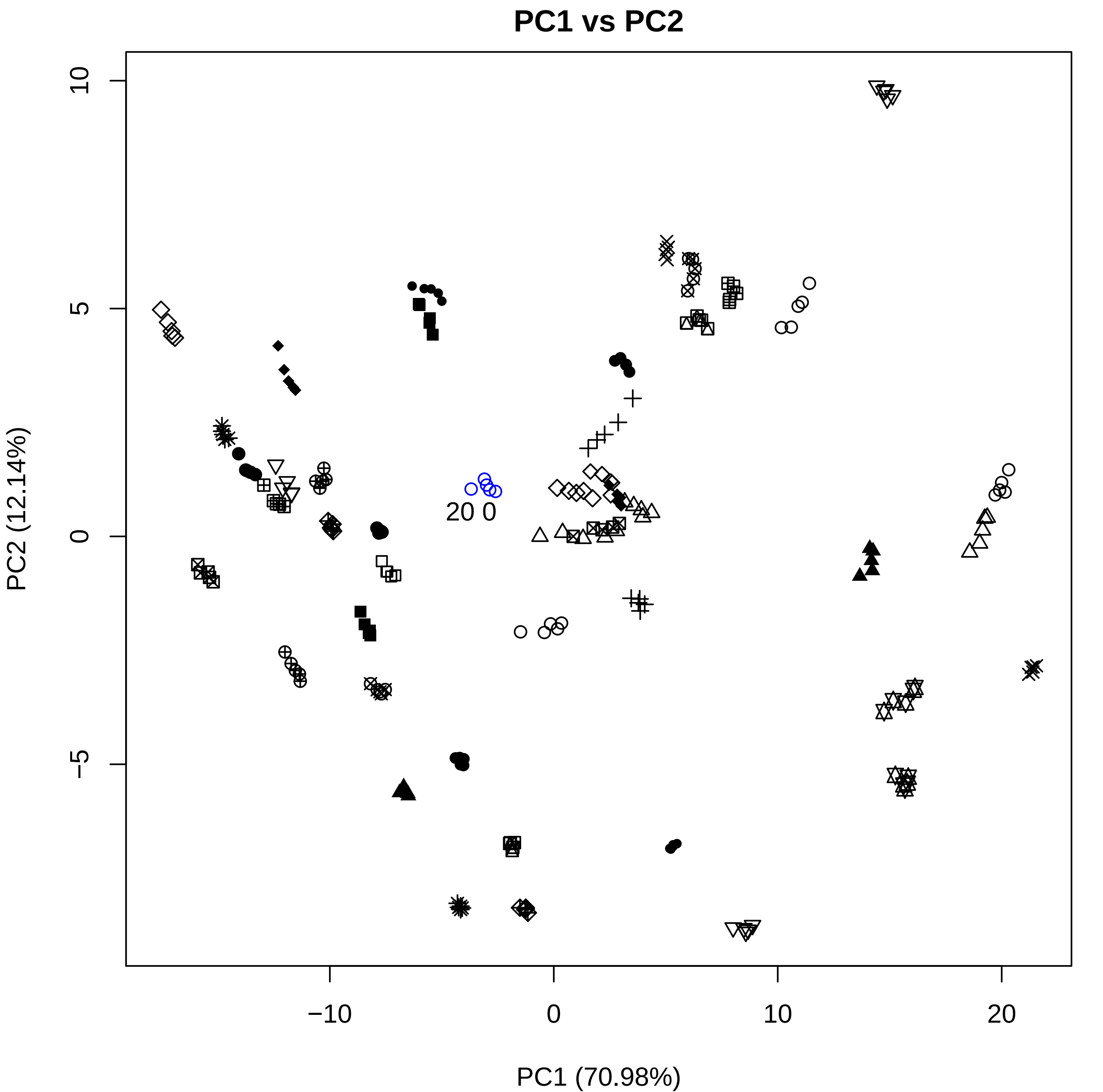
PC1 vs PC2
PC2 (12.14%)
PC1 (70.98%)
10
5
0
−5
−10
0
10
20
20 0

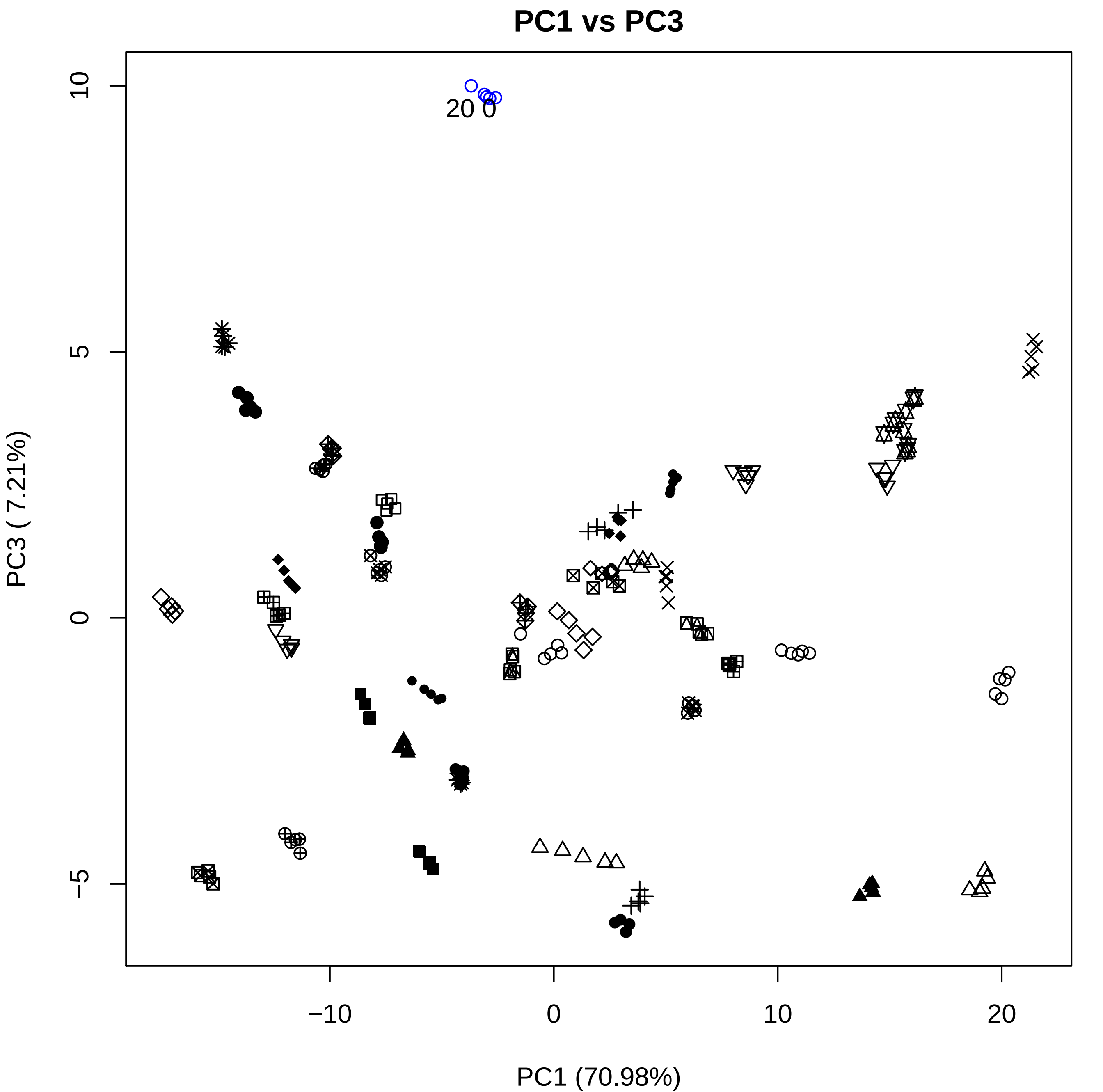
PC1 vs PC3
PC3 ( 7.21%)
PC1 (70.98%)
20 0
10
5
0
−5
−10
0
10
20

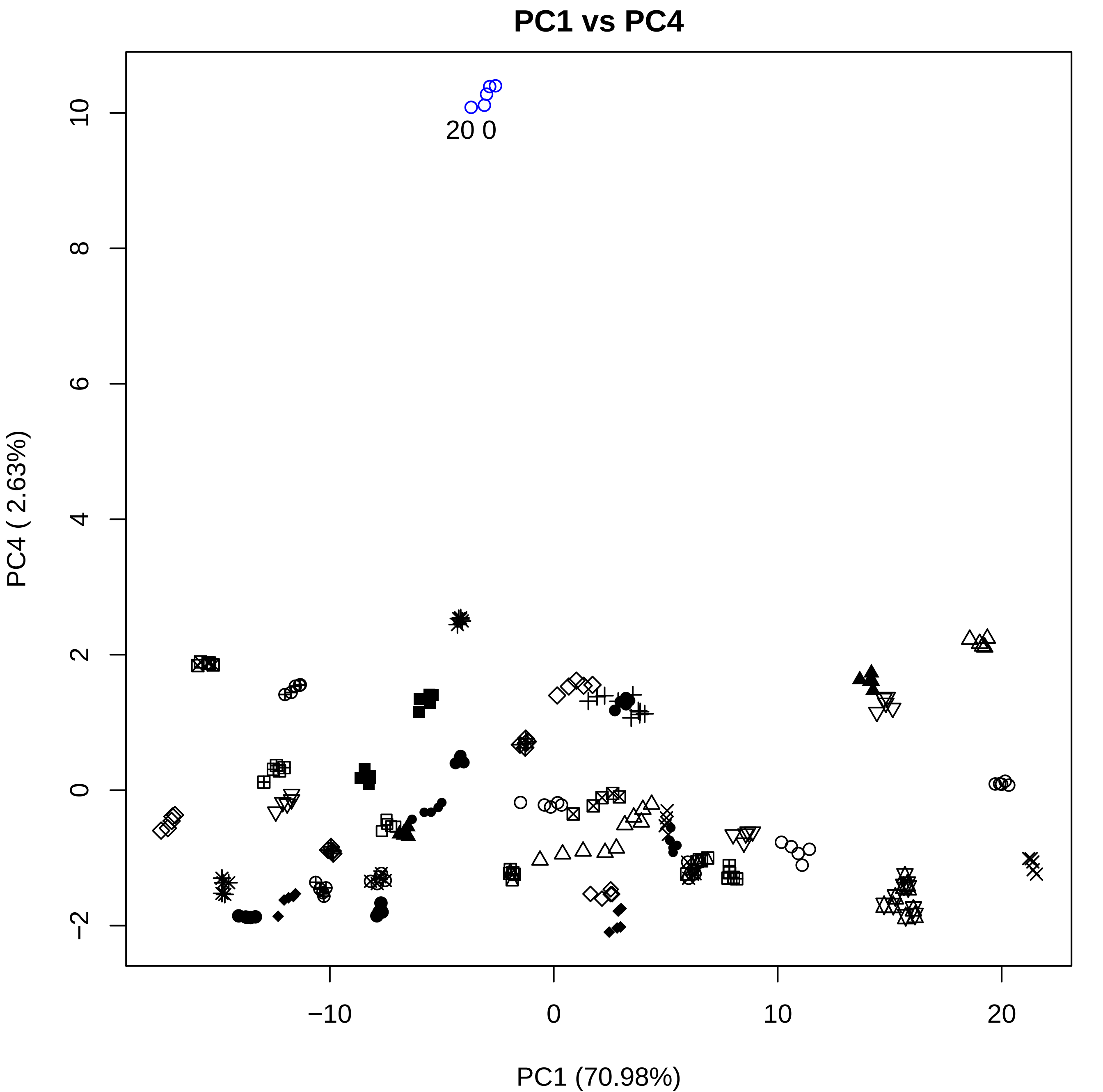
PC1 vs PC4
20 0
PC4 ( 2.63%)
PC1 (70.98%)
10
8
6
4
2
0
−2
−10
0
10
20

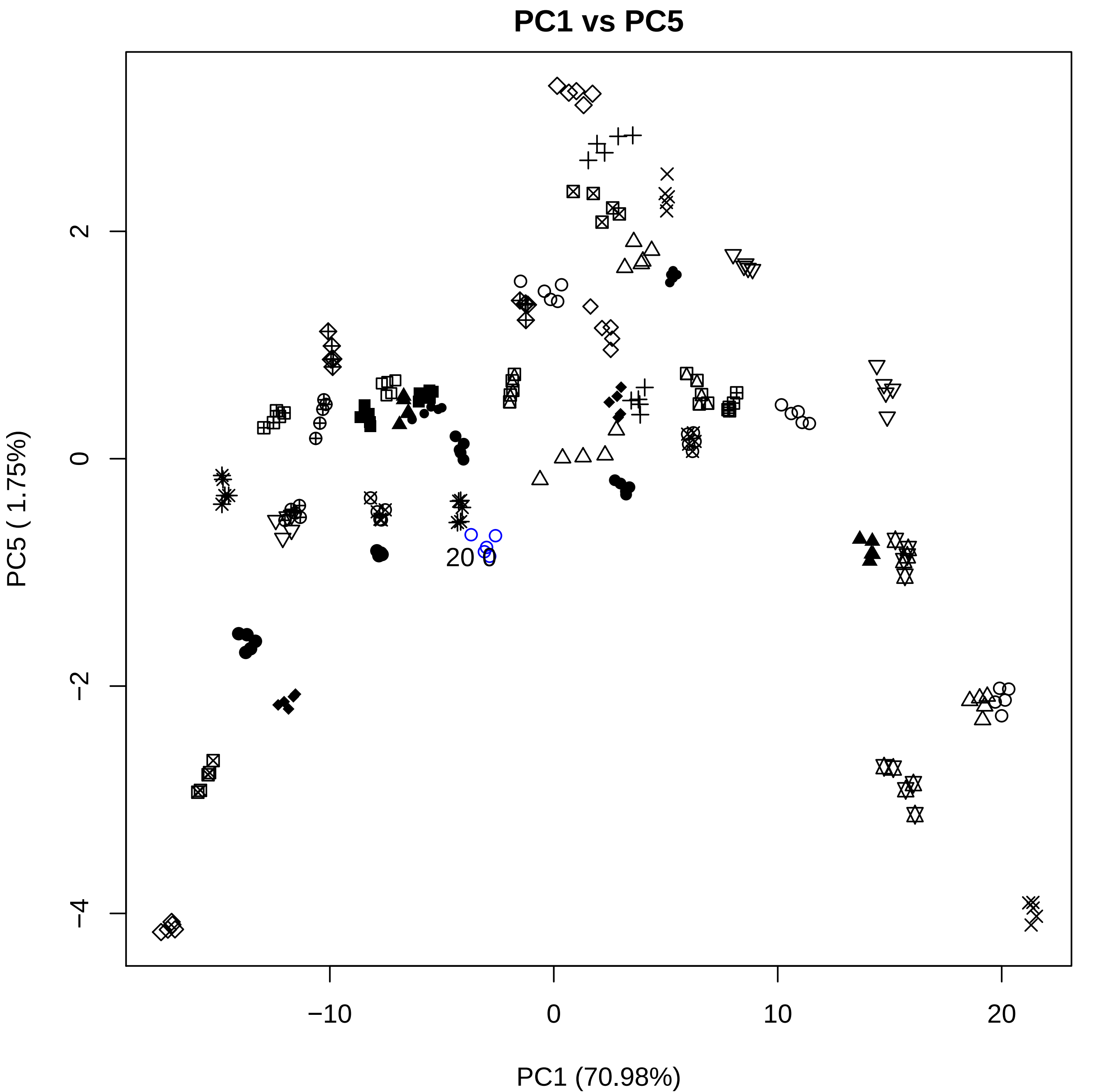
PC1 vs PC5
PC5 ( 1.75%)
PC1 (70.98%)
2
0
−2
−4
−10
0
10
20
20 0

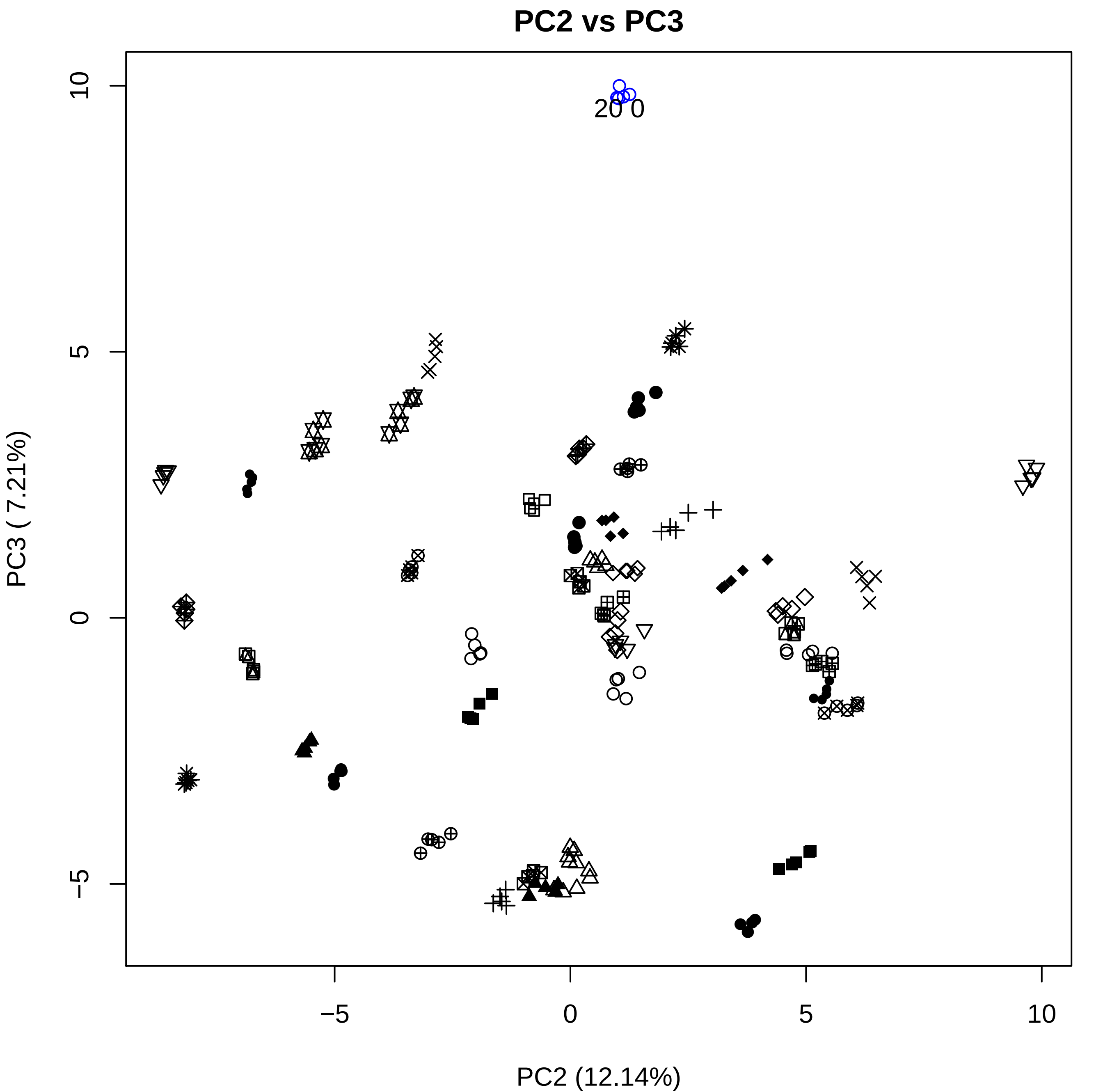
PC2 vs PC3
20 0
10
5
0
−5
−5
0
5
10
PC3 ( 7.21%)
PC2 (12.14%)

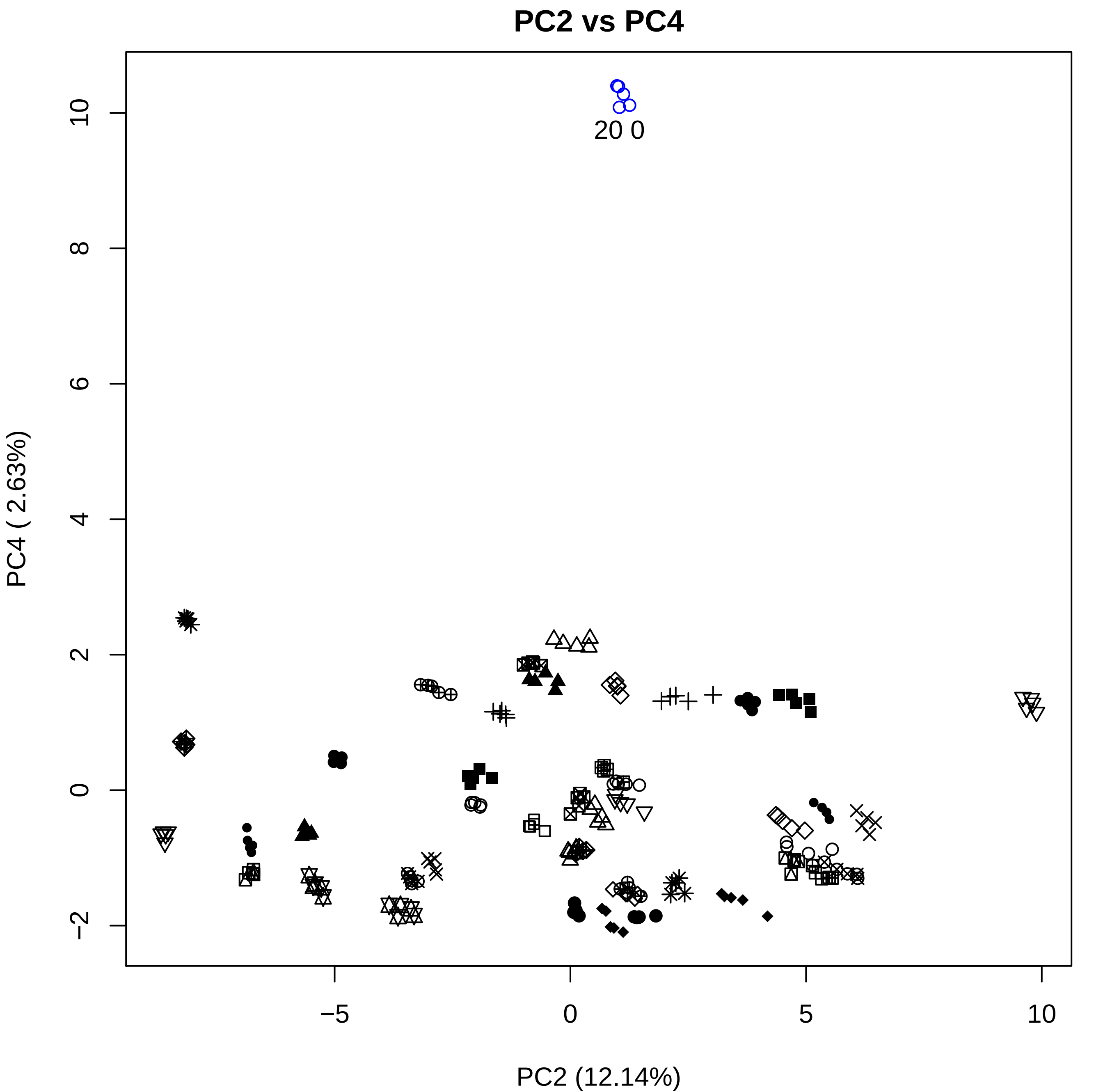
PC2 vs PC4
20 0
PC4 ( 2.63%)
PC2 (12.14%)
−2
0
2
4
6
8
10
−5
0
5
10

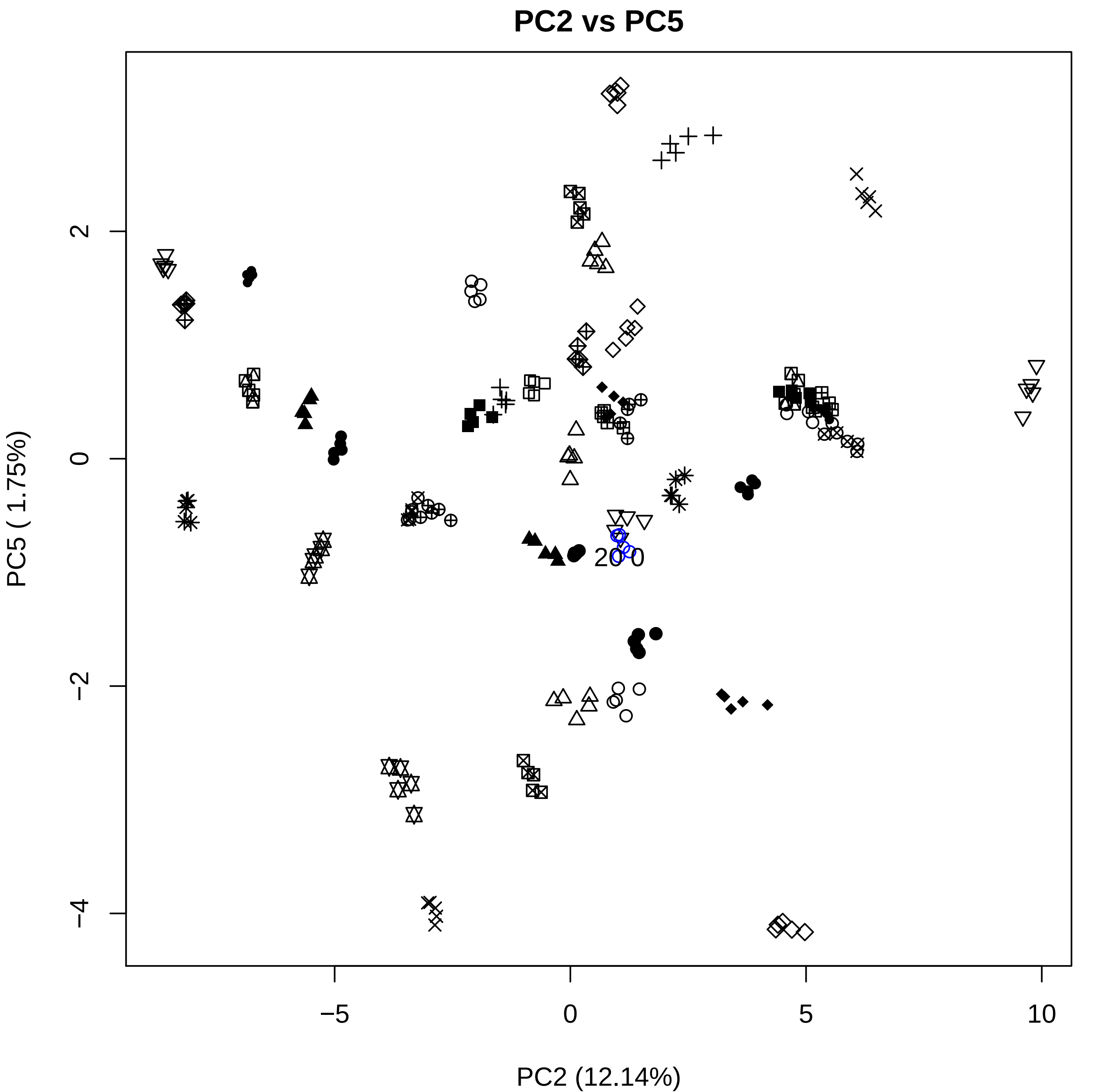
PC2 vs PC5
PC5 ( 1.75%)
PC2 (12.14%)
2
0
−2
−4
−5
0
5
10
20 0

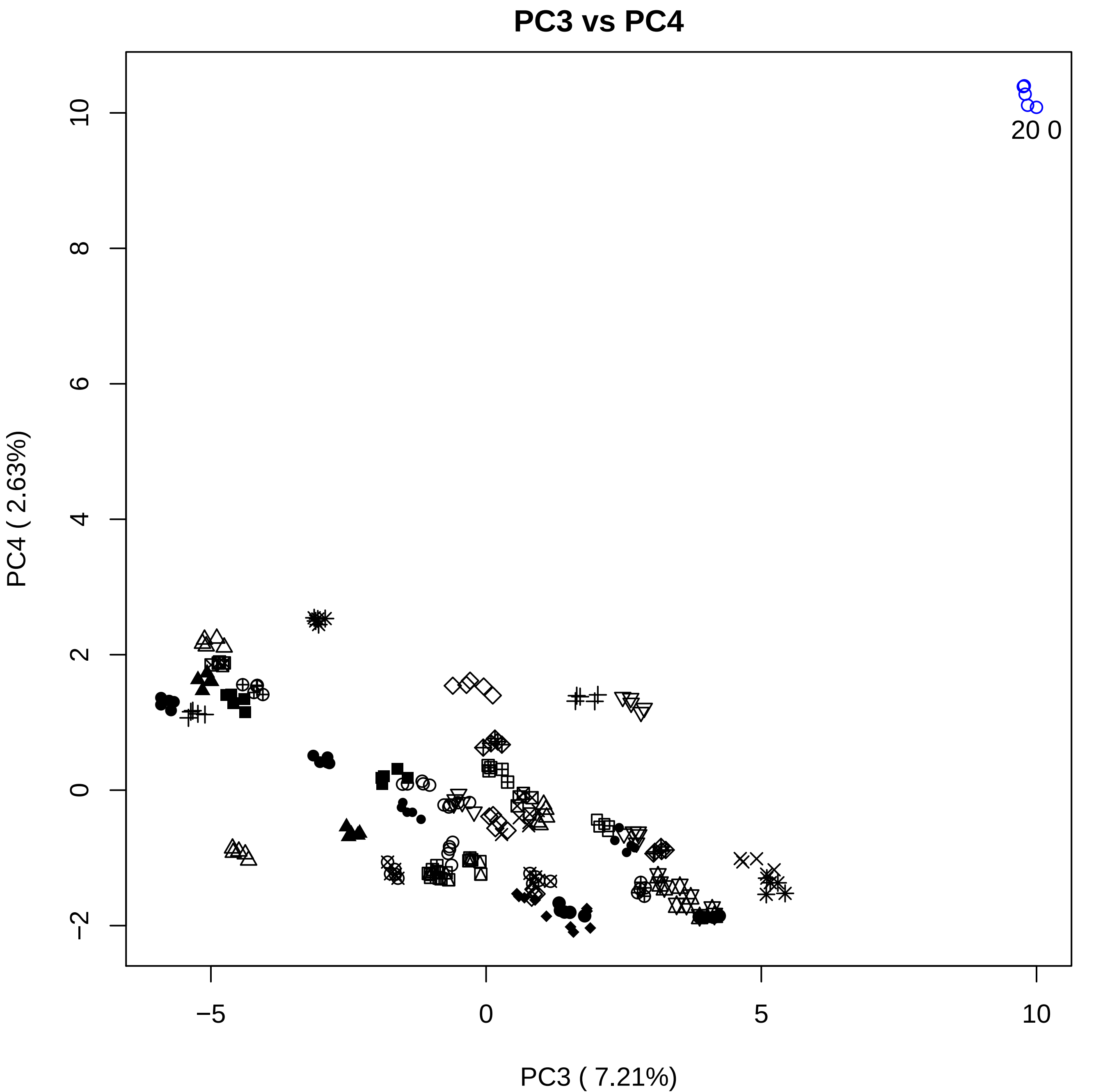
PC3 vs PC4
PC4 ( 2.63%)
PC3 ( 7.21%)
20 0
−5
0
5
10
−2
0
2
4
6
8
10

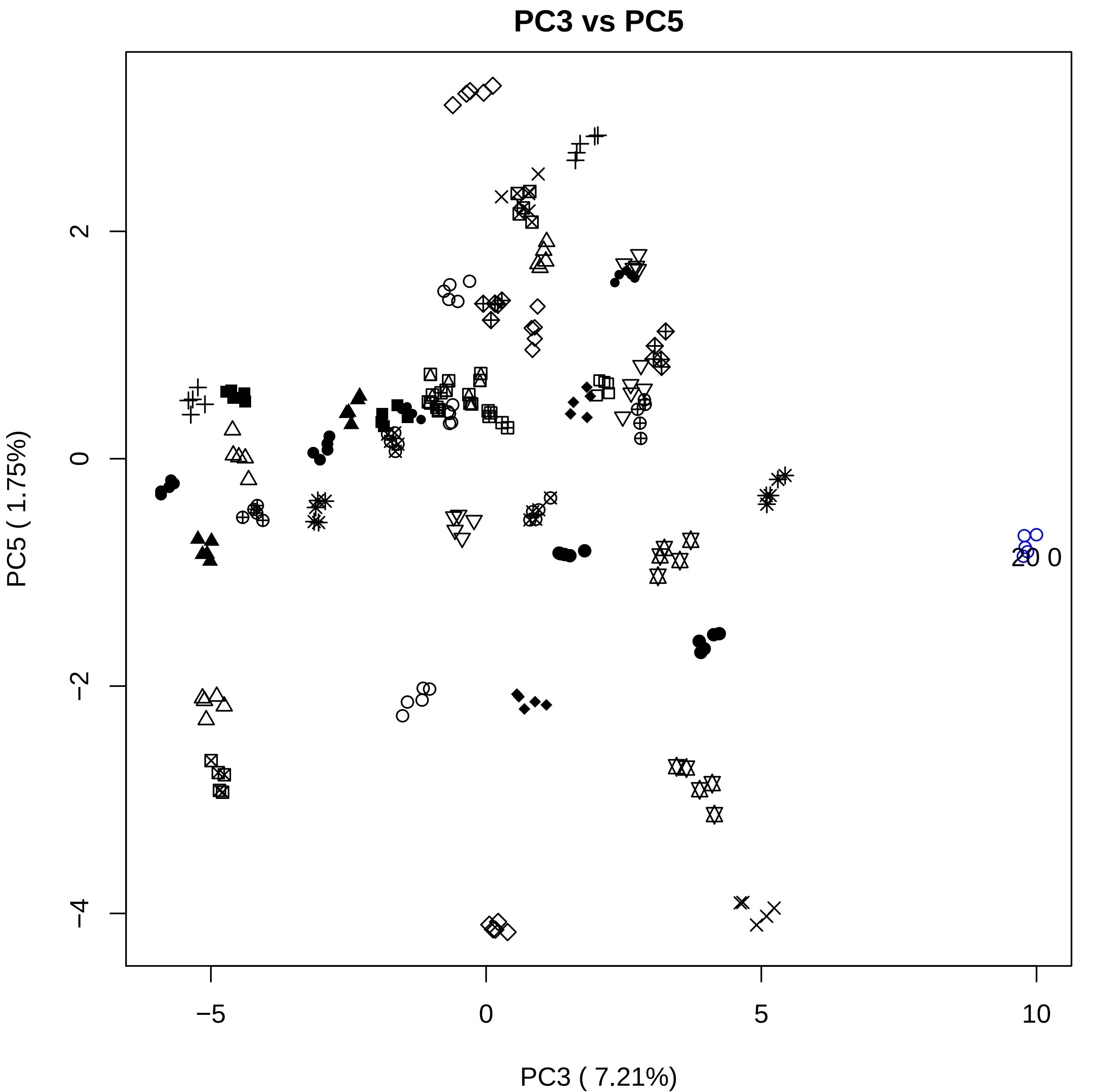
PC3 vs PC5
PC5 ( 1.75%)
PC3 ( 7.21%)
2
0
−2
−4
−5
0
5
10
20 0

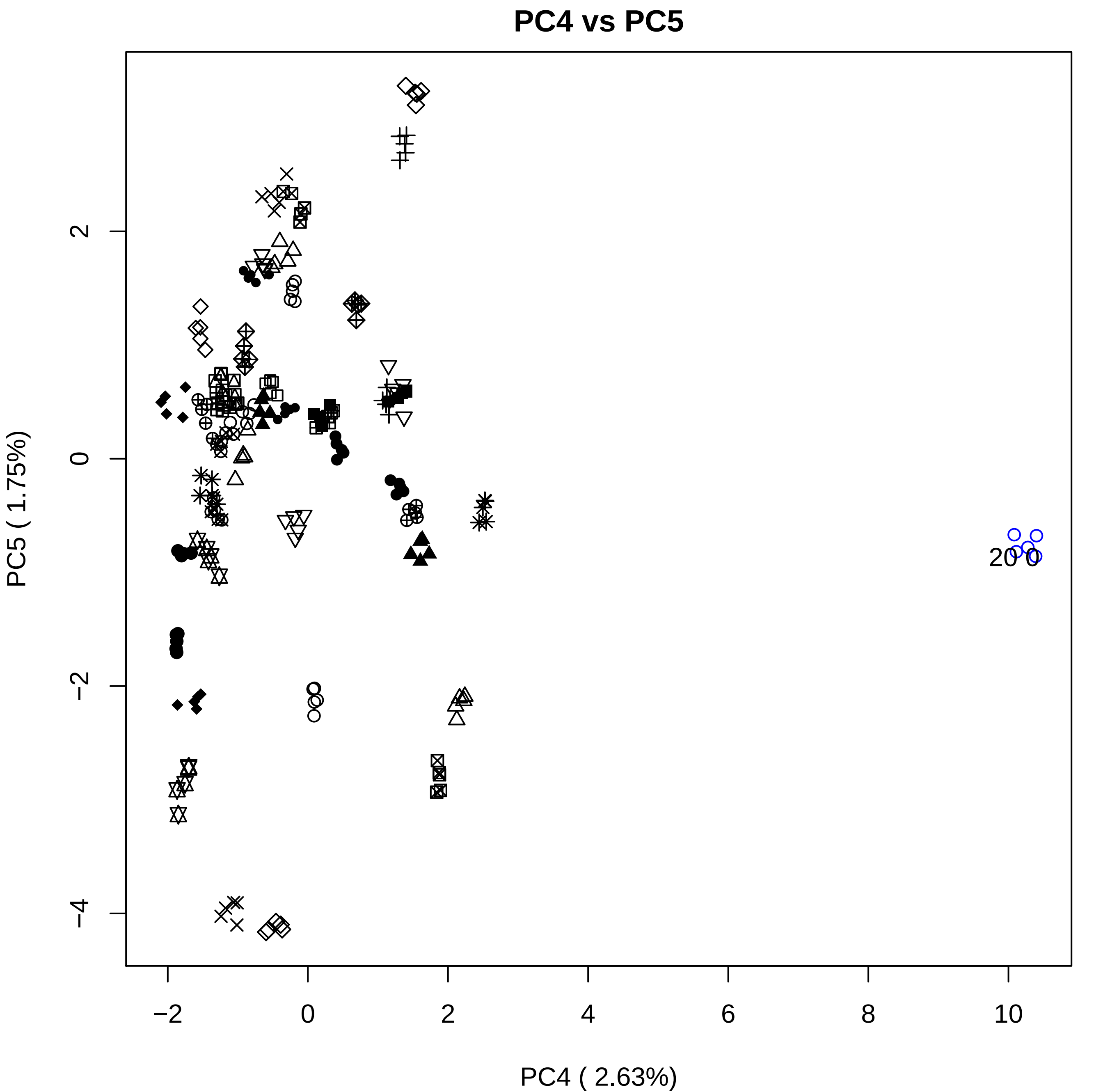
PC4 vs PC5
PC5 ( 1.75%)
PC4 ( 2.63%)
2
0
−2
−4
−2
0
2
4
6
8
10
20 0

### PCA of 40%  OSAS vs PMH

Scaled Matrix PCAed

7th order polynomial baseline correction performed

Raw spectra smoothed to 21 Neighbours, 2nd order PN, Savtisky Golay

Normalisation between 0 and 100

D2 spectra smoothed to 41 Neighbours, 2nd order PN, Savtisky Golay

Regions Analysed: 3601−2501 and 2001−700cm−1

Samples Analysed: 1 Contam's 0 OSAS's 45 PMH's

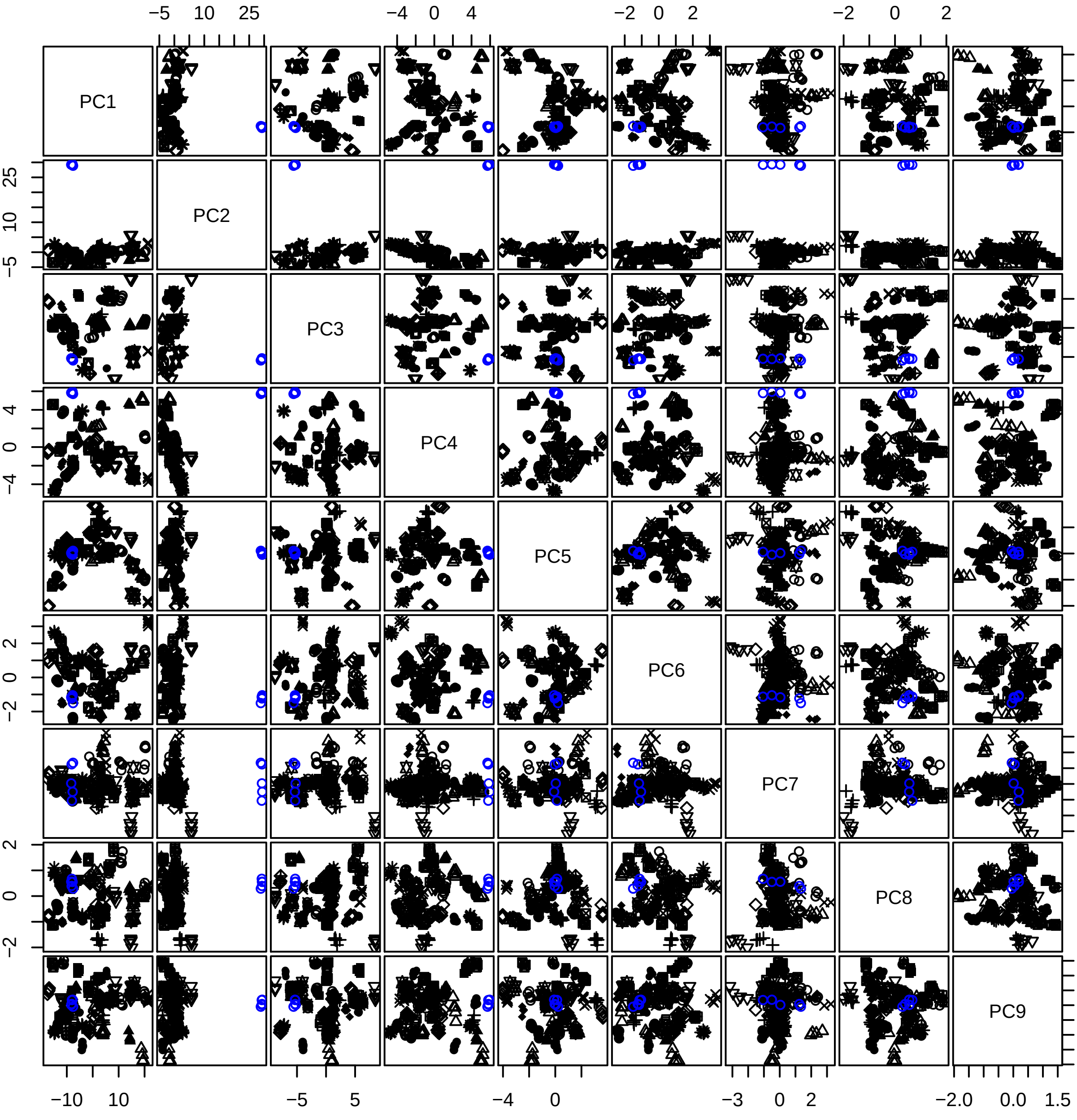
PC1
PC2
PC3
PC4
PC5
PC6
PC7
PC8
PC9

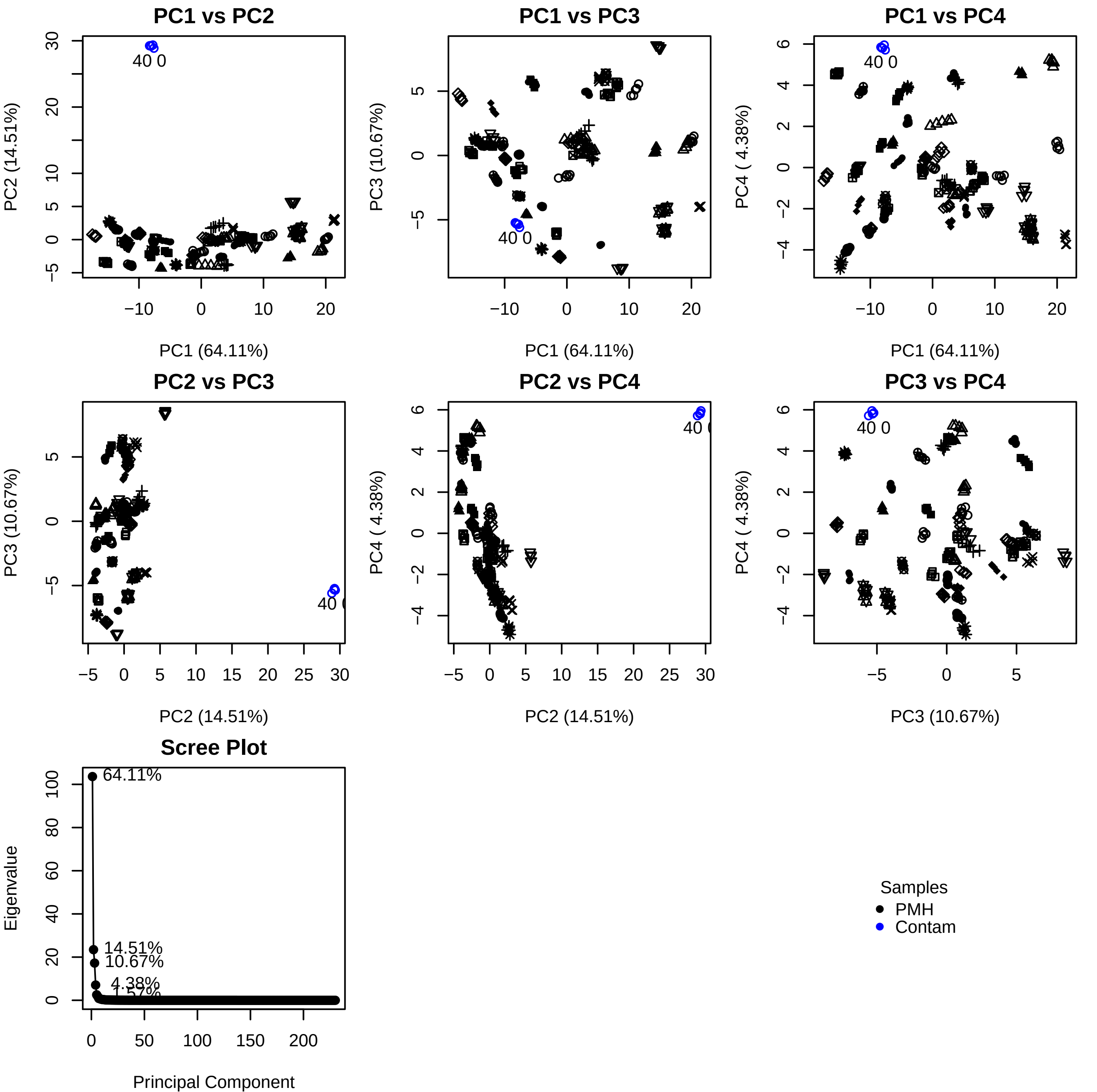
PC1 vs PC2
PC1 vs PC3
PC1 vs PC4
PC2 vs PC3
PC2 vs PC4
PC3 vs PC4
Scree Plot
PC1 (64.11%)
PC2 (14.51%)
PC3 (10.67%)
PC4 ( 4.38%)
40 0
Eigenvalue
Principal Component
64.11%
14.51%
10.67%
4.38%
1.57%
Samples
PMH
Contam

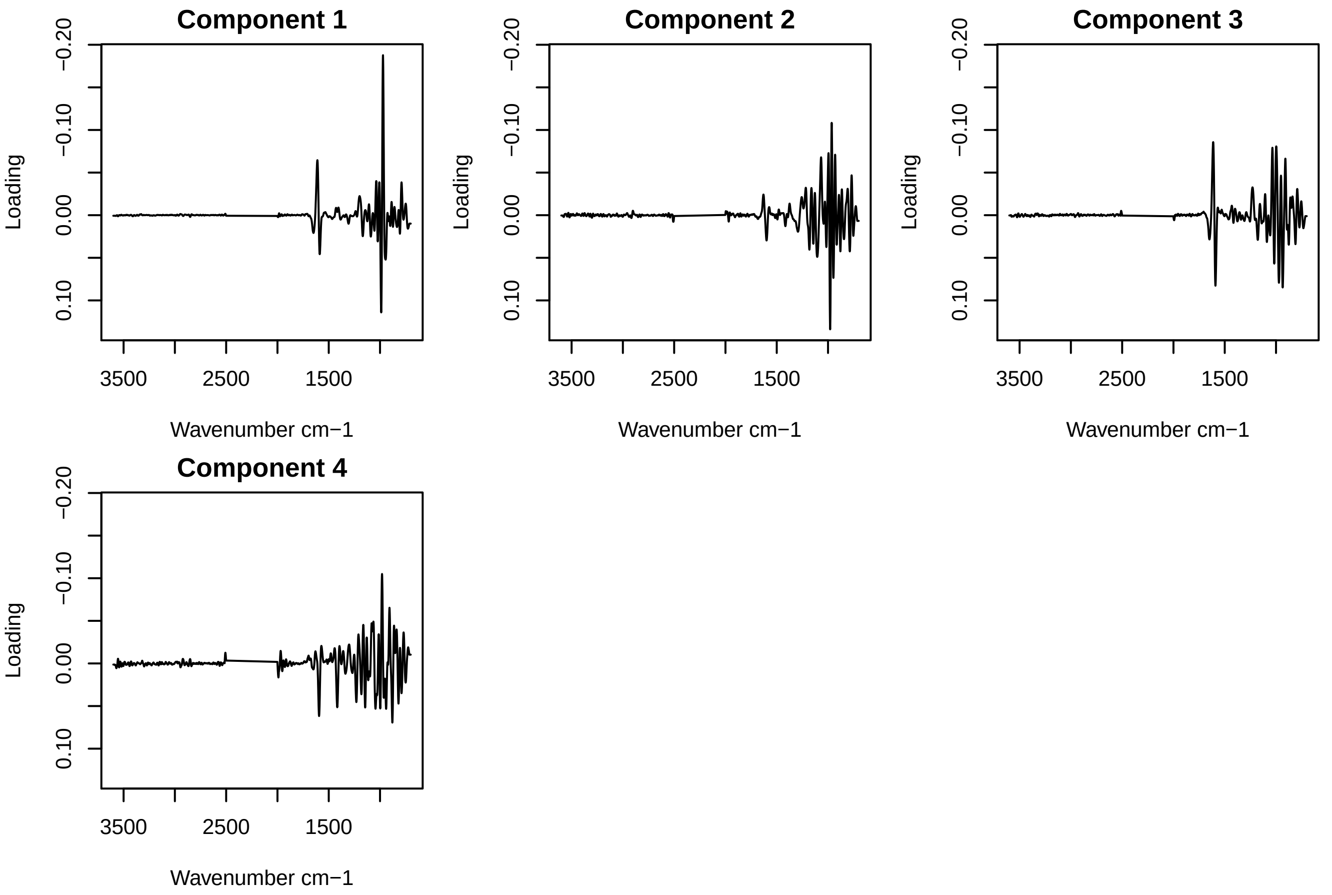
Component 1
Component 2
Component 3
Component 4
Loading
−0.20
−0.10
0.00
0.10
3500
2500
1500
Wavenumber cm−1

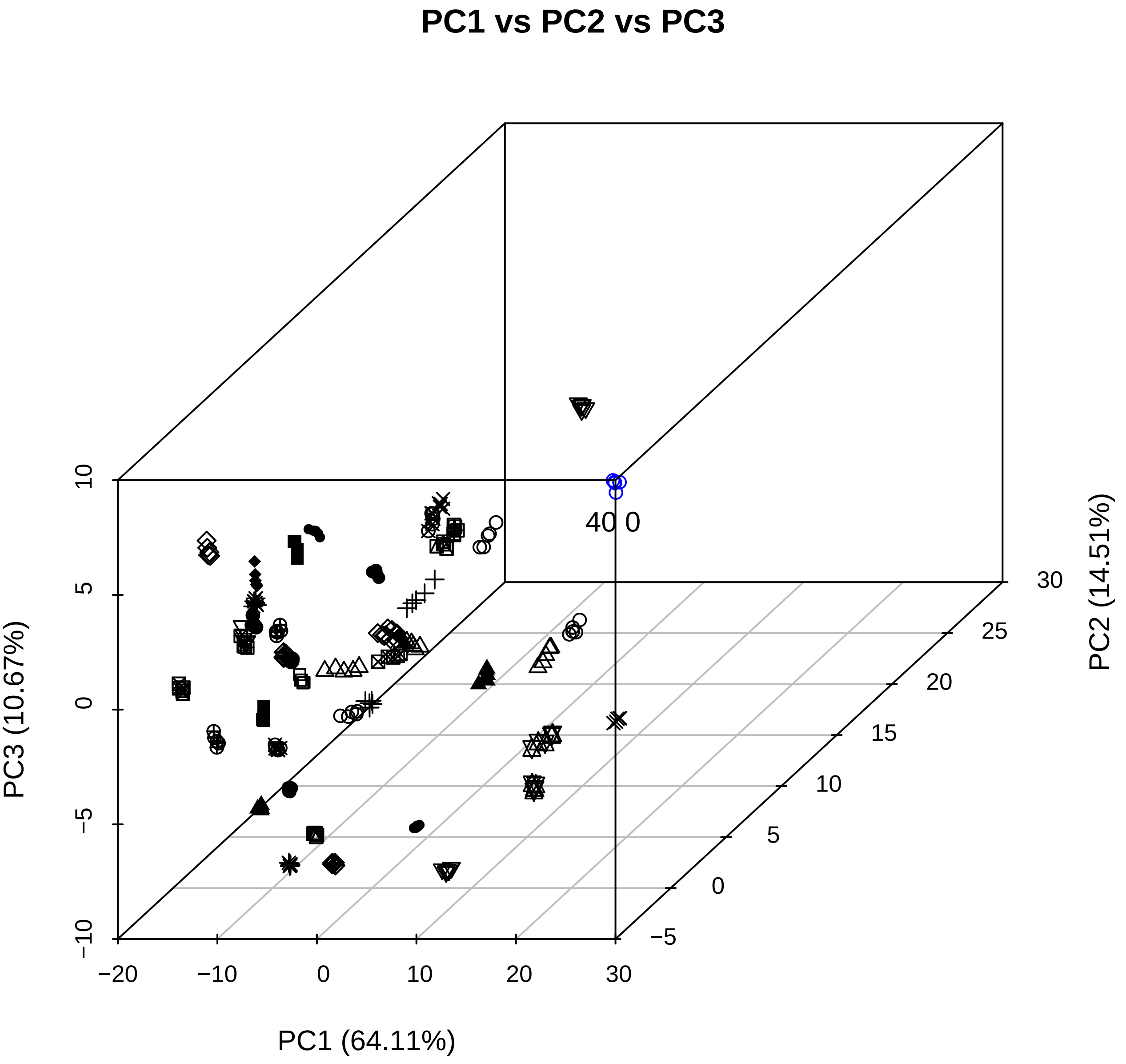
PC1 vs PC2 vs PC3
PC1 (64.11%)
PC2 (14.51%)
PC3 (10.67%)
40 0
−20
−10
0
10
20
30
−5
0
5
10
15
20
25
30
−10
−5
0
5
10

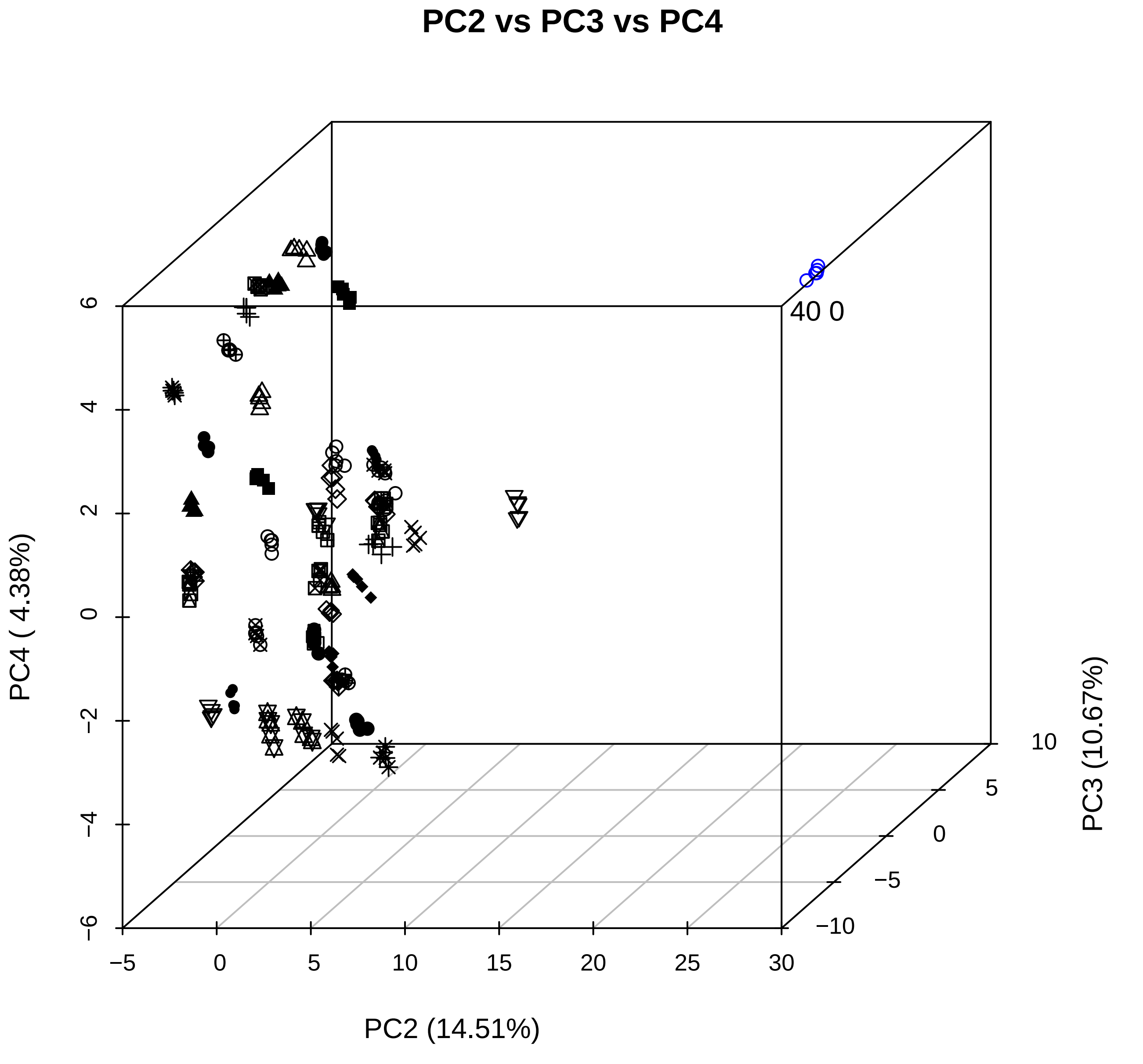
PC2 vs PC3 vs PC4
PC4 ( 4.38%)
6
4
2
0
−2
−4
−6
−5
0
5
10
15
20
25
30
PC2 (14.51%)
PC3 (10.67%)
10
5
0
−5
−10
40 0

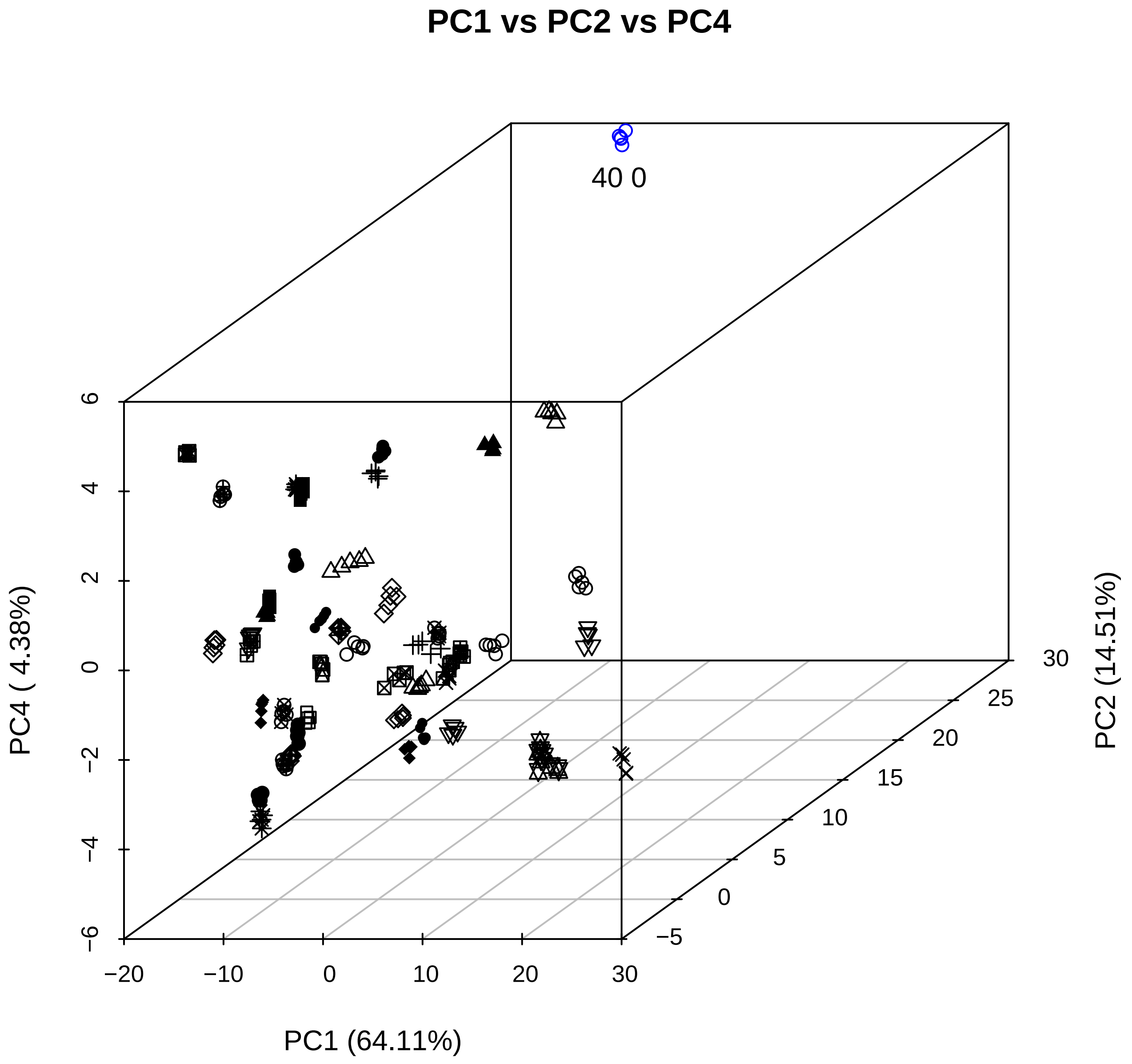
PC1 vs PC2 vs PC4
40 0
PC1 (64.11%)
PC2 (14.51%)
PC4 ( 4.38%)
−20
−10
0
10
20
30
−5
0
5
10
15
20
25
30
−6
−4
−2
0
2
4
6

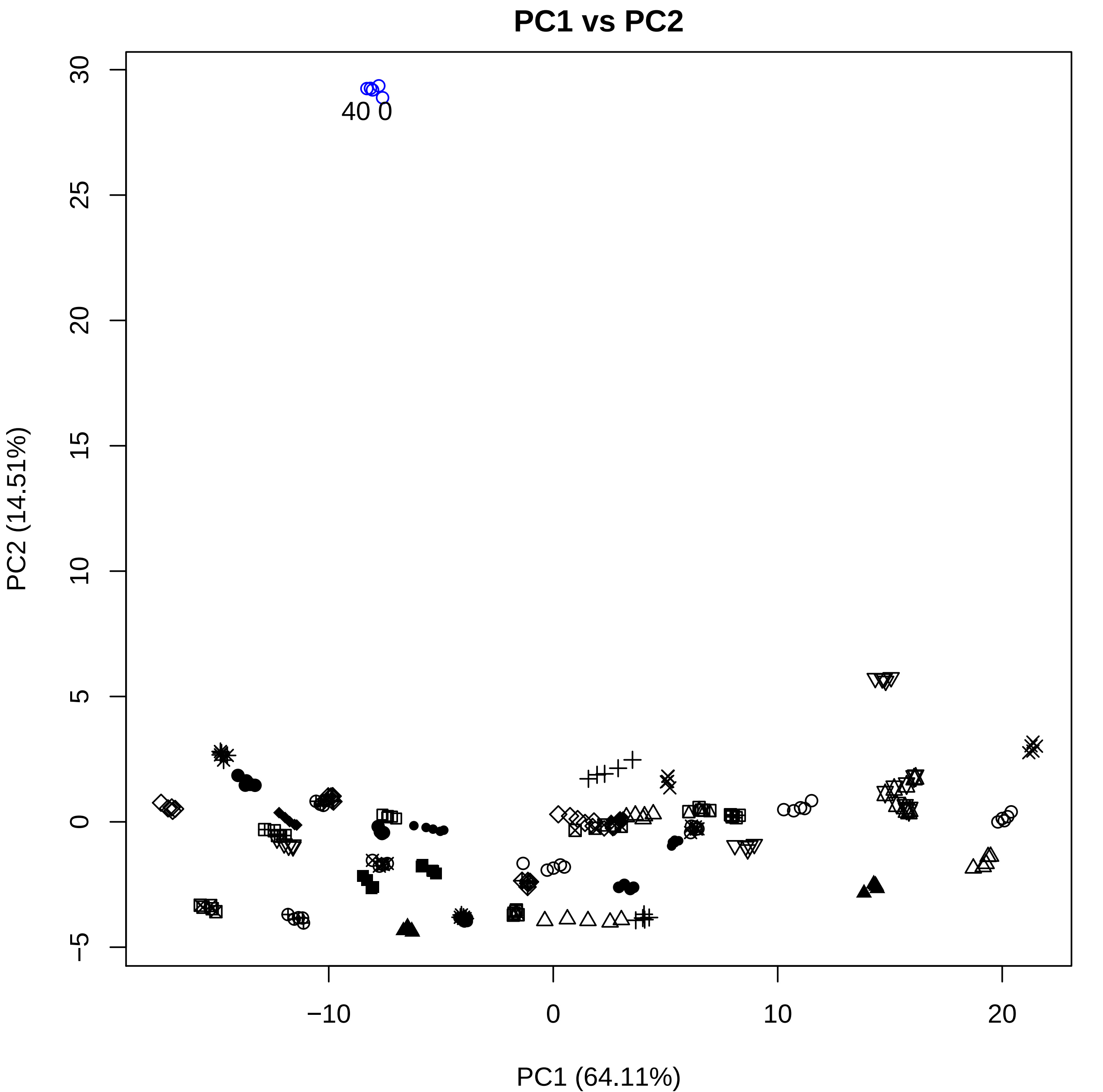
PC1 vs PC2
40 0
PC2 (14.51%)
PC1 (64.11%)
30
25
20
15
10
5
0
−5
−10
0
10
20

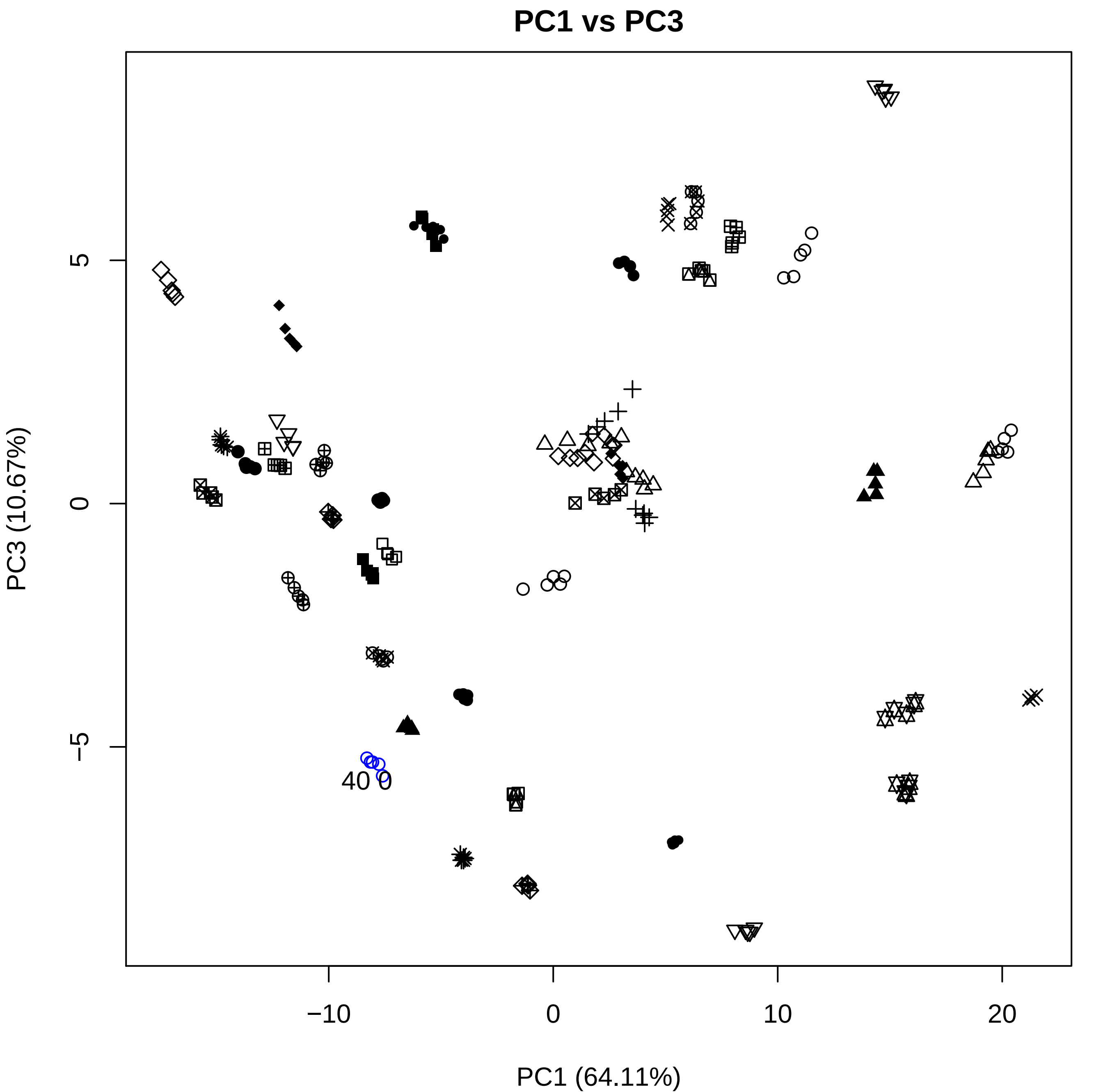
PC1 vs PC3
PC3 (10.67%)
PC1 (64.11%)
5
0
−5
−10
0
10
20
40 0

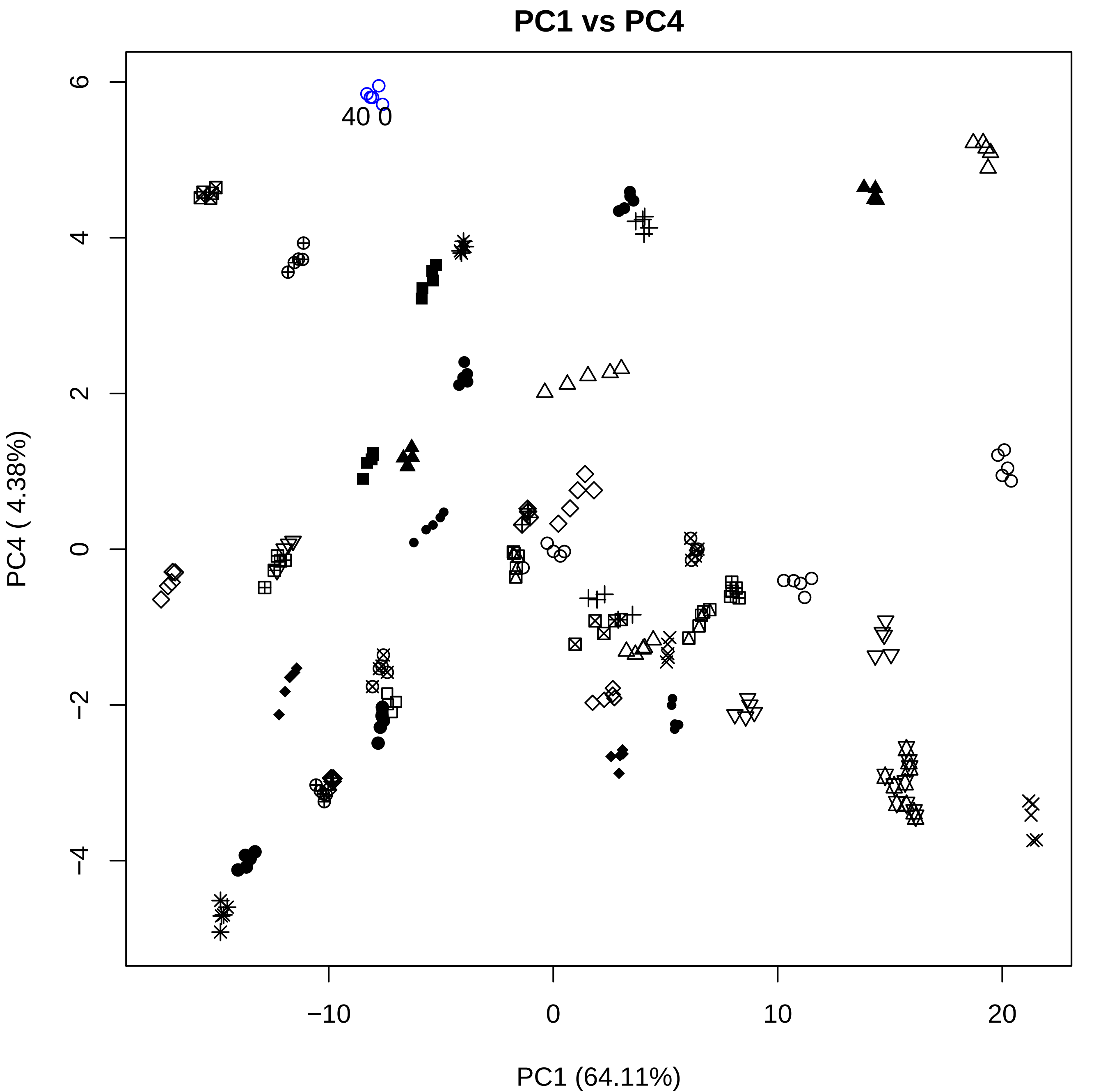
PC1 vs PC4
40 0
PC4 ( 4.38%)
PC1 (64.11%)
6
4
2
0
−2
−4
−10
0
10
20

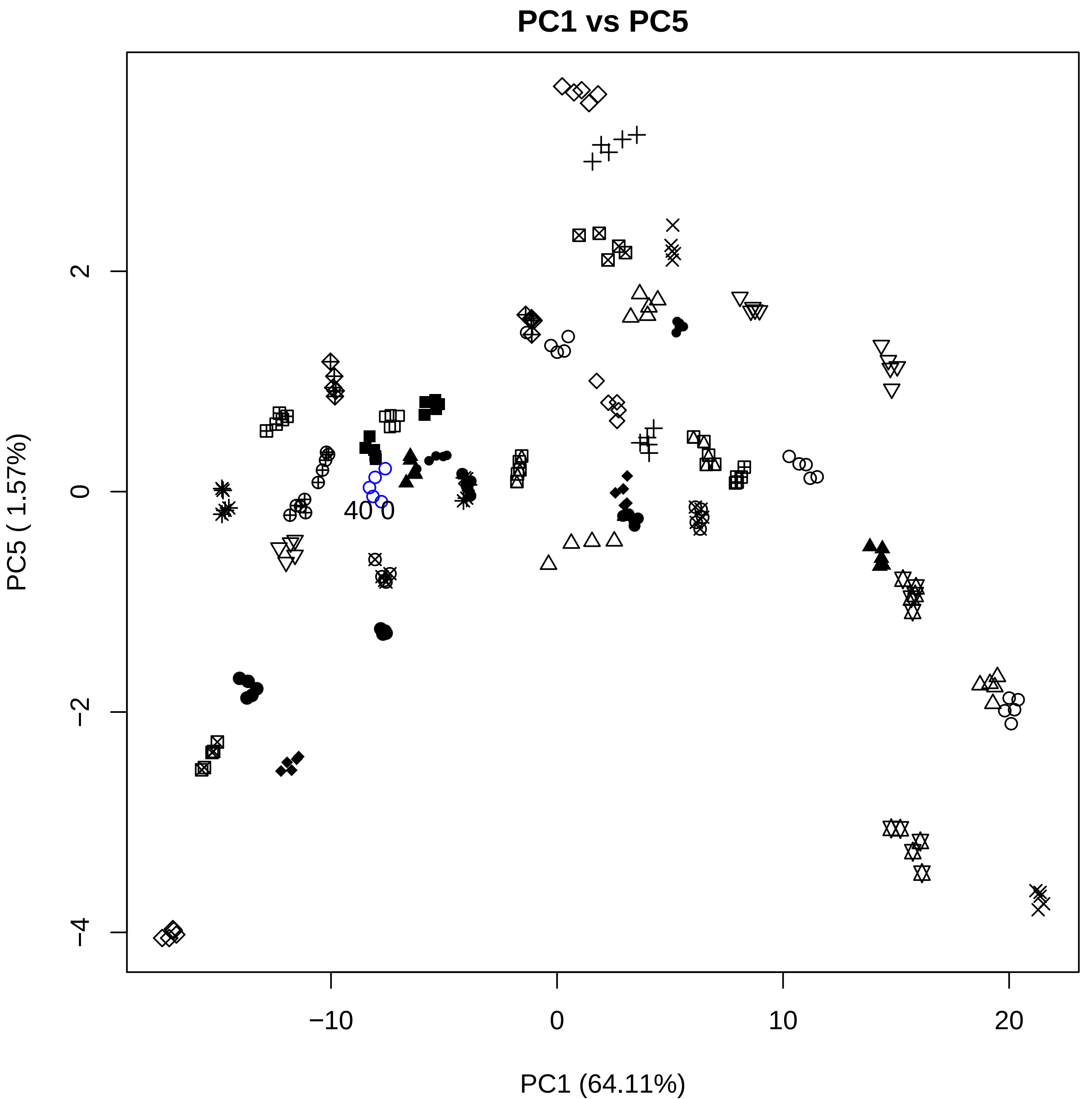

PC1 vs PC5
PC5 ( 1.57%)
PC1 (64.11%)
−10
0
10
20
−4
−2
0
2
40 0

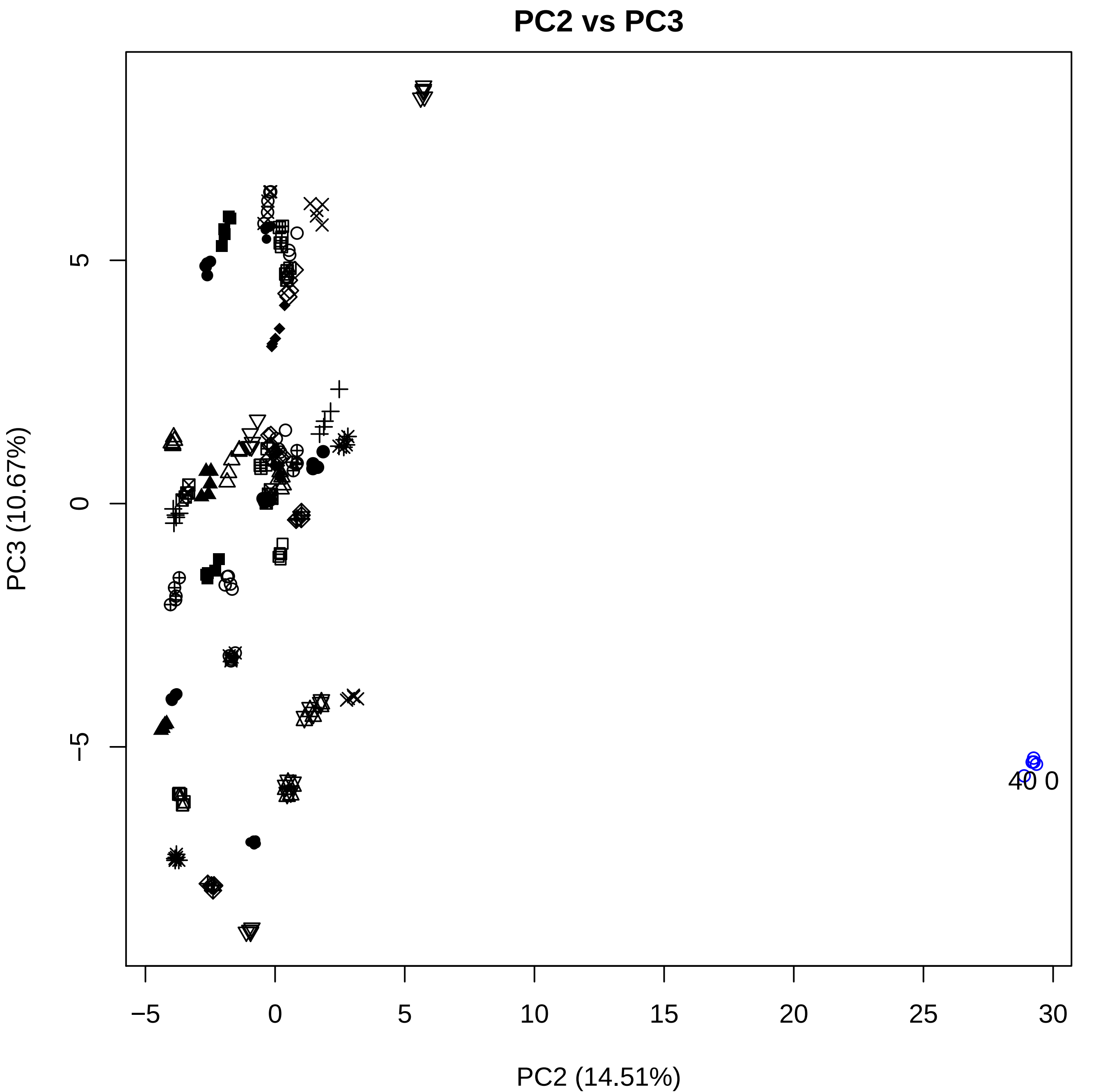

PC2 vs PC3
PC3 (10.67%)
PC2 (14.51%)
5
0
−5
−5
0
5
10
15
20
25
30
40 0

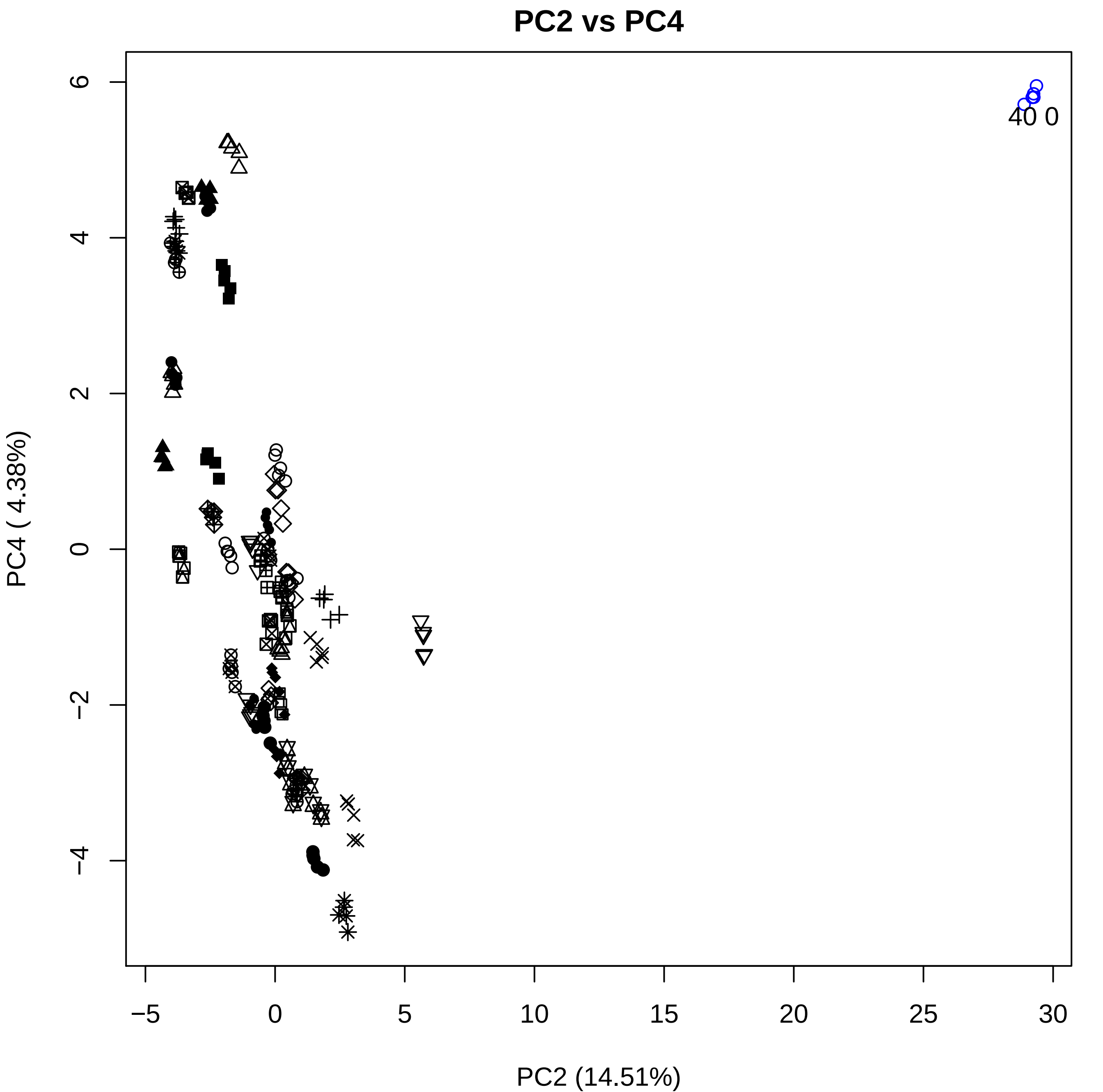
PC2 vs PC4
40 0
PC4 ( 4.38%)
PC2 (14.51%)
6
4
2
0
−2
−4
−5
0
5
10
15
20
25
30

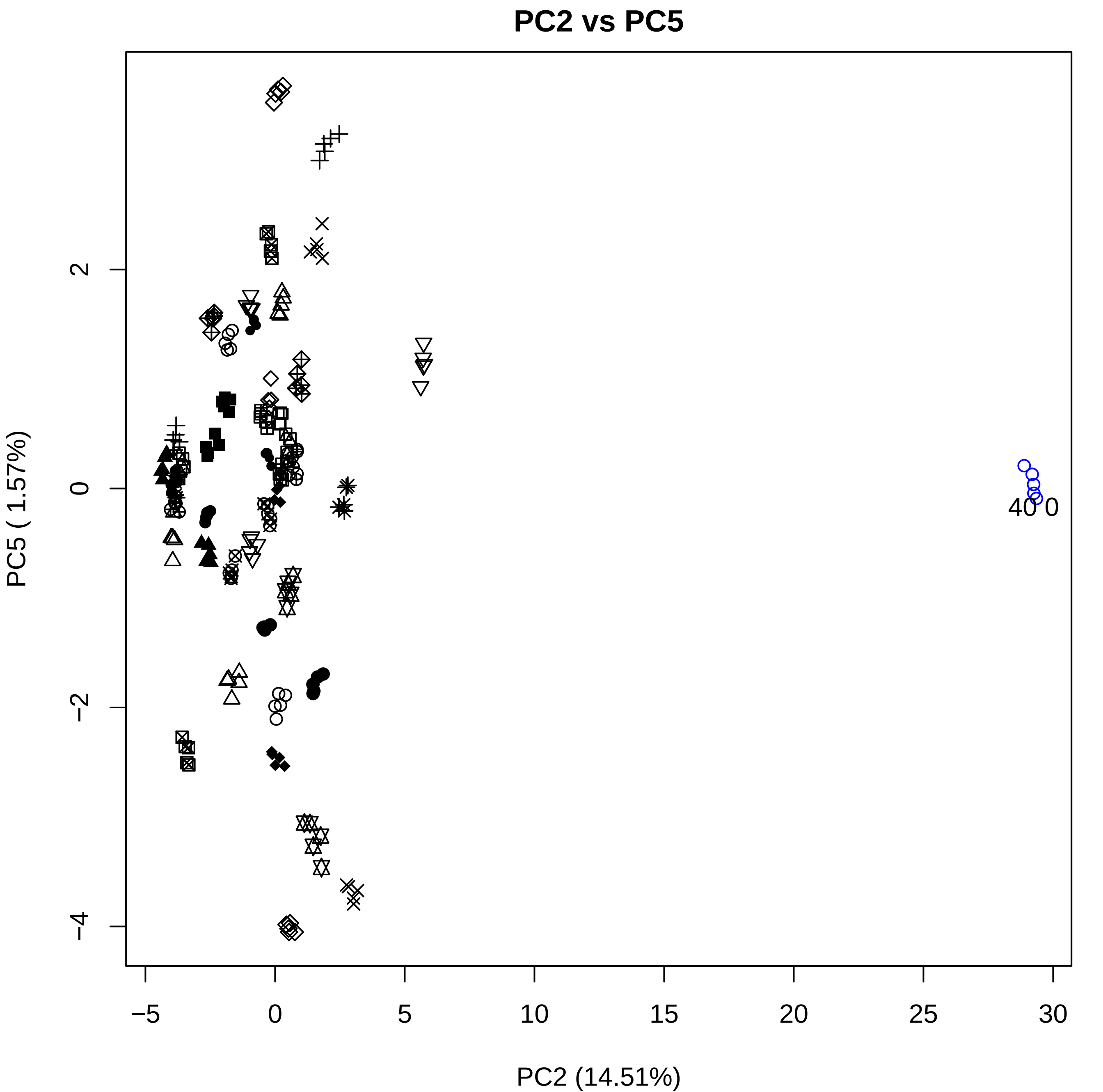

PC2 vs PC5
PC5 ( 1.57%)
PC2 (14.51%)
2
0
−2
−4
−5
0
5
10
15
20
25
30
40 0

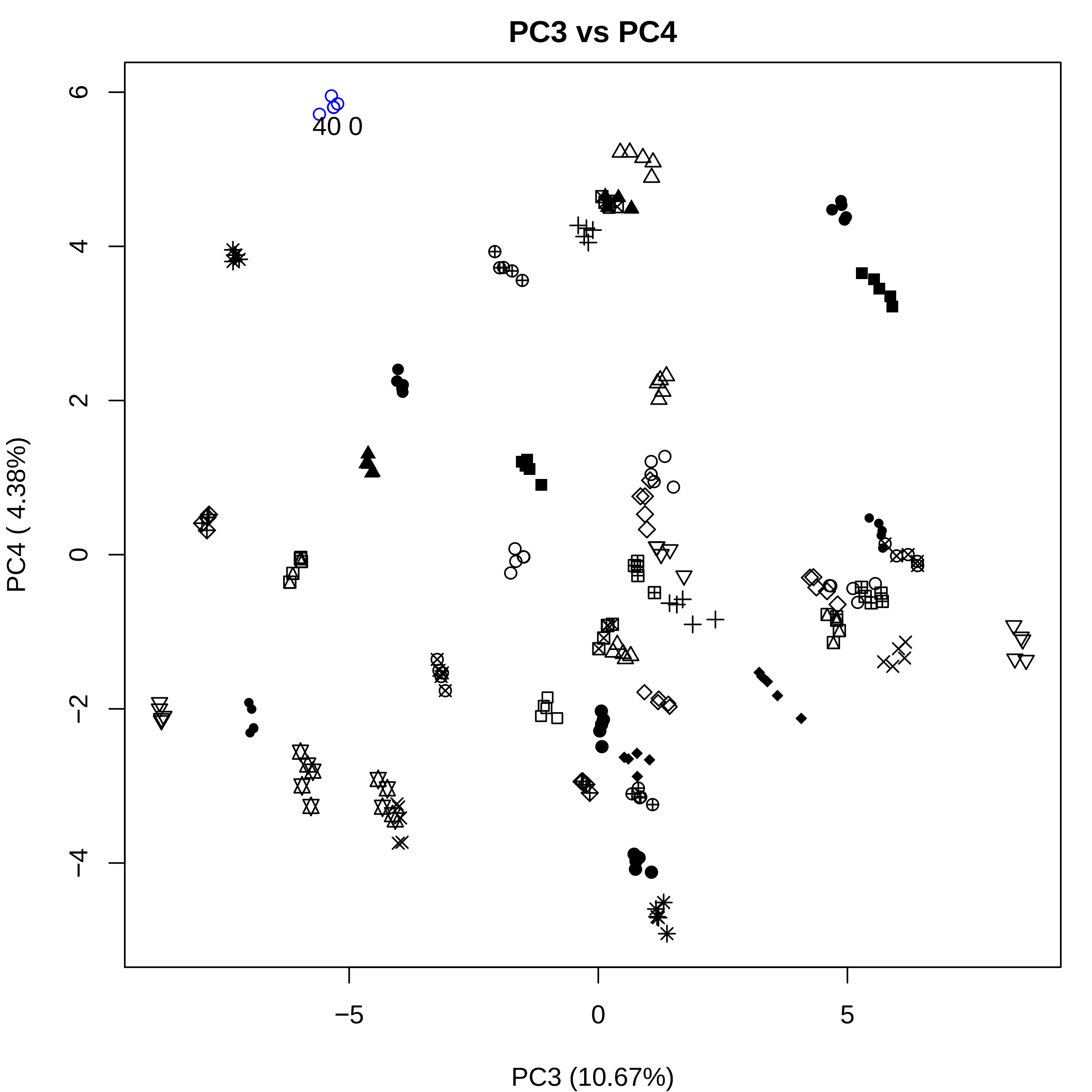
PC3 vs PC4
40 0
PC4 ( 4.38%)
PC3 (10.67%)
6
4
2
0
−2
−4
−5
0
5

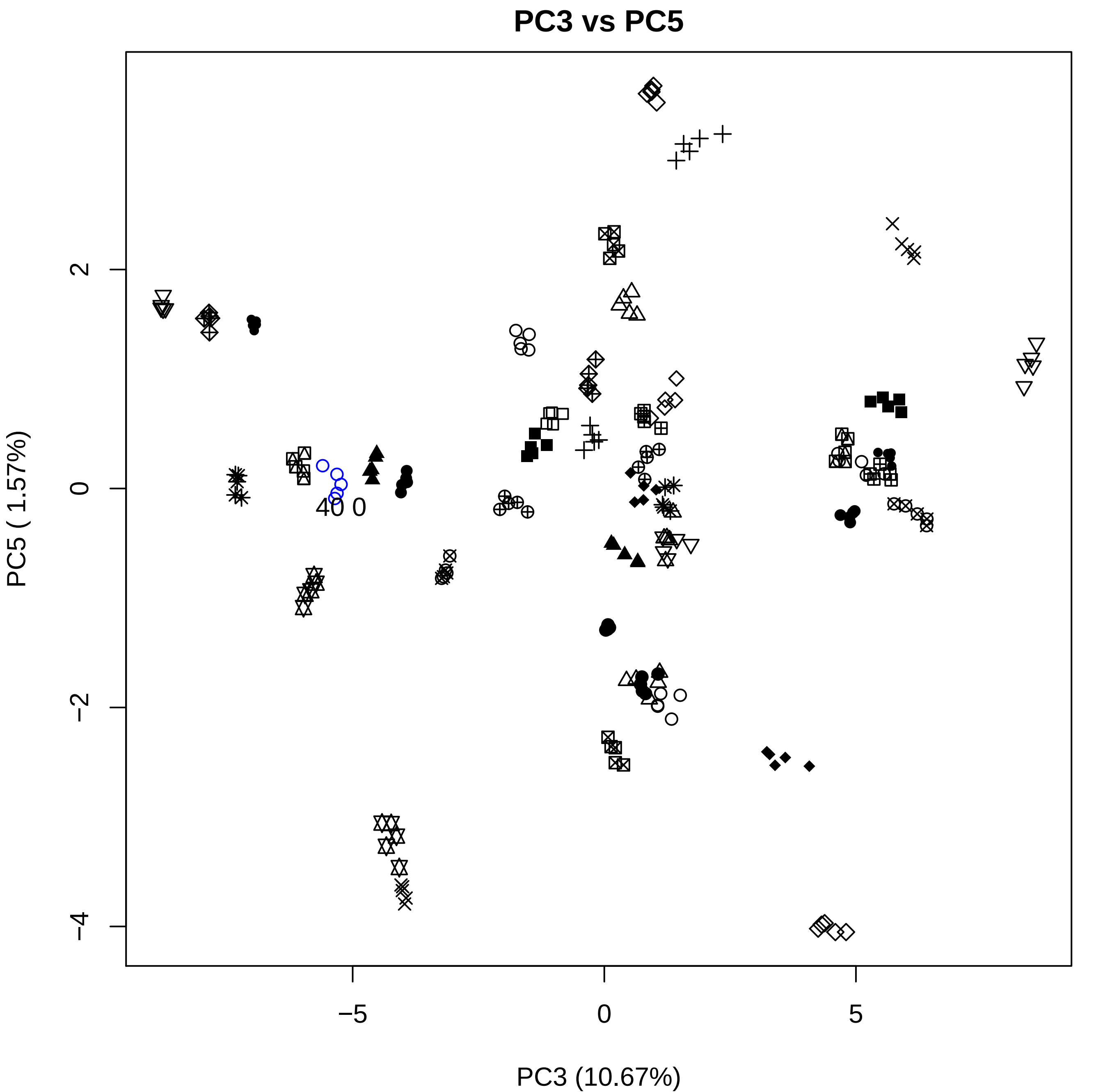
PC3 vs PC5
PC5 ( 1.57%)
PC3 (10.67%)
2
0
−2
−4
−5
0
5
40 0

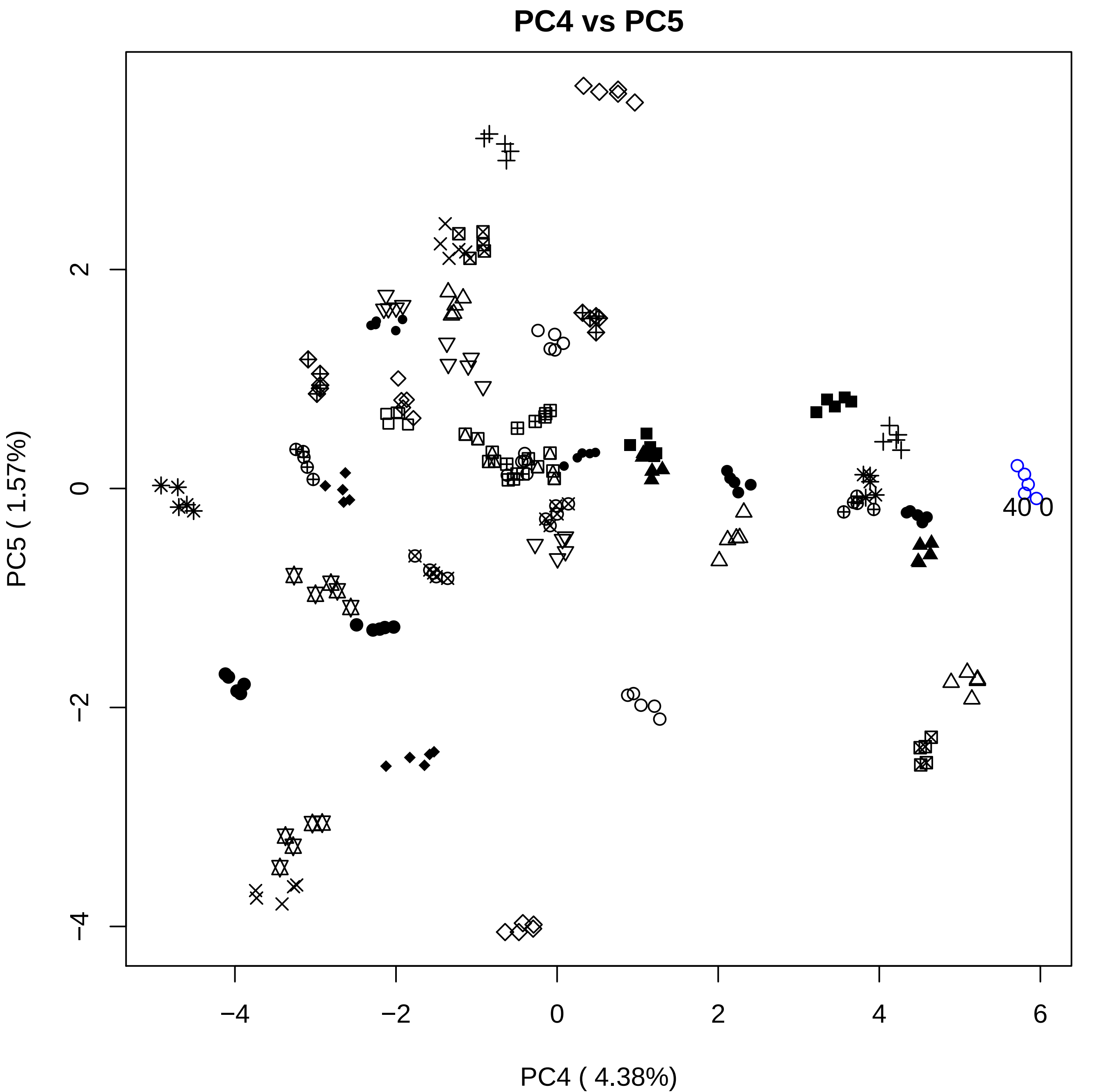
PC4 vs PC5
PC5 ( 1.57%)
PC4 ( 4.38%)
2
0
−2
−4
−4
−2
0
2
4
6
40 0

### PCA of 100%  OSAS vs PMH

Scaled Matrix PCAed

7th order polynomial baseline correction performed

Raw spectra smoothed to 21 Neighbours, 2nd order PN, Savtisky Golay

Normalisation between 0 and 100

D2 spectra smoothed to 41 Neighbours, 2nd order PN, Savtisky Golay

Regions Analysed: 3601−2501 and 2001−700cm−1

Samples Analysed: 0 Contam's 1 OSAS's 45 PMH's

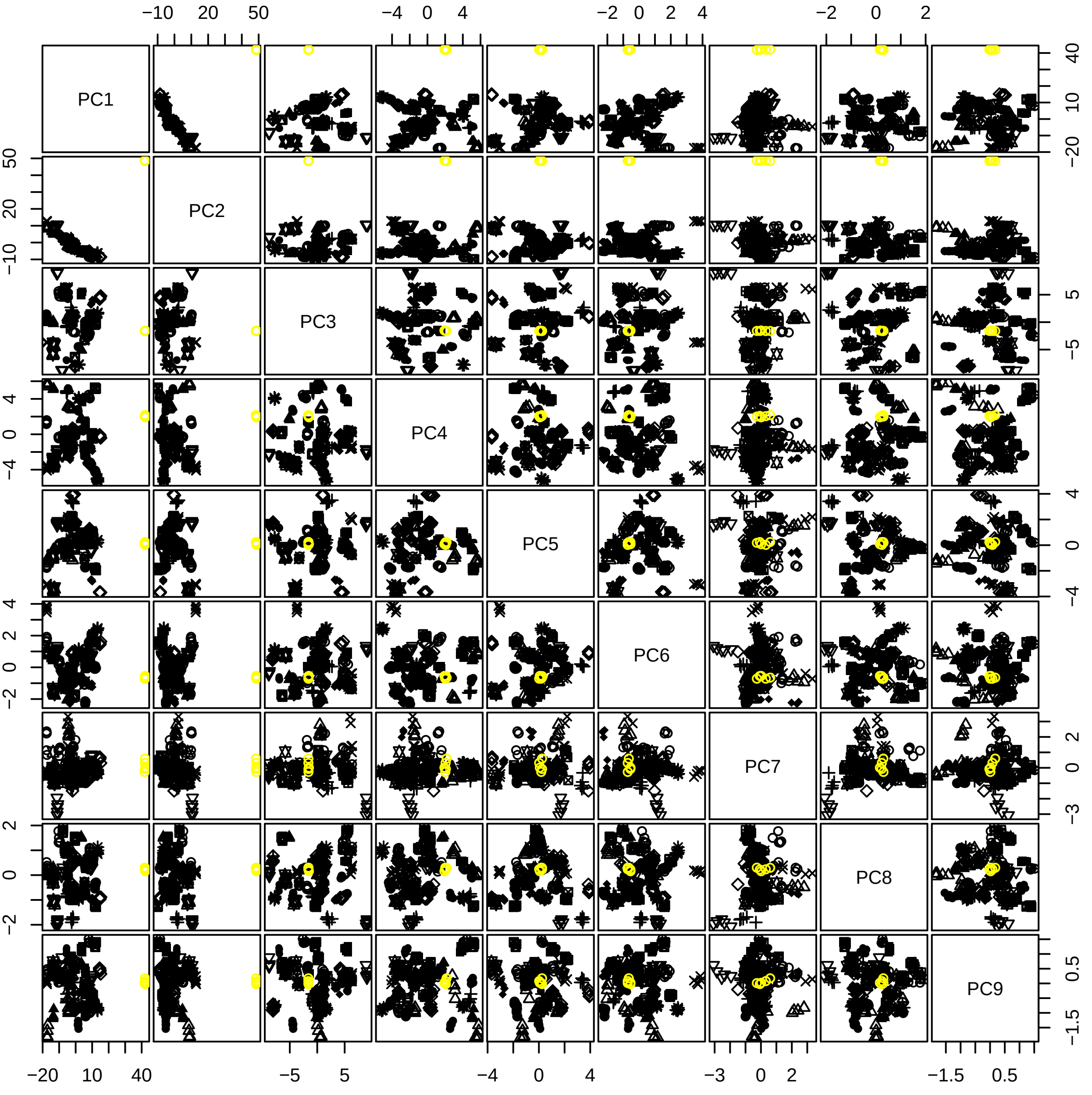
PC1
PC2
PC3
PC4
PC5
PC6
PC7
PC8
PC9

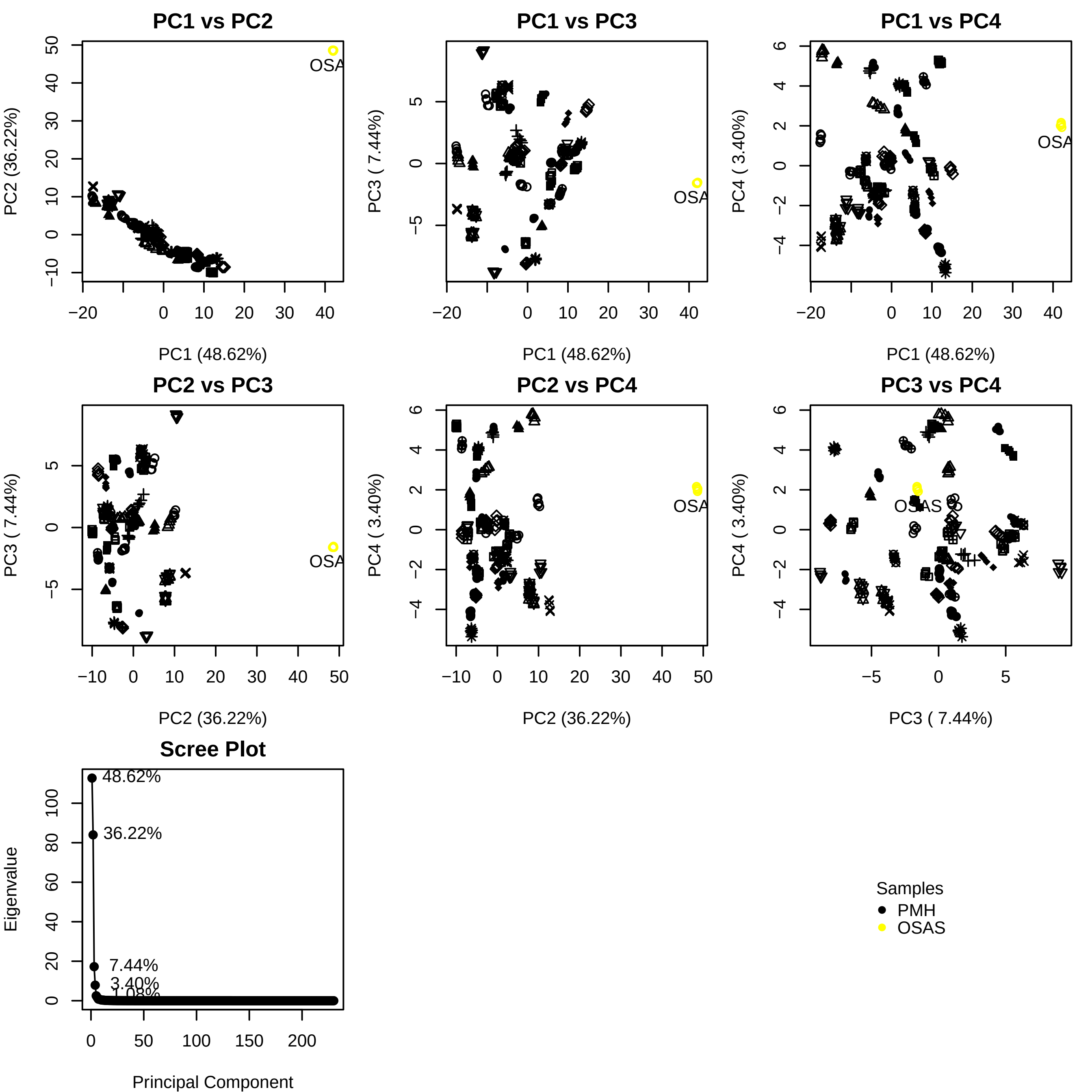

PC1 vs PC2
PC1 vs PC3
PC1 vs PC4
PC2 vs PC3
PC2 vs PC4
PC3 vs PC4
Scree Plot
PC1 (48.62%)
PC2 (36.22%)
PC3 ( 7.44%)
PC4 ( 3.40%)
OSA
OSAS
Eigenvalue
Principal Component
48.62%
36.22%
7.44%
3.40%
Samples
PMH
OSAS

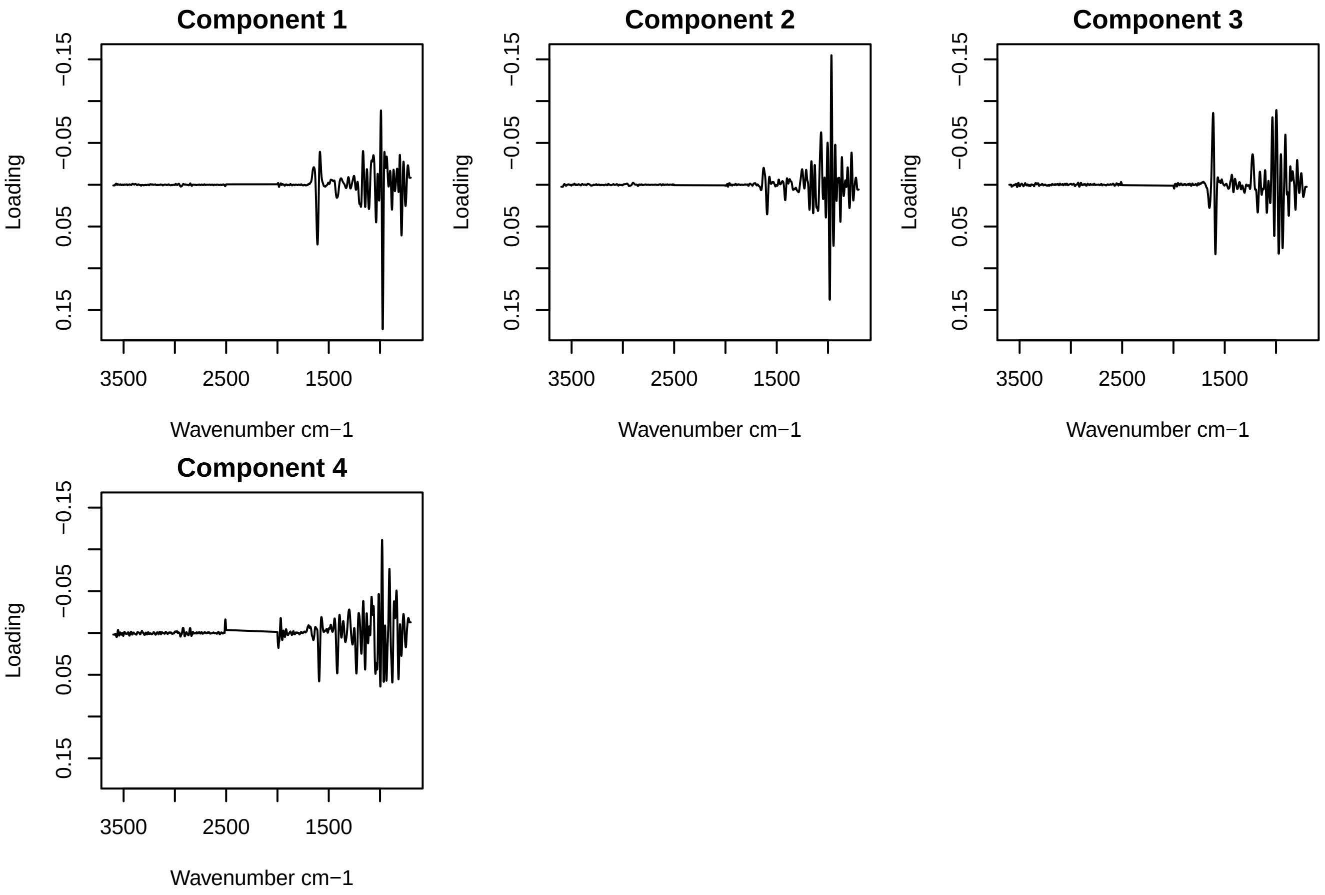
Component 1
Component 2
Component 3
Component 4
Loading
−0.15
−0.05
0.05
0.15
3500
2500
1500
Wavenumber cm−1

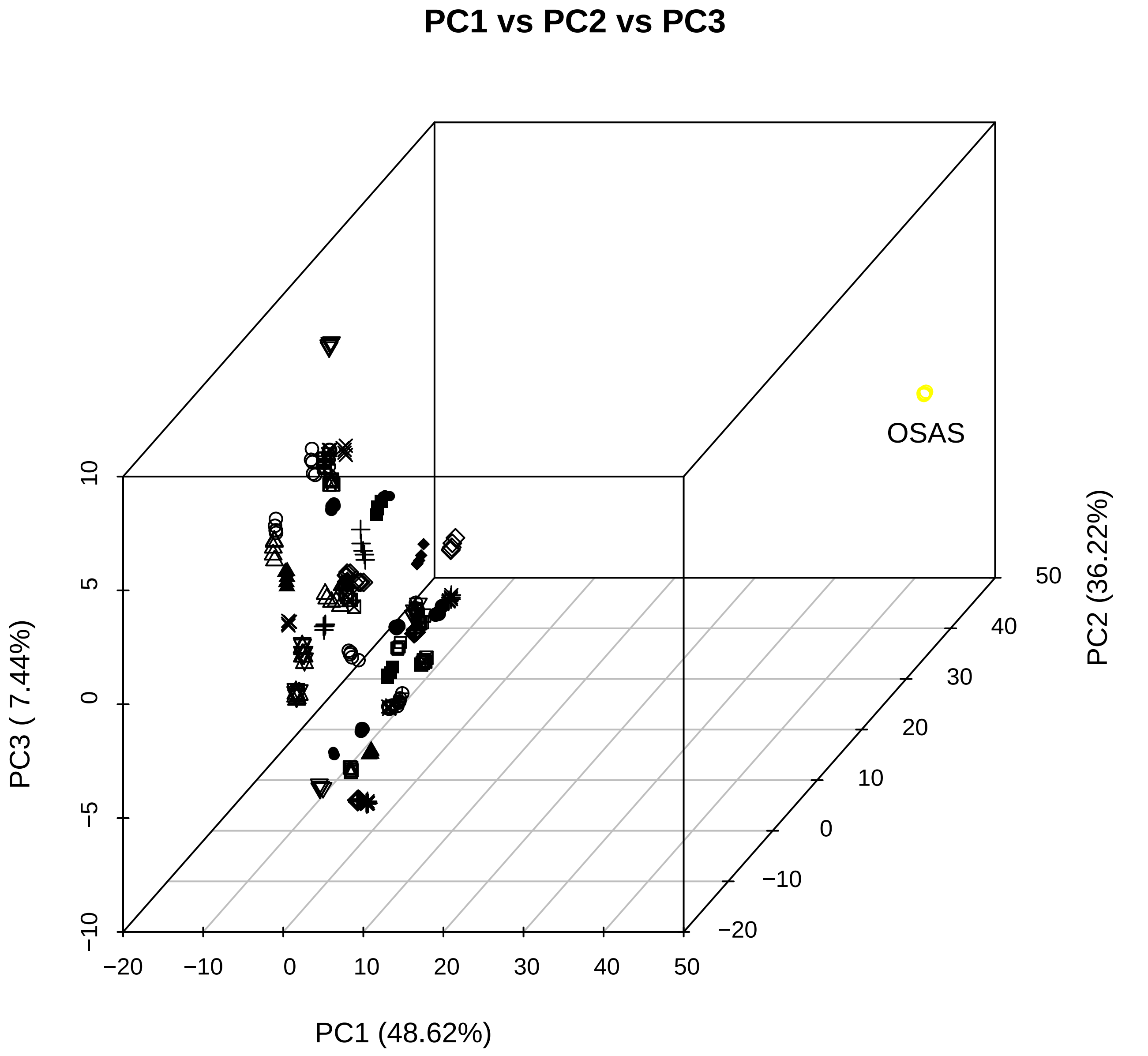
PC1 vs PC2 vs PC3
OSAS
PC1 (48.62%)
PC2 (36.22%)
PC3 ( 7.44%)
−20
−10
0
10
20
30
40
50
−20
−10
0
10
20
30
40
50
−10
−5
0
5
10

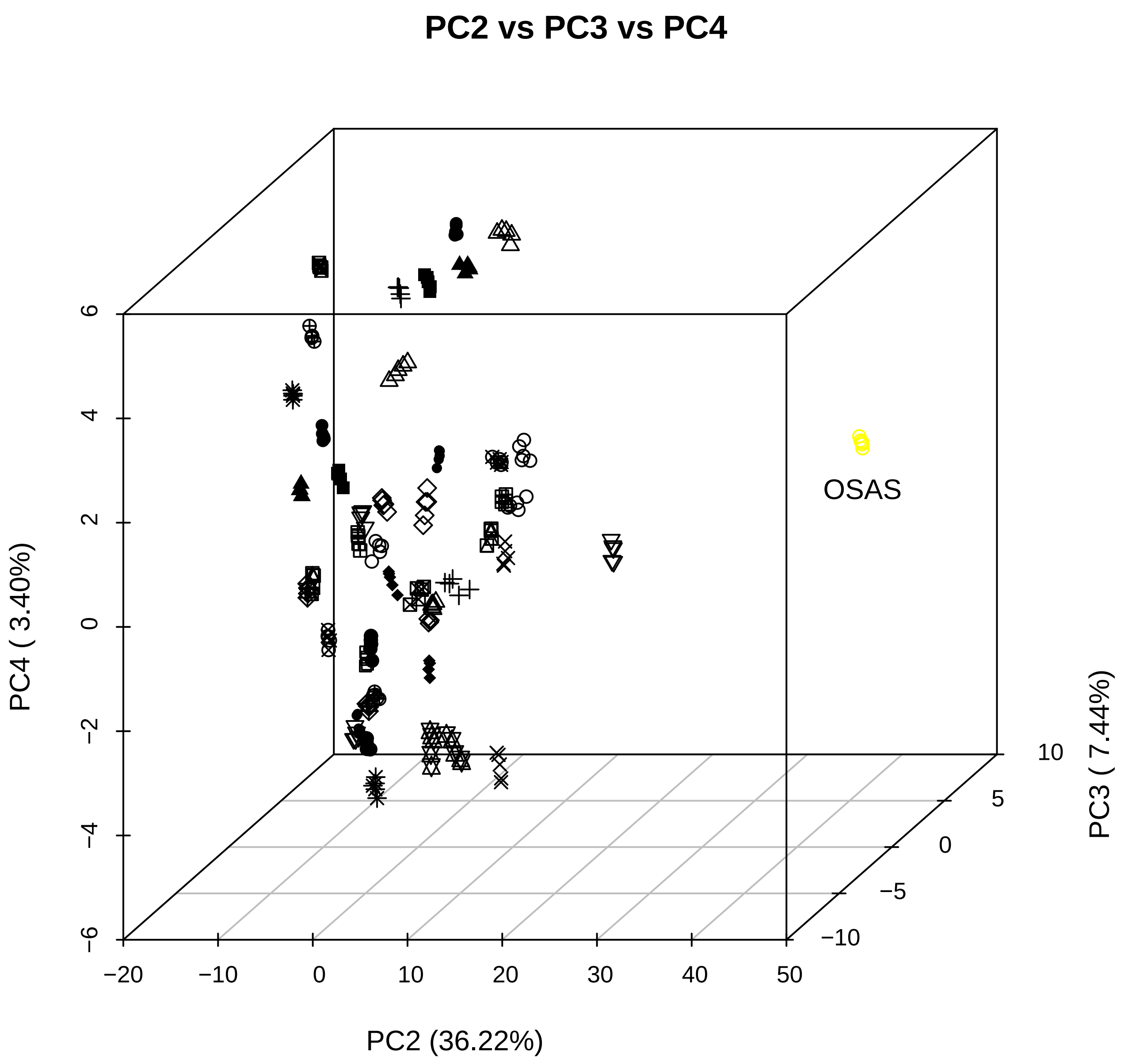

PC2 vs PC3 vs PC4
PC4 ( 3.40%)
6
4
2
0
−2
−4
−6
−20
−10
0
10
20
30
40
50
PC2 (36.22%)
PC3 ( 7.44%)
10
5
0
−5
−10
OSAS

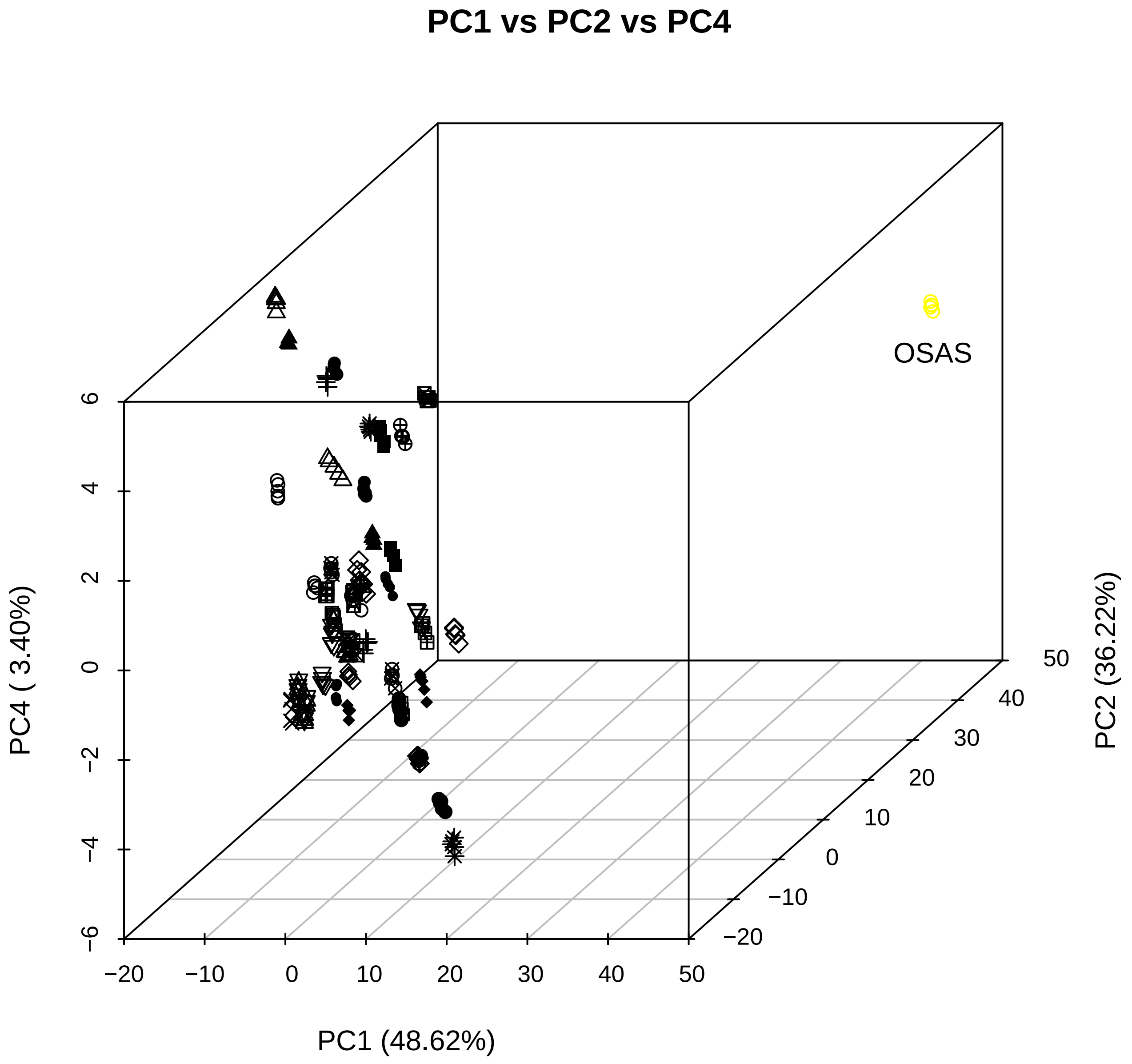
PC1 vs PC2 vs PC4
OSAS
PC1 (48.62%)
PC2 (36.22%)
PC4 ( 3.40%)
−20
−10
0
10
20
30
40
50
−6
−4
−2
0
2
4
6

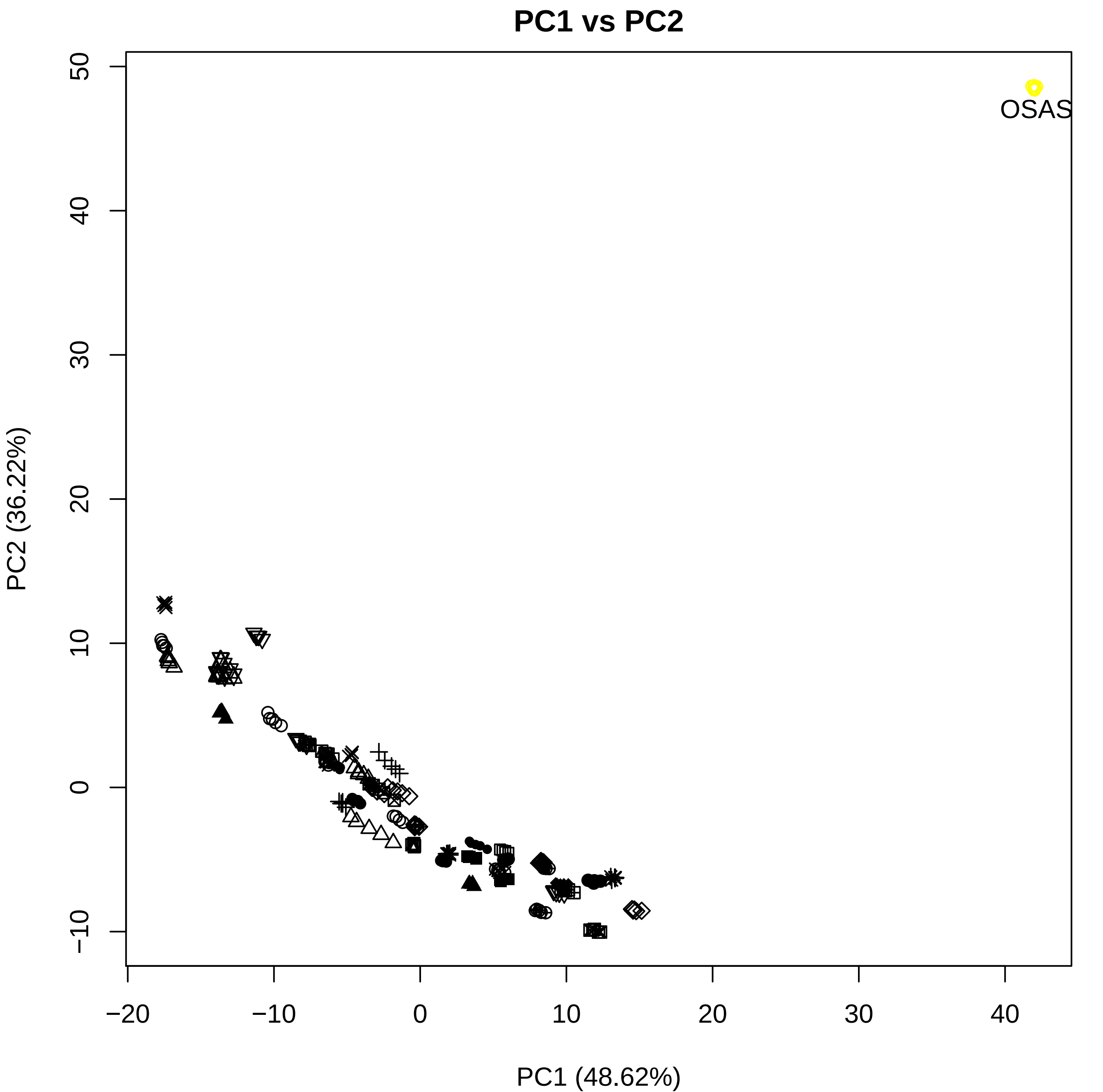

PC1 vs PC2
OSAS
PC2 (36.22%)
PC1 (48.62%)
50
40
30
20
10
0
−10
−20
−10
0
10
20
30
40

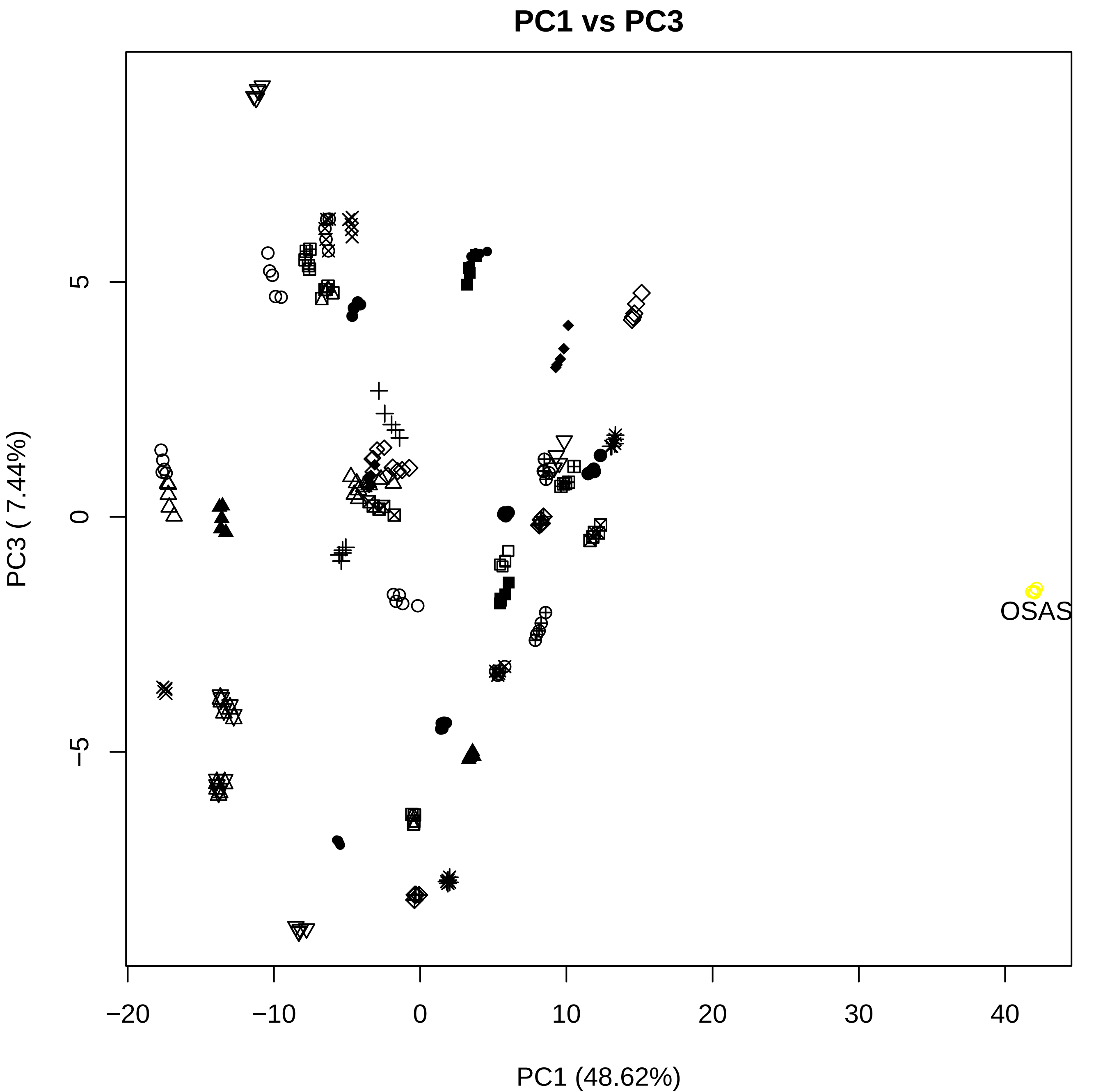
PC1 vs PC3
PC3 ( 7.44%)
PC1 (48.62%)
5
0
−5
−20
−10
0
10
20
30
40
OSAS

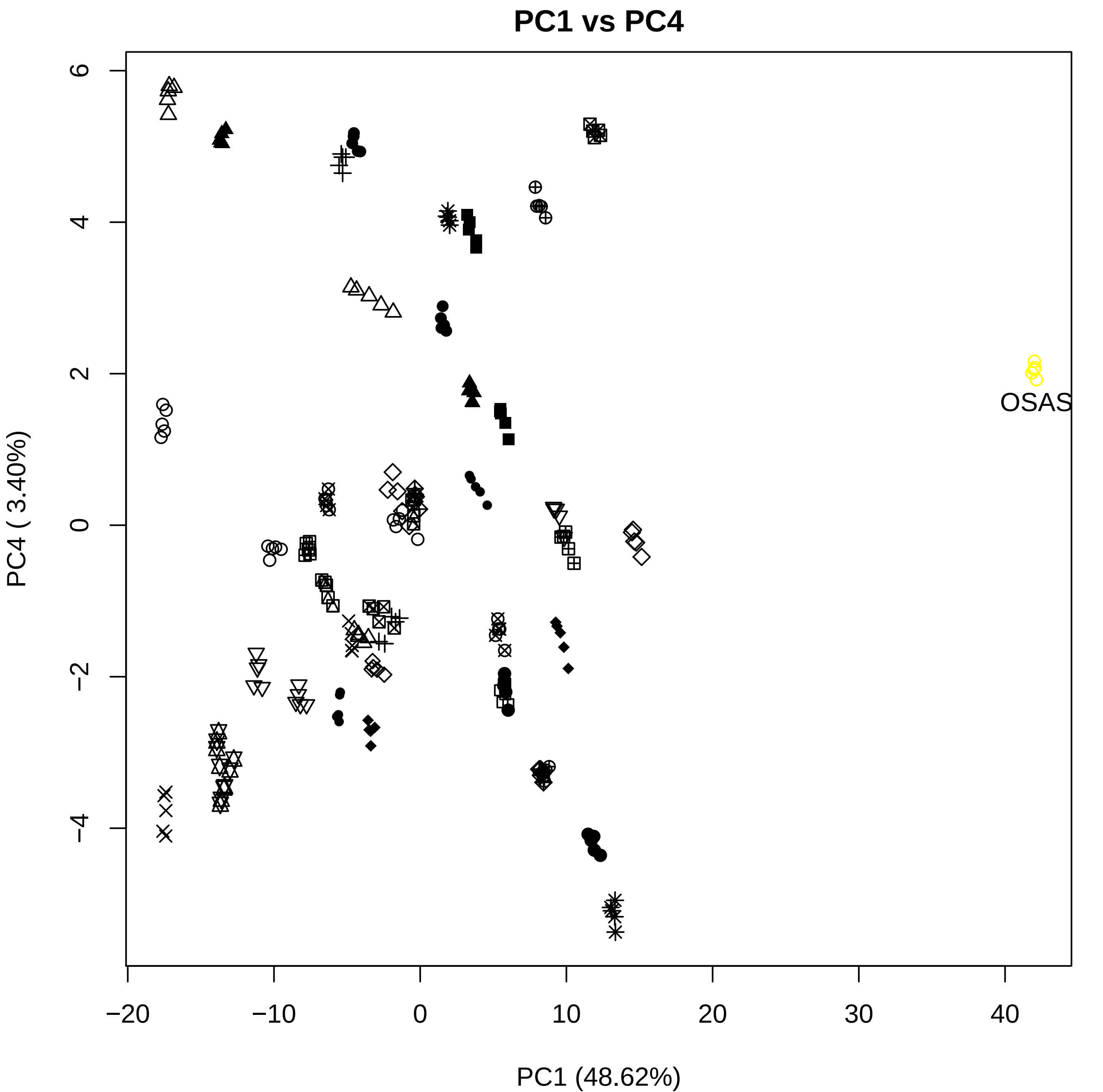
PC1 vs PC4
PC4 ( 3.40%)
PC1 (48.62%)
OSAS
6
4
2
0
−2
−4
−20
−10
0
10
20
30
40

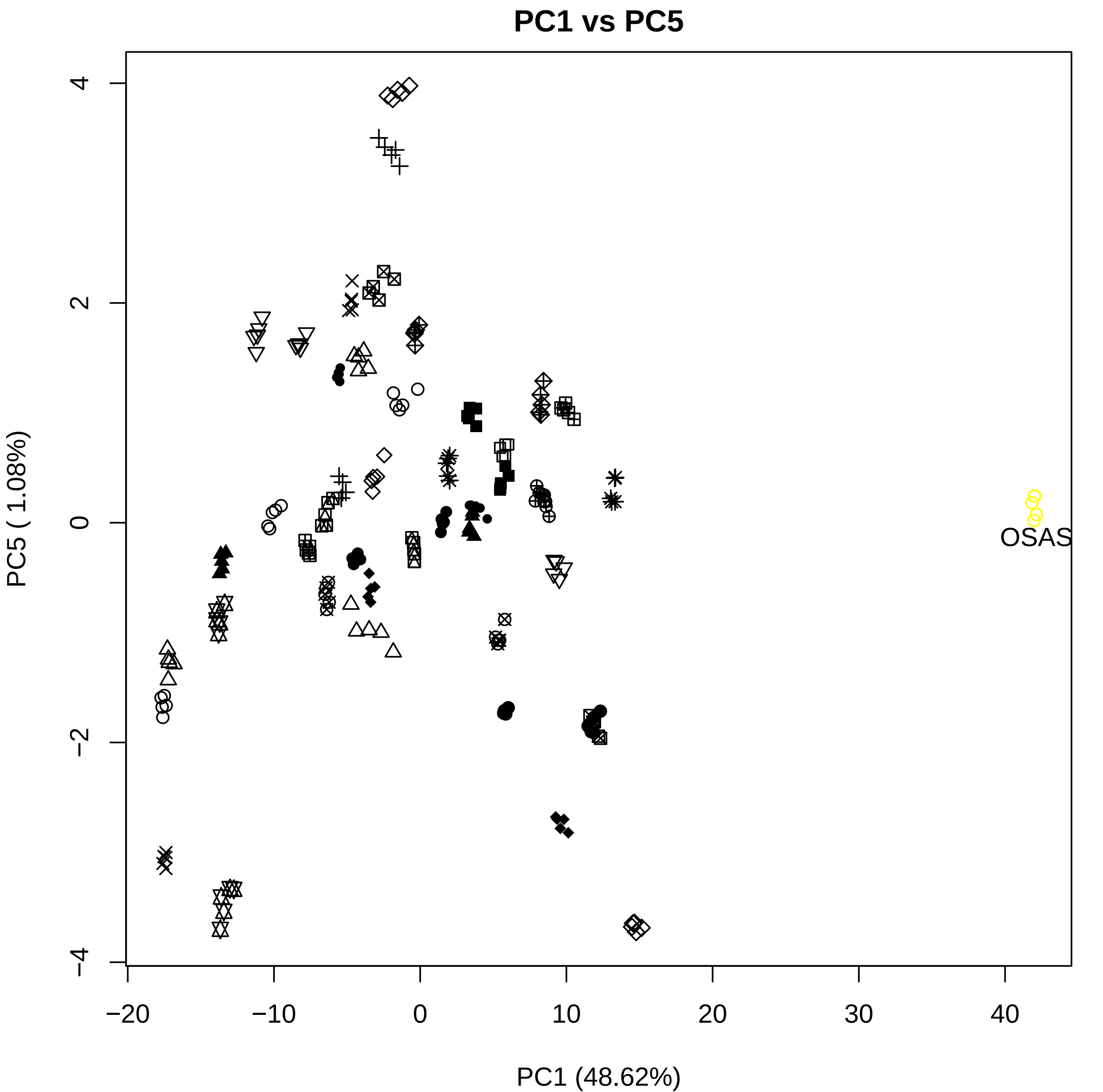

PC1 vs PC5
4
2
0
−2
−4
PC5 ( 1.08%)
−20
−10
0
10
20
30
40
PC1 (48.62%)
OSAS

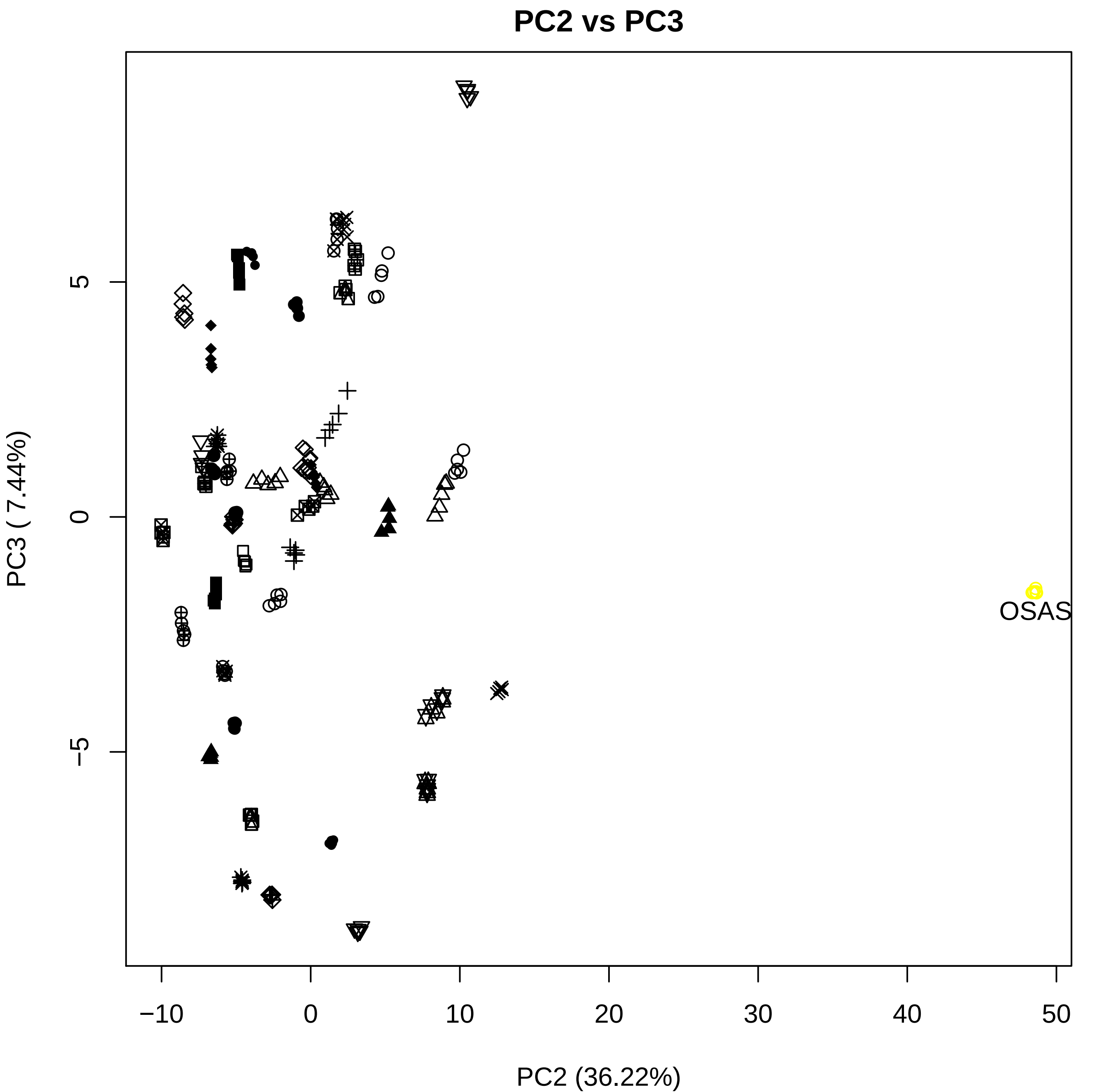

PC2 vs PC3
PC3 ( 7.44%)
PC2 (36.22%)
OSAS
−10
0
10
20
30
40
50
−5
0
5

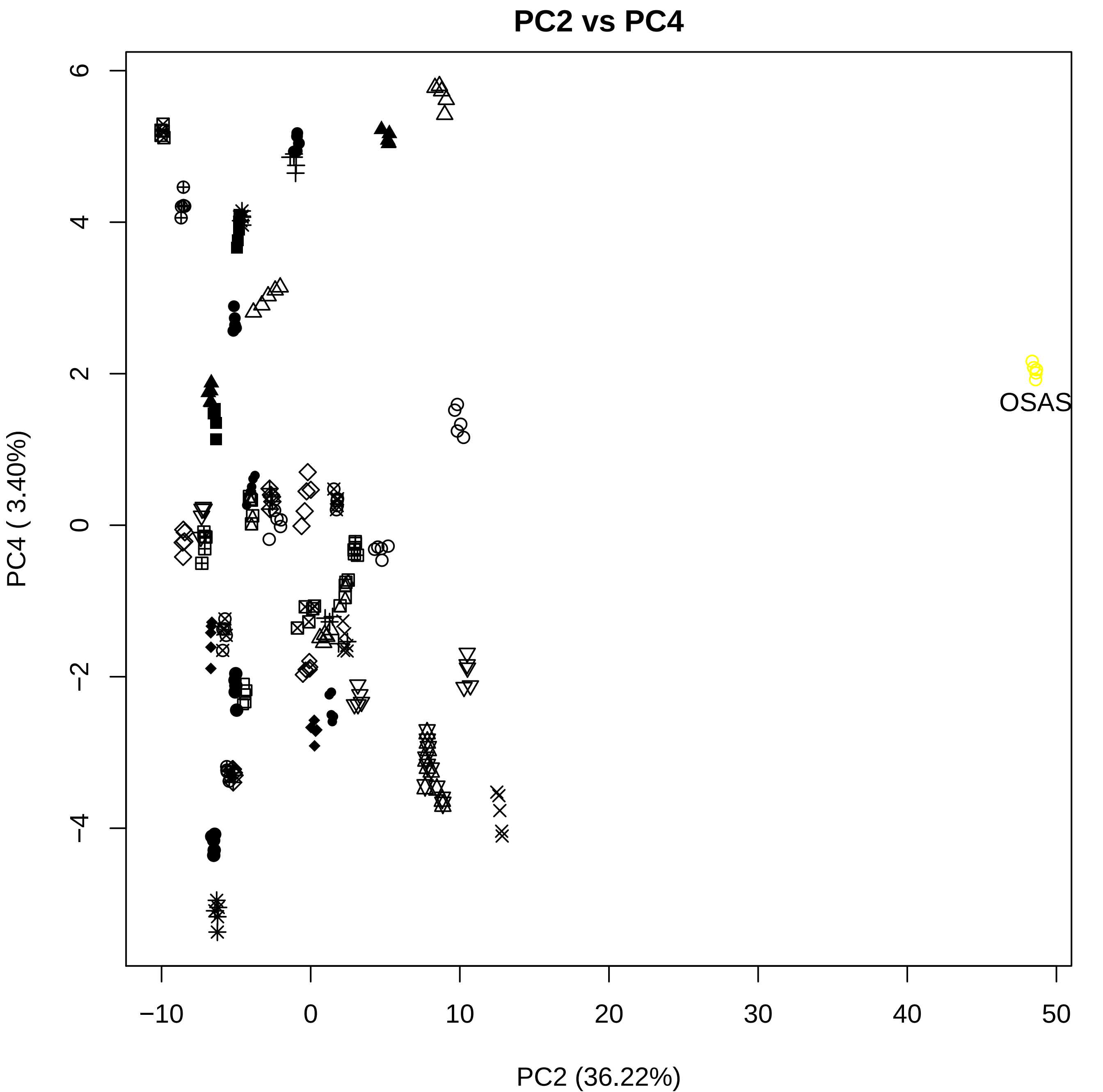

PC2 vs PC4
PC4 ( 3.40%)
PC2 (36.22%)
6
4
2
0
−2
−4
−10
0
10
20
30
40
50
OSAS

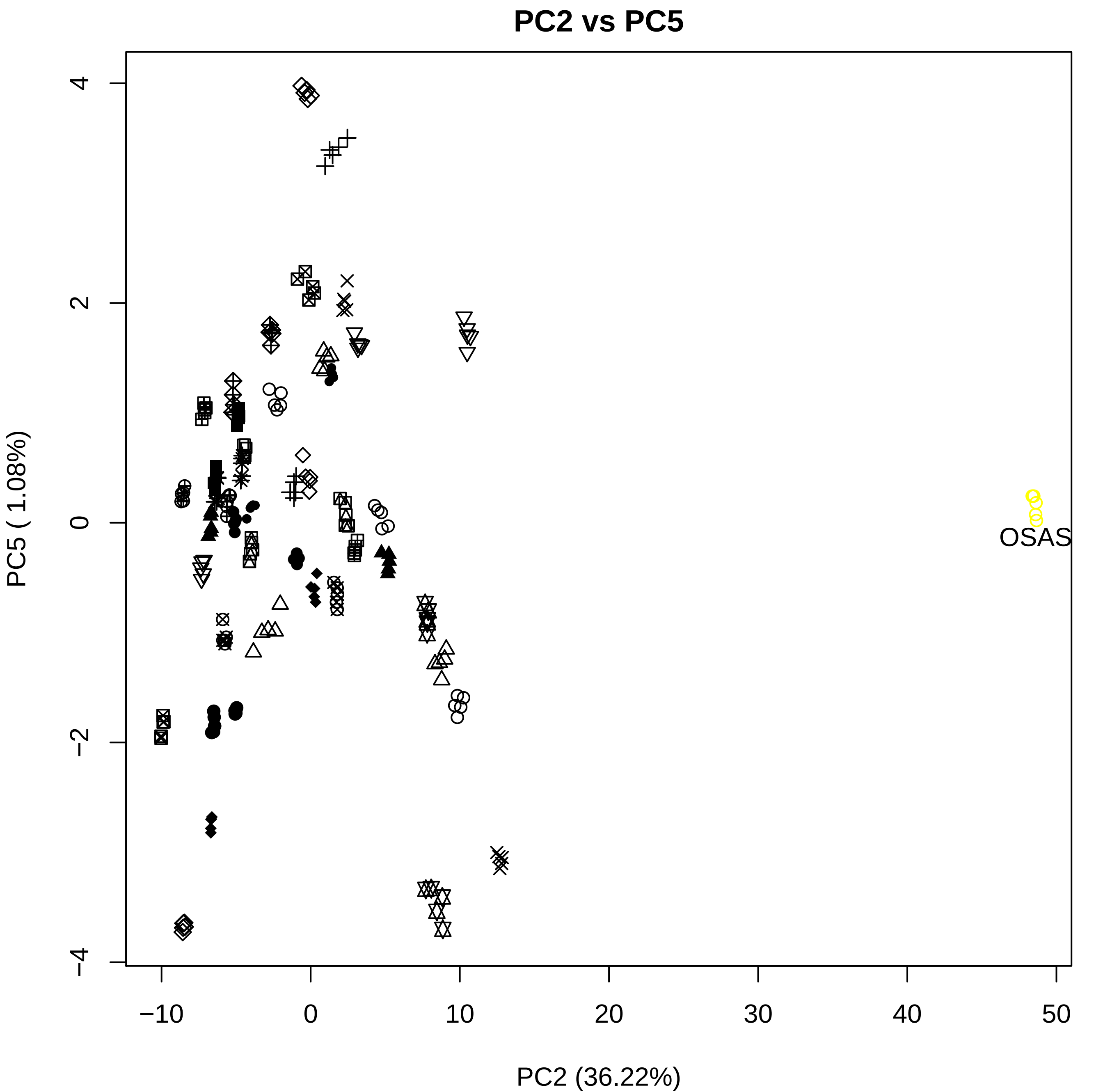
PC2 vs PC5
PC5 ( 1.08%)
PC2 (36.22%)
OSAS
4
2
0
−2
−4
−10
0
10
20
30
40
50

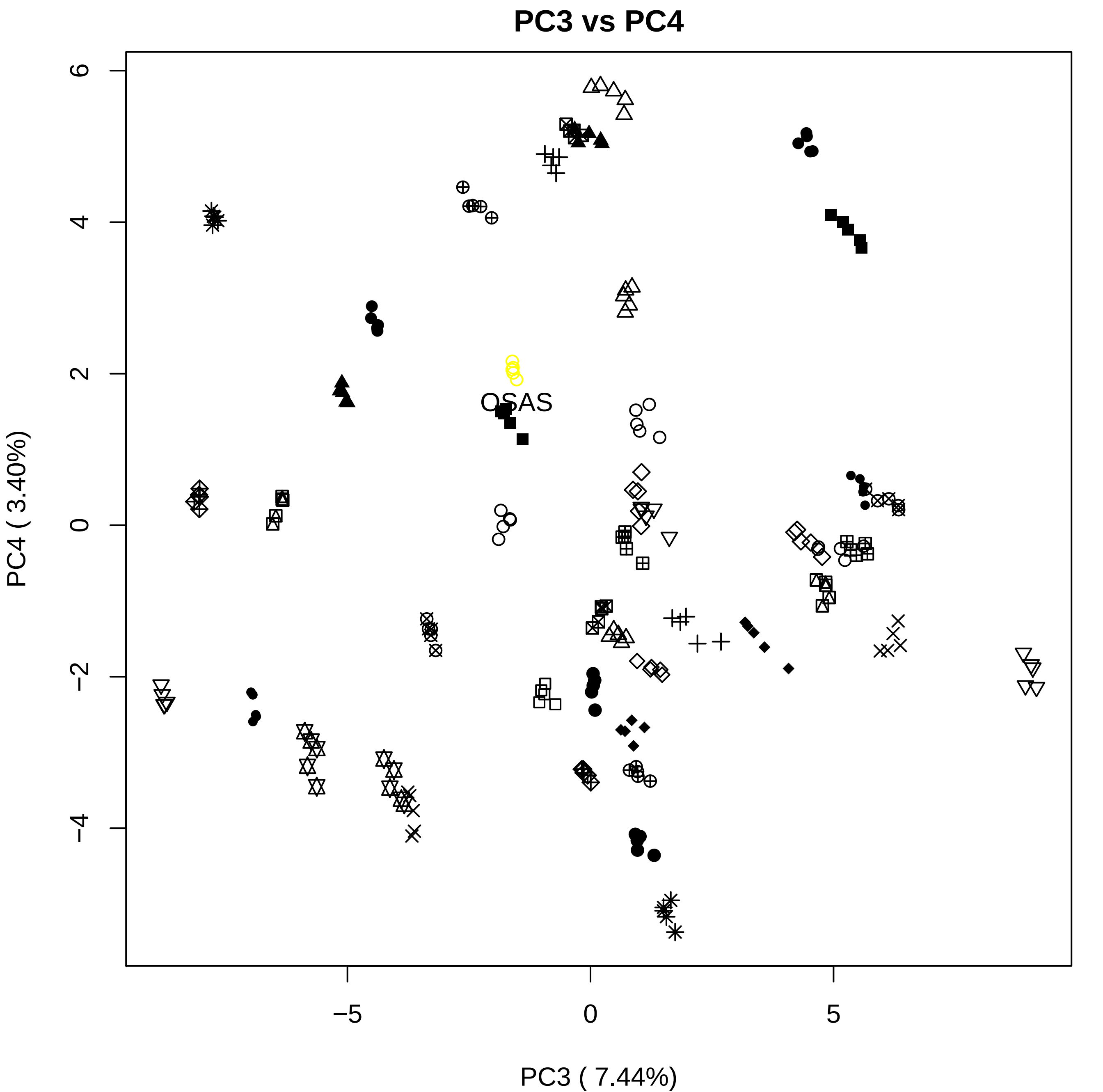
PC3 vs PC4
PC4 ( 3.40%)
PC3 ( 7.44%)
OSAS
6
4
2
0
−2
−4
−5
0
5

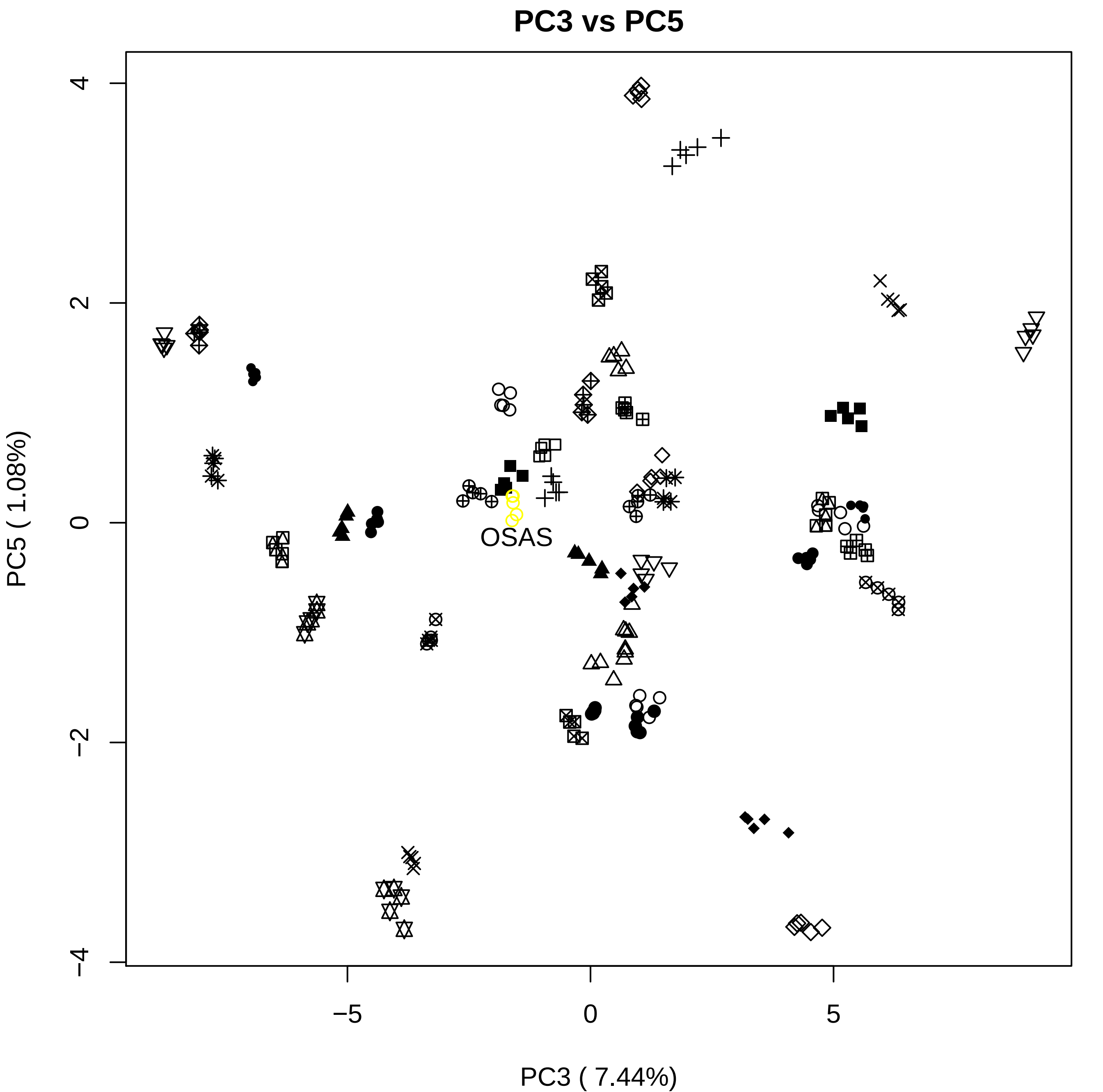
PC3 vs PC5
OSAS
PC5 ( 1.08%)
PC3 ( 7.44%)
4
2
0
−2
−4
−5
0
5

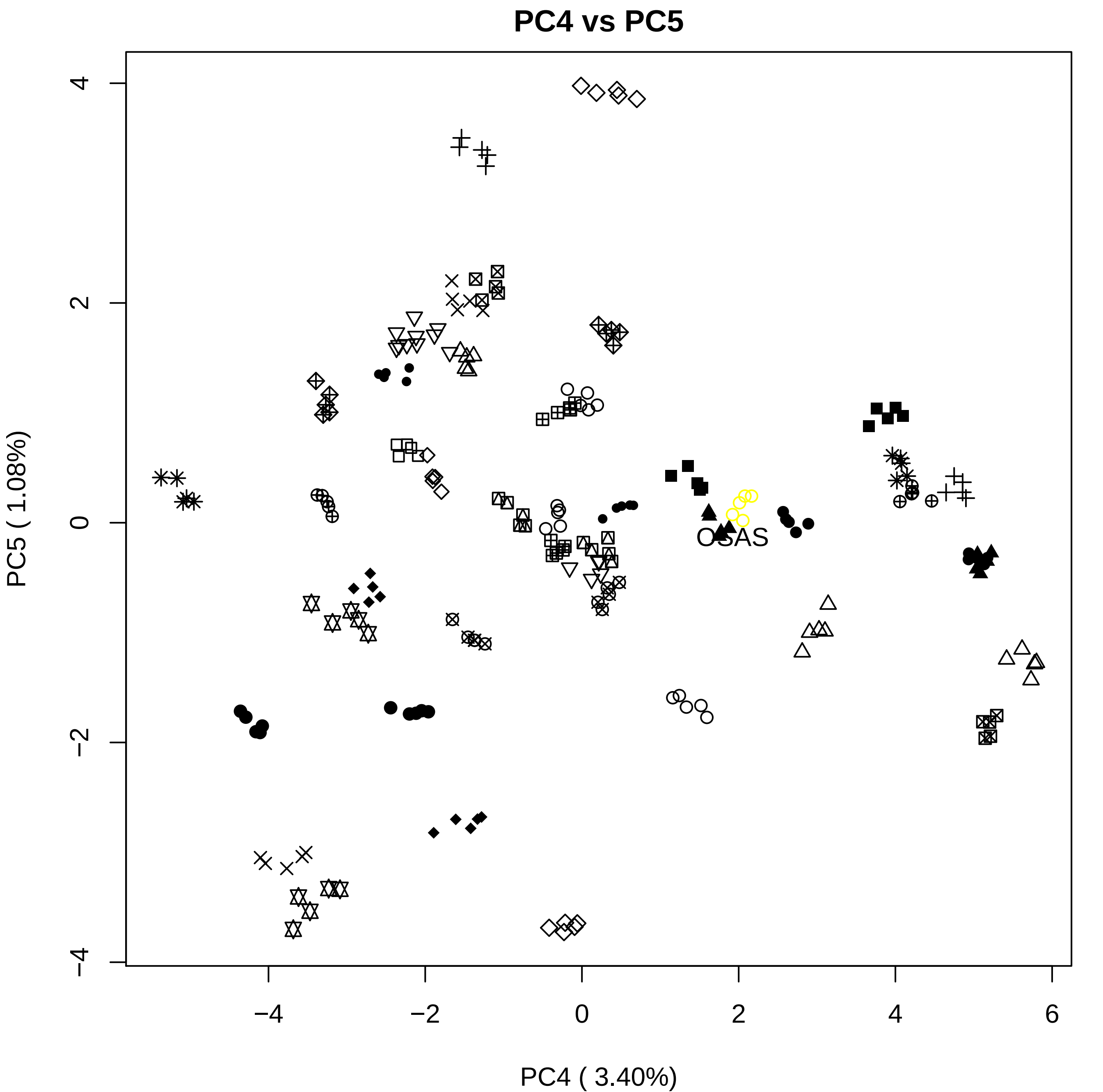
PC4 vs PC5
PC5 ( 1.08%)
PC4 ( 3.40%)
OSAS
4
2
0
−2
−4
6

### DeS Contam 7th

Anthony Devlin, Keele University

### PCA of All  DeS vs PMH

Scaled Matrix PCAed

7th order polynomial baseline correction performed

Raw spectra smoothed to 21 Neighbours, 2nd order PN, Savtisky Golay

Normalisation between 0 and 100

D2 spectra smoothed to 41 Neighbours, 2nd order PN, Savtisky Golay

Regions Analysed: 3601−2501 and 2001−700cm−1

Samples Analysed: 12 Contam's 1 DeS's 45 PMH's

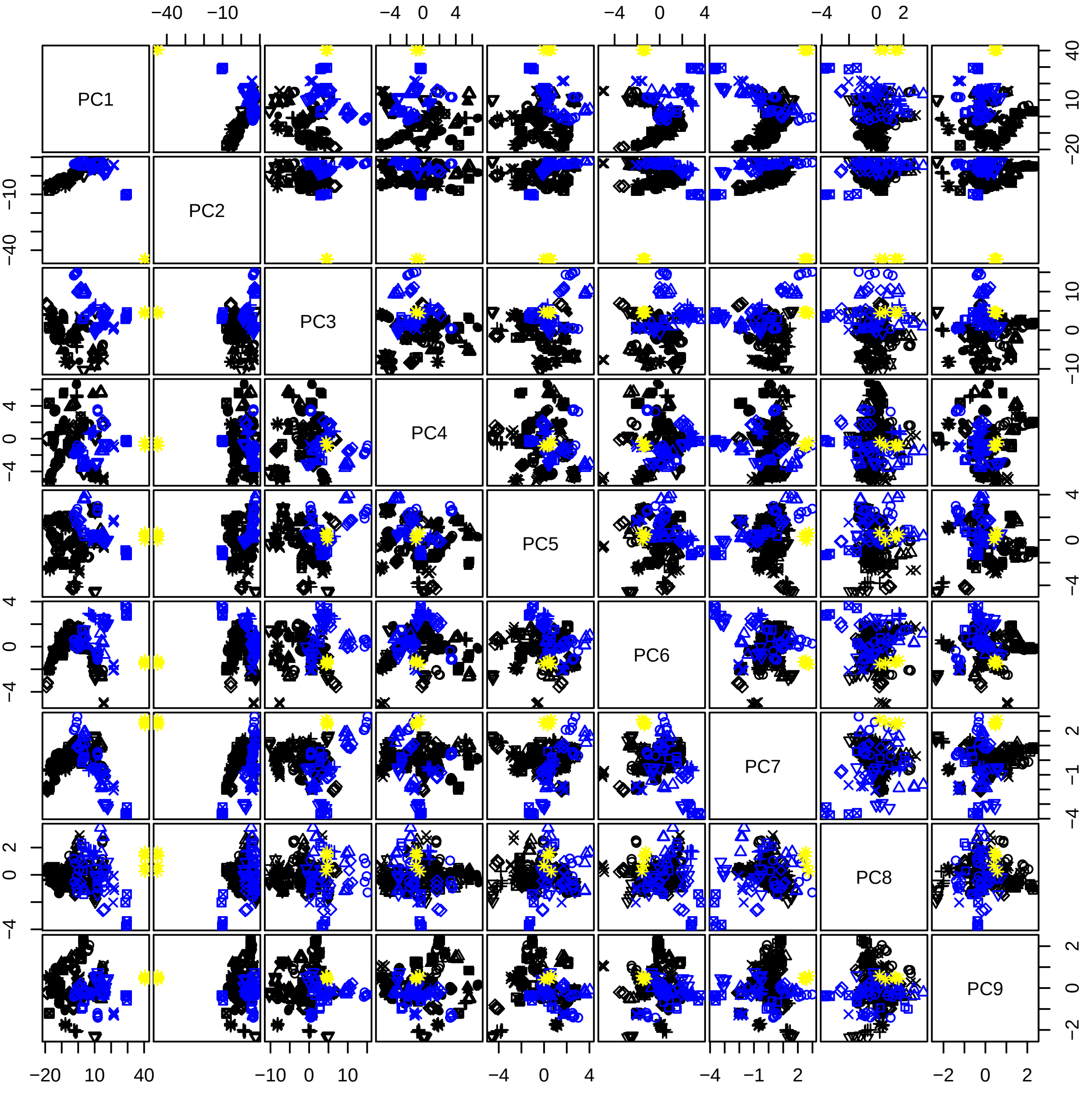
PC1
PC2
PC3
PC4
PC5
PC6
PC7
PC8
PC9
−40
−10
−4
0
4
2
−2
−1
10
40
−20

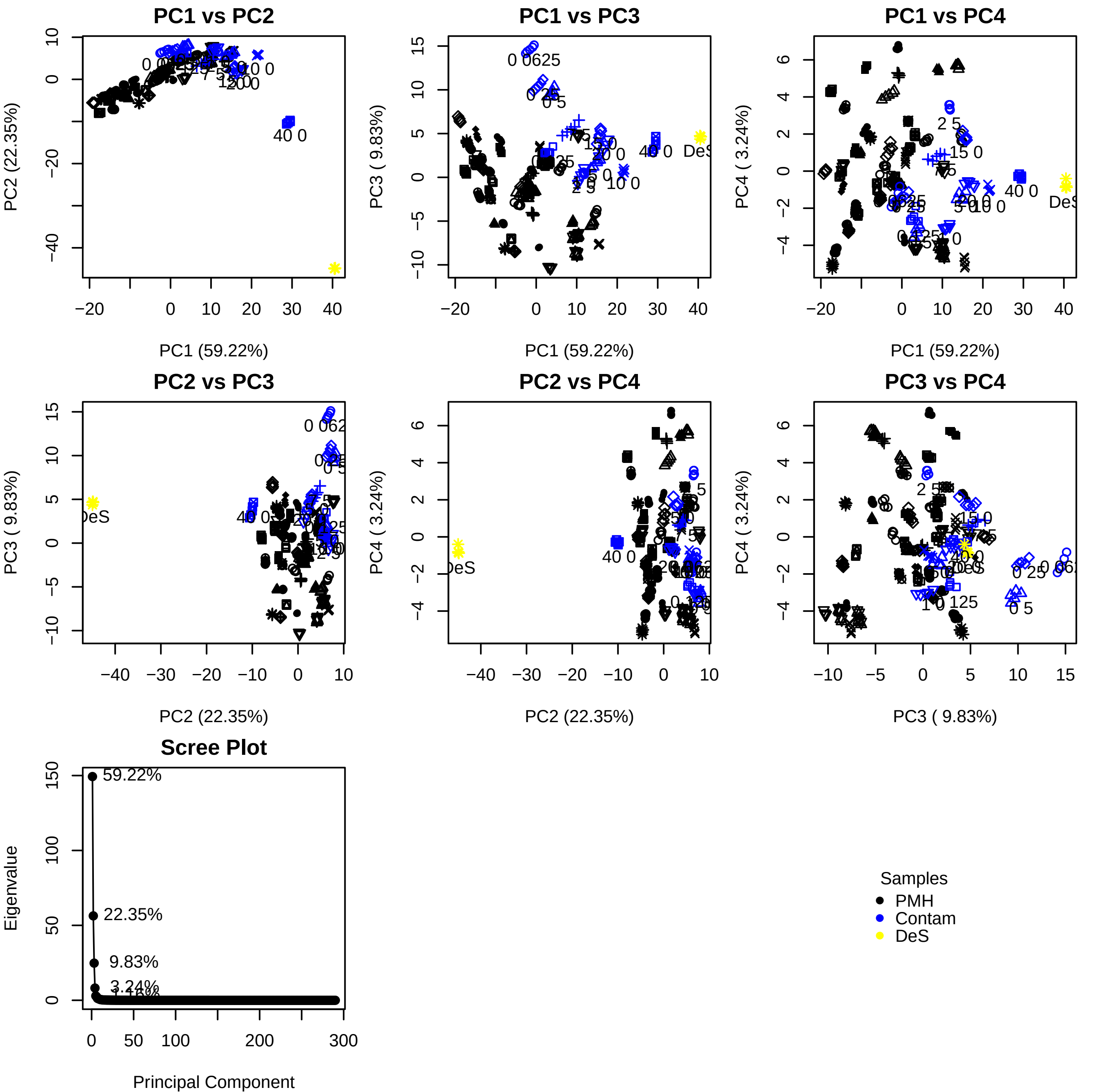
PC1 vs PC2
PC1 vs PC3
PC1 vs PC4
PC2 vs PC3
PC2 vs PC4
PC3 vs PC4
Scree Plot
PC1 (59.22%)
PC2 (22.35%)
PC3 ( 9.83%)
PC4 ( 3.24%)
Eigenvalue
Principal Component
59.22%
22.35%
9.83%
3.24%
Samples
PMH
Contam
DeS

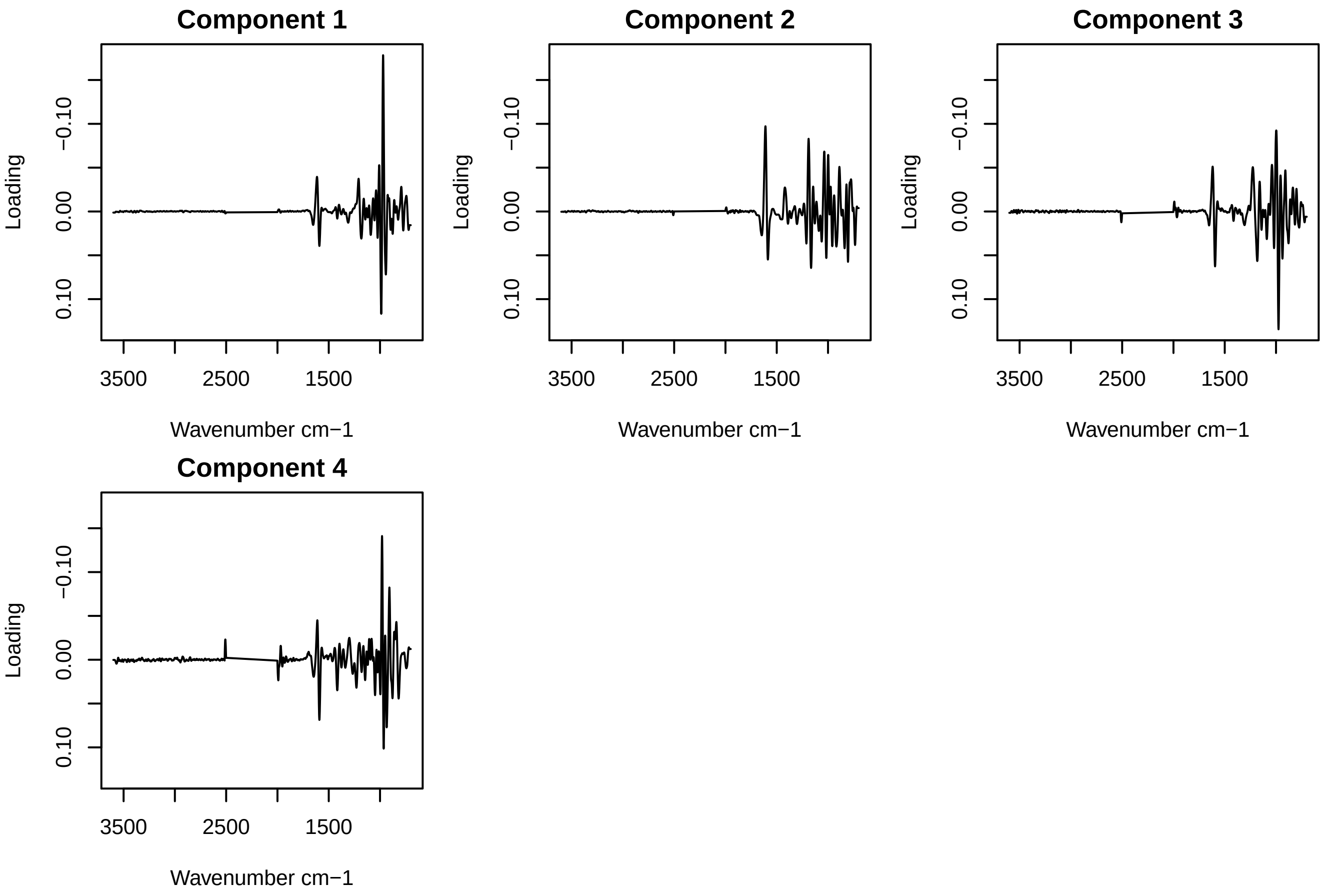
Component 1
Component 2
Component 3
Component 4
Loading
−0.10
0.00
0.10
3500
2500
1500
Wavenumber cm−1

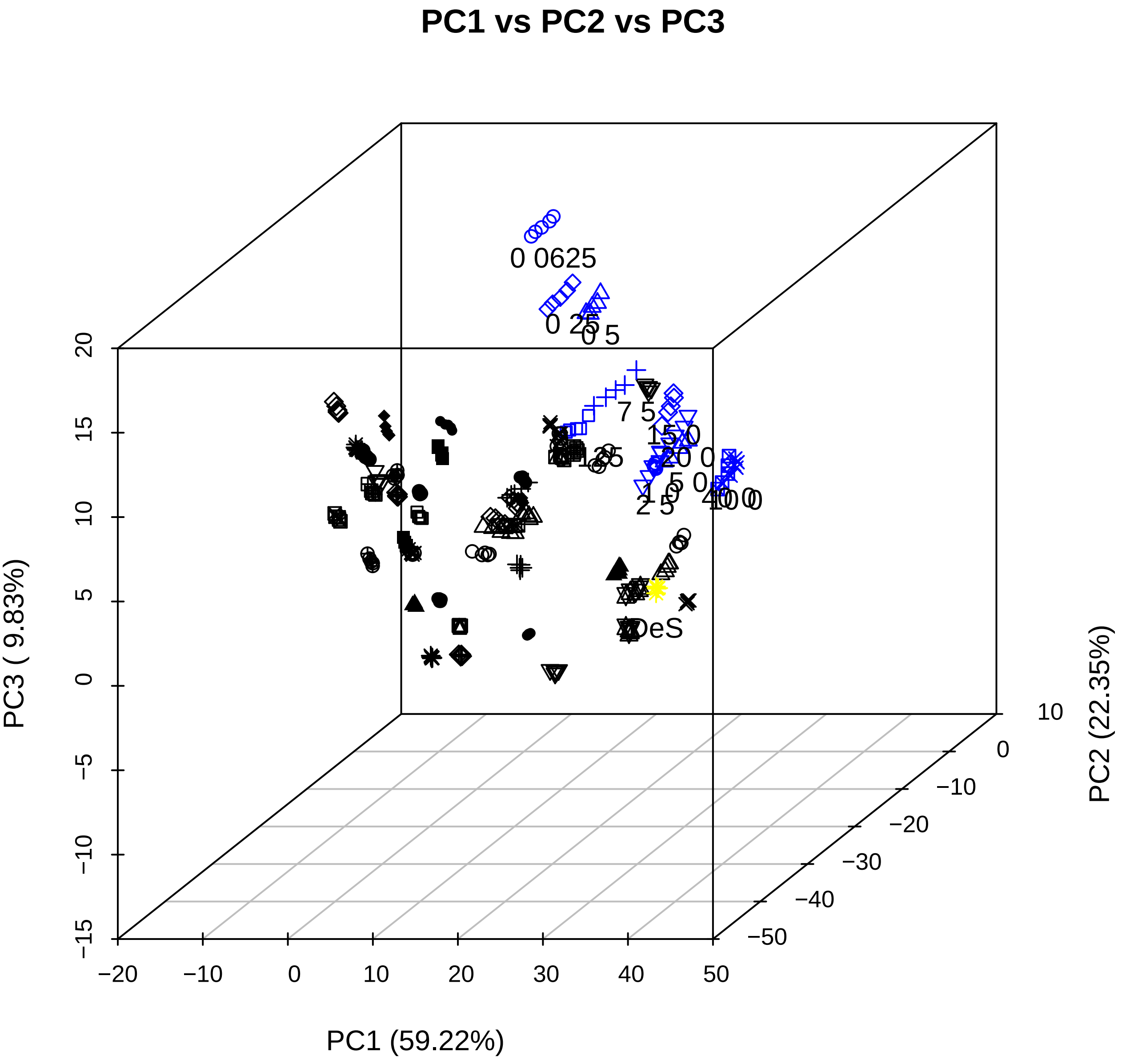
PC1 vs PC2 vs PC3
PC1 (59.22%)
PC2 (22.35%)
PC3 ( 9.83%)
−20
−10
0
10
20
30
40
50
−50
−40
−30
−20
−10
0
10
−15
−10
−5
0
5
10
15
20
0 0625
0 25
0 5
7 5
DeS

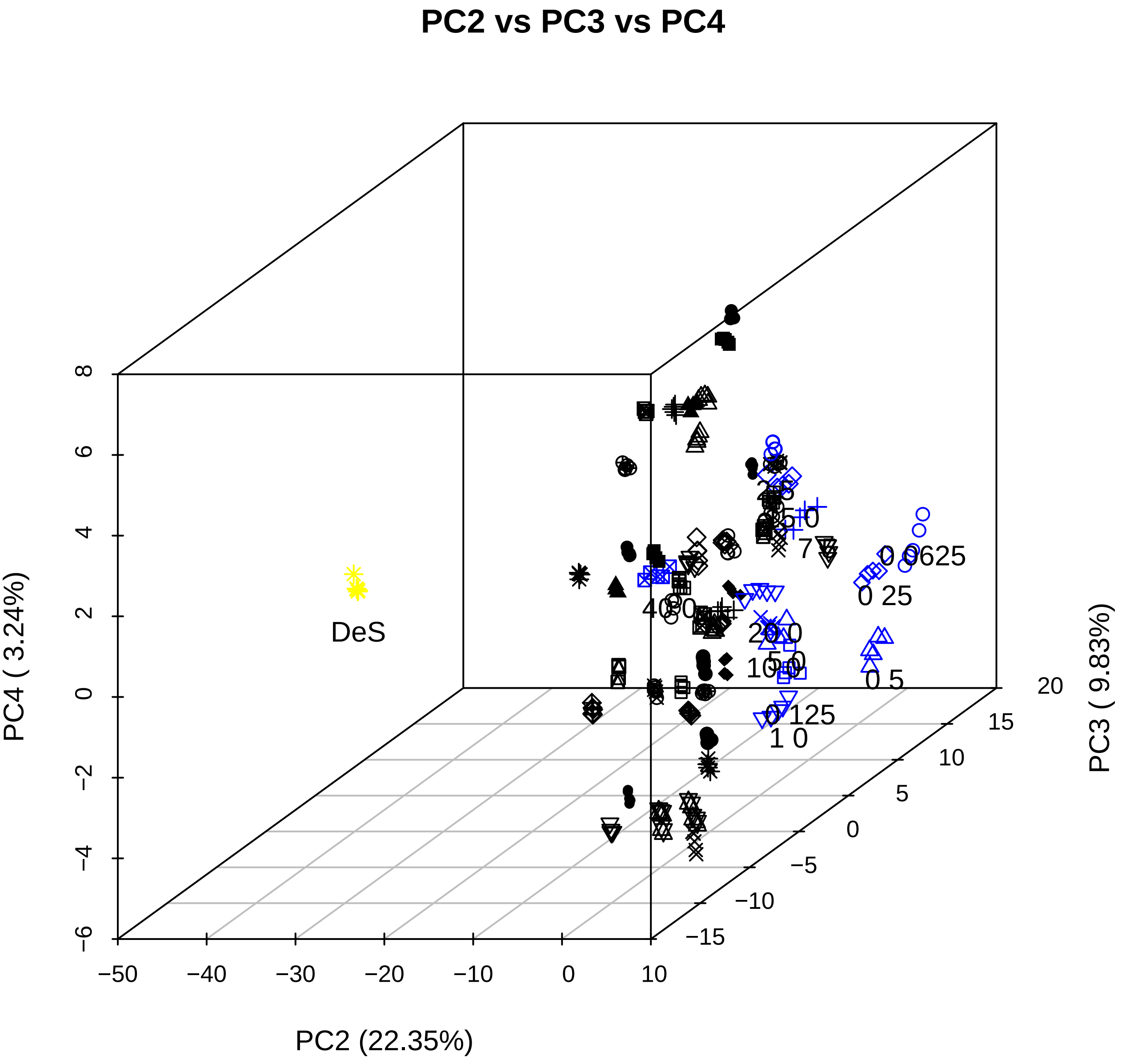

PC2 vs PC3 vs PC4
PC2 (22.35%)
PC3 ( 9.83%)
PC4 ( 3.24%)
−50
−40
−30
−20
−10
0
10
−15
−10
−5
0
5
10
15
20
−6
−4
−2
0
2
4
6
8
DeS
0 0625
0 25
0 5
0 125
1 0

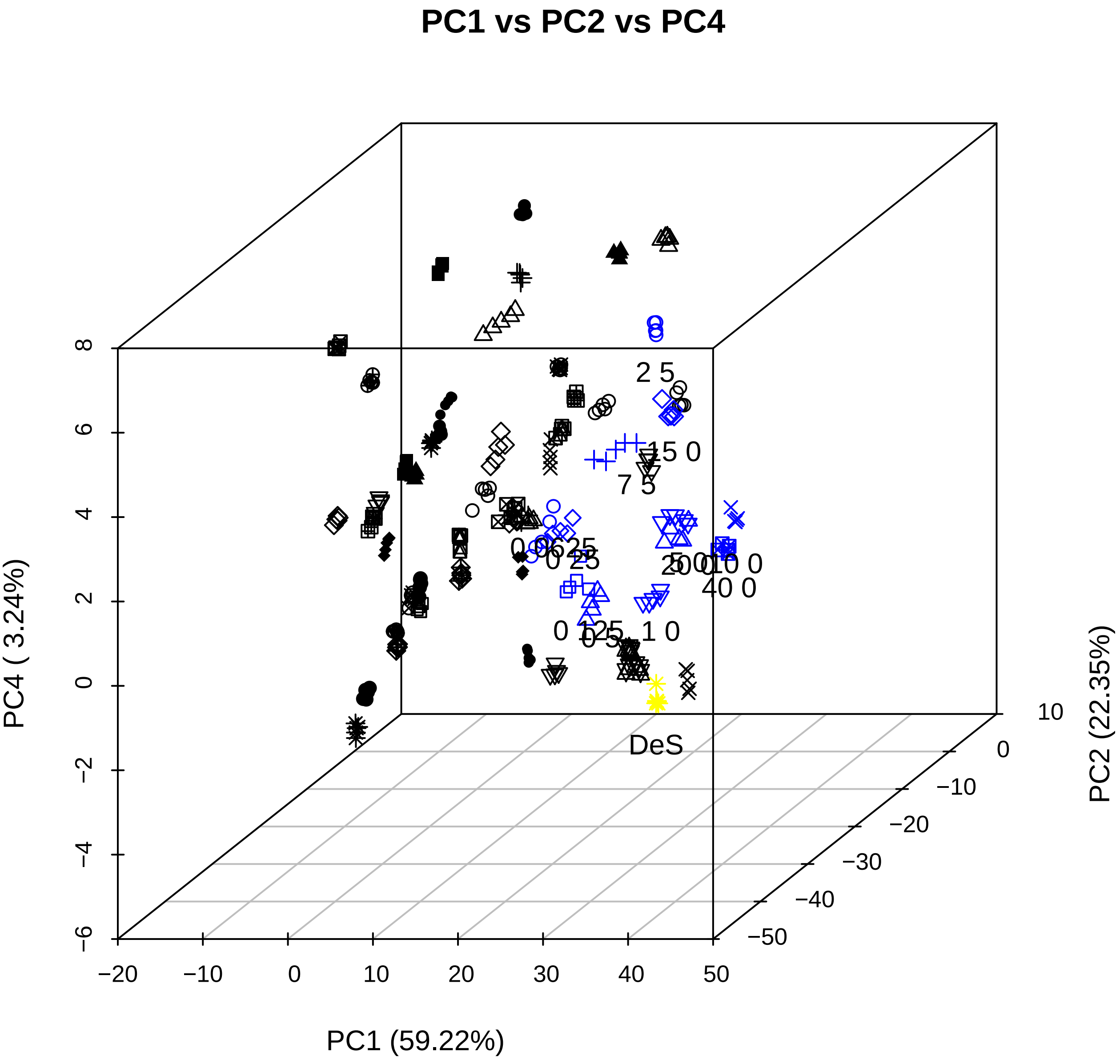
PC1 vs PC2 vs PC4
PC1 (59.22%)
PC2 (22.35%)
PC4 ( 3.24%)
−20
−10
0
10
20
30
40
50
−50
−40
−30
−20
−10
0
10
−6
−4
−2
0
2
4
6
8
2 5
15 0
7 5
0 25
40 0
1 0
DeS

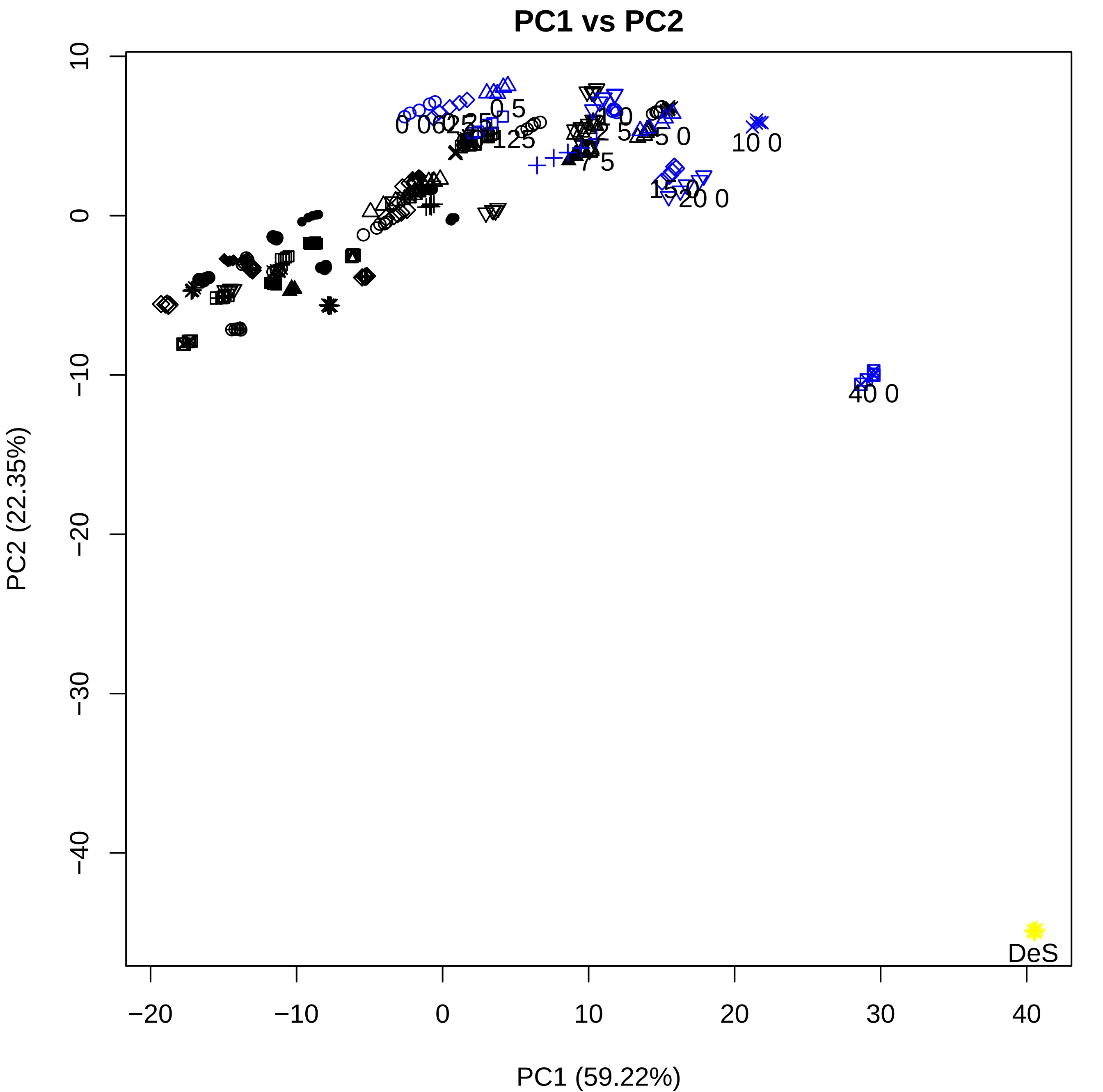

PC1 vs PC2
PC1 (59.22%)
PC2 (22.35%)
−20
−10
0
10
20
30
40
10
0
−10
−20
−30
−40
0 5
0 0625
0 125
10 0
5 0
2 5
7 5
10 0
15 0
20 0
40 0
DeS

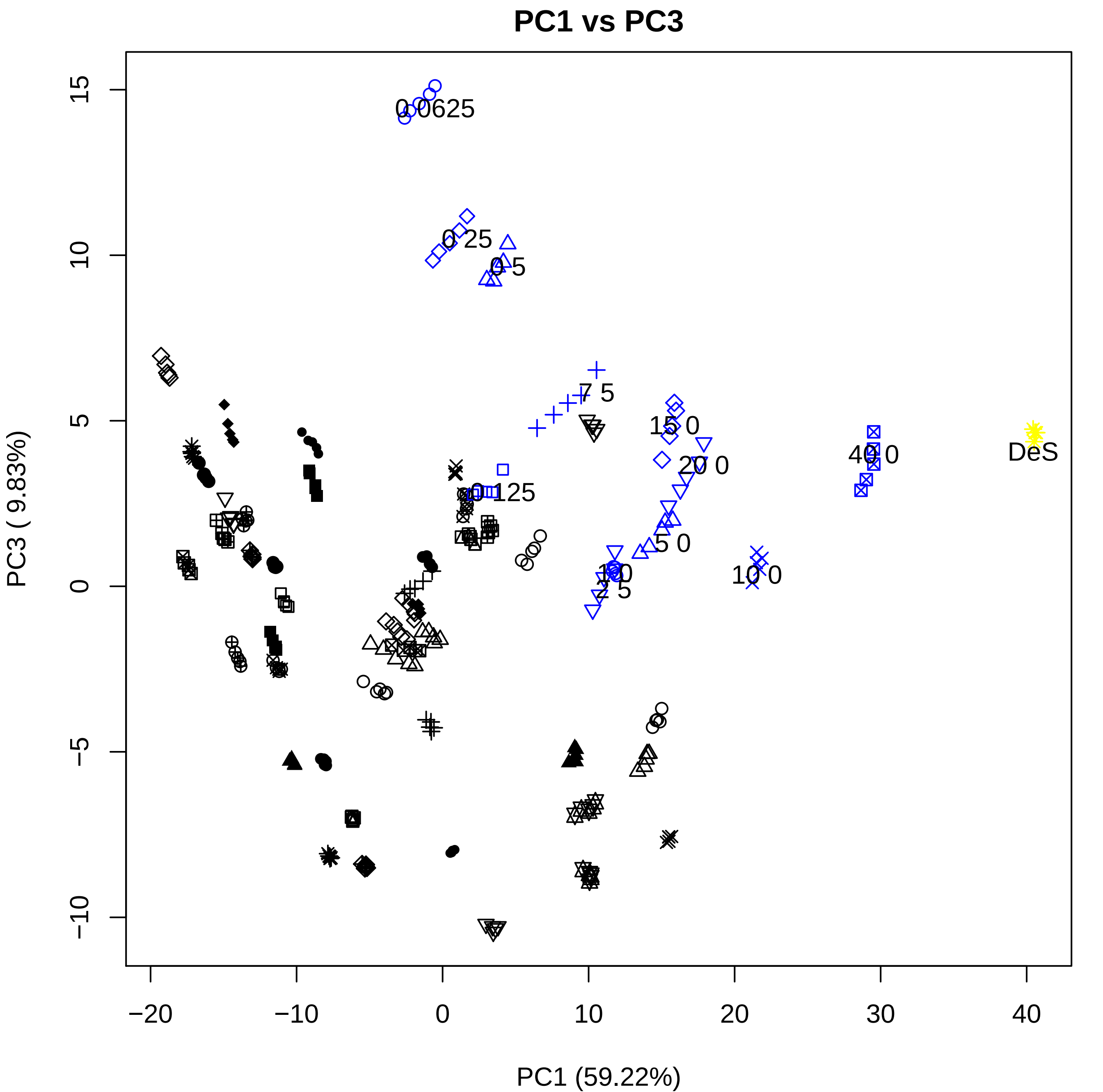
PC1 vs PC3
PC3 ( 9.83%)
PC1 (59.22%)
15
10
5
0
−5
−10
−20
−10
0
10
20
30
40
0.0625
0.25
0.5
7.5
15.0
20.0
40.0
DeS
0.125
5.0
1.0
2.5
10.0

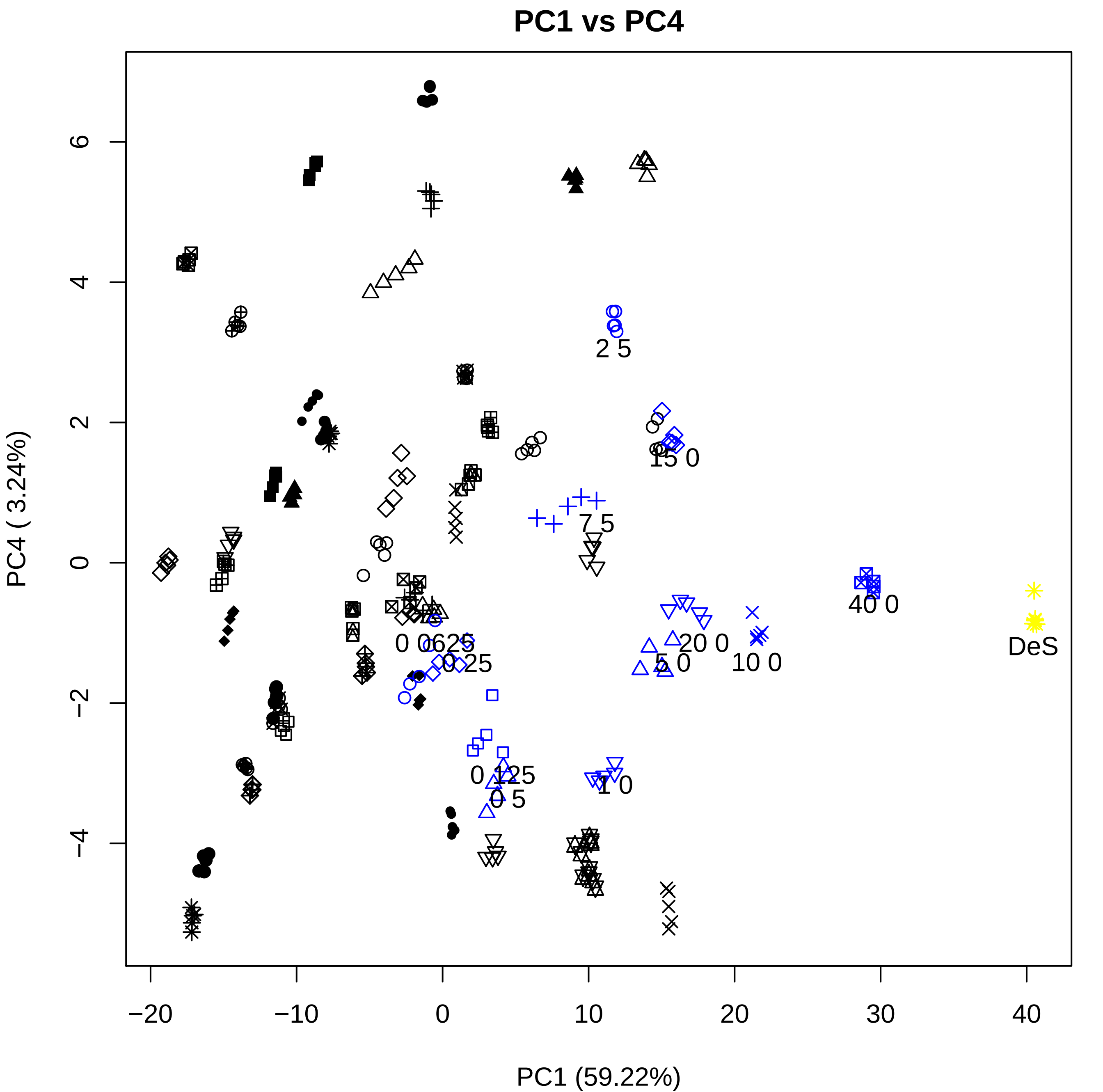
PC1 vs PC4
PC4 ( 3.24%)
PC1 (59.22%)
6
4
2
0
−2
−4
−20
−10
0
10
20
30
40
2 5
15 0
7 5
40 0
DeS
0 0625
0 25
20 0
5 0
10 0
0 125
0 5
1 0

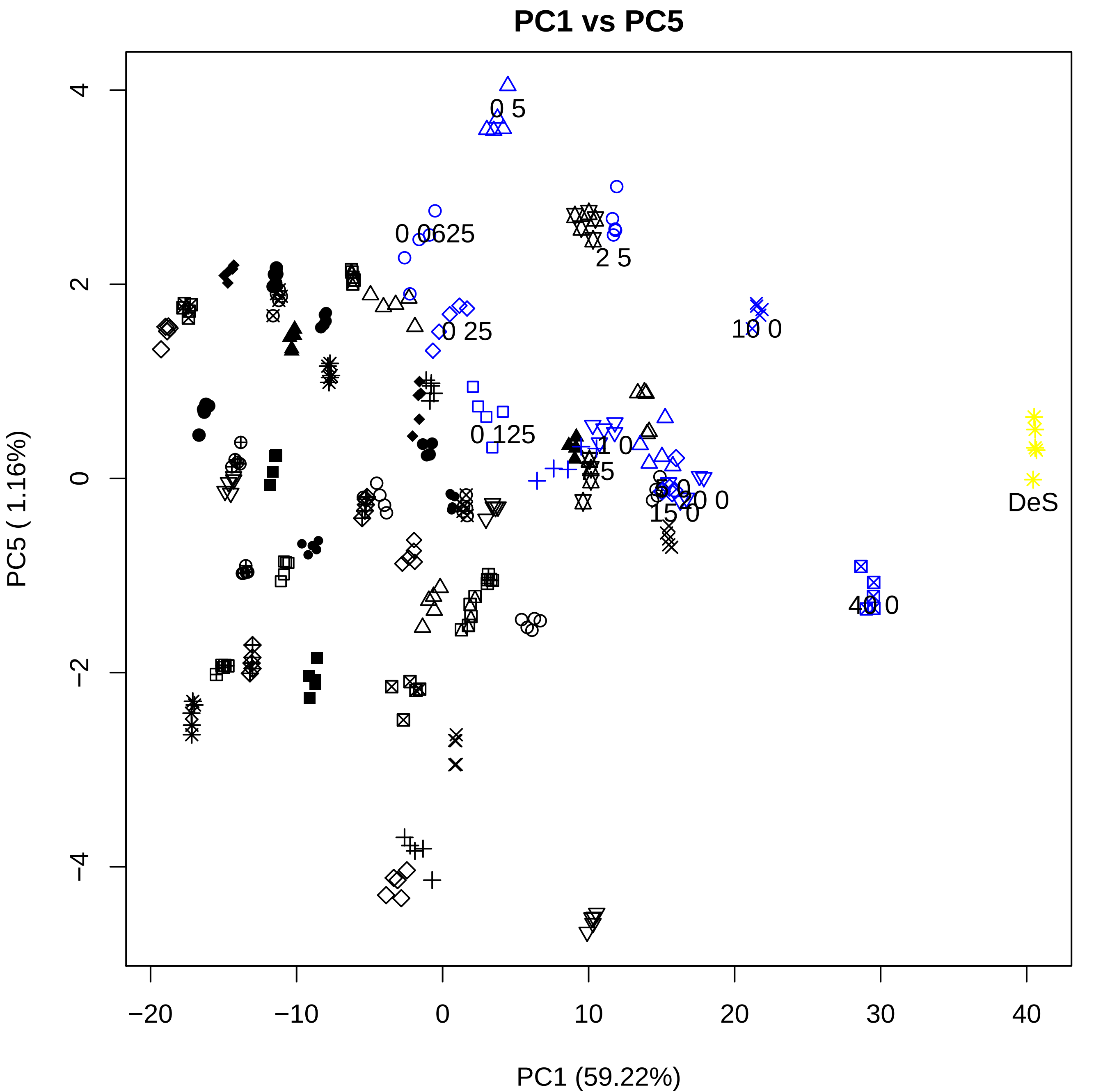
PC1 vs PC5
PC5 ( 1.16%)
PC1 (59.22%)
4
2
0
−2
−4
−20
−10
0
10
20
30
40
0 5
0 0625
2 5
0 25
10 0
0 125
1 0
5
5 0
20 0
15 0
DeS
40 0

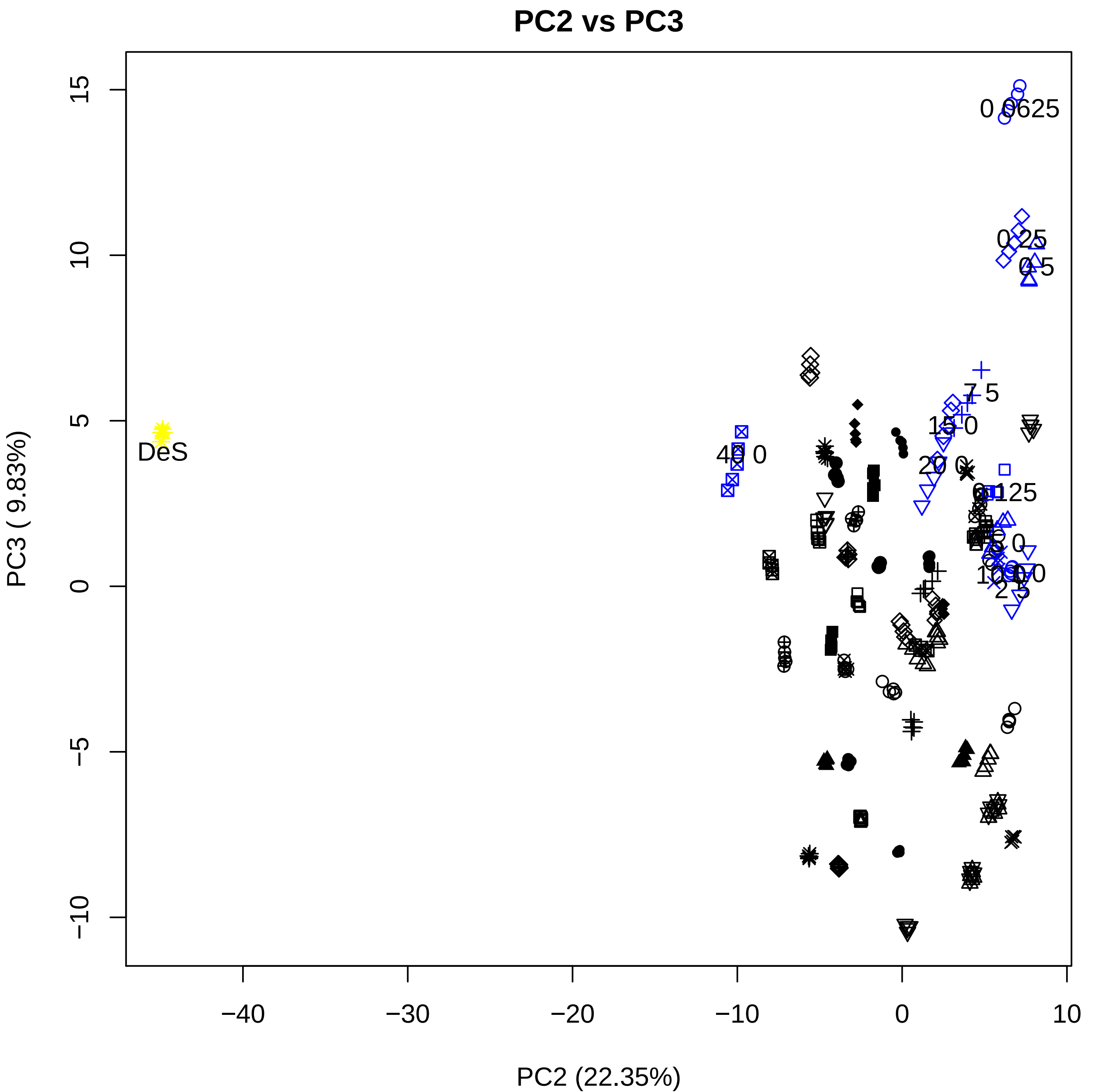

PC2 vs PC3
0.0625
0.25
0.5
7.5
15.0
40.0
20.0
0.125
5.0
10.0
2.5
DeS
PC3 ( 9.83%)
PC2 (22.35%)
15
10
5
0
−5
−10
−40
−30
−20
−10
0
10

**PC2 vs PC4**

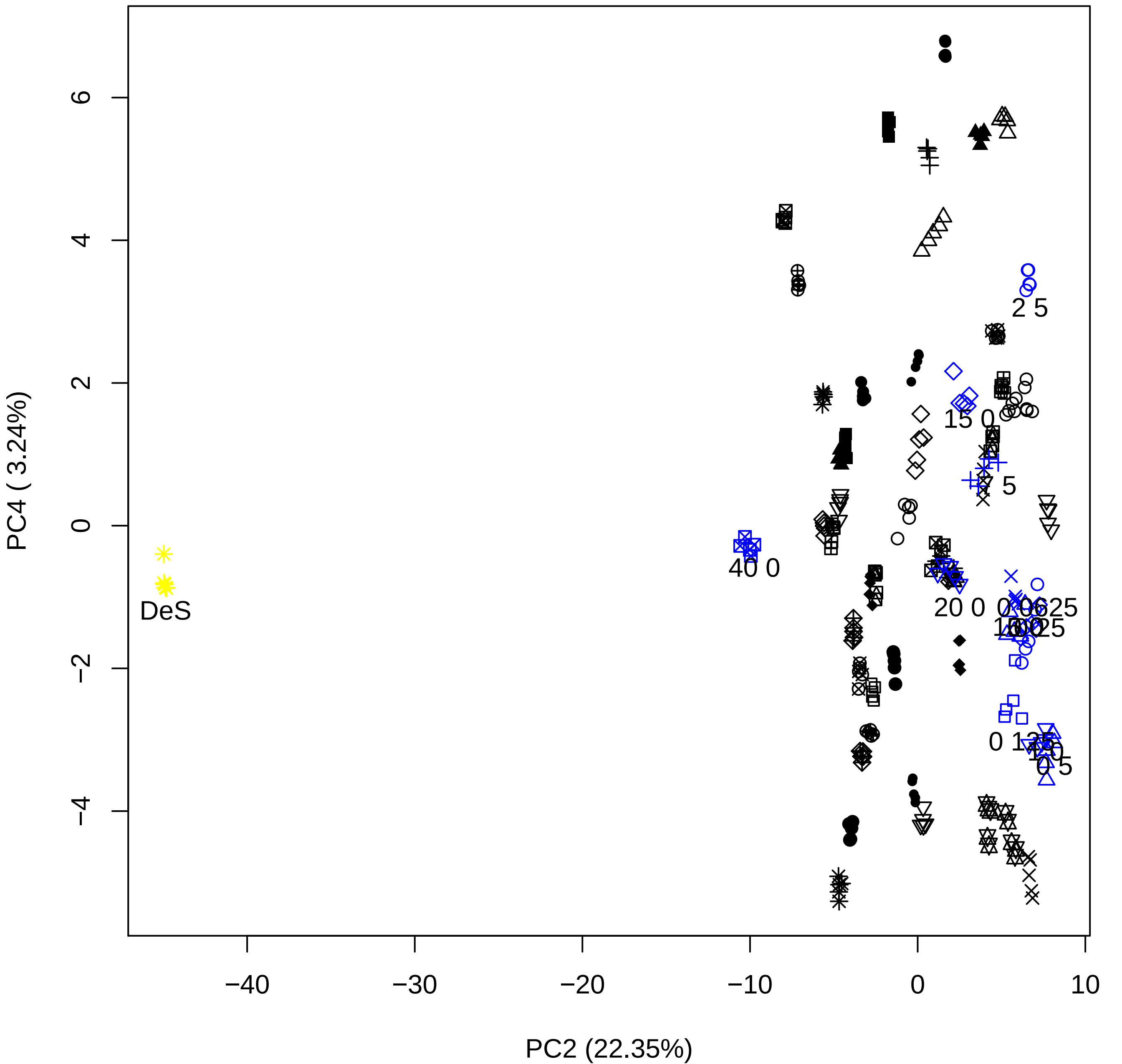
PC4 ( 3.24%)
PC2 (22.35%)
6
4
2
0
−2
−4
−40
−30
−20
−10
0
10
DeS
40 0
2 5
15 0
7 5
20 0
0 125
0 5

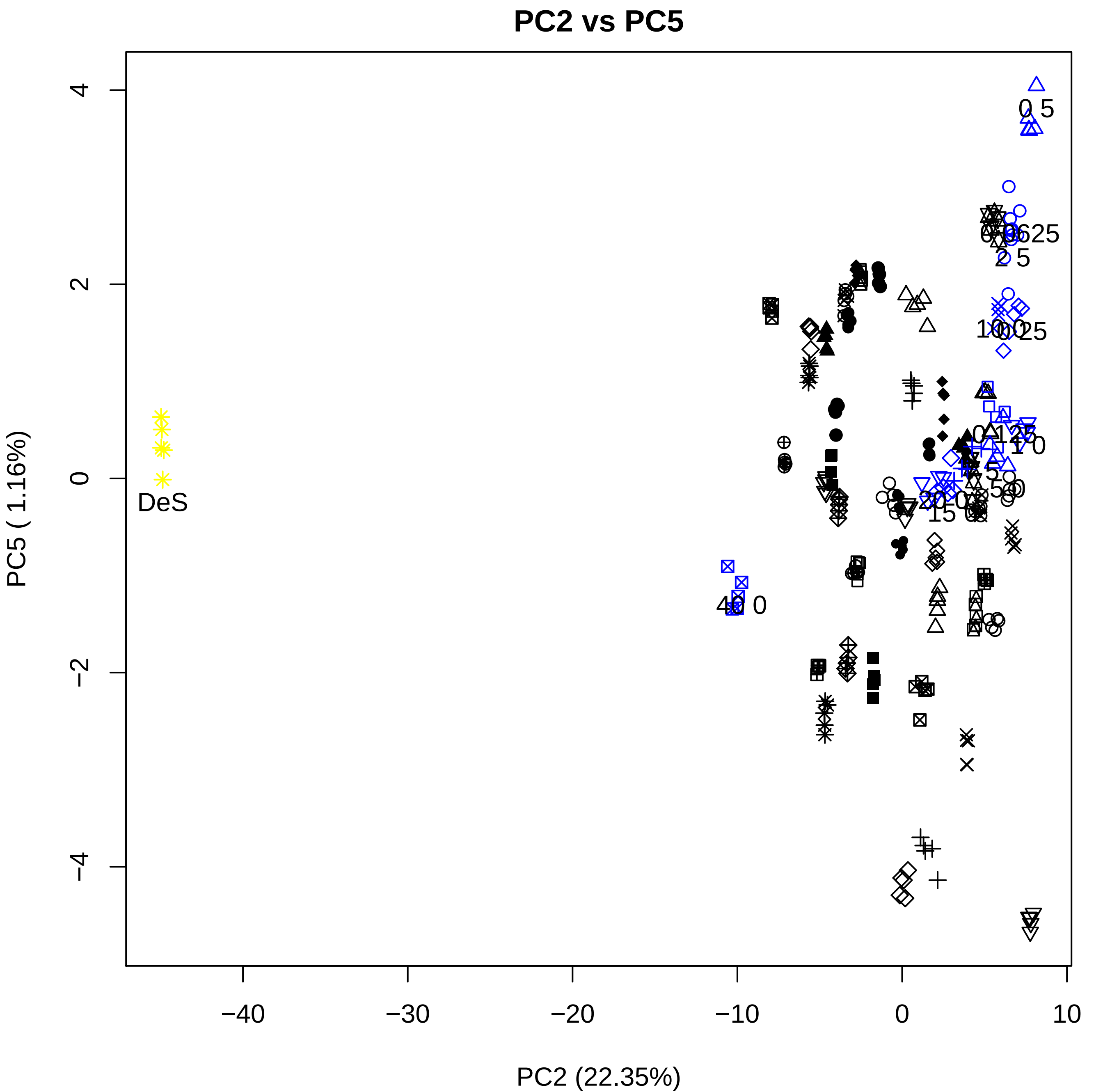
PC2 vs PC5
PC5 ( 1.16%)
PC2 (22.35%)
4
2
0
−2
−4
−40
−30
−20
−10
0
10
DeS
0 5
2 5

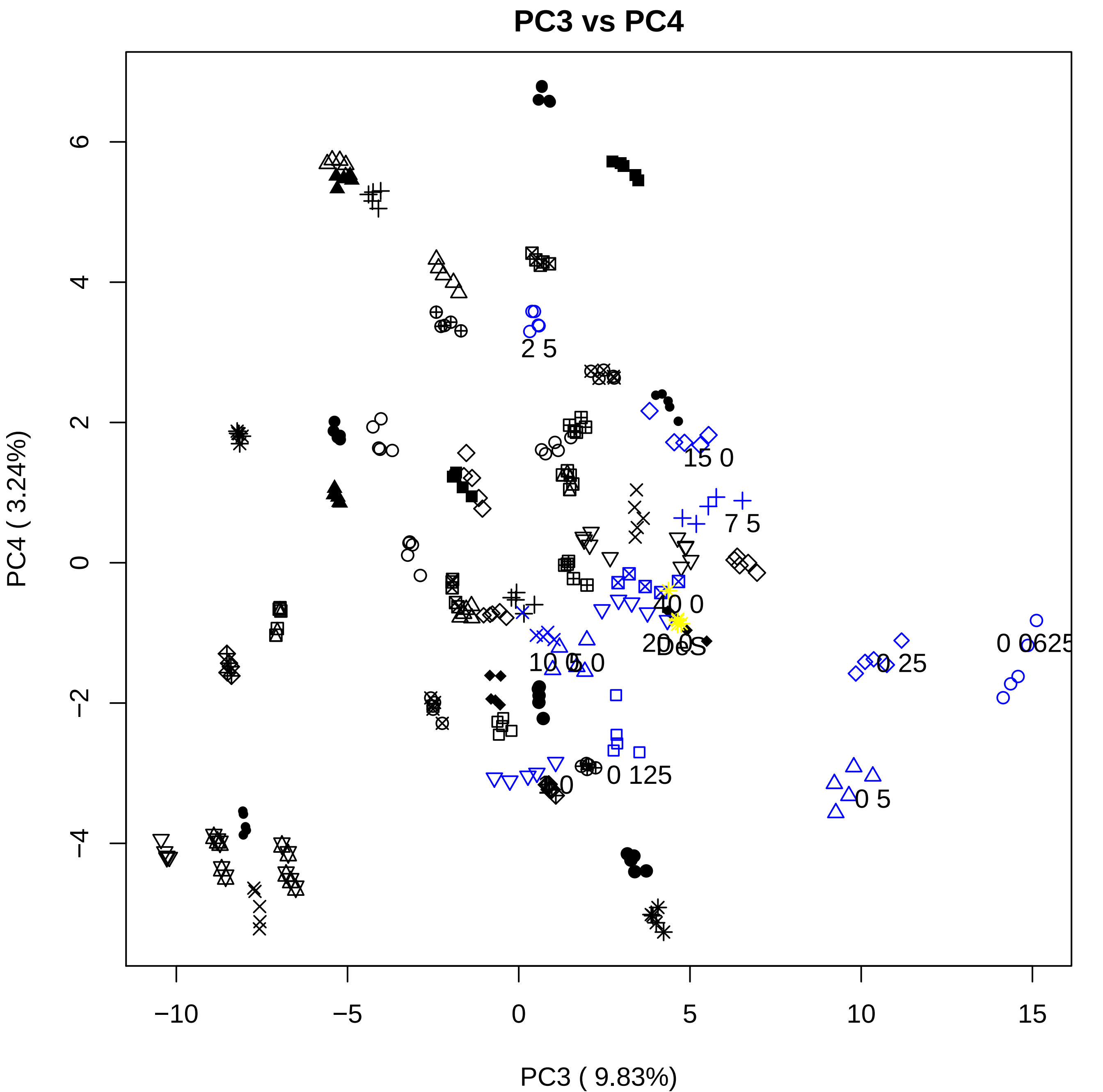
PC3 vs PC4
PC4 ( 3.24%)
PC3 ( 9.83%)
6
4
2
0
−2
−4
−10
−5
0
5
10
15
2 5
15 0
7 5
40 0
20 0
DeS
0 0625
0 25
0 125
0 5

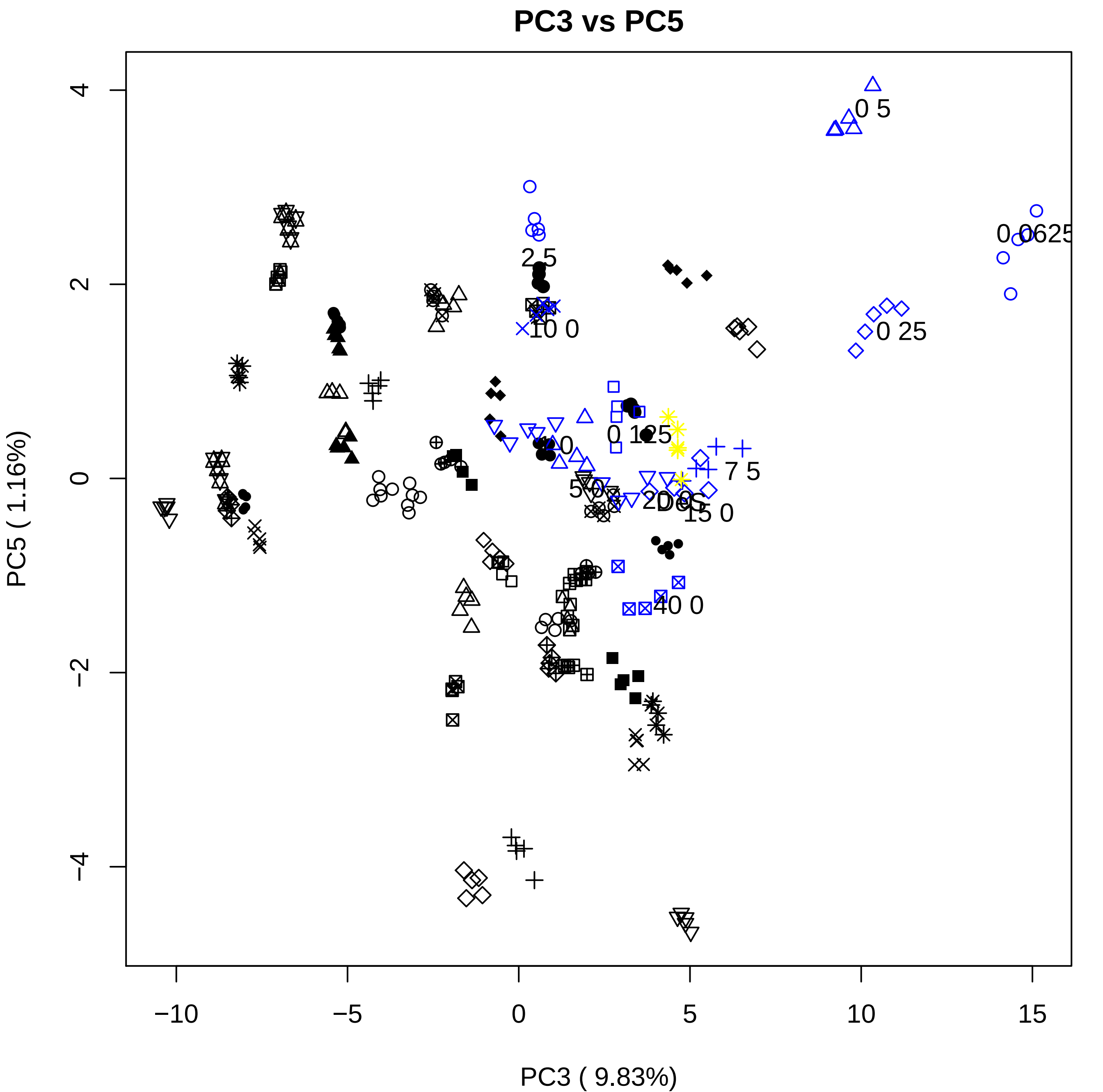
PC3 vs PC5
PC5 ( 1.16%)
PC3 ( 9.83%)
4
2
0
−2
−4
−10
−5
0
5
10
15
0 5
0.0625
2 5
0 25
10 0
0 125
7 5
5 0
15 0
40 0

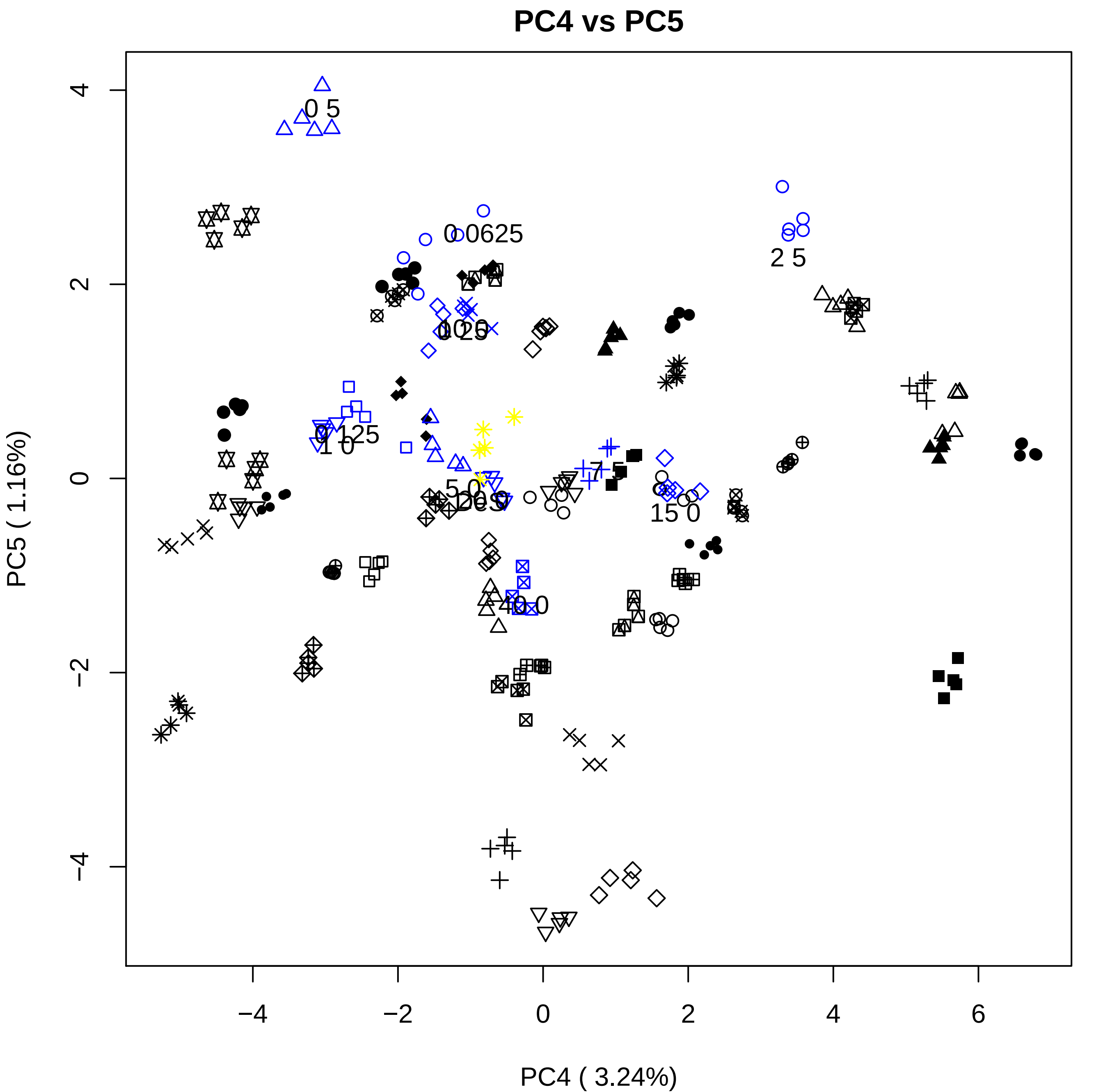

PC4 vs PC5
PC5 ( 1.16%)
PC4 ( 3.24%)
4
2
0
−2
−4
−4
−2
0
2
4
6
0 5
0 0625
2 5
1 0
7 5
15 0
40 0

### PCA of 0.06%  DeS vs PMH

Scaled Matrix PCAed

7th order polynomial baseline correction performed

Raw spectra smoothed to 21 Neighbours, 2nd order PN, Savtisky Golay

Normalisation between 0 and 100

D2 spectra smoothed to 41 Neighbours, 2nd order PN, Savtisky Golay

Regions Analysed: 3601−2501 and 2001−700cm−1

Samples Analysed: 1 Contam's 0 DeS's 45 PMH's

Keele University AD                                    Sun Dec 09 17:03:40 2018

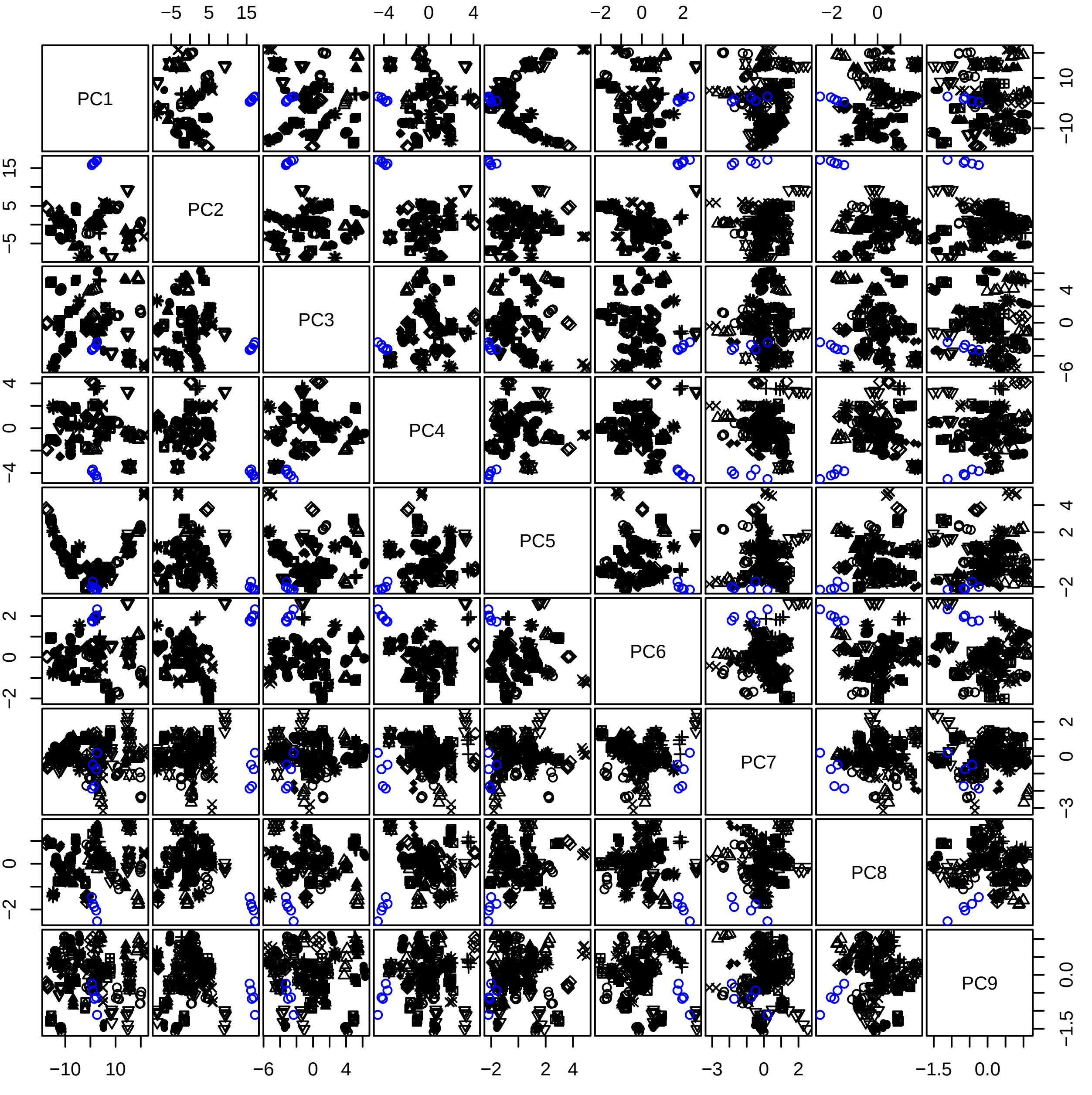
PC1
PC2
PC3
PC4
PC5
PC6
PC7
PC8
PC9

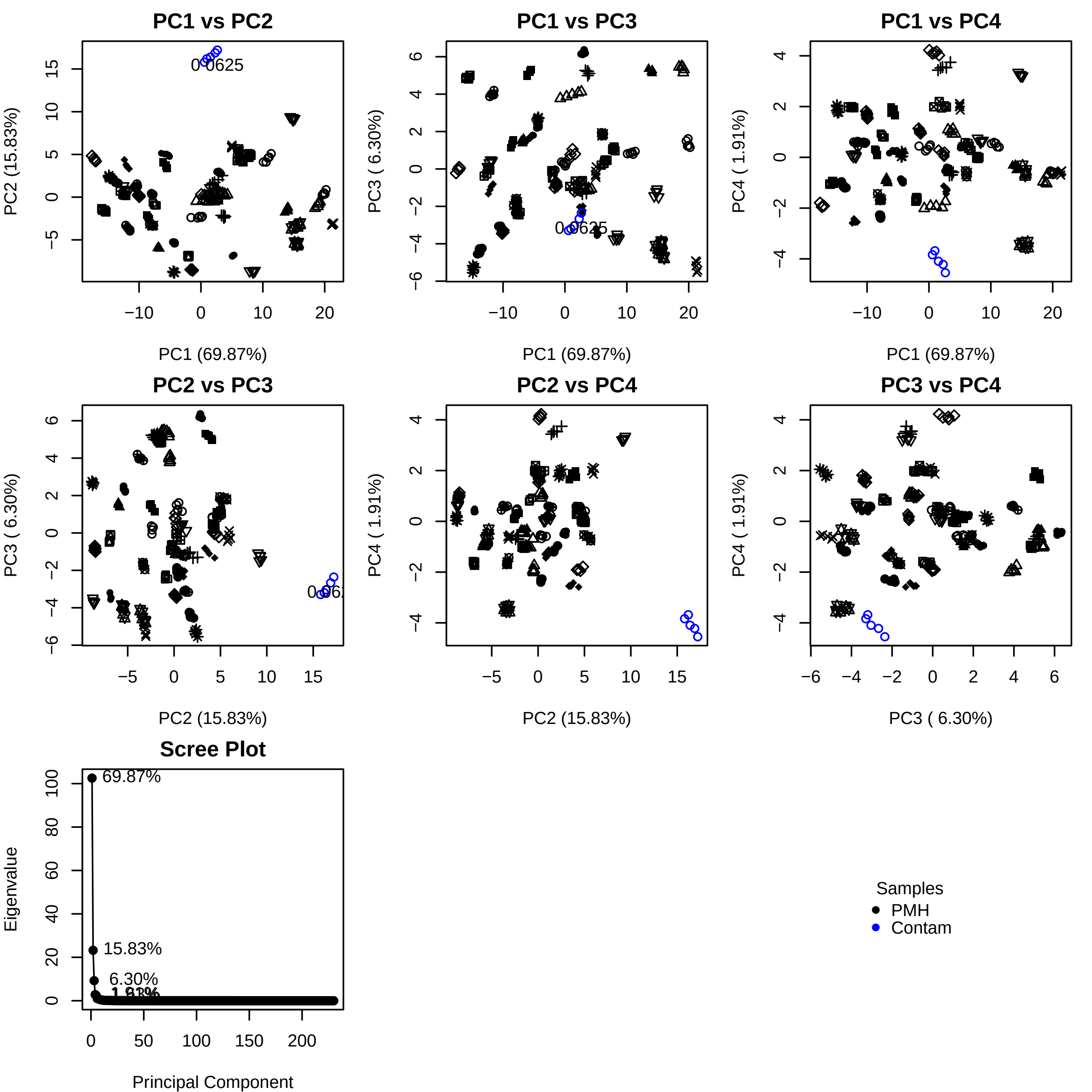

PC1 vs PC2
PC1 vs PC3
PC1 vs PC4
PC2 vs PC3
PC2 vs PC4
PC3 vs PC4
Scree Plot
PC1 (69.87%)
PC2 (15.83%)
PC3 ( 6.30%)
PC4 ( 1.91%)
0.0625
Eigenvalue
Principal Component
69.87%
15.83%
6.30%
Samples
PMH
Contam

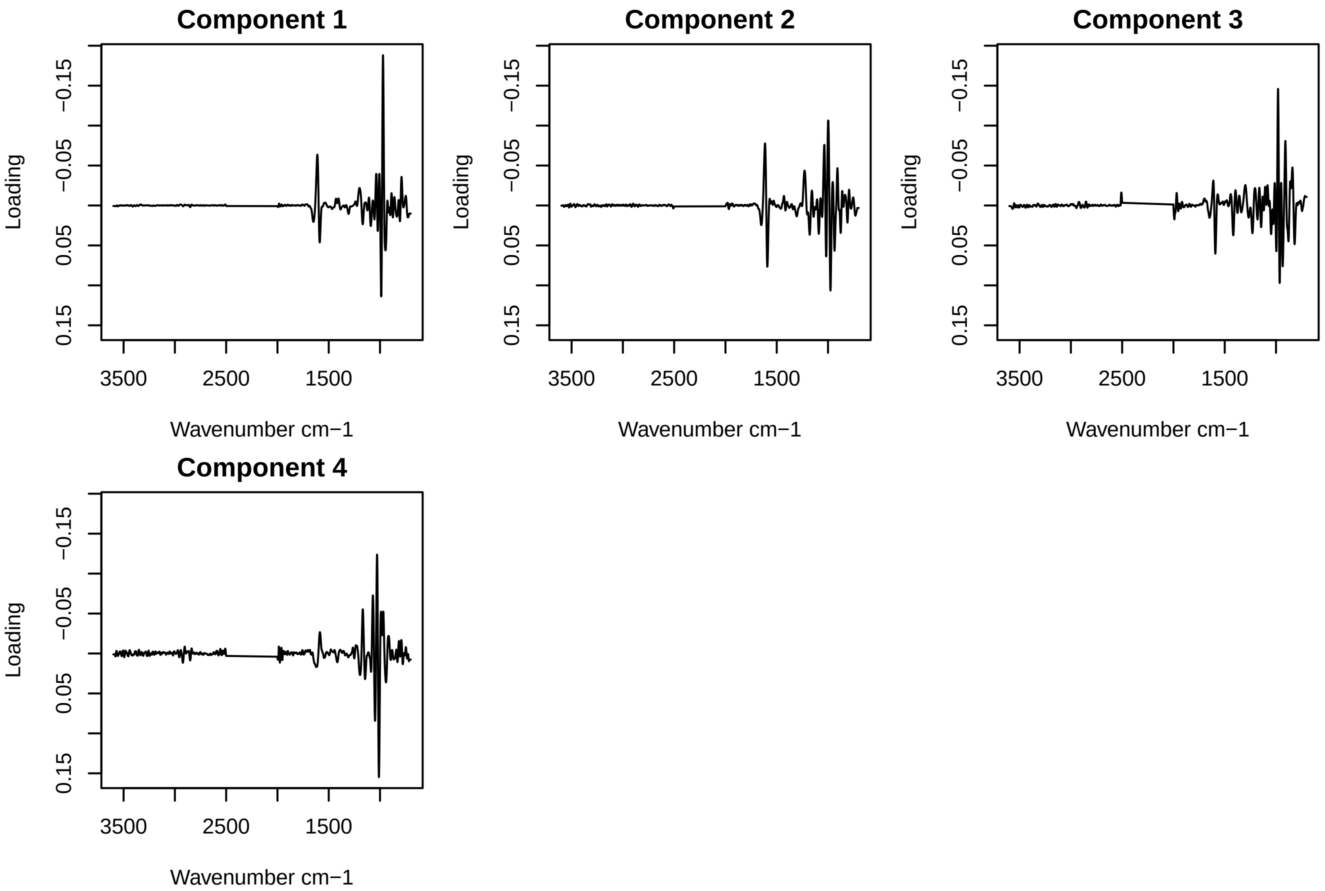
Component 1
Component 2
Component 3
Component 4
Loading
−0.15
−0.05
0.05
0.15
3500
2500
1500
Wavenumber cm−1

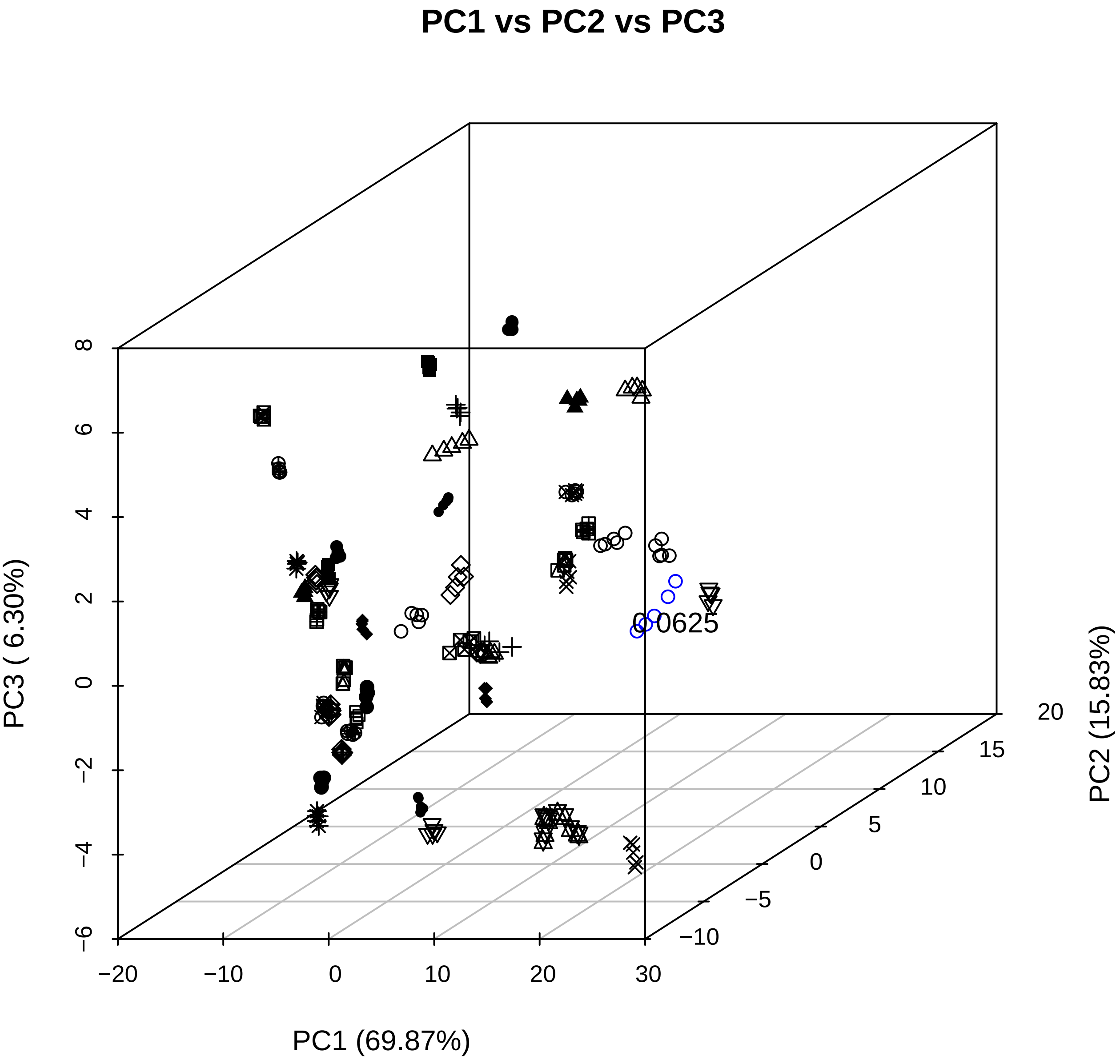
PC1 vs PC2 vs PC3
PC1 (69.87%)
PC2 (15.83%)
PC3 ( 6.30%)
−20
−10
0
10
20
30
−10
−5
0
5
10
15
20
−6
−4
−2
0
2
4
6
8
0.0625

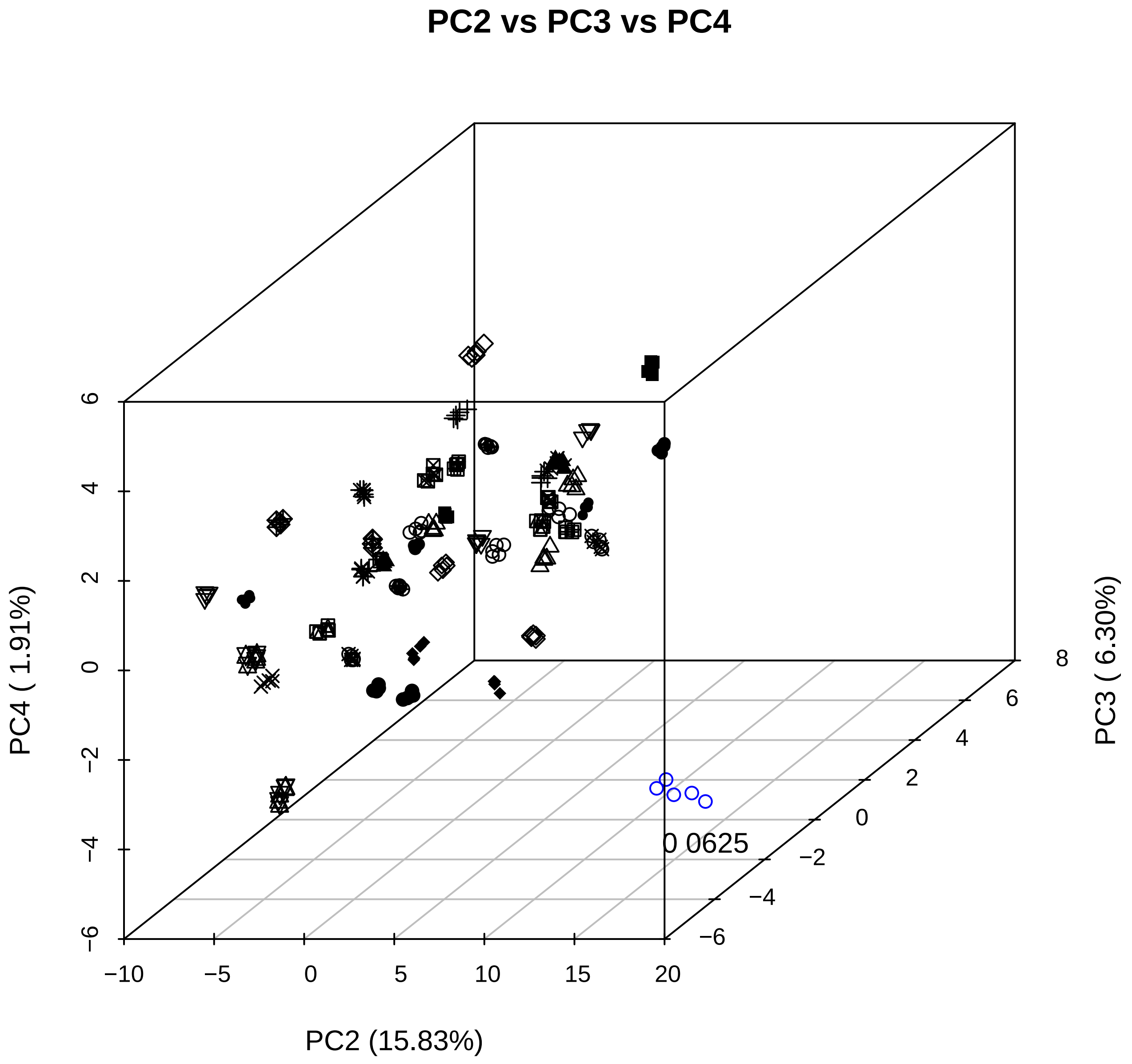
PC2 vs PC3 vs PC4
PC2 (15.83%)
PC3 ( 6.30%)
PC4 ( 1.91%)
−10
−5
0
5
10
15
20
−6
−4
−2
0
2
4
6
8
0 0625

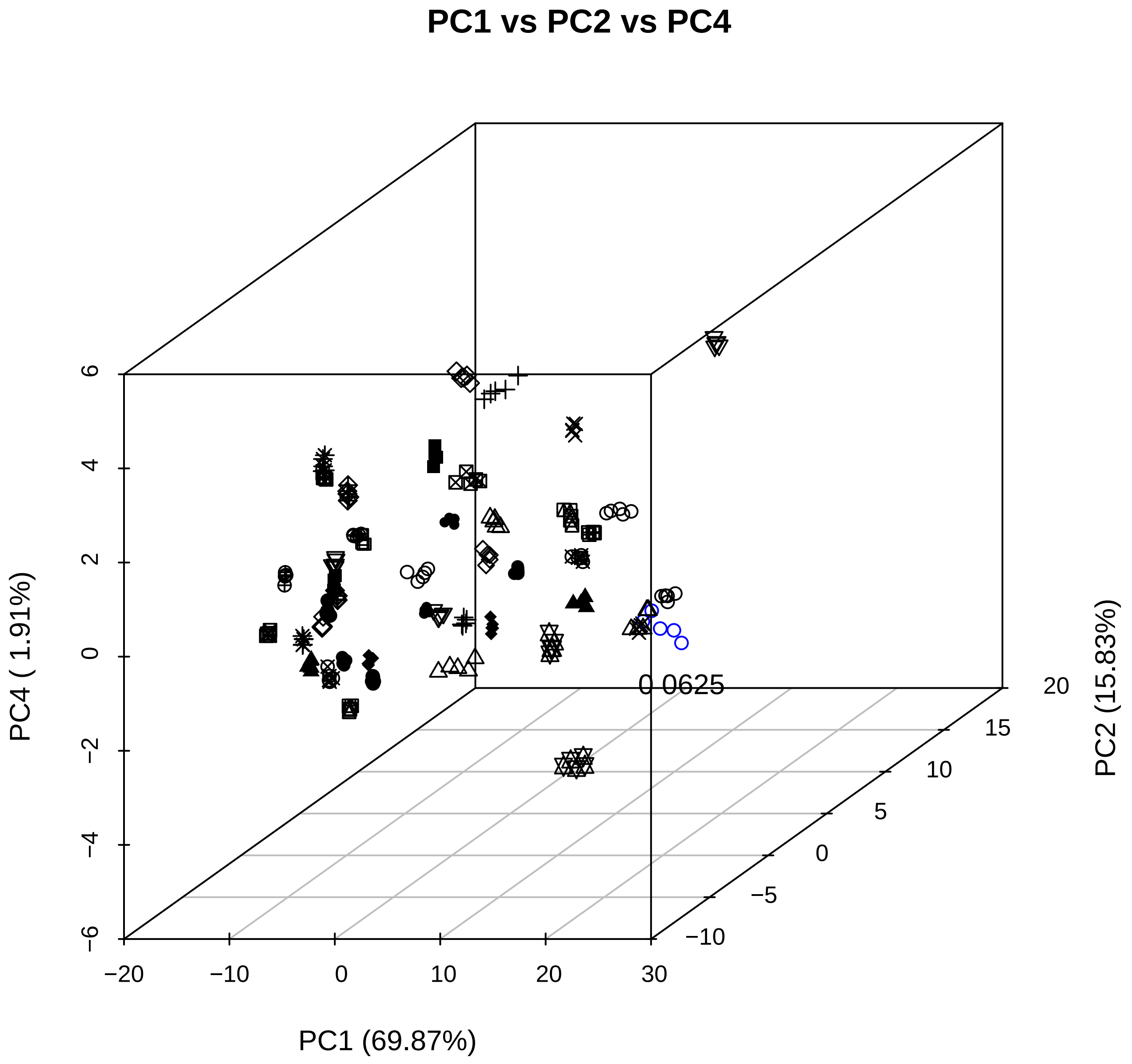

PC1 vs PC2 vs PC4
PC1 (69.87%)
PC2 (15.83%)
PC4 ( 1.91%)
0.0625
−20
−10
0
10
20
30
−10
−5
0
5
10
15
20
−6
−4
−2
0
2
4
6

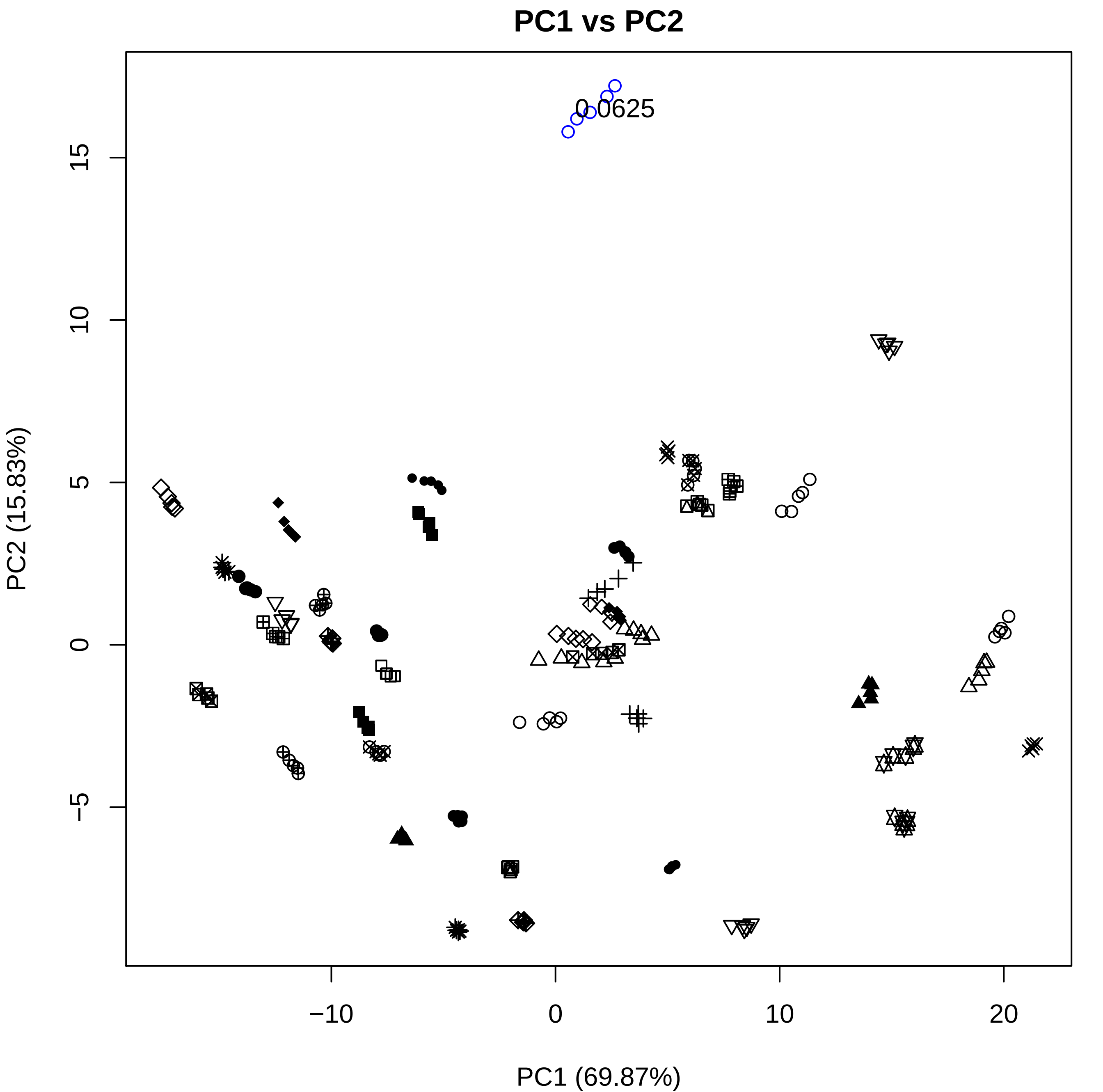

PC1 vs PC2
0.0625
15
10
5
0
−5
PC2 (15.83%)
−10
0
10
20
PC1 (69.87%)

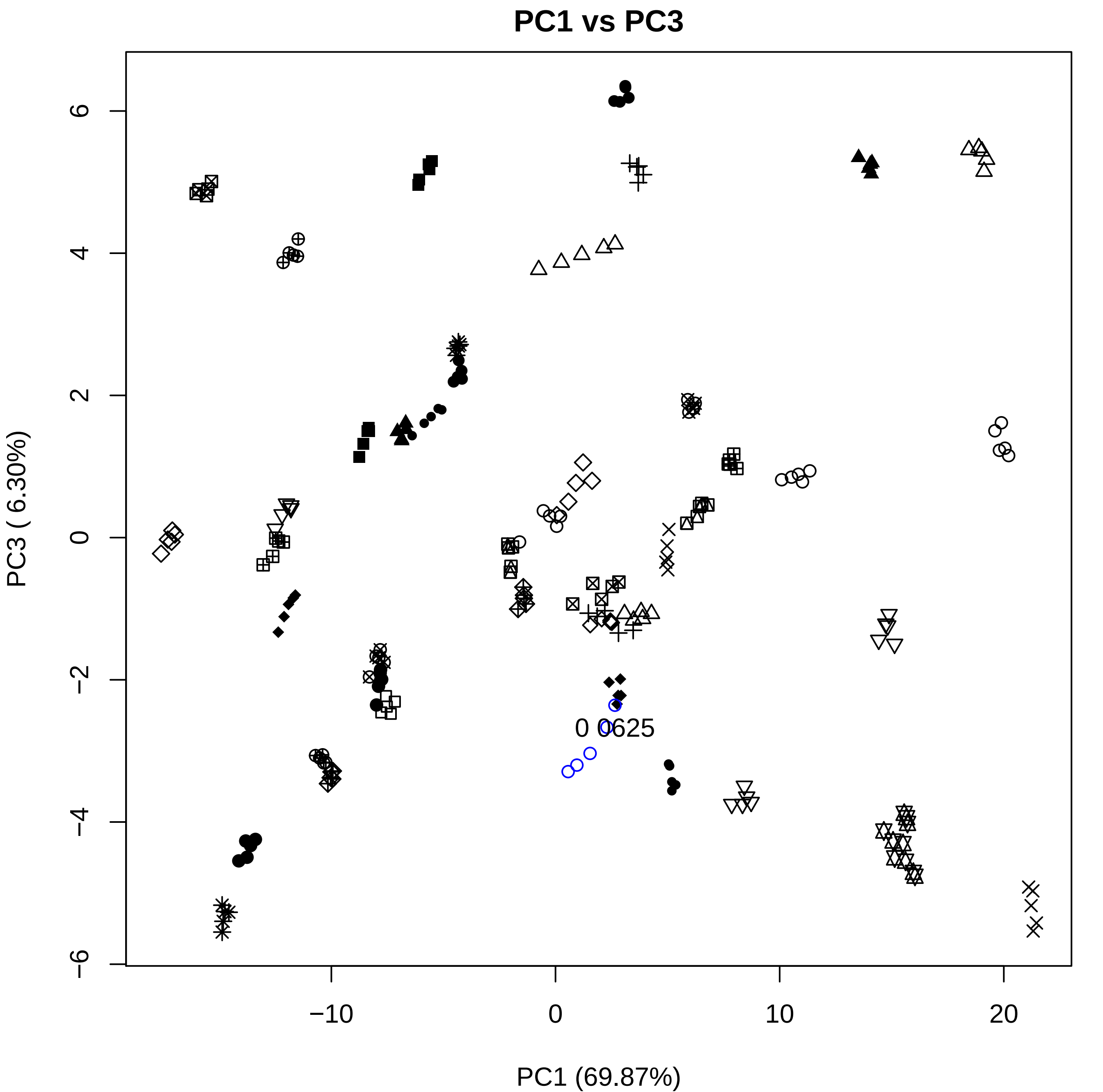
PC1 vs PC3
PC3 ( 6.30%)
PC1 (69.87%)
6
4
2
0
−2
−4
−6
−10
0
10
20
0 0625

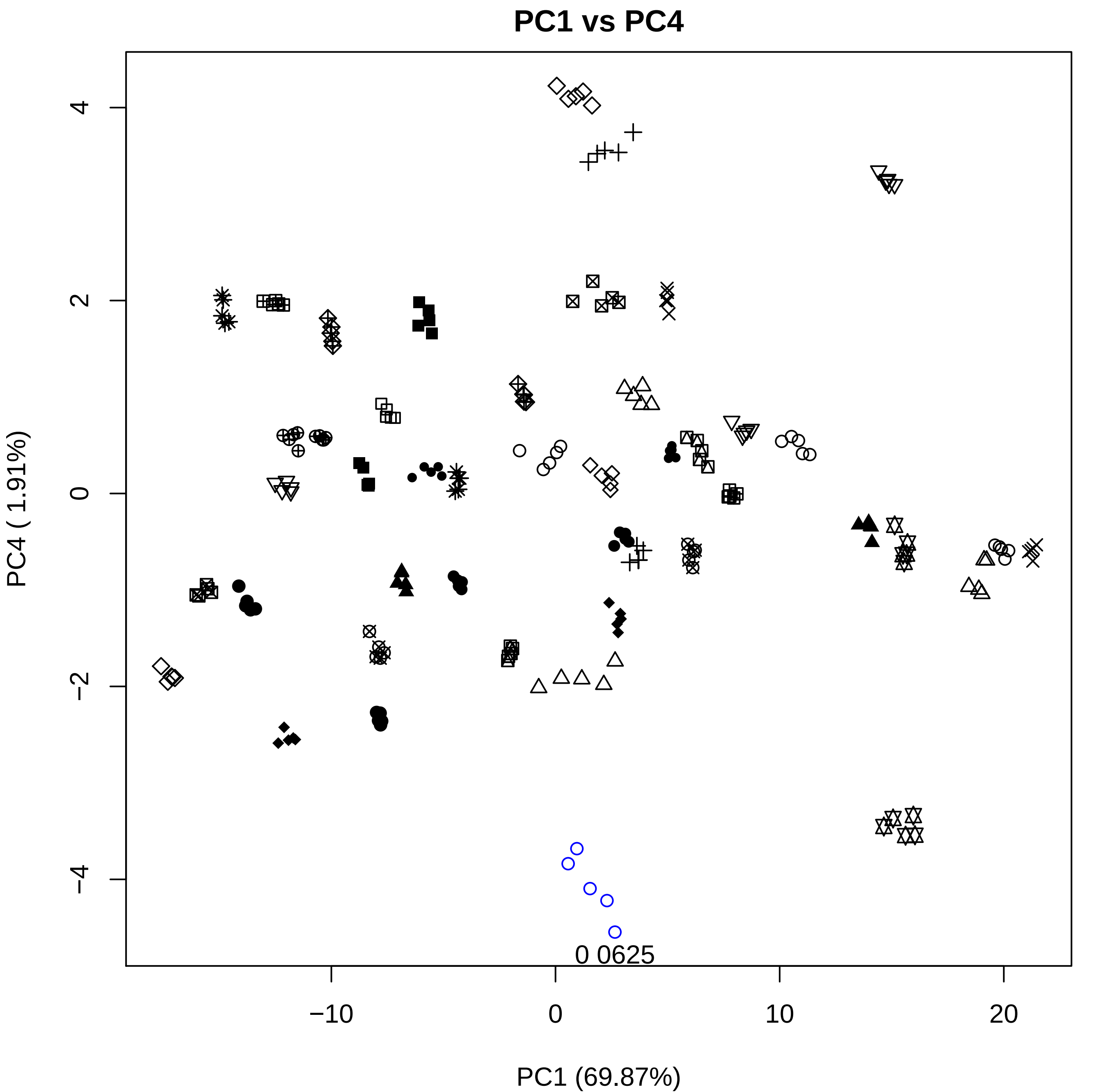

PC1 vs PC4
PC4 ( 1.91%)
PC1 (69.87%)
4
2
0
−2
−4
−10
0
10
20
0 0625

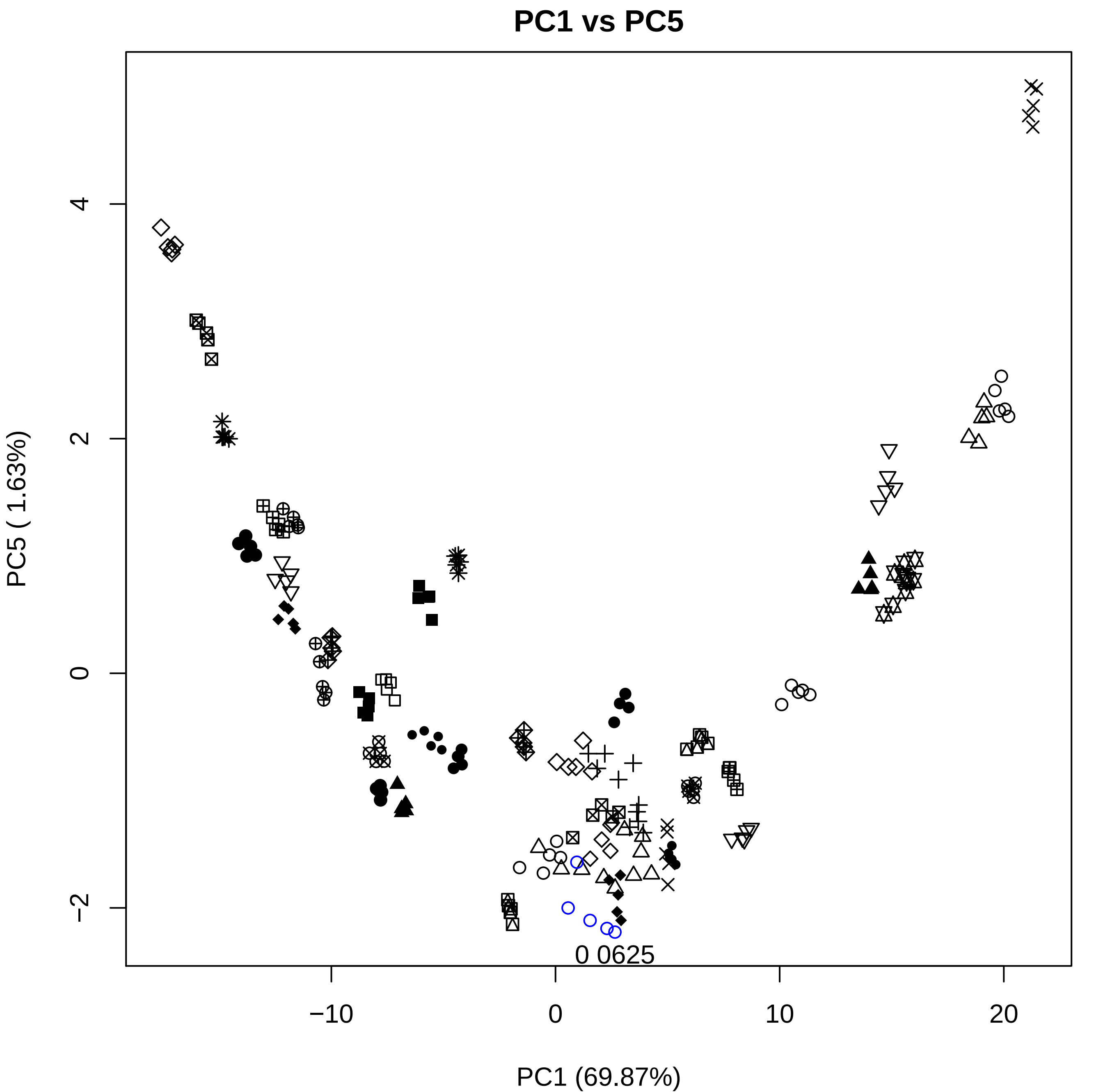
PC1 vs PC5
PC5 ( 1.63%)
PC1 (69.87%)
4
2
0
−2
−10
0
10
20
0 0625

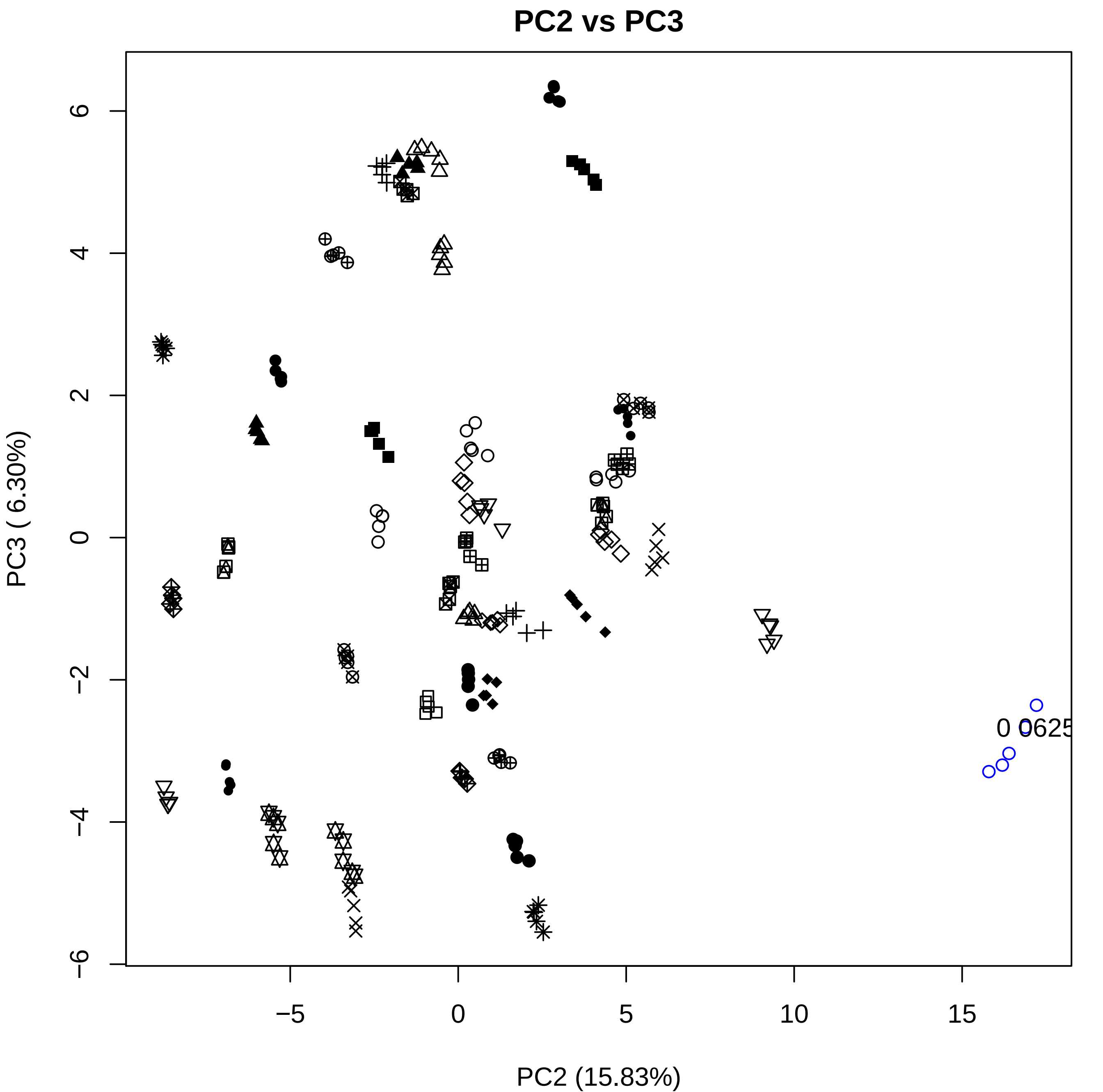
PC2 vs PC3
PC3 ( 6.30%)
PC2 (15.83%)
6
4
2
0
−2
−4
−6
−5
0
5
10
15
0 0625

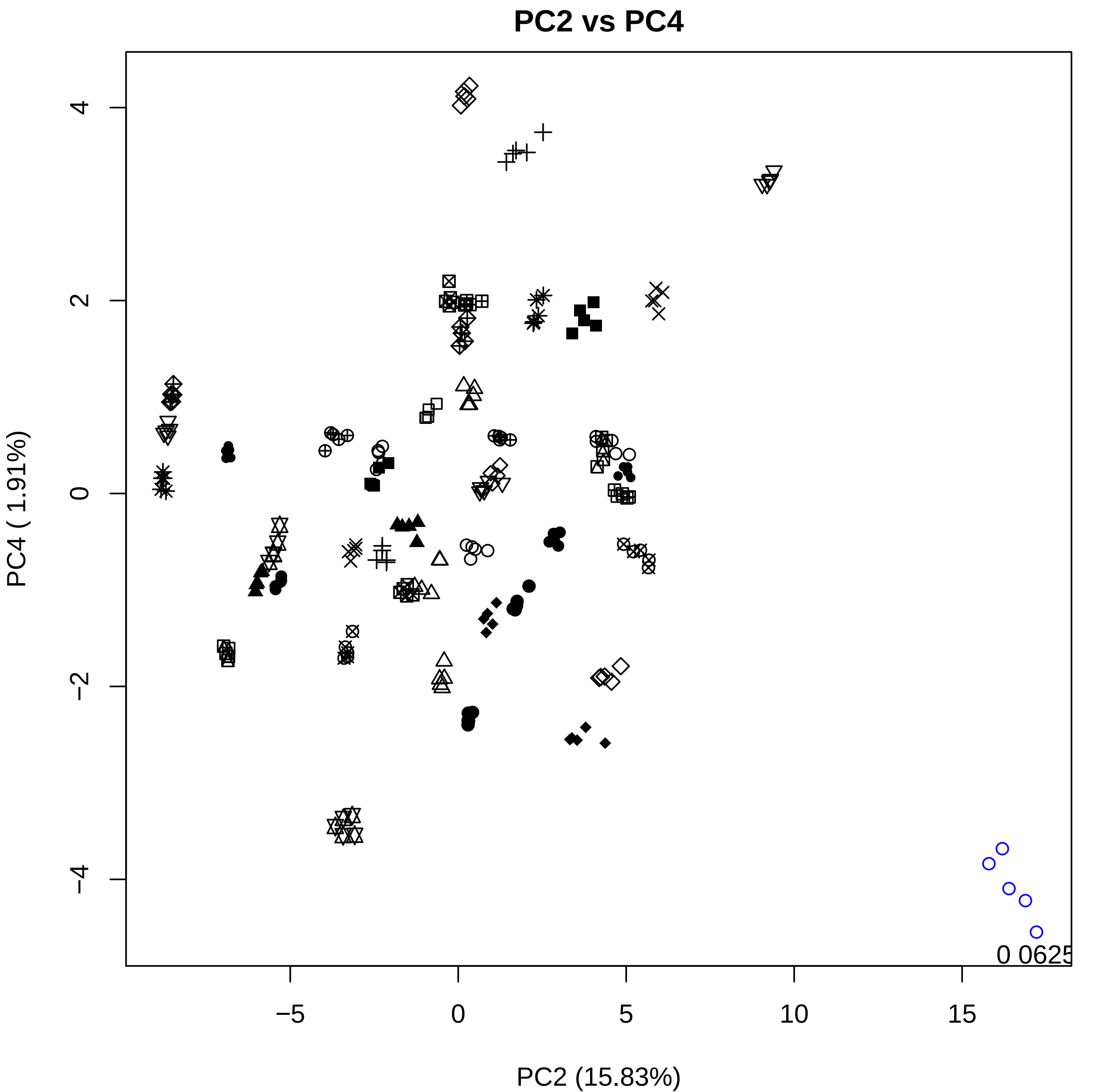
PC2 vs PC4
PC4 ( 1.91%)
PC2 (15.83%)
4
2
0
−2
−4
−5
0
5
10
15
0 0625

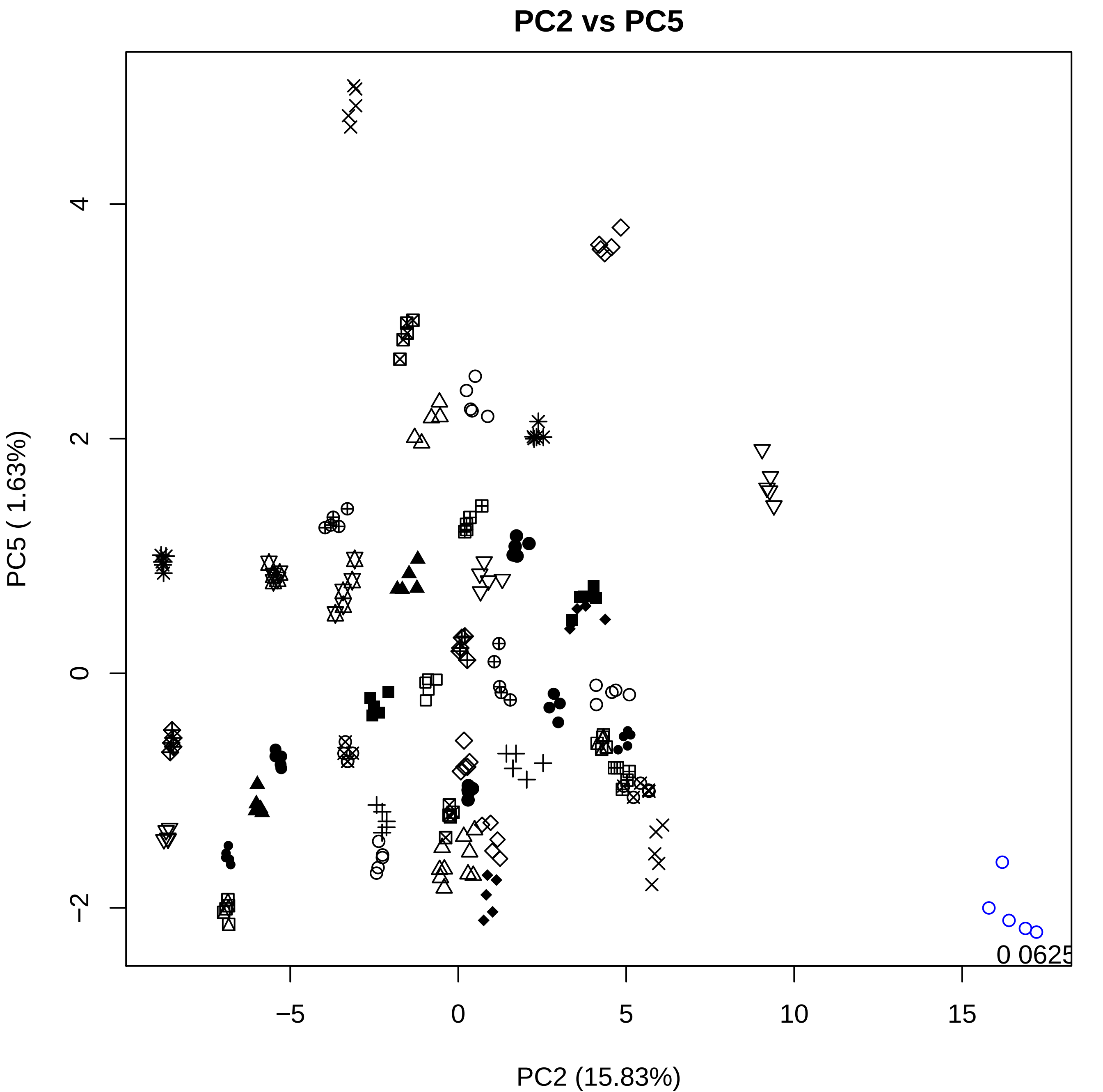
PC2 vs PC5
PC5 ( 1.63%)
PC2 (15.83%)
4
2
0
−2
−5
0
5
10
15
0 0625

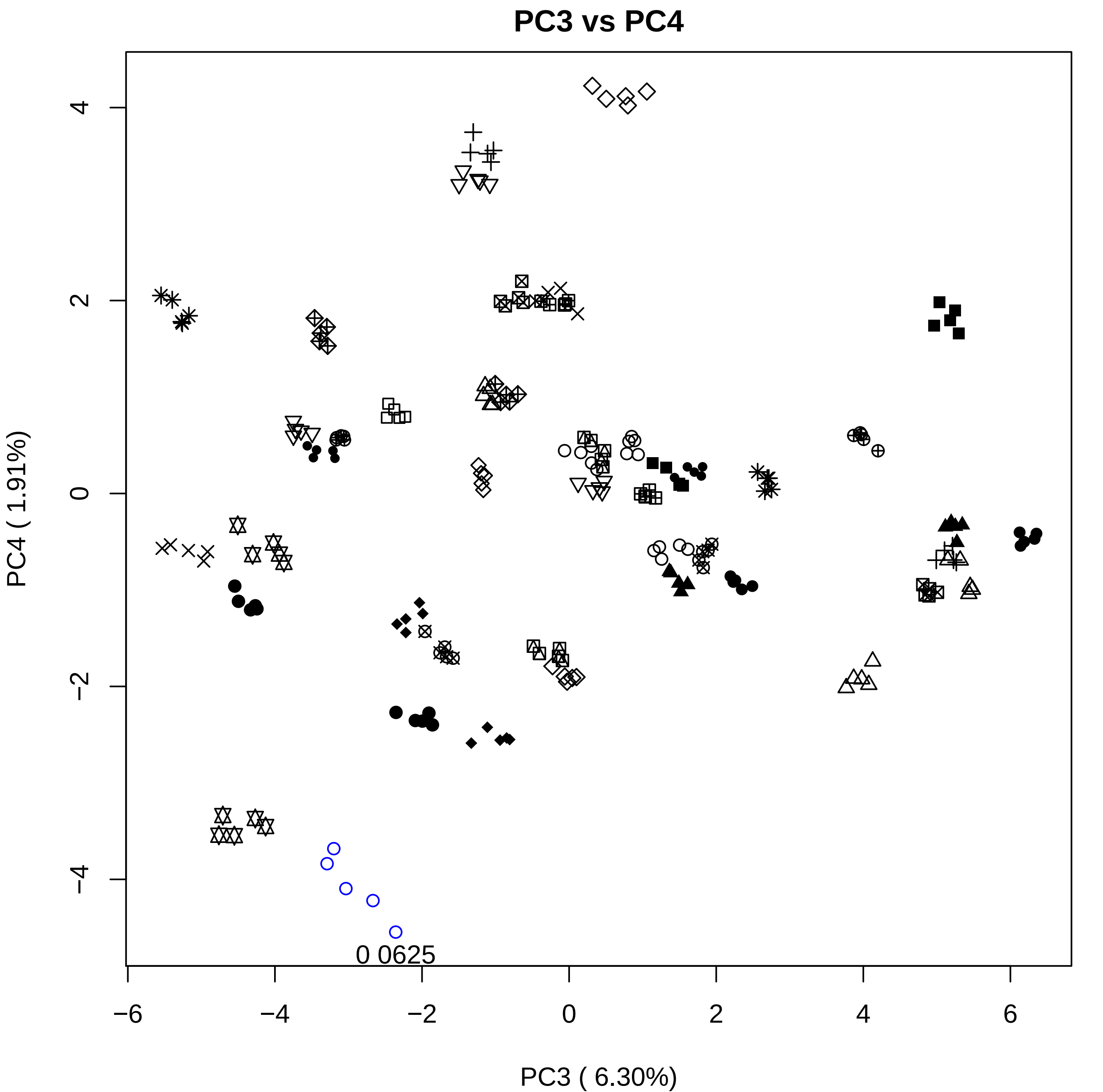

PC3 vs PC4
4
2
0
−2
−4
PC4 ( 1.91%)
0 0625
−6
−4
−2
0
2
4
6
PC3 ( 6.30%)

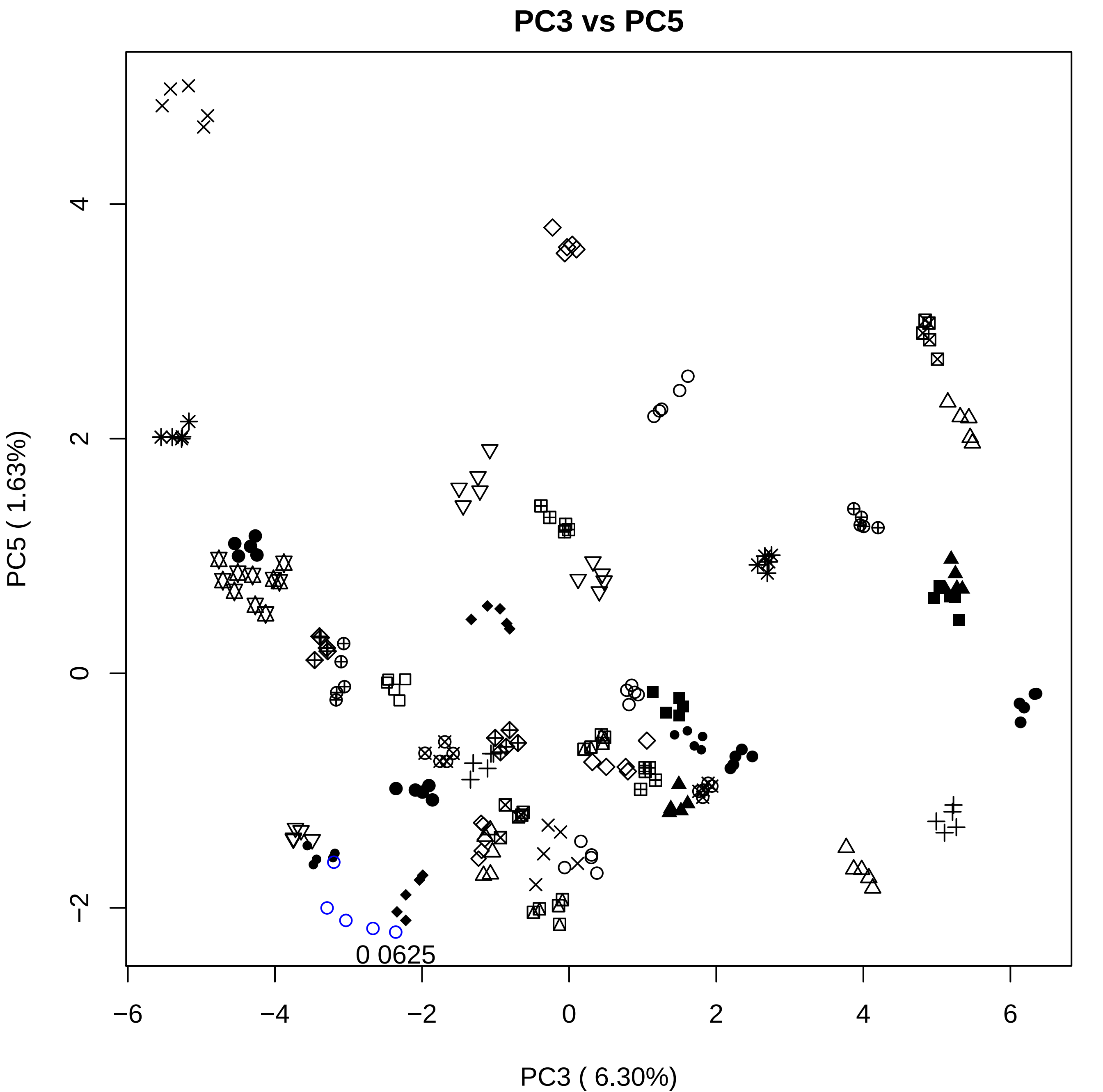

PC3 vs PC5
PC5 ( 1.63%)
PC3 ( 6.30%)
−6
−4
−2
0
2
4
6
4
2
0
−2
0 0625

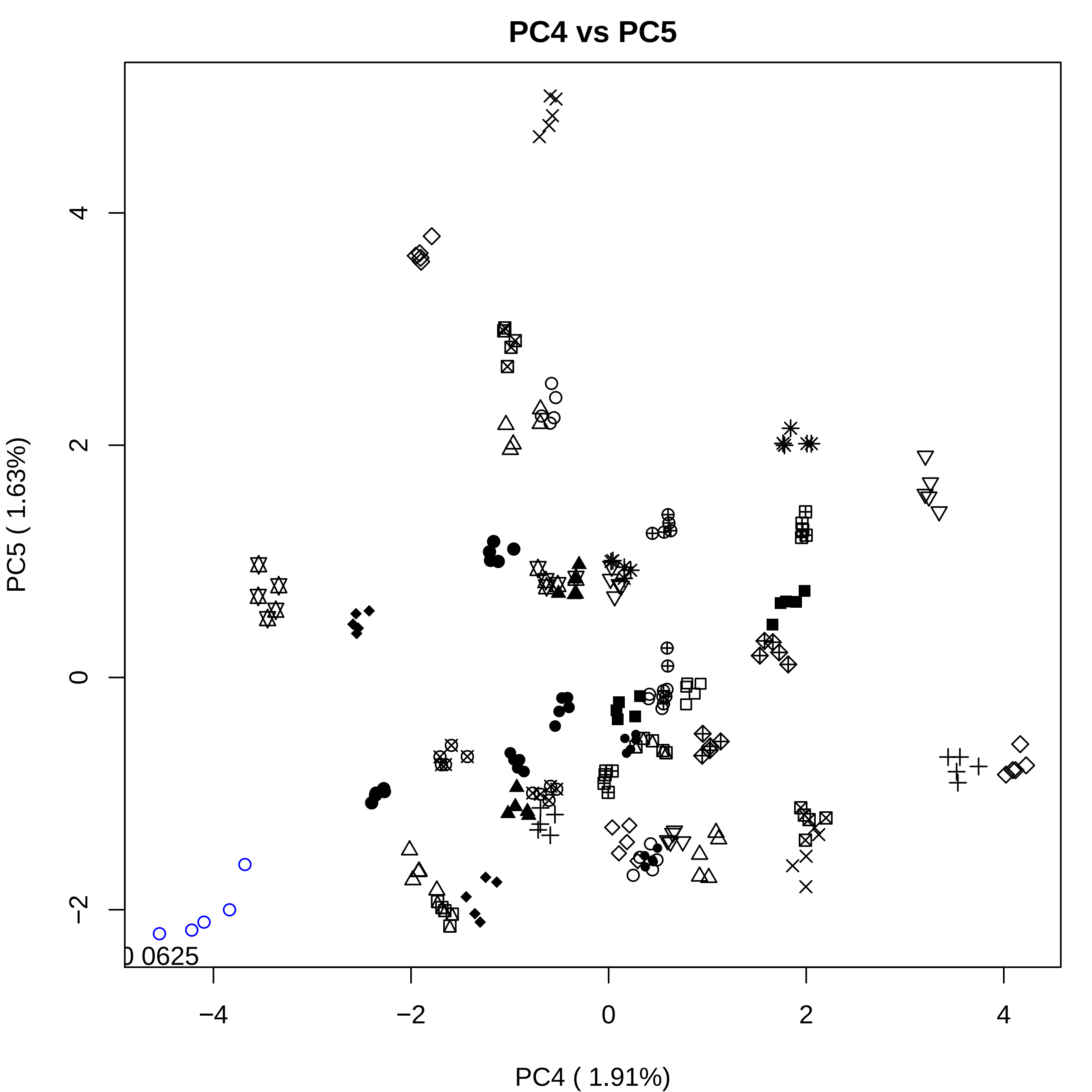

PC4 vs PC5
PC5 ( 1.63%)
PC4 ( 1.91%)
4
2
0
−2
−4
−2
0
2
4
0 0625

### PCA of 0.12%  DeS vs PMH

Scaled Matrix PCAed

7th order polynomial baseline correction performed

Raw spectra smoothed to 21 Neighbours, 2nd order PN, Savtisky Golay

Normalisation between 0 and 100

D2 spectra smoothed to 41 Neighbours, 2nd order PN, Savtisky Golay

Regions Analysed: 3601−2501 and 2001−700cm−1

Samples Analysed: 1 Contam's 0 DeS's 45 PMH's

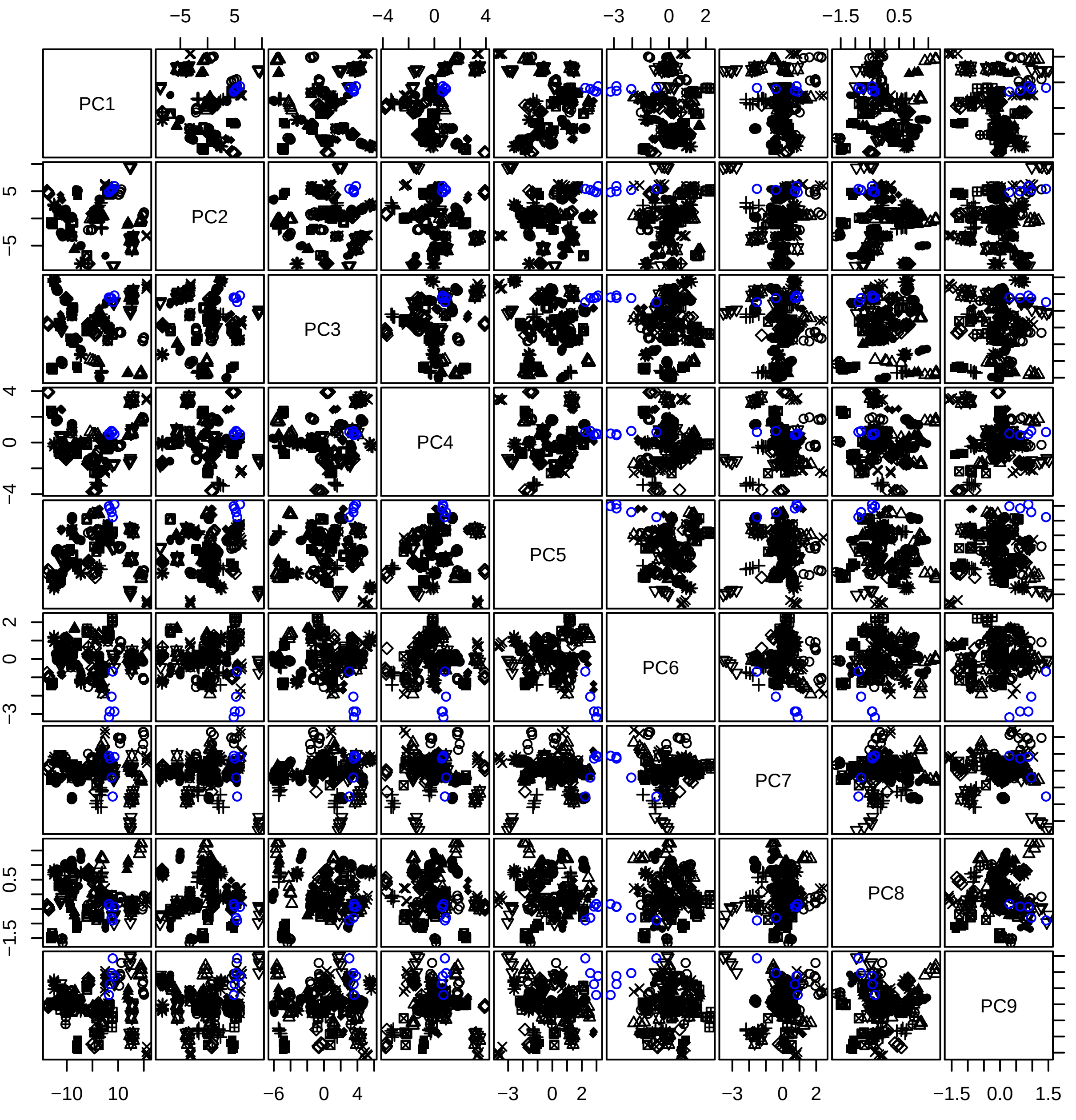

PC1
PC2
PC3
PC4
PC5
PC6
PC7
PC8
PC9
−5
5
−4
0
4
−3
0
2
−1.5
0.5
10
−10
5
−5
4
0
−6
4
0
−4
2
0
−3
2
0
−3
0.5
−1.5
1.5
0.0
−1.5
−10
10
−6
0
4
−3
0
2
−3
0
2
−1.5
0.0
1.5

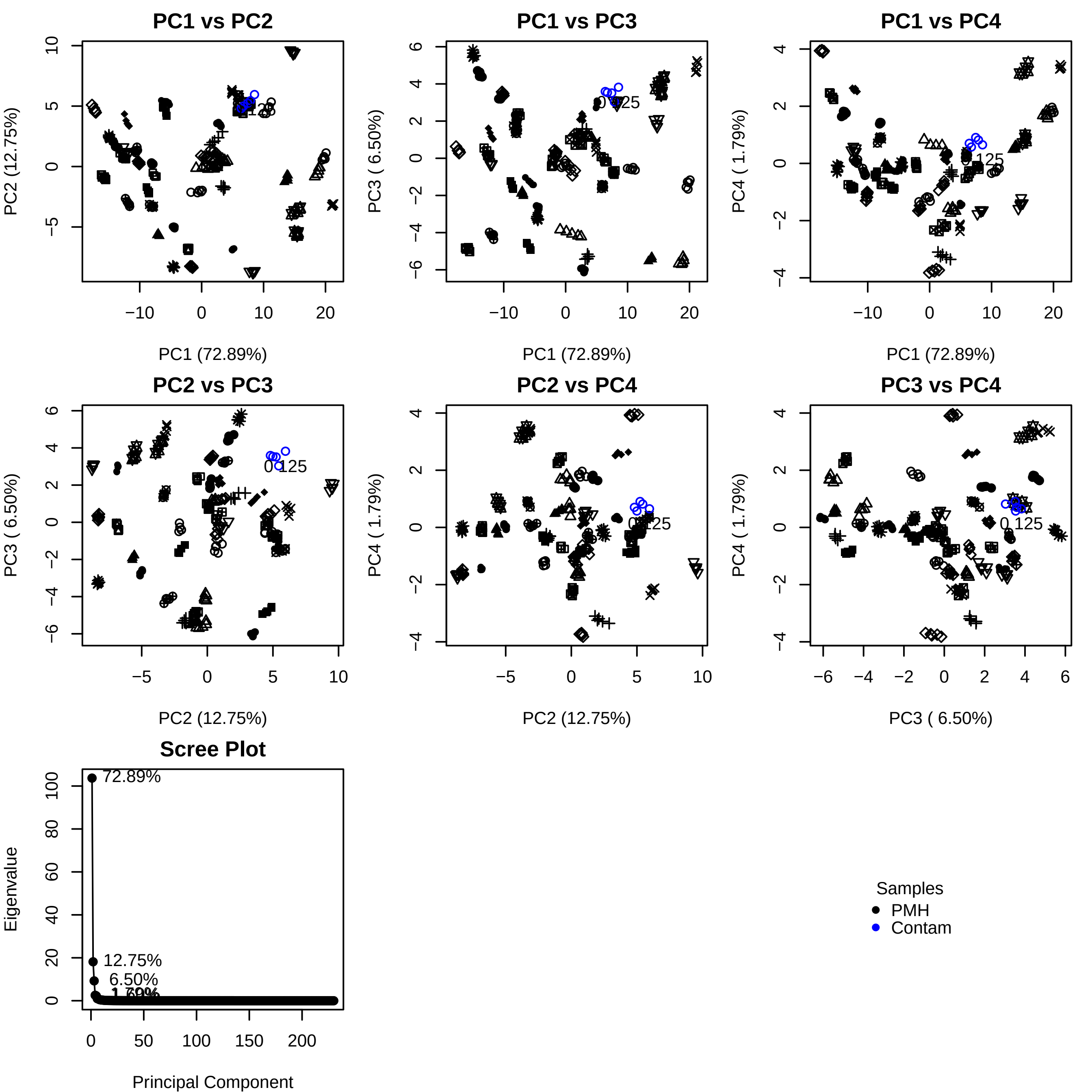
PC1 vs PC2
PC1 vs PC3
PC1 vs PC4
PC2 vs PC3
PC2 vs PC4
PC3 vs PC4
Scree Plot
PC1 (72.89%)
PC2 (12.75%)
PC3 ( 6.50%)
PC4 ( 1.79%)
72.89%
12.75%
6.50%
Eigenvalue
Principal Component
Samples
PMH
Contam

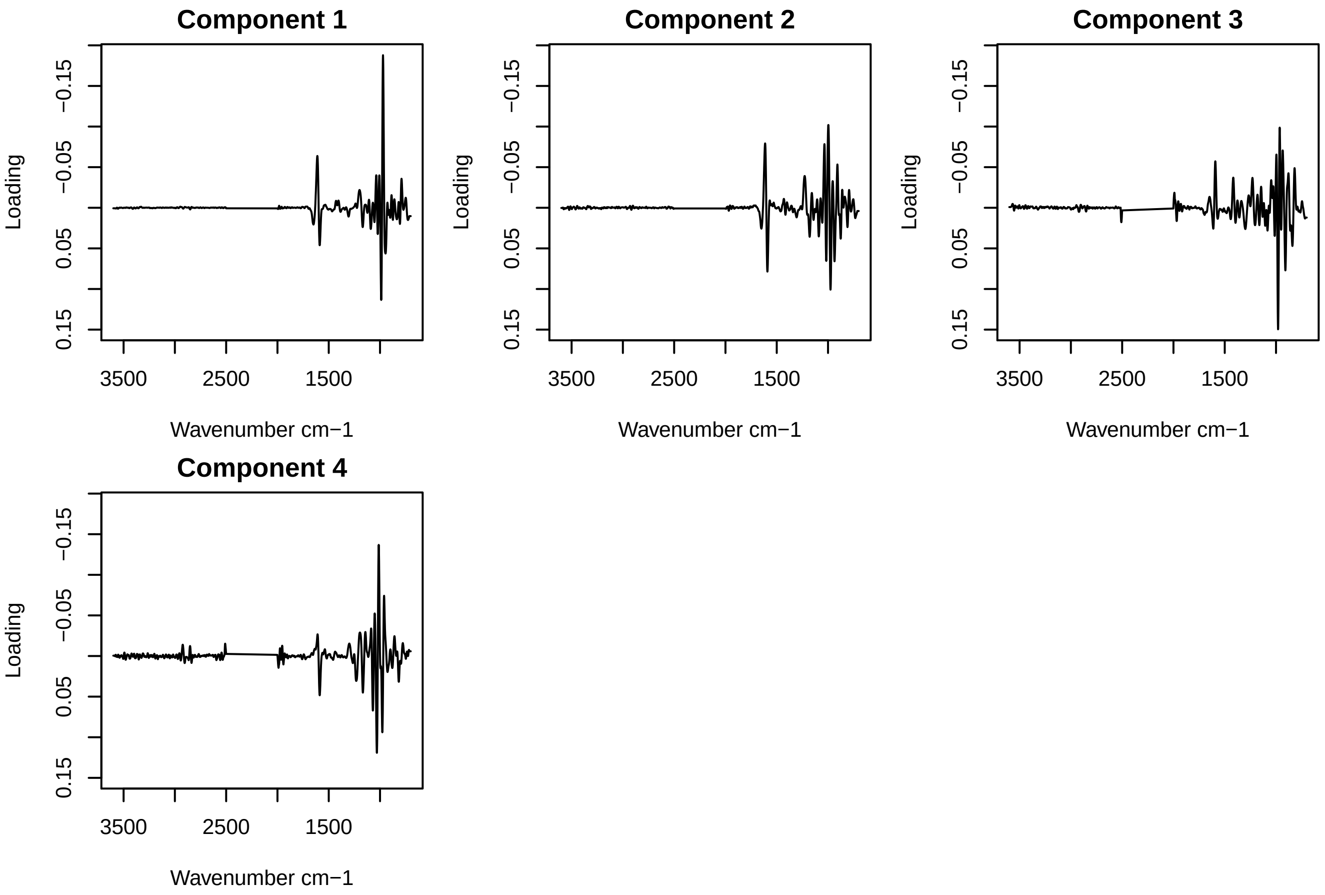

Component 1
Loading
−0.15
−0.05
0.05
0.15
3500
2500
1500
Wavenumber cm−1
Component 2
Loading
−0.15
−0.05
0.05
0.15
3500
2500
1500
Wavenumber cm−1
Component 3
Loading
−0.15
−0.05
0.05
0.15
3500
2500
1500
Wavenumber cm−1
Component 4
Loading
−0.15
−0.05
0.05
0.15
3500
2500
1500
Wavenumber cm−1

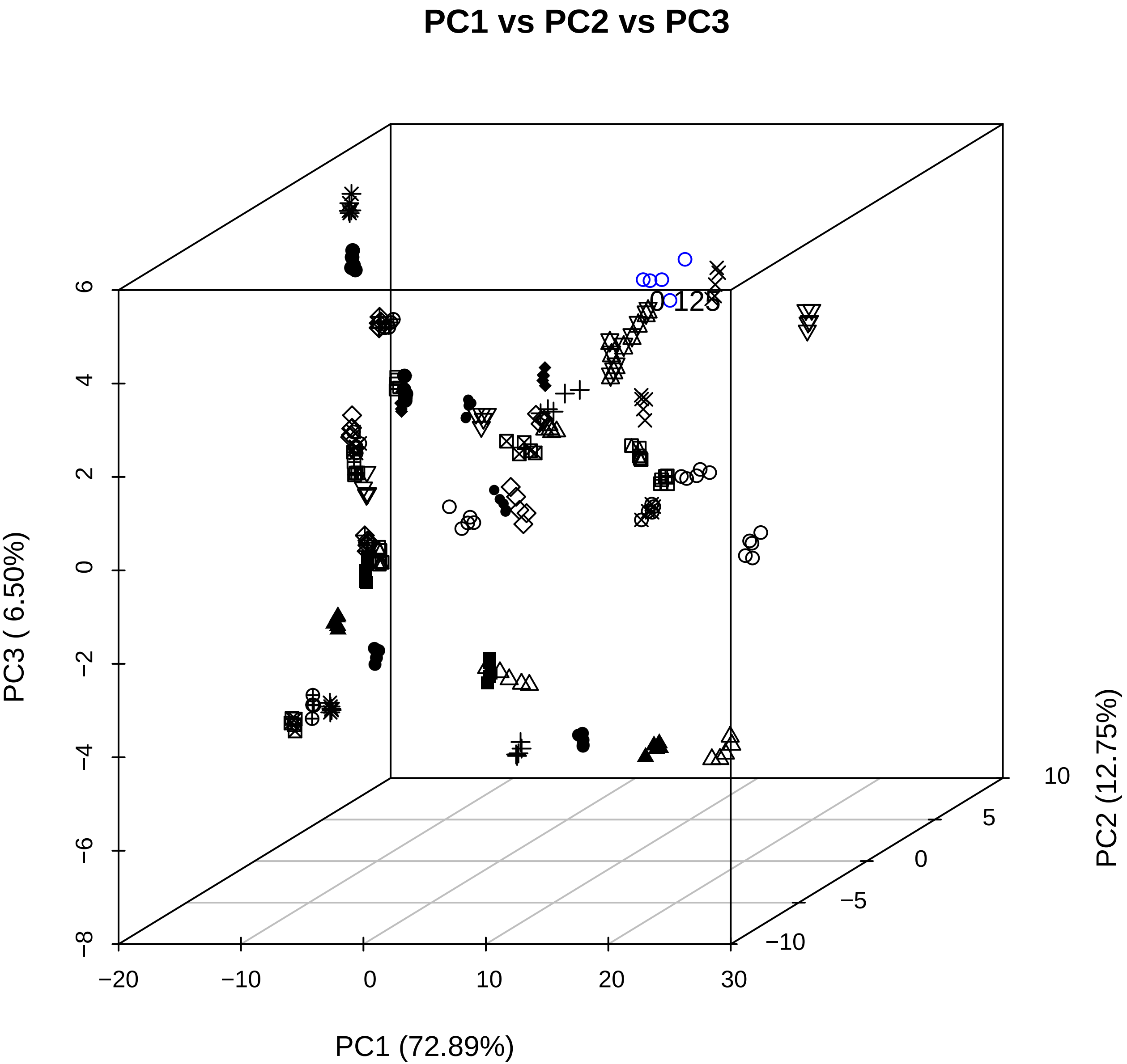
PC1 vs PC2 vs PC3
PC1 (72.89%)
PC2 (12.75%)
PC3 ( 6.50%)
−20
−10
0
10
20
30
−10
−5
0
5
10
−8
−6
−4
−2
0
2
4
6
0125

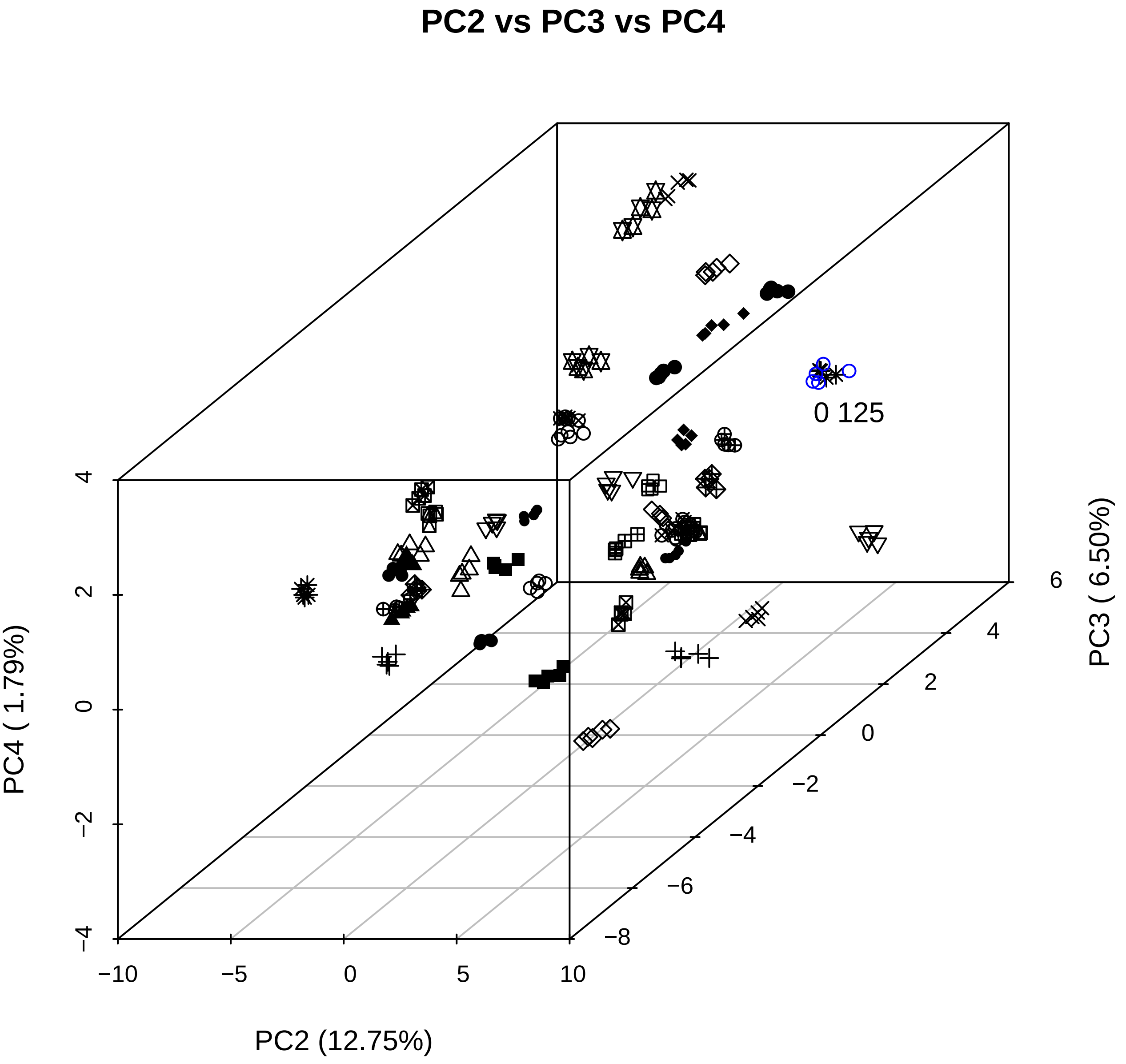
PC2 vs PC3 vs PC4
0 125
PC2 (12.75%)
PC3 ( 6.50%)
PC4 ( 1.79%)
−10
−5
0
5
10
−8
−6
−4
−2
0
2
4
6
−4
−2
0
2
4

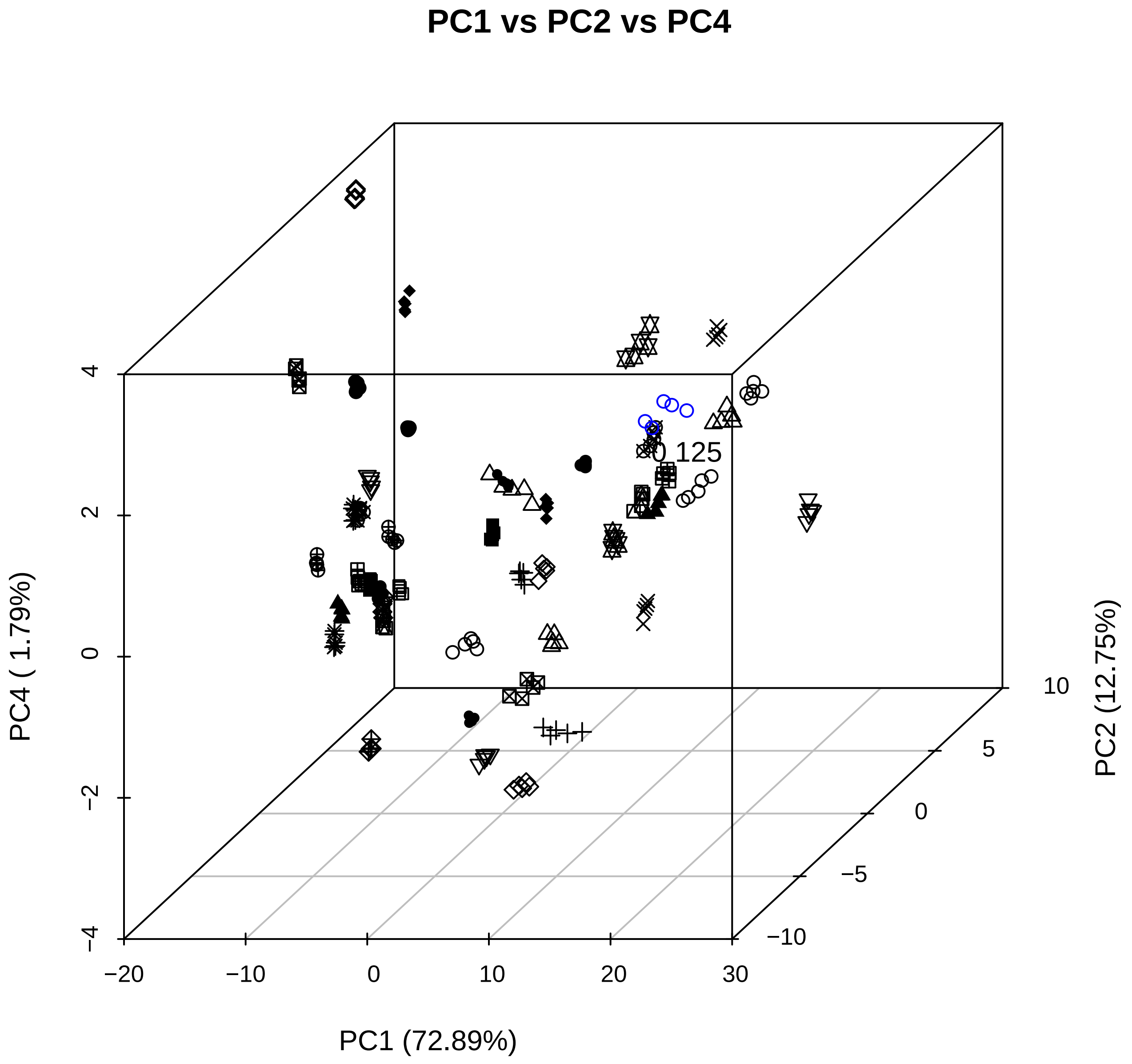

PC1 vs PC2 vs PC4
0.125
PC1 (72.89%)
PC2 (12.75%)
PC4 ( 1.79%)
−20
−10
0
10
20
30
−10
−5
0
5
10
−4
−2
0
2
4

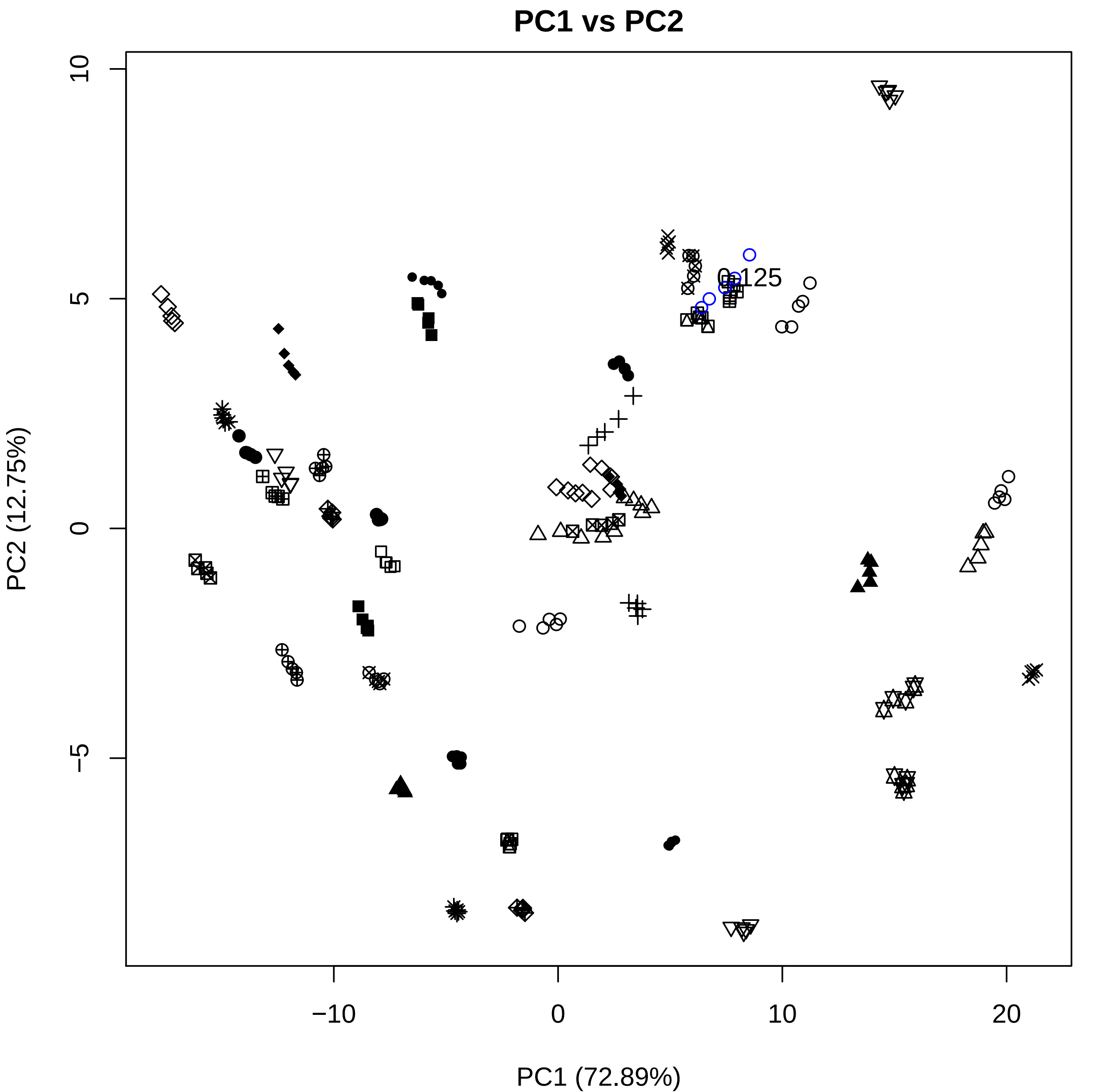
PC1 vs PC2
PC2 (12.75%)
PC1 (72.89%)
10
5
0
−5
−10
0
10
20
0.125

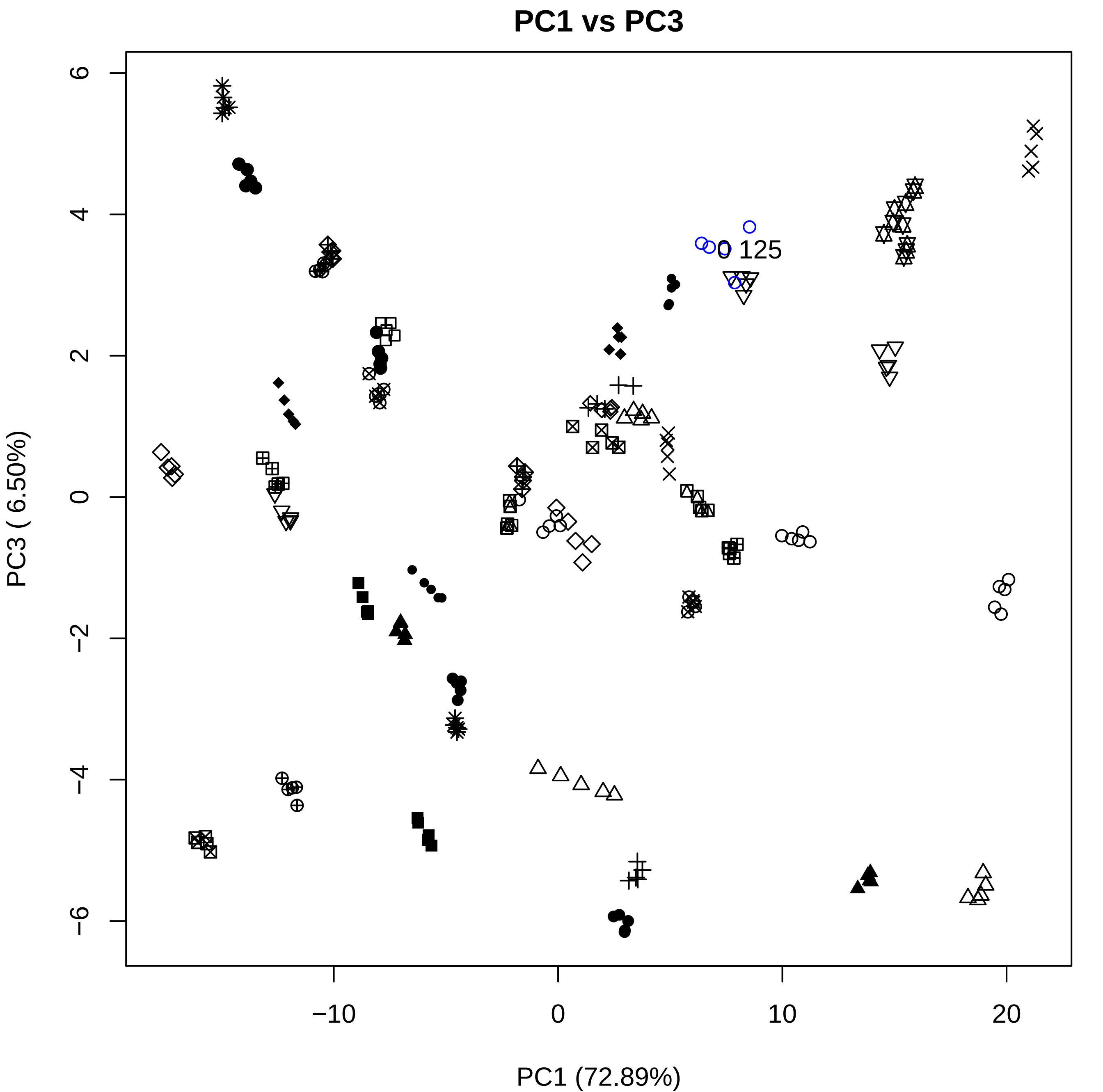
PC1 vs PC3
0 125
PC3 ( 6.50%)
PC1 (72.89%)
6
4
2
0
−2
−4
−6
−10
0
10
20

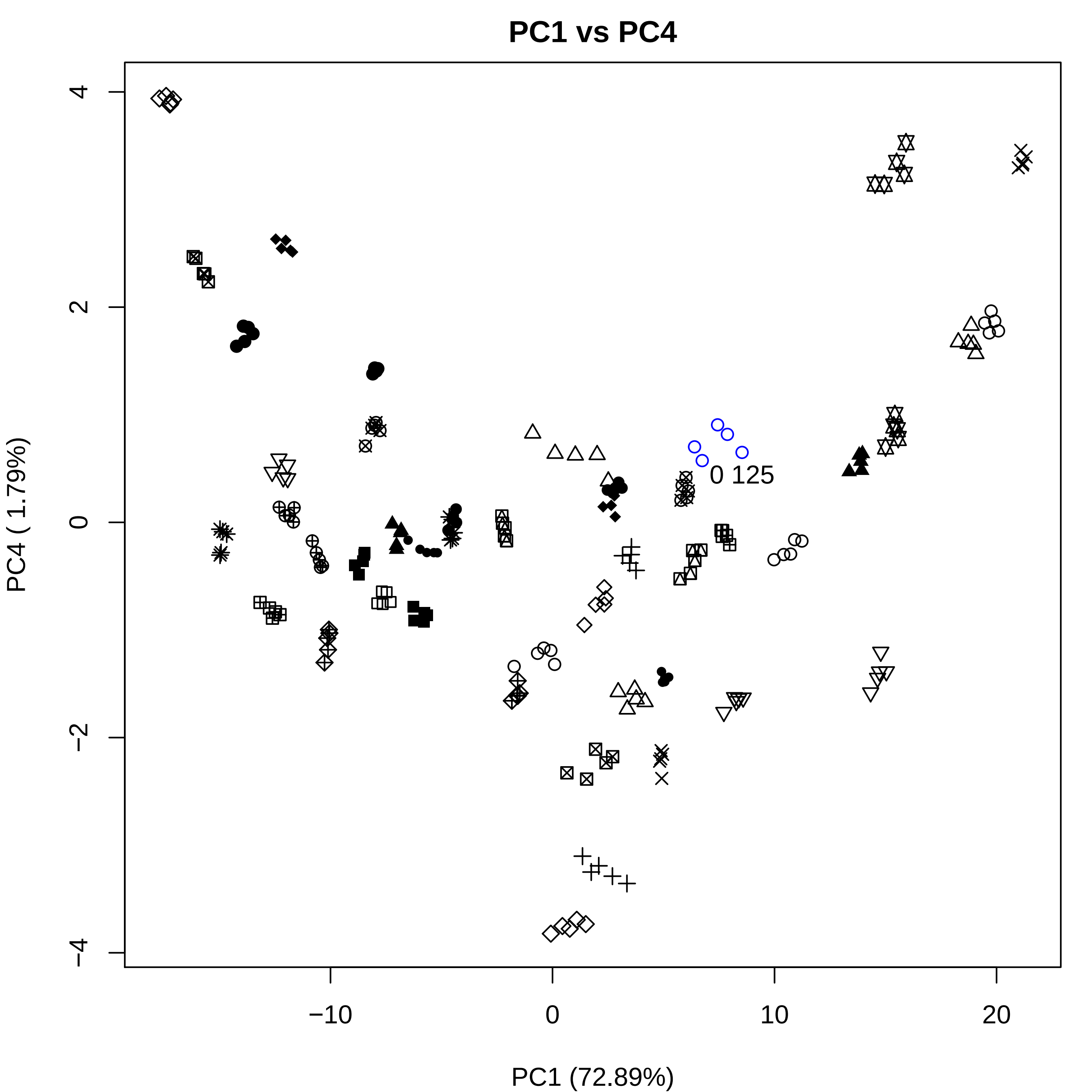
PC1 vs PC4
PC4 ( 1.79%)
PC1 (72.89%)
4
2
0
−2
−4
−10
0
10
20
0 125

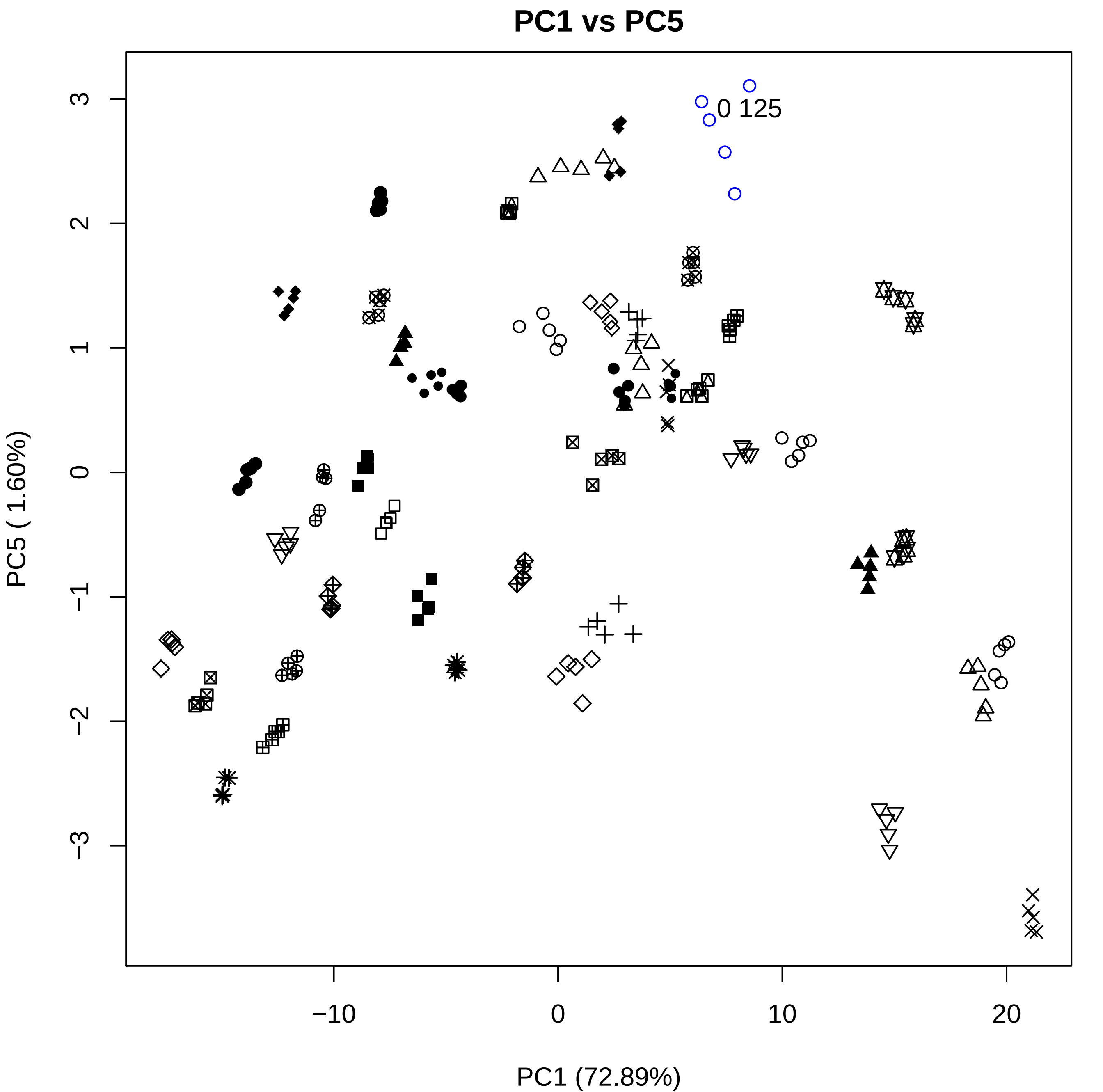
PC1 vs PC5
0 125
PC5 ( 1.60%)
PC1 (72.89%)
3
2
1
0
−1
−2
−3
−10
0
10
20

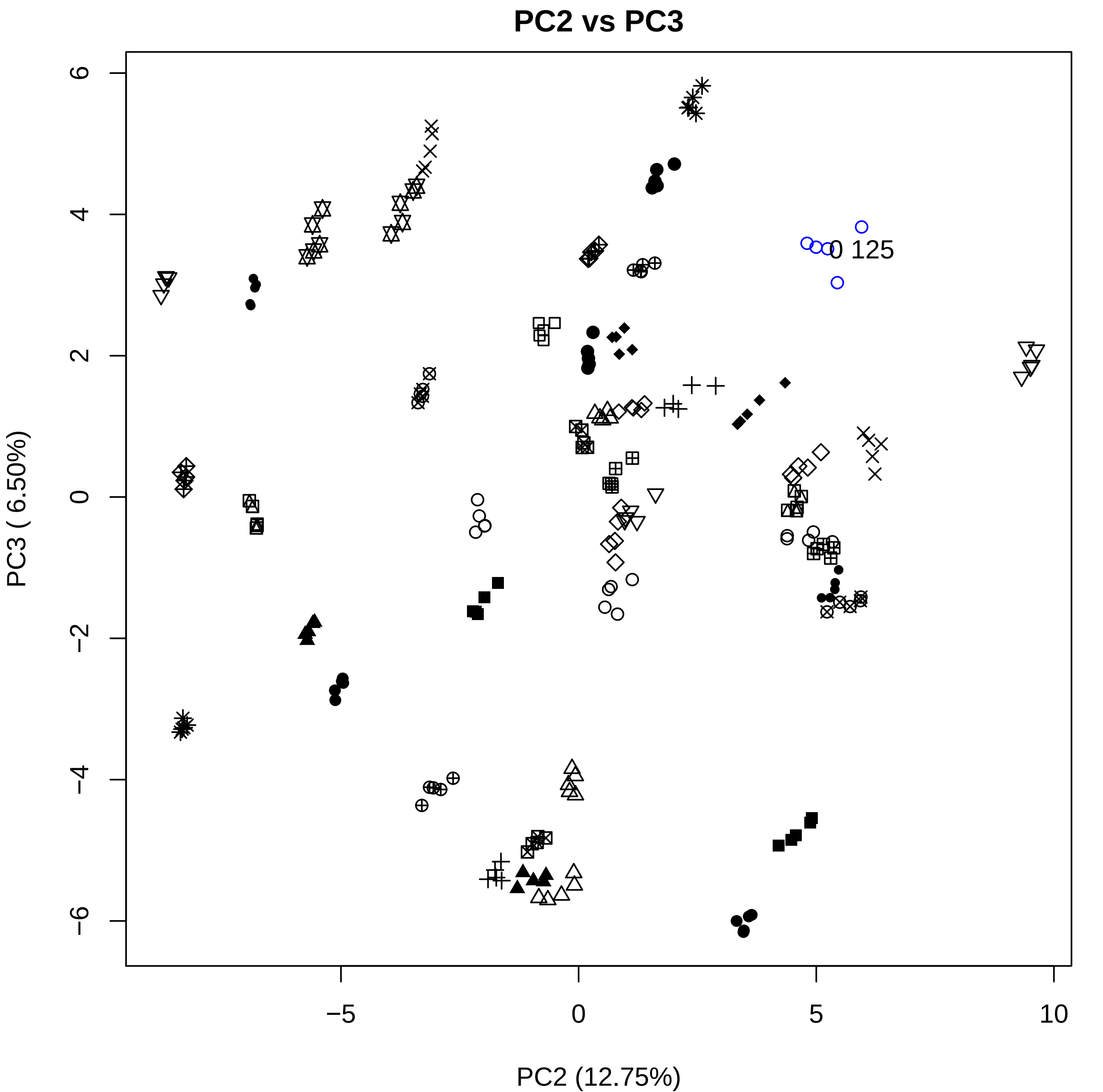
PC2 vs PC3
0 125
PC3 ( 6.50%)
PC2 (12.75%)
−6
−4
−2
0
2
4
6
−5
0
5
10

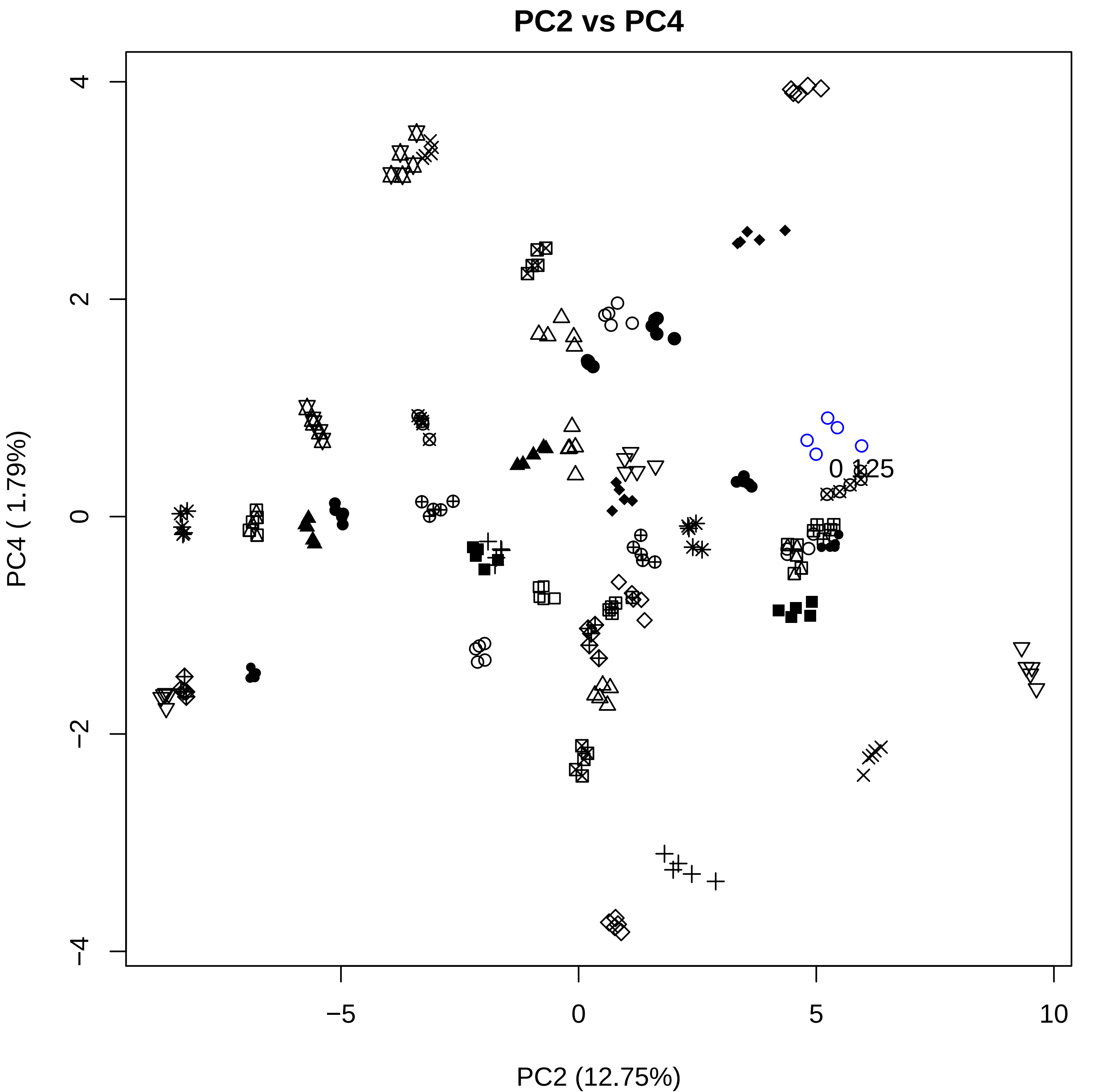
PC2 vs PC4
PC4 ( 1.79%)
PC2 (12.75%)
4
2
0
−2
−4
−5
0
5
10
0.125

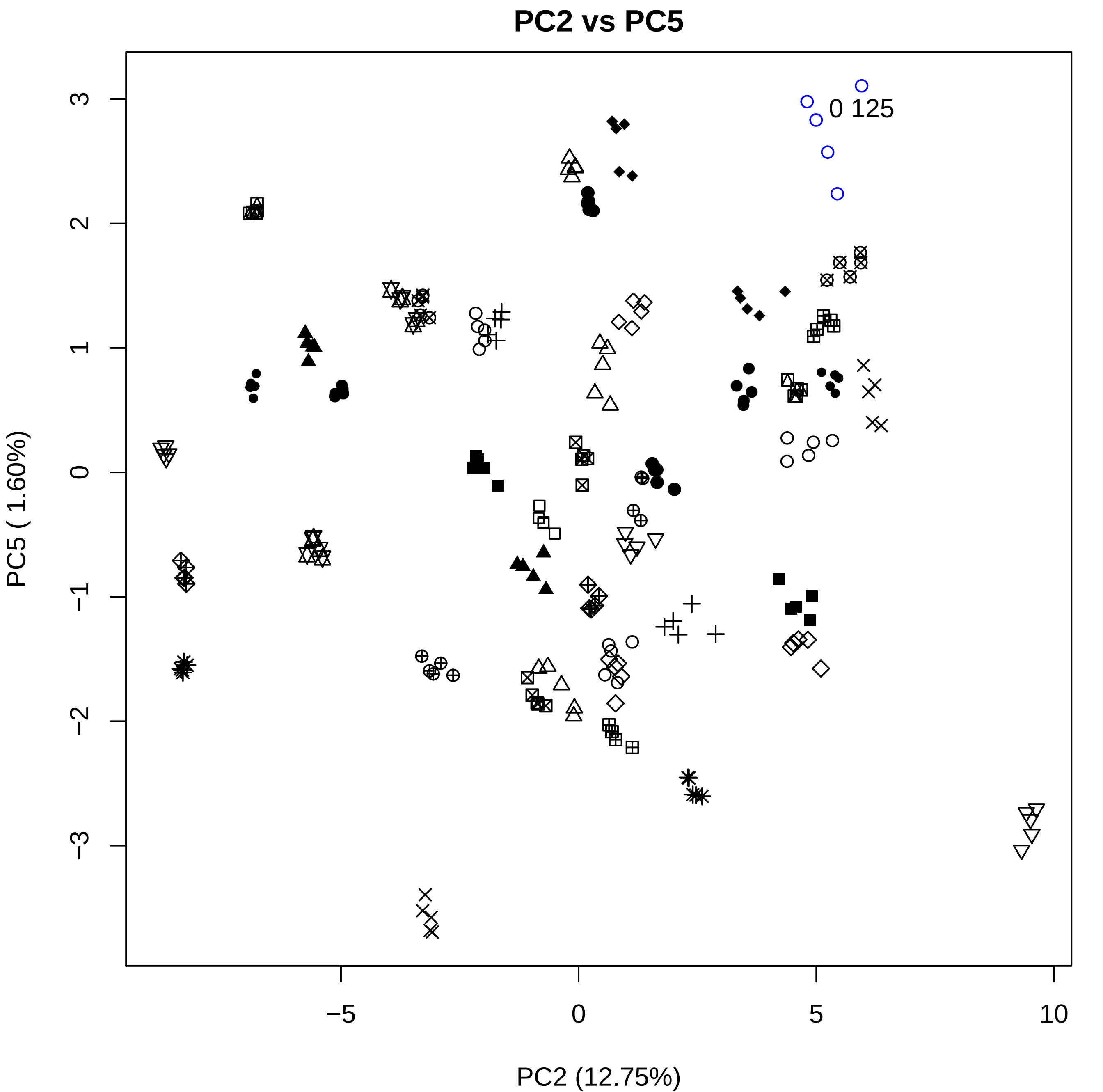
PC2 vs PC5
0 125
PC5 ( 1.60%)
PC2 (12.75%)
−5
0
5
10
−3
−2
−1
0
1
2
3

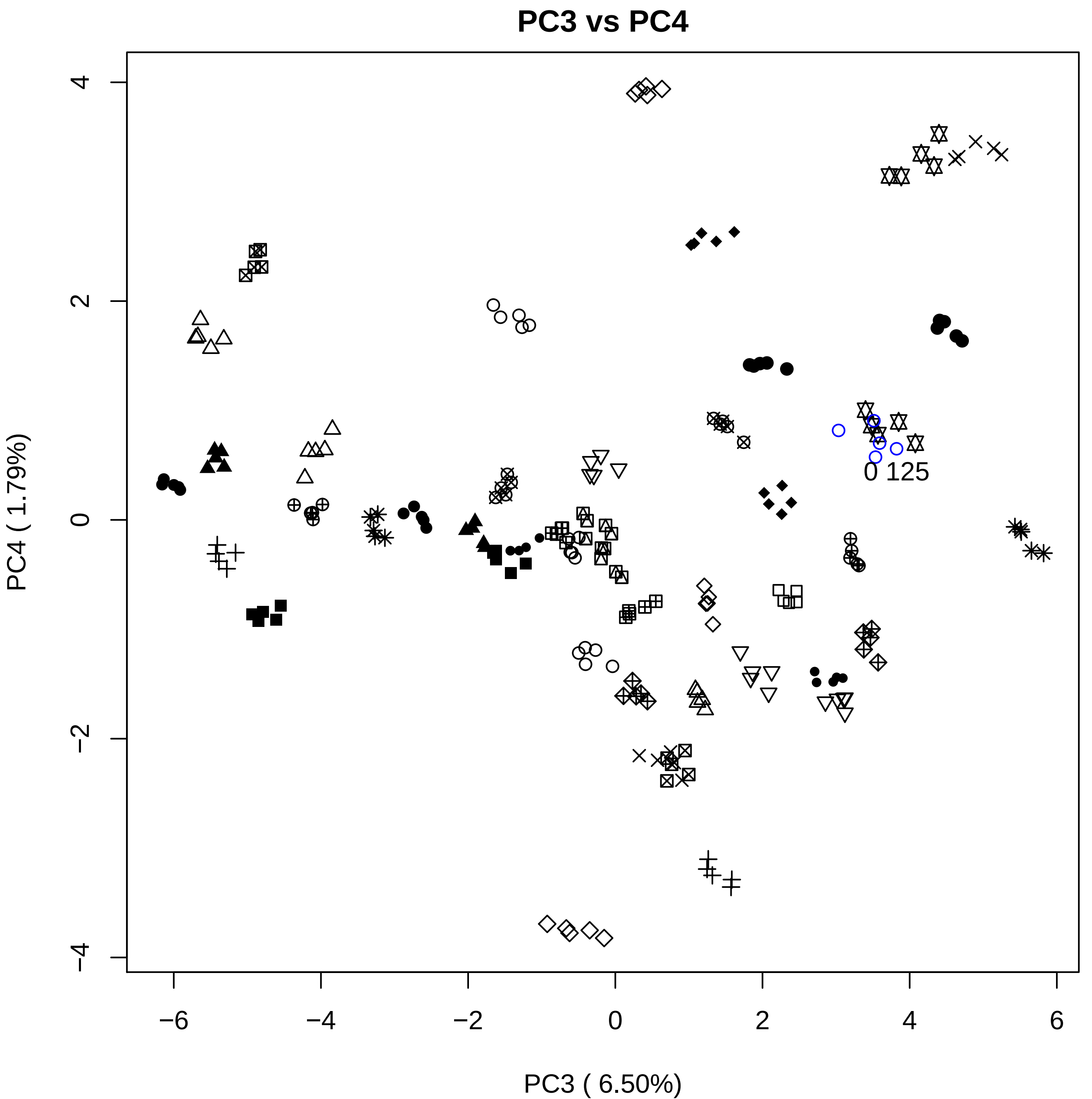
PC3 vs PC4
PC4 ( 1.79%)
PC3 ( 6.50%)
0 125
4
2
0
−2
−4
−6
−4
−2
0
2
4
6

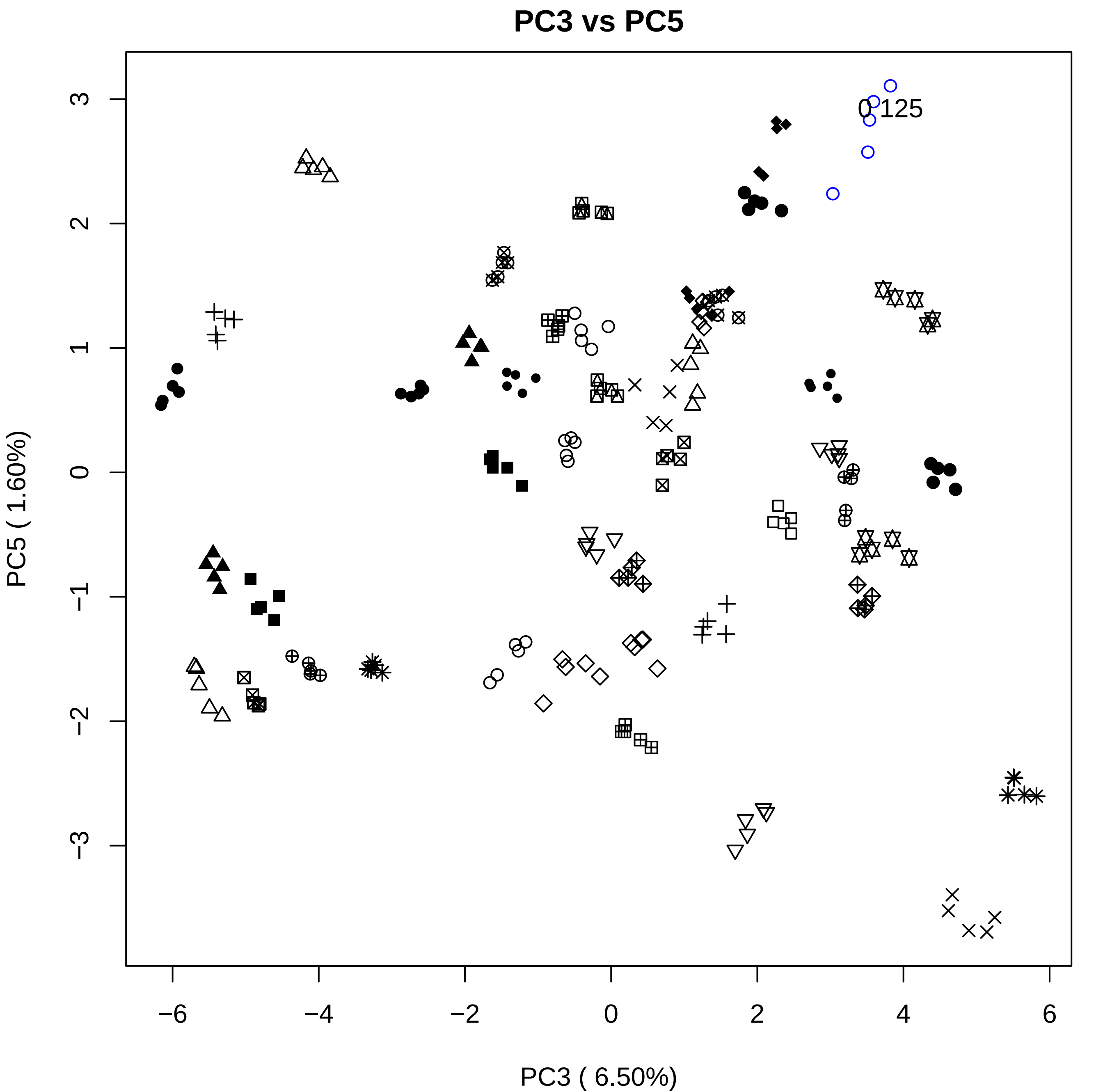
PC3 vs PC5
0.125
PC5 ( 1.60%)
PC3 ( 6.50%)
−6
−4
−2
0
2
4
6
3
2
1
0
−1
−2
−3

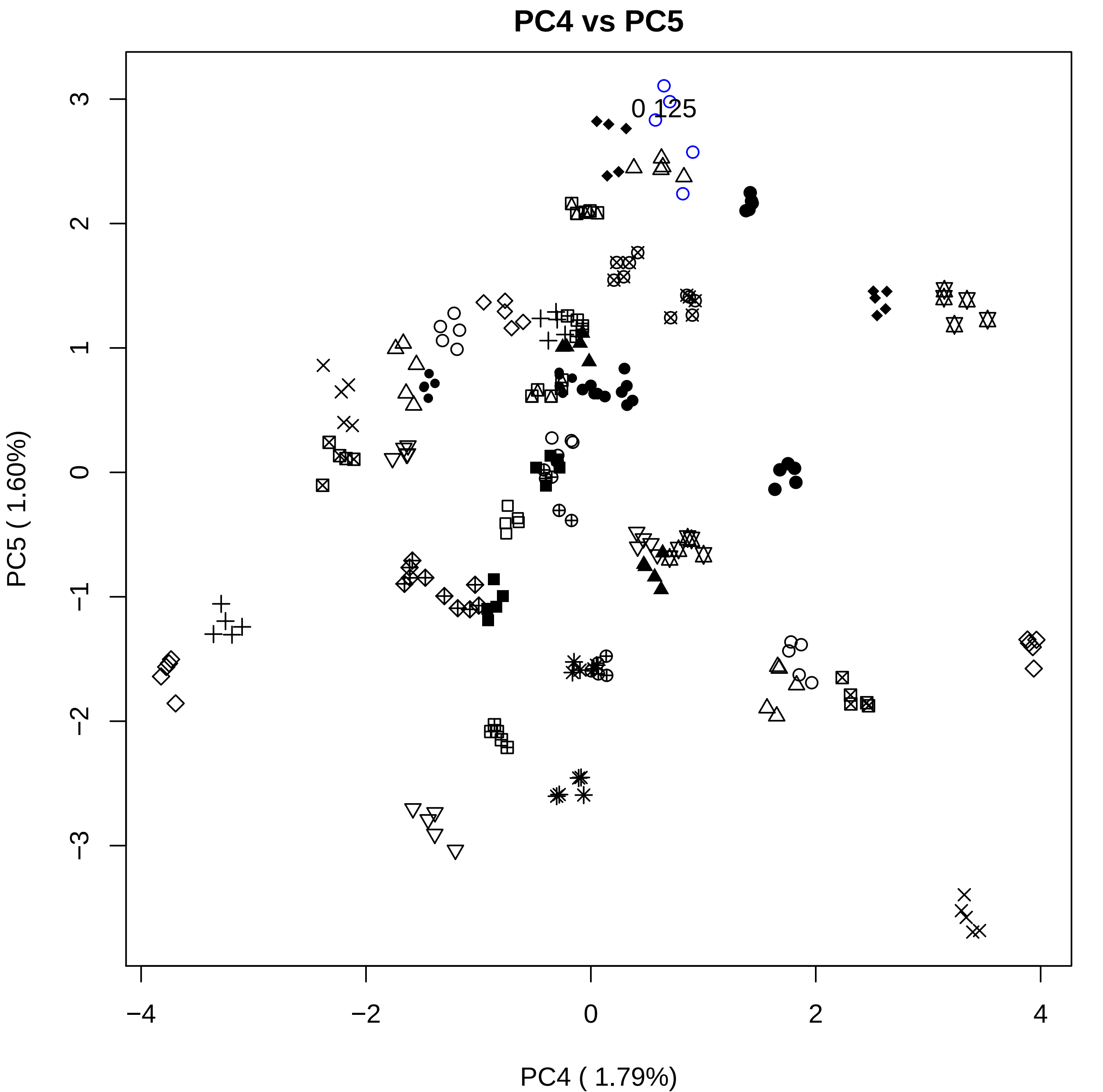
PC4 vs PC5
0.125
PC4 ( 1.79%)
PC5 ( 1.60%)
−4
−2
0
2
4
3
2
1
0
−1
−2
−3

### PCA of 0.25%  DeS vs PMH

Scaled Matrix PCAed

7th order polynomial baseline correction performed

Raw spectra smoothed to 21 Neighbours, 2nd order PN, Savtisky Golay

Normalisation between 0 and 100

D2 spectra smoothed to 41 Neighbours, 2nd order PN, Savtisky Golay

Regions Analysed: 3601−2501 and 2001−700cm−1

Samples Analysed: 1 Contam's 0 DeS's 45 PMH's

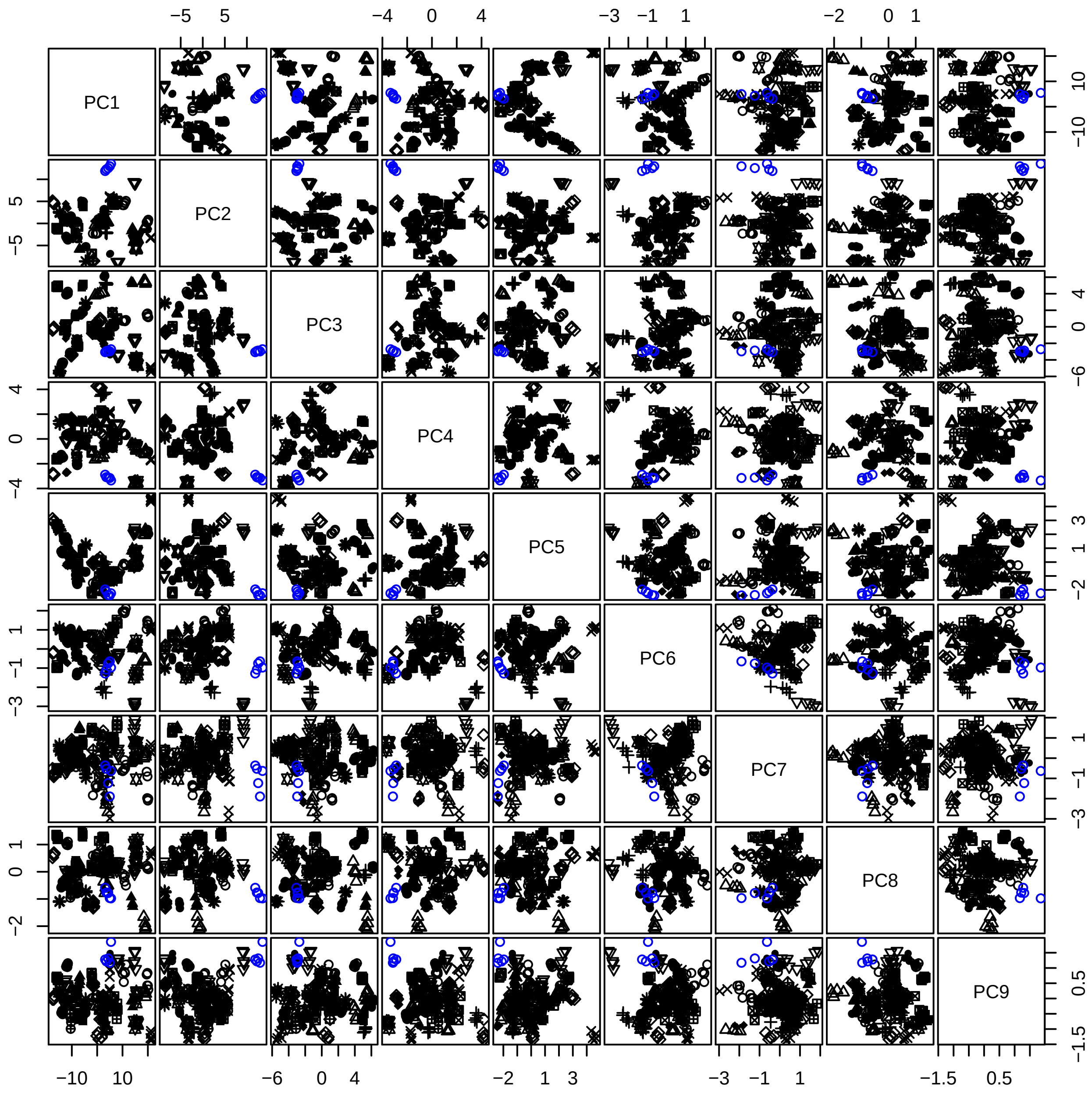
PC1
PC2
PC3
PC4
PC5
PC6
PC7
PC8
PC9

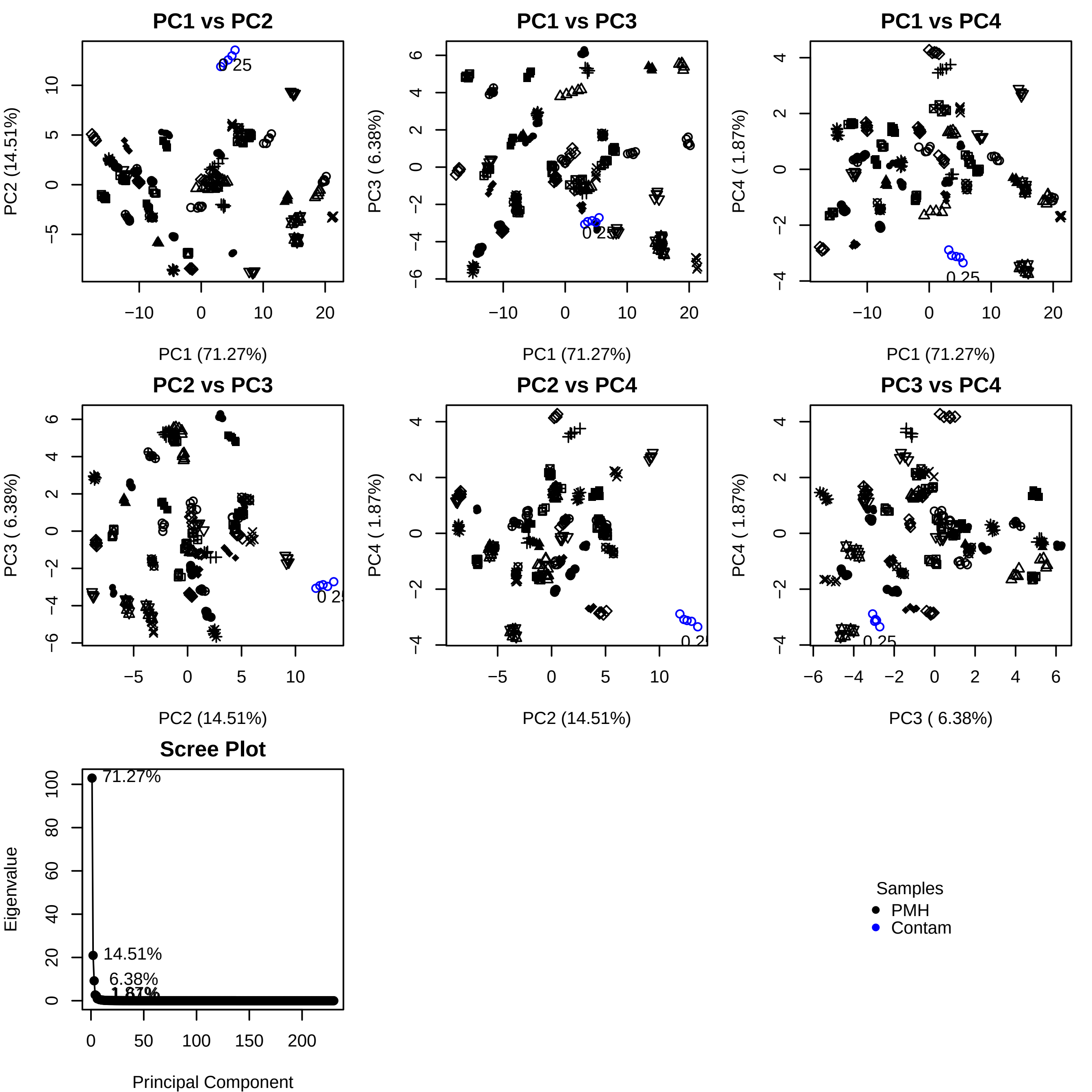
PC1 vs PC2
PC1 vs PC3
PC1 vs PC4
PC2 vs PC3
PC2 vs PC4
PC3 vs PC4
Scree Plot
PC1 (71.27%)
PC2 (14.51%)
PC3 ( 6.38%)
PC4 ( 1.87%)
Eigenvalue
Principal Component
71.27%
14.51%
6.38%
Samples
PMH
Contam

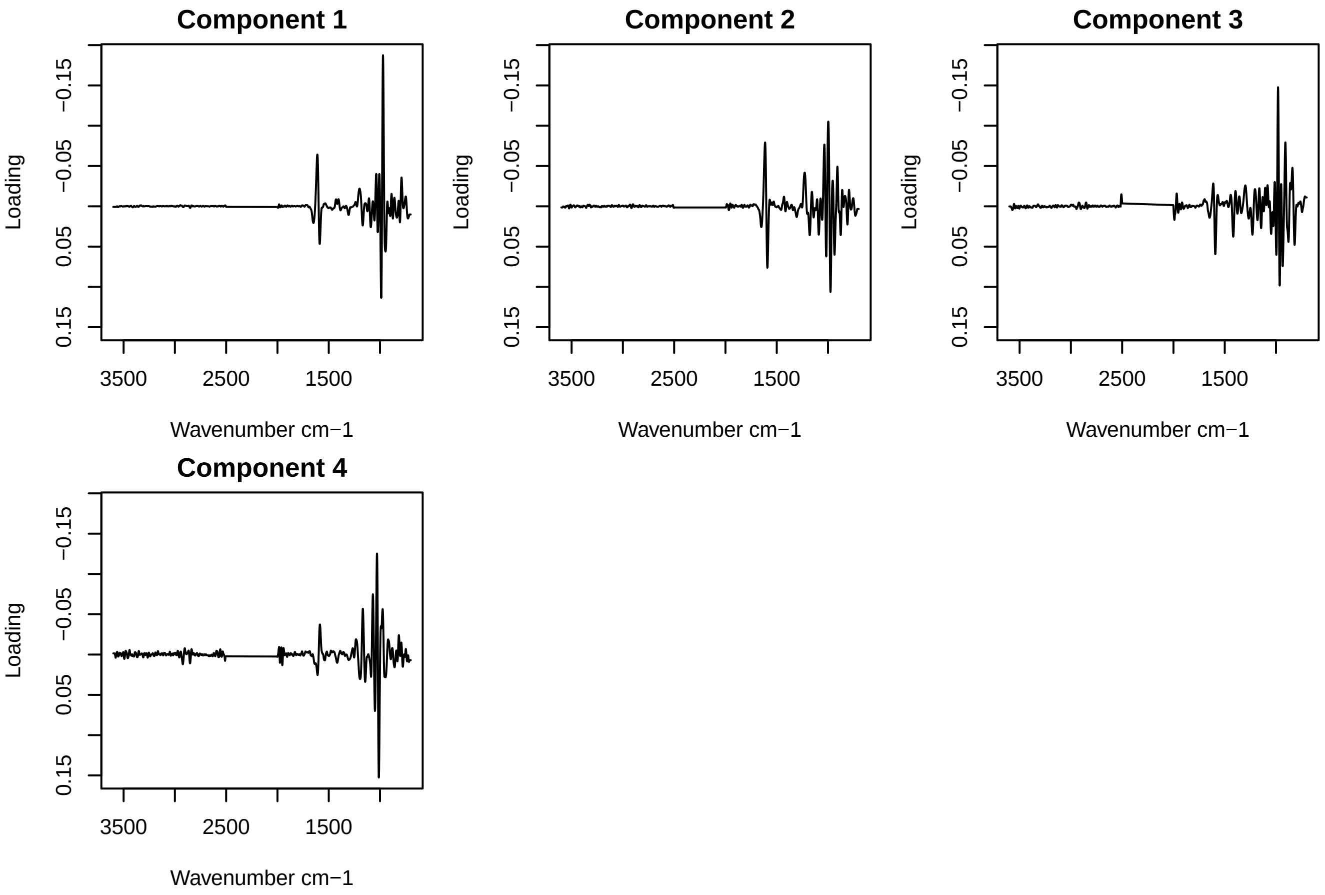
Component 1
Component 2
Component 3
Component 4
Loading
−0.15
−0.05
0.05
0.15
3500
2500
1500
Wavenumber cm−1

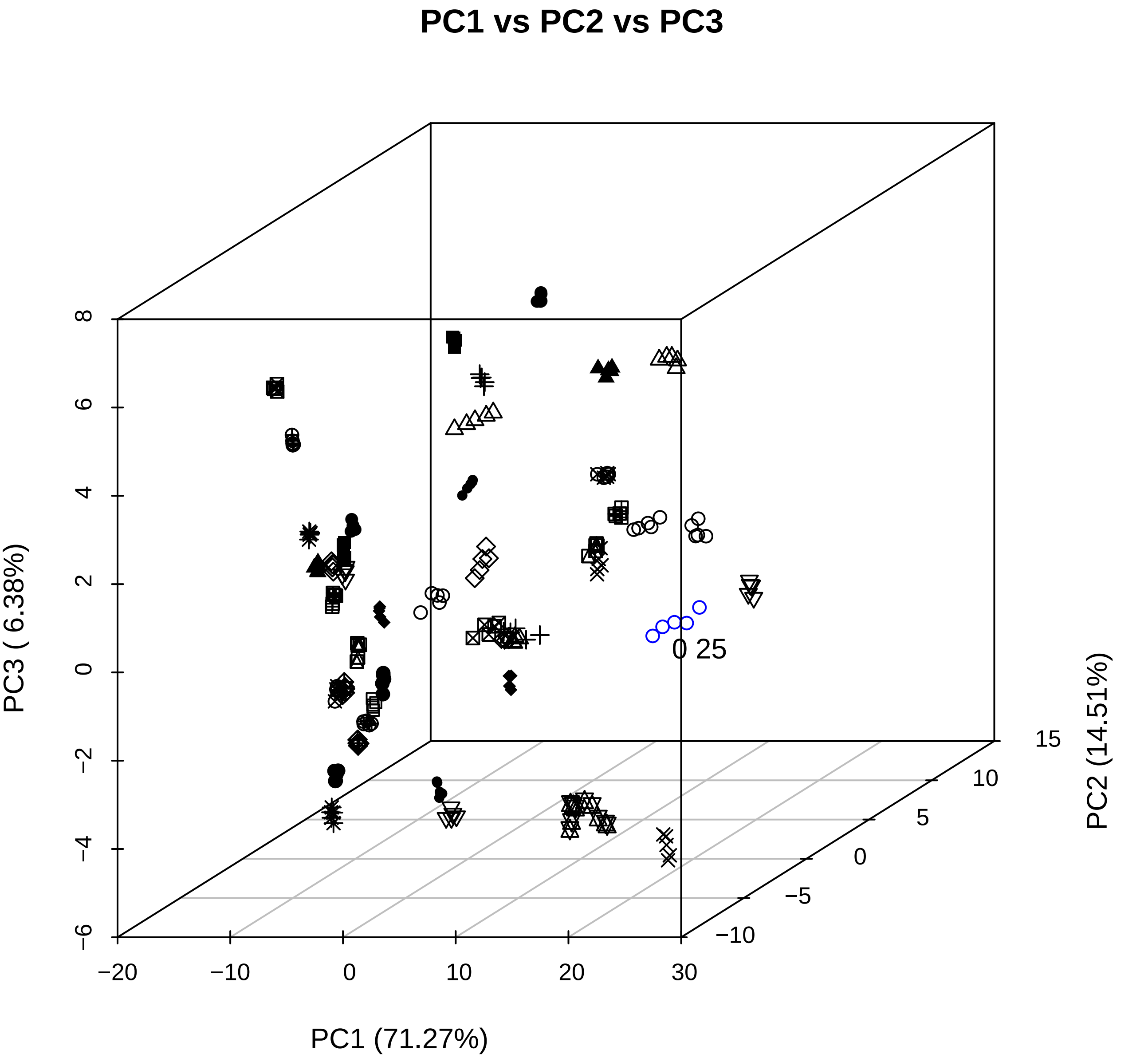

PC1 vs PC2 vs PC3
PC1 (71.27%)
PC2 (14.51%)
PC3 ( 6.38%)
−20
−10
0
10
20
30
−10
−5
0
5
10
15
−6
−4
−2
0
2
4
6
8
0 25

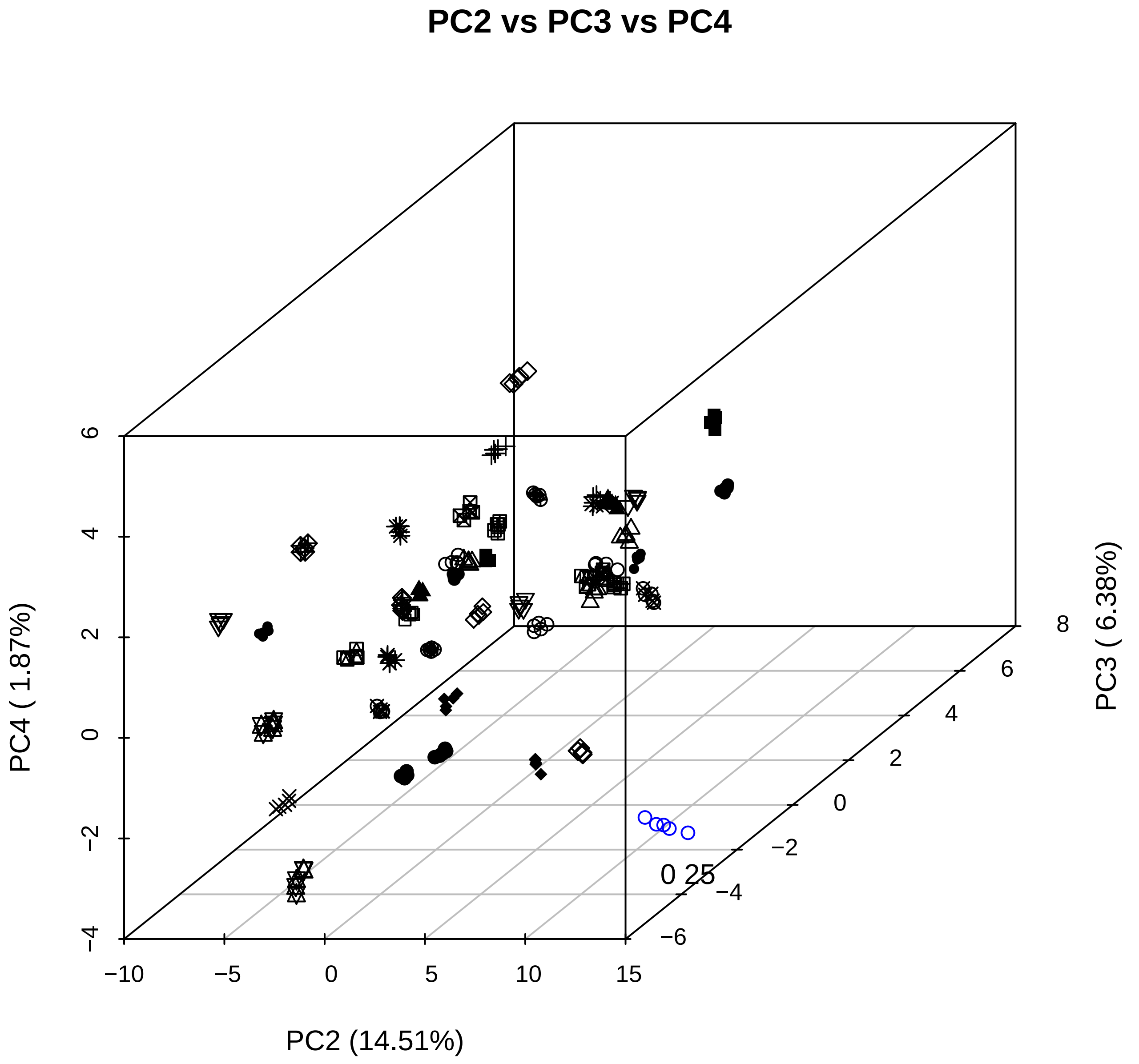
PC2 vs PC3 vs PC4
PC4 ( 1.87%)
PC2 (14.51%)
PC3 ( 6.38%)
0 25
−10
−5
0
5
10
15
−4
−2
0
2
4
6
−6
−4
−2
0
2
4
6
8

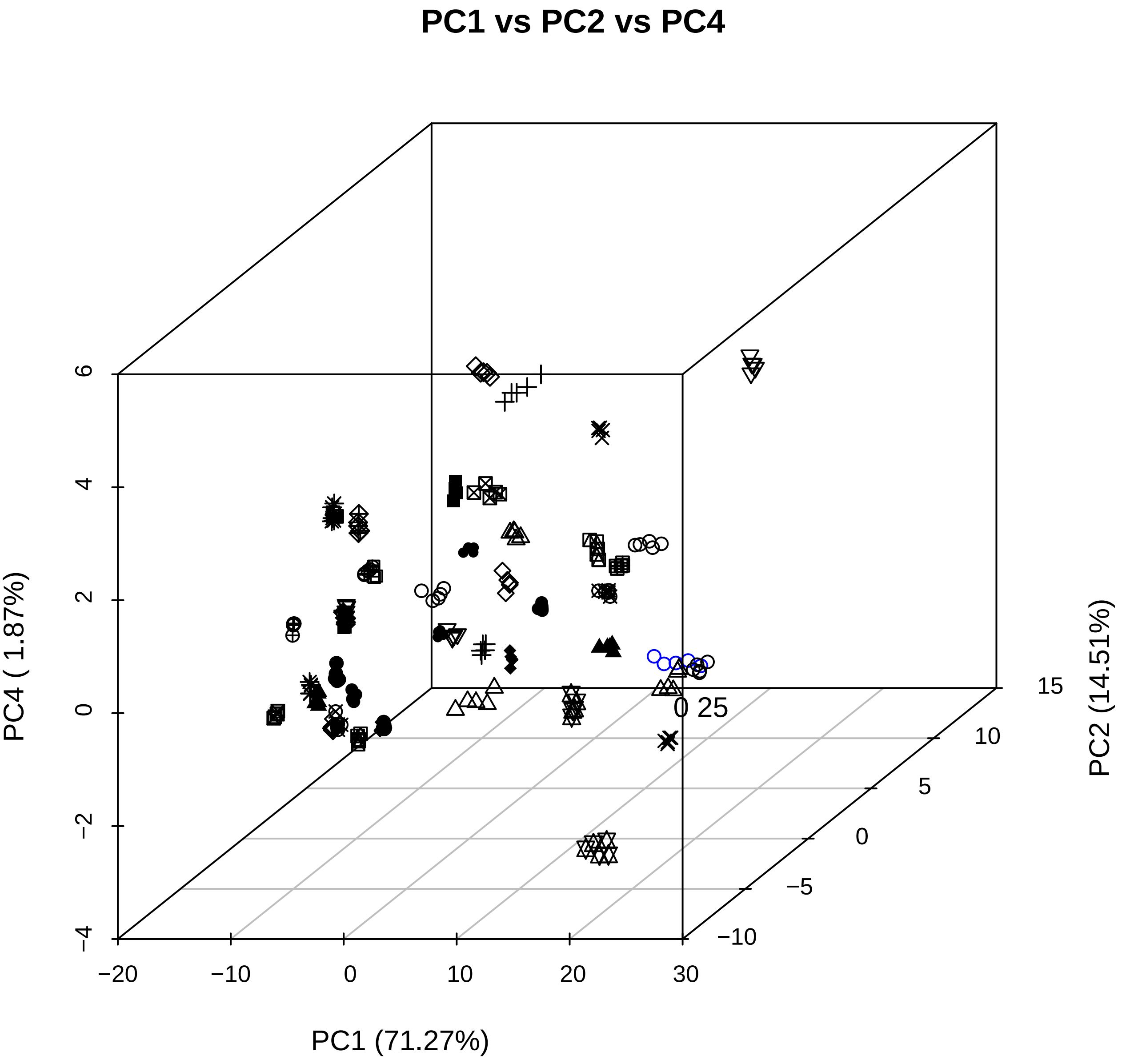

PC1 vs PC2 vs PC4
PC1 (71.27%)
PC2 (14.51%)
PC4 ( 1.87%)
−20
−10
0
10
20
30
−10
−5
0
5
10
15
−4
−2
0
2
4
6
0 25

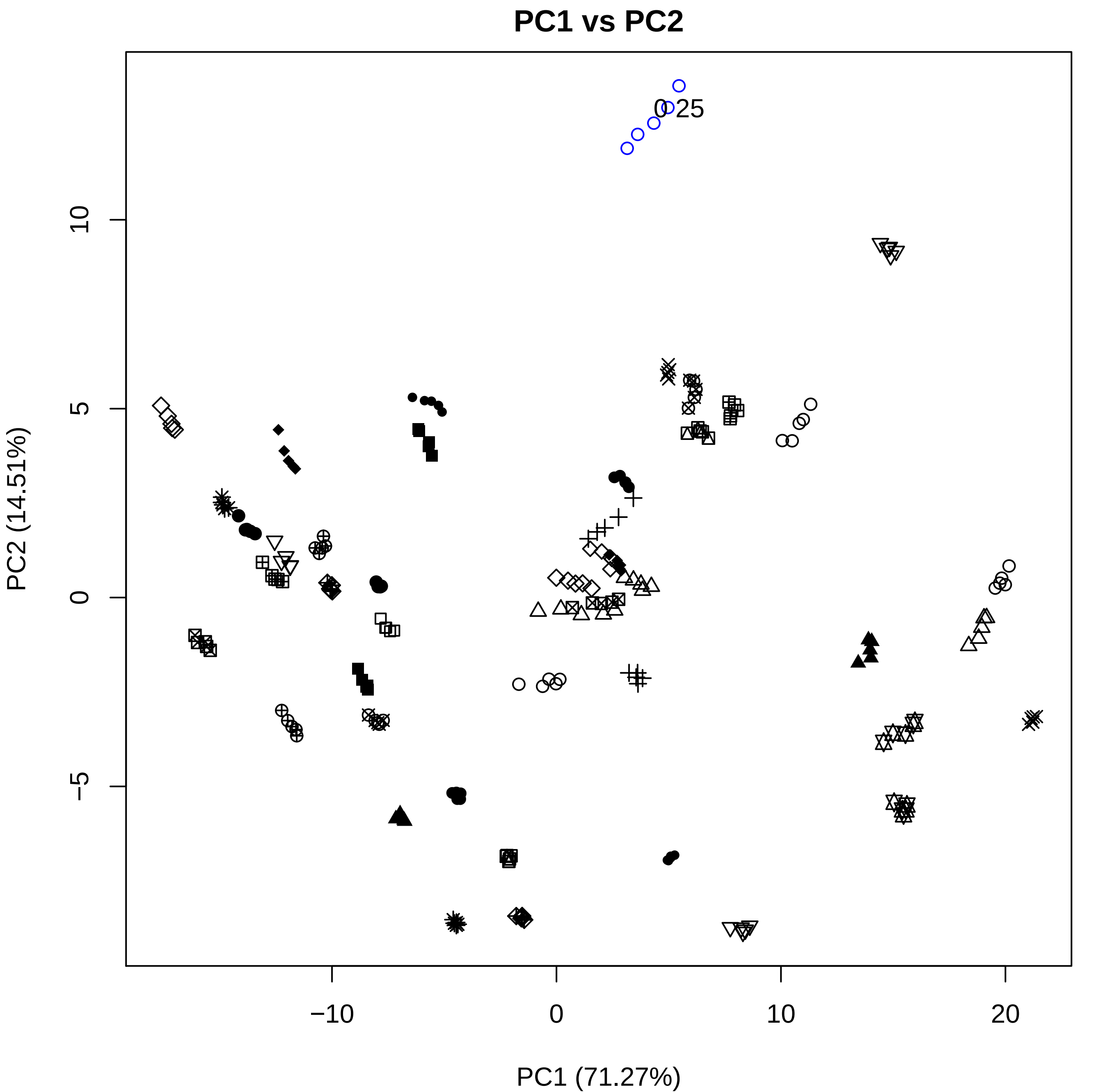

PC1 vs PC2
0.25
PC2 (14.51%)
PC1 (71.27%)
−10
0
10
20
−5
0
5
10

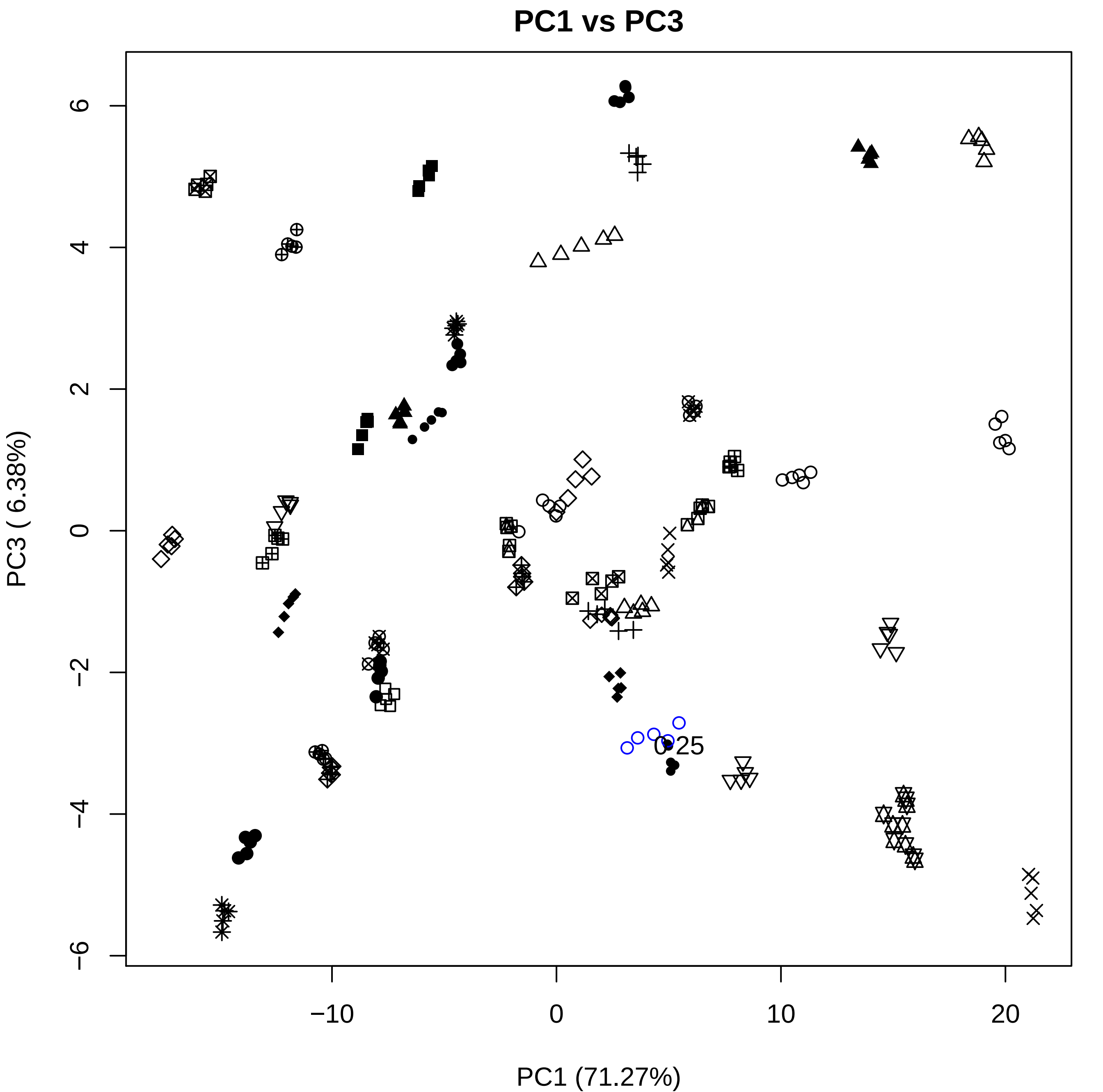

PC1 vs PC3
PC3 ( 6.38%)
PC1 (71.27%)
6
4
2
0
−2
−4
−6
−10
0
10
20
0.25

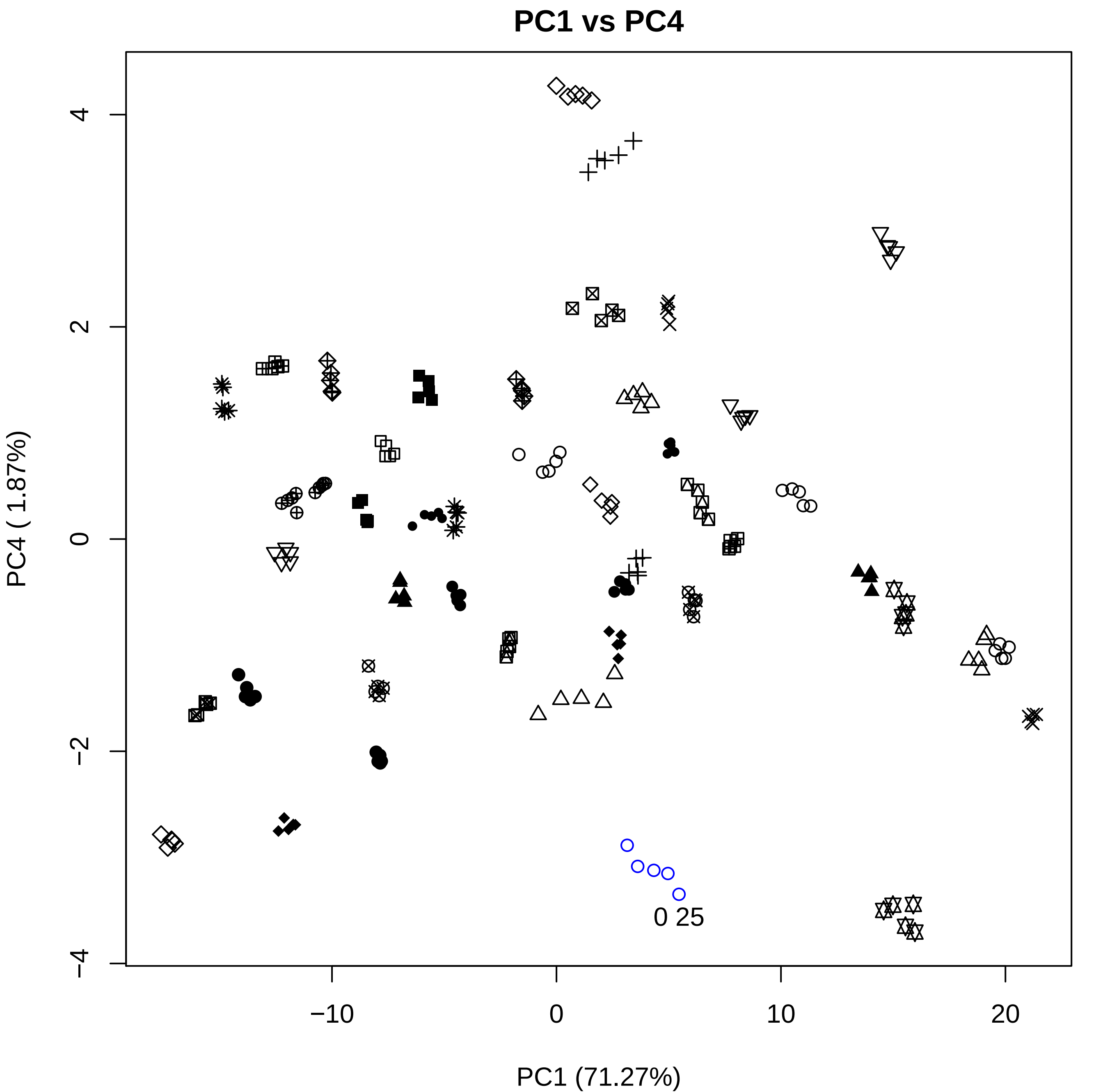

PC1 vs PC4
PC4 ( 1.87%)
PC1 (71.27%)
4
2
0
−2
−4
−10
0
10
20
0 25

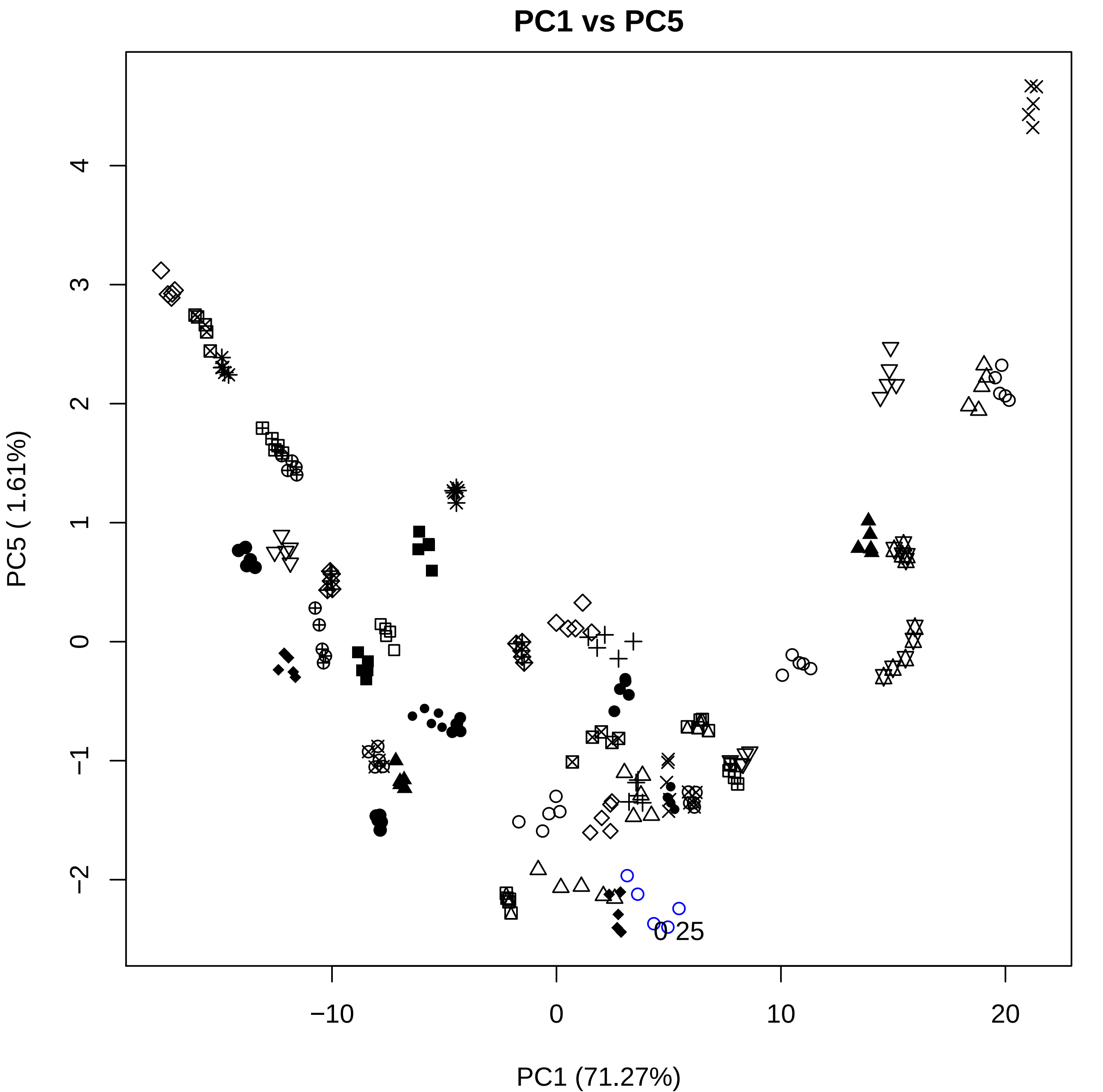
PC1 vs PC5
PC5 ( 1.61%)
PC1 (71.27%)
−10
0
10
20
−2
−1
0
1
2
3
4
0.25

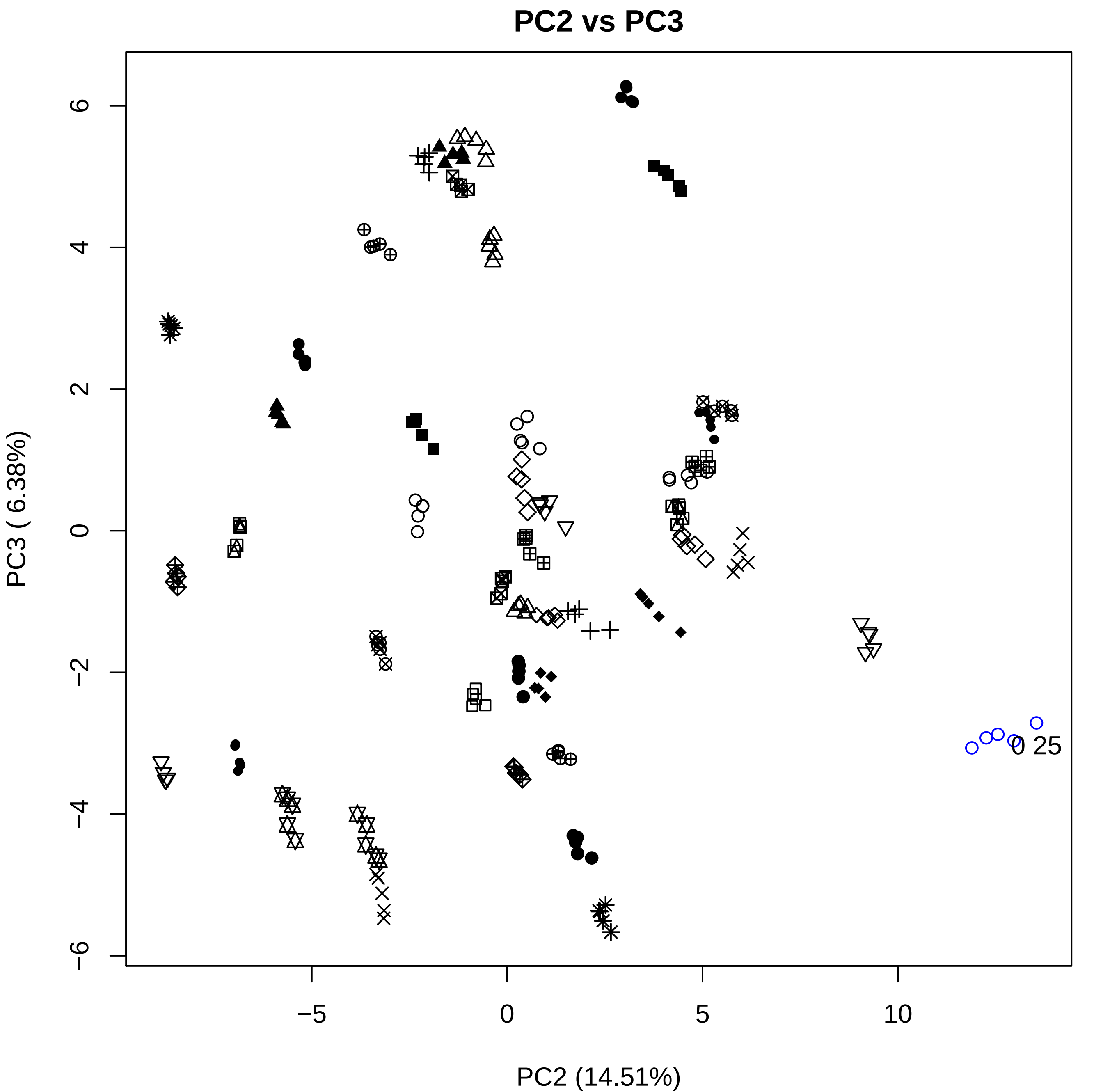
PC2 vs PC3
PC3 ( 6.38%)
PC2 (14.51%)
6
4
2
0
−2
−4
−6
−5
0
5
10
0 25

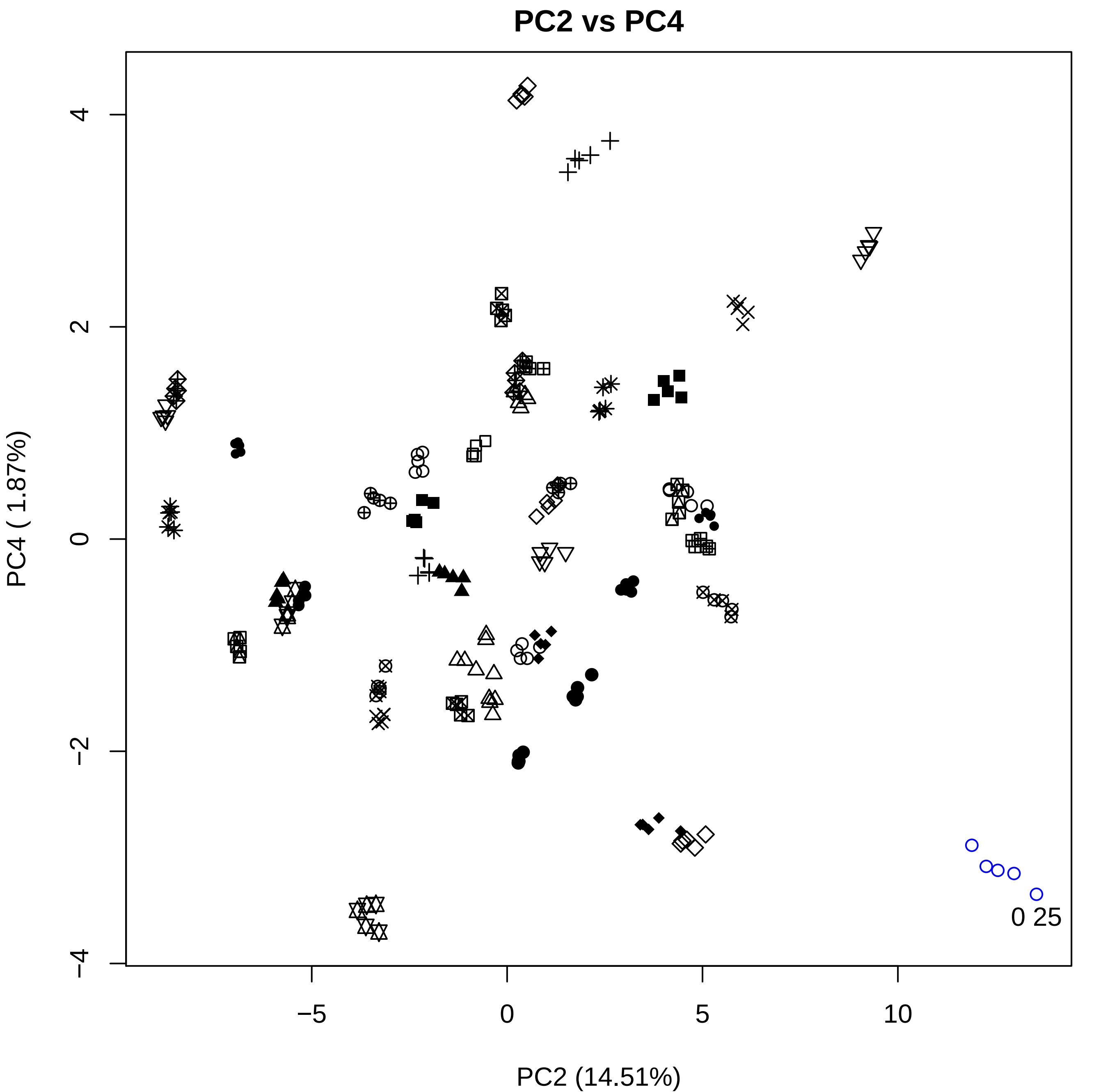

PC2 vs PC4
PC4 ( 1.87%)
PC2 (14.51%)
4
2
0
−2
−4
−5
0
5
10
0 25

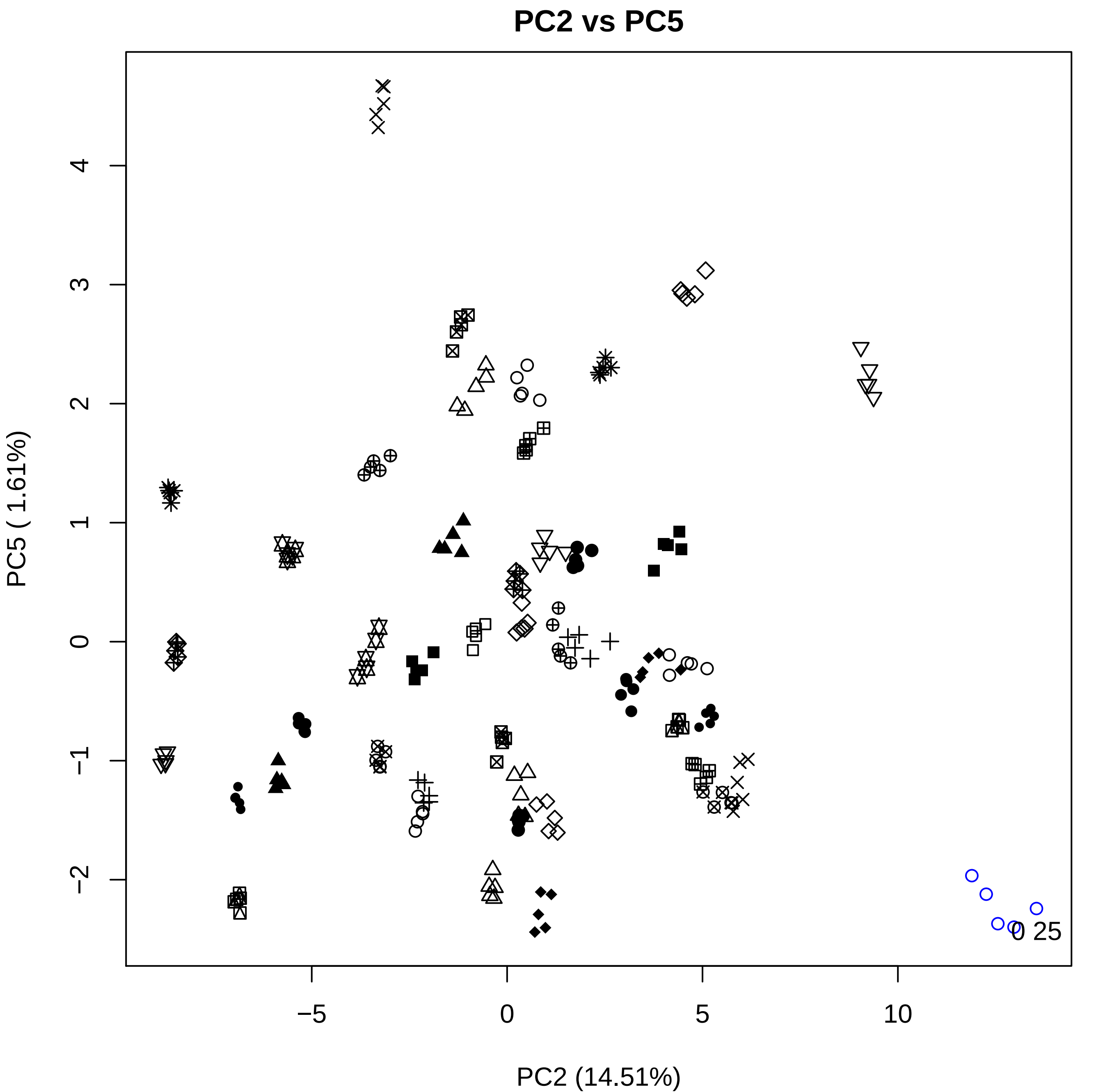
PC2 vs PC5
PC5 ( 1.61%)
PC2 (14.51%)
4
3
2
1
0
−1
−2
−5
0
5
10
0 25

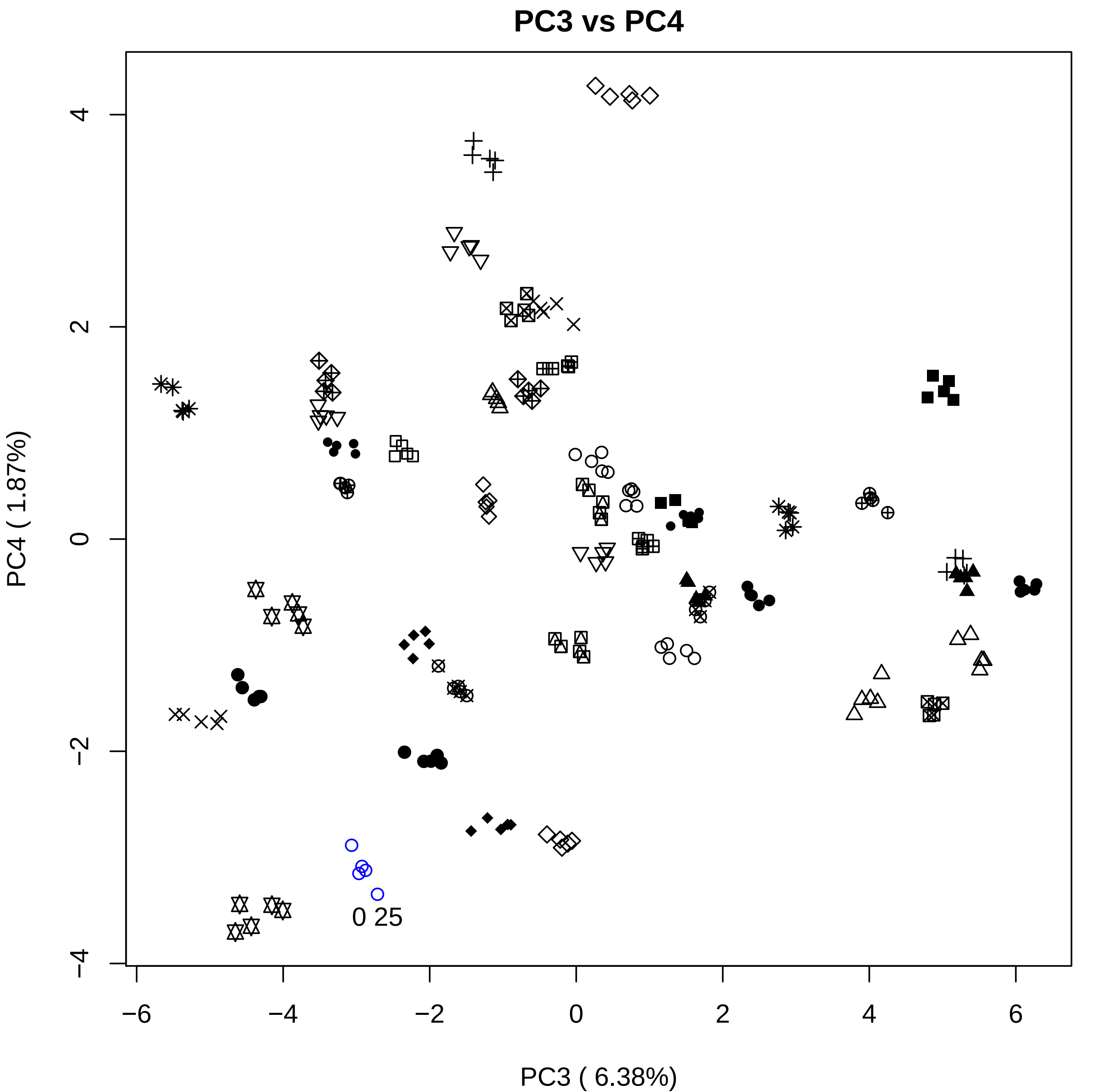

PC3 vs PC4
PC4 ( 1.87%)
PC3 ( 6.38%)
4
2
0
−2
−4
−6
−4
−2
0
2
4
6
0 25

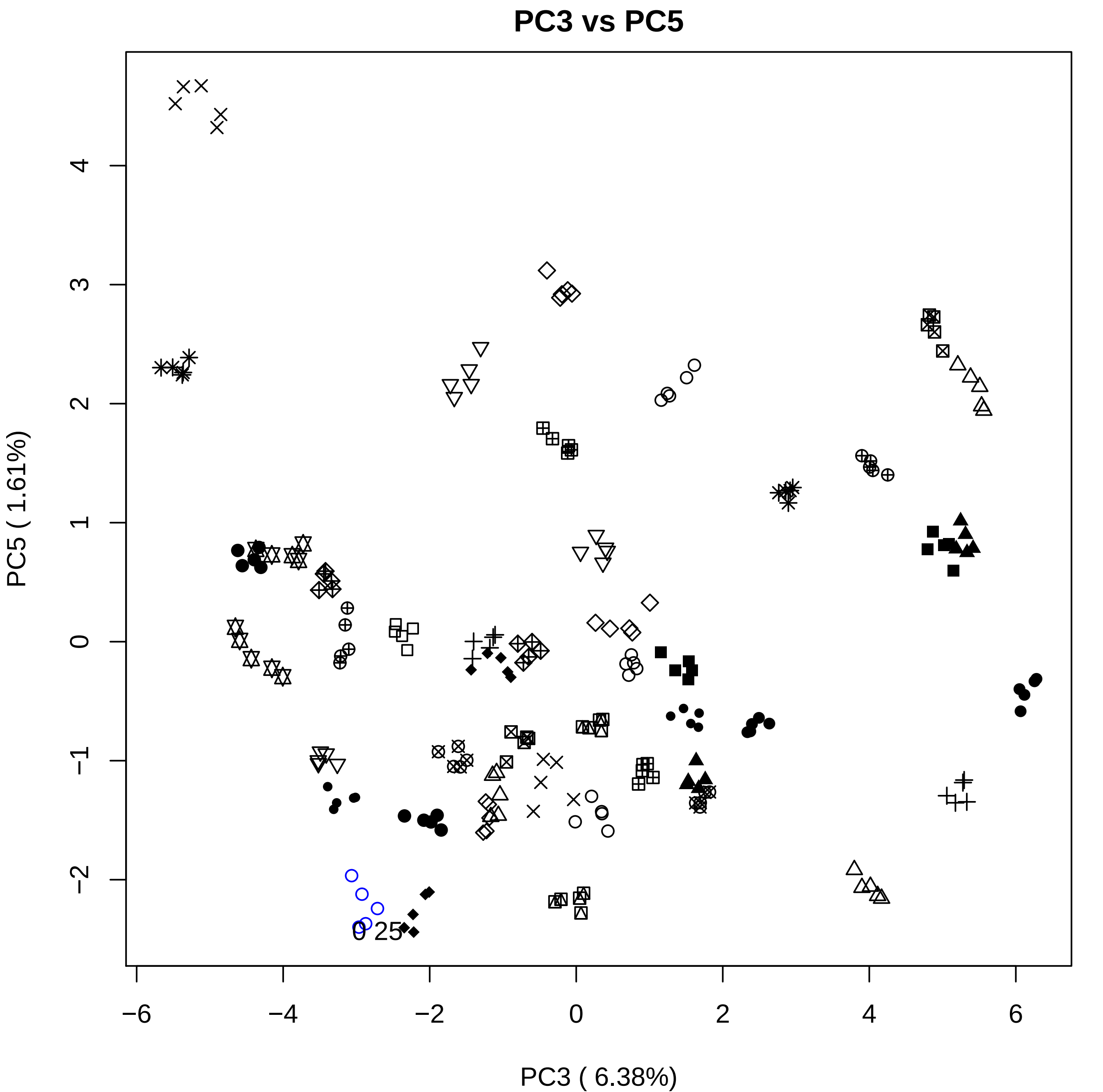
PC3 vs PC5
PC5 ( 1.61%)
PC3 ( 6.38%)
−6
−4
−2
0
2
4
6
−2
−1
0
1
2
3
4
0 25

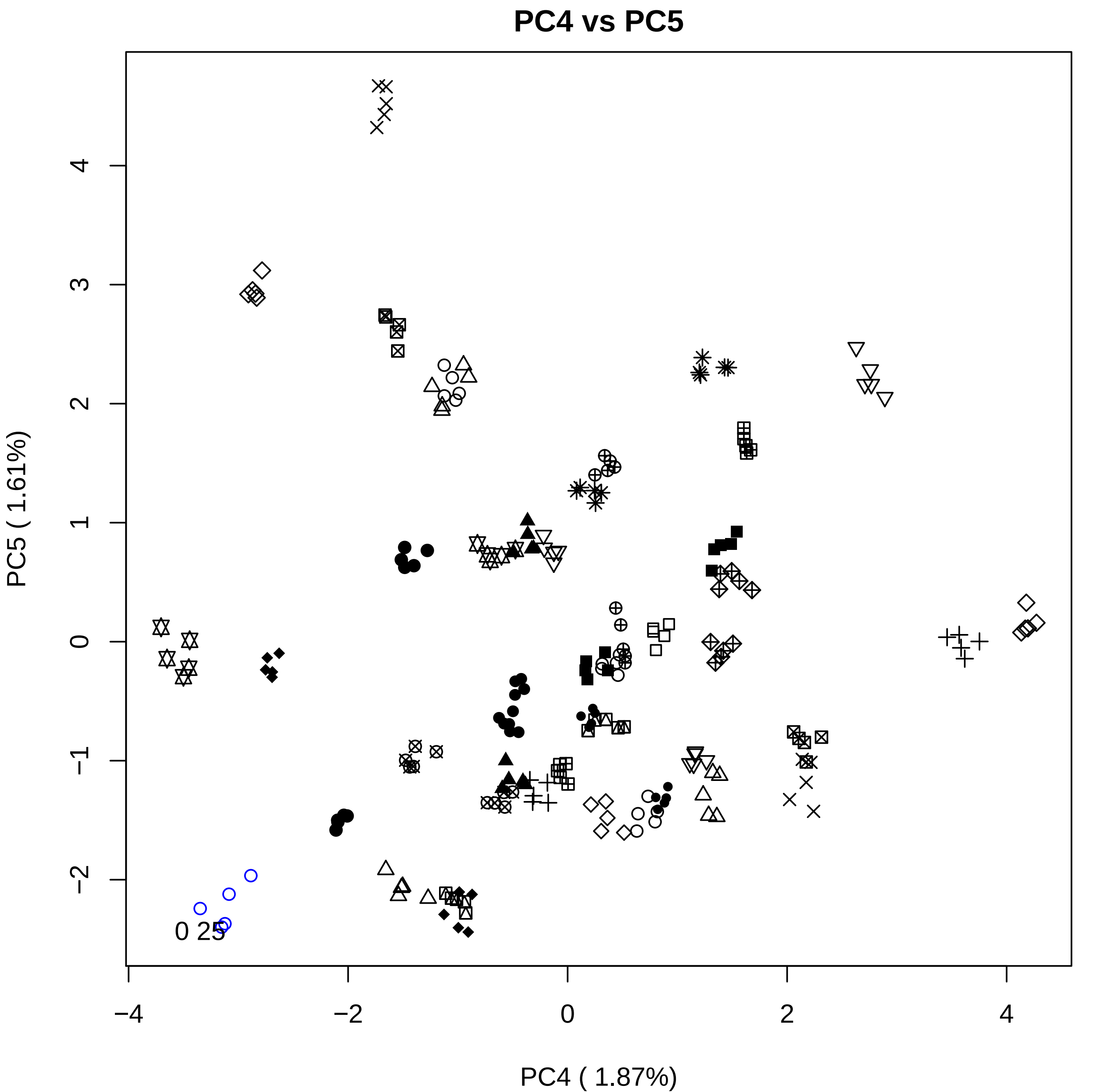

PC4 vs PC5
PC5 ( 1.61%)
PC4 ( 1.87%)
−4
−2
0
2
4
−2
−1
0
1
2
3
4
0 25

### PCA of 0.5%  DeS vs PMH

Scaled Matrix PCAed

7th order polynomial baseline correction performed

Raw spectra smoothed to 21 Neighbours, 2nd order PN, Savtisky Golay

Normalisation between 0 and 100

D2 spectra smoothed to 41 Neighbours, 2nd order PN, Savtisky Golay

Regions Analysed: 3601−2501 and 2001−700cm−1

Samples Analysed: 1 Contam's 0 DeS's 45 PMH's

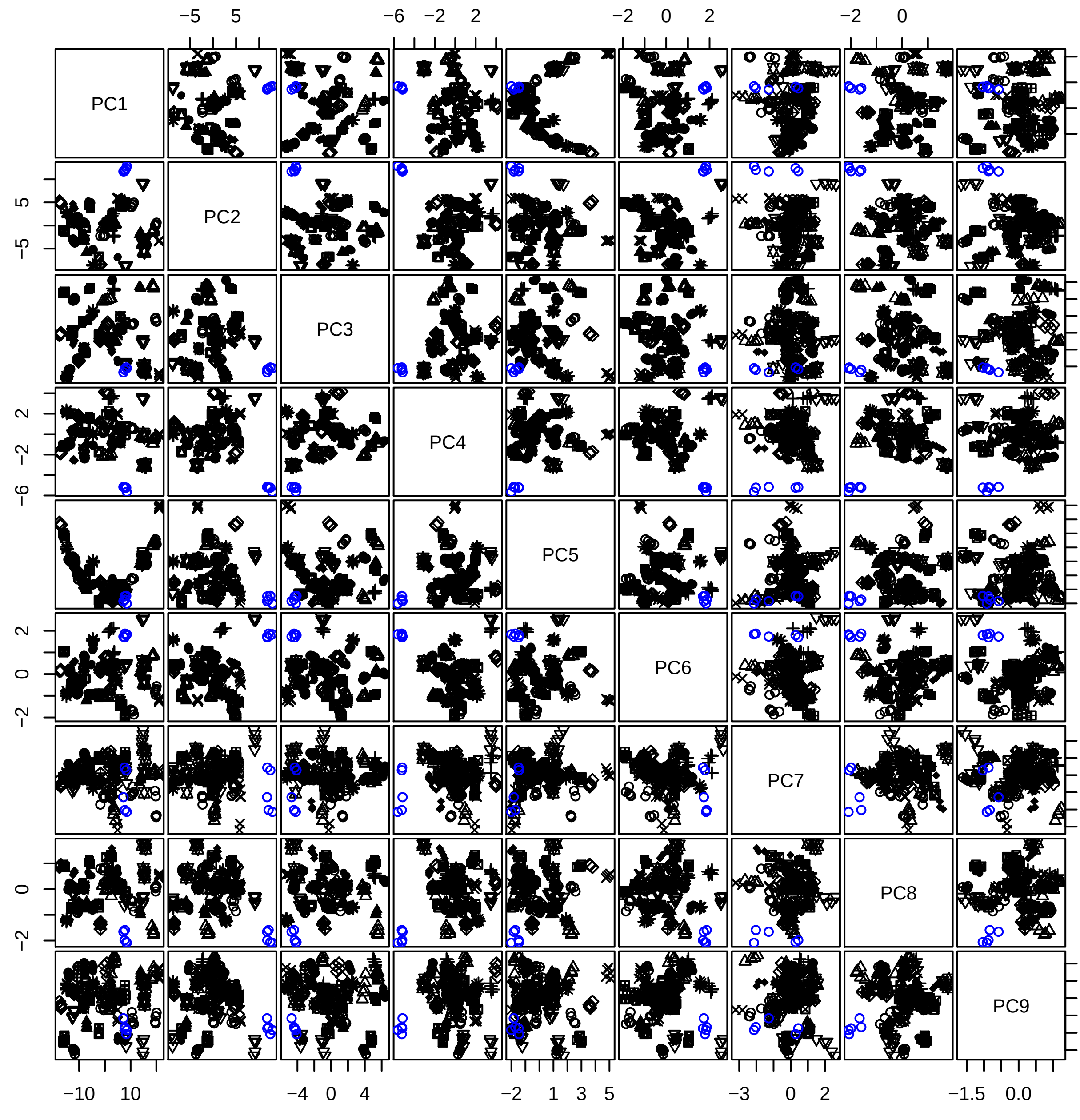
PC1
PC2
PC3
PC4
PC5
PC6
PC7
PC8
PC9

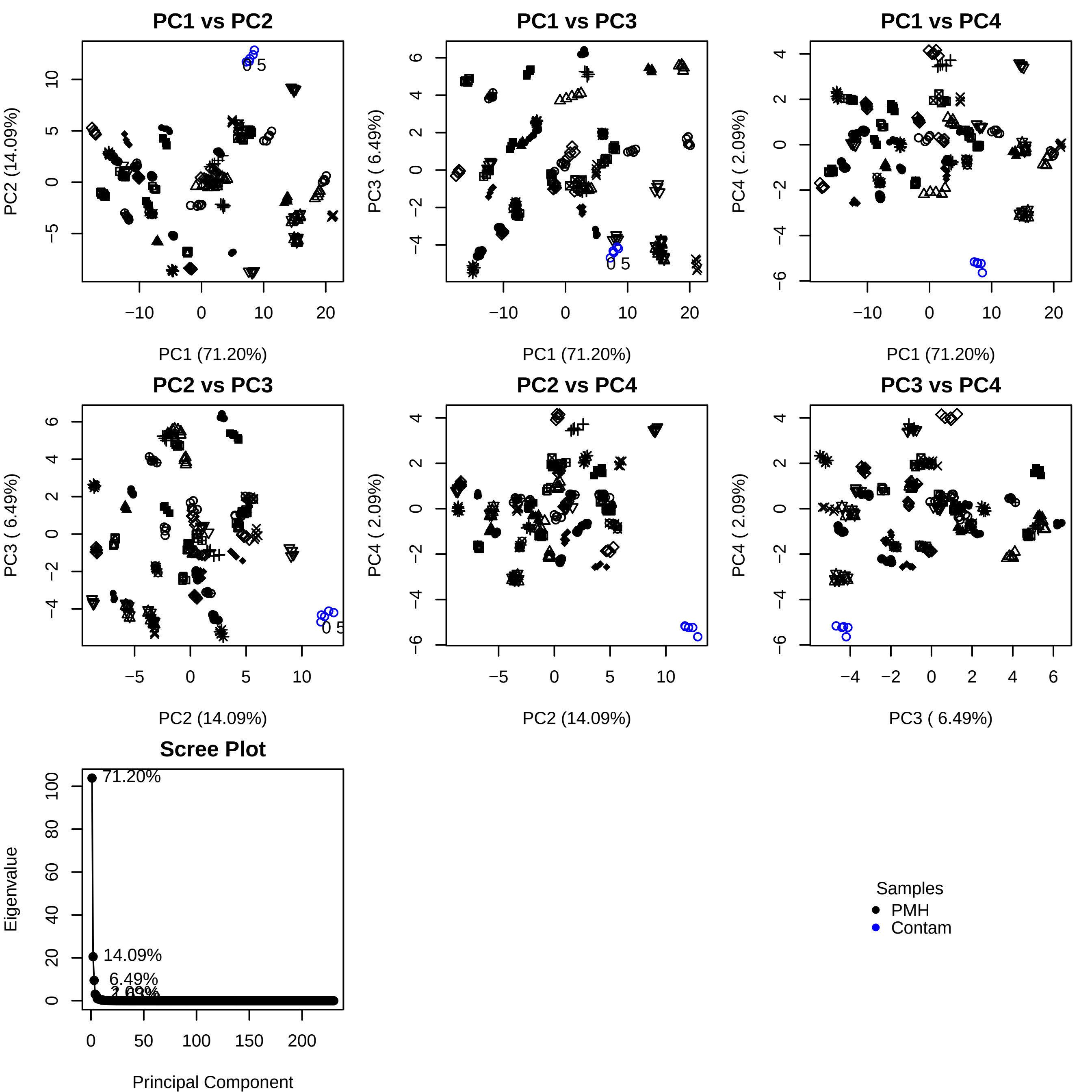
PC1 vs PC2
PC1 vs PC3
PC1 vs PC4
PC2 vs PC3
PC2 vs PC4
PC3 vs PC4
Scree Plot
PC1 (71.20%)
PC2 (14.09%)
PC3 ( 6.49%)
PC4 ( 2.09%)
Eigenvalue
Principal Component
71.20%
14.09%
6.49%
Samples
PMH
Contam

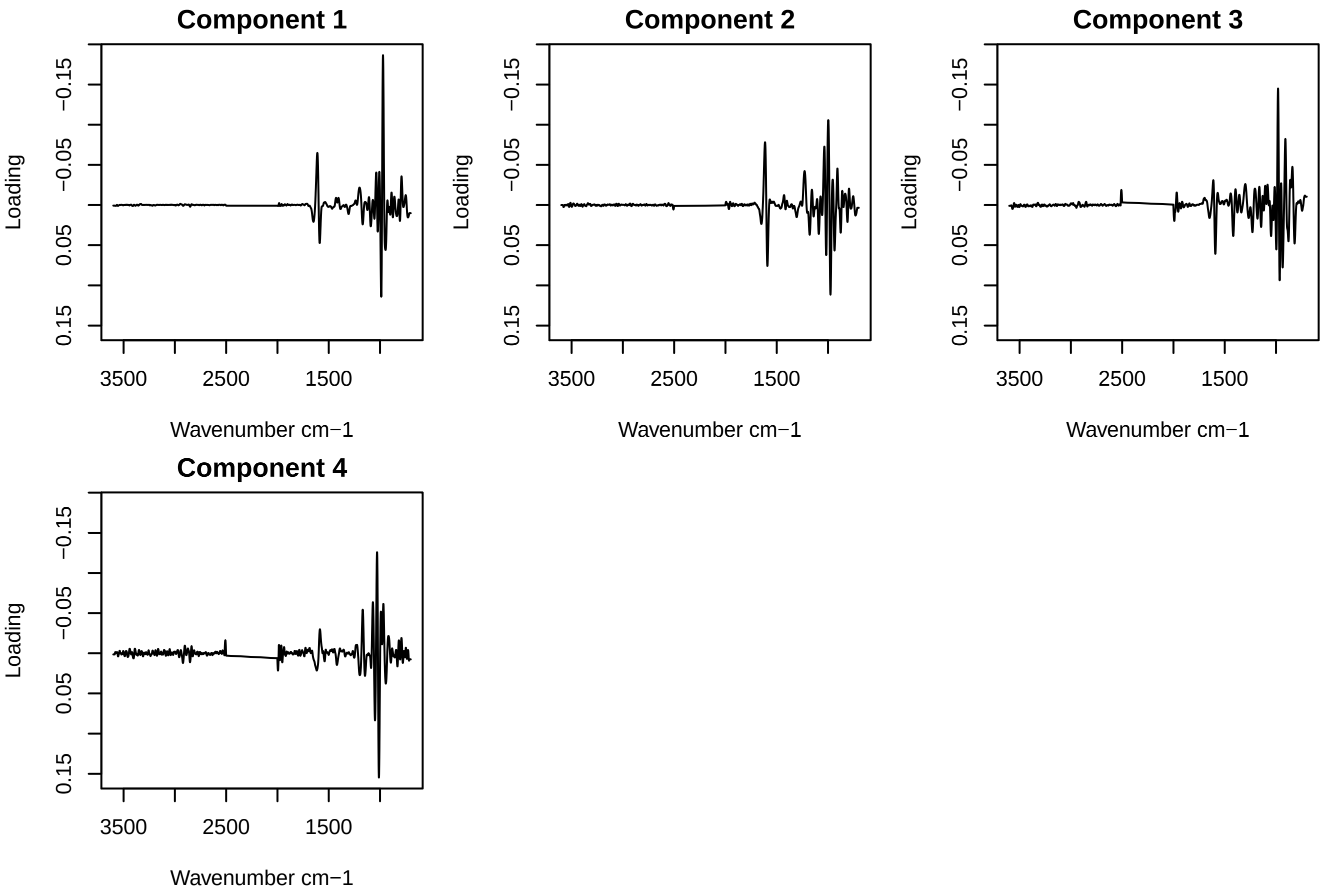
Component 1
Component 2
Component 3
Component 4
Loading
−0.15
−0.05
0.05
0.15
3500
2500
1500
Wavenumber cm−1

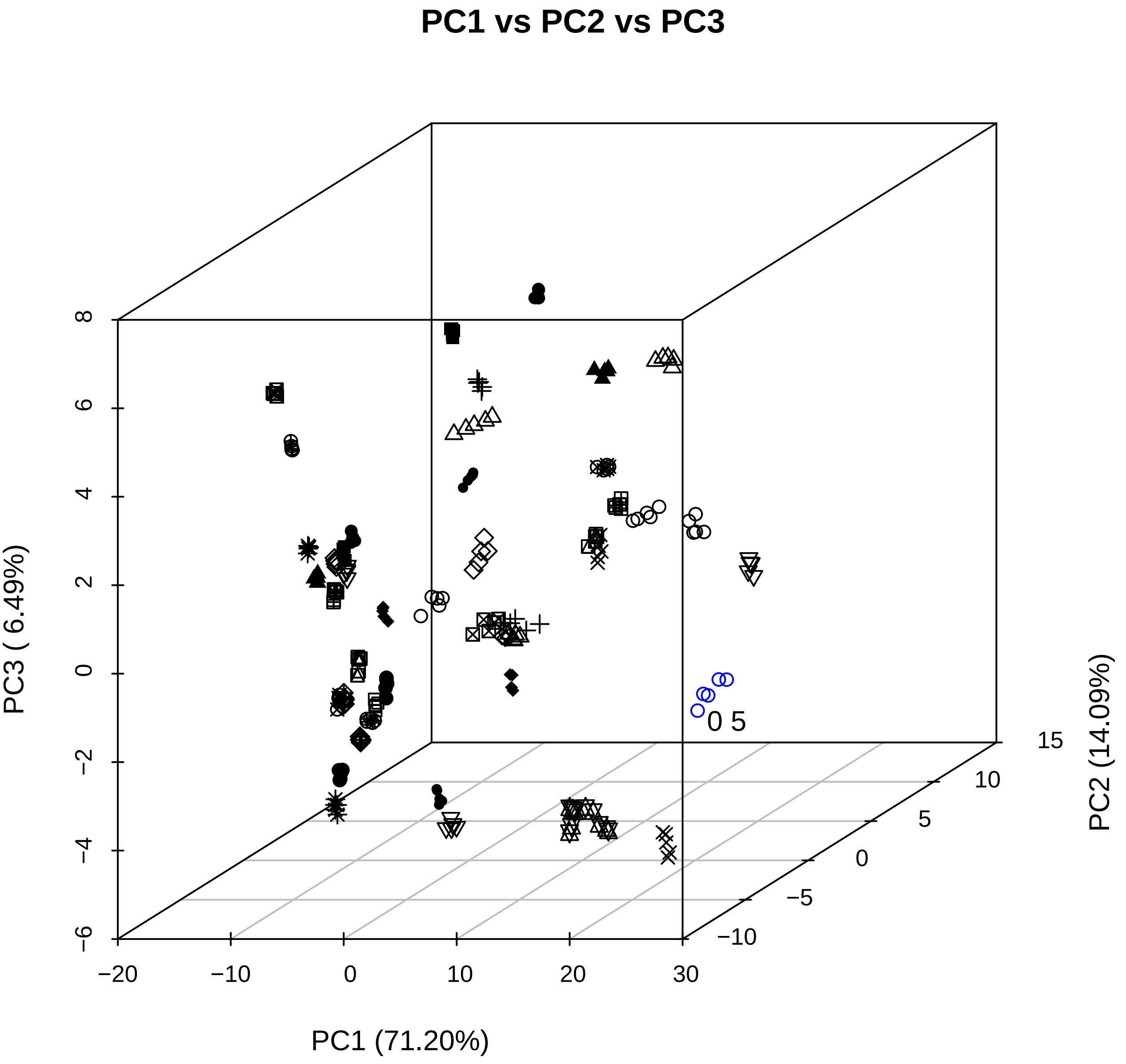

PC1 vs PC2 vs PC3
PC1 (71.20%)
PC2 (14.09%)
PC3 ( 6.49%)
−20
−10
0
10
20
30
−10
−5
0
5
10
15
−6
−4
−2
0
2
4
6
8
0 5

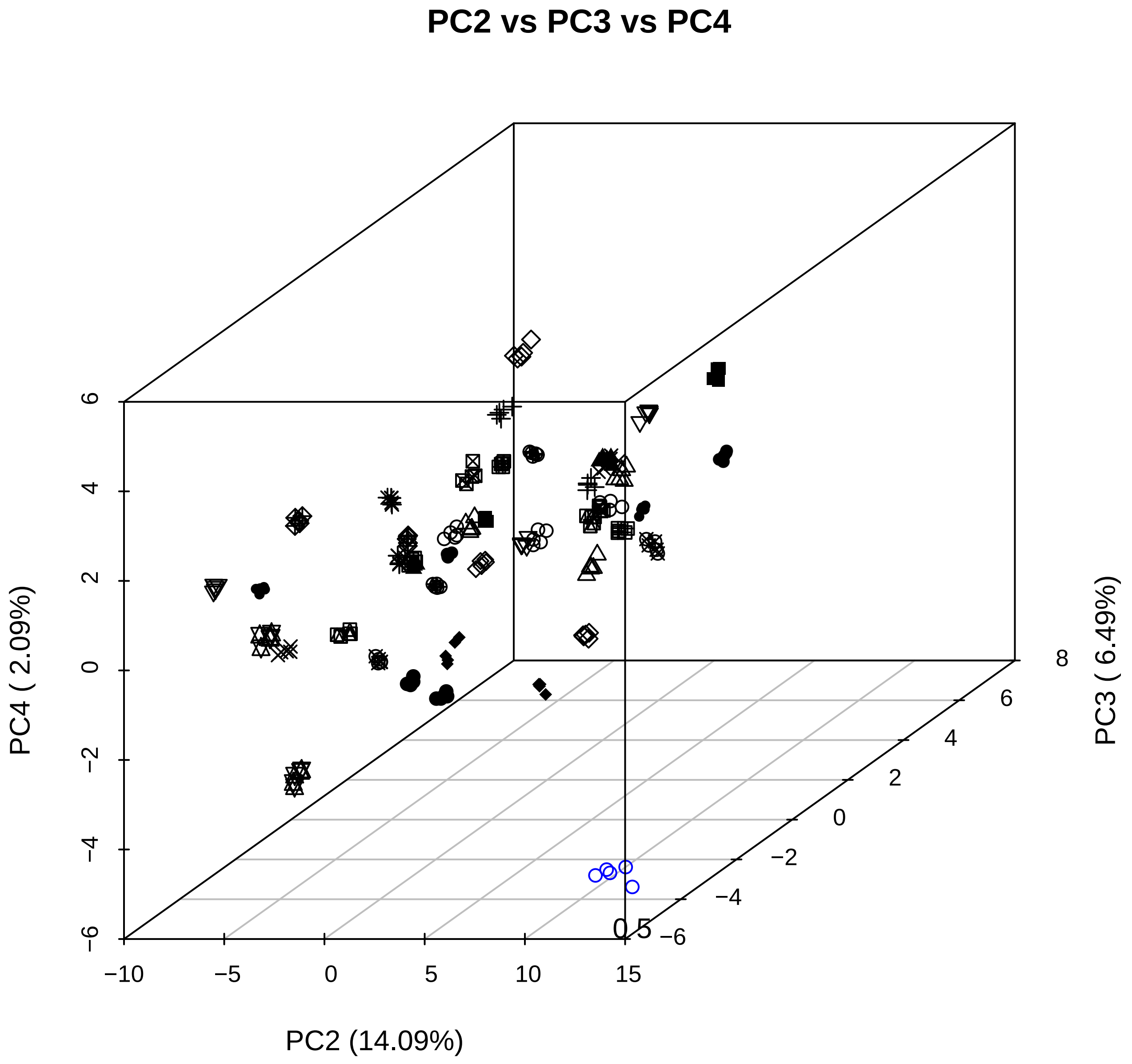
PC2 vs PC3 vs PC4
PC2 (14.09%)
PC4 ( 2.09%)
PC3 ( 6.49%)
−10
−5
0
5
10
15
−6
−4
−2
0
2
4
6
8
0.5

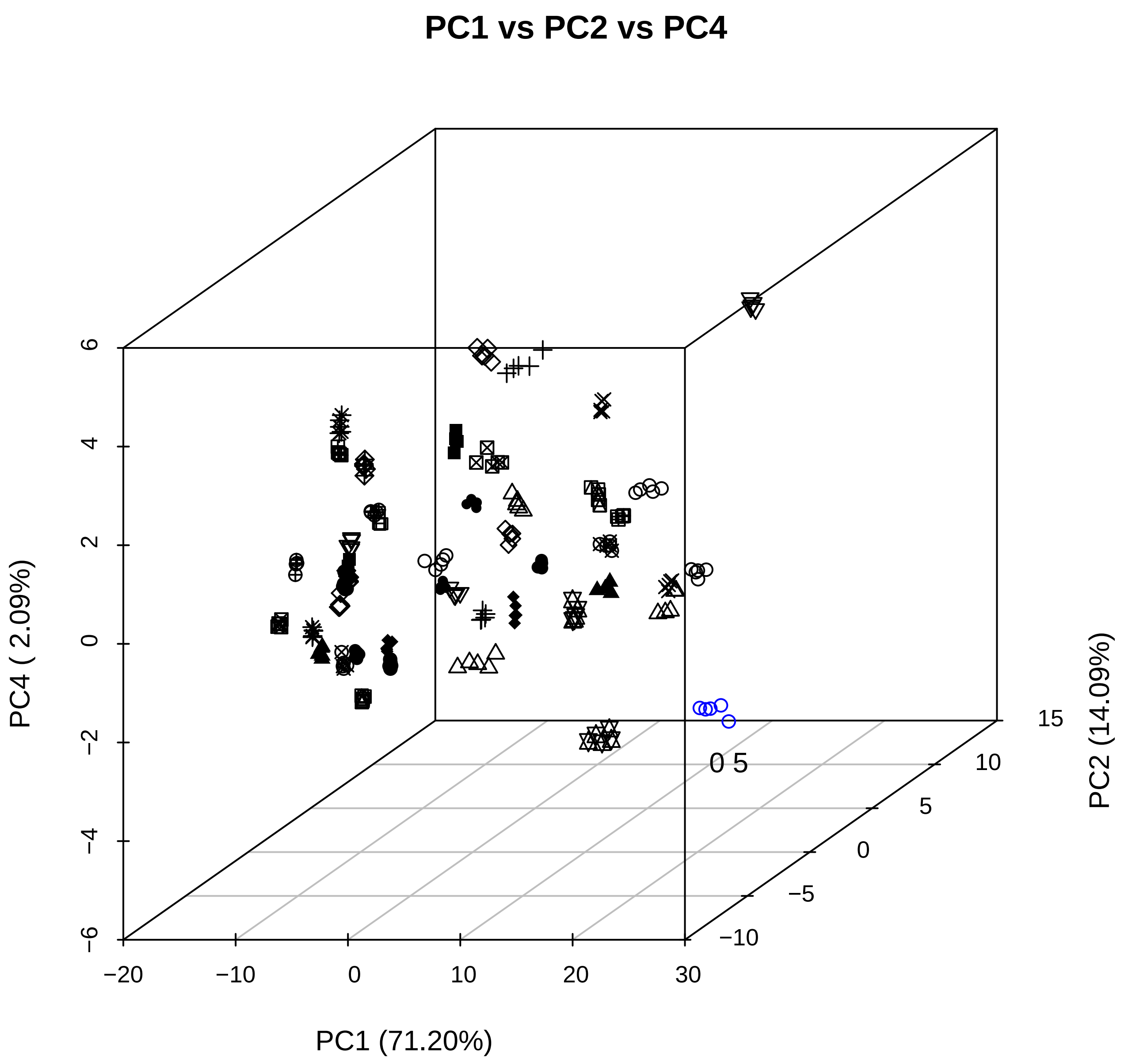
PC1 vs PC2 vs PC4
PC4 ( 2.09%)
6
4
2
0
−2
−4
−6
−20
−10
0
10
20
30
PC1 (71.20%)
PC2 (14.09%)
15
10
5
0
−5
−10
0 5

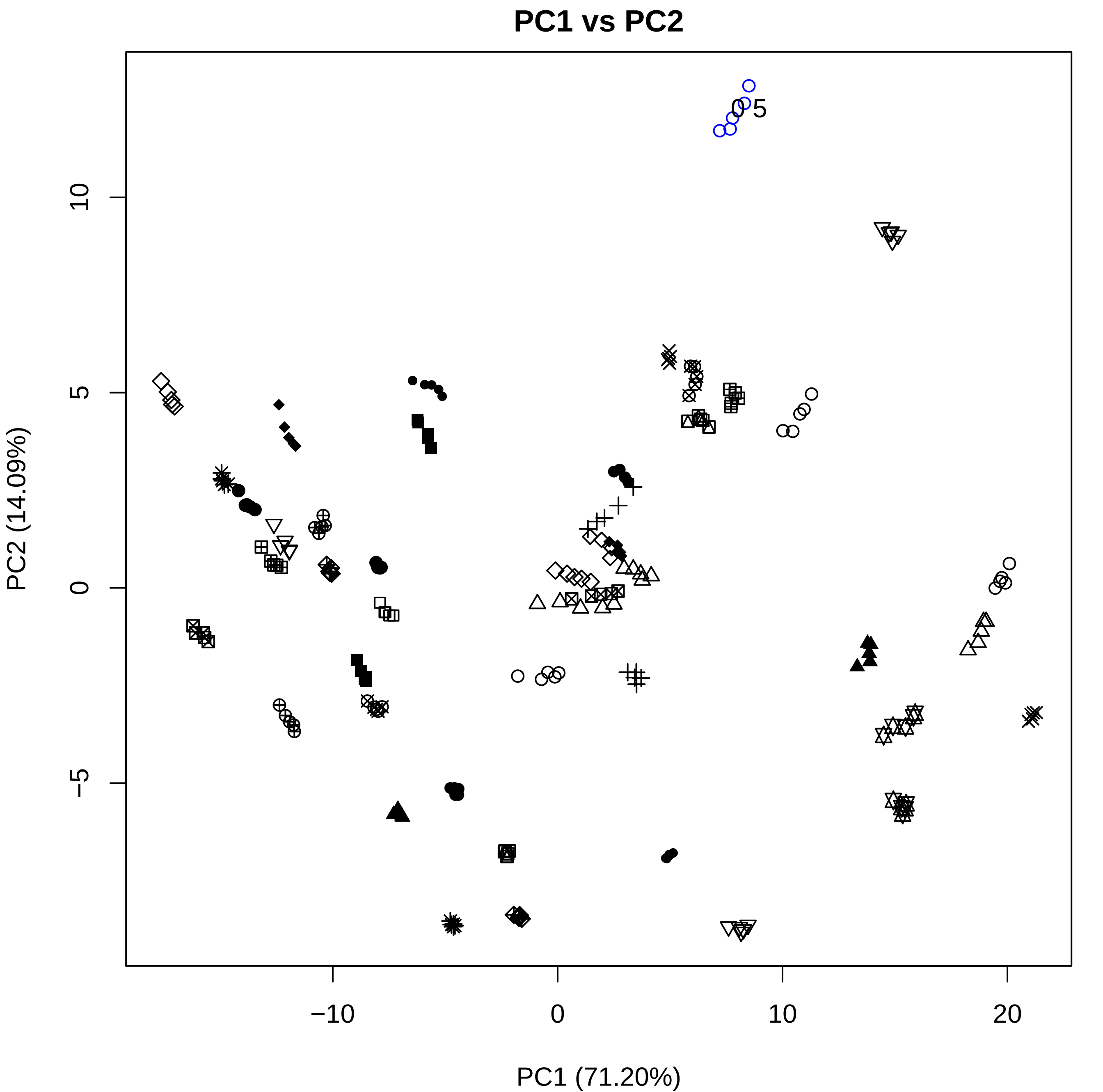
PC1 vs PC2
05
PC1 (71.20%)
PC2 (14.09%)
−10
0
10
20
−5
0
5
10

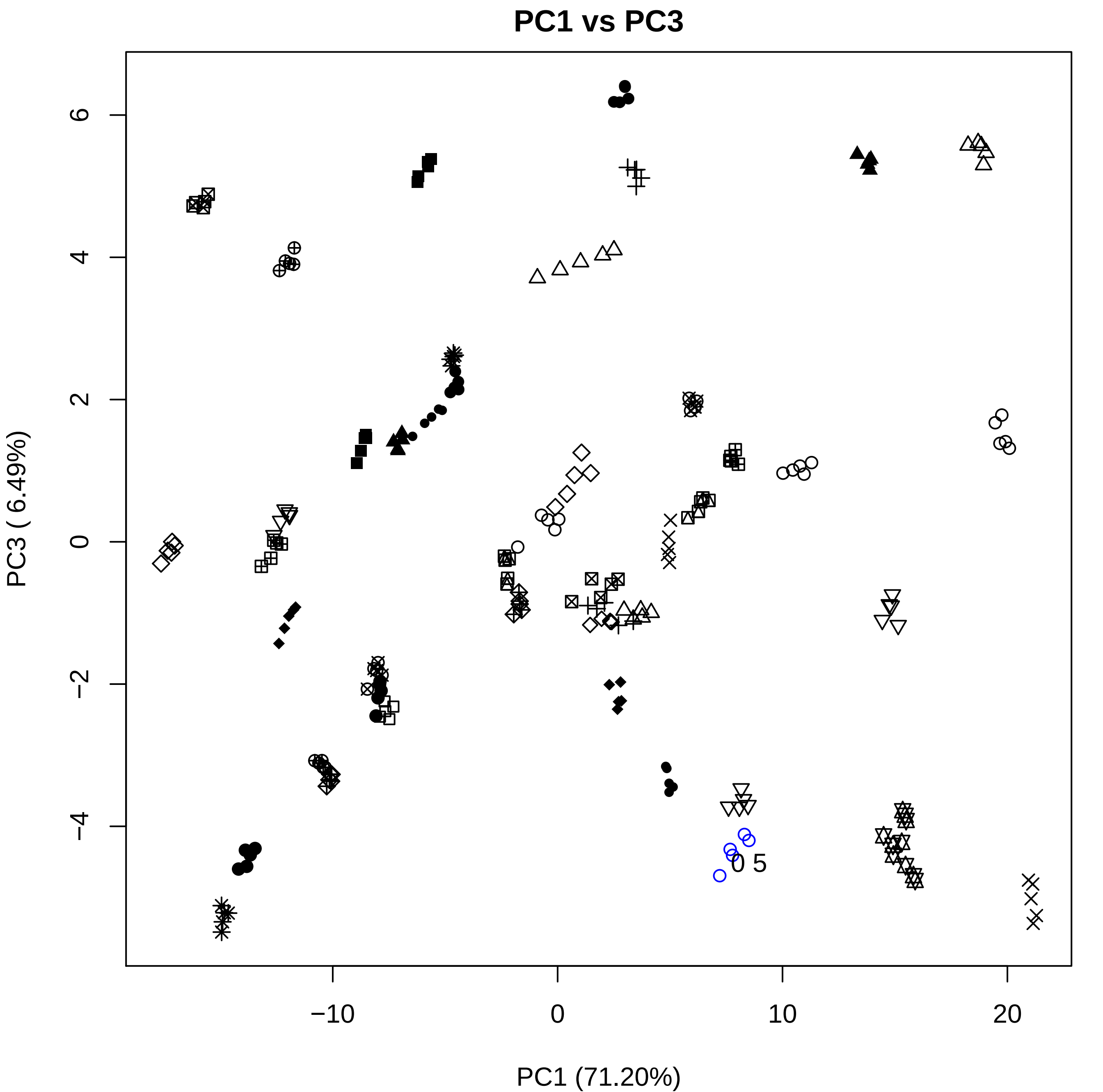
PC1 vs PC3
PC3 ( 6.49%)
PC1 (71.20%)
6
4
2
0
−2
−4
−10
0
10
20
0 5

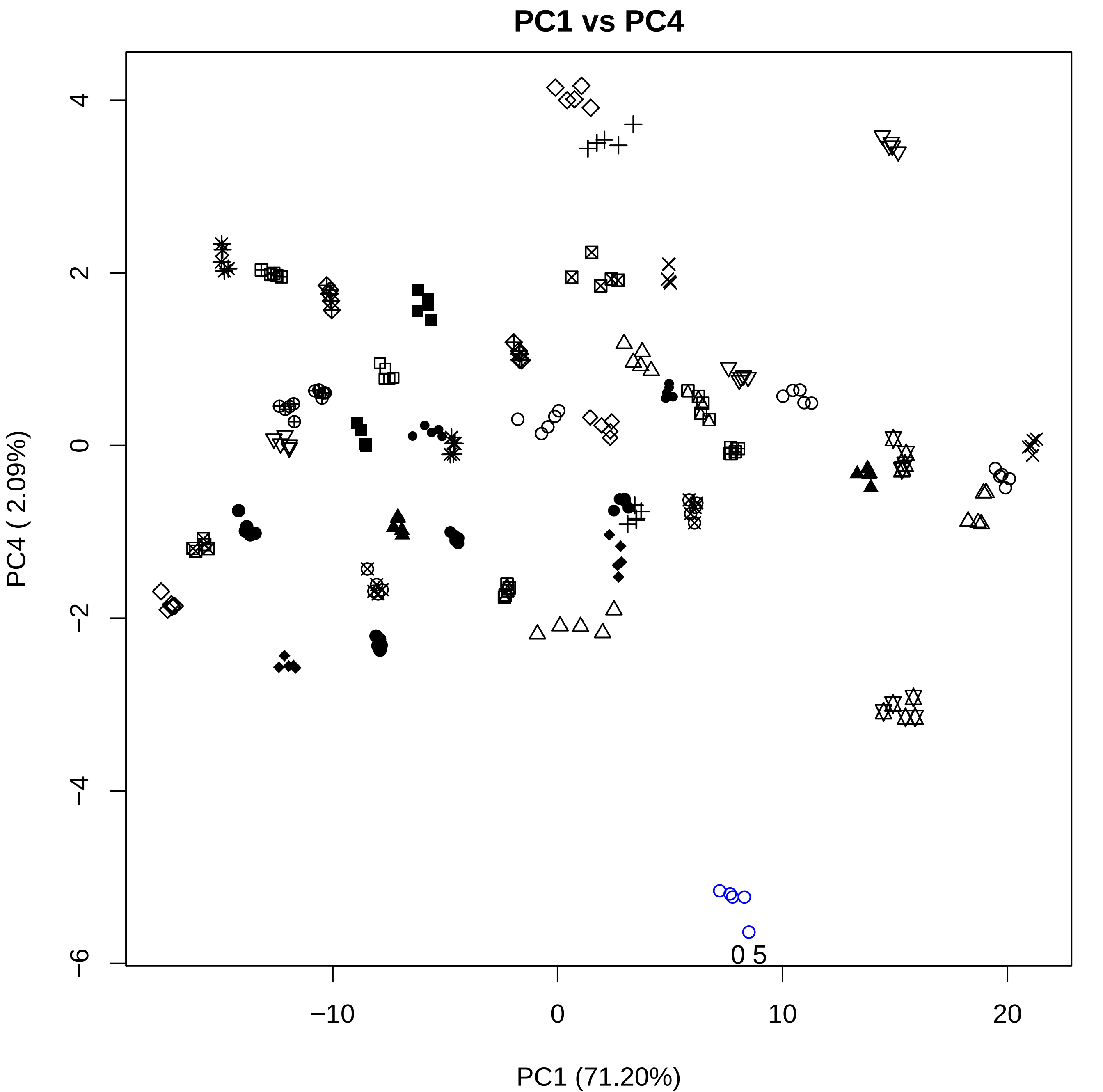
PC1 vs PC4
4
2
0
−2
−4
−6
PC4 ( 2.09%)
−10
0
10
20
PC1 (71.20%)
0 5

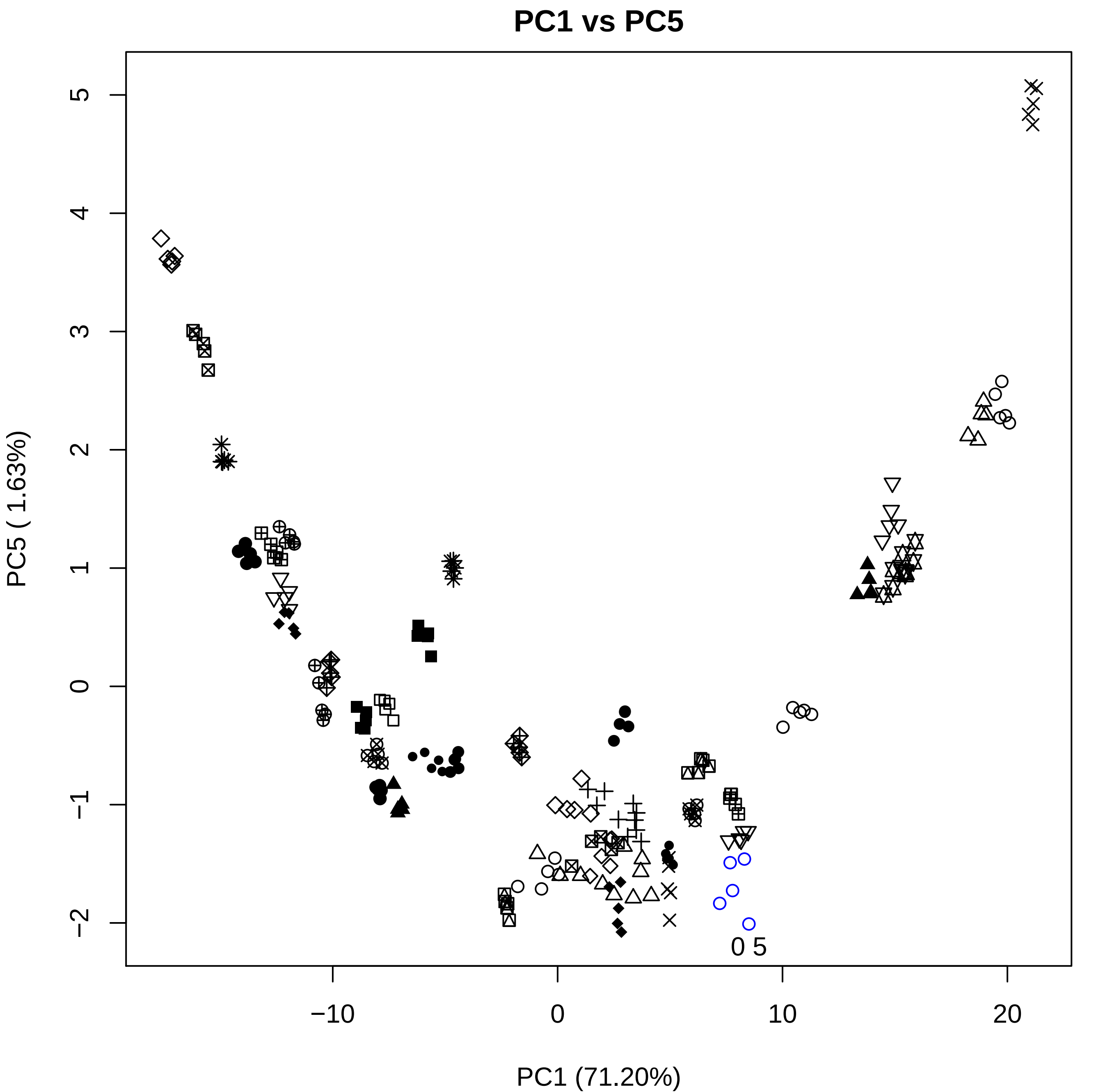

PC1 vs PC5
PC5 ( 1.63%)
PC1 (71.20%)
−10
0
10
20
−2
−1
0
1
2
3
4
5
0 5

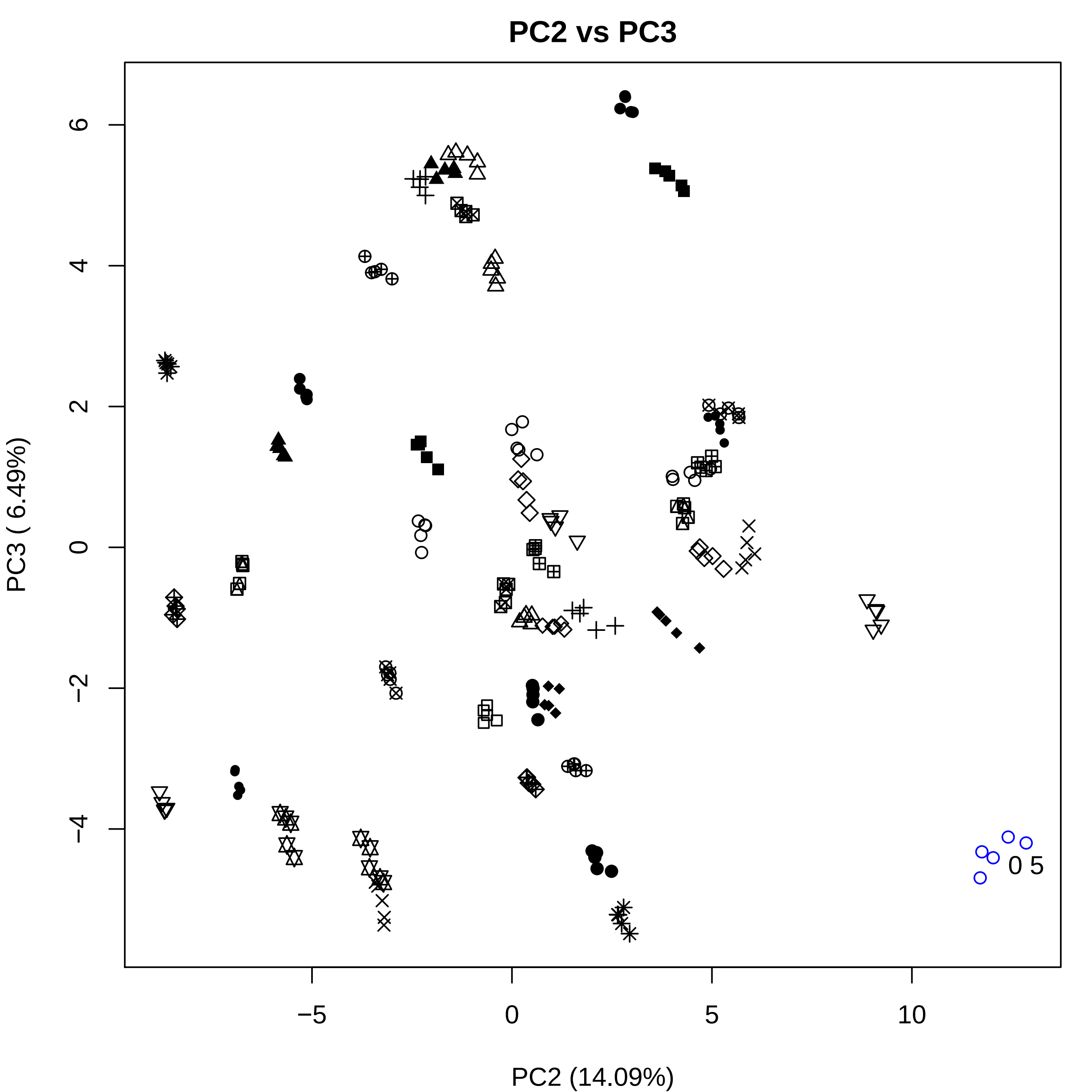
PC2 vs PC3
PC3 ( 6.49%)
PC2 (14.09%)
6
4
2
0
−2
−4
−5
0
5
10
0 5

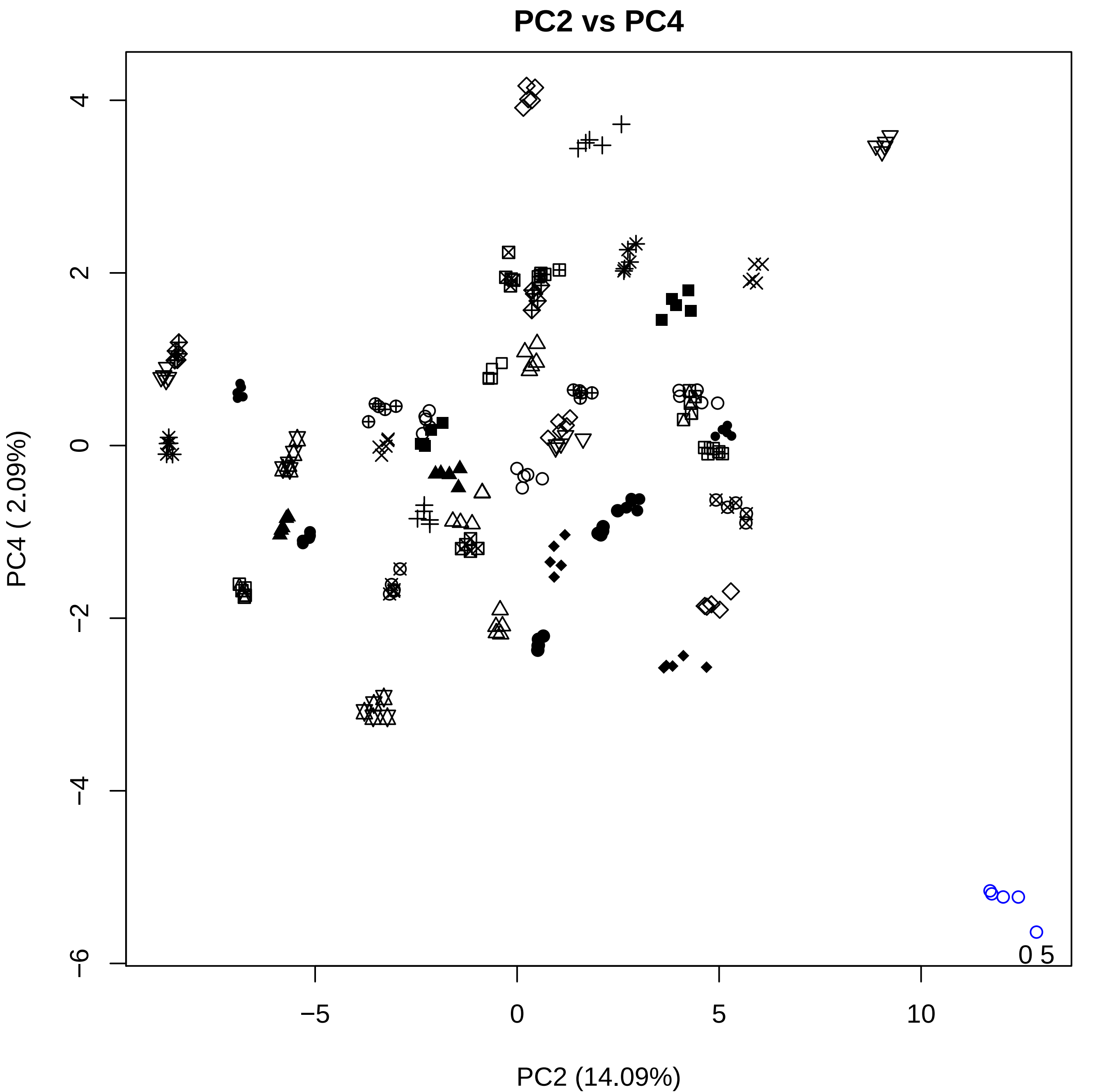

PC2 vs PC4
PC4 ( 2.09%)
PC2 (14.09%)
4
2
0
−2
−4
−6
−5
0
5
10
0 5

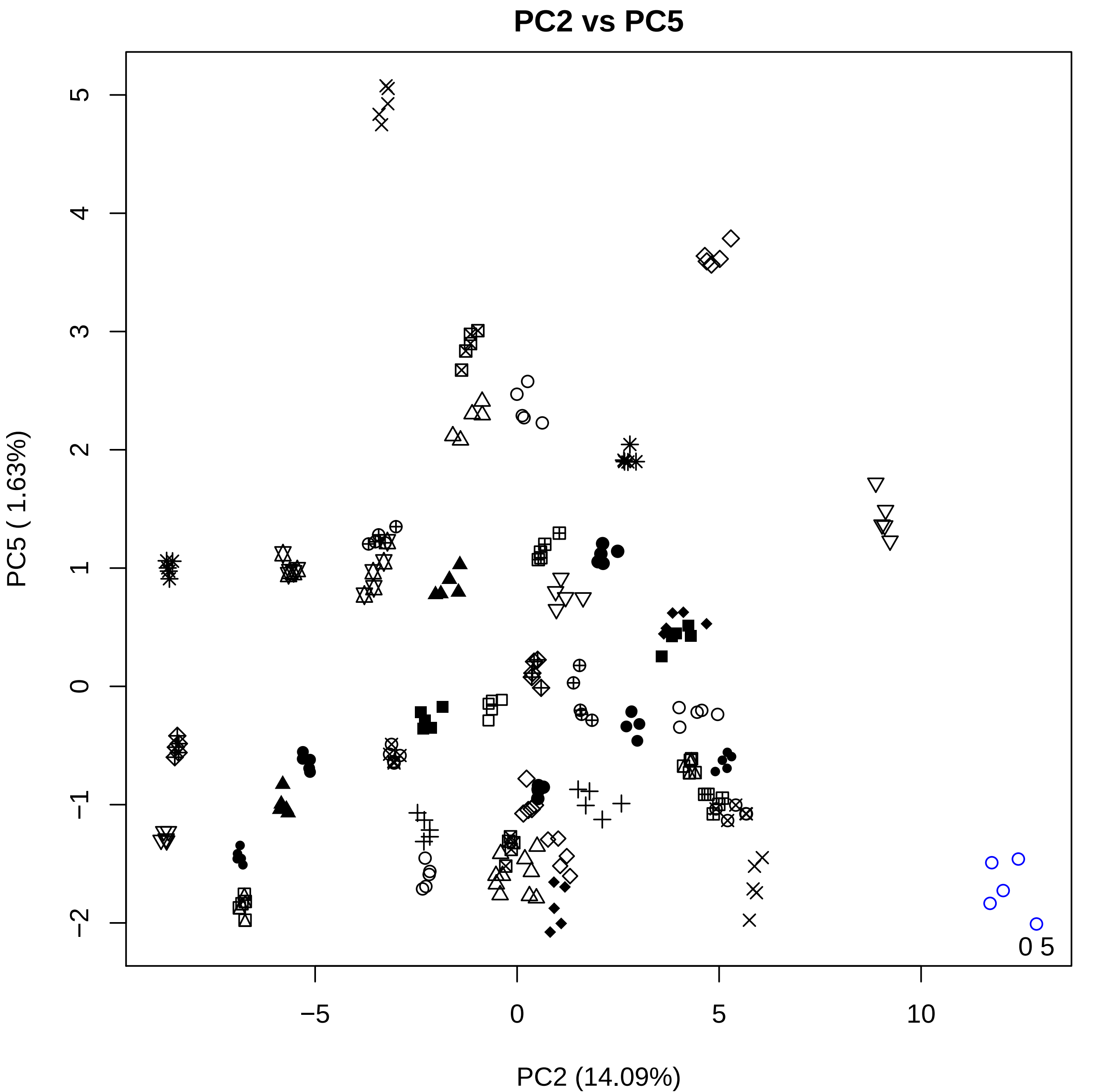
PC2 vs PC5
PC5 ( 1.63%)
PC2 (14.09%)
5
4
3
2
1
0
−1
−2
−5
0
5
10
0 5

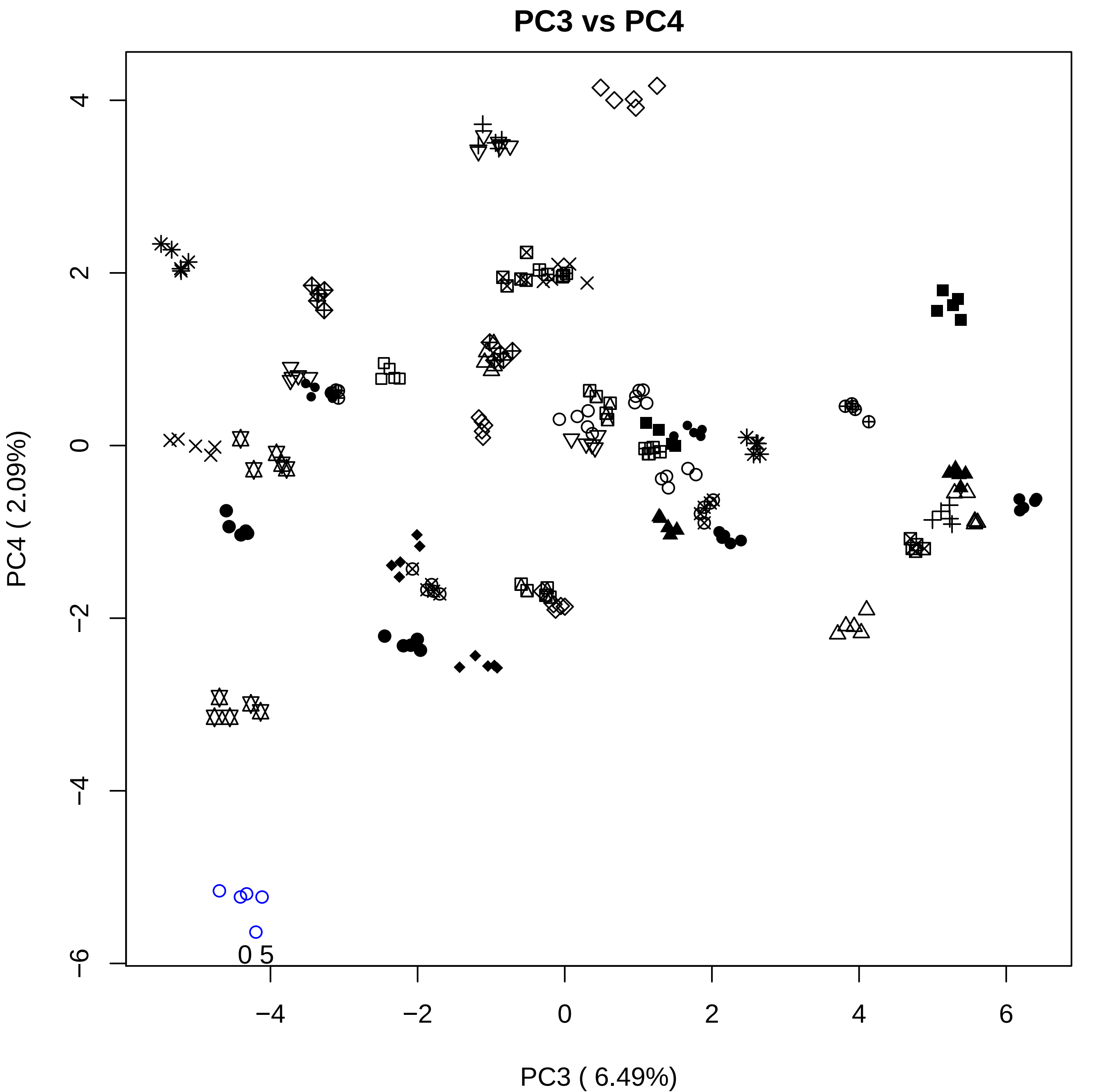
PC3 vs PC4
PC4 ( 2.09%)
PC3 ( 6.49%)
4
2
0
−2
−4
−6
−4
−2
0
2
4
6
0 5

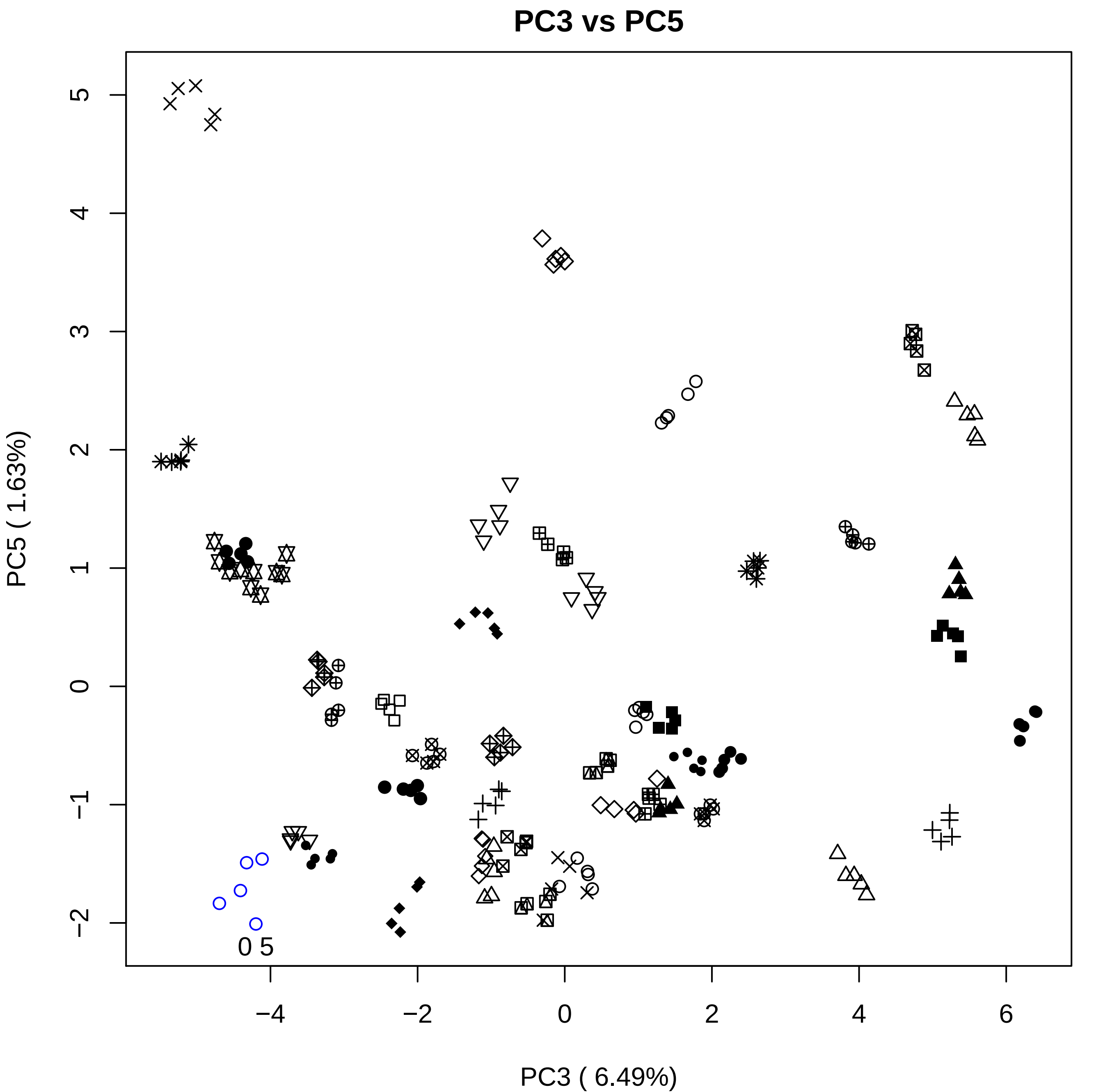
PC3 vs PC5
PC5 ( 1.63%)
PC3 ( 6.49%)
0 5
−4
−2
0
2
4
6
5
4
3
2
1
0
−1
−2

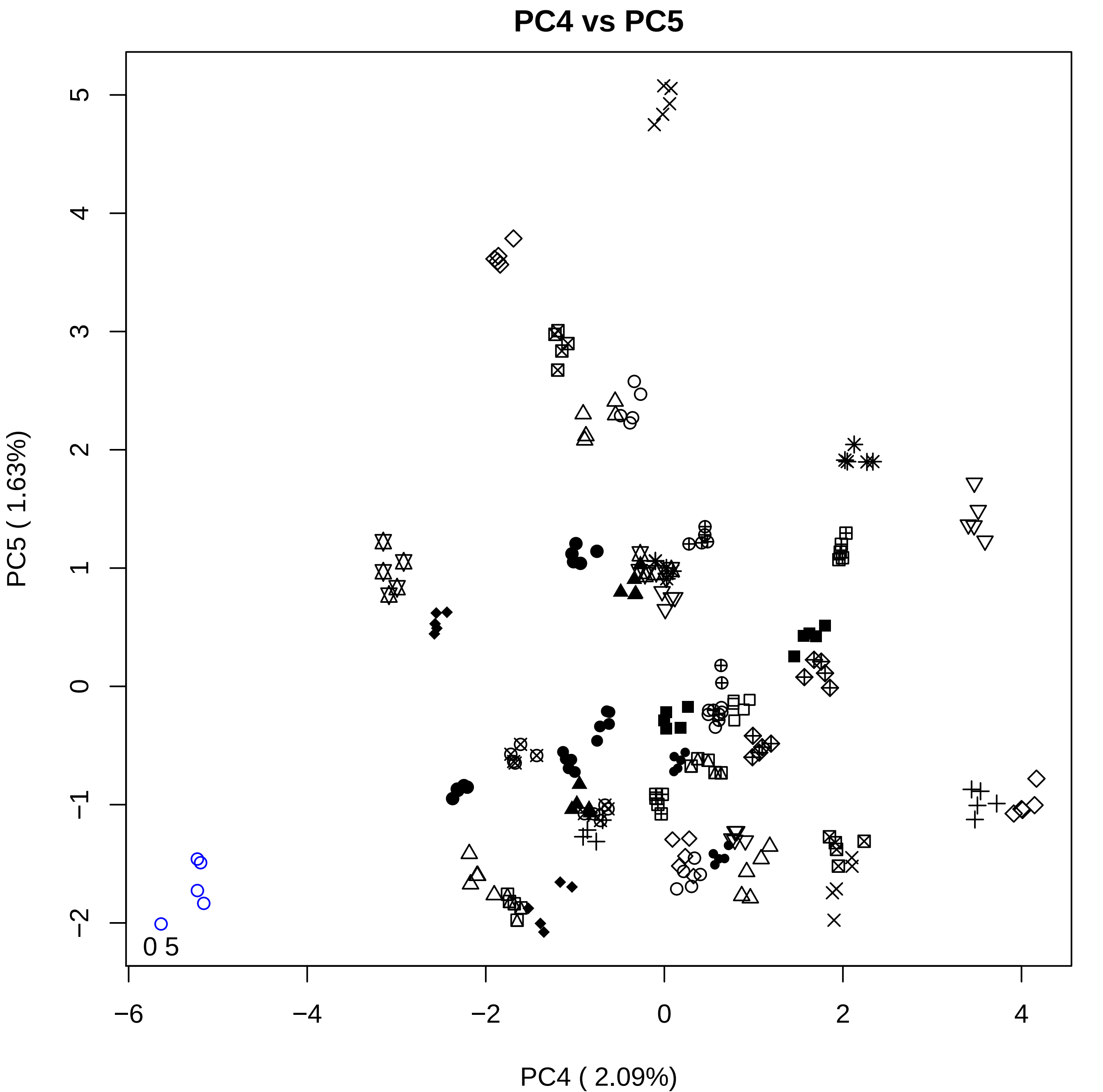

PC4 vs PC5
PC5 ( 1.63%)
PC4 ( 2.09%)
5
4
3
2
1
0
−1
−2
−6
−4
−2
0
2
4
0 5

### PCA of 1%  DeS vs PMH

Scaled Matrix PCAed

7th order polynomial baseline correction performed

Raw spectra smoothed to 21 Neighbours, 2nd order PN, Savtisky Golay

Normalisation between 0 and 100

D2 spectra smoothed to 41 Neighbours, 2nd order PN, Savtisky Golay

Regions Analysed: 3601−2501 and 2001−700cm−1

Samples Analysed: 1 Contam's 0 DeS's 45 PMH's

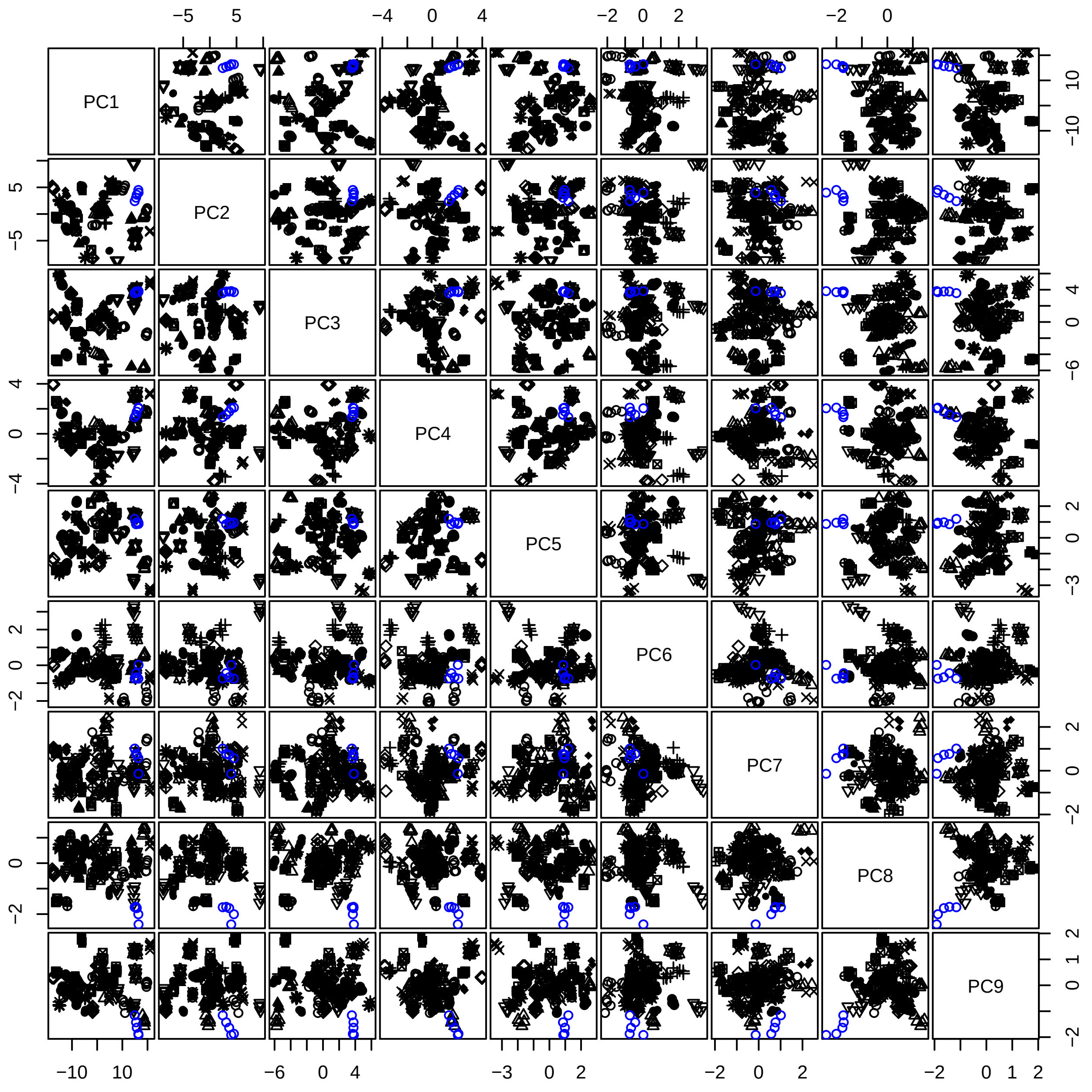

PC1
PC2
PC3
PC4
PC5
PC6
PC7
PC8
PC9
−5
5
−4
0
4
−2
0
2
−2
0
10
−10
5
−5
4
0
−6
4
0
−4
2
0
−3
2
0
−2
2
0
−2
0
−2
2
1
0
−2
−10
10
−6
0
4
−3
0
2
−2
0
2
−2
0
1
2

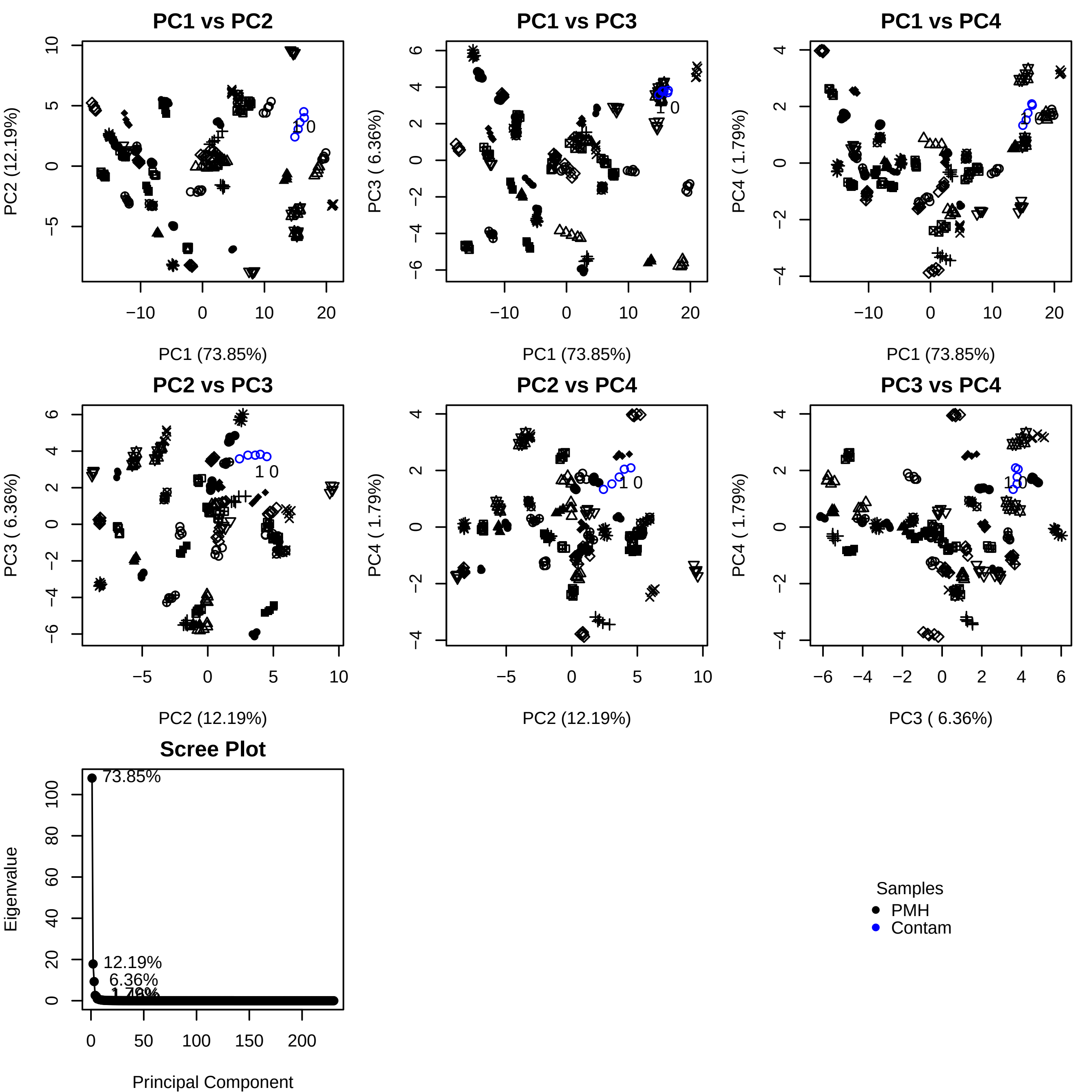
PC1 vs PC2
PC1 vs PC3
PC1 vs PC4
PC2 vs PC3
PC2 vs PC4
PC3 vs PC4
Scree Plot
PC1 (73.85%)
PC2 (12.19%)
PC3 ( 6.36%)
PC4 ( 1.79%)
73.85%
12.19%
6.36%
Eigenvalue
Principal Component
Samples
PMH
Contam

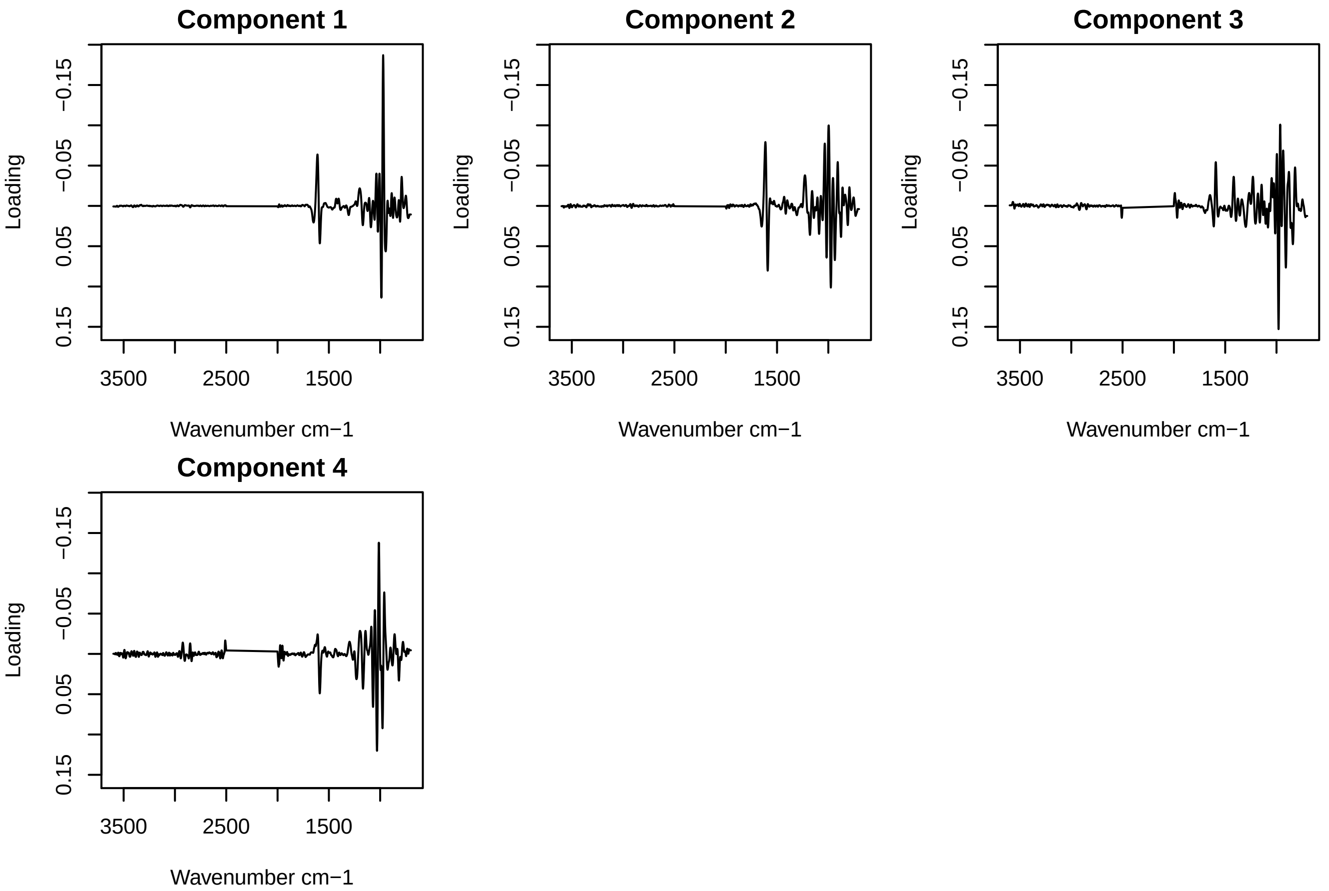

Component 1
Component 2
Component 3
Component 4
Loading
−0.15
−0.05
0.05
0.15
3500
2500
1500
Wavenumber cm−1

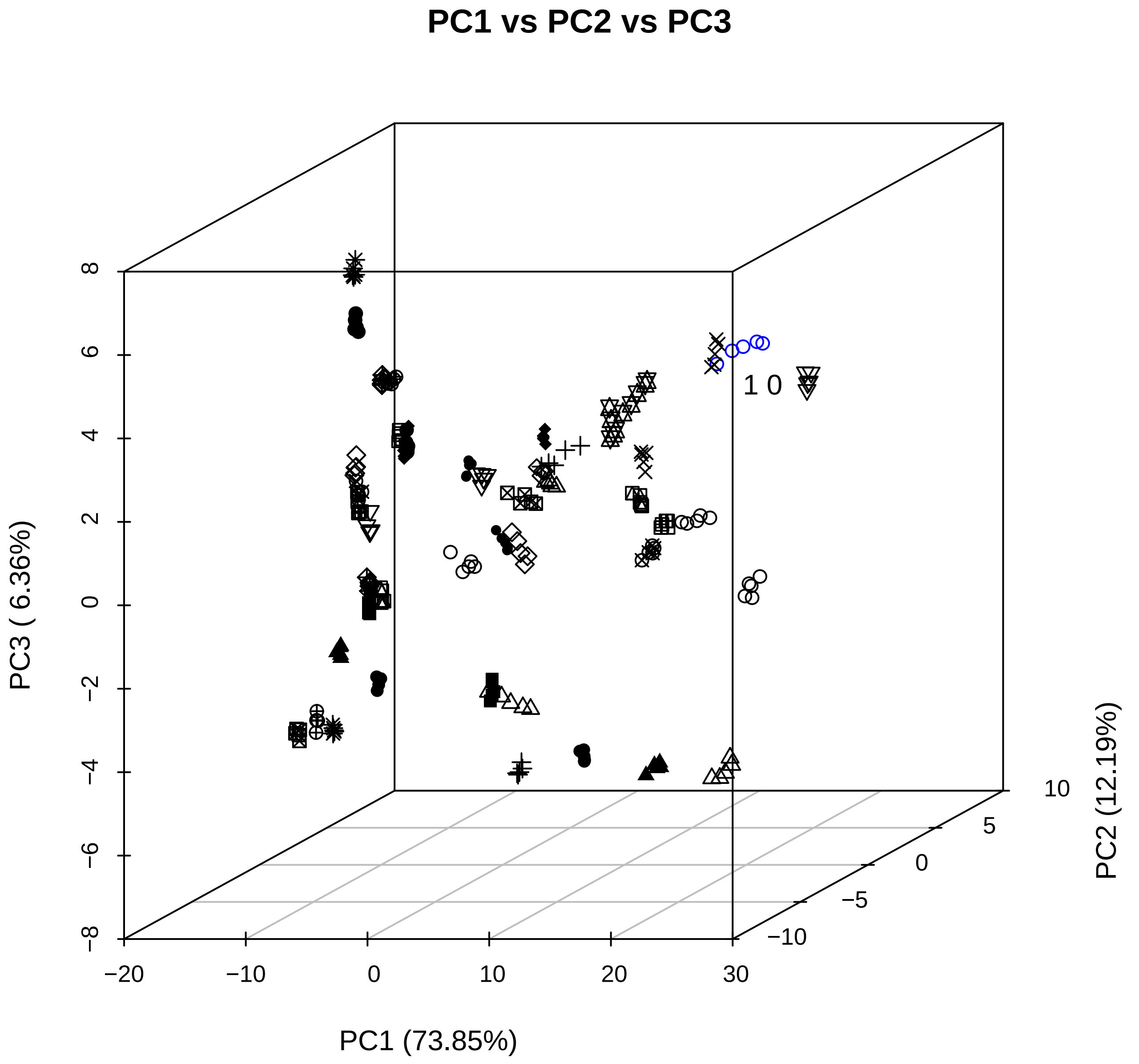
PC1 vs PC2 vs PC3
PC3 ( 6.36%)
8
6
4
2
0
−2
−4
−6
−8
−20
−10
0
10
20
30
PC1 (73.85%)
PC2 (12.19%)
10
5
0
−5
−10
1 0

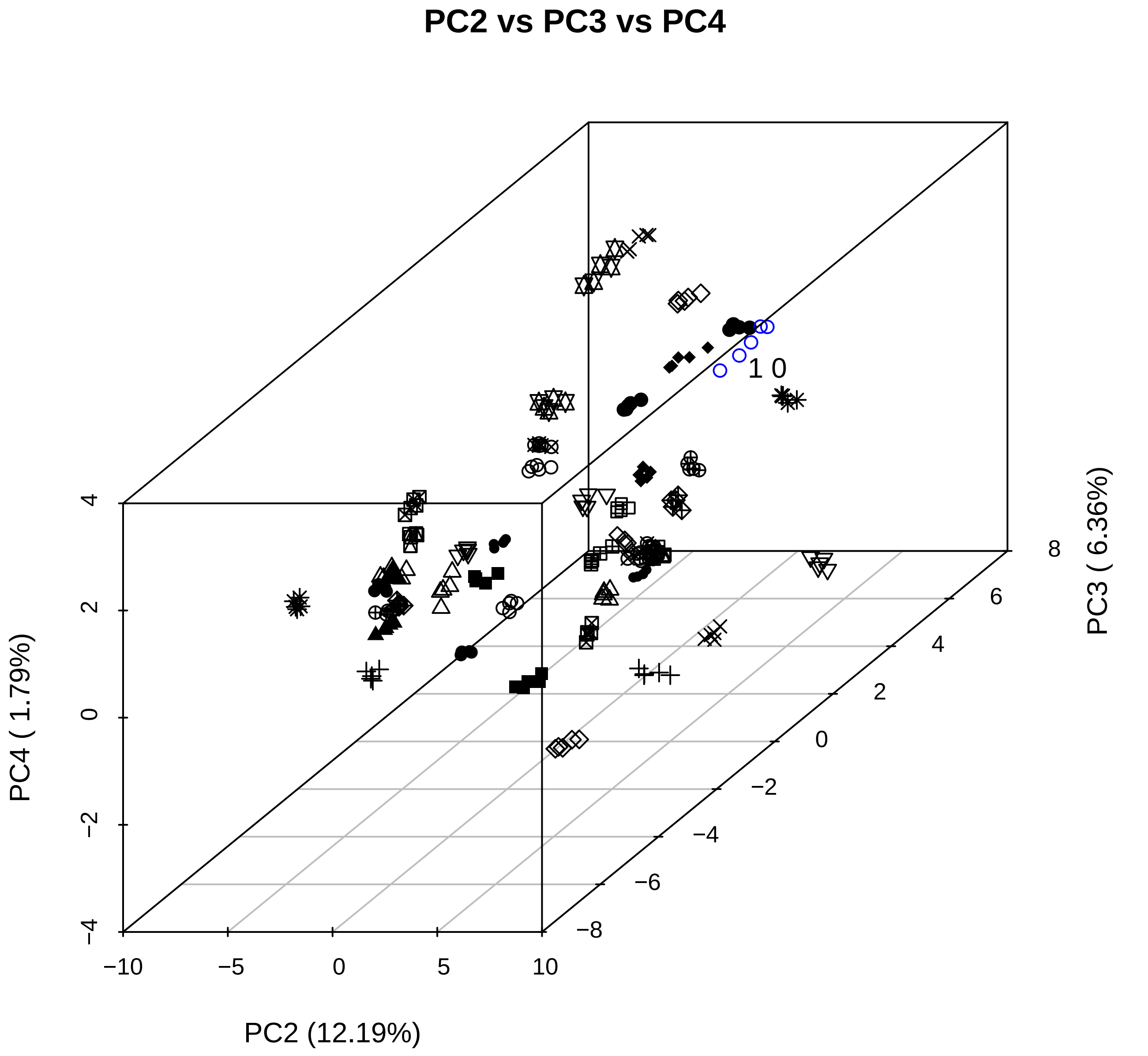

PC2 vs PC3 vs PC4
10
PC2 (12.19%)
PC3 ( 6.36%)
PC4 ( 1.79%)
−10
−5
0
5
10
−8
−6
−4
−2
0
2
4
6
8
−4
−2
0
2
4

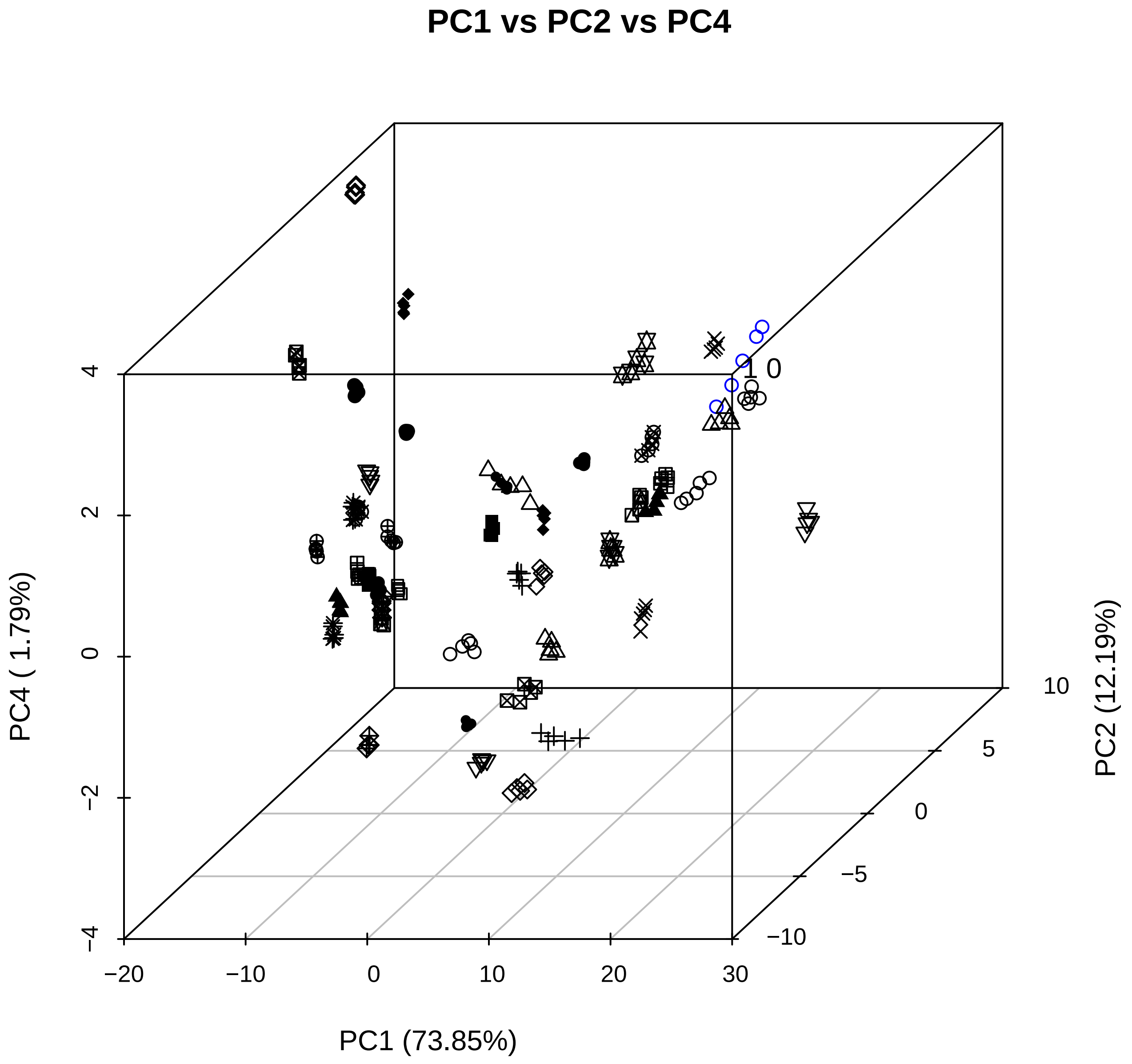
PC1 vs PC2 vs PC4
PC1 (73.85%)
PC2 (12.19%)
PC4 ( 1.79%)
−20
−10
0
10
20
30
−10
−5
0
5
10
−4
−2
0
2
4
1 0

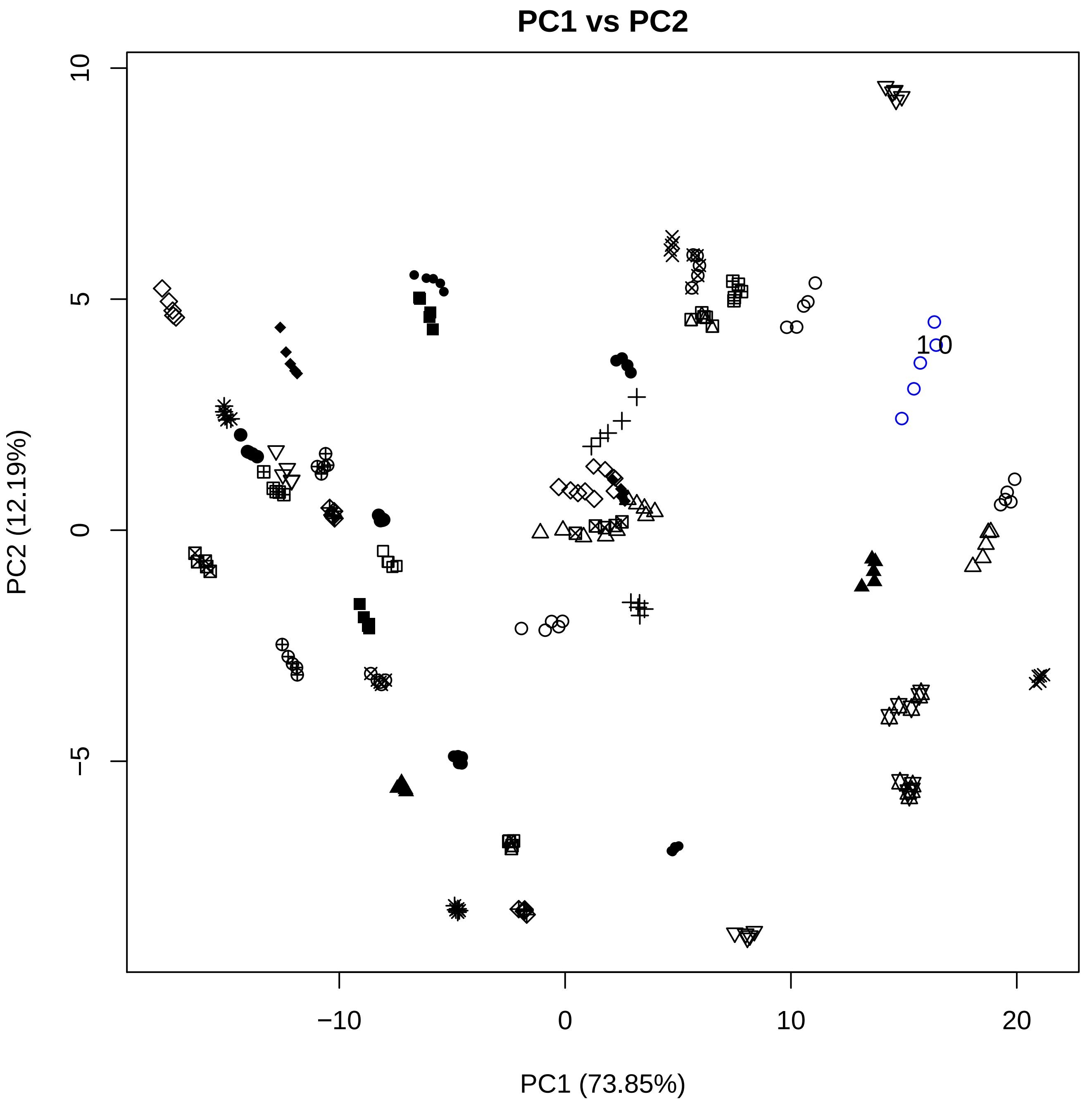

PC1 vs PC2
10
5
0
−5
PC2 (12.19%)
−10
0
10
20
PC1 (73.85%)
10

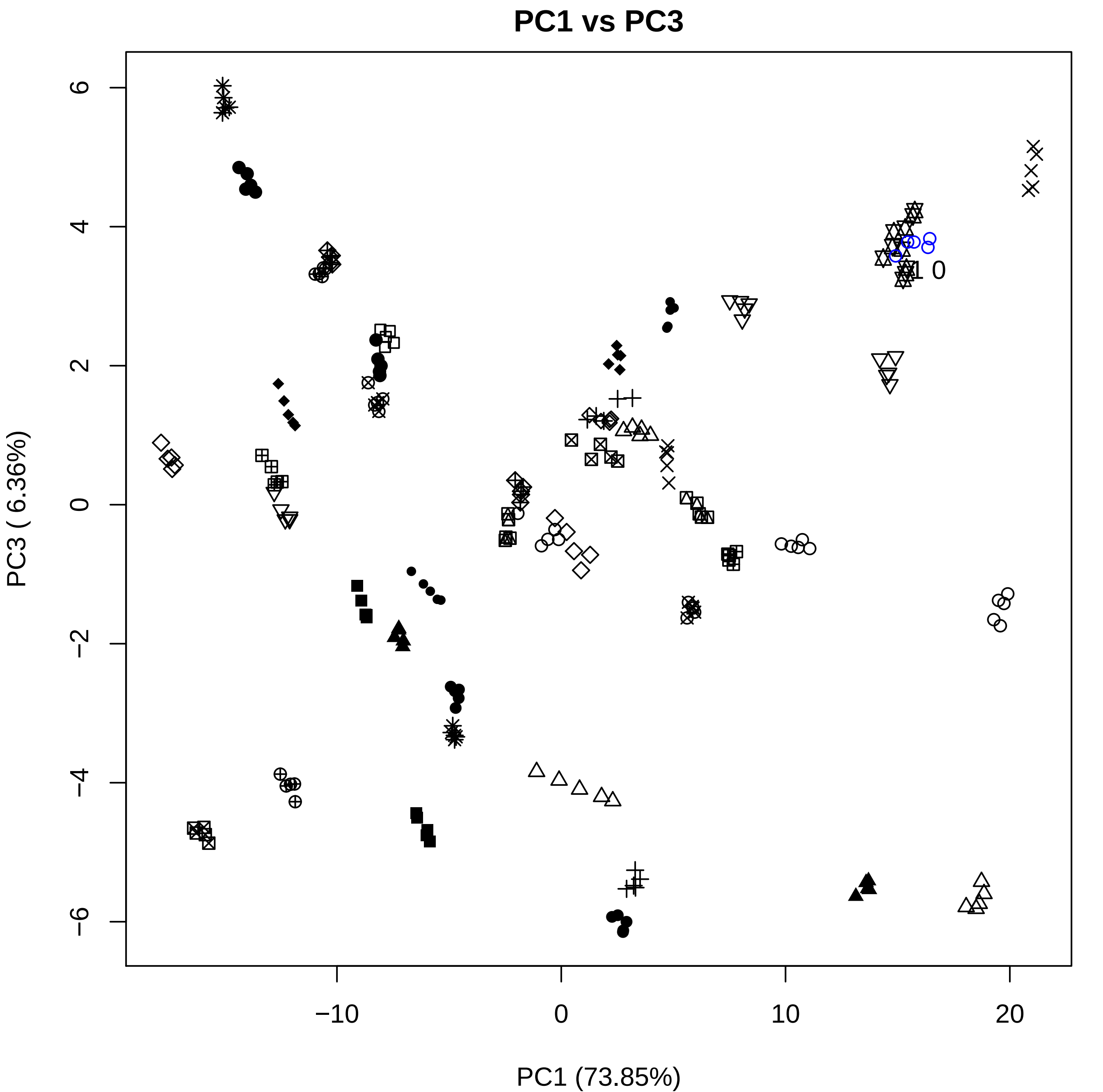
PC1 vs PC3
PC3 ( 6.36%)
PC1 (73.85%)
6
4
2
0
−2
−4
−6
−10
0
10
20
1 0

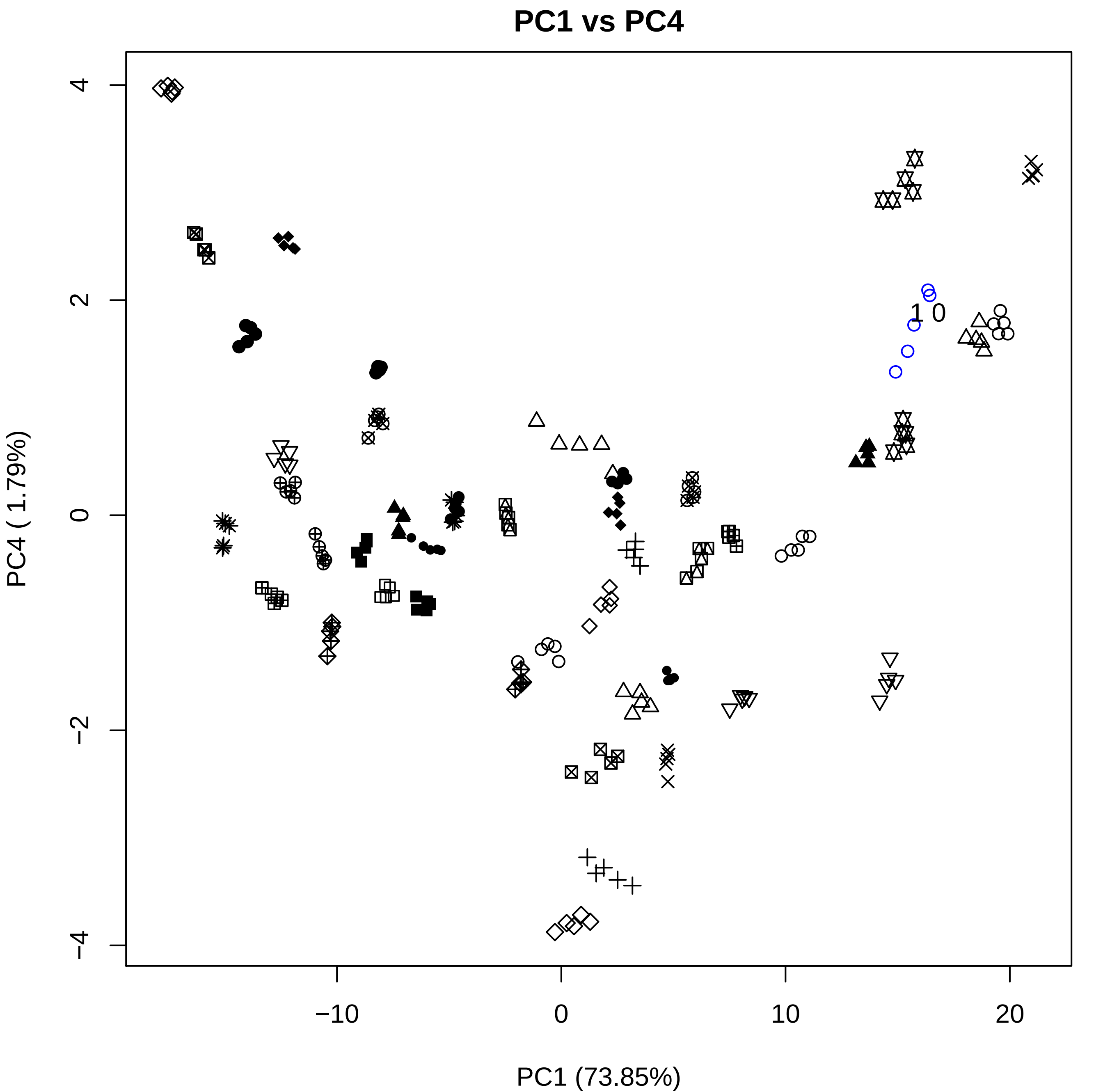
PC1 vs PC4
PC4 ( 1.79%)
PC1 (73.85%)
4
2
0
−2
−4
−10
0
10
20
1 0

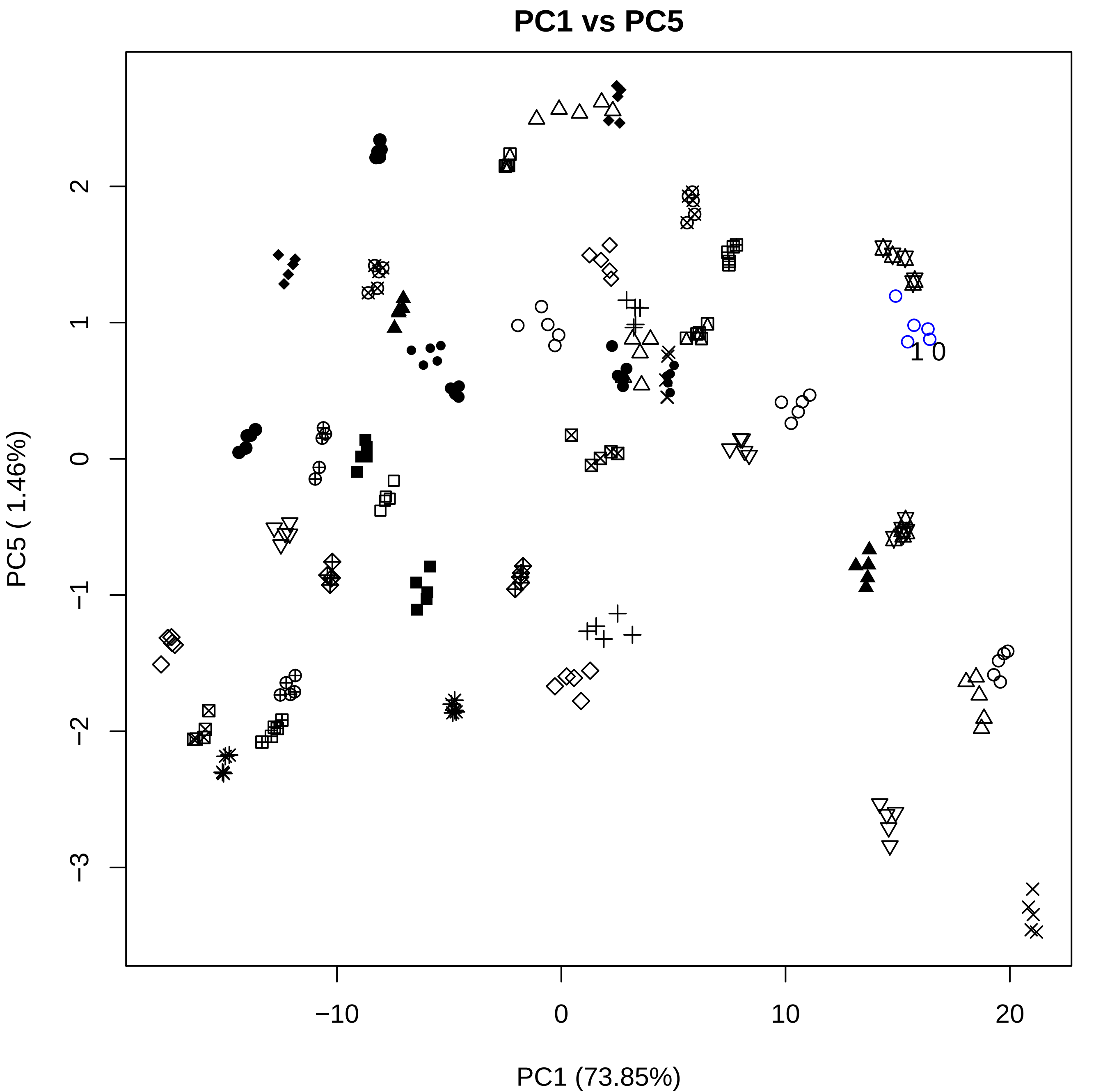
PC1 vs PC5
PC5 ( 1.46%)
PC1 (73.85%)
−10
0
10
20
−3
−2
−1
0
1
2
10

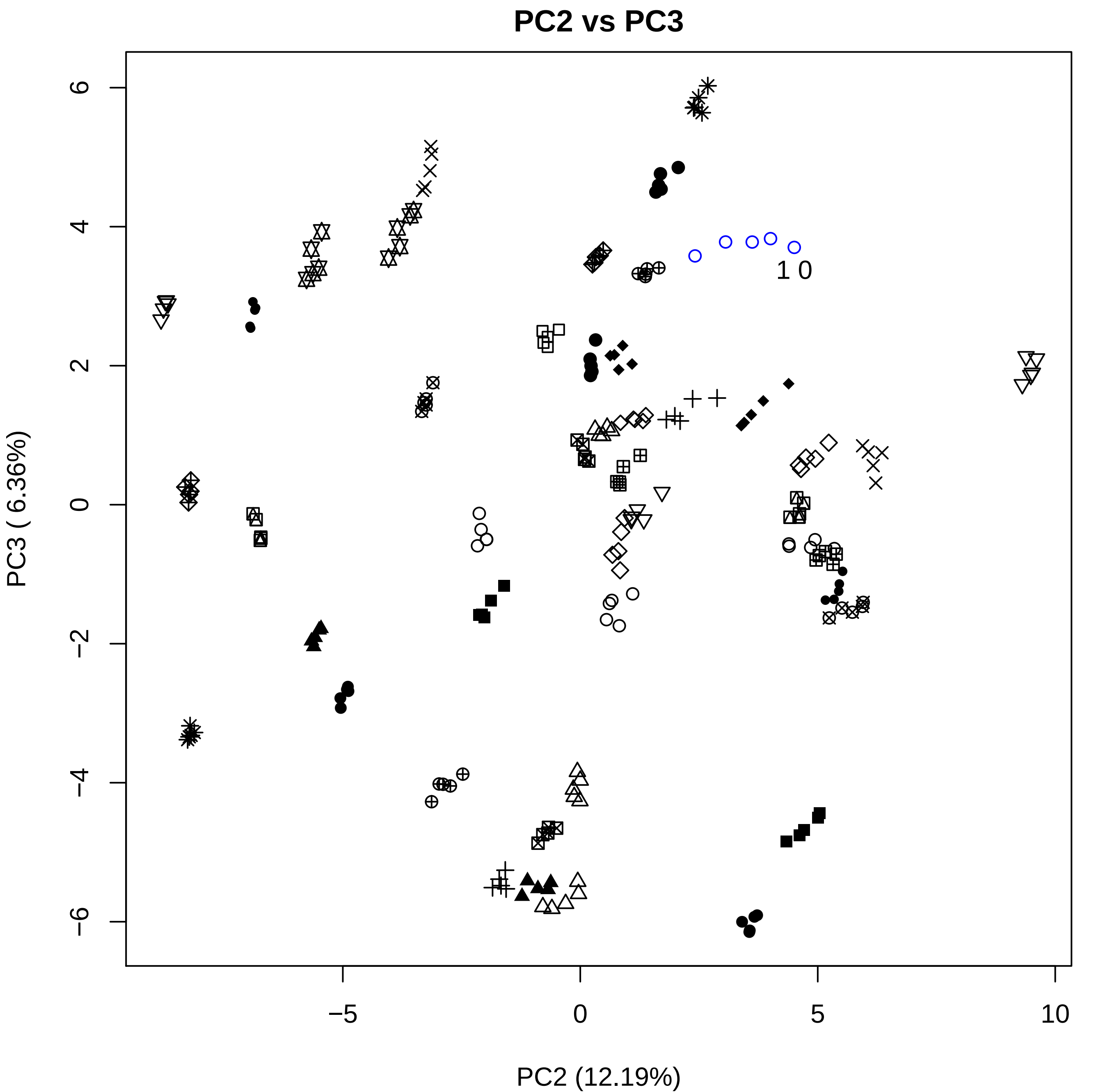
PC2 vs PC3
PC3 ( 6.36%)
PC2 (12.19%)
6
4
2
0
−2
−4
−6
−5
0
5
10
1 0

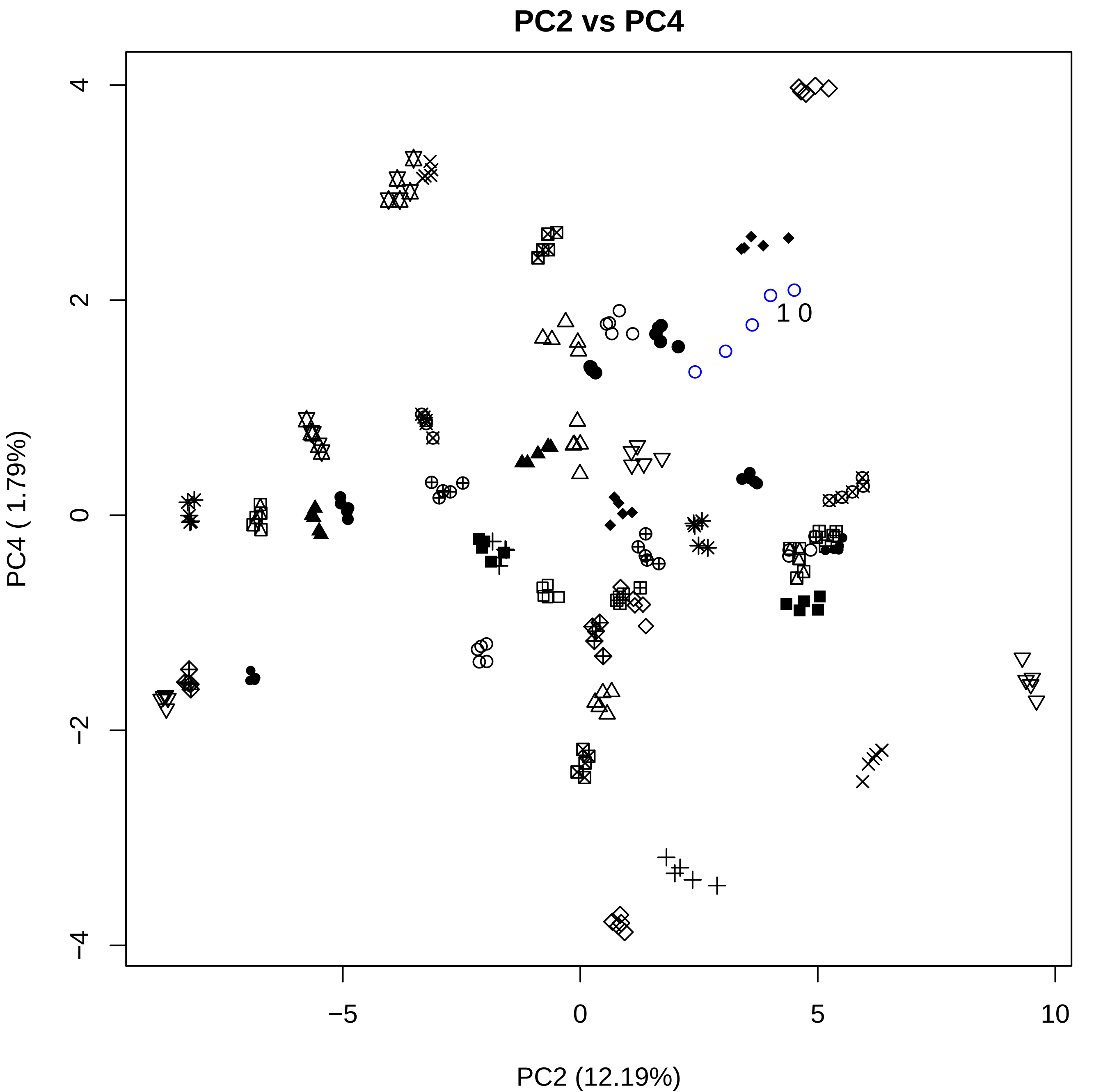

PC2 vs PC4
4
2
0
−2
−4
−5
0
5
10
PC4 ( 1.79%)
PC2 (12.19%)
1 0

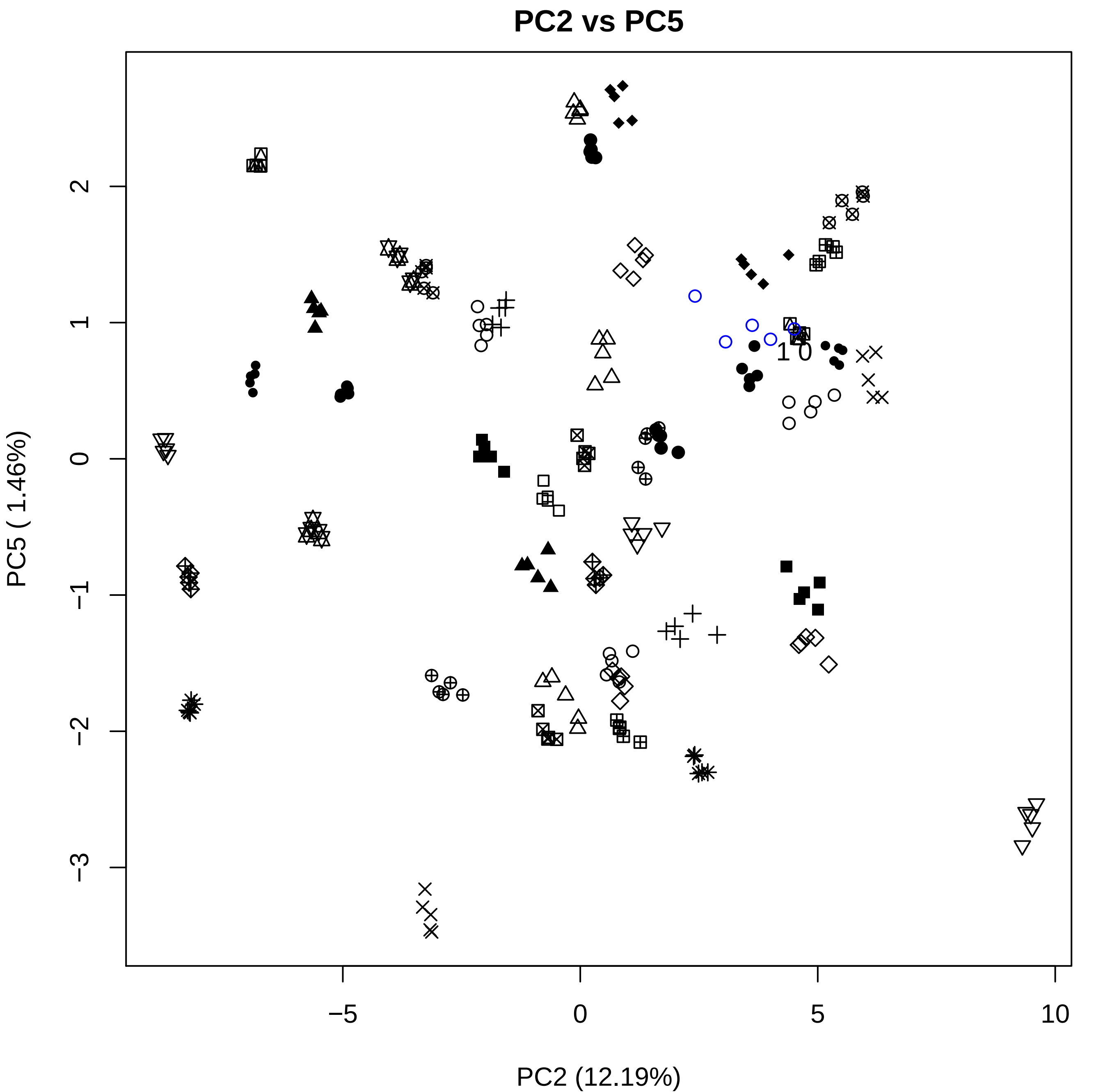
PC2 vs PC5
PC5 ( 1.46%)
PC2 (12.19%)
−5
0
5
10
−3
−2
−1
0
1
2
10

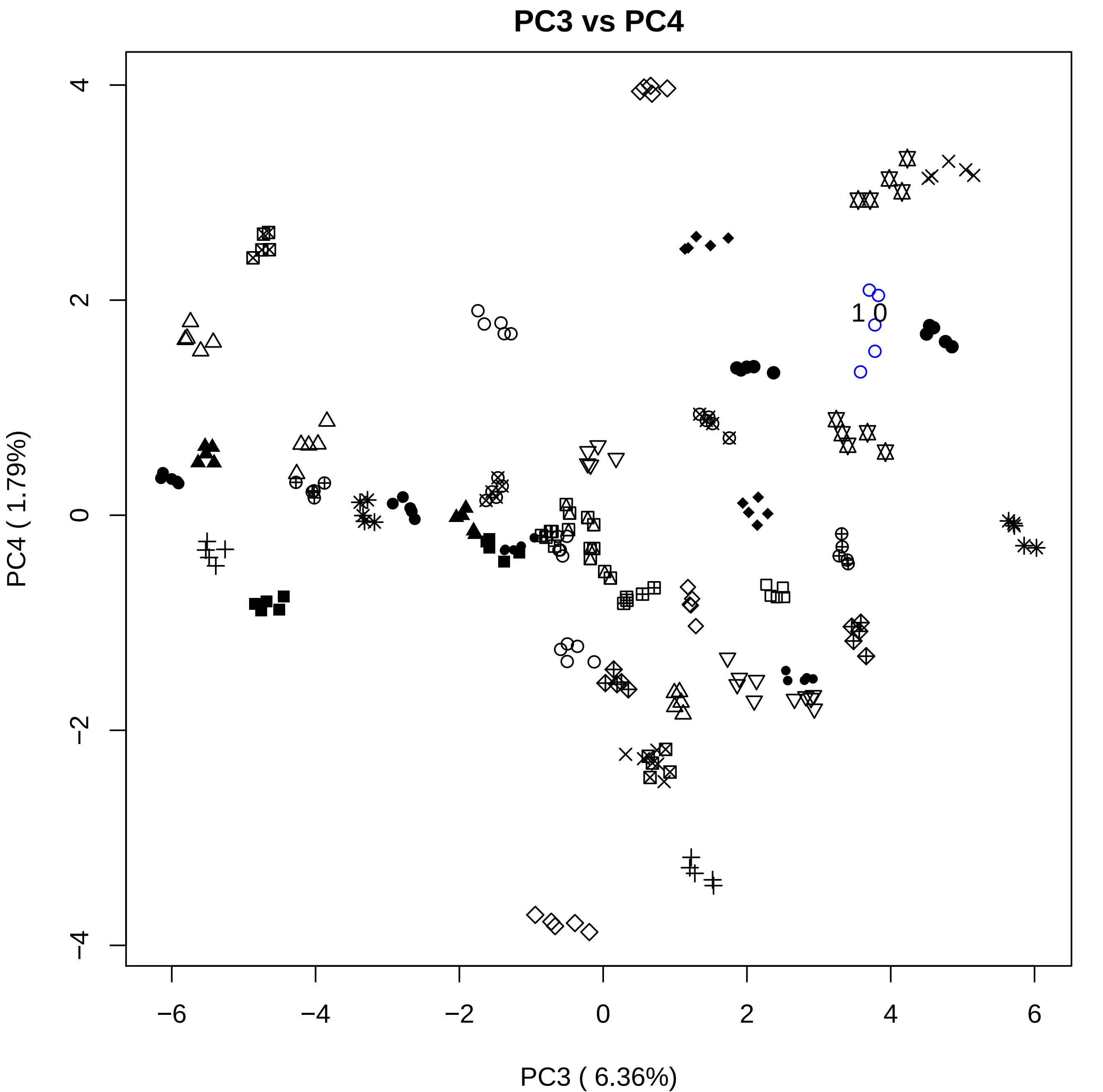
PC3 vs PC4
PC4 ( 1.79%)
PC3 ( 6.36%)
4
2
0
−2
−4
−6
−4
−2
0
2
4
6
1 0

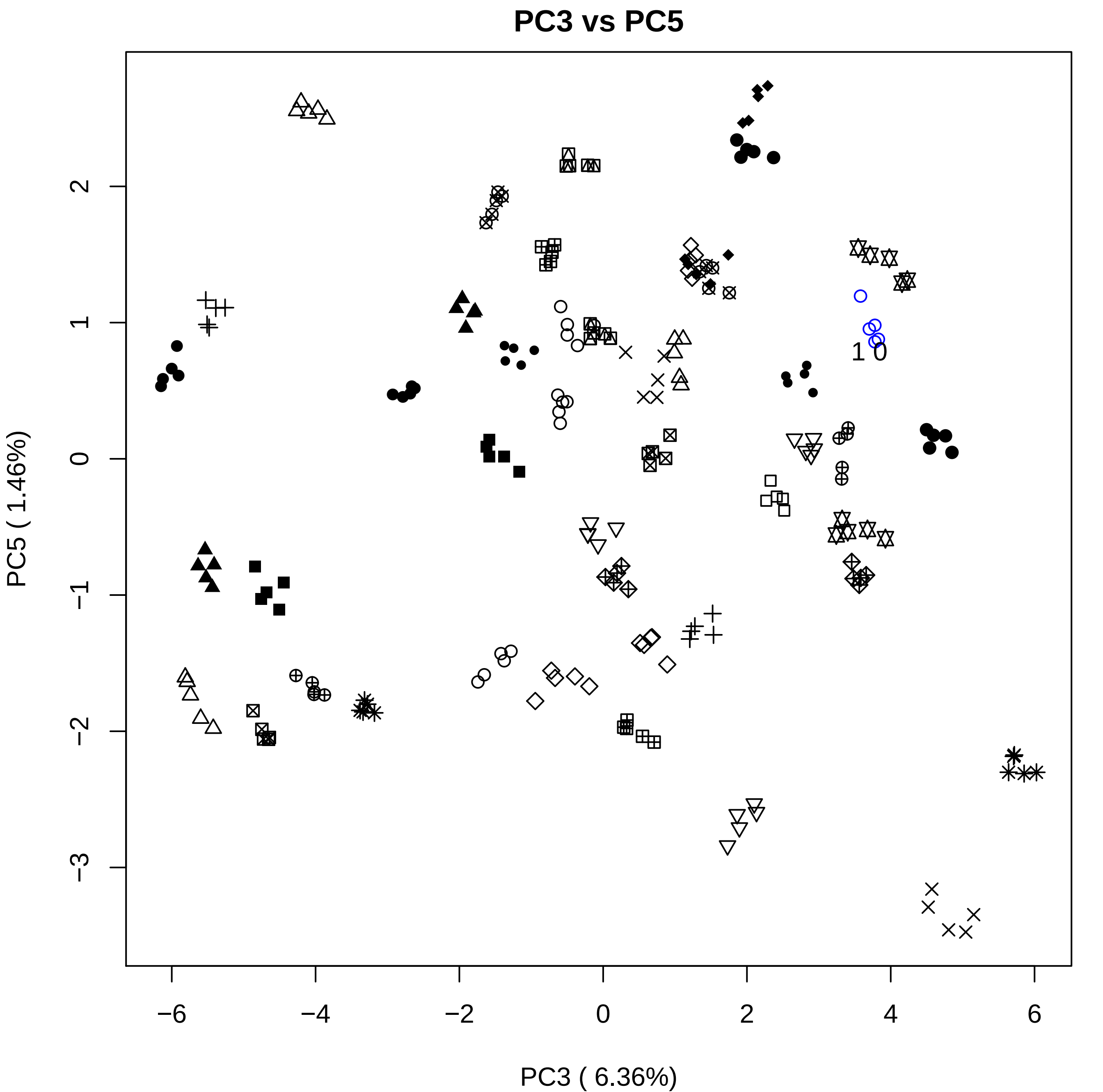
PC3 vs PC5
PC5 ( 1.46%)
PC3 ( 6.36%)
10
−6
−4
−2
0
2
4
6
−3
−2
−1
0
1
2

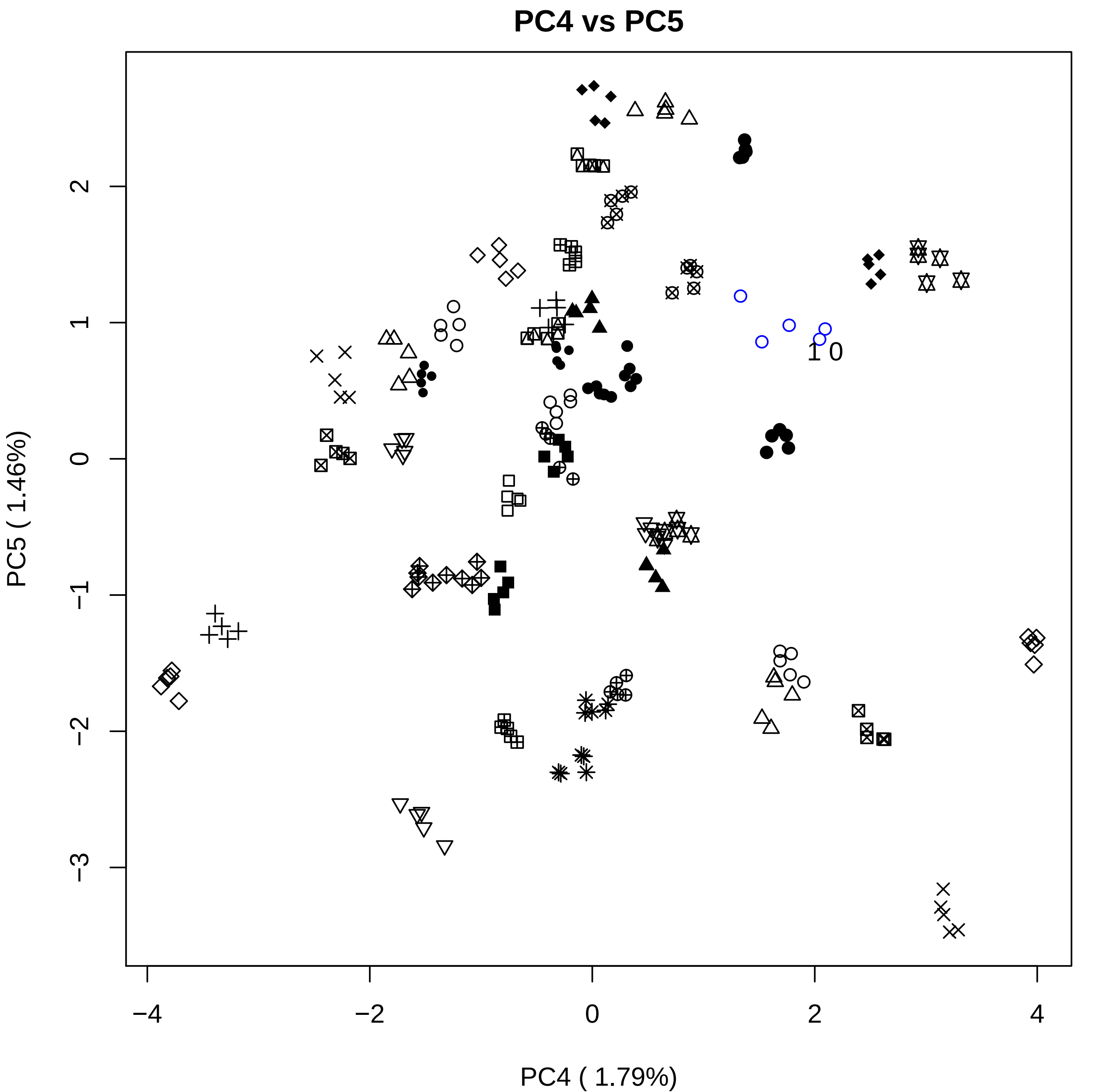
PC4 vs PC5
PC5 ( 1.46%)
PC4 ( 1.79%)
10
−4
−2
0
2
4
−3
−2
−1
0
1
2

### PCA of 2.5%  DeS vs PMH

Scaled Matrix PCAed

7th order polynomial baseline correction performed

Raw spectra smoothed to 21 Neighbours, 2nd order PN, Savtisky Golay

Normalisation between 0 and 100

D2 spectra smoothed to 41 Neighbours, 2nd order PN, Savtisky Golay

Regions Analysed: 3601−2501 and 2001−700cm−1

Samples Analysed: 1 Contam's 0 DeS's 45 PMH's

Keele University AD

Sun Dec 09 17:04:06 2018

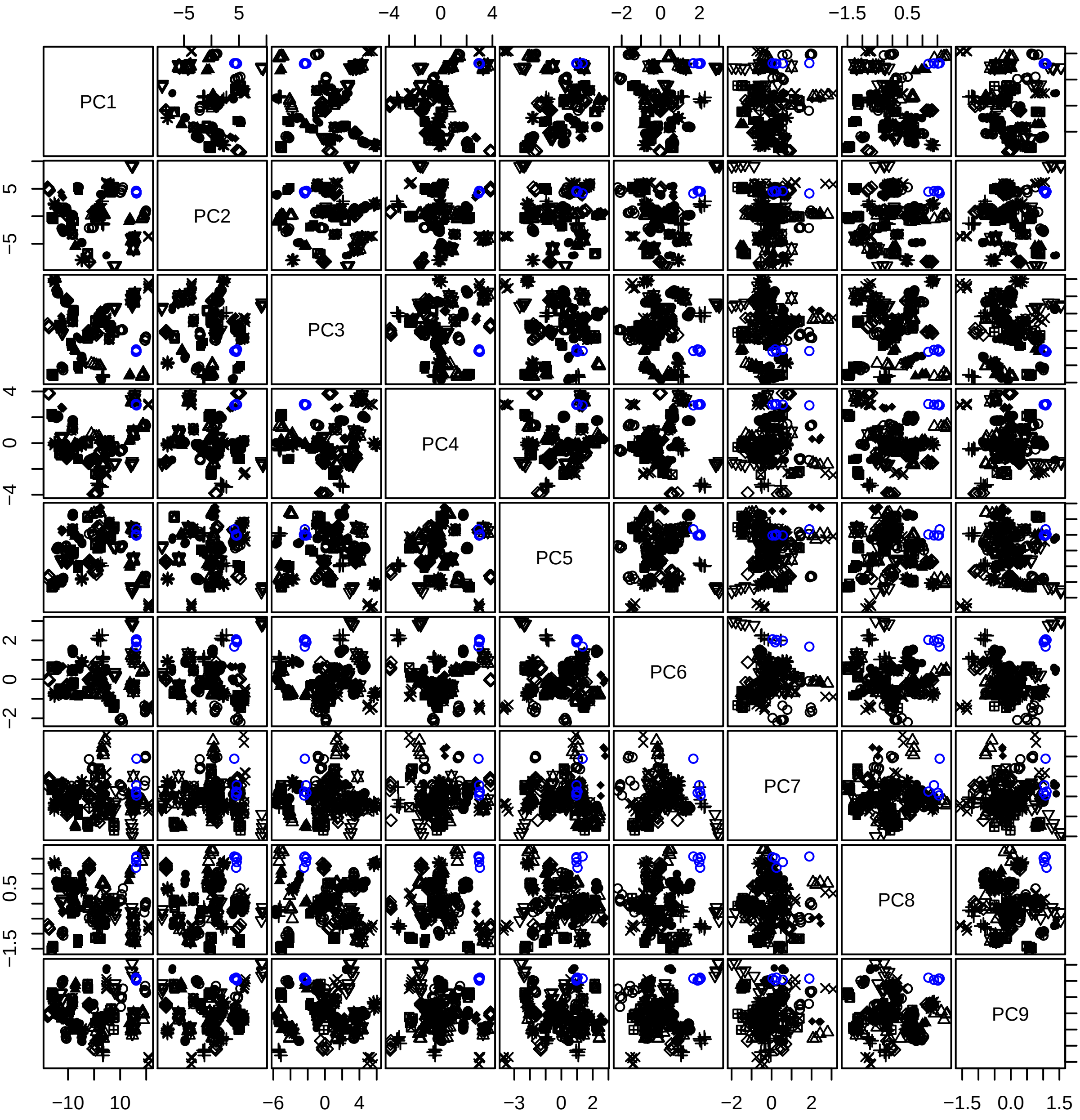
PC1
PC2
PC3
PC4
PC5
PC6
PC7
PC8
PC9

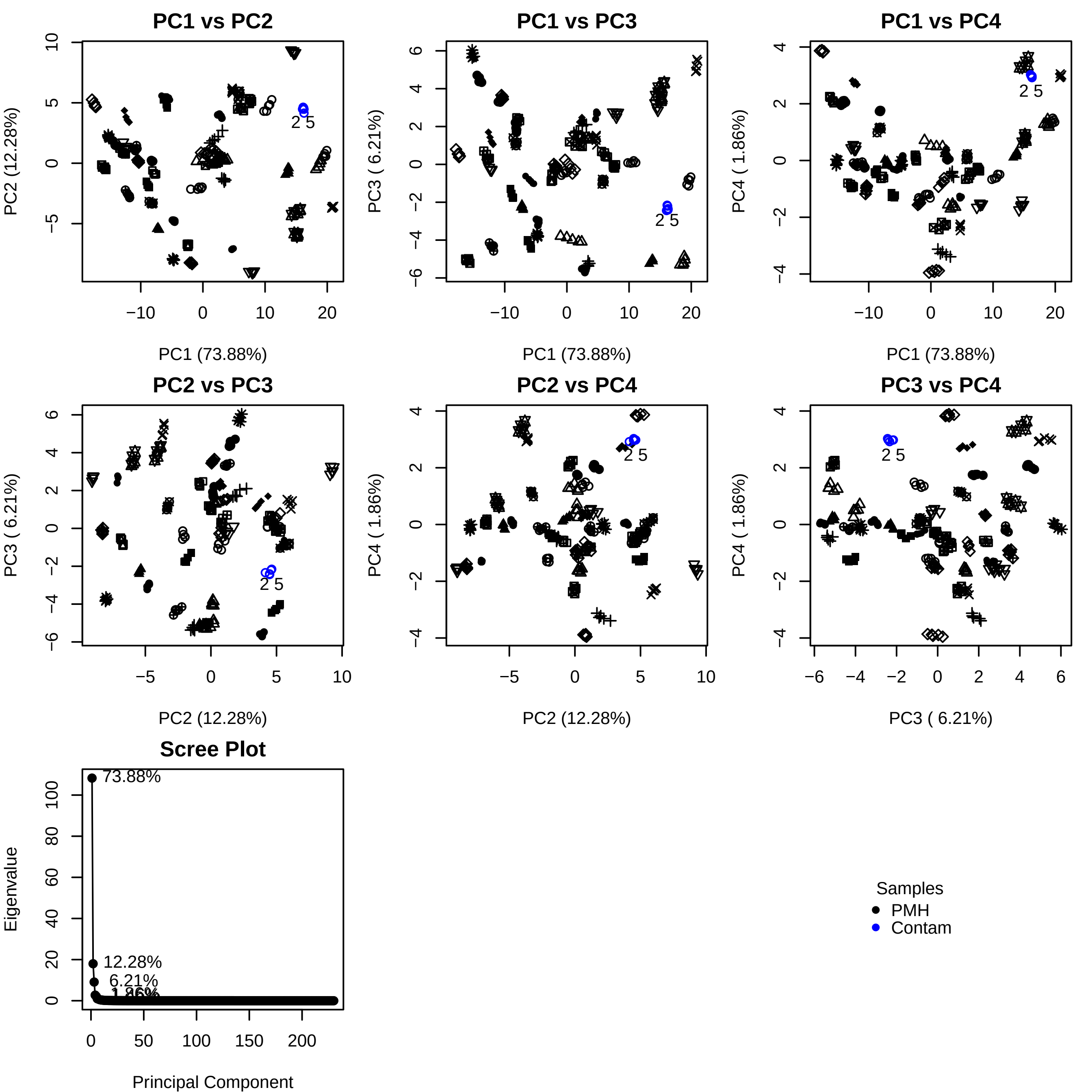
PC1 vs PC2
PC1 vs PC3
PC1 vs PC4
PC1 (73.88%)
PC2 (12.28%)
PC3 ( 6.21%)
PC4 ( 1.86%)
PC2 vs PC3
PC2 vs PC4
PC3 vs PC4
2 5
Scree Plot
73.88%
12.28%
6.21%
Eigenvalue
Principal Component
Samples
PMH
Contam

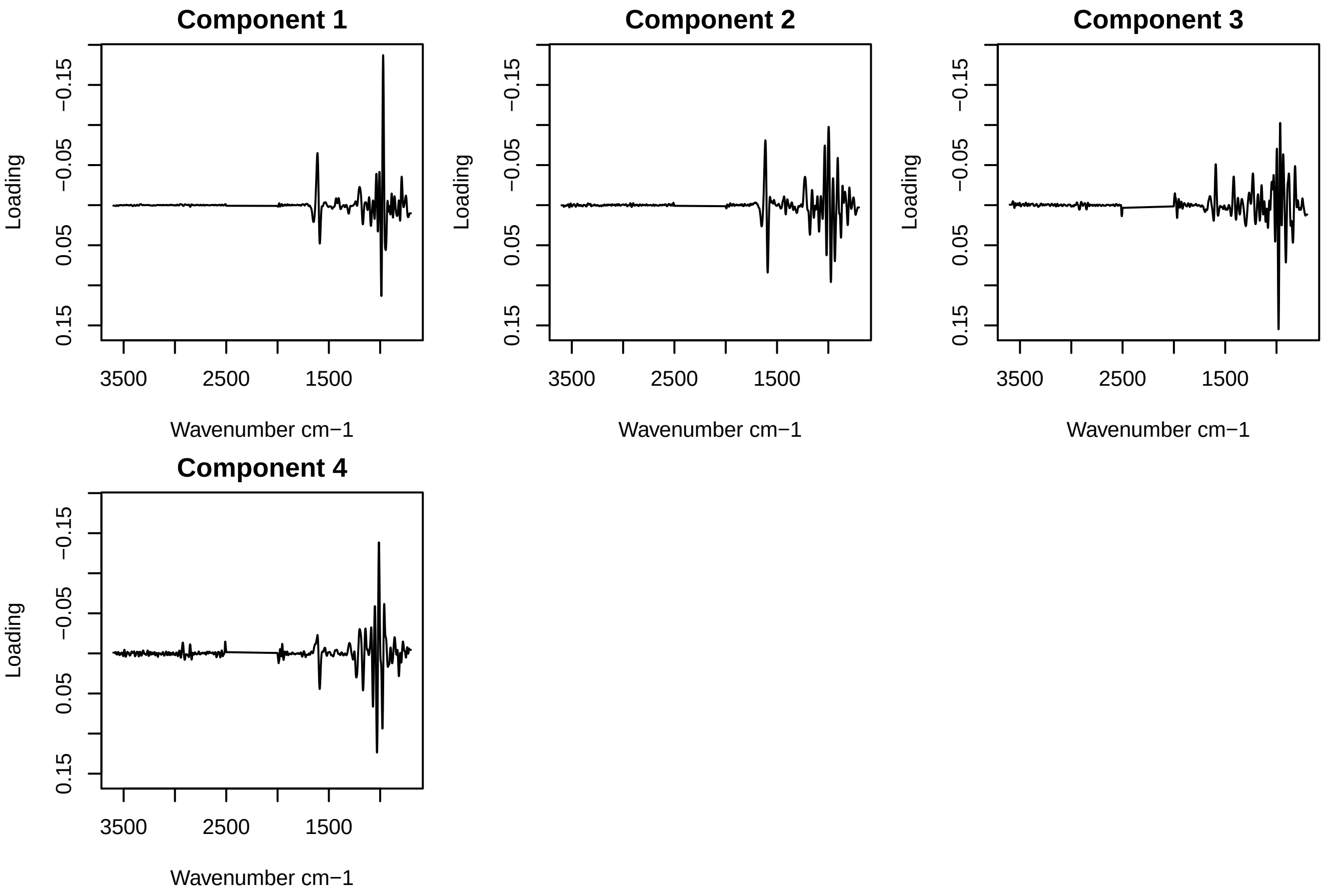

Component 1
Component 2
Component 3
Component 4
Loading
−0.15
−0.05
0.05
0.15
3500
2500
1500
Wavenumber cm−1

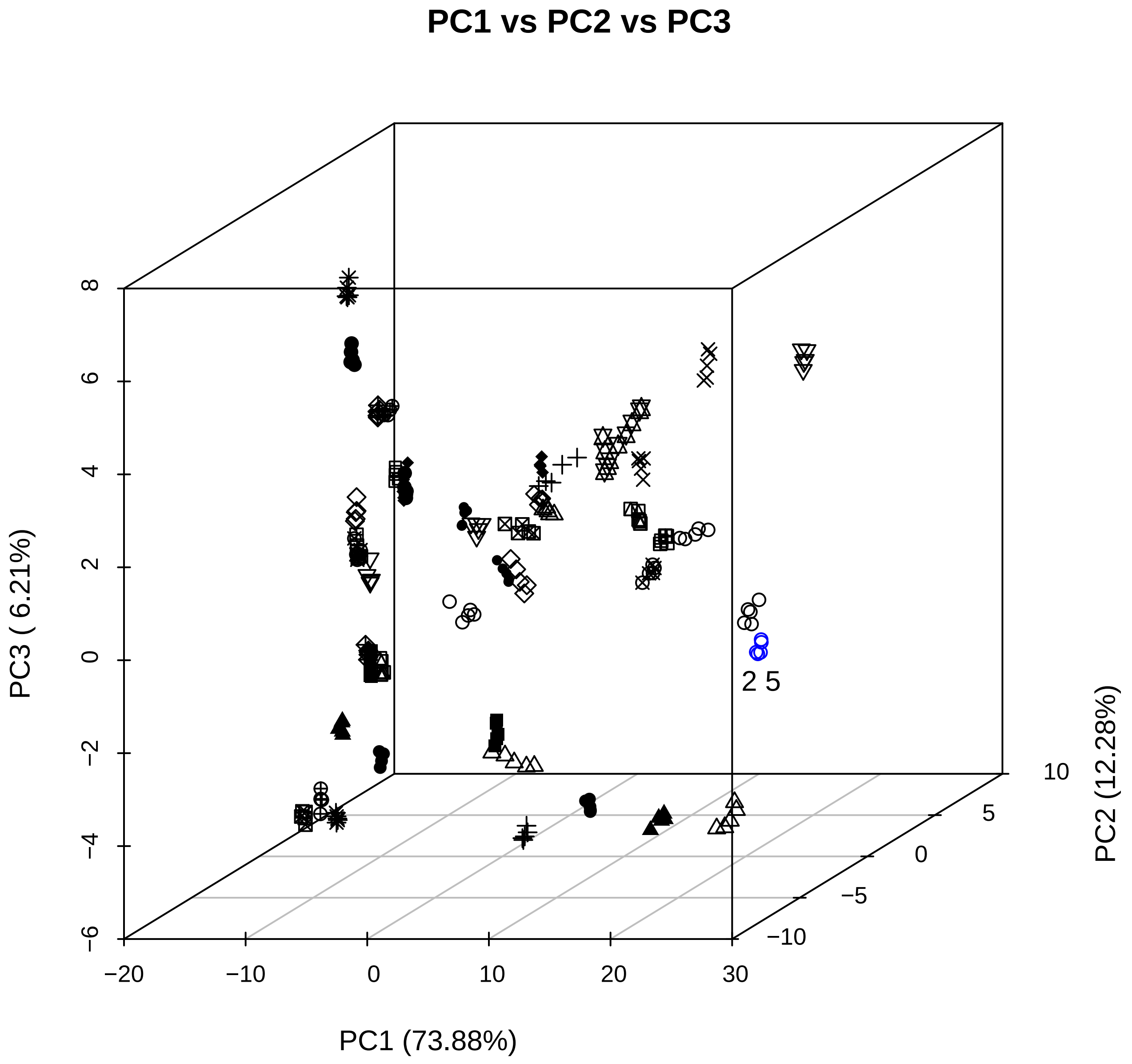
PC1 vs PC2 vs PC3
PC1 (73.88%)
PC2 (12.28%)
PC3 ( 6.21%)
−20
−10
0
10
20
30
−10
−5
0
5
10
−6
−4
−2
0
2
4
6
8
2 5

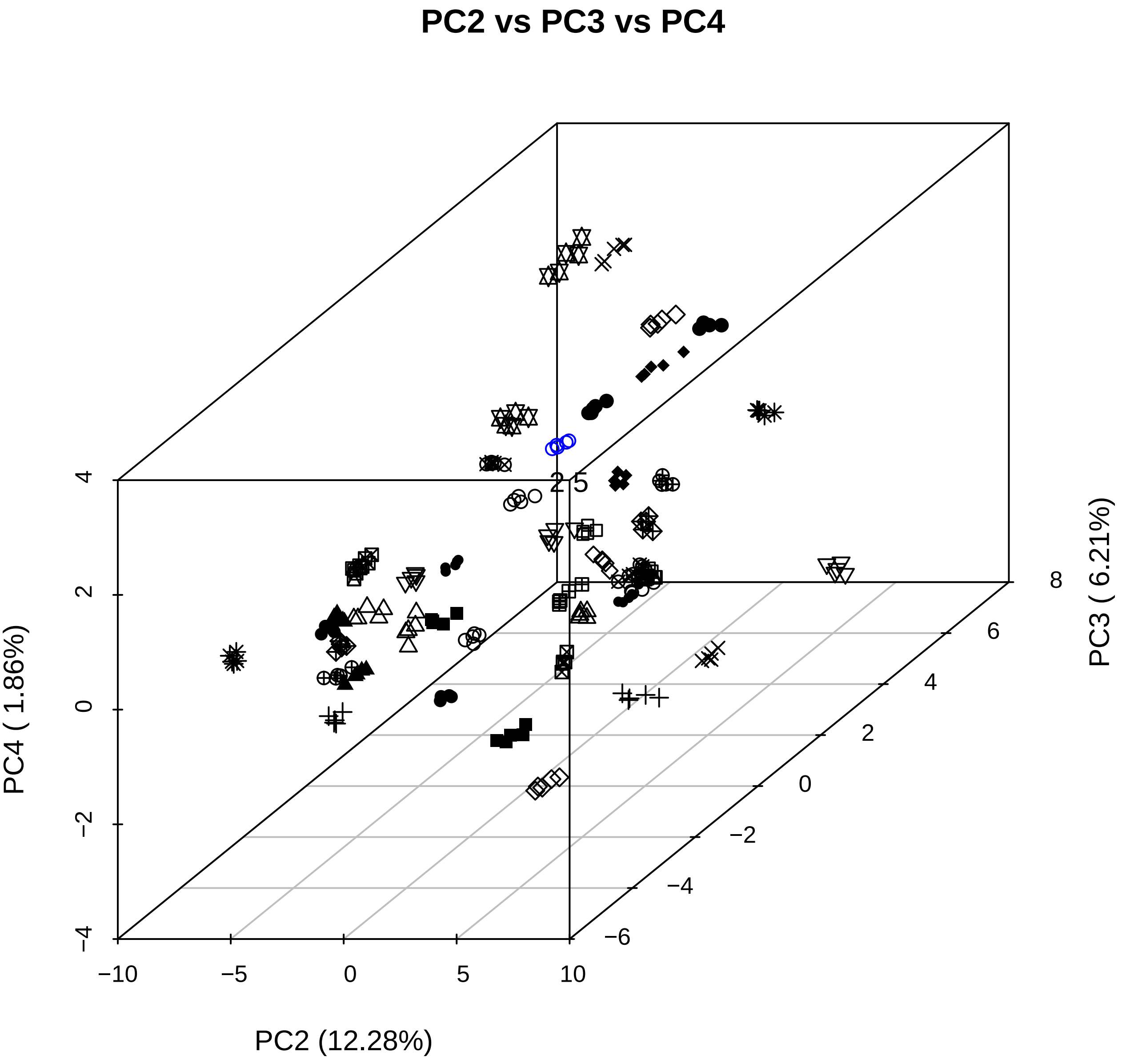
PC2 vs PC3 vs PC4
PC2 (12.28%)
PC4 ( 1.86%)
PC3 ( 6.21%)
−10
−5
0
5
10
−4
−2
0
2
4
−6
−4
−2
0
2
4
6
8
2.5

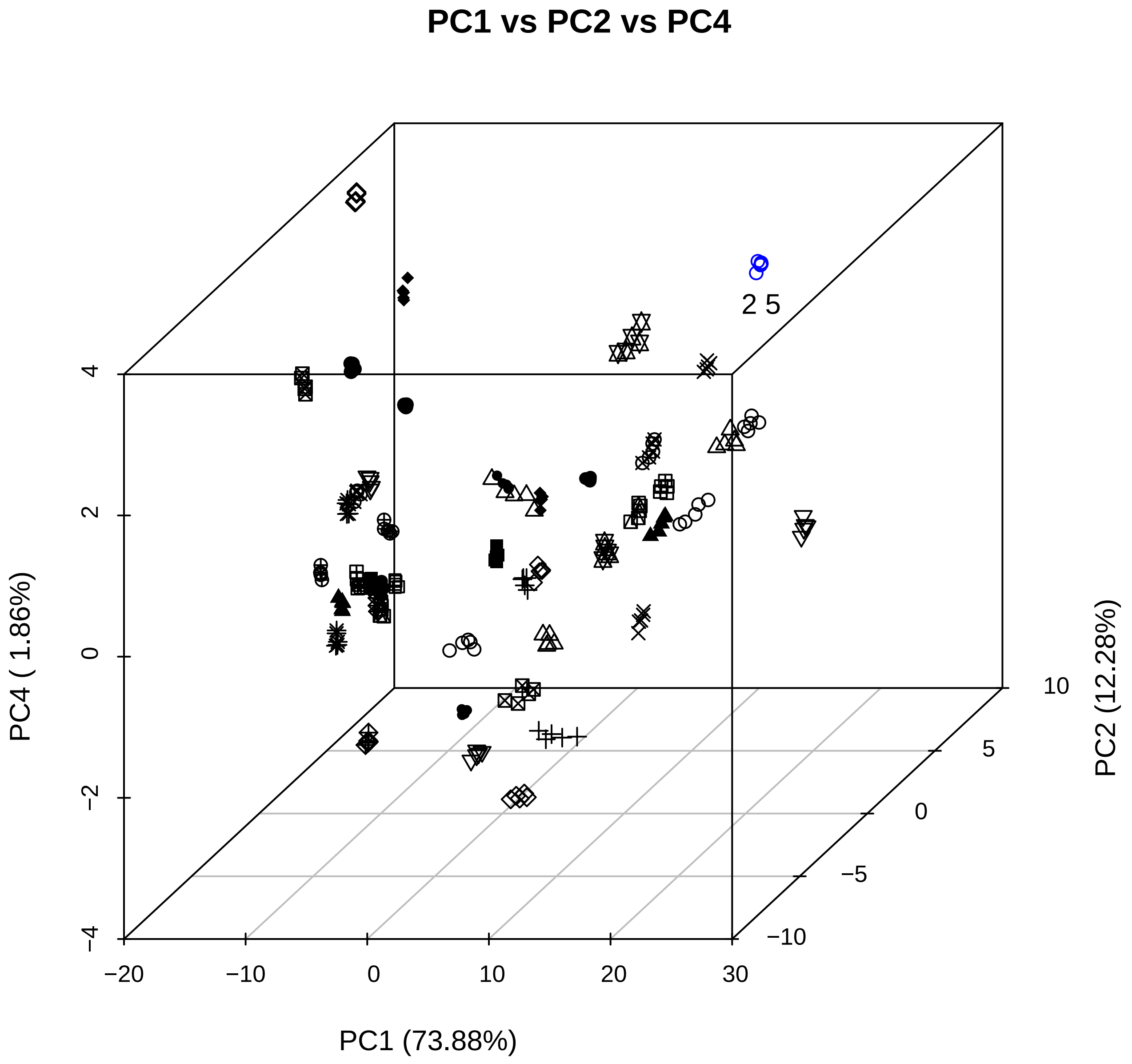
PC1 vs PC2 vs PC4
2 5
PC1 (73.88%)
PC2 (12.28%)
PC4 ( 1.86%)
−20
−10
0
10
20
30
−10
−5
0
5
10
−4
−2
0
2
4

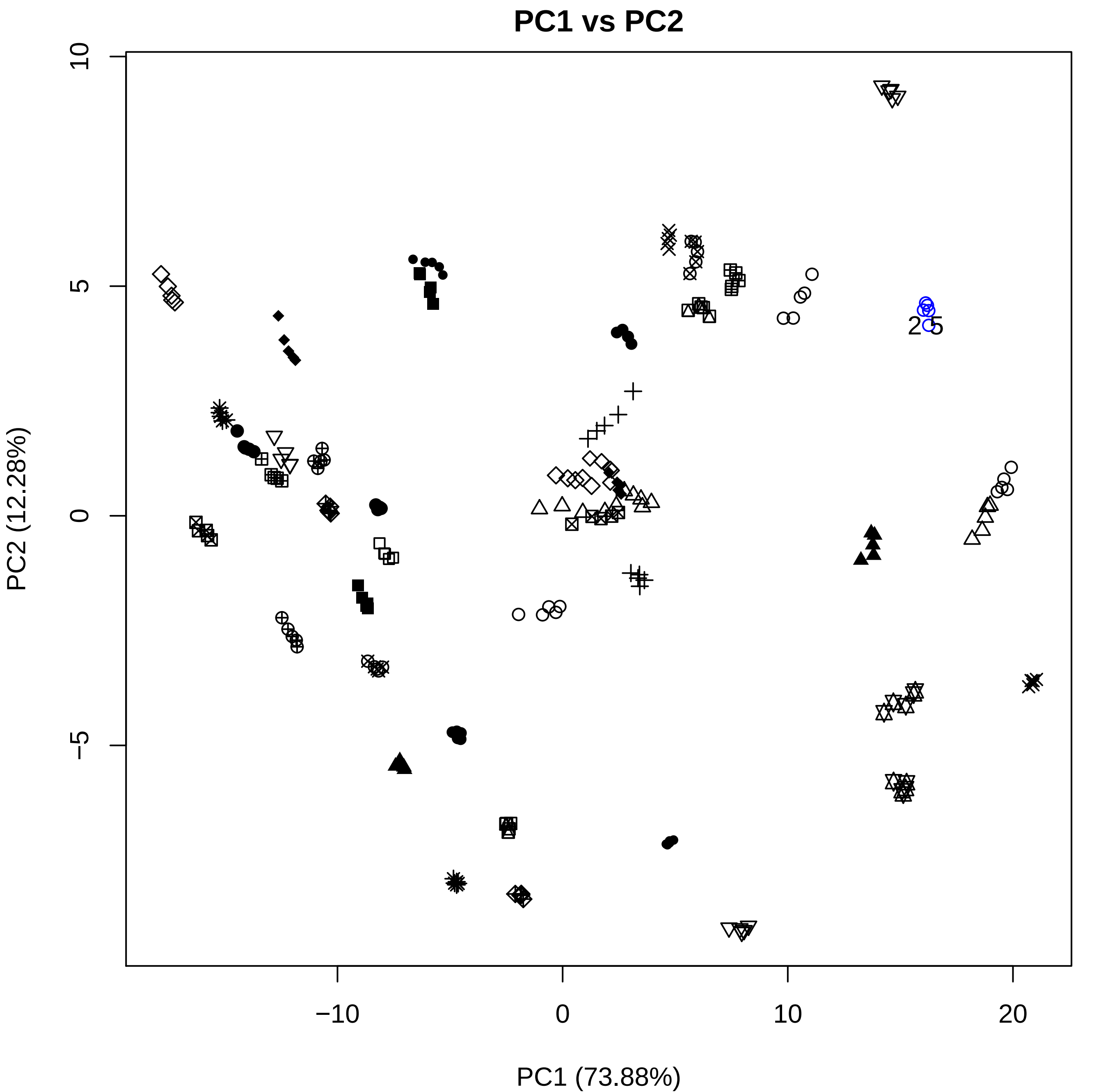
PC1 vs PC2
PC2 (12.28%)
PC1 (73.88%)
10
5
0
−5
−10
0
10
20
25

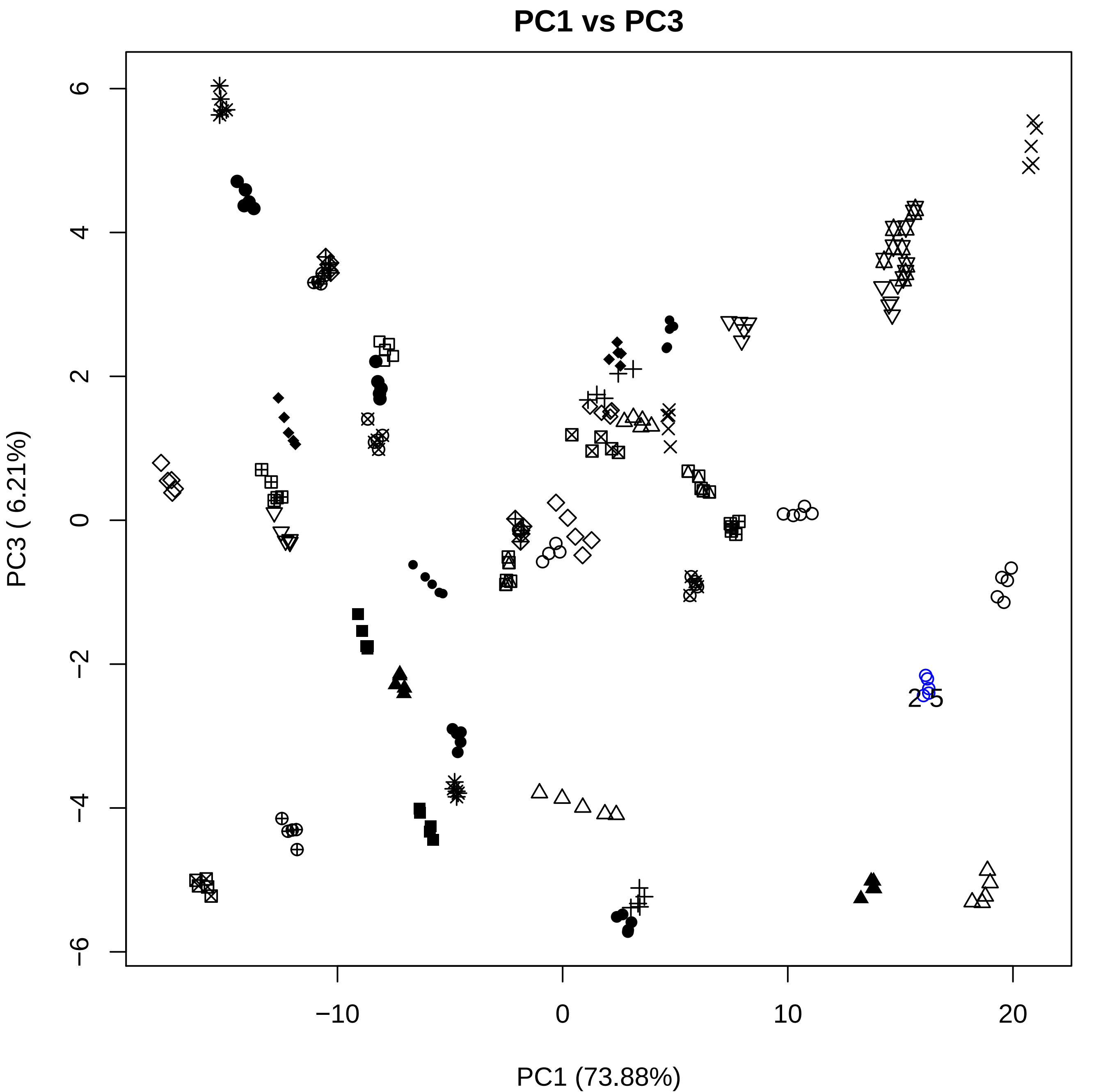
PC1 vs PC3
PC3 ( 6.21%)
PC1 (73.88%)
6
4
2
0
−2
−4
−6
−10
0
10
20
25

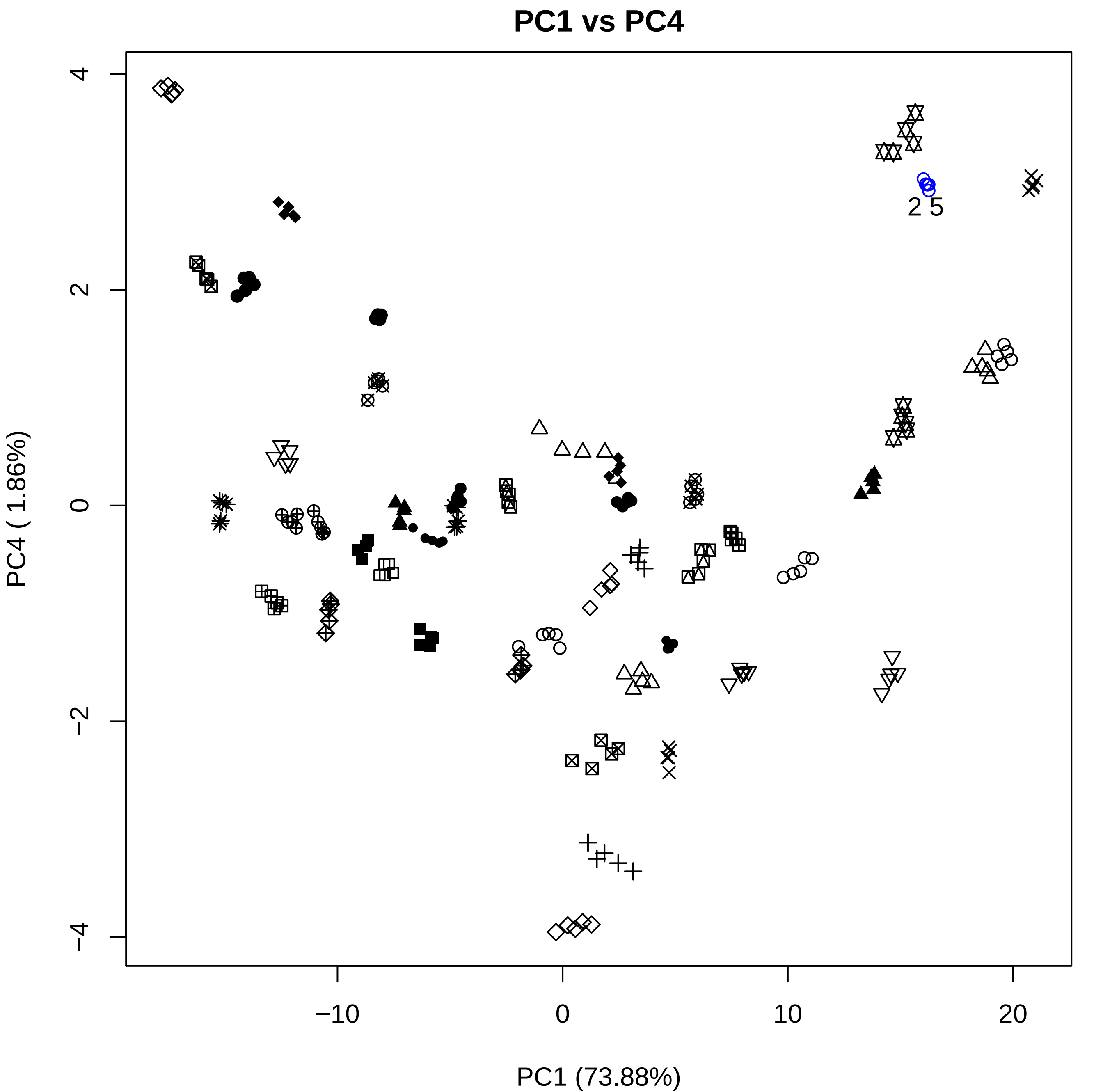
PC1 vs PC4
PC4 ( 1.86%)
PC1 (73.88%)
4
2
0
−2
−4
−10
0
10
20
2 5

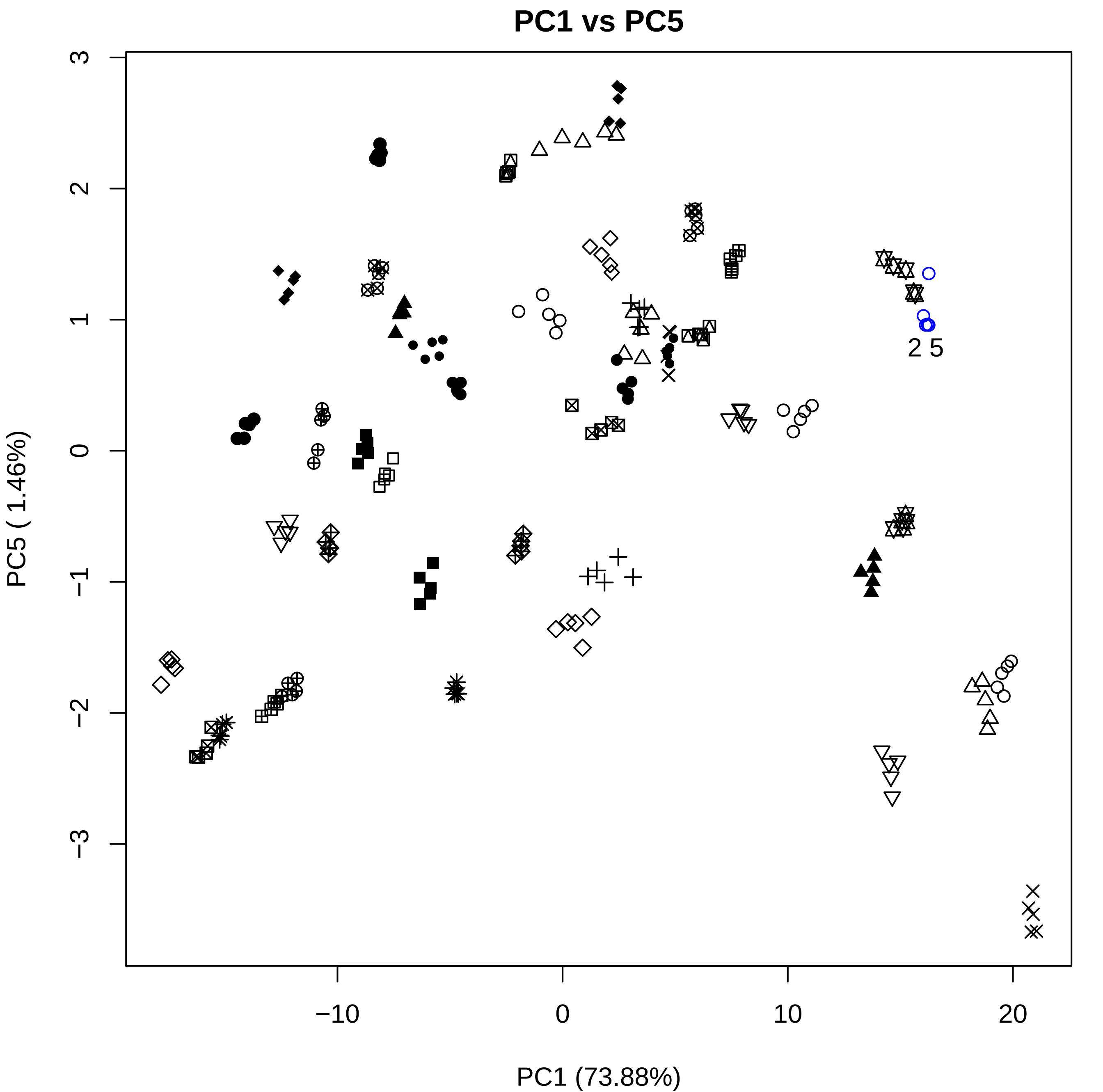

PC1 vs PC5
3
2
1
0
−1
−2
−3
PC5 ( 1.46%)
−10
0
10
20
PC1 (73.88%)
2 5

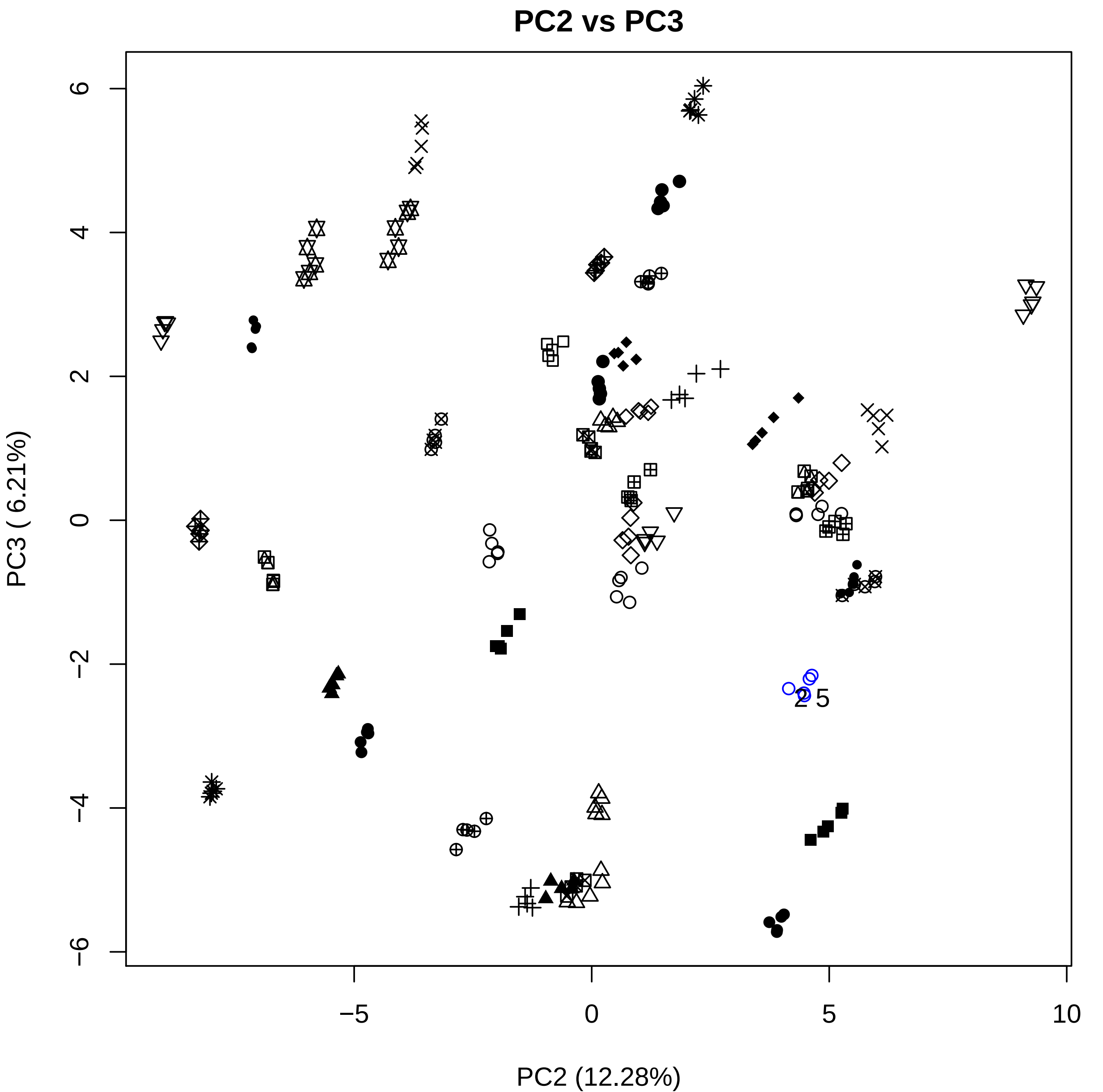
PC2 vs PC3
PC3 ( 6.21%)
PC2 (12.28%)
6
4
2
0
−2
−4
−6
−5
0
5
10
25

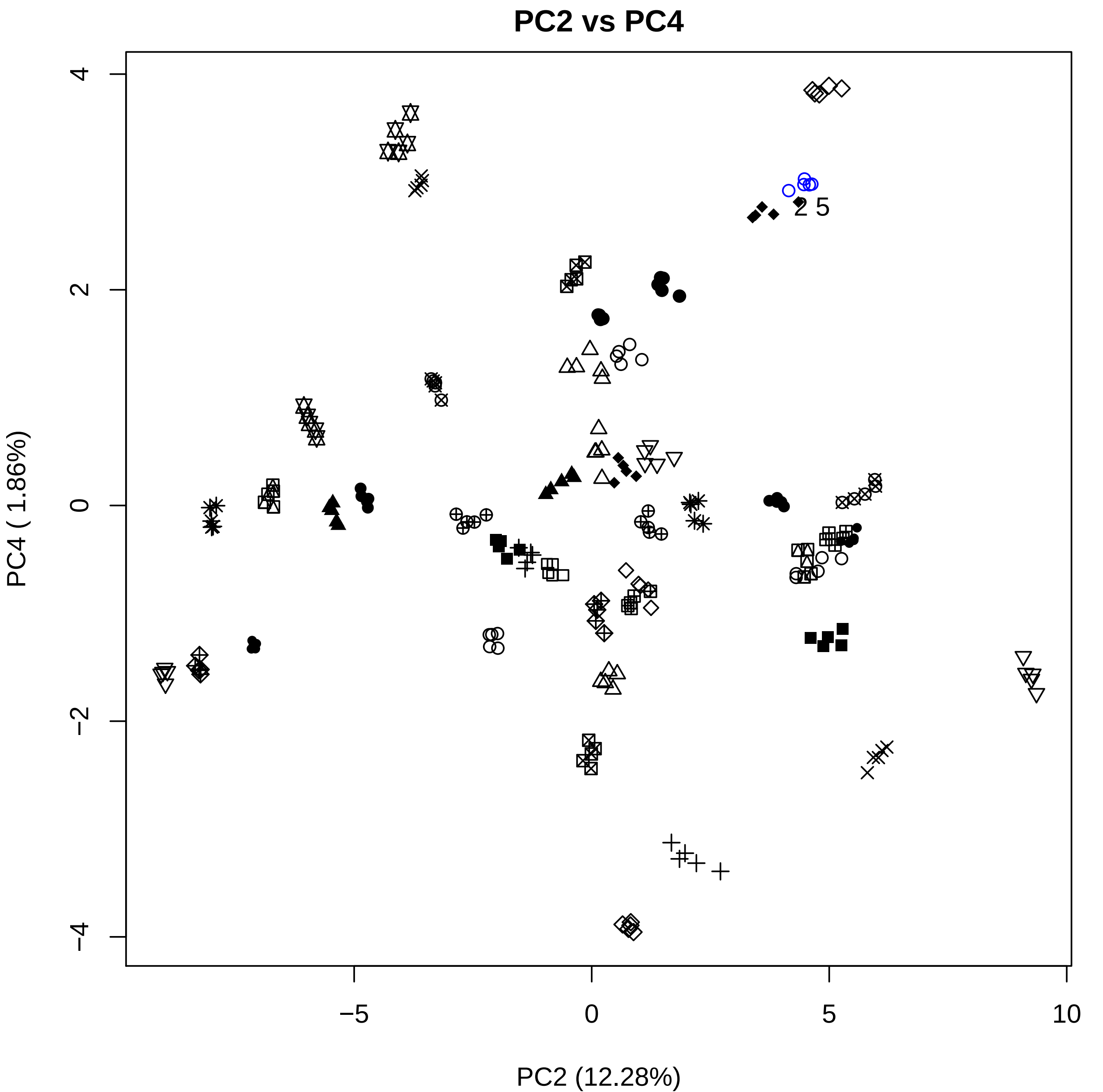
PC2 vs PC4
PC4 ( 1.86%)
PC2 (12.28%)
4
2
0
−2
−4
−5
0
5
10
2 5

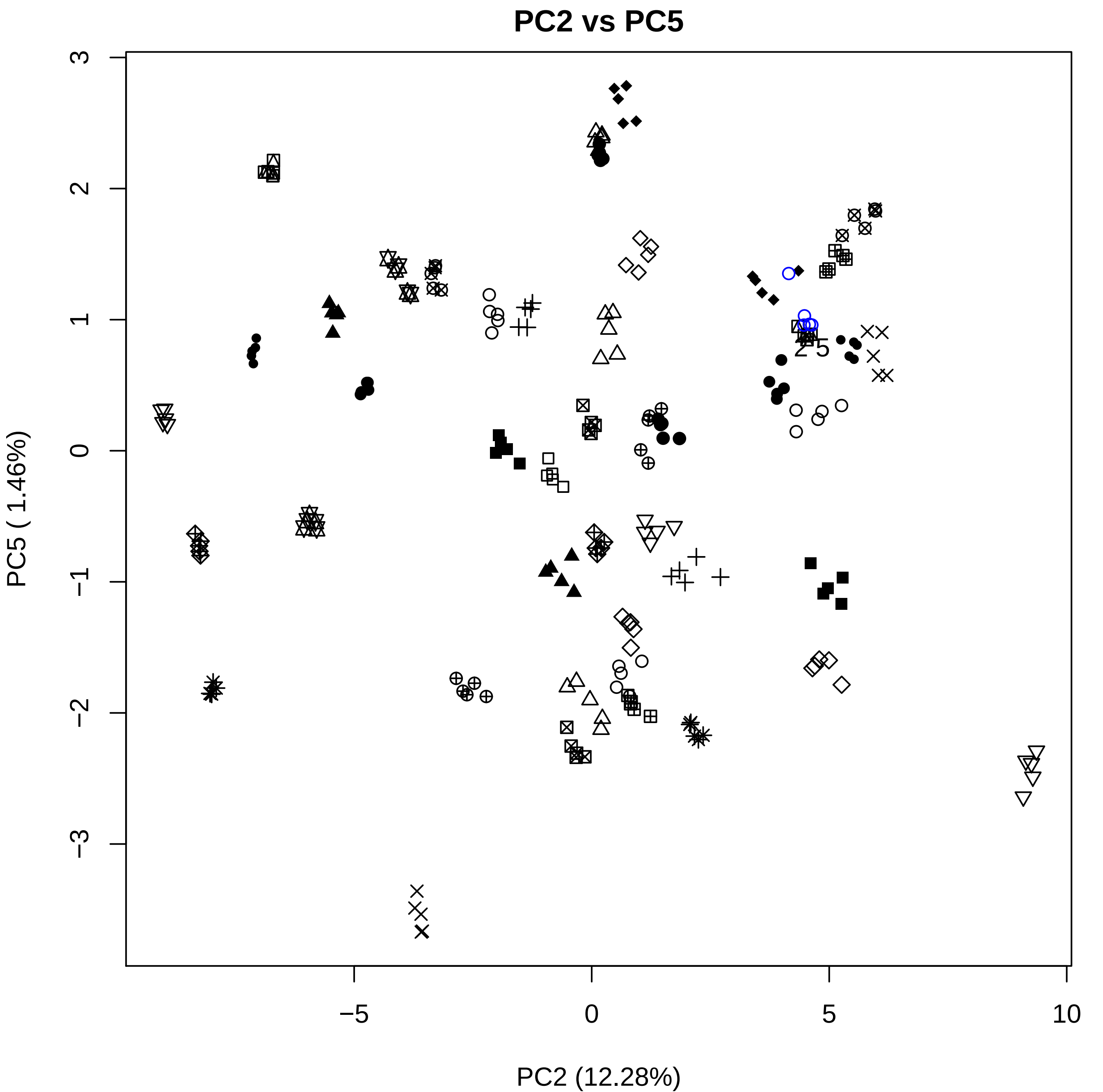
PC2 vs PC5
PC5 ( 1.46%)
PC2 (12.28%)
3
2
1
0
−1
−2
−3
−5
0
5
10

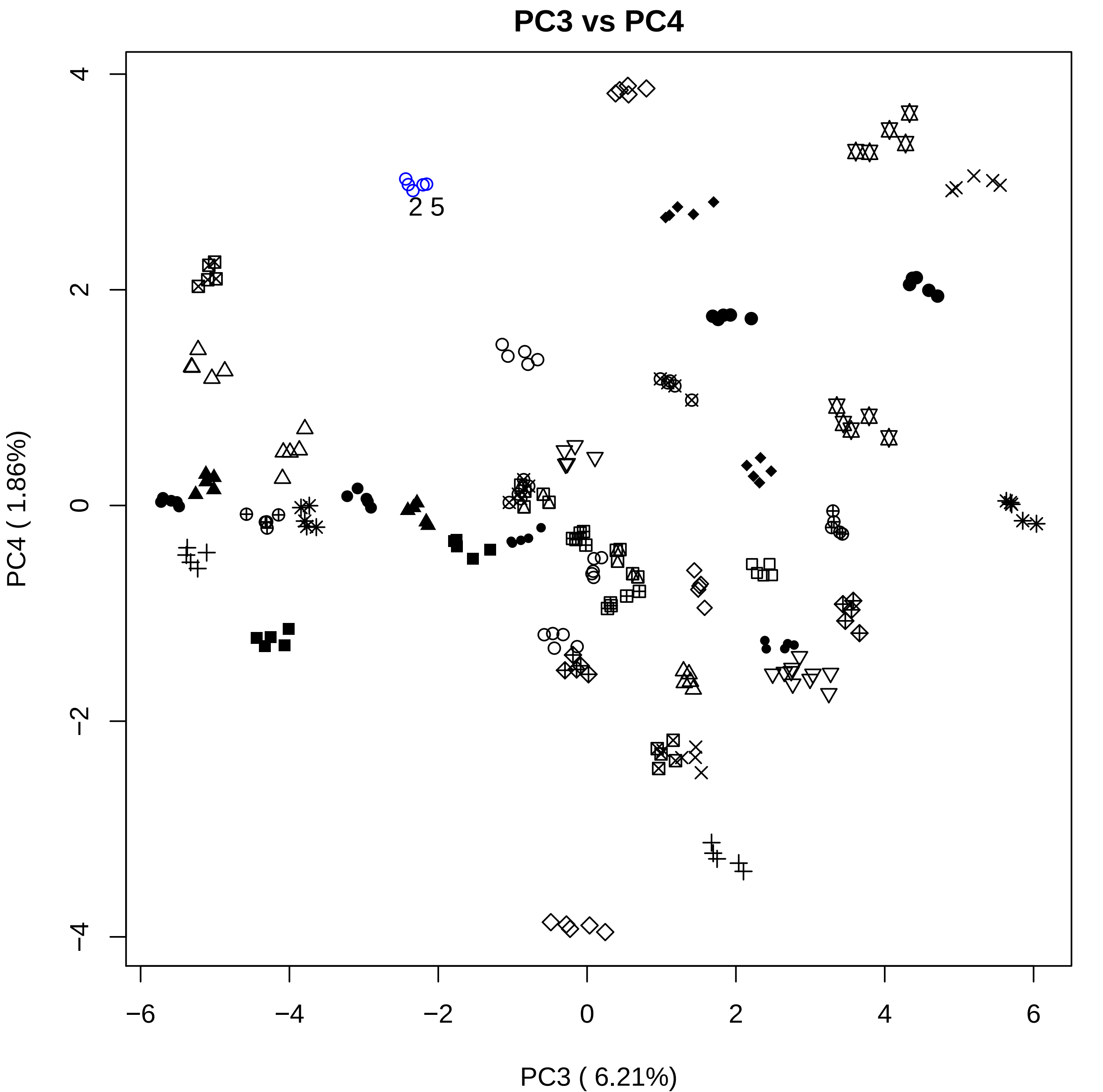

PC3 vs PC4
PC4 ( 1.86%)
PC3 ( 6.21%)
4
2
0
−2
−4
−6
−4
−2
0
2
4
6
2 5

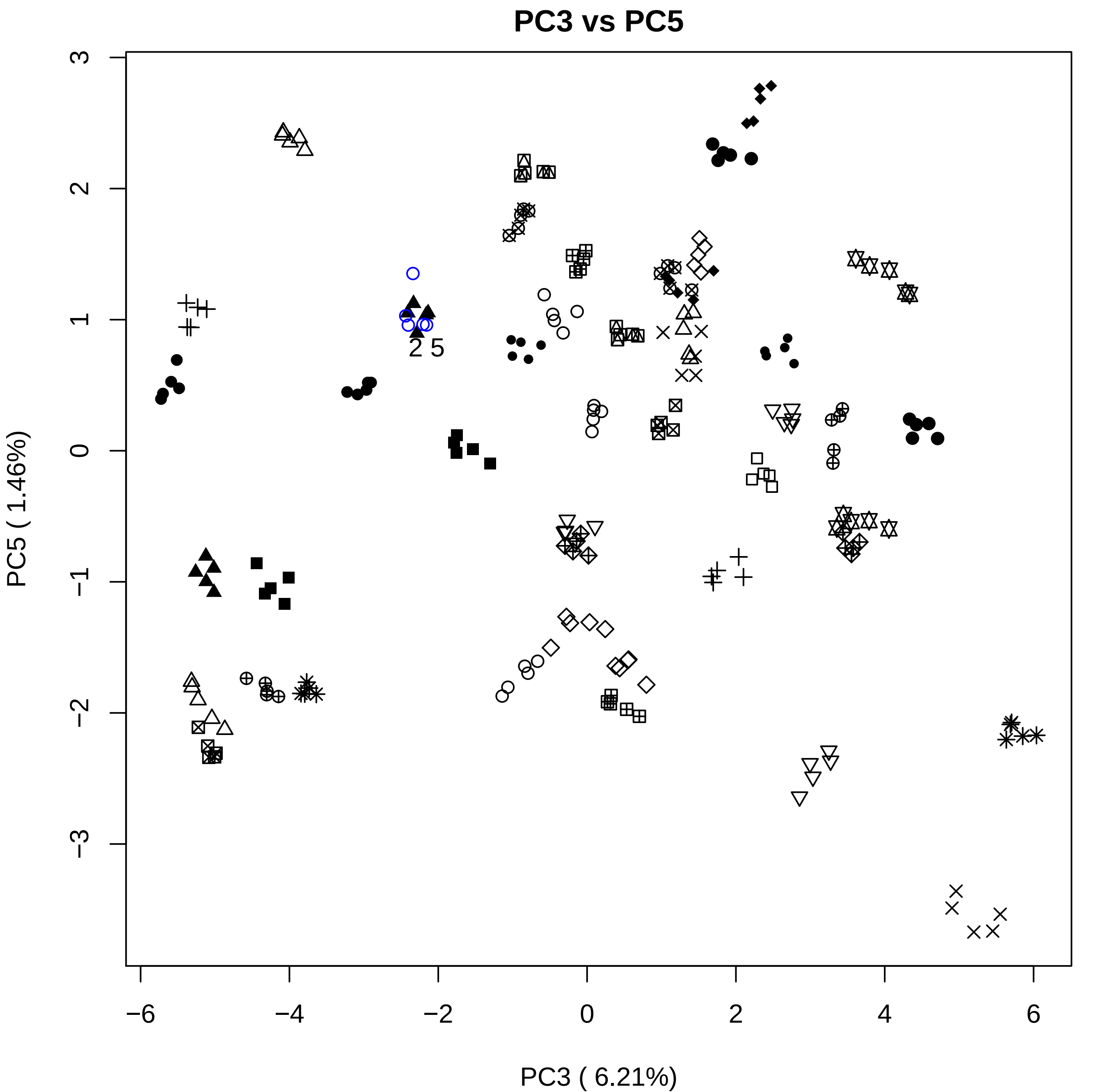
PC3 vs PC5
PC5 ( 1.46%)
PC3 ( 6.21%)
−6
−4
−2
0
2
4
6
−3
−2
−1
0
1
2
3
2
5

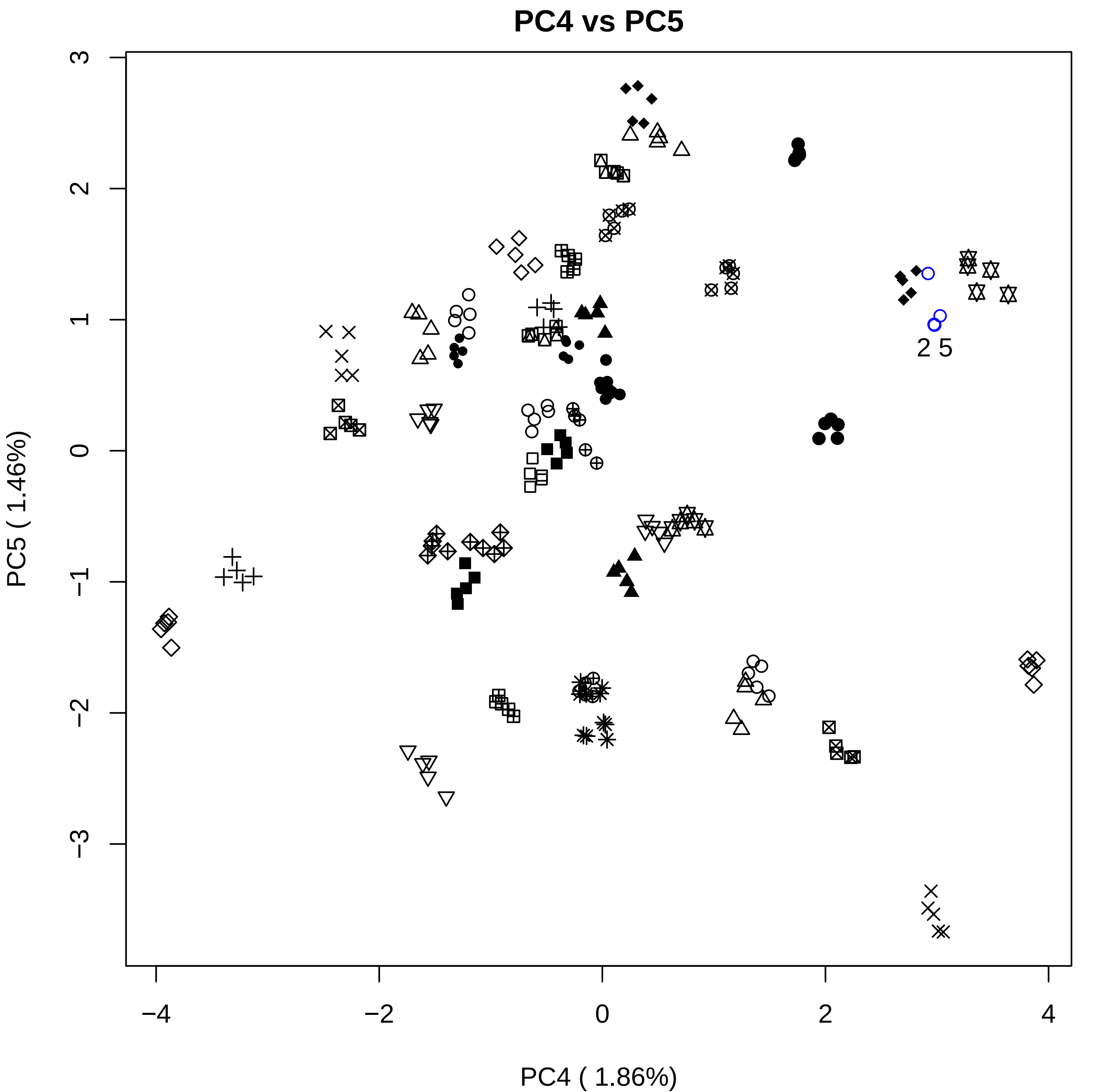

PC4 vs PC5
PC5 ( 1.46%)
PC4 ( 1.86%)
2 5
3
2
1
0
−1
−2
−3
−4
−2
0
2
4

### PCA of 5%  DeS vs PMH

Scaled Matrix PCAed

7th order polynomial baseline correction performed

Raw spectra smoothed to 21 Neighbours, 2nd order PN, Savtisky Golay

Normalisation between 0 and 100

D2 spectra smoothed to 41 Neighbours, 2nd order PN, Savtisky Golay

Regions Analysed: 3601−2501 and 2001−700cm−1

Samples Analysed: 1 Contam's 0 DeS's 45 PMH's

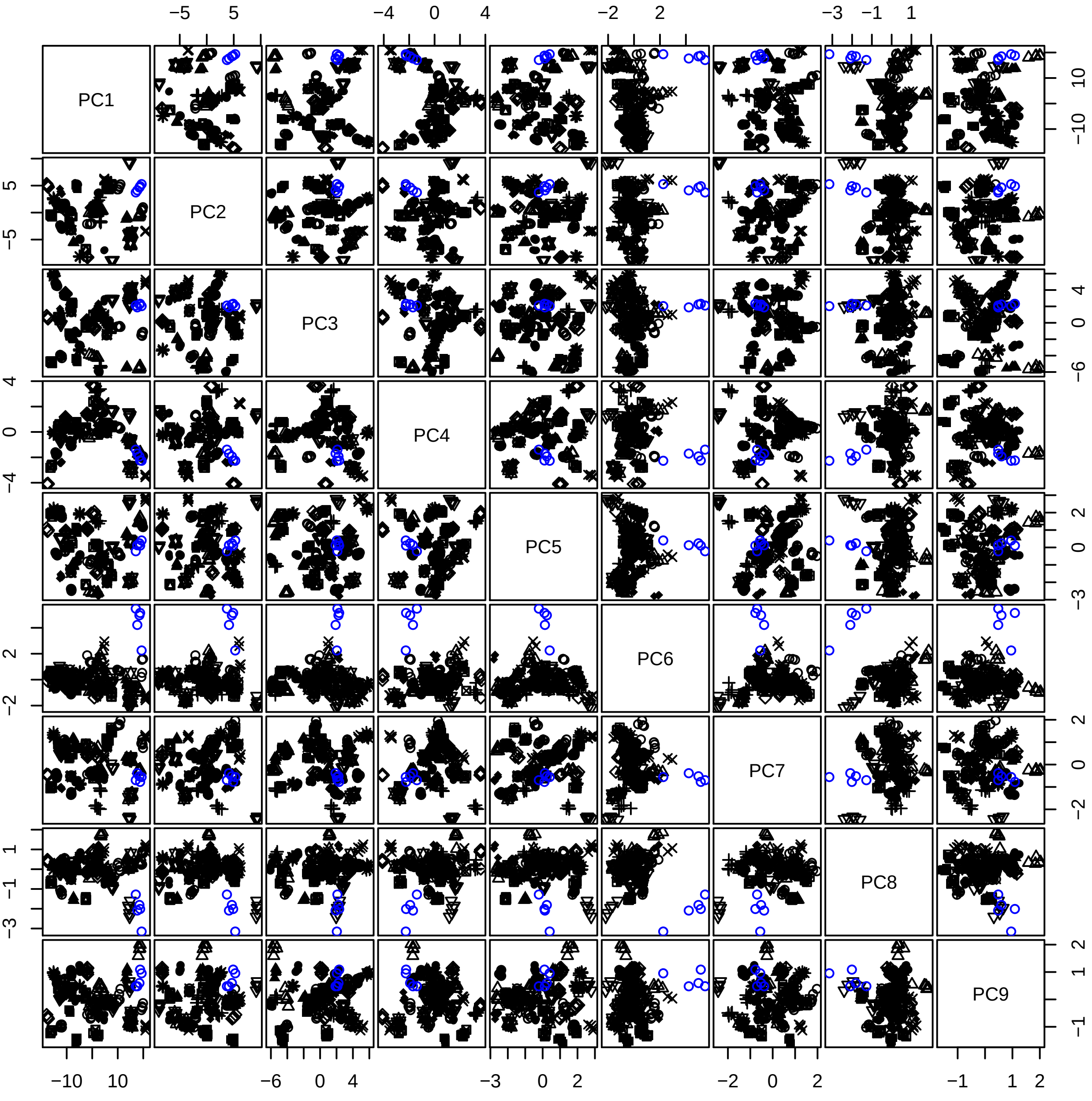
PC1
PC2
PC3
PC4
PC5
PC6
PC7
PC8
PC9

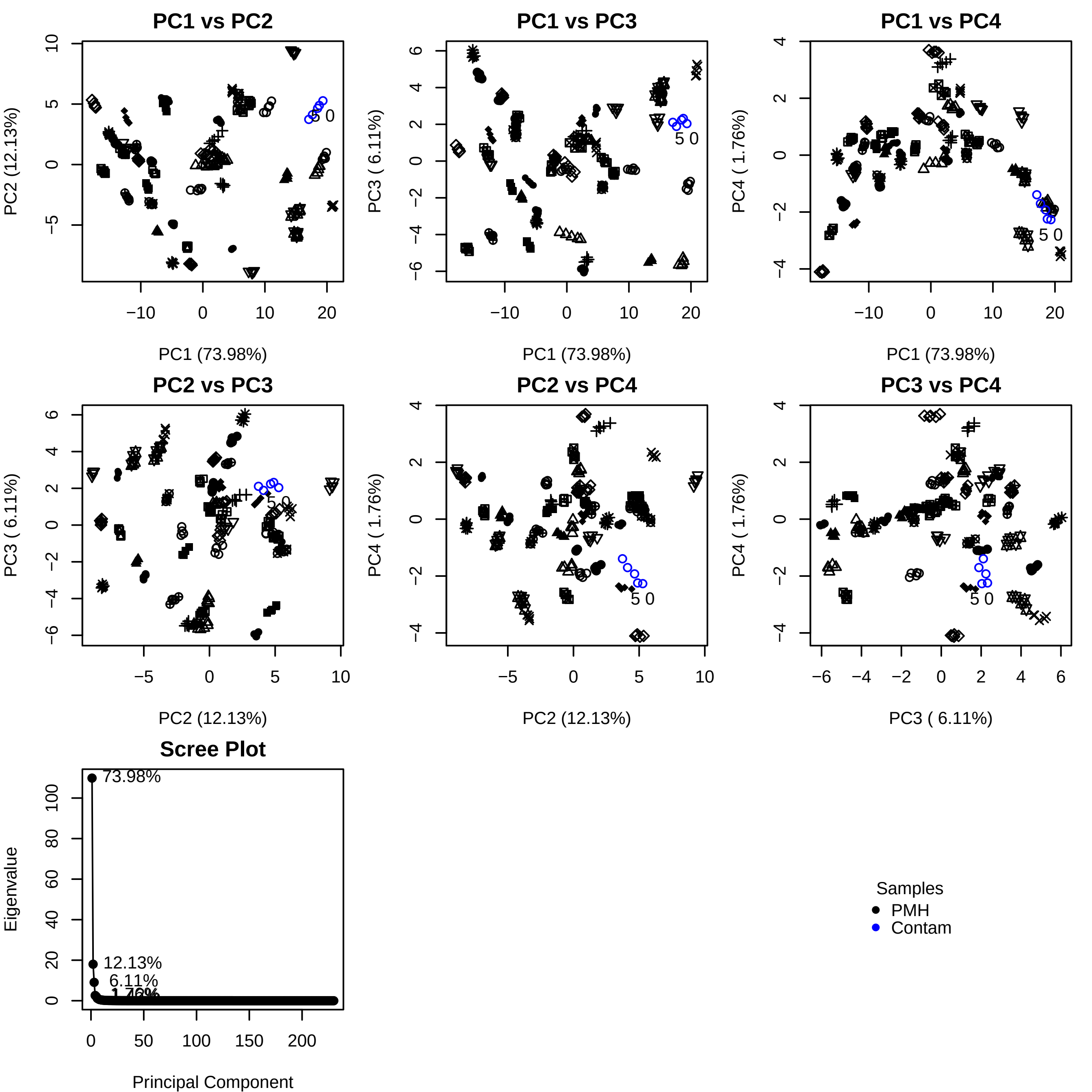

PC1 vs PC2
PC1 vs PC3
PC1 vs PC4
PC2 vs PC3
PC2 vs PC4
PC3 vs PC4
Scree Plot
PC1 (73.98%)
PC2 (12.13%)
PC3 ( 6.11%)
PC4 ( 1.76%)
Eigenvalue
Principal Component
73.98%
12.13%
6.11%
Samples
PMH
Contam

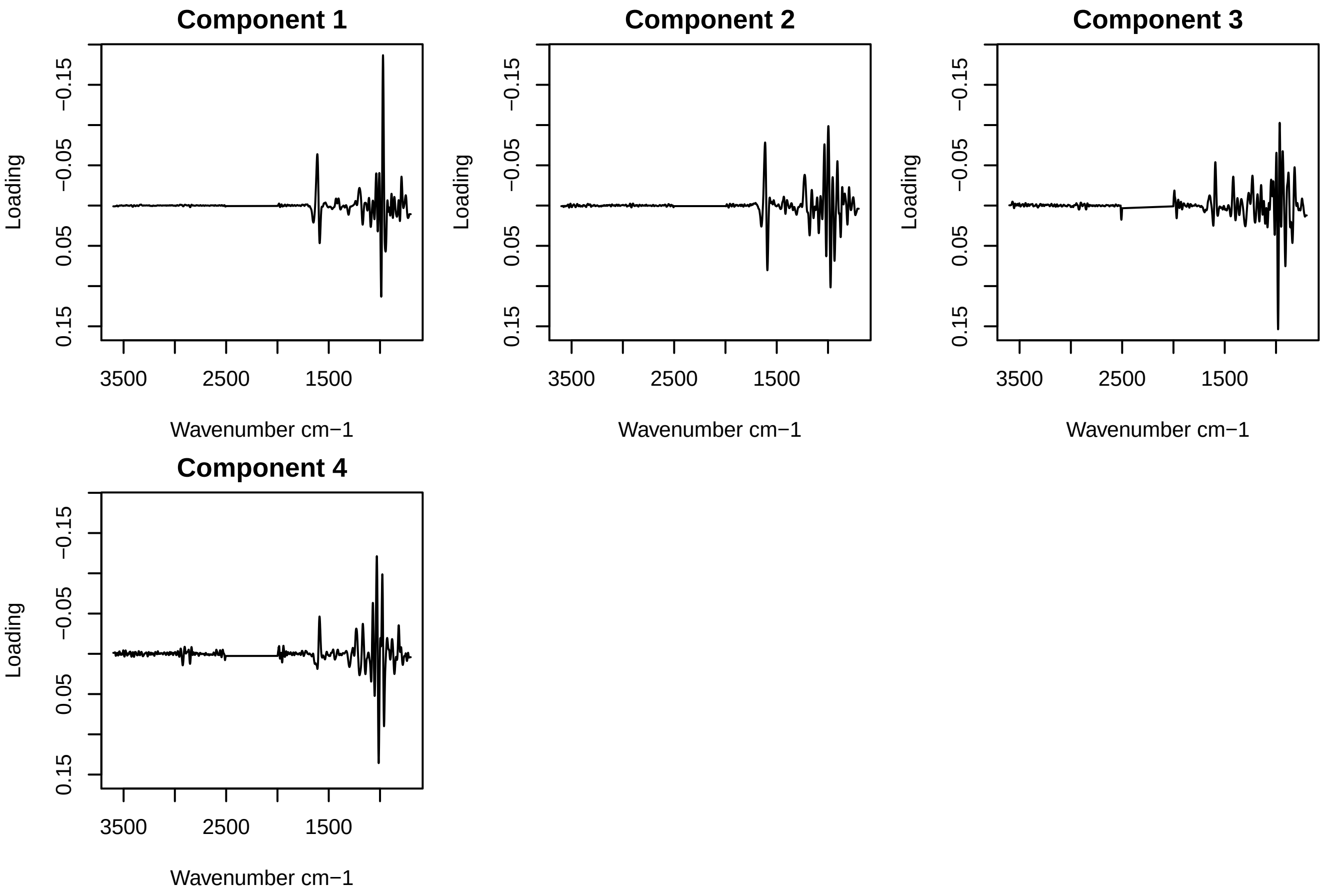
Component 1
Component 2
Component 3
Component 4
Loading
Wavenumber cm−1
−0.15
−0.05
0.05
0.15
3500
2500
1500

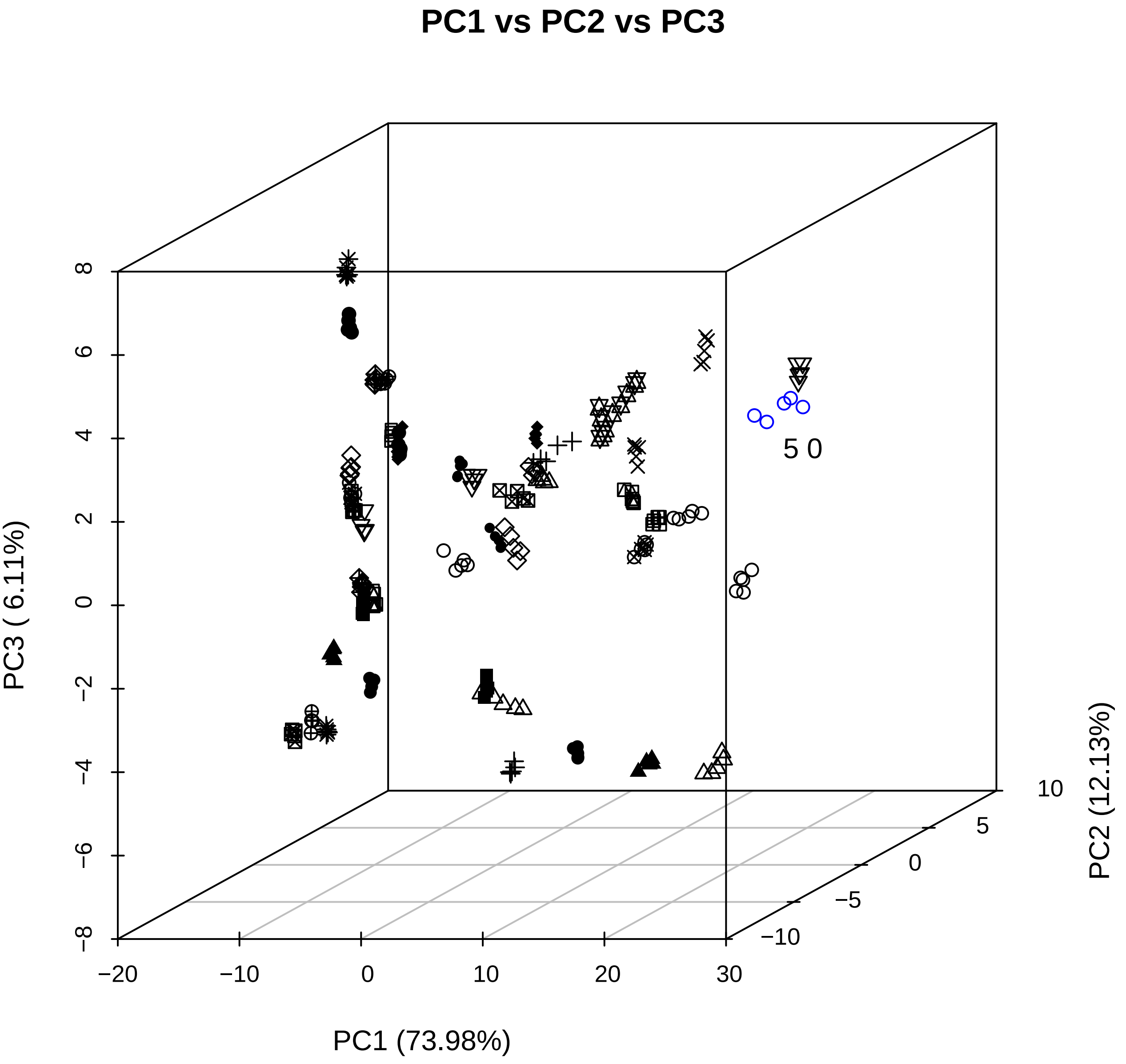
PC1 vs PC2 vs PC3
PC3 ( 6.11%)
8
6
4
2
0
−2
−4
−6
−8
−20
−10
0
10
20
30
PC1 (73.98%)
PC2 (12.13%)
10
5
0
−5
−10
5 0

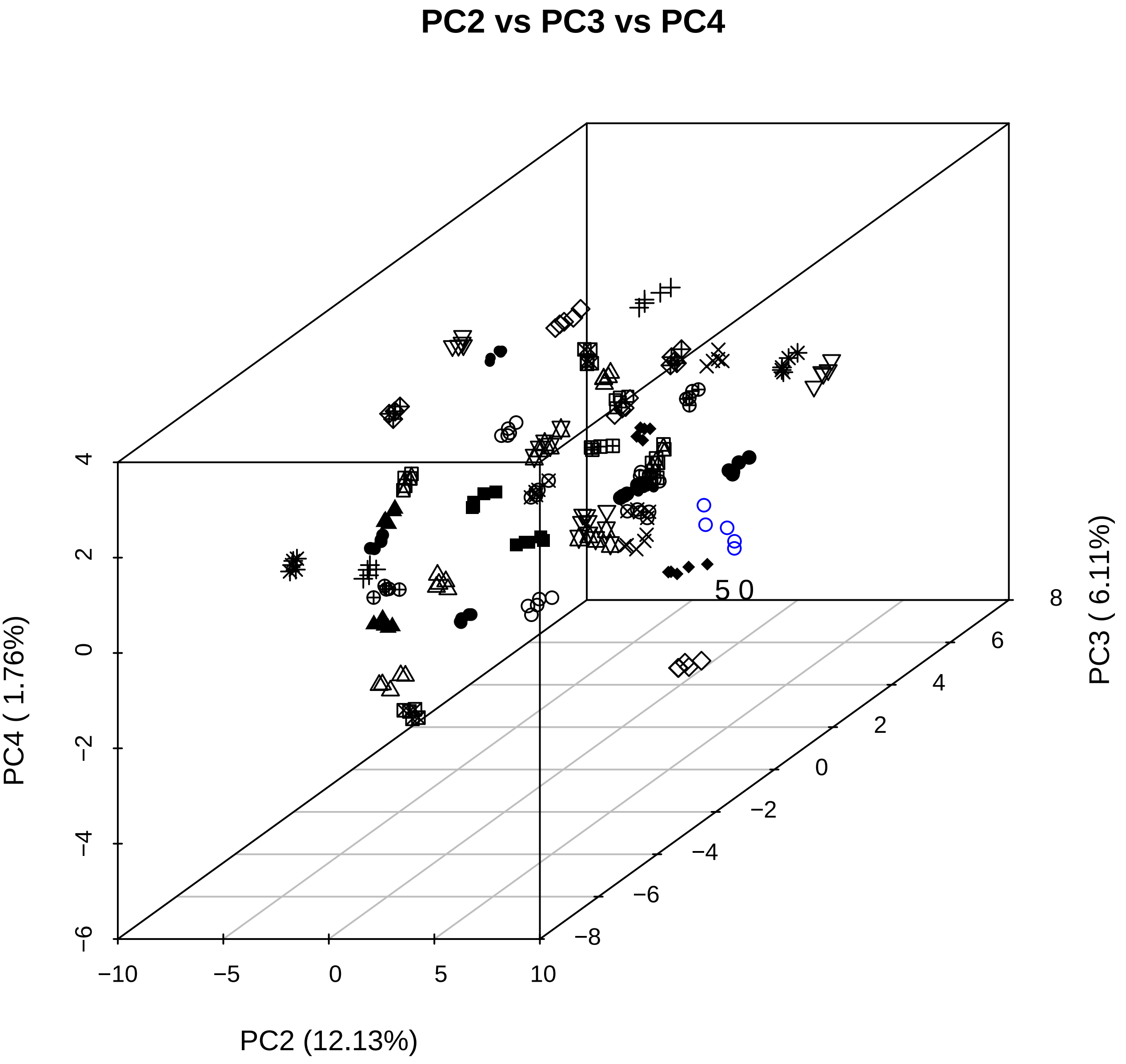
PC2 vs PC3 vs PC4
50
PC2 (12.13%)
PC4 ( 1.76%)
PC3 ( 6.11%)
−10
−5
0
5
10
4
2
0
−2
−4
−6
−8
−6
−4
−2
0
2
4
6
8

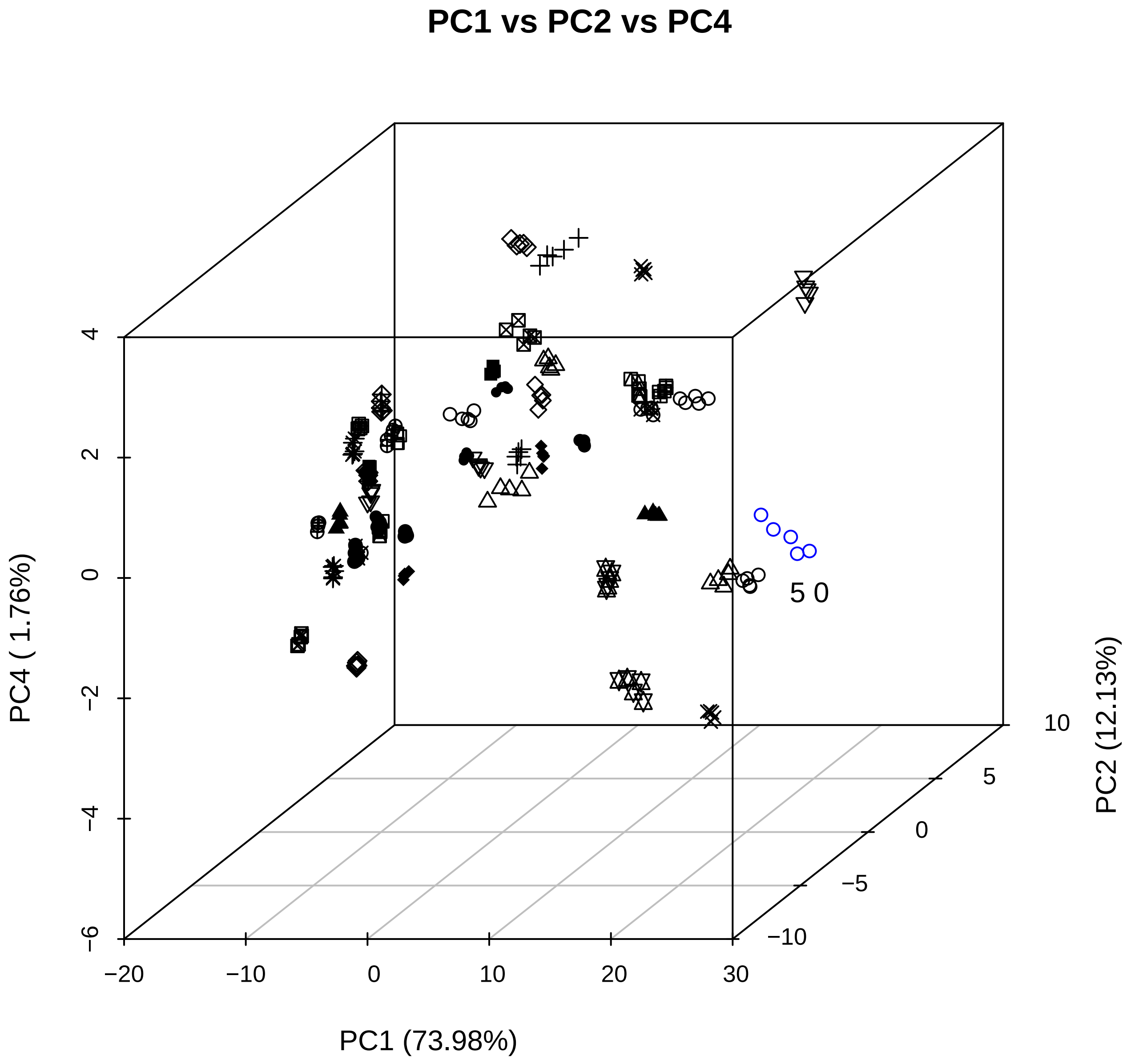

PC1 vs PC2 vs PC4
PC4 ( 1.76%)
4
2
0
−2
−4
−6
5 0
PC1 (73.98%)
−20
−10
0
10
20
30
PC2 (12.13%)
10
5
0
−5
−10

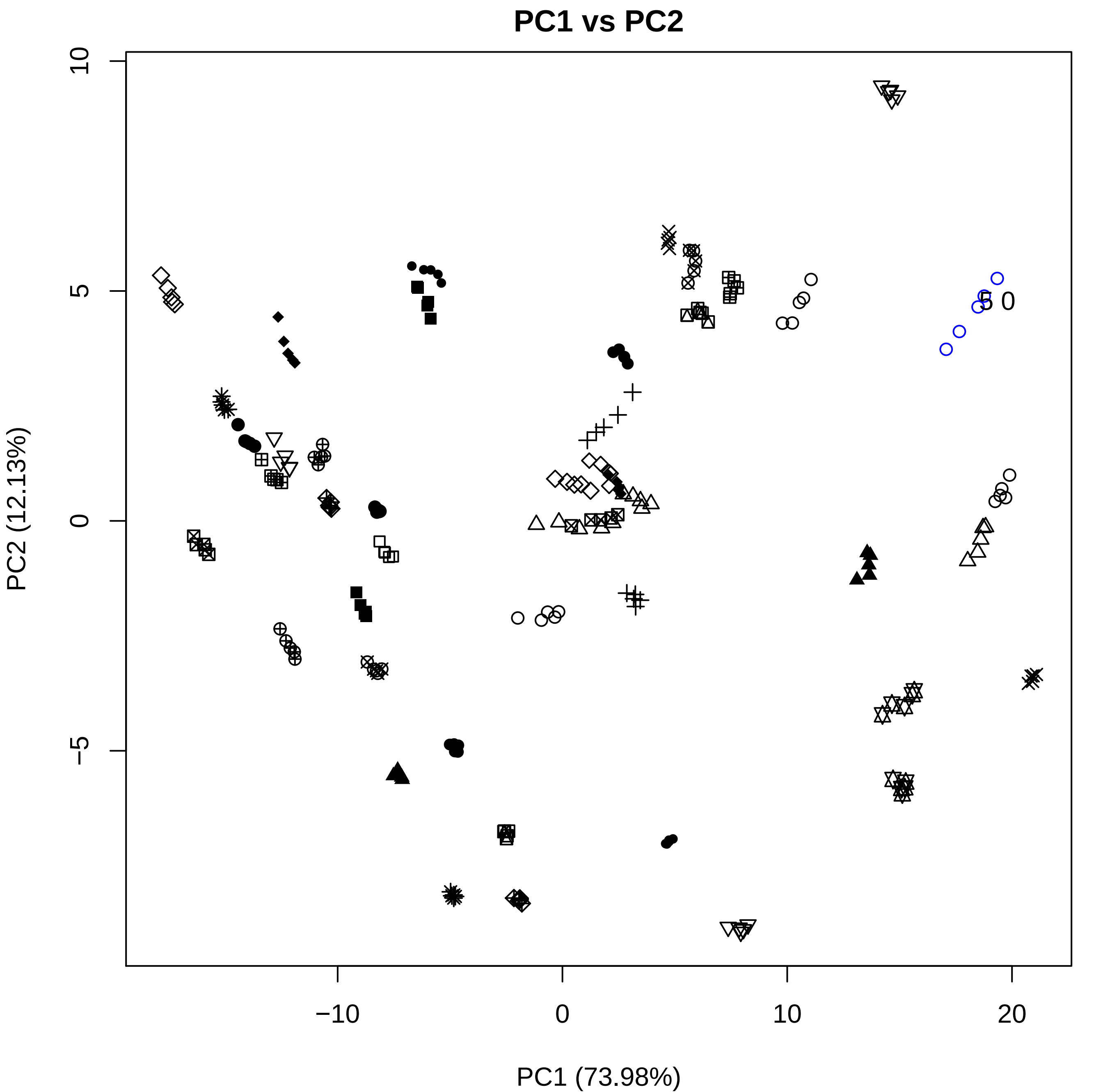
PC1 vs PC2
PC2 (12.13%)
PC1 (73.98%)
10
5
0
−5
−10
0
10
20
5 0

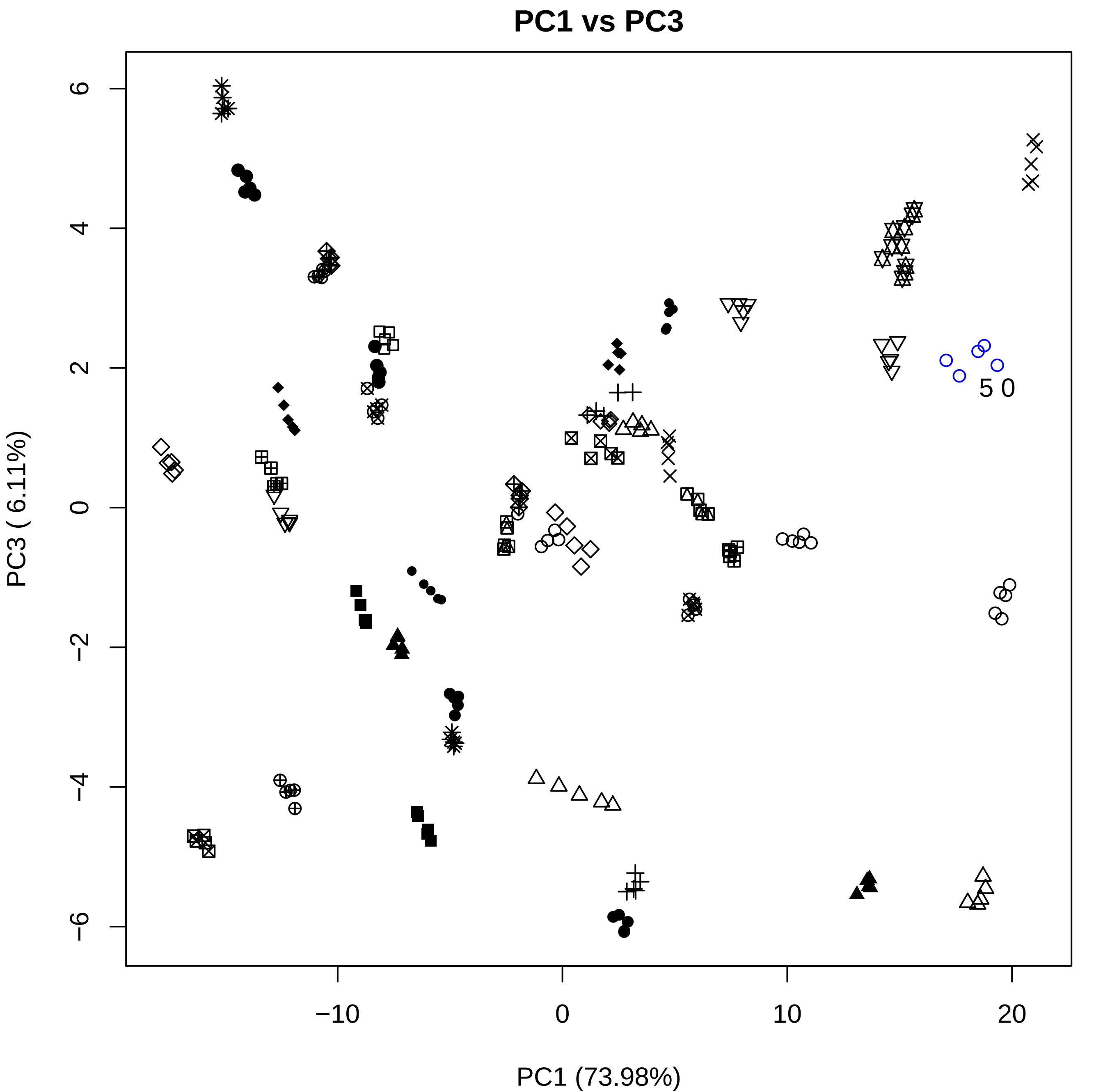
PC1 vs PC3
PC1 (73.98%)
PC3 ( 6.11%)
−10
0
10
20
−6
−4
−2
0
2
4
6
5 0

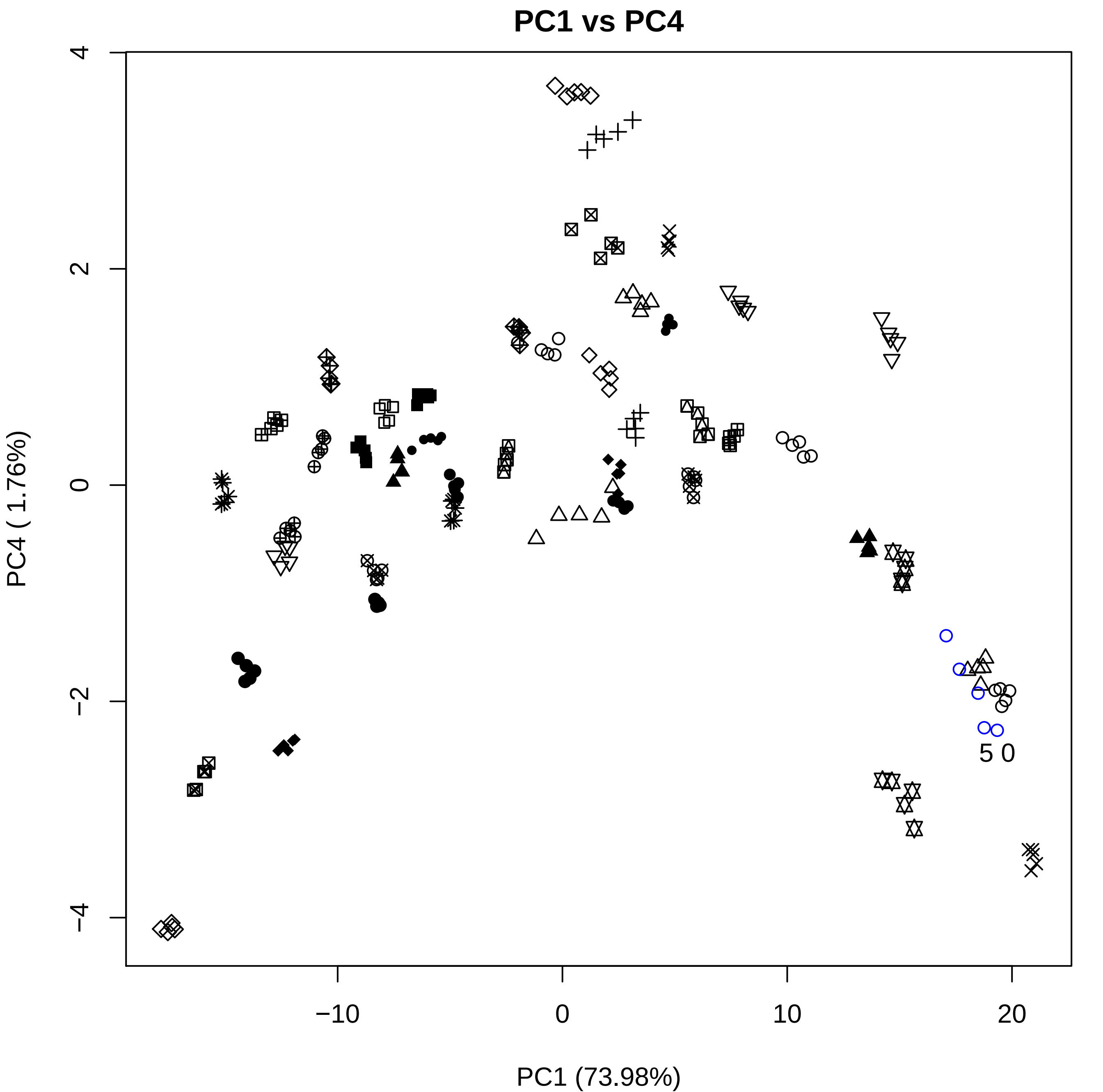

PC1 vs PC4
PC4 ( 1.76%)
PC1 (73.98%)
4
2
0
−2
−4
−10
0
10
20
5 0

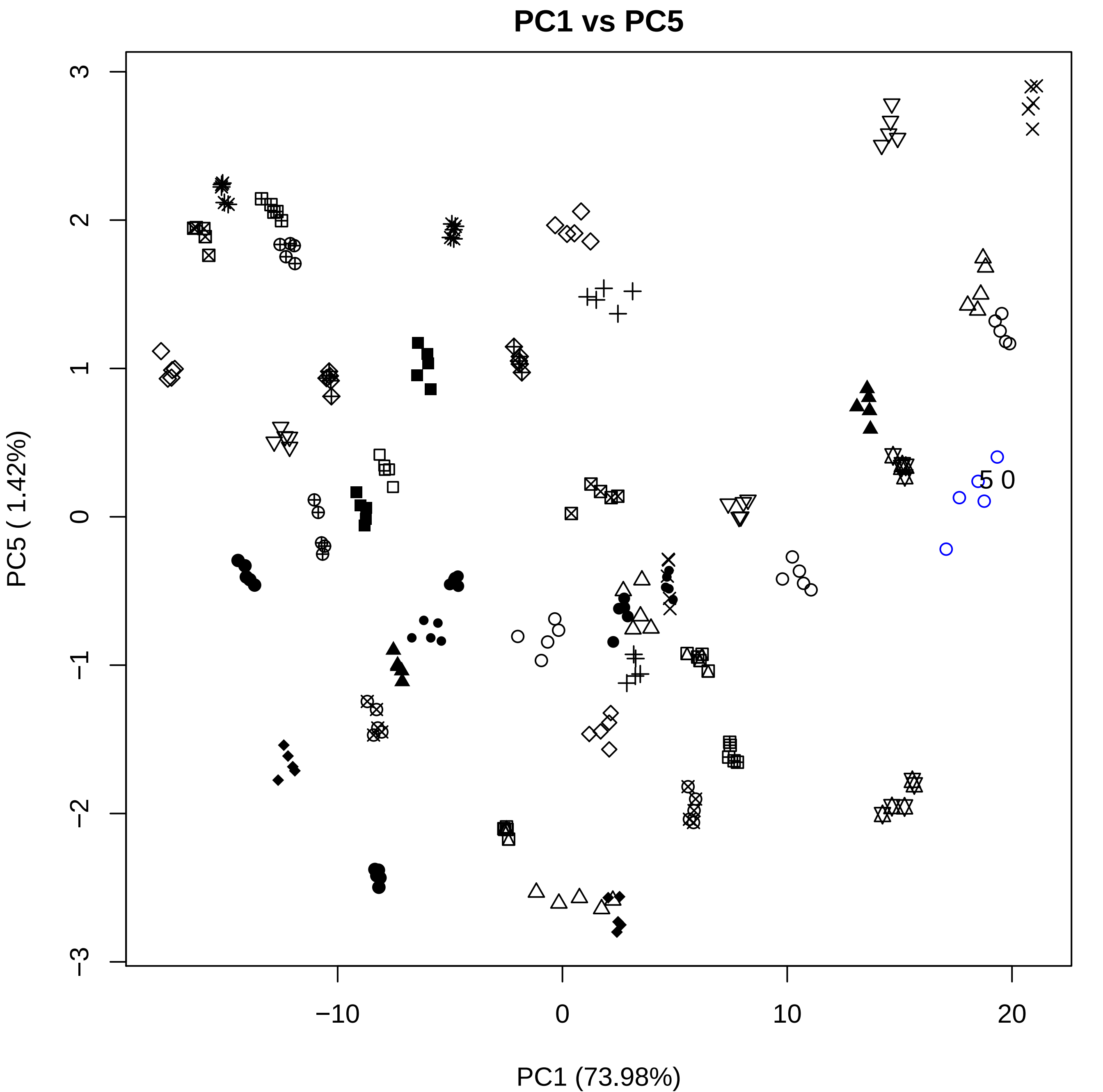

PC1 vs PC5
PC5 ( 1.42%)
PC1 (73.98%)
3
2
1
0
−1
−2
−3
−10
0
10
20
5 0

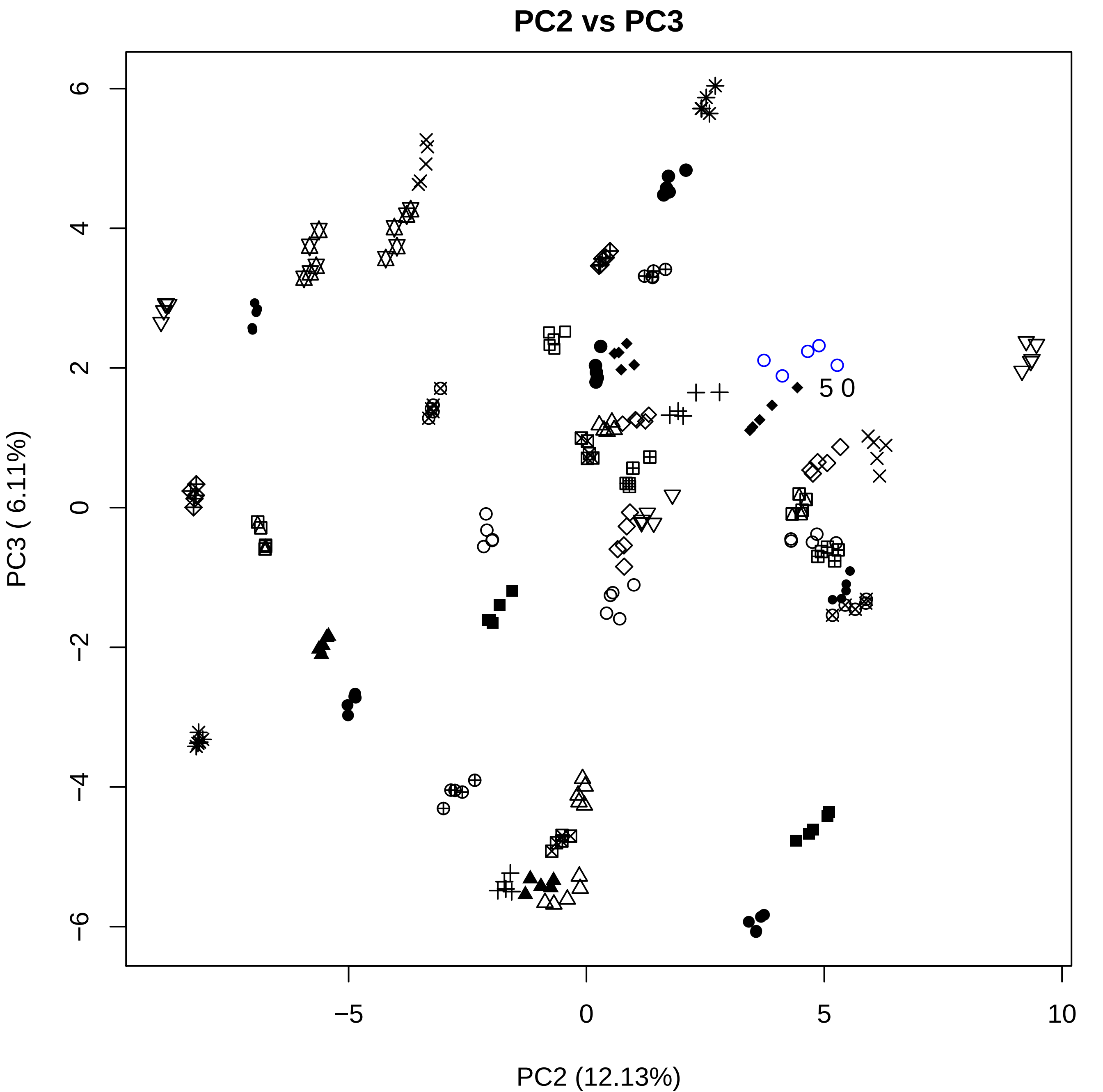
PC2 vs PC3
PC3 ( 6.11%)
PC2 (12.13%)
6
4
2
0
−2
−4
−6
−5
0
5
10
5 0

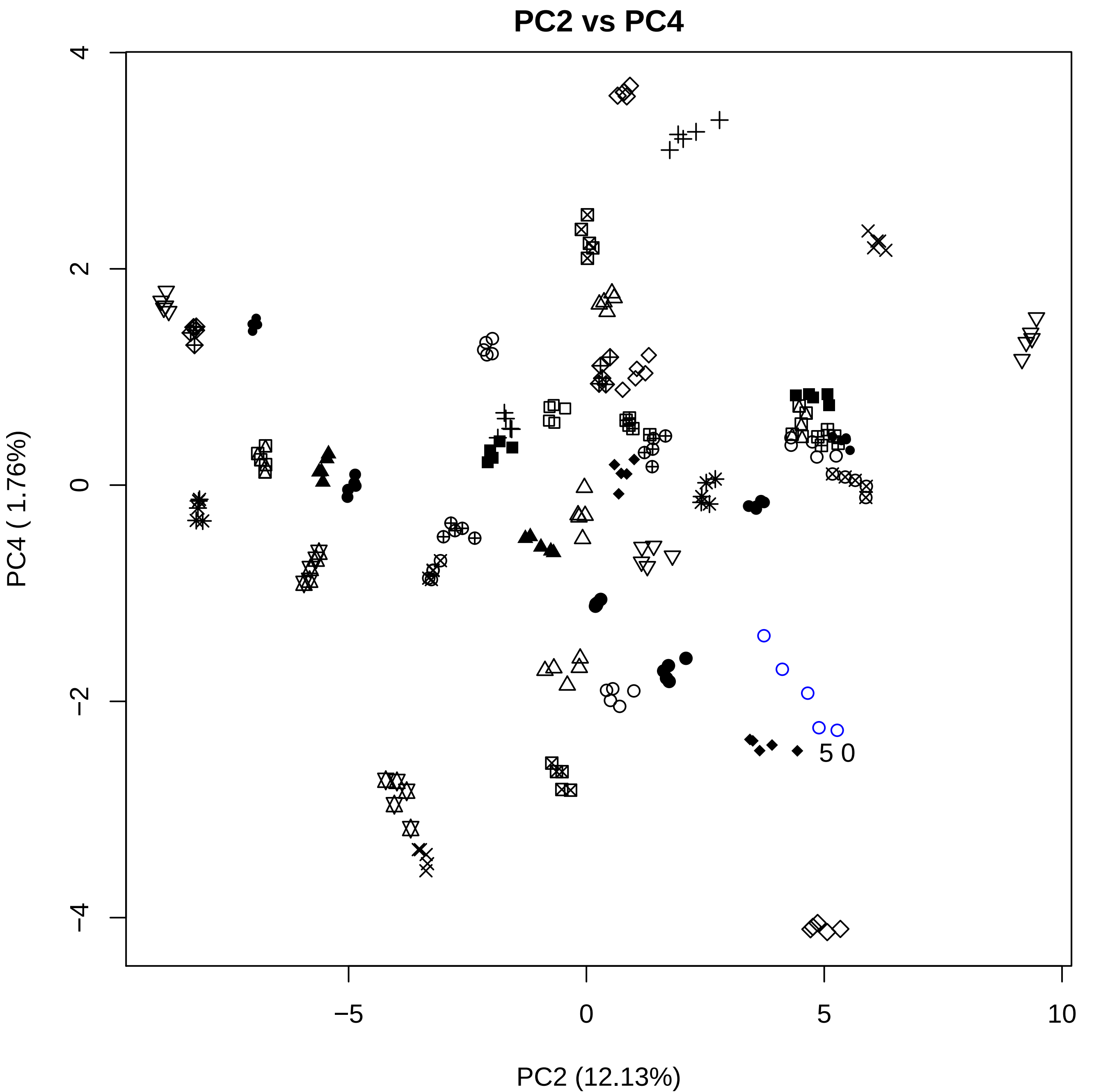
PC2 vs PC4
PC4 ( 1.76%)
PC2 (12.13%)
4
2
0
−2
−4
−5
0
5
10
5 0

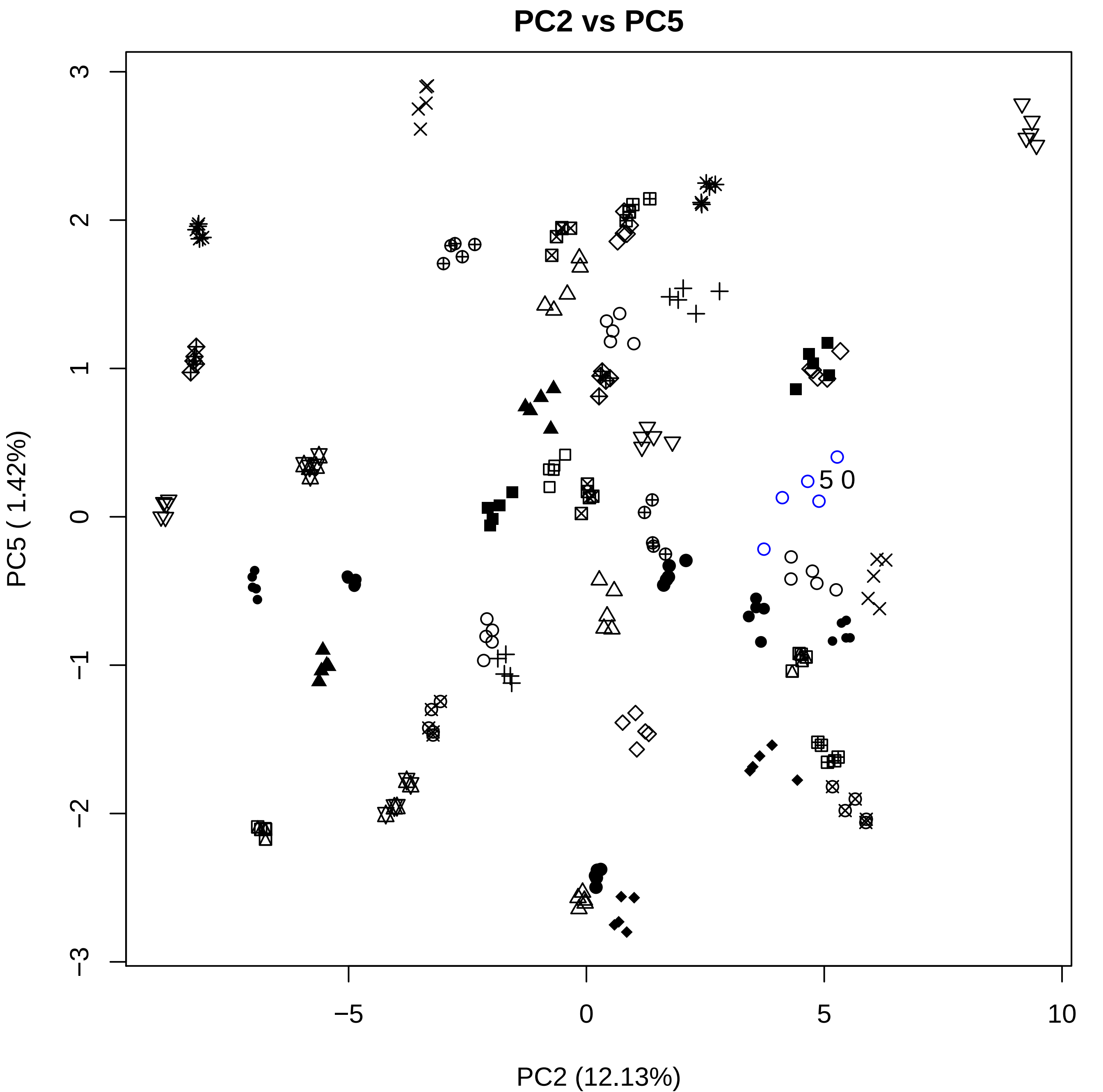

PC2 vs PC5
PC5 ( 1.42%)
PC2 (12.13%)
3
2
1
0
−1
−2
−3
−5
0
5
10
5 0

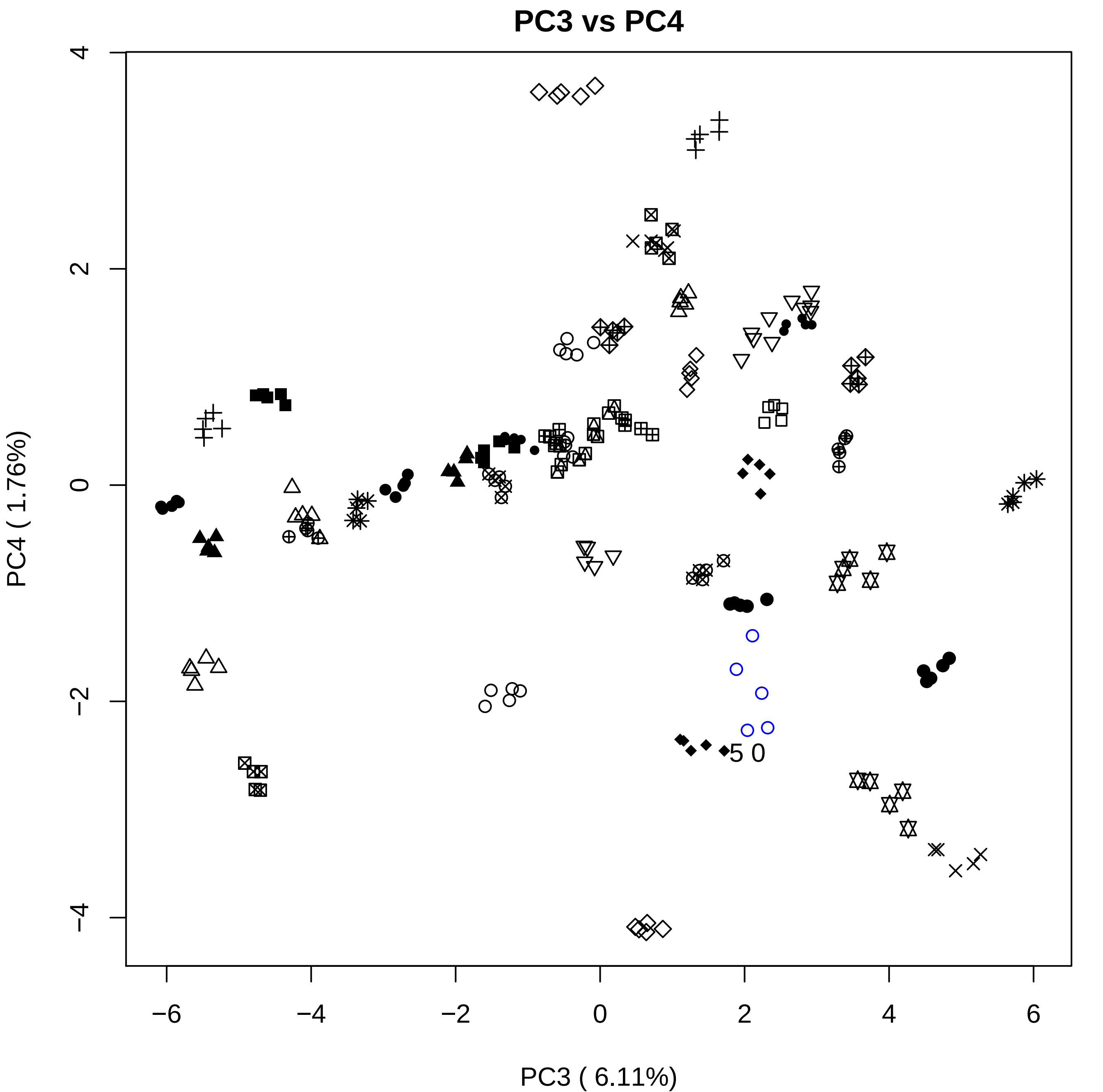
PC3 vs PC4
PC4 ( 1.76%)
PC3 ( 6.11%)
−6
−4
−2
0
2
4
6
4
2
0
−2
−4
5 0

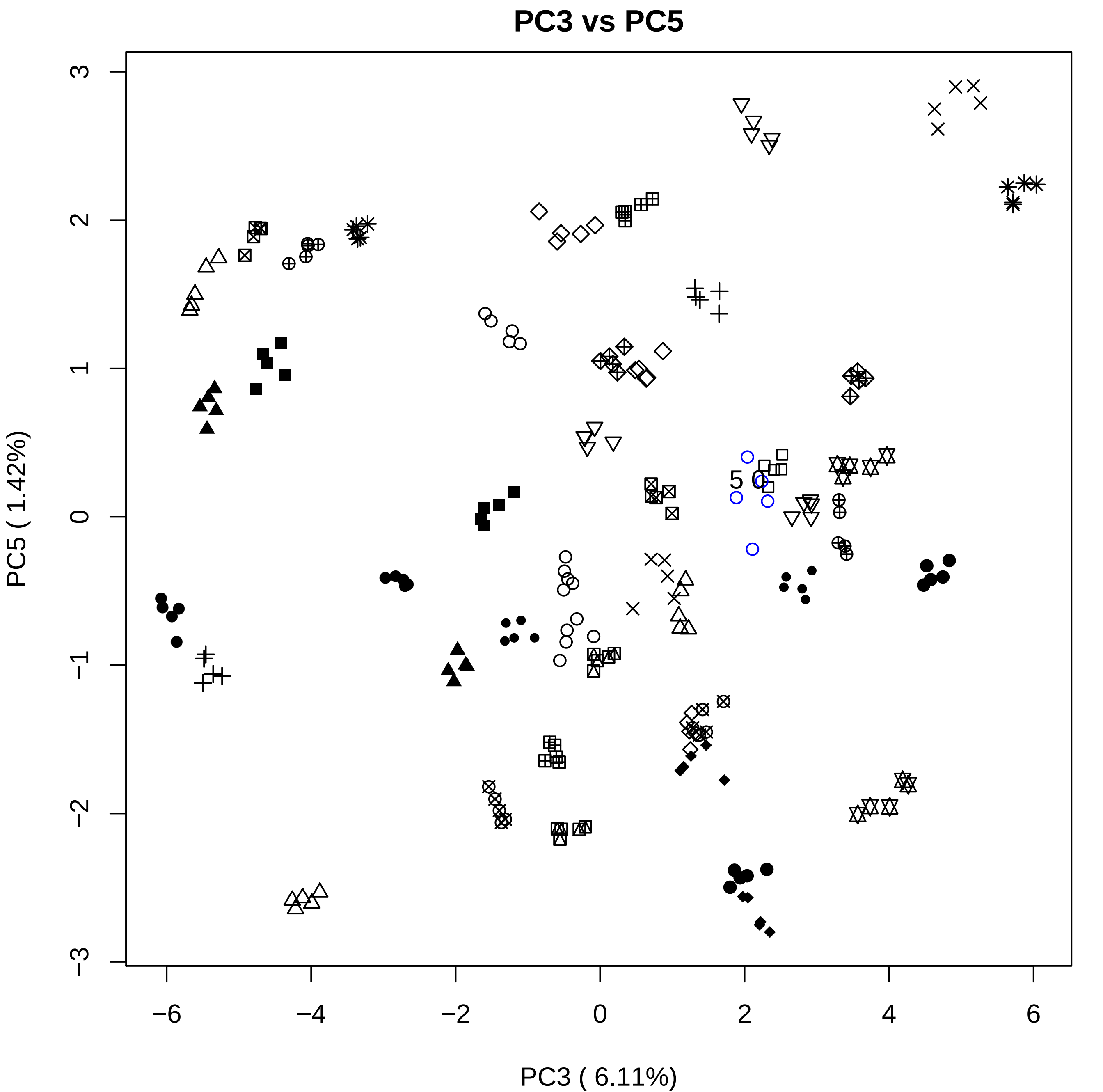
PC3 vs PC5
PC5 ( 1.42%)
PC3 ( 6.11%)
3
2
1
0
−1
−2
−3
−6
−4
−2
0
2
4
6
50

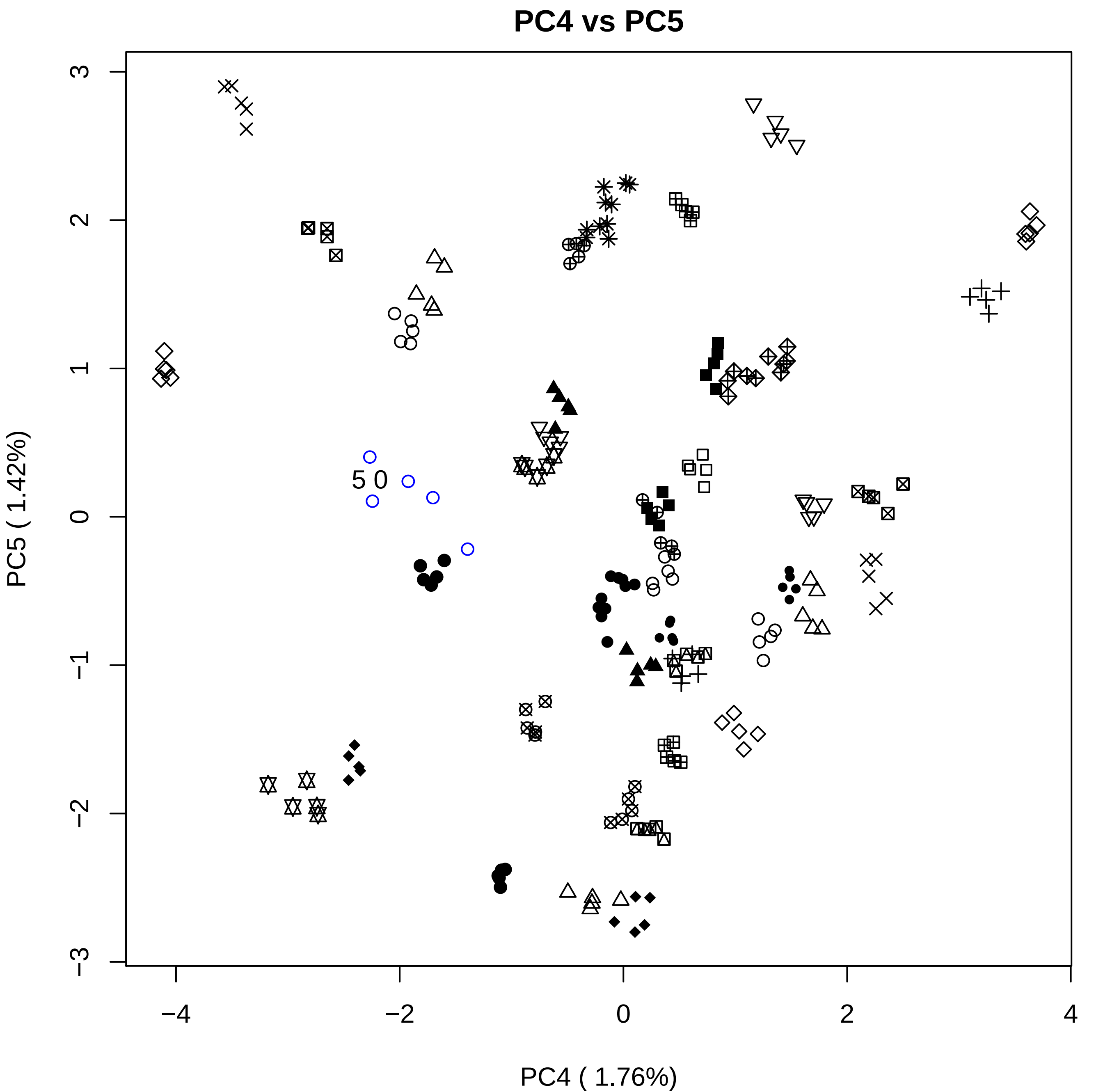
PC4 vs PC5
PC4 ( 1.76%)
PC5 ( 1.42%)
−4
−2
0
2
4
−3
−2
−1
0
1
2
3
5 0

### PCA of 7.5%  DeS vs PMH

Scaled Matrix PCAed

7th order polynomial baseline correction performed

Raw spectra smoothed to 21 Neighbours, 2nd order PN, Savtisky Golay

Normalisation between 0 and 100

D2 spectra smoothed to 41 Neighbours, 2nd order PN, Savtisky Golay

Regions Analysed: 3601−2501 and 2001−700cm−1

Samples Analysed: 1 Contam's 0 DeS's 45 PMH's

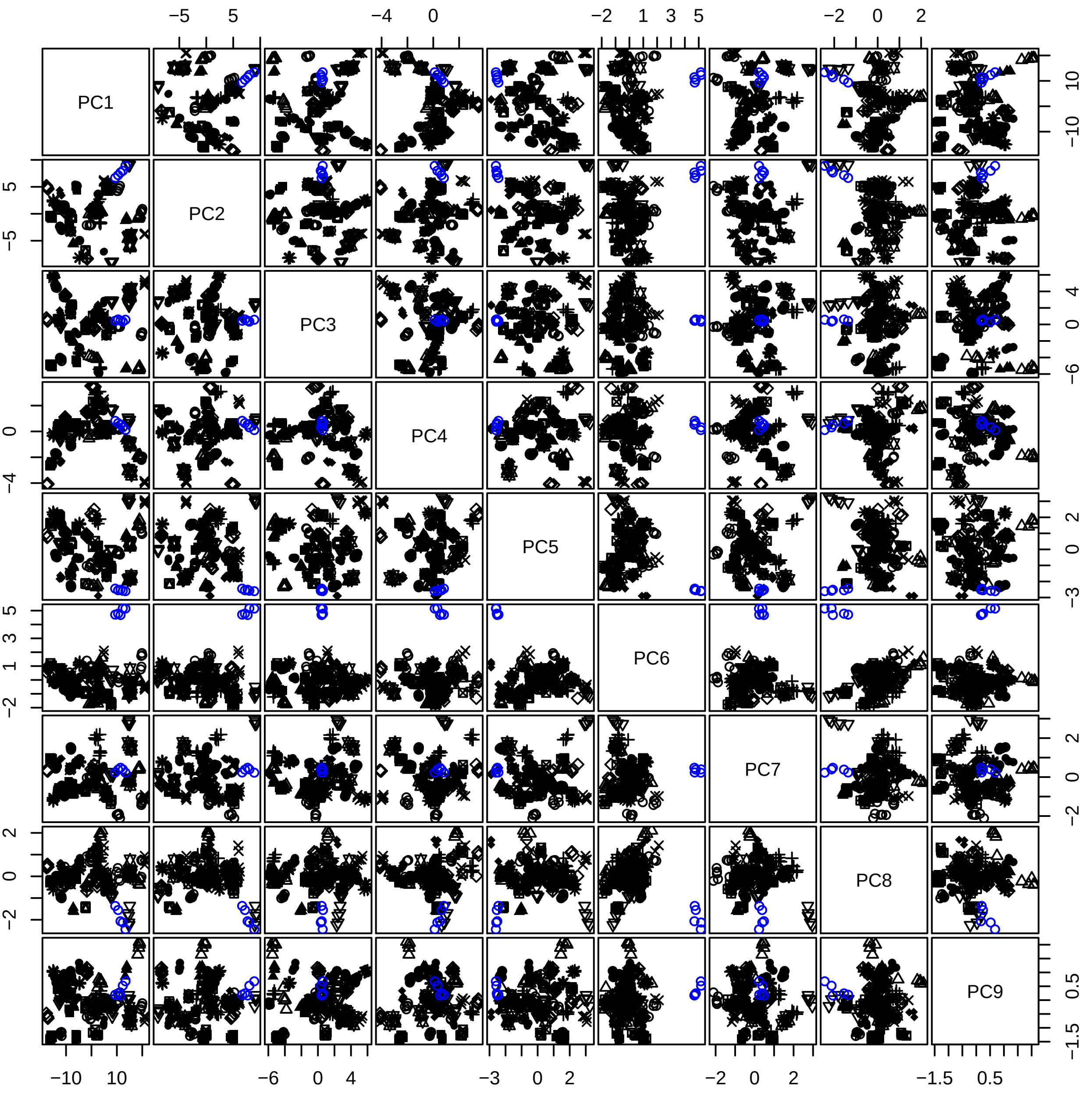
PC1
PC2
PC3
PC4
PC5
PC6
PC7
PC8
PC9
−5 5
−4 0
−2 1 3 5
−2 0 2
10 −10
5 −5
4 0 −6
0 −4
2 0 −3
5 3 1 −2
2 0 −2
2 0 −2
0.5 −1.5
−10 10
−6 0 4
−3 0 2
−2 0 2
−1.5 0.5

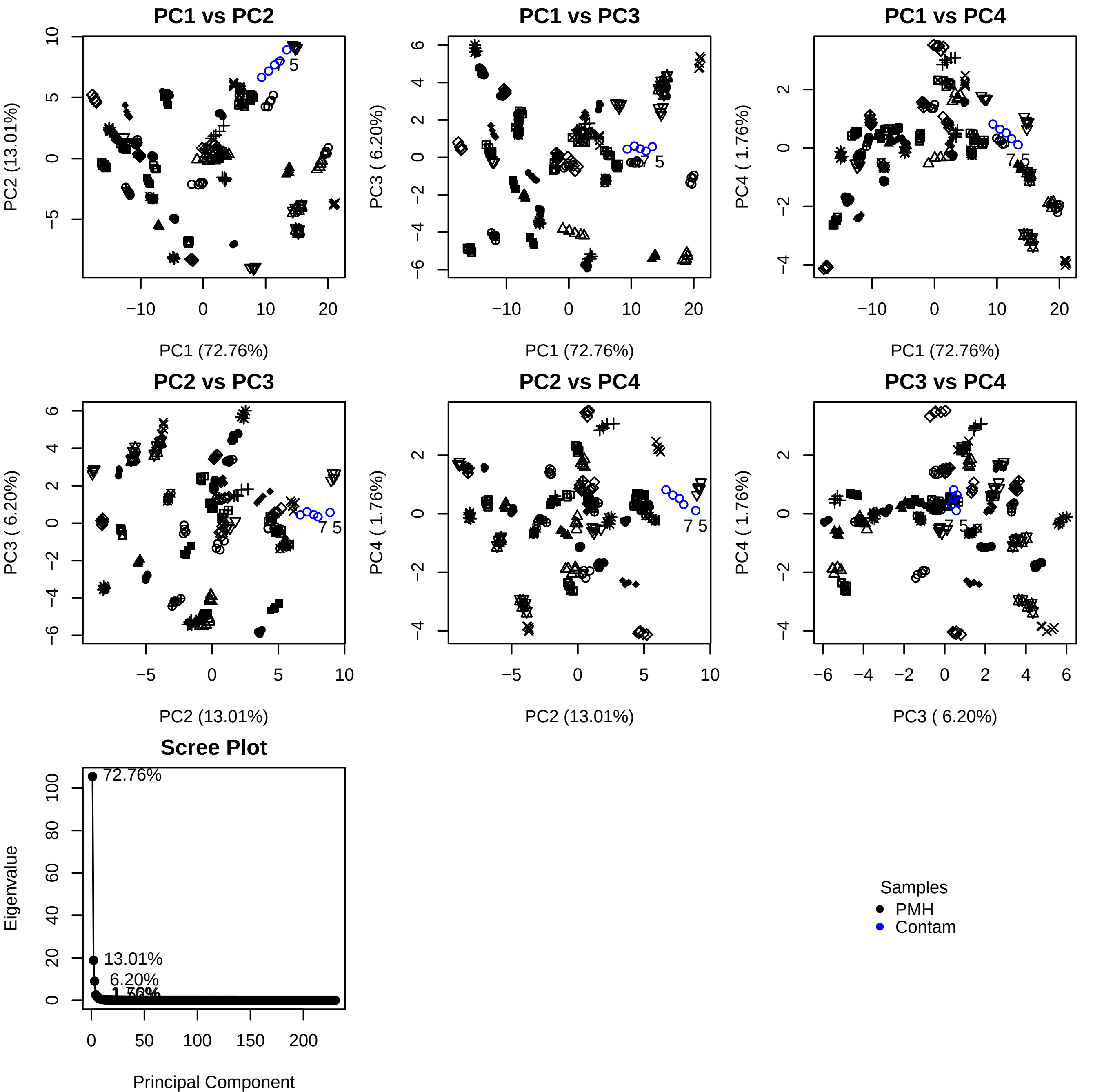
PC1 vs PC2
PC1 vs PC3
PC1 vs PC4
PC2 vs PC3
PC2 vs PC4
PC3 vs PC4
Scree Plot
PC1 (72.76%)
PC2 (13.01%)
PC3 ( 6.20%)
PC4 ( 1.76%)
7 5
Eigenvalue
Principal Component
72.76%
13.01%
6.20%
Samples
PMH
Contam

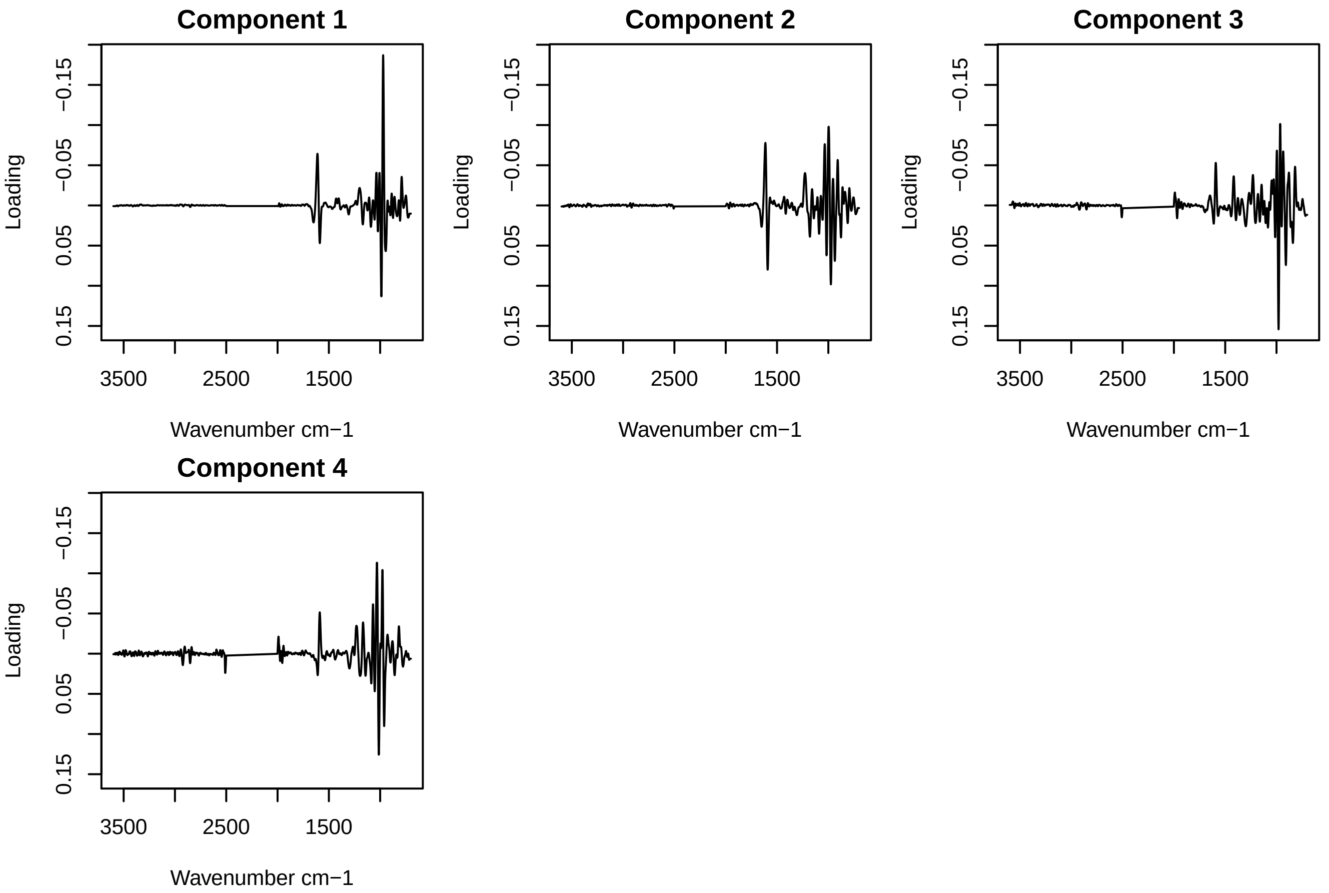
Component 1
Component 2
Component 3
Component 4
Loading
−0.15
−0.05
0.05
0.15
3500
2500
1500
Wavenumber cm−1

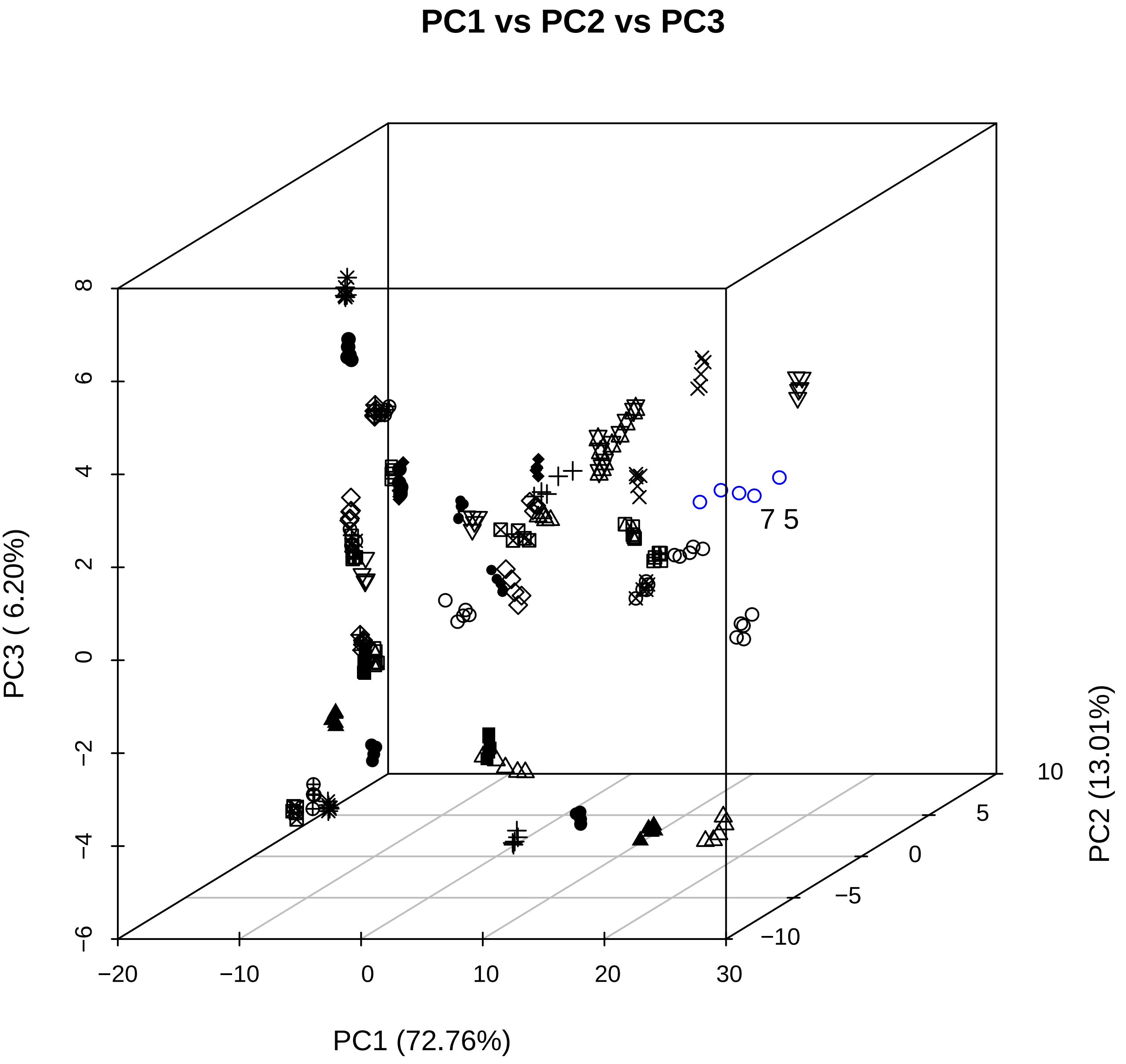
PC1 vs PC2 vs PC3
PC3 ( 6.20%)
PC1 (72.76%)
PC2 (13.01%)
−6
−4
−2
0
2
4
6
8
−20
−10
0
10
20
30
−10
−5
0
5
10
7 5

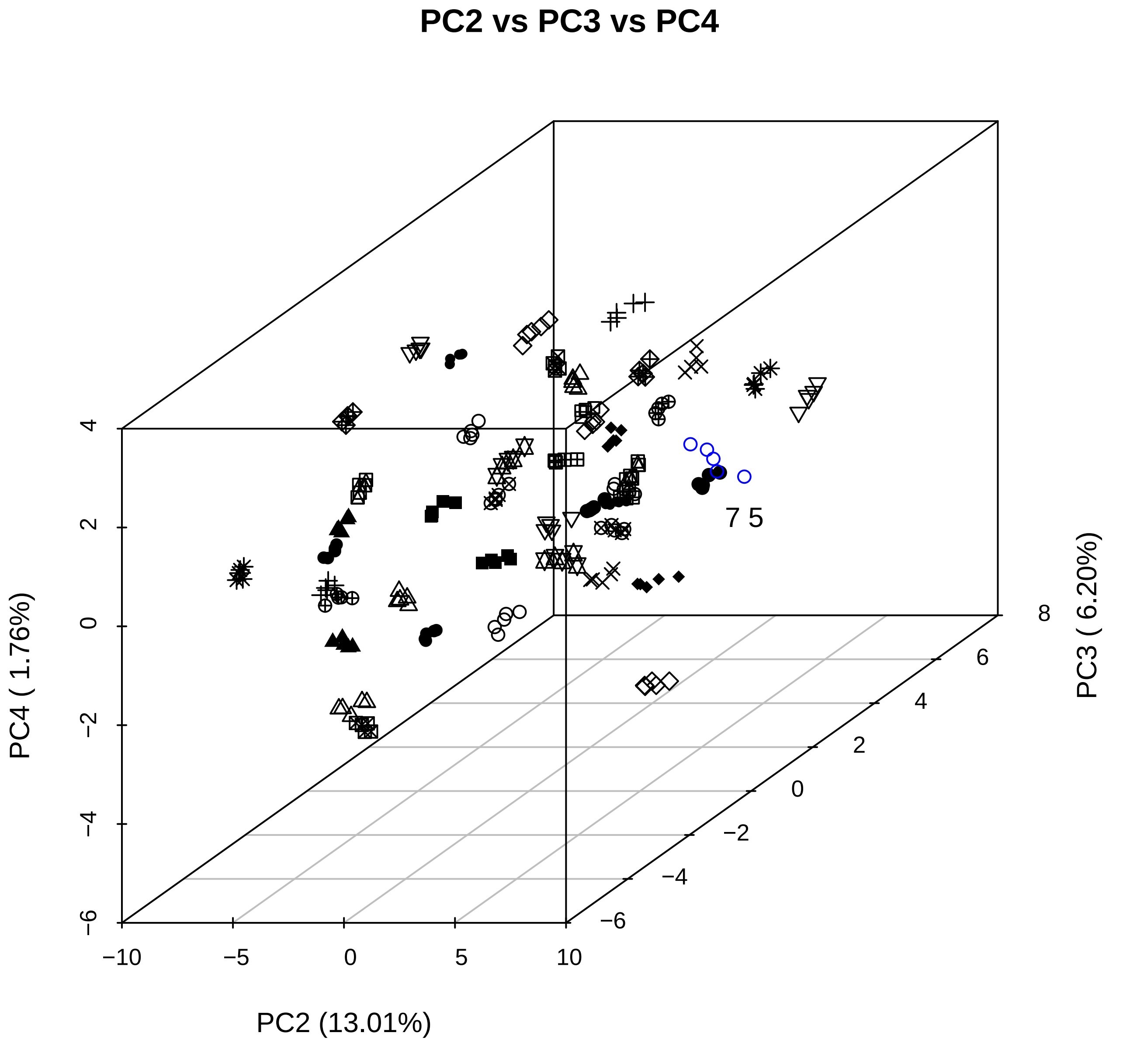
PC2 vs PC3 vs PC4
PC2 (13.01%)
PC3 ( 6.20%)
PC4 ( 1.76%)
−10
−5
0
5
10
−6
−4
−2
0
2
4
6
8
7 5

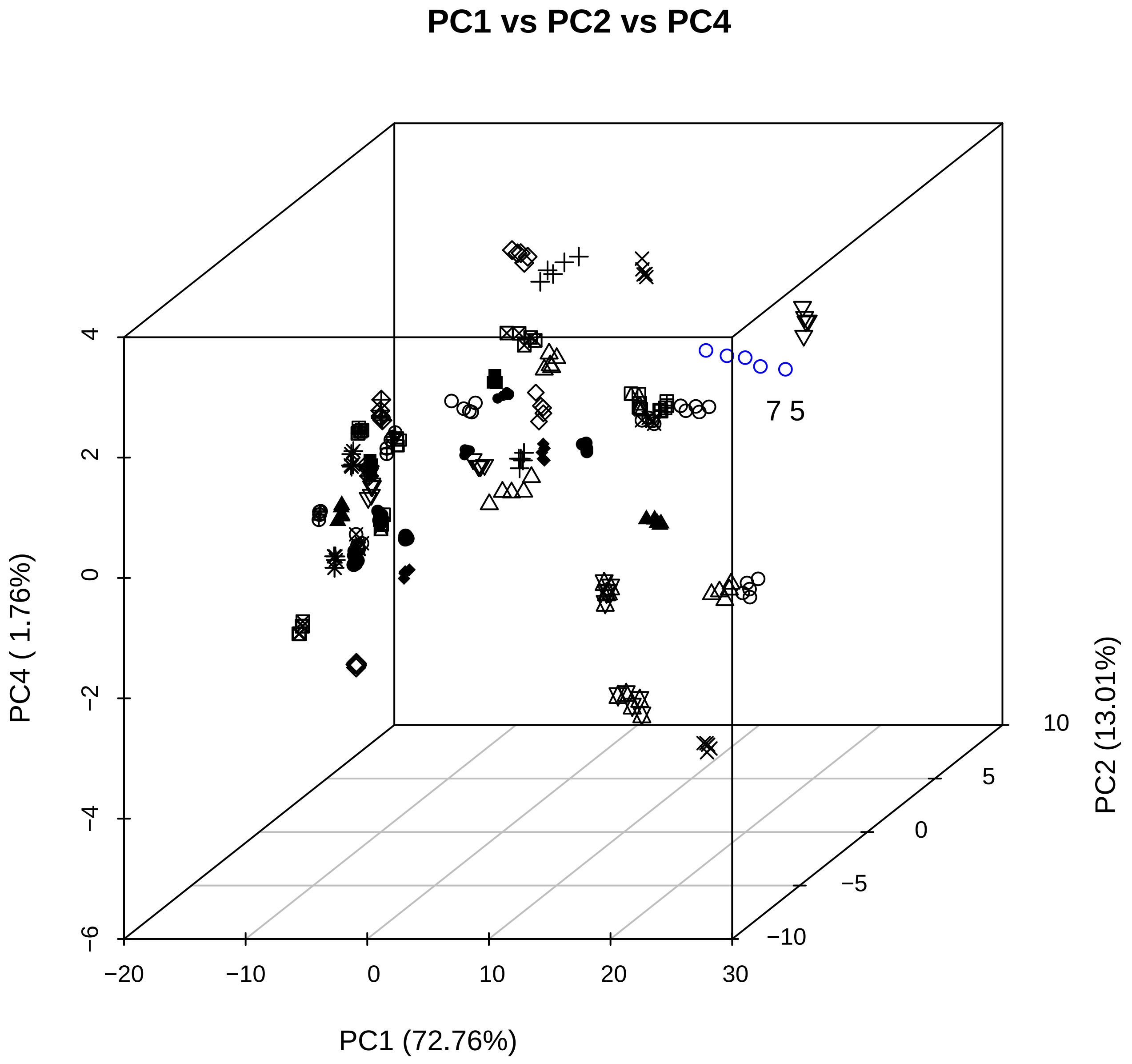

PC1 vs PC2 vs PC4
PC1 (72.76%)
PC2 (13.01%)
PC4 ( 1.76%)
−20
−10
0
10
20
30
−10
−5
0
5
10
−6
−4
−2
0
2
4
7 5

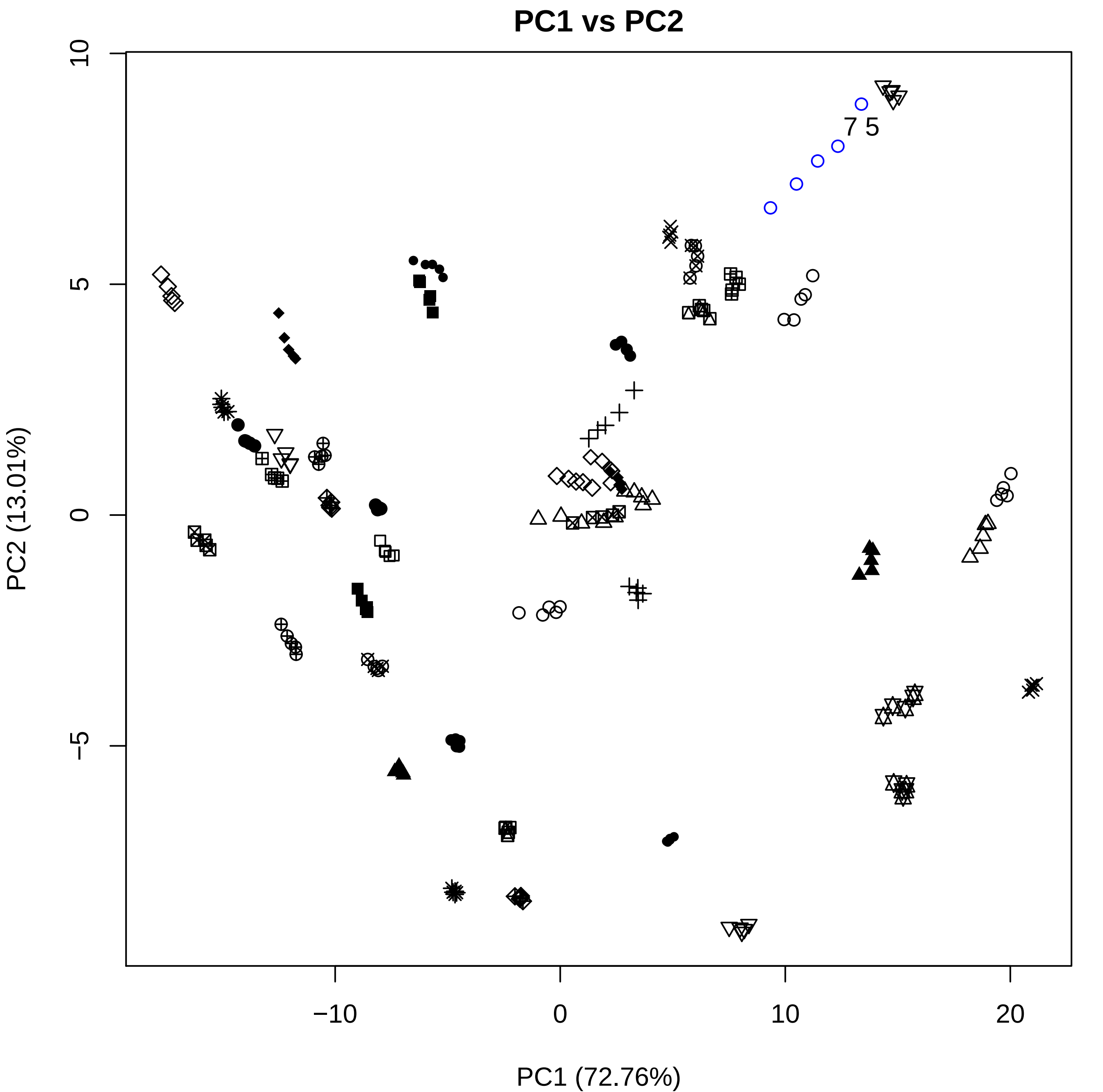
PC1 vs PC2
PC2 (13.01%)
PC1 (72.76%)
10
5
0
−5
−10
0
10
20
7 5

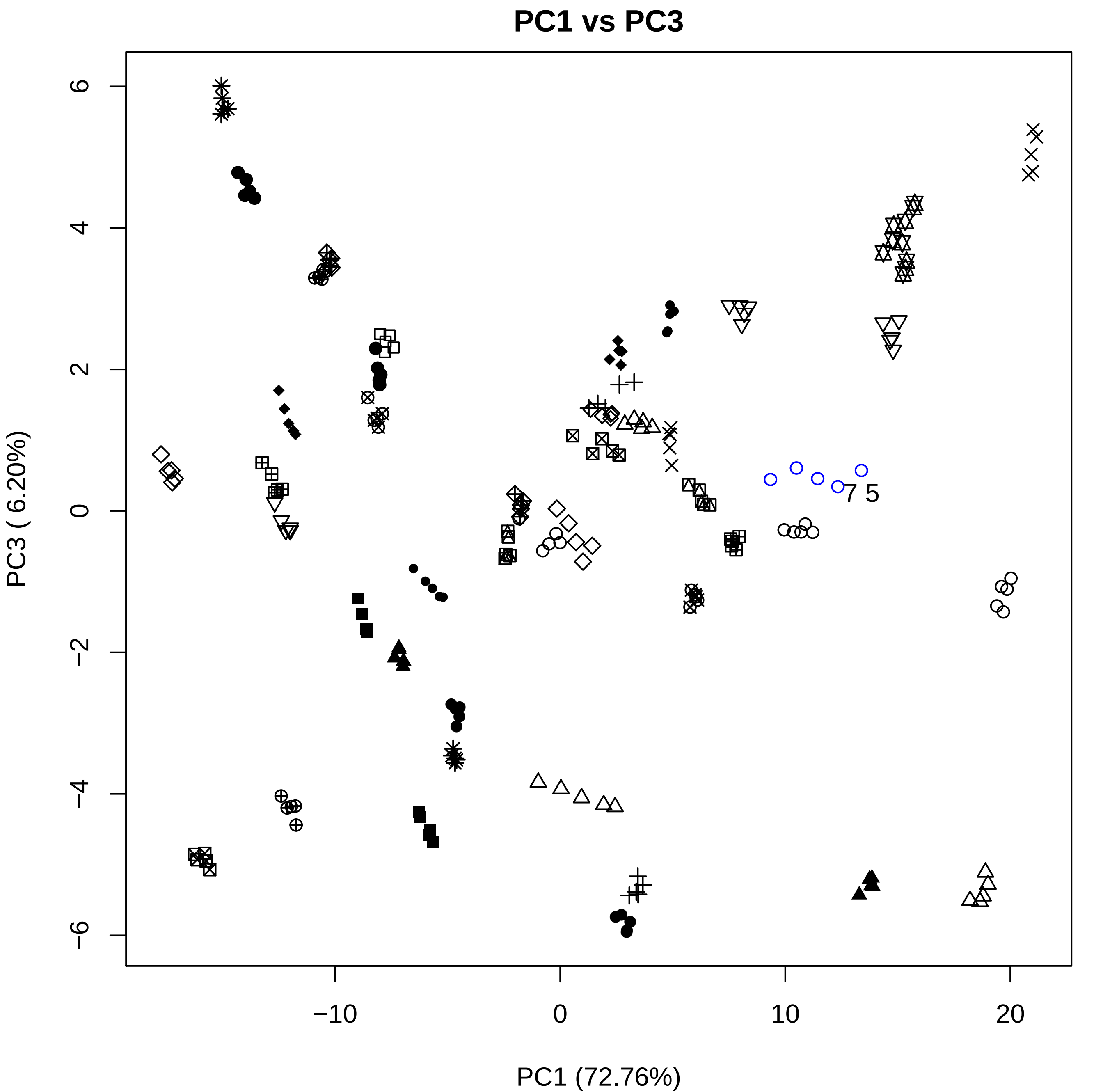

PC1 vs PC3
PC3 ( 6.20%)
PC1 (72.76%)
6
4
2
0
−2
−4
−6
−10
0
10
20
7 5

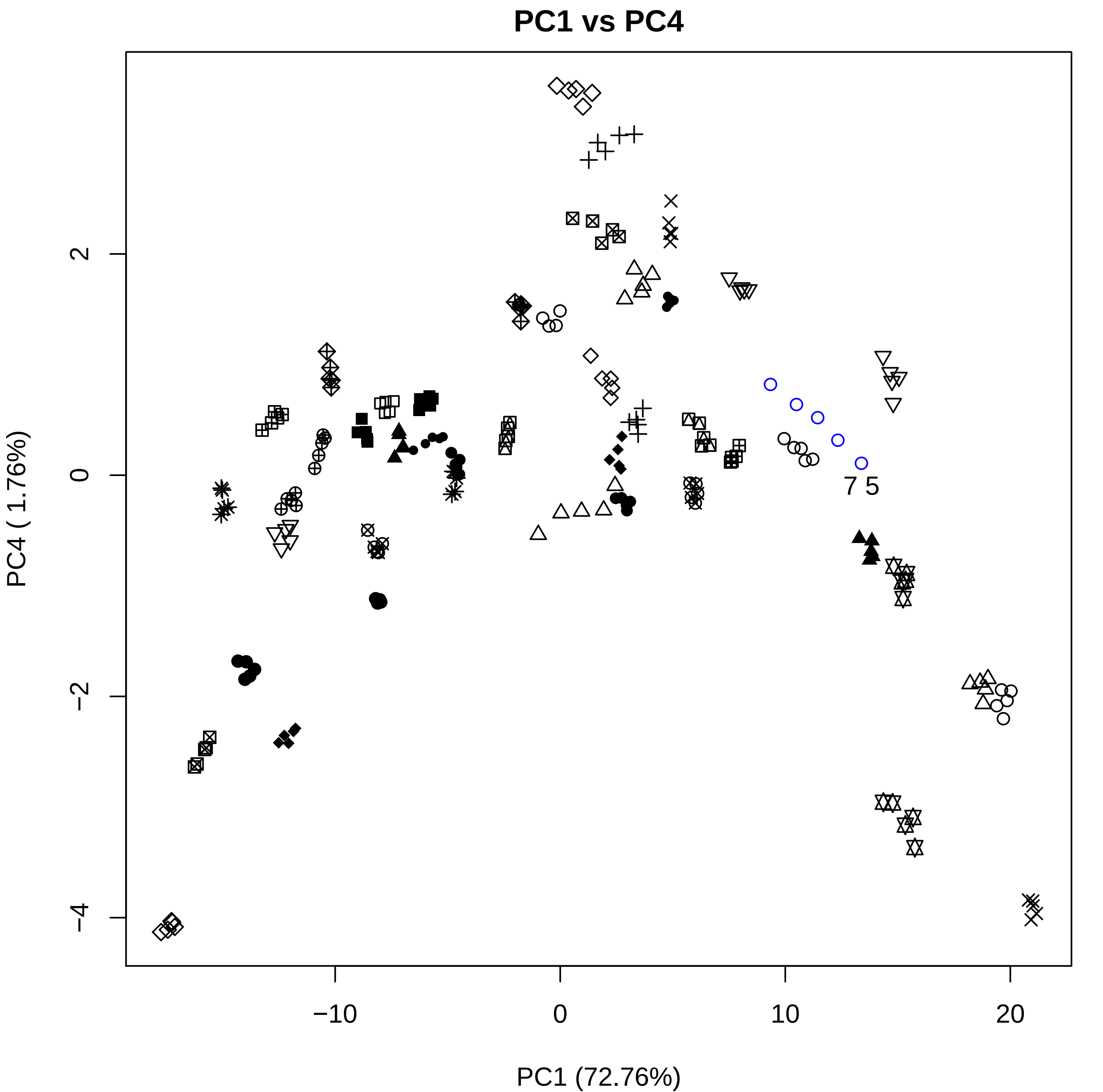
PC1 vs PC4
PC4 ( 1.76%)
PC1 (72.76%)
−4
−2
0
2
−10
0
10
20
7 5

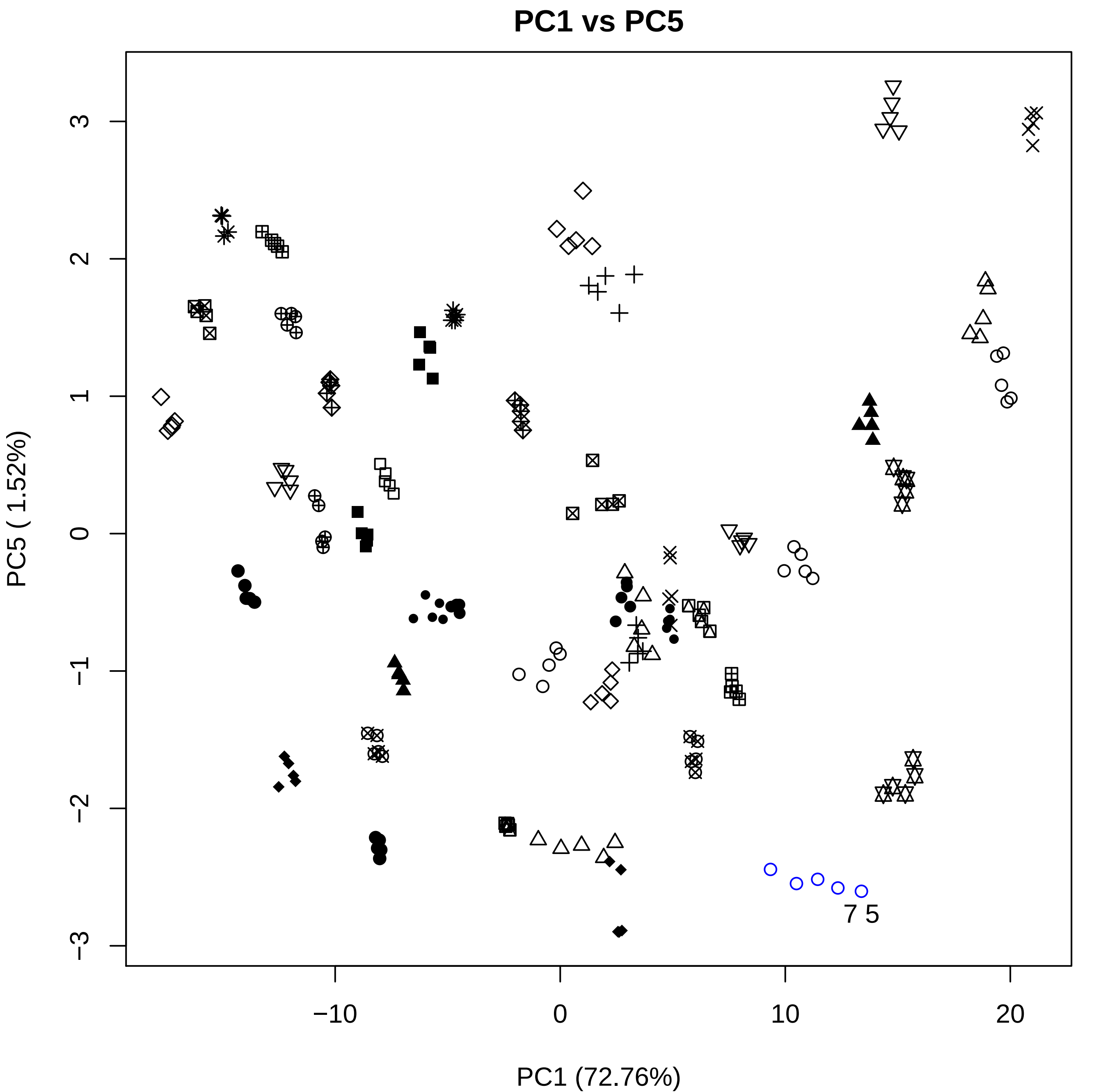
PC1 vs PC5
PC5 ( 1.52%)
PC1 (72.76%)
3
2
1
0
−1
−2
−3
−10
0
10
20
7 5

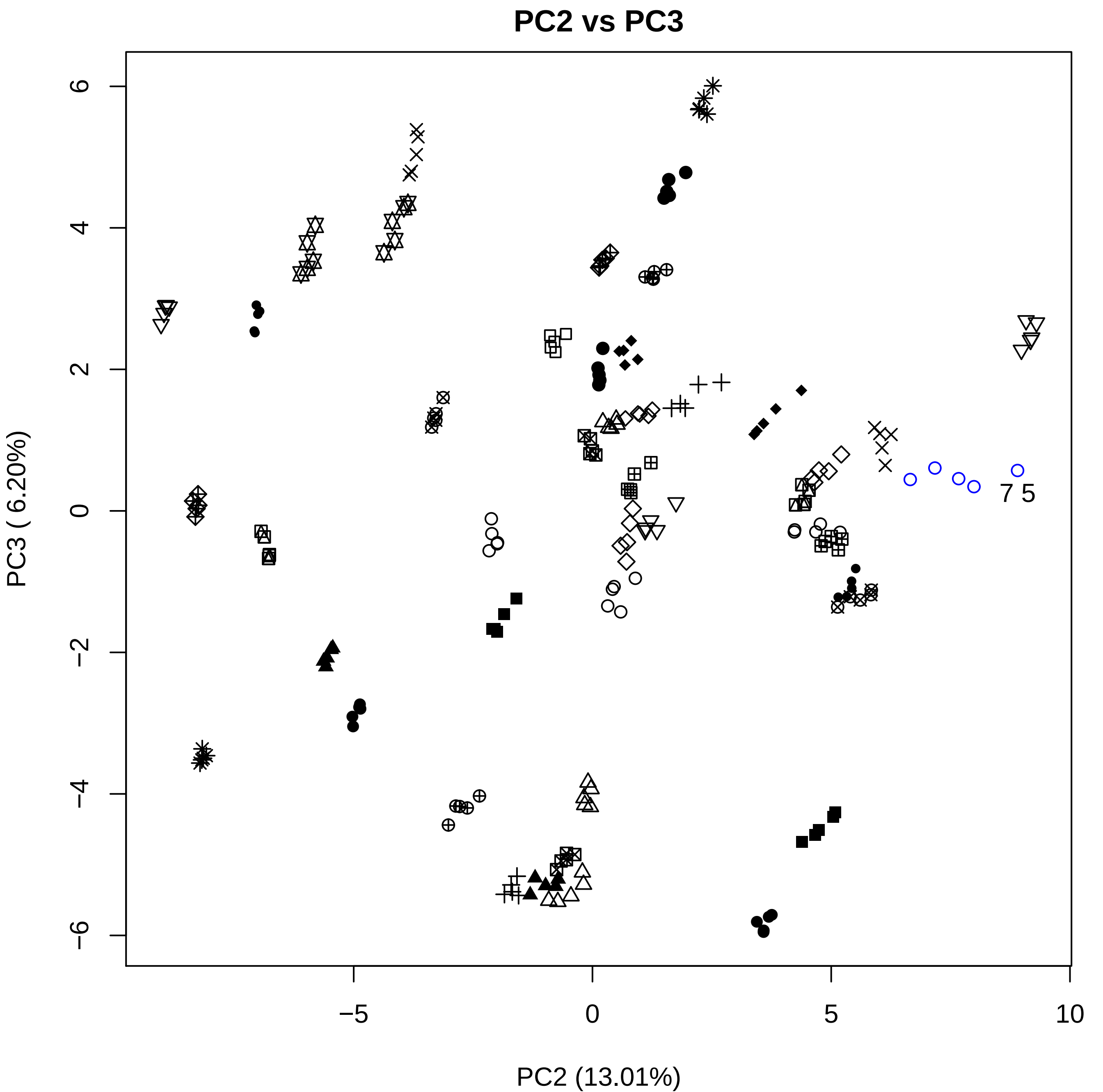
PC2 vs PC3
PC3 ( 6.20%)
PC2 (13.01%)
6
4
2
0
−2
−4
−6
−5
0
5
10
7 5

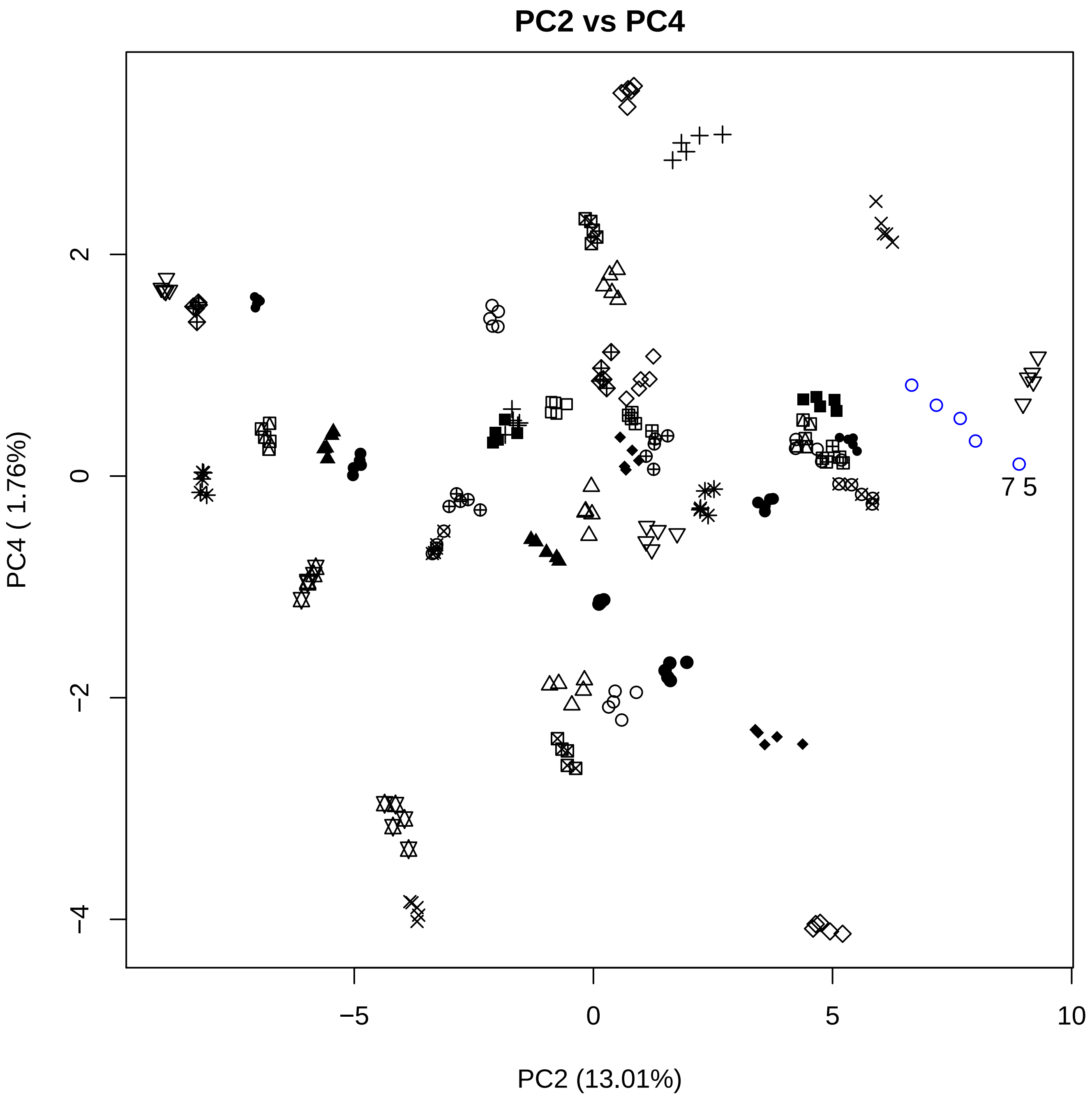
PC2 vs PC4
PC4 ( 1.76%)
PC2 (13.01%)
2
0
−2
−4
−5
0
5
10
7 5

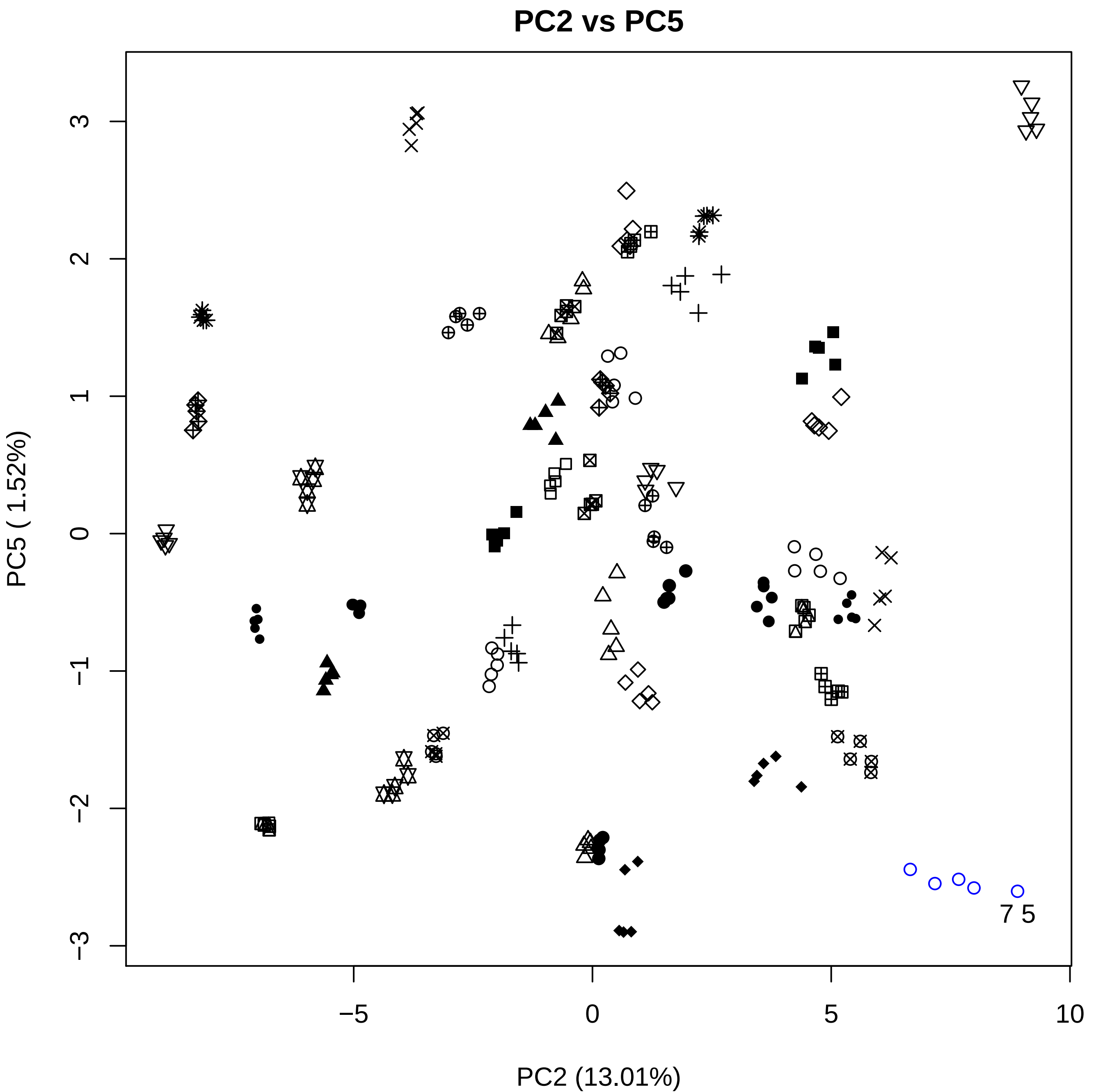
PC2 vs PC5
PC5 ( 1.52%)
PC2 (13.01%)
3
2
1
0
−1
−2
−3
−5
0
5
10
7 5

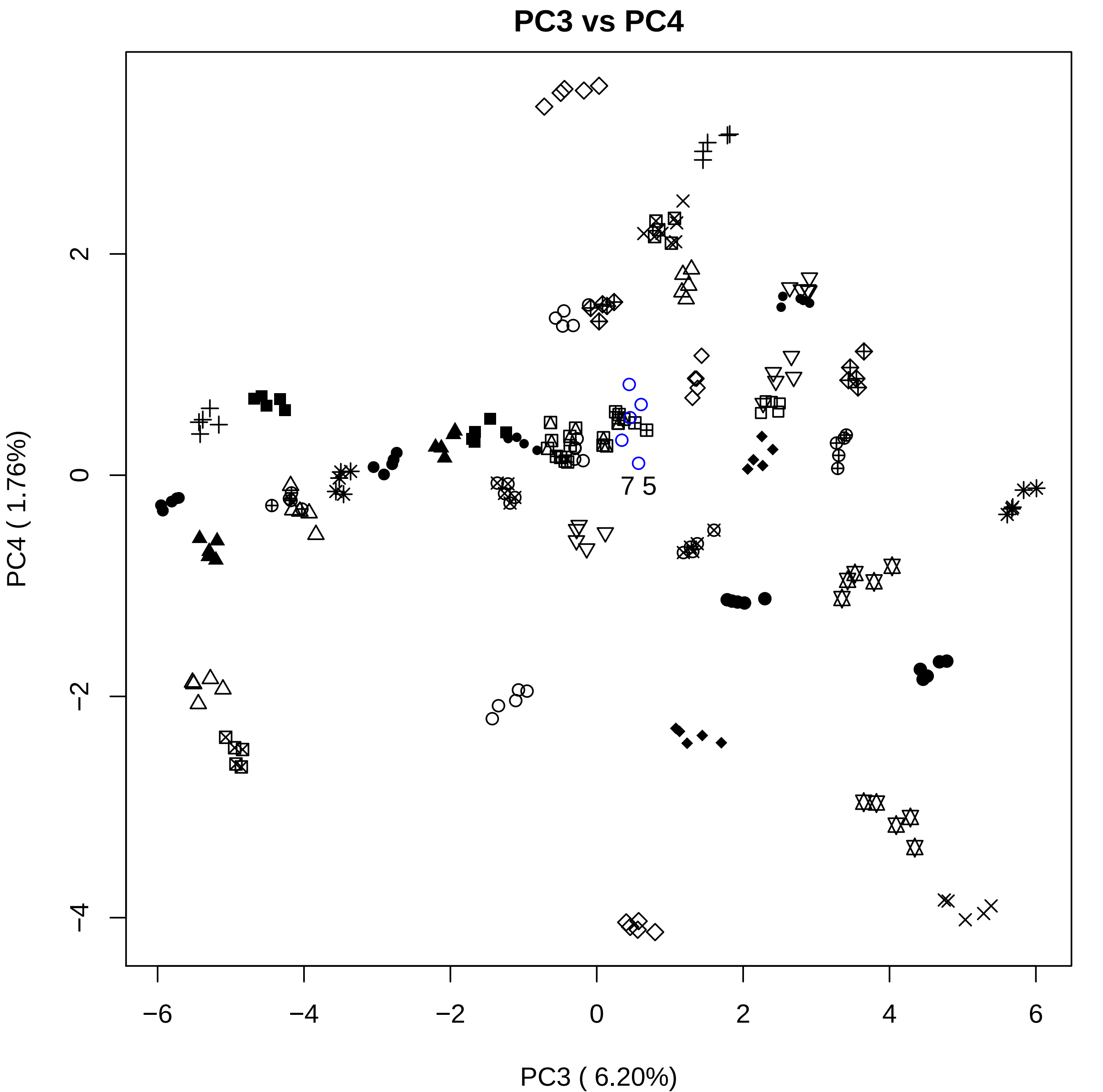
PC3 vs PC4
PC4 ( 1.76%)
PC3 ( 6.20%)
−6
−4
−2
0
2
4
6
−4
−2
0
2
7 5

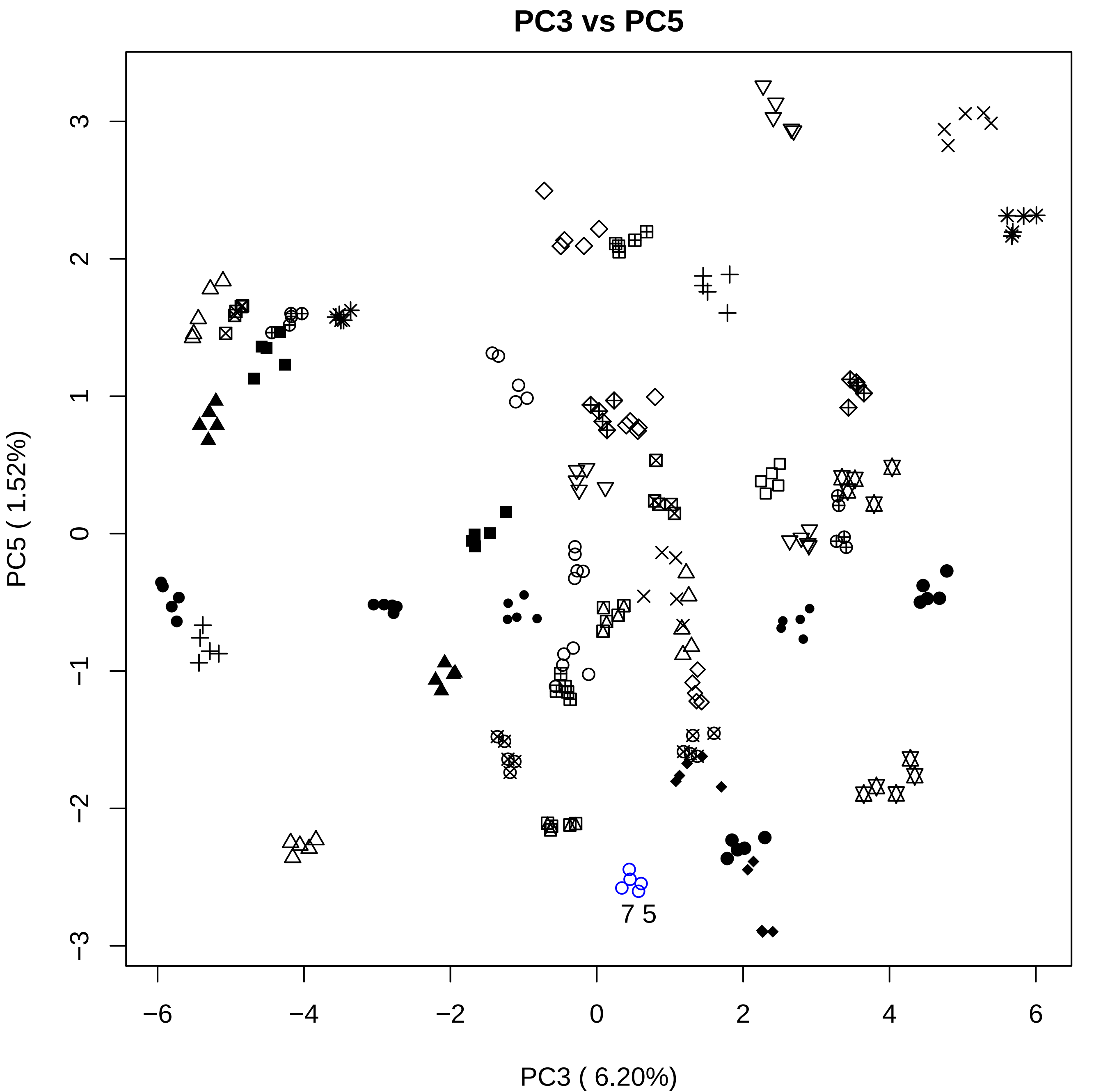

PC3 vs PC5
PC5 ( 1.52%)
PC3 ( 6.20%)
7 5
−6
−4
−2
0
2
4
6
−3
−2
−1
0
1
2
3

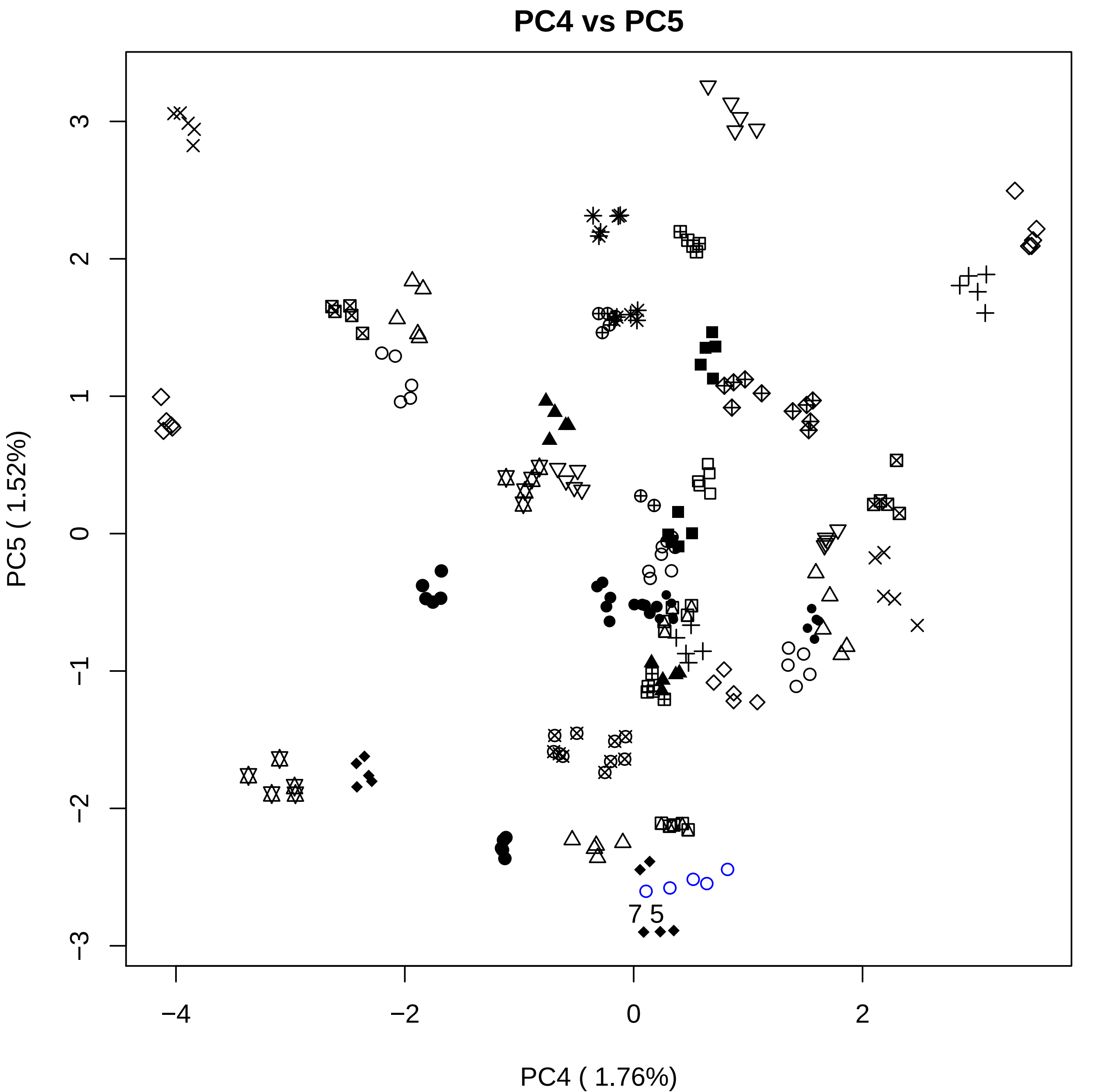
PC4 vs PC5
PC5 ( 1.52%)
PC4 ( 1.76%)
3
2
1
0
−1
−2
−3
−4
−2
0
2
7 5

### PCA of 10%  DeS vs PMH

Scaled Matrix PCAed

7th order polynomial baseline correction performed

Raw spectra smoothed to 21 Neighbours, 2nd order PN, Savtisky Golay

Normalisation between 0 and 100

D2 spectra smoothed to 41 Neighbours, 2nd order PN, Savtisky Golay

Regions Analysed: 3601−2501 and 2001−700cm−1

Samples Analysed: 1 Contam's 0 DeS's 45 PMH's

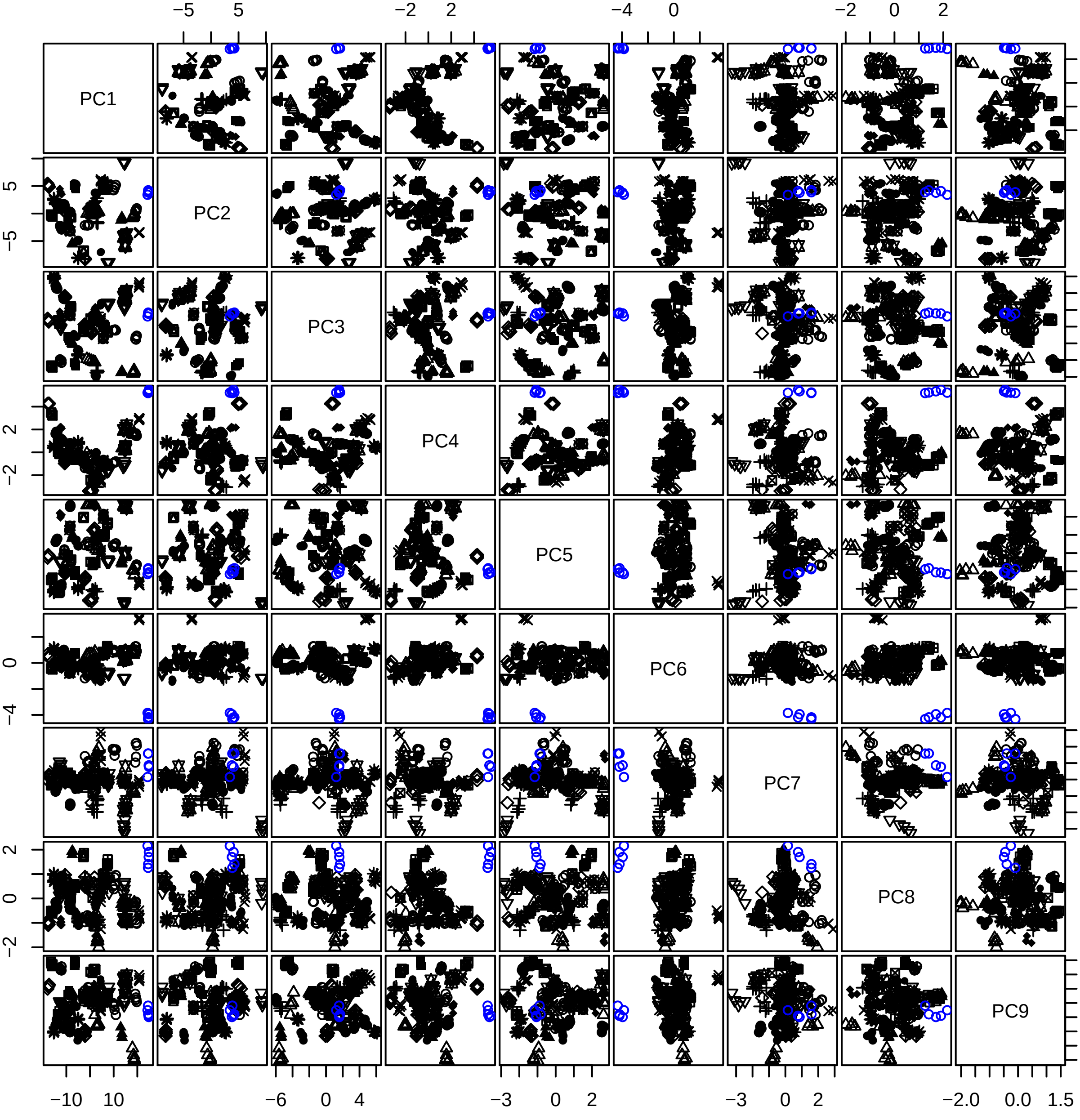

PC1
PC2
PC3
PC4
PC5
PC6
PC7
PC8
PC9
−5
5
−2
2
−4
0
−2
0
2
10
−10
5
−5
4
0
−6
2
−2
2
0
−3
0
−4
2
0
−3
2
0
−2
1.5
0.0
−2.0
−10
10
−6
0
4
−3
0
2
−3
0
2
−2.0
0.0
1.5

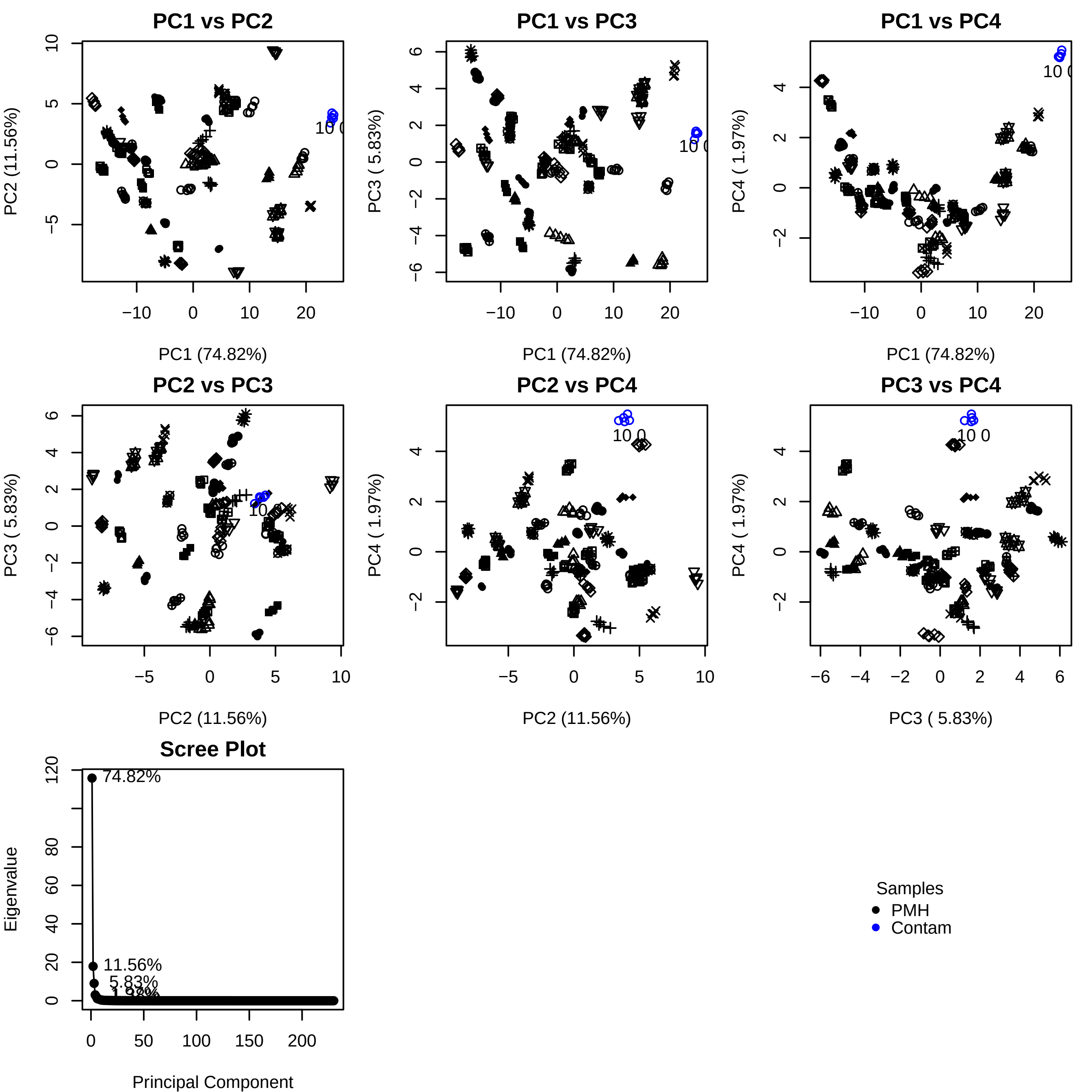
PC1 vs PC2
PC1 vs PC3
PC1 vs PC4
PC2 vs PC3
PC2 vs PC4
PC3 vs PC4
Scree Plot
PC1 (74.82%)
PC2 (11.56%)
PC3 ( 5.83%)
PC4 ( 1.97%)
74.82%
11.56%
5.83%
Eigenvalue
Principal Component
Samples
PMH
Contam

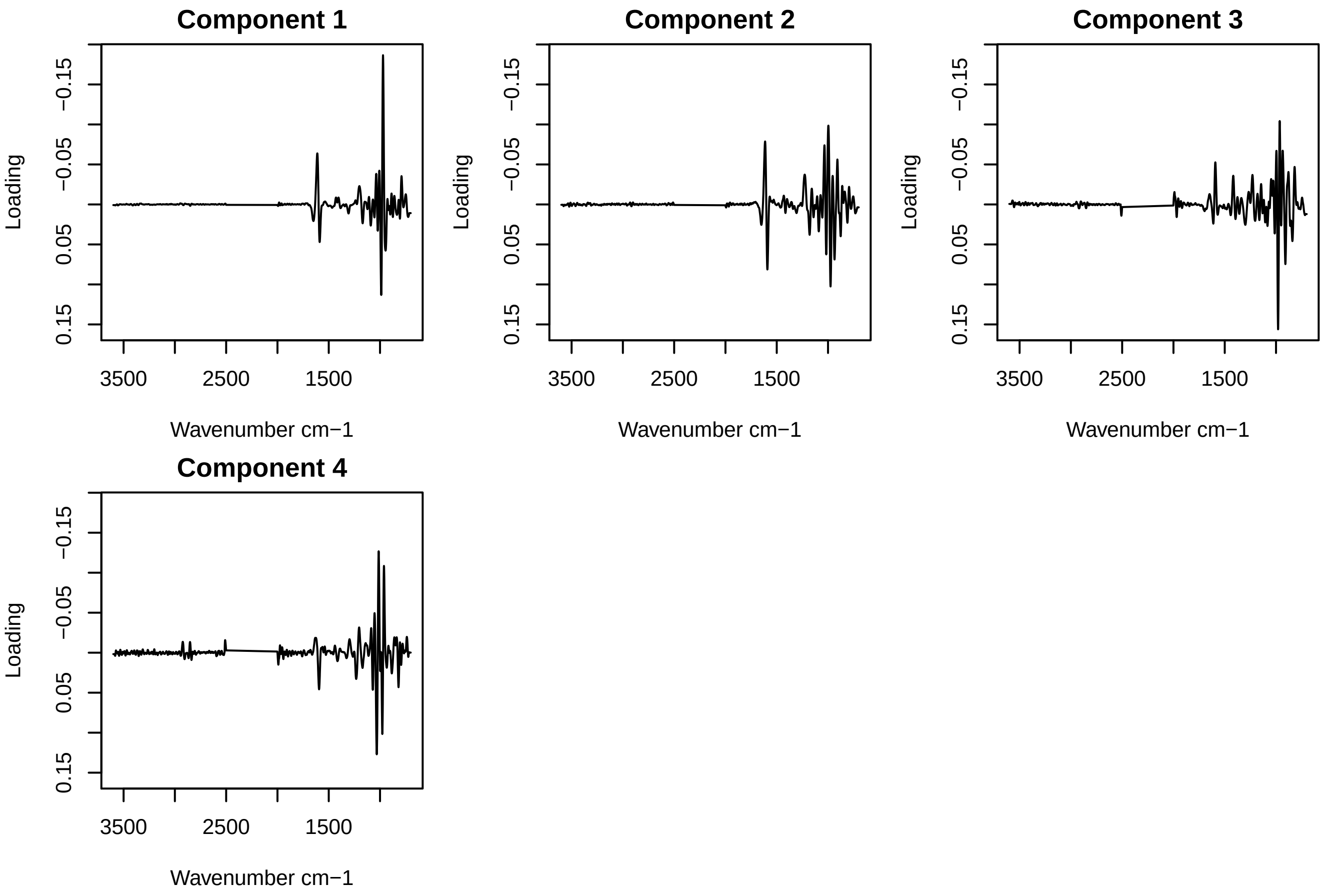
Component 1
Component 2
Component 3
Component 4
Loading
−0.15
−0.05
0.05
0.15
3500
2500
1500
Wavenumber cm−1

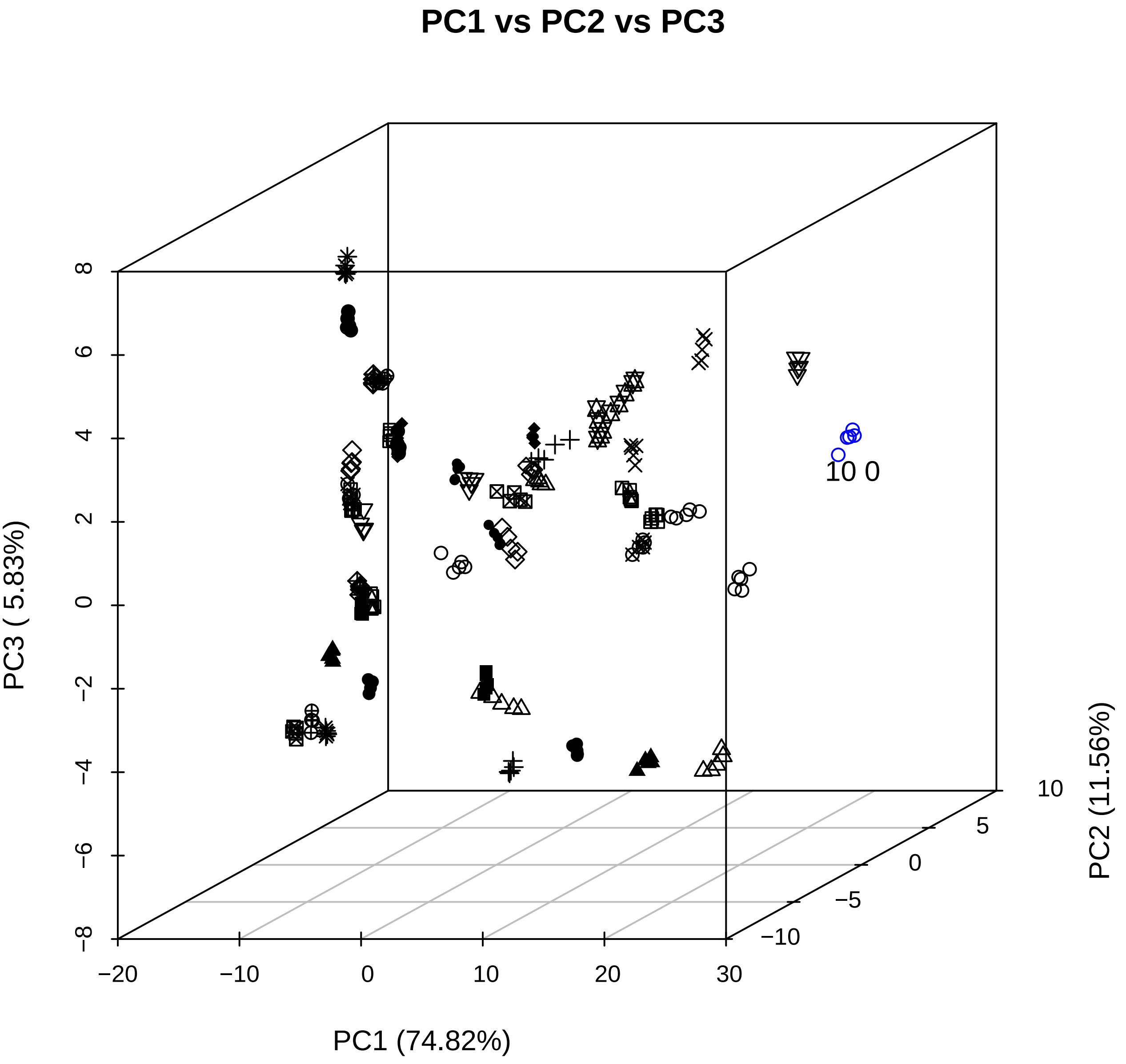

PC1 vs PC2 vs PC3
PC1 (74.82%)
PC2 (11.56%)
PC3 ( 5.83%)
−20
−10
0
10
20
30
−10
−5
0
5
10
−8
−6
−4
−2
0
2
4
6
8
10 0

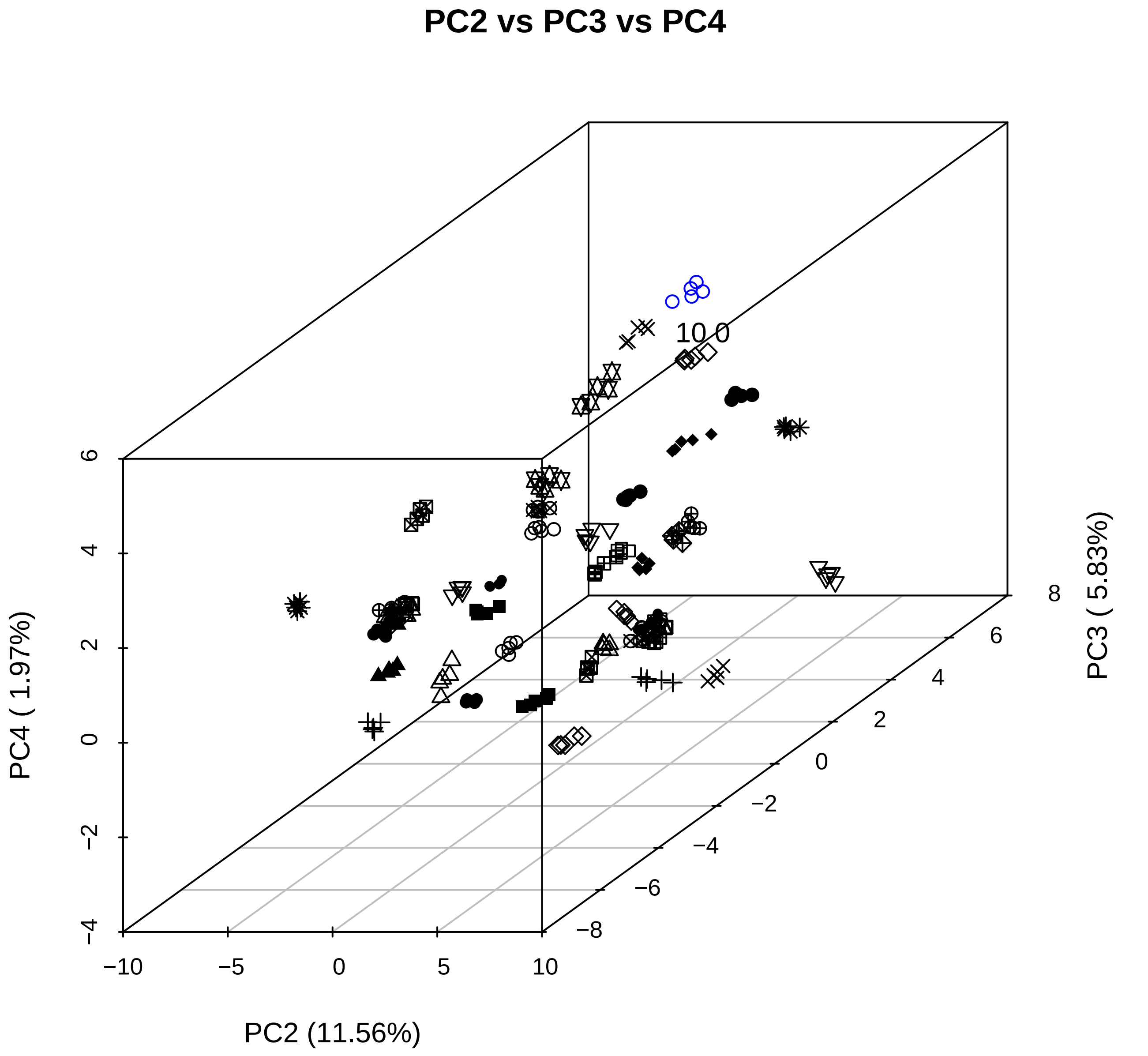

PC2 vs PC3 vs PC4
10.0
PC4 ( 1.97%)
6
4
2
0
−2
−4
−10
−5
0
5
10
PC2 (11.56%)
PC3 ( 5.83%)
8
6
4
2
0
−2
−4
−6
−8

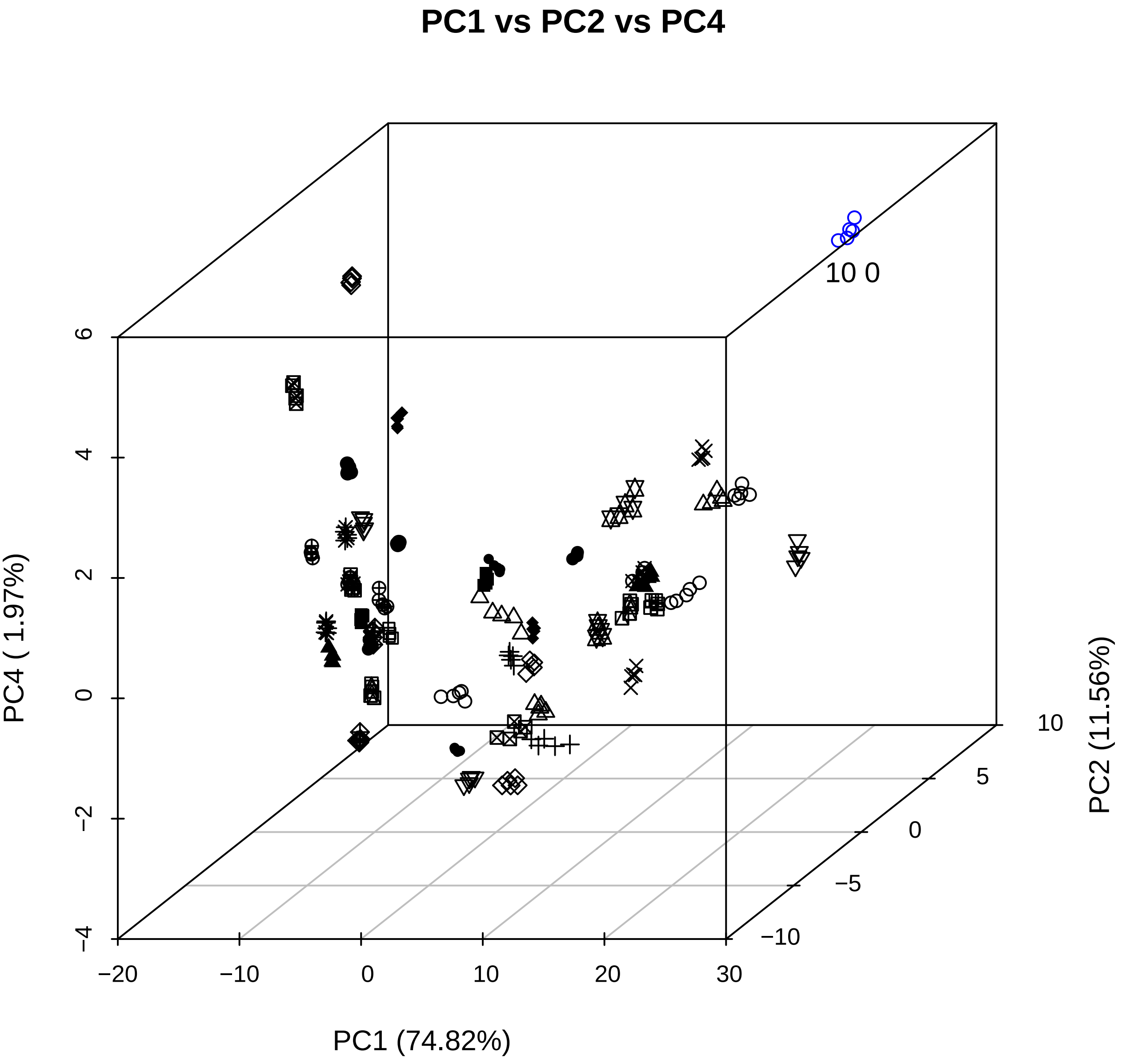
PC1 vs PC2 vs PC4
PC4 ( 1.97%)
PC1 (74.82%)
PC2 (11.56%)
10 0
−20
−10
0
10
20
30
−4
−2
0
2
4
6
−10
−5
0
5
10

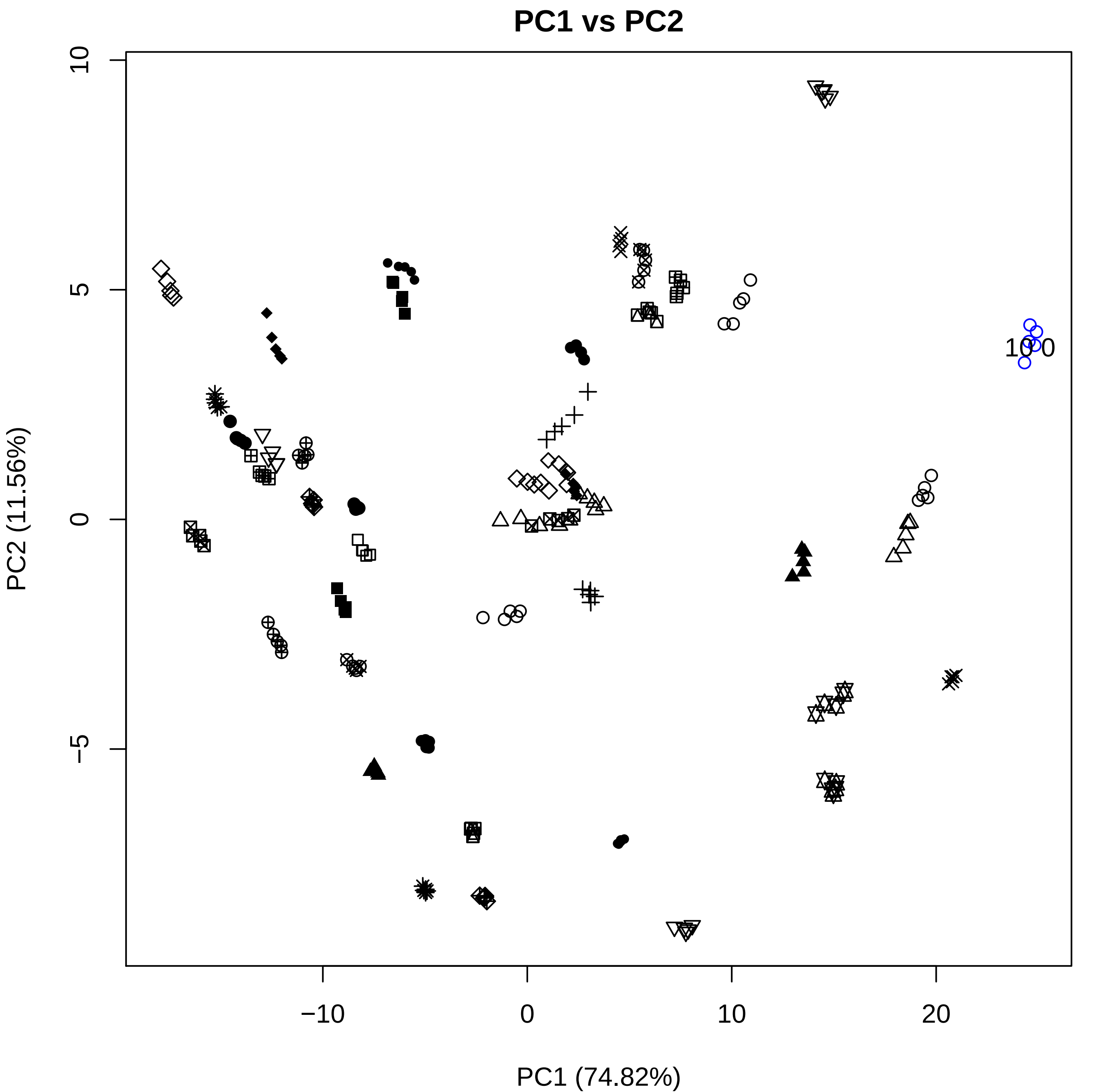
PC1 vs PC2
PC1 (74.82%)
PC2 (11.56%)
−10
0
10
20
−5
0
5
10
100

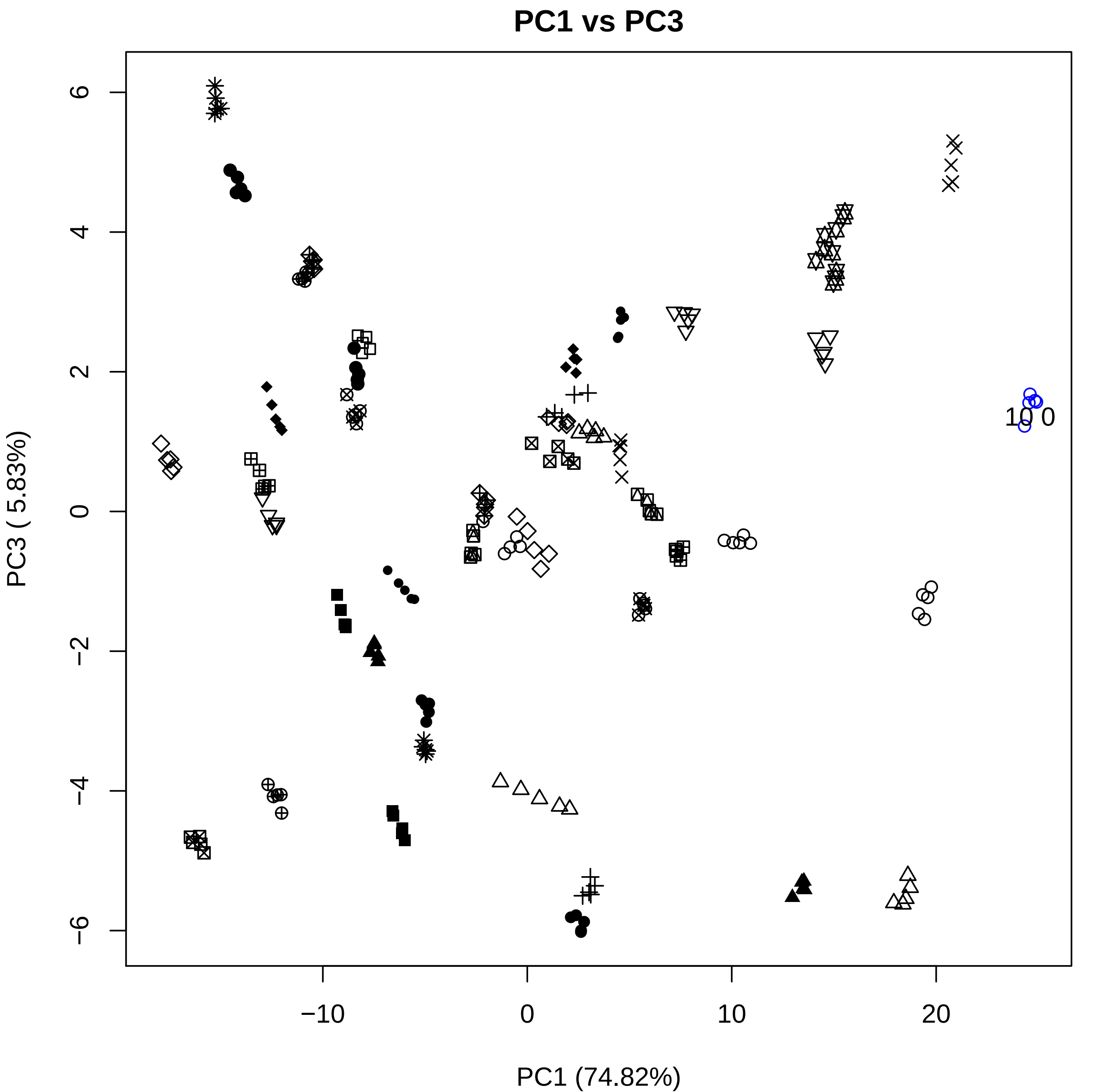
PC1 vs PC3
PC3 ( 5.83%)
PC1 (74.82%)
6
4
2
0
−2
−4
−6
−10
0
10
20
10 0

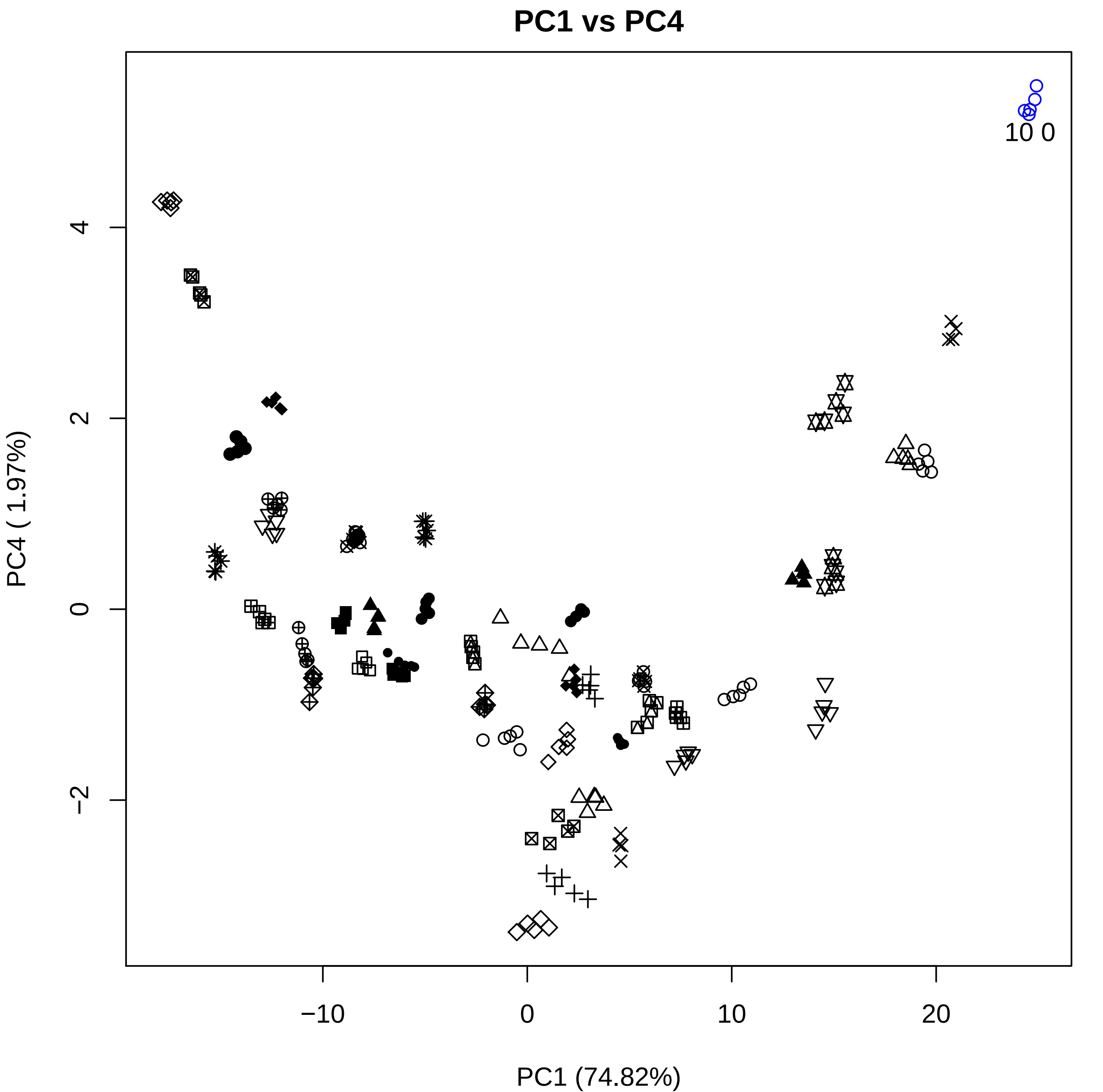

PC1 vs PC4
10 0
4
2
0
−2
PC4 ( 1.97%)
−10
0
10
20
PC1 (74.82%)

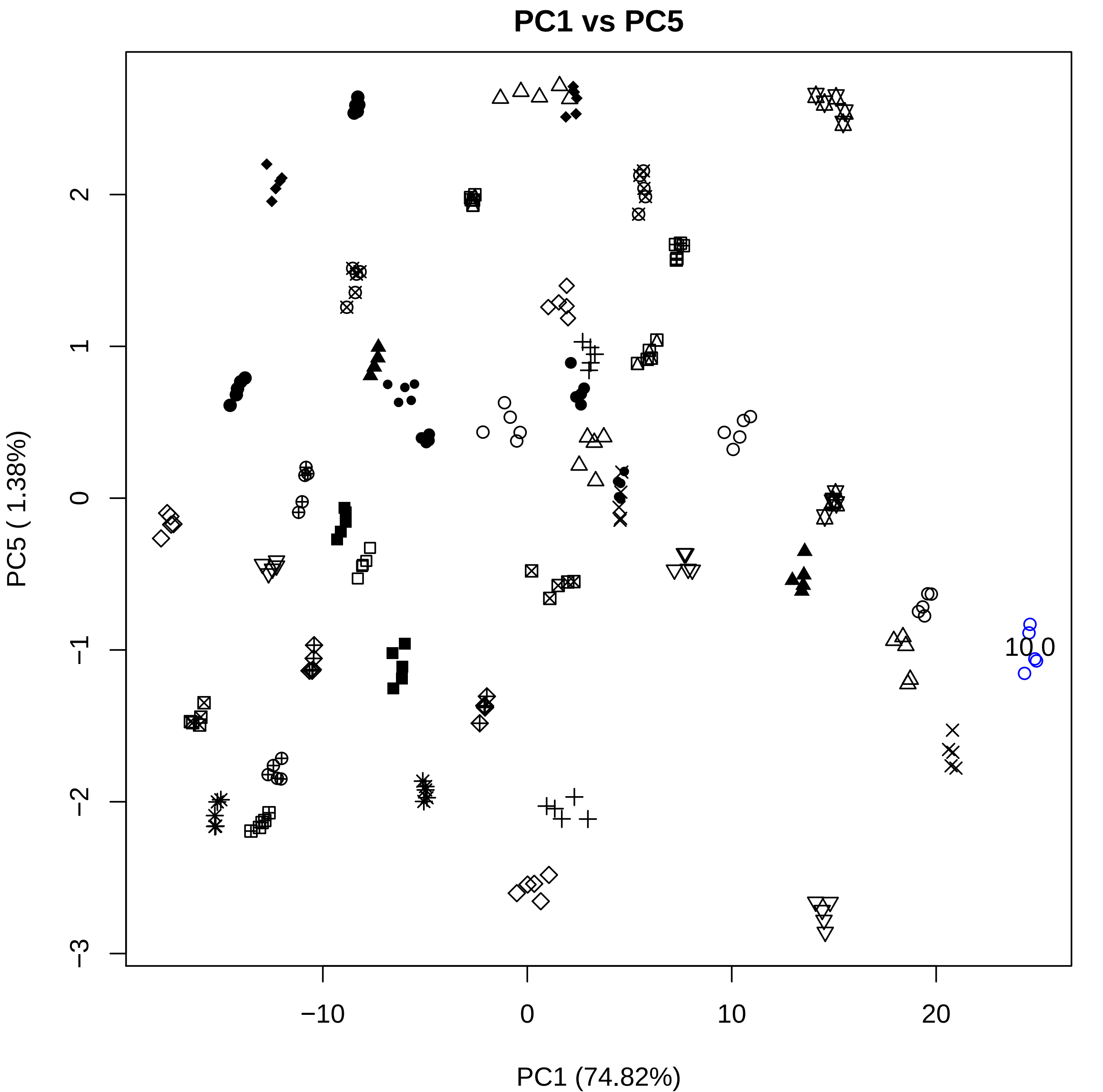

PC1 vs PC5
PC5 ( 1.38%)
PC1 (74.82%)
−3
−2
−1
0
1
2
−10
0
10
20
10.0

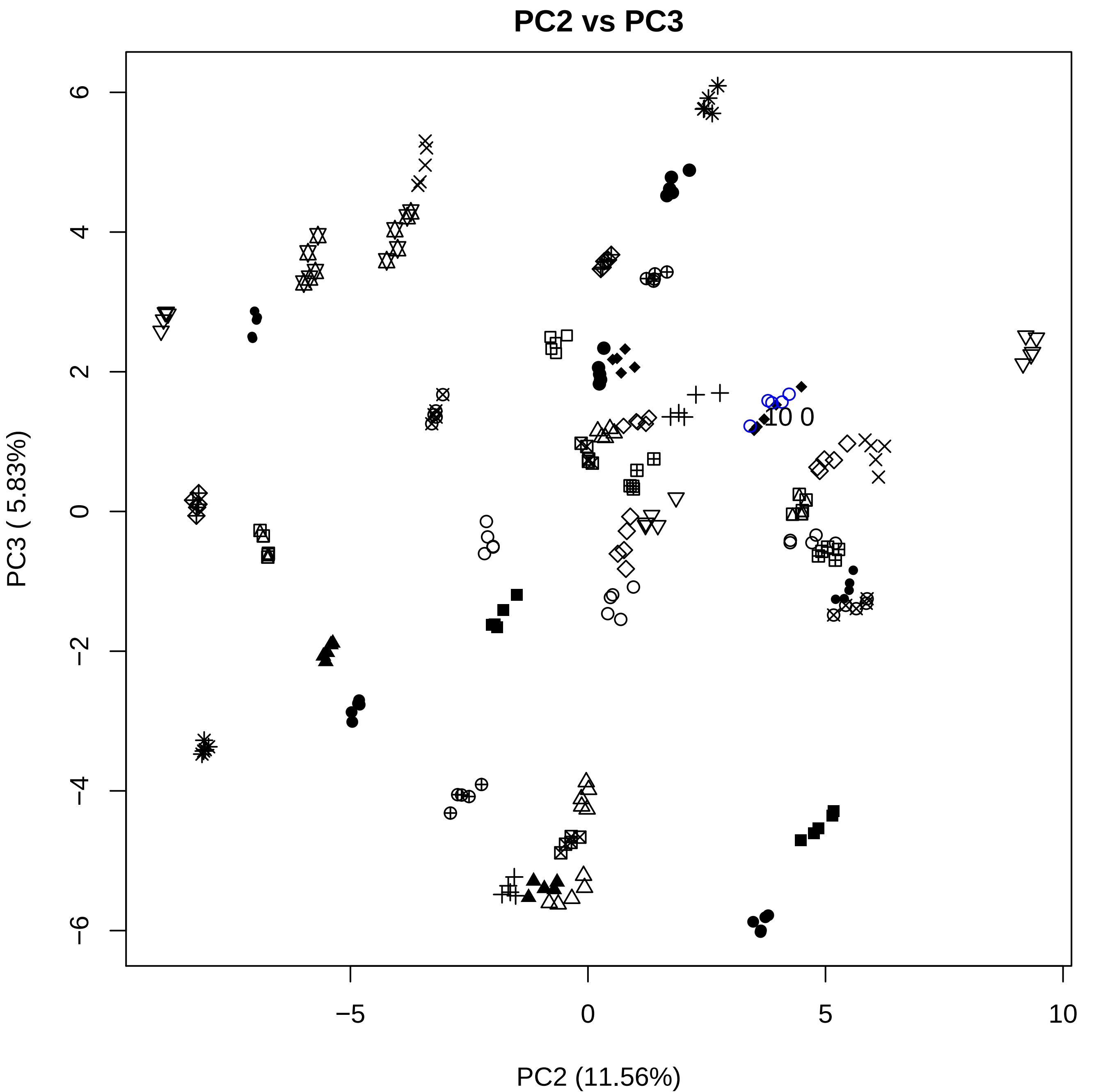
PC2 vs PC3
PC3 ( 5.83%)
PC2 (11.56%)
6
4
2
0
−2
−4
−6
−5
0
5
10
10 0

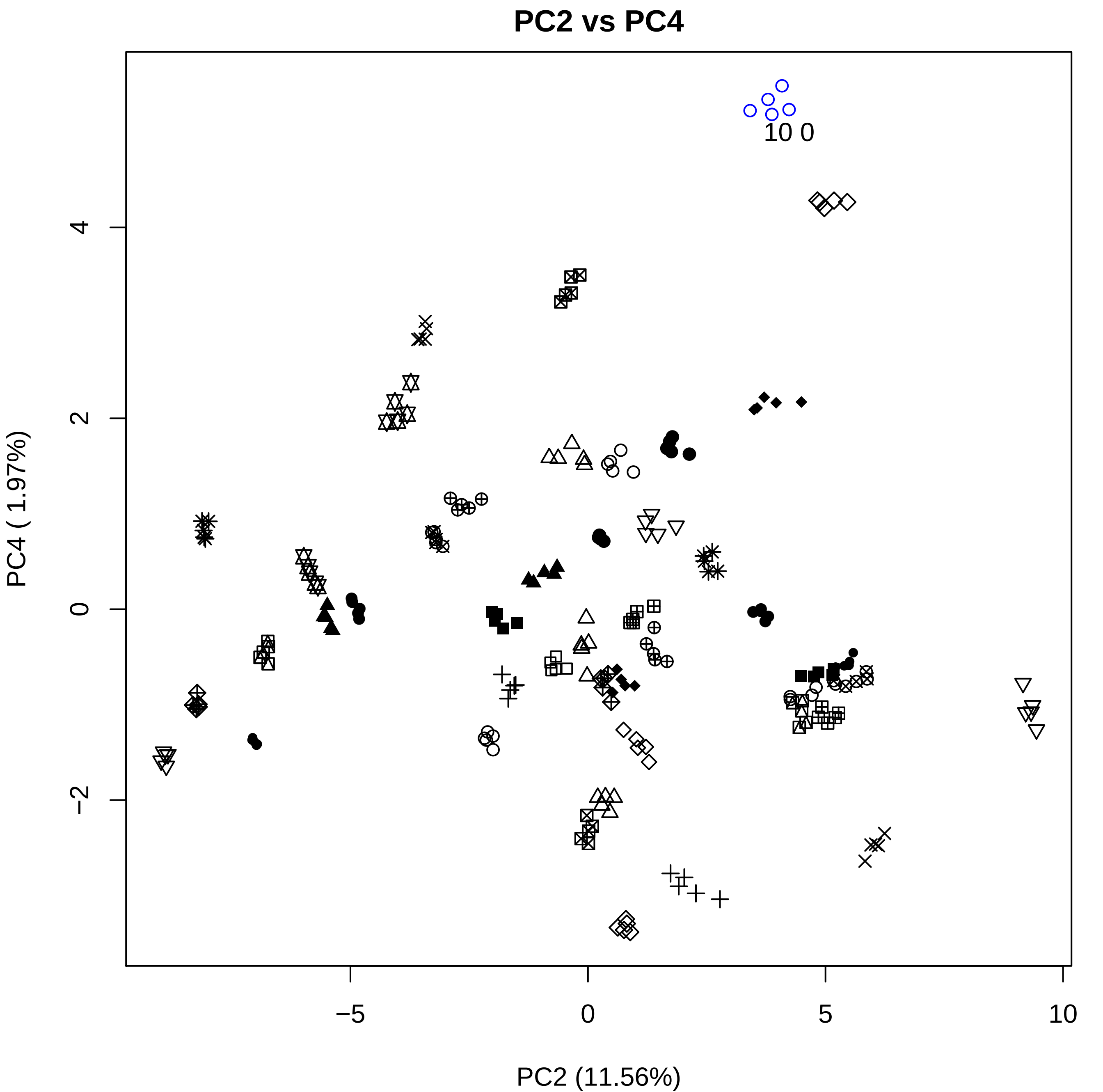
PC2 vs PC4
10 0
4
2
0
−2
−5
0
5
10
PC4 ( 1.97%)
PC2 (11.56%)

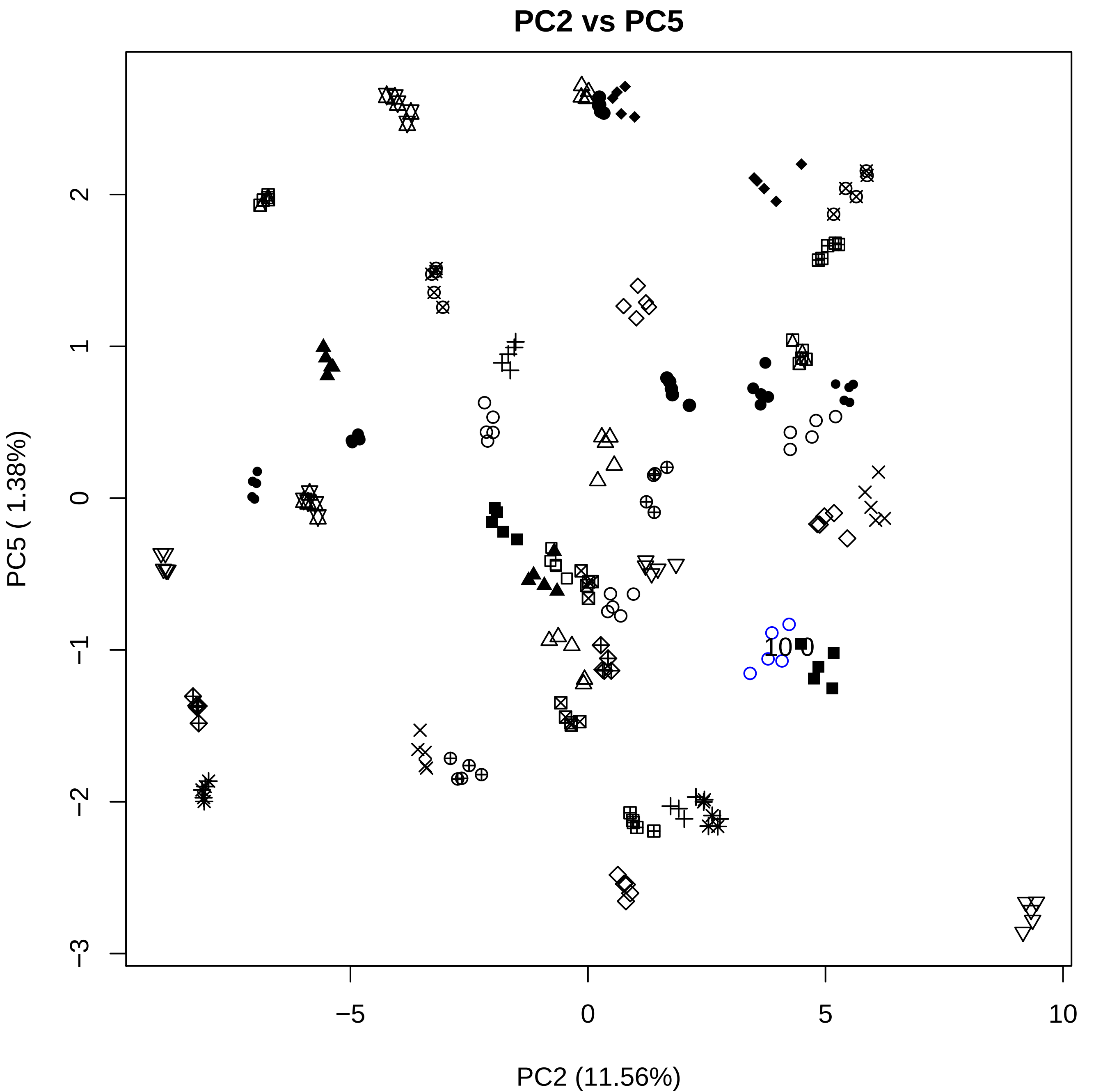

PC2 vs PC5
PC5 ( 1.38%)
PC2 (11.56%)
2
1
0
−1
−2
−3
−5
0
5
10
100

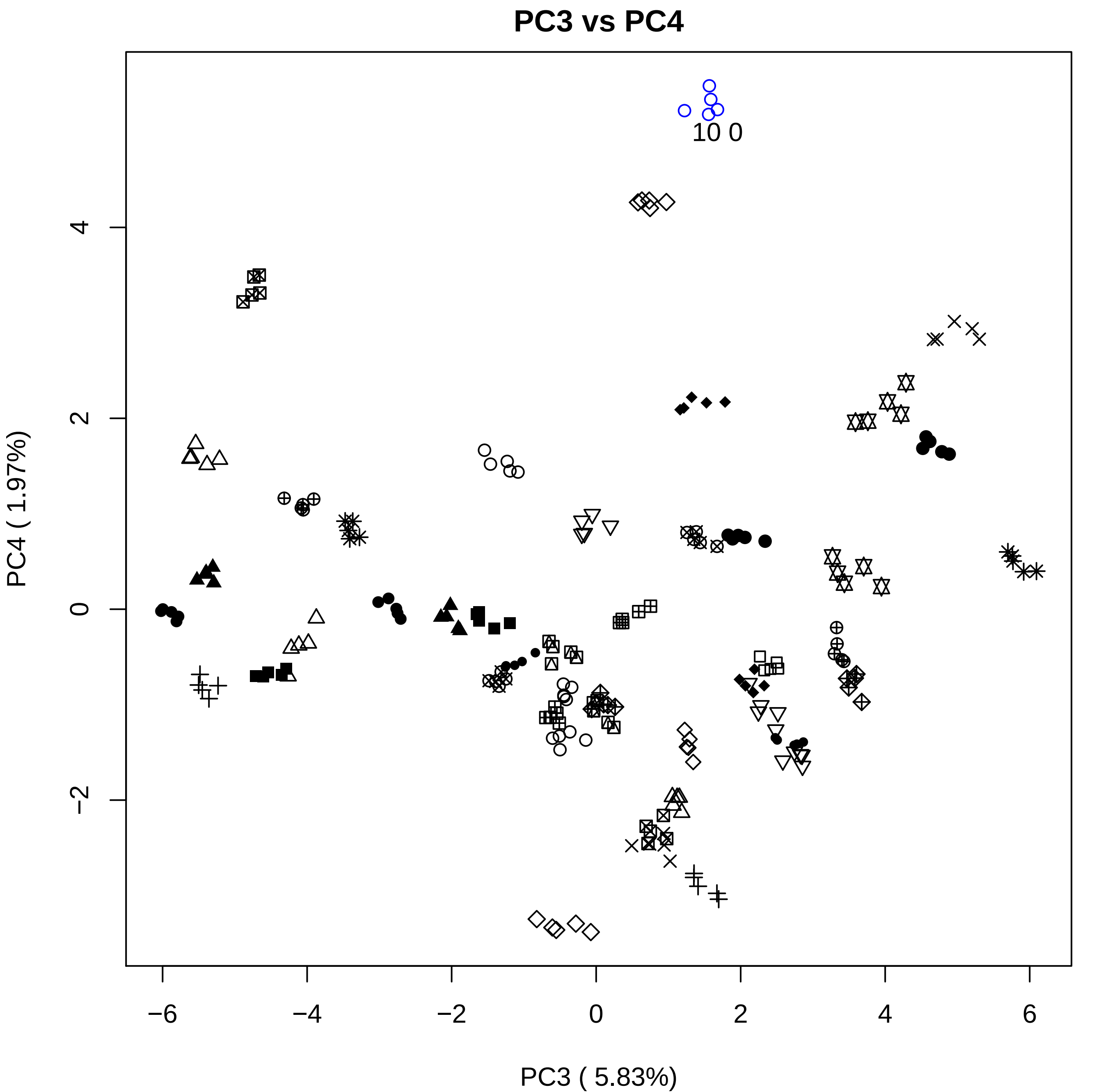
PC3 vs PC4
10 0
PC4 ( 1.97%)
PC3 ( 5.83%)
4
2
0
−2
−6
−4
−2
0
2
4
6

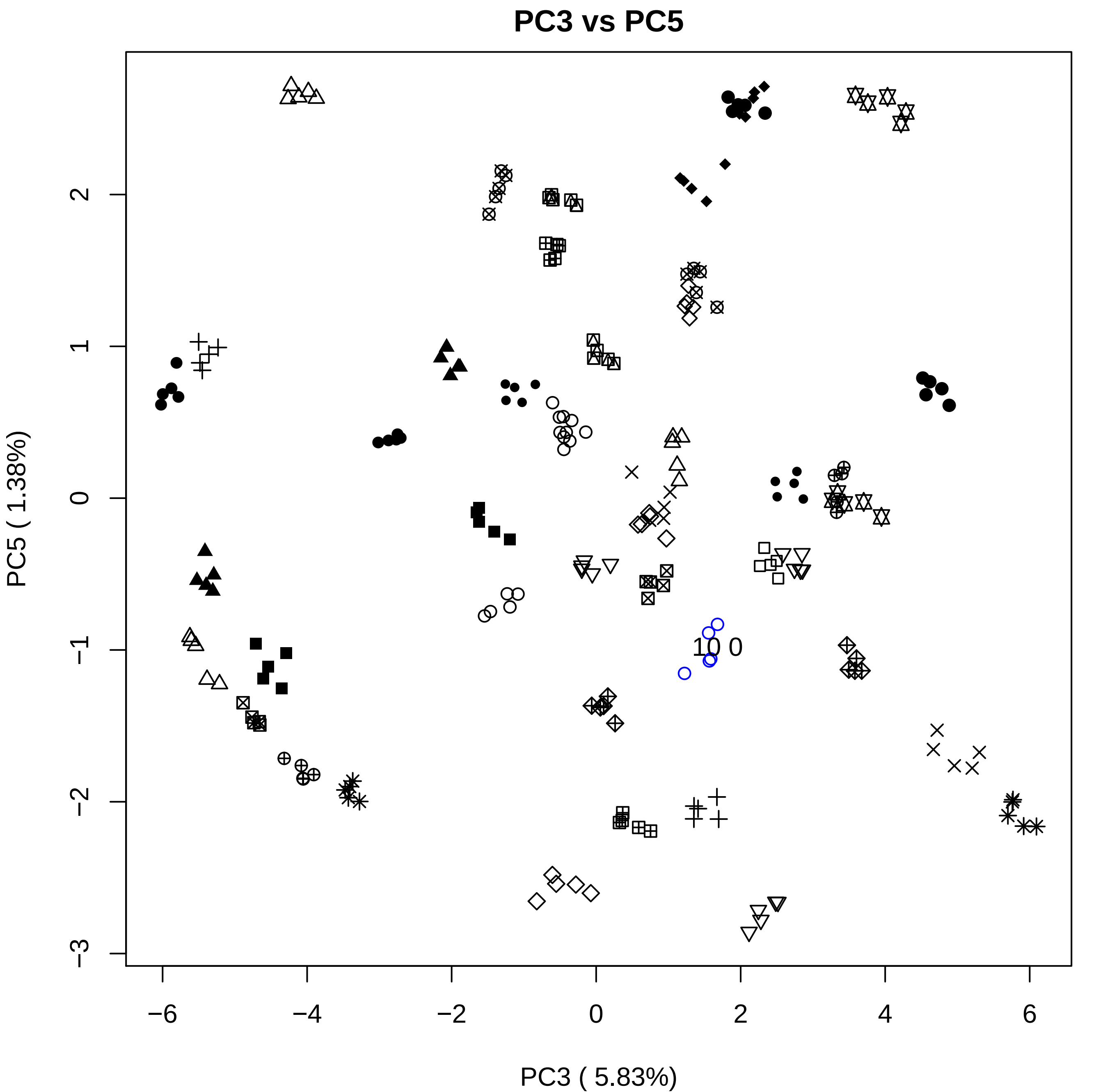
PC3 vs PC5
PC5 ( 1.38%)
PC3 ( 5.83%)
10 0
2
1
0
−1
−2
−3
−6
−4
−2
0
2
4
6

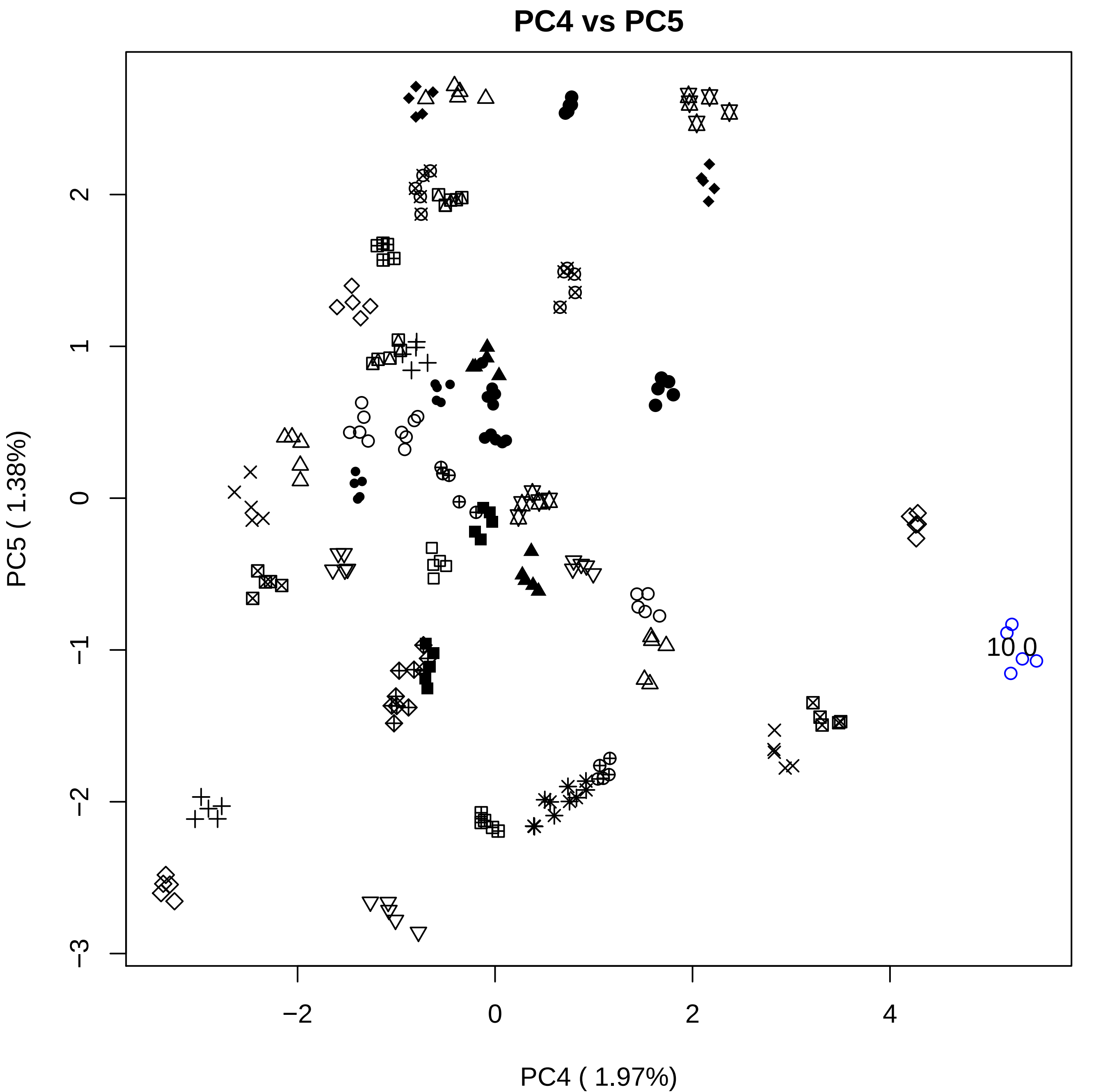

PC4 vs PC5
PC5 ( 1.38%)
PC4 ( 1.97%)
−2
0
2
4
−3
−2
−1
0
1
2
10.0

### PCA of 15%  DeS vs PMH

Scaled Matrix PCAed

7th order polynomial baseline correction performed

Raw spectra smoothed to 21 Neighbours, 2nd order PN, Savtisky Golay

Normalisation between 0 and 100

D2 spectra smoothed to 41 Neighbours, 2nd order PN, Savtisky Golay

Regions Analysed: 3601−2501 and 2001−700cm−1

Samples Analysed: 1 Contam's 0 DeS's 45 PMH's

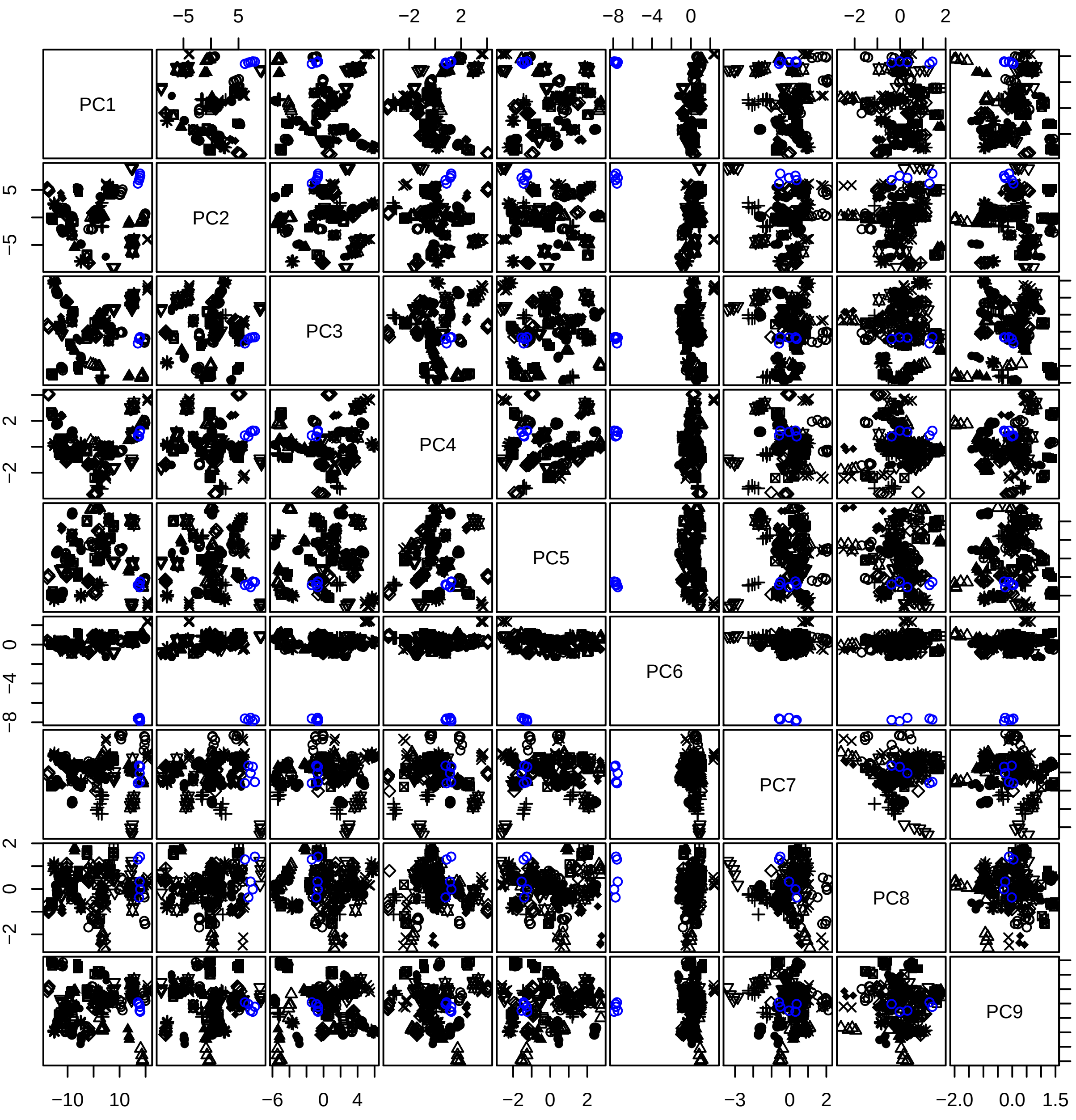
PC1
PC2
PC3
PC4
PC5
PC6
PC7
PC8
PC9

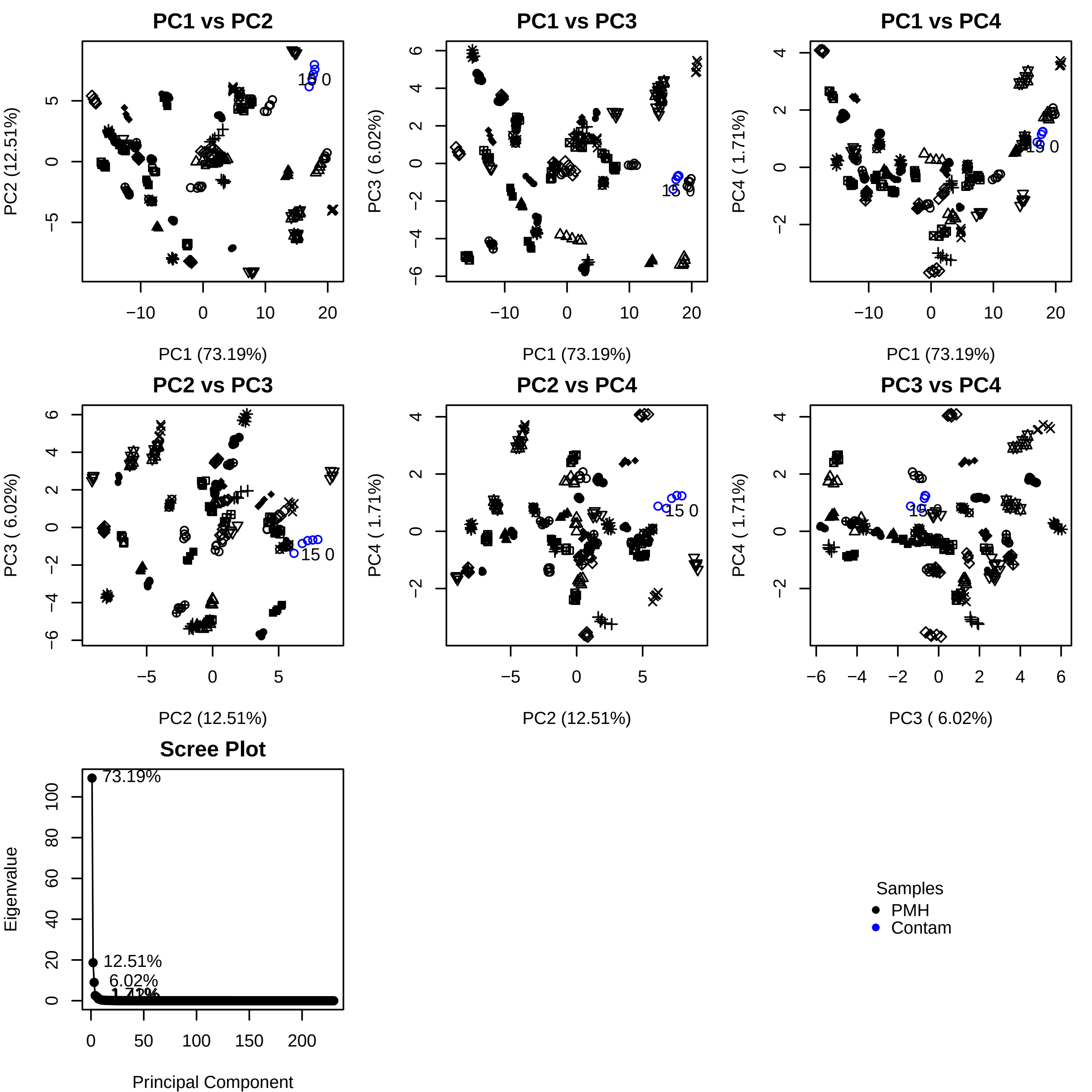

PC1 vs PC2
PC1 vs PC3
PC1 vs PC4
PC1 (73.19%)
PC2 (12.51%)
PC3 ( 6.02%)
PC4 ( 1.71%)
PC2 vs PC3
PC2 vs PC4
PC3 vs PC4
Scree Plot
73.19%
12.51%
6.02%
Eigenvalue
Principal Component
Samples
PMH
Contam

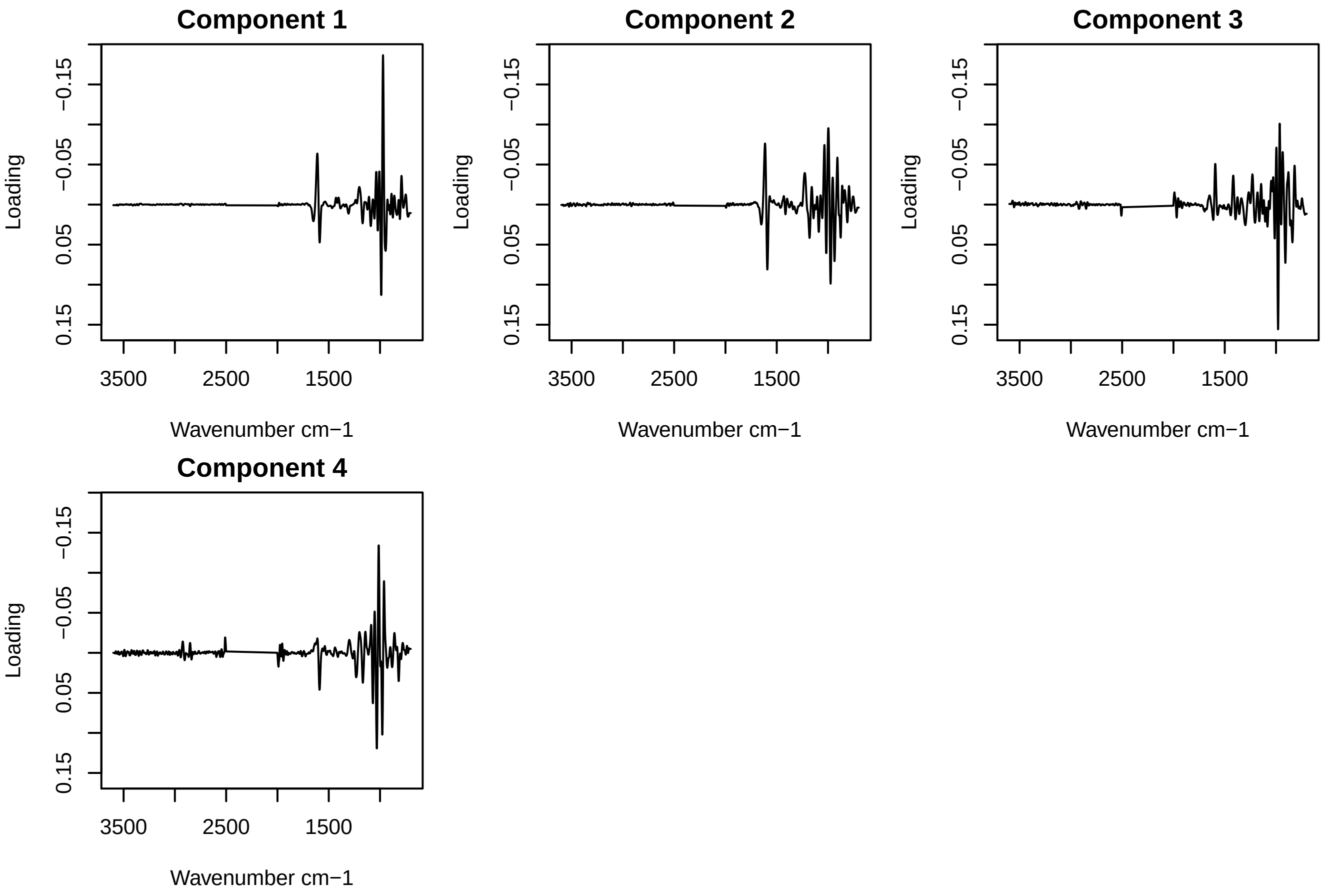
Component 1
Component 2
Component 3
Component 4
Loading
−0.15
−0.05
0.05
0.15
3500
2500
1500
Wavenumber cm−1

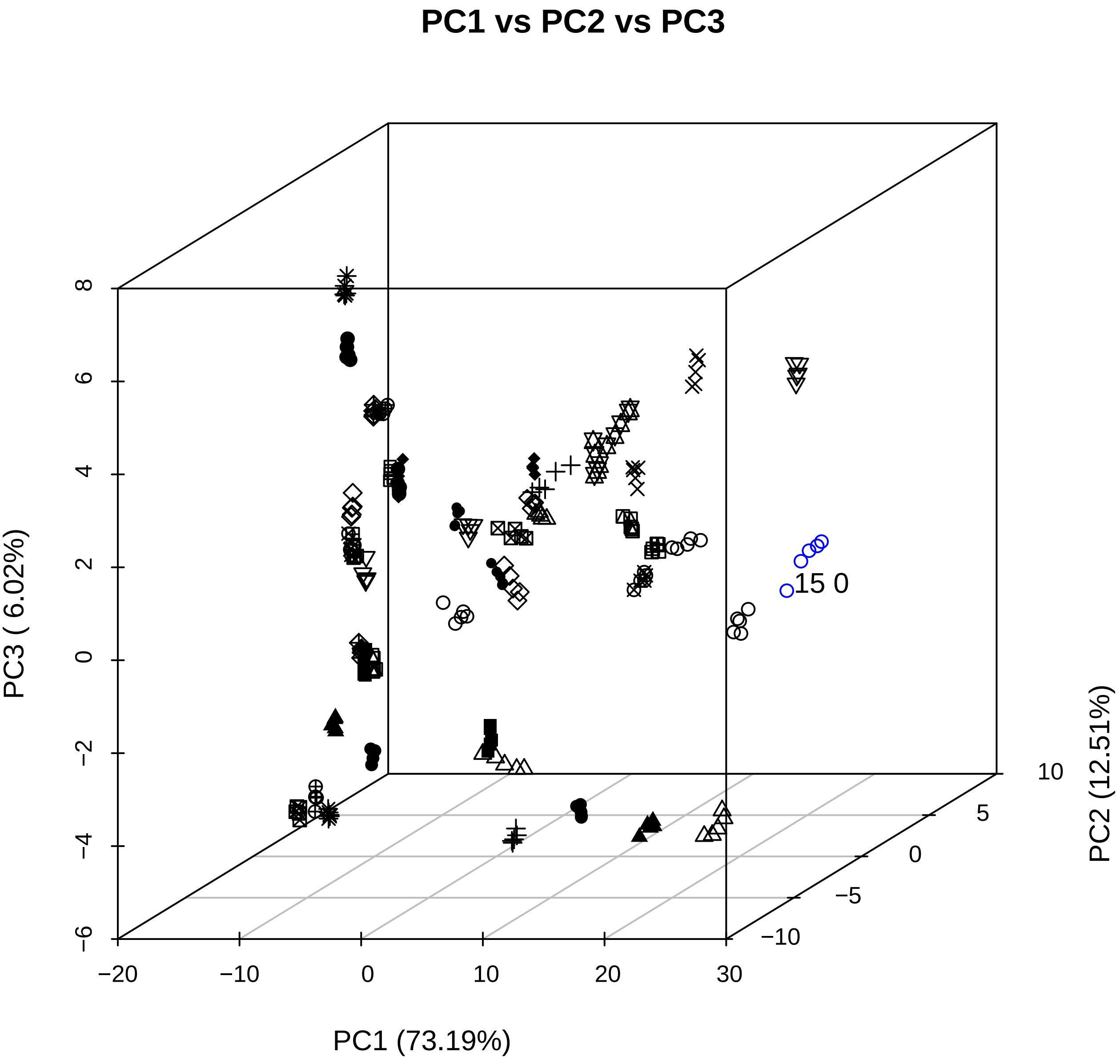
PC1 vs PC2 vs PC3
PC1 (73.19%)
PC2 (12.51%)
PC3 ( 6.02%)
−20
−10
0
10
20
30
−10
−5
0
5
10
−6
−4
−2
0
2
4
6
8
15 0

# PC2 vs PC3 vs PC4

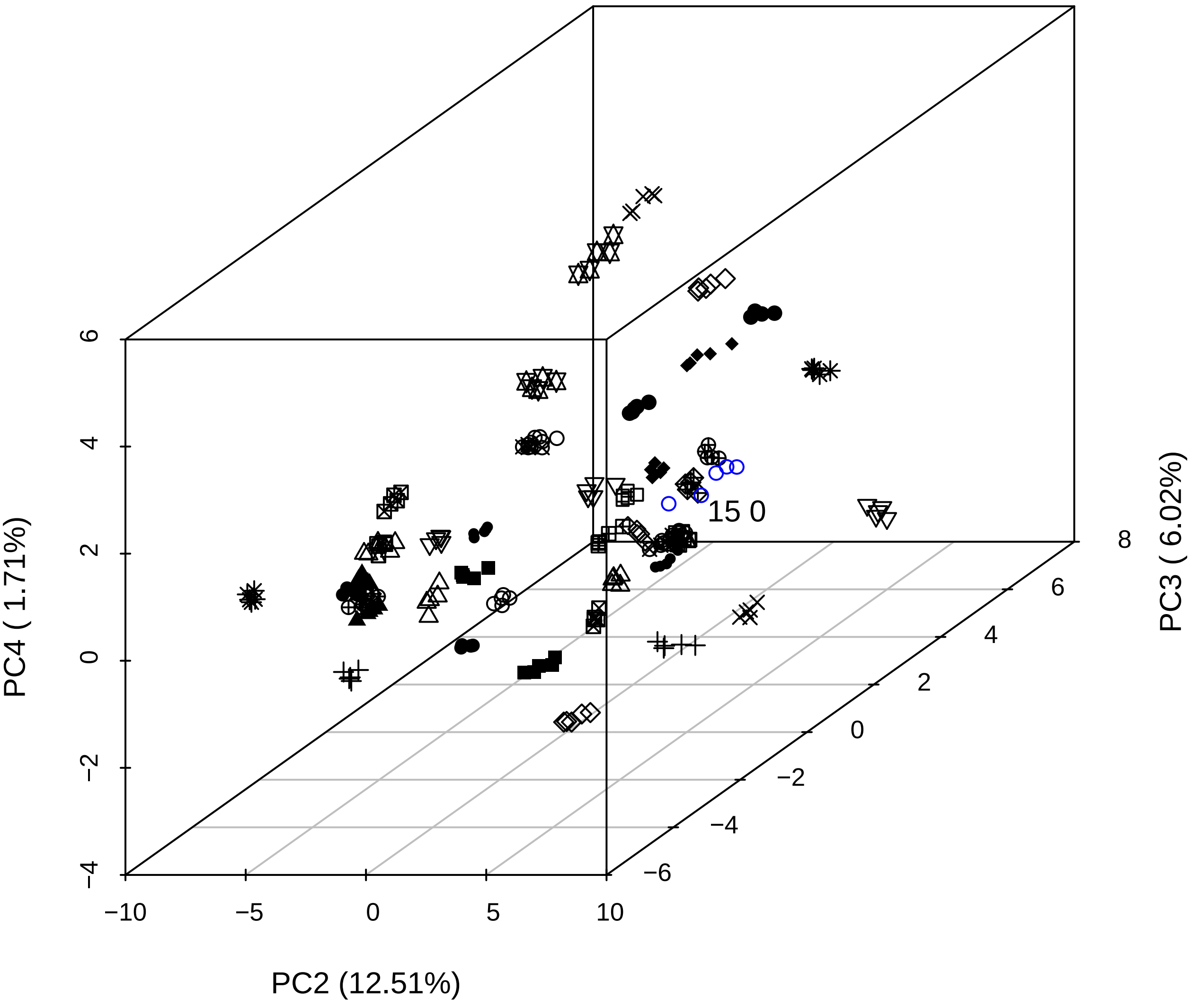

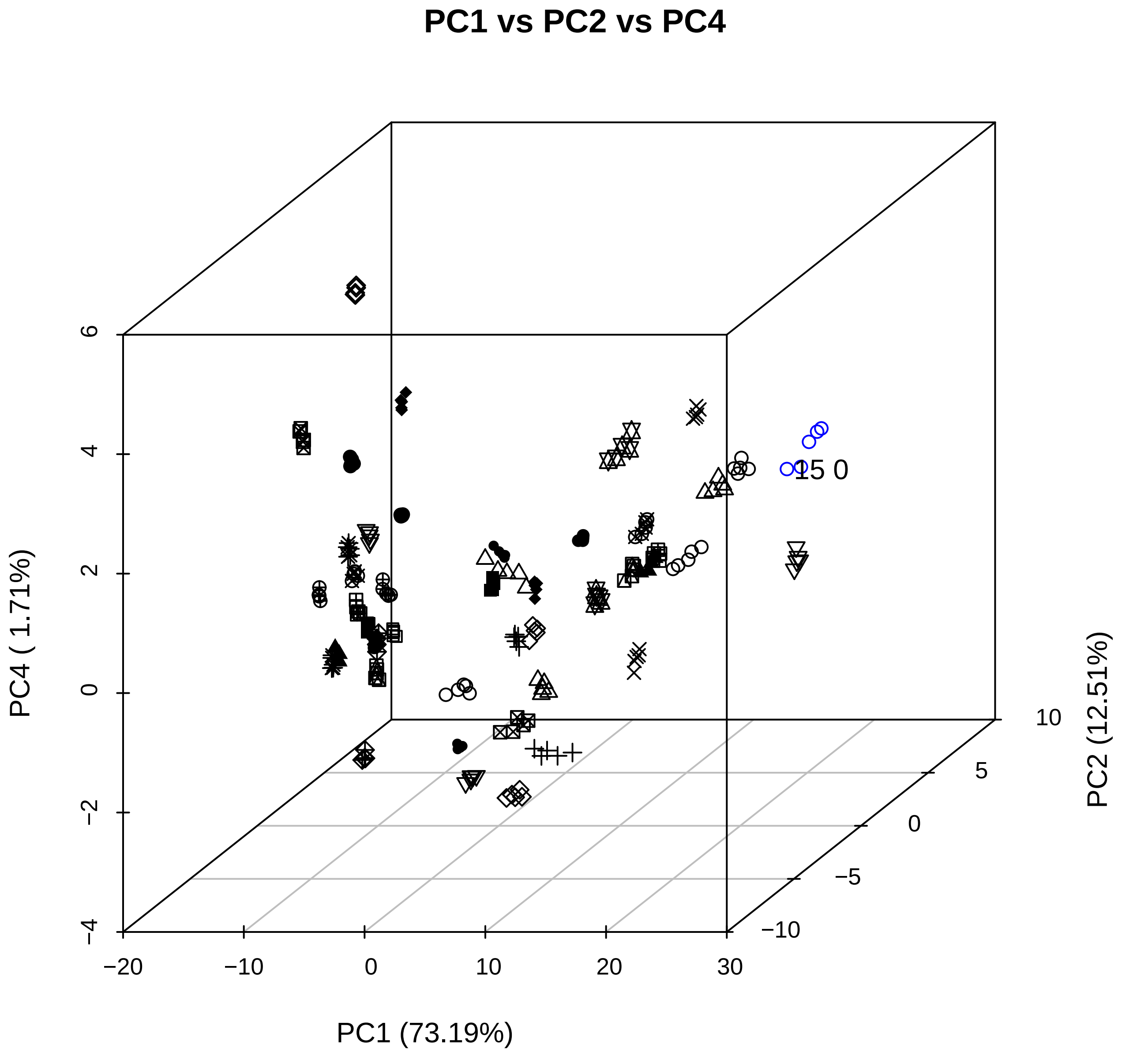

PC1 vs PC2 vs PC4
PC4 ( 1.71%)
PC1 (73.19%)
PC2 (12.51%)
−20
−10
0
10
20
30
−4
−2
0
2
4
6
−10
−5
0
5
10
15 0

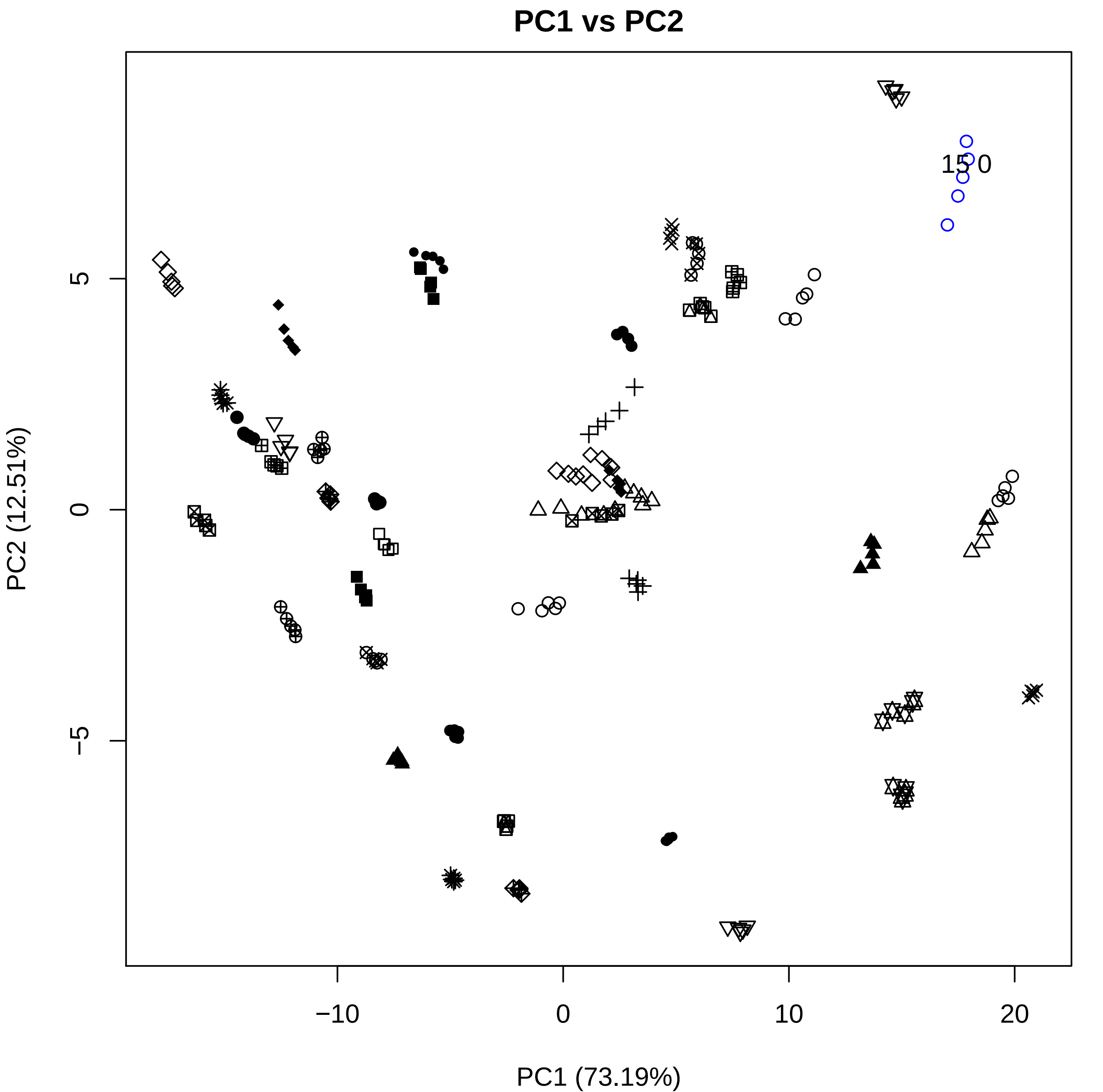
PC1 vs PC2
PC2 (12.51%)
PC1 (73.19%)
15 0
5
0
−5
−10
0
10
20

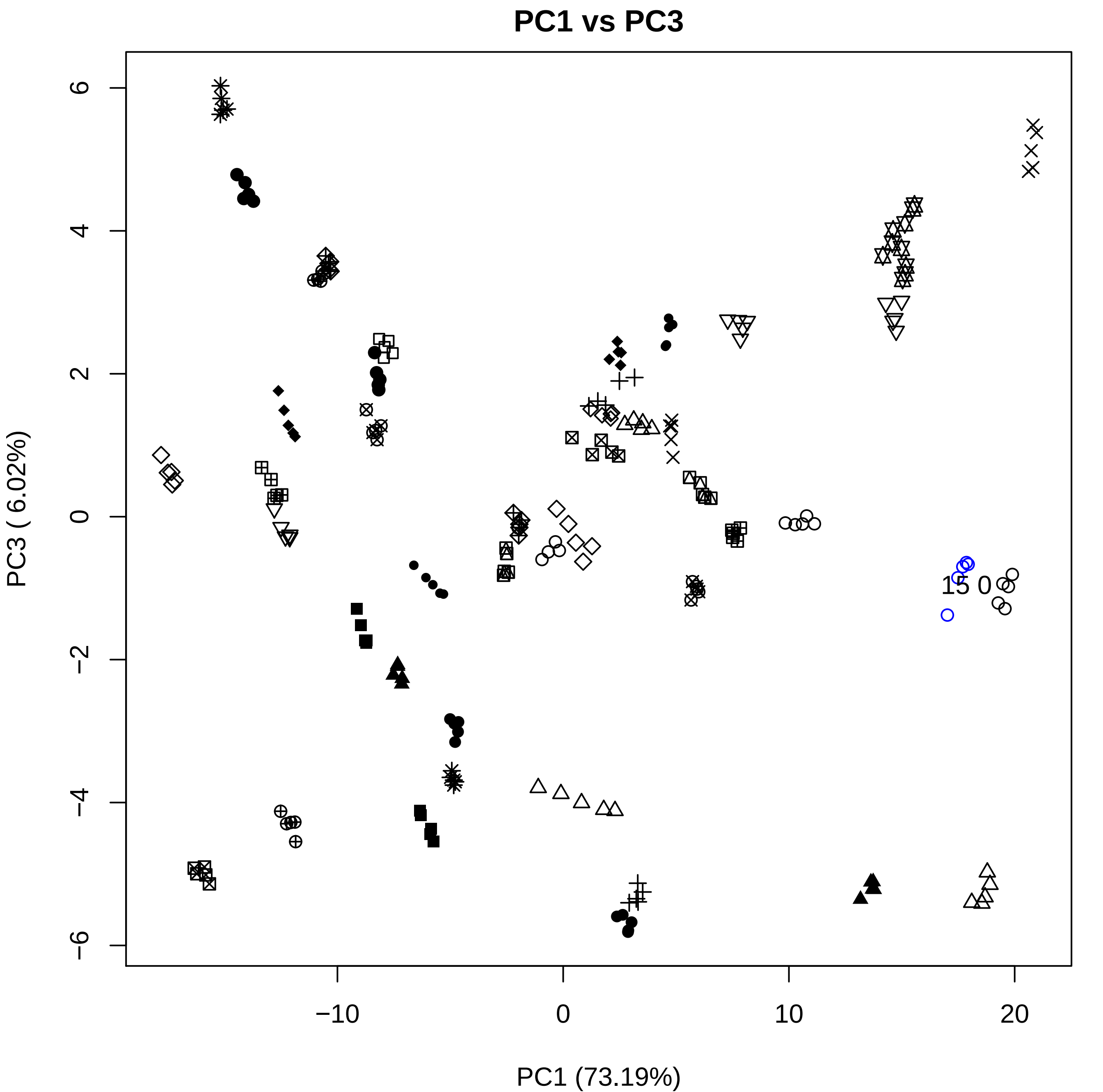

PC1 vs PC3
PC3 ( 6.02%)
PC1 (73.19%)
6
4
2
0
−2
−4
−6
−10
0
10
20
15 0

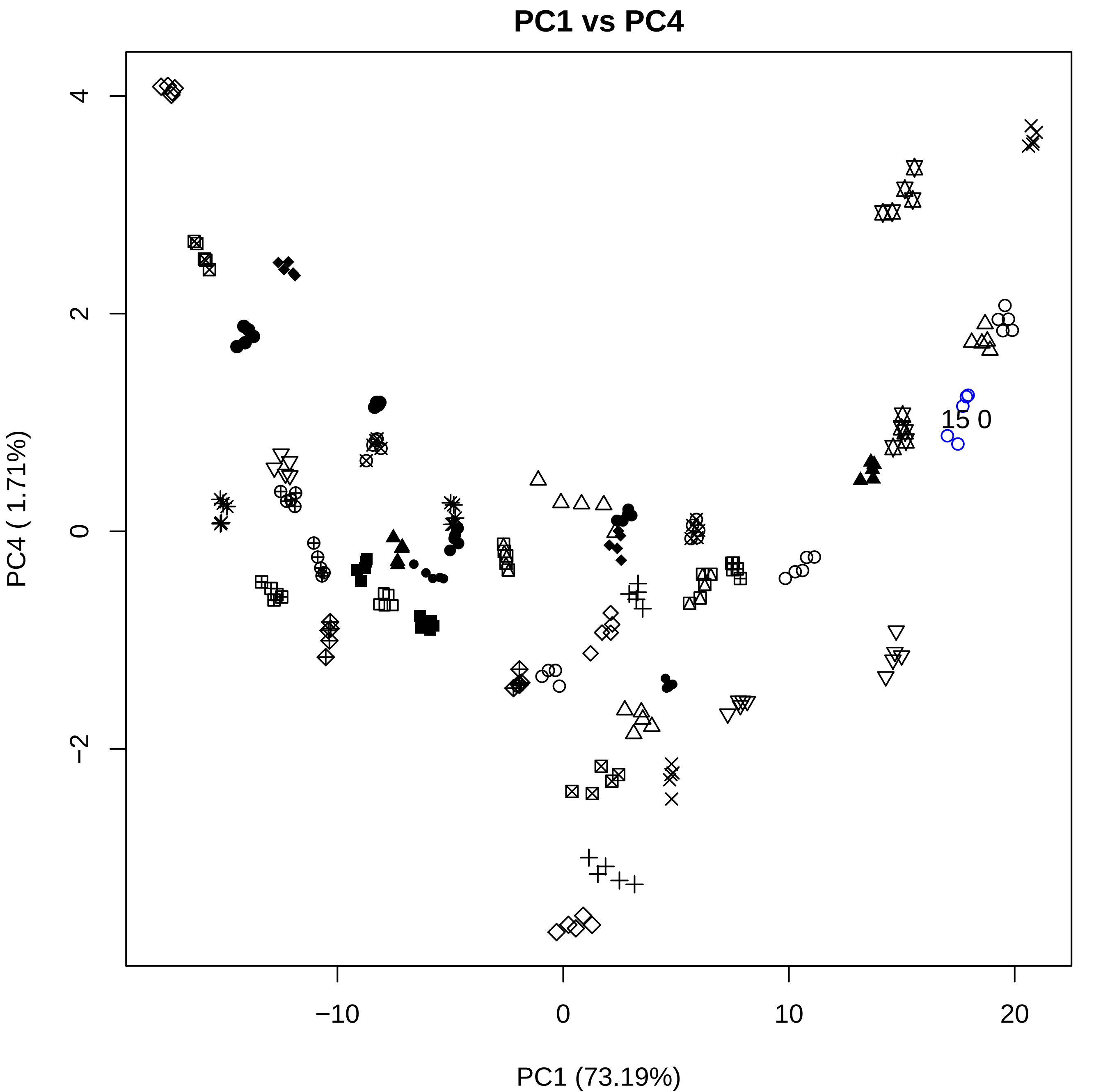
PC1 vs PC4
PC4 ( 1.71%)
PC1 (73.19%)
4
2
0
−2
−10
0
10
20
15 0

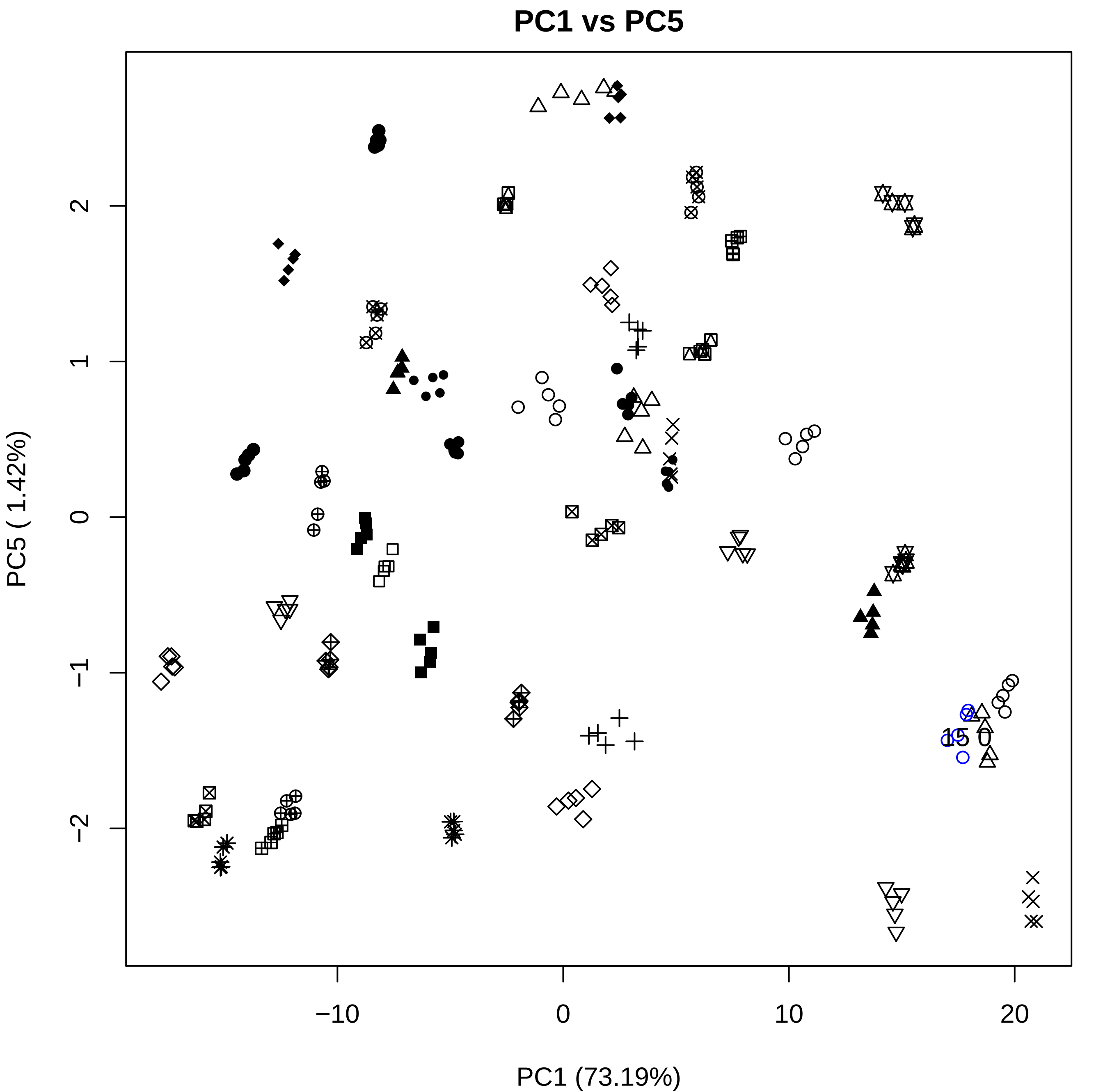
PC1 vs PC5
PC5 ( 1.42%)
PC1 (73.19%)
−10
0
10
20
−2
−1
0
1
2
15 0

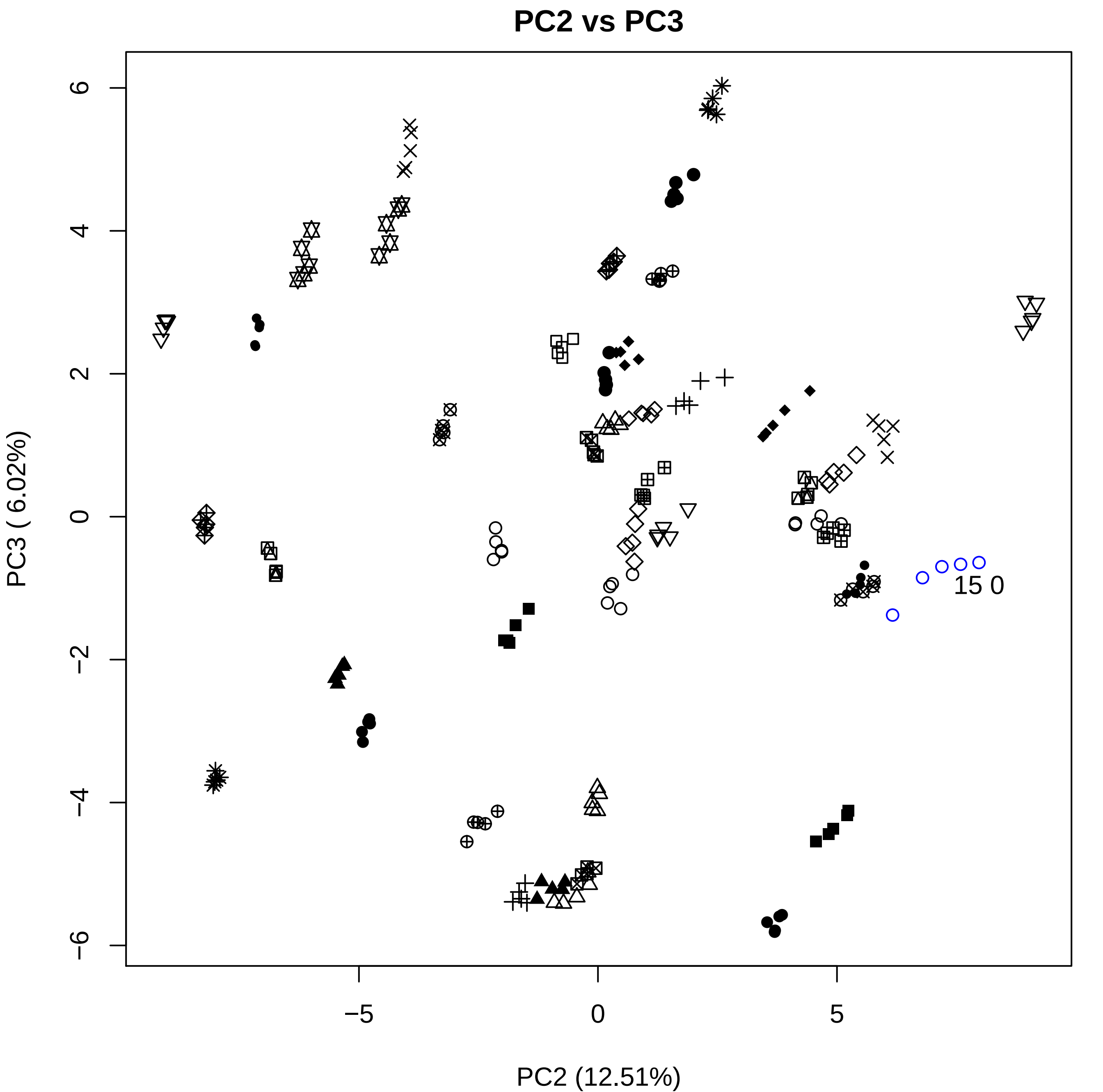
PC2 vs PC3
PC3 ( 6.02%)
PC2 (12.51%)
6
4
2
0
−2
−4
−6
−5
0
5
15 0

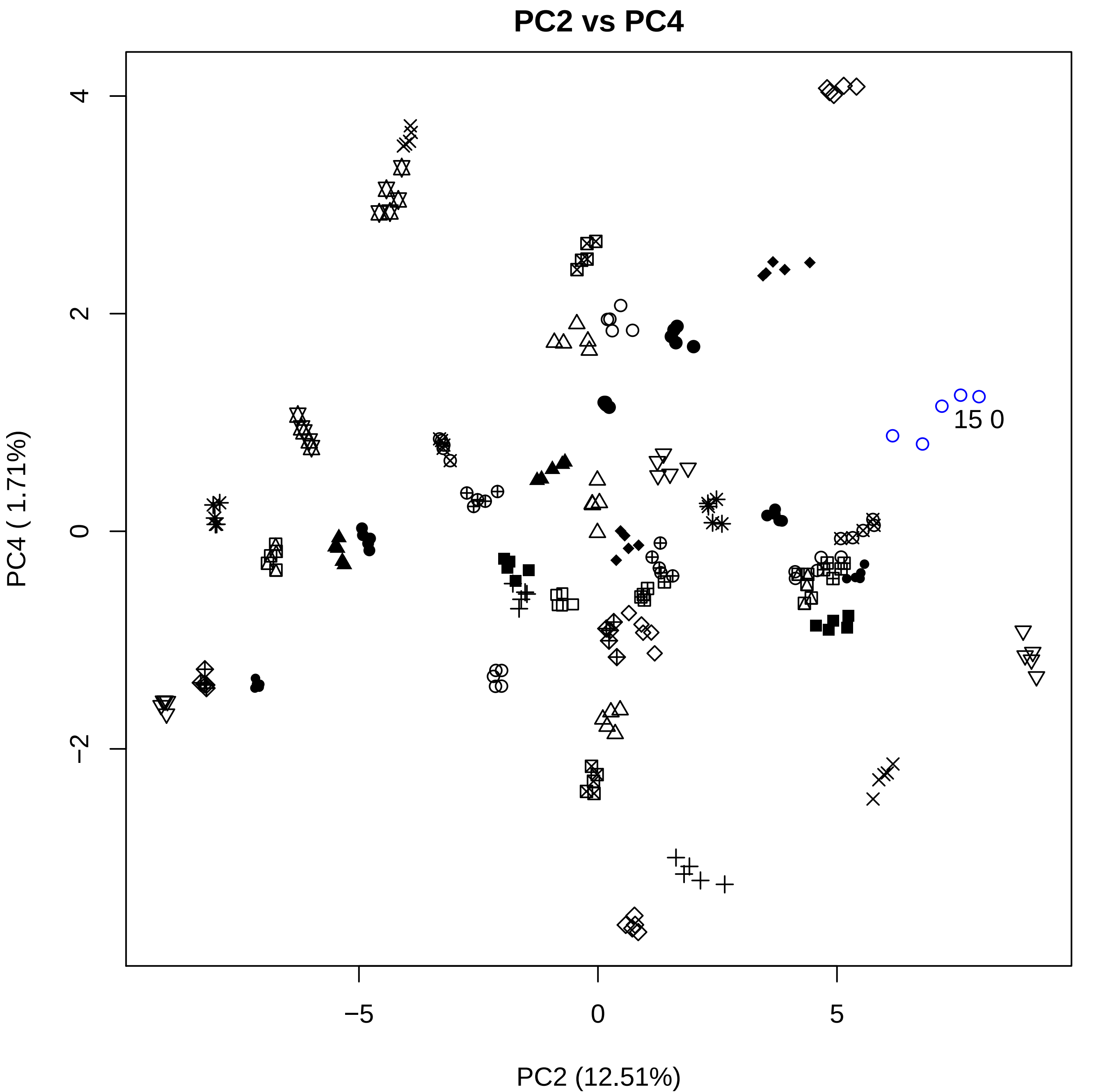

PC2 vs PC4
PC4 ( 1.71%)
PC2 (12.51%)
4
2
0
−2
−5
0
5
15 0

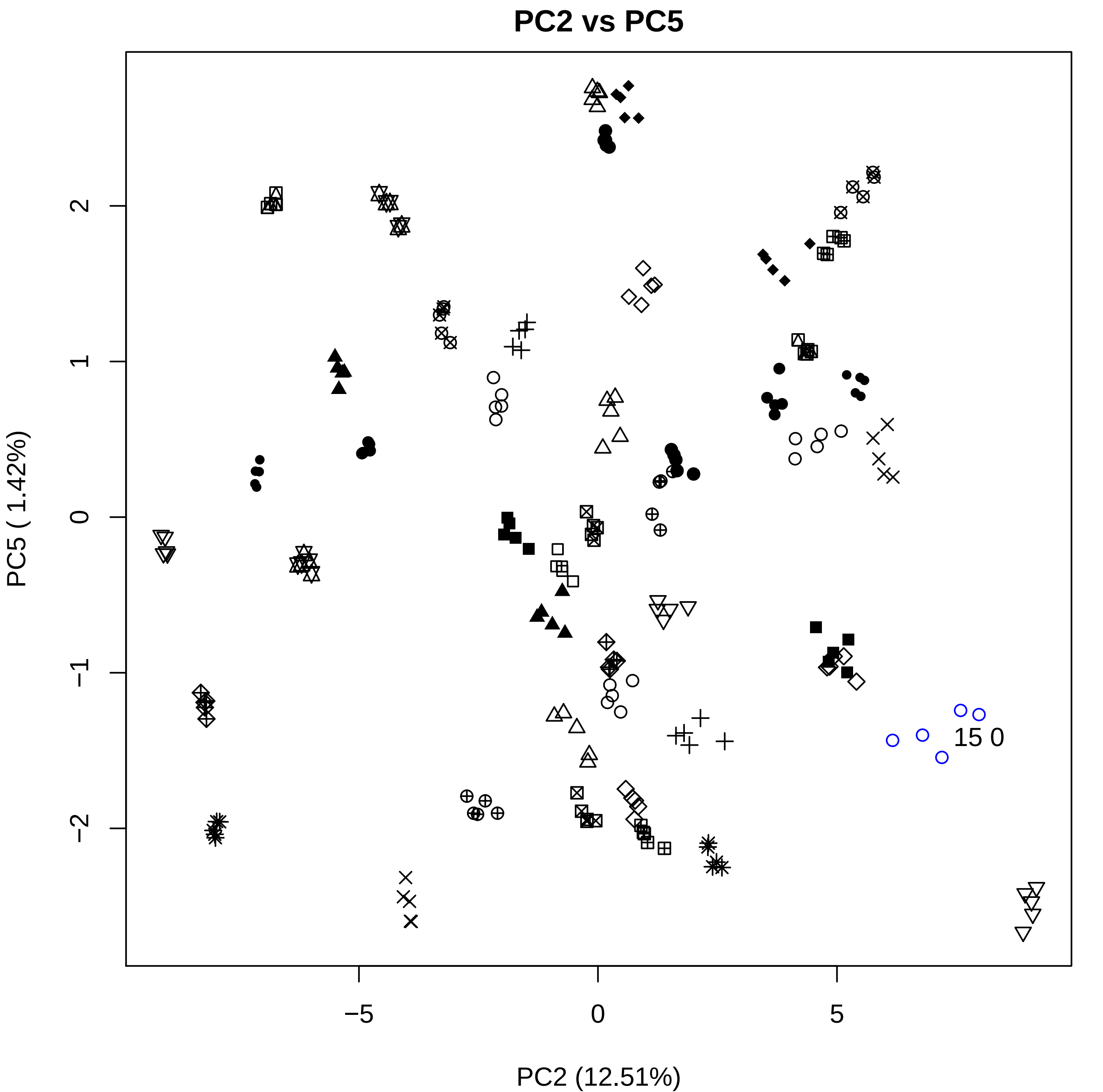
PC2 vs PC5
PC5 ( 1.42%)
PC2 (12.51%)
2
1
0
−1
−2
−5
0
5
15 0

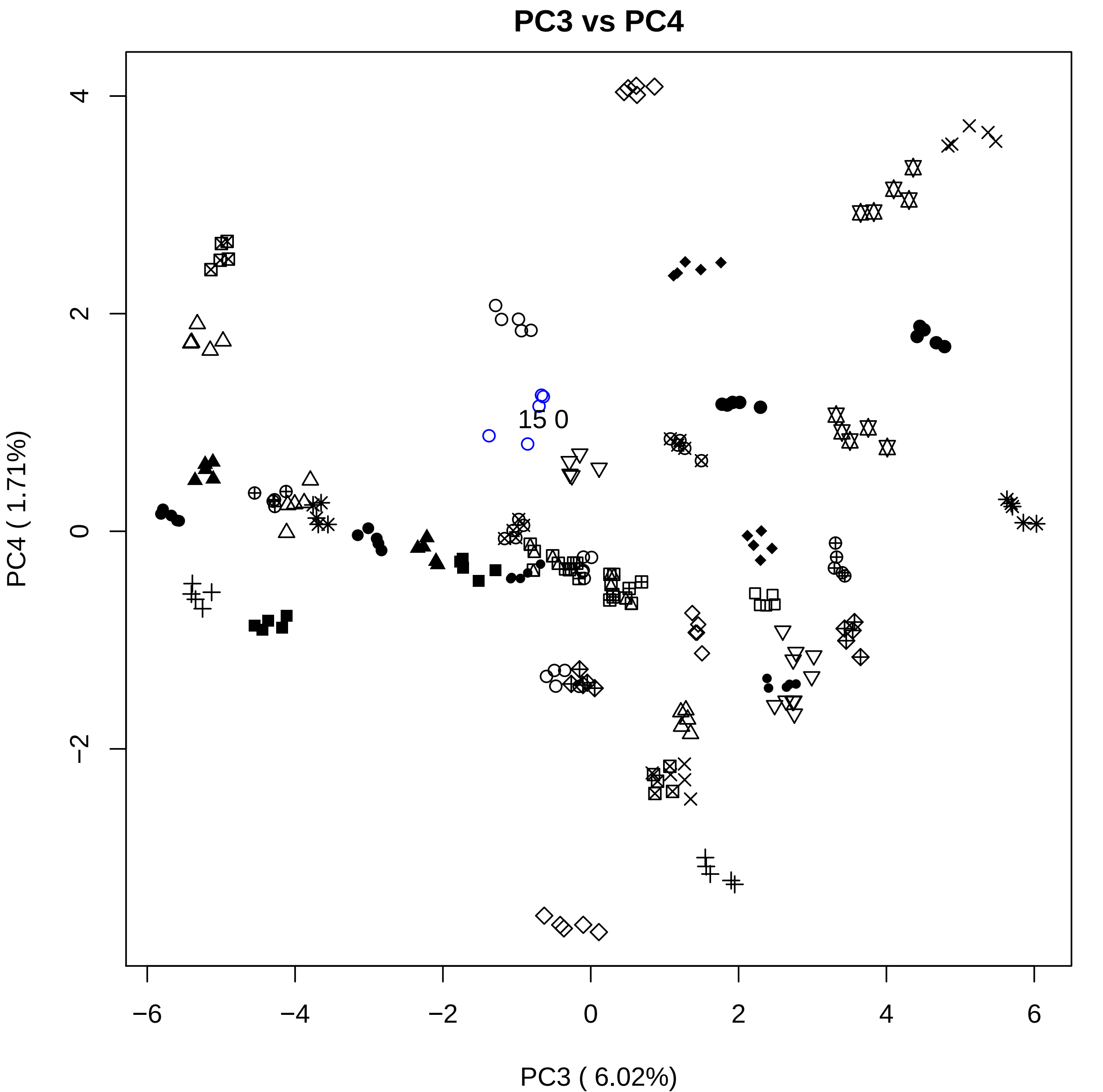
PC3 vs PC4
15 0
PC4 ( 1.71%)
PC3 ( 6.02%)
−6
−4
−2
0
2
4
6
−2
0
2
4

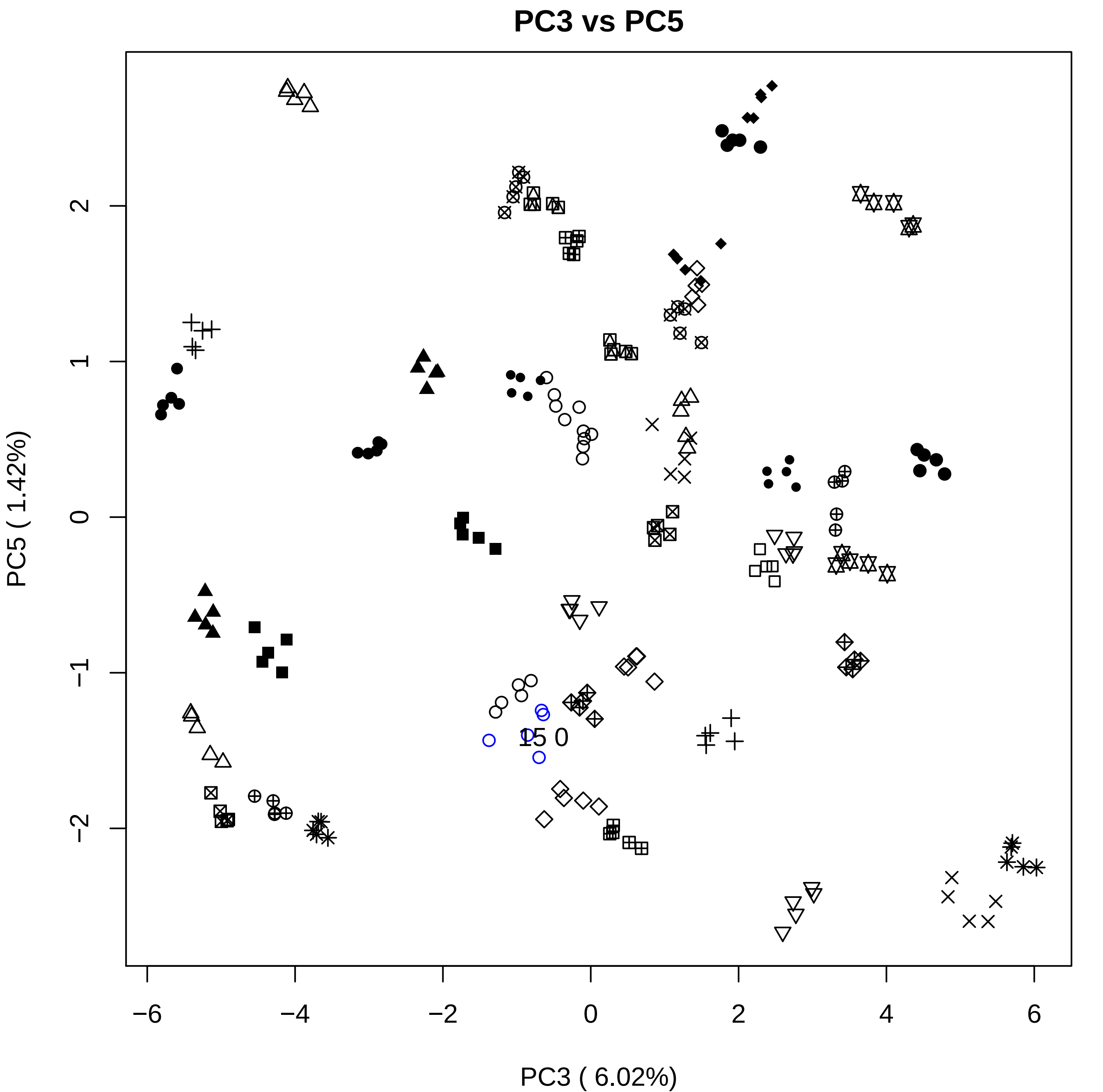
PC3 vs PC5
PC5 ( 1.42%)
PC3 ( 6.02%)
2
1
0
−1
−2
−6
−4
−2
0
2
4
6
15 0

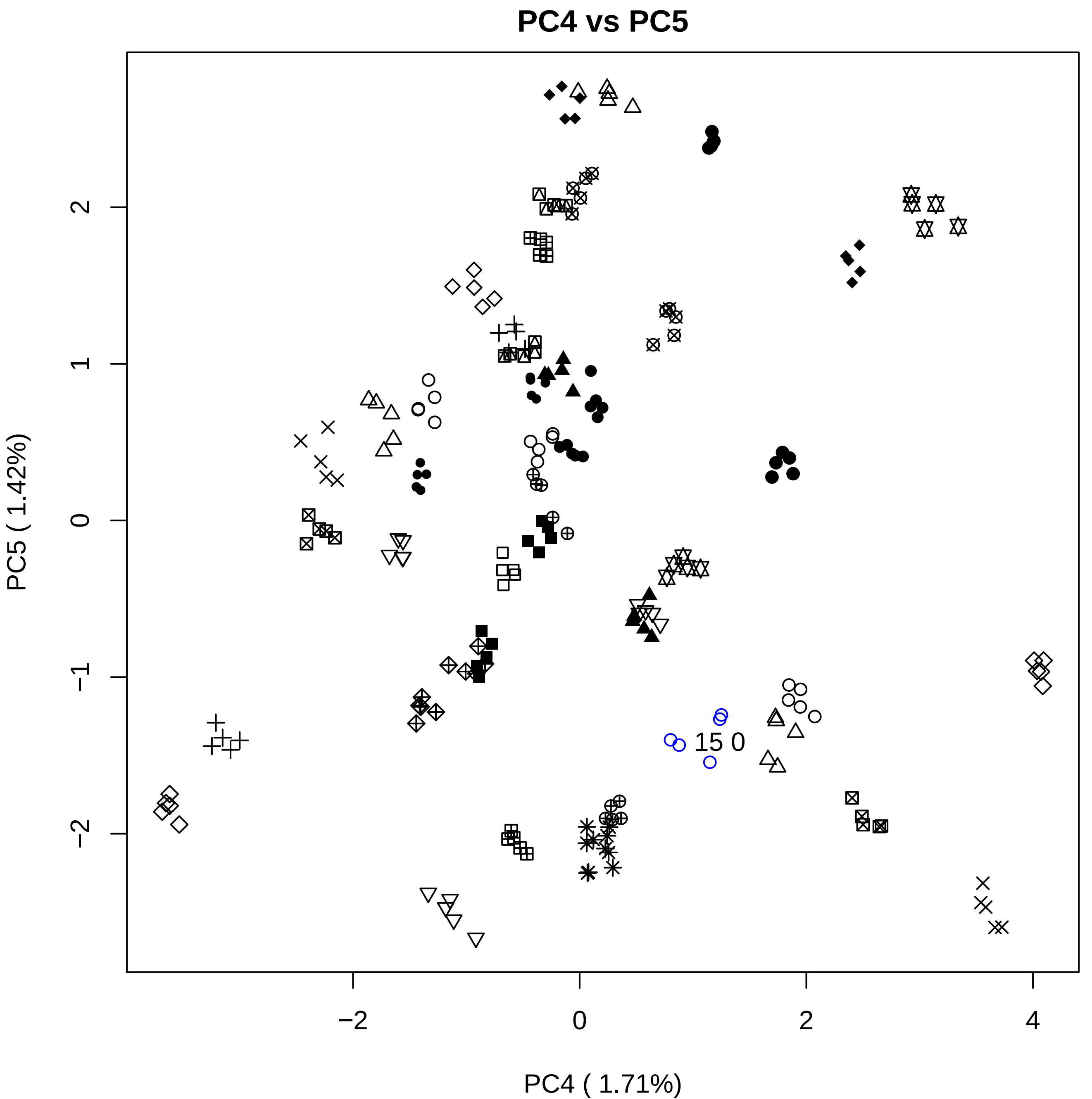
PC4 vs PC5
PC5 ( 1.42%)
PC4 ( 1.71%)
−2
0
2
4
−2
−1
0
1
2
15 0

### PCA of 20%  DeS vs PMH

Scaled Matrix PCAed

7th order polynomial baseline correction performed

Raw spectra smoothed to 21 Neighbours, 2nd order PN, Savtisky Golay

Normalisation between 0 and 100

D2 spectra smoothed to 41 Neighbours, 2nd order PN, Savtisky Golay

Regions Analysed: 3601−2501 and 2001−700cm−1

Samples Analysed: 1 Contam's 0 DeS's 45 PMH's

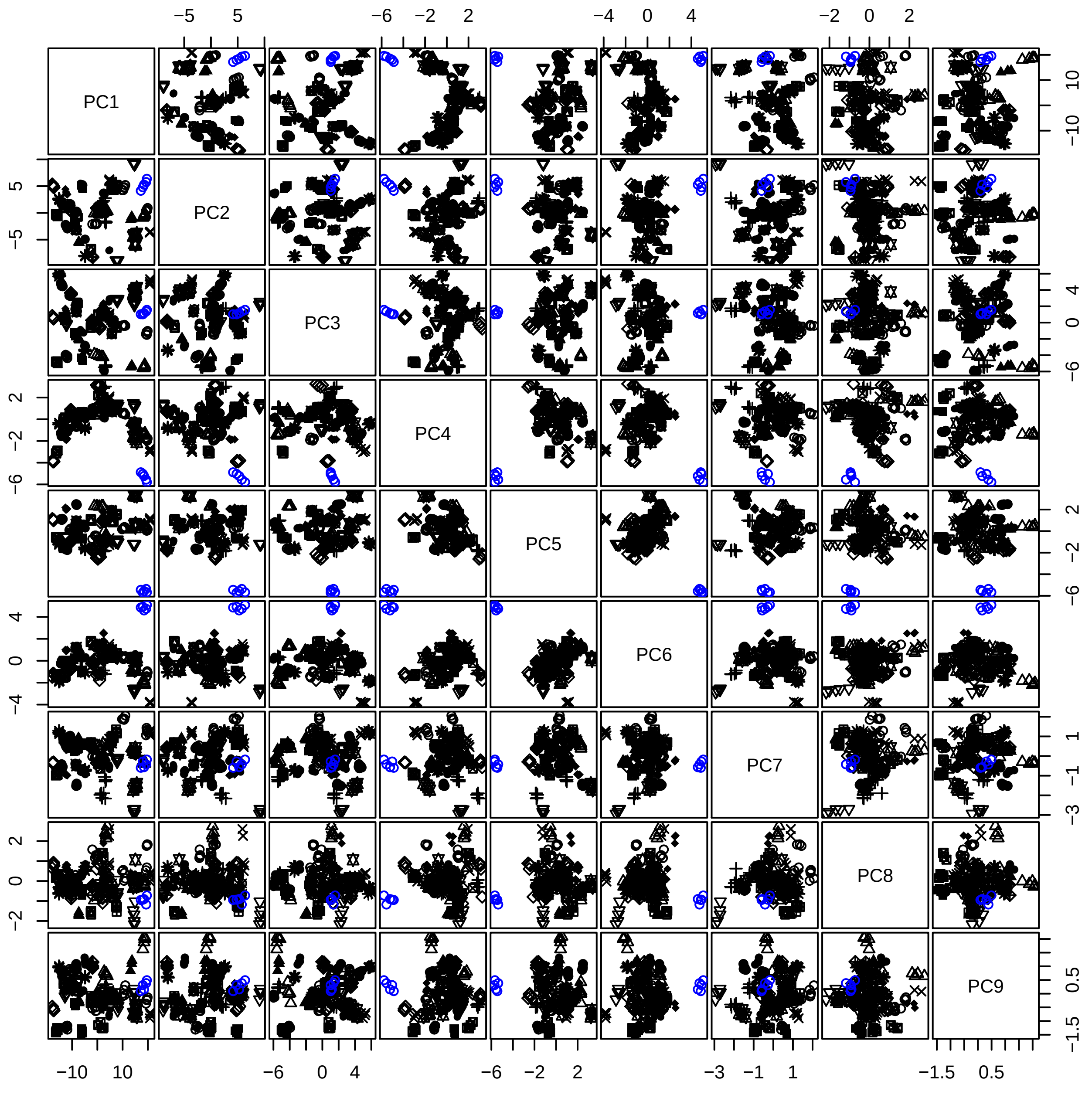
PC1
PC2
PC3
PC4
PC5
PC6
PC7
PC8
PC9

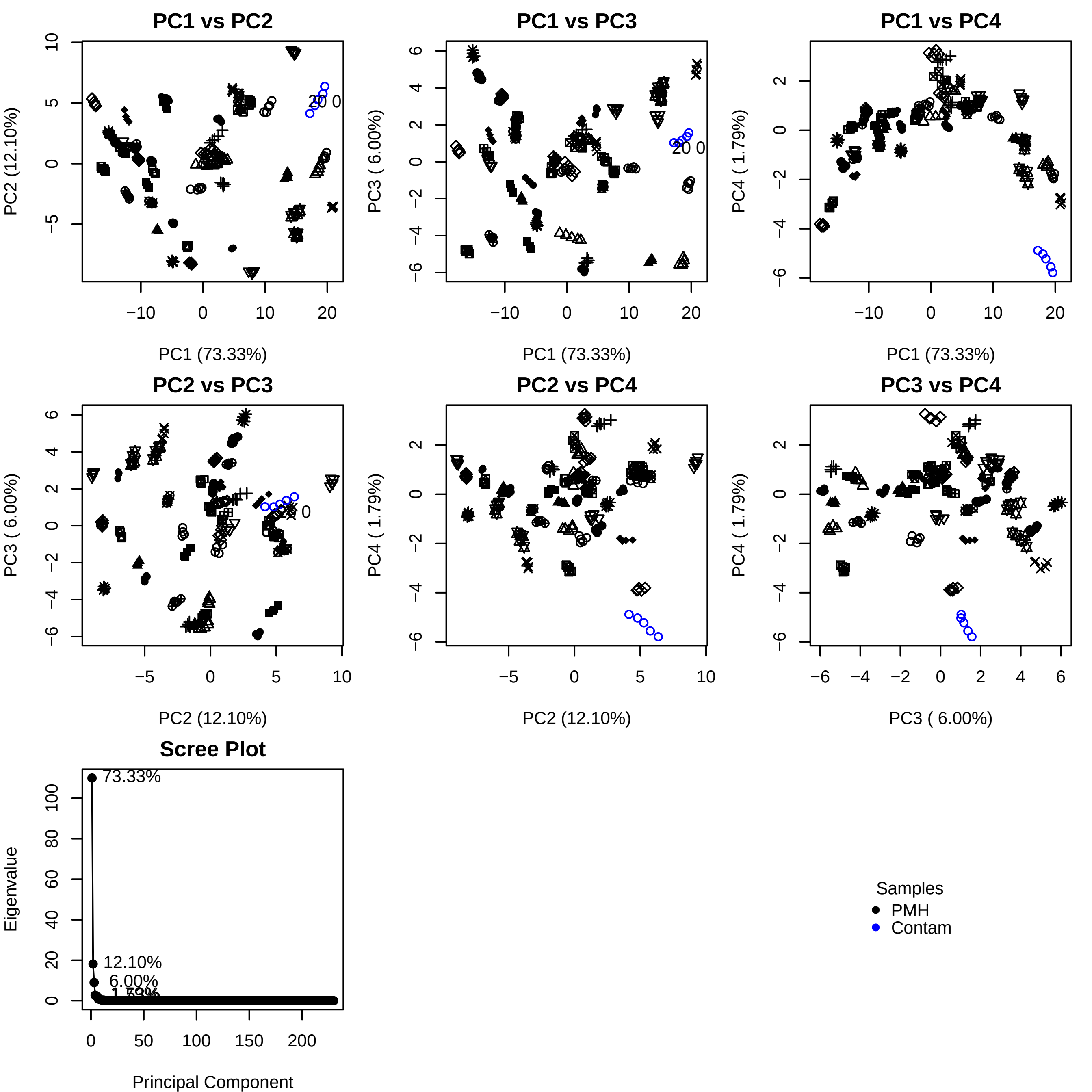
PC1 vs PC2
PC1 vs PC3
PC1 vs PC4
PC2 vs PC3
PC2 vs PC4
PC3 vs PC4
Scree Plot
PC1 (73.33%)
PC2 (12.10%)
PC3 ( 6.00%)
PC4 ( 1.79%)
Eigenvalue
Principal Component
73.33%
12.10%
6.00%
Samples
PMH
Contam

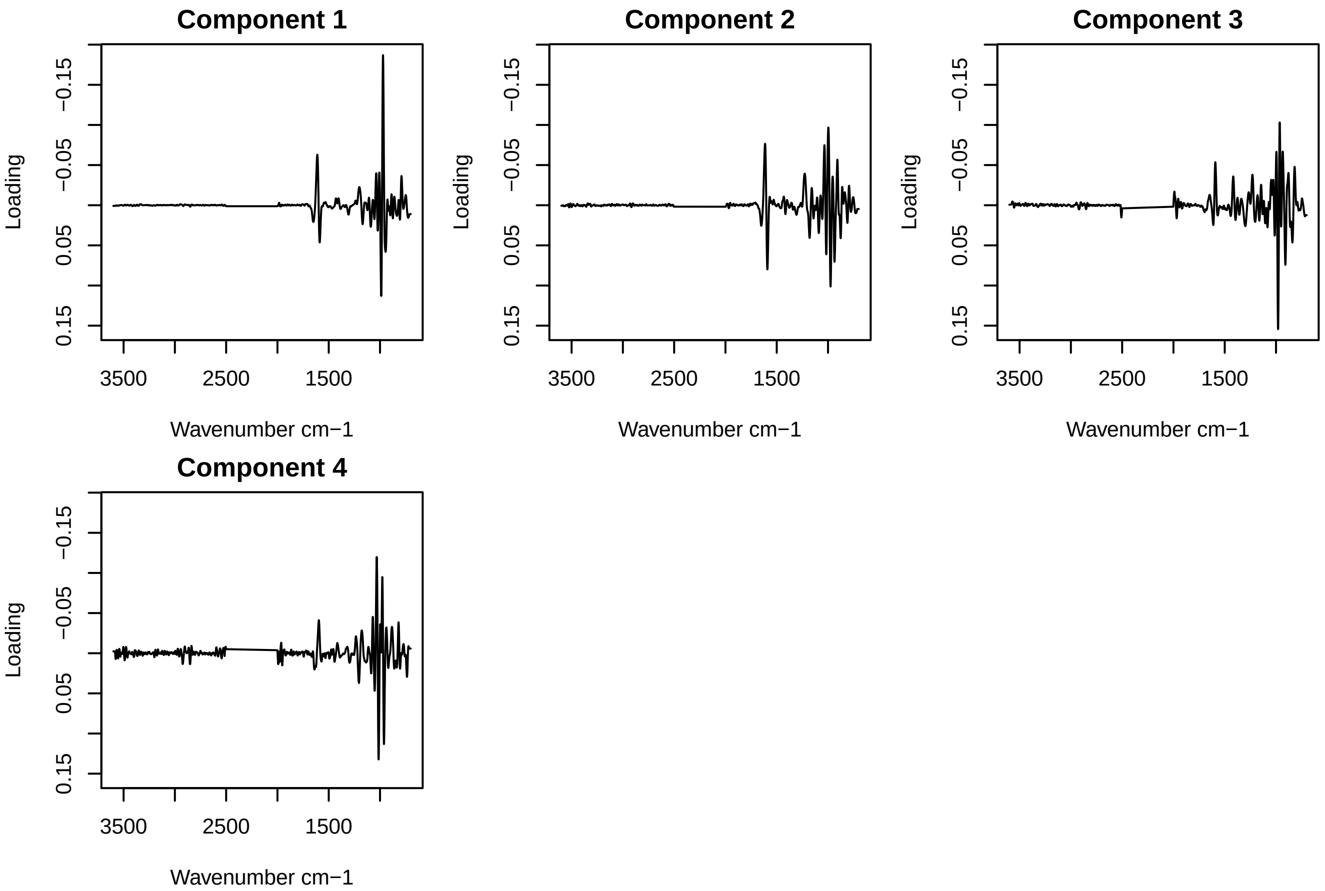
Component 1
Component 2
Component 3
Component 4
Loading
Wavenumber cm−1
−0.15
−0.05
0.05
0.15
3500
2500
1500

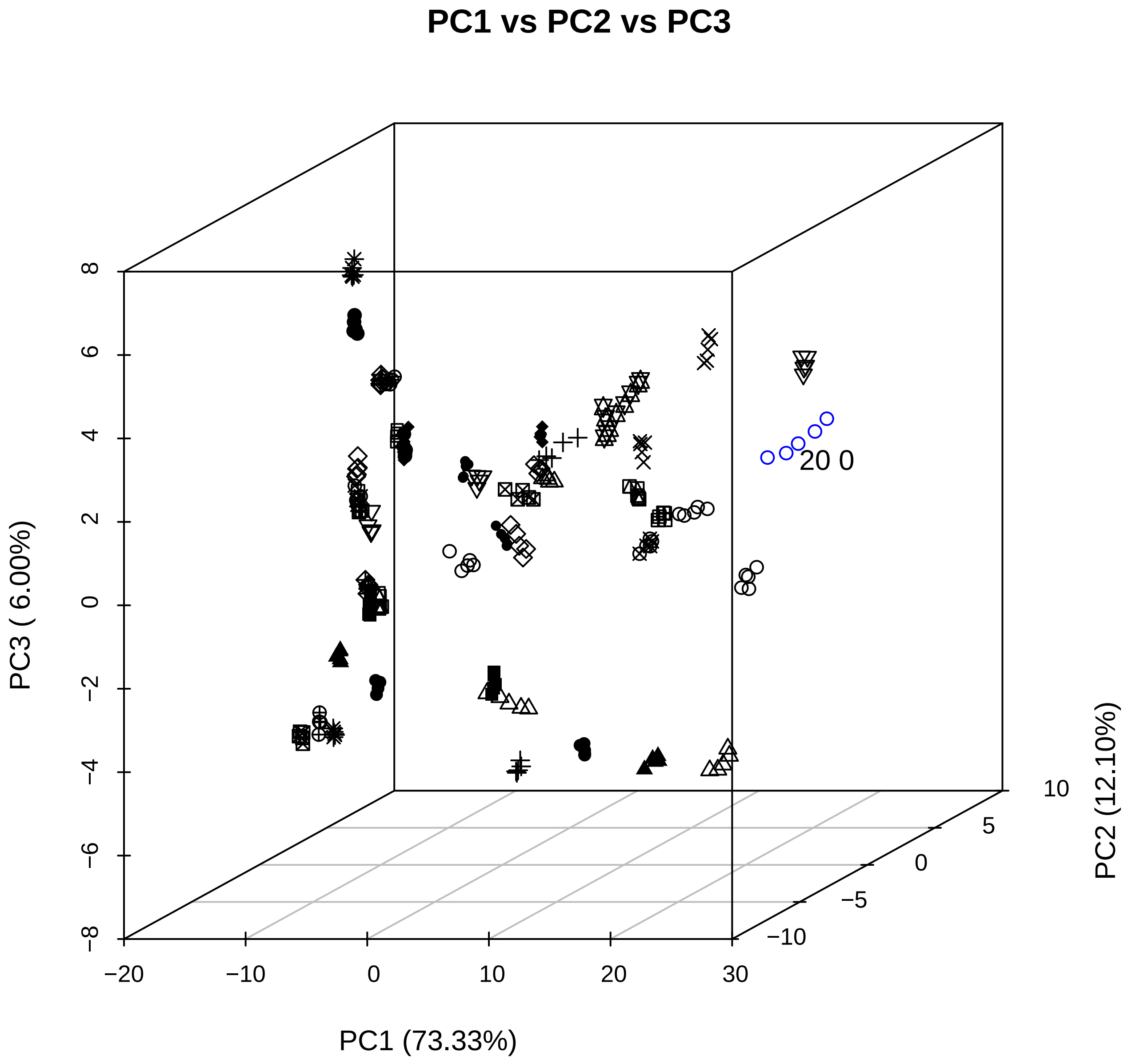

PC1 vs PC2 vs PC3
PC1 (73.33%)
PC2 (12.10%)
PC3 ( 6.00%)
−20
−10
0
10
20
30
−10
−5
0
5
10
−8
−6
−4
−2
0
2
4
6
8
20 0

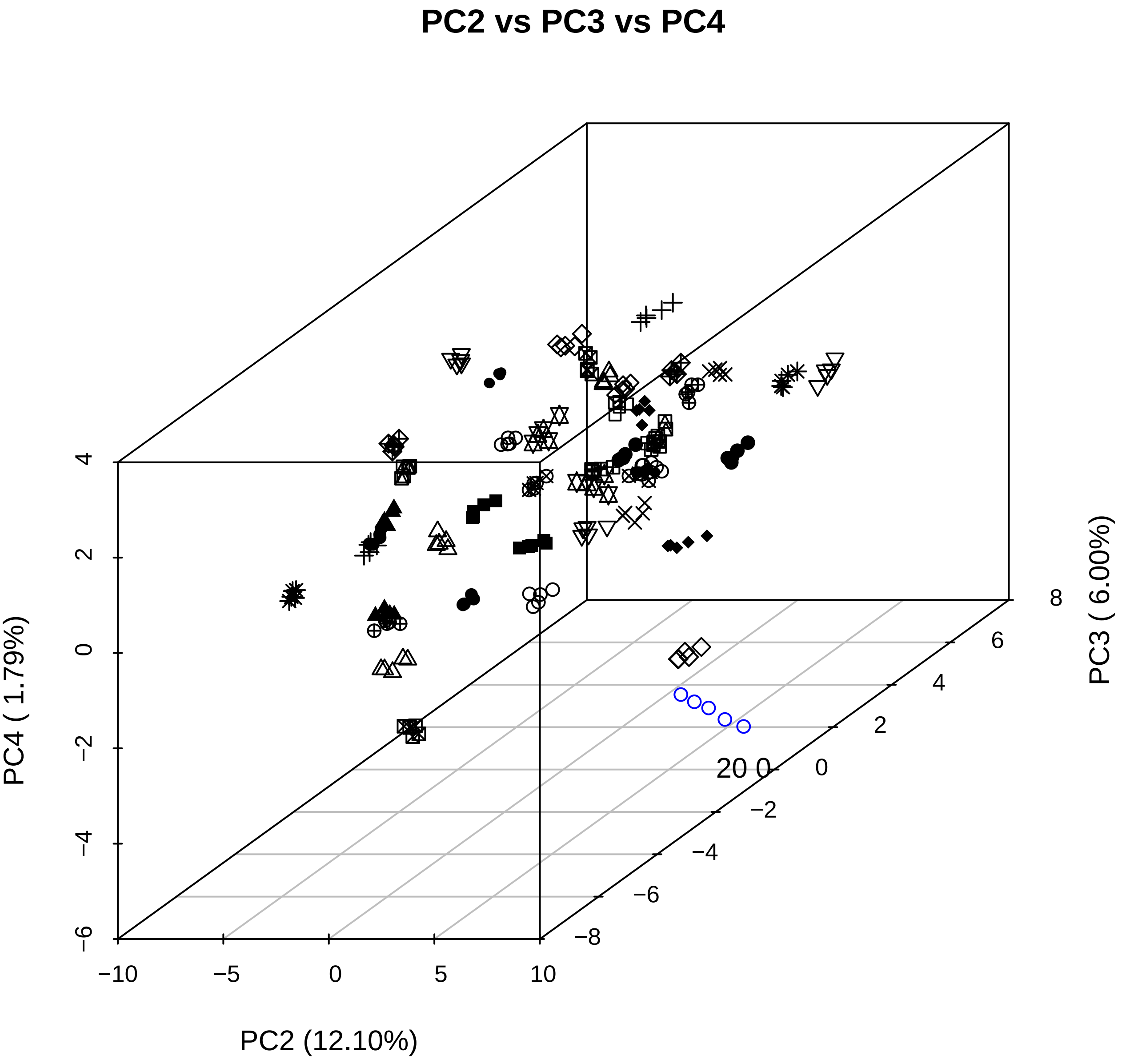
PC2 vs PC3 vs PC4
PC2 (12.10%)
PC3 ( 6.00%)
PC4 ( 1.79%)
−10
−5
0
5
10
−8
−6
−4
−2
0
2
4
6
8
−6
−4
−2
0
2
4
20 0

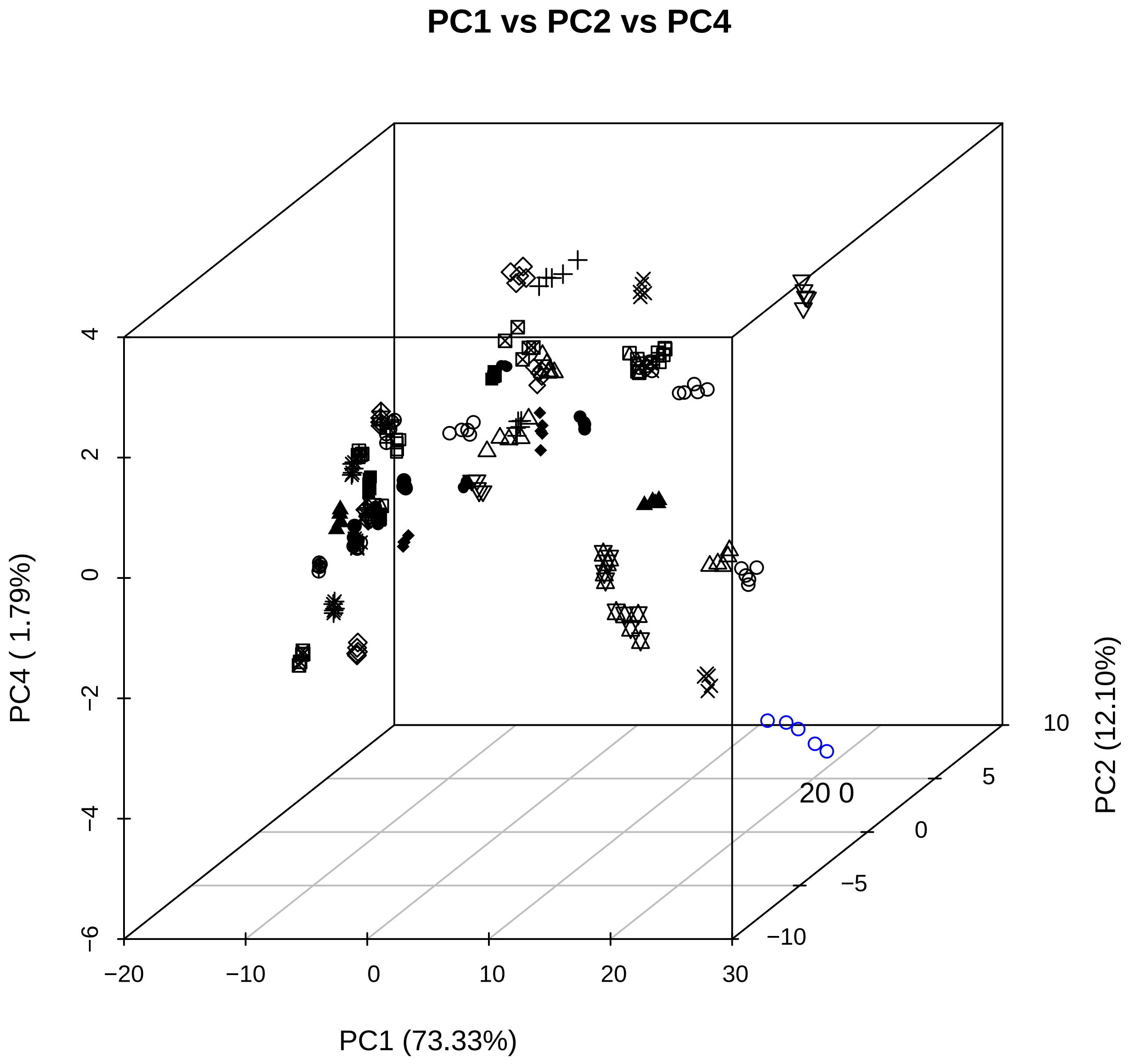

PC1 vs PC2 vs PC4
PC1 (73.33%)
PC2 (12.10%)
PC4 ( 1.79%)
−20
−10
0
10
20
30
−10
−5
0
5
10
−6
−4
−2
0
2
4
20 0

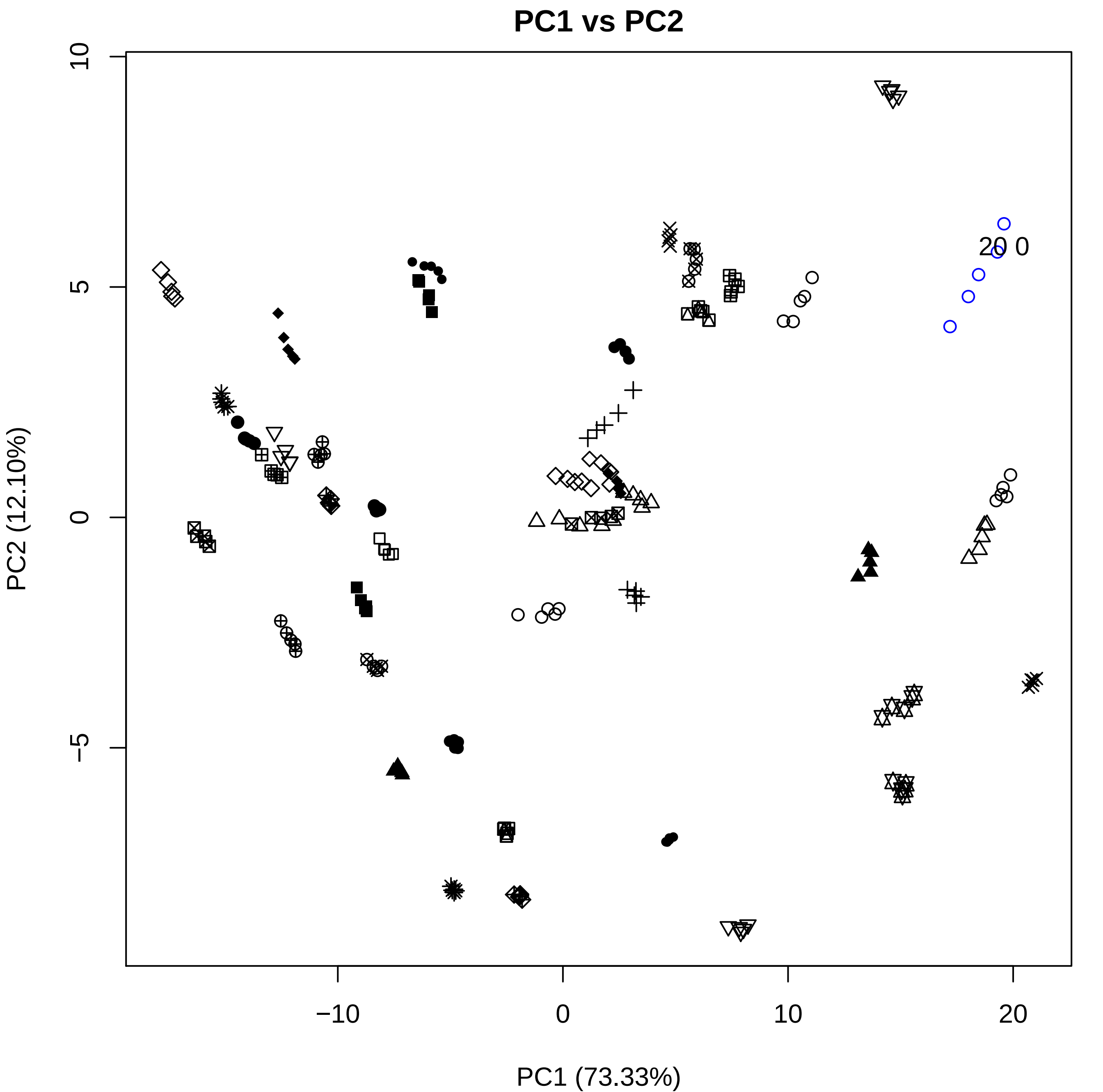
PC1 vs PC2
PC2 (12.10%)
PC1 (73.33%)
10
5
0
−5
−10
0
10
20
20 0

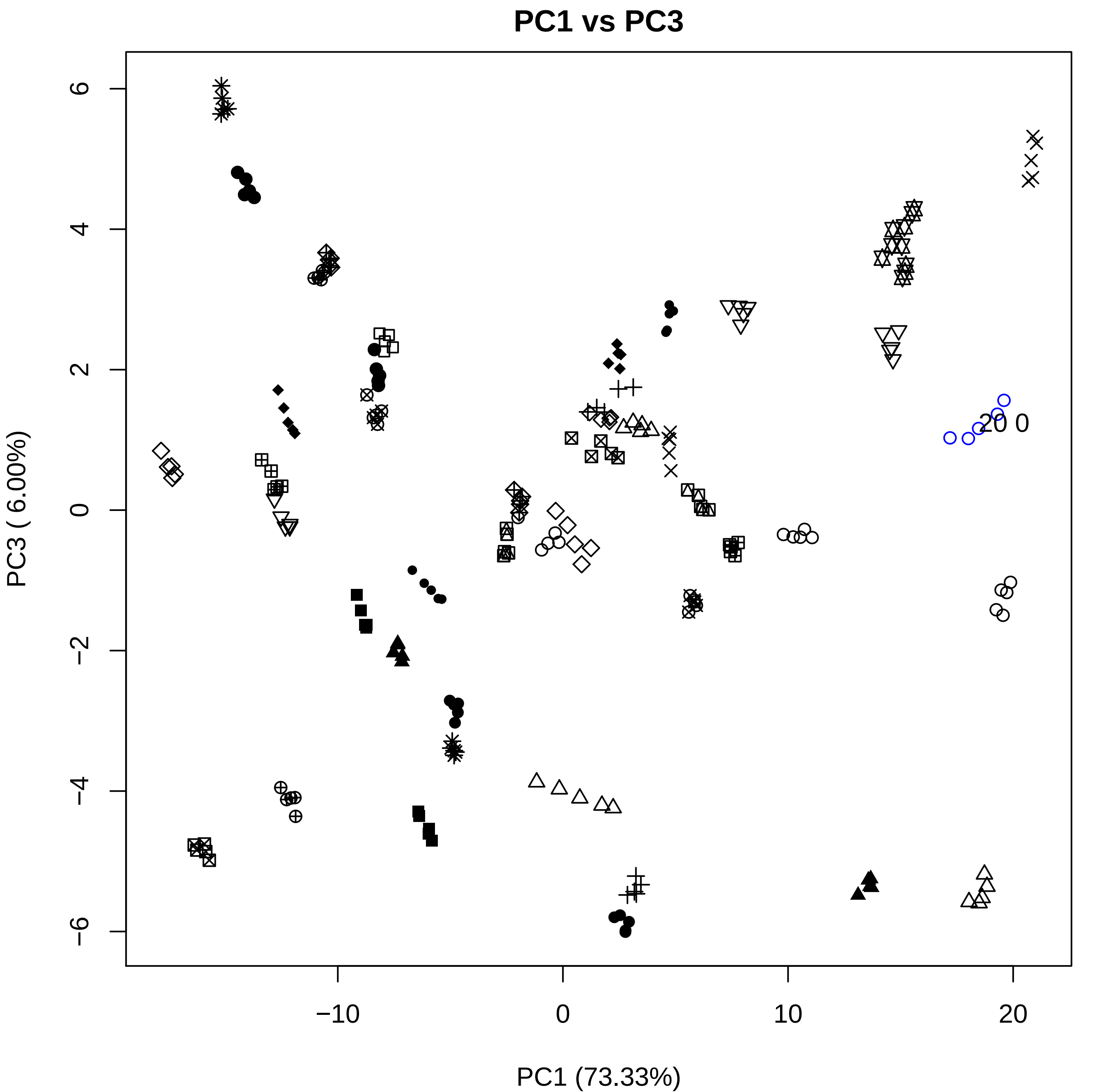
PC1 vs PC3
PC3 ( 6.00%)
PC1 (73.33%)
6
4
2
0
−2
−4
−6
−10
0
10
20
20 0

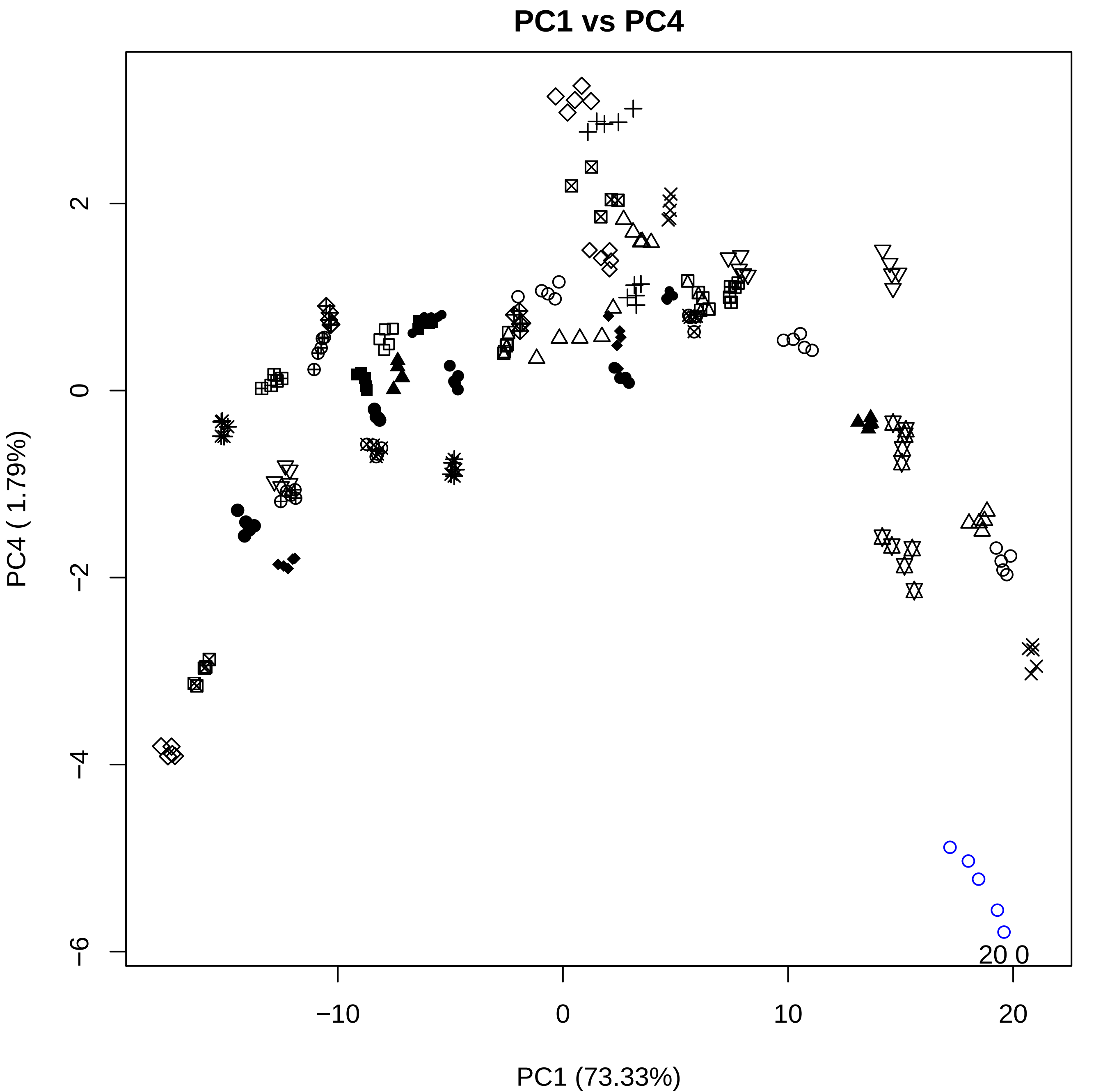
PC1 vs PC4
PC4 ( 1.79%)
PC1 (73.33%)
2
0
−2
−4
−6
−10
0
10
20
20 0

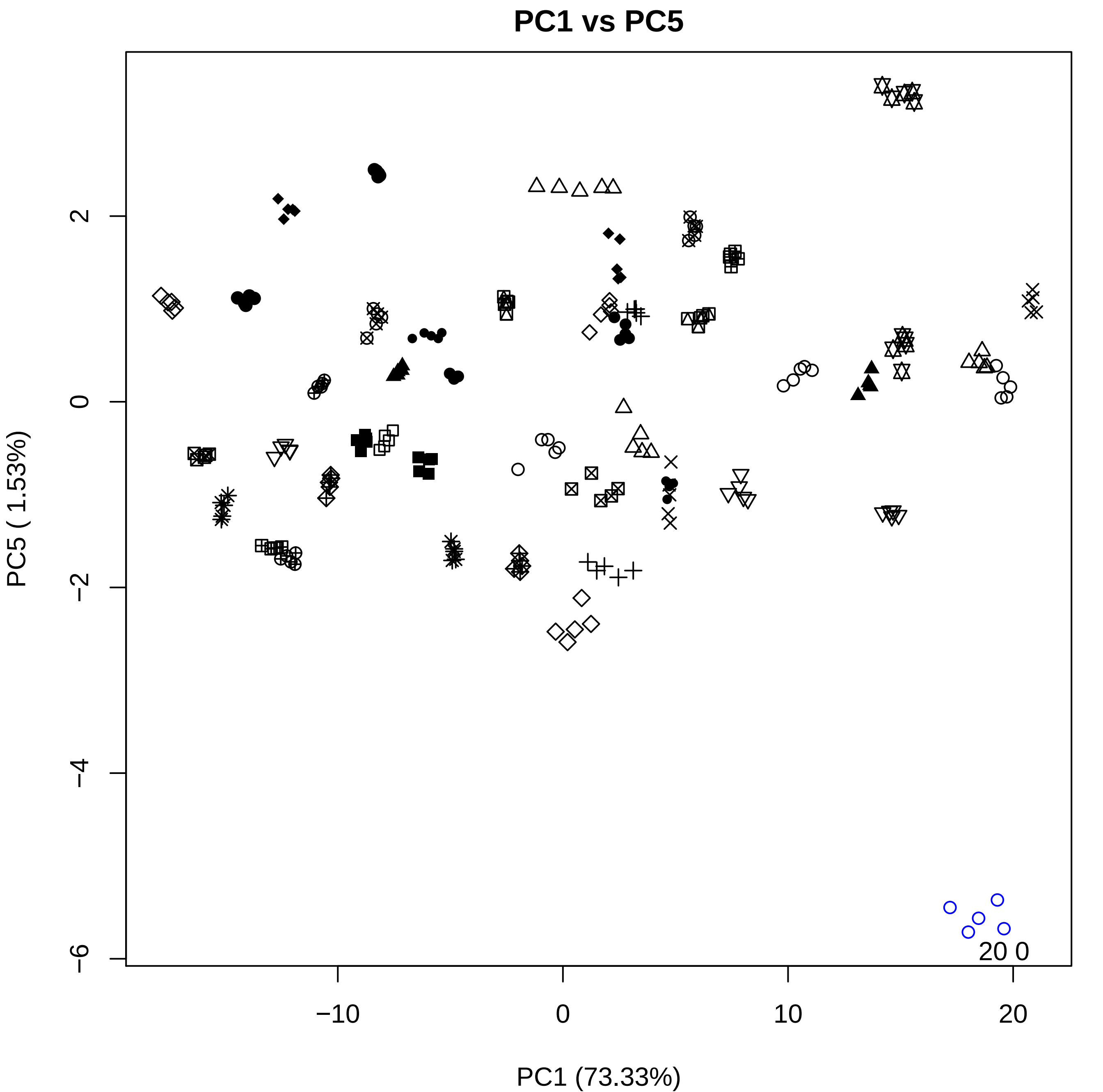

PC1 vs PC5
PC5 ( 1.53%)
PC1 (73.33%)
2
0
−2
−4
−6
−10
0
10
20
20 0

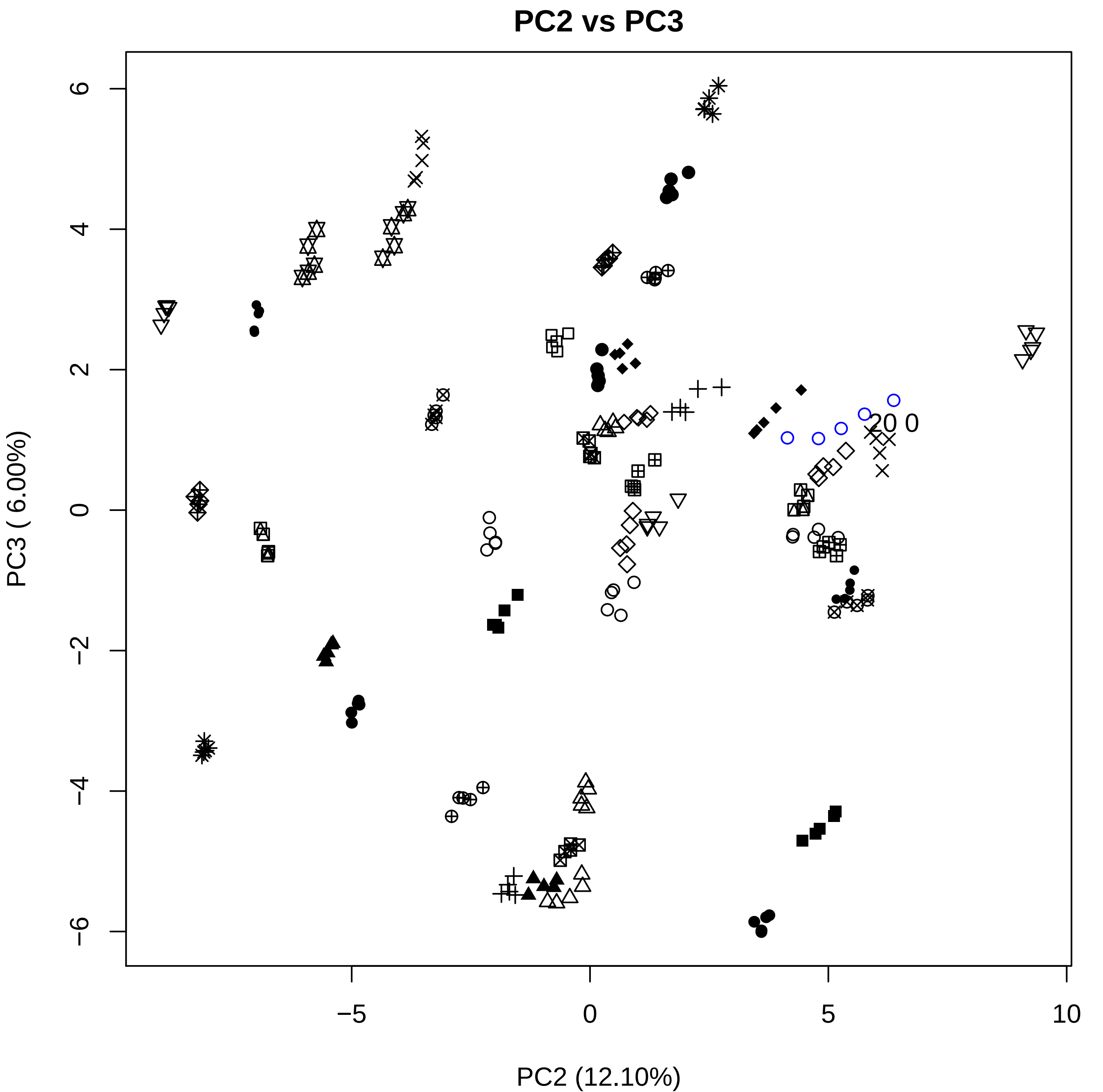
PC2 vs PC3
PC3 ( 6.00%)
PC2 (12.10%)
6
4
2
0
−2
−4
−6
−5
0
5
10
20 0

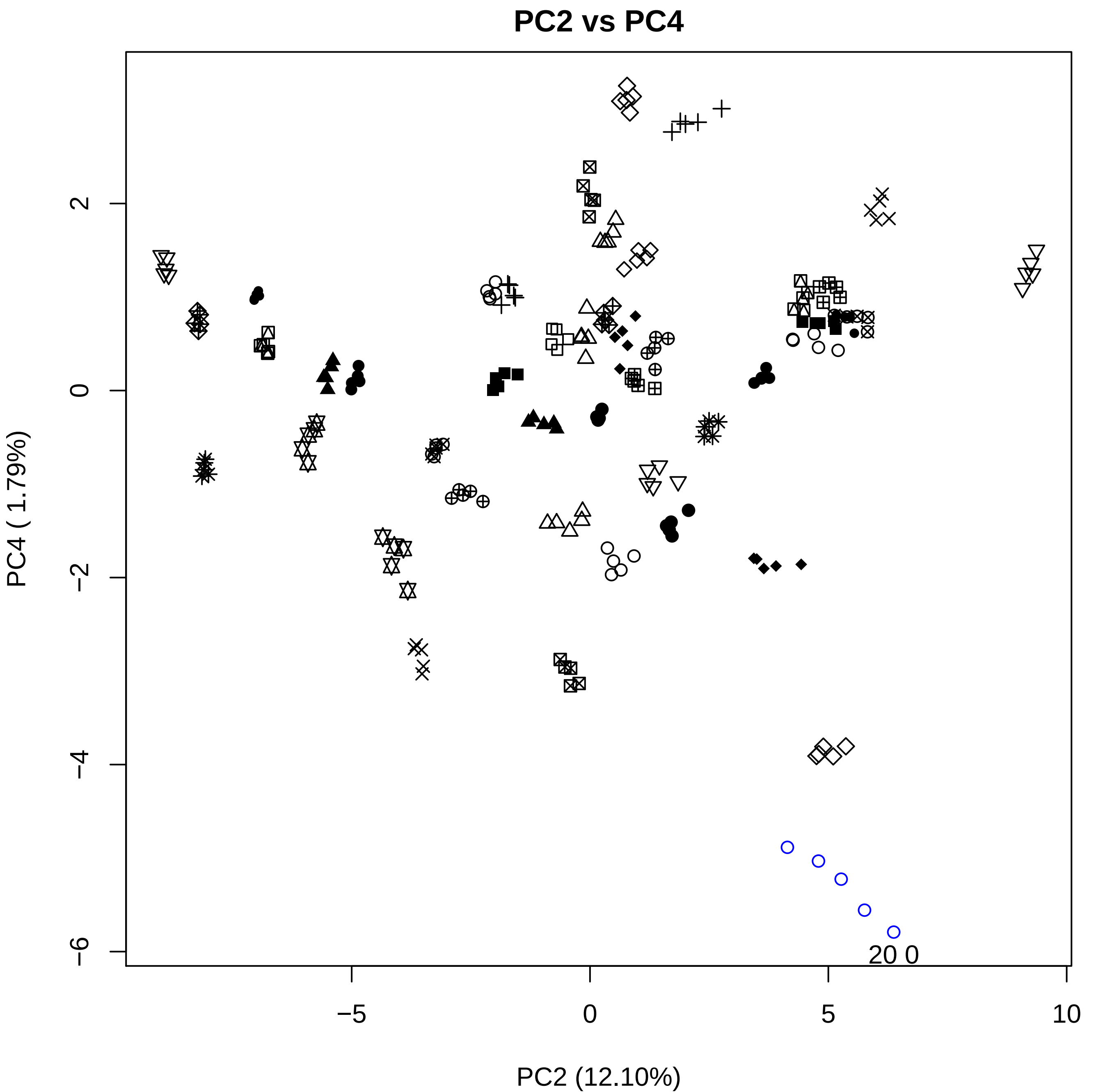
PC2 vs PC4
PC4 ( 1.79%)
PC2 (12.10%)
2
0
−2
−4
−6
−5
0
5
10
20 0

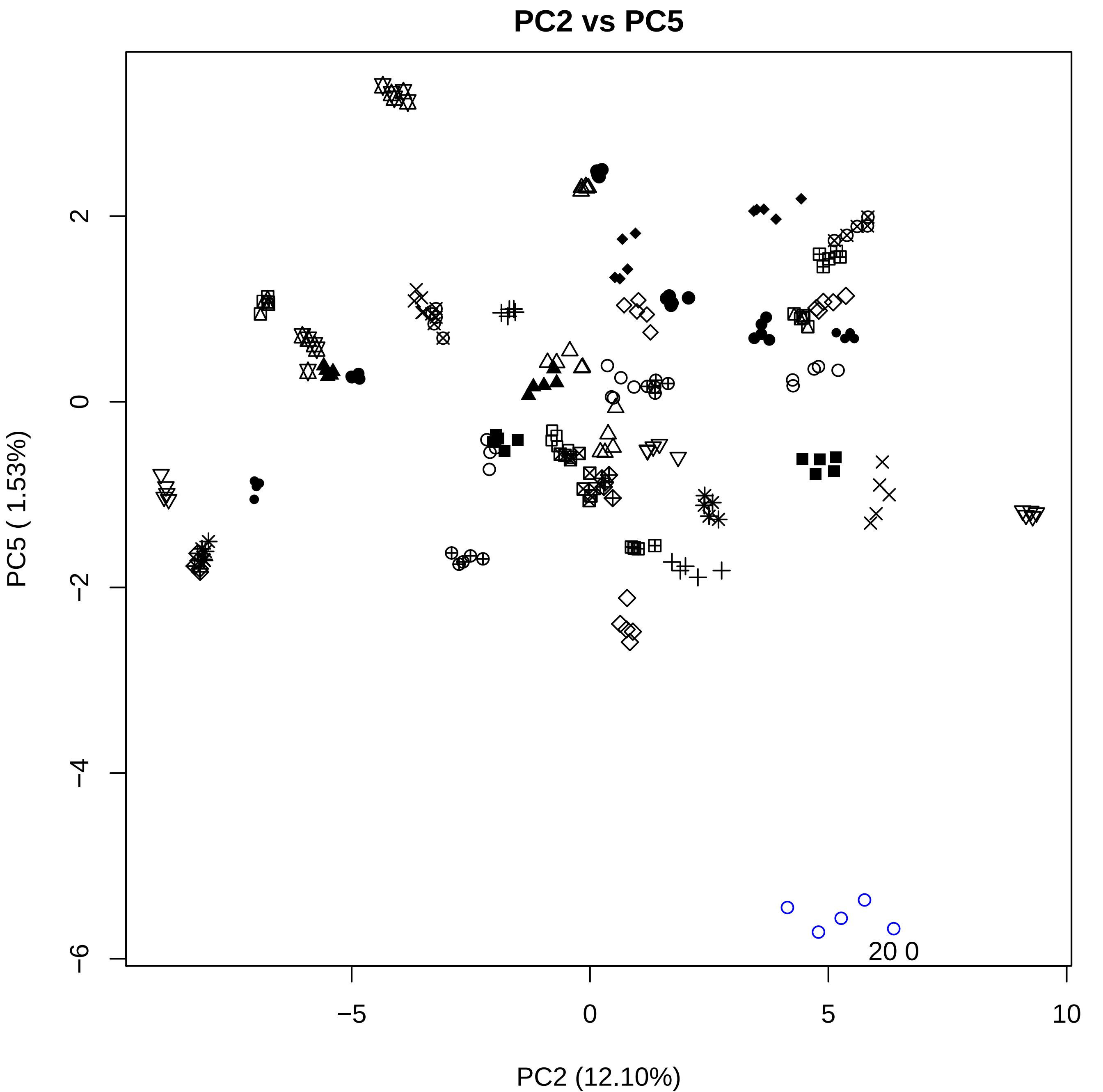
PC2 vs PC5
PC5 ( 1.53%)
PC2 (12.10%)
2
0
−2
−4
−6
−5
0
5
10
20 0

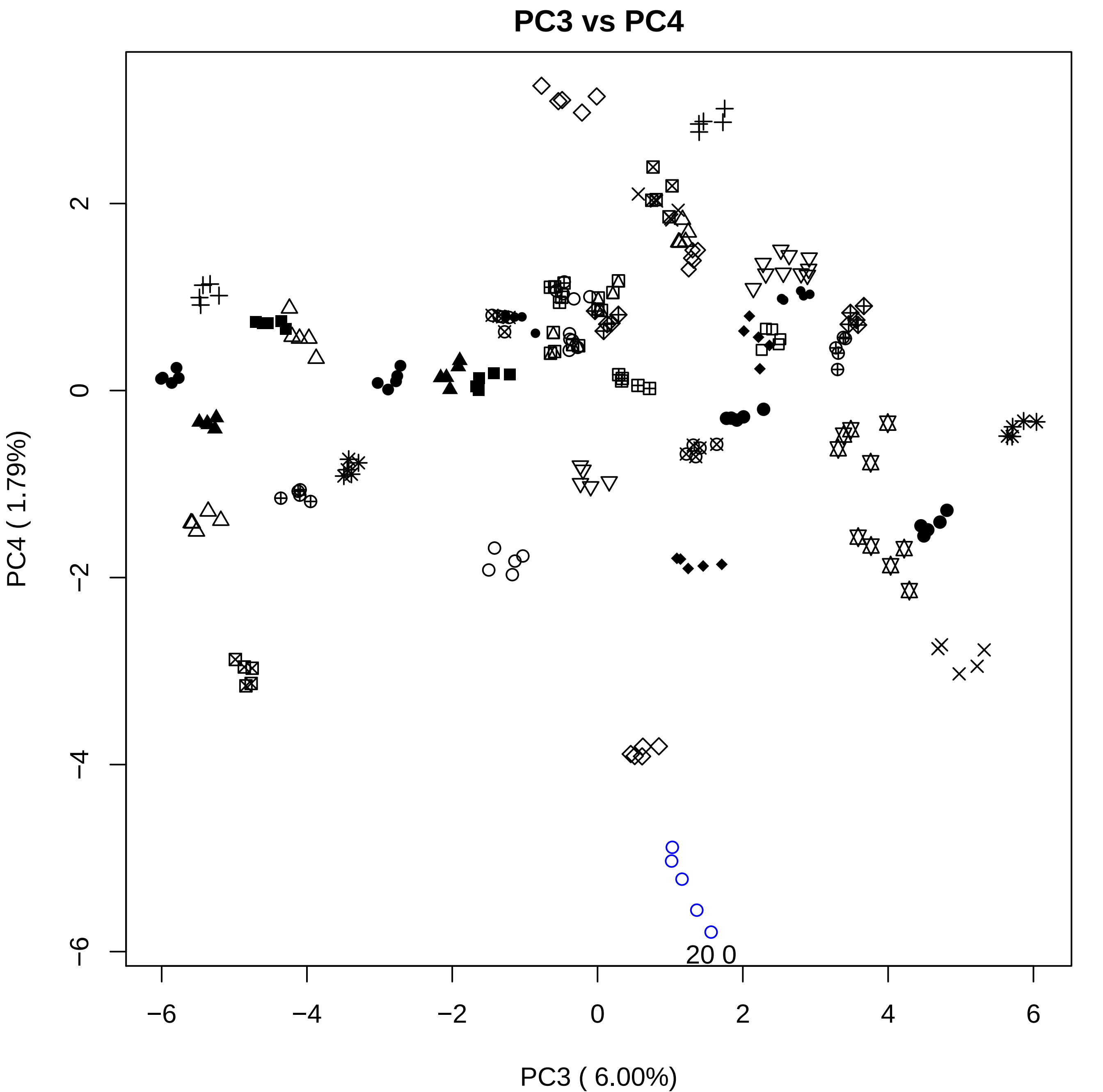
PC3 vs PC4
PC4 ( 1.79%)
PC3 ( 6.00%)
2
0
−2
−4
−6
−6
−4
−2
0
2
4
6
20 0

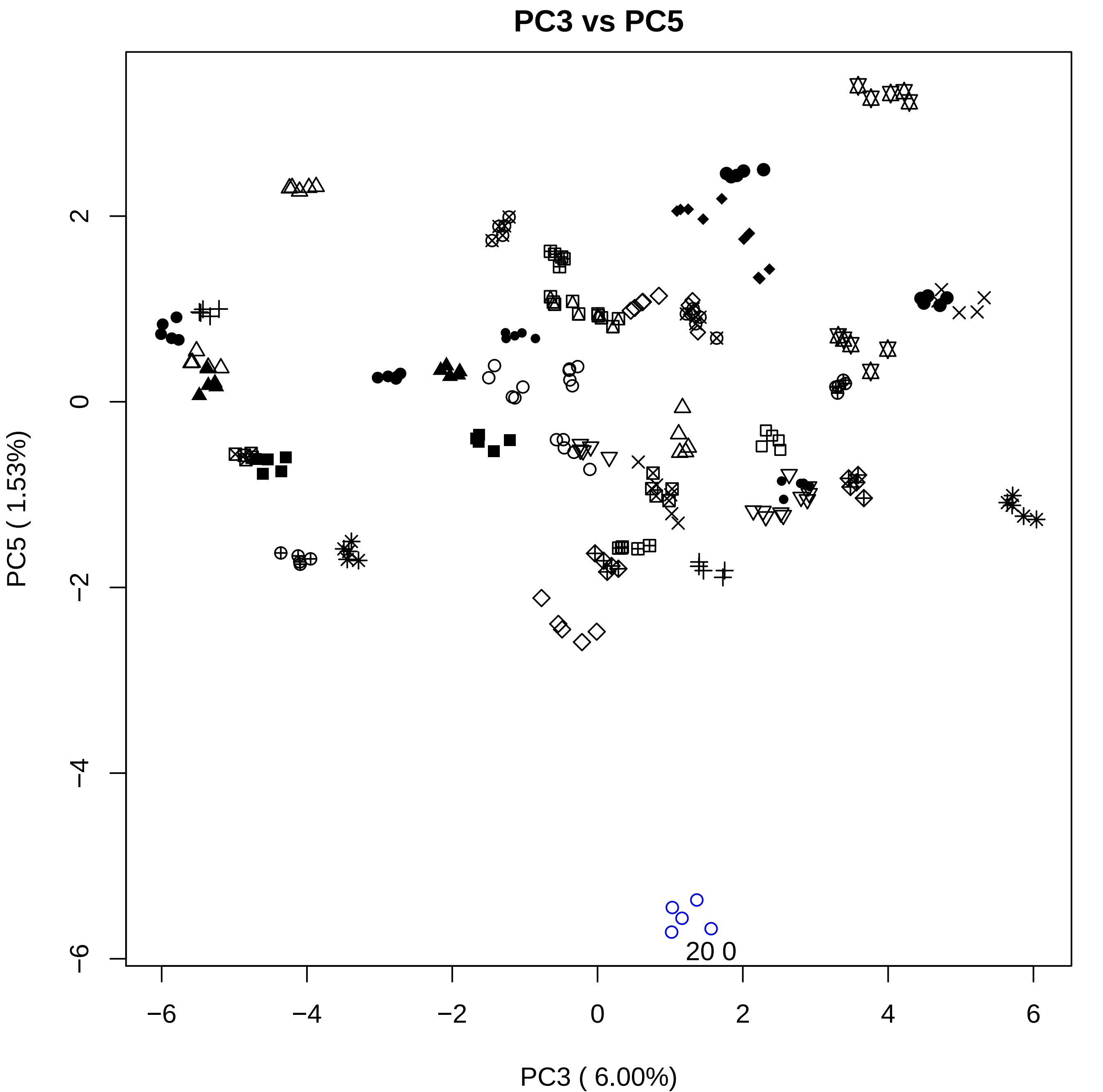

PC3 vs PC5
PC5 ( 1.53%)
PC3 ( 6.00%)
2
0
−2
−4
−6
−6
−4
−2
0
2
4
6
20 0

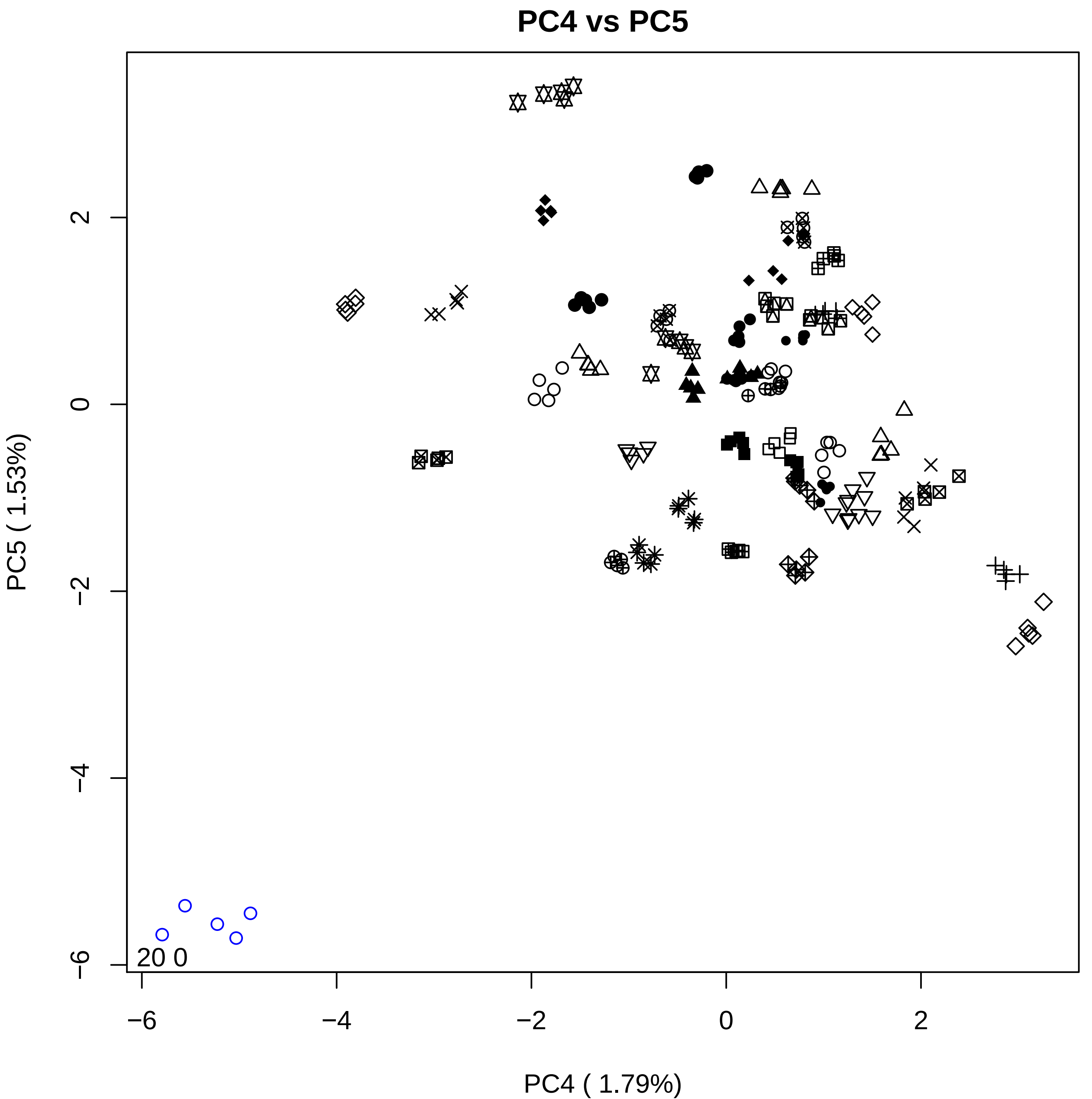
PC4 vs PC5
PC5 ( 1.53%)
PC4 ( 1.79%)
2
0
−2
−4
−6
−6
−4
−2
0
2
20 0

### PCA of 40%  DeS vs PMH

Scaled Matrix PCAed

7th order polynomial baseline correction performed

Raw spectra smoothed to 21 Neighbours, 2nd order PN, Savtisky Golay

Normalisation between 0 and 100

D2 spectra smoothed to 41 Neighbours, 2nd order PN, Savtisky Golay

Regions Analysed: 3601−2501 and 2001−700cm−1

Samples Analysed: 1 Contam's 0 DeS's 45 PMH's

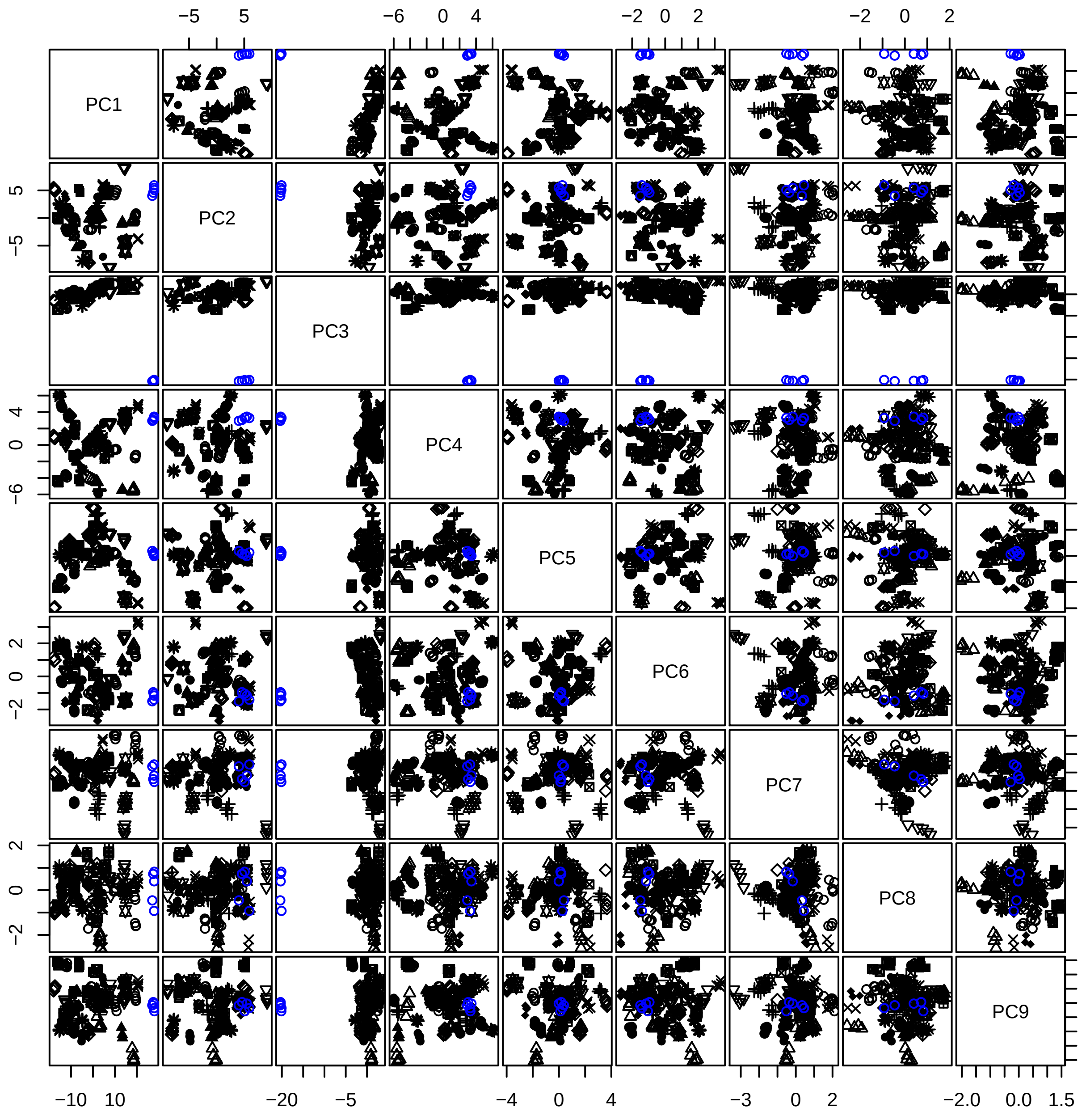

PC1
PC2
PC3
PC4
PC5
PC6
PC7
PC8
PC9

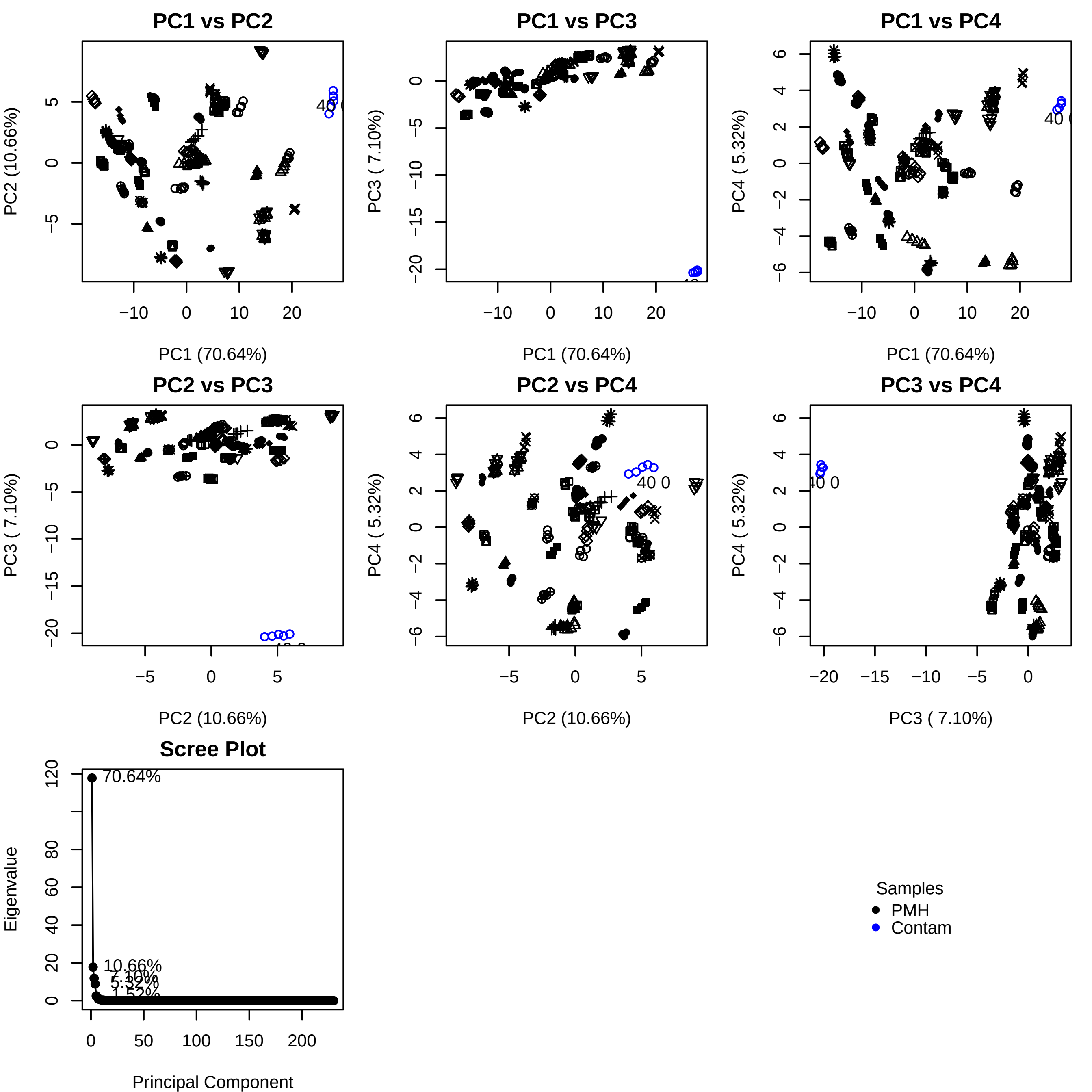
PC1 vs PC2
PC1 (70.64%)
PC2 (10.66%)
PC1 vs PC3
PC1 (70.64%)
PC3 ( 7.10%)
PC1 vs PC4
PC1 (70.64%)
PC4 ( 5.32%)
PC2 vs PC3
PC2 (10.66%)
PC3 ( 7.10%)
PC2 vs PC4
PC2 (10.66%)
PC4 ( 5.32%)
PC3 vs PC4
PC3 ( 7.10%)
PC4 ( 5.32%)
Scree Plot
Principal Component
Eigenvalue
70.64%
10.66%
5.32%
Samples
PMH
Contam

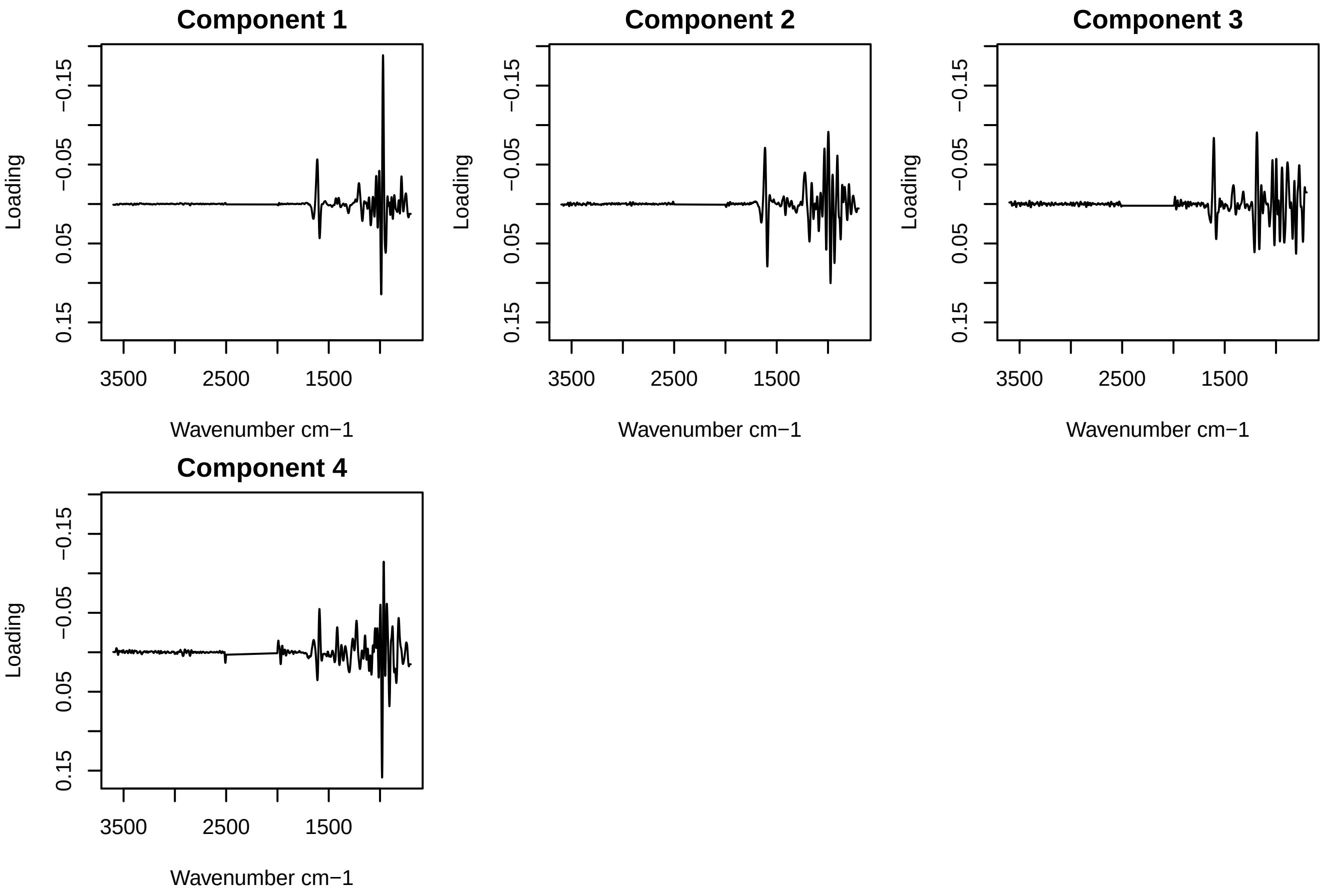
Component 1
Component 2
Component 3
Component 4
Loading
−0.15
−0.05
0.05
0.15
3500
2500
1500
Wavenumber cm−1

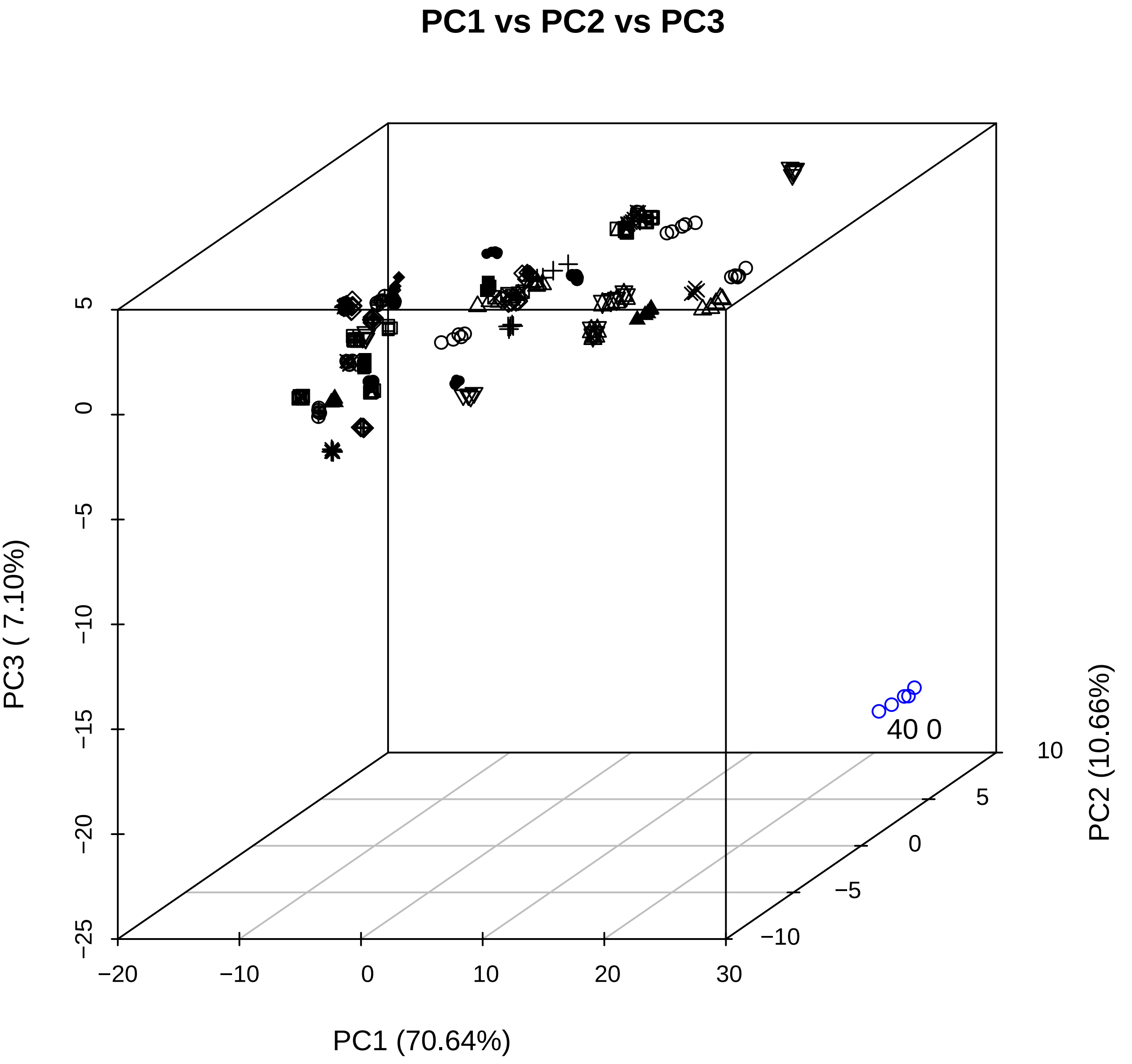

PC1 vs PC2 vs PC3
PC1 (70.64%)
PC2 (10.66%)
PC3 ( 7.10%)
−20
−10
0
10
20
30
−10
−5
0
5
10
−25
−20
−15
−10
−5
0
5
40 0

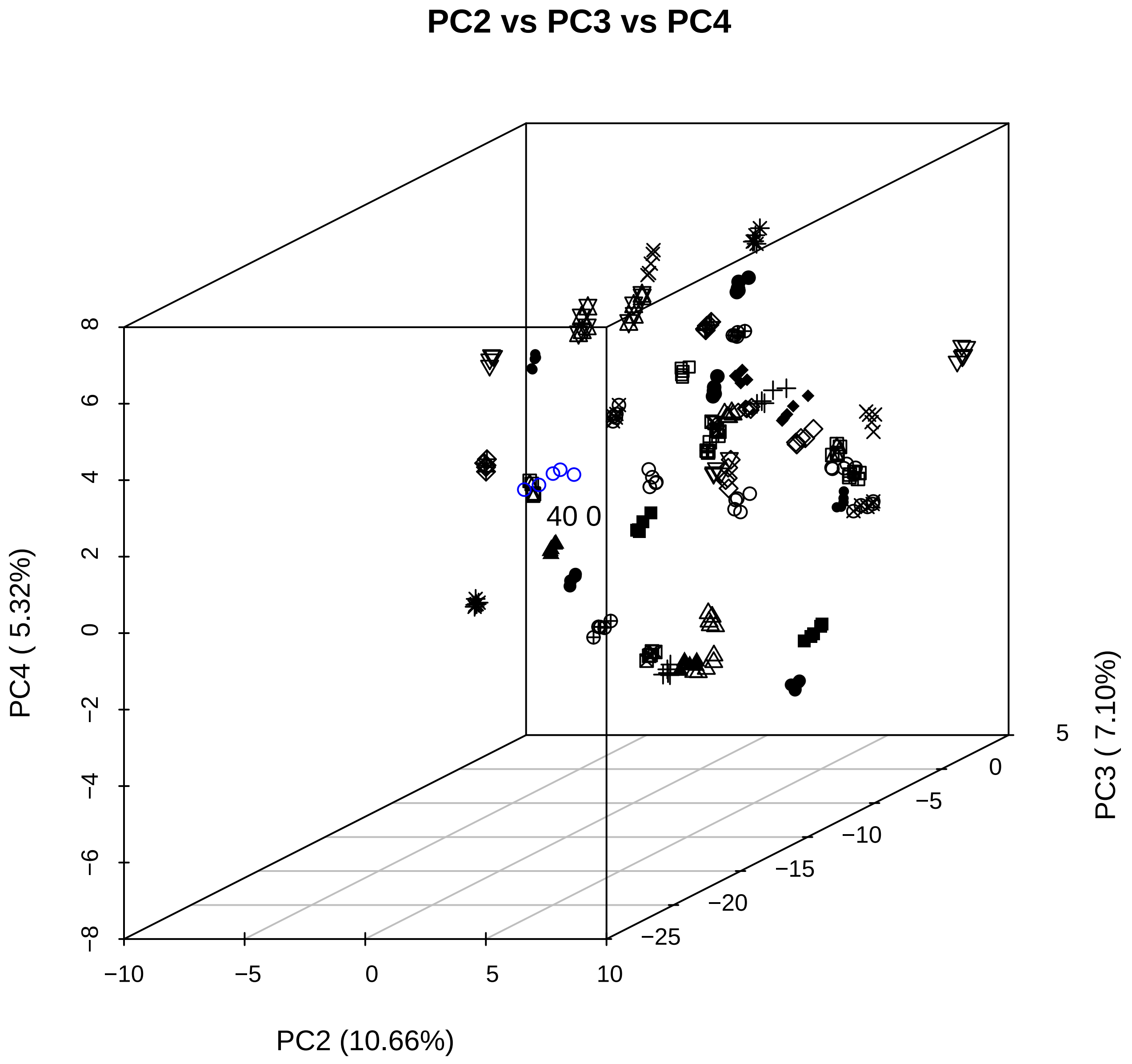
PC2 vs PC3 vs PC4
PC4 ( 5.32%)
−8
−6
−4
−2
0
2
4
6
8
PC2 (10.66%)
−10
−5
0
5
10
PC3 ( 7.10%)
−25
−20
−15
−10
−5
0
5
40 0

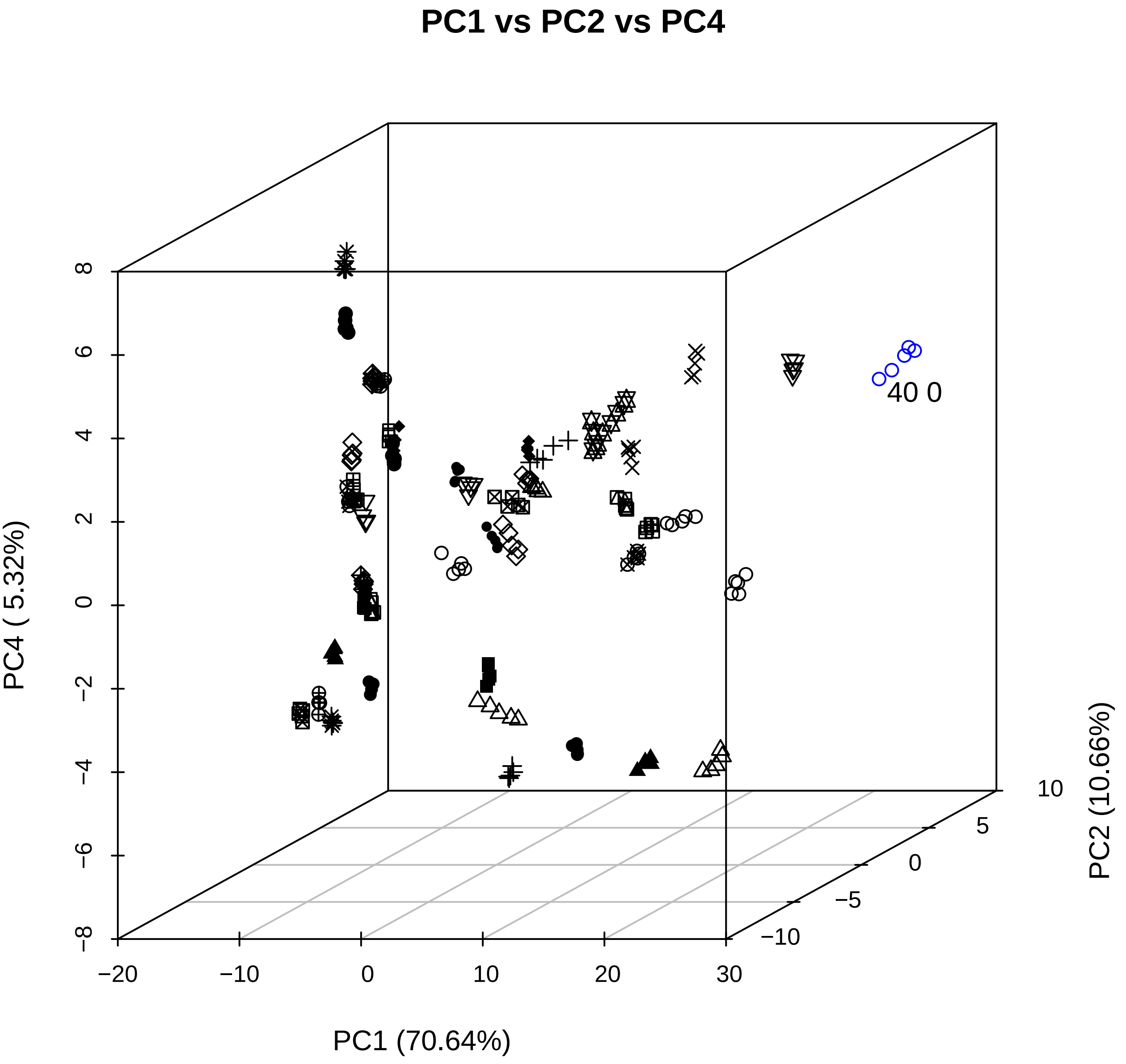
PC1 vs PC2 vs PC4
PC1 (70.64%)
PC2 (10.66%)
PC4 ( 5.32%)
−20
−10
0
10
20
30
−10
−5
0
5
10
−8
−6
−4
−2
0
2
4
6
8
40 0

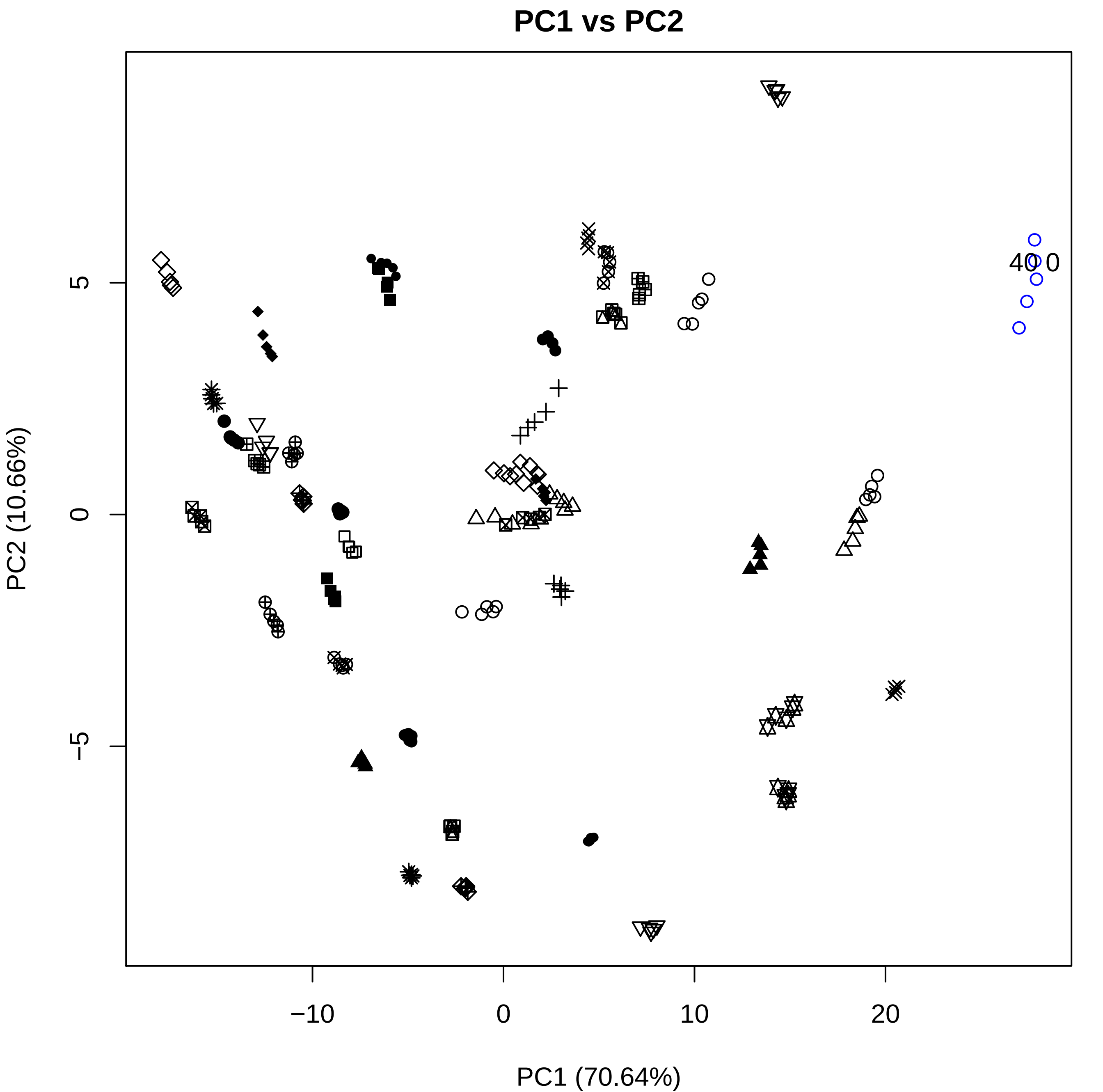

PC1 vs PC2
PC2 (10.66%)
PC1 (70.64%)
−10
0
10
20
5
0
−5
40
0

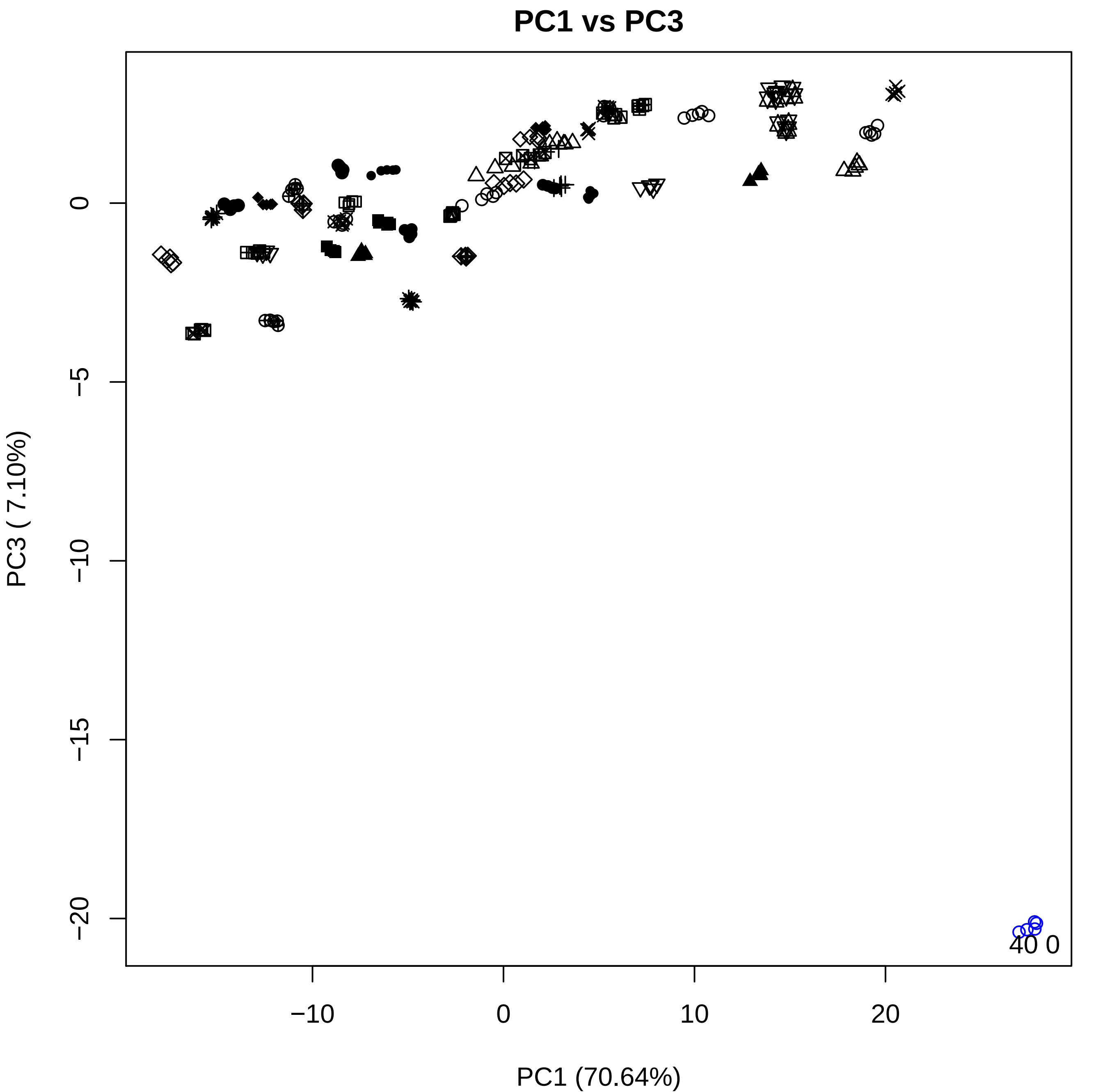
PC1 vs PC3
PC3 ( 7.10%)
PC1 (70.64%)
0
−5
−10
−15
−20
−10
0
10
20
40 0

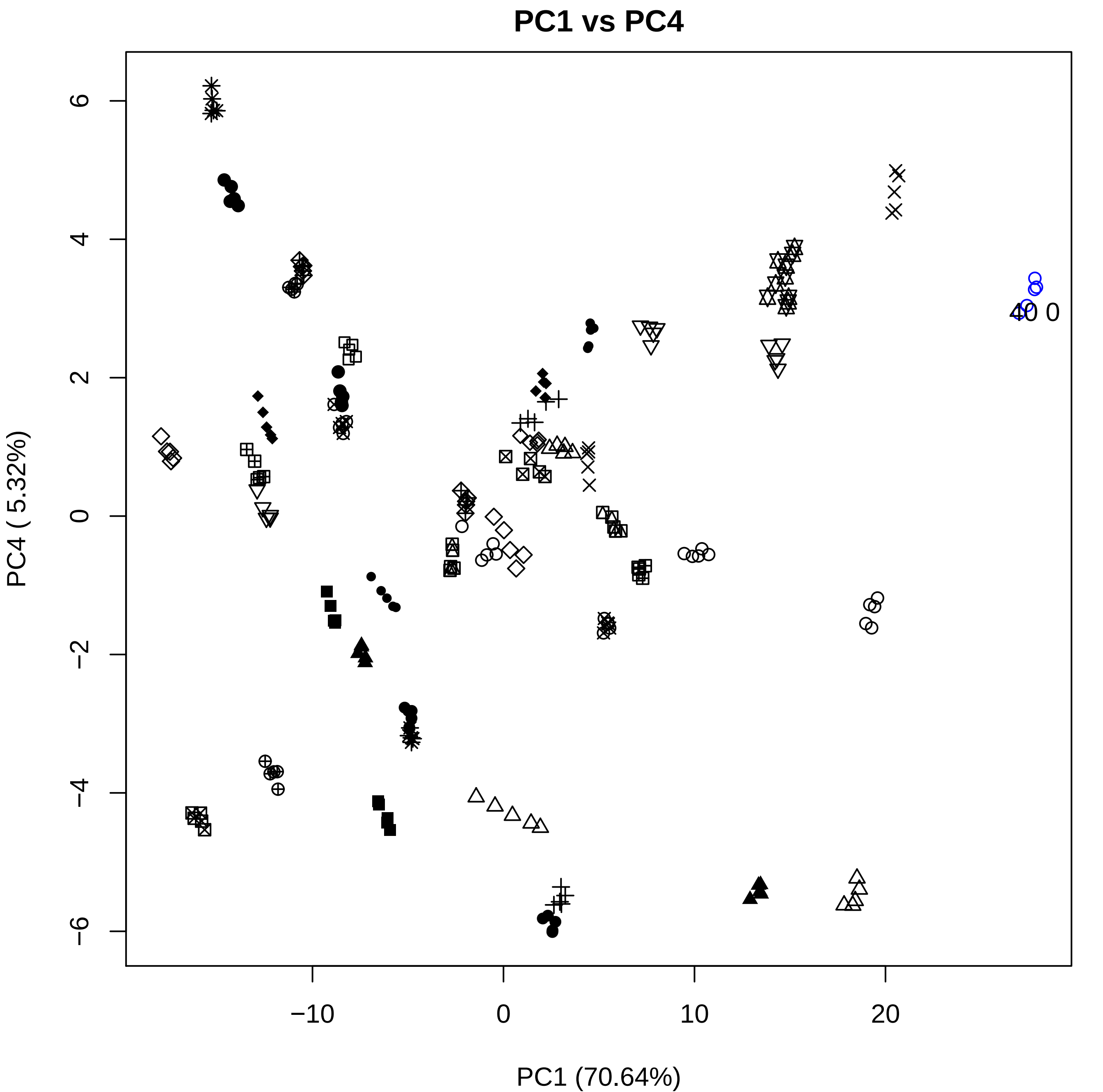
PC1 vs PC4
PC4 ( 5.32%)
PC1 (70.64%)
6
4
2
0
−2
−4
−6
−10
0
10
20
40 0

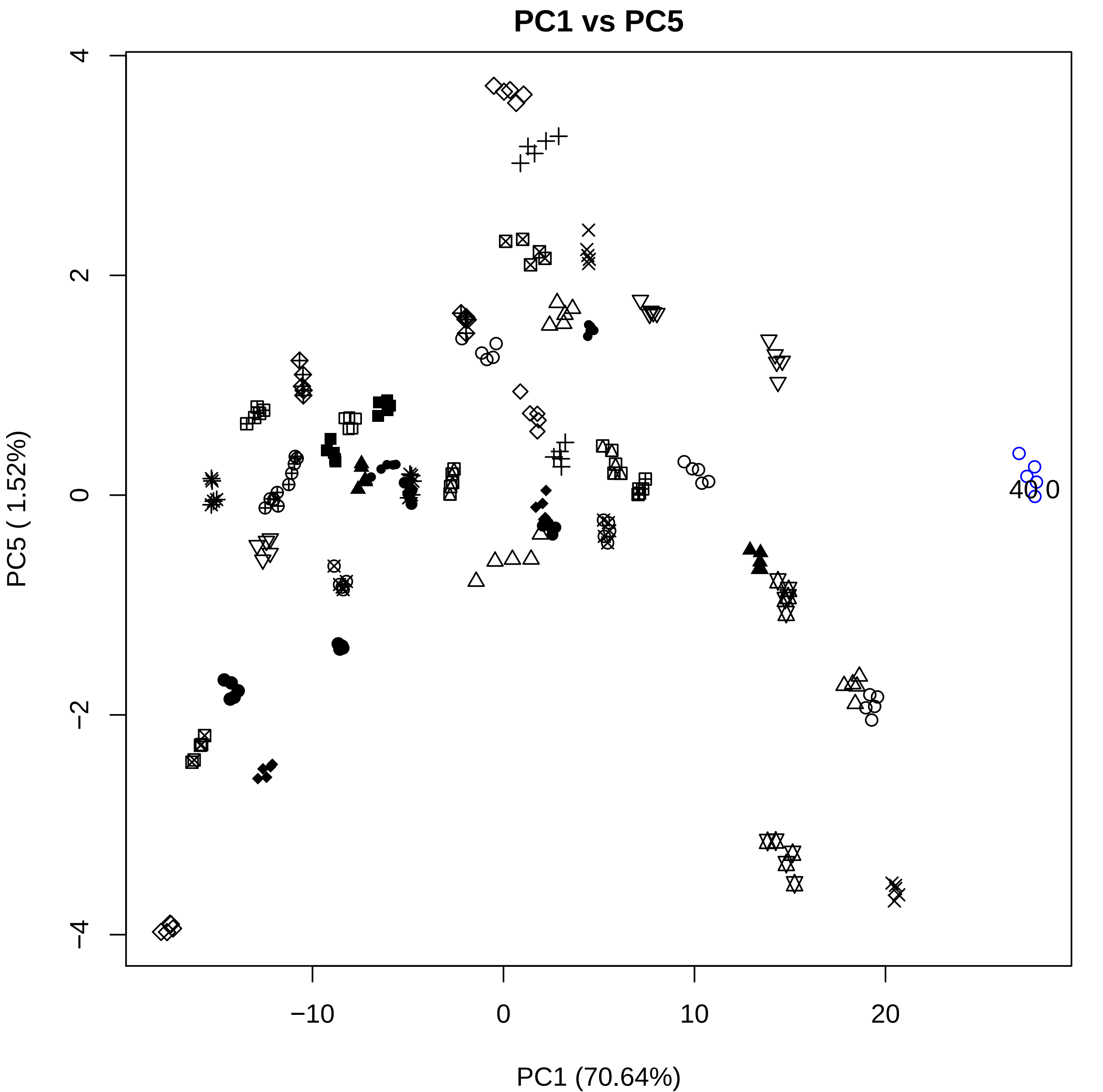
PC1 vs PC5
PC5 ( 1.52%)
PC1 (70.64%)
4
2
0
−2
−4
−10
0
10
20
400

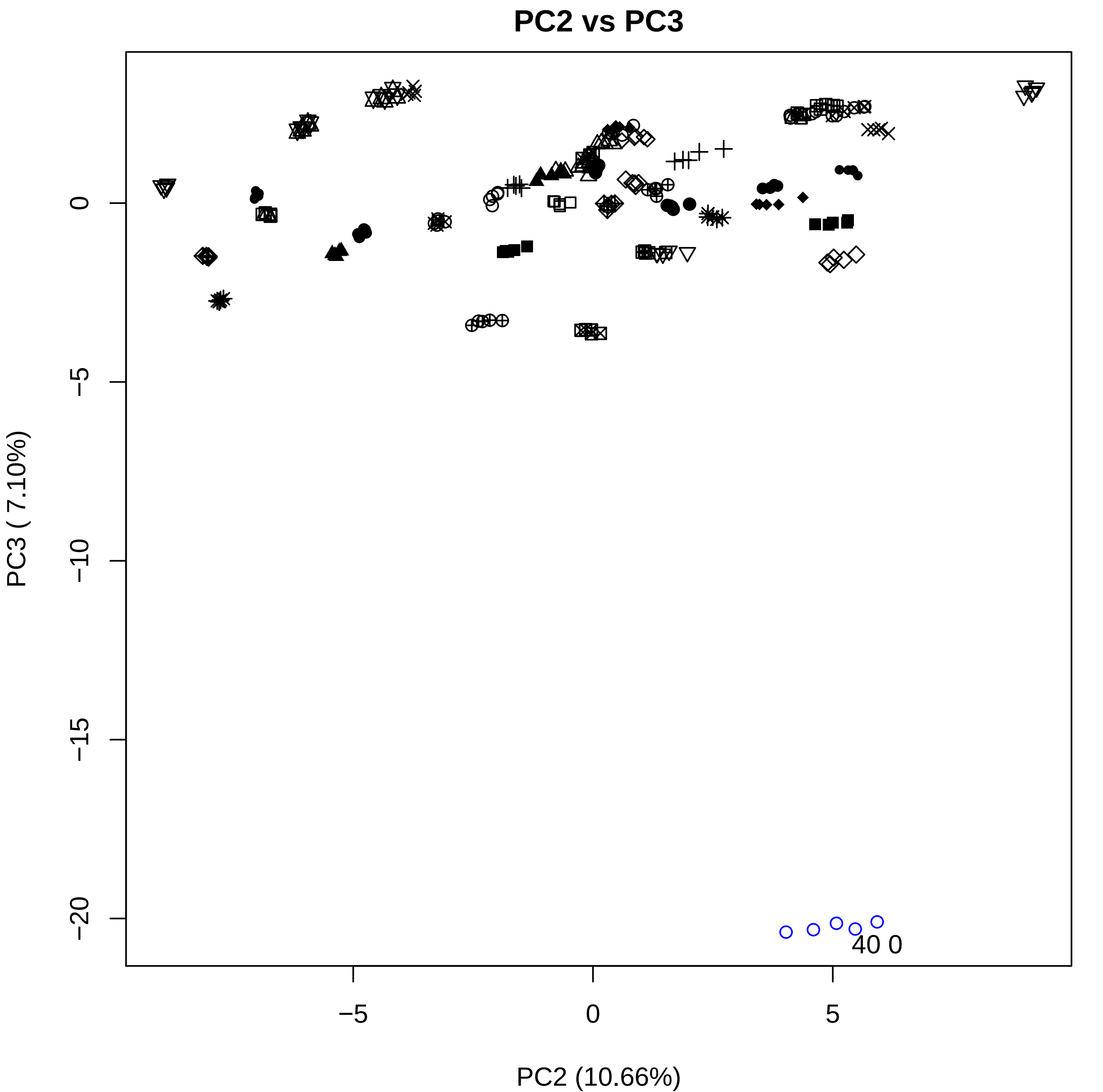
PC2 vs PC3
PC3 ( 7.10%)
PC2 (10.66%)
0
−5
−10
−15
−20
−5
0
5
40 0

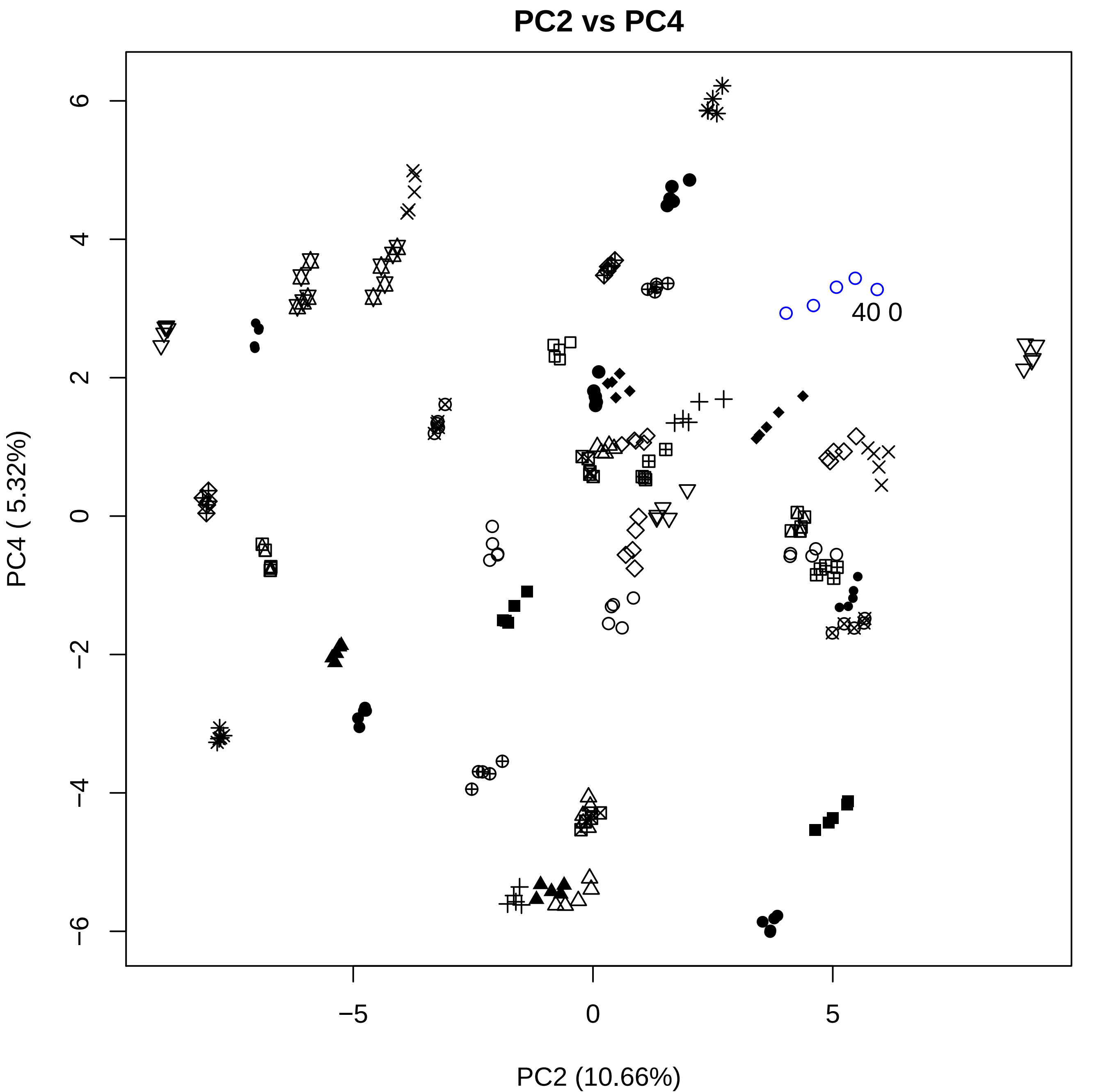
PC2 vs PC4
40 0
PC4 ( 5.32%)
PC2 (10.66%)
6
4
2
0
−2
−4
−6
−5
0
5

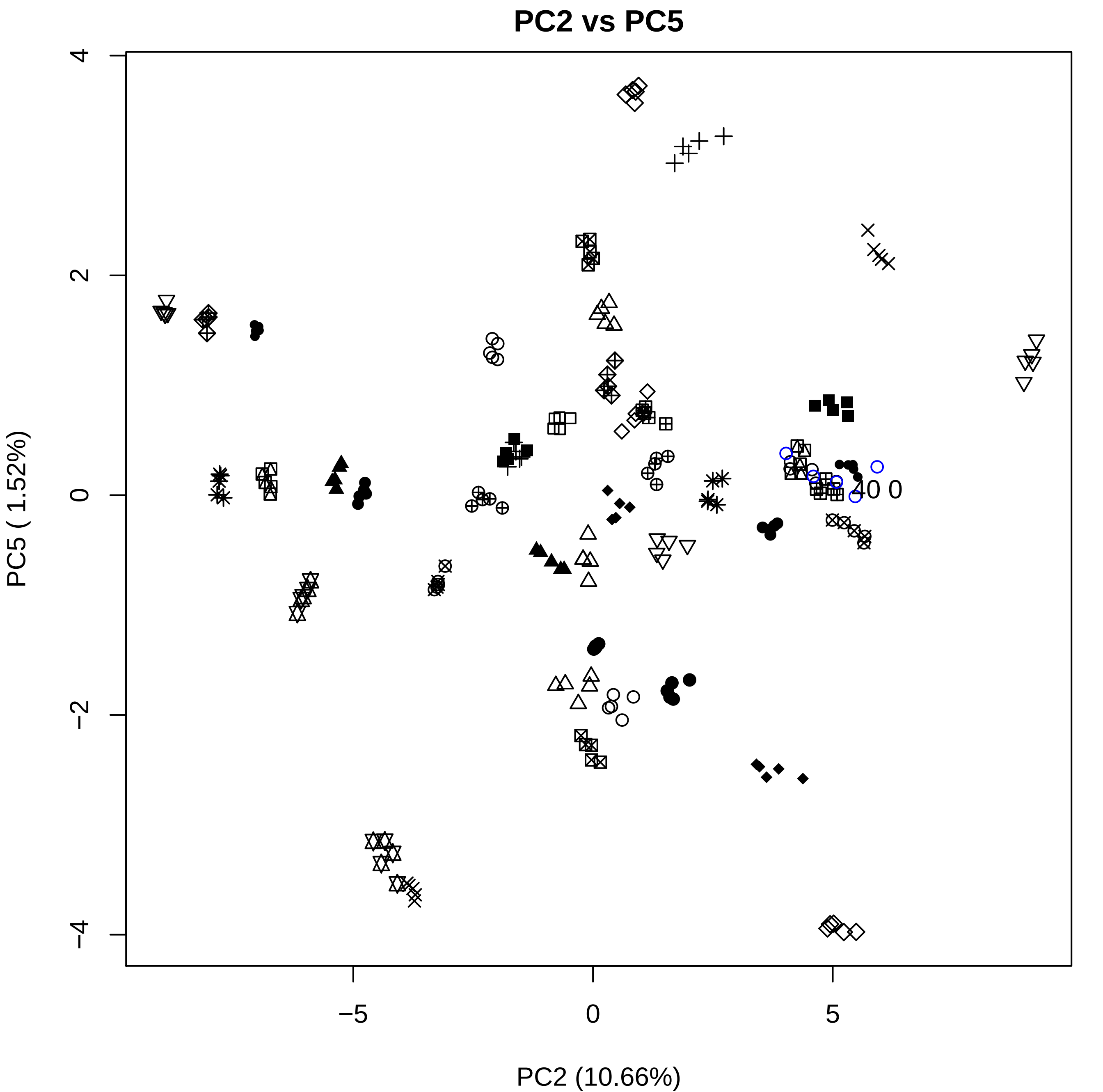
PC2 vs PC5
PC5 ( 1.52%)
PC2 (10.66%)
4
2
0
−2
−4
−5
0
5
40 0

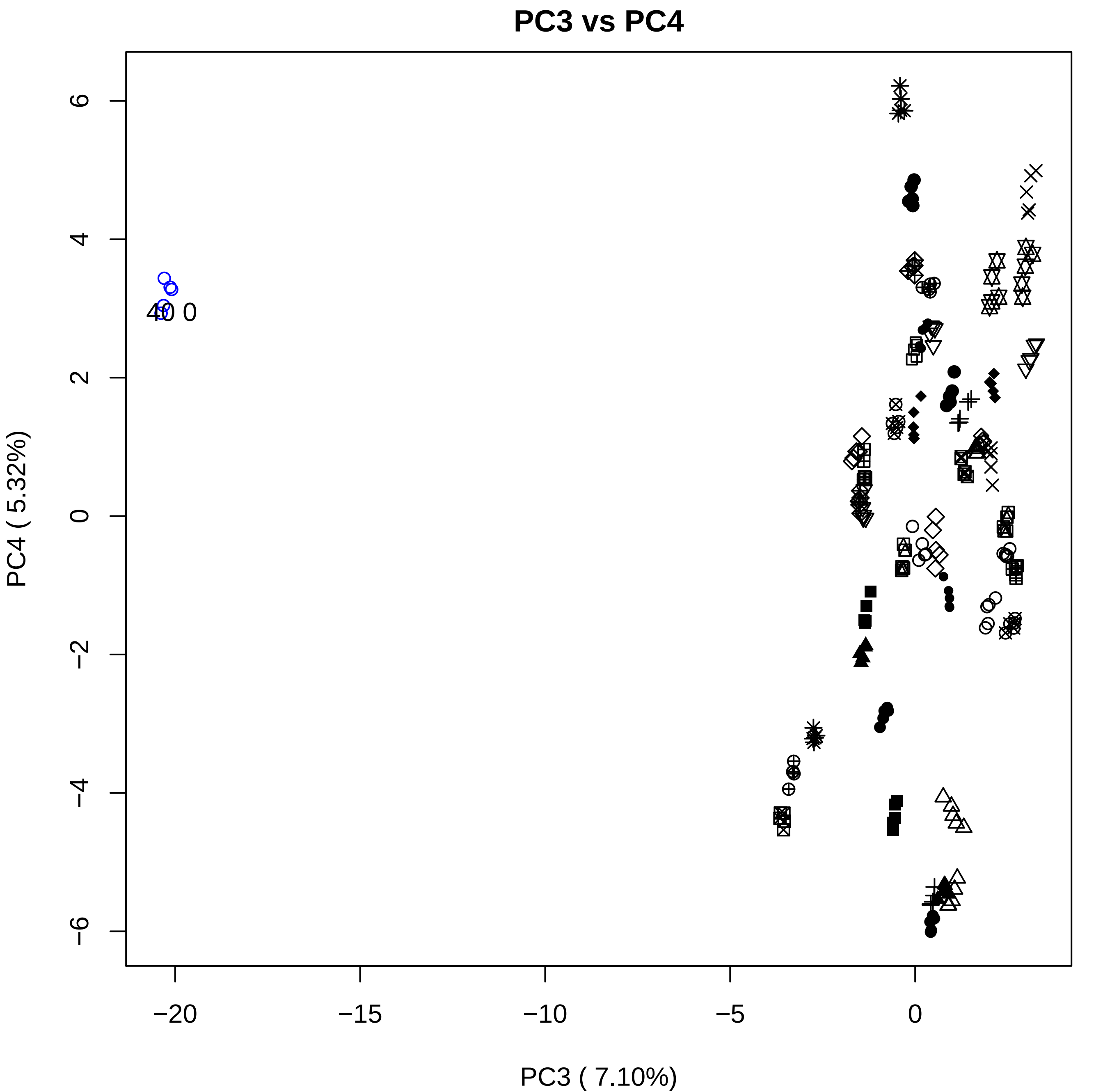
PC3 vs PC4
6
4
2
0
−2
−4
−6
PC4 ( 5.32%)
40 0
−20
−15
−10
−5
0
PC3 ( 7.10%)

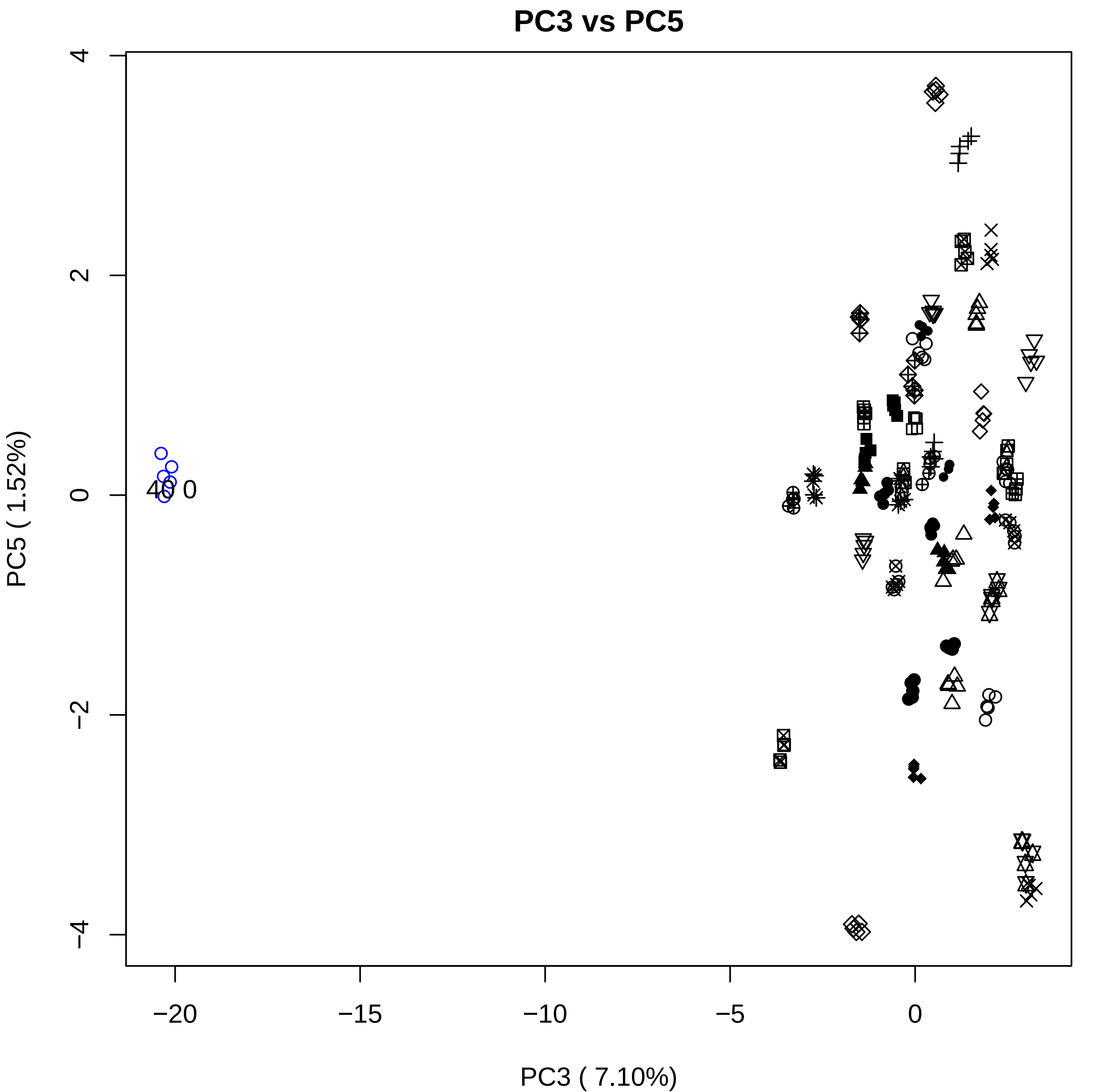
PC3 vs PC5
PC5 ( 1.52%)
PC3 ( 7.10%)
4
2
0
−2
−4
−20
−15
−10
−5
0
40 0

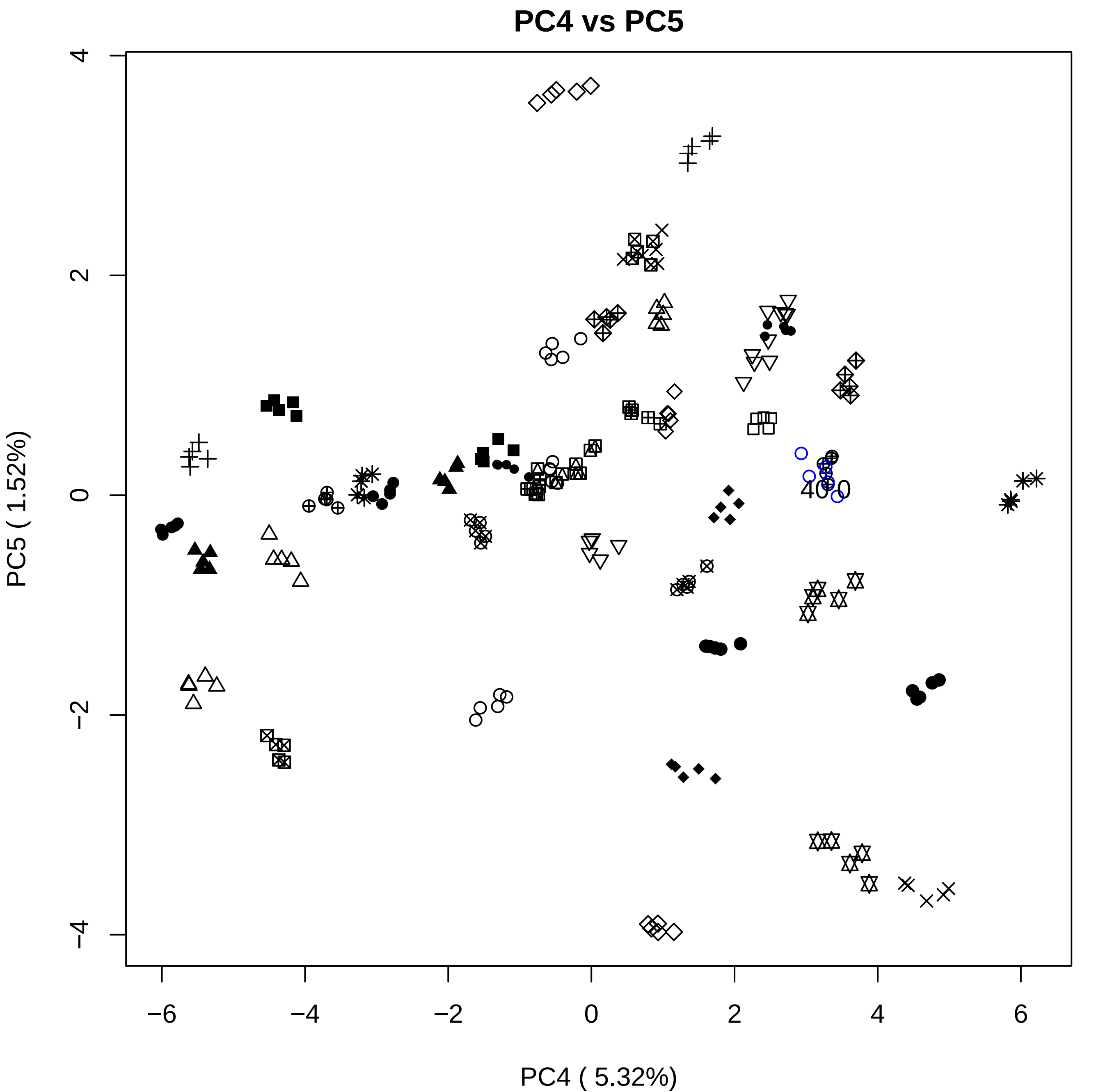
PC4 vs PC5
PC5 ( 1.52%)
PC4 ( 5.32%)
4
2
0
−2
−4
−6
−4
−2
0
2
4
6
400

### PCA of 100%  DeS vs PMH

Scaled Matrix PCAed

7th order polynomial baseline correction performed

Raw spectra smoothed to 21 Neighbours, 2nd order PN, Savtisky Golay

Normalisation between 0 and 100

D2 spectra smoothed to 41 Neighbours, 2nd order PN, Savtisky Golay

Regions Analysed: 3601−2501 and 2001−700cm−1

Samples Analysed: 0 Contam's 1 DeS's 45 PMH's

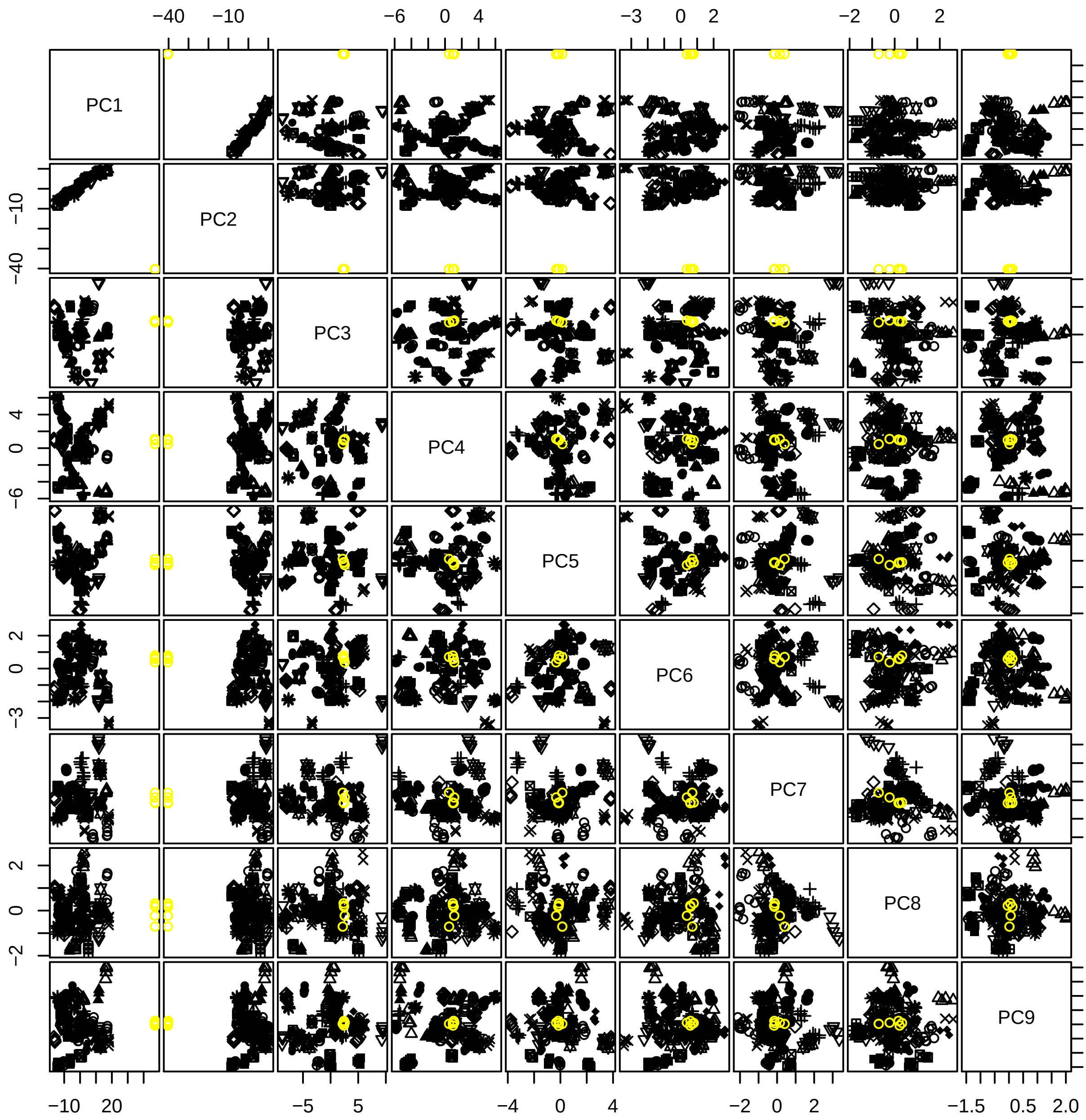

PC1
PC2
PC3
PC4
PC5
PC6
PC7
PC8
PC9
−40
−10
−6
0
4
−3
0
2
−2
0
2
−10
20
−5
5
−4
0
4
−2
0
2
−1.5
0.5
2.0

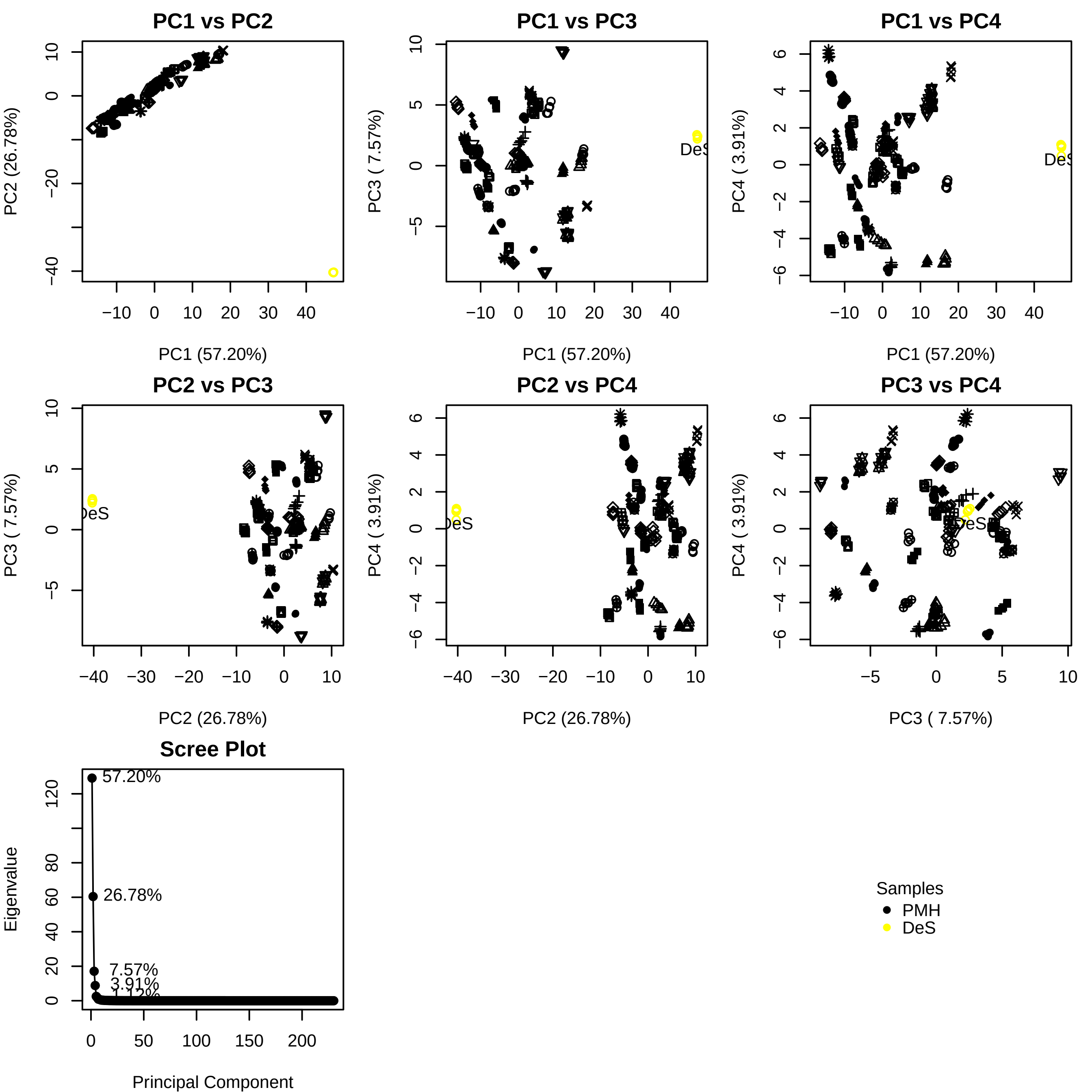
PC1 vs PC2
PC1 vs PC3
PC1 vs PC4
PC2 vs PC3
PC2 vs PC4
PC3 vs PC4
Scree Plot
PC1 (57.20%)
PC2 (26.78%)
PC3 ( 7.57%)
PC4 ( 3.91%)
DeS
57.20%
26.78%
7.57%
3.91%
1.12%
Eigenvalue
Principal Component
Samples
PMH
DeS

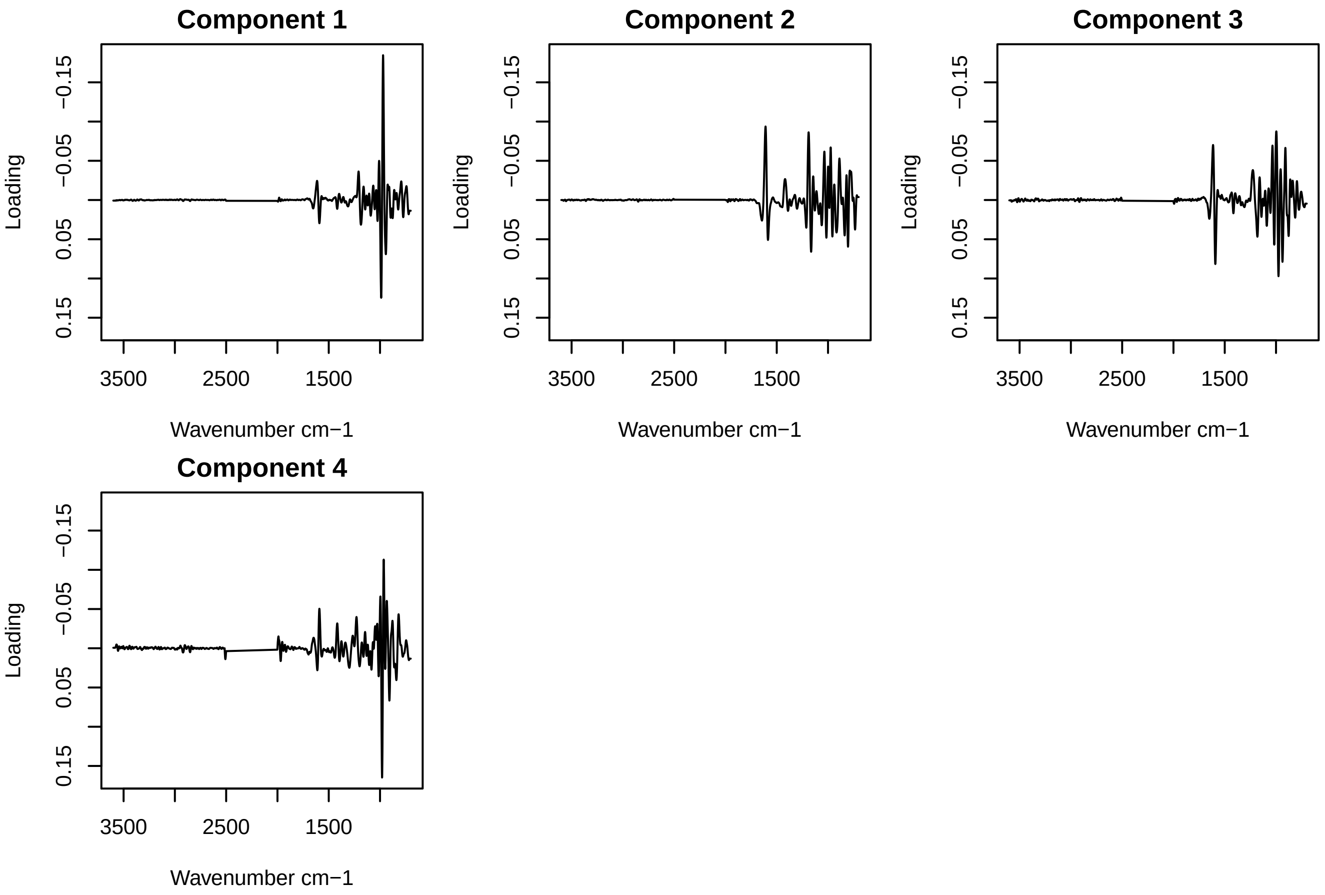
Component 1
Component 2
Component 3
Component 4
Loading
−0.15
−0.05
0.05
0.15
3500
2500
1500
Wavenumber cm−1

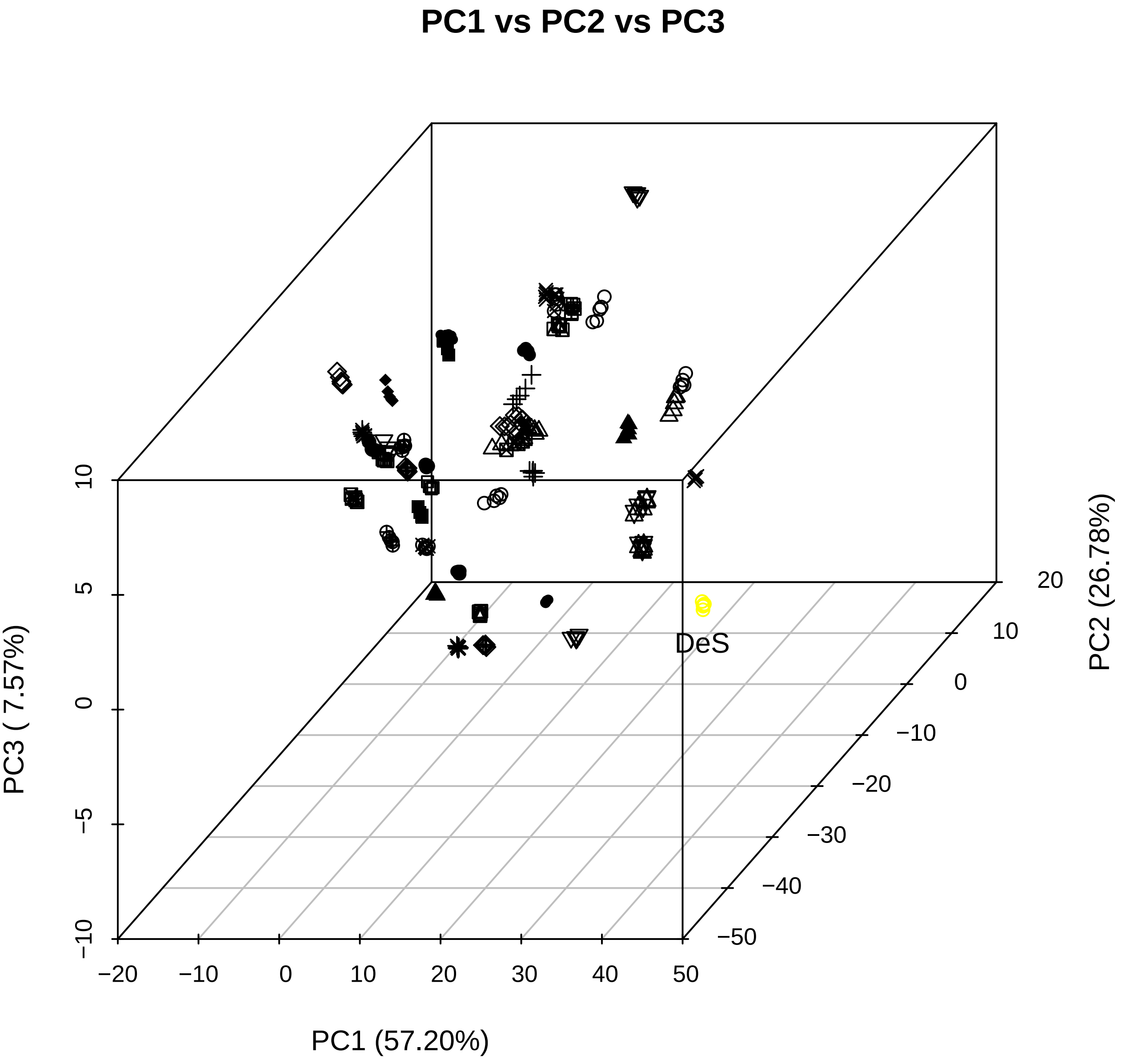
PC1 vs PC2 vs PC3
PC1 (57.20%)
PC2 (26.78%)
PC3 ( 7.57%)
−20
−10
0
10
20
30
40
50
20
10
0
−10
−20
−30
−40
−50
10
5
0
−5
−10
DeS

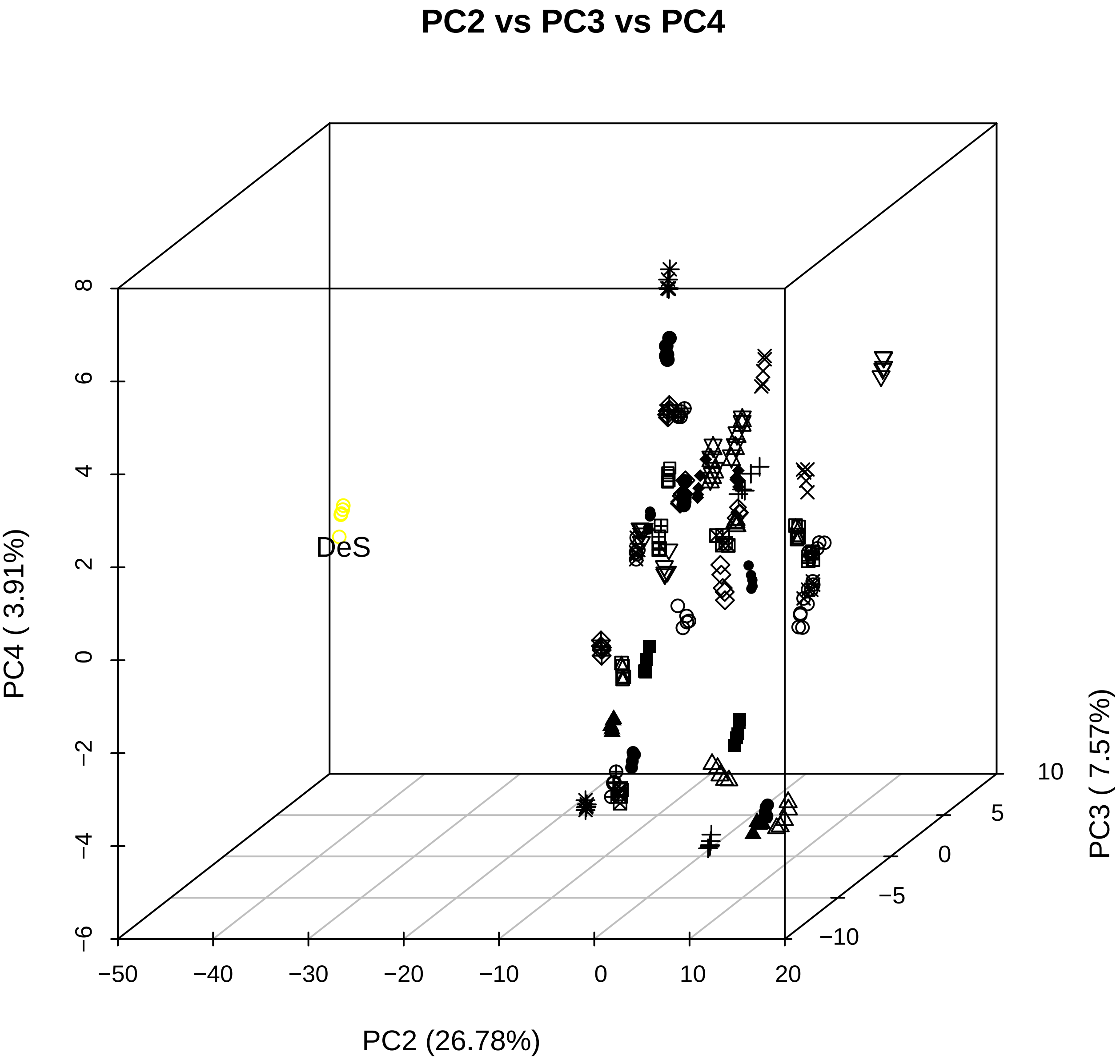

PC2 vs PC3 vs PC4
PC4 ( 3.91%)
8
6
4
2
0
−2
−4
−6
DeS
PC3 ( 7.57%)
10
5
0
−5
−10
−50
−40
−30
−20
−10
0
10
20
PC2 (26.78%)

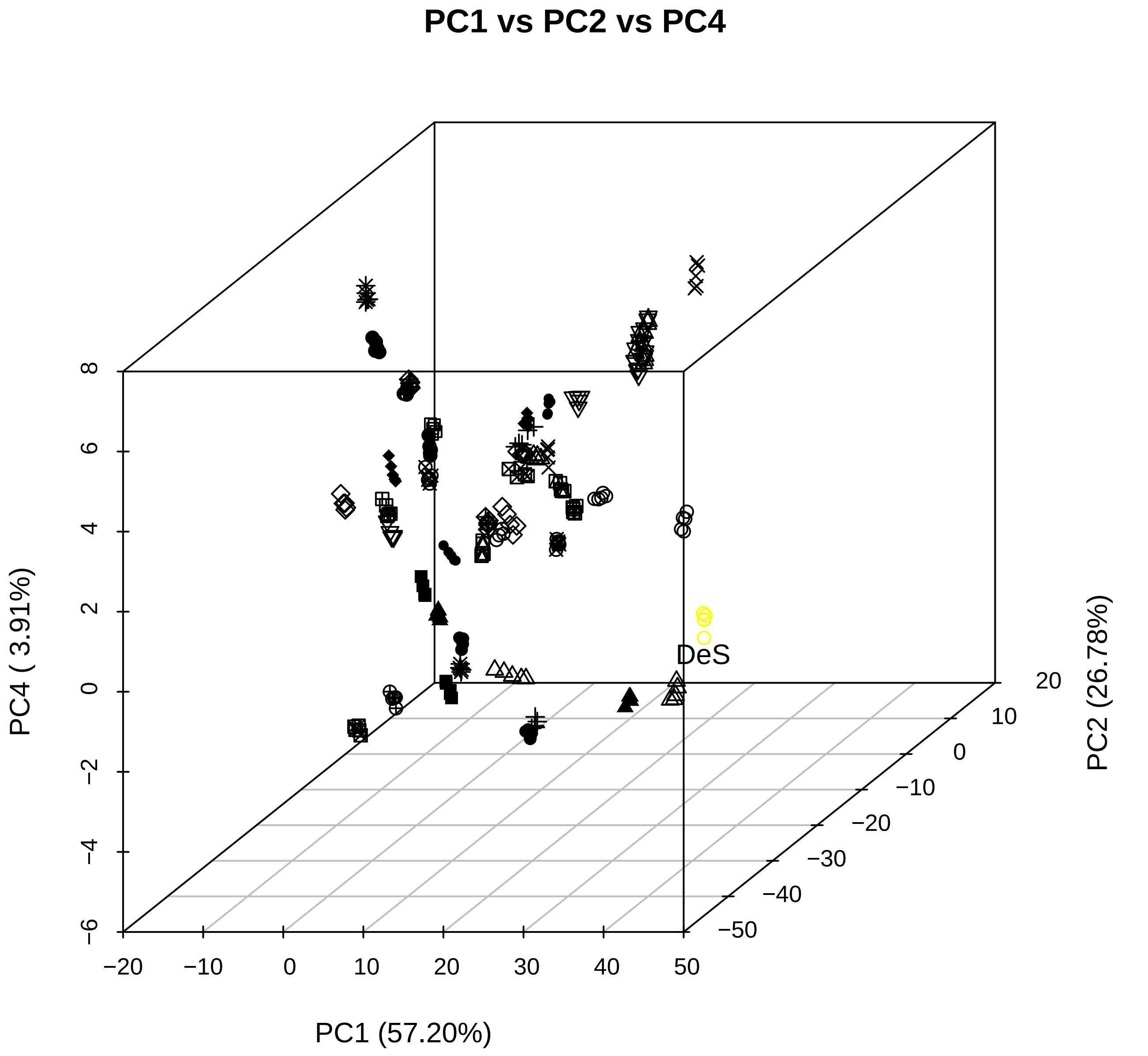
PC1 vs PC2 vs PC4
PC1 (57.20%)
PC2 (26.78%)
PC4 ( 3.91%)
DeS
−20
−10
0
10
20
30
40
50
−50
−40
−30
−20
−10
0
10
20
−6
−4
−2
0
2
4
6
8

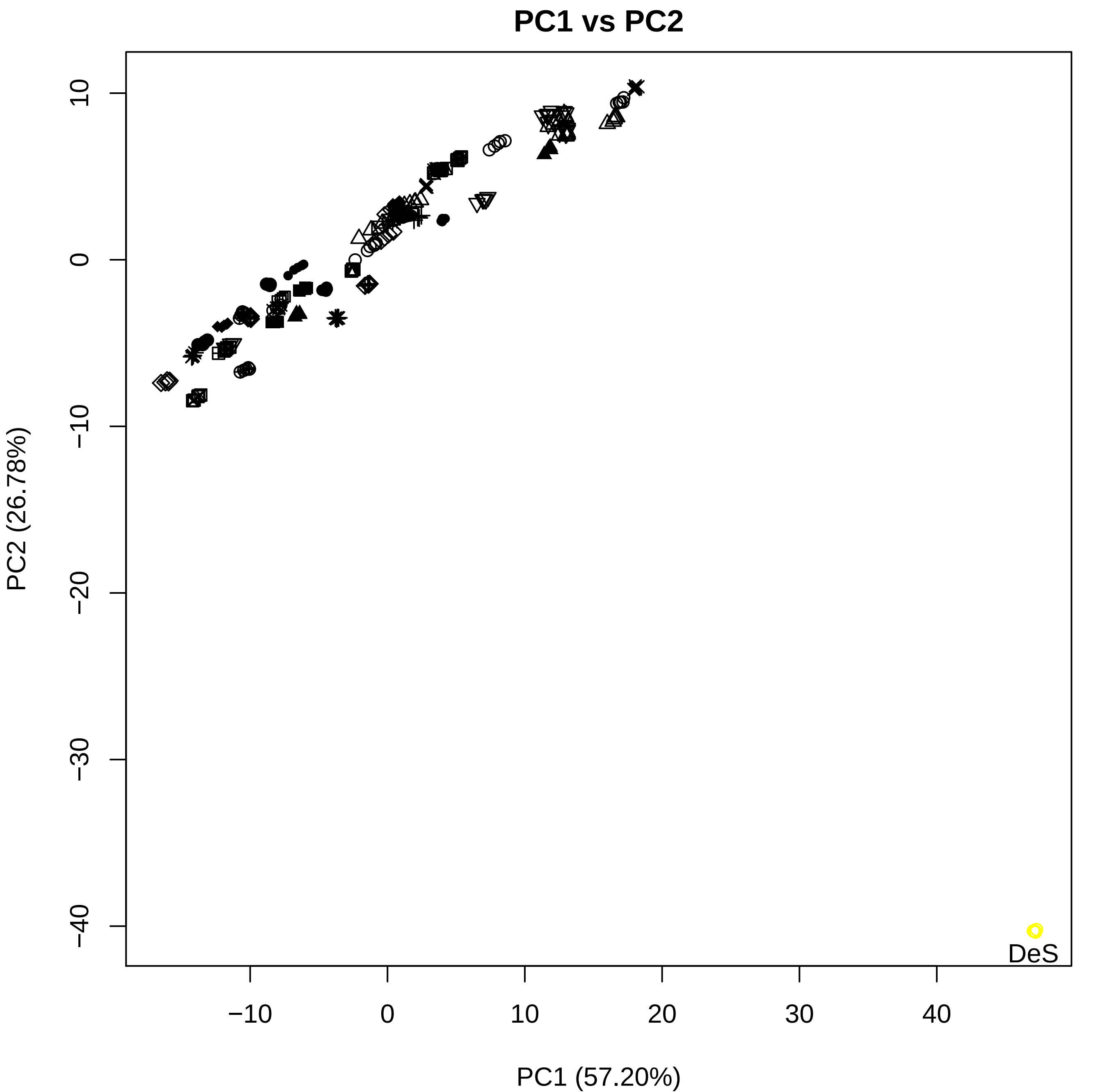
PC1 vs PC2
PC1 (57.20%)
PC2 (26.78%)
−10
0
10
20
30
40
10
0
−10
−20
−30
−40
DeS

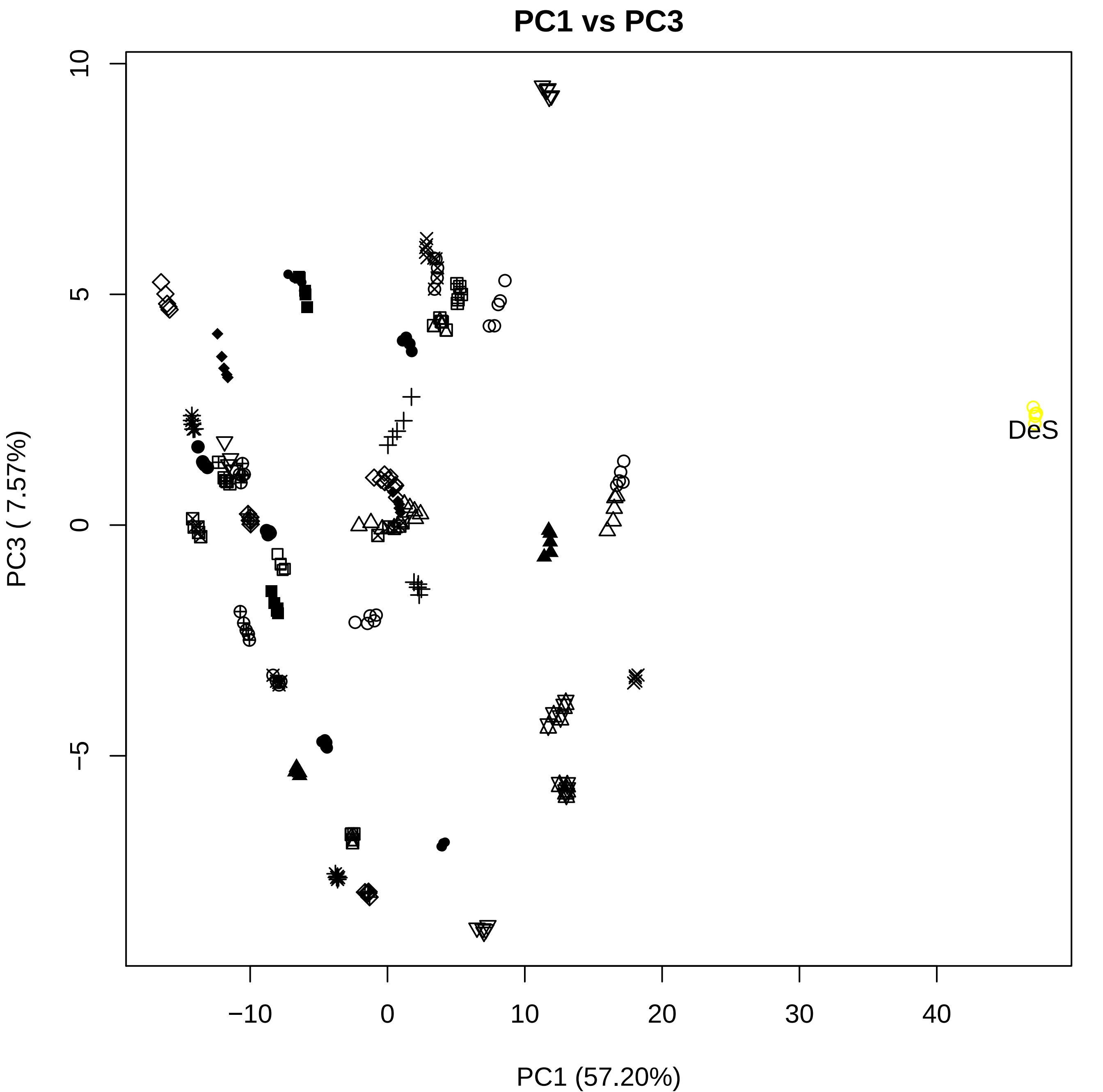

PC1 vs PC3
10
5
0
−5
PC3 ( 7.57%)
−10
0
10
20
30
40
PC1 (57.20%)
DeS

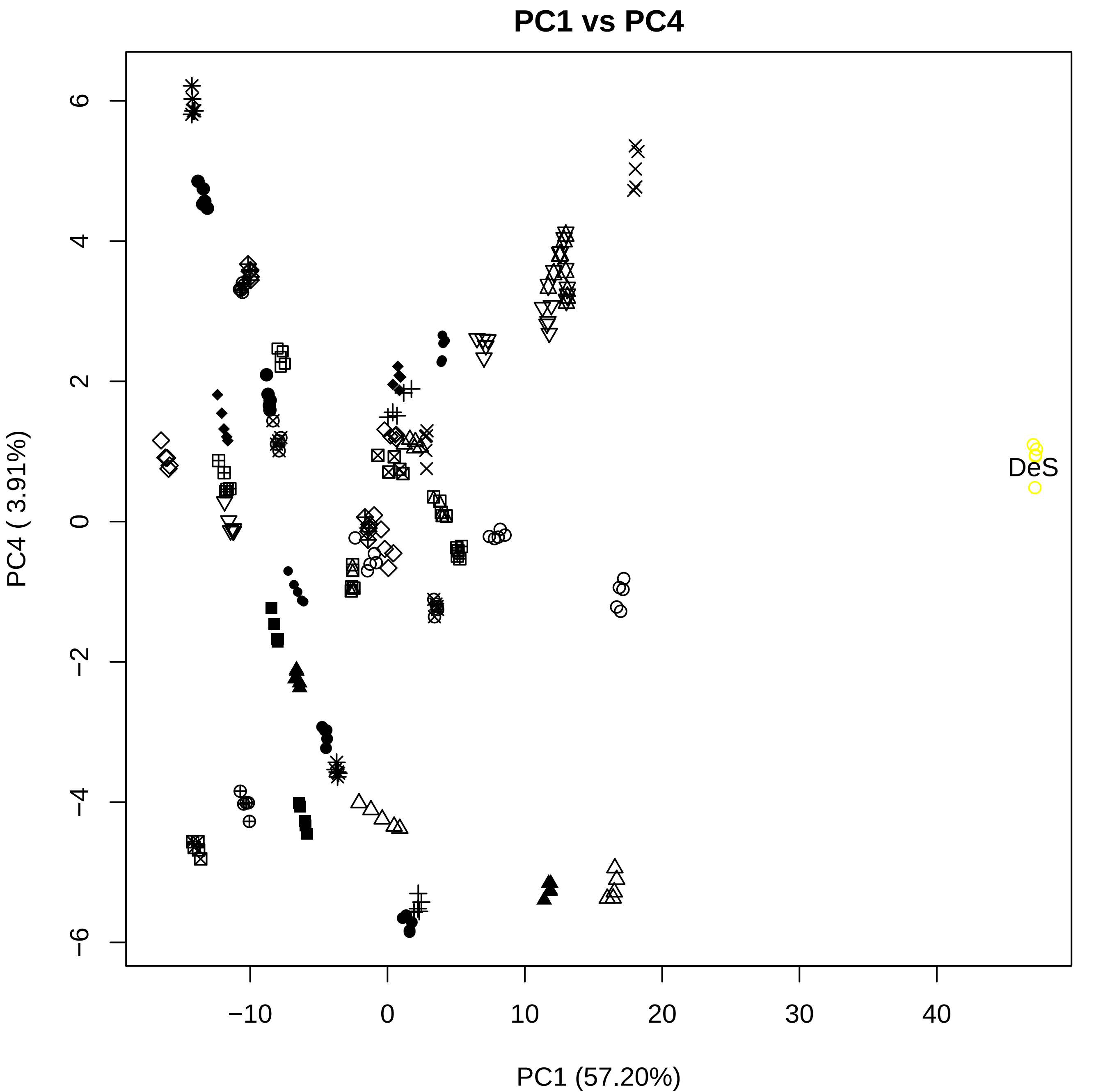

PC1 vs PC4
DeS
PC4 ( 3.91%)
PC1 (57.20%)
6
4
2
0
−2
−4
−6
−10
0
10
20
30
40

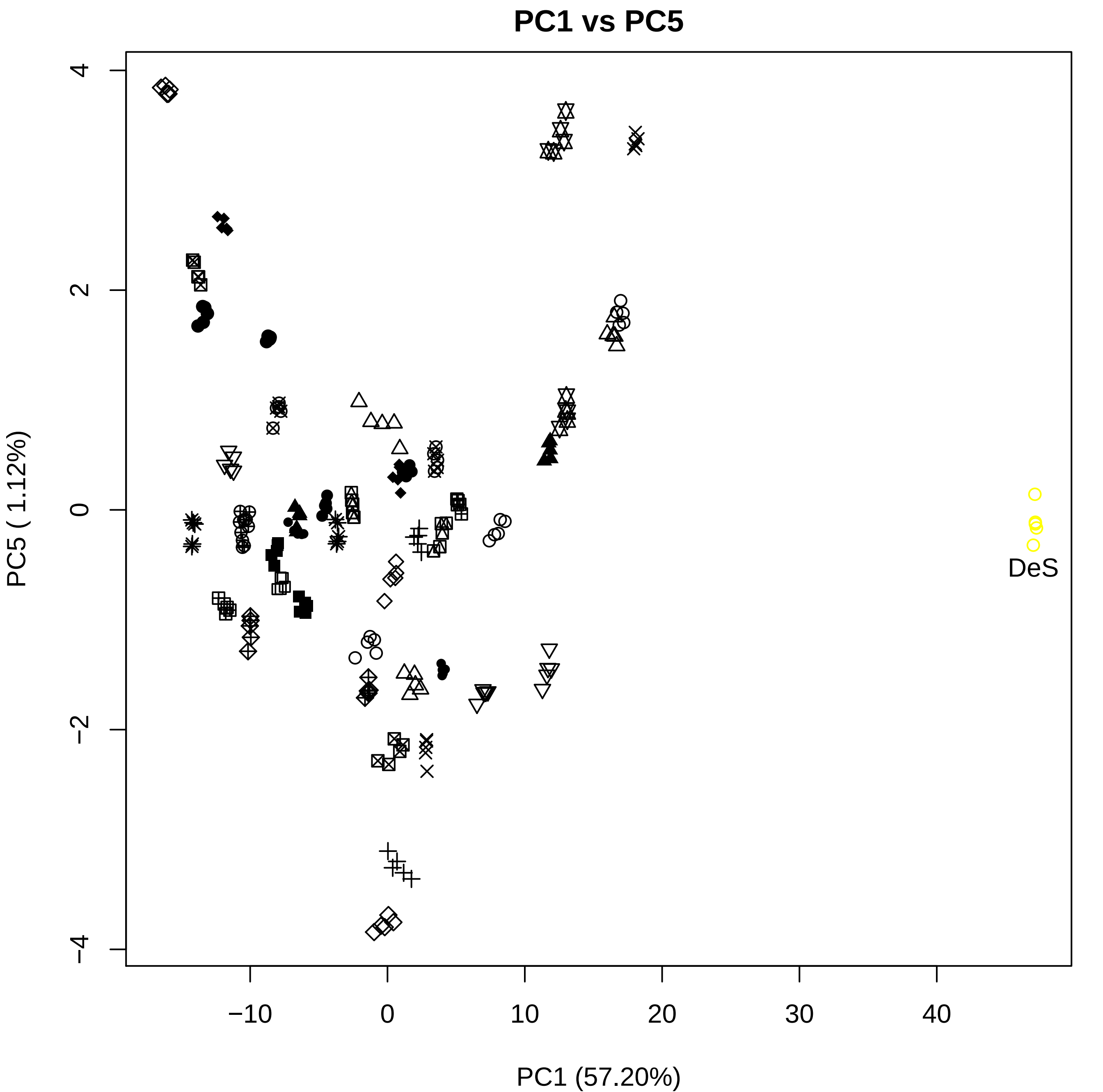
PC1 vs PC5
PC5 ( 1.12%)
PC1 (57.20%)
DeS
4
2
0
−2
−4
−10
0
10
20
30
40

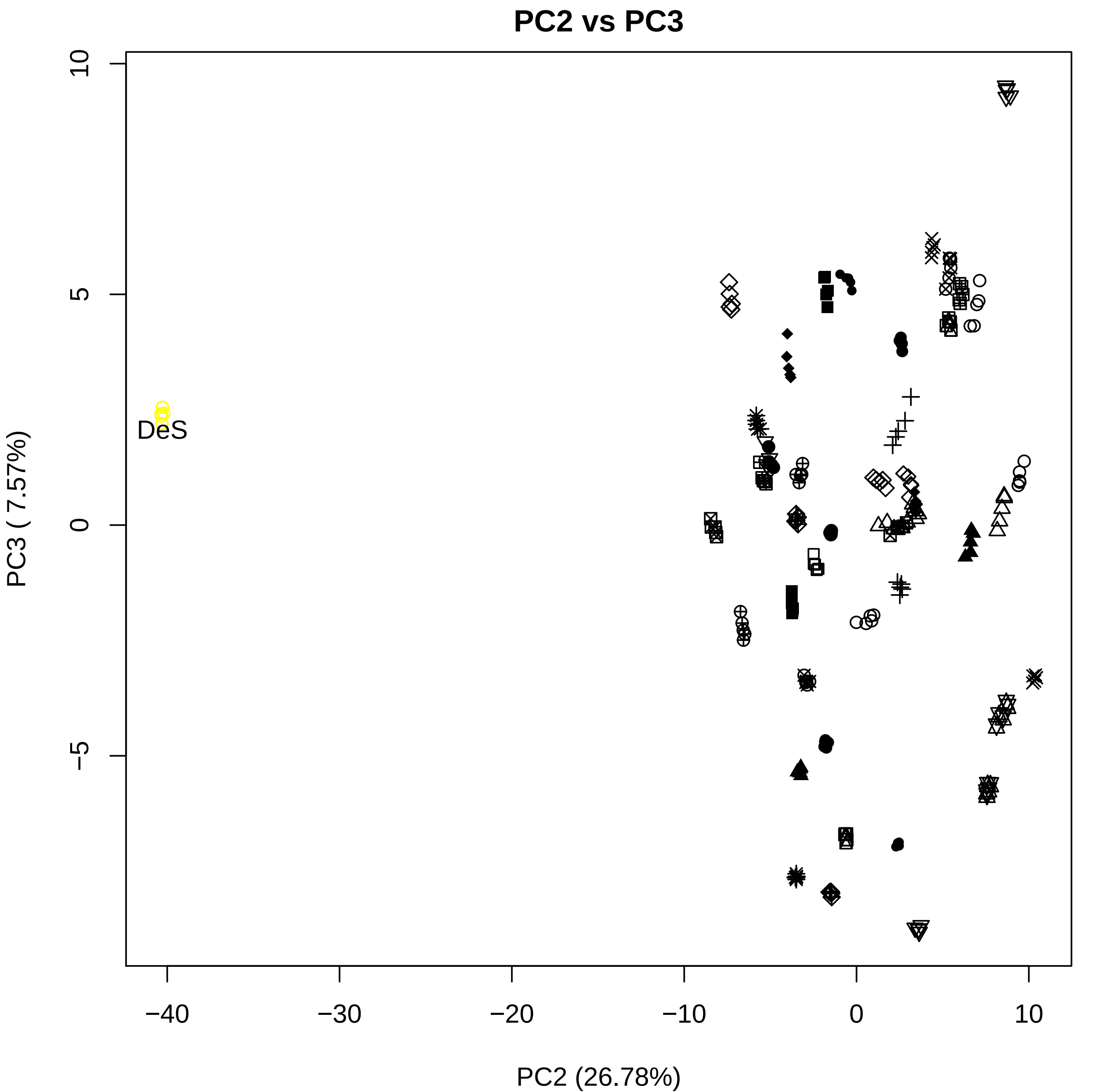

PC2 vs PC3
DeS
PC3 ( 7.57%)
PC2 (26.78%)
10
5
0
−5
−40
−30
−20
−10
0
10

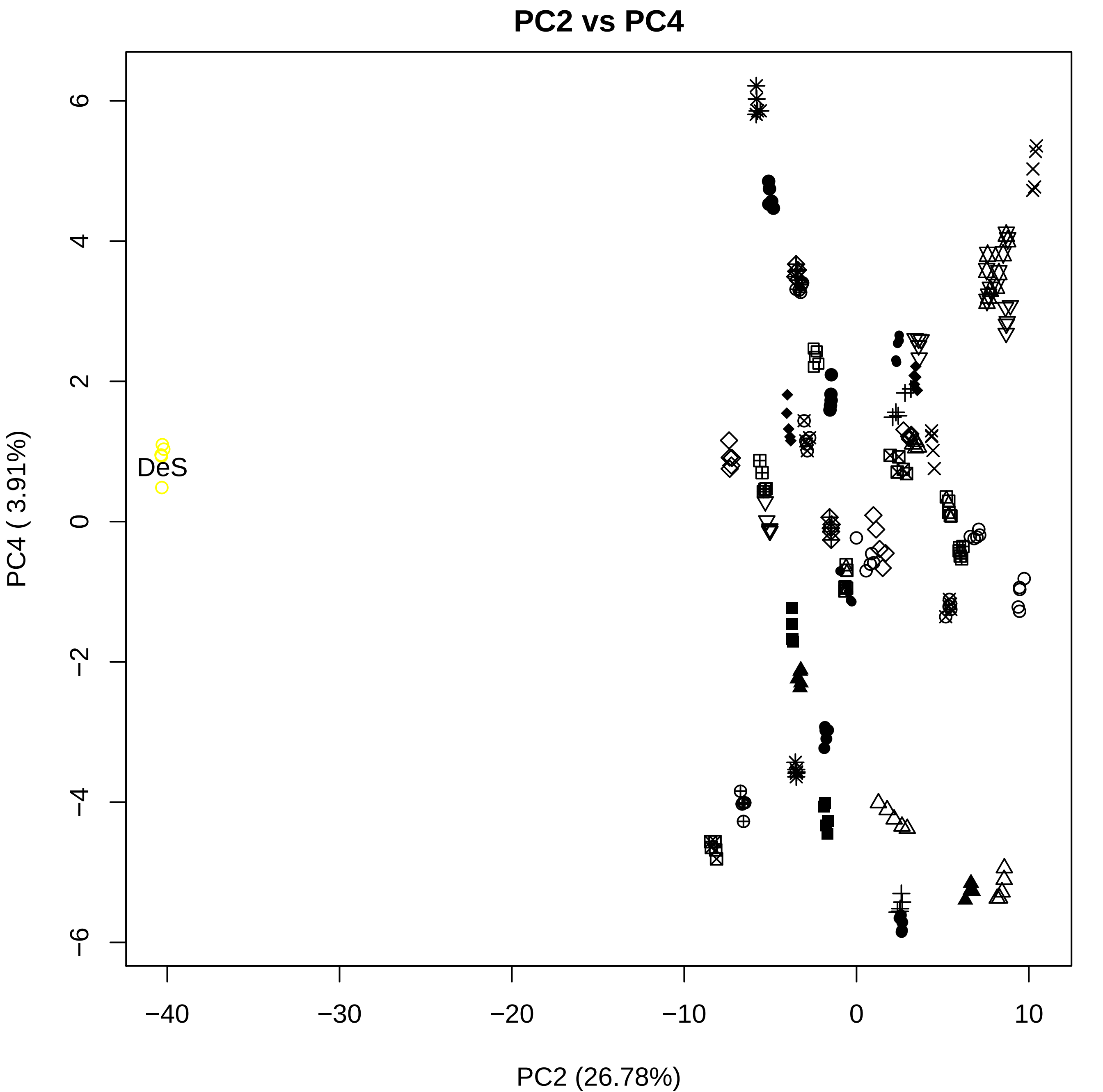
PC2 vs PC4
PC4 ( 3.91%)
PC2 (26.78%)
DeS
6
4
2
0
−2
−4
−6
−40
−30
−20
−10
0
10

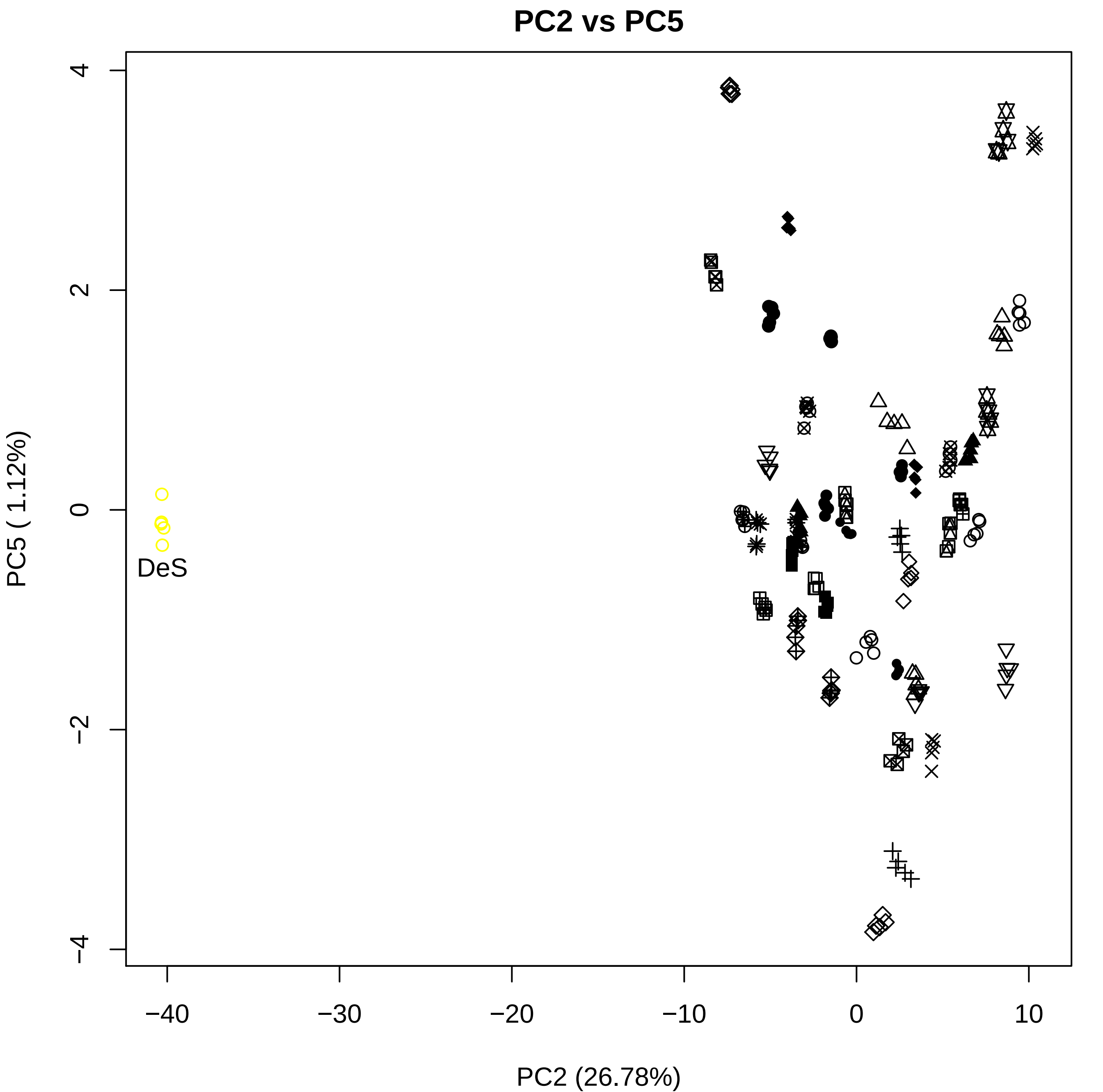
PC2 vs PC5
PC5 ( 1.12%)
PC2 (26.78%)
DeS
4
2
0
−2
−4
−40
−30
−20
−10
0
10

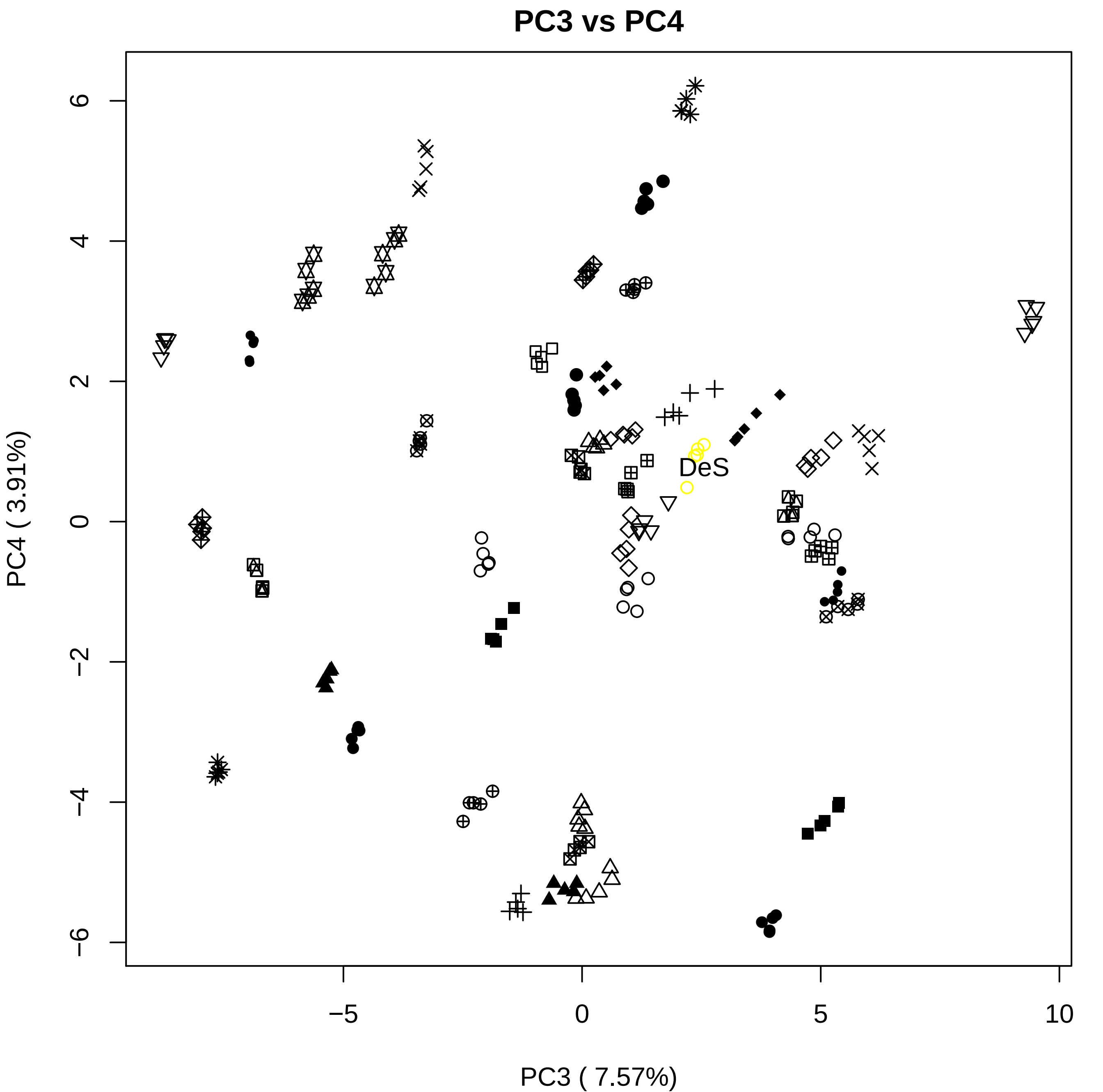
PC3 vs PC4
PC4 ( 3.91%)
PC3 ( 7.57%)
DeS
6
4
2
0
−2
−4
−6
−5
0
5
10

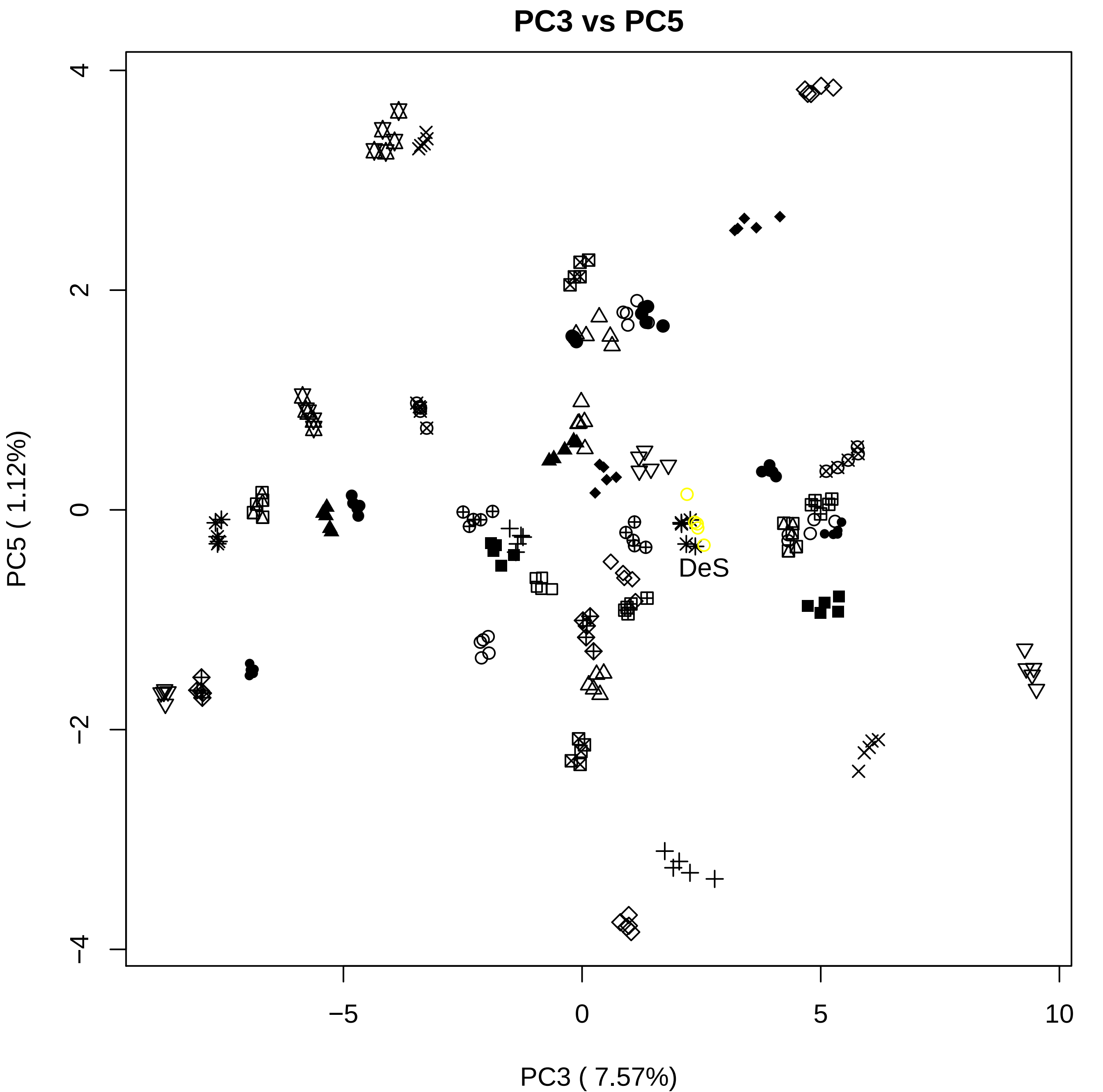

PC3 vs PC5
PC5 ( 1.12%)
PC3 ( 7.57%)
DeS
4
2
0
−2
−4
−5
0
5
10

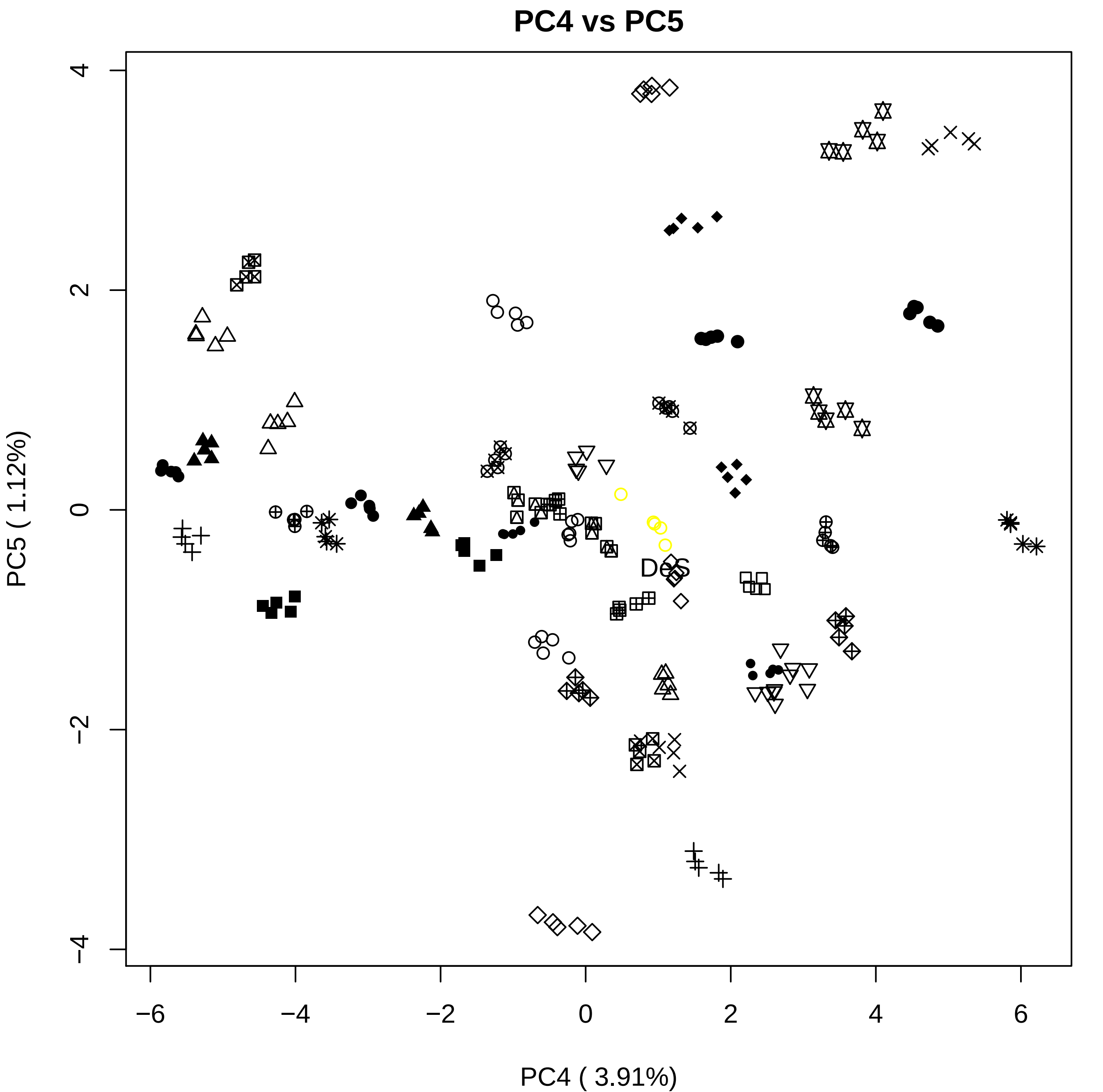
PC4 vs PC5
PC5 ( 1.12%)
PC4 ( 3.91%)
4
2
0
−2
−4
−6
−4
−2
0
2
4
6
DeS

### HpSp for paper new Prep

Anthony Devlin, Keele University

### PCA of All

Scaled Matrix PCAed

7th order polynomial baseline correction performed

Raw spectra smoothed to 21 Neighbours, 2nd order PN, Savtisky Golay

Normalisation between 0 and 100

D2 spectra smoothed to 41 Neighbours, 2nd order PN, Savtisky Golay

Regions Analysed: 3601−2501 and 2001−700cm−1

Samples Analysed: 0 bContam's 16 BLH's 50 BMH's 30 OMH's 0 pContam's
0 PLH's 60 PMH's

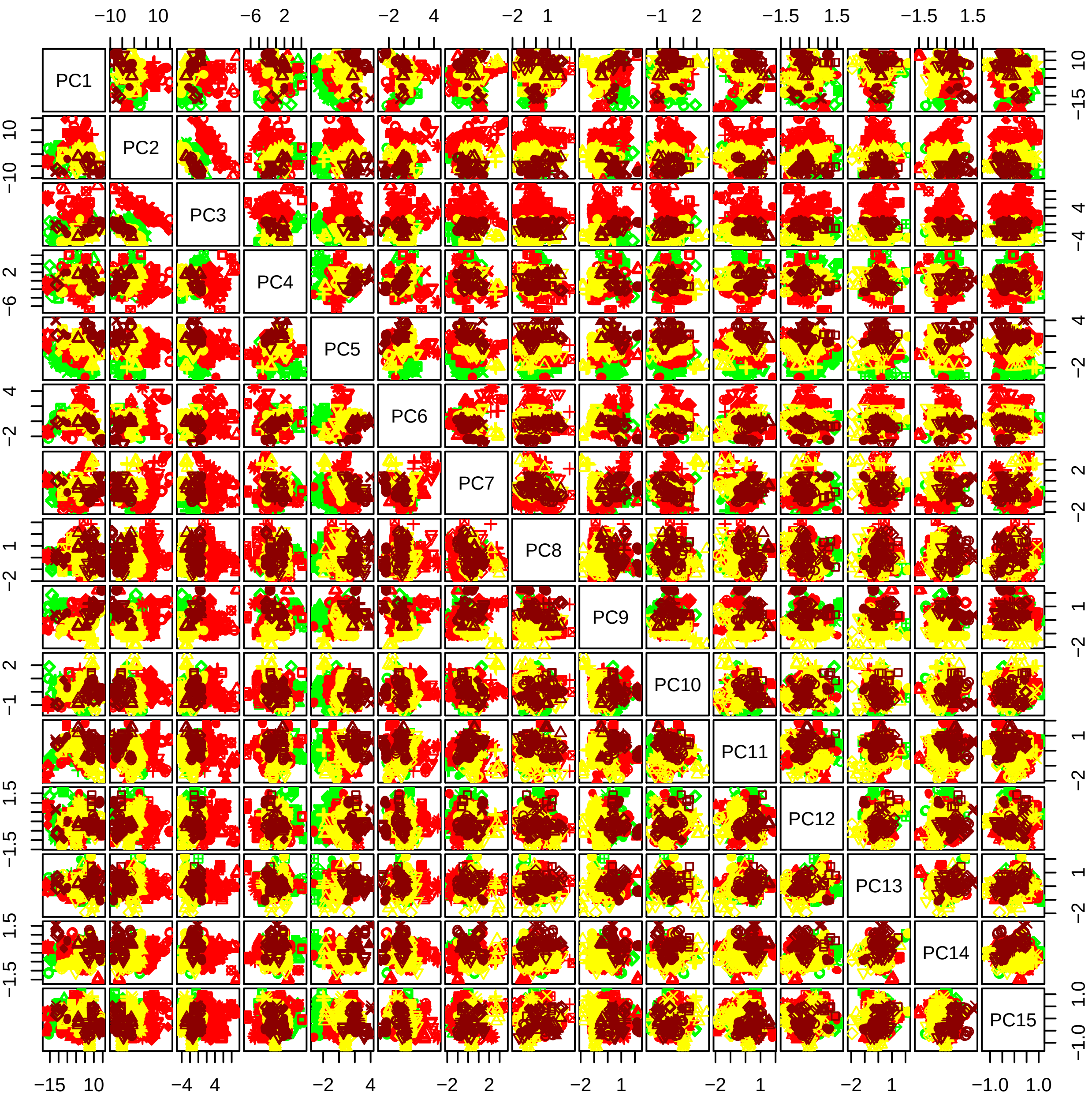

PC1
PC2
PC3
PC4
PC5
PC6
PC7
PC8
PC9
PC10
PC11
PC12
PC13
PC14
PC15

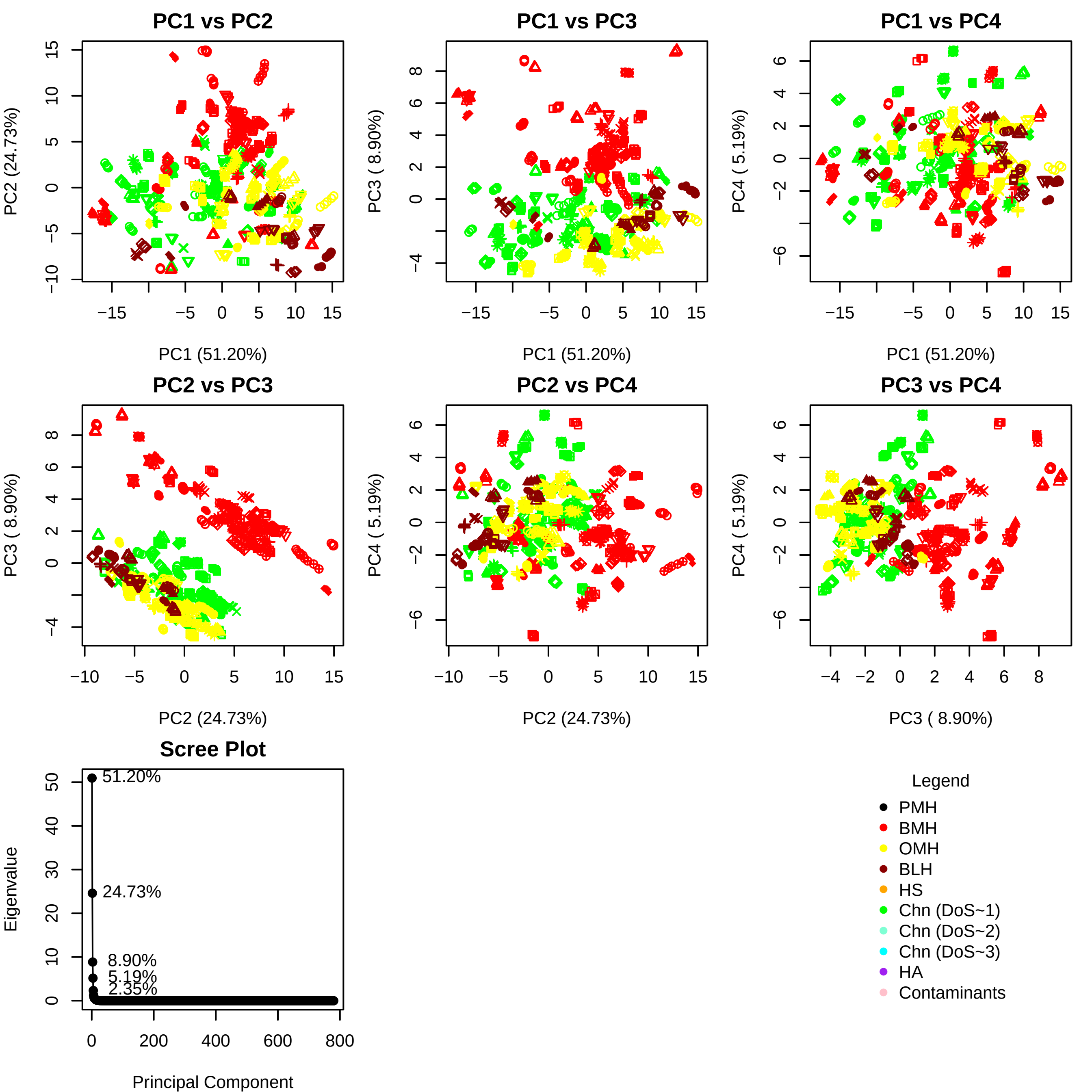

PC1 vs PC2
PC1 vs PC3
PC1 vs PC4
PC2 vs PC3
PC2 vs PC4
PC3 vs PC4
Scree Plot
PC1 (51.20%)
PC2 (24.73%)
PC3 ( 8.90%)
PC4 ( 5.19%)
Eigenvalue
Principal Component
51.20%
24.73%
8.90%
5.19%
2.35%
Legend
PMH
BMH
OMH
BLH
HS
Chn (DoS~1)
Chn (DoS~2)
Chn (DoS~3)
HA
Contaminants

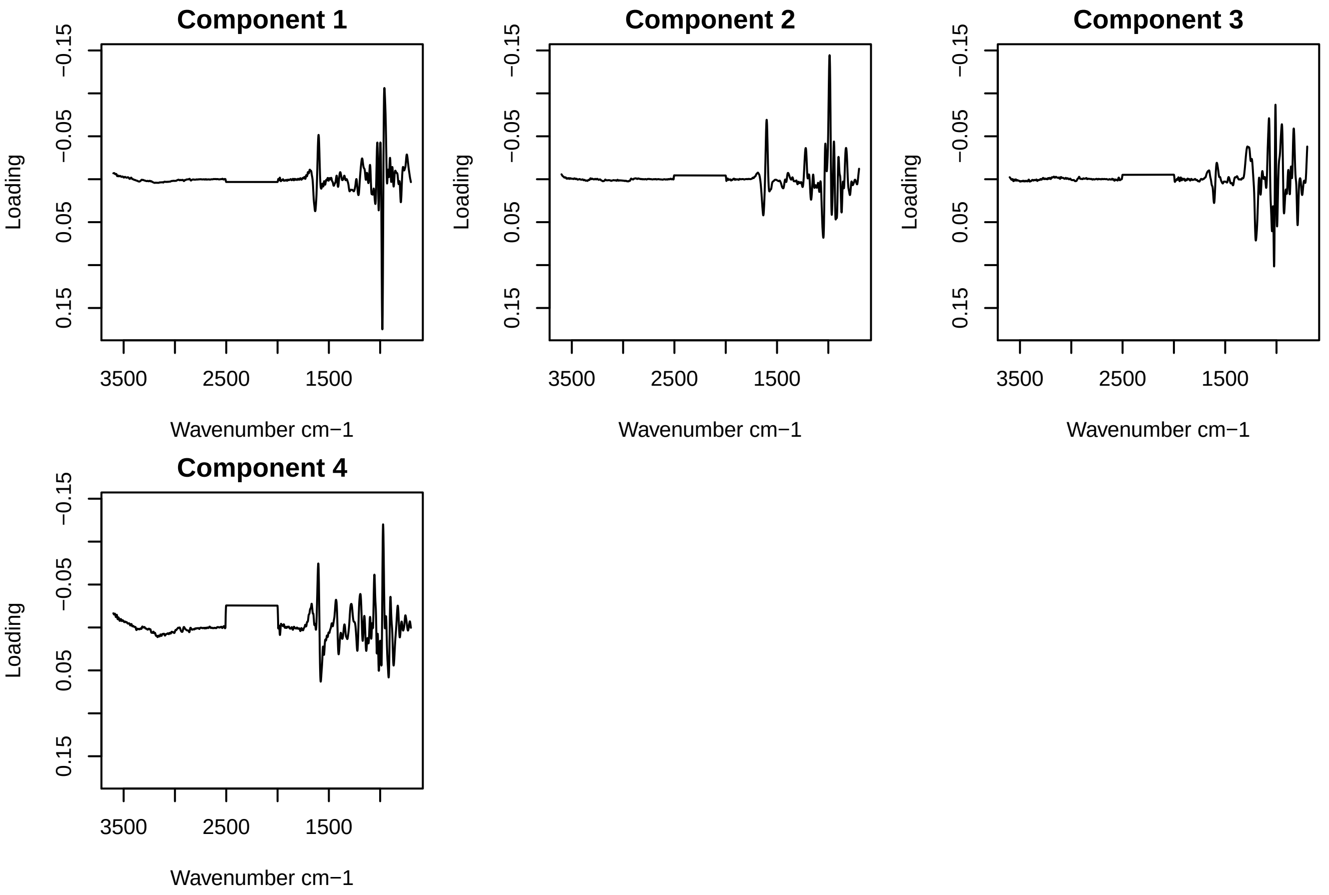

Component 1
Component 2
Component 3
Component 4
Loading
−0.15
−0.05
0.05
0.15
3500
2500
1500
Wavenumber cm−1

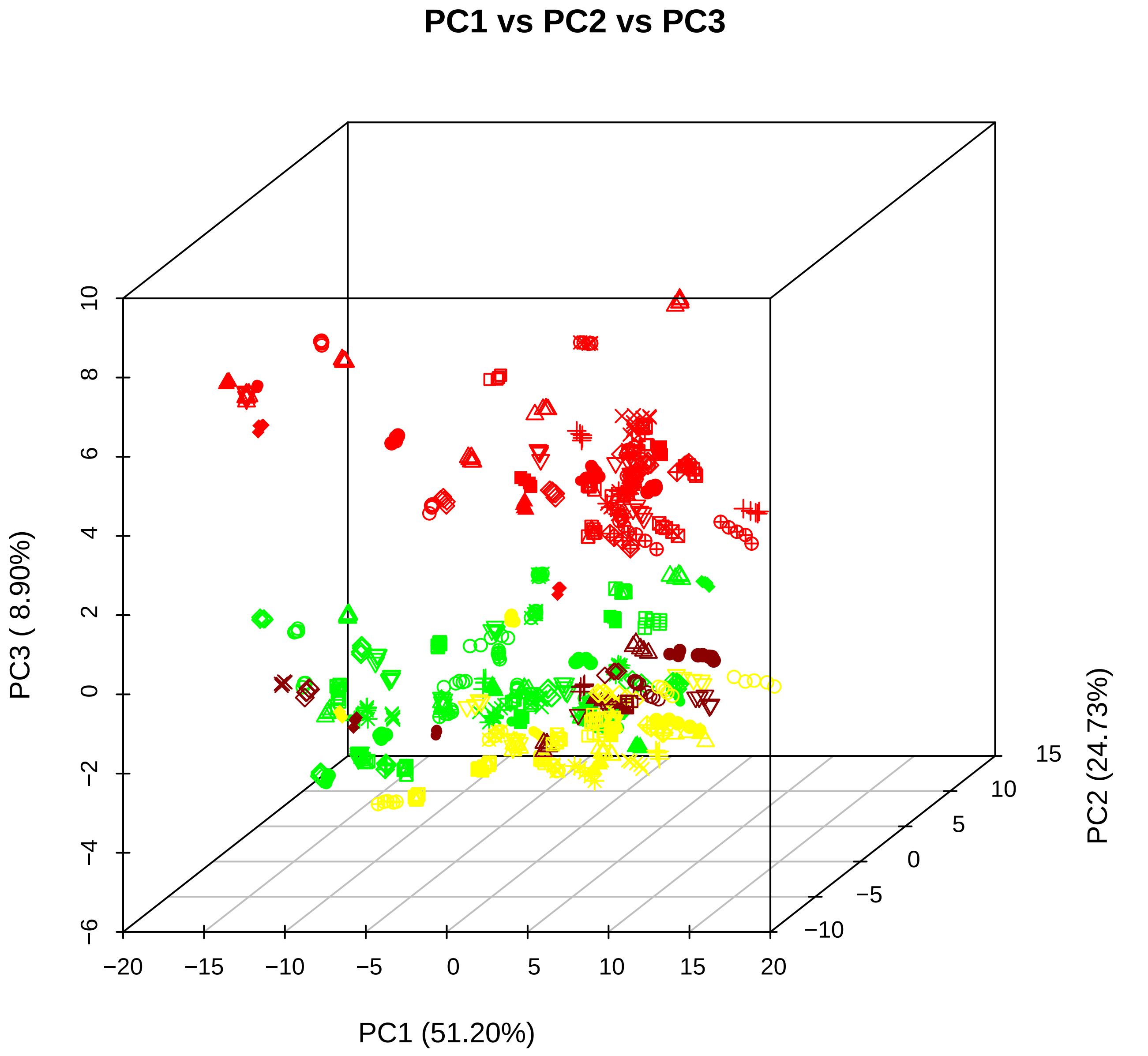
PC1 vs PC2 vs PC3
PC1 (51.20%)
PC2 (24.73%)
PC3 ( 8.90%)
−20 −15 −10 −5 0 5 10 15 20
−10 −5 0 5 10 15
−6 −4 −2 0 2 4 6 8 10

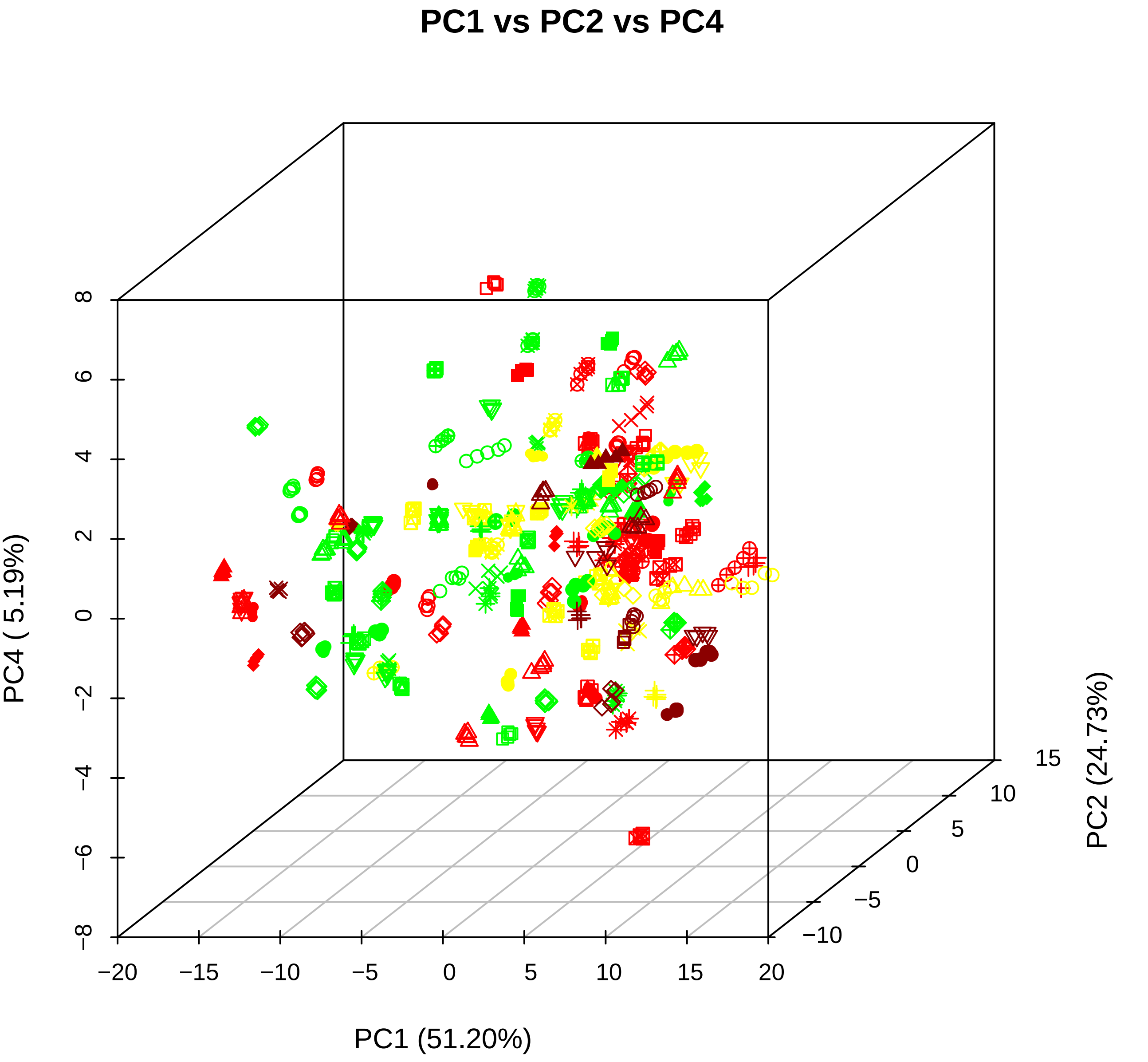
PC1 vs PC2 vs PC4
PC1 (51.20%)
PC2 (24.73%)
PC4 ( 5.19%)
−20
−15
−10
−5
0
5
10
15
20
8
6
4
2
0
−2
−4
−6
−8
15
10
5
0
−5
−10

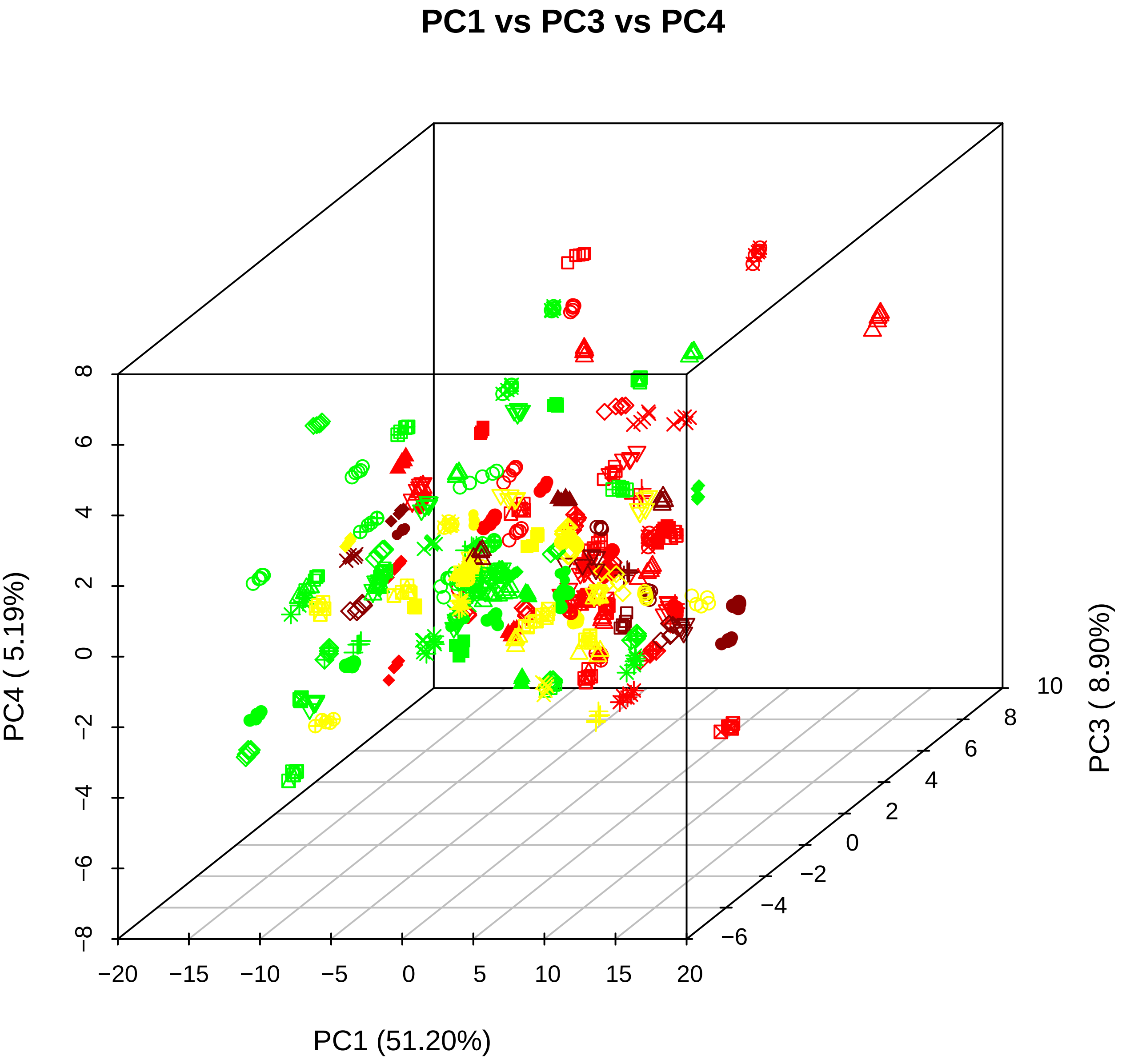

PC1 vs PC3 vs PC4
PC1 (51.20%)
PC4 ( 5.19%)
PC3 ( 8.90%)
−20 −15 −10 −5 0 5 10 15 20
−8 −6 −4 −2 0 2 4 6 8
−6 −4 −2 0 2 4 6 8 10

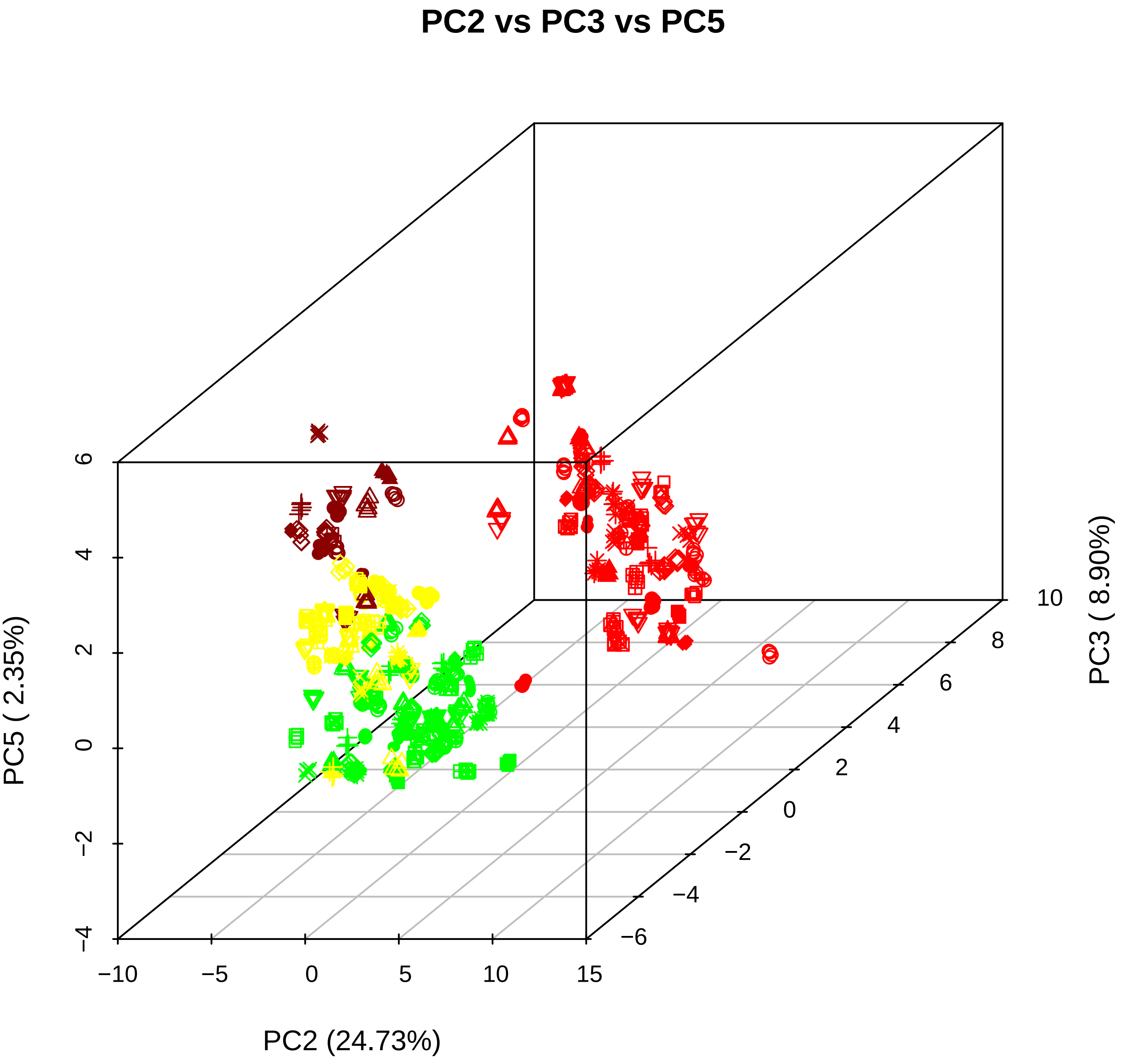
PC2 vs PC3 vs PC5
PC2 (24.73%)
PC3 ( 8.90%)
PC5 ( 2.35%)
−10
−5
0
5
10
15
−6
−4
−2
0
2
4
6
8
10
−4
−2
0
2
4
6

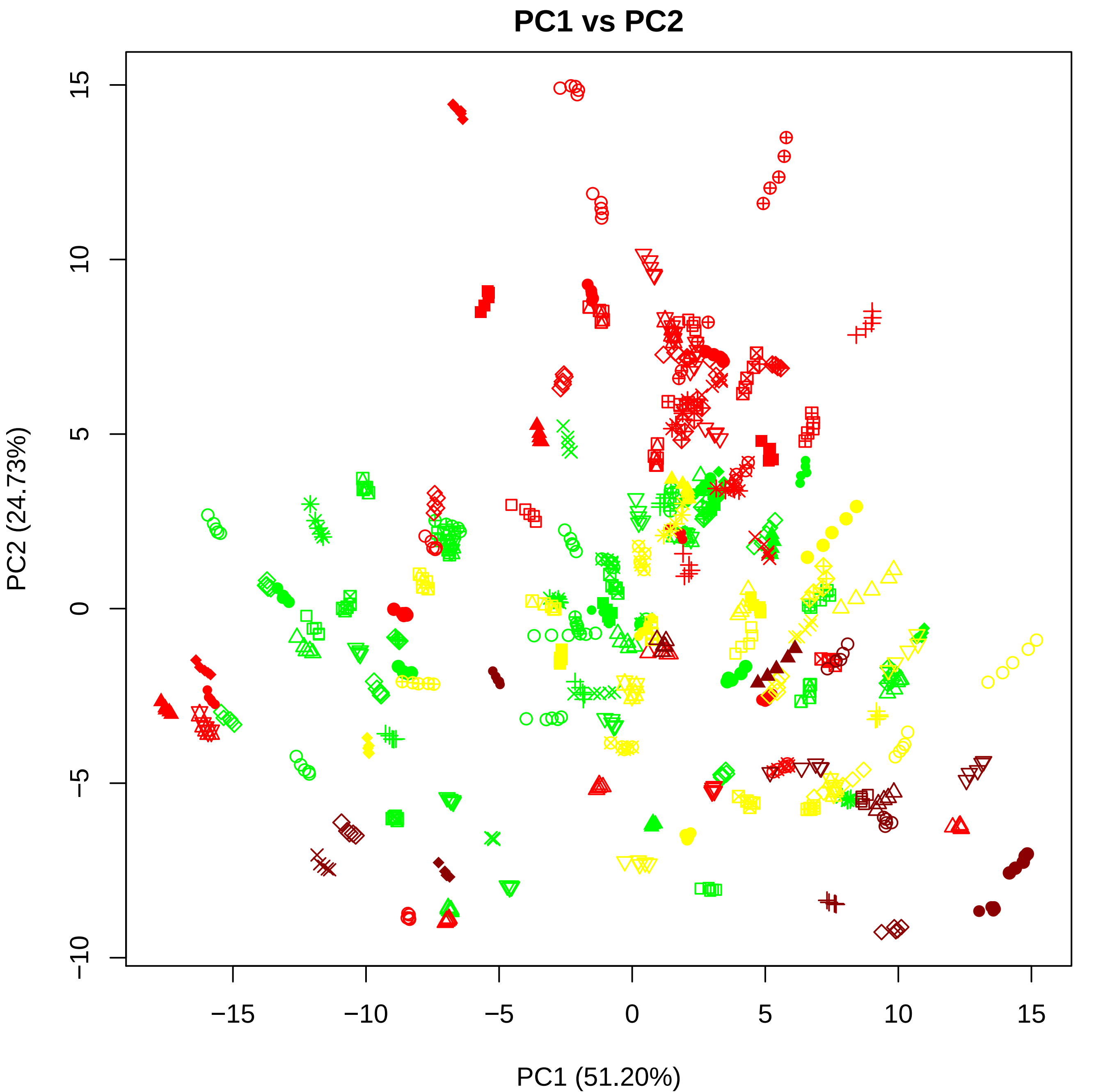
PC1 vs PC2
PC2 (24.73%)
PC1 (51.20%)
15
10
5
0
−5
−10
−15
−10
−5
0
5
10
15

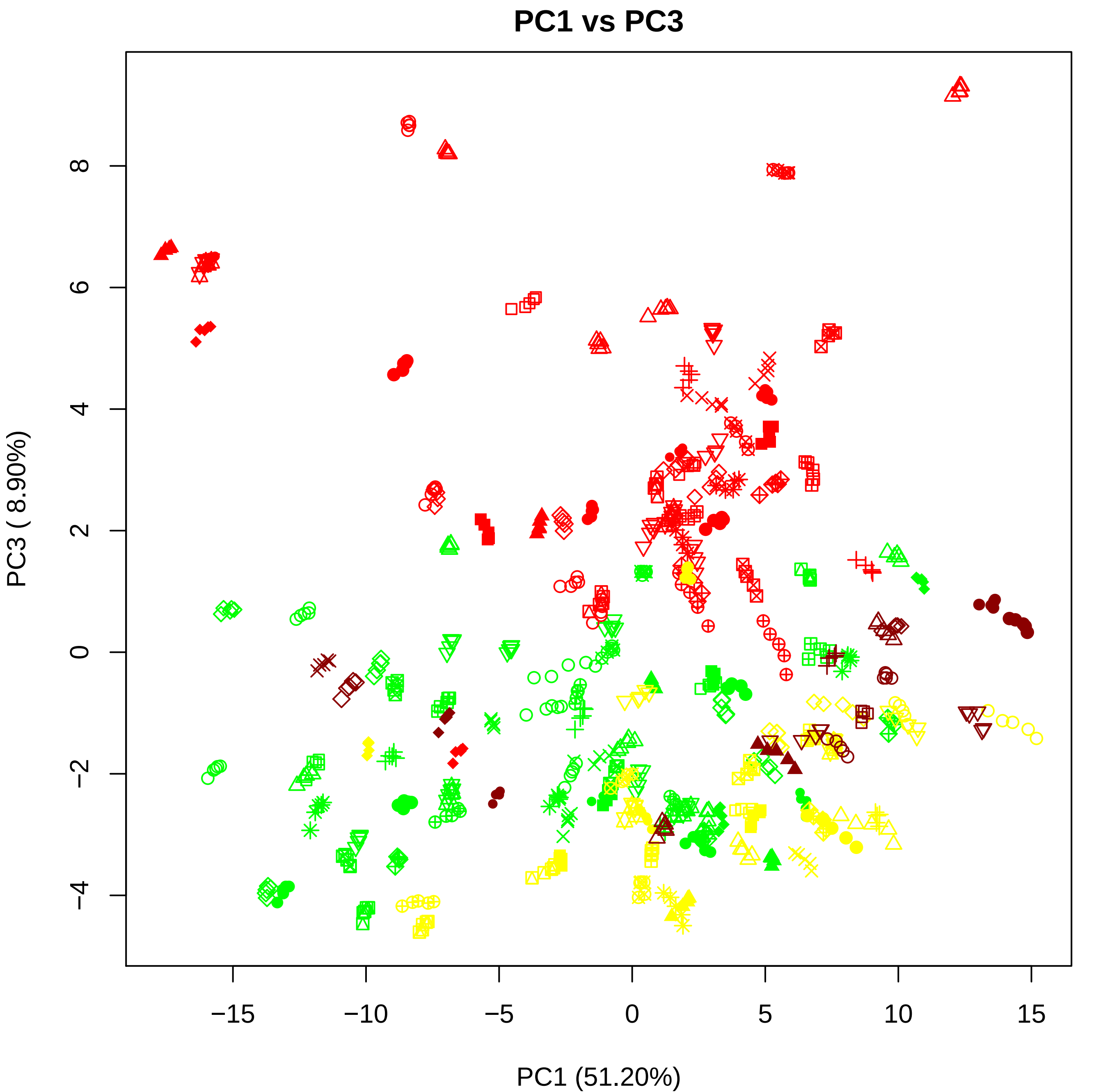
PC1 vs PC3
PC3 ( 8.90%)
PC1 (51.20%)
−15
−10
−5
0
5
10
15
−4
−2
0
2
4
6
8

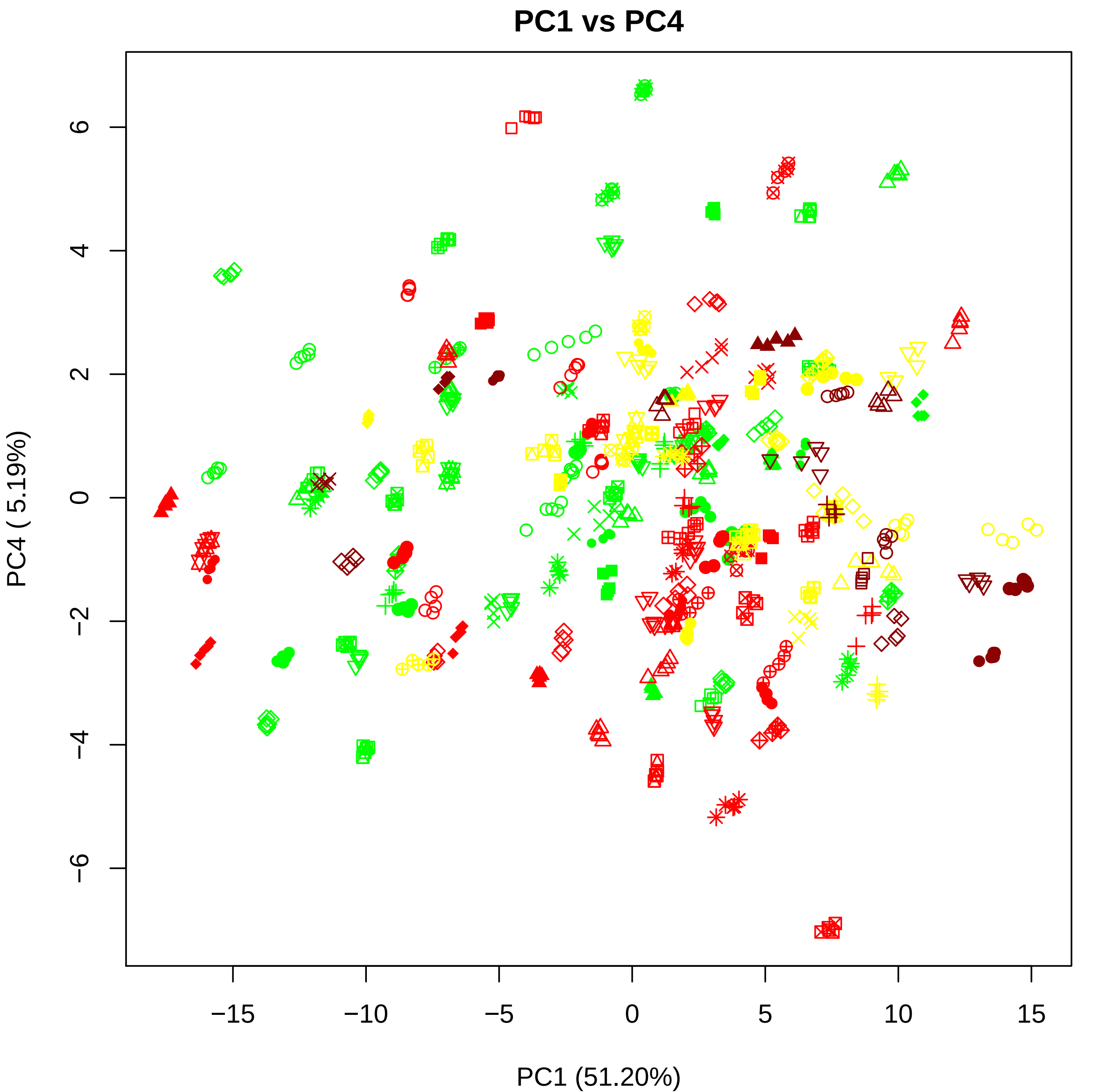
PC1 vs PC4
PC4 ( 5.19%)
PC1 (51.20%)
6
4
2
0
−2
−4
−6
−15
−10
−5
0
5
10
15

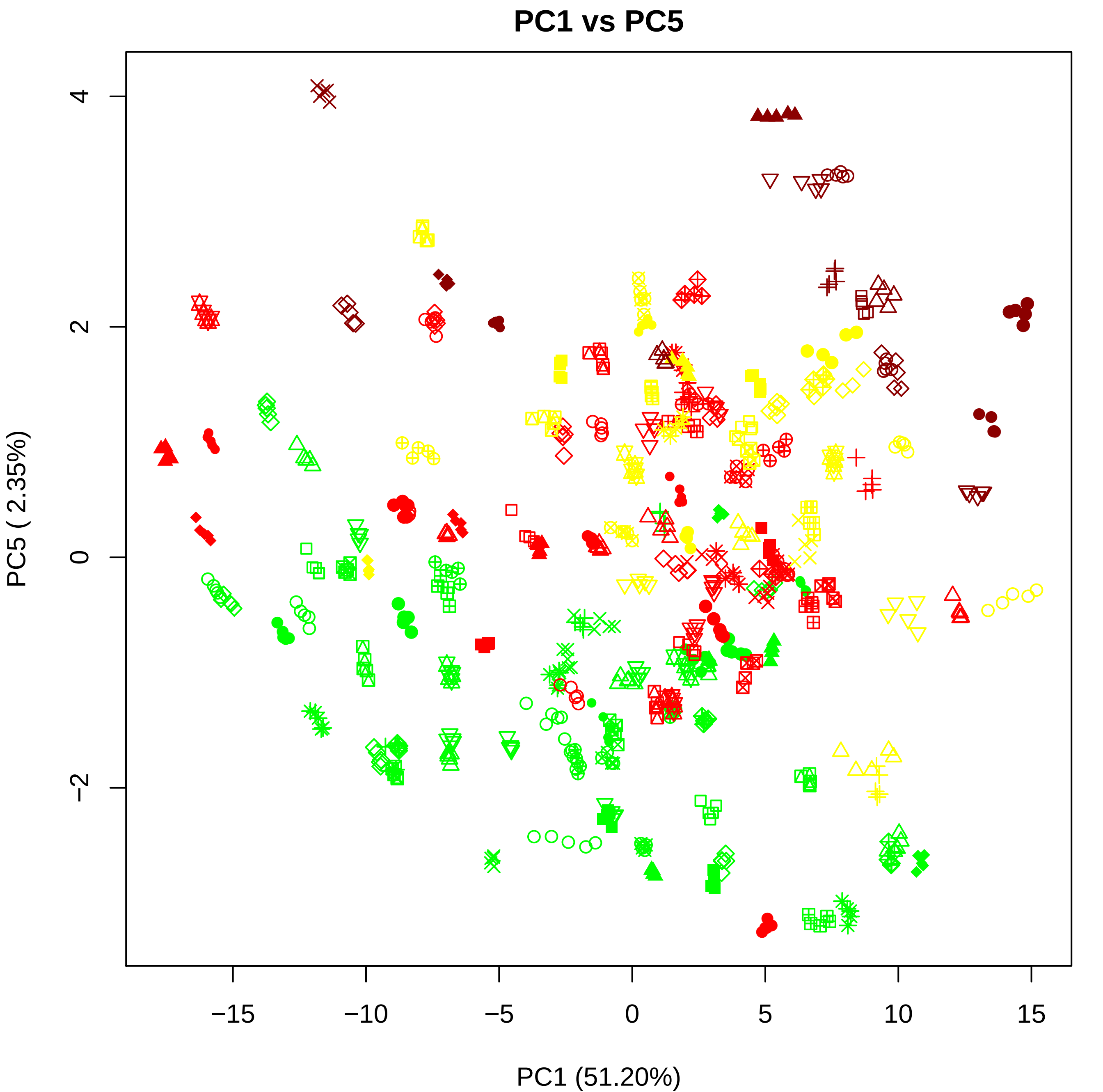
PC1 vs PC5
PC5 ( 2.35%)
PC1 (51.20%)
4
2
0
−2
−15
−10
−5
0
5
10
15

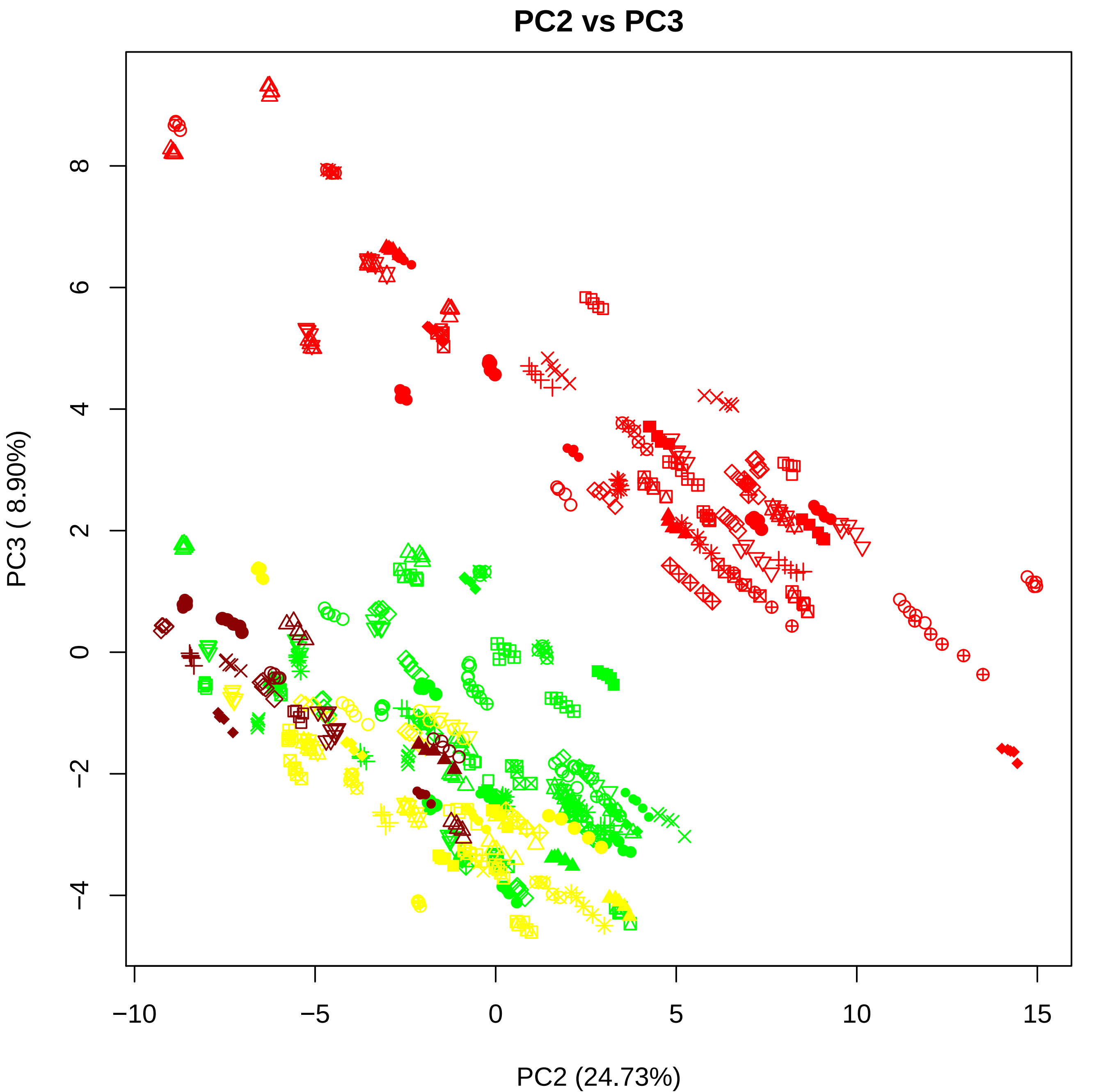

PC2 vs PC3
PC3 ( 8.90%)
PC2 (24.73%)
−10
−5
0
5
10
15
−4
−2
0
2
4
6
8

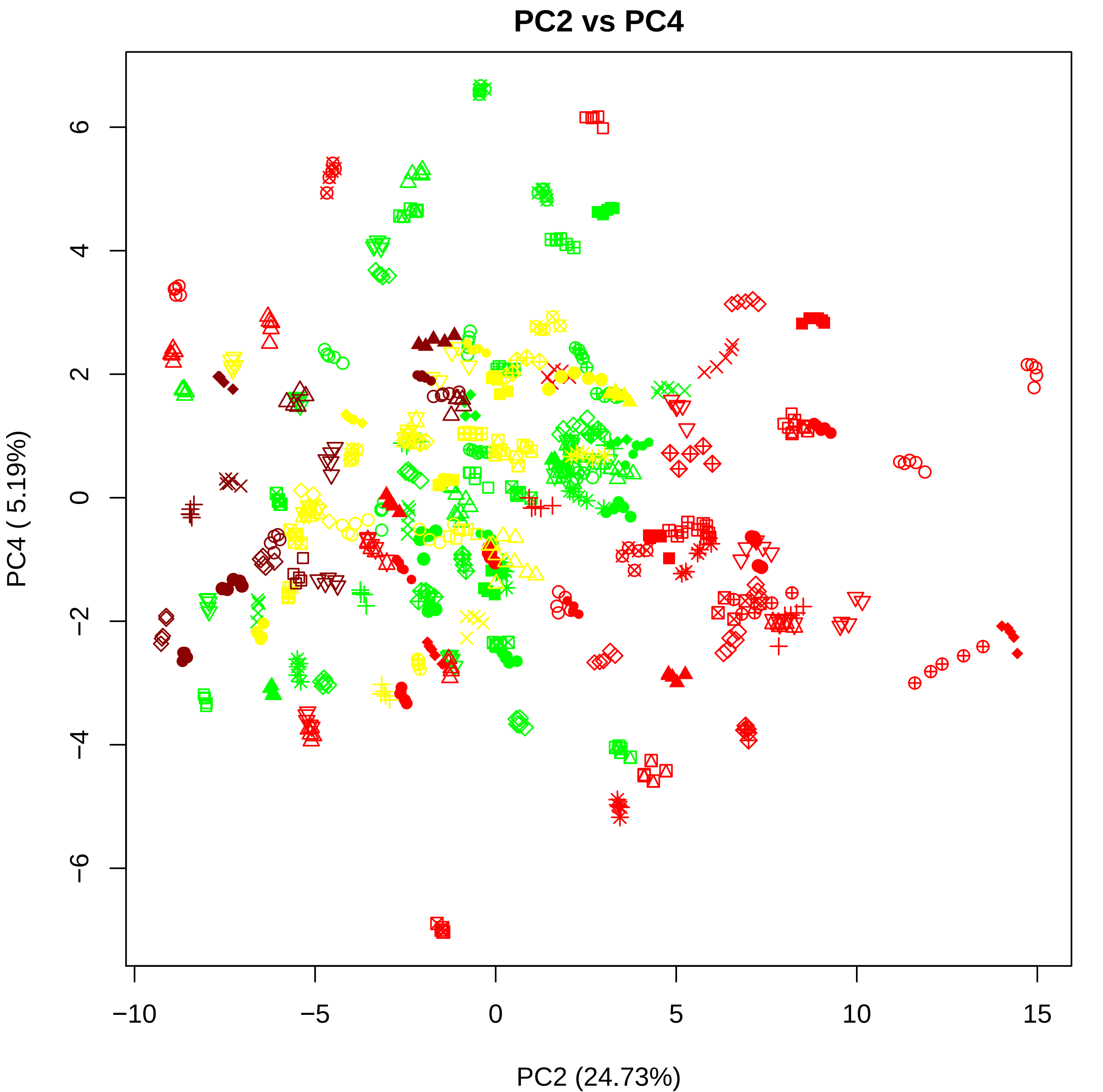

PC2 vs PC4
PC4 ( 5.19%)
PC2 (24.73%)
6
4
2
0
−2
−4
−6
−10
−5
0
5
10
15

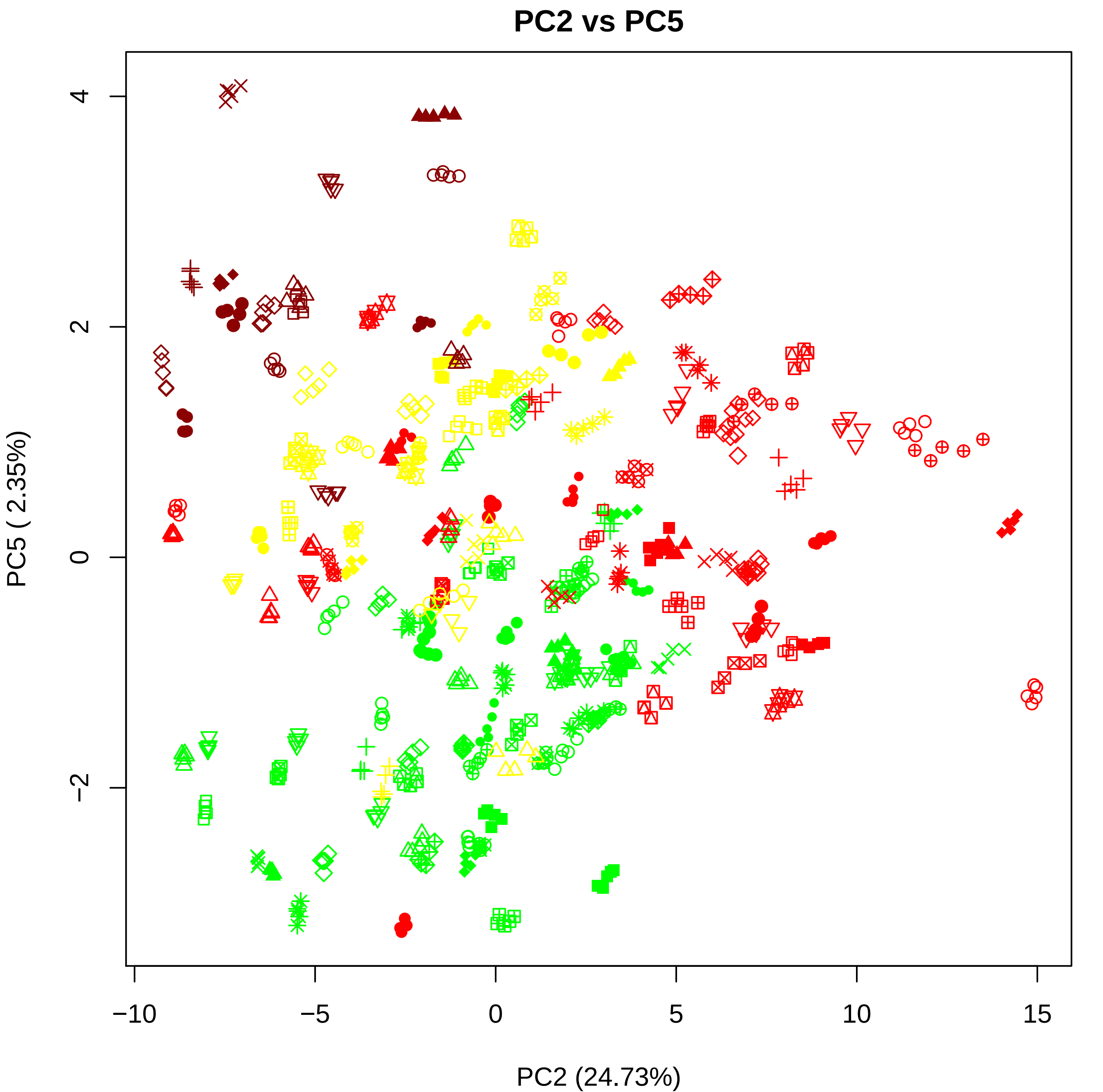
PC2 vs PC5
PC5 ( 2.35%)
PC2 (24.73%)
4
2
0
−2
−10
−5
0
5
10
15

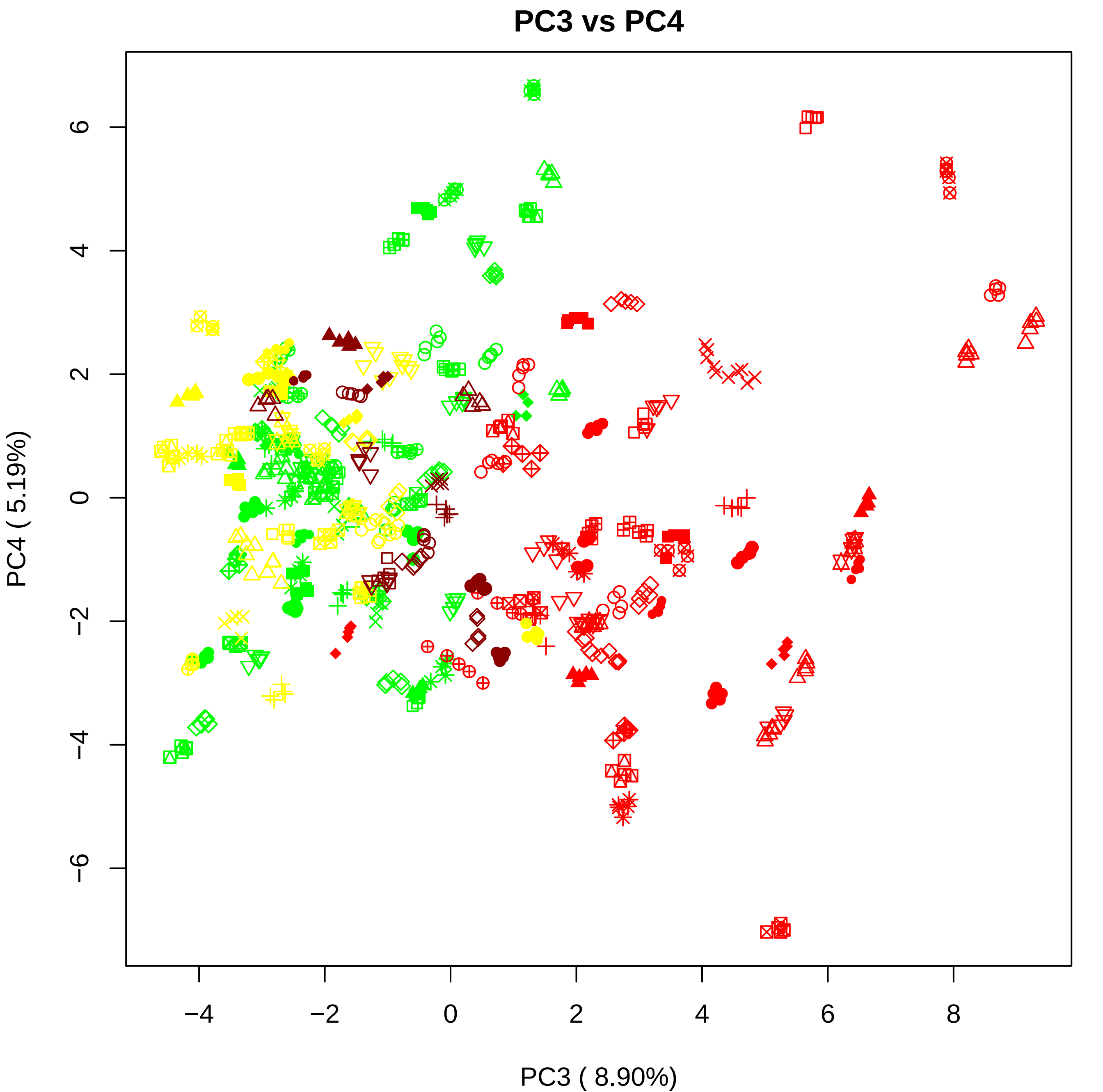
PC3 vs PC4
PC4 ( 5.19%)
PC3 ( 8.90%)
6
4
2
0
−2
−4
−6
−4
−2
0
2
4
6
8

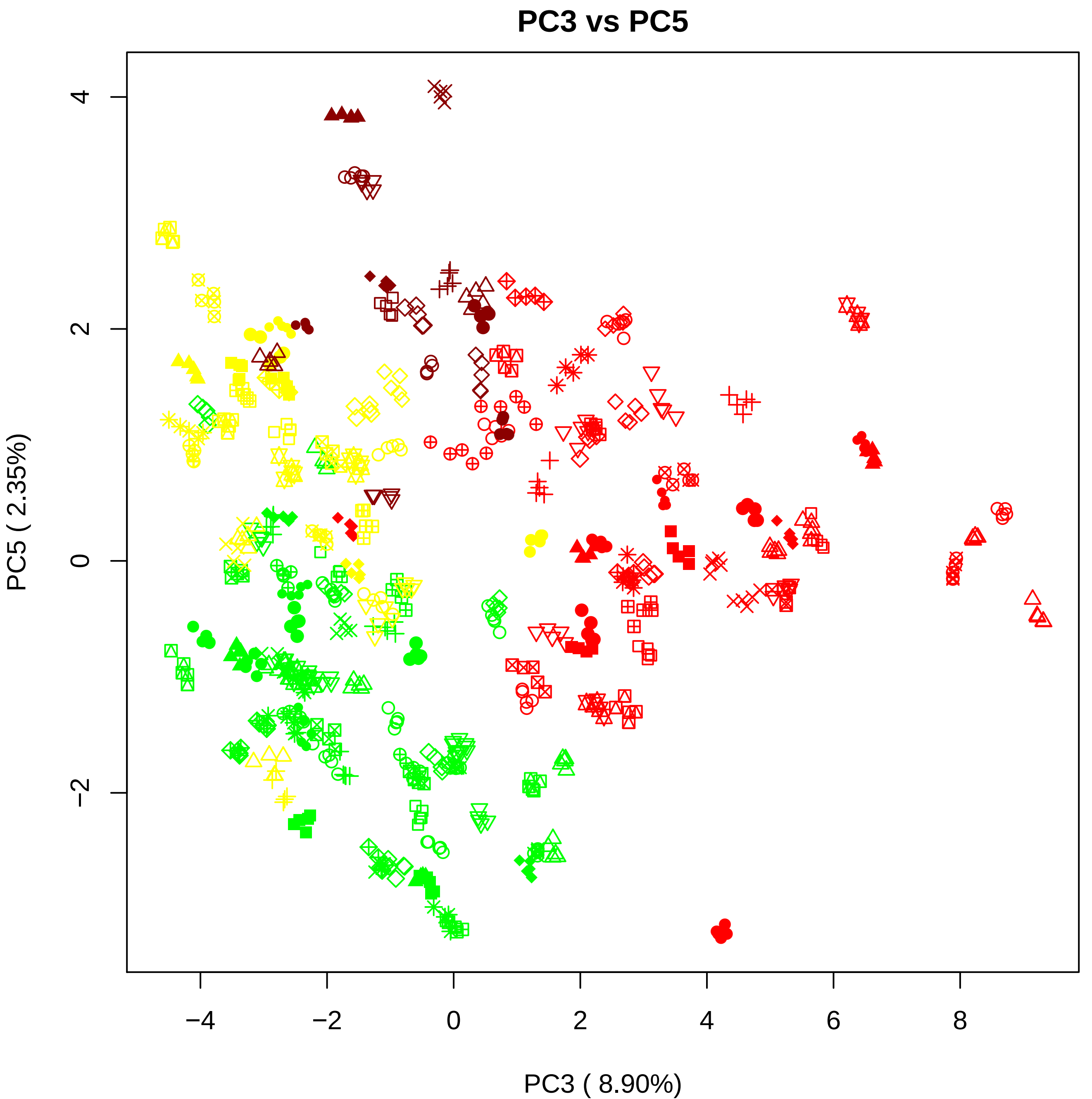
PC3 vs PC5
PC5 ( 2.35%)
PC3 ( 8.90%)
4
2
0
−2
−4
−2
0
2
4
6
8

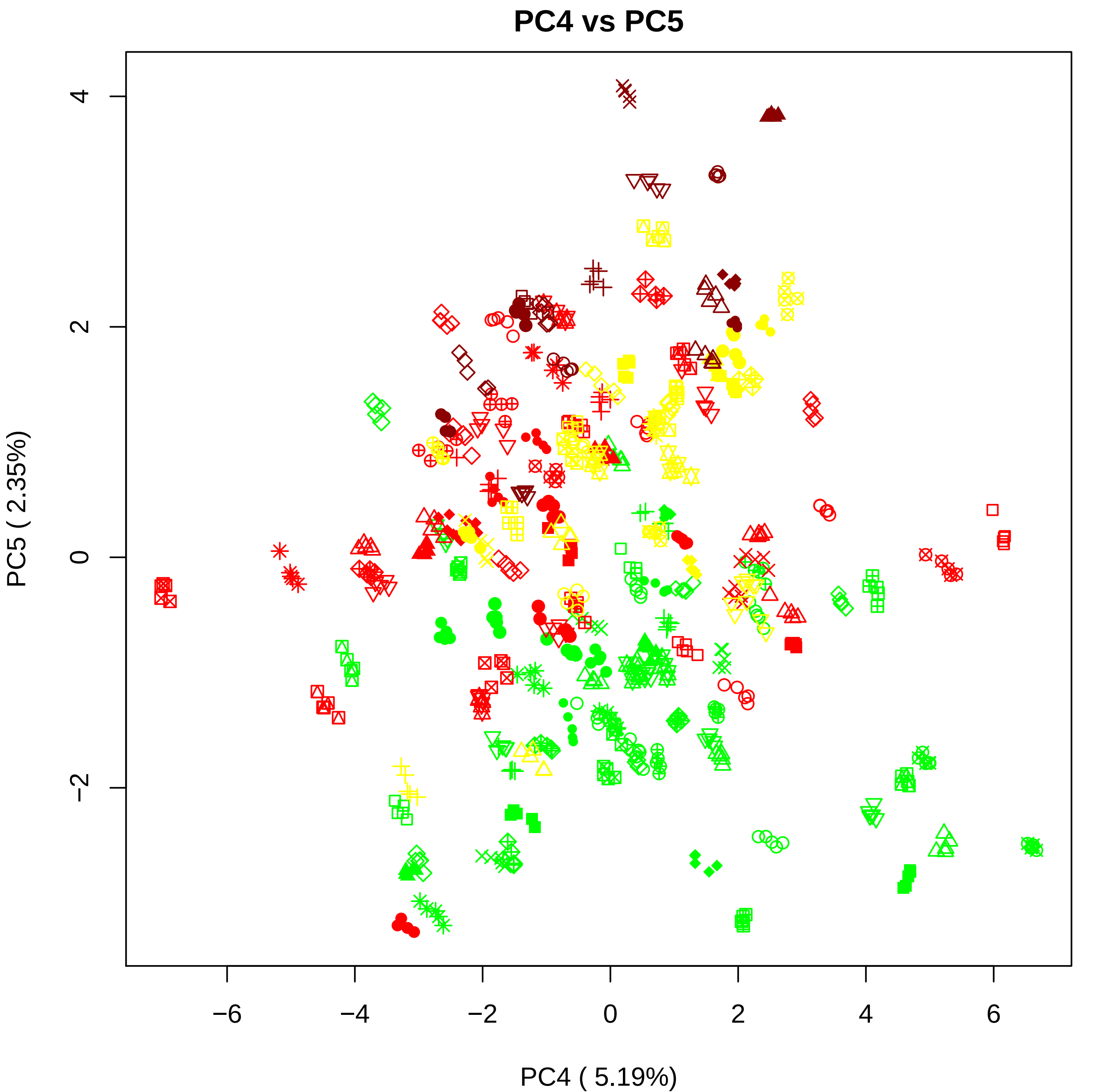

PC4 vs PC5
PC5 ( 2.35%)
PC4 ( 5.19%)
4
2
0
−2
−6
−4
−2
0
2
4
6

### PCA of PMH vs OMH

Scaled Matrix PCAed

7th order polynomial baseline correction performed

Raw spectra smoothed to 21 Neighbours, 2nd order PN, Savtisky Golay

Normalisation between 0 and 100

D2 spectra smoothed to 41 Neighbours, 2nd order PN, Savtisky Golay

Regions Analysed: 3601−2501 and 2001−700cm−1

Samples Analysed: 0 bContam's 0 BLH's 0 BMH's 30 OMH's 0 pContam's 0 PLH's 60 PMH's

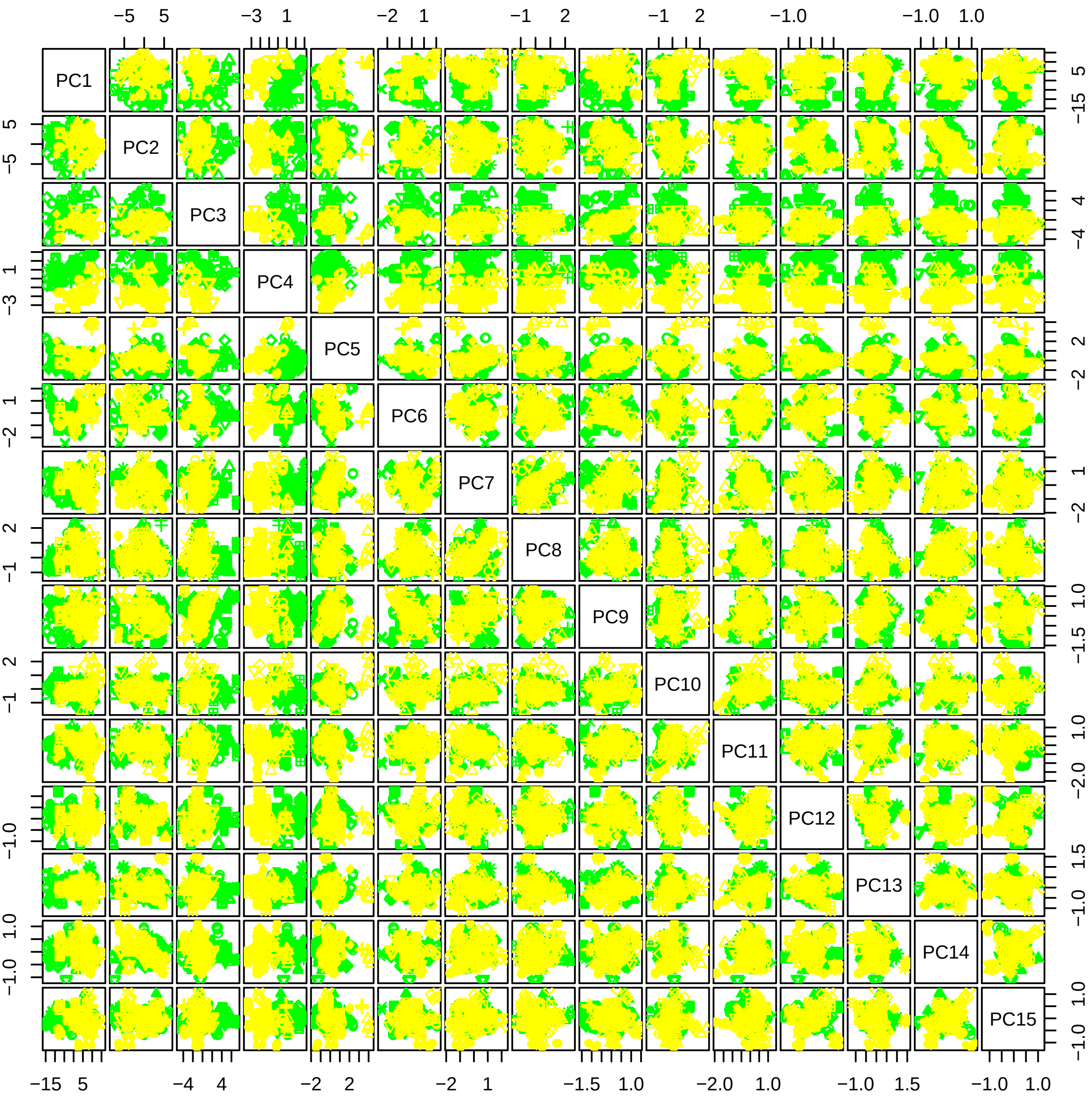

PC1
PC2
PC3
PC4
PC5
PC6
PC7
PC8
PC9
PC10
PC11
PC12
PC13
PC14
PC15

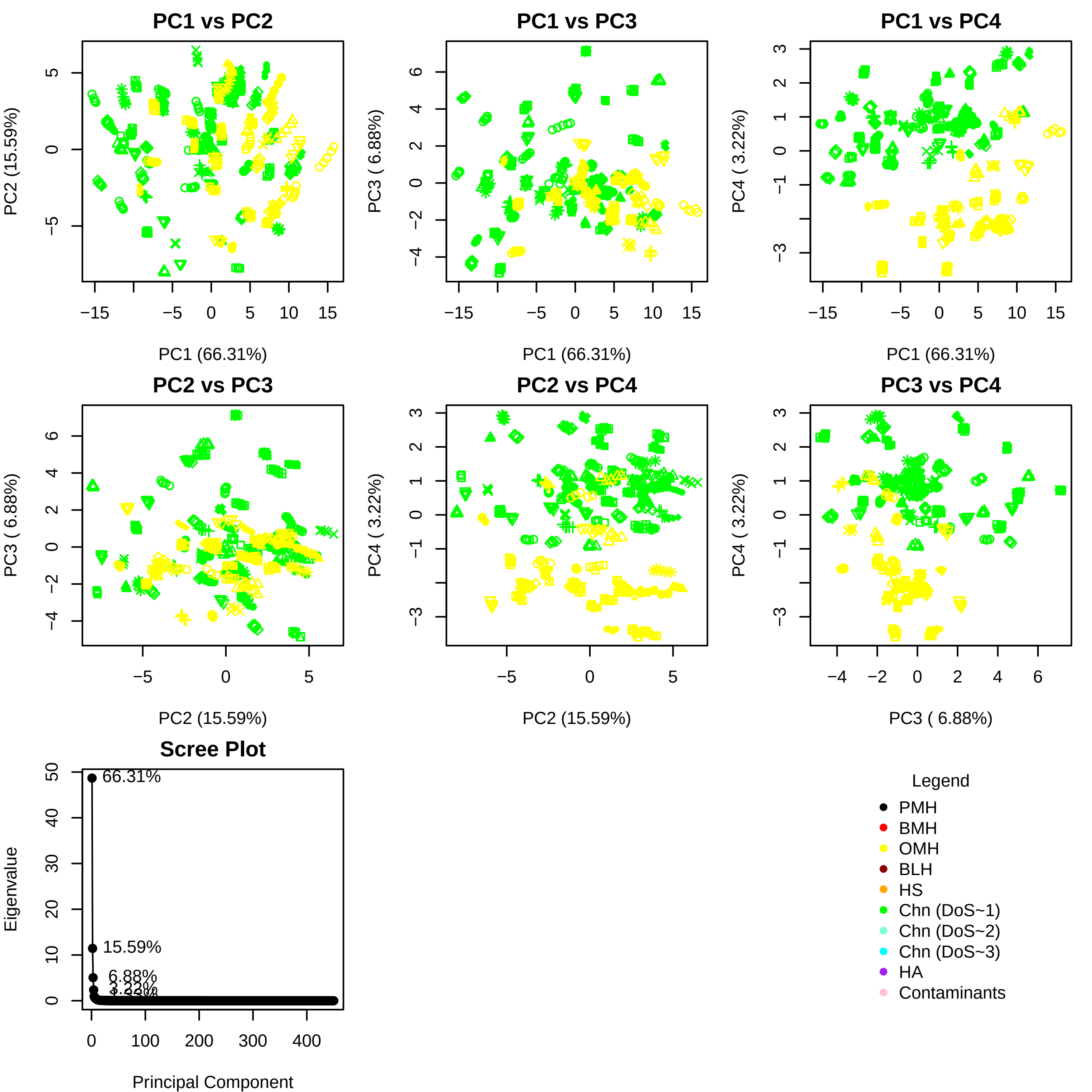
PC1 vs PC2
PC1 vs PC3
PC1 vs PC4
PC2 vs PC3
PC2 vs PC4
PC3 vs PC4
Scree Plot
PC1 (66.31%)
PC2 (15.59%)
PC3 ( 6.88%)
PC4 ( 3.22%)
Eigenvalue
Principal Component
66.31%
15.59%
6.88%
3.22%
Legend
PMH
BMH
OMH
BLH
HS
Chn (DoS~1)
Chn (DoS~2)
Chn (DoS~3)
HA
Contaminants

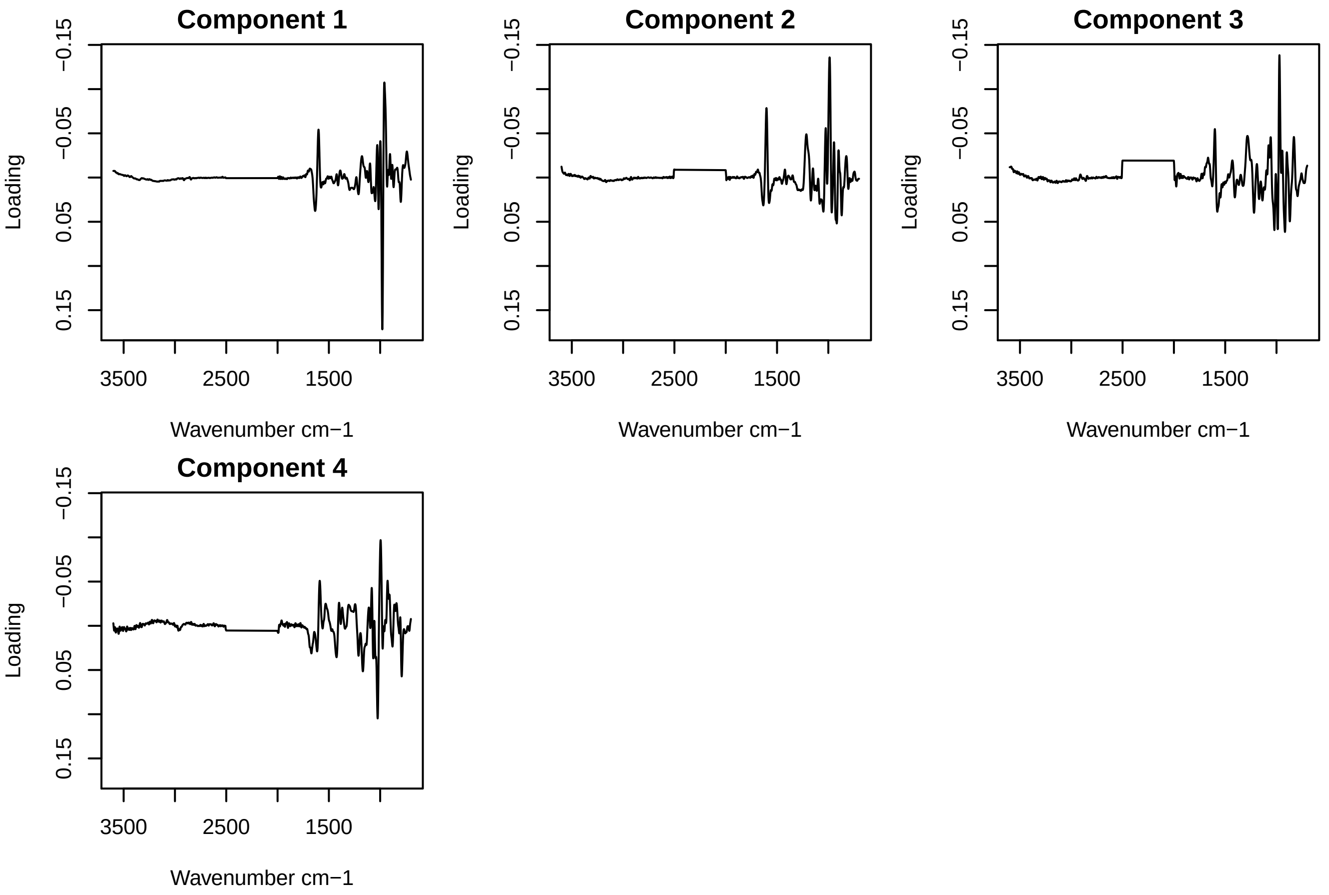

Component 1
Component 2
Component 3
Component 4
Loading
−0.15
−0.05
0.05
0.15
3500
2500
1500
Wavenumber cm−1

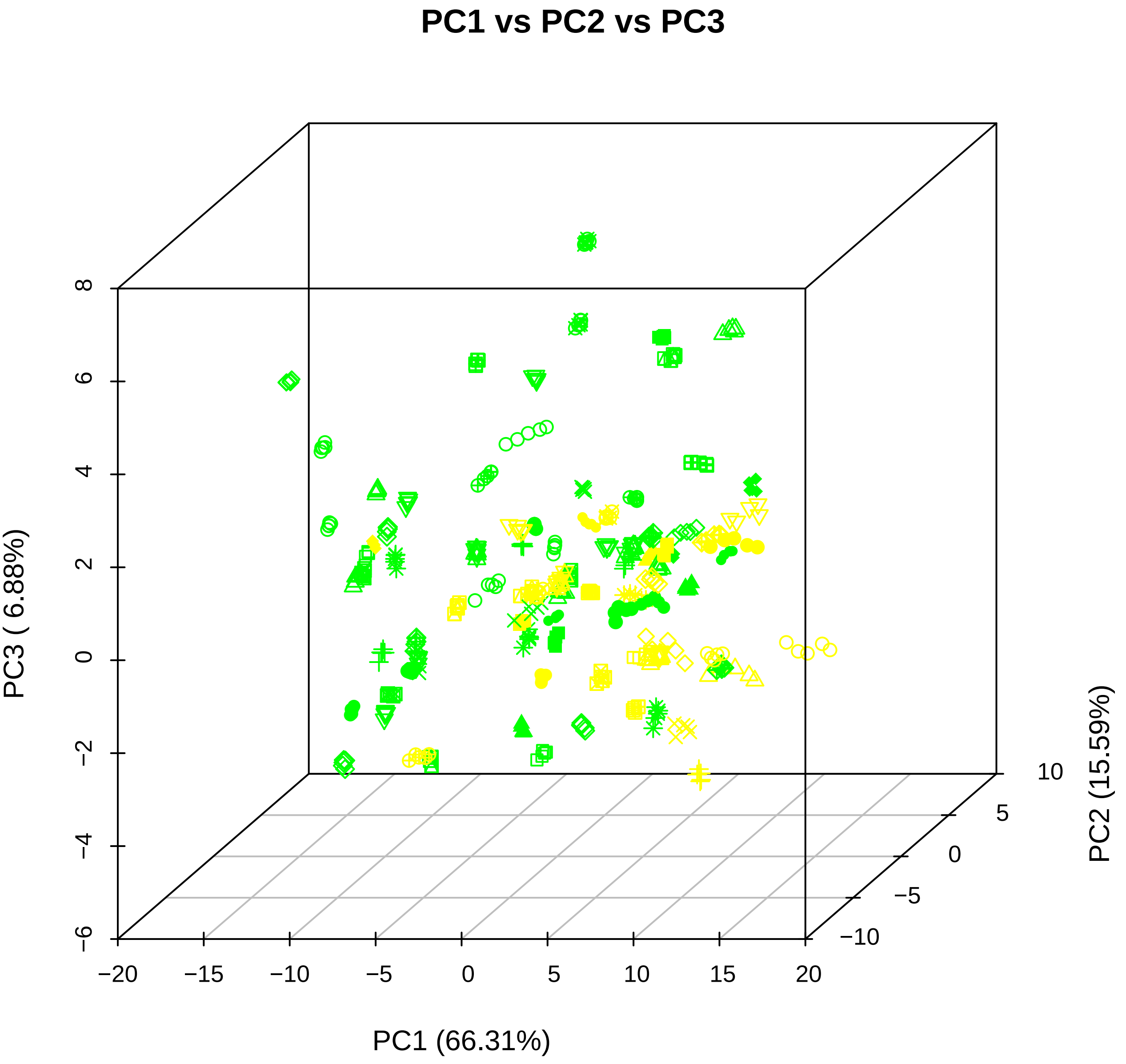

PC1 vs PC2 vs PC3
PC1 (66.31%)
PC2 (15.59%)
PC3 ( 6.88%)
−20
−15
−10
−5
0
5
10
15
20
−6
−4
−2
0
2
4
6
8
−10
−5
0
5
10

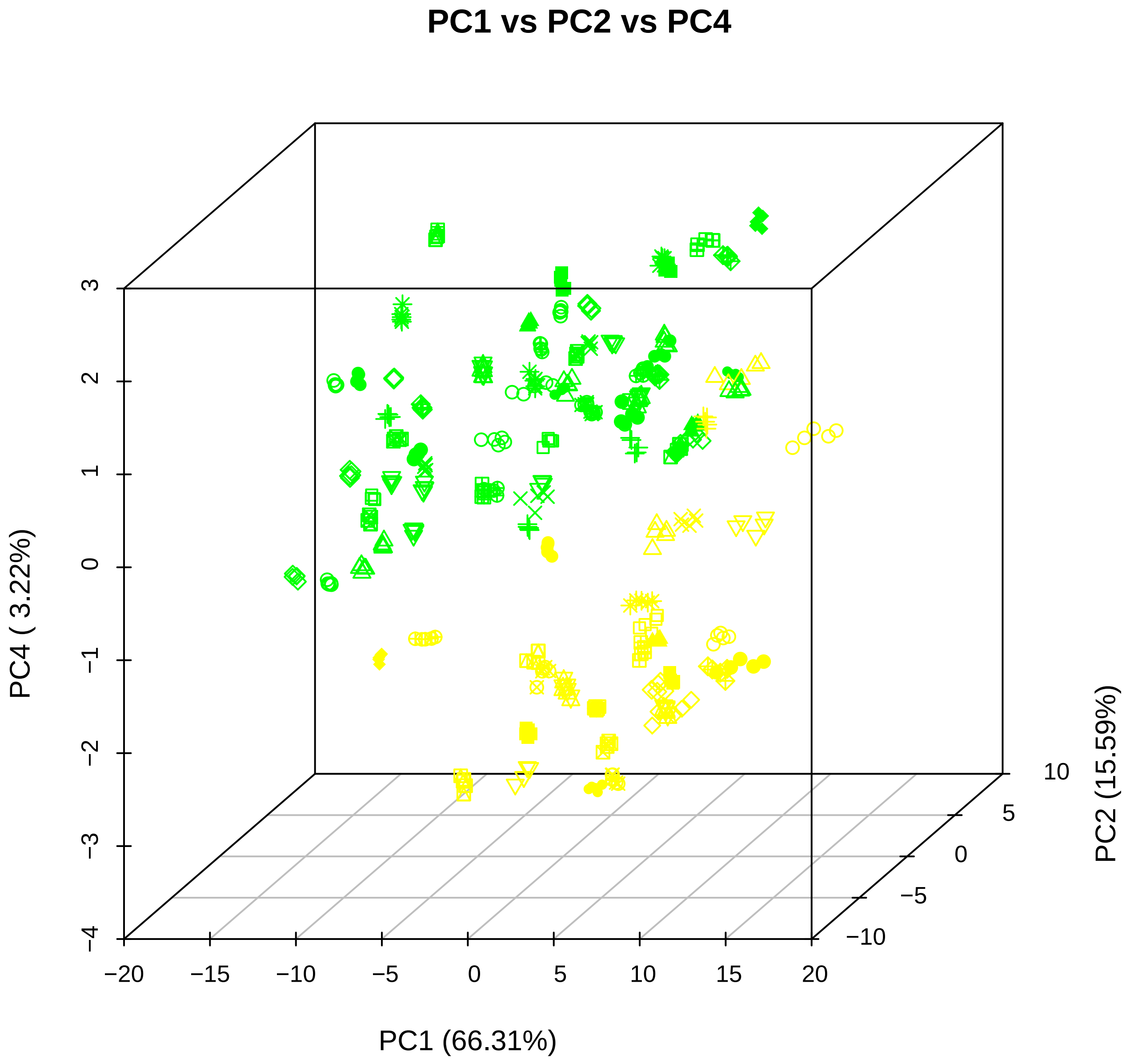
PC1 vs PC2 vs PC4
PC4 ( 3.22%)
3
2
1
0
−1
−2
−3
−4
−20
−15
−10
−5
0
5
10
15
20
PC1 (66.31%)
PC2 (15.59%)
10
5
0
−5
−10

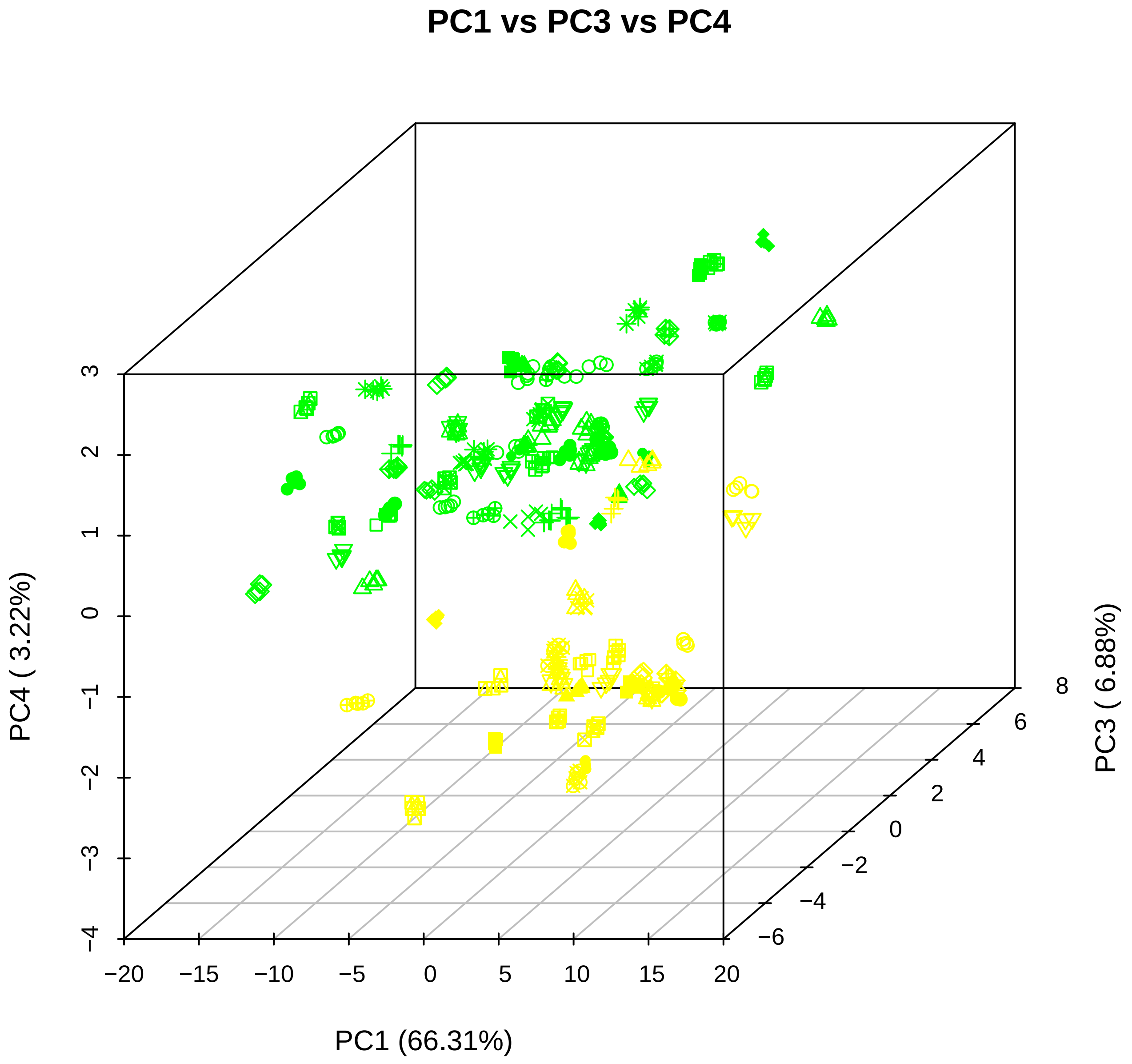
PC1 vs PC3 vs PC4
PC1 (66.31%)
PC3 ( 6.88%)
PC4 ( 3.22%)
−20
−15
−10
−5
0
5
10
15
20
−6
−4
−2
0
2
4
6
8
−4
−3
−2
−1
0
1
2
3

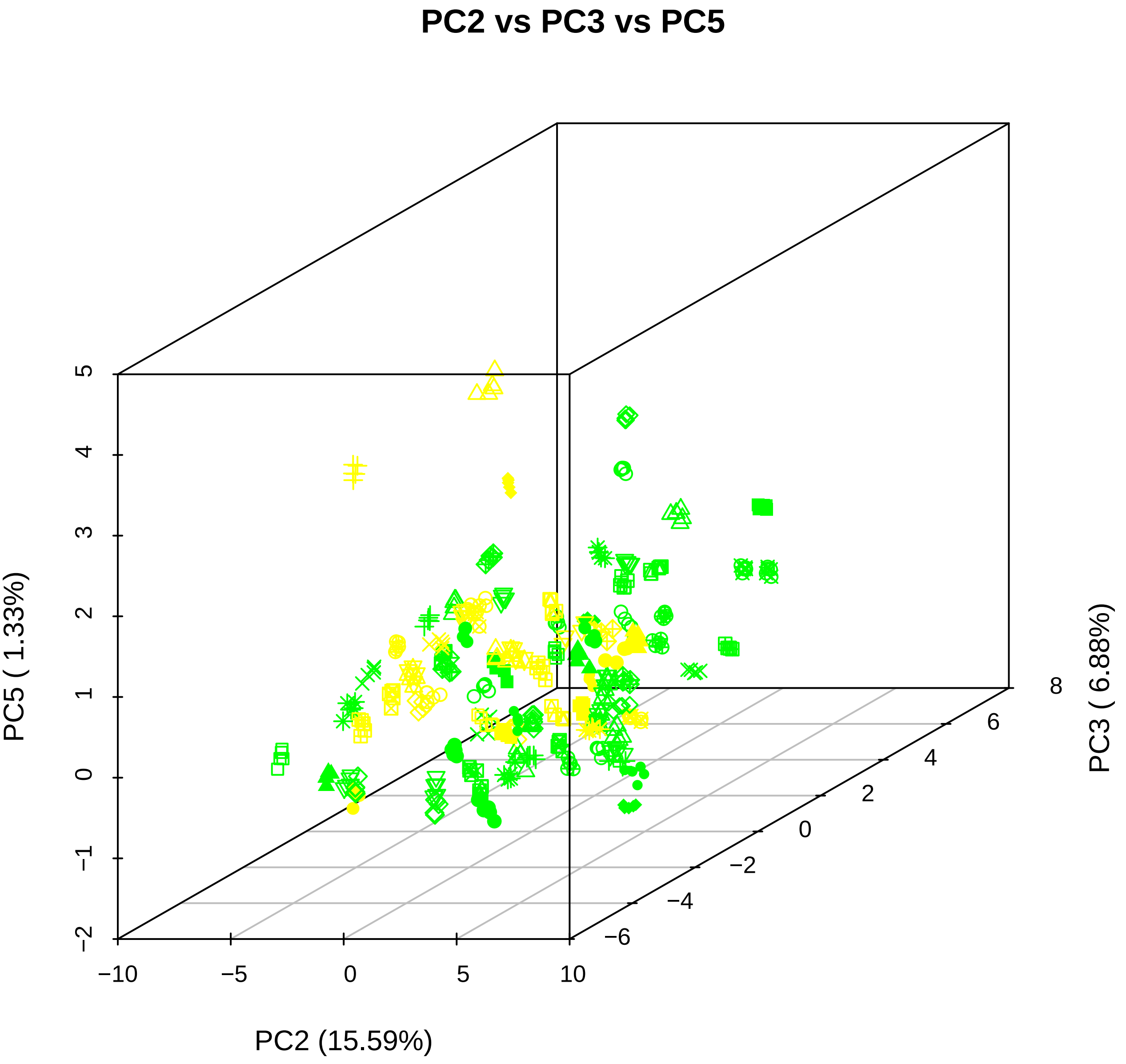
PC2 vs PC3 vs PC5
PC5 ( 1.33%)
PC2 (15.59%)
PC3 ( 6.88%)
−2
−1
0
1
2
3
4
5
−10
−5
0
5
10
−6
−4
−2
0
2
4
6
8

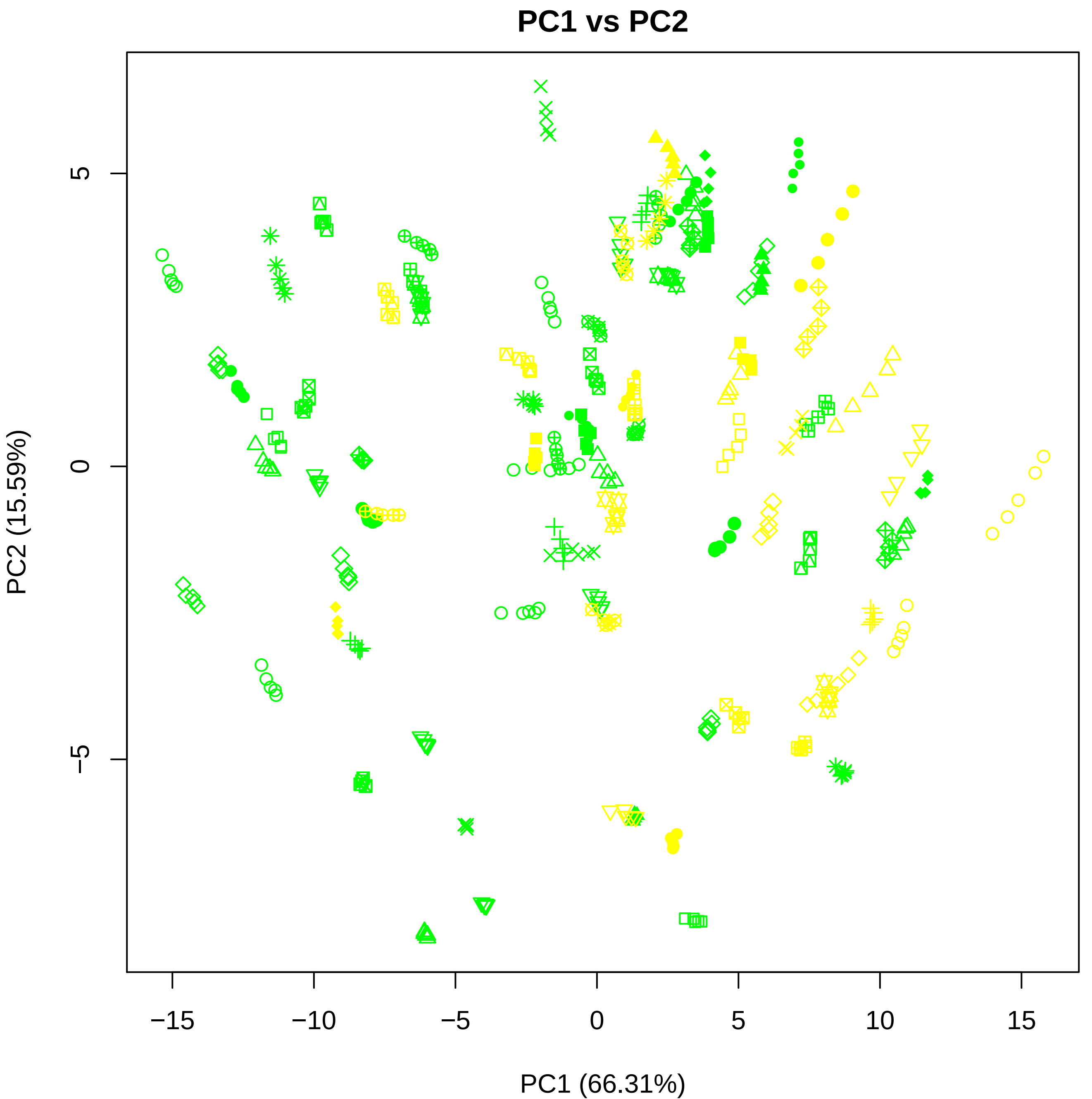
PC1 vs PC2
PC2 (15.59%)
PC1 (66.31%)
5
0
−5
−15
−10
−5
0
5
10
15

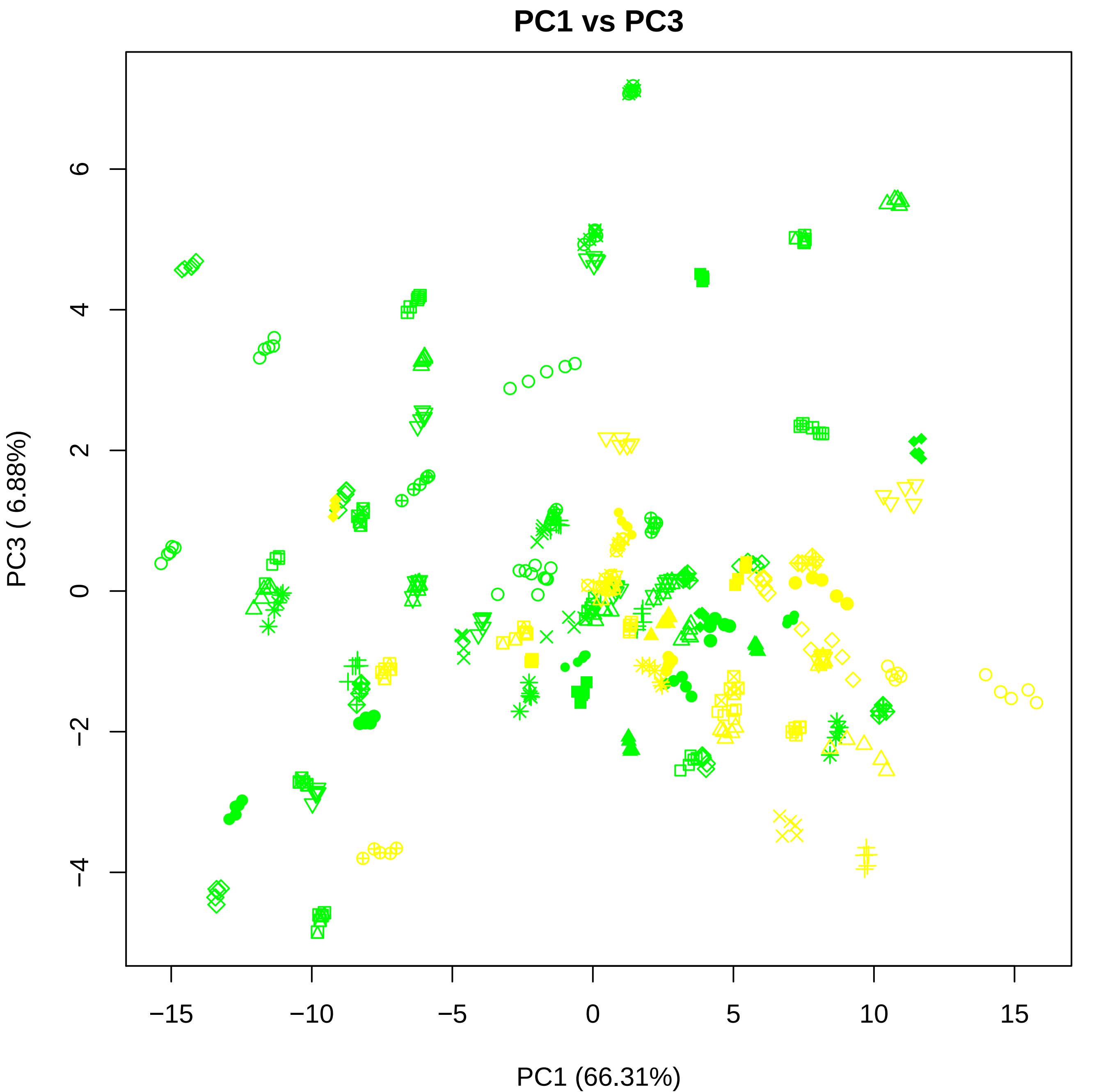
PC1 vs PC3
PC3 ( 6.88%)
PC1 (66.31%)
6
4
2
0
−2
−4
−15
−10
−5
0
5
10
15

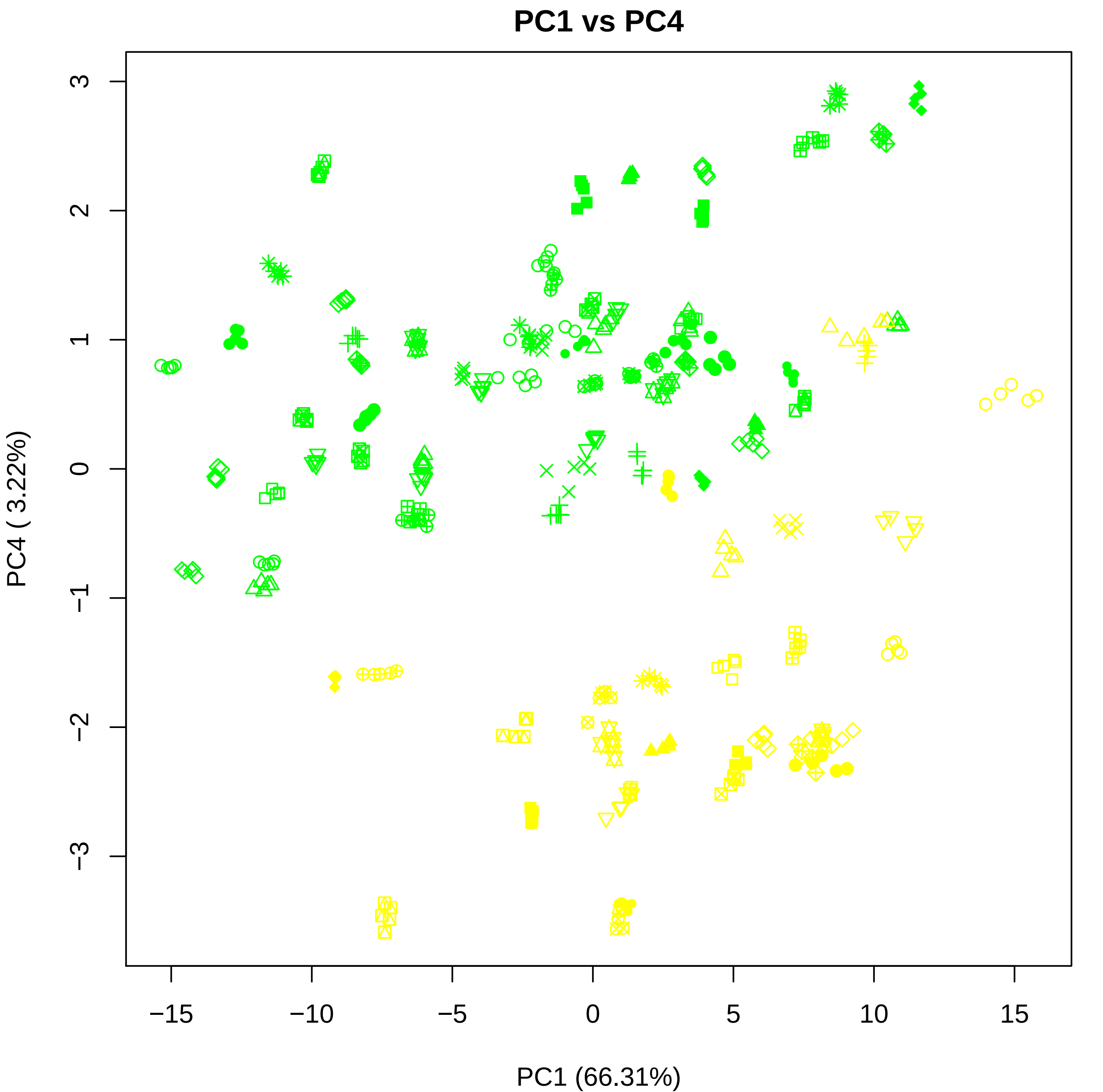
PC1 vs PC4
PC4 ( 3.22%)
PC1 (66.31%)
3
2
1
0
−1
−2
−3
−15
−10
−5
0
5
10
15

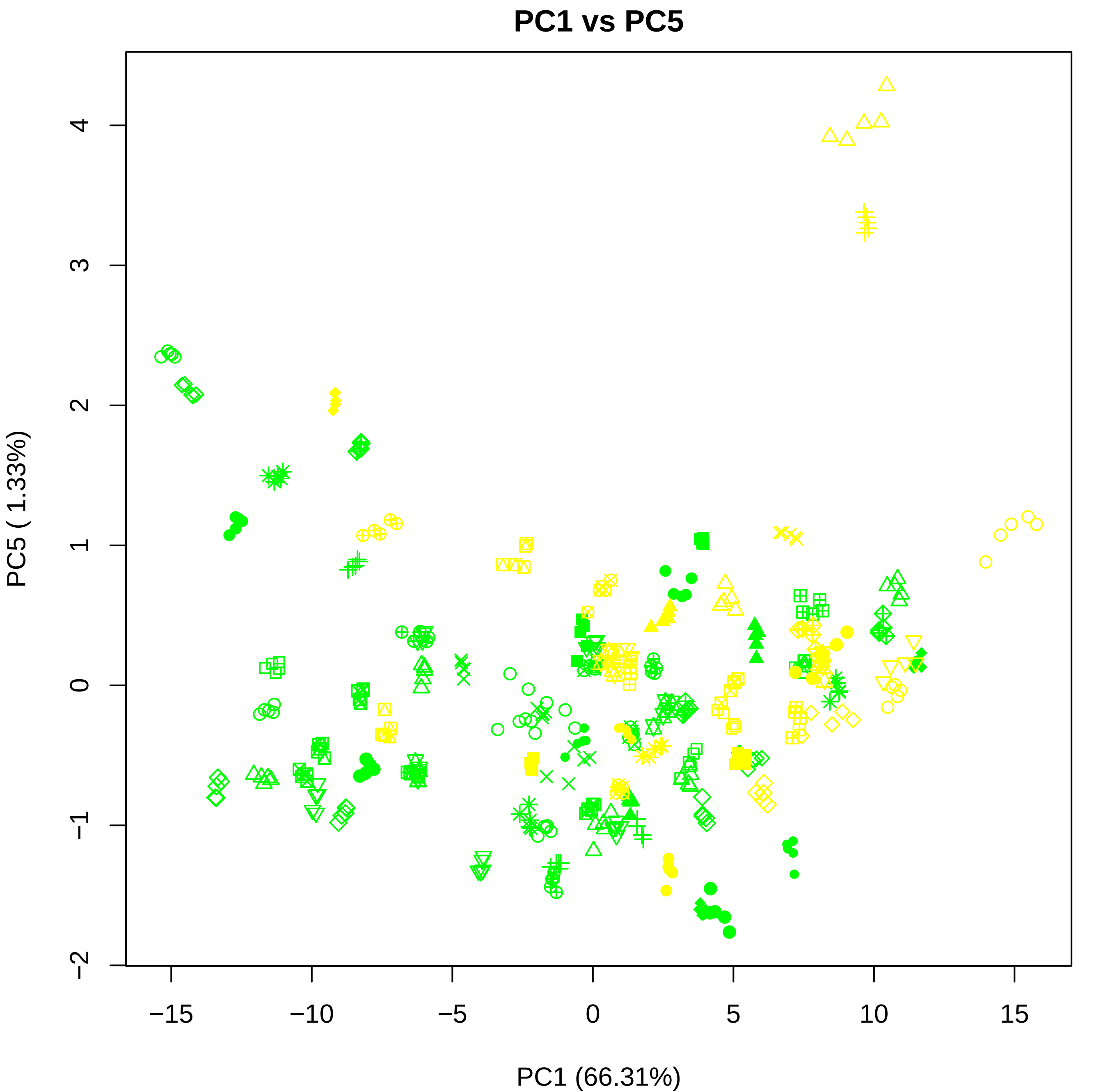
PC1 vs PC5
PC5 ( 1.33%)
PC1 (66.31%)
−2
−1
0
1
2
3
4
−15
−10
−5
0
5
10
15

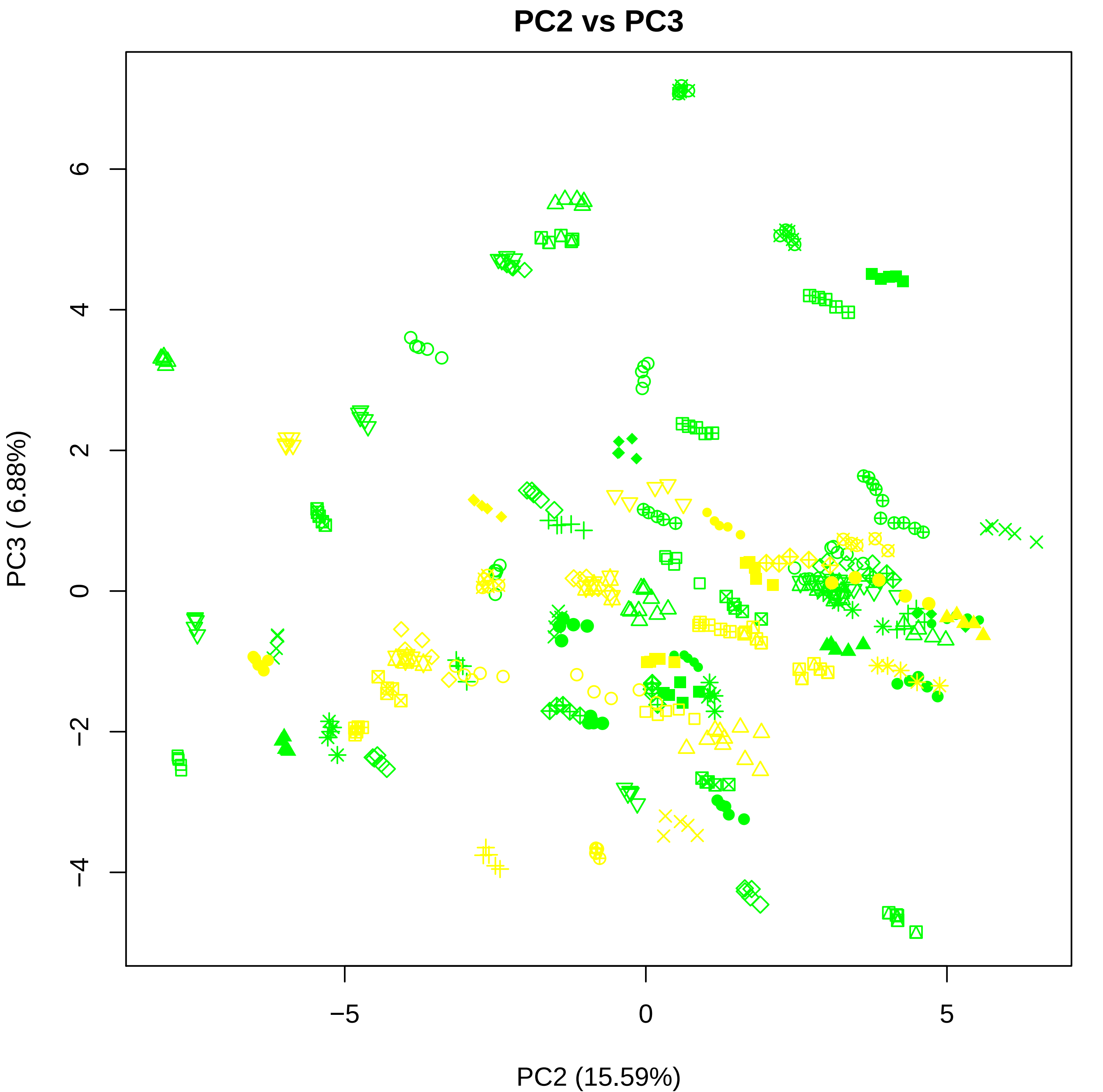

PC2 vs PC3
PC3 ( 6.88%)
PC2 (15.59%)
6
4
2
0
−2
−4
−5
0
5

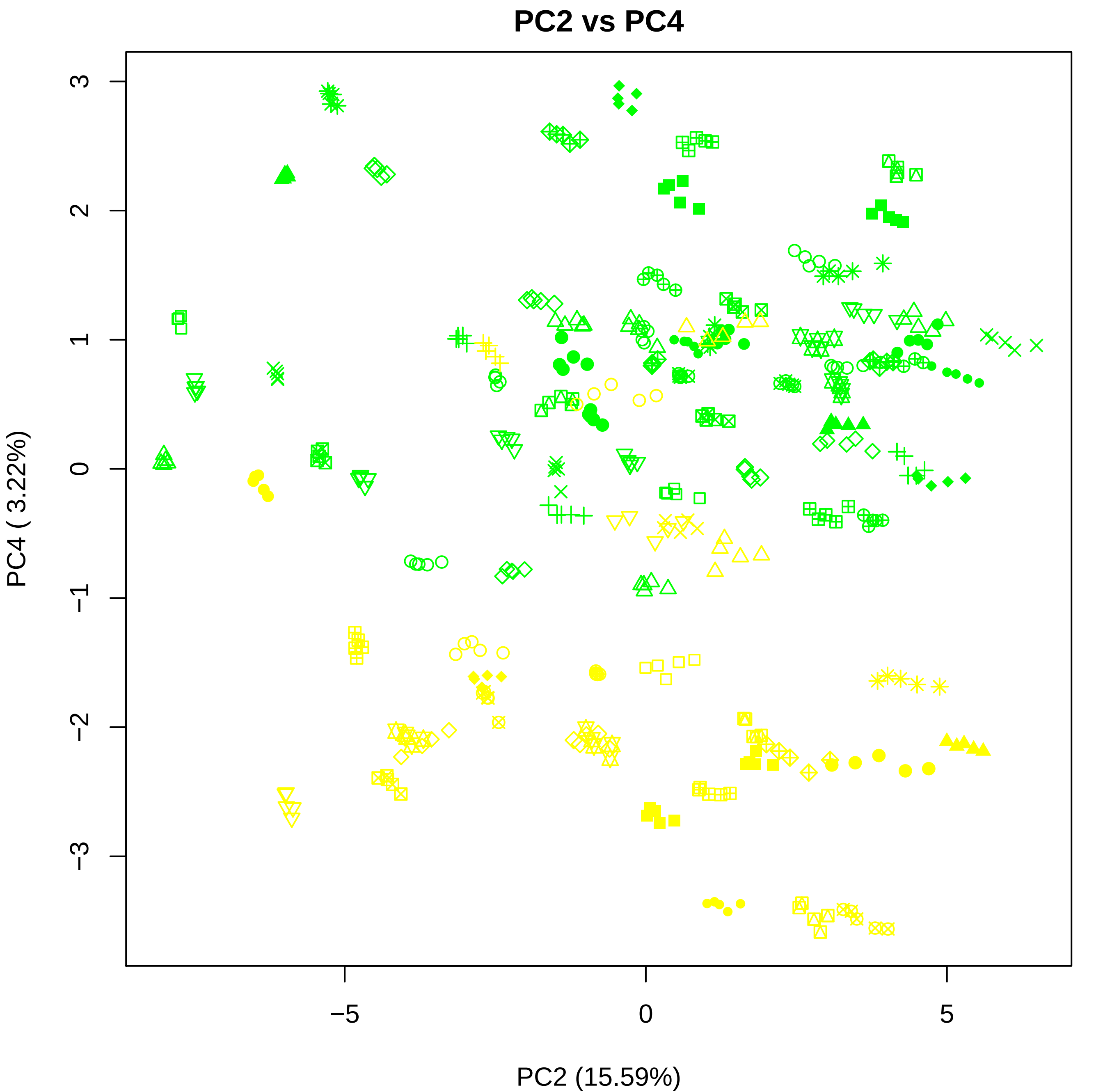
PC2 vs PC4
PC4 ( 3.22%)
PC2 (15.59%)
3
2
1
0
−1
−2
−3
−5
0
5

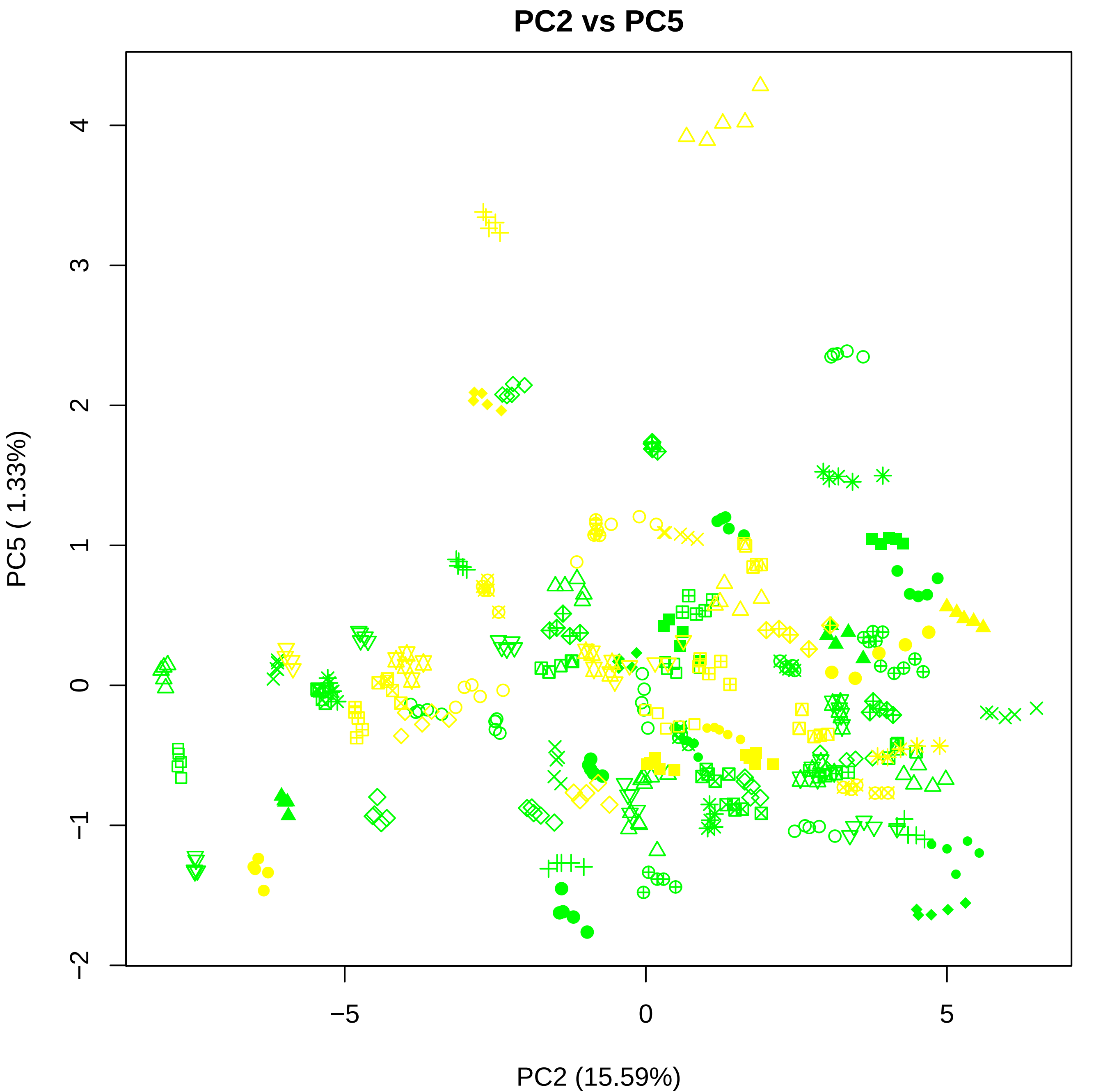
PC2 vs PC5
PC5 ( 1.33%)
PC2 (15.59%)
−2
−1
0
1
2
3
4
−5
0
5

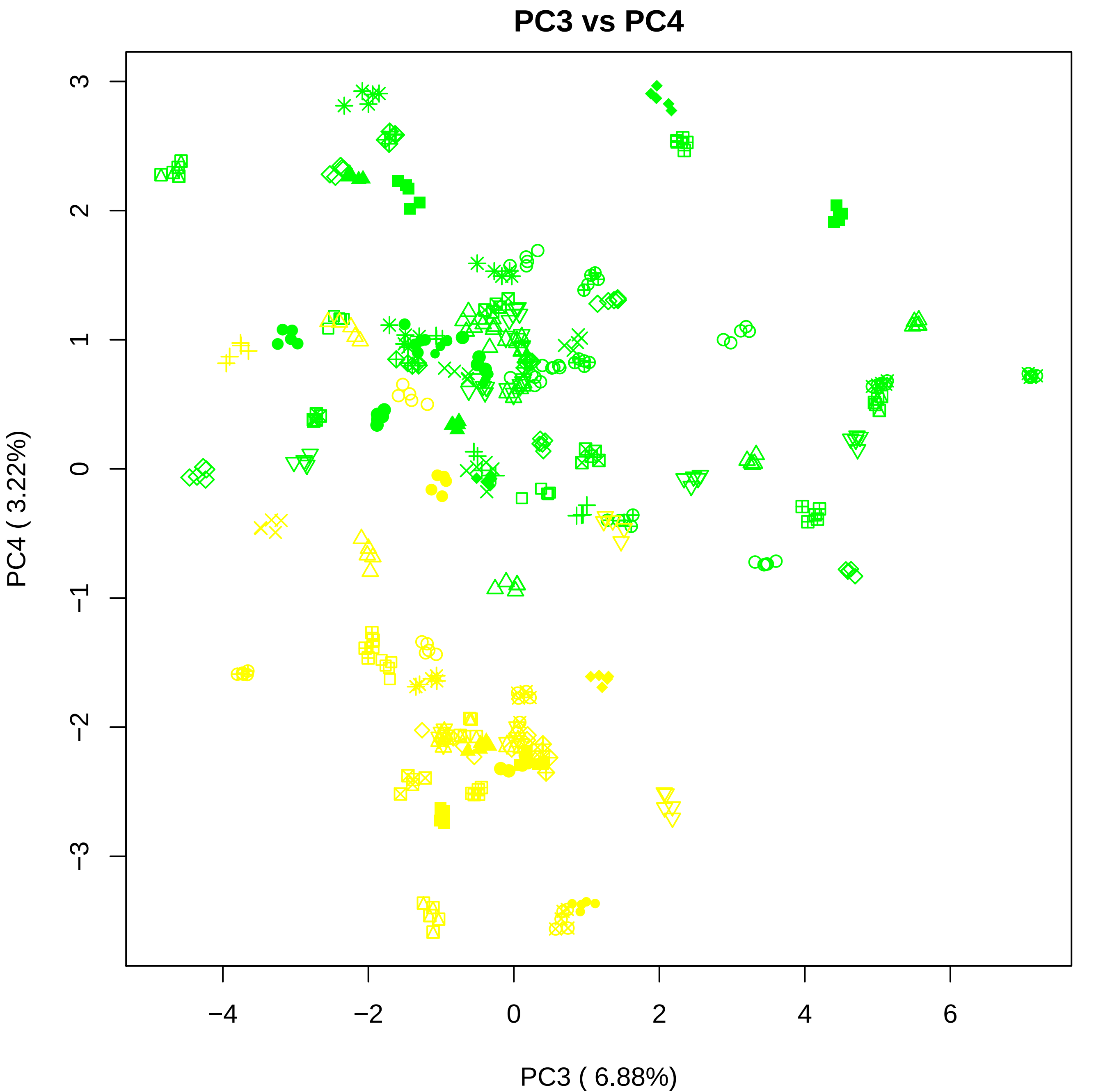
PC3 vs PC4
PC4 ( 3.22%)
PC3 ( 6.88%)
3
2
1
0
−1
−2
−3
−4
−2
0
2
4
6

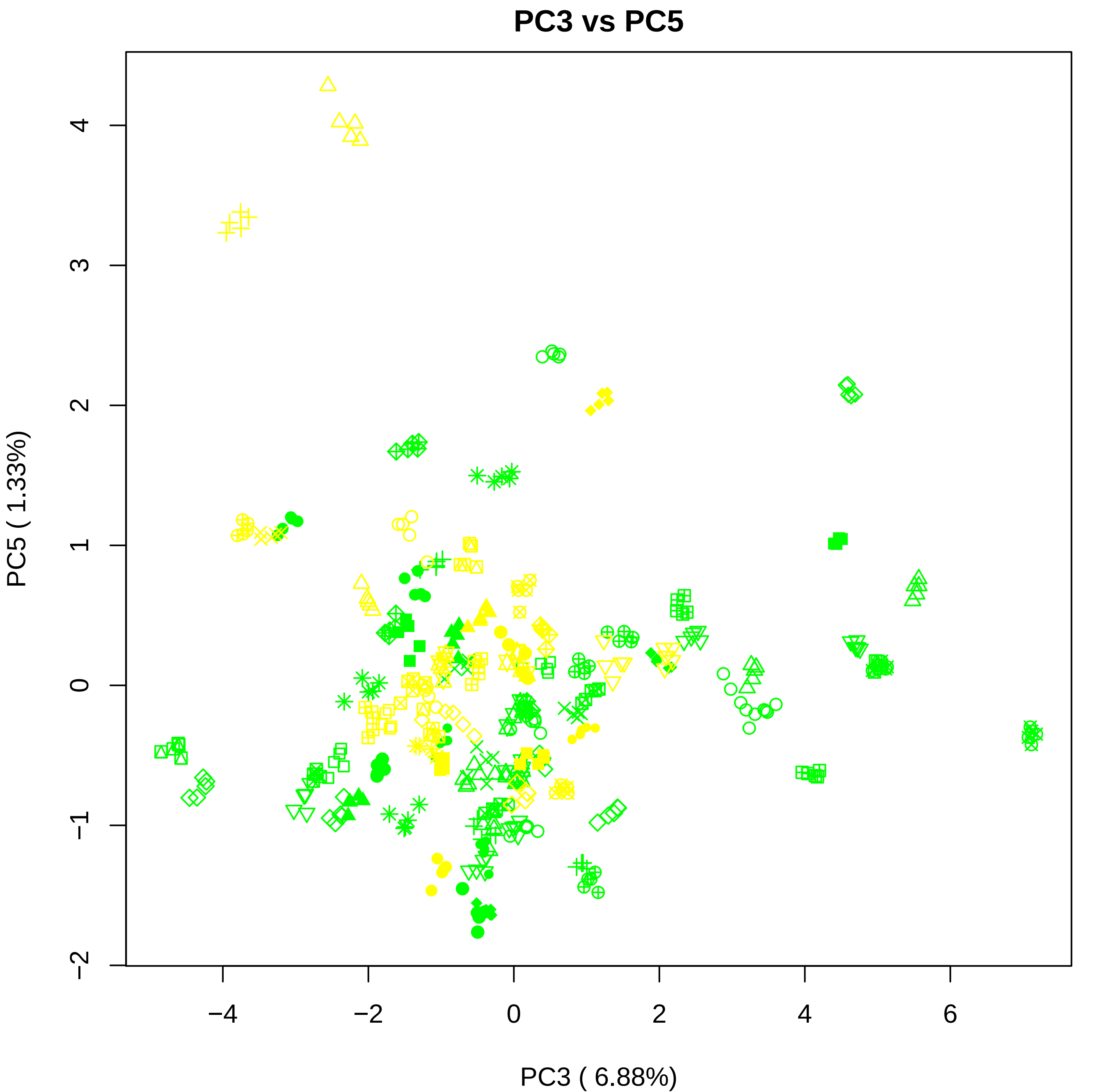
PC3 vs PC5
PC5 ( 1.33%)
PC3 ( 6.88%)
−2
−1
0
1
2
3
4
−4
−2
0
2
4
6

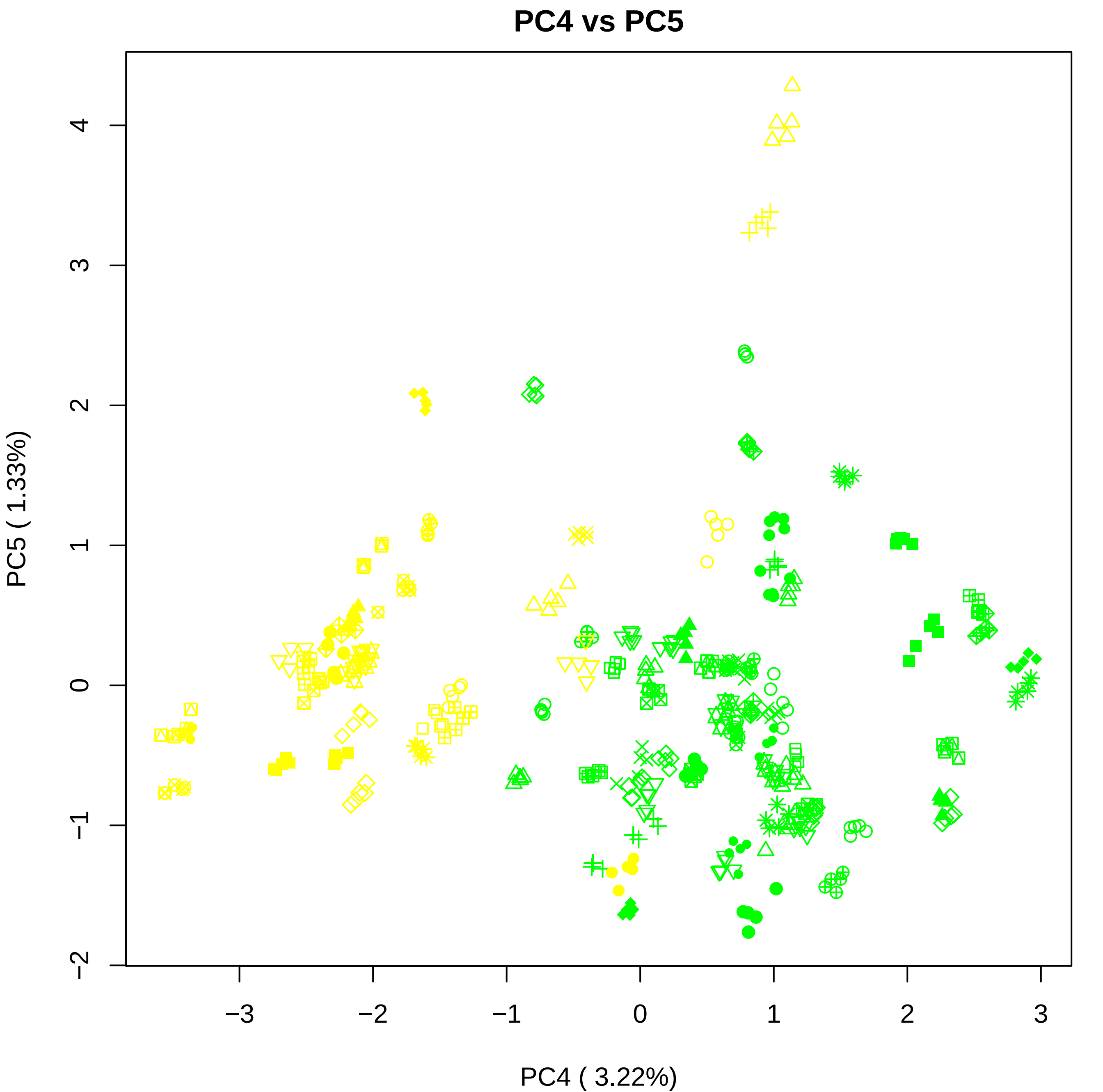
PC4 vs PC5
PC5 ( 1.33%)
PC4 ( 3.22%)
−3
−2
−1
0
1
2
3
4

### PCA of PMH vs BLH

Scaled Matrix PCAed

7th order polynomial baseline correction performed

Raw spectra smoothed to 21 Neighbours, 2nd order PN, Savtisky Golay

Normalisation between 0 and 100

D2 spectra smoothed to 41 Neighbours, 2nd order PN, Savtisky Golay

Regions Analysed: 3601−2501 and 2001−700cm−1

Samples Analysed: 0 bContam's 16 BLH's 0 BMH's 0 OMH's 0 pContam's 0 PLH's 60 PMH's

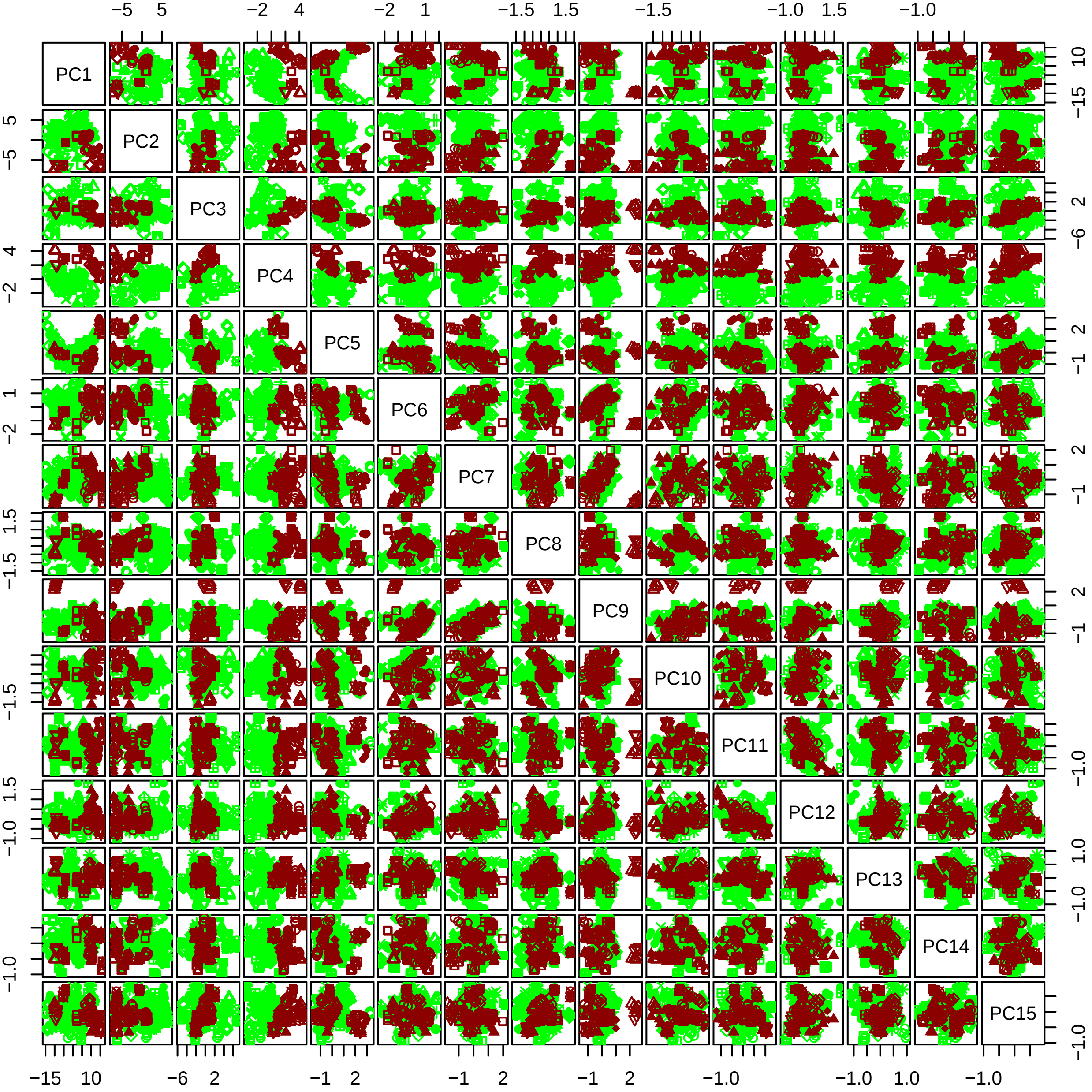
PC1
PC2
PC3
PC4
PC5
PC6
PC7
PC8
PC9
PC10
PC11
PC12
PC13
PC14
PC15
−5 5
−2 4
−2 1
−1.5 1.5
−1.5
−1.0 1.5
−1.0
10
−15
−5
2
−6
4
−2
−1
1.5
−1.5
−1.0
1.0
−15 10
−6 2
−1 2
−1.0 1.0

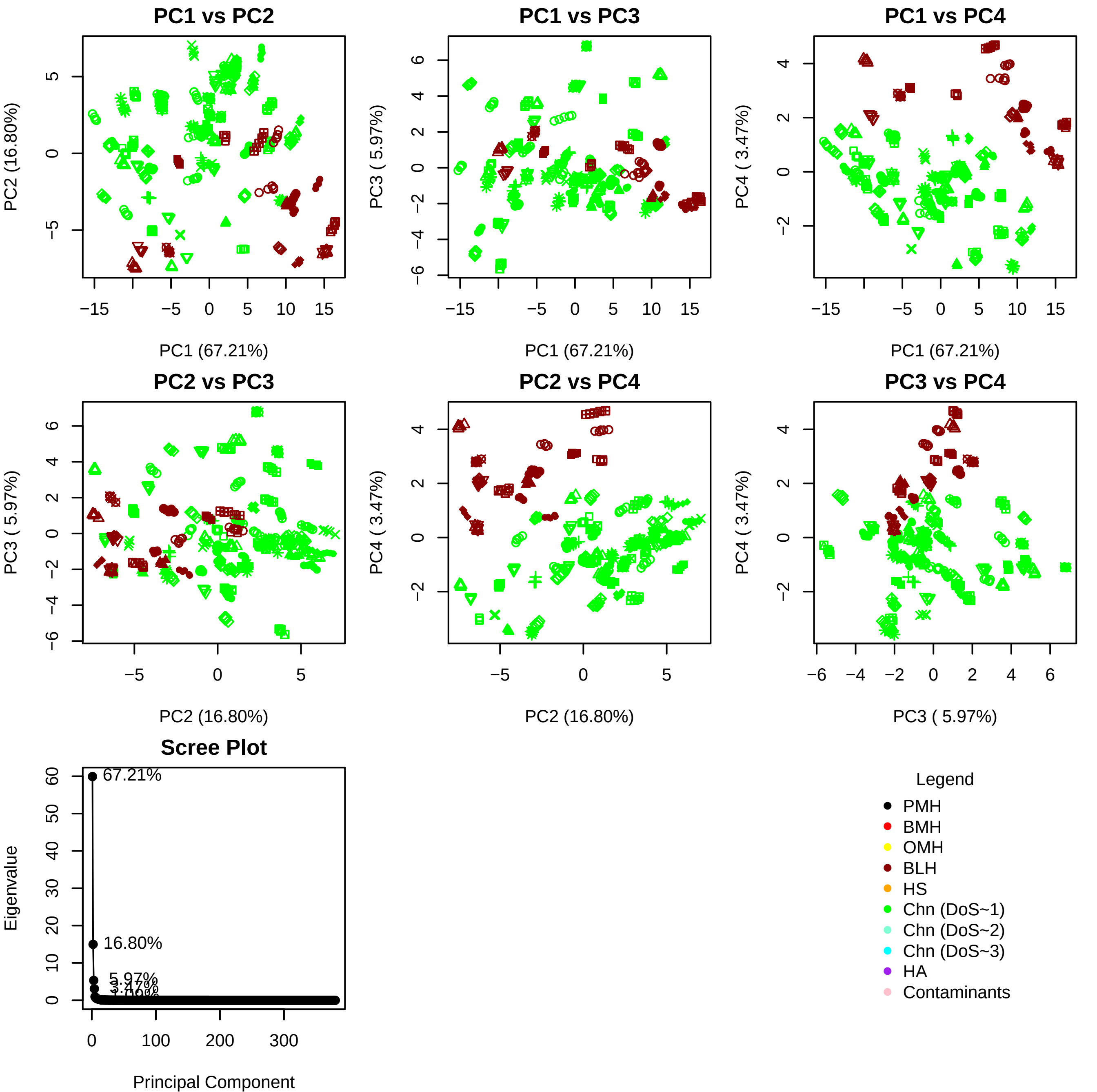
PC1 vs PC2
PC2 (16.80%)
PC1 (67.21%)
PC1 vs PC3
PC3 ( 5.97%)
PC1 (67.21%)
PC1 vs PC4
PC4 ( 3.47%)
PC1 (67.21%)
PC2 vs PC3
PC3 ( 5.97%)
PC2 (16.80%)
PC2 vs PC4
PC4 ( 3.47%)
PC2 (16.80%)
PC3 vs PC4
PC4 ( 3.47%)
PC3 ( 5.97%)
Scree Plot
67.21%
16.80%
5.97%
Eigenvalue
Principal Component
Legend
PMH
BMH
OMH
BLH
HS
Chn (DoS~1)
Chn (DoS~2)
Chn (DoS~3)
HA
Contaminants

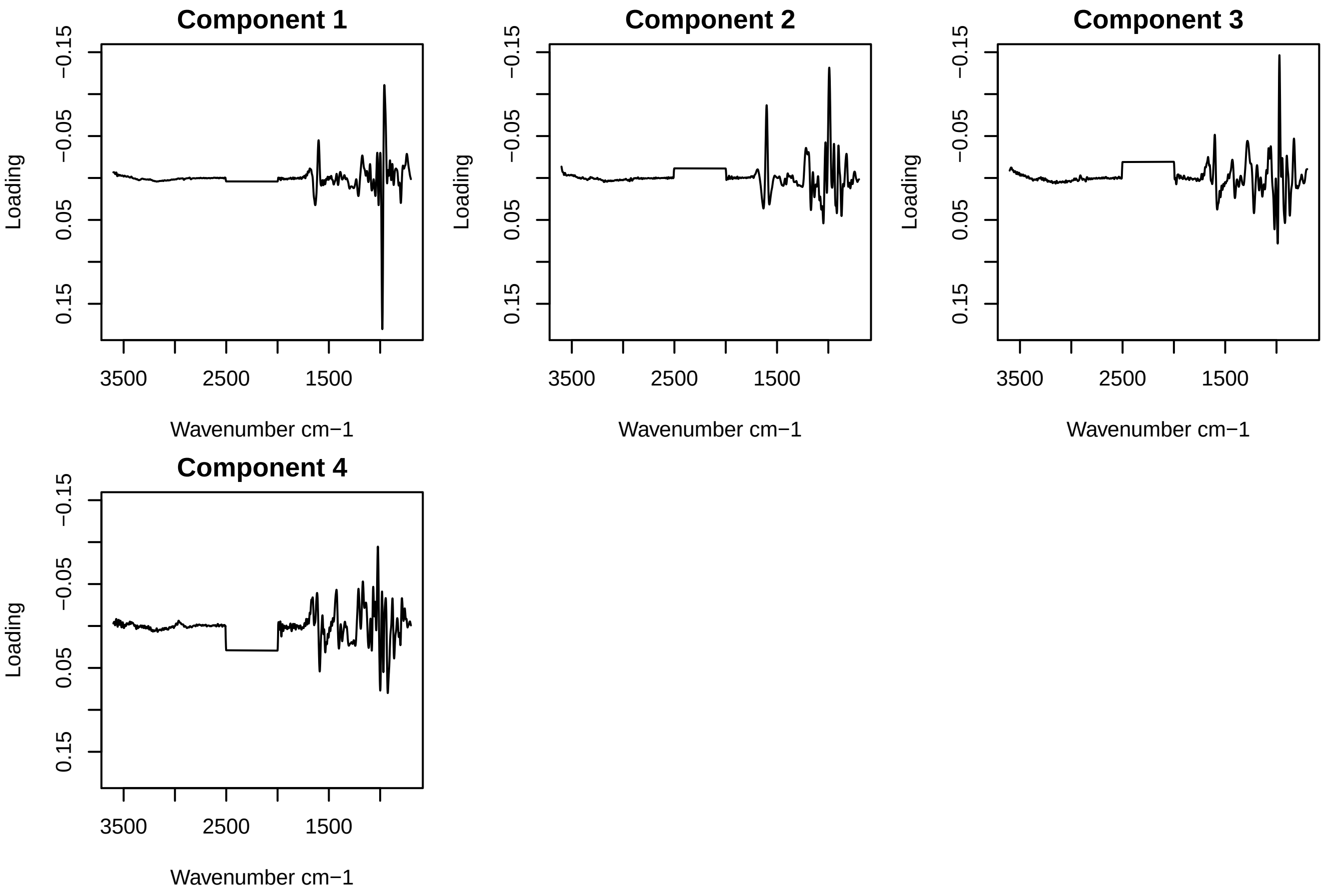
Component 1
Component 2
Component 3
Component 4
Loading
−0.15
−0.05
0.05
0.15
3500
2500
1500
Wavenumber cm−1

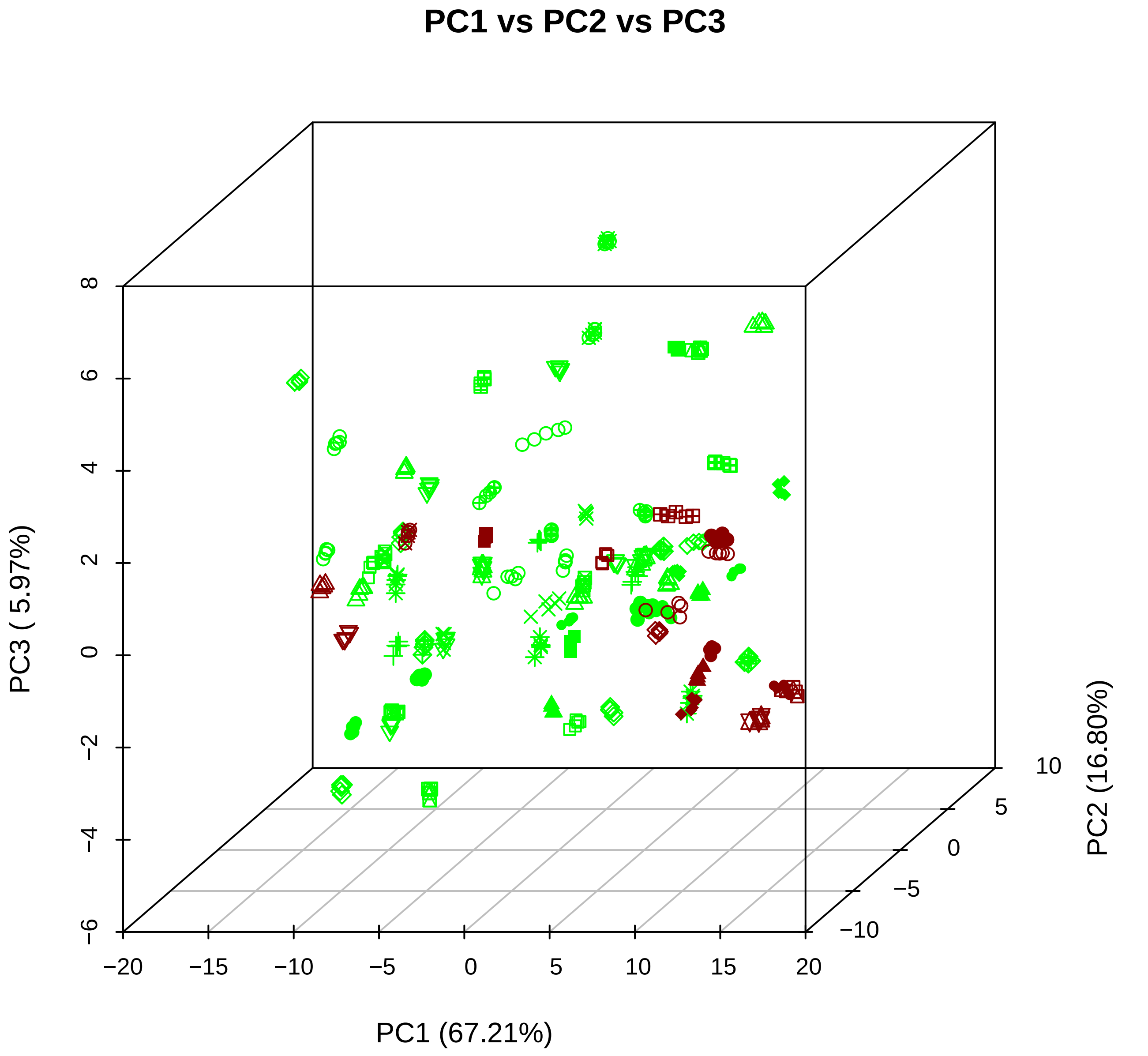
PC1 vs PC2 vs PC3
PC1 (67.21%)
PC2 (16.80%)
PC3 ( 5.97%)
−20
−15
−10
−5
0
5
10
15
20
−10
−5
0
5
10
−6
−4
−2
0
2
4
6
8

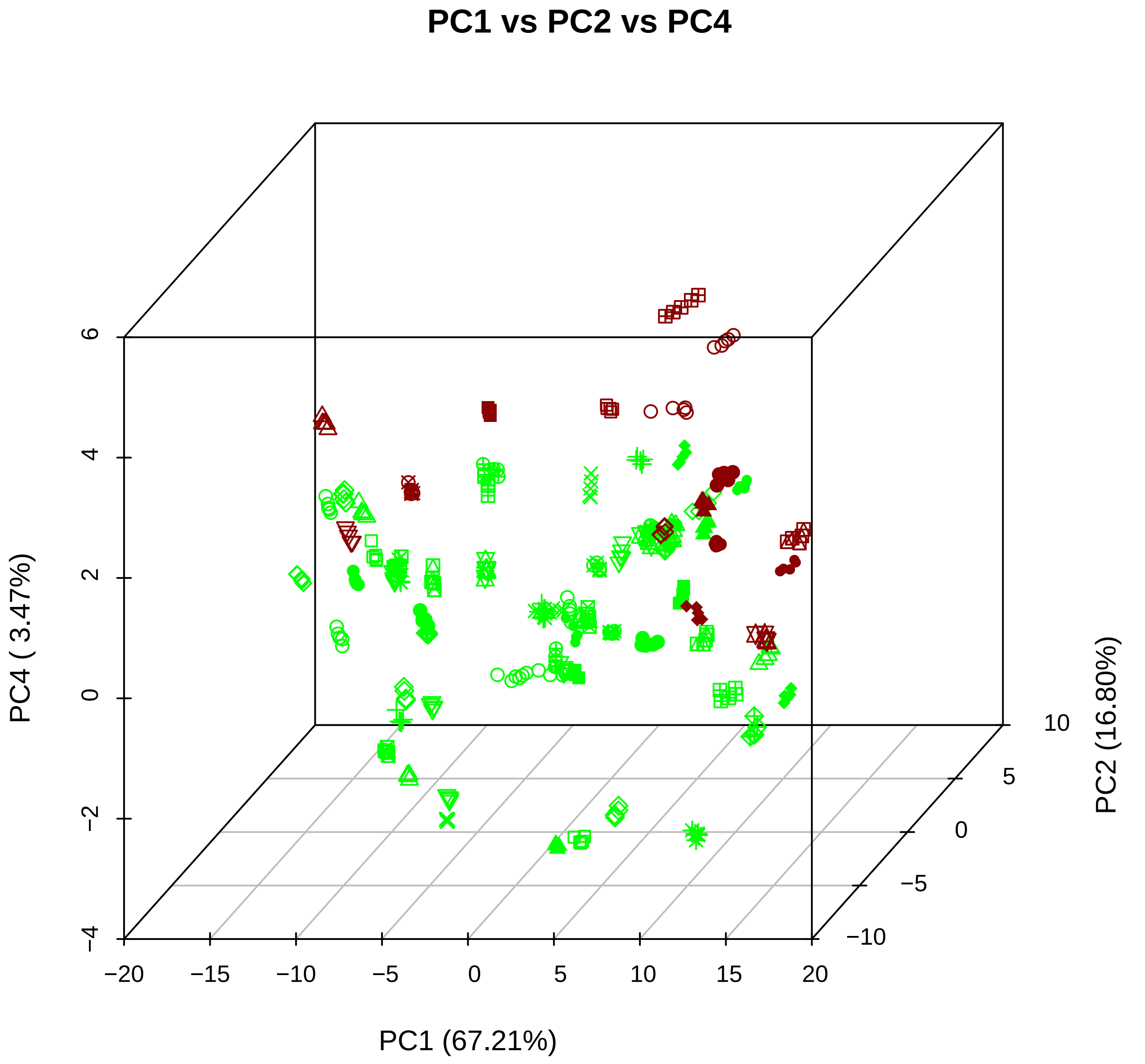
PC1 vs PC2 vs PC4
PC4 ( 3.47%)
PC1 (67.21%)
PC2 (16.80%)
−4
−2
0
2
4
6
−20
−15
−10
−5
0
5
10
15
20
−10
−5
0
5
10

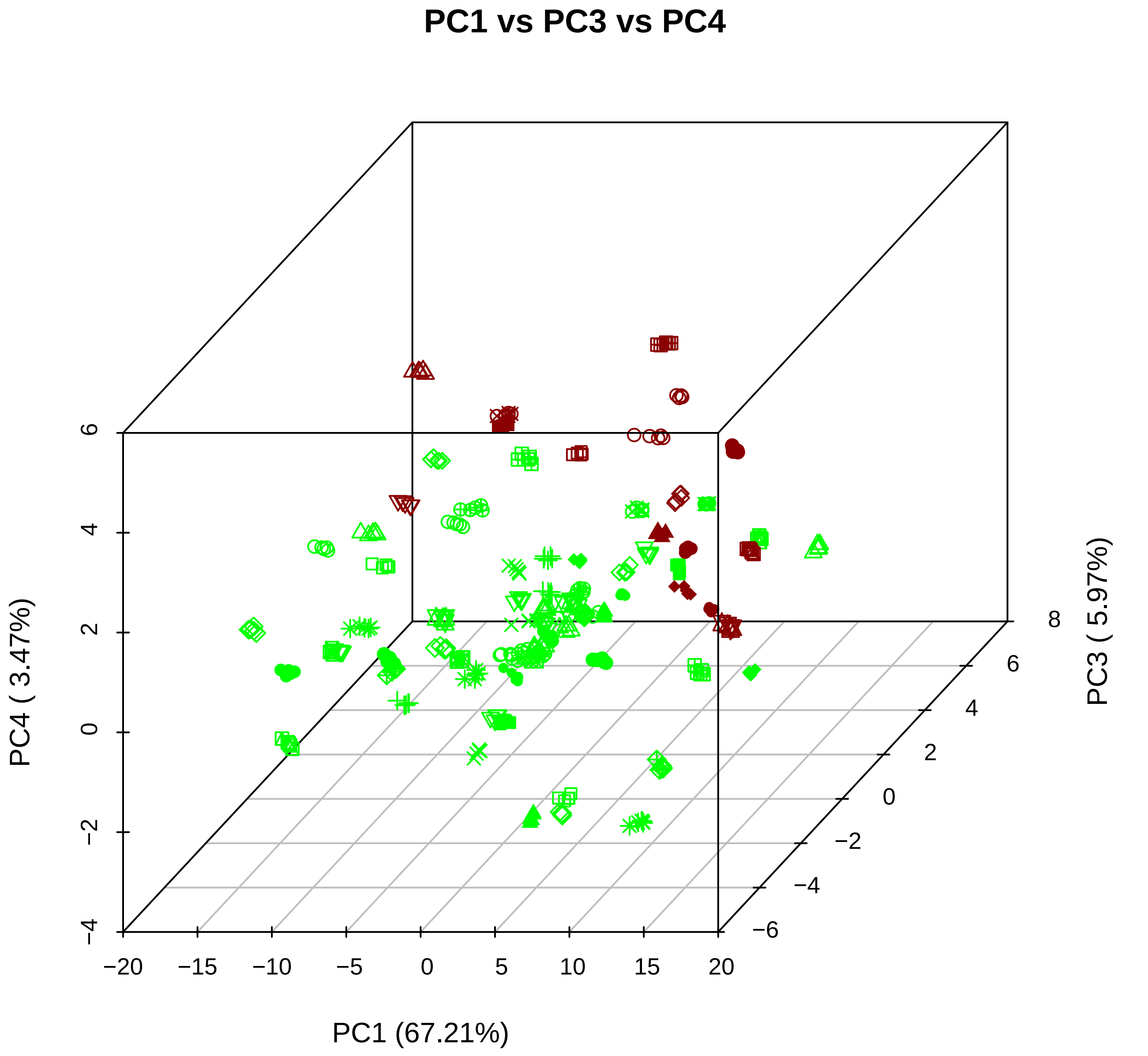
PC1 vs PC3 vs PC4
PC1 (67.21%)
PC4 ( 3.47%)
PC3 ( 5.97%)
−20
−15
−10
−5
0
5
10
15
20
−4
−2
0
2
4
6
−6
−4
−2
0
2
4
6
8

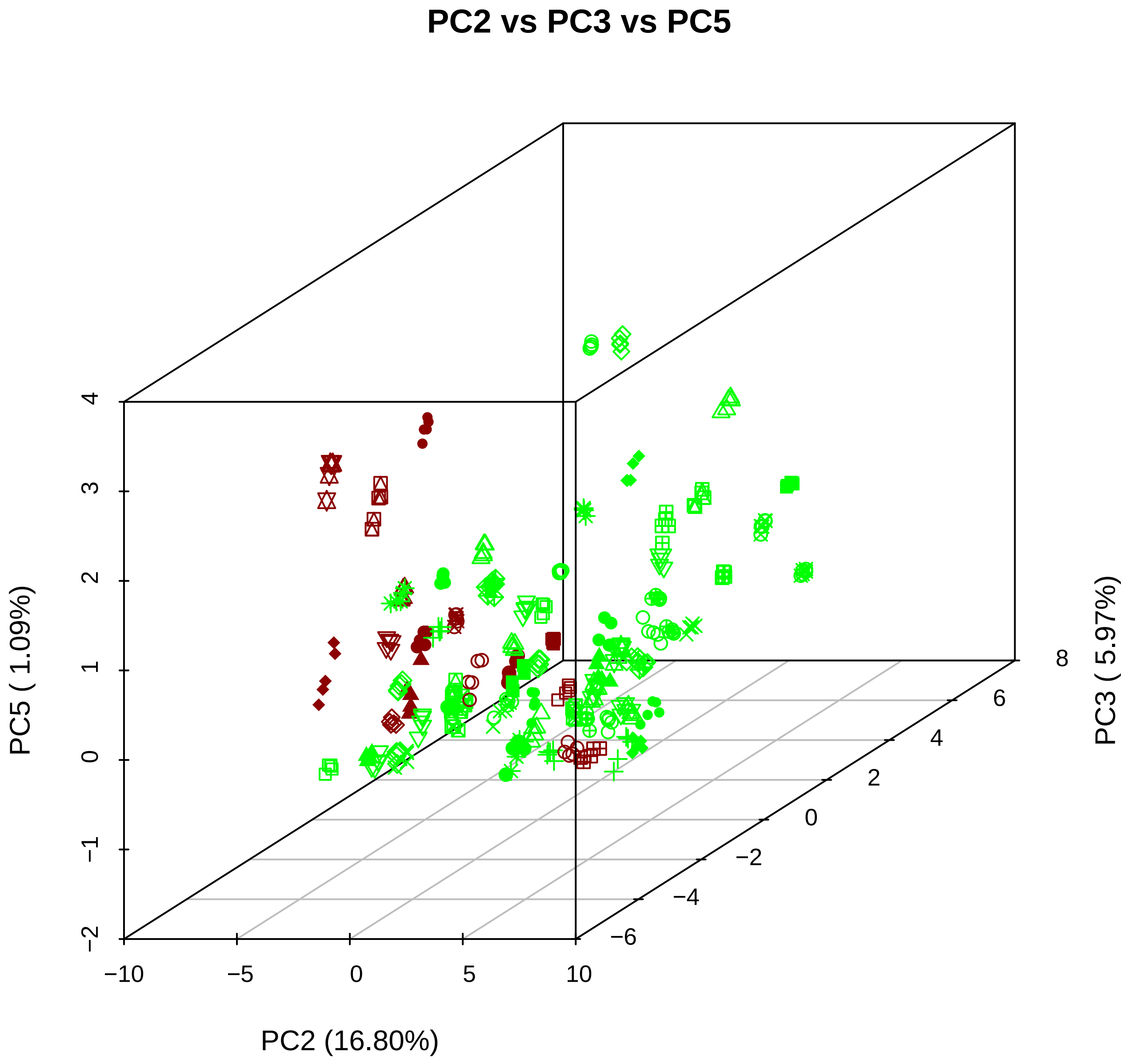

PC2 vs PC3 vs PC5
PC5 ( 1.09%)
PC2 (16.80%)
PC3 ( 5.97%)
−2
−1
0
1
2
3
4
−10
−5
0
5
10
−6
−4
−2
0
2
4
6
8

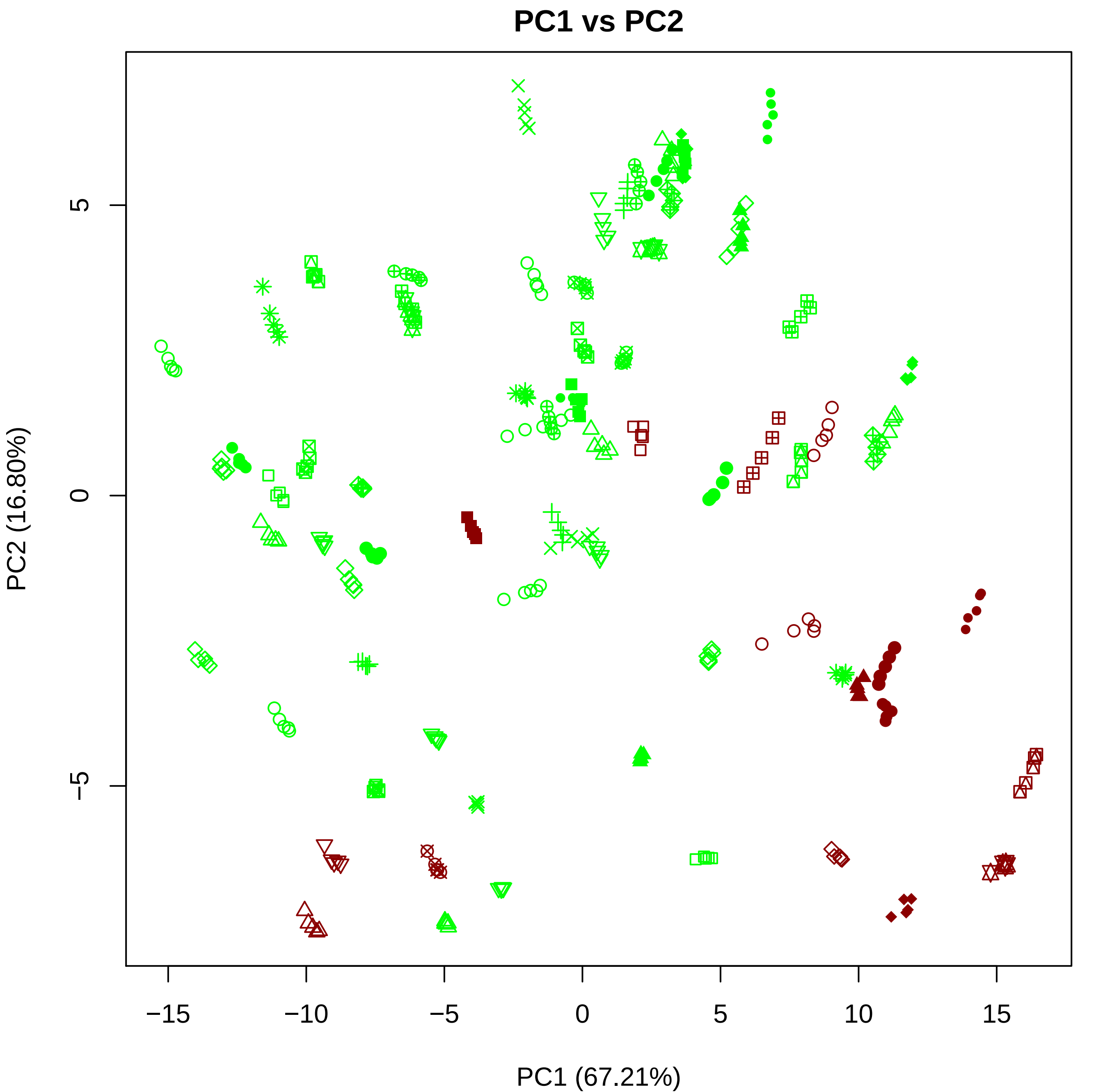

PC1 vs PC2
PC2 (16.80%)
PC1 (67.21%)
−15
−10
−5
0
5
10
15
−5
0
5

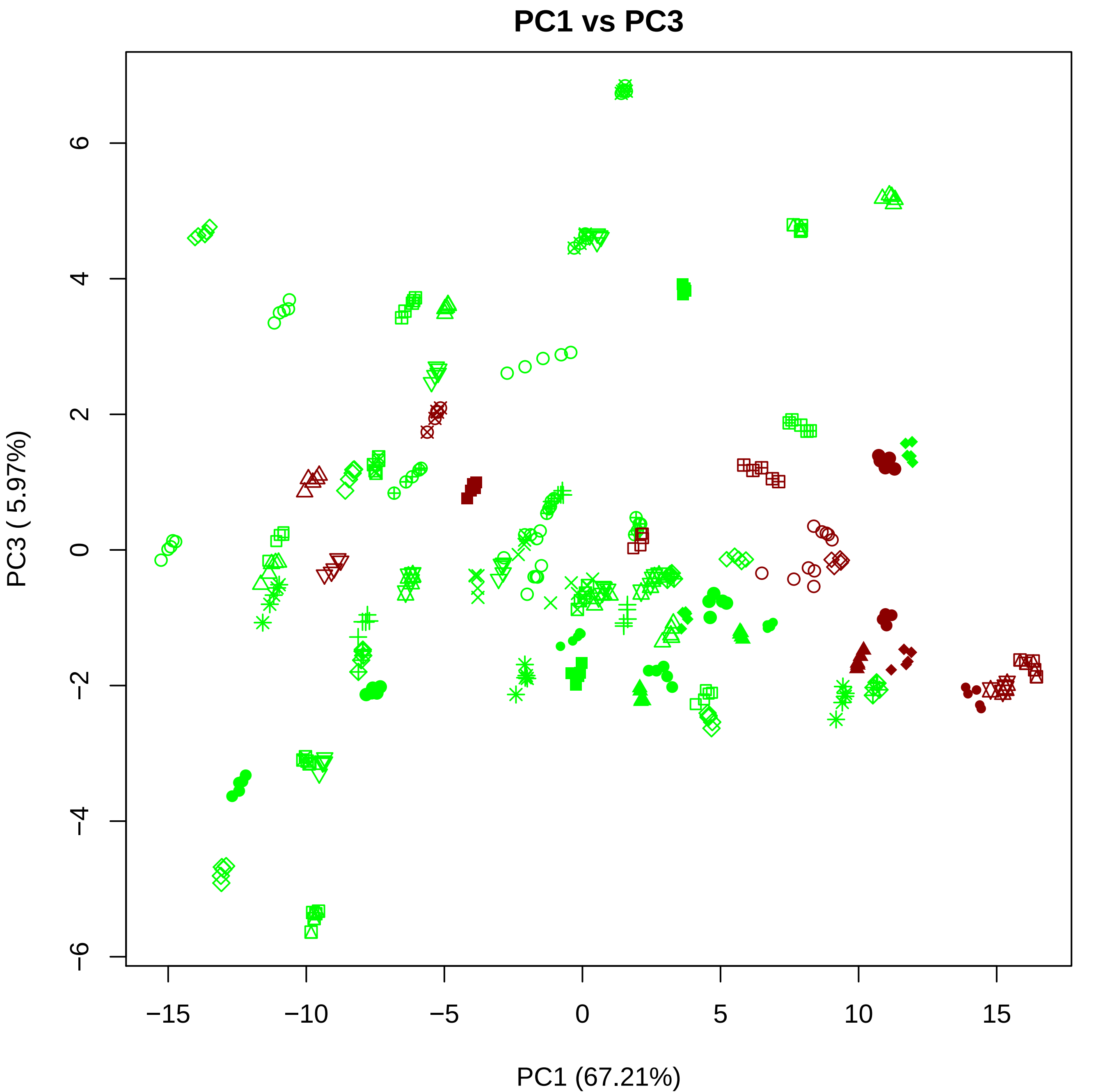
PC1 vs PC3
PC3 ( 5.97%)
PC1 (67.21%)
−15
−10
−5
0
5
10
15
−6
−4
−2
0
2
4
6

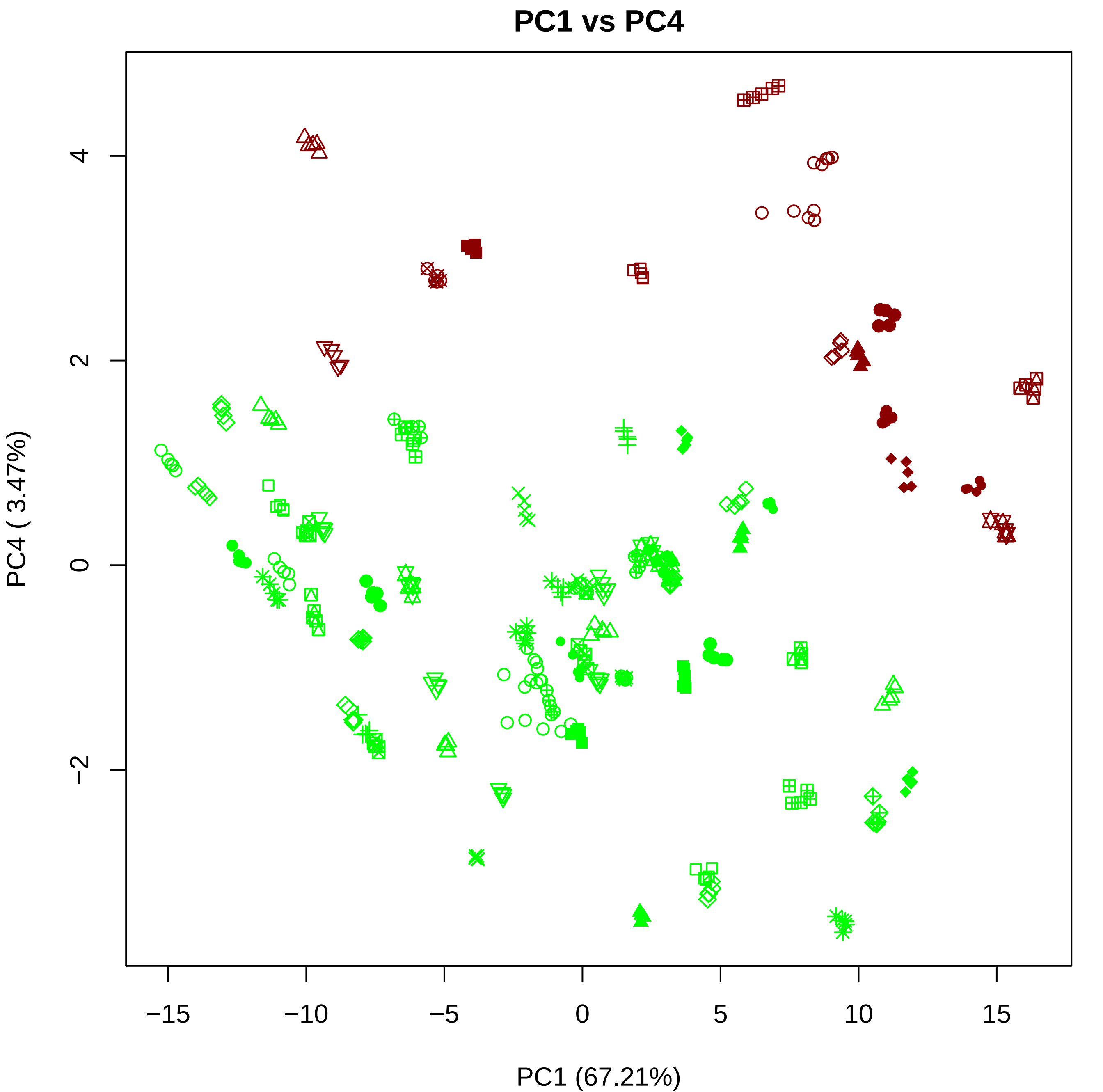
PC1 vs PC4
PC4 ( 3.47%)
PC1 (67.21%)
−15
−10
−5
0
5
10
15
−2
0
2
4

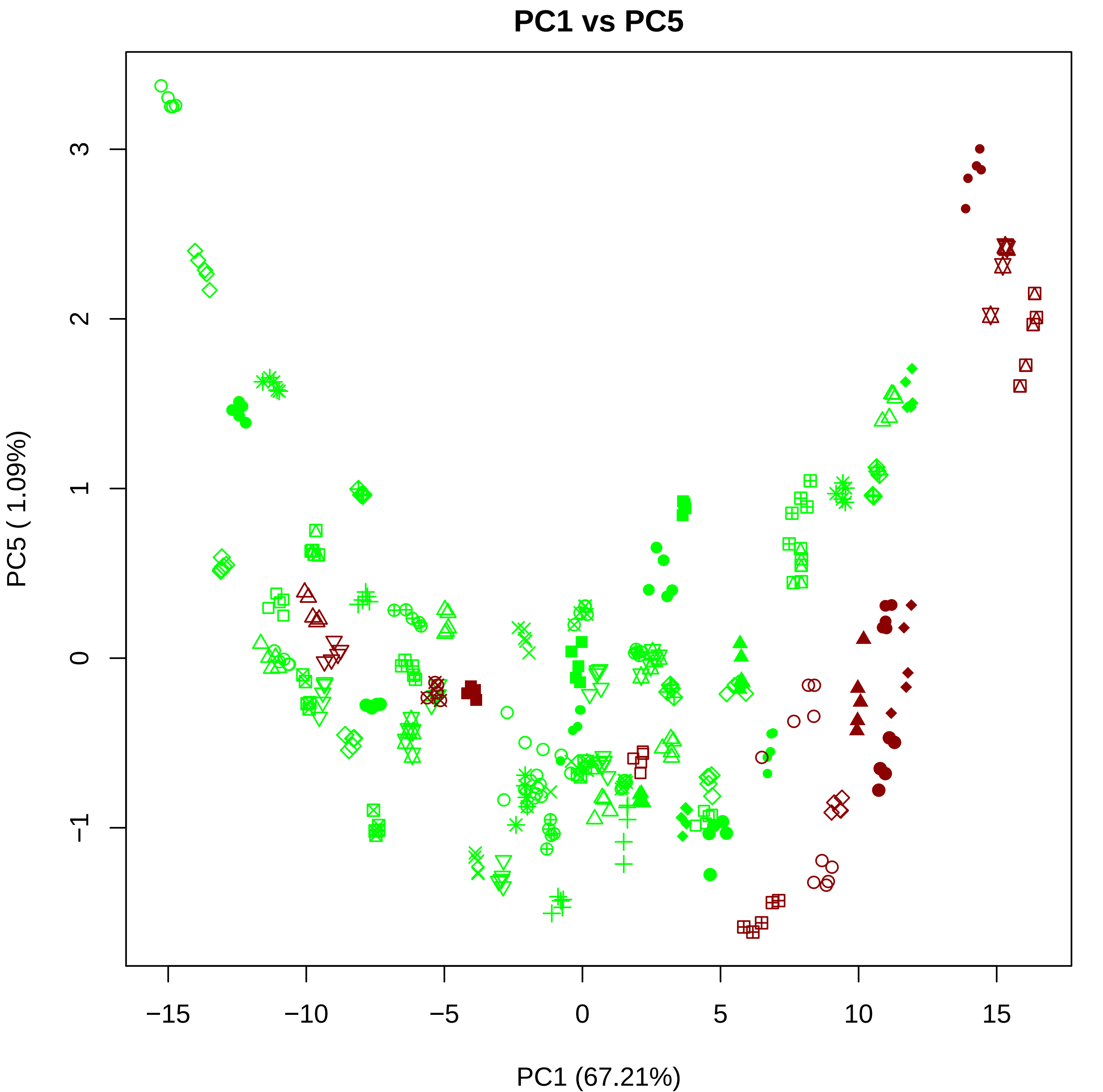

PC1 vs PC5
PC5 ( 1.09%)
PC1 (67.21%)
−15
−10
−5
0
5
10
15
−1
0
1
2
3

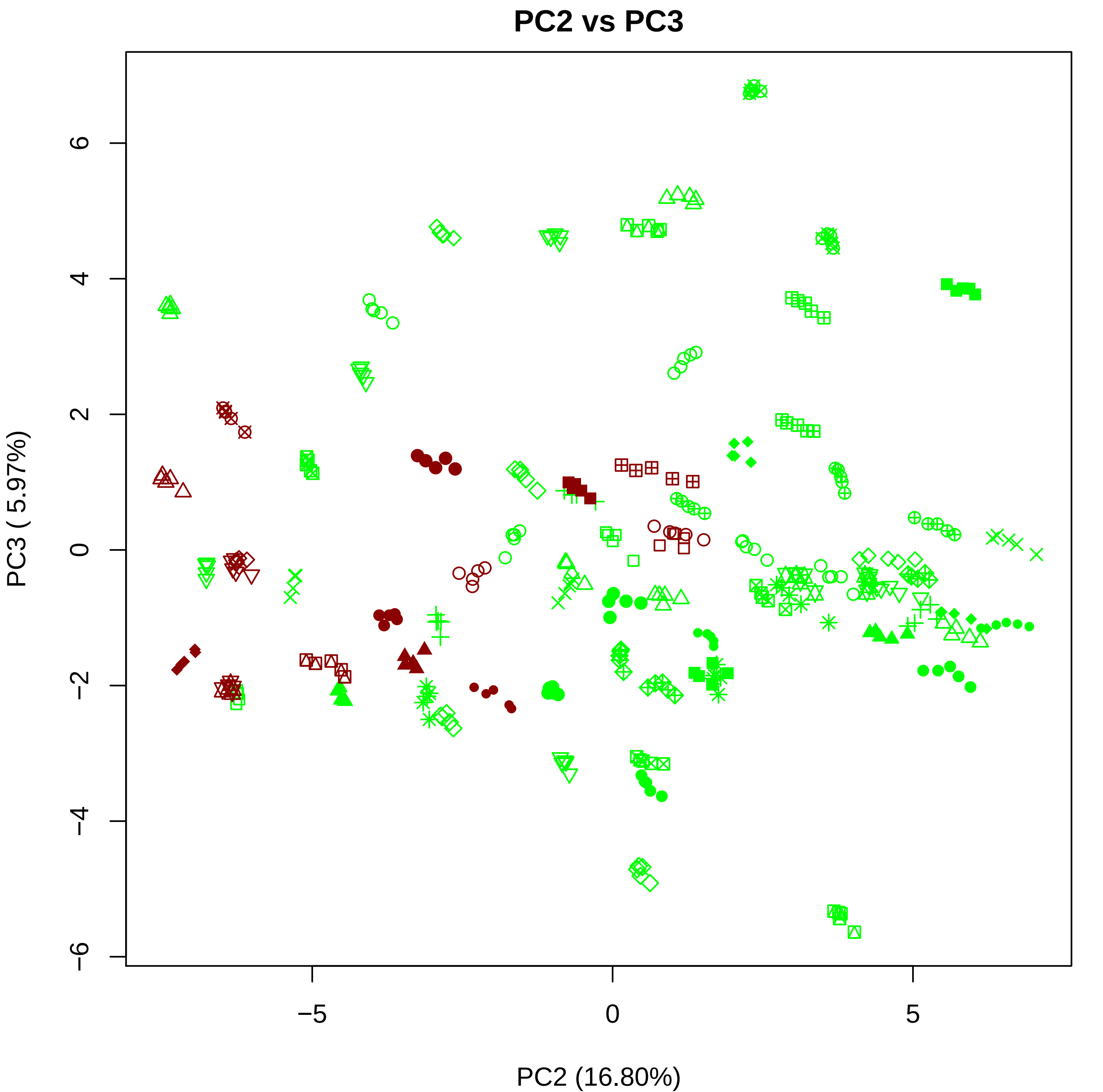

PC2 vs PC3
PC3 ( 5.97%)
PC2 (16.80%)
6
4
2
0
−2
−4
−6
−5
0
5

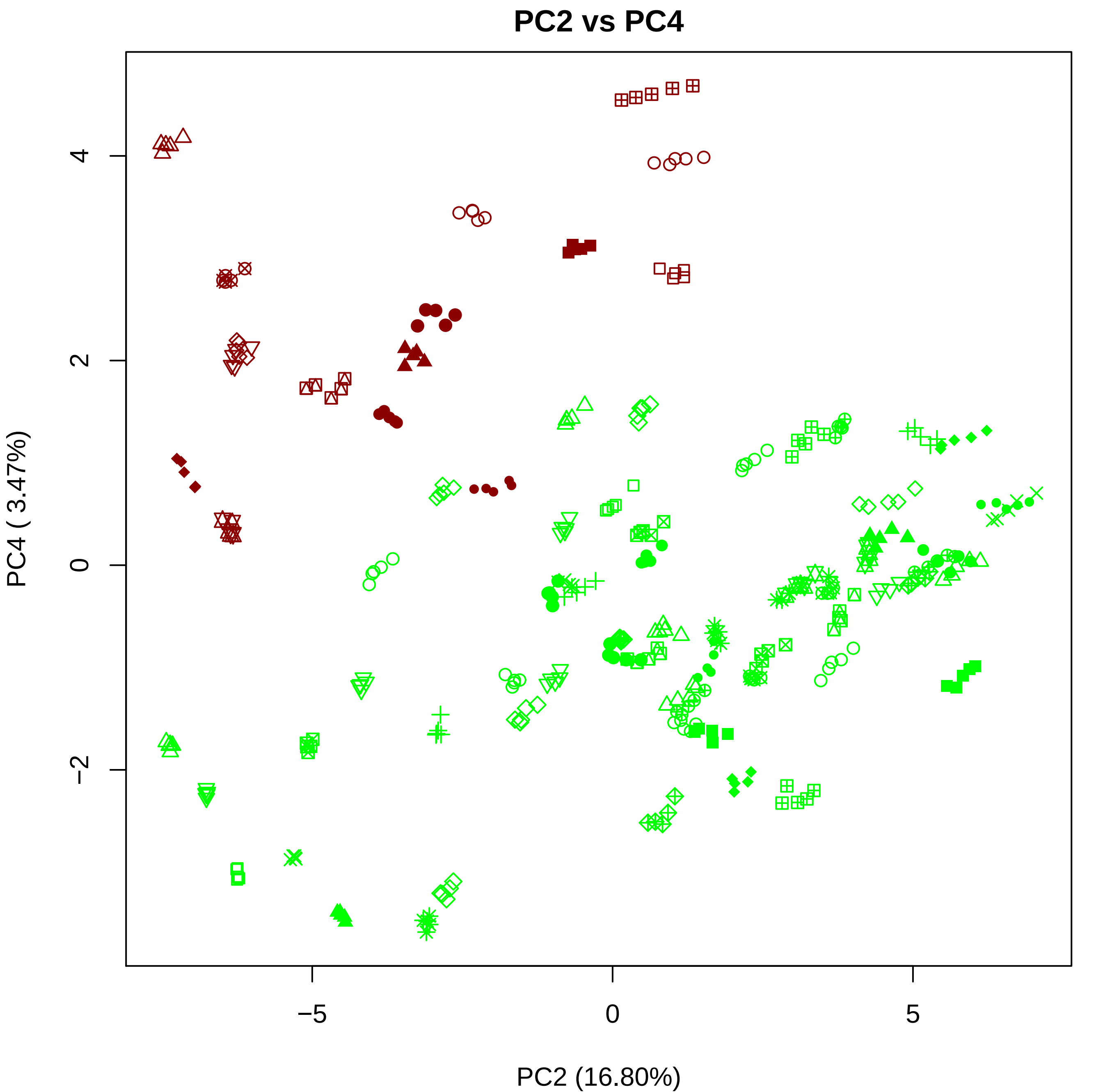
PC2 vs PC4
PC4 ( 3.47%)
4
2
0
−2
−5
0
5
PC2 (16.80%)

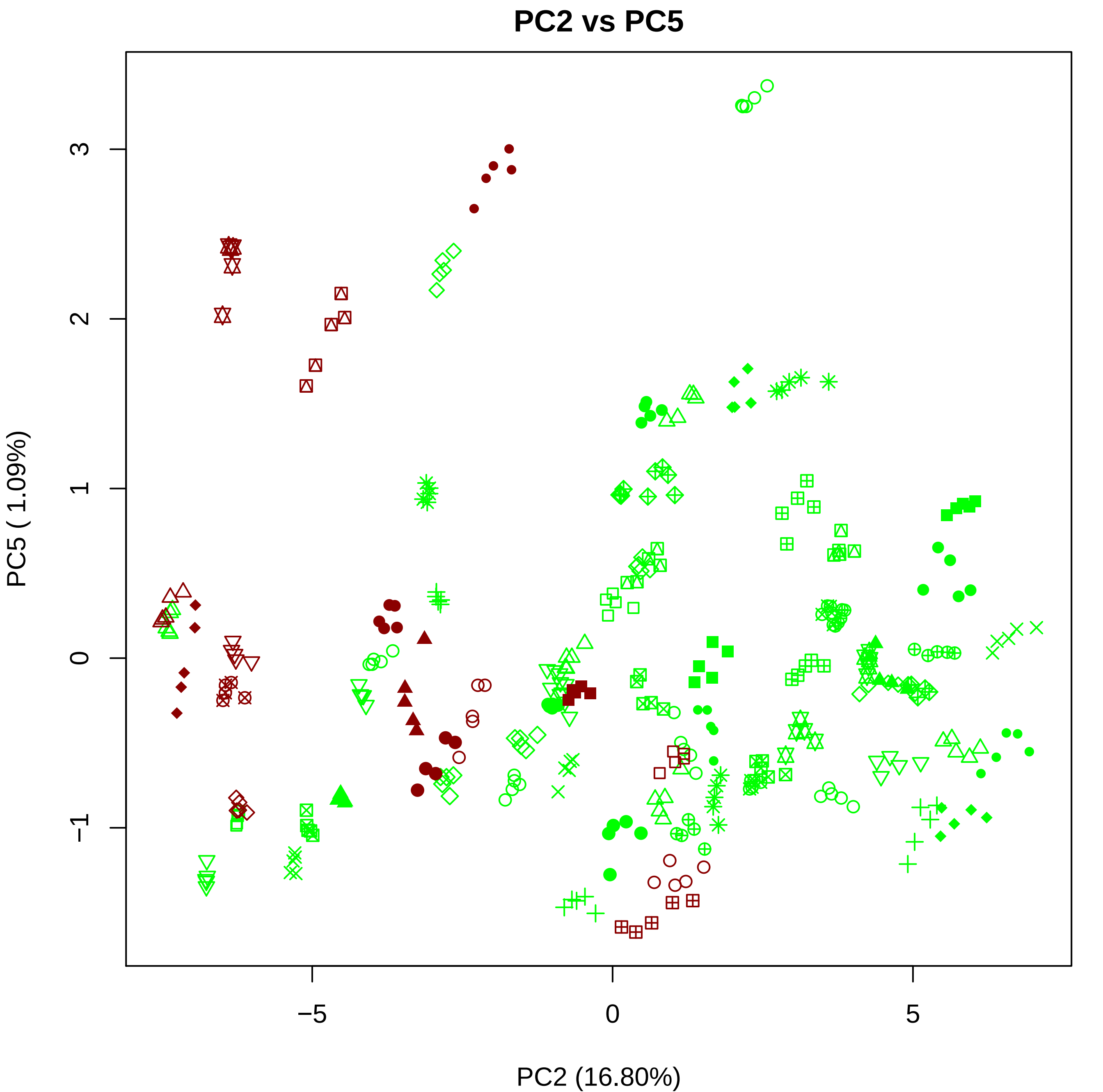
PC2 vs PC5
PC5 ( 1.09%)
PC2 (16.80%)
−5
0
5
−1
0
1
2
3

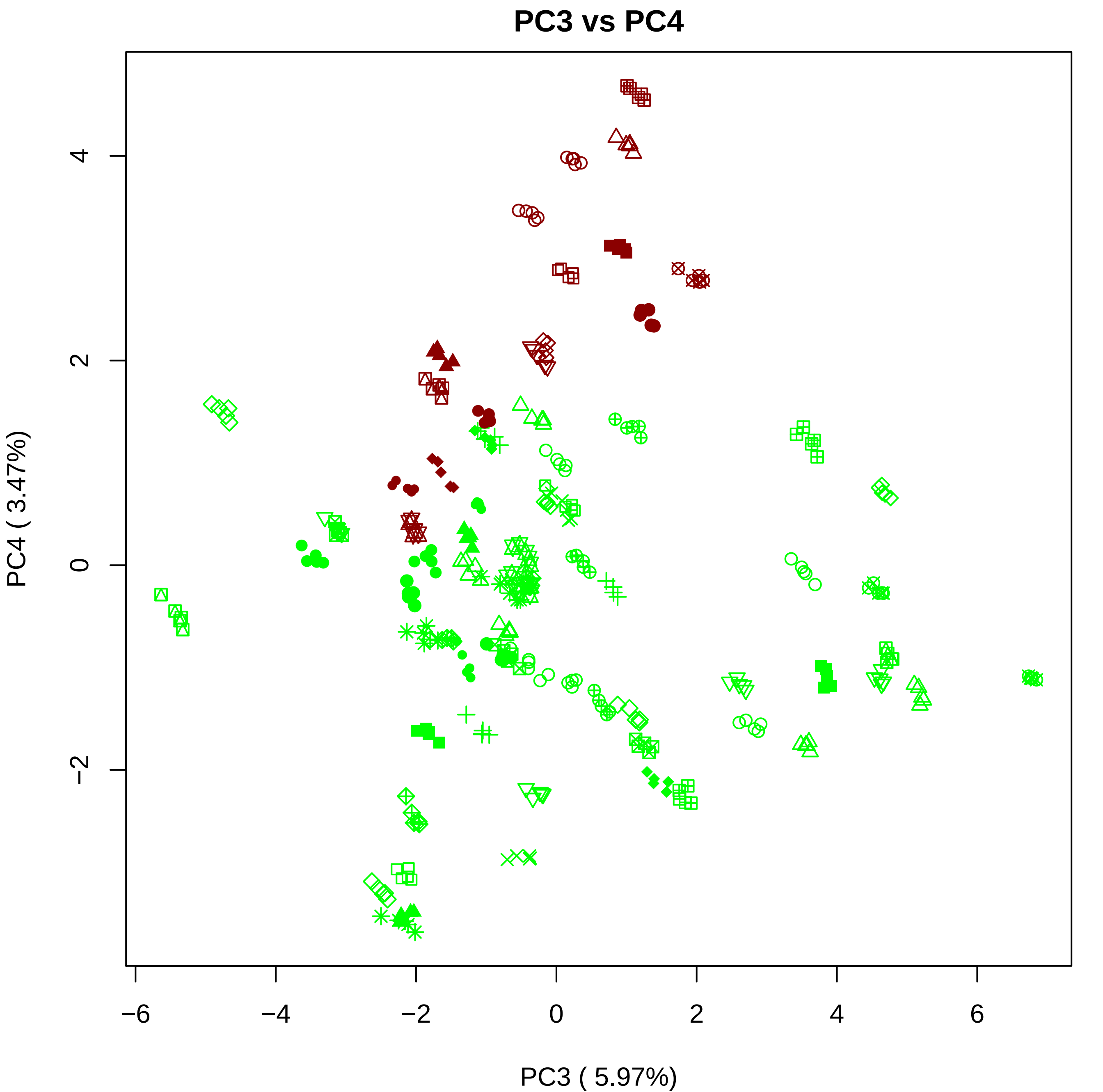

PC3 vs PC4
PC4 ( 3.47%)
PC3 ( 5.97%)
−6
−4
−2
0
2
4
6
−2
0
2
4

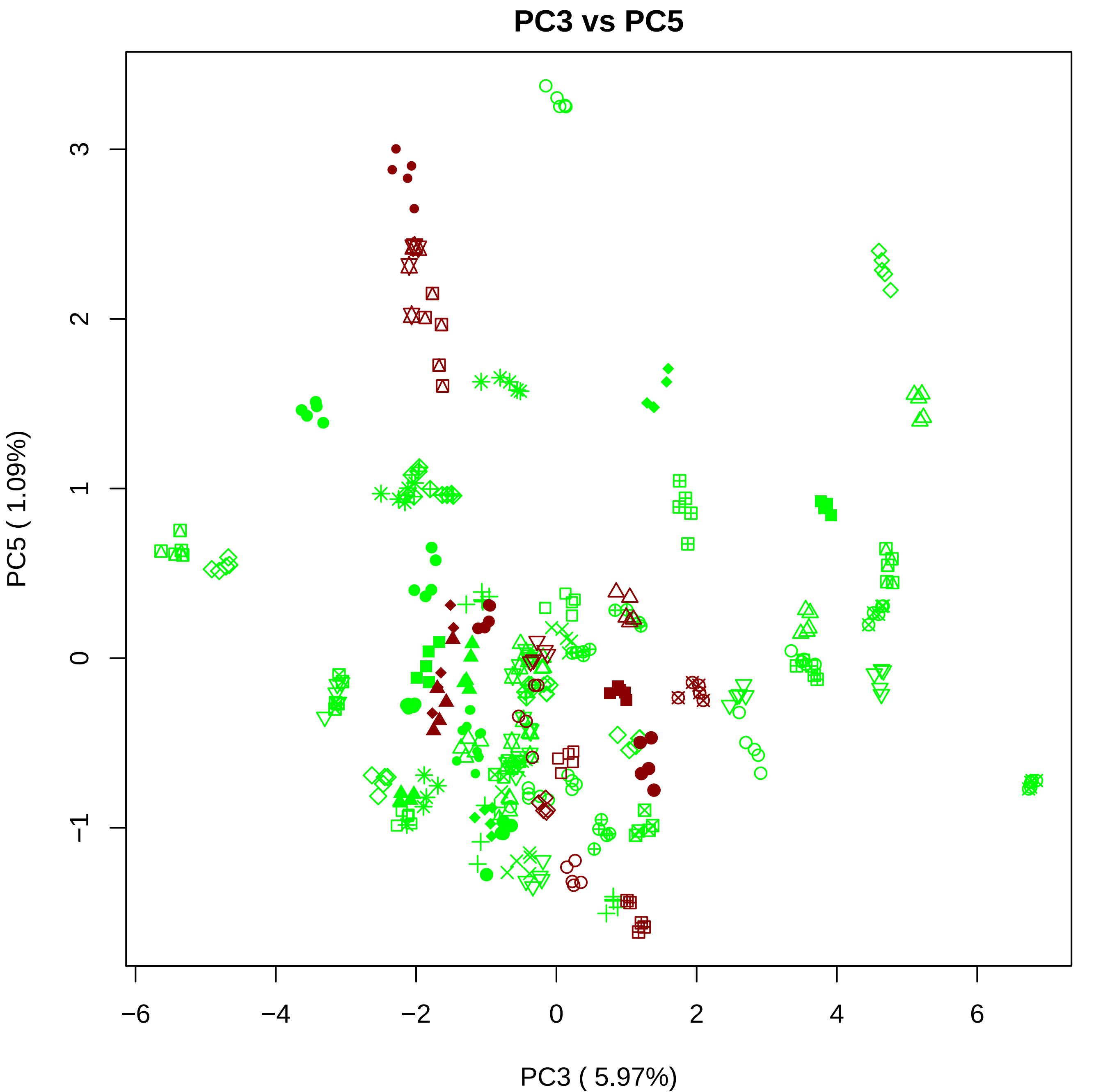

PC3 vs PC5
PC5 ( 1.09%)
PC3 ( 5.97%)
−6
−4
−2
0
2
4
6
−1
0
1
2
3

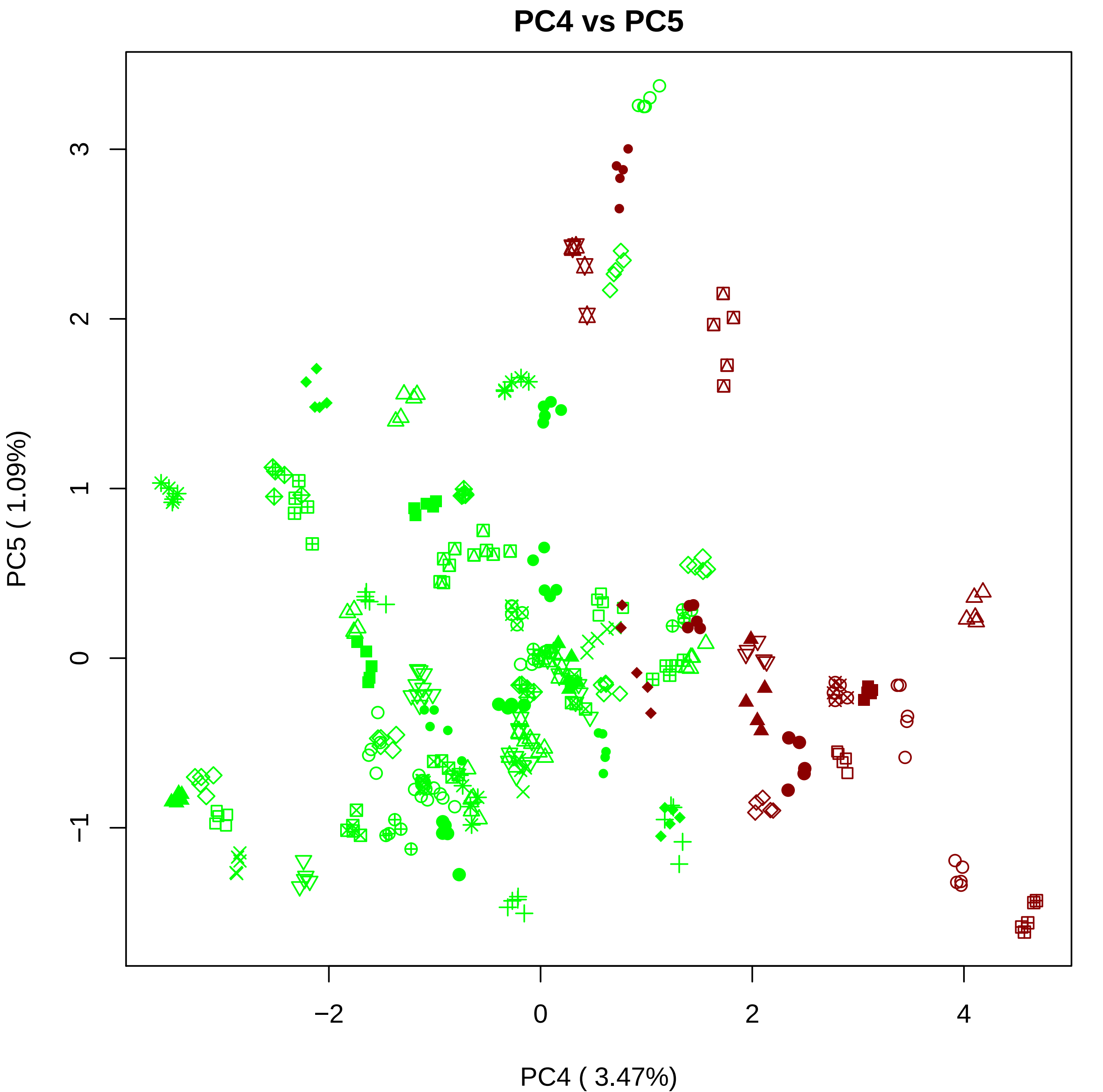

PC4 vs PC5
PC5 ( 1.09%)
PC4 ( 3.47%)
−2
0
2
4
−1
0
1
2
3

### PCA of All  BMH(a) vs PMH(a)

Scaled Matrix PCAed

7th order polynomial baseline correction performed

Raw spectra smoothed to 21 Neighbours, 2nd order PN, Savtisky Golay

Normalisation between 0 and 100

D2 spectra smoothed to 41 Neighbours, 2nd order PN, Savtisky Golay

Regions Analysed: 3601−2501 and 2001−700cm−1

Samples Analysed: 8 bContam's 0 BLH's 0 BMH's 0 OMH's 0 pContam's 0 PLH's 60 PMH's

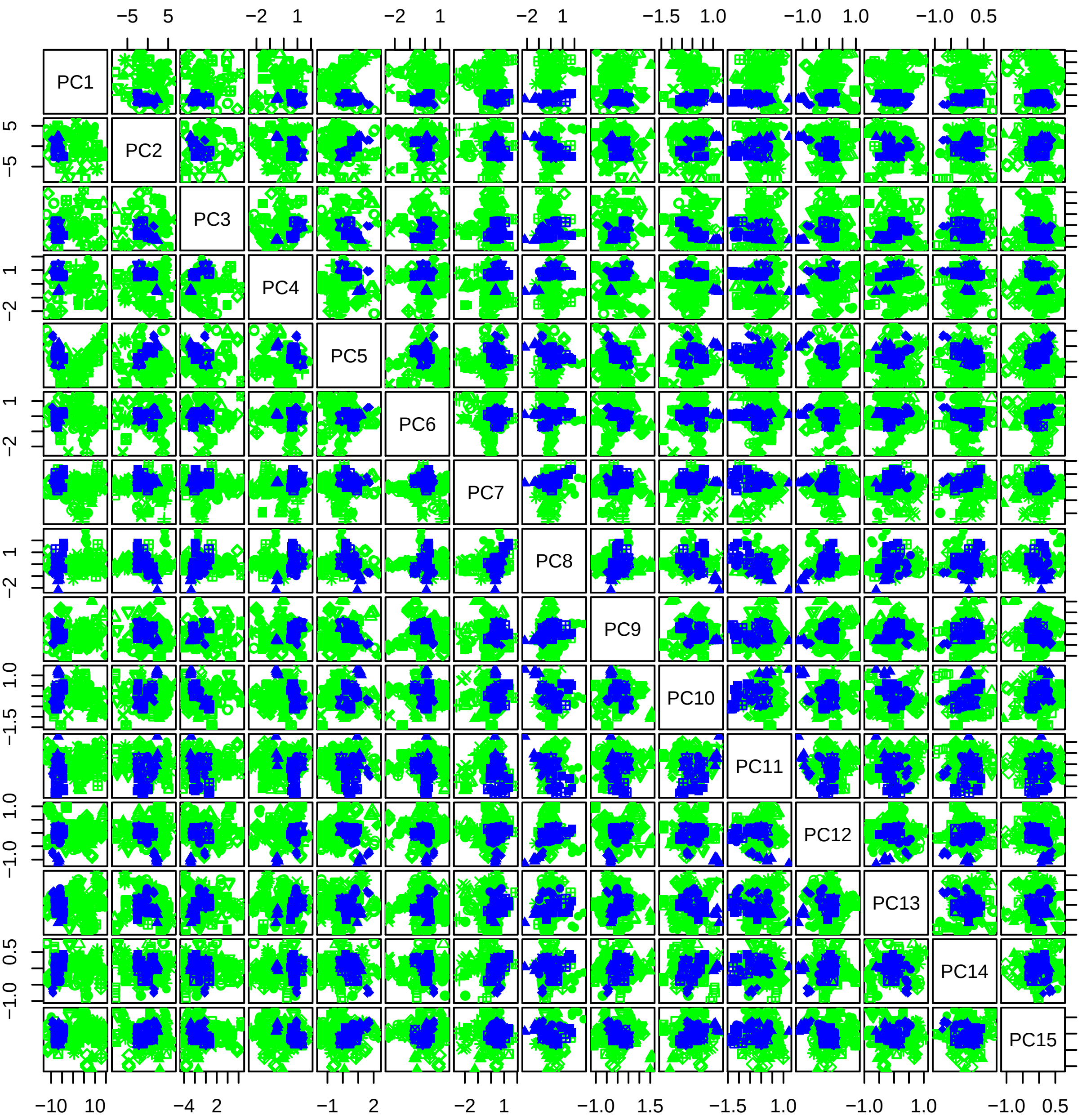
PC1
PC2
PC3
PC4
PC5
PC6
PC7
PC8
PC9
PC10
PC11
PC12
PC13
PC14
PC15

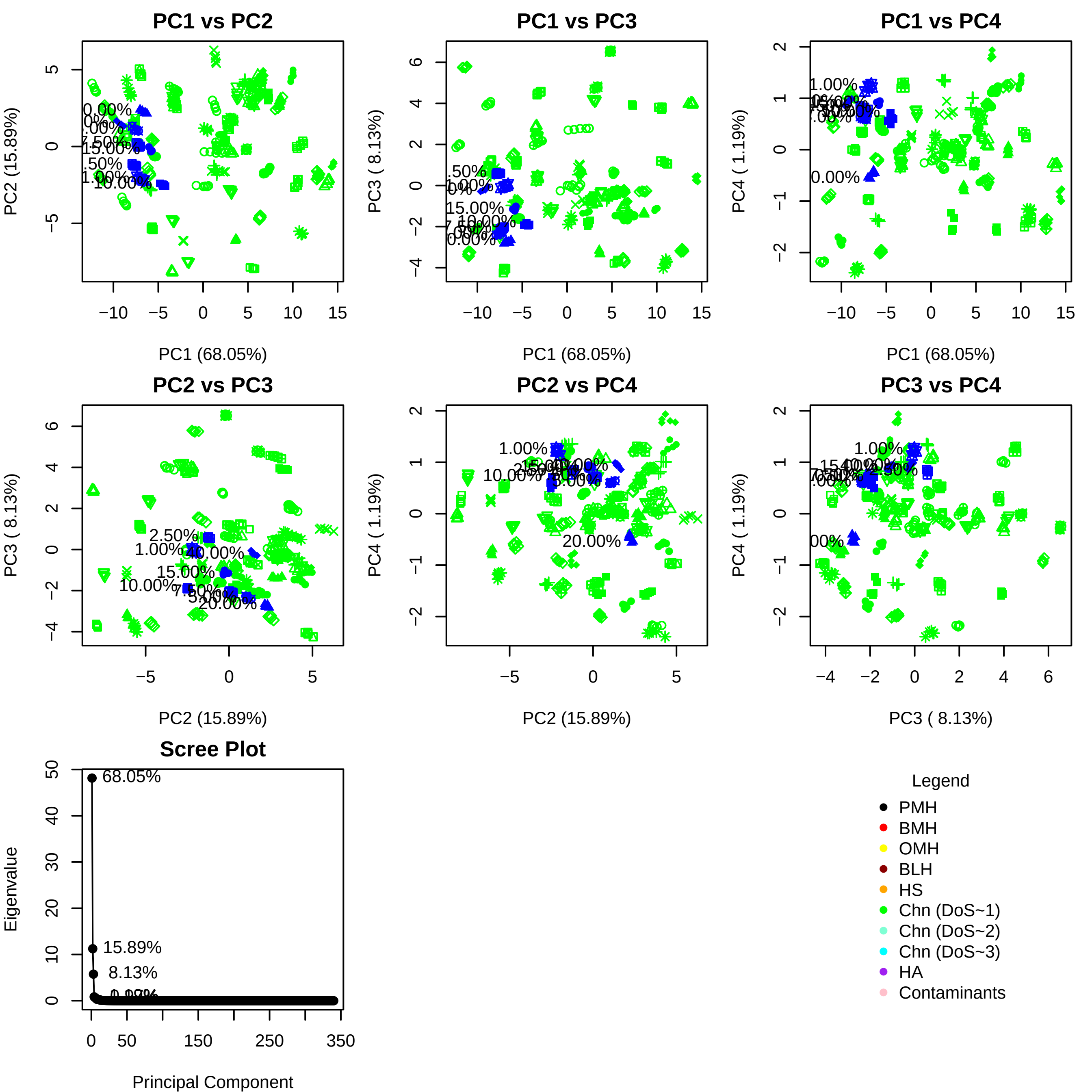

PC1 vs PC2
PC1 vs PC3
PC1 vs PC4
PC2 vs PC3
PC2 vs PC4
PC3 vs PC4
PC1 (68.05%)
PC2 (15.89%)
PC3 ( 8.13%)
PC4 ( 1.19%)
Scree Plot
Eigenvalue
Principal Component
68.05%
15.89%
8.13%
Legend
PMH
BMH
OMH
BLH
HS
Chn (DoS~1)
Chn (DoS~2)
Chn (DoS~3)
HA
Contaminants

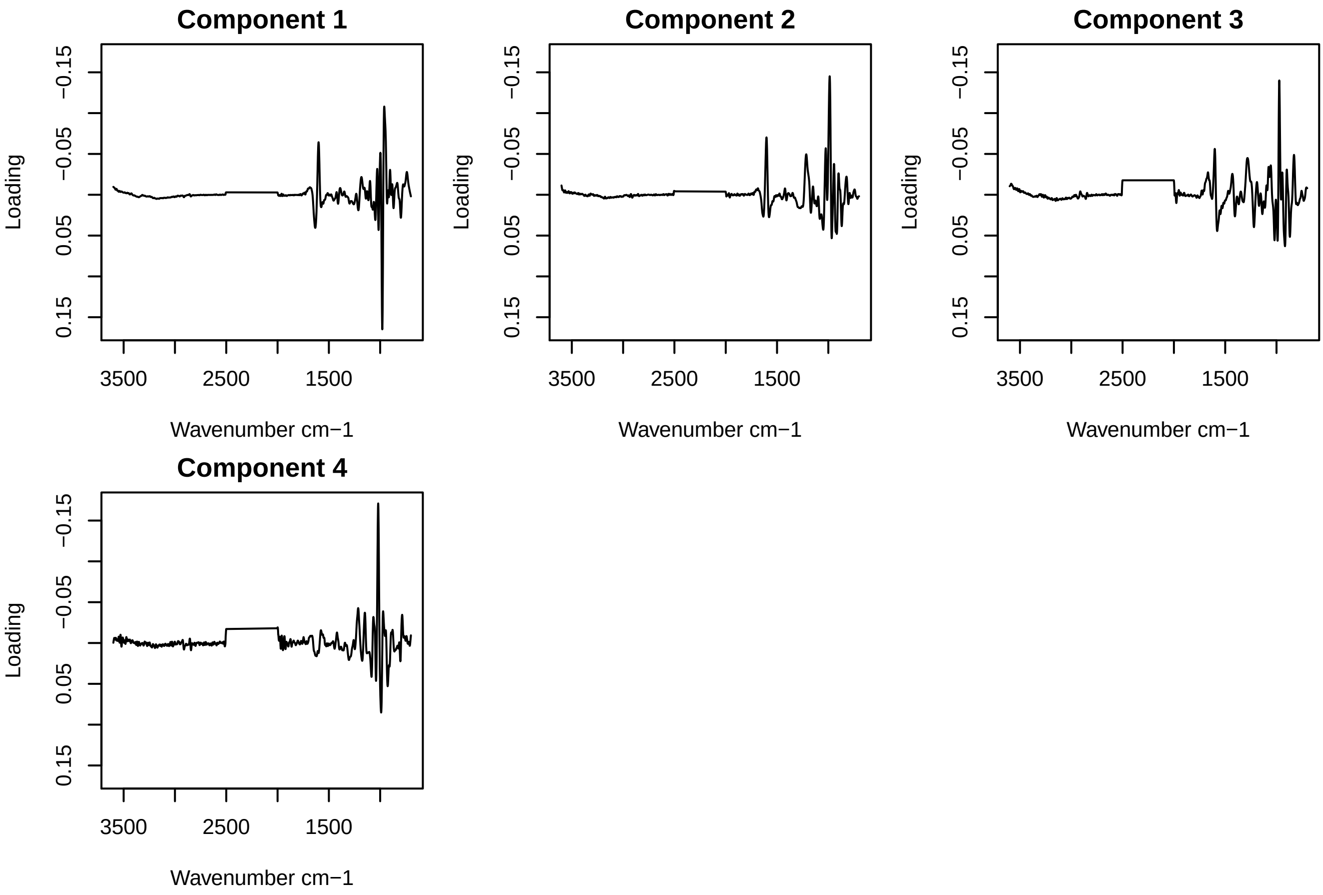
Component 1
Component 2
Component 3
Component 4
Loading
−0.15
−0.05
0.05
0.15
3500
2500
1500
Wavenumber cm−1

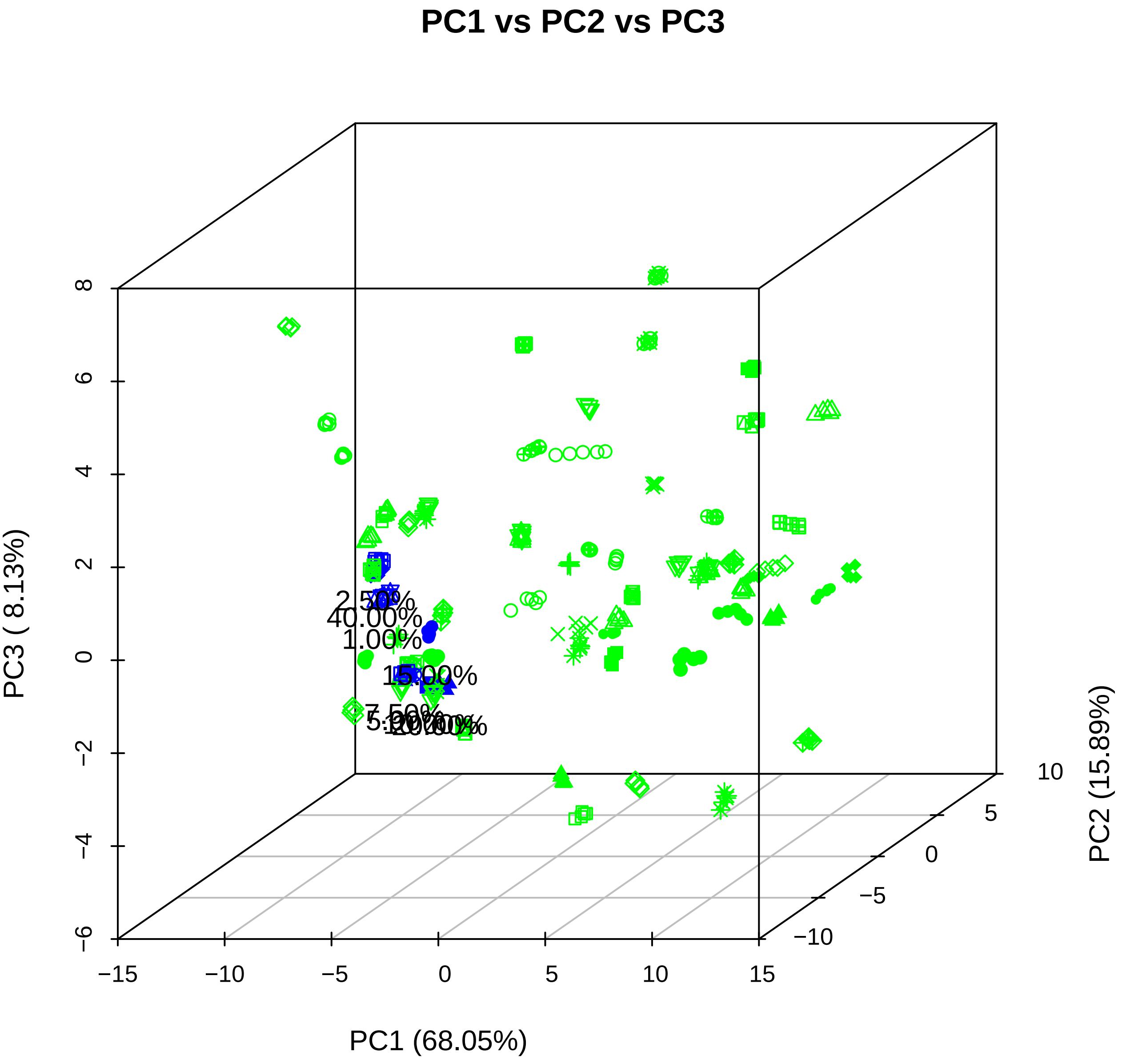

PC1 vs PC2 vs PC3
PC1 (68.05%)
PC2 (15.89%)
PC3 ( 8.13%)
−15
−10
−5
0
5
10
15
−6
−4
−2
0
2
4
6
8
2.50%
40.00%
1.00%
15.00%
7.50%

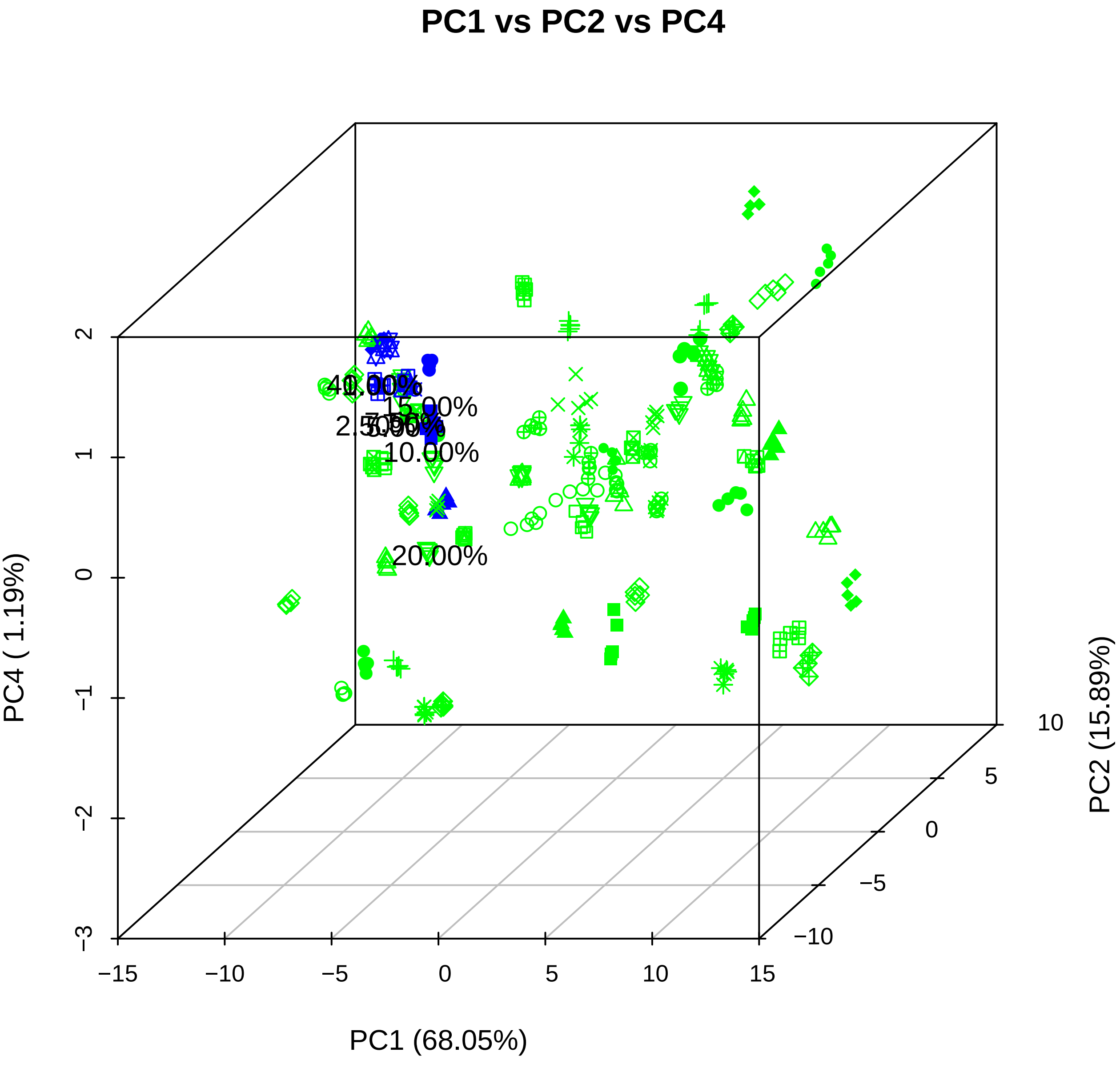
PC1 vs PC2 vs PC4
PC1 (68.05%)
PC2 (15.89%)
PC4 ( 1.19%)
−15
−10
−5
0
5
10
15
10
5
0
−5
−10
2
1
0
−1
−2
−3
15.00%
10.00%
20.00%

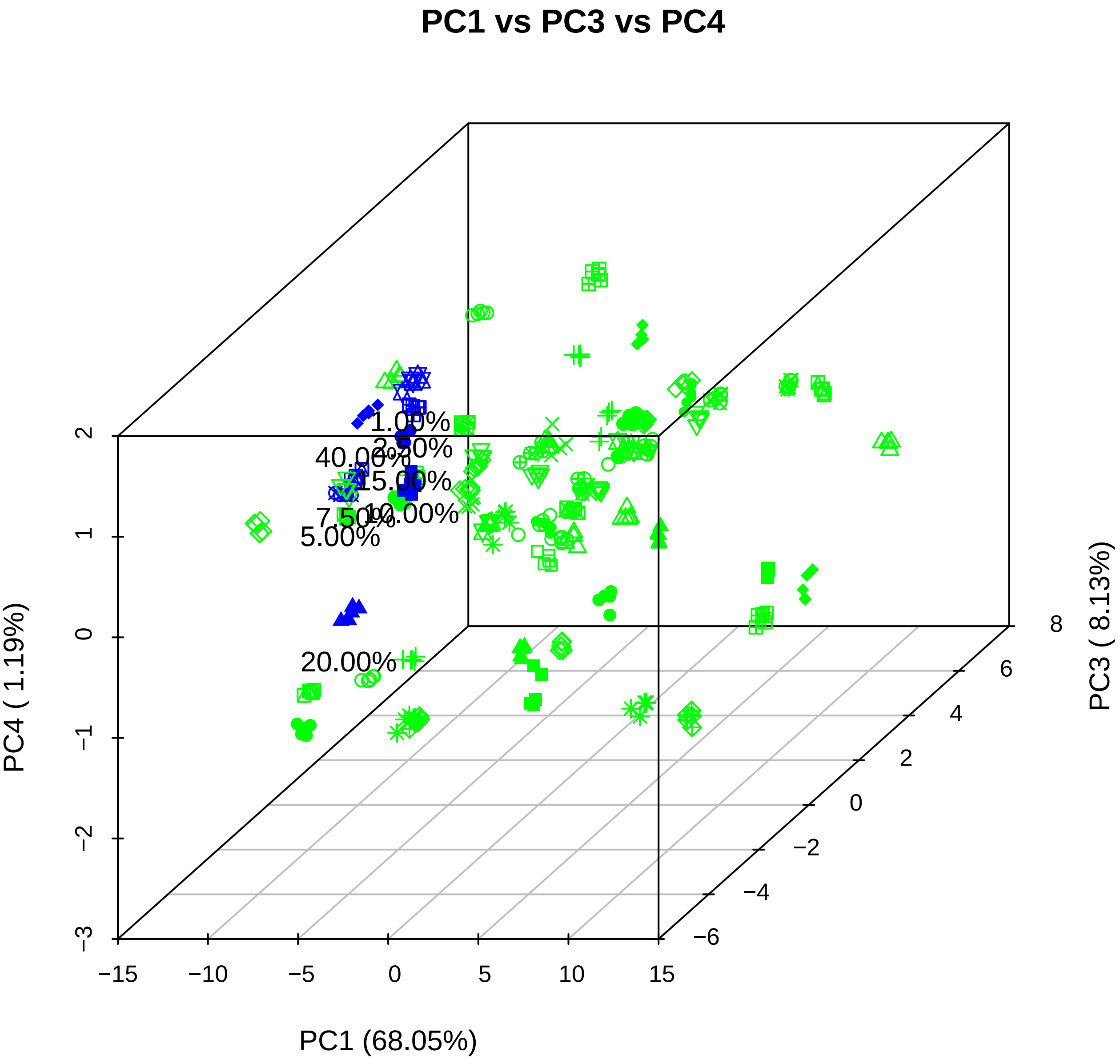
PC1 vs PC3 vs PC4
1.00%
2.50%
40.00%
15.00%
10.00%
7.50%
5.00%
20.00%
PC4 ( 1.19%)
PC1 (68.05%)
PC3 ( 8.13%)
−15
−10
−5
0
5
10
15
−3
−2
−1
0
1
2
−6
−4
−2
0
2
4
6
8

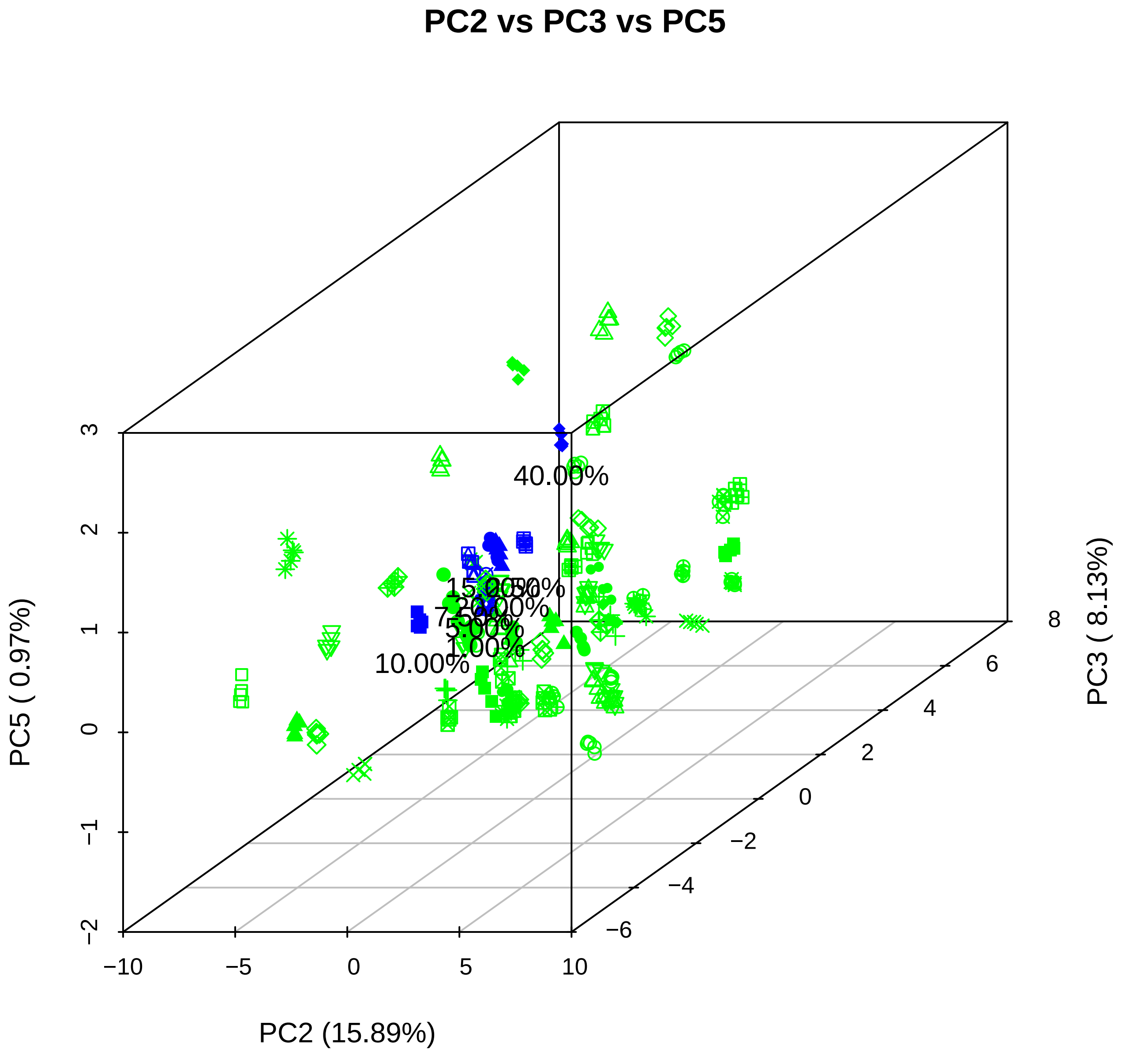
PC2 vs PC3 vs PC5
40.00%
15.00%
25.00%
20.00%
7.50%
5.00%
1.00%
10.00%
PC2 (15.89%)
PC5 ( 0.97%)
PC3 ( 8.13%)
−10
−5
0
5
10
−2
−1
0
1
2
3
−6
−4
−2
0
2
4
6
8

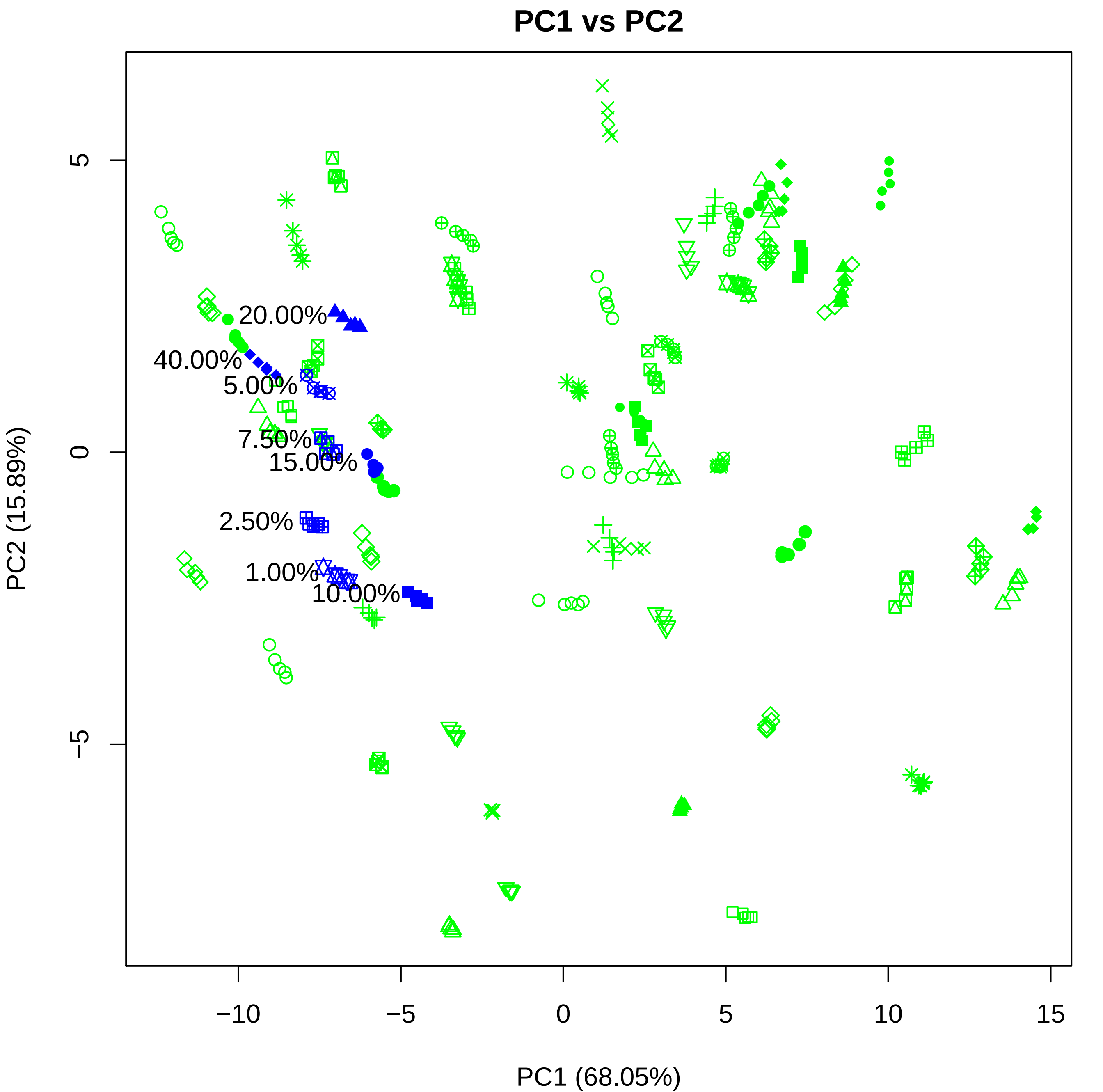

PC1 vs PC2
20.00%
40.00%
5.00%
7.50%
15.00%
2.50%
1.00%
10.00%
PC1 (68.05%)
PC2 (15.89%)
−10
−5
0
5
10
15

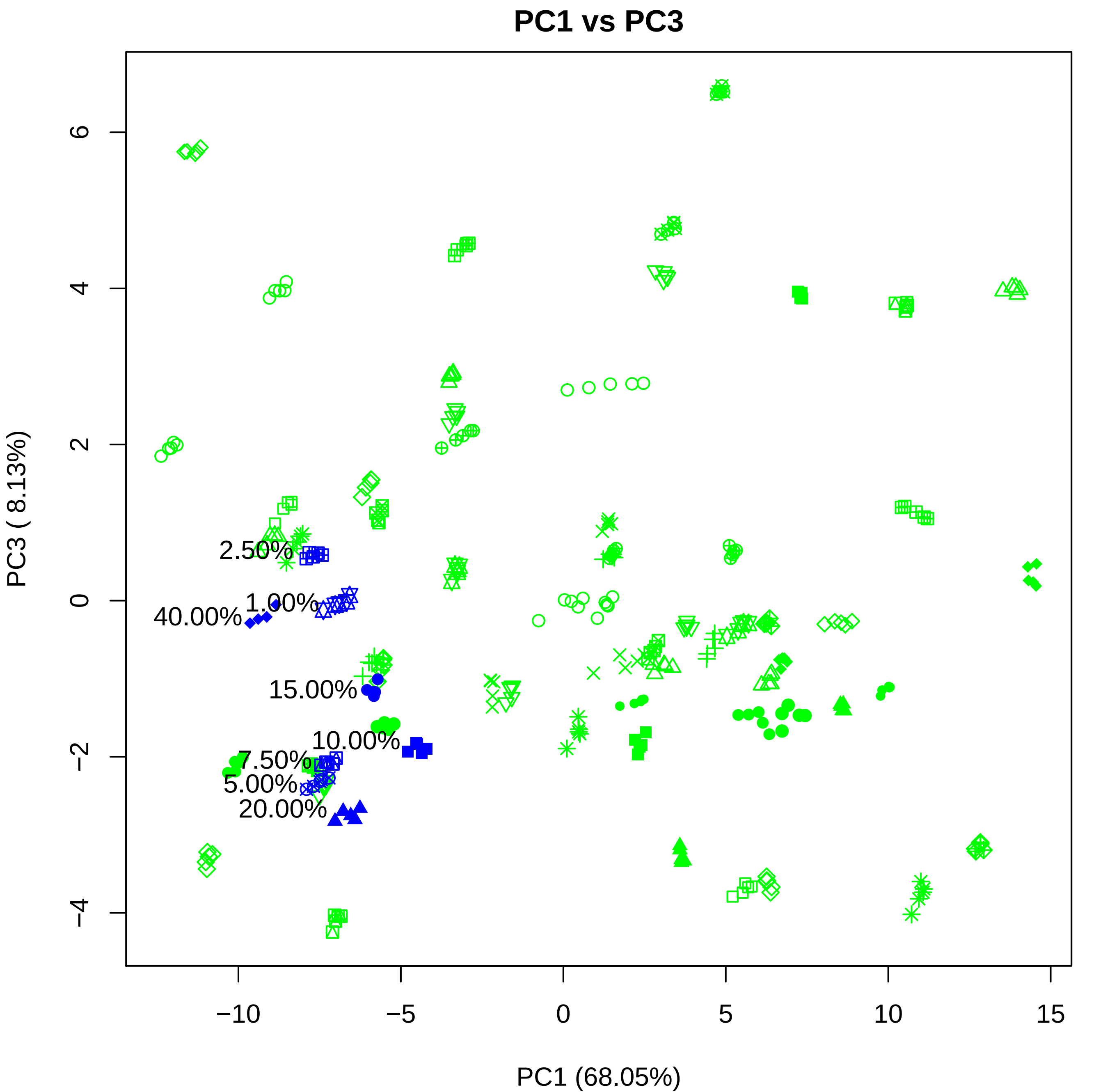
PC1 vs PC3
PC3 ( 8.13%)
PC1 (68.05%)
6
4
2
0
−2
−4
−10
−5
0
5
10
15
2.50%
1.00%
40.00%
15.00%
10.00%
7.50%
5.00%
20.00%

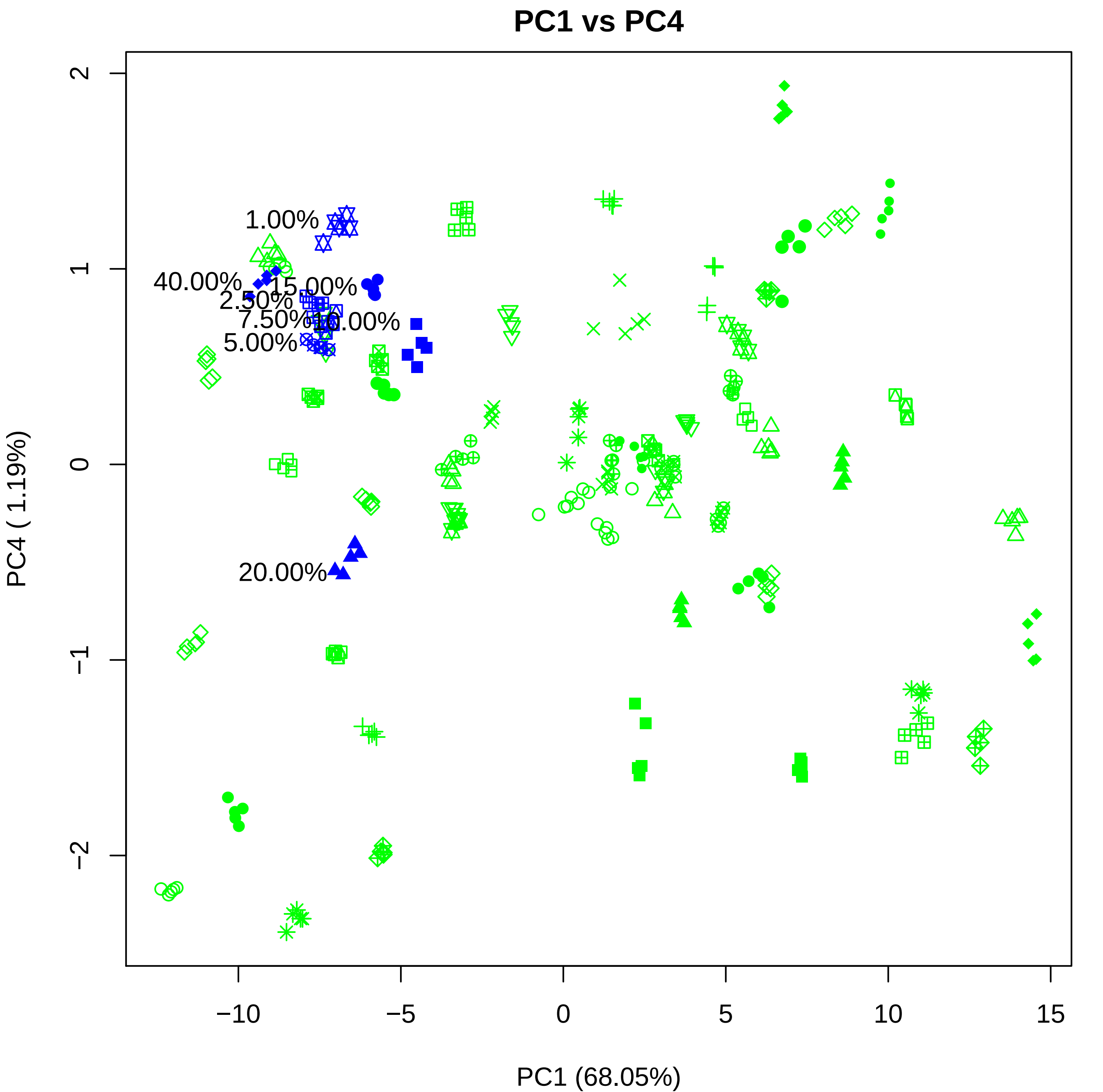

PC1 vs PC4
1.00%
40.00%
15.00%
2.50%
7.50%
10.00%
5.00%
20.00%
PC4 ( 1.19%)
PC1 (68.05%)
2
1
0
−1
−2
−10
−5
0
5
10
15

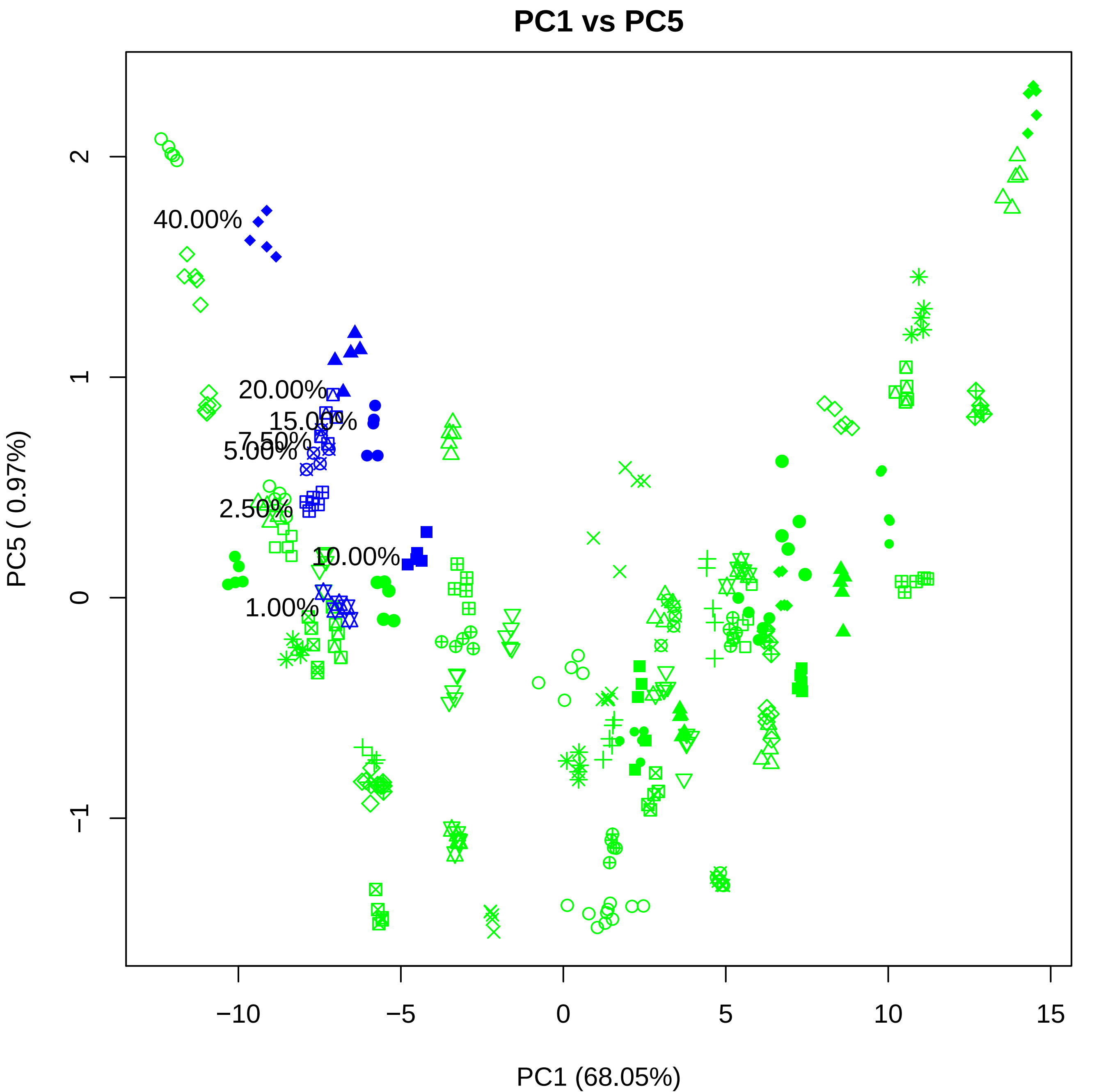
PC1 vs PC5
PC5 ( 0.97%)
PC1 (68.05%)
−1
0
1
2
−10
−5
0
5
10
15
40.00%
20.00%
15.00%
7.50%
5.00%
2.50%
10.00%
1.00%

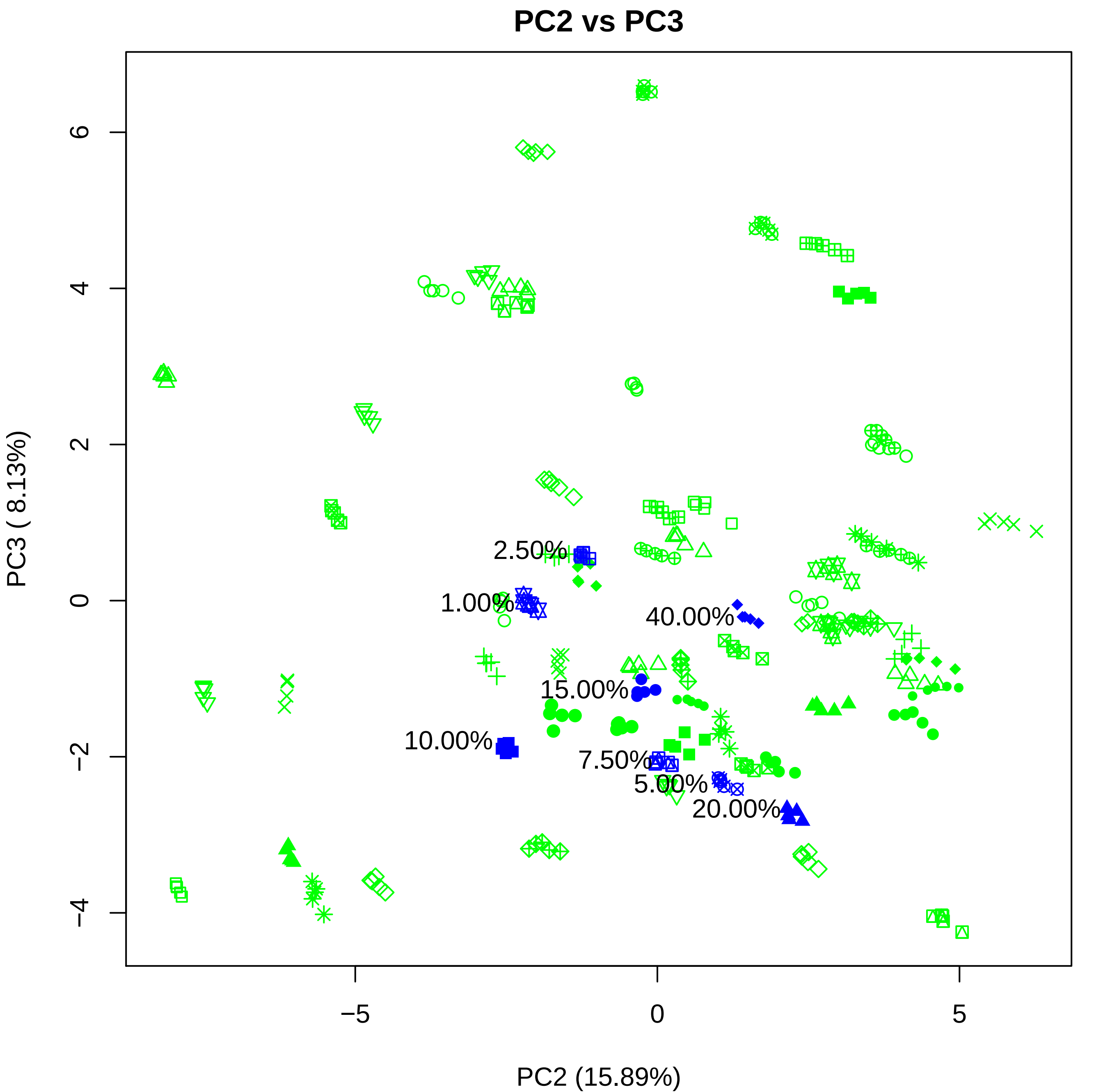

PC2 vs PC3
2.50%
1.00%
40.00%
15.00%
10.00%
7.50%
5.00%
20.00%
6
4
2
0
−2
−4
−5
0
5
PC3 ( 8.13%)
PC2 (15.89%)

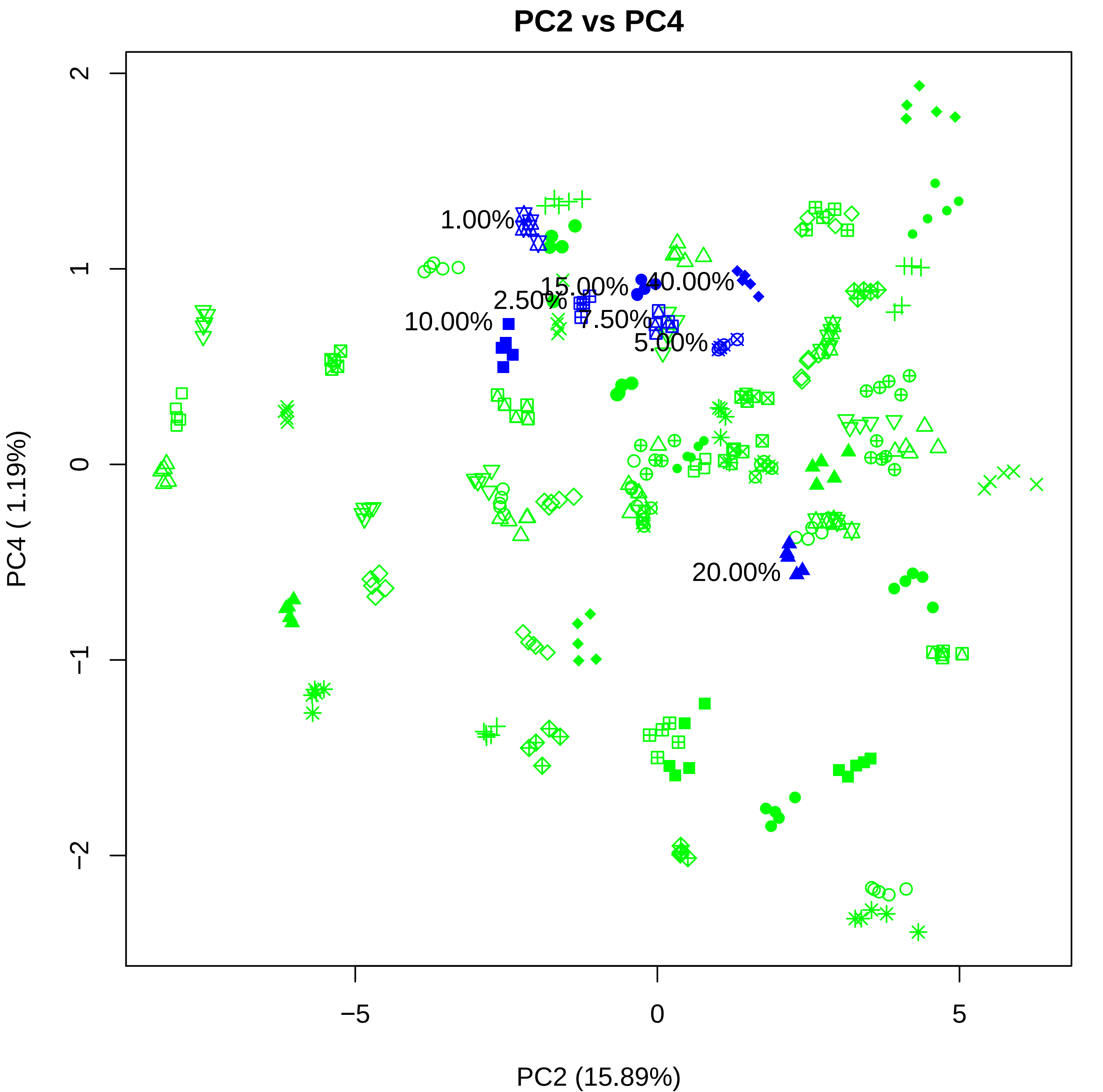
PC2 vs PC4
PC4 ( 1.19%)
PC2 (15.89%)
2
1
0
−1
−2
−5
0
5
1.00%
15.00%
40.00%
2.50%
10.00%
7.50%
5.00%
20.00%

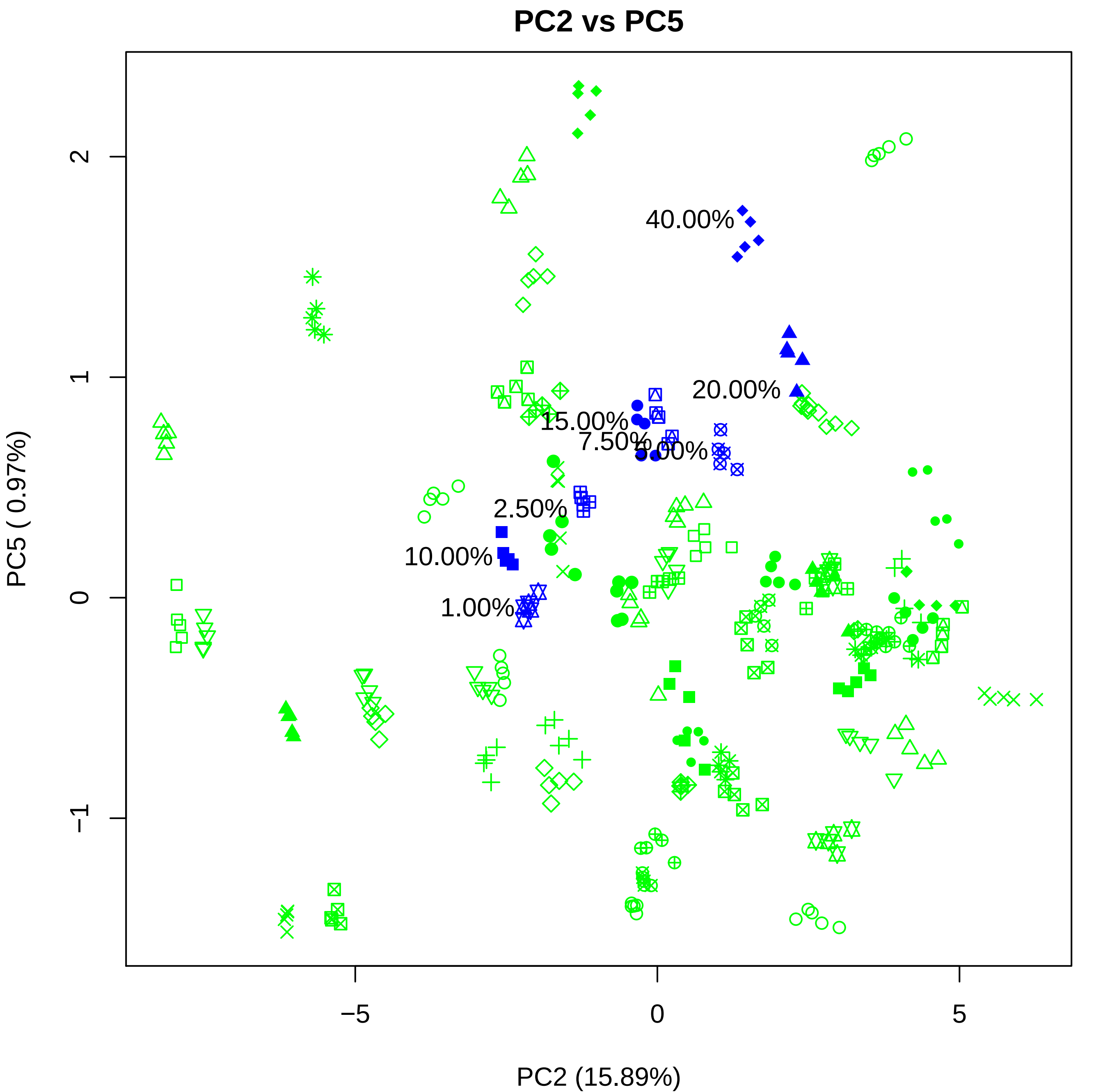

PC2 vs PC5
40.00%
20.00%
15.00%
7.50%
5.00%
2.50%
10.00%
1.00%
2
1
0
−1
−5
0
5
PC5 ( 0.97%)
PC2 (15.89%)

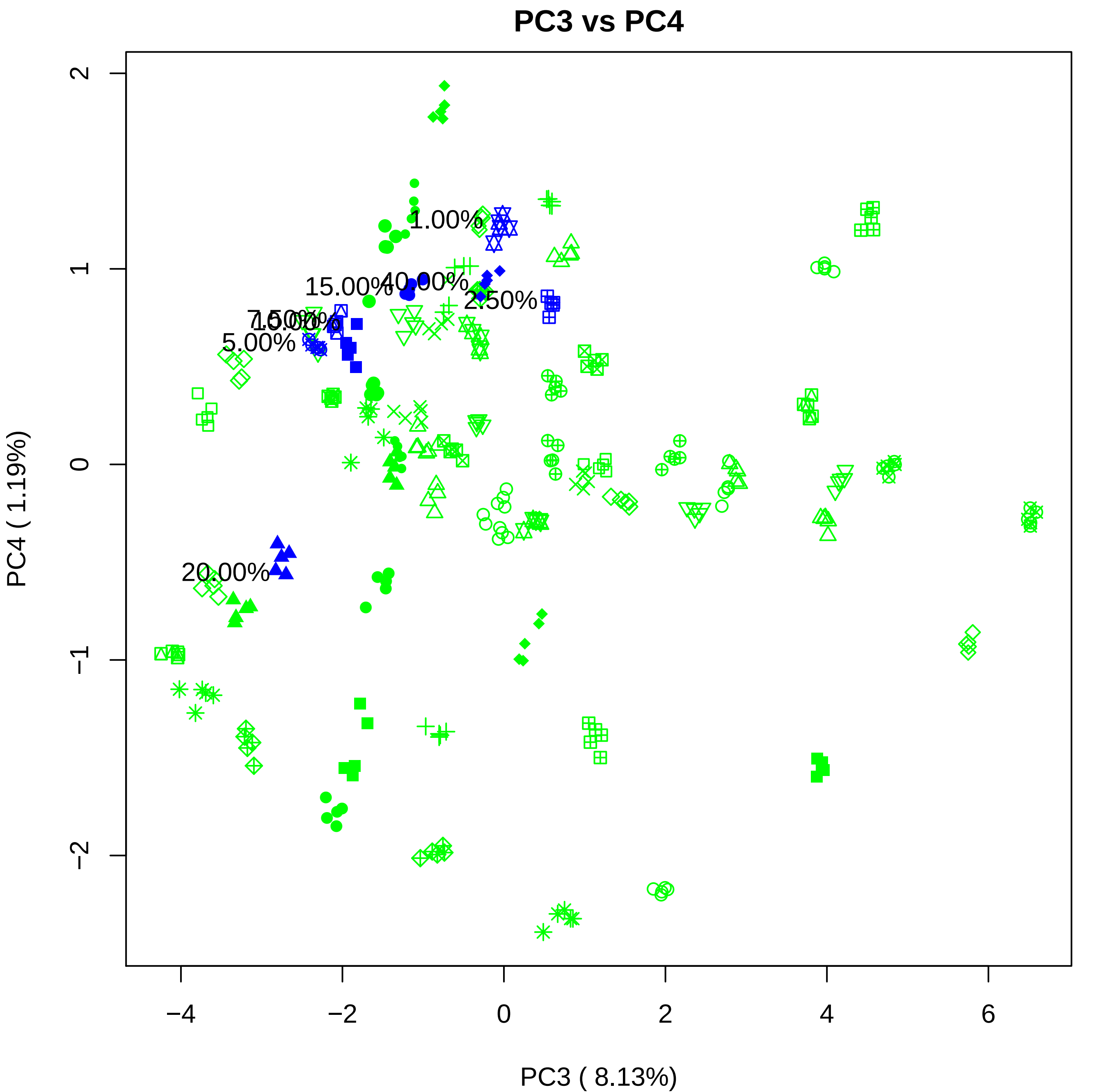
PC3 vs PC4
PC4 ( 1.19%)
PC3 ( 8.13%)
2
1
0
−1
−2
−4
−2
0
2
4
6
1.00%
15.00%
40.00%
2.50%
7.50%
10.00%
5.00%
20.00%

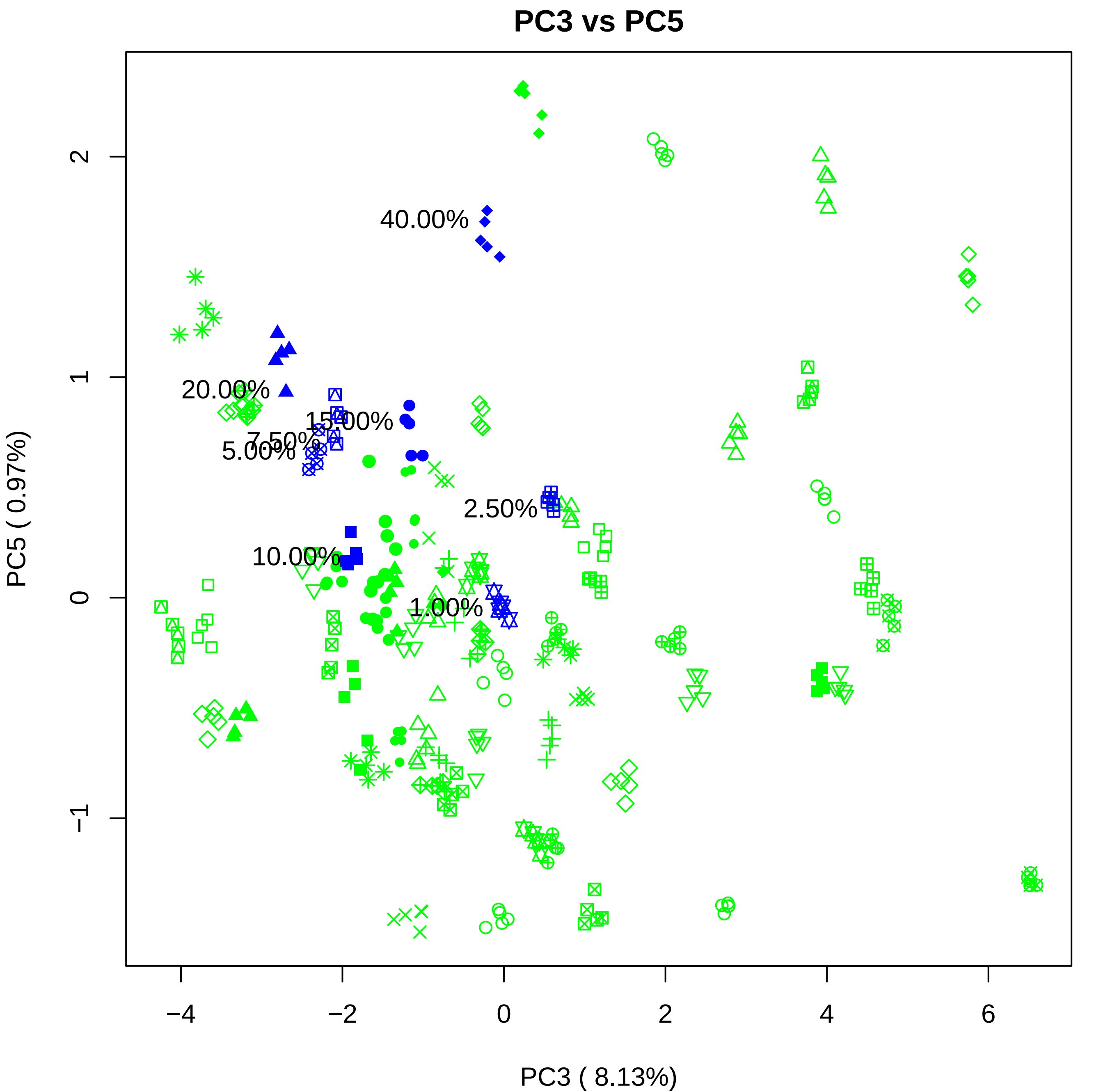
PC3 vs PC5
40.00%
20.00%
15.00%
7.50%
5.00%
2.50%
10.00%
1.00%
PC5 ( 0.97%)
PC3 ( 8.13%)
−4
−2
0
2
4
6
2
1
0
−1

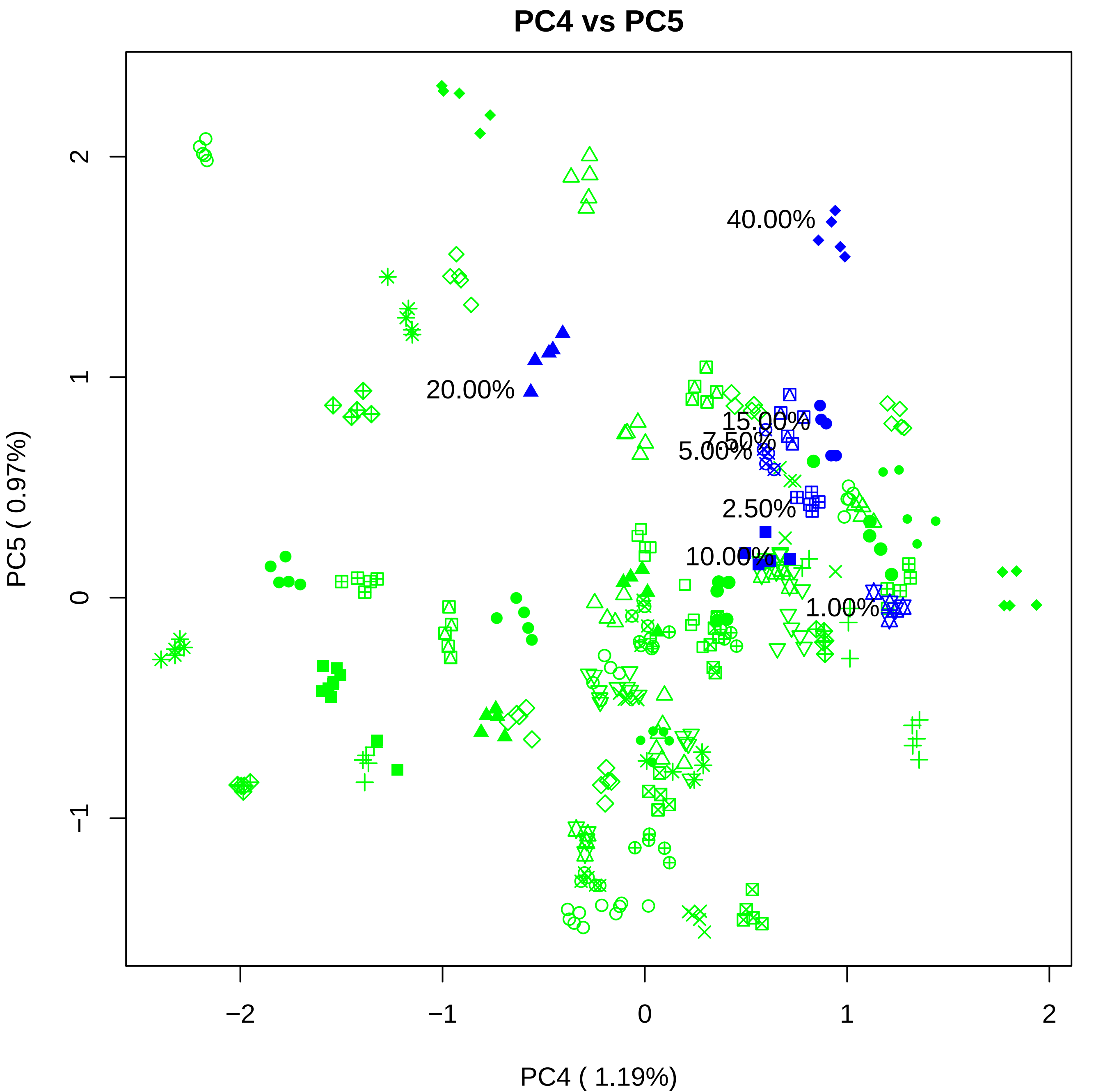
PC4 vs PC5
PC5 ( 0.97%)
PC4 ( 1.19%)
40.00%
20.00%
15.00%
7.50%
5.00%
2.50%
10.00%
1.00%
−2
−1
0
1
2

### PCA of 1%  BMH(a) vs PMH(a)

Scaled Matrix PCAed

7th order polynomial baseline correction performed

Raw spectra smoothed to 21 Neighbours, 2nd order PN, Savtisky Golay

Normalisation between 0 and 100

D2 spectra smoothed to 41 Neighbours, 2nd order PN, Savtisky Golay

Regions Analysed: 3601−2501 and 2001−700cm−1

Samples Analysed: 1 bContam's 0 BLH's 0 BMH's 0 OMH's 0 pContam's 0 PLH's 60 PMH's

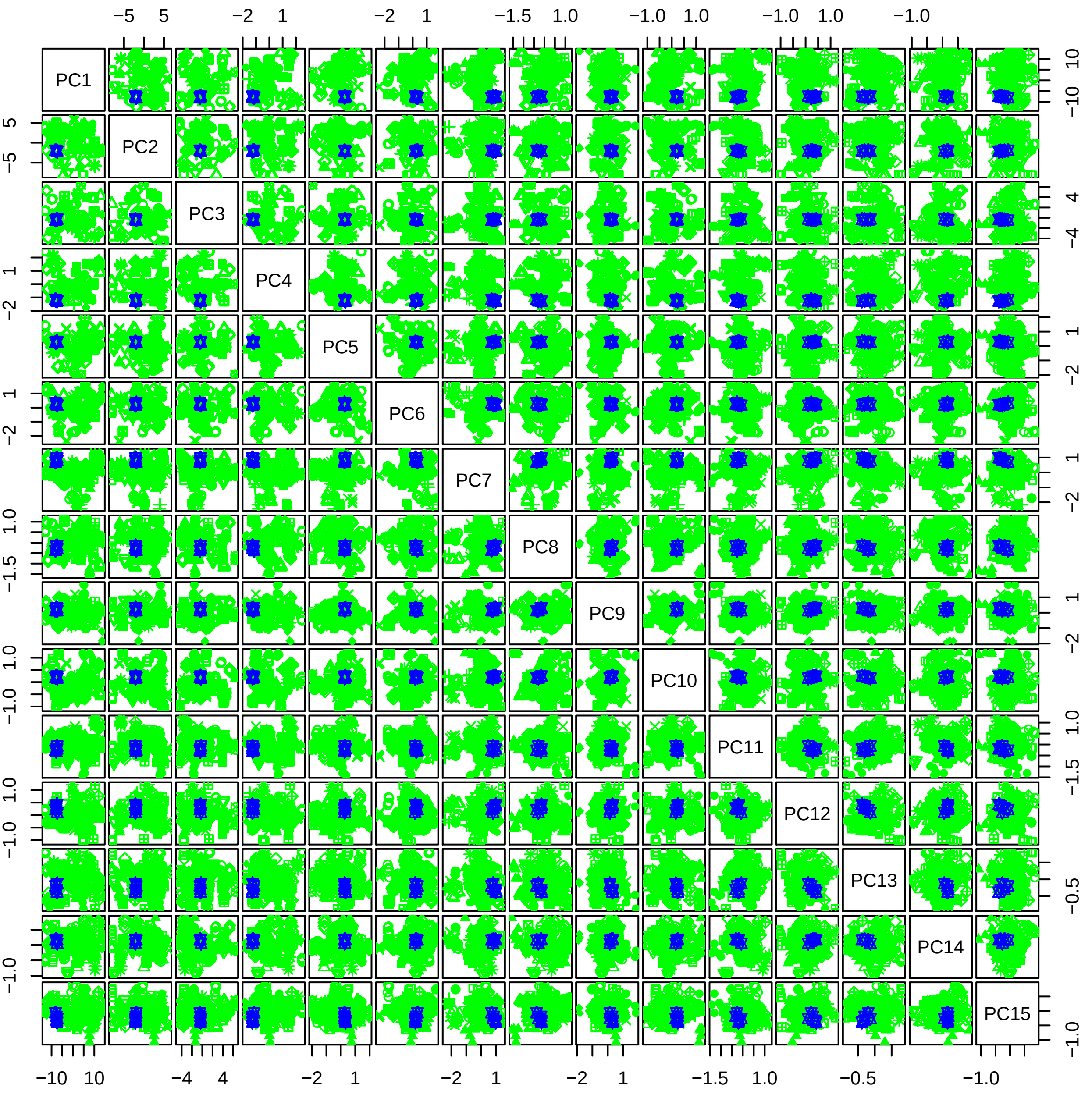
PC1
PC2
PC3
PC4
PC5
PC6
PC7
PC8
PC9
PC10
PC11
PC12
PC13
PC14
PC15

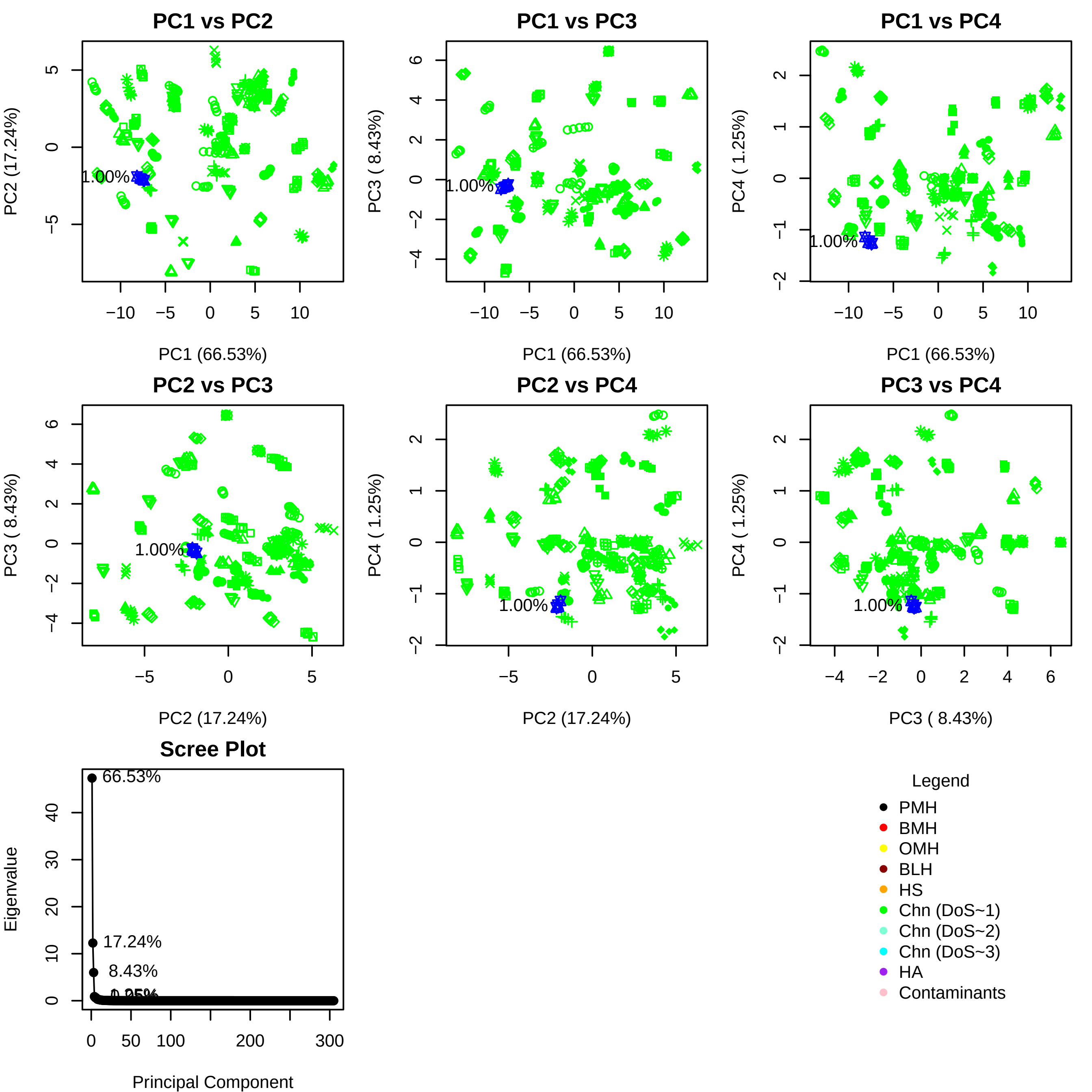
PC1 vs PC2
PC1 vs PC3
PC1 vs PC4
PC2 vs PC3
PC2 vs PC4
PC3 vs PC4
Scree Plot
PC1 (66.53%)
PC2 (17.24%)
PC3 ( 8.43%)
PC4 ( 1.25%)
1.00%
66.53%
17.24%
8.43%
Eigenvalue
Principal Component
Legend
PMH
BMH
OMH
BLH
HS
Chn (DoS~1)
Chn (DoS~2)
Chn (DoS~3)
HA
Contaminants

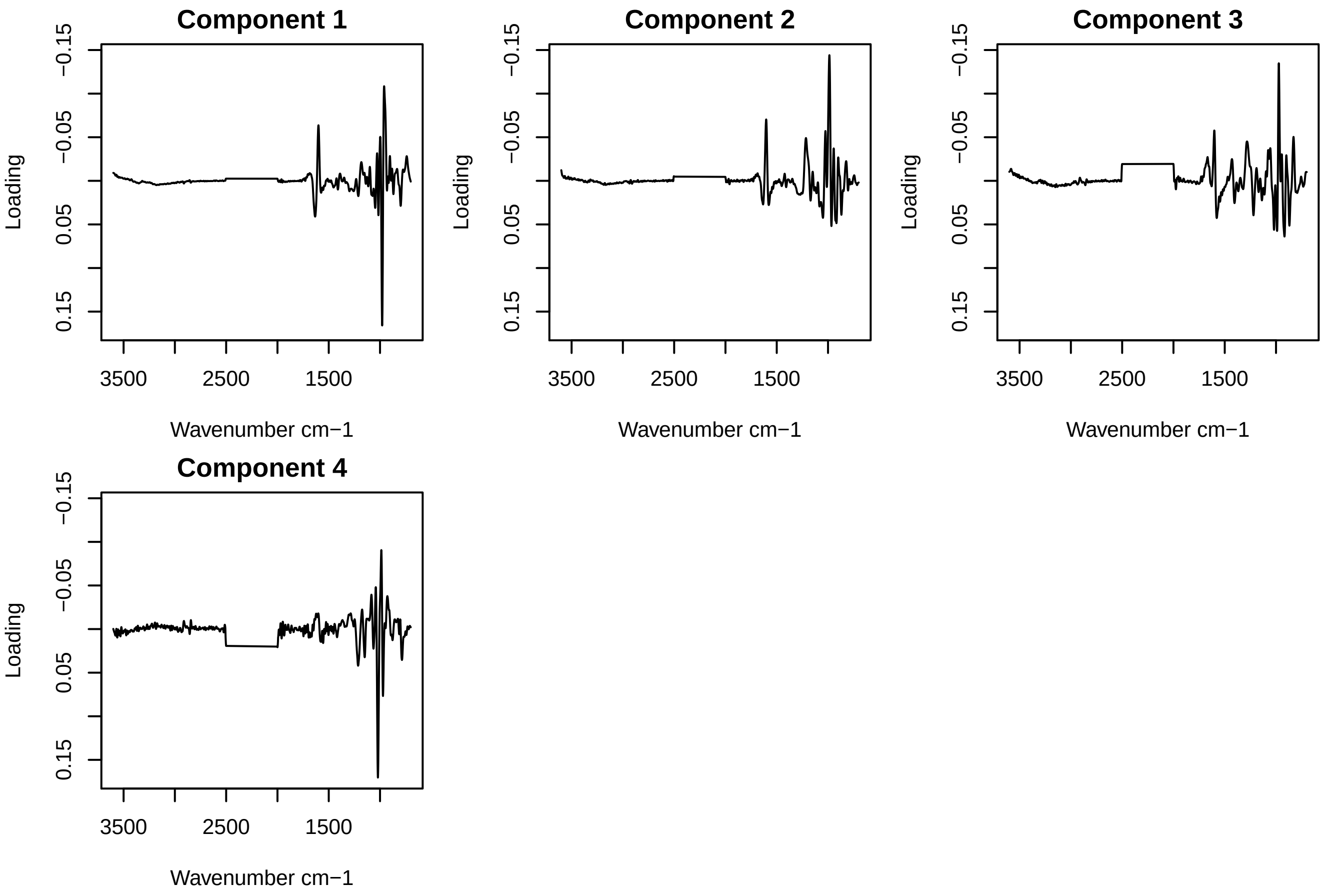
Component 1
Component 2
Component 3
Component 4
Loading
−0.15
−0.05
0.05
0.15
3500
2500
1500
Wavenumber cm−1

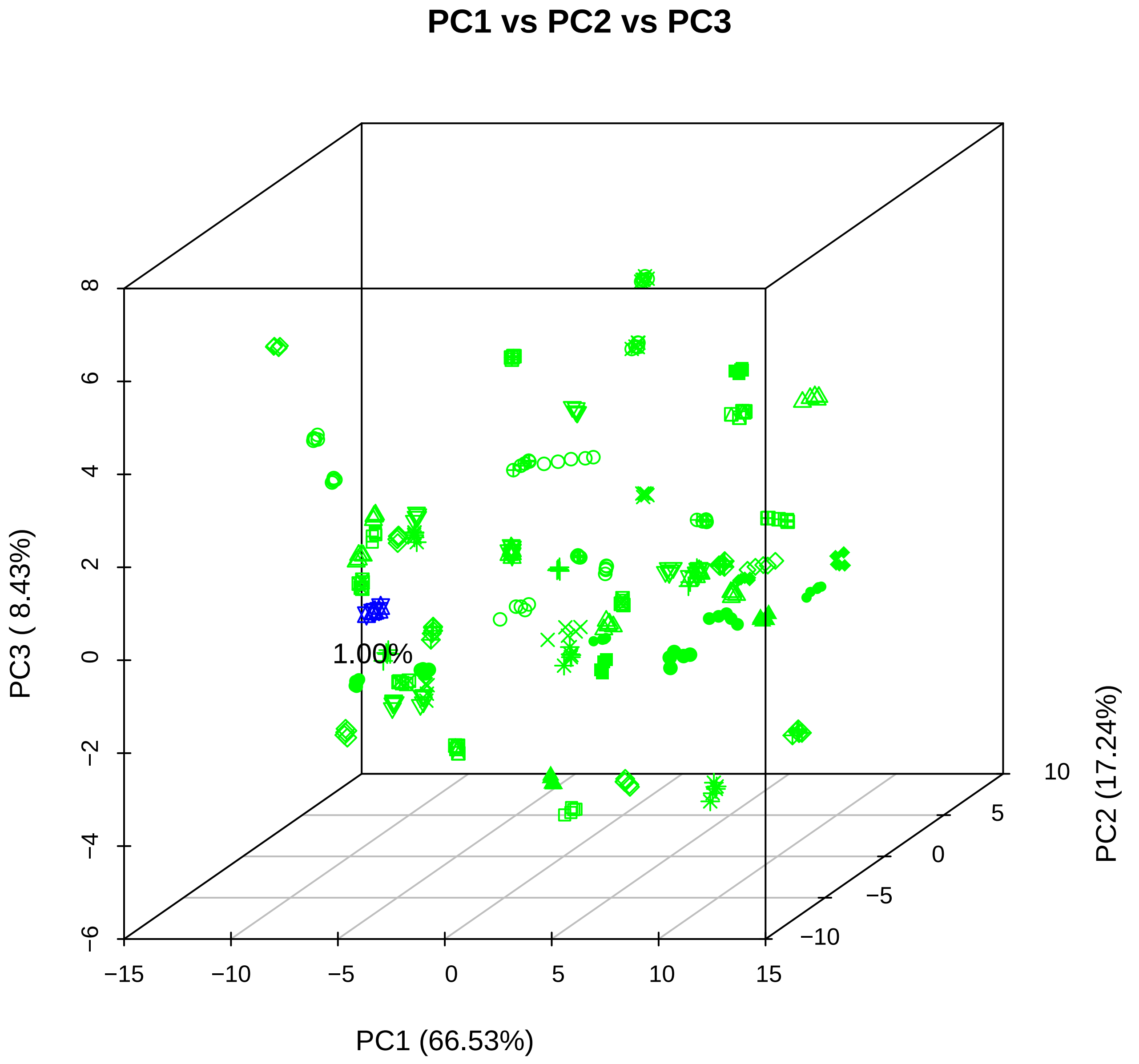
PC1 vs PC2 vs PC3
PC1 (66.53%)
PC2 (17.24%)
PC3 ( 8.43%)
1.00%
−15
−10
−5
0
5
10
15
−10
−5
0
5
10
−6
−4
−2
0
2
4
6
8

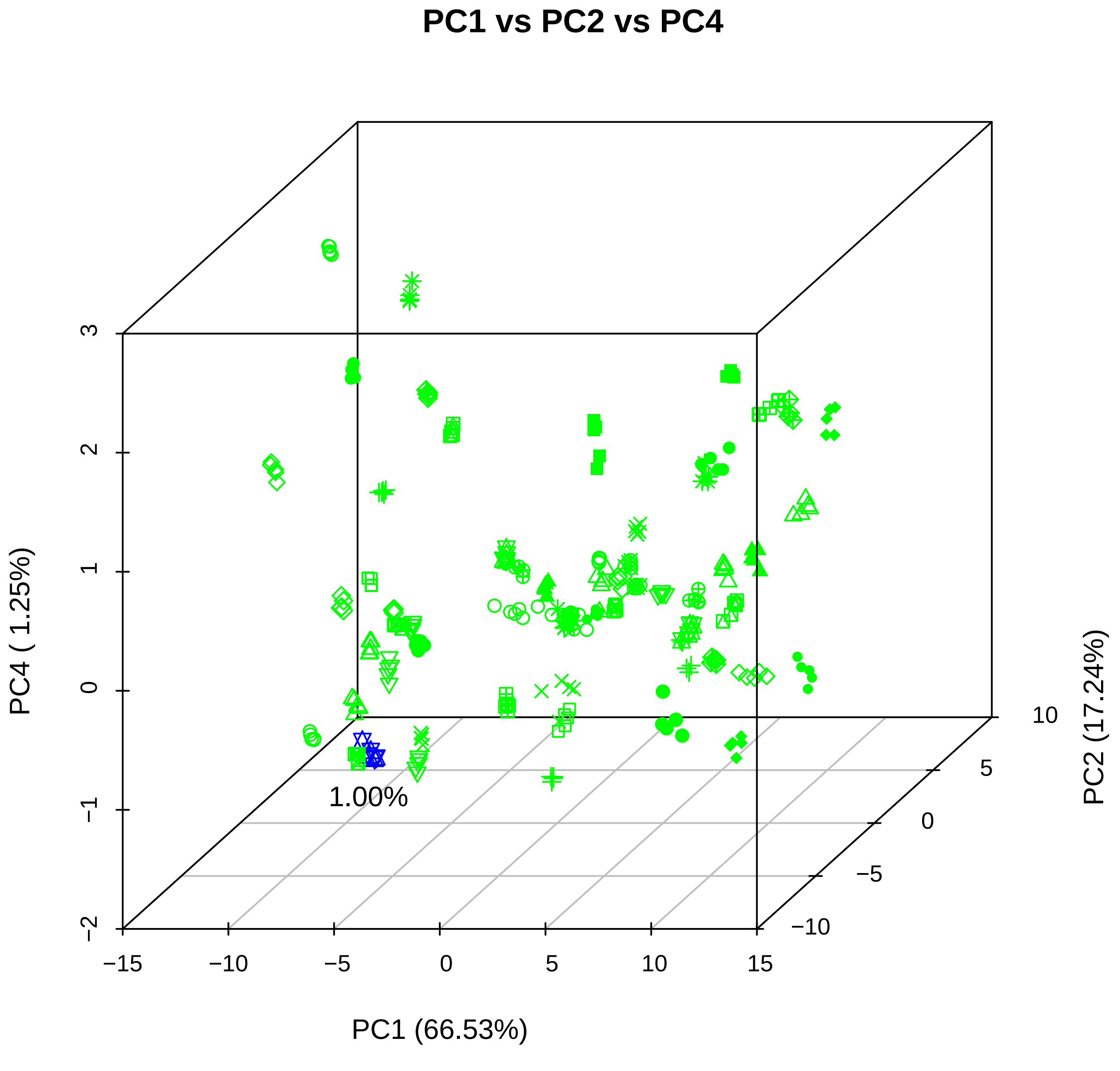

PC1 vs PC2 vs PC4
PC1 (66.53%)
PC2 (17.24%)
PC4 ( 1.25%)
1.00%
−15
−10
−5
0
5
10
15
−2
−1
0
1
2
3

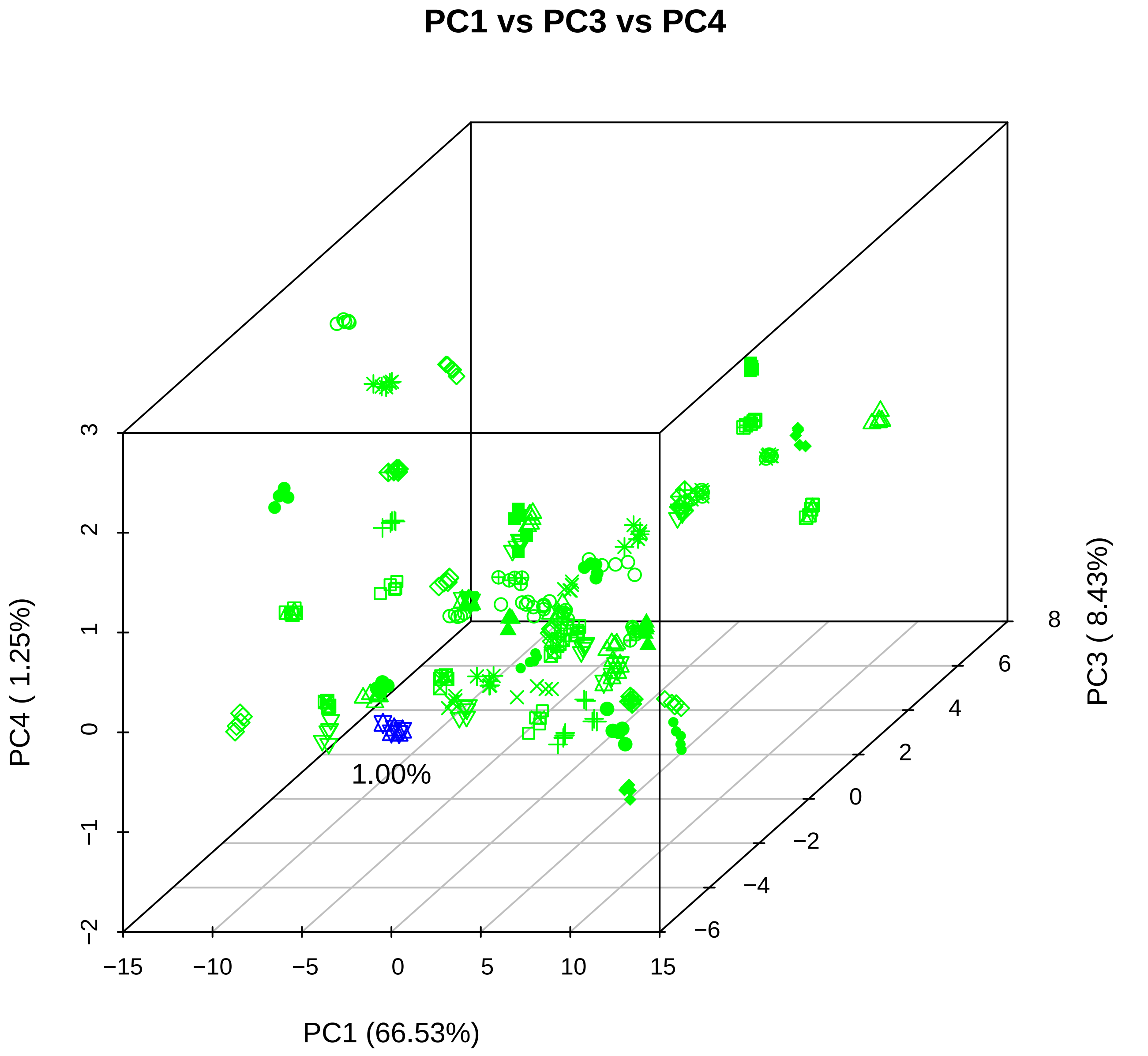
PC1 vs PC3 vs PC4
PC1 (66.53%)
PC4 ( 1.25%)
PC3 ( 8.43%)
1.00%
−15
−10
−5
0
5
10
15
−2
−1
0
1
2
3
−6
−4
−2
0
2
4
6
8

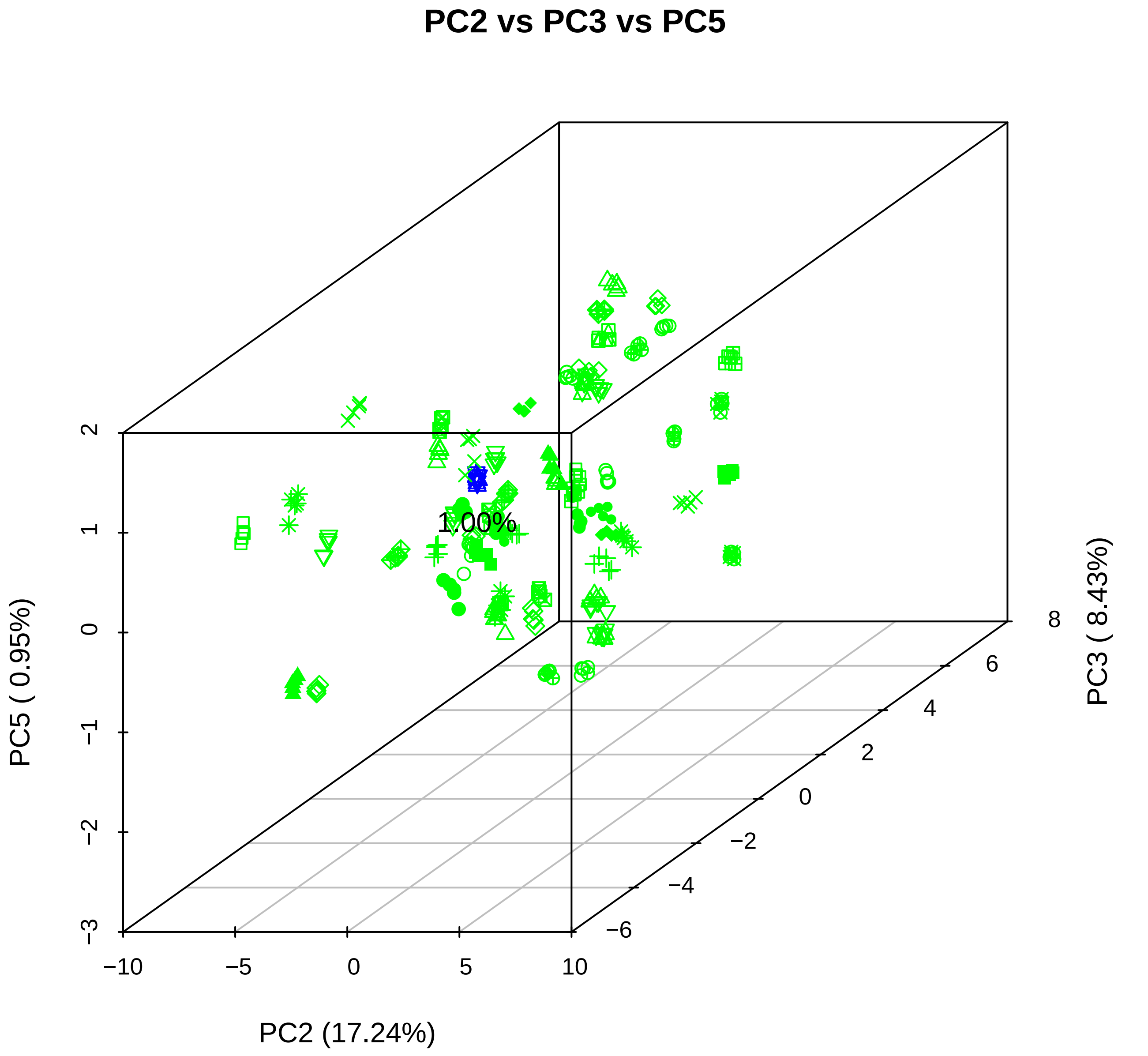

PC2 vs PC3 vs PC5
1.00%
PC2 (17.24%)
PC3 ( 8.43%)
PC5 ( 0.95%)
−10
−5
0
5
10
−6
−4
−2
0
2
4
6
8
−3
−2
−1
0
1
2

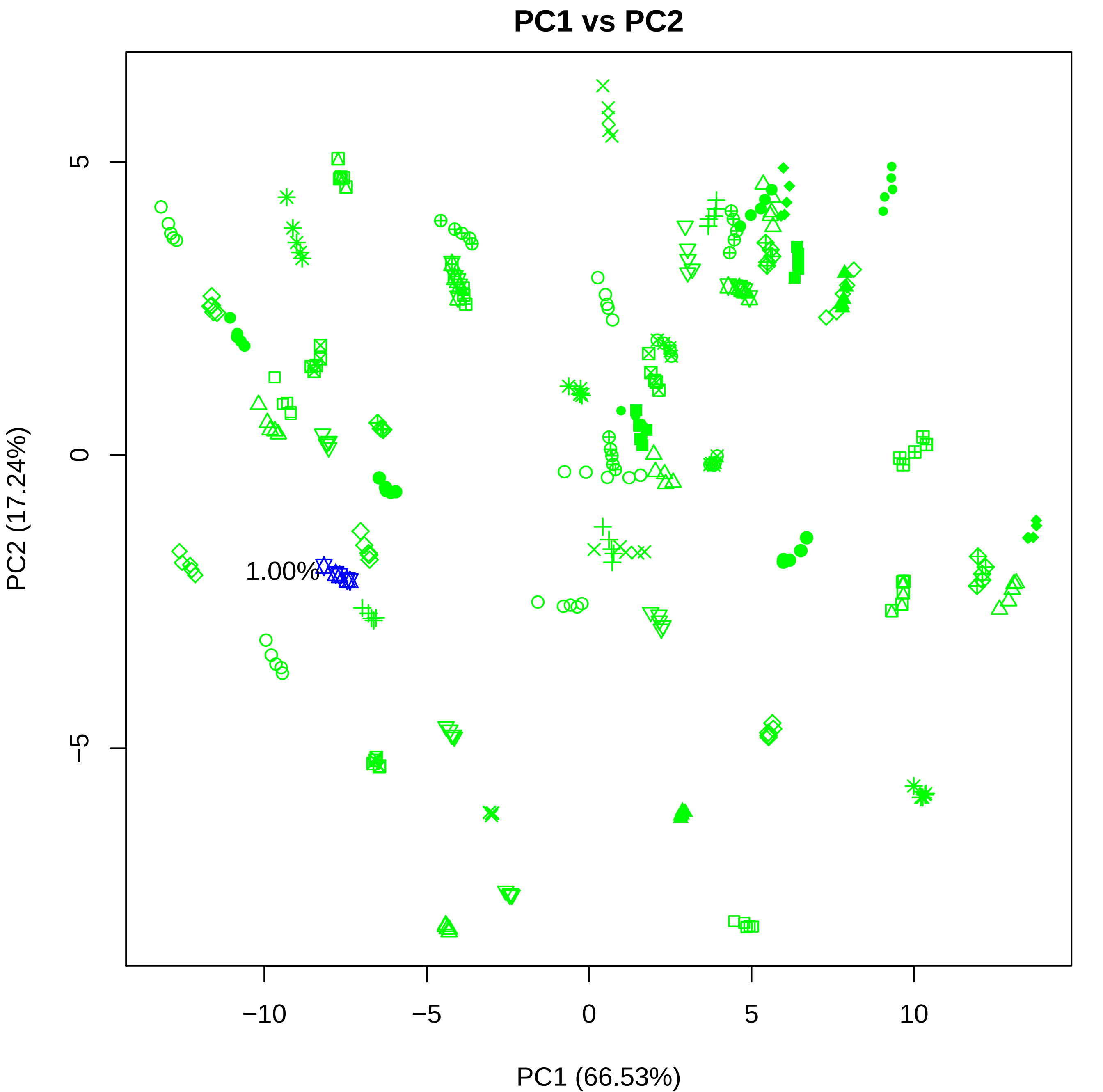

PC1 vs PC2
1.00%
5
0
−5
−10
−5
0
5
10
PC1 (66.53%)
PC2 (17.24%)

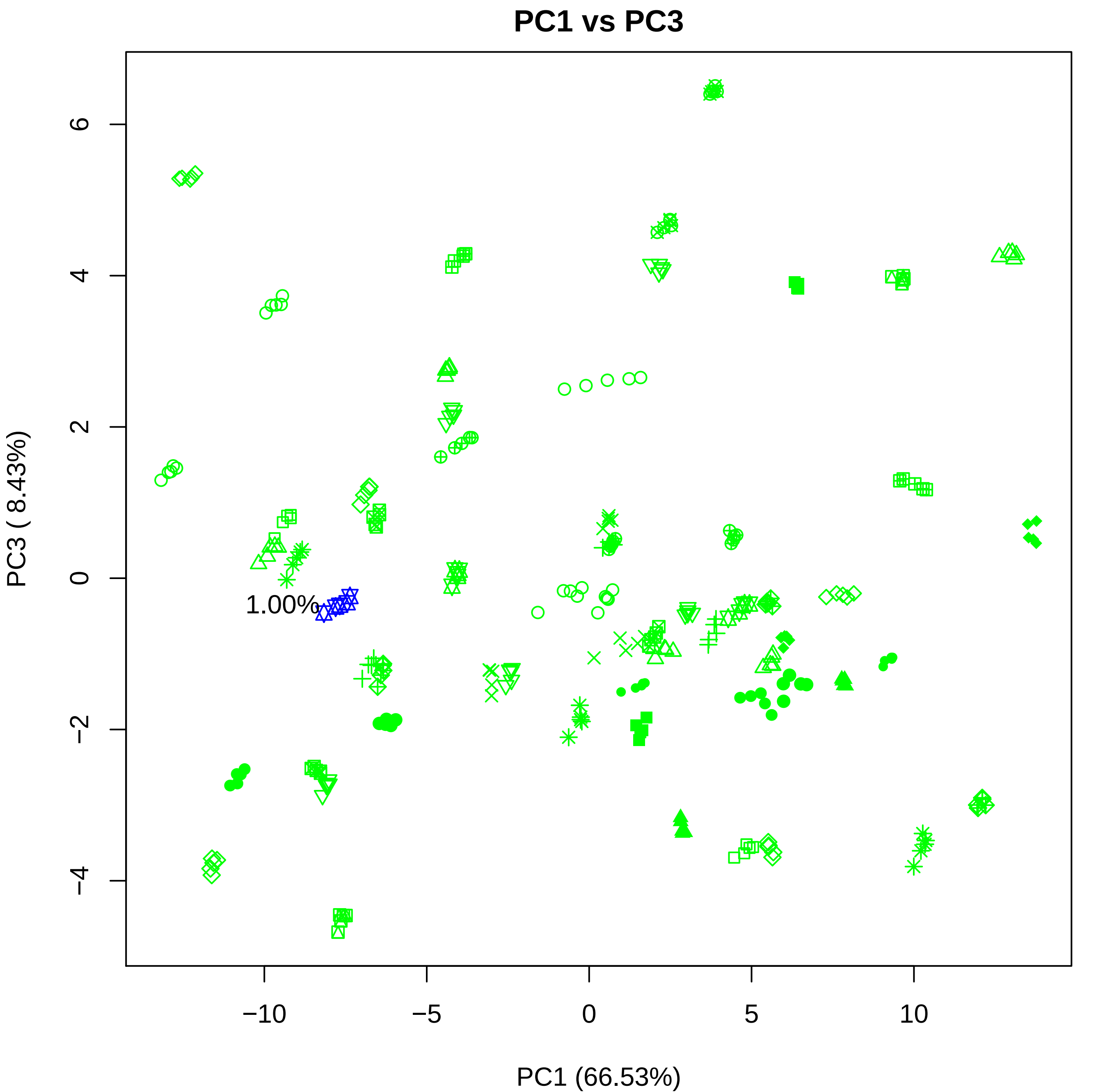

PC1 vs PC3
1.00%
PC3 ( 8.43%)
PC1 (66.53%)
6
4
2
0
−2
−4
−10
−5
0
5
10

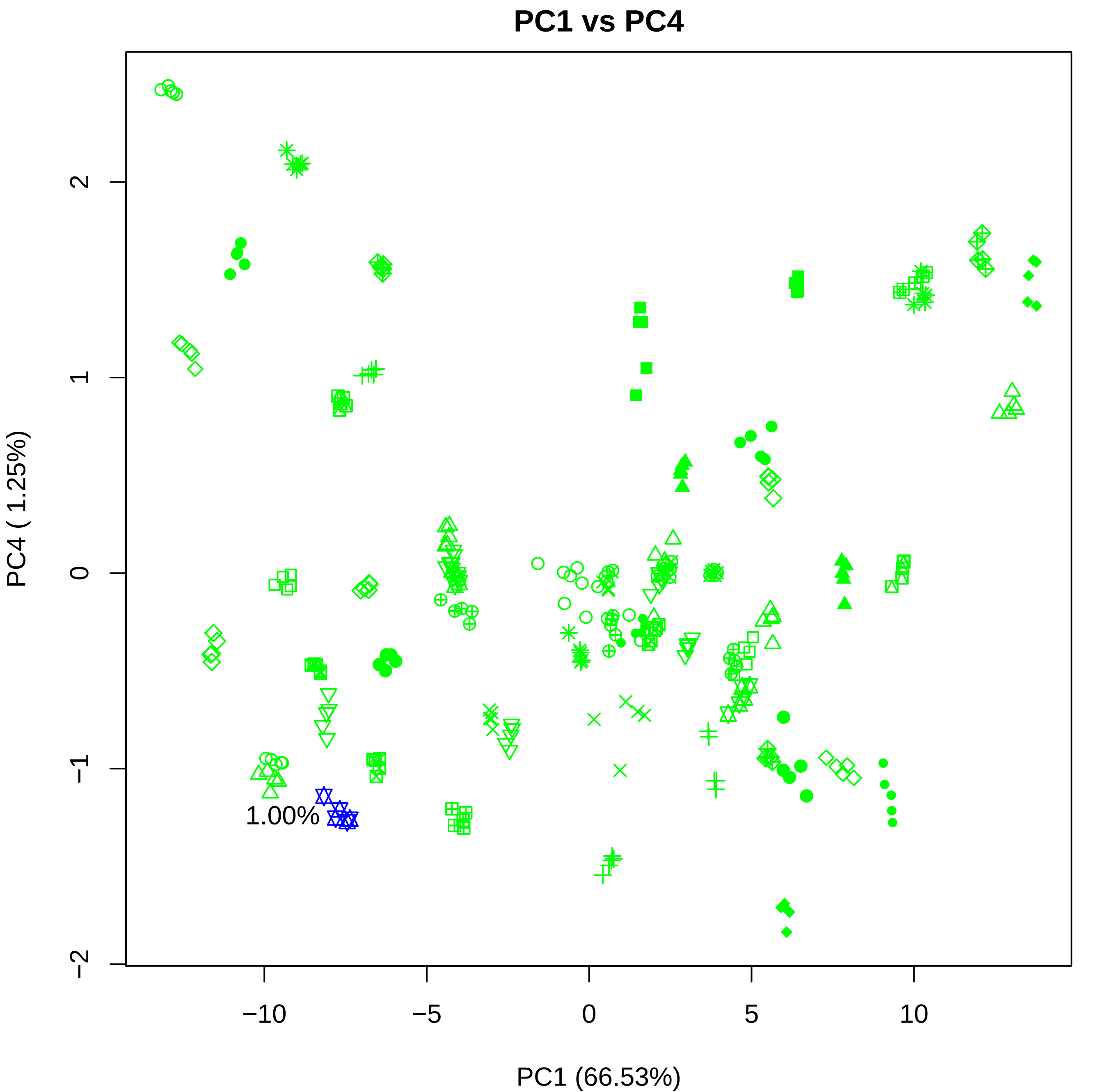
PC1 vs PC4
PC4 ( 1.25%)
PC1 (66.53%)
1.00%
2
1
0
−1
−2
−10
−5
0
5
10

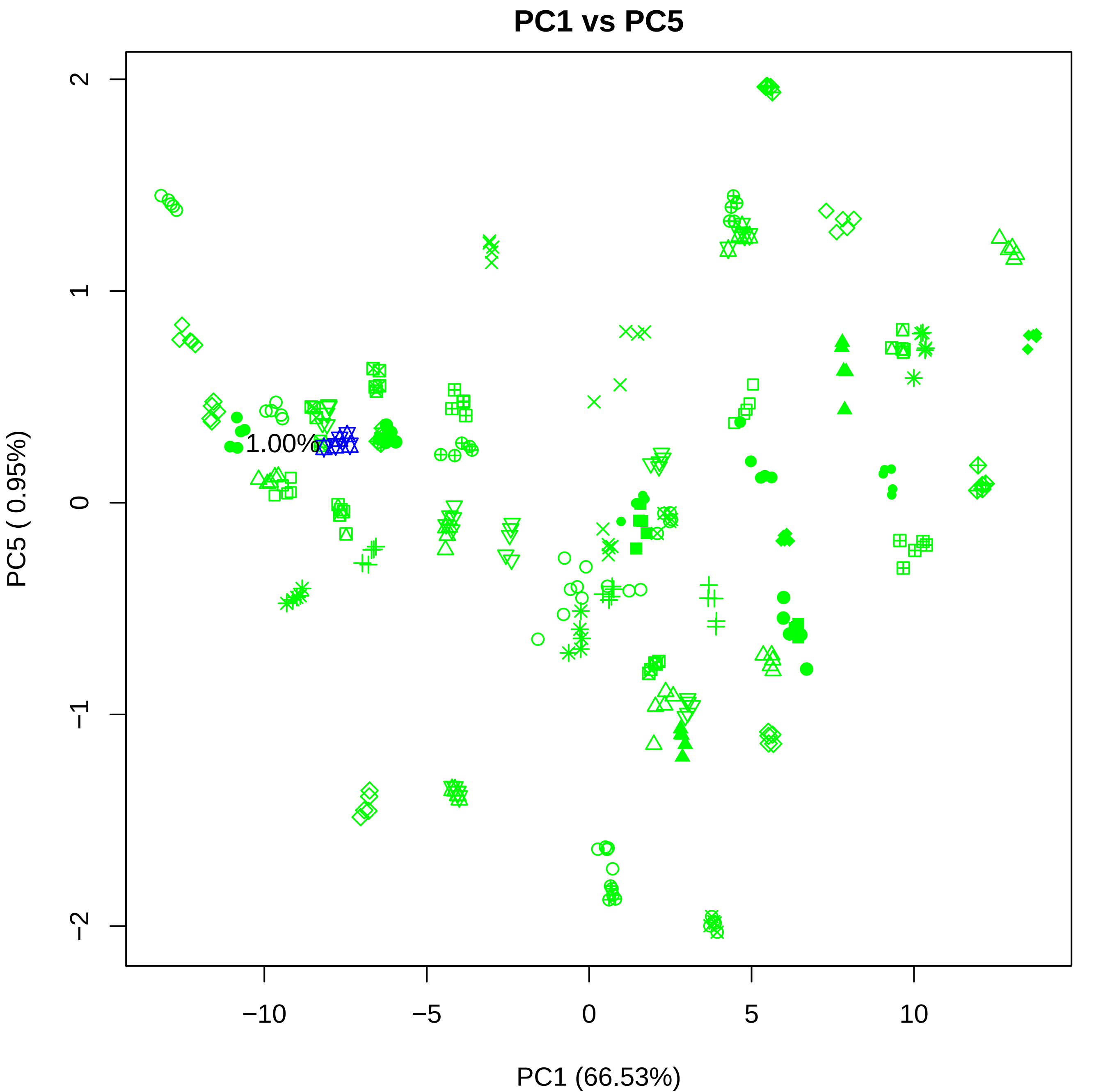
PC1 vs PC5
1.00%
PC5 ( 0.95%)
PC1 (66.53%)
2
1
0
−1
−2
−10
−5
0
5
10

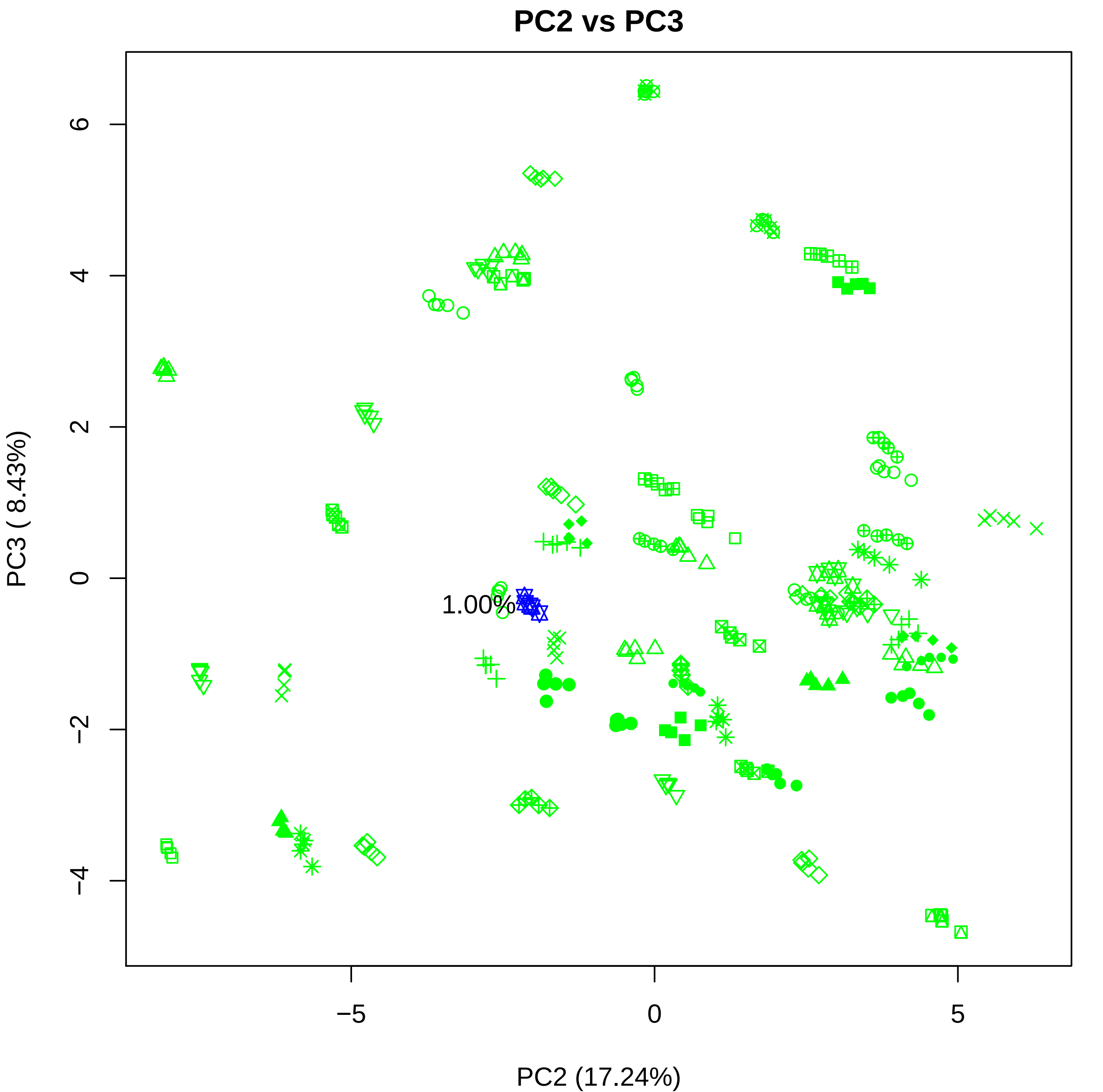
PC2 vs PC3
PC2 (17.24%)
PC3 ( 8.43%)
1.00%
−5
0
5
−4
−2
0
2
4
6

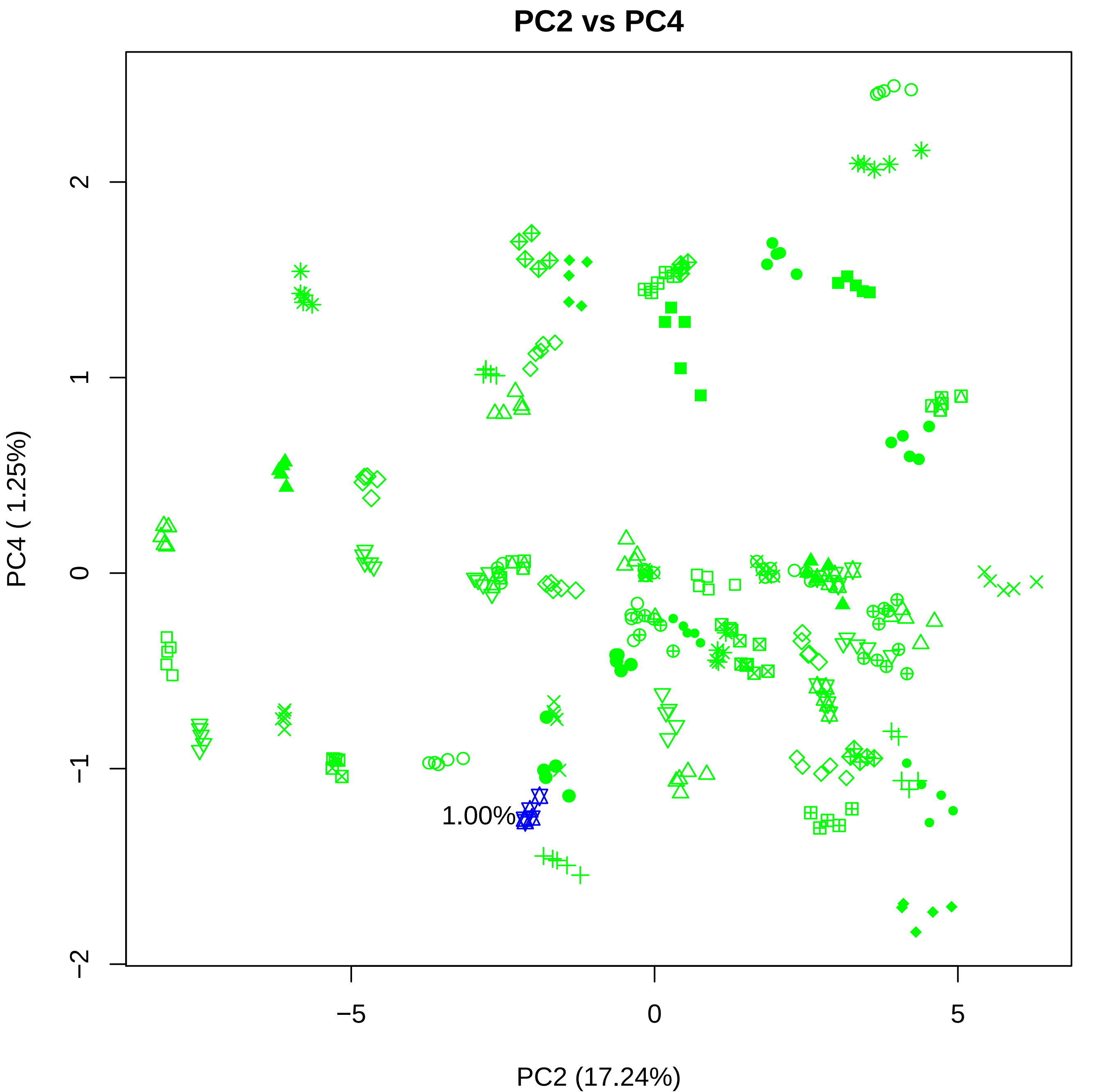
PC2 vs PC4
PC4 ( 1.25%)
PC2 (17.24%)
1.00%
2
1
0
−1
−2
−5
0
5

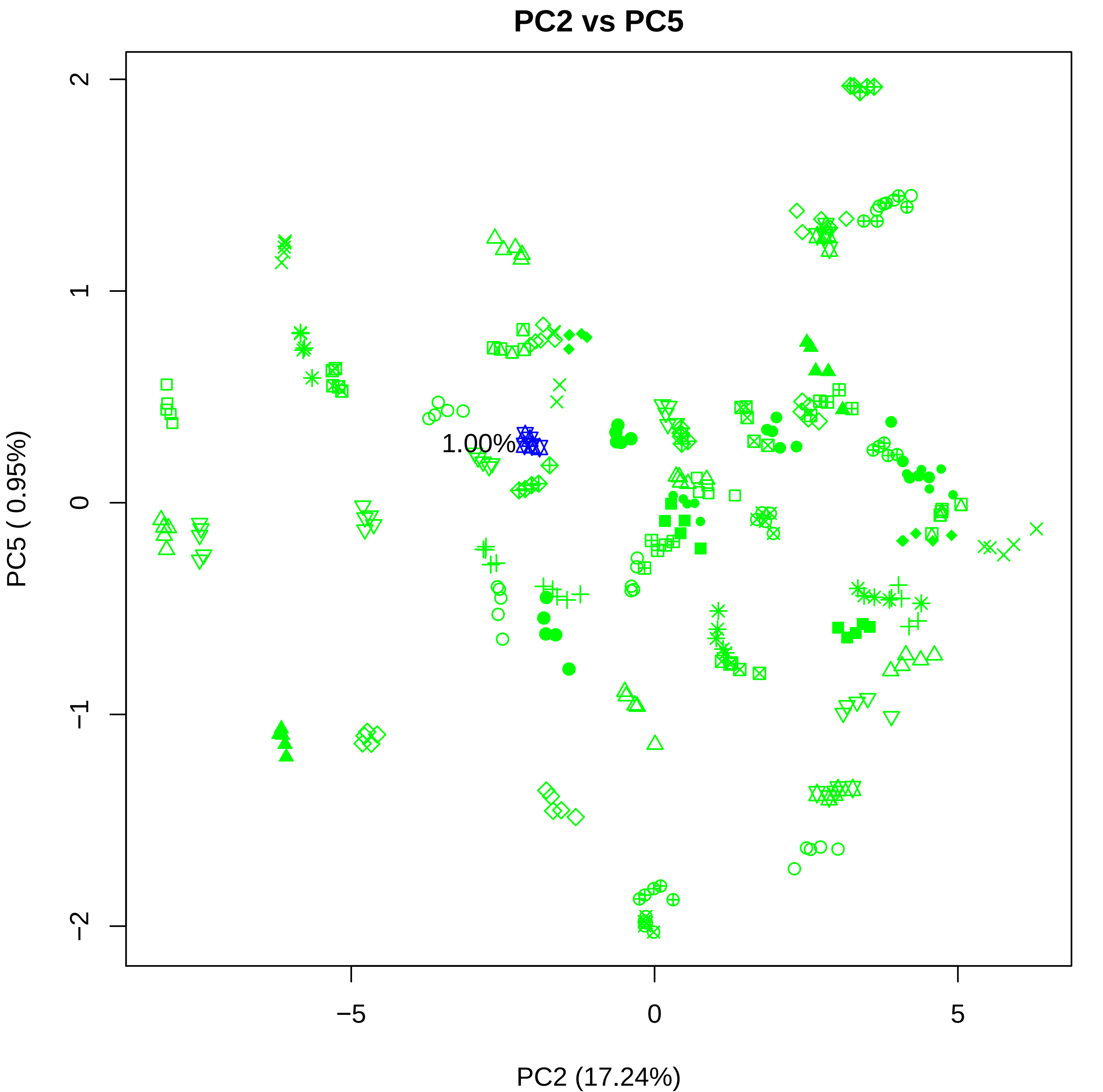

PC2 vs PC5
1.00%
PC5 ( 0.95%)
PC2 (17.24%)
2
1
0
−1
−2
−5
0
5

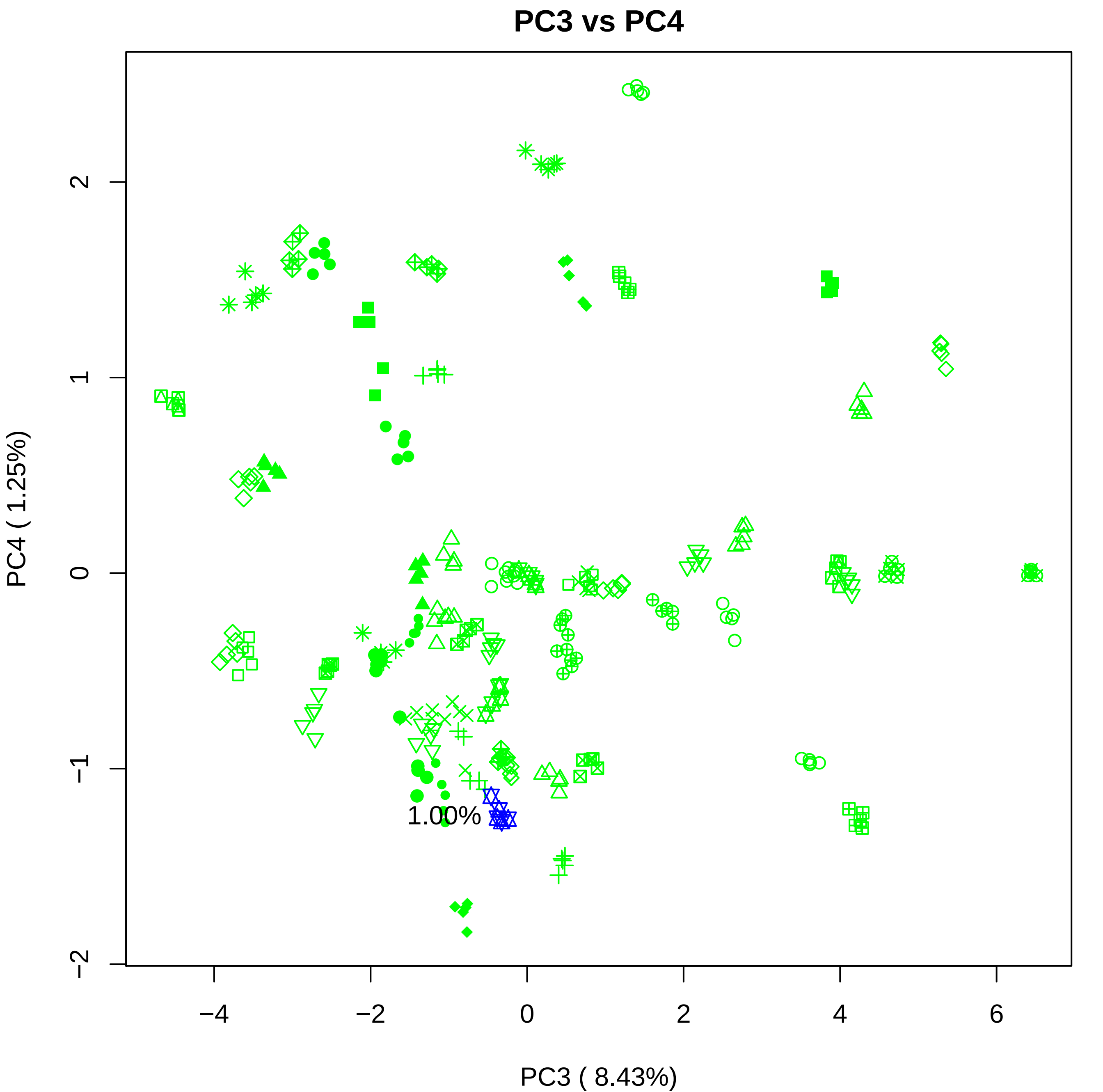

PC3 vs PC4
PC4 ( 1.25%)
PC3 ( 8.43%)
1.00%
2
1
0
−1
−2
−4
−2
0
2
4
6

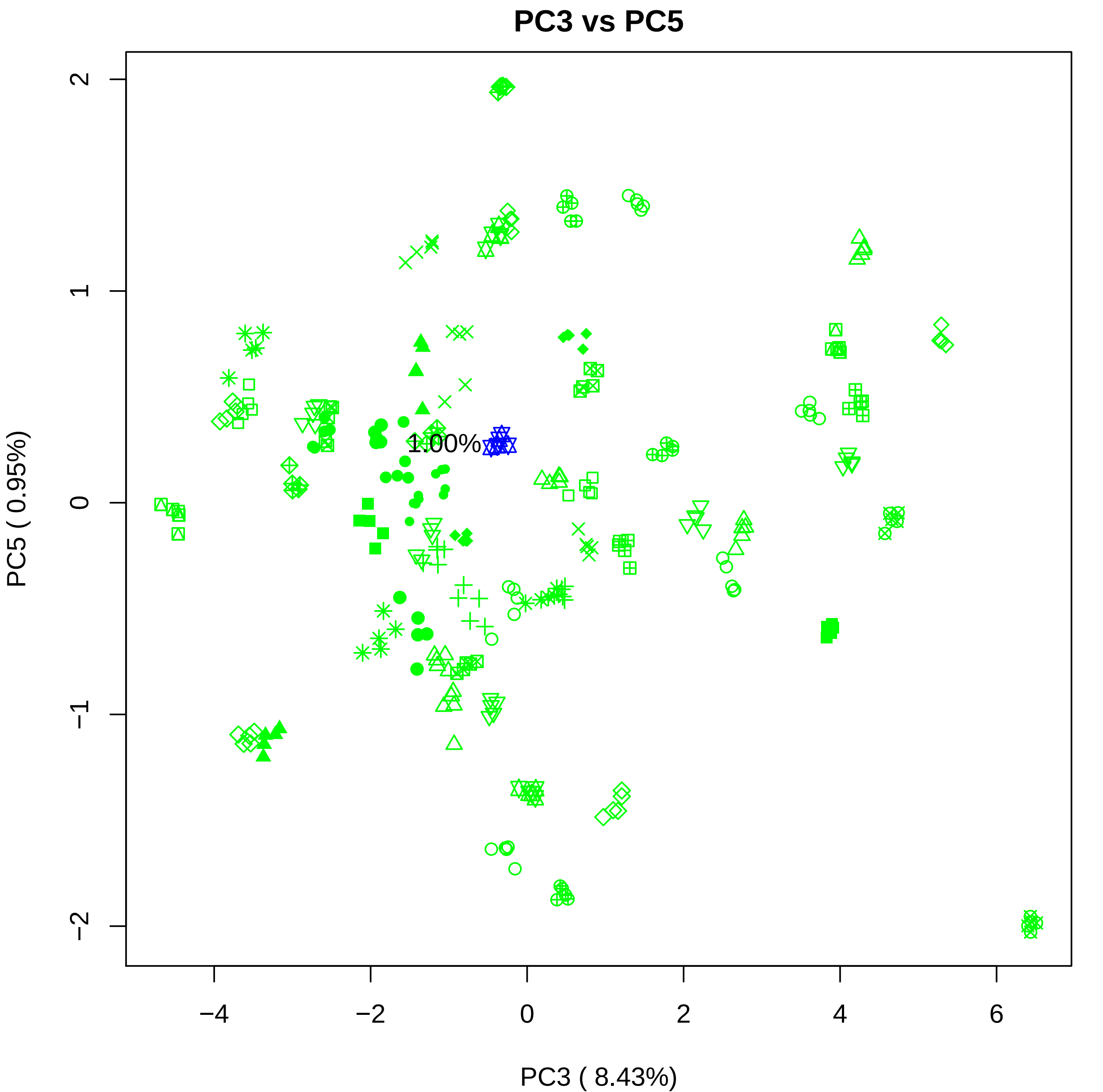
PC3 vs PC5
1.00%
PC5 ( 0.95%)
PC3 ( 8.43%)
−4
−2
0
2
4
6
−2
−1
0
1
2

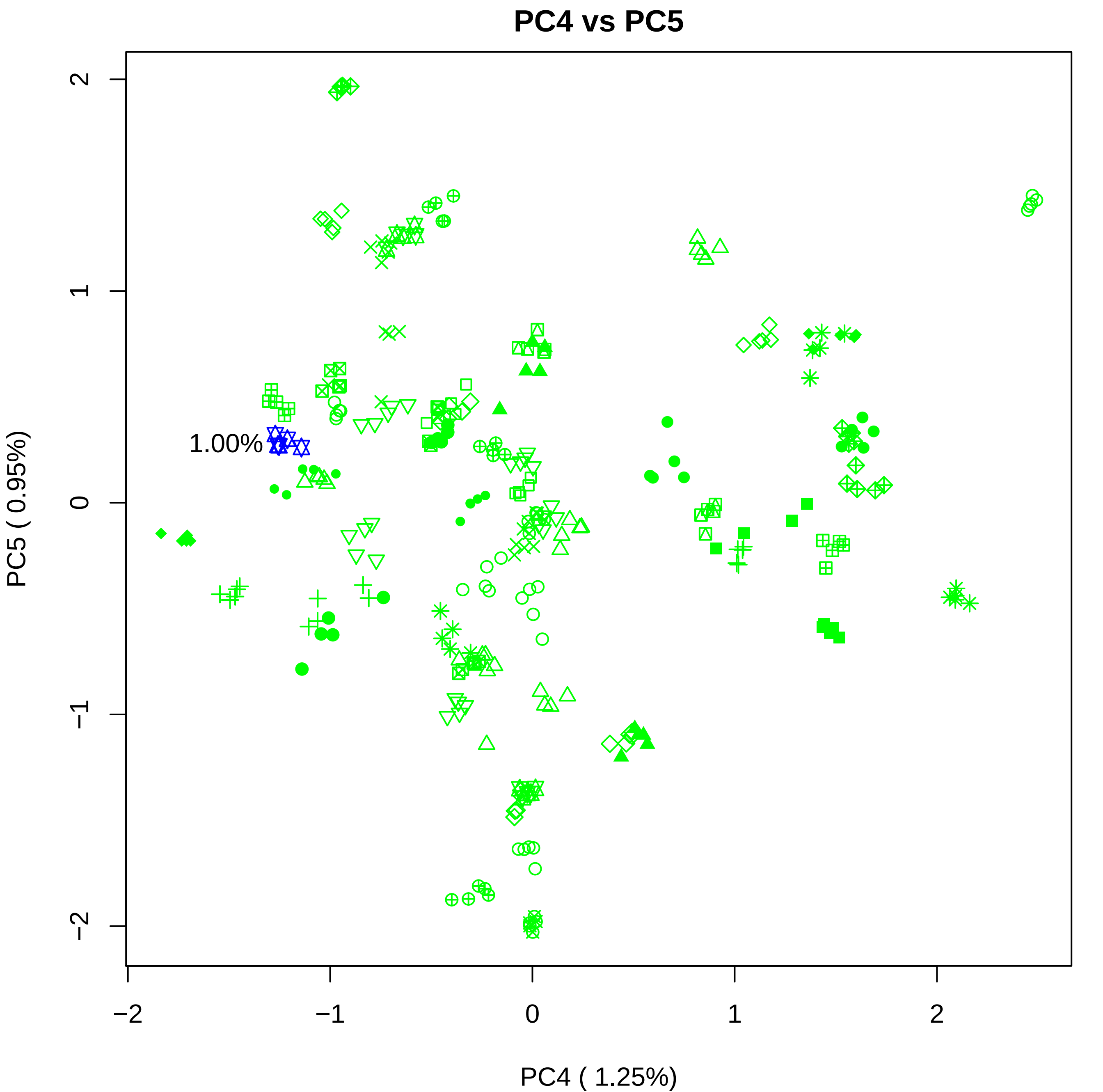
PC4 vs PC5
1.00%
PC5 ( 0.95%)
PC4 ( 1.25%)
2
1
0
−1
−2
−2
−1
0
1
2

### PCA of 2.5% BMH(a) vs PMH (a)

Scaled Matrix PCAed

7th order polynomial baseline correction performed

Raw spectra smoothed to 21 Neighbours, 2nd order PN, Savtisky Golay

Normalisation between 0 and 100

D2 spectra smoothed to 41 Neighbours, 2nd order PN, Savtisky Golay

Regions Analysed: 3601−2501 and 2001−700cm−1

Samples Analysed: 1 bContam's 0 BLH's 0 BMH's 0 OMH's 0 pContam's 0 PLH's 60 PMH's

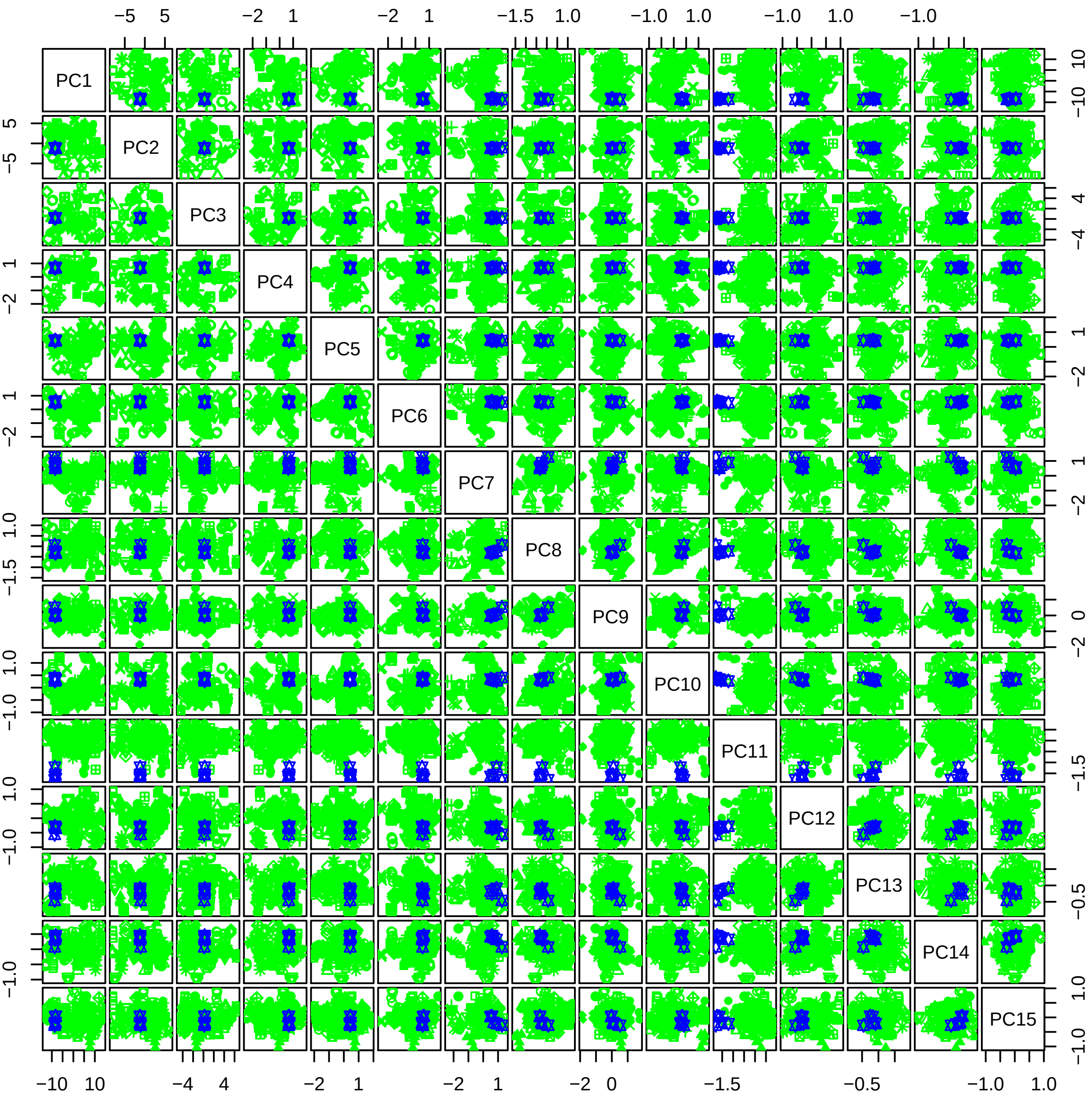

PC1
PC2
PC3
PC4
PC5
PC6
PC7
PC8
PC9
PC10
PC11
PC12
PC13
PC14
PC15

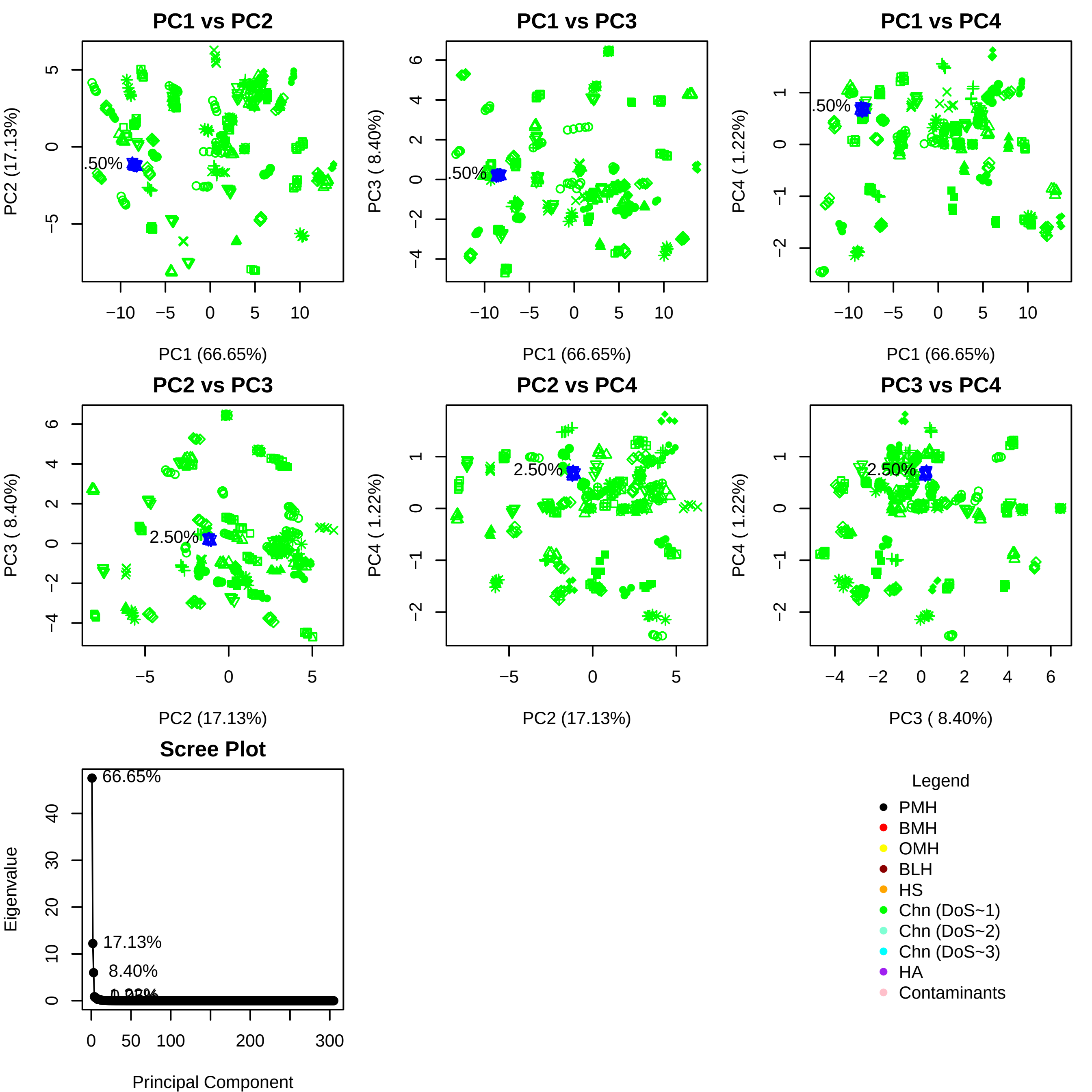

PC1 vs PC2
PC1 vs PC3
PC1 vs PC4
PC2 vs PC3
PC2 vs PC4
PC3 vs PC4
Scree Plot
PC1 (66.65%)
PC2 (17.13%)
PC3 ( 8.40%)
PC4 ( 1.22%)
Eigenvalue
Principal Component
66.65%
17.13%
8.40%
.50%
2.50%
Legend
PMH
BMH
OMH
BLH
HS
Chn (DoS~1)
Chn (DoS~2)
Chn (DoS~3)
HA
Contaminants

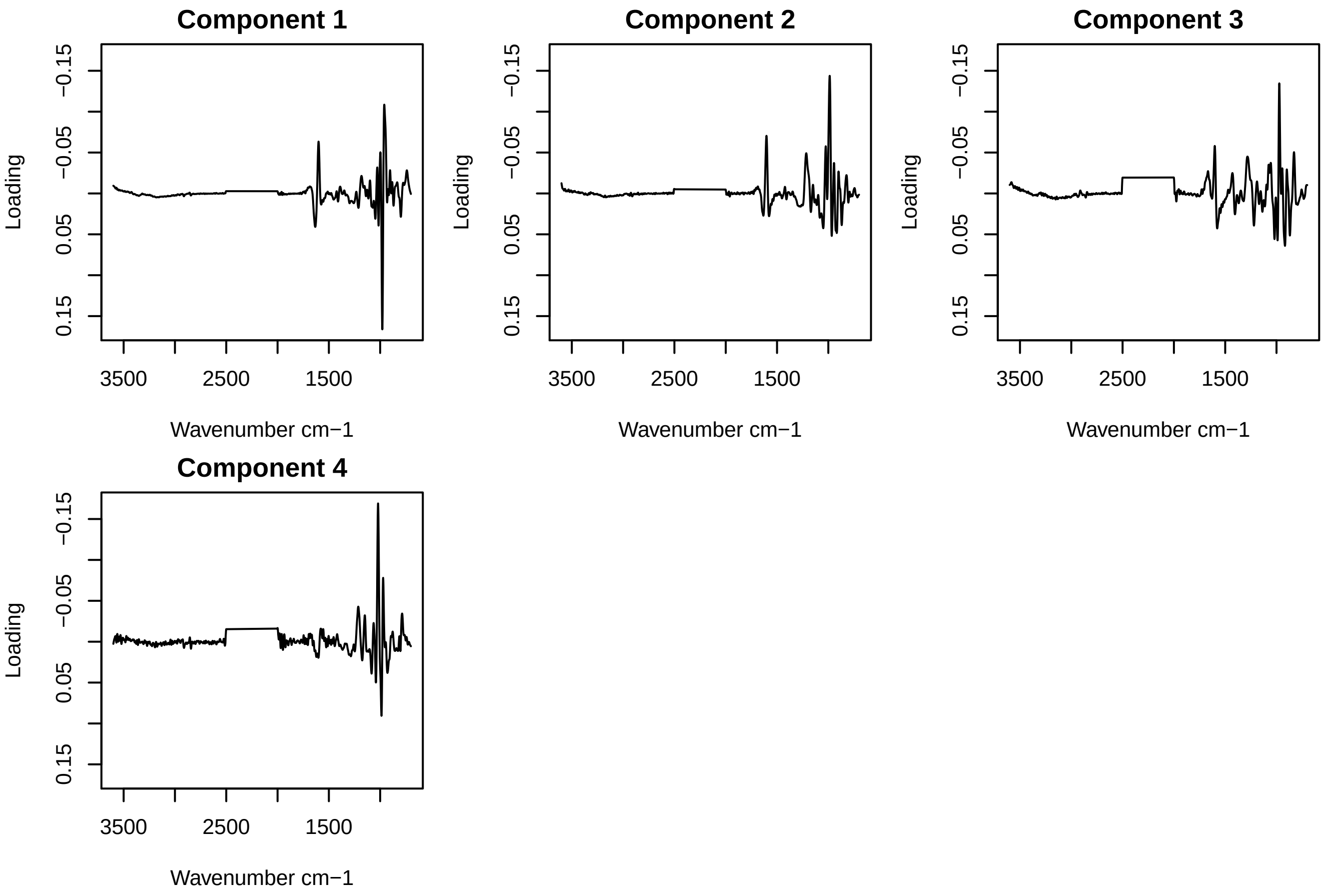
Component 1
Component 2
Component 3
Component 4
Loading
−0.15
−0.05
0.05
0.15
3500
2500
1500
Wavenumber cm−1

# PC1 vs PC2 vs PC3

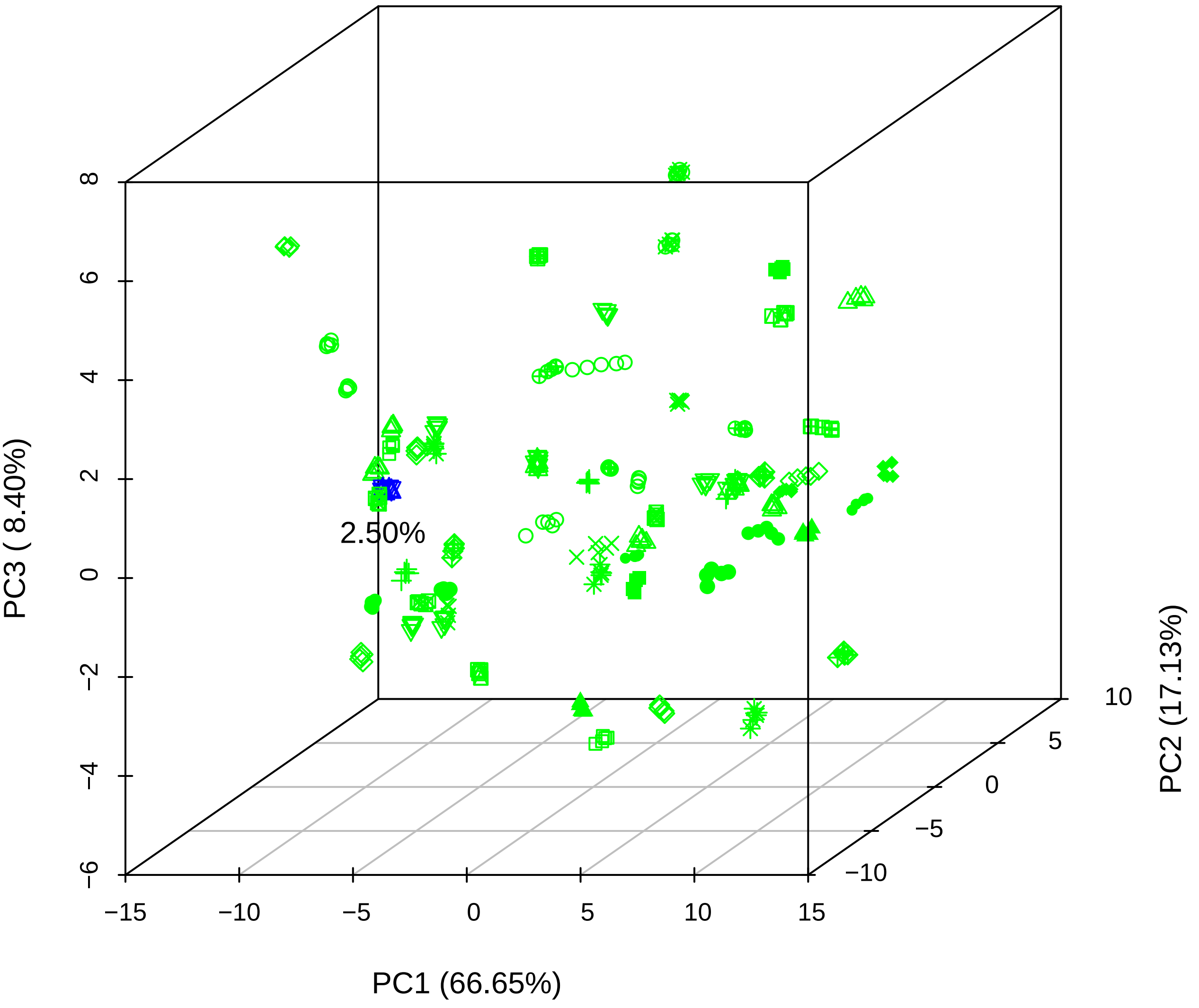

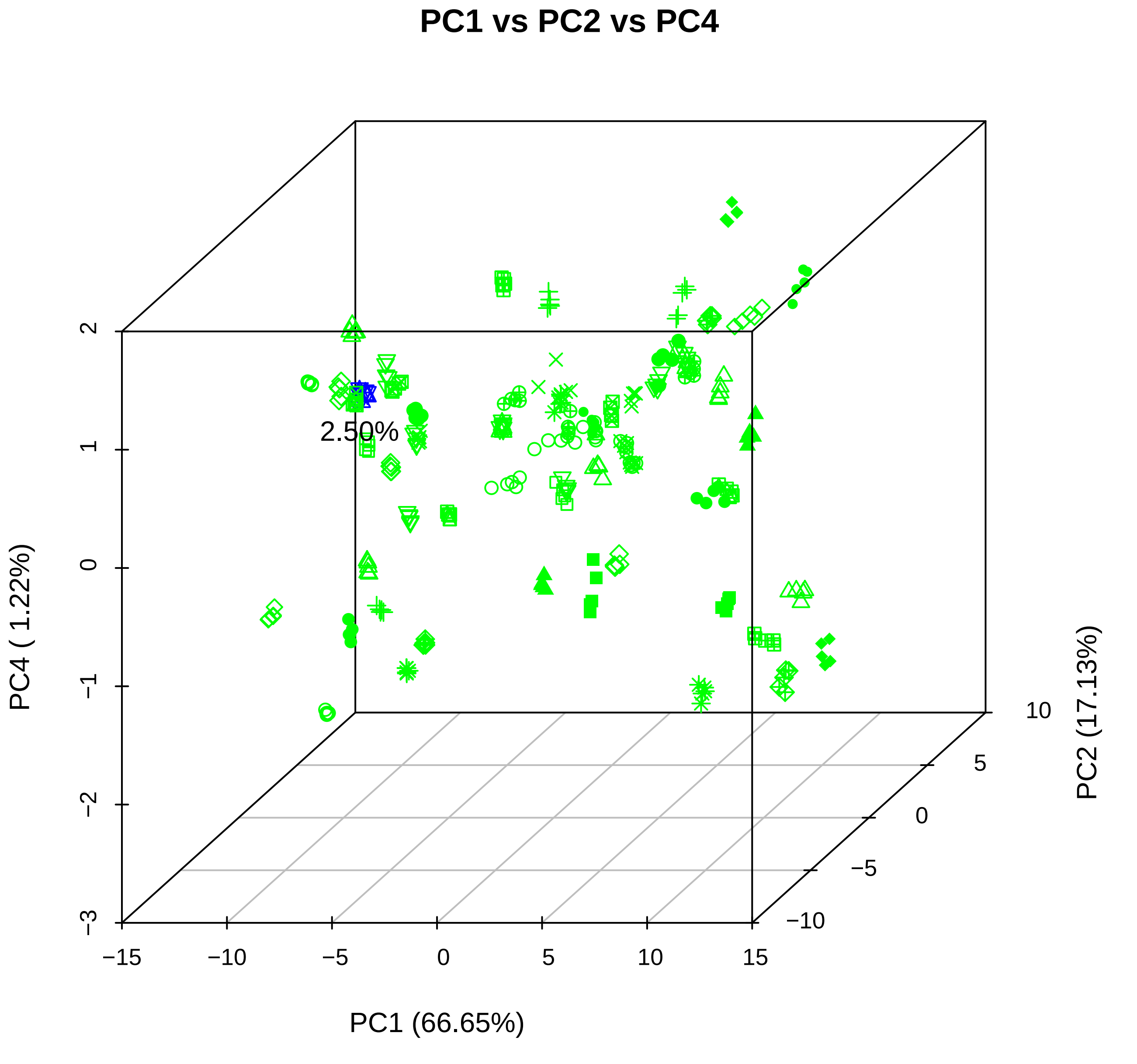
PC1 vs PC2 vs PC4
2.50%
PC1 (66.65%)
PC2 (17.13%)
PC4 ( 1.22%)
−15
−10
−5
0
5
10
15
−10
−5
0
5
10
−3
−2
−1
0
1
2

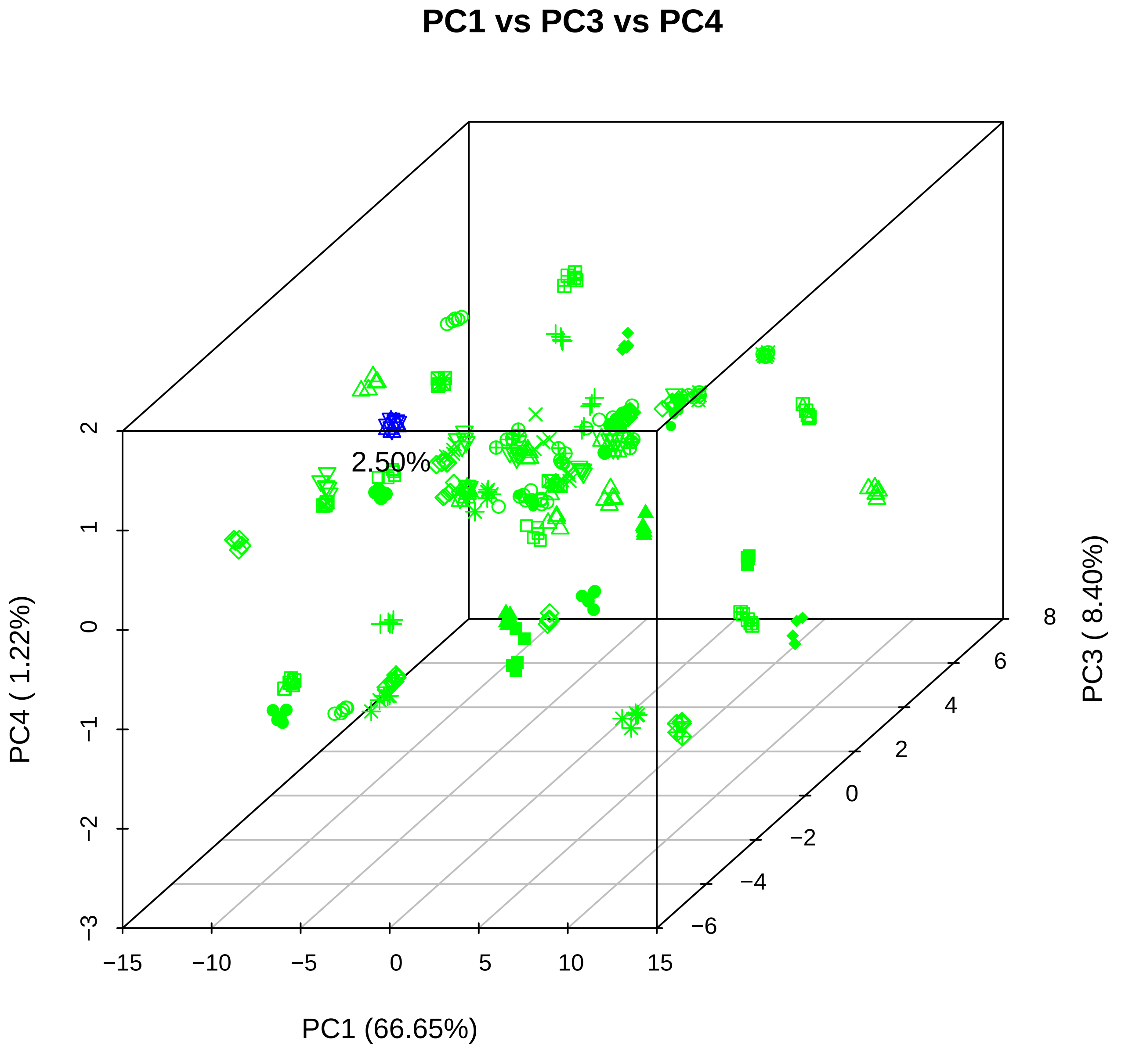

PC1 vs PC3 vs PC4
2.50%
PC1 (66.65%)
PC3 ( 8.40%)
PC4 ( 1.22%)
−15
−10
−5
0
5
10
15
−6
−4
−2
0
2
4
6
8
−3
−2
−1
0
1
2

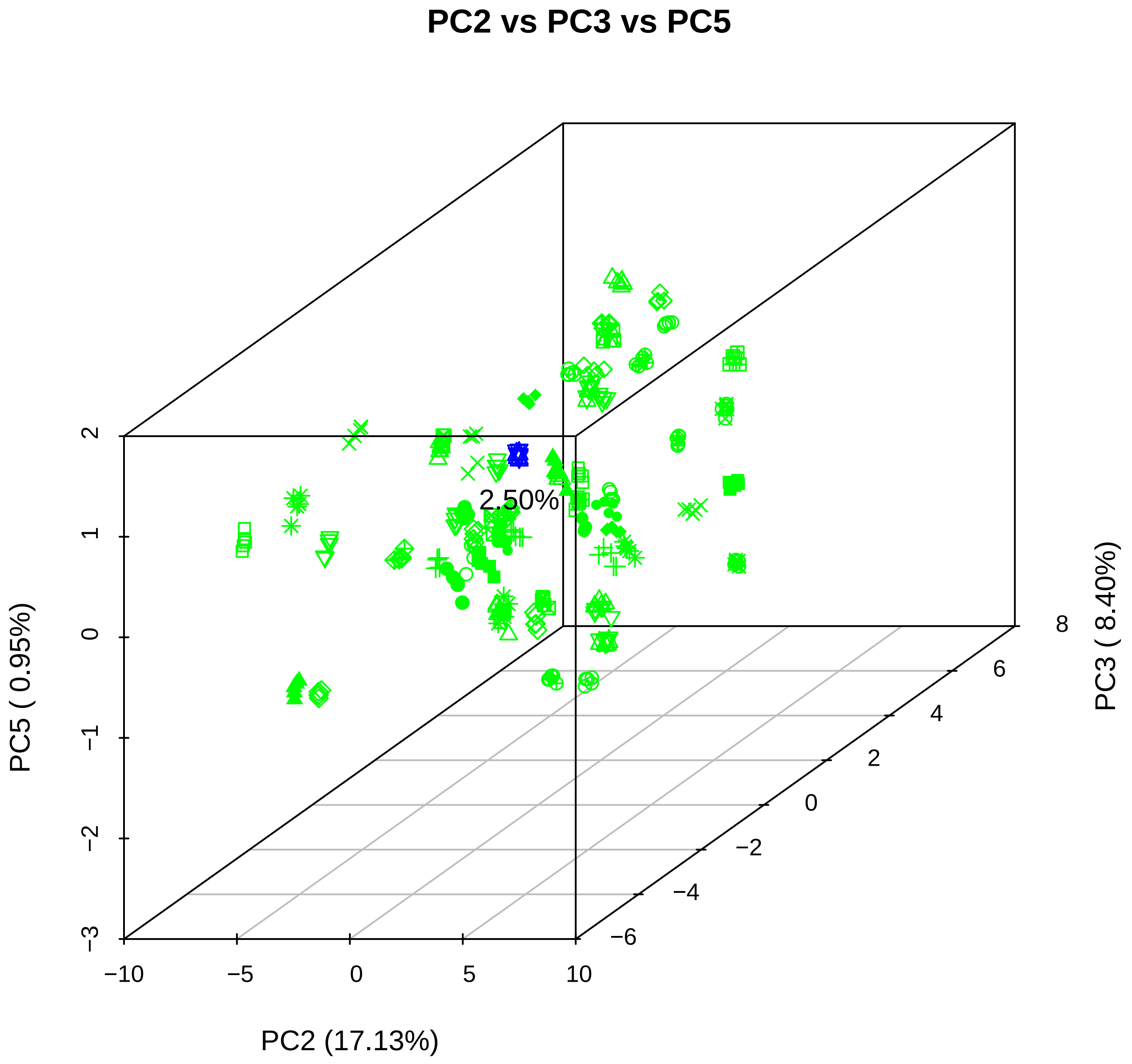

PC2 vs PC3 vs PC5
2.50%
PC2 (17.13%)
PC5 ( 0.95%)
PC3 ( 8.40%)
−10
−5
0
5
10
−3
−2
−1
0
1
2
−6
−4
−2
0
2
4
6
8

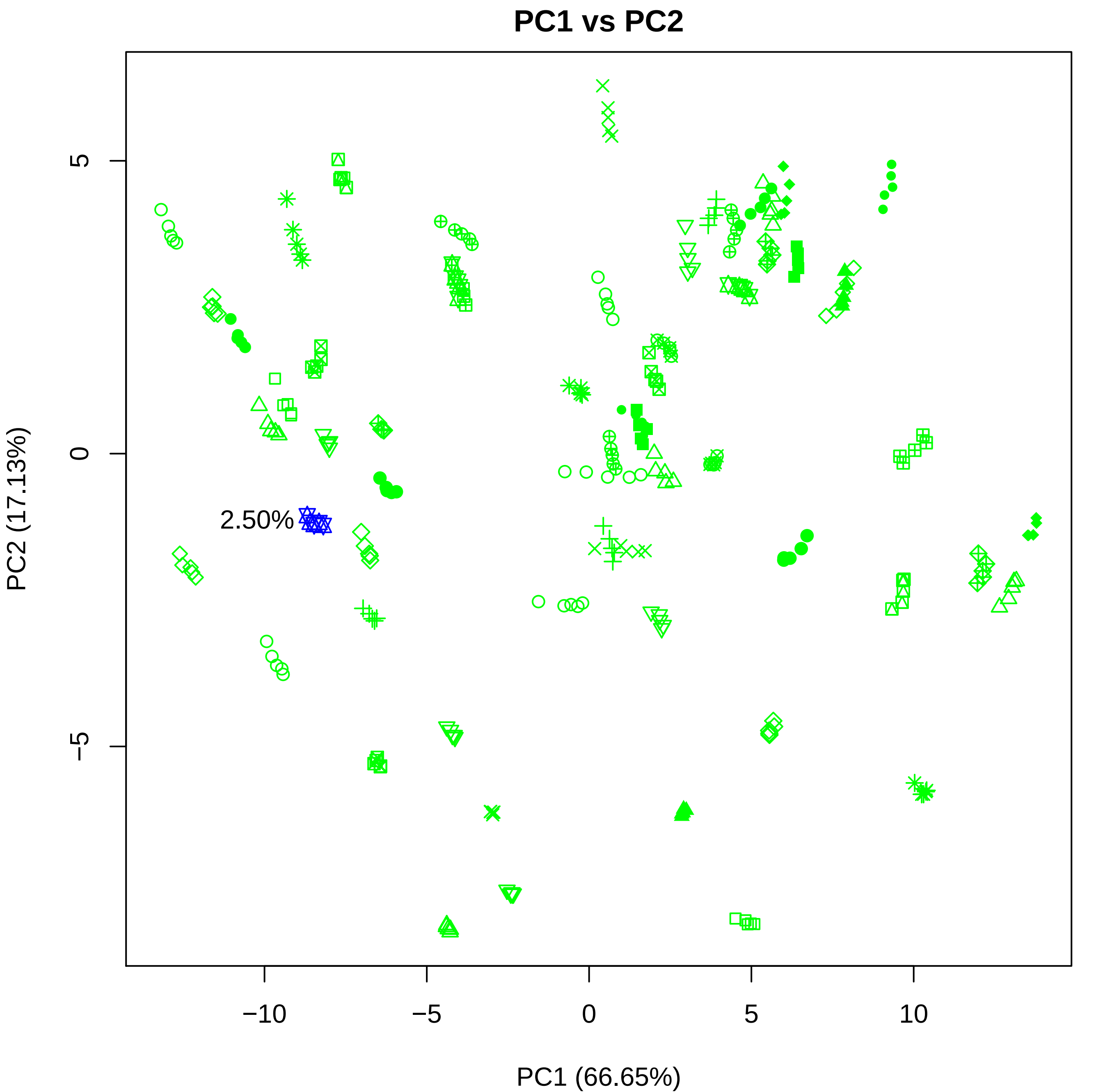
PC1 vs PC2
2.50%
5
0
−5
−10
−5
0
5
10
PC2 (17.13%)
PC1 (66.65%)

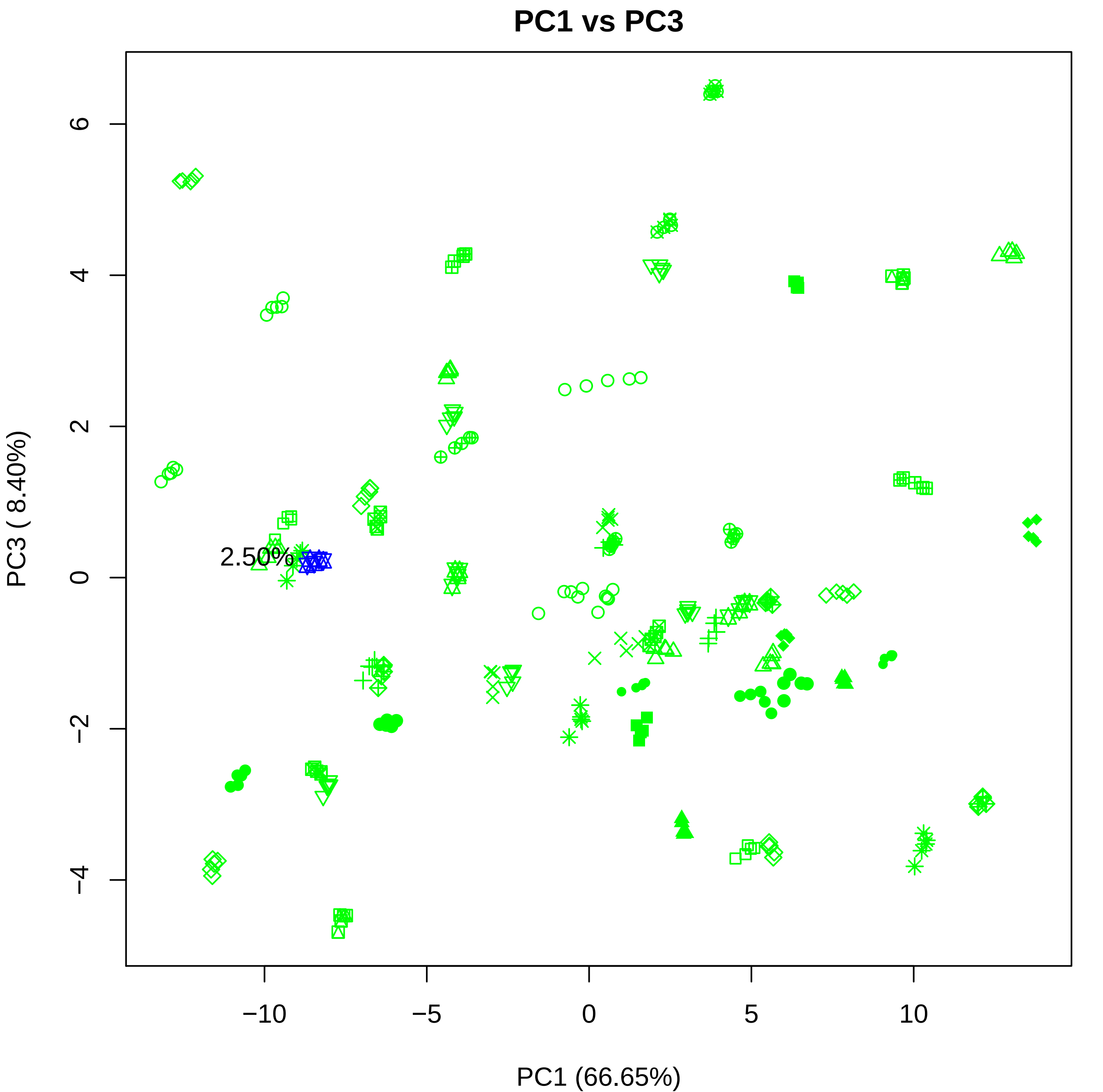
PC1 vs PC3
PC3 ( 8.40%)
PC1 (66.65%)
2.50%
6
4
2
0
−2
−4
−10
−5
0
5
10

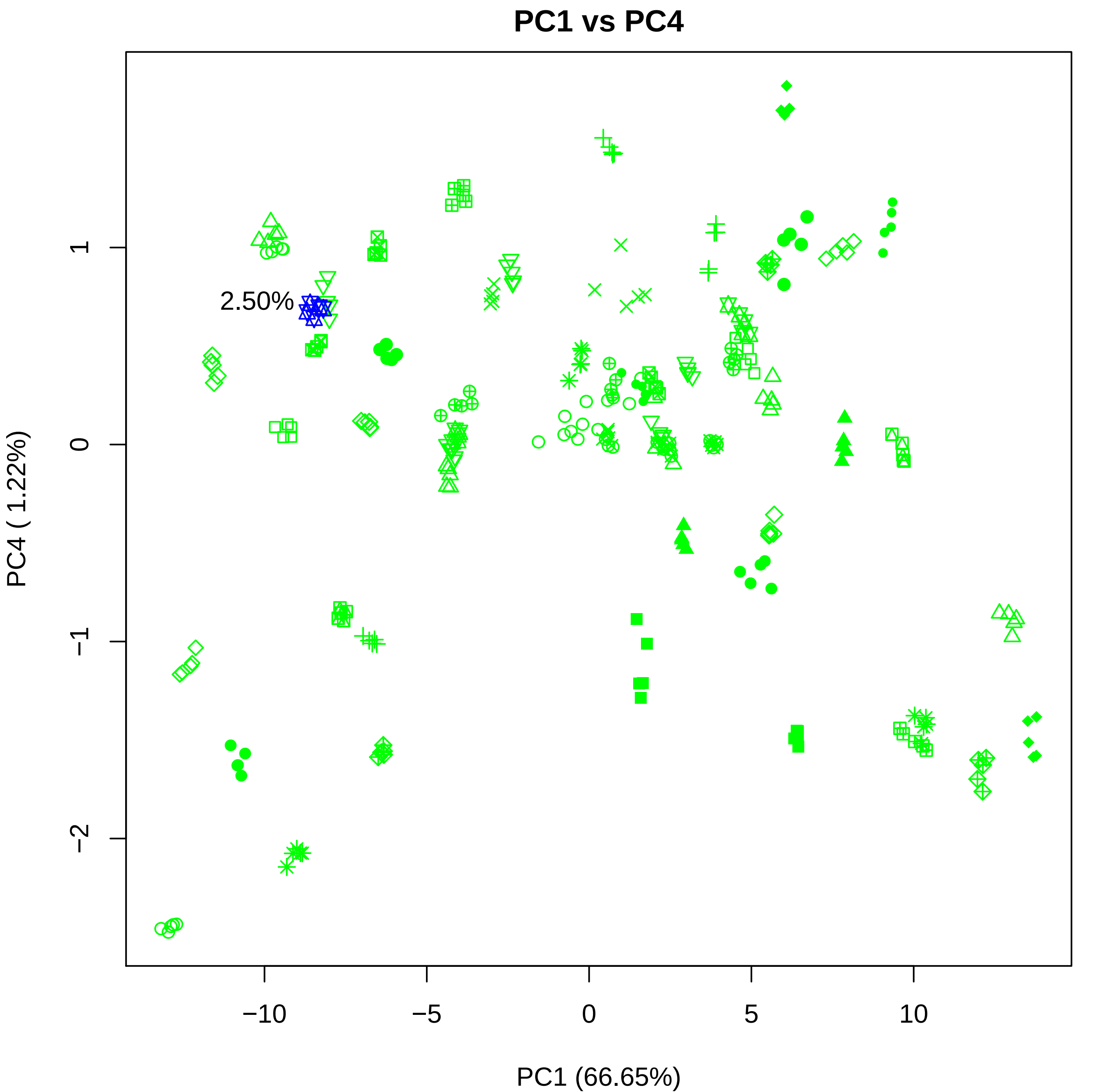

PC1 vs PC4
2.50%
1
0
−1
−2
−10
−5
0
5
10
PC1 (66.65%)
PC4 ( 1.22%)

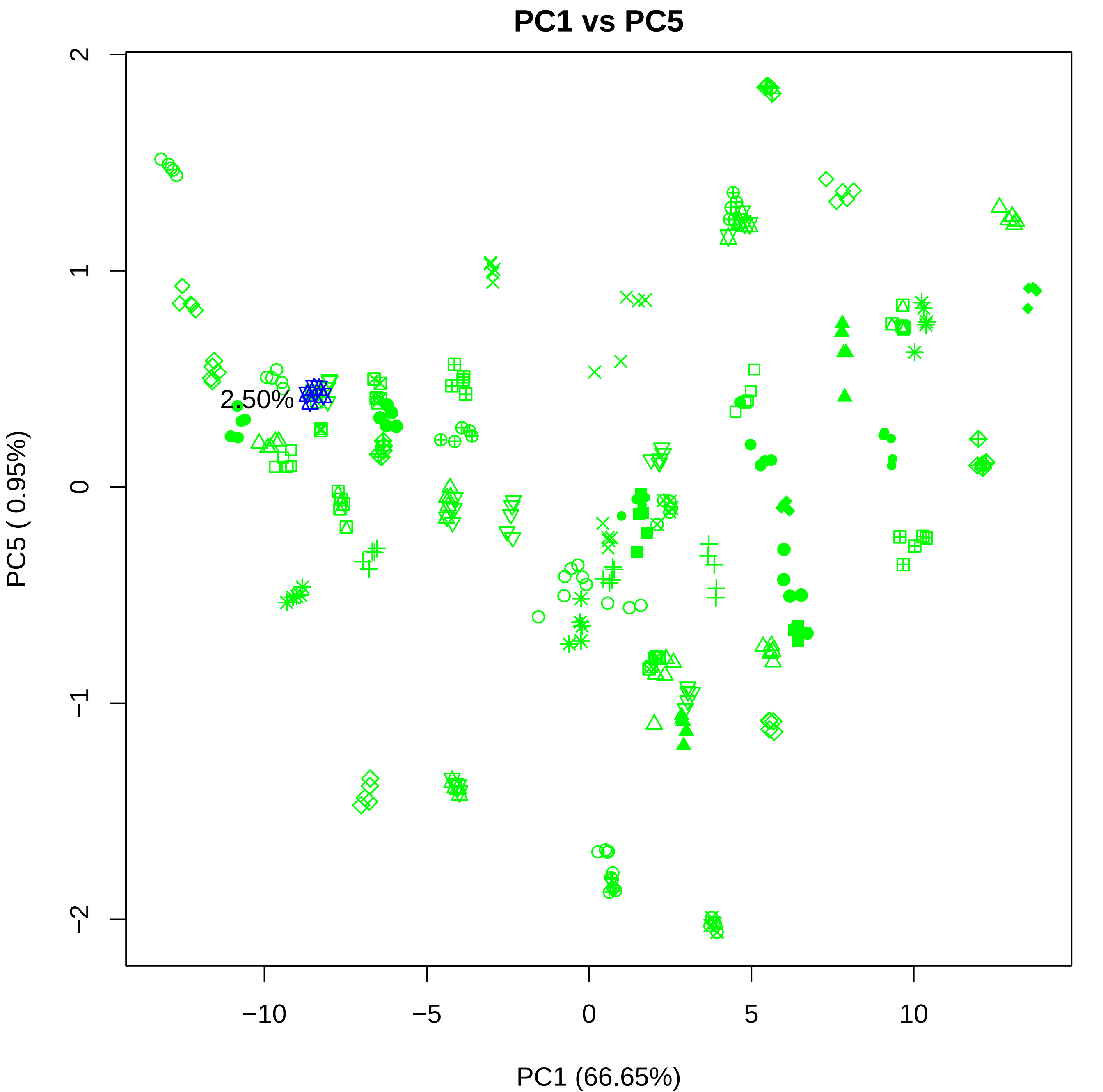

PC1 vs PC5
2.50%
PC5 ( 0.95%)
PC1 (66.65%)
2
1
0
−1
−2
−10
−5
0
5
10

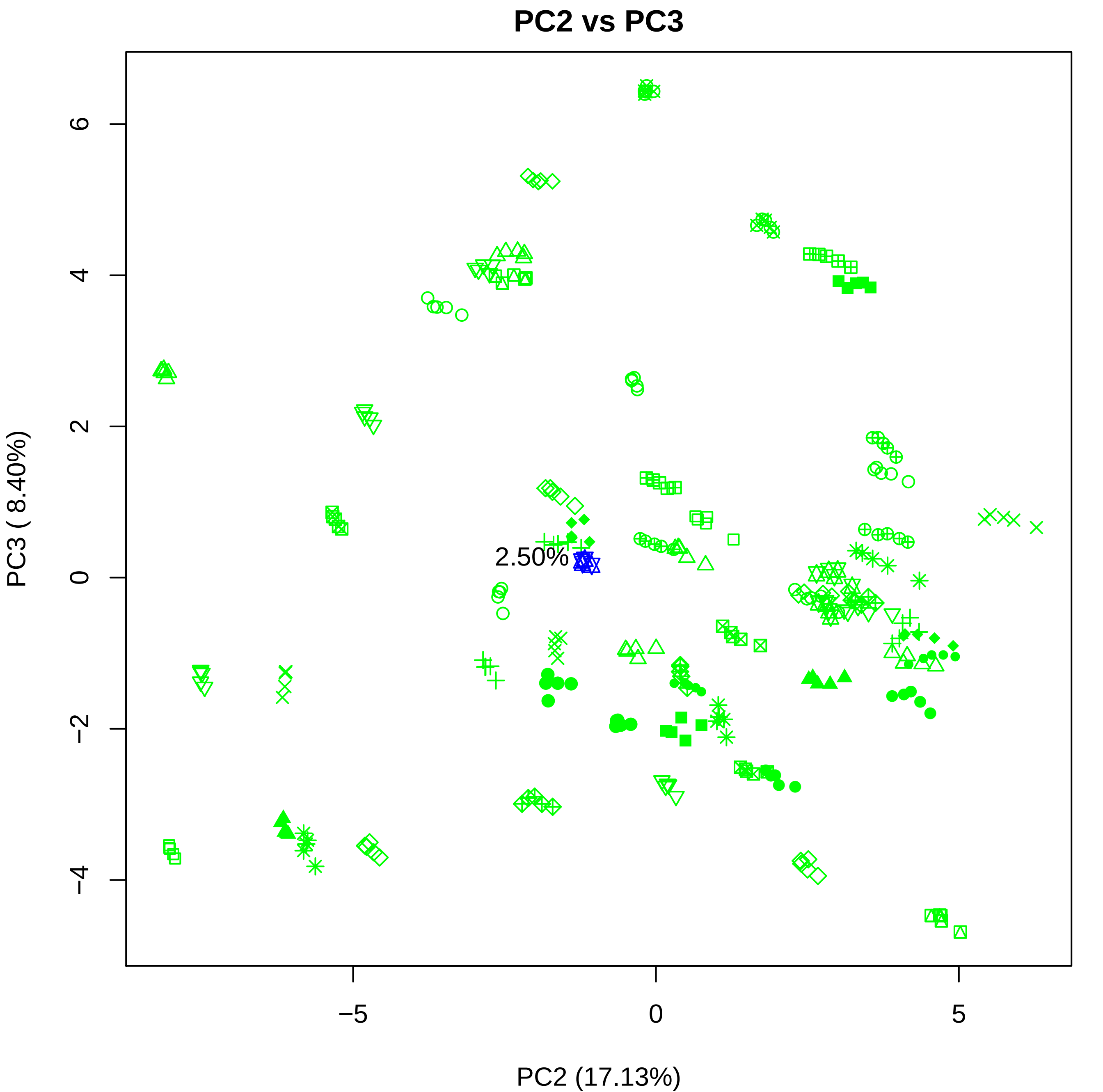

PC2 vs PC3
PC3 ( 8.40%)
PC2 (17.13%)
2.50%
6
4
2
0
−2
−4
−5
0
5

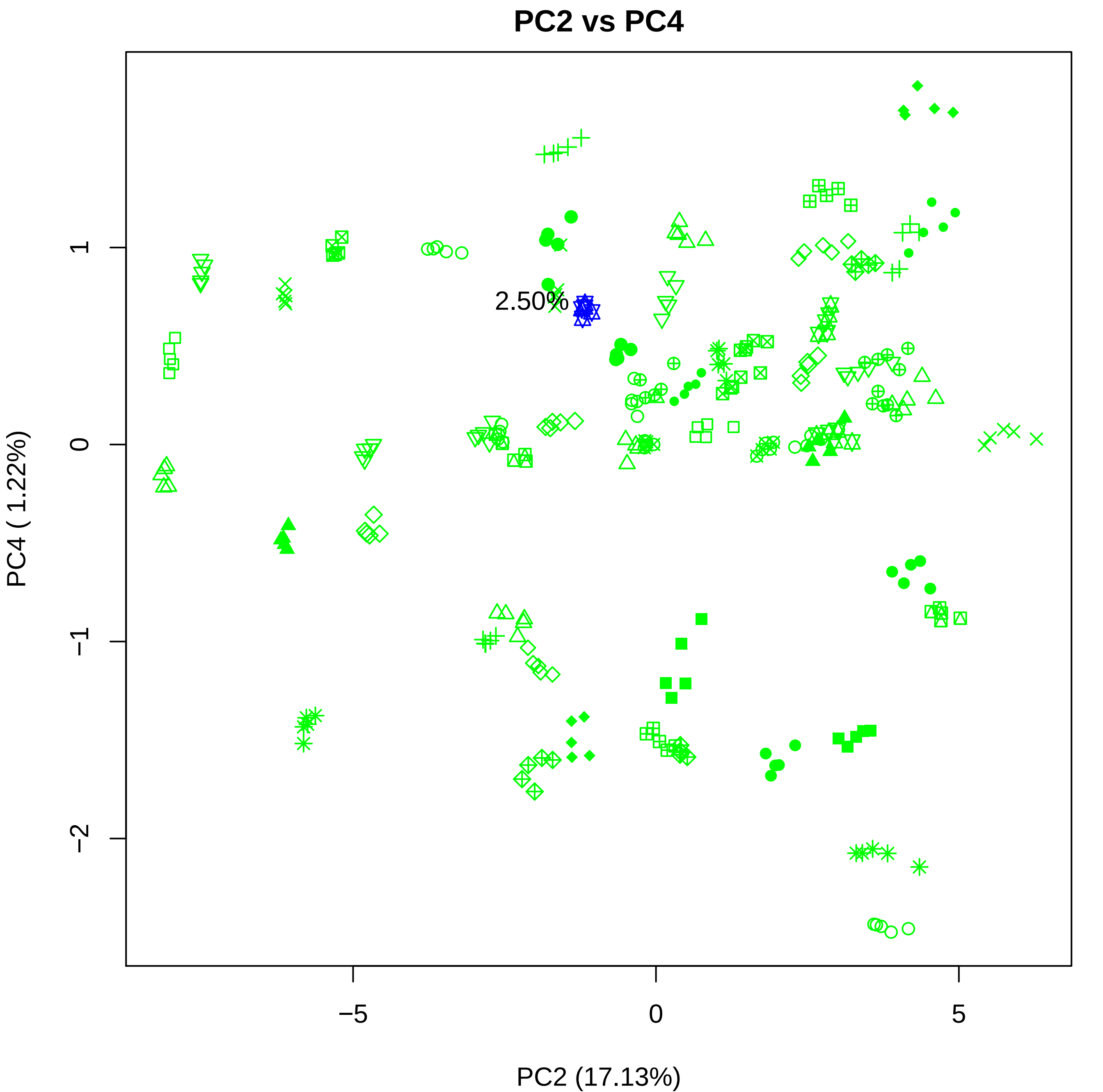

PC2 vs PC4
2.50%
PC4 ( 1.22%)
PC2 (17.13%)
1
0
−1
−2
−5
0
5

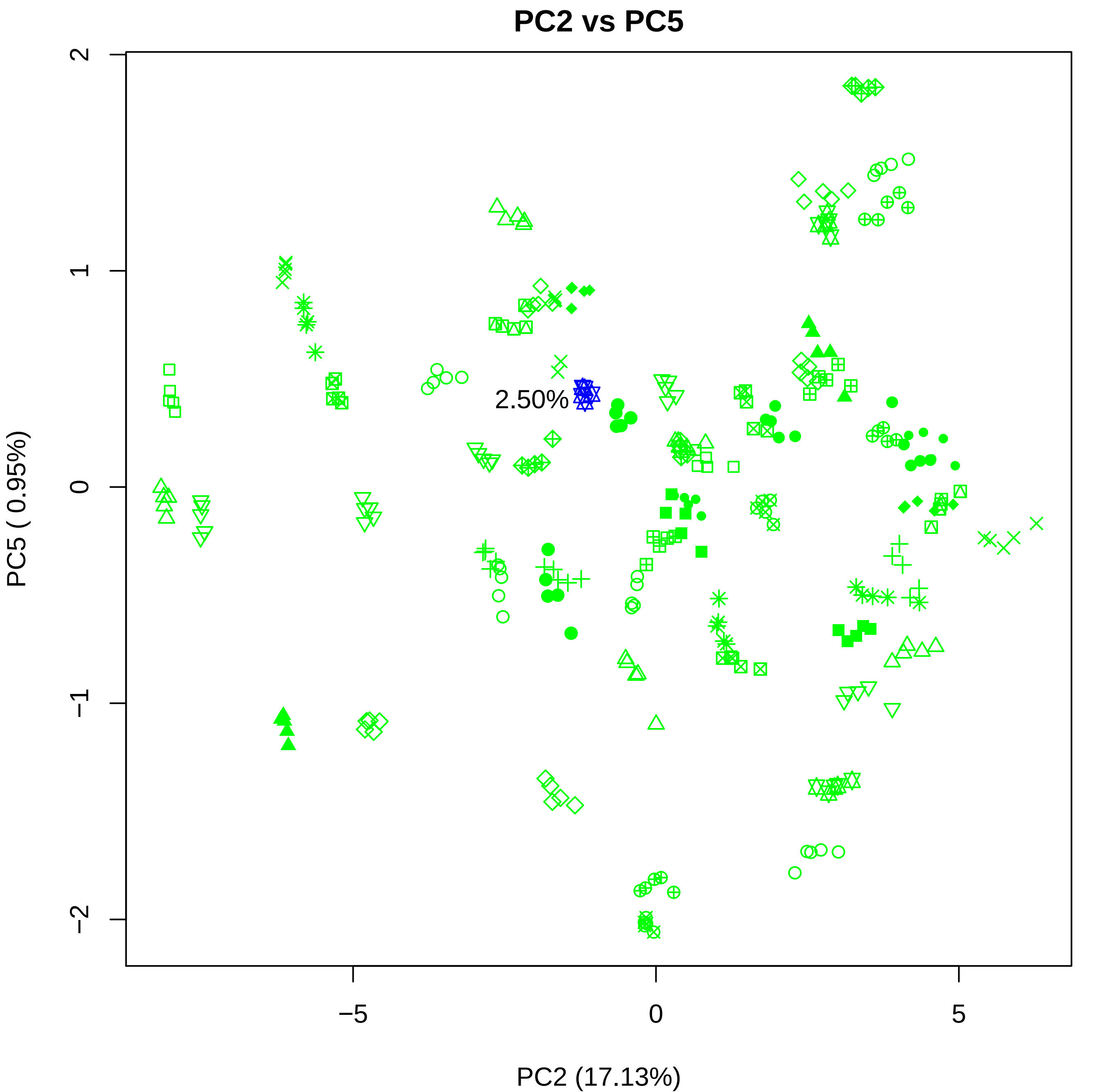
PC2 vs PC5
2.50%
PC5 ( 0.95%)
PC2 (17.13%)
2
1
0
−1
−2
−5
0
5

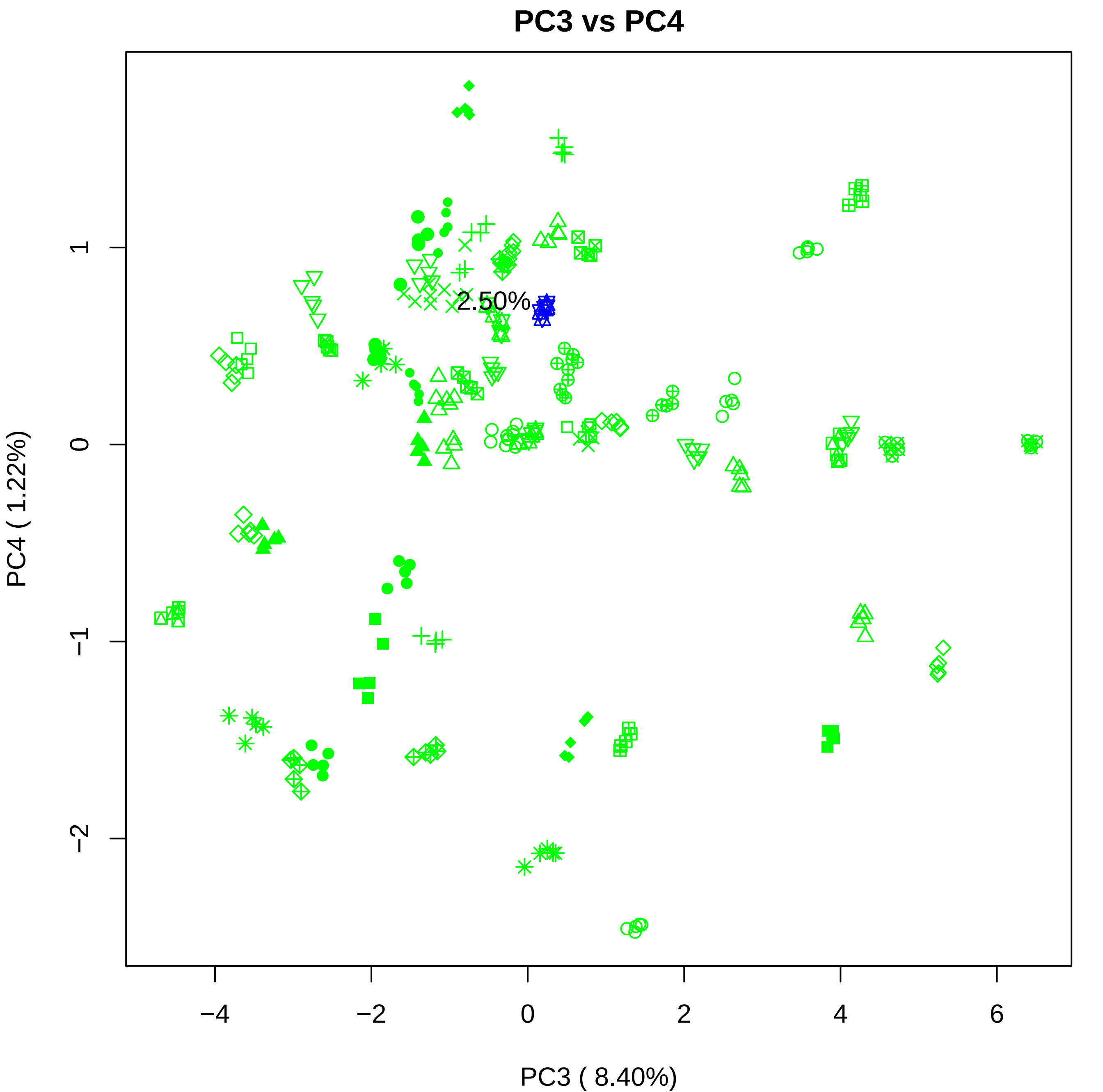

PC3 vs PC4
2.50%
PC4 ( 1.22%)
PC3 ( 8.40%)
−4
−2
0
2
4
6
1
0
−1
−2

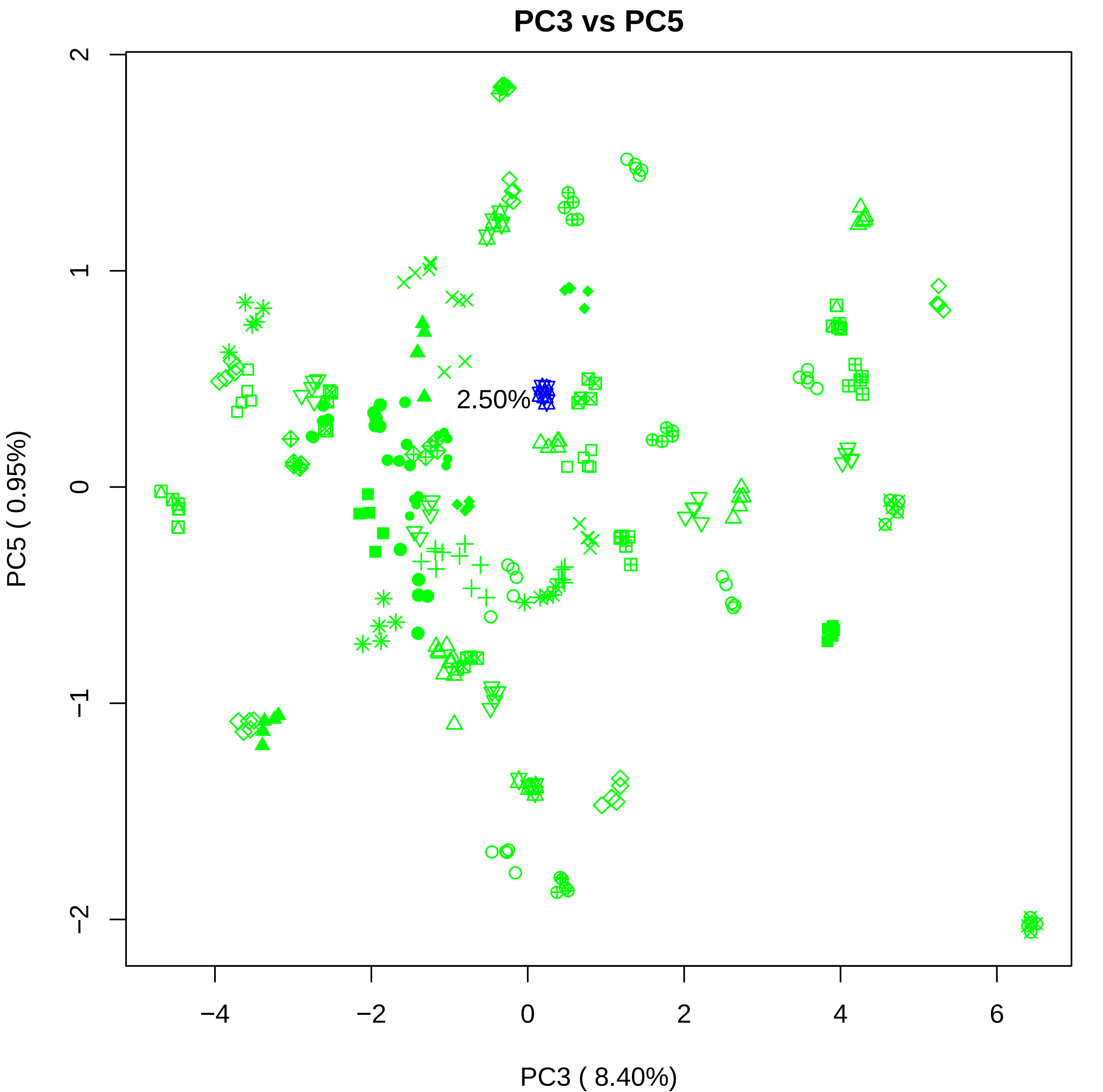
PC3 vs PC5
2.50%
PC5 ( 0.95%)
PC3 ( 8.40%)
−4
−2
0
2
4
6
−2
−1
0
1
2

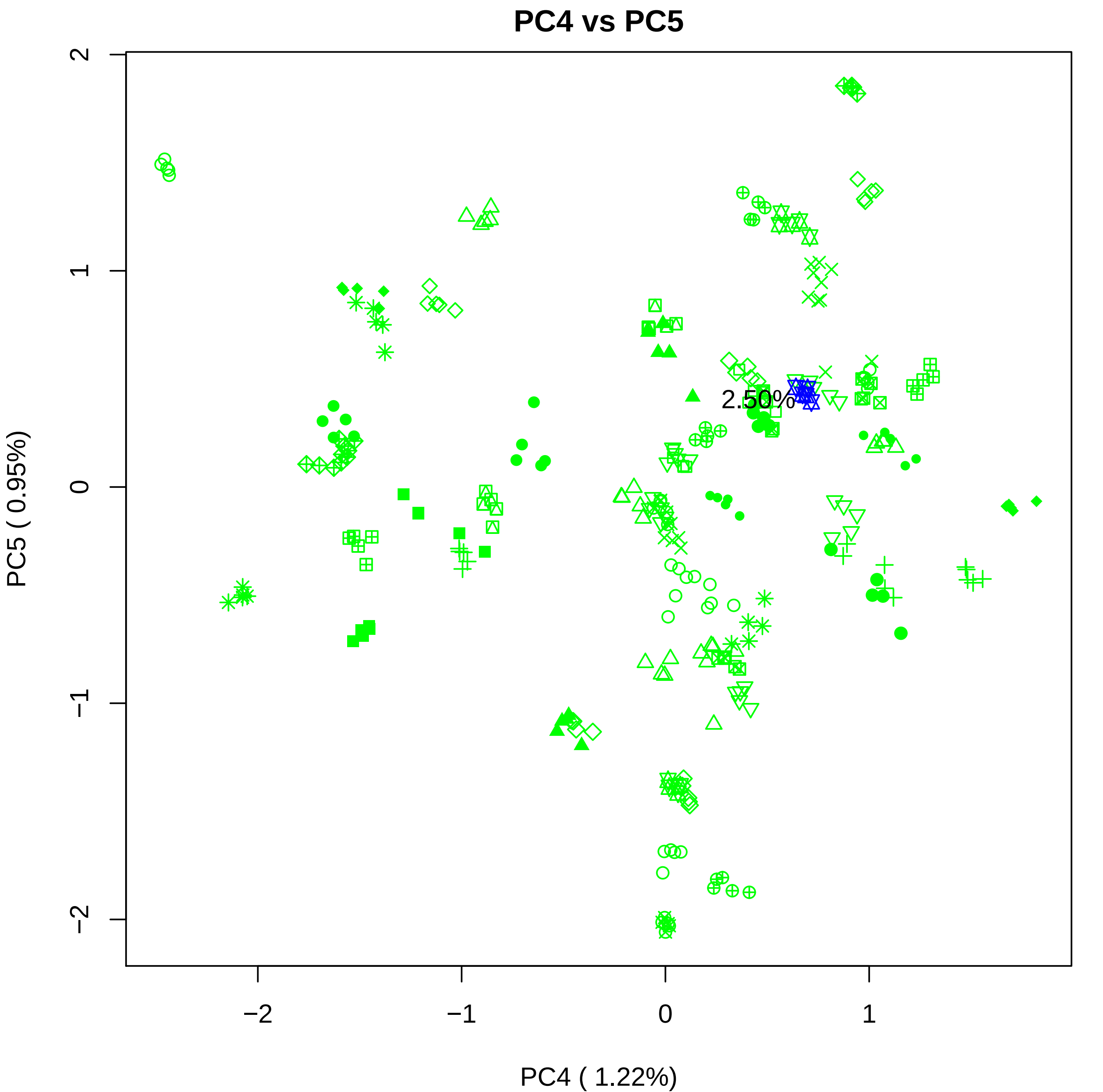

PC4 vs PC5
2.50%
PC4 ( 1.22%)
PC5 ( 0.95%)
−2
−1
0
1
2

### PCA of 5% BMH(a) vs PMH(a)

Scaled Matrix PCAed

7th order polynomial baseline correction performed

Raw spectra smoothed to 21 Neighbours, 2nd order PN, Savtisky Golay

Normalisation between 0 and 100

D2 spectra smoothed to 41 Neighbours, 2nd order PN, Savtisky Golay

Regions Analysed: 3601−2501 and 2001−700cm−1

Samples Analysed: 1 bContam's 0 BLH's 0 BMH's 0 OMH's 0 pContam's 0 PLH's 60 PMH's

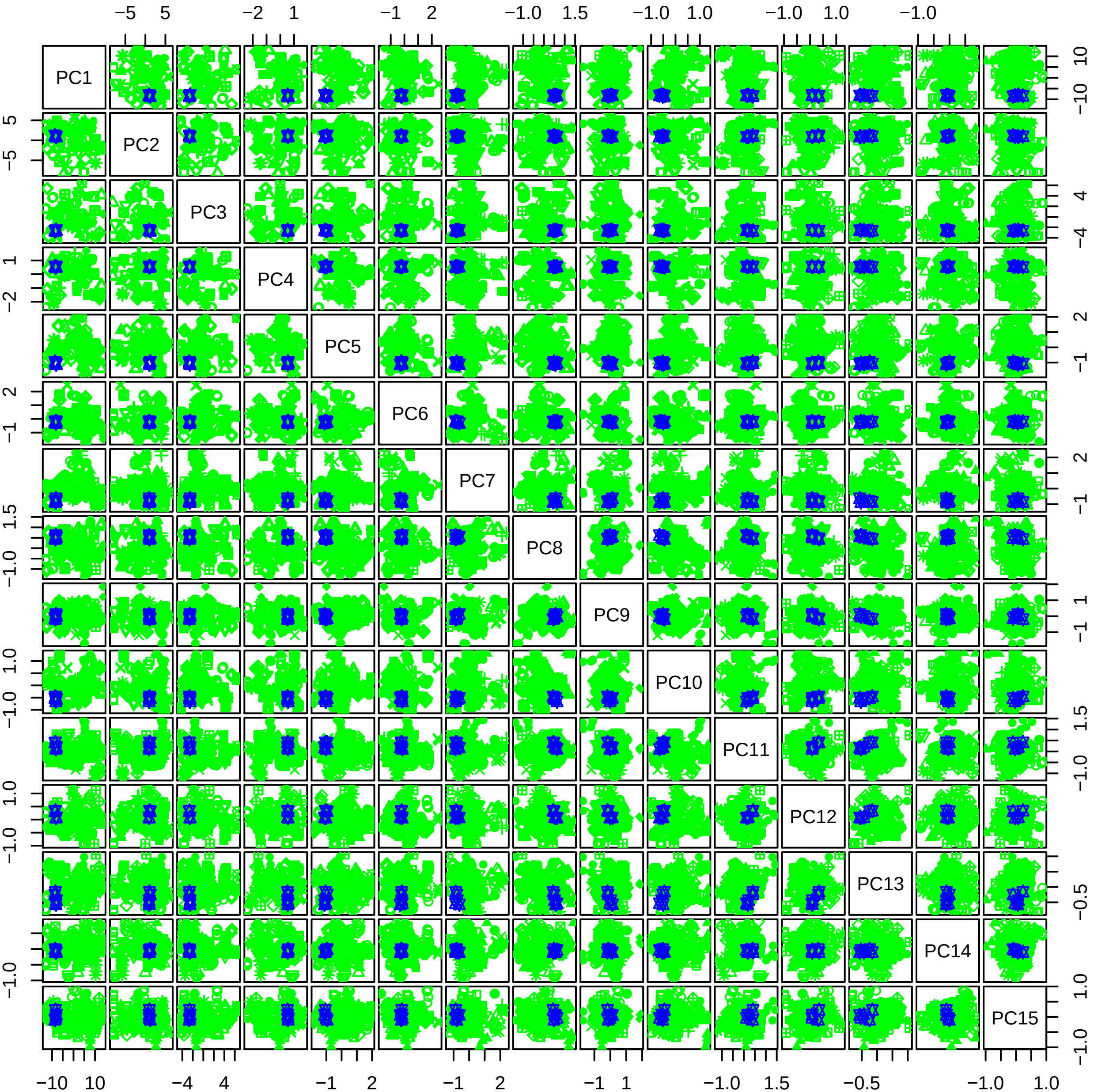
PC1
PC2
PC3
PC4
PC5
PC6
PC7
PC8
PC9
PC10
PC11
PC12
PC13
PC14
PC15

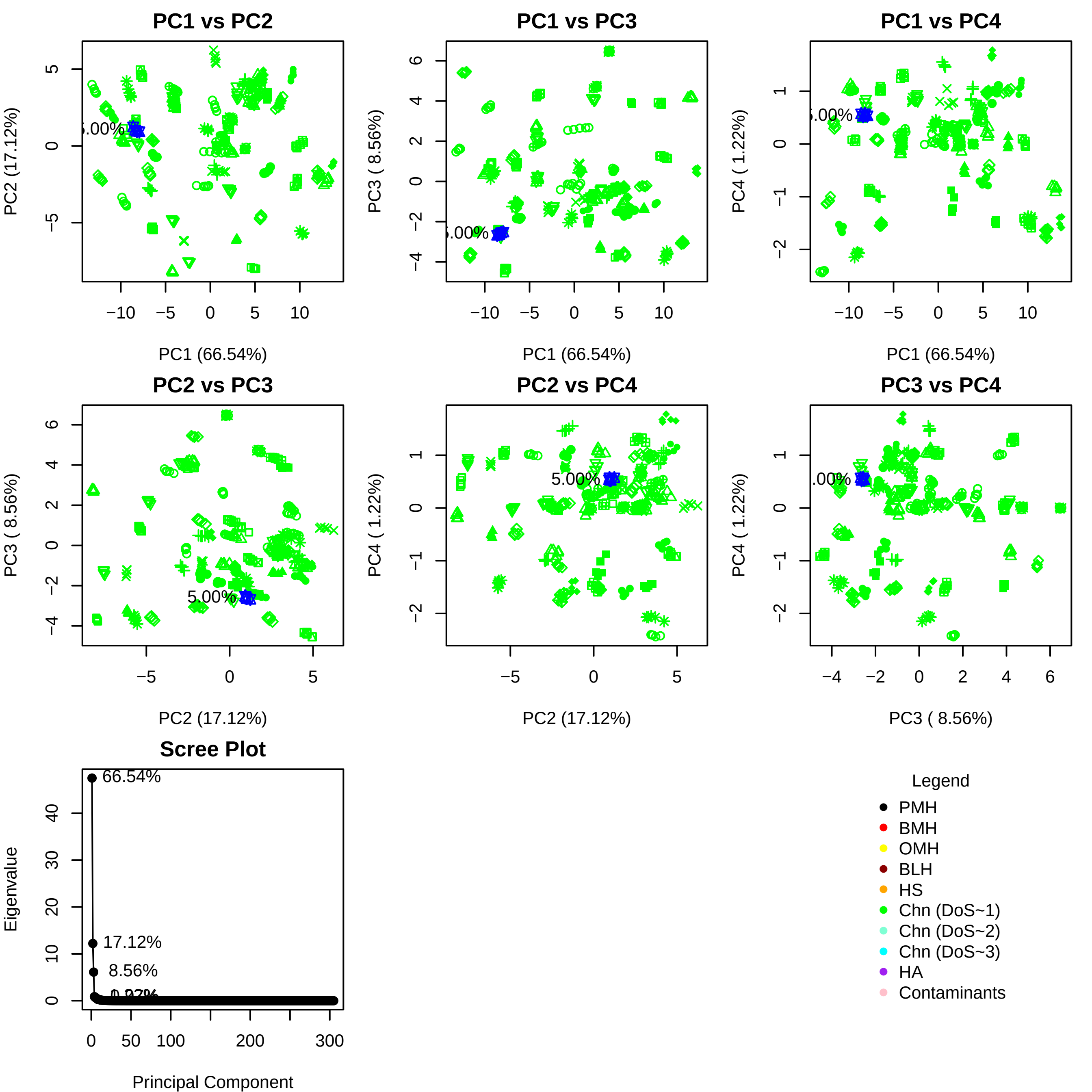

PC1 vs PC2
PC1 vs PC3
PC1 vs PC4
PC2 vs PC3
PC2 vs PC4
PC3 vs PC4
PC1 (66.54%)
PC2 (17.12%)
PC3 ( 8.56%)
PC4 ( 1.22%)
5.00%
Scree Plot
66.54%
17.12%
8.56%
Eigenvalue
Principal Component
Legend
PMH
BMH
OMH
BLH
HS
Chn (DoS~1)
Chn (DoS~2)
Chn (DoS~3)
HA
Contaminants

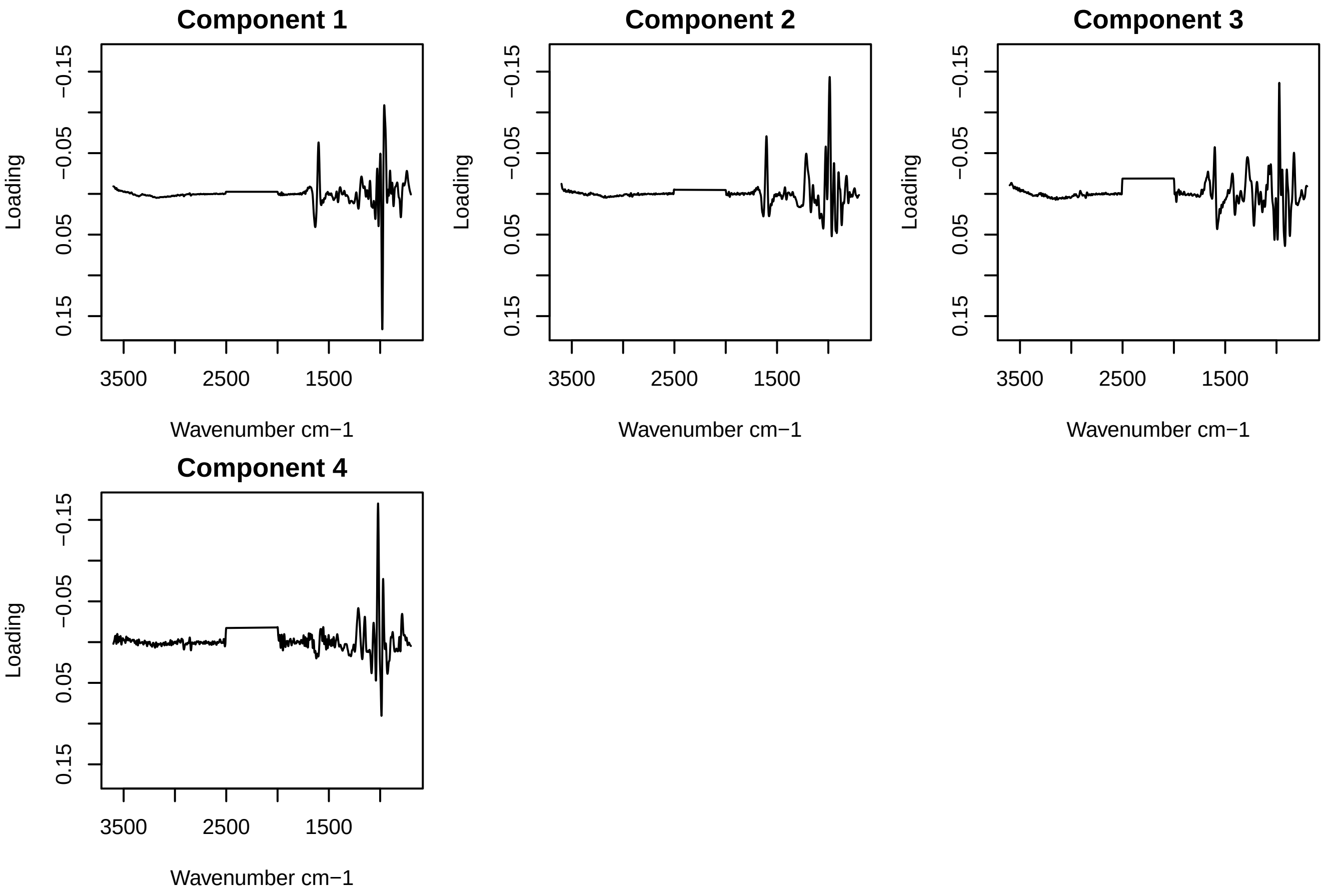
Component 1
Component 2
Component 3
Component 4
Loading
−0.15
−0.05
0.05
0.15
3500
2500
1500
Wavenumber cm−1

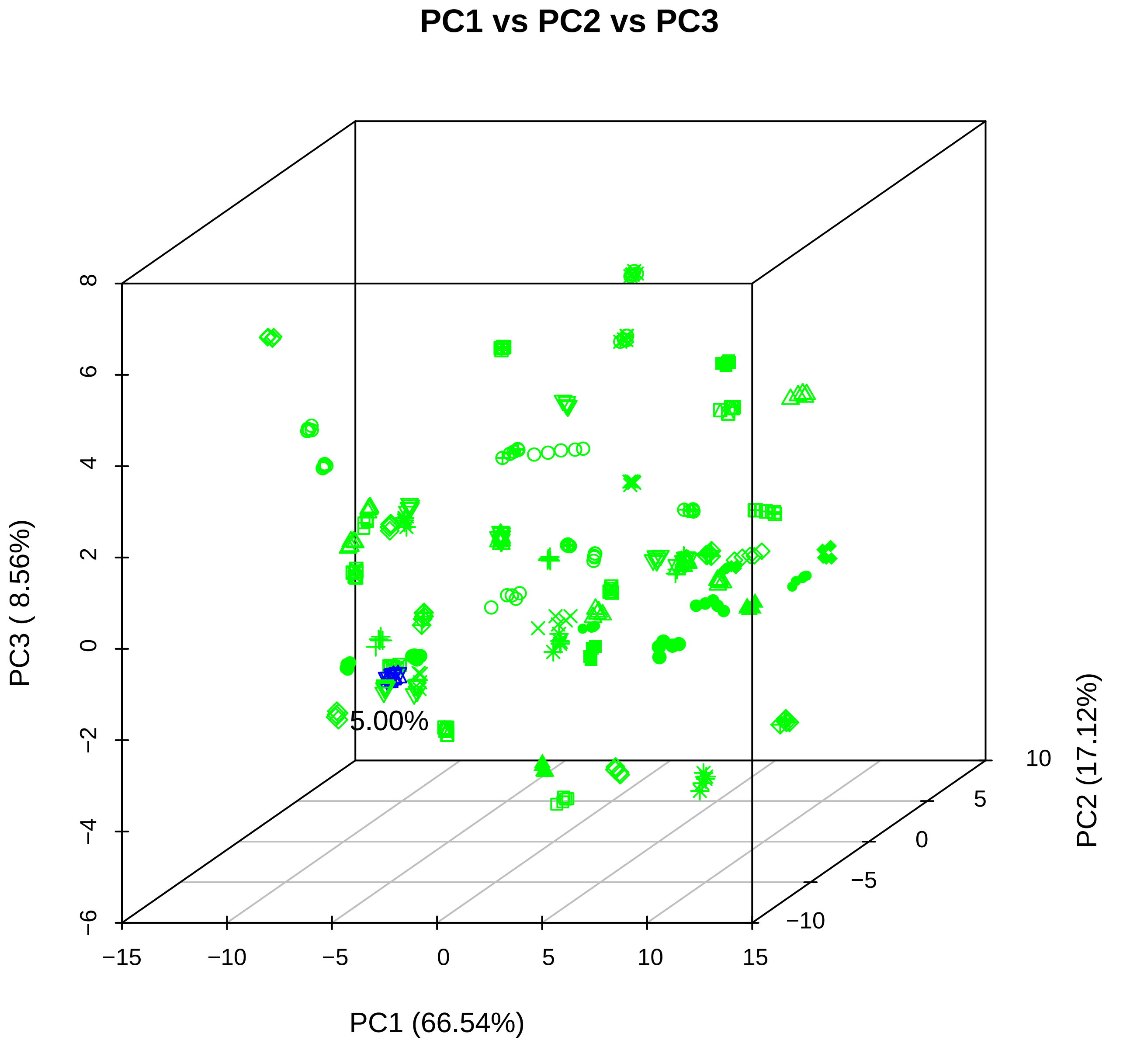
PC1 vs PC2 vs PC3
PC1 (66.54%)
PC2 (17.12%)
PC3 ( 8.56%)
5.00%
−15
−10
−5
0
5
10
15
−6
−4
−2
0
2
4
6
8
−10
−5
0
5
10

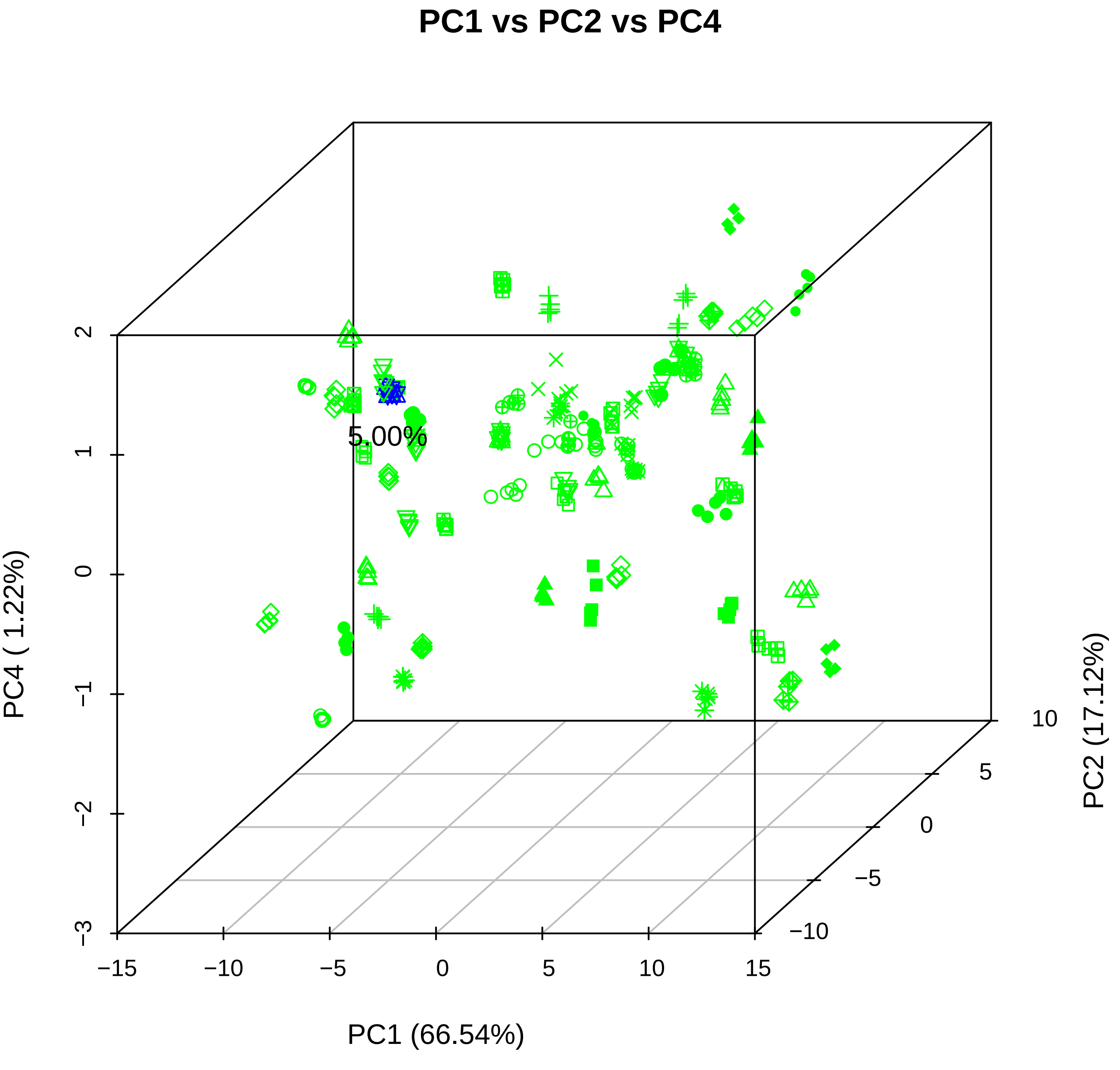
PC1 vs PC2 vs PC4
5.00%
PC1 (66.54%)
PC2 (17.12%)
PC4 ( 1.22%)
−15
−10
−5
0
5
10
15
−3
−2
−1
0
1
2
−10
−5
0
5
10

# PC1 vs PC3 vs PC4

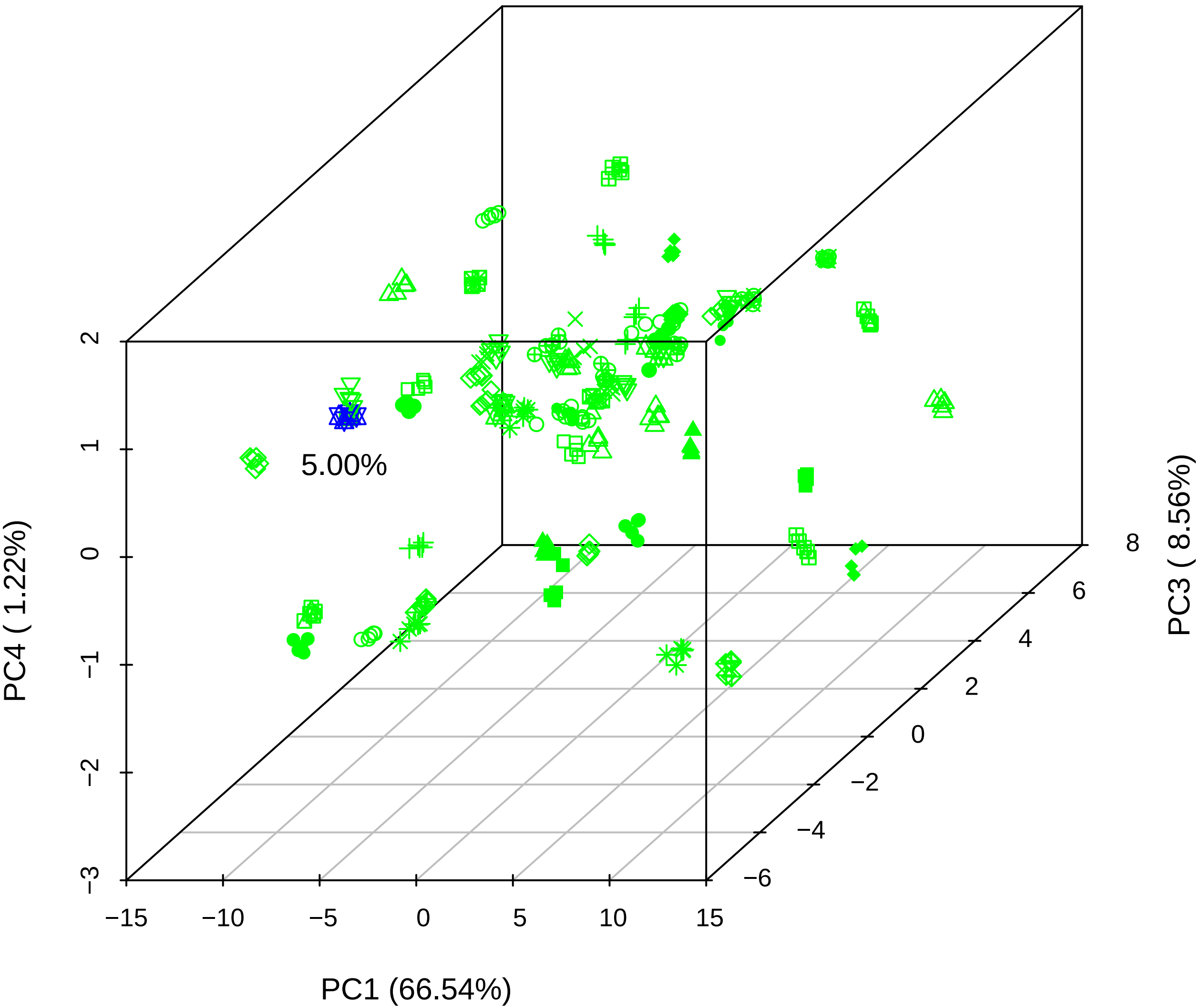

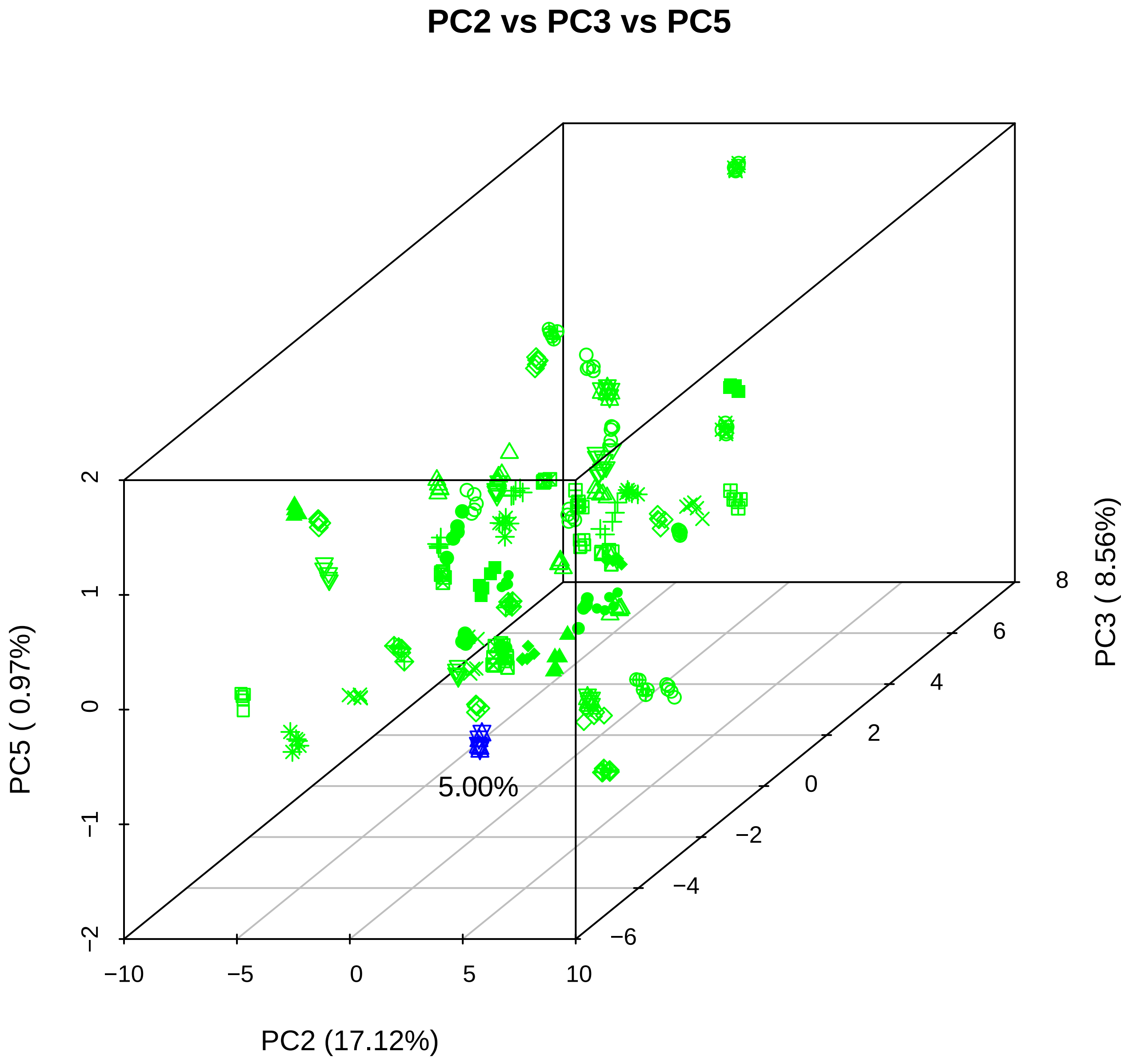
PC2 vs PC3 vs PC5
PC2 (17.12%)
PC3 ( 8.56%)
PC5 ( 0.97%)
5.00%
−10
−5
0
5
10
−6
−4
−2
0
2
4
6
8
−2
−1
0
1
2

PC1 vs PC2

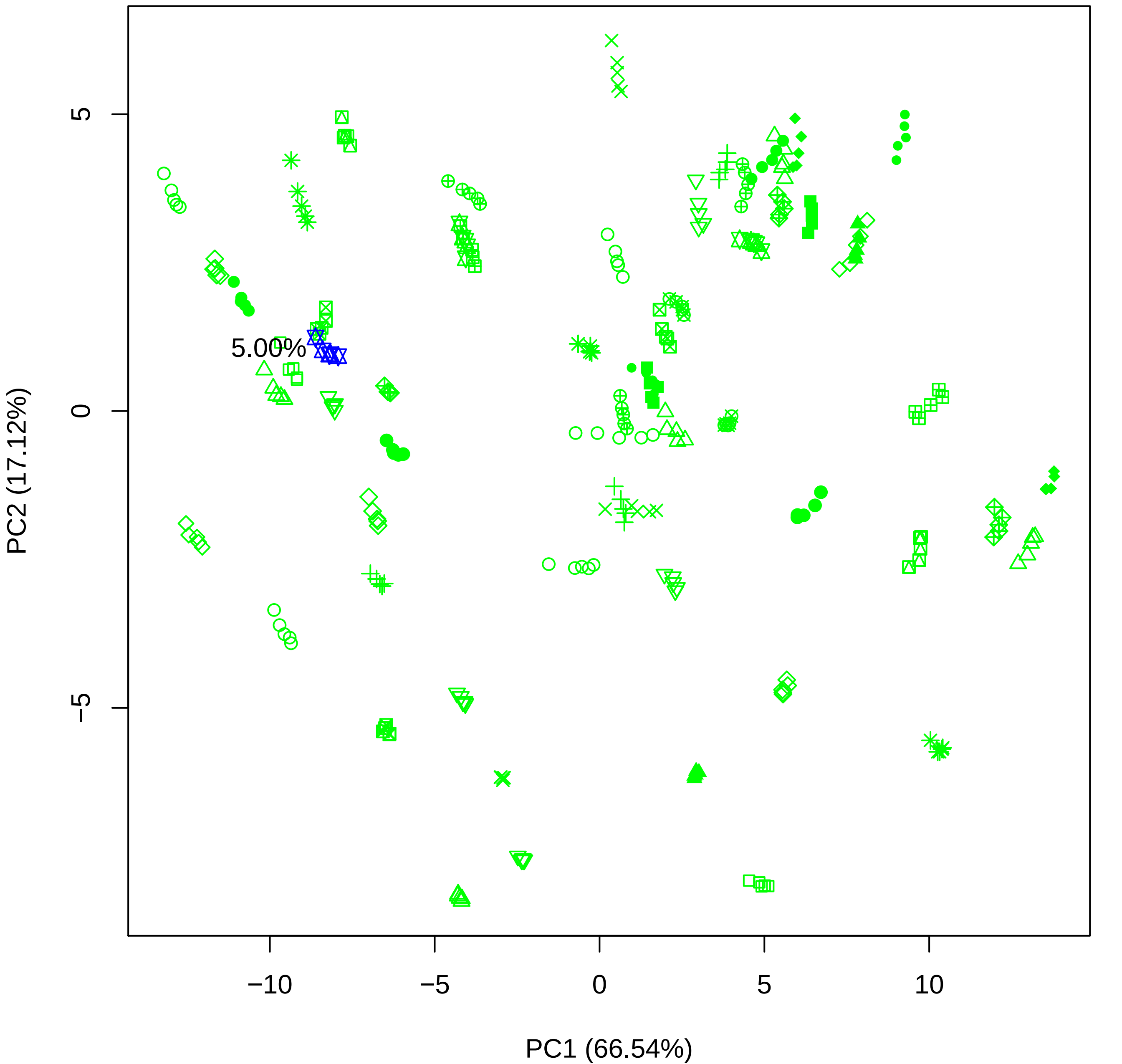

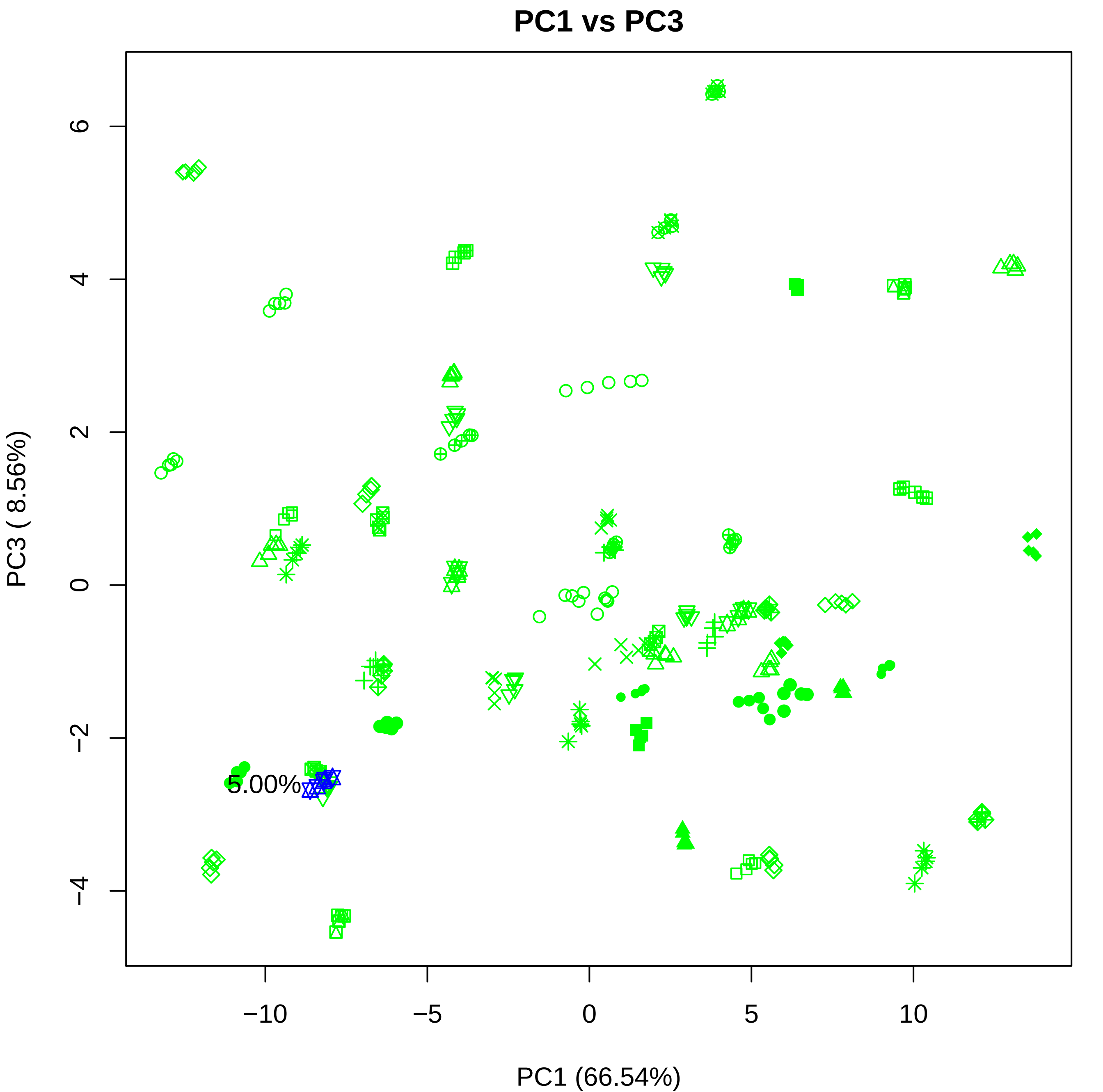
PC1 vs PC3
PC3 ( 8.56%)
PC1 (66.54%)
6
4
2
0
−2
−4
−10
−5
0
5
10
5.00%

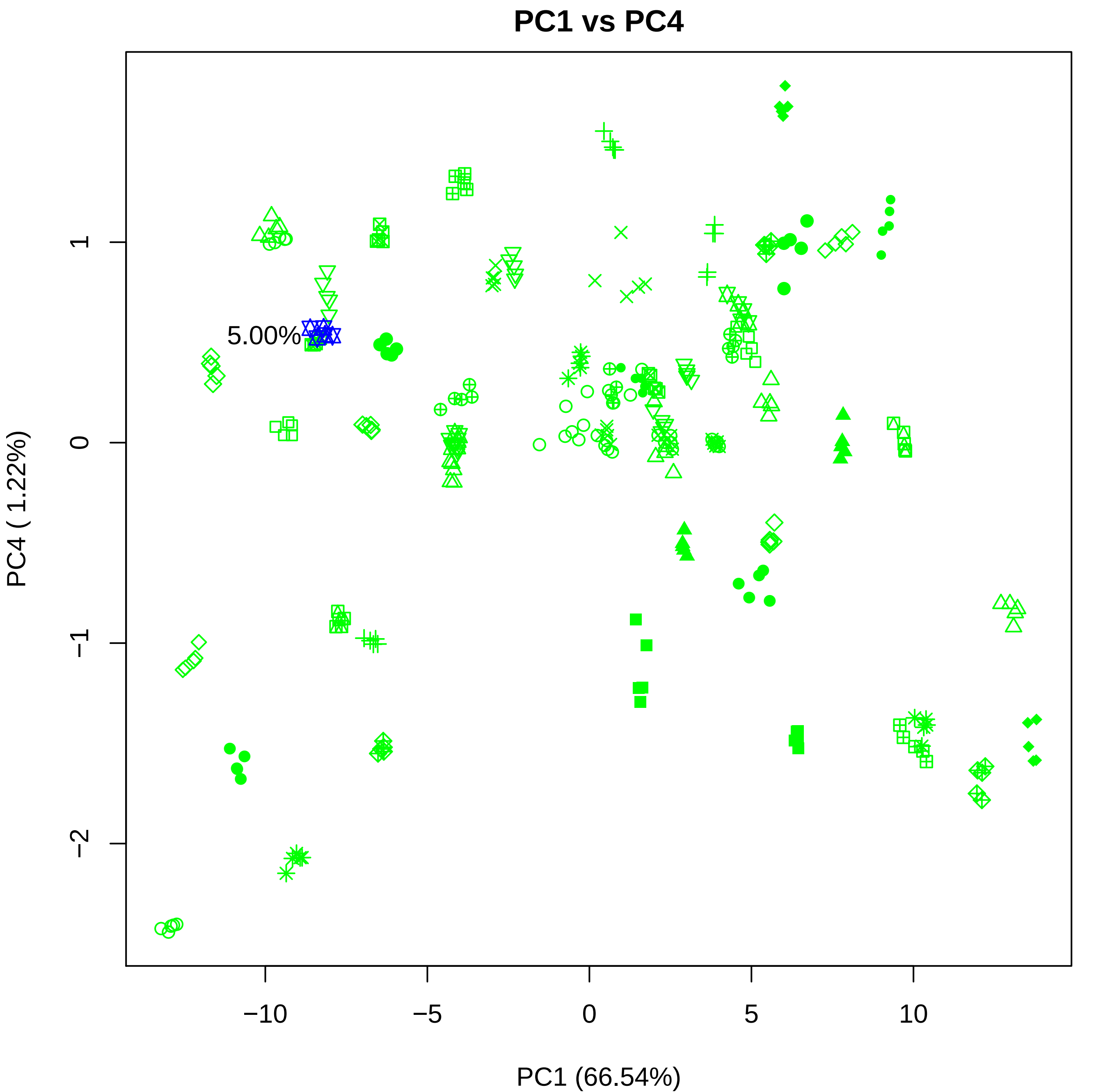
PC1 vs PC4
5.00%
PC4 ( 1.22%)
PC1 (66.54%)
1
0
−1
−2
−10
−5
0
5
10

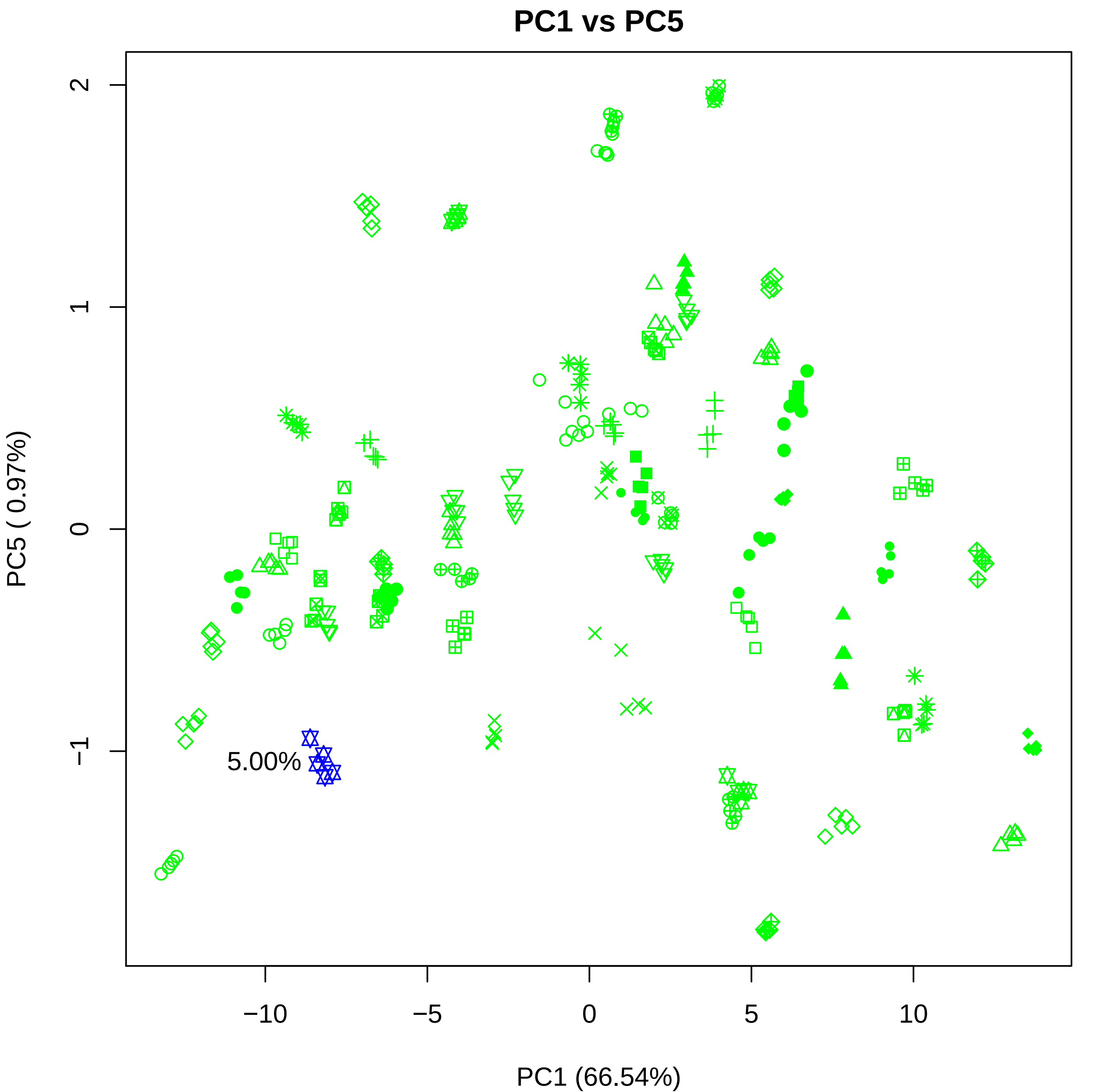
PC1 vs PC5
PC5 ( 0.97%)
PC1 (66.54%)
5.00%
2
1
0
−1
−10
−5
0
5
10

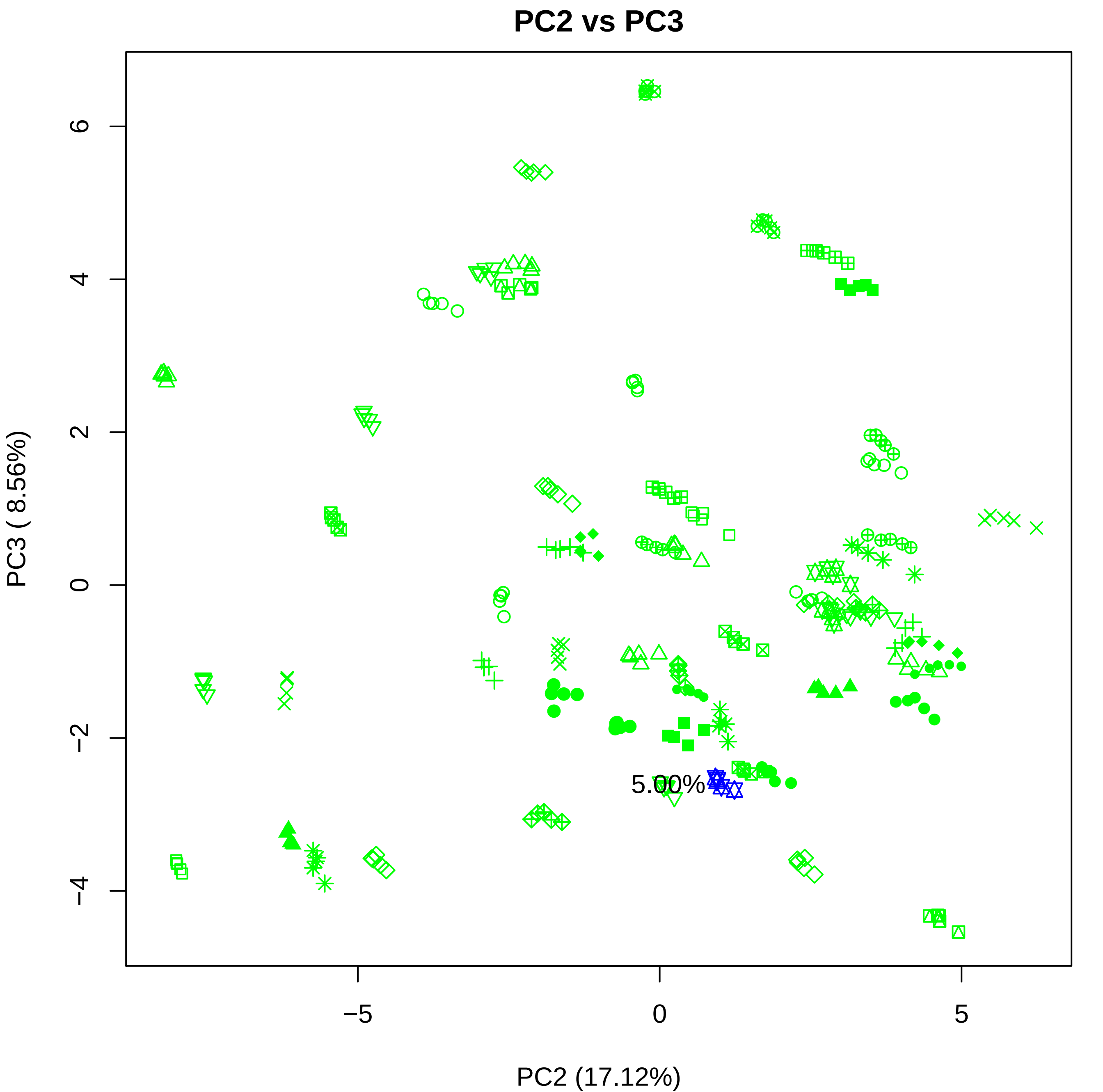
PC2 vs PC3
PC3 ( 8.56%)
PC2 (17.12%)
6
4
2
0
−2
−4
−5
0
5
5.00%

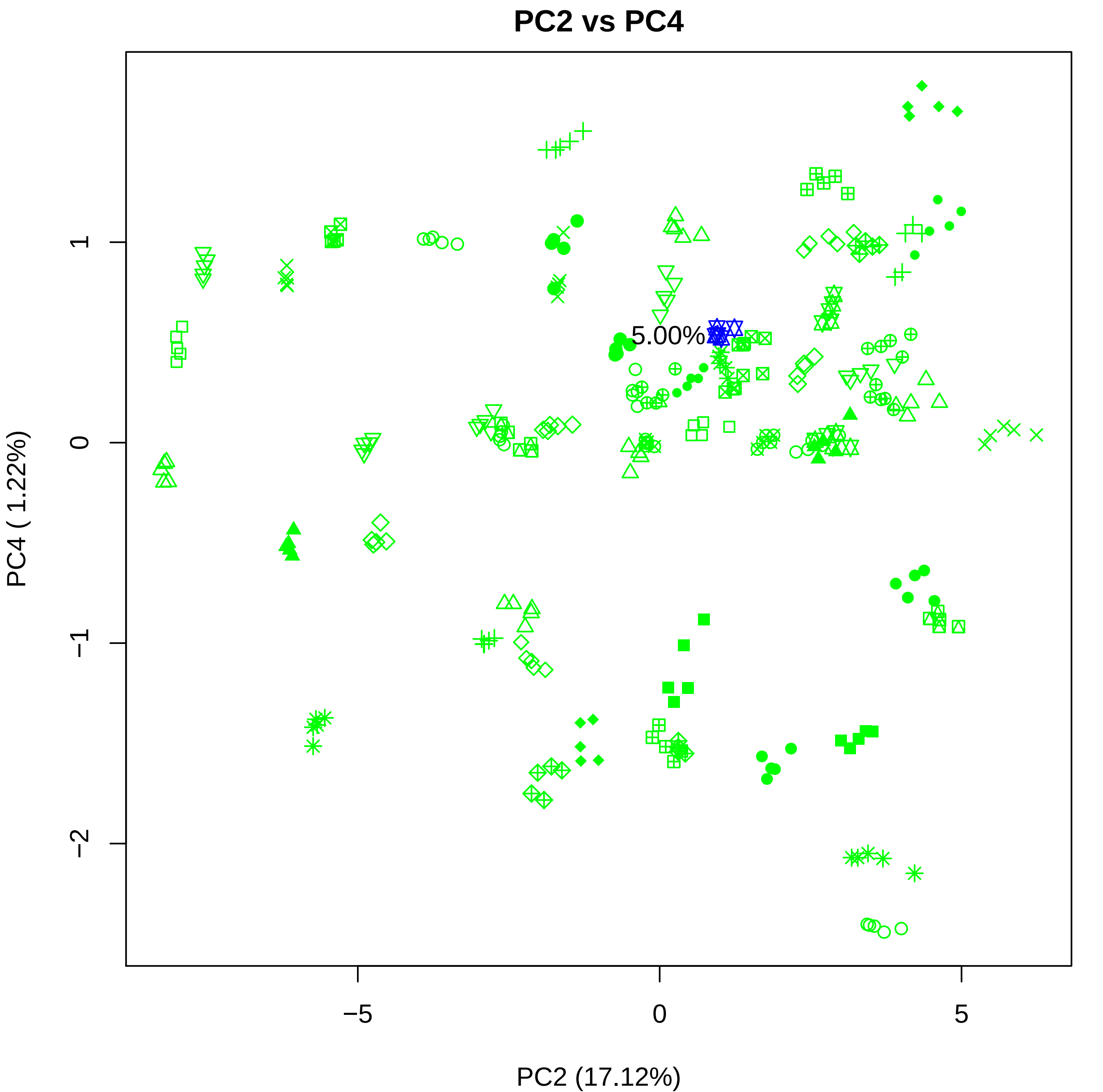
PC2 vs PC4
5.00%
PC4 ( 1.22%)
PC2 (17.12%)
1
0
−1
−2
−5
0
5

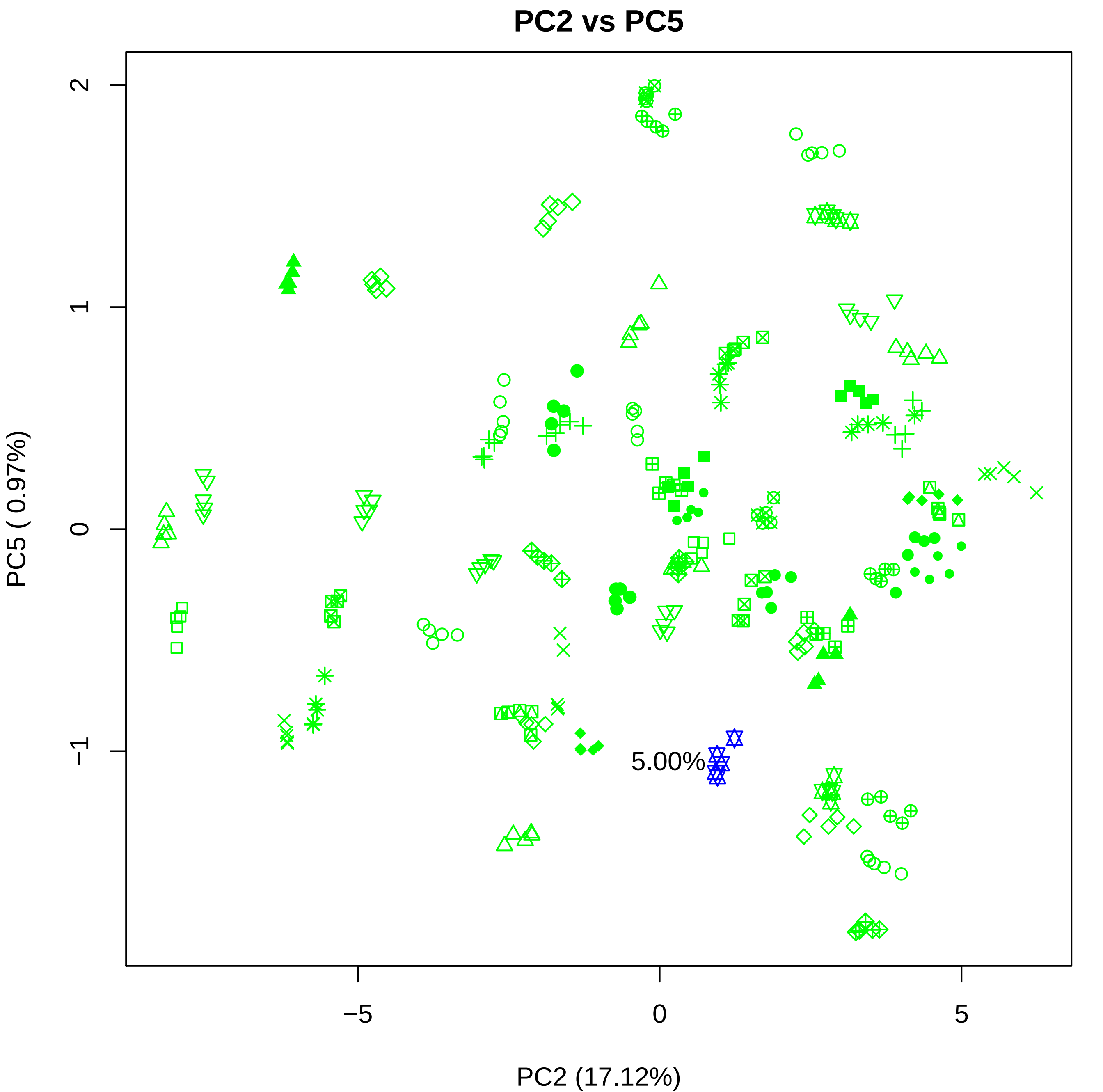
PC2 vs PC5
PC5 ( 0.97%)
PC2 (17.12%)
5.00%
2
1
0
−1
−5
0
5

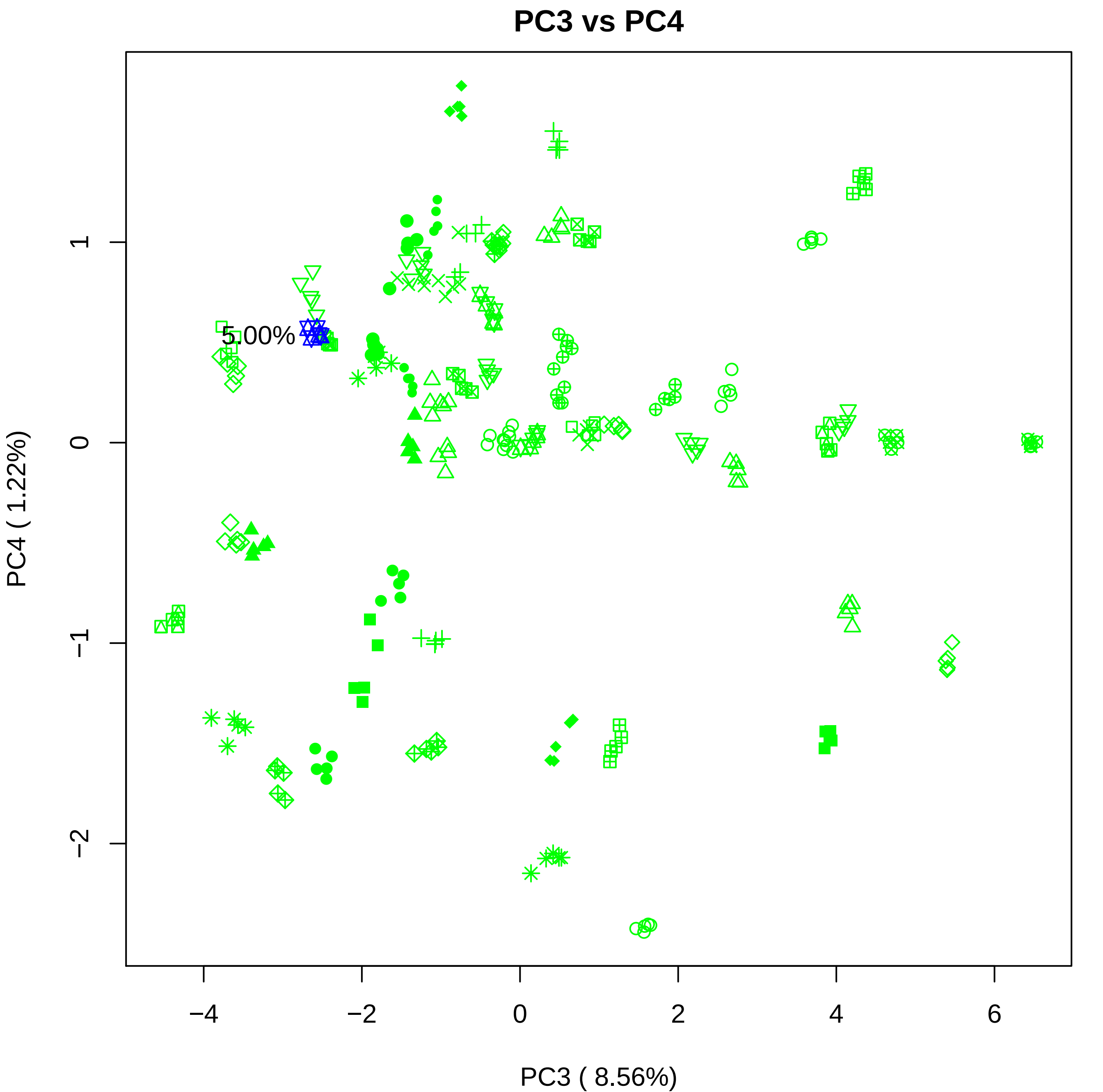
PC3 vs PC4
5.00%
PC3 ( 8.56%)
PC4 ( 1.22%)
−4
−2
0
2
4
6
−2
−1
0
1

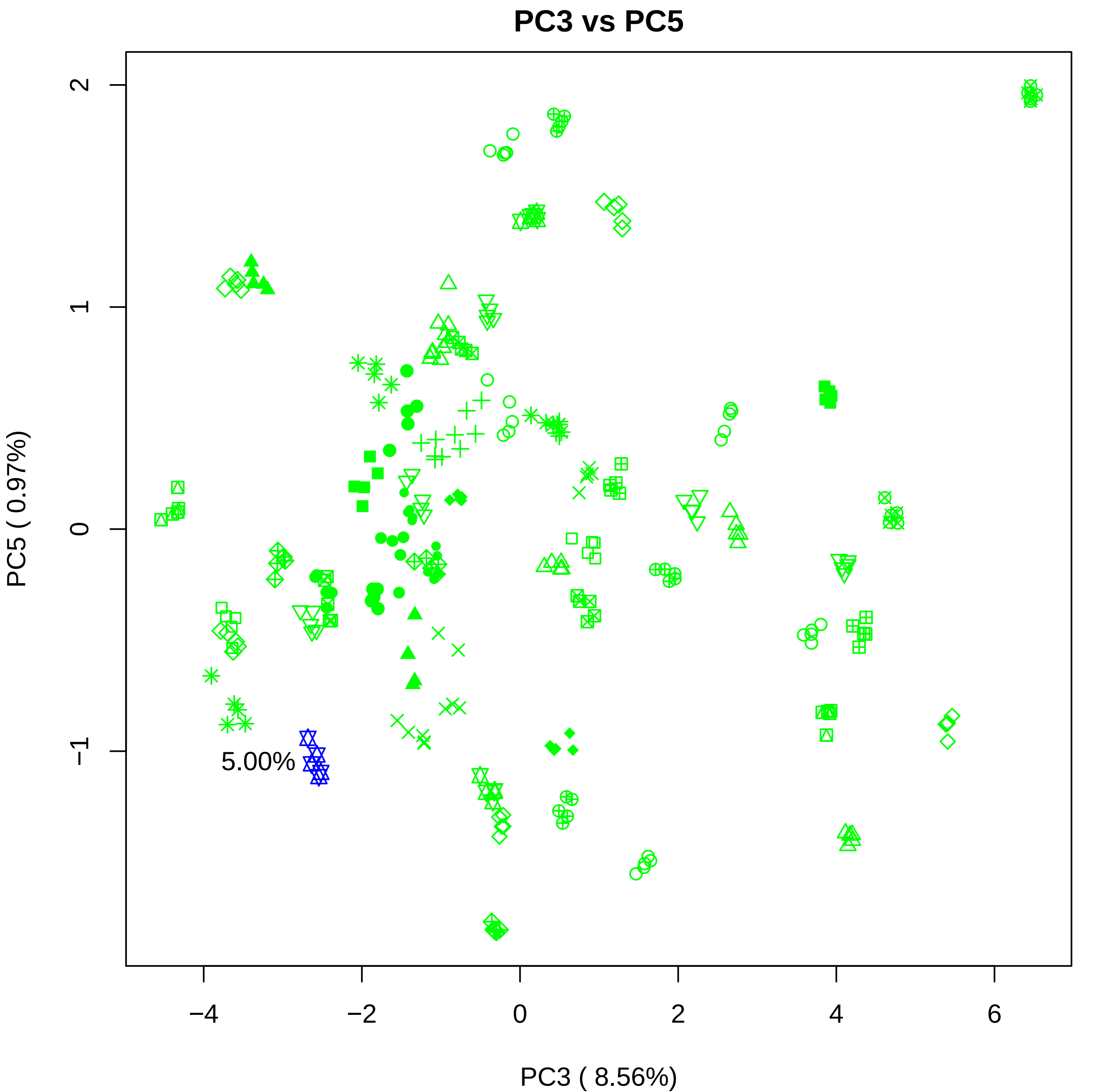
PC3 vs PC5
PC5 ( 0.97%)
PC3 ( 8.56%)
5.00%
2
1
0
−1
−4
−2
0
2
4
6

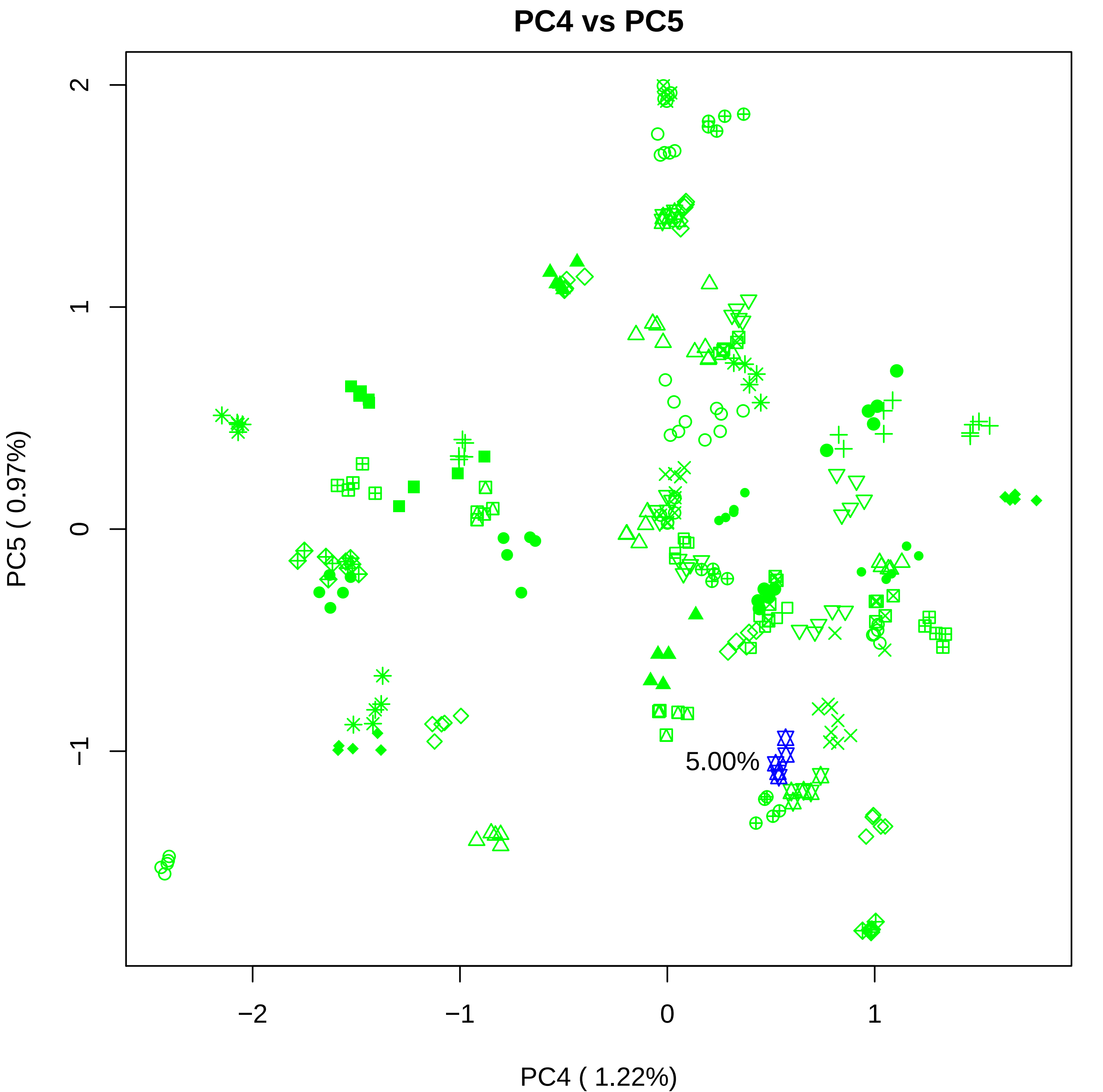

PC4 vs PC5
PC5 ( 0.97%)
PC4 ( 1.22%)
5.00%
−2
−1
0
1
2

### PCA of 7.5% BMH(a) vs PMH(a)

Scaled Matrix PCAed

7th order polynomial baseline correction performed

Raw spectra smoothed to 21 Neighbours, 2nd order PN, Savtisky Golay

Normalisation between 0 and 100

D2 spectra smoothed to 41 Neighbours, 2nd order PN, Savtisky Golay

Regions Analysed: 3601−2501 and 2001−700cm−1

Samples Analysed: 1 bContam's 0 BLH's 0 BMH's 0 OMH's 0 pContam's 0 PLH's 60 PMH's

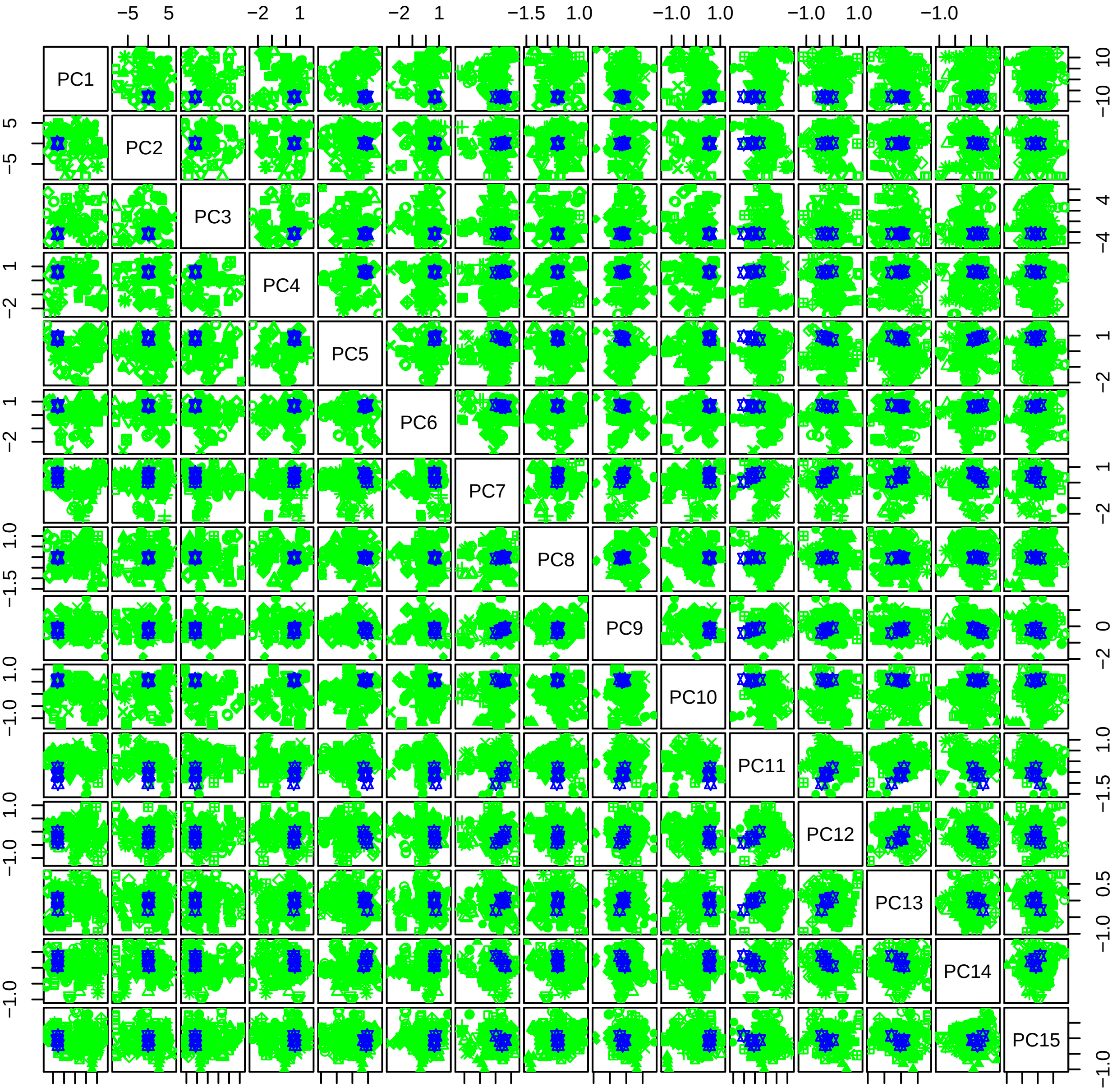
PC1
PC2
PC3
PC4
PC5
PC6
PC7
PC8
PC9
PC10
PC11
PC12
PC13
PC14
PC15

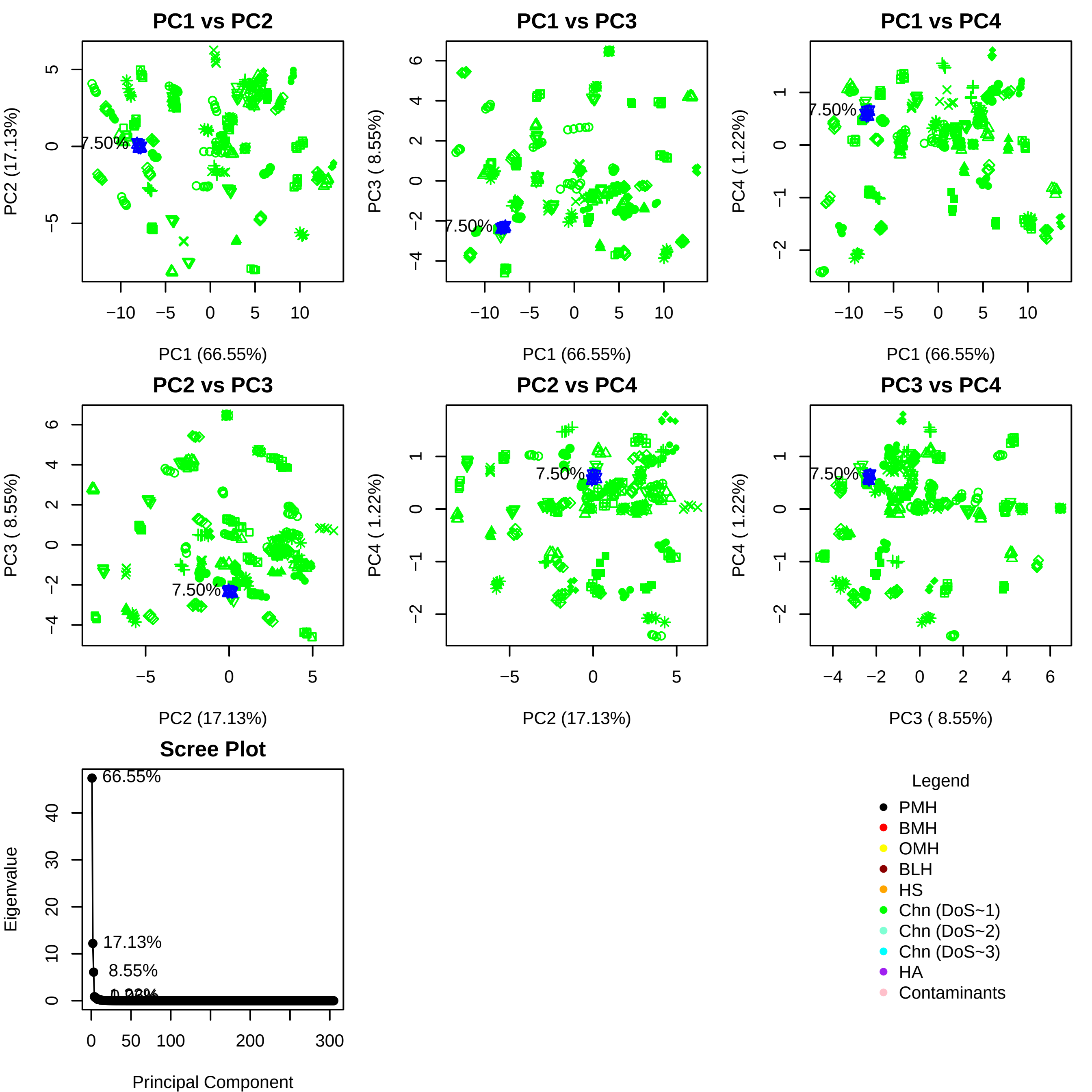

PC1 vs PC2
PC1 vs PC3
PC1 vs PC4
PC2 vs PC3
PC2 vs PC4
PC3 vs PC4
Scree Plot
PC1 (66.55%)
PC2 (17.13%)
PC3 ( 8.55%)
PC4 ( 1.22%)
7.50%
Eigenvalue
Principal Component
66.55%
17.13%
8.55%
Legend
PMH
BMH
OMH
BLH
HS
Chn (DoS~1)
Chn (DoS~2)
Chn (DoS~3)
HA
Contaminants

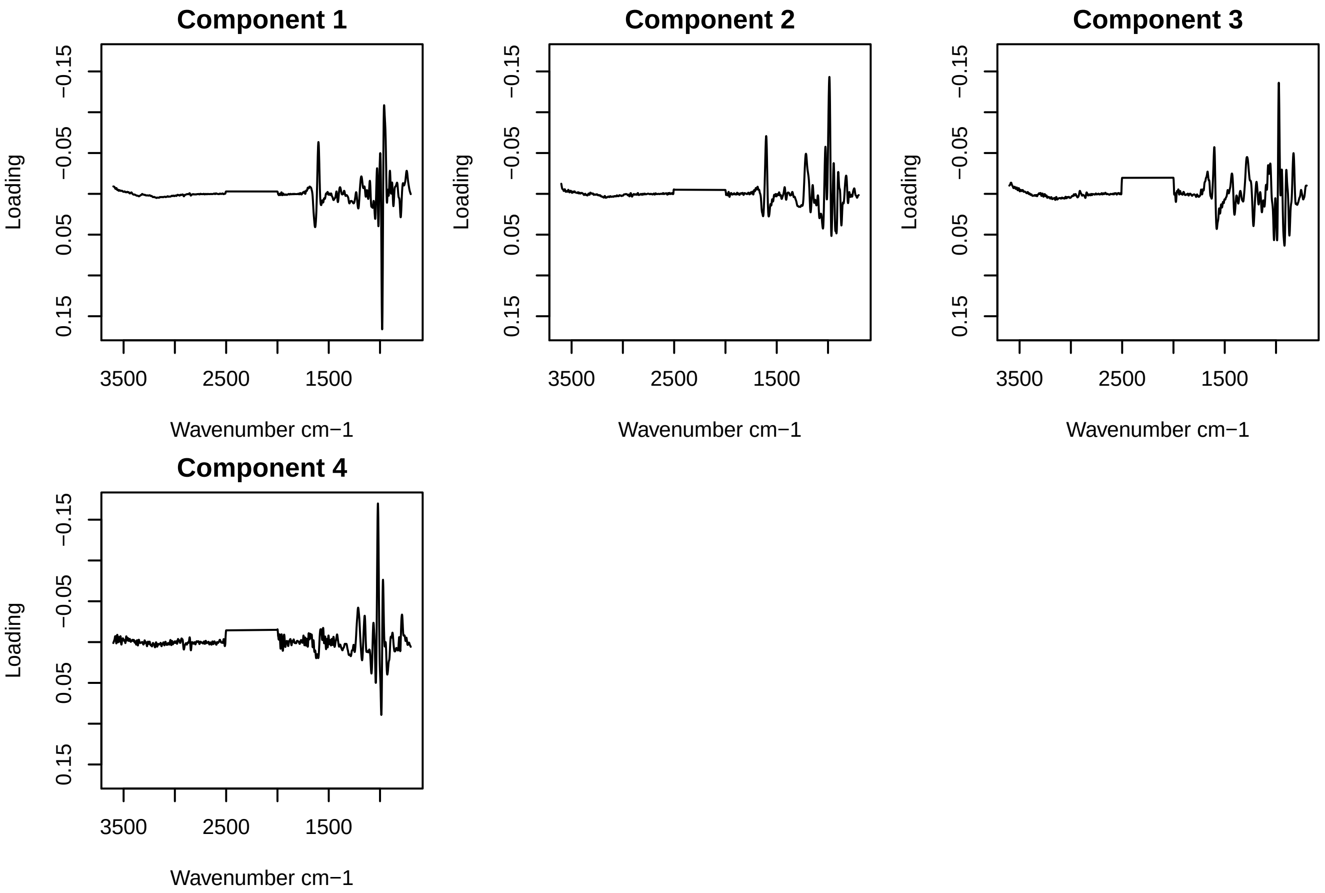
Component 1
Component 2
Component 3
Component 4
Loading
−0.15
−0.05
0.05
0.15
3500
2500
1500
Wavenumber cm−1

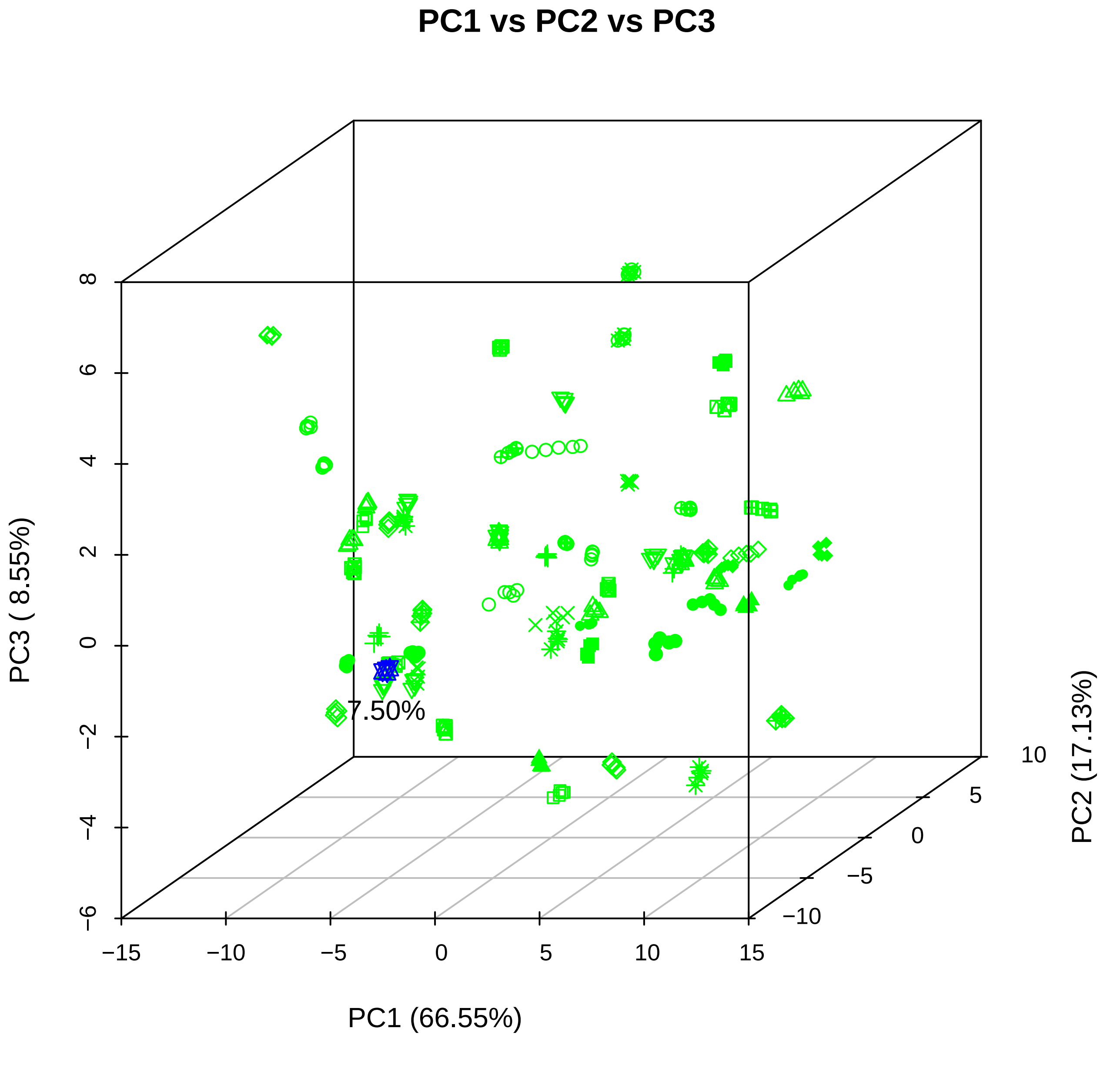
PC1 vs PC2 vs PC3
PC1 (66.55%)
PC2 (17.13%)
PC3 ( 8.55%)
7.50%
−15
−10
−5
0
5
10
15
−6
−4
−2
0
2
4
6
8
−10
−5
0
5
10

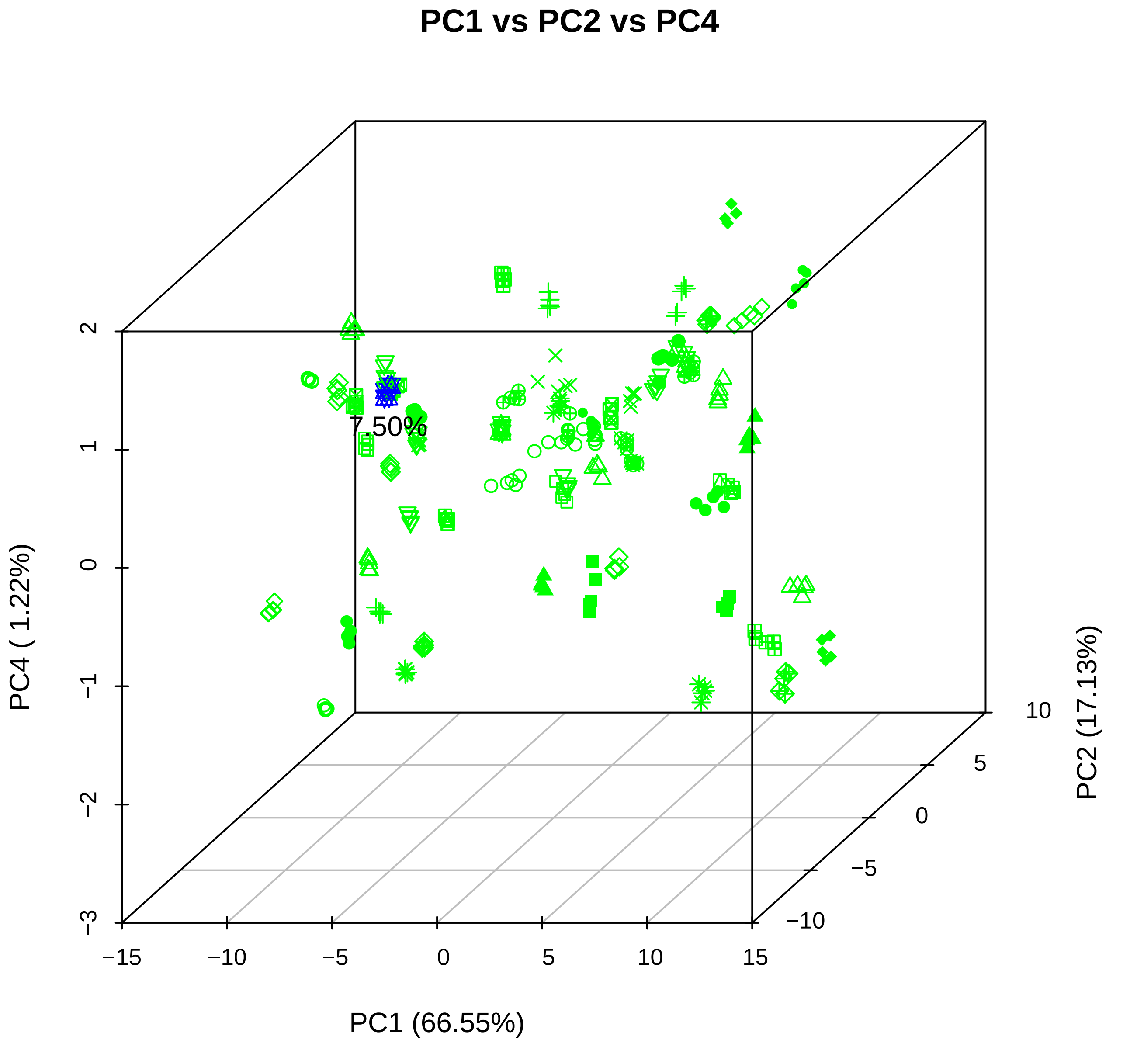

PC1 vs PC2 vs PC4
7.50%
PC1 (66.55%)
PC2 (17.13%)
PC4 ( 1.22%)
−15
−10
−5
0
5
10
15
−10
−5
0
5
10
−3
−2
−1
0
1
2

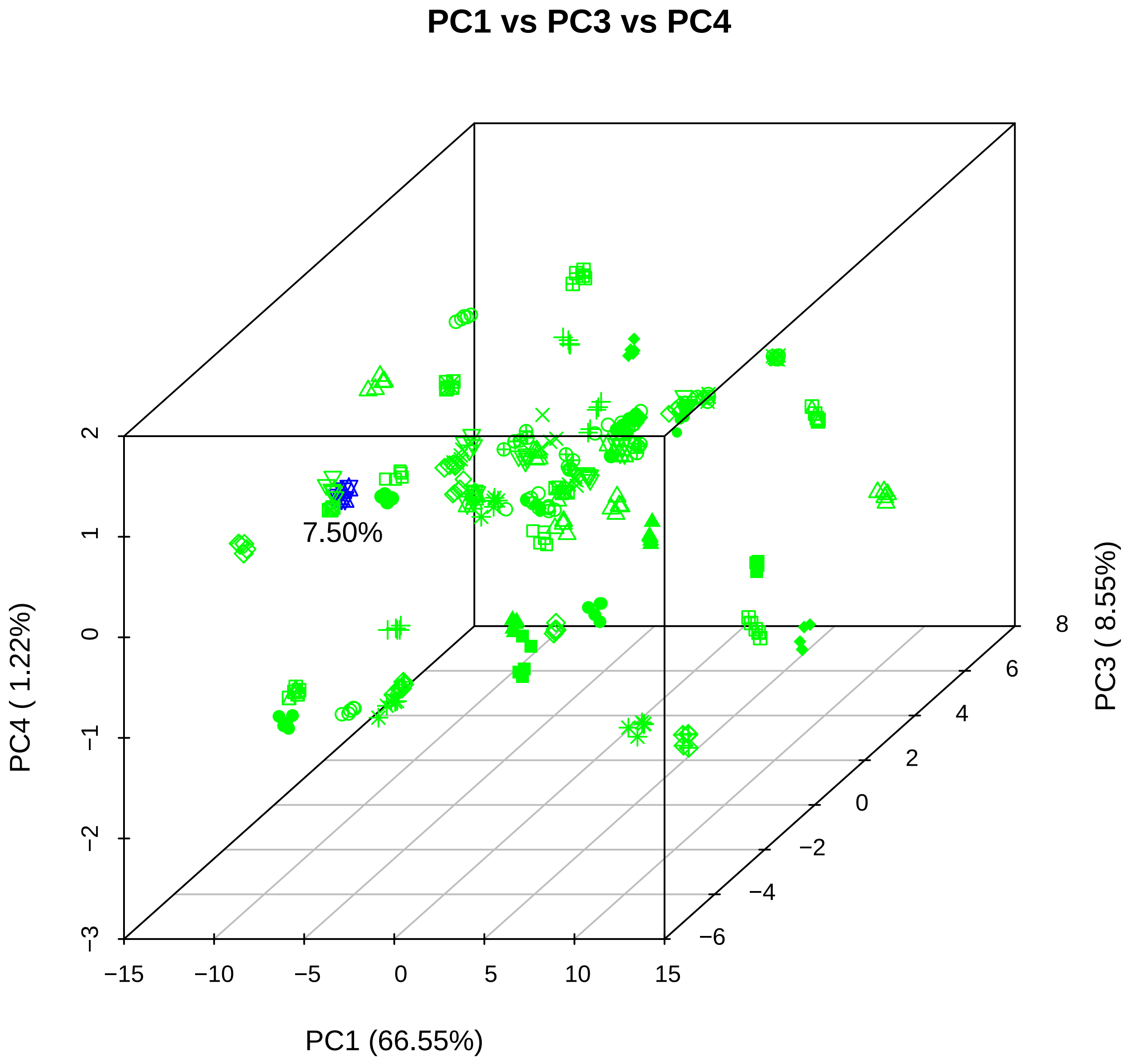

PC1 vs PC3 vs PC4
7.50%
PC1 (66.55%)
PC4 ( 1.22%)
PC3 ( 8.55%)
−15
−10
−5
0
5
10
15
−3
−2
−1
0
1
2
−6
−4
−2
0
2
4
6
8

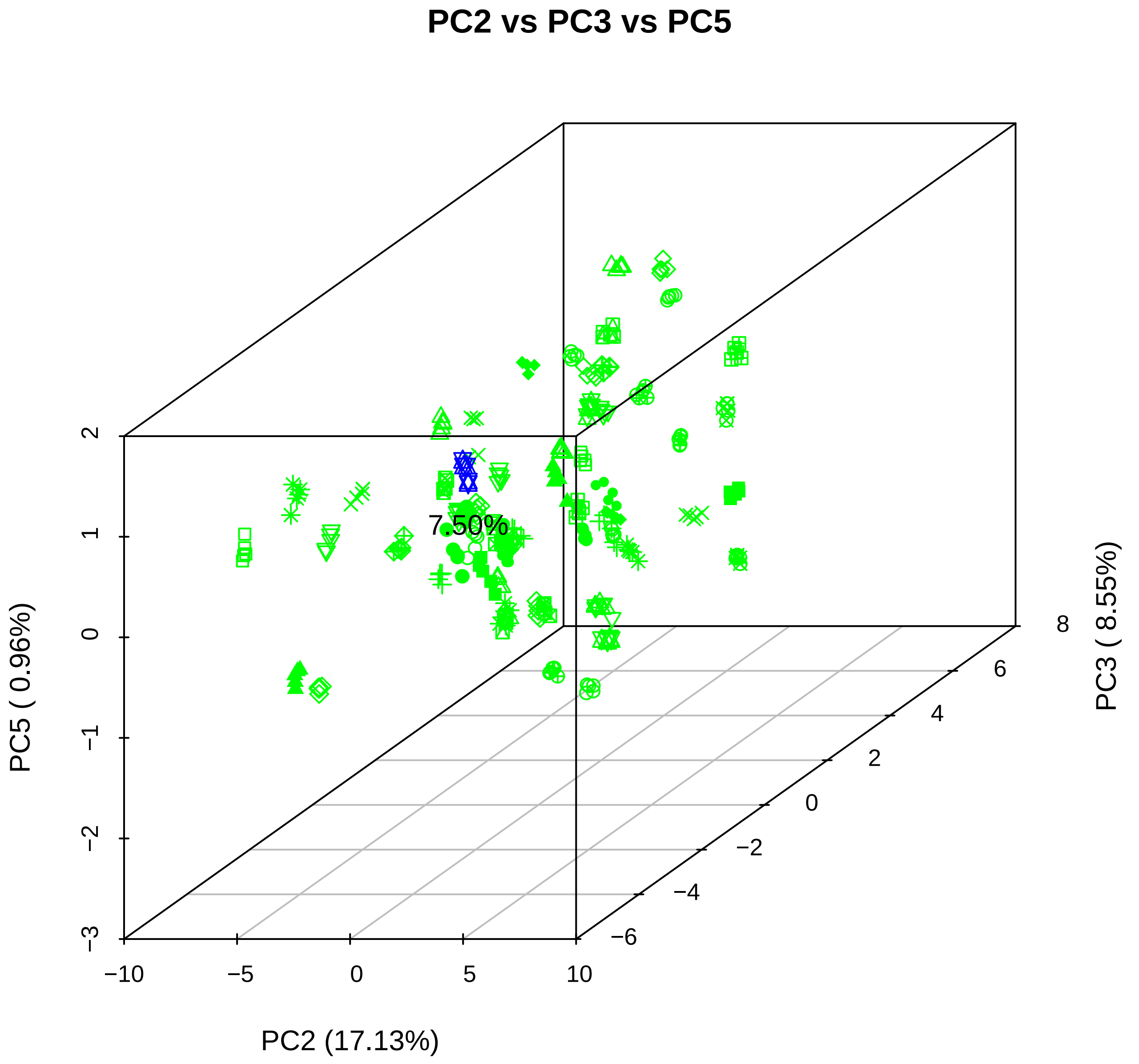
PC2 vs PC3 vs PC5
7.50%
PC5 ( 0.96%)
PC2 (17.13%)
PC3 ( 8.55%)
2
1
0
−1
−2
−3
−10
−5
0
5
10
−6
−4
−2
0
2
4
6
8

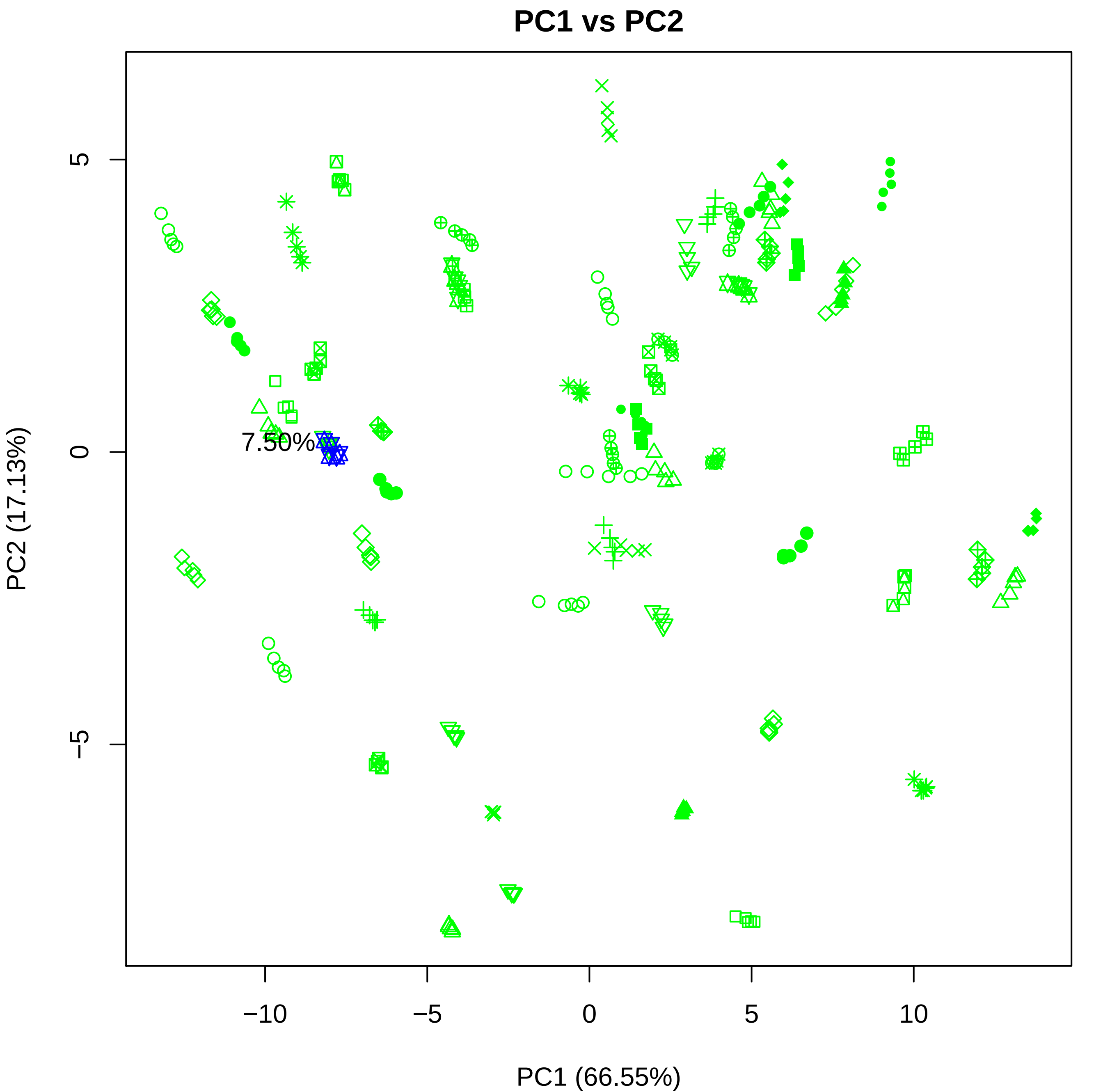
PC1 vs PC2
7.50%
PC1 (66.55%)
PC2 (17.13%)
−10
−5
0
5
10

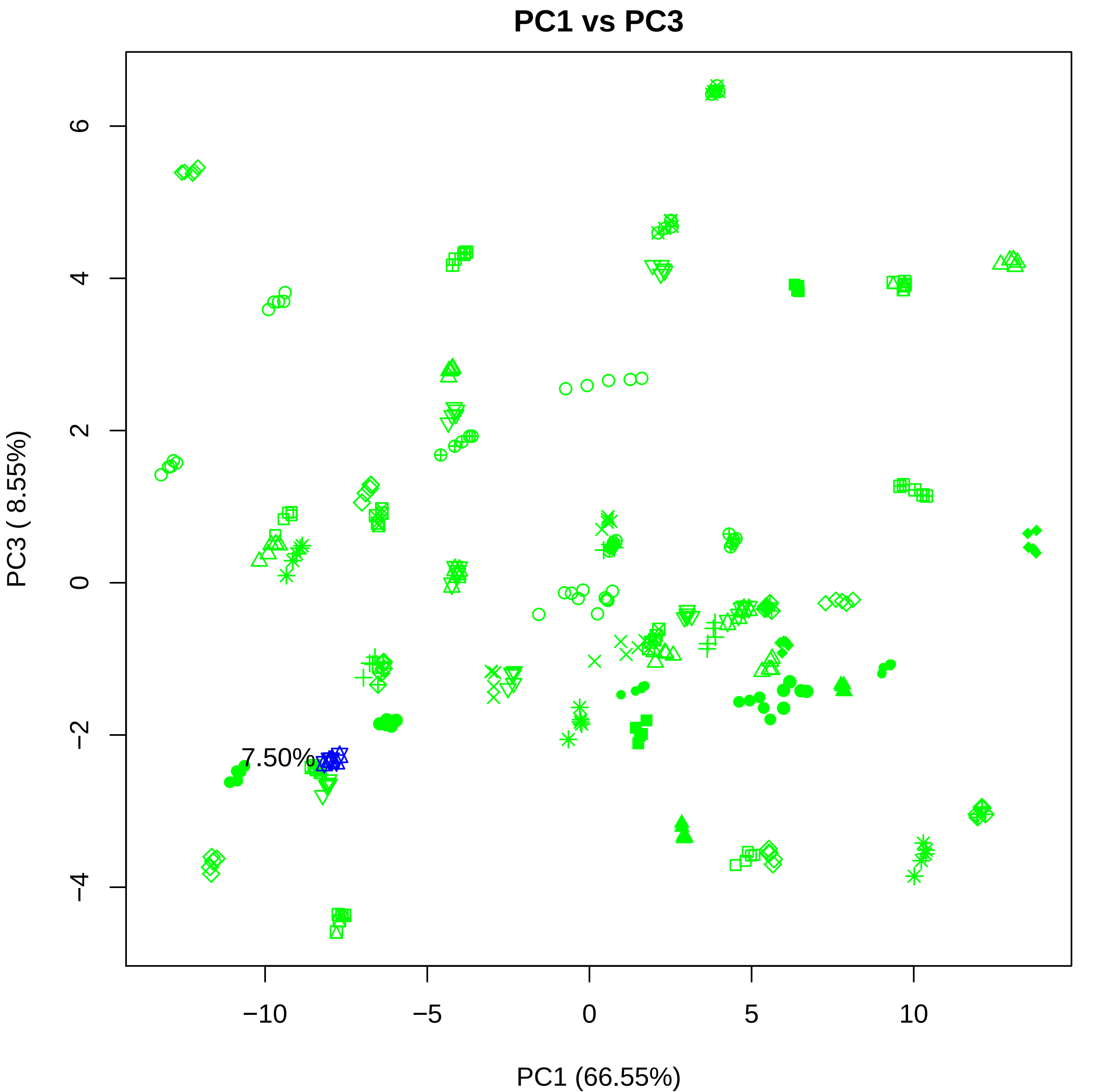
PC1 vs PC3
PC3 ( 8.55%)
PC1 (66.55%)
7.50%
6
4
2
0
−2
−4
−10
−5
0
5
10

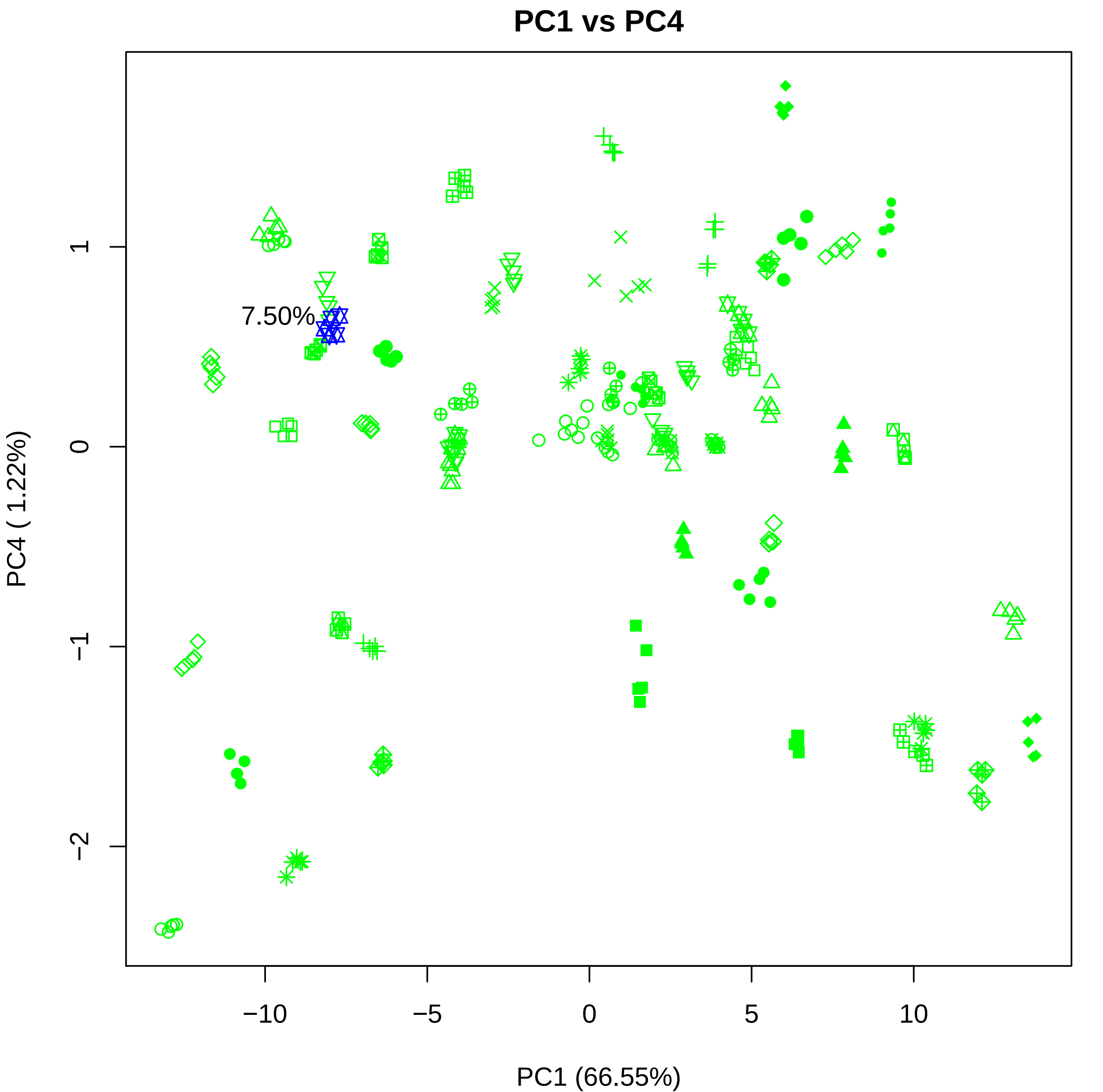
PC1 vs PC4
7.50%
PC4 ( 1.22%)
PC1 (66.55%)
1
0
−1
−2
−10
−5
0
5
10

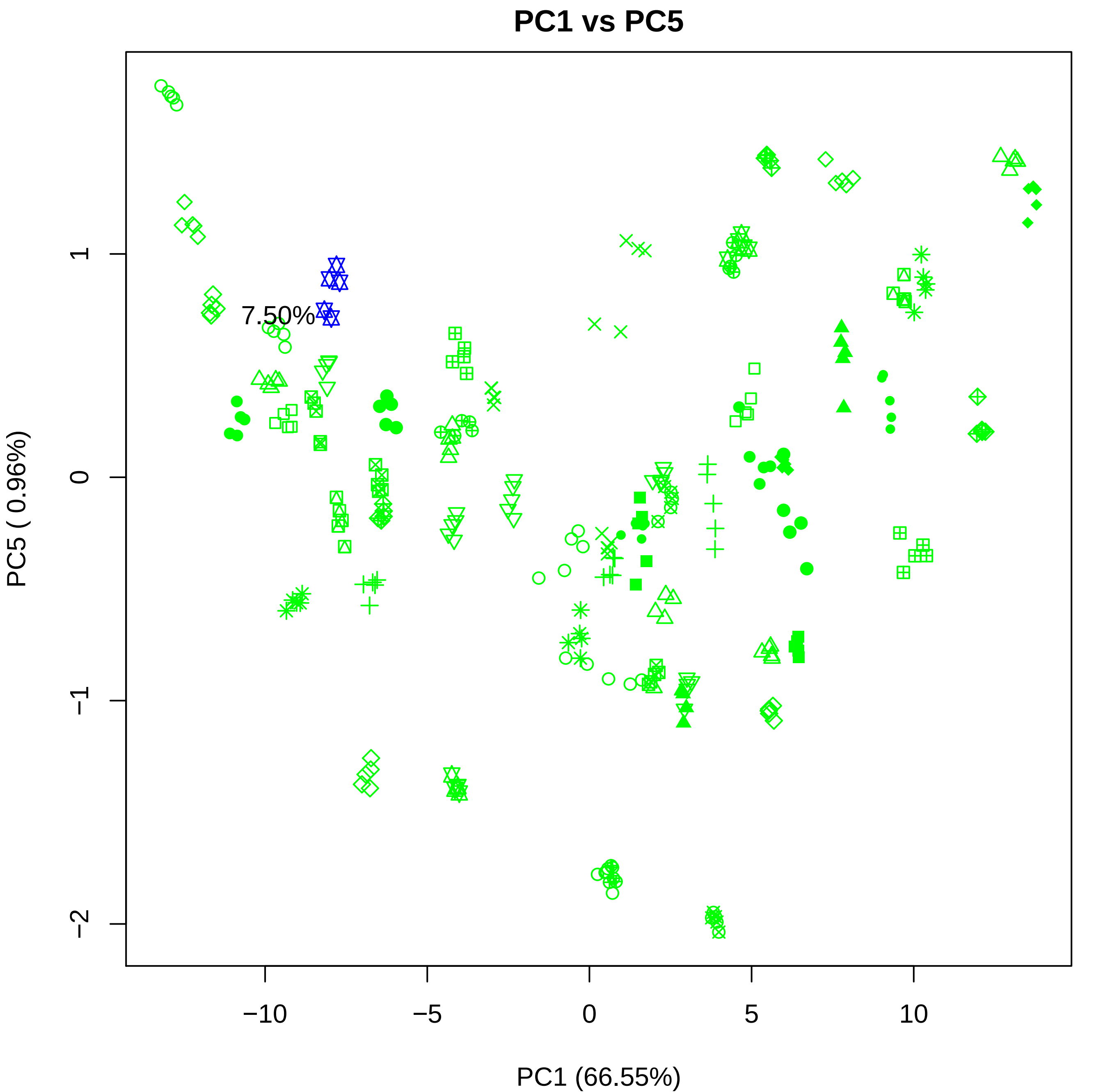

PC1 vs PC5
7.50%
PC5 ( 0.96%)
PC1 (66.55%)
−10
−5
0
5
10
−2
−1
0
1

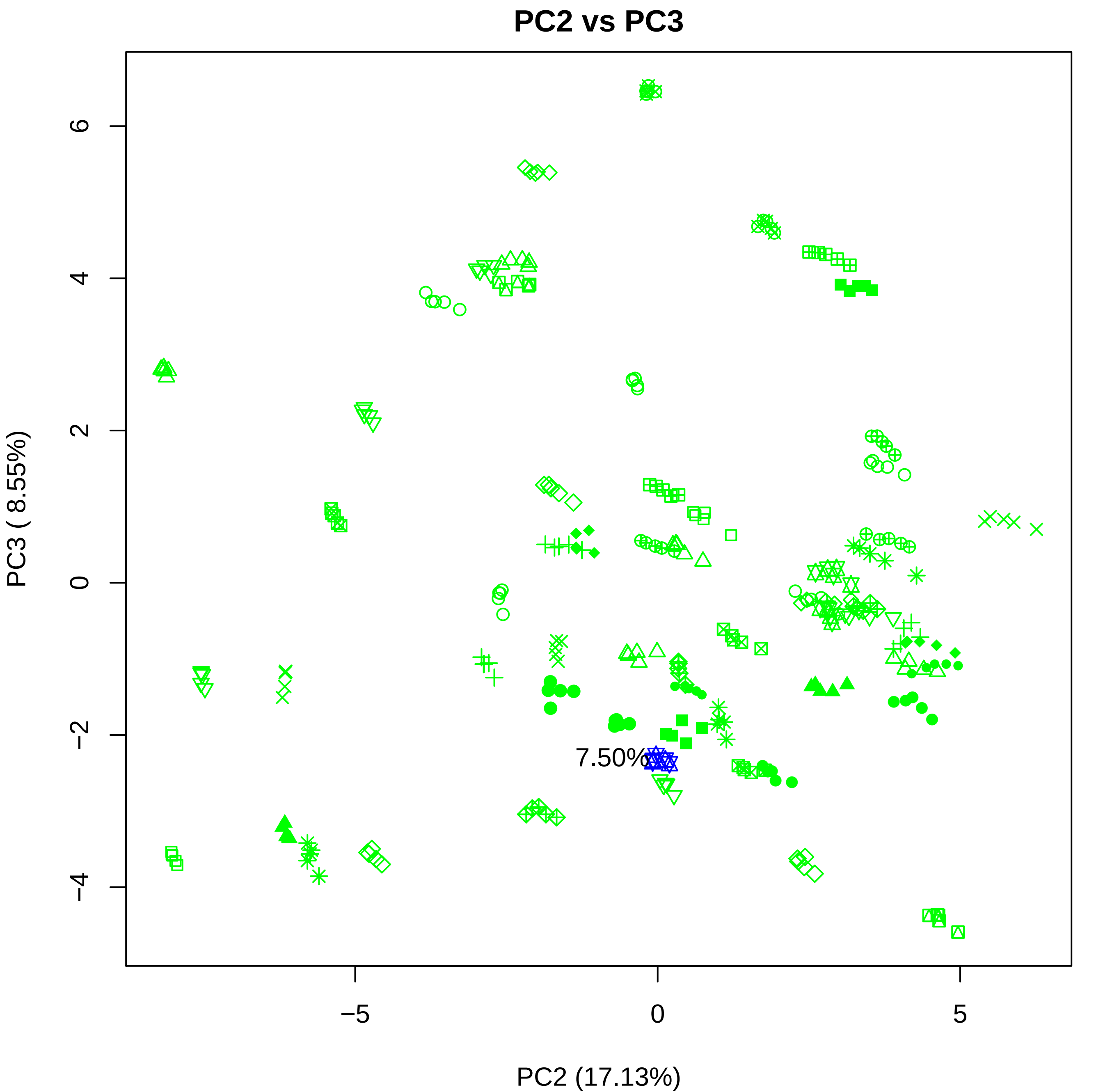
PC2 vs PC3
PC3 ( 8.55%)
PC2 (17.13%)
6
4
2
0
−2
−4
−5
0
5
7.50%

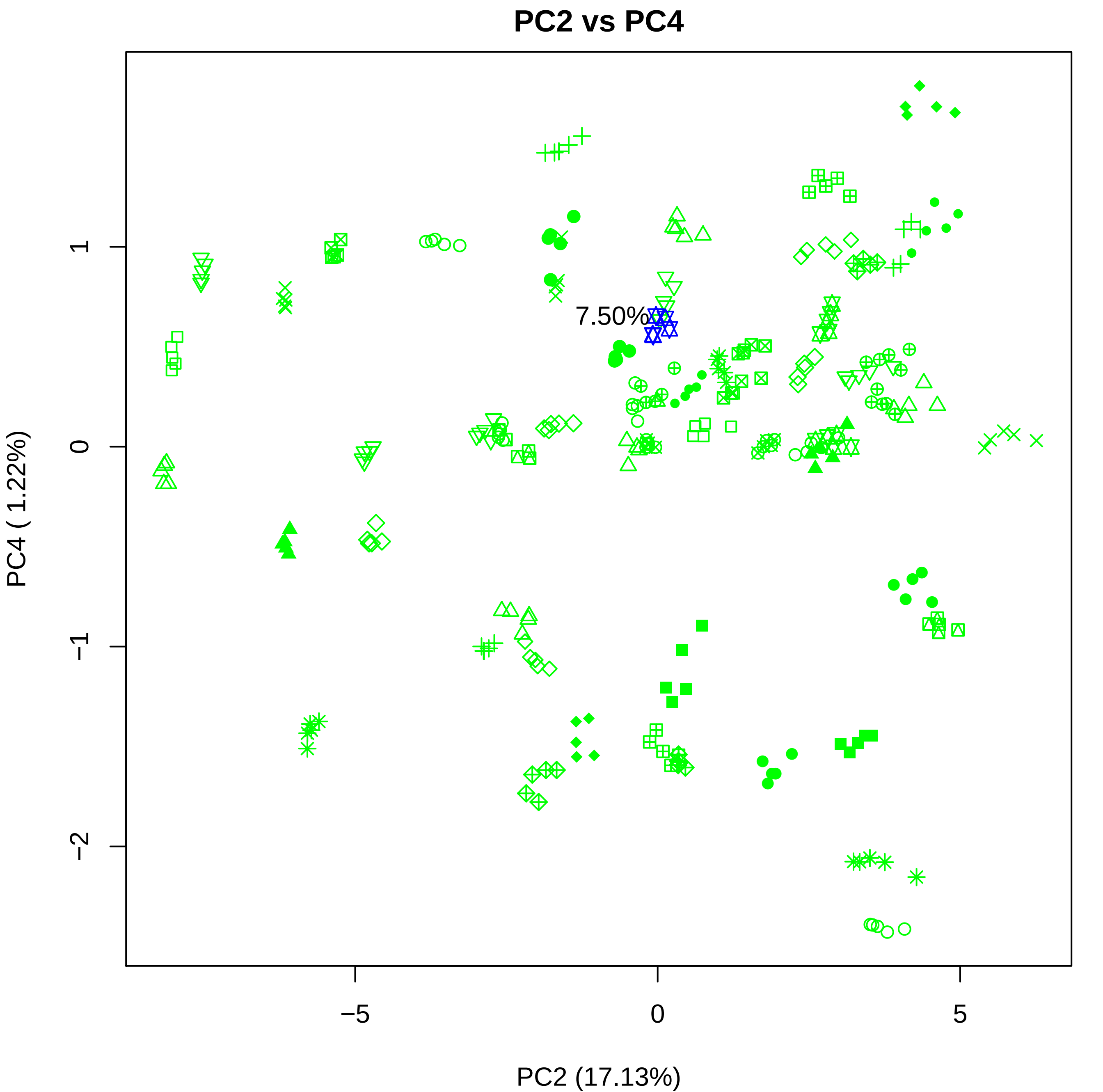

PC2 vs PC4
7.50%
PC4 ( 1.22%)
PC2 (17.13%)
−5
0
5
1
0
−1
−2

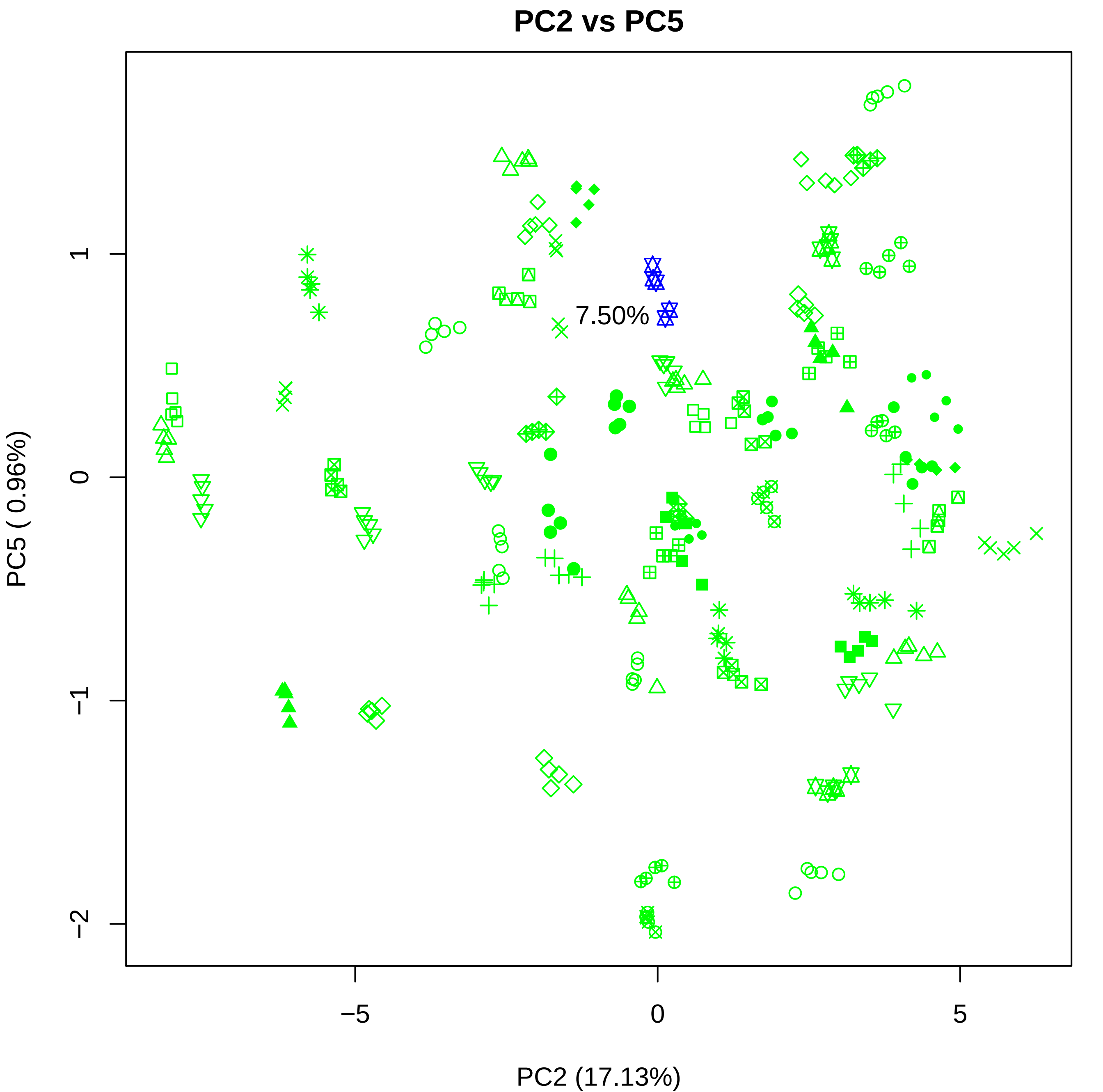

PC2 vs PC5
7.50%
PC5 ( 0.96%)
PC2 (17.13%)
1
0
−1
−2
−5
0
5

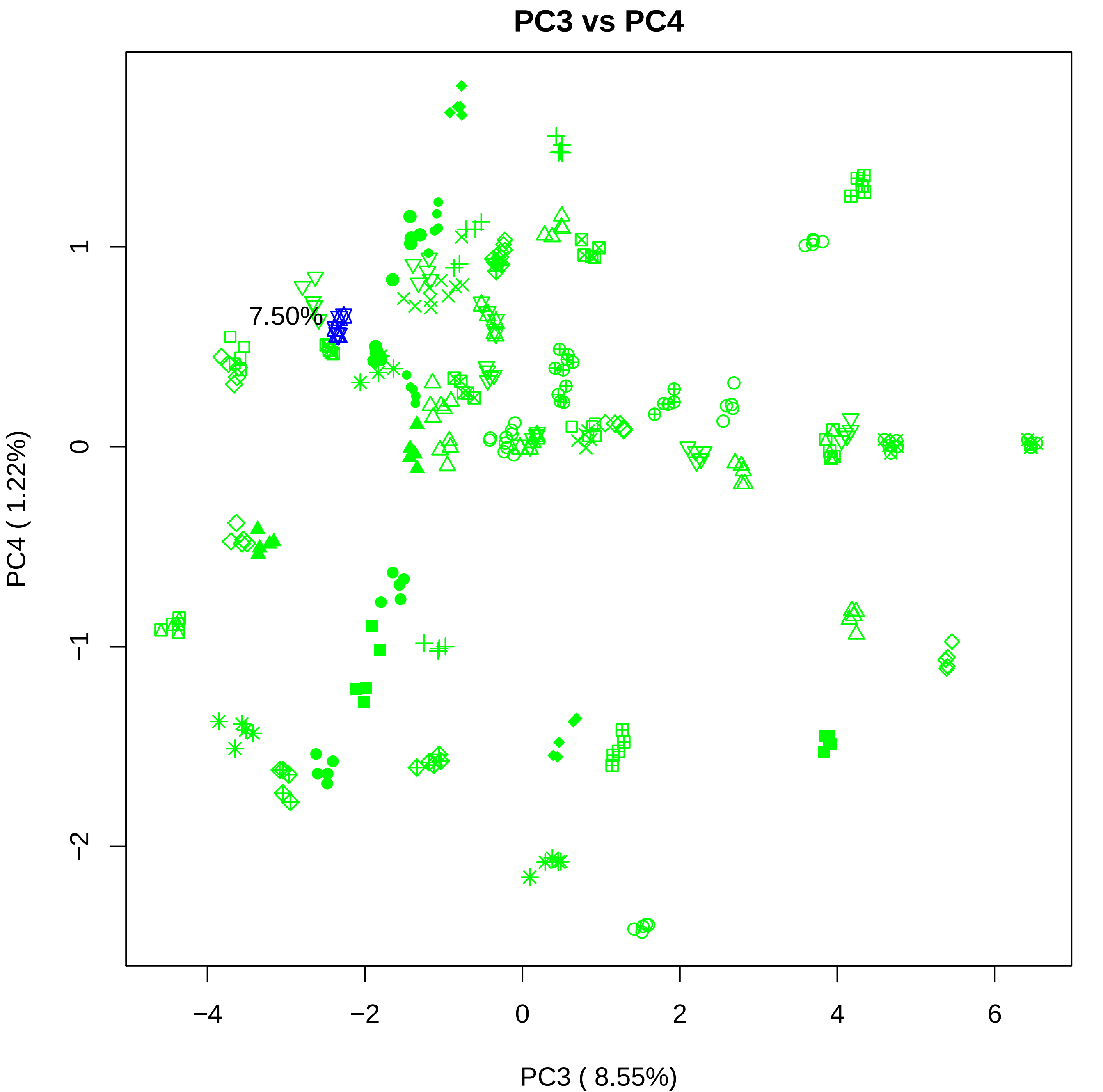
PC3 vs PC4
7.50%
PC4 ( 1.22%)
PC3 ( 8.55%)
−4
−2
0
2
4
6
−2
−1
0
1

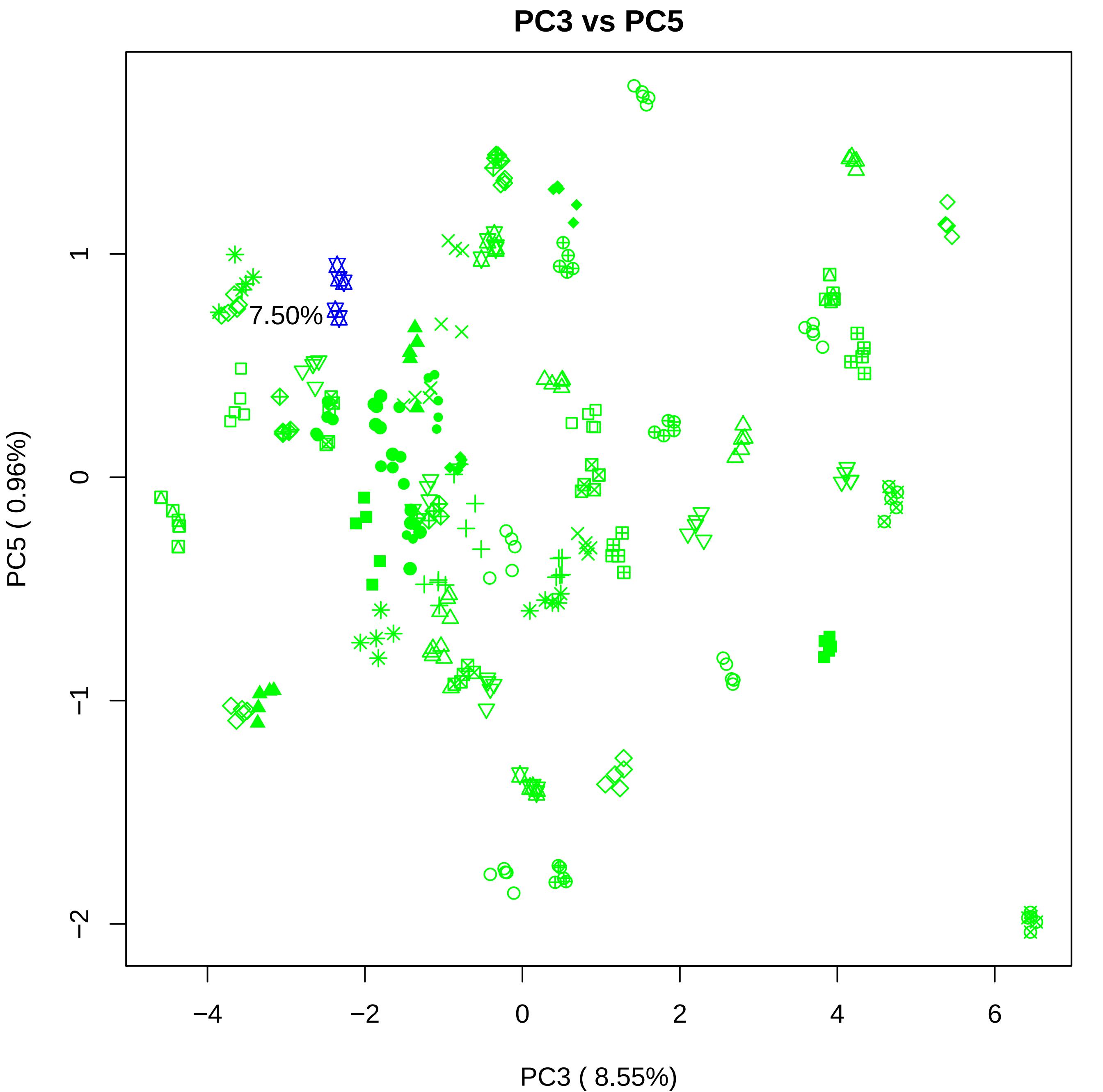

PC3 vs PC5
7.50%
1
0
−1
−2
PC5 ( 0.96%)
−4
−2
0
2
4
6
PC3 ( 8.55%)

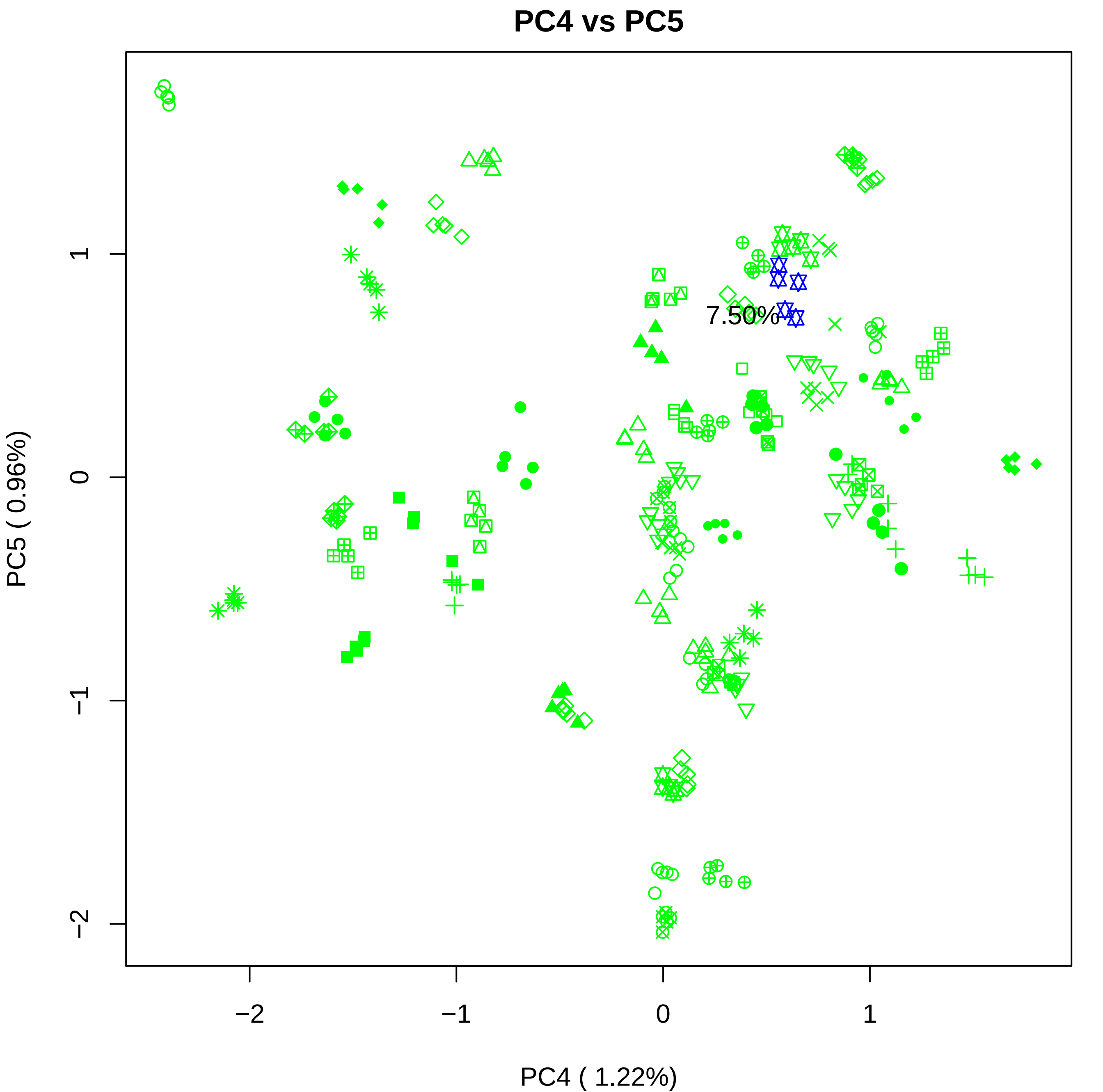
PC4 vs PC5
7.50%
PC4 ( 1.22%)
PC5 ( 0.96%)
−2
−1
0
1
−2
−1
0
1

### PCA of 10% BMH(a) vs PMH(a)

Scaled Matrix PCAed

7th order polynomial baseline correction performed

Raw spectra smoothed to 21 Neighbours, 2nd order PN, Savtisky Golay

Normalisation between 0 and 100

D2 spectra smoothed to 41 Neighbours, 2nd order PN, Savtisky Golay

Regions Analysed: 3601−2501 and 2001−700cm−1

Samples Analysed: 1 bContam's 0 BLH's 0 BMH's 0 OMH's 0 pContam's 0 PLH's 60 PMH's

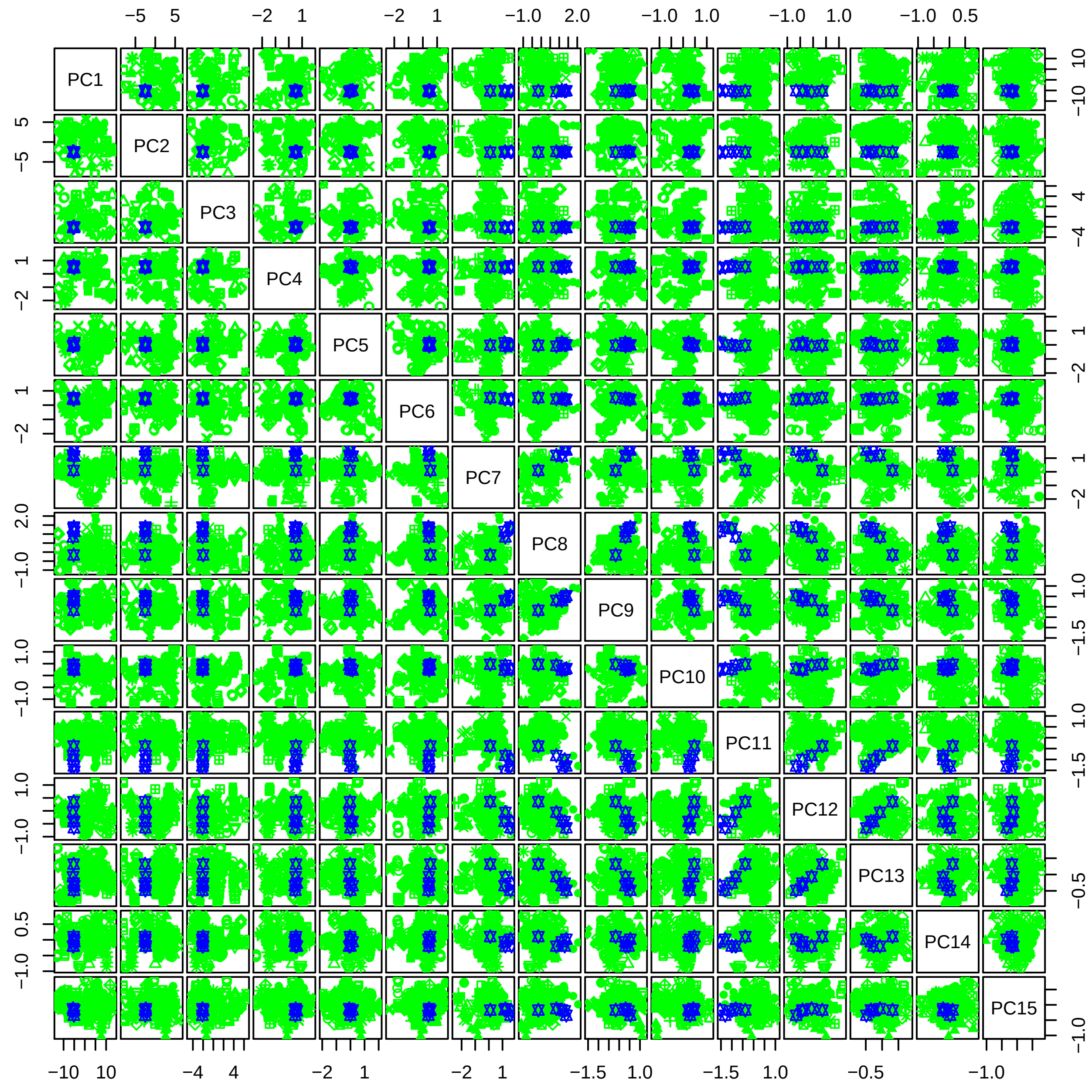
PC1
PC2
PC3
PC4
PC5
PC6
PC7
PC8
PC9
PC10
PC11
PC12
PC13
PC14
PC15

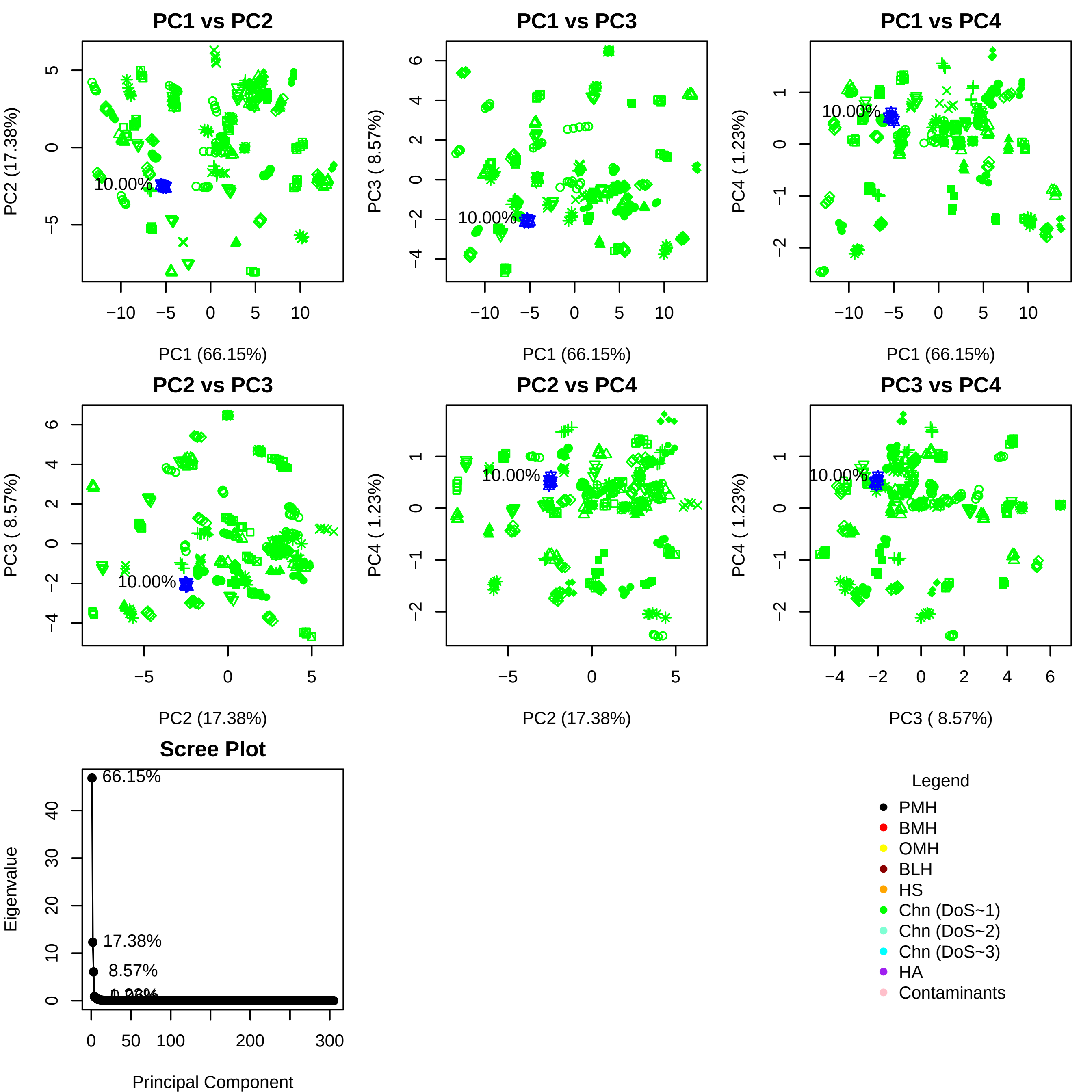

PC1 vs PC2
PC1 vs PC3
PC1 vs PC4
PC2 vs PC3
PC2 vs PC4
PC3 vs PC4
Scree Plot
PC1 (66.15%)
PC2 (17.38%)
PC3 ( 8.57%)
PC4 ( 1.23%)
10.00%
Eigenvalue
Principal Component
66.15%
17.38%
8.57%
Legend
PMH
BMH
OMH
BLH
HS
Chn (DoS~1)
Chn (DoS~2)
Chn (DoS~3)
HA
Contaminants

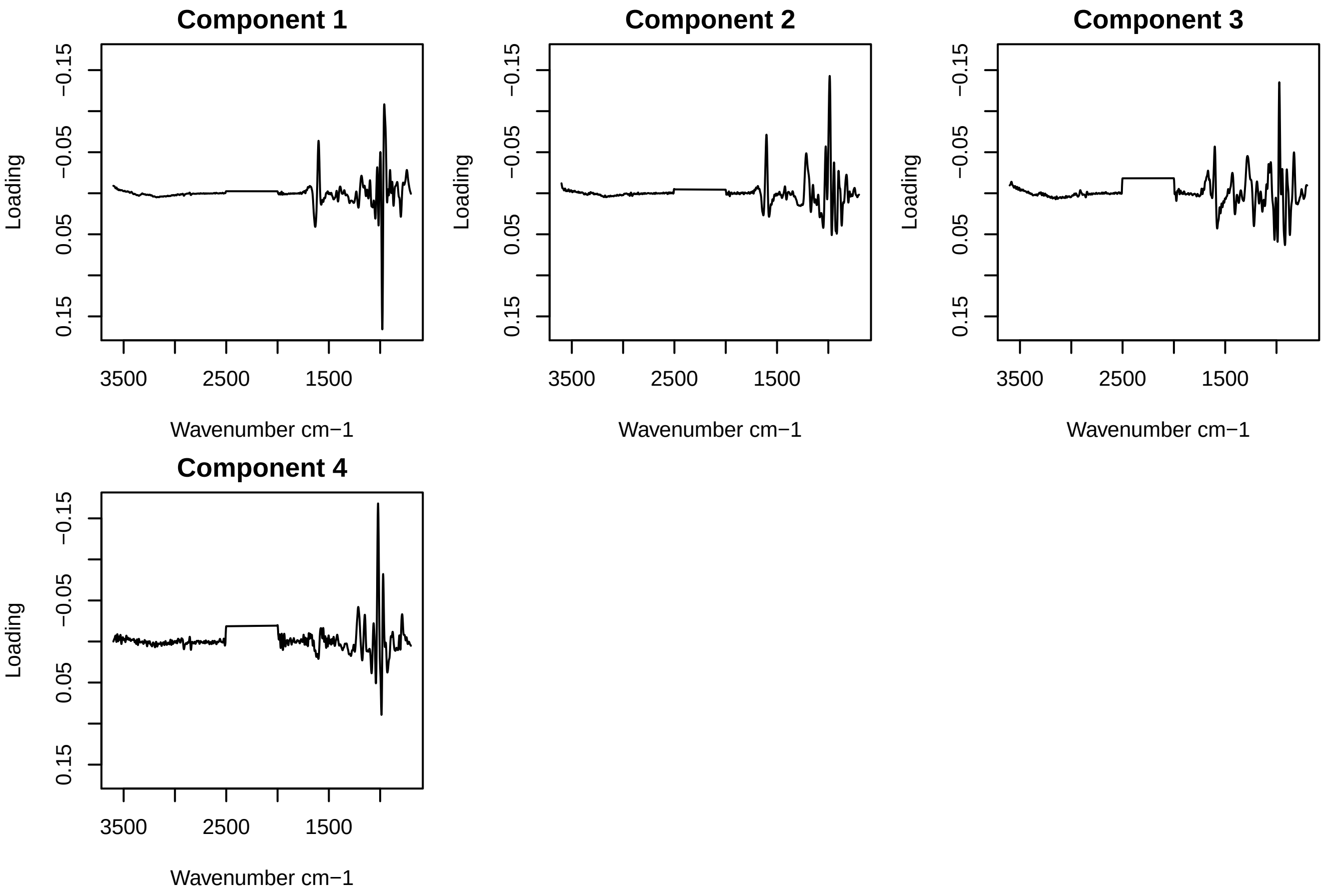
Component 1
Component 2
Component 3
Component 4
Loading
−0.15
−0.05
0.05
0.15
3500
2500
1500
Wavenumber cm−1

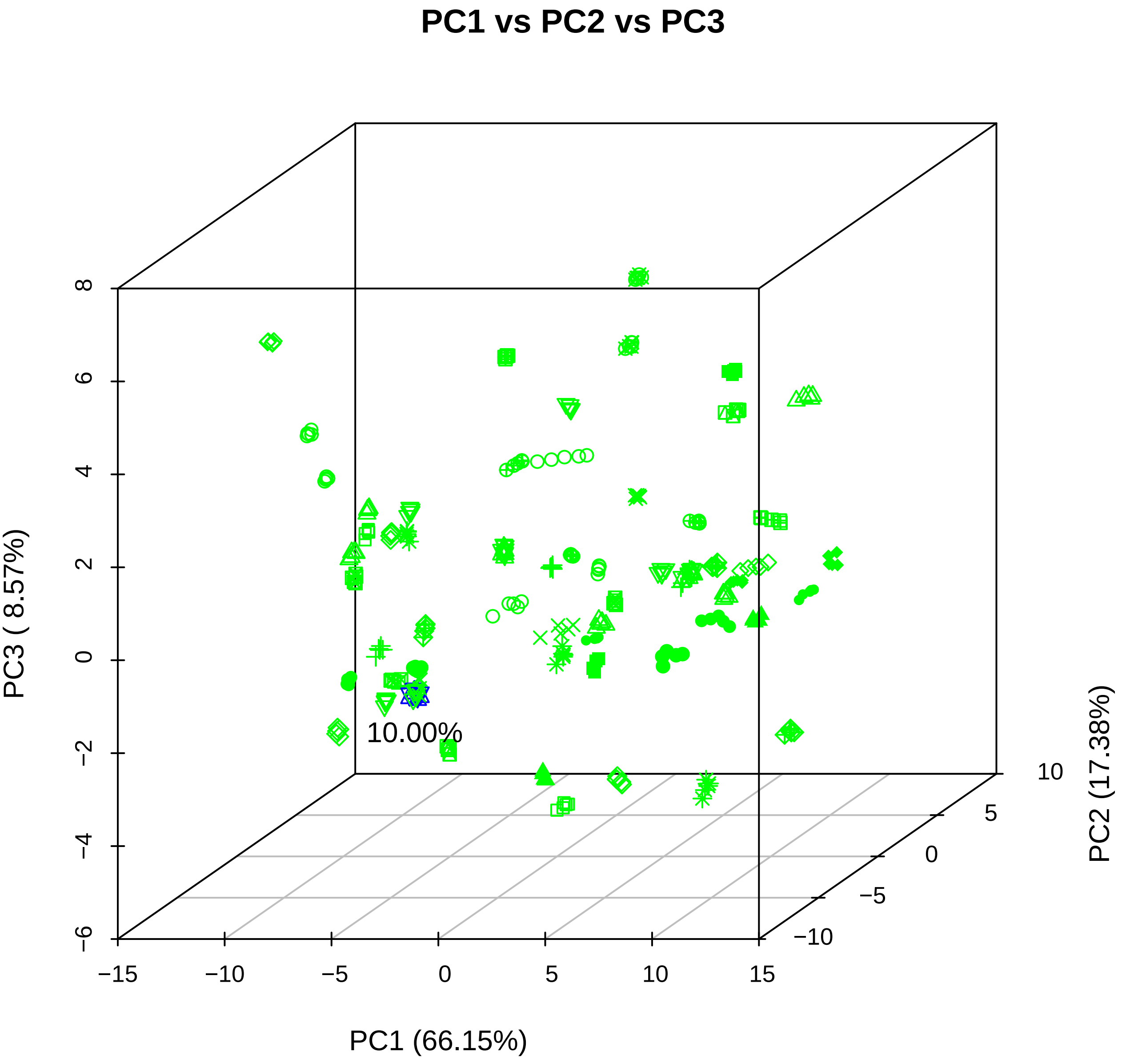
PC1 vs PC2 vs PC3
PC3 ( 8.57%)
8
6
4
2
0
−2
−4
−6
10.00%
−15
−10
−5
0
5
10
15
PC1 (66.15%)
10
5
0
−5
−10
PC2 (17.38%)

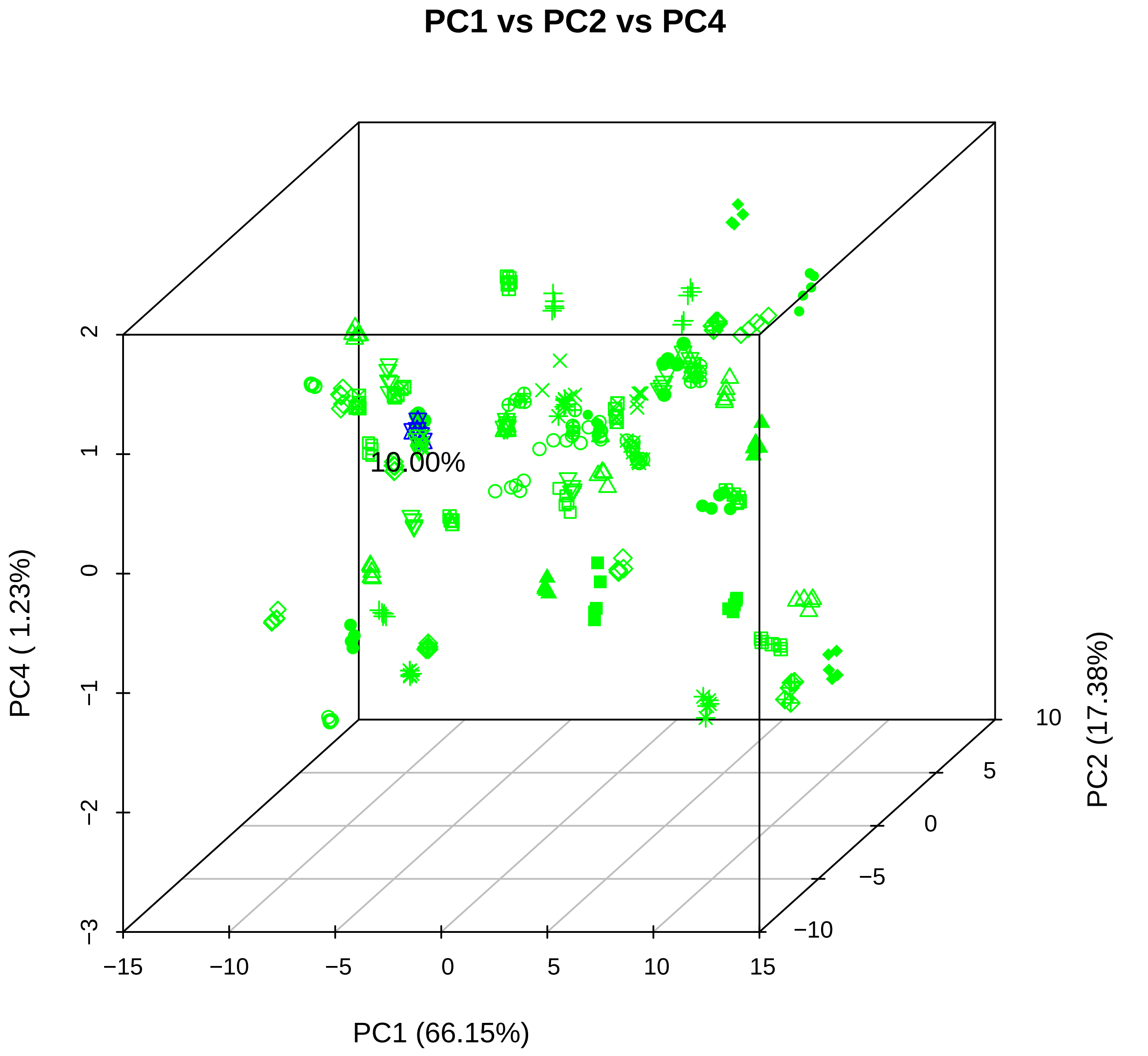
PC1 vs PC2 vs PC4
10.00%
PC1 (66.15%)
PC4 ( 1.23%)
PC2 (17.38%)
−15
−10
−5
0
5
10
15
−3
−2
−1
0
1
2
−10
−5
0
5
10

## PC1 vs PC3 vs PC4

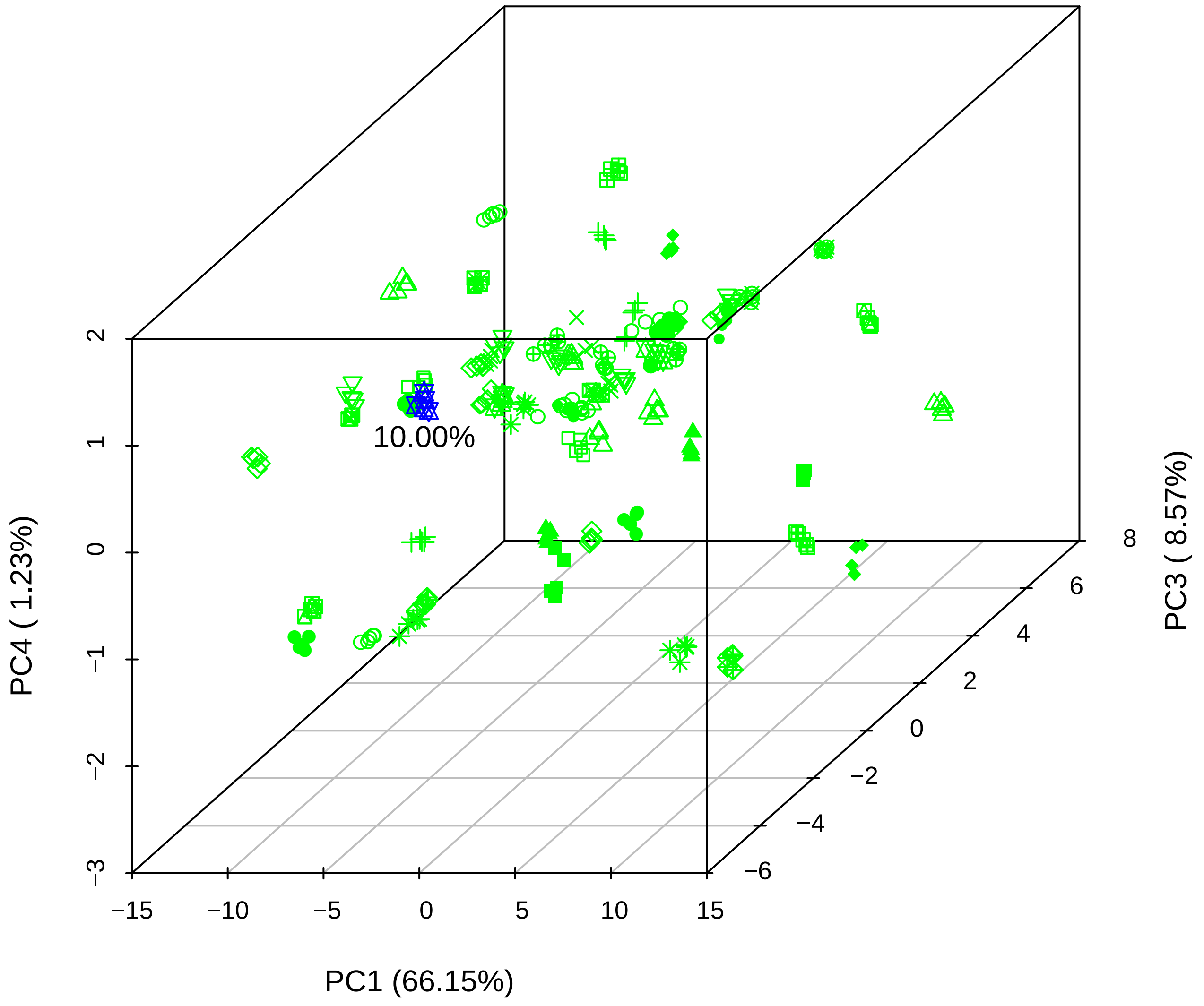

# PC2 vs PC3 vs PC5

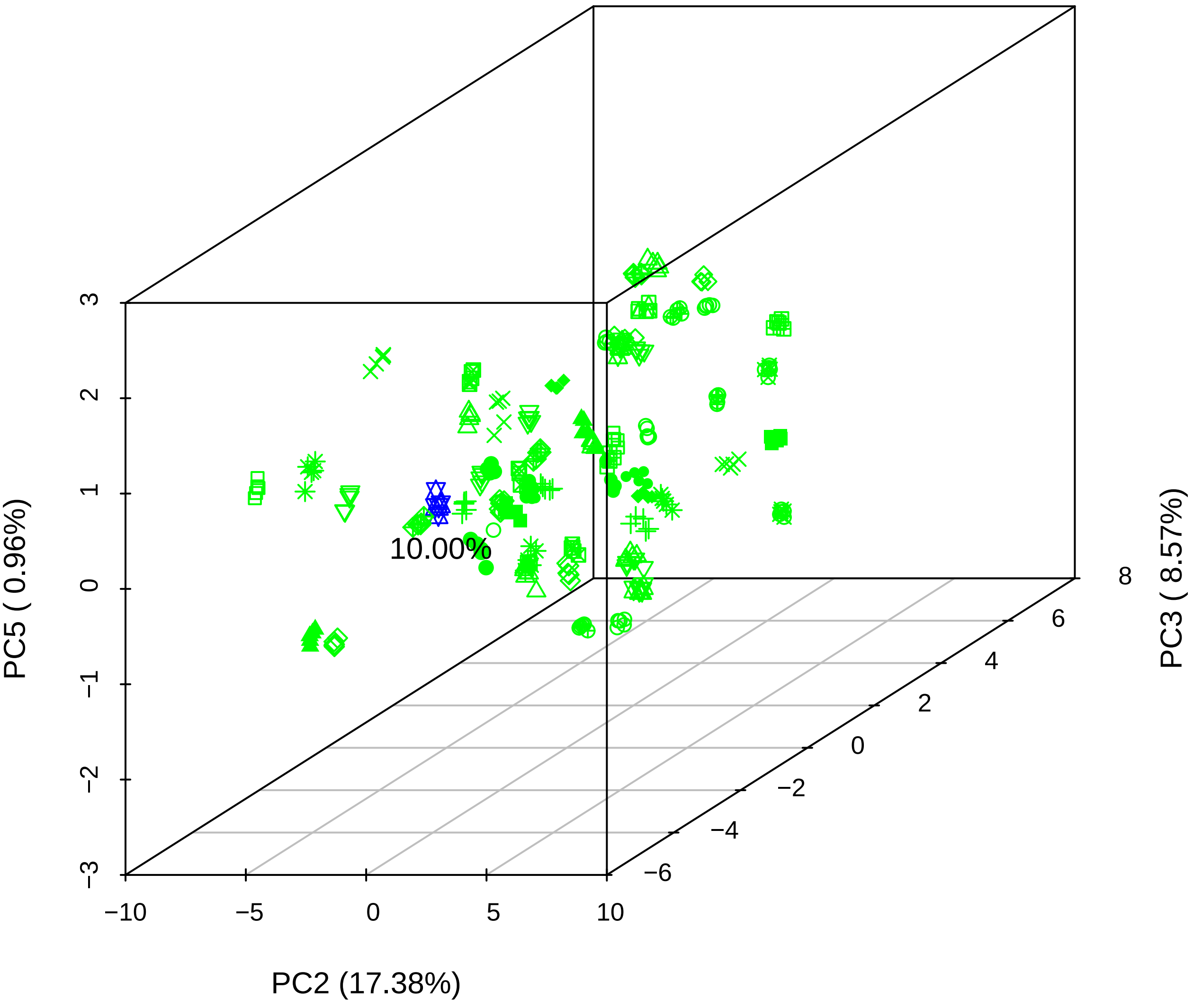

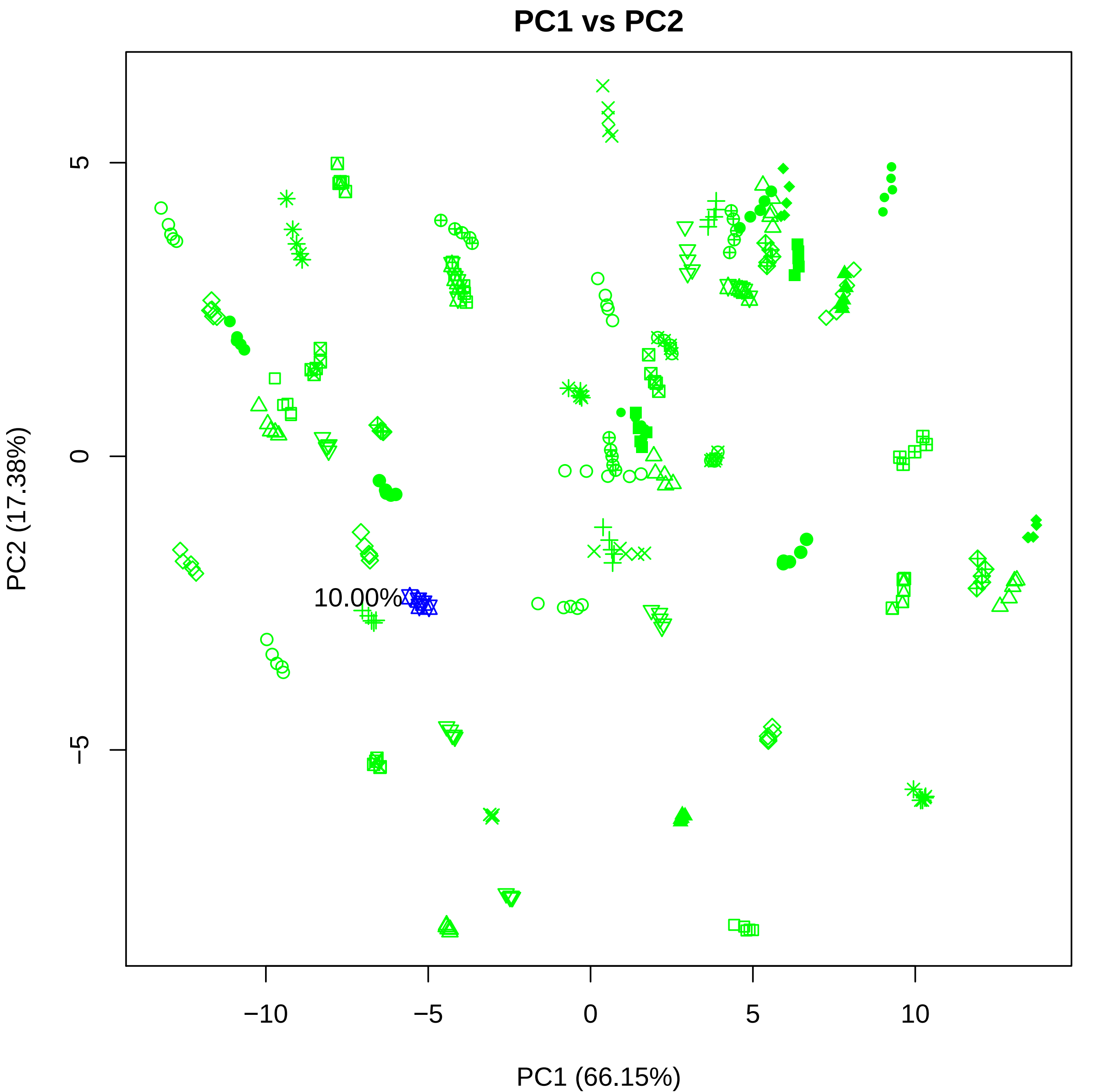
PC1 vs PC2
10.00%
PC2 (17.38%)
PC1 (66.15%)
5
0
−5
−10
−5
0
5
10

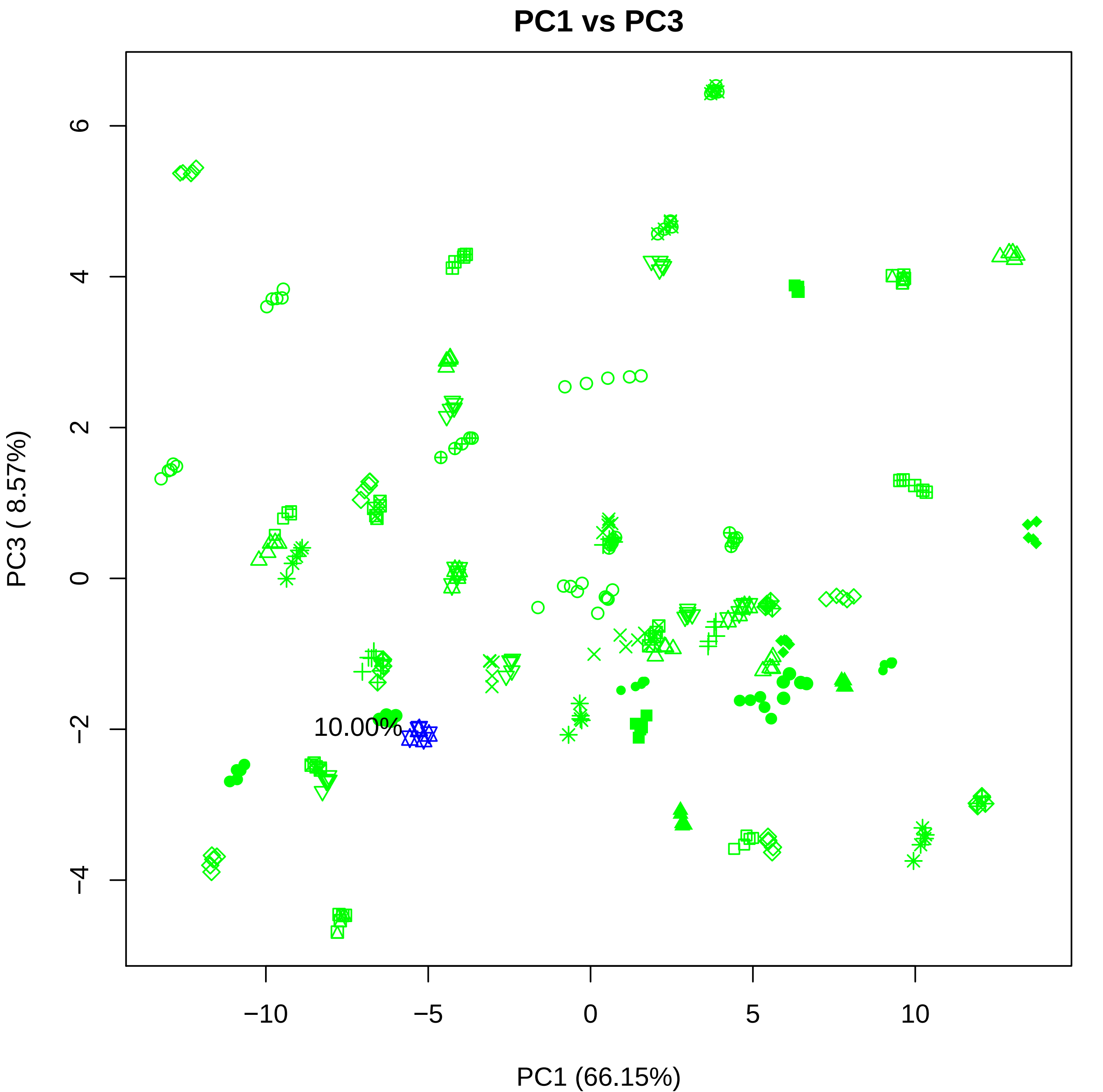
PC1 vs PC3
PC3 ( 8.57%)
PC1 (66.15%)
10.00%
6
4
2
0
−2
−4
−10
−5
0
5
10

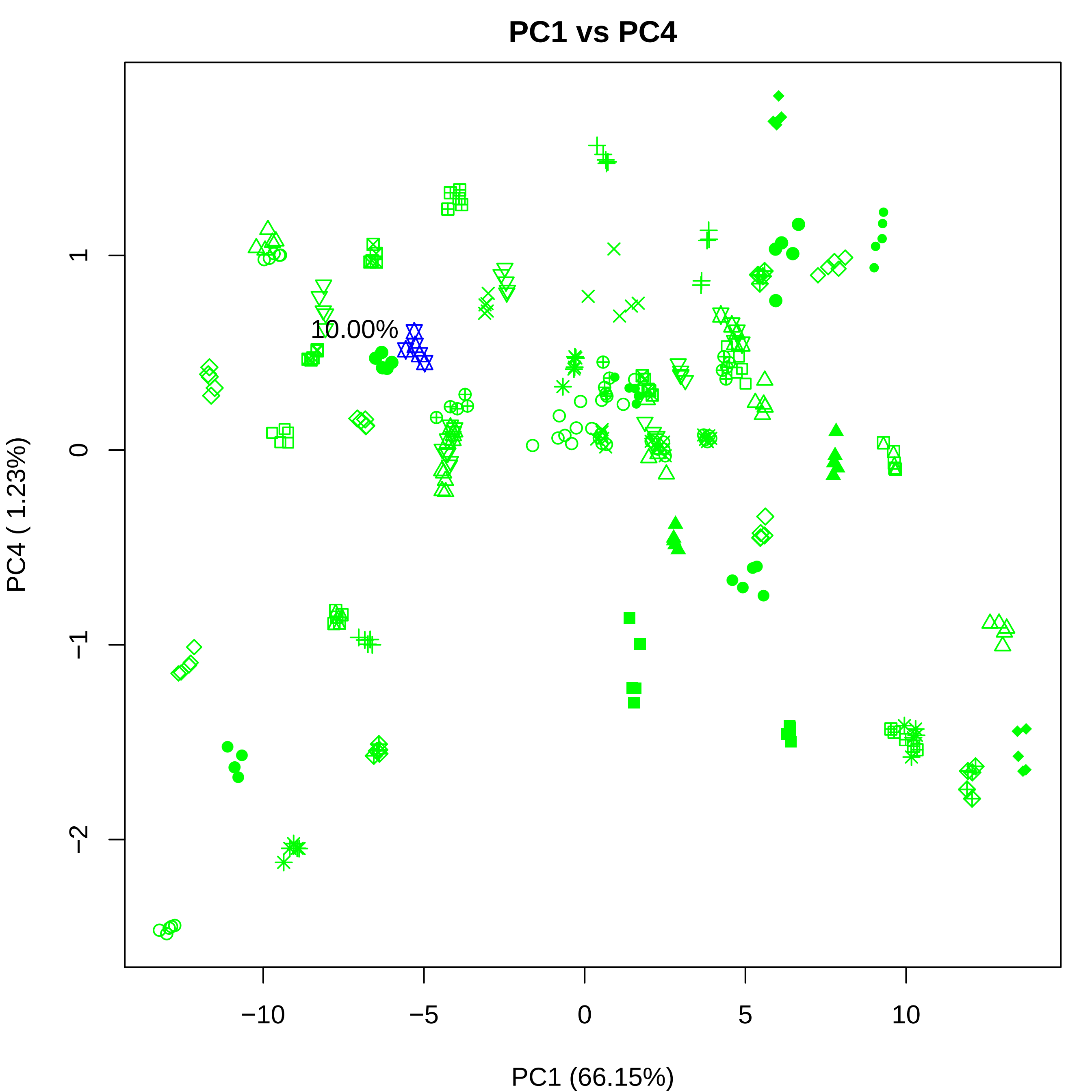

PC1 vs PC4
10.00%
PC4 ( 1.23%)
PC1 (66.15%)
−10
−5
0
5
10
−2
−1
0
1

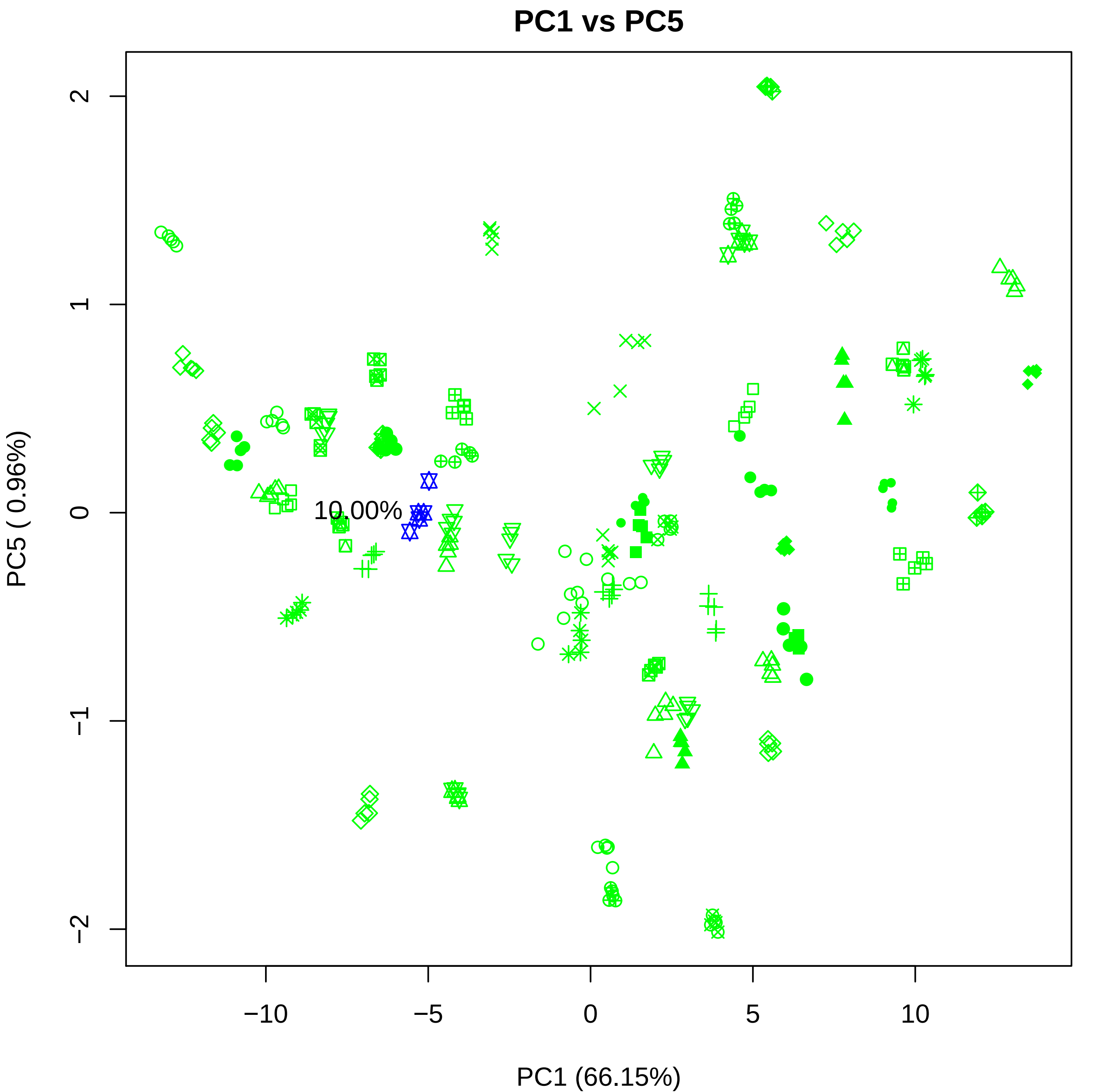
PC1 vs PC5
10.00%
PC5 ( 0.96%)
PC1 (66.15%)
2
1
0
−1
−2
−10
−5
0
5
10

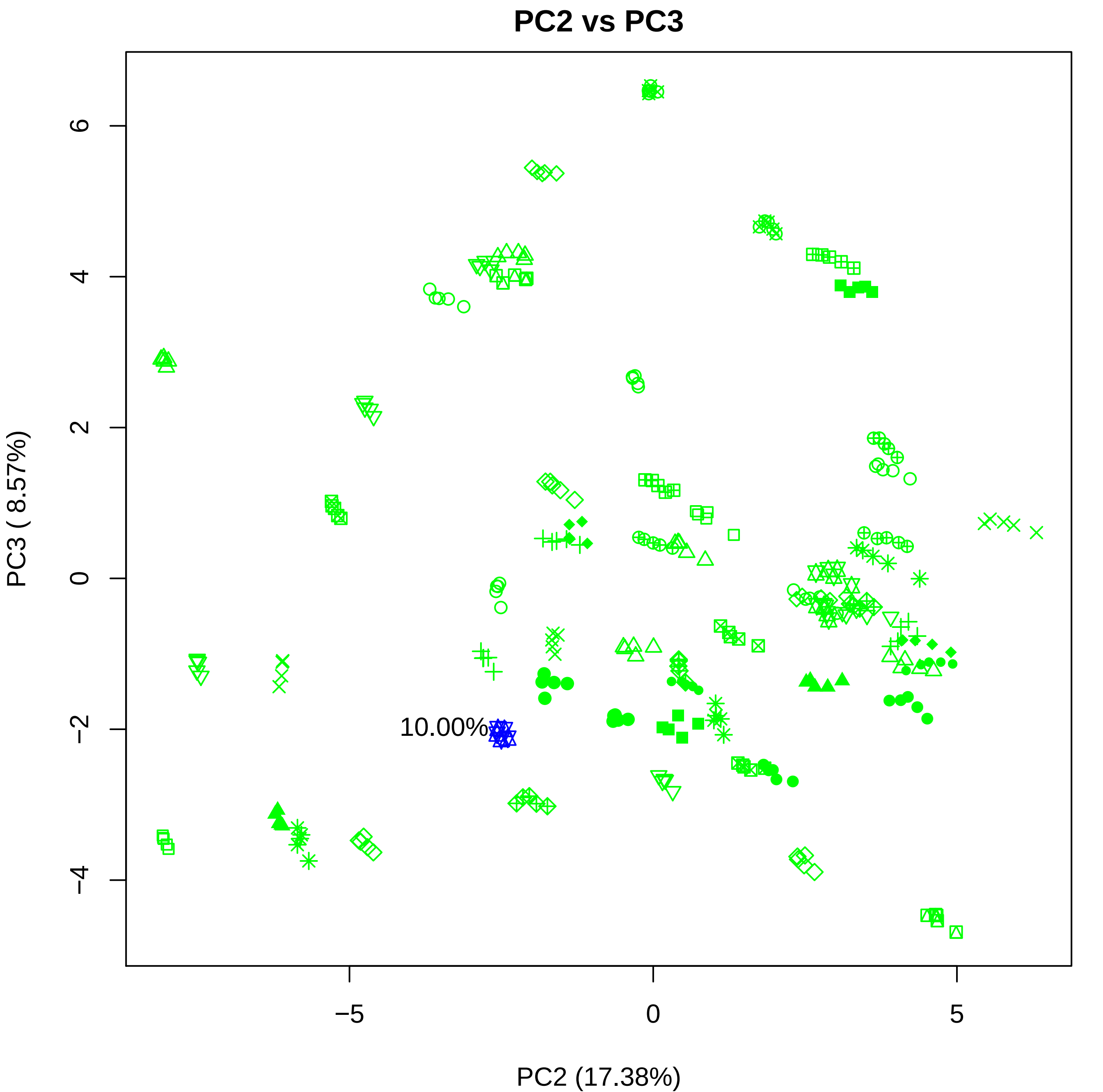

PC2 vs PC3
10.00%
PC3 ( 8.57%)
PC2 (17.38%)
6
4
2
0
−2
−4
−5
0
5

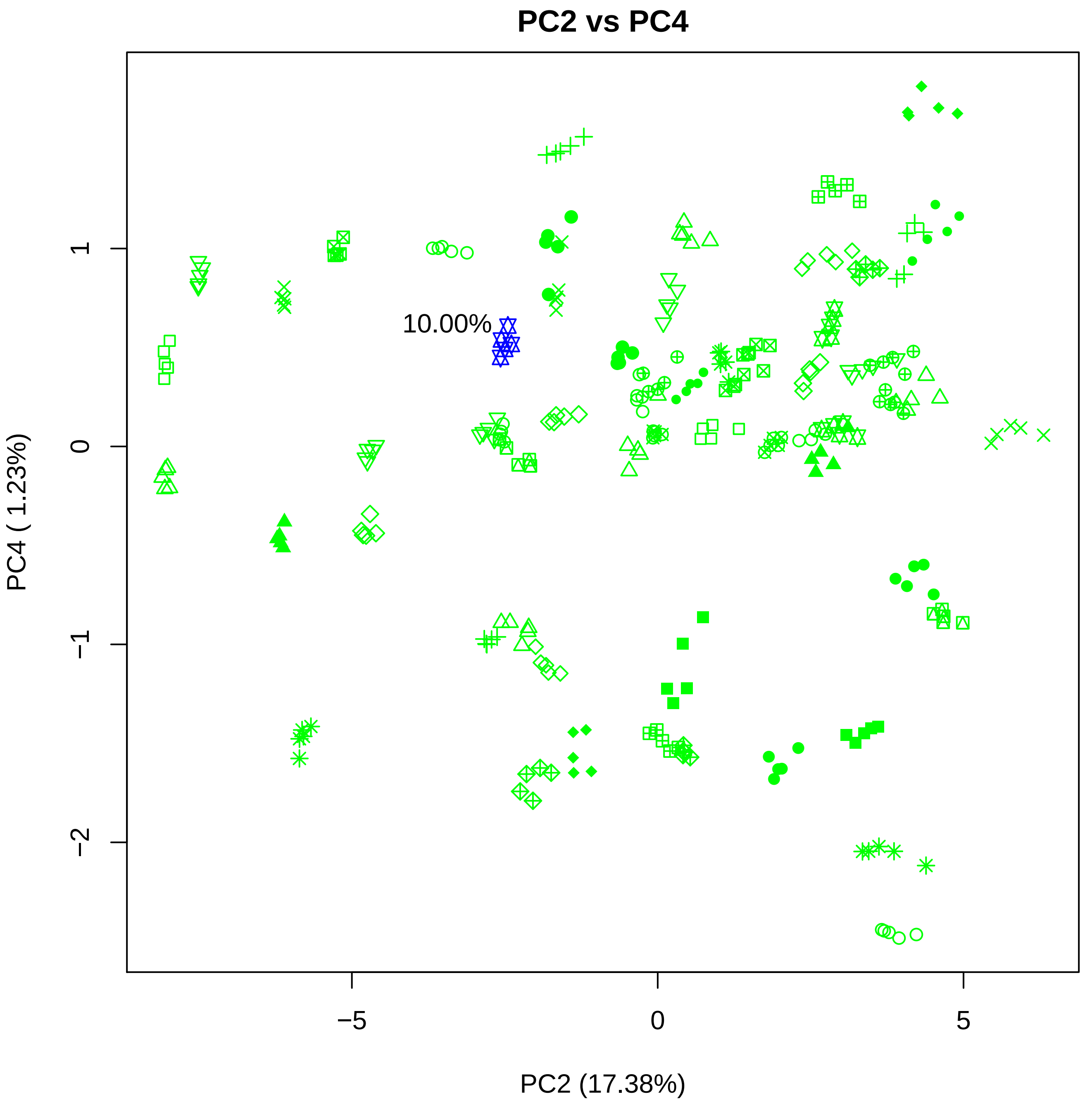
PC2 vs PC4
10.00%
PC4 ( 1.23%)
PC2 (17.38%)
−5
0
5
−2
−1
0
1

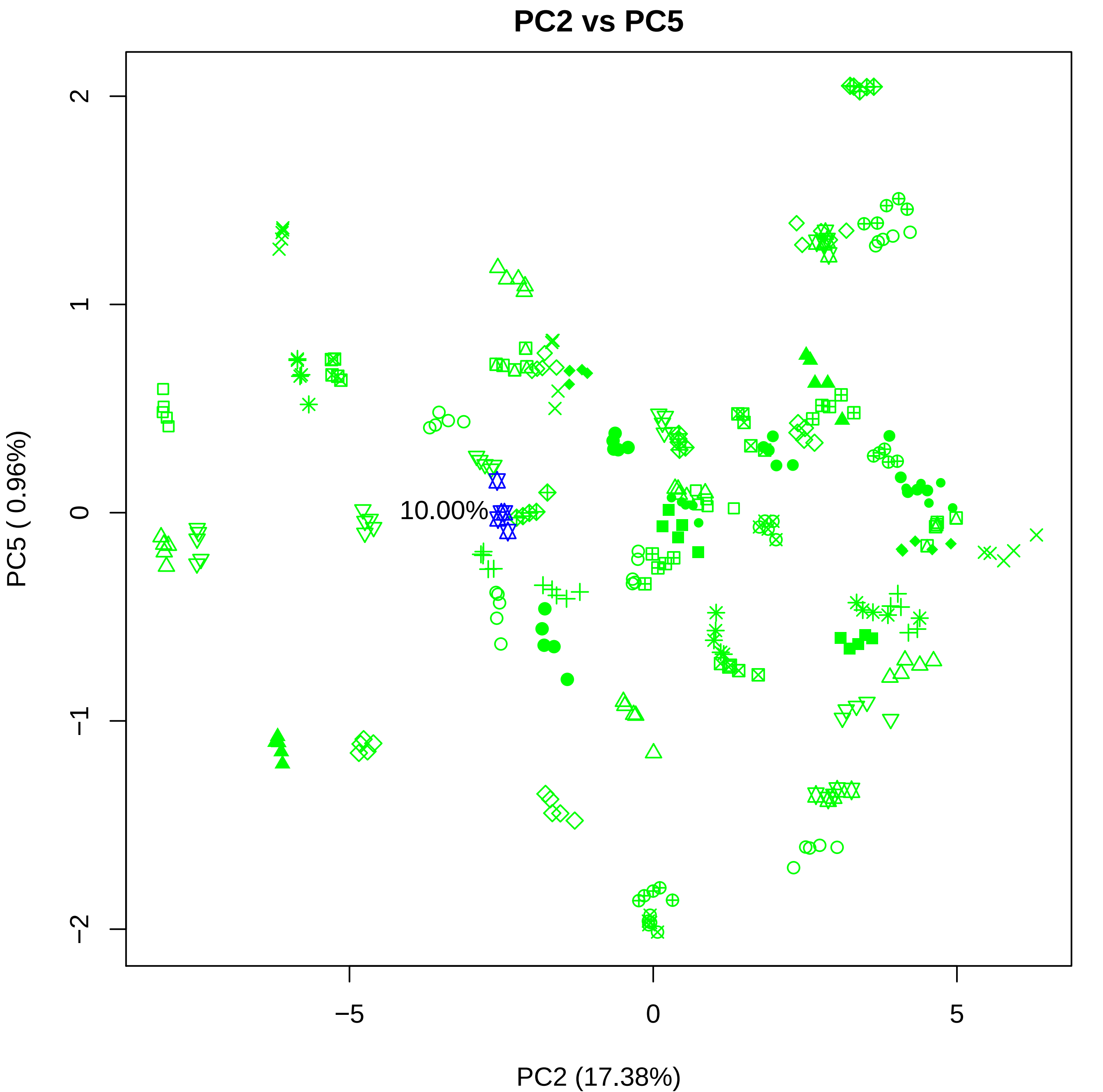
PC2 vs PC5
PC5 ( 0.96%)
PC2 (17.38%)
10.00%
2
1
0
−1
−2
−5
0
5

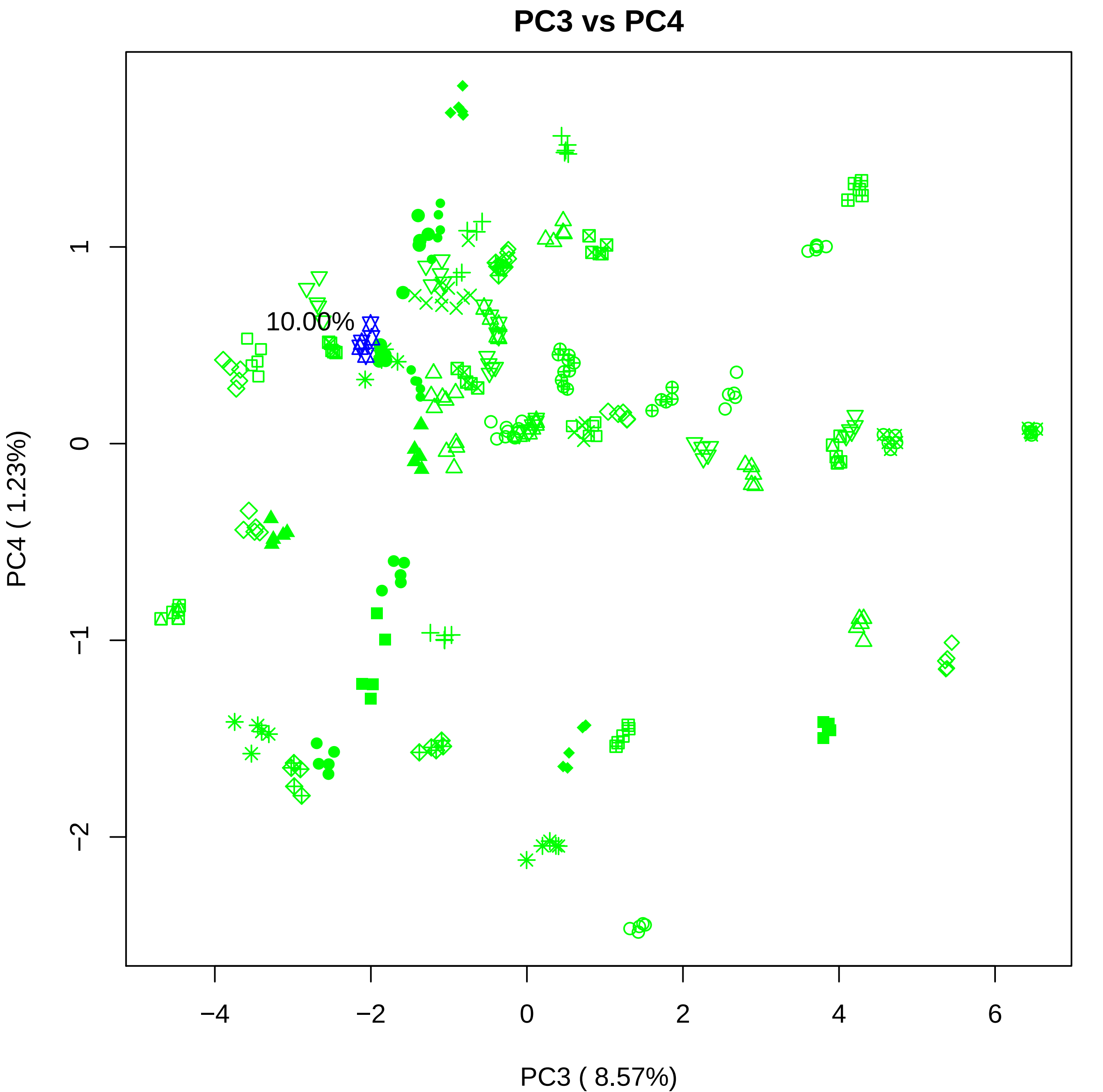

PC3 vs PC4
10.00%
PC4 ( 1.23%)
PC3 ( 8.57%)
1
0
−1
−2
−4
−2
0
2
4
6

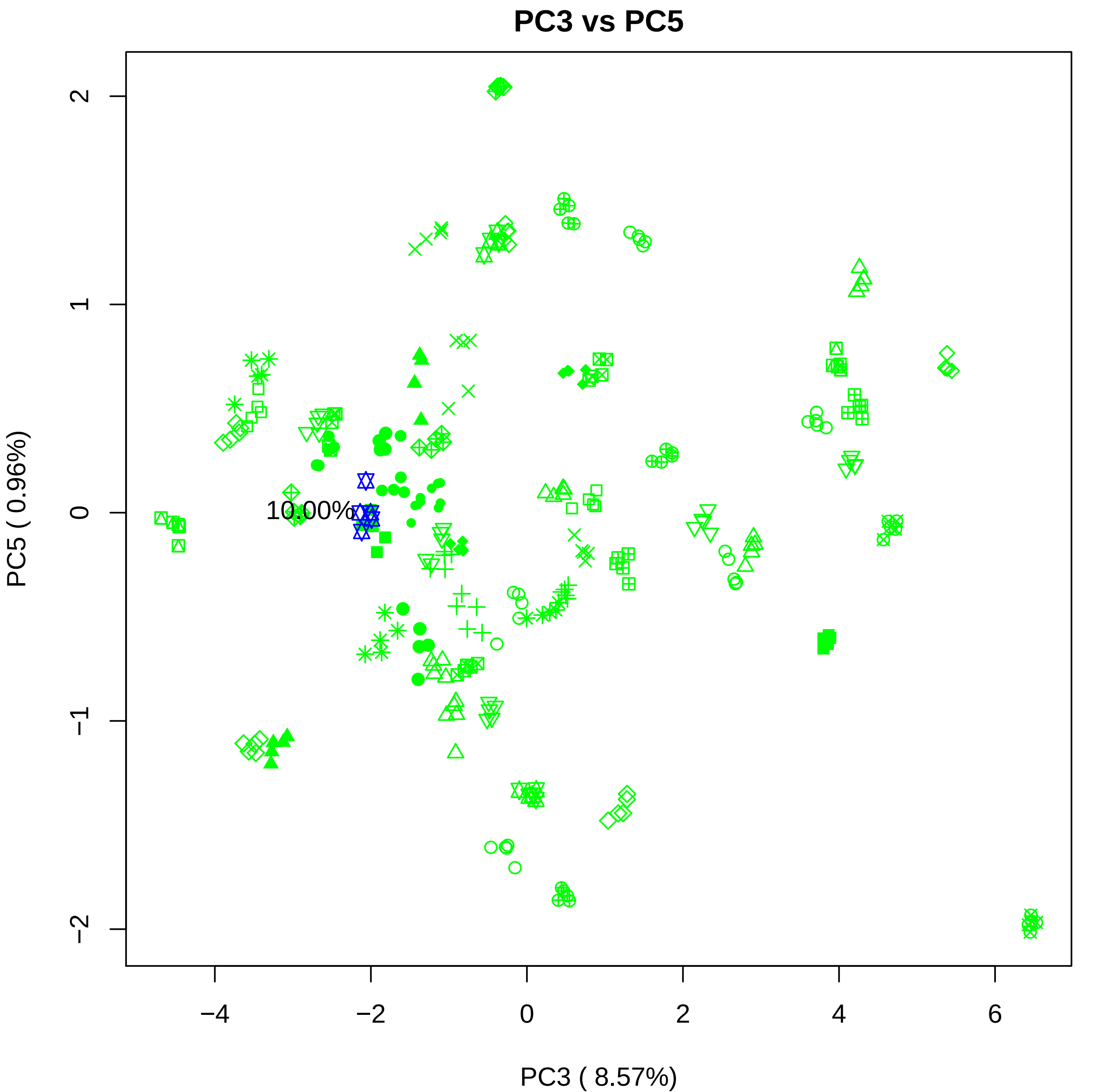
PC3 vs PC5
PC5 ( 0.96%)
PC3 ( 8.57%)
10.00%
2
1
0
−1
−2
−4
−2
0
2
4
6

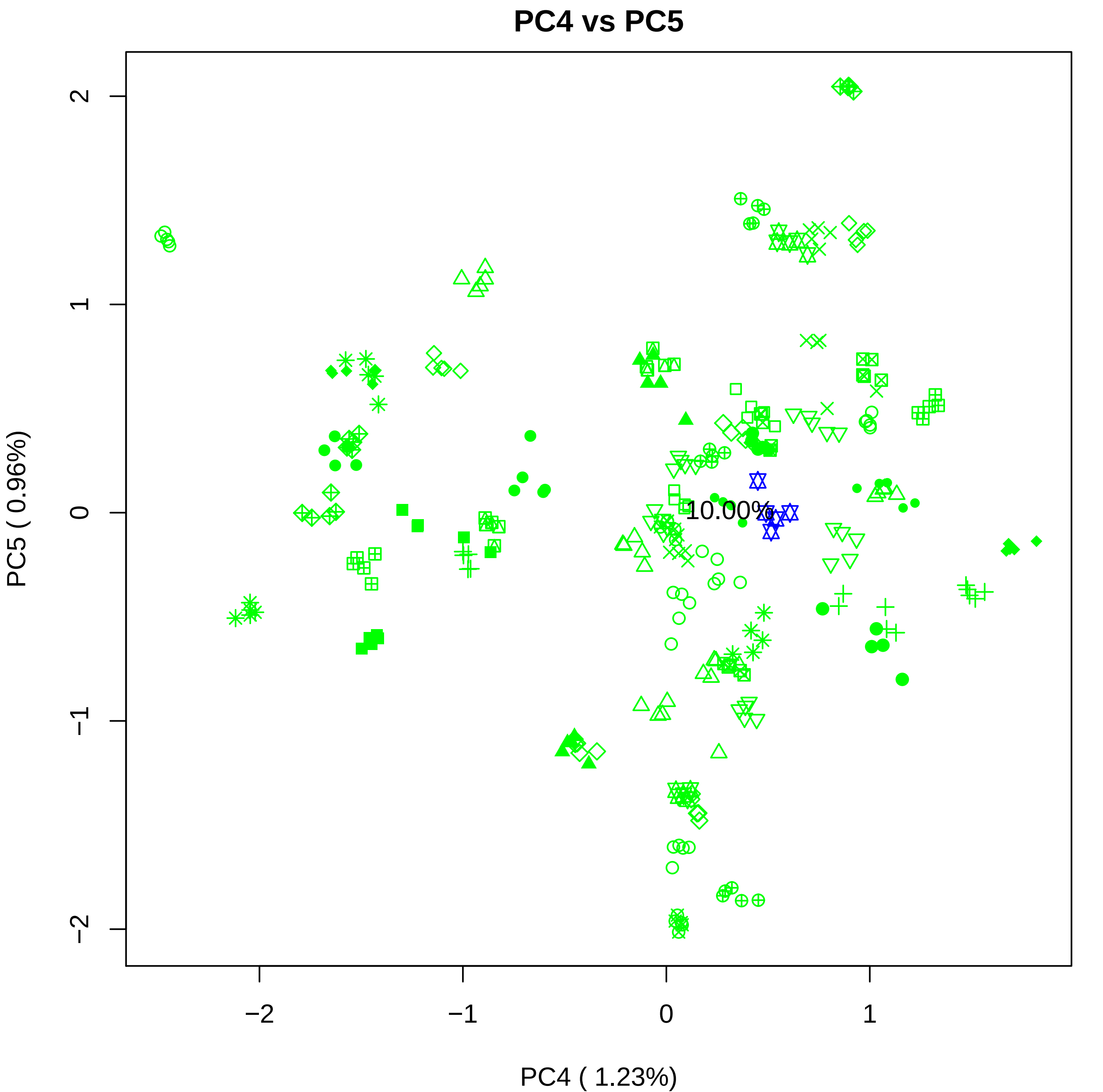
PC4 vs PC5
PC5 ( 0.96%)
PC4 ( 1.23%)
10.00%
2
1
0
−1
−2
−2
−1
0
1

### PCA of 15%  BMH (a)  vs PMH(a)

Scaled Matrix PCAed

7th order polynomial baseline correction performed

Raw spectra smoothed to 21 Neighbours, 2nd order PN, Savtisky Golay

Normalisation between 0 and 100

D2 spectra smoothed to 41 Neighbours, 2nd order PN, Savtisky Golay

Regions Analysed: 3601−2501 and 2001−700cm−1

Samples Analysed: 1 bContam's 0 BLH's 0 BMH's 0 OMH's 0 pContam's 0 PLH's 60 PMH's

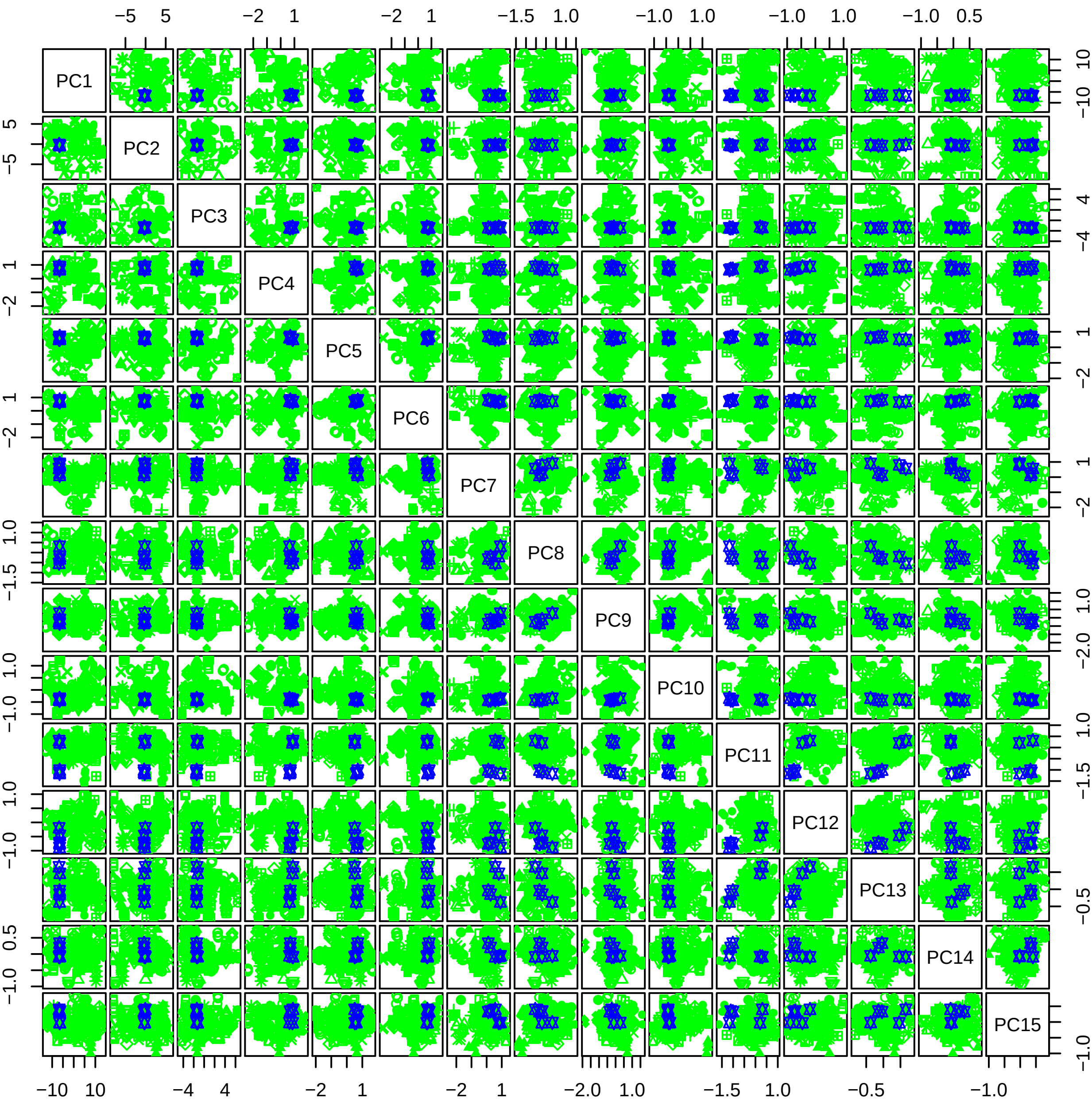
PC1
PC2
PC3
PC4
PC5
PC6
PC7
PC8
PC9
PC10
PC11
PC12
PC13
PC14
PC15
−5 5
−2 1
−2 1
−1.5 1.0
−1.0 1.0
−1.0 1.0
−1.0 0.5
10
−10
5
−5
4
−4
1
−2
1
−2
1
−2
1
−2
1.0
−1.5
1.0
−2.0
1.0
−1.0
1.0
−1.5
1.0
−1.0
−0.5
0.5
−1.0
−1.0
−10 10
−4 4
−2 1
−2 1
−2.0 1.0
−1.5 1.0
−0.5
−1.0

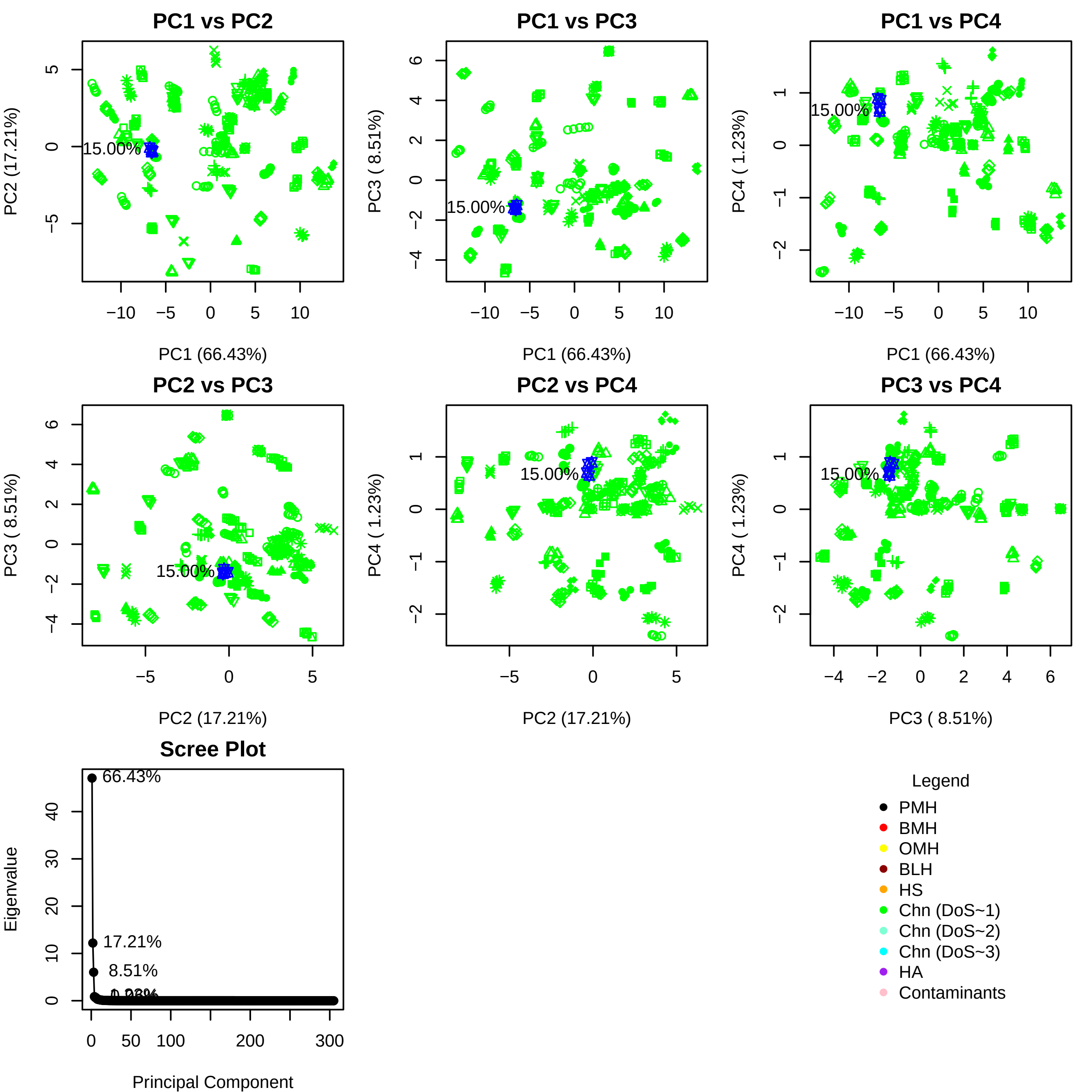

PC1 vs PC2
PC1 vs PC3
PC1 vs PC4
PC2 vs PC3
PC2 vs PC4
PC3 vs PC4
Scree Plot
PC1 (66.43%)
PC2 (17.21%)
PC3 ( 8.51%)
PC4 ( 1.23%)
15.00%
Eigenvalue
Principal Component
66.43%
17.21%
8.51%
Legend
PMH
BMH
OMH
BLH
HS
Chn (DoS~1)
Chn (DoS~2)
Chn (DoS~3)
HA
Contaminants

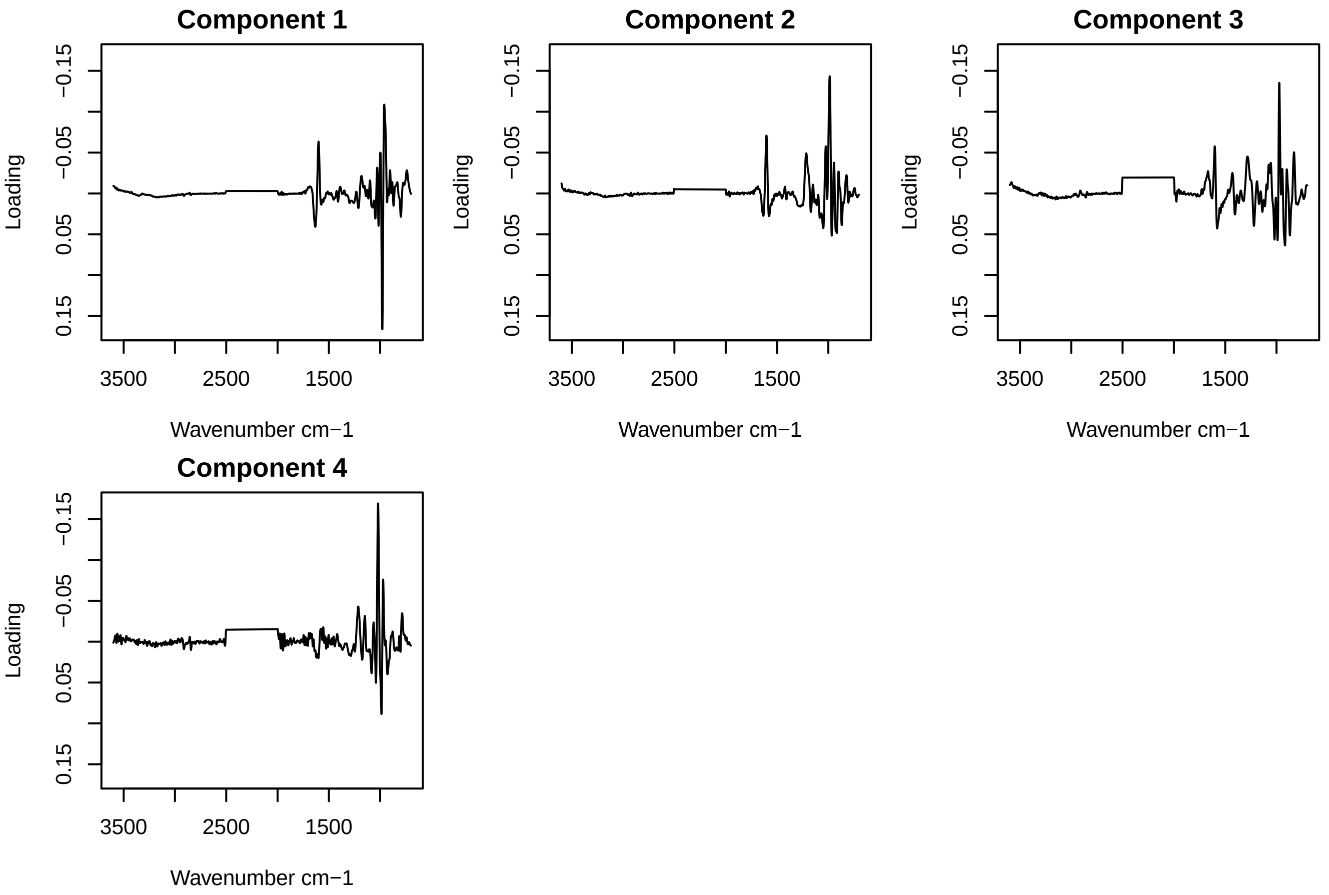

Component 1
Component 2
Component 3
Component 4
Loading
Wavenumber cm−1
3500
2500
1500
−0.15
−0.05
0.05
0.15

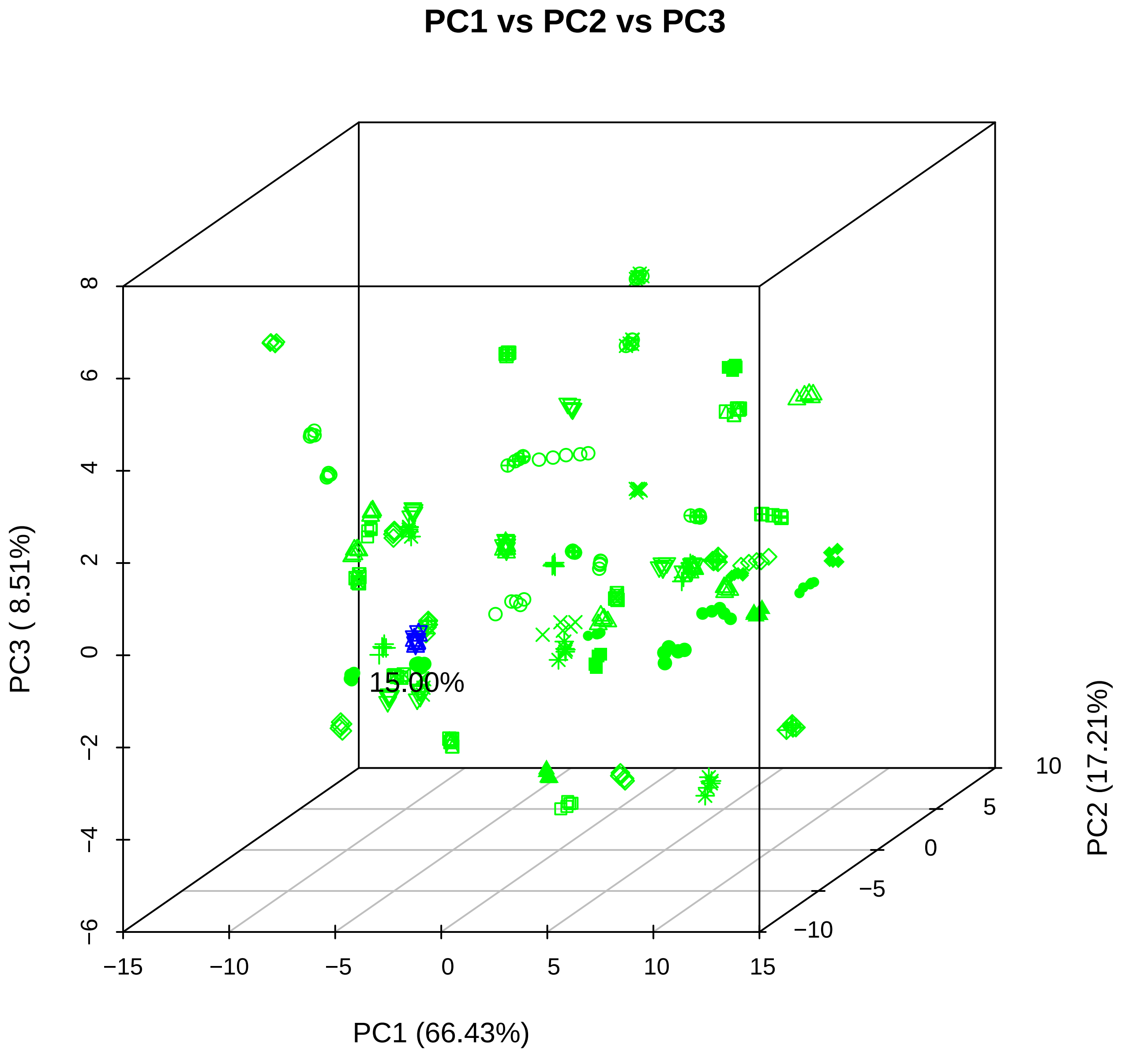
PC1 vs PC2 vs PC3
PC1 (66.43%)
PC2 (17.21%)
PC3 ( 8.51%)
15.00%
−15
−10
−5
0
5
10
15
−6
−4
−2
0
2
4
6
8
−10
−5
0
5
10

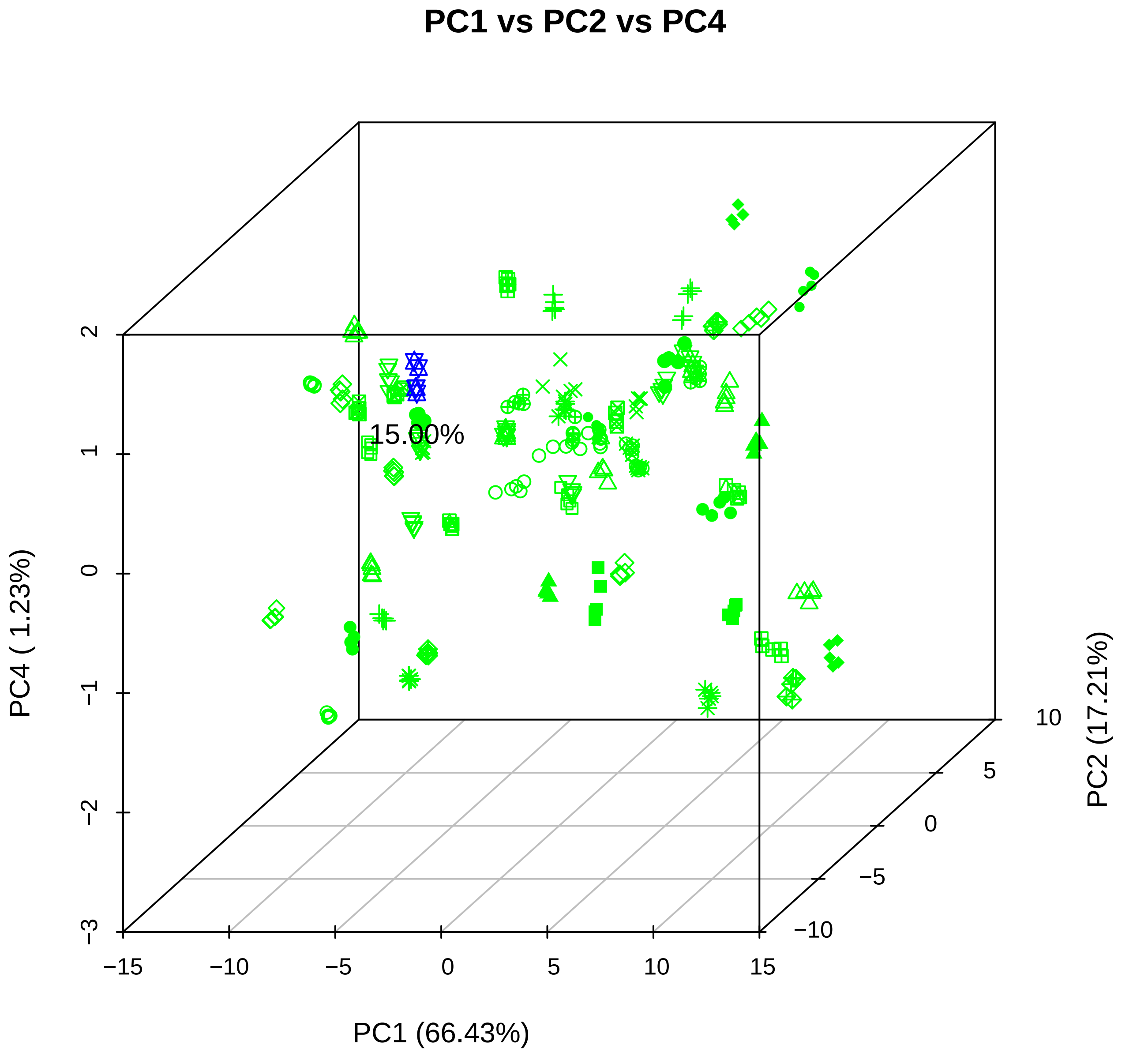
PC1 vs PC2 vs PC4
15.00%
PC1 (66.43%)
PC2 (17.21%)
PC4 ( 1.23%)
−15
−10
−5
0
5
10
15
−3
−2
−1
0
1
2
−10
−5
0
5
10

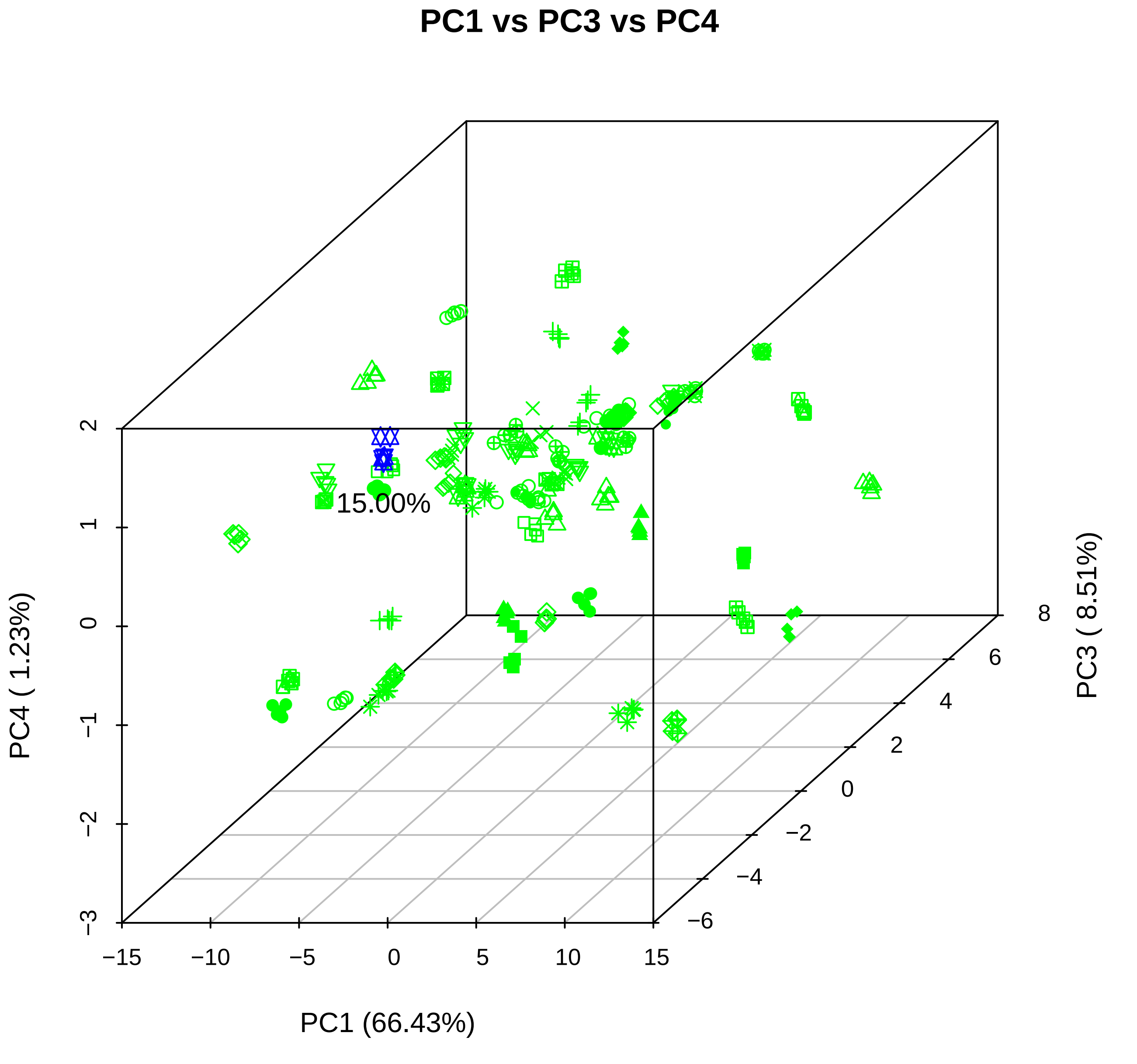
PC1 vs PC3 vs PC4
15.00%
PC1 (66.43%)
PC3 ( 8.51%)
PC4 ( 1.23%)
−15 −10 −5 0 5 10 15
−6 −4 −2 0 2 4 6 8
−3 −2 −1 0 1 2

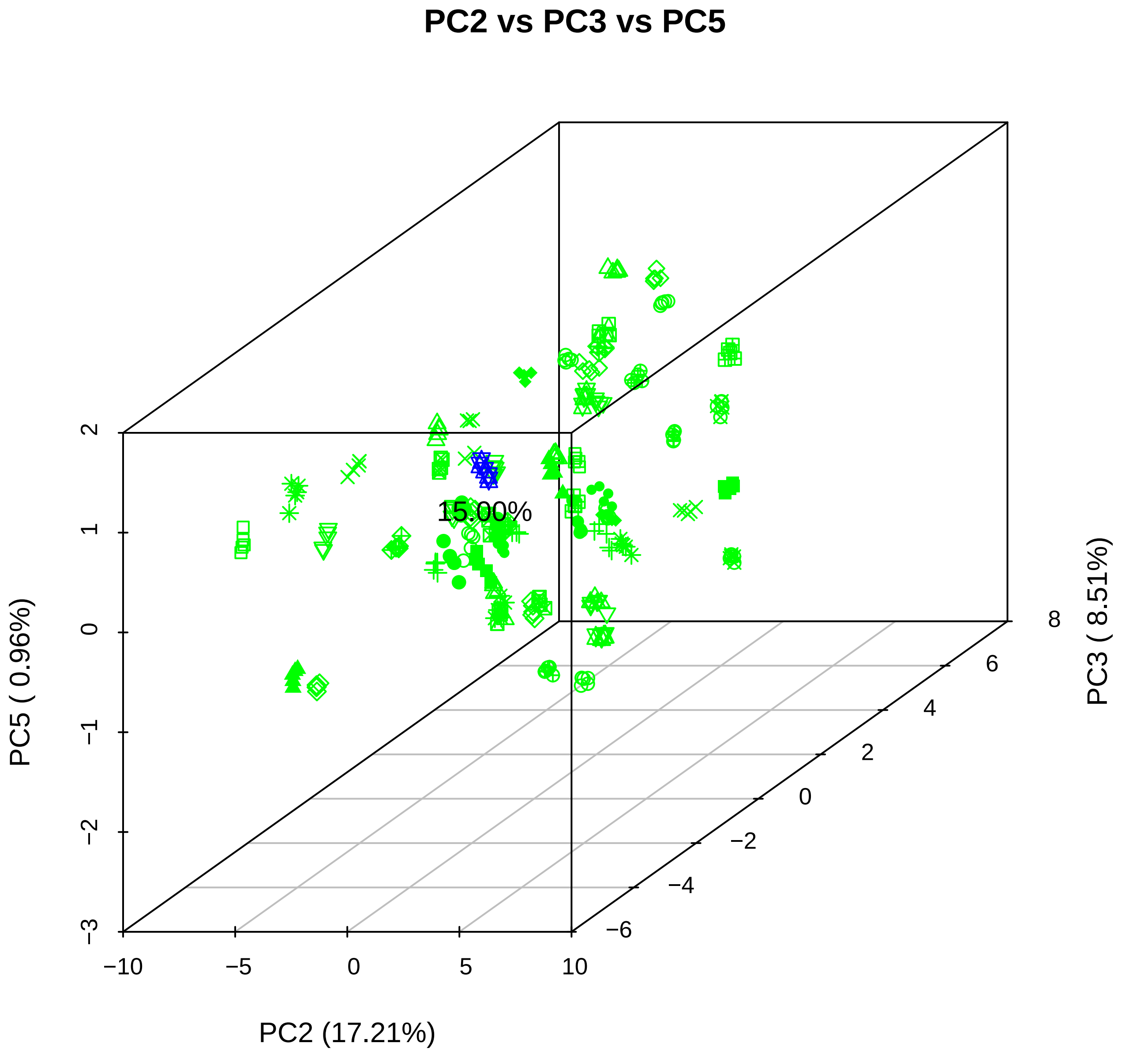

PC2 vs PC3 vs PC5
15.00%
PC2 (17.21%)
PC3 ( 8.51%)
PC5 ( 0.96%)
−10
−5
0
5
10
−6
−4
−2
0
2
4
6
8
−3
−2
−1
0
1
2

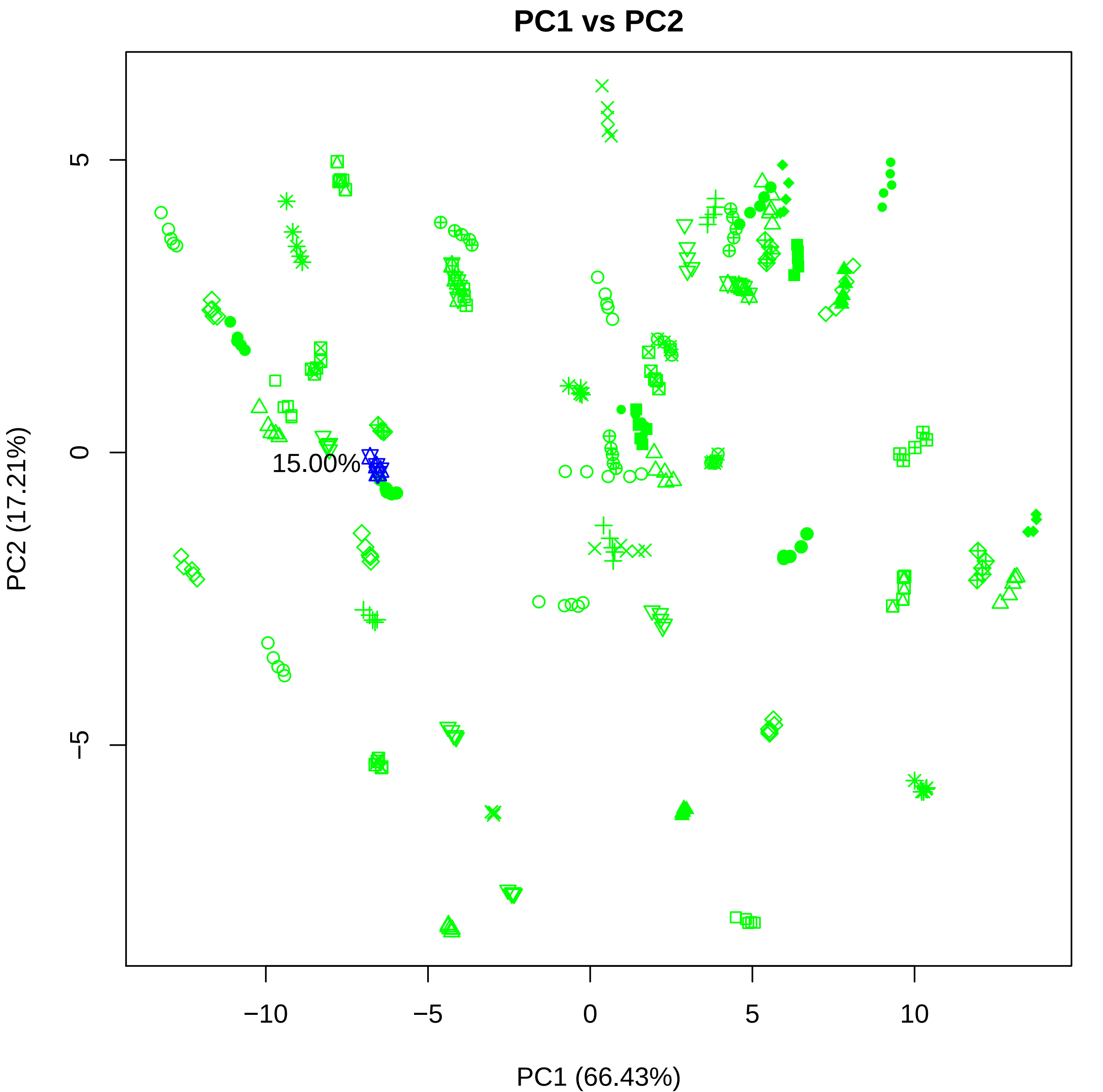

PC1 vs PC2
PC2 (17.21%)
PC1 (66.43%)
15.00%
5
0
−5
−10
−5
0
5
10

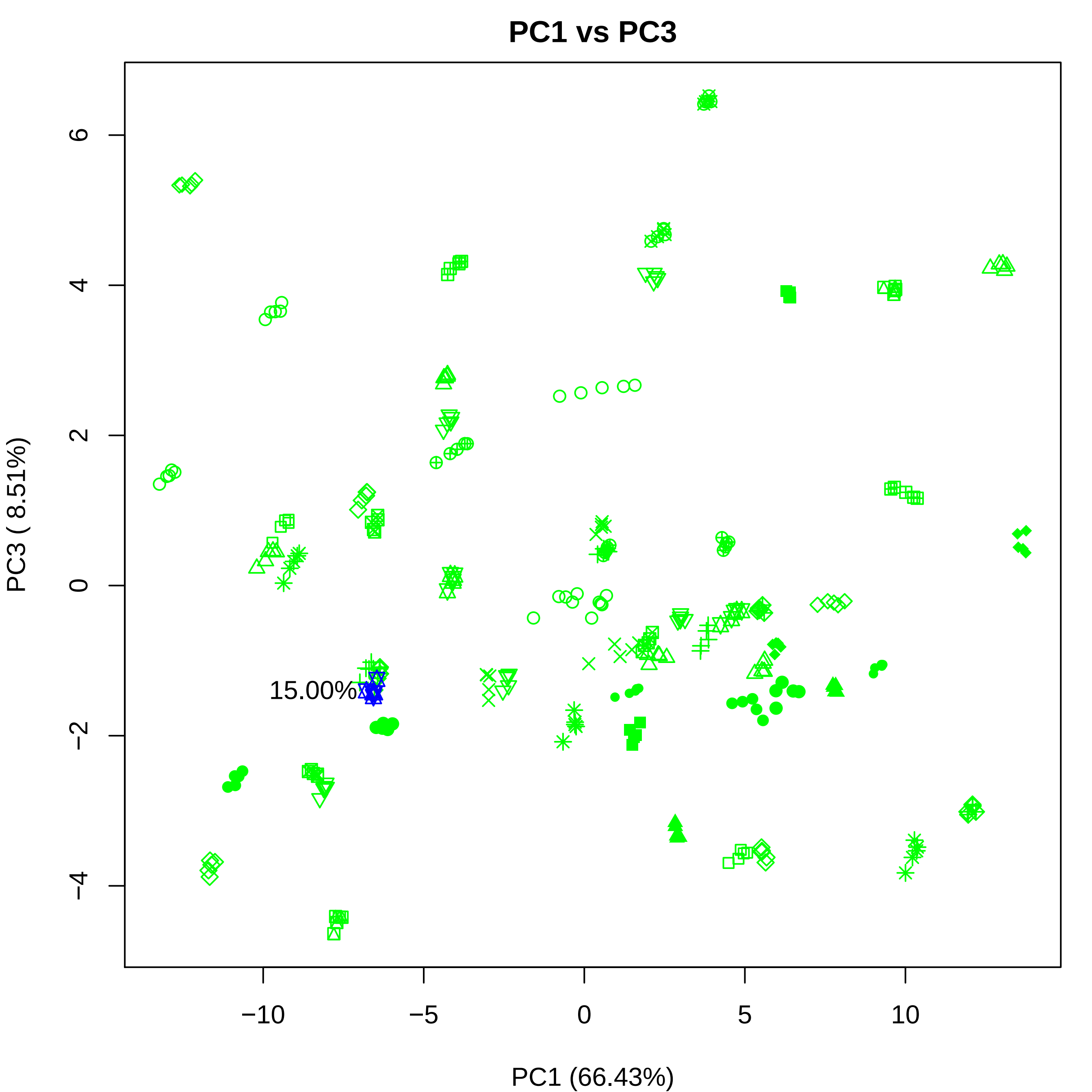

PC1 vs PC3
15.00%
PC3 ( 8.51%)
PC1 (66.43%)
6
4
2
0
−2
−4
−10
−5
0
5
10

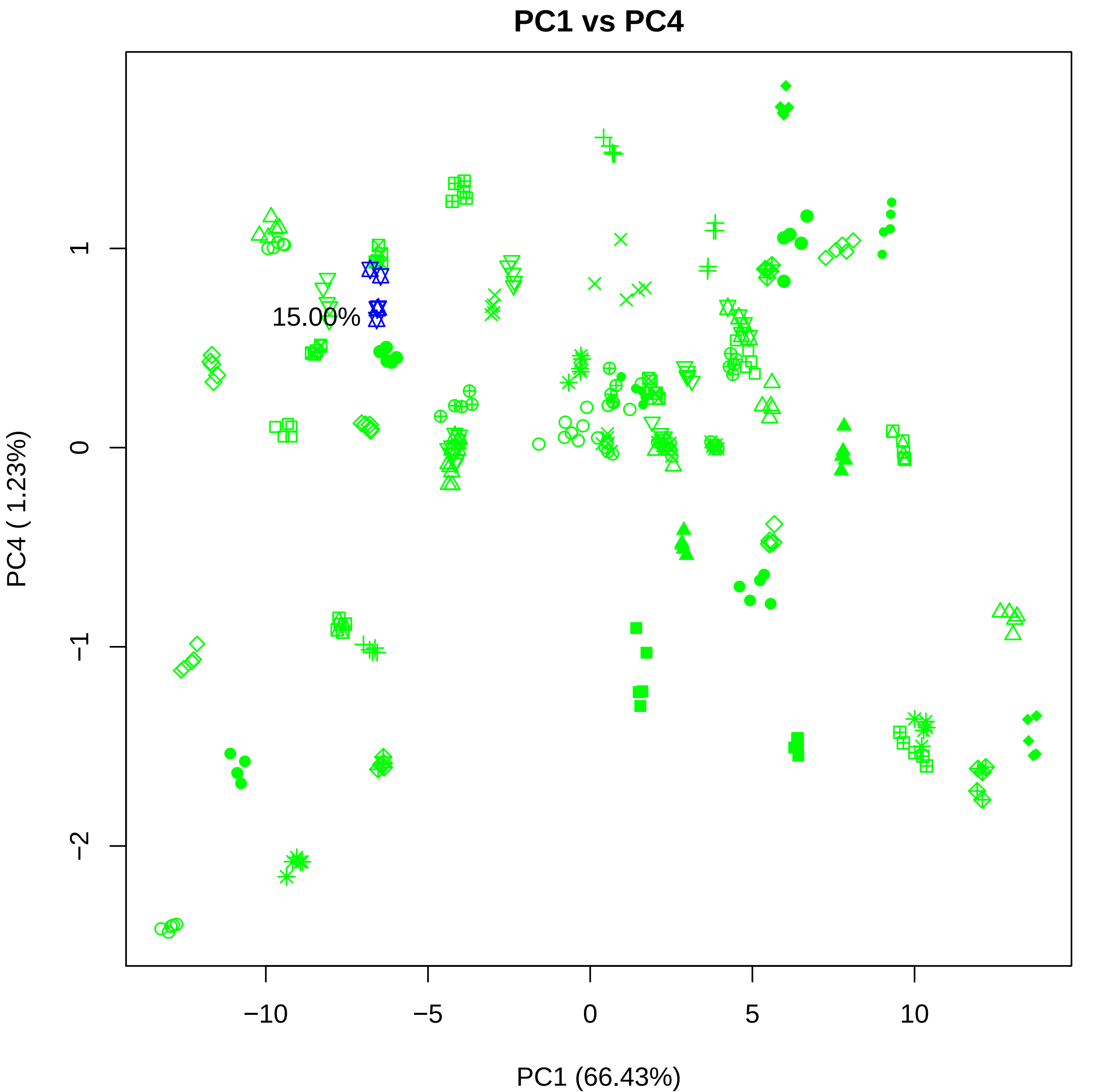

PC1 vs PC4
15.00%
1
0
−1
−2
−10
−5
0
5
10
PC1 (66.43%)
PC4 ( 1.23%)

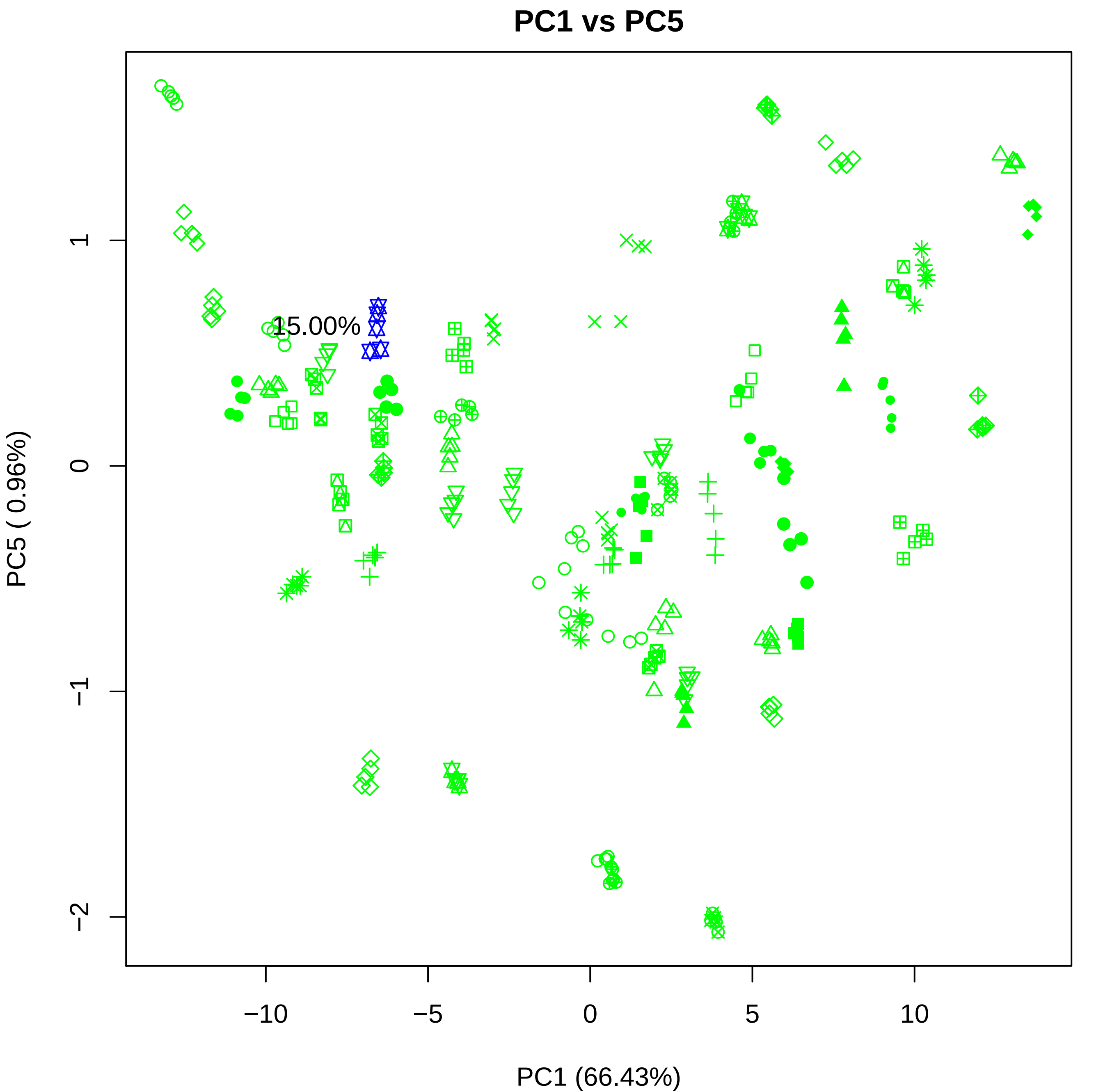

PC1 vs PC5
15.00%
1
0
−1
−2
PC5 ( 0.96%)
−10
−5
0
5
10
PC1 (66.43%)

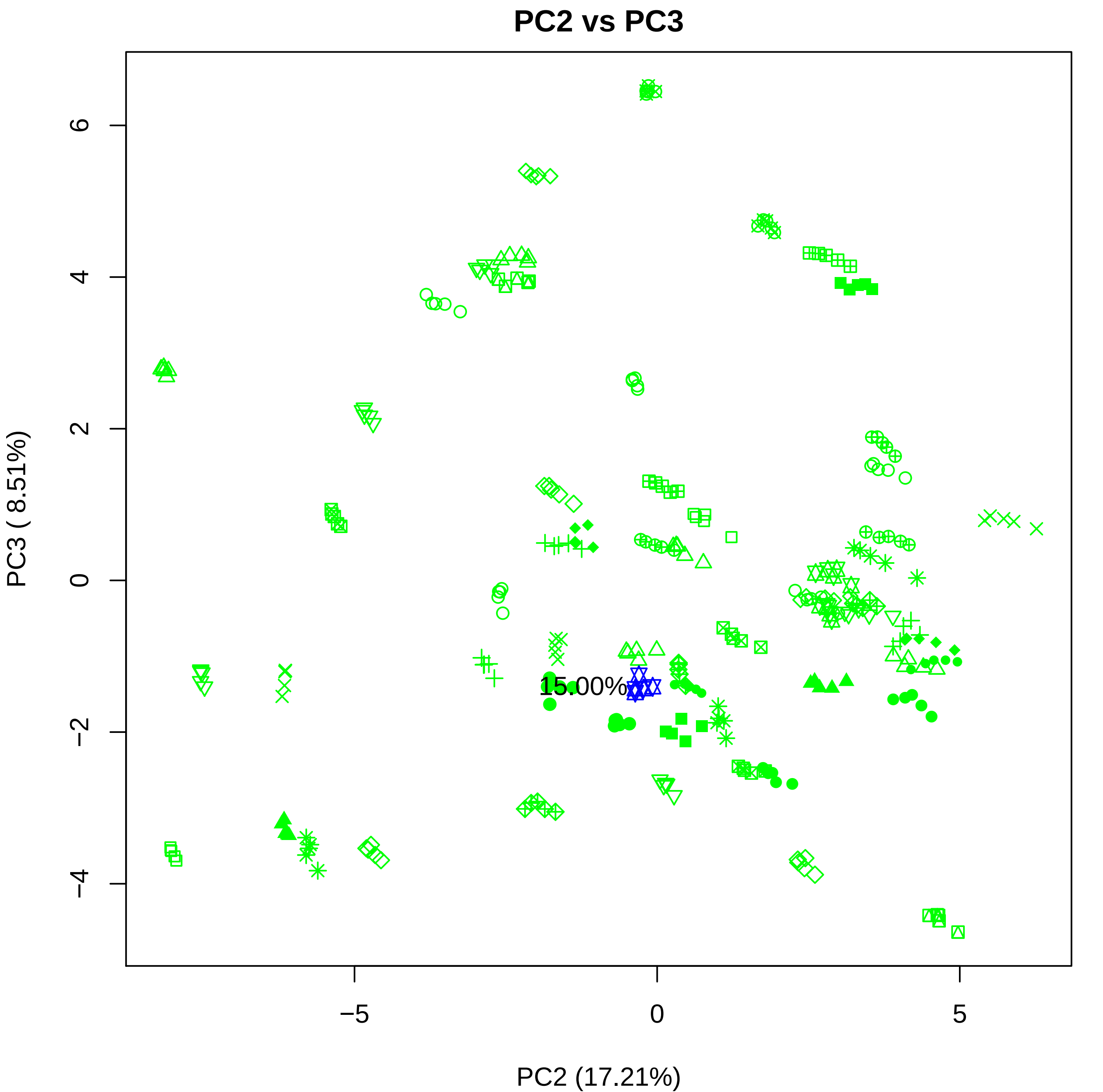
PC2 vs PC3
PC3 ( 8.51%)
PC2 (17.21%)
15.00%
6
4
2
0
−2
−4
−5
0
5

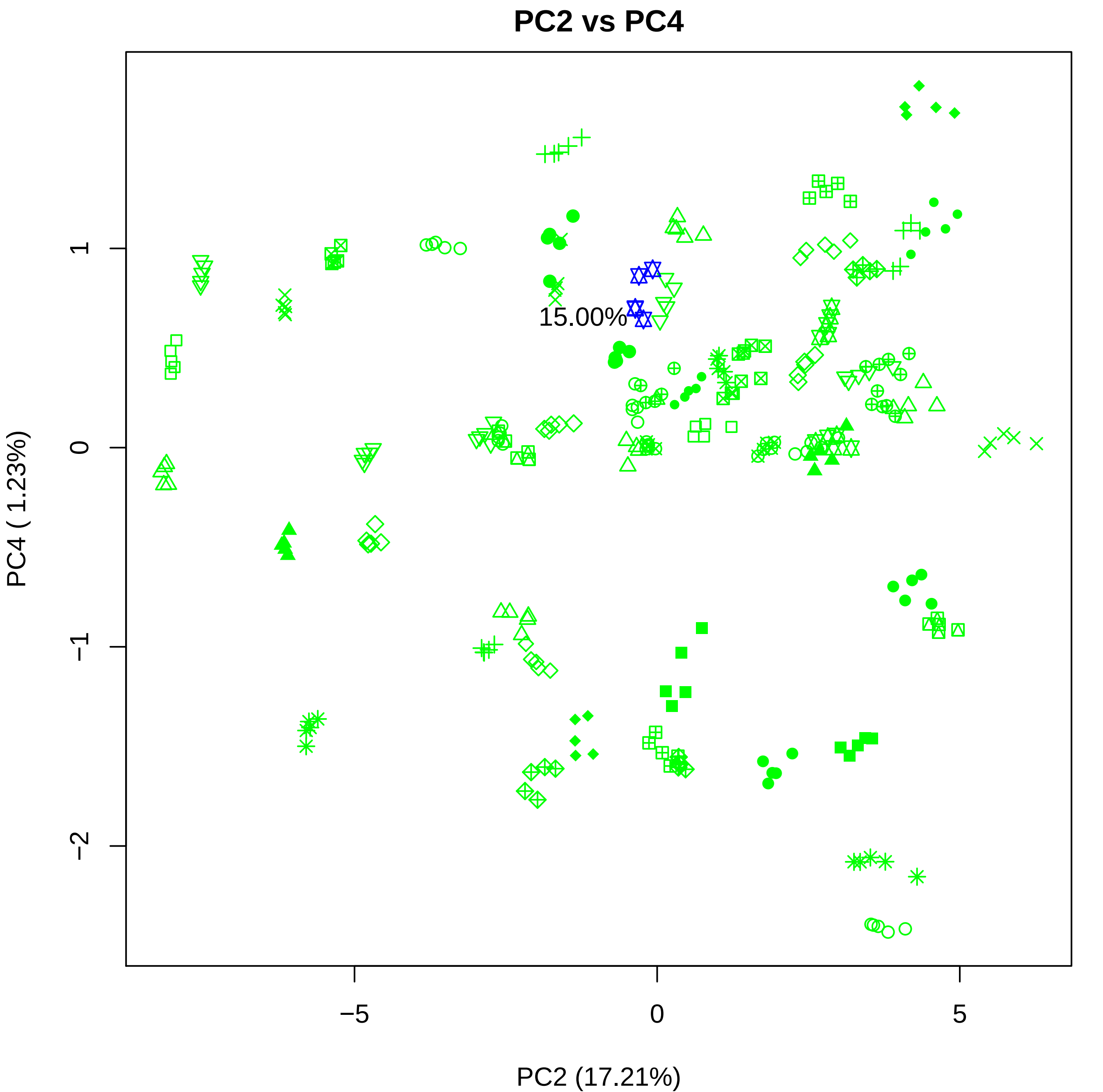

PC2 vs PC4
15.00%
PC4 ( 1.23%)
PC2 (17.21%)
−2
−1
0
1
−5
0
5

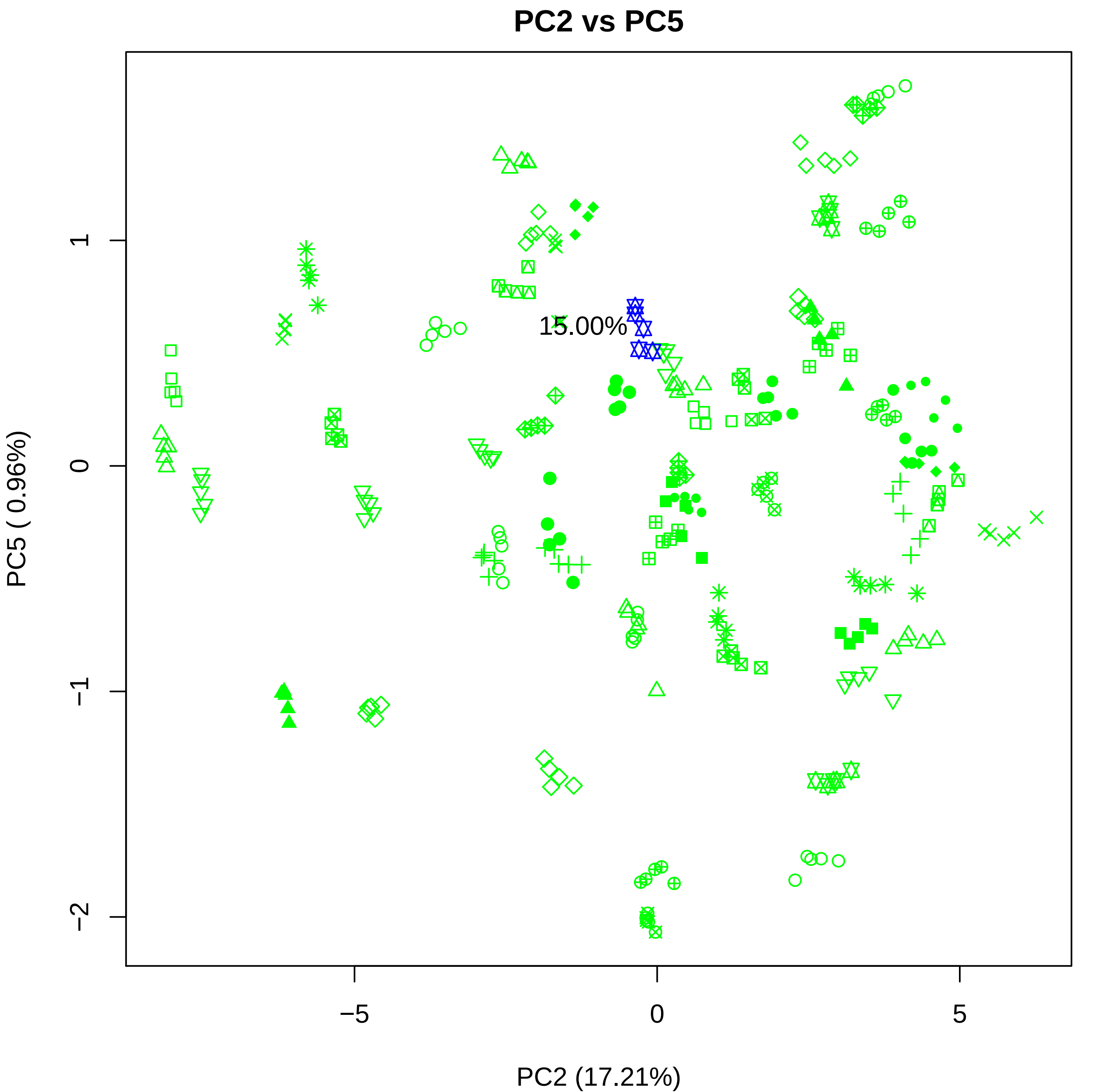
PC2 vs PC5
15.00%
PC5 ( 0.96%)
PC2 (17.21%)
1
0
−1
−2
−5
0
5

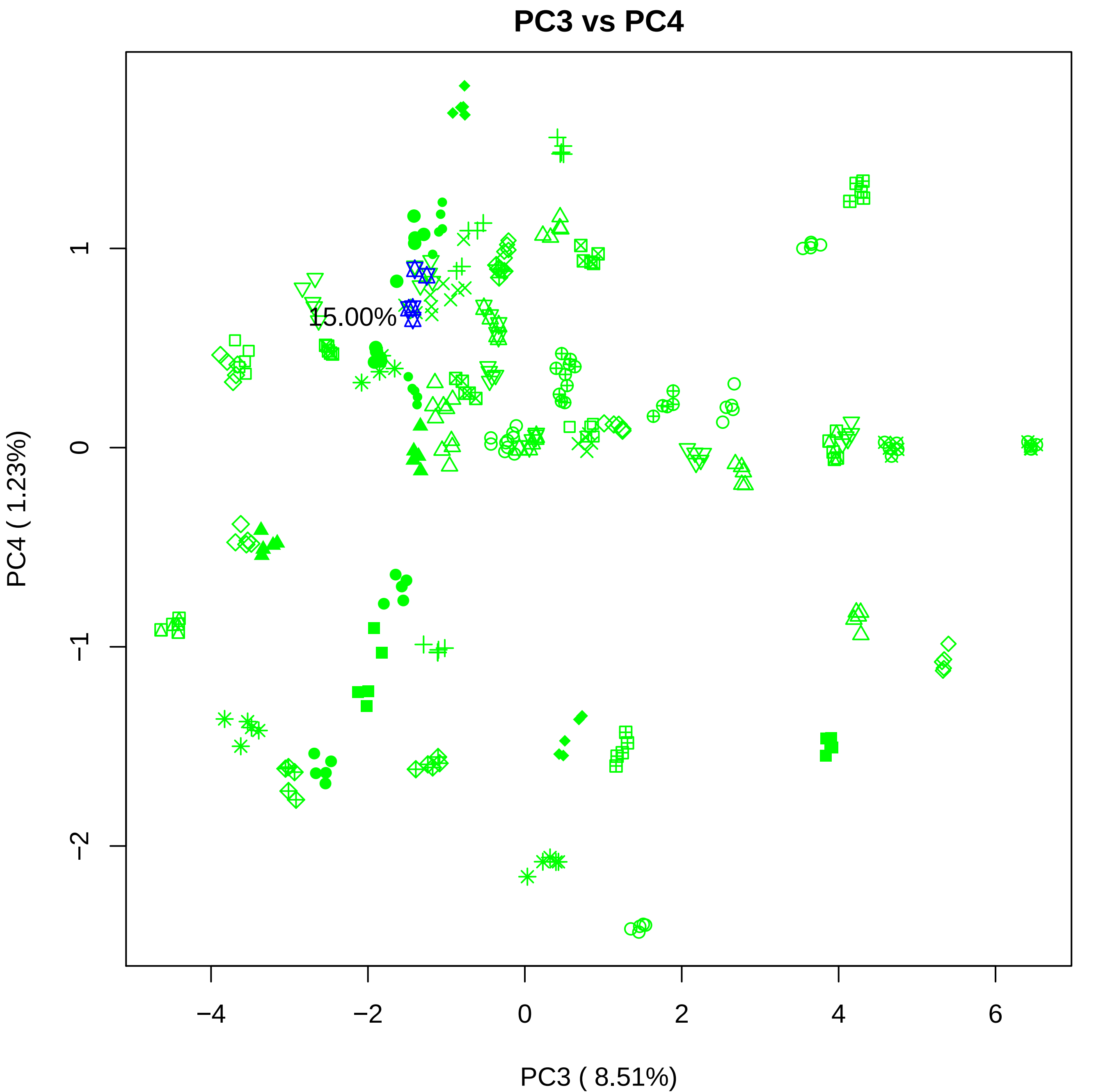
PC3 vs PC4
15.00%
PC4 ( 1.23%)
PC3 ( 8.51%)
−4
−2
0
2
4
6
−2
−1
0
1

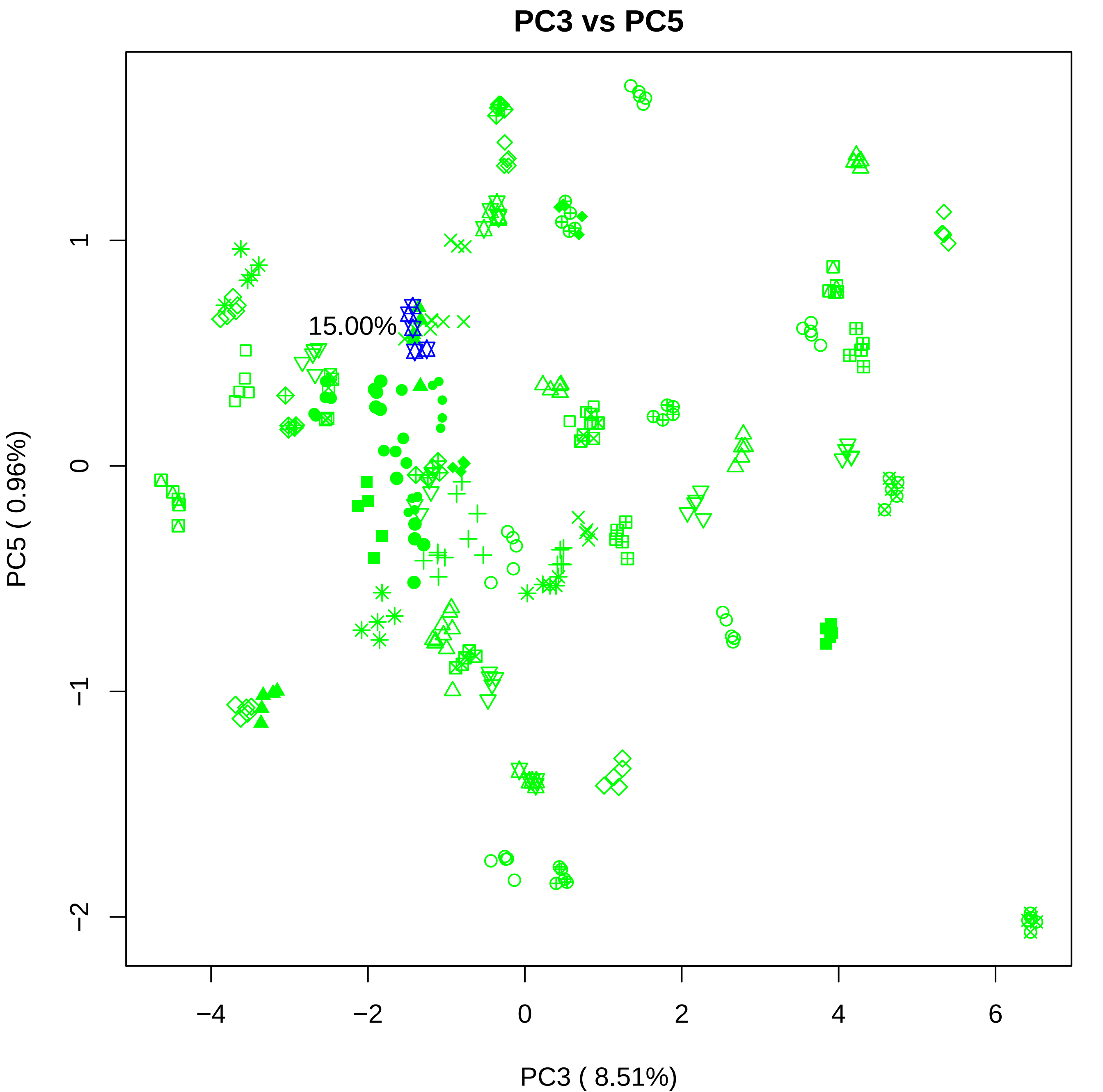
PC3 vs PC5
15.00%
PC5 ( 0.96%)
PC3 ( 8.51%)
−4
−2
0
2
4
6
1
0
−1
−2

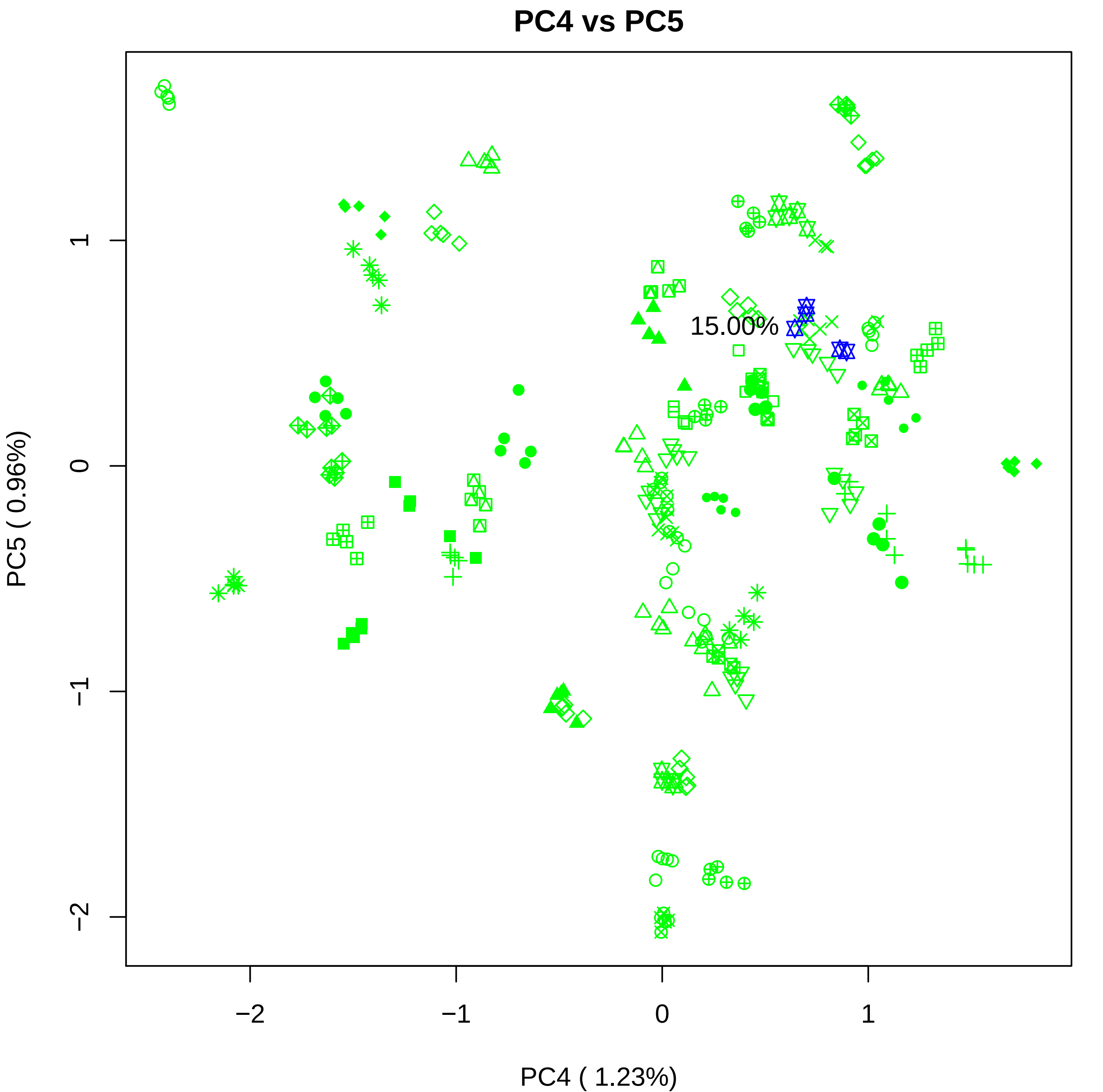
PC4 vs PC5
15.00%
PC5 ( 0.96%)
1
0
−1
−2
PC4 ( 1.23%)
−2
−1
0
1

### PCA of 20%  BMH(a)  vs PMH(a)

Scaled Matrix PCAed

7th order polynomial baseline correction performed

Raw spectra smoothed to 21 Neighbours, 2nd order PN, Savtisky Golay

Normalisation between 0 and 100

D2 spectra smoothed to 41 Neighbours, 2nd order PN, Savtisky Golay

Regions Analysed: 3601−2501 and 2001−700cm−1

Samples Analysed: 1 bContam's 0 BLH's 0 BMH's 0 OMH's 0 pContam's 0 PLH's 60 PMH's

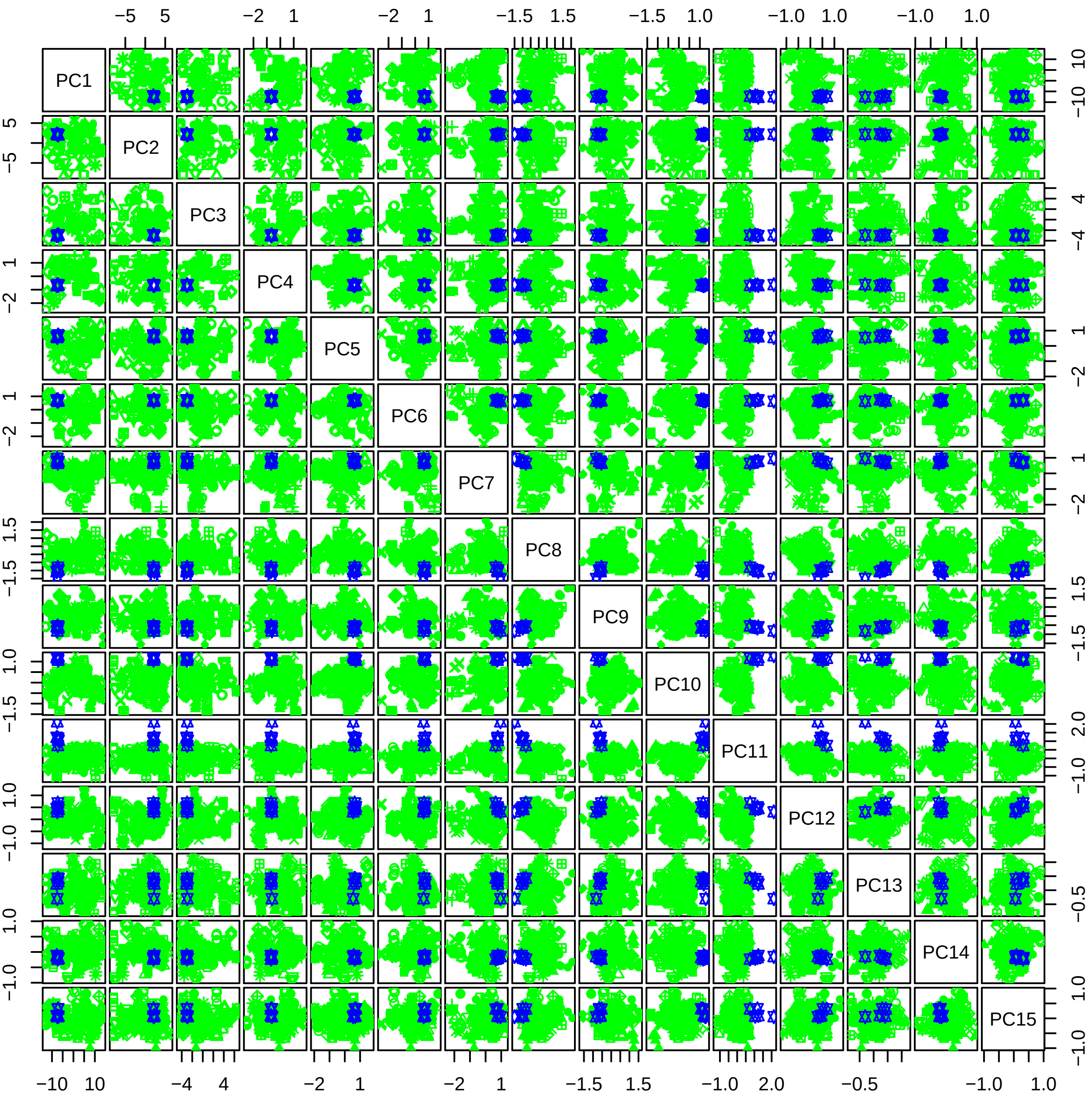
PC1
PC2
PC3
PC4
PC5
PC6
PC7
PC8
PC9
PC10
PC11
PC12
PC13
PC14
PC15

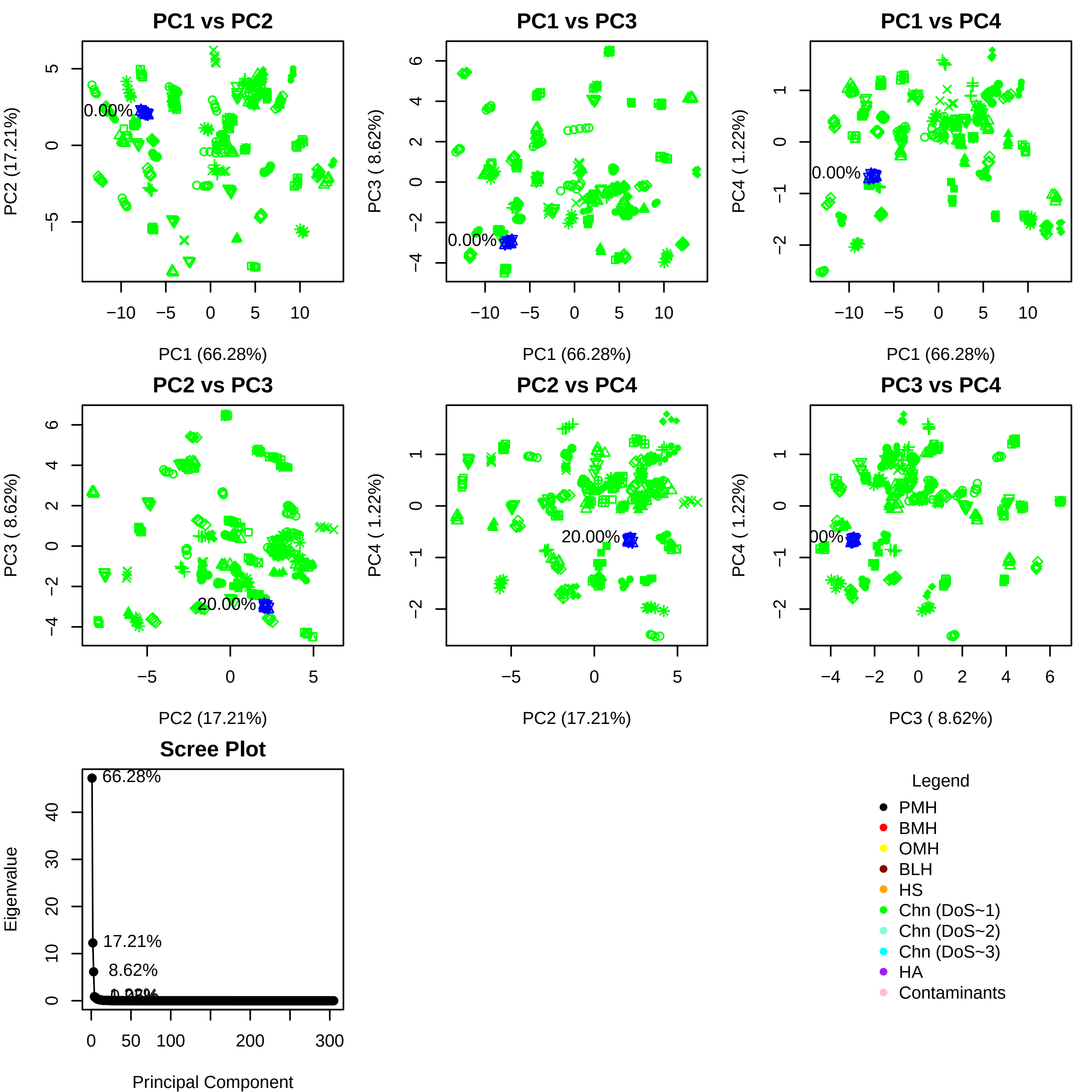
PC1 vs PC2
PC2 (17.21%)
PC1 (66.28%)
0.00%
PC1 vs PC3
PC3 ( 8.62%)
PC1 (66.28%)
0.00%
PC1 vs PC4
PC4 ( 1.22%)
PC1 (66.28%)
0.00%
PC2 vs PC3
PC3 ( 8.62%)
PC2 (17.21%)
20.00%
PC2 vs PC4
PC4 ( 1.22%)
PC2 (17.21%)
20.00%
PC3 vs PC4
PC4 ( 1.22%)
PC3 ( 8.62%)
Scree Plot
Eigenvalue
Principal Component
66.28%
17.21%
8.62%
Legend
PMH
BMH
OMH
BLH
HS
Chn (DoS~1)
Chn (DoS~2)
Chn (DoS~3)
HA
Contaminants

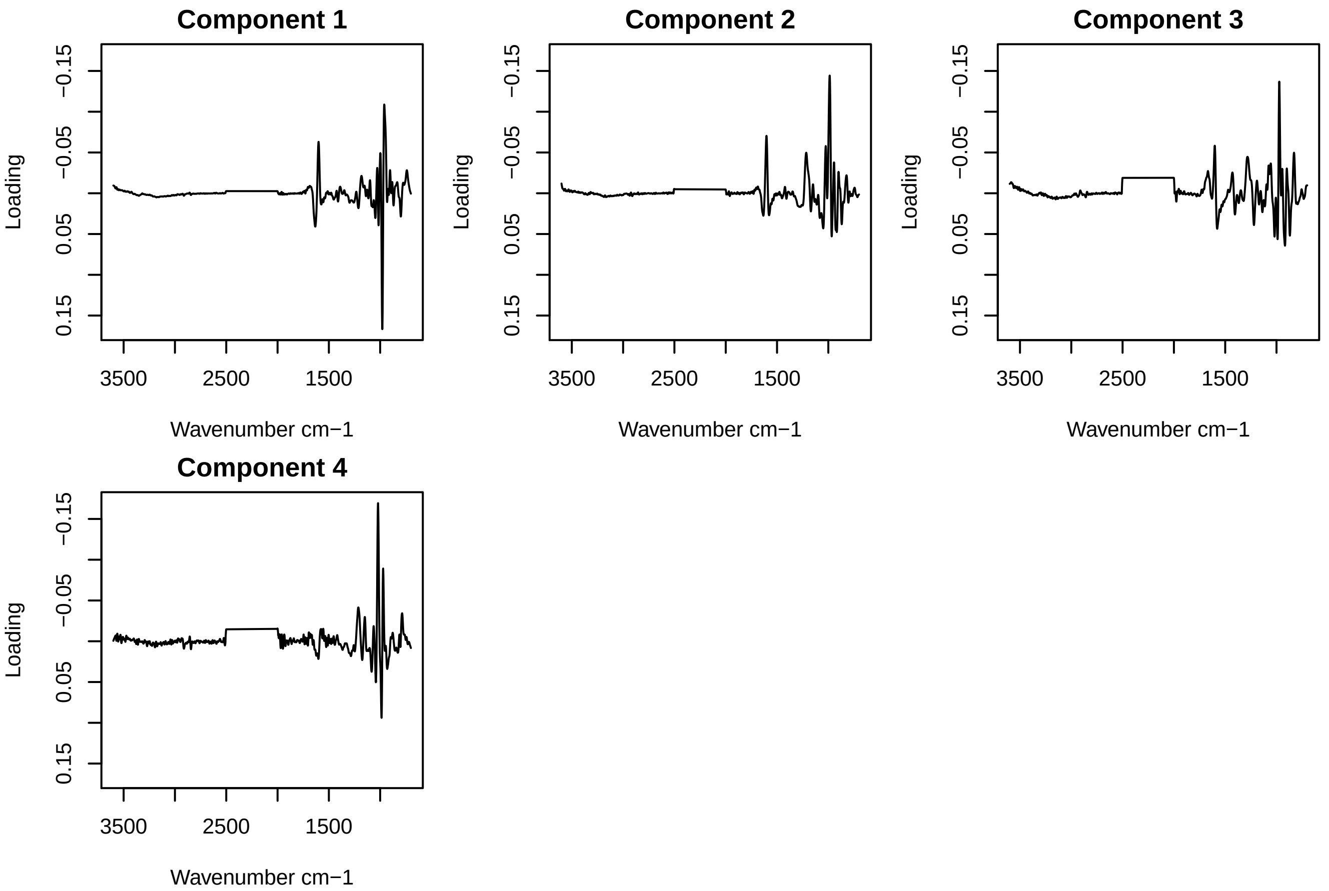
Component 1
Component 2
Component 3
Component 4
Loading
−0.15
−0.05
0.05
0.15
3500
2500
1500
Wavenumber cm−1

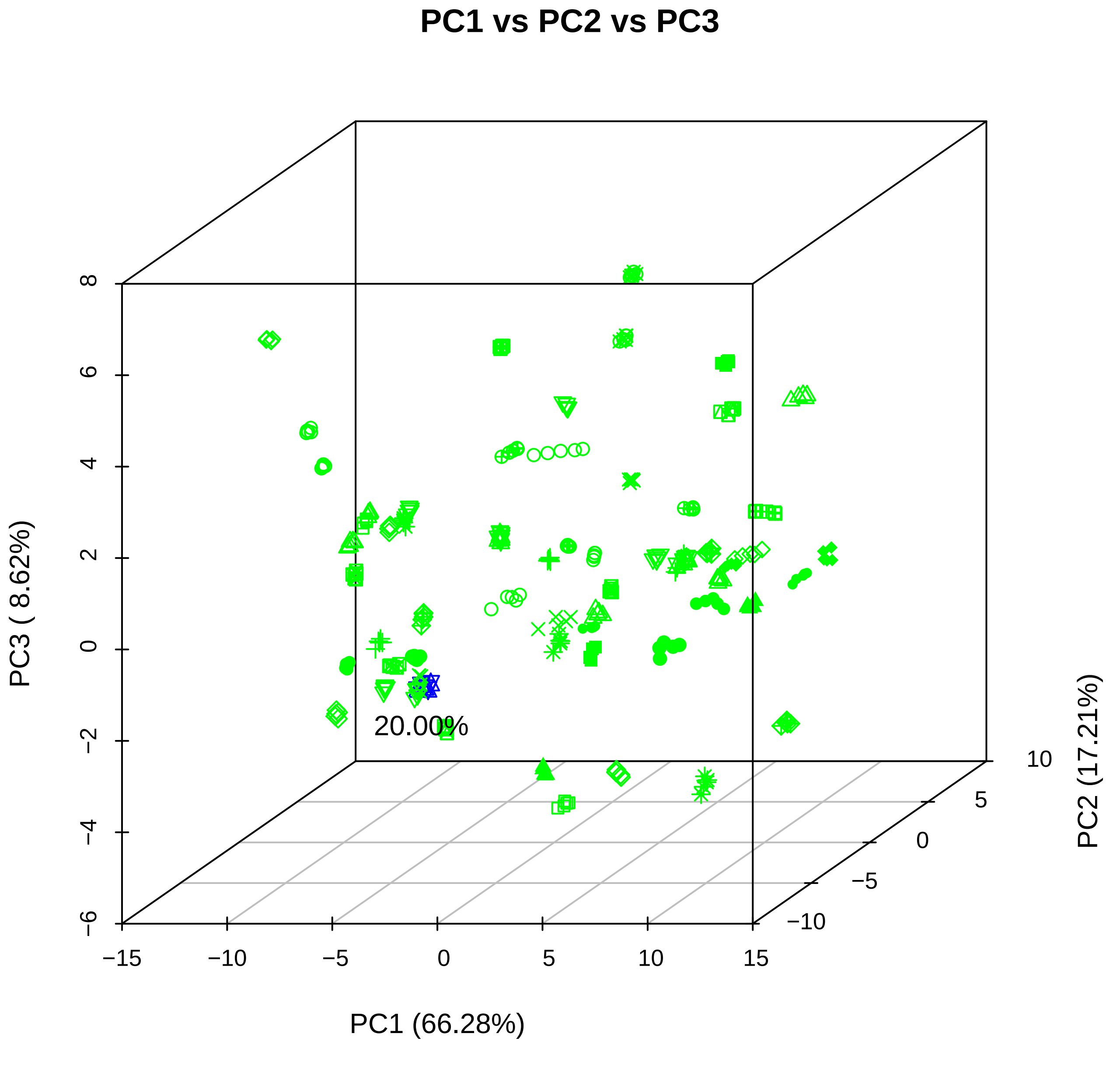

PC1 vs PC2 vs PC3
PC1 (66.28%)
PC2 (17.21%)
PC3 ( 8.62%)
20.00%
−15
−10
−5
0
5
10
15
−6
−4
−2
0
2
4
6
8
−10
−5
0
5
10

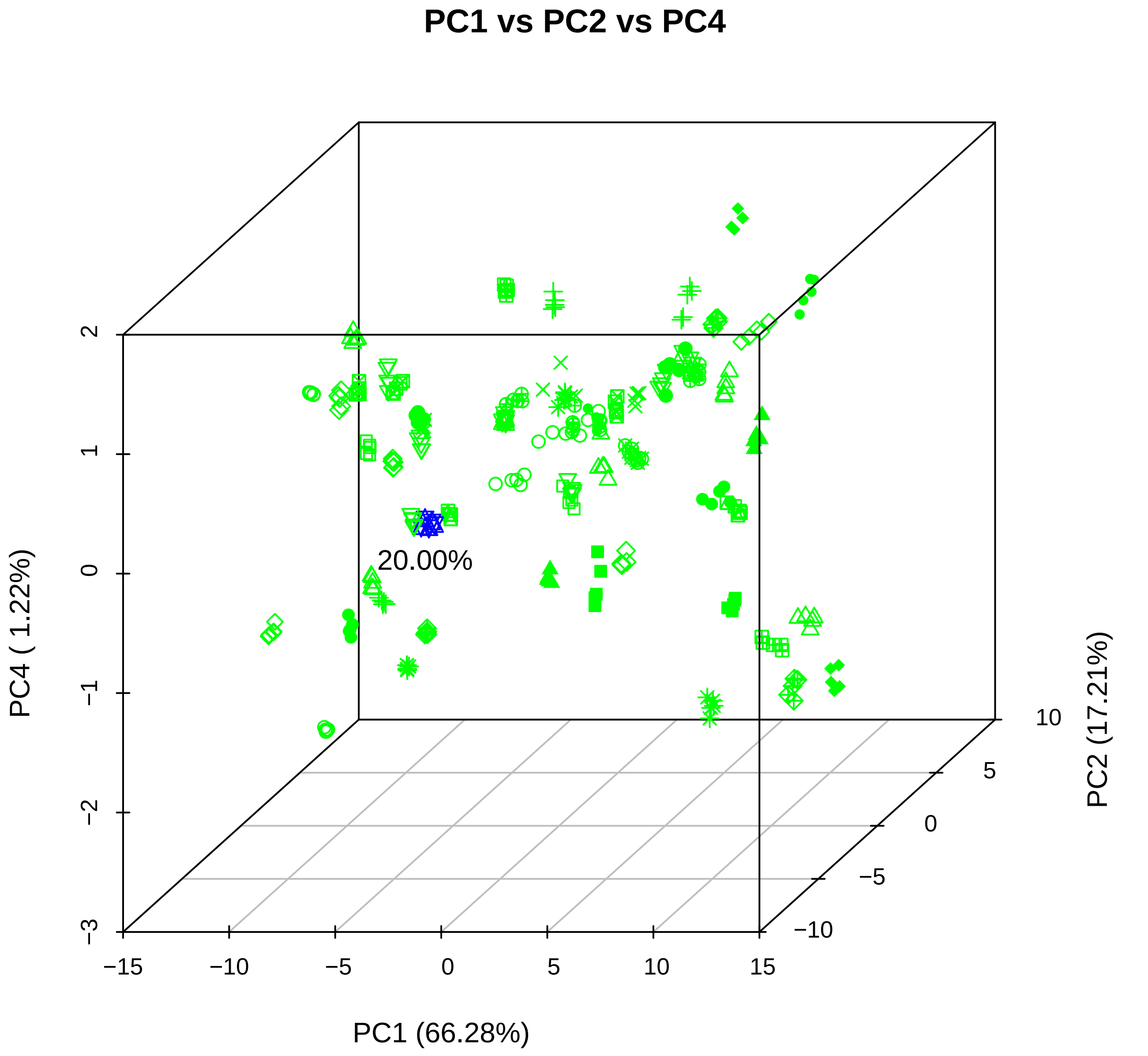
PC1 vs PC2 vs PC4
20.00%
PC1 (66.28%)
PC2 (17.21%)
PC4 ( 1.22%)
−15
−10
−5
0
5
10
15
−10
−5
0
5
10
−3
−2
−1
0
1
2

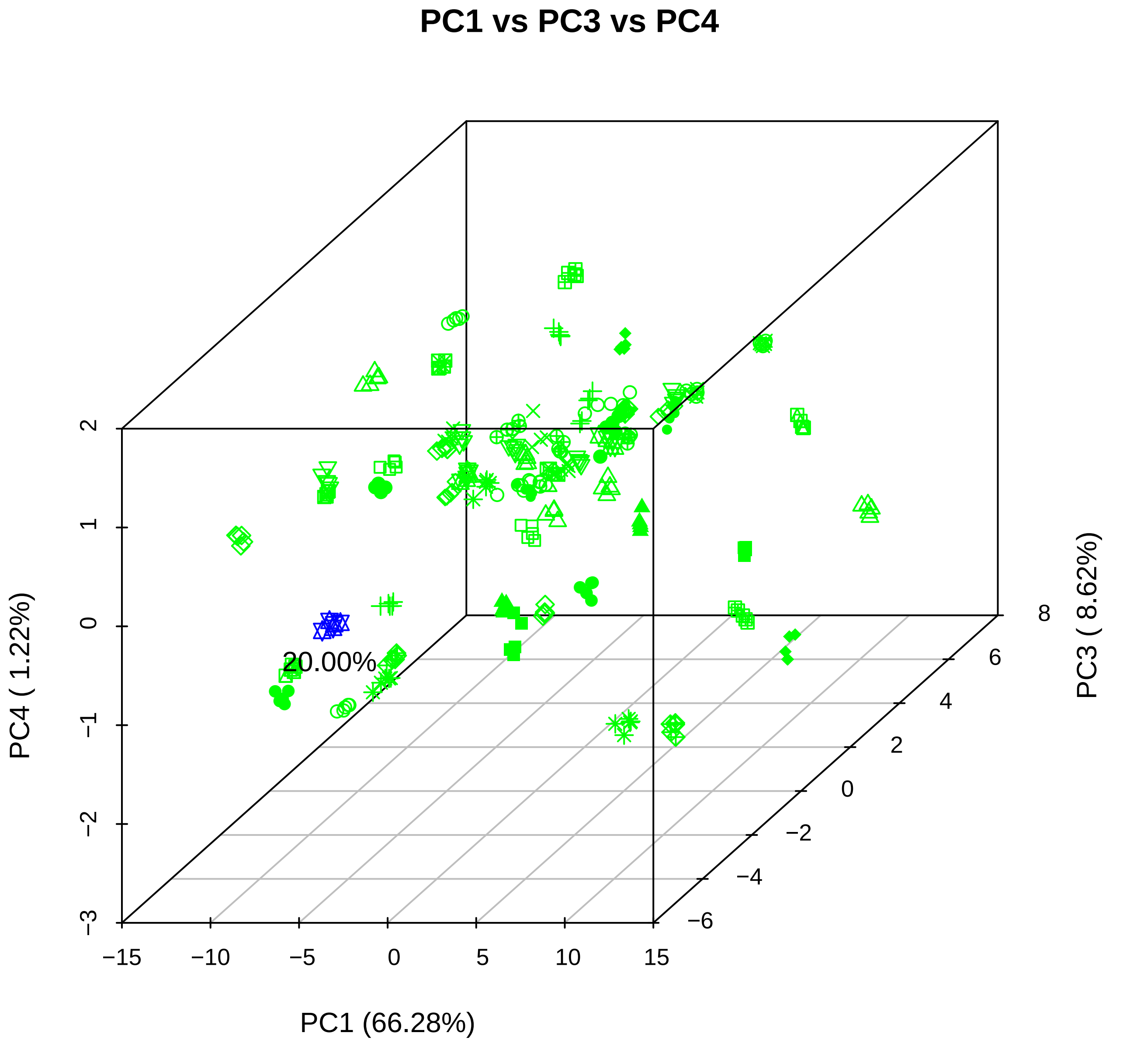

PC1 vs PC3 vs PC4
PC1 (66.28%)
PC4 ( 1.22%)
PC3 ( 8.62%)
20.00%
−15
−10
−5
0
5
10
15
−3
−2
−1
0
1
2
−6
−4
−2
0
2
4
6
8

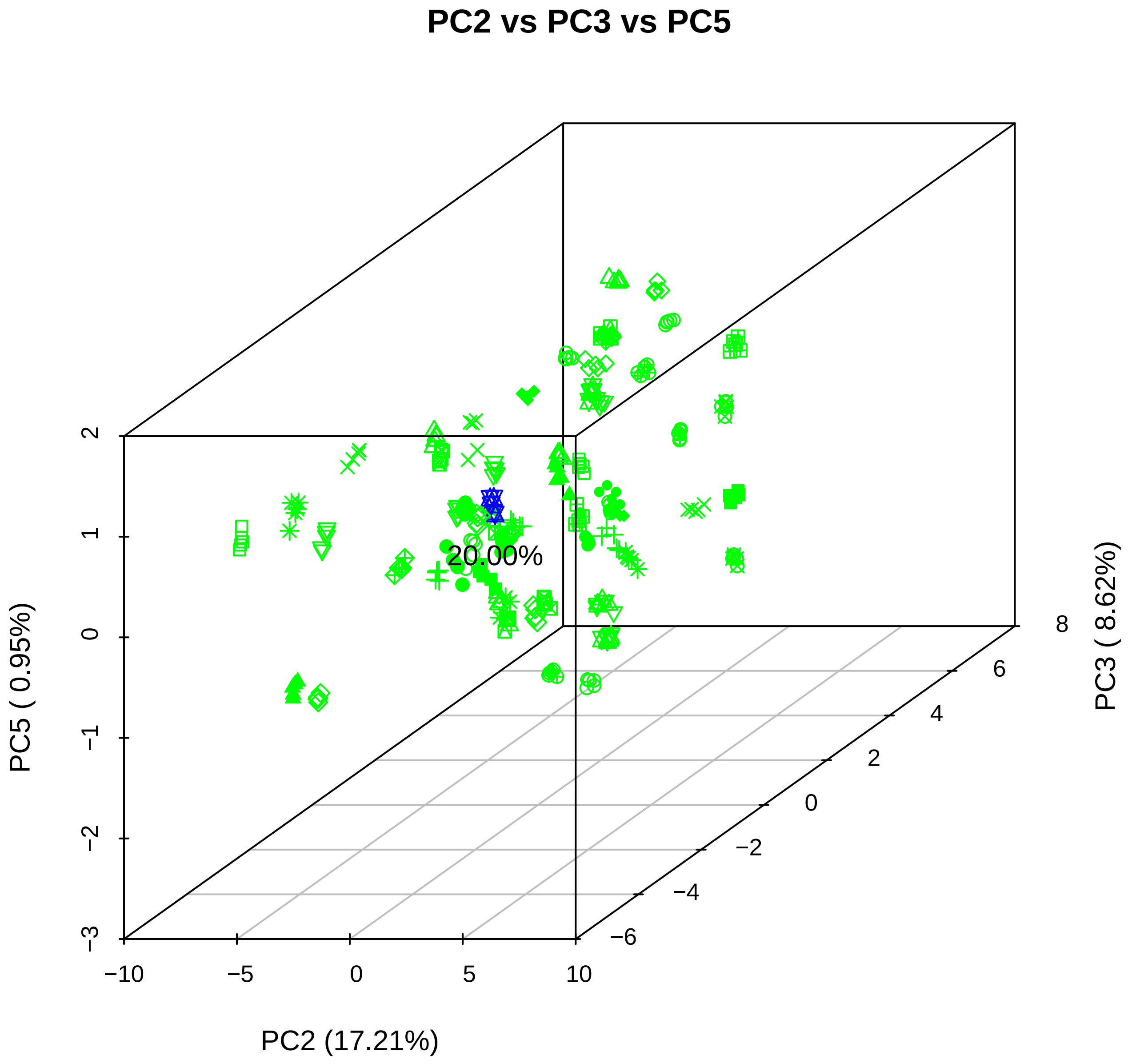

PC2 vs PC3 vs PC5
20.00%
PC2 (17.21%)
PC5 ( 0.95%)
PC3 ( 8.62%)
−10
−5
0
5
10
−3
−2
−1
0
1
2
−6
−4
−2
0
2
4
6
8

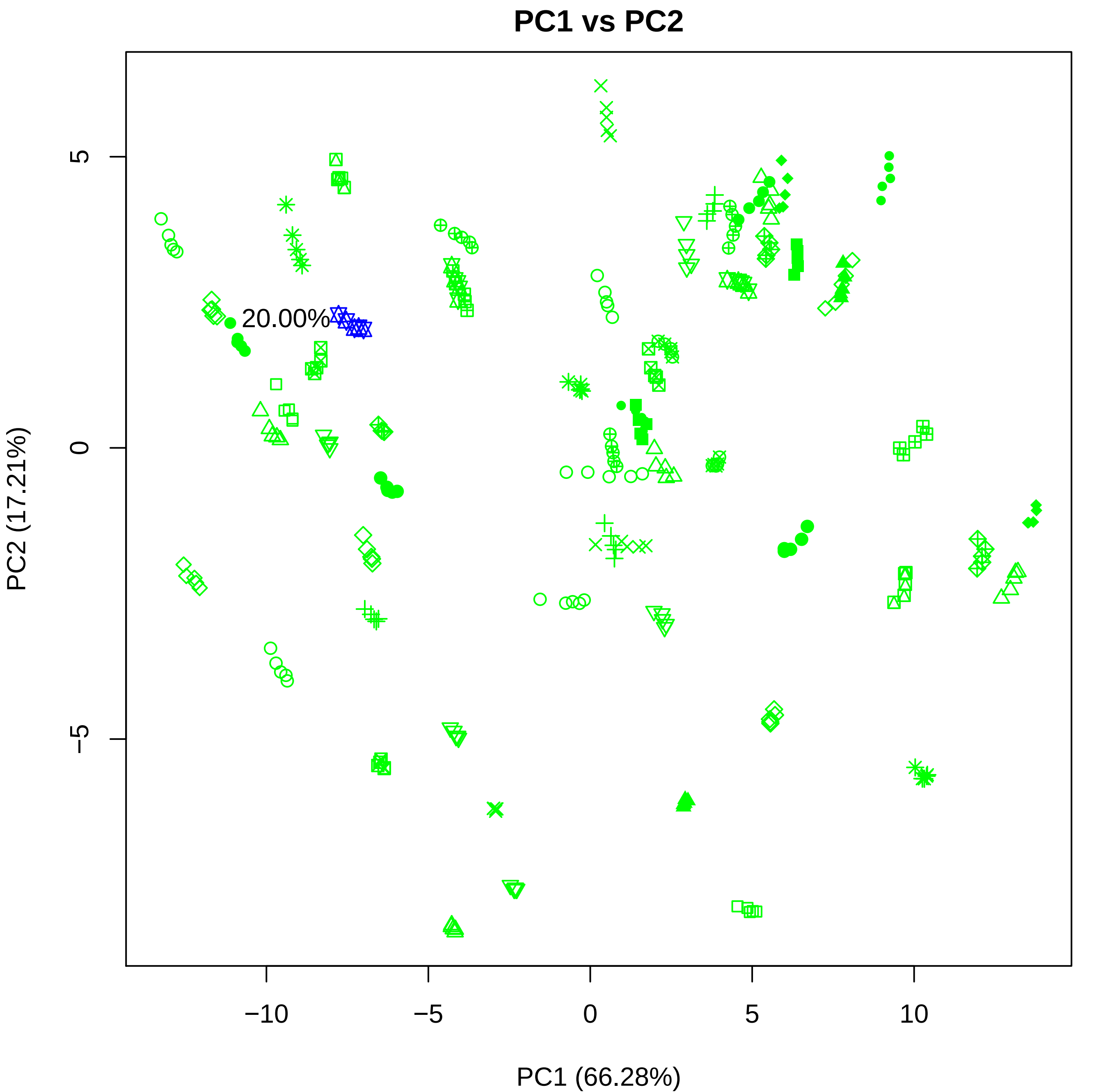

PC1 vs PC2
20.00%
PC2 (17.21%)
PC1 (66.28%)
5
0
−5
−10
−5
0
5
10

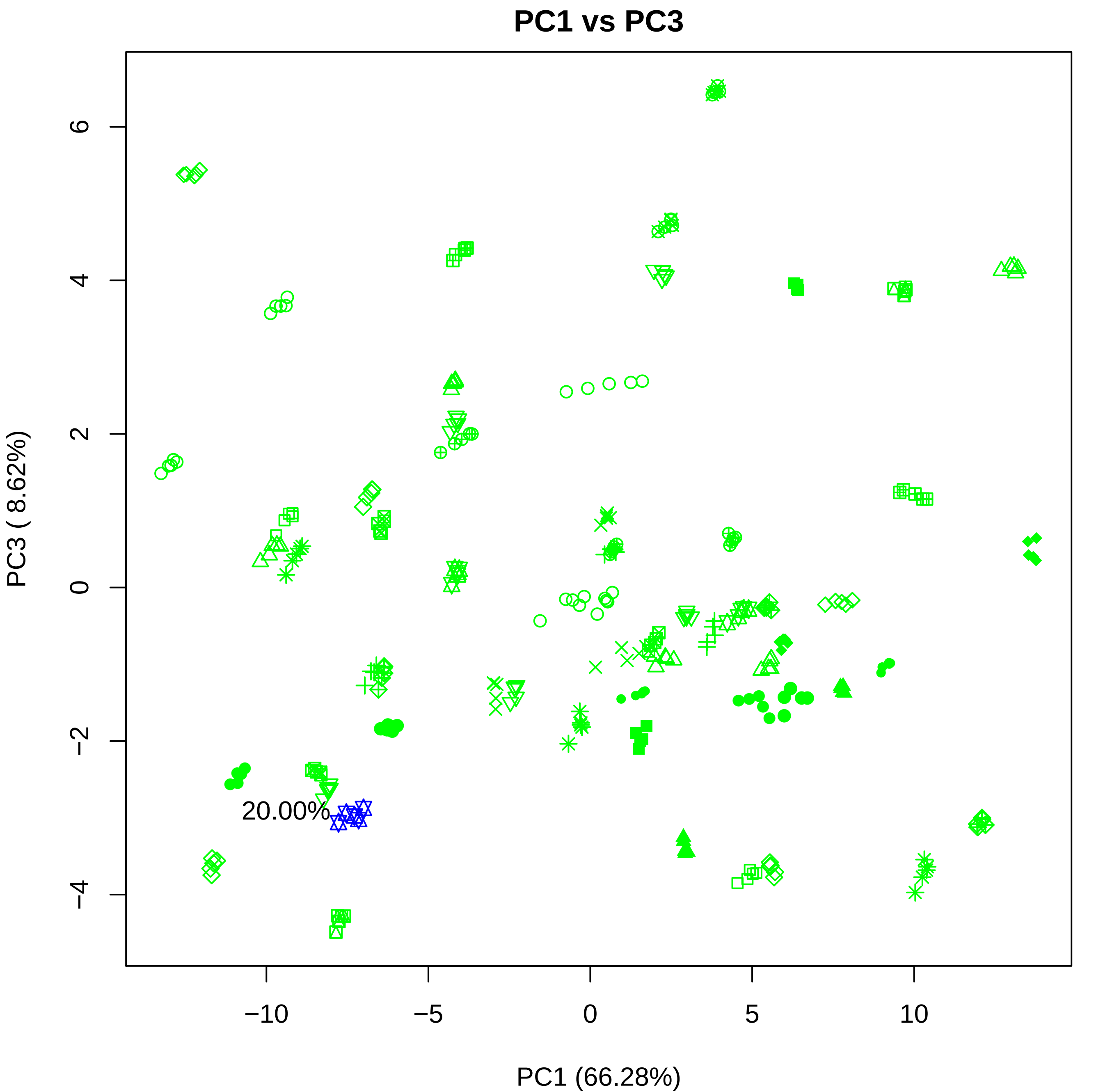
PC1 vs PC3
PC3 ( 8.62%)
PC1 (66.28%)
20.00%
−10
−5
0
5
10
−4
−2
0
2
4
6

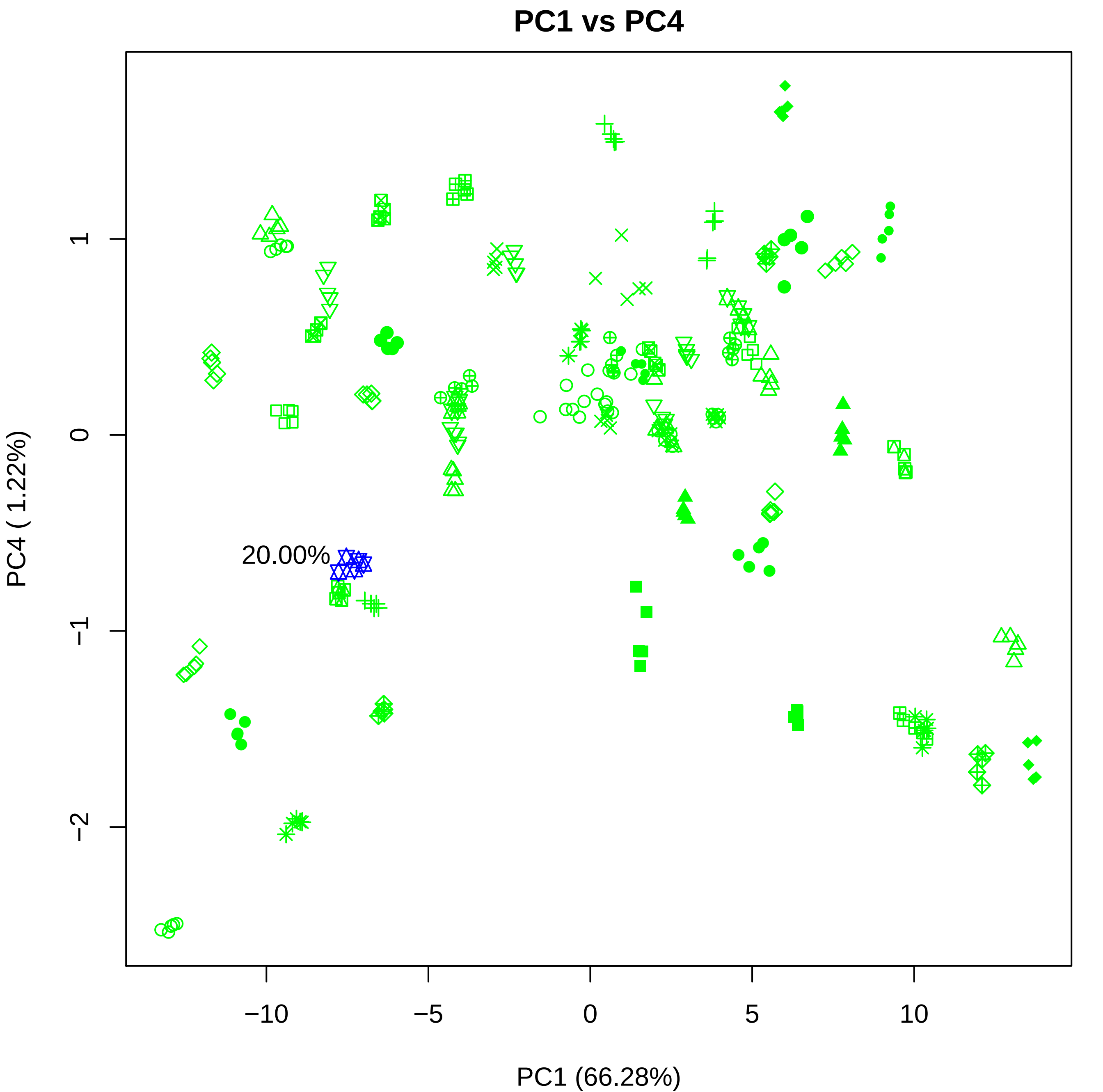
PC1 vs PC4
20.00%
1
0
−1
−2
PC4 ( 1.22%)
−10
−5
0
5
10
PC1 (66.28%)

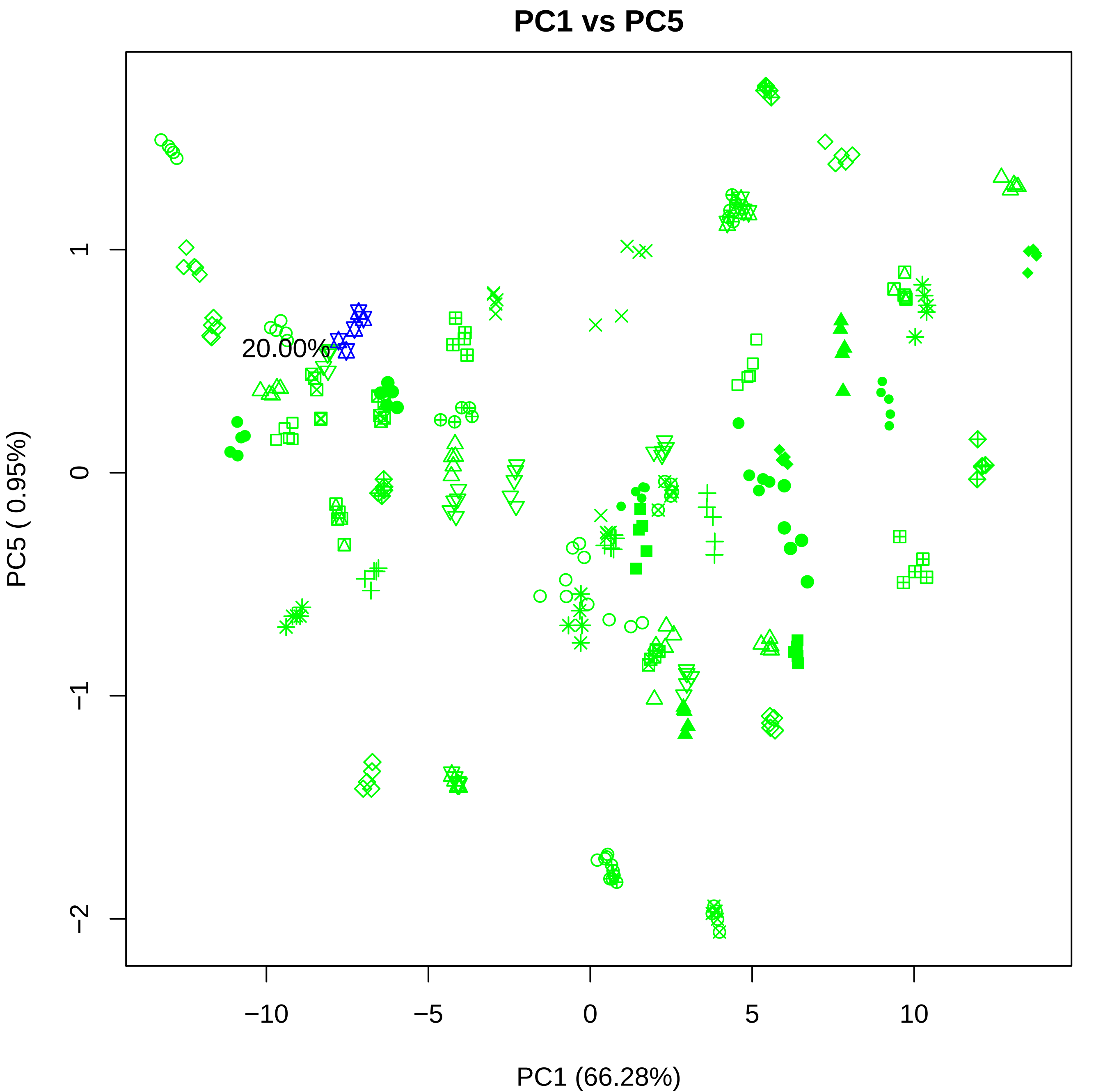

PC1 vs PC5
20.00%
1
0
−1
−2
PC5 ( 0.95%)
−10
−5
0
5
10
PC1 (66.28%)

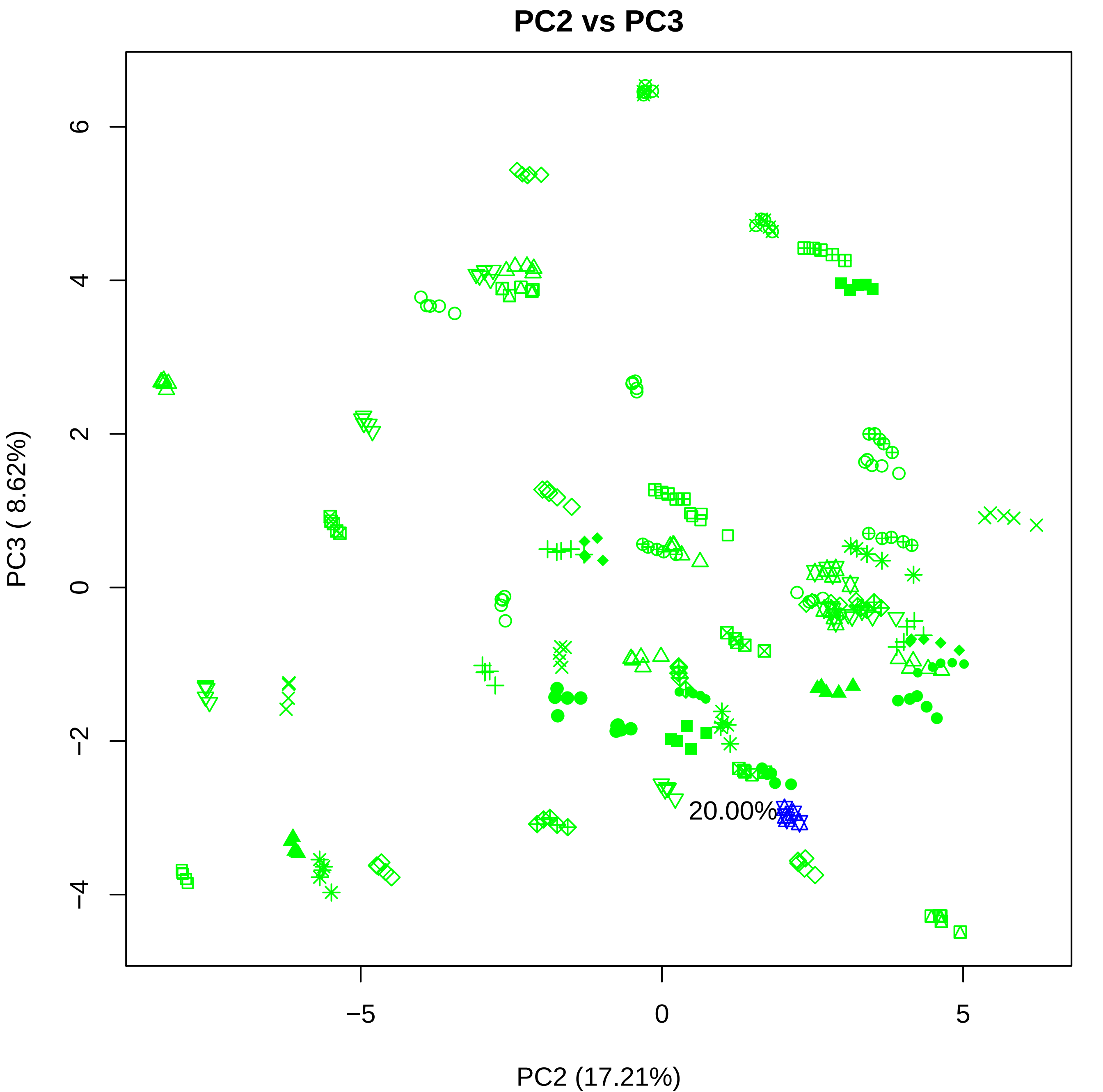
PC2 vs PC3
PC3 ( 8.62%)
6
4
2
0
−2
−4
−5
0
5
20.00%
PC2 (17.21%)

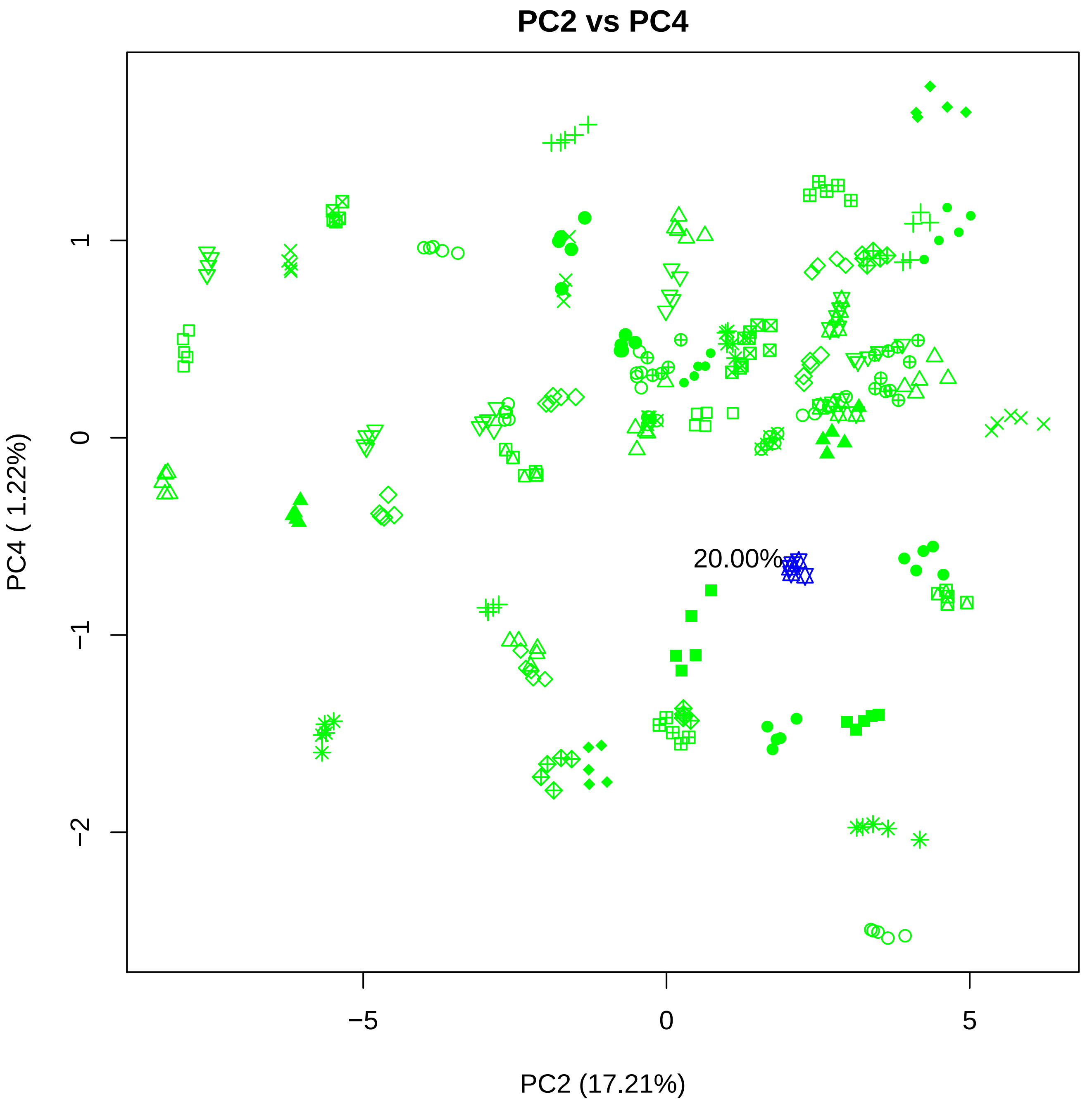

PC2 vs PC4
20.00%
PC4 ( 1.22%)
PC2 (17.21%)
1
0
−1
−2
−5
0
5

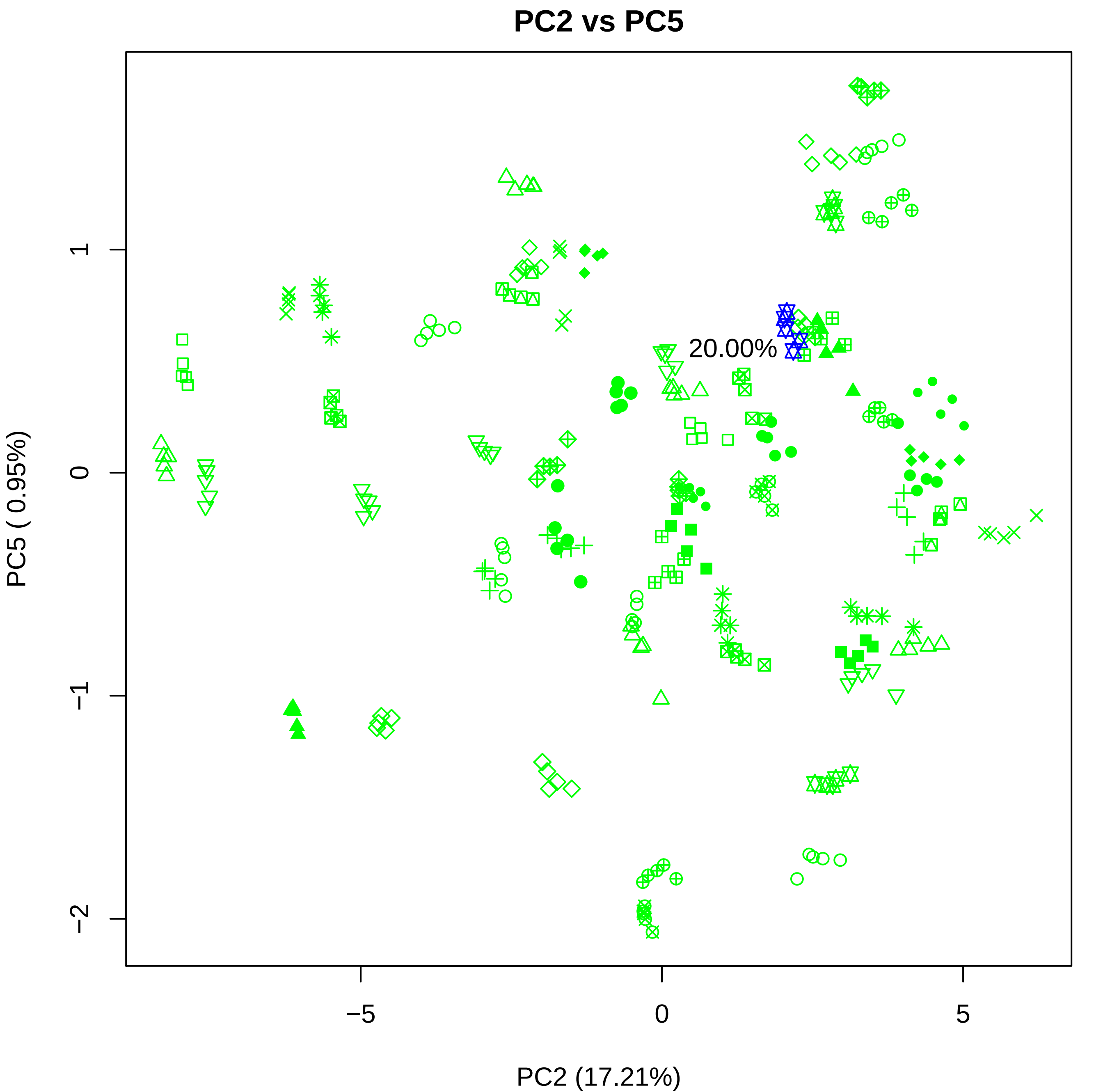

PC2 vs PC5
20.00%
PC5 ( 0.95%)
PC2 (17.21%)
1
0
−1
−2
−5
0
5

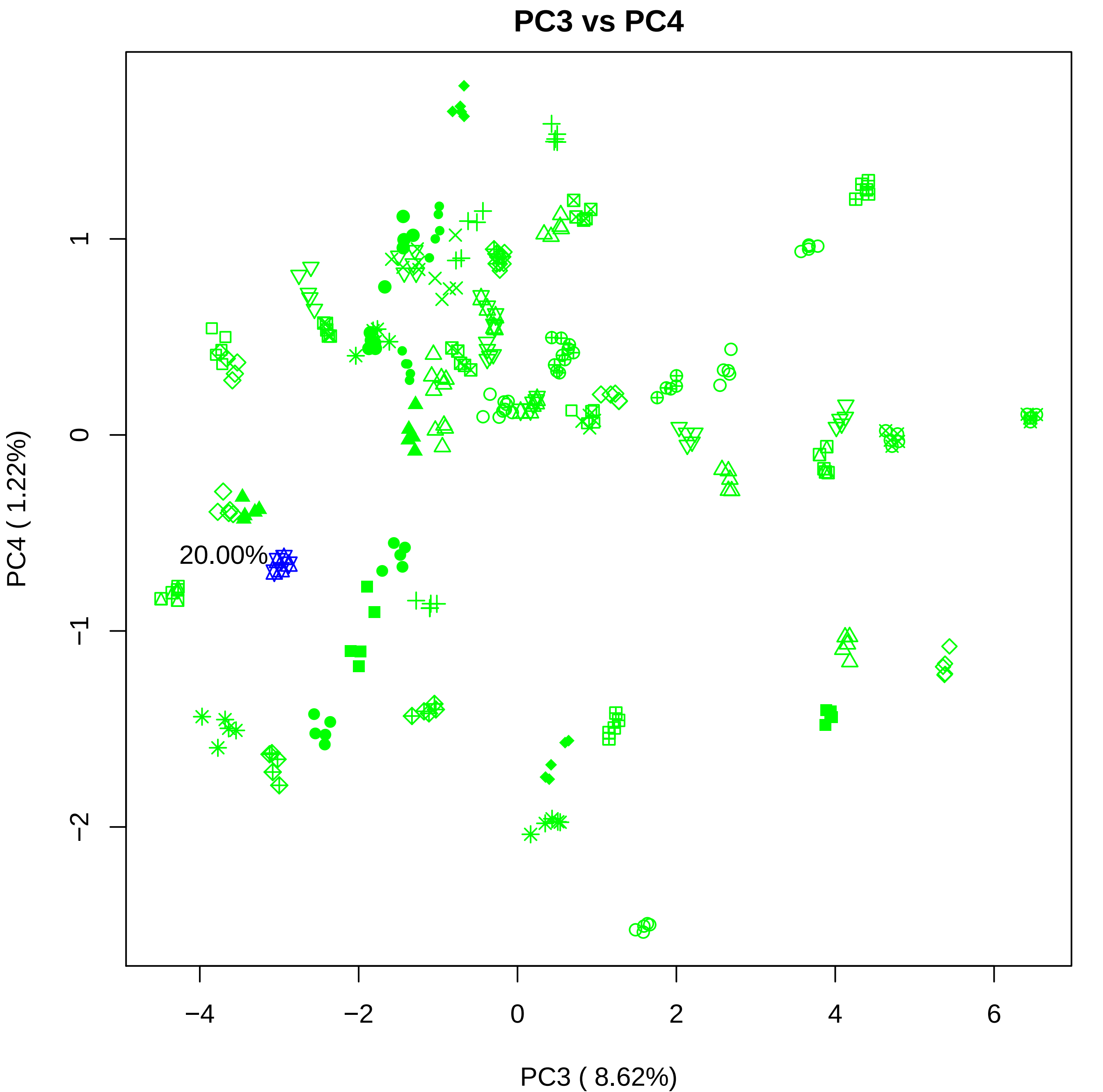
PC3 vs PC4
20.00%
PC4 ( 1.22%)
PC3 ( 8.62%)
1
0
−1
−2
−4
−2
0
2
4
6

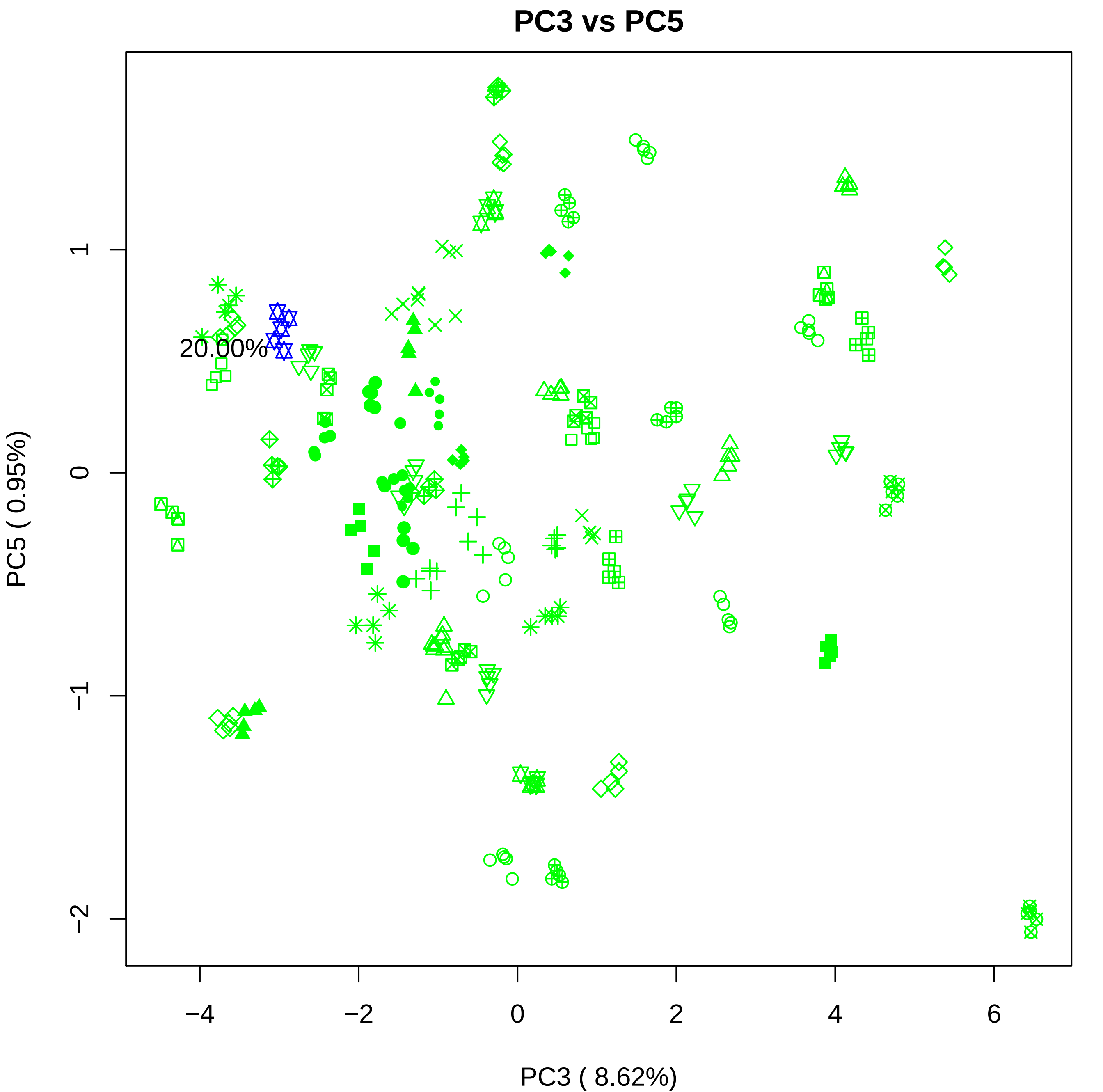
PC3 vs PC5
20.00%
PC5 ( 0.95%)
PC3 ( 8.62%)
−4
−2
0
2
4
6
−2
−1
0
1

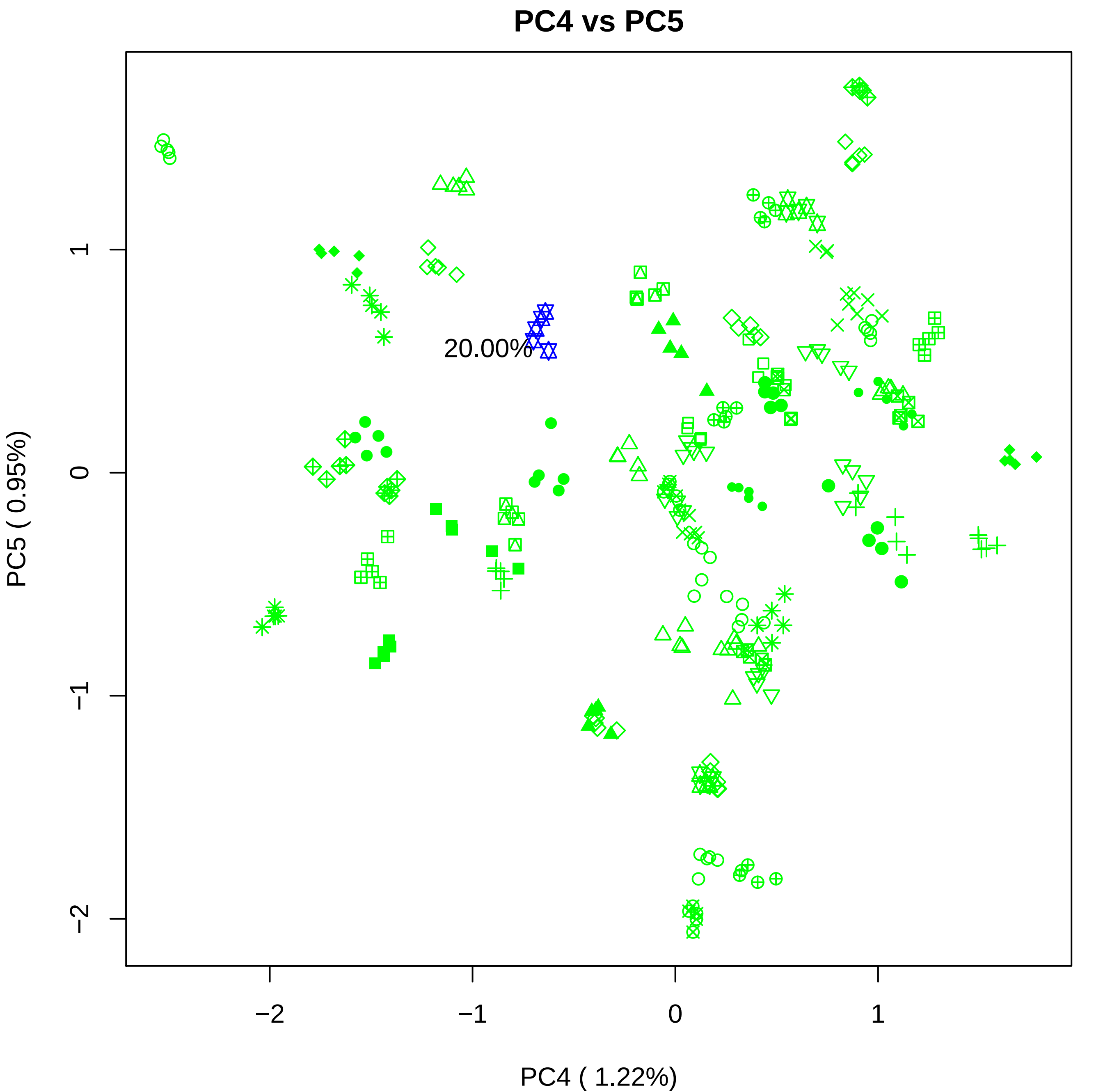
PC4 vs PC5
20.00%
PC5 ( 0.95%)
1
0
−1
−2
−2
−1
0
1
PC4 ( 1.22%)

### PCA of 40% BMH(a) vs PMH(a)

Scaled Matrix PCAed

7th order polynomial baseline correction performed

Raw spectra smoothed to 21 Neighbours, 2nd order PN, Savtisky Golay

Normalisation between 0 and 100

D2 spectra smoothed to 41 Neighbours, 2nd order PN, Savtisky Golay

Regions Analysed: 3601−2501 and 2001−700cm−1

Samples Analysed: 1 bContam's 0 BLH's 0 BMH's 0 OMH's 0 pContam's 0 PLH's 60 PMH's

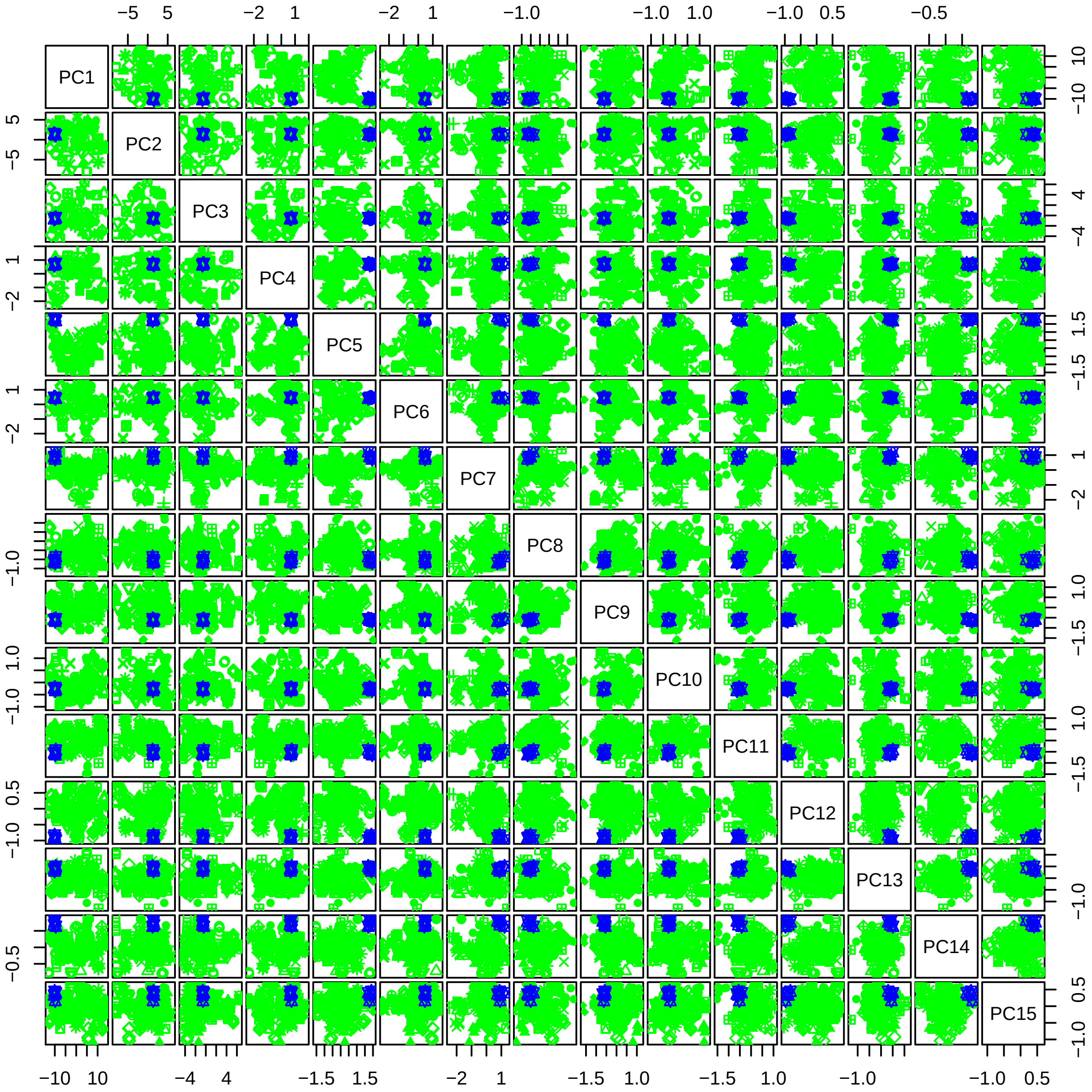
PC1
PC2
PC3
PC4
PC5
PC6
PC7
PC8
PC9
PC10
PC11
PC12
PC13
PC14
PC15

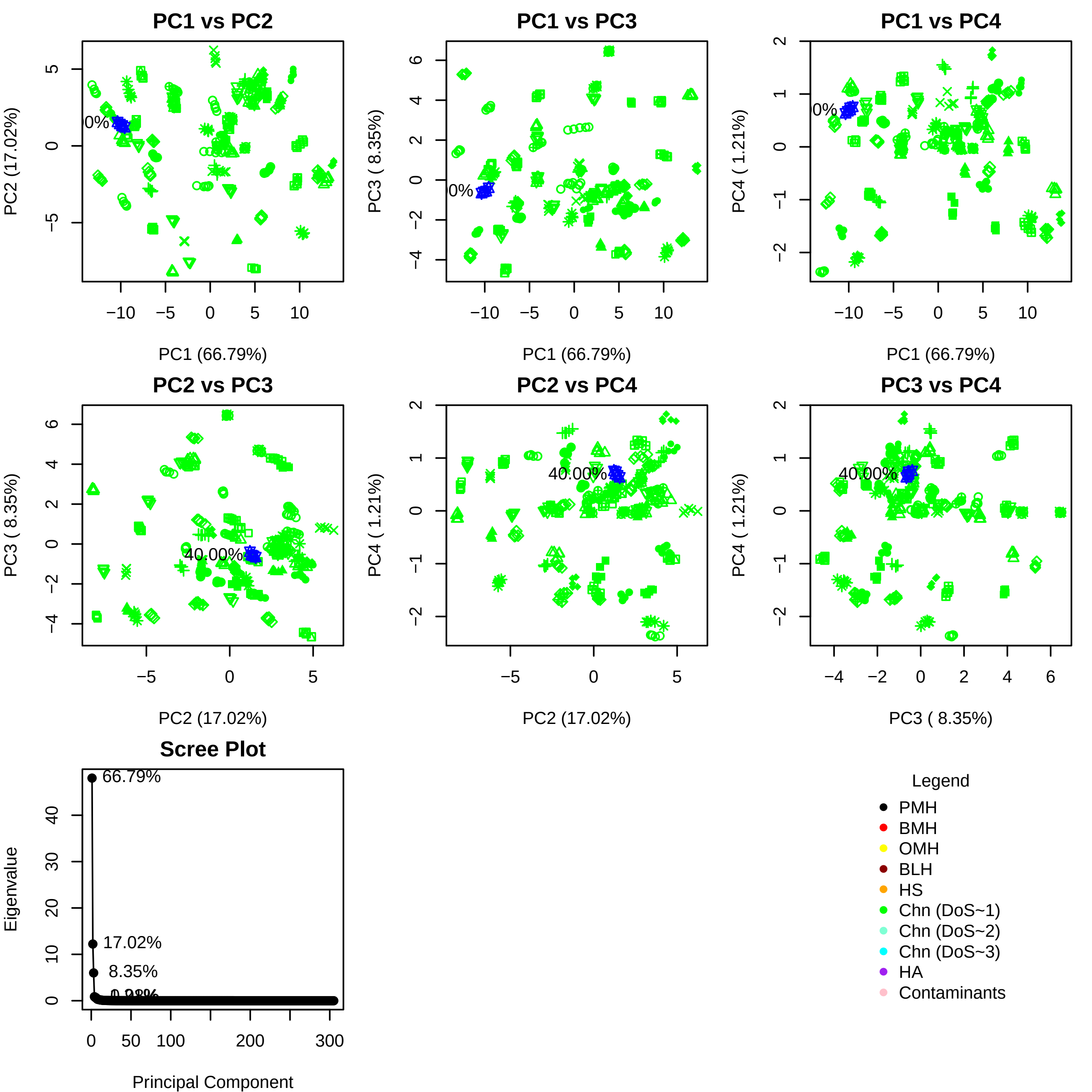

PC1 vs PC2
PC1 vs PC3
PC1 vs PC4
PC2 vs PC3
PC2 vs PC4
PC3 vs PC4
Scree Plot
PC1 (66.79%)
PC2 (17.02%)
PC3 ( 8.35%)
PC4 ( 1.21%)
0%
40.00%
66.79%
17.02%
8.35%
Eigenvalue
Principal Component
Legend
PMH
BMH
OMH
BLH
HS
Chn (DoS~1)
Chn (DoS~2)
Chn (DoS~3)
HA
Contaminants

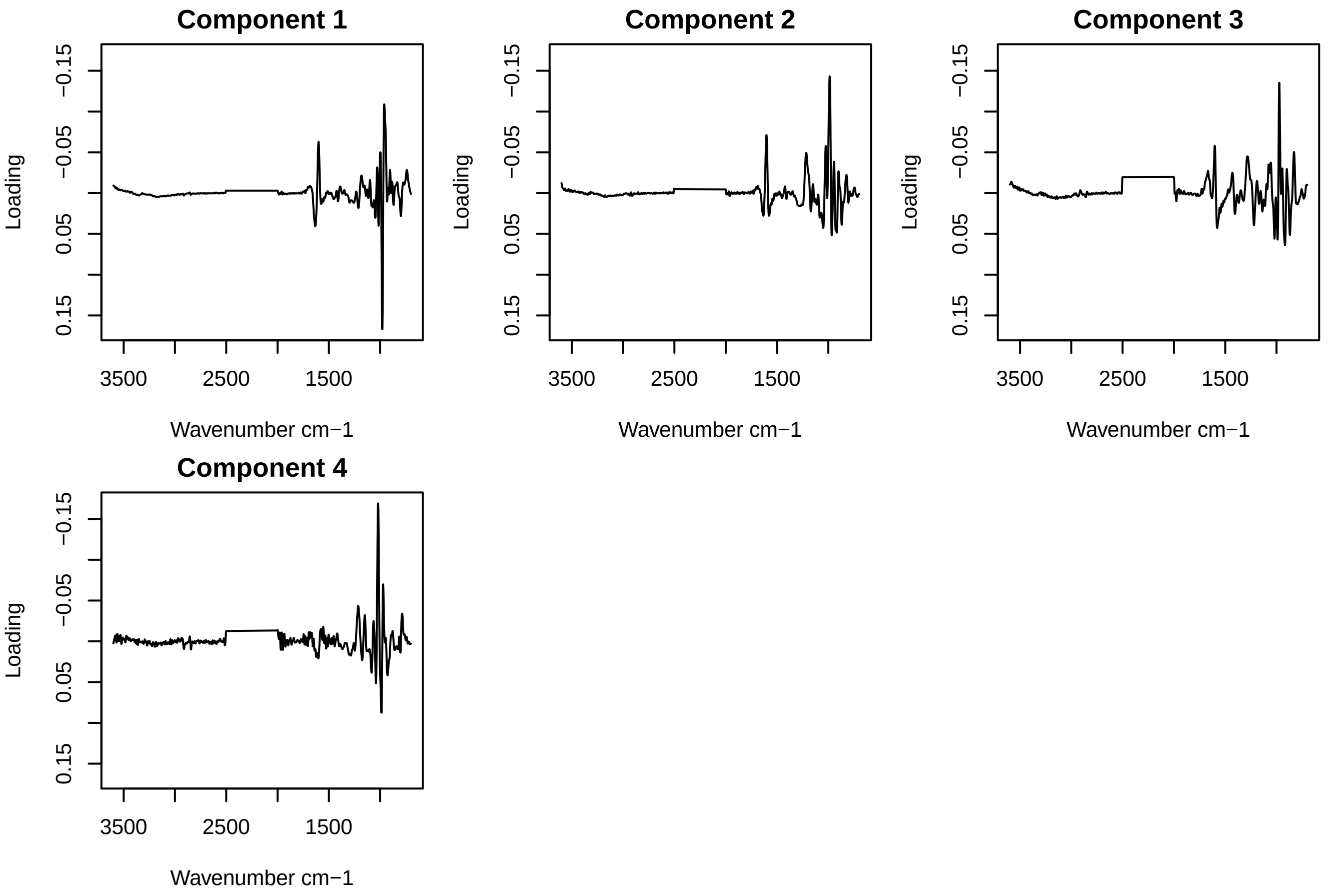
Component 1
Component 2
Component 3
Component 4
Loading
−0.15
−0.05
0.05
0.15
3500
2500
1500
Wavenumber cm−1

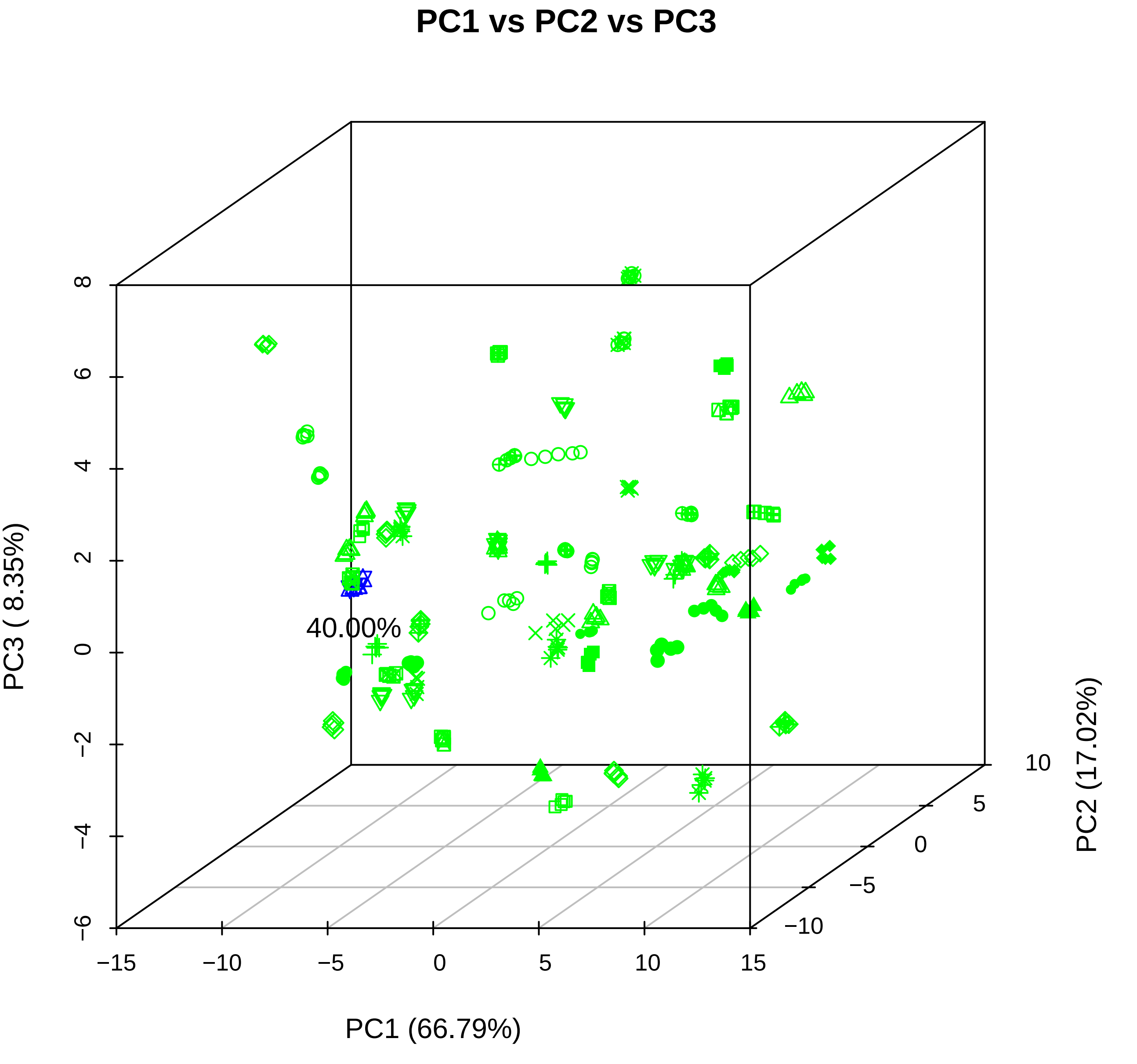

PC1 vs PC2 vs PC3
PC1 (66.79%)
PC2 (17.02%)
PC3 ( 8.35%)
40.00%
−15
−10
−5
0
5
10
15
−10
−5
0
5
10
−6
−4
−2
0
2
4
6
8

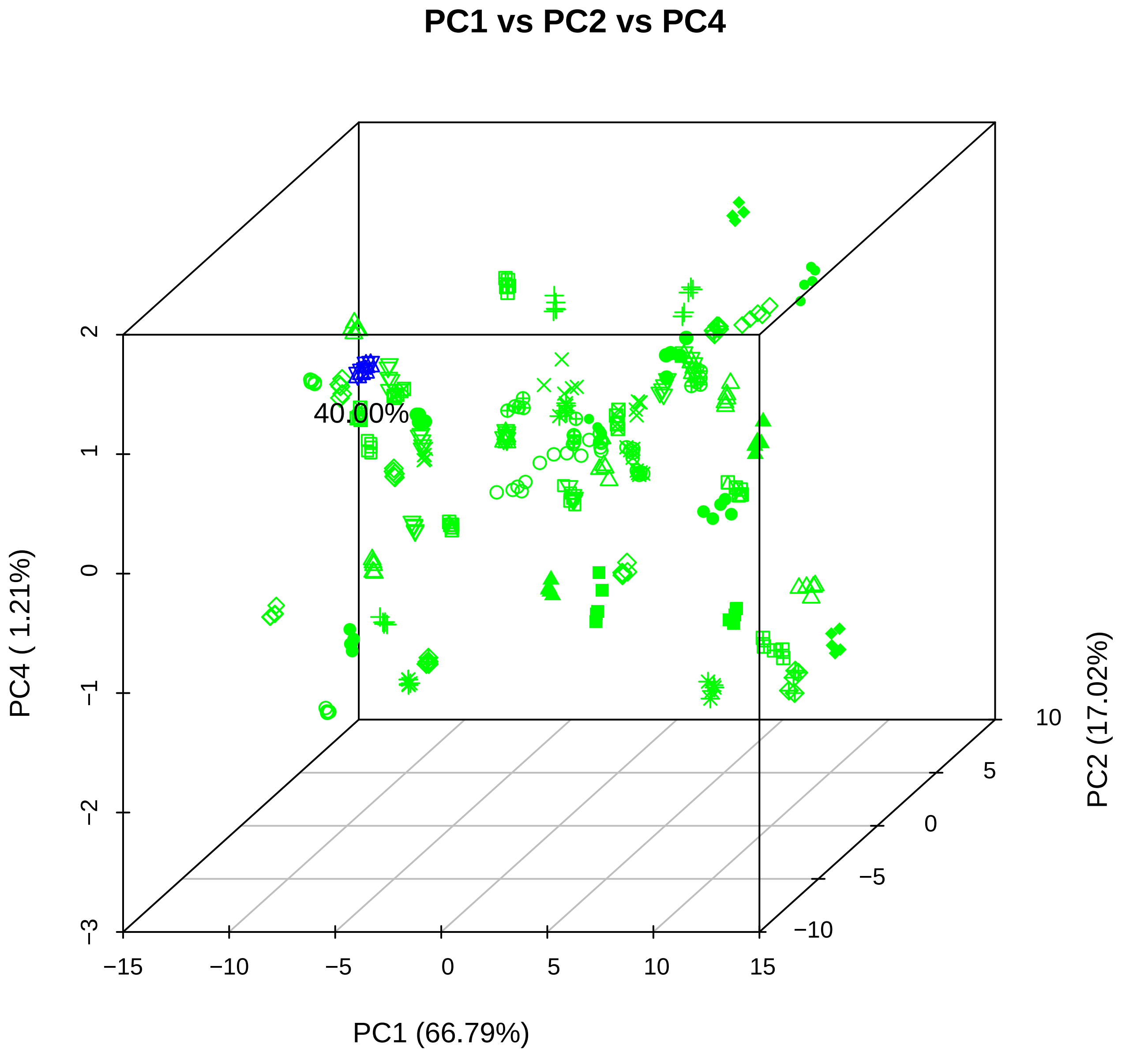
PC1 vs PC2 vs PC4
40.00%
PC1 (66.79%)
PC2 (17.02%)
PC4 ( 1.21%)
−15
−10
−5
0
5
10
15
10
5
0
−5
−10
2
1
0
−1
−2
−3

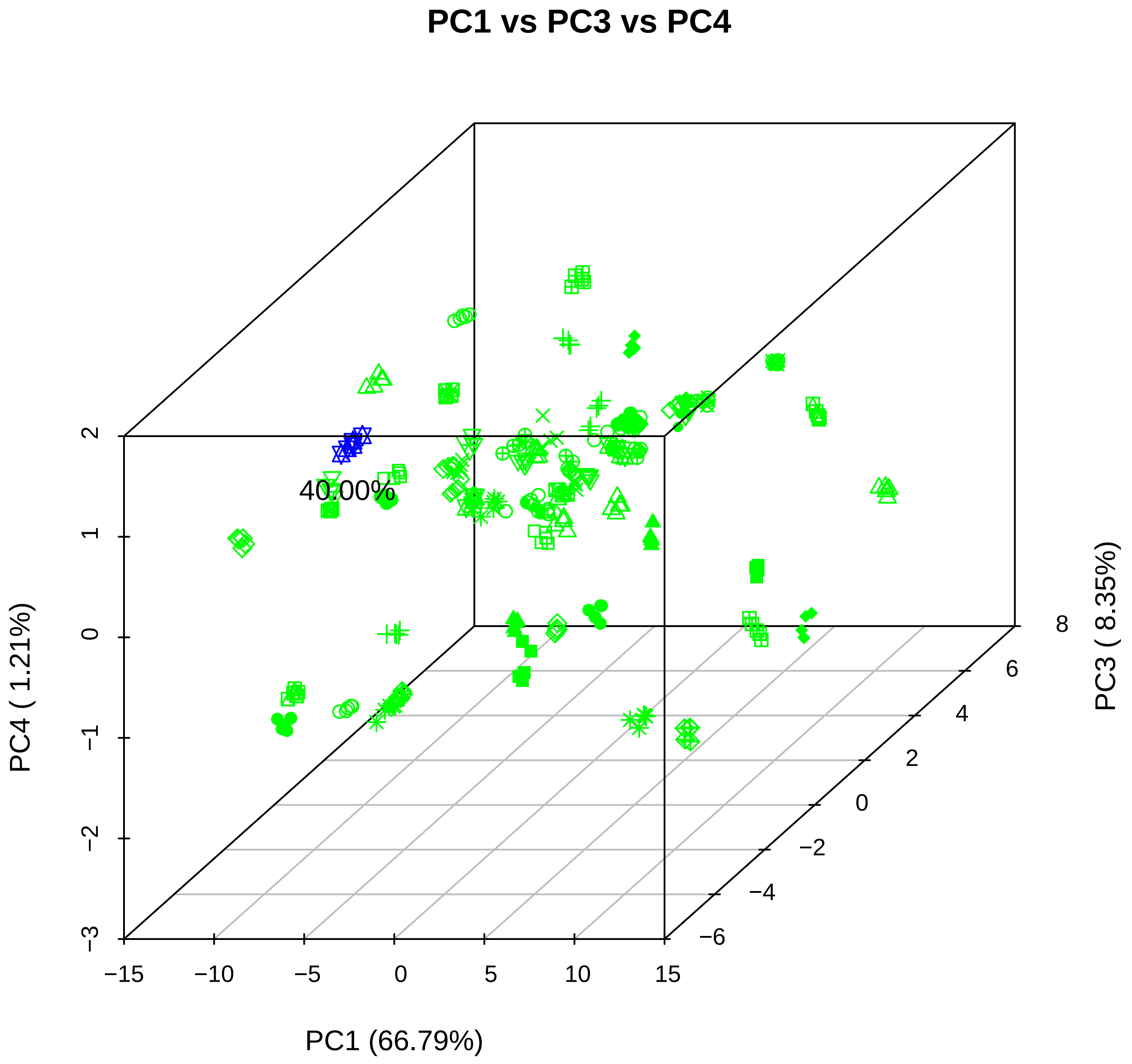
PC1 vs PC3 vs PC4
40.00%
PC1 (66.79%)
PC3 ( 8.35%)
PC4 ( 1.21%)
−15
−10
−5
0
5
10
15
−6
−4
−2
0
2
4
6
8
−3
−2
−1
0
1
2

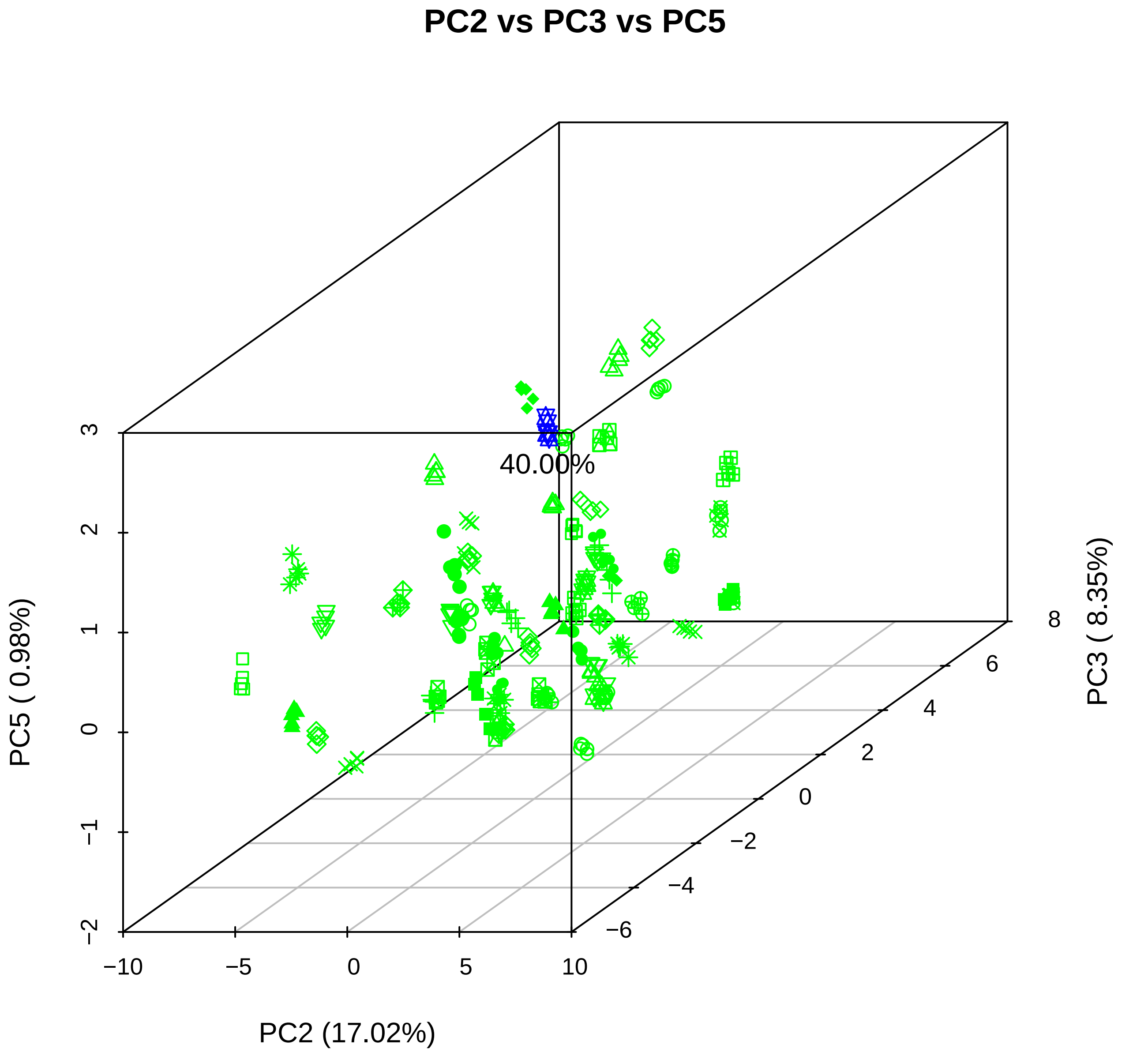
PC2 vs PC3 vs PC5
40.00%
PC2 (17.02%)
PC3 ( 8.35%)
PC5 ( 0.98%)
−10
−5
0
5
10
−6
−4
−2
0
2
4
6
8
−2
−1
0
1
2
3

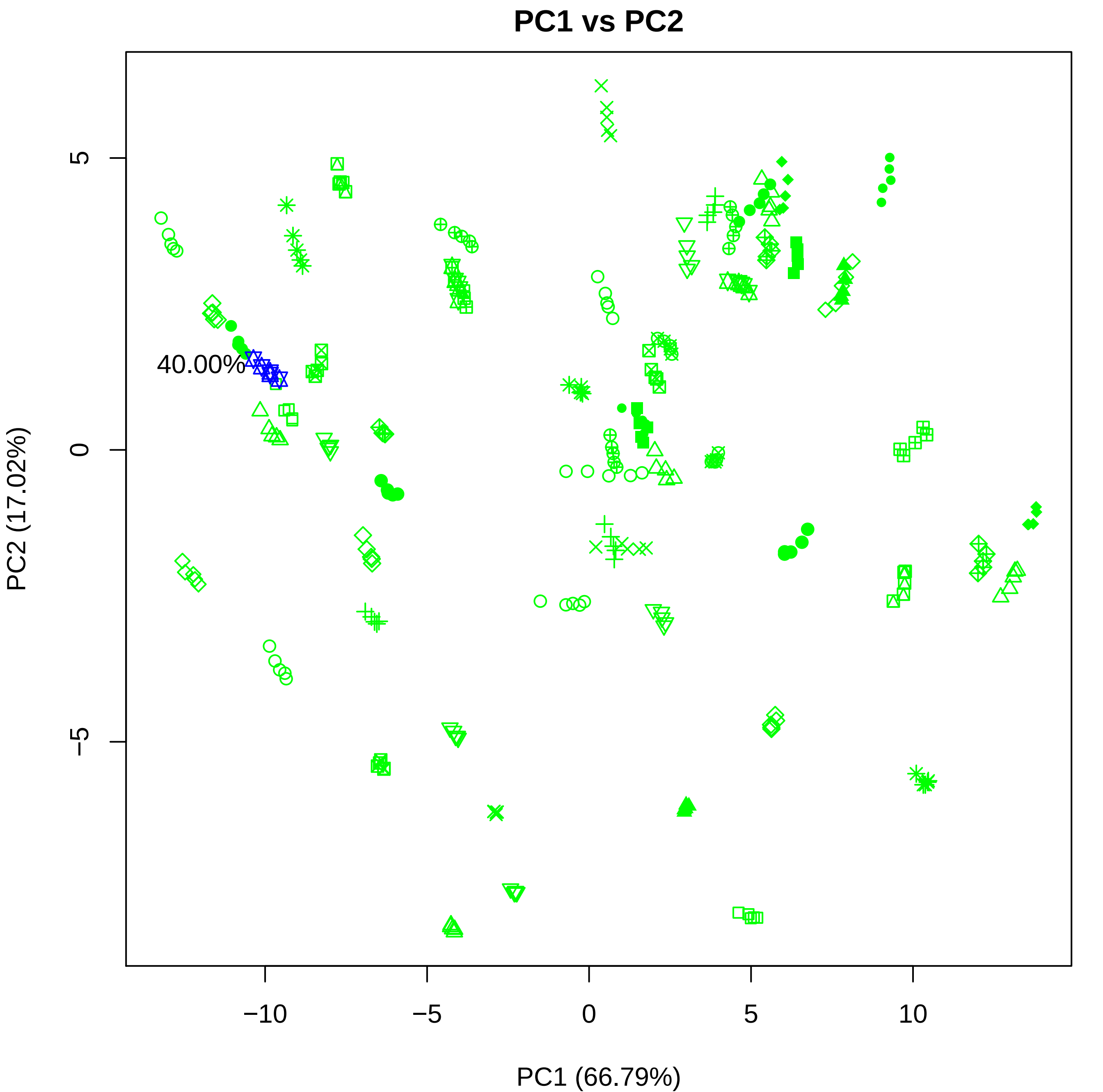
PC1 vs PC2
40.00%
PC1 (66.79%)
PC2 (17.02%)
−10
−5
0
5
10
5
0
−5

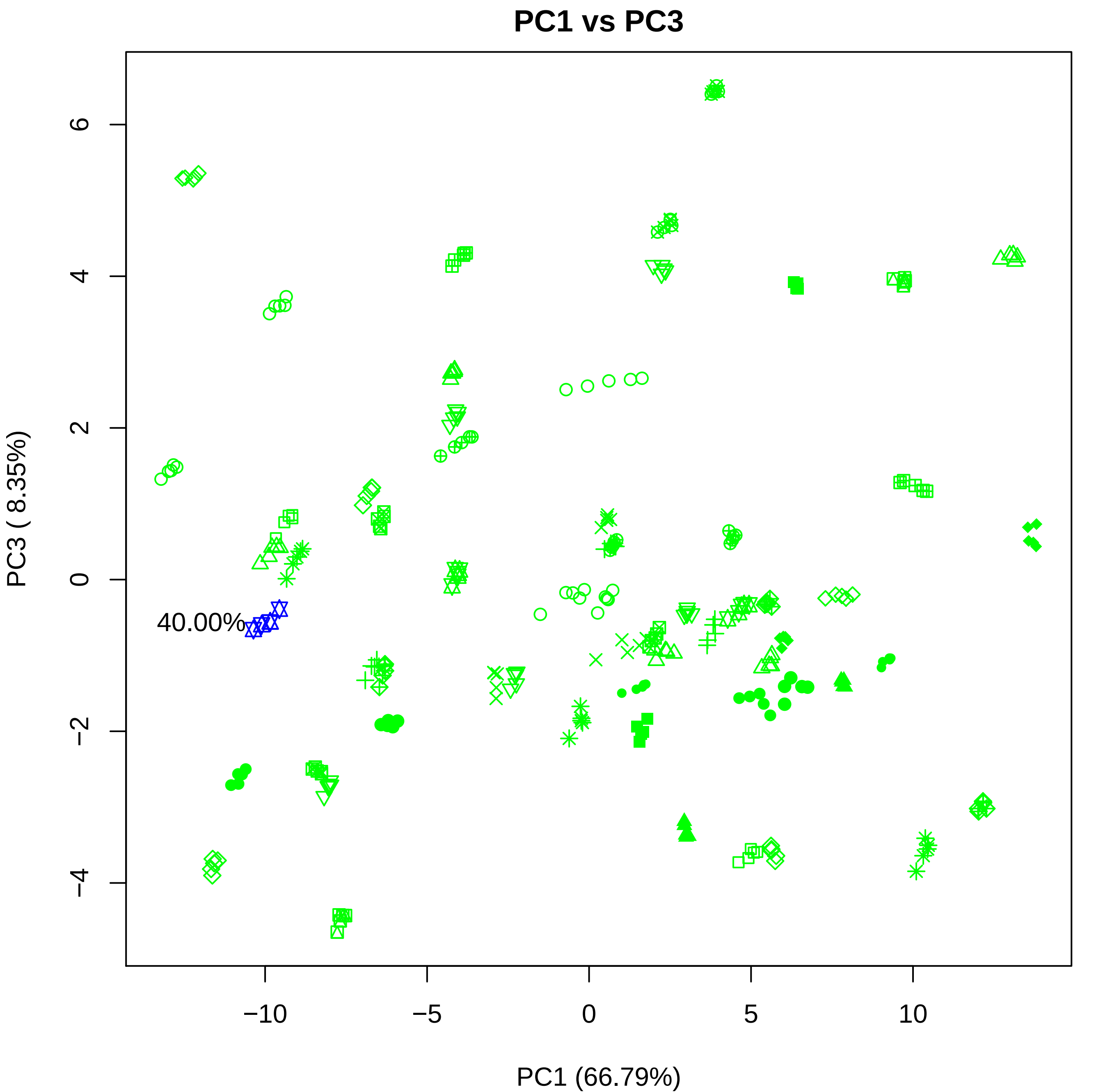
PC1 vs PC3
40.00%
PC3 ( 8.35%)
6
4
2
0
−2
−4
−10
−5
0
5
10
PC1 (66.79%)

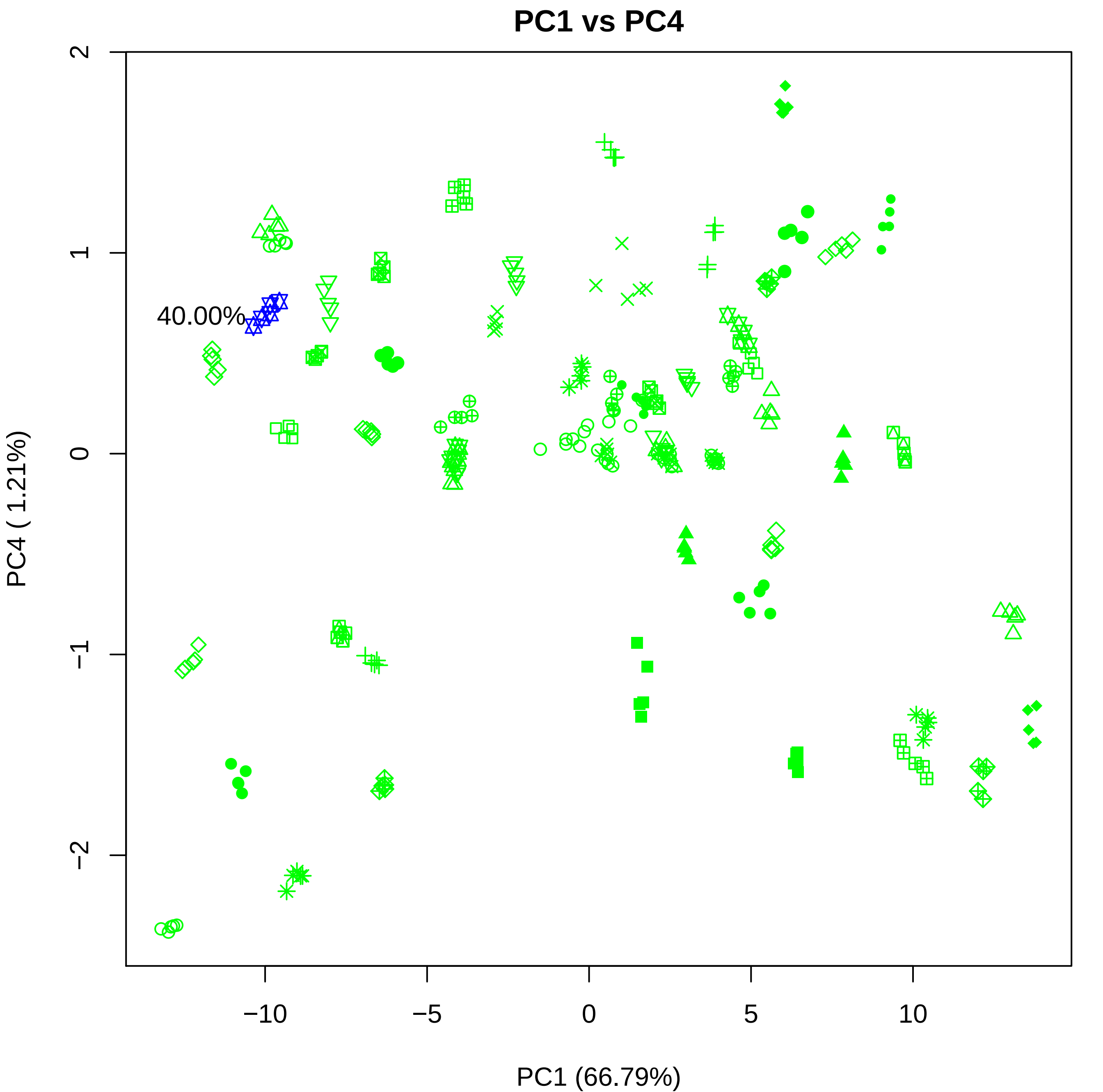
PC1 vs PC4
40.00%
PC4 ( 1.21%)
PC1 (66.79%)
2
1
0
−1
−2
−10
−5
0
5
10

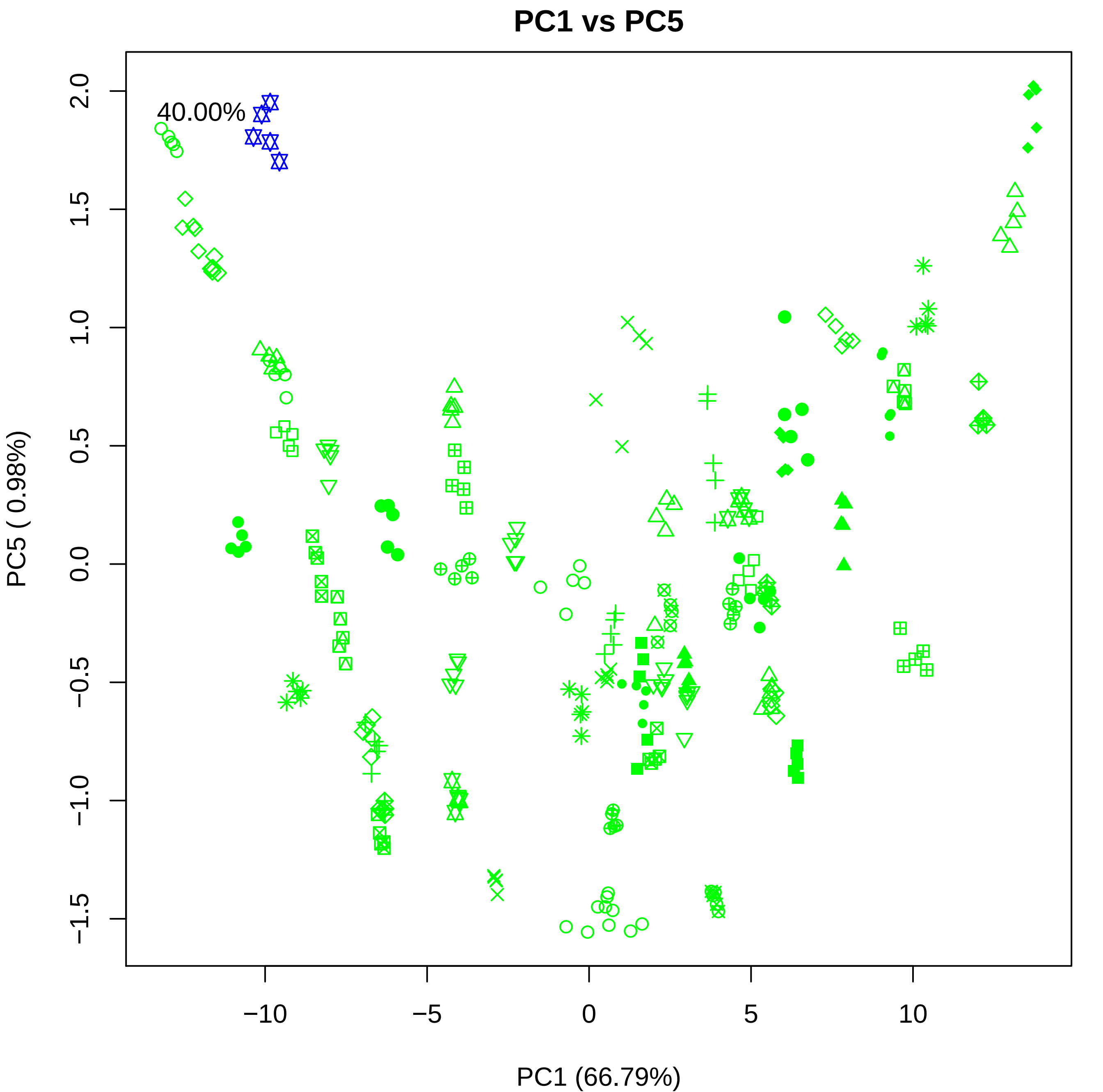
PC1 vs PC5
40.00%
PC5 ( 0.98%)
PC1 (66.79%)
2.0
1.5
1.0
0.5
0.0
−0.5
−1.0
−1.5
−10
−5
0
5
10

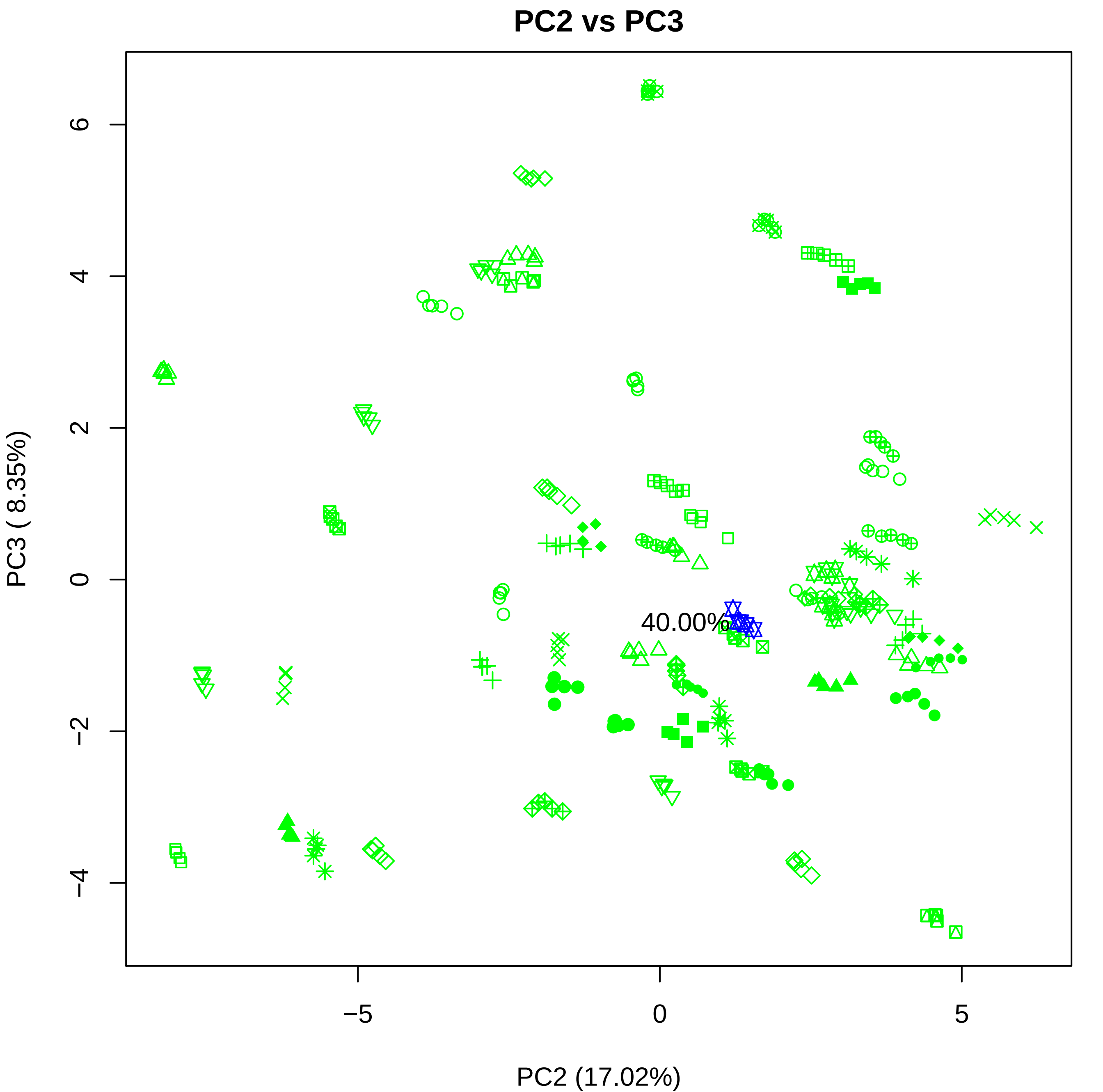

PC2 vs PC3
PC3 ( 8.35%)
PC2 (17.02%)
40.00%
6
4
2
0
−2
−4
−5
0
5

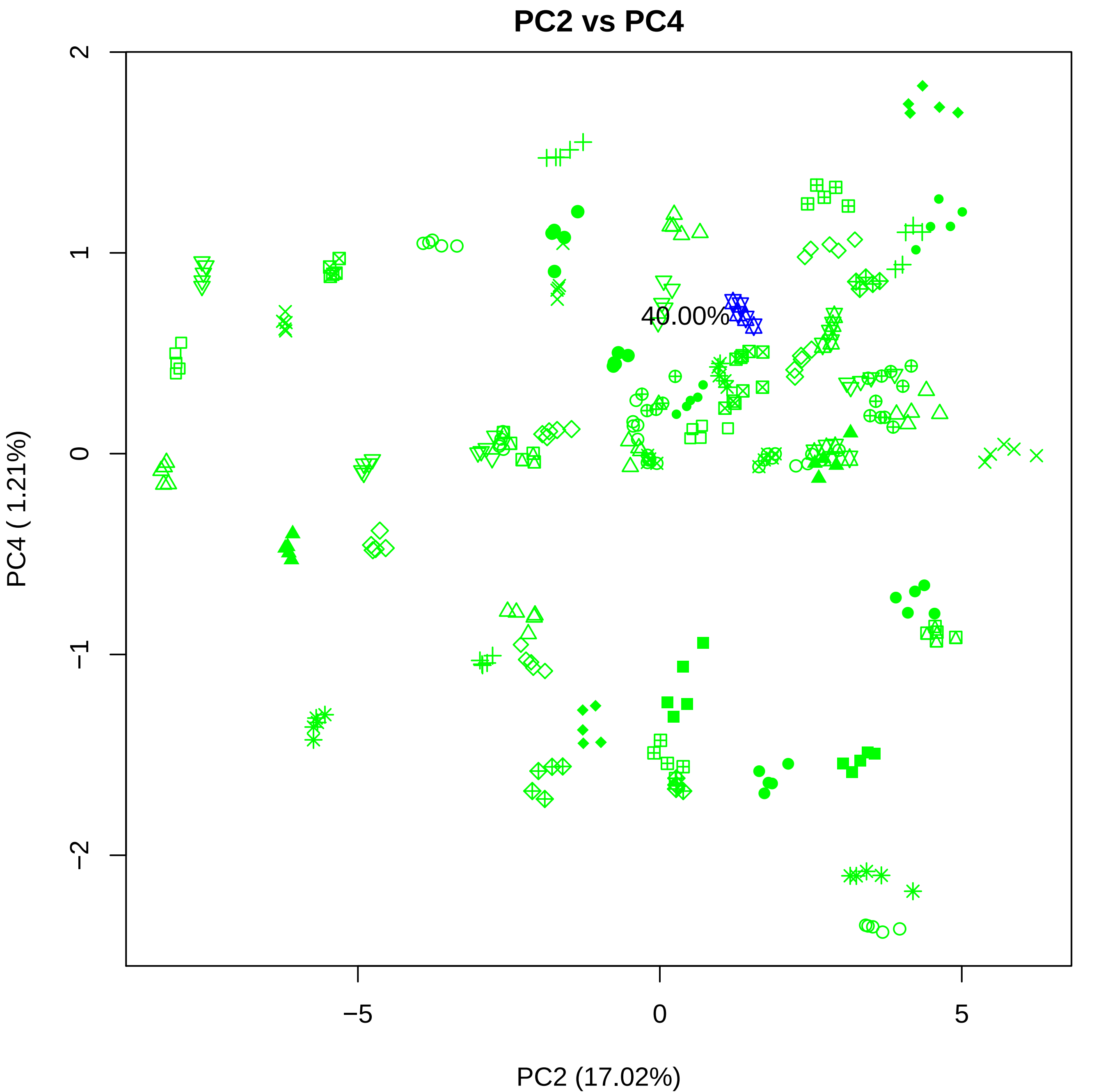
PC2 vs PC4
PC4 ( 1.21%)
PC2 (17.02%)
40.00%
2
1
0
−1
−2
−5
0
5

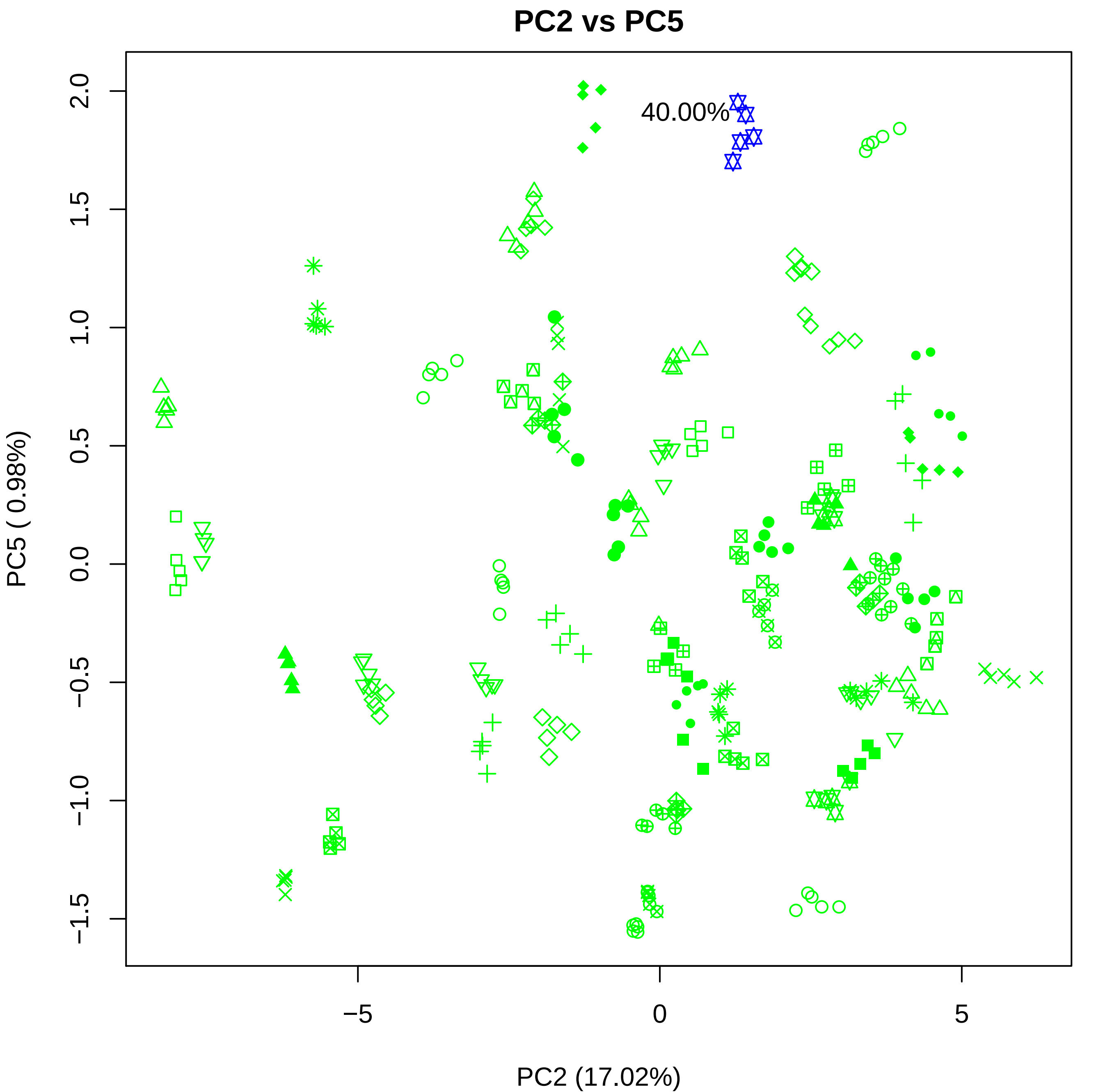
PC2 vs PC5
40.00%
PC5 ( 0.98%)
PC2 (17.02%)
2.0
1.5
1.0
0.5
0.0
−0.5
−1.0
−1.5
−5
0
5

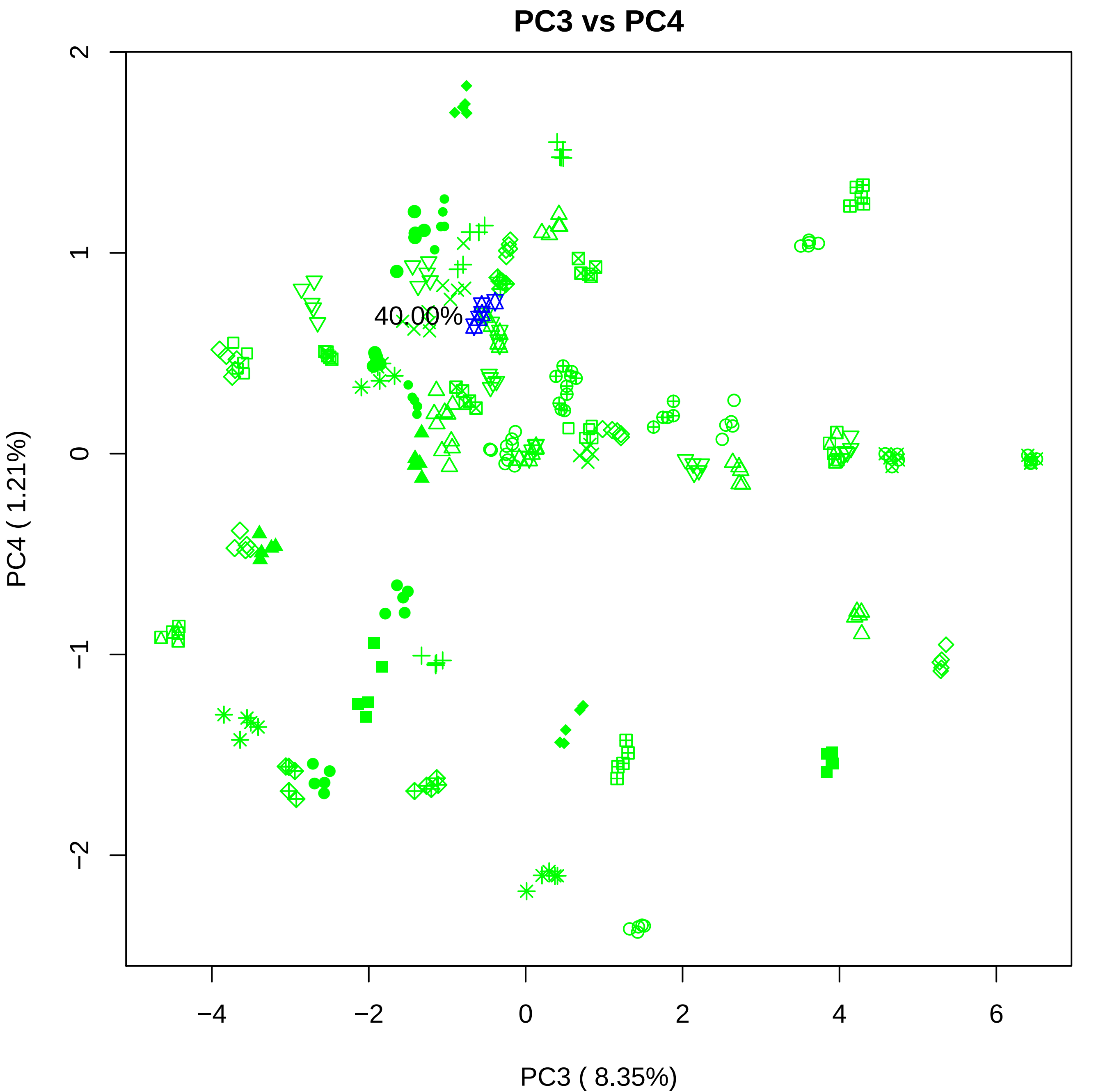
PC3 vs PC4
40.00%
PC3 ( 8.35%)
PC4 ( 1.21%)
−4
−2
0
2
4
6
−2
−1
0
1
2

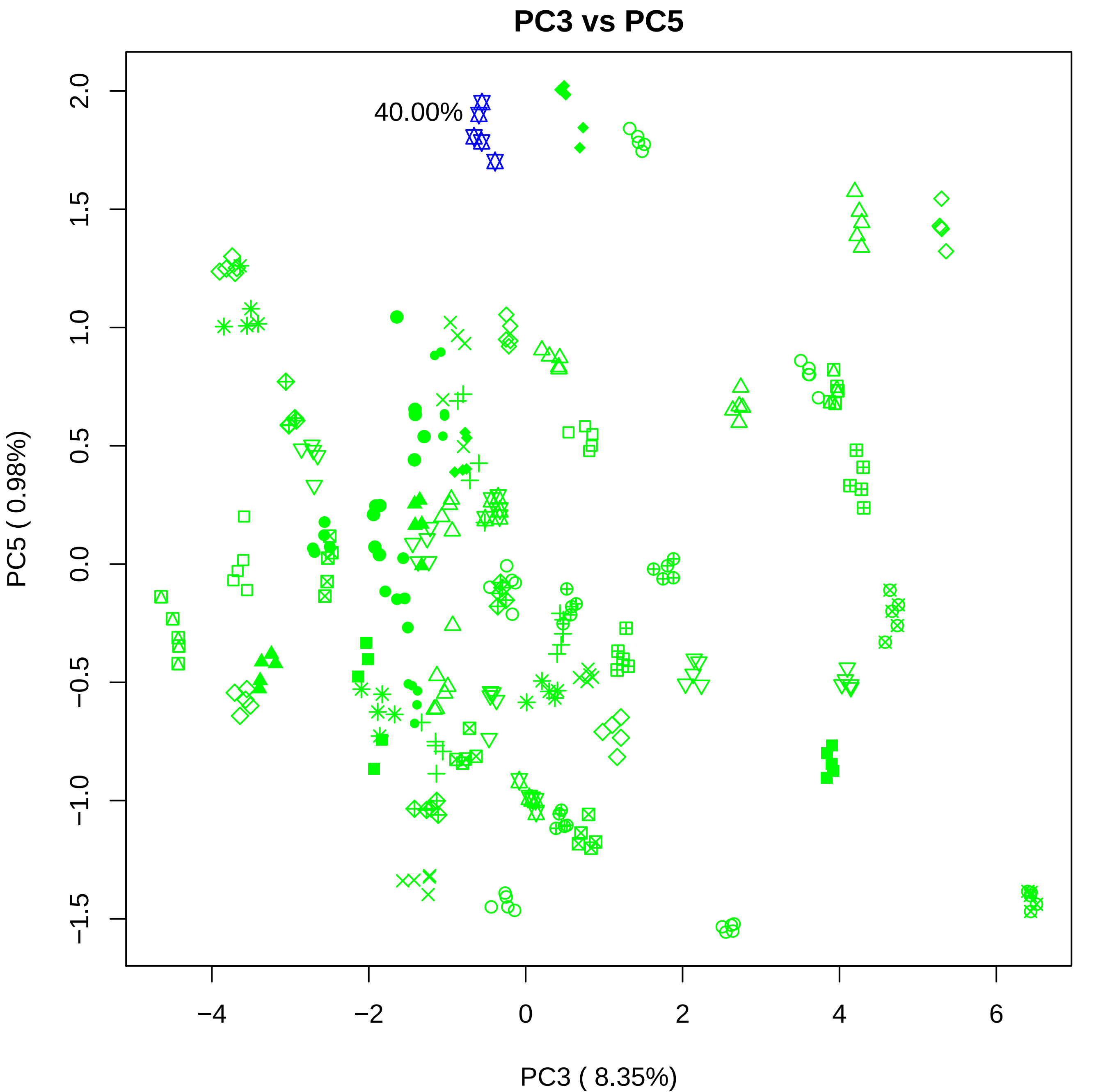
PC3 vs PC5
40.00%
2.0
1.5
1.0
0.5
0.0
−0.5
−1.0
−1.5
−4
−2
0
2
4
6
PC5 ( 0.98%)
PC3 ( 8.35%)

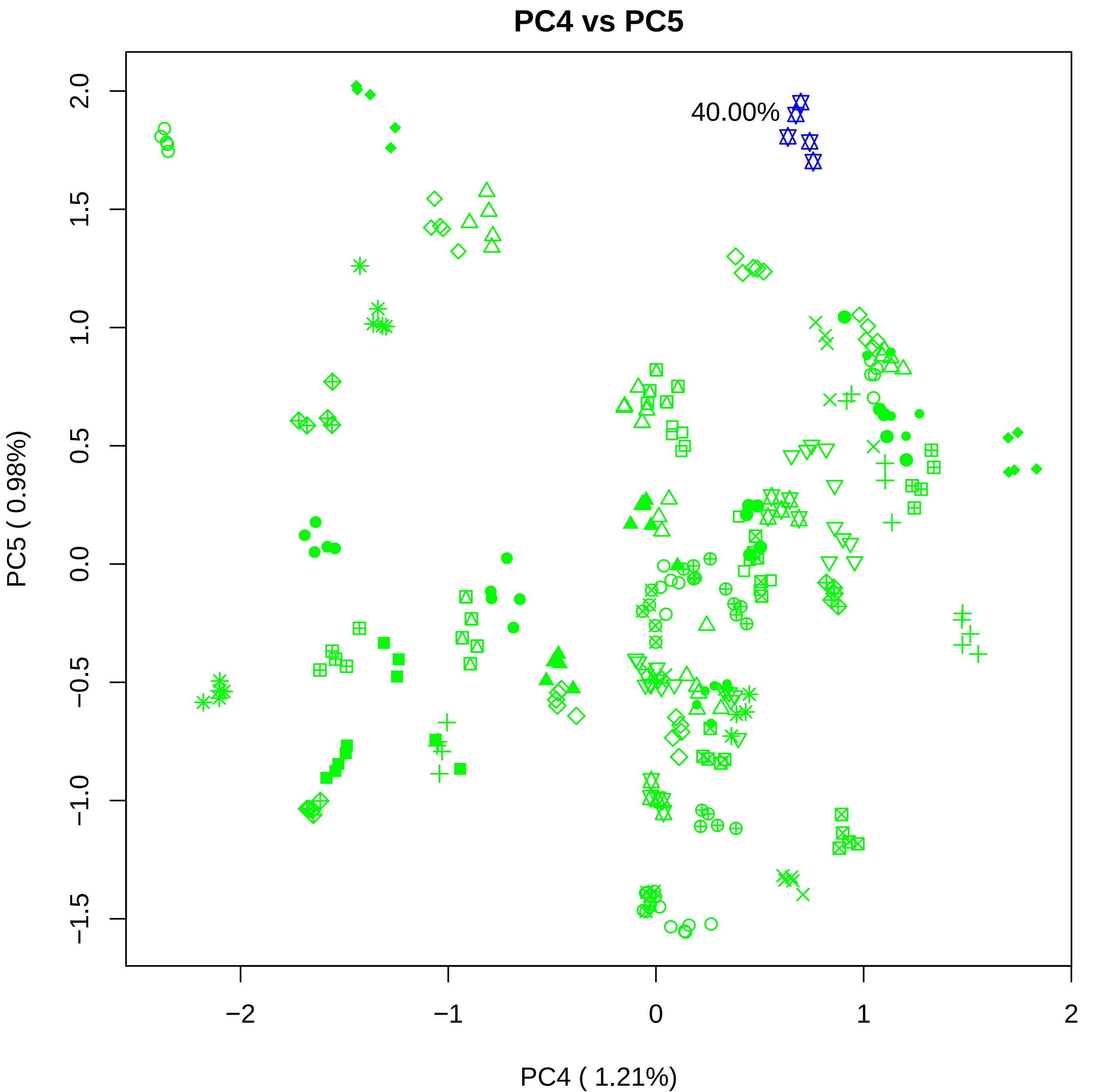

PC4 vs PC5
40.00%
PC5 ( 0.98%)
PC4 ( 1.21%)
2.0
1.5
1.0
0.5
0.0
−0.5
−1.0
−1.5
−2
−1
0
1
2

### PCA of All  BMH(b) vs PMH(b)

Scaled Matrix PCAed

7th order polynomial baseline correction performed

Raw spectra smoothed to 21 Neighbours, 2nd order PN, Savtisky Golay

Normalisation between 0 and 100

D2 spectra smoothed to 41 Neighbours, 2nd order PN, Savtisky Golay

Regions Analysed: 3601−2501 and 2001−700cm−1

Samples Analysed: 8 bContam's 0 BLH's 0 BMH's 0 OMH's 0 pContam's 0 PLH's 60 PMH's

Keele University AD

Fri Dec 14 14:21:00 2018

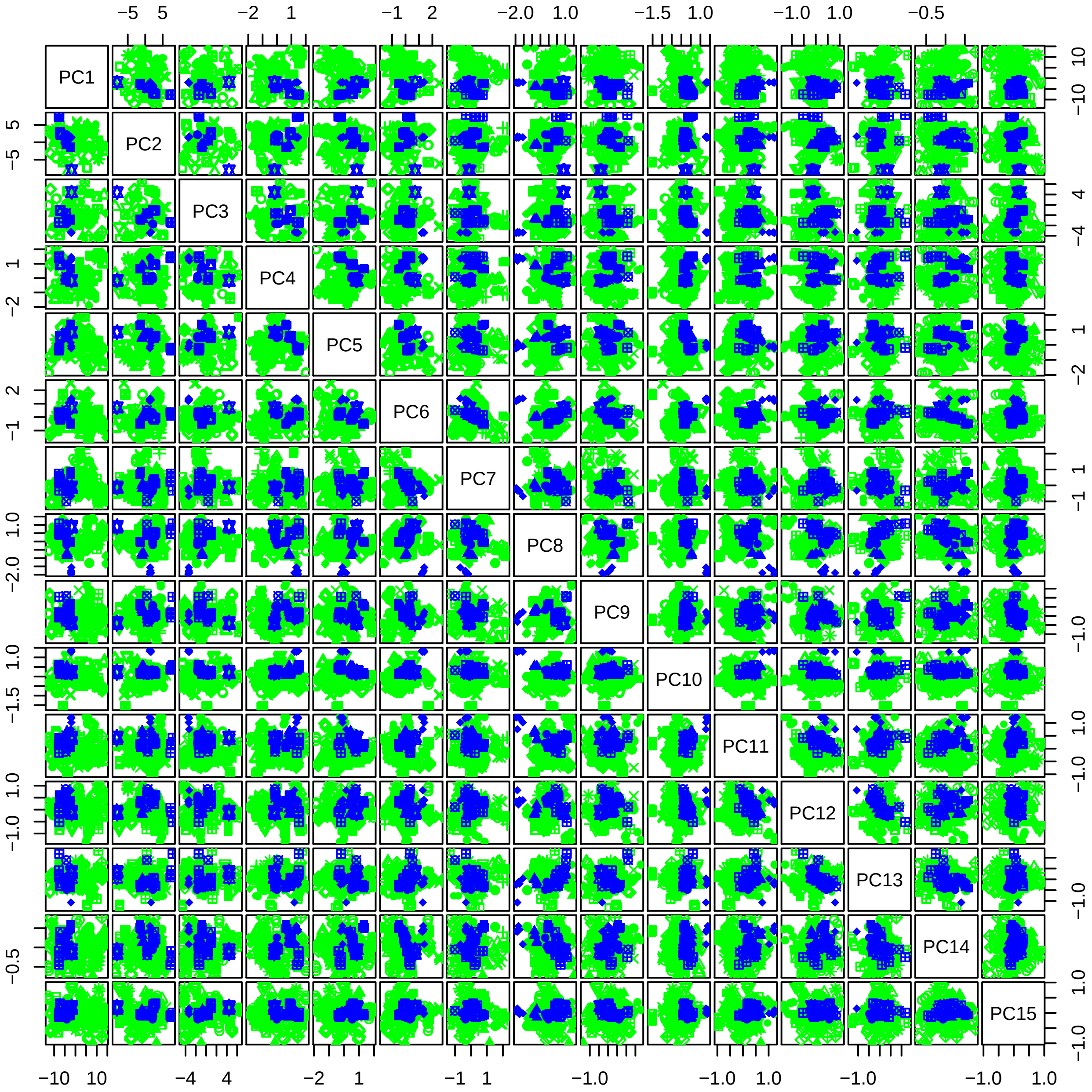
PC1
PC2
PC3
PC4
PC5
PC6
PC7
PC8
PC9
PC10
PC11
PC12
PC13
PC14
PC15

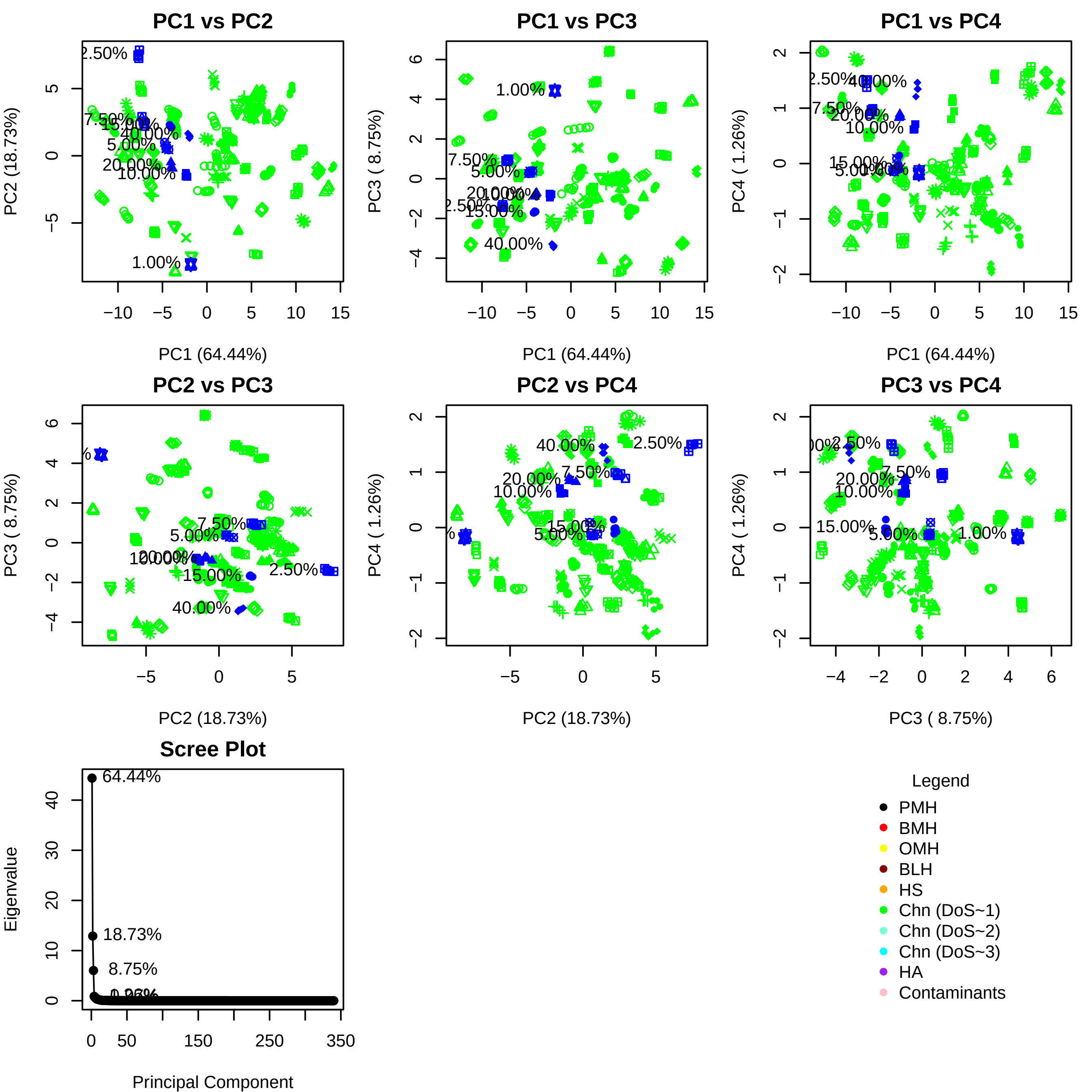
PC1 vs PC2
PC1 vs PC3
PC1 vs PC4
PC2 vs PC3
PC2 vs PC4
PC3 vs PC4
Scree Plot
PC1 (64.44%)
PC2 (18.73%)
PC3 ( 8.75%)
PC4 ( 1.26%)
Eigenvalue
Principal Component
64.44%
18.73%
8.75%
Legend
PMH
BMH
OMH
BLH
HS
Chn (DoS~1)
Chn (DoS~2)
Chn (DoS~3)
HA
Contaminants

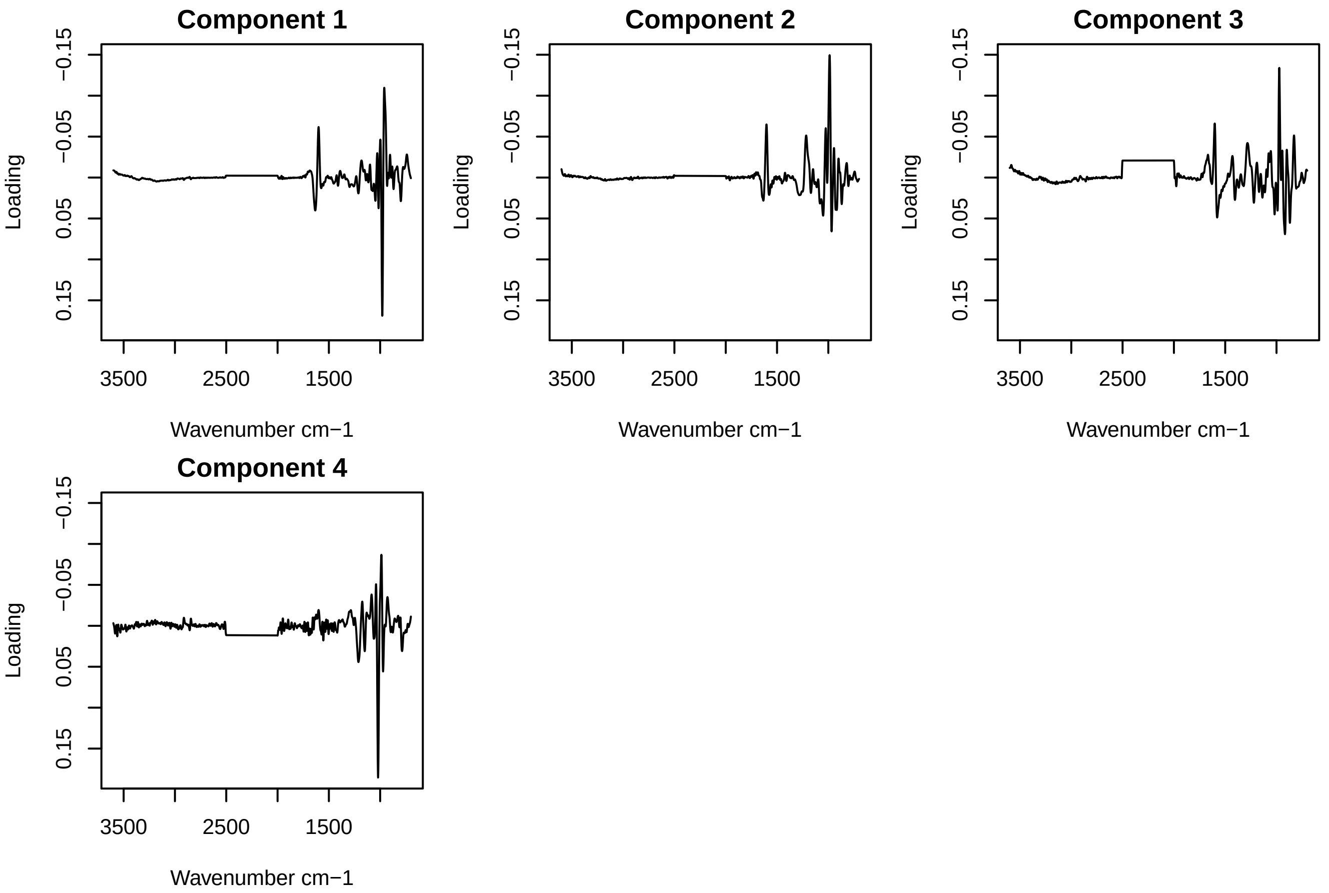
Component 1
Component 2
Component 3
Component 4
Loading
−0.15
−0.05
0.05
0.15
3500
2500
1500
Wavenumber cm−1

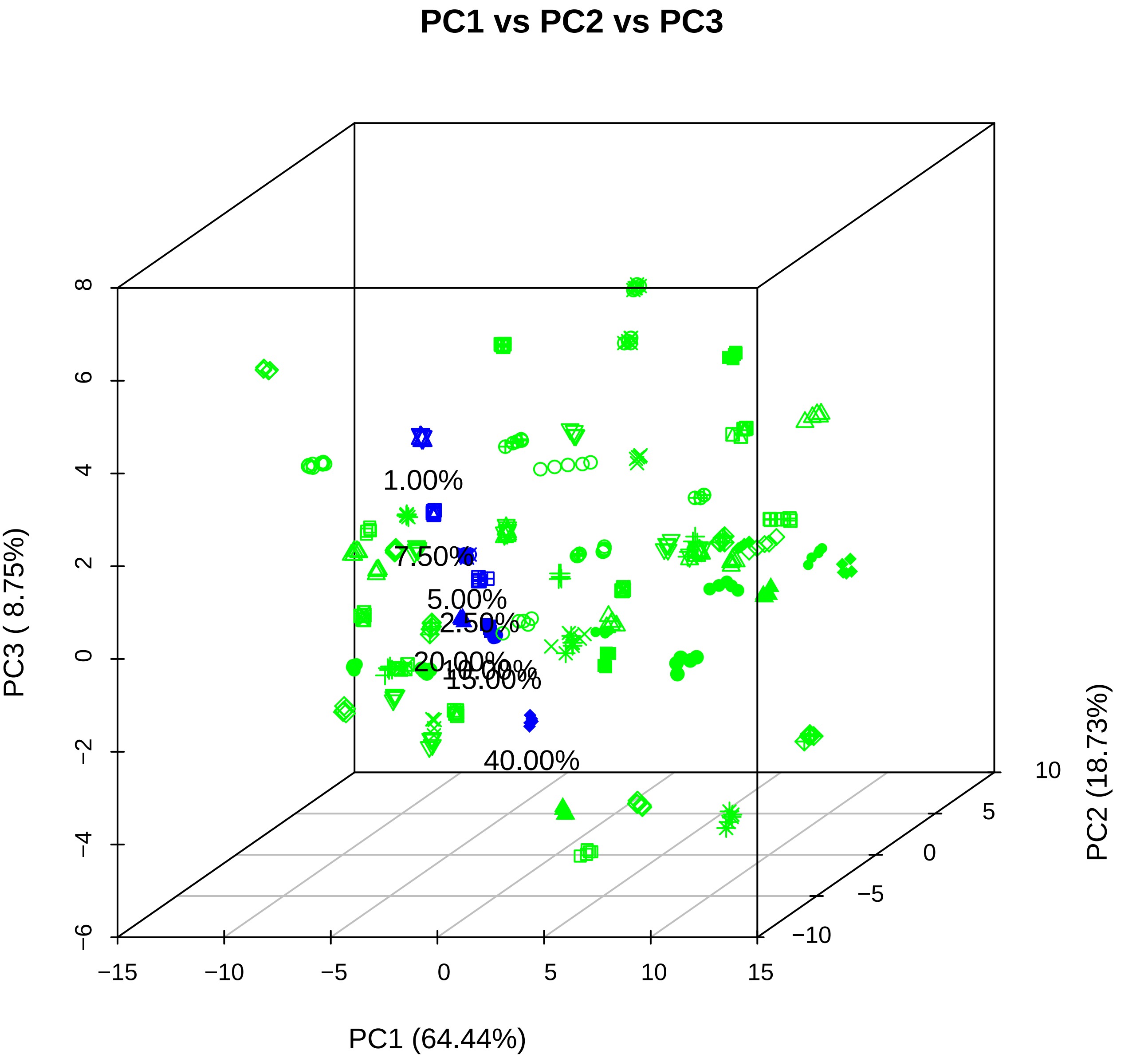
PC1 vs PC2 vs PC3
1.00%
7.50%
5.00%
2.50%
20.00%
10.00%
15.00%
40.00%
PC1 (64.44%)
PC2 (18.73%)
PC3 ( 8.75%)
−15
−10
−5
0
5
10
15
−6
−4
−2
0
2
4
6
8

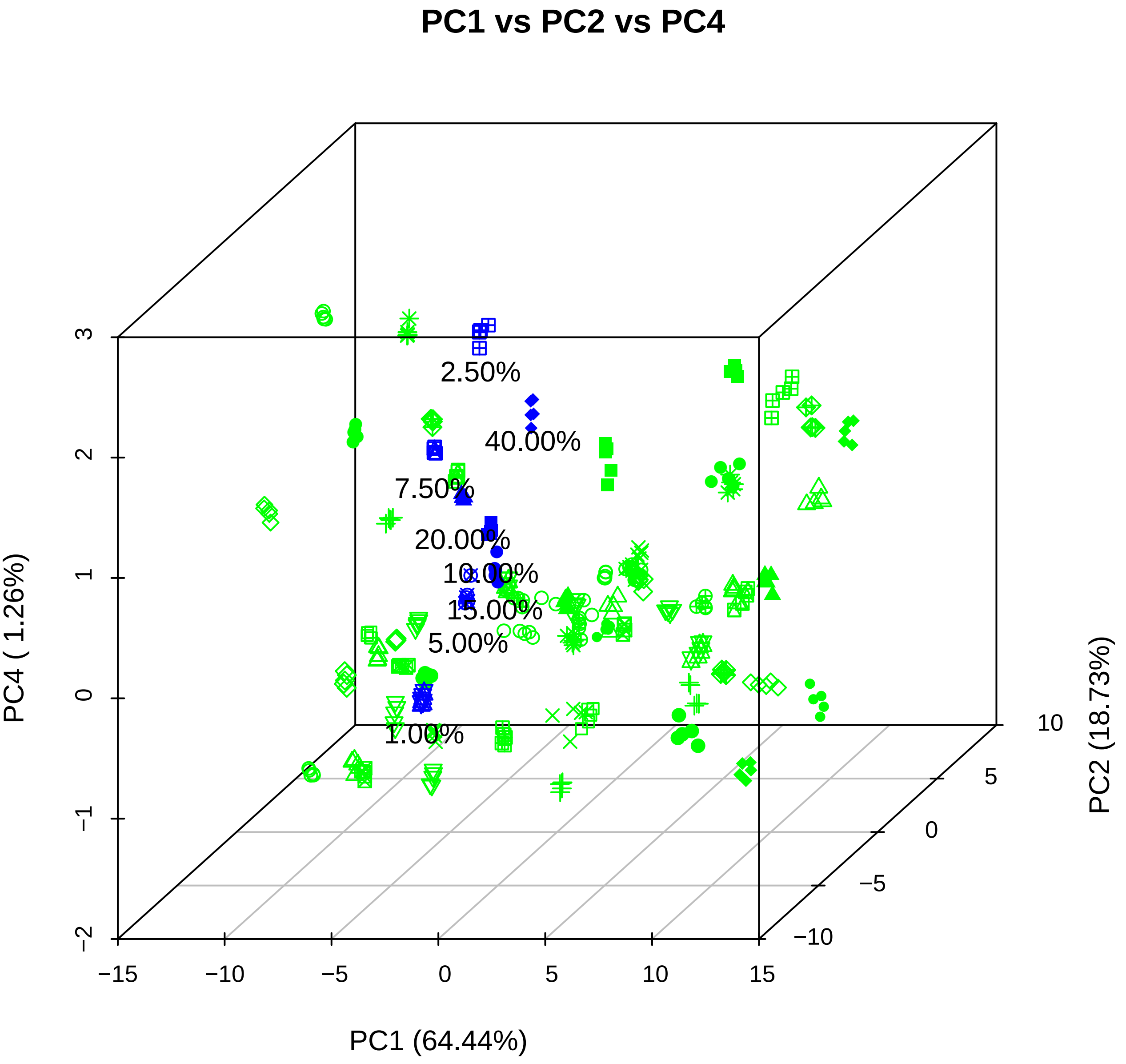

PC1 vs PC2 vs PC4
2.50%
40.00%
7.50%
20.00%
10.00%
15.00%
5.00%
1.00%
PC4 ( 1.26%)
PC1 (64.44%)
PC2 (18.73%)
−15
−10
−5
0
5
10
15
−2
−1
0
1
2
3
−10
−5
0
5
10

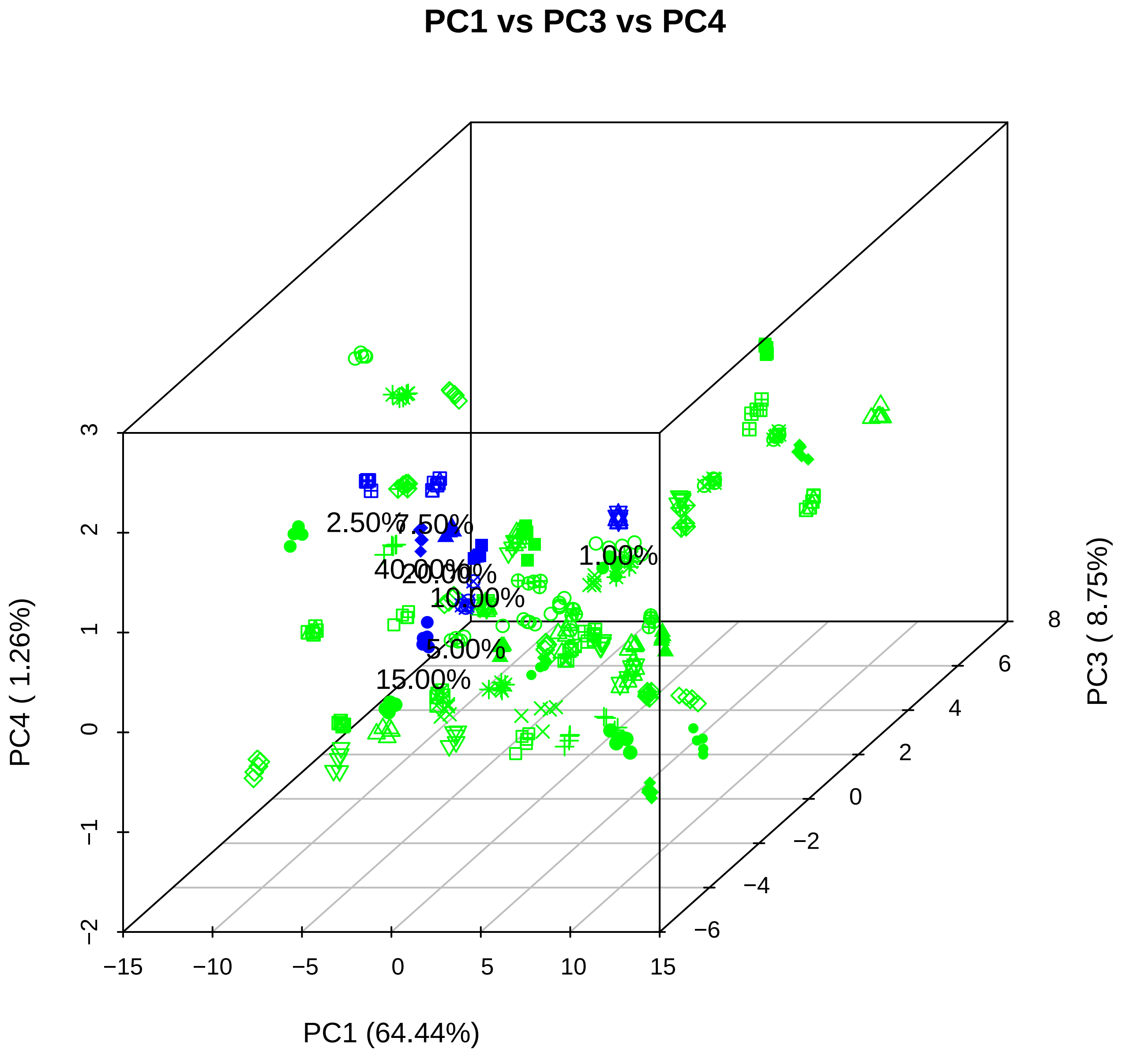

PC1 vs PC3 vs PC4
PC1 (64.44%)
PC4 ( 1.26%)
PC3 ( 8.75%)
−15
−10
−5
0
5
10
15
−2
−1
0
1
2
3
−6
−4
−2
0
2
4
6
8
2.50%
7.50%
40.00%
20.00%
10.00%
1.00%
5.00%
15.00%

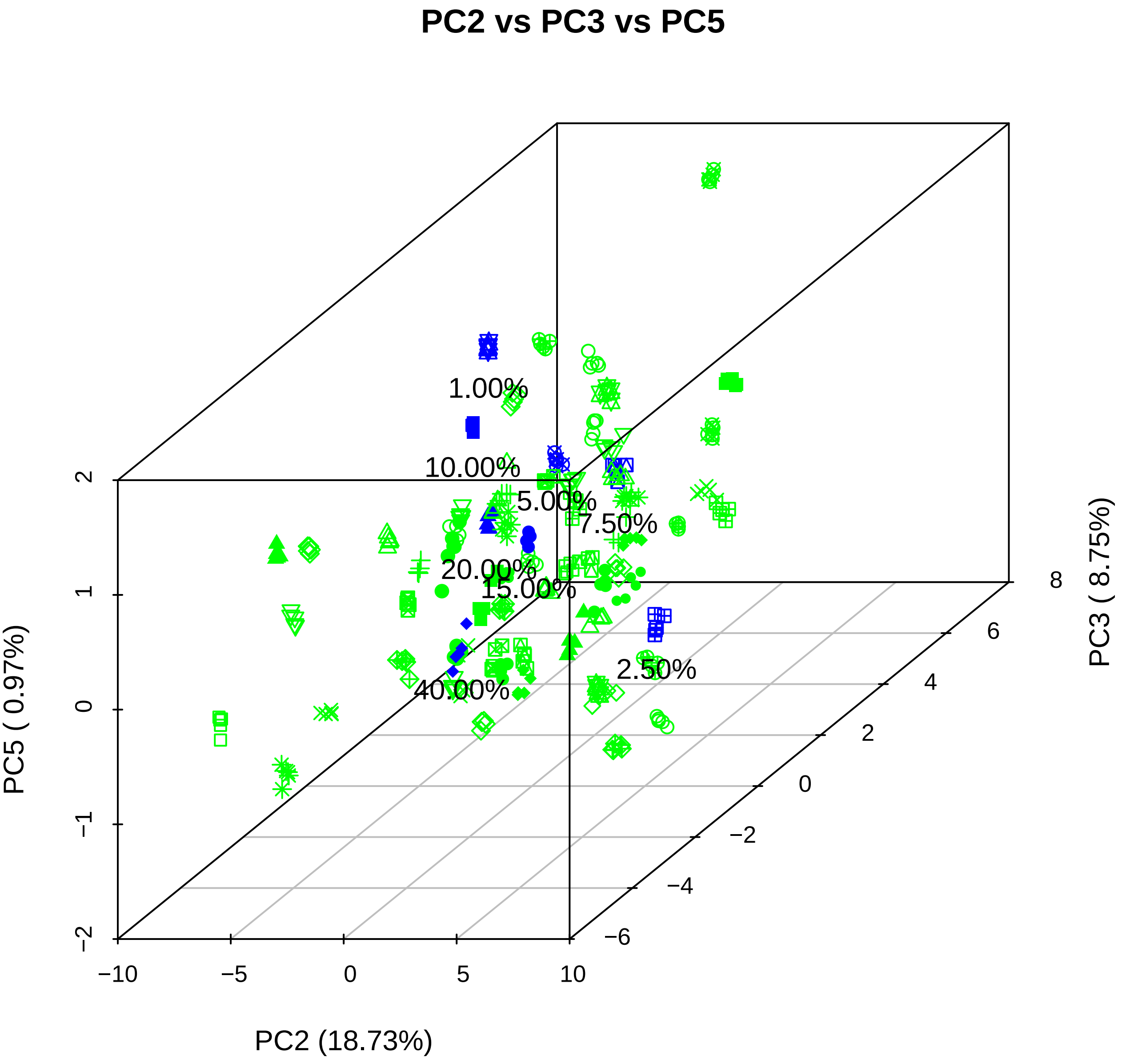

PC2 vs PC3 vs PC5
PC2 (18.73%)
PC5 ( 0.97%)
PC3 ( 8.75%)
−10
−5
0
5
10
−2
−1
0
1
2
−6
−4
−2
0
2
4
6
8
1.00%
10.00%
5.00%
7.50%
20.00%
15.00%
2.50%
40.00%

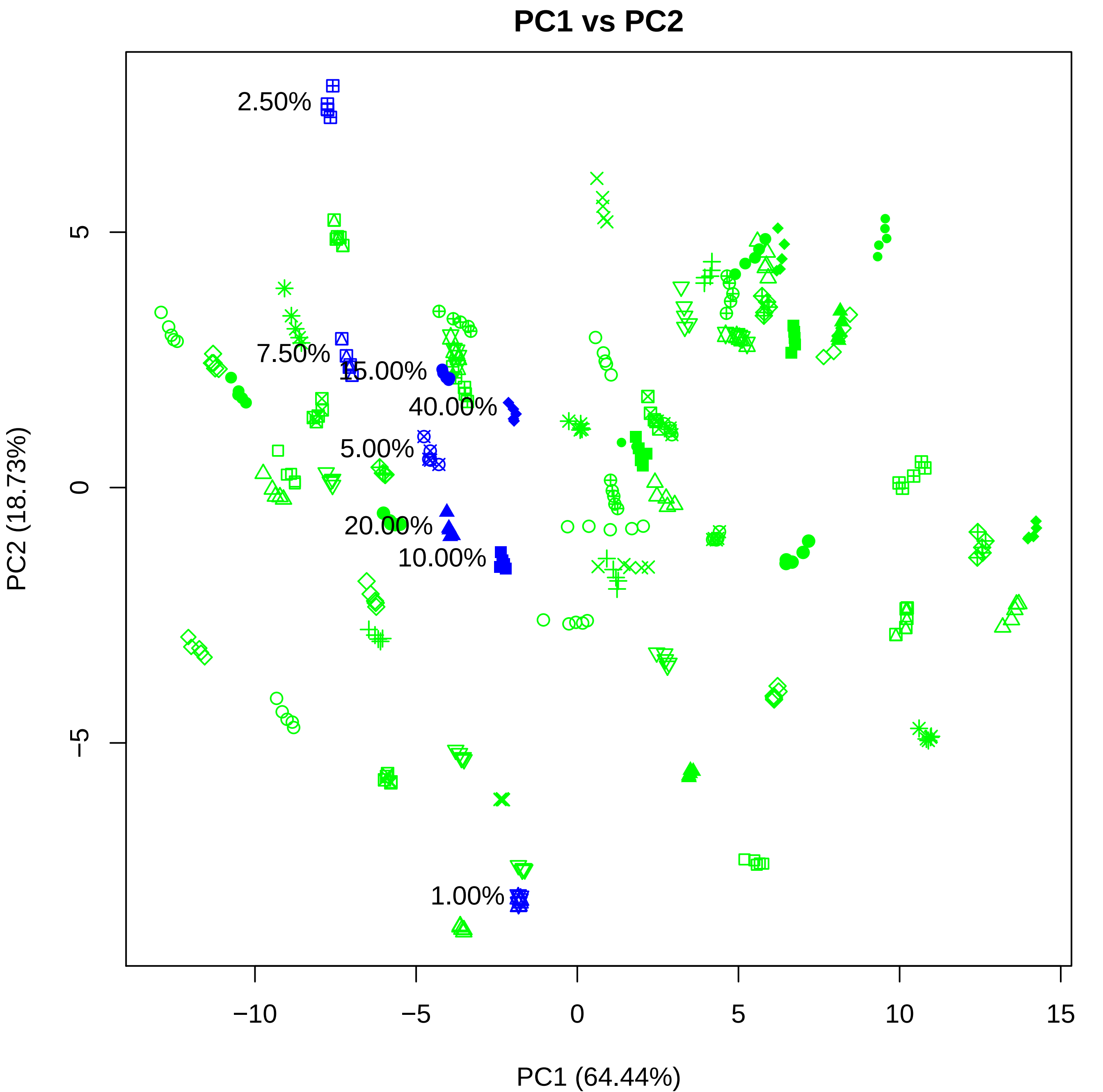

PC1 vs PC2
2.50%
7.50%
15.00%
40.00%
5.00%
20.00%
10.00%
1.00%
5
0
−5
−10
−5
0
5
10
15
PC2 (18.73%)
PC1 (64.44%)

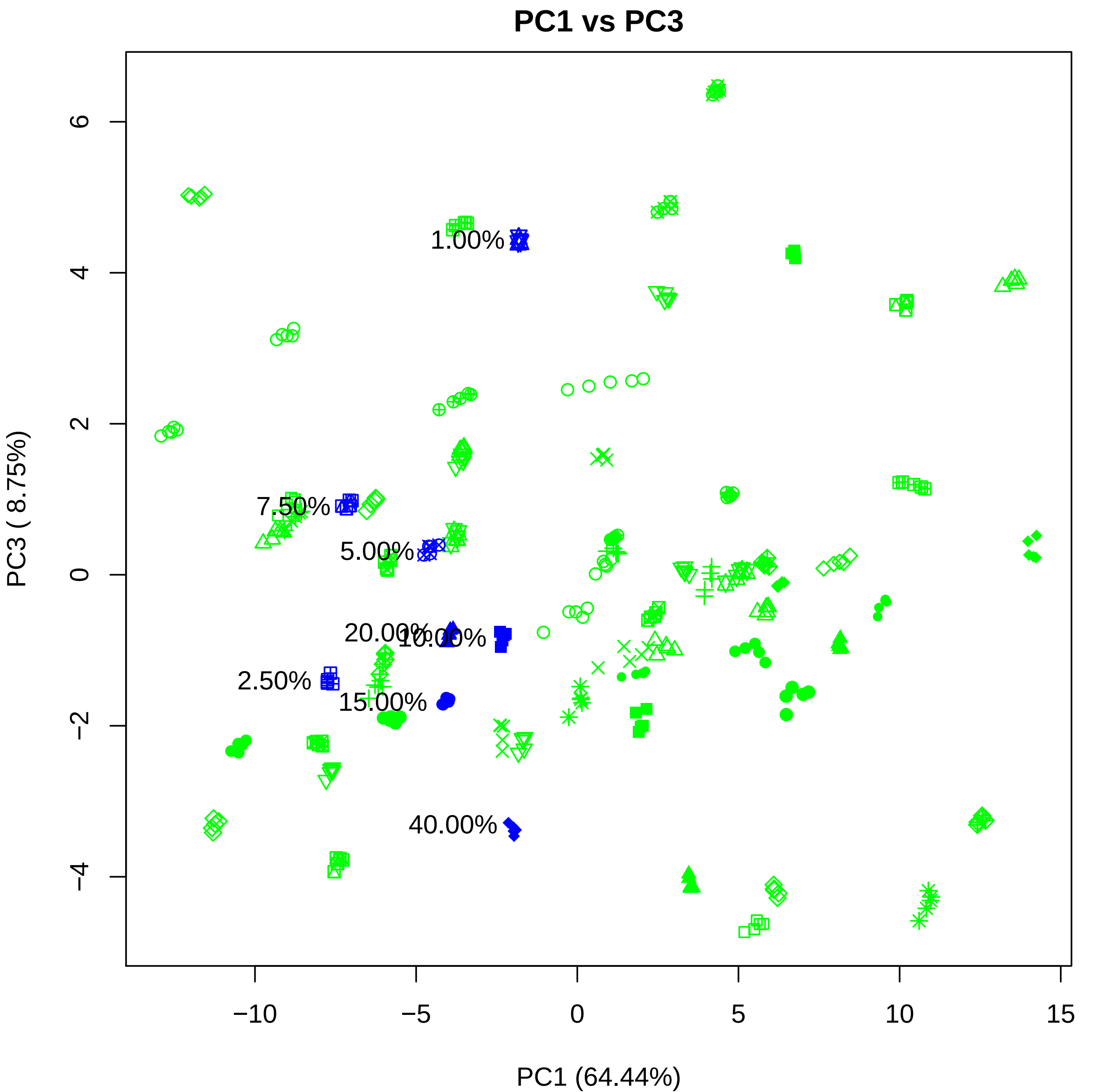
PC1 vs PC3
PC3 ( 8.75%)
PC1 (64.44%)
6
4
2
0
−2
−4
−10
−5
0
5
10
15
1.00%
7.50%
5.00%
20.00%
10.00%
2.50%
15.00%
40.00%

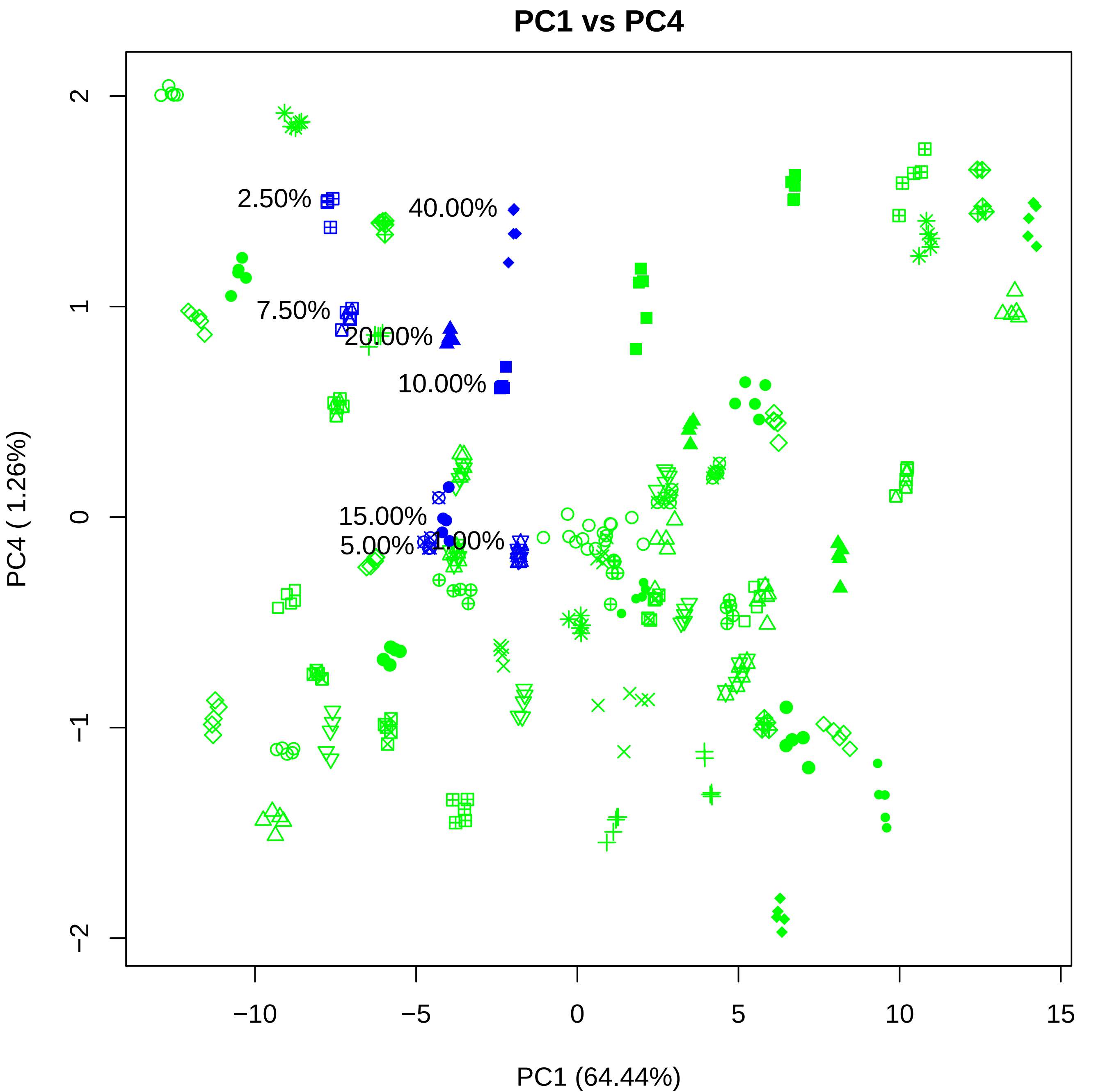
PC1 vs PC4
2.50%
40.00%
7.50%
20.00%
10.00%
15.00%
5.00%
1.00%
PC4 ( 1.26%)
PC1 (64.44%)
2
1
0
−1
−2
−10
−5
0
5
10
15

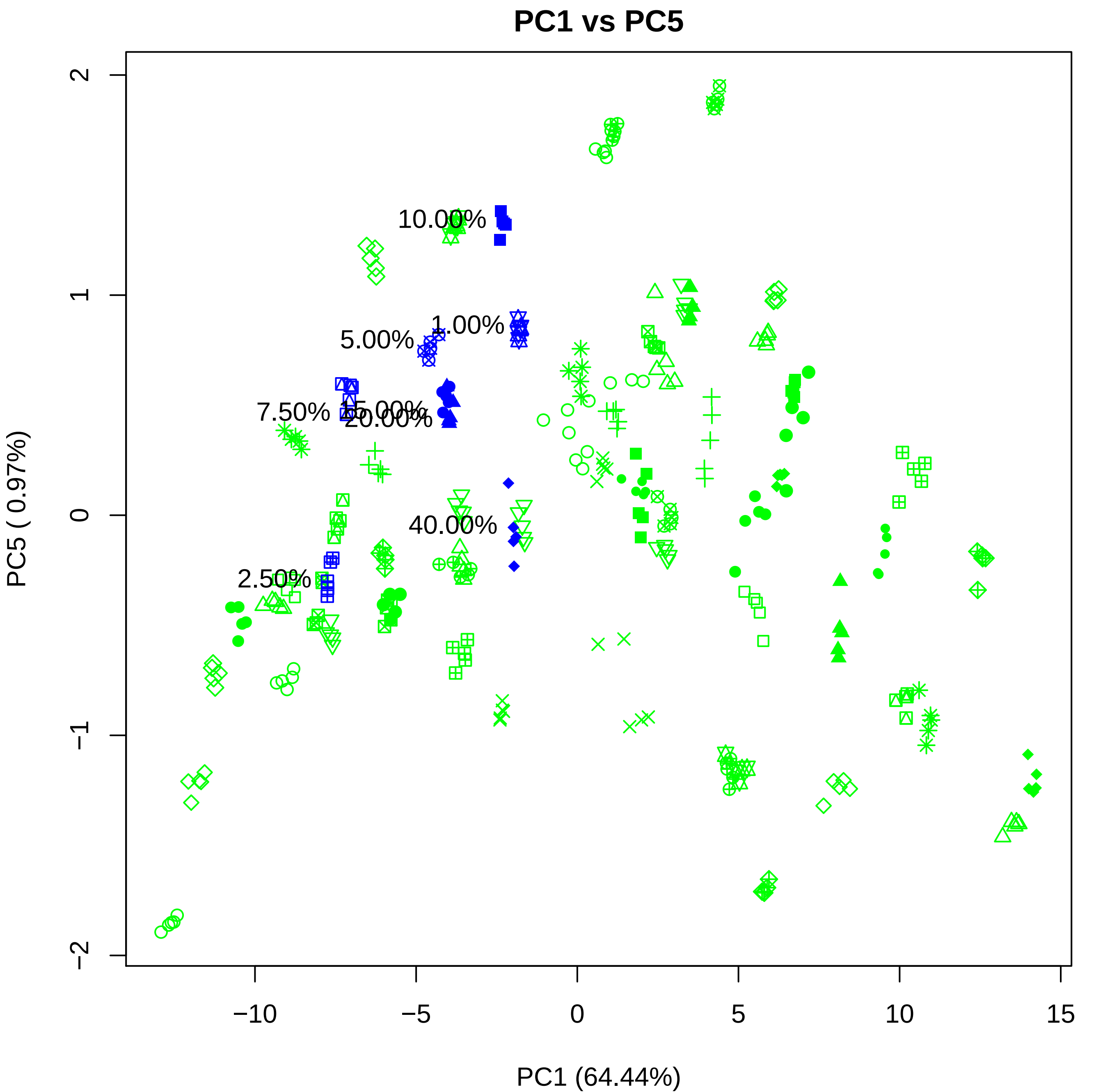

PC1 vs PC5
PC5 ( 0.97%)
PC1 (64.44%)
2
1
0
−1
−2
−10
−5
0
5
10
15
10.00%
1.00%
5.00%
7.50%
15.00%
20.00%
40.00%
2.50%

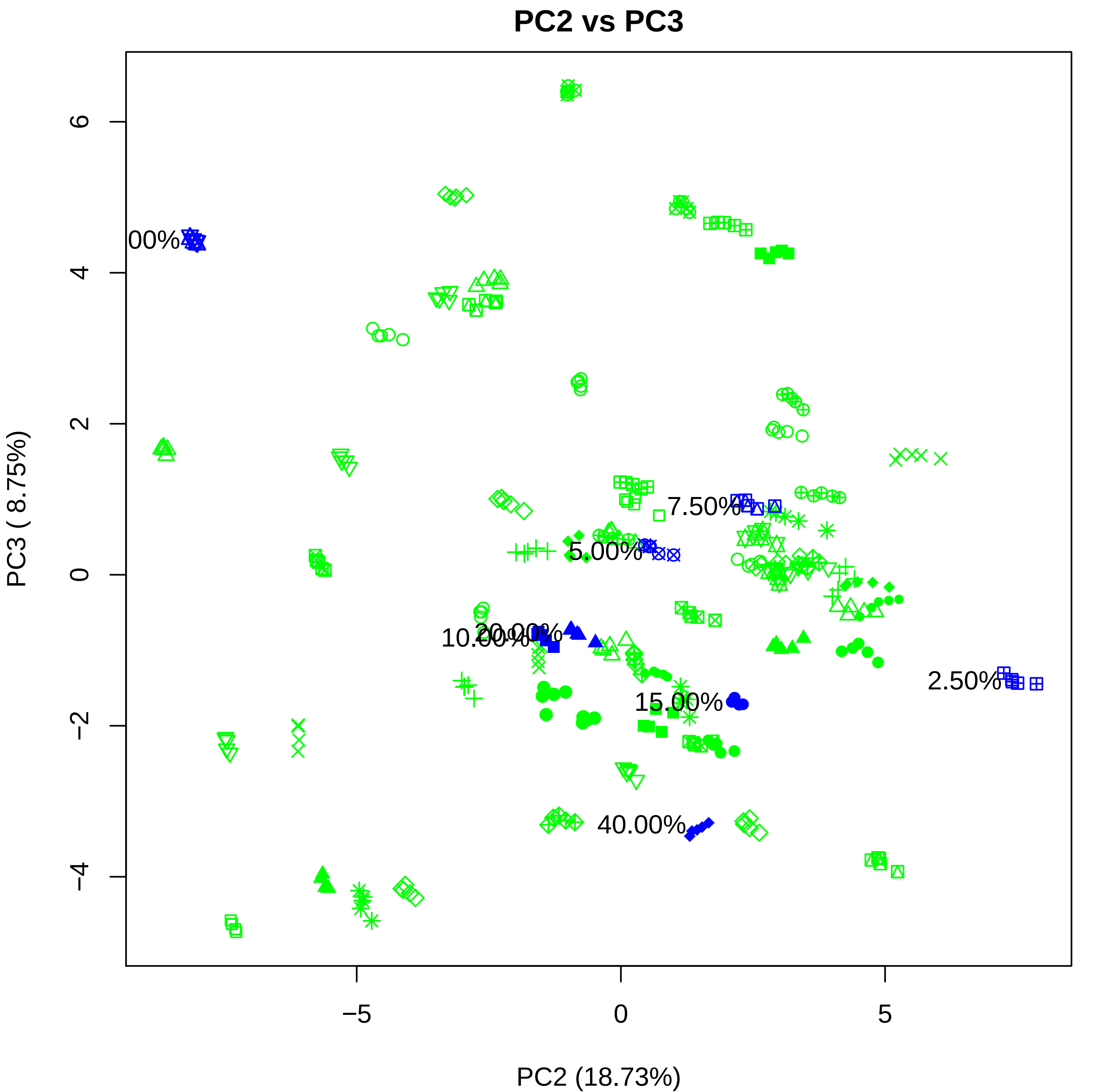

PC2 vs PC3
PC3 ( 8.75%)
PC2 (18.73%)
6
4
2
0
−2
−4
−5
0
5
00%
7.50%
5.00%
20.00%
10.00%
15.00%
2.50%
40.00%

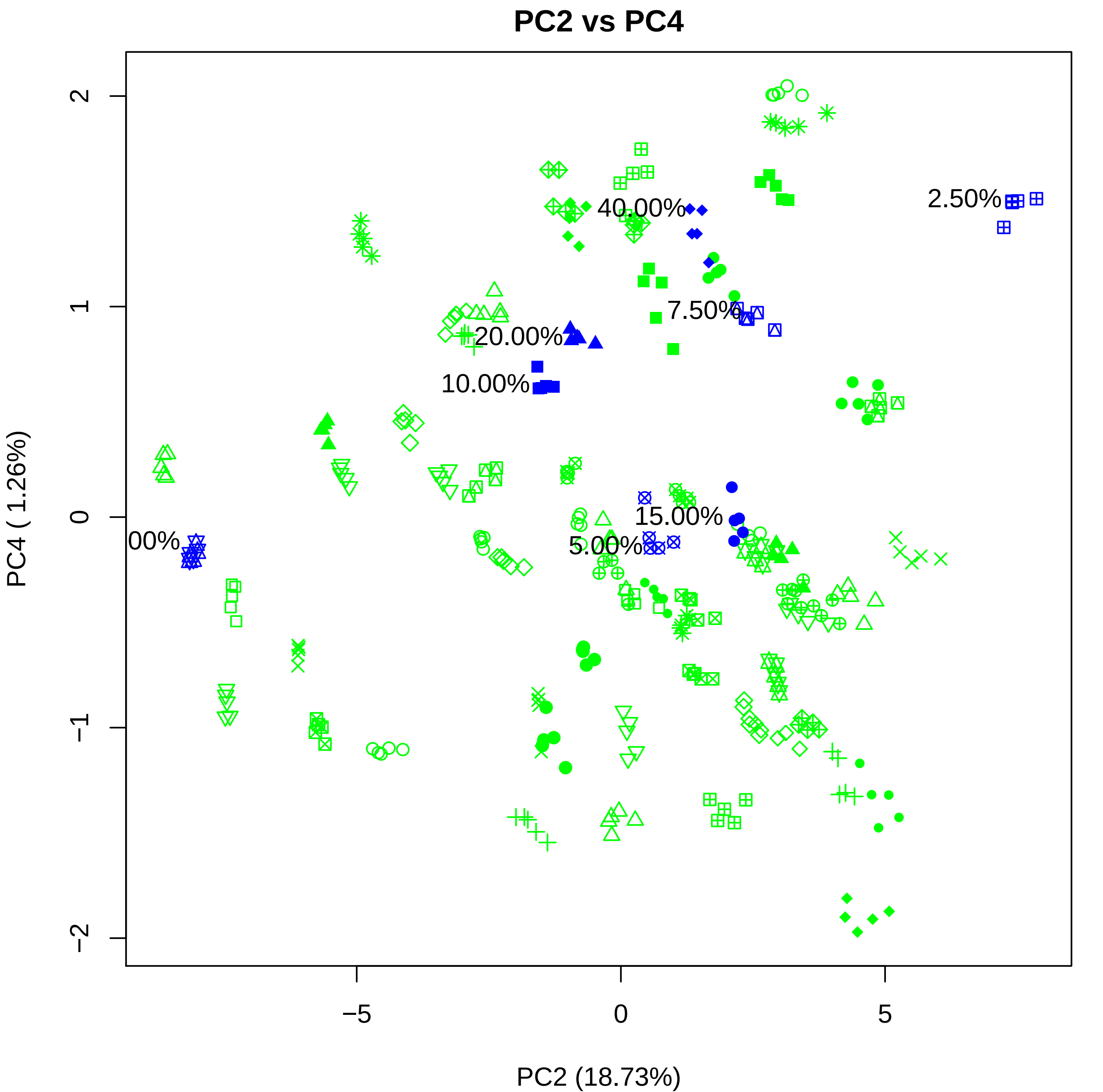
PC2 vs PC4
PC4 ( 1.26%)
PC2 (18.73%)
2
1
0
−1
−2
−5
0
5
2.50%
40.00%
7.50%
20.00%
10.00%
15.00%
00%
5.00%

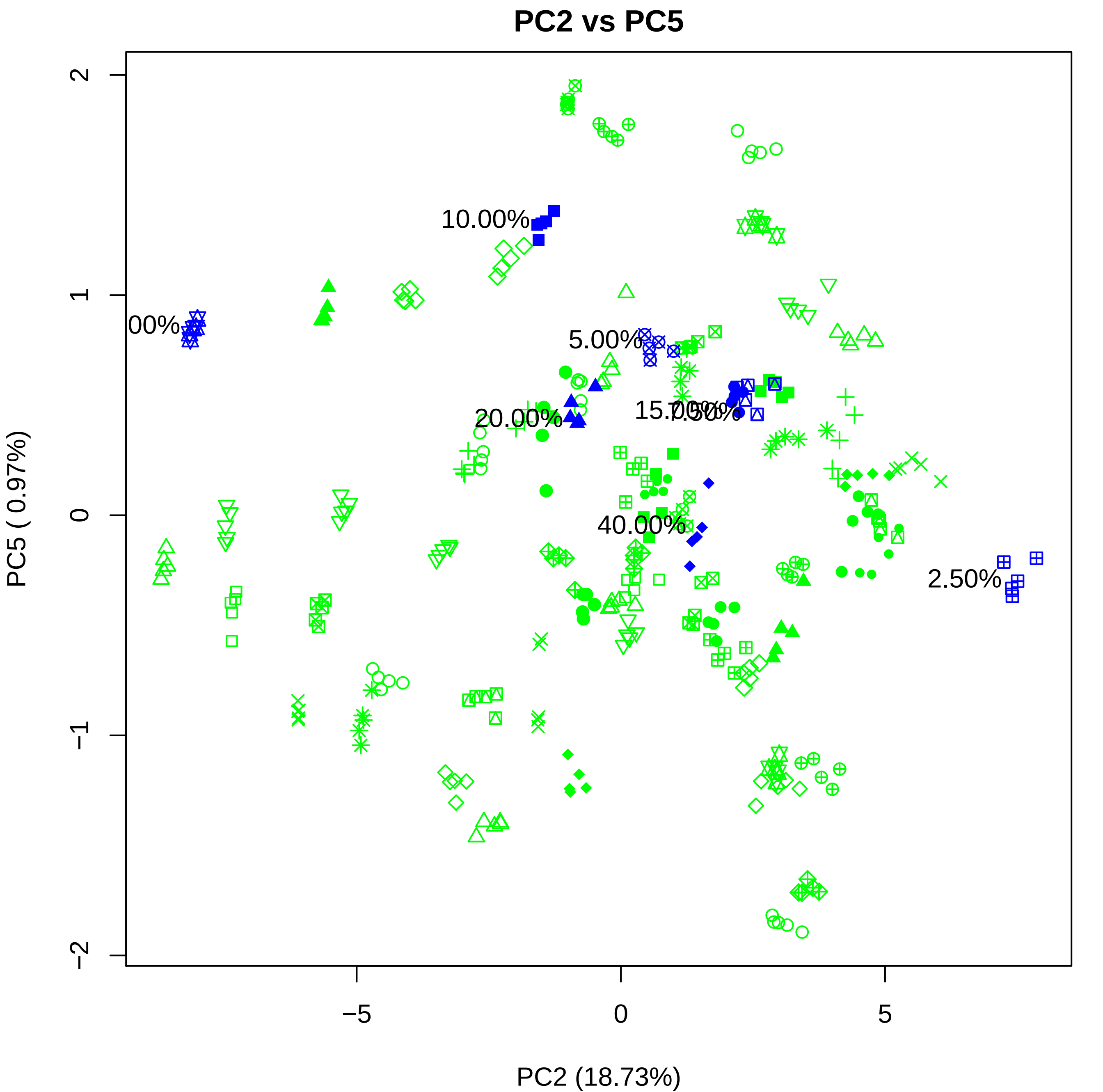
PC2 vs PC5
10.00%
00%
5.00%
20.00%
40.00%
2.50%
2
1
0
−1
−2
−5
0
5
PC5 ( 0.97%)
PC2 (18.73%)

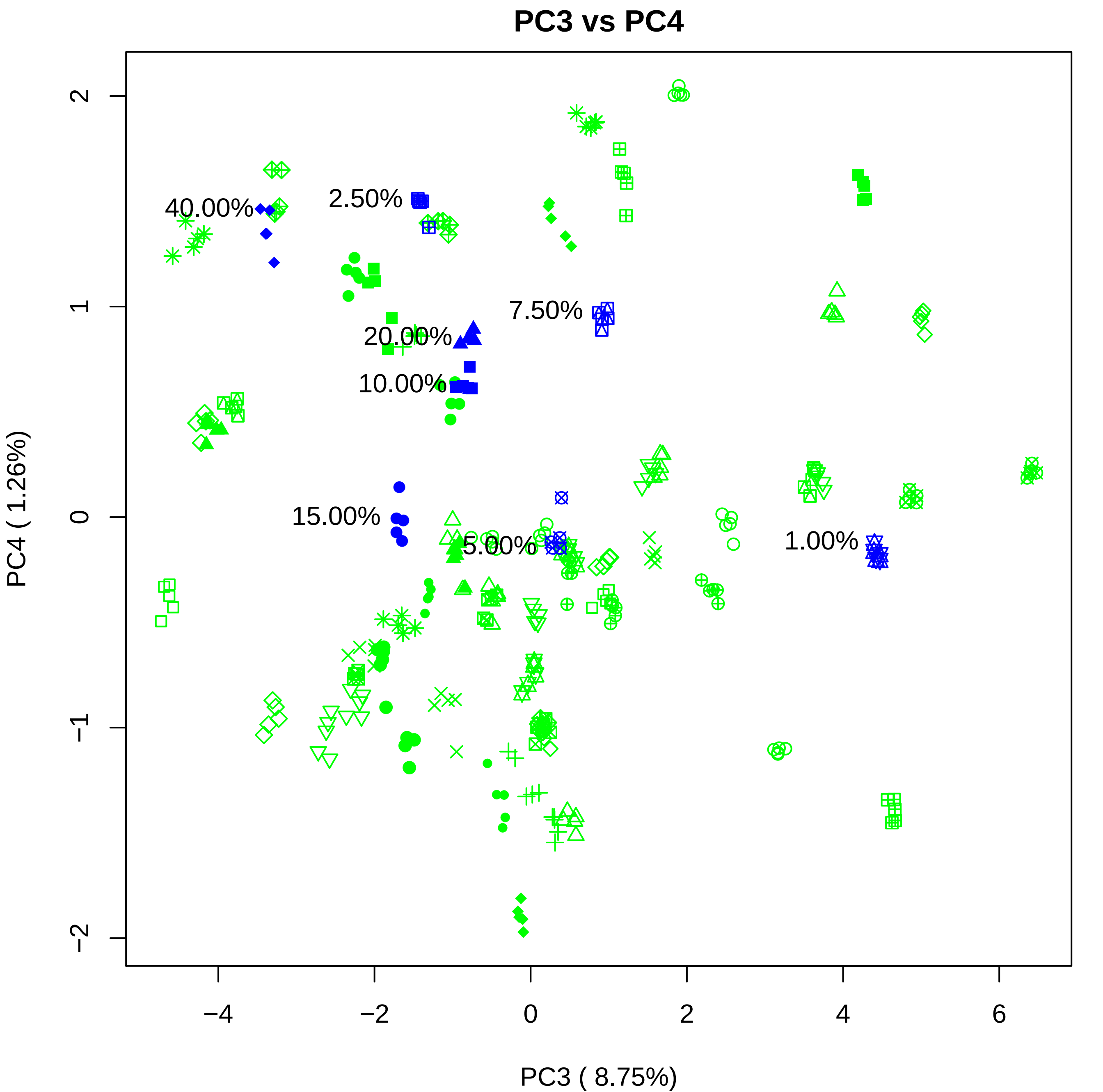
PC3 vs PC4
40.00%
2.50%
7.50%
20.00%
10.00%
15.00%
5.00%
1.00%
PC4 ( 1.26%)
PC3 ( 8.75%)
2
1
0
−1
−2
−4
−2
0
2
4
6

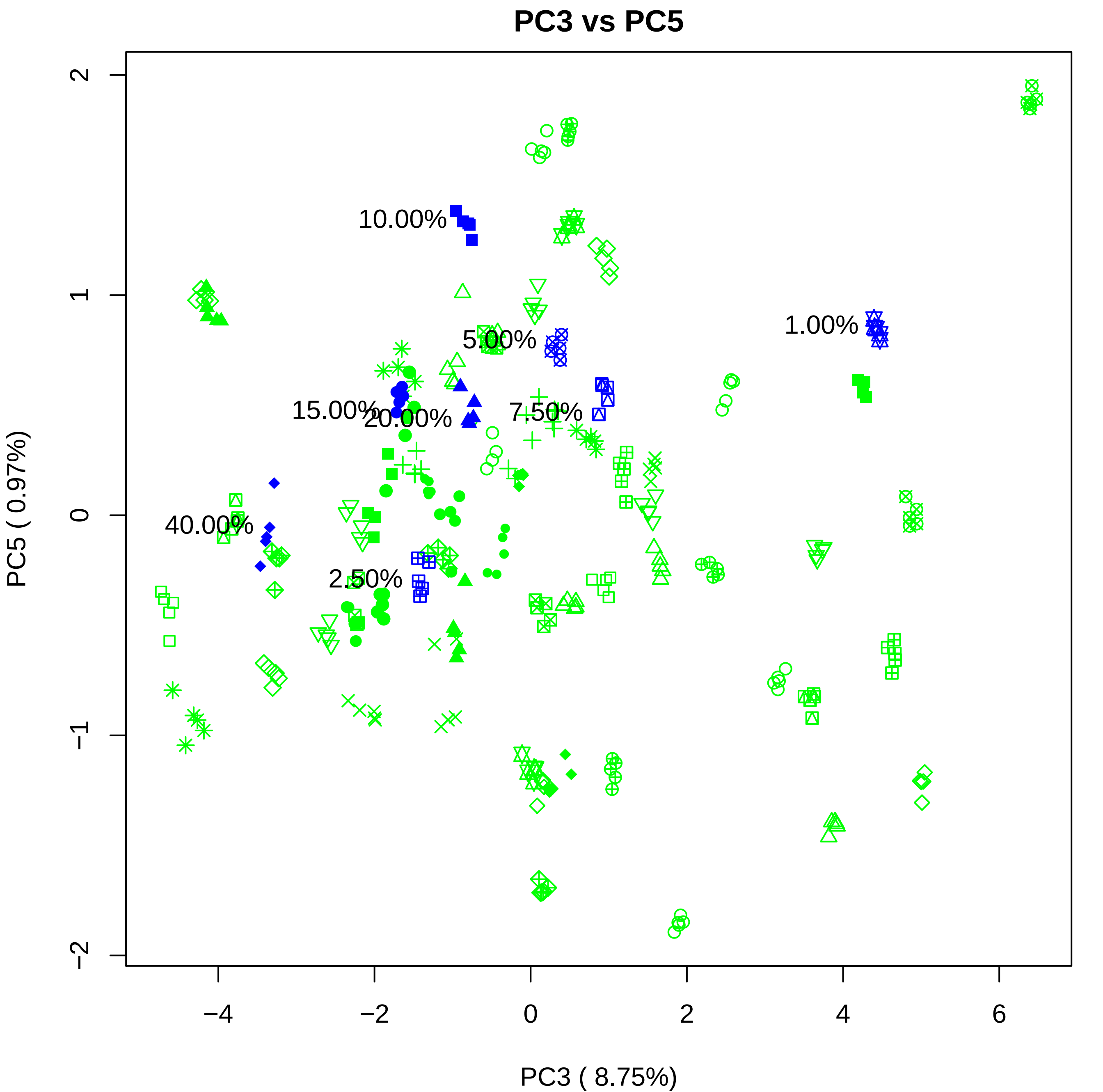

PC3 vs PC5
10.00%
1.00%
5.00%
15.00%
20.00%
7.50%
40.00%
2.50%
PC5 ( 0.97%)
PC3 ( 8.75%)
2
1
0
−1
−2
−4
−2
0
2
4
6

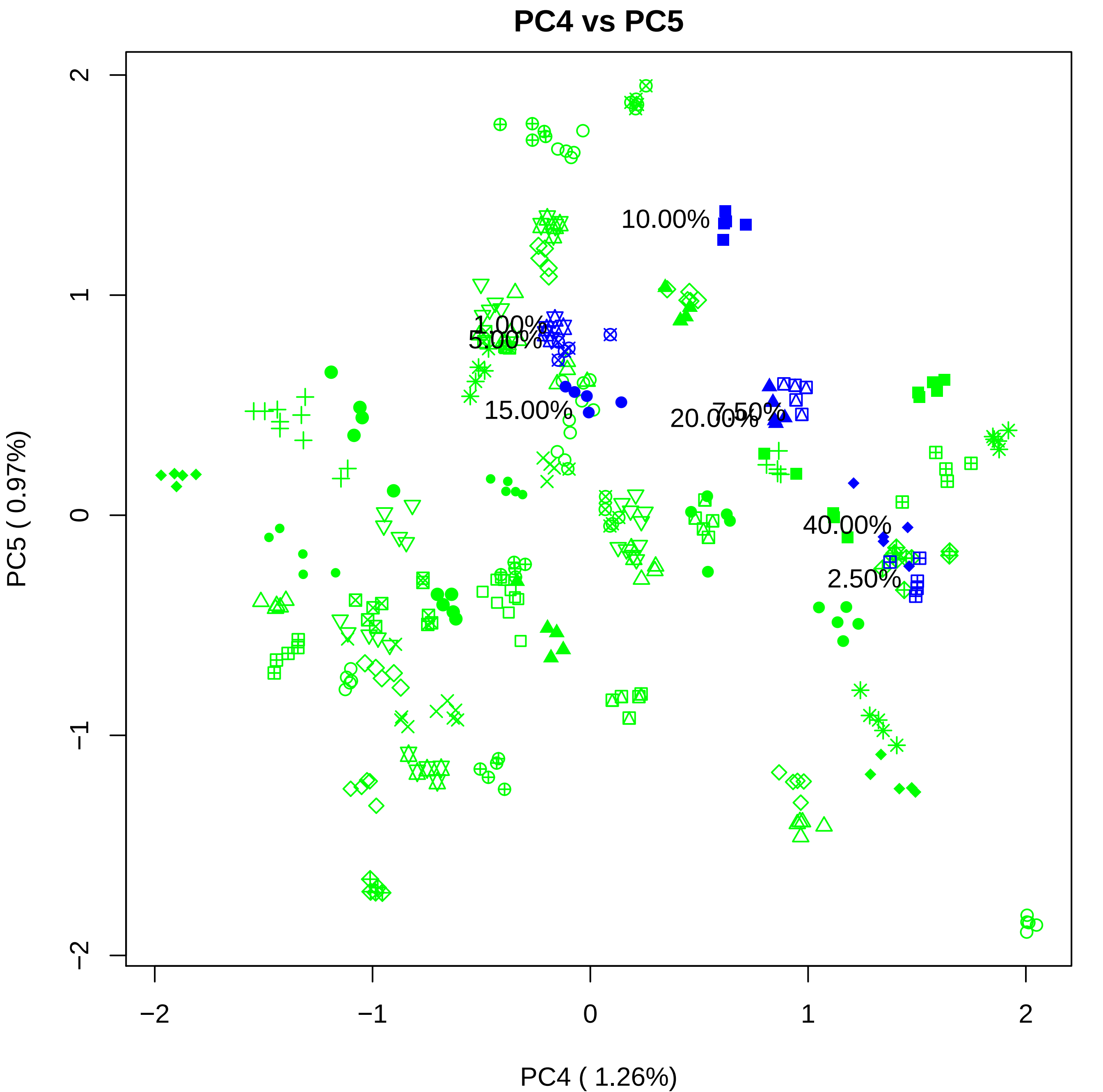
PC4 vs PC5
10.00%
1.00%
5.00%
15.00%
20.00%
7.50%
40.00%
2.50%
PC5 ( 0.97%)
PC4 ( 1.26%)
2
1
0
−1
−2
−2
−1
0
1
2

### PCA of 1%  BMH(b) vs PMH(b)

Scaled Matrix PCAed

7th order polynomial baseline correction performed

Raw spectra smoothed to 21 Neighbours, 2nd order PN, Savtisky Golay

Normalisation between 0 and 100

D2 spectra smoothed to 41 Neighbours, 2nd order PN, Savtisky Golay

Regions Analysed: 3601−2501 and 2001−700cm−1

Samples Analysed: 1 bContam's 0 BLH's 0 BMH's 0 OMH's 0 pContam's 0 PLH's 60 PMH's

Keele University AD

Fri Dec 14 14:21:09 2018

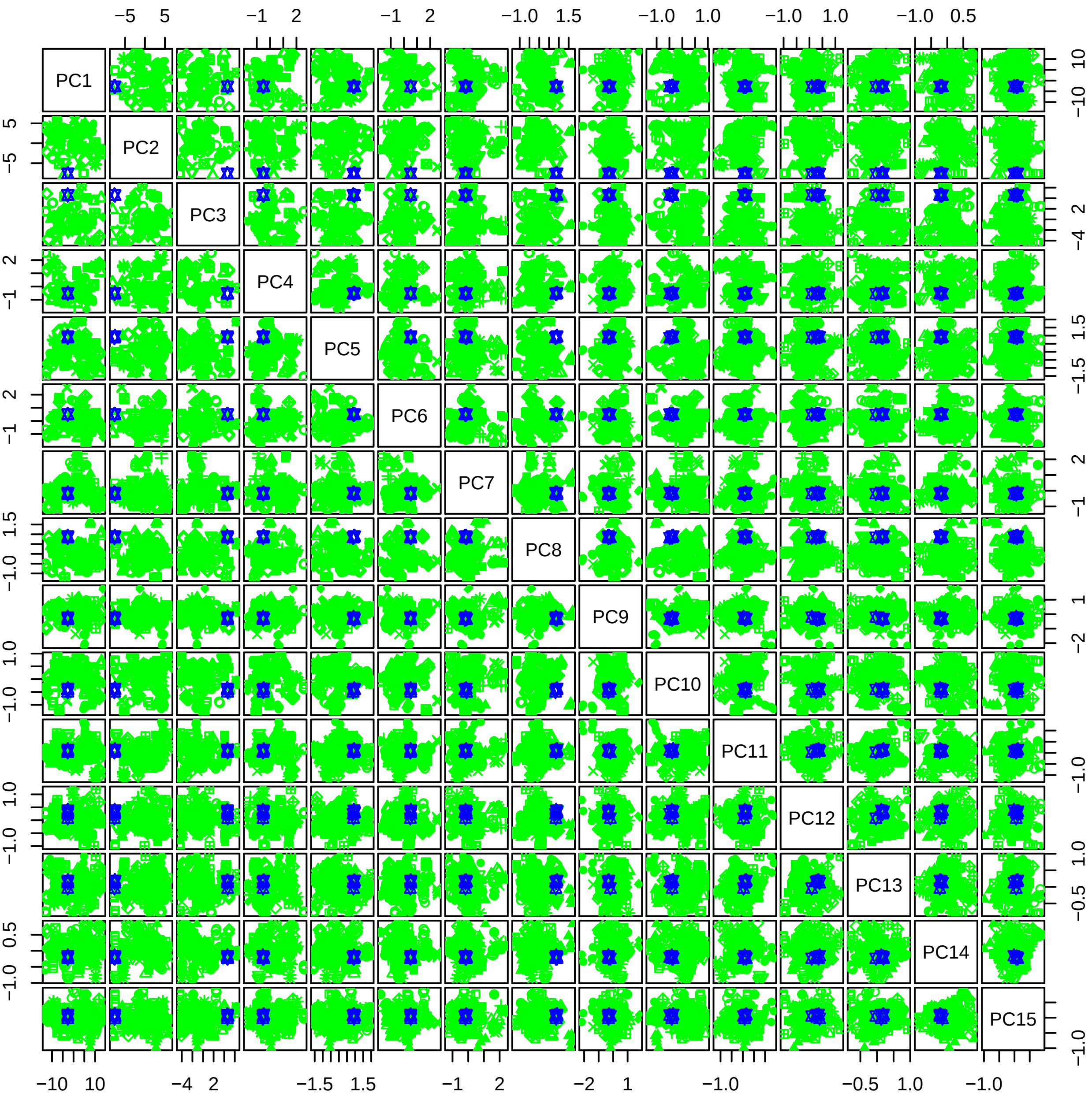
PC1
PC2
PC3
PC4
PC5
PC6
PC7
PC8
PC9
PC10
PC11
PC12
PC13
PC14
PC15

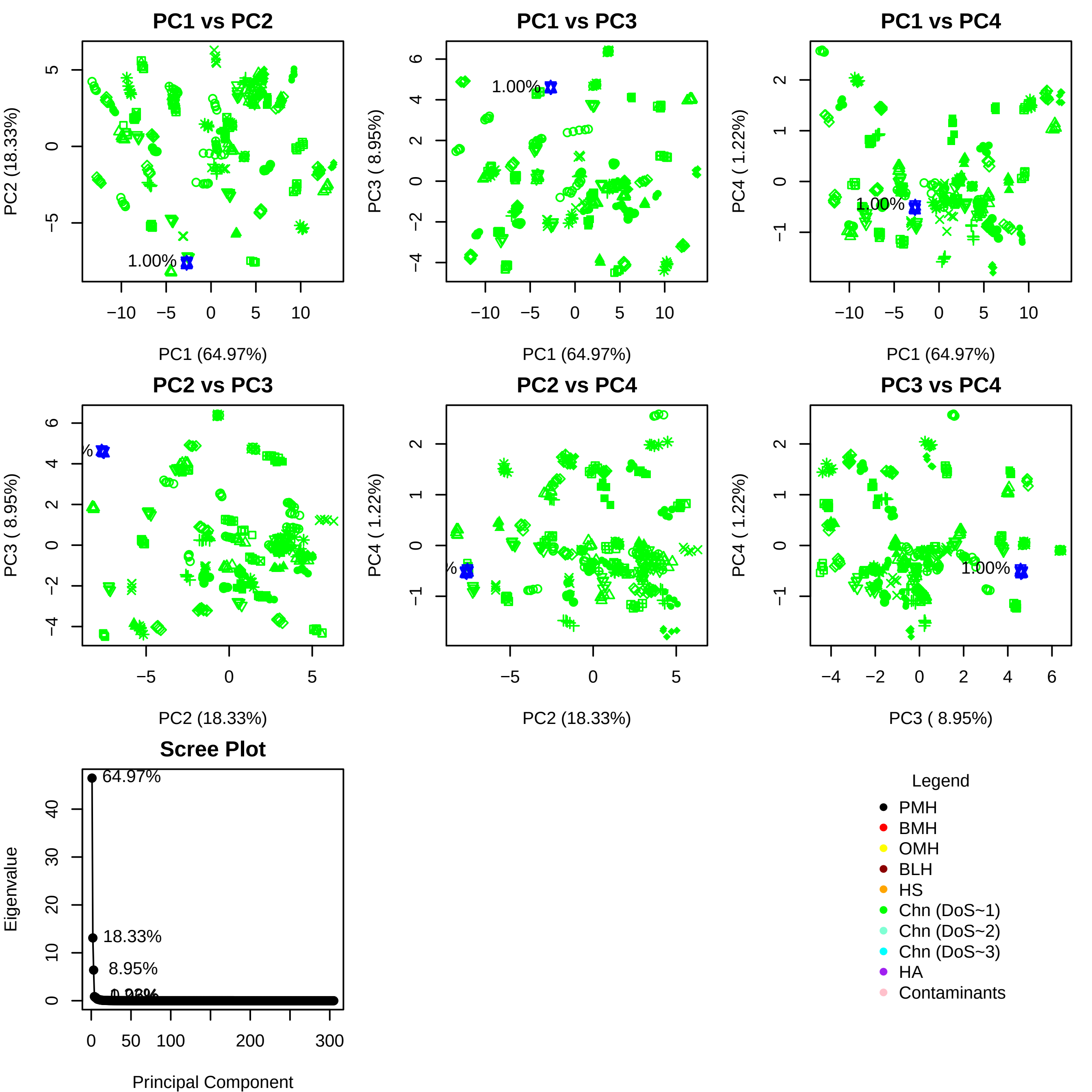
PC1 vs PC2
PC1 vs PC3
PC1 vs PC4
PC2 vs PC3
PC2 vs PC4
PC3 vs PC4
Scree Plot
PC1 (64.97%)
PC2 (18.33%)
PC3 ( 8.95%)
PC4 ( 1.22%)
1.00%
64.97%
18.33%
8.95%
Eigenvalue
Principal Component
Legend
PMH
BMH
OMH
BLH
HS
Chn (DoS~1)
Chn (DoS~2)
Chn (DoS~3)
HA
Contaminants

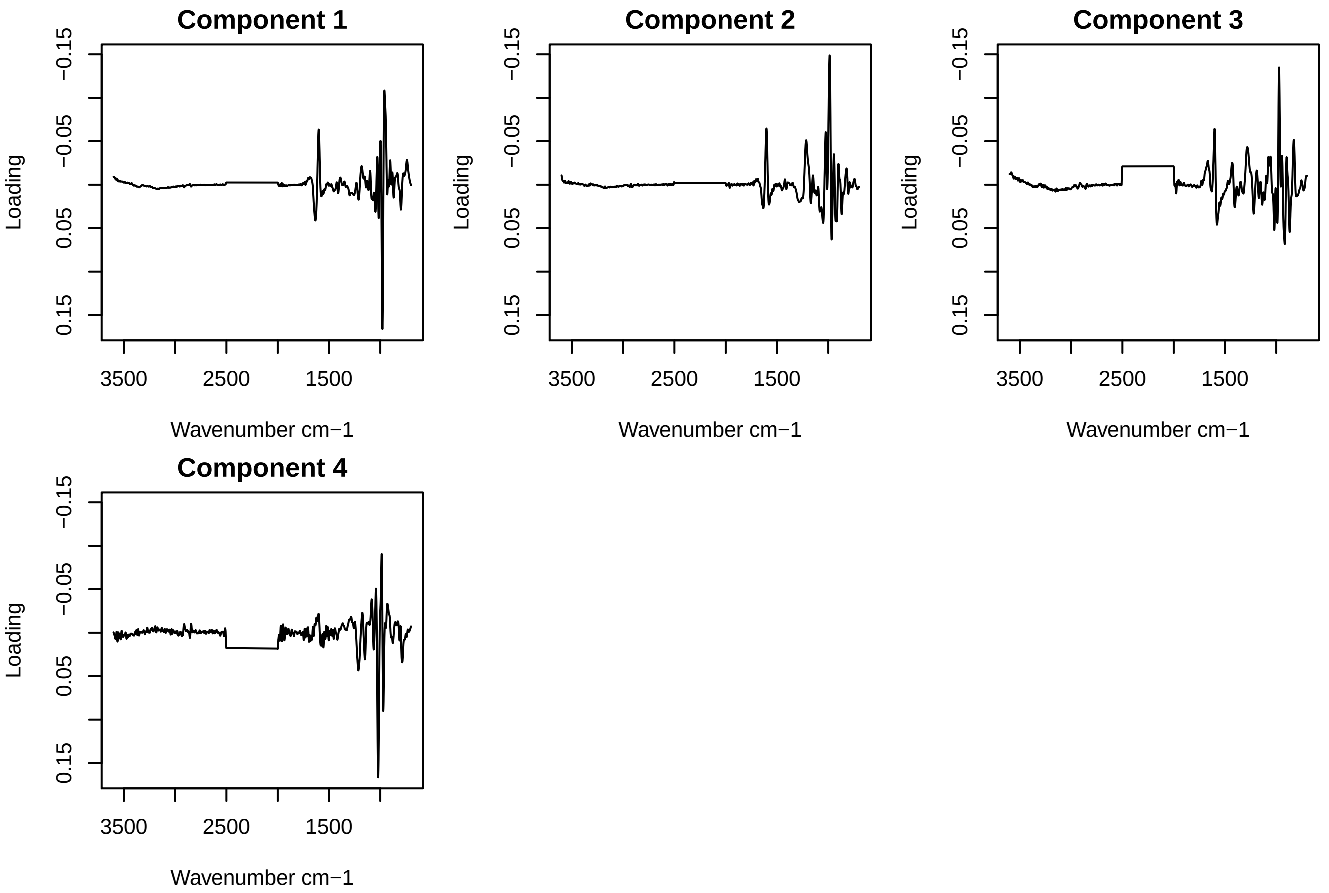
Component 1
Component 2
Component 3
Component 4
Loading
−0.15
−0.05
0.05
0.15
3500
2500
1500
Wavenumber cm−1

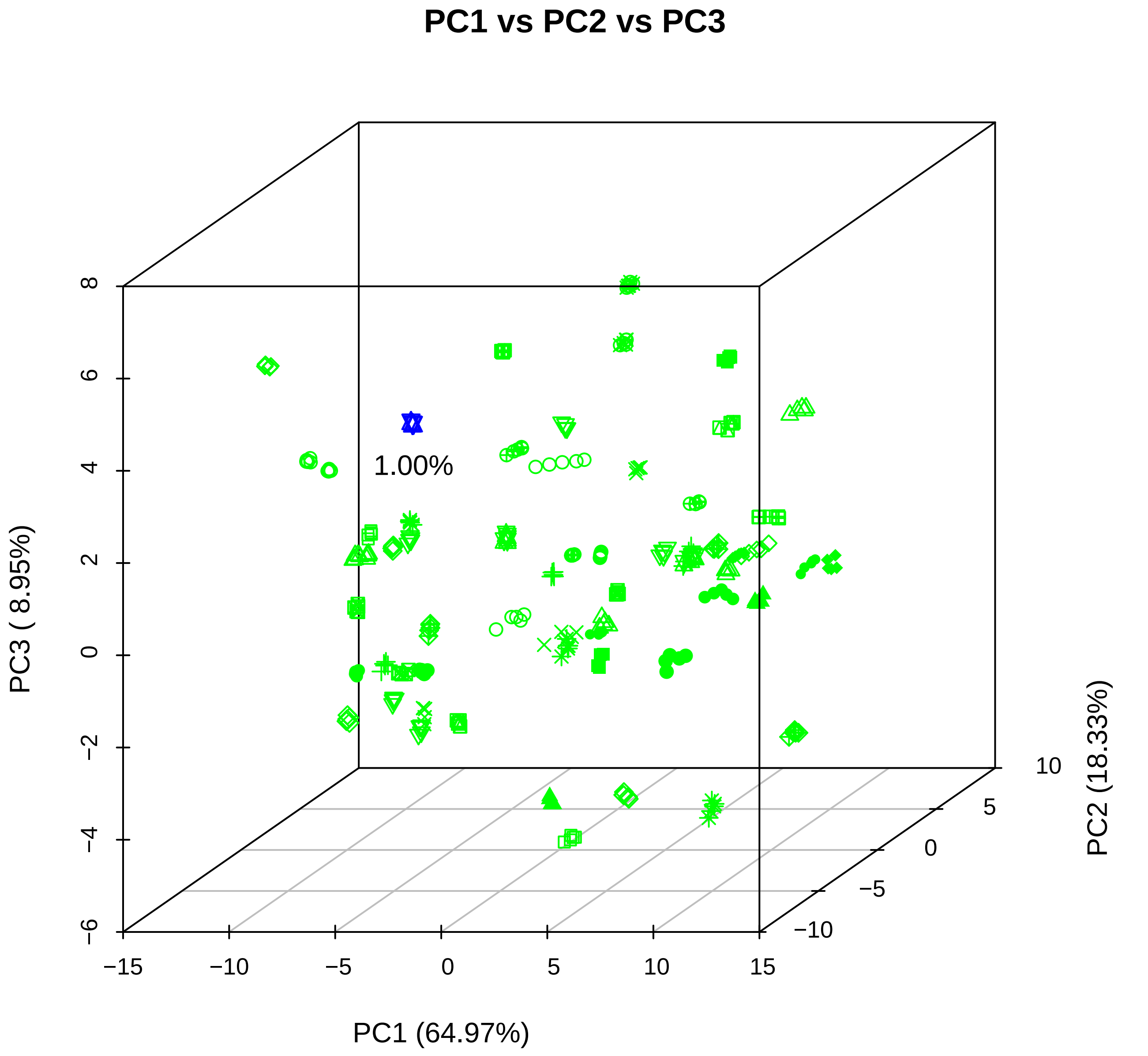
PC1 vs PC2 vs PC3
1.00%
PC1 (64.97%)
PC2 (18.33%)
PC3 ( 8.95%)
−15
−10
−5
0
5
10
15
−10
−5
0
5
10
−6
−4
−2
0
2
4
6
8

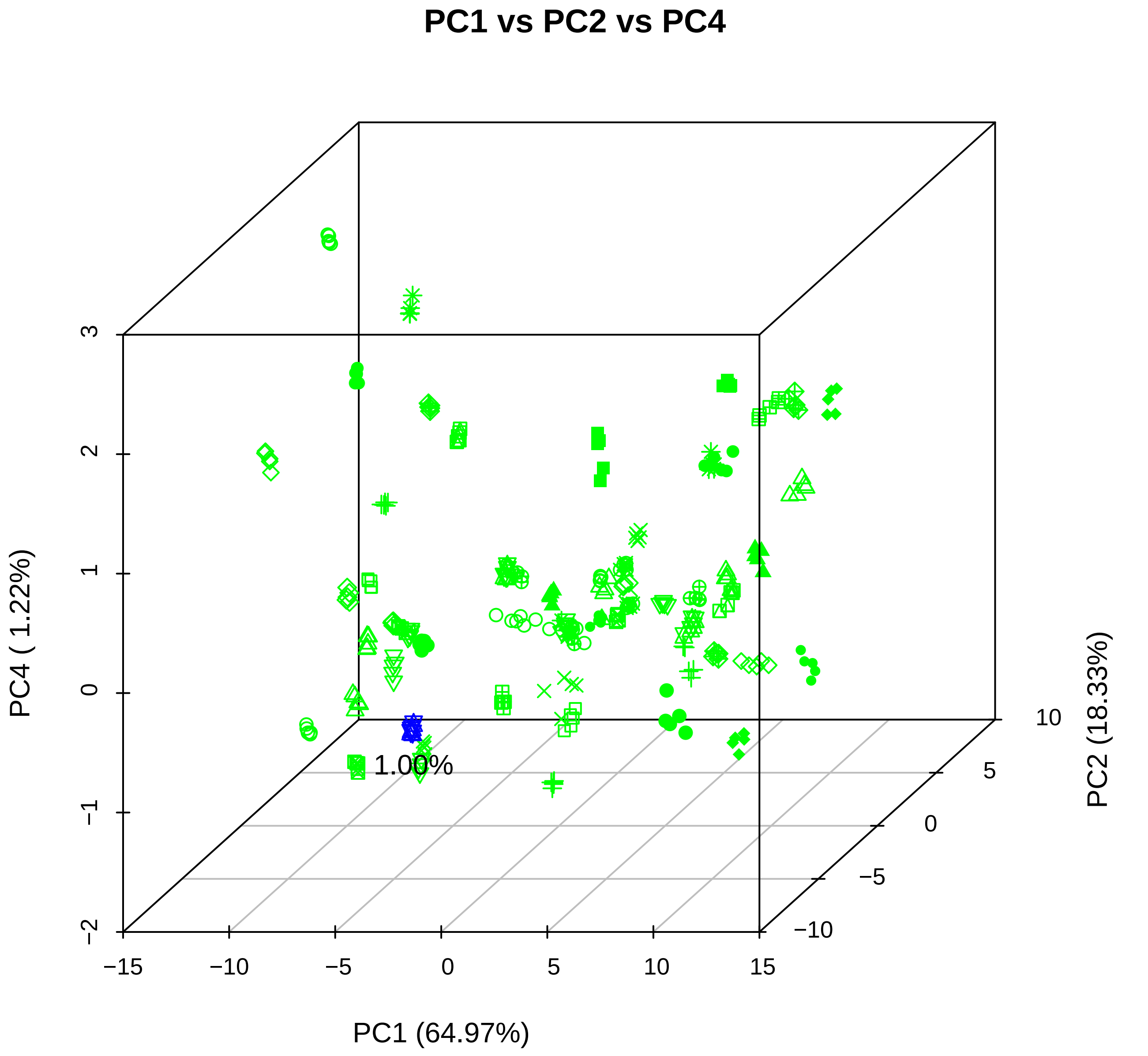

PC1 vs PC2 vs PC4
PC1 (64.97%)
PC2 (18.33%)
PC4 ( 1.22%)
1.00%
−15
−10
−5
0
5
10
15
−2
−1
0
1
2
3
−10
−5
0
5
10

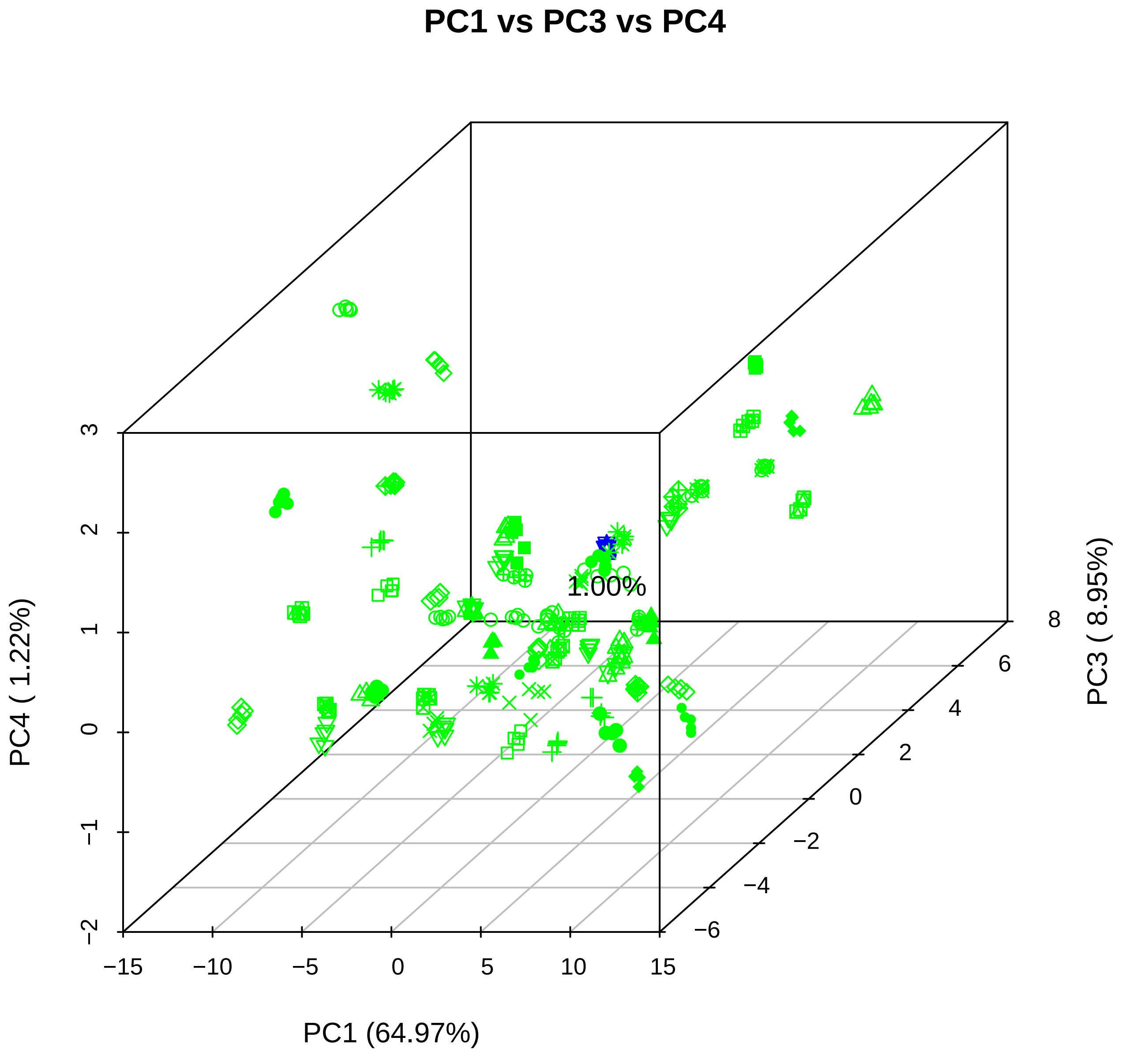
PC1 vs PC3 vs PC4
PC1 (64.97%)
PC3 ( 8.95%)
PC4 ( 1.22%)
1.00%
−15
−10
−5
0
5
10
15
−6
−4
−2
0
2
4
6
8
−2
−1
0
1
2
3

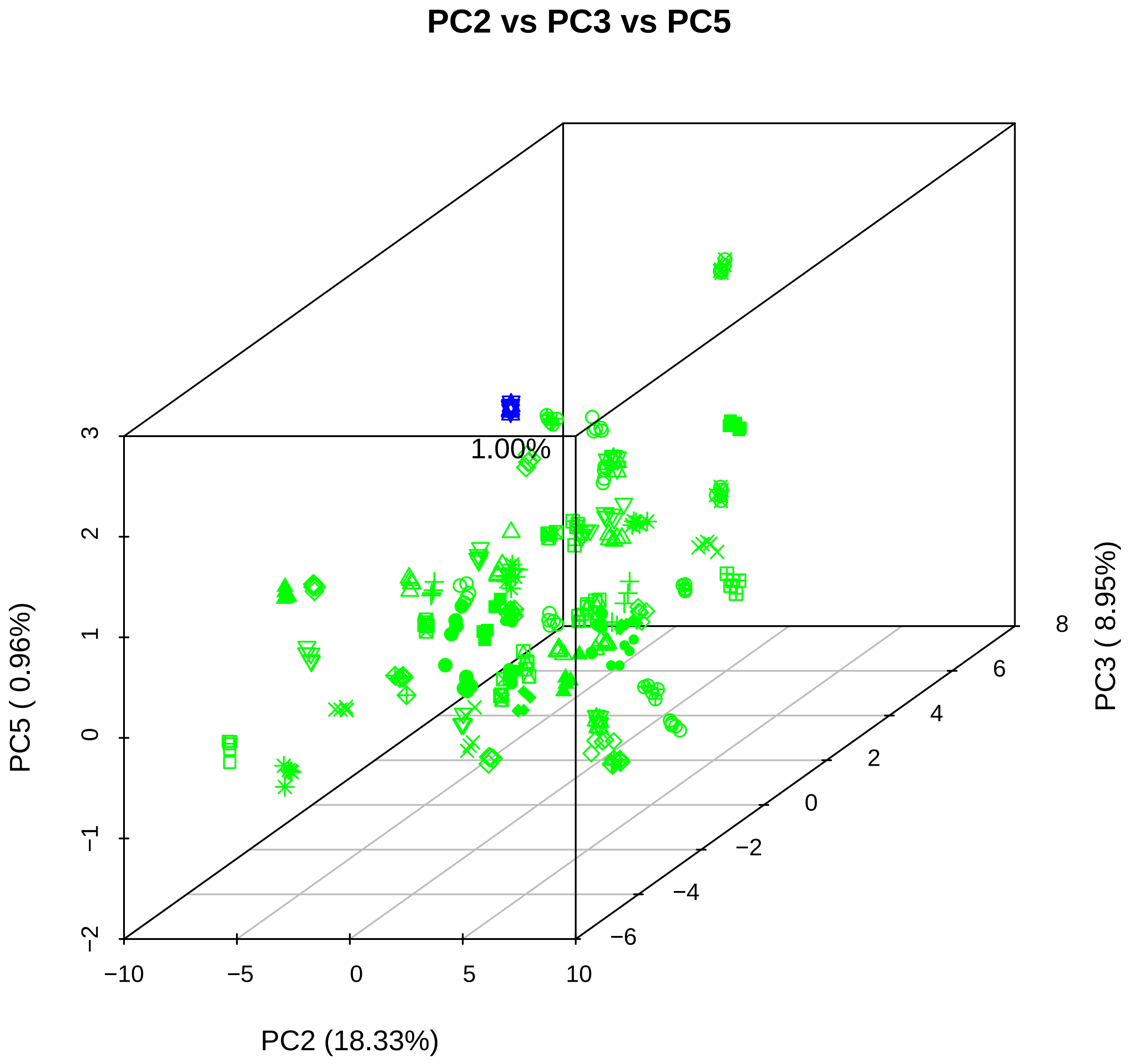

PC2 vs PC3 vs PC5
1.00%
PC5 ( 0.96%)
PC2 (18.33%)
PC3 ( 8.95%)
3
2
1
0
−1
−2
−10
−5
0
5
10
−6
−4
−2
0
2
4
6
8

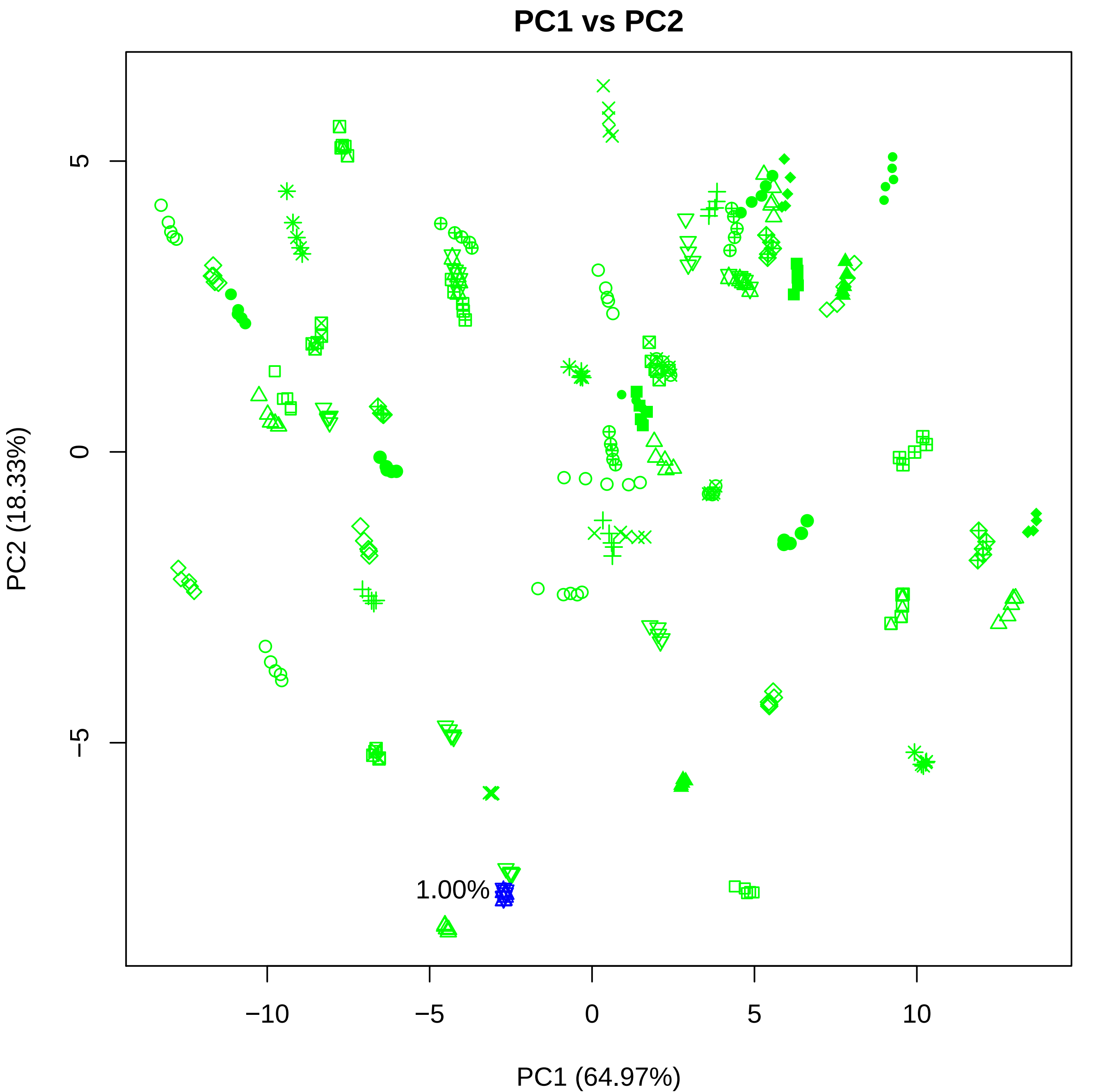
PC1 vs PC2
PC2 (18.33%)
5
0
−5
1.00%
−10
−5
0
5
10
PC1 (64.97%)

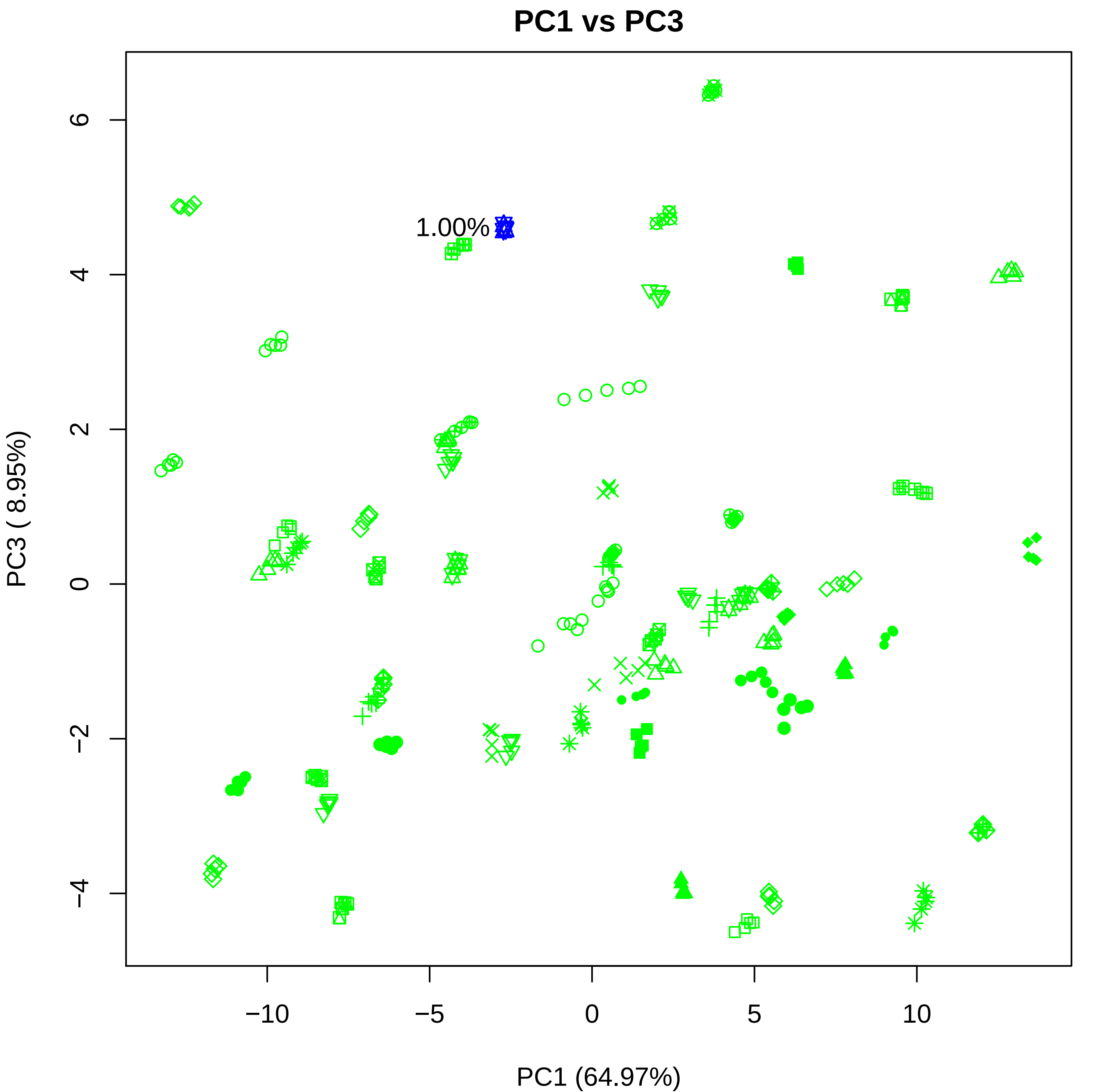

PC1 vs PC3
1.00%
PC3 ( 8.95%)
PC1 (64.97%)
6
4
2
0
−2
−4
−10
−5
0
5
10

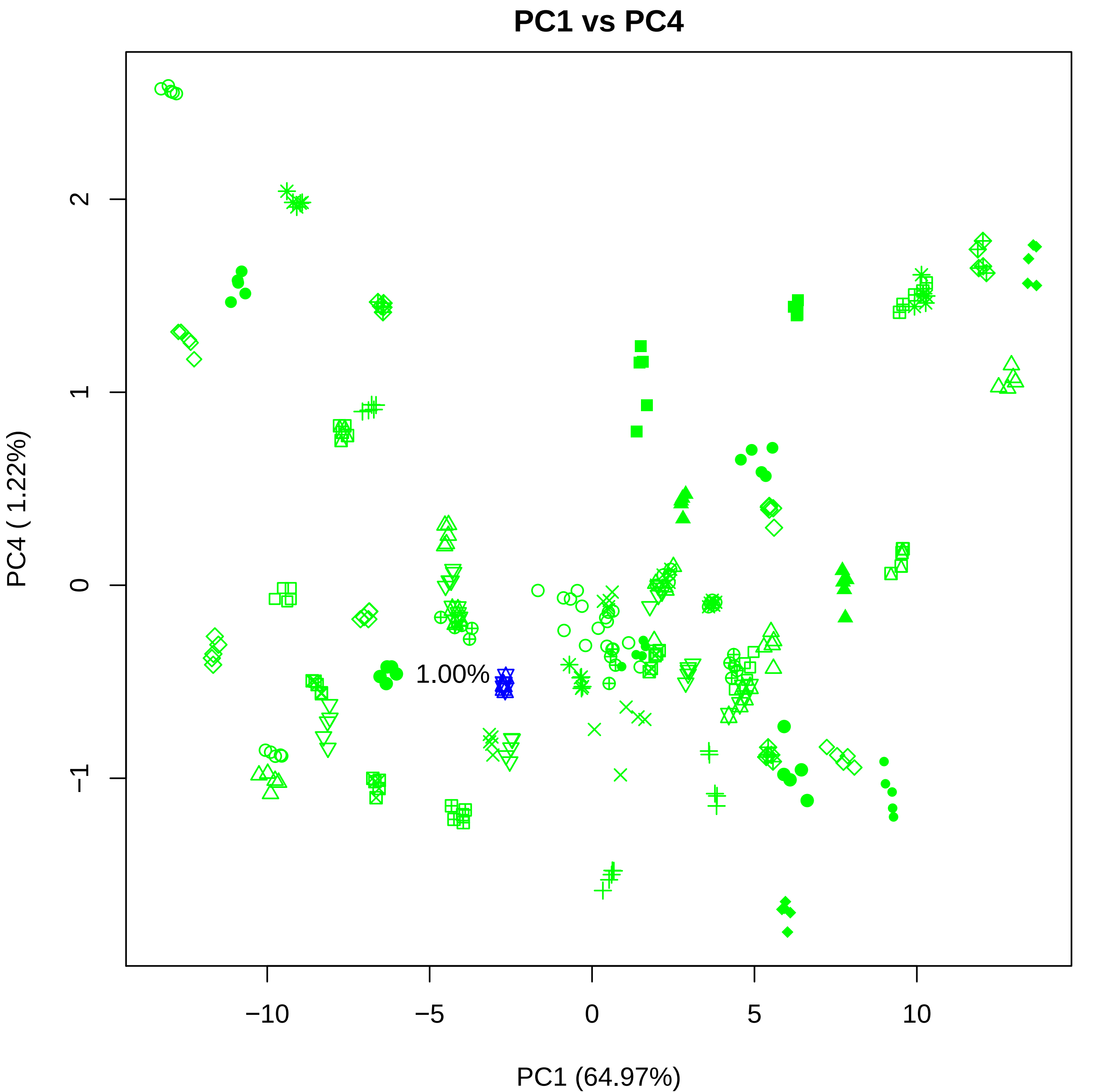

PC1 vs PC4
1.00%
PC4 ( 1.22%)
PC1 (64.97%)
−10
−5
0
5
10
2
1
0
−1

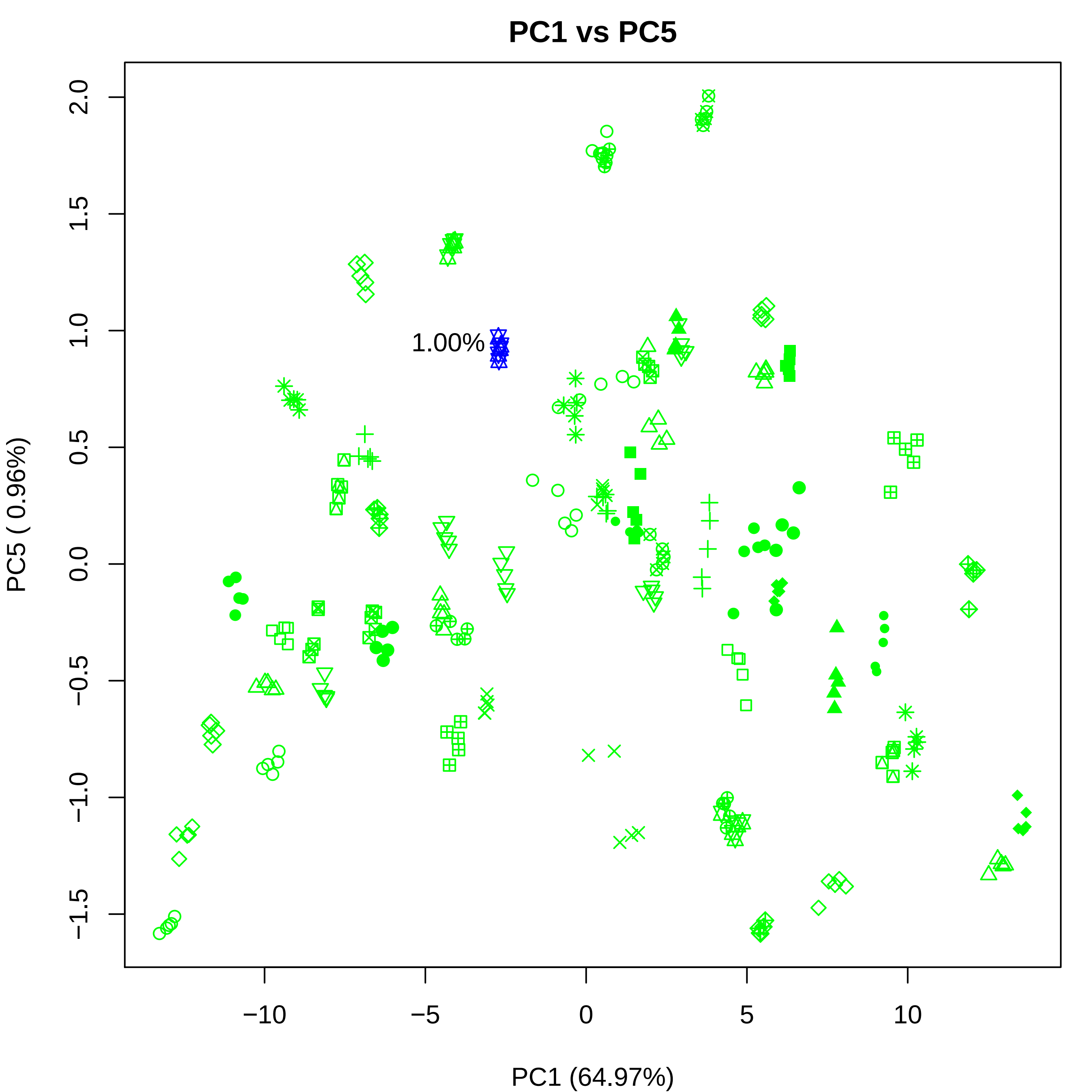
PC1 vs PC5
1.00%
PC5 ( 0.96%)
PC1 (64.97%)
2.0
1.5
1.0
0.5
0.0
−0.5
−1.0
−1.5
−10
−5
0
5
10

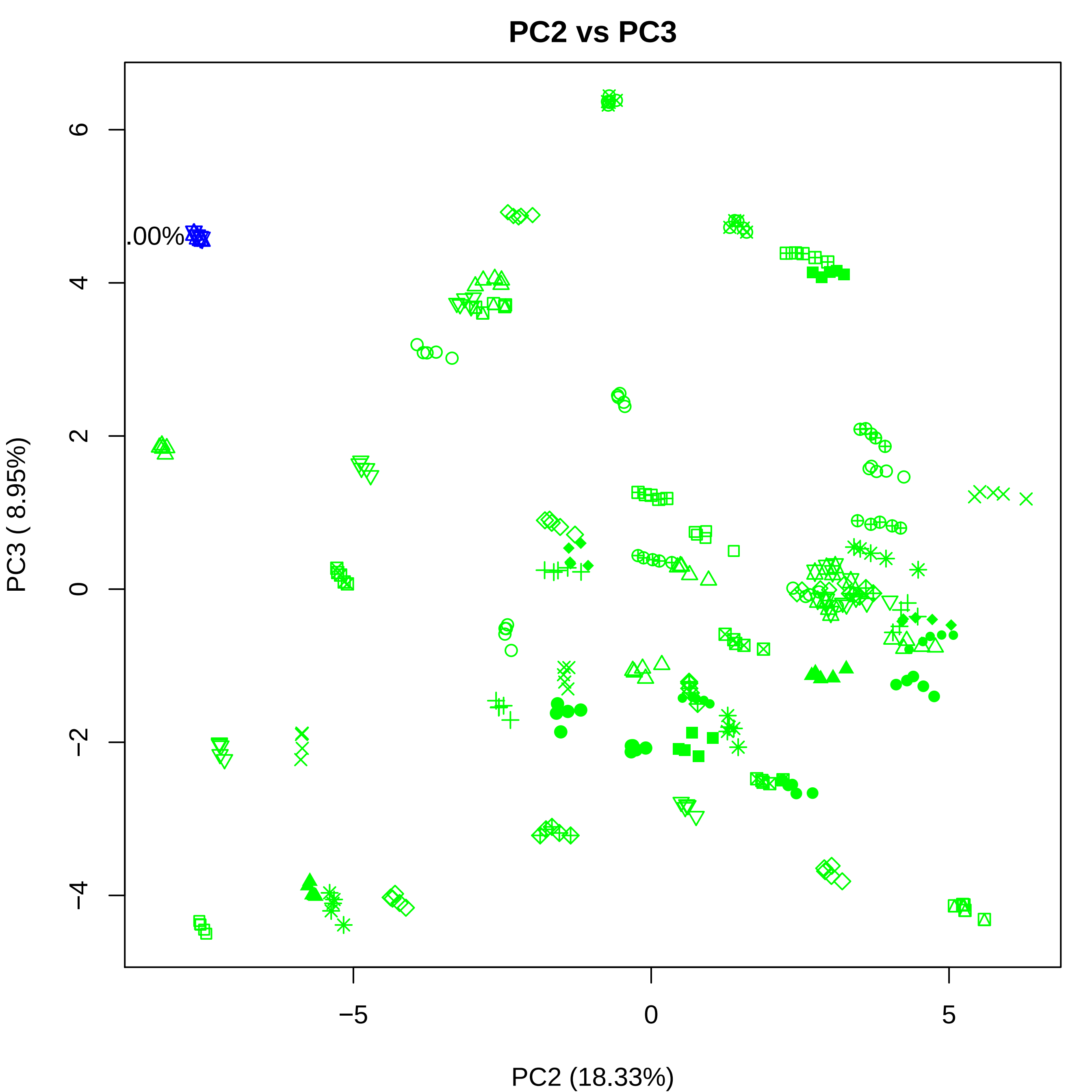

PC2 vs PC3
PC3 ( 8.95%)
PC2 (18.33%)
6
4
2
0
−2
−4
−5
0
5
.00%

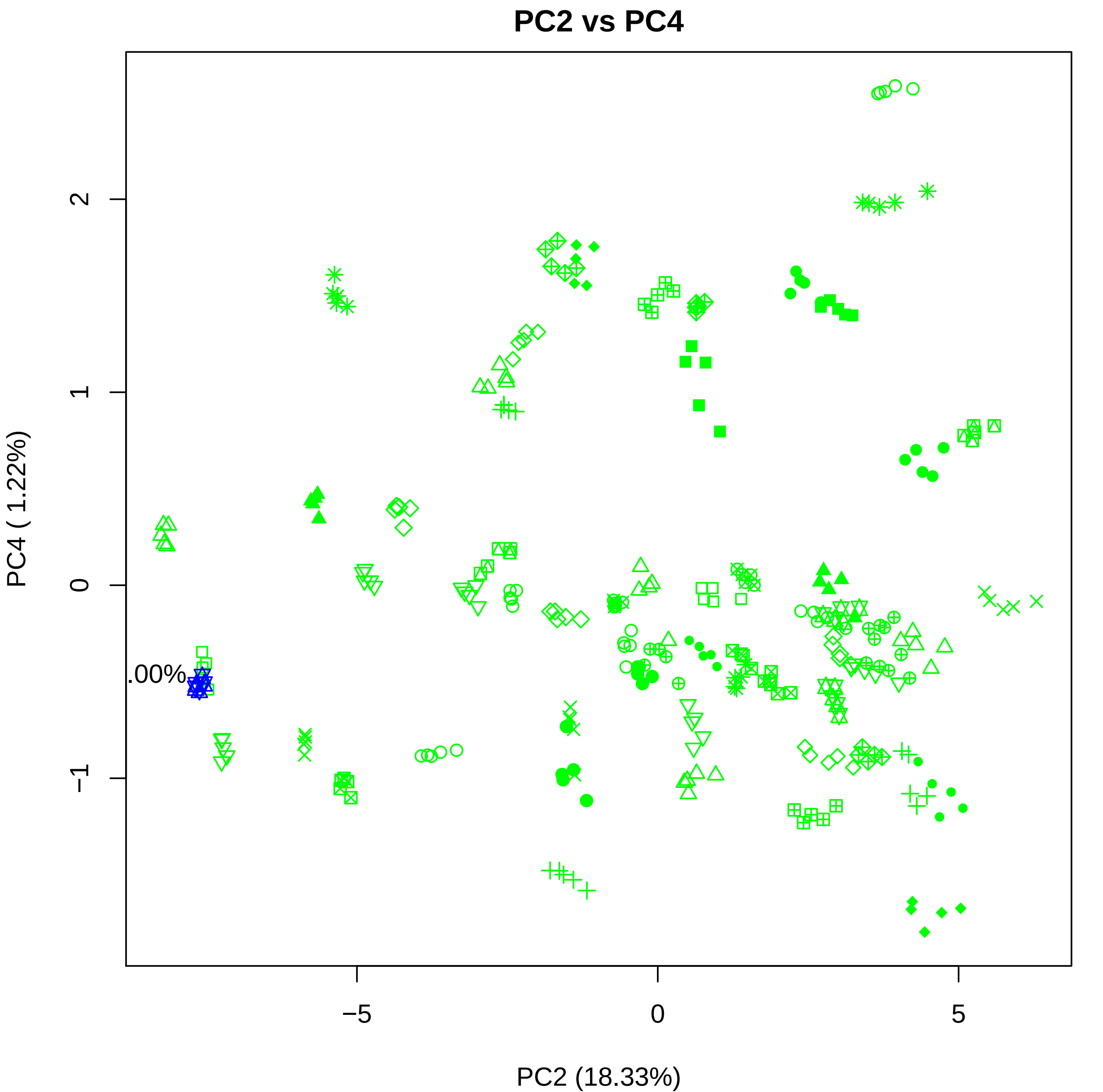
PC2 vs PC4
PC4 ( 1.22%)
PC2 (18.33%)
.00%
2
1
0
−1
−5
0
5

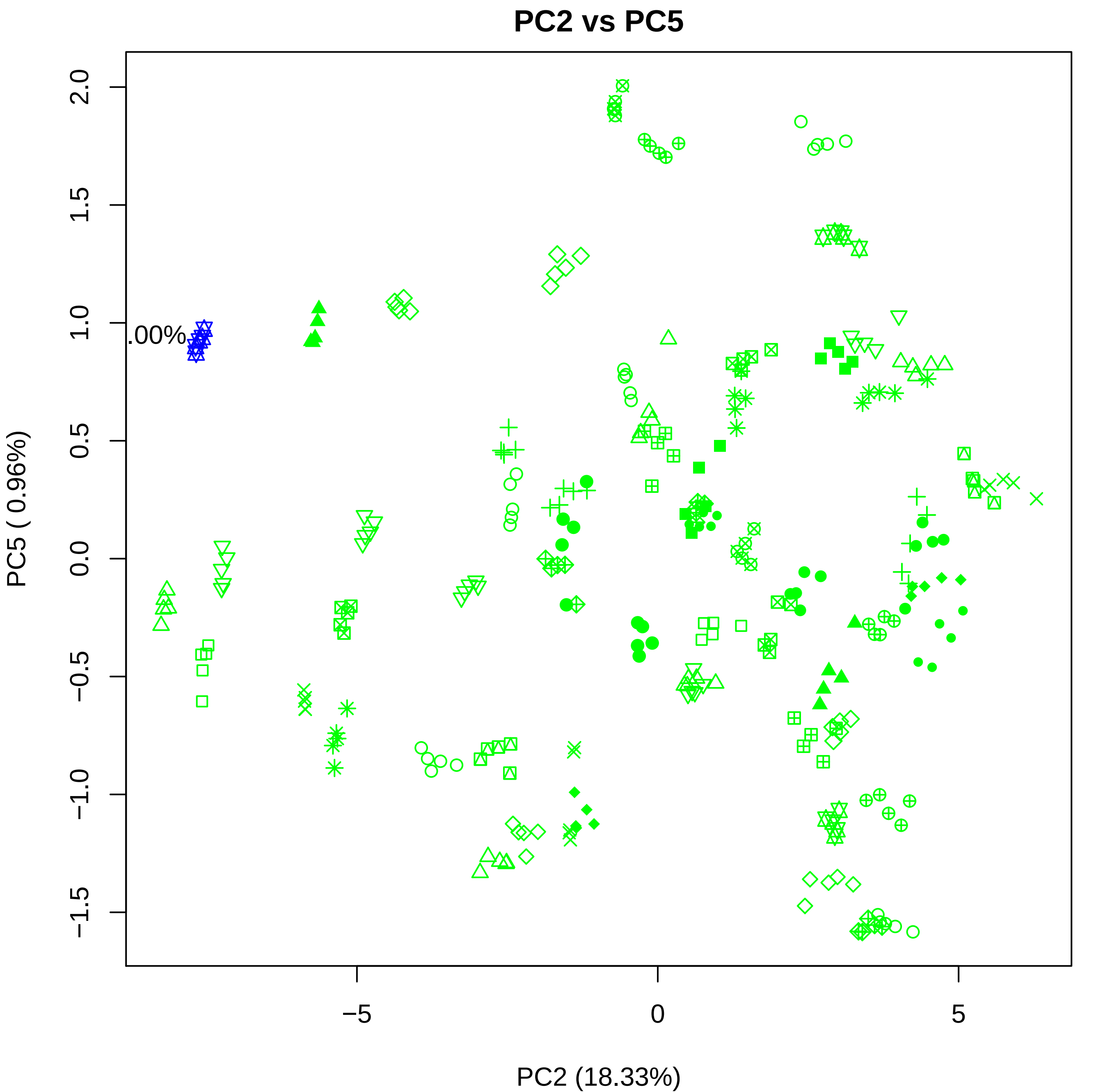

PC2 vs PC5
.00%
2.0
1.5
1.0
0.5
0.0
−0.5
−1.0
−1.5
PC5 ( 0.96%)
−5
0
5
PC2 (18.33%)

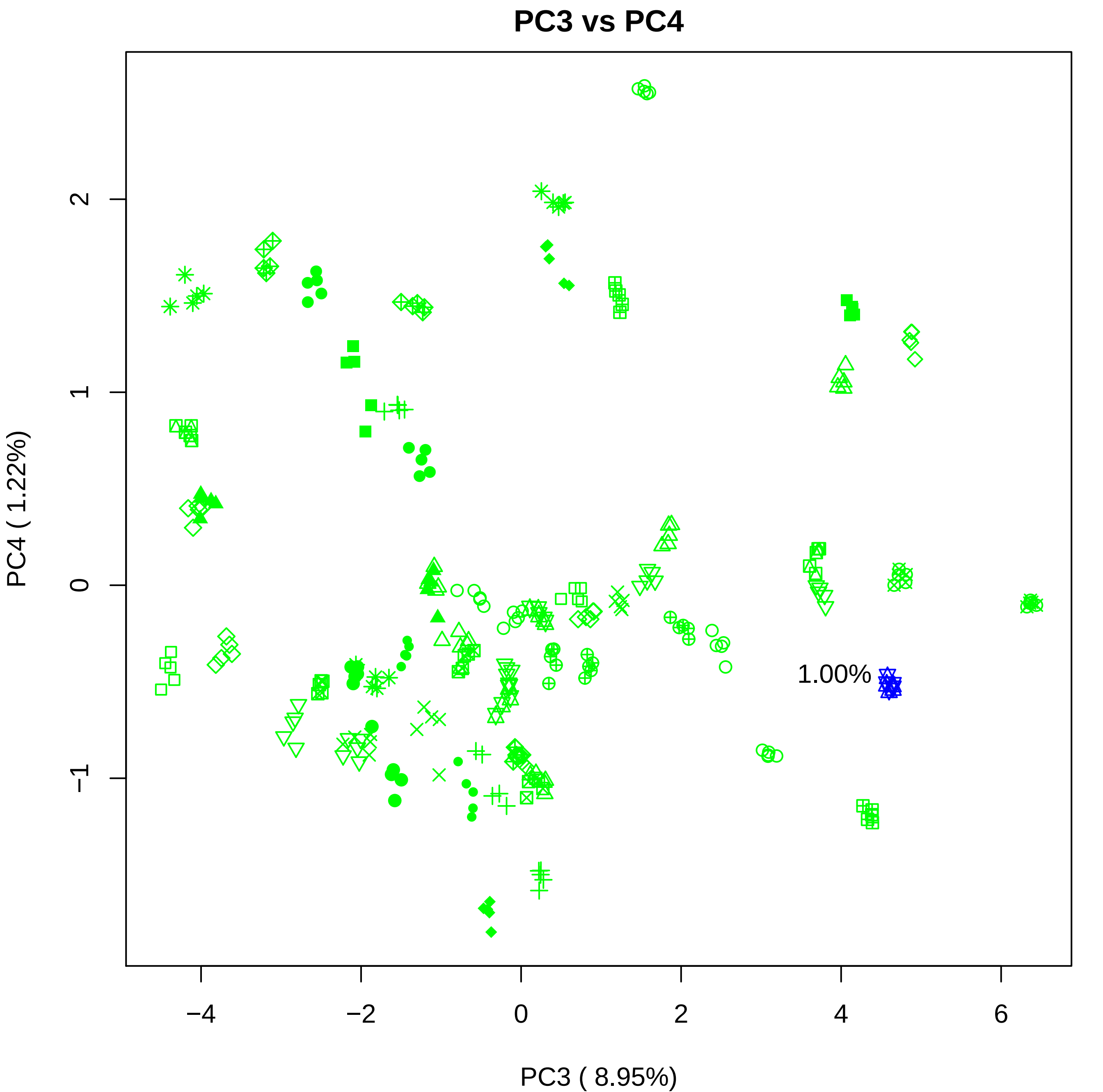
PC3 vs PC4
PC4 ( 1.22%)
PC3 ( 8.95%)
1.00%
2
1
0
−1
−4
−2
0
2
4
6

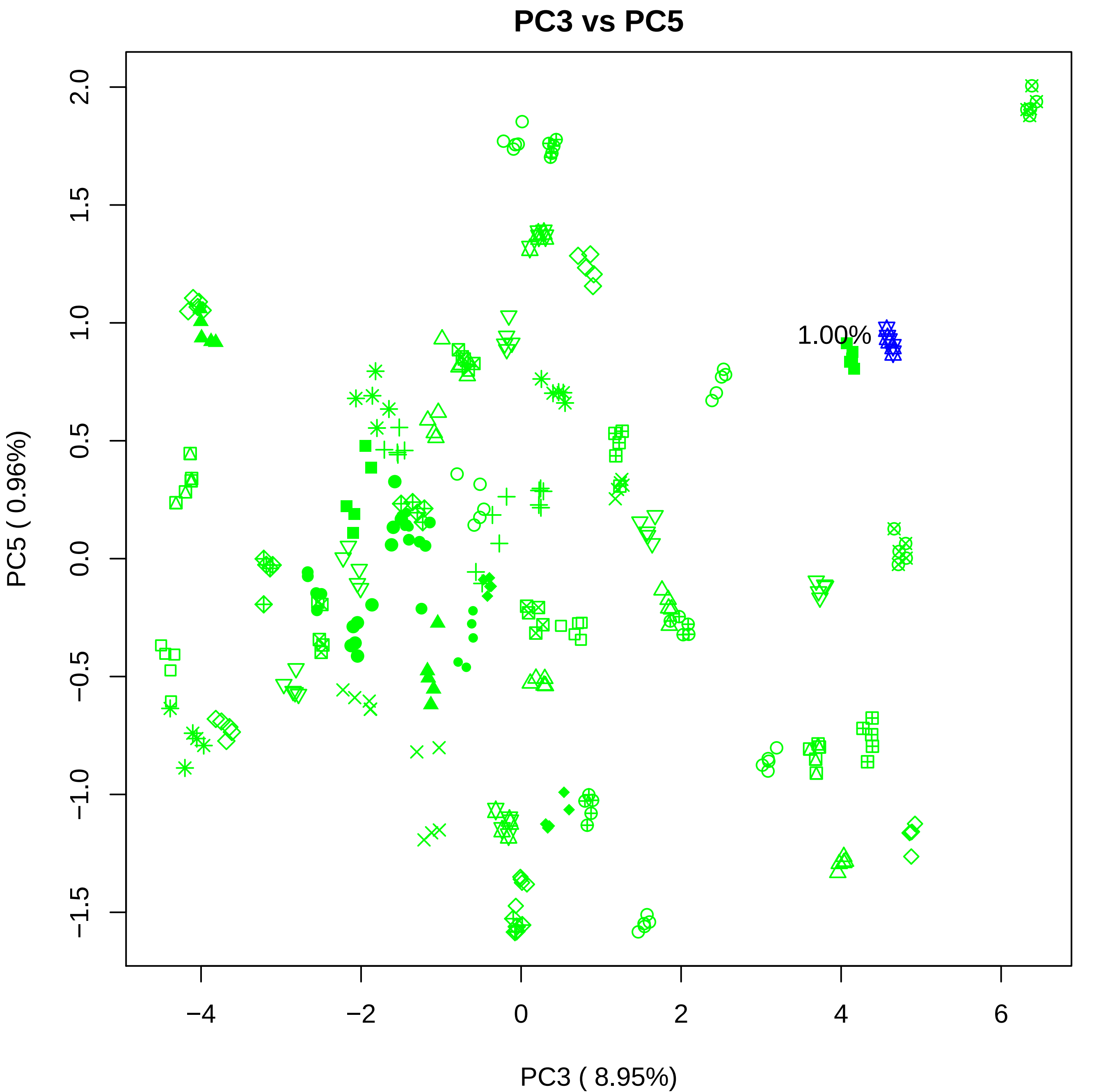
PC3 vs PC5
1.00%
PC5 ( 0.96%)
PC3 ( 8.95%)
2.0
1.5
1.0
0.5
0.0
−0.5
−1.0
−1.5
−4
−2
0
2
4
6

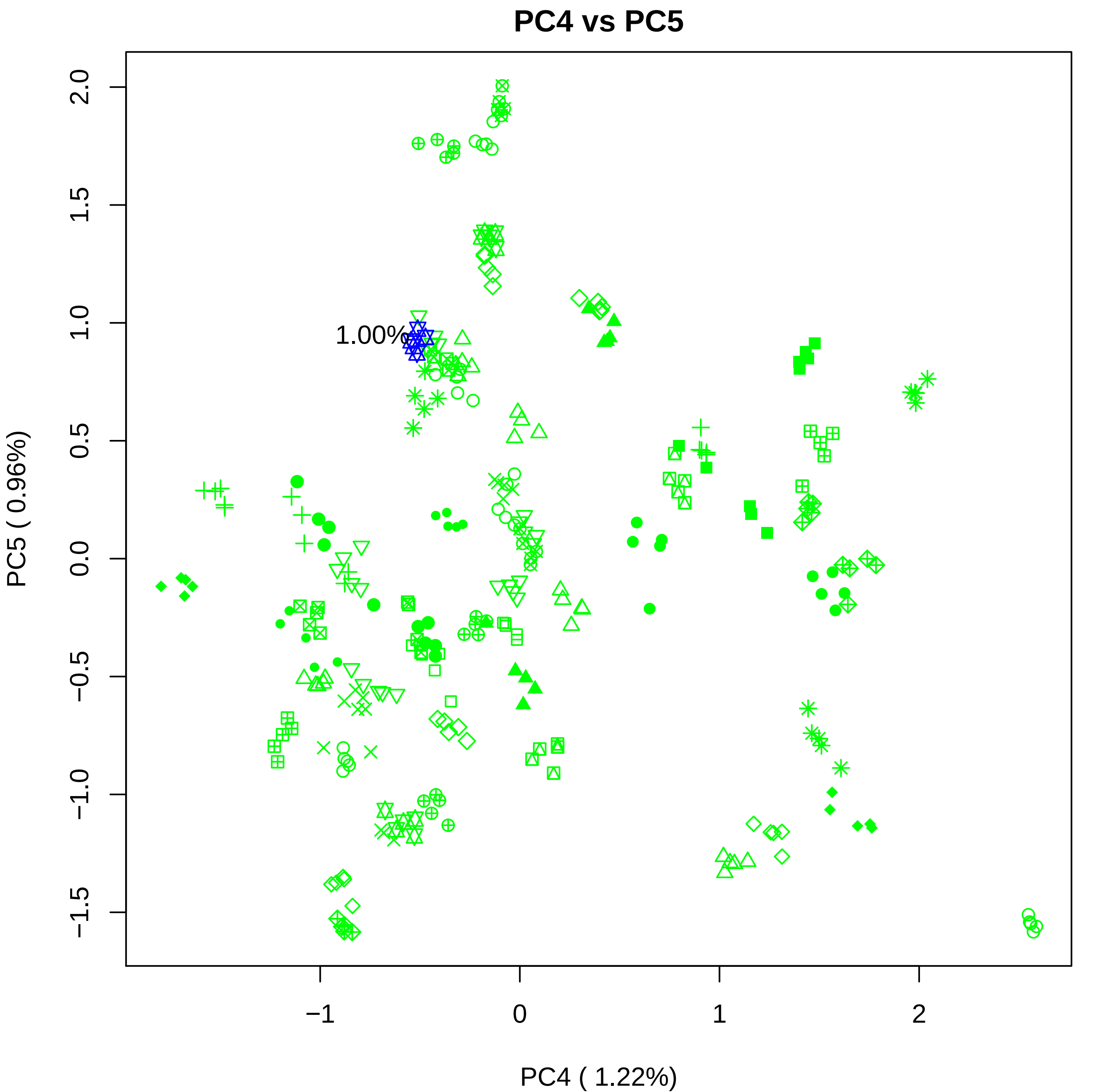
PC4 vs PC5
1.00%
PC5 ( 0.96%)
PC4 ( 1.22%)
2.0
1.5
1.0
0.5
0.0
−0.5
−1.0
−1.5
−1
0
1
2

### PCA of 2.5% BMH (b) vs PMH (b)

Scaled Matrix PCAed

7th order polynomial baseline correction performed

Raw spectra smoothed to 21 Neighbours, 2nd order PN, Savtisky Golay

Normalisation between 0 and 100

D2 spectra smoothed to 41 Neighbours, 2nd order PN, Savtisky Golay

Regions Analysed: 3601−2501 and 2001−700cm−1

Samples Analysed: 1 bContam's 0 BLH's 0 BMH's 0 OMH's 0 pContam's 0 PLH's 60 PMH's

Keele University AD

Fri Dec 14 14:21:17 2018

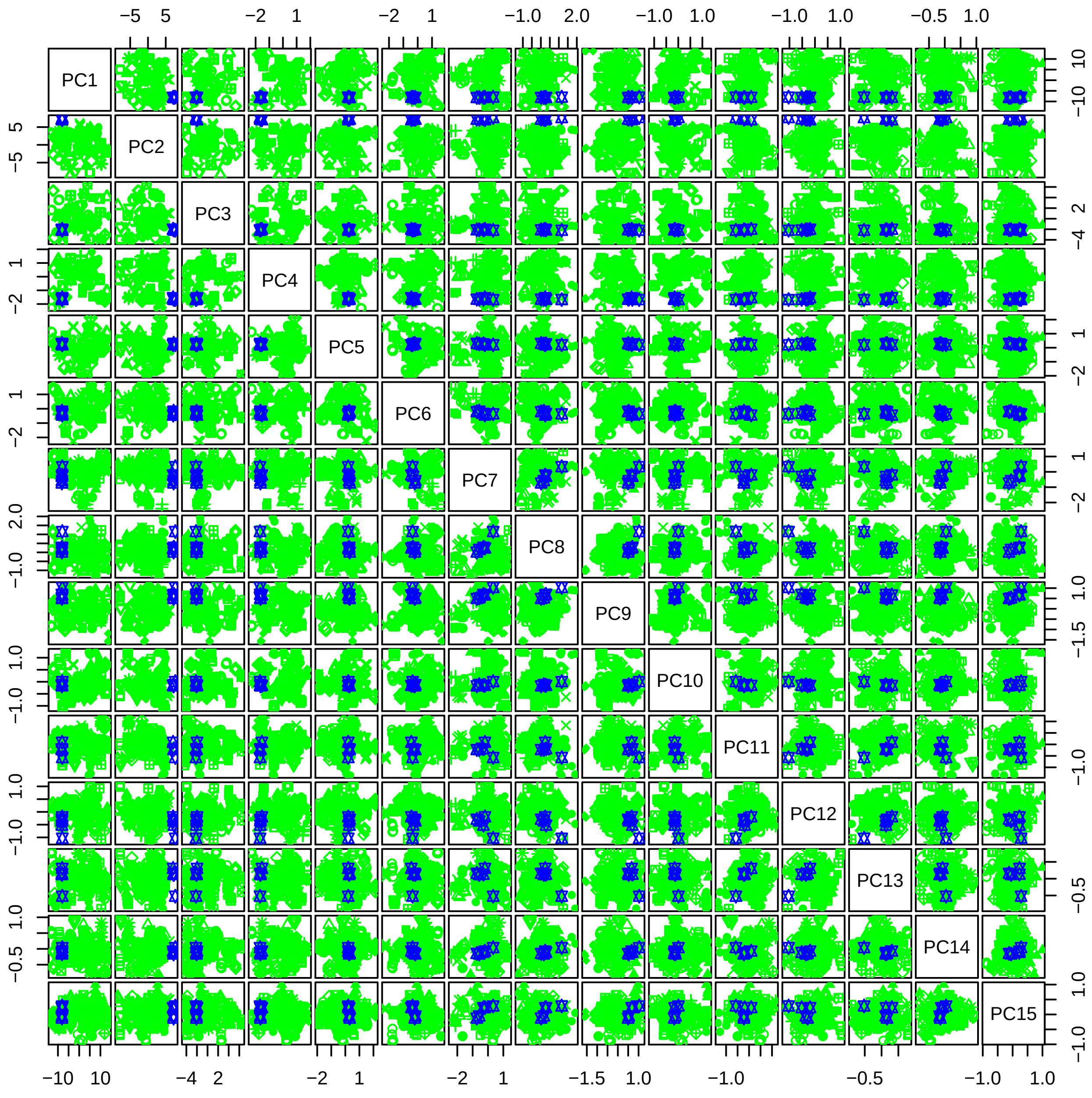
PC1
PC2
PC3
PC4
PC5
PC6
PC7
PC8
PC9
PC10
PC11
PC12
PC13
PC14
PC15

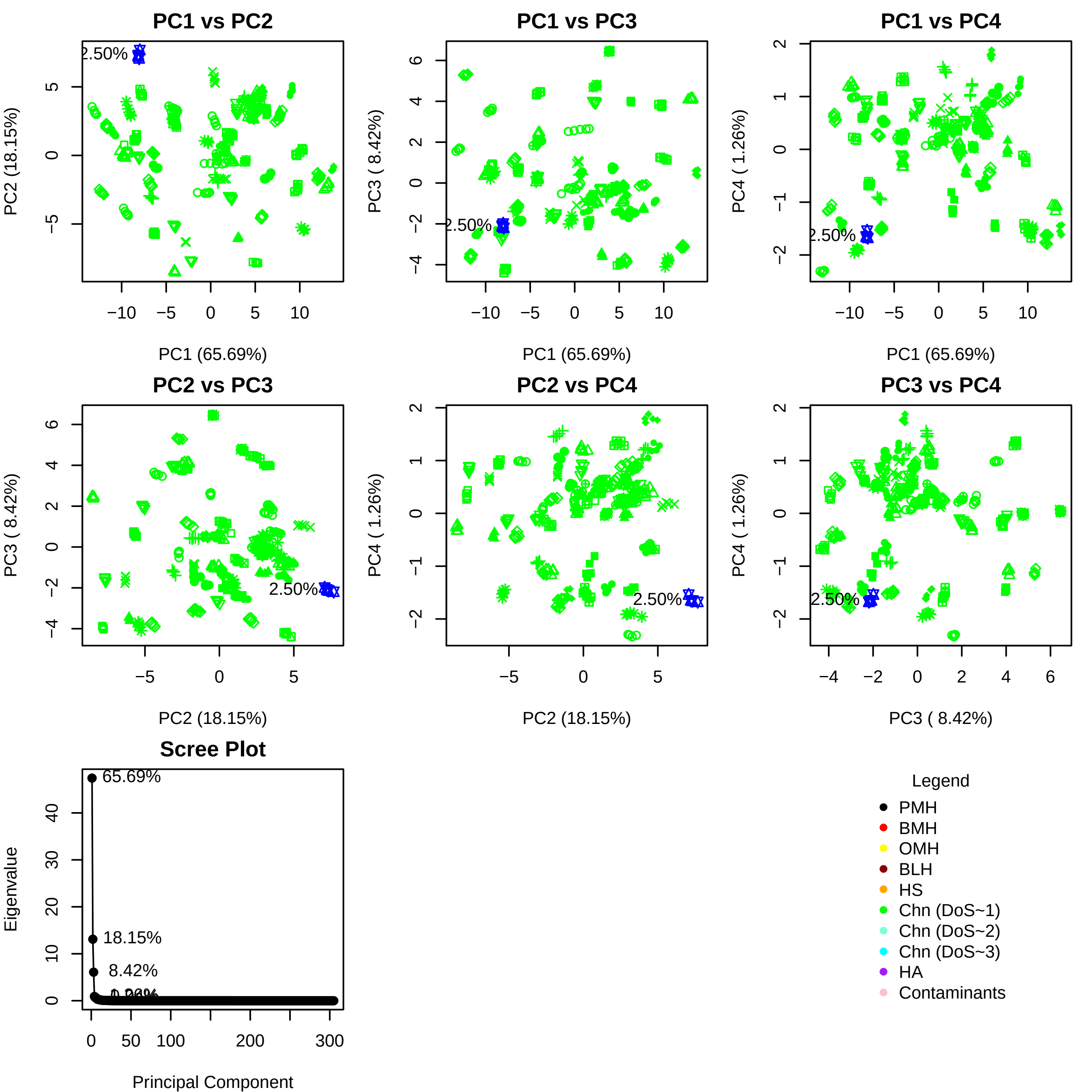
PC1 vs PC2
PC2 (18.15%)
PC1 (65.69%)
2.50%
PC1 vs PC3
PC3 ( 8.42%)
PC1 (65.69%)
2.50%
PC1 vs PC4
PC4 ( 1.26%)
PC1 (65.69%)
2.50%
PC2 vs PC3
PC3 ( 8.42%)
PC2 (18.15%)
2.50%
PC2 vs PC4
PC4 ( 1.26%)
PC2 (18.15%)
2.50%
PC3 vs PC4
PC4 ( 1.26%)
PC3 ( 8.42%)
2.50%
Scree Plot
Eigenvalue
Principal Component
65.69%
18.15%
8.42%
Legend
PMH
BMH
OMH
BLH
HS
Chn (DoS~1)
Chn (DoS~2)
Chn (DoS~3)
HA
Contaminants

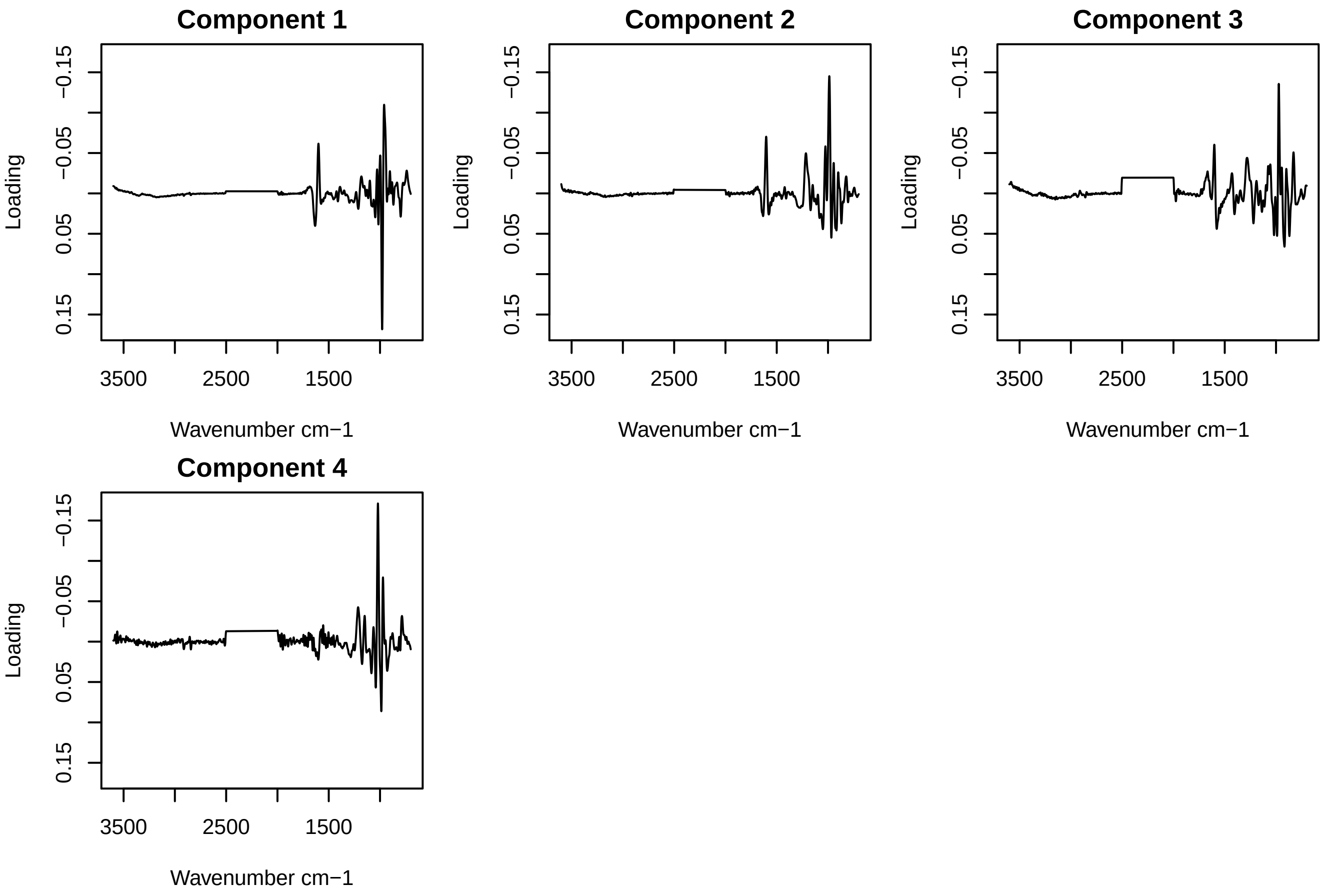
Component 1
Component 2
Component 3
Component 4
Loading
−0.15
−0.05
0.05
0.15
3500
2500
1500
Wavenumber cm−1

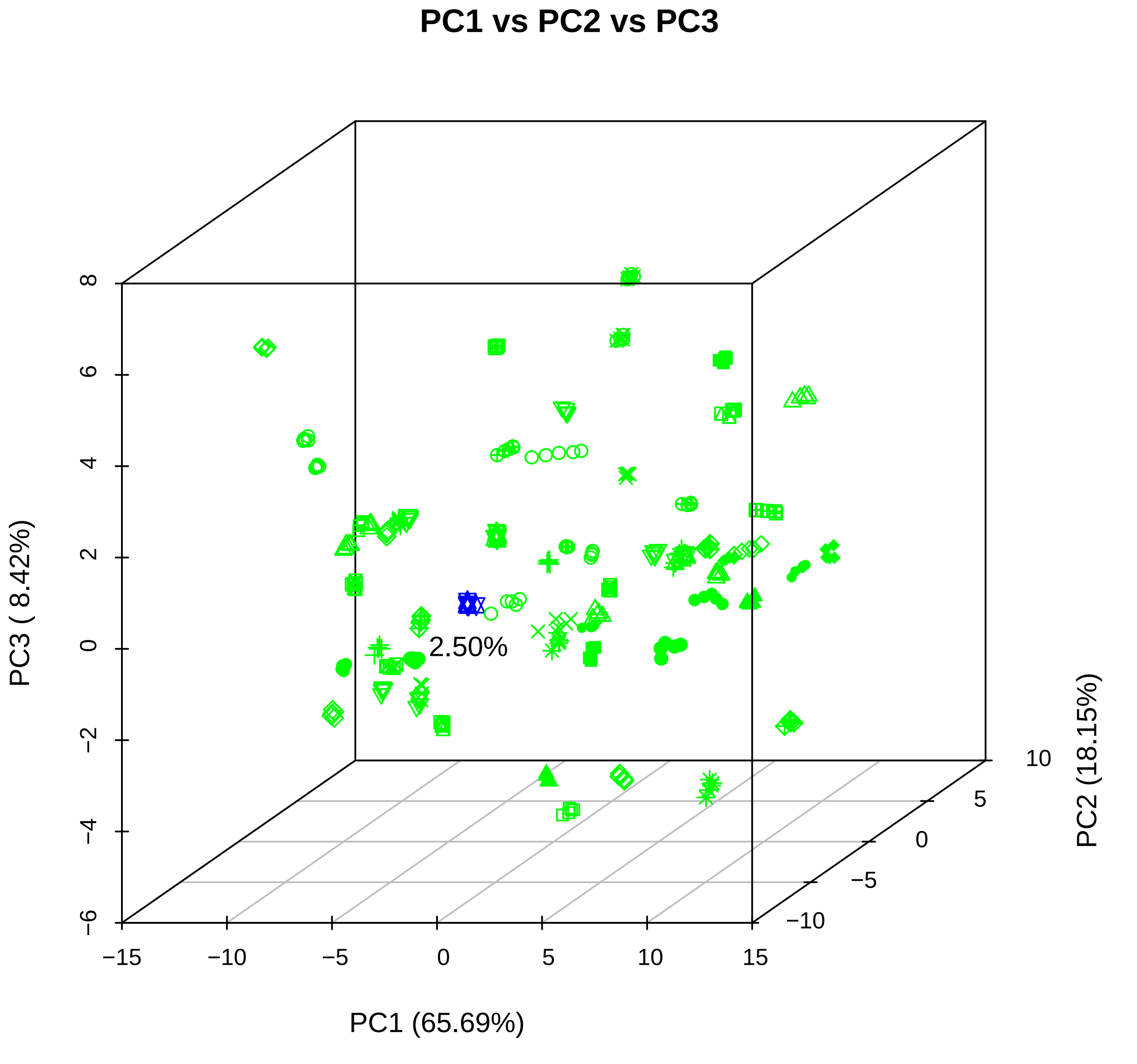
PC1 vs PC2 vs PC3
PC1 (65.69%)
PC2 (18.15%)
PC3 ( 8.42%)
2.50%
−15
−10
−5
0
5
10
15
−6
−4
−2
0
2
4
6
8
−10
−5
0
5
10

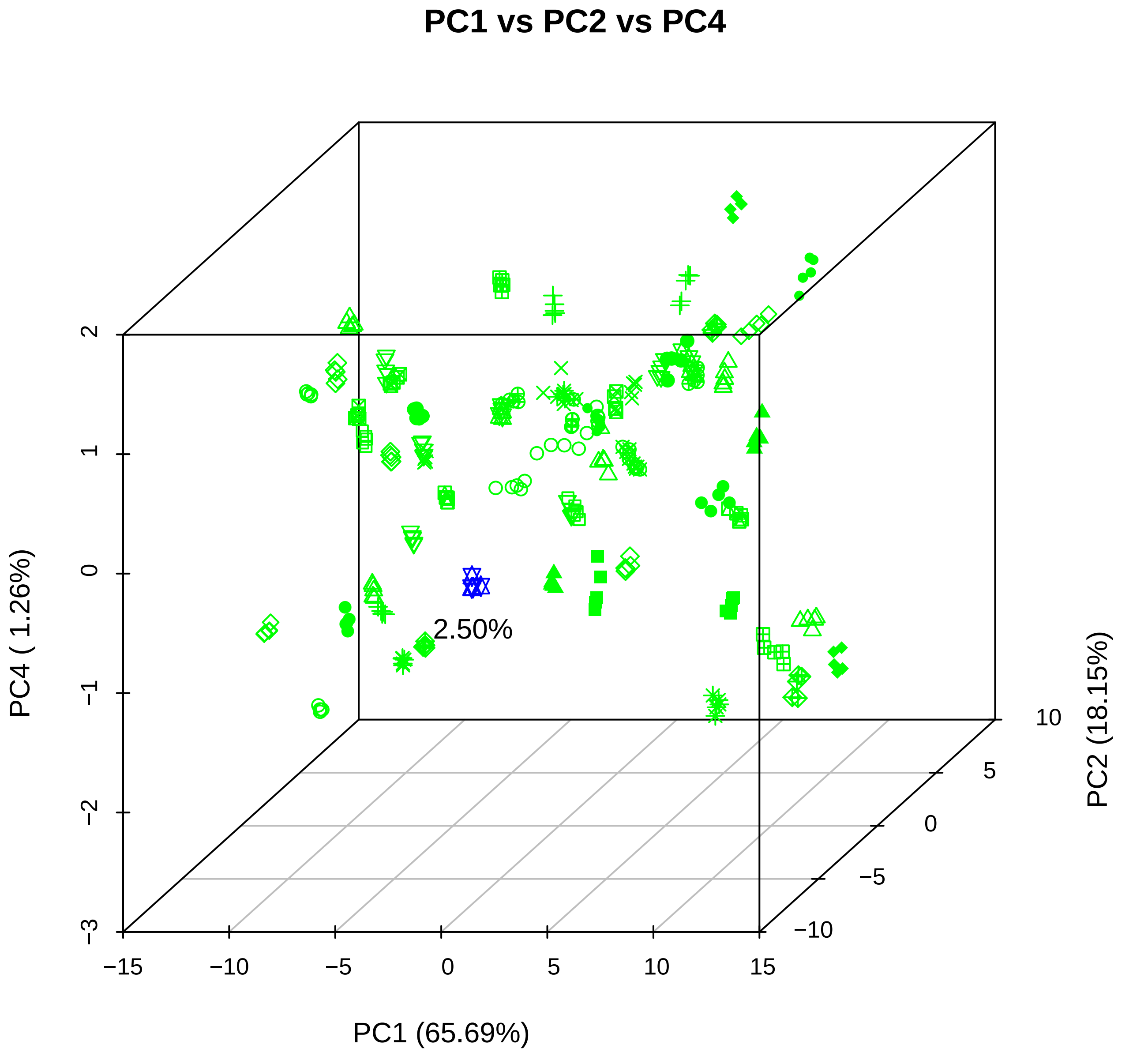
PC1 vs PC2 vs PC4
2.50%
PC1 (65.69%)
PC2 (18.15%)
PC4 ( 1.26%)
−15
−10
−5
0
5
10
15
10
5
0
−5
−10
2
1
0
−1
−2
−3

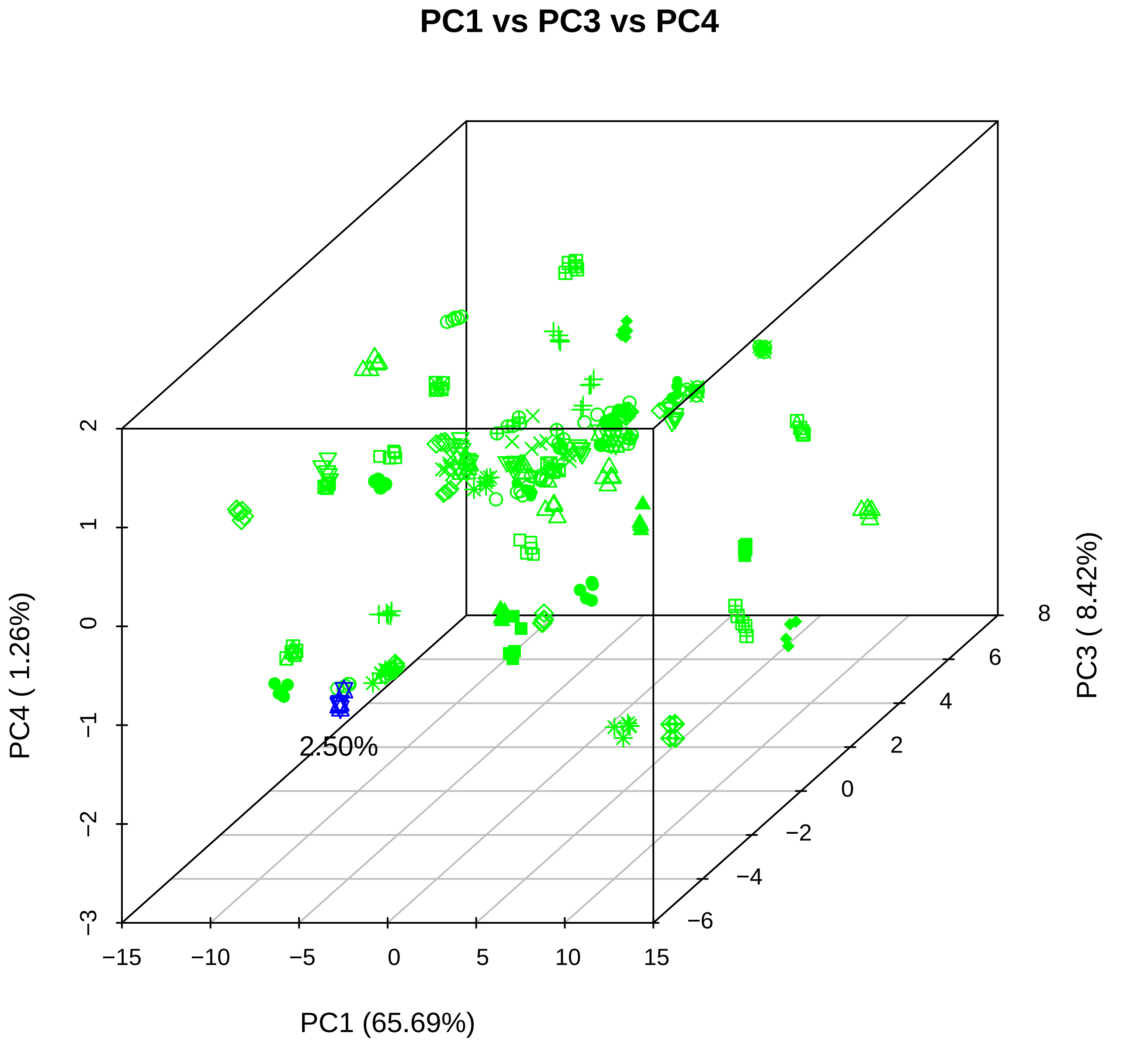

PC1 vs PC3 vs PC4
PC4 ( 1.26%)
2
1
0
−1
−2
−3
−15
−10
−5
0
5
10
15
PC1 (65.69%)
−6
−4
−2
0
2
4
6
8
PC3 ( 8.42%)
2.50%

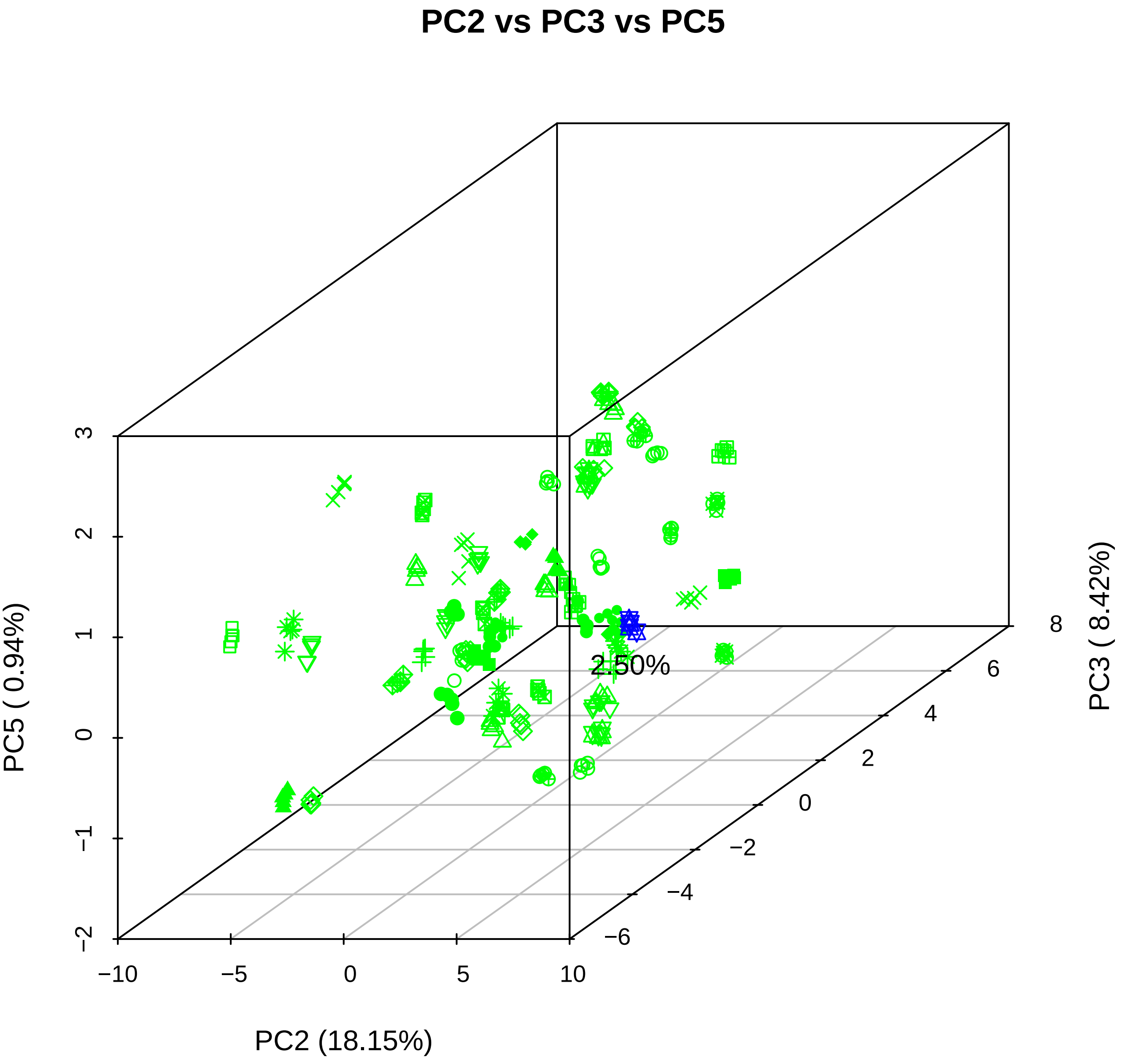

PC2 vs PC3 vs PC5
PC2 (18.15%)
PC5 ( 0.94%)
PC3 ( 8.42%)
2.50%
−10
−5
0
5
10
−2
−1
0
1
2
3
−6
−4
−2
0
2
4
6
8

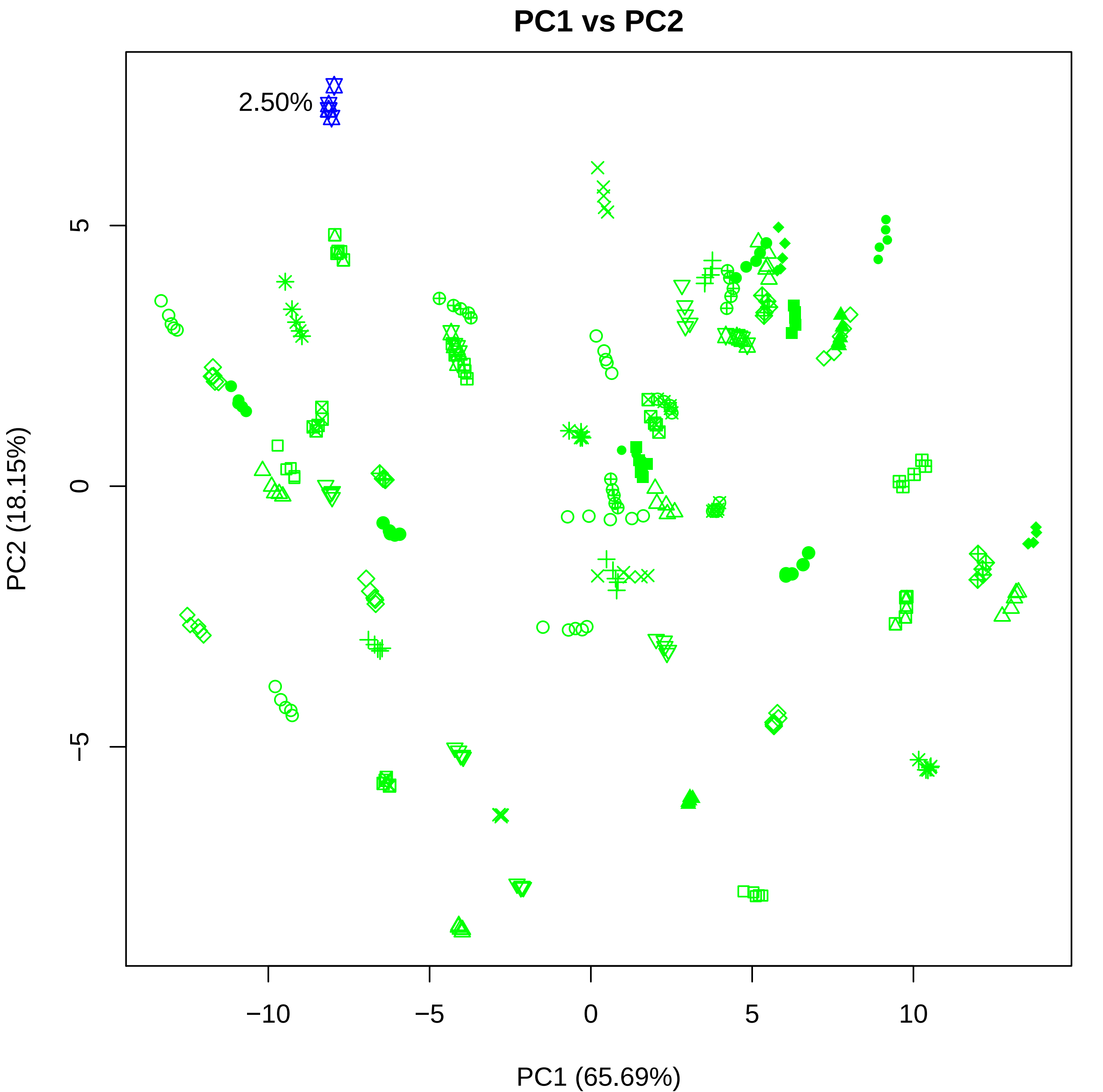

PC1 vs PC2
2.50%
5
0
−5
PC2 (18.15%)
−10
−5
0
5
10
PC1 (65.69%)

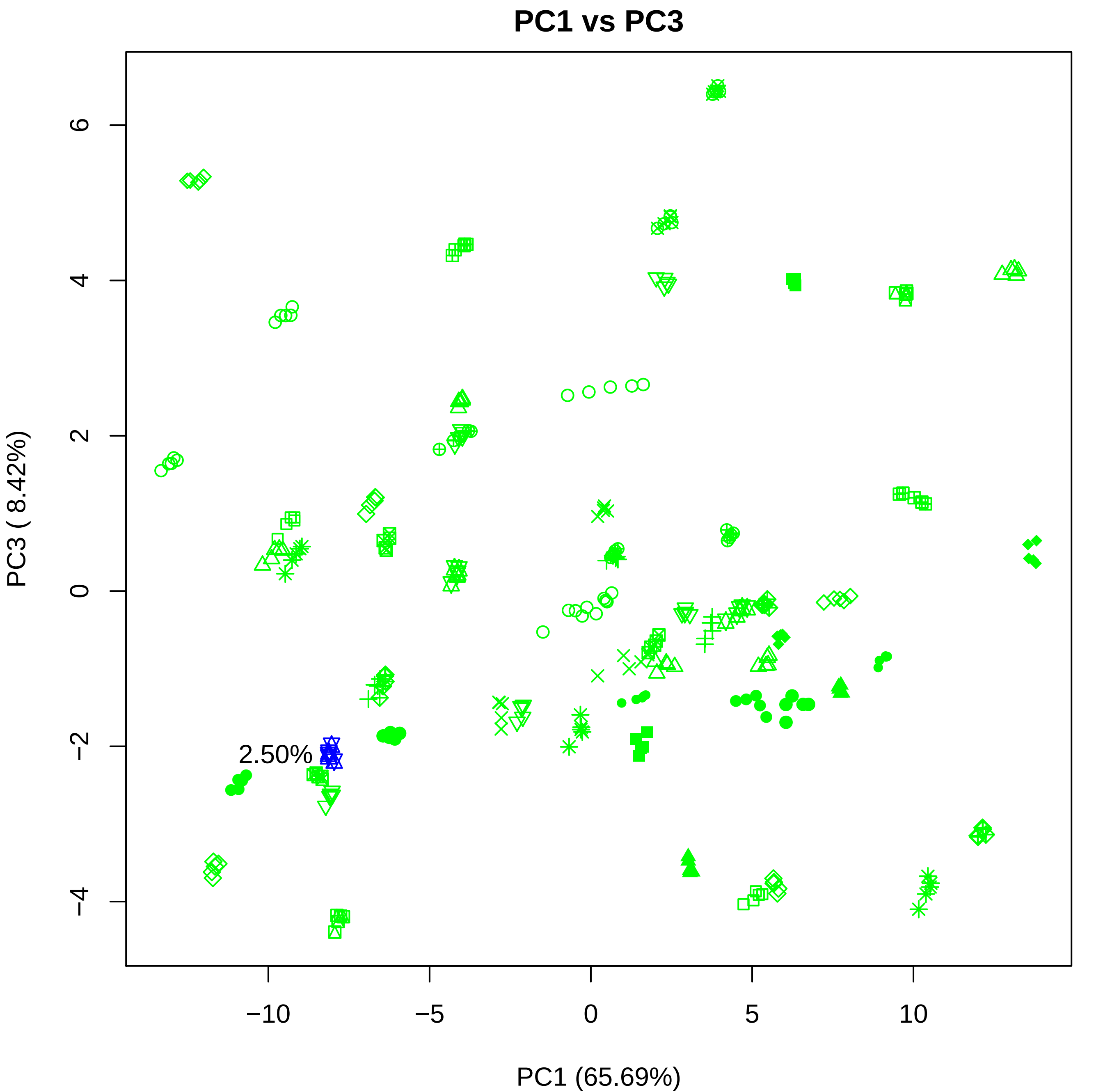
PC1 vs PC3
PC3 ( 8.42%)
PC1 (65.69%)
2.50%
6
4
2
0
−2
−4
−10
−5
0
5
10

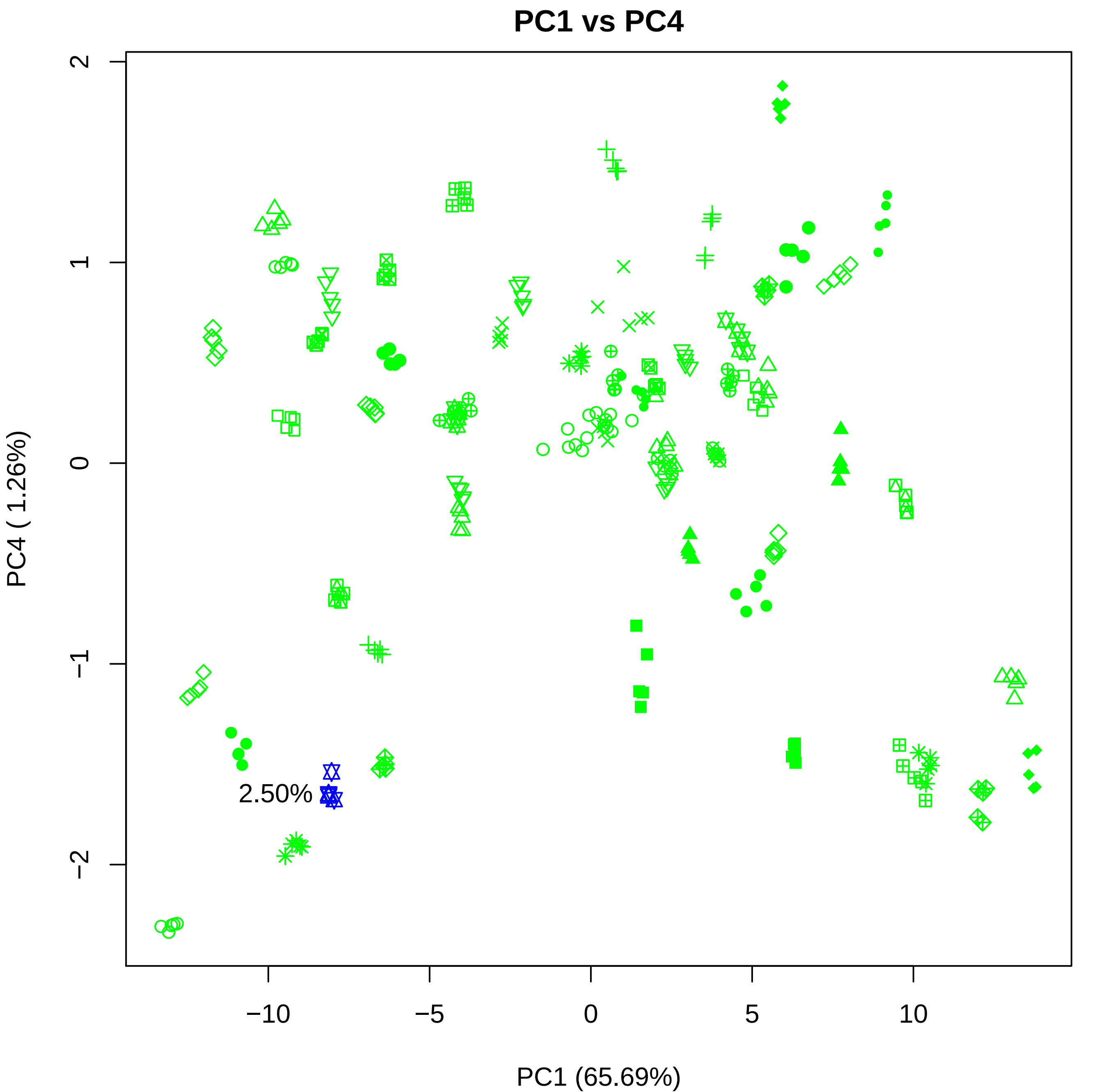
PC1 vs PC4
2.50%
PC4 ( 1.26%)
PC1 (65.69%)
2
1
0
−1
−2
−10
−5
0
5
10

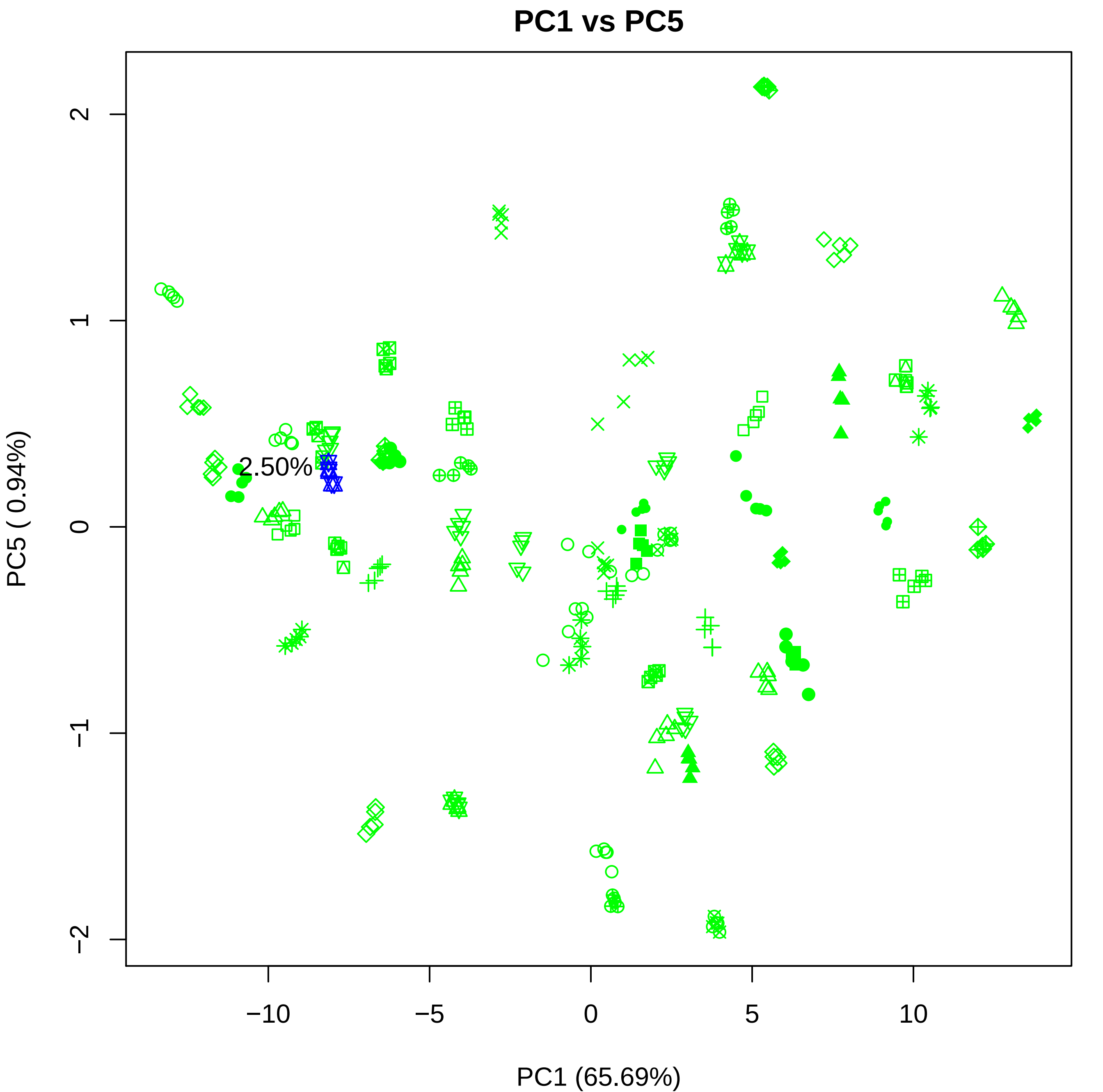

PC1 vs PC5
2.50%
PC5 ( 0.94%)
PC1 (65.69%)
−10
−5
0
5
10
−2
−1
0
1
2

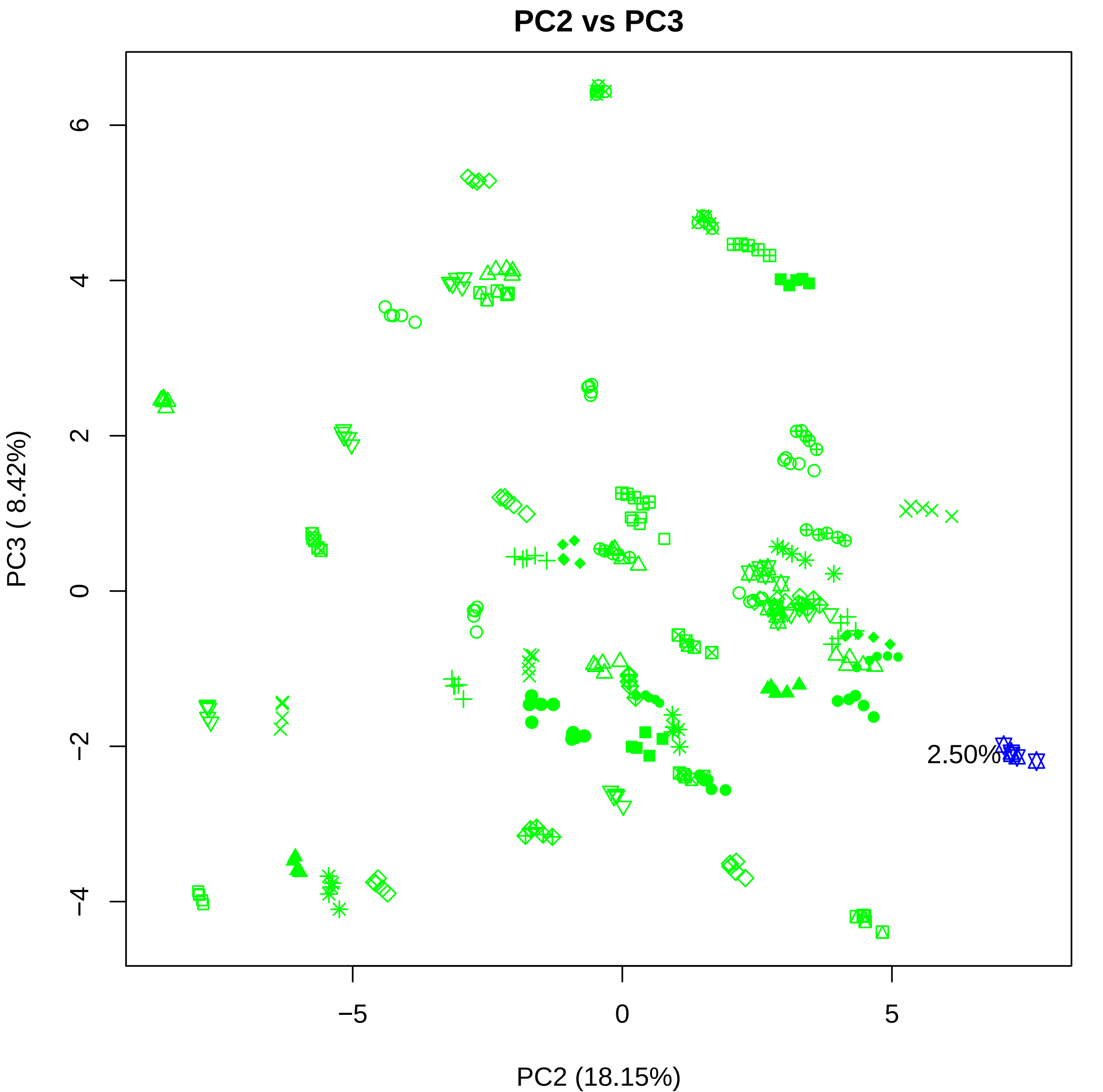

PC2 vs PC3
PC3 ( 8.42%)
PC2 (18.15%)
6
4
2
0
−2
−4
−5
0
5
2.50%

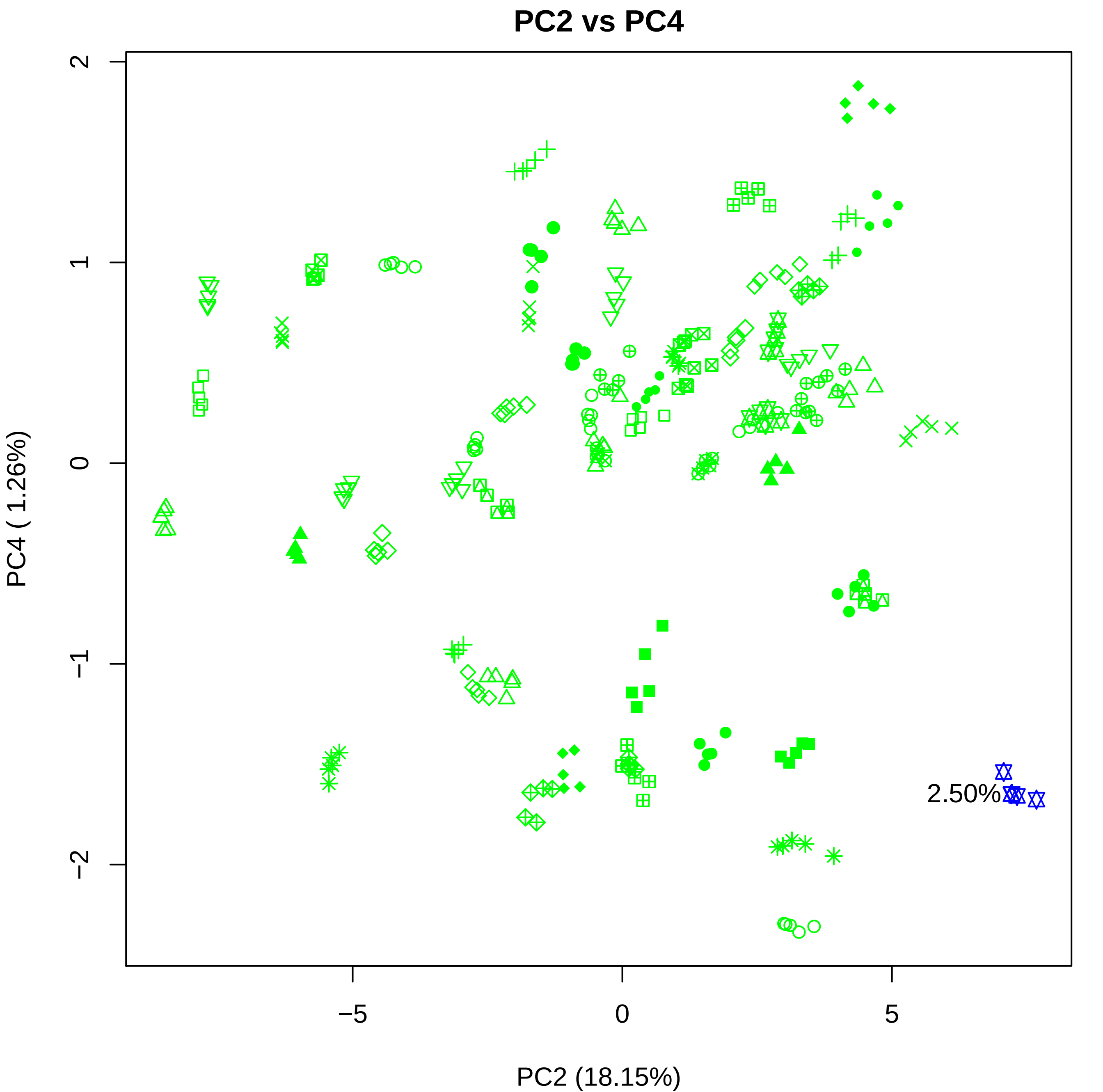
PC2 vs PC4
PC4 ( 1.26%)
PC2 (18.15%)
2.50%
2
1
0
−1
−2
−5
0
5

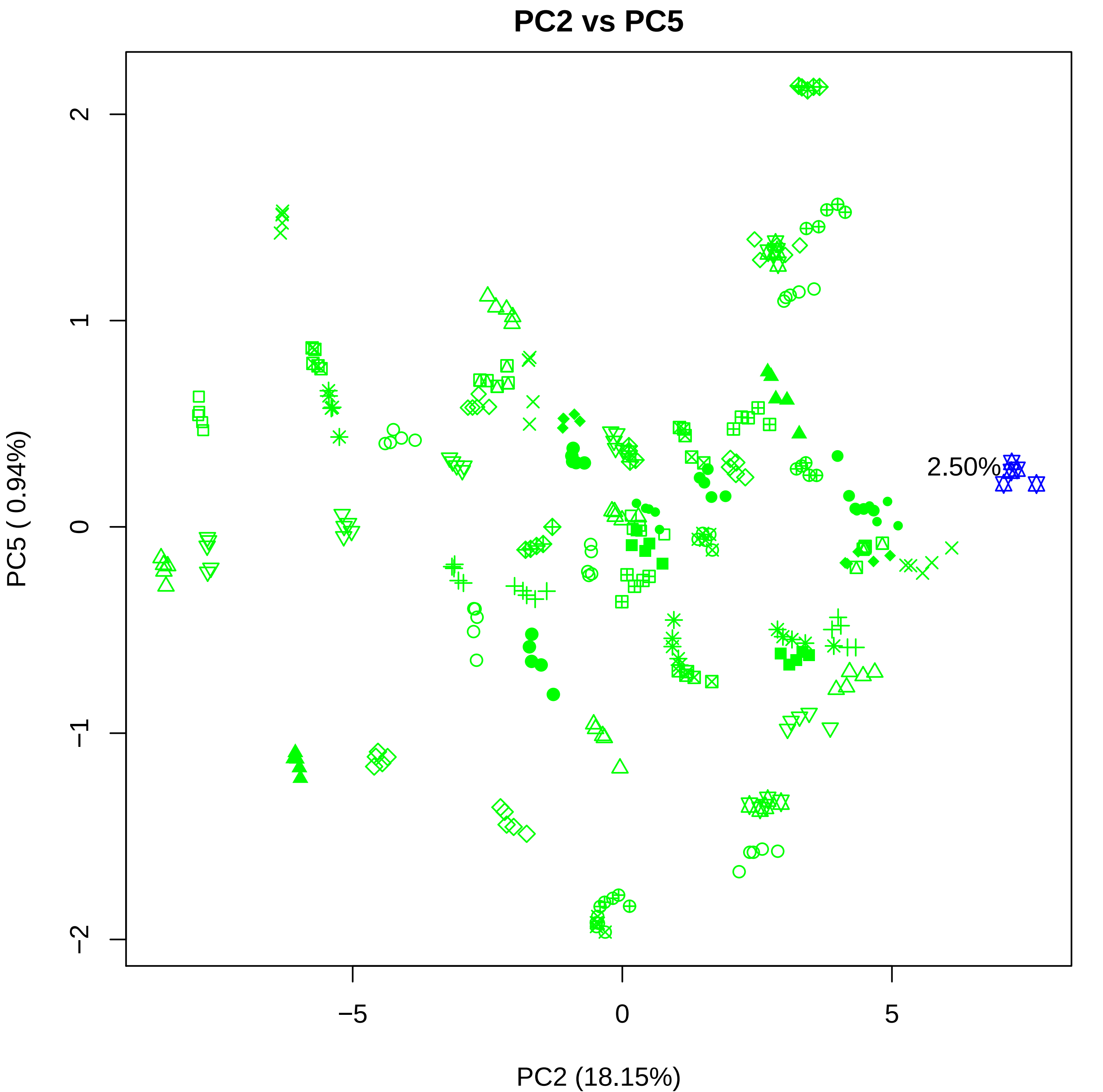
PC2 vs PC5
2.50%
PC5 ( 0.94%)
PC2 (18.15%)
2
1
0
−1
−2
−5
0
5

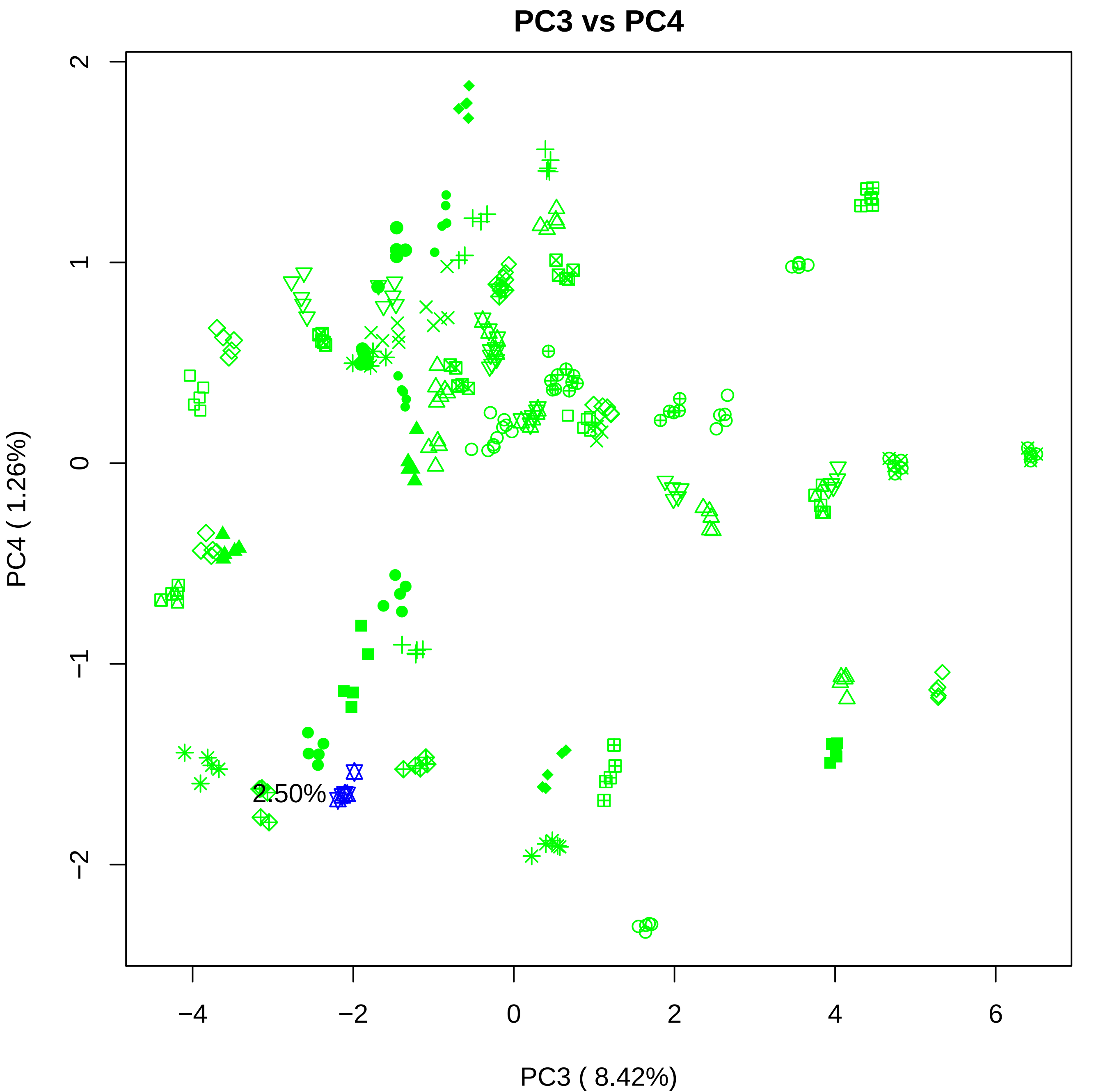

PC3 vs PC4
PC4 ( 1.26%)
PC3 ( 8.42%)
2.50%
2
1
0
−1
−2
−4
−2
0
2
4
6

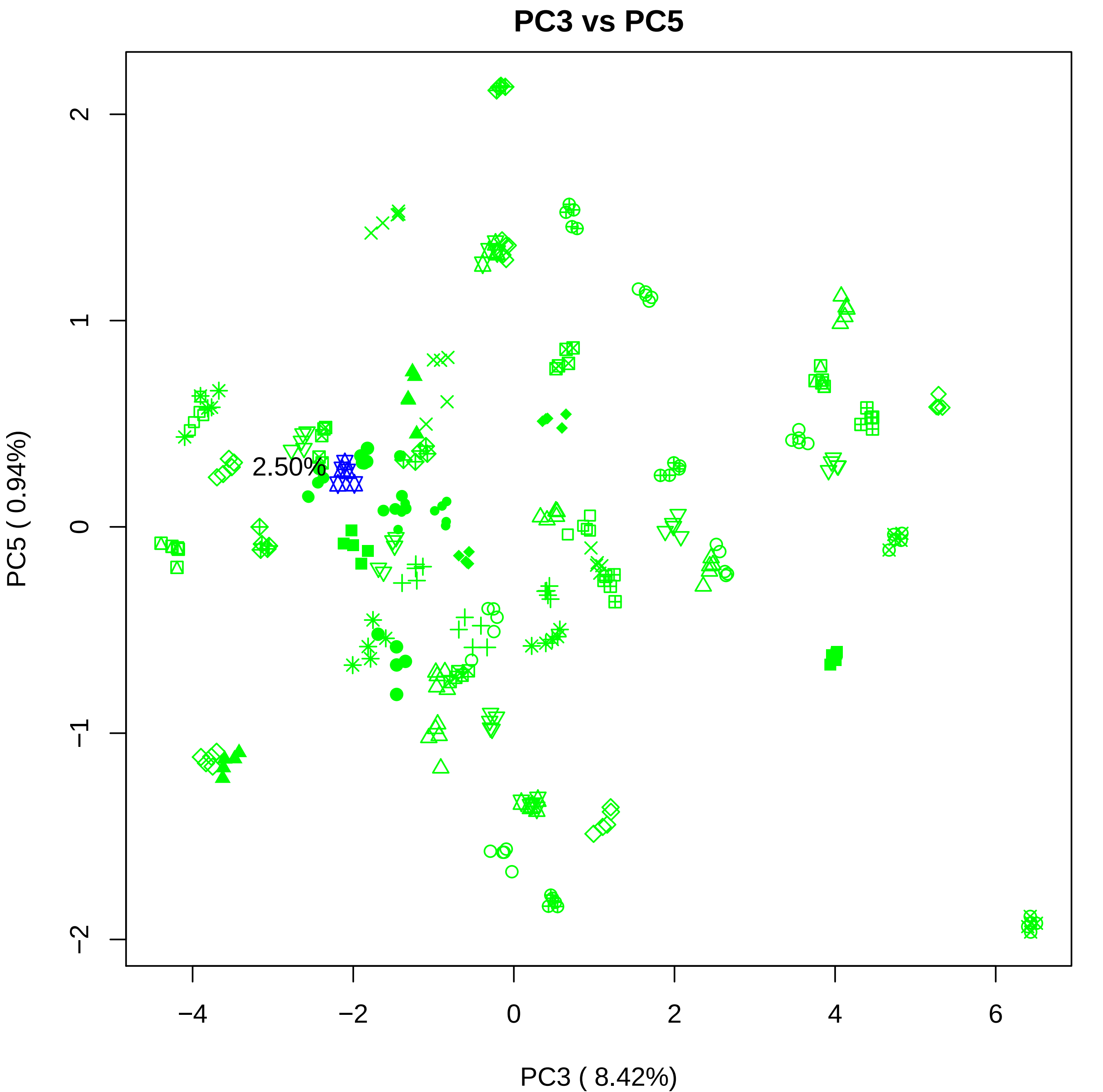
PC3 vs PC5
2.50%
PC5 ( 0.94%)
PC3 ( 8.42%)
2
1
0
−1
−2
−4
−2
0
2
4
6

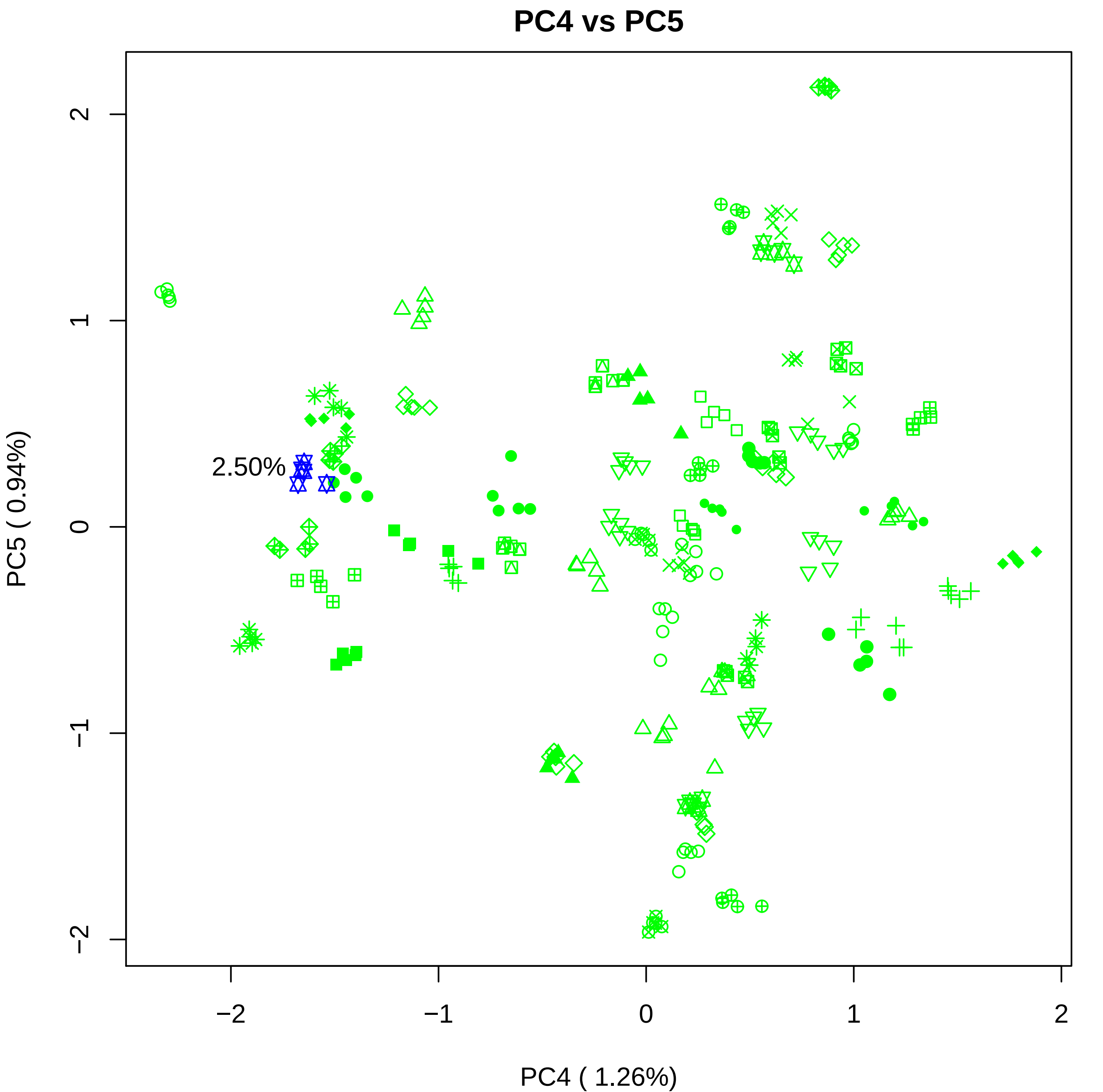
PC4 vs PC5
2
1
0
−1
−2
PC5 ( 0.94%)
2.50%
−2
−1
0
1
2
PC4 ( 1.26%)

### PCA of 5% BMH(b) vs PMH(b)

Scaled Matrix PCAed

7th order polynomial baseline correction performed

Raw spectra smoothed to 21 Neighbours, 2nd order PN, Savtisky Golay

Normalisation between 0 and 100

D2 spectra smoothed to 41 Neighbours, 2nd order PN, Savtisky Golay

Regions Analysed: 3601−2501 and 2001−700cm−1

Samples Analysed: 1 bContam's 0 BLH's 0 BMH's 0 OMH's 0 pContam's 0 PLH's 60 PMH's

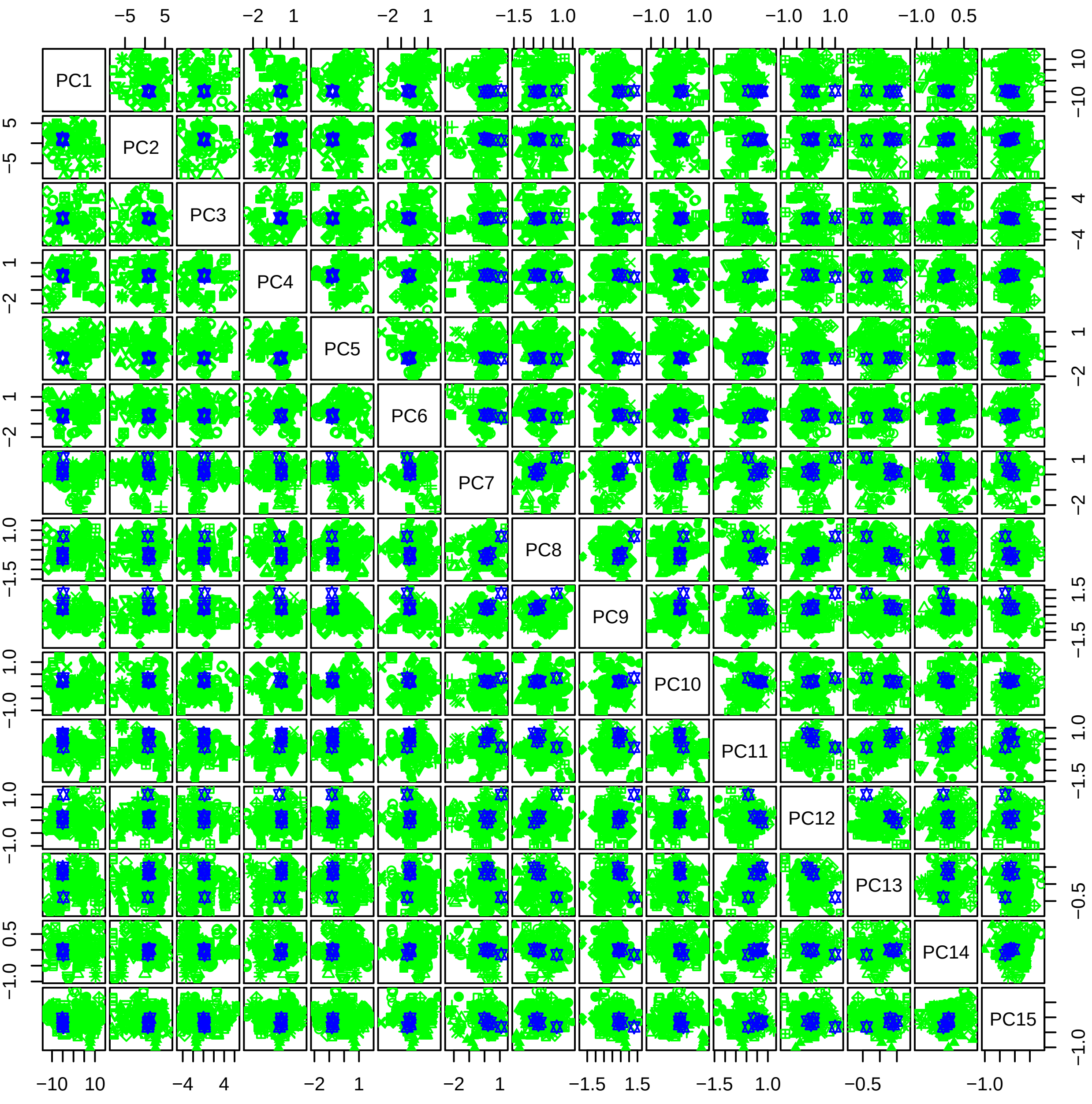

PC1
PC2
PC3
PC4
PC5
PC6
PC7
PC8
PC9
PC10
PC11
PC12
PC13
PC14
PC15

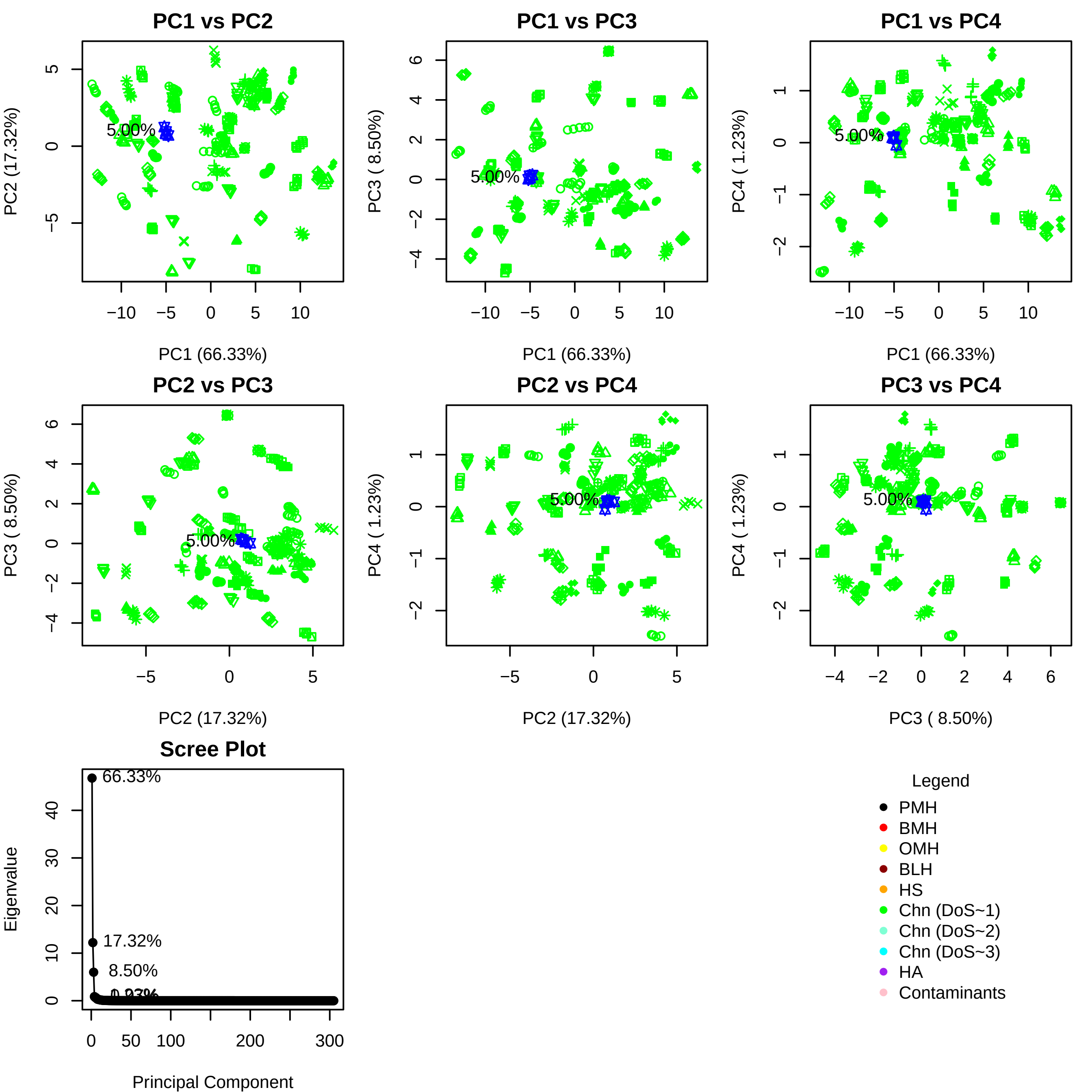
PC1 vs PC2
PC1 vs PC3
PC1 vs PC4
PC2 vs PC3
PC2 vs PC4
PC3 vs PC4
Scree Plot
PC1 (66.33%)
PC2 (17.32%)
PC3 ( 8.50%)
PC4 ( 1.23%)
5.00%
Eigenvalue
Principal Component
66.33%
17.32%
8.50%
Legend
PMH
BMH
OMH
BLH
HS
Chn (DoS~1)
Chn (DoS~2)
Chn (DoS~3)
HA
Contaminants

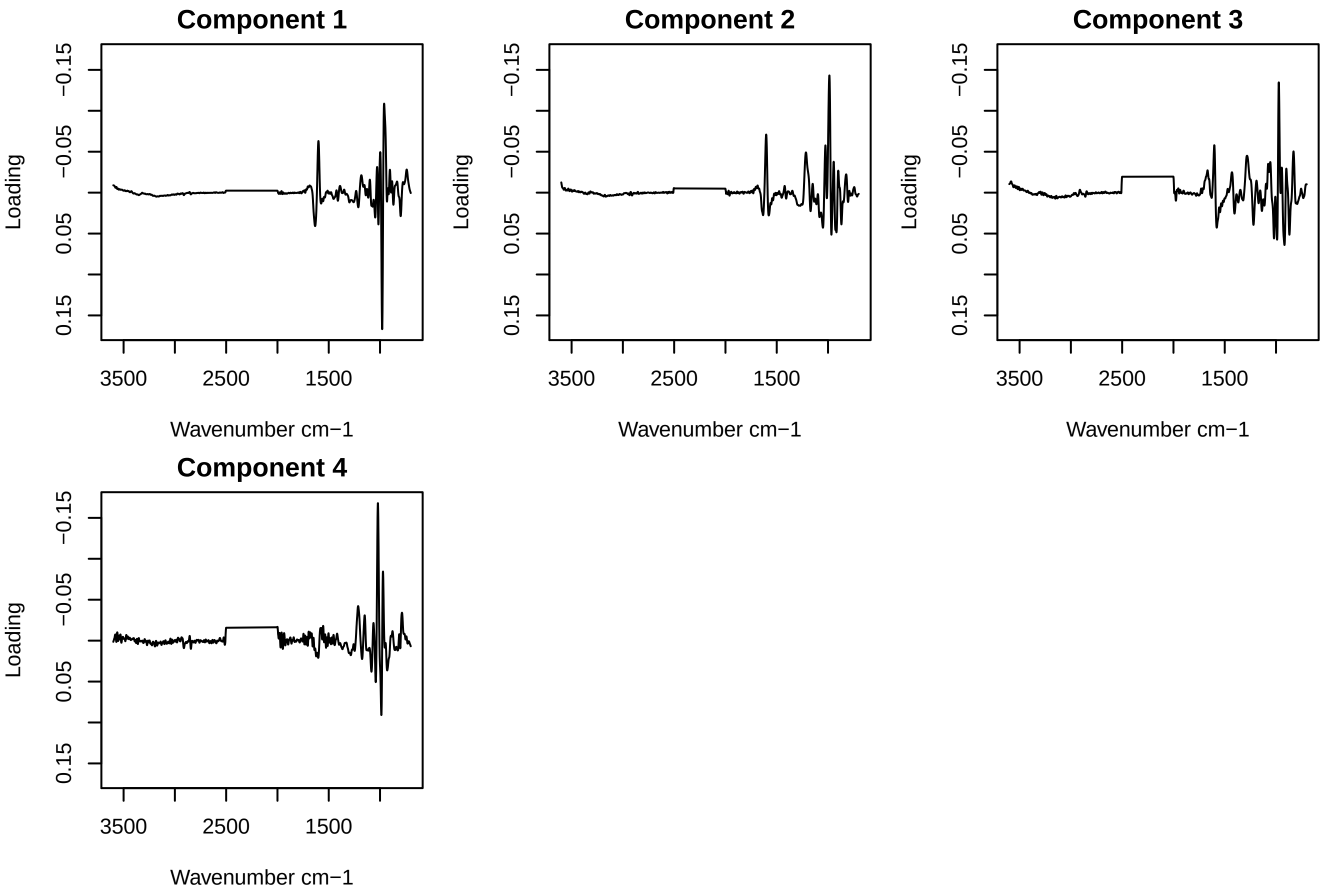
Component 1
Component 2
Component 3
Component 4
Loading
−0.15
−0.05
0.05
0.15
3500
2500
1500
Wavenumber cm−1

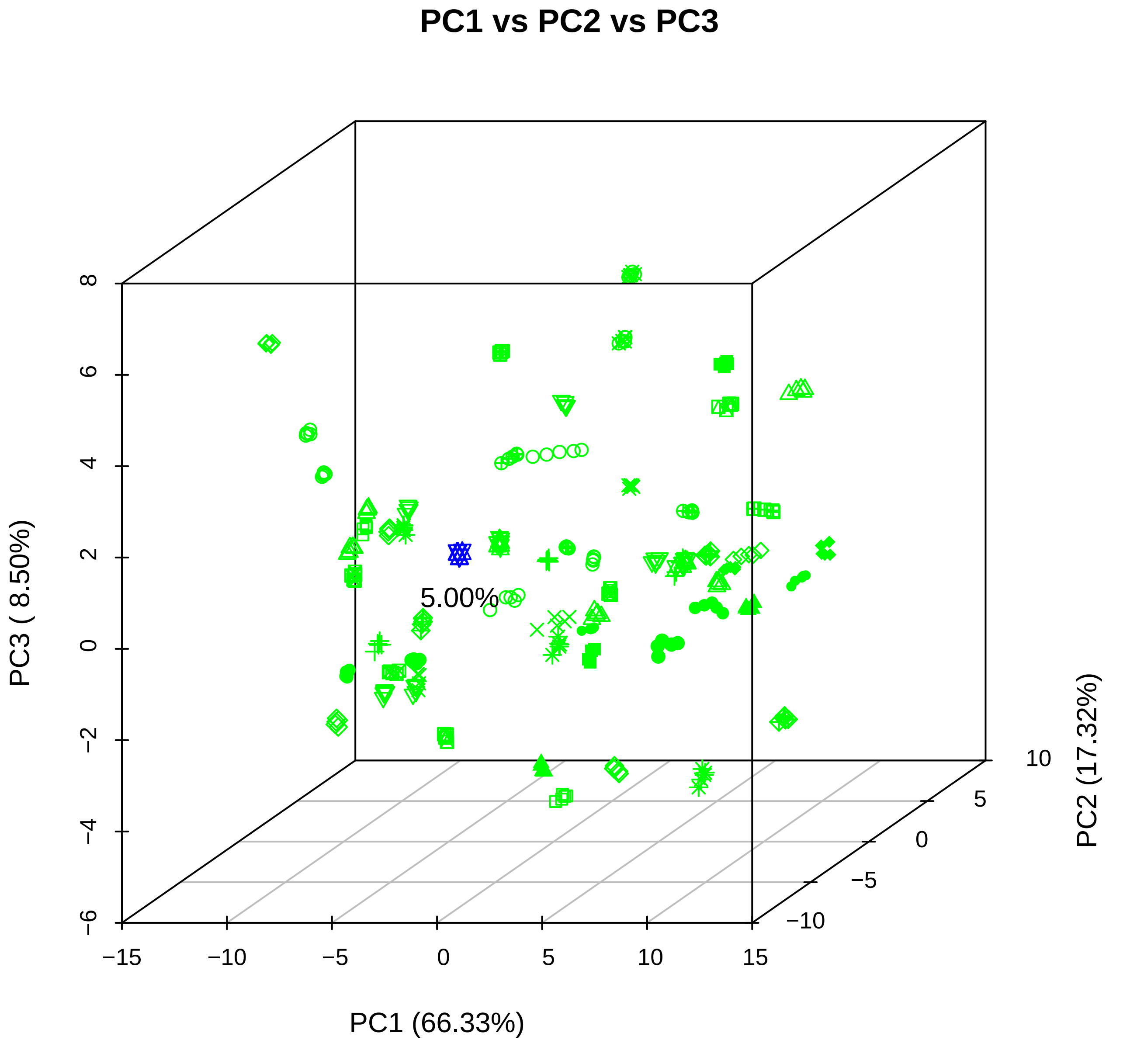
PC1 vs PC2 vs PC3
5.00%
PC1 (66.33%)
PC2 (17.32%)
PC3 ( 8.50%)
−15
−10
−5
0
5
10
15
−10
−5
0
5
10
−6
−4
−2
0
2
4
6
8

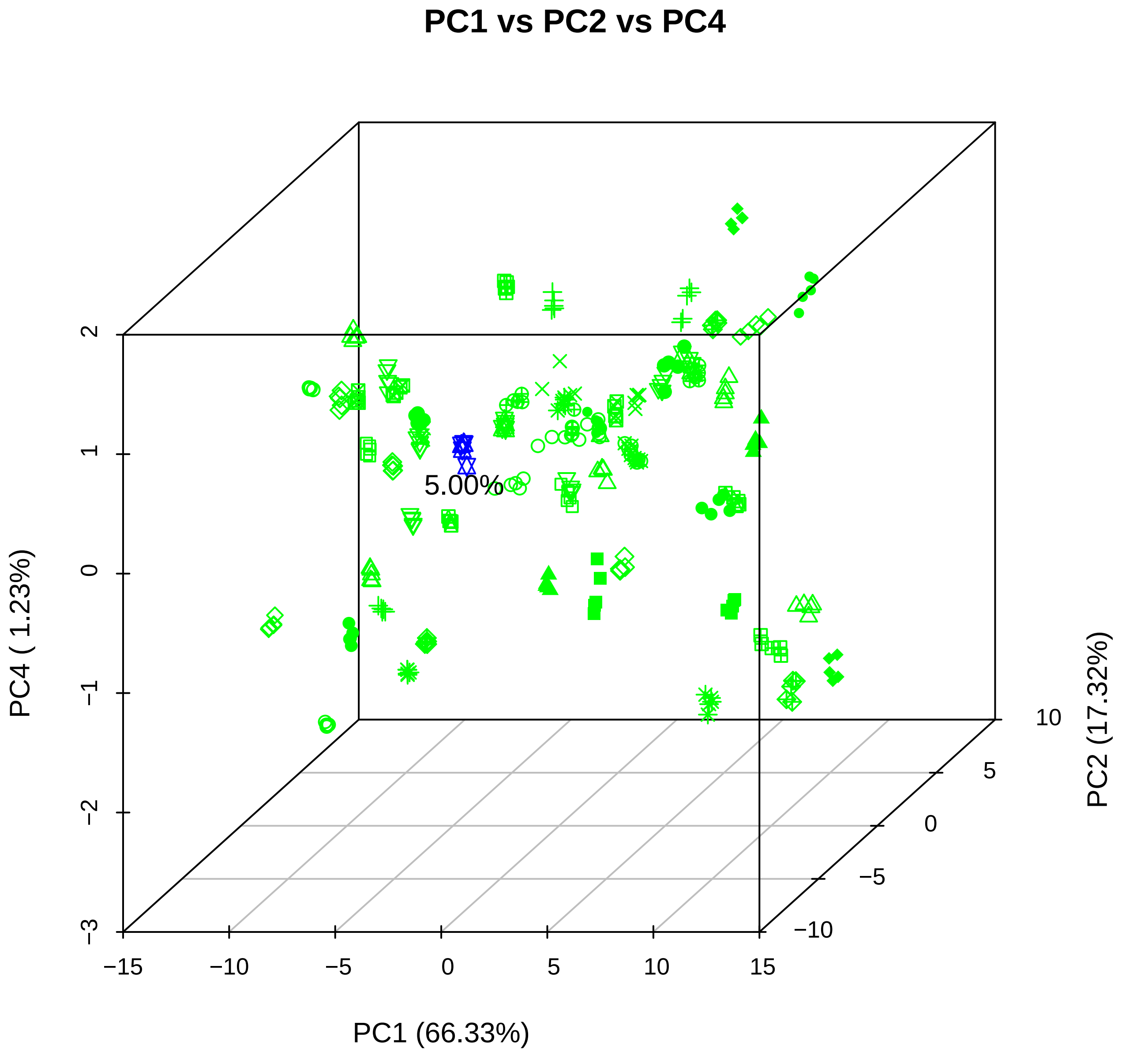
PC1 vs PC2 vs PC4
5.00%
PC1 (66.33%)
PC2 (17.32%)
PC4 ( 1.23%)
−15
−10
−5
0
5
10
15
−10
−5
0
5
10
−3
−2
−1
0
1
2

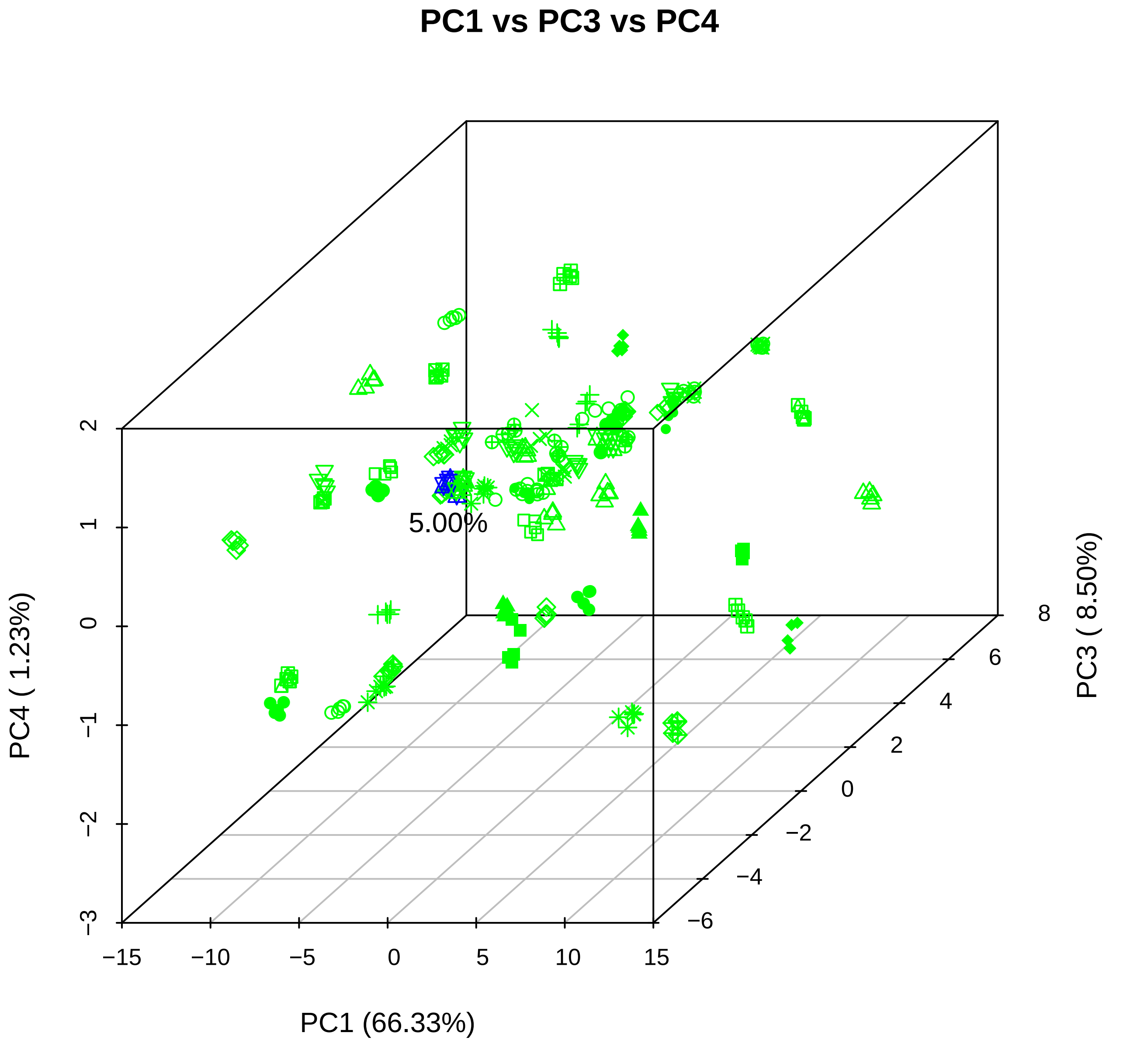

PC1 vs PC3 vs PC4
5.00%
PC1 (66.33%)
PC4 ( 1.23%)
PC3 ( 8.50%)
−15
−10
−5
0
5
10
15
−3
−2
−1
0
1
2
−6
−4
−2
0
2
4
6
8

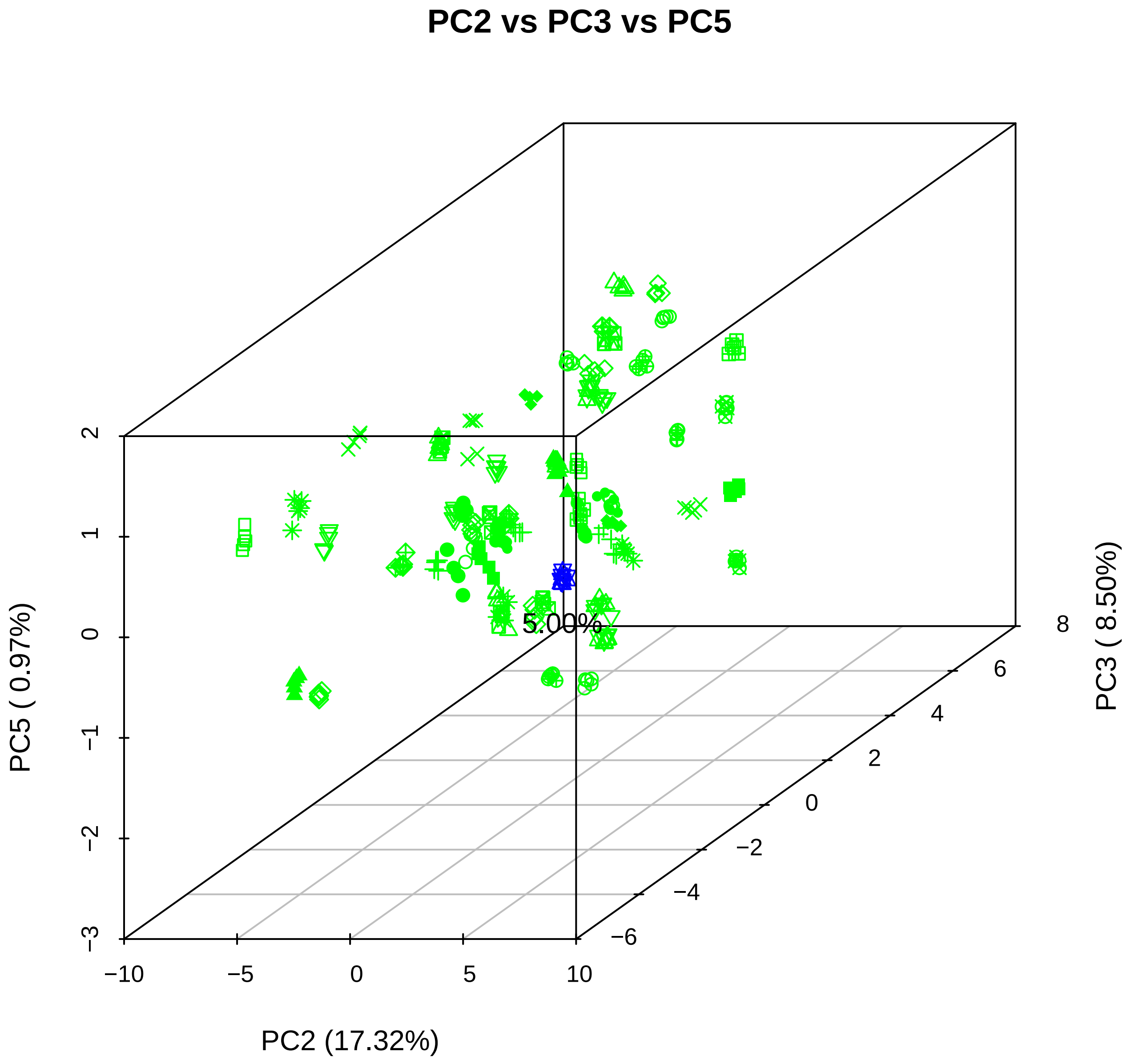

PC2 vs PC3 vs PC5
5.00%
PC2 (17.32%)
PC5 ( 0.97%)
PC3 ( 8.50%)
−10
−5
0
5
10
−3
−2
−1
0
1
2
−6
−4
−2
0
2
4
6
8

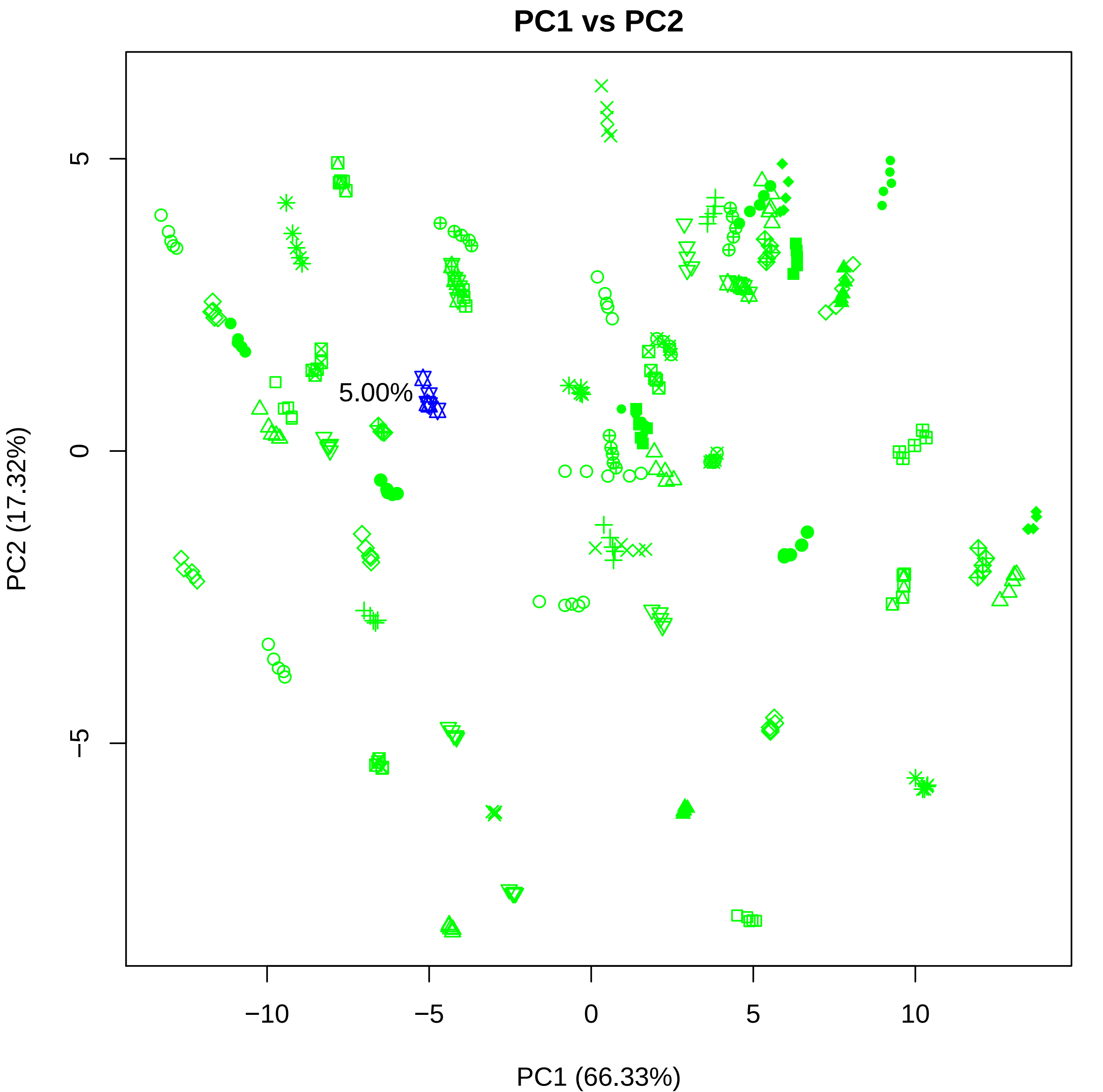
PC1 vs PC2
5.00%
PC2 (17.32%)
PC1 (66.33%)
5
0
−5
−10
−5
0
5
10

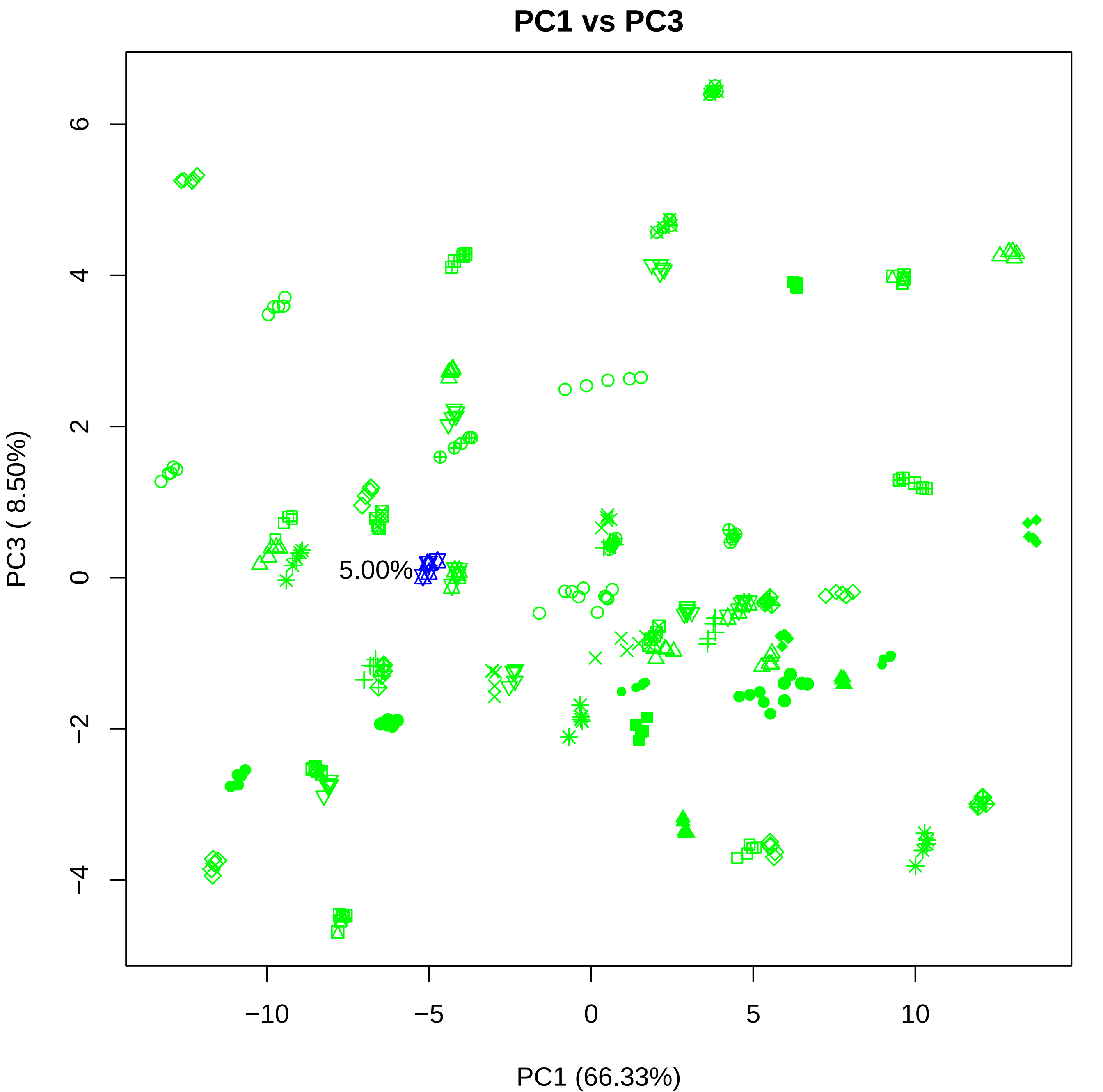
PC1 vs PC3
PC3 ( 8.50%)
PC1 (66.33%)
5.00%
−10
−5
0
5
10
−4
−2
0
2
4
6

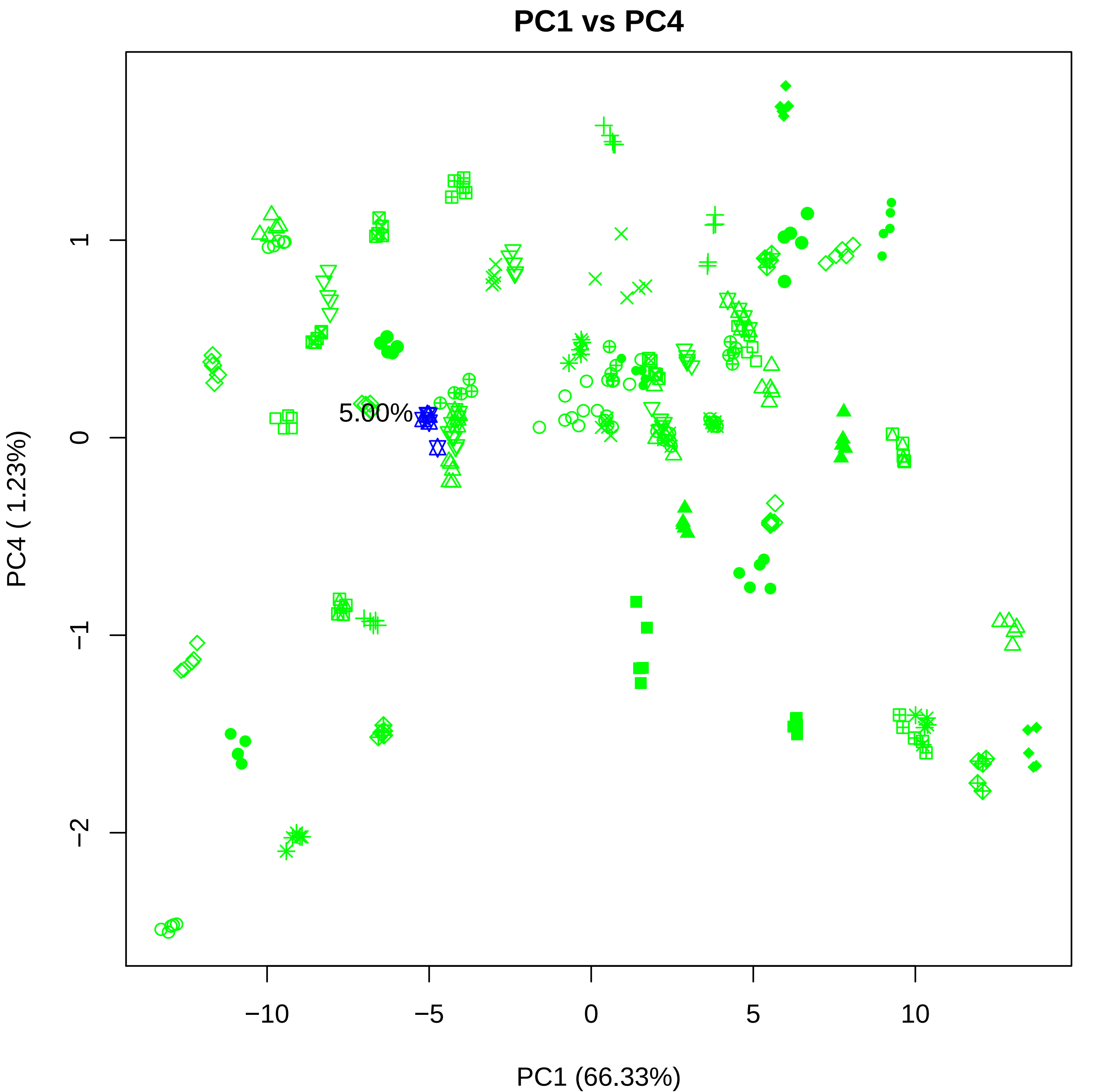
PC1 vs PC4
5.00%
PC4 ( 1.23%)
PC1 (66.33%)
−10
−5
0
5
10
1
0
−1
−2

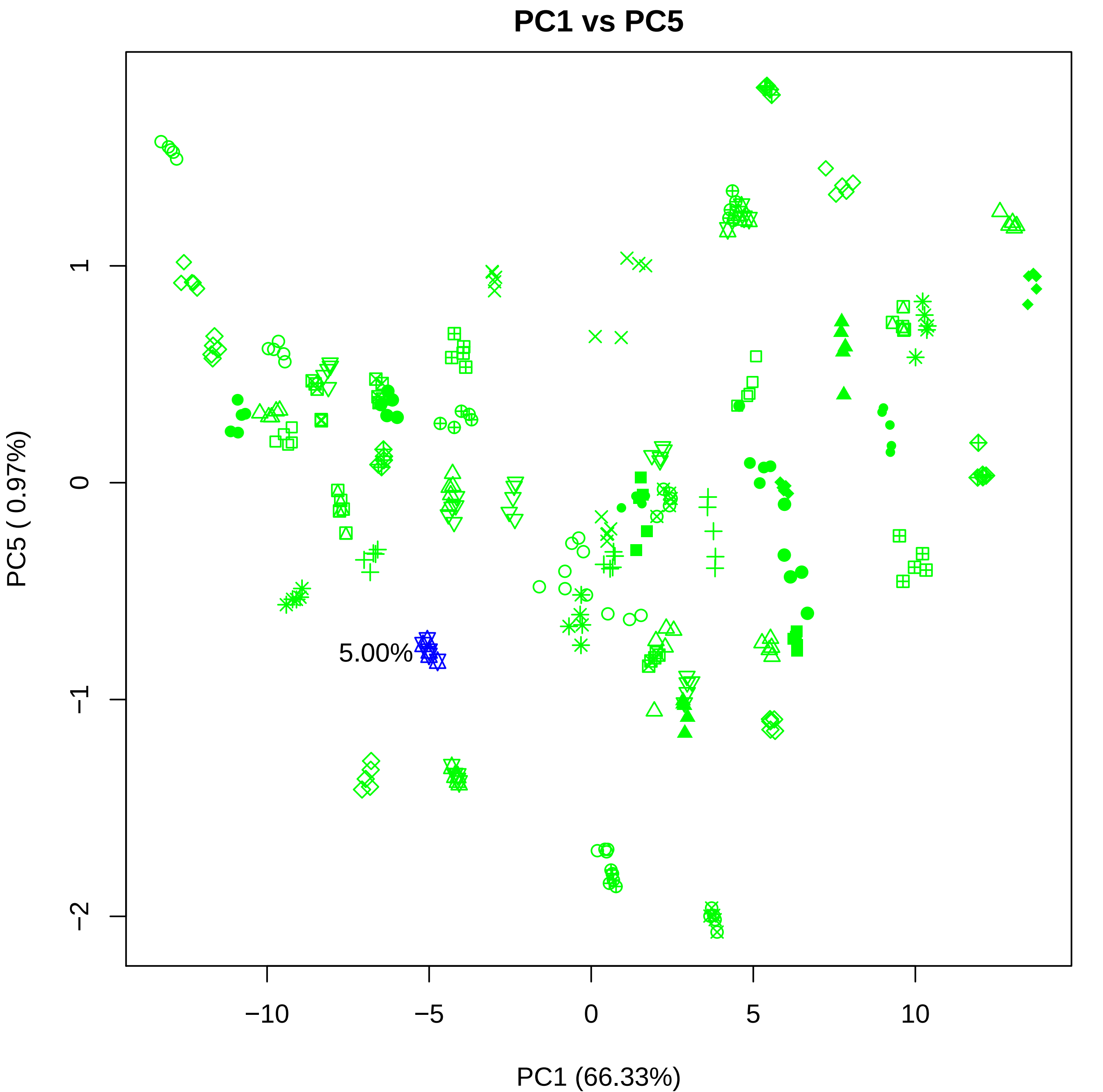
PC1 vs PC5
PC5 ( 0.97%)
PC1 (66.33%)
5.00%
1
0
−1
−2
−10
−5
0
5
10

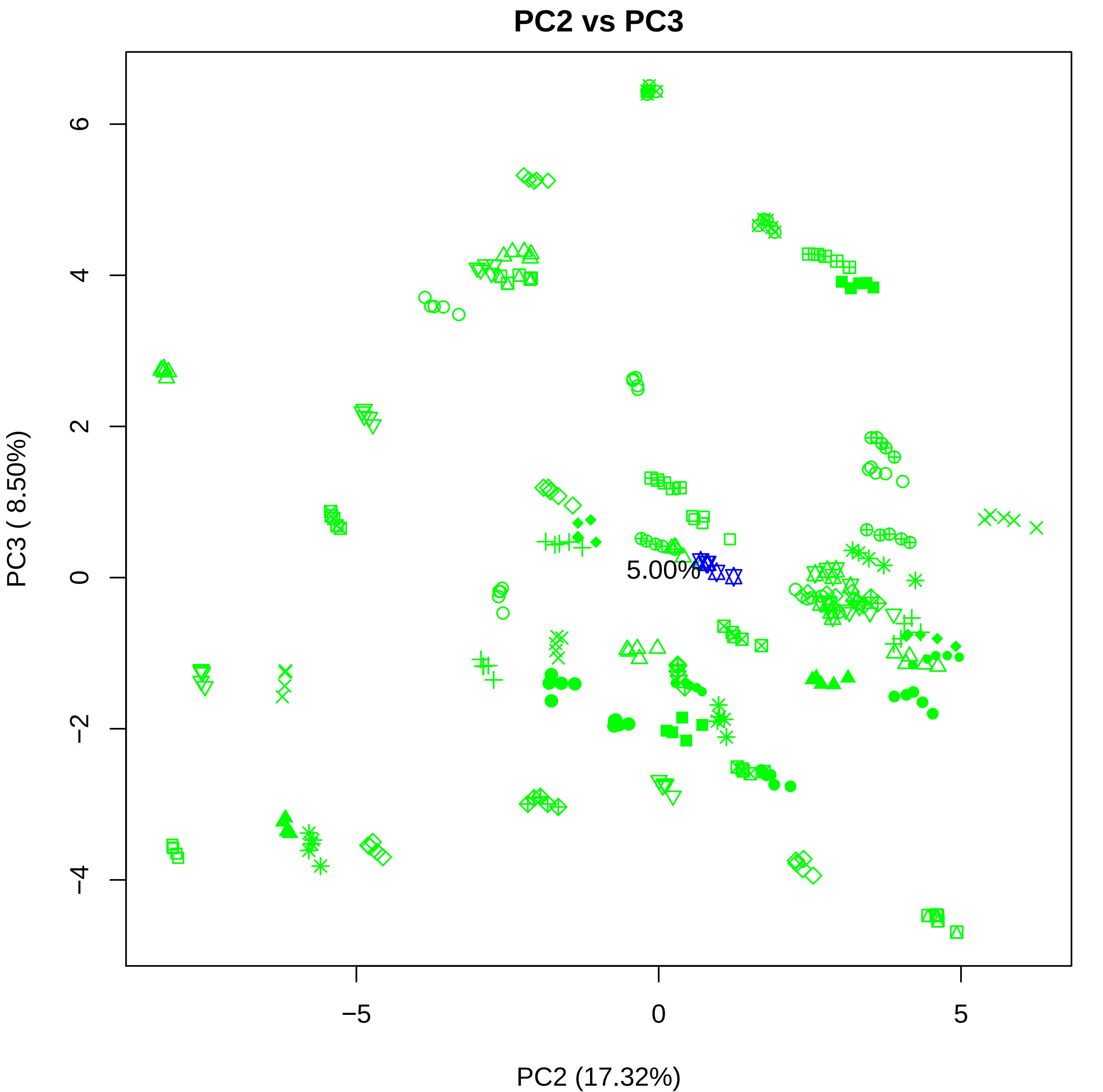

PC2 vs PC3
PC3 ( 8.50%)
PC2 (17.32%)
6
4
2
0
−2
−4
−5
0
5
5.00%

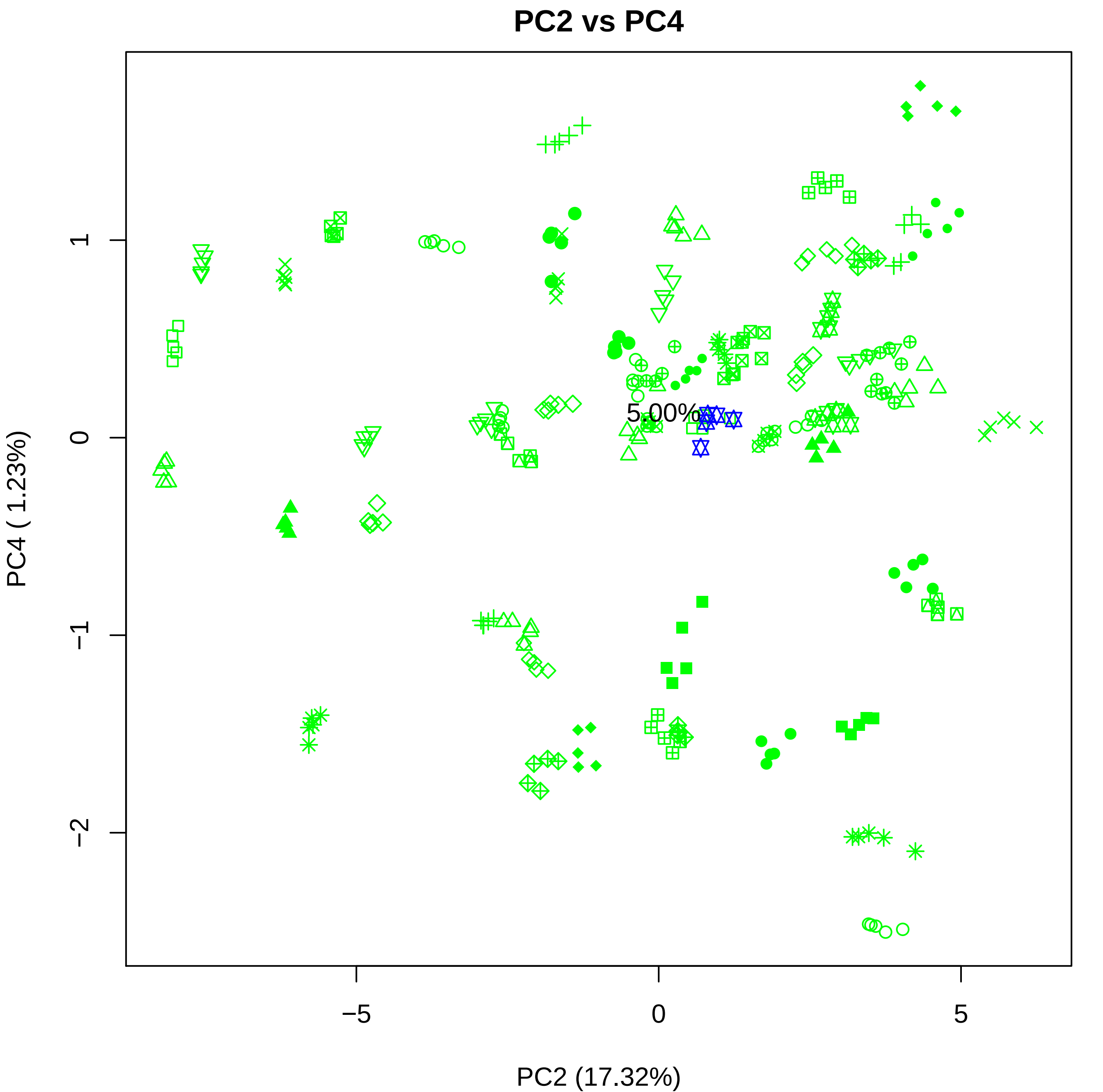
PC2 vs PC4
5.00%
PC4 ( 1.23%)
PC2 (17.32%)
1
0
−1
−2
−5
0
5

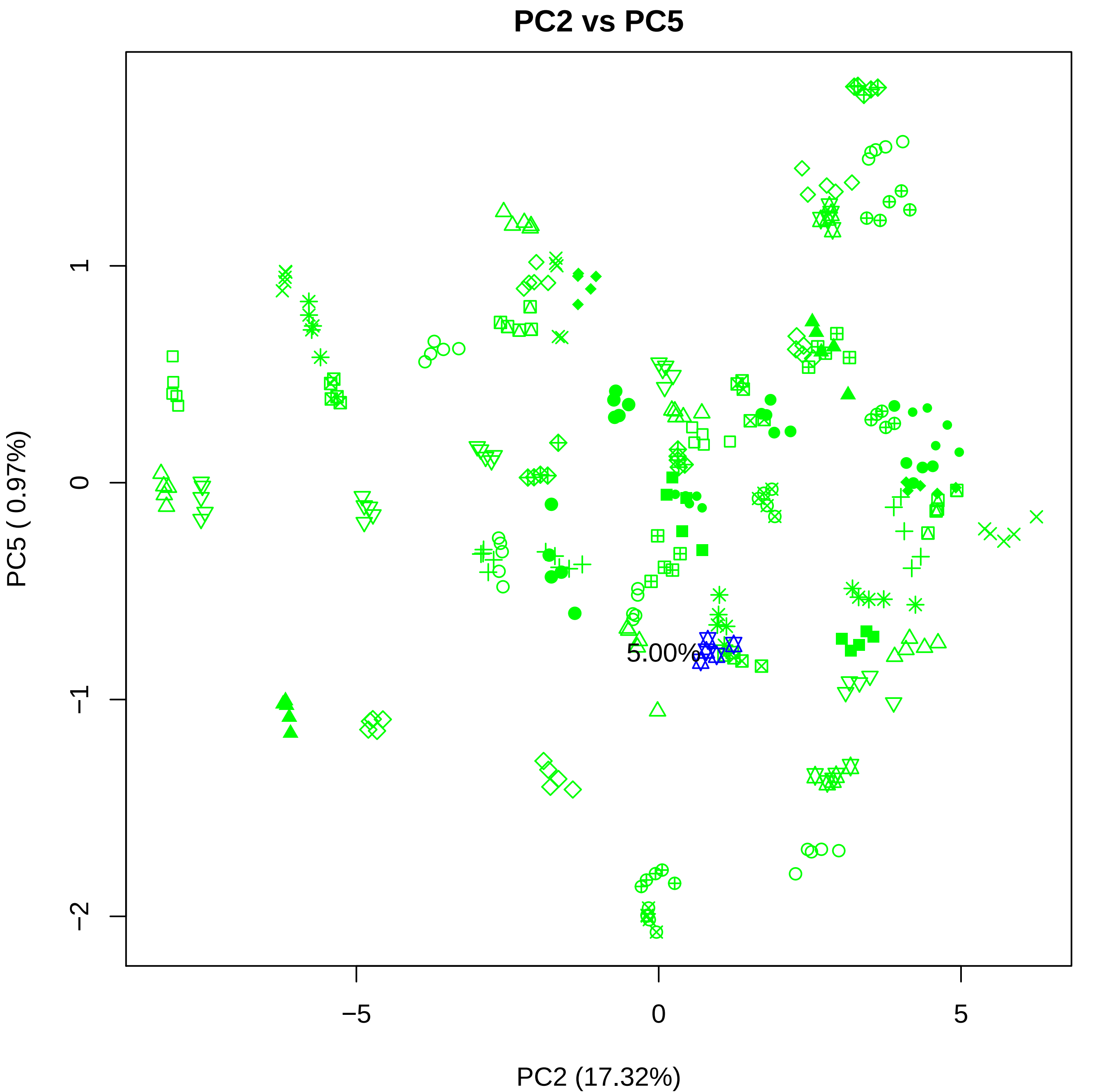
PC2 vs PC5
1
0
−1
−2
PC5 ( 0.97%)
−5
0
5
PC2 (17.32%)
5.00%

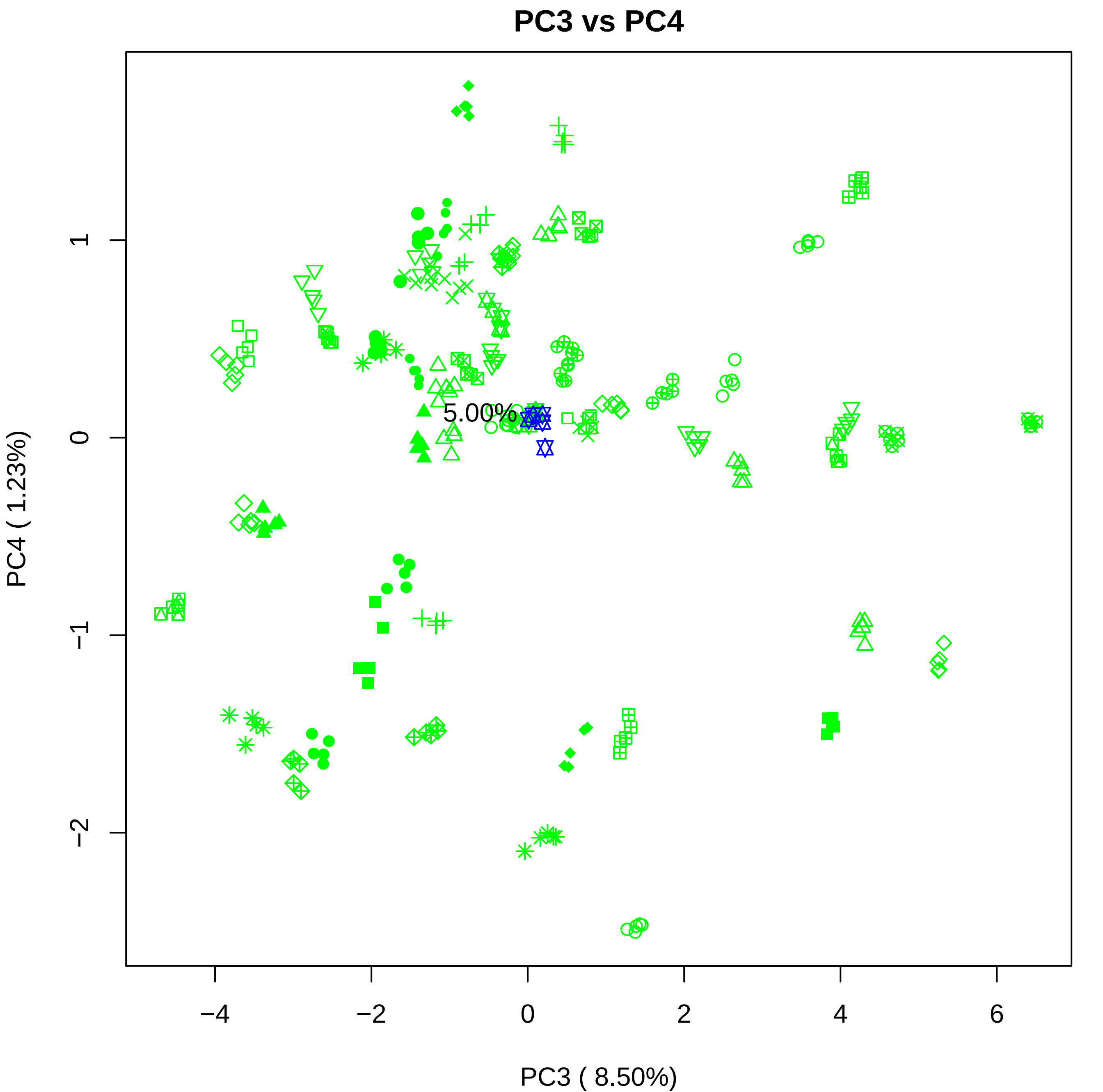
PC3 vs PC4
5.00%
PC4 ( 1.23%)
1
0
−1
−2
−4
−2
0
2
4
6
PC3 ( 8.50%)

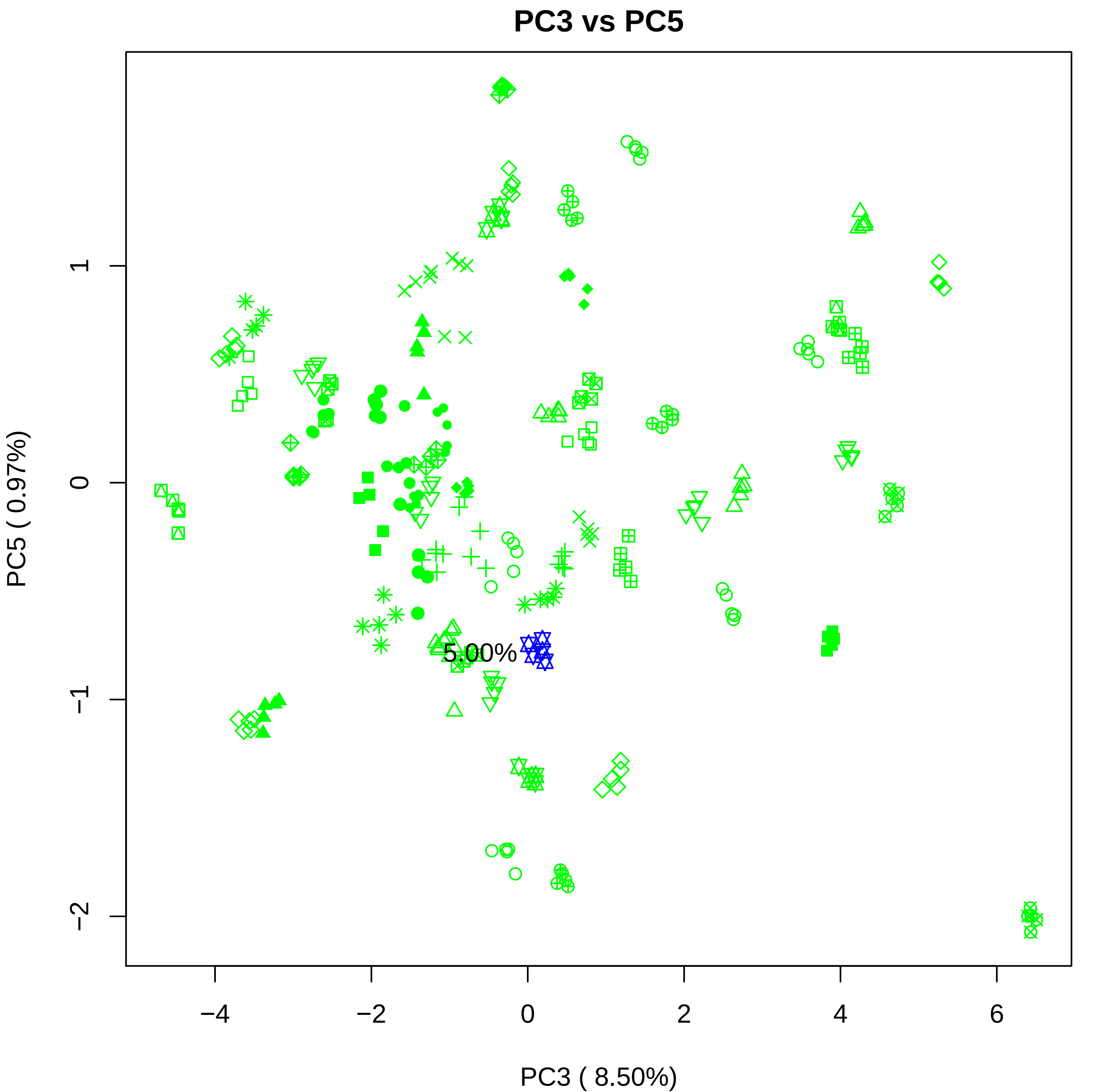
PC3 vs PC5
PC5 ( 0.97%)
PC3 ( 8.50%)
5.00%
−4
−2
0
2
4
6
1
0
−1
−2

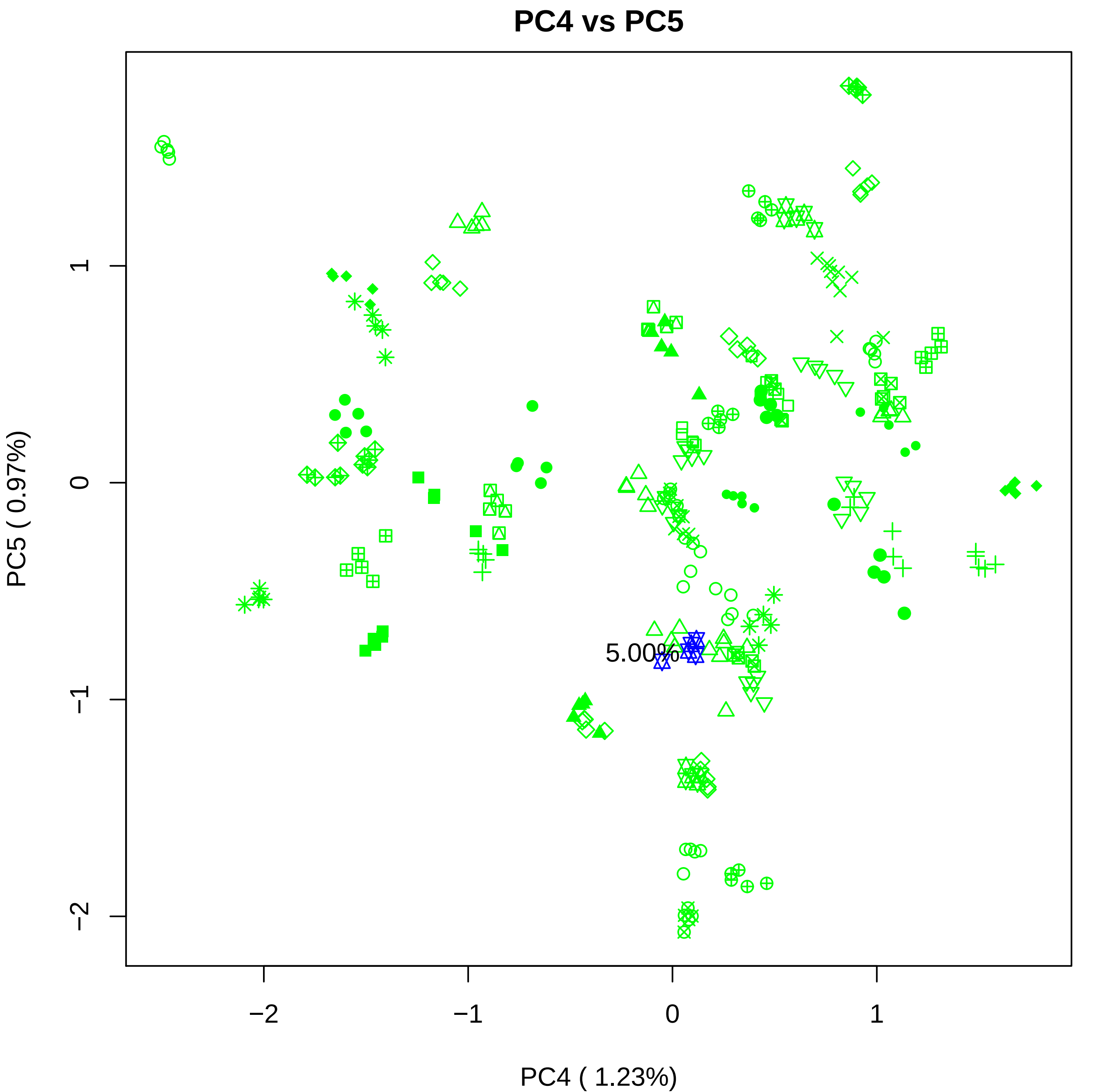

PC4 vs PC5
PC5 ( 0.97%)
PC4 ( 1.23%)
5.00%
1
0
−1
−2
−2
−1
0
1

### PCA of 7.5%  BMH(b)  vs PMH(b)

Scaled Matrix PCAed

7th order polynomial baseline correction performed

Raw spectra smoothed to 21 Neighbours, 2nd order PN, Savtisky Golay

Normalisation between 0 and 100

D2 spectra smoothed to 41 Neighbours, 2nd order PN, Savtisky Golay

Regions Analysed: 3601−2501 and 2001−700cm−1

Samples Analysed: 1 bContam's 0 BLH's 0 BMH's 0 OMH's 0 pContam's 0 PLH's 60 PMH's

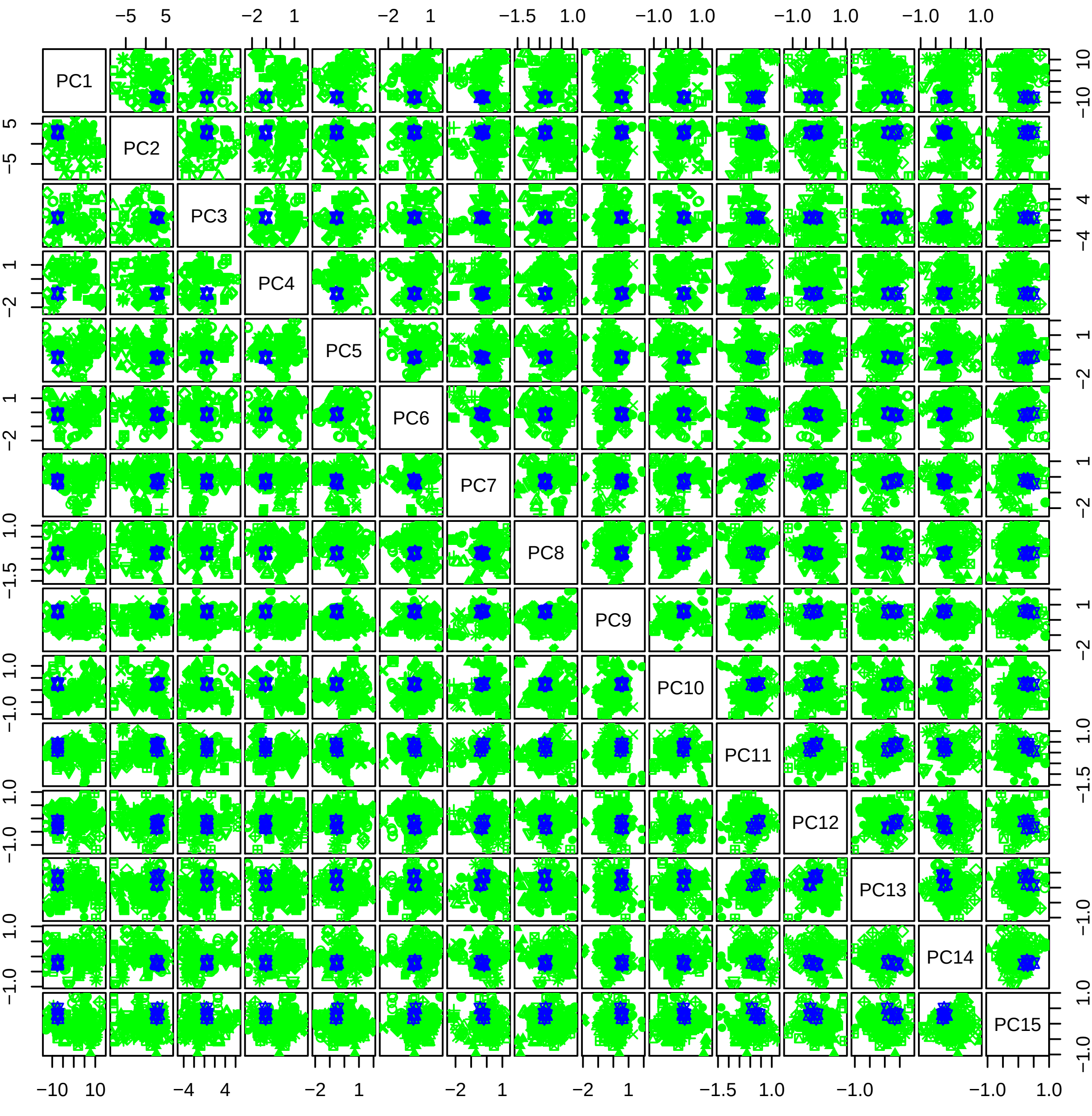
PC1
PC2
PC3
PC4
PC5
PC6
PC7
PC8
PC9
PC10
PC11
PC12
PC13
PC14
PC15

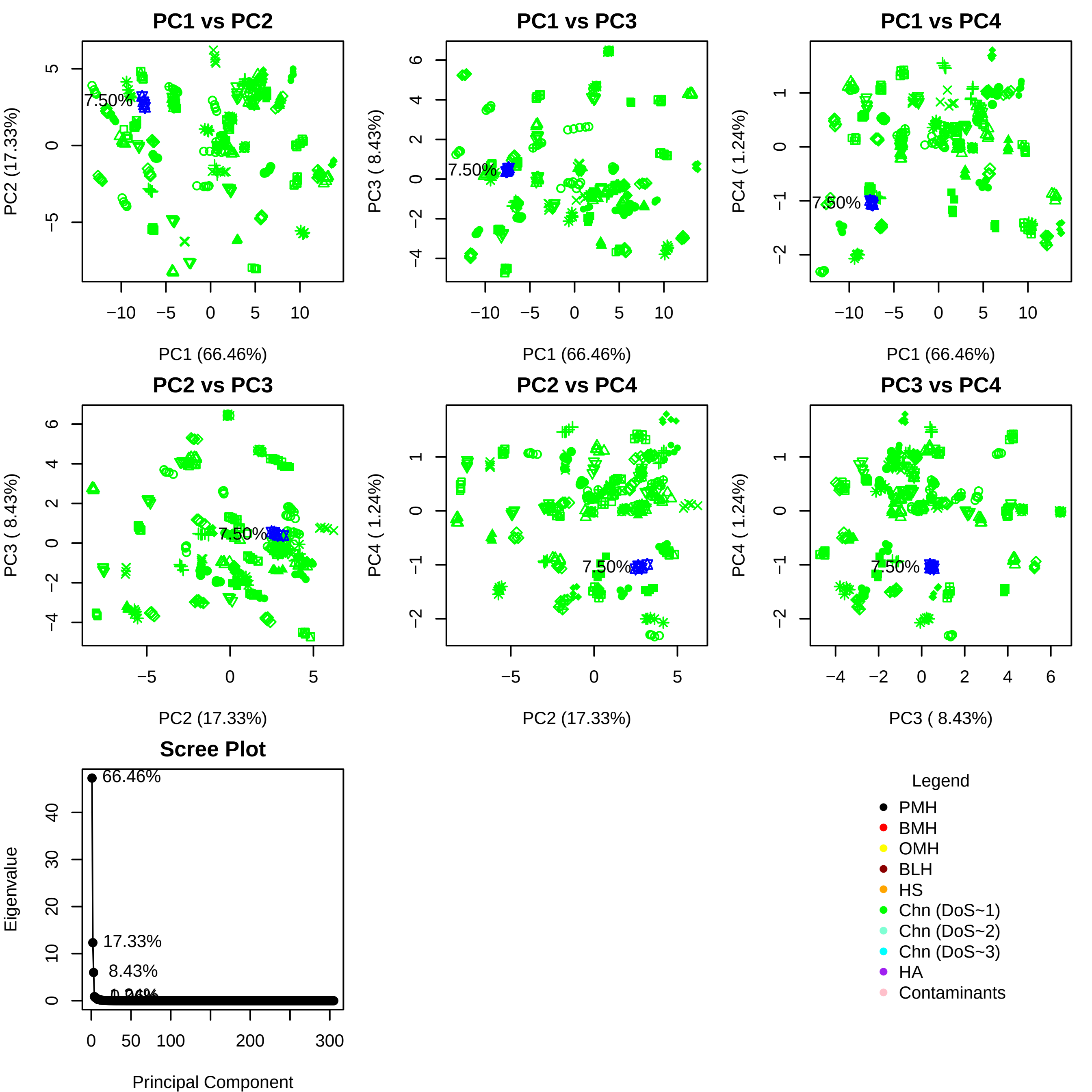

PC1 vs PC2
PC1 vs PC3
PC1 vs PC4
PC2 vs PC3
PC2 vs PC4
PC3 vs PC4
Scree Plot
PC1 (66.46%)
PC2 (17.33%)
PC3 ( 8.43%)
PC4 ( 1.24%)
7.50%
Eigenvalue
Principal Component
66.46%
17.33%
8.43%
Legend
PMH
BMH
OMH
BLH
HS
Chn (DoS~1)
Chn (DoS~2)
Chn (DoS~3)
HA
Contaminants

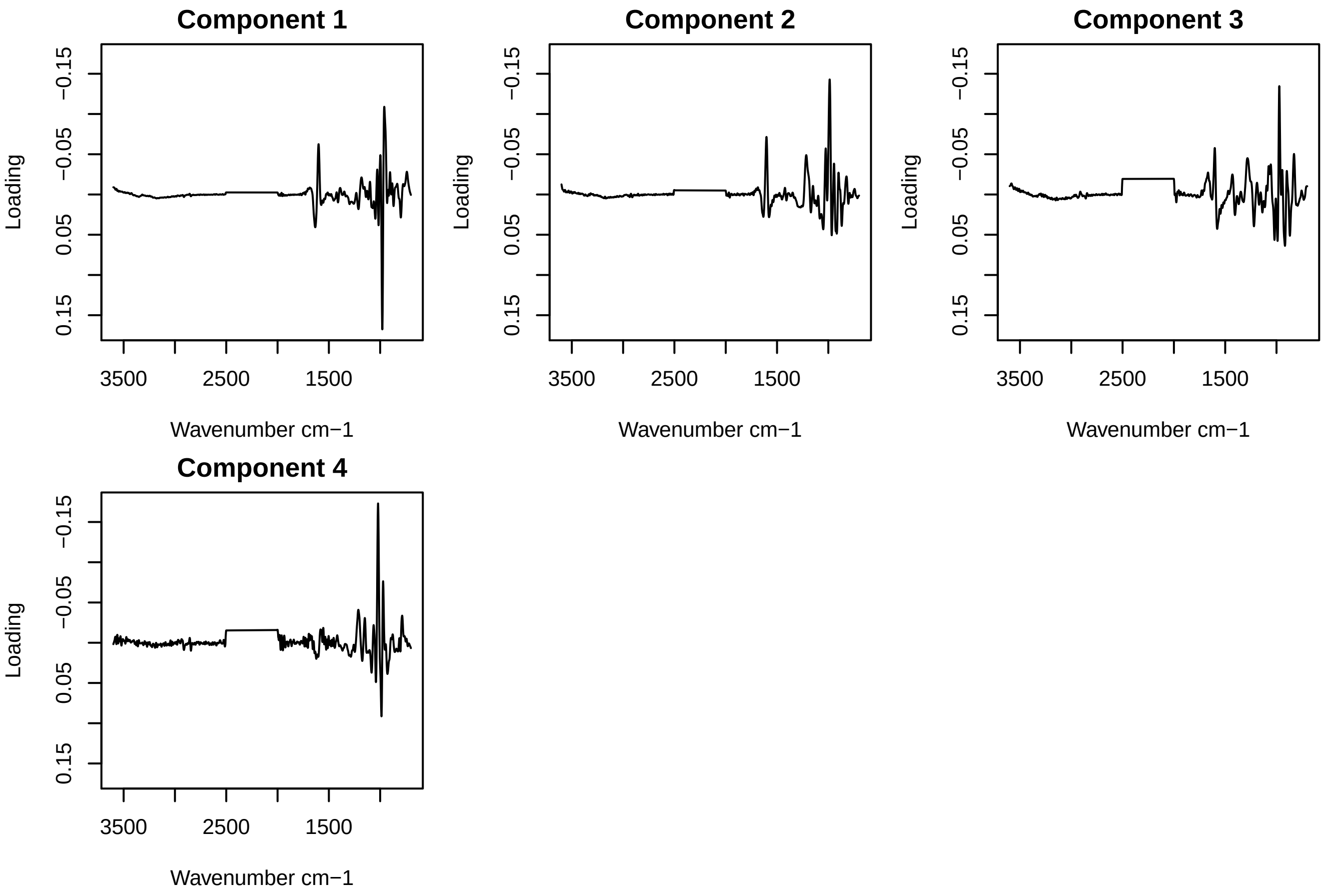

Component 1
Component 2
Component 3
Component 4
Loading
−0.15
−0.05
0.05
0.15
3500
2500
1500
Wavenumber cm−1

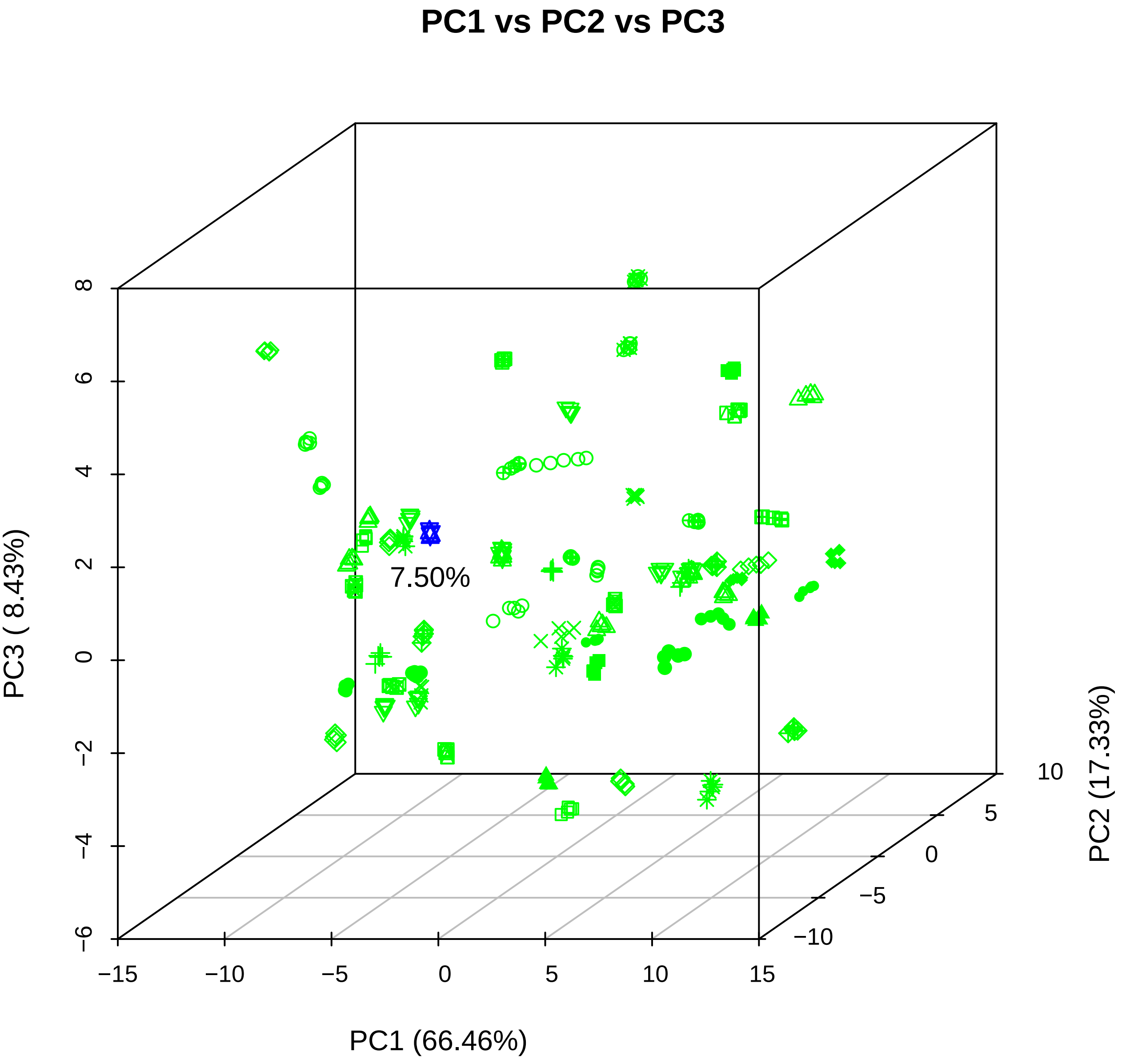

PC1 vs PC2 vs PC3
7.50%
PC1 (66.46%)
PC2 (17.33%)
PC3 ( 8.43%)
−15
−10
−5
0
5
10
15
−6
−4
−2
0
2
4
6
8

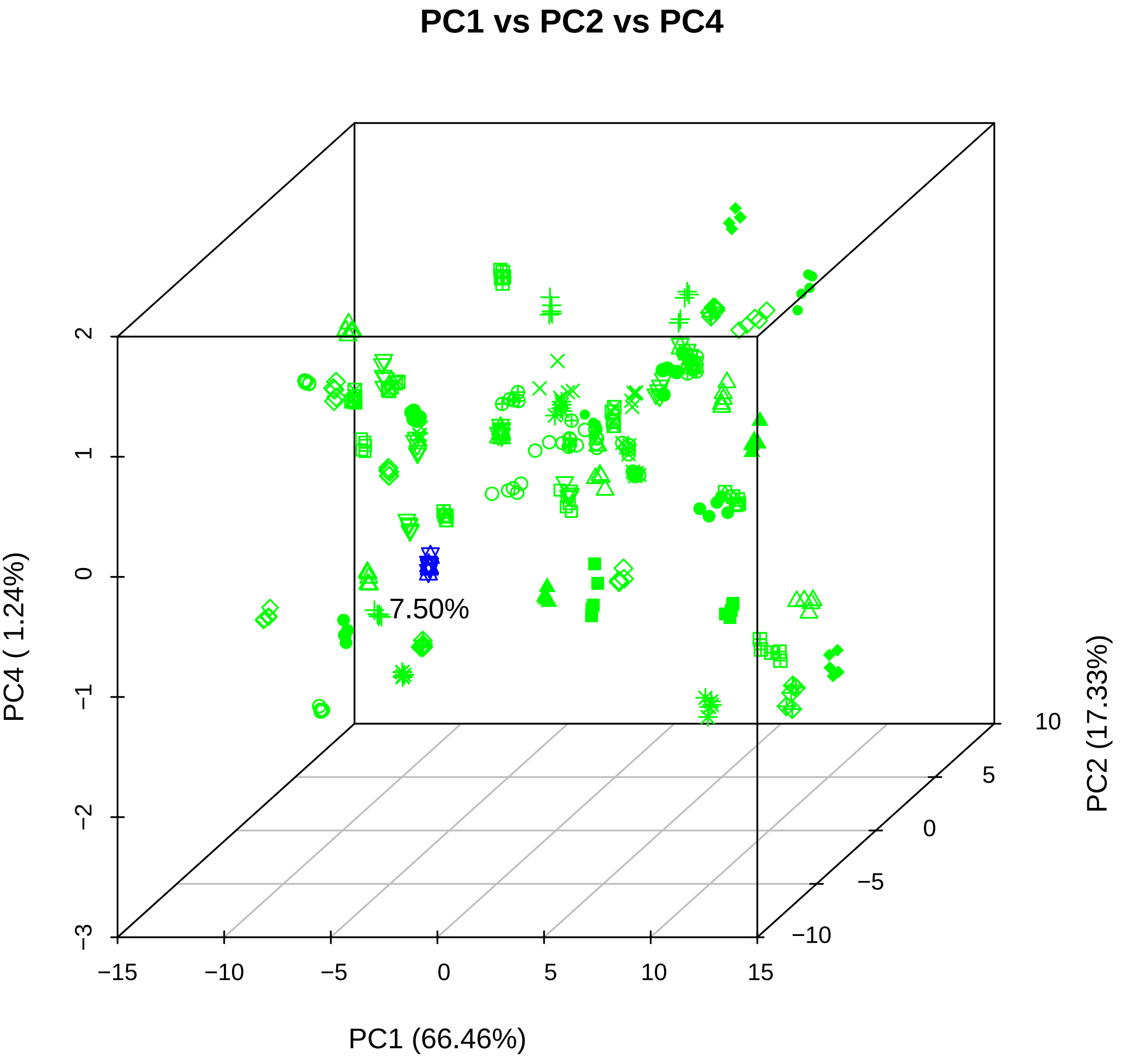

PC1 vs PC2 vs PC4
PC4 ( 1.24%)
PC1 (66.46%)
PC2 (17.33%)
7.50%
2
1
0
−1
−2
−3
−15
−10
−5
0
5
10
15
10
5
0
−5
−10

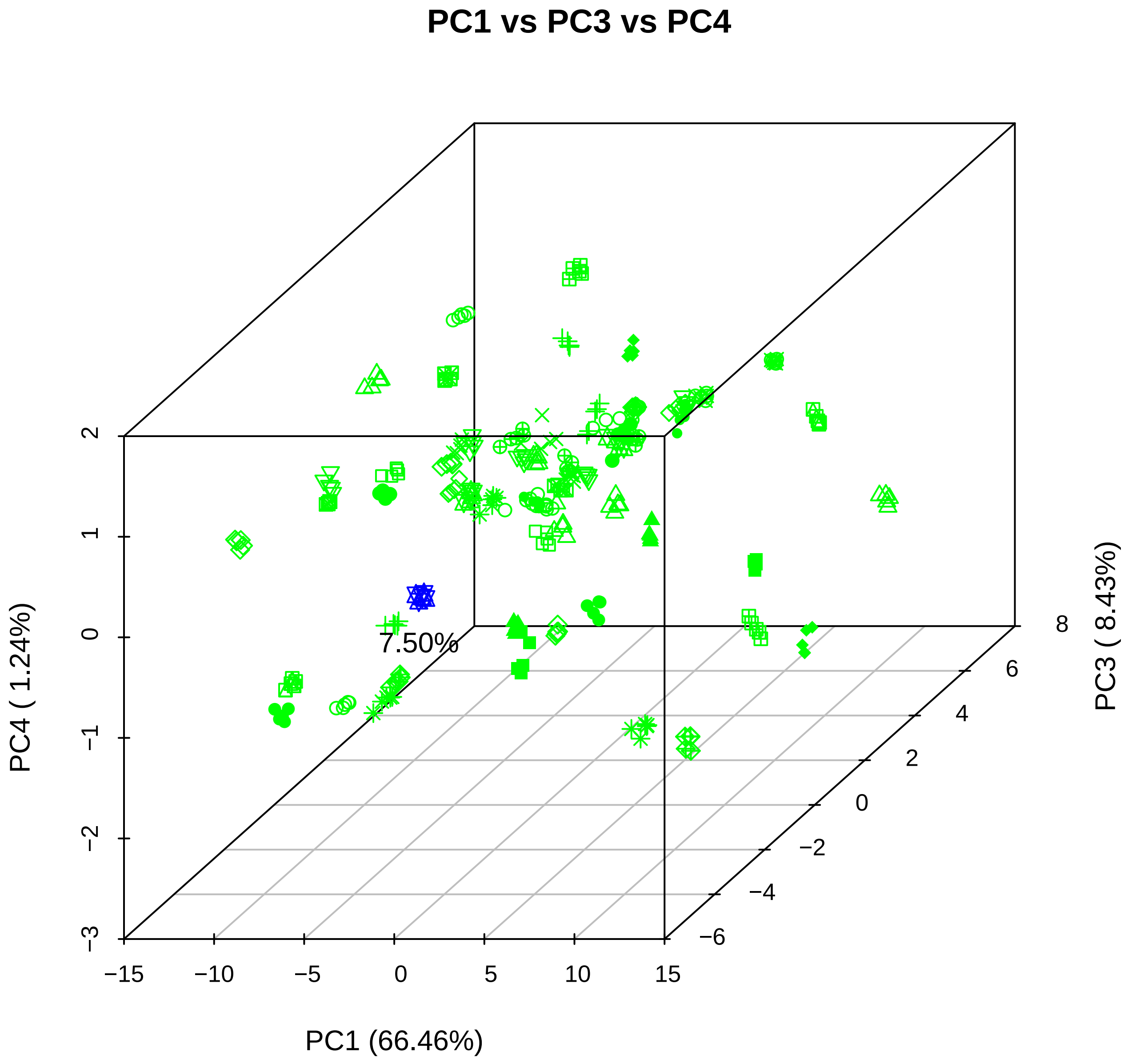

PC1 vs PC3 vs PC4
PC1 (66.46%)
PC4 ( 1.24%)
PC3 ( 8.43%)
7.50%
−15
−10
−5
0
5
10
15
−3
−2
−1
0
1
2
−6
−4
−2
0
2
4
6
8

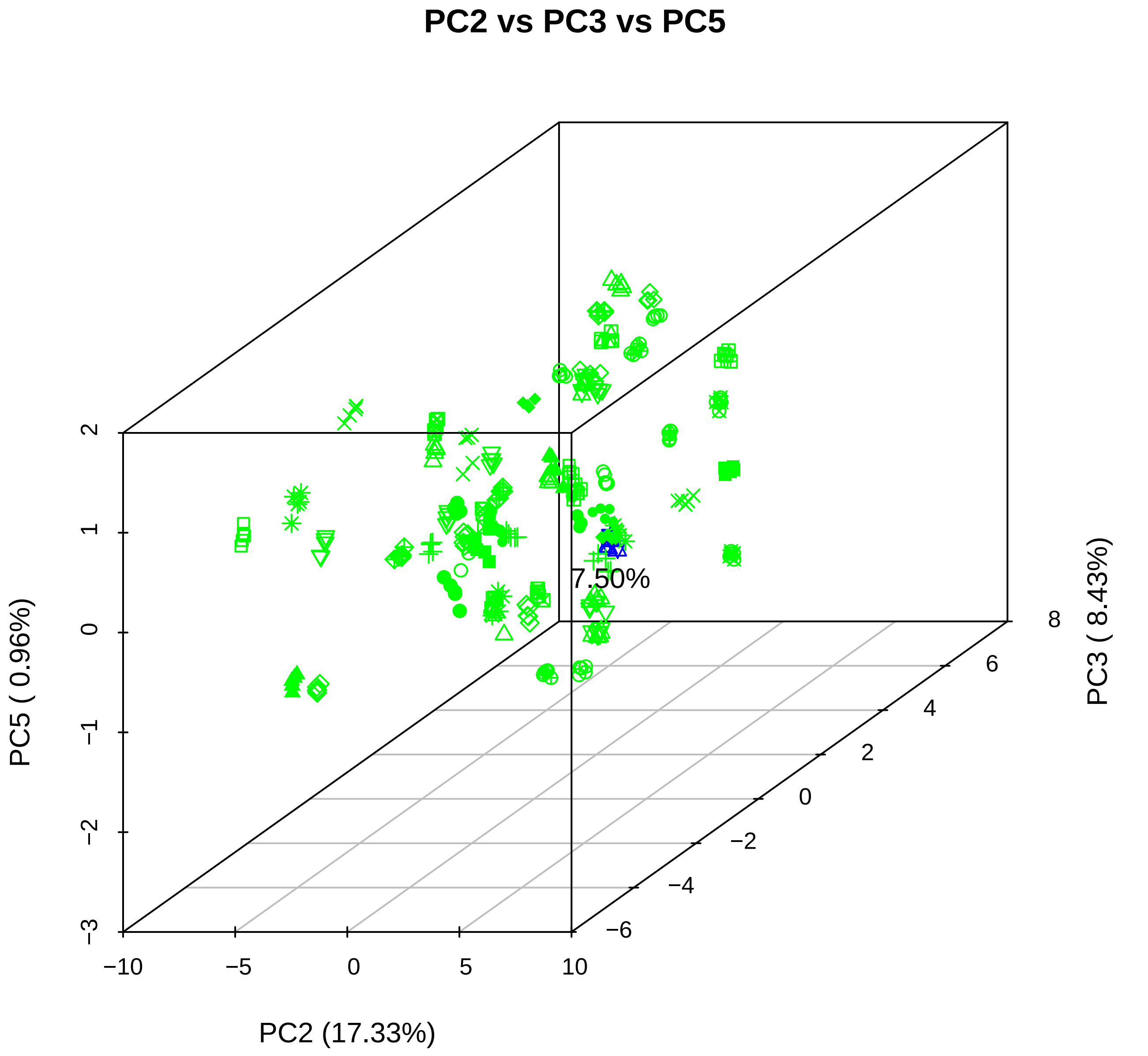
PC2 vs PC3 vs PC5
7.50%
PC2 (17.33%)
PC3 ( 8.43%)
PC5 ( 0.96%)
−10
−5
0
5
10
−6
−4
−2
0
2
4
6
8
−3
−2
−1
0
1
2

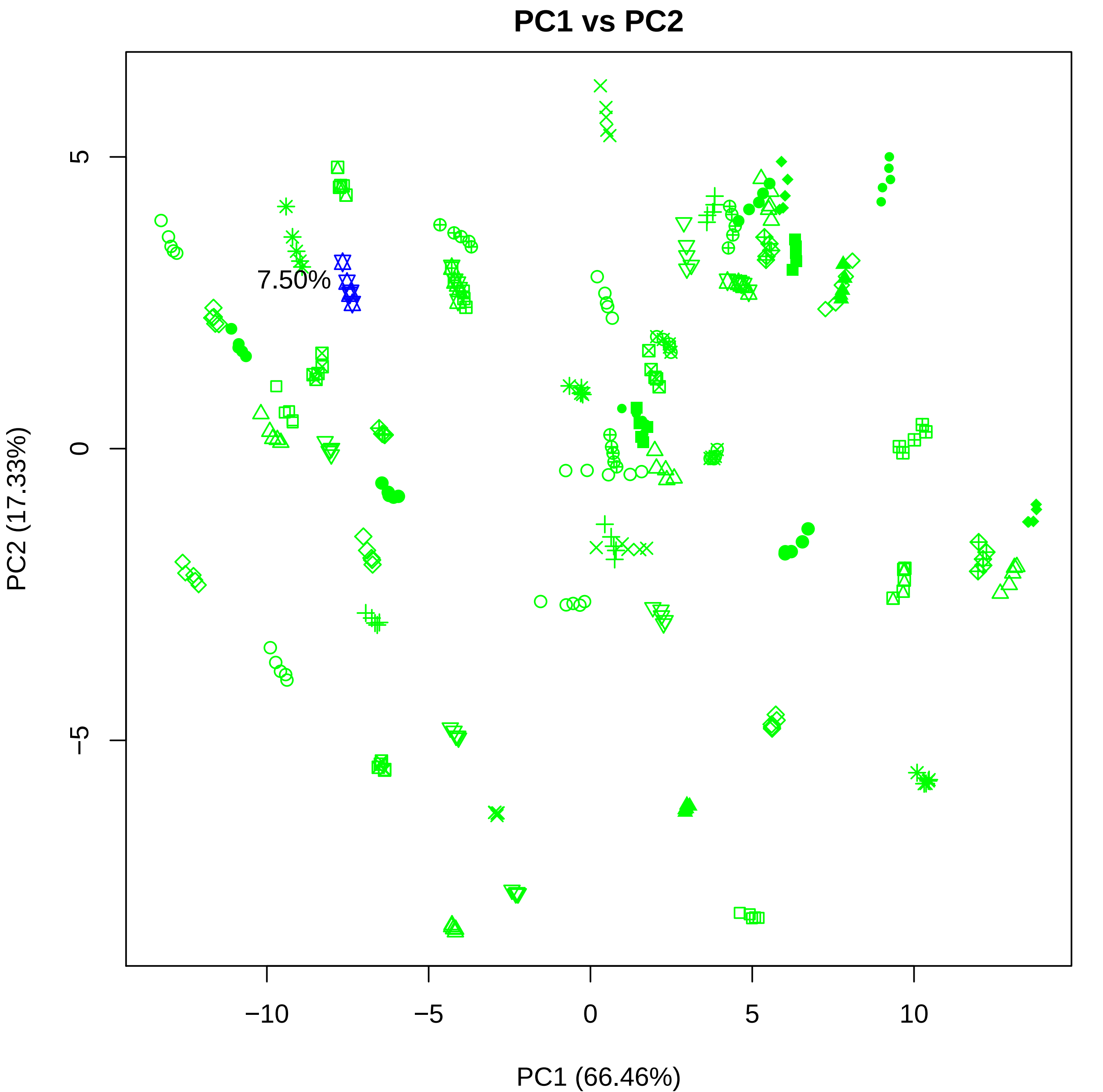
PC1 vs PC2
7.50%
PC2 (17.33%)
PC1 (66.46%)
5
0
−5
−10
−5
0
5
10

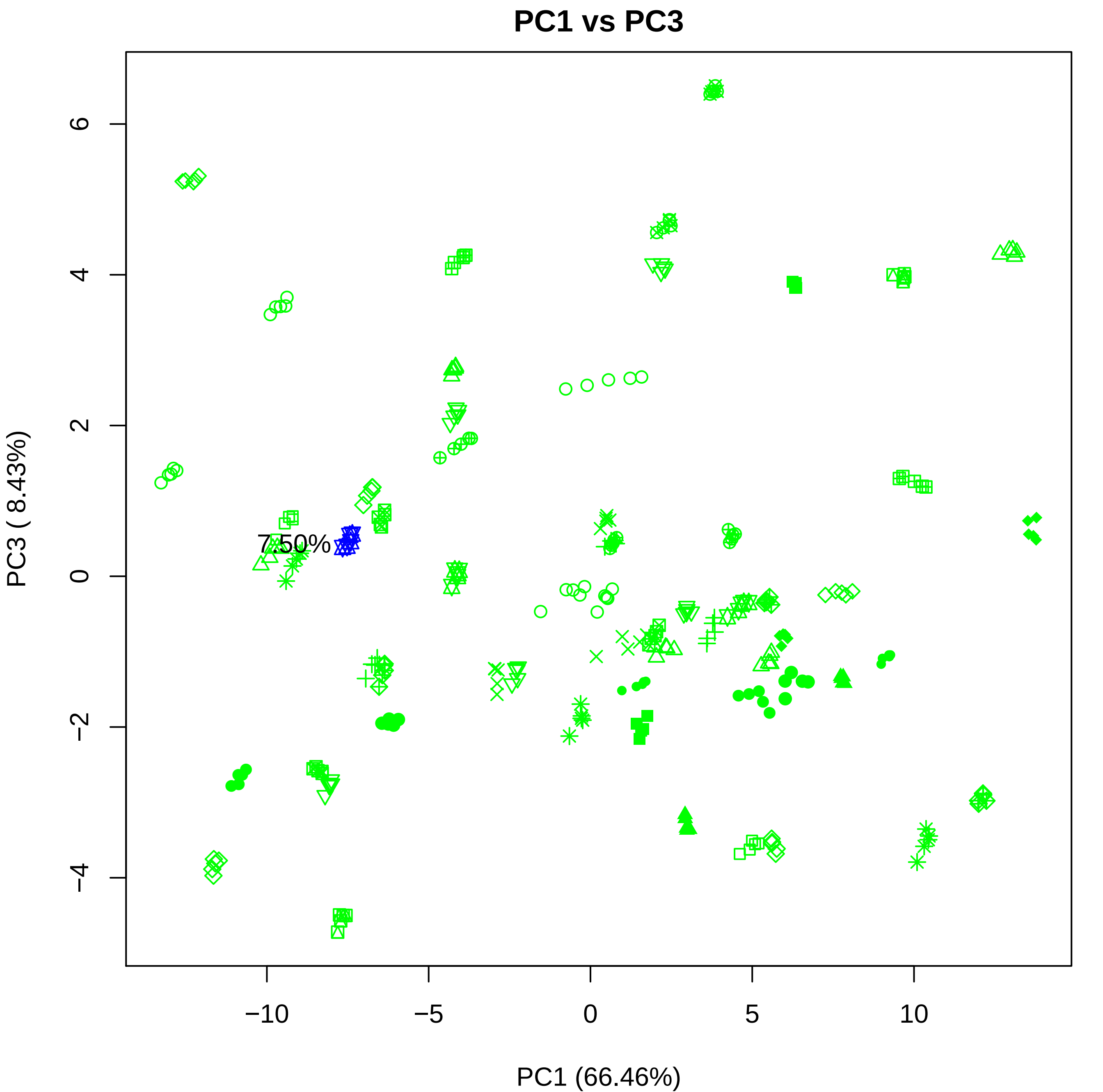
PC1 vs PC3
PC3 ( 8.43%)
PC1 (66.46%)
7.50%
6
4
2
0
−2
−4
−10
−5
0
5
10

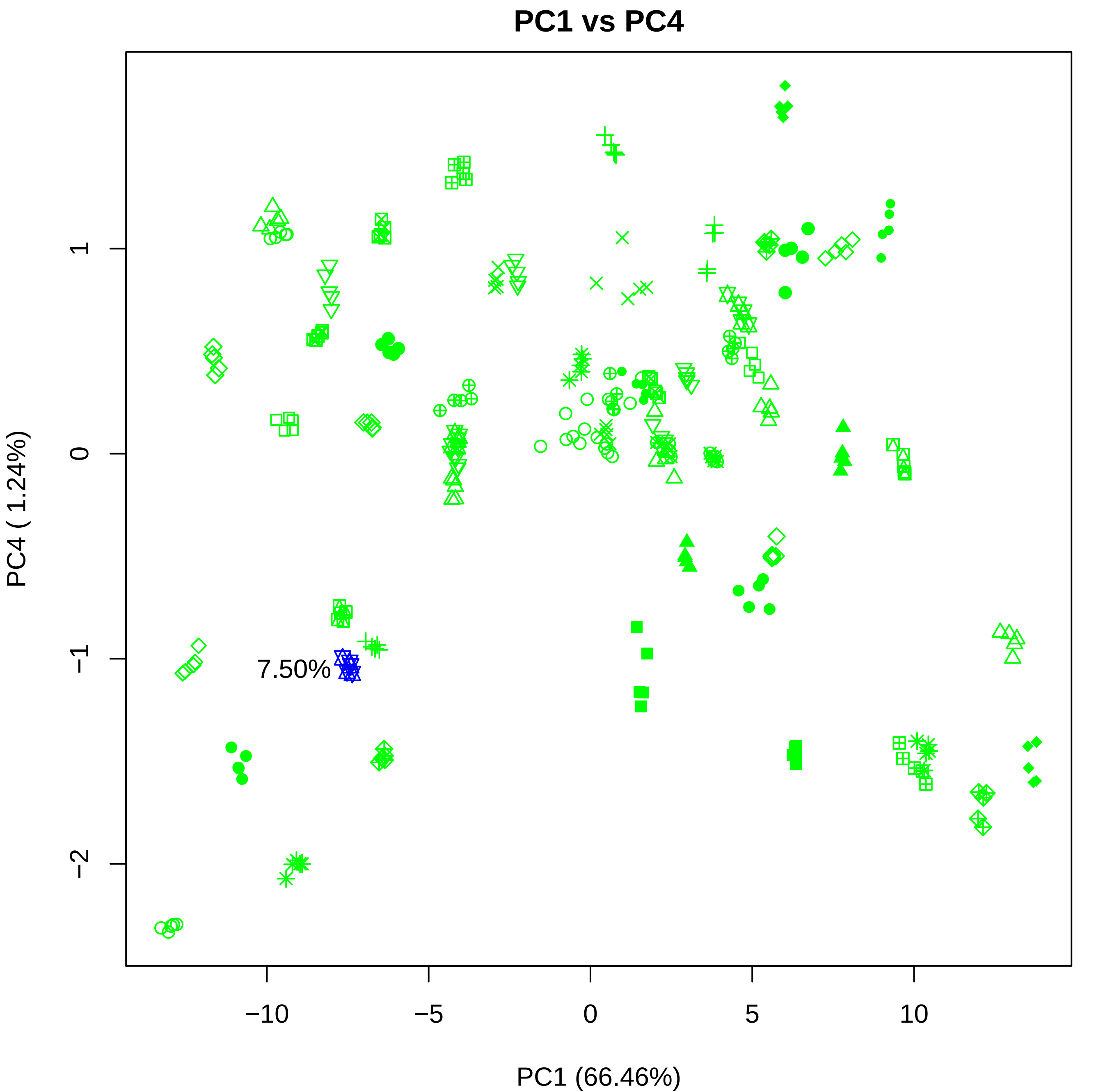
PC1 vs PC4
7.50%
PC4 ( 1.24%)
PC1 (66.46%)
1
0
−1
−2
−10
−5
0
5
10

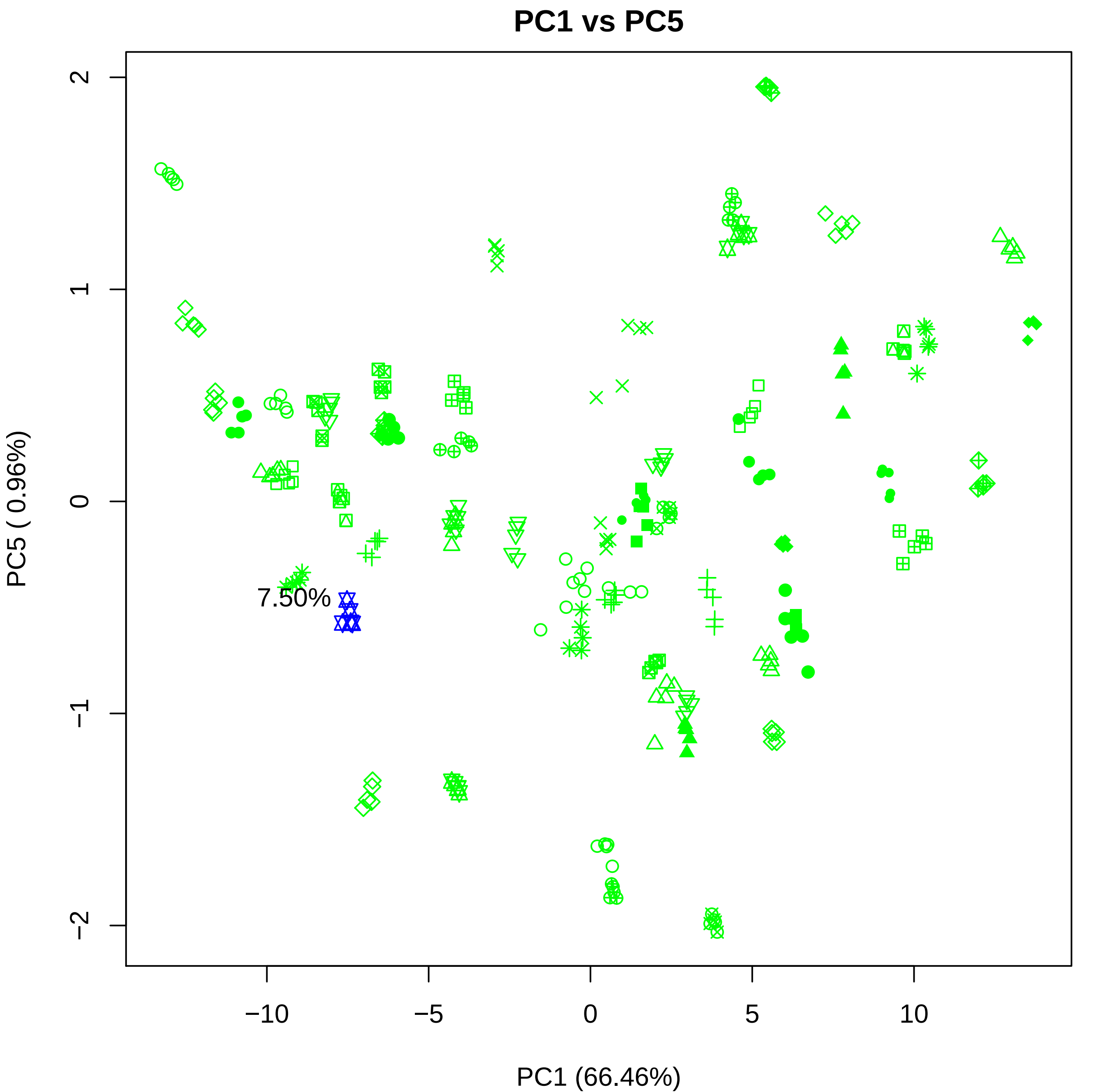
PC1 vs PC5
7.50%
PC5 ( 0.96%)
PC1 (66.46%)
2
1
0
−1
−2
−10
−5
0
5
10

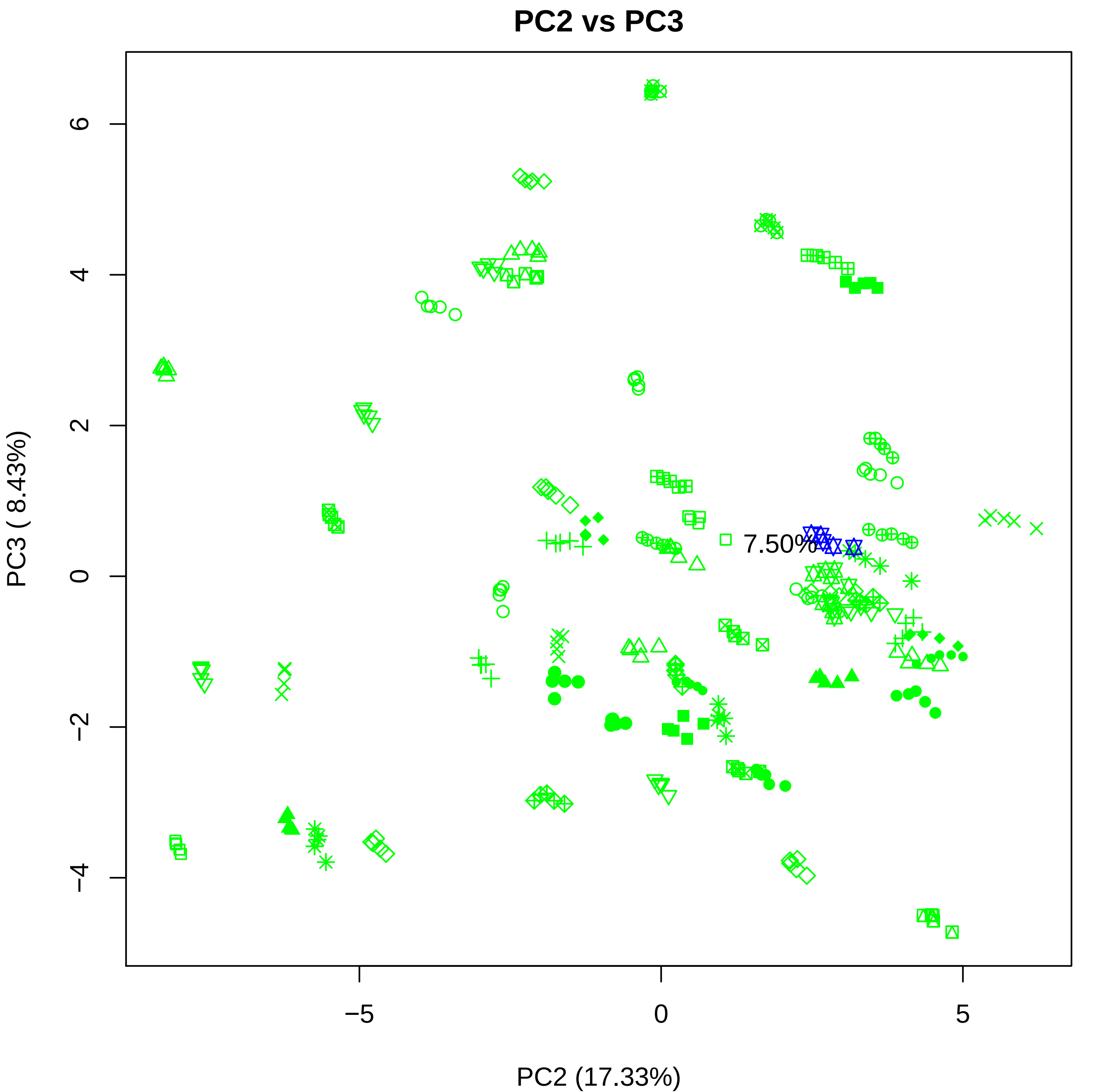
PC2 vs PC3
PC3 ( 8.43%)
PC2 (17.33%)
7.50%
6
4
2
0
−2
−4
−5
0
5

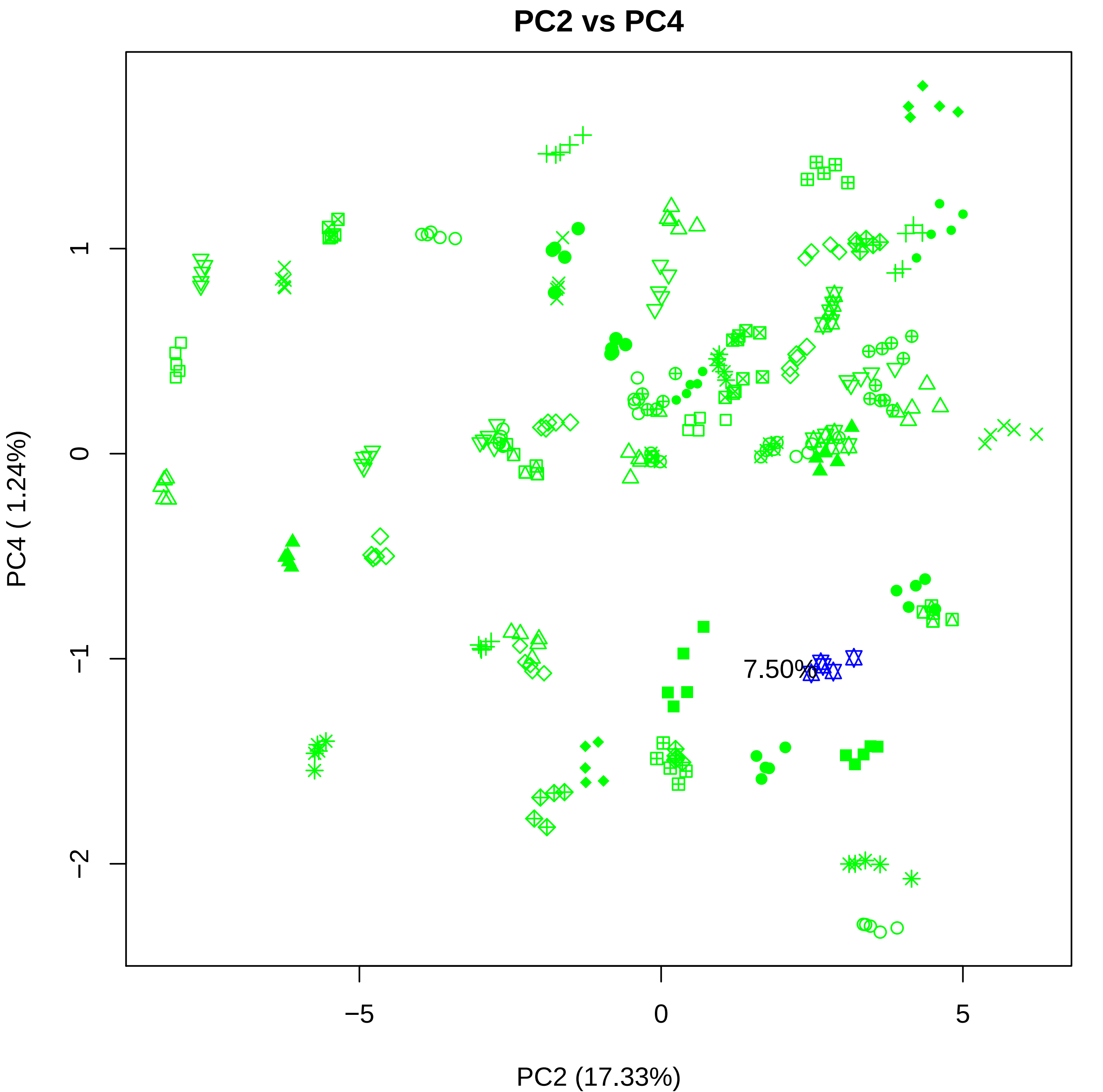
PC2 vs PC4
PC4 ( 1.24%)
PC2 (17.33%)
7.50%
1
0
−1
−2
−5
0
5

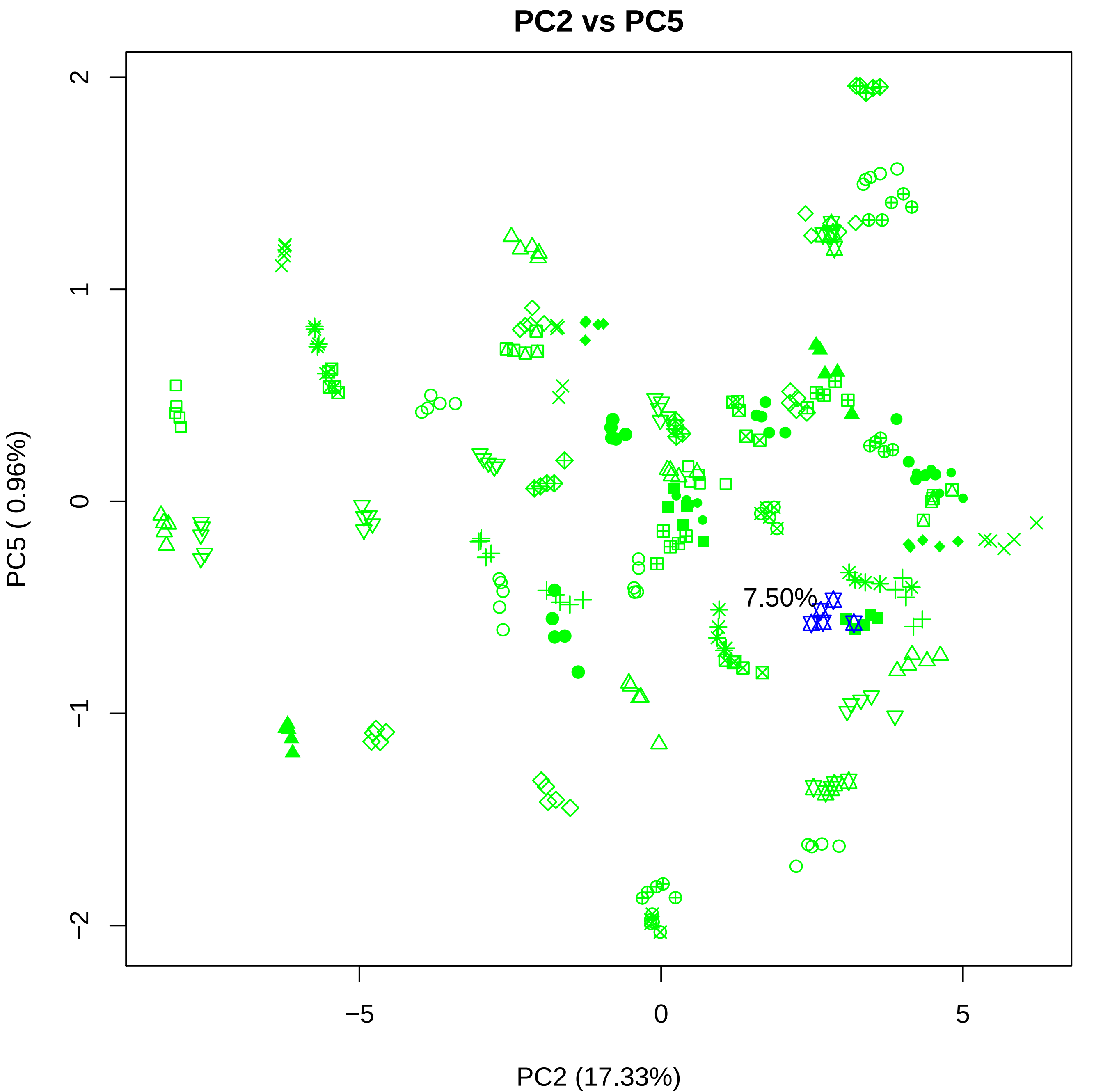
PC2 vs PC5
PC5 ( 0.96%)
PC2 (17.33%)
7.50%
2
1
0
−1
−2
−5
0
5

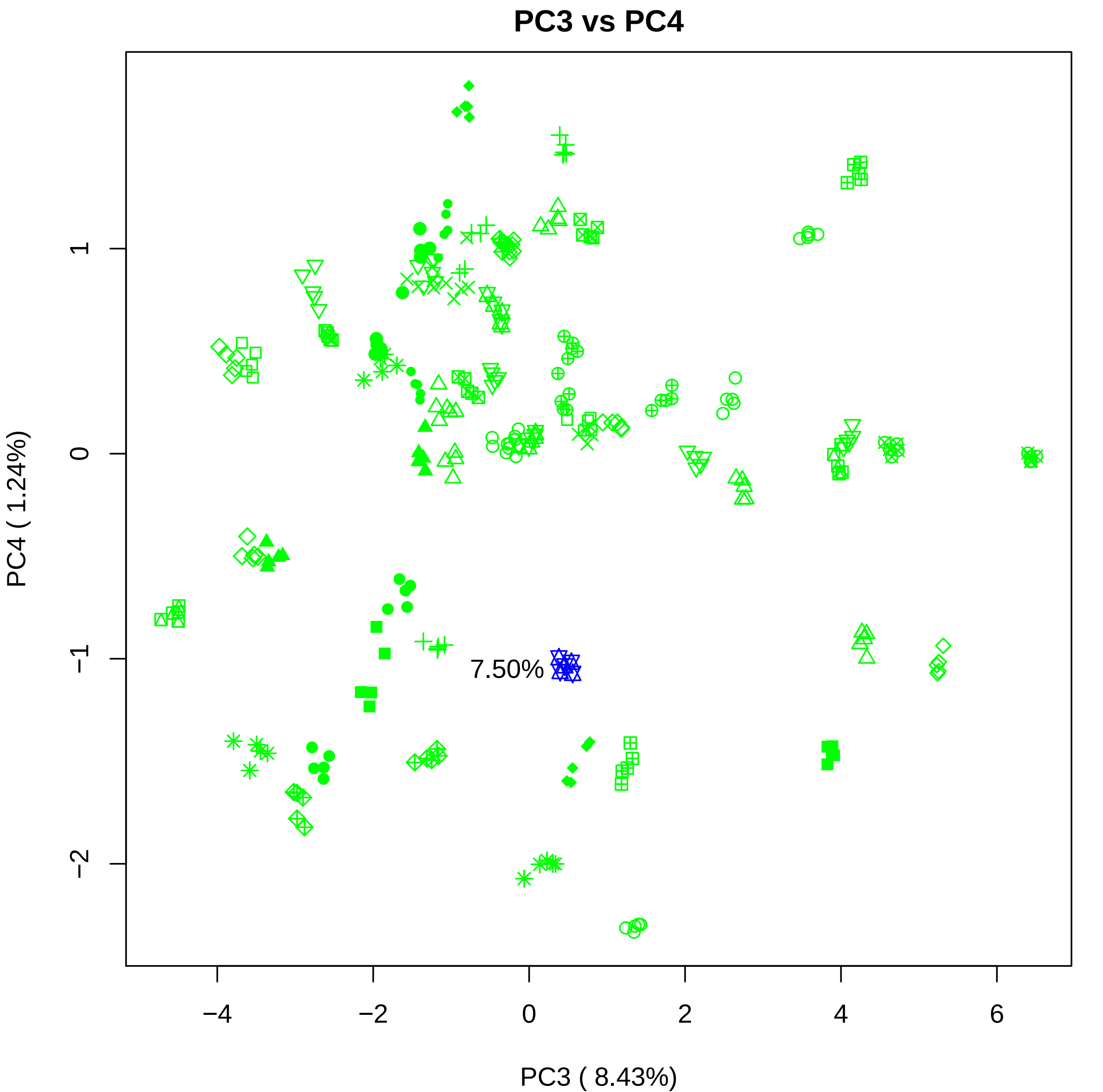

PC3 vs PC4
PC4 ( 1.24%)
PC3 ( 8.43%)
7.50%
1
0
−1
−2
−4
−2
0
2
4
6

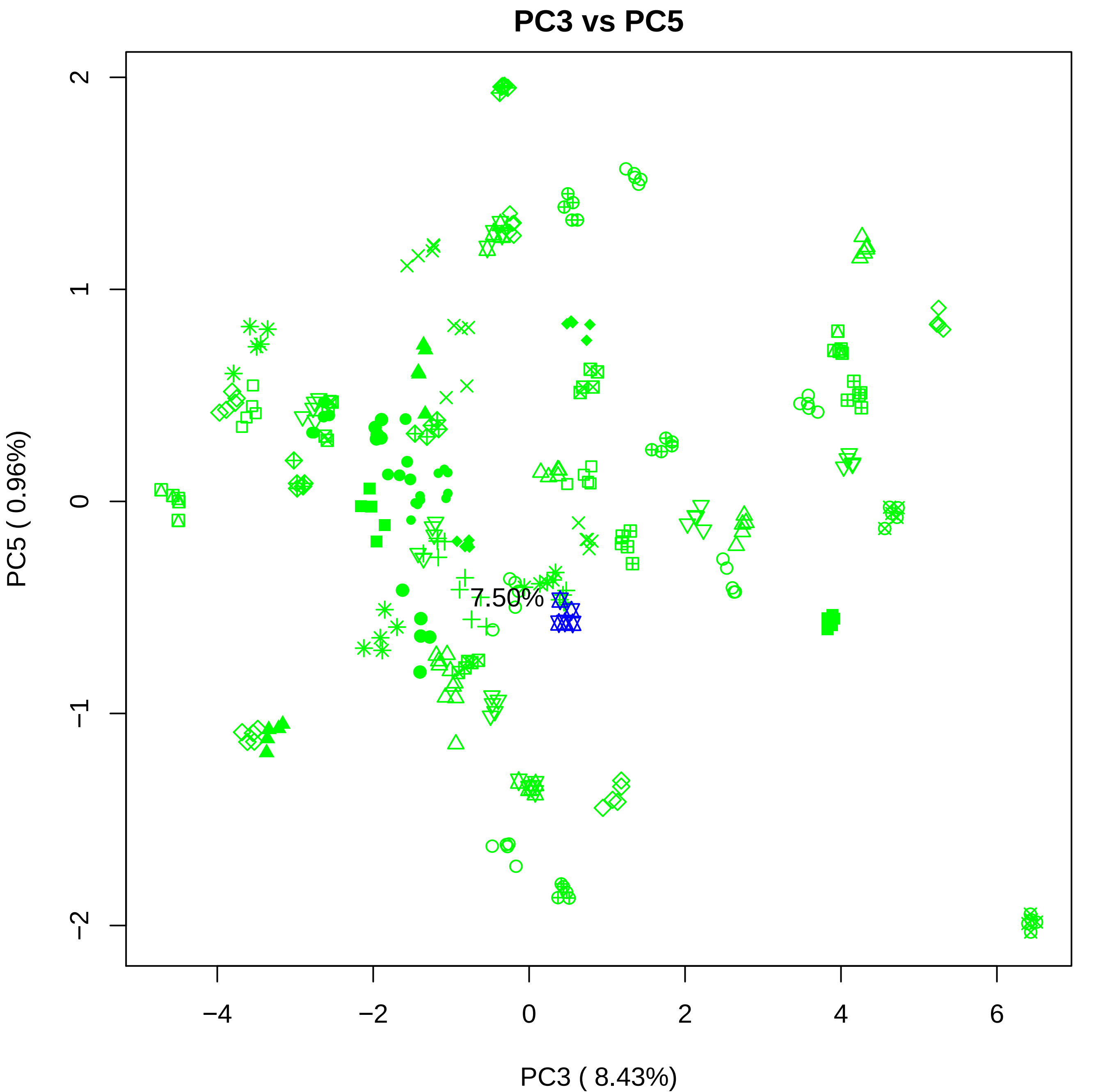

PC3 vs PC5
PC5 ( 0.96%)
PC3 ( 8.43%)
7.50%
2
1
0
−1
−2
−4
−2
0
2
4
6

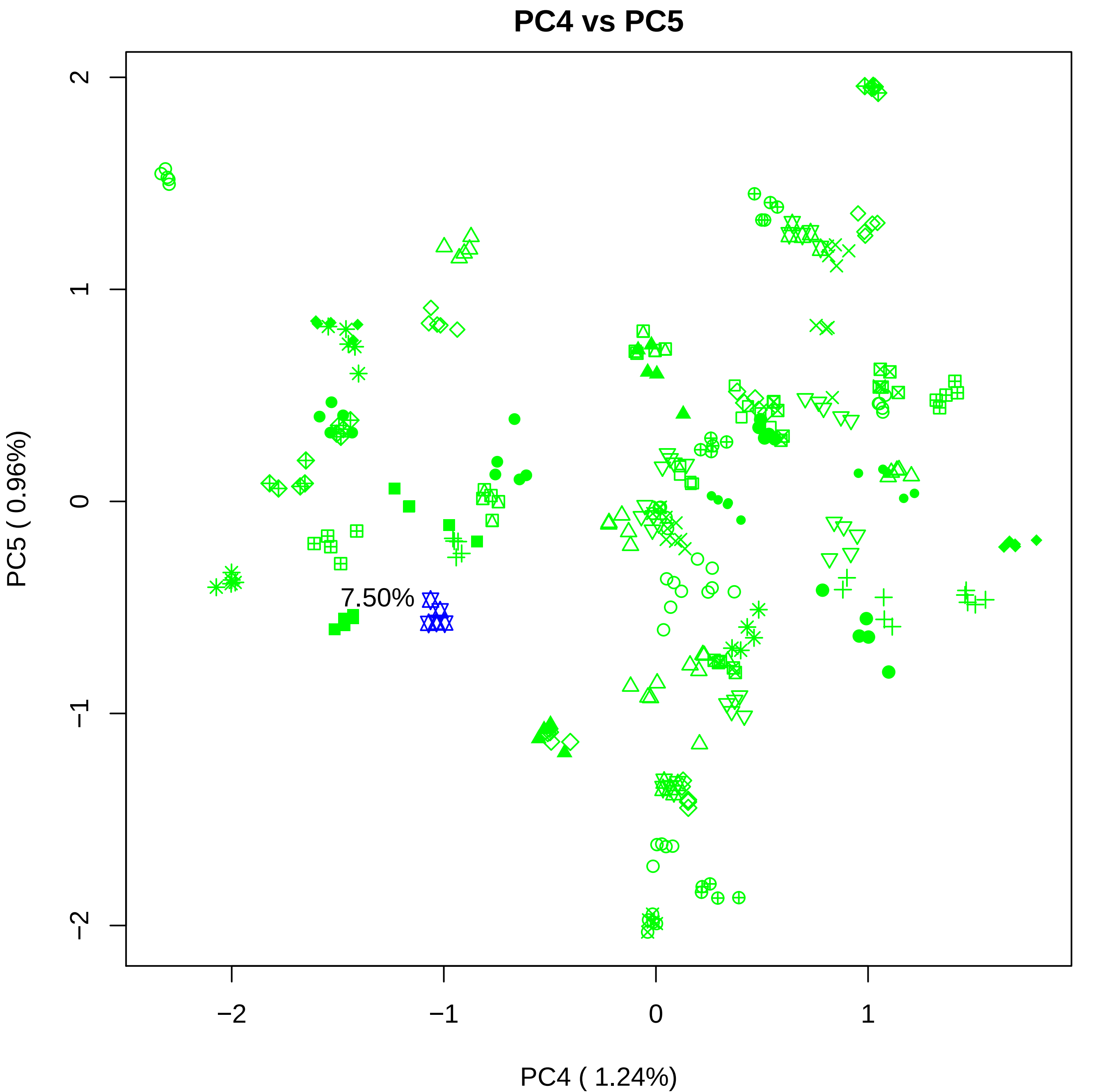

PC4 vs PC5
2
1
0
−1
−2
PC5 ( 0.96%)
7.50%
−2
−1
0
1
PC4 ( 1.24%)

### PCA of 10%  BMH(b) vs PMH(b)

Scaled Matrix PCAed

7th order polynomial baseline correction performed

Raw spectra smoothed to 21 Neighbours, 2nd order PN, Savtisky Golay

Normalisation between 0 and 100

D2 spectra smoothed to 41 Neighbours, 2nd order PN, Savtisky Golay

Regions Analysed: 3601−2501 and 2001−700cm−1

Samples Analysed: 1 bContam's 0 BLH's 0 BMH's 0 OMH's 0 pContam's 0 PLH's 60 PMH's

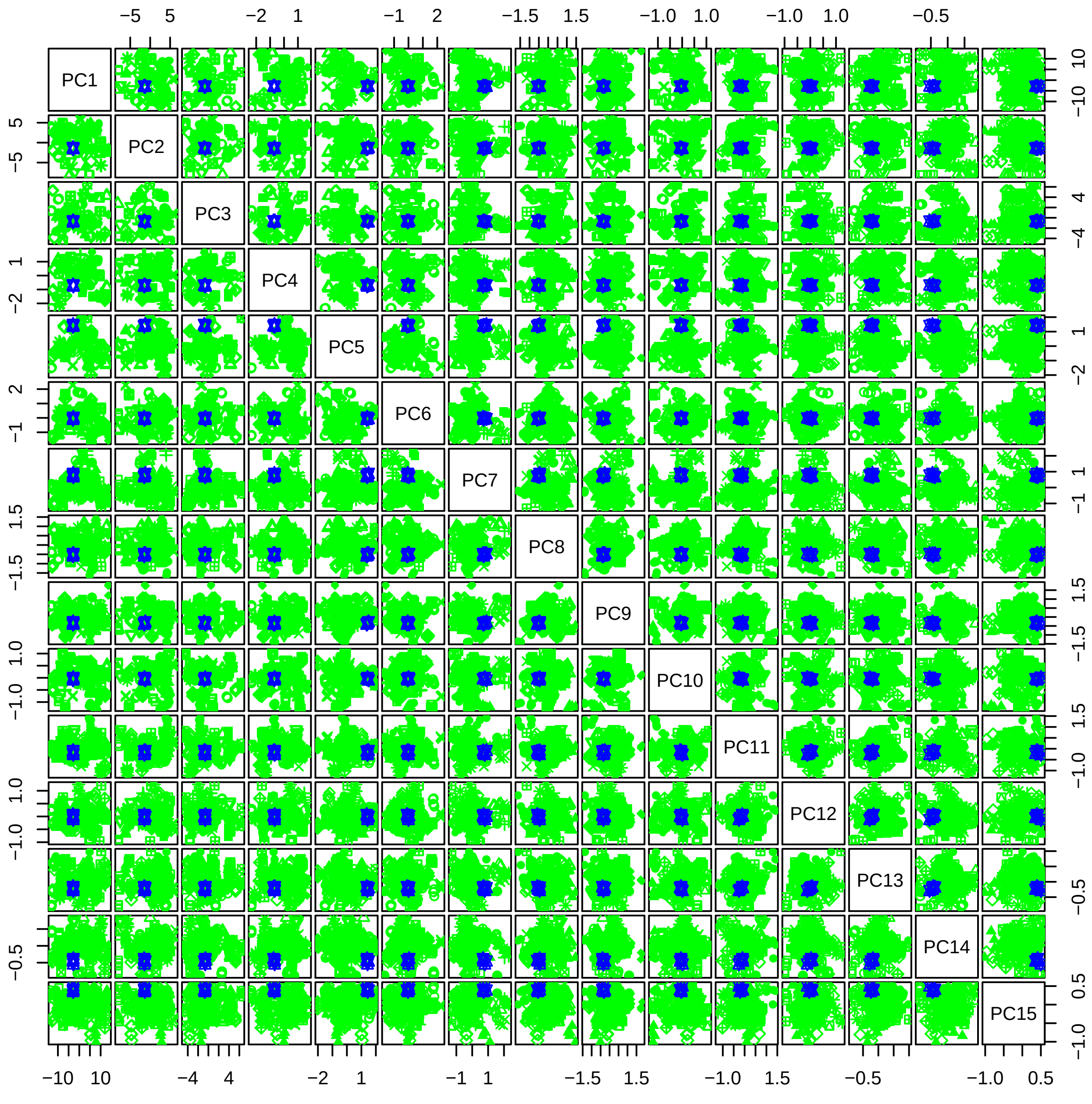
PC1
PC2
PC3
PC4
PC5
PC6
PC7
PC8
PC9
PC10
PC11
PC12
PC13
PC14
PC15
−5
5
−2
1
−1
2
−1.5
1.5
−1.0
1.0
−1.0
1.0
−0.5
10
−10
5
−5
4
−4
1
−2
1
−2
2
−1
1
−1
1.5
−1.5
1.5
−1.5
1.0
−1.0
1.5
−1.0
1.0
−1.0
−0.5
−0.5
0.5
−1.0
−10
10
−4
4
−2
1
−1
1
−1.5
1.5
−1.0
1.5
−0.5
−1.0
0.5

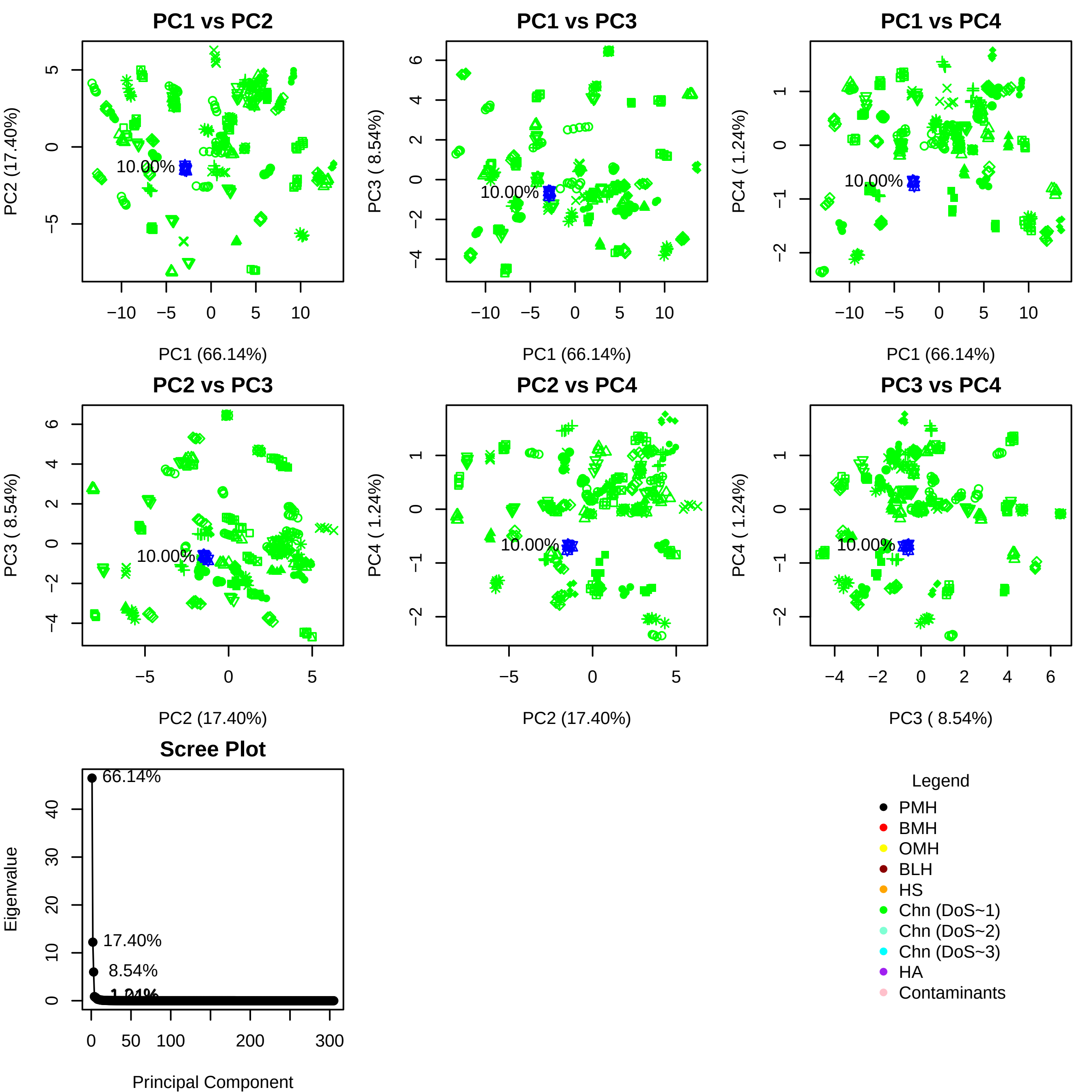
PC1 vs PC2
PC1 vs PC3
PC1 vs PC4
PC2 vs PC3
PC2 vs PC4
PC3 vs PC4
Scree Plot
PC1 (66.14%)
PC2 (17.40%)
PC3 ( 8.54%)
PC4 ( 1.24%)
10.00%
Eigenvalue
Principal Component
66.14%
17.40%
8.54%
Legend
PMH
BMH
OMH
BLH
HS
Chn (DoS~1)
Chn (DoS~2)
Chn (DoS~3)
HA
Contaminants

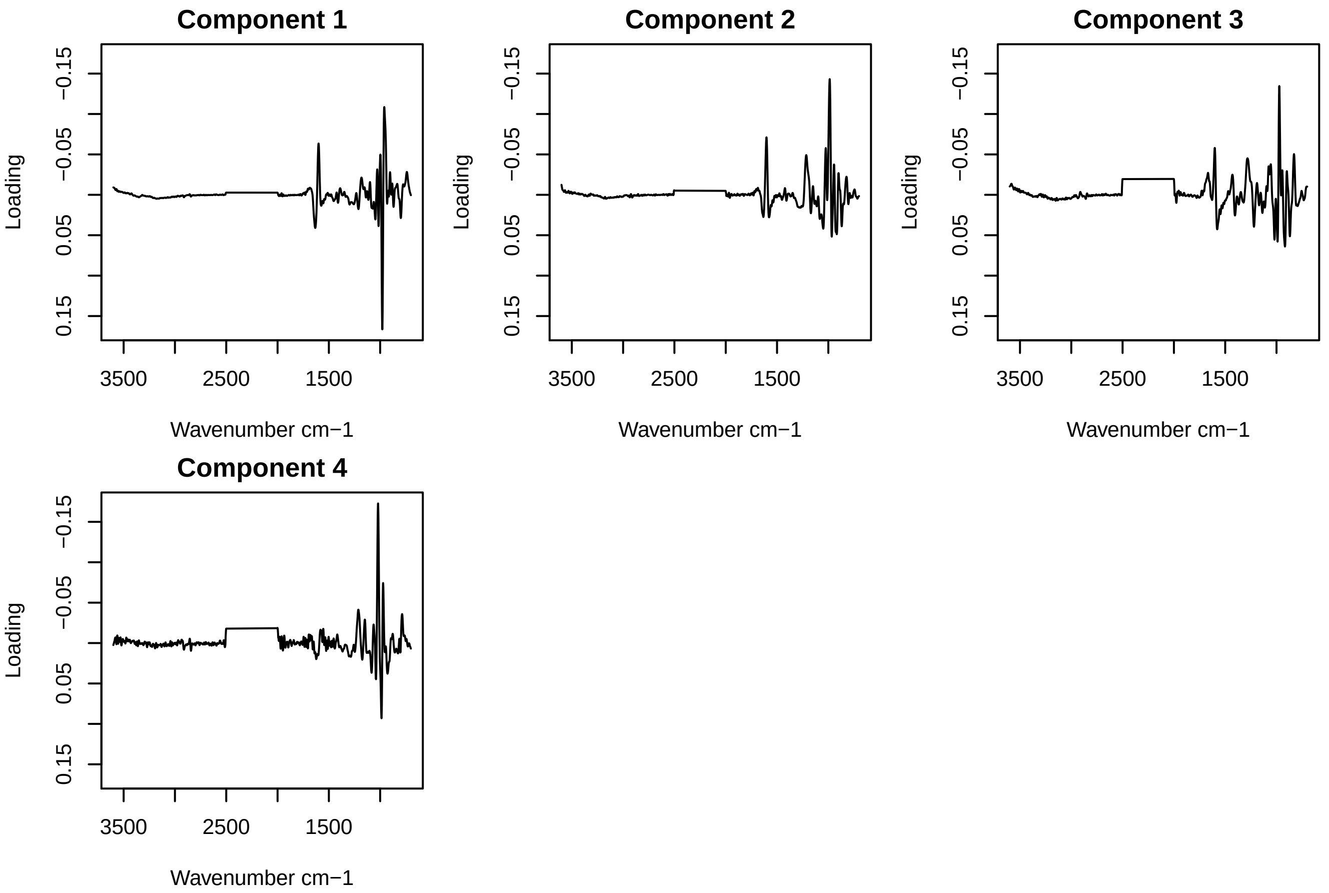
Component 1
Component 2
Component 3
Component 4
Loading
−0.15
−0.05
0.05
0.15
3500
2500
1500
Wavenumber cm−1

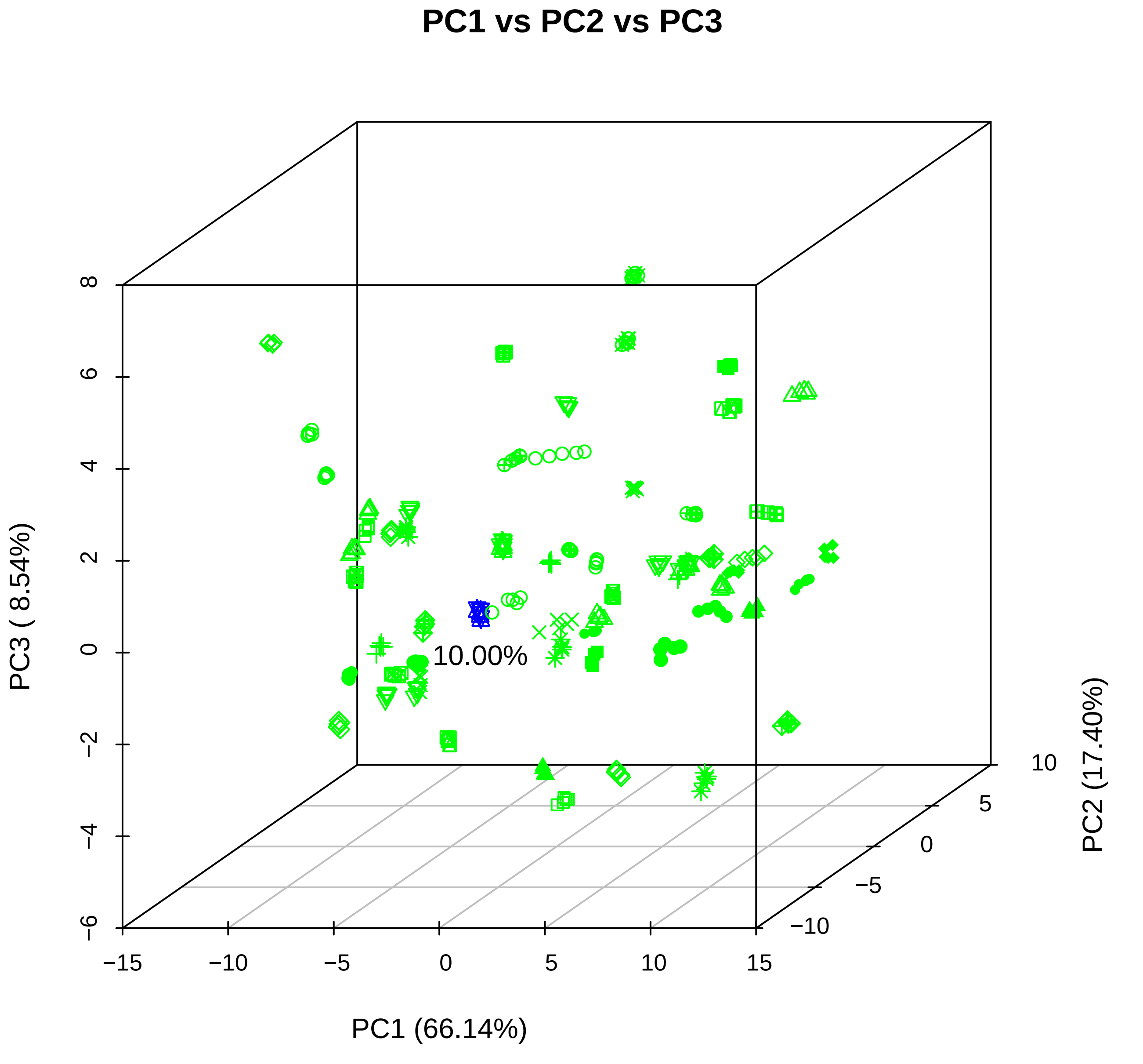
PC1 vs PC2 vs PC3
PC1 (66.14%)
PC2 (17.40%)
PC3 ( 8.54%)
10.00%
−15
−10
−5
0
5
10
15
−10
−5
0
5
10
−6
−4
−2
0
2
4
6
8

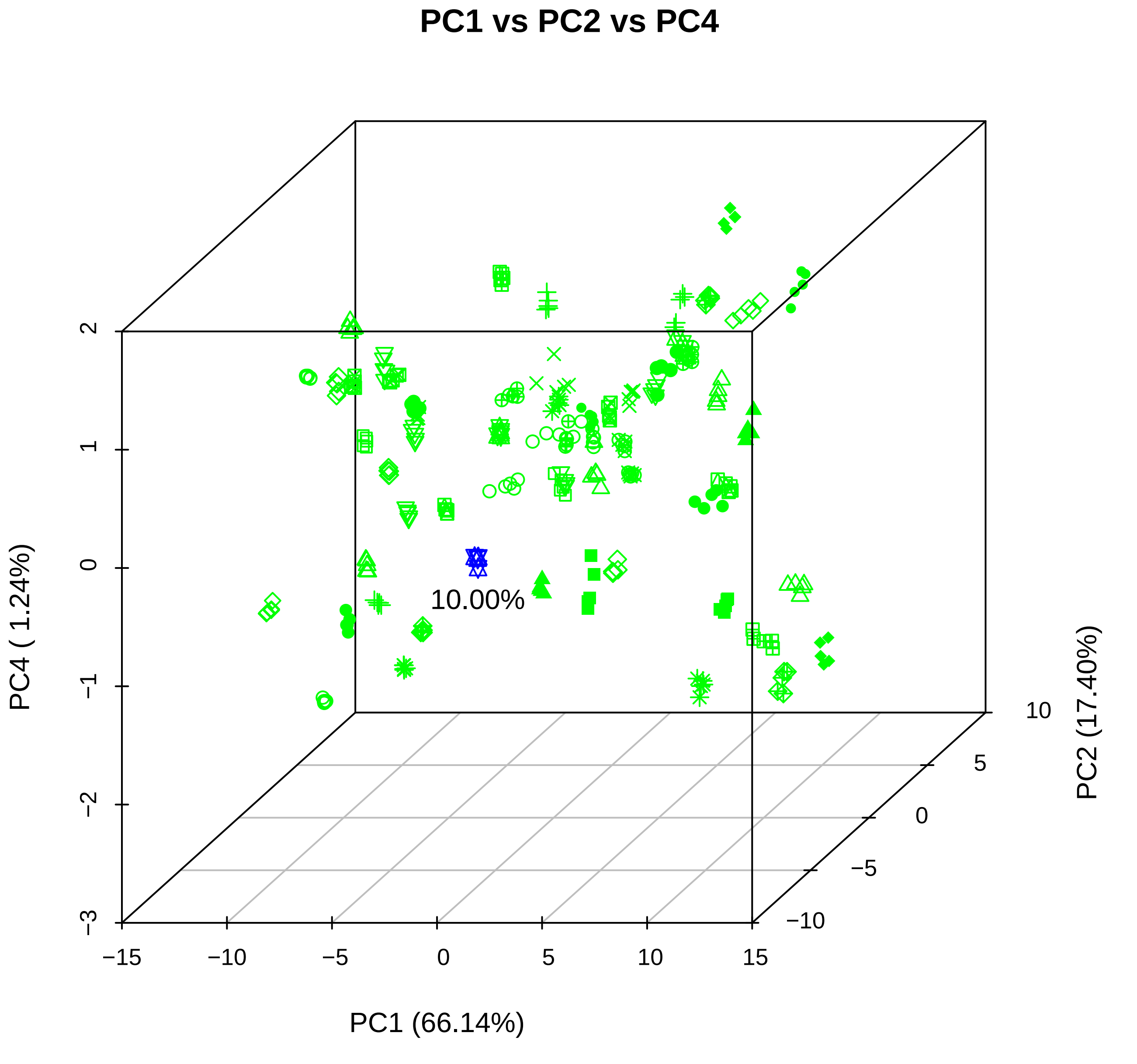
PC1 vs PC2 vs PC4
10.00%
PC1 (66.14%)
PC2 (17.40%)
PC4 ( 1.24%)
−15
−10
−5
0
5
10
15
−3
−2
−1
0
1
2
−10
−5
0
5
10

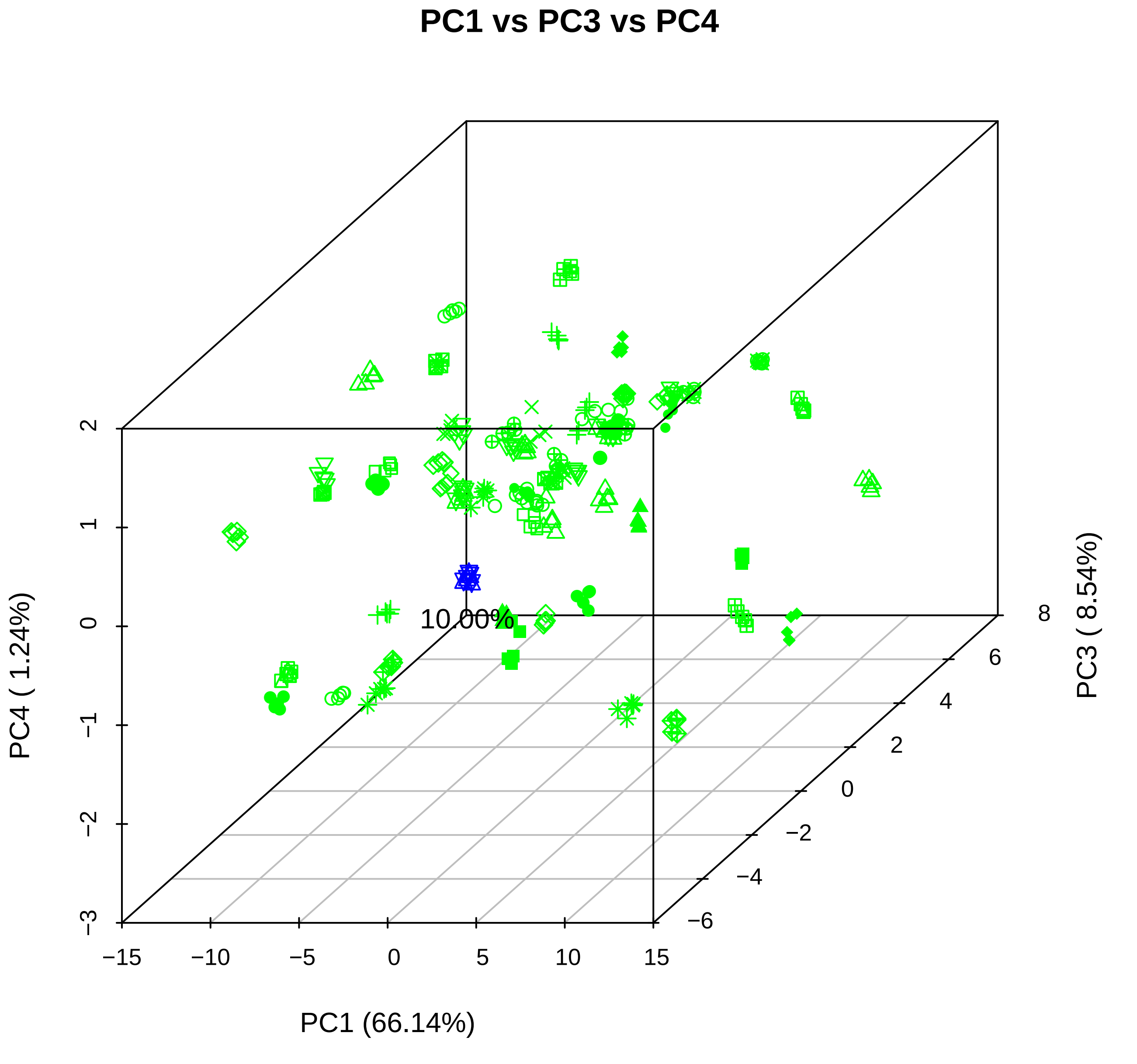

PC1 vs PC3 vs PC4
PC1 (66.14%)
PC4 ( 1.24%)
PC3 ( 8.54%)
10.00%
−15
−10
−5
0
5
10
15
−3
−2
−1
0
1
2
−6
−4
−2
0
2
4
6
8

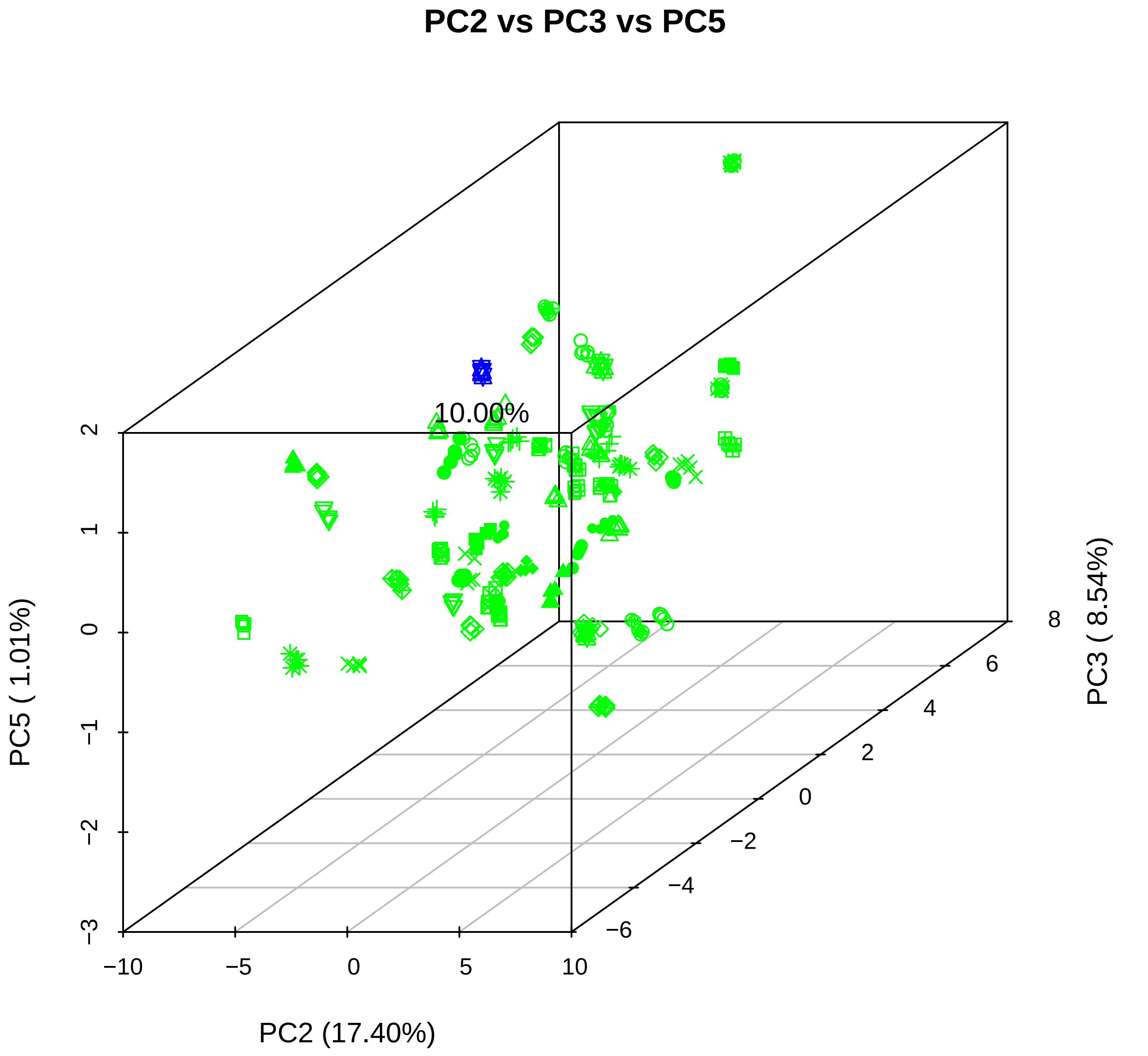

PC2 vs PC3 vs PC5
10.00%
PC2 (17.40%)
PC5 ( 1.01%)
PC3 ( 8.54%)
−10
−5
0
5
10
−3
−2
−1
0
1
2
−6
−4
−2
0
2
4
6
8

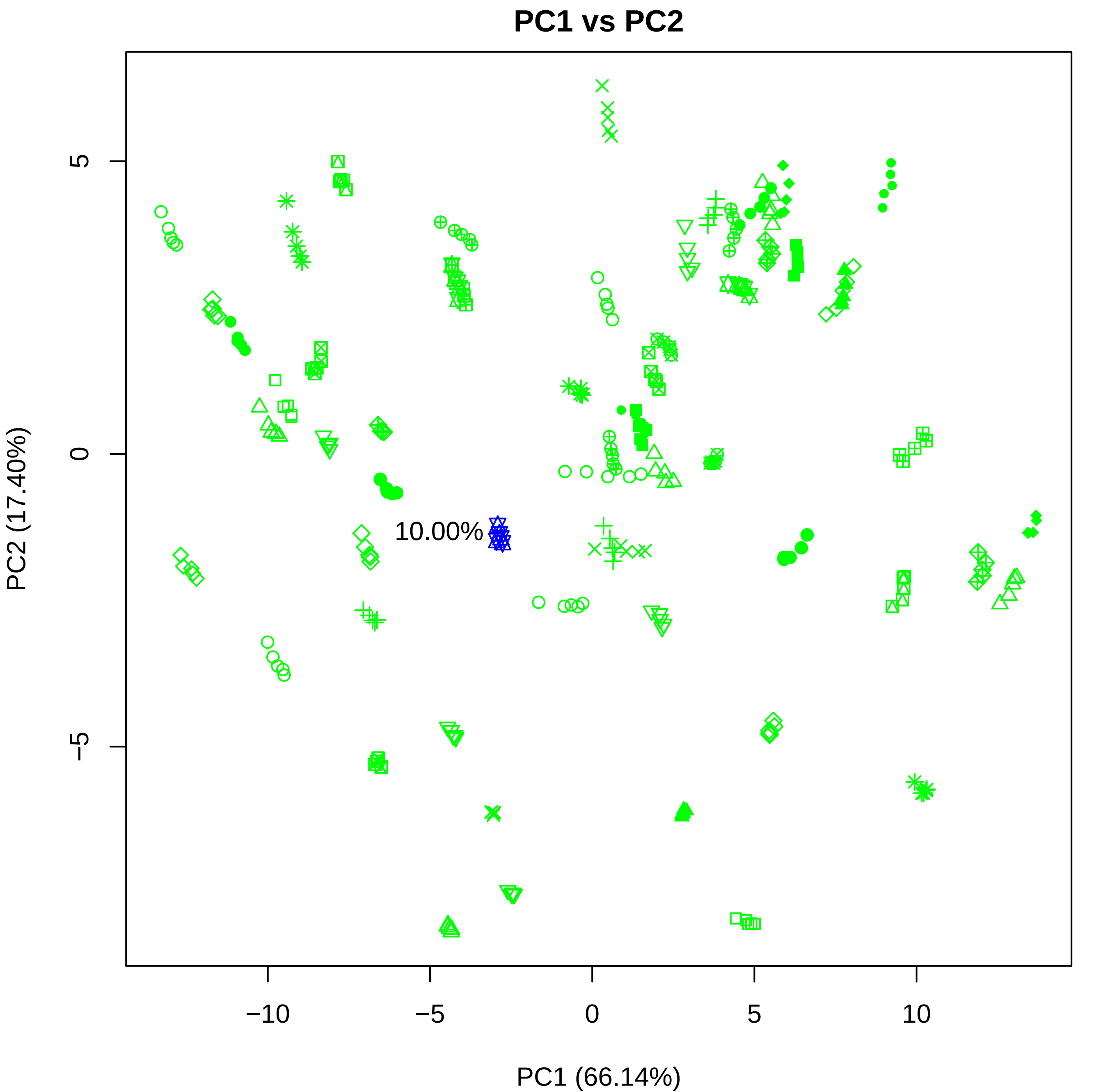

PC1 vs PC2
PC2 (17.40%)
PC1 (66.14%)
10.00%
5
0
−5
−10
−5
0
5
10

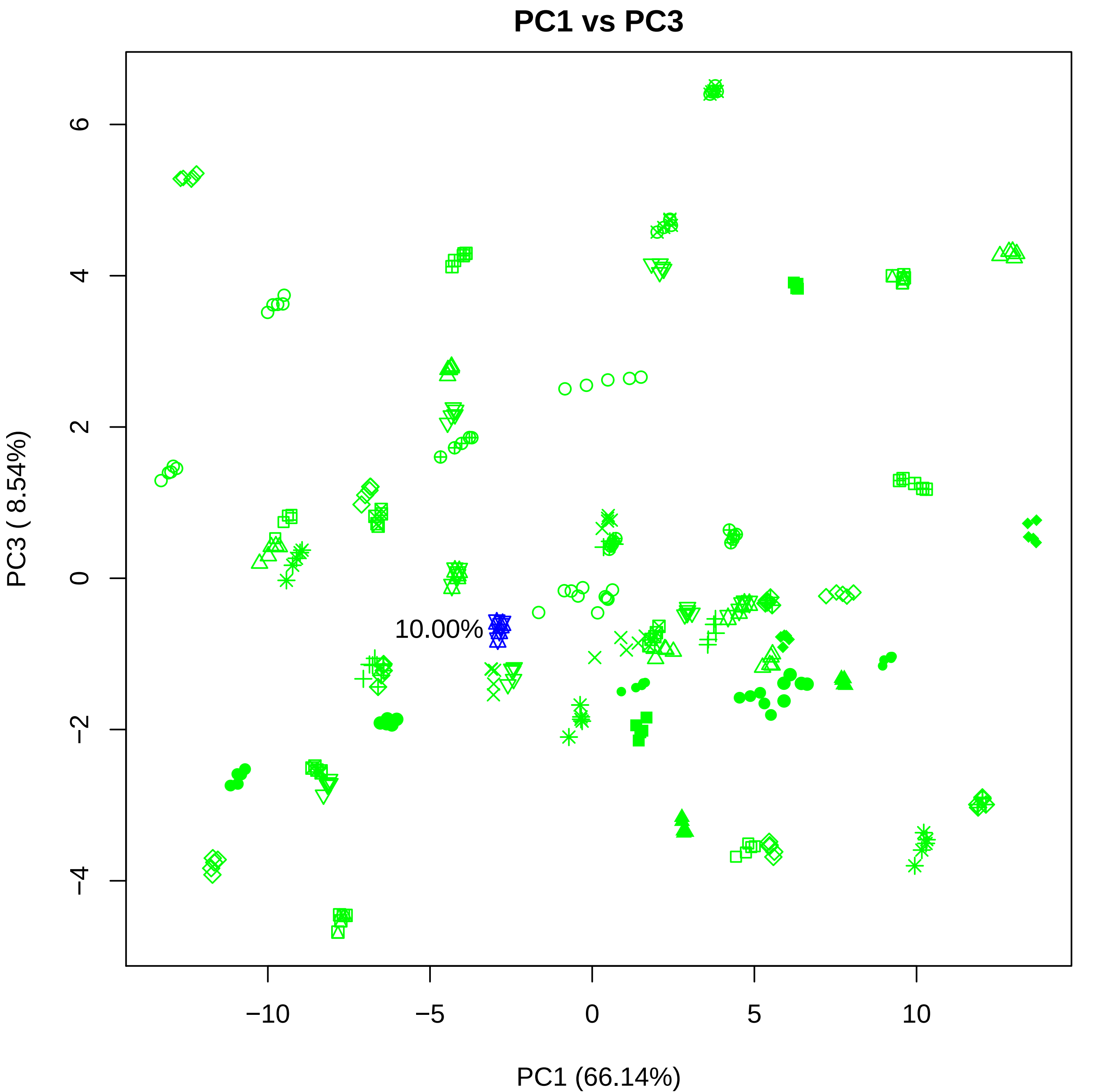

PC1 vs PC3
PC3 ( 8.54%)
PC1 (66.14%)
10.00%
6
4
2
0
−2
−4
−10
−5
0
5
10

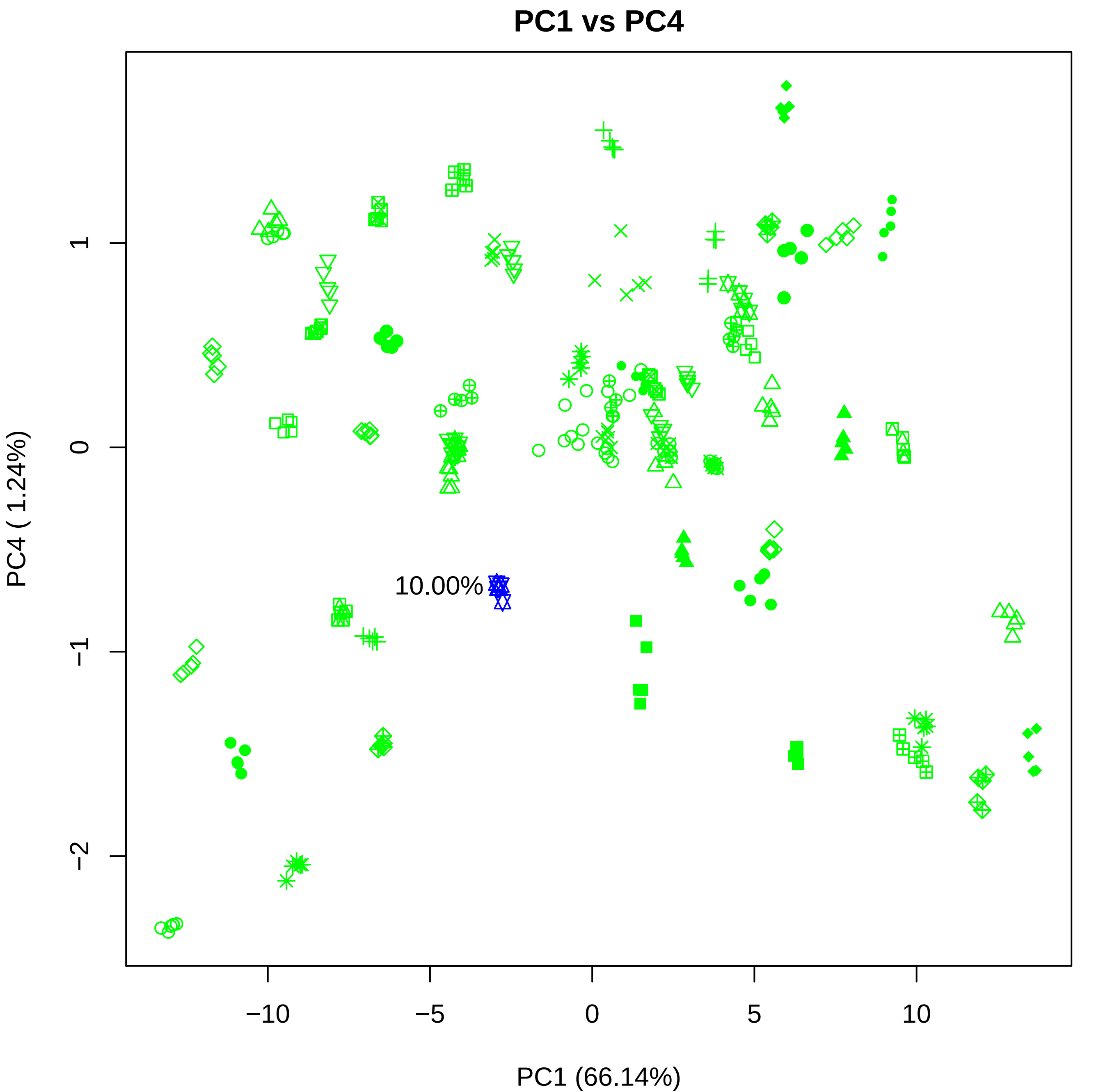
PC1 vs PC4
10.00%
PC4 ( 1.24%)
PC1 (66.14%)
−10
−5
0
5
10
1
0
−1
−2

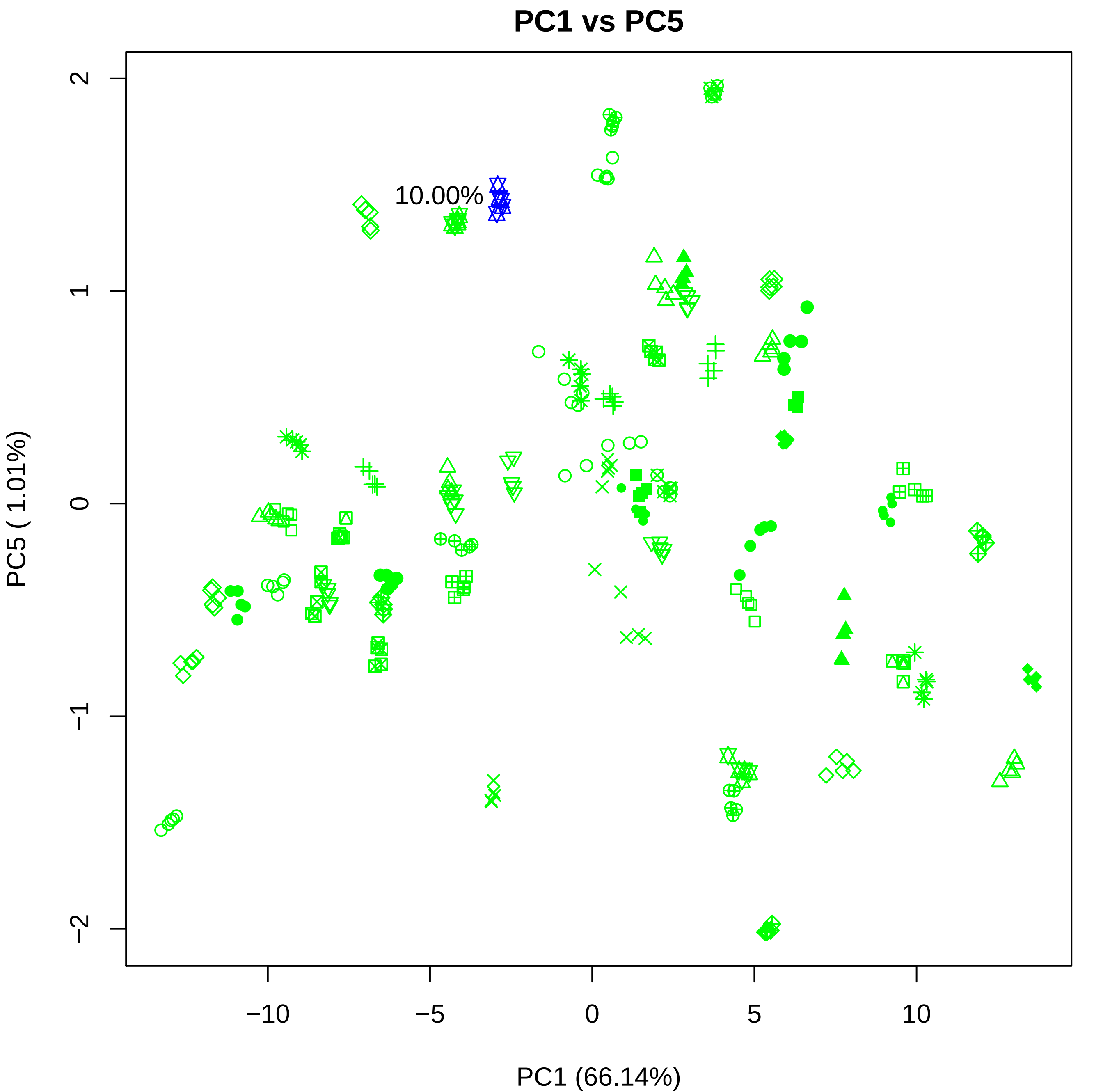
PC1 vs PC5
10.00%
PC5 ( 1.01%)
PC1 (66.14%)
2
1
0
−1
−2
−10
−5
0
5
10

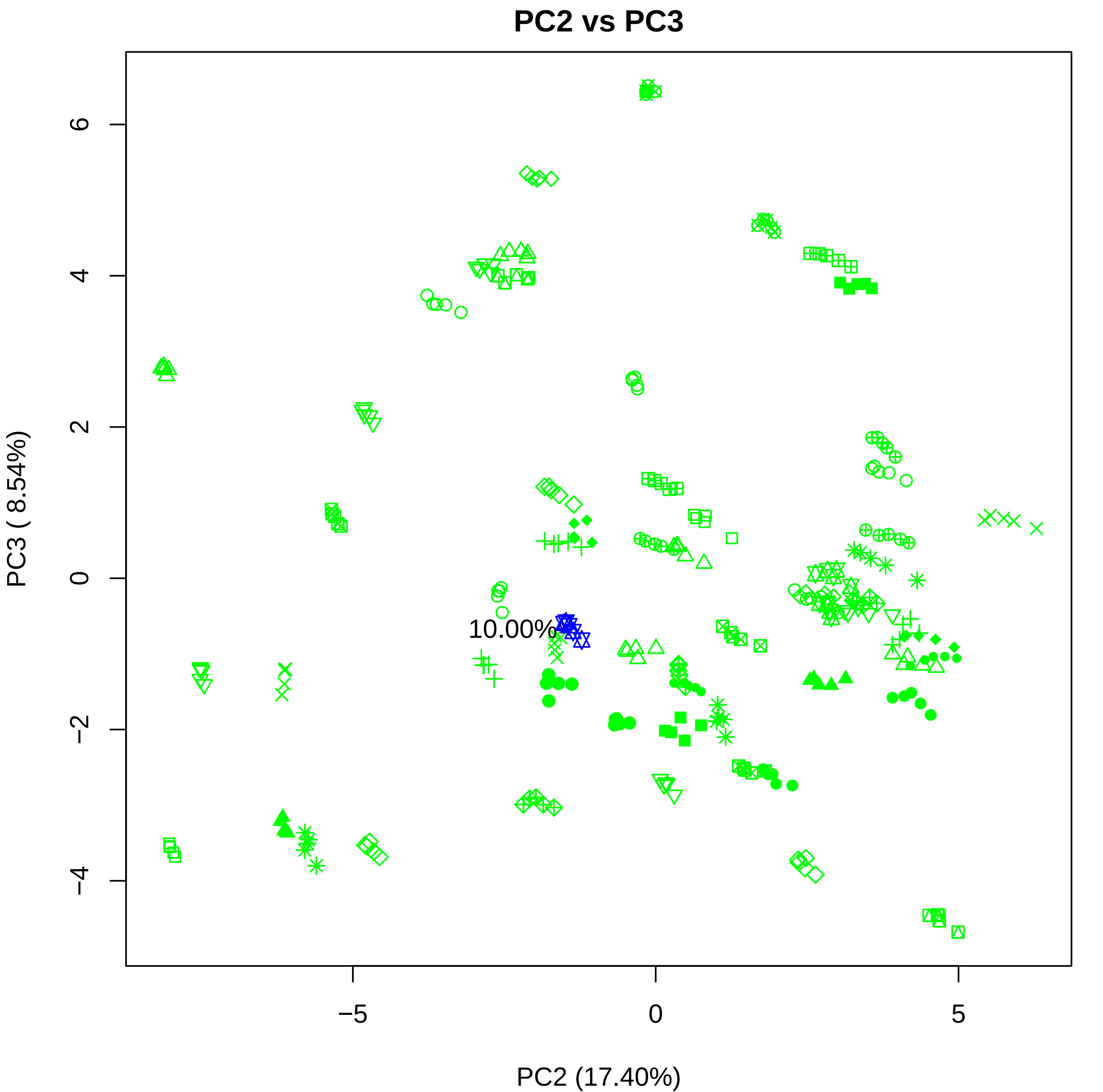
PC2 vs PC3
10.00%
PC3 ( 8.54%)
PC2 (17.40%)
6
4
2
0
−2
−4
−5
0
5

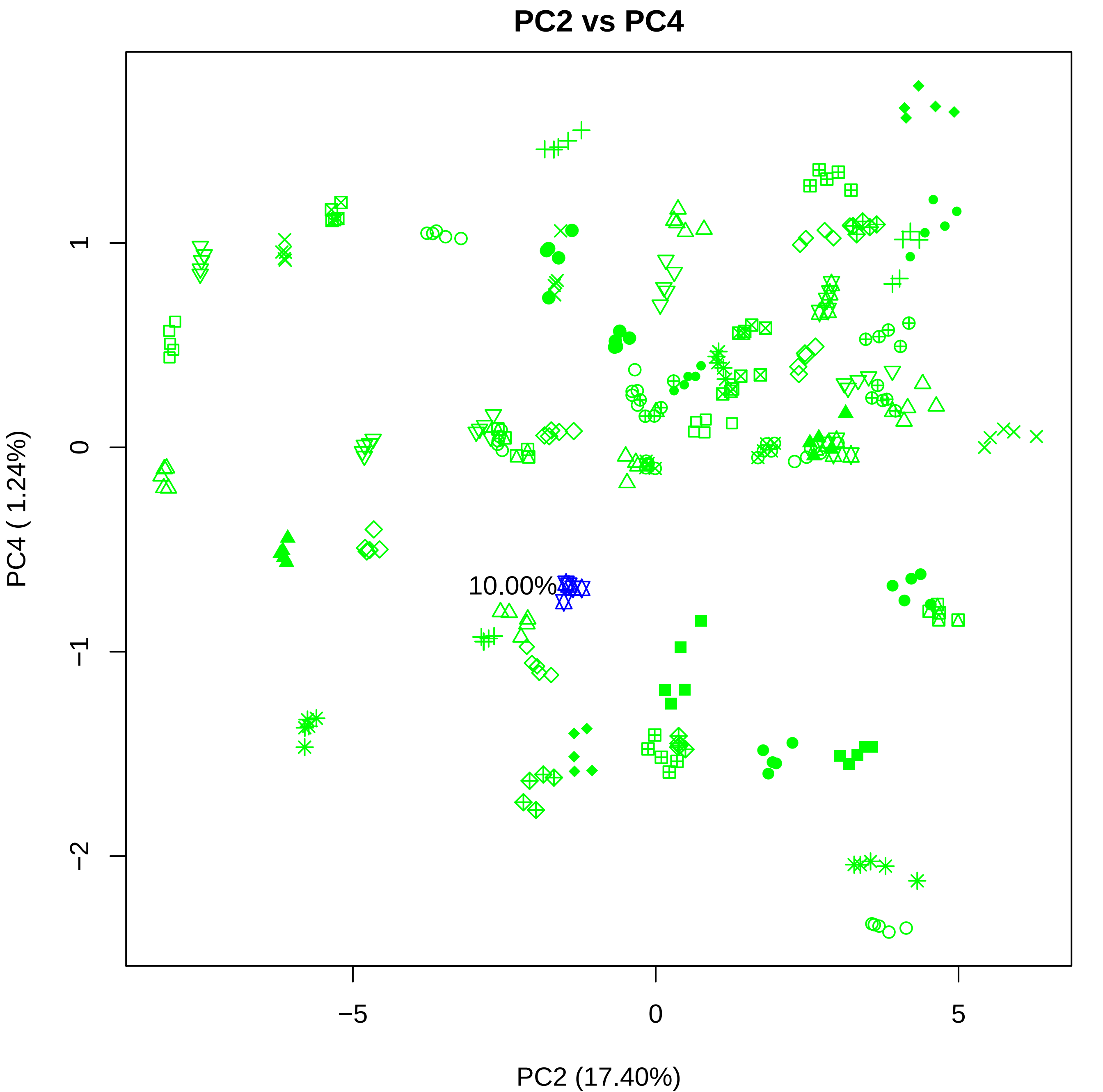

PC2 vs PC4
10.00%
1
0
−1
−2
−5
0
5
PC4 ( 1.24%)
PC2 (17.40%)

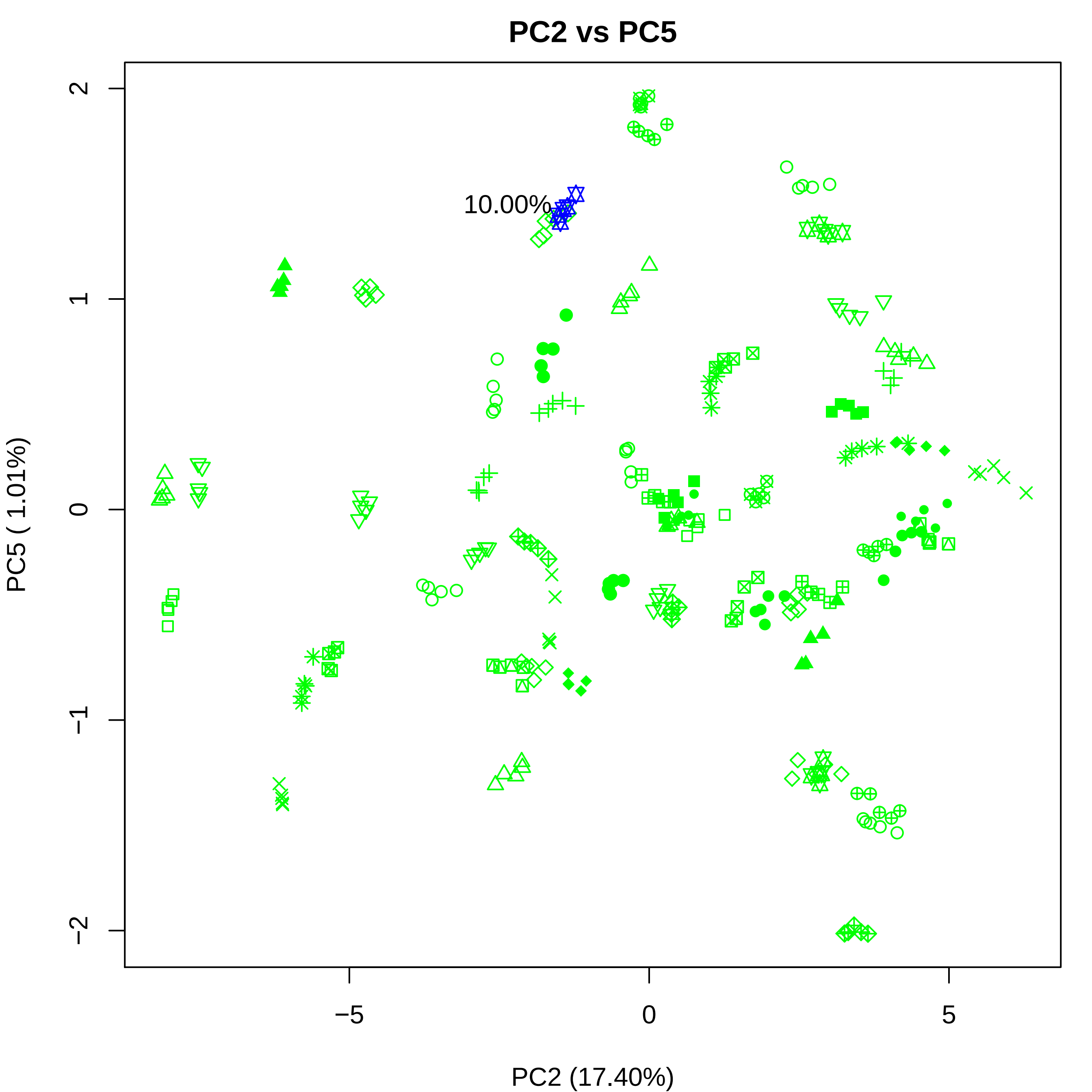
PC2 vs PC5
10.00%
PC5 ( 1.01%)
PC2 (17.40%)
2
1
0
−1
−2
−5
0
5

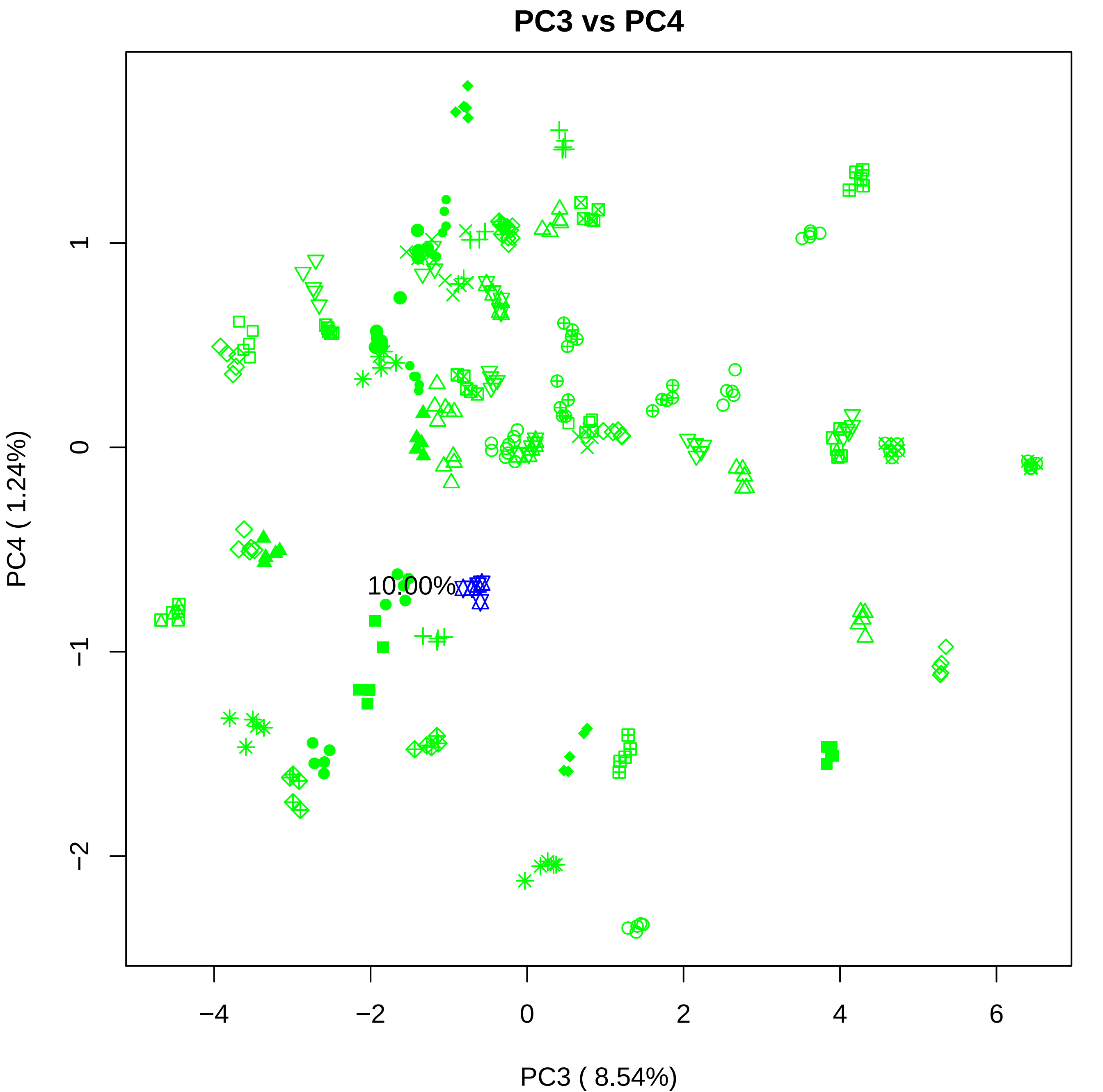

PC3 vs PC4
10.00%
PC4 ( 1.24%)
PC3 ( 8.54%)
−4
−2
0
2
4
6
1
0
−1
−2

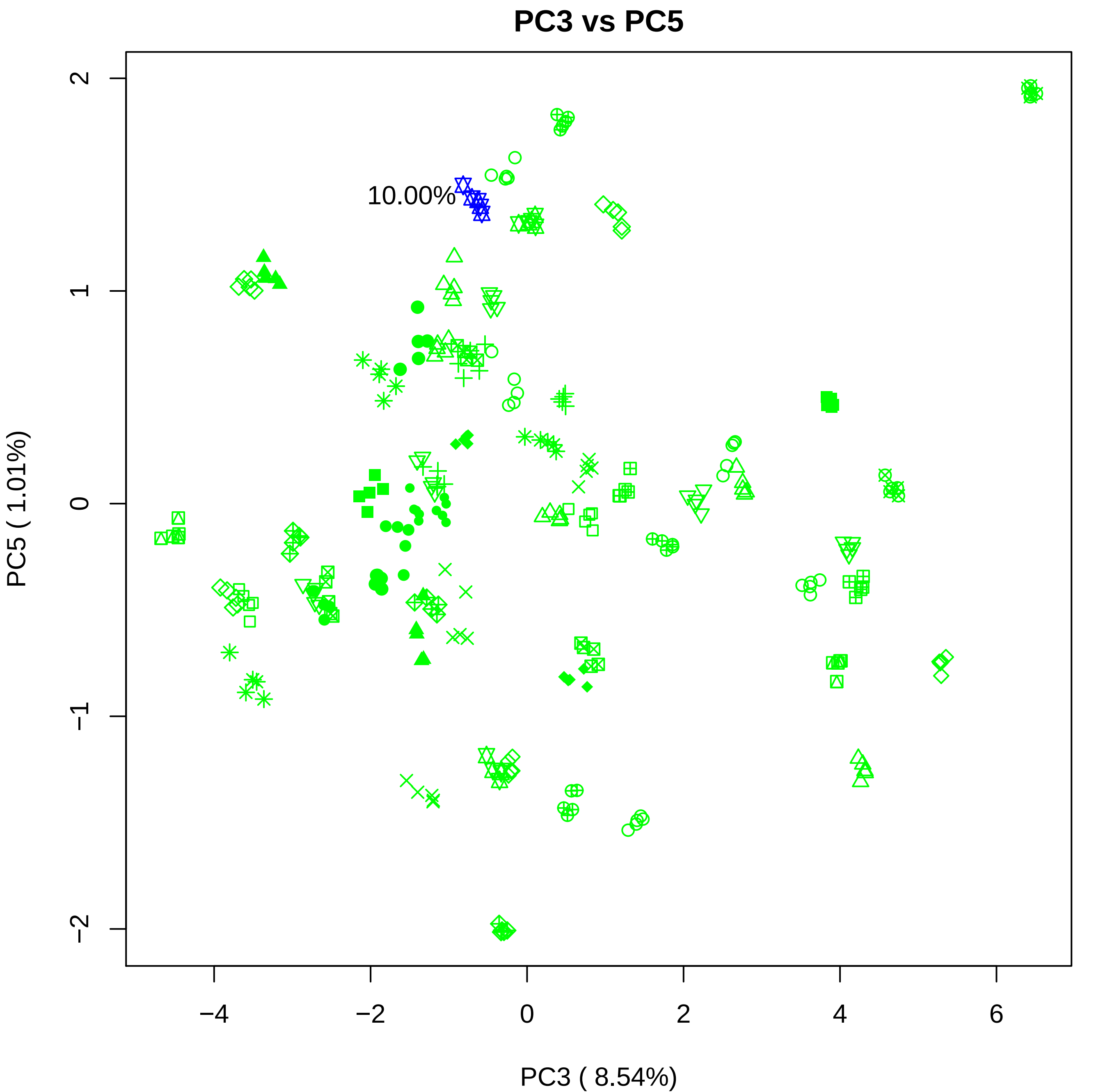

PC3 vs PC5
10.00%
PC5 ( 1.01%)
PC3 ( 8.54%)
2
1
0
−1
−2
−4
−2
0
2
4
6

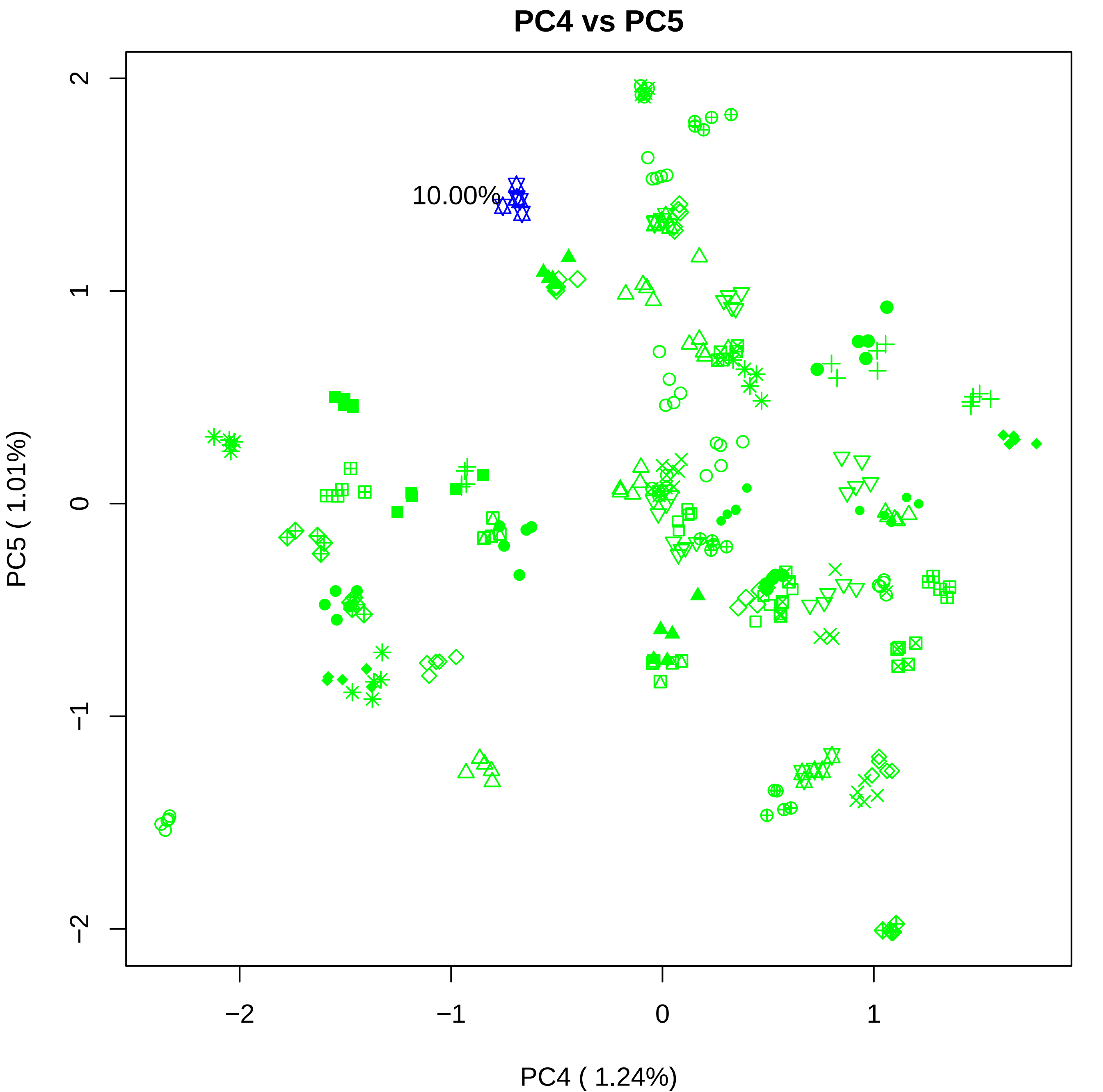
PC4 vs PC5
10.00%
PC5 ( 1.01%)
PC4 ( 1.24%)
2
1
0
−1
−2
−2
−1
0
1

### PCA of 15%  BMH(b)  vs PMH(b)

Scaled Matrix PCAed

7th order polynomial baseline correction performed

Raw spectra smoothed to 21 Neighbours, 2nd order PN, Savtisky Golay

Normalisation between 0 and 100

D2 spectra smoothed to 41 Neighbours, 2nd order PN, Savtisky Golay

Regions Analysed: 3601−2501 and 2001−700cm−1

Samples Analysed: 1 bContam's 0 BLH's 0 BMH's 0 OMH's 0 pContam's 0 PLH's 60 PMH's

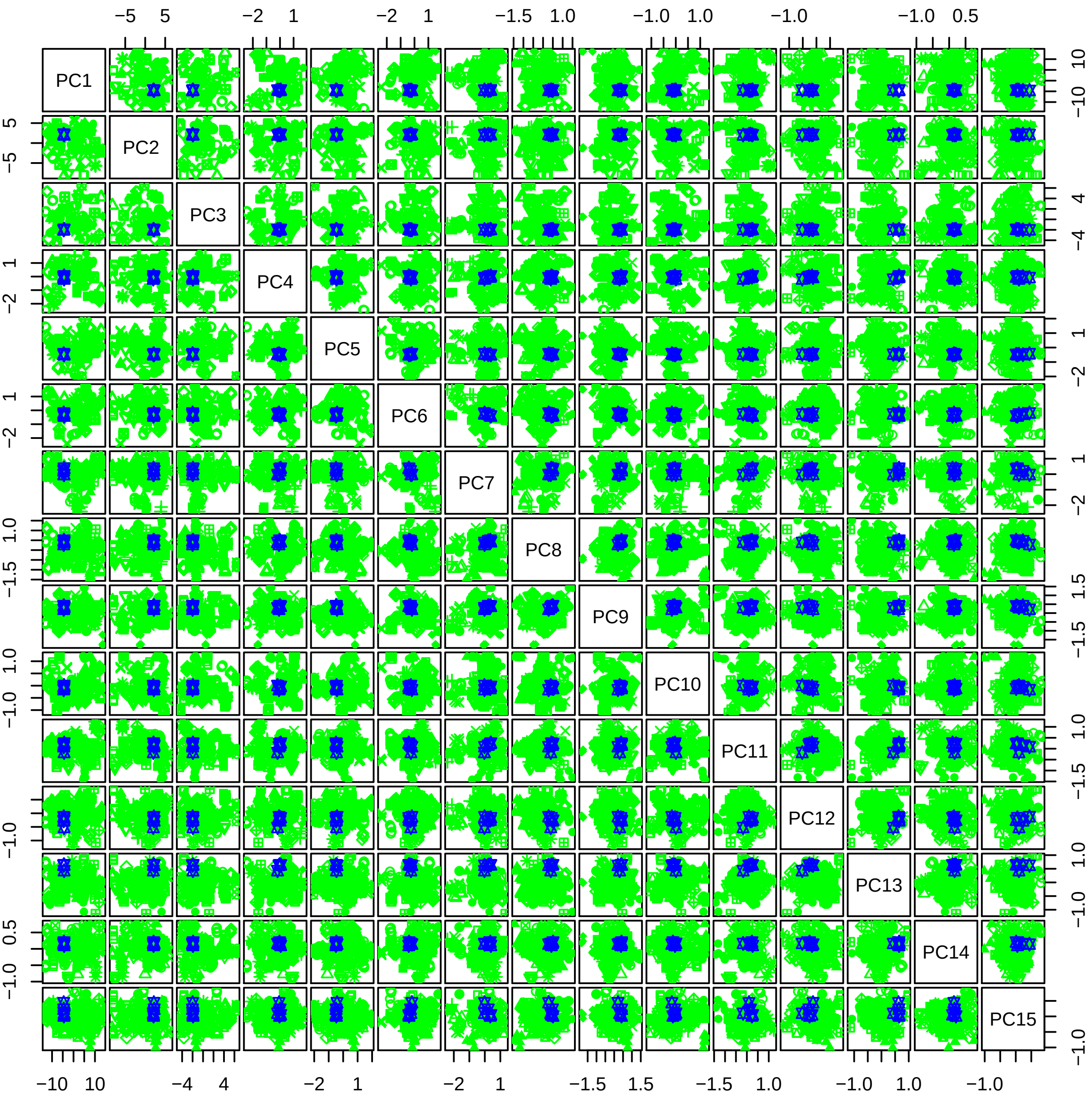
PC1
PC2
PC3
PC4
PC5
PC6
PC7
PC8
PC9
PC10
PC11
PC12
PC13
PC14
PC15

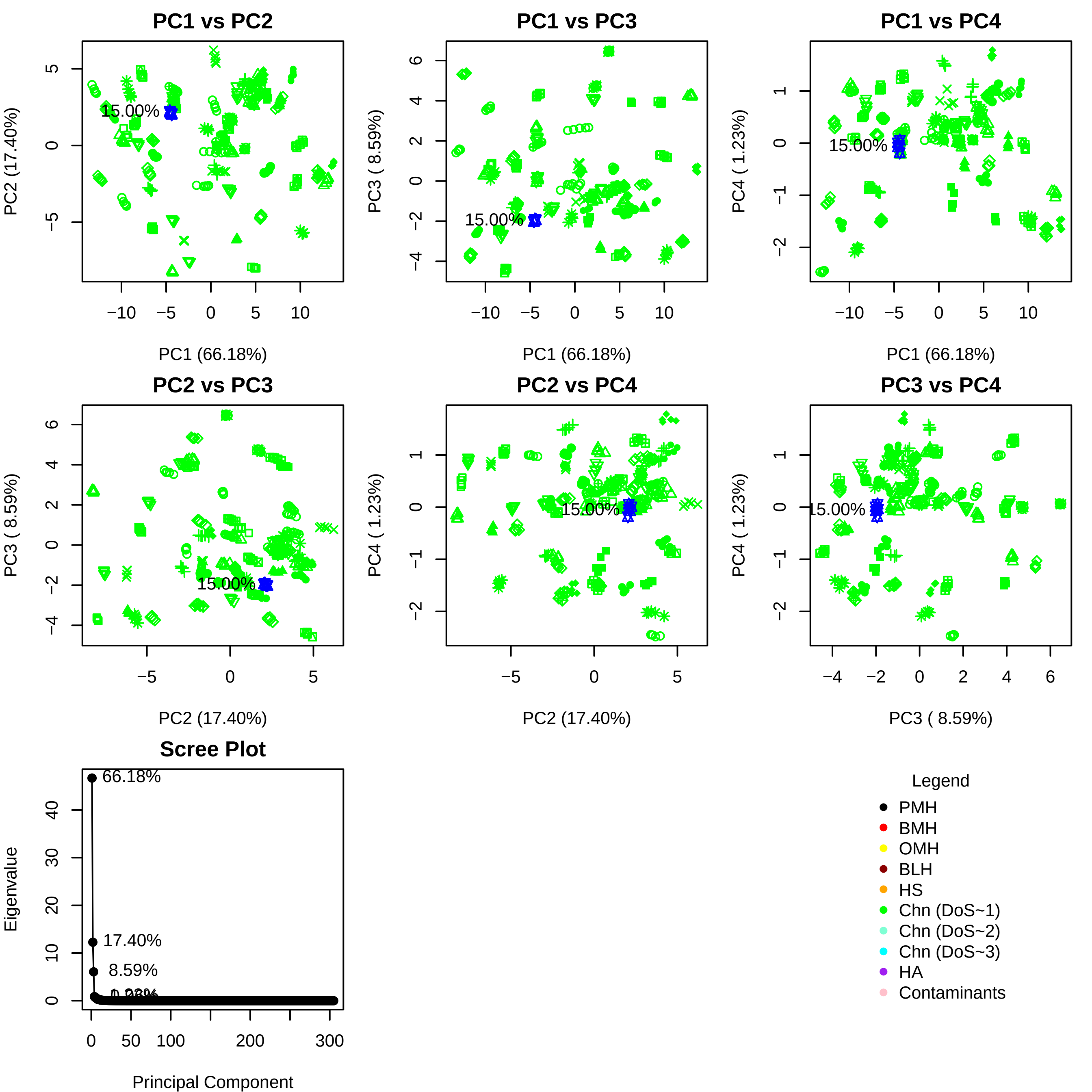

PC1 vs PC2
PC1 vs PC3
PC1 vs PC4
PC2 vs PC3
PC2 vs PC4
PC3 vs PC4
Scree Plot
PC1 (66.18%)
PC2 (17.40%)
PC3 ( 8.59%)
PC4 ( 1.23%)
15.00%
Eigenvalue
Principal Component
66.18%
17.40%
8.59%
Legend
PMH
BMH
OMH
BLH
HS
Chn (DoS~1)
Chn (DoS~2)
Chn (DoS~3)
HA
Contaminants

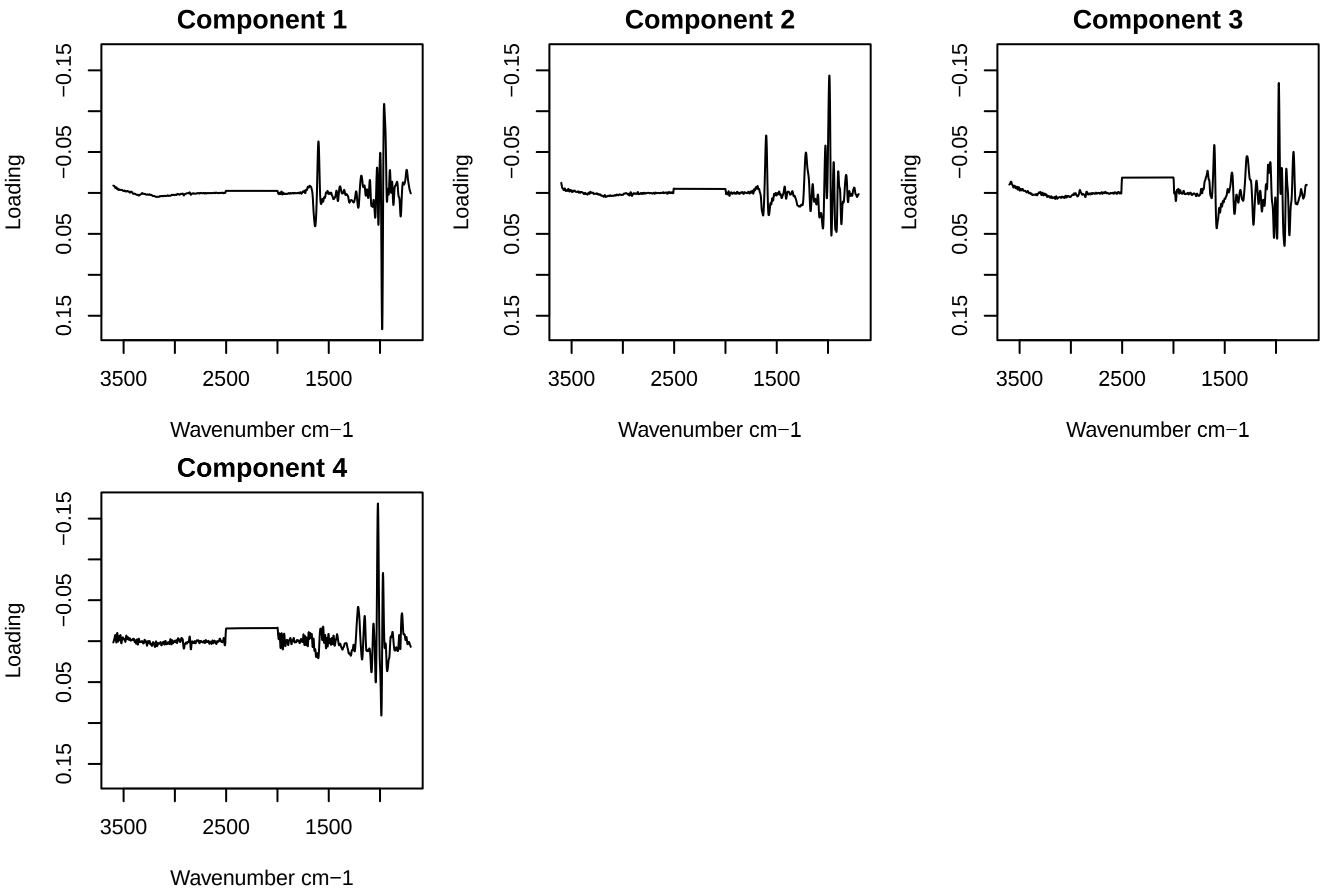
Component 1
Component 2
Component 3
Component 4
Loading
−0.15
−0.05
0.05
0.15
3500
2500
1500
Wavenumber cm−1

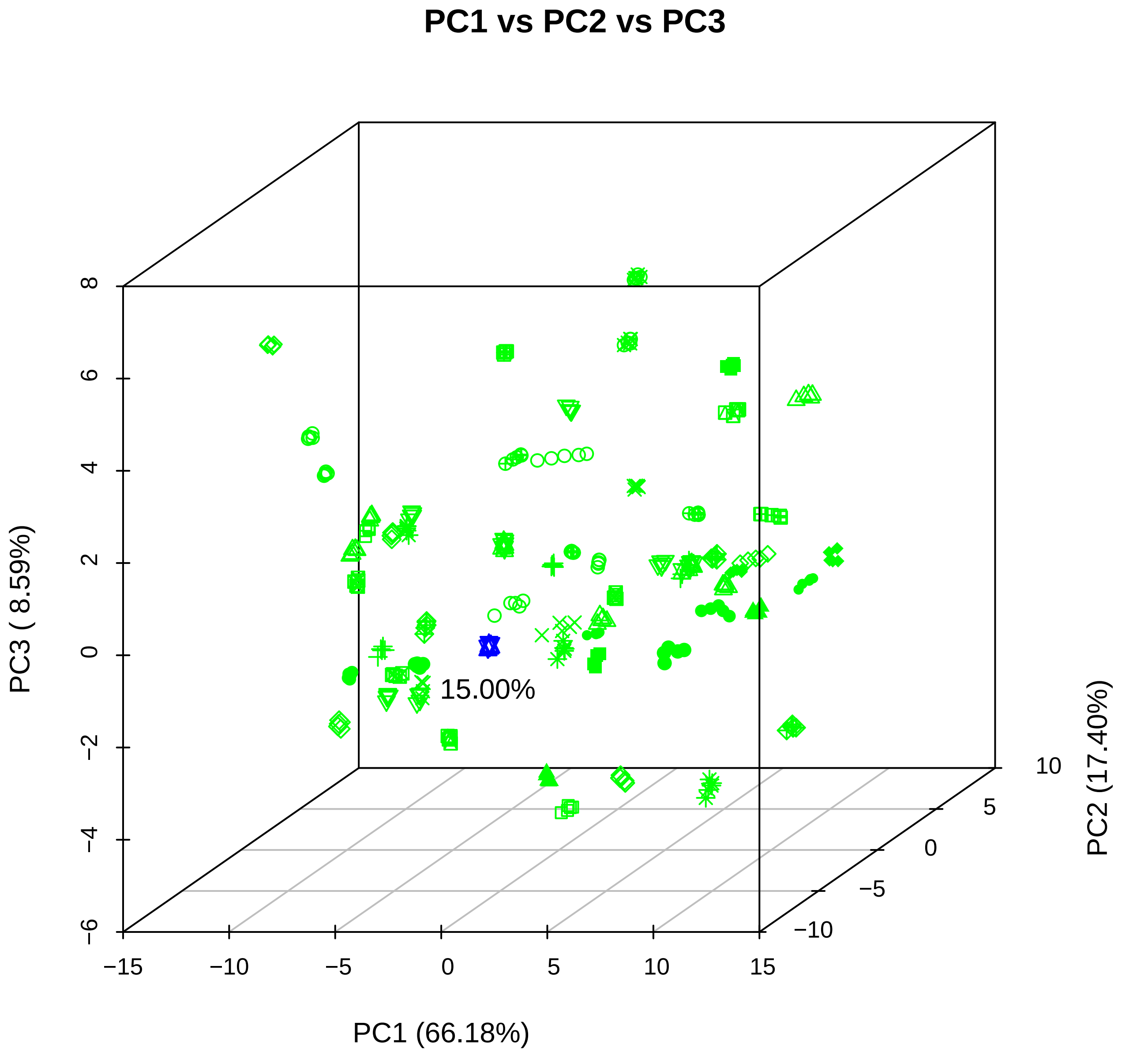
PC1 vs PC2 vs PC3
15.00%
PC1 (66.18%)
PC2 (17.40%)
PC3 ( 8.59%)
−15
−10
−5
0
5
10
15
−10
−5
0
5
10
−6
−4
−2
0
2
4
6
8

## PC1 vs PC2 vs PC4

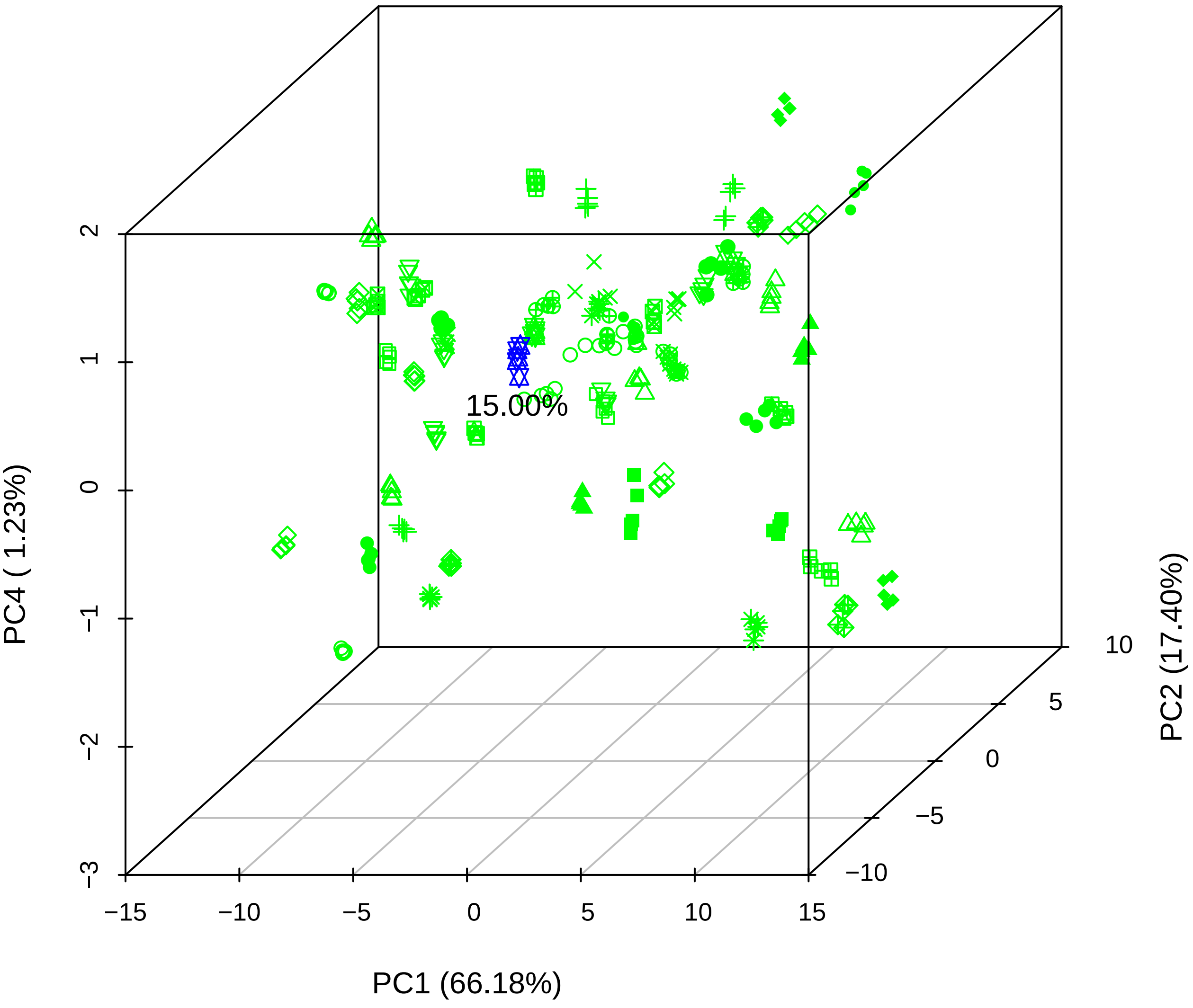

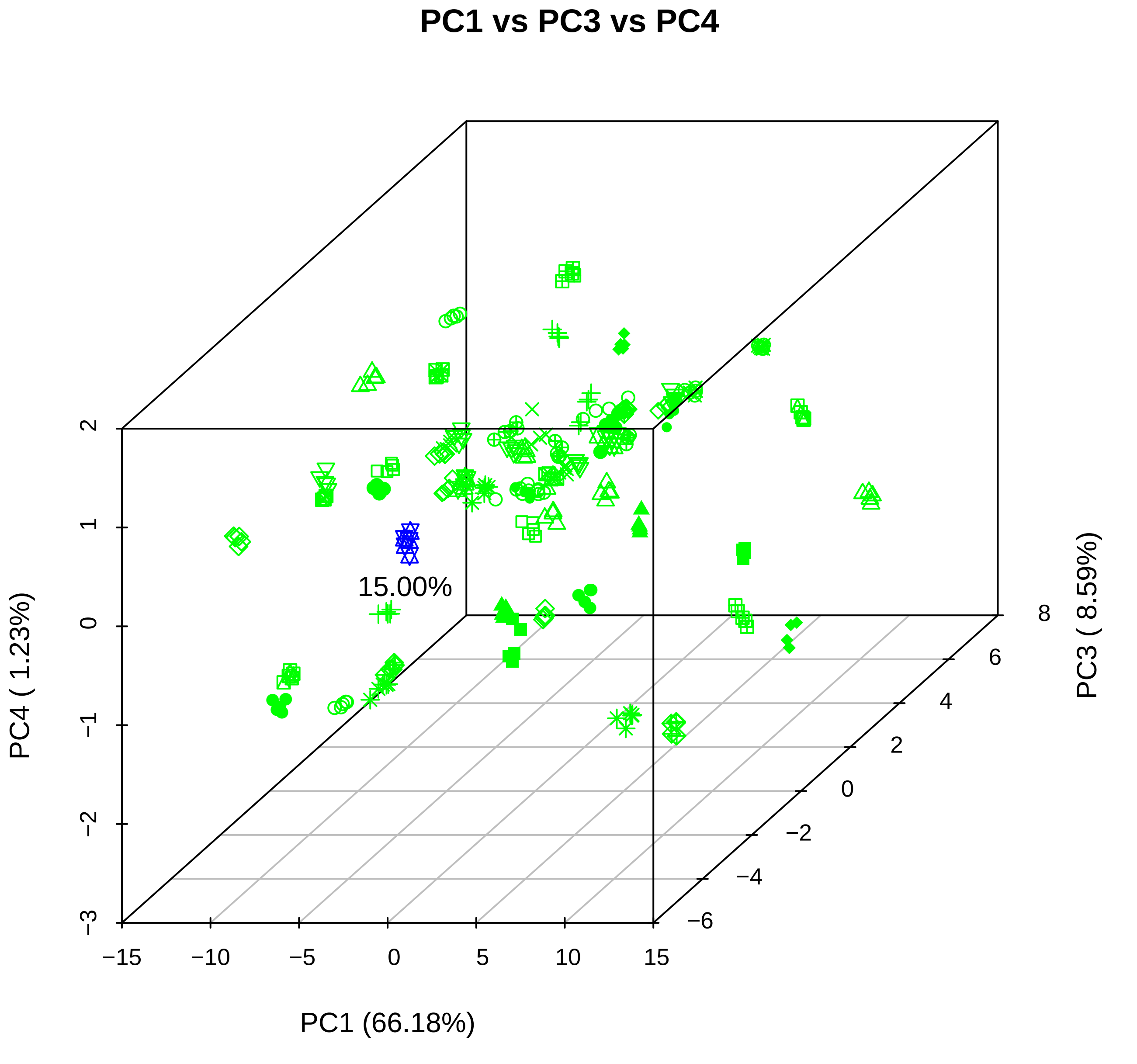
PC1 vs PC3 vs PC4
PC1 (66.18%)
PC4 ( 1.23%)
PC3 ( 8.59%)
15.00%
−15
−10
−5
0
5
10
15
−3
−2
−1
0
1
2
−6
−4
−2
0
2
4
6
8

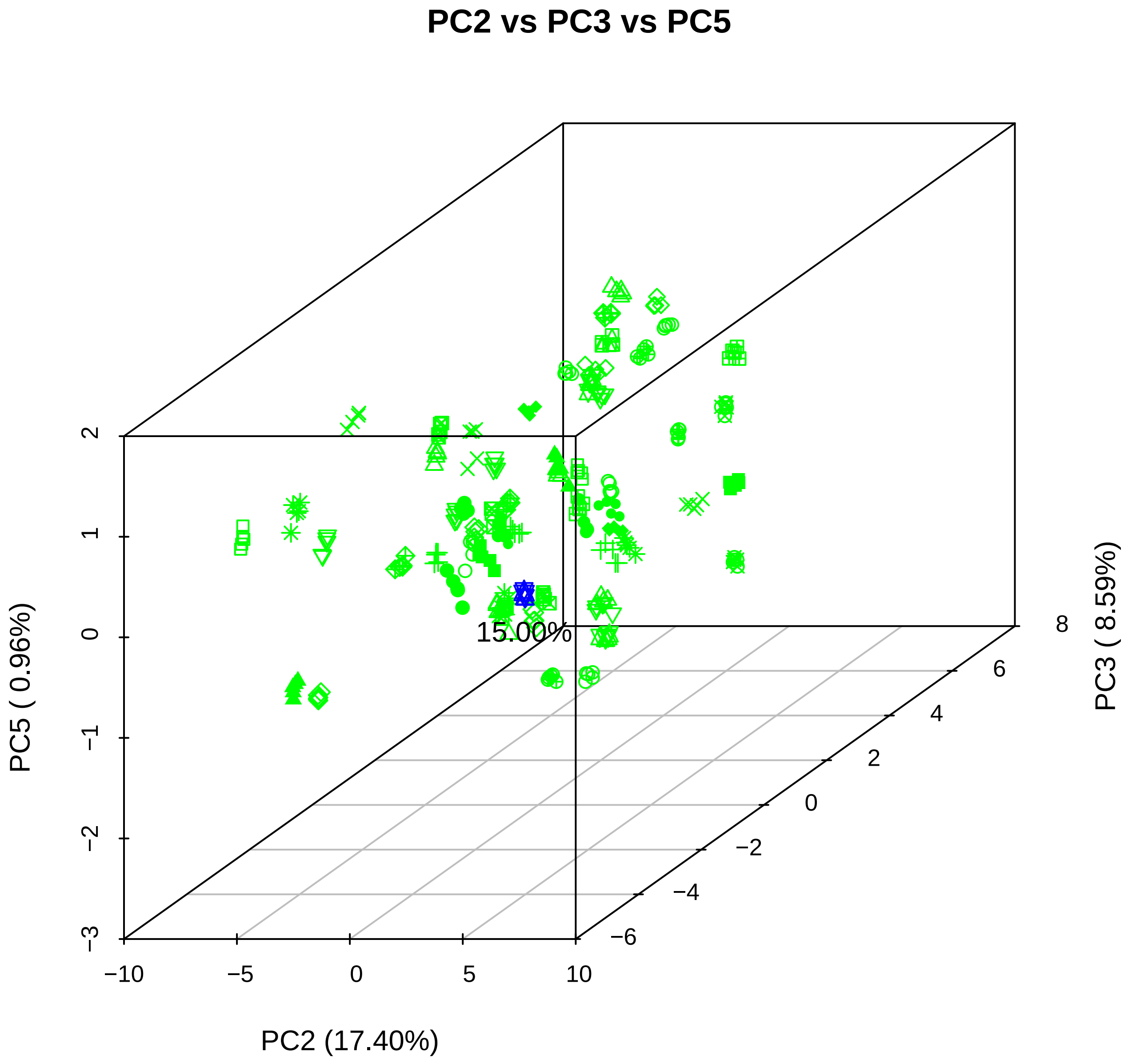
PC2 vs PC3 vs PC5
15.00%
PC2 (17.40%)
PC5 ( 0.96%)
PC3 ( 8.59%)
−10
−5
0
5
10
−3
−2
−1
0
1
2
−6
−4
−2
0
2
4
6
8

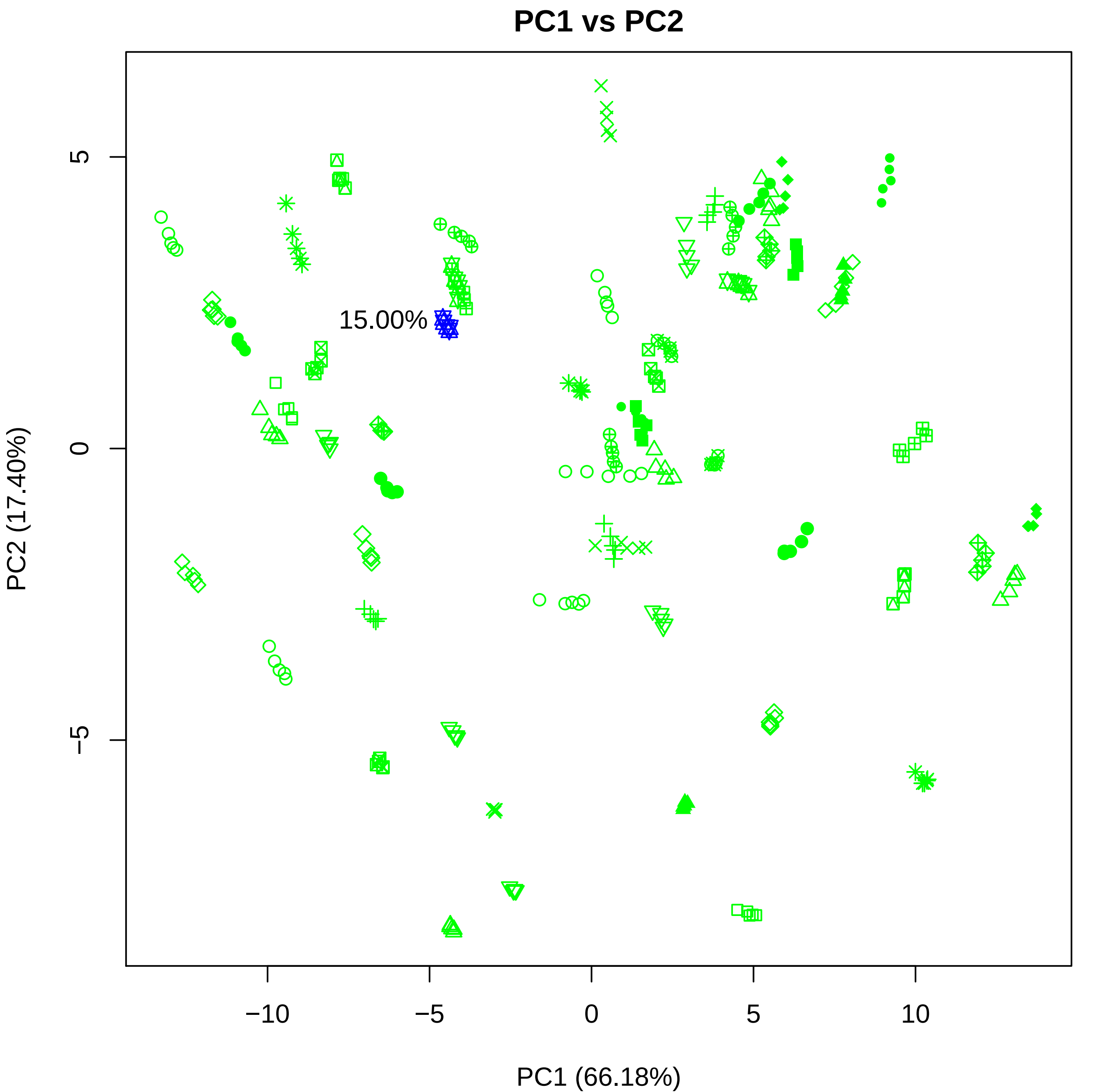

PC1 vs PC2
15.00%
PC2 (17.40%)
5
0
−5
−10
−5
0
5
10
PC1 (66.18%)

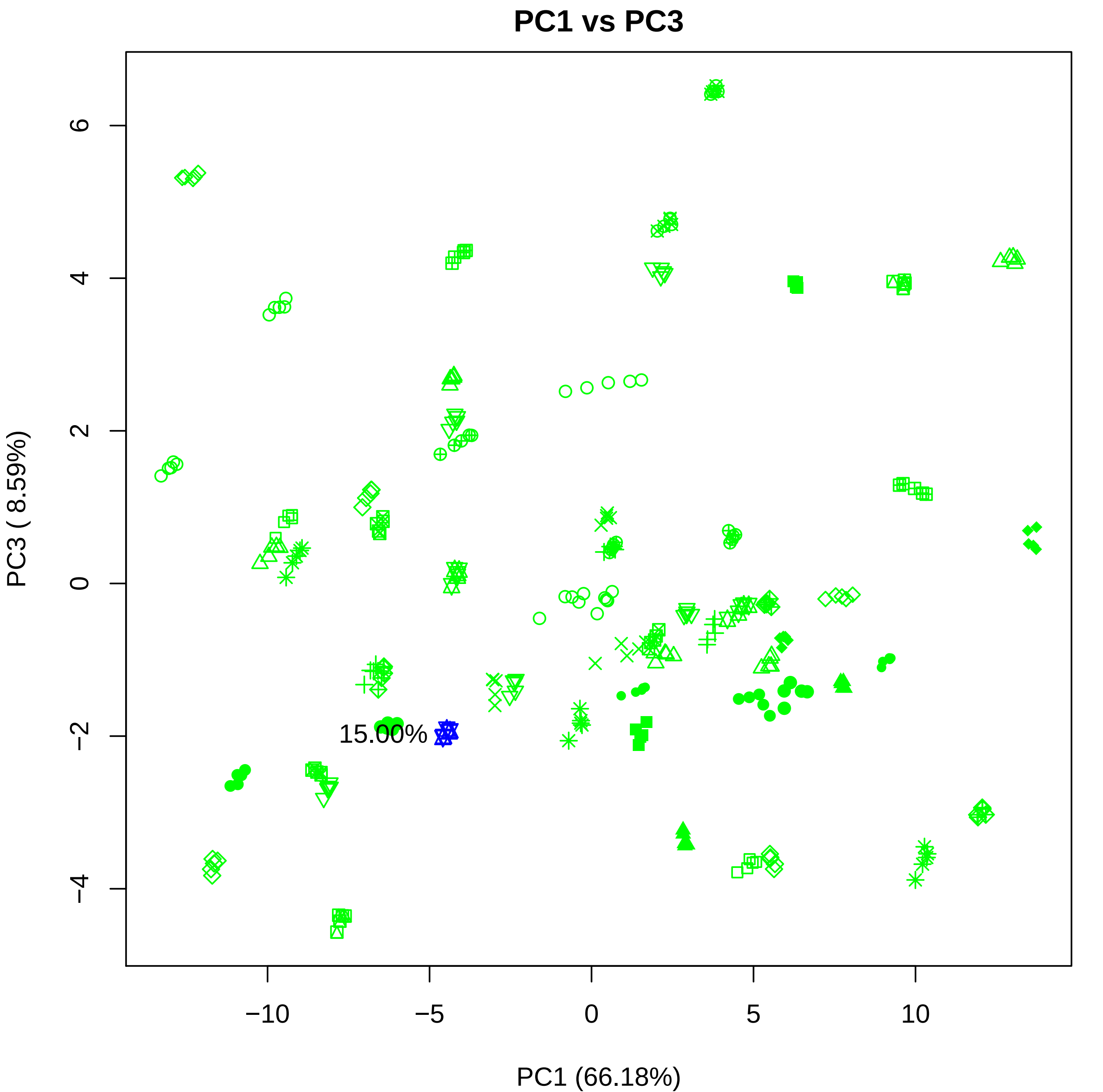

PC1 vs PC3
PC3 ( 8.59%)
PC1 (66.18%)
15.00%
−10
−5
0
5
10
−4
−2
0
2
4
6

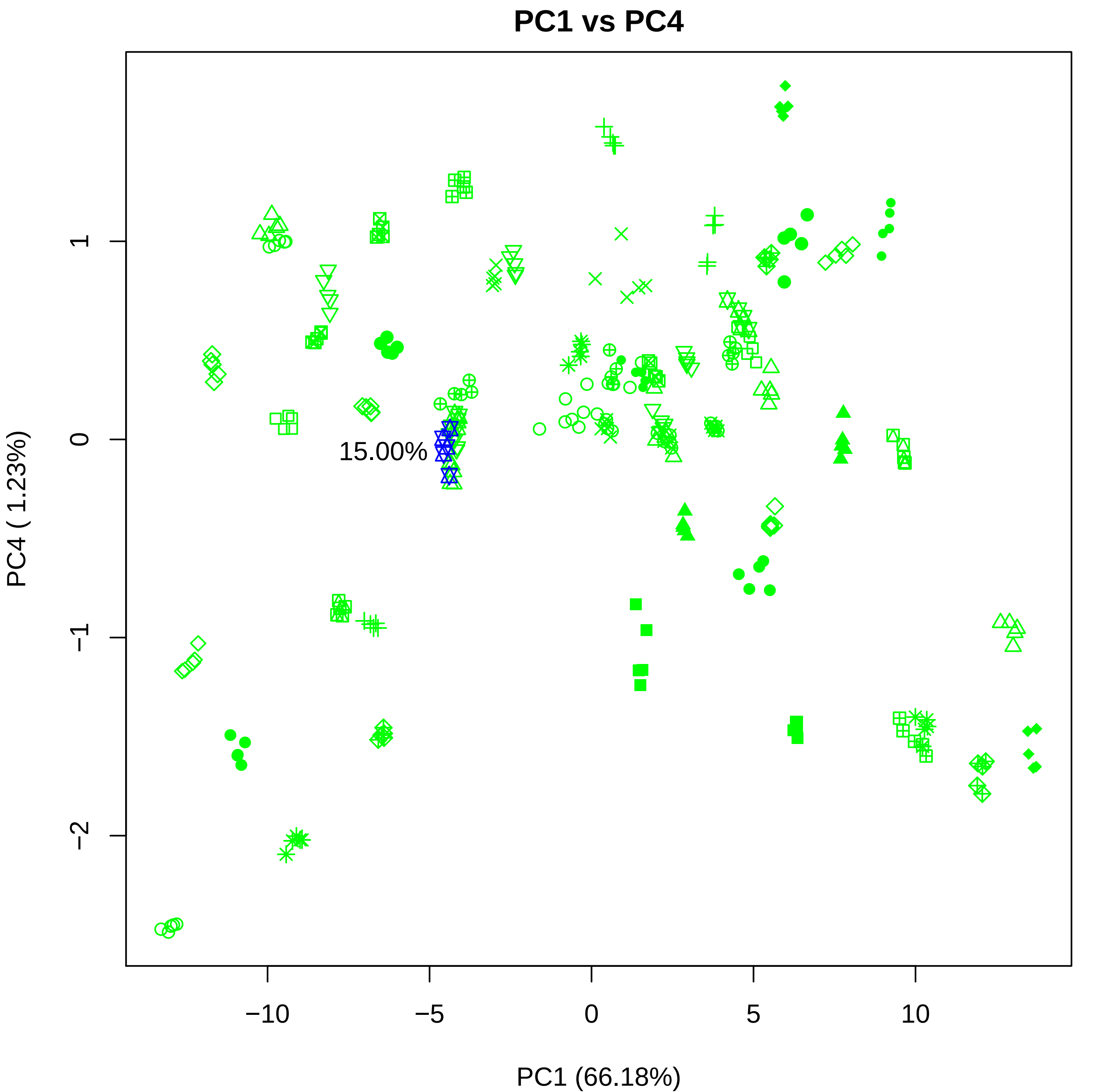

PC1 vs PC4
15.00%
PC1 (66.18%)
PC4 ( 1.23%)
−10
−5
0
5
10
−2
−1
0
1

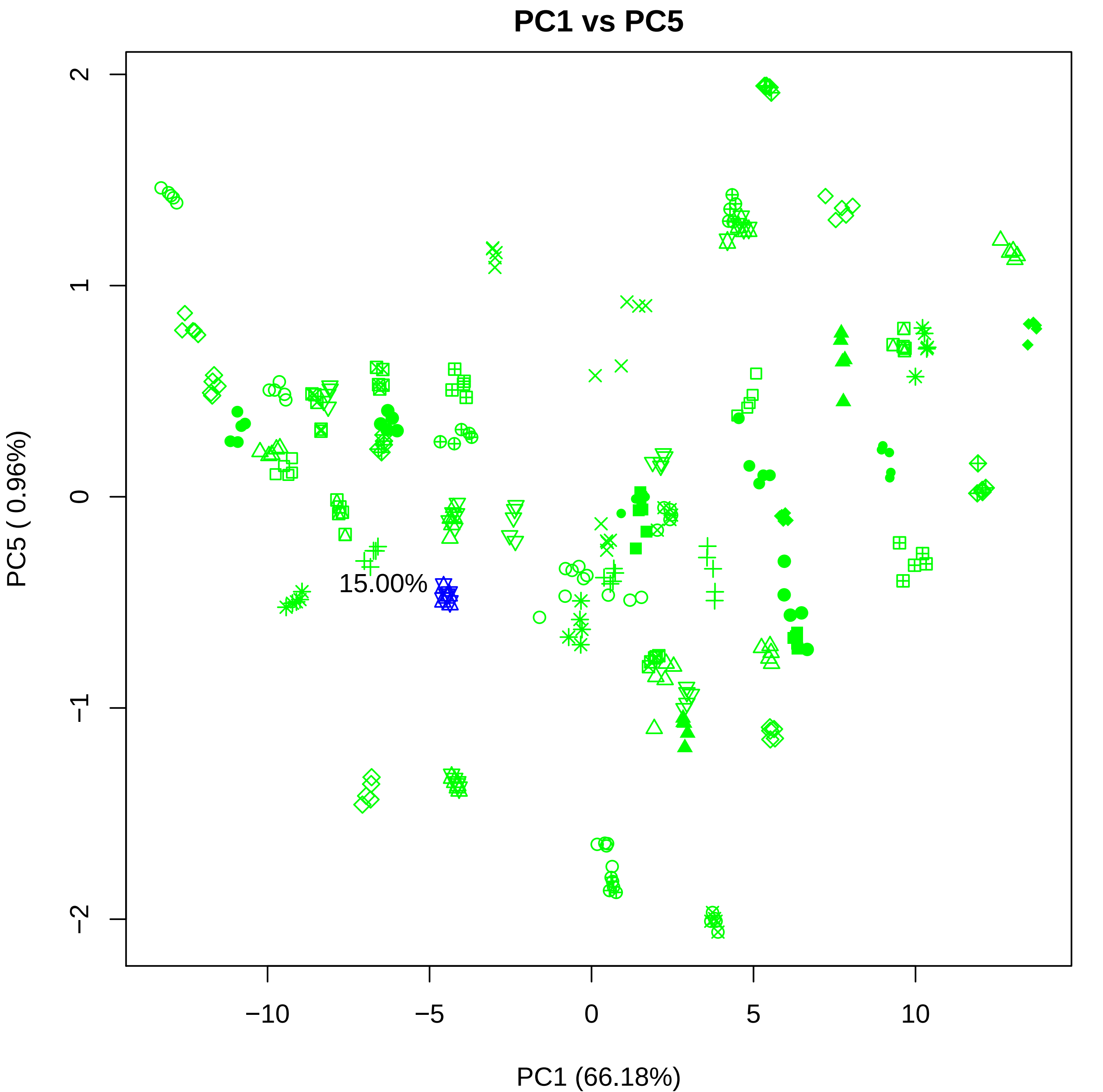

PC1 vs PC5
2
1
0
−1
−2
PC5 ( 0.96%)
15.00%
−10
−5
0
5
10
PC1 (66.18%)

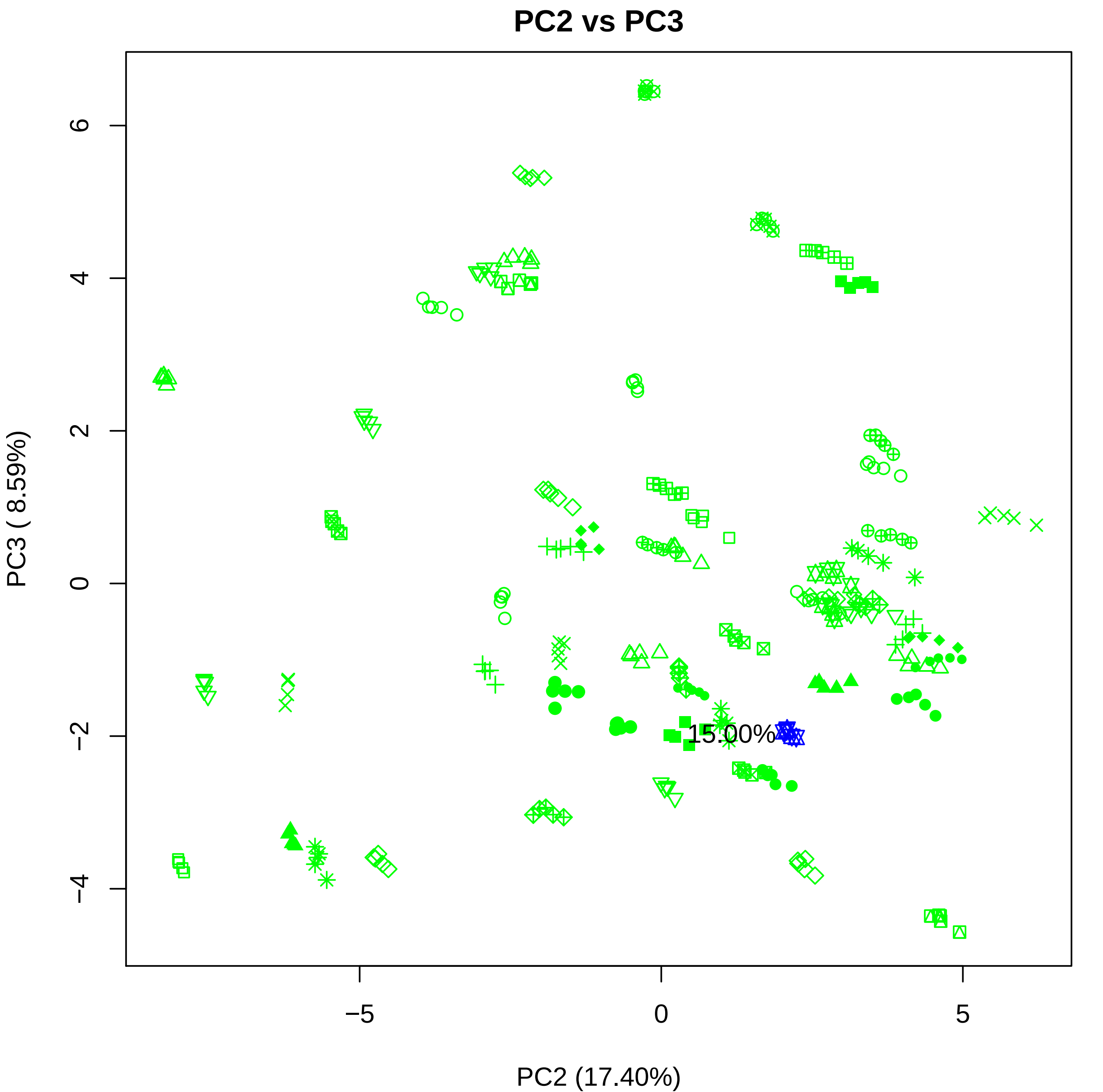

PC2 vs PC3
PC3 ( 8.59%)
PC2 (17.40%)
6
4
2
0
−2
−4
−5
0
5
15.00%

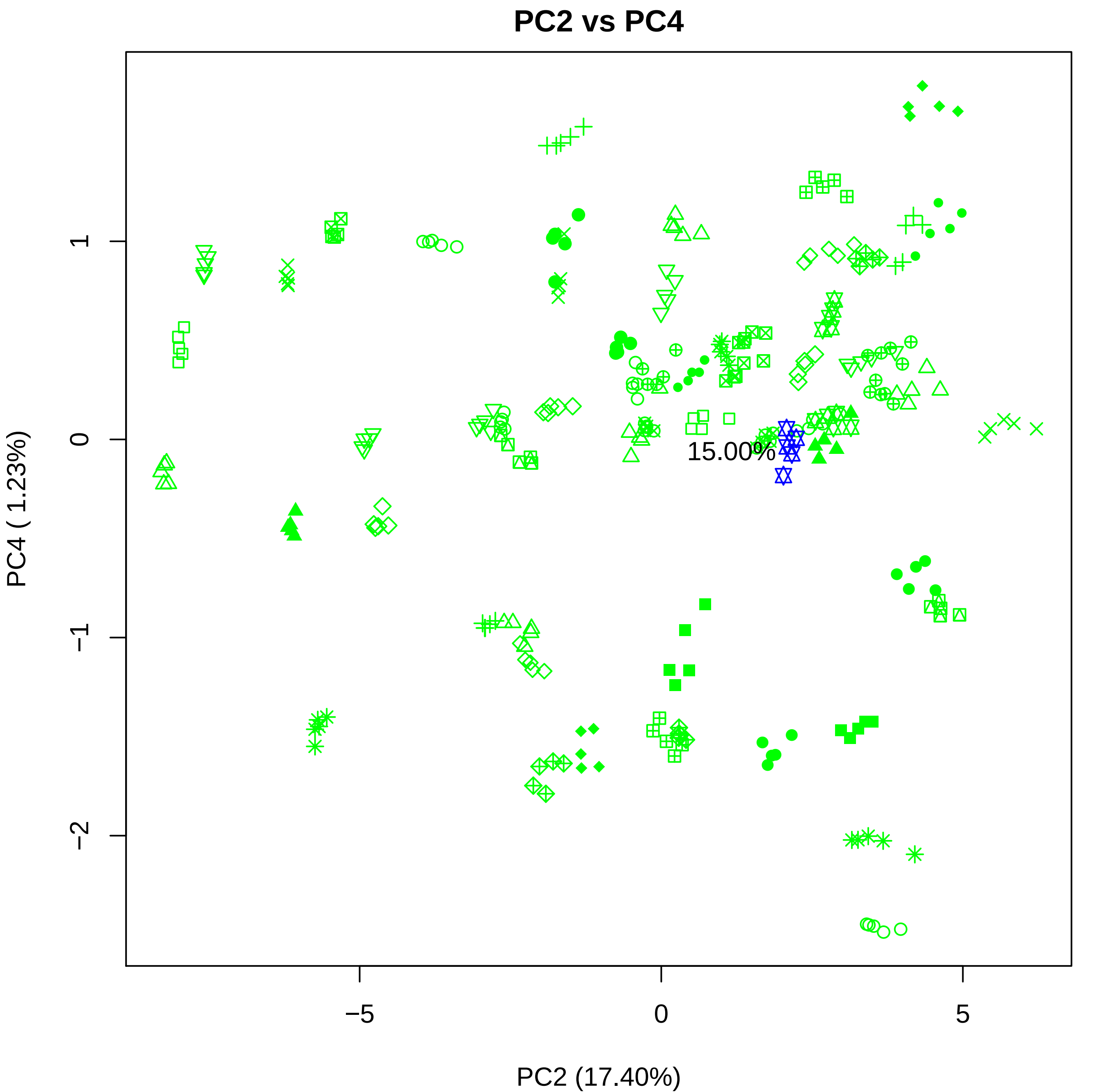

PC2 vs PC4
15.00%
PC4 ( 1.23%)
PC2 (17.40%)
−5
0
5
−2
−1
0
1

## PC2 vs PC5

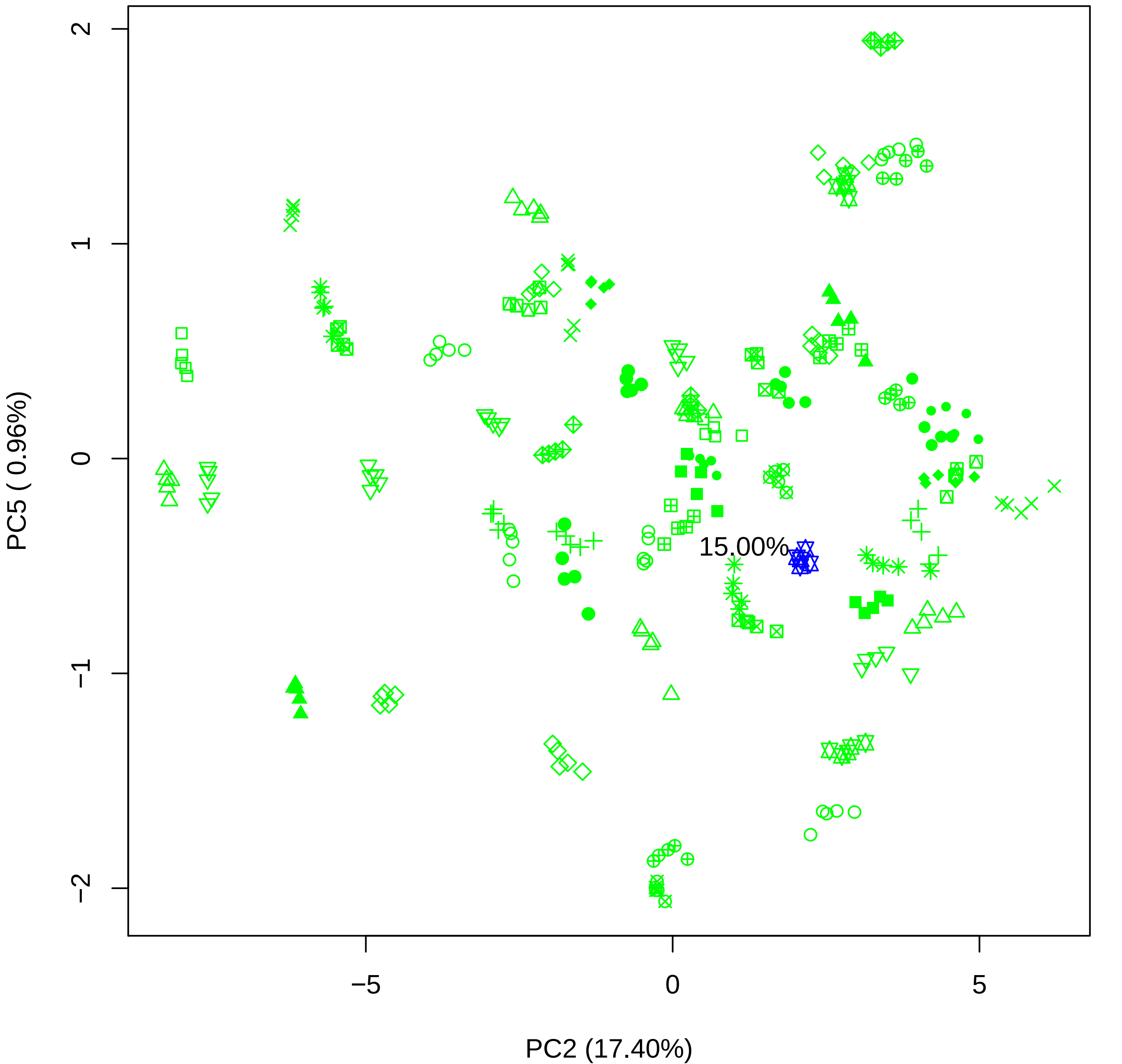

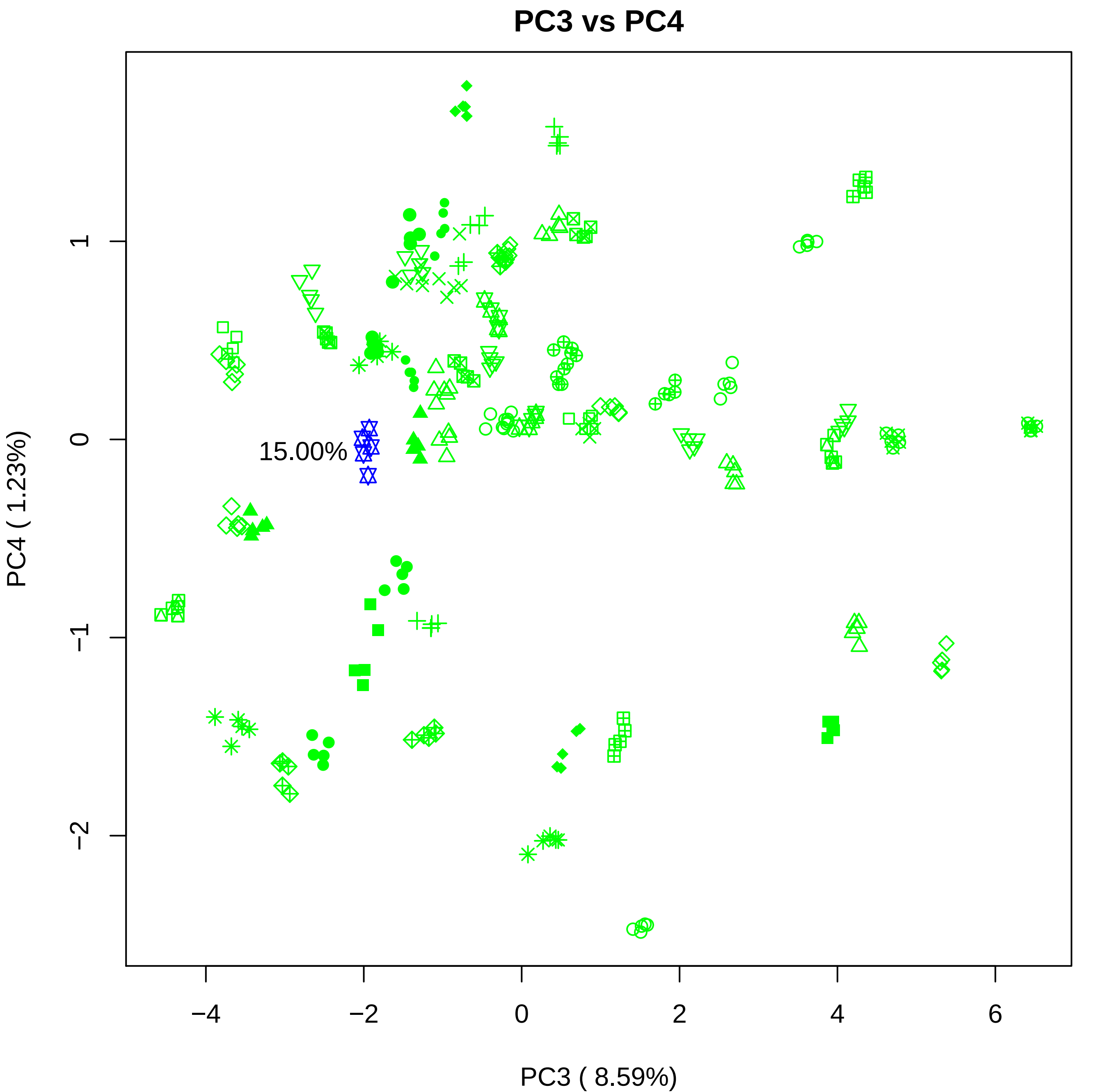
PC3 vs PC4
15.00%
PC4 ( 1.23%)
PC3 ( 8.59%)
1
0
−1
−2
−4
−2
0
2
4
6

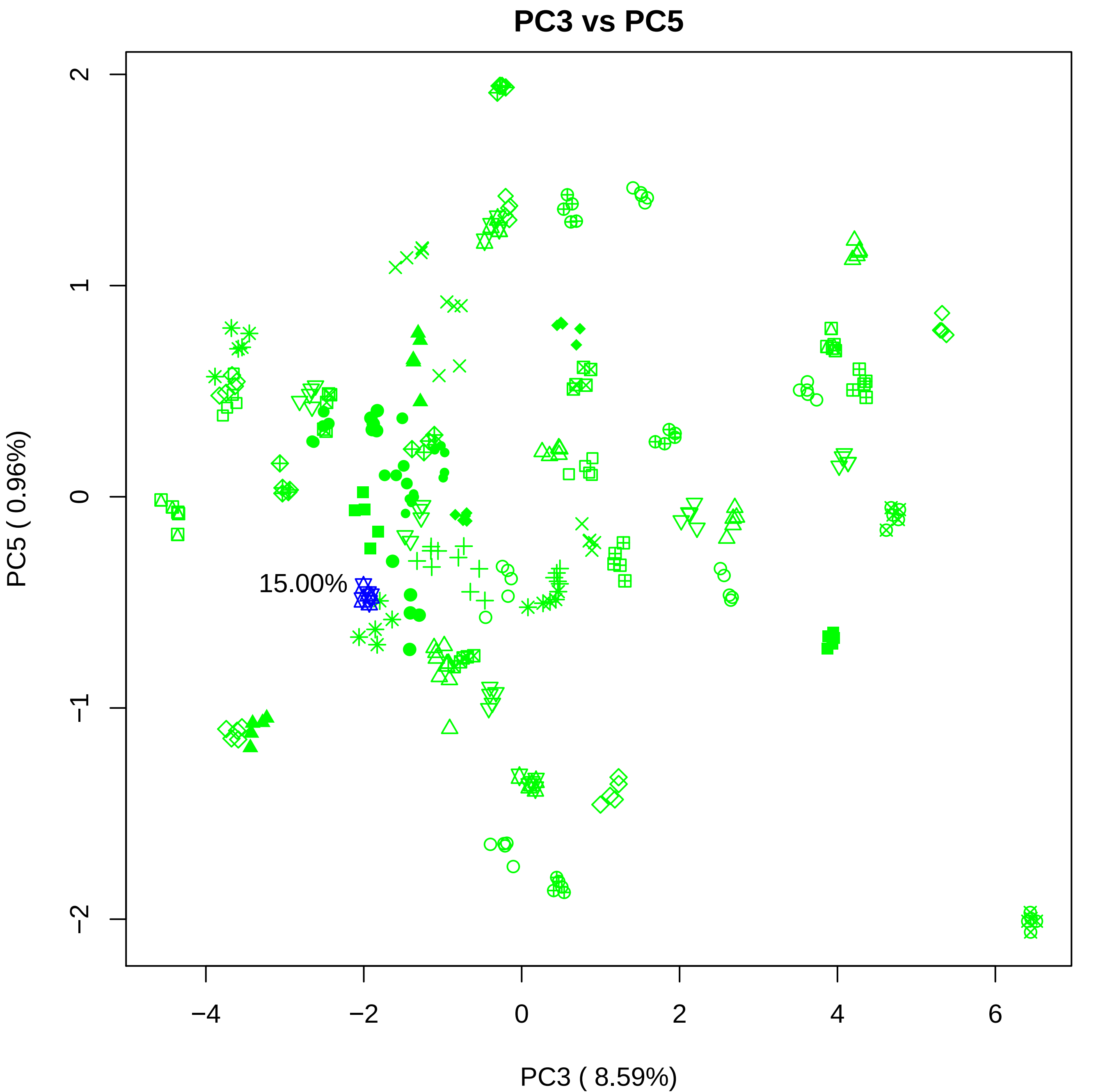

PC3 vs PC5
15.00%
PC5 ( 0.96%)
PC3 ( 8.59%)
2
1
0
−1
−2
−4
−2
0
2
4
6

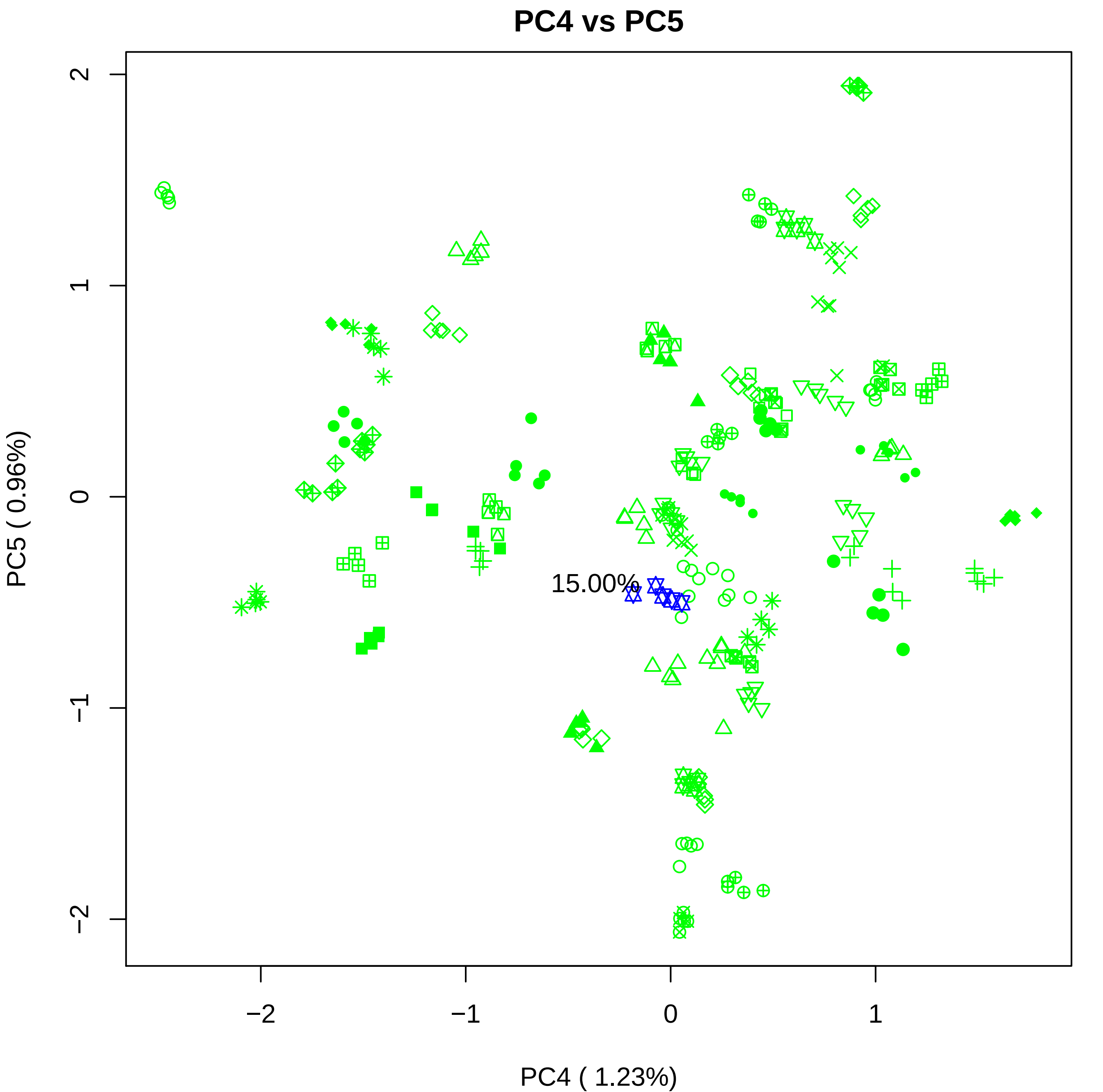
PC4 vs PC5
PC5 ( 0.96%)
PC4 ( 1.23%)
15.00%
2
1
0
−1
−2
−2
−1
0
1

### PCA of 20%  BMH(b) vs PMH(b)

Scaled Matrix PCAed

7th order polynomial baseline correction performed

Raw spectra smoothed to 21 Neighbours, 2nd order PN, Savtisky Golay

Normalisation between 0 and 100

D2 spectra smoothed to 41 Neighbours, 2nd order PN, Savtisky Golay

Regions Analysed: 3601−2501 and 2001−700cm−1

Samples Analysed: 1 bContam's 0 BLH's 0 BMH's 0 OMH's 0 pContam's 0 PLH's 60 PMH's

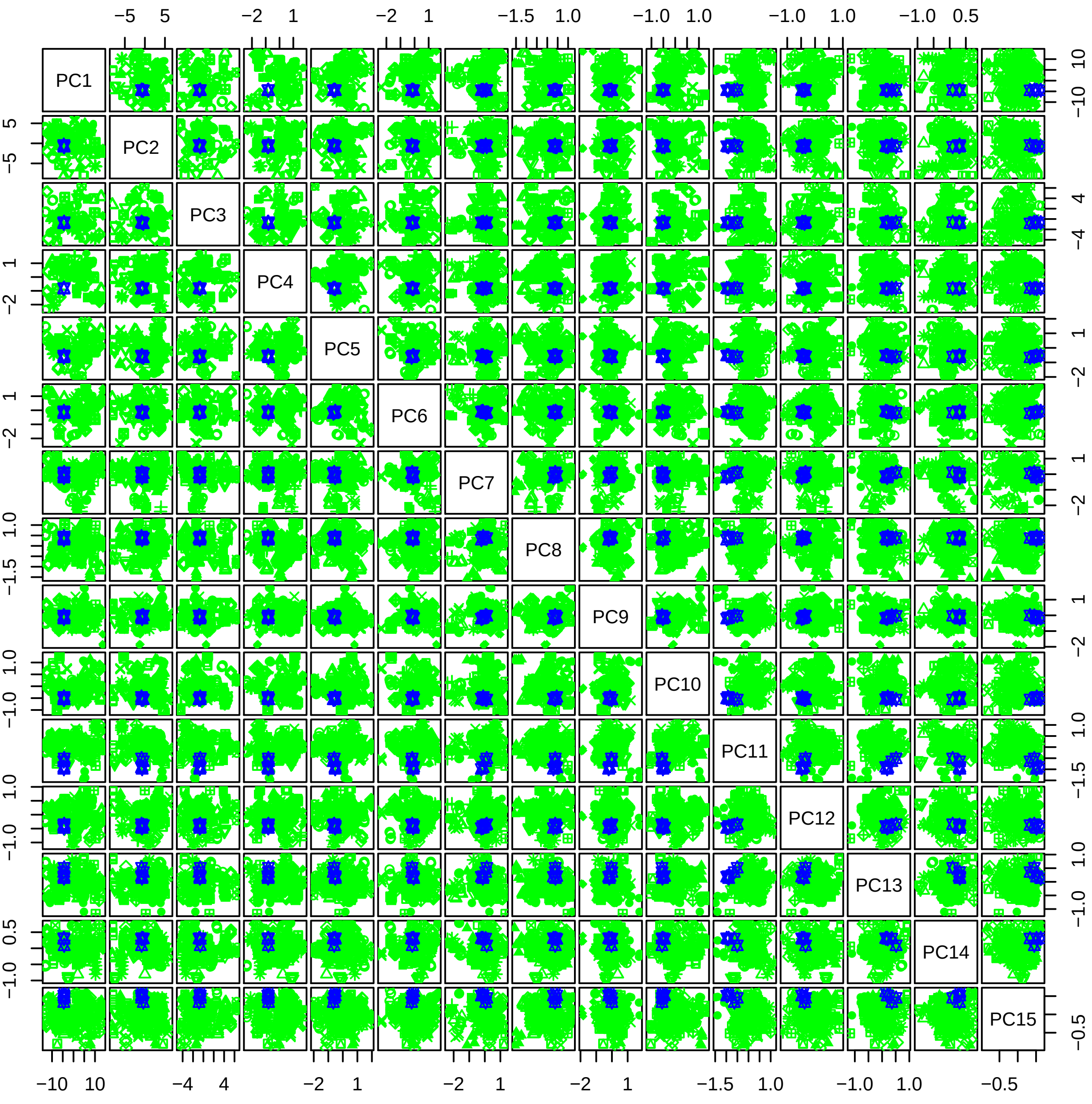
PC1
PC2
PC3
PC4
PC5
PC6
PC7
PC8
PC9
PC10
PC11
PC12
PC13
PC14
PC15
−5 5
−2 1
−2 1
−1.5 1.0
−1.0 1.0
−1.0 1.0
−1.0 0.5
10 −10
4 −4
1 −2
1 −2
1.0 −1.5
1.0 −1.0
−0.5
5 −5
1 −2
1 −2
1.0 −1.5
1.0 −1.0
0.5 −1.0
−10 10
−4 4
−2 1
−2 1
−2 1
−1.5 1.0
−1.0 1.0
−0.5

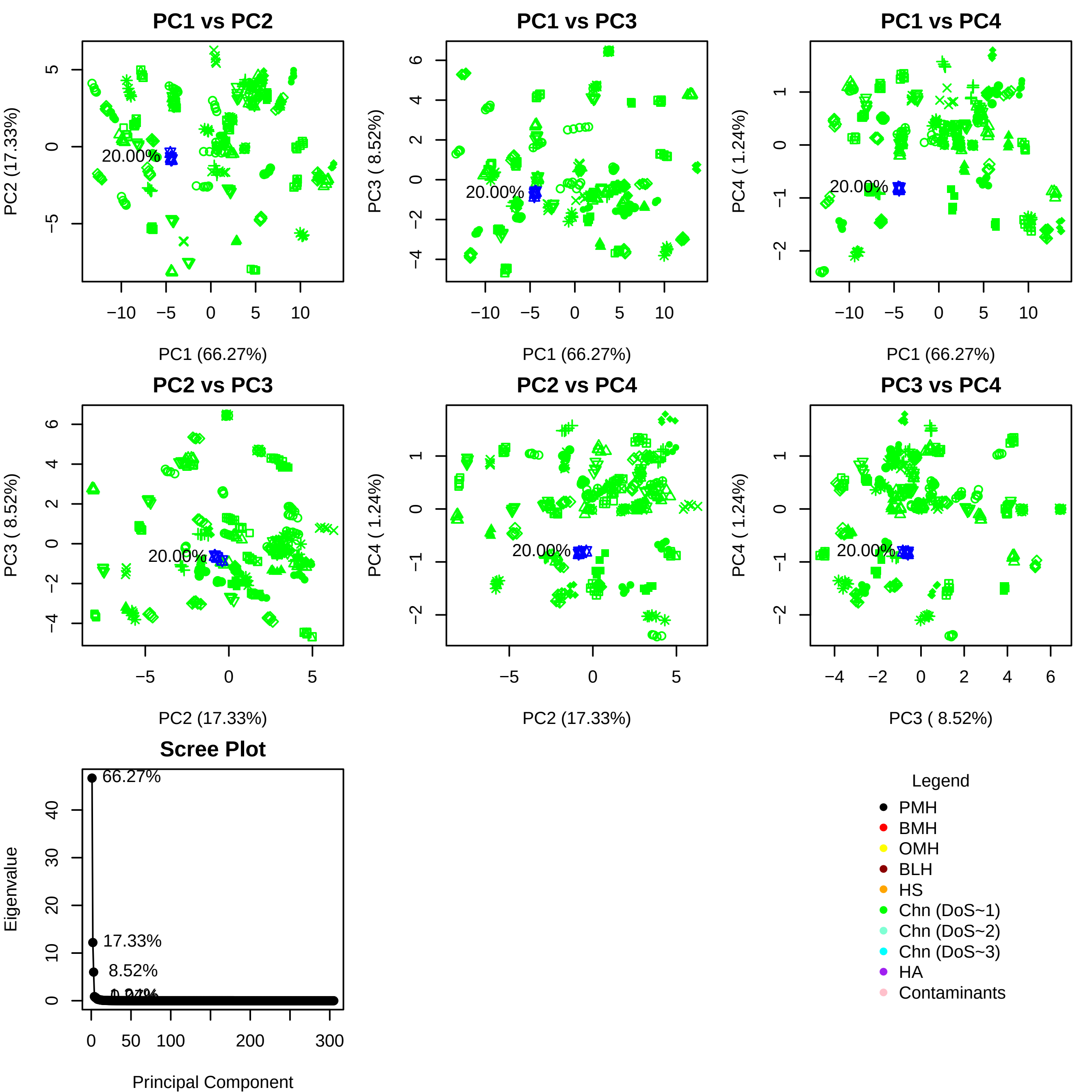
PC1 vs PC2
PC1 vs PC3
PC1 vs PC4
PC2 vs PC3
PC2 vs PC4
PC3 vs PC4
Scree Plot
PC1 (66.27%)
PC2 (17.33%)
PC3 ( 8.52%)
PC4 ( 1.24%)
20.00%
Eigenvalue
Principal Component
66.27%
17.33%
8.52%
Legend
PMH
BMH
OMH
BLH
HS
Chn (DoS~1)
Chn (DoS~2)
Chn (DoS~3)
HA
Contaminants

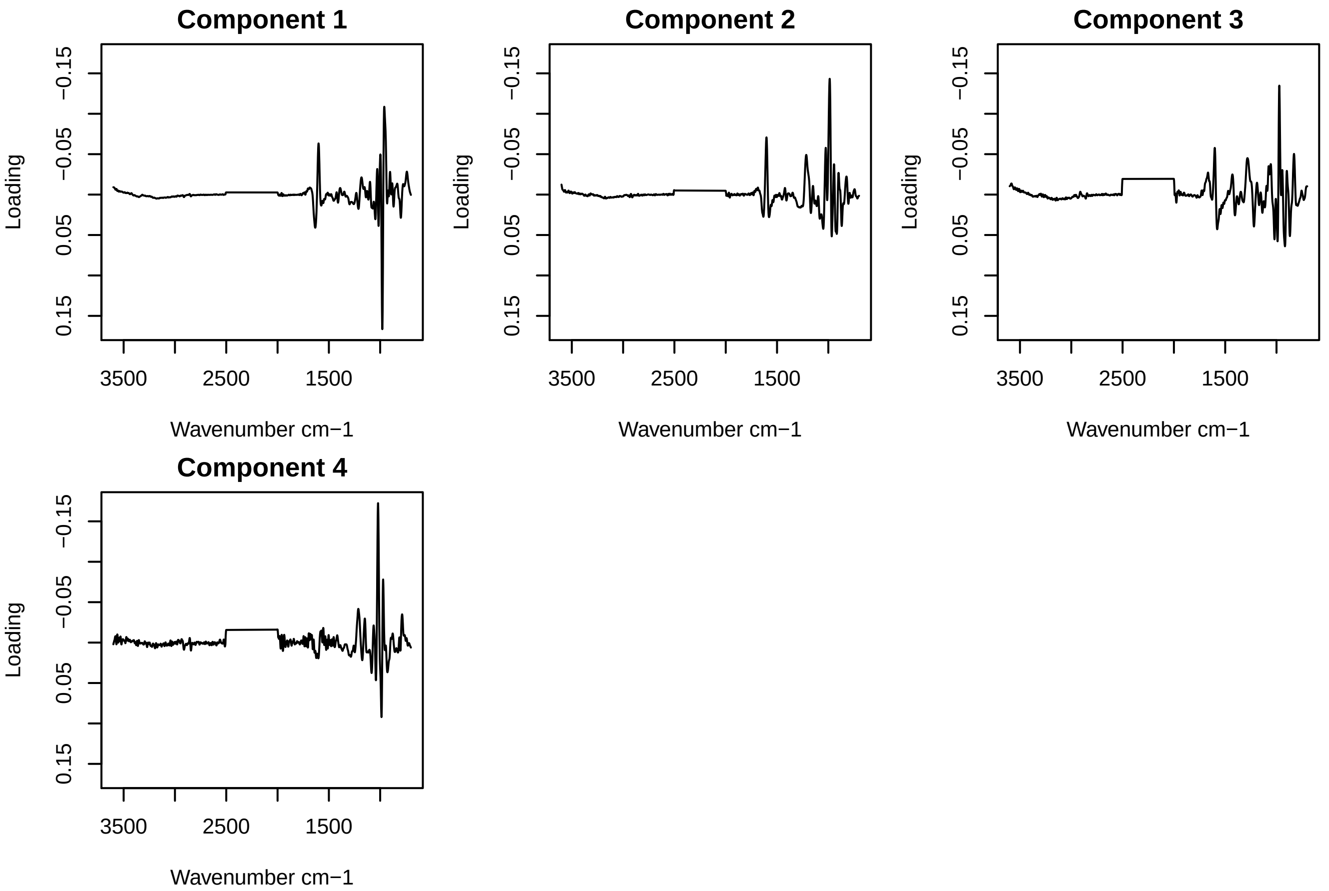

Component 1
Component 2
Component 3
Component 4
Loading
−0.15
−0.05
0.05
0.15
3500
2500
1500
Wavenumber cm−1

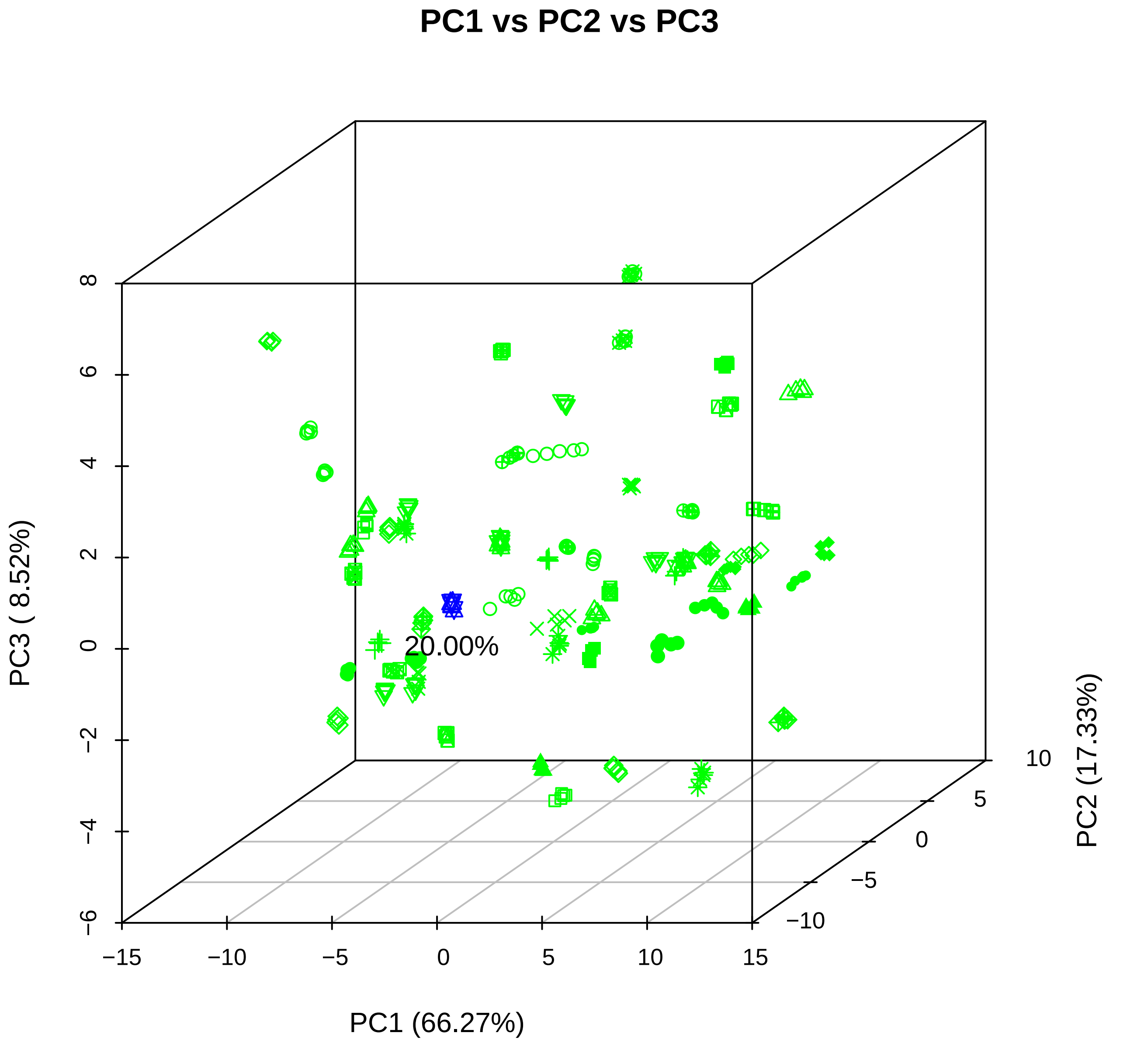
PC1 vs PC2 vs PC3
PC1 (66.27%)
PC2 (17.33%)
PC3 ( 8.52%)
20.00%
−15
−10
−5
0
5
10
15
−10
−5
0
5
10
−6
−4
−2
0
2
4
6
8

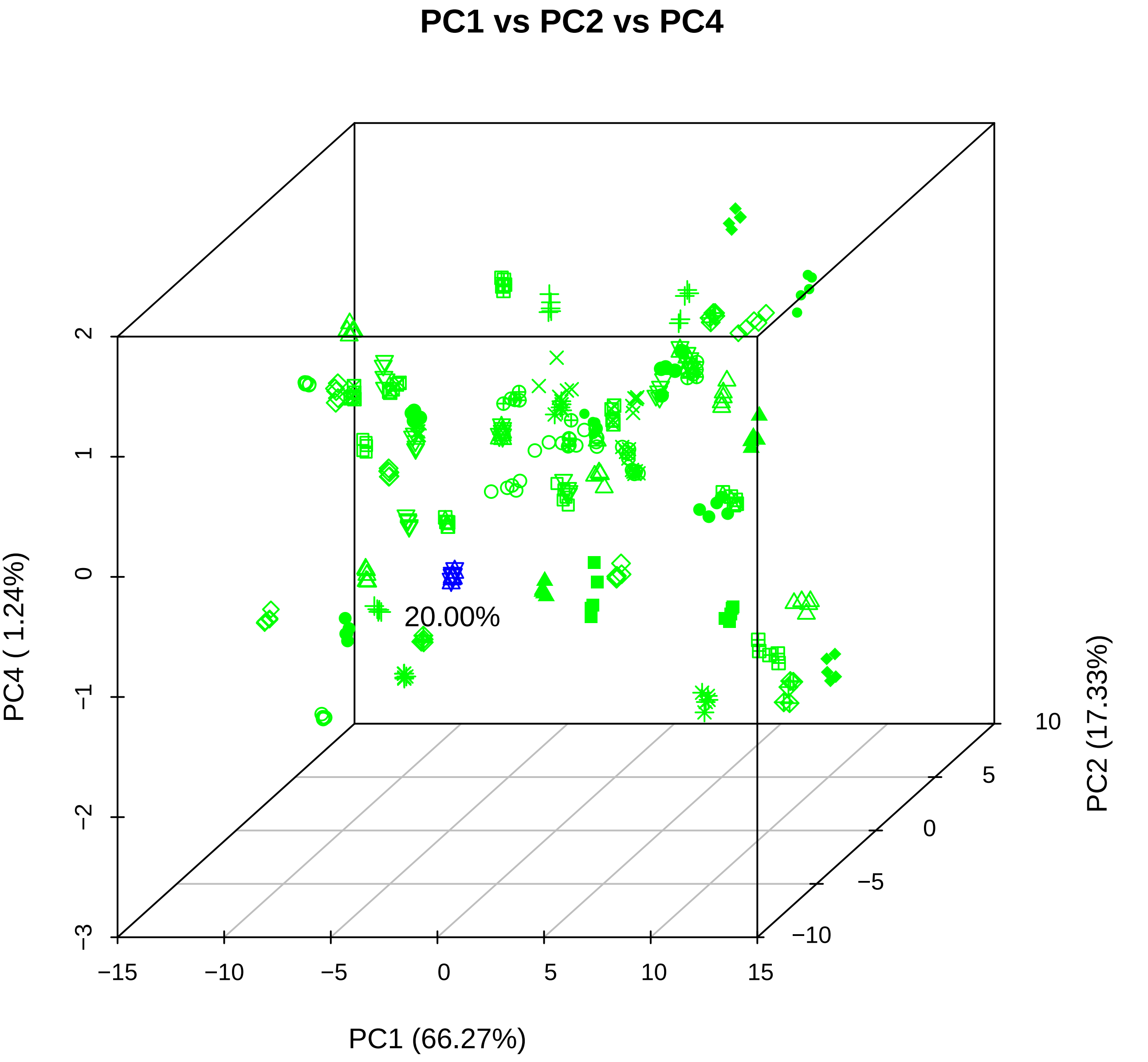

PC1 vs PC2 vs PC4
20.00%
PC1 (66.27%)
PC2 (17.33%)
PC4 ( 1.24%)
−15
−10
−5
0
5
10
15
−3
−2
−1
0
1
2
−10
−5
0
5
10

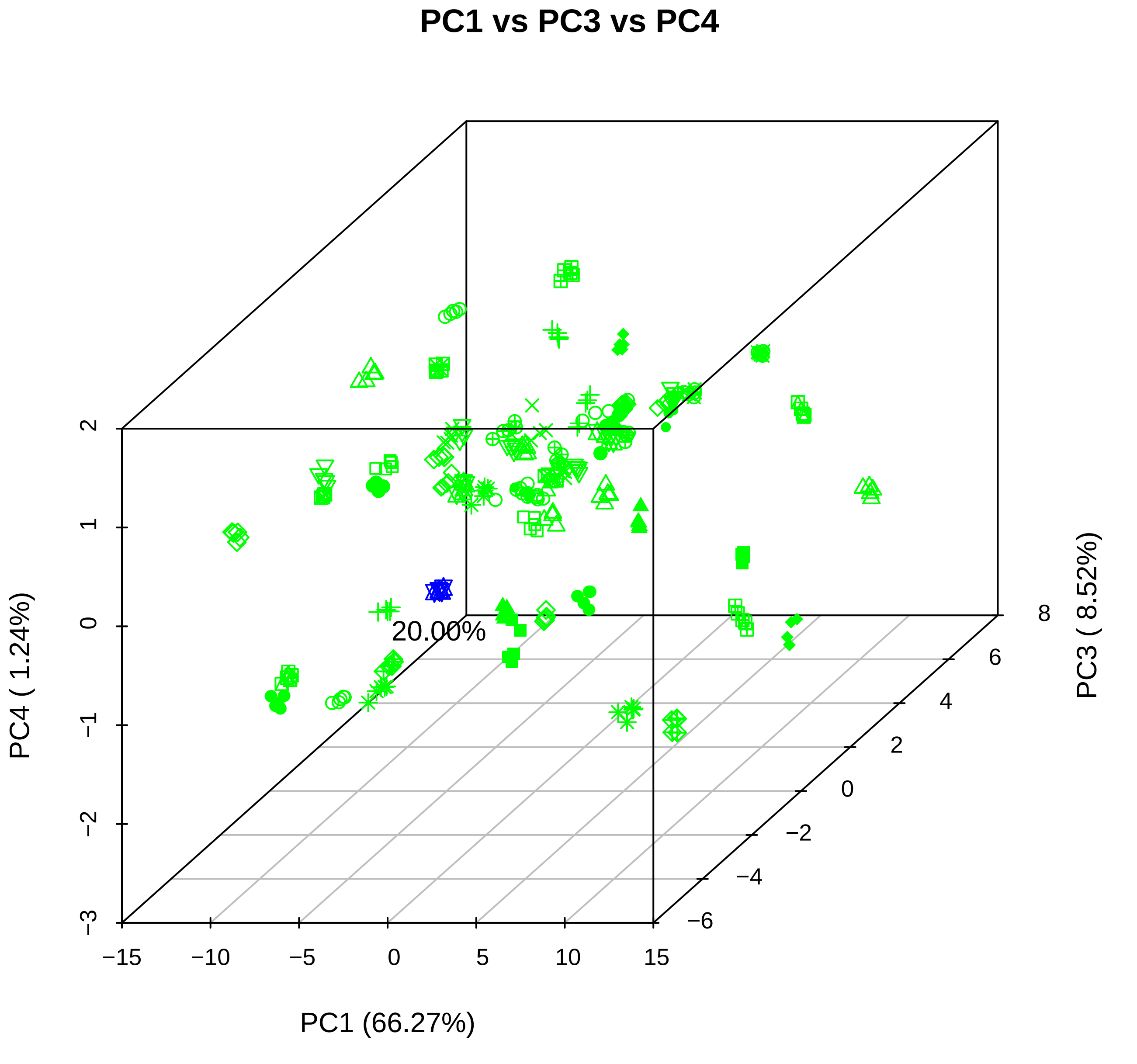

PC1 vs PC3 vs PC4
20.00%
PC1 (66.27%)
PC3 ( 8.52%)
PC4 ( 1.24%)
−15
−10
−5
0
5
10
15
−6
−4
−2
0
2
4
6
8
−3
−2
−1
0
1
2

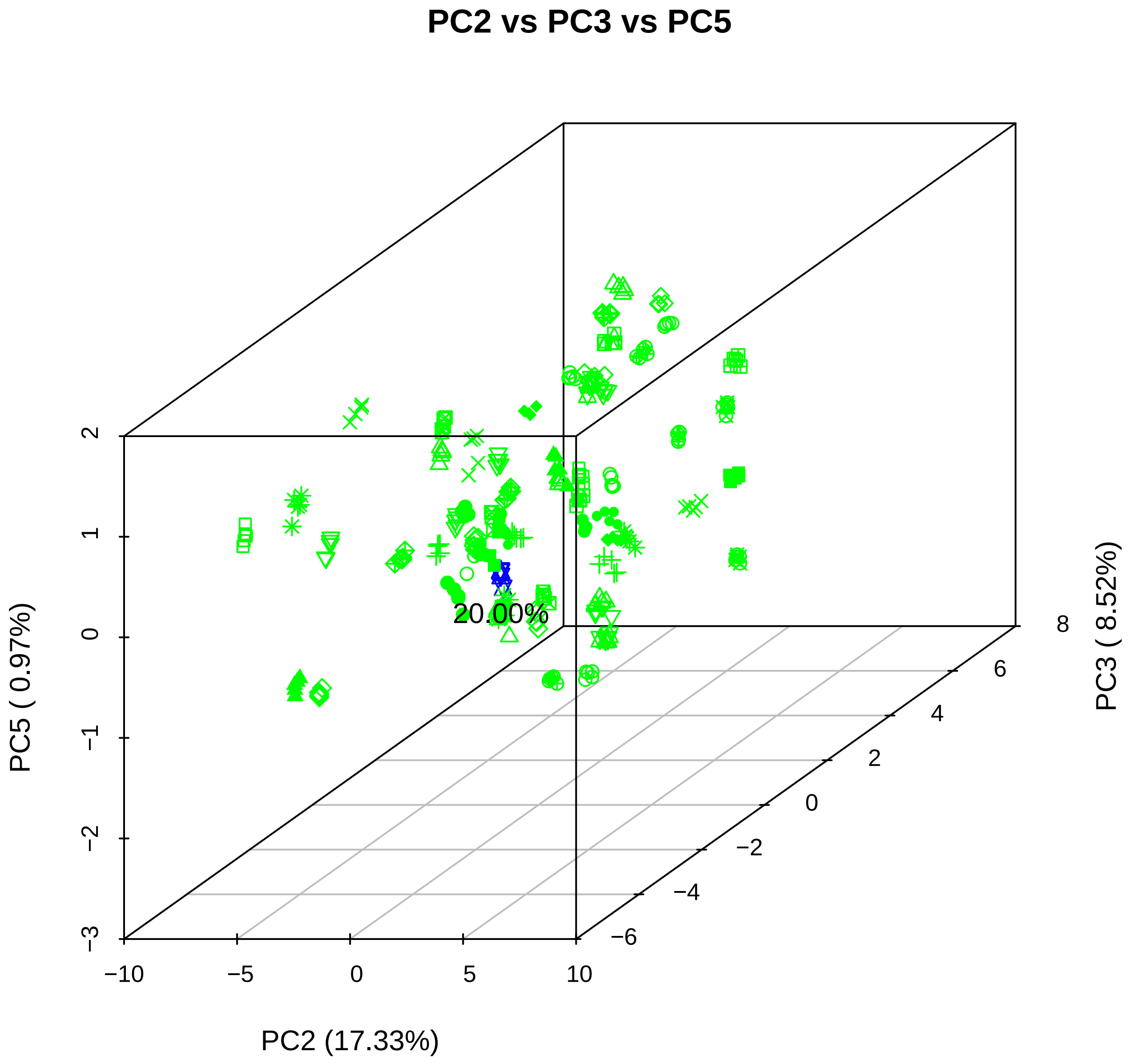

PC2 vs PC3 vs PC5
PC5 ( 0.97%)
PC2 (17.33%)
PC3 ( 8.52%)
20.00%
2
1
0
−1
−2
−3
−10
−5
0
5
10
−6
−4
−2
0
2
4
6
8

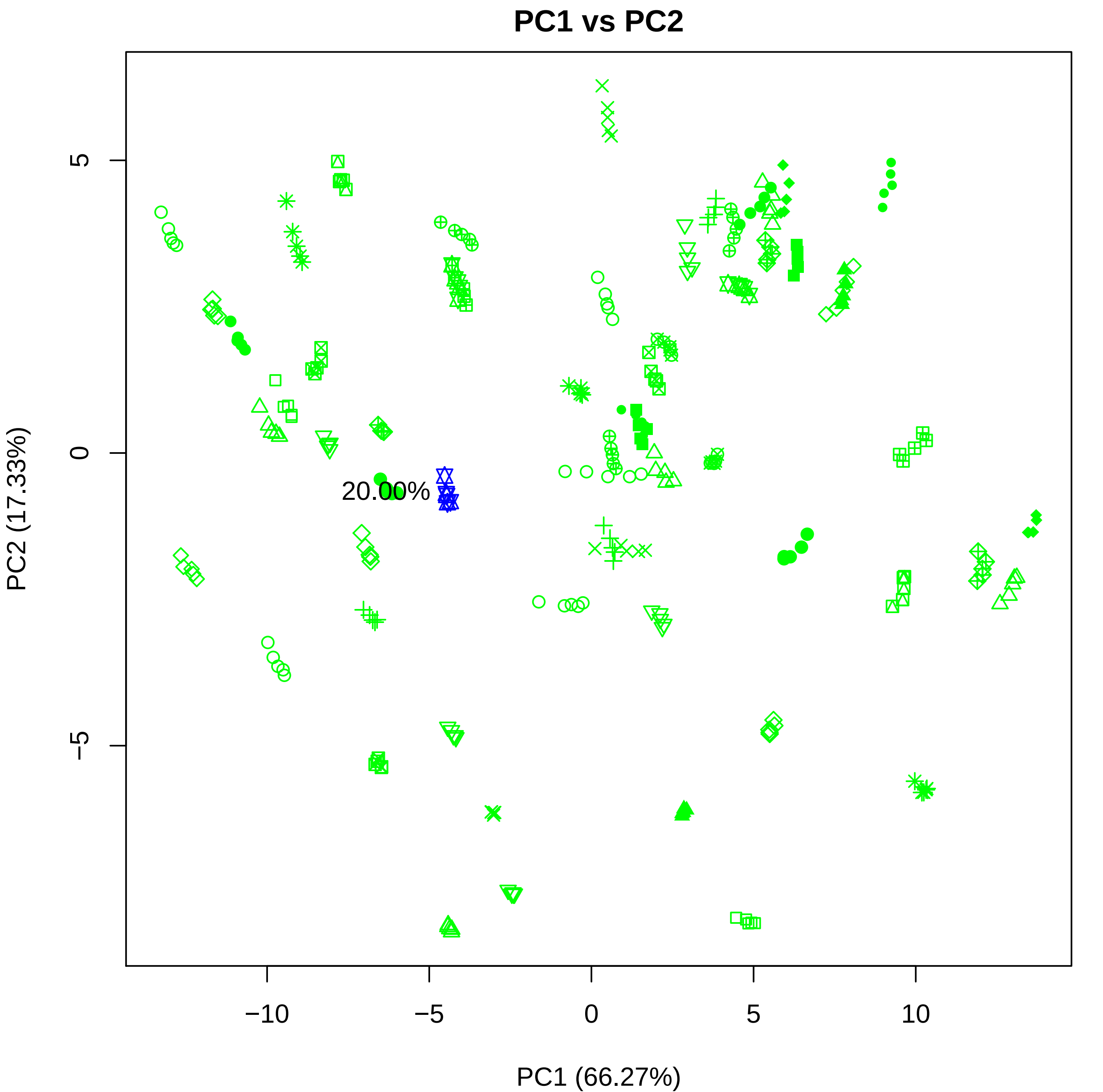
PC1 vs PC2
PC2 (17.33%)
PC1 (66.27%)
5
0
−5
−10
−5
0
5
10
20.00%

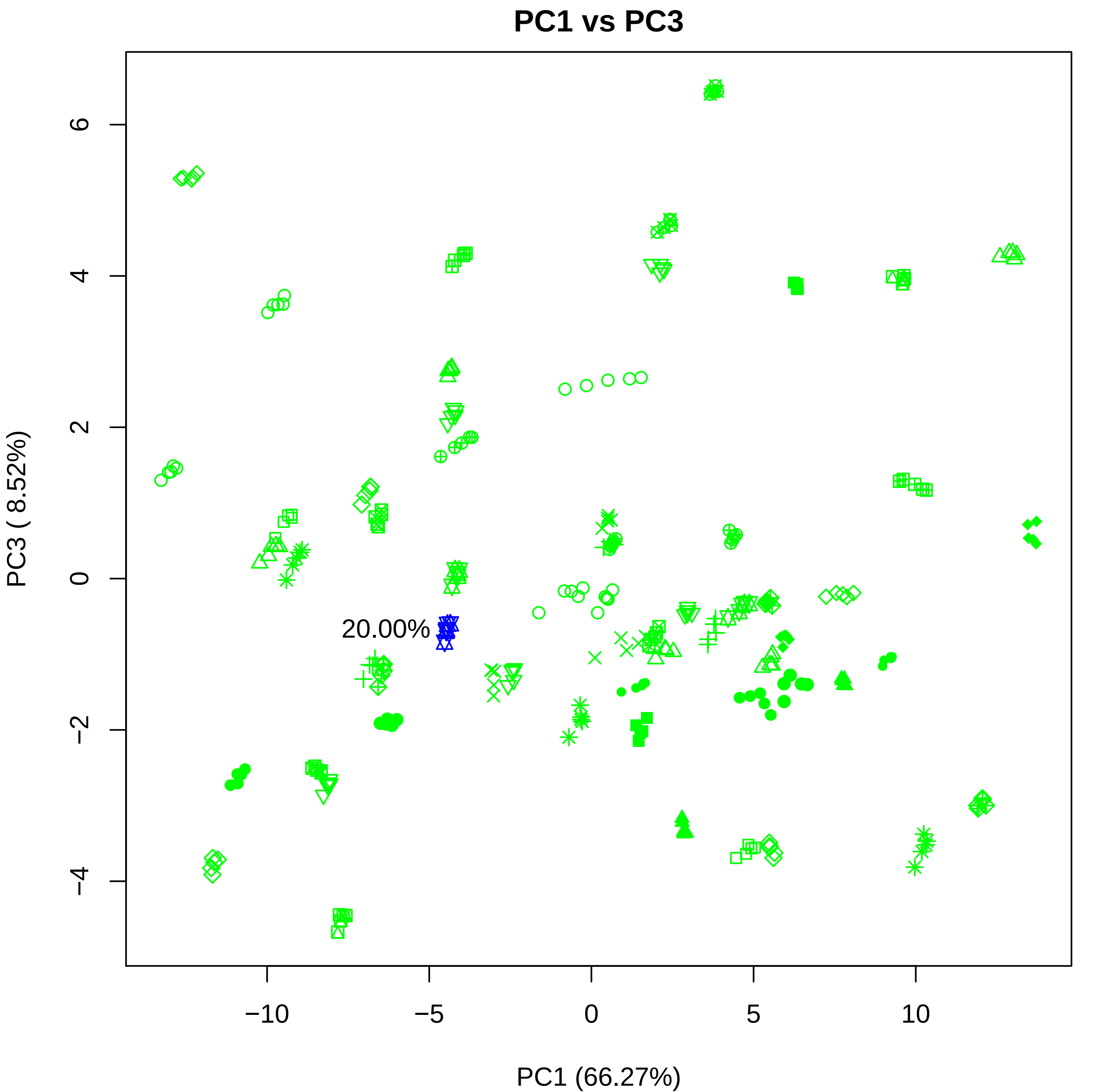

PC1 vs PC3
PC3 ( 8.52%)
PC1 (66.27%)
20.00%
6
4
2
0
−2
−4
−10
−5
0
5
10

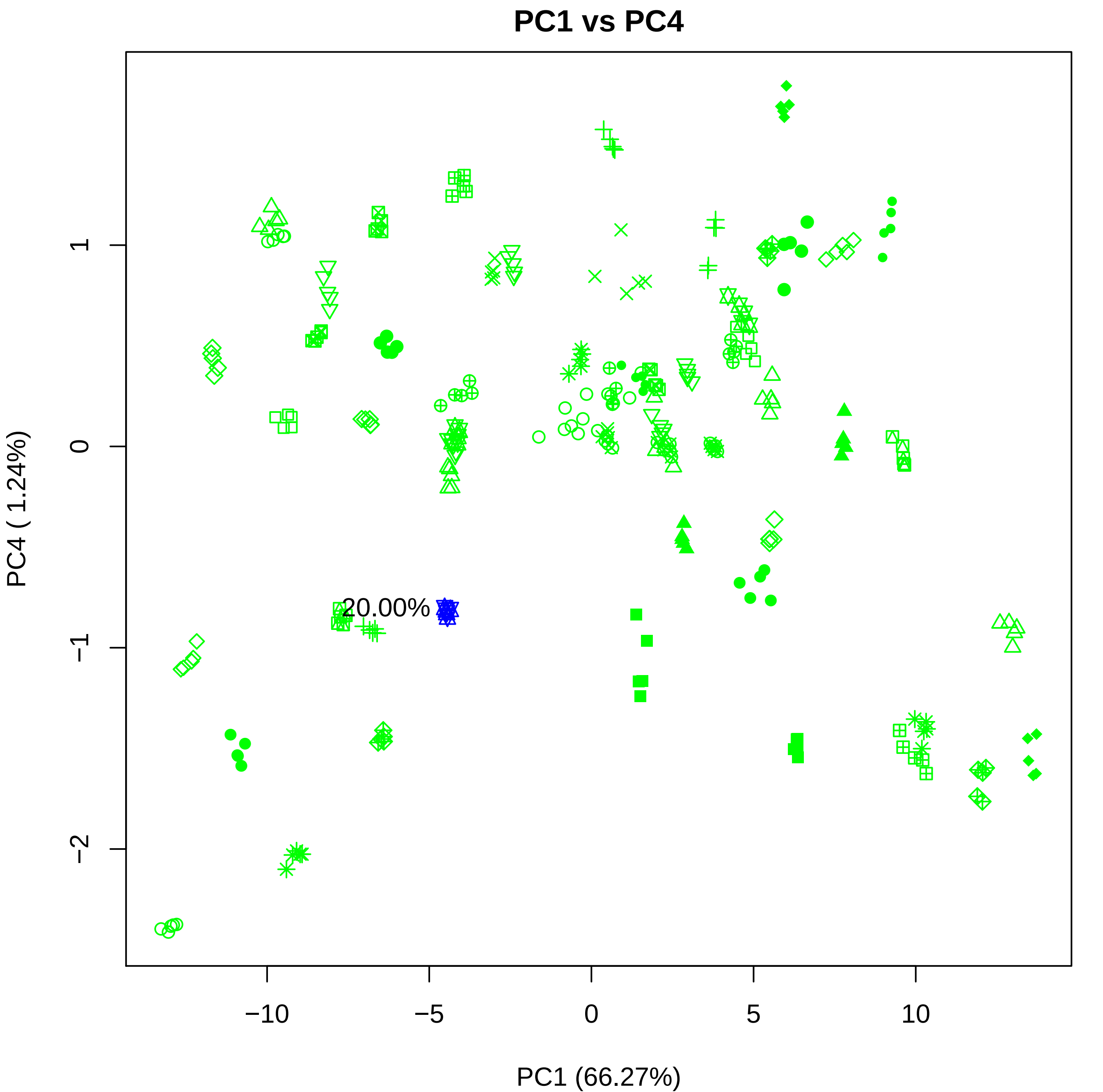
PC1 vs PC4
20.00%
PC4 ( 1.24%)
PC1 (66.27%)
1
0
−1
−2
−10
−5
0
5
10

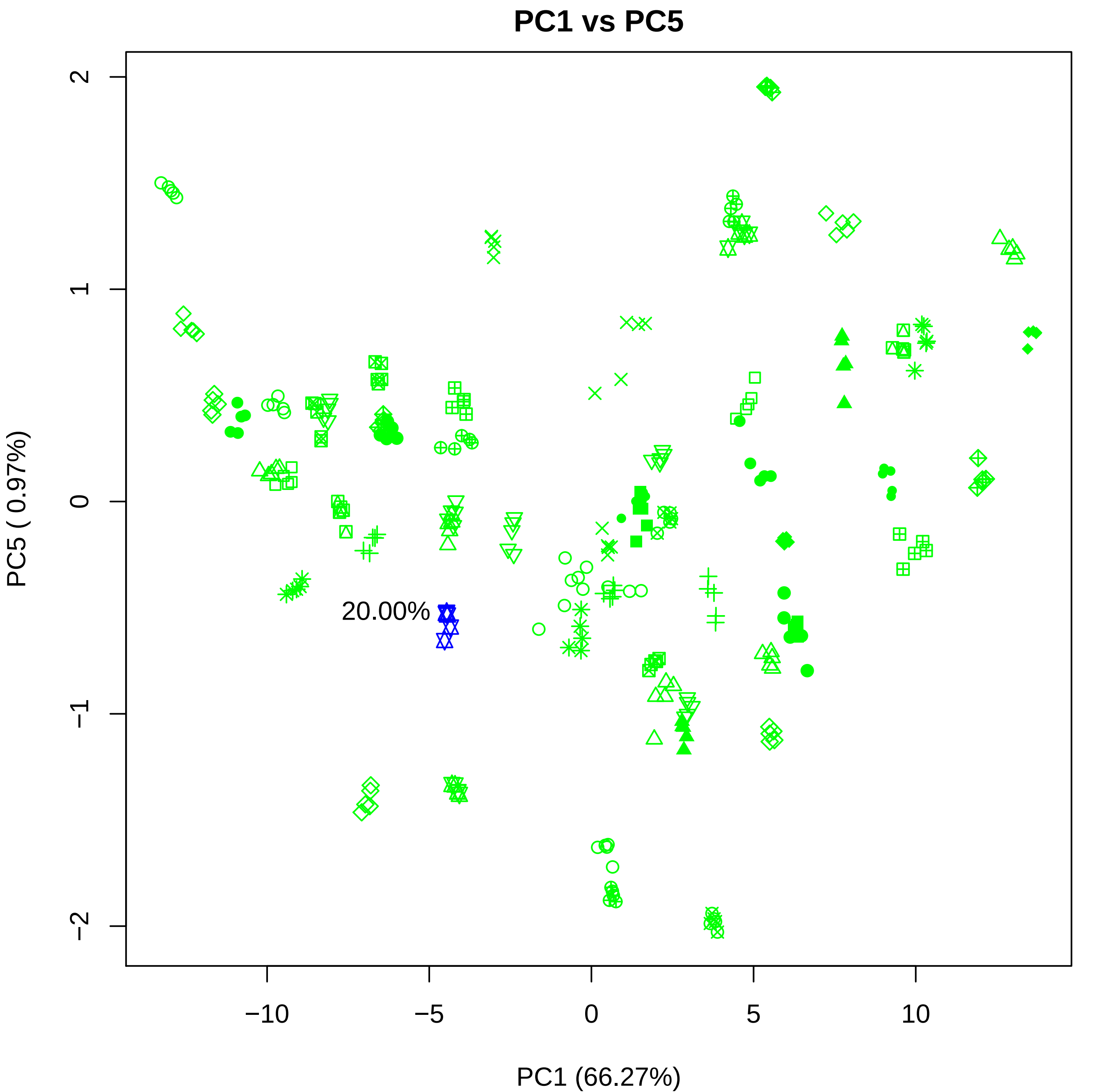
PC1 vs PC5
2
1
0
−1
−2
PC5 ( 0.97%)
20.00%
−10
−5
0
5
10
PC1 (66.27%)

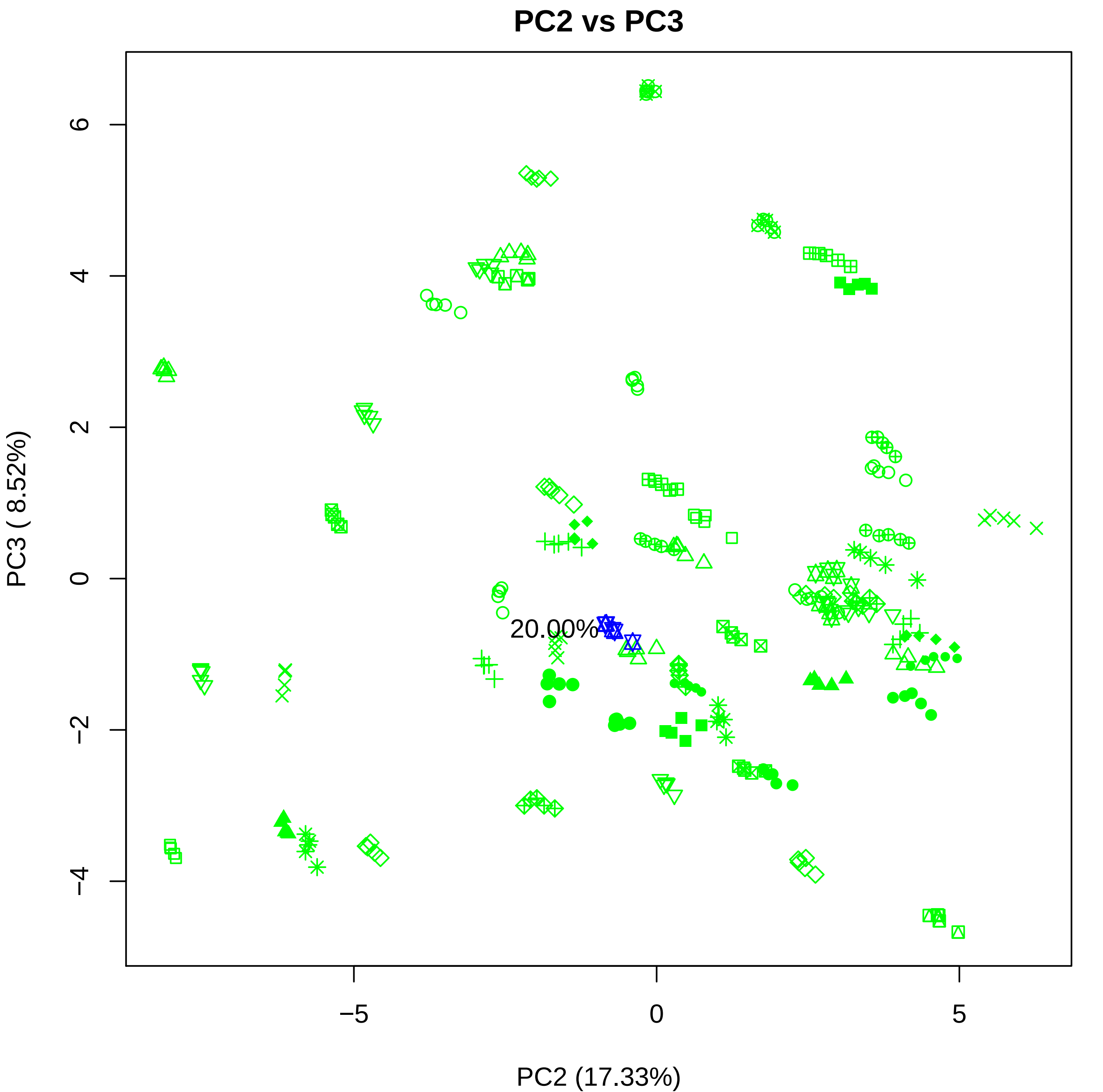

PC2 vs PC3
20.00%
PC3 ( 8.52%)
PC2 (17.33%)
6
4
2
0
−2
−4
−5
0
5

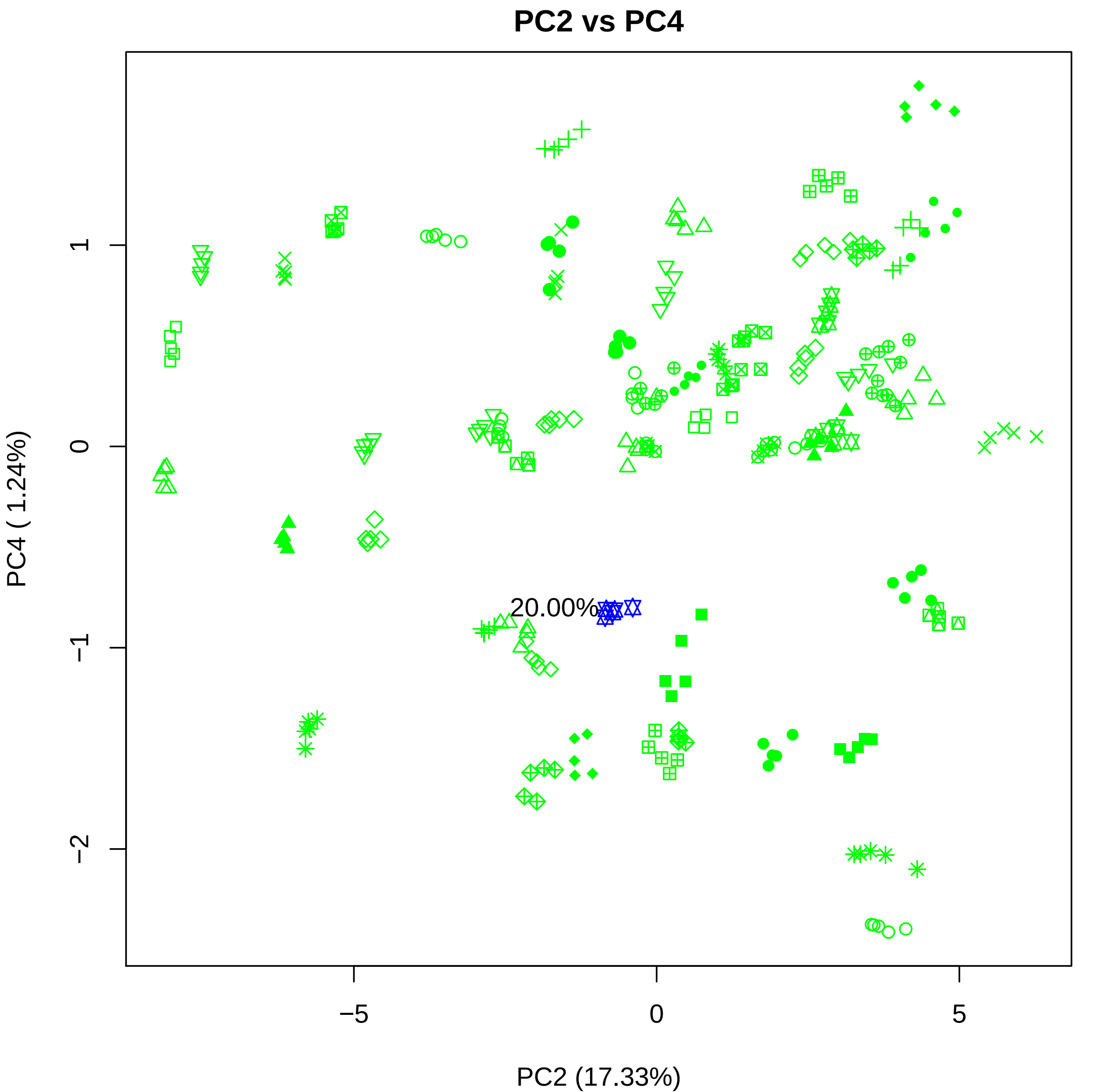
PC2 vs PC4
PC4 ( 1.24%)
PC2 (17.33%)
20.00%
1
0
−1
−2
−5
0
5

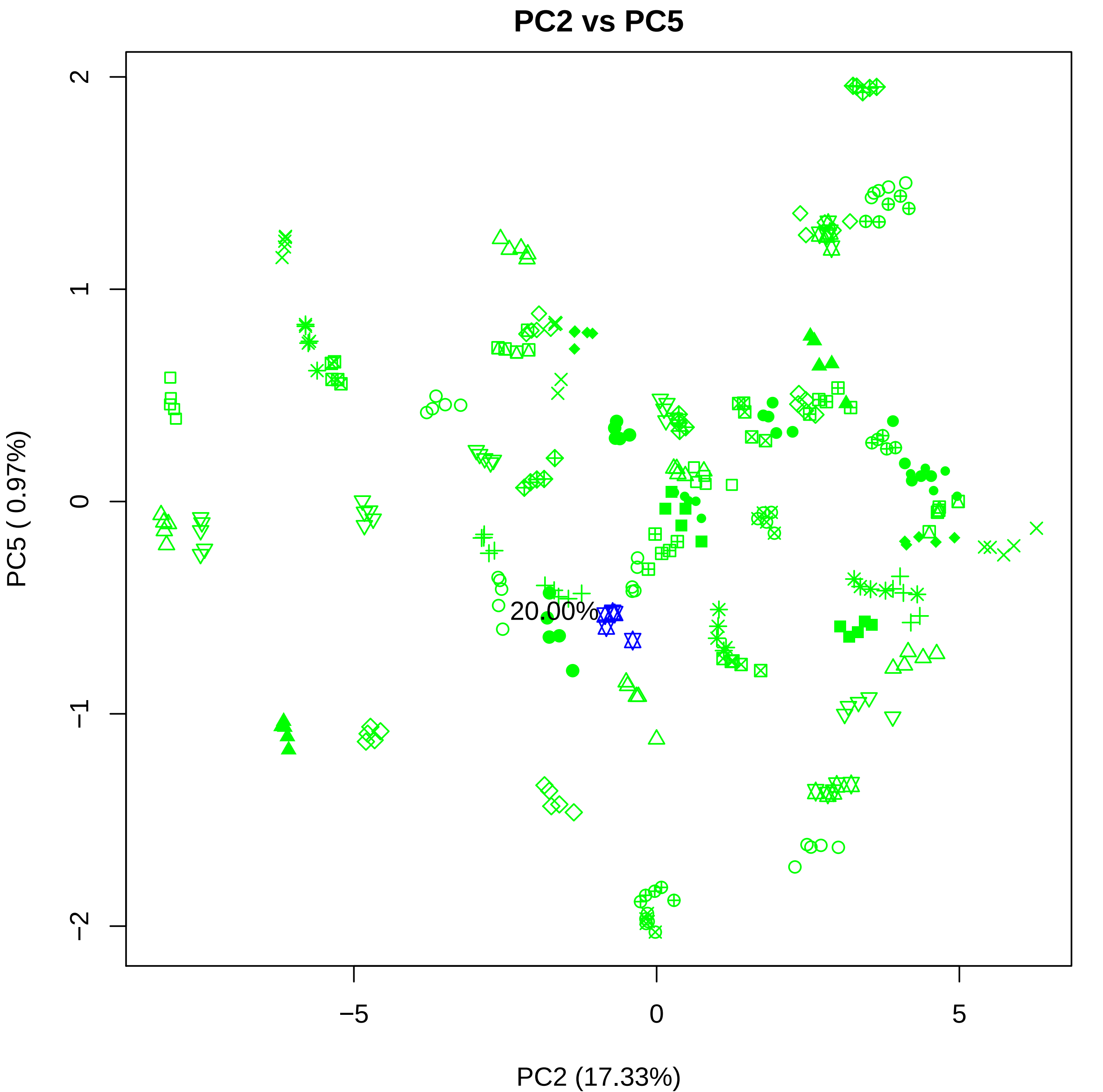
PC2 vs PC5
PC5 ( 0.97%)
PC2 (17.33%)
20.00%
2
1
0
−1
−2
−5
0
5

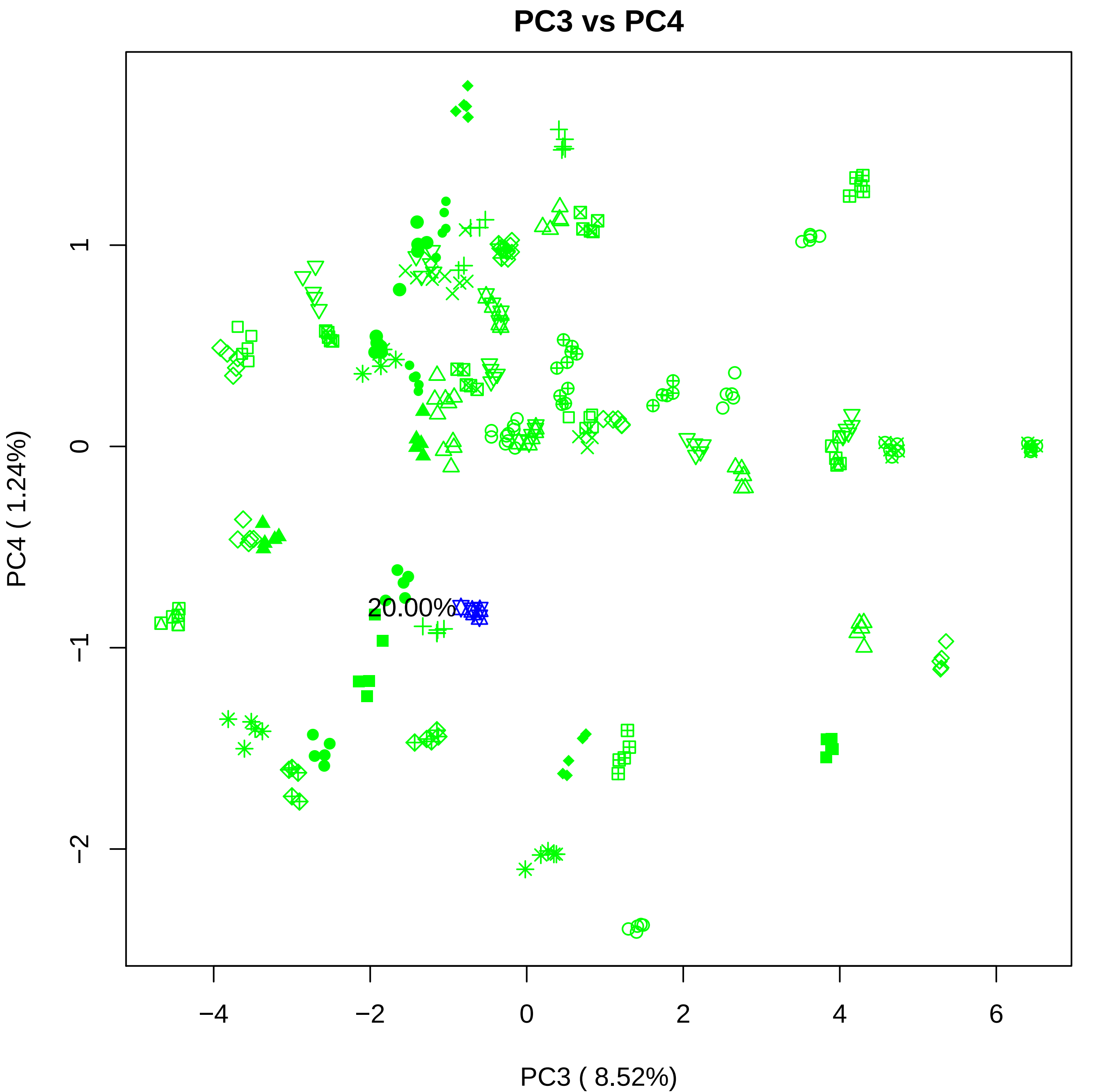
PC3 vs PC4
PC4 ( 1.24%)
PC3 ( 8.52%)
20.00%
1
0
−1
−2
−4
−2
0
2
4
6

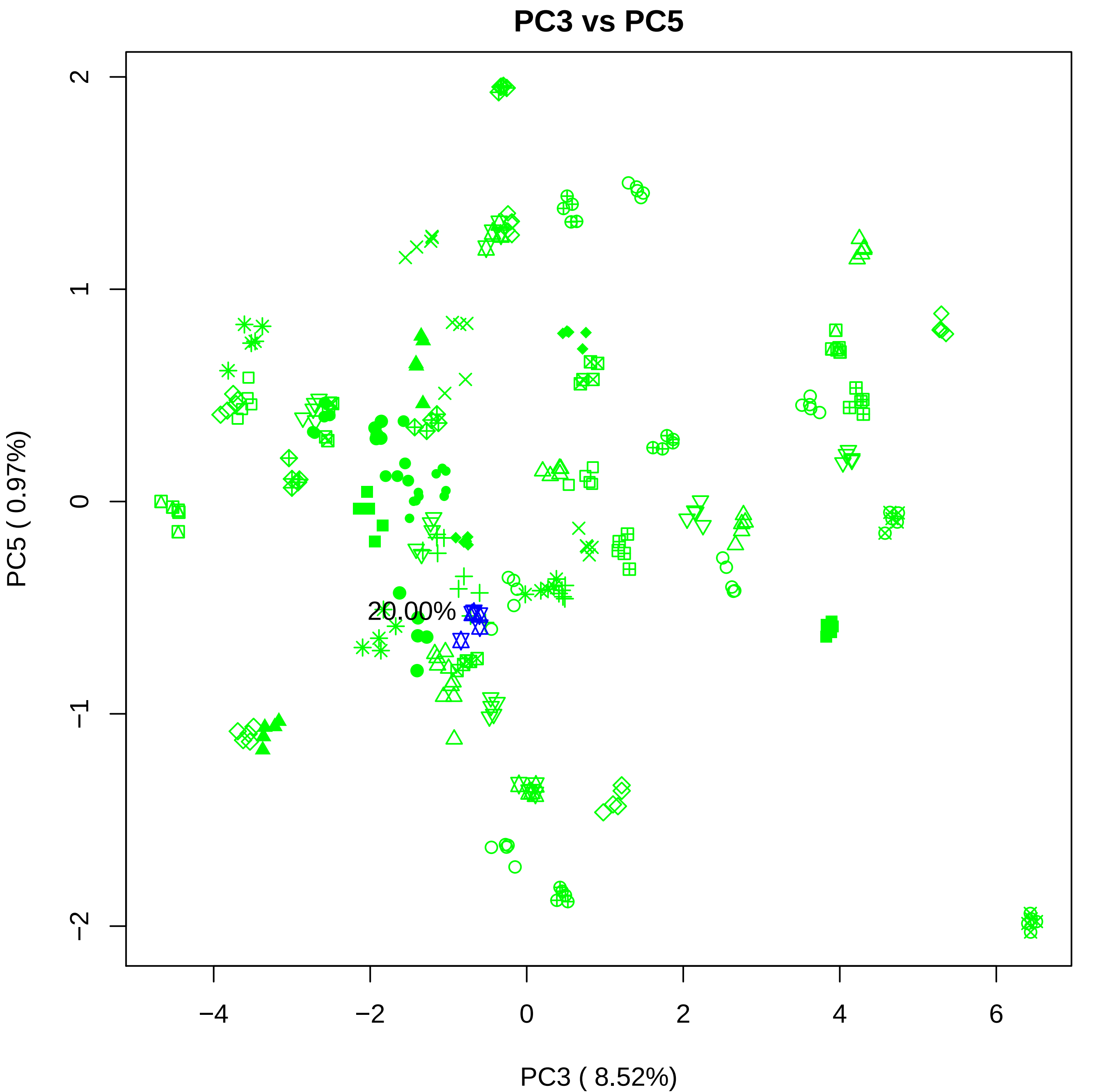

PC3 vs PC5
PC5 ( 0.97%)
PC3 ( 8.52%)
20.00%
2
1
0
−1
−2
−4
−2
0
2
4
6

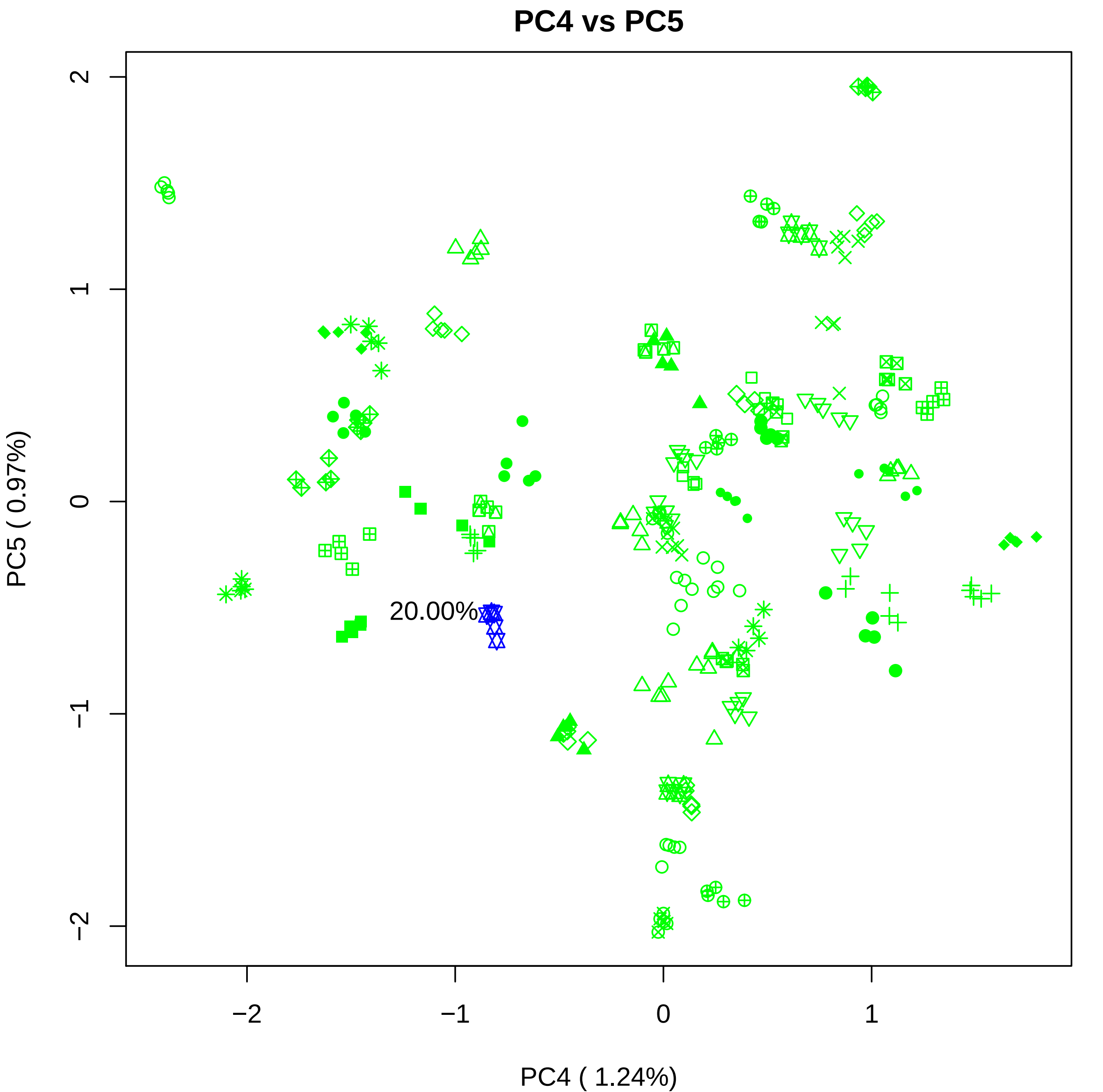

PC4 vs PC5
PC5 ( 0.97%)
PC4 ( 1.24%)
20.00%
−2
−1
0
1
2

### PCA of 40% BMH(b) vs PMH(b)

Scaled Matrix PCAed

7th order polynomial baseline correction performed

Raw spectra smoothed to 21 Neighbours, 2nd order PN, Savtisky Golay

Normalisation between 0 and 100

D2 spectra smoothed to 41 Neighbours, 2nd order PN, Savtisky Golay

Regions Analysed: 3601−2501 and 2001−700cm−1

Samples Analysed: 1 bContam's 0 BLH's 0 BMH's 0 OMH's 0 pContam's 0 PLH's 60 PMH's

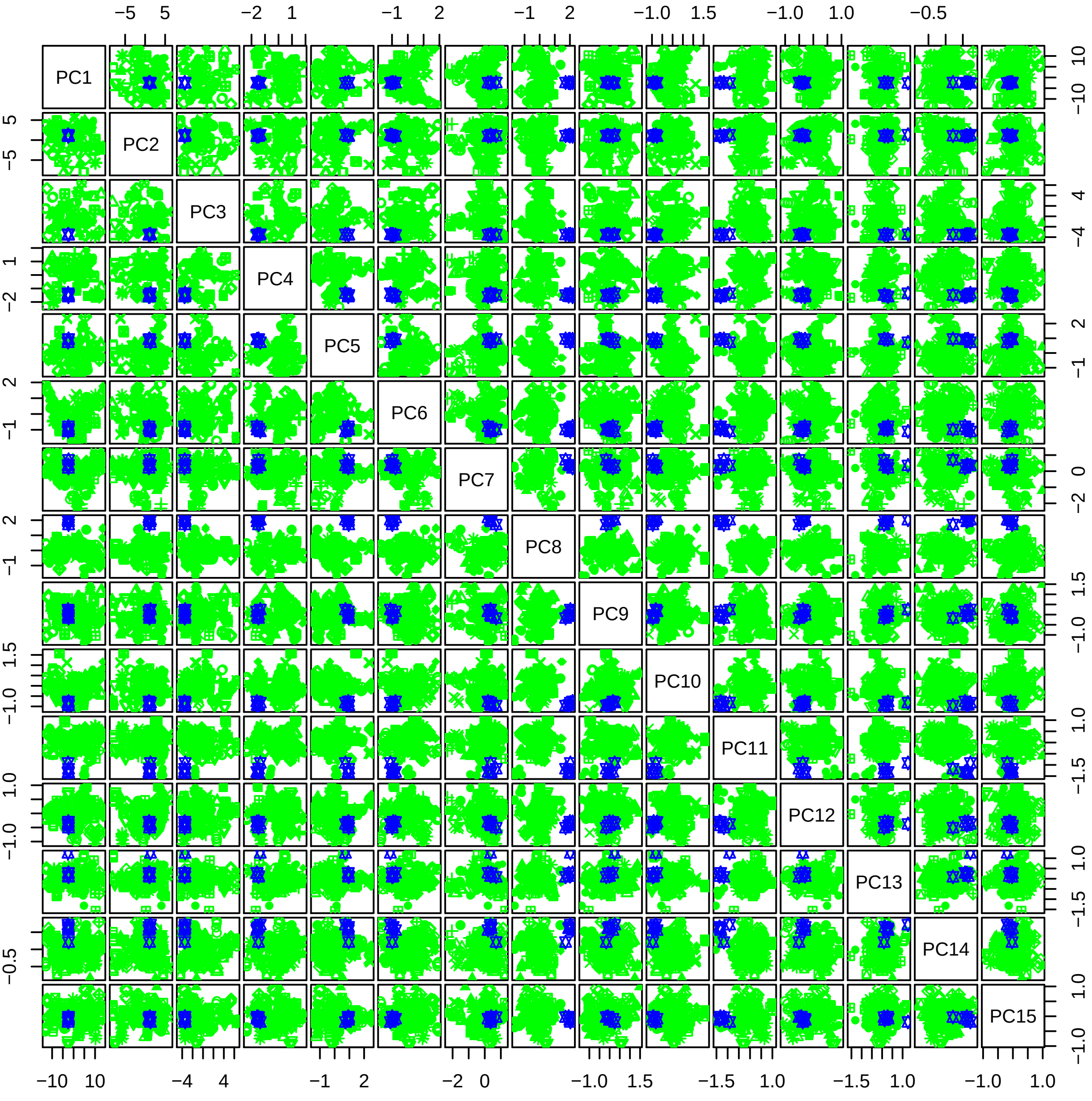
PC1
PC2
PC3
PC4
PC5
PC6
PC7
PC8
PC9
PC10
PC11
PC12
PC13
PC14
PC15

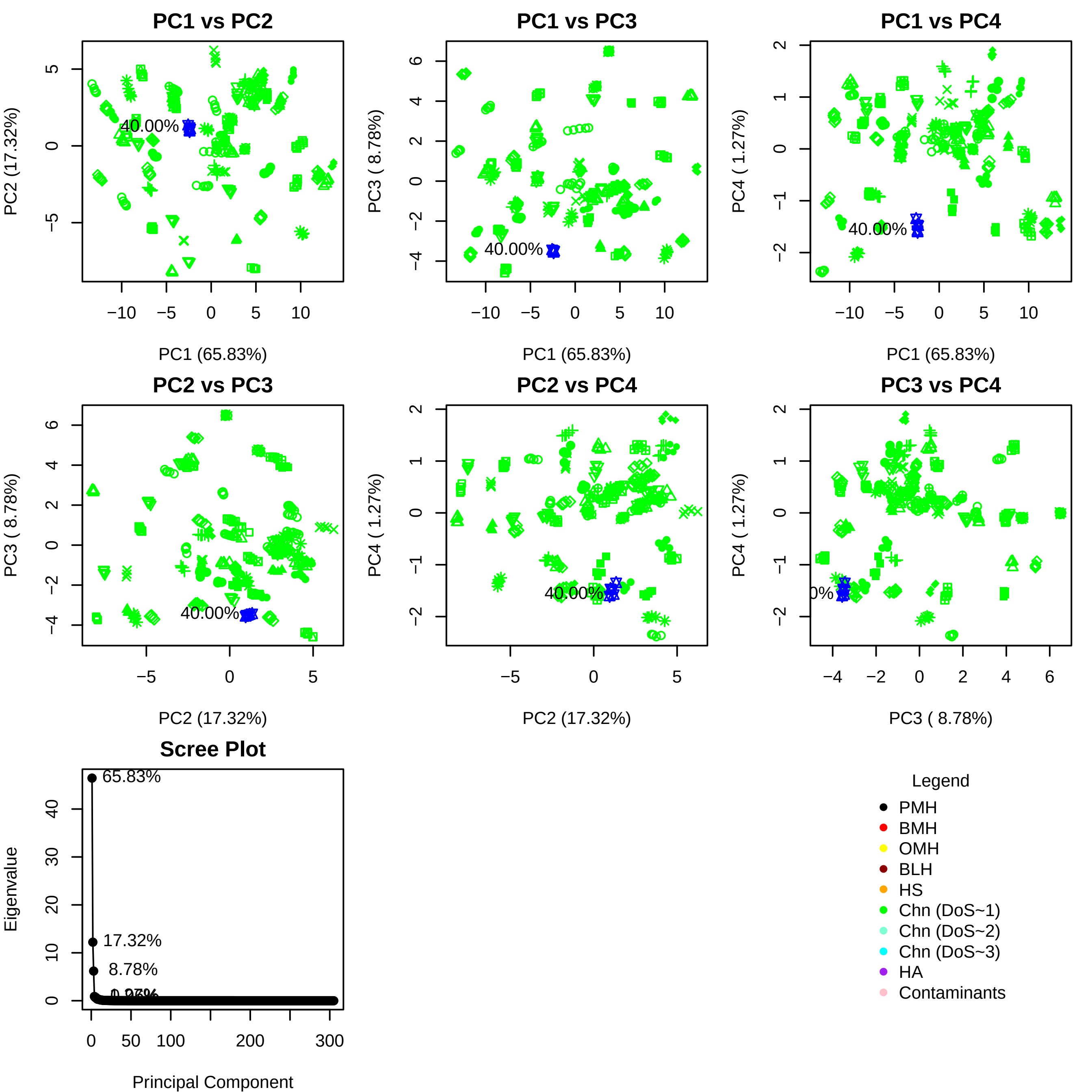
PC1 vs PC2
PC1 vs PC3
PC1 vs PC4
PC2 vs PC3
PC2 vs PC4
PC3 vs PC4
Scree Plot
PC1 (65.83%)
PC2 (17.32%)
PC3 ( 8.78%)
PC4 ( 1.27%)
40.00%
Eigenvalue
Principal Component
65.83%
17.32%
8.78%
Legend
PMH
BMH
OMH
BLH
HS
Chn (DoS~1)
Chn (DoS~2)
Chn (DoS~3)
HA
Contaminants

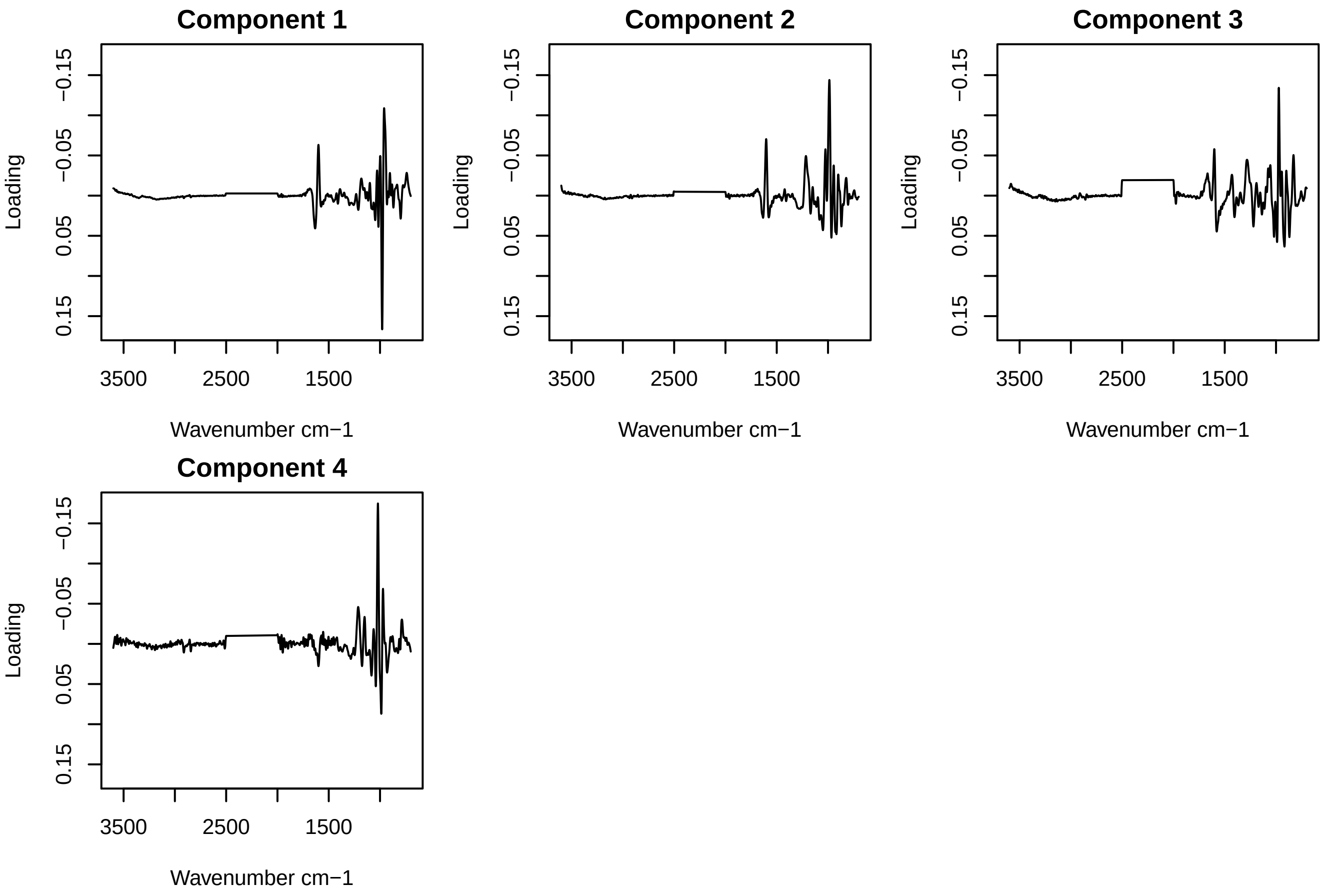
Component 1
Component 2
Component 3
Component 4
Loading
−0.15
−0.05
0.05
0.15
3500
2500
1500
Wavenumber cm−1

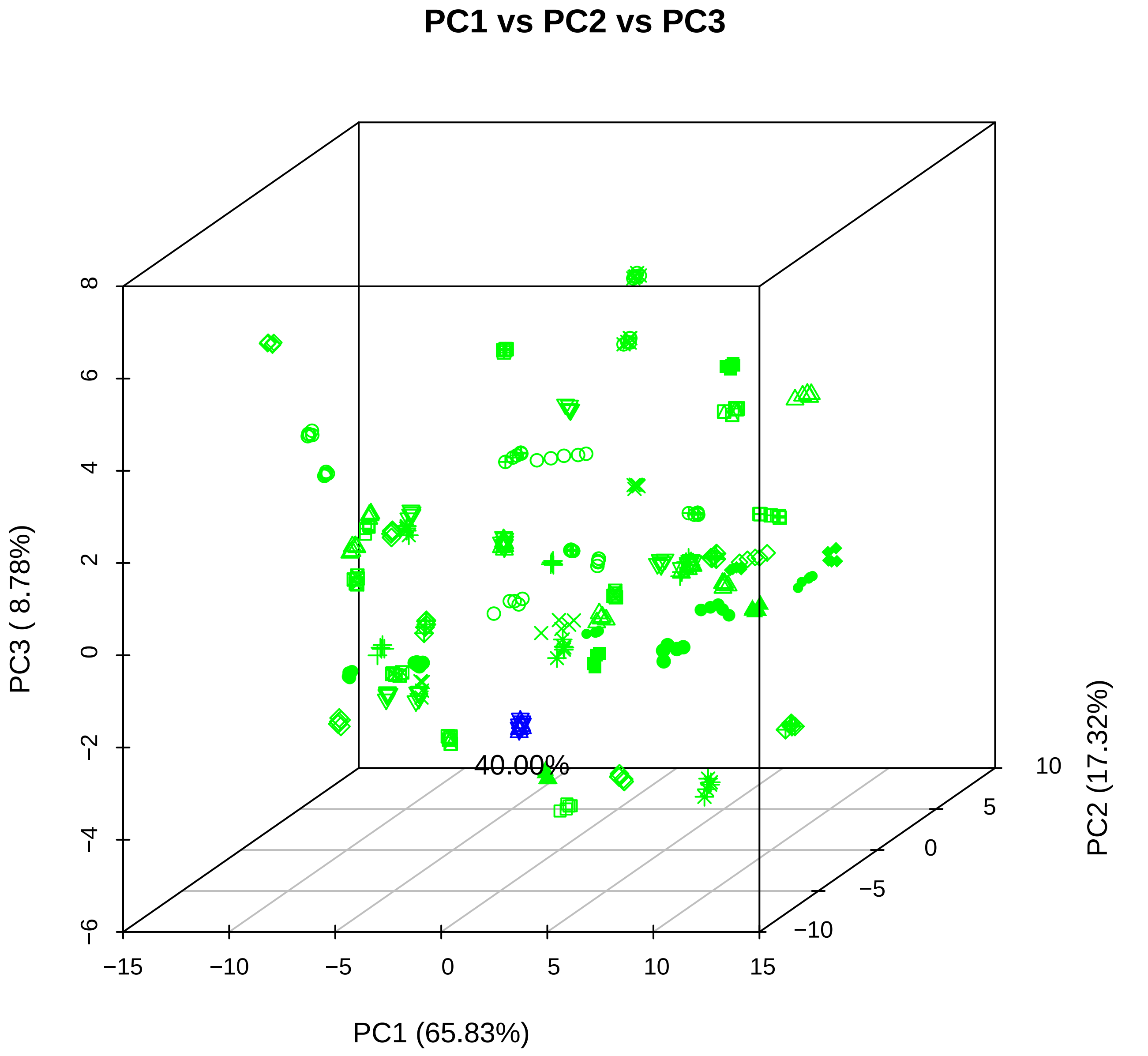

PC1 vs PC2 vs PC3
PC1 (65.83%)
PC2 (17.32%)
PC3 ( 8.78%)
40.00%
−15
−10
−5
0
5
10
15
−10
−5
0
5
10
−6
−4
−2
0
2
4
6
8

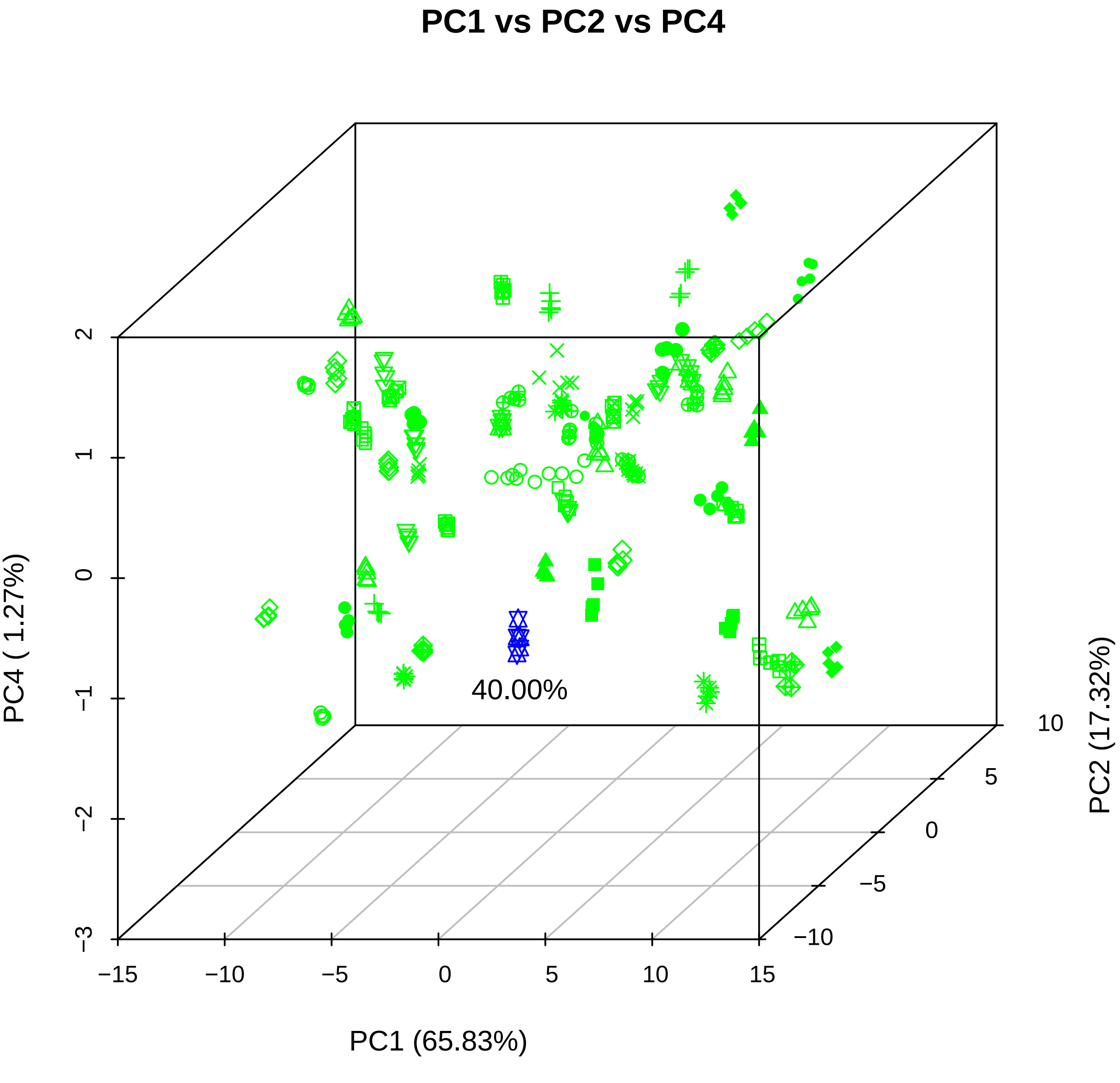

PC1 vs PC2 vs PC4
40.00%
PC1 (65.83%)
PC2 (17.32%)
PC4 ( 1.27%)
−15
−10
−5
0
5
10
15
−3
−2
−1
0
1
2
−10
−5
0
5
10

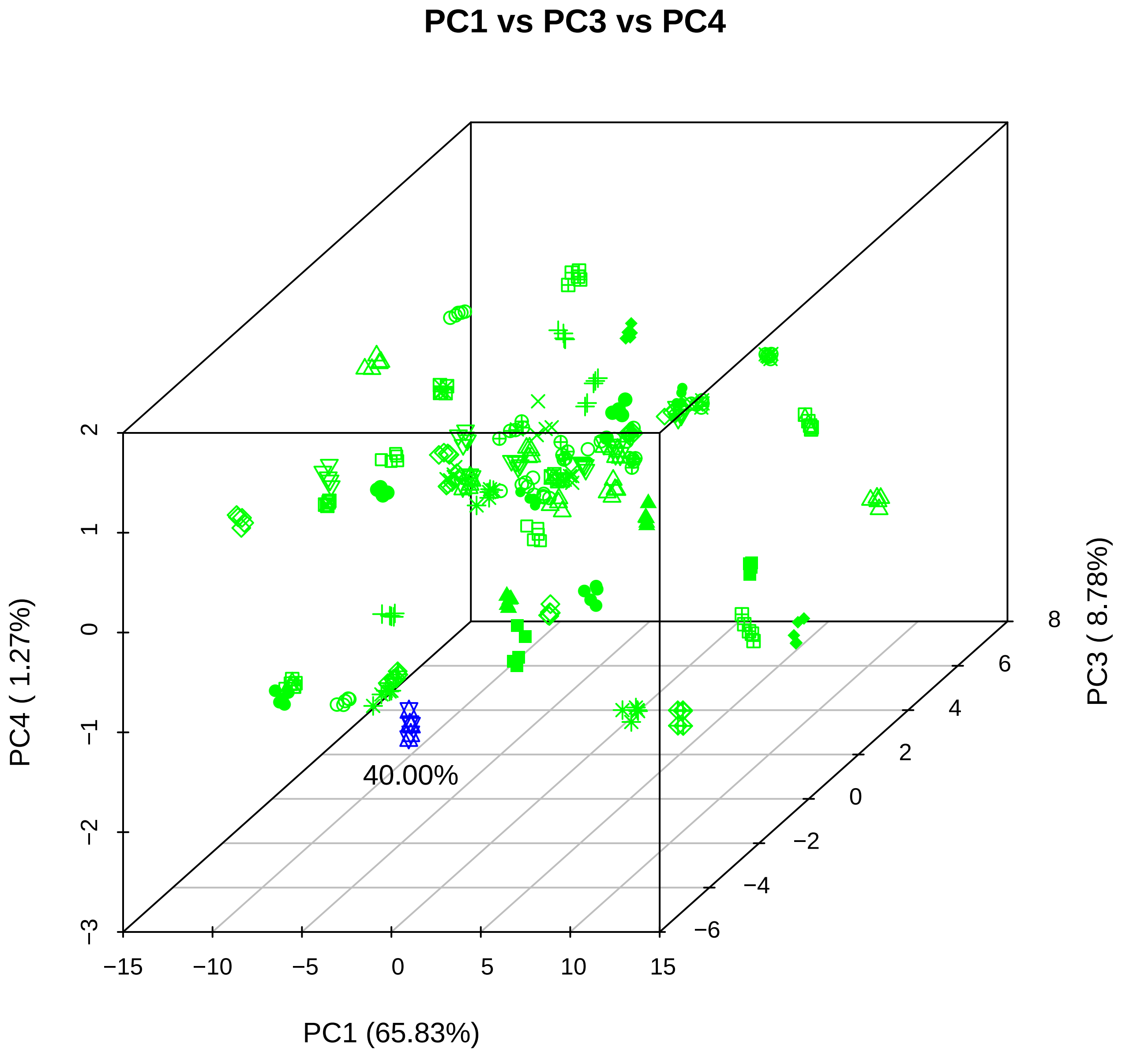

PC1 vs PC3 vs PC4
PC1 (65.83%)
PC4 ( 1.27%)
PC3 ( 8.78%)
40.00%
−15
−10
−5
0
5
10
15
−3
−2
−1
0
1
2
−6
−4
−2
0
2
4
6
8

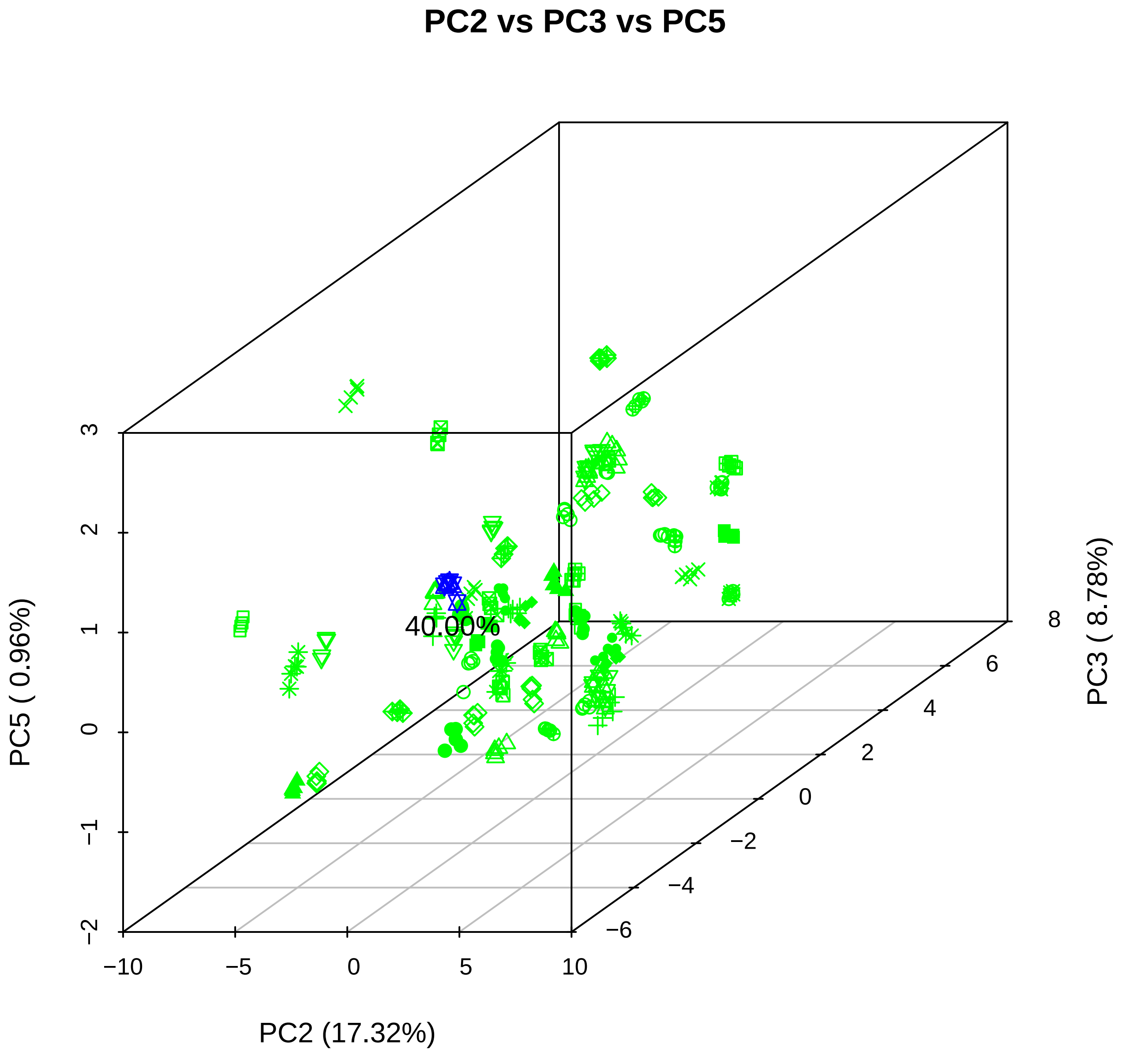

PC2 vs PC3 vs PC5
40.00%
PC5 ( 0.96%)
3
2
1
0
−1
−2
−10
−5
0
5
10
PC2 (17.32%)
−6
−4
−2
0
2
4
6
8
PC3 ( 8.78%)

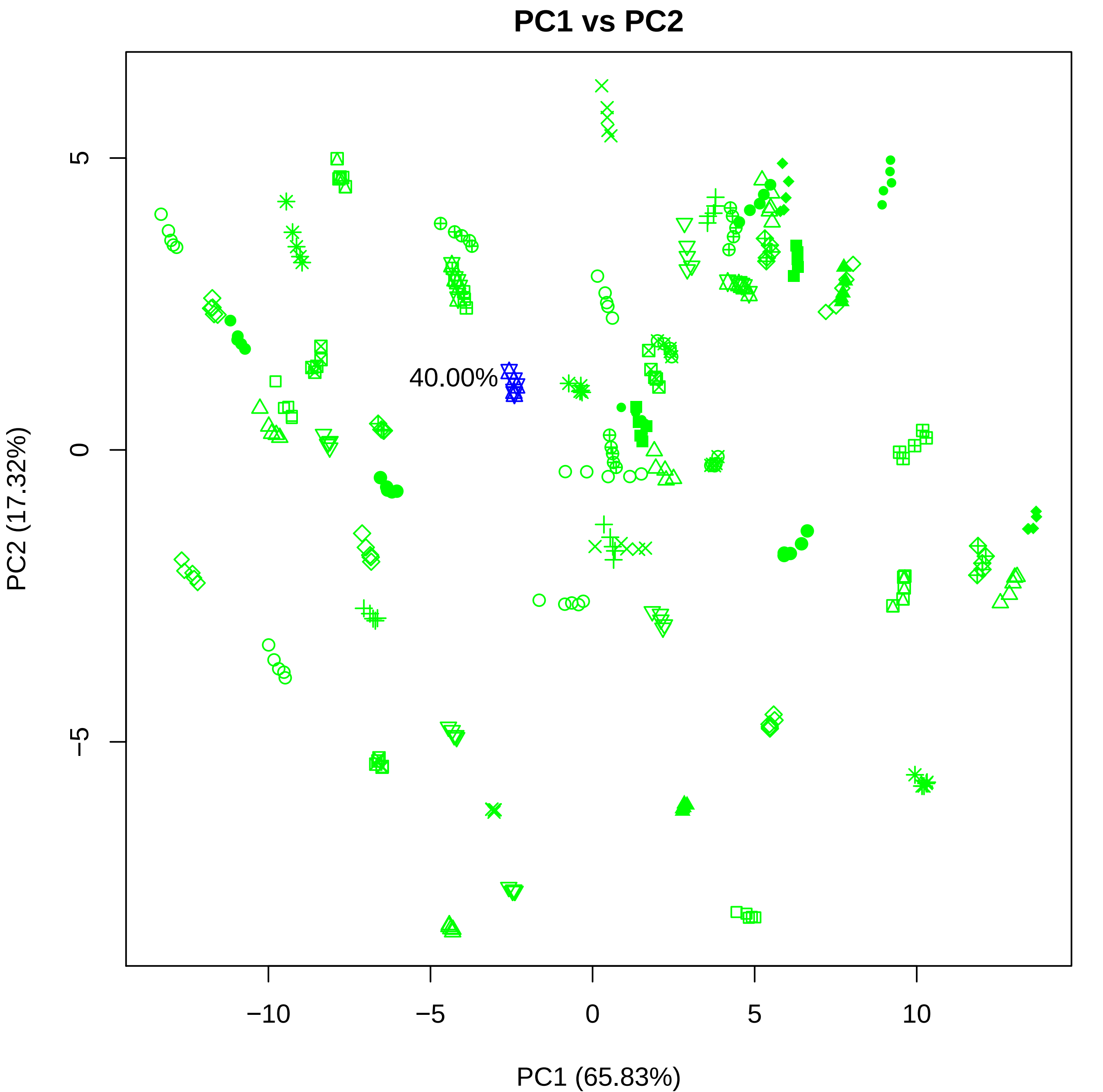
PC1 vs PC2
40.00%
PC1 (65.83%)
PC2 (17.32%)
−10
−5
0
5
10
5
0
−5

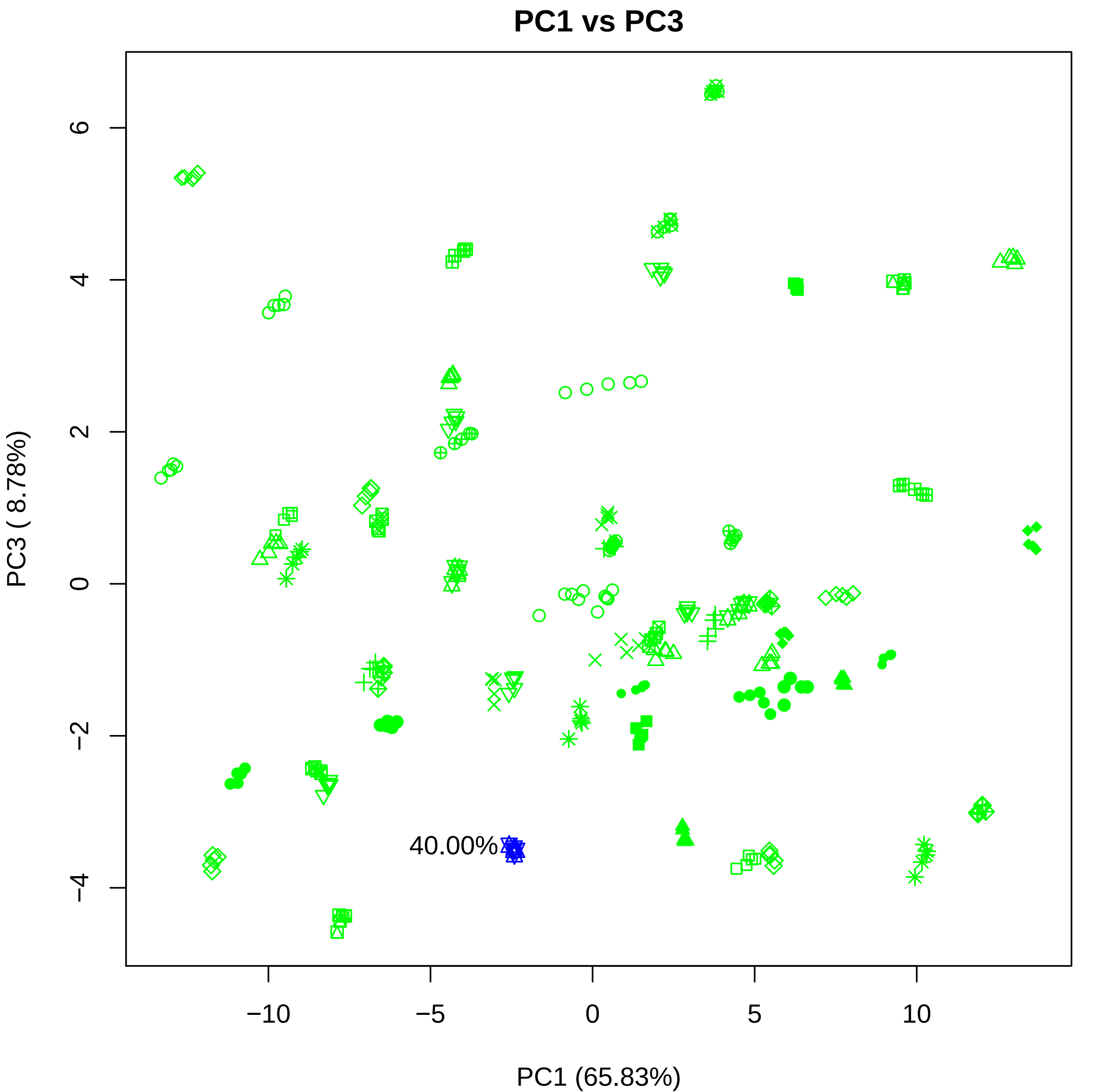

PC1 vs PC3
PC3 ( 8.78%)
PC1 (65.83%)
40.00%
−4
−2
0
2
4
6
−10
−5
0
5
10

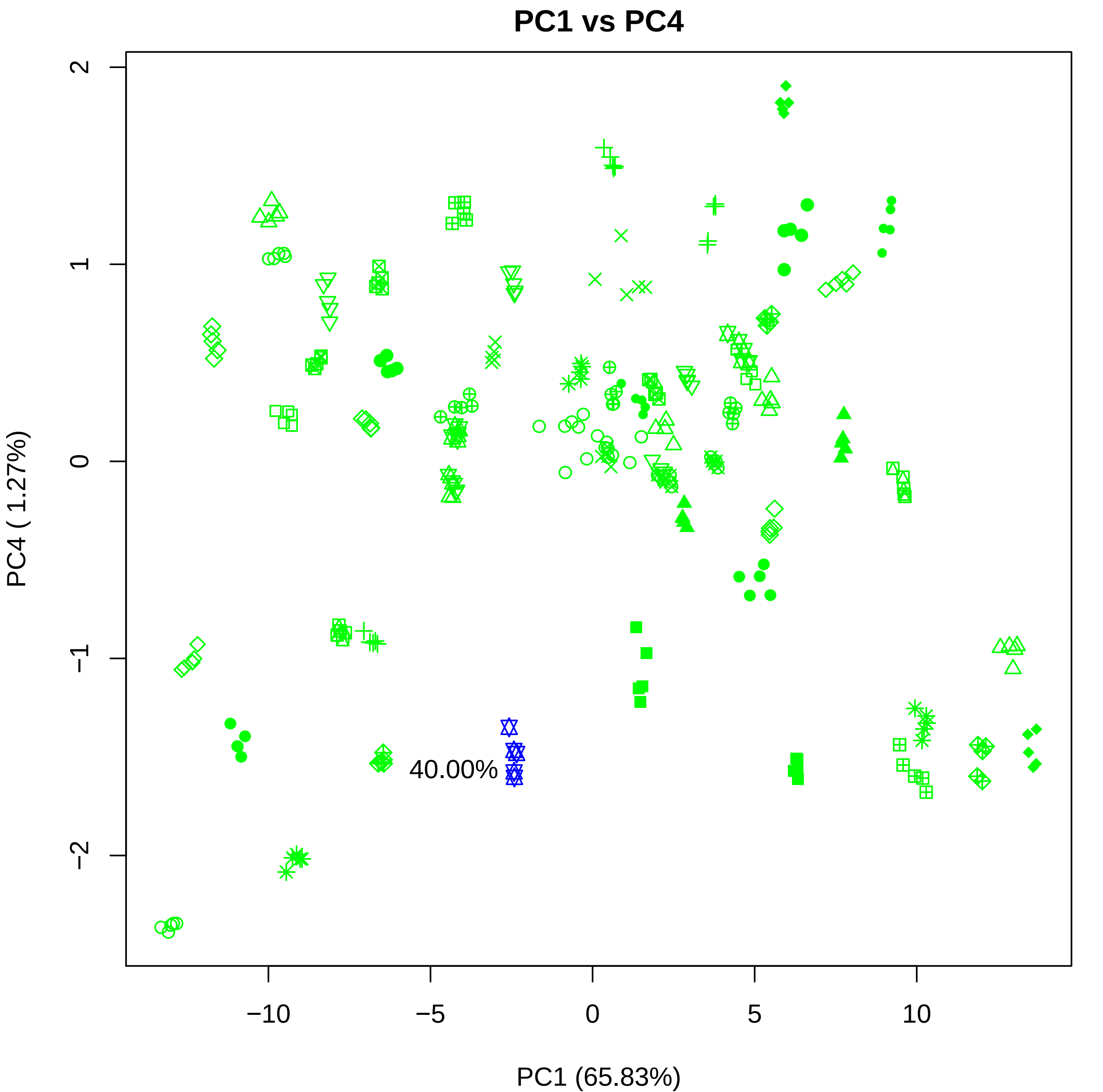

PC1 vs PC4
PC4 ( 1.27%)
PC1 (65.83%)
40.00%
2
1
0
−1
−2
−10
−5
0
5
10

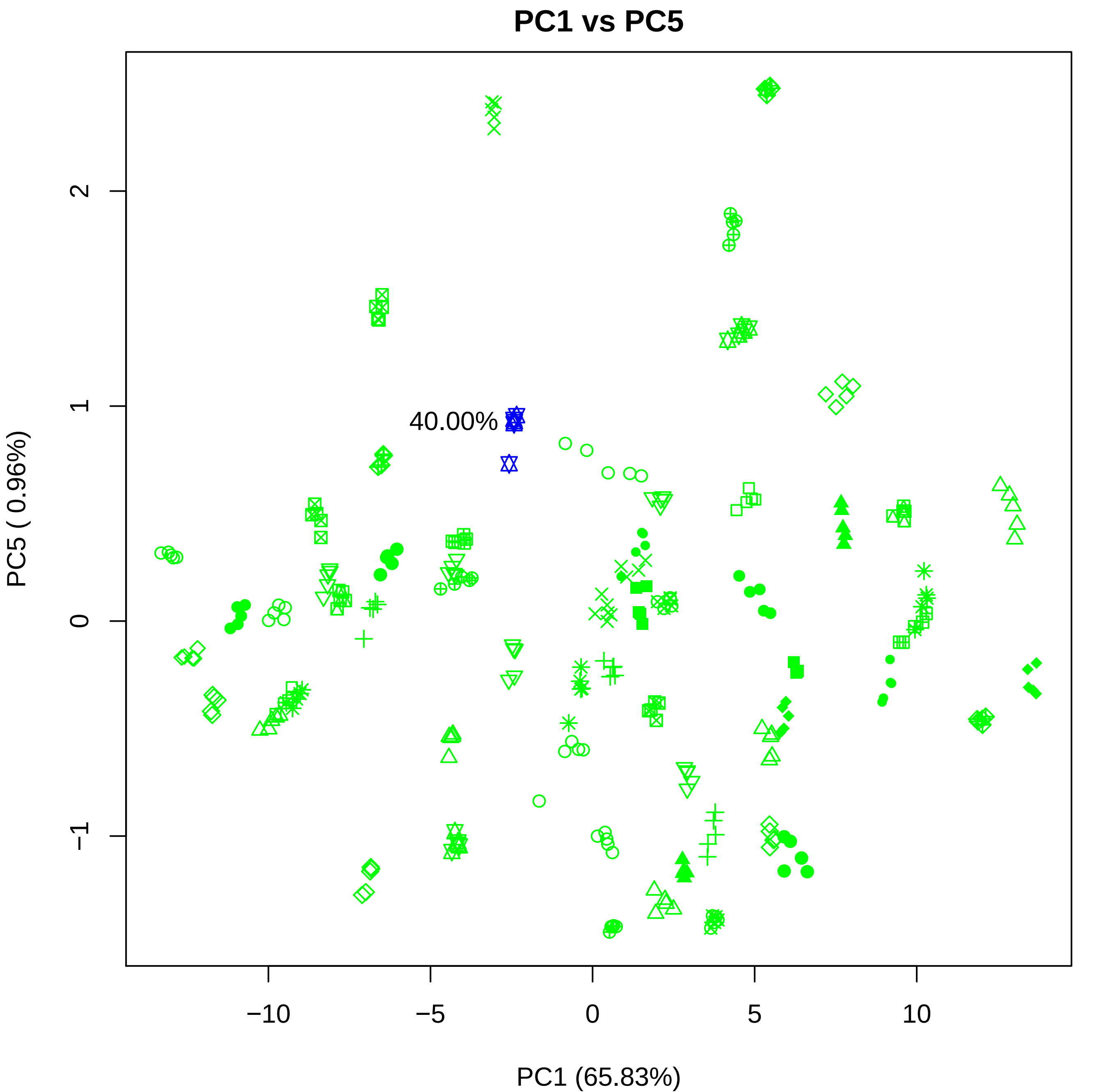
PC1 vs PC5
40.00%
2
1
0
−1
PC5 ( 0.96%)
−10
−5
0
5
10
PC1 (65.83%)

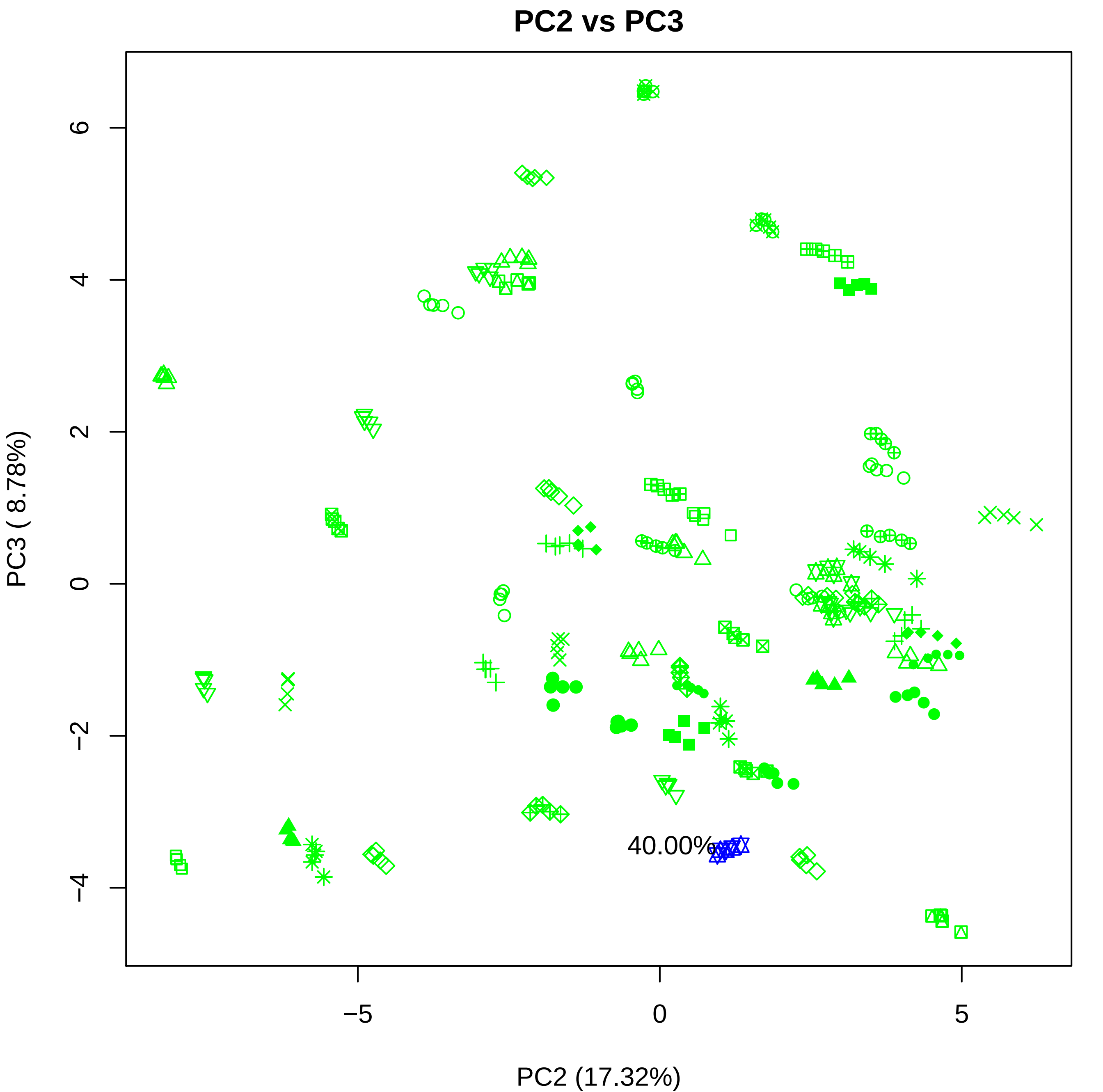
PC2 vs PC3
6
4
2
0
−2
−4
PC3 ( 8.78%)
40.00%
−5
0
5
PC2 (17.32%)

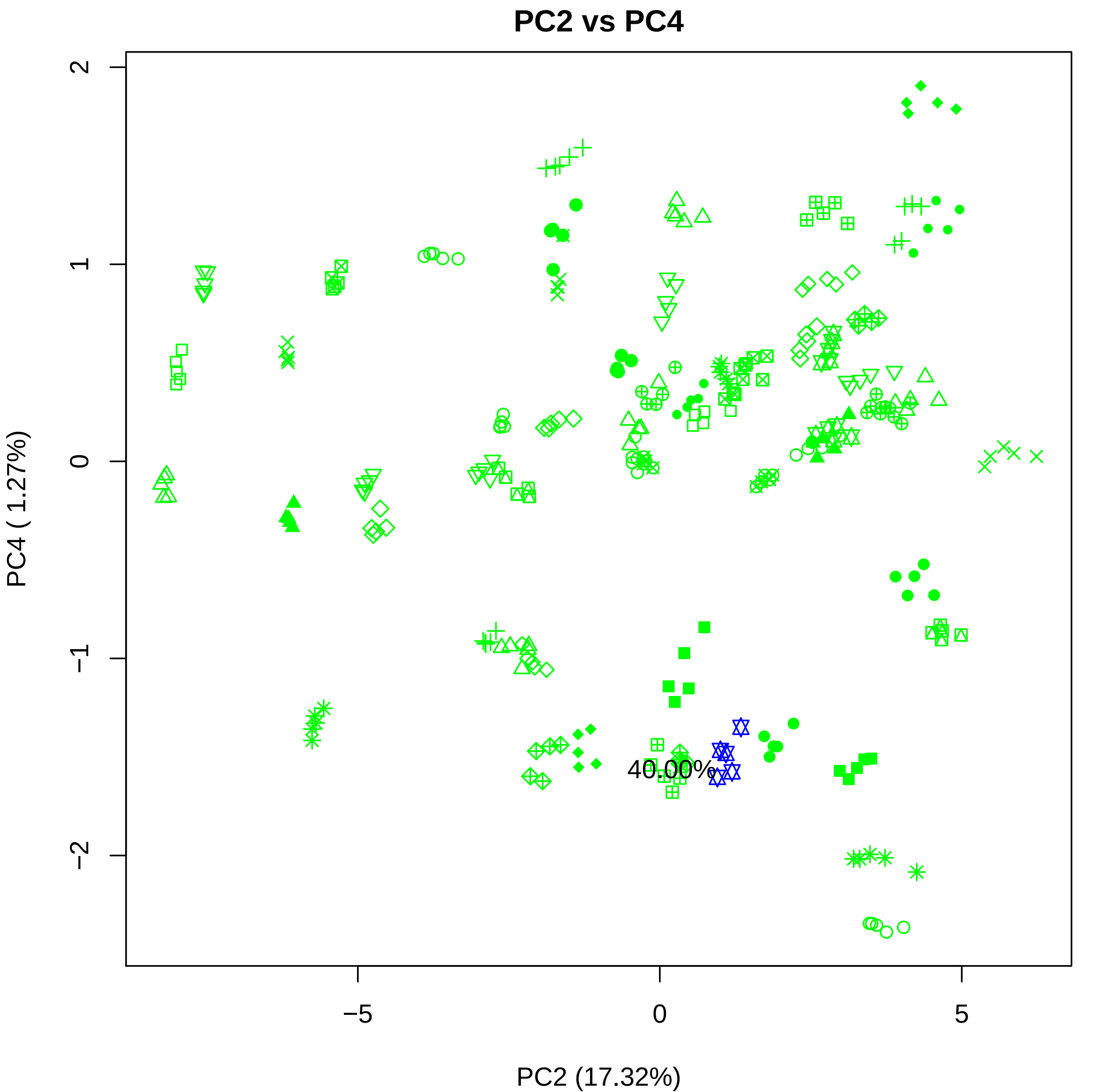
PC2 vs PC4
PC4 ( 1.27%)
PC2 (17.32%)
40.00%
2
1
0
−1
−2
−5
0
5

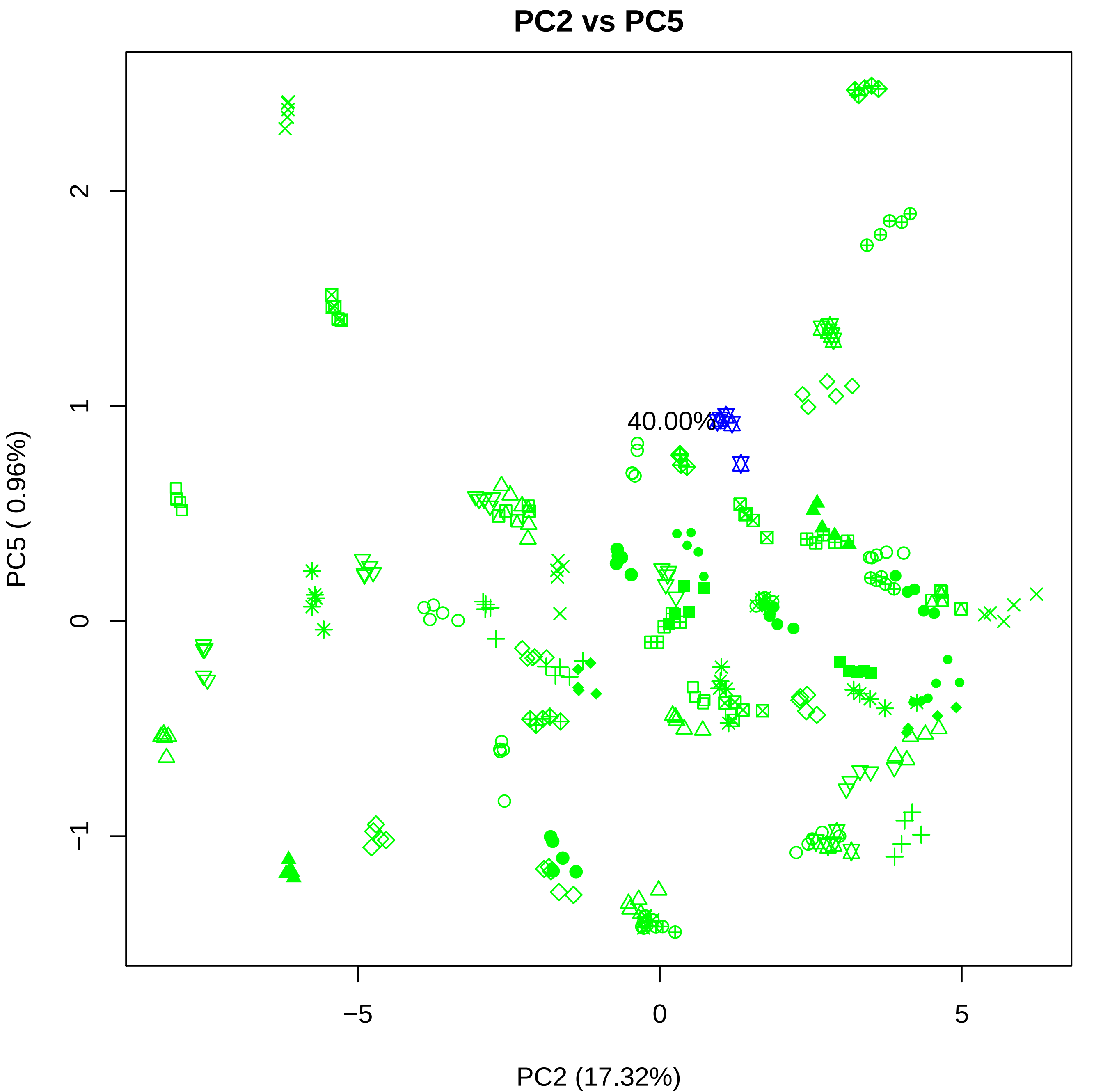
PC2 vs PC5
40.00%
PC5 ( 0.96%)
PC2 (17.32%)
2
1
0
−1
−5
0
5

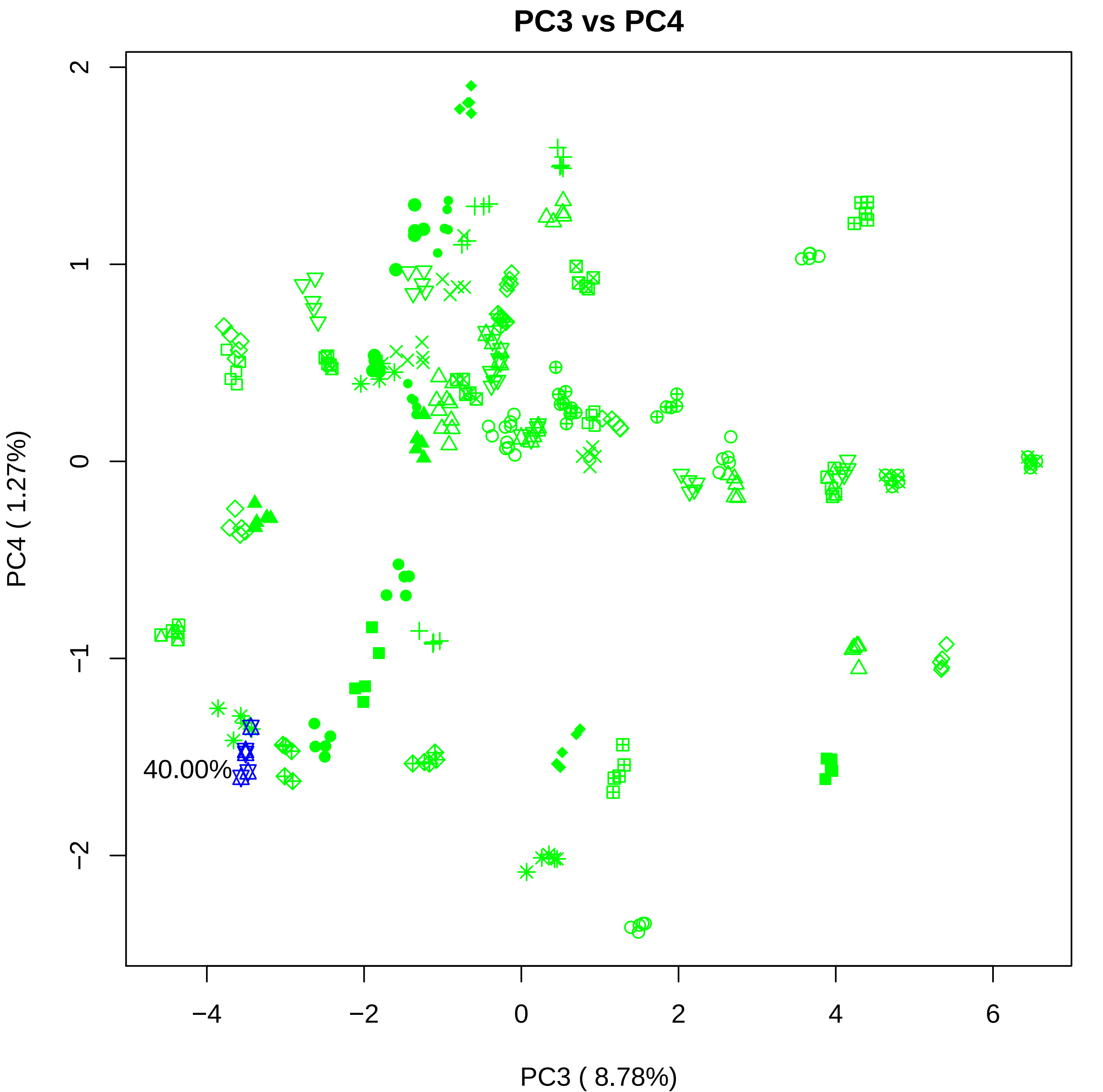
PC3 vs PC4
PC4 ( 1.27%)
PC3 ( 8.78%)
40.00%
2
1
0
−1
−2
−4
−2
0
2
4
6

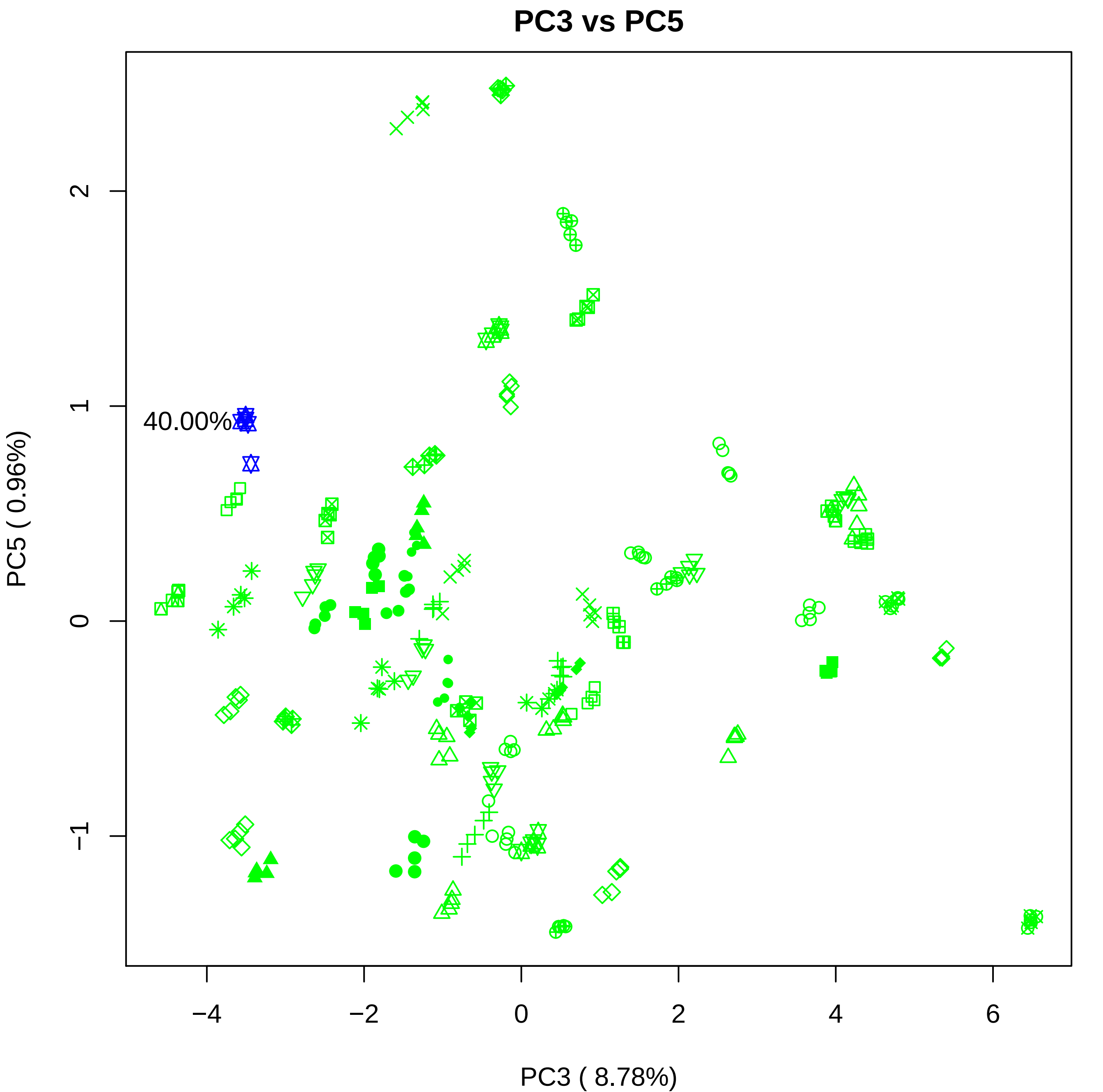
PC3 vs PC5
40.00%
PC5 ( 0.96%)
PC3 ( 8.78%)
−4
−2
0
2
4
6
−1
0
1
2

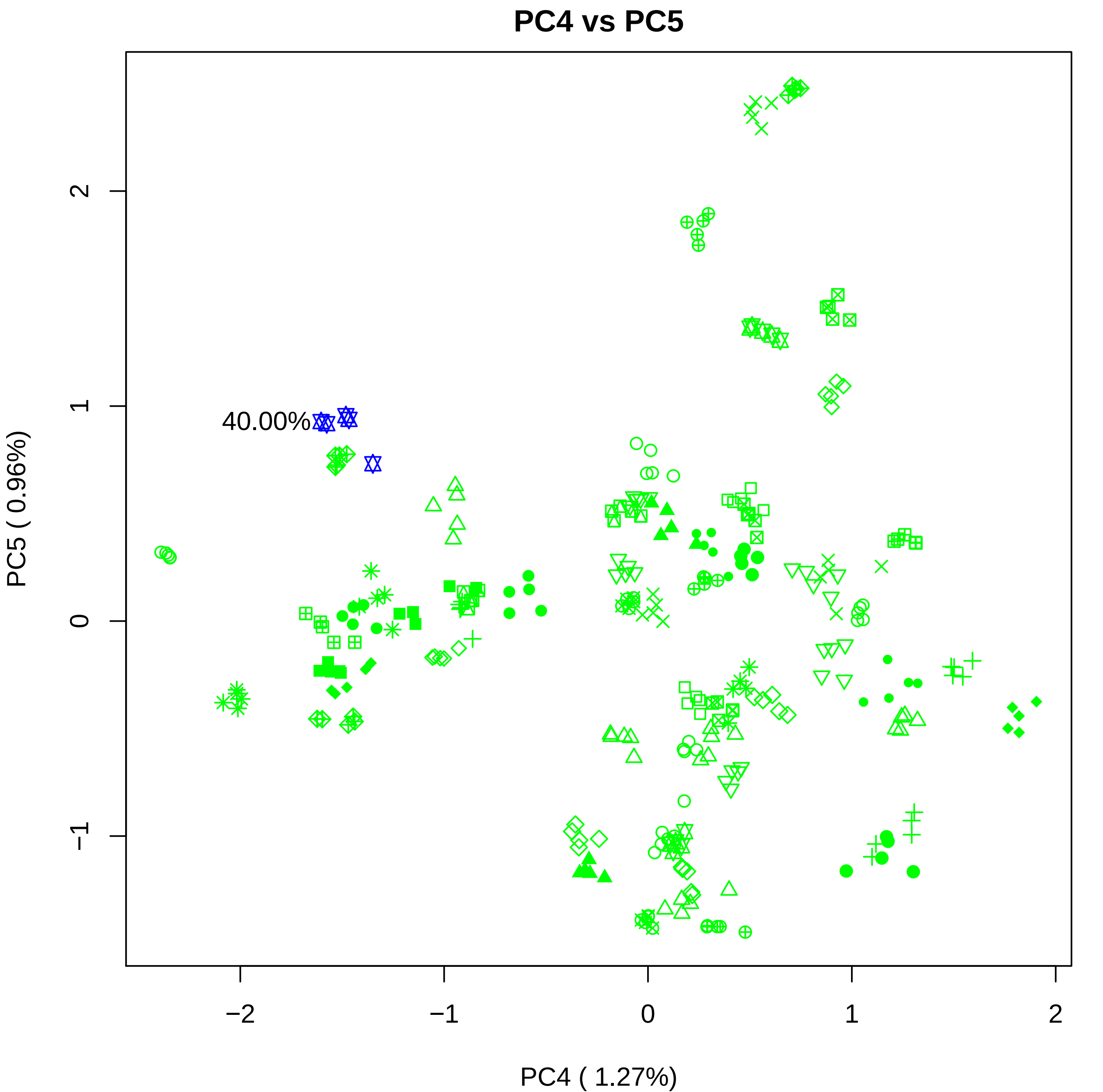

PC4 vs PC5
40.00%
PC5 ( 0.96%)
PC4 ( 1.27%)
2
1
0
−1
−2
−1
0
1
2

### PCA of PMH vs PMH(c) contaminated with BMH(c)

Scaled Matrix PCAed

7th order polynomial baseline correction performed

Raw spectra smoothed to 21 Neighbours, 2nd order PN, Savtisky Golay

Normalisation between 0 and 100

D2 spectra smoothed to 41 Neighbours, 2nd order PN, Savtisky Golay

Regions Analysed: 3601−2501 and 2001−700cm−1

Samples Analysed: 8 bContam's 0 BLH's 0 BMH's 0 OMH's 0 pContam's 0 PLH's 60 PMH's

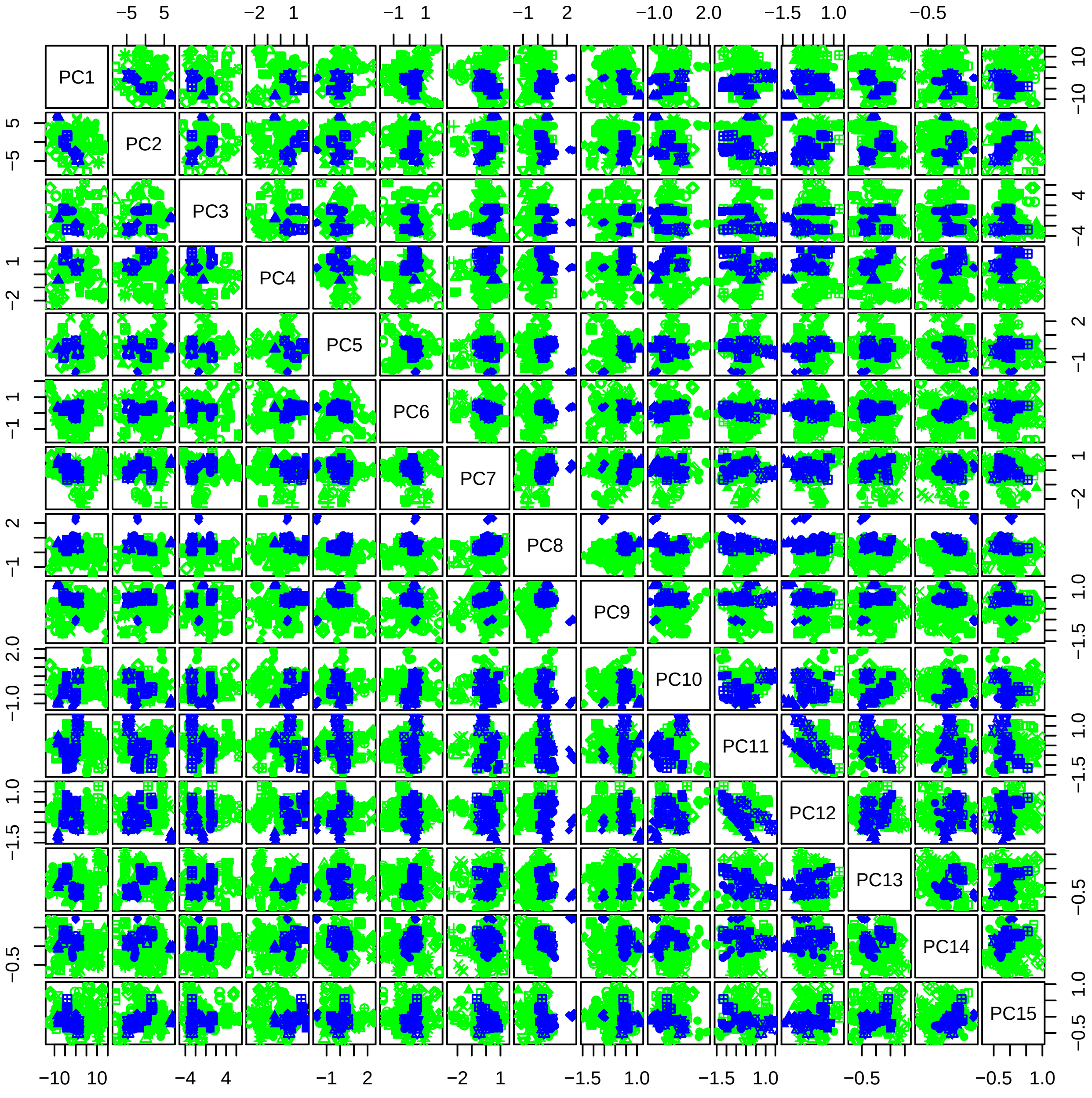
PC1
PC2
PC3
PC4
PC5
PC6
PC7
PC8
PC9
PC10
PC11
PC12
PC13
PC14
PC15
−5 5
−2 1
−1 1
−1 2
−1.0 2.0
−1.5 1.0
−0.5
10
−10
5
−5
4
−4
1
−2
2
−1
1
−1
1
−2
2
−1
1.0
−1.5
2.0
−1.0
1.0
−1.5
1.0
−1.5
−0.5
−0.5
1.0
−0.5
−10 10
−4 4
−1 2
−2 1
−1.5 1.0
−1.5 1.0
−0.5
−0.5 1.0

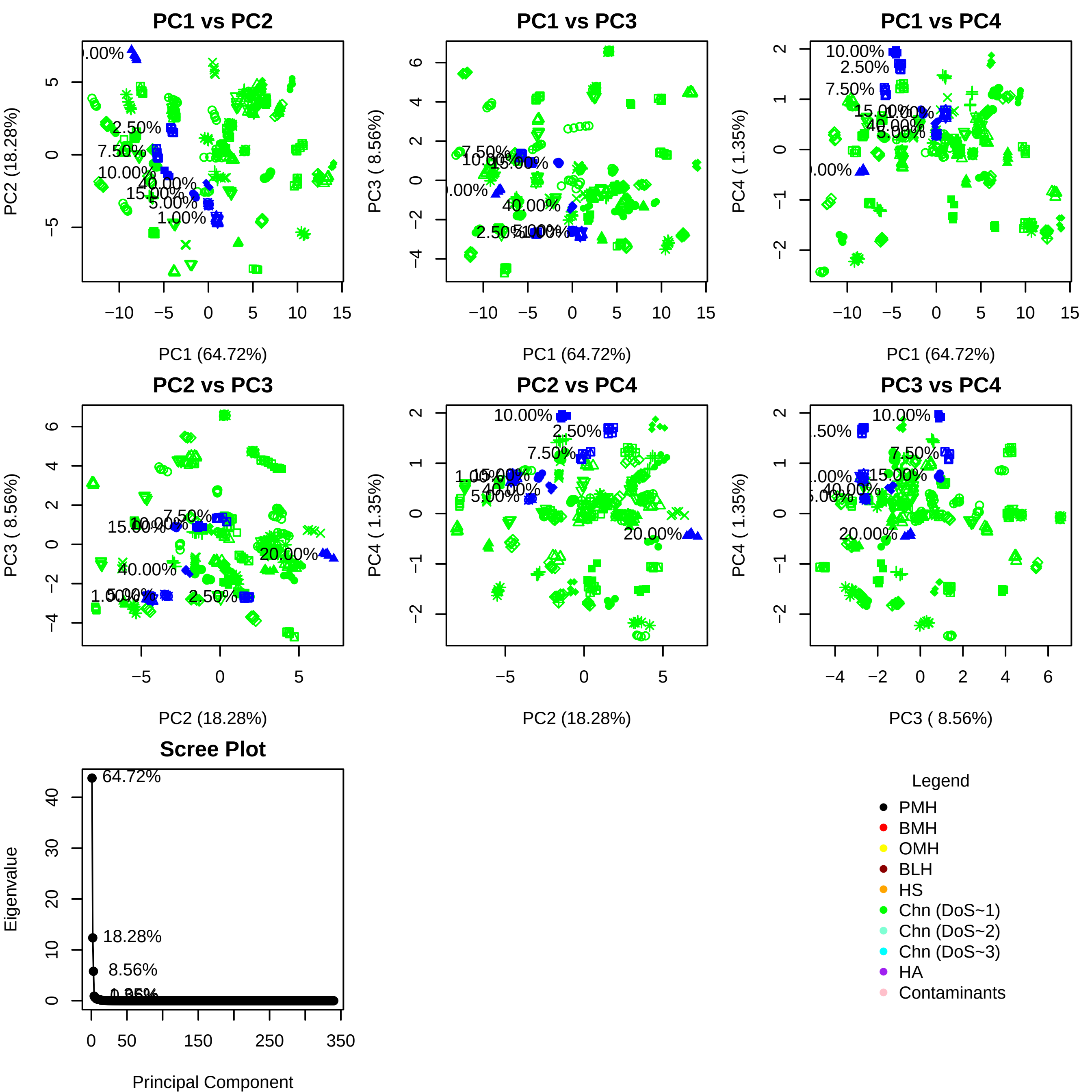
PC1 vs PC2
PC1 vs PC3
PC1 vs PC4
PC2 vs PC3
PC2 vs PC4
PC3 vs PC4
PC1 (64.72%)
PC2 (18.28%)
PC3 ( 8.56%)
PC4 ( 1.35%)
Scree Plot
Eigenvalue
Principal Component
64.72%
18.28%
8.56%
Legend
PMH
BMH
OMH
BLH
HS
Chn (DoS~1)
Chn (DoS~2)
Chn (DoS~3)
HA
Contaminants

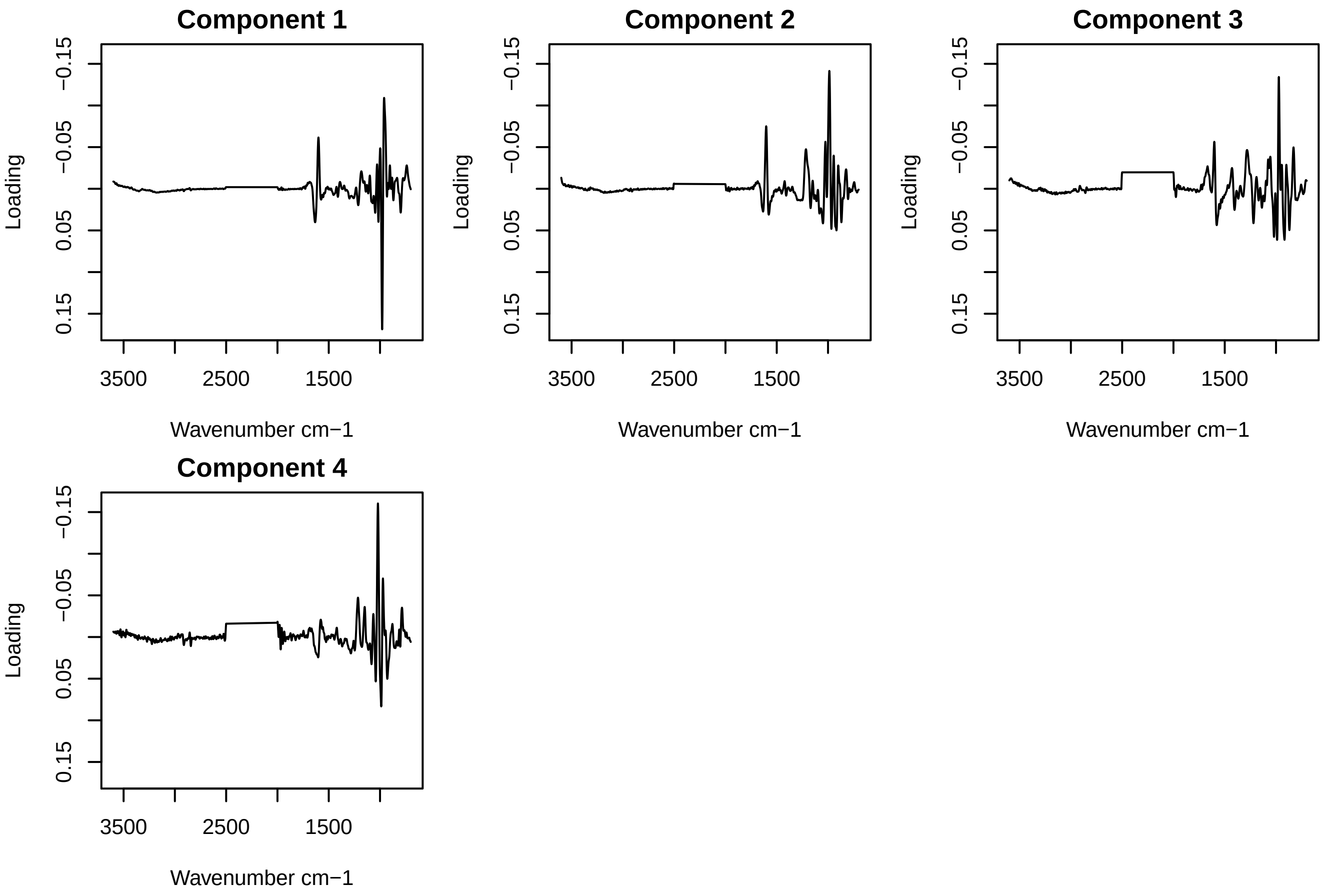
Component 1
Component 2
Component 3
Component 4
Loading
−0.15
−0.05
0.05
0.15
3500
2500
1500
Wavenumber cm−1

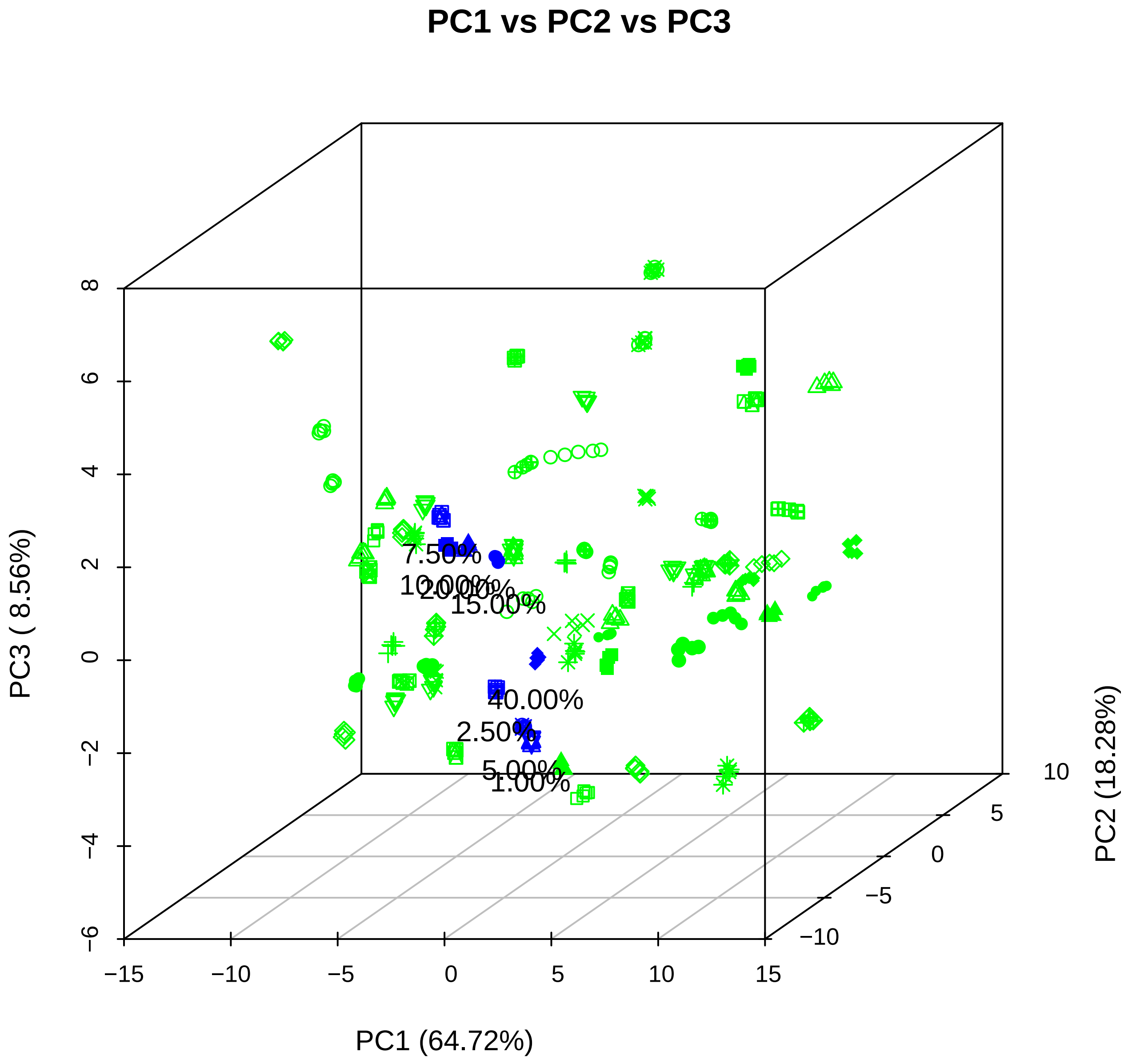

PC1 vs PC2 vs PC3
PC1 (64.72%)
PC2 (18.28%)
PC3 ( 8.56%)
−15
−10
−5
0
5
10
15
−10
−5
0
5
10
−6
−4
−2
0
2
4
6
8
7.50%
10.00%
20.00%
15.00%
40.00%
2.50%
5.00%
1.00%

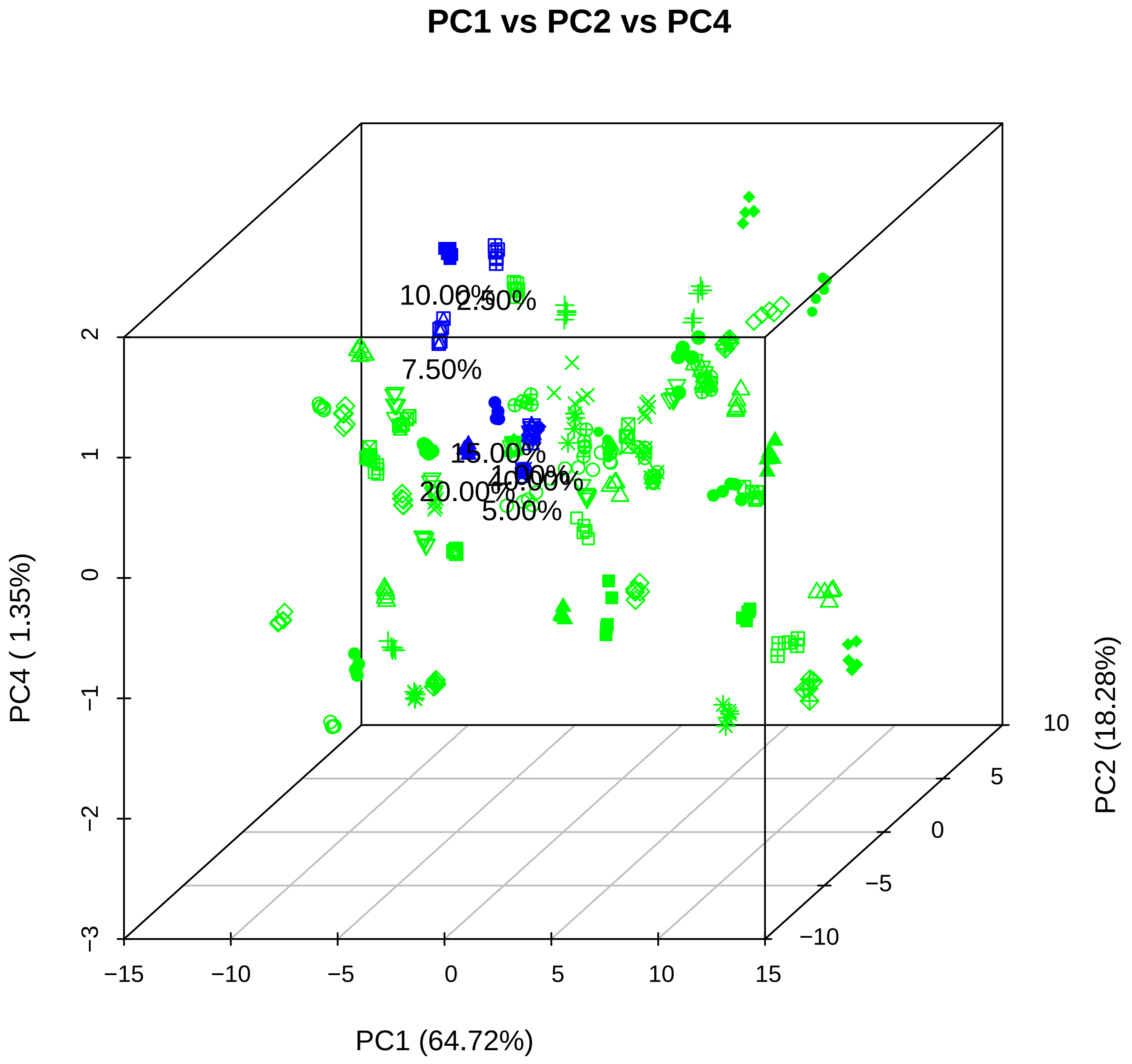

PC1 vs PC2 vs PC4
10.00%
2.50%
7.50%
15.00%
1.00%
40.00%
20.00%
5.00%
PC1 (64.72%)
PC2 (18.28%)
PC4 ( 1.35%)
−15
−10
−5
0
5
10
15
−10
−5
0
5
10
−3
−2
−1
0
1
2

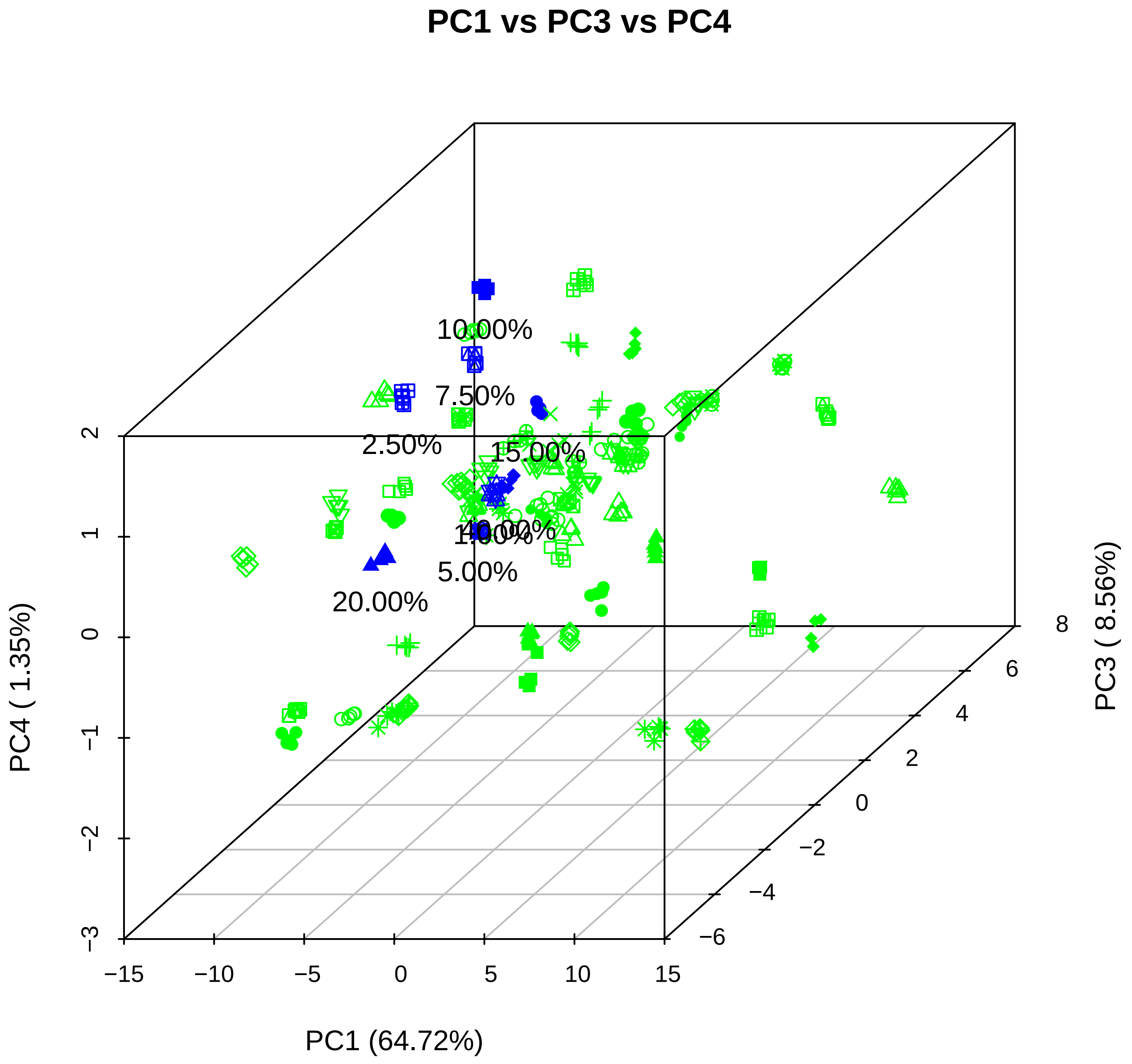

PC1 vs PC3 vs PC4
10.00%
7.50%
2.50%
15.00%
5.00%
20.00%
PC4 ( 1.35%)
PC3 ( 8.56%)
PC1 (64.72%)
−15
−10
−5
0
5
10
15
−3
−2
−1
0
1
2
−6
−4
−2
0
2
4
6
8

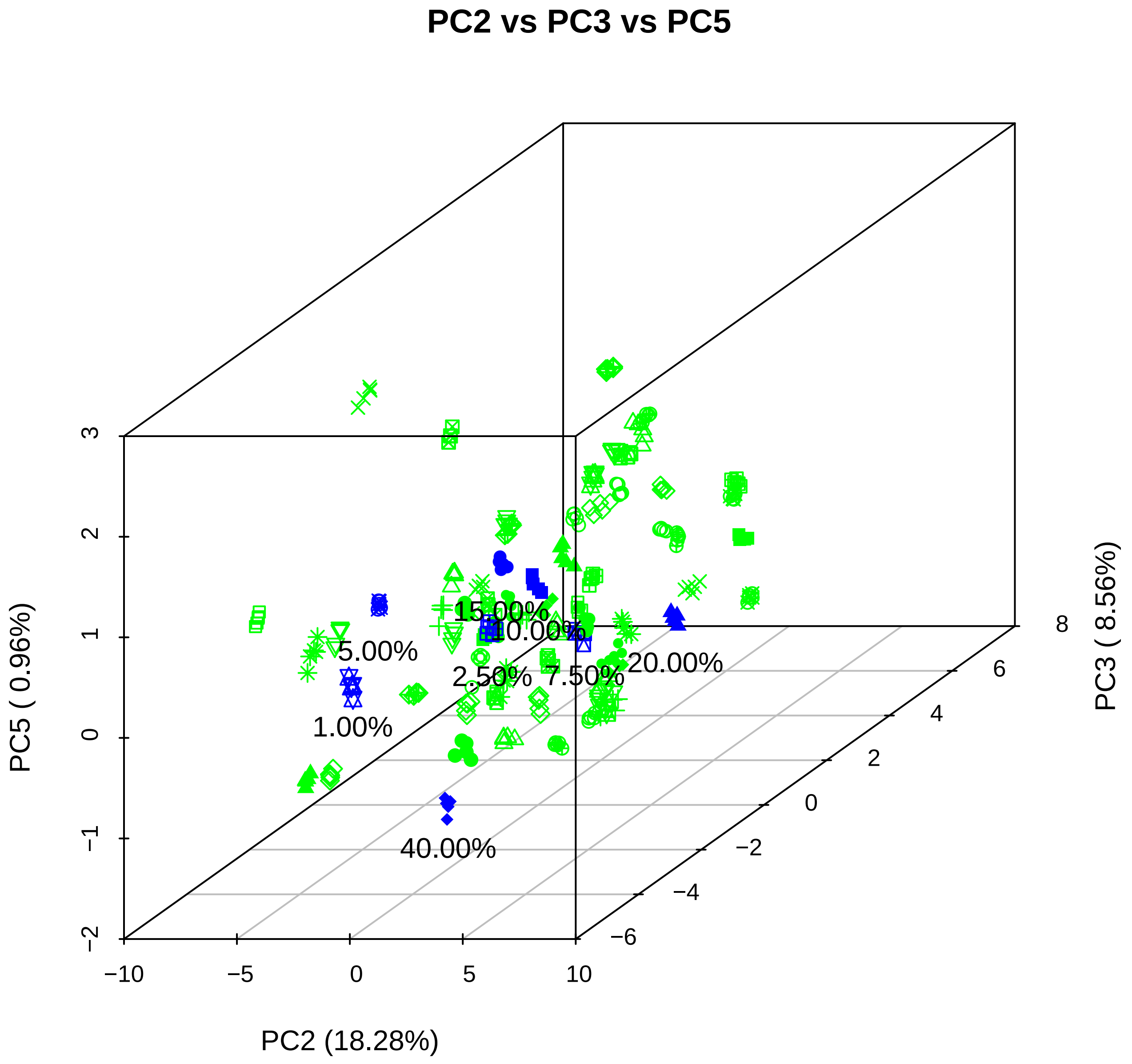

PC2 vs PC3 vs PC5
PC2 (18.28%)
PC3 ( 8.56%)
PC5 ( 0.96%)
−10
−5
0
5
10
−6
−4
−2
0
2
4
6
8
−2
−1
0
1
2
3
15.00%
10.00%
5.00%
2.50%
7.50%
20.00%
1.00%
40.00%

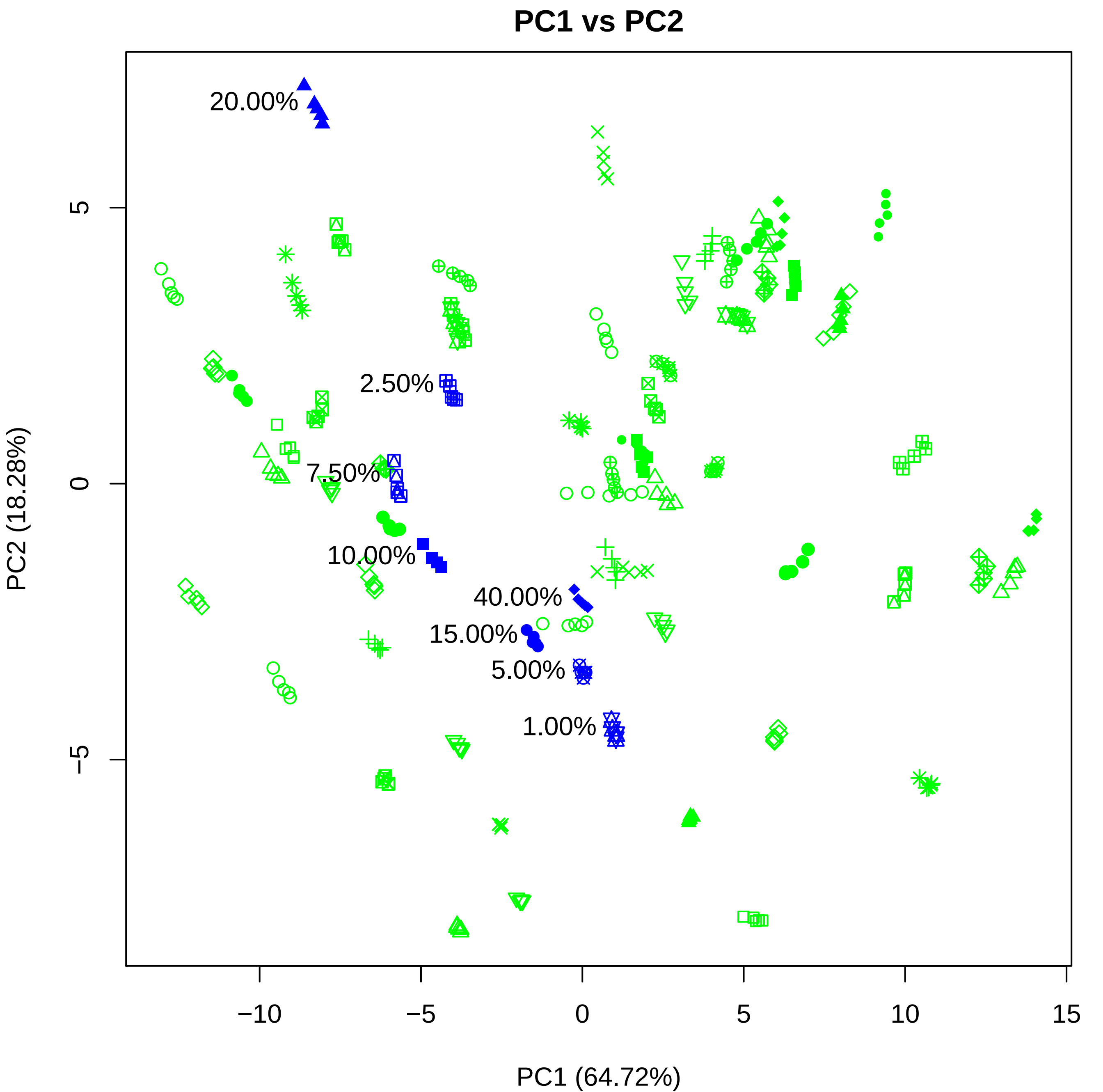
PC1 vs PC2
20.00%
2.50%
7.50%
10.00%
40.00%
15.00%
5.00%
1.00%
5
0
−5
−10
−5
0
5
10
15
PC2 (18.28%)
PC1 (64.72%)

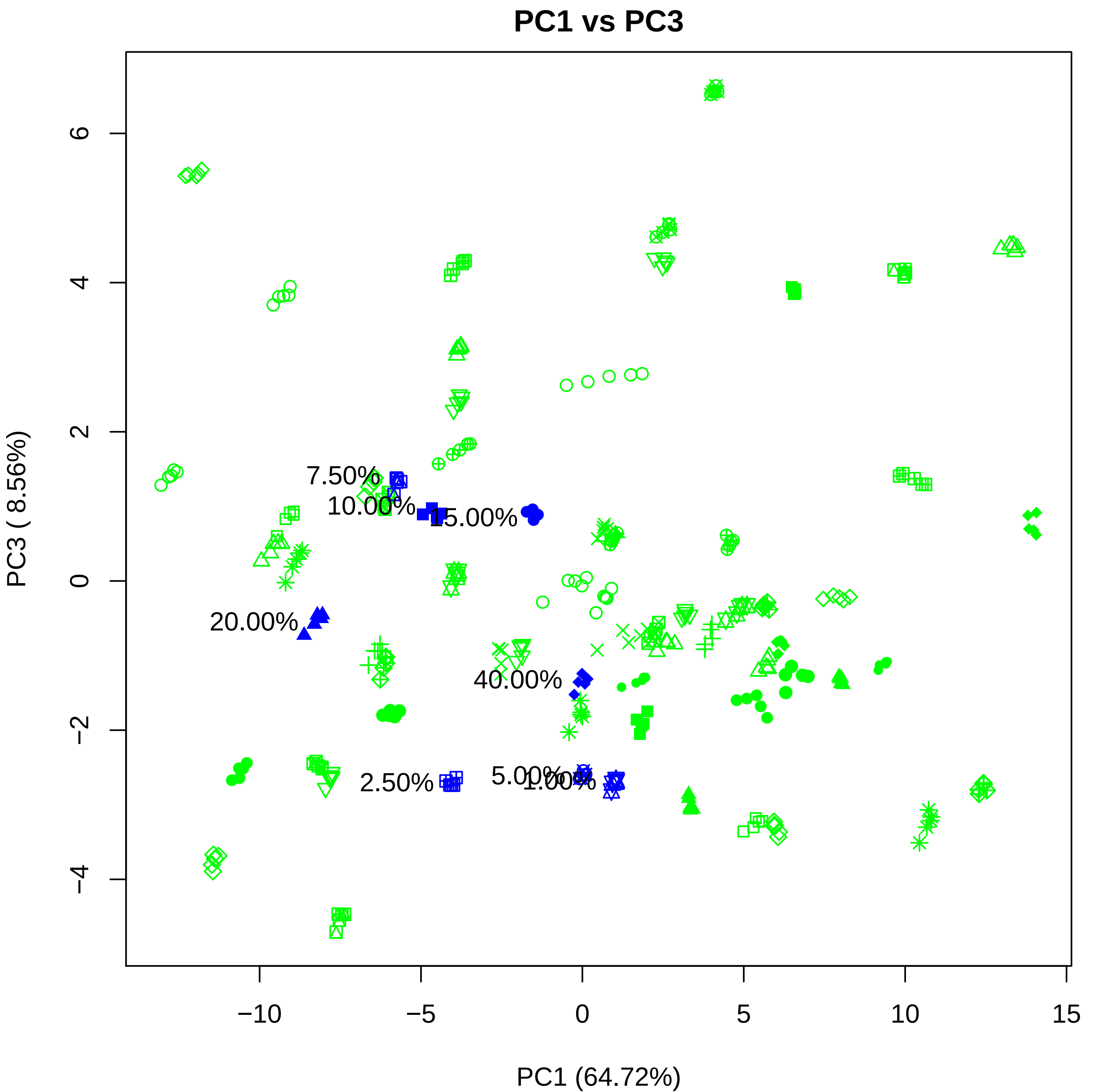

PC1 vs PC3
7.50%
10.00%
15.00%
20.00%
40.00%
2.50%
5.00%
1.00%
PC3 ( 8.56%)
PC1 (64.72%)
6
4
2
0
−2
−4
−10
−5
0
5
10
15

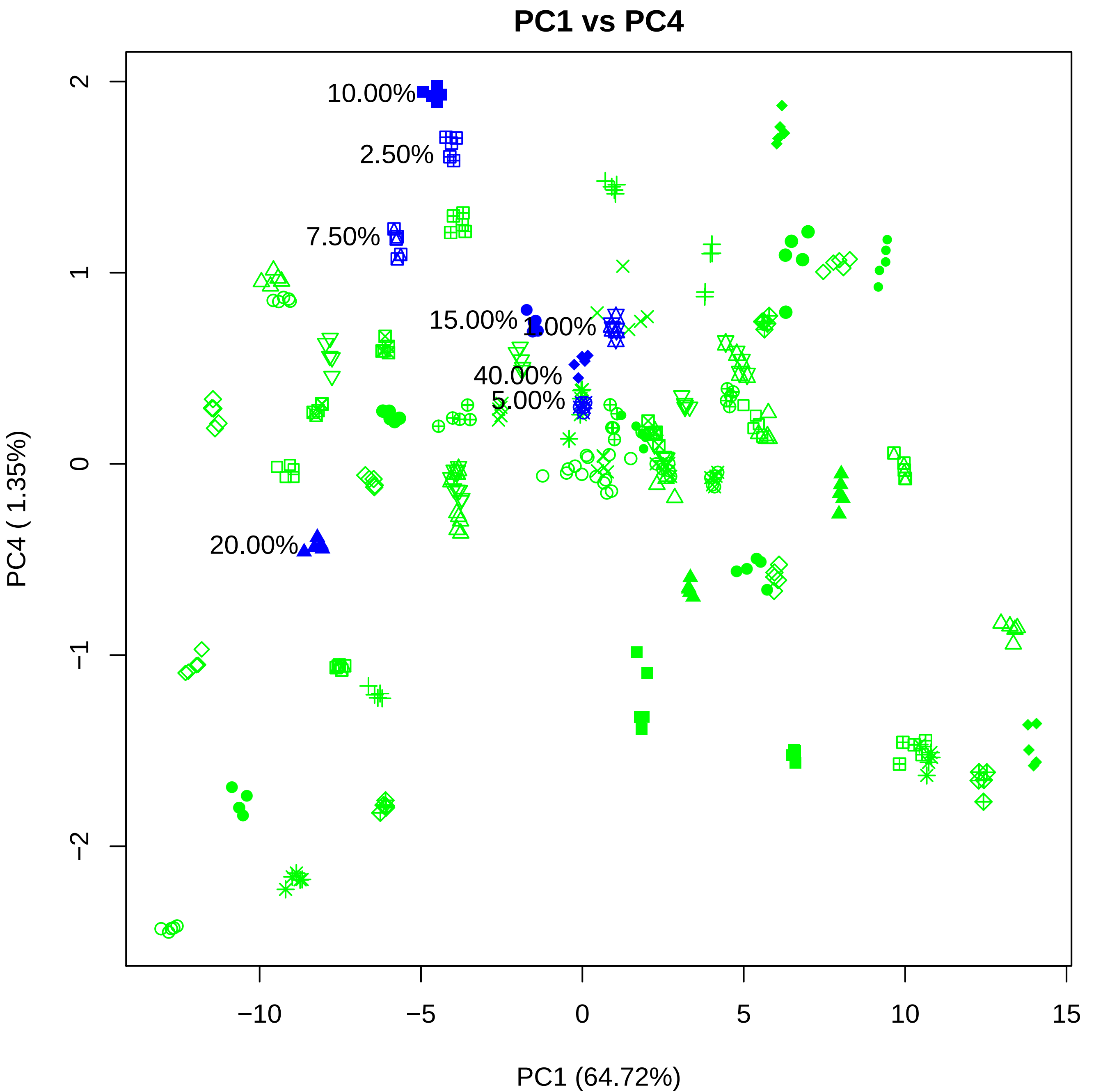

PC1 vs PC4
10.00%
2.50%
7.50%
15.00%
1.00%
40.00%
5.00%
20.00%
2
1
0
−1
−2
−10
−5
0
5
10
15
PC4 ( 1.35%)
PC1 (64.72%)

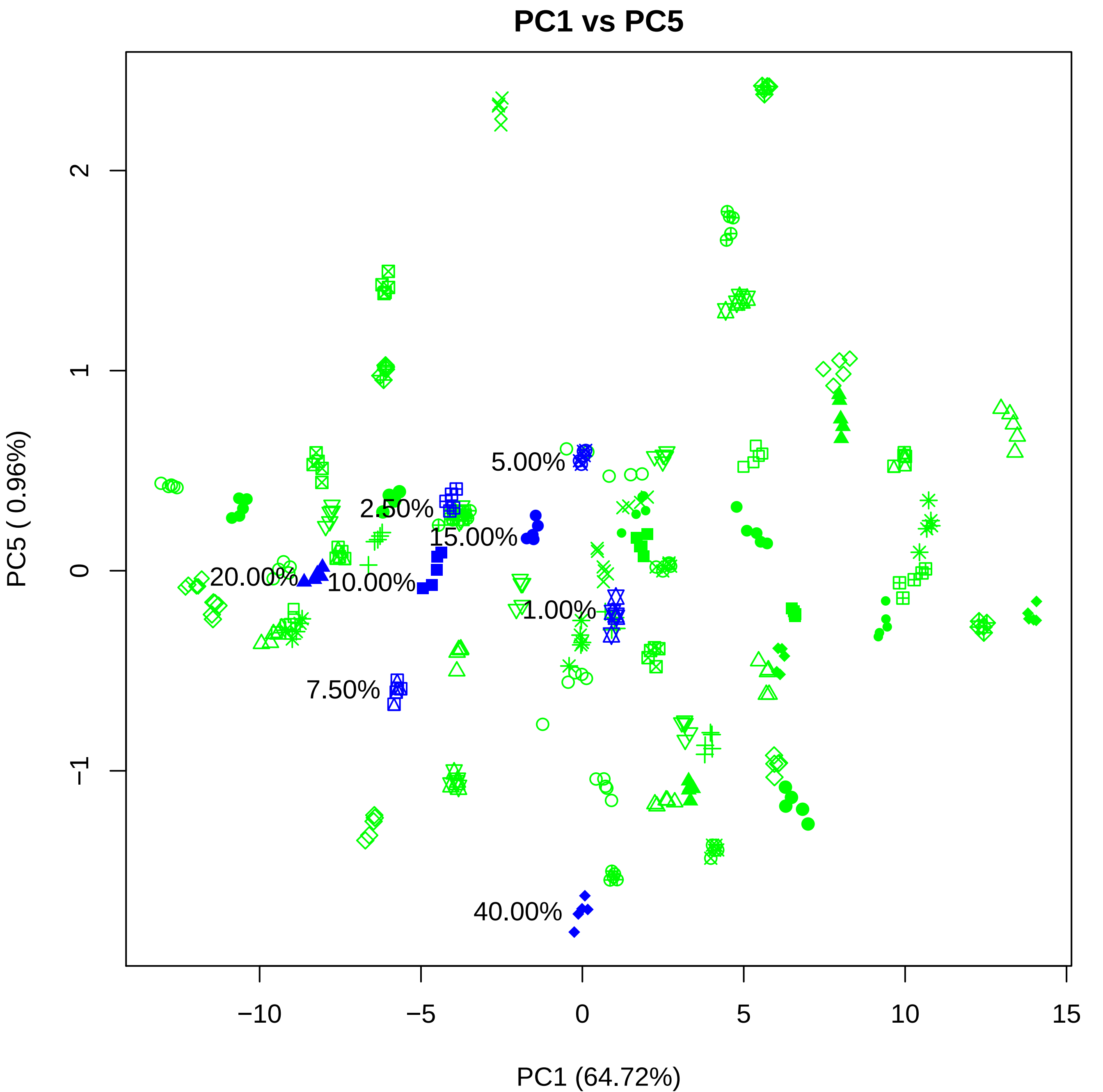

PC1 vs PC5
PC5 ( 0.96%)
PC1 (64.72%)
2
1
0
−1
−10
−5
0
5
10
15
5.00%
2.50%
15.00%
20.00%
10.00%
1.00%
7.50%
40.00%

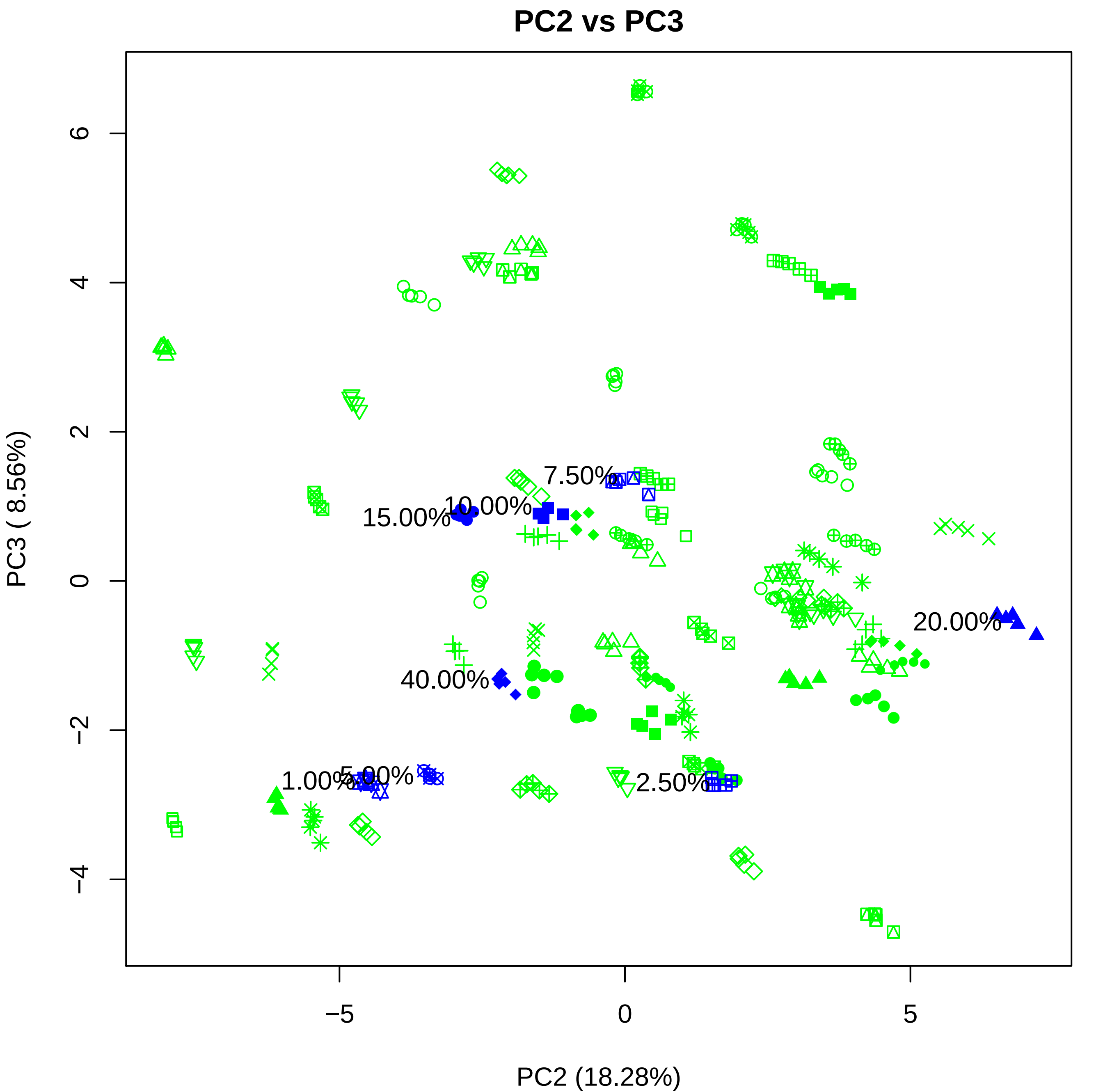

PC2 vs PC3
6
4
2
0
−2
−4
−5
0
5
PC3 ( 8.56%)
PC2 (18.28%)
7.50%
10.00%
15.00%
20.00%
40.00%
1.00%
5.00%
2.50%

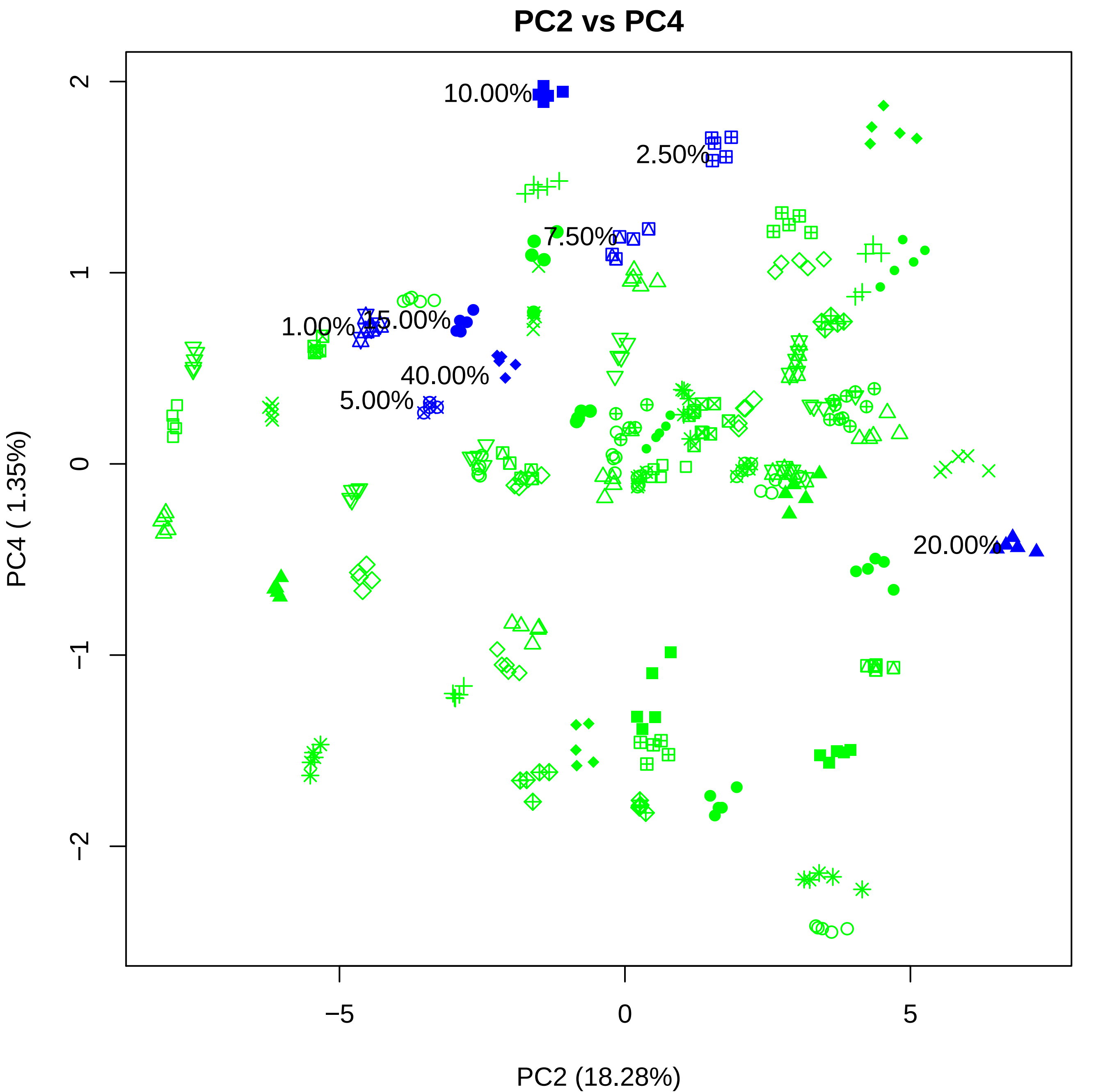
PC2 vs PC4
10.00%
2.50%
7.50%
1.00%
15.00%
40.00%
5.00%
20.00%
PC4 ( 1.35%)
PC2 (18.28%)
2
1
0
−1
−2
−5
0
5

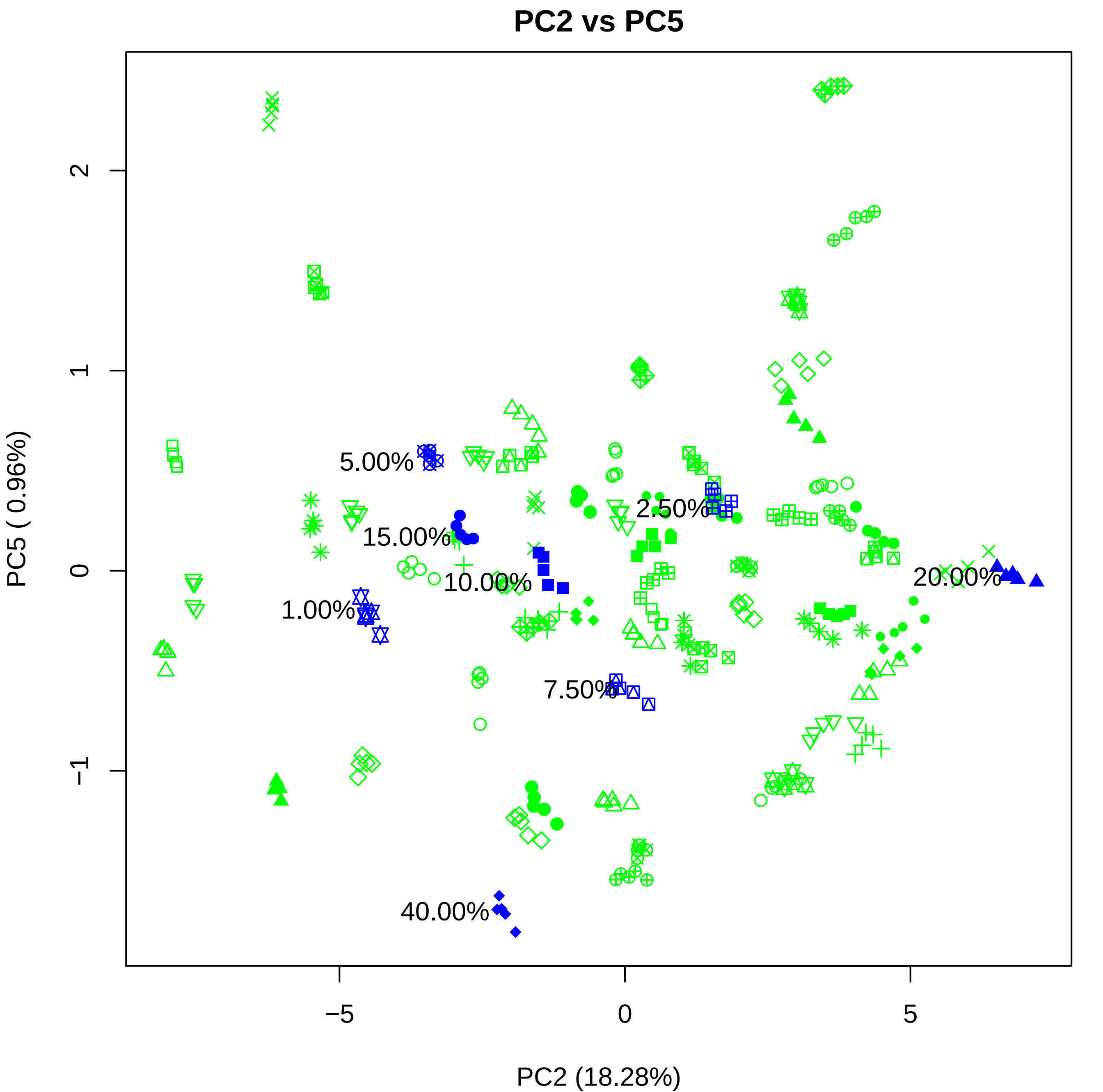
PC2 vs PC5
PC5 ( 0.96%)
PC2 (18.28%)
2
1
0
−1
−5
0
5
5.00%
15.00%
1.00%
10.00%
2.50%
7.50%
20.00%
40.00%

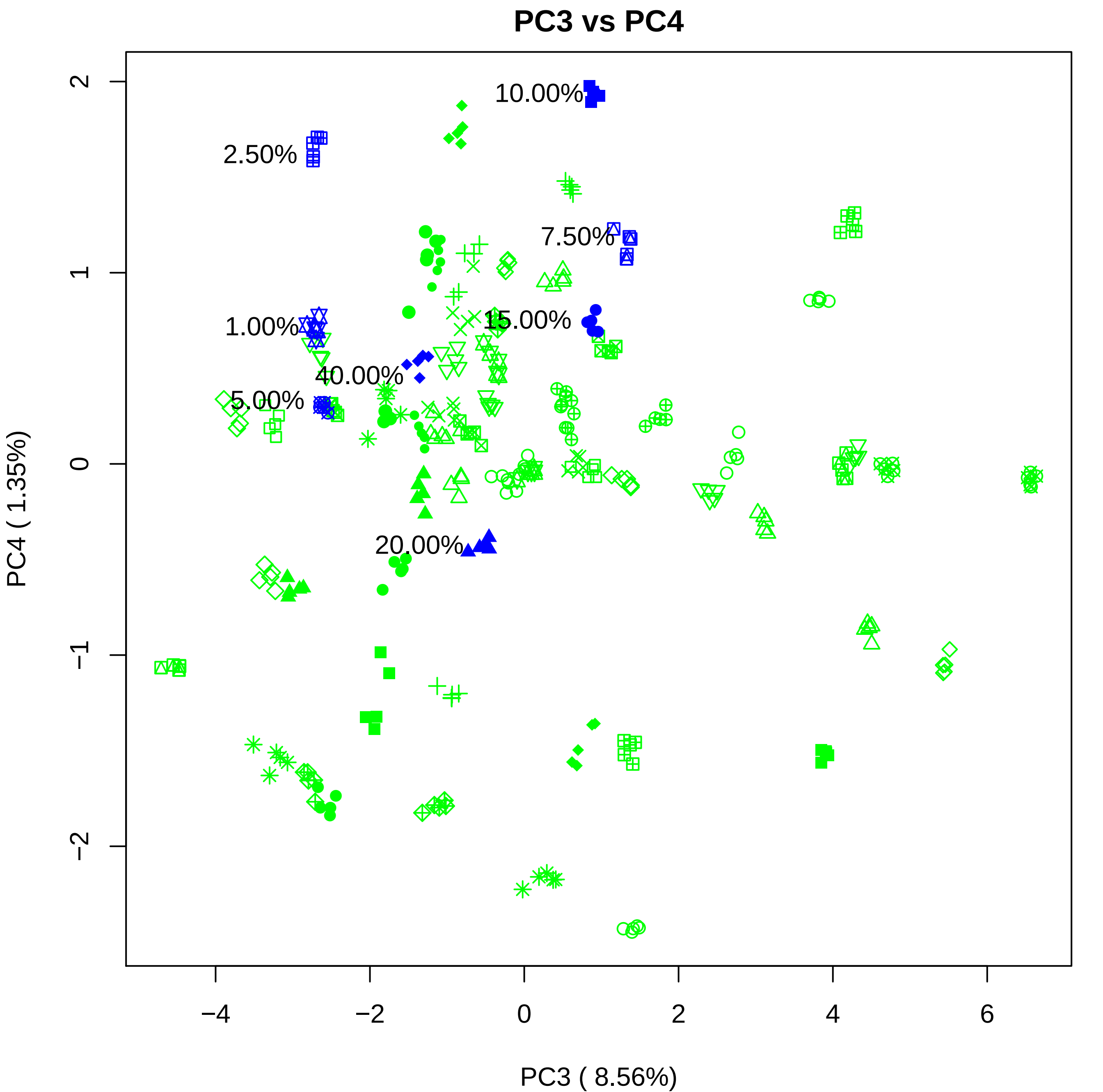
PC3 vs PC4
10.00%
2.50%
7.50%
1.00%
15.00%
40.00%
5.00%
20.00%
PC4 ( 1.35%)
PC3 ( 8.56%)
2
1
0
−1
−2
−4
−2
0
2
4
6

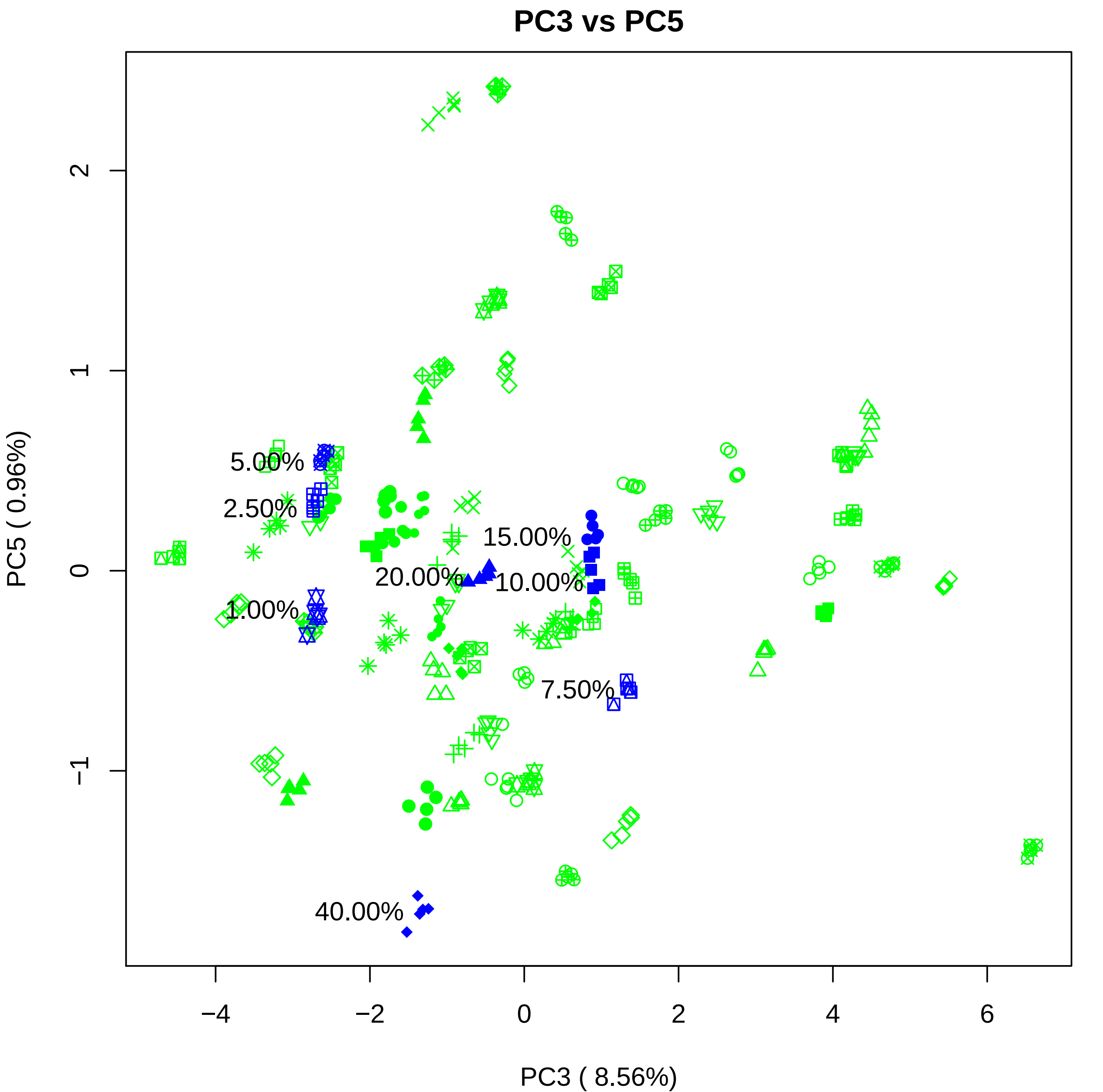
PC3 vs PC5
PC5 ( 0.96%)
PC3 ( 8.56%)
5.00%
2.50%
15.00%
20.00%
10.00%
1.00%
7.50%
40.00%
−4
−2
0
2
4
6
2
1
0
−1

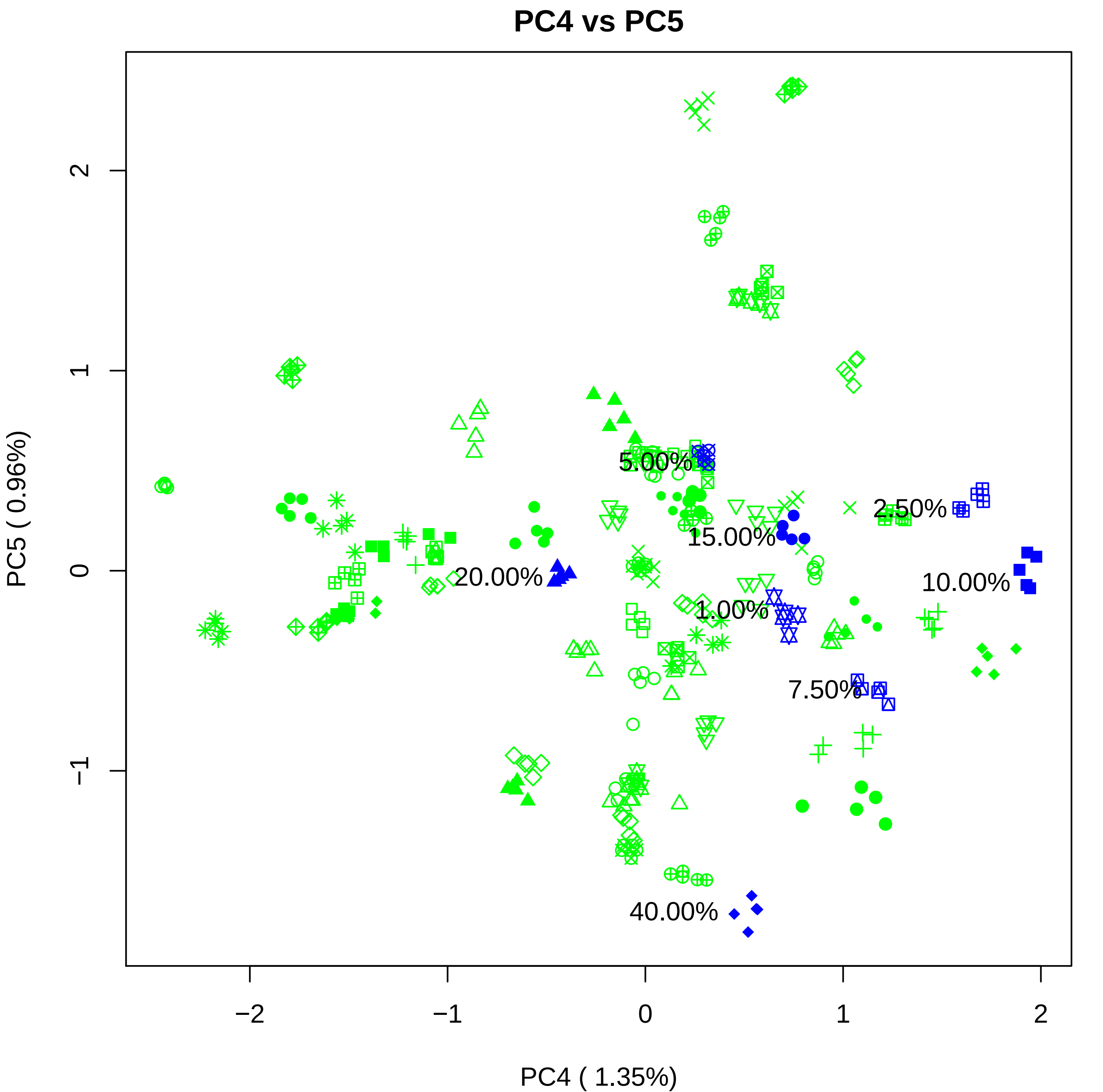
PC4 vs PC5
PC5 ( 0.96%)
PC4 ( 1.35%)
−2
−1
0
1
2
5.00%
2.50%
15.00%
20.00%
10.00%
1.00%
7.50%
40.00%

### PCA of 1%  BMH (c)  vs PMH (c)

Scaled Matrix PCAed

7th order polynomial baseline correction performed

Raw spectra smoothed to 21 Neighbours, 2nd order PN, Savtisky Golay

Normalisation between 0 and 100

D2 spectra smoothed to 41 Neighbours, 2nd order PN, Savtisky Golay

Regions Analysed: 3601−2501 and 2001−700cm−1

Samples Analysed: 1 bContam's 0 BLH's 0 BMH's 0 OMH's 0 pContam's 0 PLH's 60 PMH's

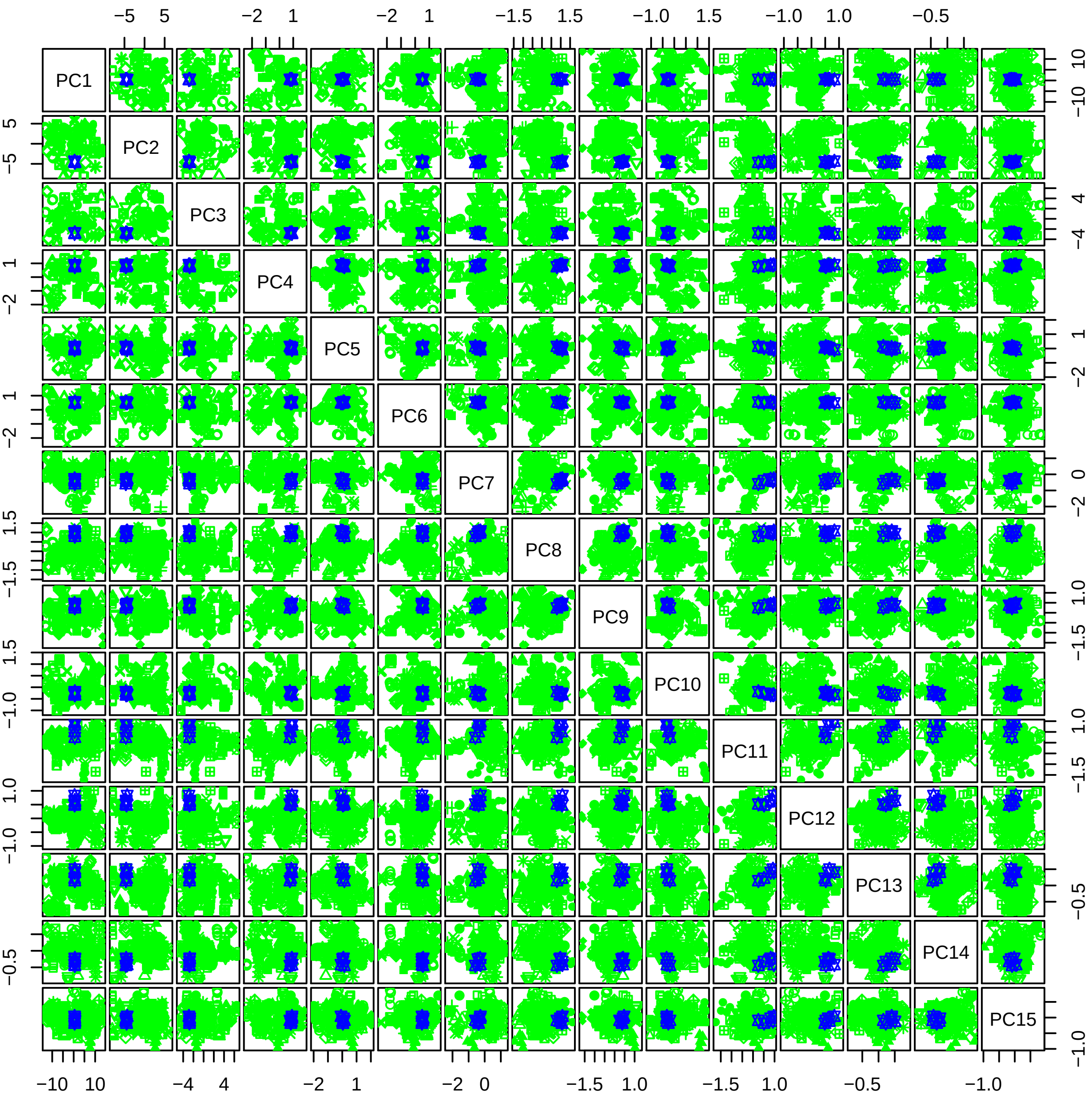
PC1
PC2
PC3
PC4
PC5
PC6
PC7
PC8
PC9
PC10
PC11
PC12
PC13
PC14
PC15

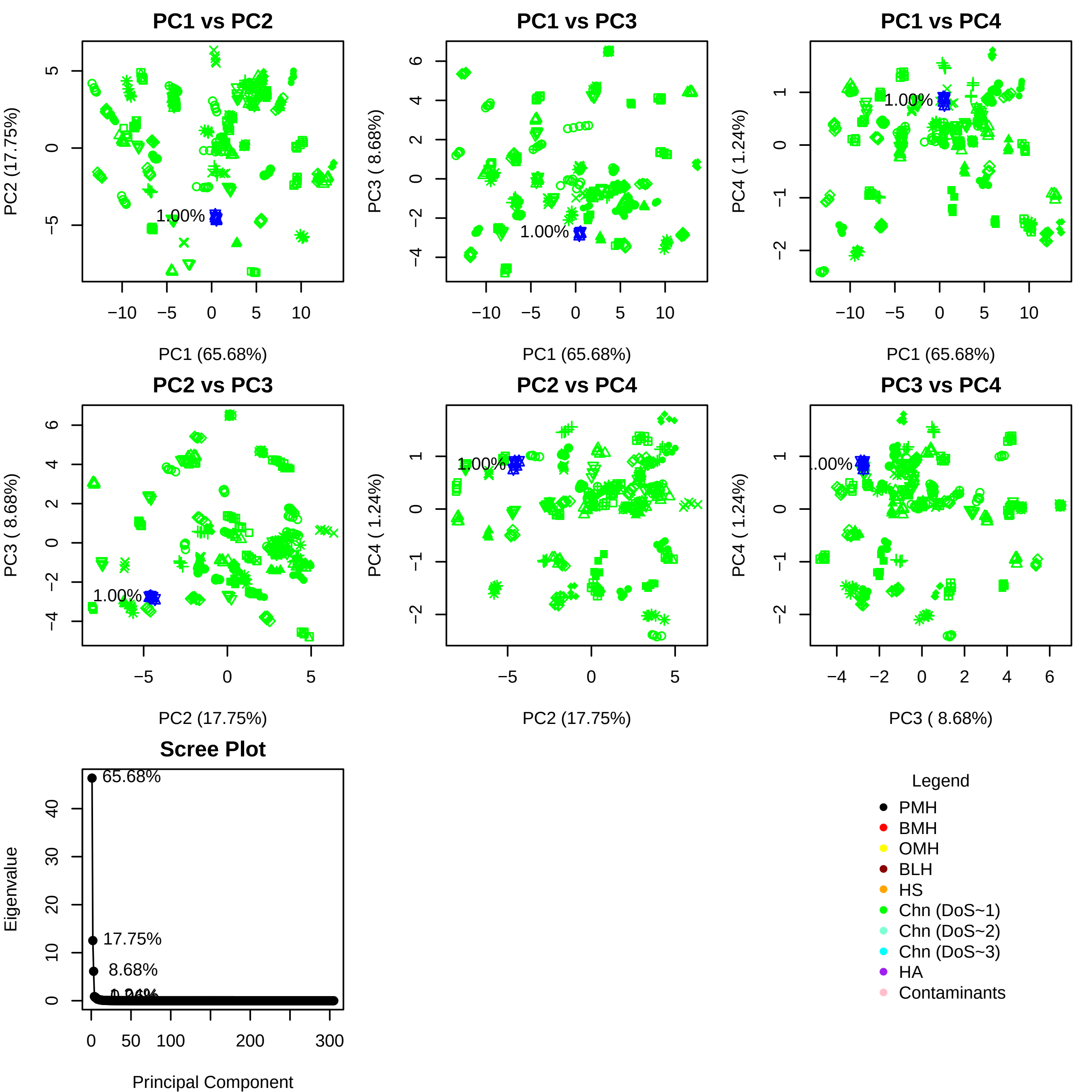

PC1 vs PC2
PC1 vs PC3
PC1 vs PC4
PC2 vs PC3
PC2 vs PC4
PC3 vs PC4
Scree Plot
PC1 (65.68%)
PC2 (17.75%)
PC3 ( 8.68%)
PC4 ( 1.24%)
1.00%
Eigenvalue
Principal Component
65.68%
17.75%
8.68%
Legend
PMH
BMH
OMH
BLH
HS
Chn (DoS~1)
Chn (DoS~2)
Chn (DoS~3)
HA
Contaminants

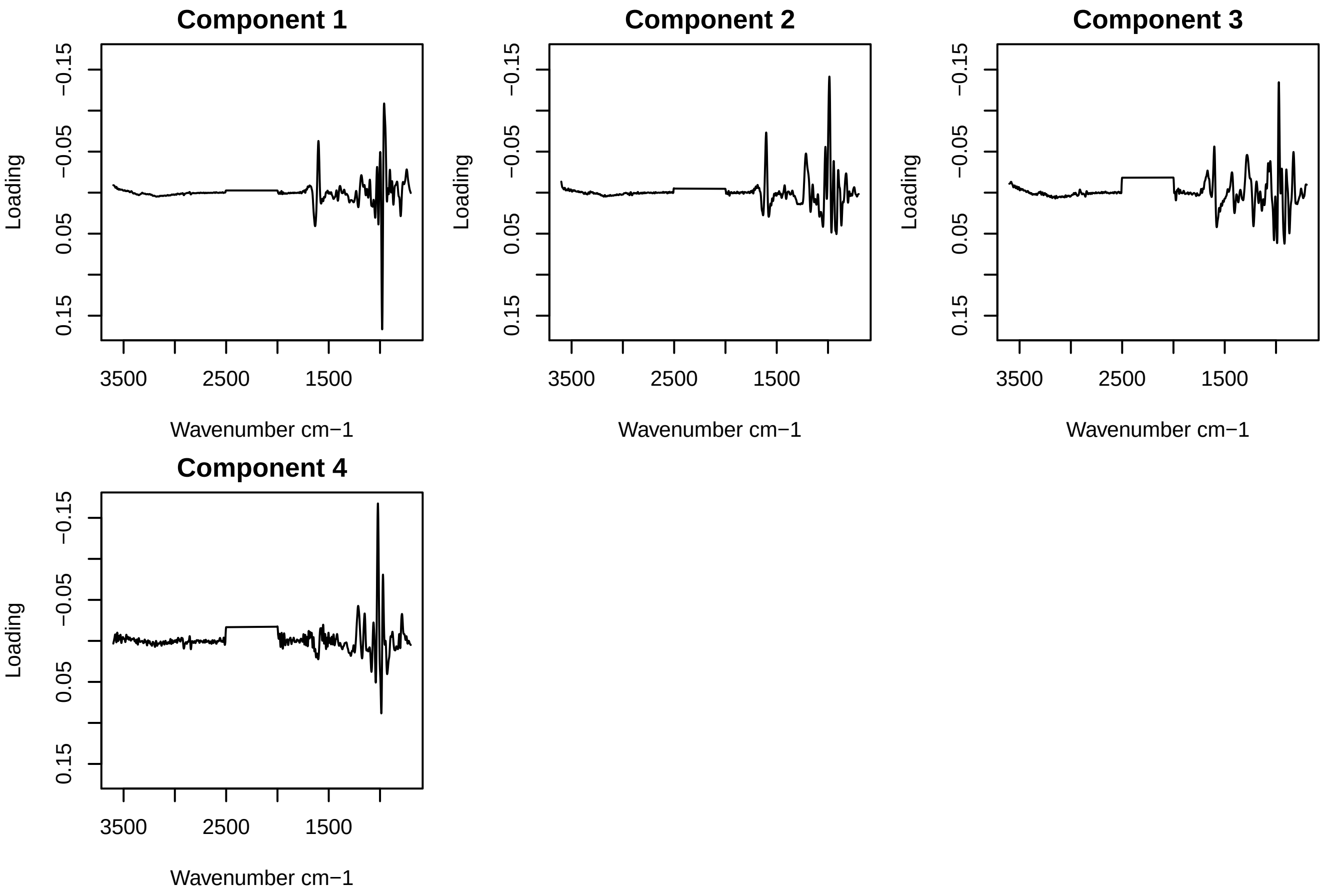
Component 1
Component 2
Component 3
Component 4
Loading
−0.15
−0.05
0.05
0.15
3500
2500
1500
Wavenumber cm−1

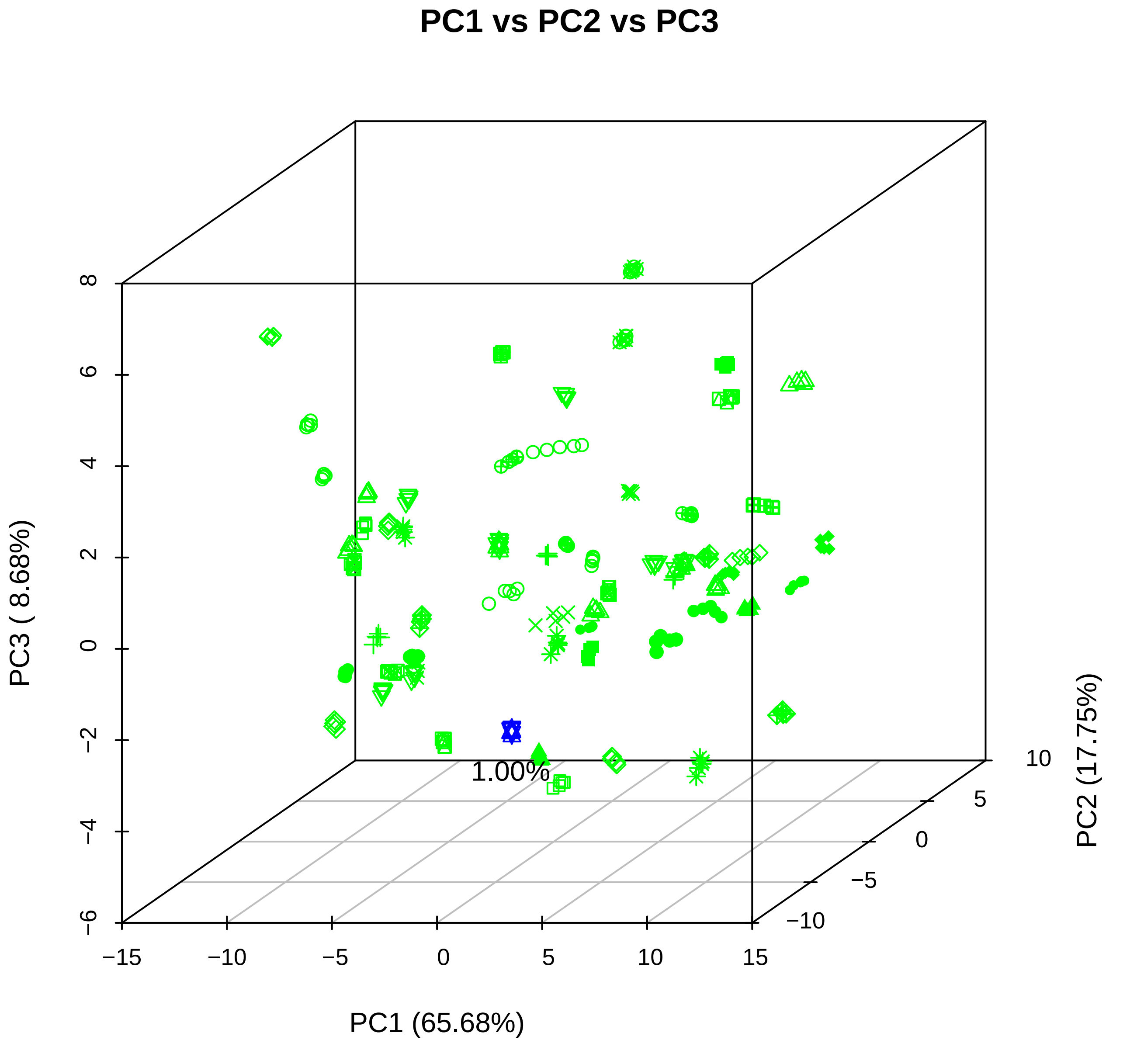
PC1 vs PC2 vs PC3
PC1 (65.68%)
PC2 (17.75%)
PC3 ( 8.68%)
1.00%
−15
−10
−5
0
5
10
15
−6
−4
−2
0
2
4
6
8
−10
−5
0
5
10

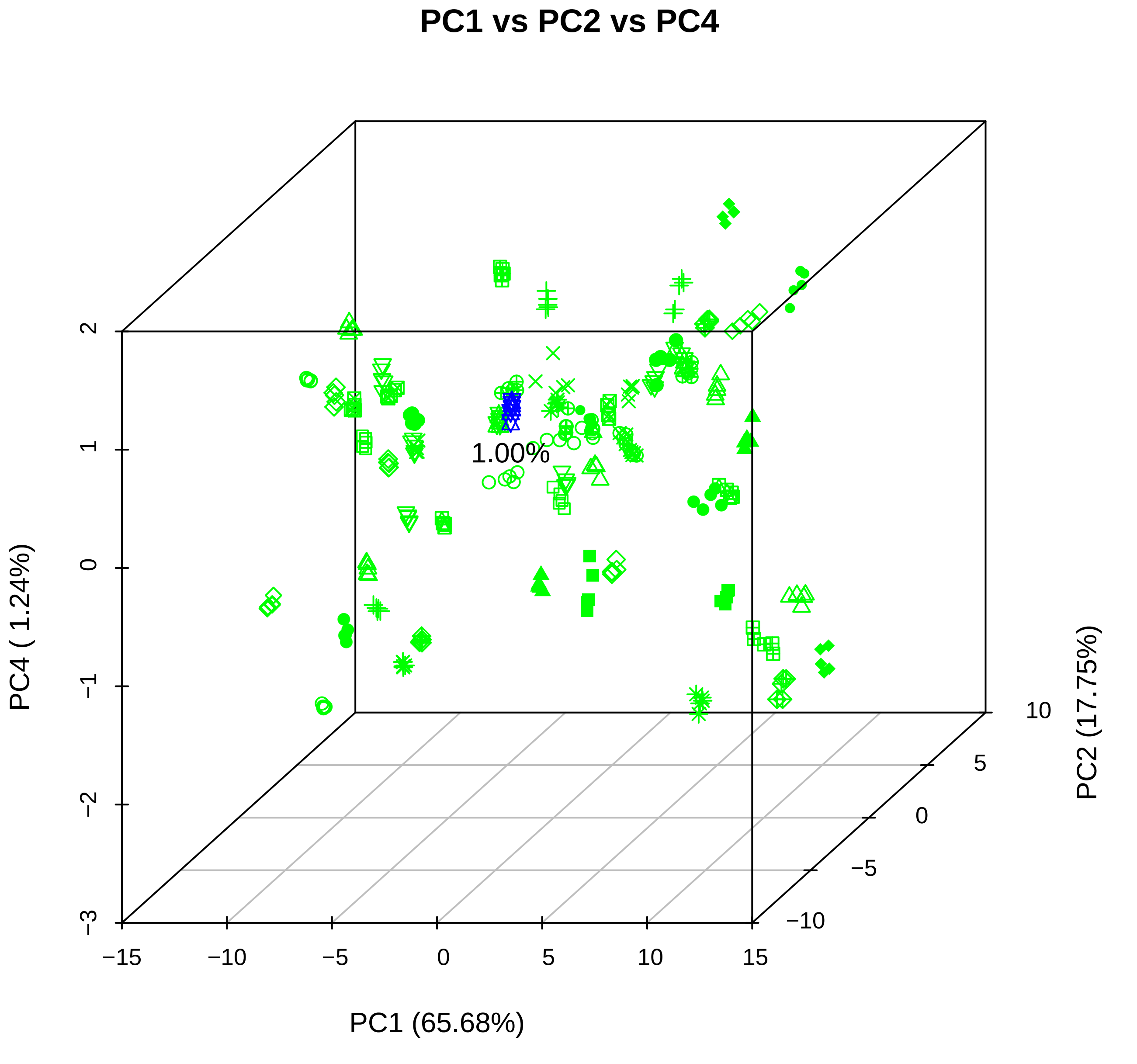

PC1 vs PC2 vs PC4
1.00%
PC4 ( 1.24%)
2
1
0
−1
−2
−3
−15
−10
−5
0
5
10
15
PC1 (65.68%)
PC2 (17.75%)
10
5
0
−5
−10

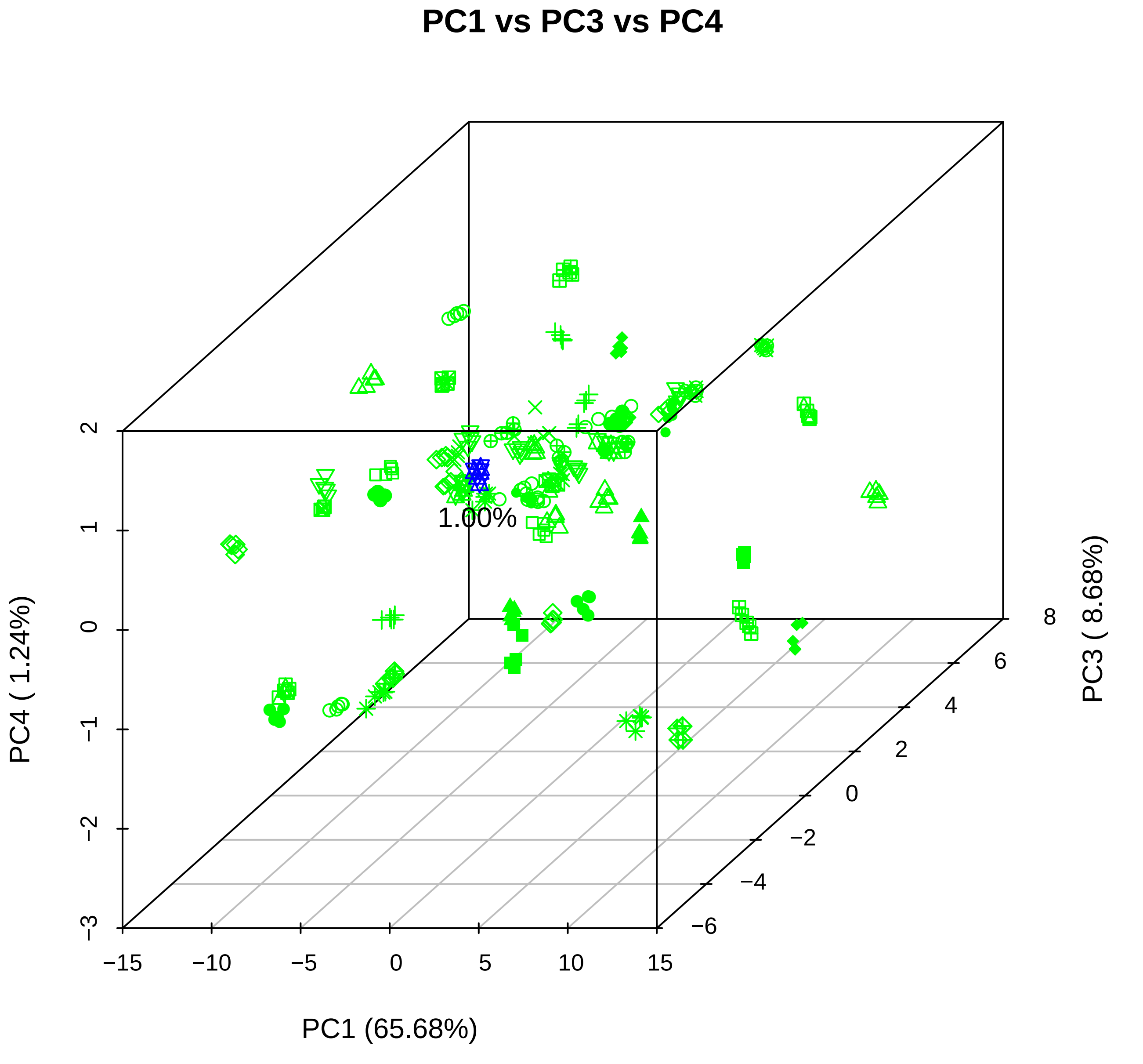

PC1 vs PC3 vs PC4
1.00%
PC1 (65.68%)
PC4 ( 1.24%)
PC3 ( 8.68%)
−15
−10
−5
0
5
10
15
−3
−2
−1
0
1
2
−6
−4
−2
0
2
4
6
8

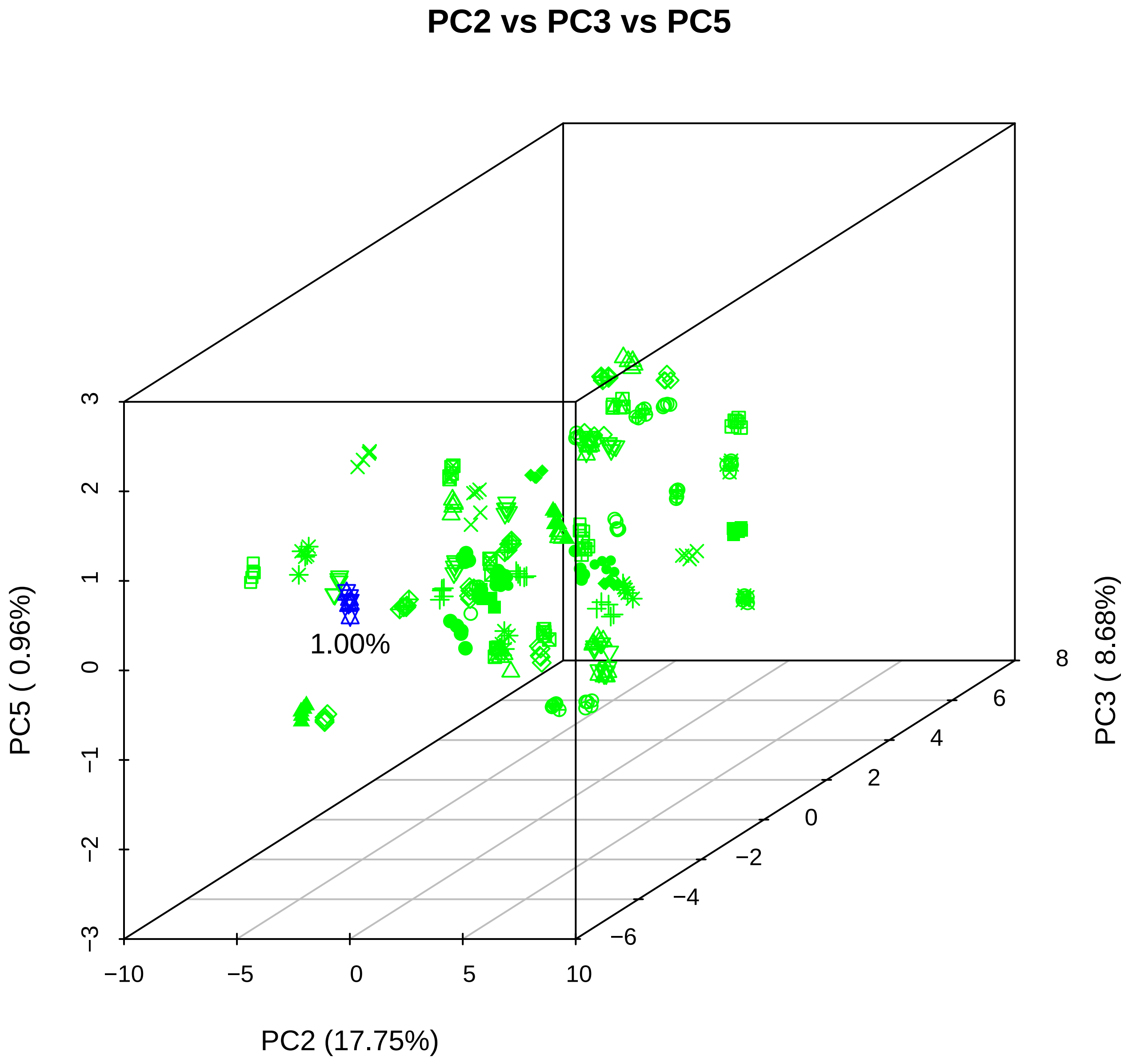

PC2 vs PC3 vs PC5
3
2
1
0
−1
−2
−3
PC5 ( 0.96%)
−10
−5
0
5
10
PC2 (17.75%)
−6
−4
−2
0
2
4
6
8
PC3 ( 8.68%)
1.00%

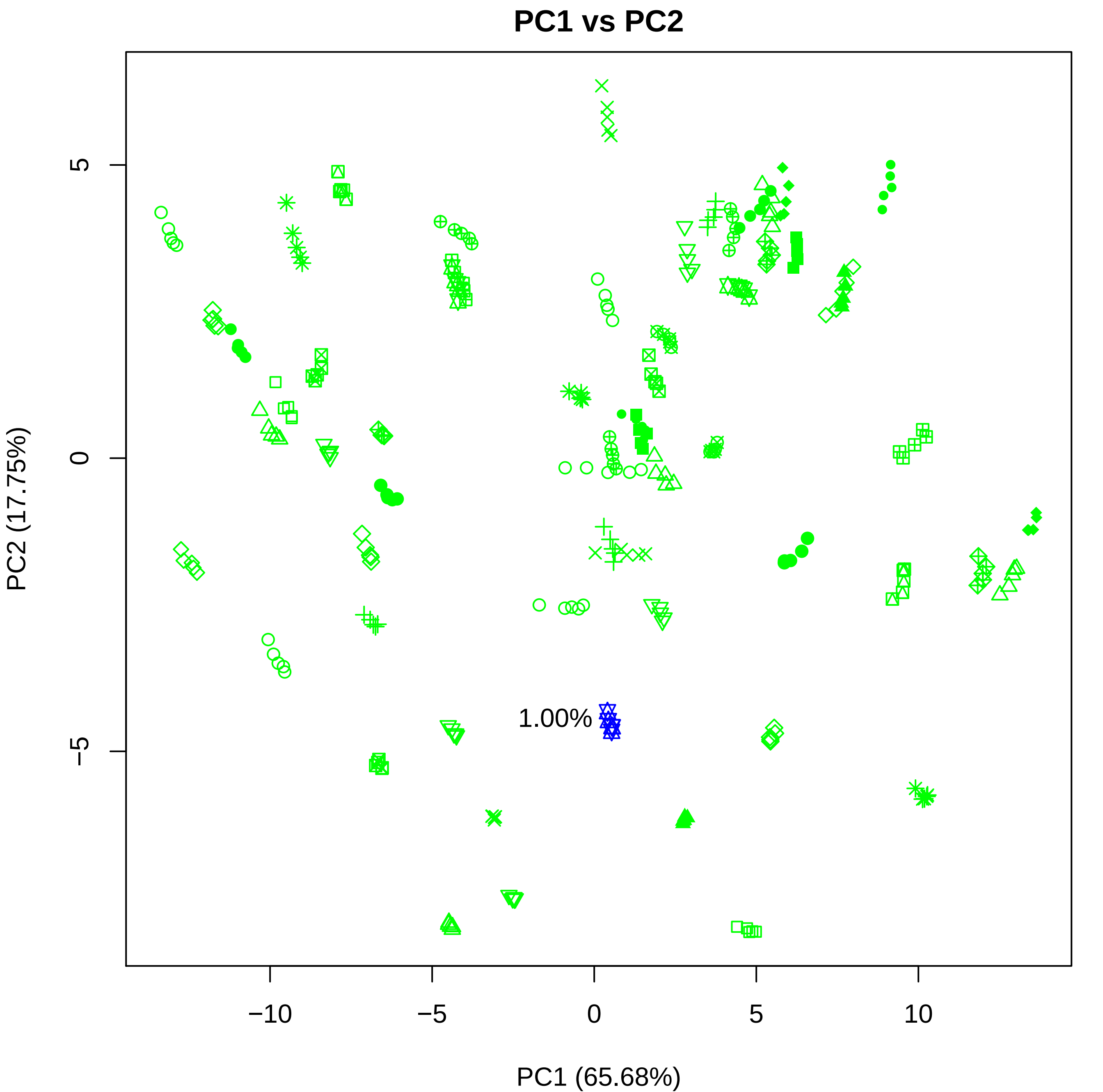

PC1 vs PC2
PC2 (17.75%)
PC1 (65.68%)
5
0
−5
−10
−5
0
5
10
1.00%

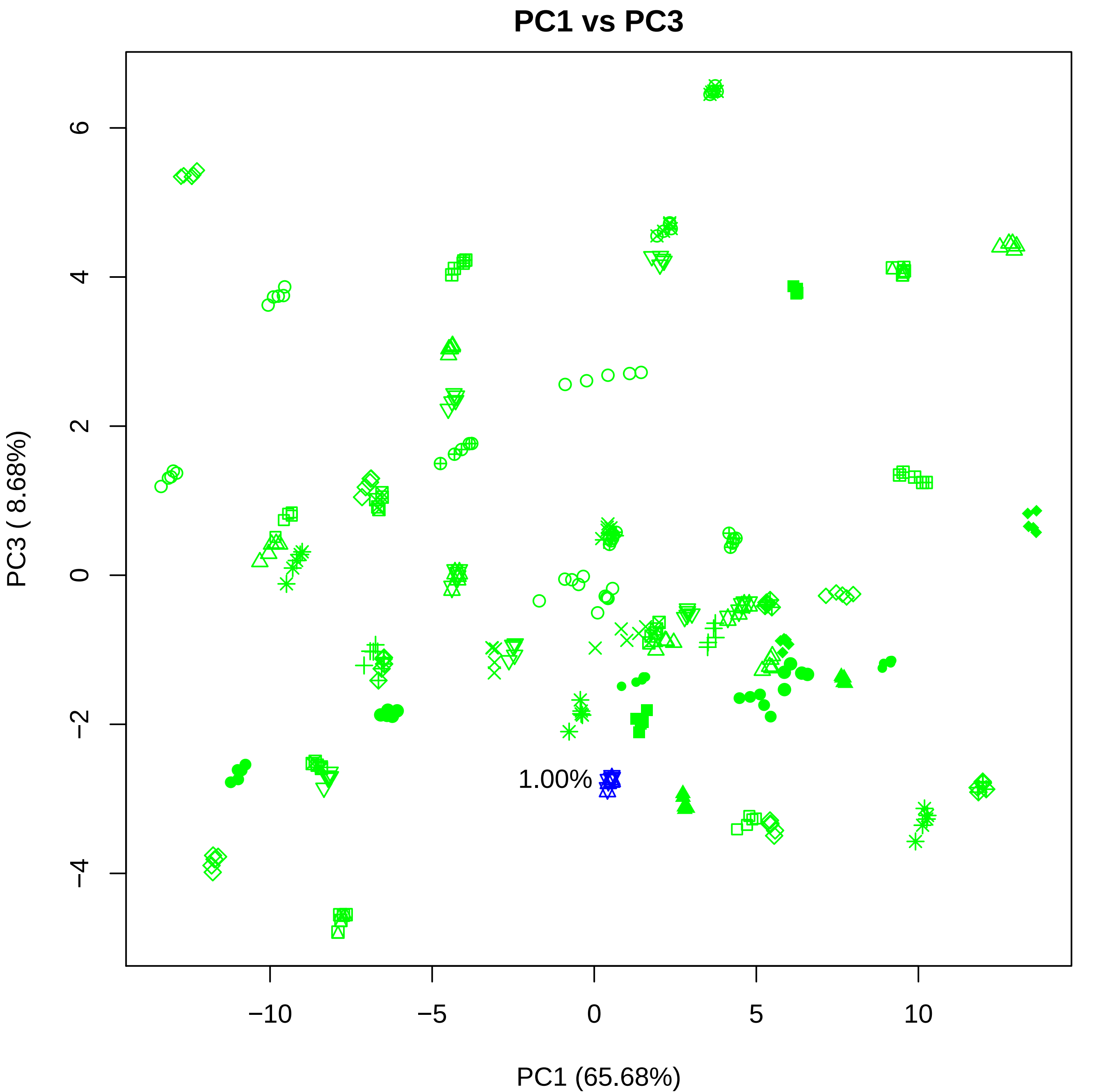

PC1 vs PC3
PC3 ( 8.68%)
PC1 (65.68%)
1.00%
−10
−5
0
5
10
−4
−2
0
2
4
6

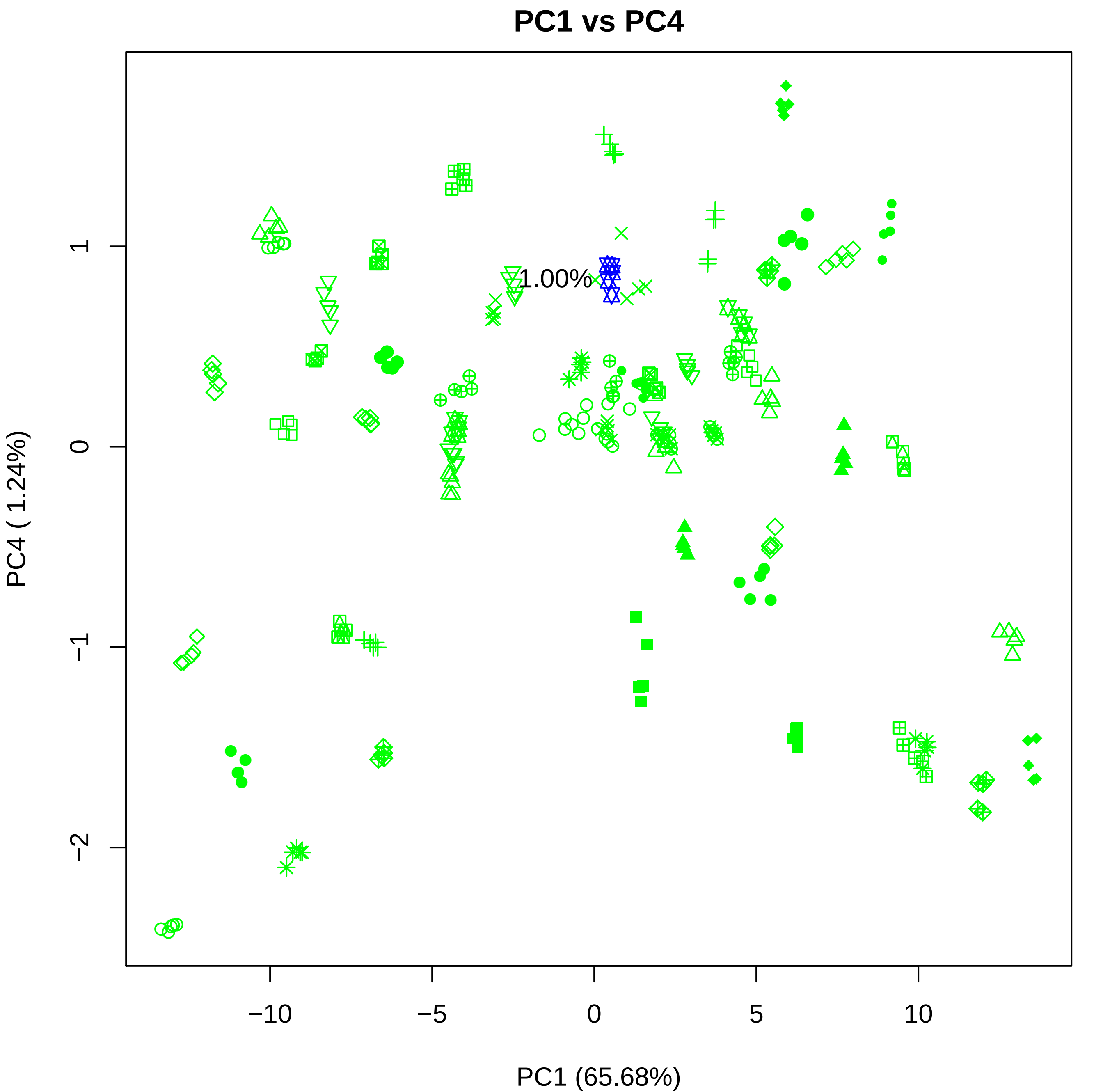
PC1 vs PC4
1.00%
PC4 ( 1.24%)
PC1 (65.68%)
−10
−5
0
5
10
1
0
−1
−2

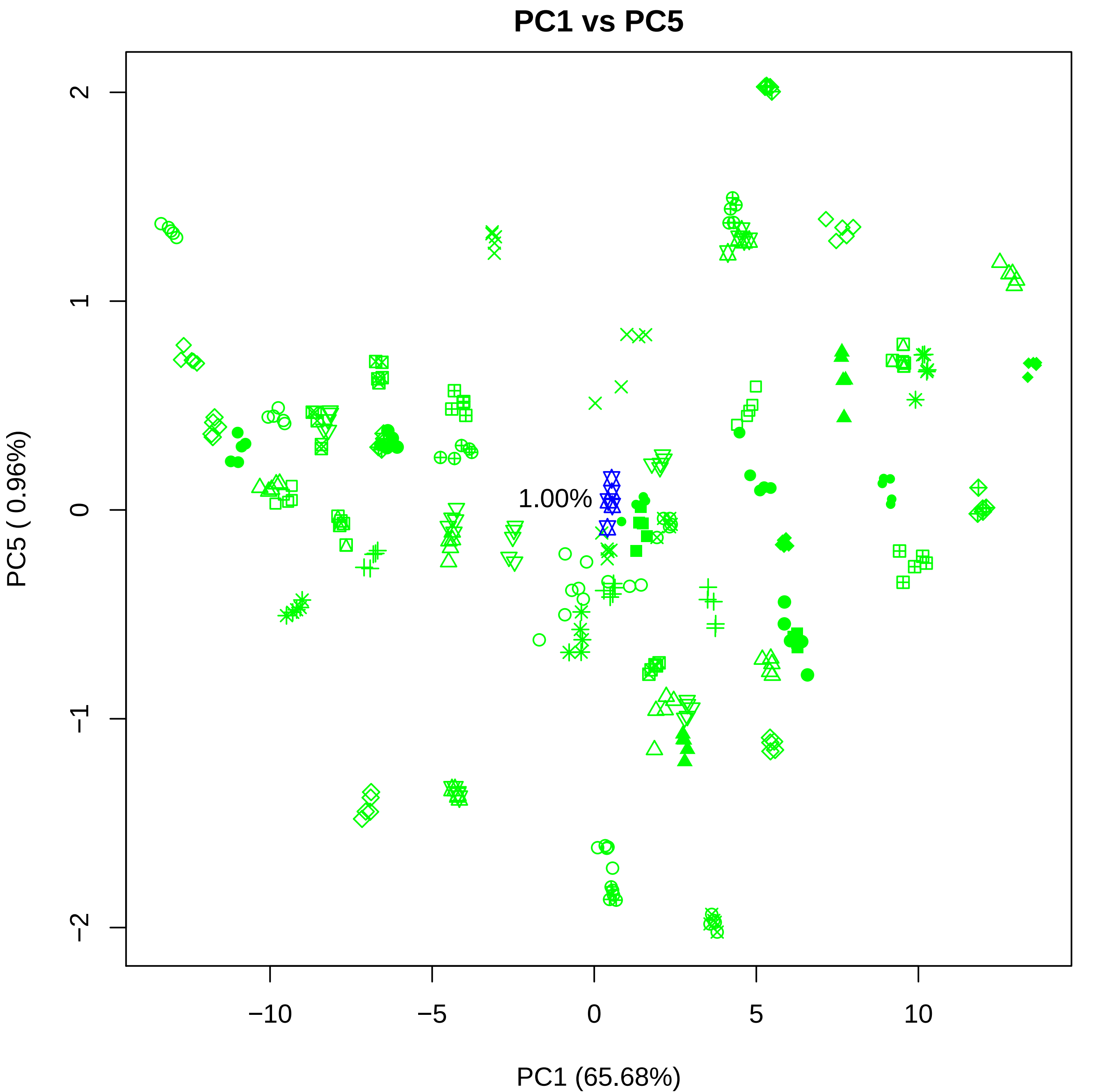
PC1 vs PC5
PC5 ( 0.96%)
PC1 (65.68%)
1.00%
2
1
0
−1
−2
−10
−5
0
5
10

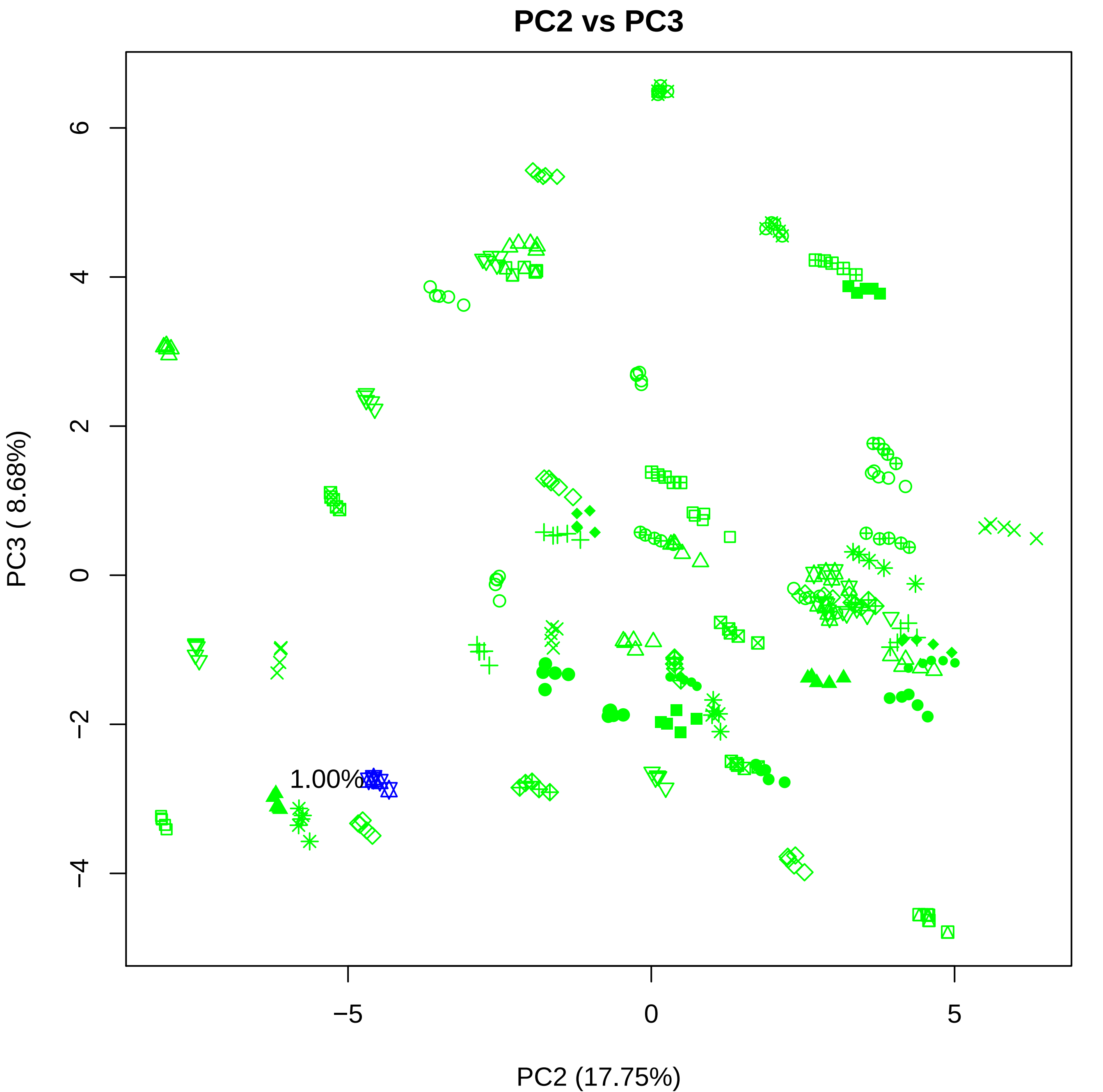
PC2 vs PC3
PC3 ( 8.68%)
PC2 (17.75%)
1.00%
6
4
2
0
−2
−4
−5
0
5

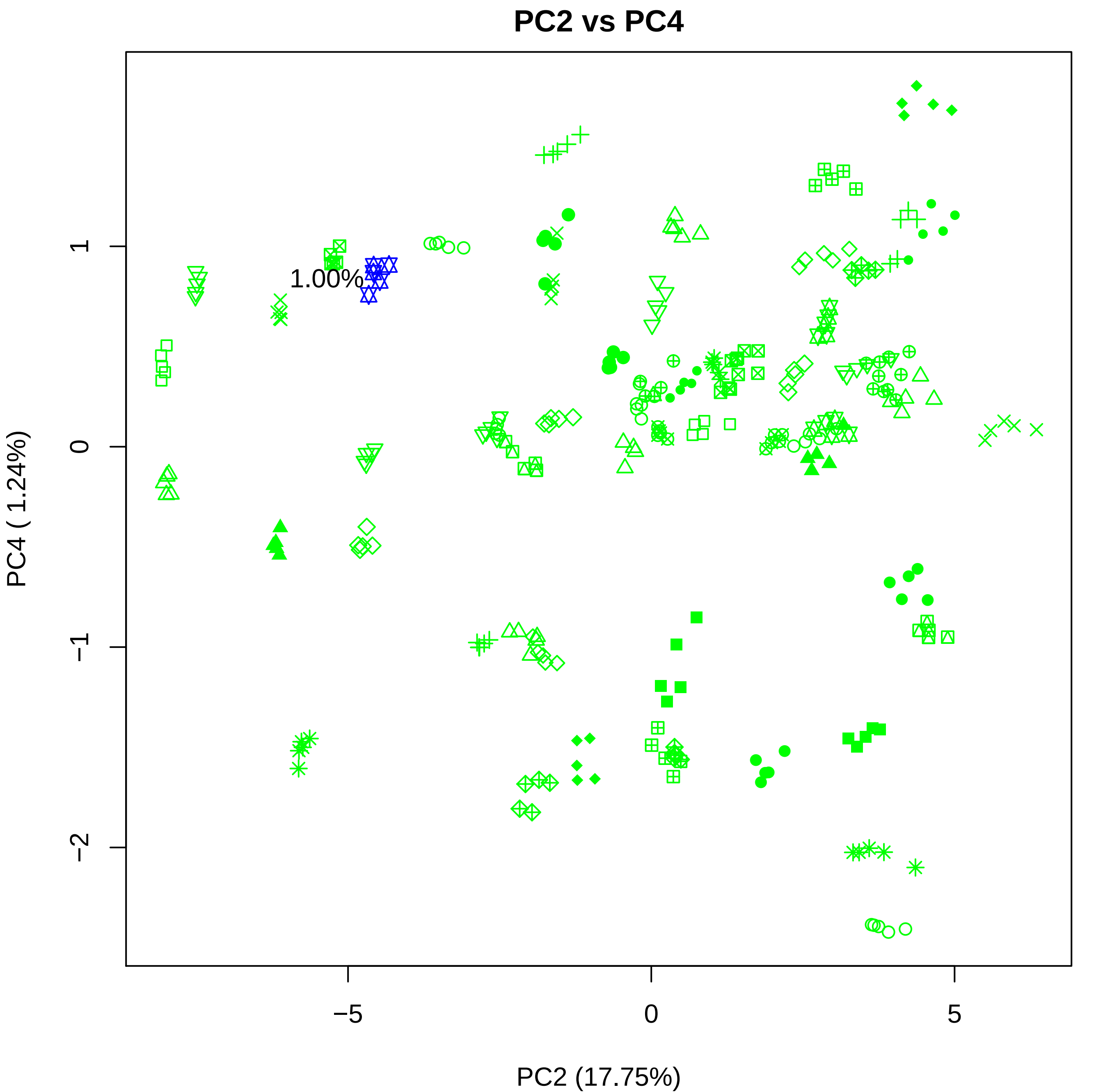
PC2 vs PC4
1.00%
PC4 ( 1.24%)
PC2 (17.75%)
−2
−1
0
1
−5
0
5

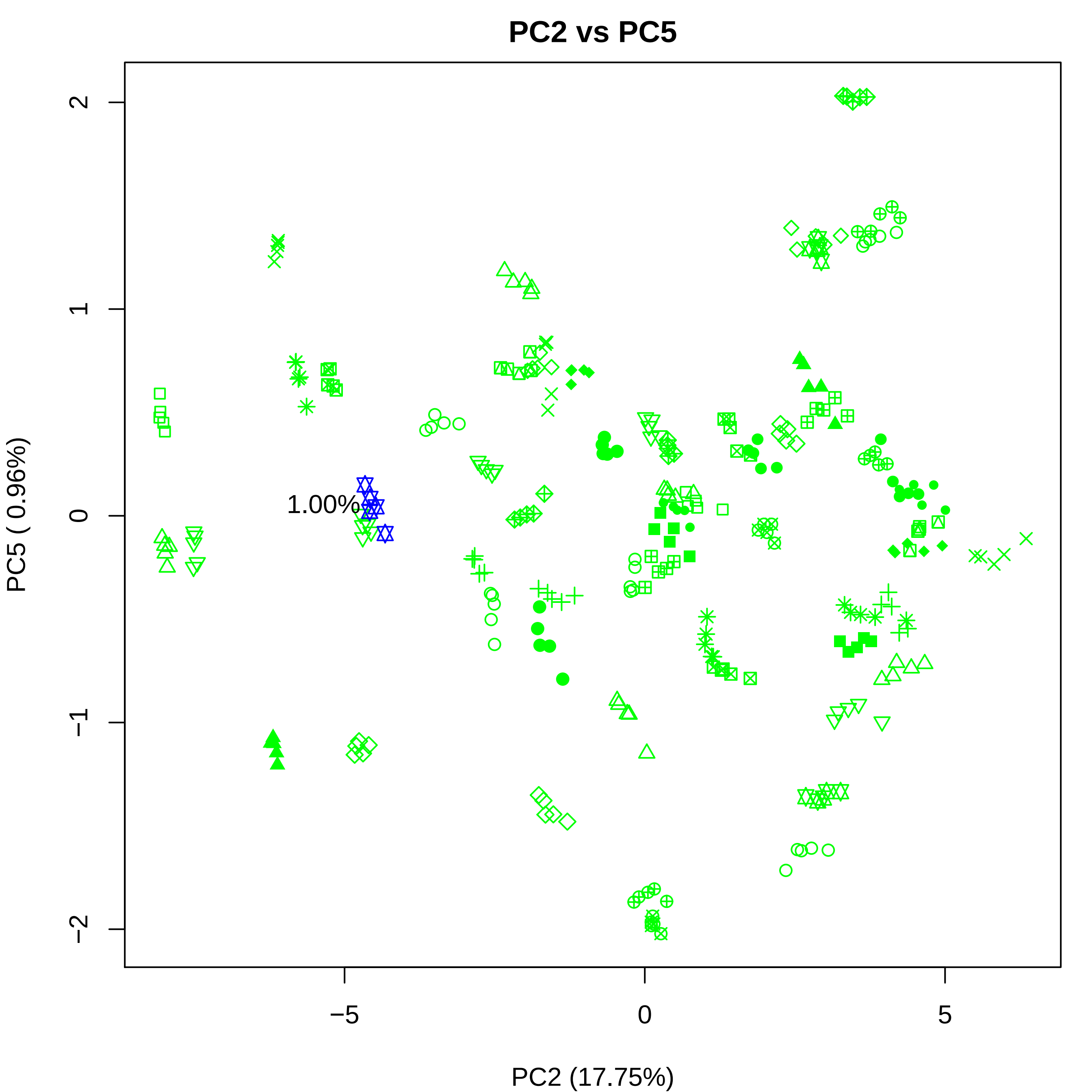

PC2 vs PC5
1.00%
PC5 ( 0.96%)
PC2 (17.75%)
2
1
0
−1
−2
−5
0
5

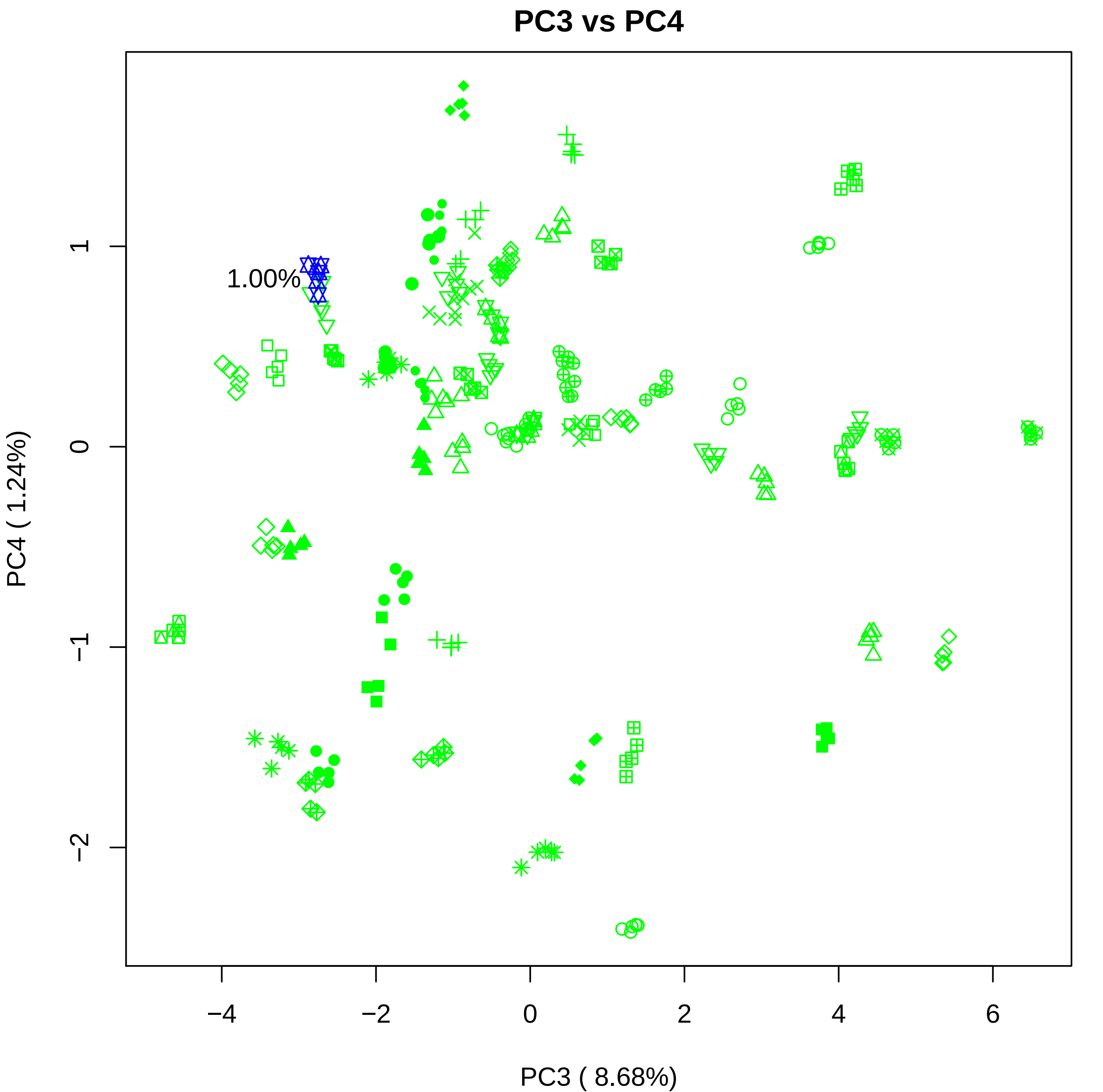
PC3 vs PC4
1.00%
PC4 ( 1.24%)
1
0
−1
−2
−4
−2
0
2
4
6
PC3 ( 8.68%)

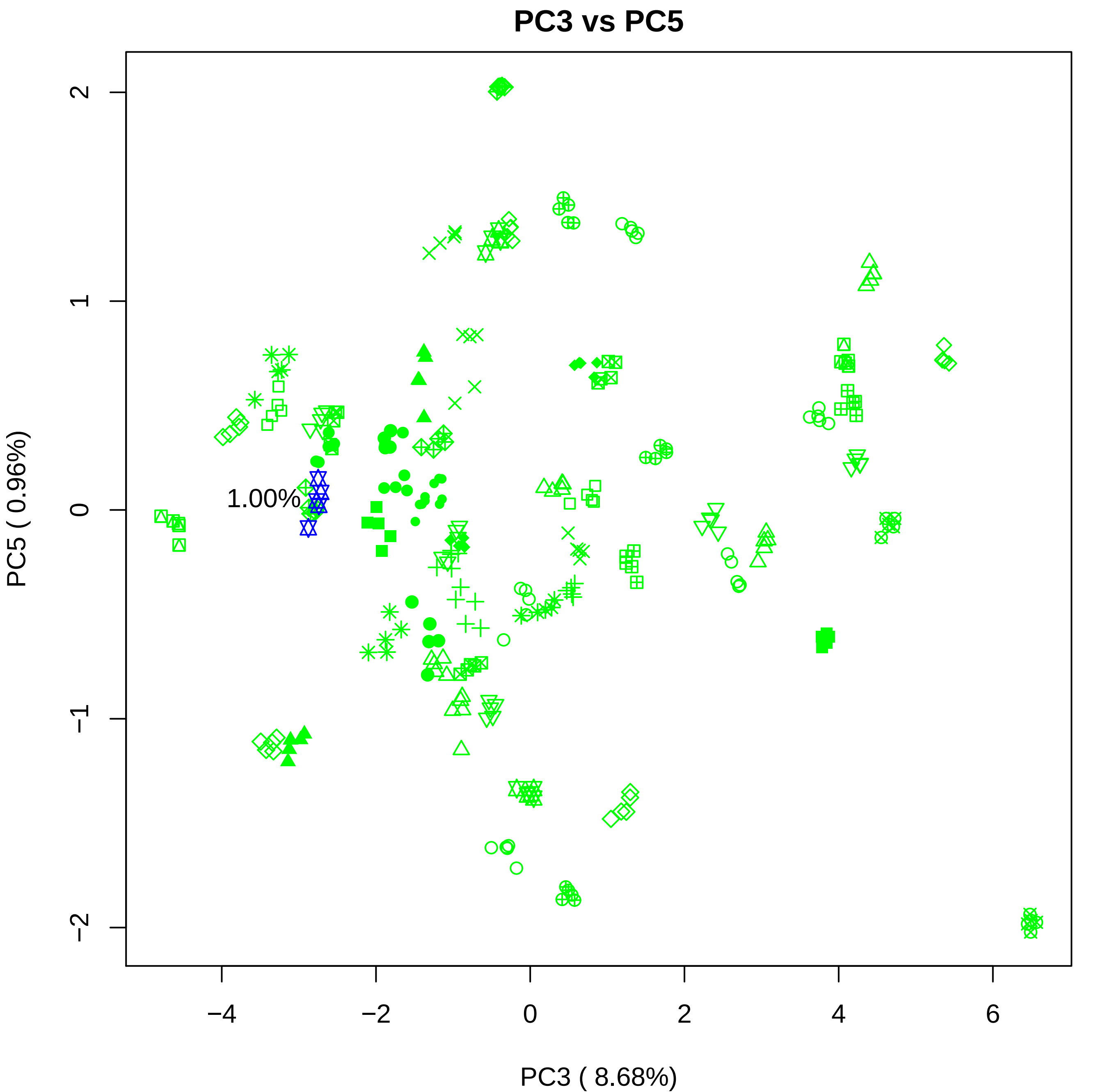
PC3 vs PC5
PC5 ( 0.96%)
PC3 ( 8.68%)
1.00%
2
1
0
−1
−2
−4
−2
0
2
4
6

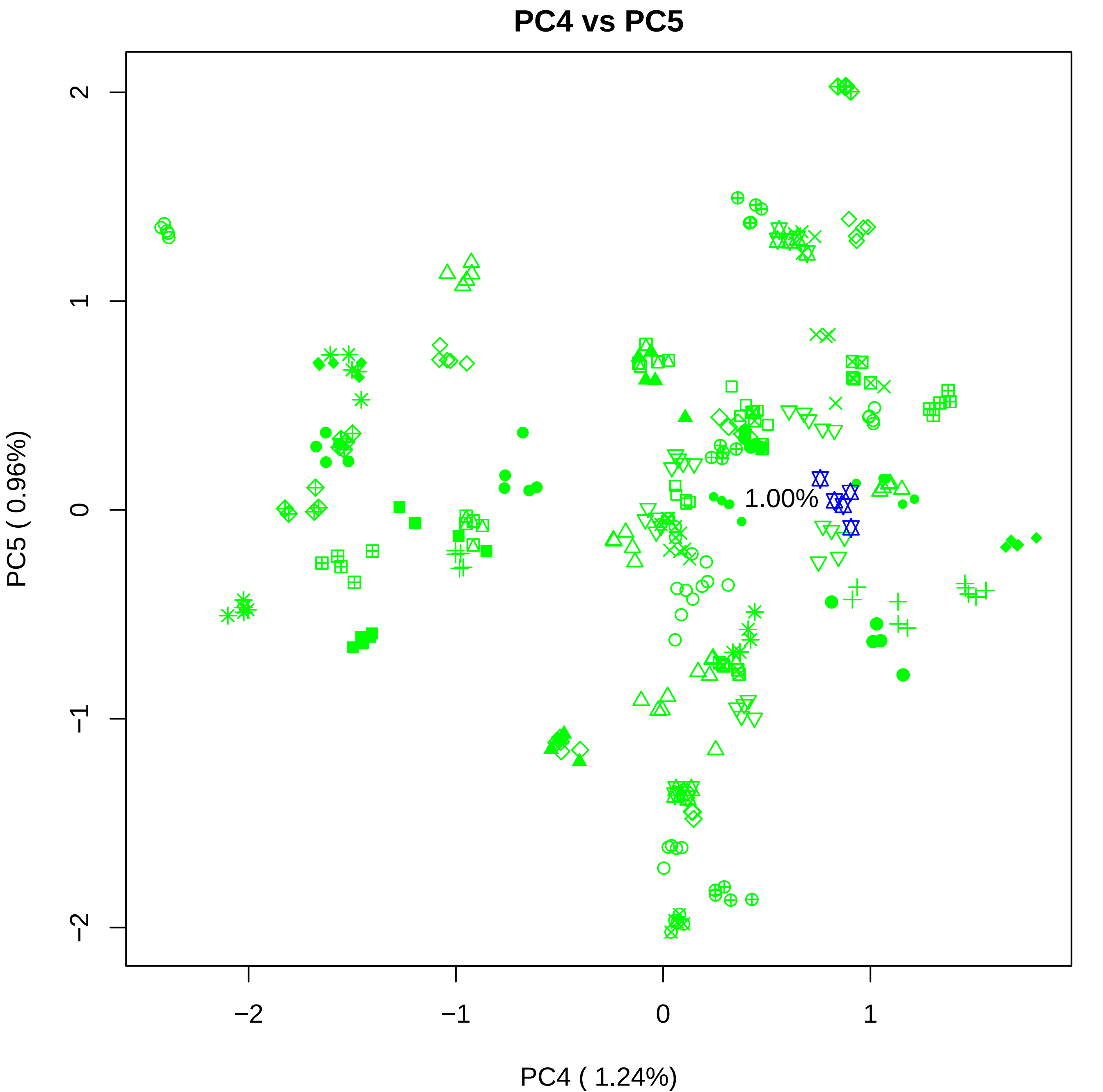

PC4 vs PC5
1.00%
PC5 ( 0.96%)
PC4 ( 1.24%)
−2
−1
0
1
2

### PCA of 2.5% BMH (c) vs PMH (c)

Scaled Matrix PCAed

7th order polynomial baseline correction performed

Raw spectra smoothed to 21 Neighbours, 2nd order PN, Savtisky Golay

Normalisation between 0 and 100

D2 spectra smoothed to 41 Neighbours, 2nd order PN, Savtisky Golay

Regions Analysed: 3601−2501 and 2001−700cm−1

Samples Analysed: 1 bContam's 0 BLH's 0 BMH's 0 OMH's 0 pContam's 0 PLH's 60 PMH's

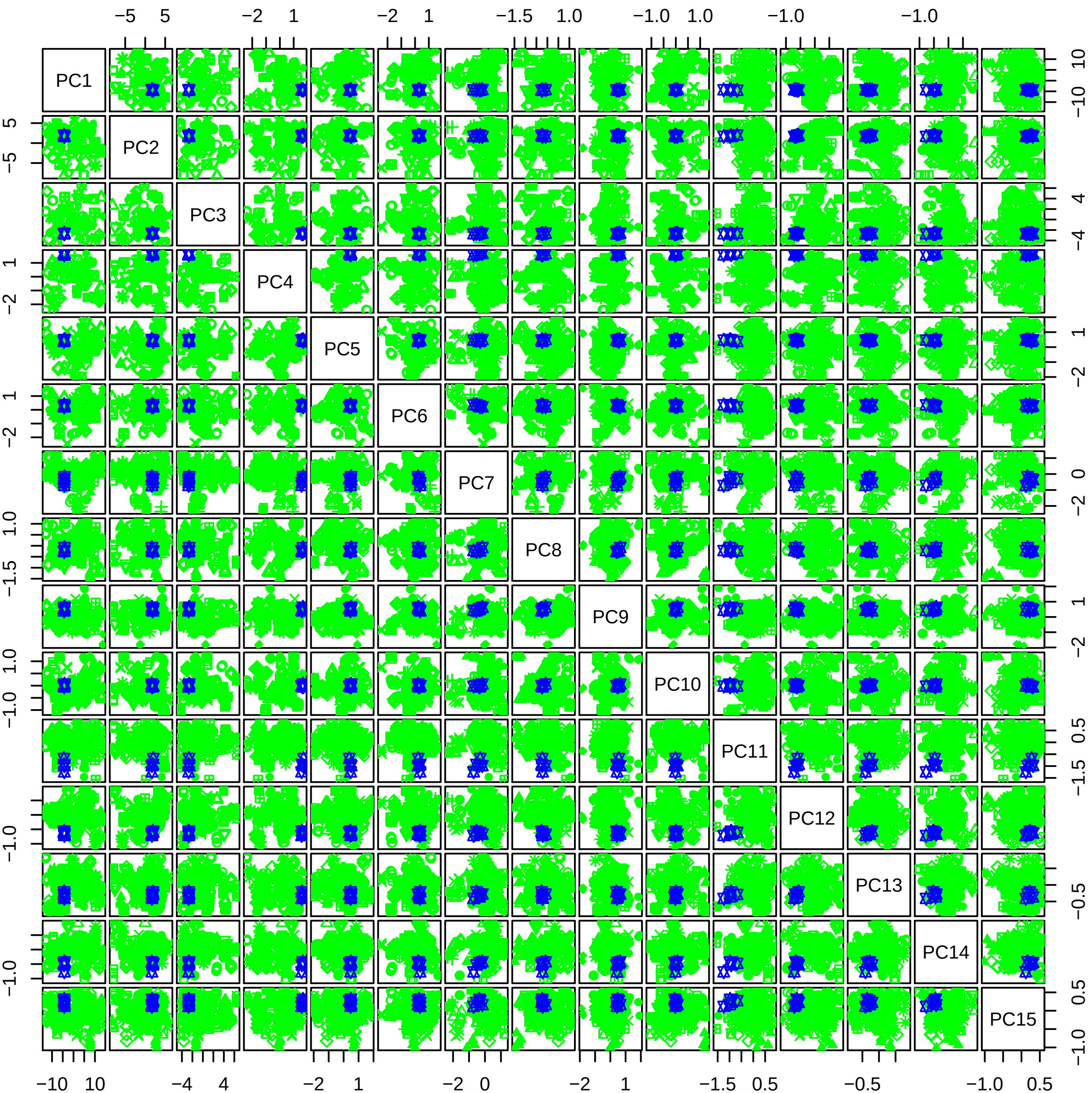
PC1
PC2
PC3
PC4
PC5
PC6
PC7
PC8
PC9
PC10
PC11
PC12
PC13
PC14
PC15

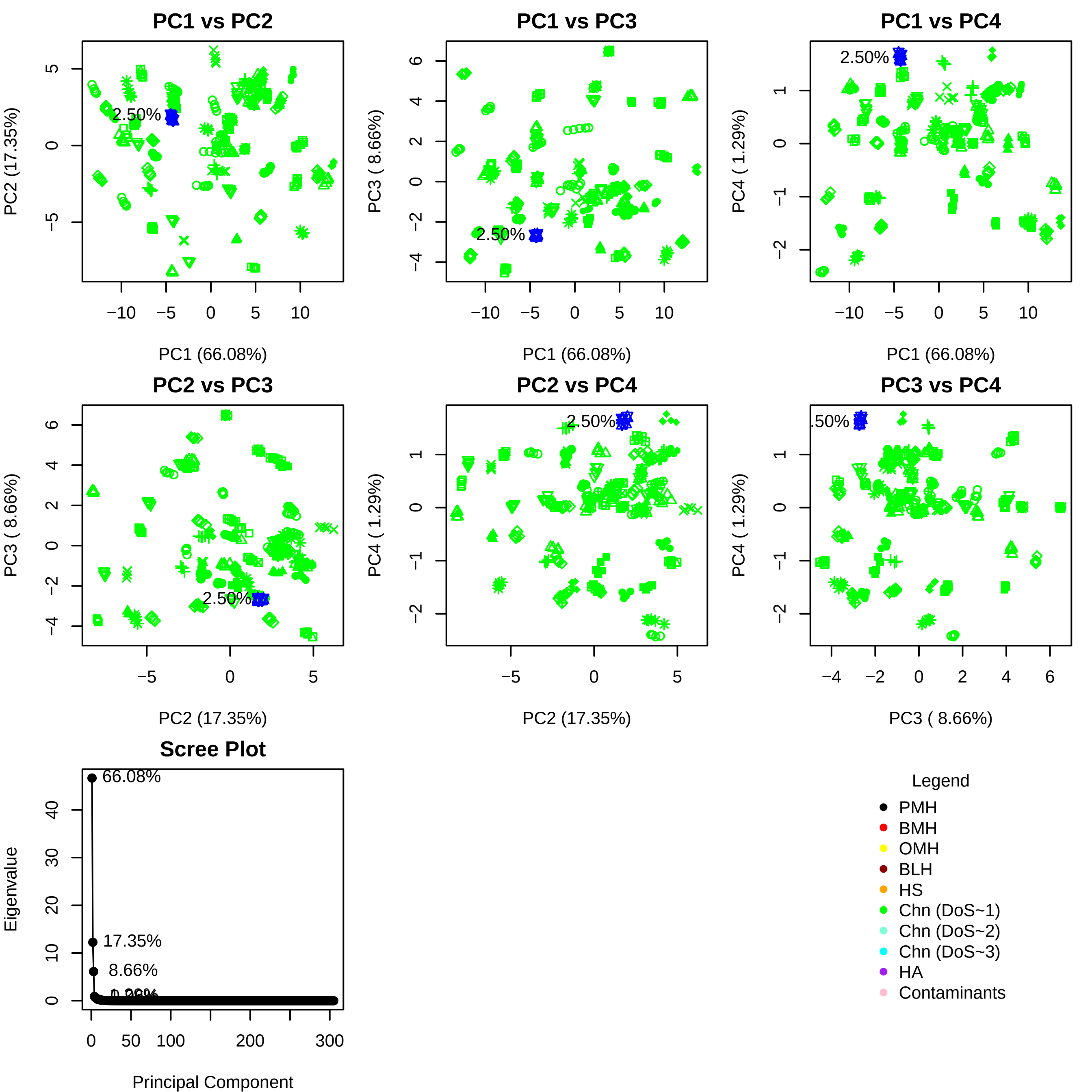
PC1 vs PC2
PC1 vs PC3
PC1 vs PC4
PC2 vs PC3
PC2 vs PC4
PC3 vs PC4
Scree Plot
PC1 (66.08%)
PC2 (17.35%)
PC3 ( 8.66%)
PC4 ( 1.29%)
2.50%
66.08%
17.35%
8.66%
Eigenvalue
Principal Component
Legend
PMH
BMH
OMH
BLH
HS
Chn (DoS~1)
Chn (DoS~2)
Chn (DoS~3)
HA
Contaminants

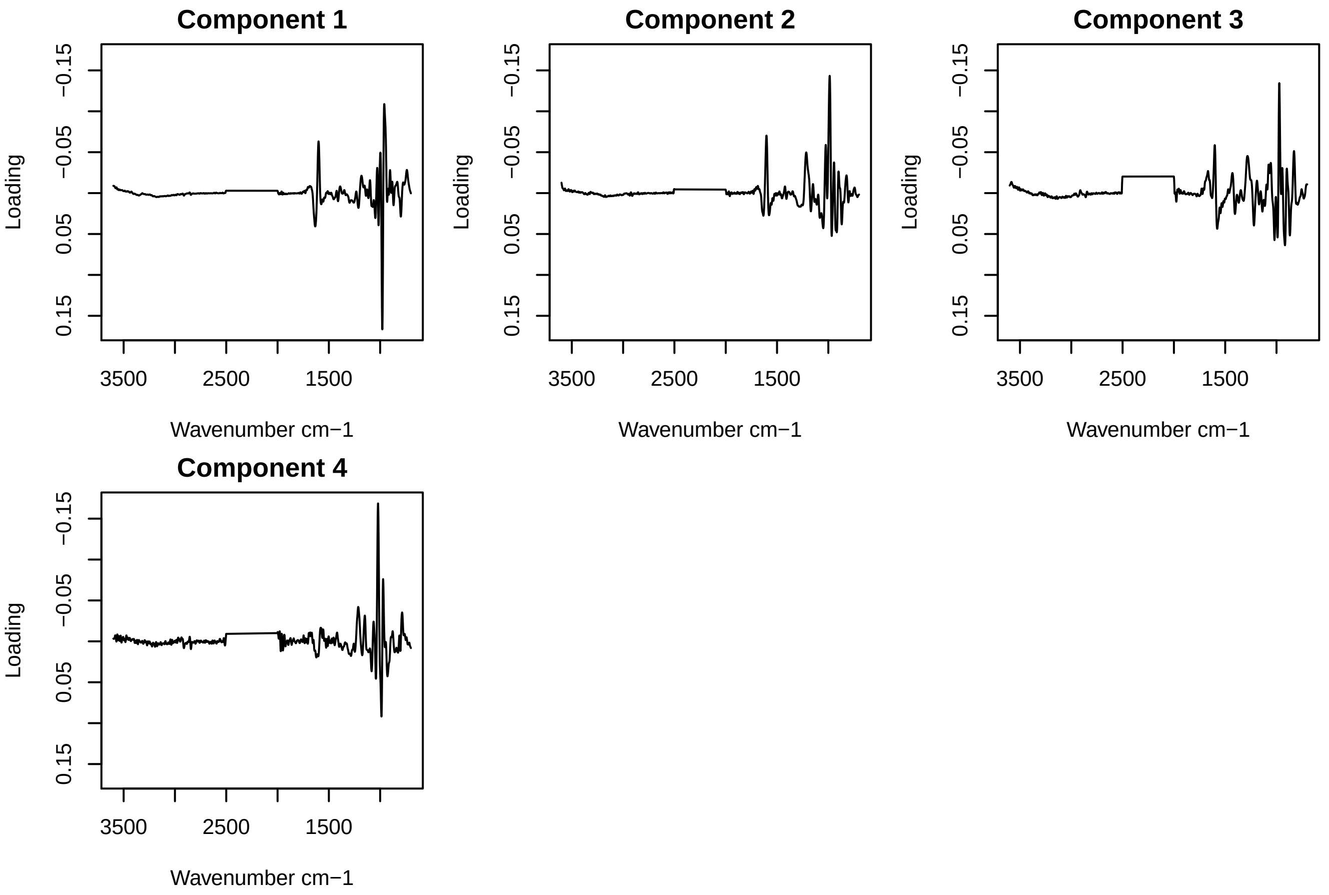
Component 1
Component 2
Component 3
Component 4
Loading
−0.15
−0.05
0.05
0.15
3500
2500
1500
Wavenumber cm−1

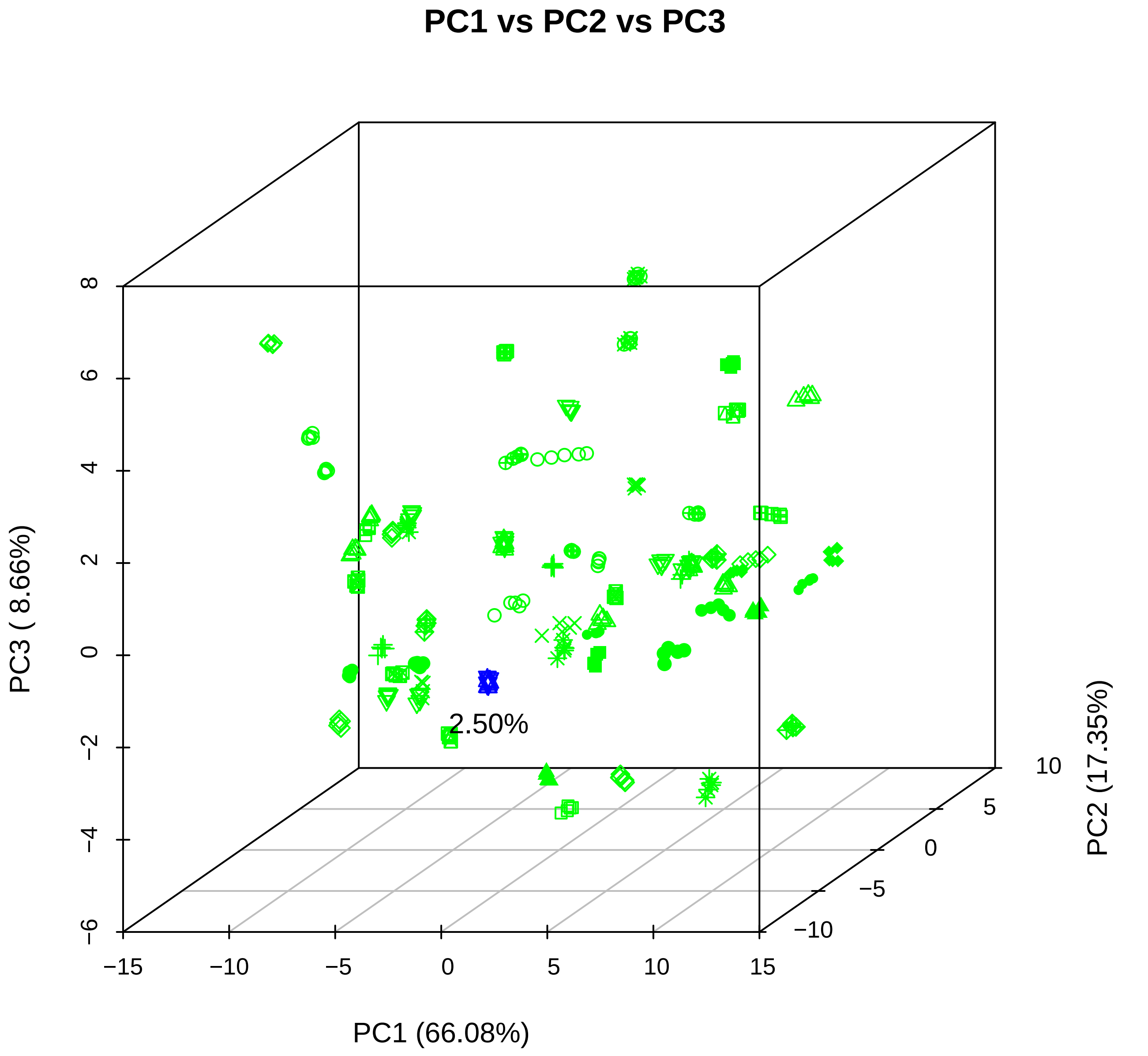
PC1 vs PC2 vs PC3
PC1 (66.08%)
PC2 (17.35%)
PC3 ( 8.66%)
2.50%
−15
−10
−5
0
5
10
15
−6
−4
−2
0
2
4
6
8
−10
−5
0
5
10

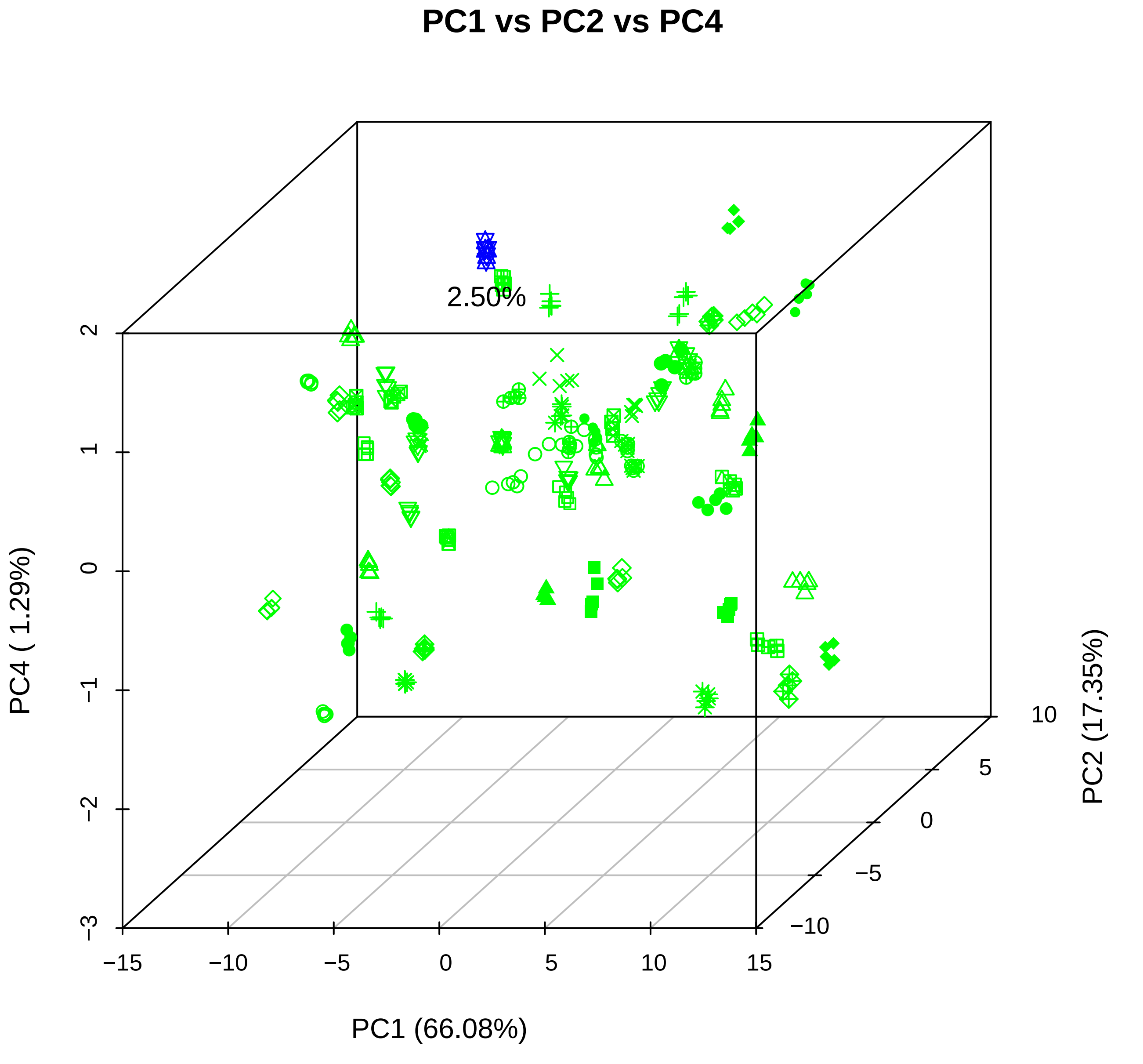

PC1 vs PC2 vs PC4
2.50%
PC4 ( 1.29%)
PC1 (66.08%)
PC2 (17.35%)
−15
−10
−5
0
5
10
15
−3
−2
−1
0
1
2
−10
−5
0
5
10

# PC1 vs PC3 vs PC4

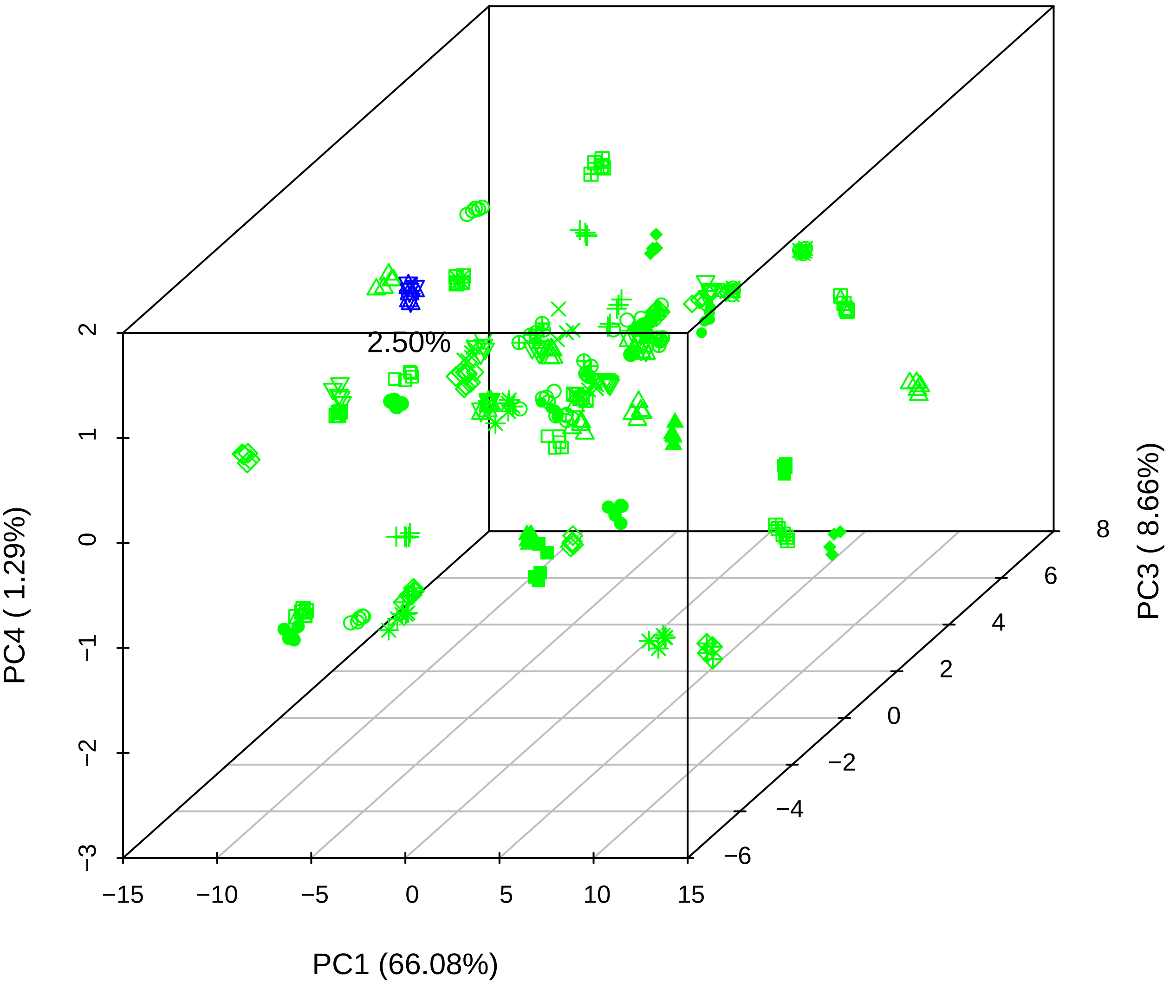

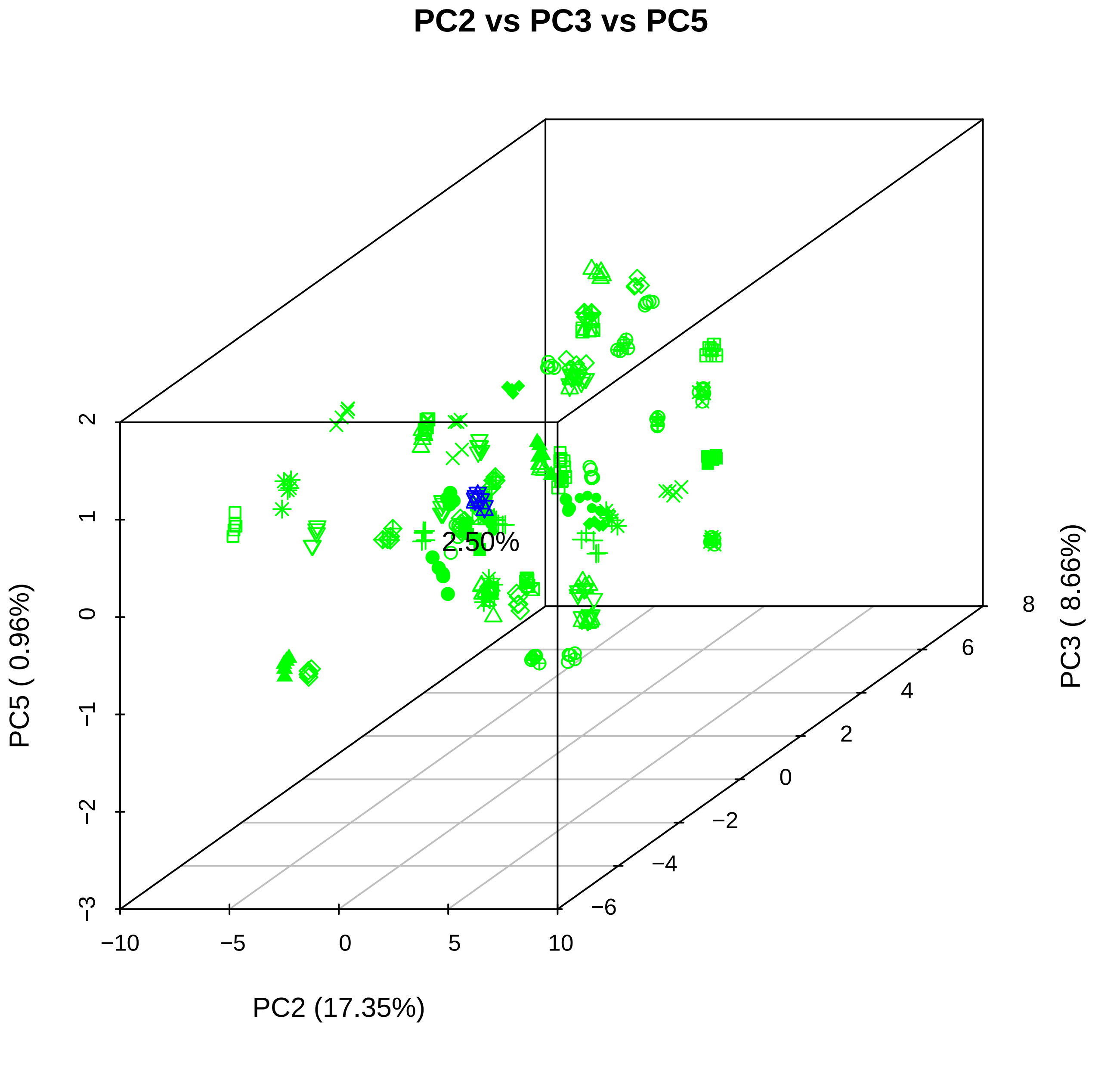

PC2 vs PC3 vs PC5
2.50%
PC2 (17.35%)
PC5 ( 0.96%)
PC3 ( 8.66%)
−10
−5
0
5
10
−3
−2
−1
0
1
2
−6
−4
−2
0
2
4
6
8

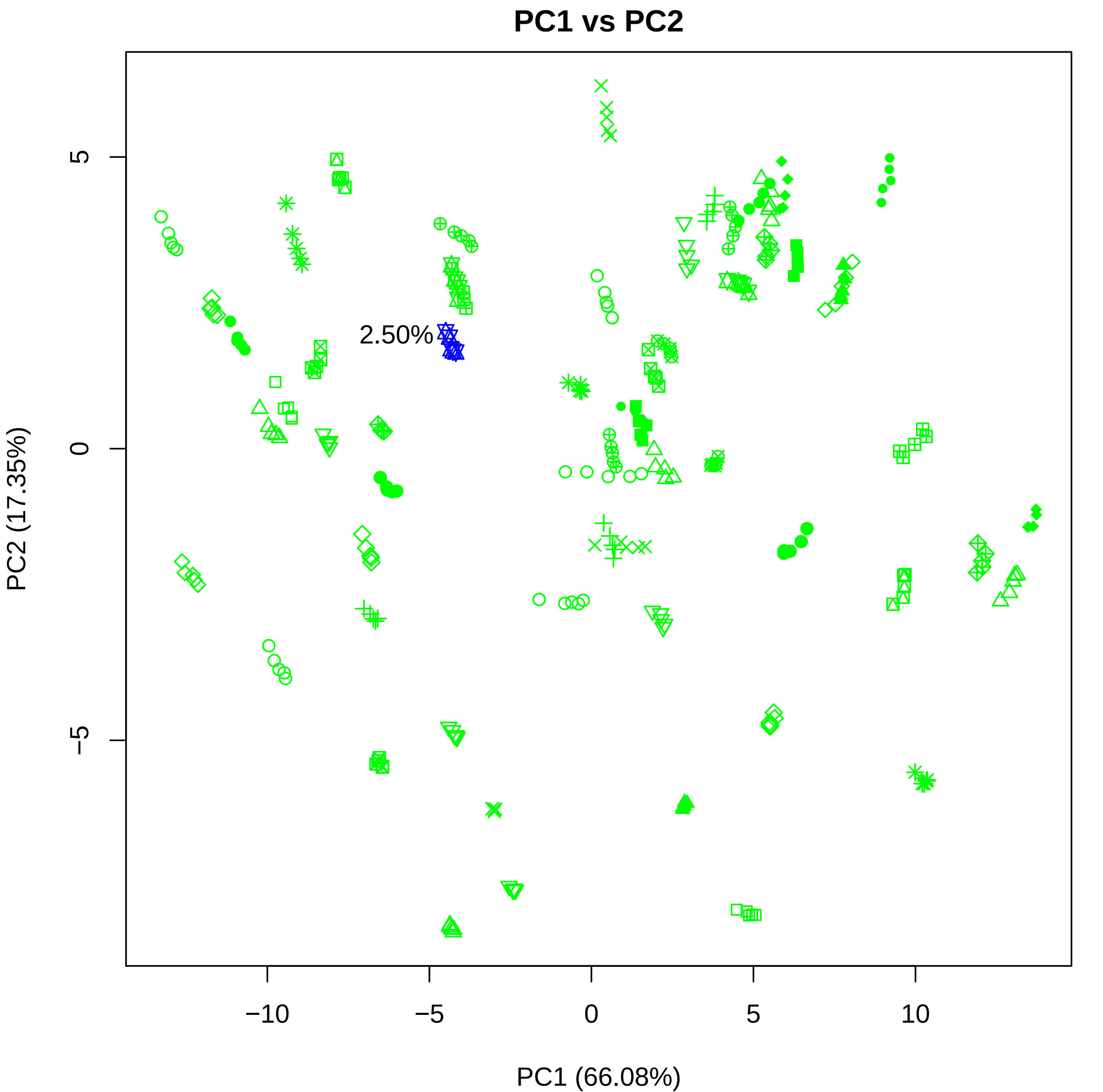
PC1 vs PC2
2.50%
PC2 (17.35%)
PC1 (66.08%)
5
0
−5
−10
−5
0
5
10

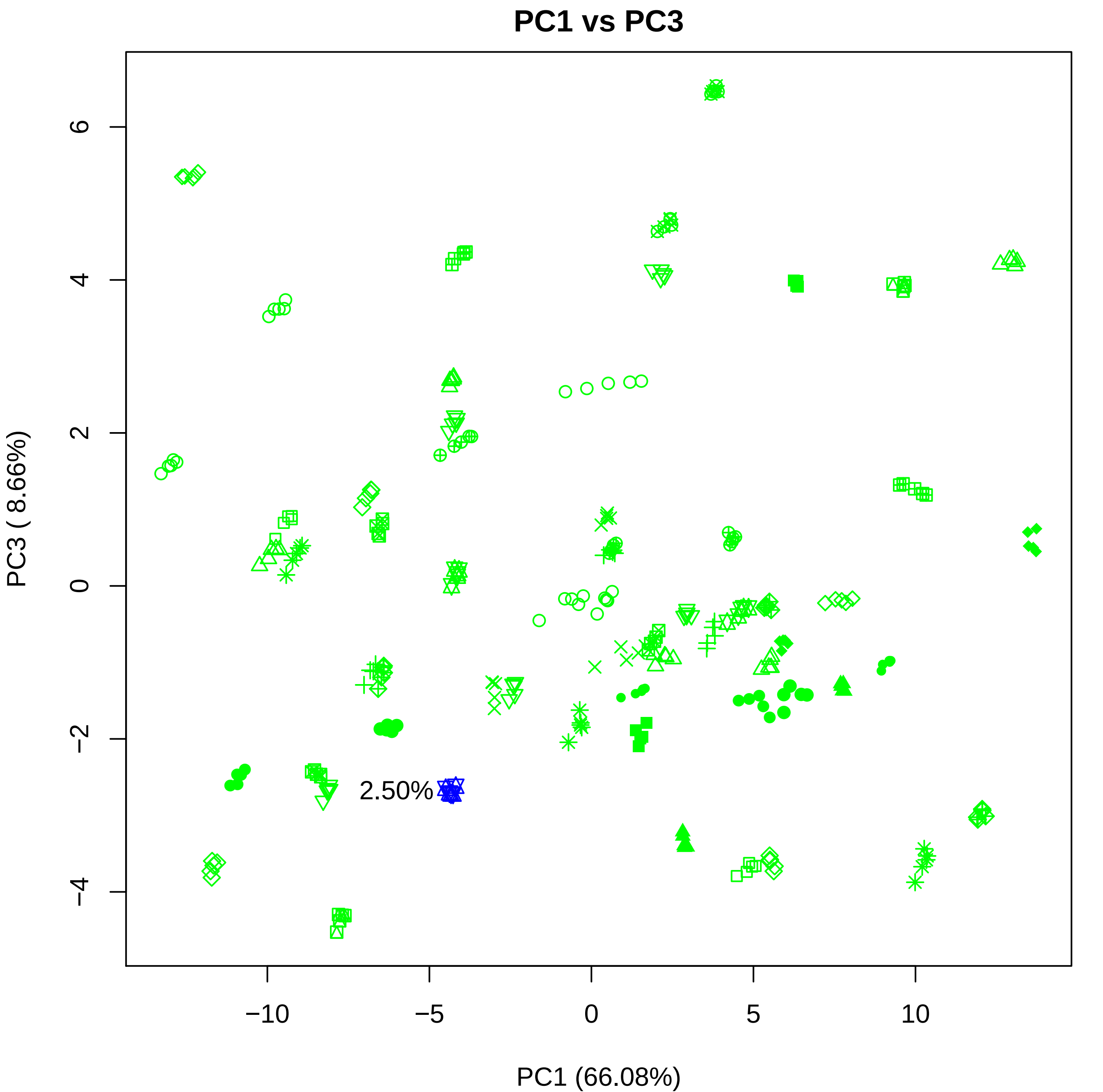
PC1 vs PC3
PC3 ( 8.66%)
PC1 (66.08%)
2.50%
6
4
2
0
−2
−4
−10
−5
0
5
10

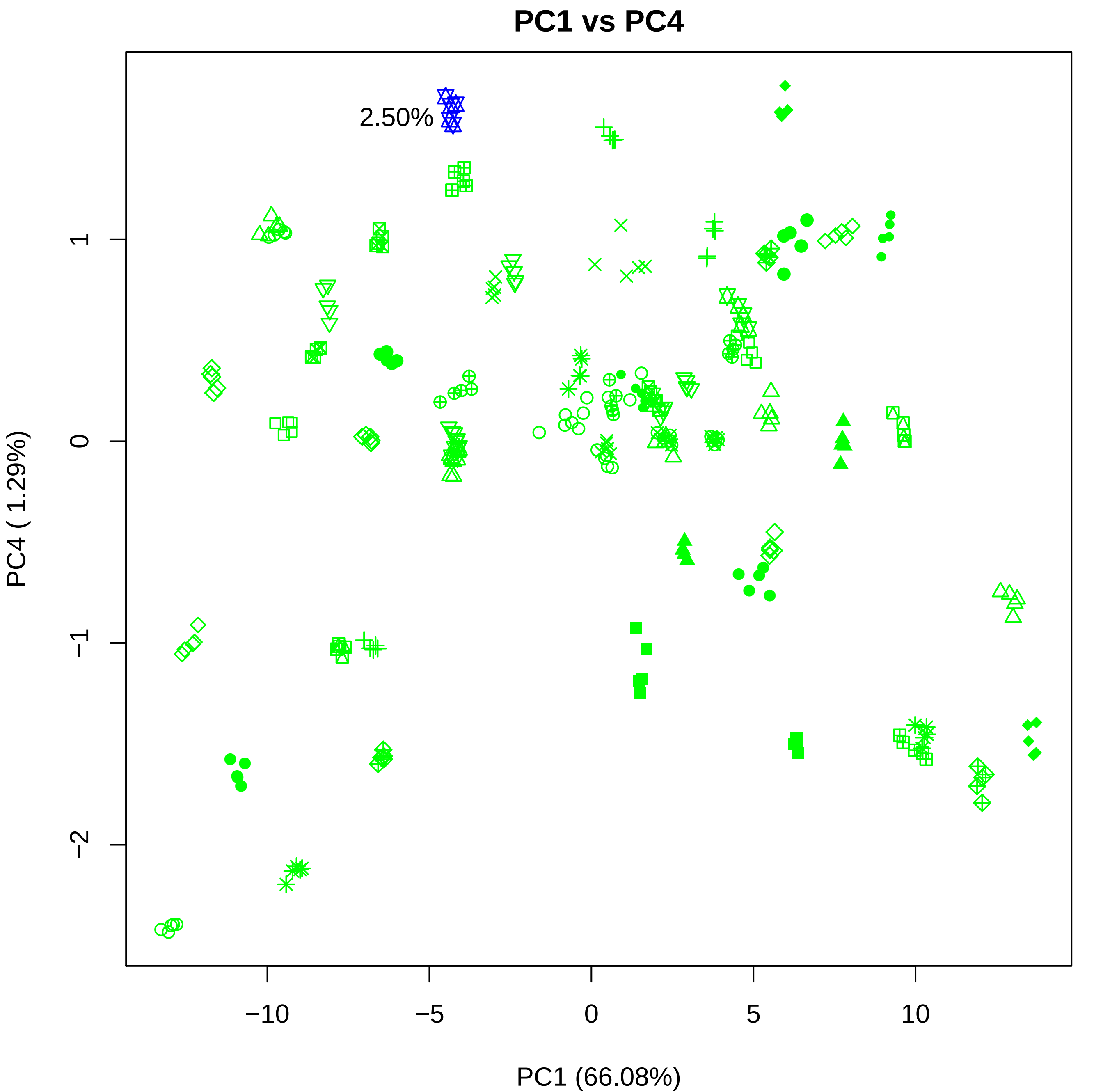

PC1 vs PC4
2.50%
PC4 ( 1.29%)
PC1 (66.08%)
−10
−5
0
5
10
−2
−1
0
1

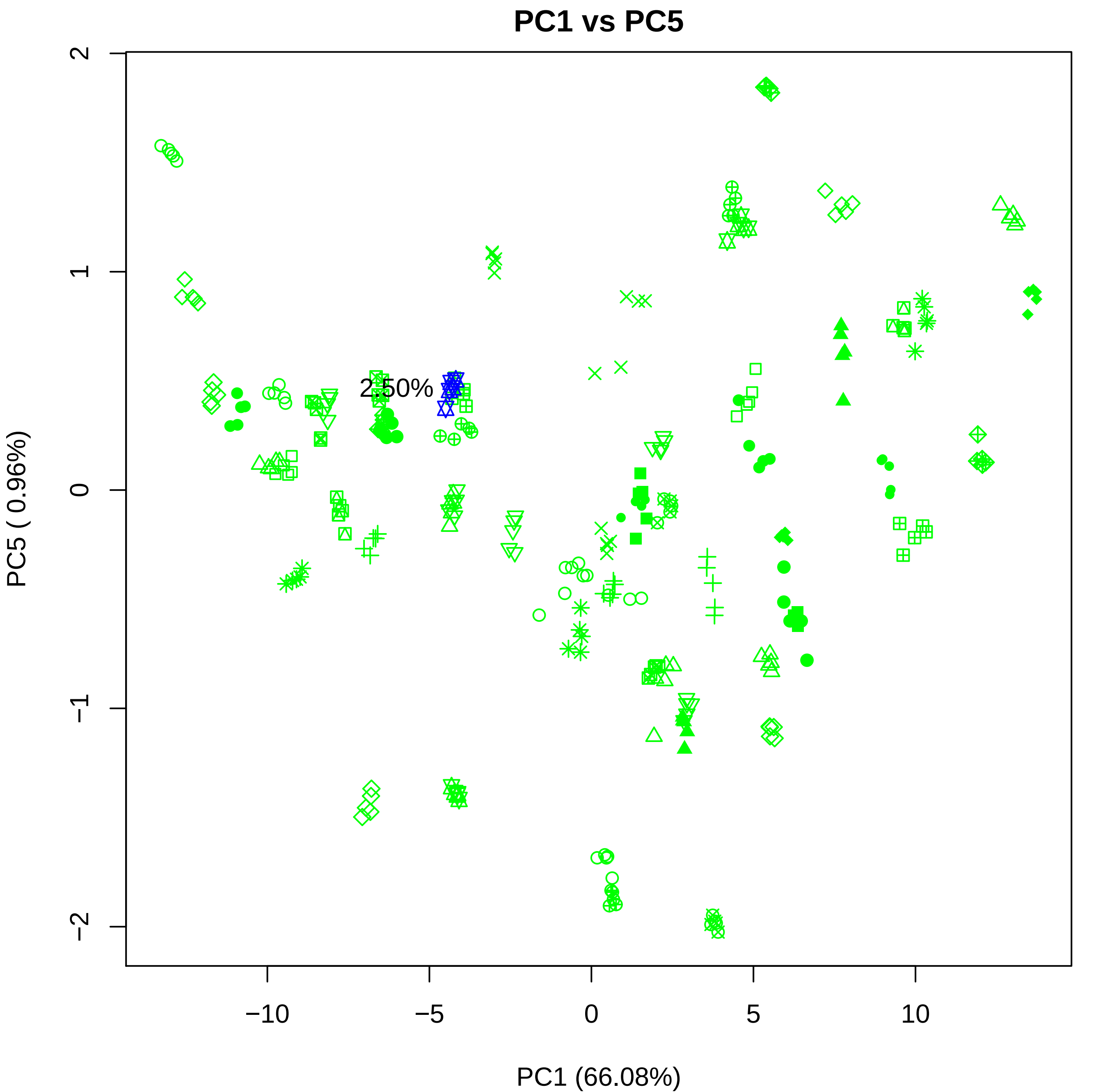

PC1 vs PC5
2.50%
PC1 (66.08%)
PC5 ( 0.96%)
−10
−5
0
5
10
−2
−1
0
1
2

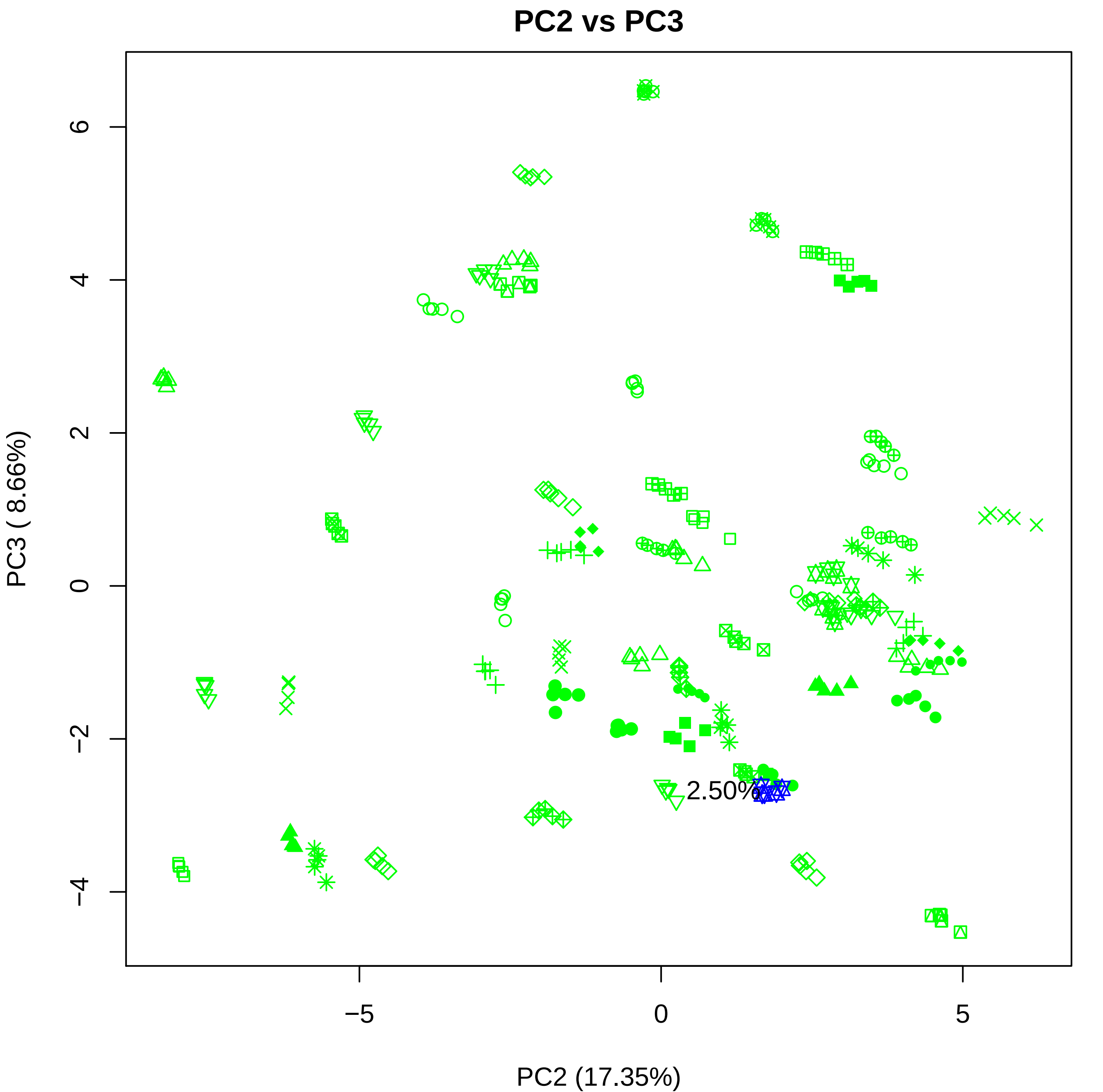

PC2 vs PC3
PC3 ( 8.66%)
PC2 (17.35%)
6
4
2
0
−2
−4
−5
0
5
2.50%

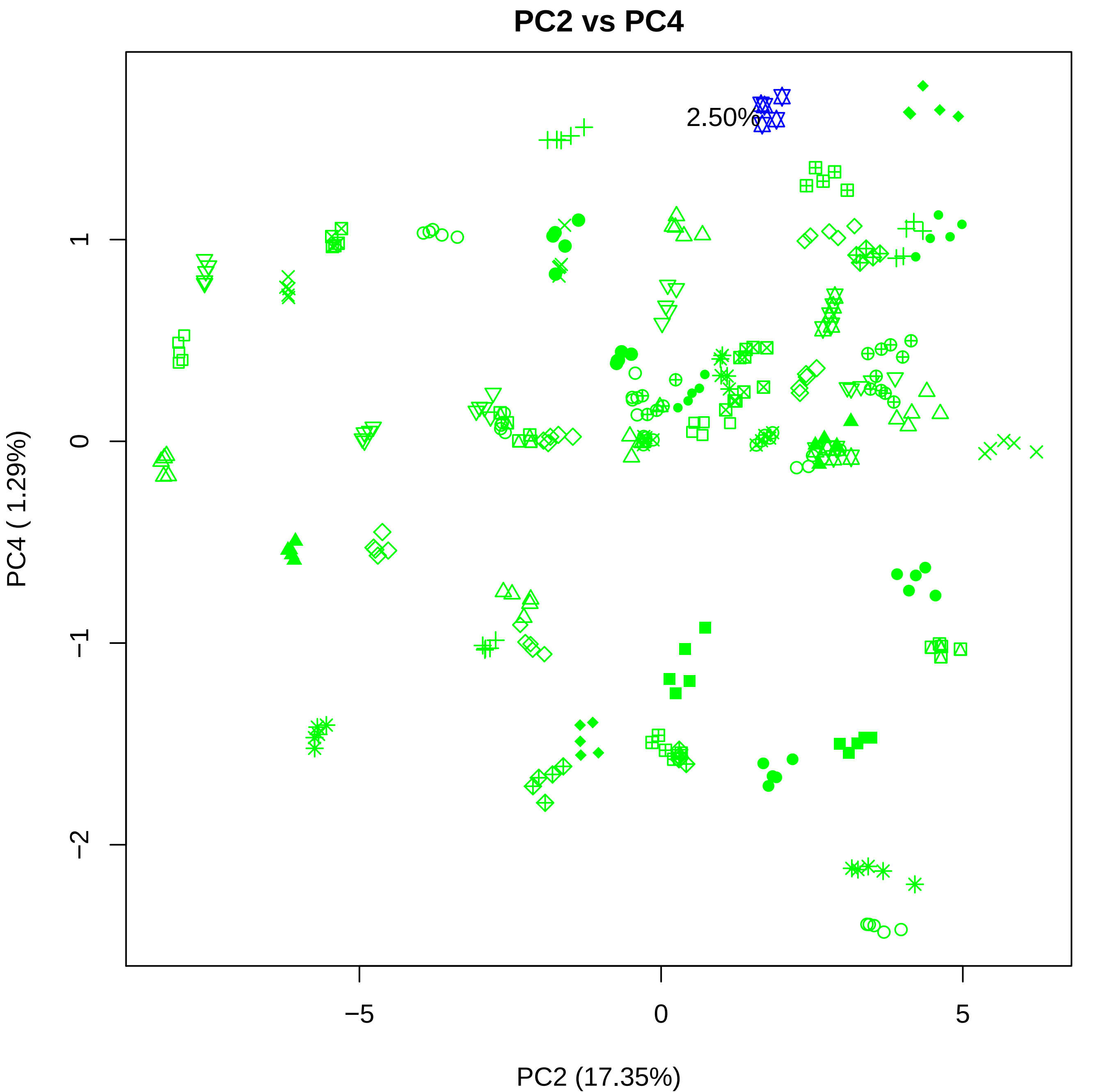

PC2 vs PC4
2.50%
1
0
−1
−2
PC4 ( 1.29%)
−5
0
5
PC2 (17.35%)

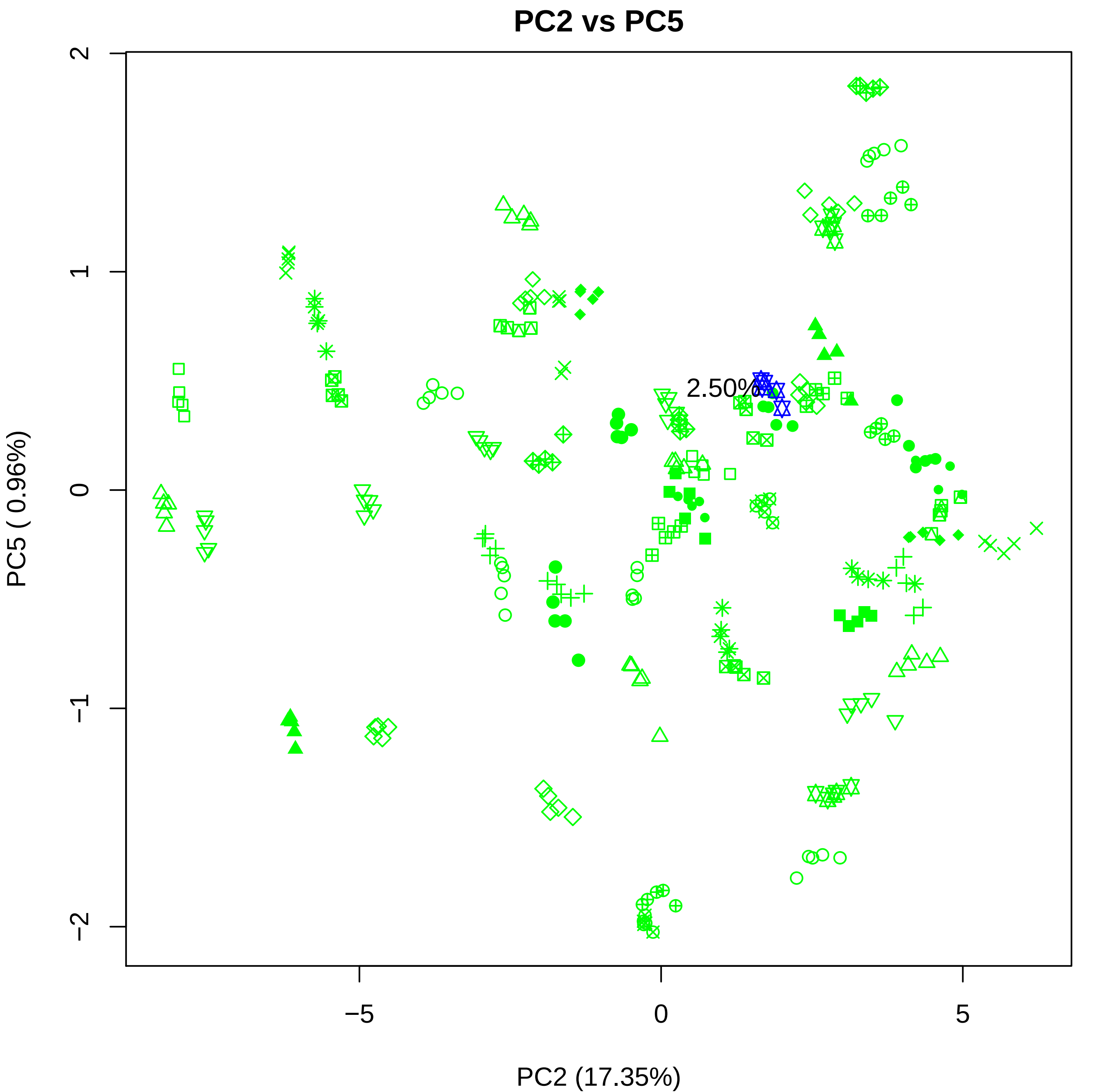
PC2 vs PC5
2.50%
PC5 ( 0.96%)
PC2 (17.35%)
2
1
0
−1
−2
−5
0
5

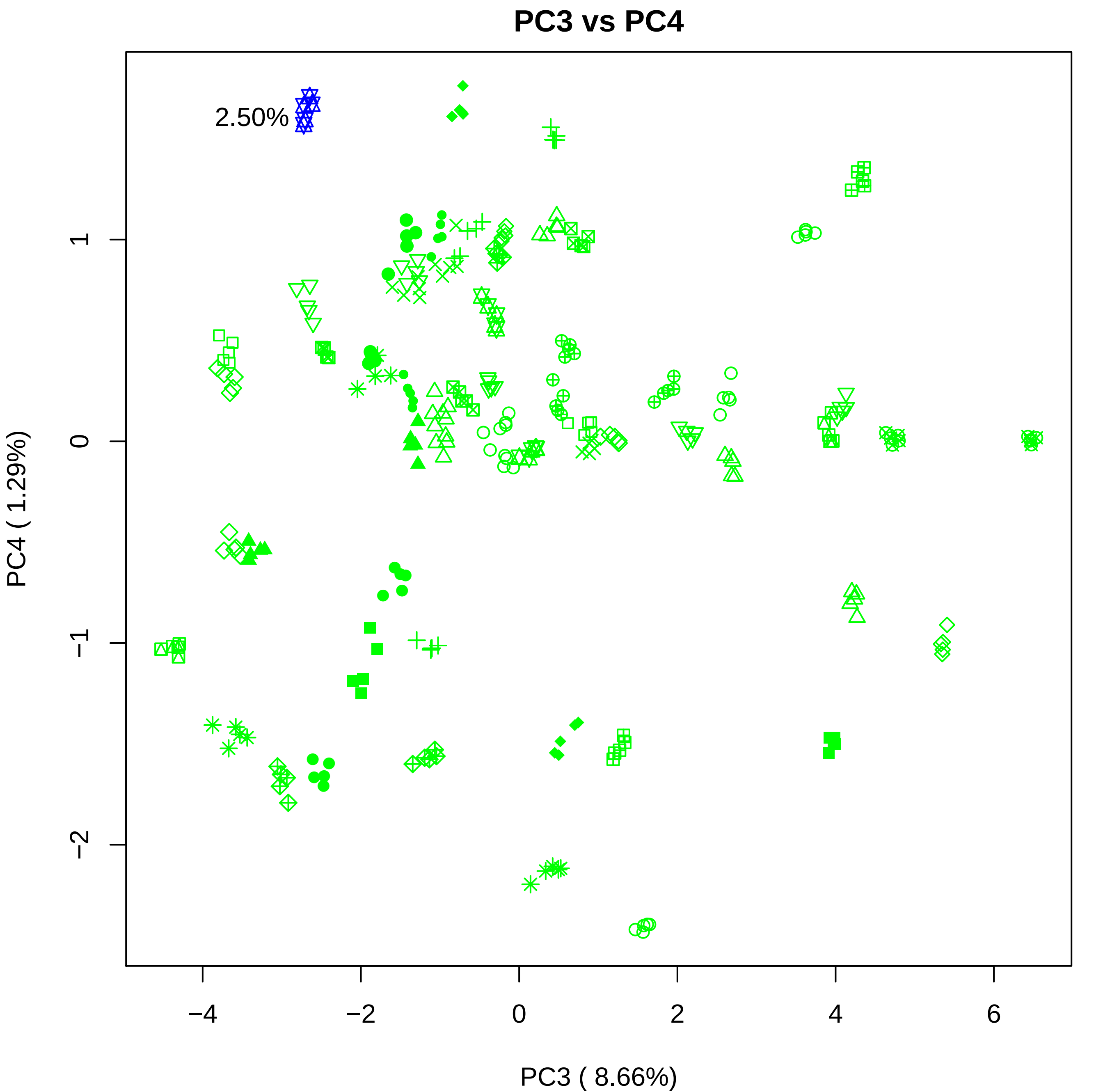
PC3 vs PC4
2.50%
PC4 ( 1.29%)
PC3 ( 8.66%)
1
0
−1
−2
−4
−2
0
2
4
6

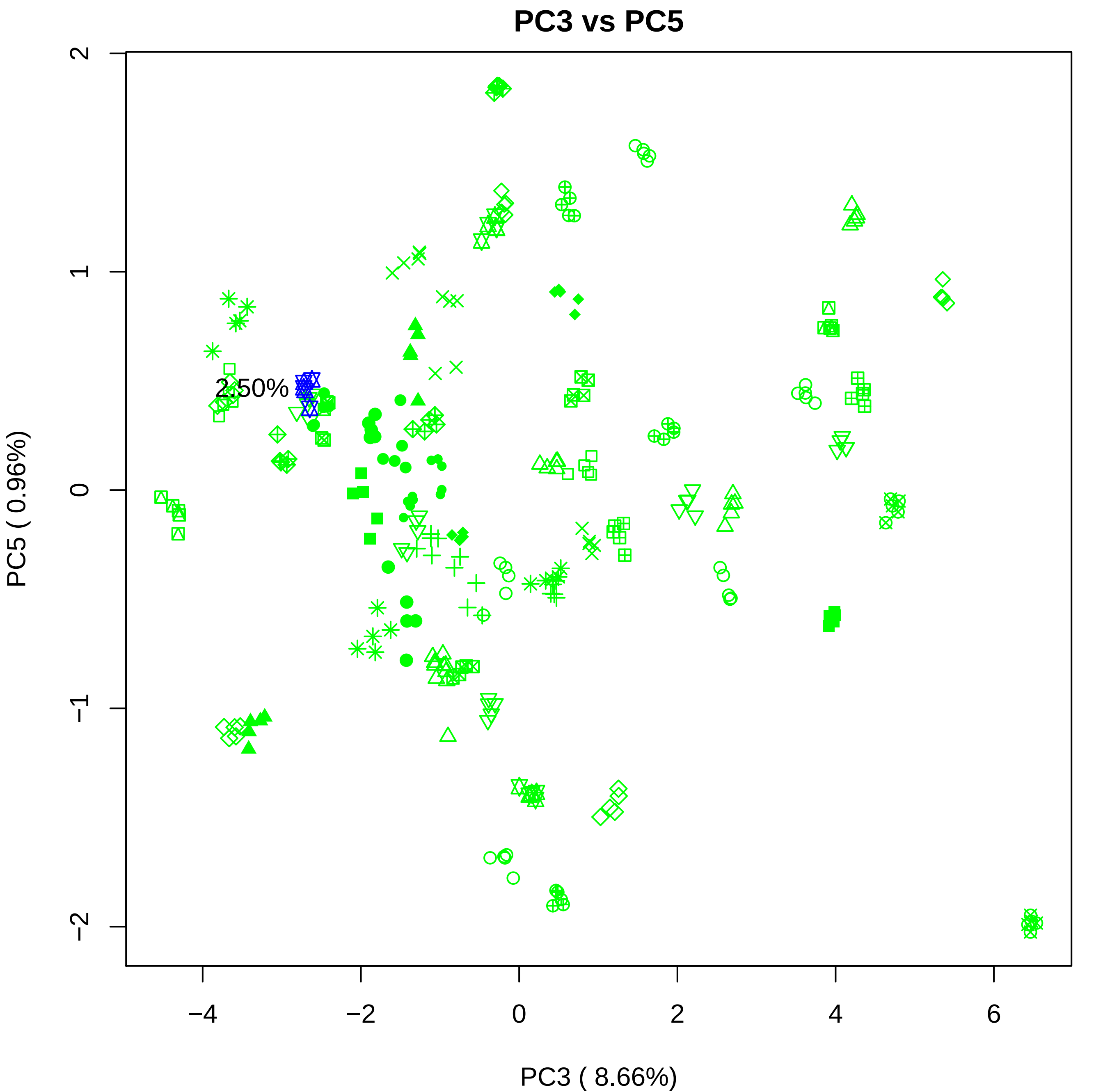
PC3 vs PC5
2.50%
PC5 ( 0.96%)
PC3 ( 8.66%)
2
1
0
−1
−2
−4
−2
0
2
4
6

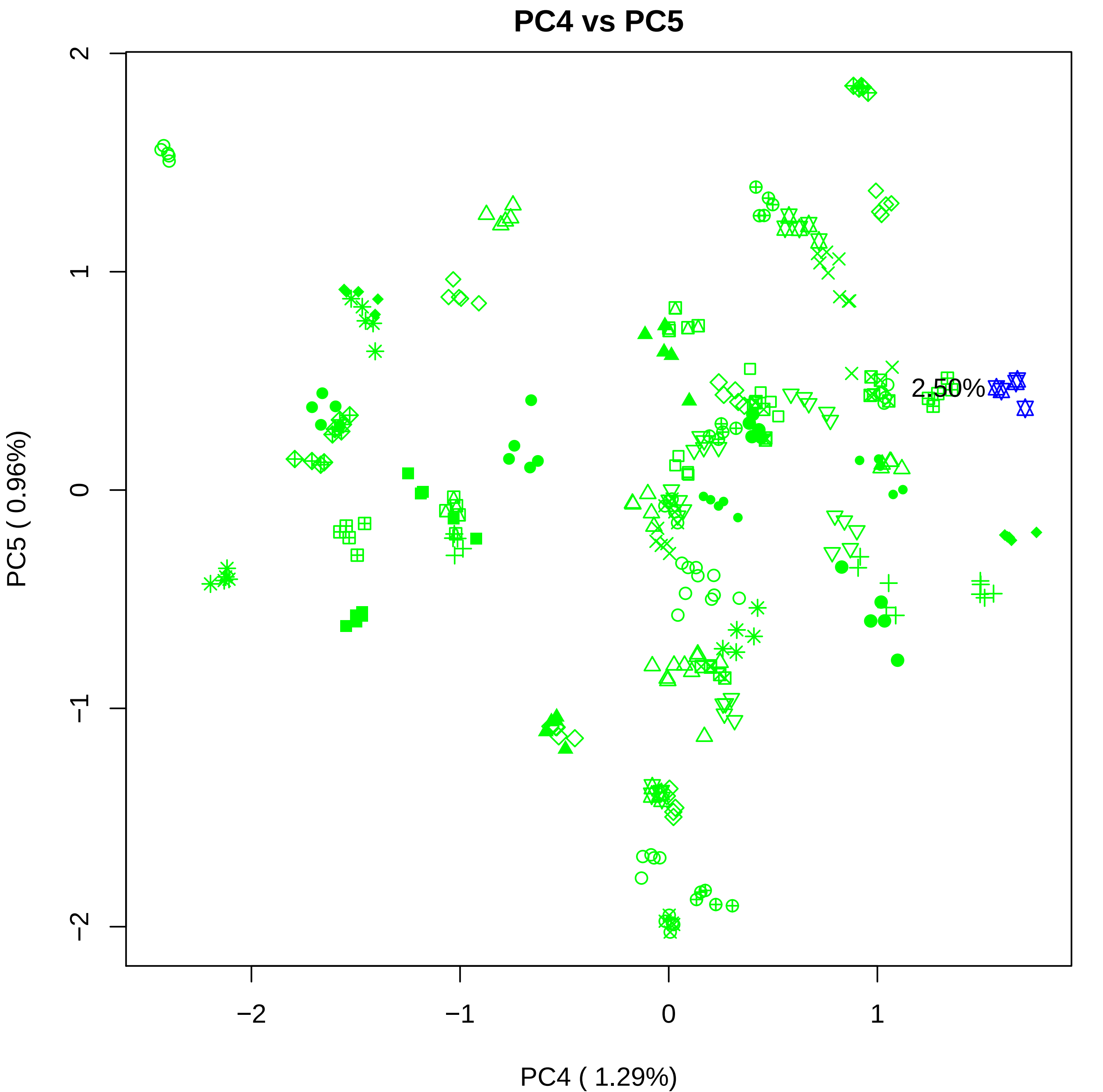

PC4 vs PC5
2
1
0
−1
−2
PC5 ( 0.96%)
−2
−1
0
1
PC4 ( 1.29%)
2.50%

### PCA of 5% BMH(c) vs PMH(c)

Scaled Matrix PCAed

7th order polynomial baseline correction performed

Raw spectra smoothed to 21 Neighbours, 2nd order PN, Savtisky Golay

Normalisation between 0 and 100

D2 spectra smoothed to 41 Neighbours, 2nd order PN, Savtisky Golay

Regions Analysed: 3601−2501 and 2001−700cm−1

Samples Analysed: 1 bContam's 0 BLH's 0 BMH's 0 OMH's 0 pContam's 0 PLH's 60 PMH's

Keele University AD

Fri Dec 14 14:18:56 2018

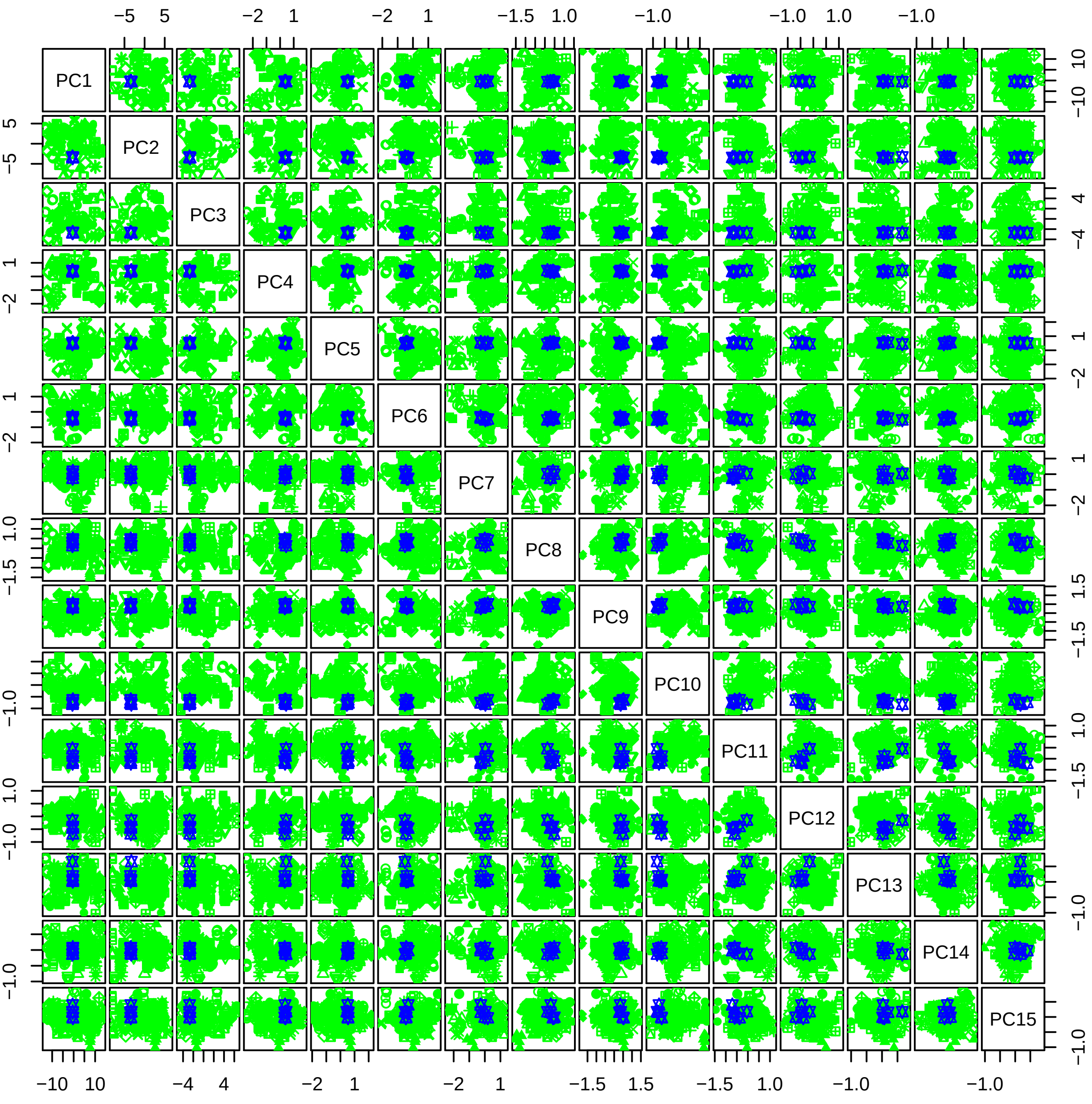

PC1
PC2
PC3
PC4
PC5
PC6
PC7
PC8
PC9
PC10
PC11
PC12
PC13
PC14
PC15

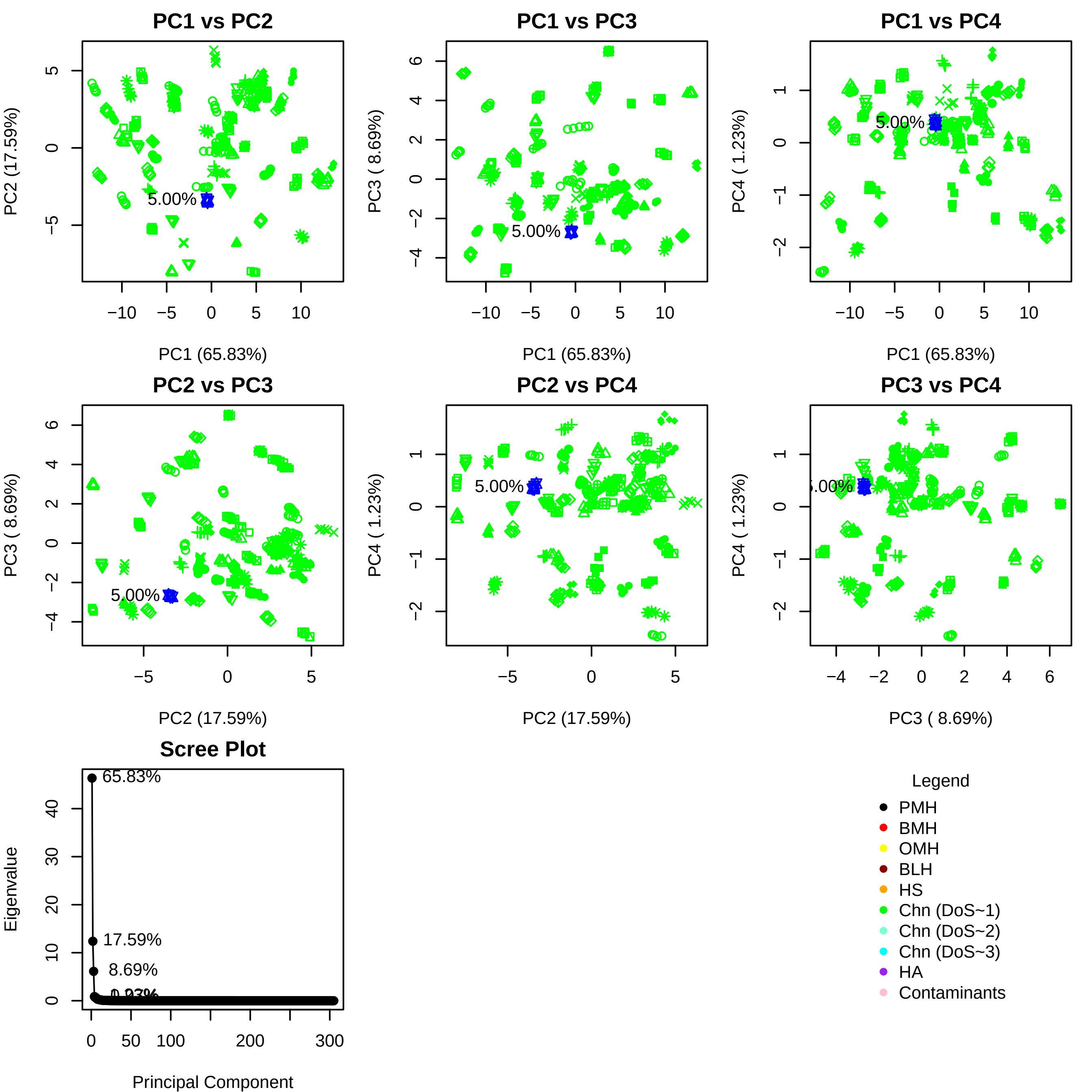

PC1 vs PC2
PC1 vs PC3
PC1 vs PC4
PC2 vs PC3
PC2 vs PC4
PC3 vs PC4
Scree Plot
PC1 (65.83%)
PC2 (17.59%)
PC3 ( 8.69%)
PC4 ( 1.23%)
5.00%
Eigenvalue
Principal Component
65.83%
17.59%
8.69%
Legend
PMH
BMH
OMH
BLH
HS
Chn (DoS~1)
Chn (DoS~2)
Chn (DoS~3)
HA
Contaminants

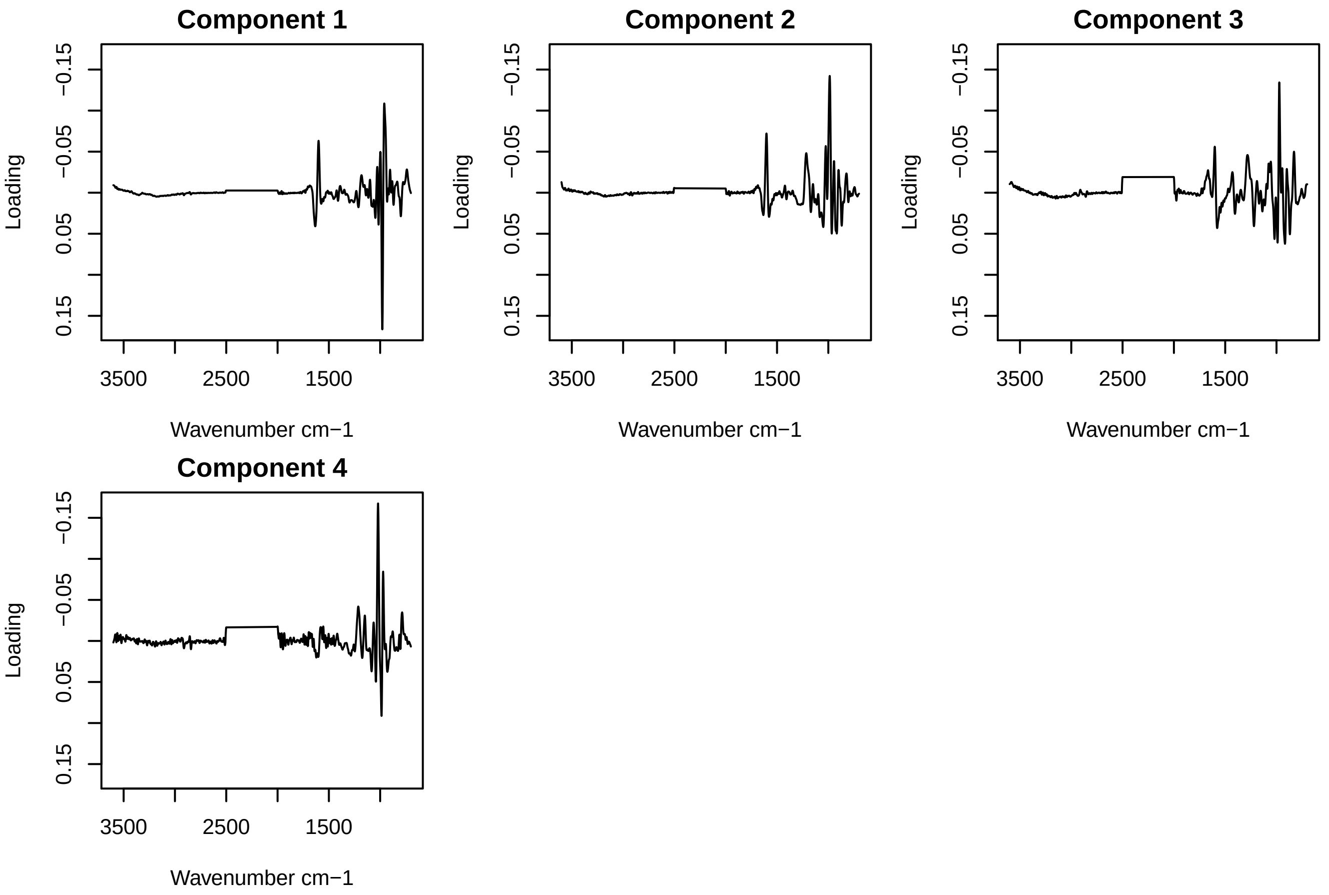
Component 1
Loading
−0.15
−0.05
0.05
0.15
3500
2500
1500
Wavenumber cm−1
Component 2
Loading
−0.15
−0.05
0.05
0.15
3500
2500
1500
Wavenumber cm−1
Component 3
Loading
−0.15
−0.05
0.05
0.15
3500
2500
1500
Wavenumber cm−1
Component 4
Loading
−0.15
−0.05
0.05
0.15
3500
2500
1500
Wavenumber cm−1

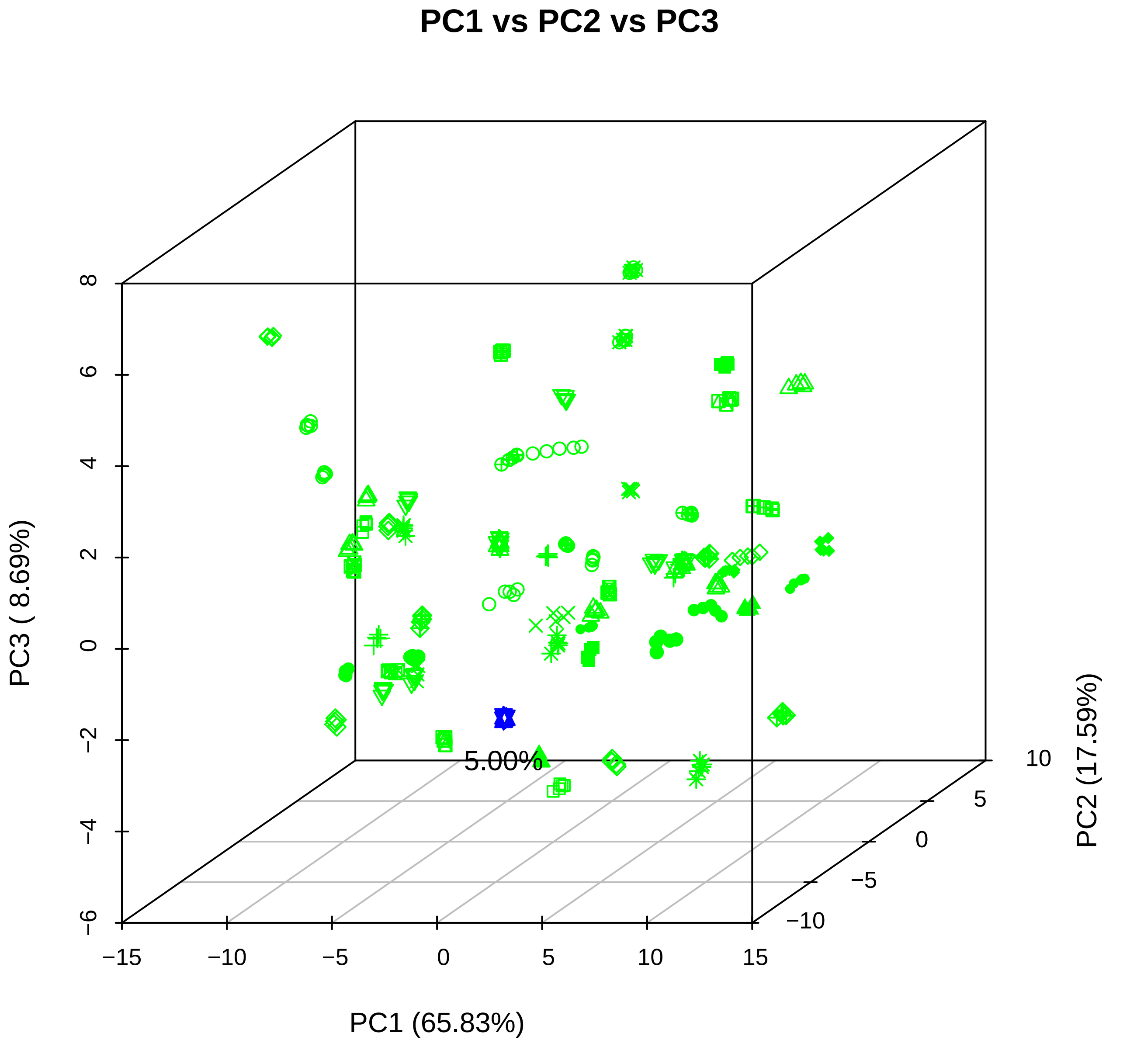
PC1 vs PC2 vs PC3
PC3 ( 8.69%)
8
6
4
2
0
−2
−4
−6
5.00%
−15
−10
−5
0
5
10
15
PC1 (65.83%)
−10
−5
0
5
10
PC2 (17.59%)

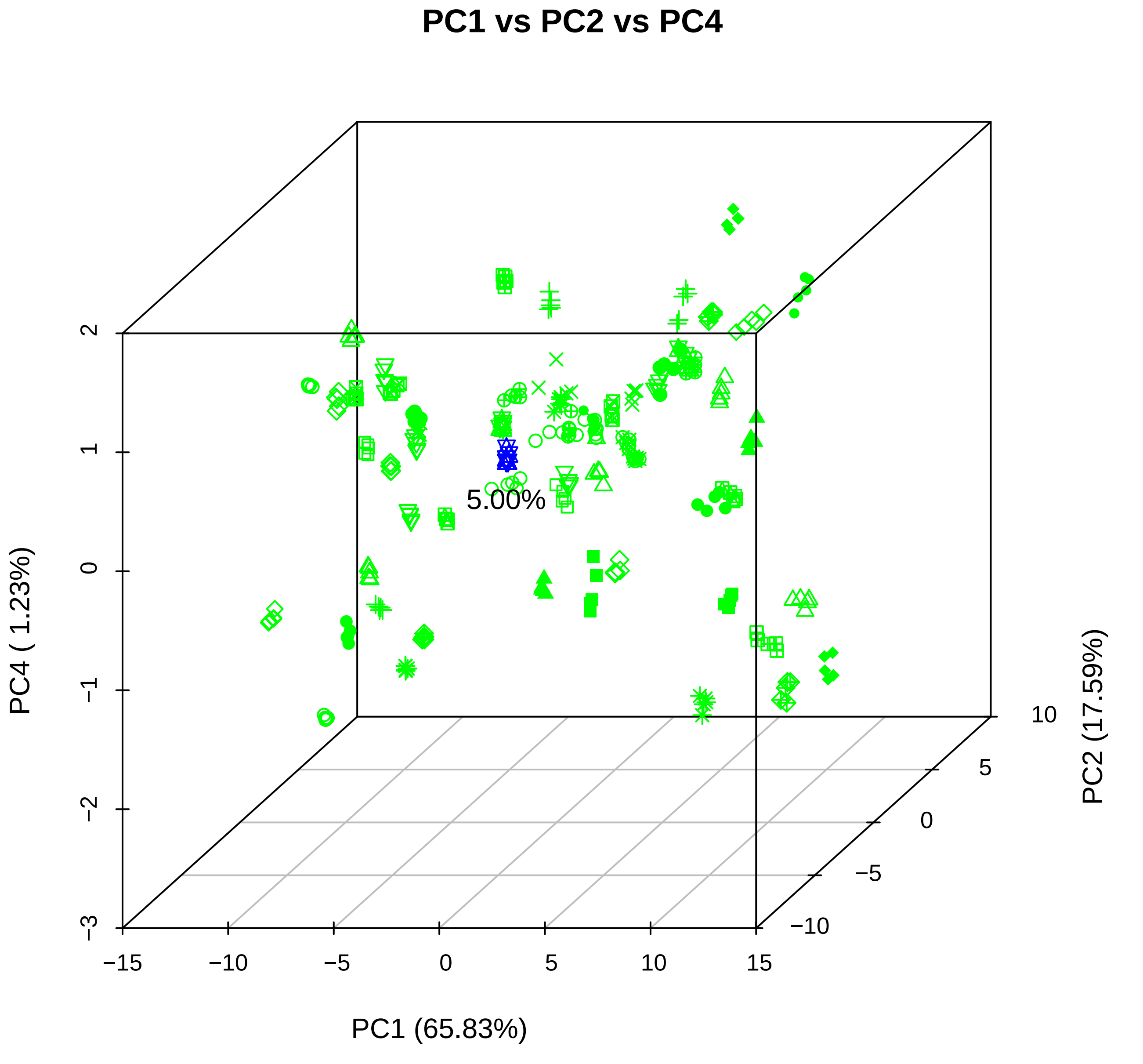

PC1 vs PC2 vs PC4
5.00%
PC1 (65.83%)
PC2 (17.59%)
PC4 ( 1.23%)
−15
−10
−5
0
5
10
15
−10
−5
0
5
10
−3
−2
−1
0
1
2

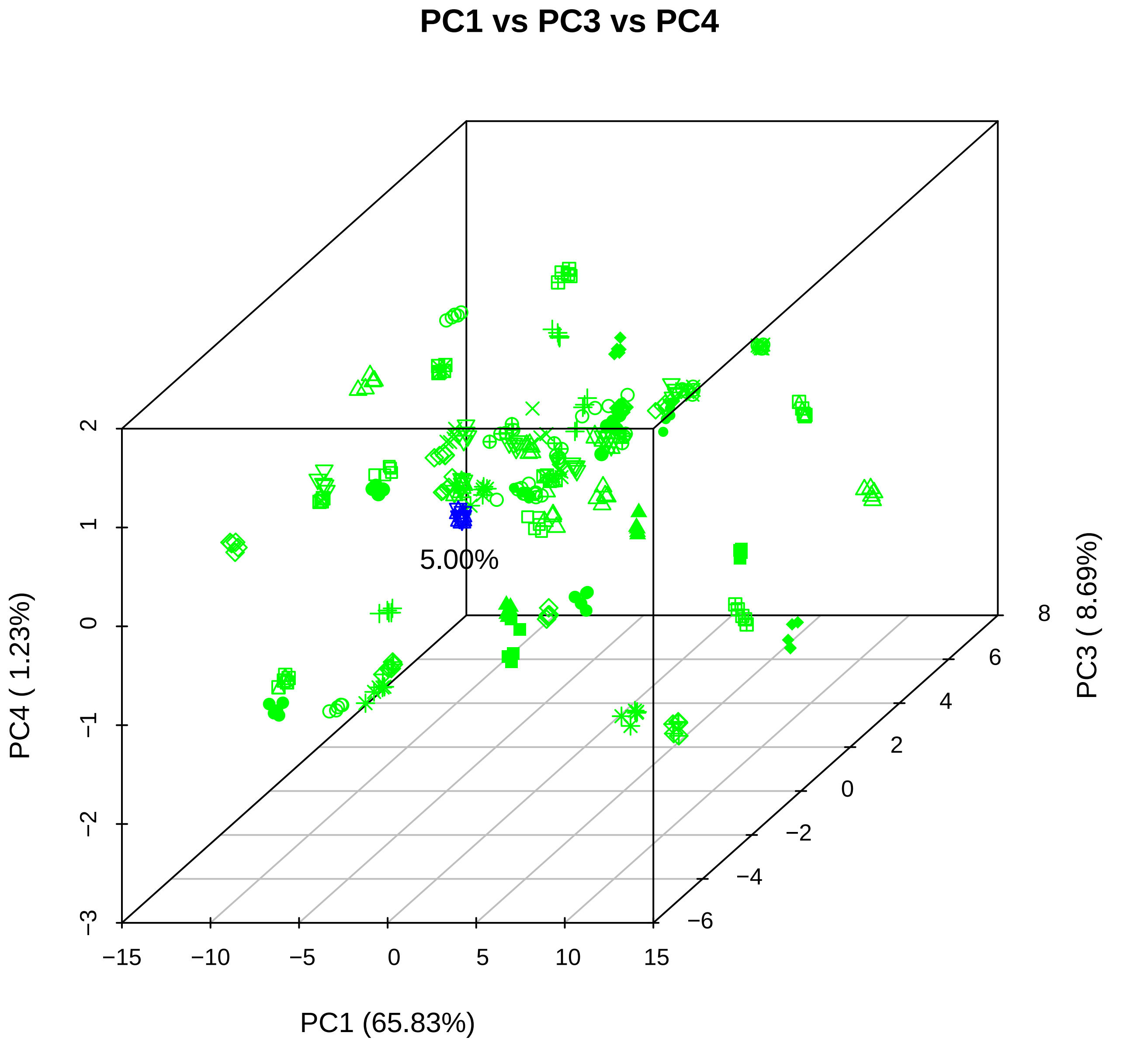
PC1 vs PC3 vs PC4
PC1 (65.83%)
PC4 ( 1.23%)
PC3 ( 8.69%)
5.00%
−15
−10
−5
0
5
10
15
−3
−2
−1
0
1
2
−6
−4
−2
0
2
4
6
8

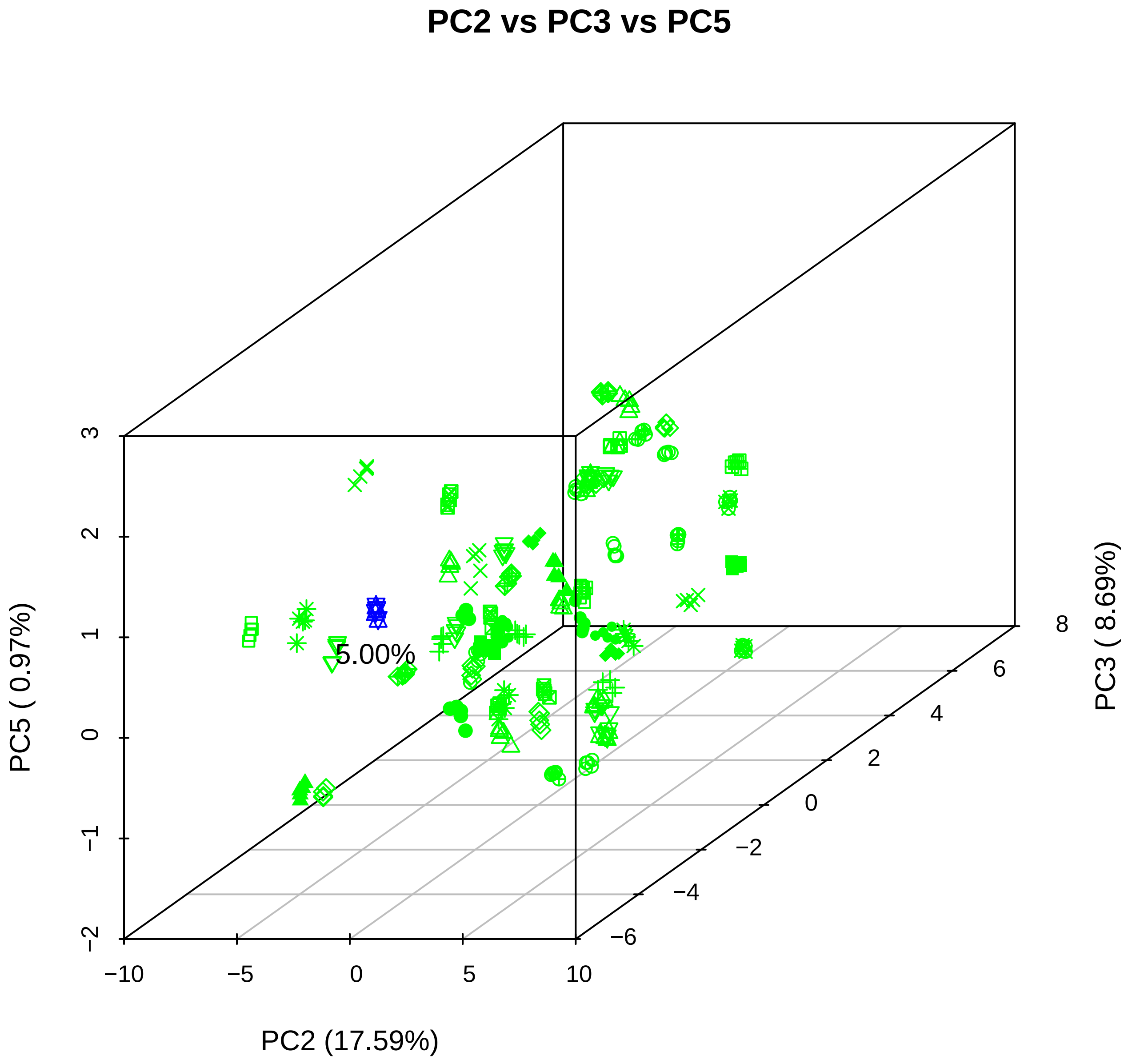

PC2 vs PC3 vs PC5
5.00%
PC5 ( 0.97%)
PC2 (17.59%)
PC3 ( 8.69%)
−10
−5
0
5
10
−2
−1
0
1
2
3
−6
−4
−2
0
2
4
6
8

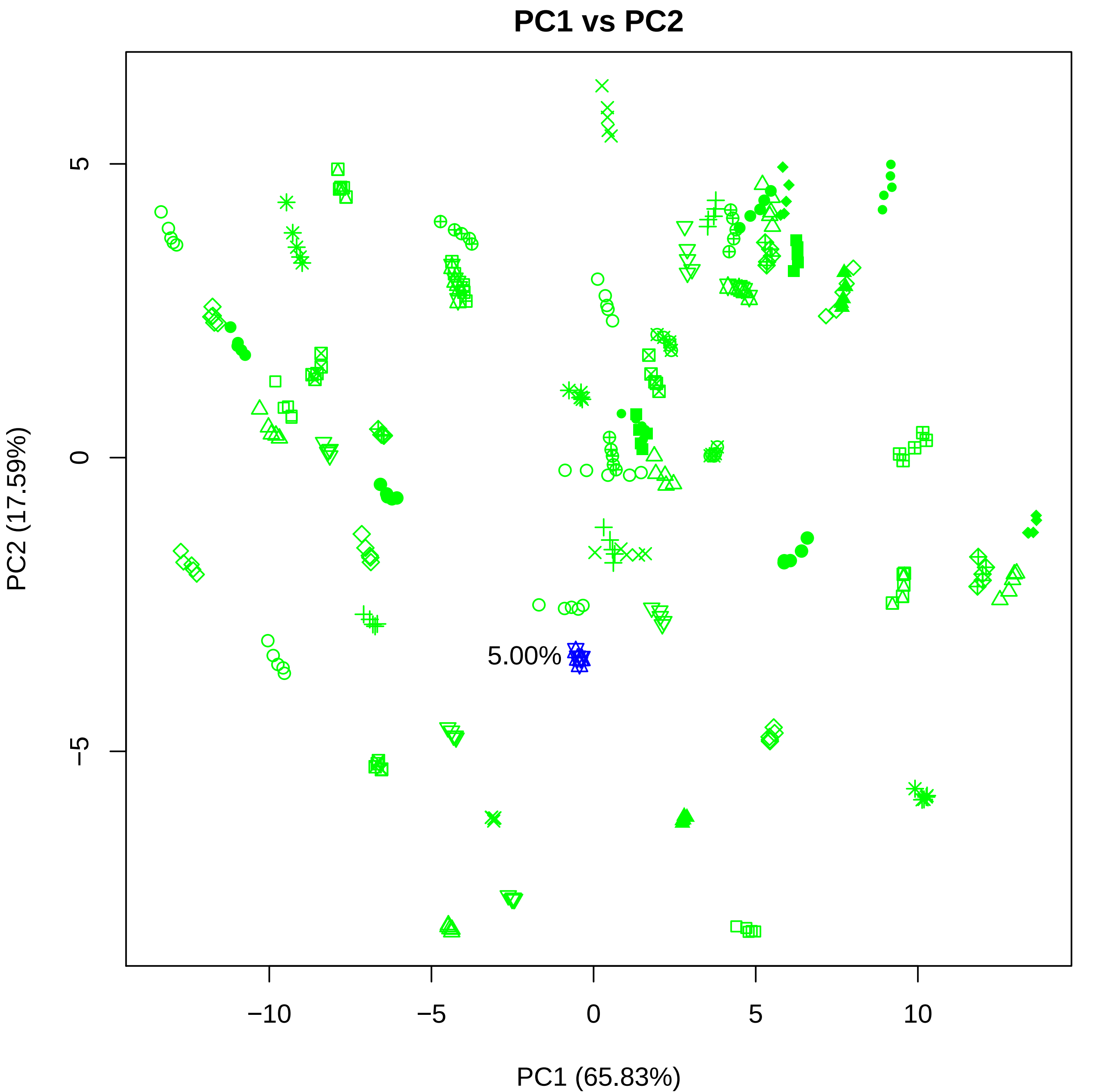
PC1 vs PC2
5
0
−5
−10
−5
0
5
10
PC2 (17.59%)
PC1 (65.83%)
5.00%

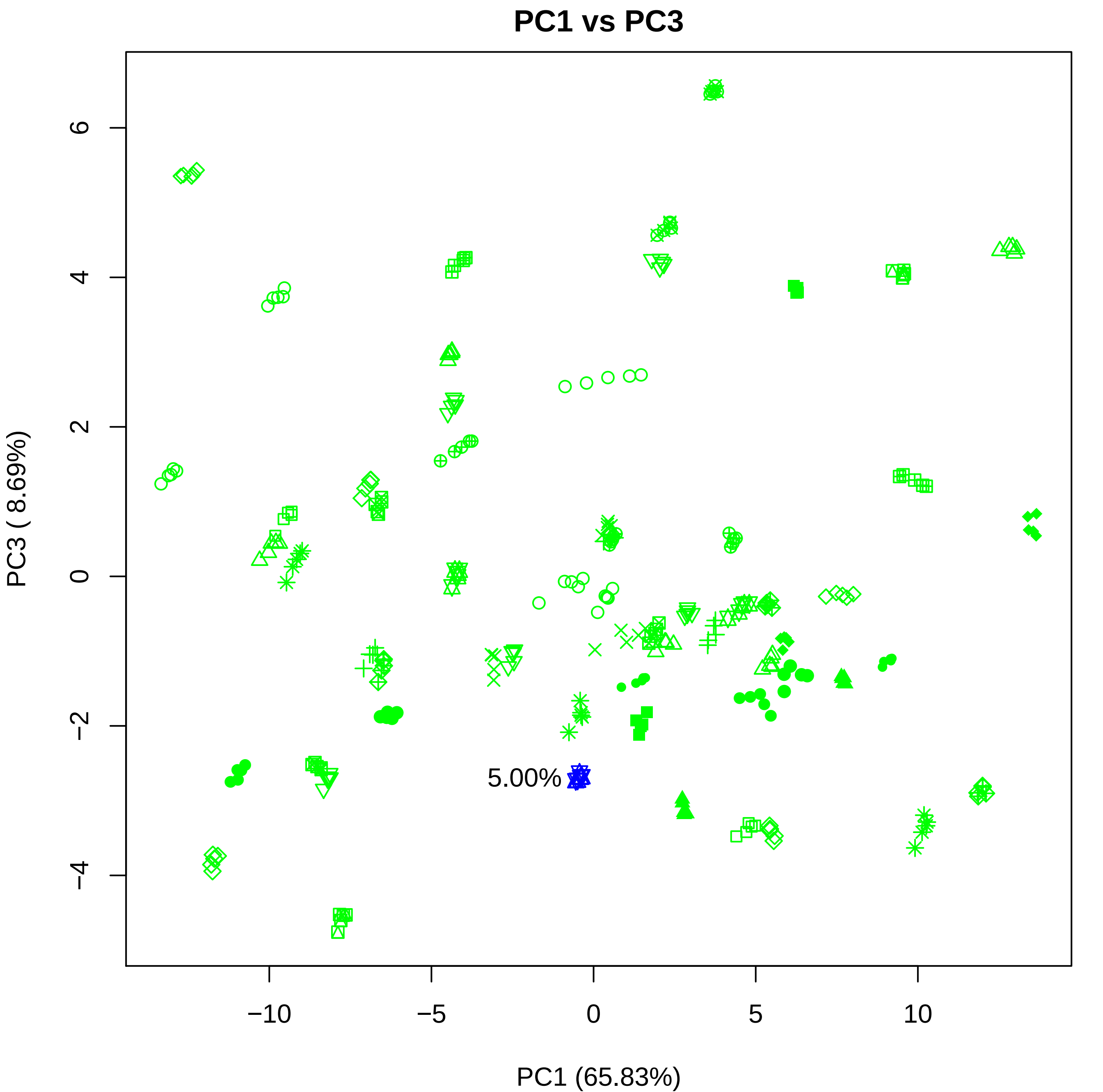
PC1 vs PC3
PC3 ( 8.69%)
PC1 (65.83%)
5.00%
6
4
2
0
−2
−4
−10
−5
0
5
10

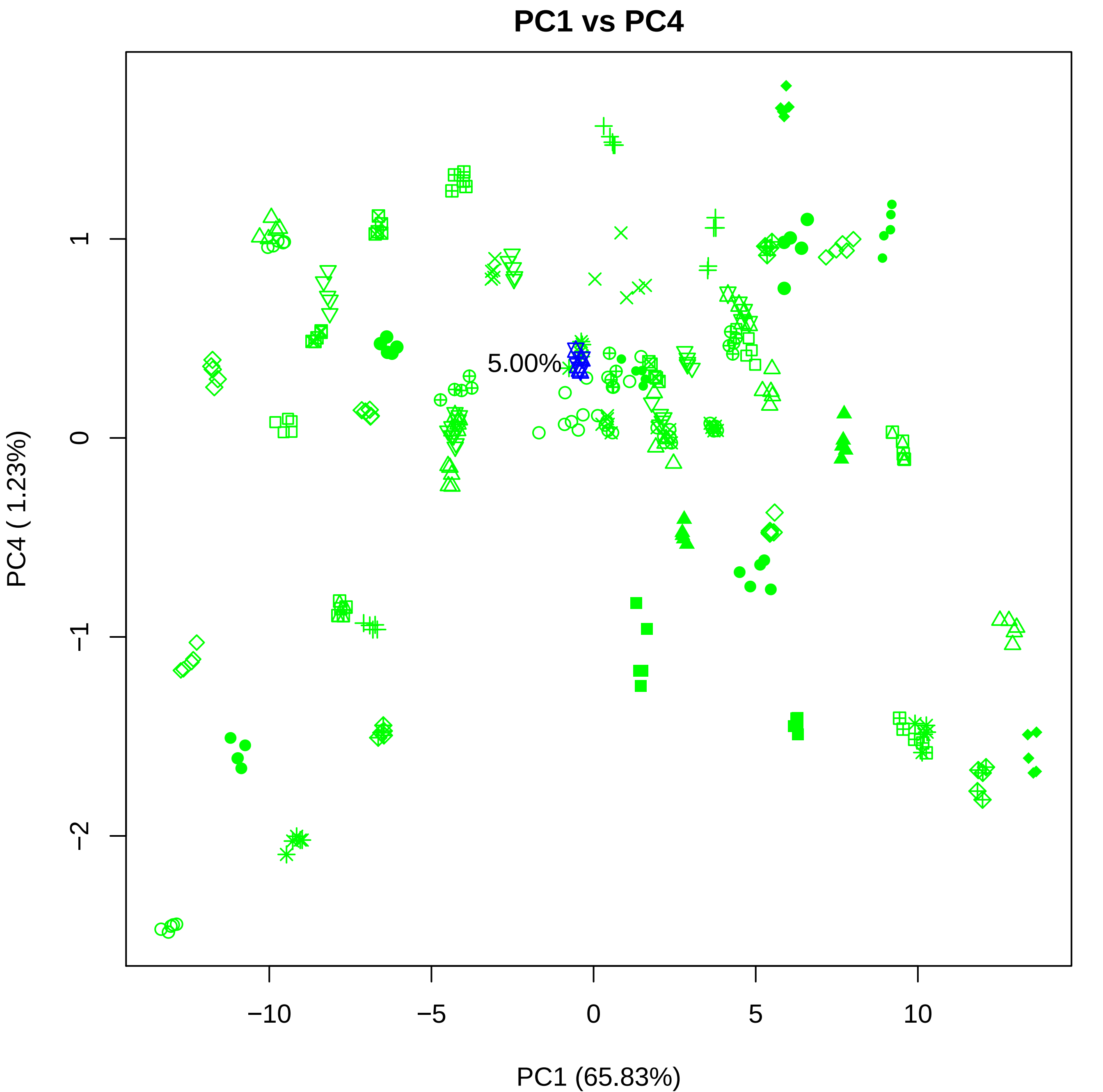

PC1 vs PC4
5.00%
PC4 ( 1.23%)
PC1 (65.83%)
−10
−5
0
5
10
−2
−1
0
1

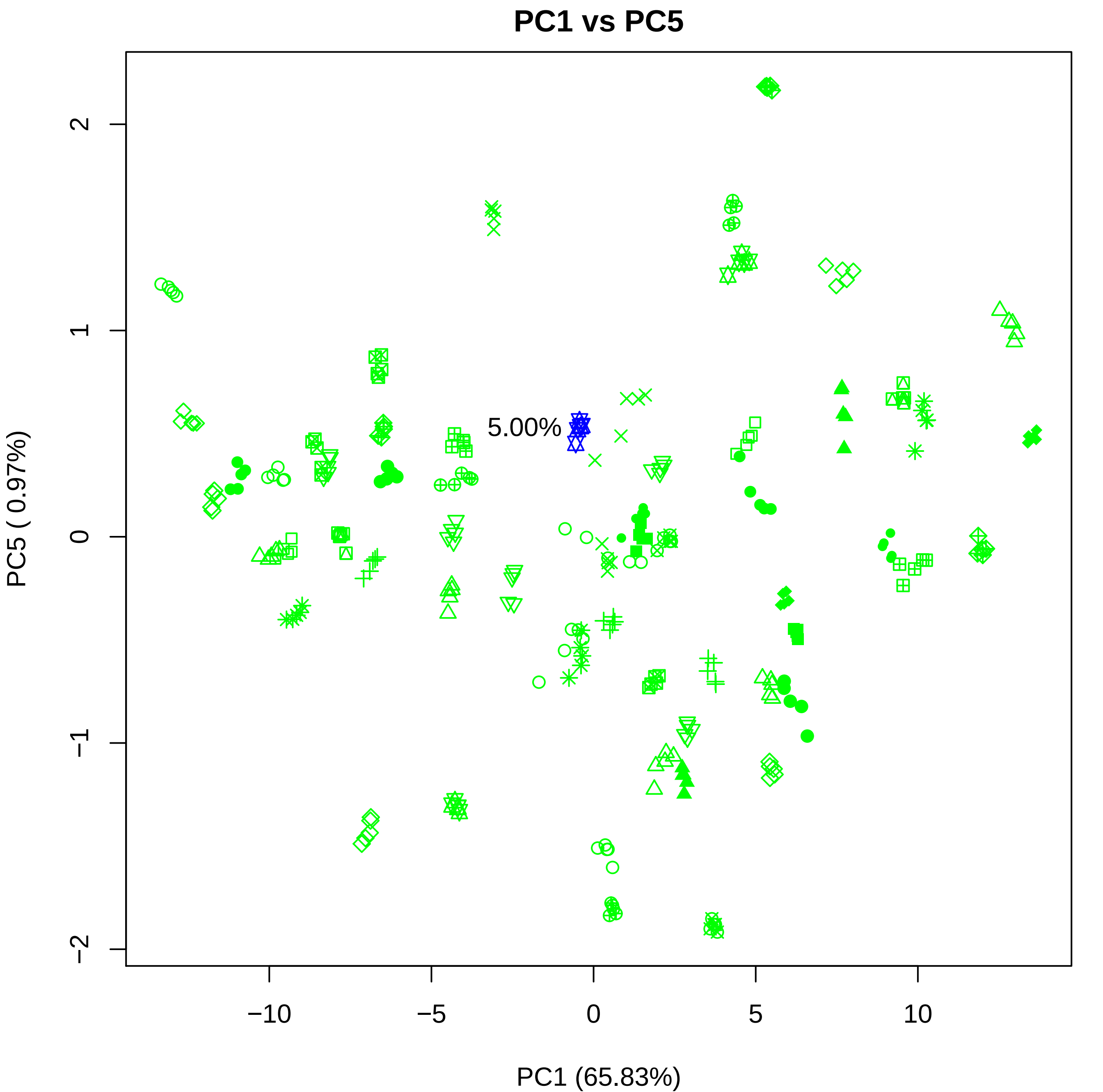

PC1 vs PC5
5.00%
PC5 ( 0.97%)
PC1 (65.83%)
2
1
0
−1
−2
−10
−5
0
5
10

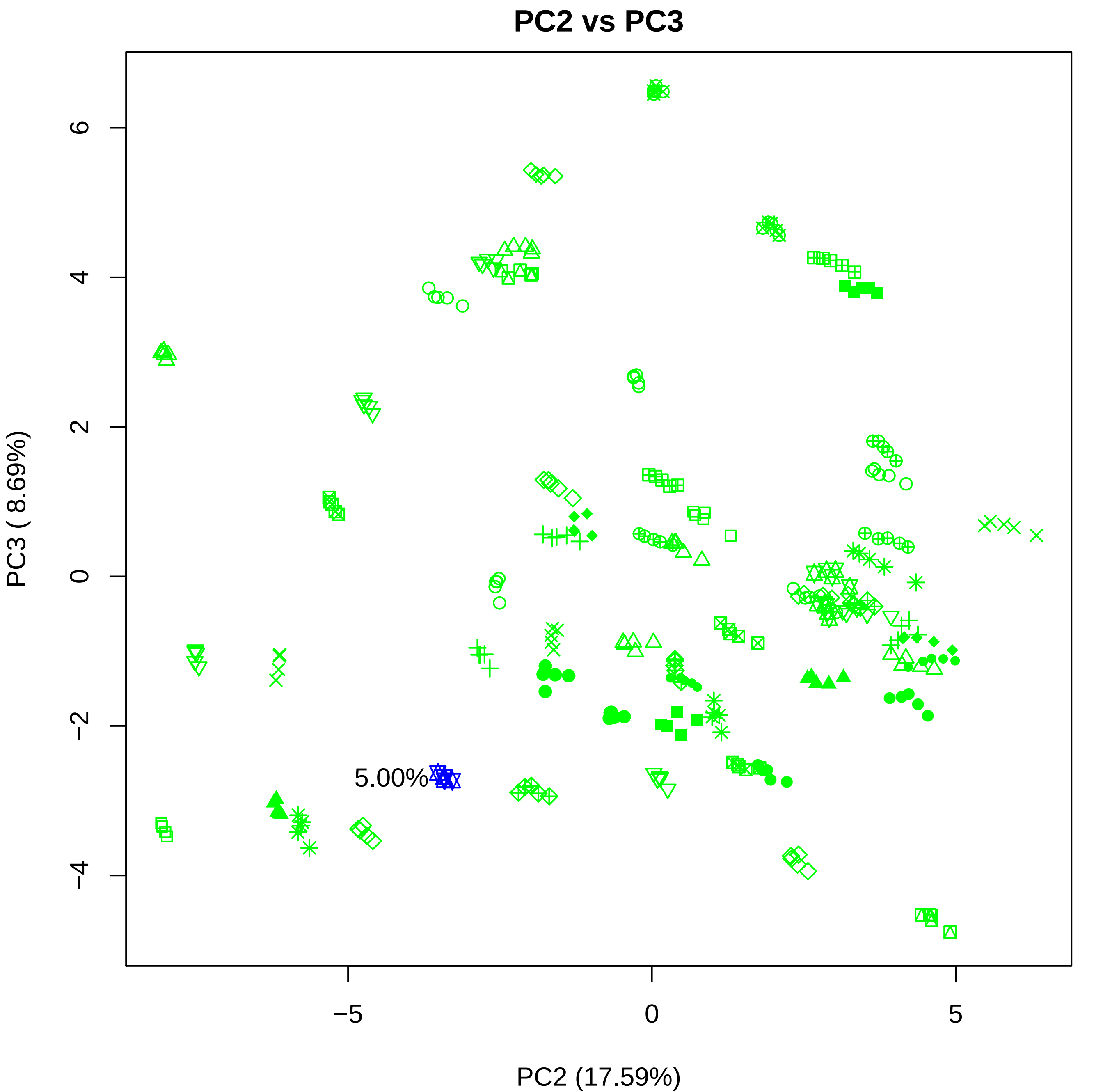
PC2 vs PC3
5.00%
PC3 ( 8.69%)
PC2 (17.59%)
−5
0
5
−4
−2
0
2
4
6

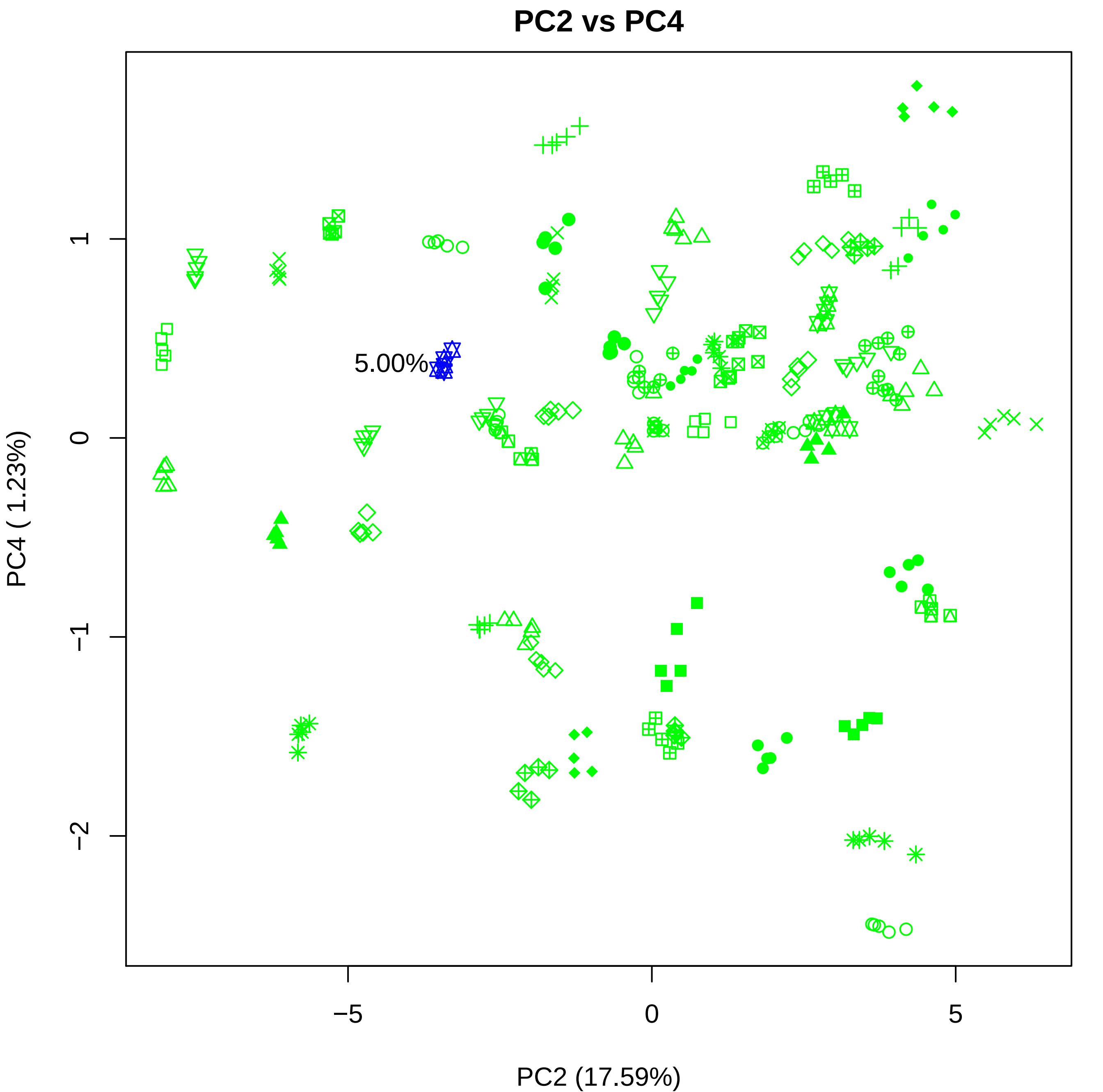
PC2 vs PC4
5.00%
PC4 ( 1.23%)
PC2 (17.59%)
−5
0
5
−2
−1
0
1

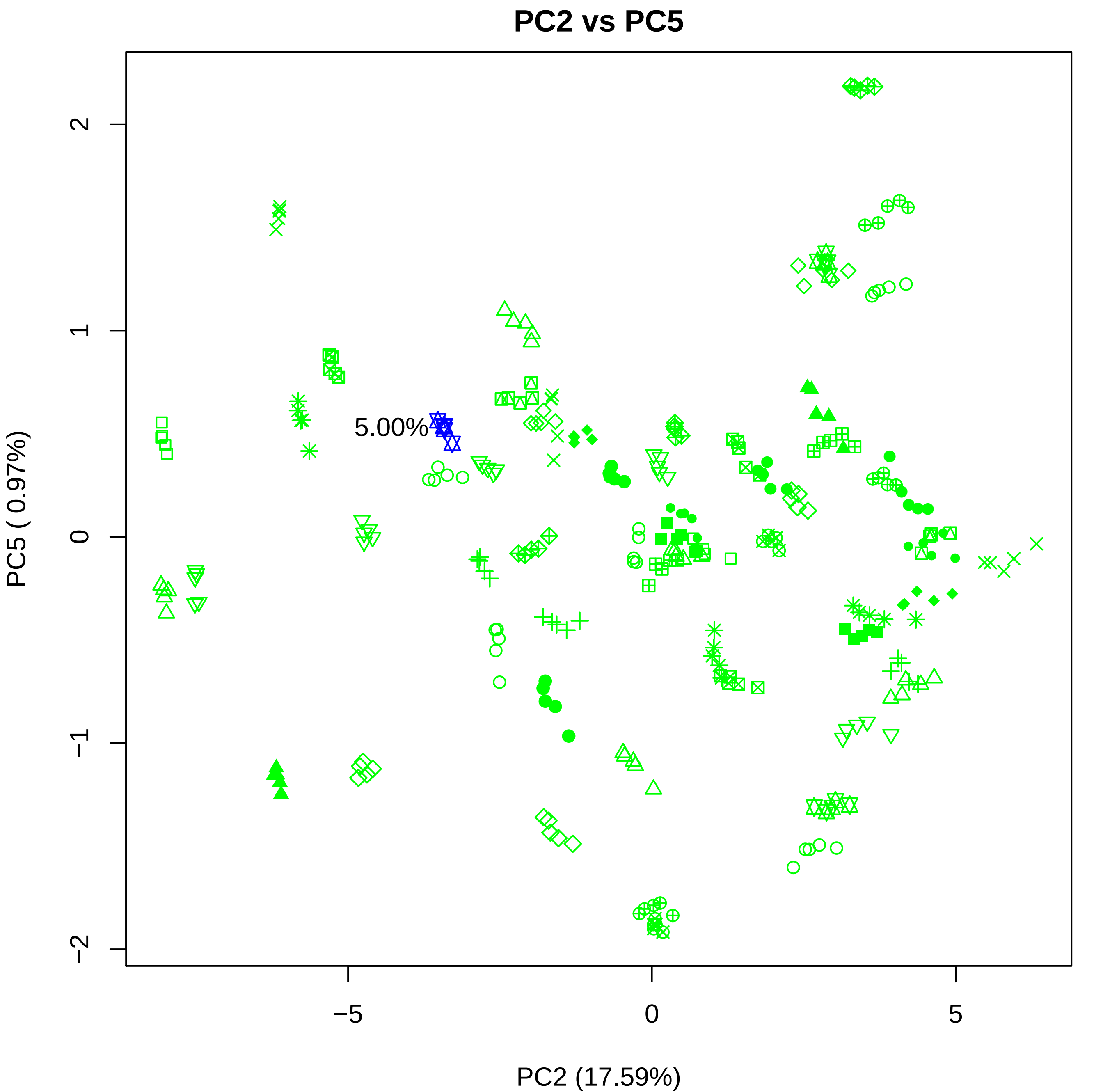
PC2 vs PC5
5.00%
PC5 ( 0.97%)
PC2 (17.59%)
2
1
0
−1
−2
−5
0
5

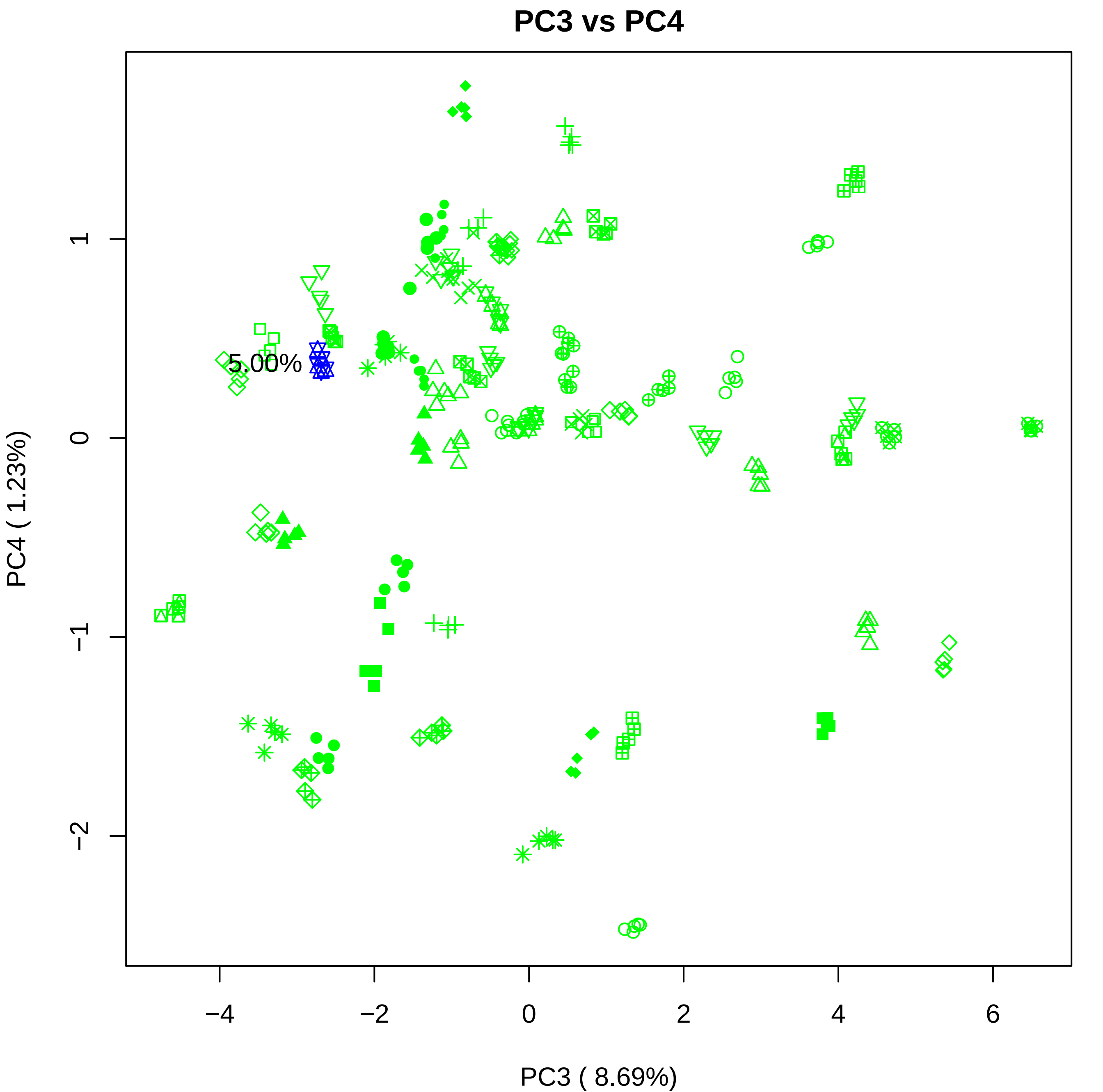
PC3 vs PC4
PC4 ( 1.23%)
PC3 ( 8.69%)
5.00%
1
0
−1
−2
−4
−2
0
2
4
6

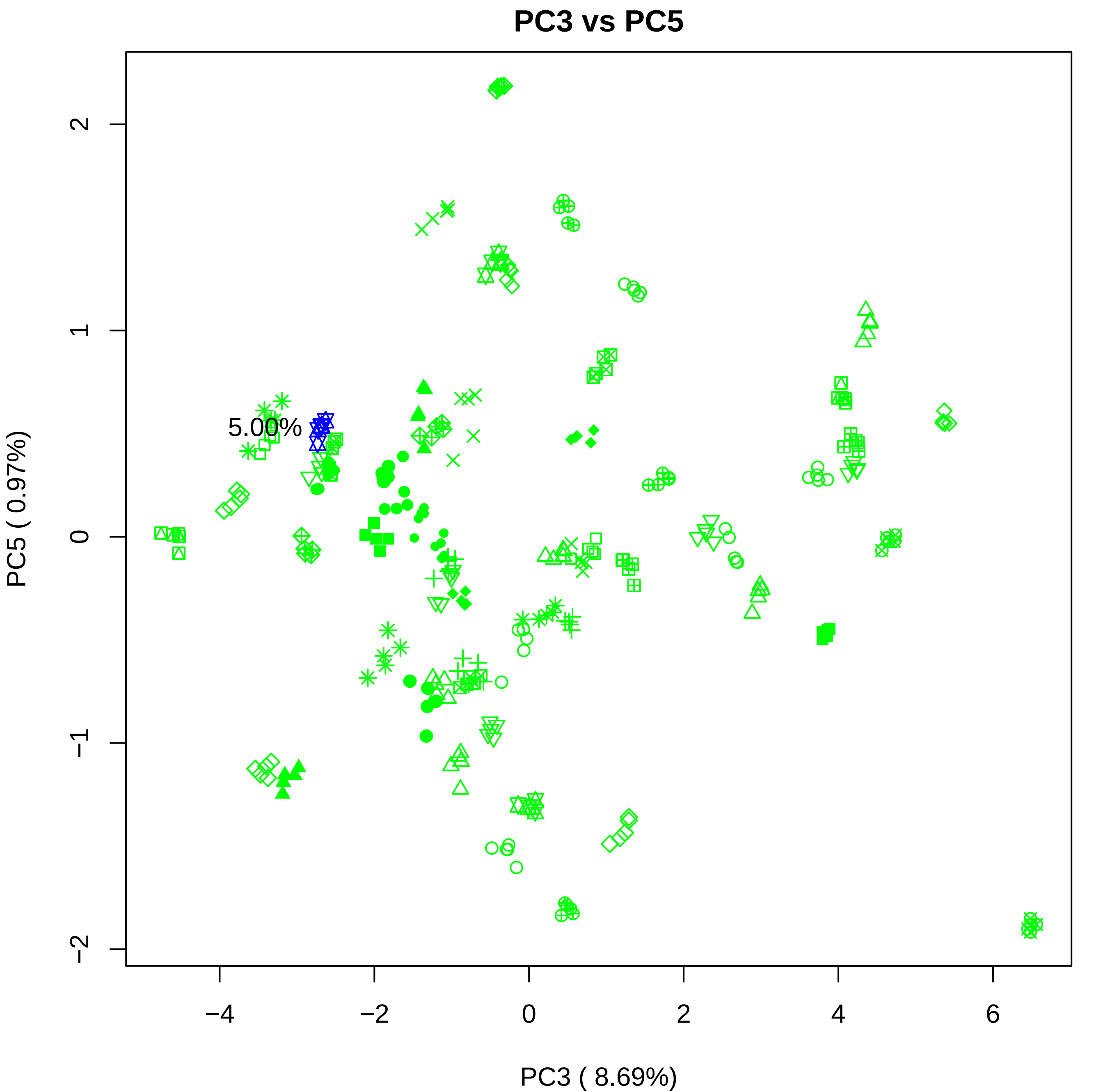
PC3 vs PC5
PC5 ( 0.97%)
PC3 ( 8.69%)
5.00%
2
1
0
−1
−2
−4
−2
0
2
4
6

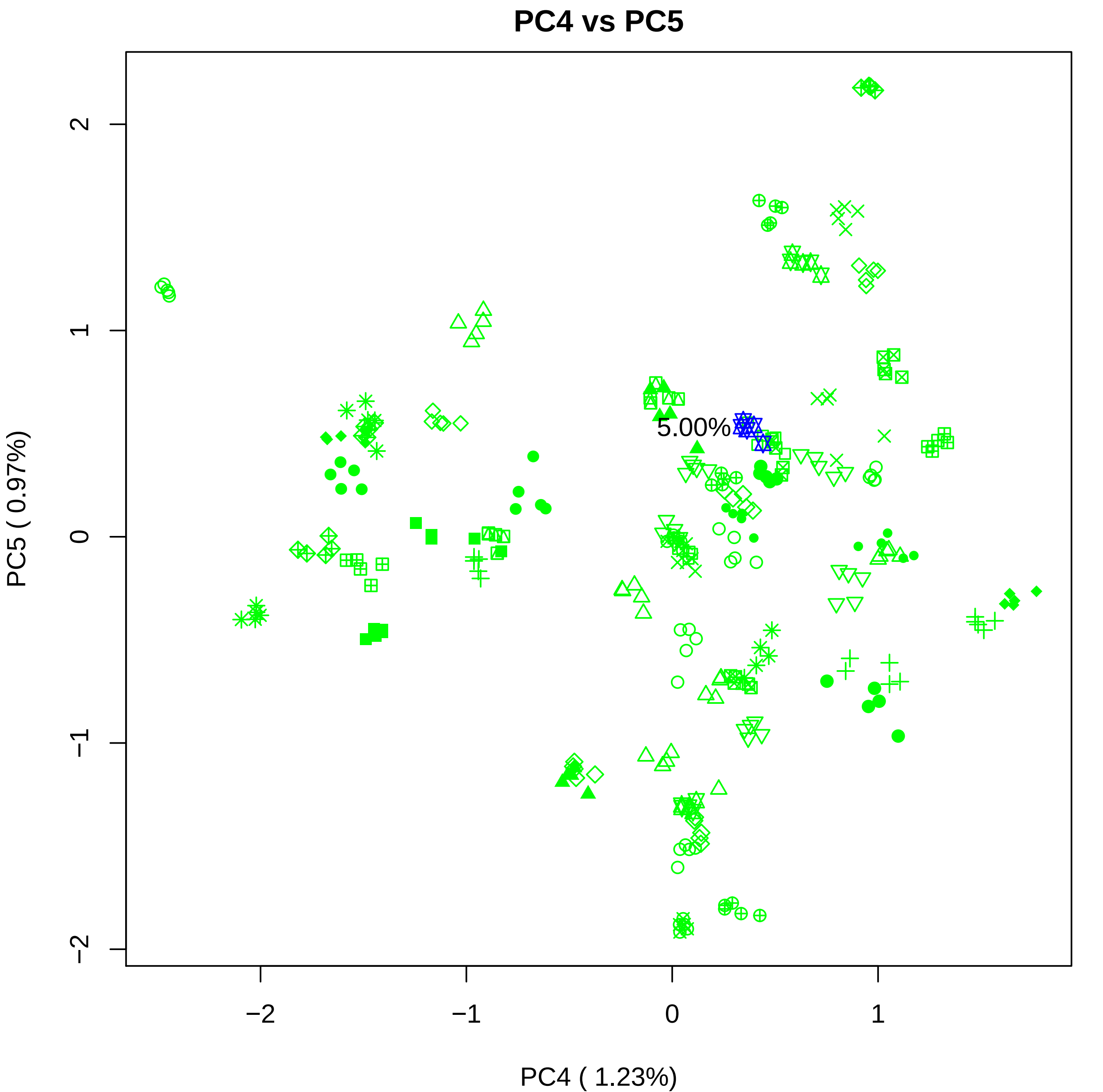
PC4 vs PC5
5.00%
PC5 ( 0.97%)
PC4 ( 1.23%)
2
1
0
−1
−2
−2
−1
0
1

### PCA of 7.5%  BMH(a) vs PMH(a)

Scaled Matrix PCAed

7th order polynomial baseline correction performed

Raw spectra smoothed to 21 Neighbours, 2nd order PN, Savtisky Golay

Normalisation between 0 and 100

D2 spectra smoothed to 41 Neighbours, 2nd order PN, Savtisky Golay

Regions Analysed: 3601−2501 and 2001−700cm−1

Samples Analysed: 1 bContam's 0 BLH's 0 BMH's 0 OMH's 0 pContam's 0 PLH's 60 PMH's

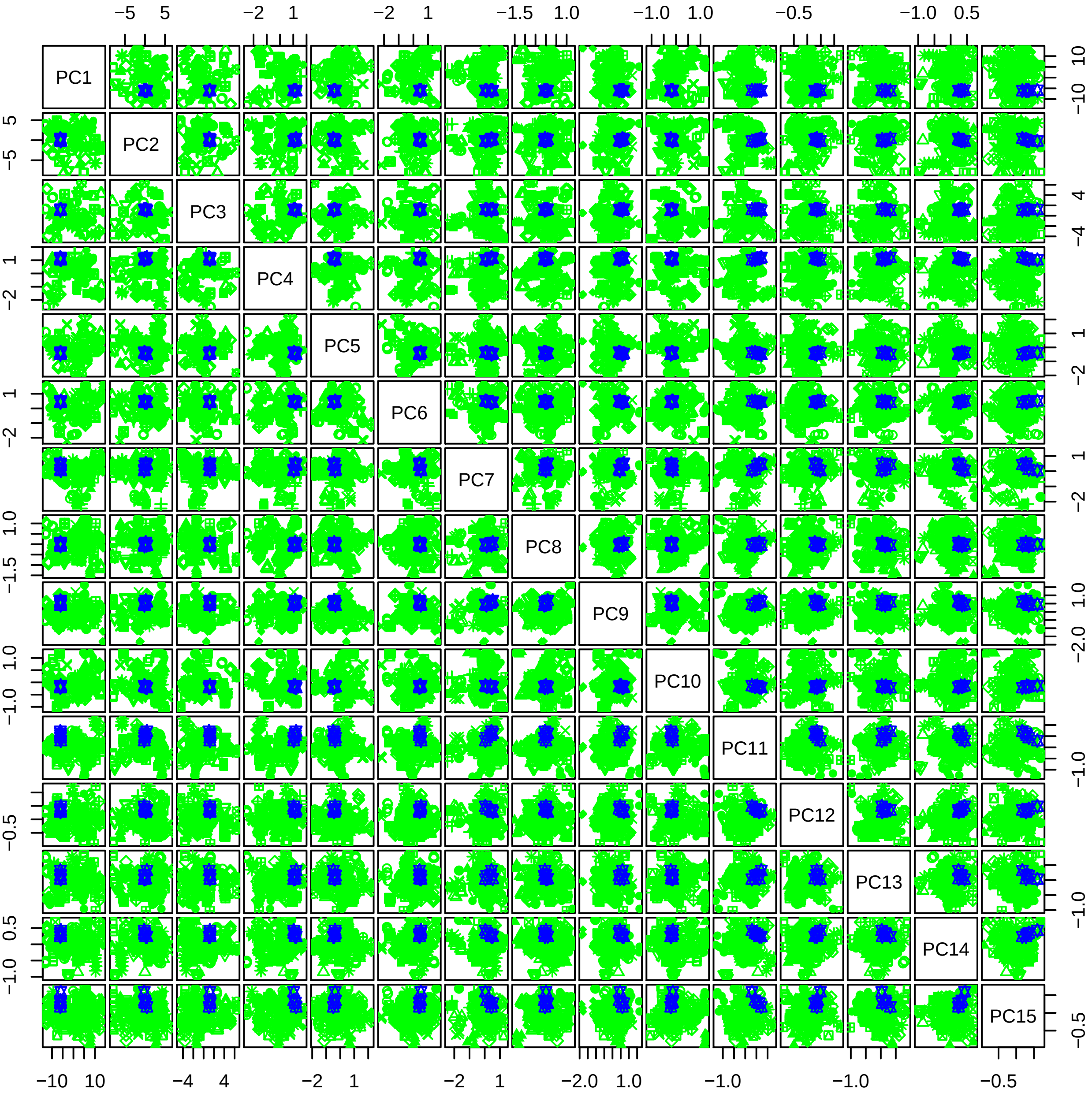
PC1
PC2
PC3
PC4
PC5
PC6
PC7
PC8
PC9
PC10
PC11
PC12
PC13
PC14
PC15

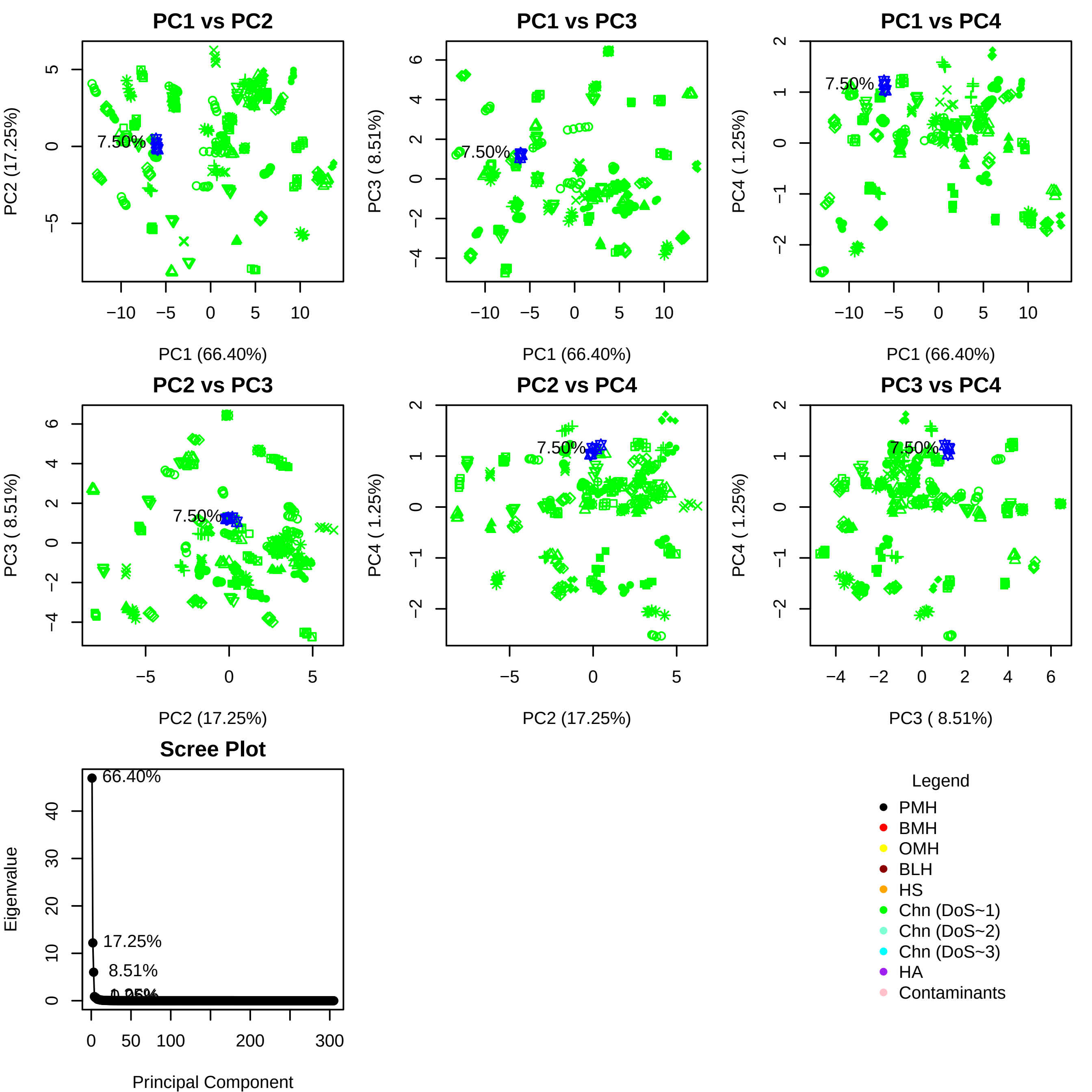

PC1 vs PC2
PC1 vs PC3
PC1 vs PC4
PC2 vs PC3
PC2 vs PC4
PC3 vs PC4
PC1 (66.40%)
PC2 (17.25%)
PC3 ( 8.51%)
PC4 ( 1.25%)
7.50%
Scree Plot
Eigenvalue
Principal Component
66.40%
17.25%
8.51%
Legend
PMH
BMH
OMH
BLH
HS
Chn (DoS~1)
Chn (DoS~2)
Chn (DoS~3)
HA
Contaminants

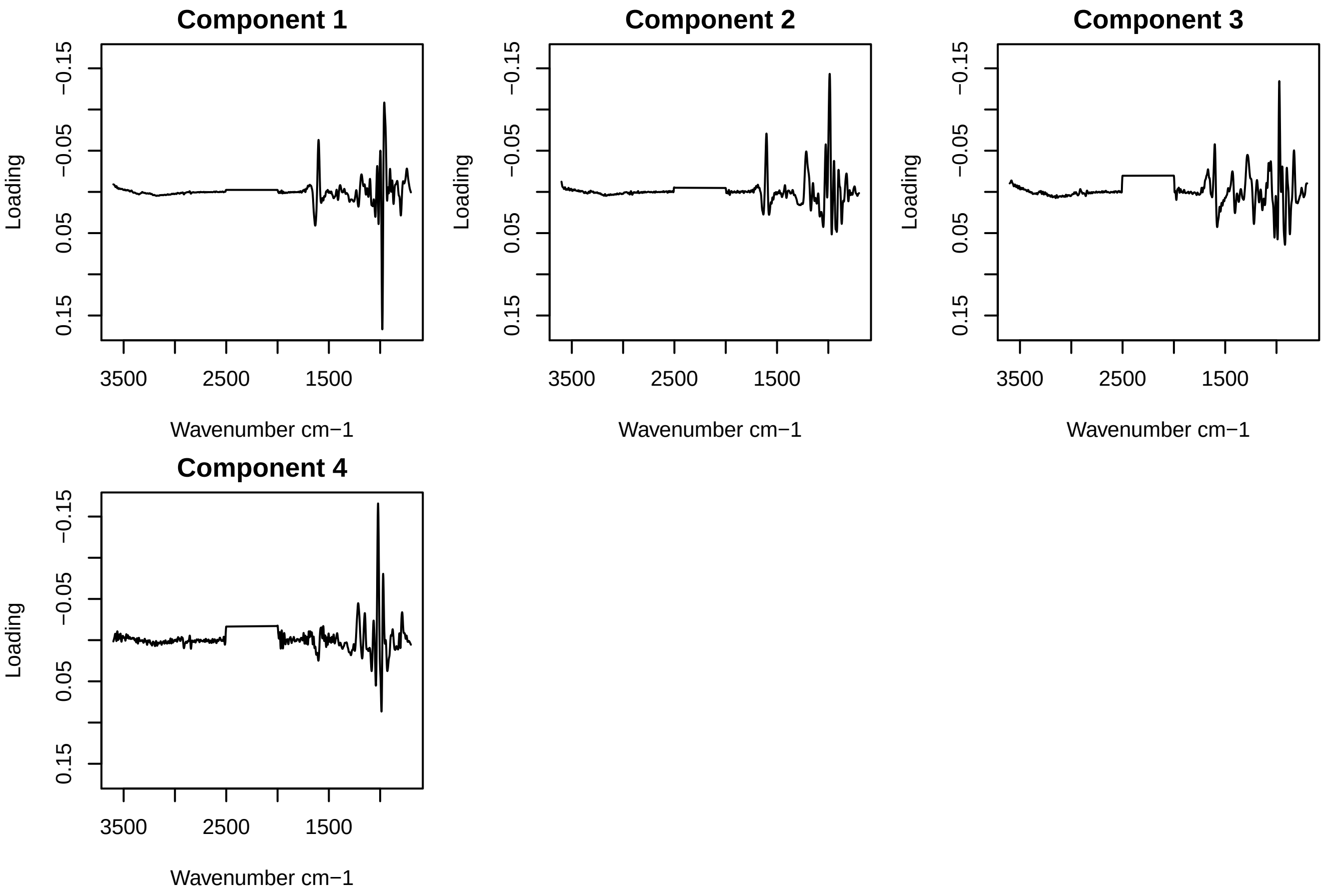
Component 1
Component 2
Component 3
Component 4
Loading
−0.15
−0.05
0.05
0.15
3500
2500
1500
Wavenumber cm−1

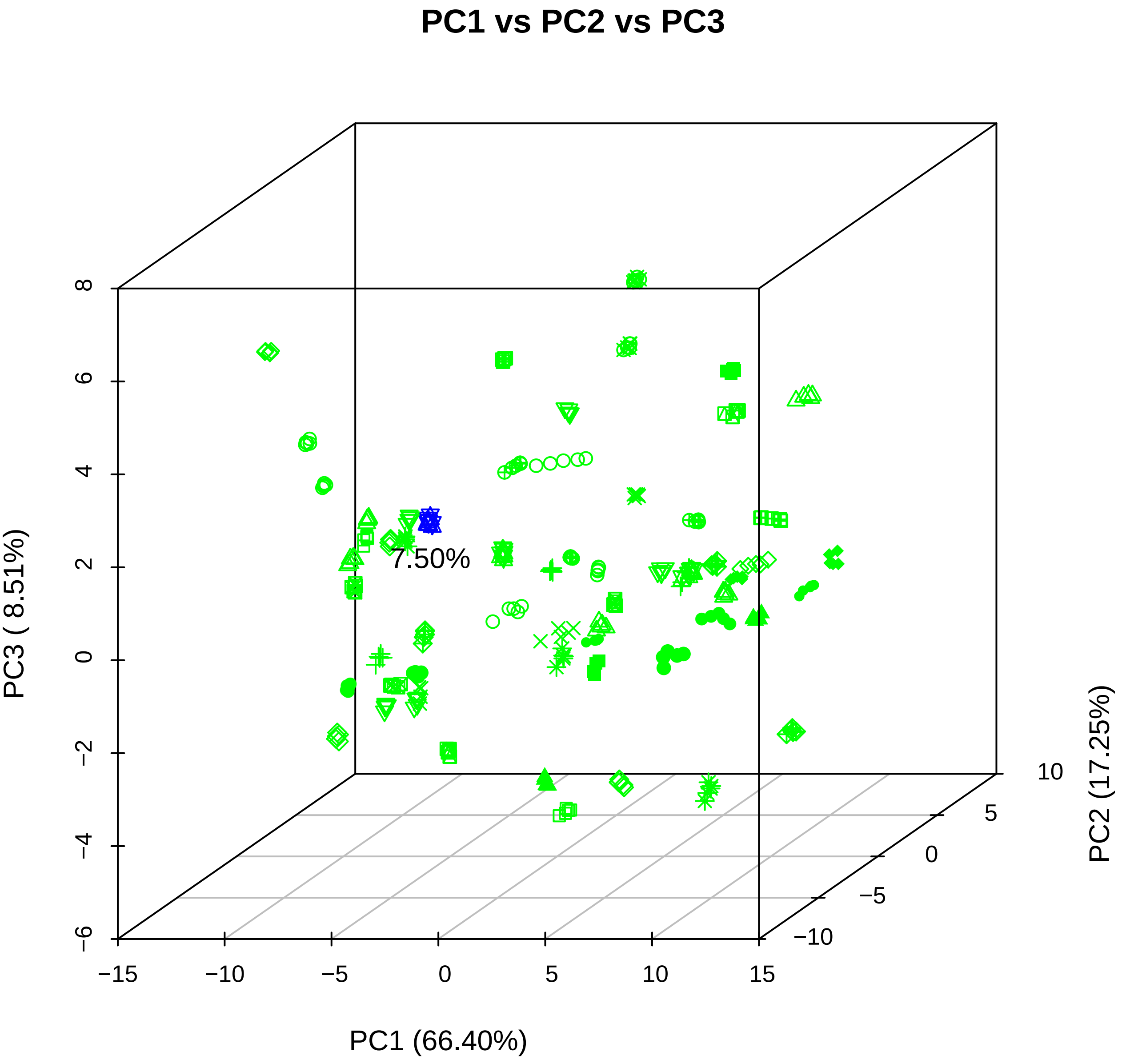
PC1 vs PC2 vs PC3
8
6
4
2
0
−2
−4
−6
PC3 ( 8.51%)
7.50%
−15
−10
−5
0
5
10
15
PC1 (66.40%)
10
5
0
−5
−10
PC2 (17.25%)

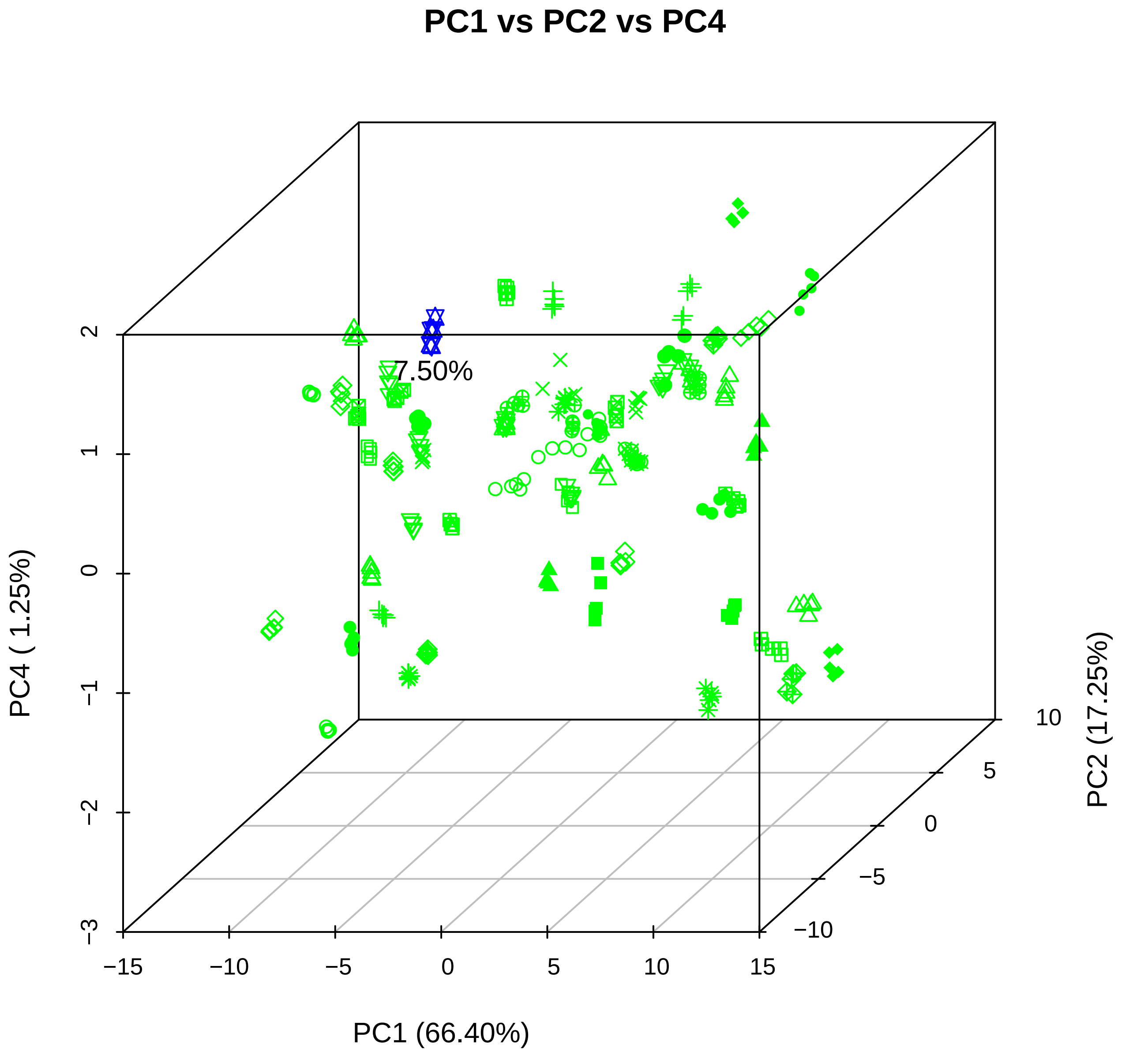
PC1 vs PC2 vs PC4
7.50%
PC1 (66.40%)
PC4 ( 1.25%)
PC2 (17.25%)
−15
−10
−5
0
5
10
15
−3
−2
−1
0
1
2
−10
−5
0
5
10

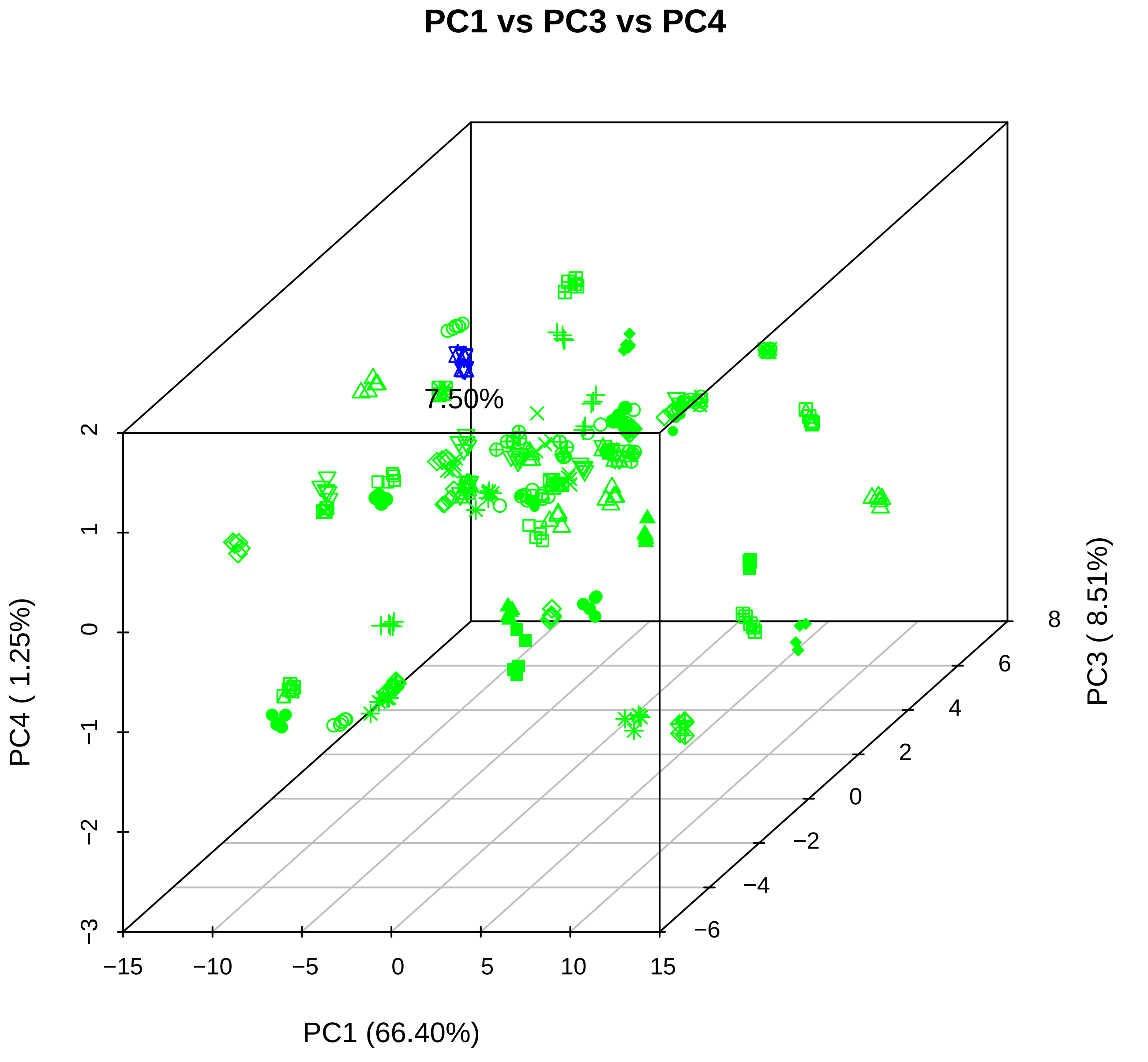
PC1 vs PC3 vs PC4
7.50%
PC1 (66.40%)
PC4 ( 1.25%)
PC3 ( 8.51%)
−15
−10
−5
0
5
10
15
−3
−2
−1
0
1
2
−6
−4
−2
0
2
4
6
8

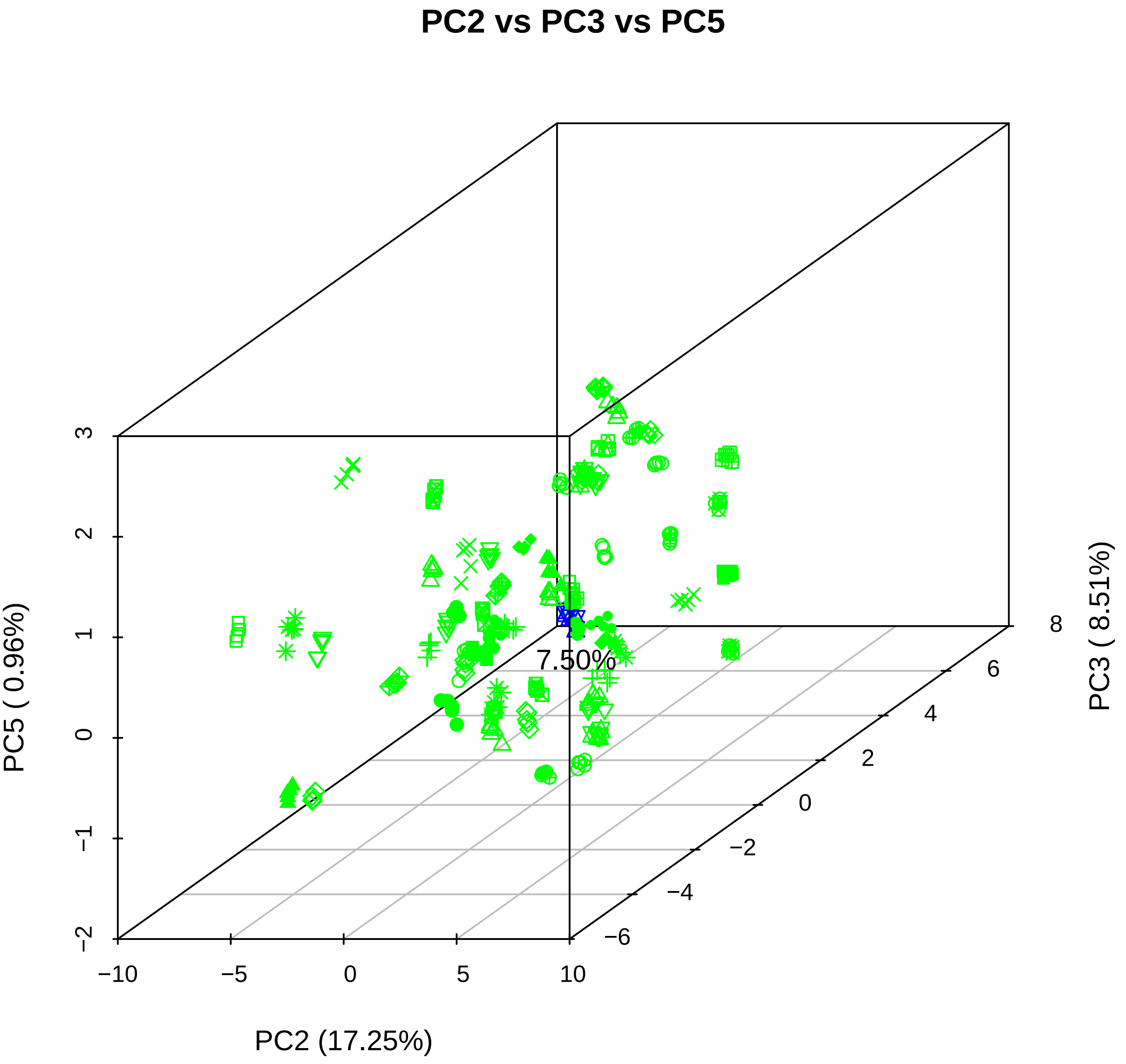

PC2 vs PC3 vs PC5
PC2 (17.25%)
PC5 ( 0.96%)
PC3 ( 8.51%)
7.50%
−10
−5
0
5
10
−2
−1
0
1
2
3
−6
−4
−2
0
2
4
6
8

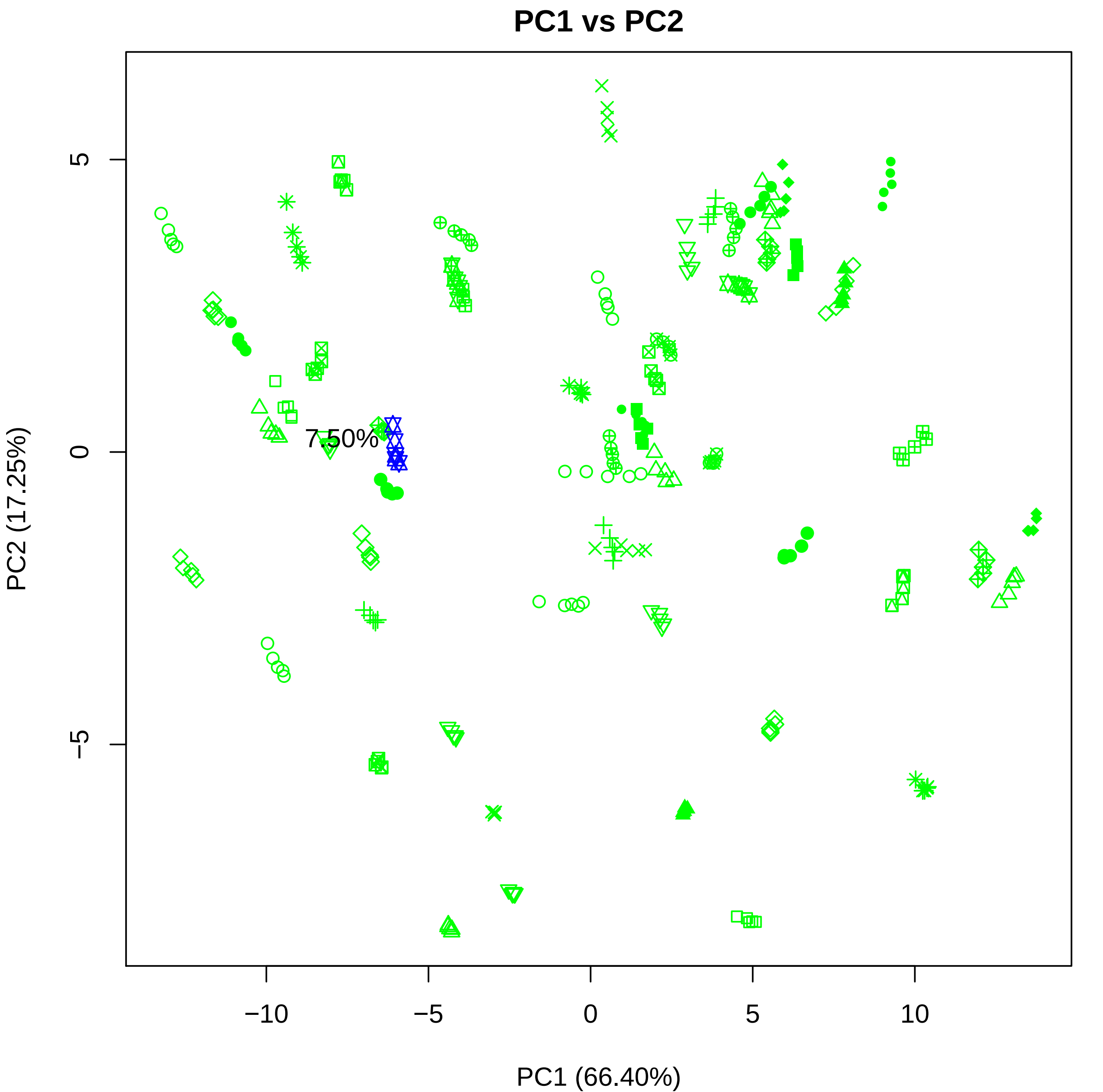

PC1 vs PC2
PC2 (17.25%)
PC1 (66.40%)
7.50%
5
0
−5
−10
−5
0
5
10

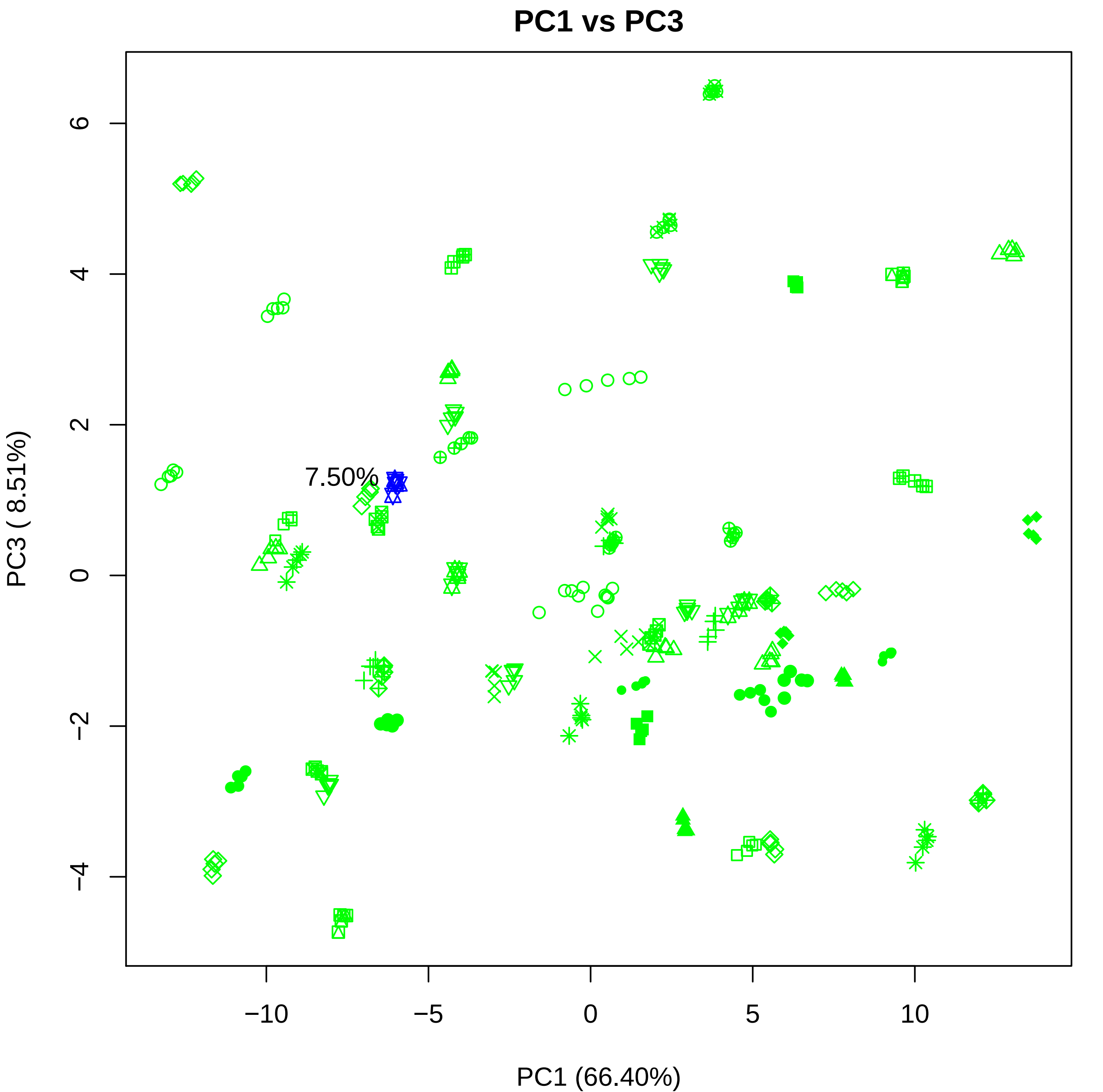
PC1 vs PC3
7.50%
PC3 ( 8.51%)
PC1 (66.40%)
−10
−5
0
5
10
−4
−2
0
2
4
6

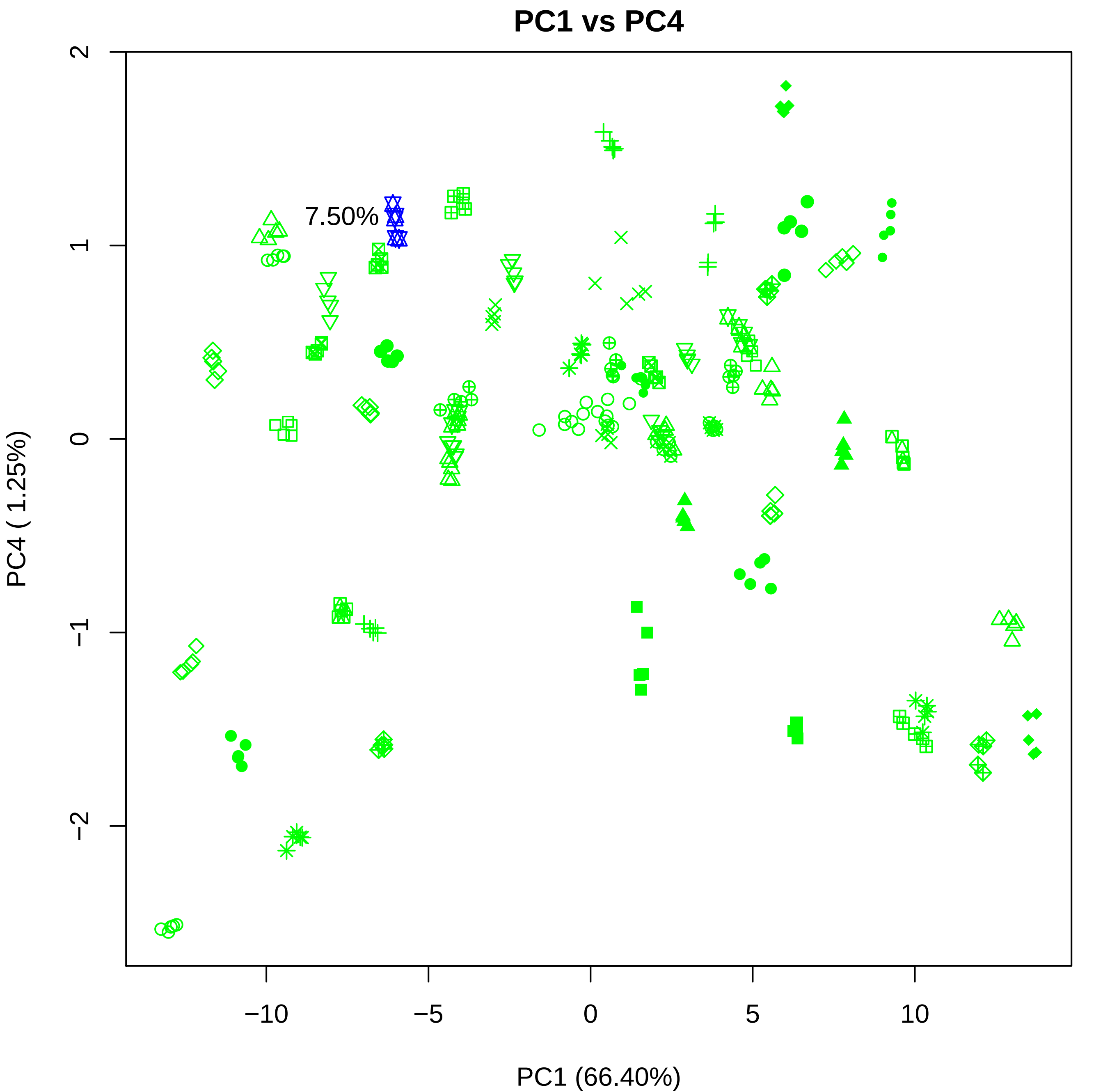
PC1 vs PC4
7.50%
PC4 ( 1.25%)
PC1 (66.40%)
2
1
0
−1
−2
−10
−5
0
5
10

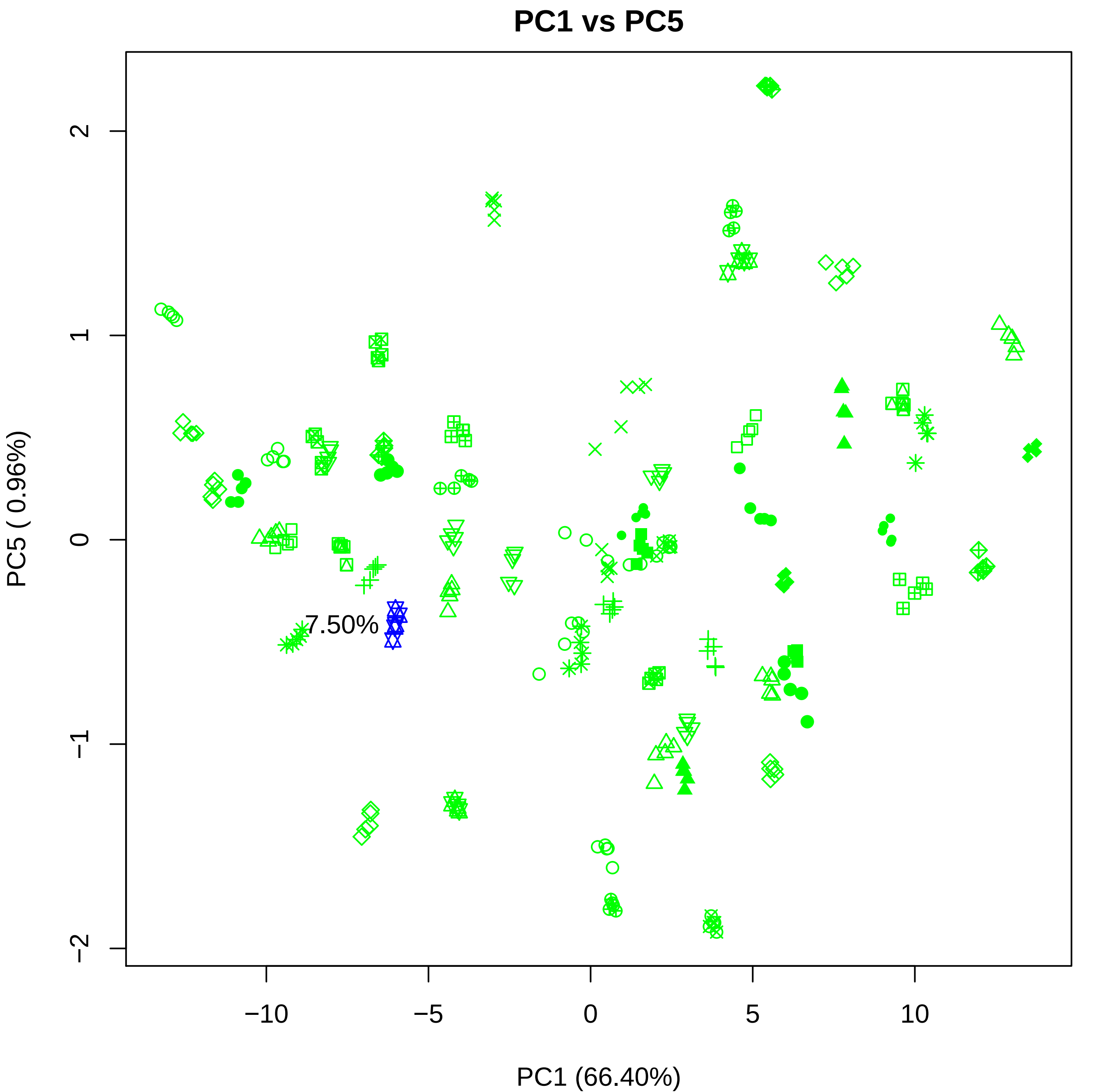
PC1 vs PC5
PC5 ( 0.96%)
PC1 (66.40%)
7.50%
2
1
0
−1
−2
−10
−5
0
5
10

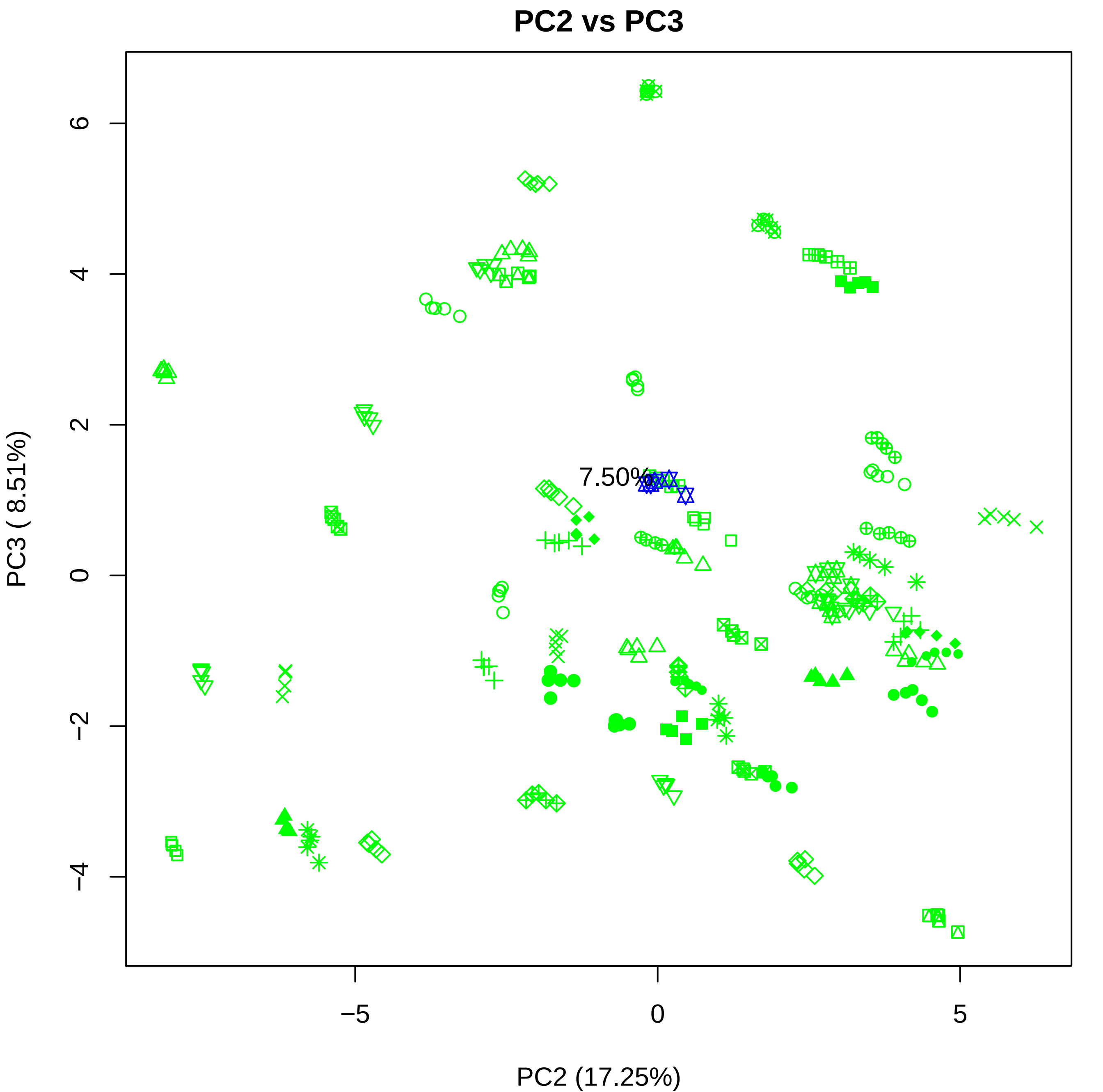
PC2 vs PC3
PC3 ( 8.51%)
PC2 (17.25%)
7.50%
6
4
2
0
−2
−4
−5
0
5

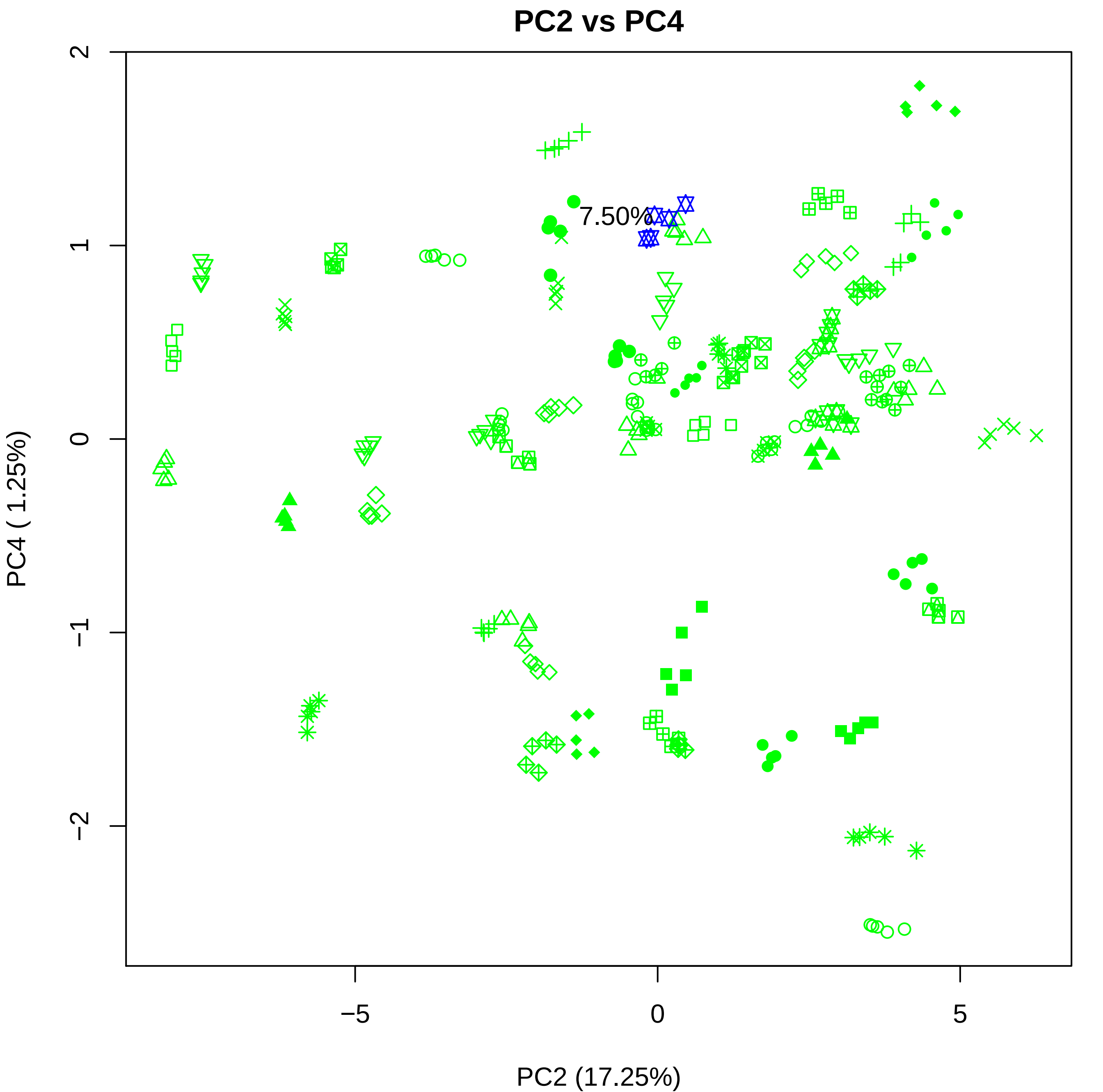
PC2 vs PC4
7.50%
2
1
0
−1
−2
PC4 ( 1.25%)
−5
0
5
PC2 (17.25%)

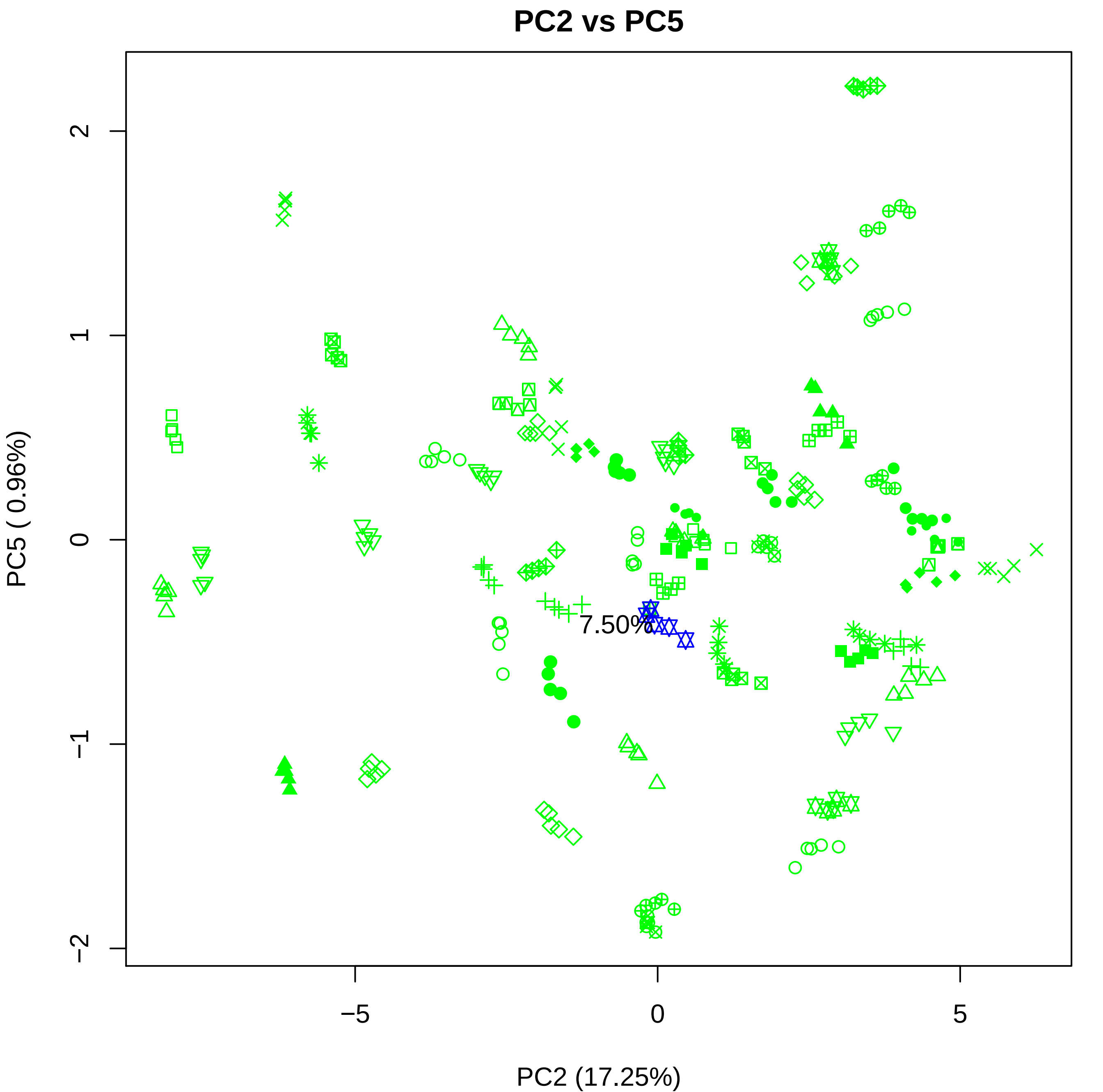
PC2 vs PC5
PC5 ( 0.96%)
PC2 (17.25%)
7.50%
2
1
0
−1
−2
−5
0
5

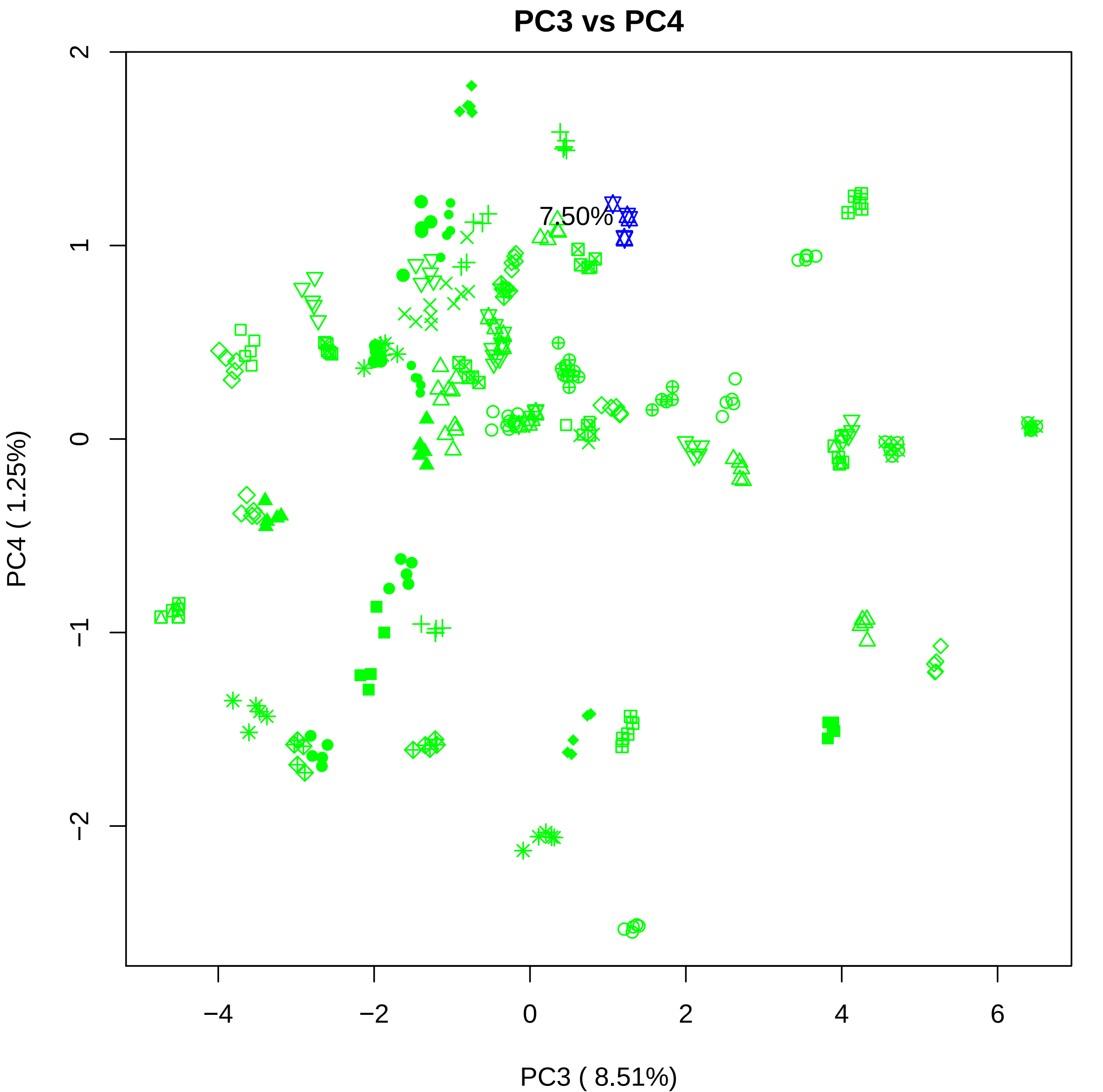
PC3 vs PC4
7.50%
PC4 ( 1.25%)
PC3 ( 8.51%)
2
1
0
−1
−2
−4
−2
0
2
4
6

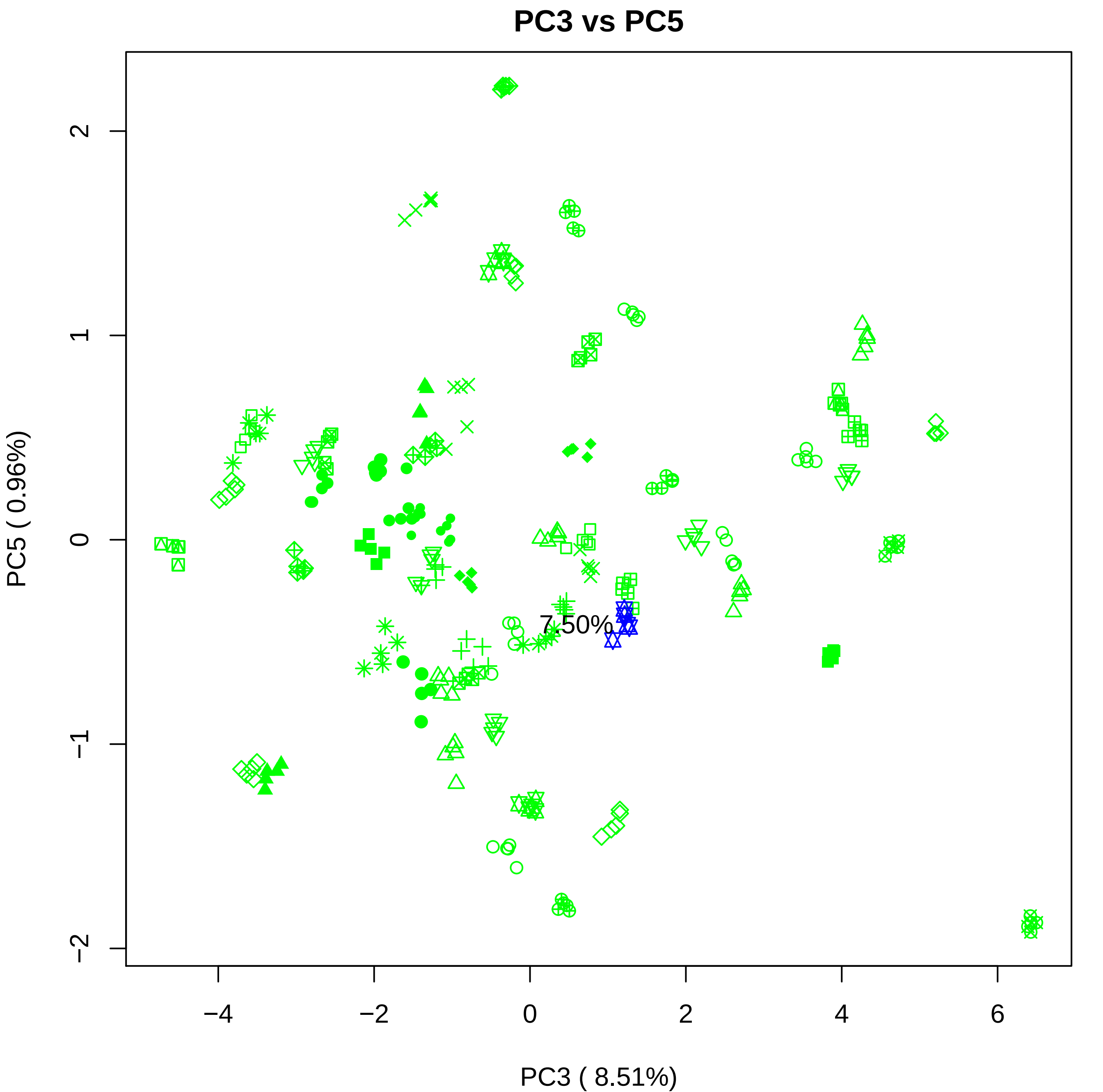
PC3 vs PC5
PC5 ( 0.96%)
PC3 ( 8.51%)
7.50%
2
1
0
−1
−2
−4
−2
0
2
4
6

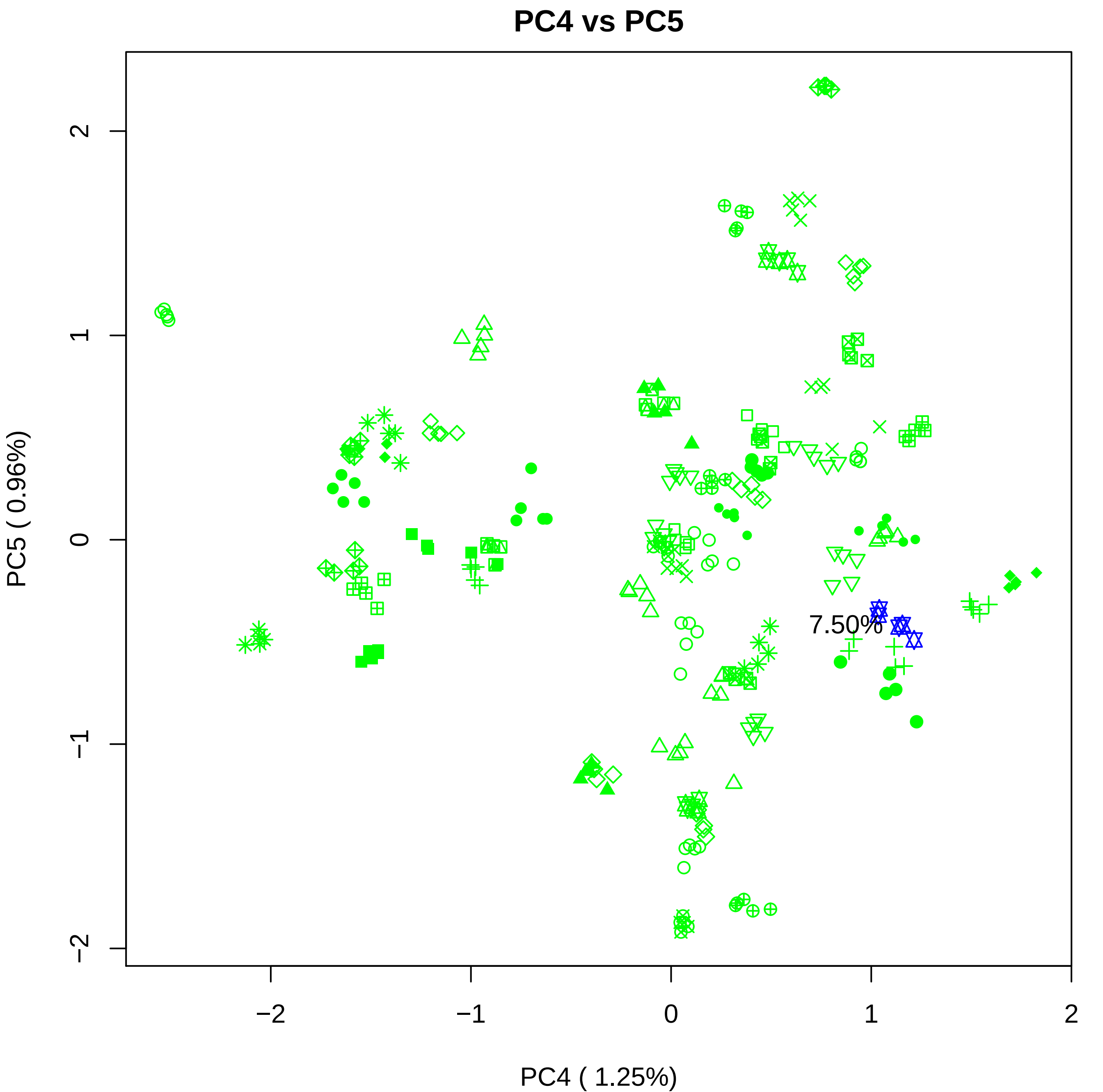
PC4 vs PC5
PC5 ( 0.96%)
PC4 ( 1.25%)
7.50%
2
1
0
−1
−2
−2
−1
0
1
2

### PCA of 10% BMH(c) vs PMH(c)

Scaled Matrix PCAed

7th order polynomial baseline correction performed

Raw spectra smoothed to 21 Neighbours, 2nd order PN, Savtisky Golay

Normalisation between 0 and 100

D2 spectra smoothed to 41 Neighbours, 2nd order PN, Savtisky Golay

Regions Analysed: 3601−2501 and 2001−700cm−1

Samples Analysed: 1 bContam's 0 BLH's 0 BMH's 0 OMH's 0 pContam's 0 PLH's 60 PMH's

Keele University AD

Fri Dec 14 14:19:13 2018

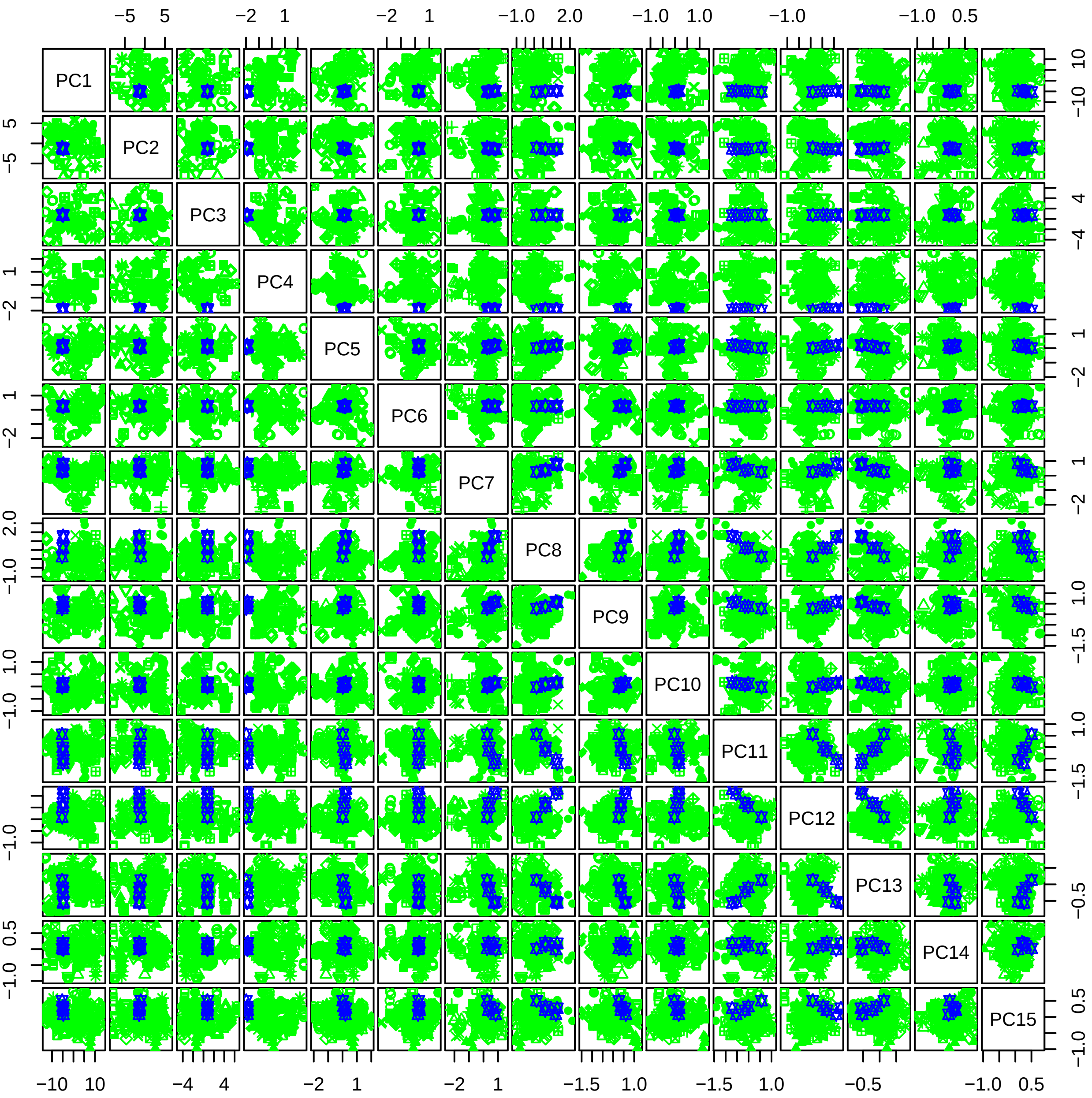
PC1
PC2
PC3
PC4
PC5
PC6
PC7
PC8
PC9
PC10
PC11
PC12
PC13
PC14
PC15

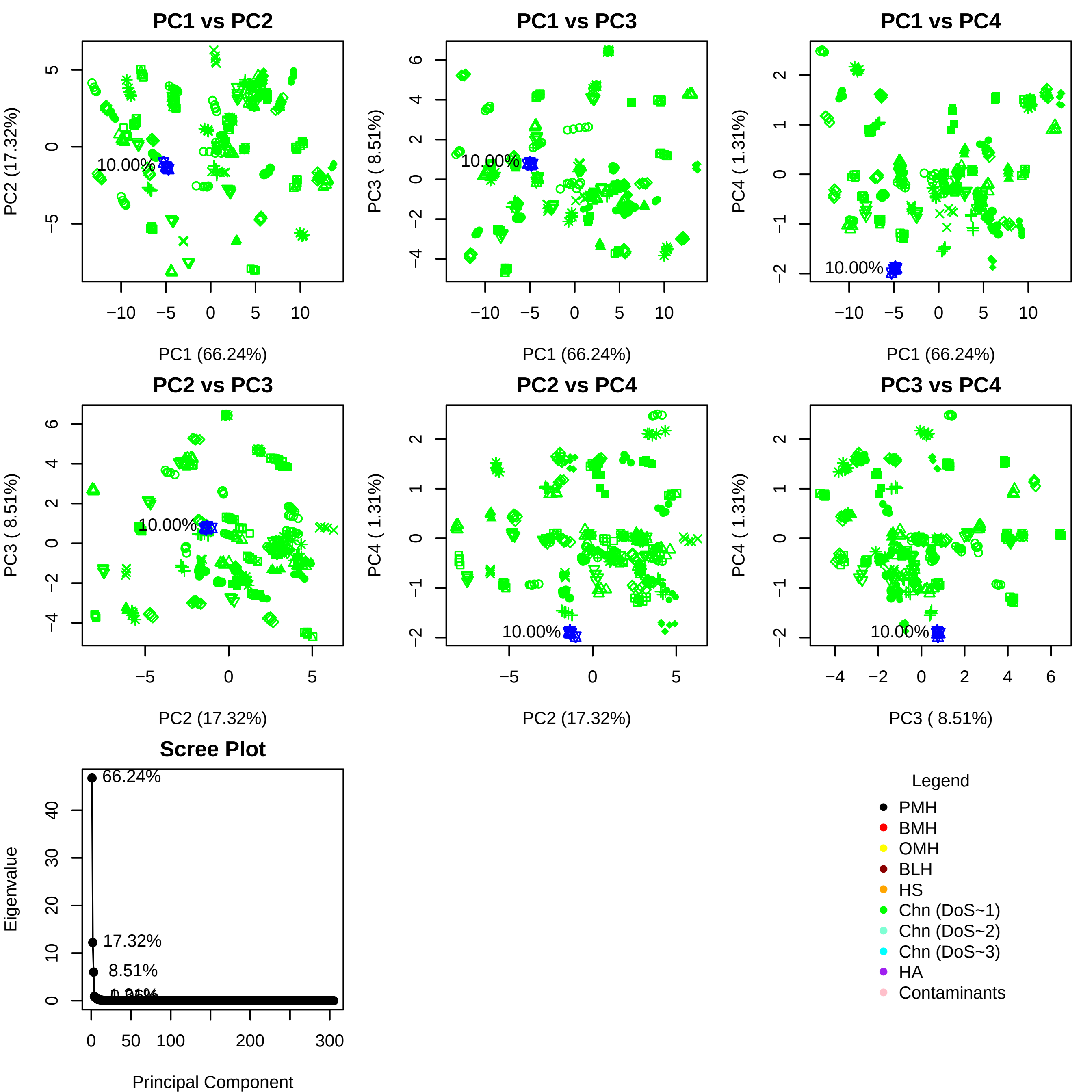

PC1 vs PC2
PC1 vs PC3
PC1 vs PC4
PC2 vs PC3
PC2 vs PC4
PC3 vs PC4
Scree Plot
PC1 (66.24%)
PC2 (17.32%)
PC3 ( 8.51%)
PC4 ( 1.31%)
10.00%
66.24%
17.32%
8.51%
Eigenvalue
Principal Component
Legend
PMH
BMH
OMH
BLH
HS
Chn (DoS~1)
Chn (DoS~2)
Chn (DoS~3)
HA
Contaminants

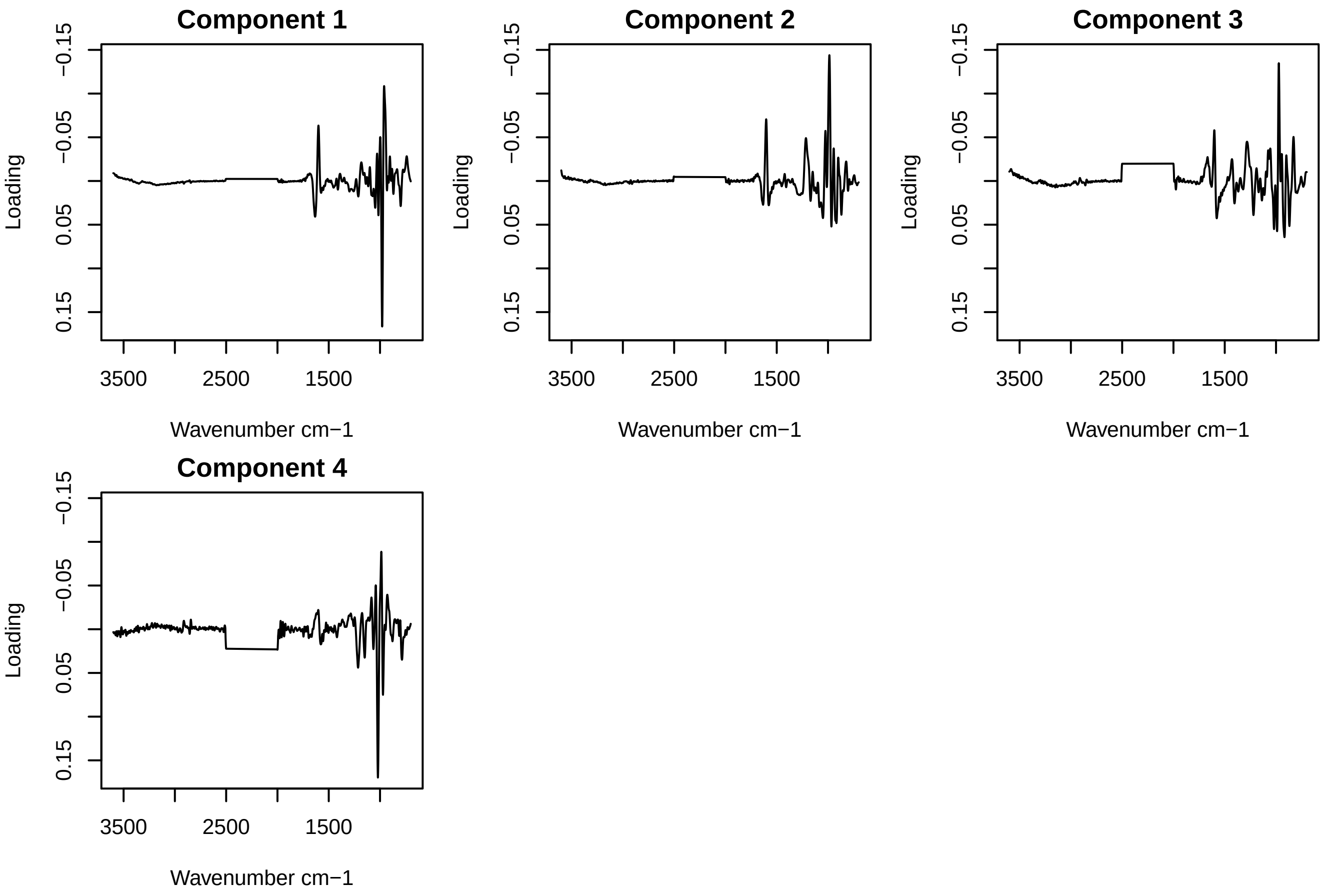
Component 1
Component 2
Component 3
Component 4
Loading
−0.15
−0.05
0.05
0.15
3500
2500
1500
Wavenumber cm−1

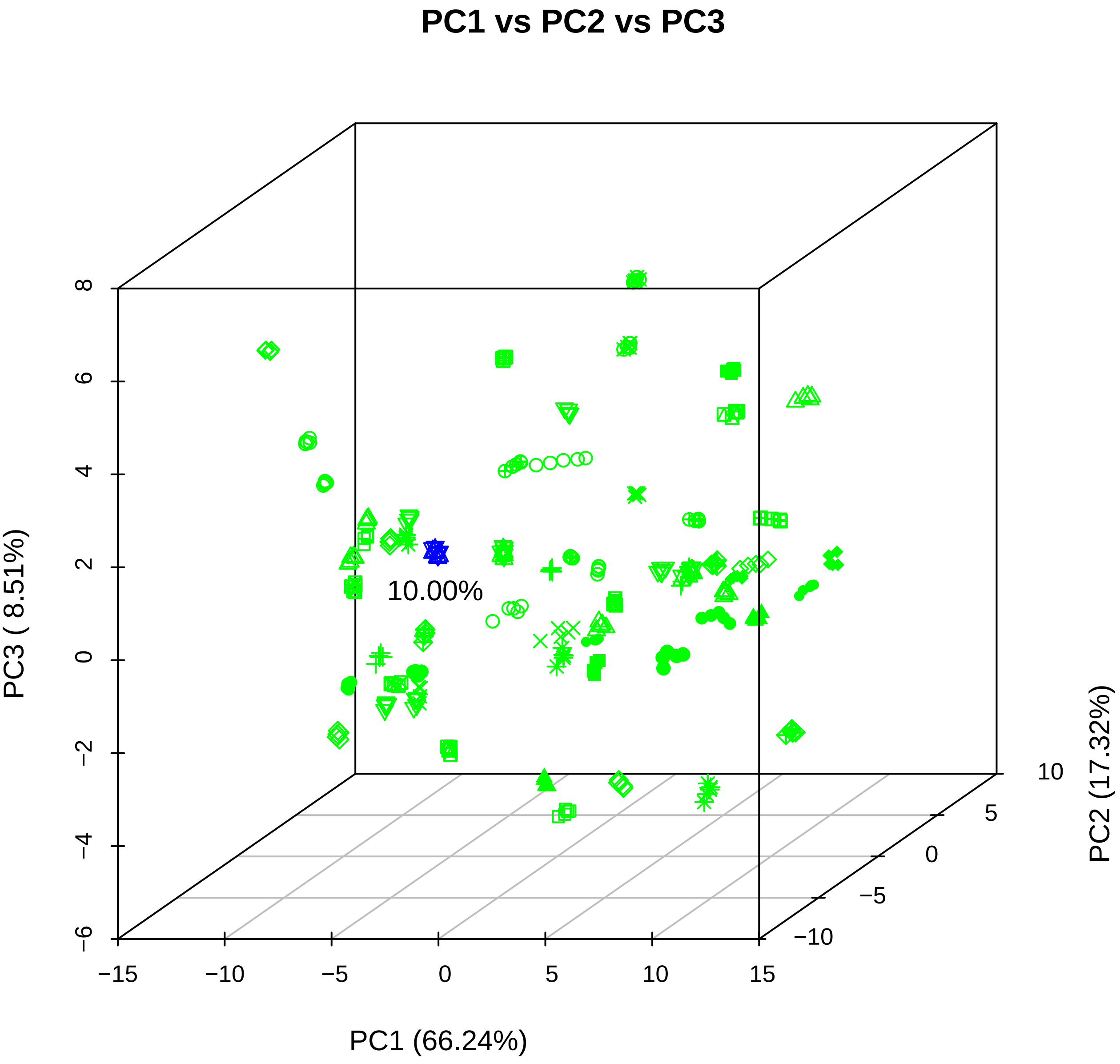

PC1 vs PC2 vs PC3
10.00%
PC1 (66.24%)
PC2 (17.32%)
PC3 ( 8.51%)
−15
−10
−5
0
5
10
15
−6
−4
−2
0
2
4
6
8
−10
−5
0
5
10

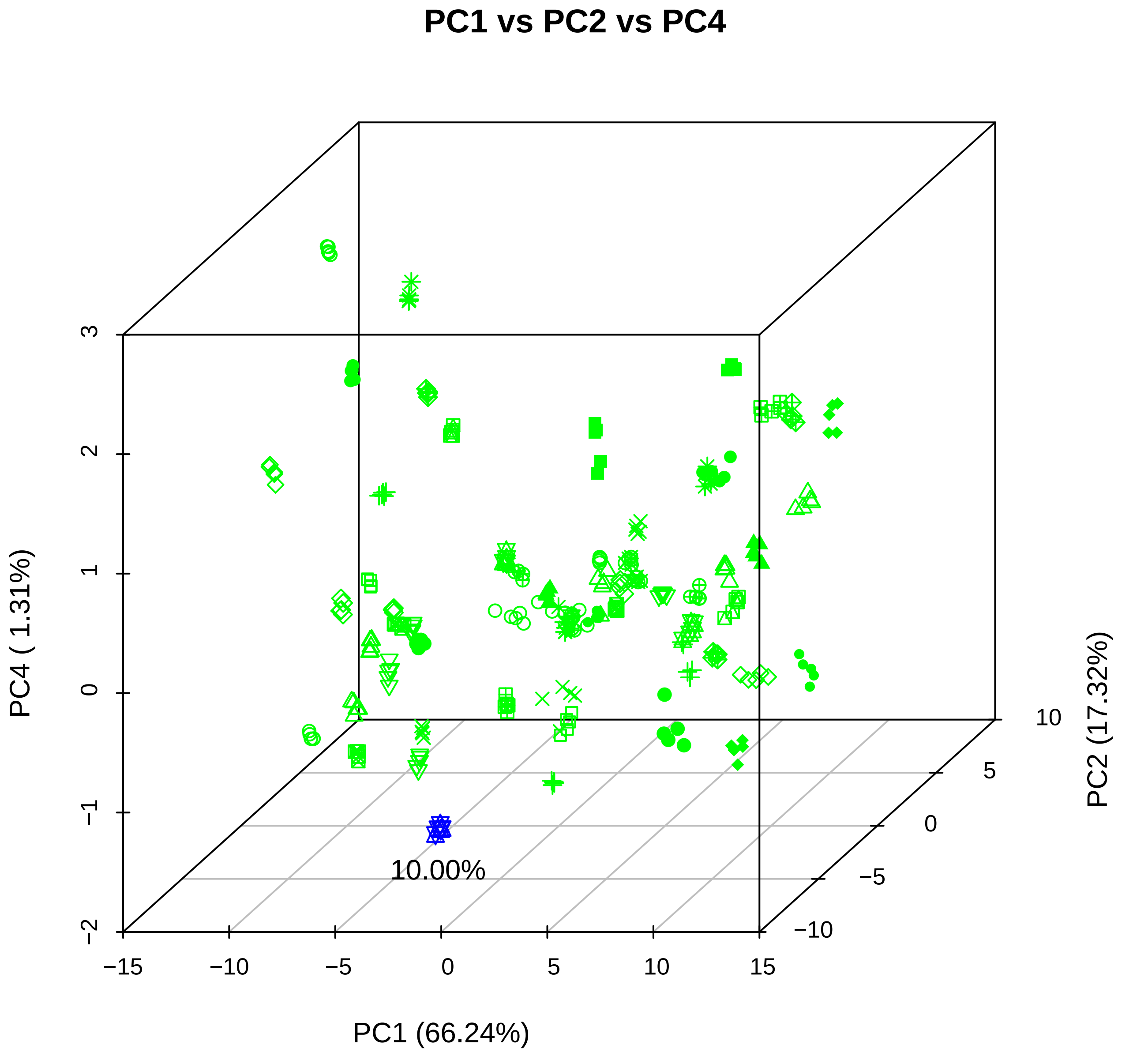
PC1 vs PC2 vs PC4
PC1 (66.24%)
PC2 (17.32%)
PC4 ( 1.31%)
10.00%
−15
−10
−5
0
5
10
15
−2
−1
0
1
2
3
−10
−5
0
5
10

## PC1 vs PC3 vs PC4

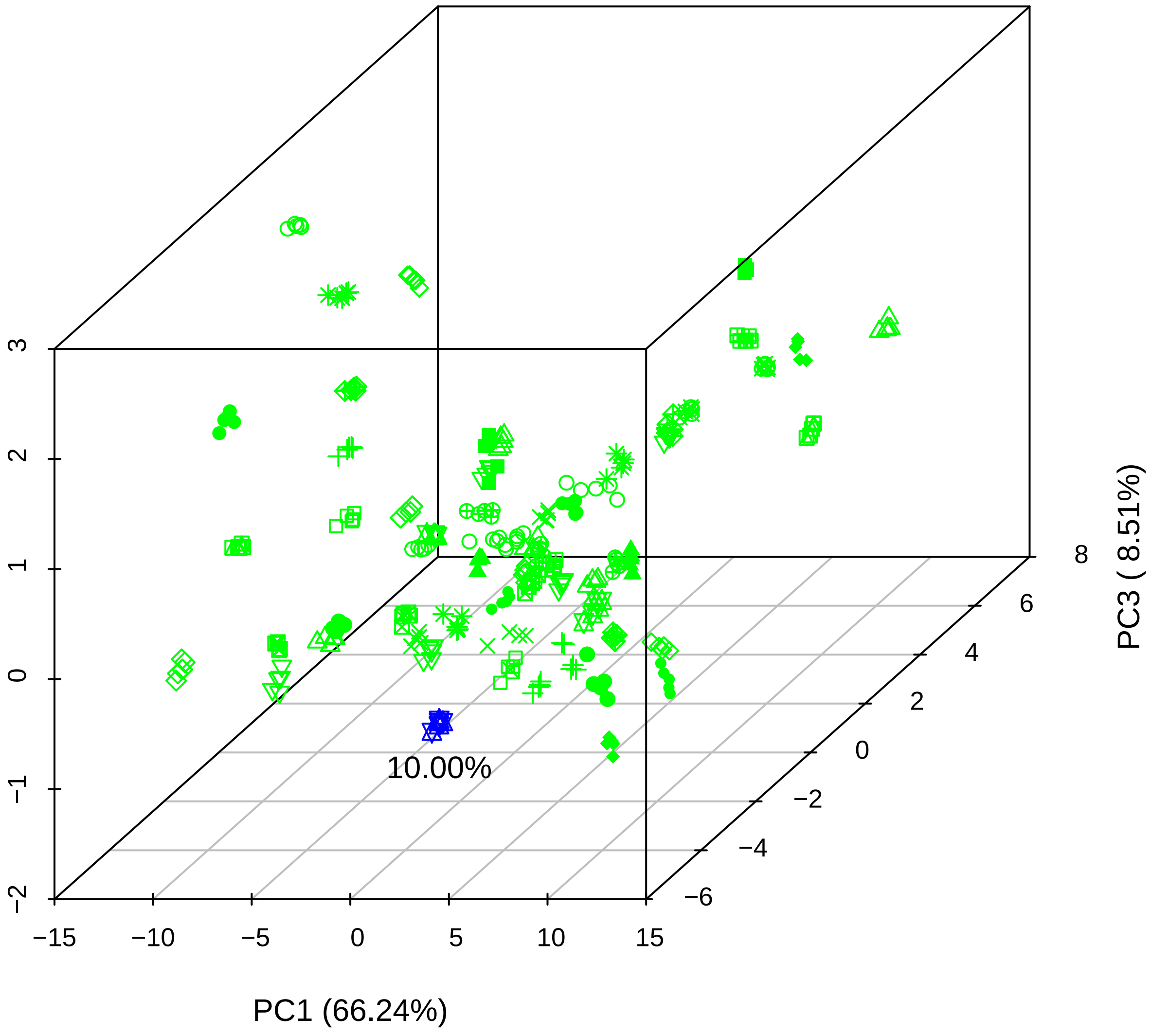

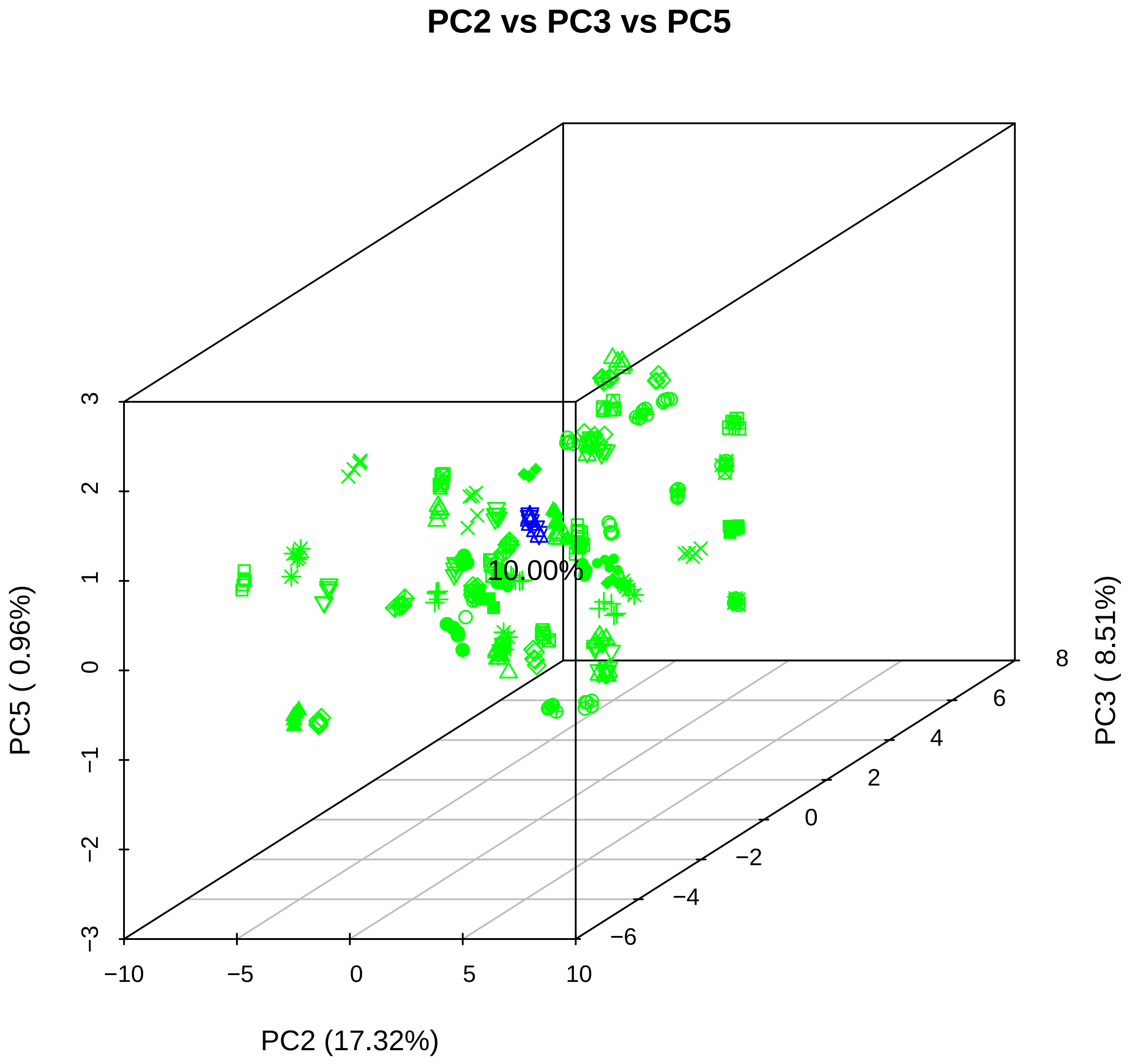
PC2 vs PC3 vs PC5
10.00%
PC2 (17.32%)
PC5 ( 0.96%)
PC3 ( 8.51%)
−10
−5
0
5
10
−3
−2
−1
0
1
2
3
−6
−4
−2
0
2
4
6
8

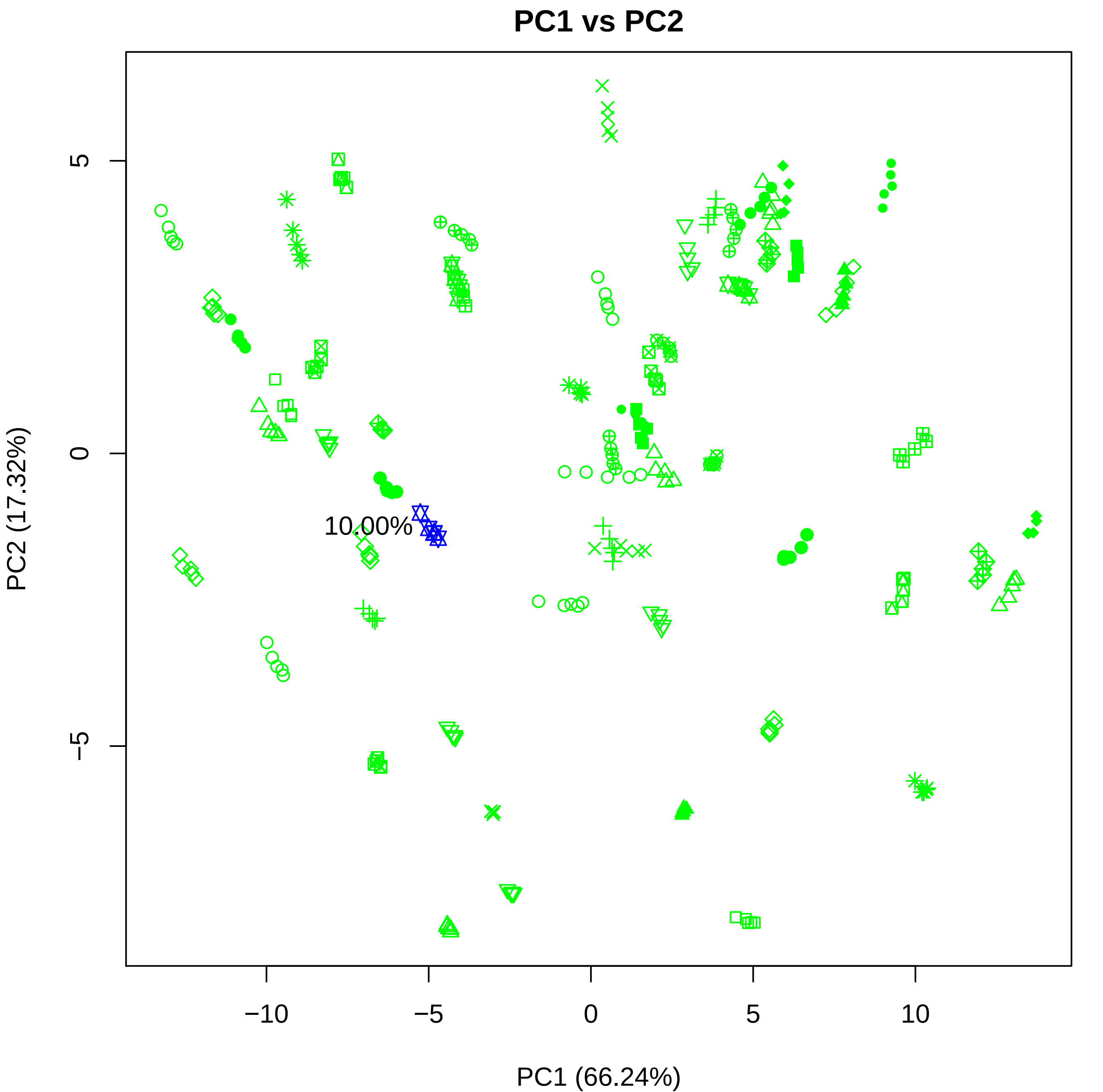
PC1 vs PC2
10.00%
PC1 (66.24%)
PC2 (17.32%)
−10
−5
0
5
10
5
0
−5

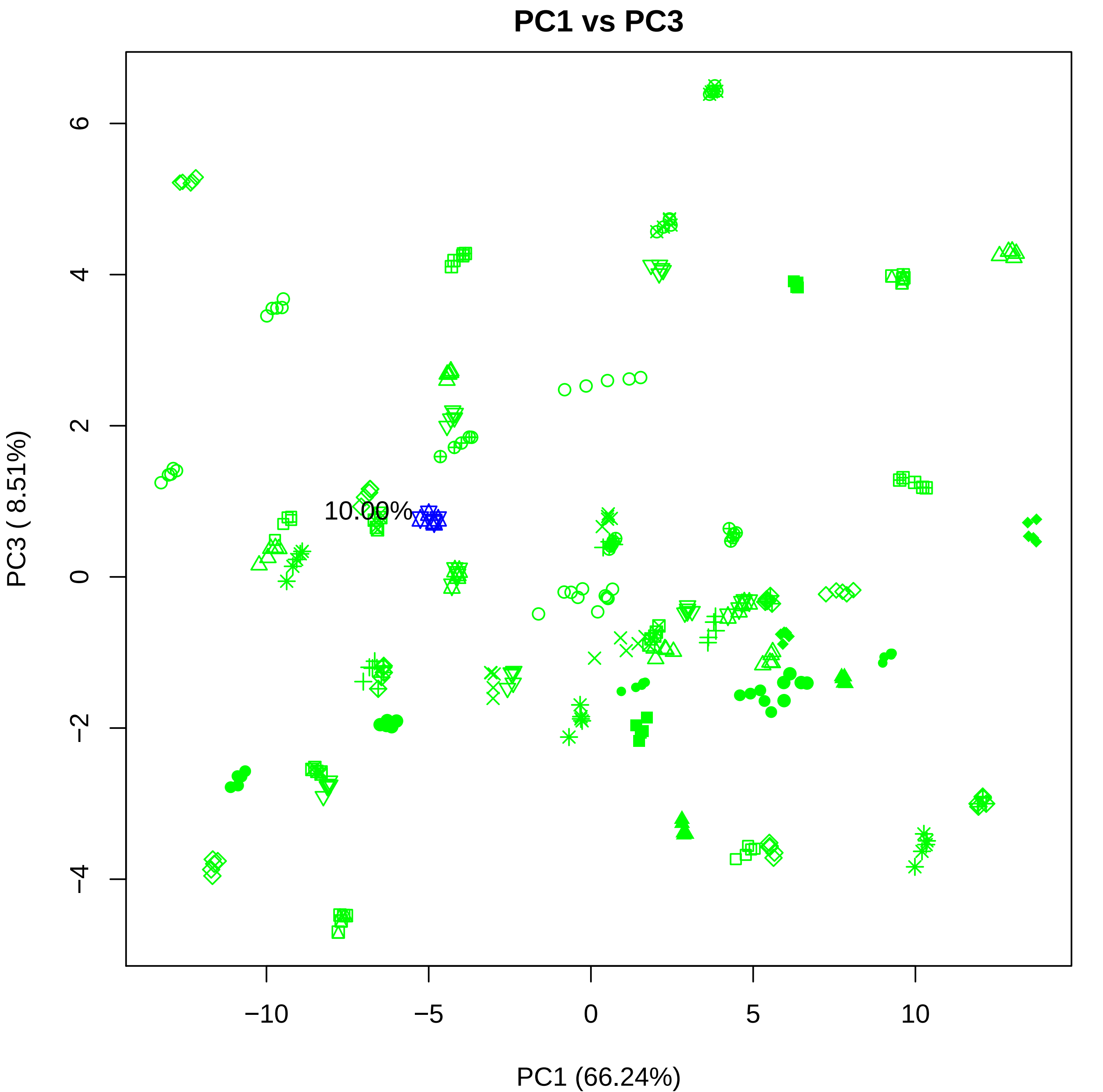

PC1 vs PC3
10.00%
PC3 ( 8.51%)
PC1 (66.24%)
6
4
2
0
−2
−4
−10
−5
0
5
10

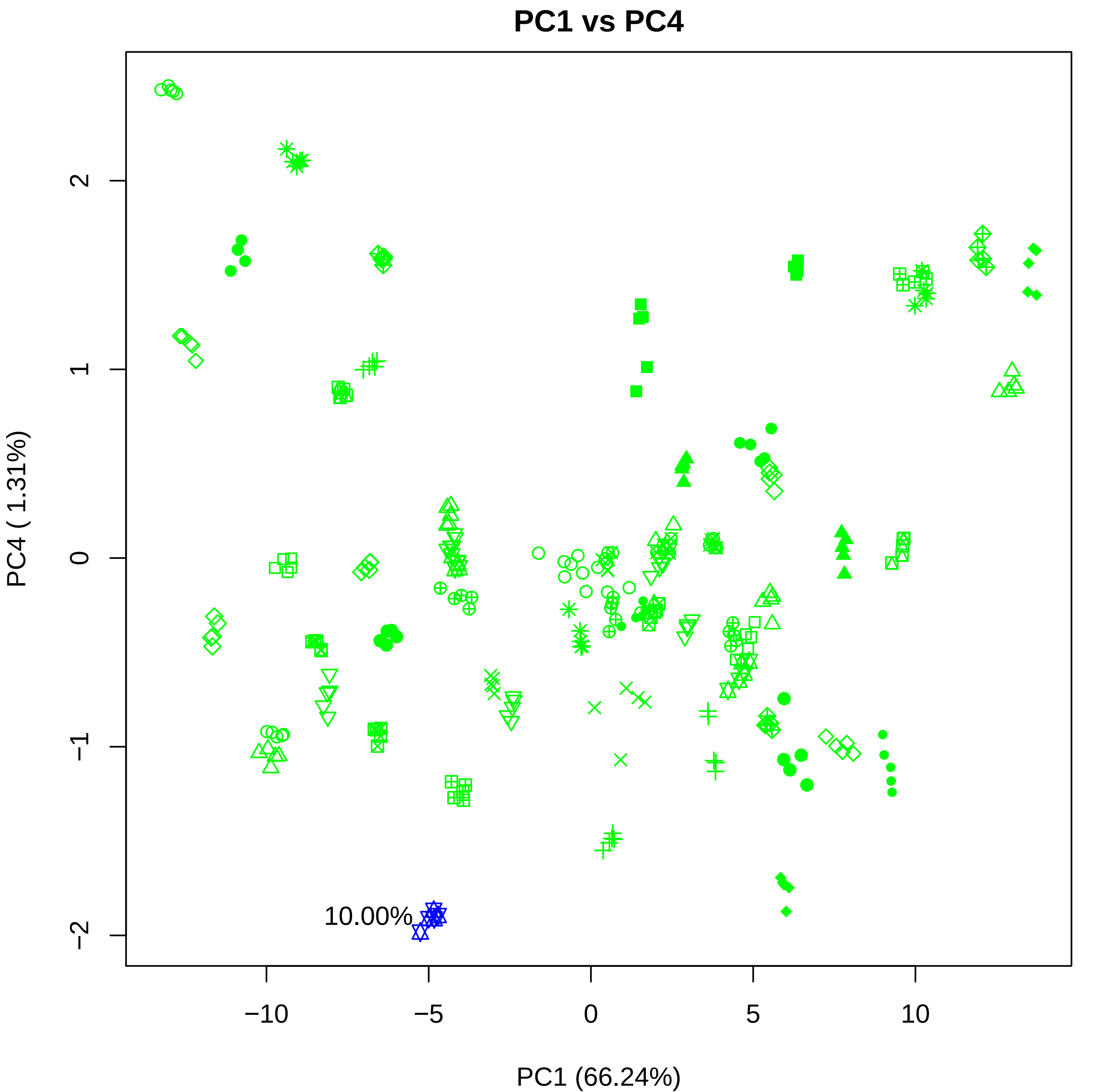
PC1 vs PC4
PC4 ( 1.31%)
PC1 (66.24%)
10.00%
2
1
0
−1
−2
−10
−5
0
5
10

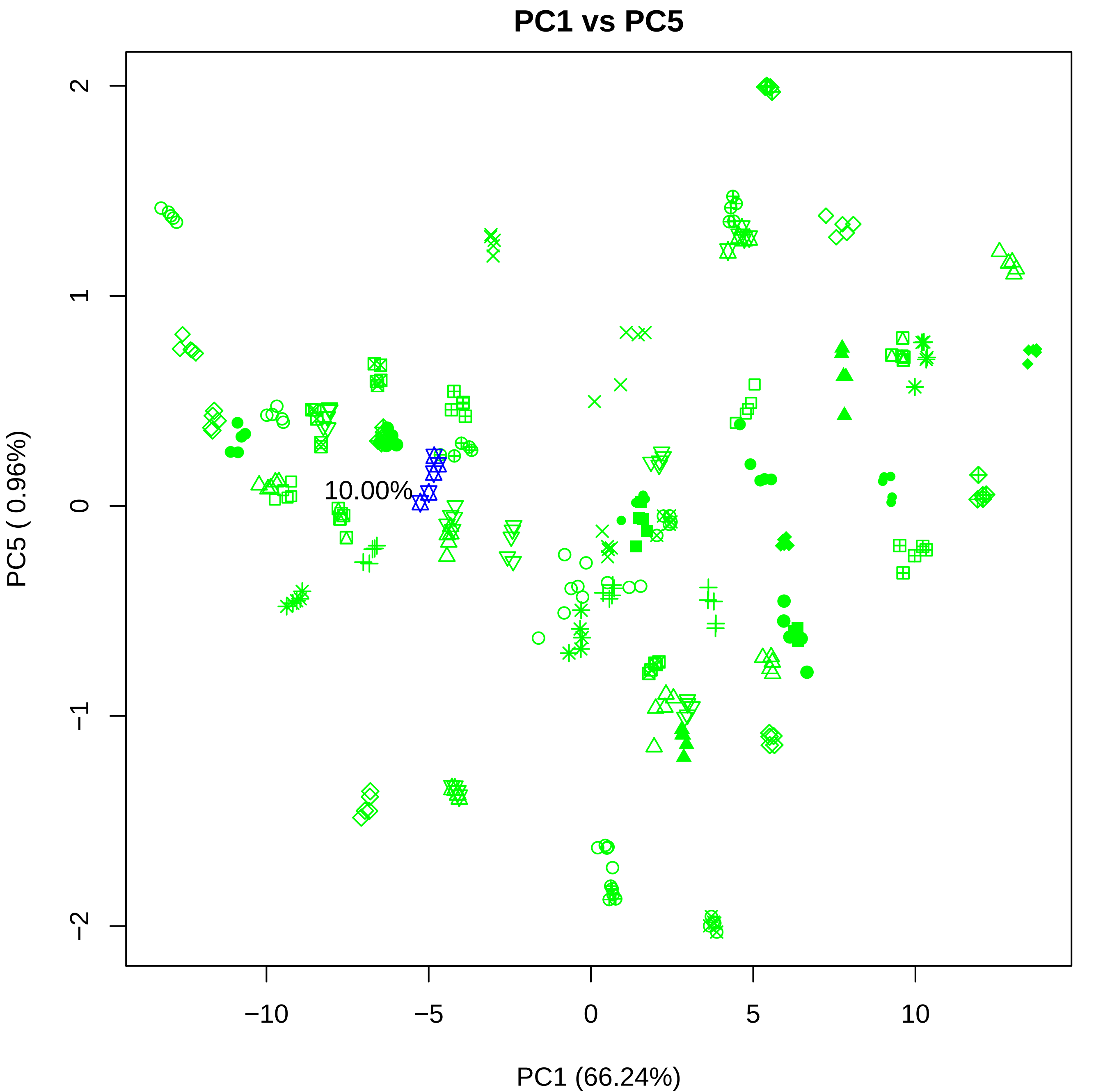

PC1 vs PC5
2
1
0
−1
−2
PC5 ( 0.96%)
10.00%
−10
−5
0
5
10
PC1 (66.24%)

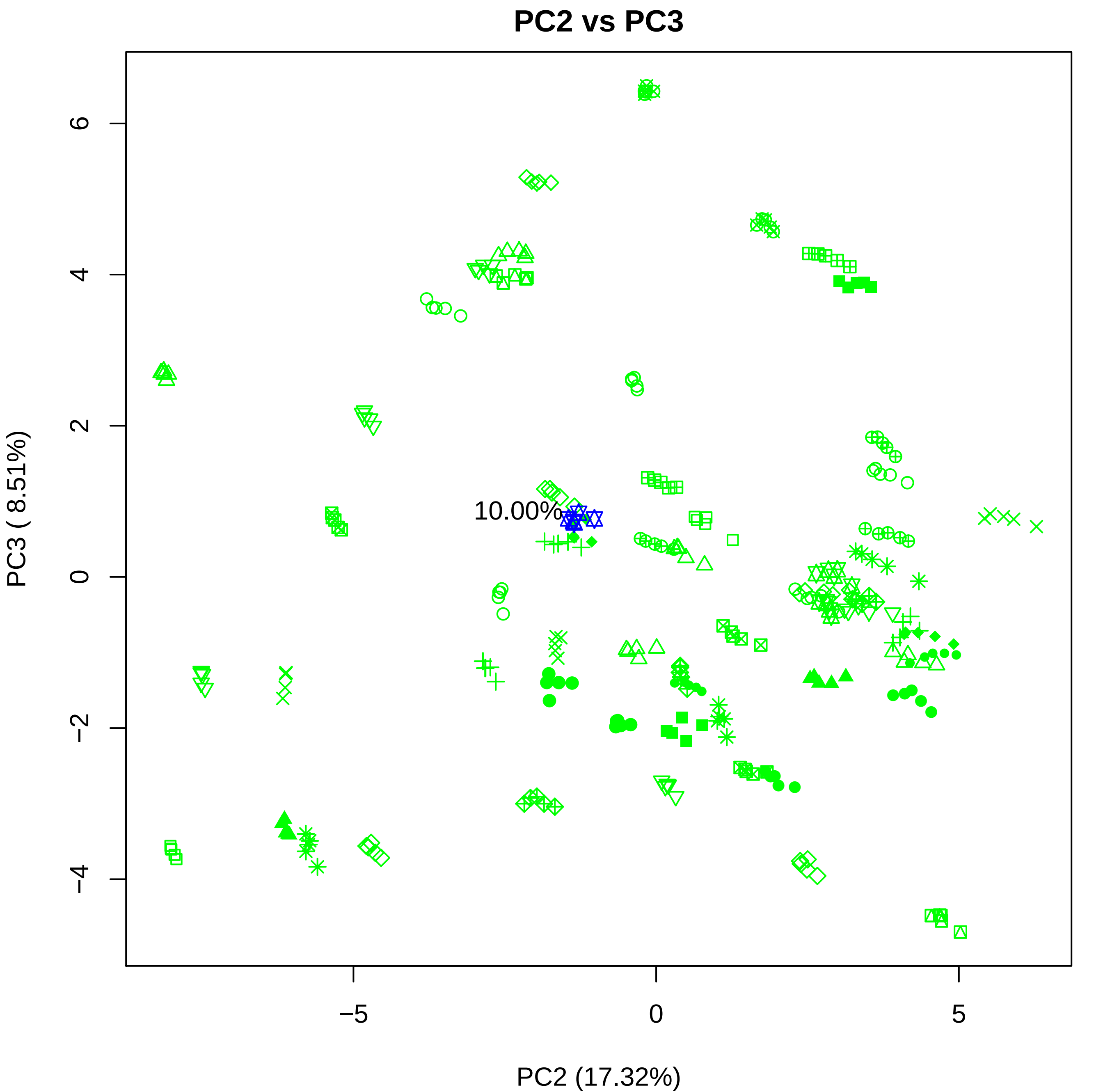

PC2 vs PC3
10.00%
PC3 ( 8.51%)
PC2 (17.32%)
−5
0
5
−4
−2
0
2
4
6

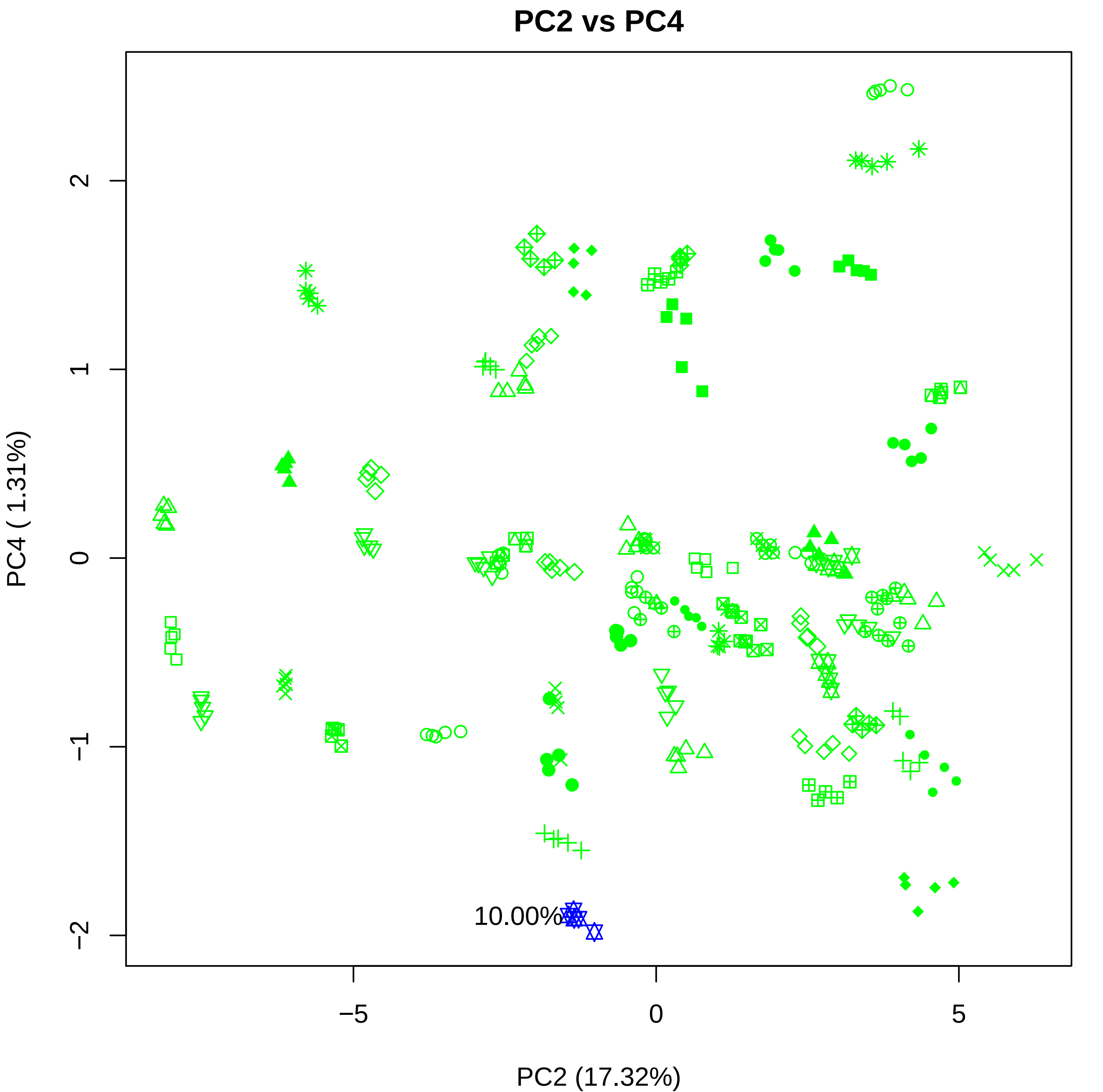

PC2 vs PC4
PC4 ( 1.31%)
PC2 (17.32%)
10.00%
−5
0
5
2
1
0
−1
−2

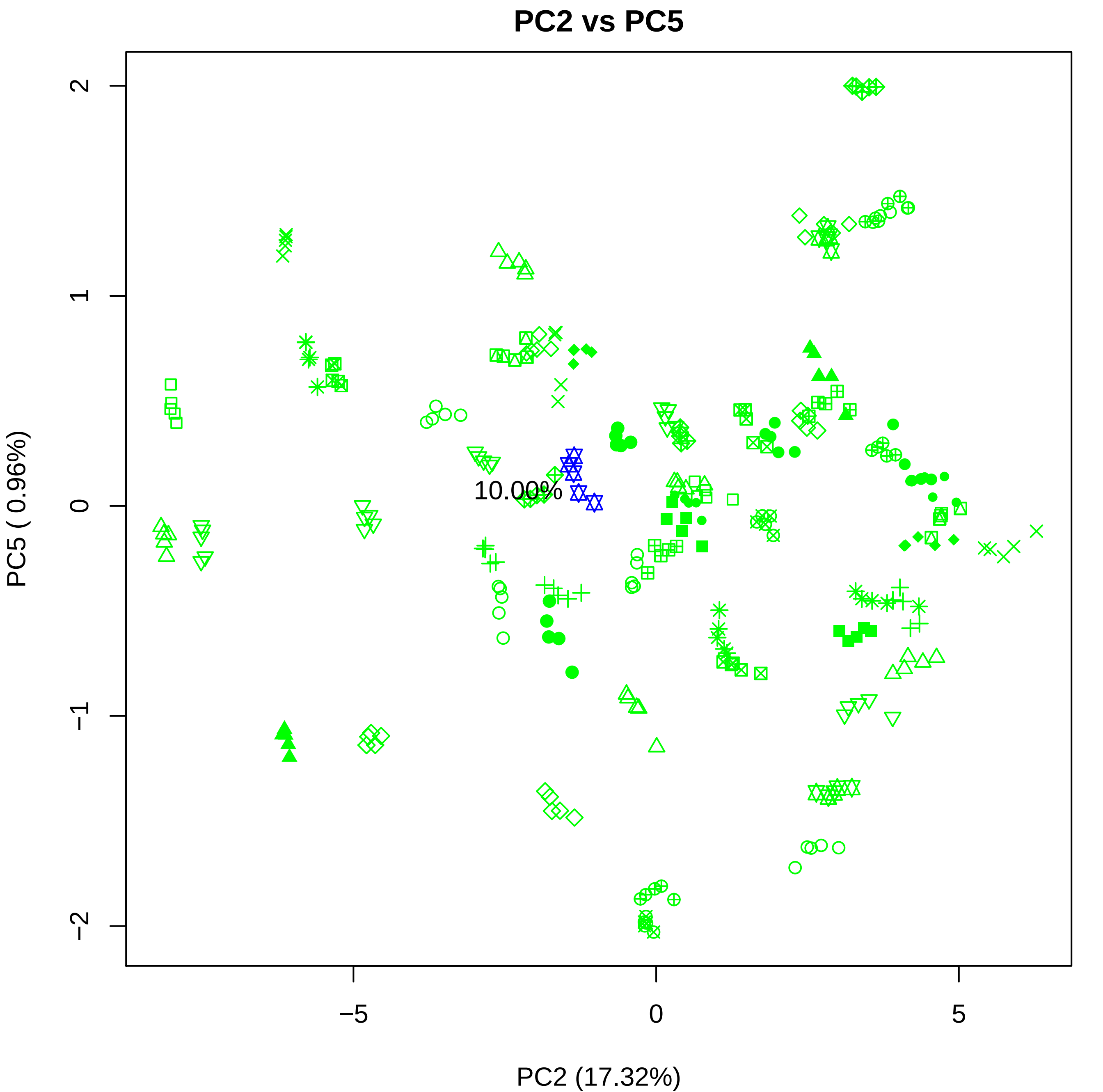

PC2 vs PC5
10.00%
PC5 ( 0.96%)
PC2 (17.32%)
2
1
0
−1
−2
−5
0
5

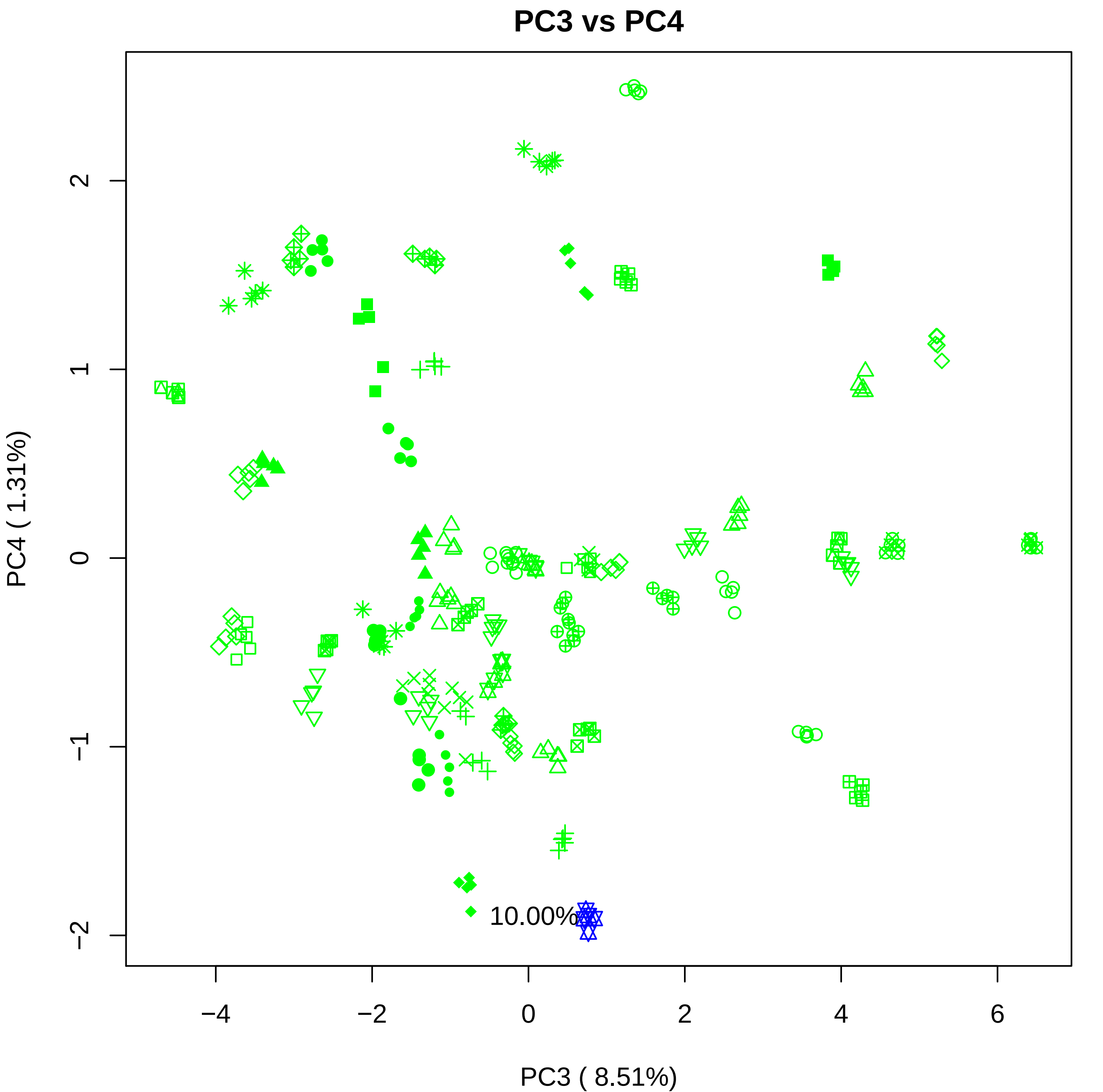

PC3 vs PC4
PC4 ( 1.31%)
PC3 ( 8.51%)
2
1
0
−1
−2
−4
−2
0
2
4
6
10.00%

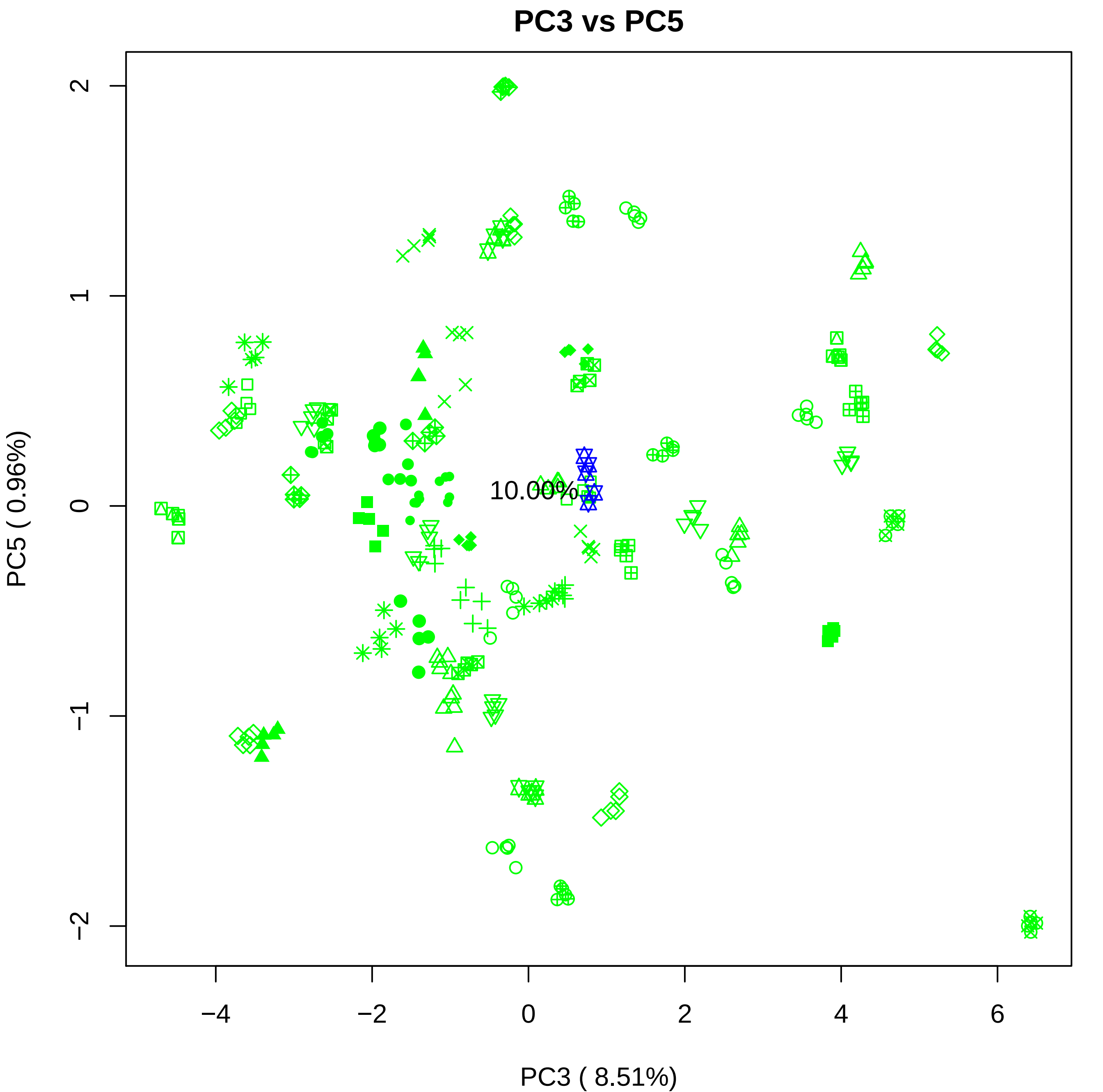
PC3 vs PC5
10.00%
PC5 ( 0.96%)
PC3 ( 8.51%)
2
1
0
−1
−2
−4
−2
0
2
4
6

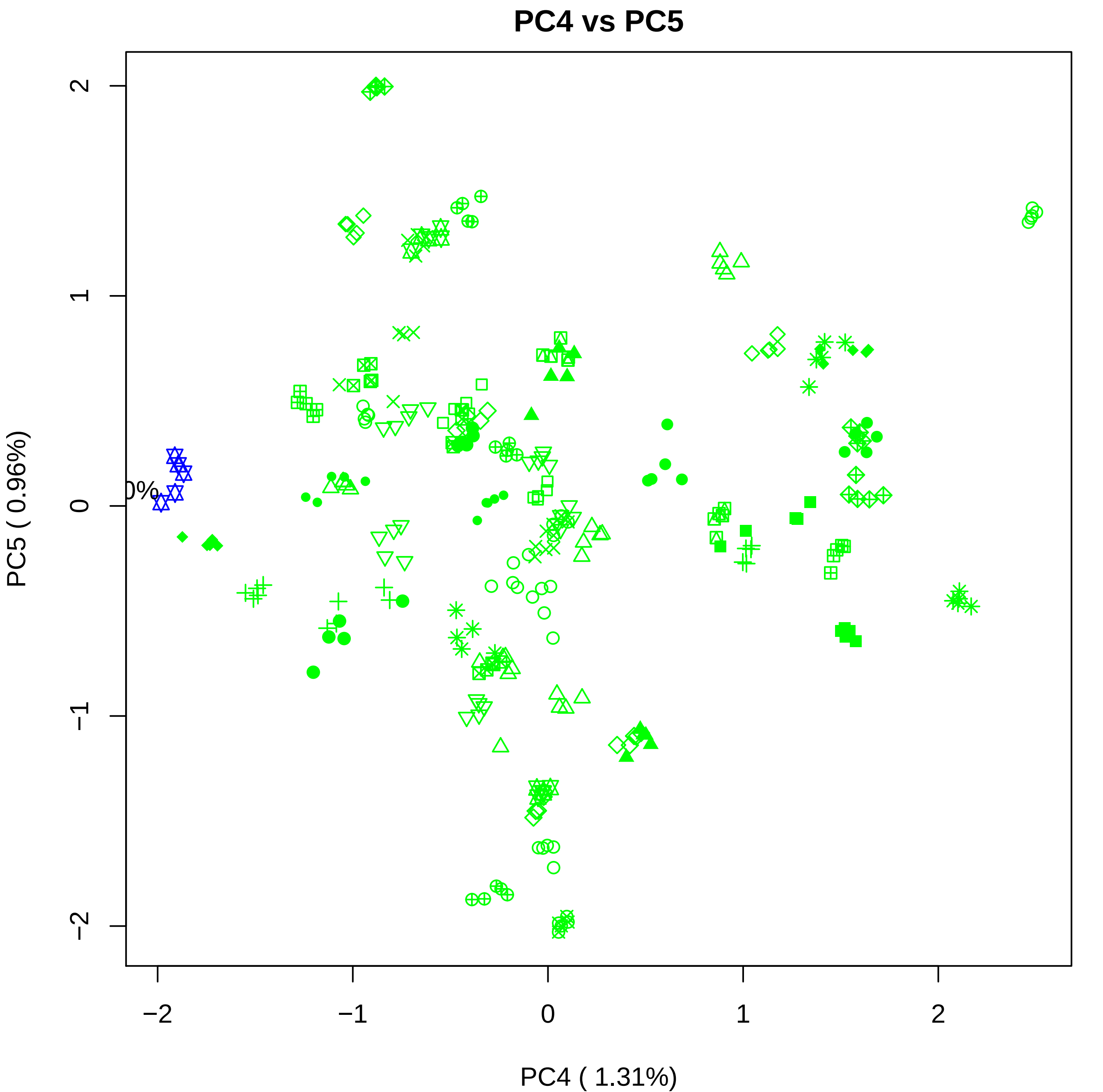
PC4 vs PC5
2
1
0
−1
−2
PC5 ( 0.96%)
0%
−2
−1
0
1
2
PC4 ( 1.31%)

### PCA of 15% BMH(c) vs PMH(c)

Scaled Matrix PCAed

7th order polynomial baseline correction performed

Raw spectra smoothed to 21 Neighbours, 2nd order PN, Savtisky Golay

Normalisation between 0 and 100

D2 spectra smoothed to 41 Neighbours, 2nd order PN, Savtisky Golay

Regions Analysed: 3601−2501 and 2001−700cm−1

Samples Analysed: 1 bContam's 0 BLH's 0 BMH's 0 OMH's 0 pContam's 0 PLH's 60 PMH's

Keele University AD

Fri Dec 14 14:19:21 2018

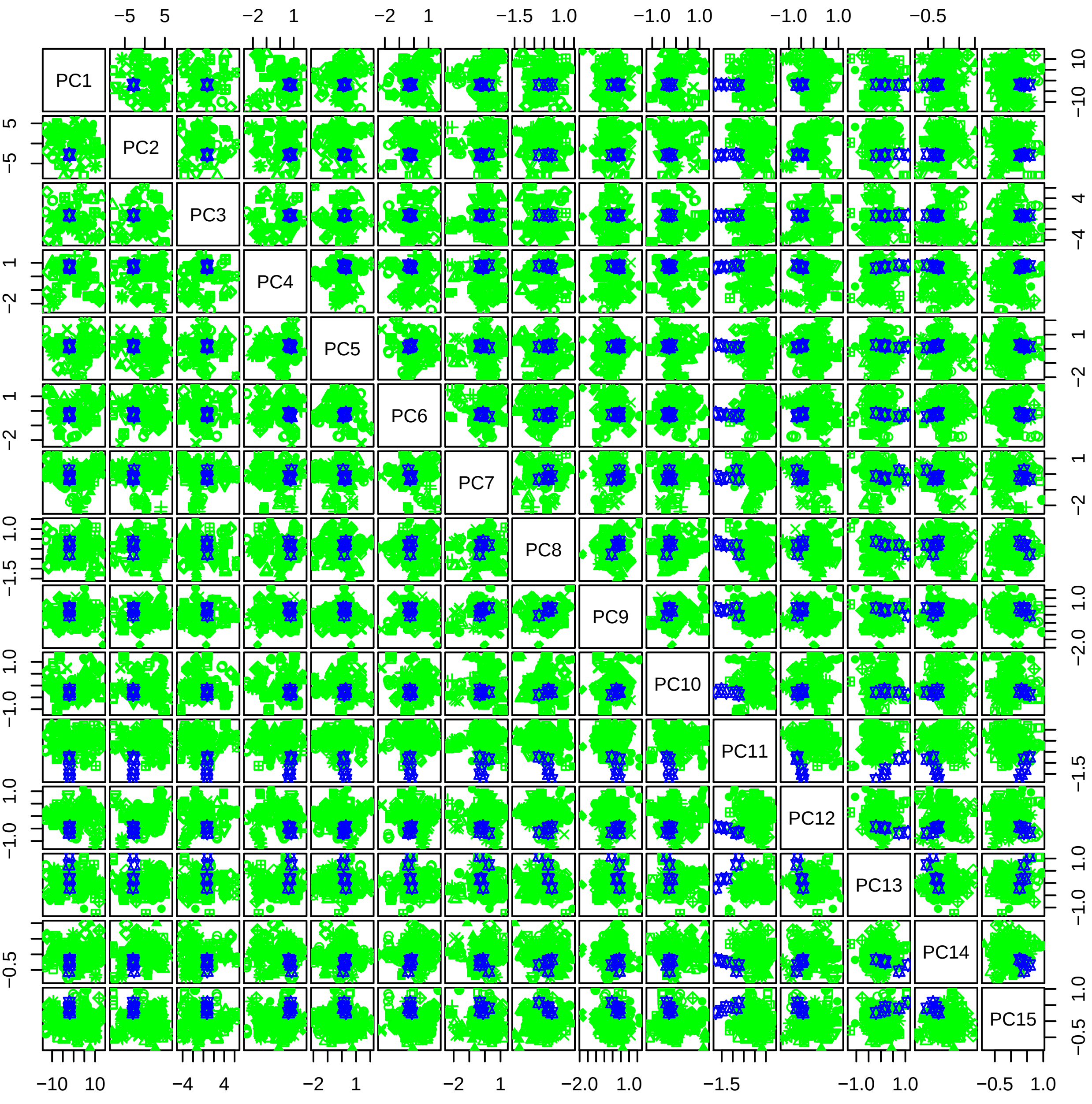
PC1
PC2
PC3
PC4
PC5
PC6
PC7
PC8
PC9
PC10
PC11
PC12
PC13
PC14
PC15

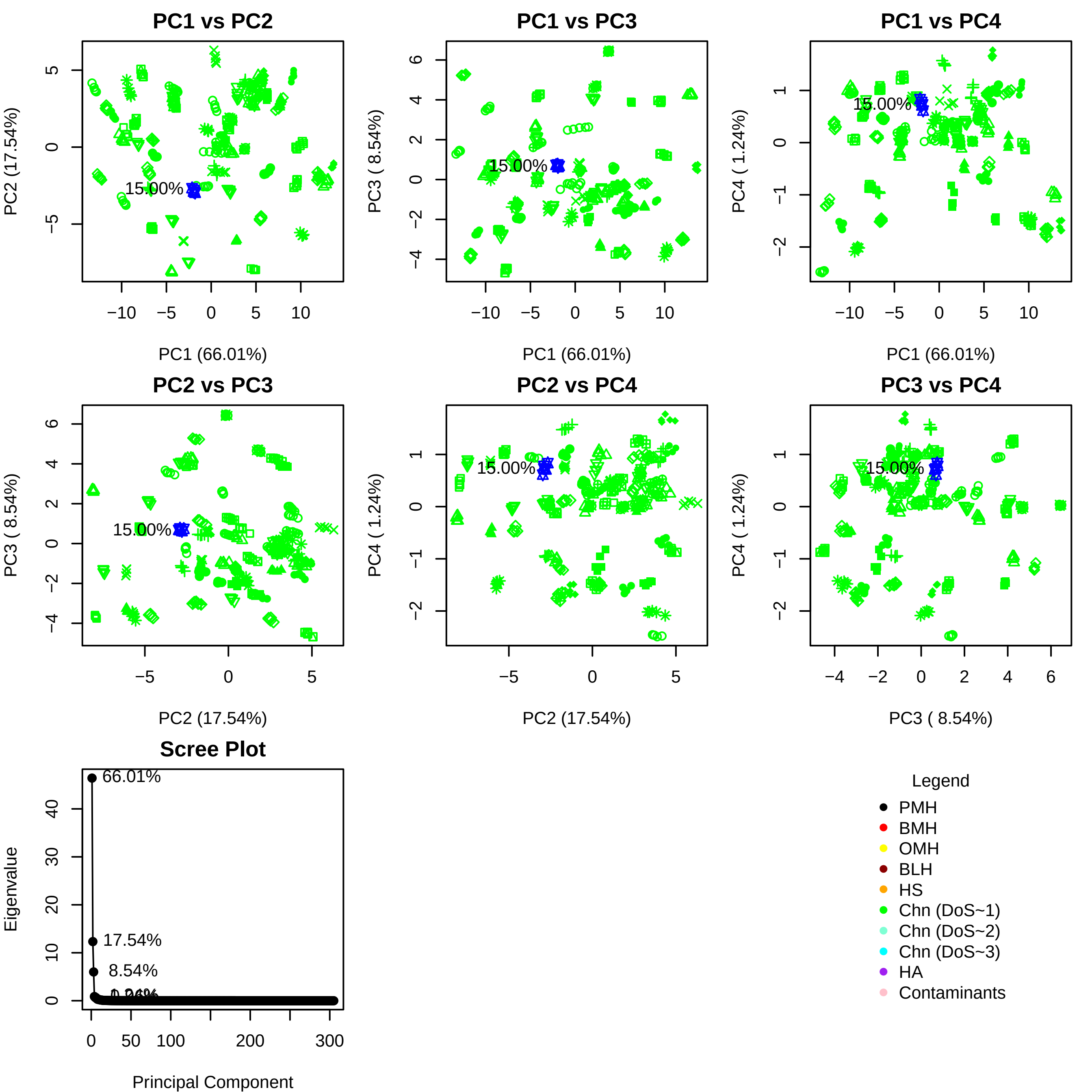

PC1 vs PC2
PC1 vs PC3
PC1 vs PC4
PC2 vs PC3
PC2 vs PC4
PC3 vs PC4
Scree Plot
PC1 (66.01%)
PC2 (17.54%)
PC3 ( 8.54%)
PC4 ( 1.24%)
15.00%
Eigenvalue
Principal Component
66.01%
17.54%
8.54%
Legend
PMH
BMH
OMH
BLH
HS
Chn (DoS~1)
Chn (DoS~2)
Chn (DoS~3)
HA
Contaminants

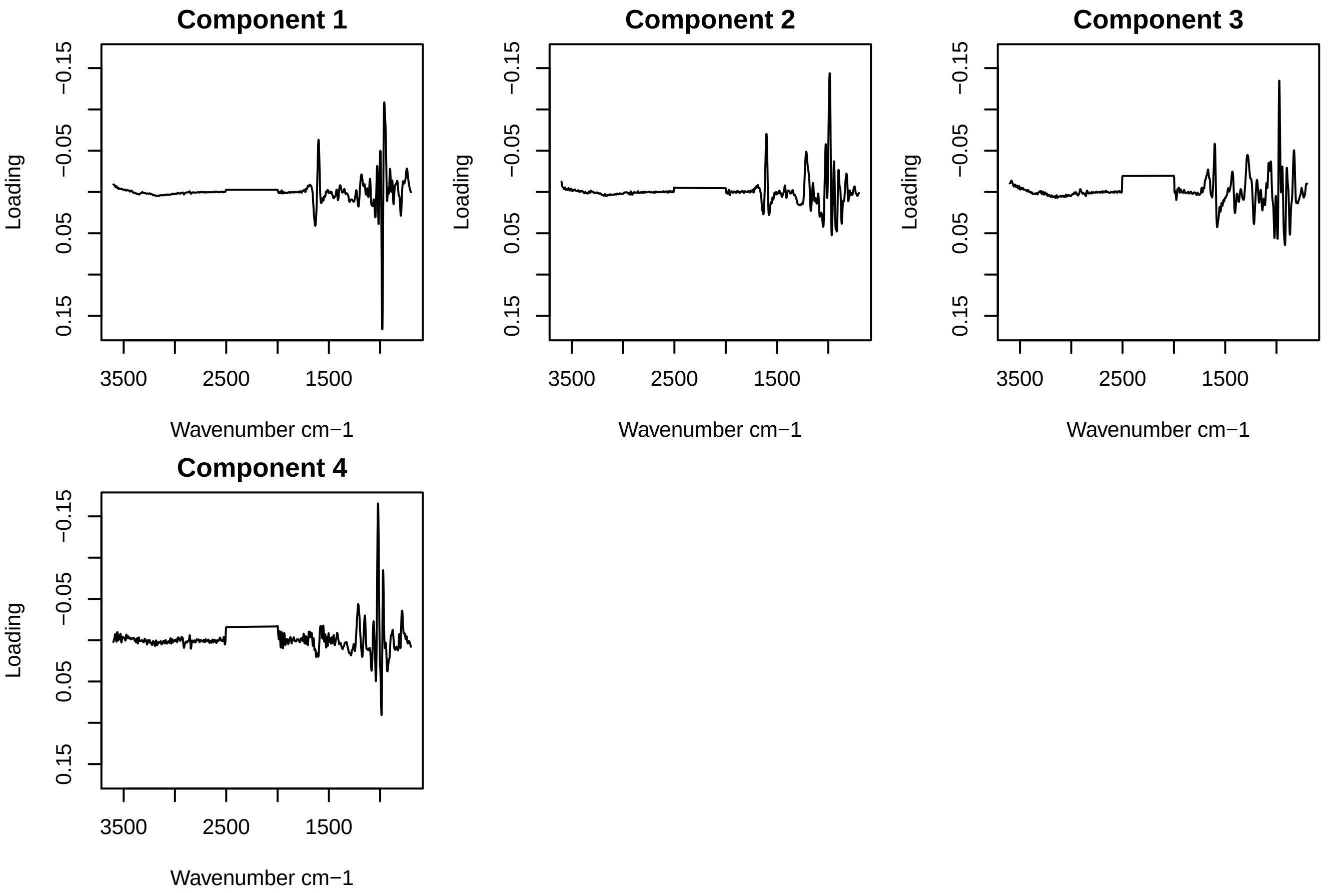
Component 1
Component 2
Component 3
Component 4
Loading
−0.15
−0.05
0.05
0.15
3500
2500
1500
Wavenumber cm−1

# PC1 vs PC2 vs PC3

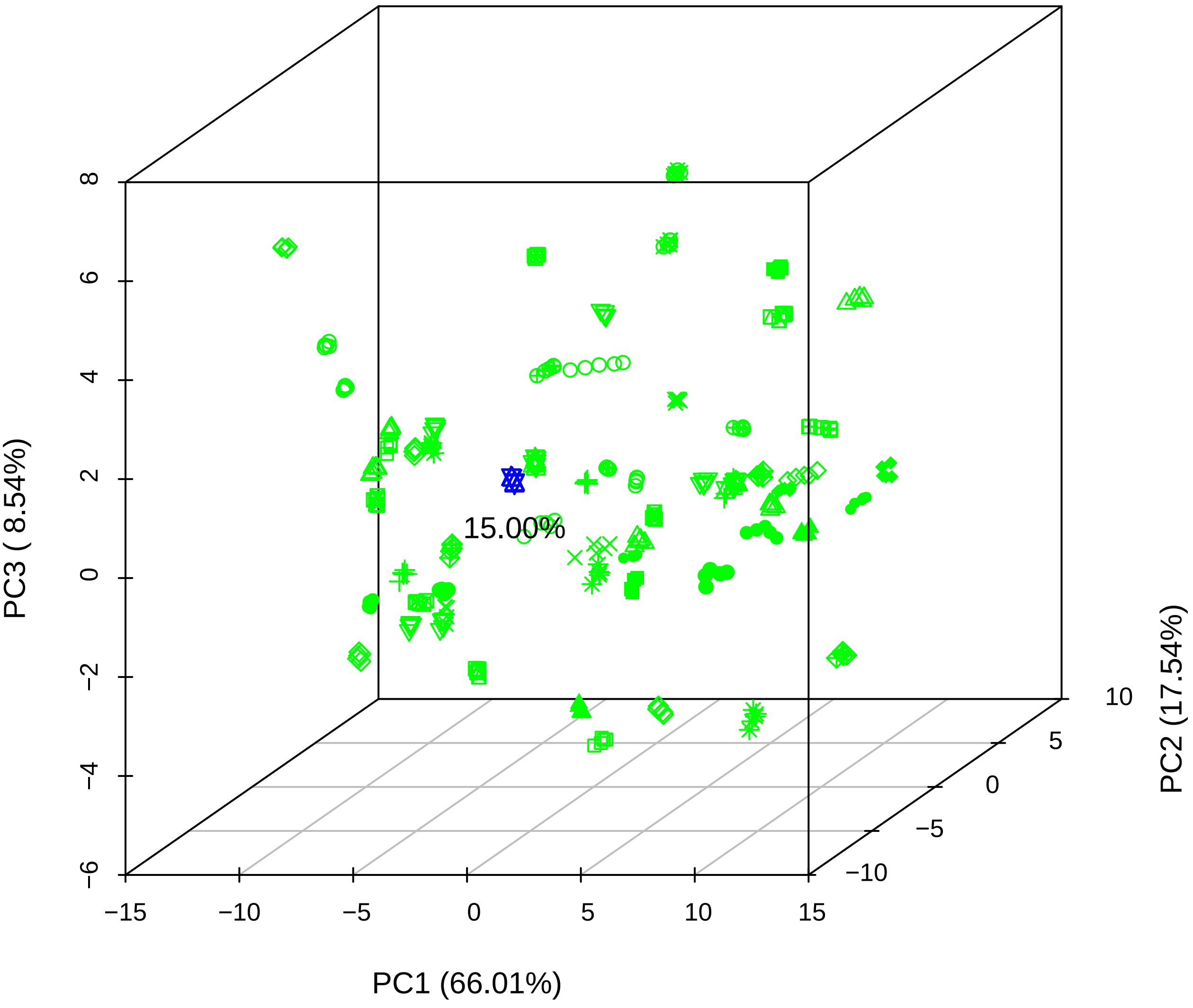

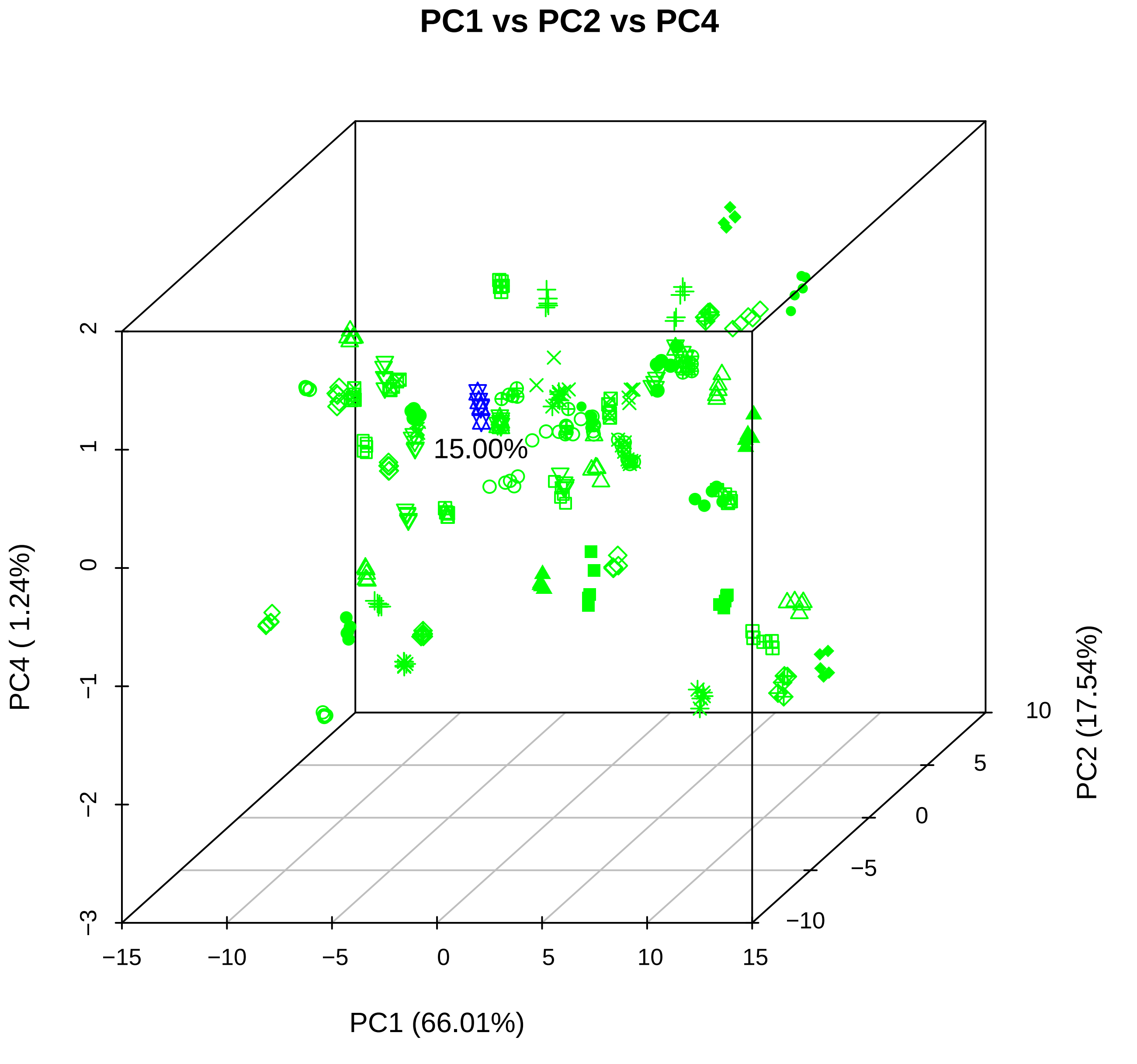

PC1 vs PC2 vs PC4
15.00%
PC1 (66.01%)
PC2 (17.54%)
PC4 ( 1.24%)
−15
−10
−5
0
5
10
15
−3
−2
−1
0
1
2
−10
−5
0
5
10

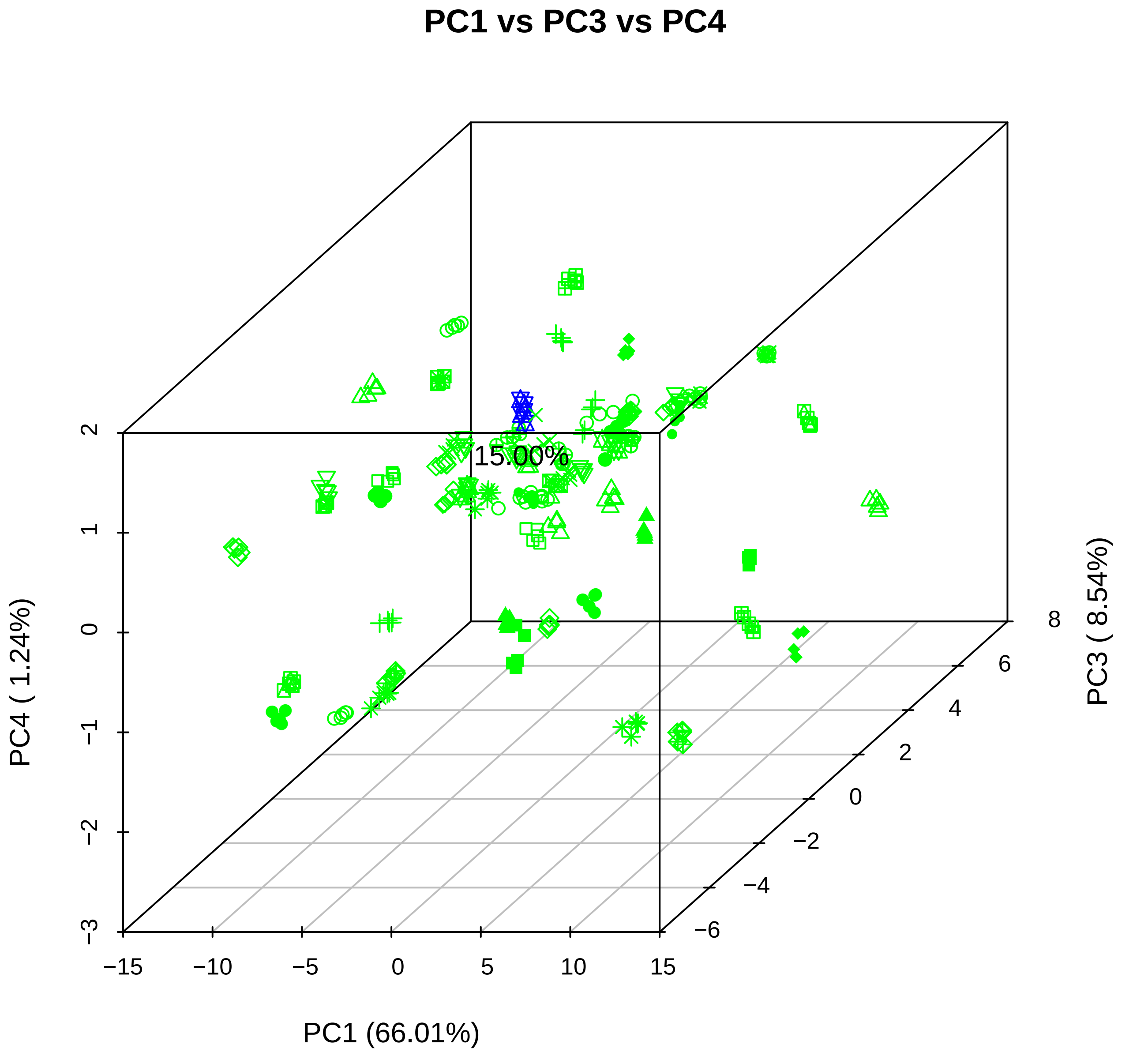
PC1 vs PC3 vs PC4
15.00%
PC1 (66.01%)
PC3 ( 8.54%)
PC4 ( 1.24%)
−15
−10
−5
0
5
10
15
−6
−4
−2
0
2
4
6
8
−3
−2
−1
0
1
2

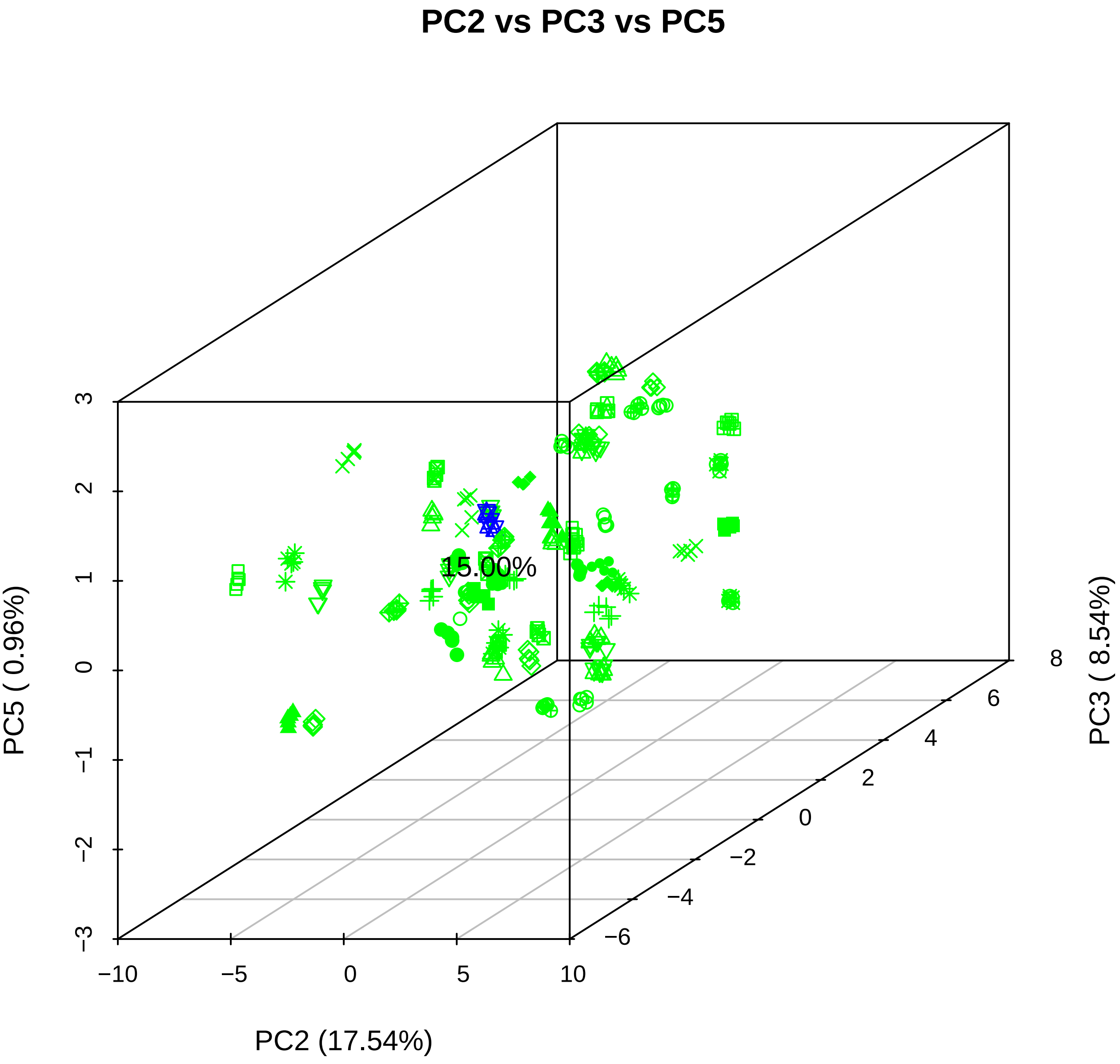

PC2 vs PC3 vs PC5
15.00%
PC2 (17.54%)
PC5 ( 0.96%)
PC3 ( 8.54%)
−10
−5
0
5
10
−3
−2
−1
0
1
2
3
−6
−4
−2
0
2
4
6
8

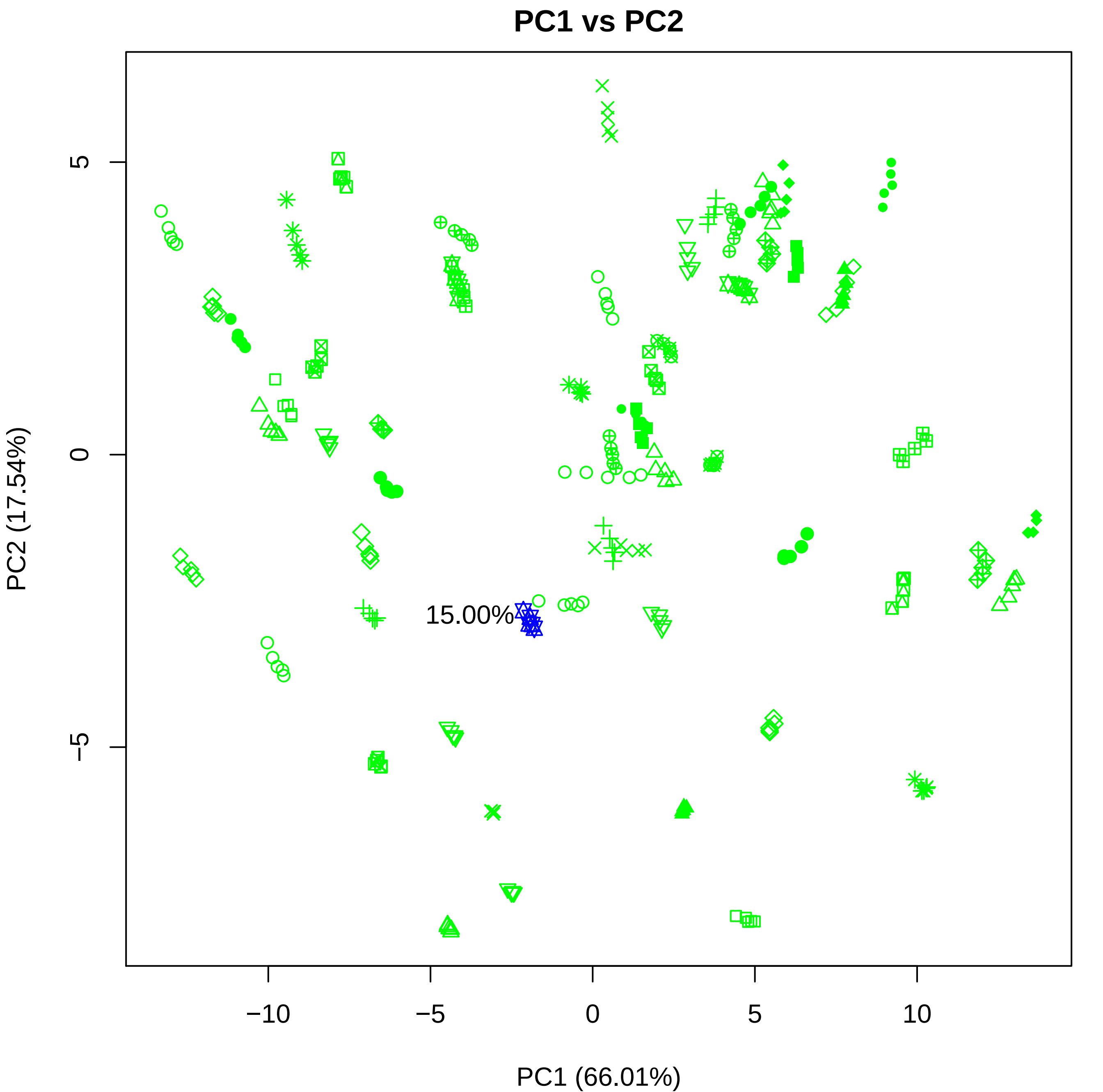
PC1 vs PC2
PC2 (17.54%)
PC1 (66.01%)
15.00%
5
0
−5
−10
−5
0
5
10

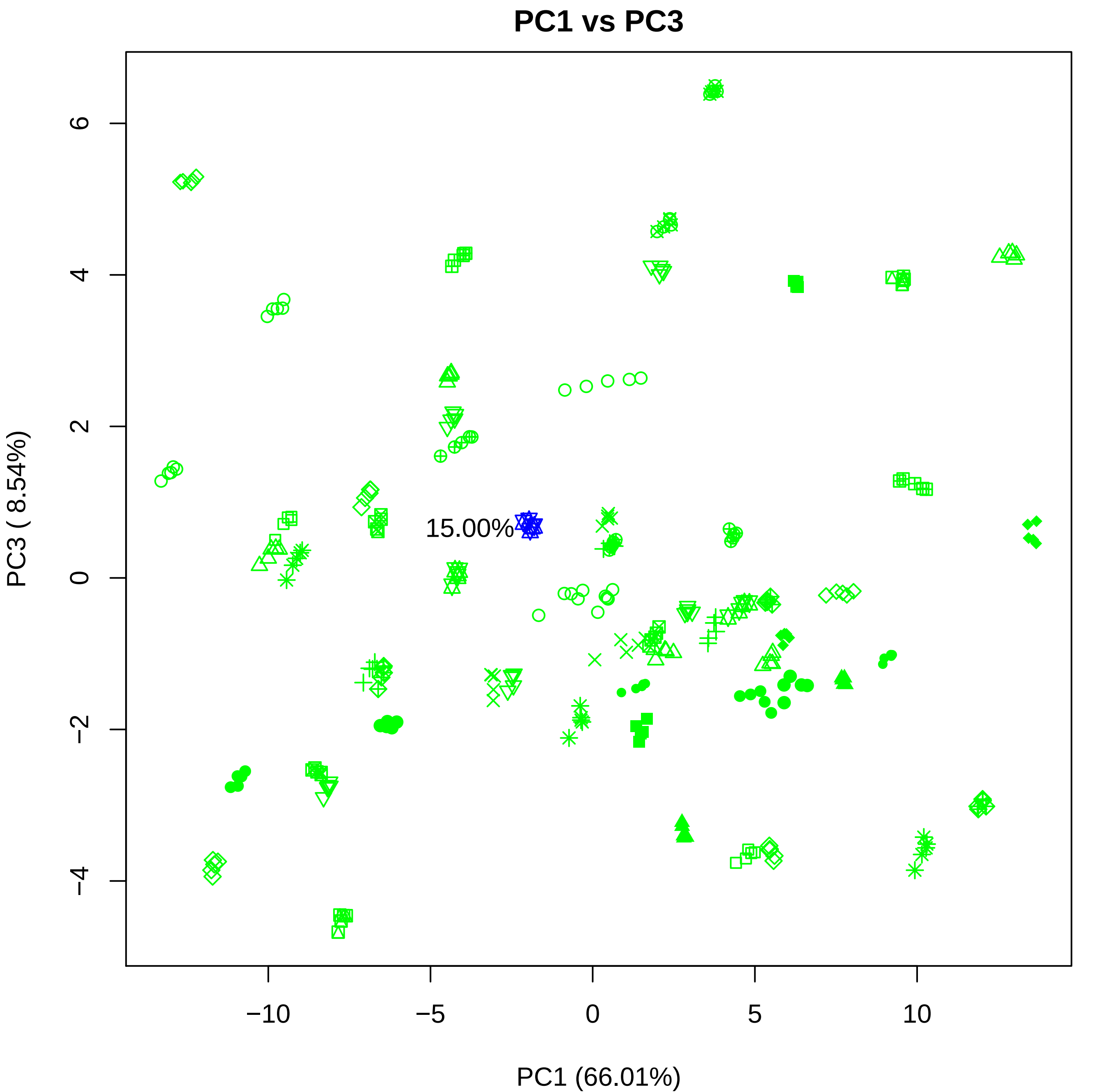

PC1 vs PC3
15.00%
PC3 ( 8.54%)
PC1 (66.01%)
6
4
2
0
−2
−4
−10
−5
0
5
10

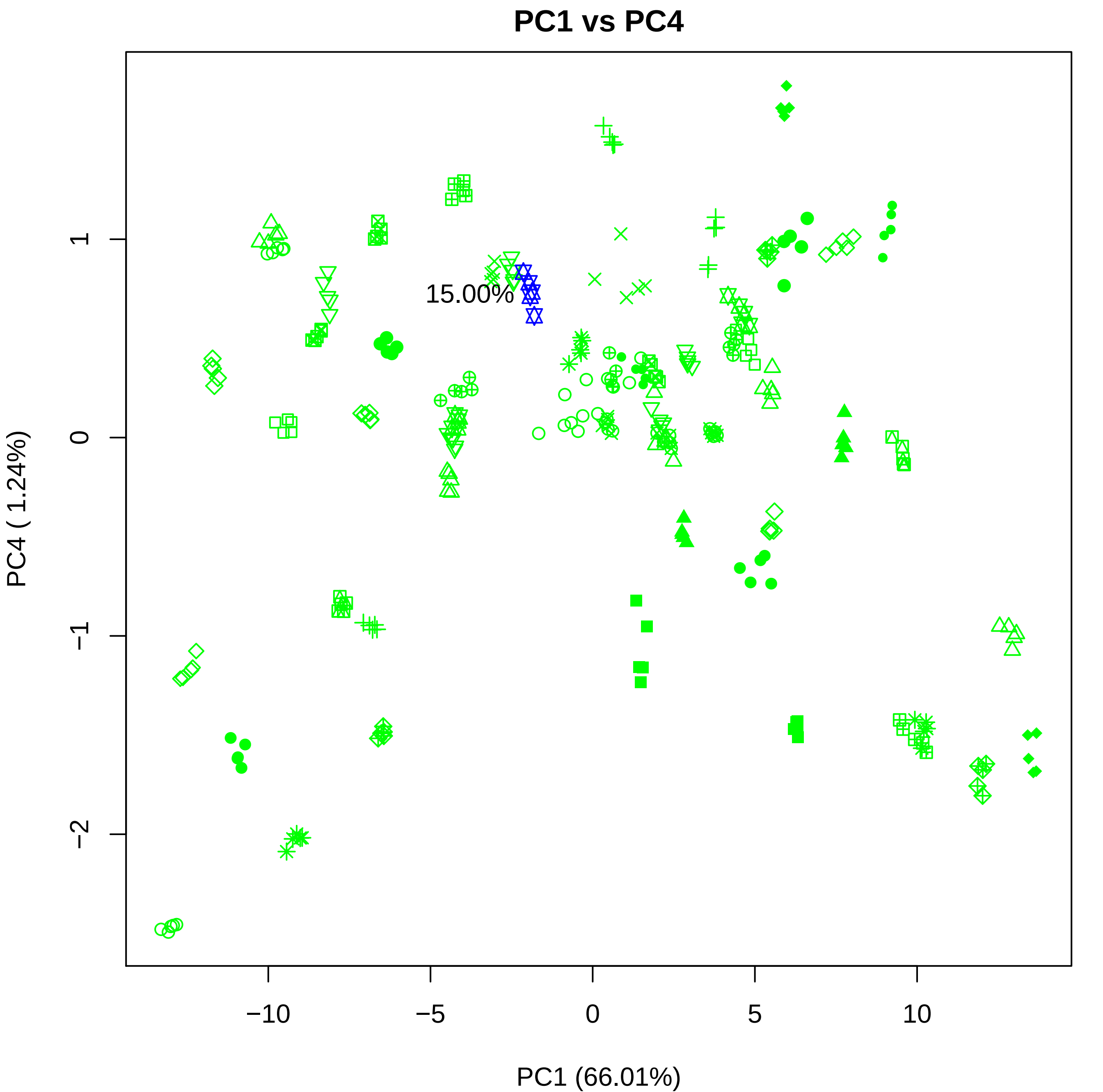
PC1 vs PC4
15.00%
PC4 ( 1.24%)
PC1 (66.01%)
1
0
−1
−2
−10
−5
0
5
10

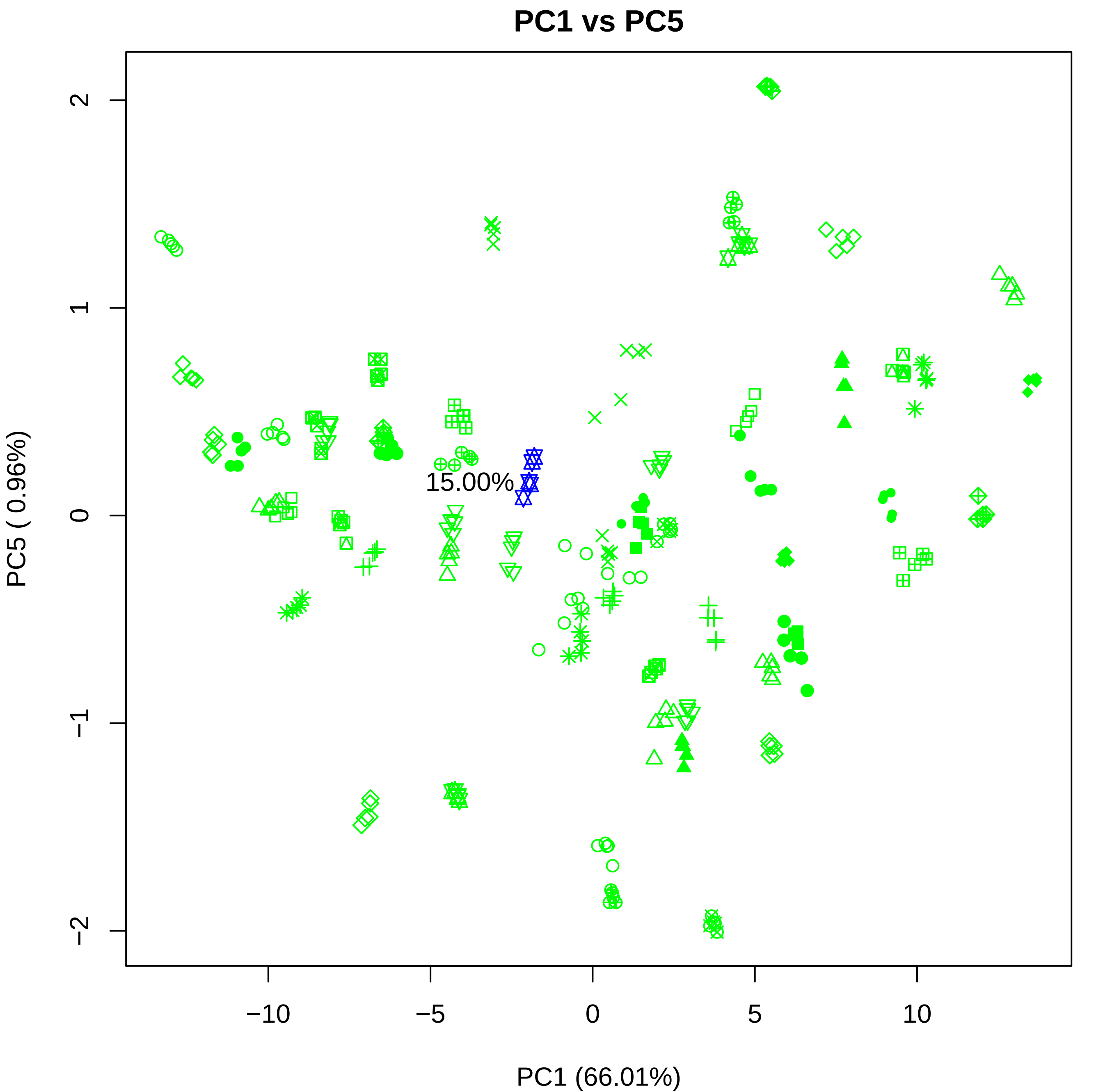
PC1 vs PC5
15.00%
PC5 ( 0.96%)
PC1 (66.01%)
2
1
0
−1
−2
−10
−5
0
5
10

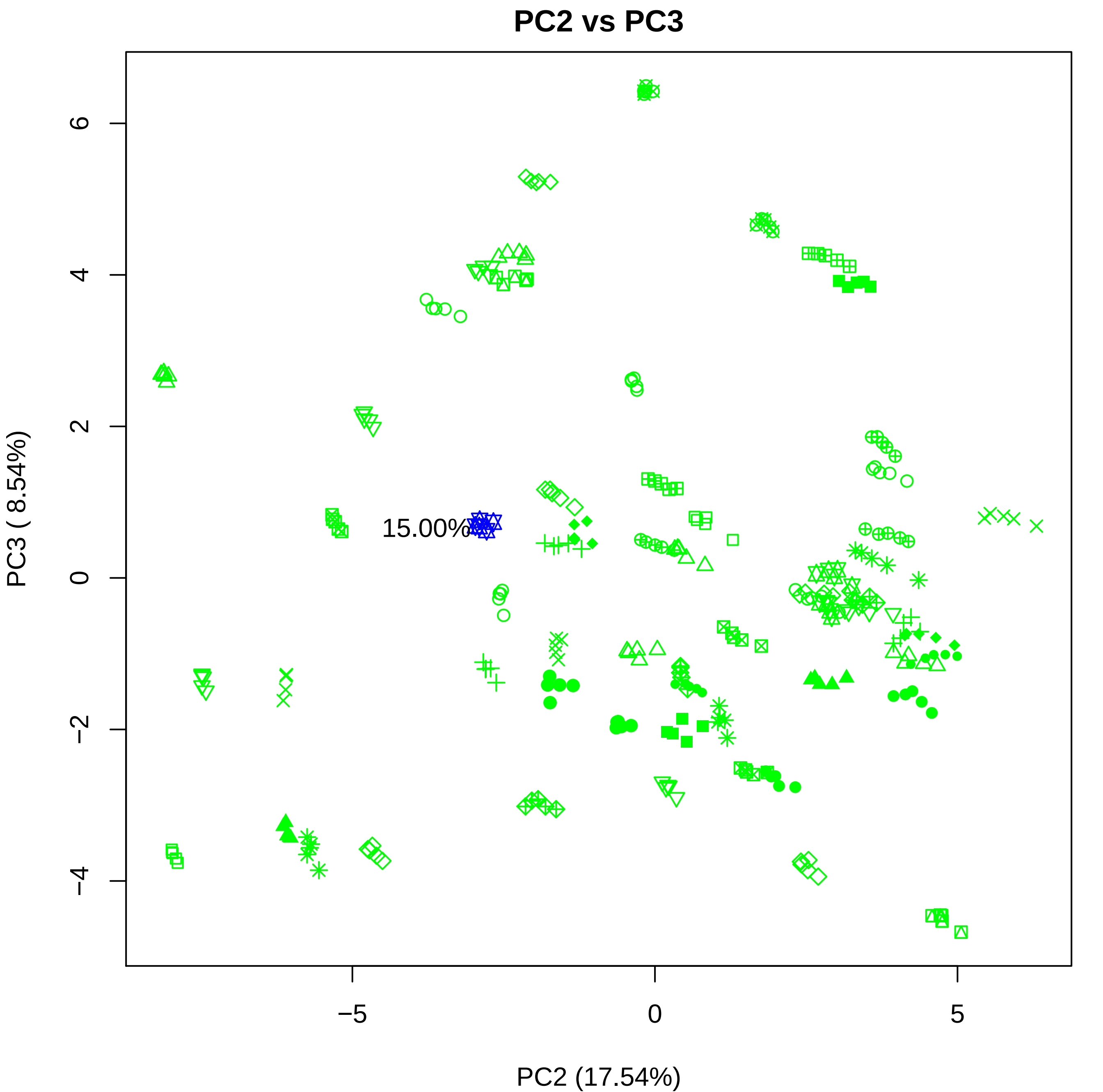

PC2 vs PC3
15.00%
PC3 ( 8.54%)
PC2 (17.54%)
6
4
2
0
−2
−4
−5
0
5

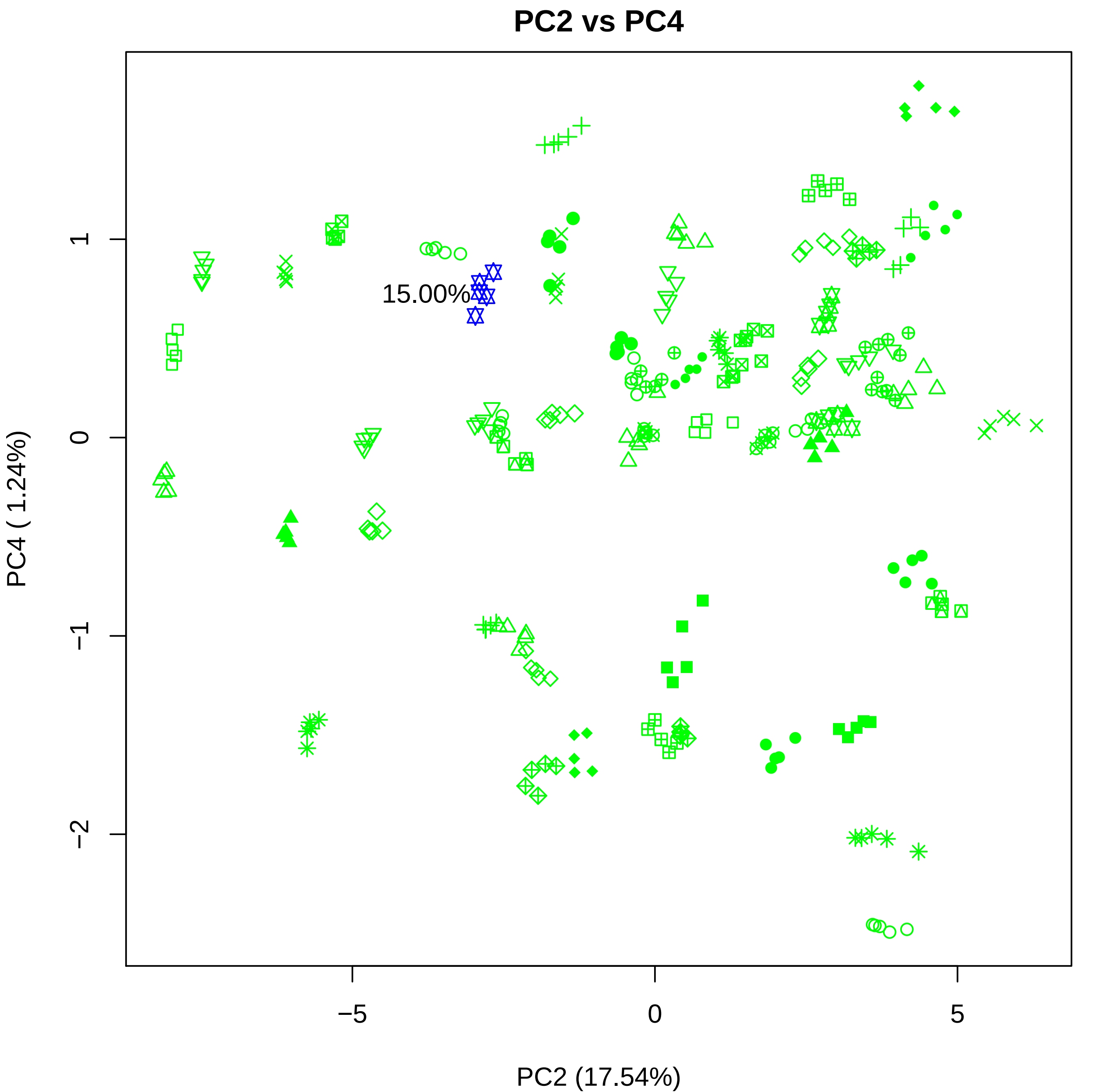
PC2 vs PC4
15.00%
PC4 ( 1.24%)
PC2 (17.54%)
1
0
−1
−2
−5
0
5

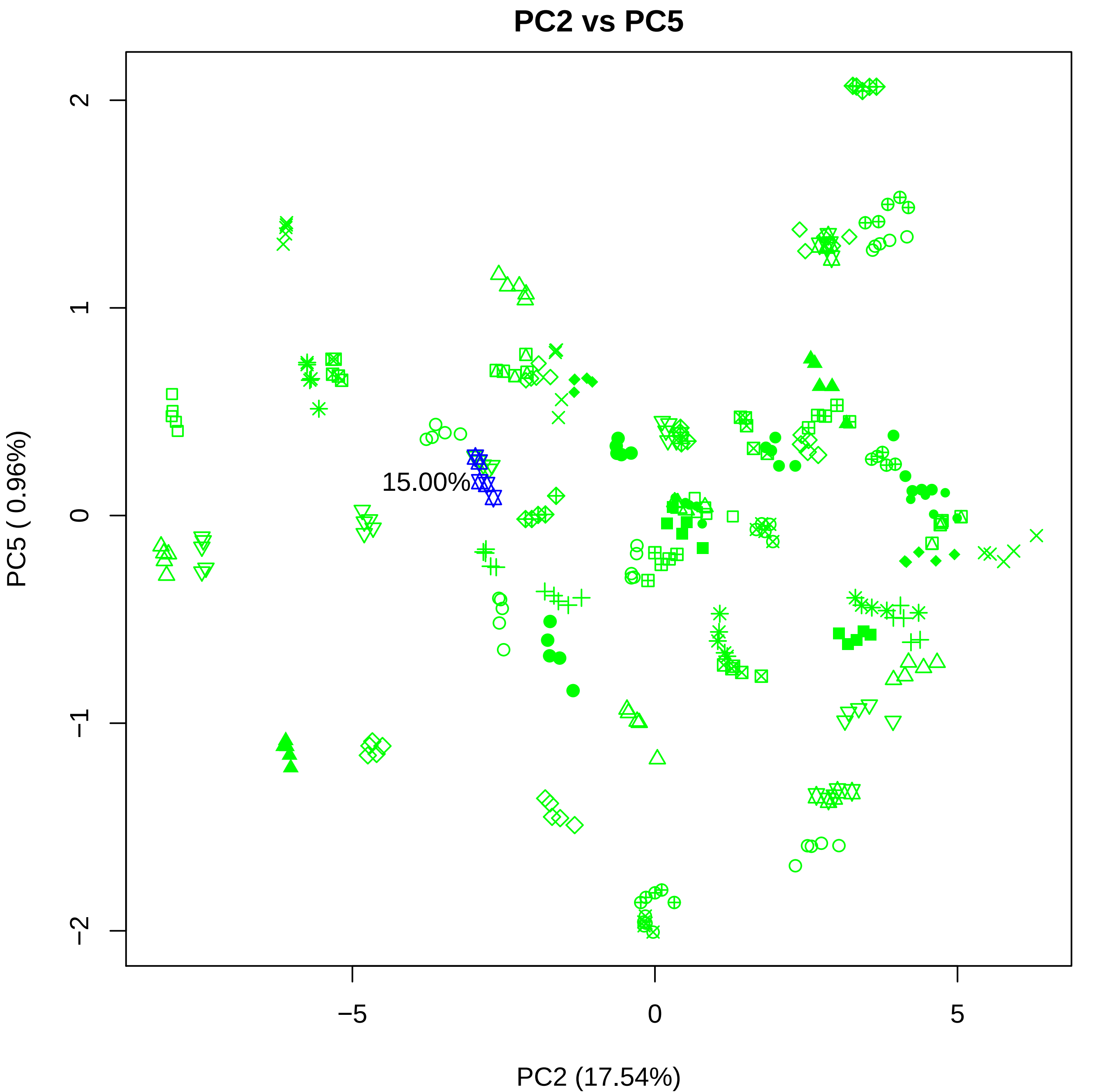

PC2 vs PC5
15.00%
PC5 ( 0.96%)
PC2 (17.54%)
−5
0
5
−2
−1
0
1
2

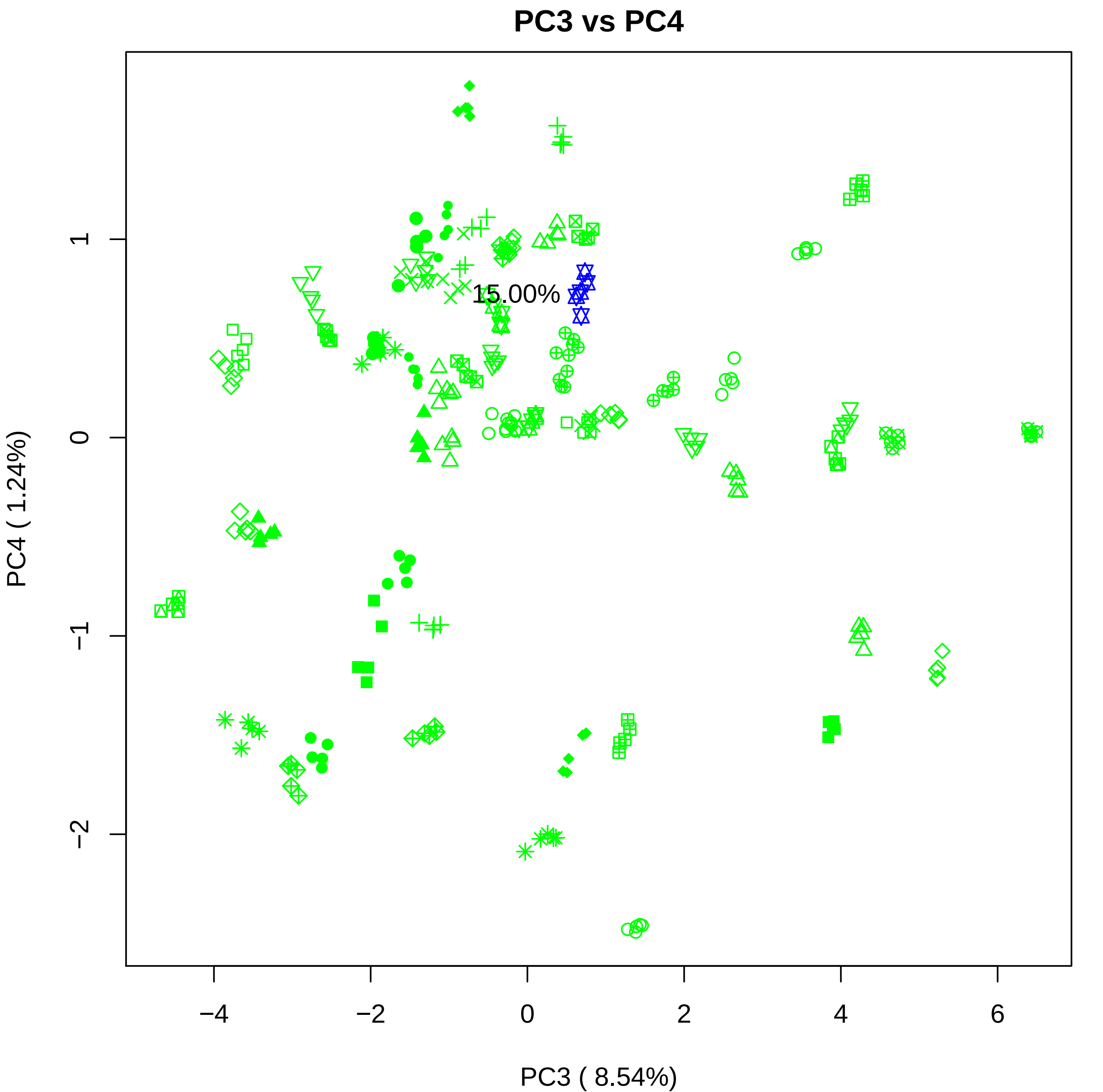

PC3 vs PC4
15.00%
PC4 ( 1.24%)
PC3 ( 8.54%)
−4
−2
0
2
4
6
1
0
−1
−2

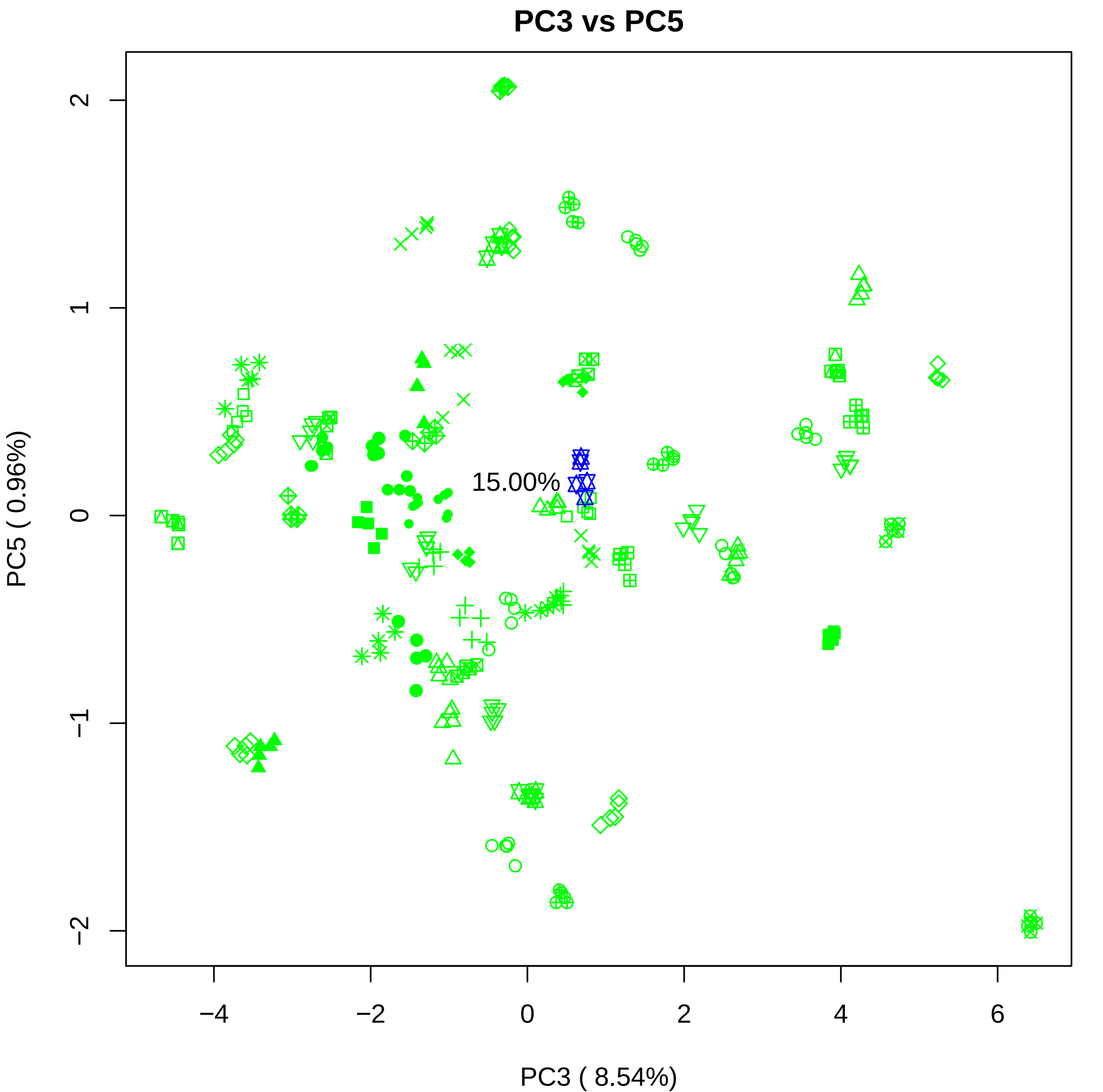
PC3 vs PC5
15.00%
PC5 ( 0.96%)
PC3 ( 8.54%)
−4
−2
0
2
4
6
−2
−1
0
1
2

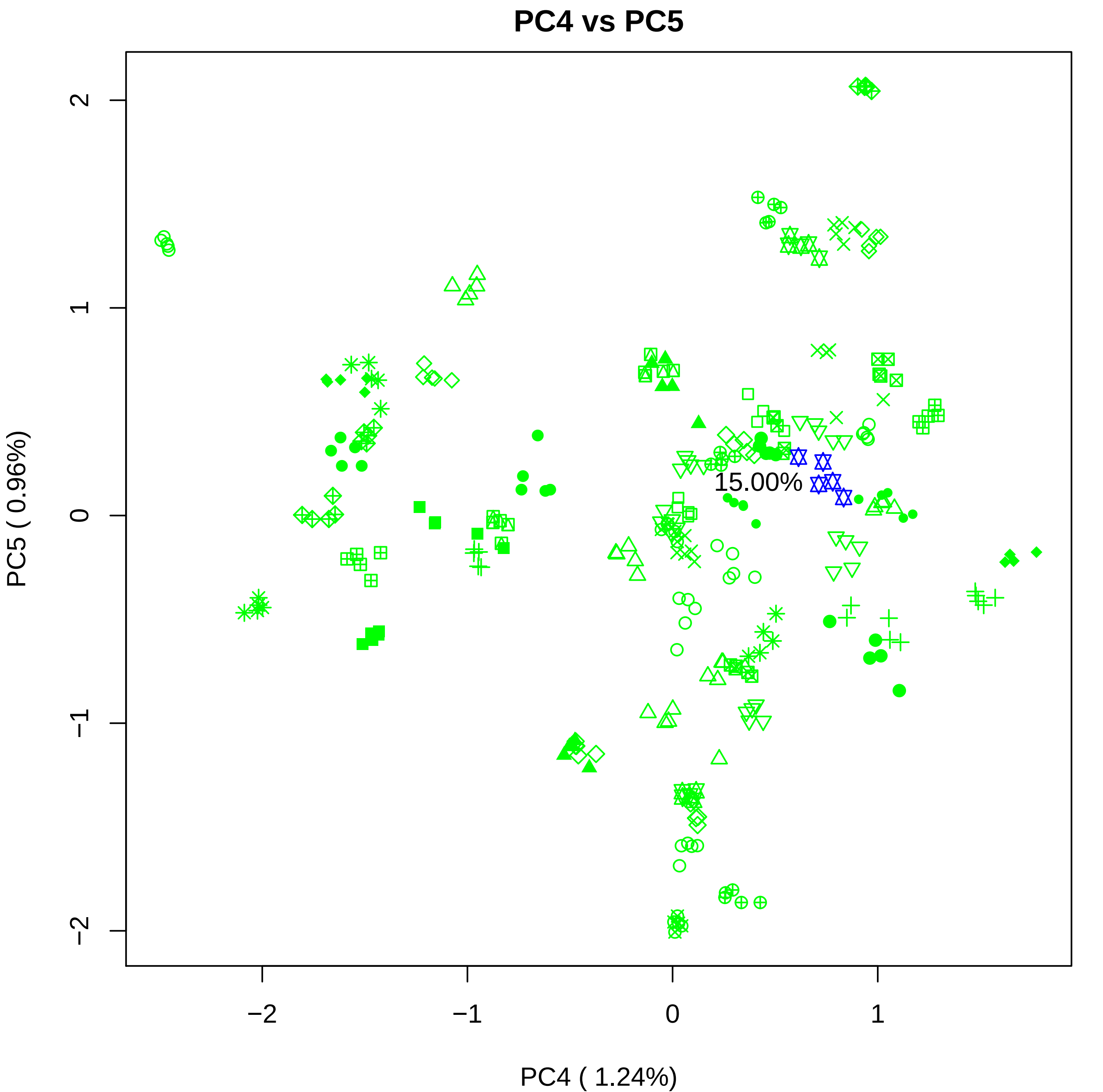

PC4 vs PC5
15.00%
PC4 ( 1.24%)
PC5 ( 0.96%)
−2
−1
0
1
2

### PCA of 20%  BMH(c)  vs PMH(c)

Scaled Matrix PCAed

7th order polynomial baseline correction performed

Raw spectra smoothed to 21 Neighbours, 2nd order PN, Savtisky Golay

Normalisation between 0 and 100

D2 spectra smoothed to 41 Neighbours, 2nd order PN, Savtisky Golay

Regions Analysed: 3601−2501 and 2001−700cm−1

Samples Analysed: 1 bContam's 0 BLH's 0 BMH's 0 OMH's 0 pContam's 0 PLH's 60 PMH's

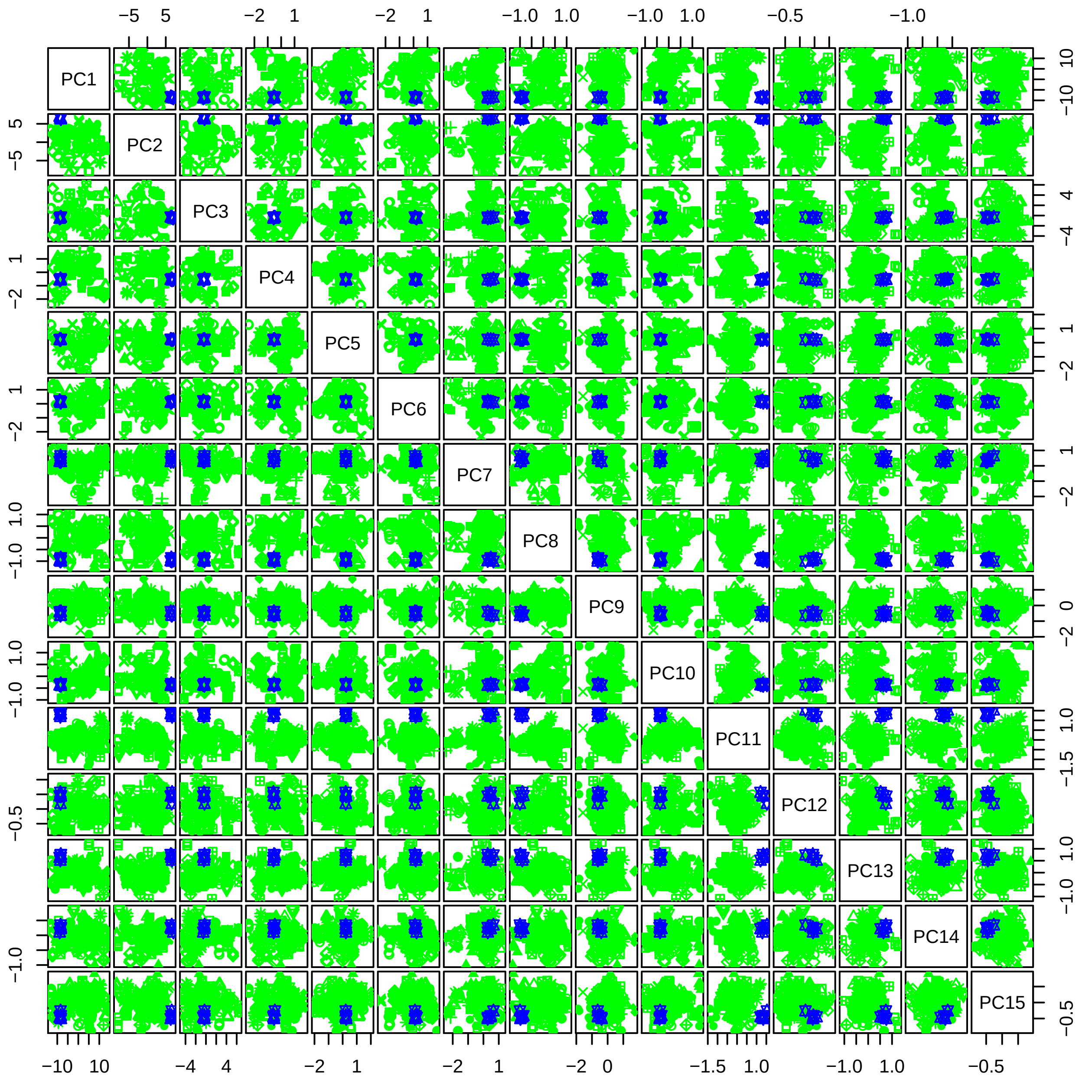
PC1
PC2
PC3
PC4
PC5
PC6
PC7
PC8
PC9
PC10
PC11
PC12
PC13
PC14
PC15

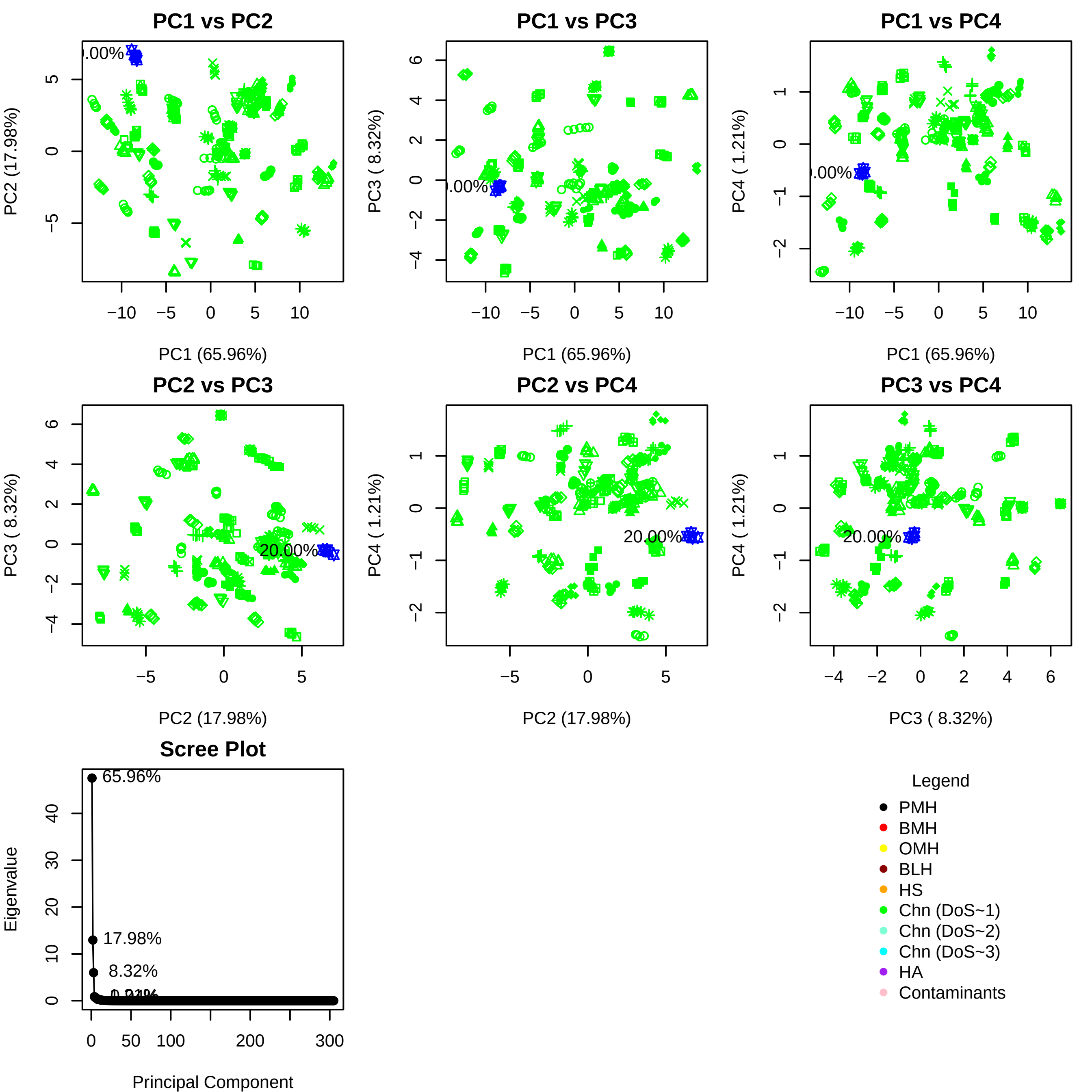
PC1 vs PC2
PC1 vs PC3
PC1 vs PC4
PC2 vs PC3
PC2 vs PC4
PC3 vs PC4
Scree Plot
PC1 (65.96%)
PC2 (17.98%)
PC3 ( 8.32%)
PC4 ( 1.21%)
20.00%
Eigenvalue
Principal Component
65.96%
17.98%
8.32%
Legend
PMH
BMH
OMH
BLH
HS
Chn (DoS~1)
Chn (DoS~2)
Chn (DoS~3)
HA
Contaminants

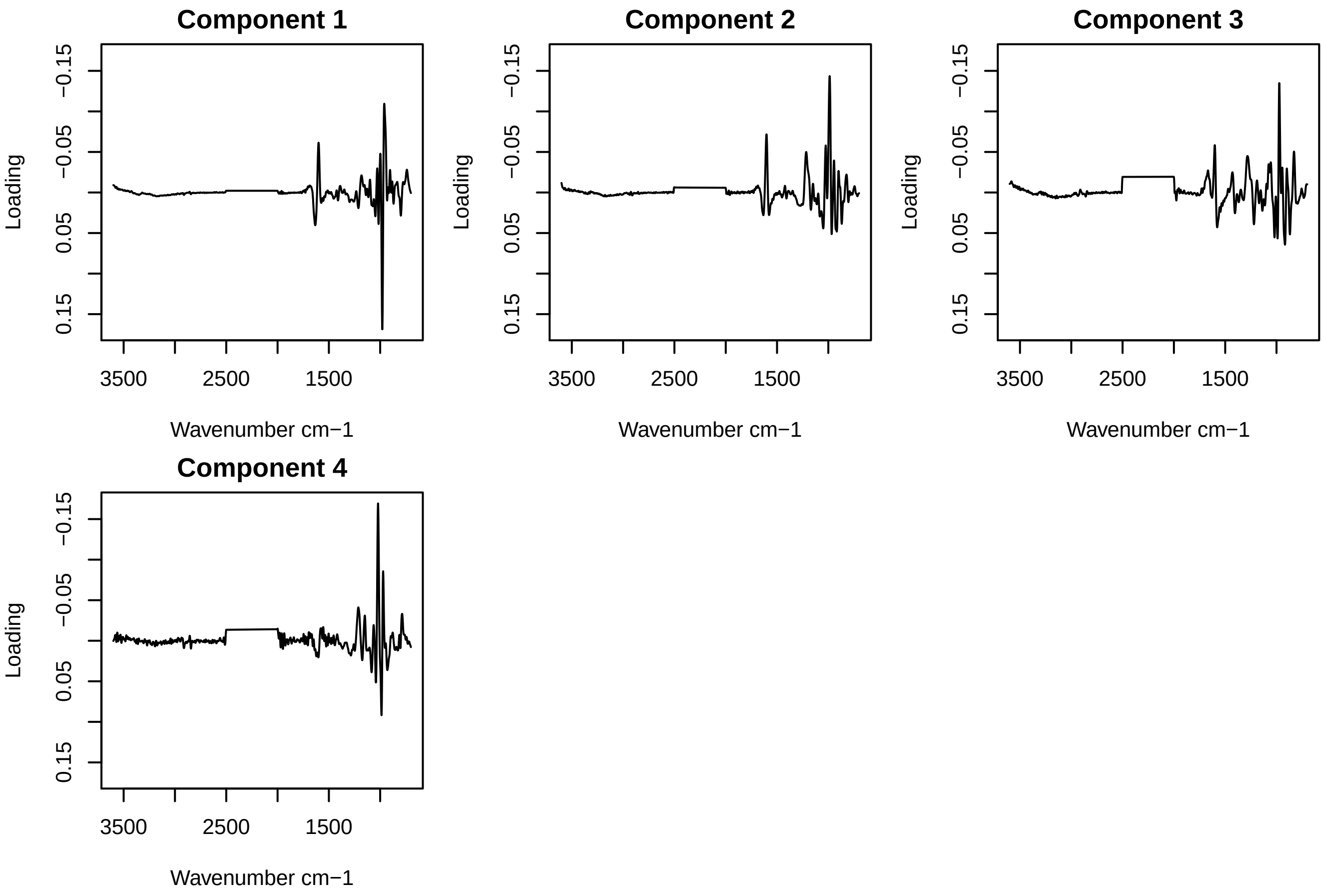
Component 1
Component 2
Component 3
Component 4
Loading
−0.15
−0.05
0.05
0.15
3500
2500
1500
Wavenumber cm−1

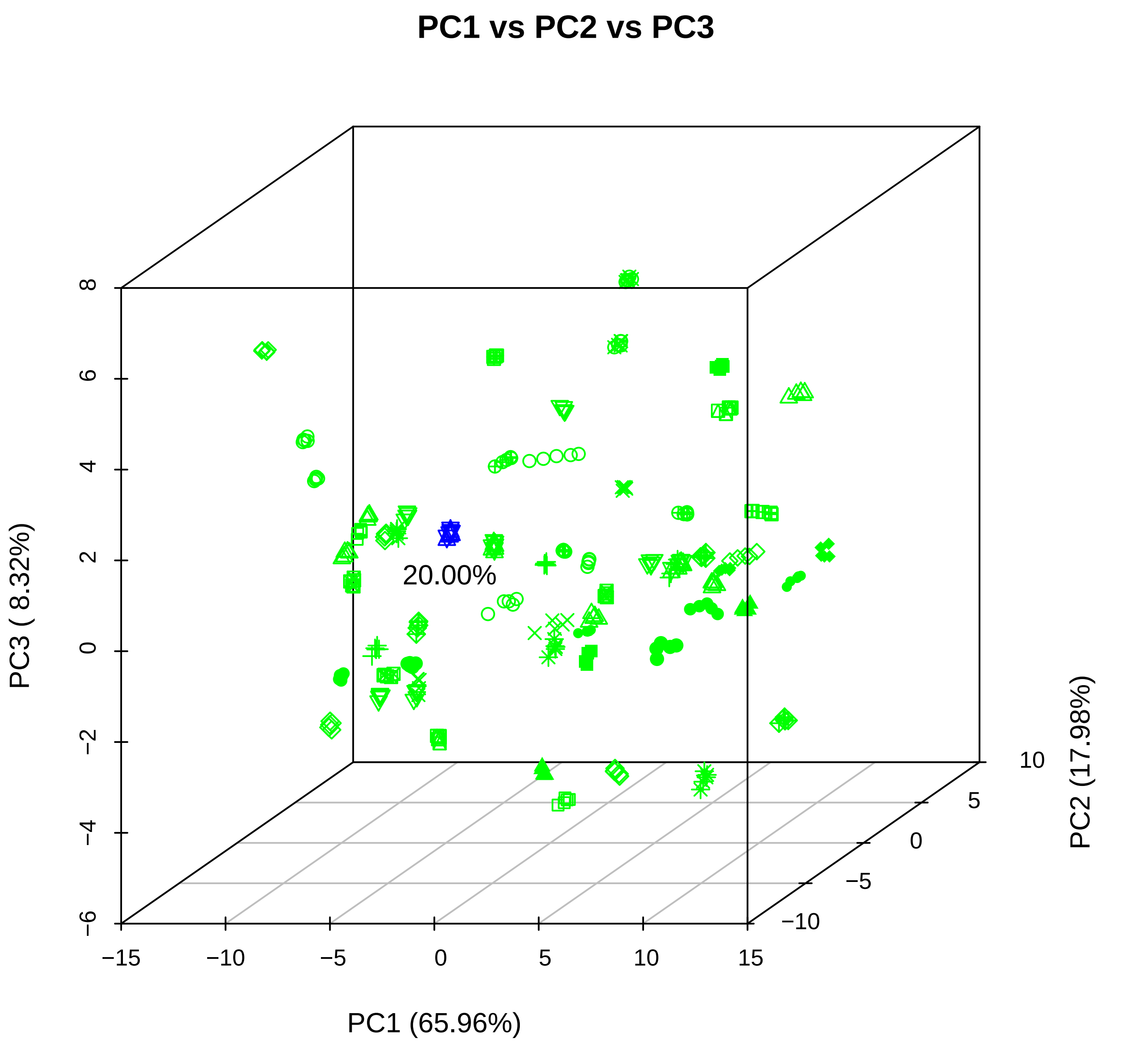
PC1 vs PC2 vs PC3
20.00%
PC1 (65.96%)
PC2 (17.98%)
PC3 ( 8.32%)
−15
−10
−5
0
5
10
15
−6
−4
−2
0
2
4
6
8

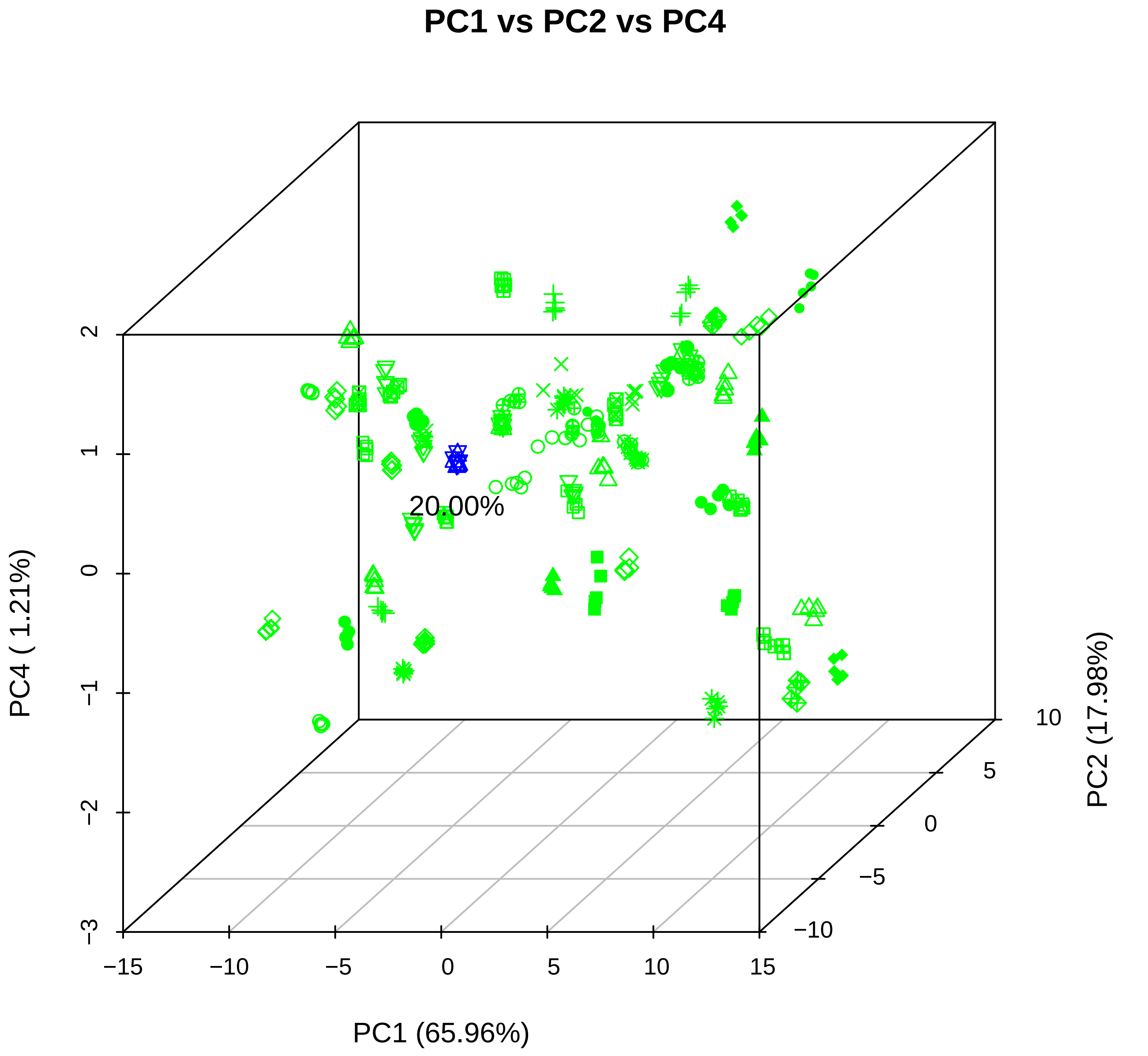

PC1 vs PC2 vs PC4
20.00%
PC1 (65.96%)
PC2 (17.98%)
PC4 ( 1.21%)
−15
−10
−5
0
5
10
15
−10
−5
0
5
10
−3
−2
−1
0
1
2

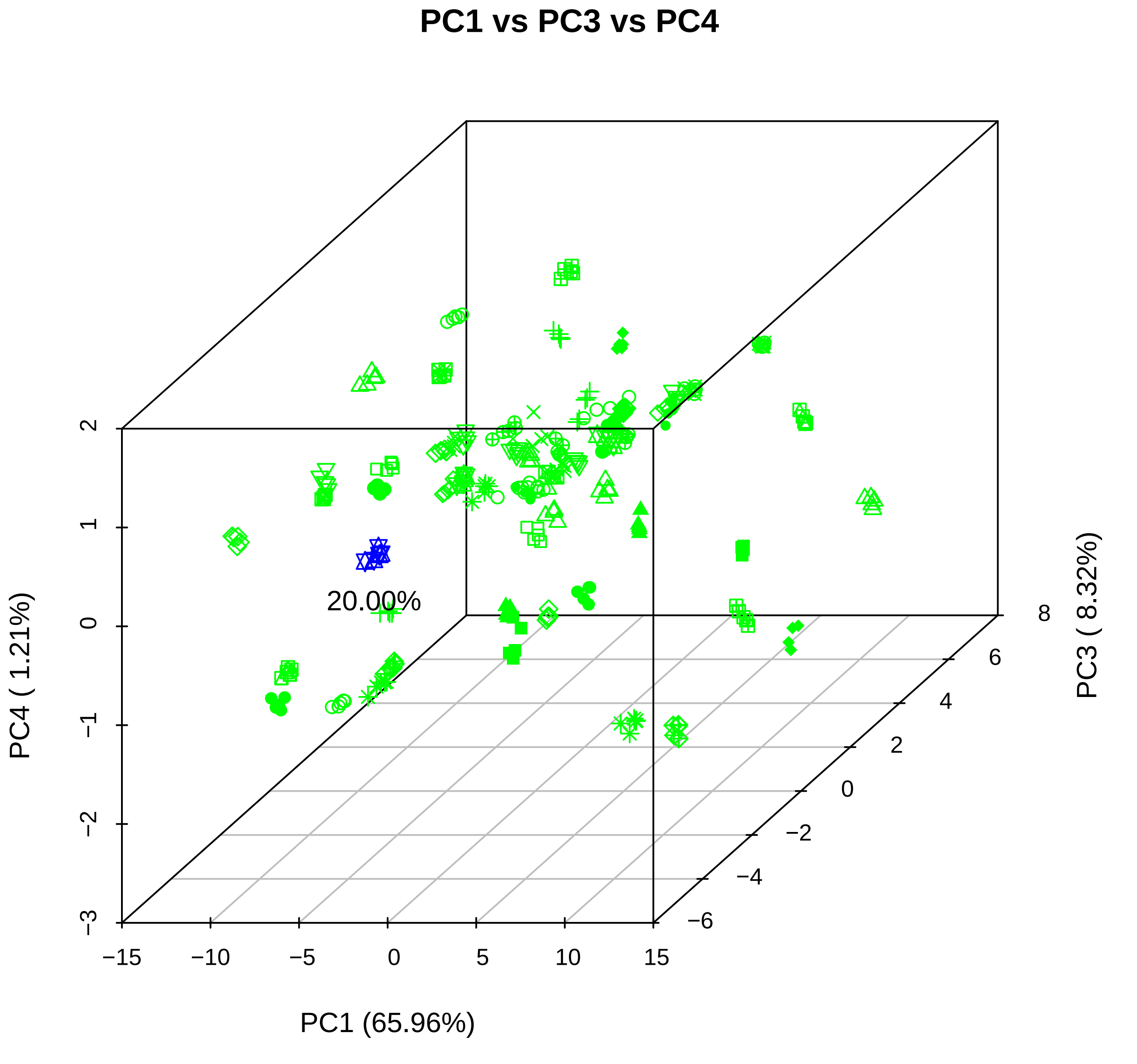

PC1 vs PC3 vs PC4
20.00%
PC1 (65.96%)
PC4 ( 1.21%)
PC3 ( 8.32%)
−15
−10
−5
0
5
10
15
−3
−2
−1
0
1
2
−6
−4
−2
0
2
4
6
8

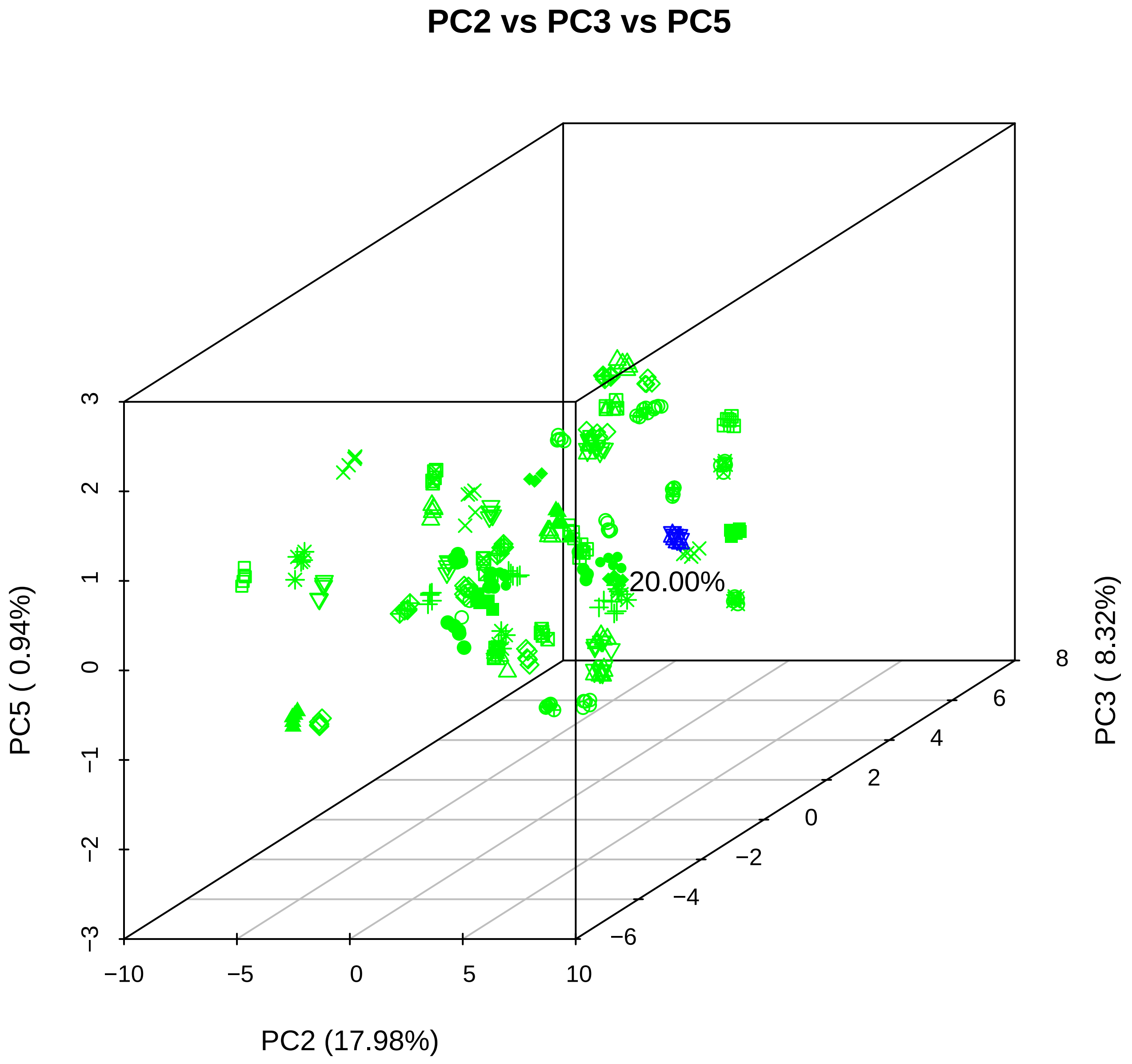
PC2 vs PC3 vs PC5
3
2
1
0
−1
−2
−3
PC5 ( 0.94%)
−10
−5
0
5
10
PC2 (17.98%)
−6
−4
−2
0
2
4
6
8
PC3 ( 8.32%)
20.00%

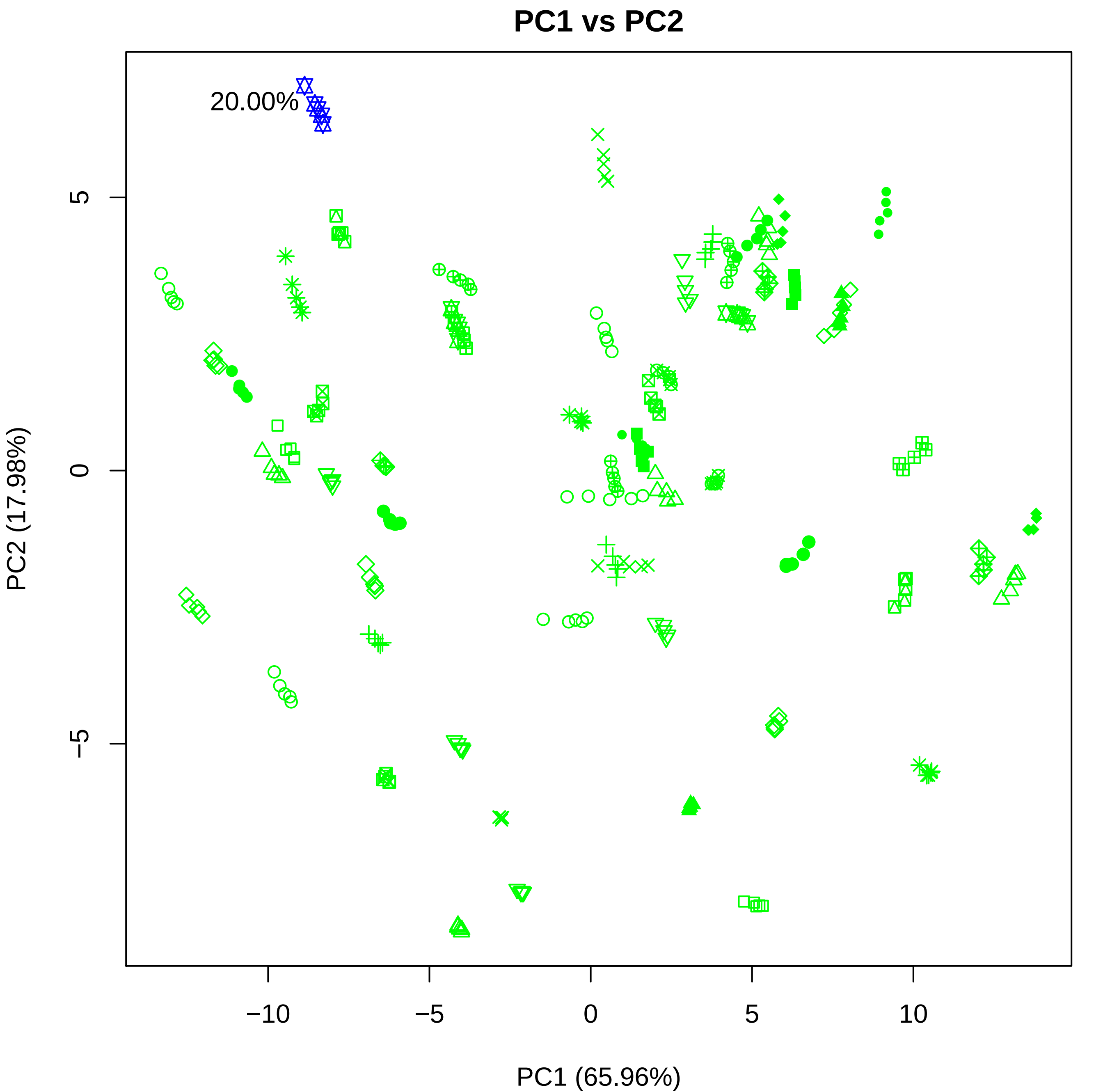
PC1 vs PC2
20.00%
PC2 (17.98%)
5
0
−5
−10
−5
0
5
10
PC1 (65.96%)

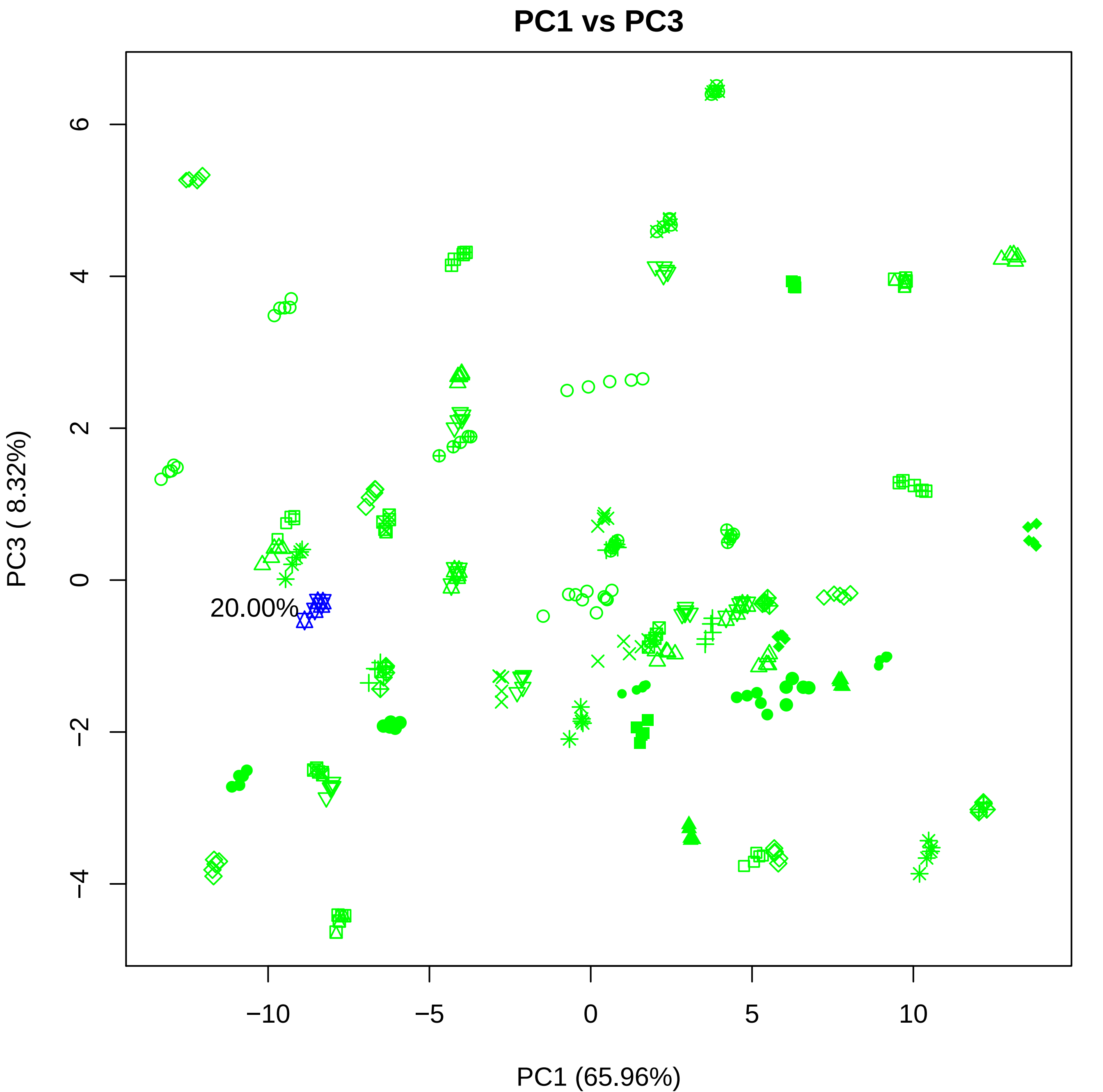
PC1 vs PC3
PC3 ( 8.32%)
PC1 (65.96%)
20.00%
6
4
2
0
−2
−4
−10
−5
0
5
10

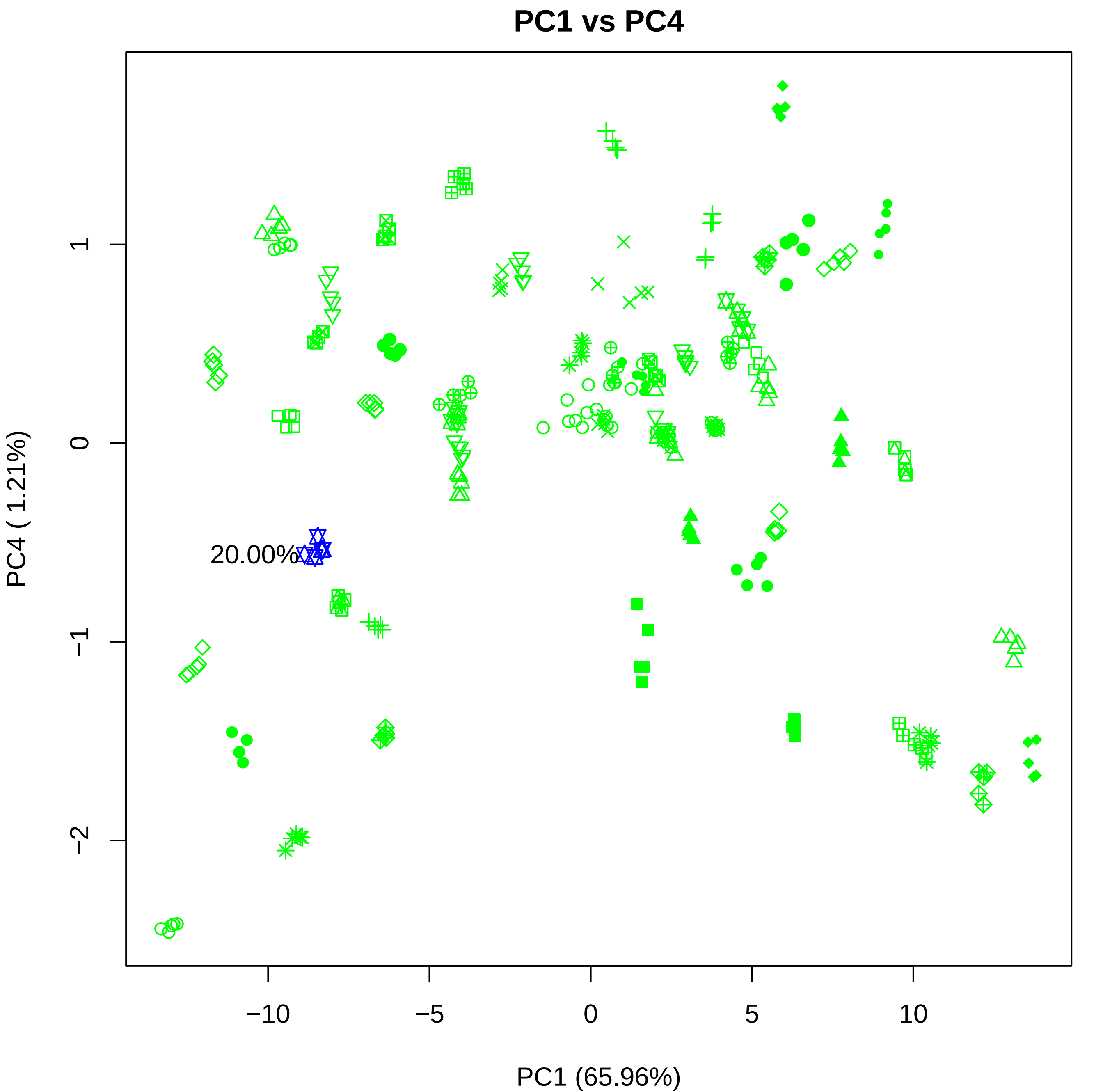
PC1 vs PC4
PC4 ( 1.21%)
PC1 (65.96%)
20.00%
1
0
−1
−2
−10
−5
0
5
10

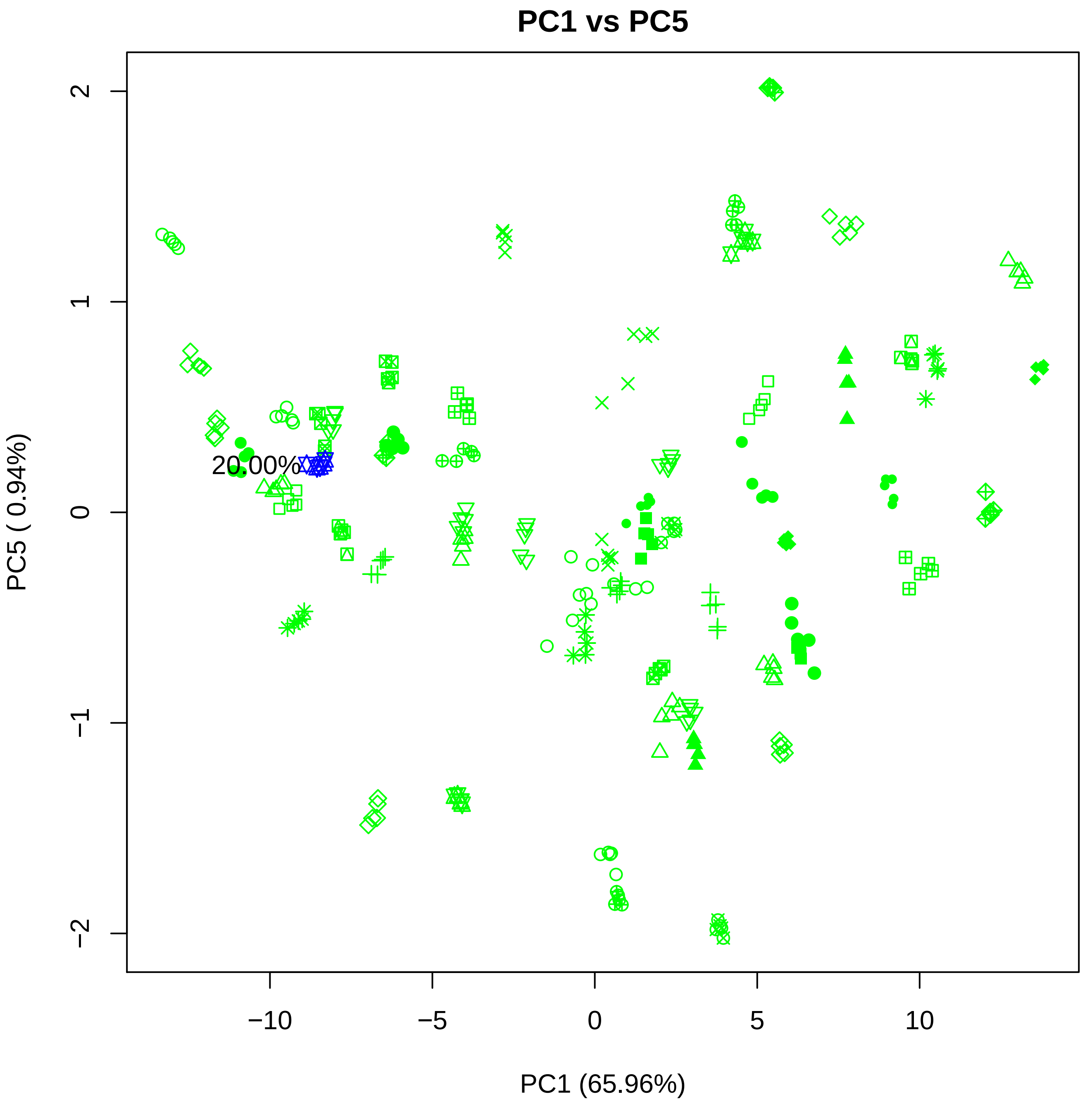
PC1 vs PC5
20.00%
PC5 ( 0.94%)
PC1 (65.96%)
2
1
0
−1
−2
−10
−5
0
5
10

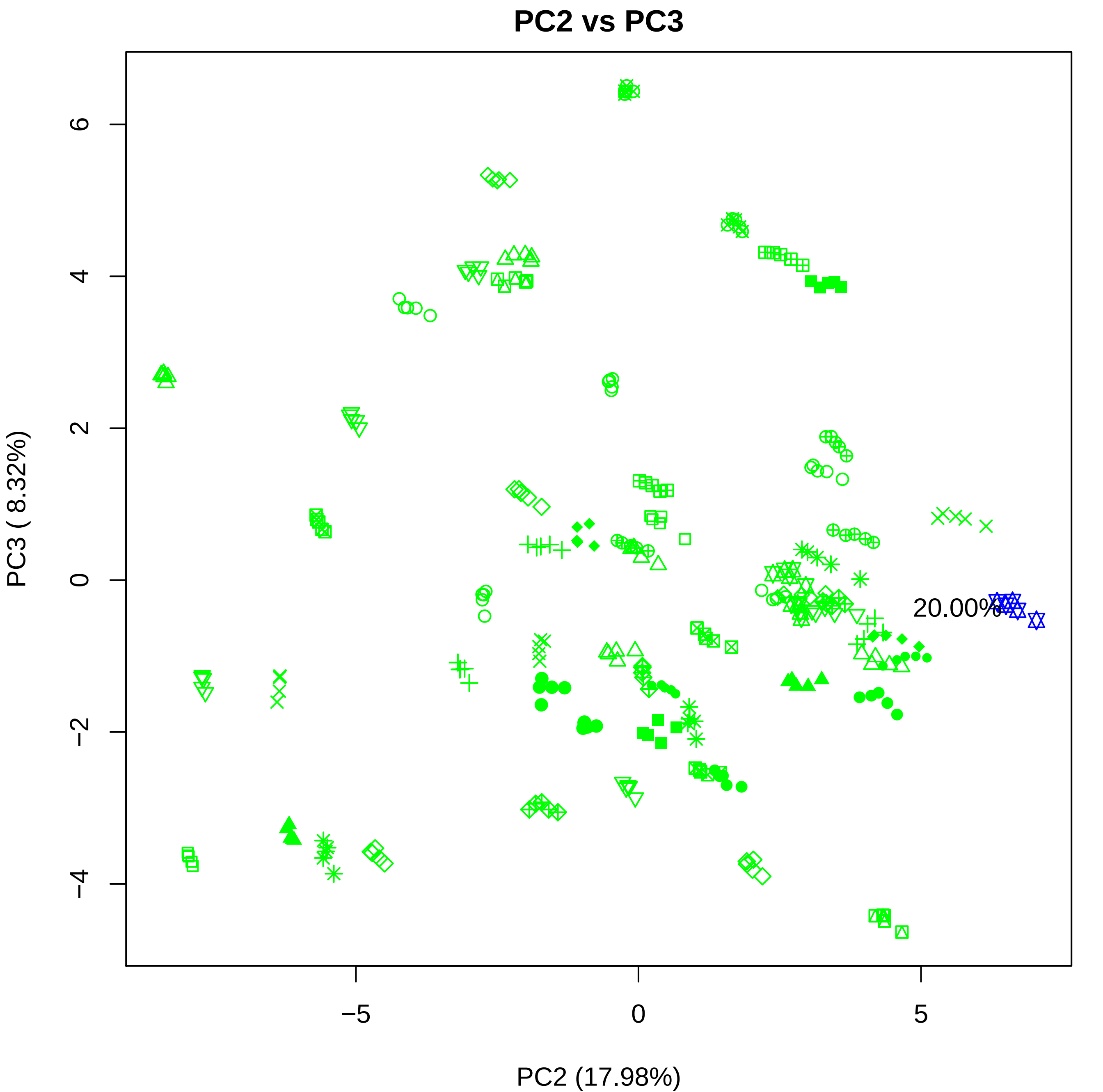
PC2 vs PC3
PC3 ( 8.32%)
PC2 (17.98%)
20.00%
6
4
2
0
−2
−4
−5
0
5

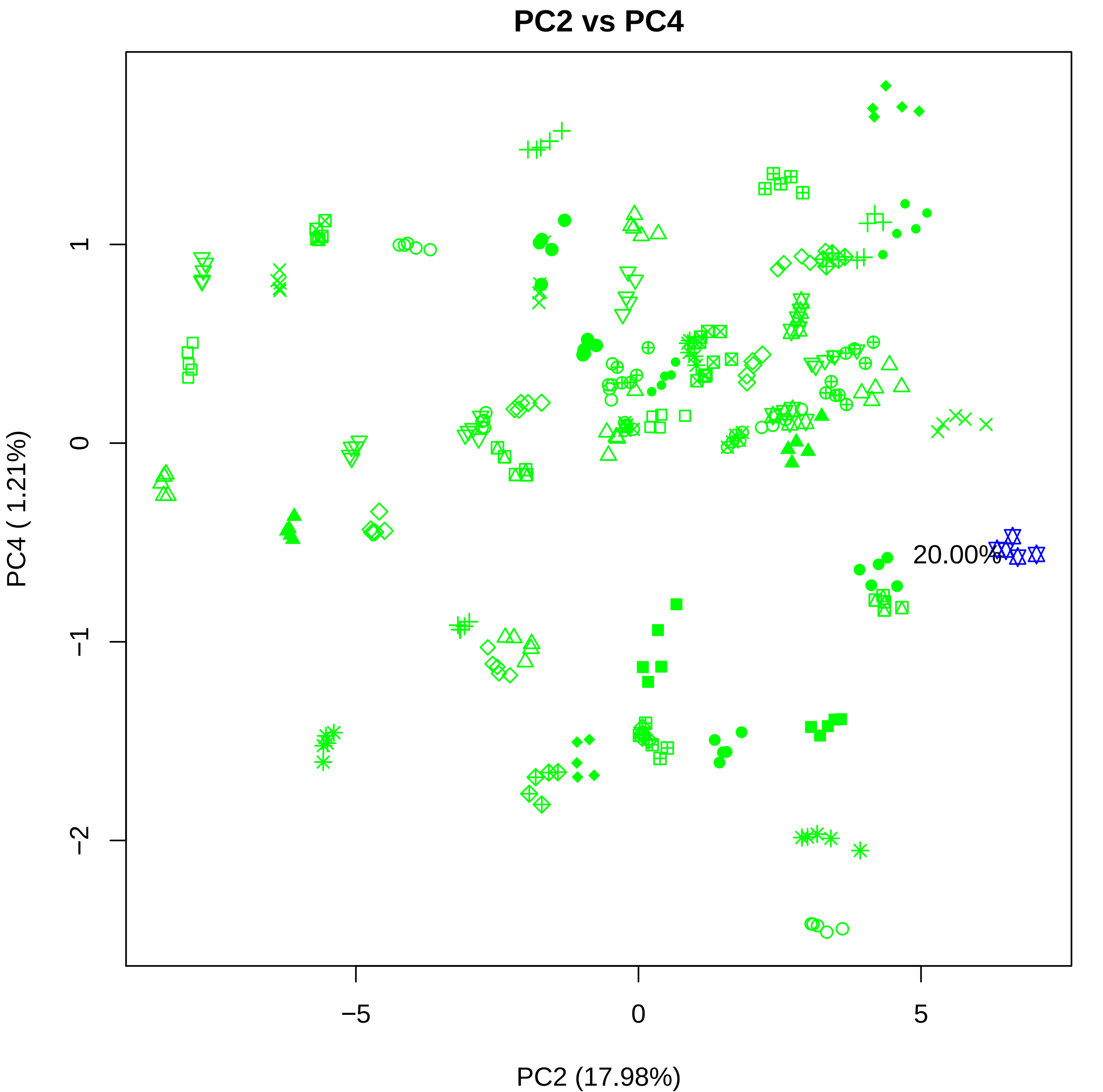

PC2 vs PC4
PC4 ( 1.21%)
PC2 (17.98%)
1
0
−1
−2
−5
0
5
20.00%

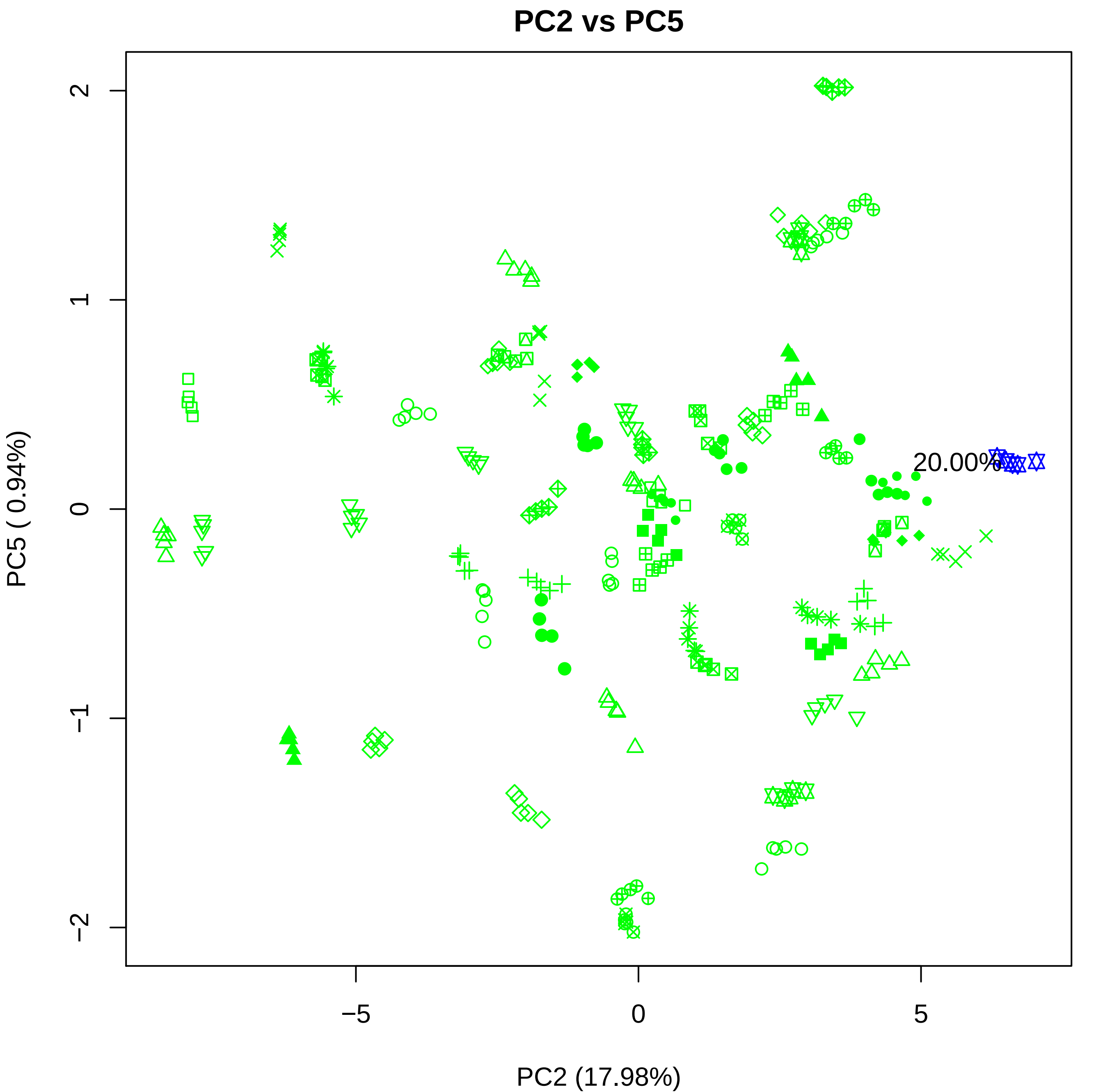
PC2 vs PC5
20.00%
PC5 ( 0.94%)
PC2 (17.98%)
−5
0
5
2
1
0
−1
−2

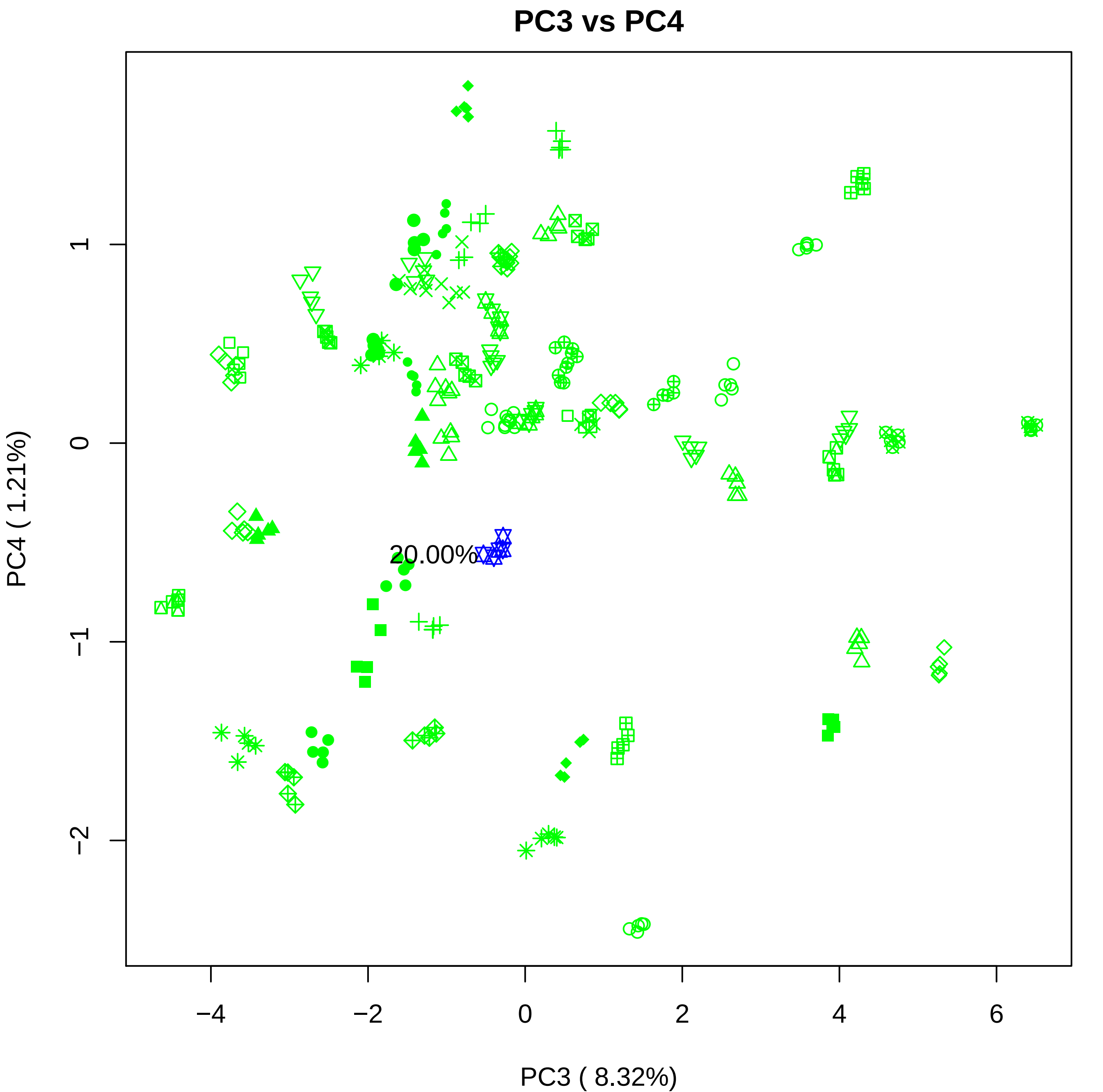
PC3 vs PC4
PC4 ( 1.21%)
PC3 ( 8.32%)
20.00%
1
0
−1
−2
−4
−2
0
2
4
6

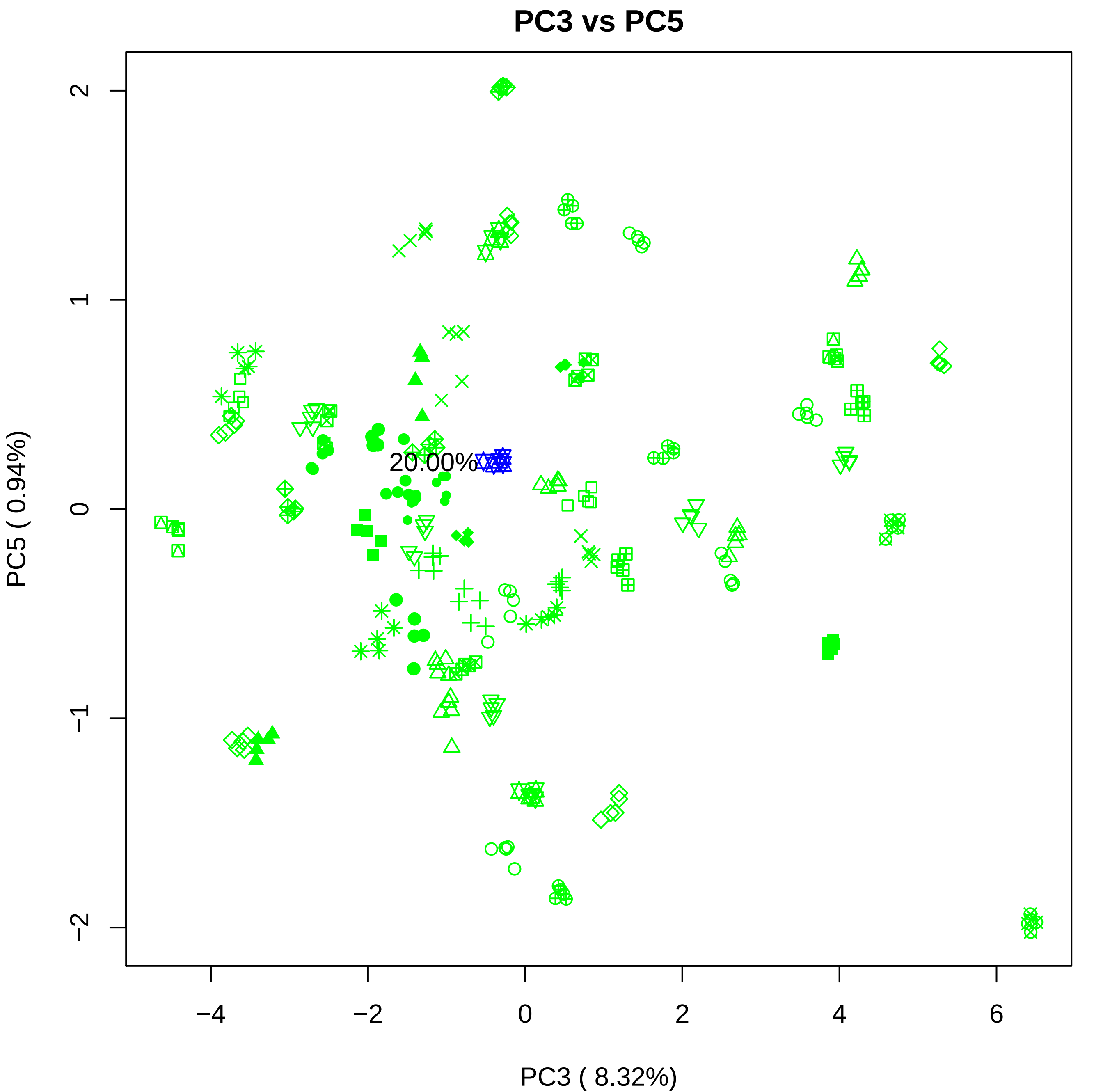

PC3 vs PC5
PC5 ( 0.94%)
PC3 ( 8.32%)
20.00%
−4
−2
0
2
4
6
−2
−1
0
1
2

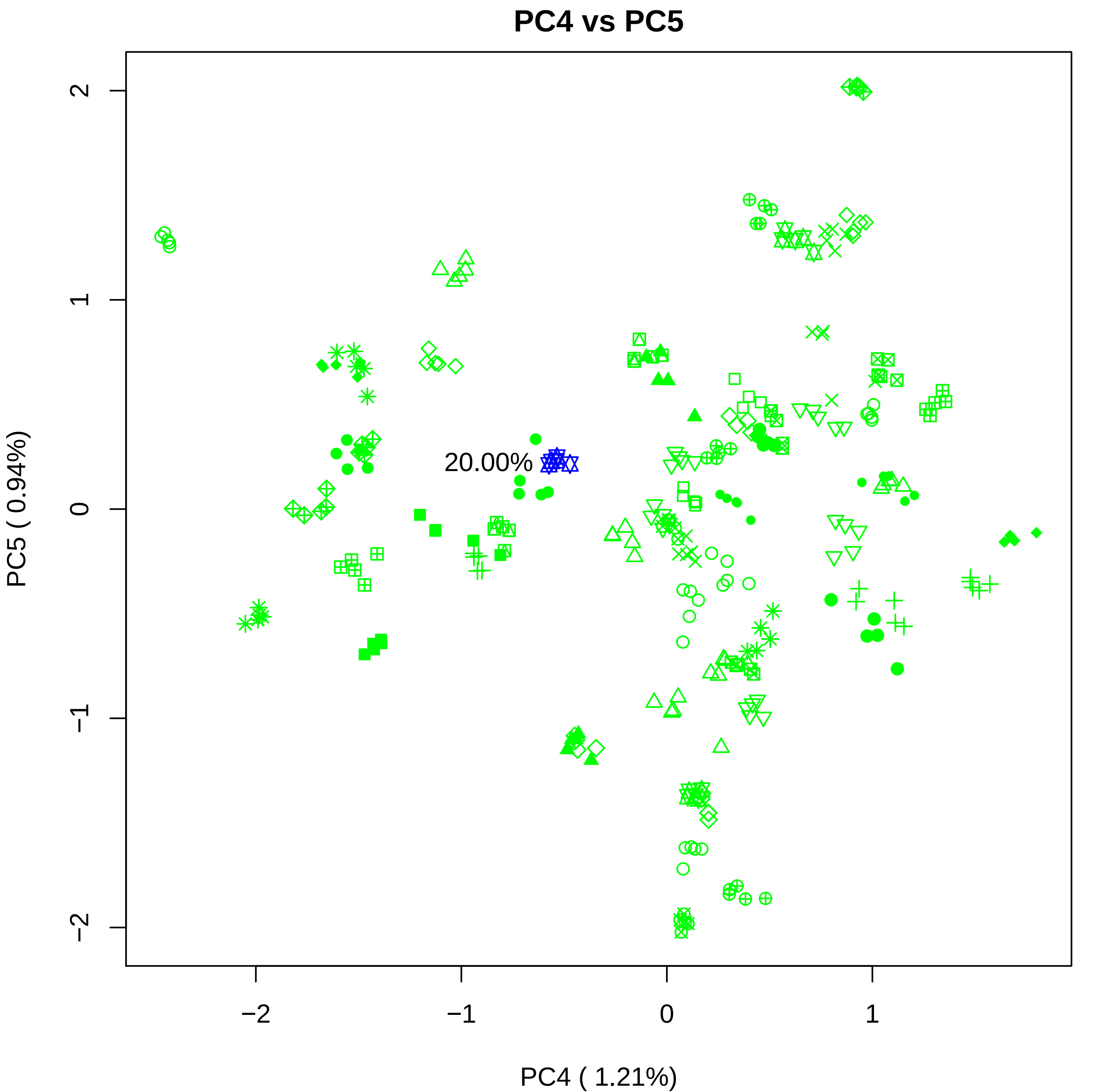
PC4 vs PC5
20.00%
PC5 ( 0.94%)
PC4 ( 1.21%)
−2
−1
0
1
2

### PCA of 40%  BMH(c)  vs PMH(c)

Scaled Matrix PCAed

7th order polynomial baseline correction performed

Raw spectra smoothed to 21 Neighbours, 2nd order PN, Savtisky Golay

Normalisation between 0 and 100

D2 spectra smoothed to 41 Neighbours, 2nd order PN, Savtisky Golay

Regions Analysed: 3601−2501 and 2001−700cm−1

Samples Analysed: 1 bContam's 0 BLH's 0 BMH's 0 OMH's 0 pContam's 0 PLH's 60 PMH's

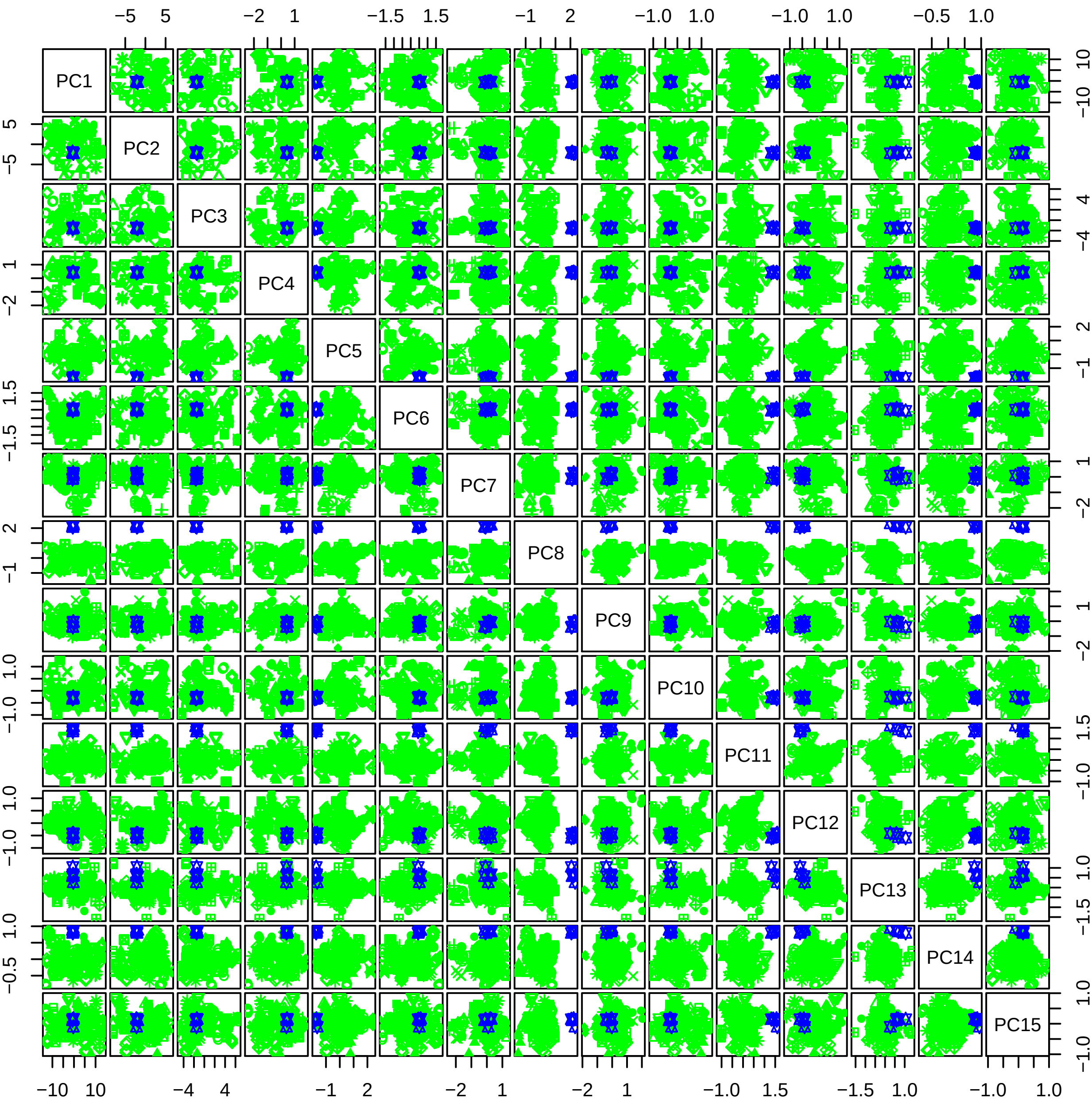
PC1
PC2
PC3
PC4
PC5
PC6
PC7
PC8
PC9
PC10
PC11
PC12
PC13
PC14
PC15

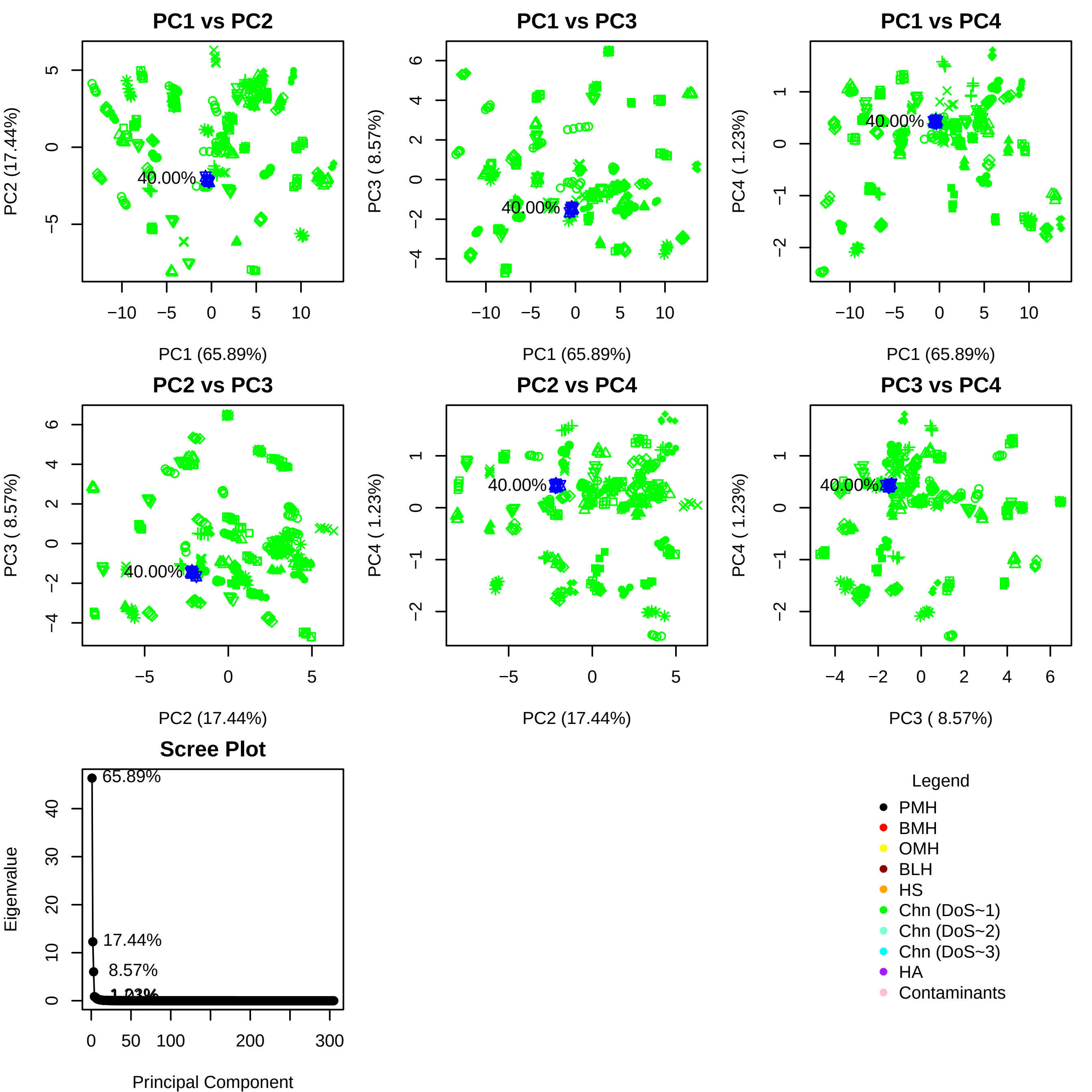
PC1 vs PC2
PC1 vs PC3
PC1 vs PC4
PC2 vs PC3
PC2 vs PC4
PC3 vs PC4
Scree Plot
PC1 (65.89%)
PC2 (17.44%)
PC3 ( 8.57%)
PC4 ( 1.23%)
40.00%
Eigenvalue
Principal Component
65.89%
17.44%
8.57%
Legend
PMH
BMH
OMH
BLH
HS
Chn (DoS~1)
Chn (DoS~2)
Chn (DoS~3)
HA
Contaminants

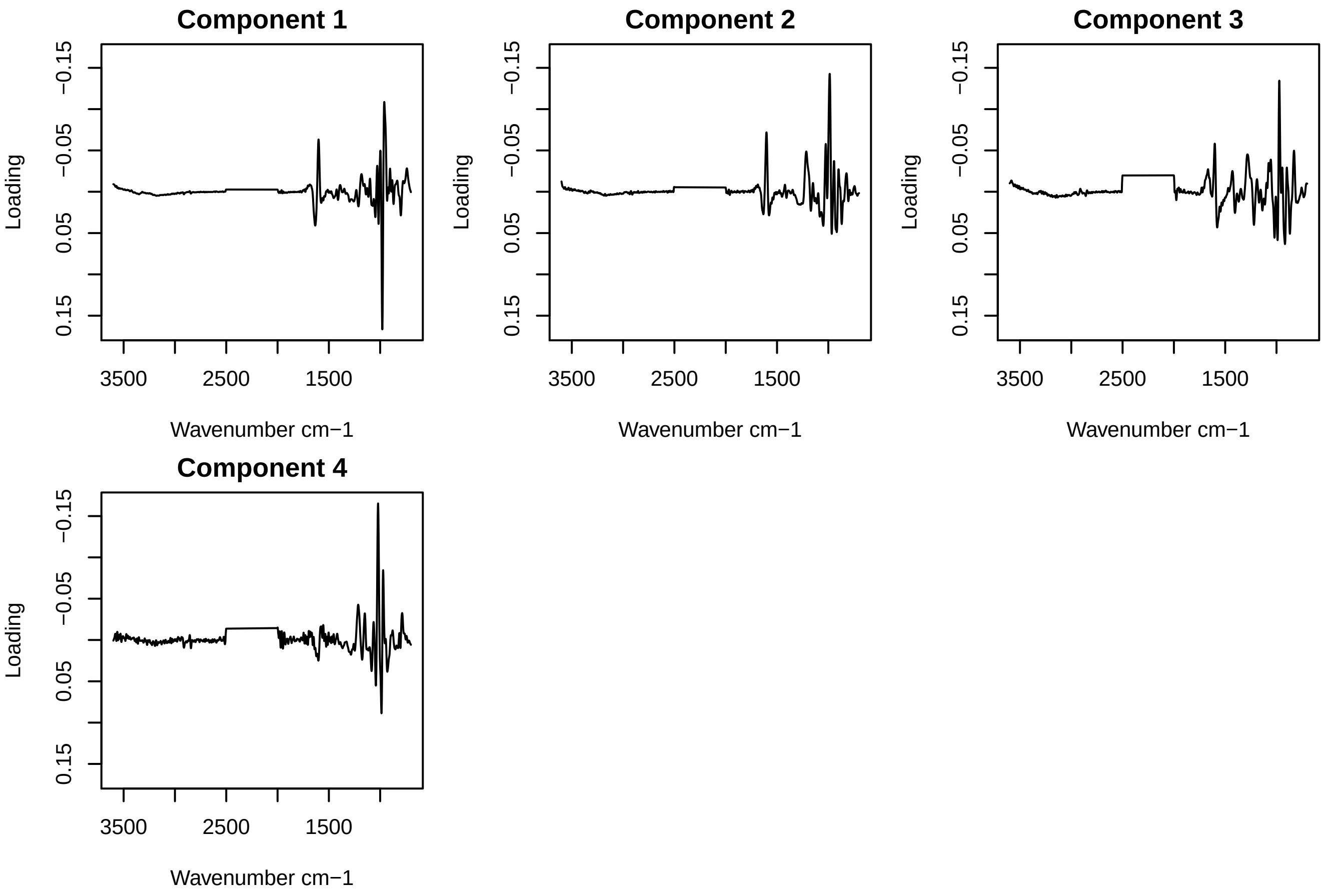
Component 1
Component 2
Component 3
Component 4
Loading
−0.15
−0.05
0.05
0.15
3500
2500
1500
Wavenumber cm−1

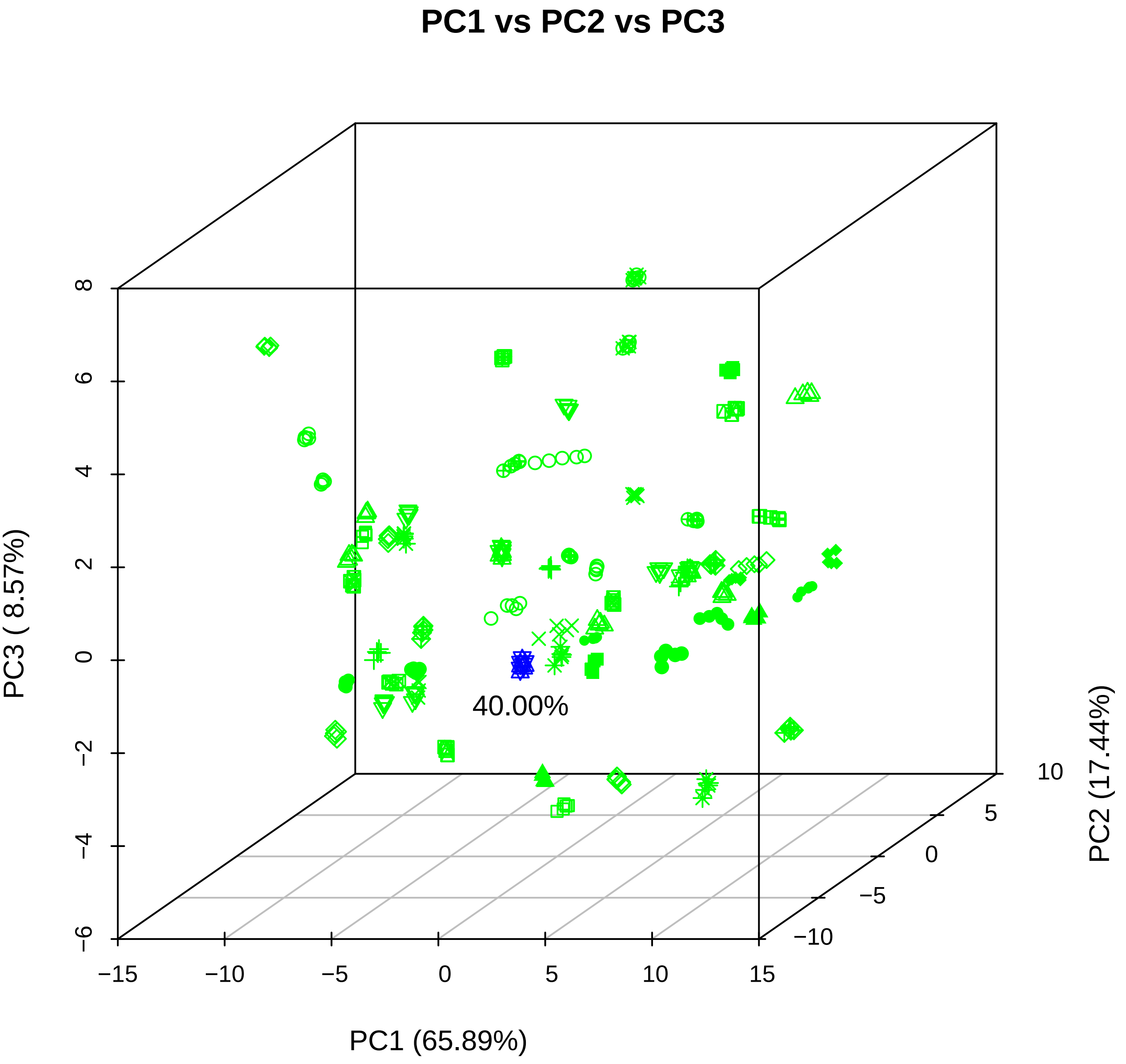

PC1 vs PC2 vs PC3
PC3 ( 8.57%)
8
6
4
2
0
−2
−4
−6
40.00%
−15
−10
−5
0
5
10
15
PC1 (65.89%)
10
5
0
−5
−10
PC2 (17.44%)

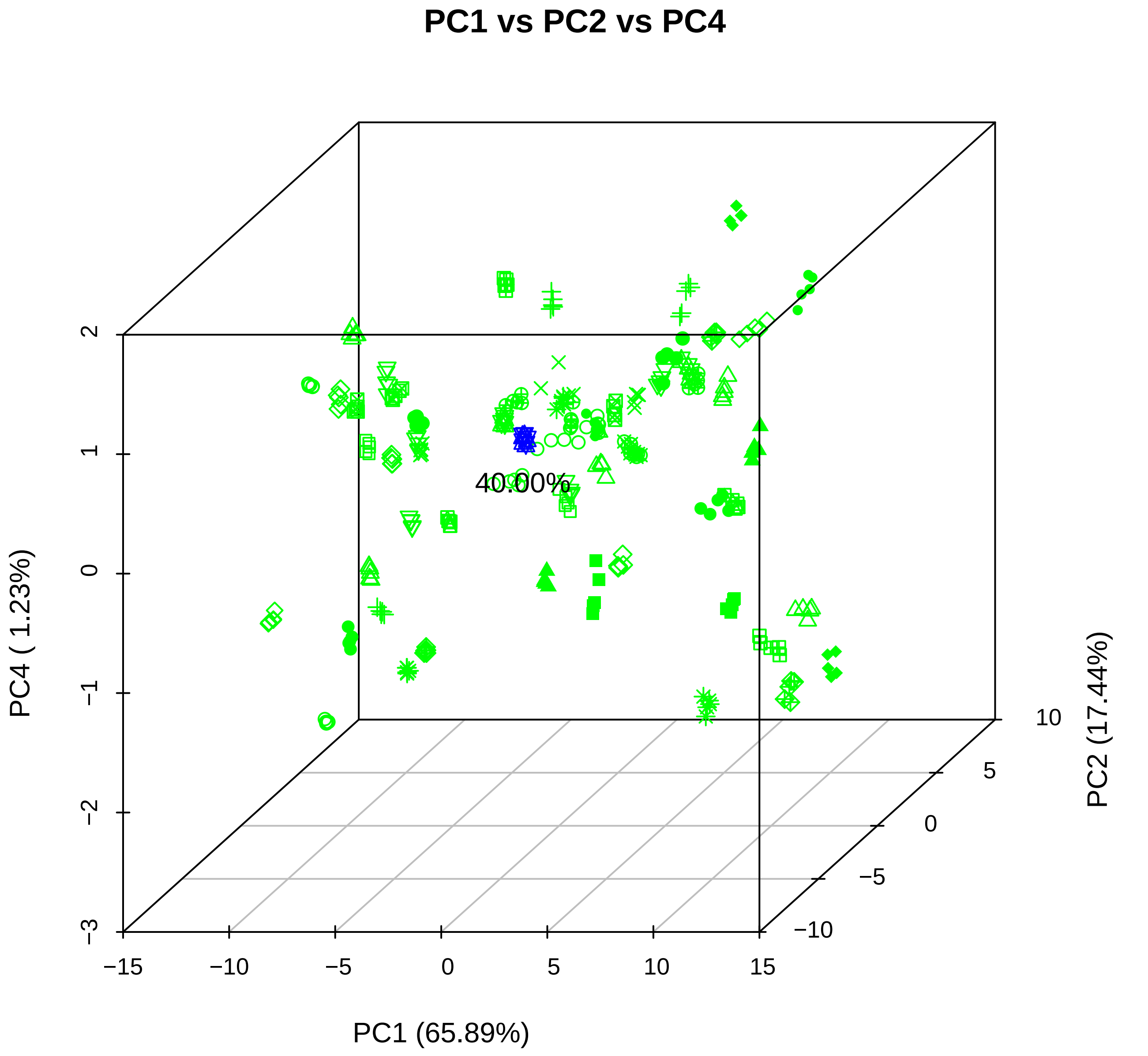
PC1 vs PC2 vs PC4
40.00%
PC1 (65.89%)
PC2 (17.44%)
PC4 ( 1.23%)
−15
−10
−5
0
5
10
15
−10
−5
0
5
10
−3
−2
−1
0
1
2

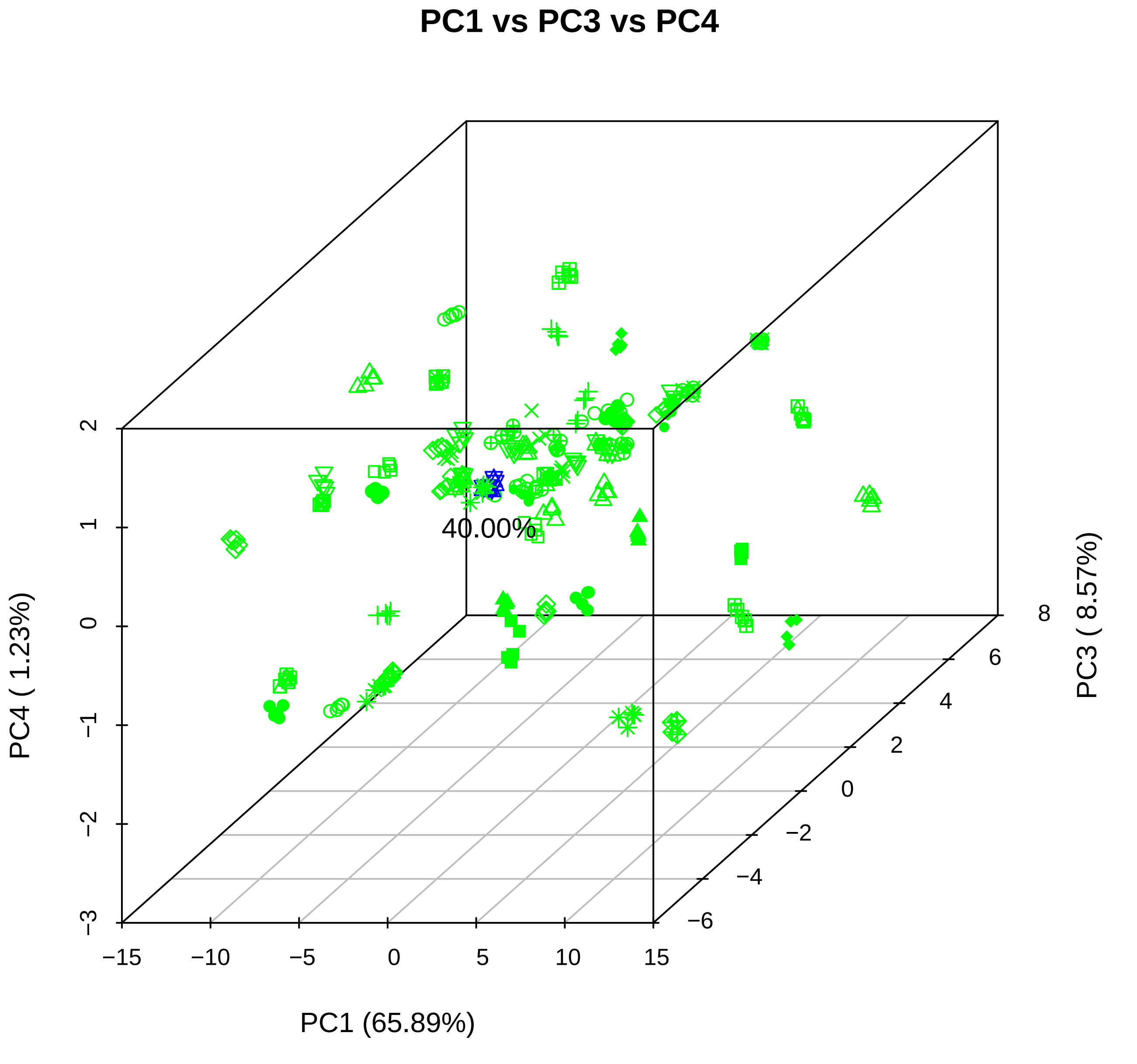
PC1 vs PC3 vs PC4
40.00%
PC1 (65.89%)
PC4 ( 1.23%)
PC3 ( 8.57%)
−15
−10
−5
0
5
10
15
−3
−2
−1
0
1
2
−6
−4
−2
0
2
4
6
8

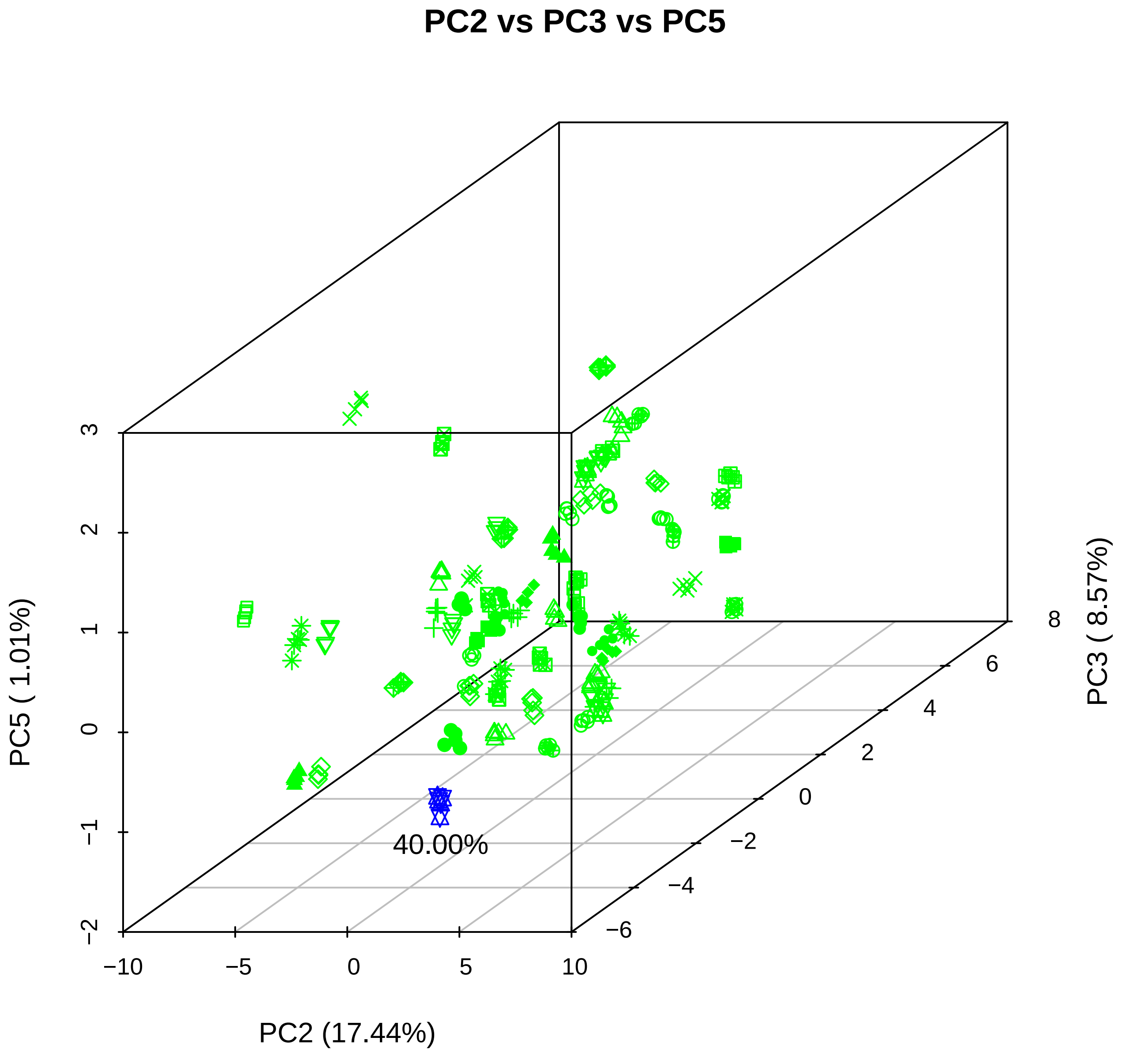
PC2 vs PC3 vs PC5
PC5 ( 1.01%)
PC3 ( 8.57%)
PC2 (17.44%)
40.00%
3
2
1
0
−1
−2
−10
−5
0
5
10
8
6
4
2
0
−2
−4
−6

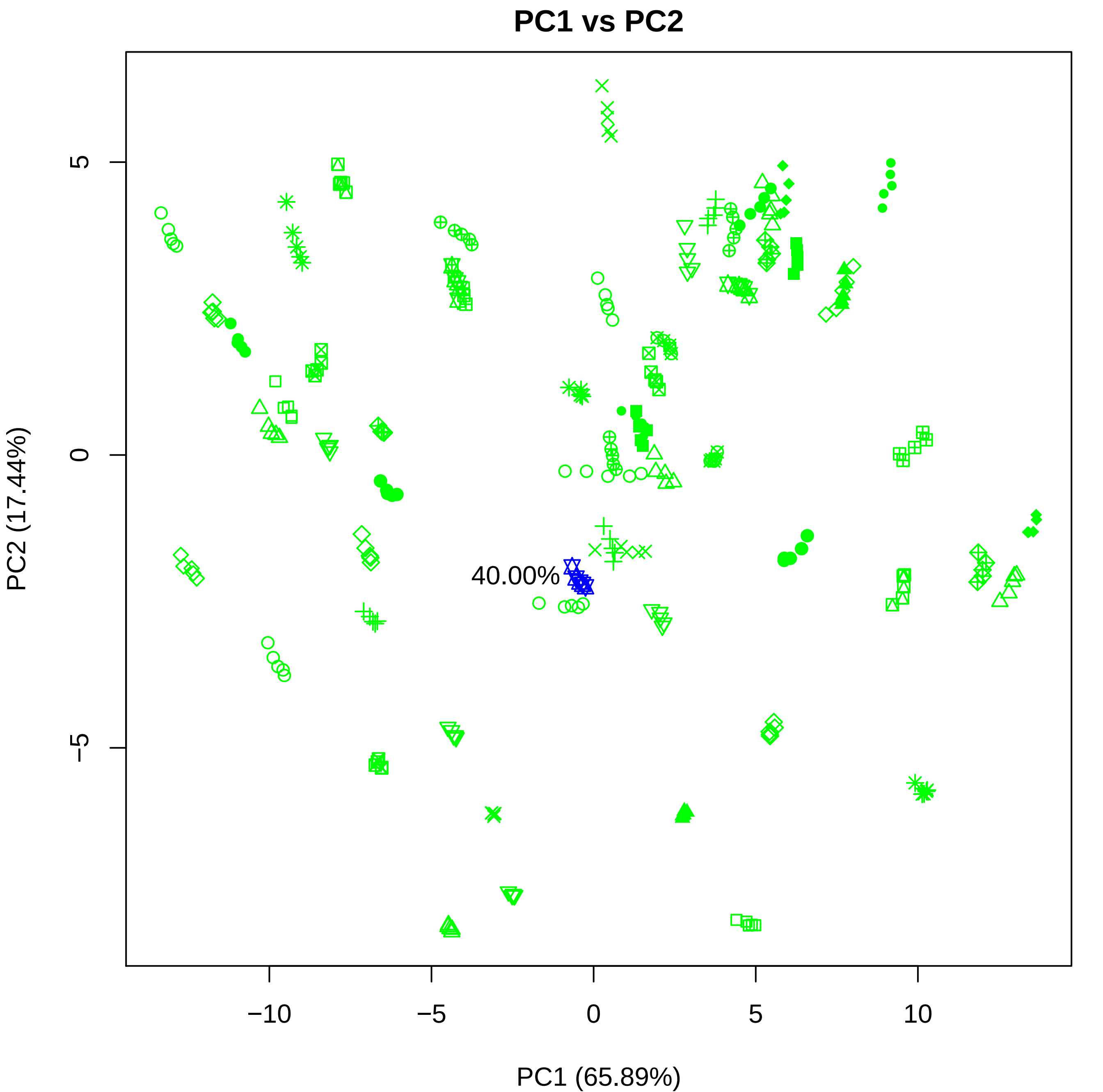

PC1 vs PC2
40.00%
5
0
−5
−10
−5
0
5
10
PC1 (65.89%)
PC2 (17.44%)

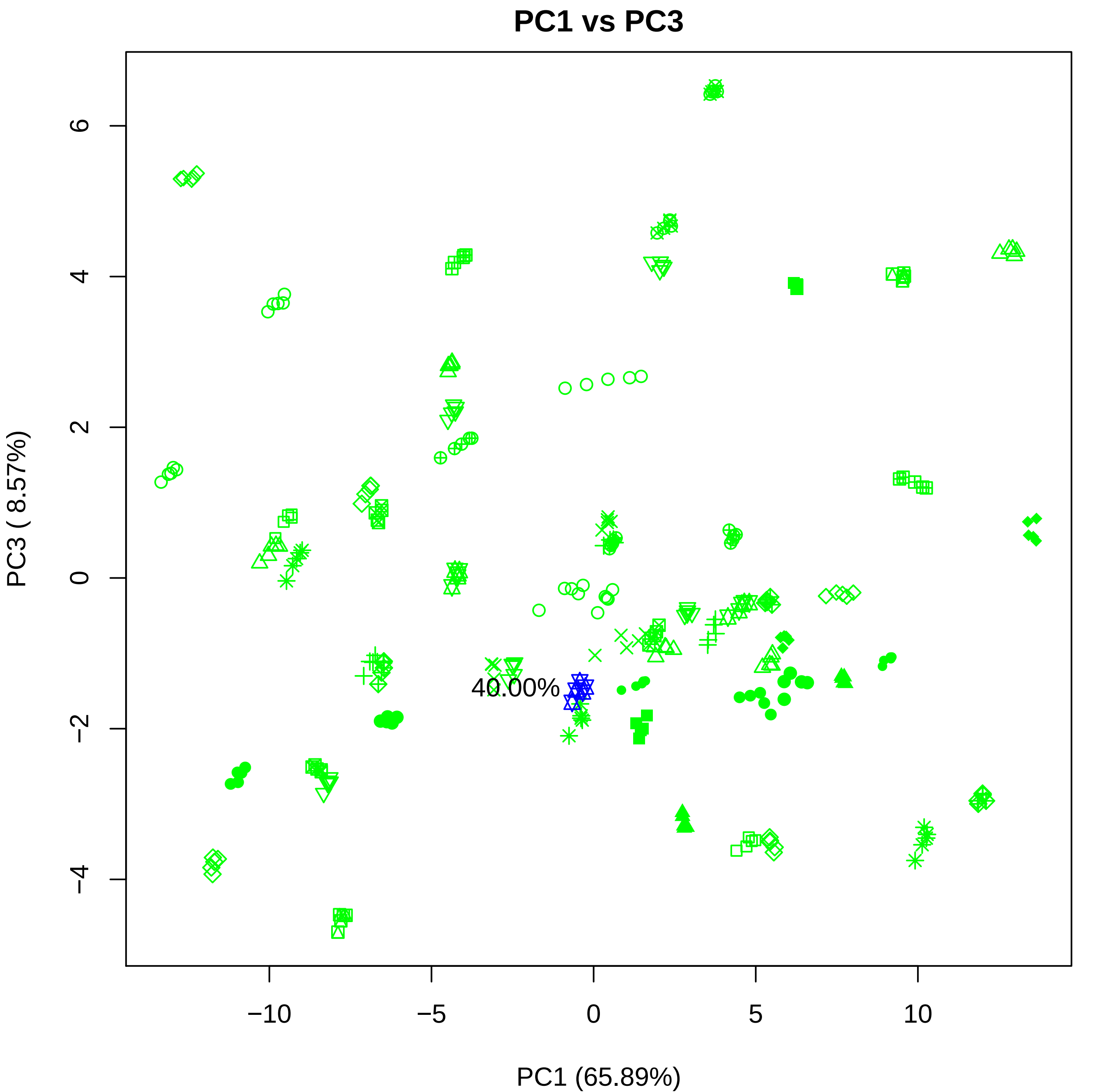
PC1 vs PC3
PC3 ( 8.57%)
PC1 (65.89%)
40.00%
−4
−2
0
2
4
6
−10
−5
0
5
10

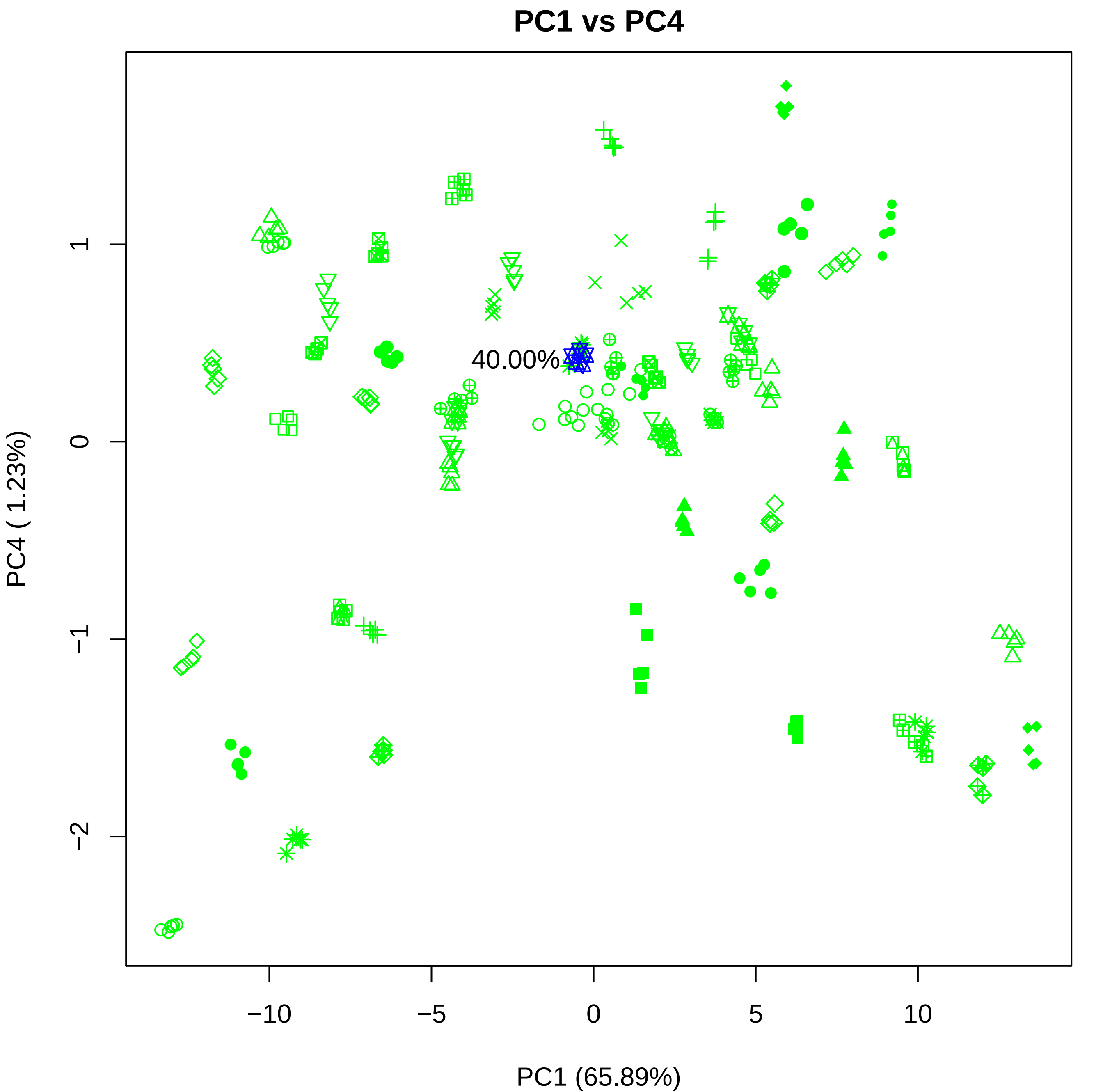
PC1 vs PC4
40.00%
PC4 ( 1.23%)
PC1 (65.89%)
−10
−5
0
5
10
1
0
−1
−2

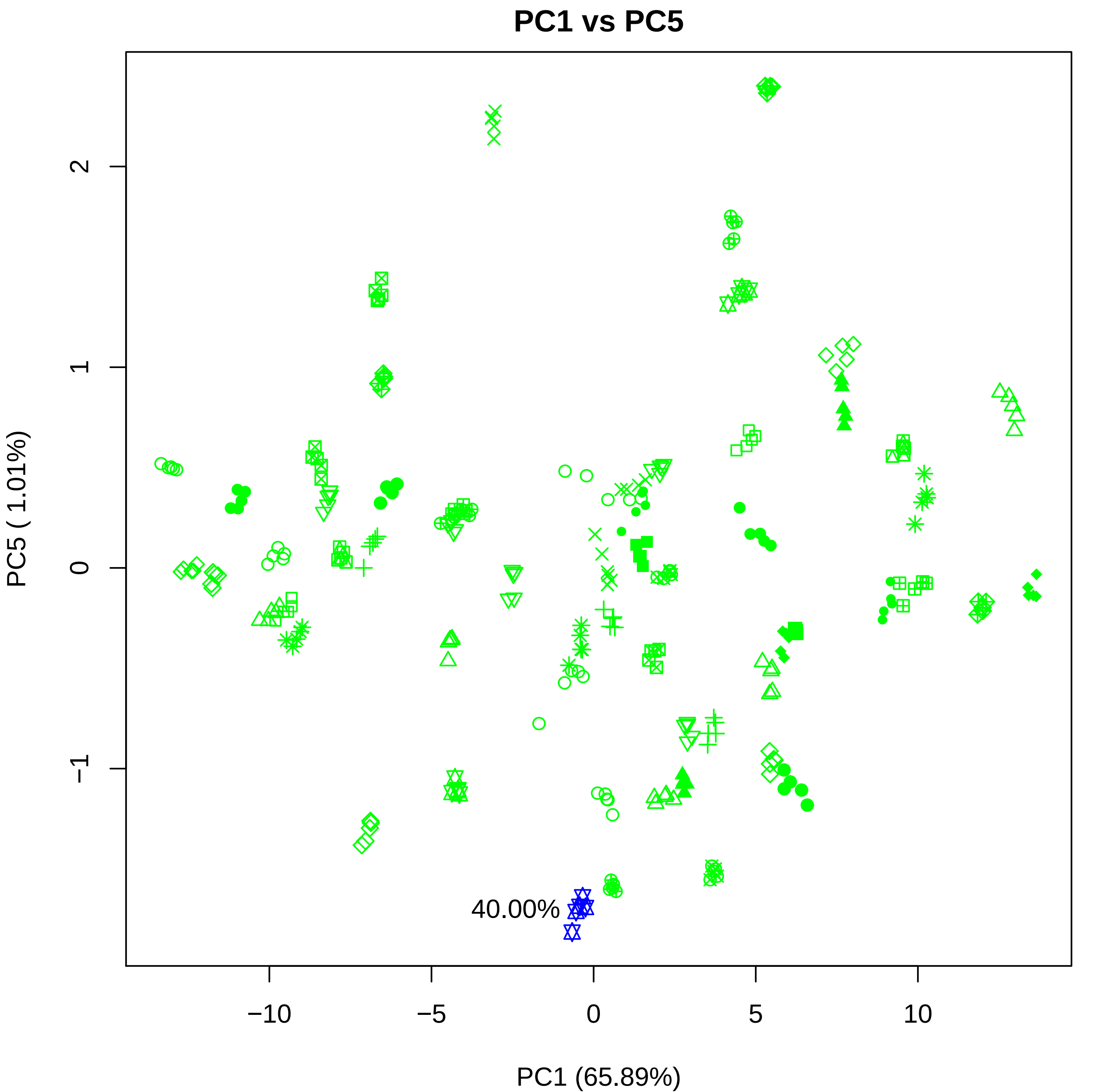
PC1 vs PC5
PC5 ( 1.01%)
PC1 (65.89%)
2
1
0
−1
−10
−5
0
5
10
40.00%

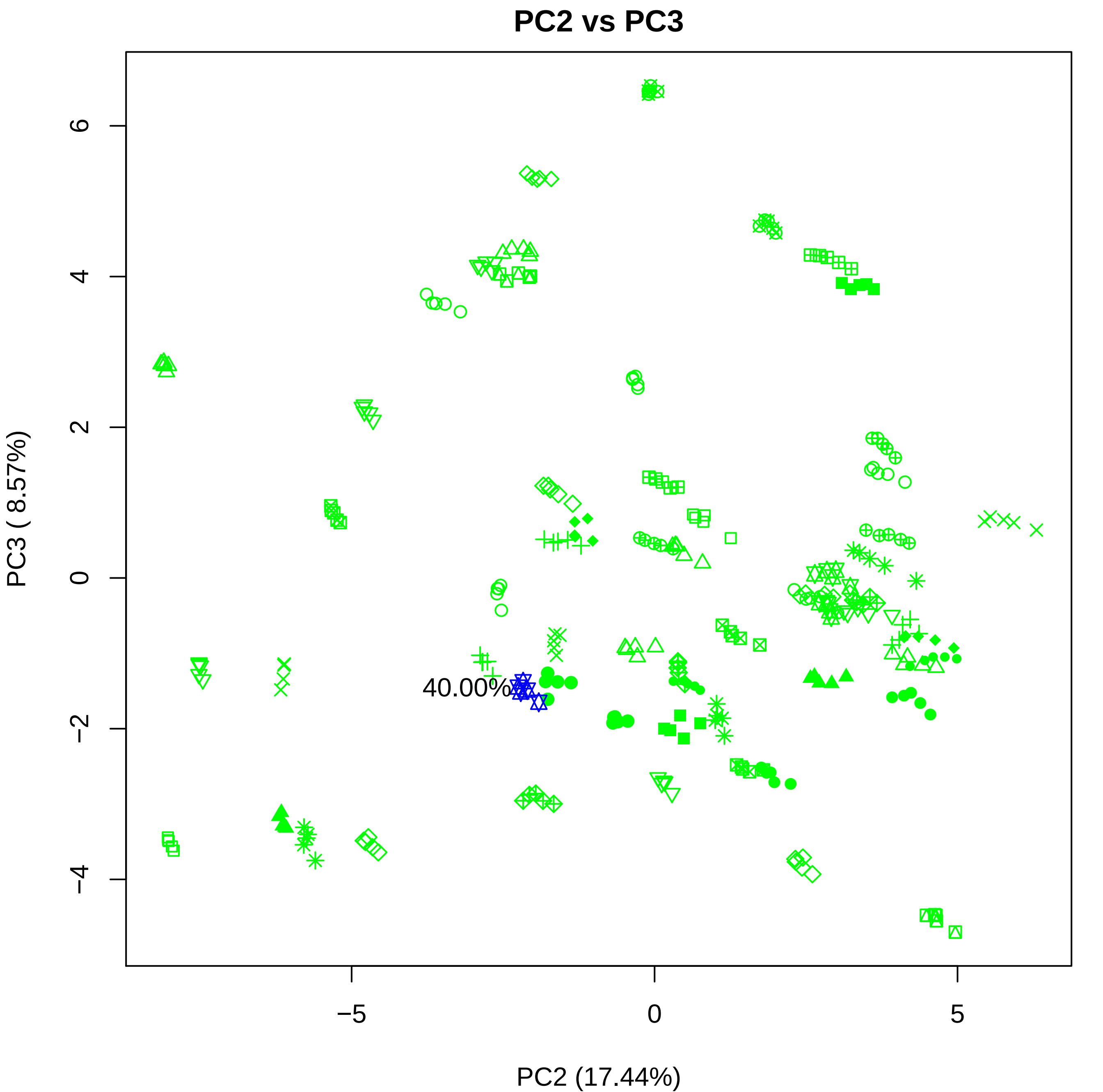

PC2 vs PC3
PC3 ( 8.57%)
PC2 (17.44%)
6
4
2
0
−2
−4
−5
0
5
40.00%

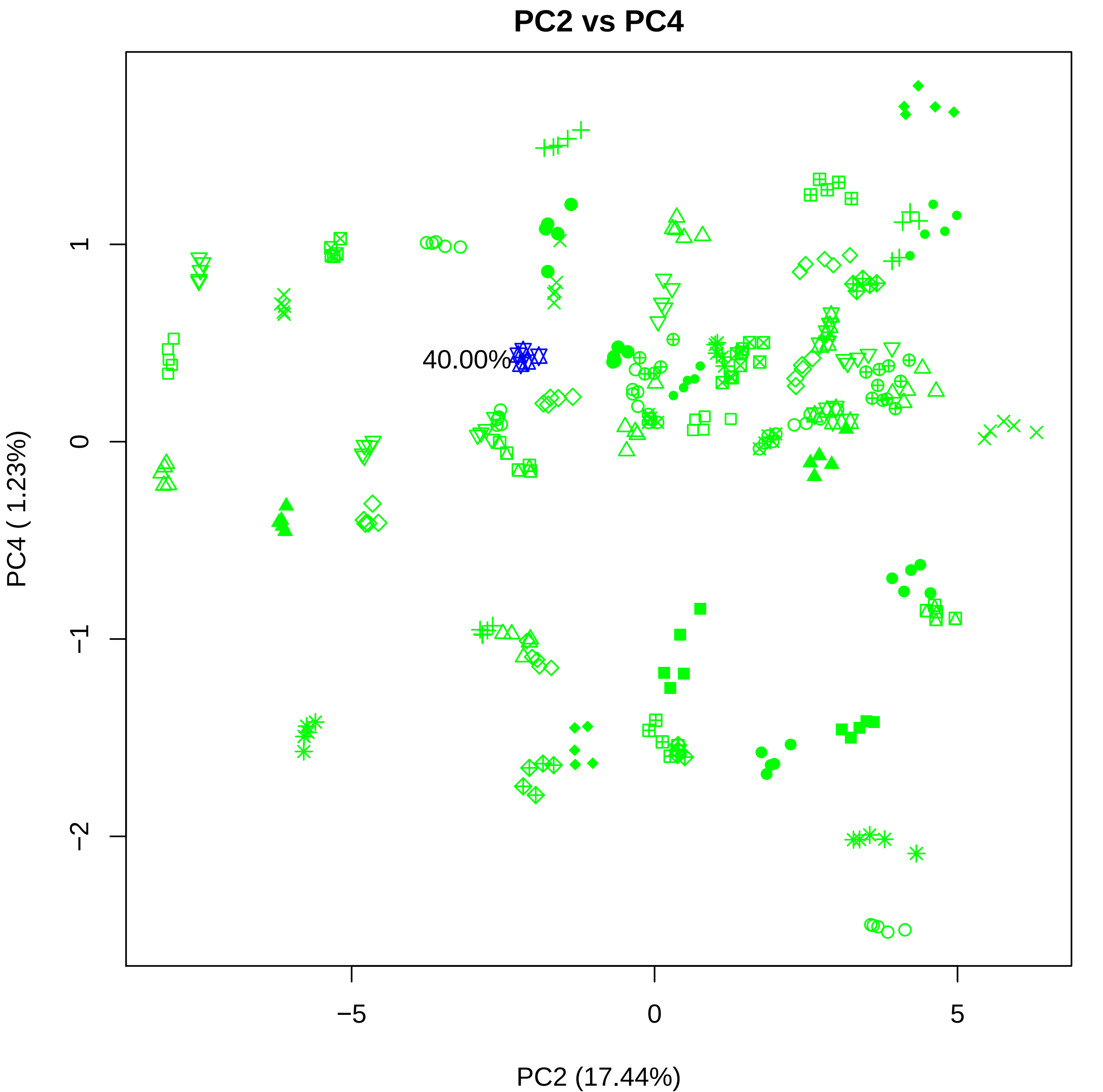

PC2 vs PC4
40.00%
PC4 ( 1.23%)
PC2 (17.44%)
1
0
−1
−2
−5
0
5

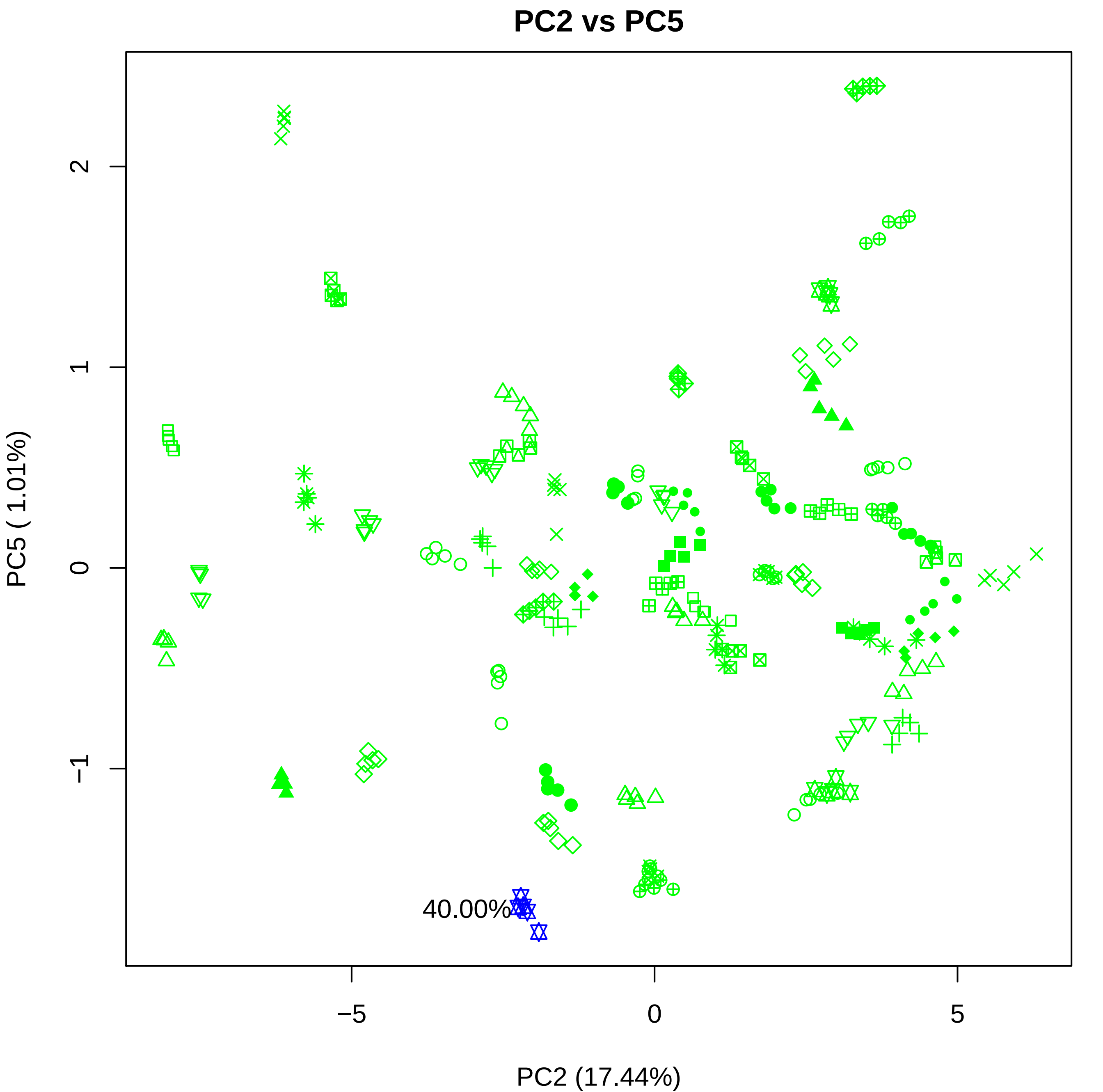
PC2 vs PC5
PC5 ( 1.01%)
PC2 (17.44%)
−5
0
5
−1
0
1
2
40.00%

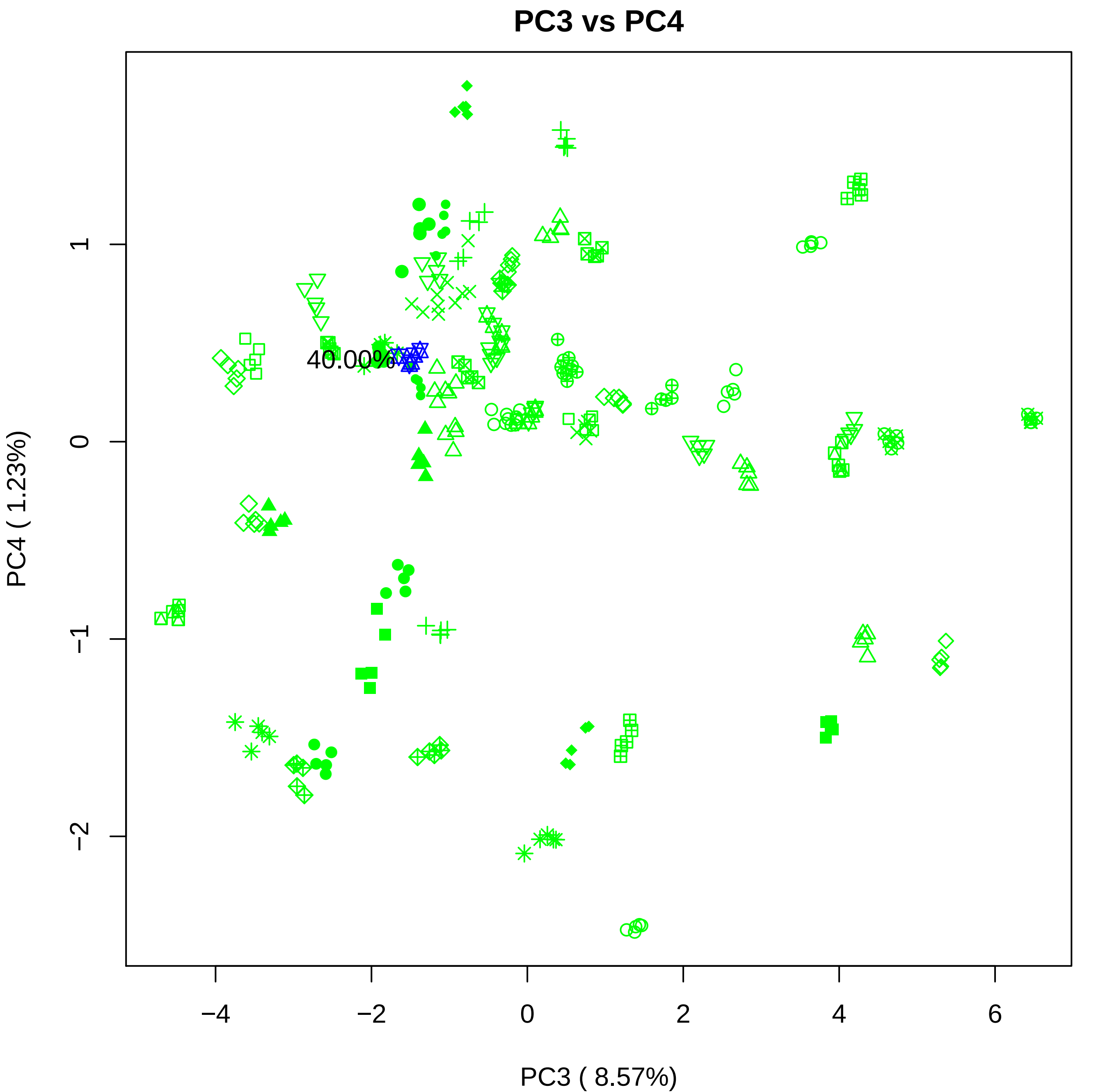

PC3 vs PC4
40.00%
1
0
−1
−2
−4
−2
0
2
4
6
PC3 ( 8.57%)
PC4 ( 1.23%)

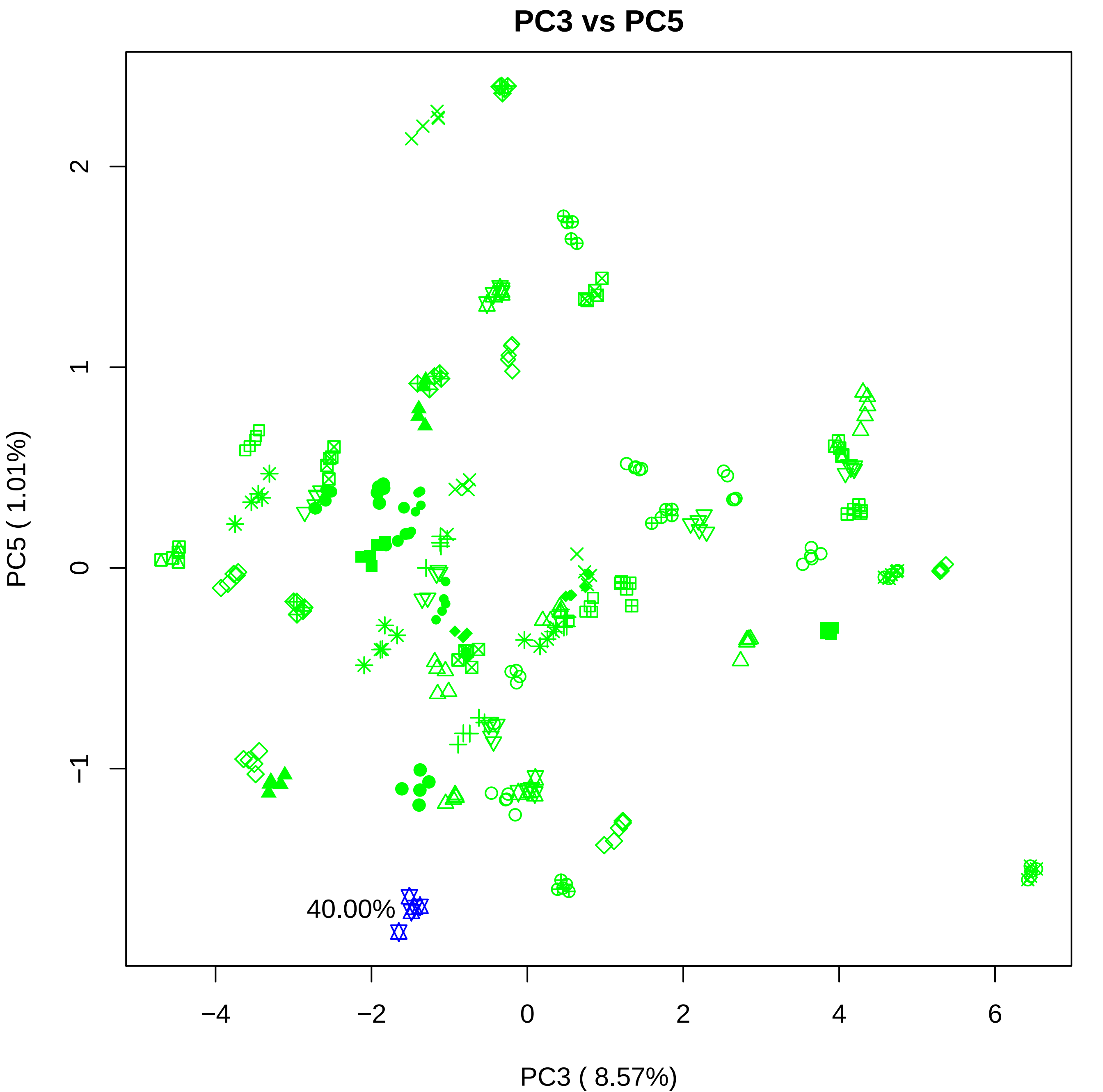
PC3 vs PC5
PC5 ( 1.01%)
PC3 ( 8.57%)
40.00%
2
1
0
−1
−4
−2
0
2
4
6

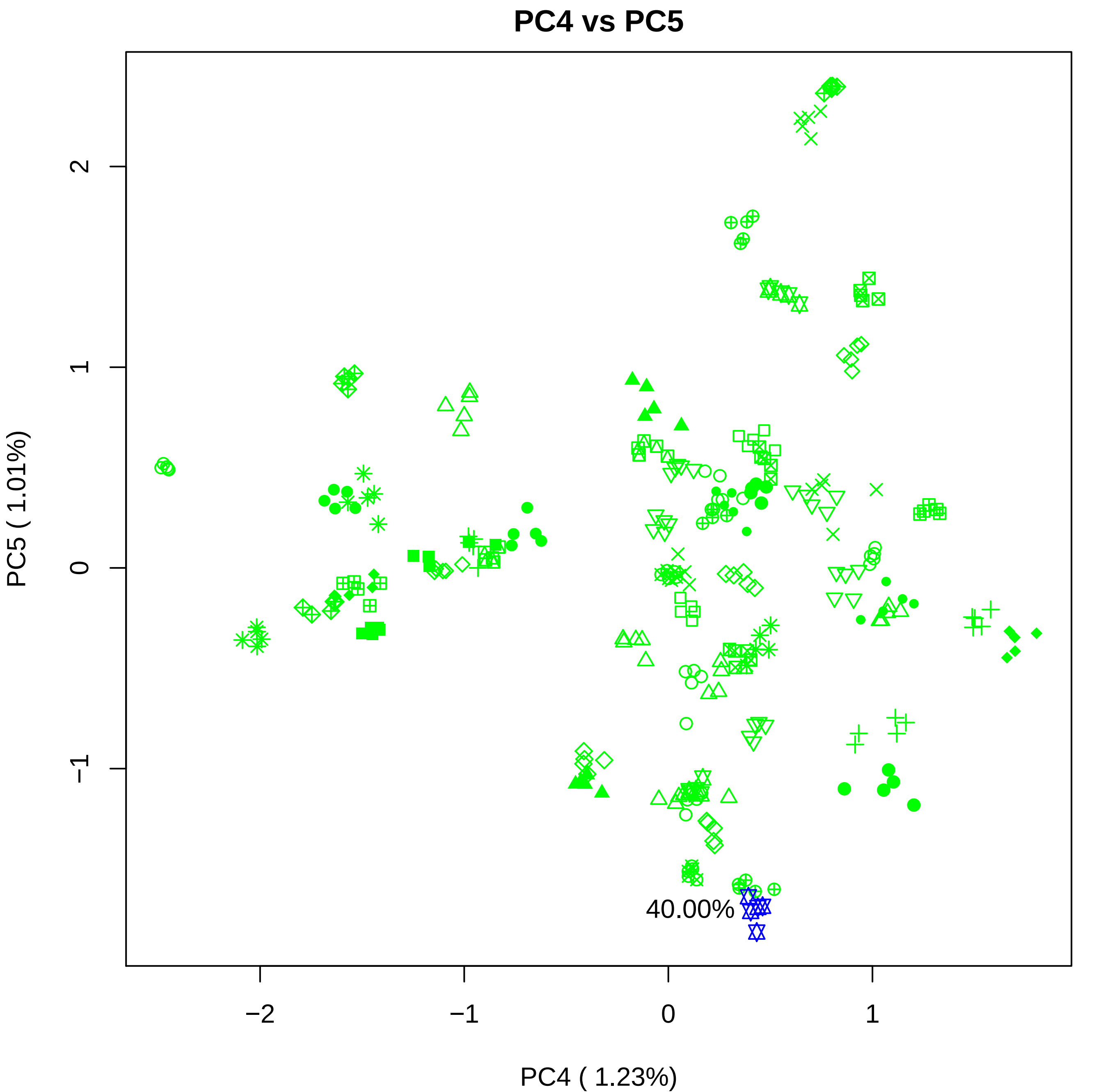
PC4 vs PC5
PC5 ( 1.01%)
PC4 ( 1.23%)
2
1
0
−1
−2
−1
0
1
40.00%

### PCA of All BMH(d) vs PMH(d)

Scaled Matrix PCAed

7th order polynomial baseline correction performed

Raw spectra smoothed to 21 Neighbours, 2nd order PN, Savtisky Golay

Normalisation between 0 and 100

D2 spectra smoothed to 41 Neighbours, 2nd order PN, Savtisky Golay

Regions Analysed: 3601−2501 and 2001−700cm−1

Samples Analysed: 8 bContam's 0 BLH's 0 BMH's 0 OMH's 0 pContam's 0 PLH's 60 PMH's

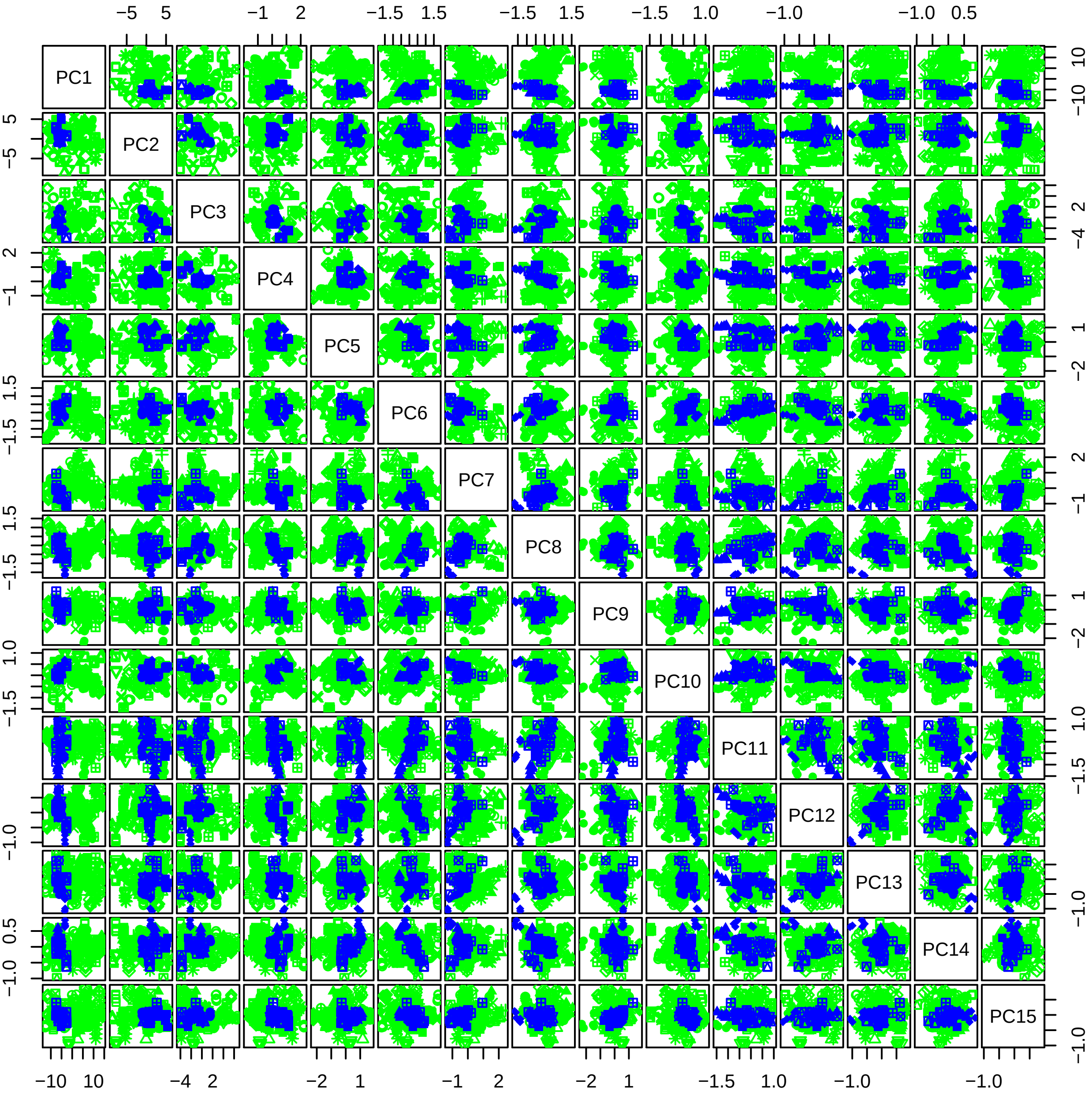

PC1
PC2
PC3
PC4
PC5
PC6
PC7
PC8
PC9
PC10
PC11
PC12
PC13
PC14
PC15

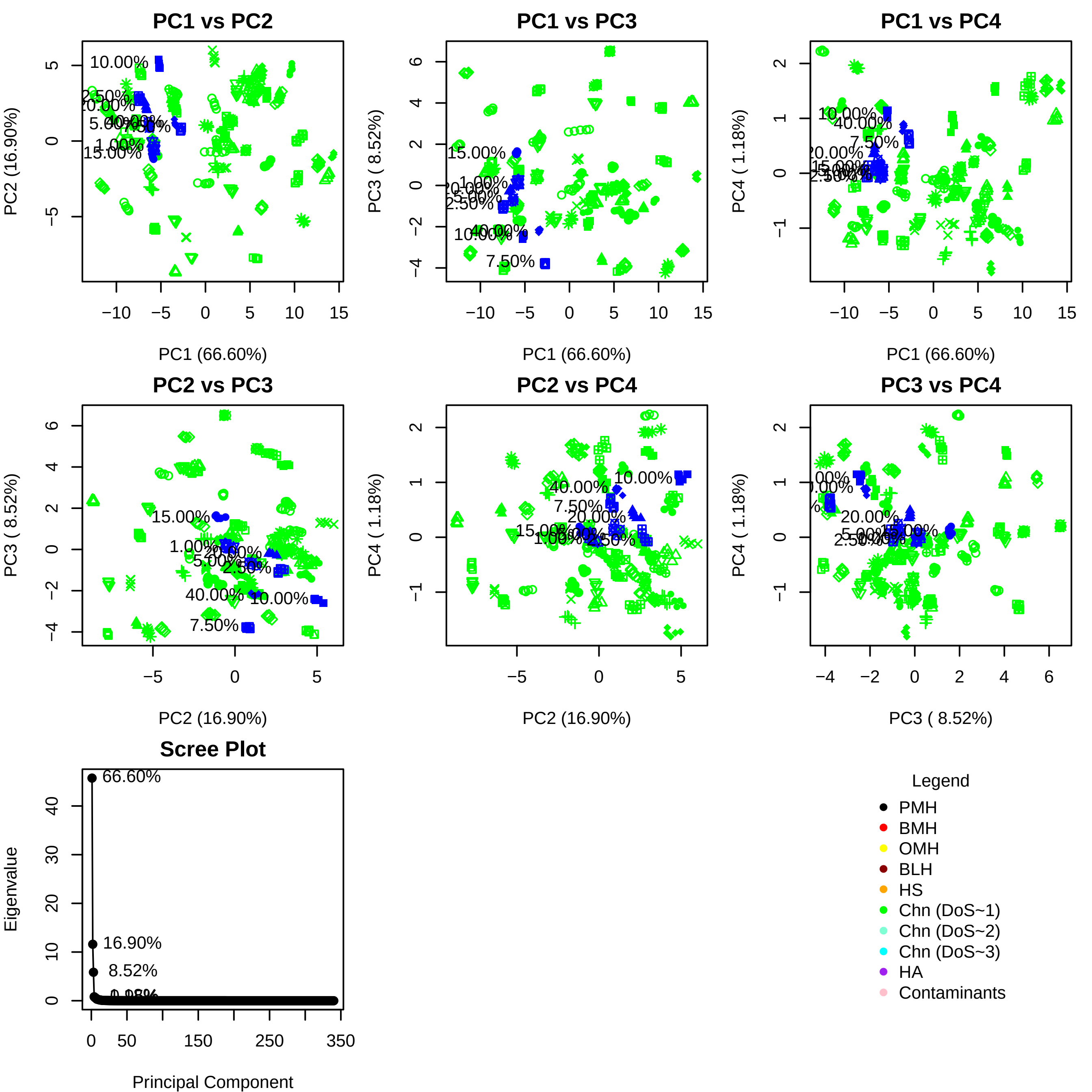

PC1 vs PC2
PC1 vs PC3
PC1 vs PC4
PC2 vs PC3
PC2 vs PC4
PC3 vs PC4
Scree Plot
PC1 (66.60%)
PC2 (16.90%)
PC3 ( 8.52%)
PC4 ( 1.18%)
Eigenvalue
Principal Component
66.60%
16.90%
8.52%
Legend
PMH
BMH
OMH
BLH
HS
Chn (DoS~1)
Chn (DoS~2)
Chn (DoS~3)
HA
Contaminants

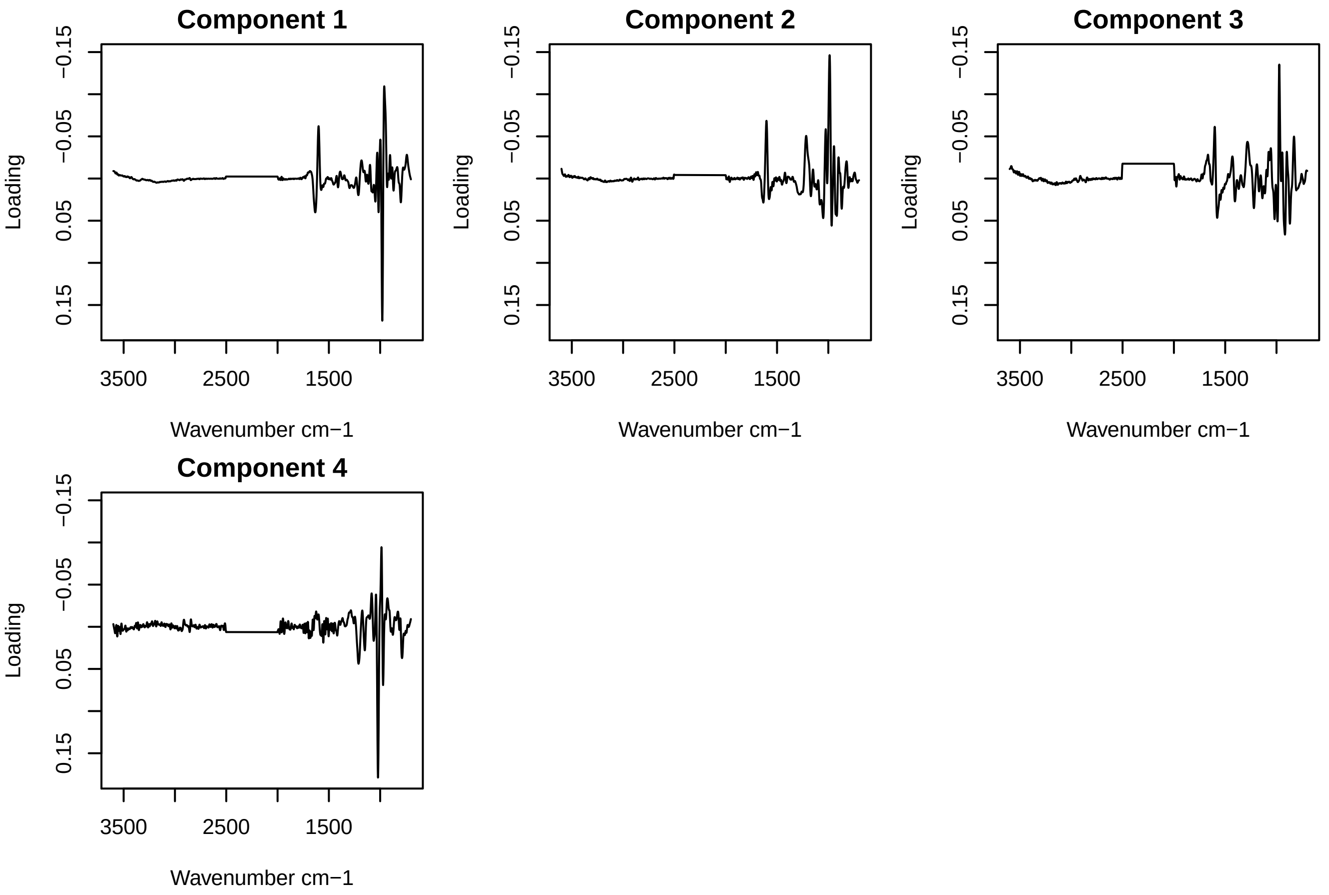
Component 1
Component 2
Component 3
Component 4
Loading
−0.15
−0.05
0.05
0.15
3500
2500
1500
Wavenumber cm−1

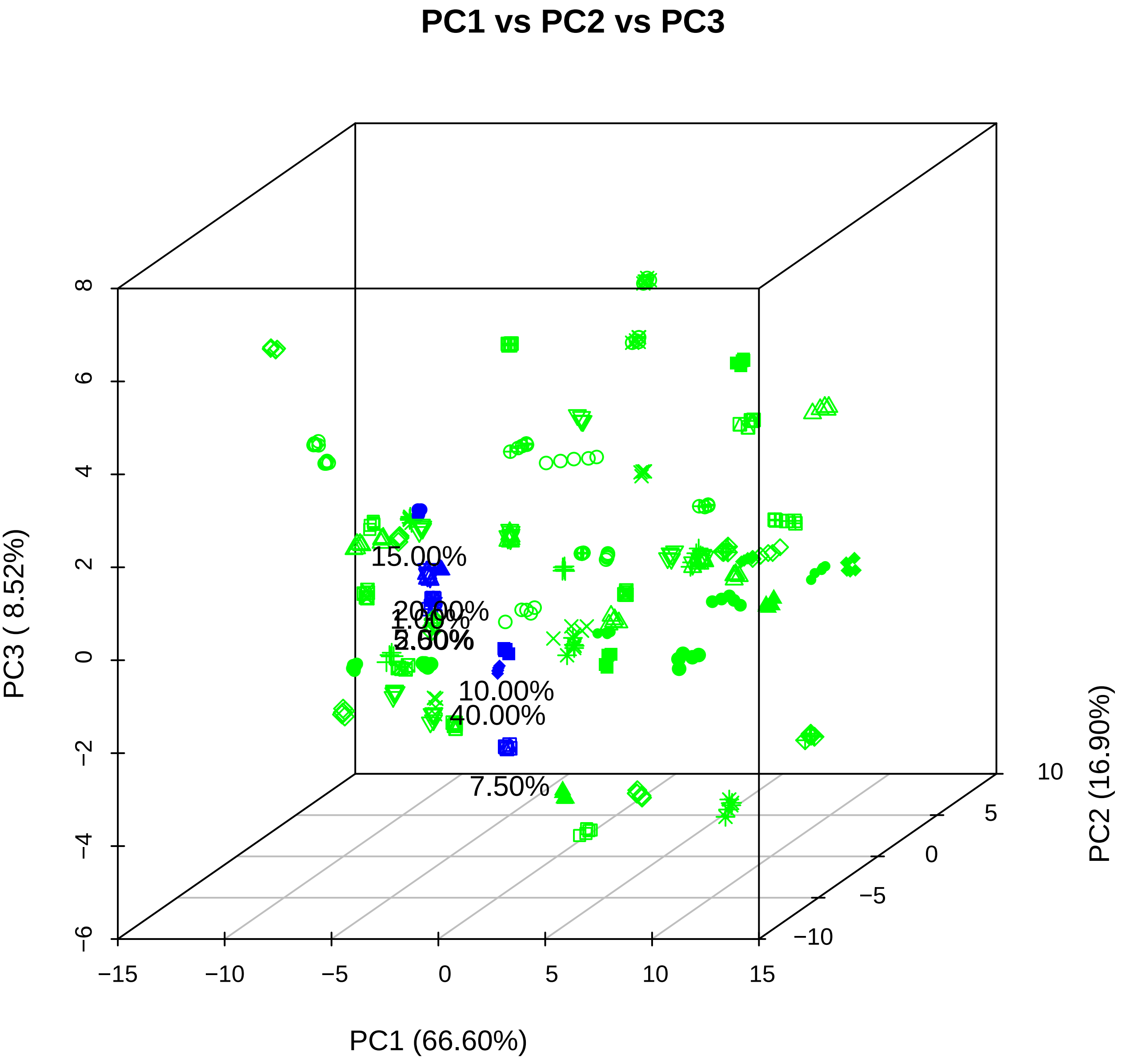

PC1 vs PC2 vs PC3
15.00%
20.00%
1.00%
5.00%
2.50%
10.00%
40.00%
7.50%
8
6
4
2
0
−2
−4
−6
PC3 ( 8.52%)
−15
−10
−5
0
5
10
15
PC1 (66.60%)
10
5
0
−5
−10
PC2 (16.90%)

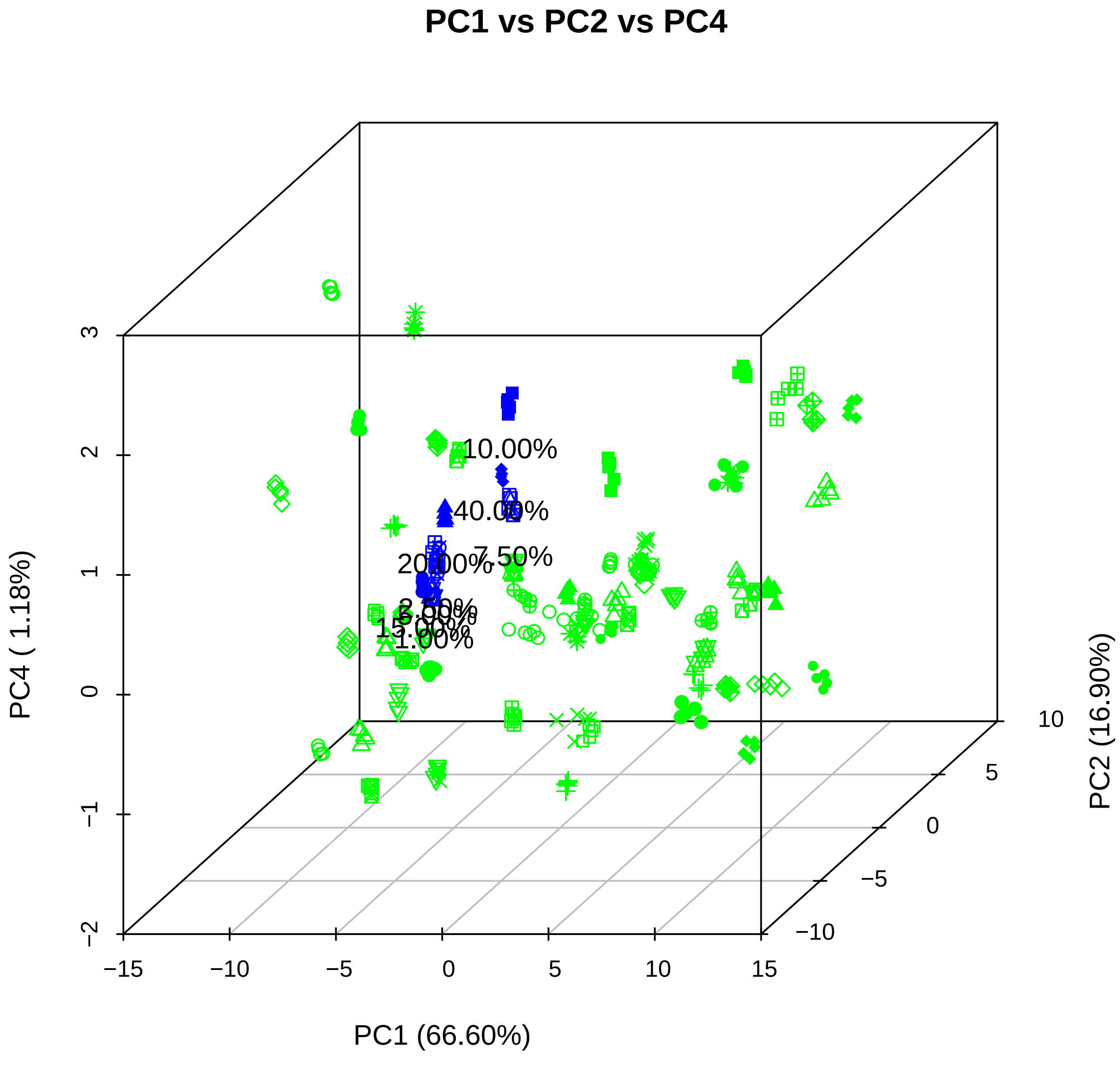
PC1 vs PC2 vs PC4
10.00%
40.00%
7.50%
20.00%
2.50%
5.00%
15.00%
1.00%
PC4 ( 1.18%)
PC1 (66.60%)
PC2 (16.90%)
−15
−10
−5
0
5
10
15
−2
−1
0
1
2
3
−10
−5
0
5
10

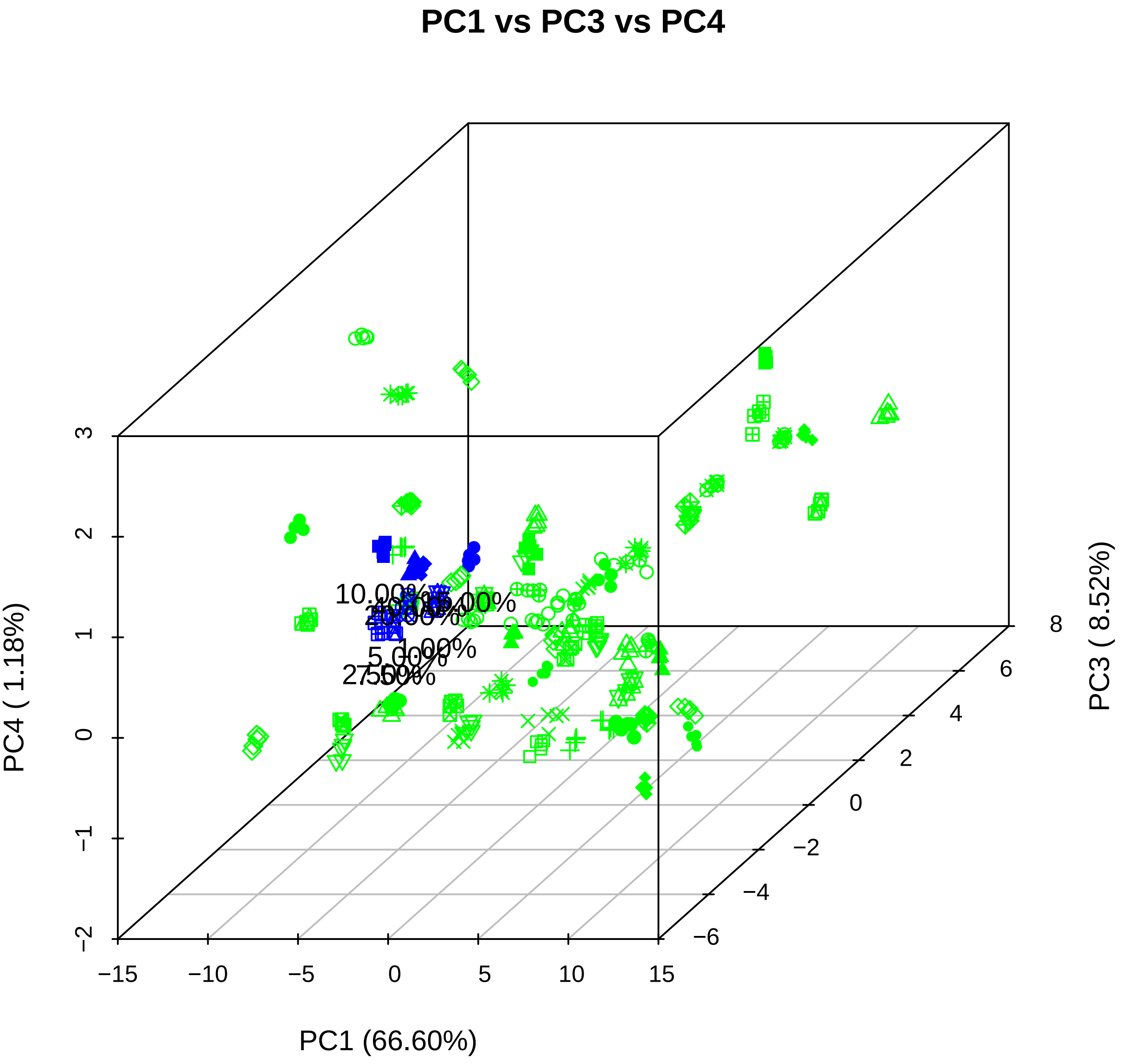

PC1 vs PC3 vs PC4
PC1 (66.60%)
PC3 ( 8.52%)
PC4 ( 1.18%)
−15
−10
−5
0
5
10
15
−6
−4
−2
0
2
4
6
8
−2
−1
0
1
2
3
10.00%
1.00%

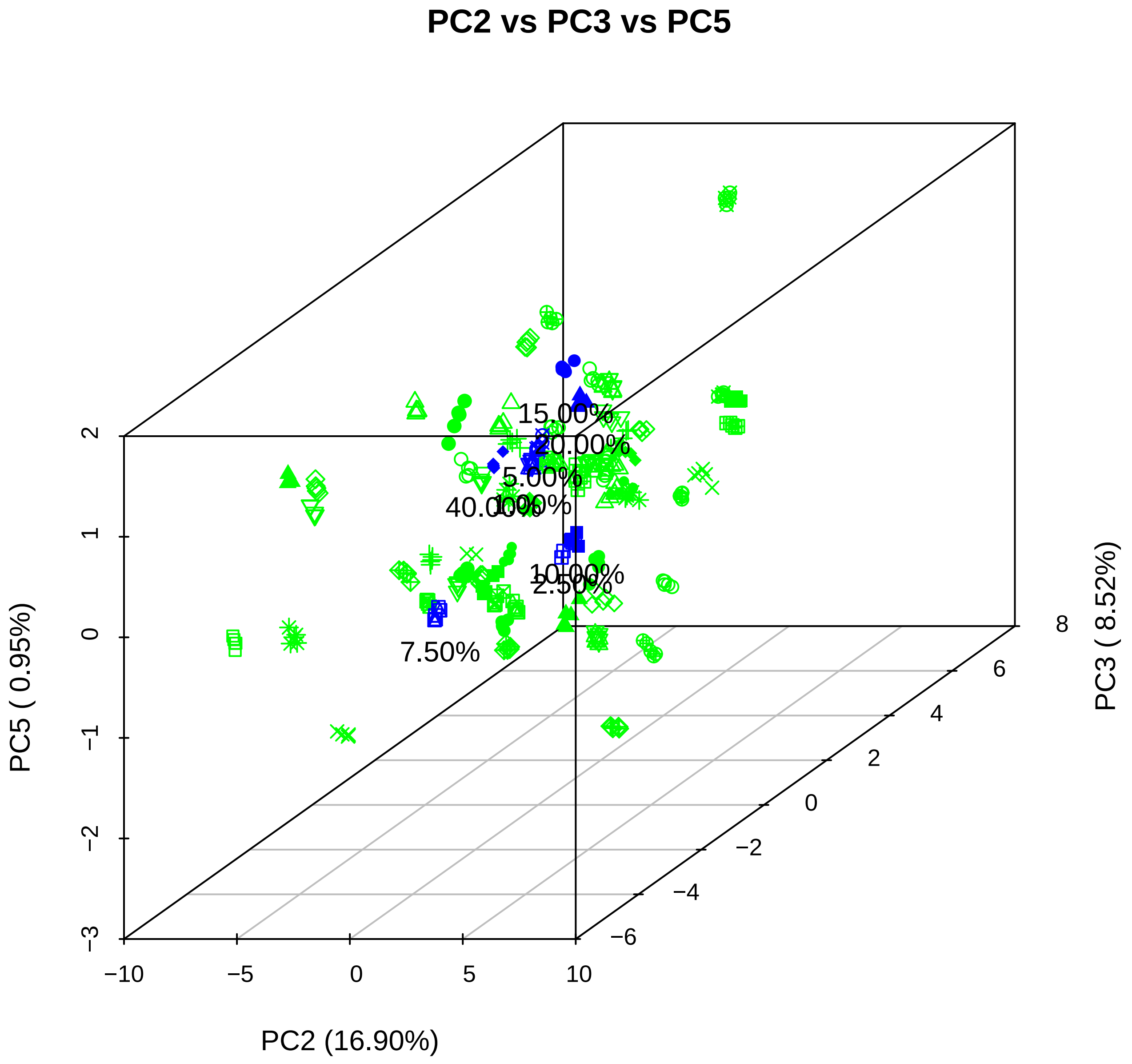
PC2 vs PC3 vs PC5
PC2 (16.90%)
PC3 ( 8.52%)
PC5 ( 0.95%)
−10
−5
0
5
10
−6
−4
−2
0
2
4
6
8
−3
−2
−1
0
1
2
15.00%
20.00%
5.00%
40.00%
10.00%
2.50%
7.50%

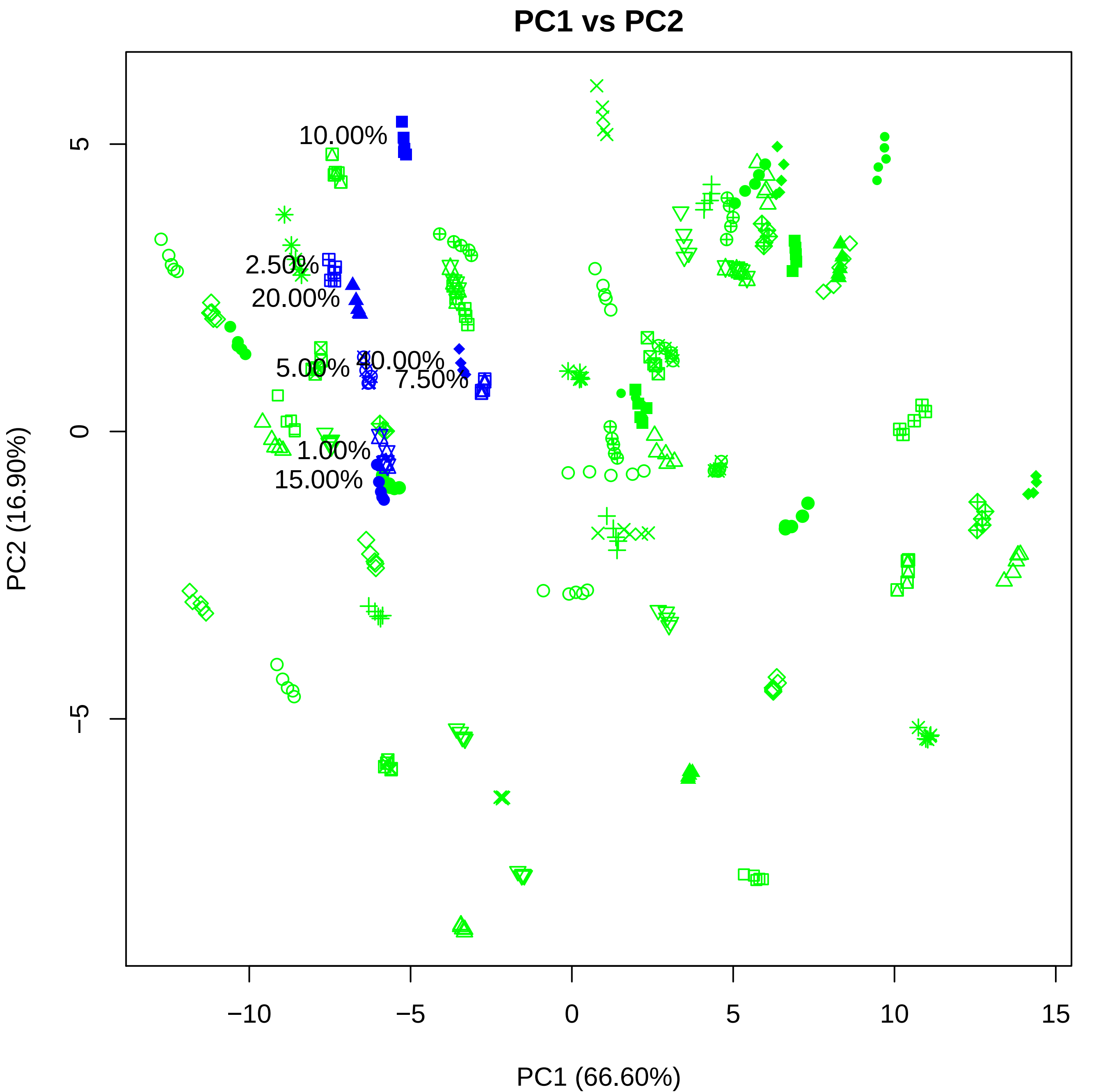

PC1 vs PC2
10.00%
2.50%
20.00%
5.00%
40.00%
7.50%
1.00%
15.00%
5
0
−5
−10
−5
0
5
10
15
PC2 (16.90%)
PC1 (66.60%)

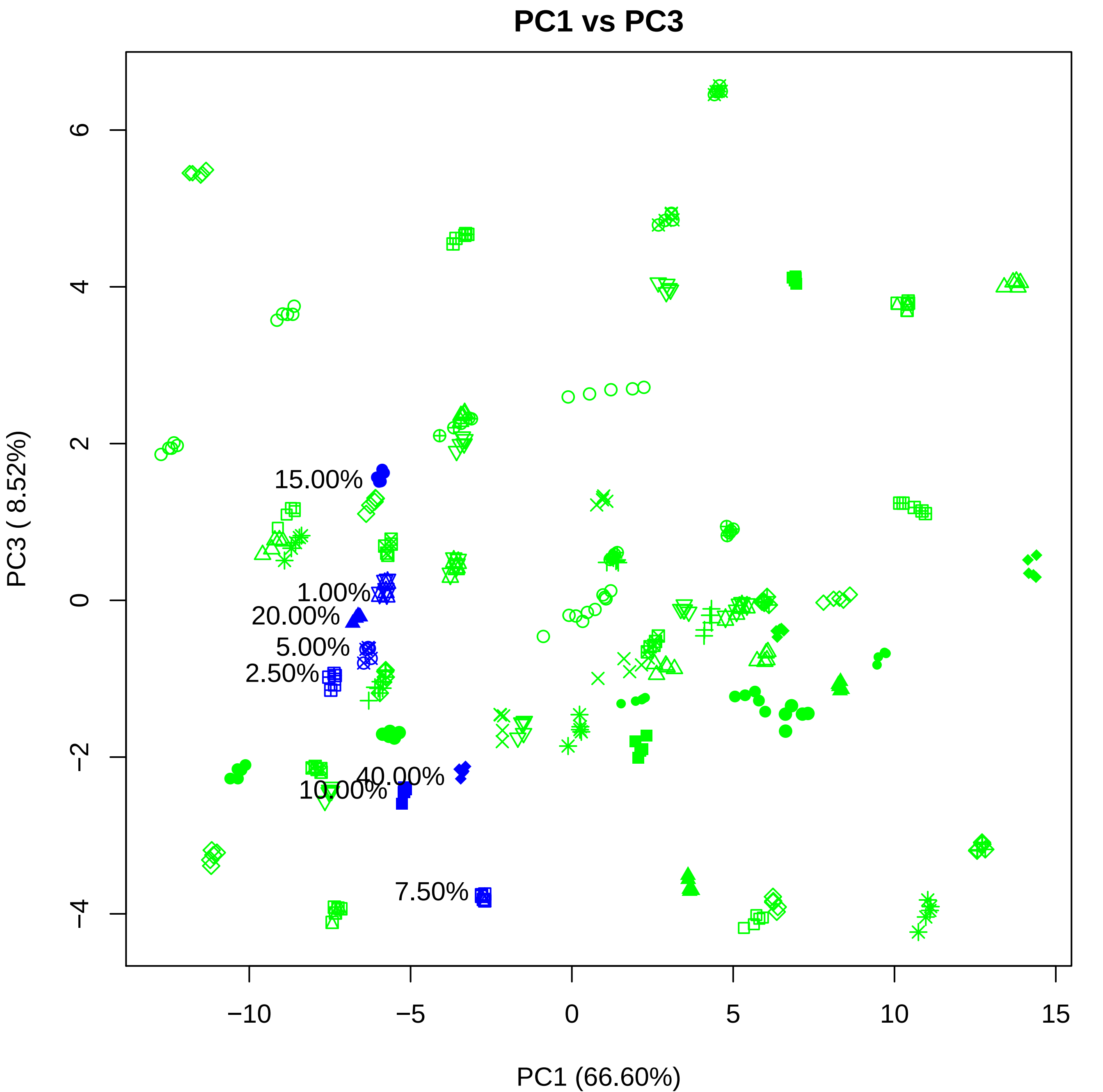

PC1 vs PC3
15.00%
1.00%
20.00%
5.00%
2.50%
40.00%
10.00%
7.50%
PC3 ( 8.52%)
PC1 (66.60%)
−10
−5
0
5
10
15
−4
−2
0
2
4
6

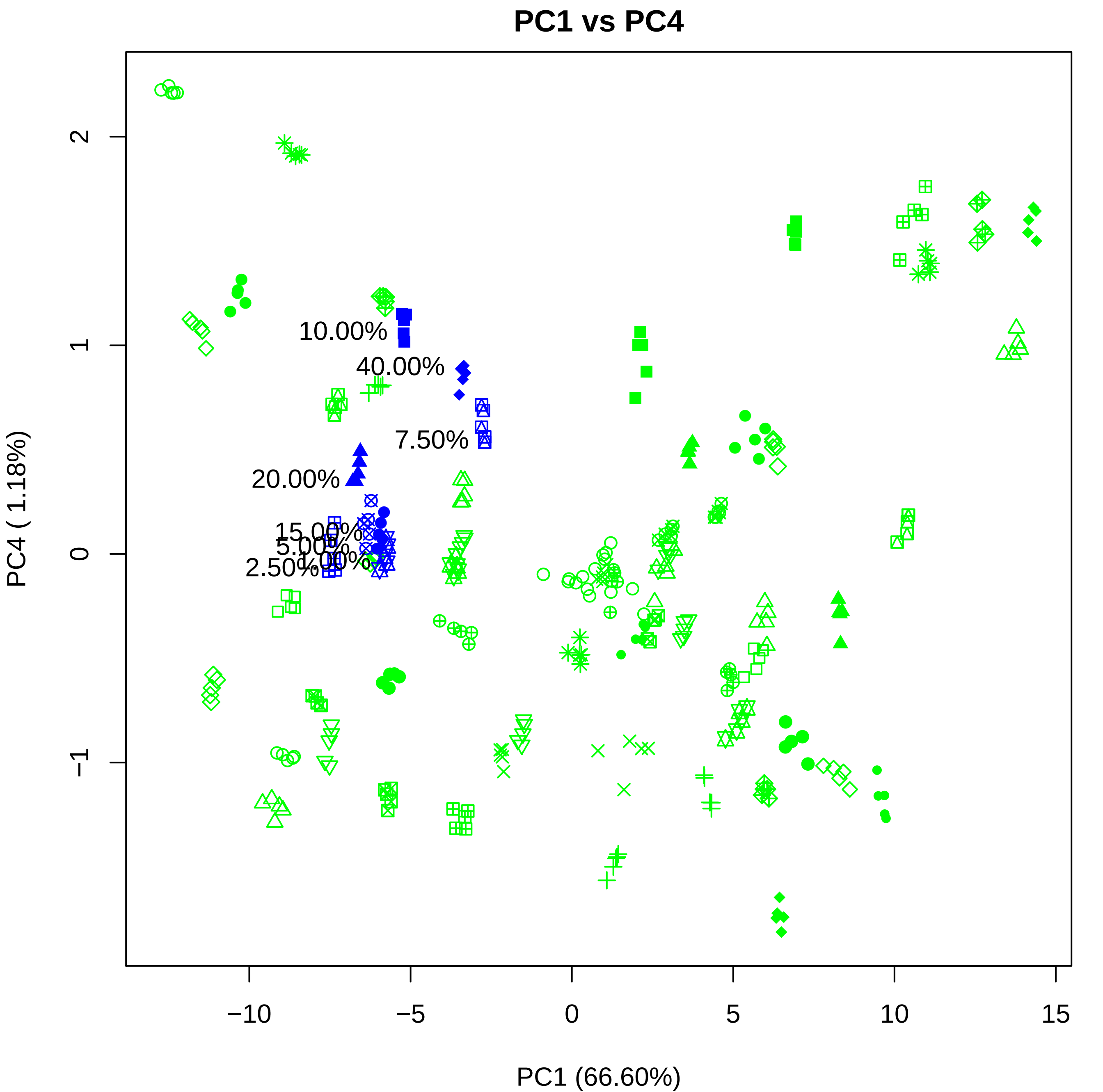
PC1 vs PC4
PC4 ( 1.18%)
PC1 (66.60%)
10.00%
40.00%
7.50%
20.00%
15.00%
5.00%
2.50%
1.00%
2
1
0
−1
−10
−5
0
5
10
15

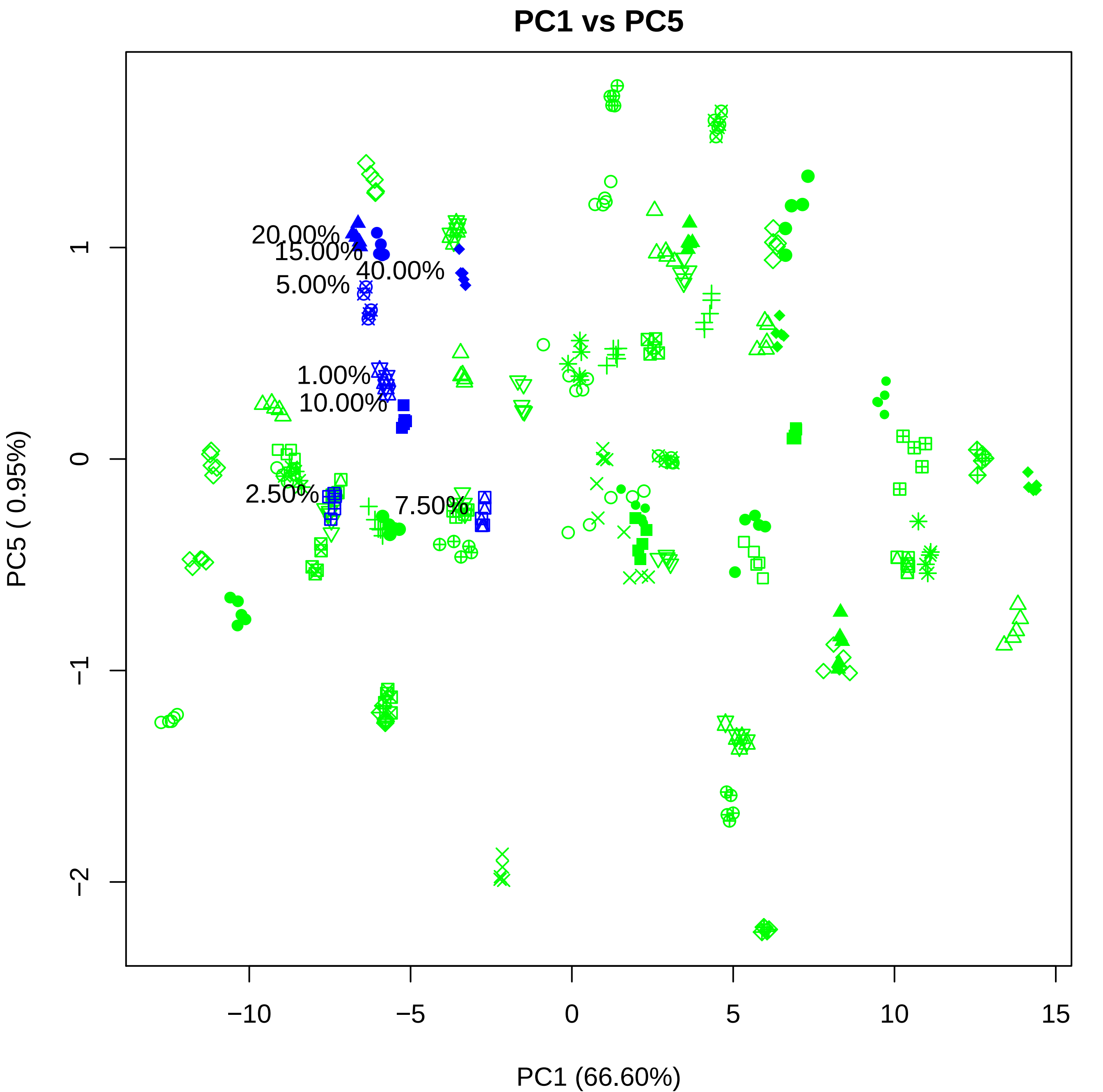
PC1 vs PC5
20.00%
15.00%
40.00%
5.00%
1.00%
10.00%
2.50%
7.50%
1
0
−1
−2
−10
−5
0
5
10
15
PC5 ( 0.95%)
PC1 (66.60%)

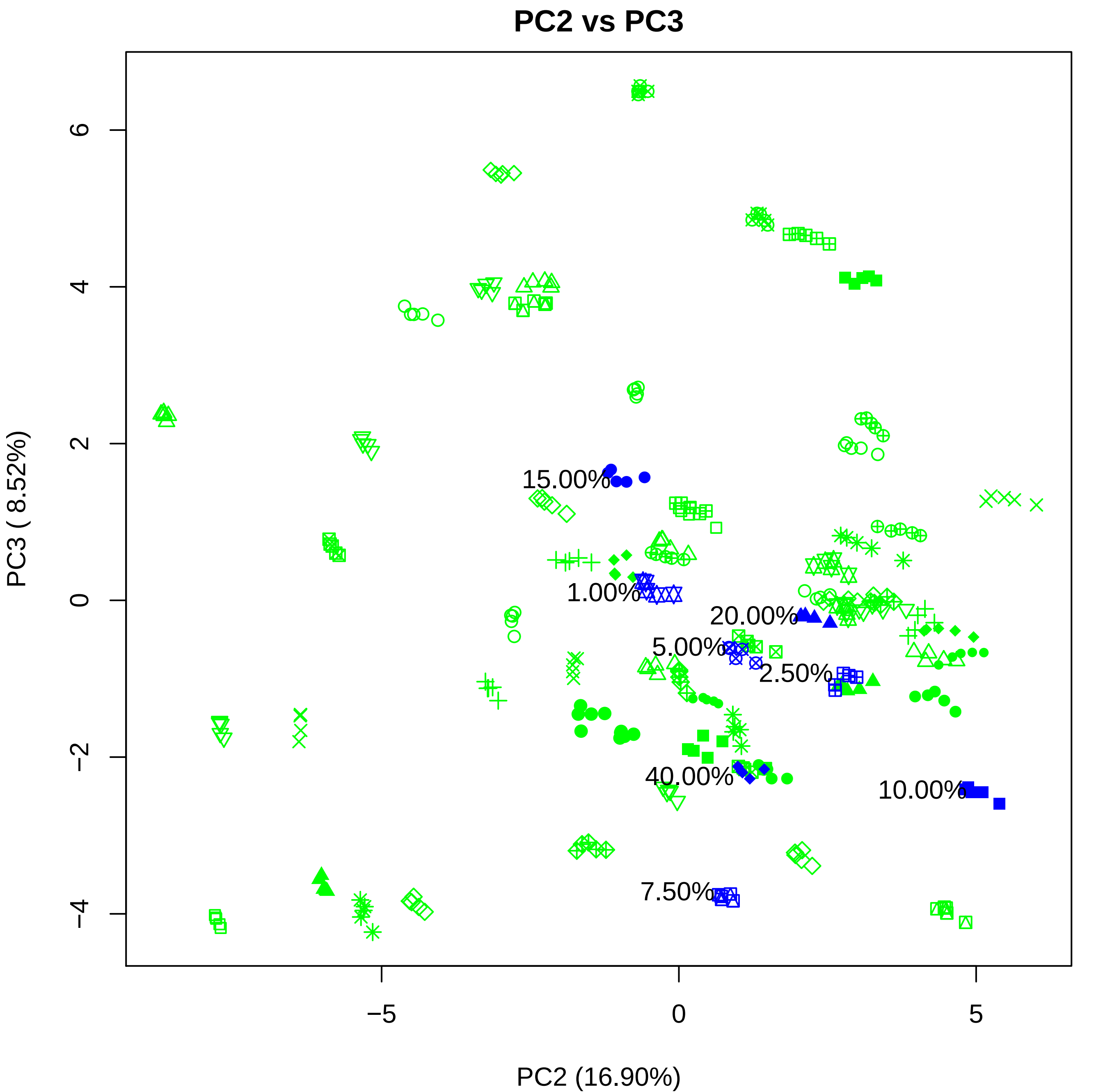
PC2 vs PC3
PC3 ( 8.52%)
PC2 (16.90%)
6
4
2
0
−2
−4
−5
0
5
15.00%
1.00%
20.00%
5.00%
2.50%
40.00%
10.00%
7.50%

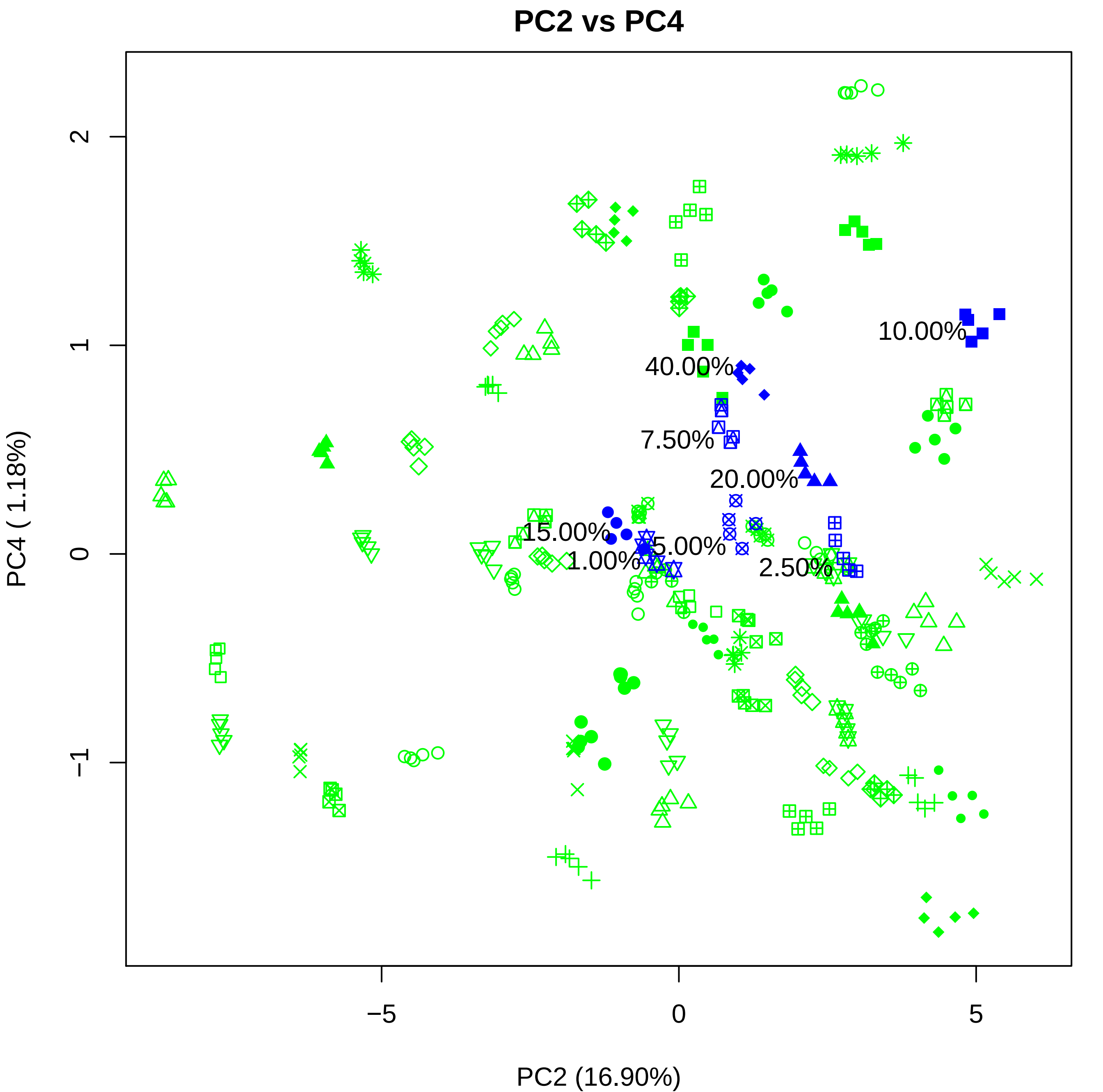

PC2 vs PC4
10.00%
40.00%
7.50%
20.00%
15.00%
5.00%
1.00%
2.50%
2
1
0
−1
−5
0
5
PC4 ( 1.18%)
PC2 (16.90%)

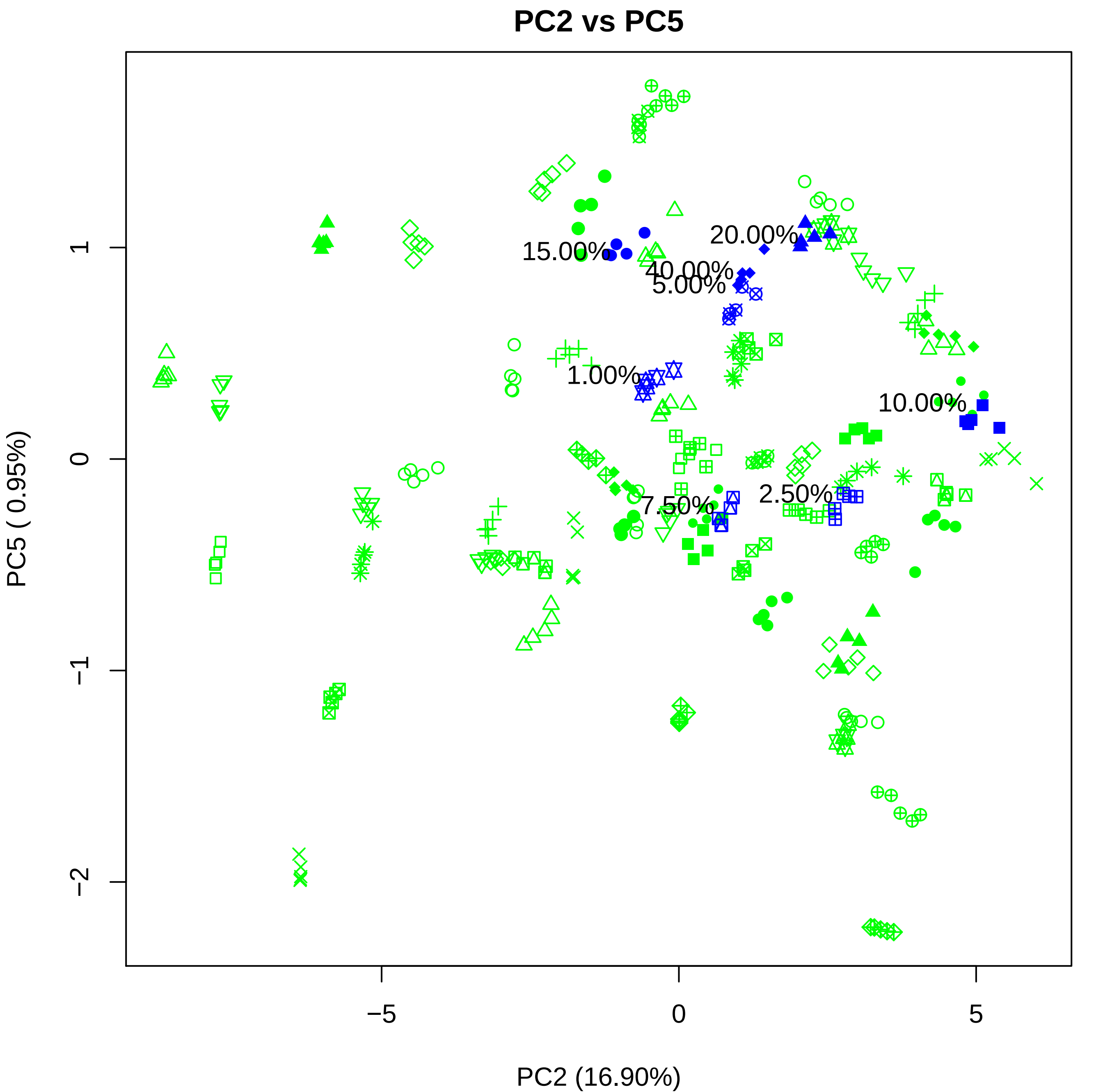

PC2 vs PC5
15.00%
20.00%
40.00%
5.00%
1.00%
10.00%
7.50%
2.50%
1
0
−1
−2
−5
0
5
PC5 ( 0.95%)
PC2 (16.90%)

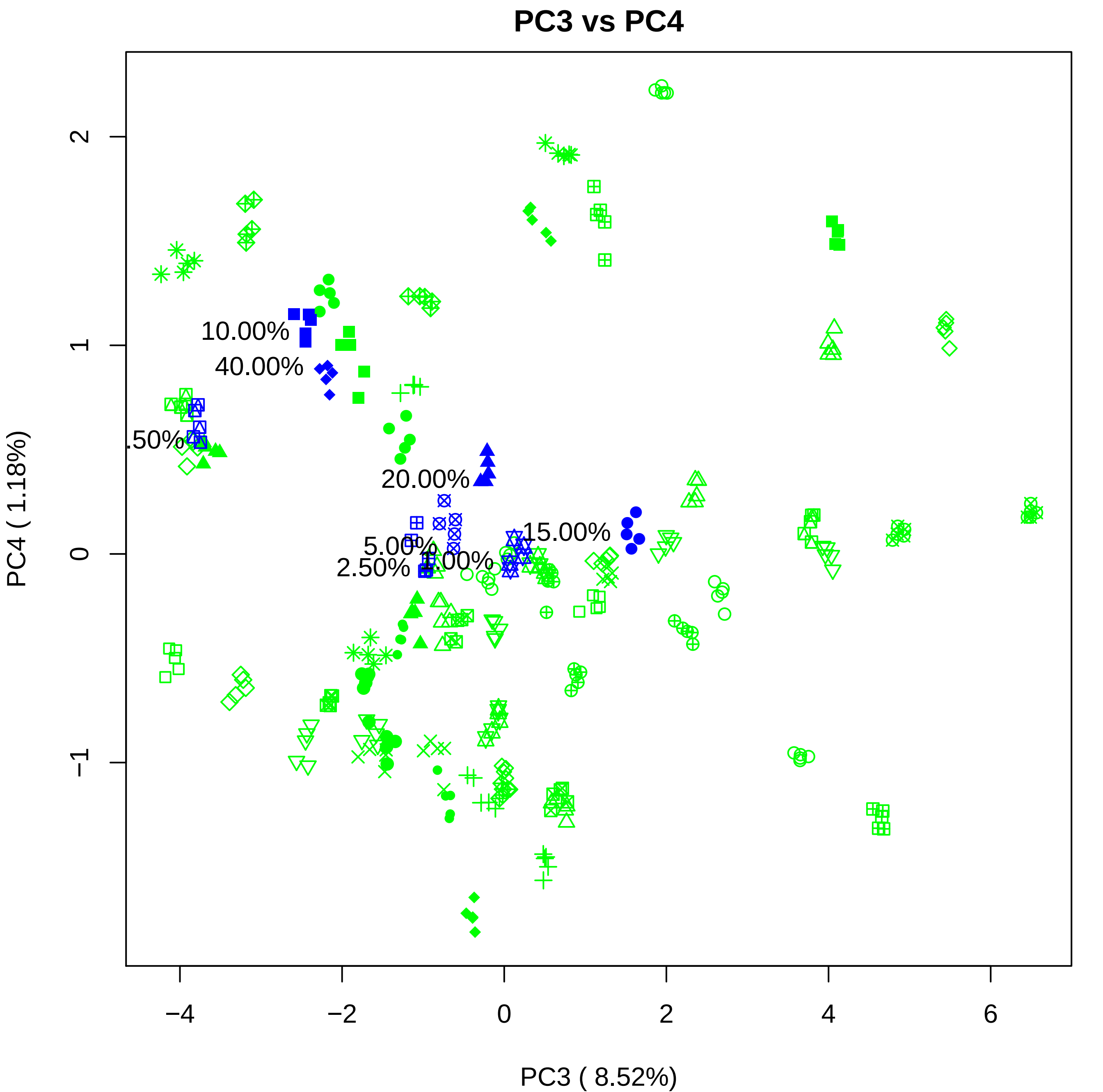

PC3 vs PC4
PC4 ( 1.18%)
PC3 ( 8.52%)
10.00%
40.00%
.50%
20.00%
15.00%
5.00%
2.50%
1.00%
2
1
0
−1
−4
−2
0
2
4
6

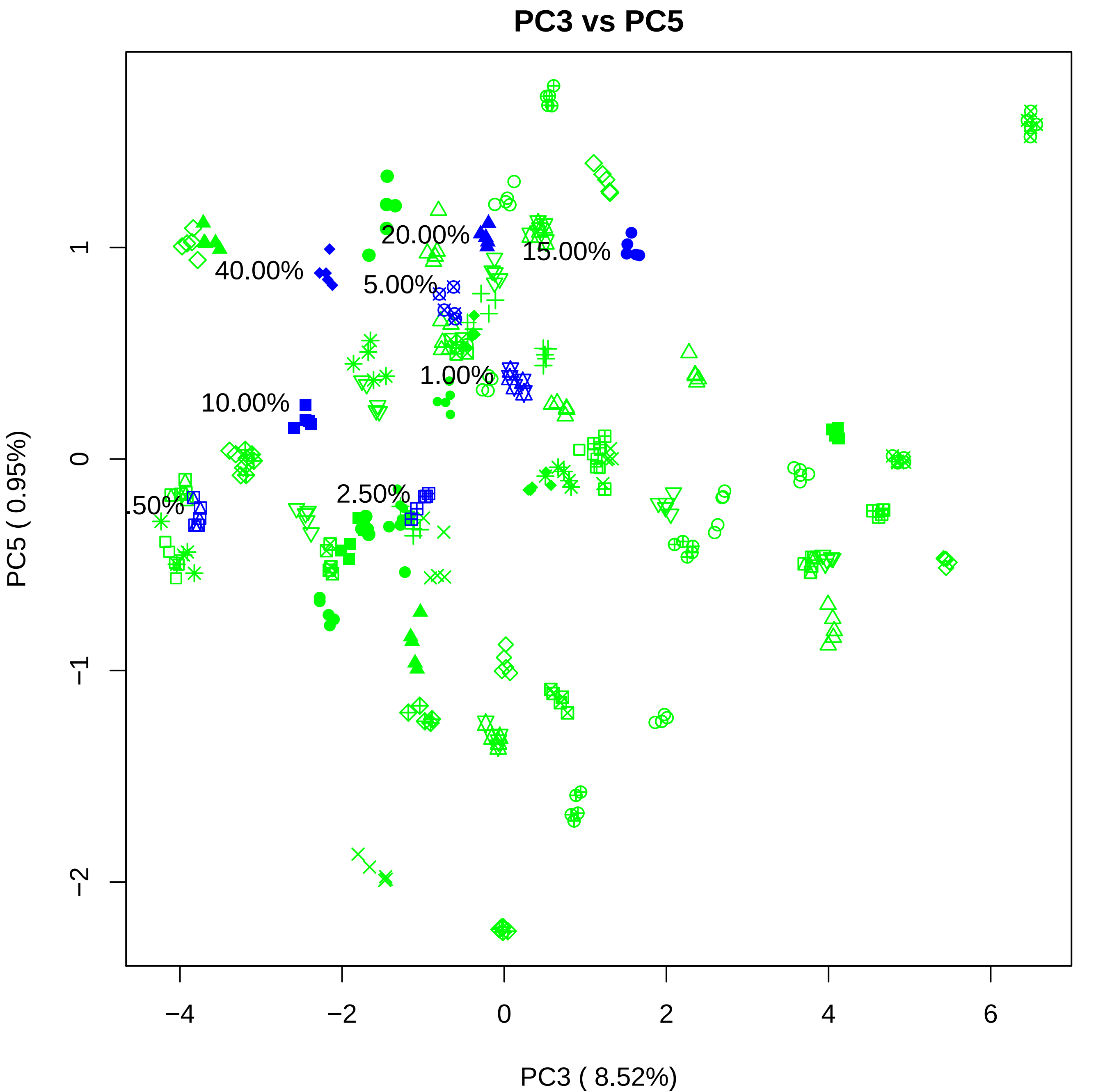
PC3 vs PC5
40.00%
20.00%
15.00%
5.00%
1.00%
10.00%
2.50%
.50%
1
0
−1
−2
−4
−2
0
2
4
6
PC5 ( 0.95%)
PC3 ( 8.52%)

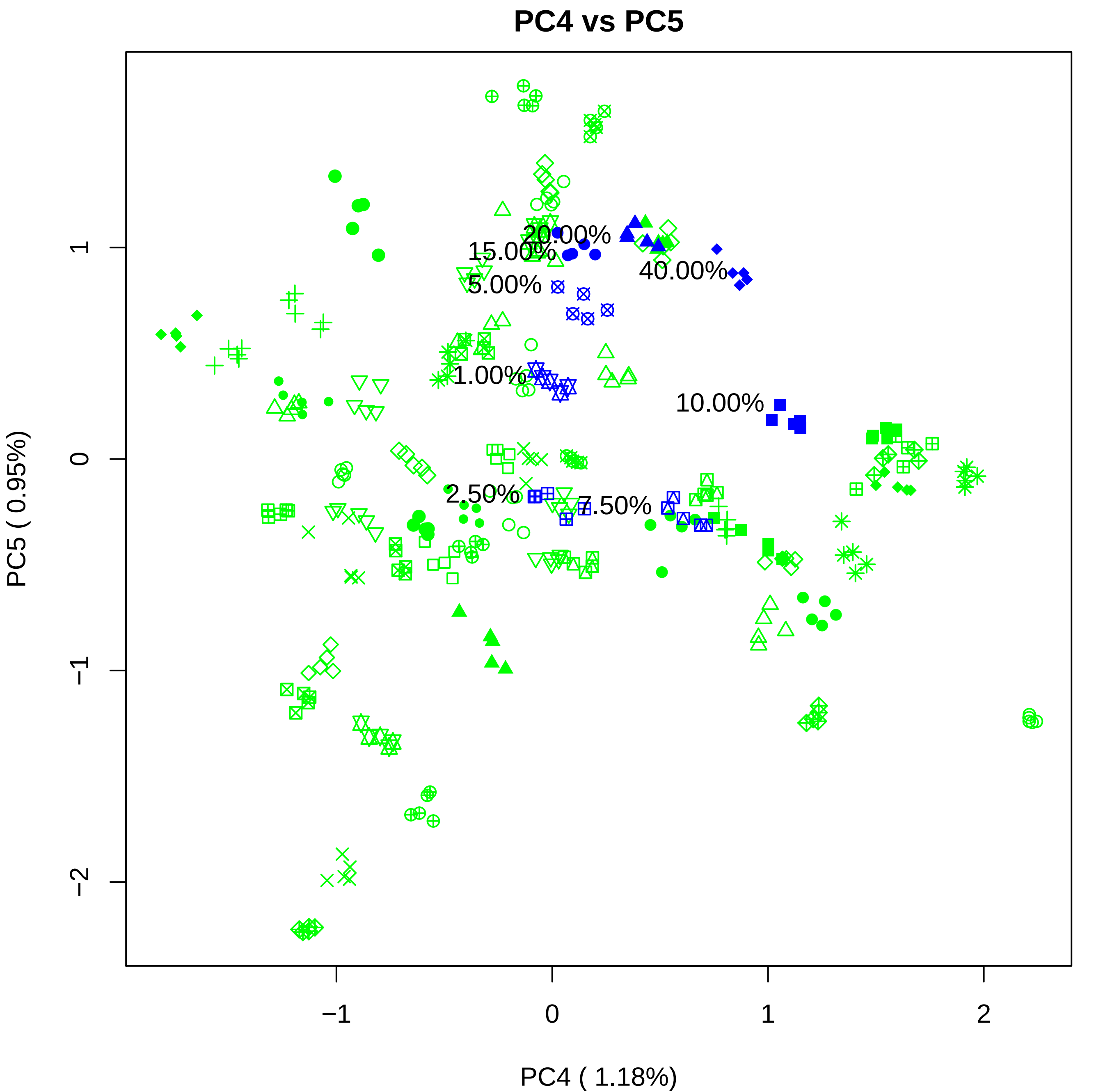

PC4 vs PC5
20.00%
15.00%
40.00%
5.00%
1.00%
10.00%
2.50%
7.50%
1
0
−1
−2
PC5 ( 0.95%)
−1
0
1
2
PC4 ( 1.18%)

### PCA of 1%  BMH(d) vs PMH(d)

Scaled Matrix PCAed

7th order polynomial baseline correction performed

Raw spectra smoothed to 21 Neighbours, 2nd order PN, Savtisky Golay

Normalisation between 0 and 100

D2 spectra smoothed to 41 Neighbours, 2nd order PN, Savtisky Golay

Regions Analysed: 3601−2501 and 2001−700cm−1

Samples Analysed: 1 bContam's 0 BLH's 0 BMH's 0 OMH's 0 pContam's 0 PLH's 60 PMH's

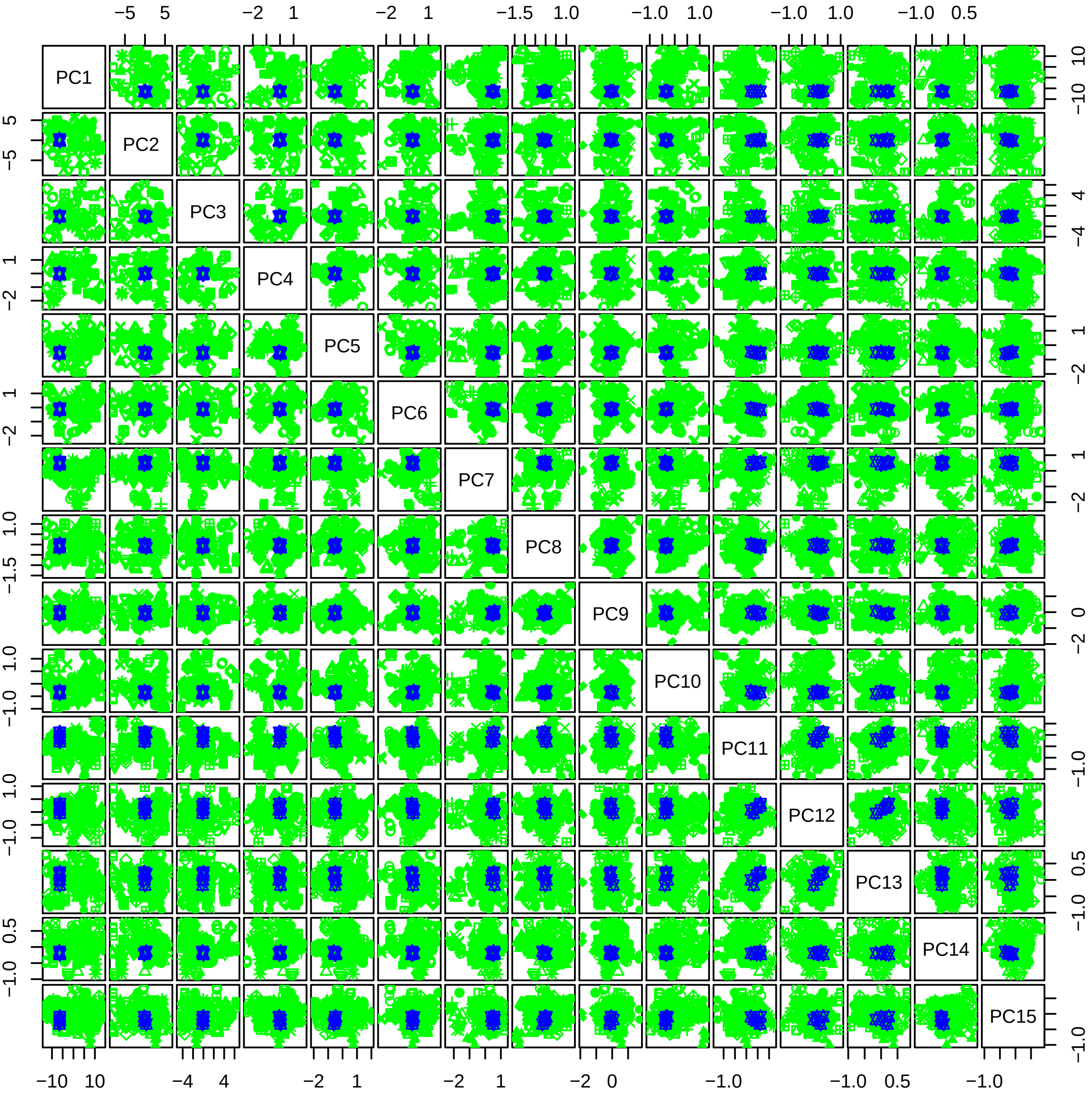

PC1
PC2
PC3
PC4
PC5
PC6
PC7
PC8
PC9
PC10
PC11
PC12
PC13
PC14
PC15

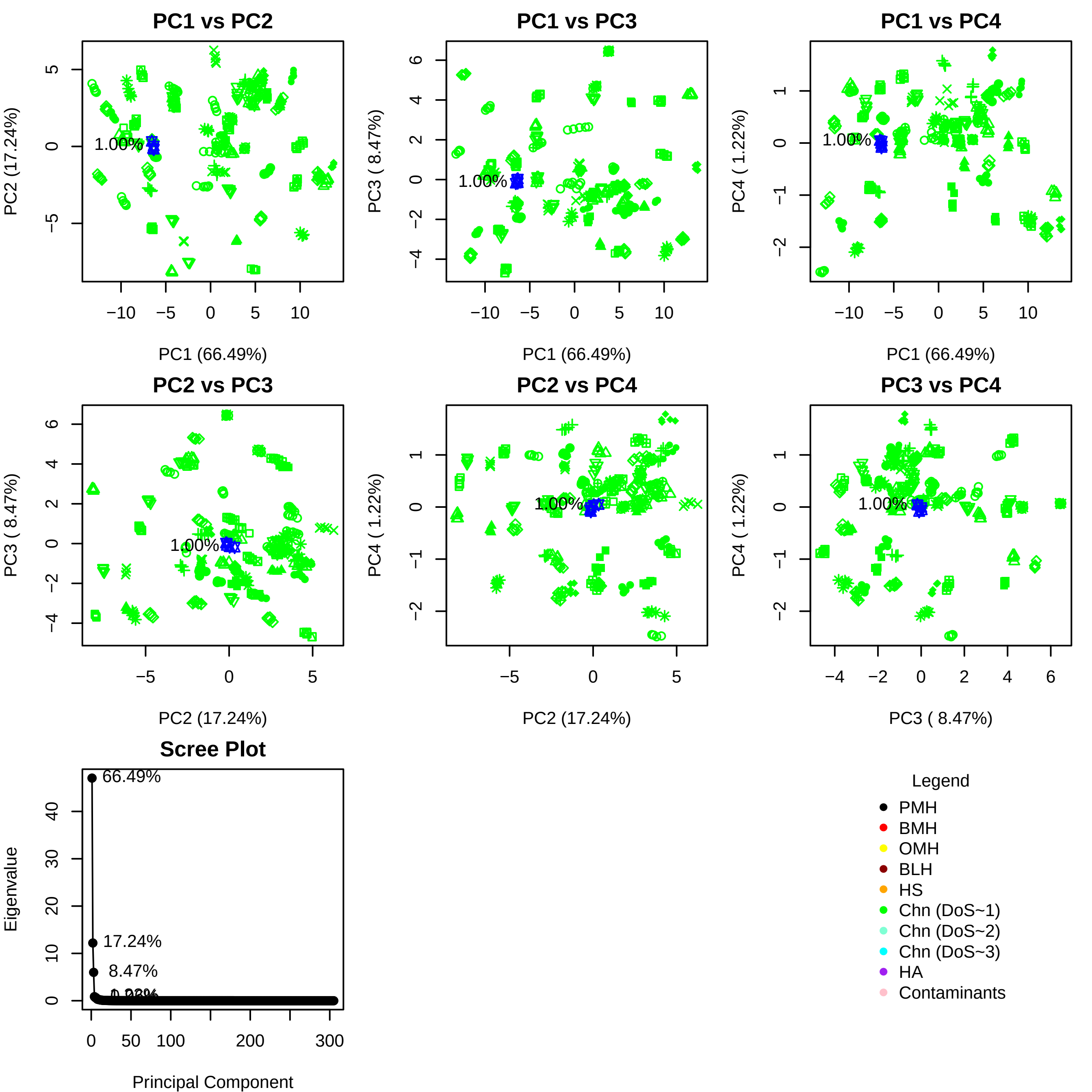

PC1 vs PC2
PC1 vs PC3
PC1 vs PC4
PC2 vs PC3
PC2 vs PC4
PC3 vs PC4
Scree Plot
PC1 (66.49%)
PC2 (17.24%)
PC3 ( 8.47%)
PC4 ( 1.22%)
1.00%
Eigenvalue
Principal Component
66.49%
17.24%
8.47%
Legend
PMH
BMH
OMH
BLH
HS
Chn (DoS~1)
Chn (DoS~2)
Chn (DoS~3)
HA
Contaminants

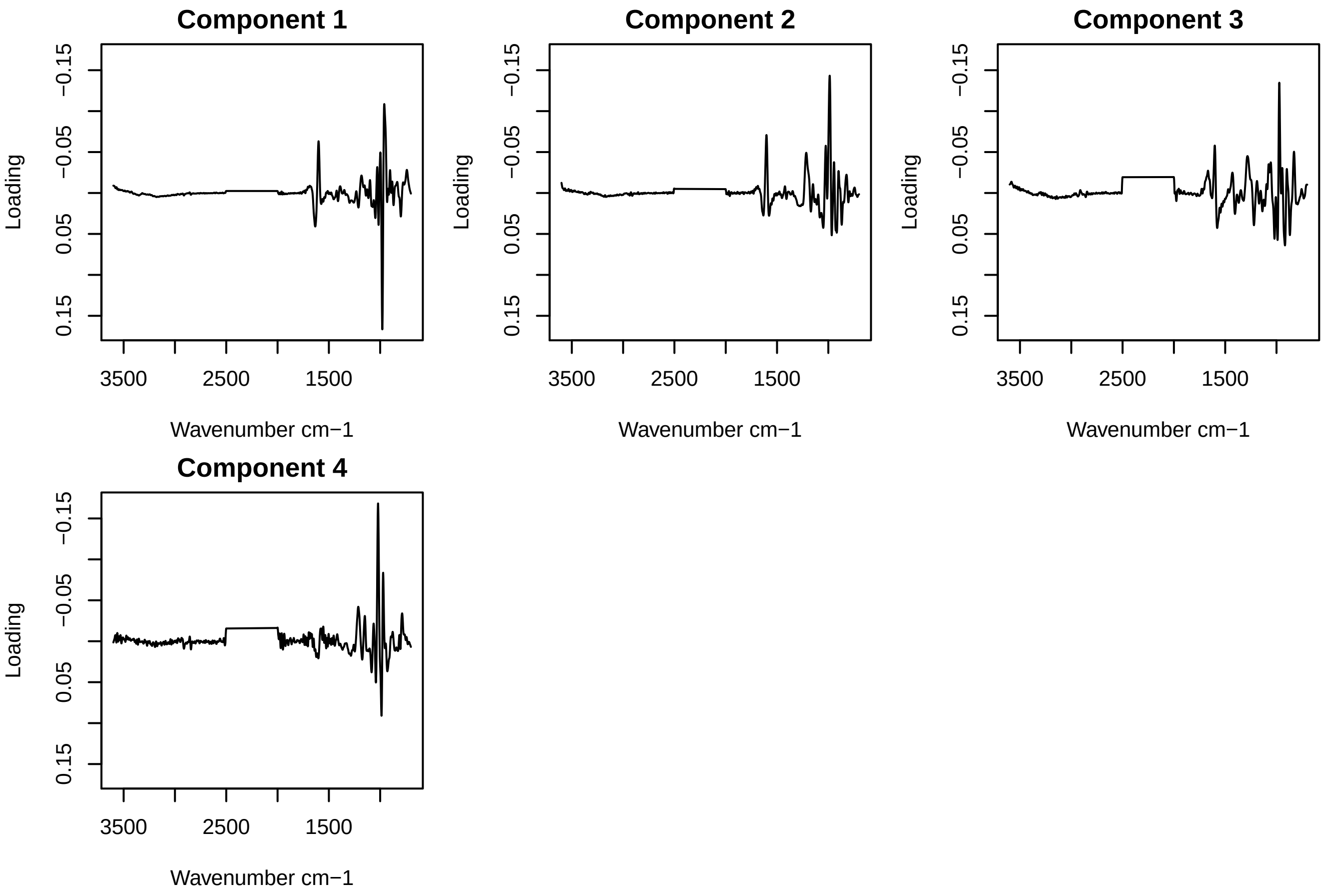
Component 1
Component 2
Component 3
Component 4
Loading
−0.15
−0.05
0.05
0.15
3500
2500
1500
Wavenumber cm−1

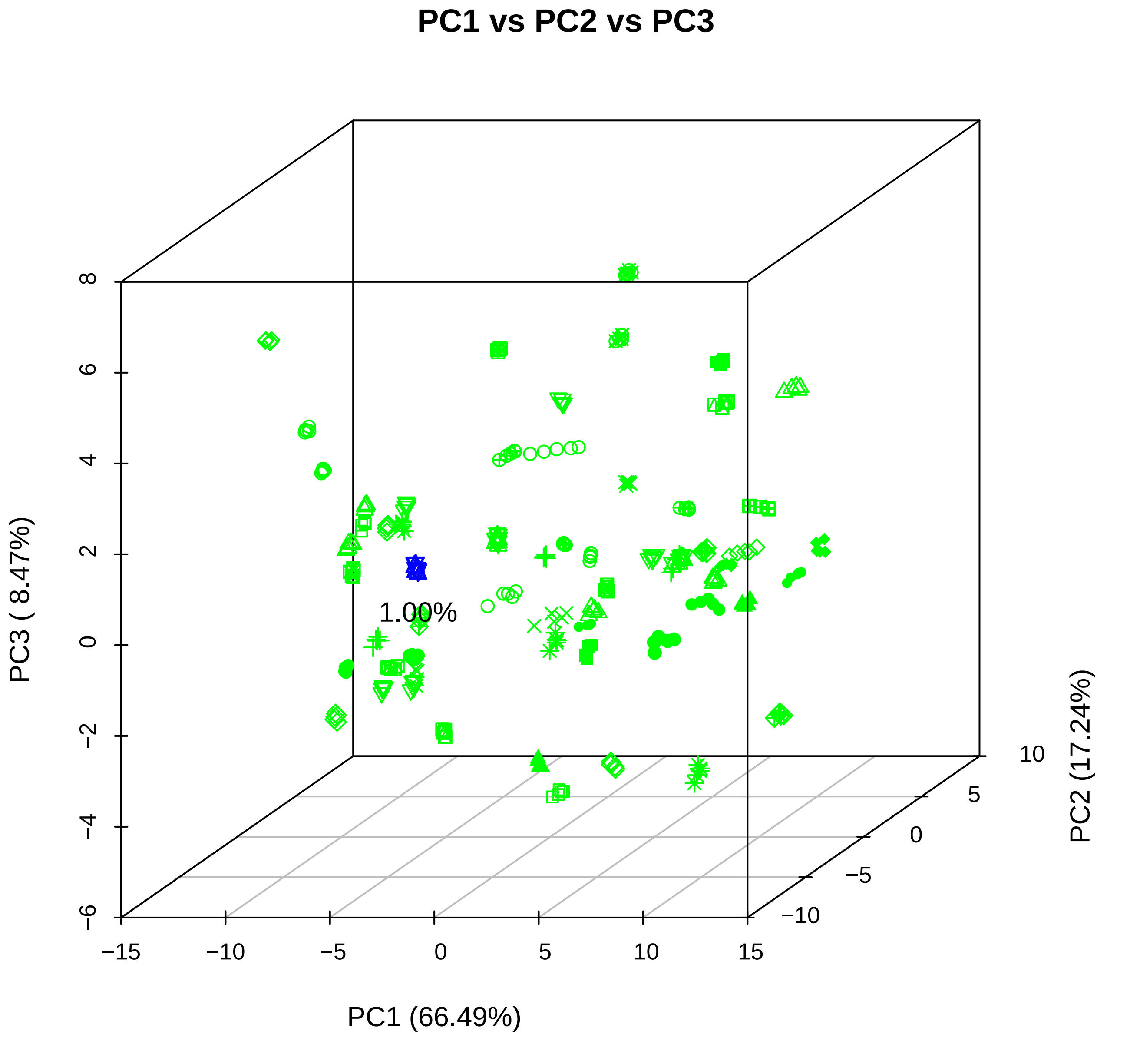
PC1 vs PC2 vs PC3
PC3 ( 8.47%)
PC2 (17.24%)
PC1 (66.49%)
1.00%
−15
−10
−5
0
5
10
15
−6
−4
−2
0
2
4
6
8
−10
−5
0
5
10

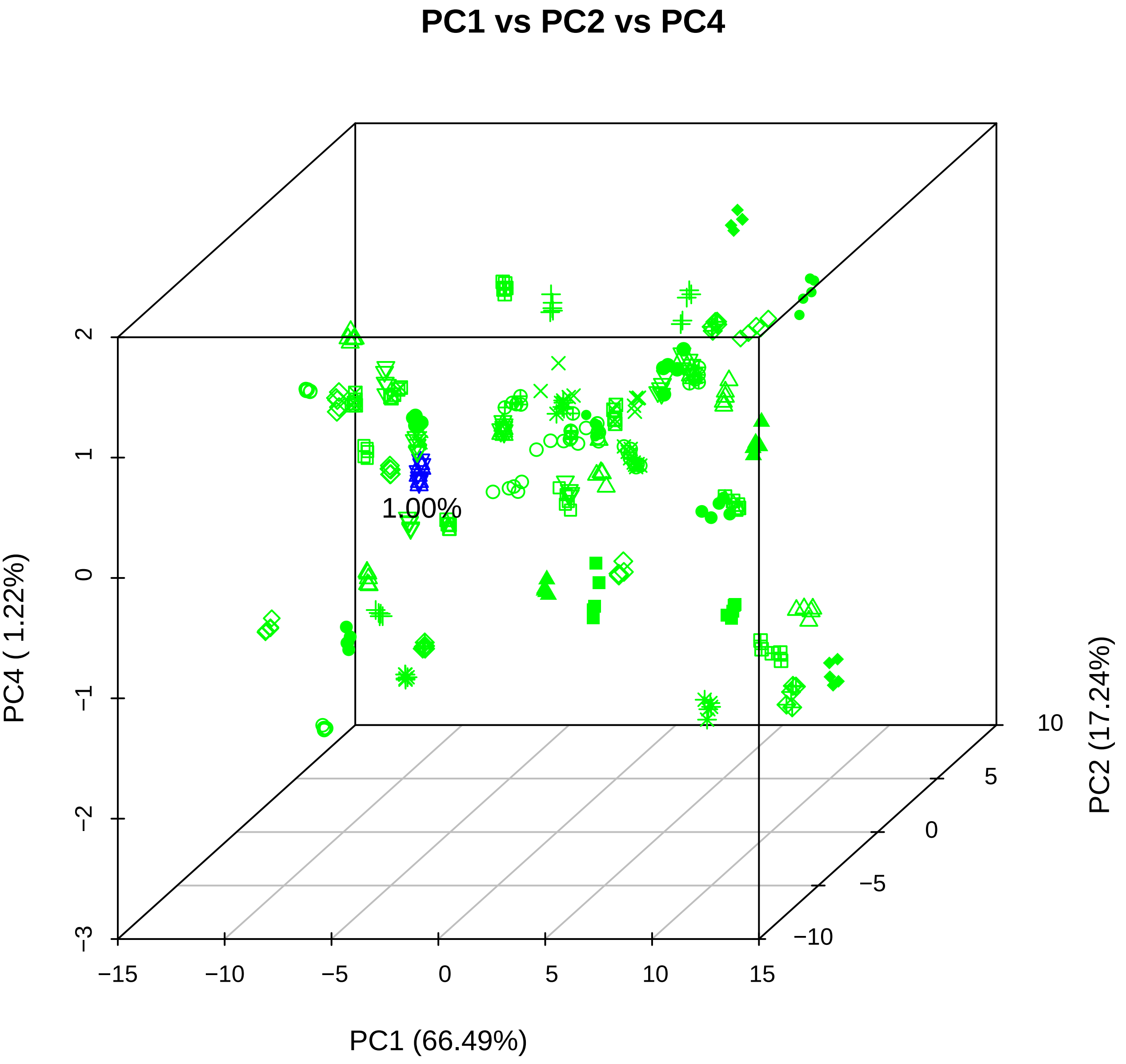
PC1 vs PC2 vs PC4
1.00%
PC1 (66.49%)
PC2 (17.24%)
PC4 ( 1.22%)
−15
−10
−5
0
5
10
15
−10
−5
0
5
10
−3
−2
−1
0
1
2

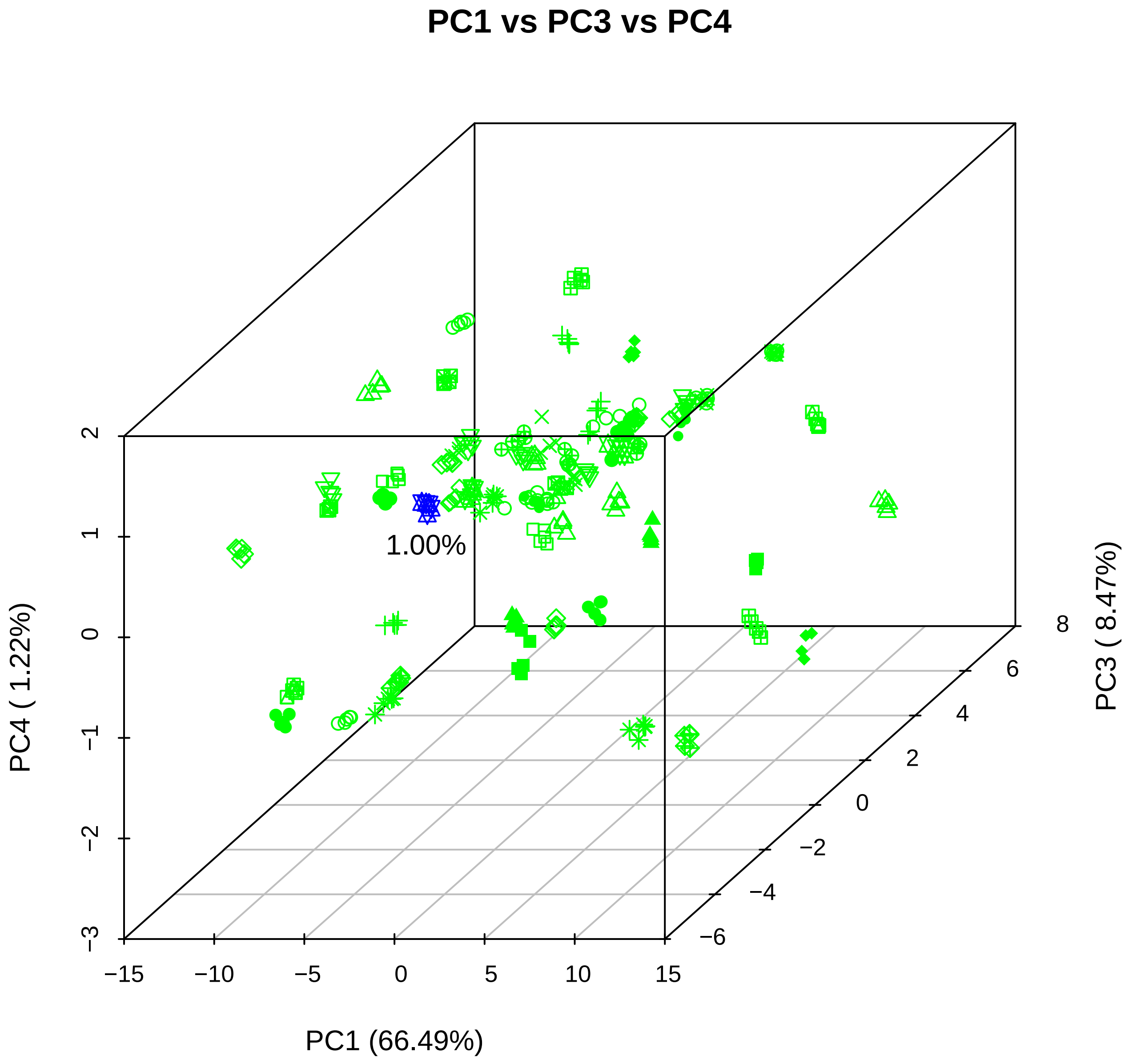

PC1 vs PC3 vs PC4
1.00%
PC1 (66.49%)
PC3 ( 8.47%)
PC4 ( 1.22%)
−15
−10
−5
0
5
10
15
−6
−4
−2
0
2
4
6
8
−3
−2
−1
0
1
2

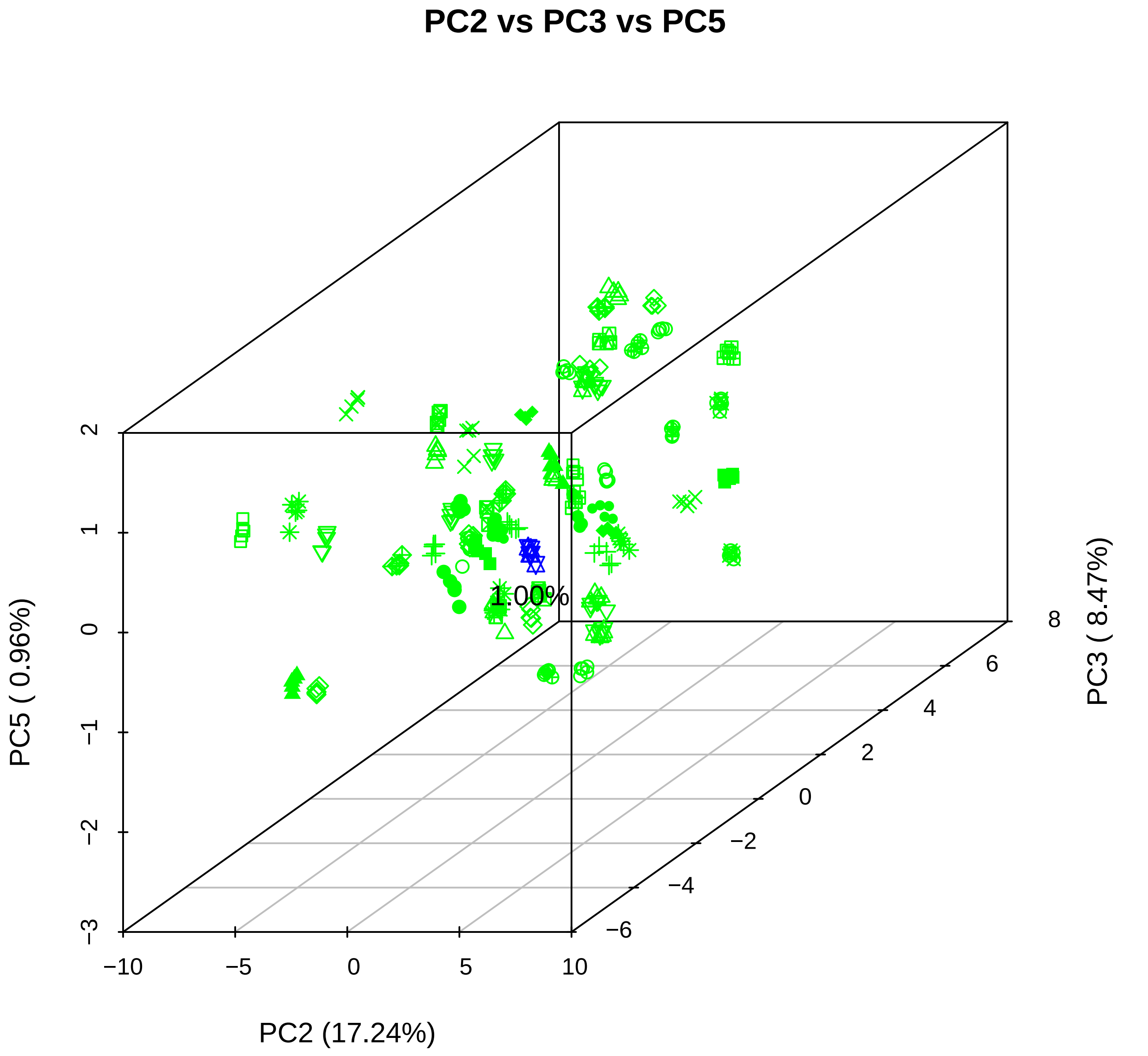
PC2 vs PC3 vs PC5
PC2 (17.24%)
PC3 ( 8.47%)
PC5 ( 0.96%)
1.00%
−10
−5
0
5
10
−6
−4
−2
0
2
4
6
8
−3
−2
−1
0
1
2

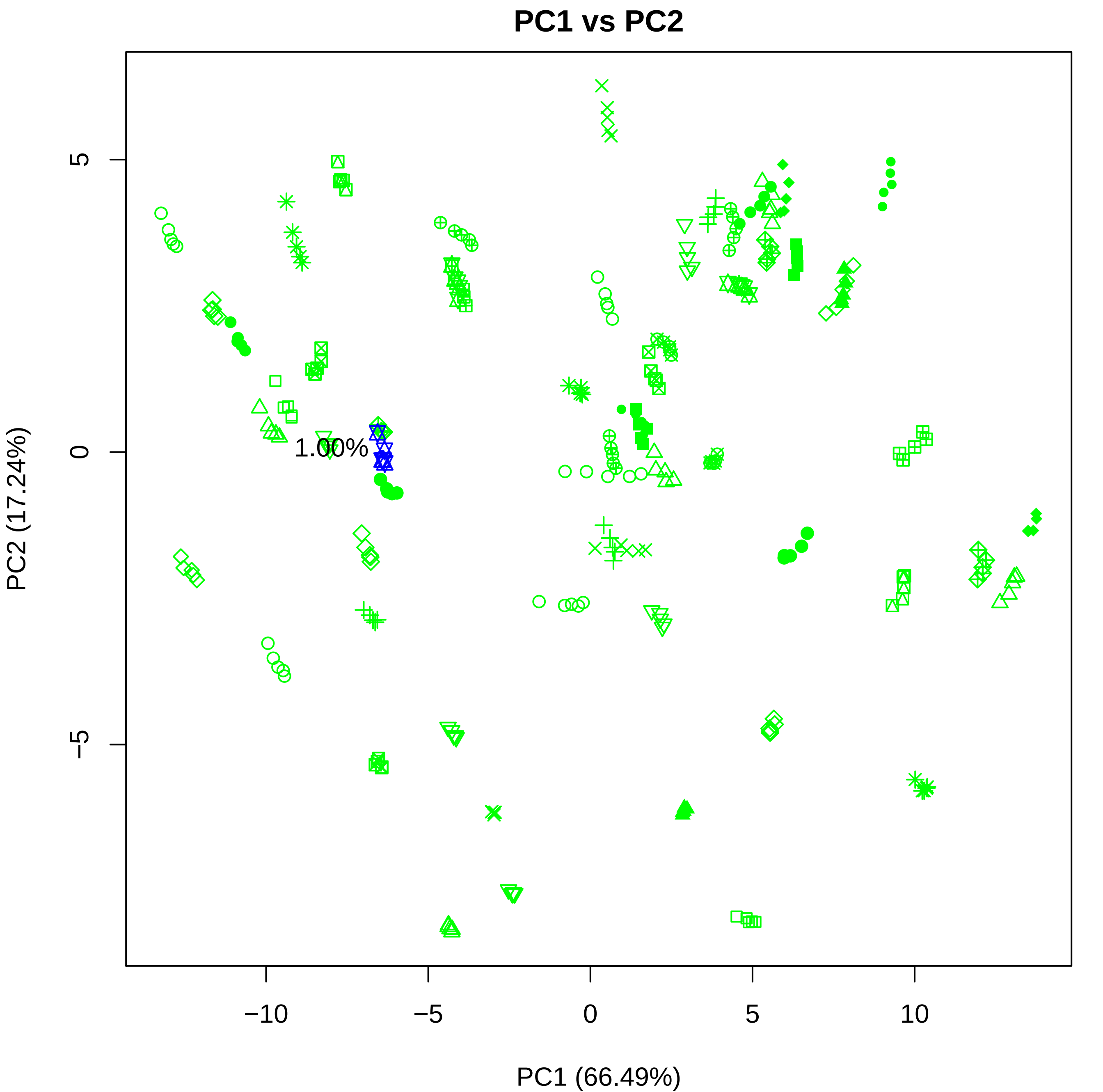

PC1 vs PC2
1.00%
PC1 (66.49%)
PC2 (17.24%)
−10
−5
0
5
10

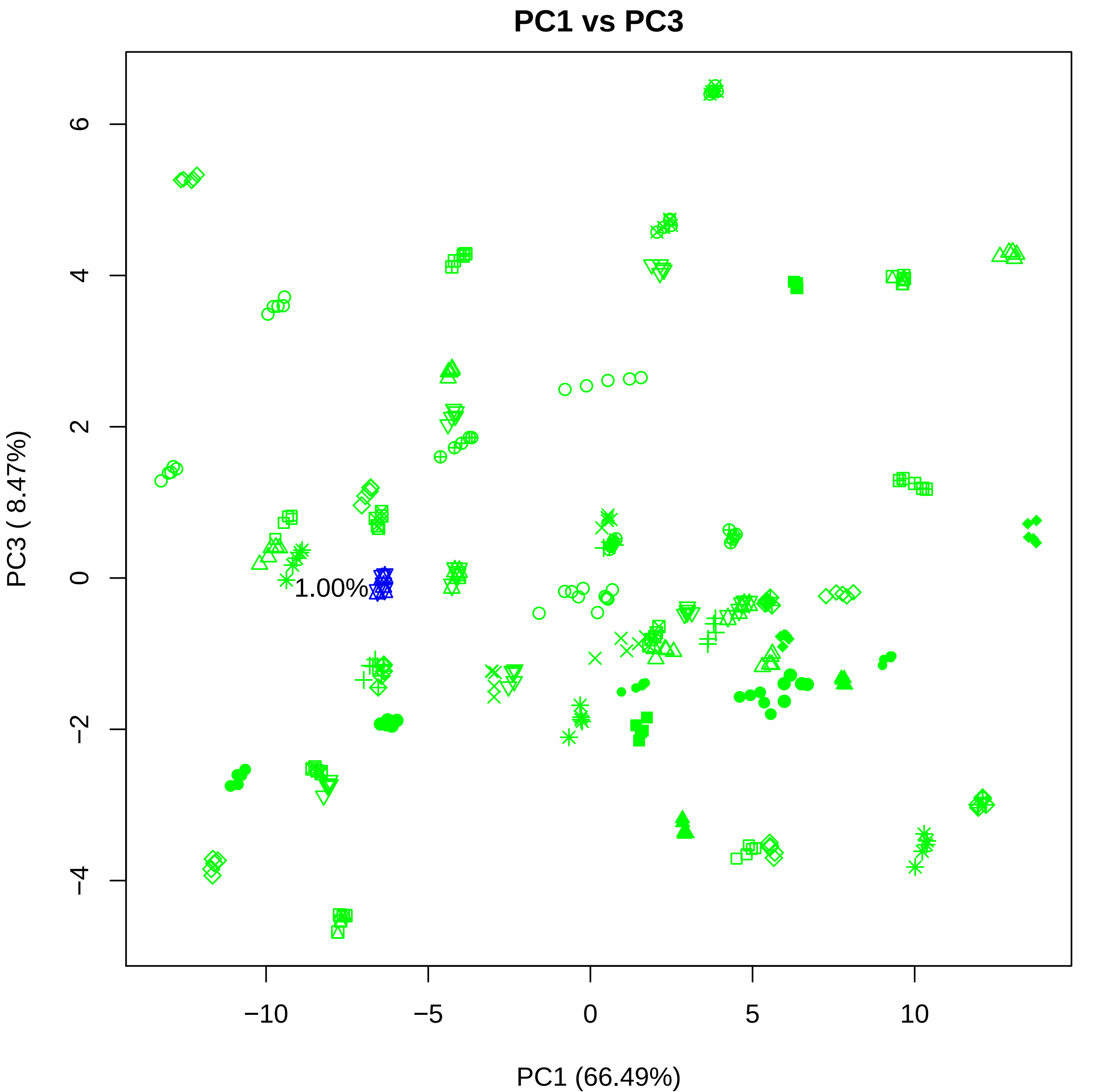
PC1 vs PC3
PC3 ( 8.47%)
PC1 (66.49%)
1.00%
6
4
2
0
−2
−4
−10
−5
0
5
10

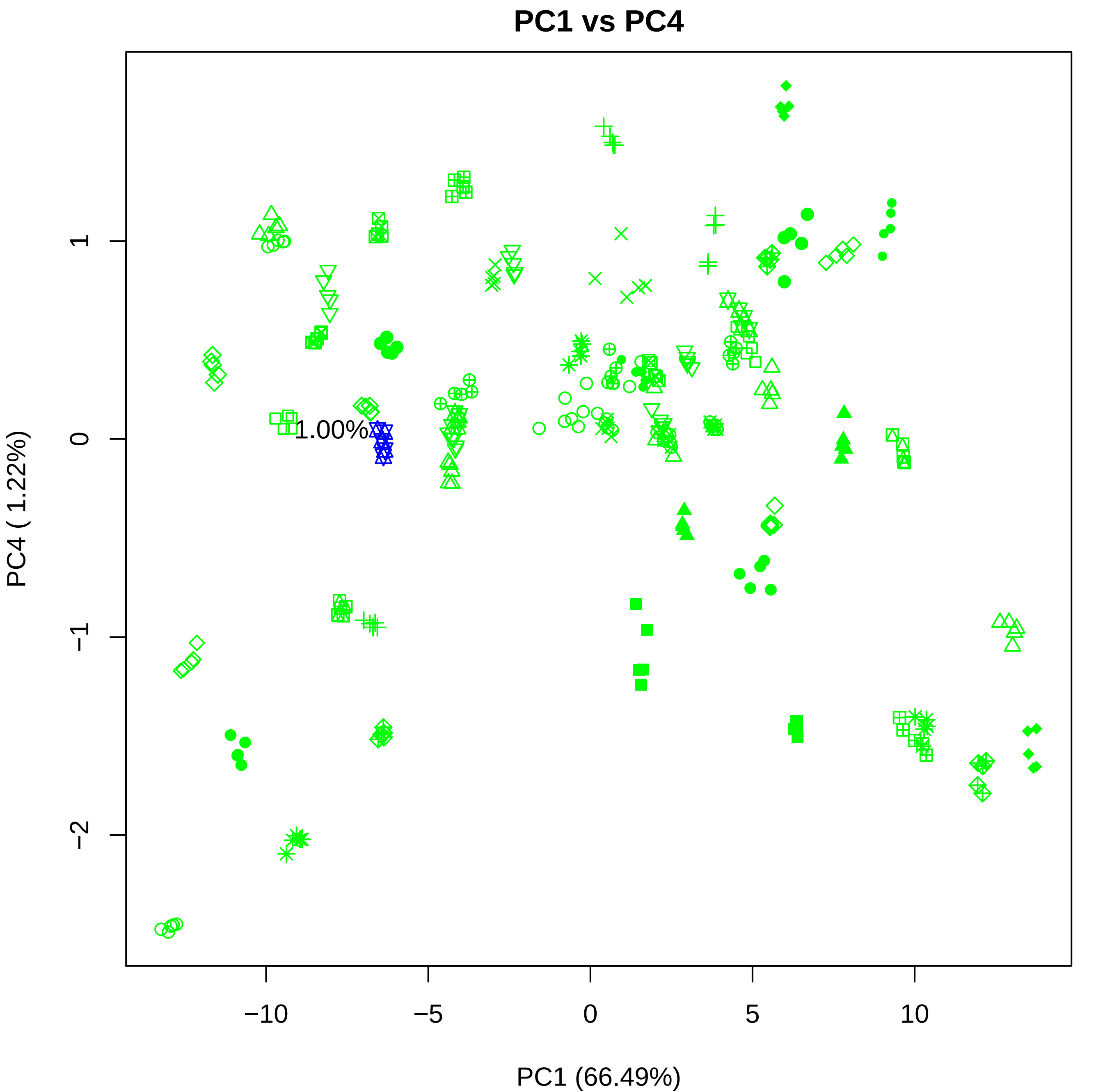

PC1 vs PC4
1.00%
PC4 ( 1.22%)
PC1 (66.49%)
−10
−5
0
5
10
1
0
−1
−2

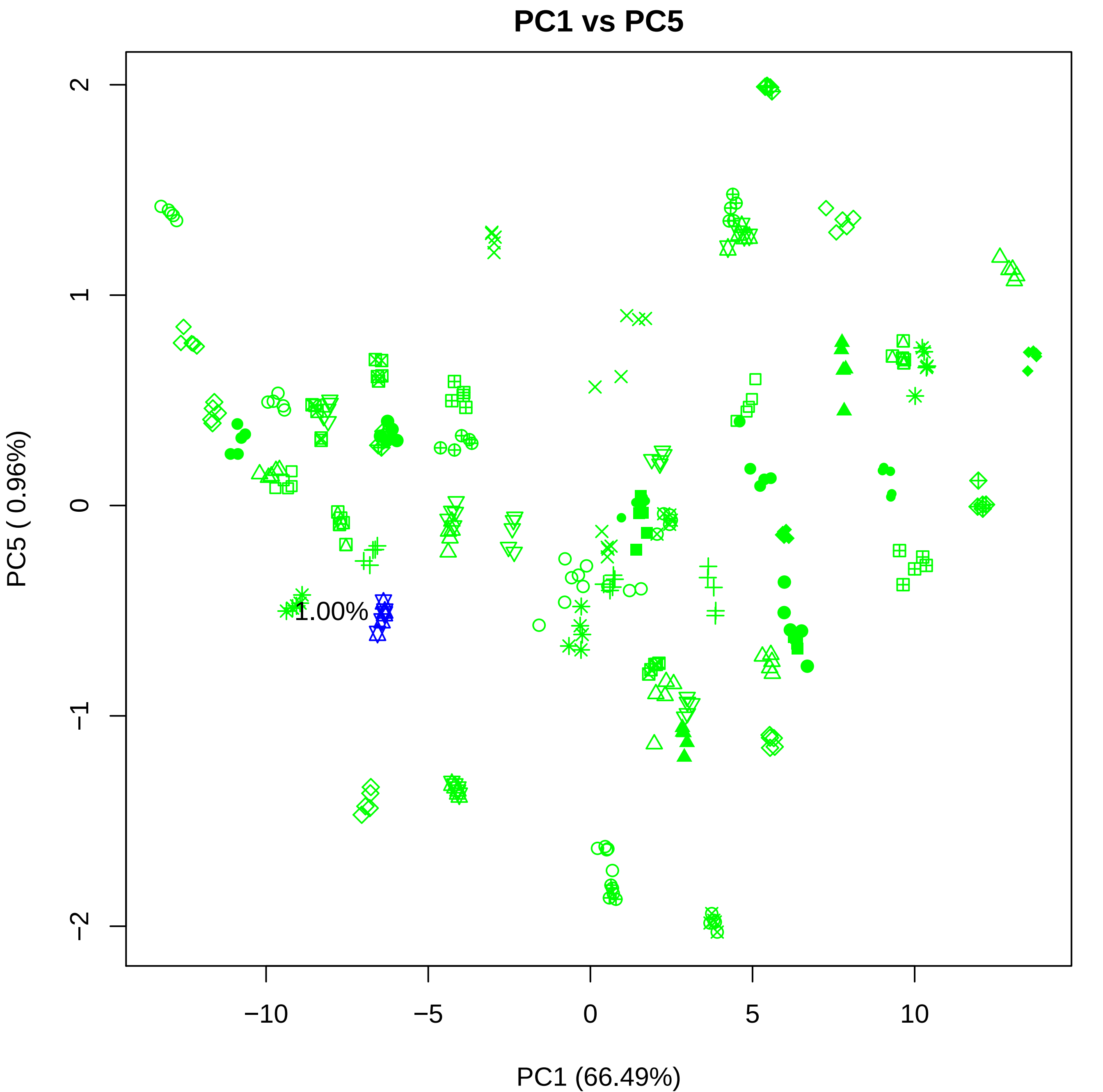
PC1 vs PC5
PC5 ( 0.96%)
PC1 (66.49%)
1.00%
2
1
0
−1
−2
−10
−5
0
5
10

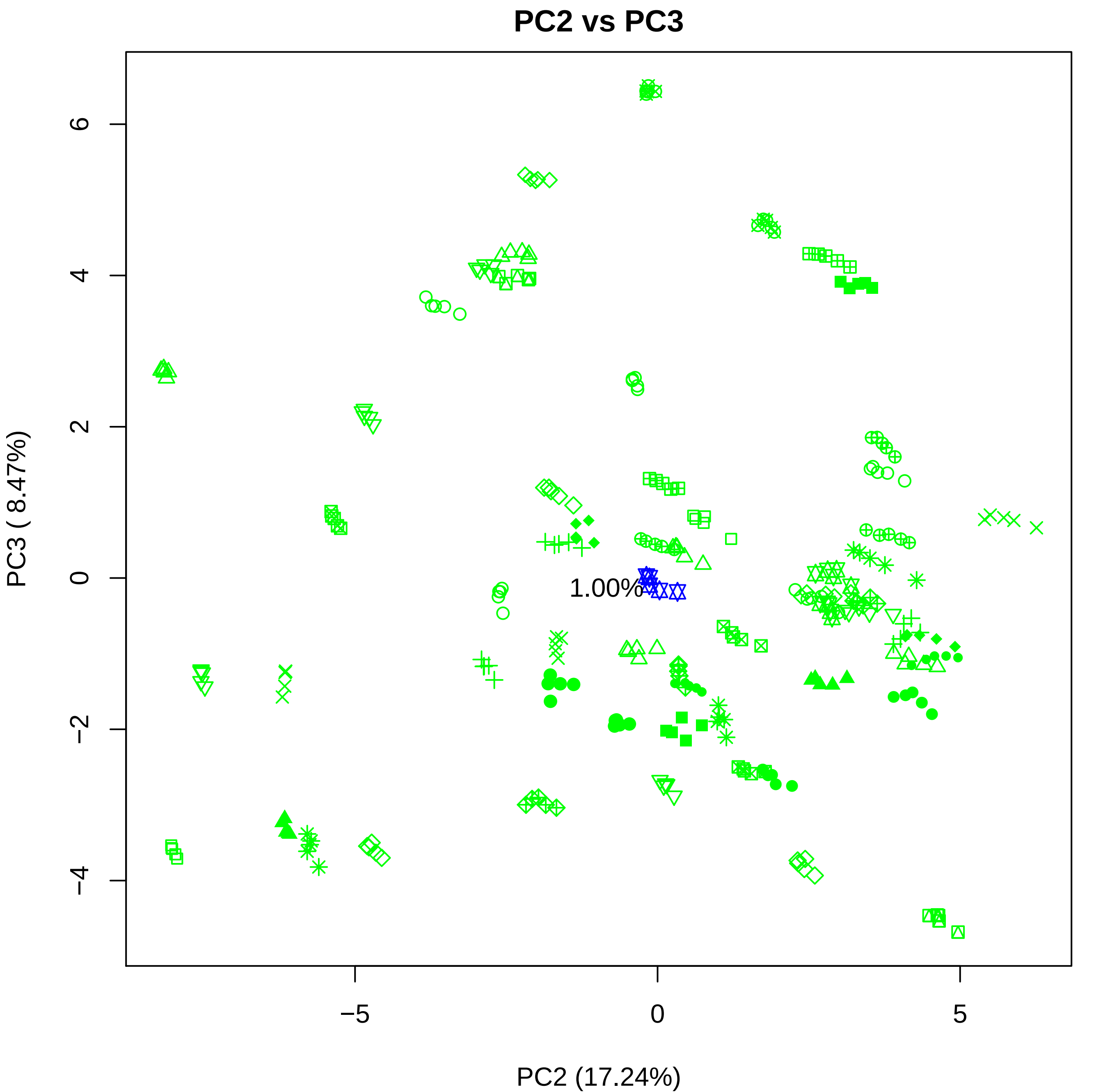

PC2 vs PC3
PC3 ( 8.47%)
PC2 (17.24%)
6
4
2
0
−2
−4
−5
0
5
1.00%

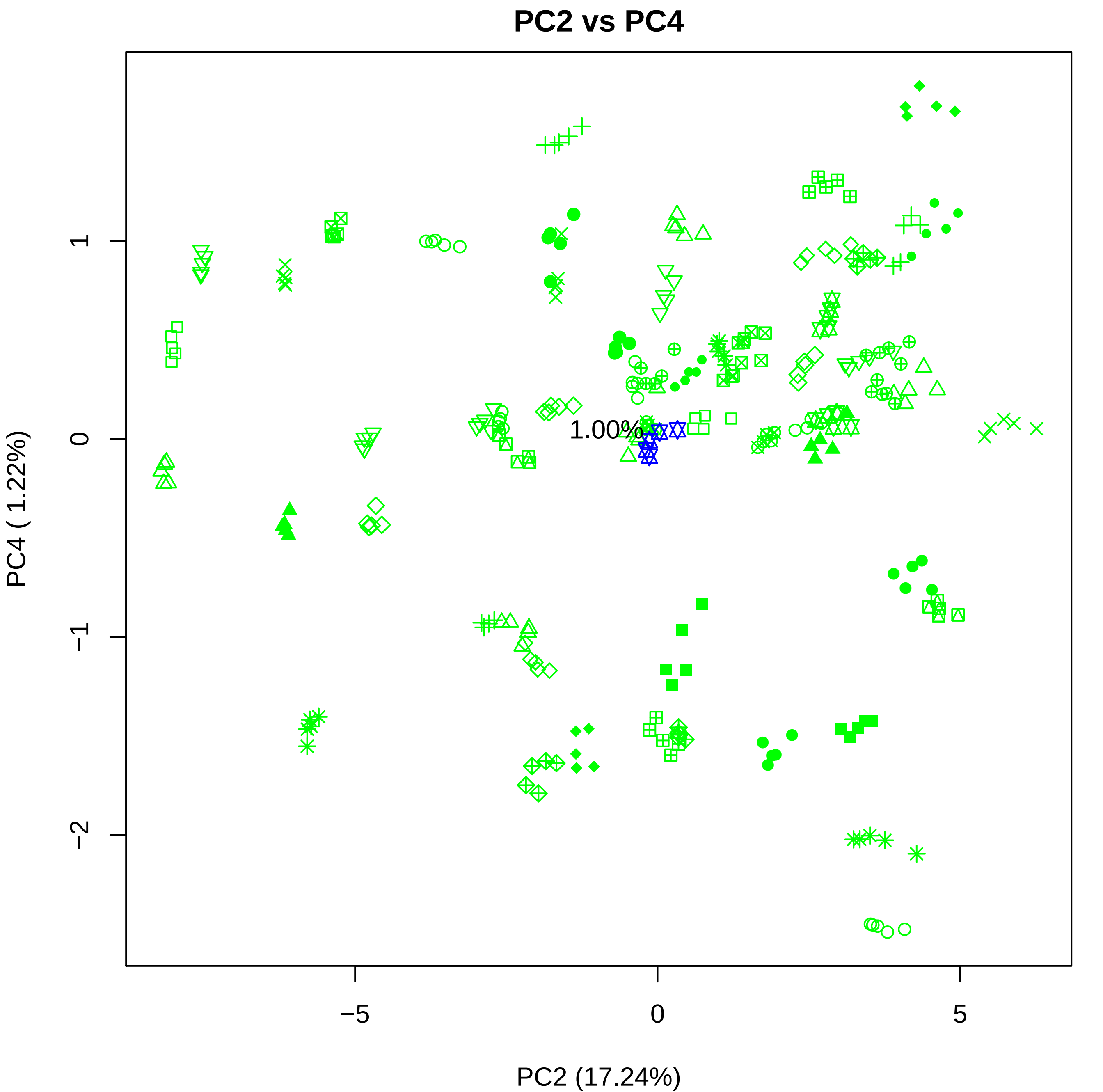

PC2 vs PC4
1.00%
1
0
−1
−2
−5
0
5
PC4 ( 1.22%)
PC2 (17.24%)

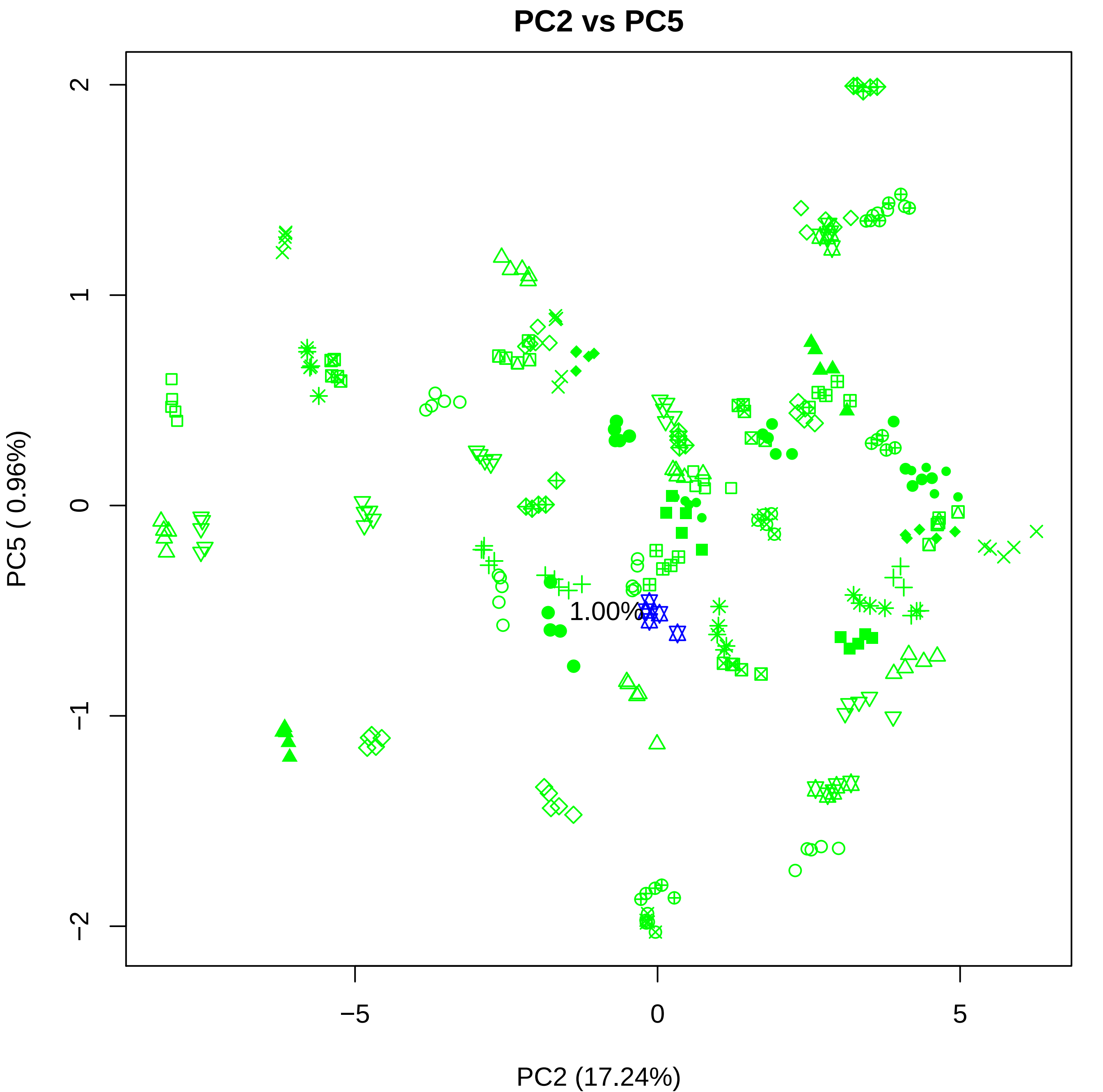
PC2 vs PC5
PC5 ( 0.96%)
PC2 (17.24%)
1.00%
2
1
0
−1
−2
−5
0
5

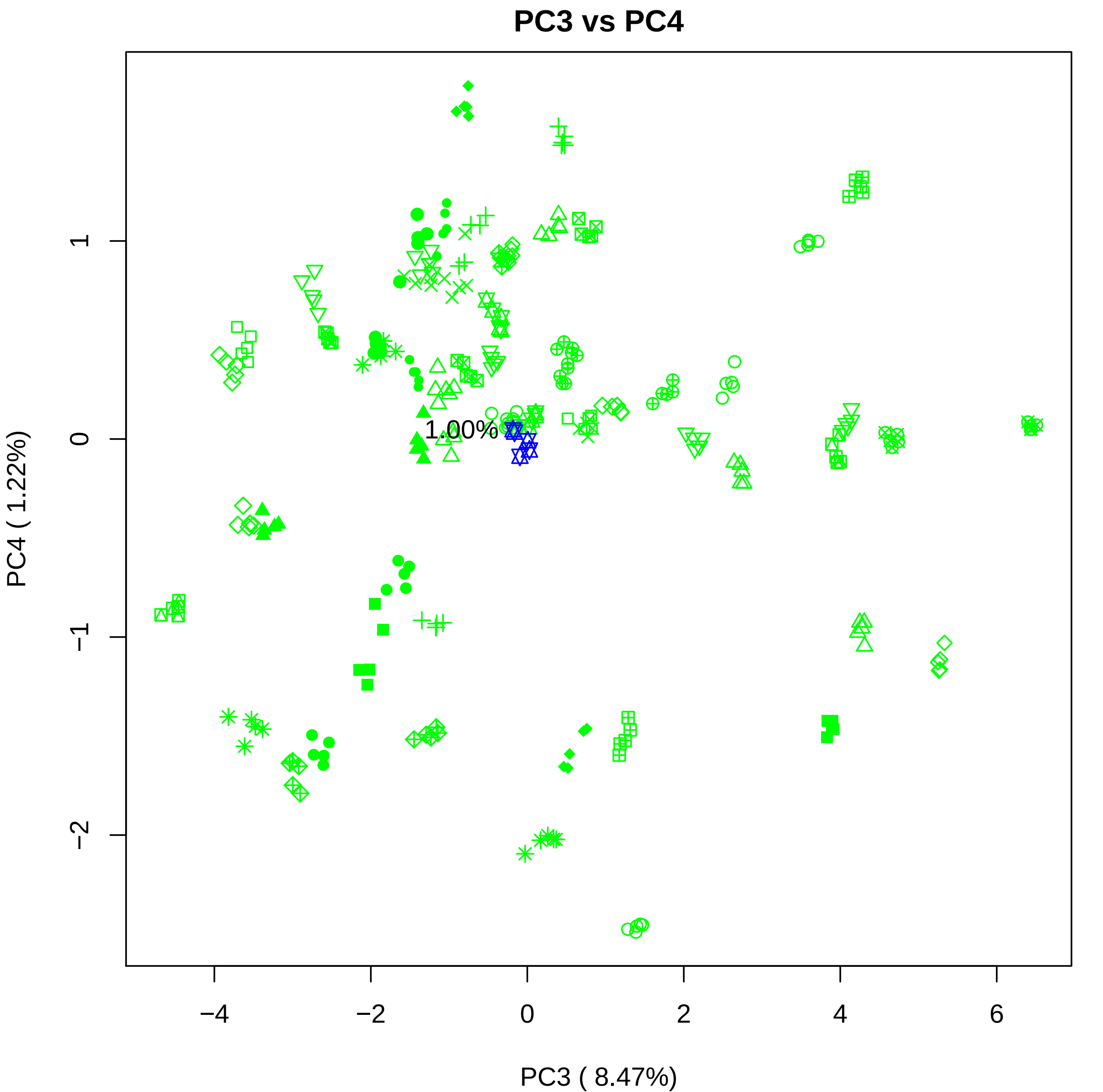
PC3 vs PC4
1.00%
PC4 ( 1.22%)
PC3 ( 8.47%)
−4
−2
0
2
4
6
1
0
−1
−2

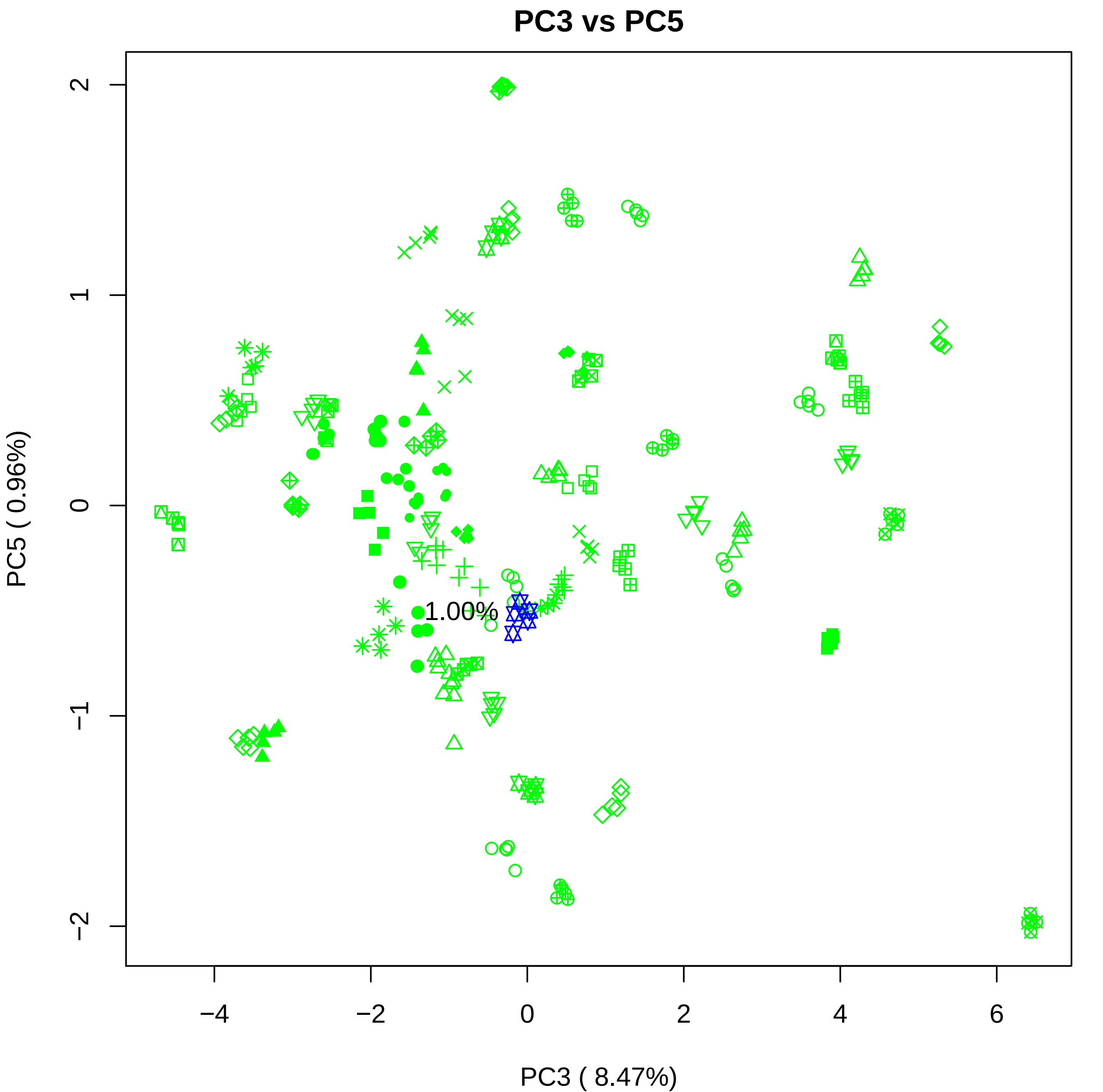
PC3 vs PC5
PC5 ( 0.96%)
PC3 ( 8.47%)
1.00%
2
1
0
−1
−2
−4
−2
0
2
4
6

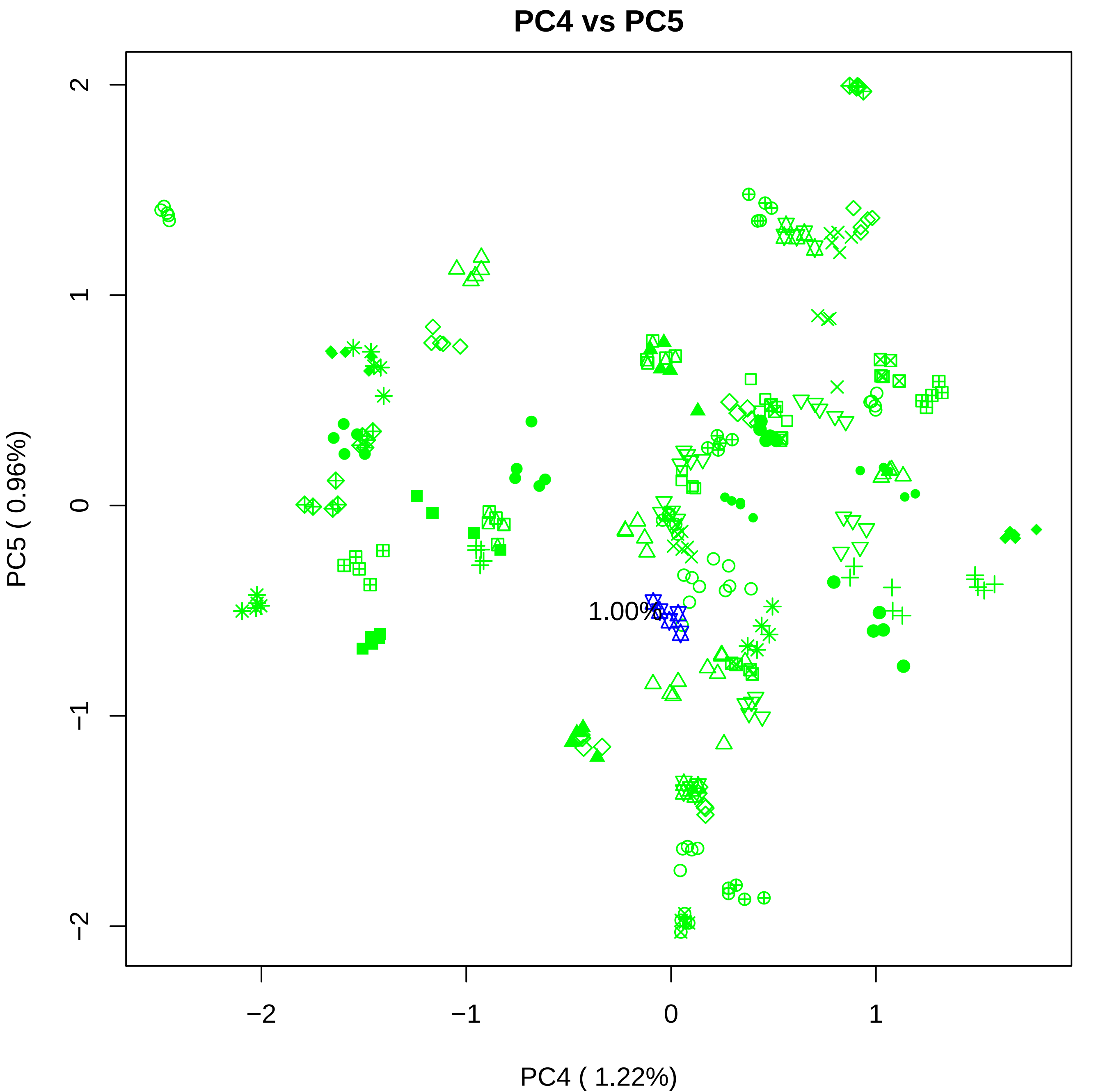
PC4 vs PC5
PC5 ( 0.96%)
PC4 ( 1.22%)
1.00%
2
1
0
−1
−2
−2
−1
0
1

### PCA of 2.5% BMH(d) vs PMH(d)

Scaled Matrix PCAed

7th order polynomial baseline correction performed

Raw spectra smoothed to 21 Neighbours, 2nd order PN, Savtisky Golay

Normalisation between 0 and 100

D2 spectra smoothed to 41 Neighbours, 2nd order PN, Savtisky Golay

Regions Analysed: 3601−2501 and 2001−700cm−1

Samples Analysed: 1 bContam's 0 BLH's 0 BMH's 0 OMH's 0 pContam's 0 PLH's 60 PMH's

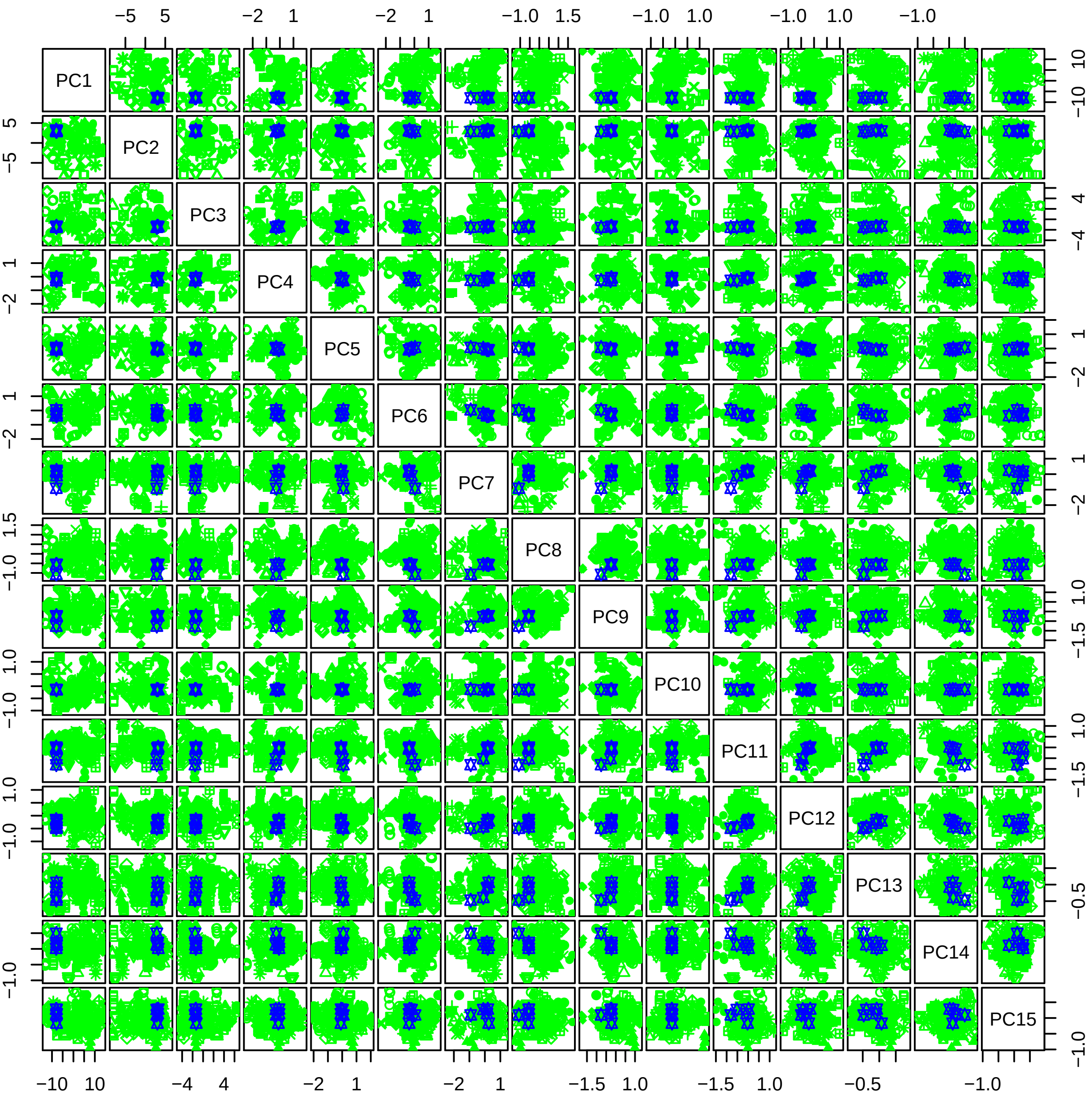
PC1
PC2
PC3
PC4
PC5
PC6
PC7
PC8
PC9
PC10
PC11
PC12
PC13
PC14
PC15

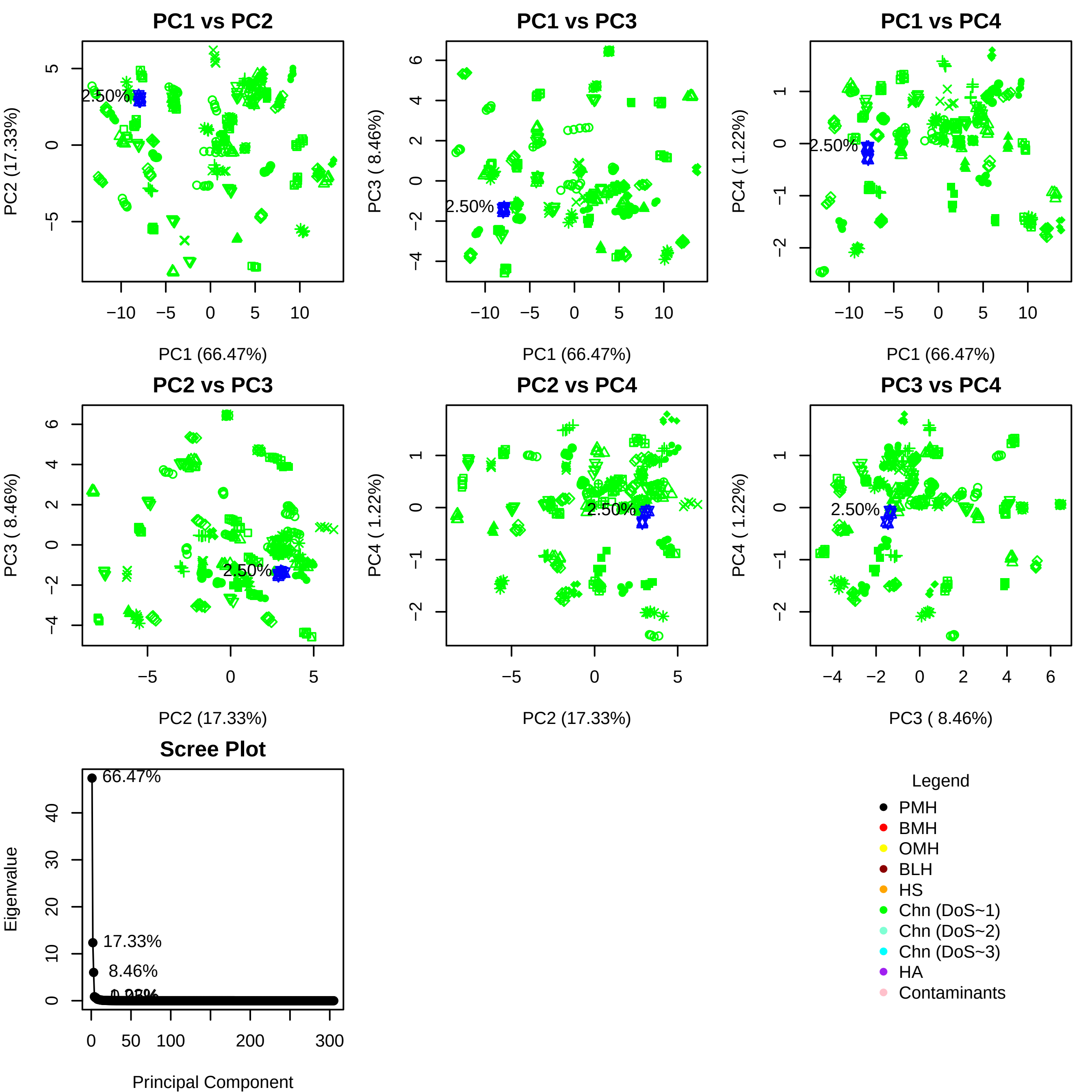

PC1 vs PC2
PC1 vs PC3
PC1 vs PC4
PC2 vs PC3
PC2 vs PC4
PC3 vs PC4
PC1 (66.47%)
PC2 (17.33%)
PC3 ( 8.46%)
PC4 ( 1.22%)
2.50%
Scree Plot
66.47%
17.33%
8.46%
Eigenvalue
Principal Component
Legend
PMH
BMH
OMH
BLH
HS
Chn (DoS~1)
Chn (DoS~2)
Chn (DoS~3)
HA
Contaminants

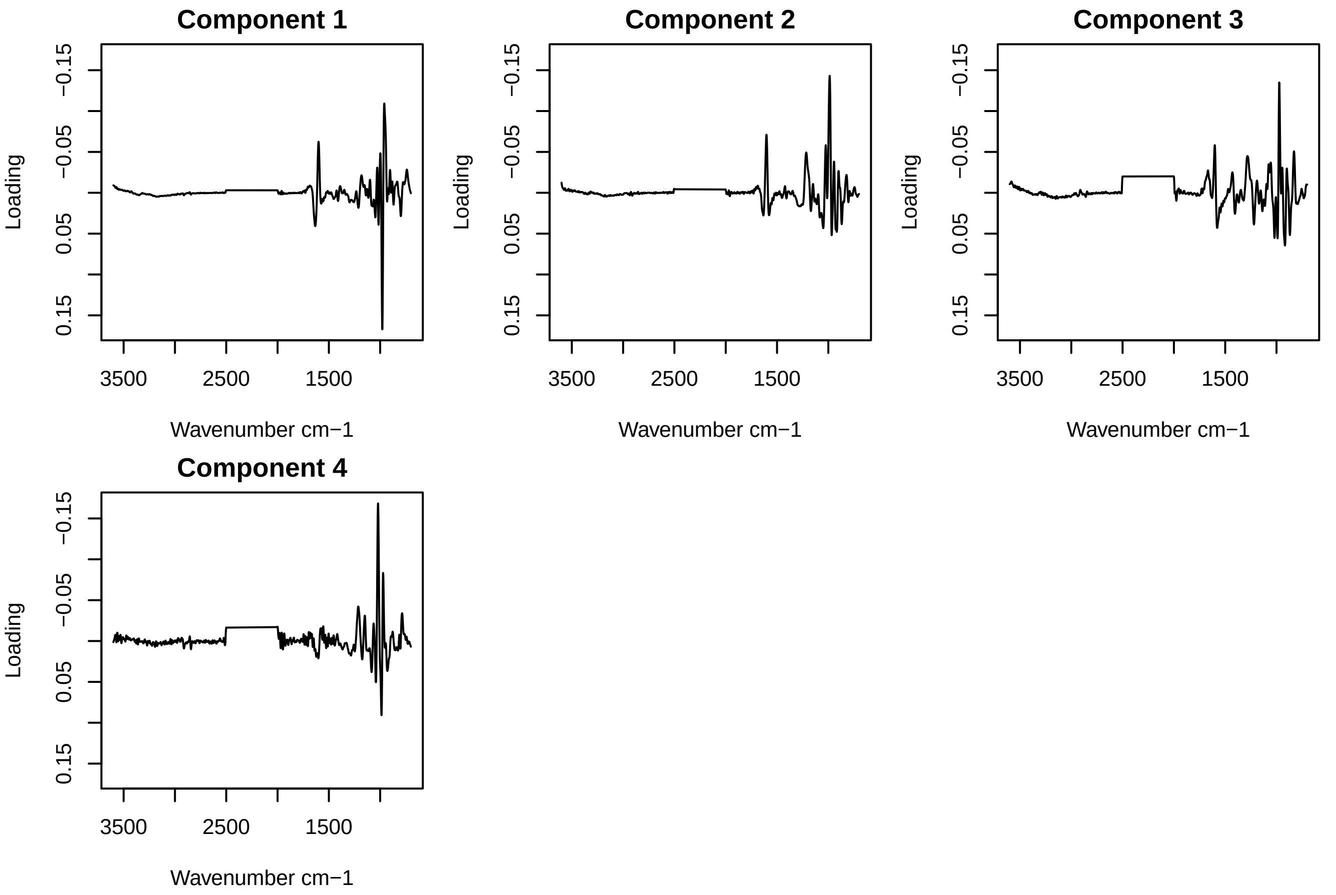
Component 1
Component 2
Component 3
Component 4
Loading
−0.15
−0.05
0.05
0.15
3500
2500
1500
Wavenumber cm−1

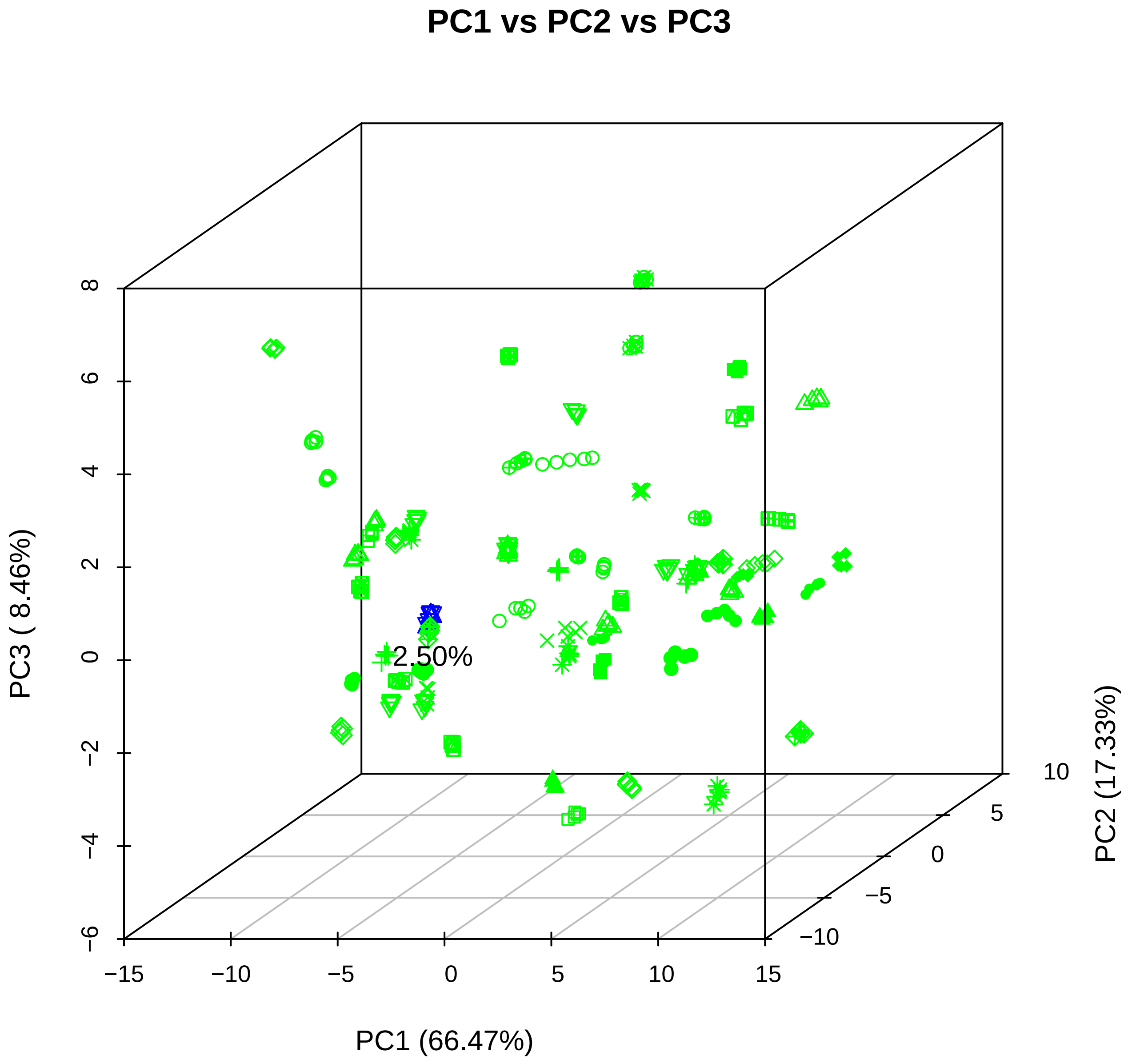
PC1 vs PC2 vs PC3
PC1 (66.47%)
PC2 (17.33%)
PC3 ( 8.46%)
2.50%
−15
−10
−5
0
5
10
15
−10
−5
0
5
10
−6
−4
−2
0
2
4
6
8

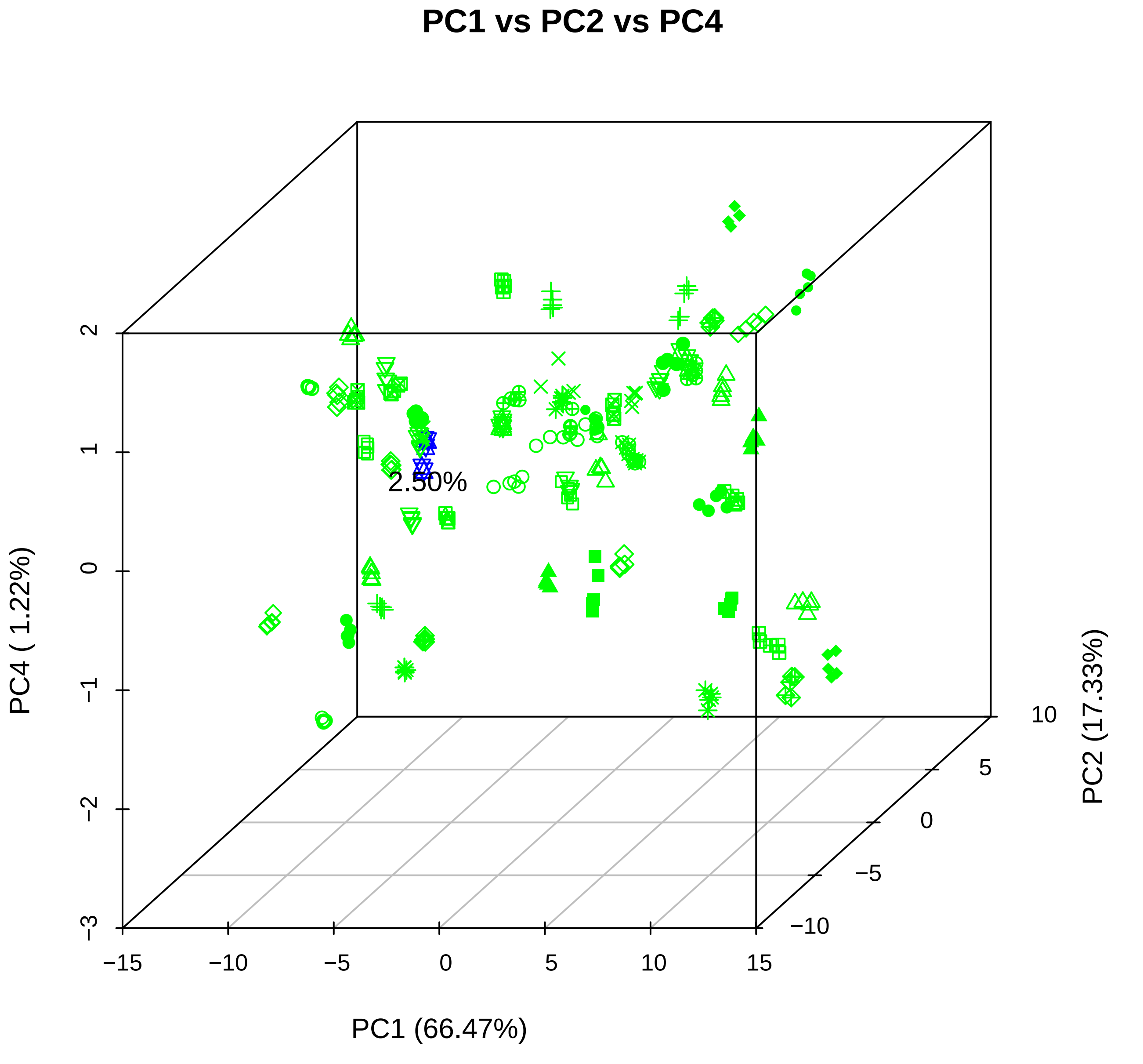

PC1 vs PC2 vs PC4
2.50%
PC1 (66.47%)
PC2 (17.33%)
PC4 ( 1.22%)
−15
−10
−5
0
5
10
15
−3
−2
−1
0
1
2

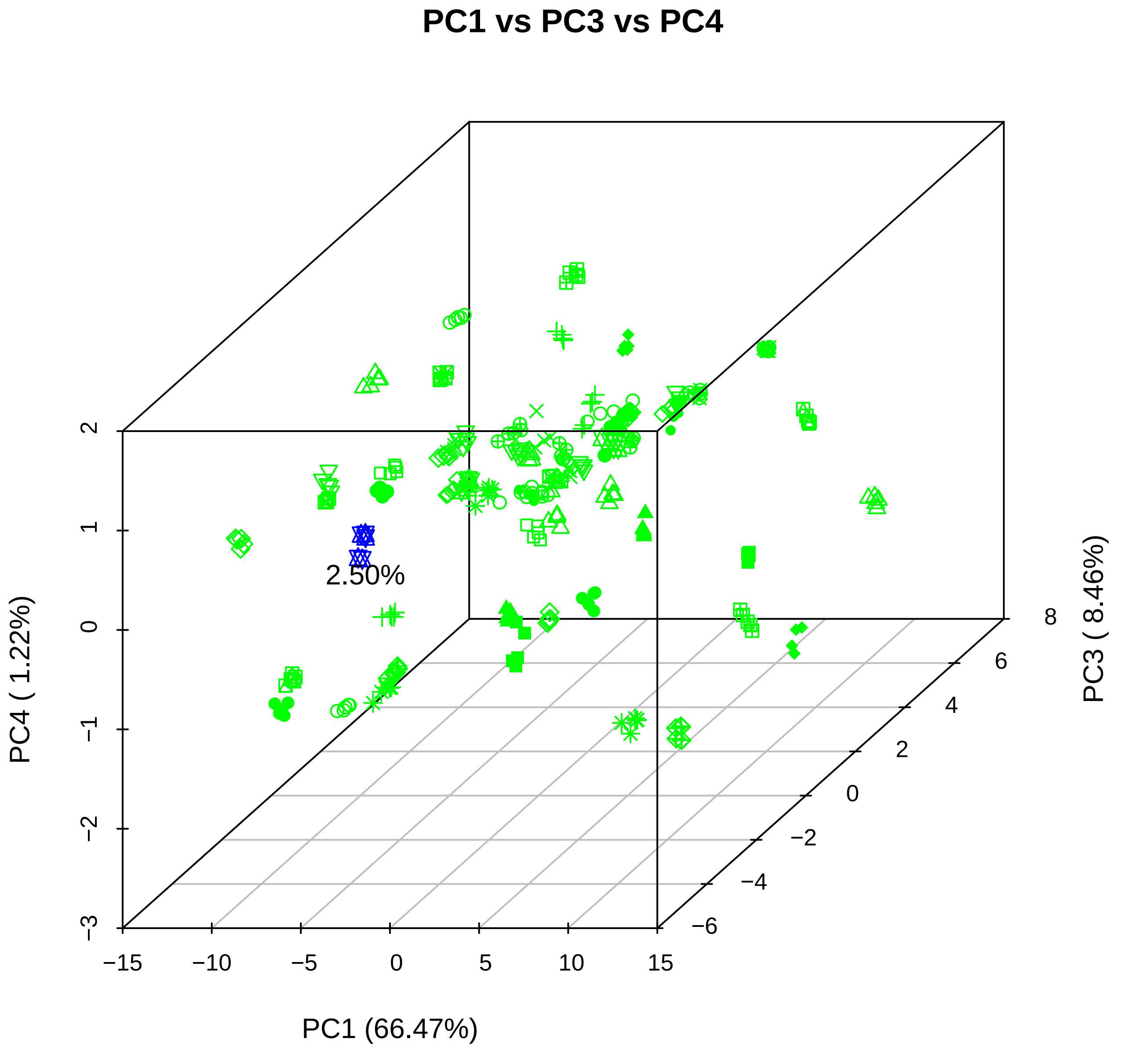

PC1 vs PC3 vs PC4
2.50%
PC1 (66.47%)
PC4 ( 1.22%)
PC3 ( 8.46%)
−15
−10
−5
0
5
10
15
−3
−2
−1
0
1
2
−6
−4
−2
0
2
4
6
8

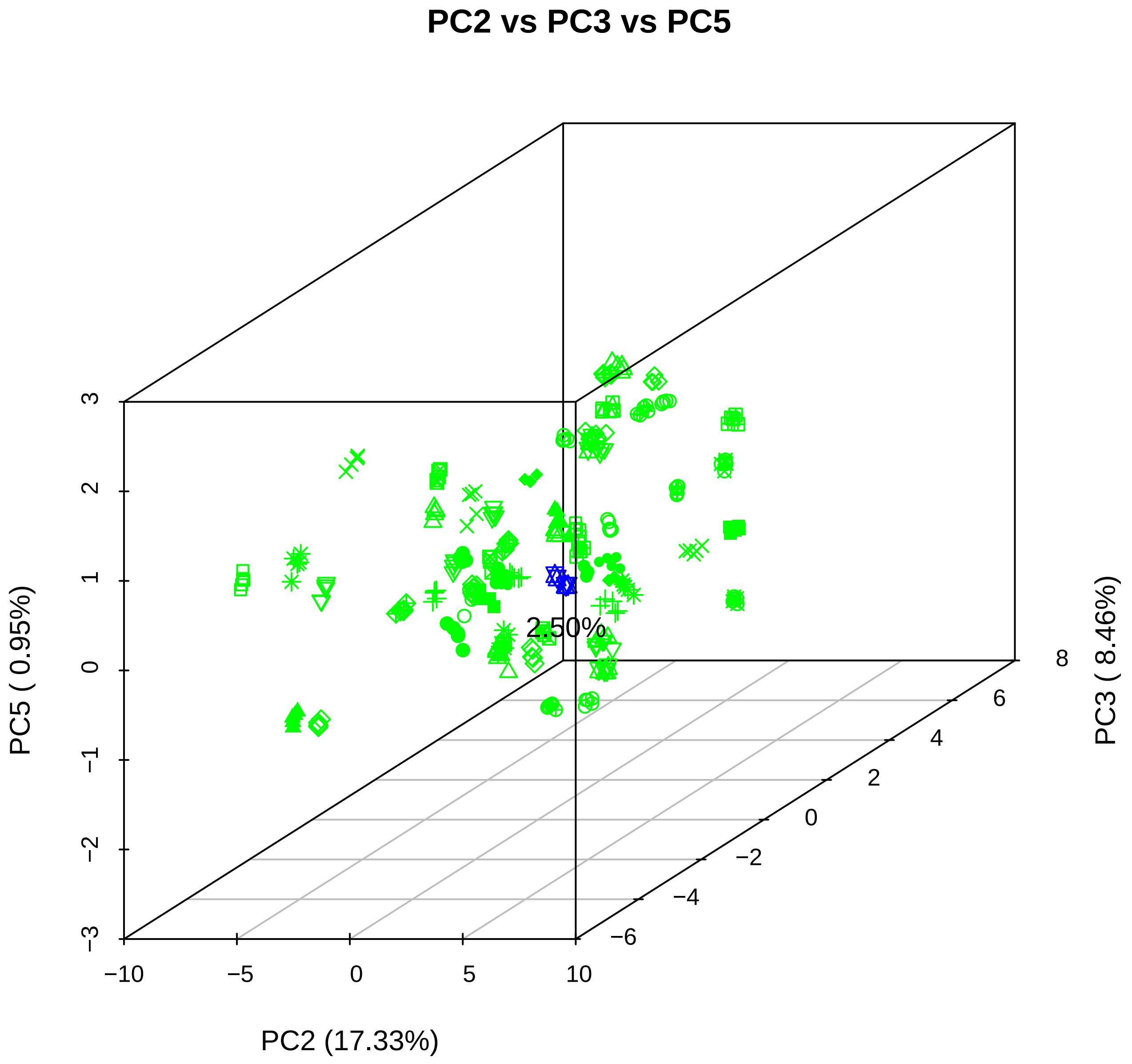

PC2 vs PC3 vs PC5
PC2 (17.33%)
PC5 ( 0.95%)
PC3 ( 8.46%)
2.50%
−10
−5
0
5
10
−3
−2
−1
0
1
2
3
−6
−4
−2
0
2
4
6
8

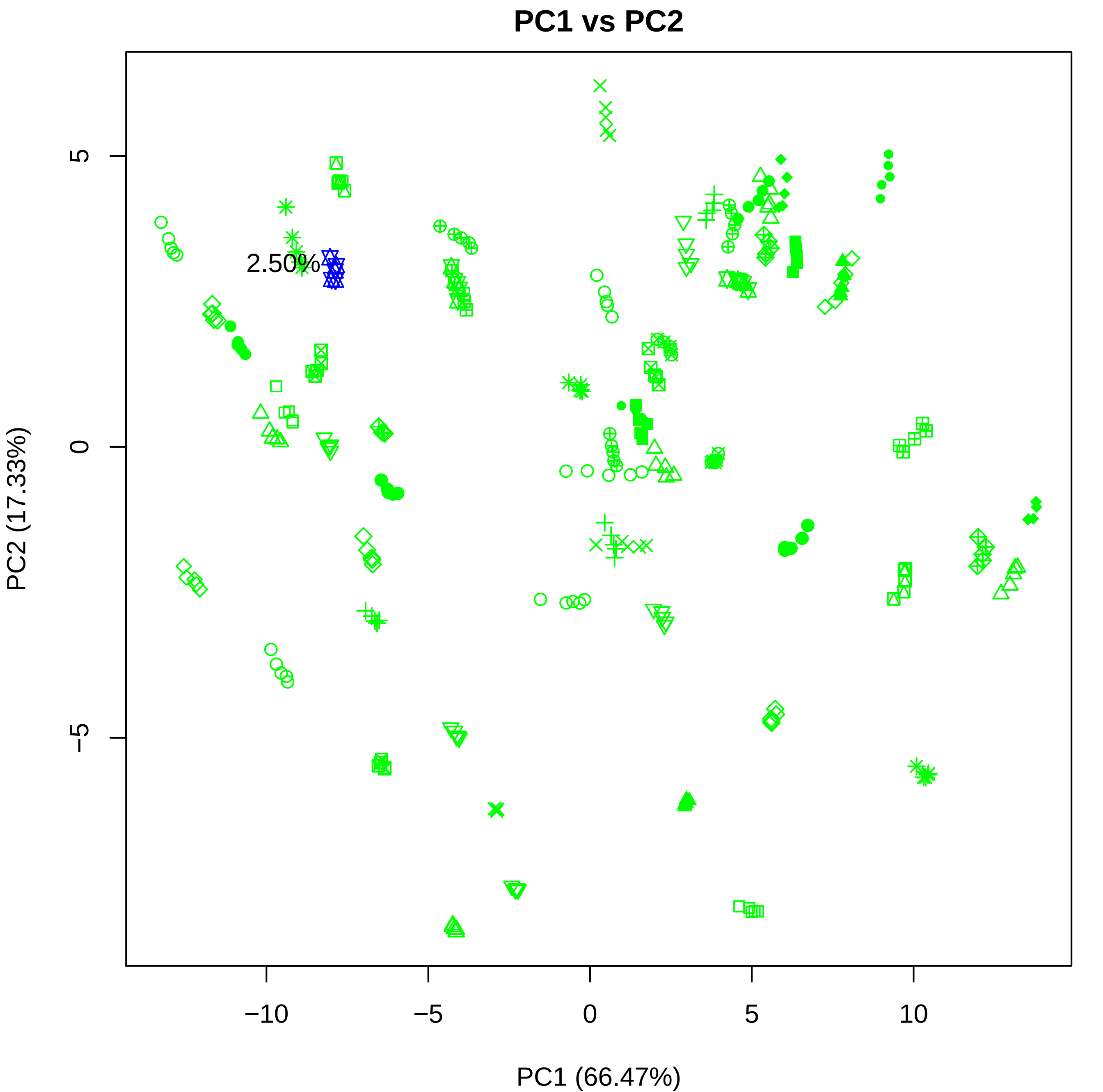

PC1 vs PC2
2.50%
PC2 (17.33%)
PC1 (66.47%)
5
0
−5
−10
−5
0
5
10

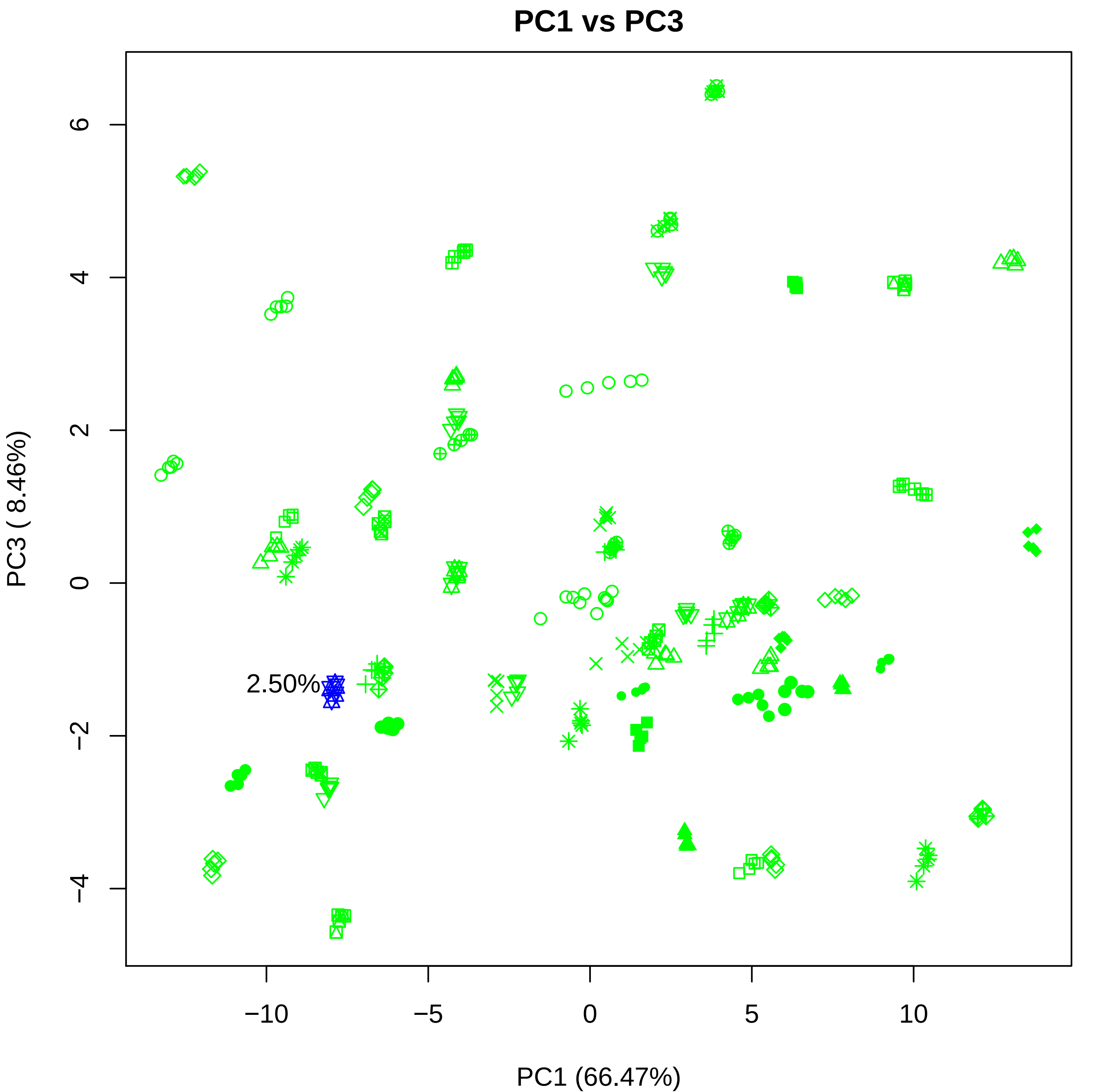
PC1 vs PC3
PC3 ( 8.46%)
PC1 (66.47%)
2.50%
6
4
2
0
−2
−4
−10
−5
0
5
10

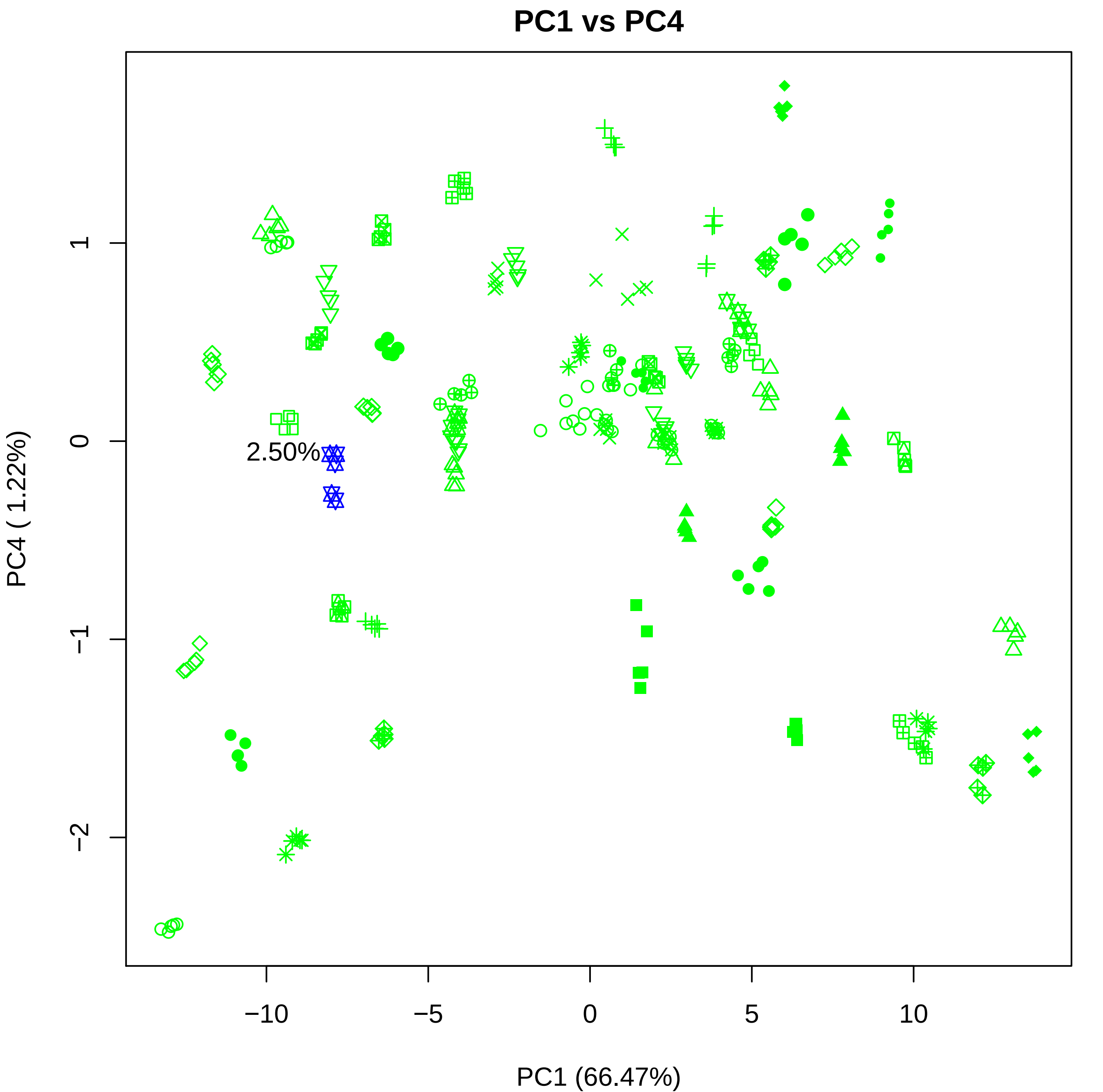
PC1 vs PC4
PC4 ( 1.22%)
PC1 (66.47%)
2.50%
1
0
−1
−2
−10
−5
0
5
10

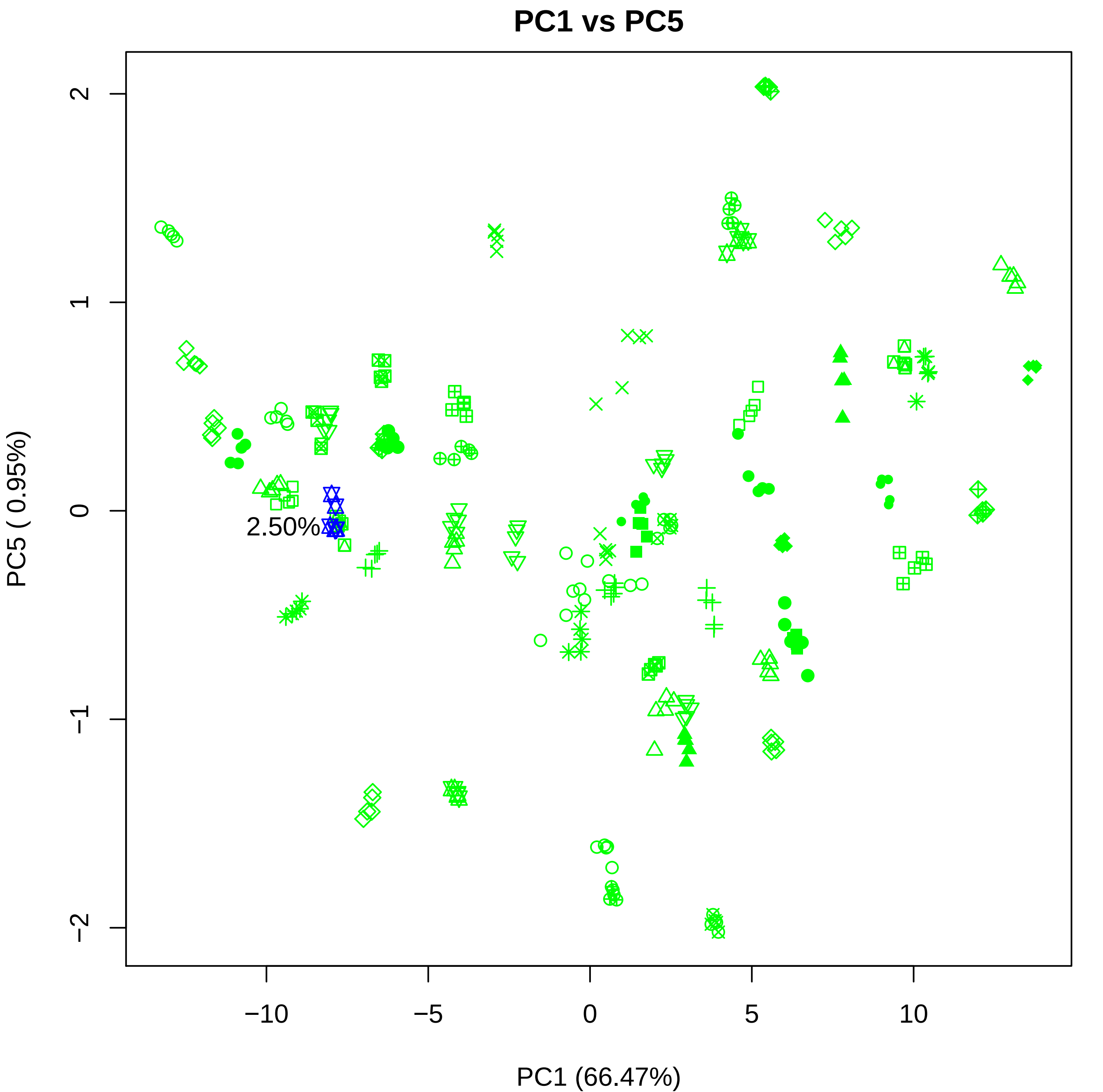
PC1 vs PC5
PC1 (66.47%)
PC5 ( 0.95%)
2.50%
−10
−5
0
5
10
−2
−1
0
1
2

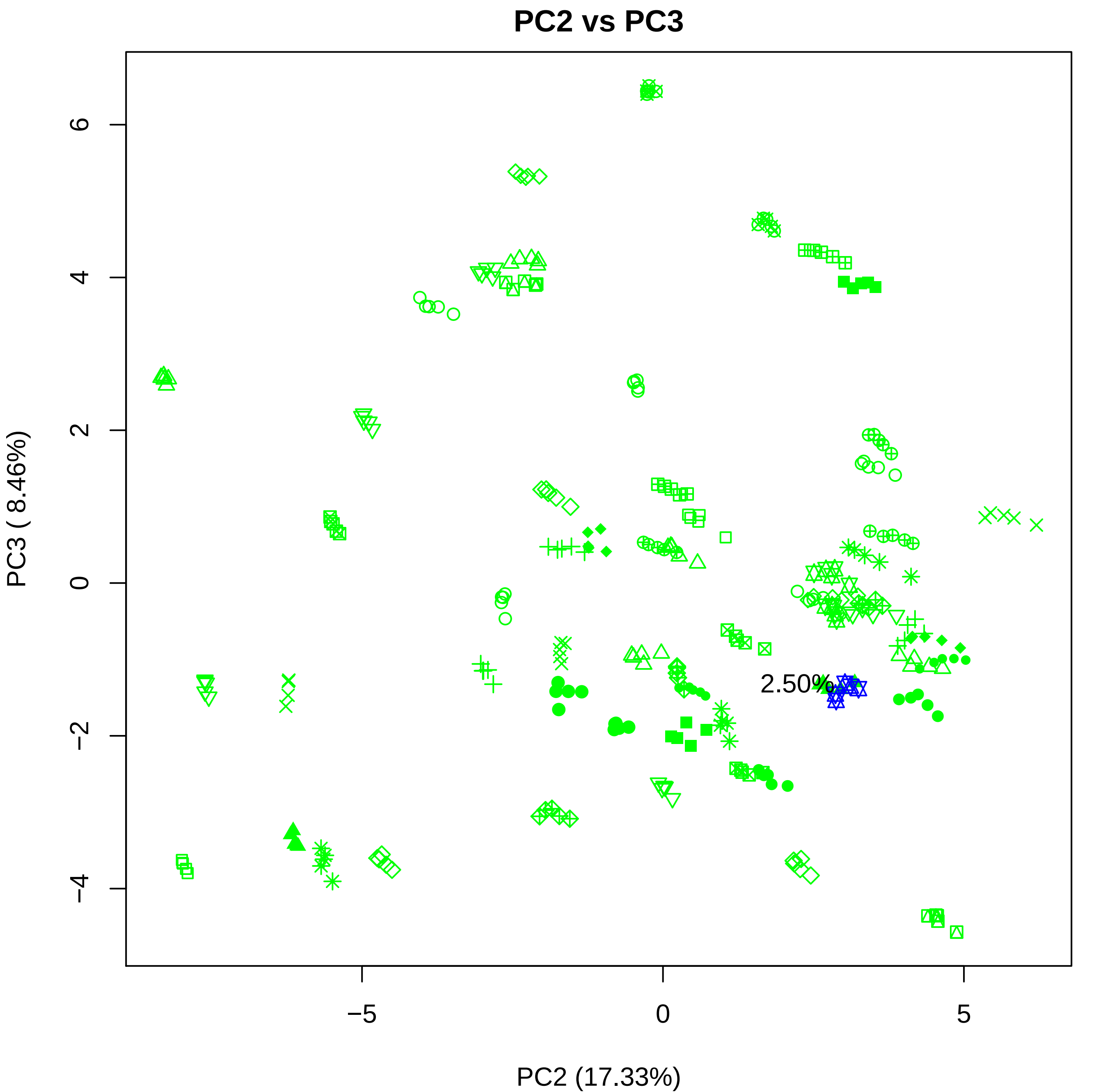

PC2 vs PC3
PC3 ( 8.46%)
PC2 (17.33%)
2.50%
6
4
2
0
−2
−4
−5
0
5

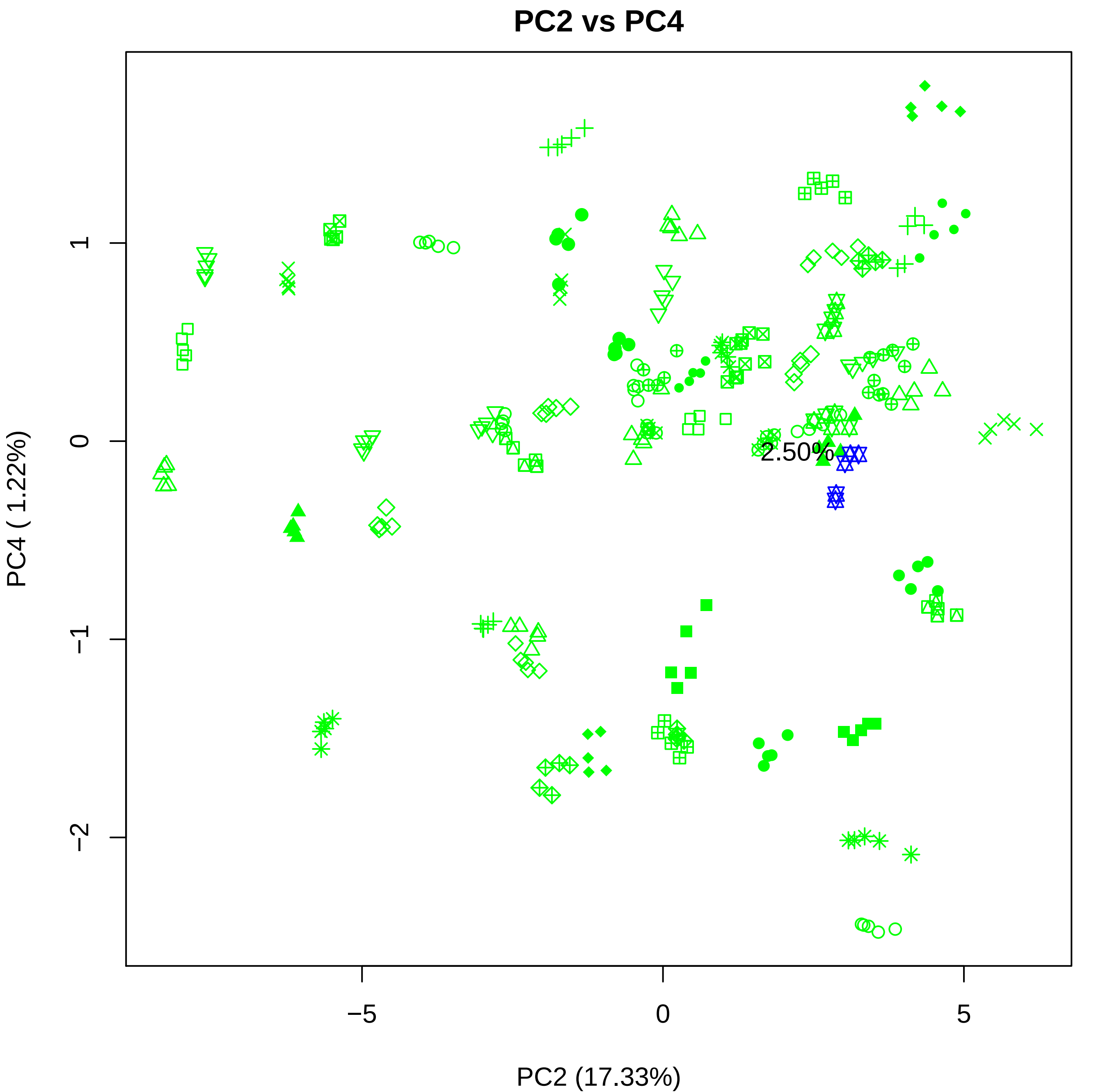

PC2 vs PC4
2.50%
1
0
−1
−2
PC4 ( 1.22%)
−5
0
5
PC2 (17.33%)

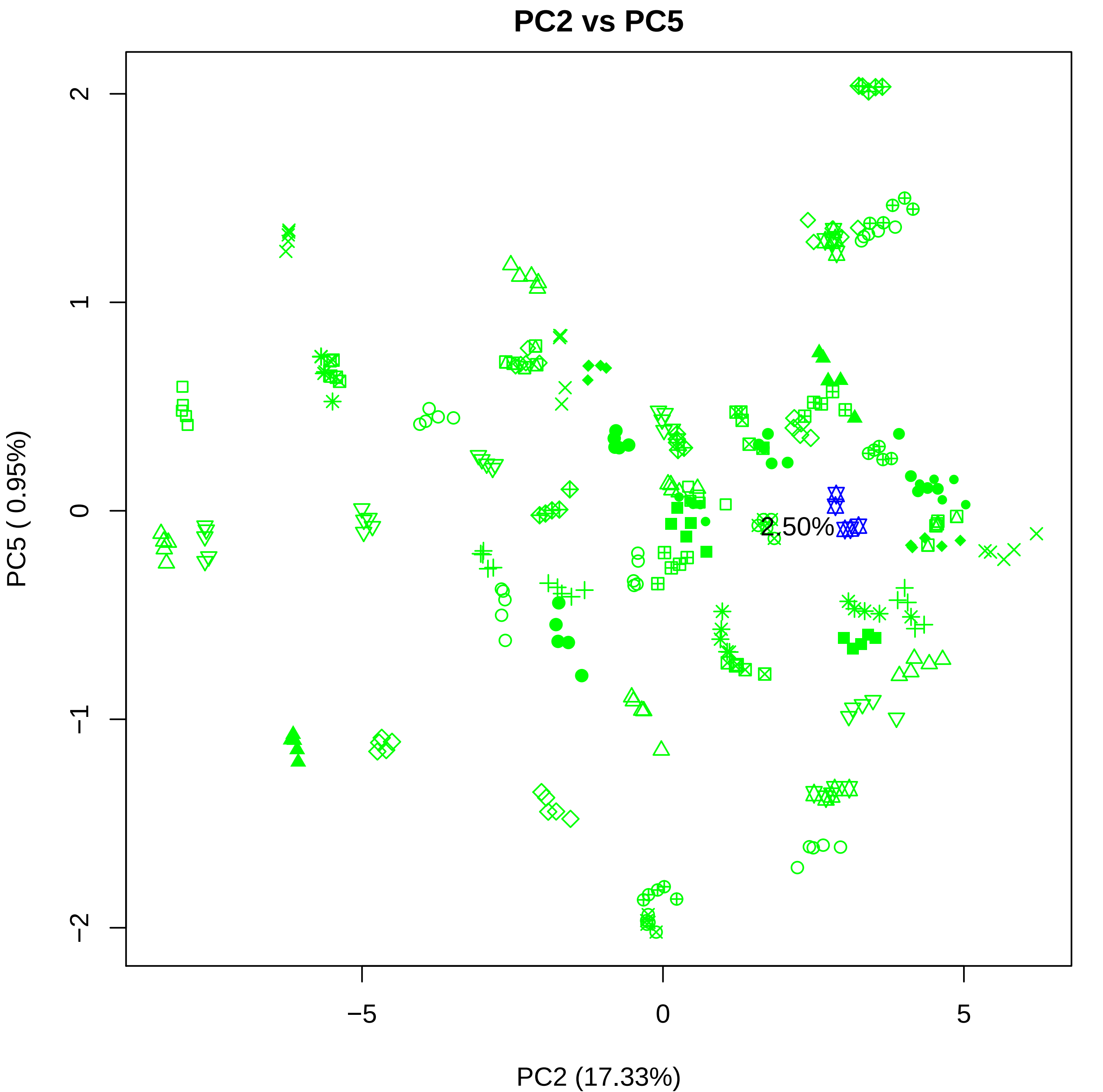
PC2 vs PC5
PC5 ( 0.95%)
PC2 (17.33%)
2.50%
2
1
0
−1
−2
−5
0
5

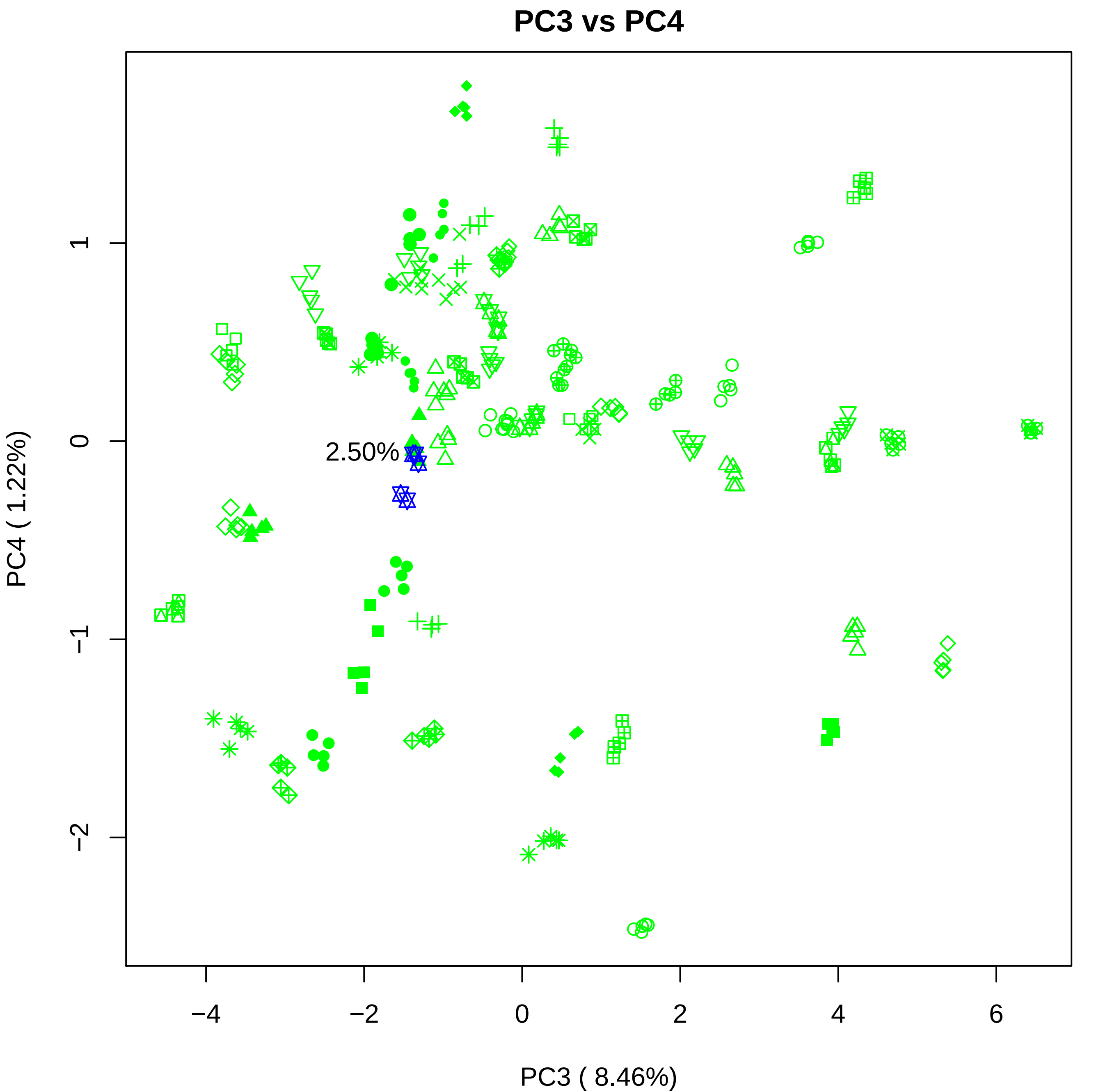
PC3 vs PC4
PC4 ( 1.22%)
PC3 ( 8.46%)
2.50%
1
0
−1
−2
−4
−2
0
2
4
6

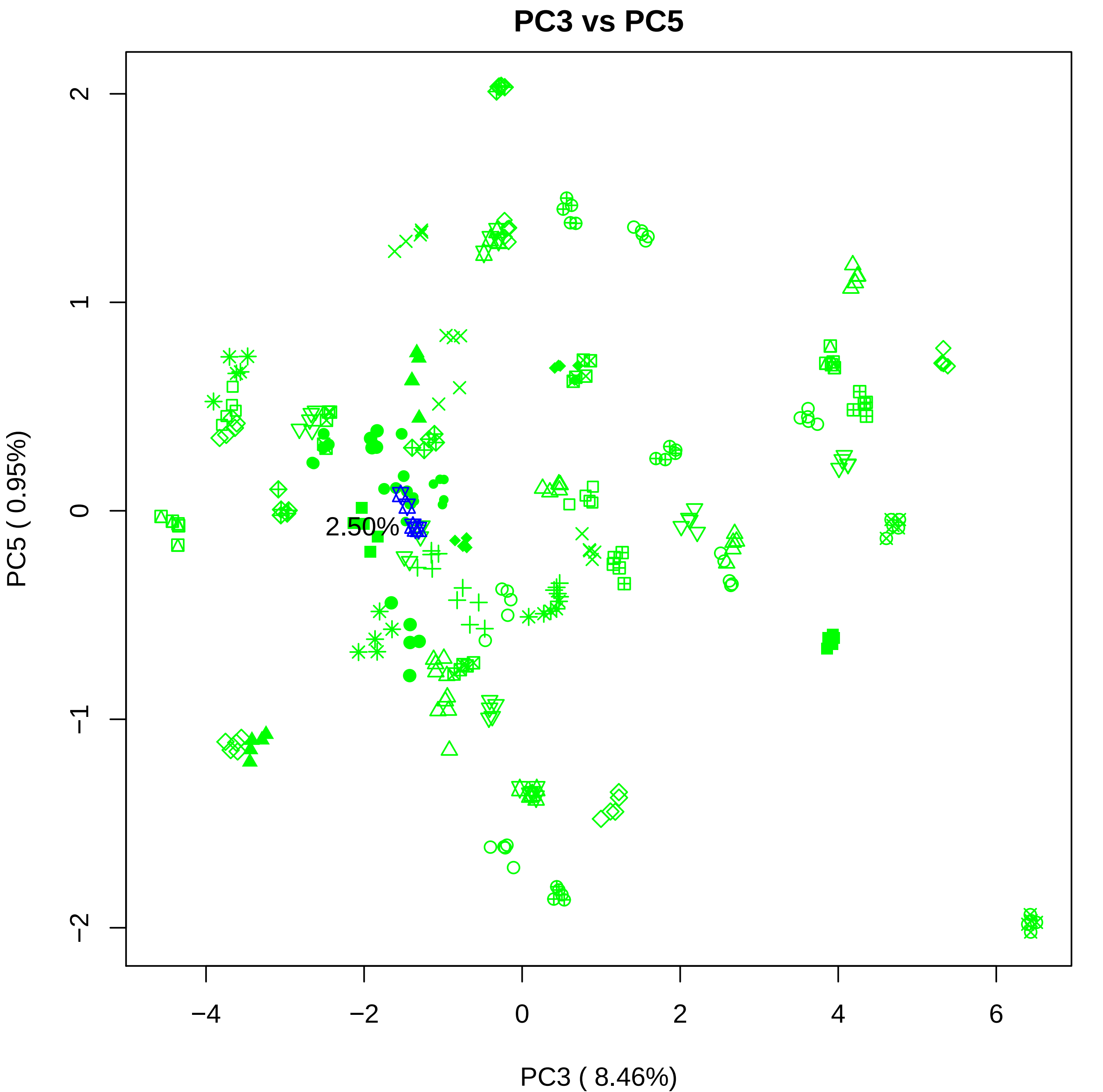
PC3 vs PC5
PC5 ( 0.95%)
PC3 ( 8.46%)
2.50%
2
1
0
−1
−2
−4
−2
0
2
4
6

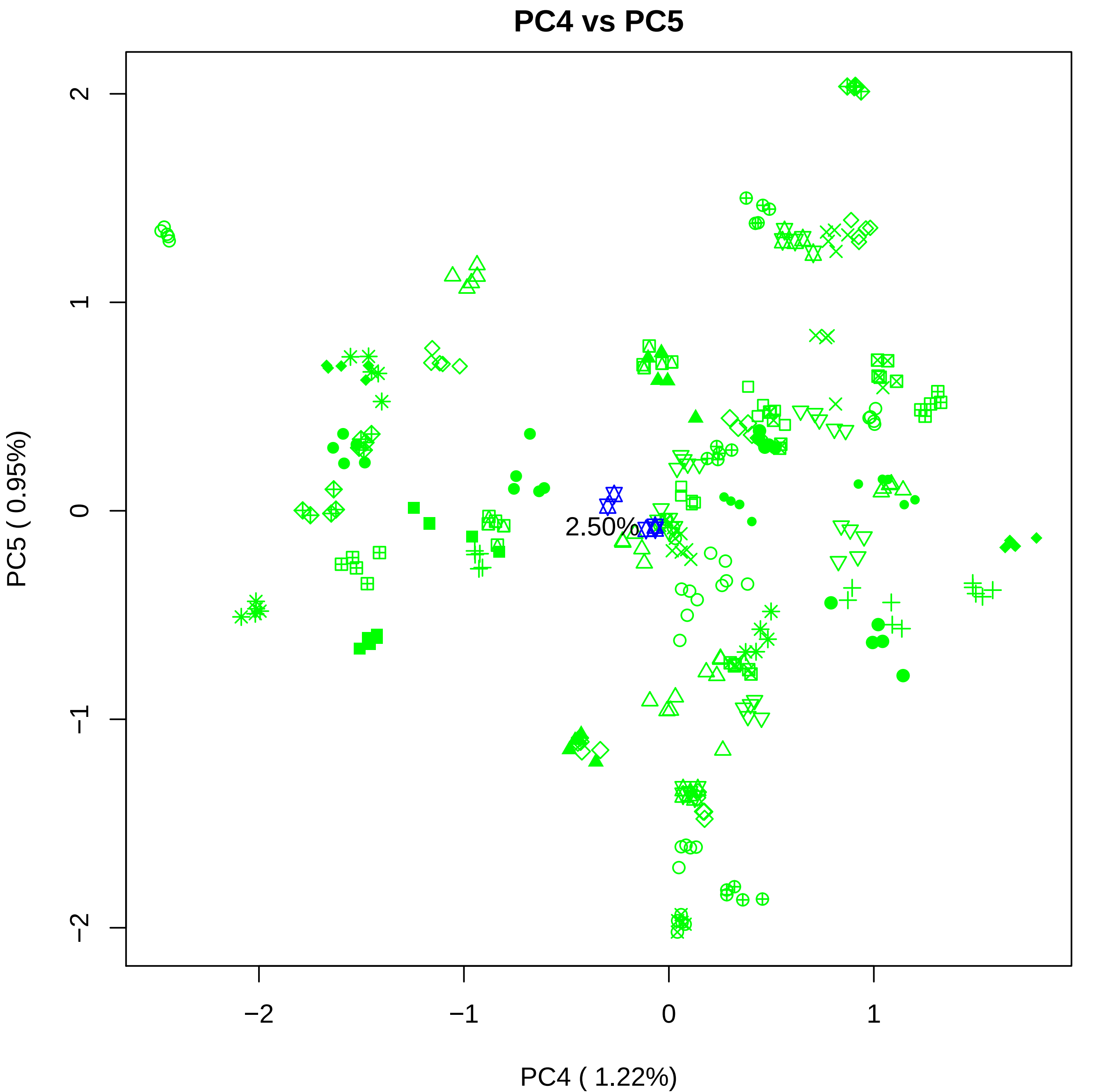

PC4 vs PC5
2.50%
2
1
0
−1
−2
−2
−1
0
1
PC4 ( 1.22%)
PC5 ( 0.95%)

### PCA of 5% BMH(d) vs PMH(d)

Scaled Matrix PCAed

7th order polynomial baseline correction performed

Raw spectra smoothed to 21 Neighbours, 2nd order PN, Savtisky Golay

Normalisation between 0 and 100

D2 spectra smoothed to 41 Neighbours, 2nd order PN, Savtisky Golay

Regions Analysed: 3601−2501 and 2001−700cm−1

Samples Analysed: 1 bContam's 0 BLH's 0 BMH's 0 OMH's 0 pContam's 0 PLH's 60 PMH's

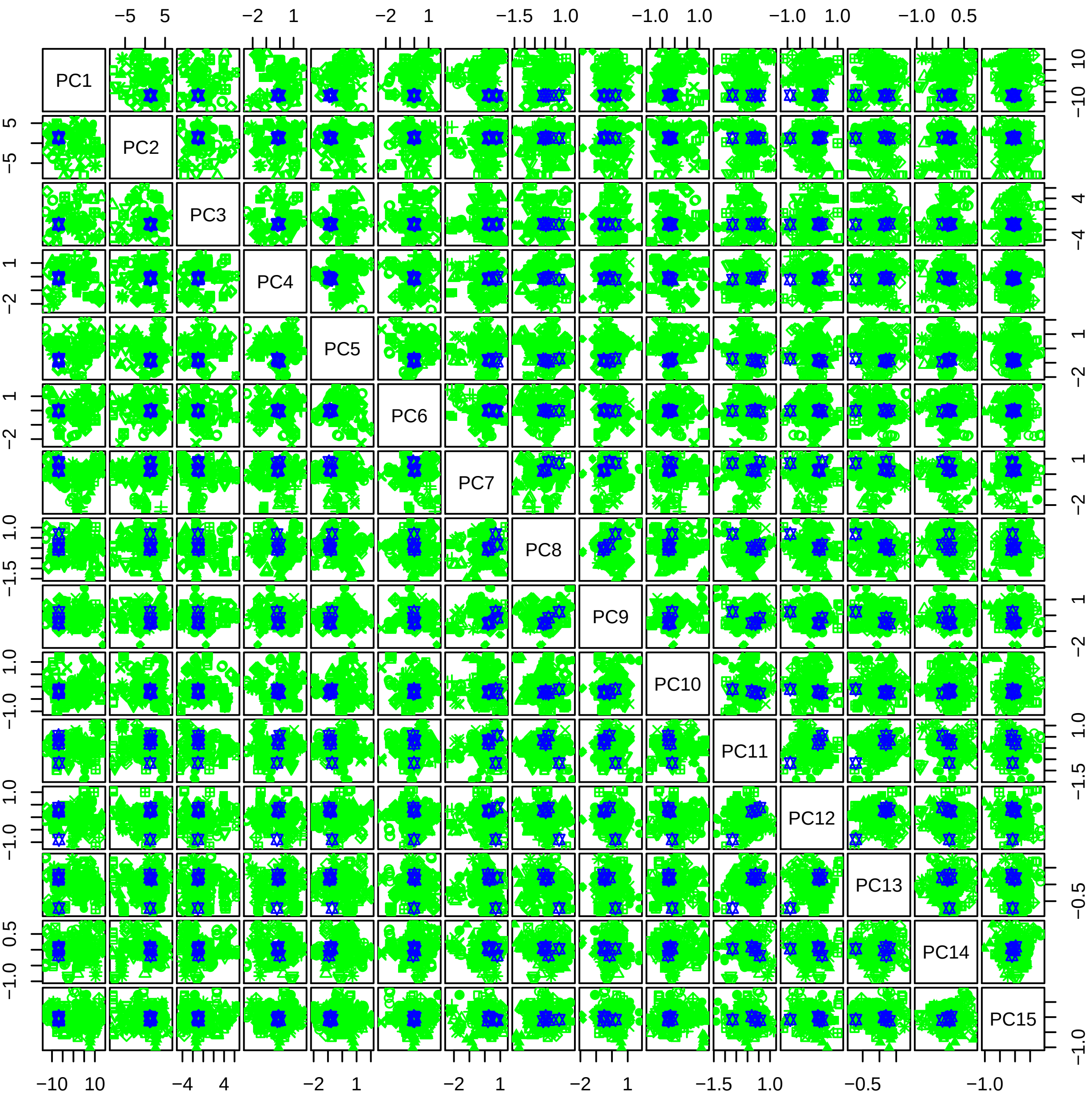

PC1
PC2
PC3
PC4
PC5
PC6
PC7
PC8
PC9
PC10
PC11
PC12
PC13
PC14
PC15

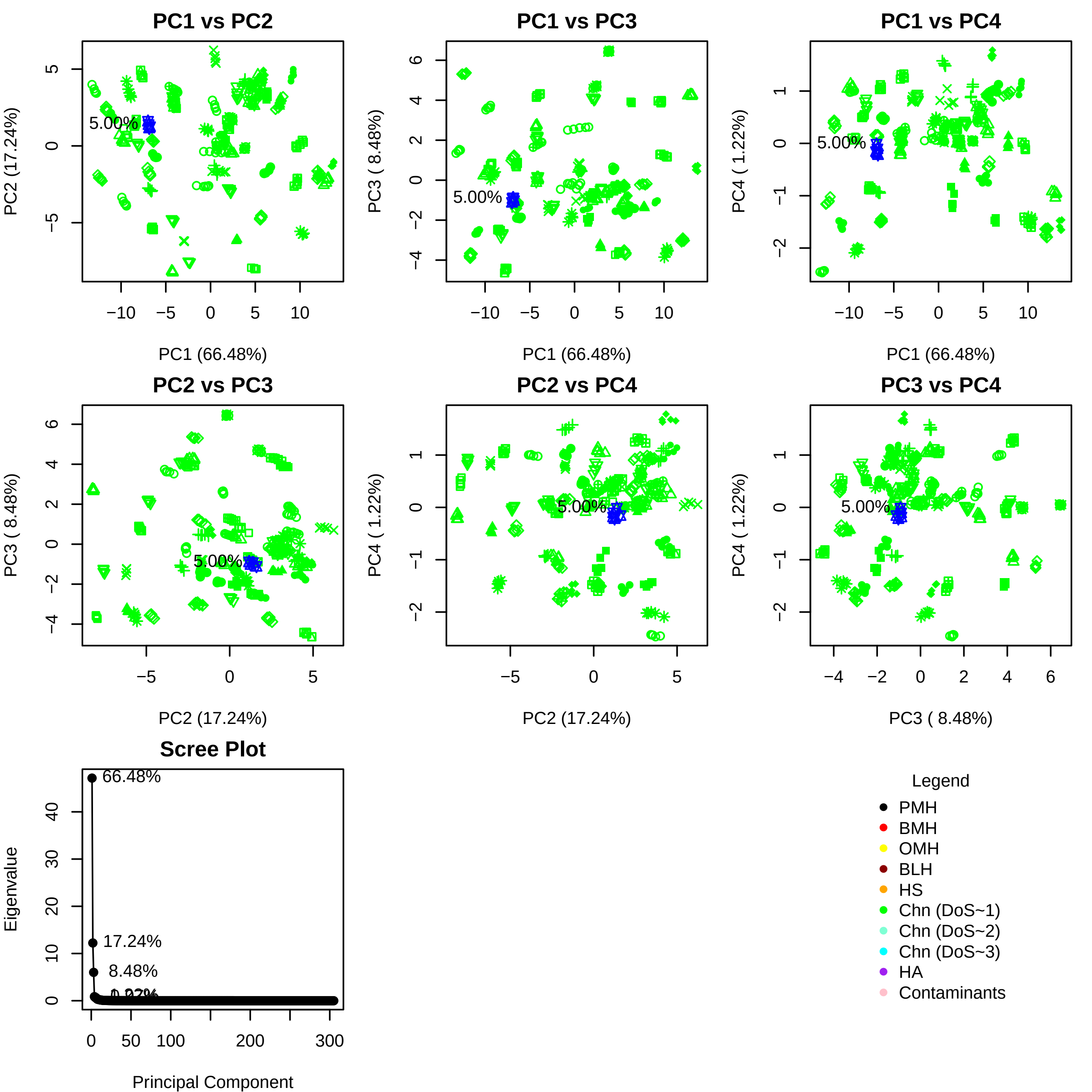
PC1 vs PC2
PC1 vs PC3
PC1 vs PC4
PC2 vs PC3
PC2 vs PC4
PC3 vs PC4
Scree Plot
PC1 (66.48%)
PC2 (17.24%)
PC3 ( 8.48%)
PC4 ( 1.22%)
5.00%
Eigenvalue
Principal Component
66.48%
17.24%
8.48%
Legend
PMH
BMH
OMH
BLH
HS
Chn (DoS~1)
Chn (DoS~2)
Chn (DoS~3)
HA
Contaminants

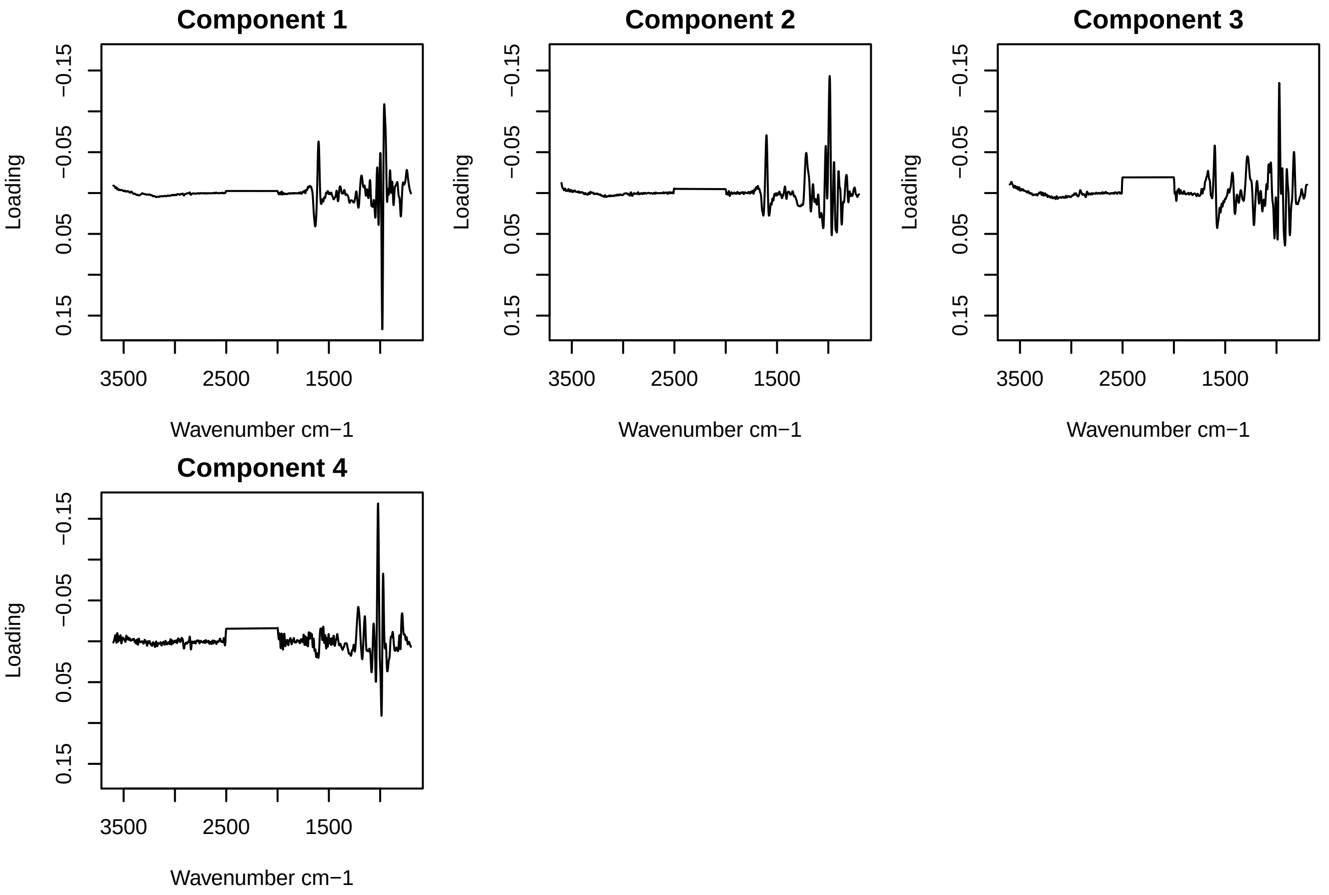
Component 1
Component 2
Component 3
Component 4
Loading
−0.15
−0.05
0.05
0.15
3500
2500
1500
Wavenumber cm−1

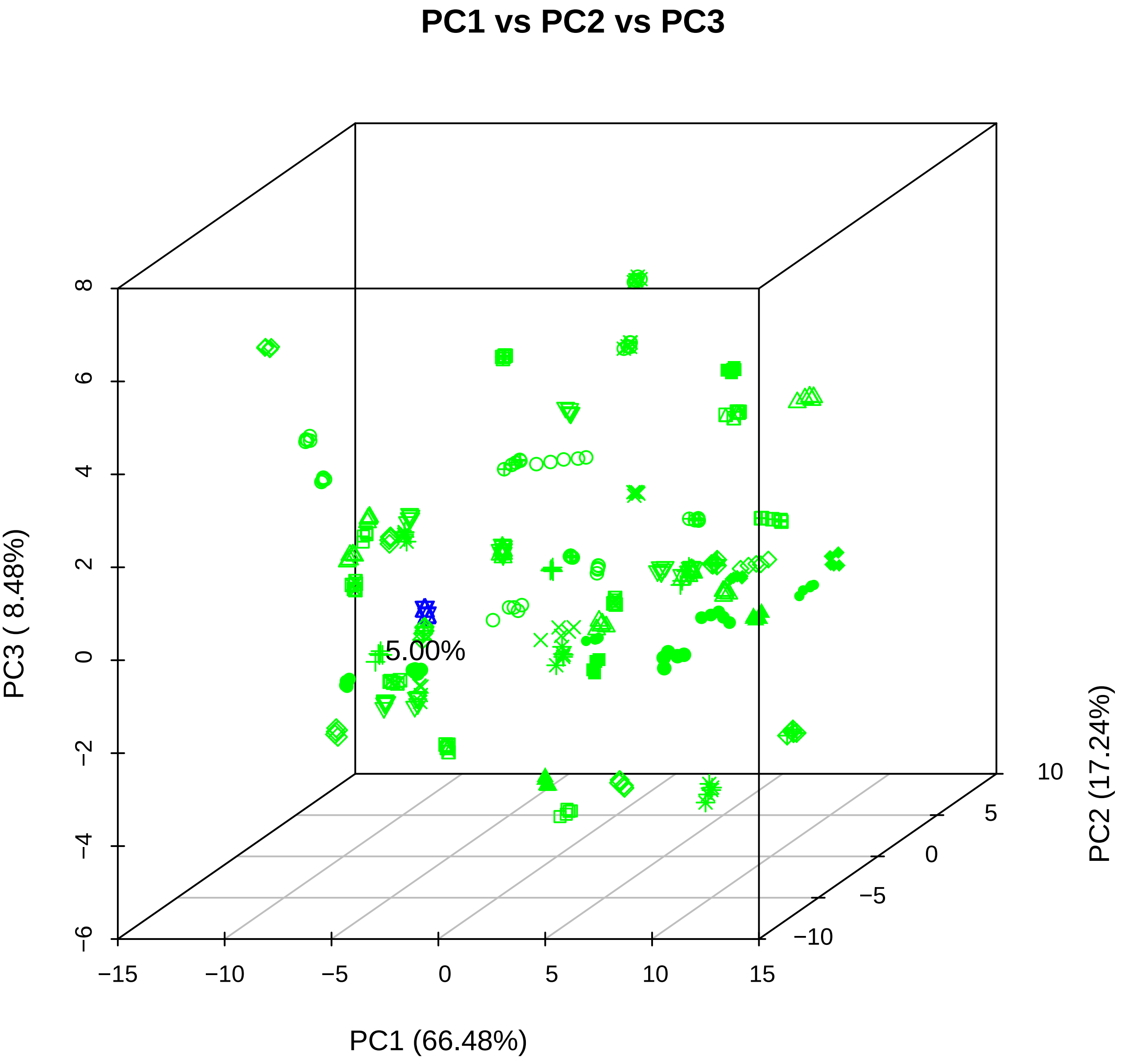

PC1 vs PC2 vs PC3
PC3 ( 8.48%)
PC1 (66.48%)
PC2 (17.24%)
5.00%
8
6
4
2
0
−2
−4
−6
−15
−10
−5
0
5
10
15
10
5
0
−5
−10

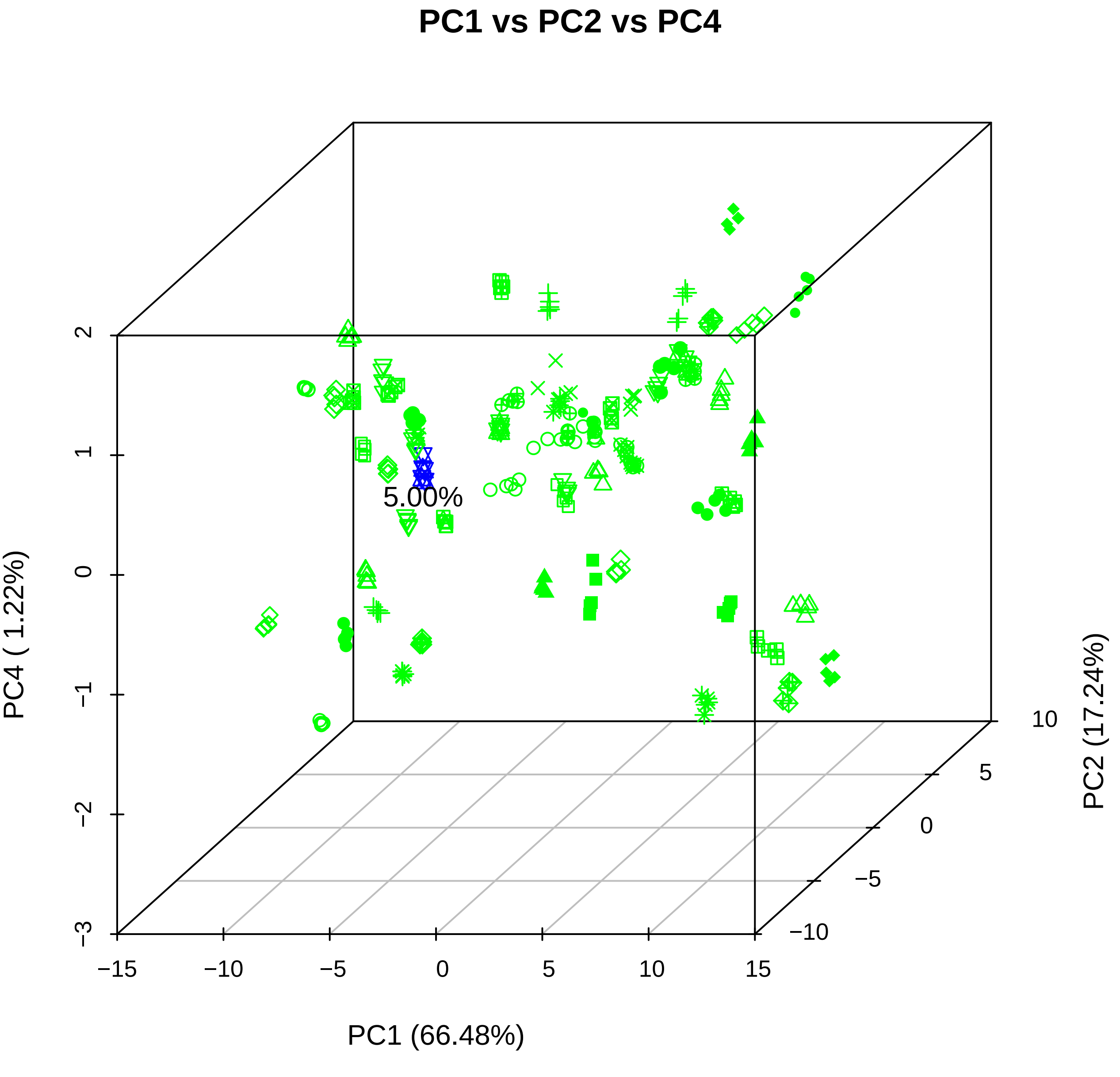
PC1 vs PC2 vs PC4
5.00%
PC1 (66.48%)
PC2 (17.24%)
PC4 ( 1.22%)
−15
−10
−5
0
5
10
15
−10
−5
0
5
10
−3
−2
−1
0
1
2

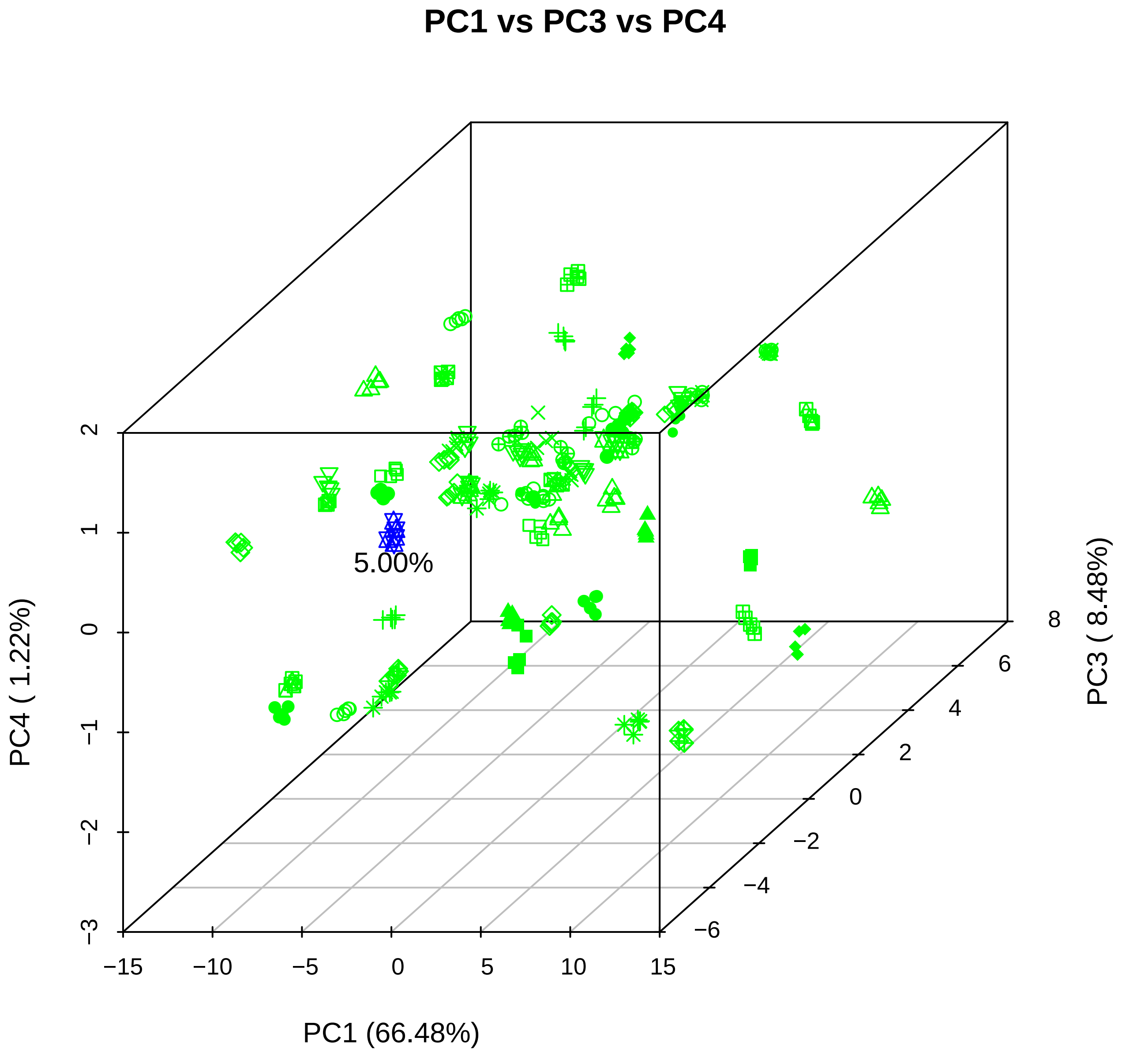
PC1 vs PC3 vs PC4
5.00%
PC1 (66.48%)
PC4 ( 1.22%)
PC3 ( 8.48%)
−15
−10
−5
0
5
10
15
−3
−2
−1
0
1
2
−6
−4
−2
0
2
4
6
8

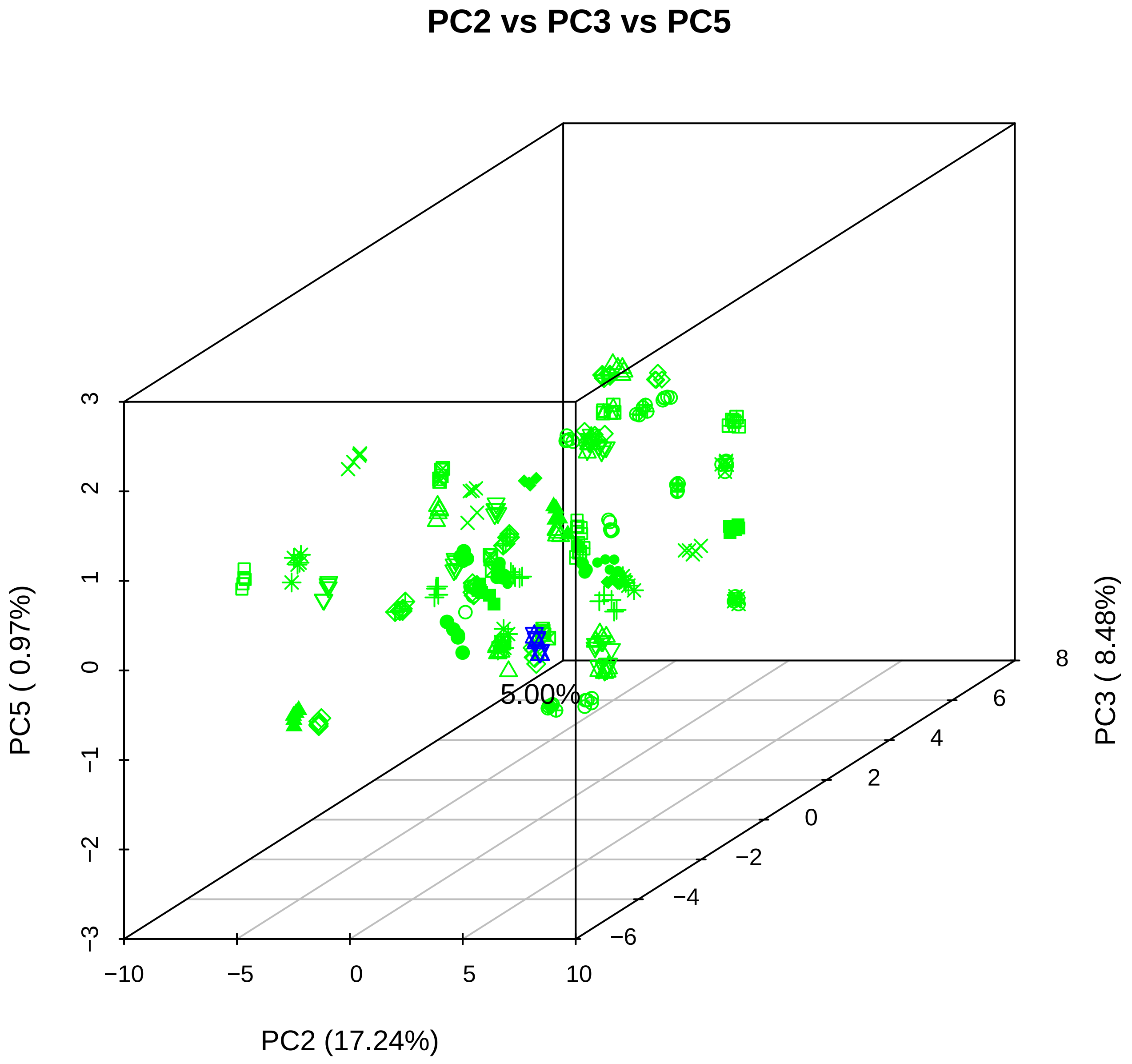

PC2 vs PC3 vs PC5
PC5 ( 0.97%)
PC2 (17.24%)
PC3 ( 8.48%)
5.00%
−10
−5
0
5
10
−3
−2
−1
0
1
2
3
−6
−4
−2
0
2
4
6
8

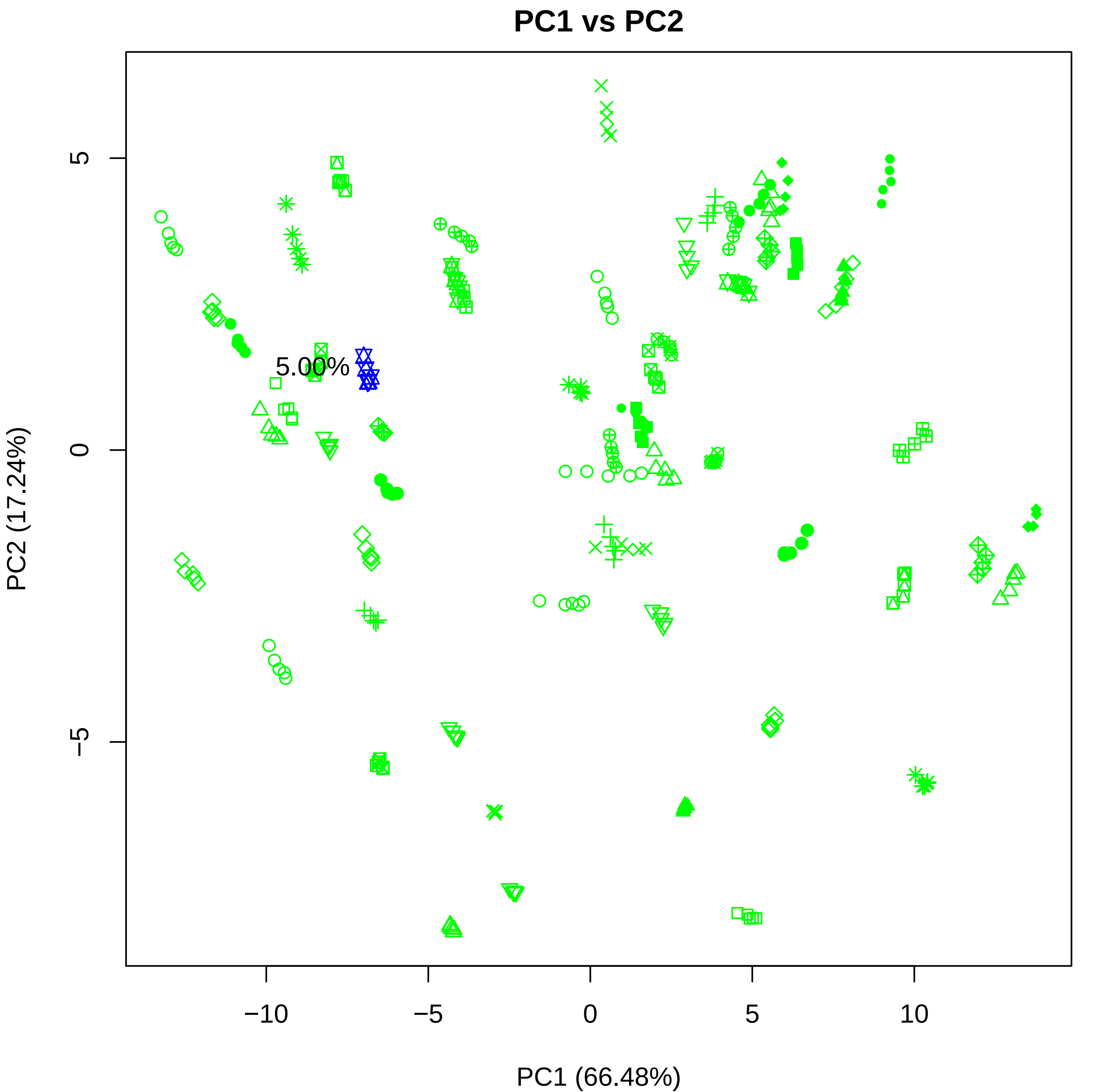
PC1 vs PC2
5.00%
PC2 (17.24%)
PC1 (66.48%)
5
0
−5
−10
−5
0
5
10

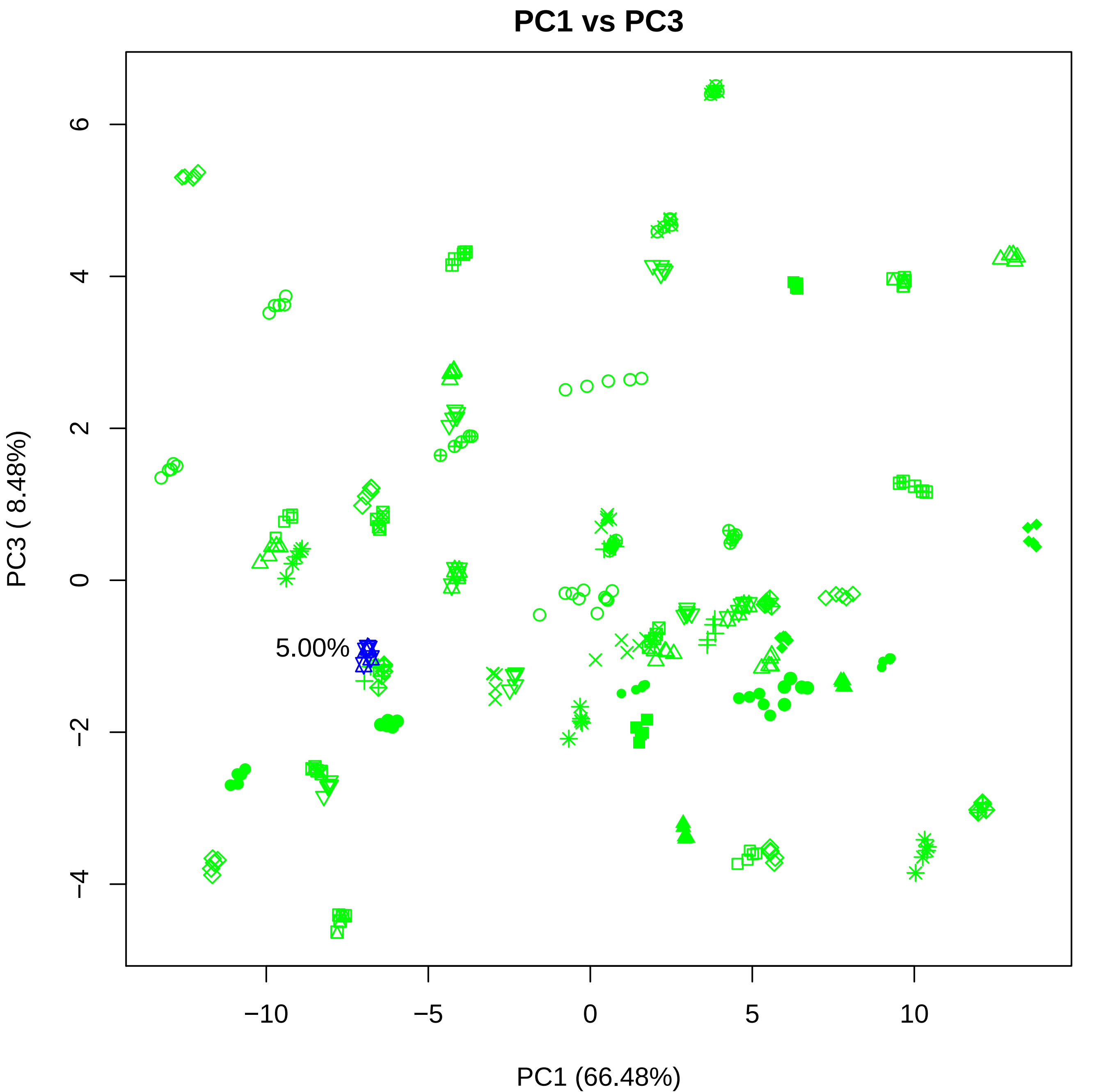

PC1 vs PC3
5.00%
PC3 ( 8.48%)
PC1 (66.48%)
6
4
2
0
−2
−4
−10
−5
0
5
10

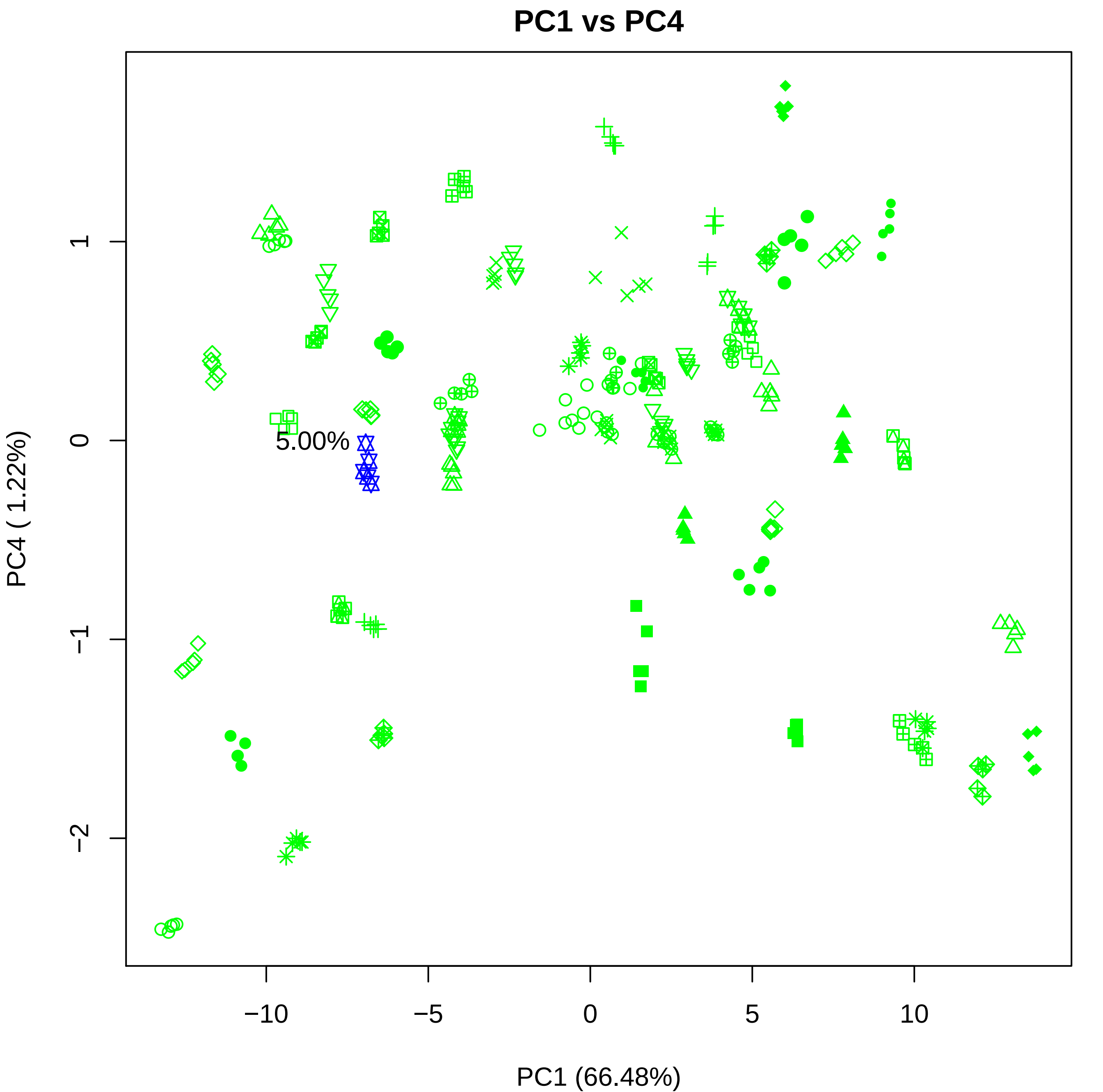

PC1 vs PC4
PC4 ( 1.22%)
PC1 (66.48%)
5.00%
−10
−5
0
5
10
1
0
−1
−2

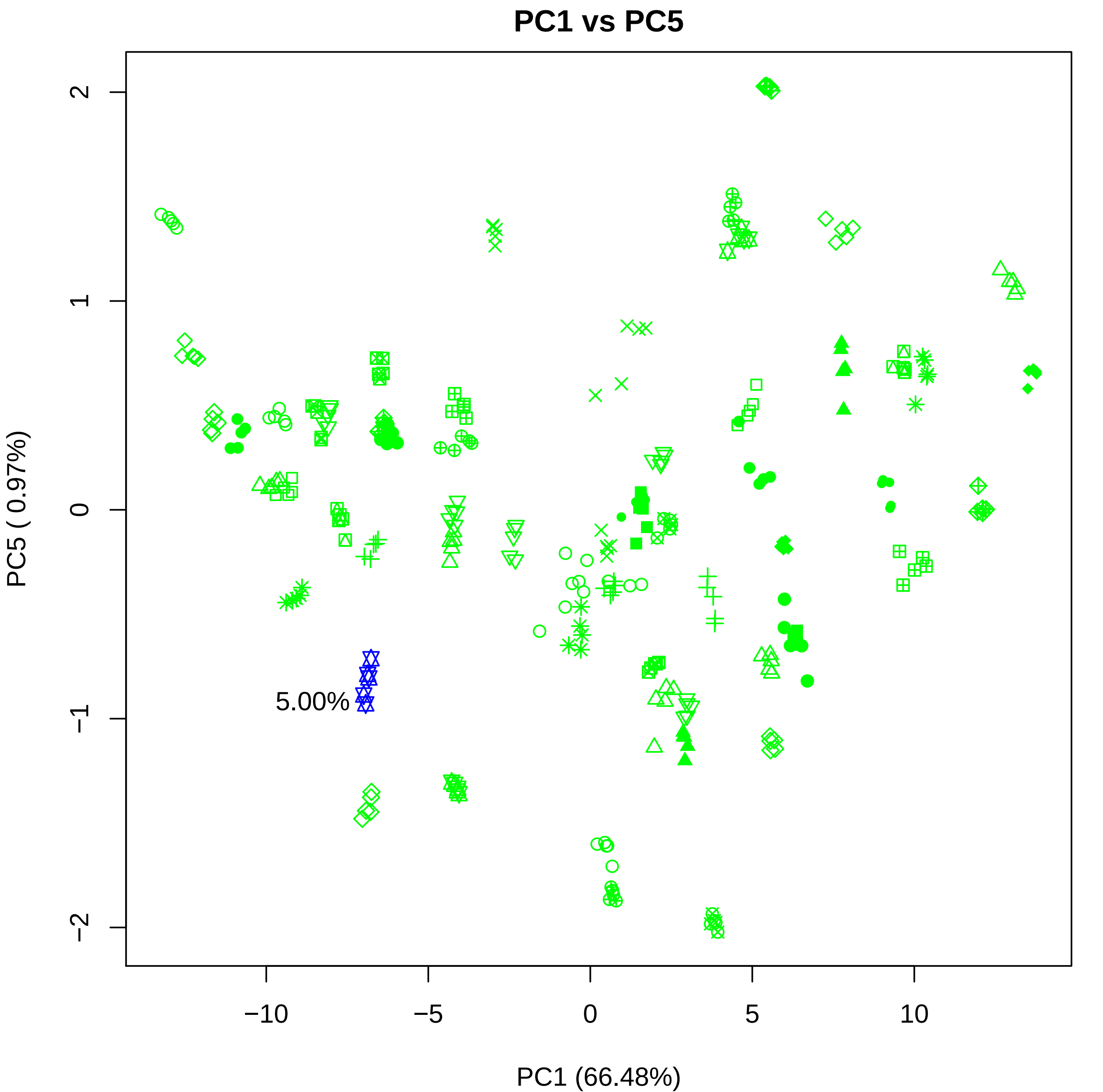
PC1 vs PC5
PC5 ( 0.97%)
PC1 (66.48%)
5.00%
2
1
0
−1
−2
−10
−5
0
5
10

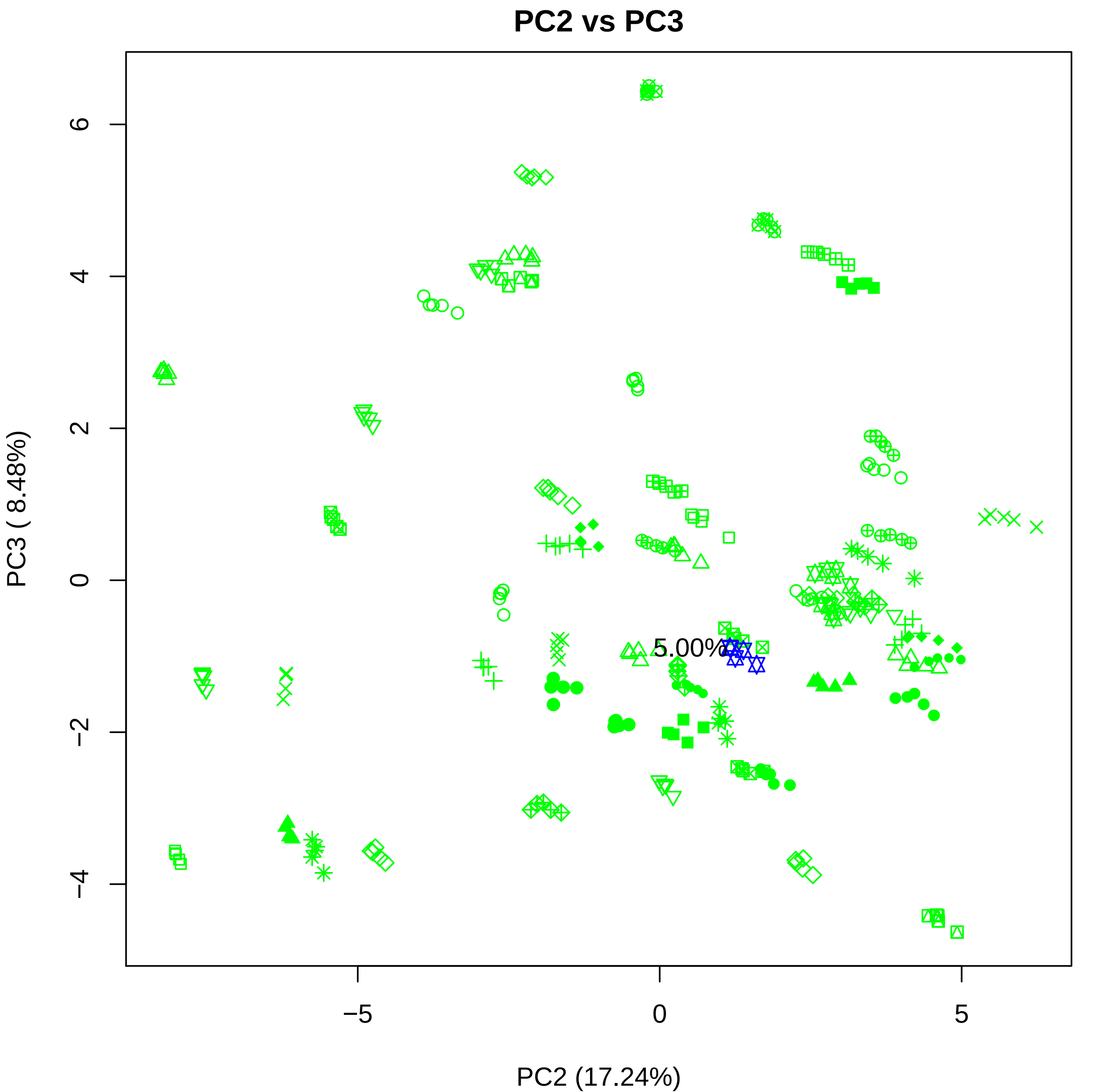

PC2 vs PC3
PC3 ( 8.48%)
PC2 (17.24%)
6
4
2
0
−2
−4
−5
0
5
5.00%

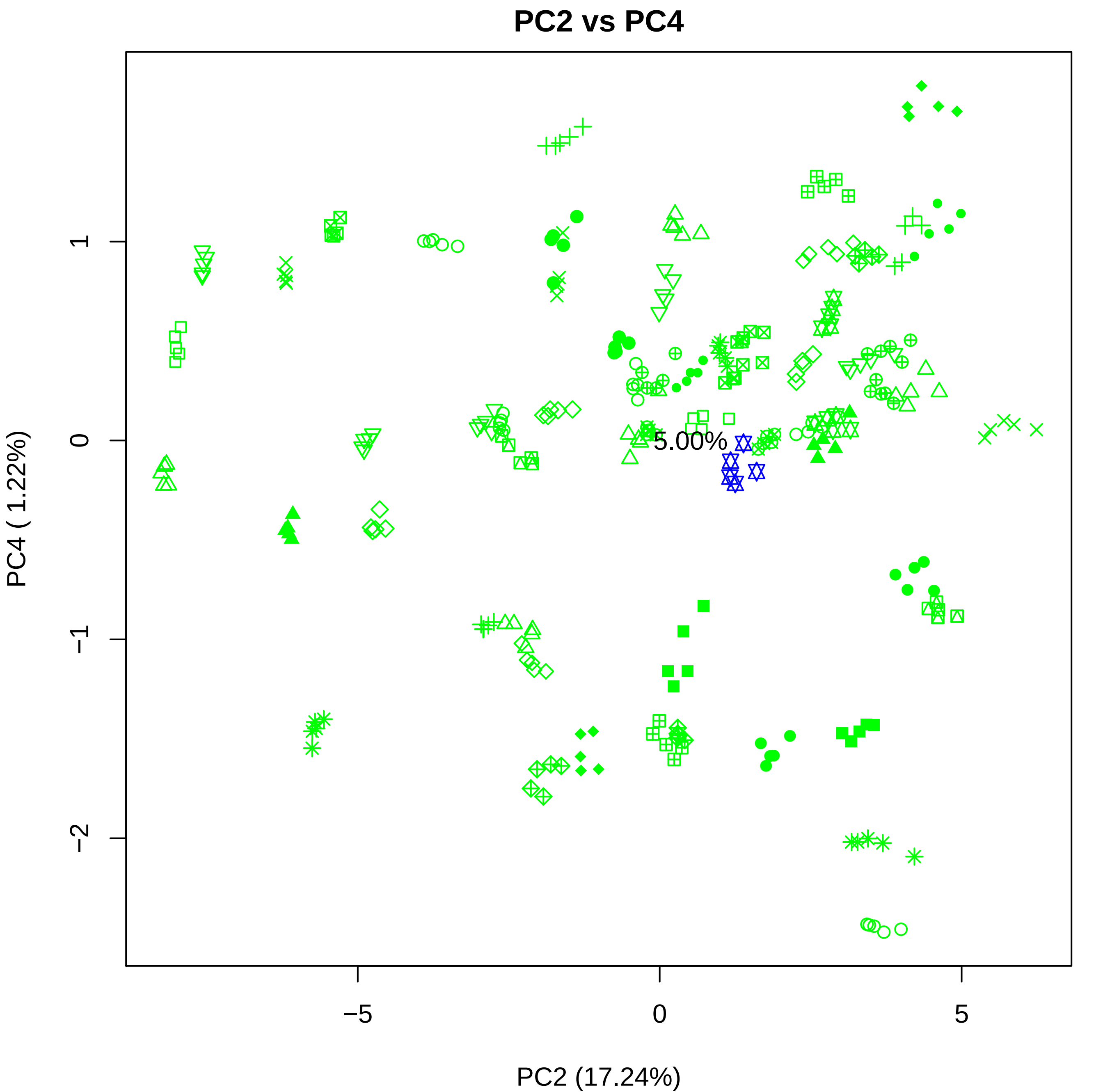

PC2 vs PC4
5.00%
1
0
−1
−2
−5
0
5
PC4 ( 1.22%)
PC2 (17.24%)

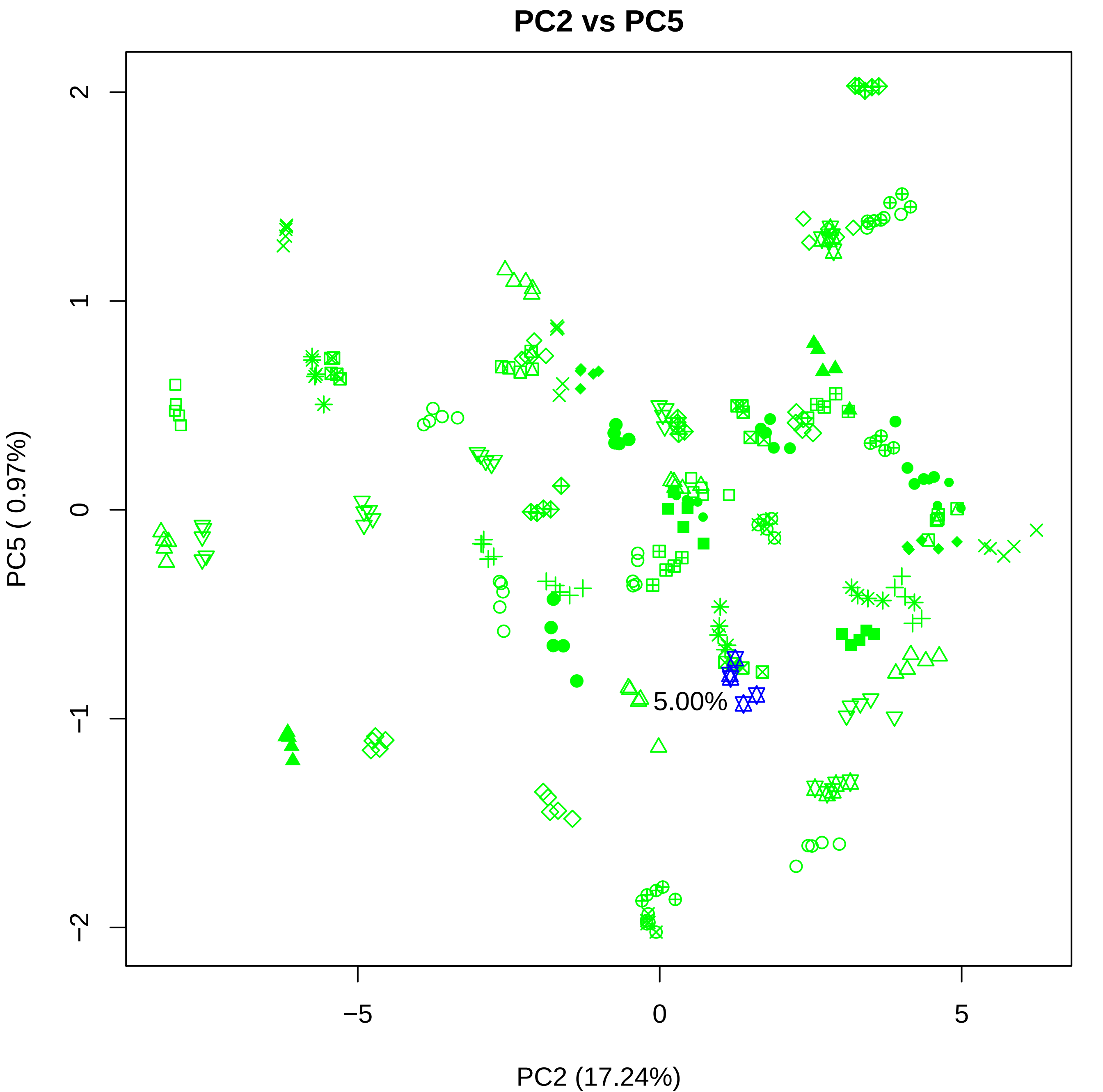
PC2 vs PC5
PC5 ( 0.97%)
PC2 (17.24%)
5.00%
−5
0
5
2
1
0
−1
−2

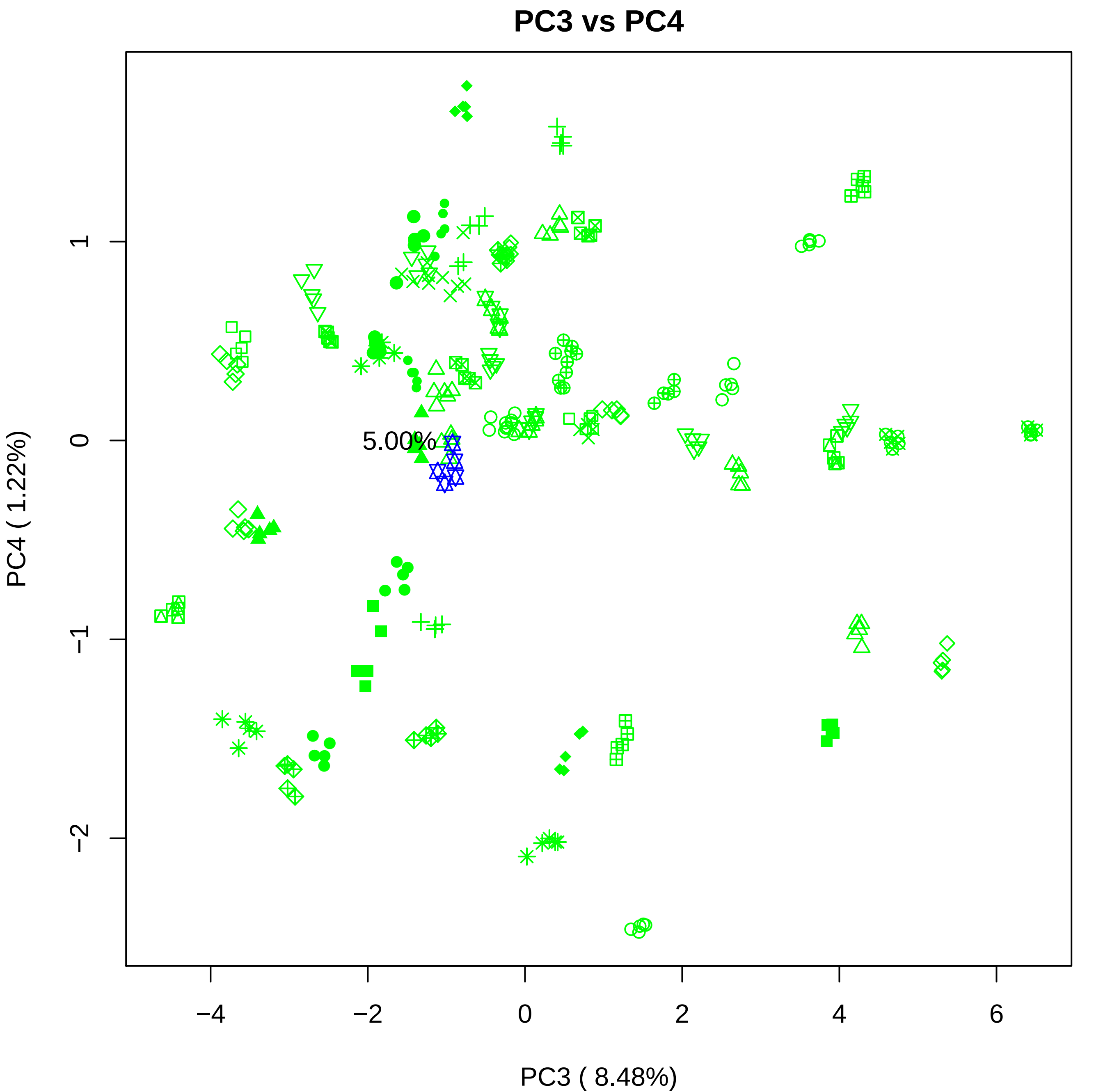
PC3 vs PC4
PC4 ( 1.22%)
PC3 ( 8.48%)
5.00%
1
0
−1
−2
−4
−2
0
2
4
6

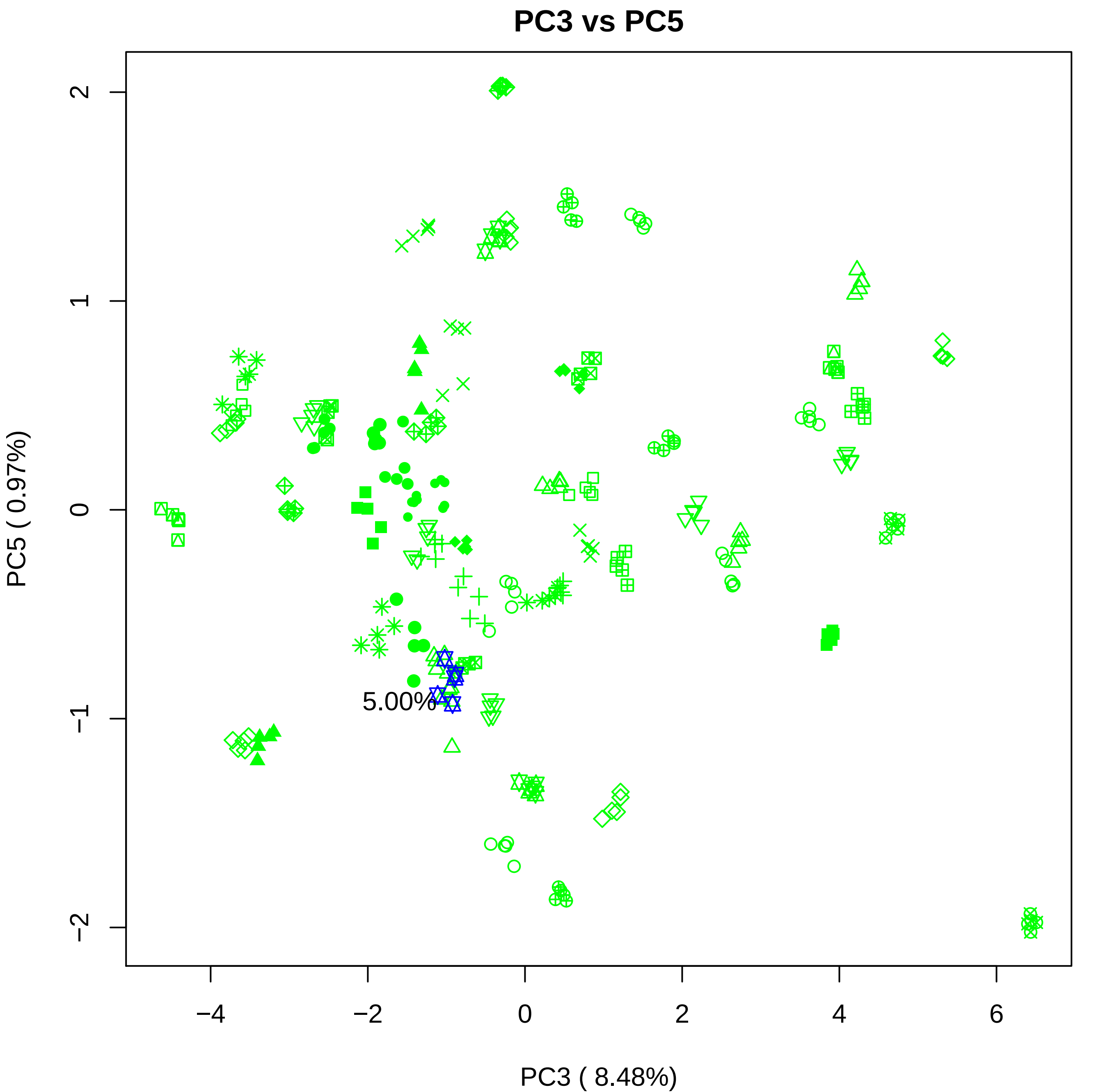
PC3 vs PC5
5.00%
PC5 ( 0.97%)
PC3 ( 8.48%)
2
1
0
−1
−2
−4
−2
0
2
4
6

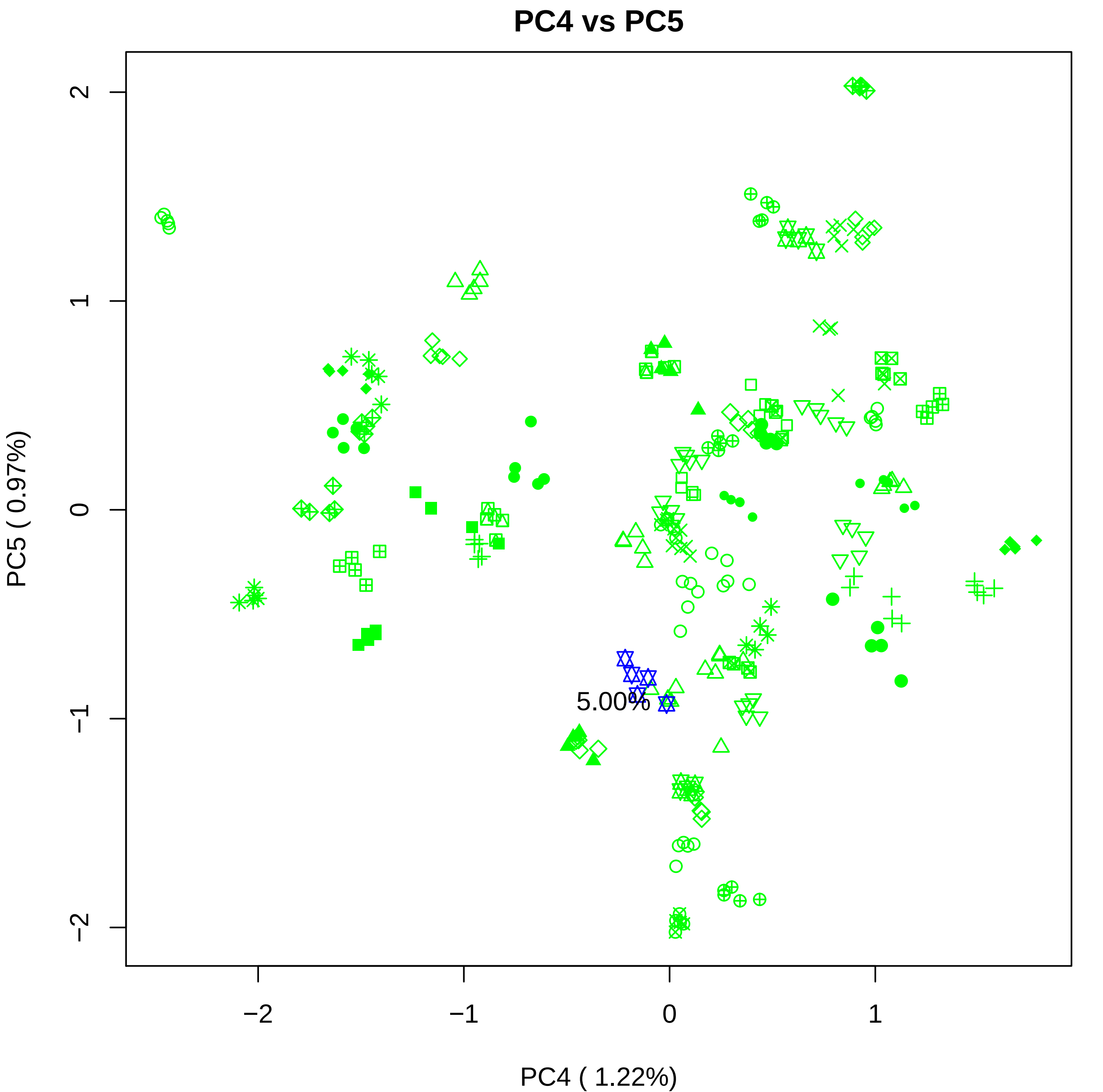

PC4 vs PC5
PC5 ( 0.97%)
PC4 ( 1.22%)
2
1
0
−1
−2
−2
−1
0
1
5.00%

### PCA of 7.5% BMH(d) vs PMH(d)

Scaled Matrix PCAed

7th order polynomial baseline correction performed

Raw spectra smoothed to 21 Neighbours, 2nd order PN, Savtisky Golay

Normalisation between 0 and 100

D2 spectra smoothed to 41 Neighbours, 2nd order PN, Savtisky Golay

Regions Analysed: 3601−2501 and 2001−700cm−1

Samples Analysed: 1 bContam's 0 BLH's 0 BMH's 0 OMH's 0 pContam's 0 PLH's 60 PMH's

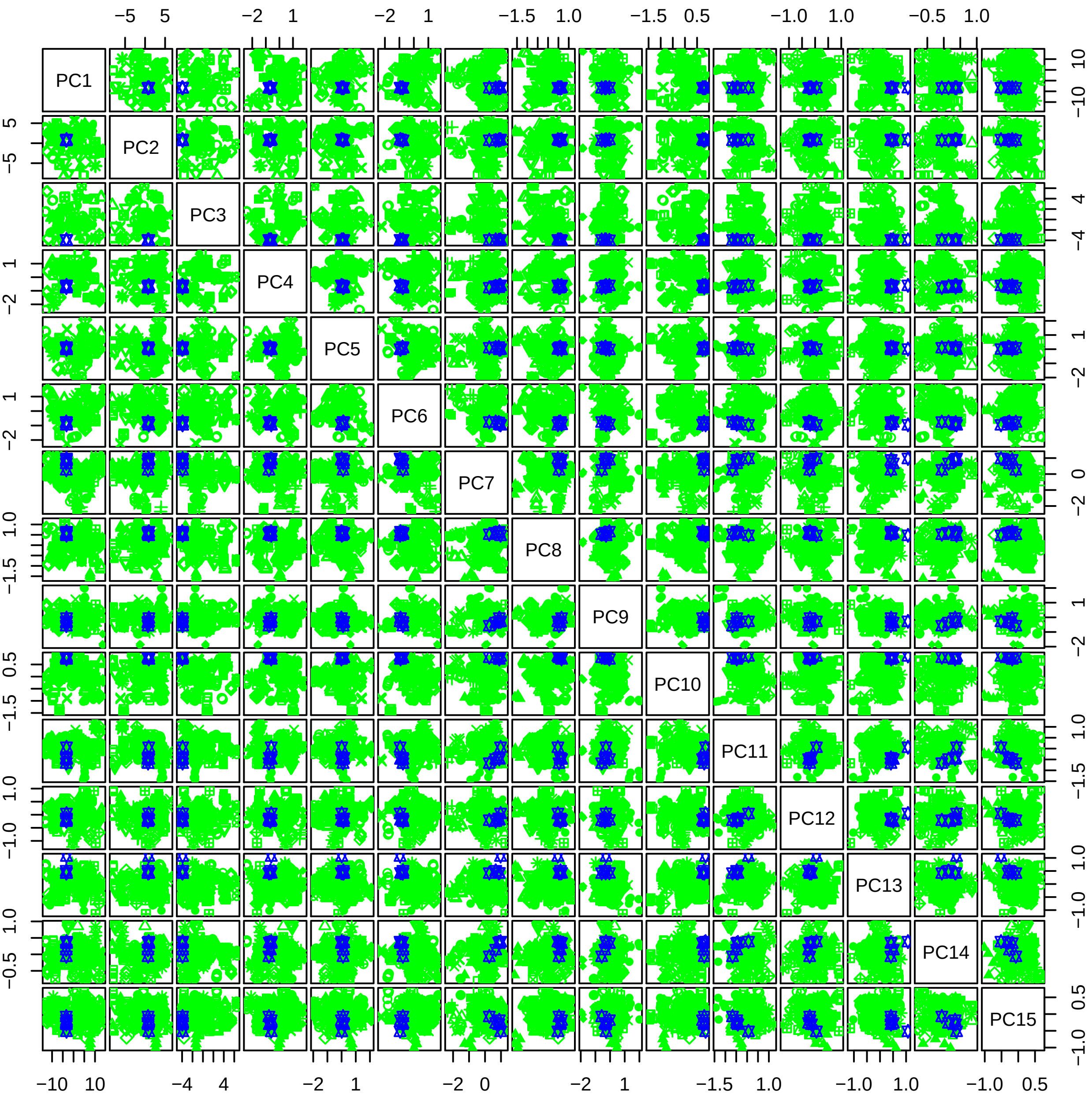
PC1
PC2
PC3
PC4
PC5
PC6
PC7
PC8
PC9
PC10
PC11
PC12
PC13
PC14
PC15

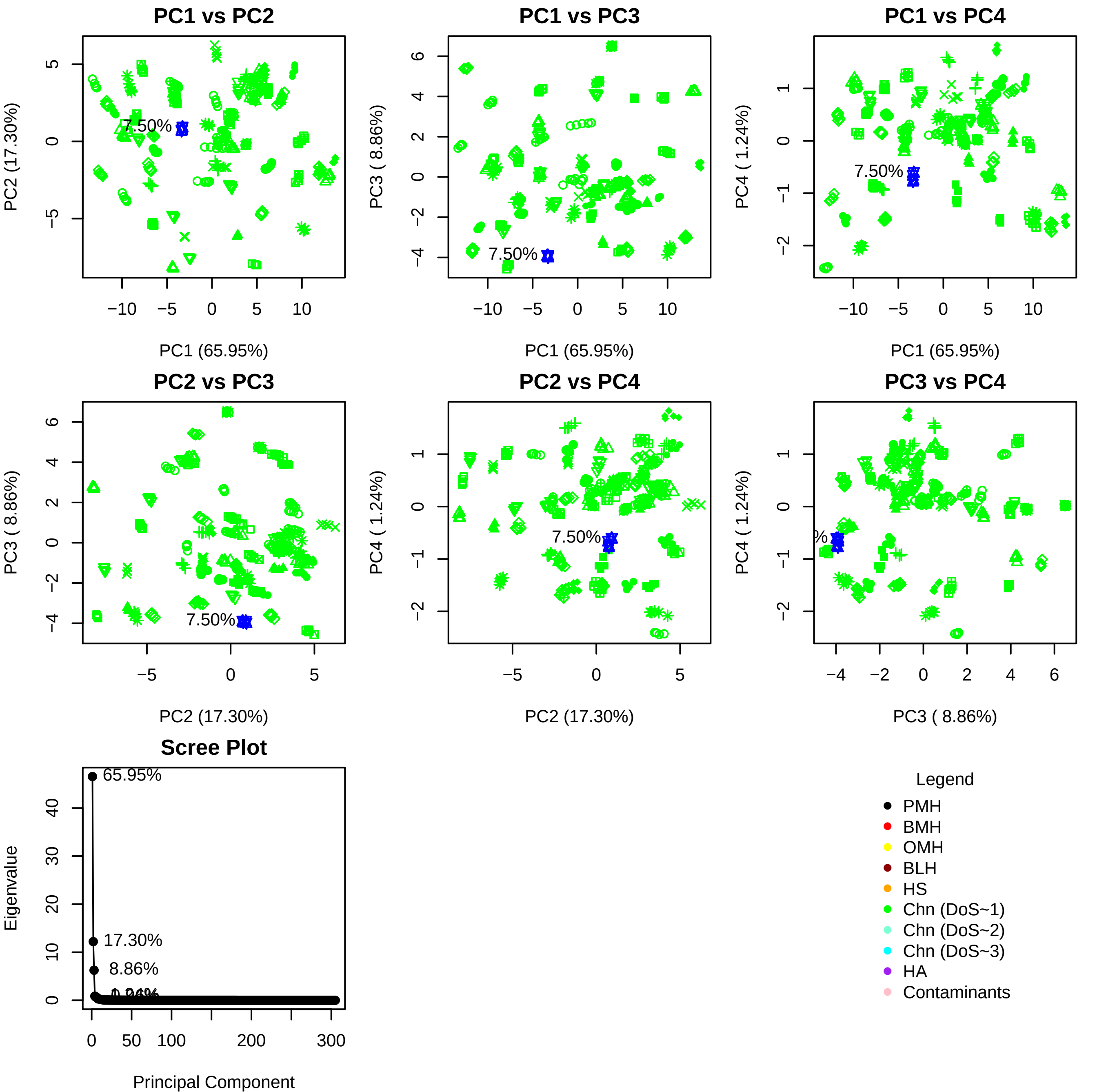
PC1 vs PC2
PC1 vs PC3
PC1 vs PC4
PC2 vs PC3
PC2 vs PC4
PC3 vs PC4
Scree Plot
PC1 (65.95%)
PC2 (17.30%)
PC3 ( 8.86%)
PC4 ( 1.24%)
7.50%
Eigenvalue
Principal Component
65.95%
17.30%
8.86%
Legend
PMH
BMH
OMH
BLH
HS
Chn (DoS~1)
Chn (DoS~2)
Chn (DoS~3)
HA
Contaminants

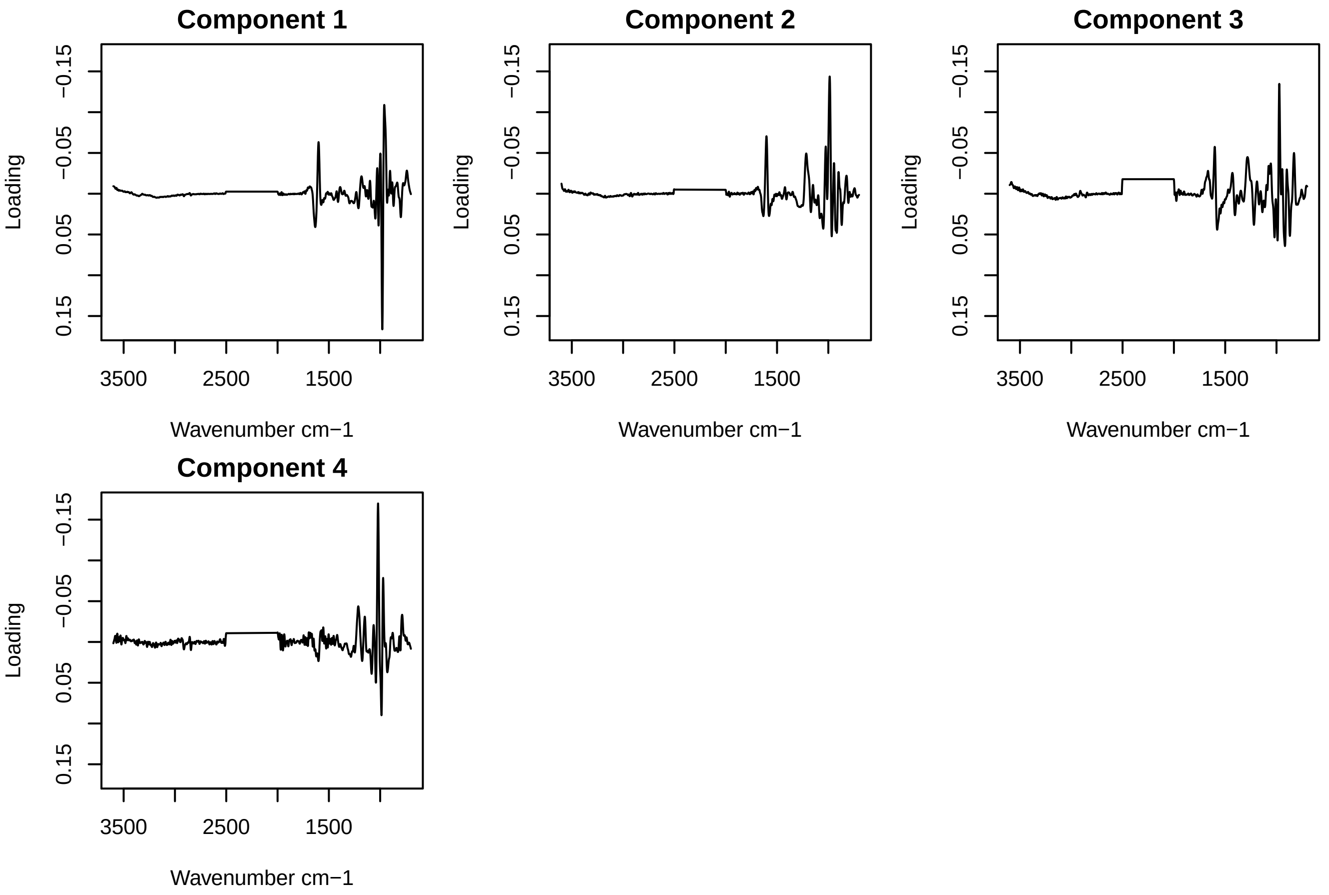
Component 1
Loading
−0.15
−0.05
0.05
0.15
3500
2500
1500
Wavenumber cm−1
Component 2
Loading
−0.15
−0.05
0.05
0.15
3500
2500
1500
Wavenumber cm−1
Component 3
Loading
−0.15
−0.05
0.05
0.15
3500
2500
1500
Wavenumber cm−1
Component 4
Loading
−0.15
−0.05
0.05
0.15
3500
2500
1500
Wavenumber cm−1

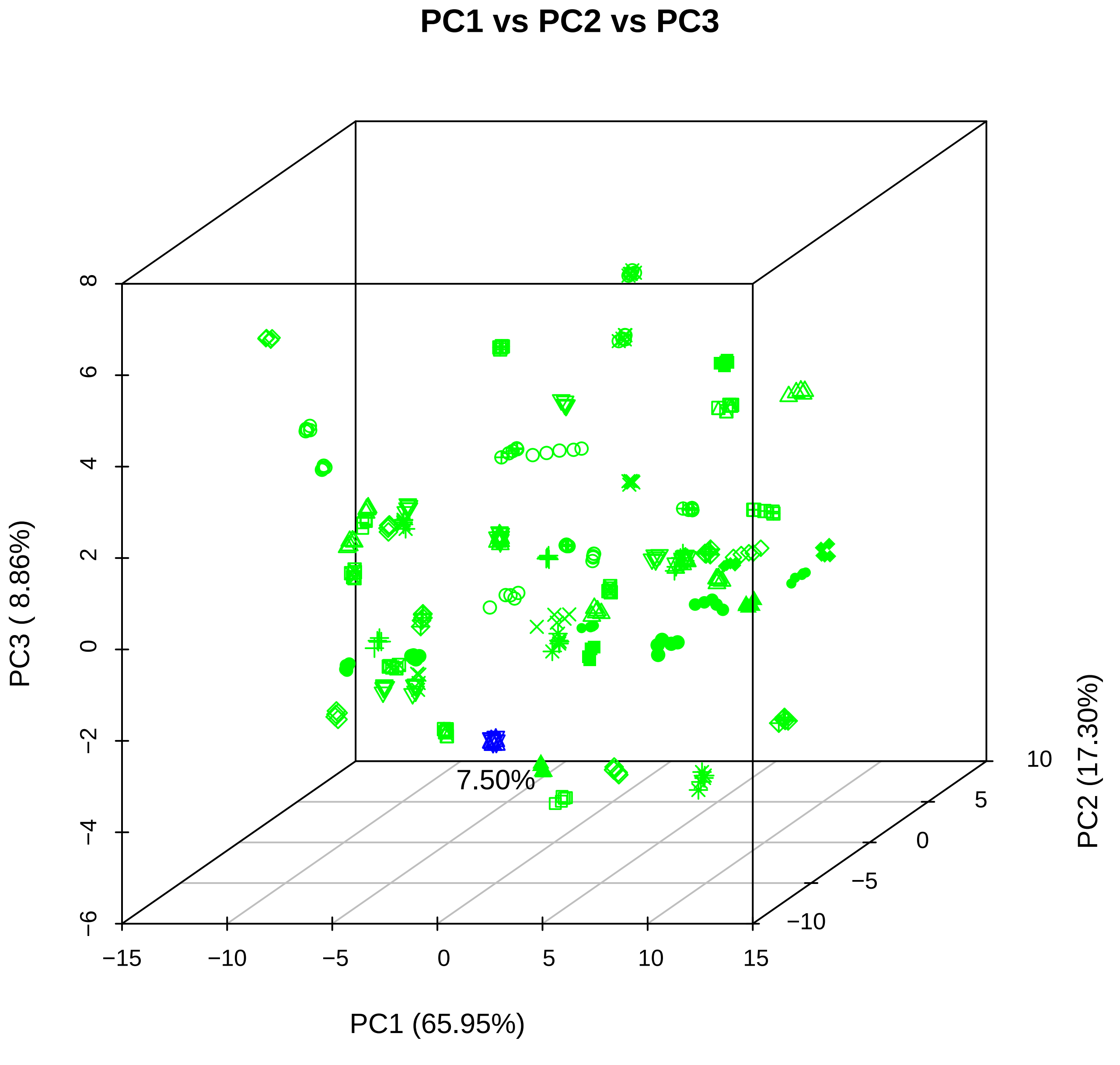

PC1 vs PC2 vs PC3
PC3 ( 8.86%)
8
6
4
2
0
−2
−4
−6
7.50%
−15
−10
−5
0
5
10
15
PC1 (65.95%)
10
5
0
−5
−10
PC2 (17.30%)

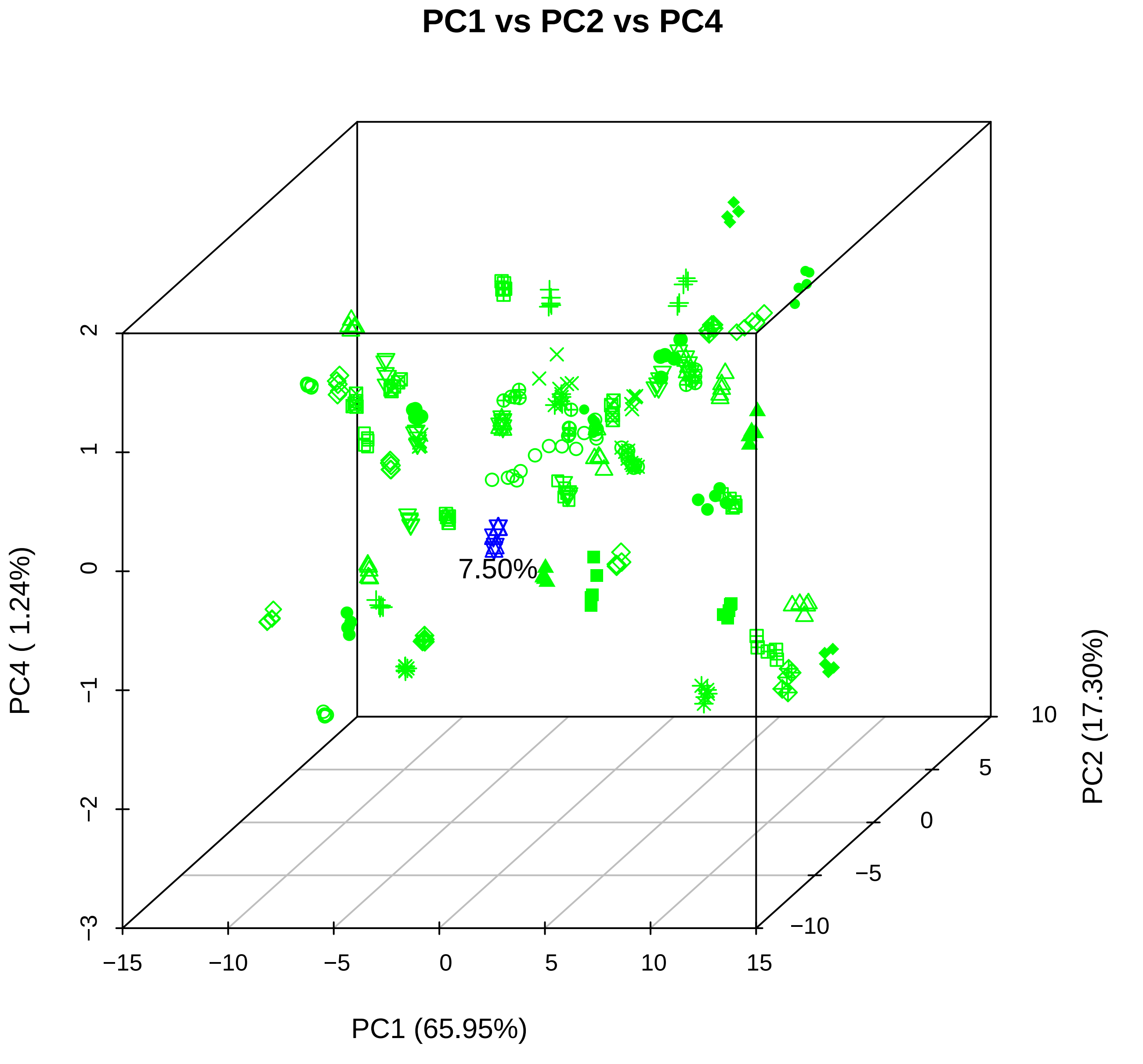

PC1 vs PC2 vs PC4
7.50%
PC1 (65.95%)
PC2 (17.30%)
PC4 ( 1.24%)
−15
−10
−5
0
5
10
15
−3
−2
−1
0
1
2
−10
−5
0
5
10

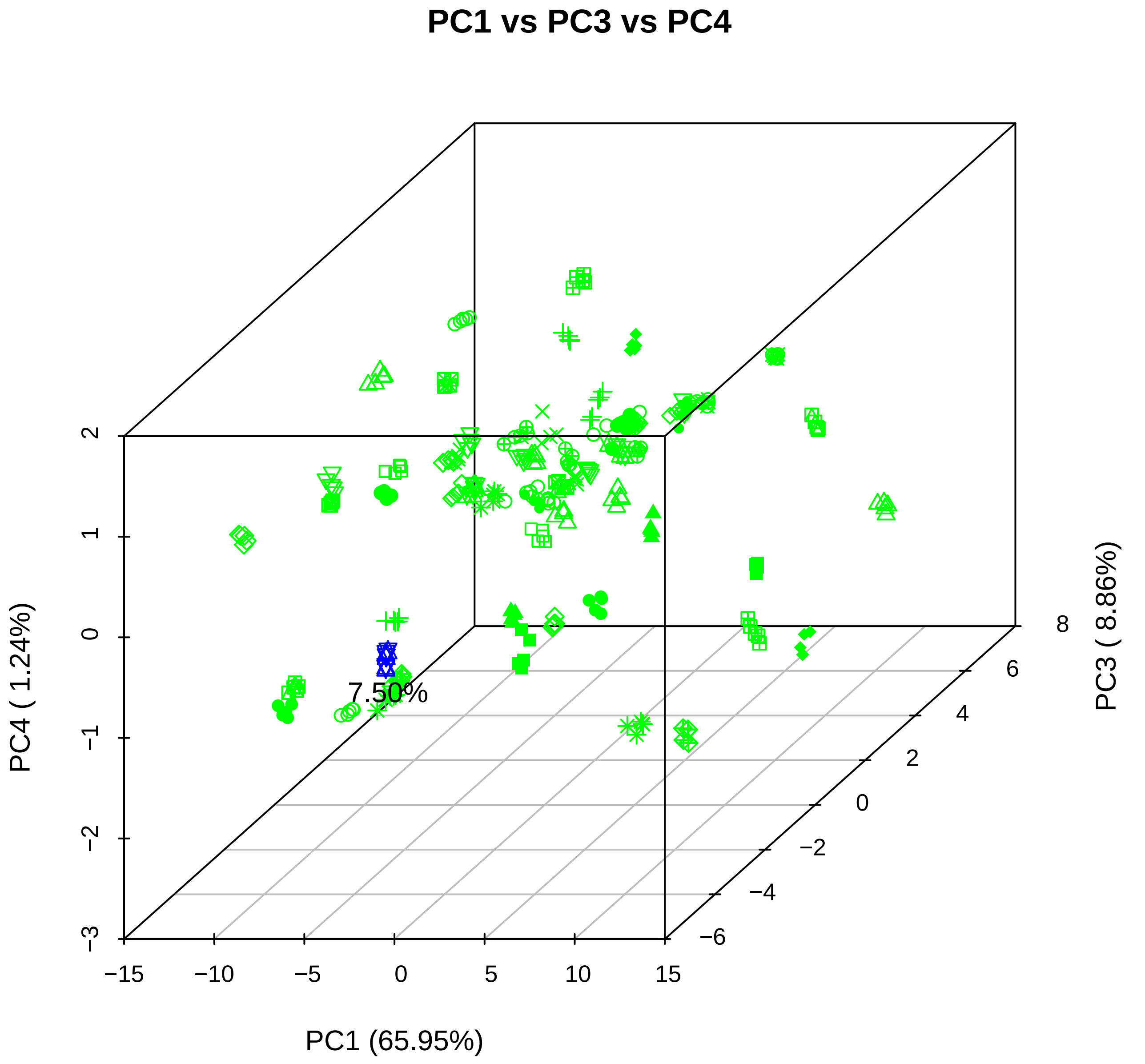

PC1 vs PC3 vs PC4
PC1 (65.95%)
PC4 ( 1.24%)
PC3 ( 8.86%)
7.50%
−15
−10
−5
0
5
10
15
−3
−2
−1
0
1
2
−6
−4
−2
0
2
4
6
8

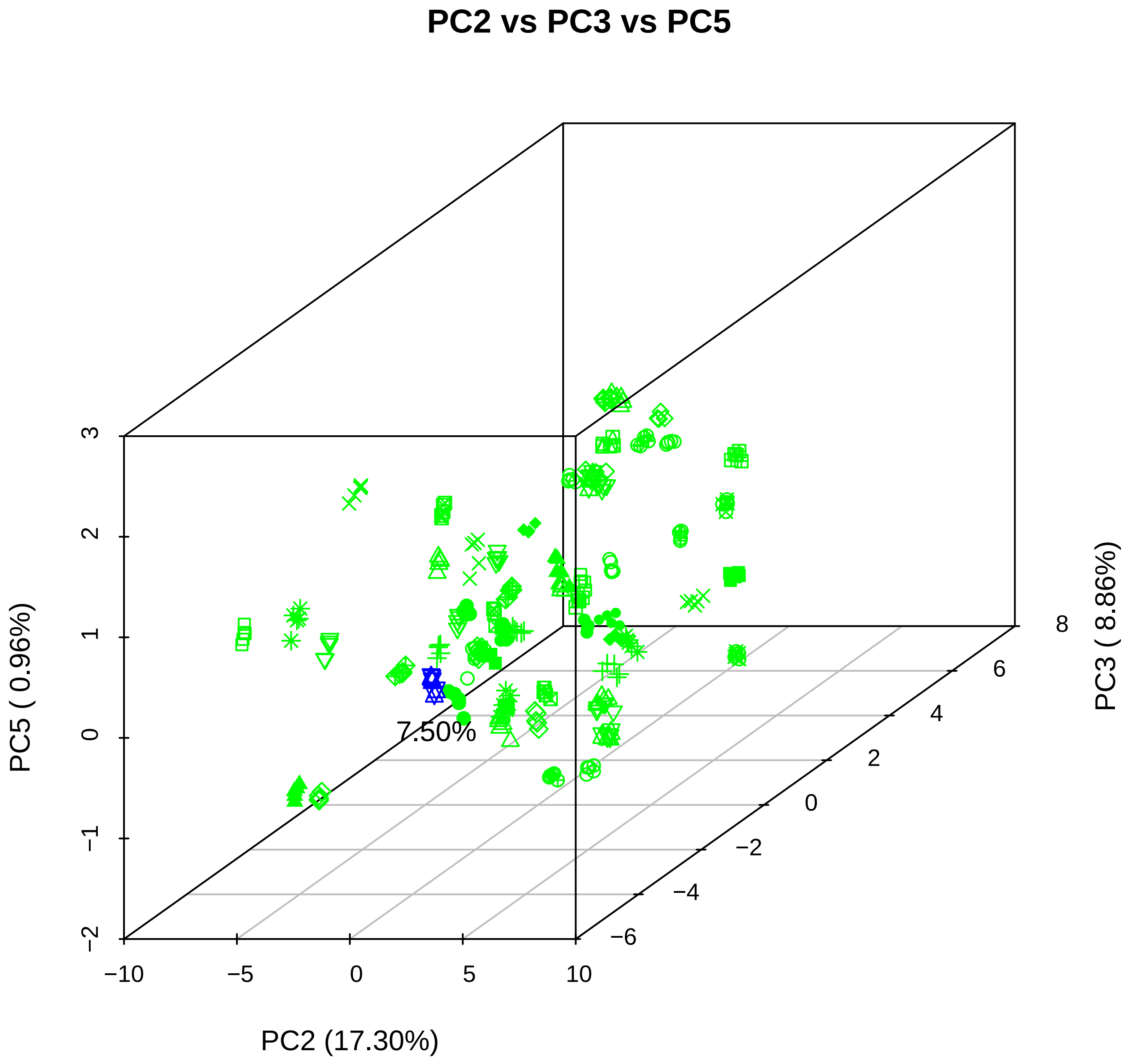

PC2 vs PC3 vs PC5
PC5 ( 0.96%)
PC2 (17.30%)
PC3 ( 8.86%)
7.50%
3
2
1
0
−1
−2
−10
−5
0
5
10
8
6
4
2
0
−2
−4
−6

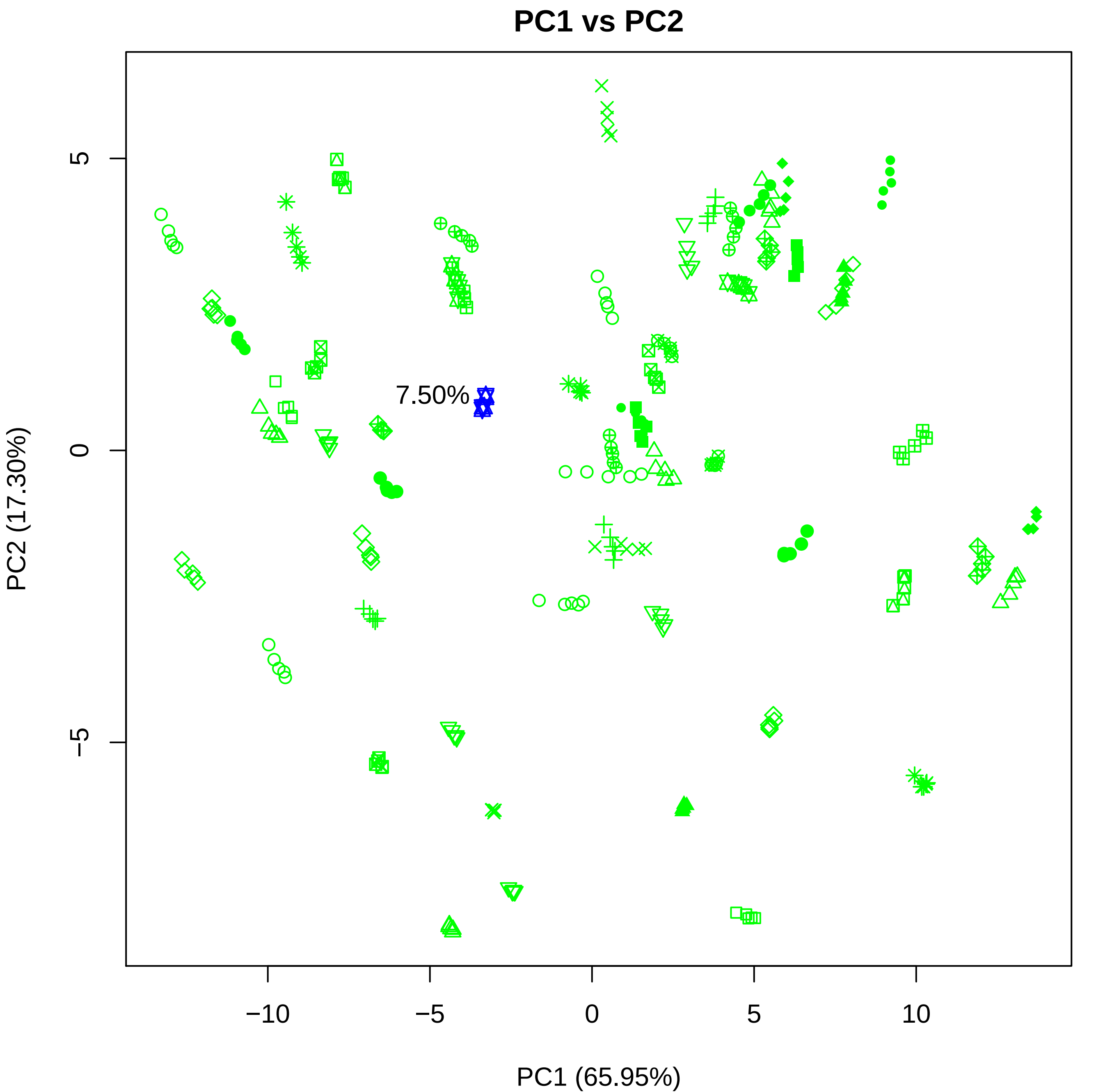

PC1 vs PC2
7.50%
5
0
−5
−10
−5
0
5
10
PC2 (17.30%)
PC1 (65.95%)

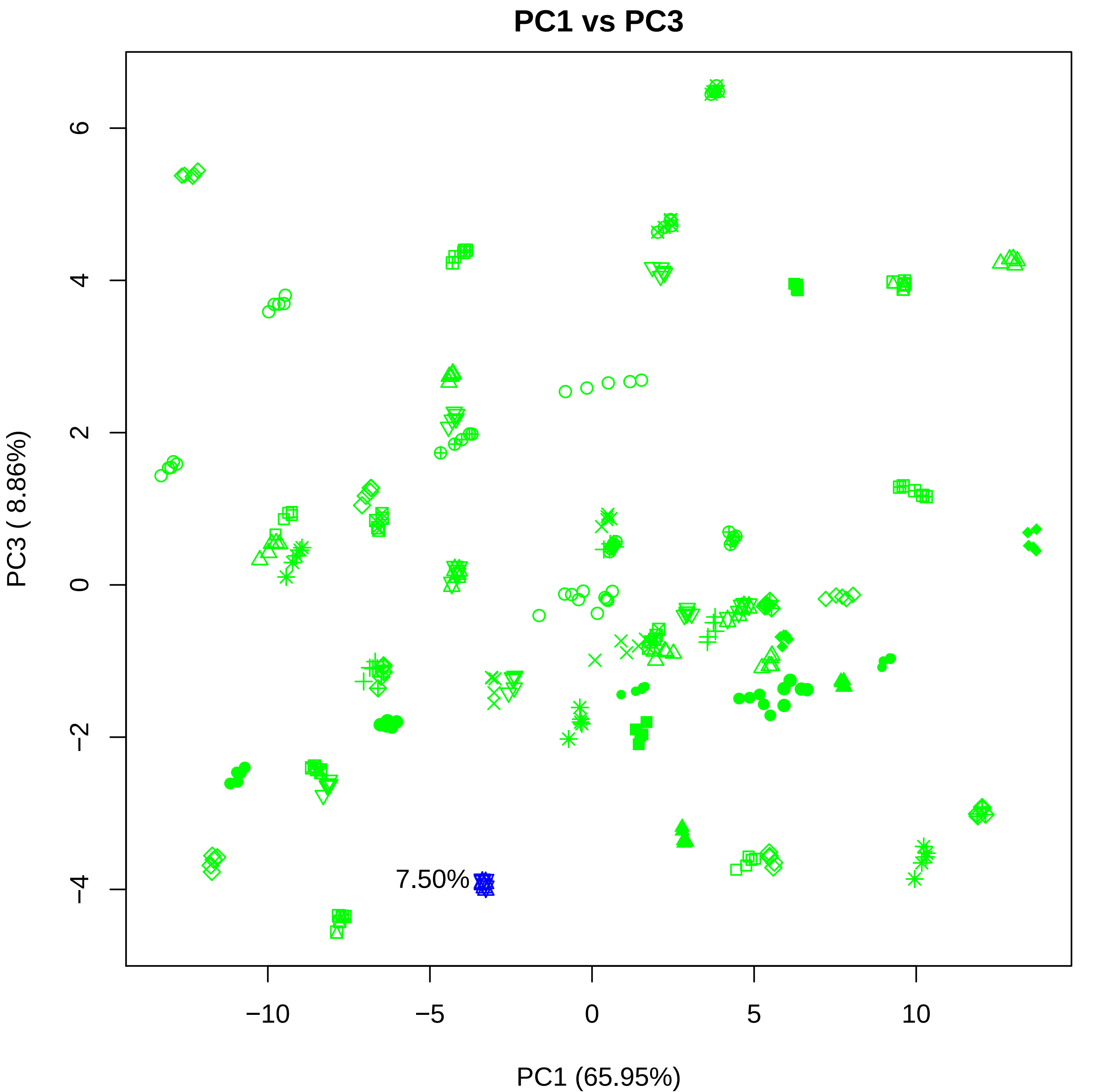
PC1 vs PC3
PC3 ( 8.86%)
PC1 (65.95%)
7.50%
6
4
2
0
−2
−4
−10
−5
0
5
10

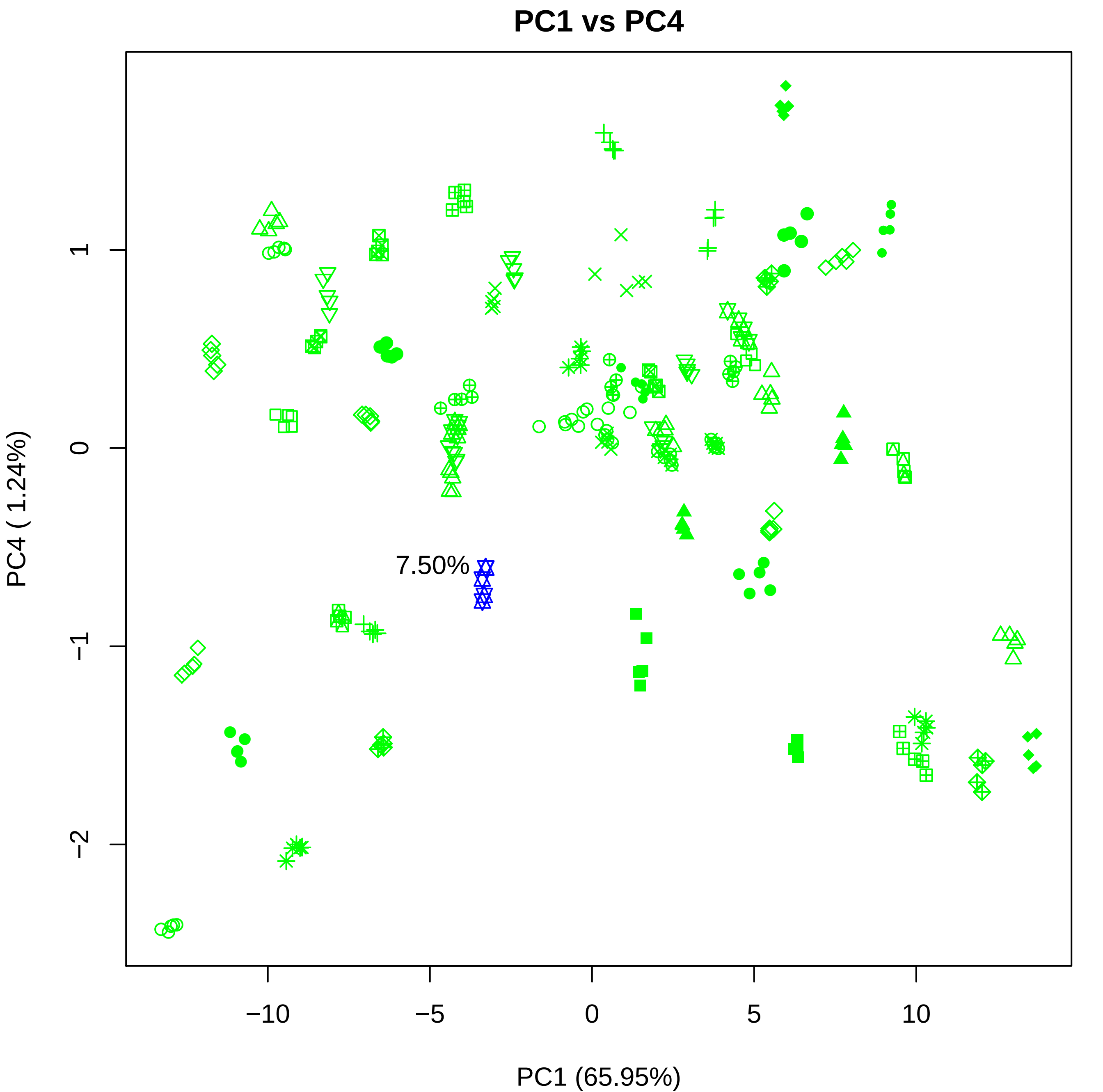

PC1 vs PC4
7.50%
PC4 ( 1.24%)
PC1 (65.95%)
1
0
−1
−2
−10
−5
0
5
10

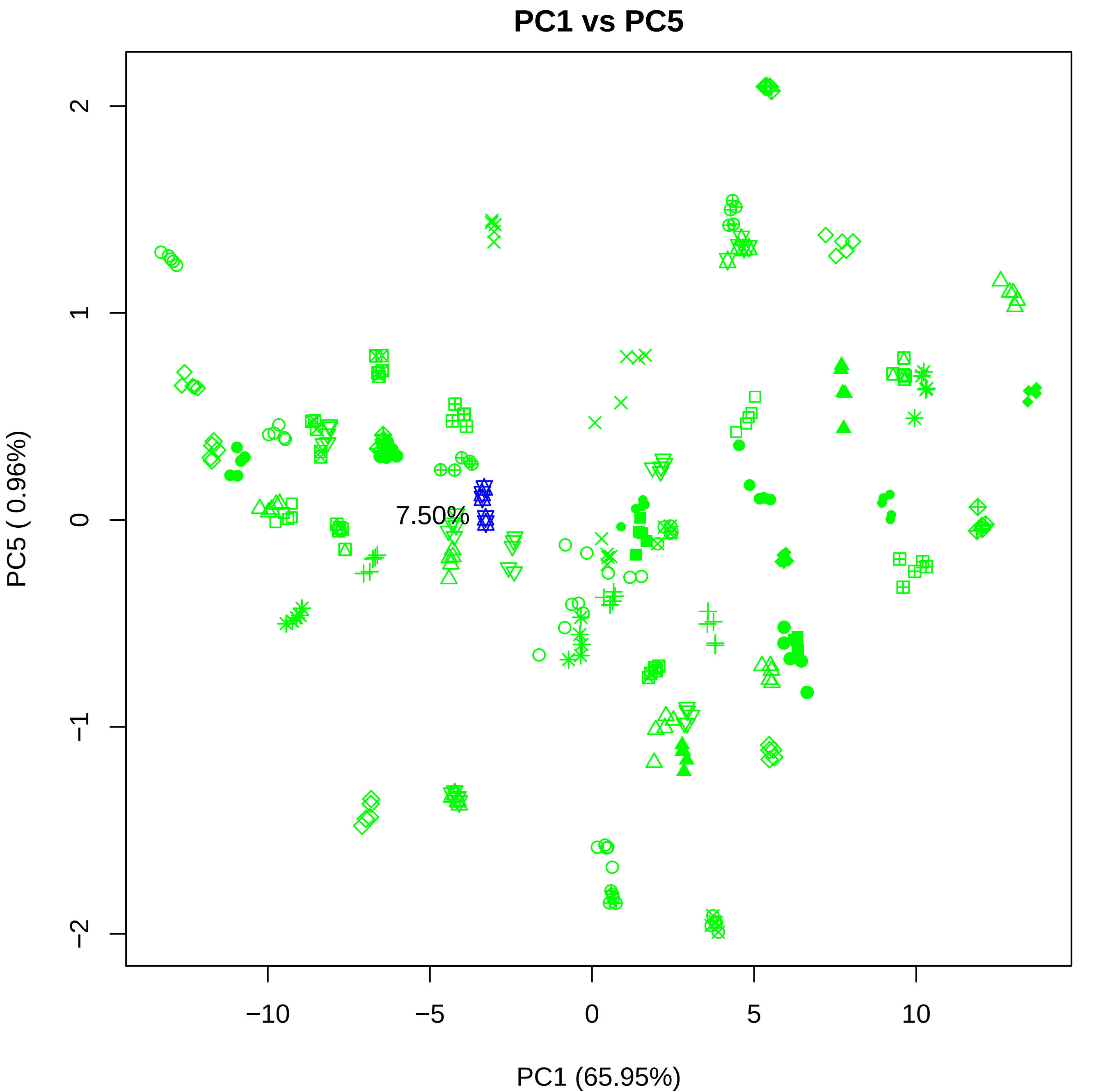

PC1 vs PC5
7.50%
PC1 (65.95%)
PC5 ( 0.96%)
−10
−5
0
5
10
2
1
0
−1
−2

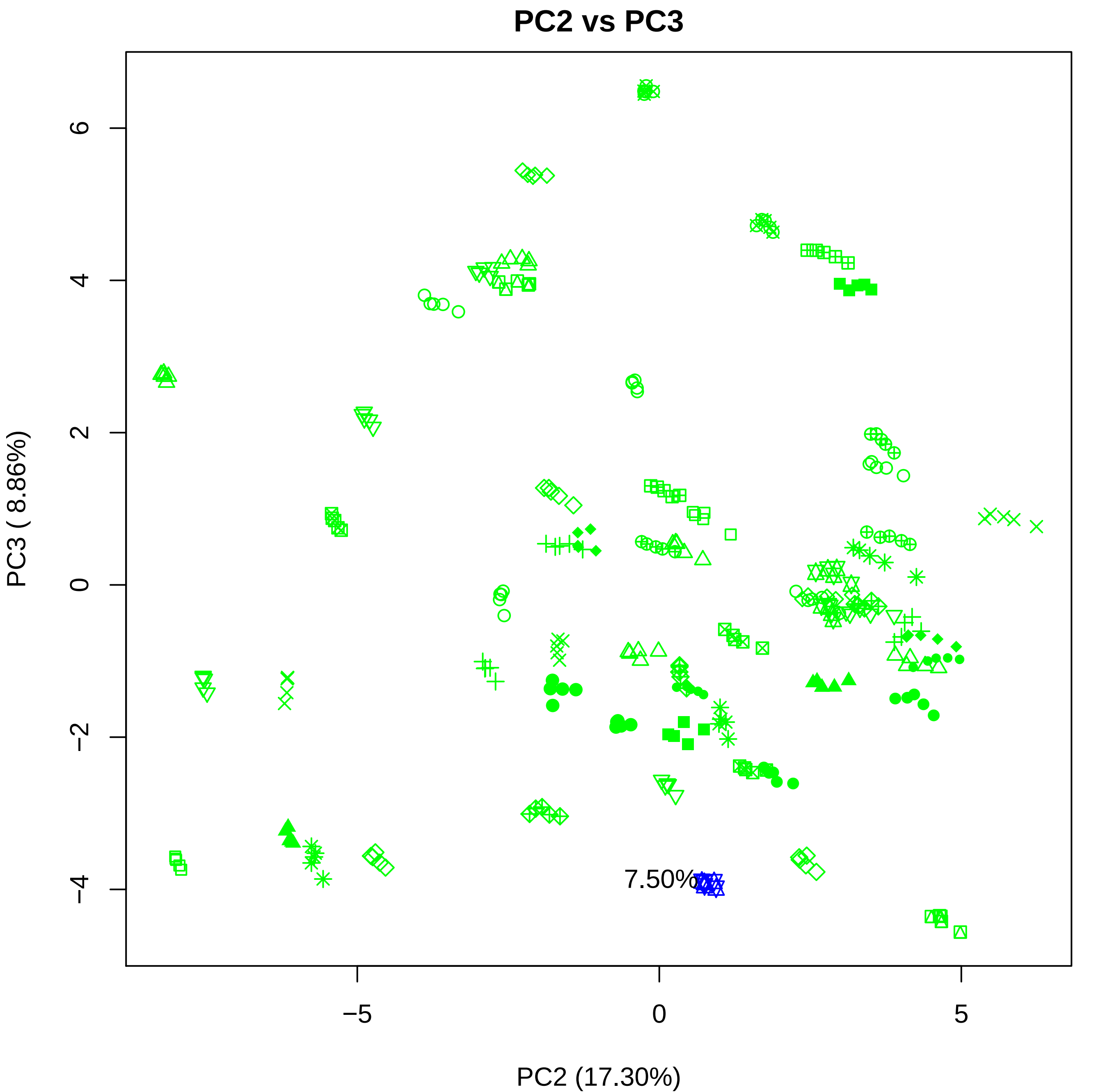
PC2 vs PC3
PC3 ( 8.86%)
PC2 (17.30%)
6
4
2
0
−2
−4
−5
0
5
7.50%

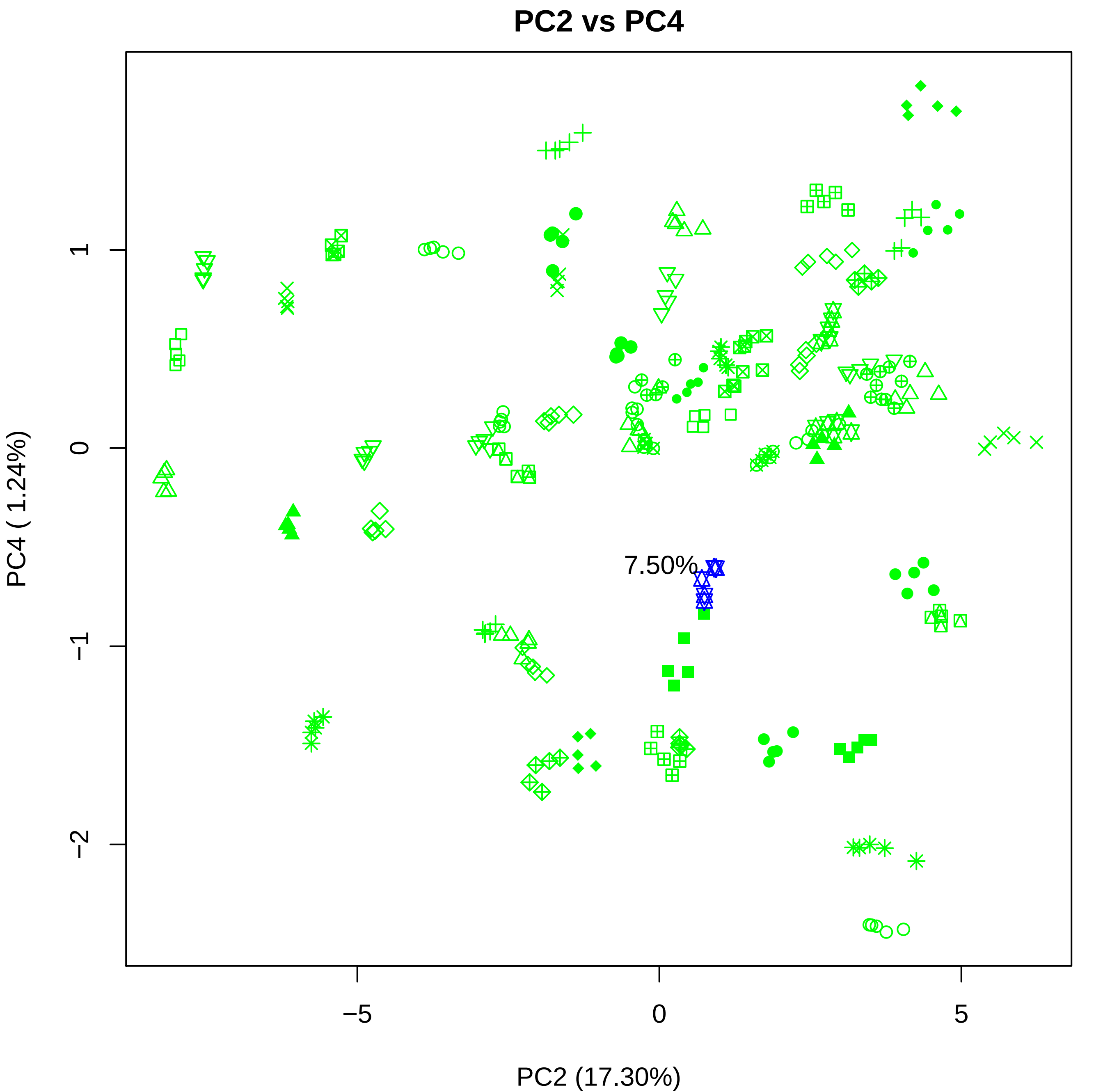

PC2 vs PC4
PC4 ( 1.24%)
PC2 (17.30%)
7.50%
1
0
−1
−2
−5
0
5

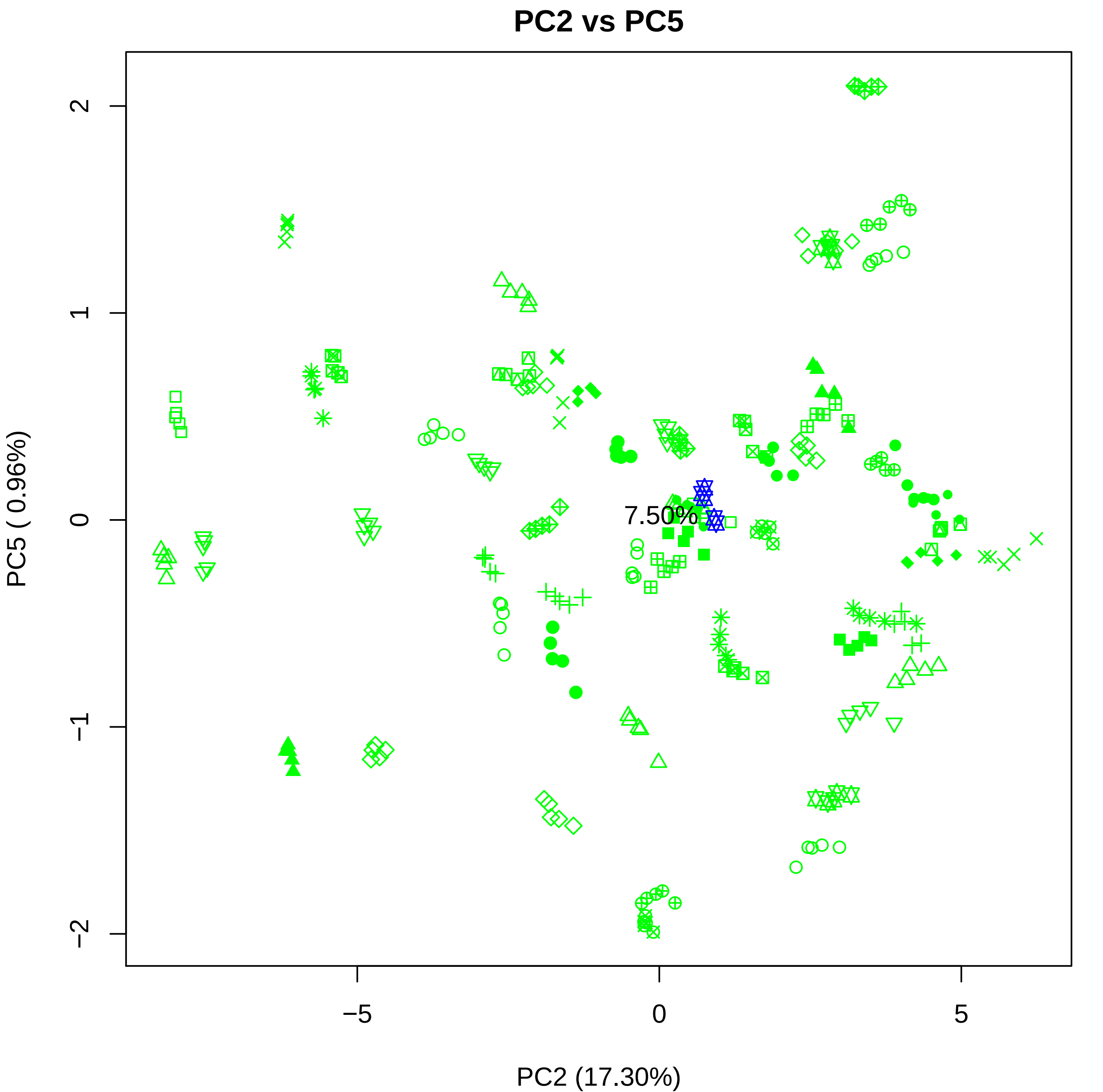
PC2 vs PC5
2
1
0
−1
−2
PC5 ( 0.96%)
7.50%
−5
0
5
PC2 (17.30%)

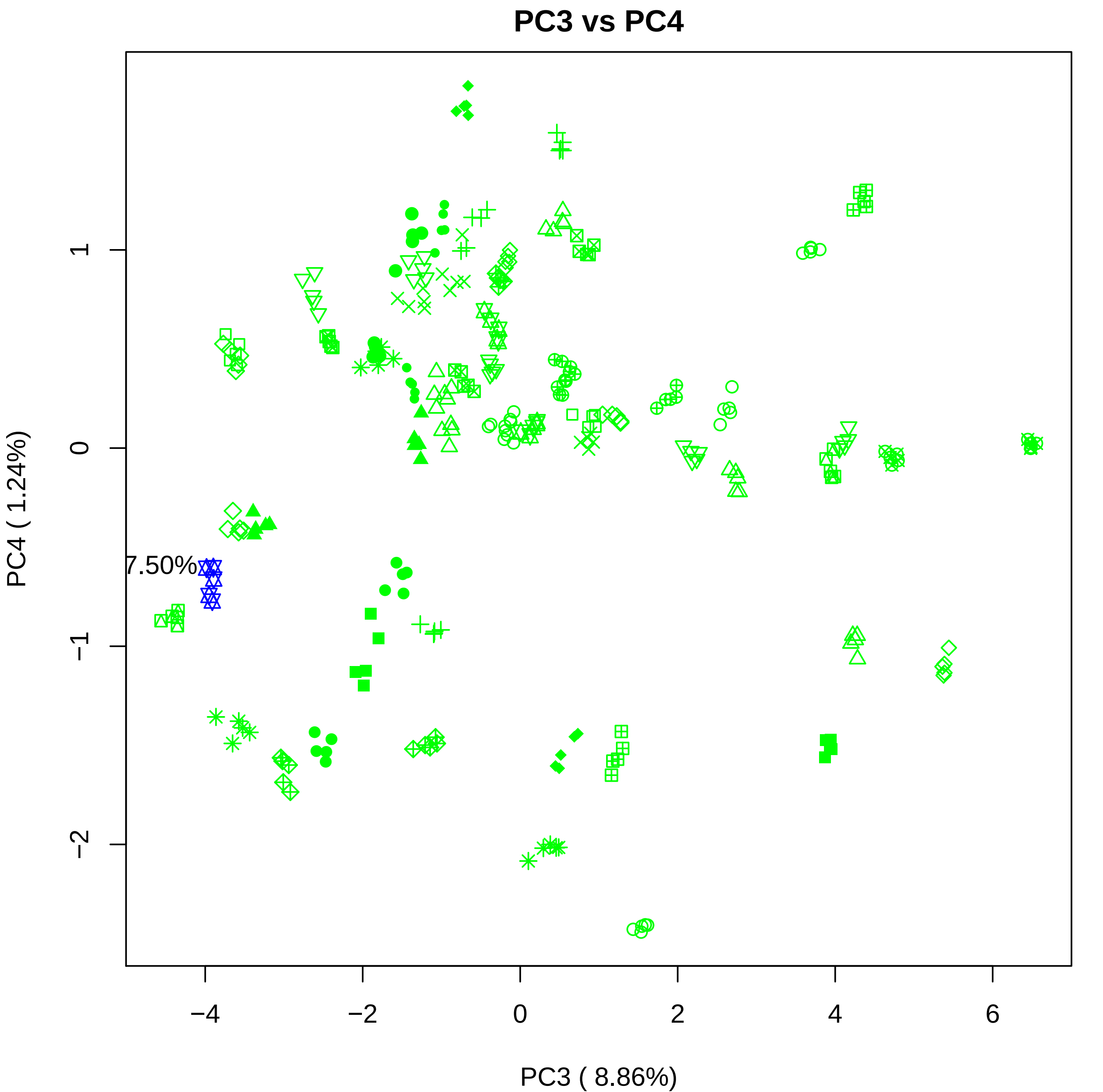
PC3 vs PC4
7.50%
PC4 ( 1.24%)
PC3 ( 8.86%)
1
0
−1
−2
−4
−2
0
2
4
6

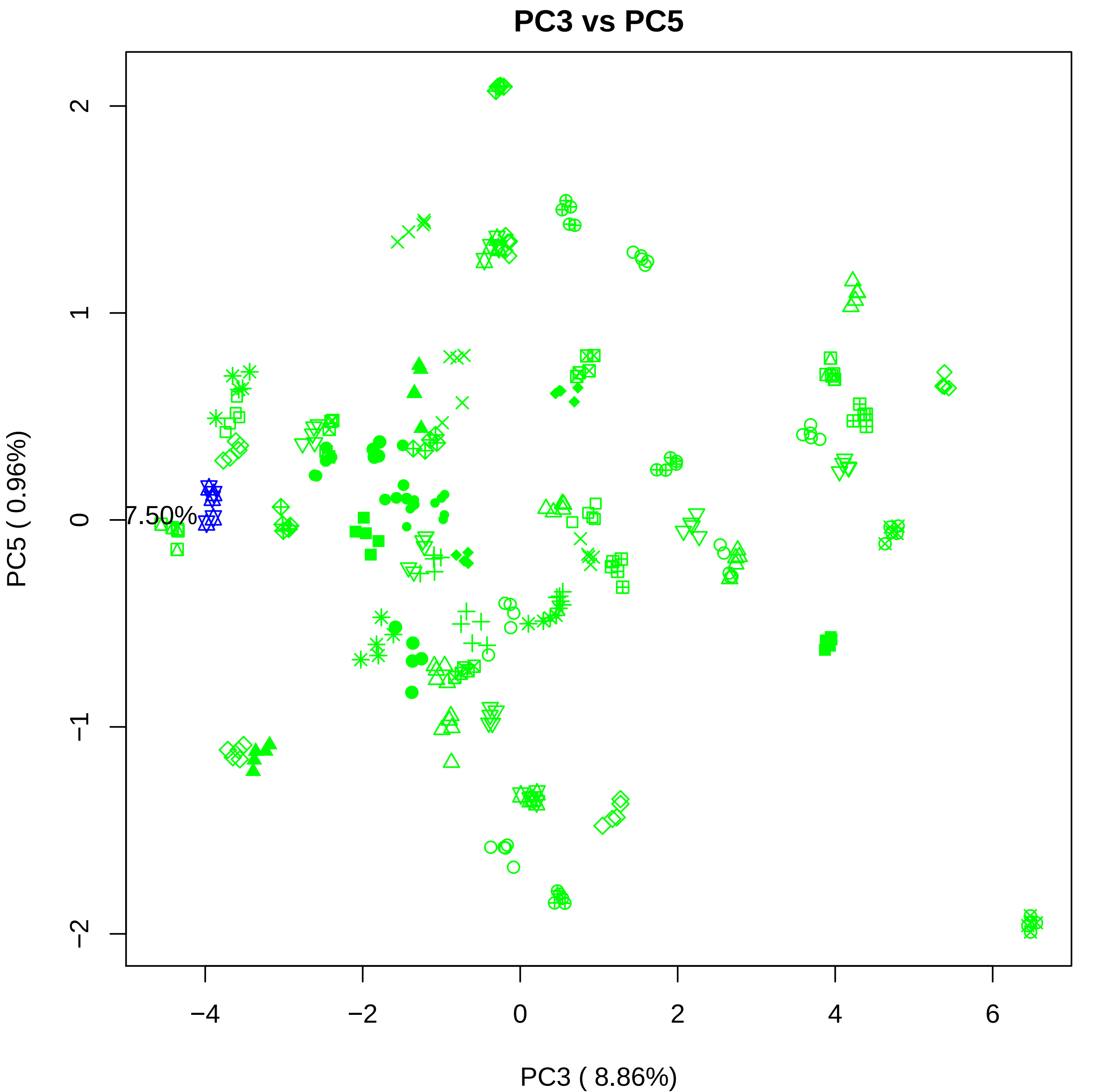
PC3 vs PC5
7.50%
PC5 ( 0.96%)
PC3 ( 8.86%)
−4
−2
0
2
4
6
2
1
0
−1
−2

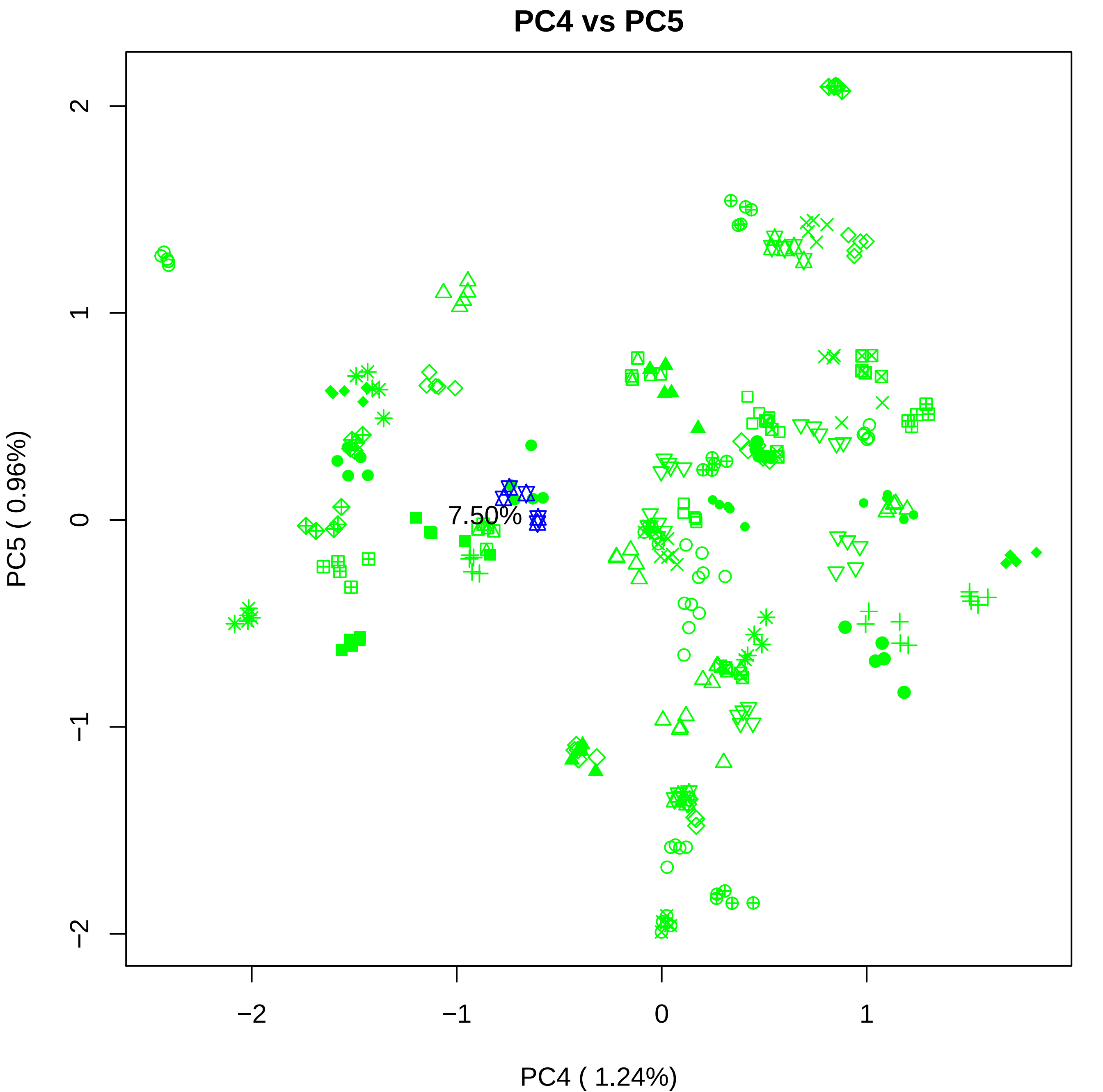
PC4 vs PC5
7.50%
PC5 ( 0.96%)
PC4 ( 1.24%)
2
1
0
−1
−2
−2
−1
0
1

### PCA of 10%  BMH(d) vs PMH(d)

Scaled Matrix PCAed

7th order polynomial baseline correction performed

Raw spectra smoothed to 21 Neighbours, 2nd order PN, Savtisky Golay

Normalisation between 0 and 100

D2 spectra smoothed to 41 Neighbours, 2nd order PN, Savtisky Golay

Regions Analysed: 3601−2501 and 2001−700cm−1

Samples Analysed: 1 bContam's 0 BLH's 0 BMH's 0 OMH's 0 pContam's 0 PLH's 60 PMH's

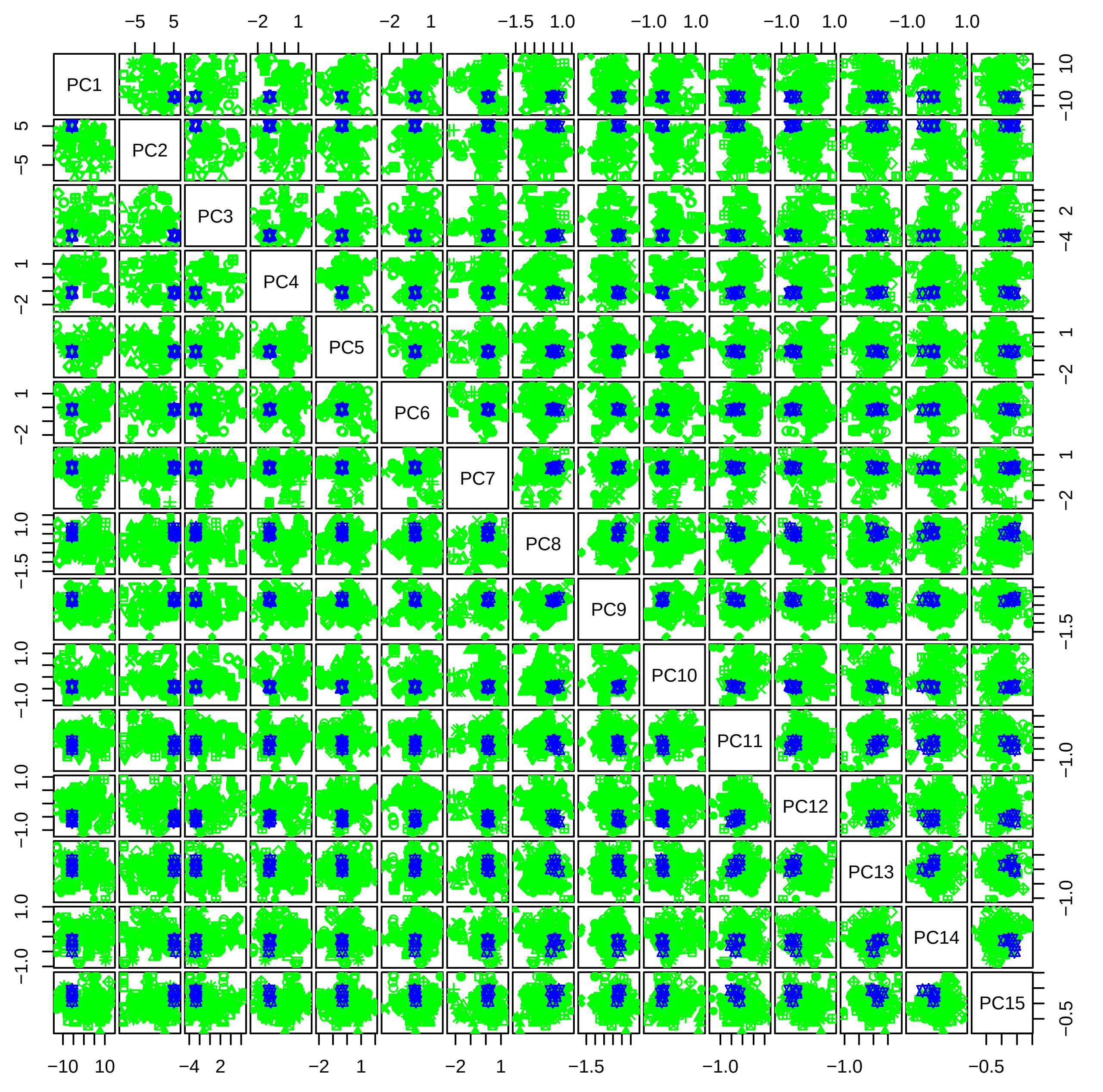

PC1
PC2
PC3
PC4
PC5
PC6
PC7
PC8
PC9
PC10
PC11
PC12
PC13
PC14
PC15
−5 5
−2 1
−2 1
−1.5 1.0
−1.0 1.0
−1.0 1.0
−1.0 1.0
−10 10
−4 2
−2 1
−2 1
−1.5
−1.0
−1.0
−0.5

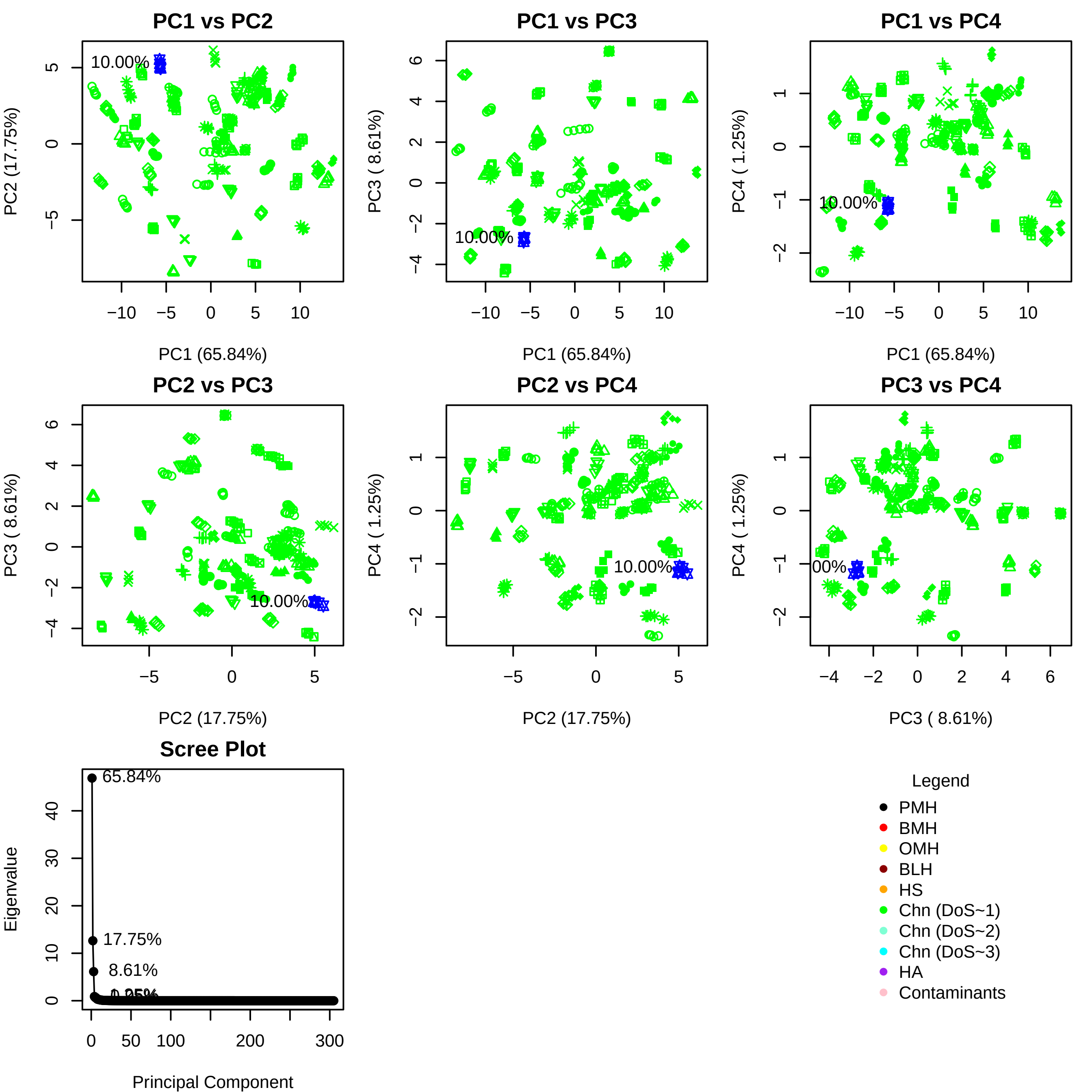
PC1 vs PC2
10.00%
PC2 (17.75%)
PC1 (65.84%)
PC1 vs PC3
10.00%
PC3 ( 8.61%)
PC1 (65.84%)
PC1 vs PC4
10.00%
PC4 ( 1.25%)
PC1 (65.84%)
PC2 vs PC3
10.00%
PC3 ( 8.61%)
PC2 (17.75%)
PC2 vs PC4
10.00%
PC4 ( 1.25%)
PC2 (17.75%)
PC3 vs PC4
00%
PC4 ( 1.25%)
PC3 ( 8.61%)
Scree Plot
65.84%
17.75%
8.61%
Eigenvalue
Principal Component
Legend
PMH
BMH
OMH
BLH
HS
Chn (DoS~1)
Chn (DoS~2)
Chn (DoS~3)
HA
Contaminants

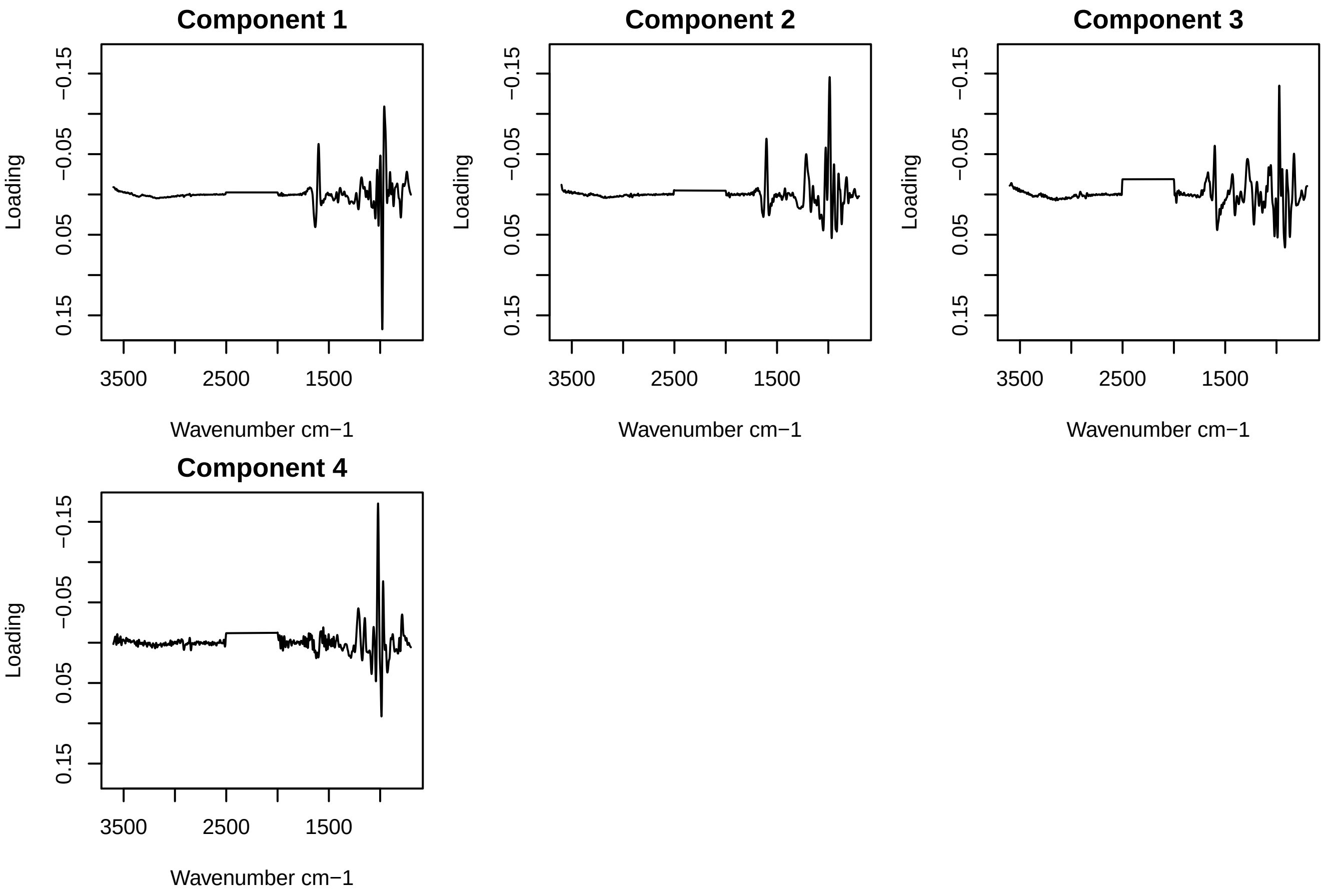
Component 1
Component 2
Component 3
Component 4
Loading
−0.15
−0.05
0.05
0.15
3500
2500
1500
Wavenumber cm−1

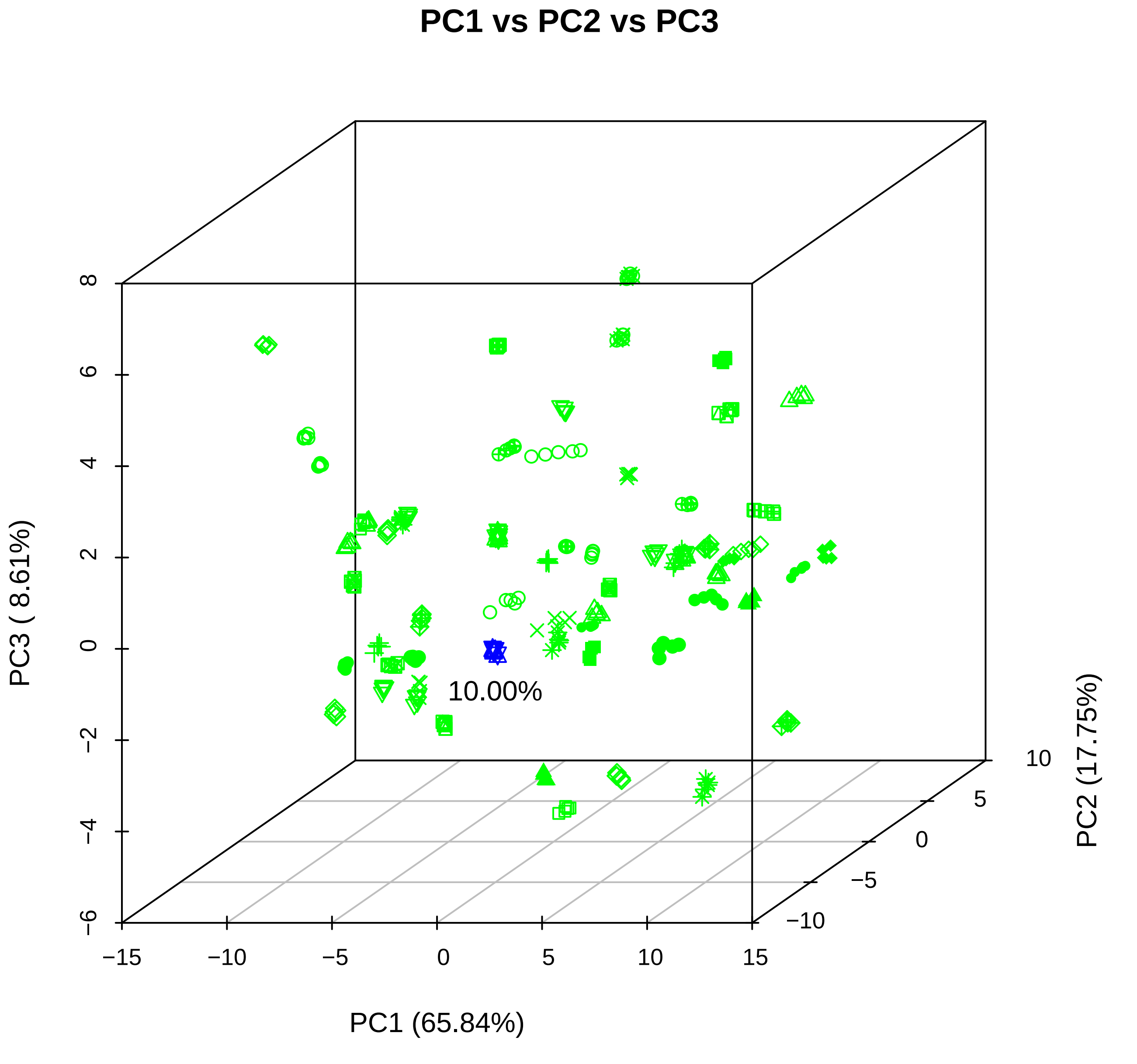
PC1 vs PC2 vs PC3
PC1 (65.84%)
PC2 (17.75%)
PC3 ( 8.61%)
10.00%
−15
−10
−5
0
5
10
15
−6
−4
−2
0
2
4
6
8
−10
−5
0
5
10

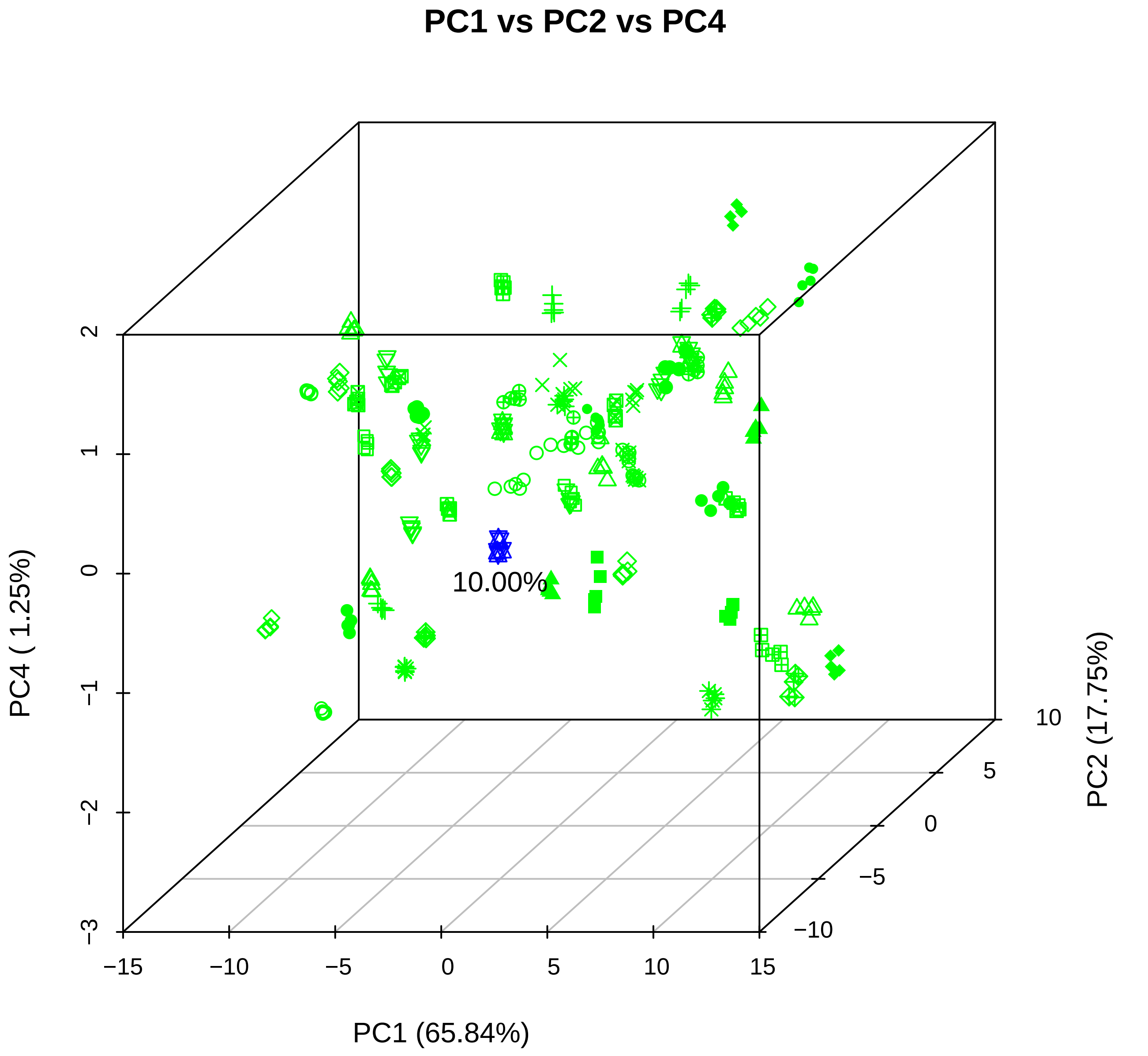
PC1 vs PC2 vs PC4
10.00%
PC1 (65.84%)
PC2 (17.75%)
PC4 ( 1.25%)
−15
−10
−5
0
5
10
15
−3
−2
−1
0
1
2
−10
−5
0
5
10

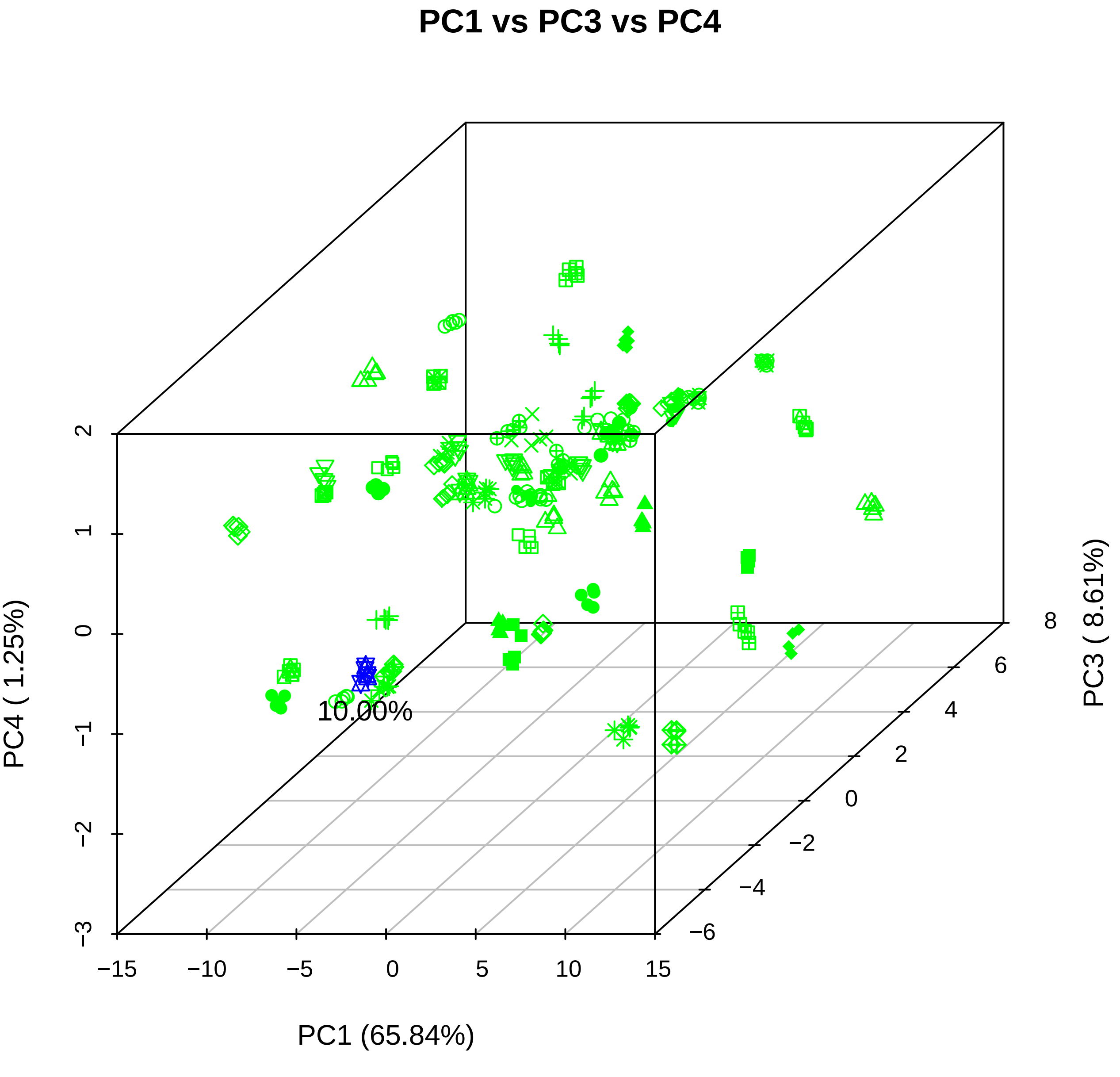
PC1 vs PC3 vs PC4
PC1 (65.84%)
PC3 ( 8.61%)
PC4 ( 1.25%)
10.00%
−15
−10
−5
0
5
10
15
−6
−4
−2
0
2
4
6
8
−3
−2
−1
0
1
2

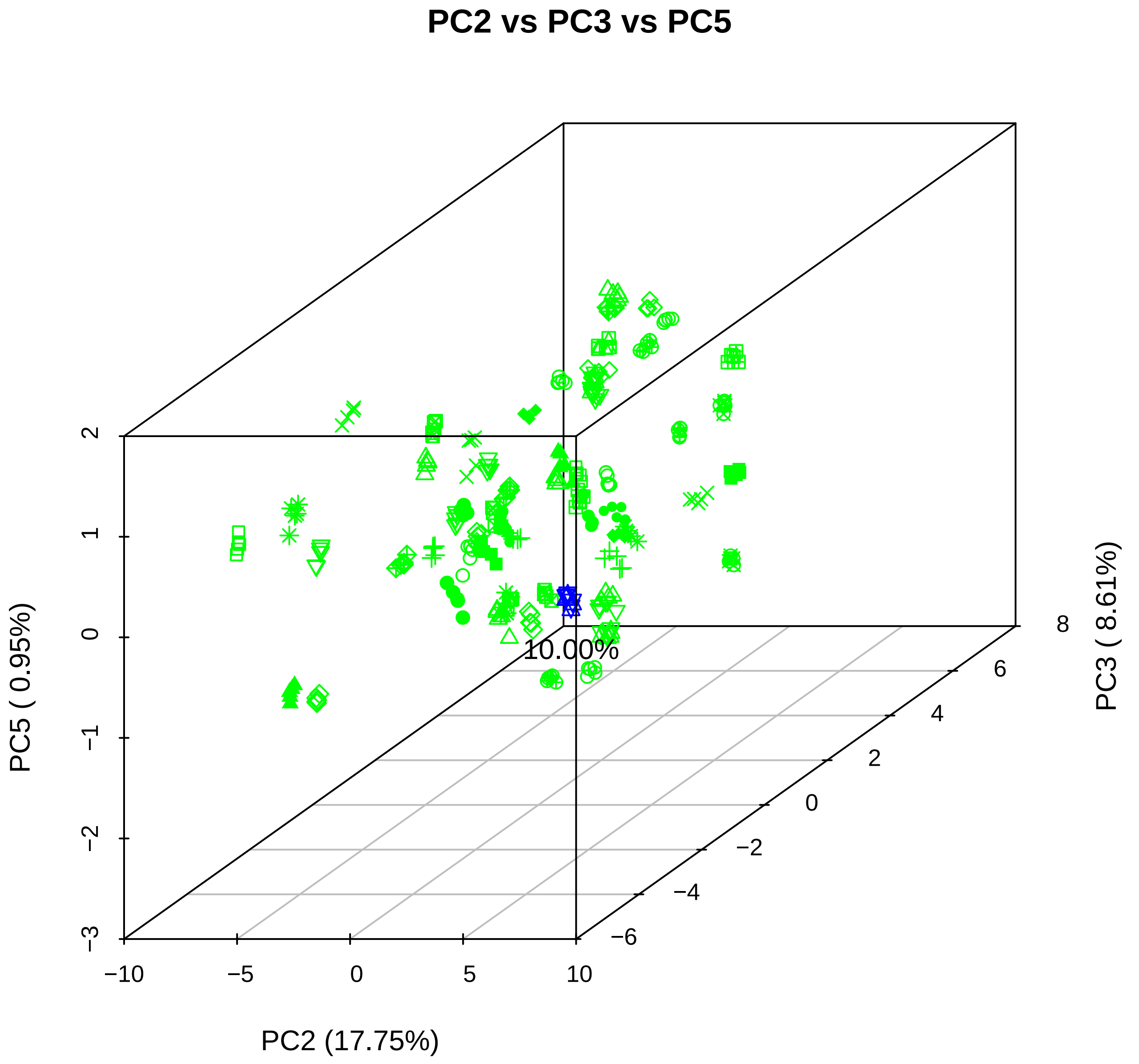

PC2 vs PC3 vs PC5
PC5 ( 0.95%)
2
1
0
−1
−2
−3
−10
−5
0
5
10
PC2 (17.75%)
PC3 ( 8.61%)
8
6
4
2
0
−2
−4
−6
10.00%

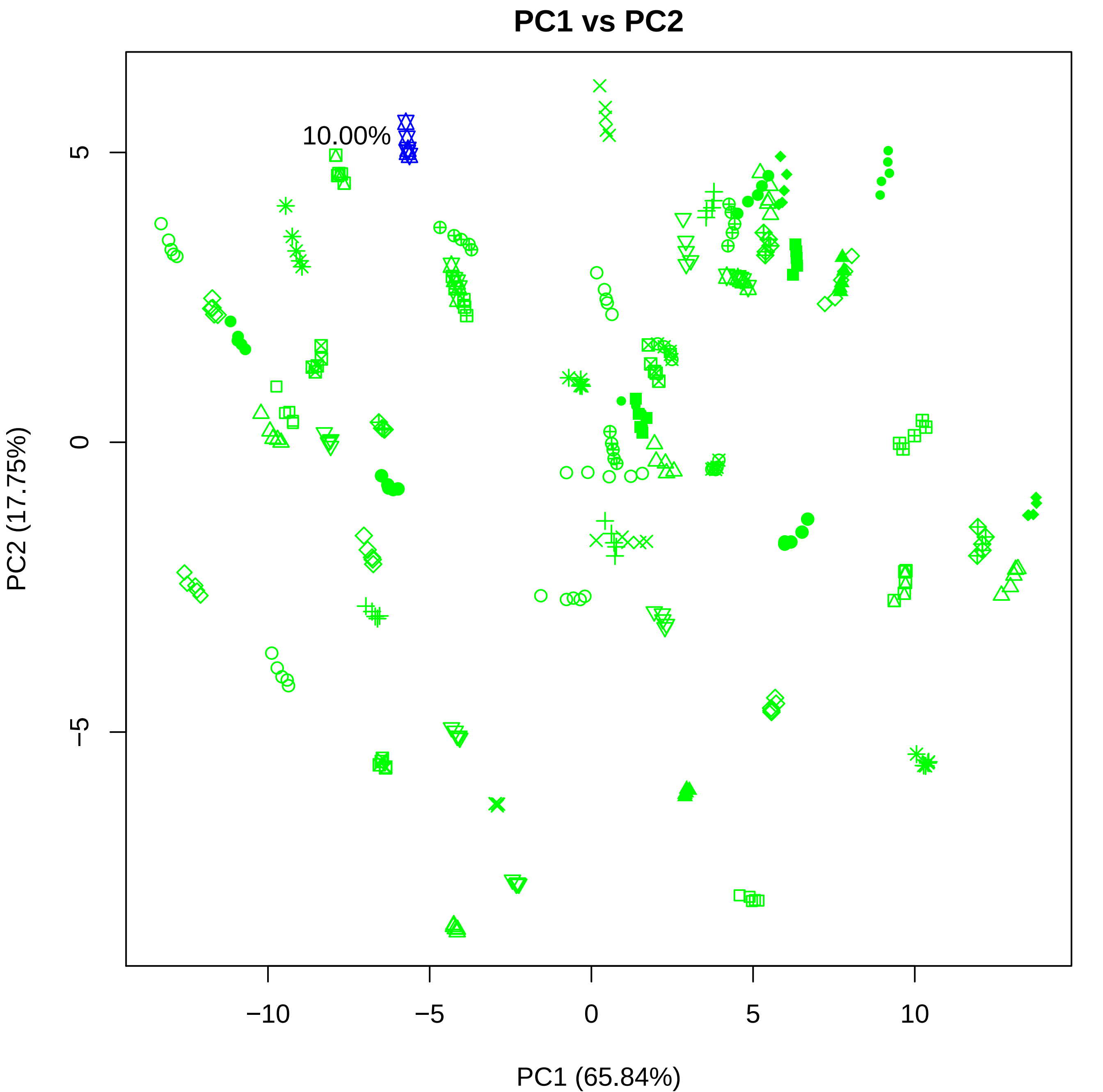
PC1 vs PC2
10.00%
5
0
−5
PC2 (17.75%)
−10
−5
0
5
10
PC1 (65.84%)

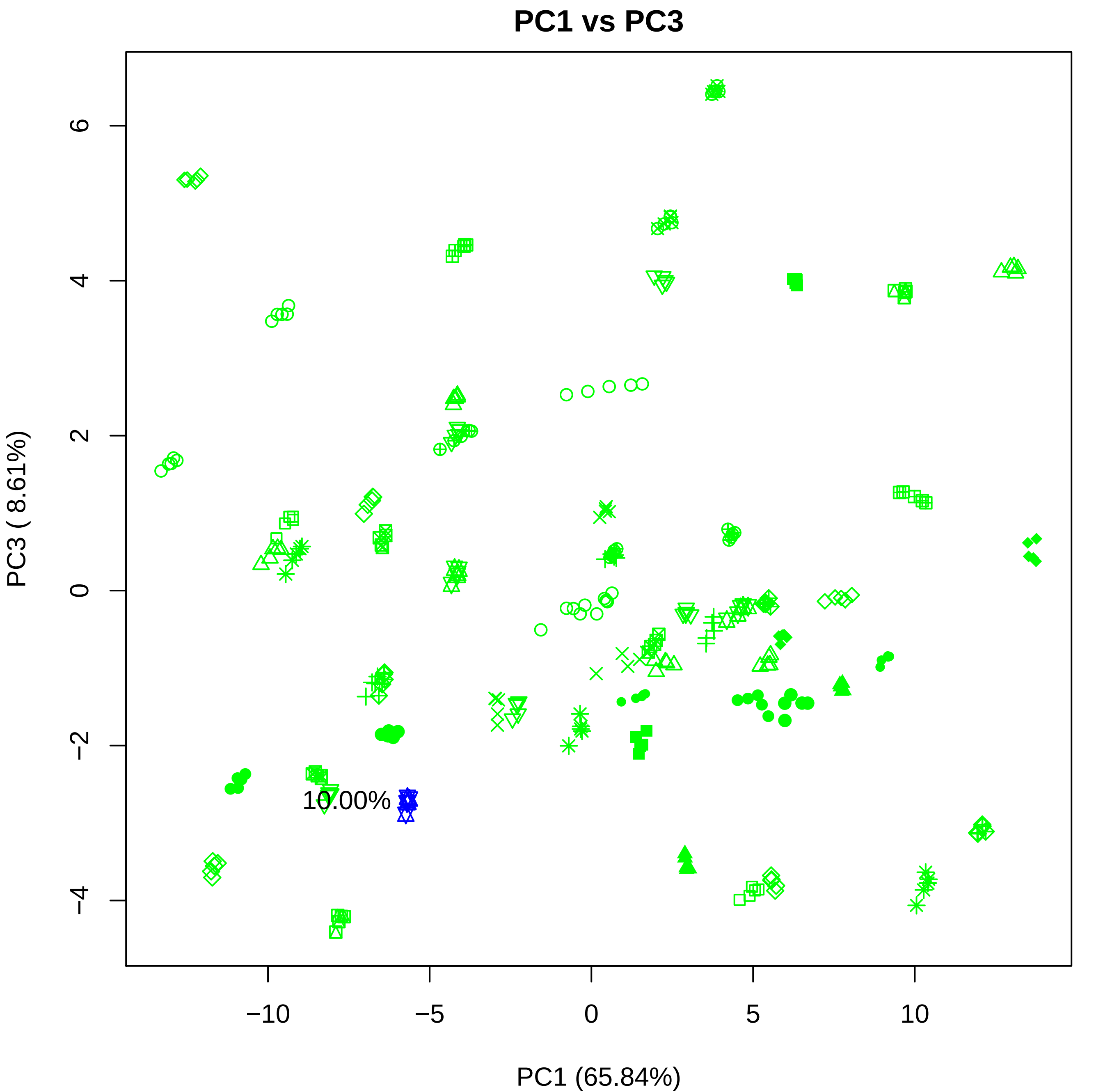

PC1 vs PC3
PC3 ( 8.61%)
PC1 (65.84%)
6
4
2
0
−2
−4
−10
−5
0
5
10
10.00%

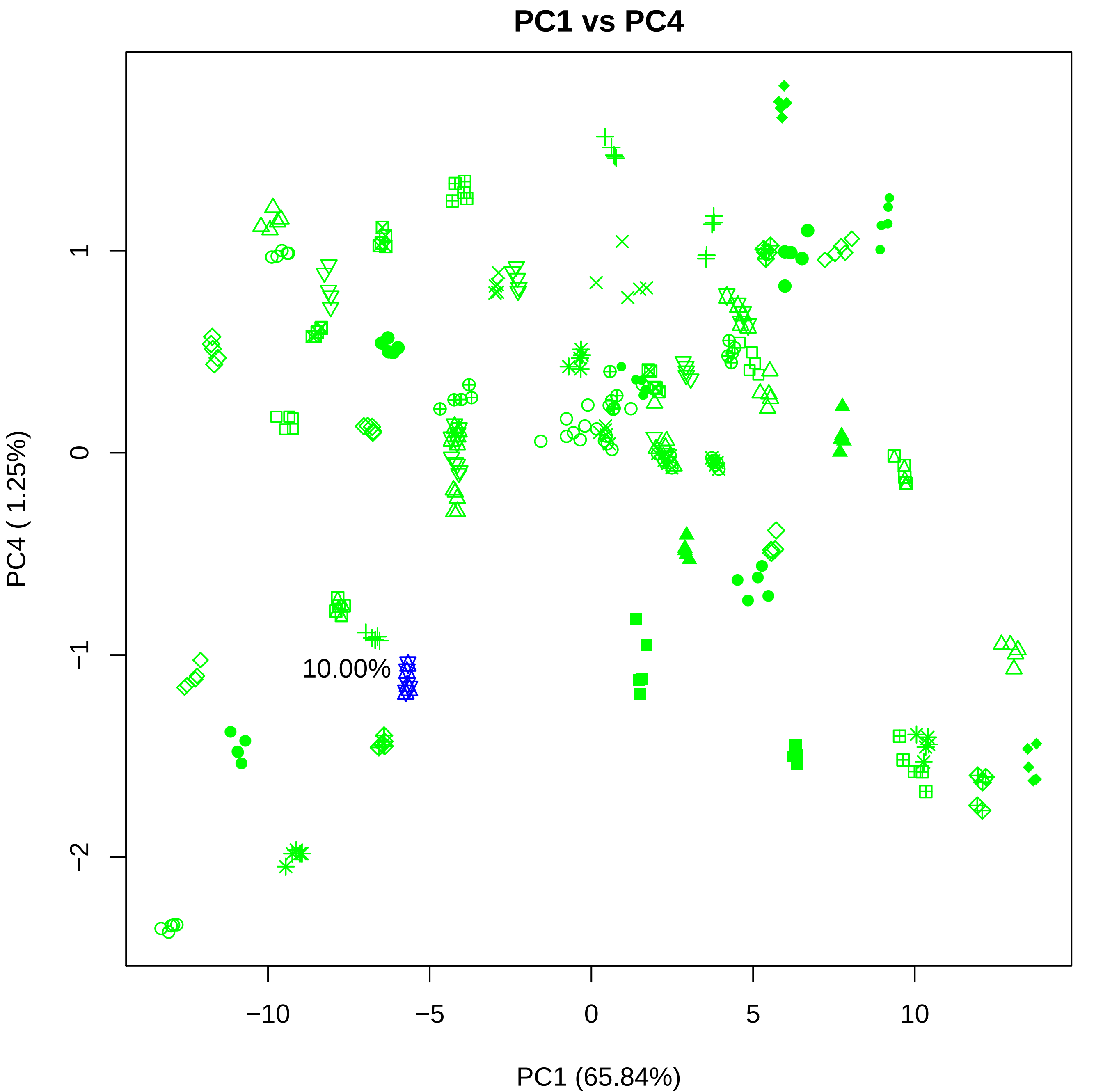
PC1 vs PC4
PC4 ( 1.25%)
PC1 (65.84%)
10.00%
1
0
−1
−2
−10
−5
0
5
10

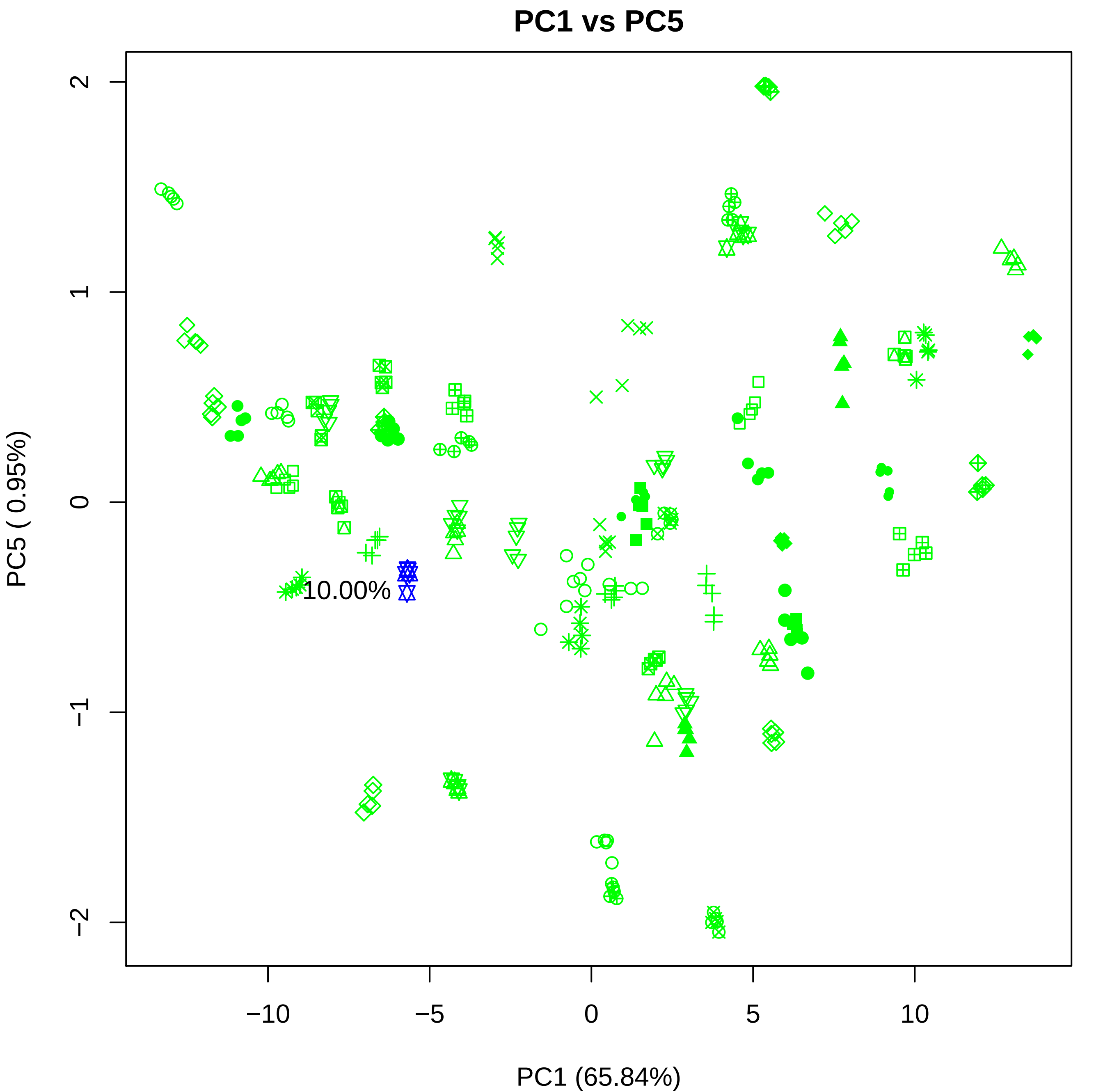

PC1 vs PC5
10.00%
PC5 ( 0.95%)
PC1 (65.84%)
2
1
0
−1
−2
−10
−5
0
5
10

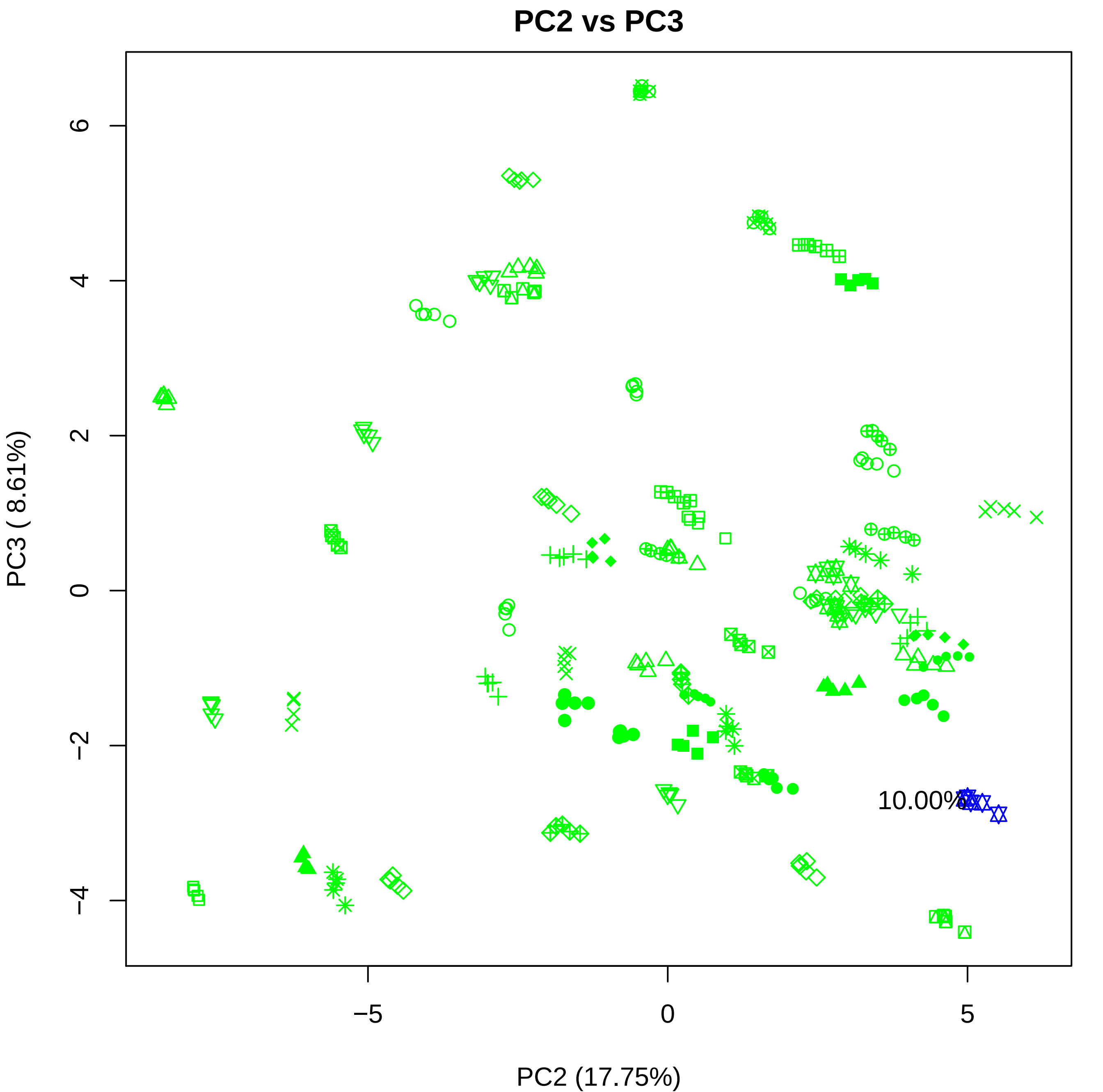
PC2 vs PC3
PC3 ( 8.61%)
PC2 (17.75%)
6
4
2
0
−2
−4
−5
0
5
10.00%

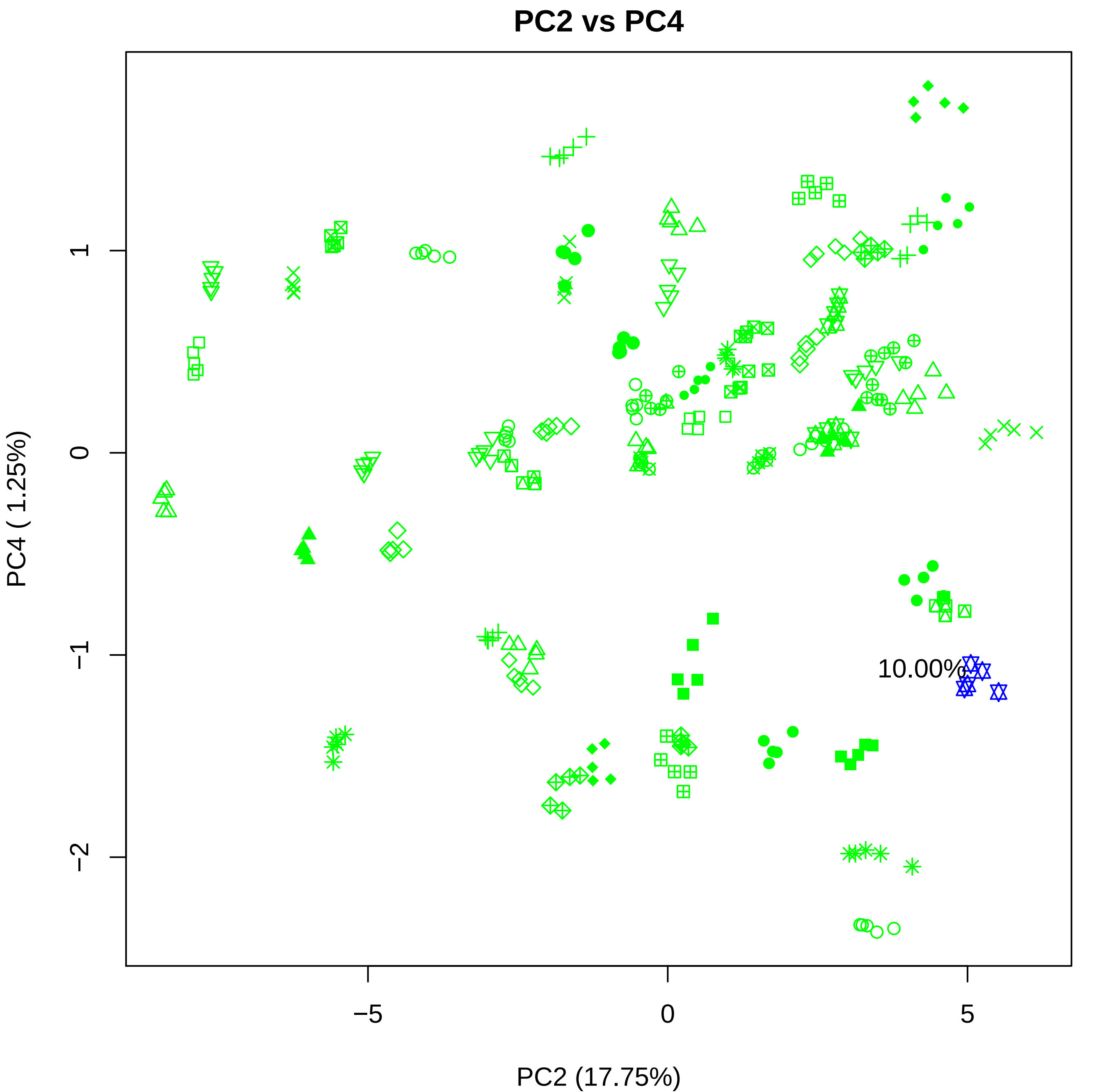

PC2 vs PC4
10.00%
PC4 ( 1.25%)
−2
−1
0
1
−5
0
5
PC2 (17.75%)

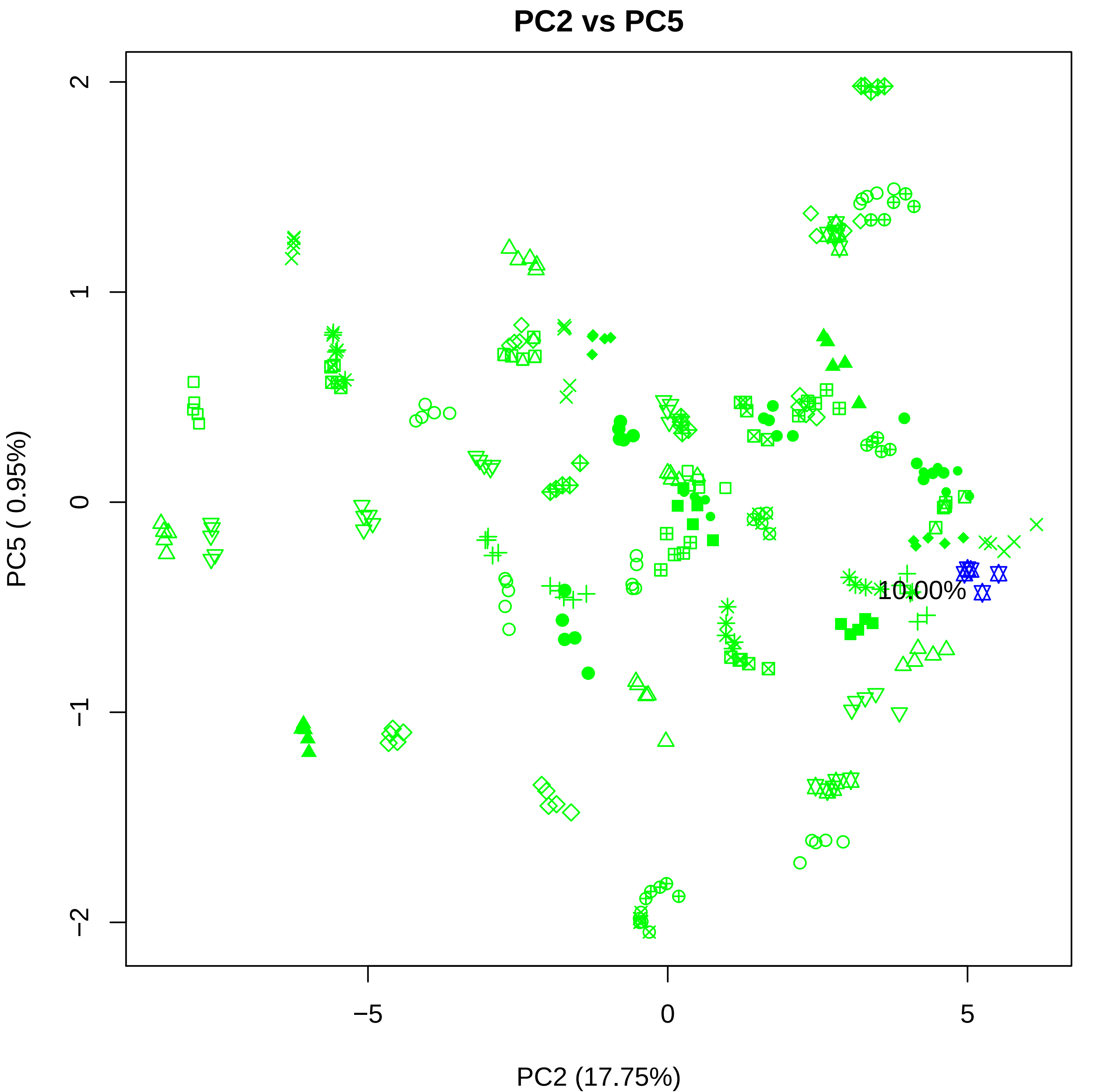
PC2 vs PC5
PC5 ( 0.95%)
PC2 (17.75%)
2
1
0
−1
−2
−5
0
5
10.00%

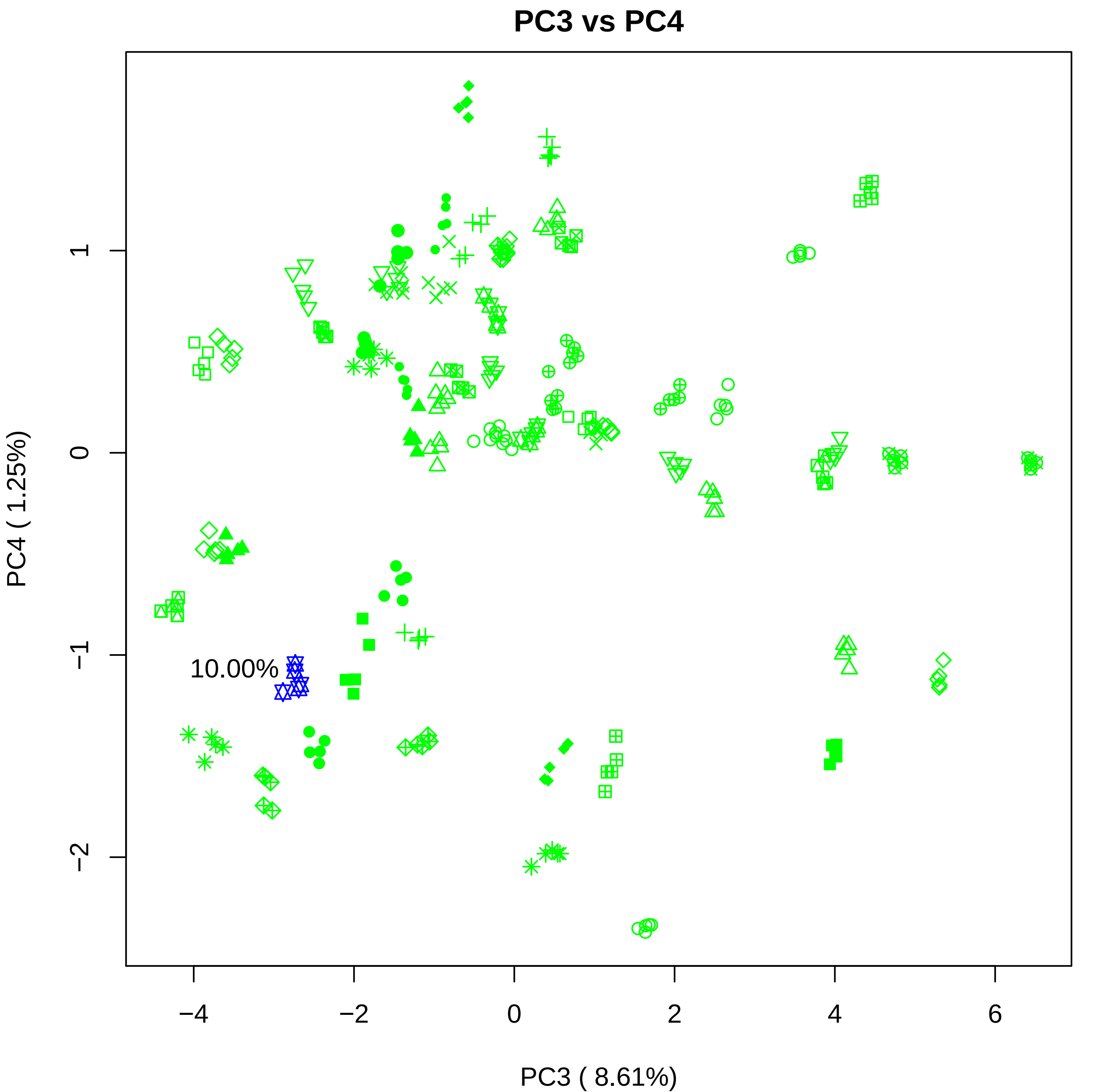
PC3 vs PC4
PC4 ( 1.25%)
PC3 ( 8.61%)
10.00%
1
0
−1
−2
−4
−2
0
2
4
6

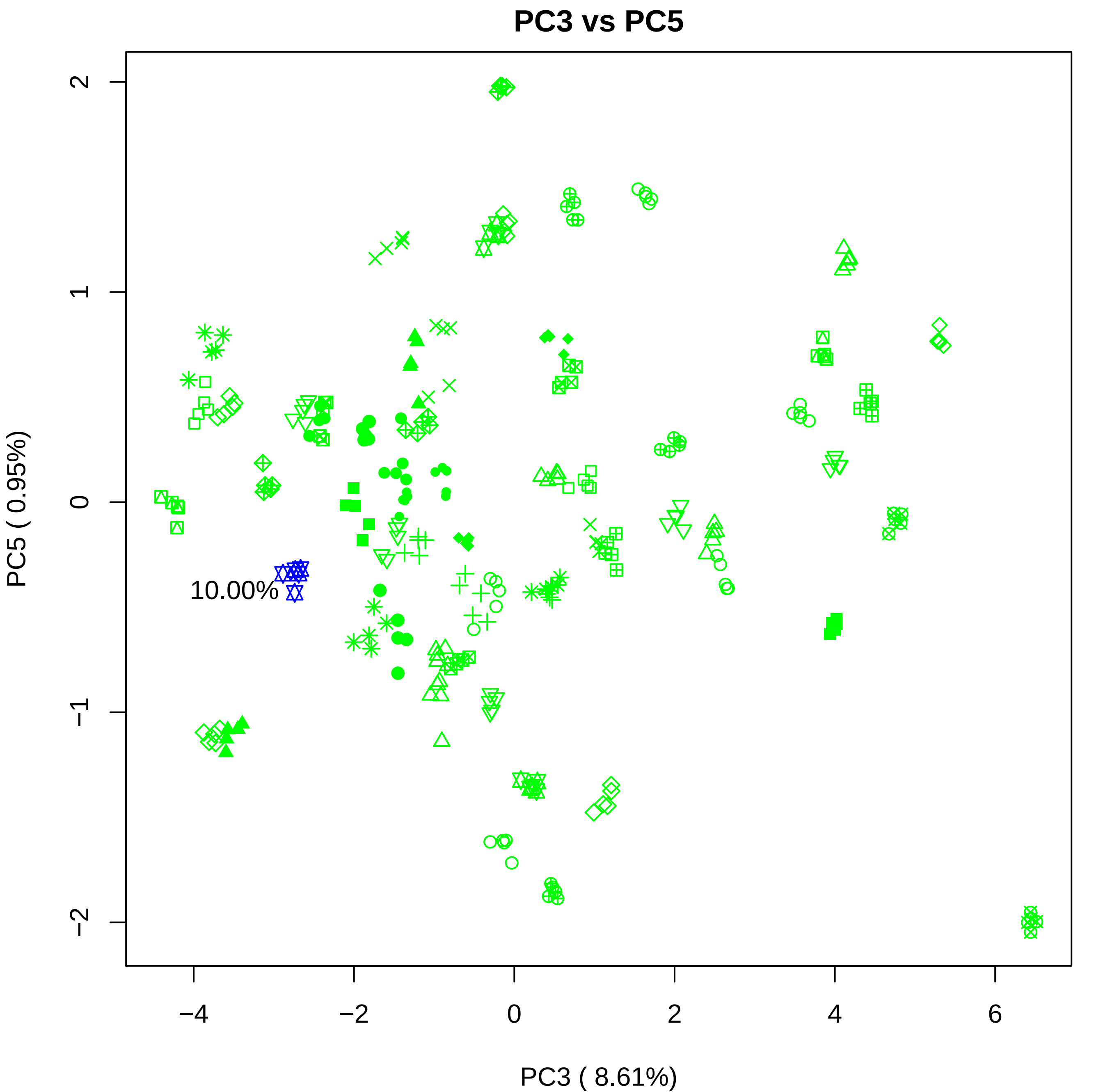

PC3 vs PC5
2
1
0
−1
−2
PC5 ( 0.95%)
10.00%
−4
−2
0
2
4
6
PC3 ( 8.61%)

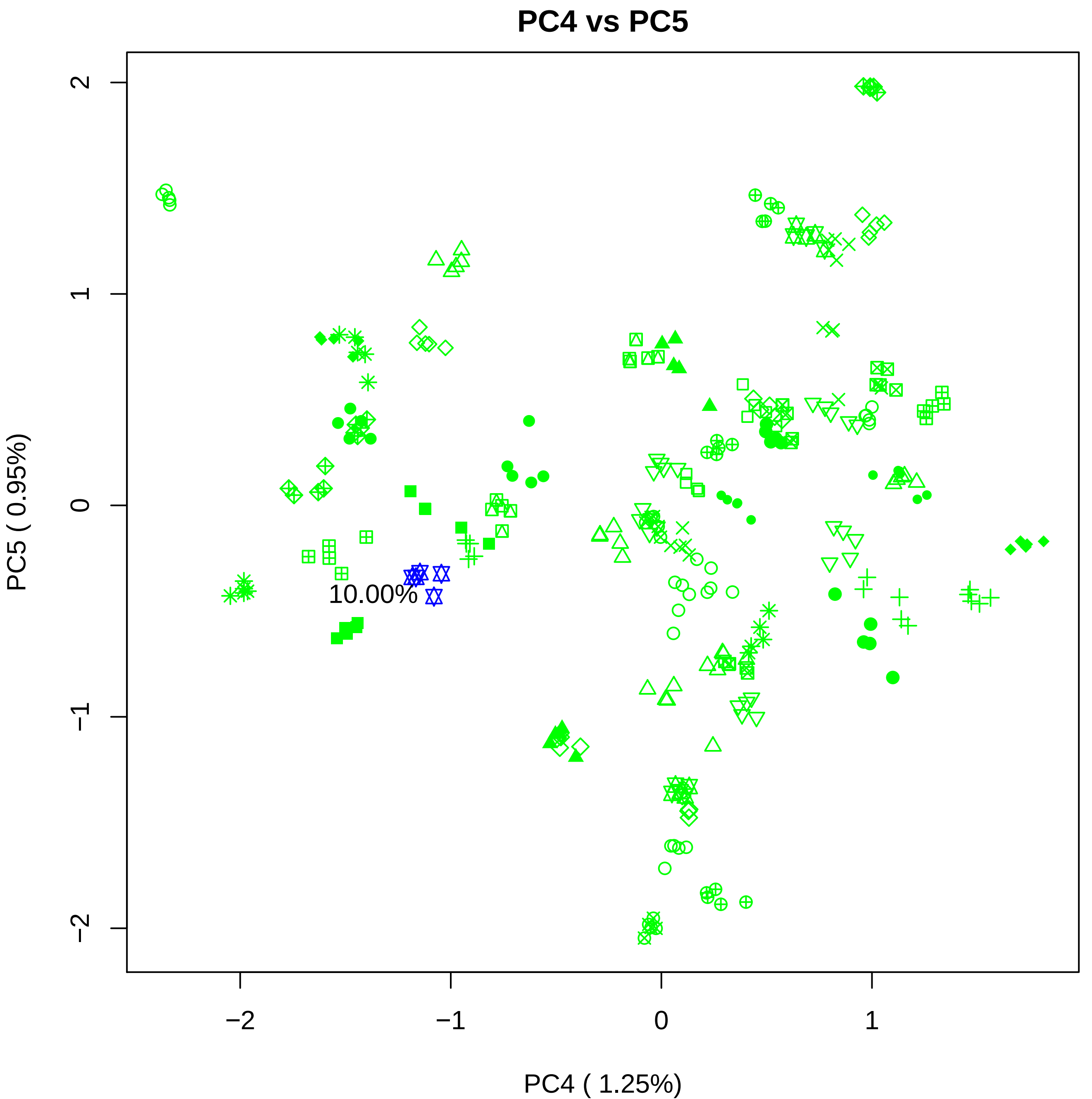

PC4 vs PC5
PC5 ( 0.95%)
PC4 ( 1.25%)
2
1
0
−1
−2
−2
−1
0
1
10.00%

### PCA of 15%  BMH(d) vs PMH(d)

Scaled Matrix PCAed

7th order polynomial baseline correction performed

Raw spectra smoothed to 21 Neighbours, 2nd order PN, Savtisky Golay

Normalisation between 0 and 100

D2 spectra smoothed to 41 Neighbours, 2nd order PN, Savtisky Golay

Regions Analysed: 3601−2501 and 2001−700cm−1

Samples Analysed: 1 bContam's 0 BLH's 0 BMH's 0 OMH's 0 pContam's 0 PLH's 60 PMH's

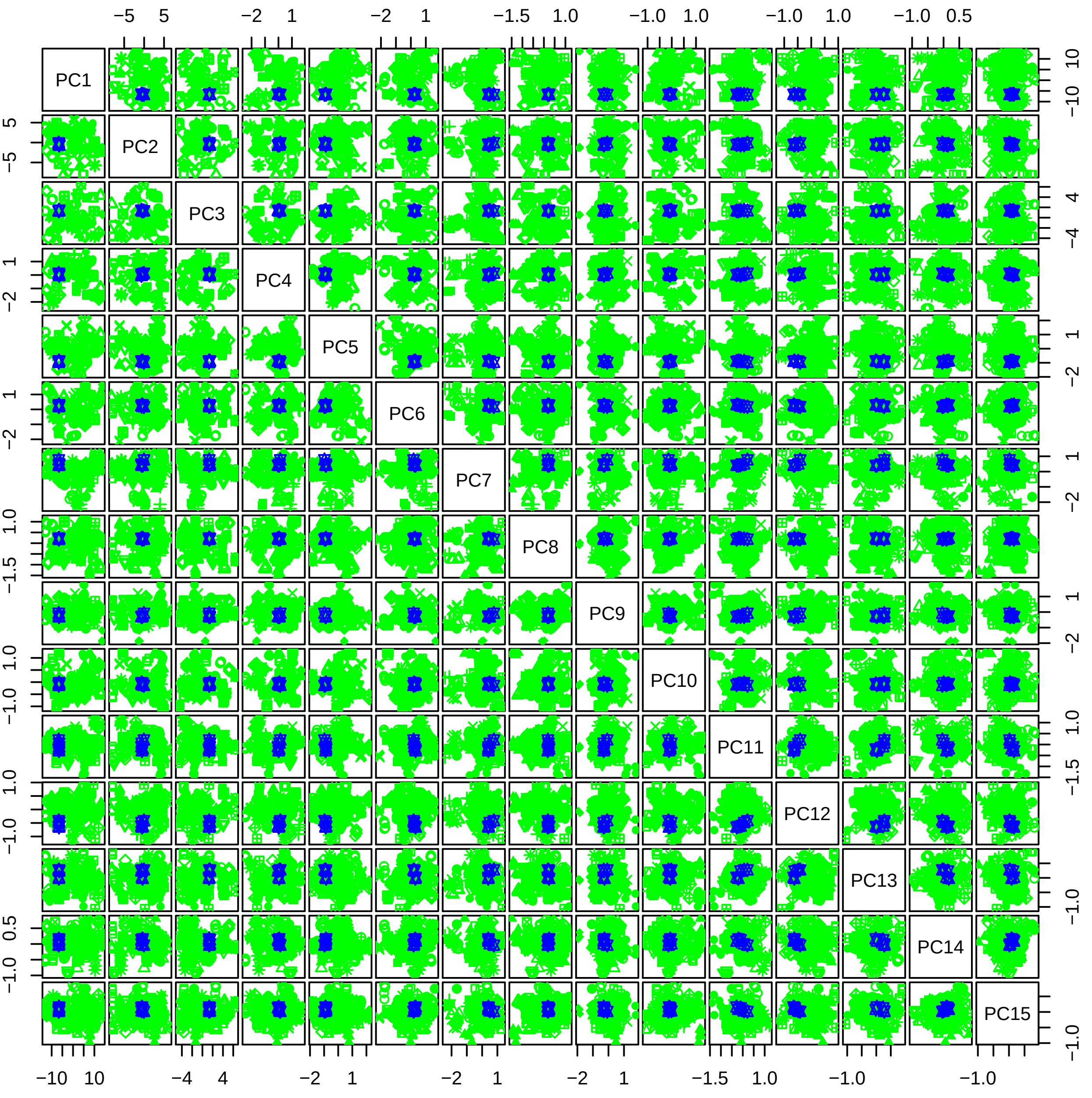
PC1
PC2
PC3
PC4
PC5
PC6
PC7
PC8
PC9
PC10
PC11
PC12
PC13
PC14
PC15

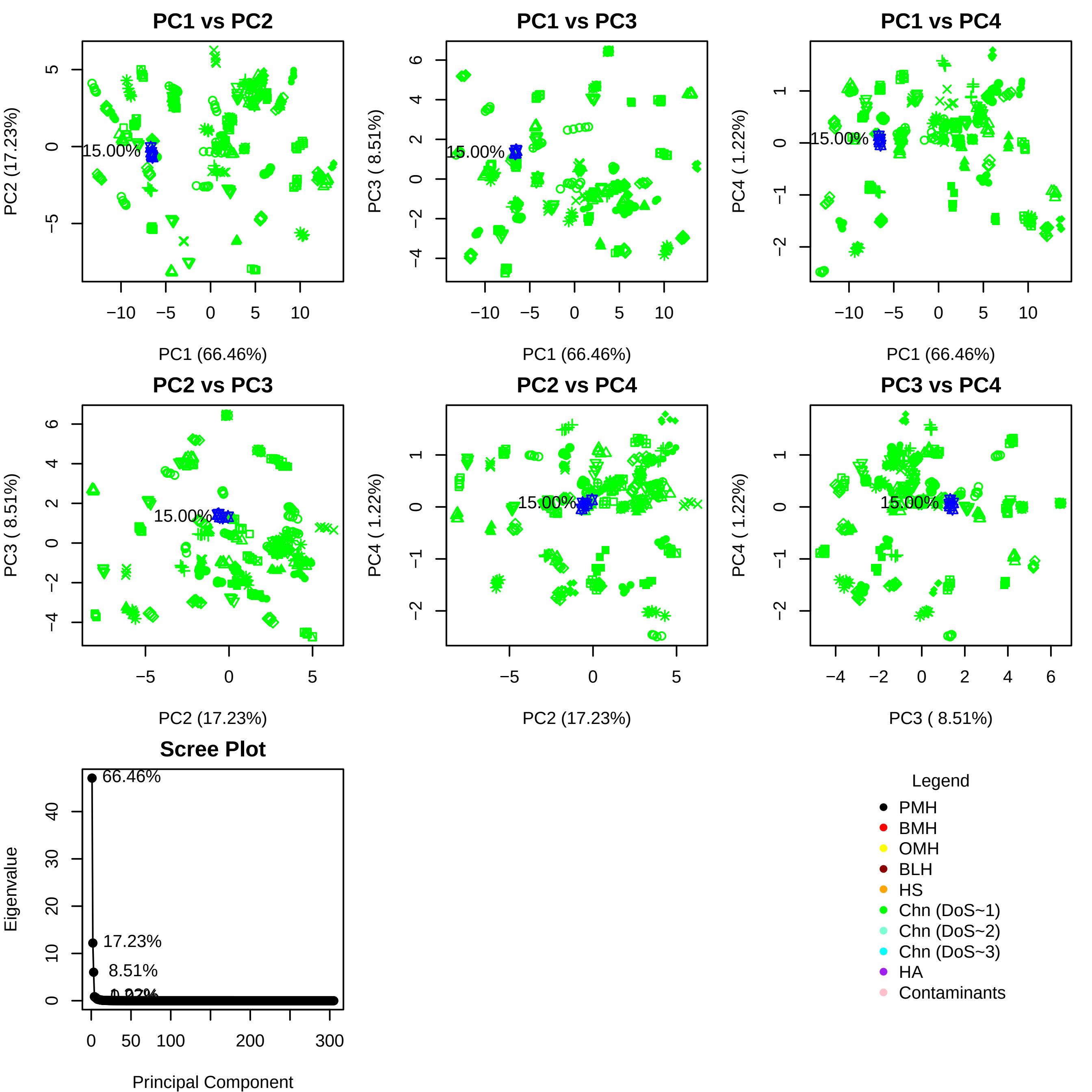

PC1 vs PC2
PC1 vs PC3
PC1 vs PC4
PC2 vs PC3
PC2 vs PC4
PC3 vs PC4
Scree Plot
PC1 (66.46%)
PC2 (17.23%)
PC3 ( 8.51%)
PC4 ( 1.22%)
15.00%
Eigenvalue
Principal Component
66.46%
17.23%
8.51%
Legend
PMH
BMH
OMH
BLH
HS
Chn (DoS~1)
Chn (DoS~2)
Chn (DoS~3)
HA
Contaminants

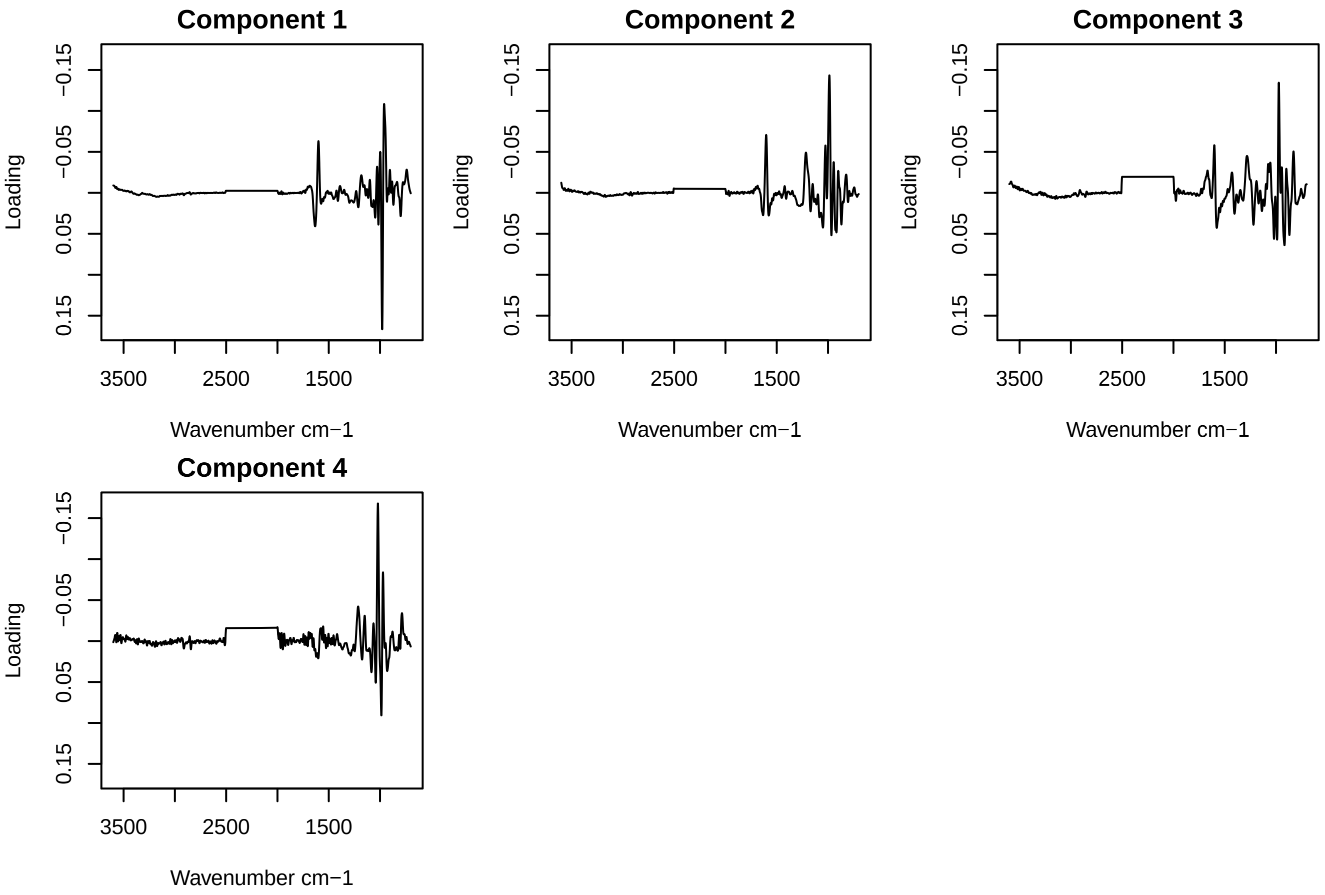
Component 1
Component 2
Component 3
Component 4
Loading
−0.15
−0.05
0.05
0.15
3500
2500
1500
Wavenumber cm−1

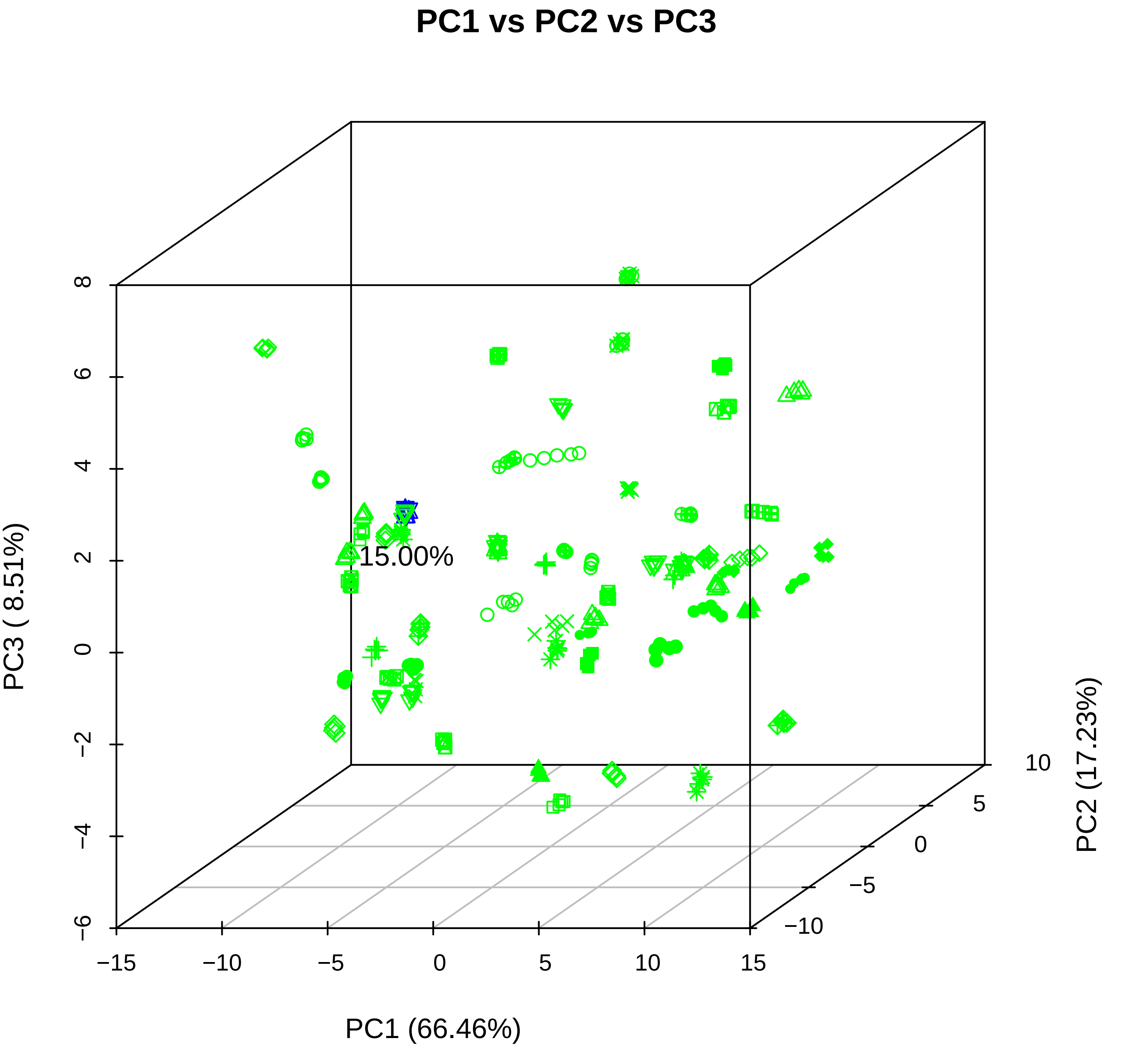
PC1 vs PC2 vs PC3
PC1 (66.46%)
PC2 (17.23%)
PC3 ( 8.51%)
15.00%
−15
−10
−5
0
5
10
15
−10
−5
0
5
10
−6
−4
−2
0
2
4
6
8

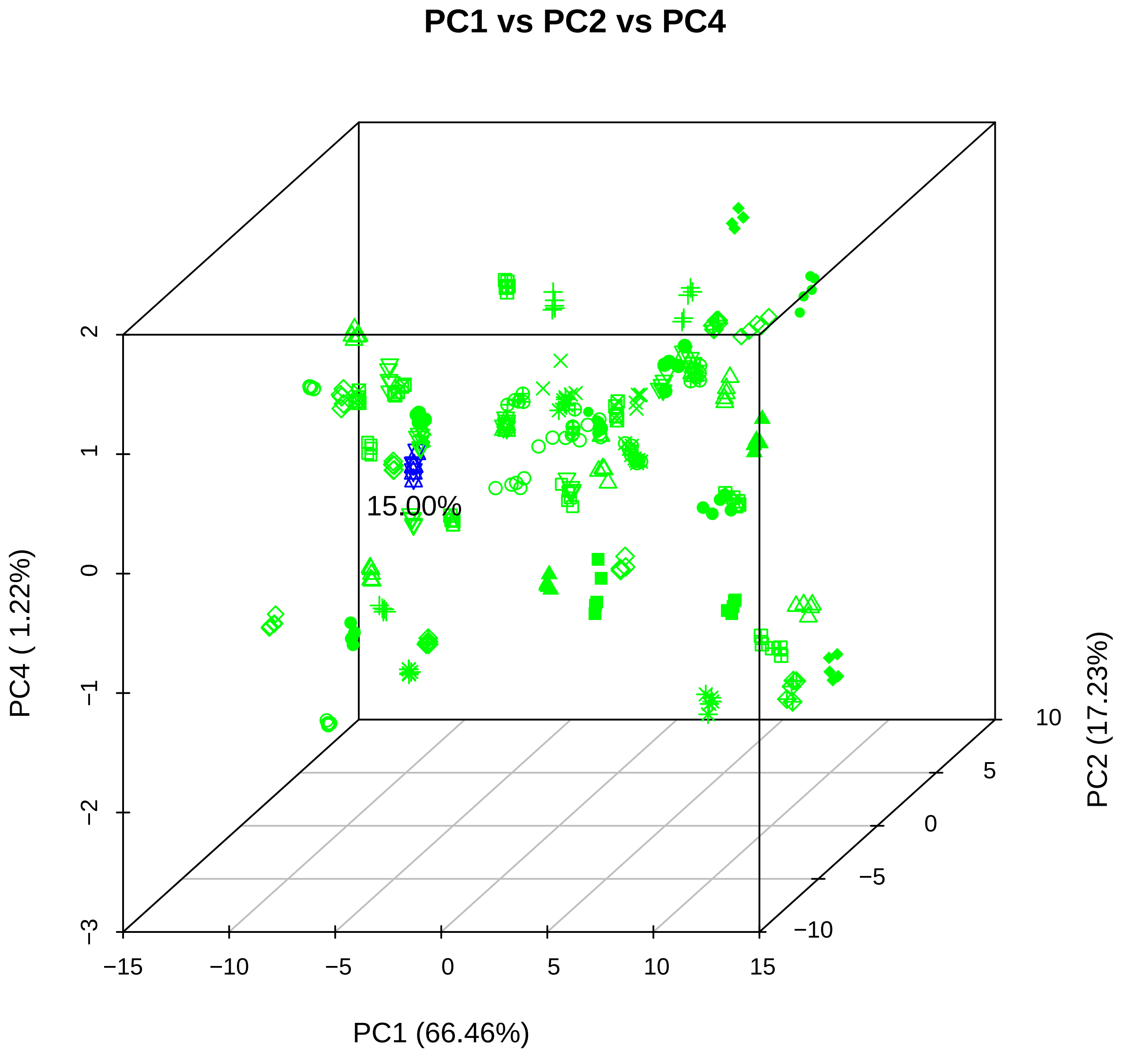
PC1 vs PC2 vs PC4
15.00%
PC1 (66.46%)
PC2 (17.23%)
PC4 ( 1.22%)
−15
−10
−5
0
5
10
15
−10
−5
0
5
10
−3
−2
−1
0
1
2

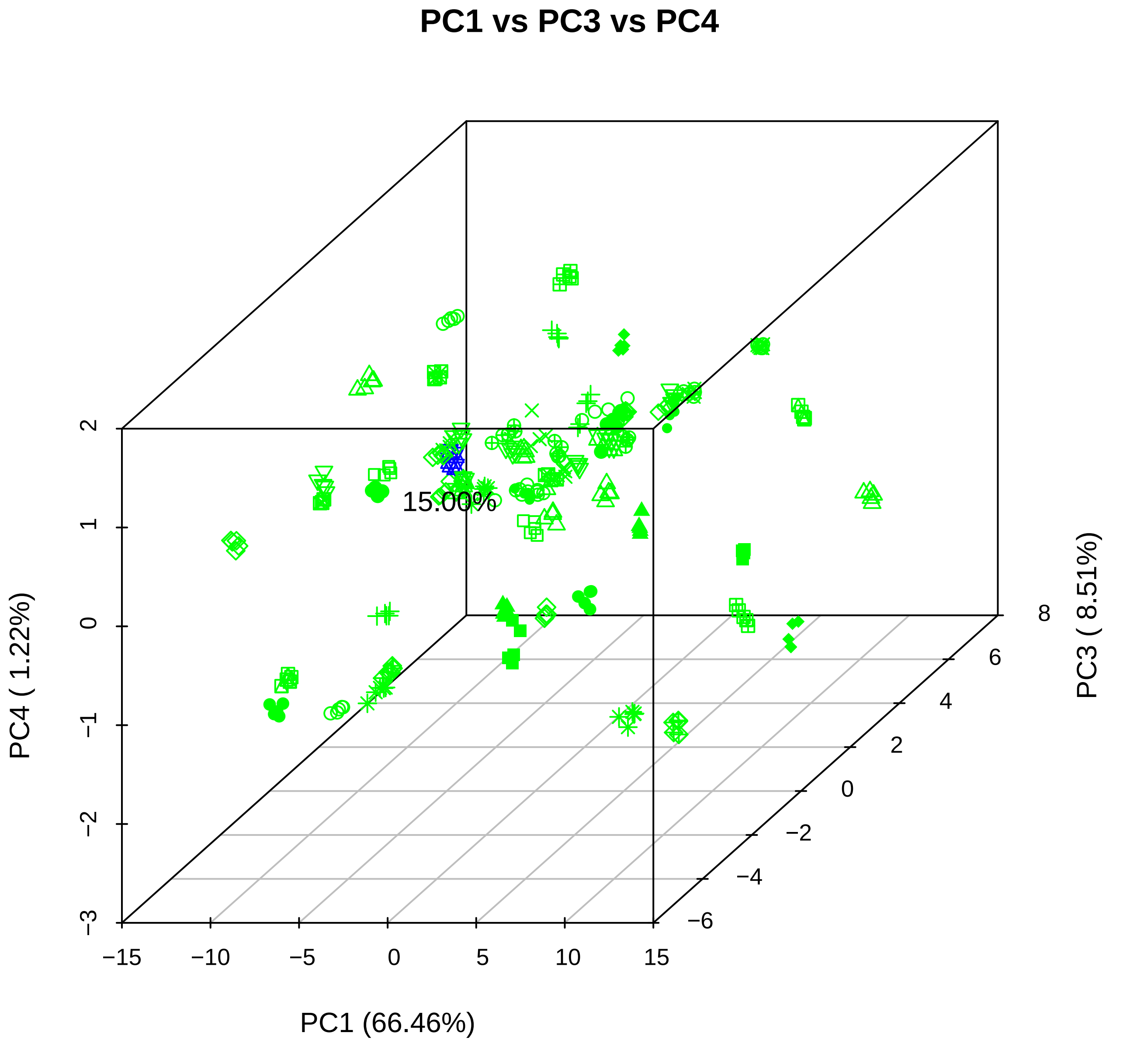

PC1 vs PC3 vs PC4
15.00%
PC1 (66.46%)
PC4 ( 1.22%)
PC3 ( 8.51%)
−15
−10
−5
0
5
10
15
−3
−2
−1
0
1
2
−6
−4
−2
0
2
4
6
8

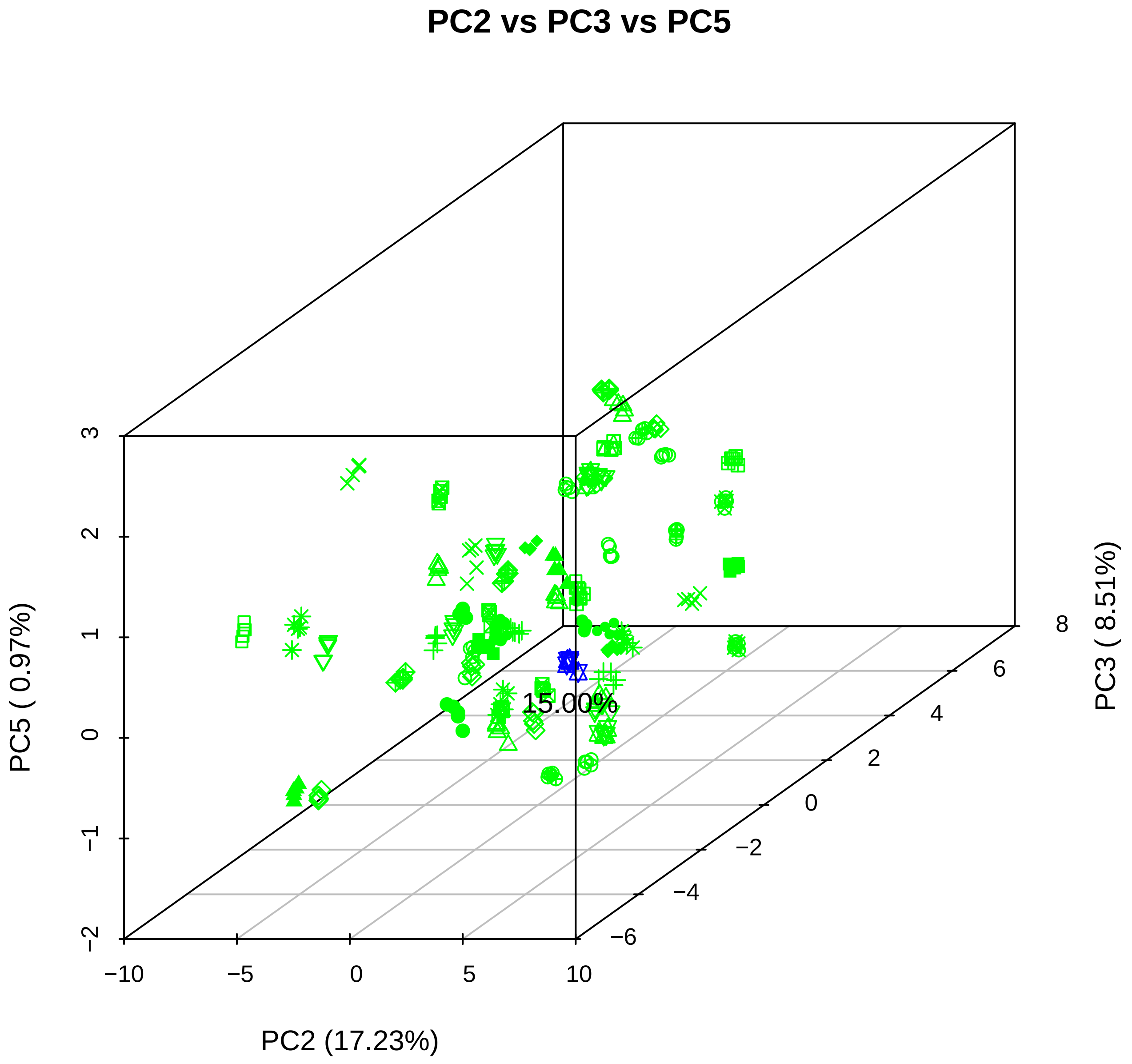
PC2 vs PC3 vs PC5
PC5 ( 0.97%)
PC2 (17.23%)
PC3 ( 8.51%)
15.00%
−2
−1
0
1
2
3
−10
−5
0
5
10
−6
−4
−2
0
2
4
6
8

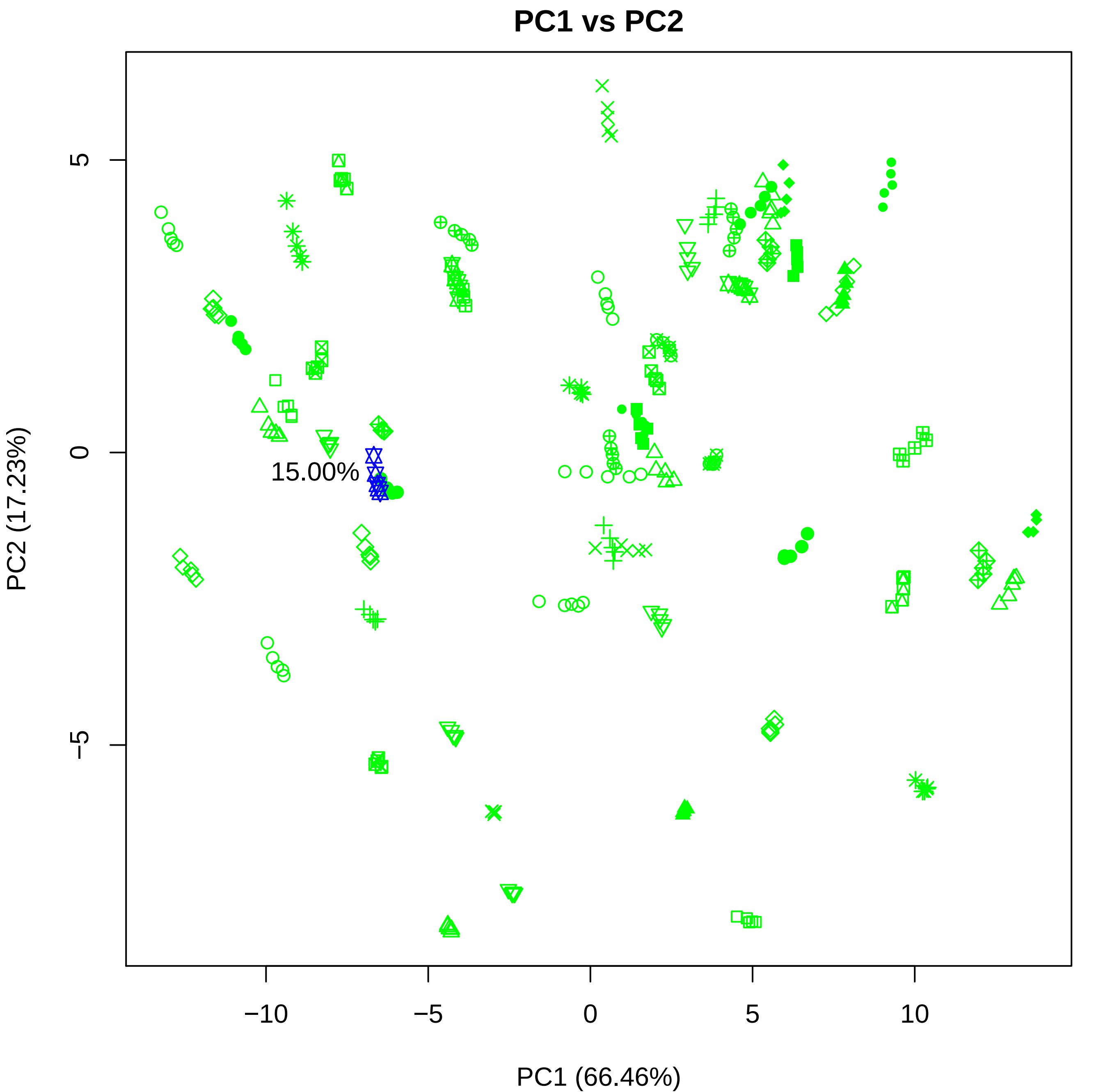
PC1 vs PC2
15.00%
5
0
−5
PC2 (17.23%)
−10
−5
0
5
10
PC1 (66.46%)

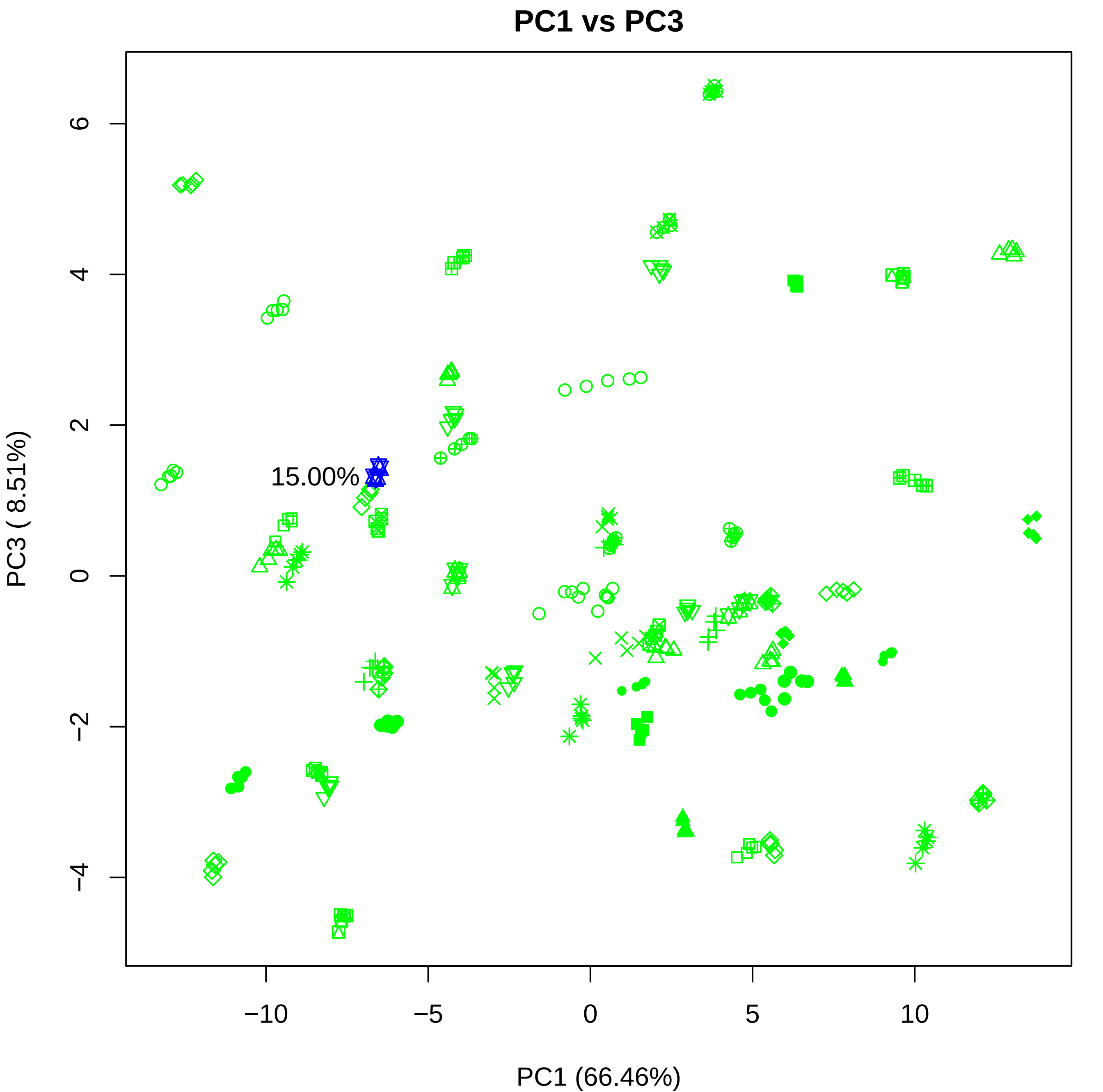
PC1 vs PC3
PC3 ( 8.51%)
PC1 (66.46%)
15.00%
6
4
2
0
−2
−4
−10
−5
0
5
10

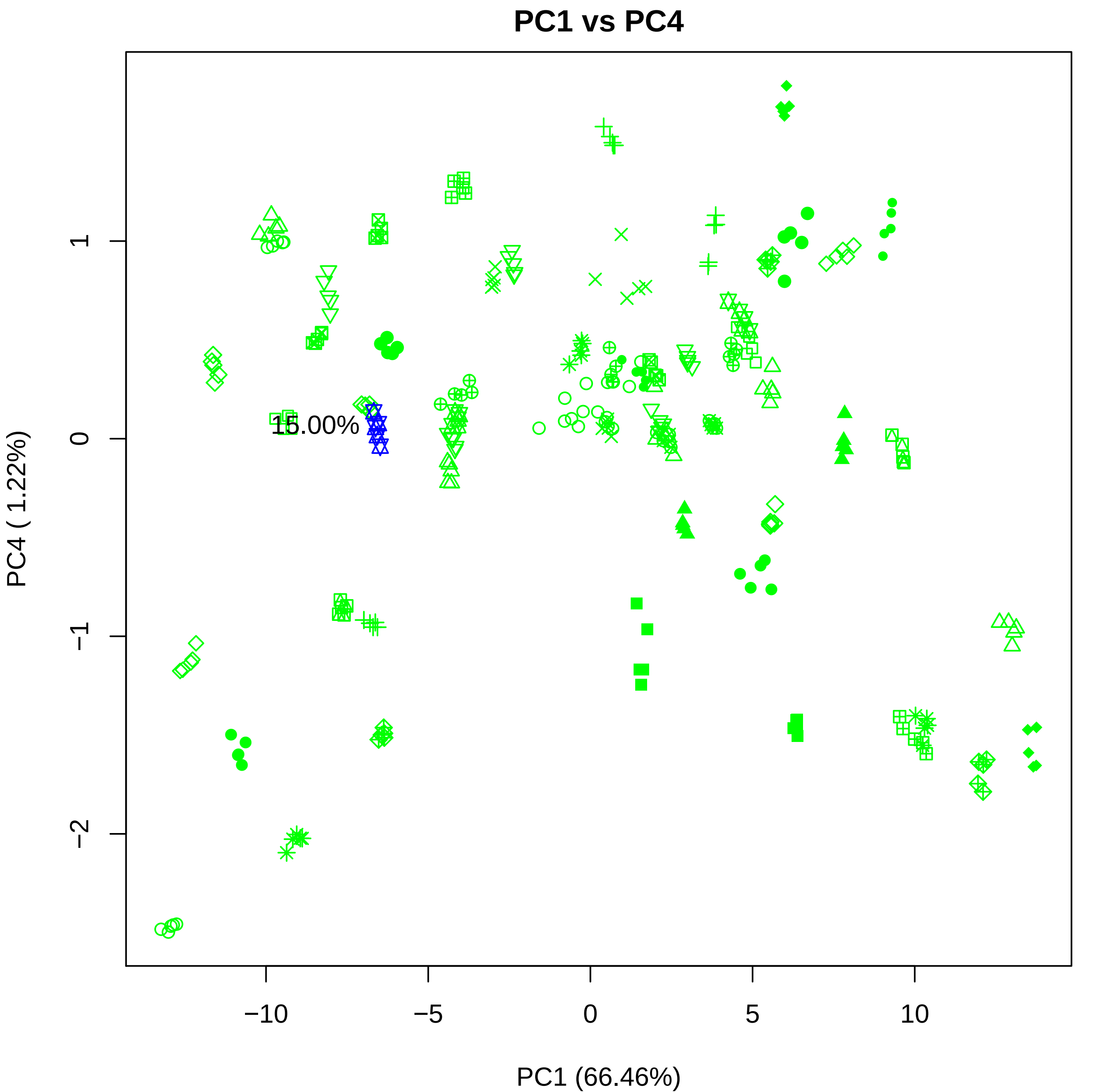
PC1 vs PC4
15.00%
PC4 ( 1.22%)
PC1 (66.46%)
−10
−5
0
5
10
−2
−1
0
1

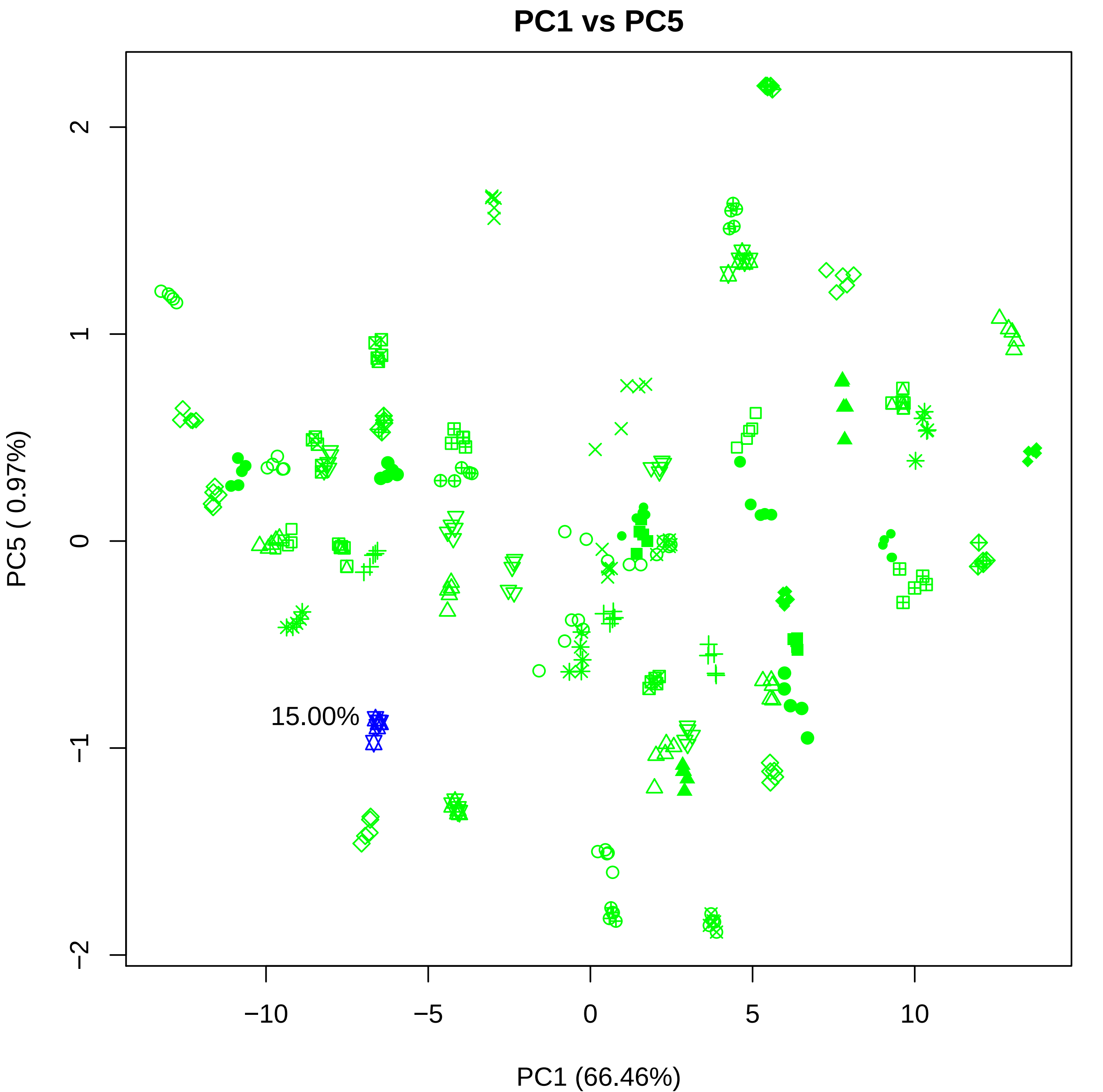
PC1 vs PC5
15.00%
PC5 ( 0.97%)
PC1 (66.46%)
2
1
0
−1
−2
−10
−5
0
5
10

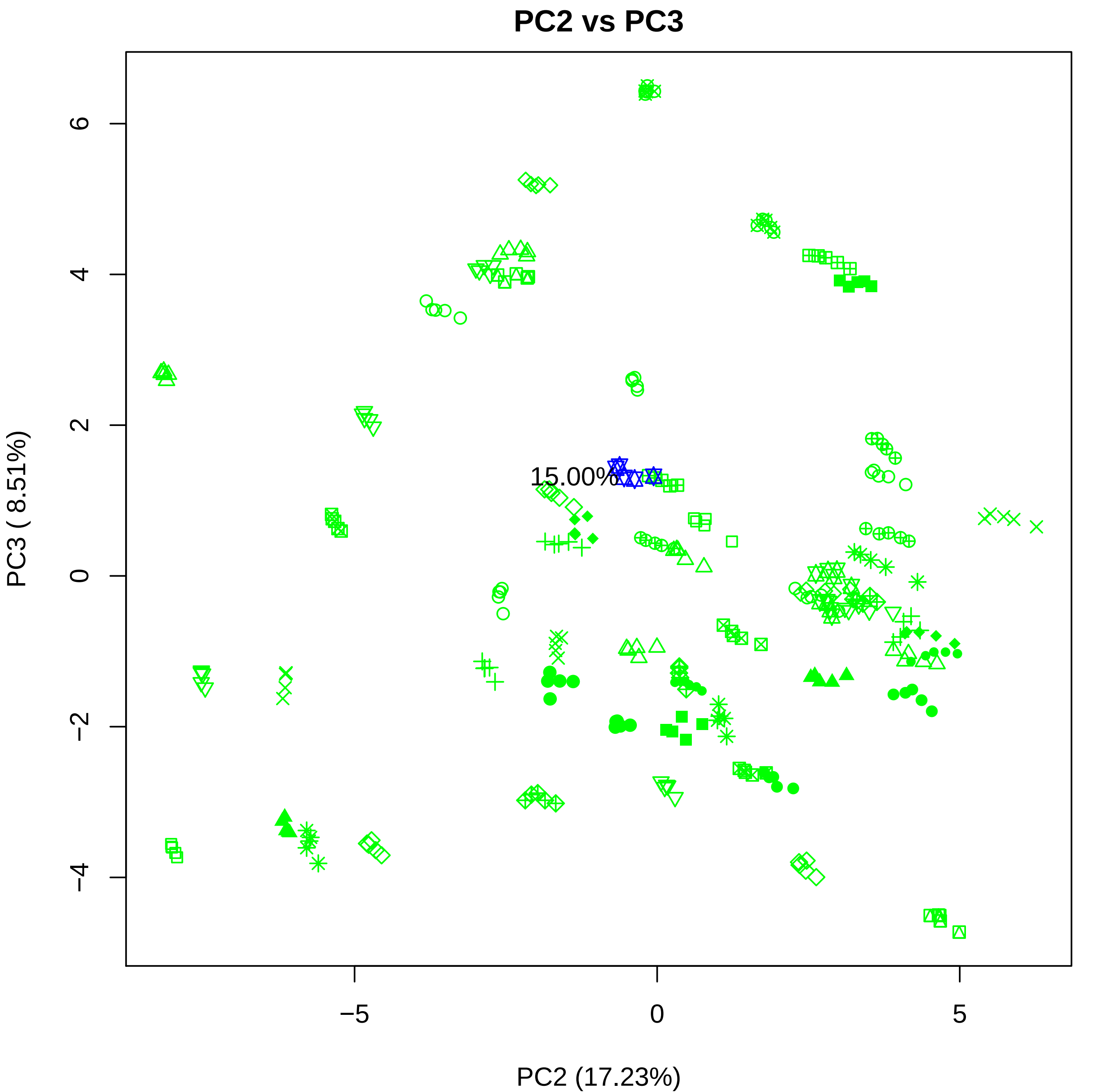
PC2 vs PC3
PC3 ( 8.51%)
PC2 (17.23%)
15.00%
6
4
2
0
−2
−4
−5
0
5

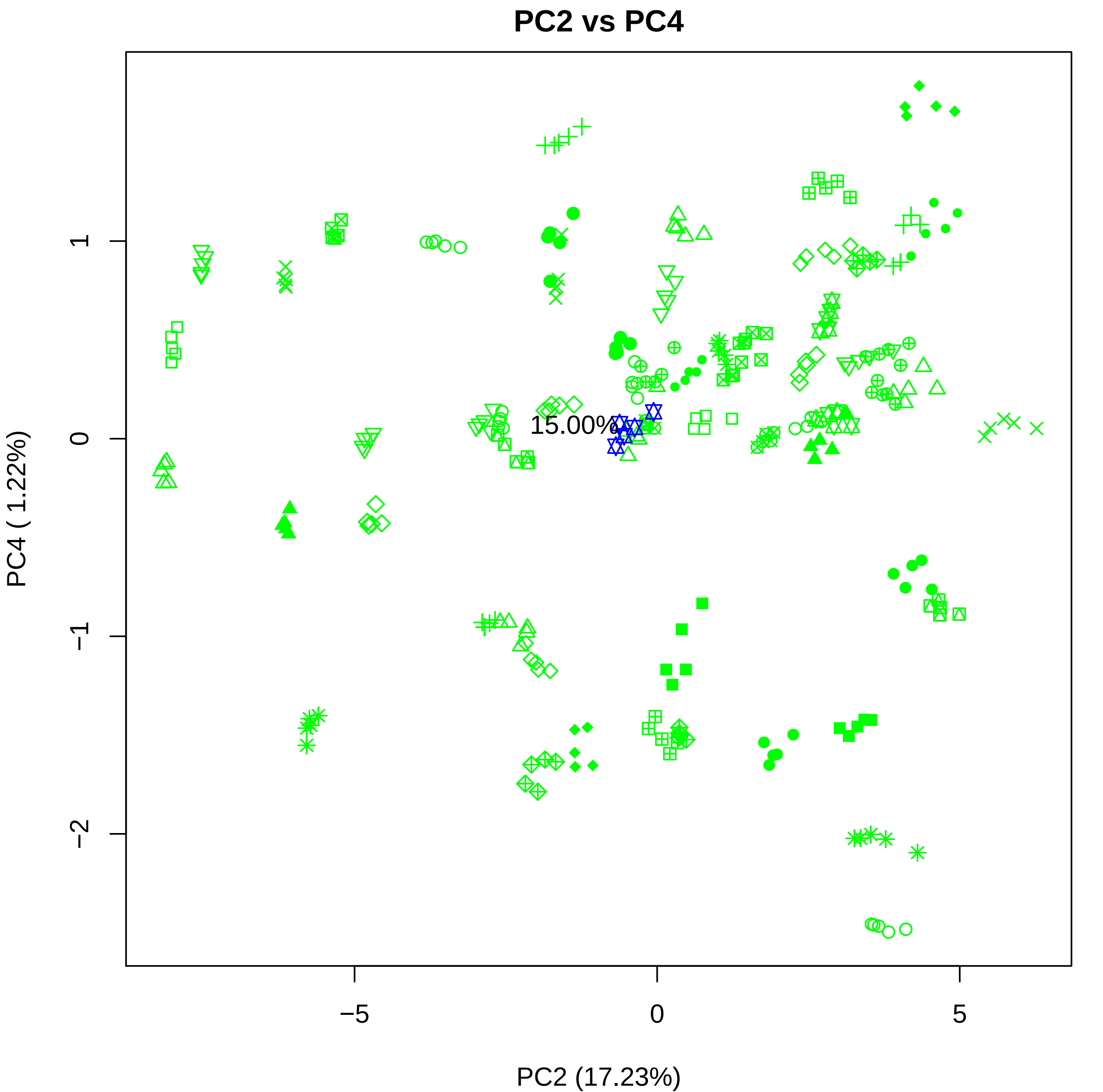

PC2 vs PC4
15.00%
PC4 ( 1.22%)
PC2 (17.23%)
1
0
−1
−2
−5
0
5

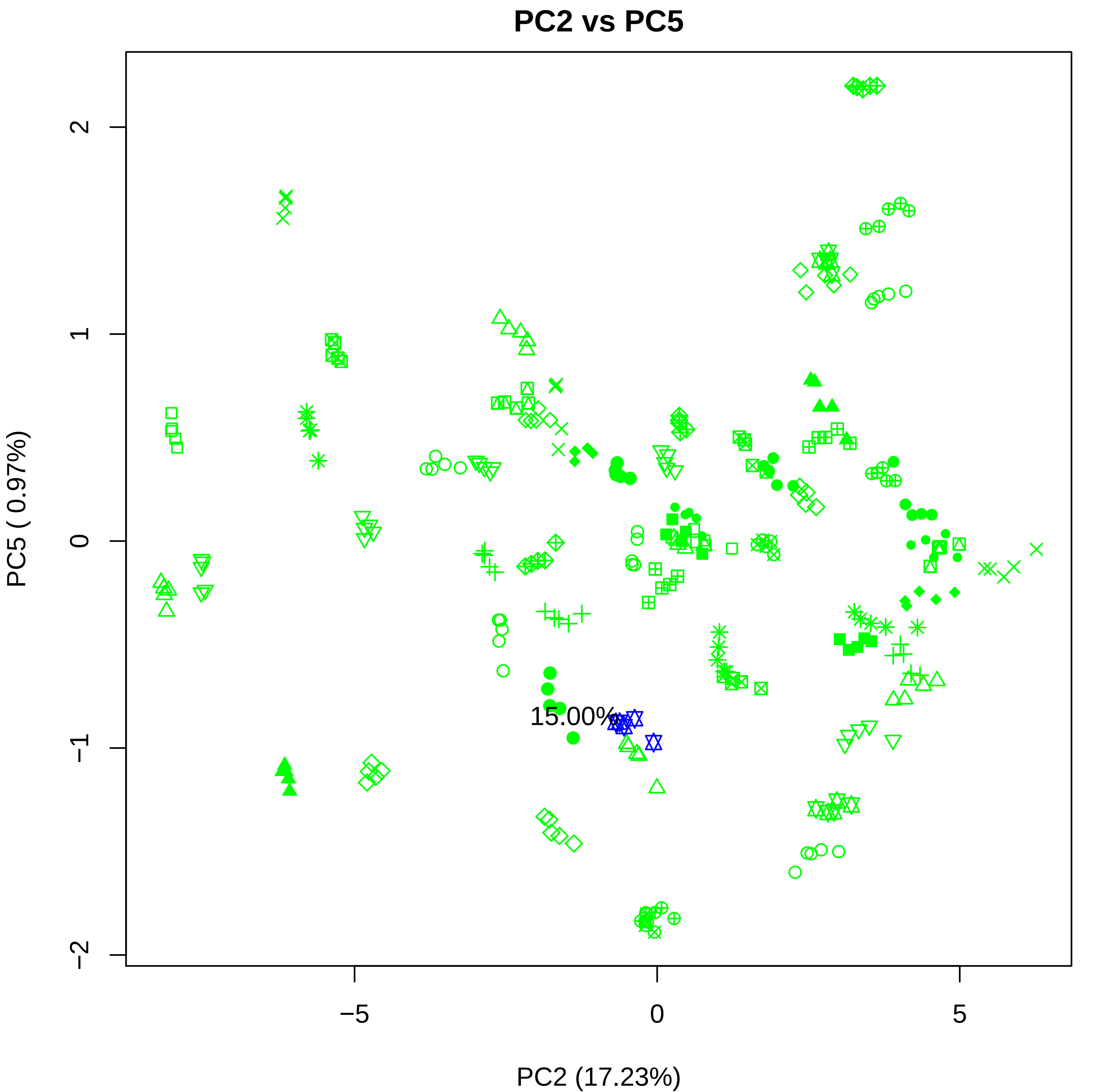
PC2 vs PC5
PC5 ( 0.97%)
PC2 (17.23%)
2
1
0
−1
−2
−5
0
5
15.00%

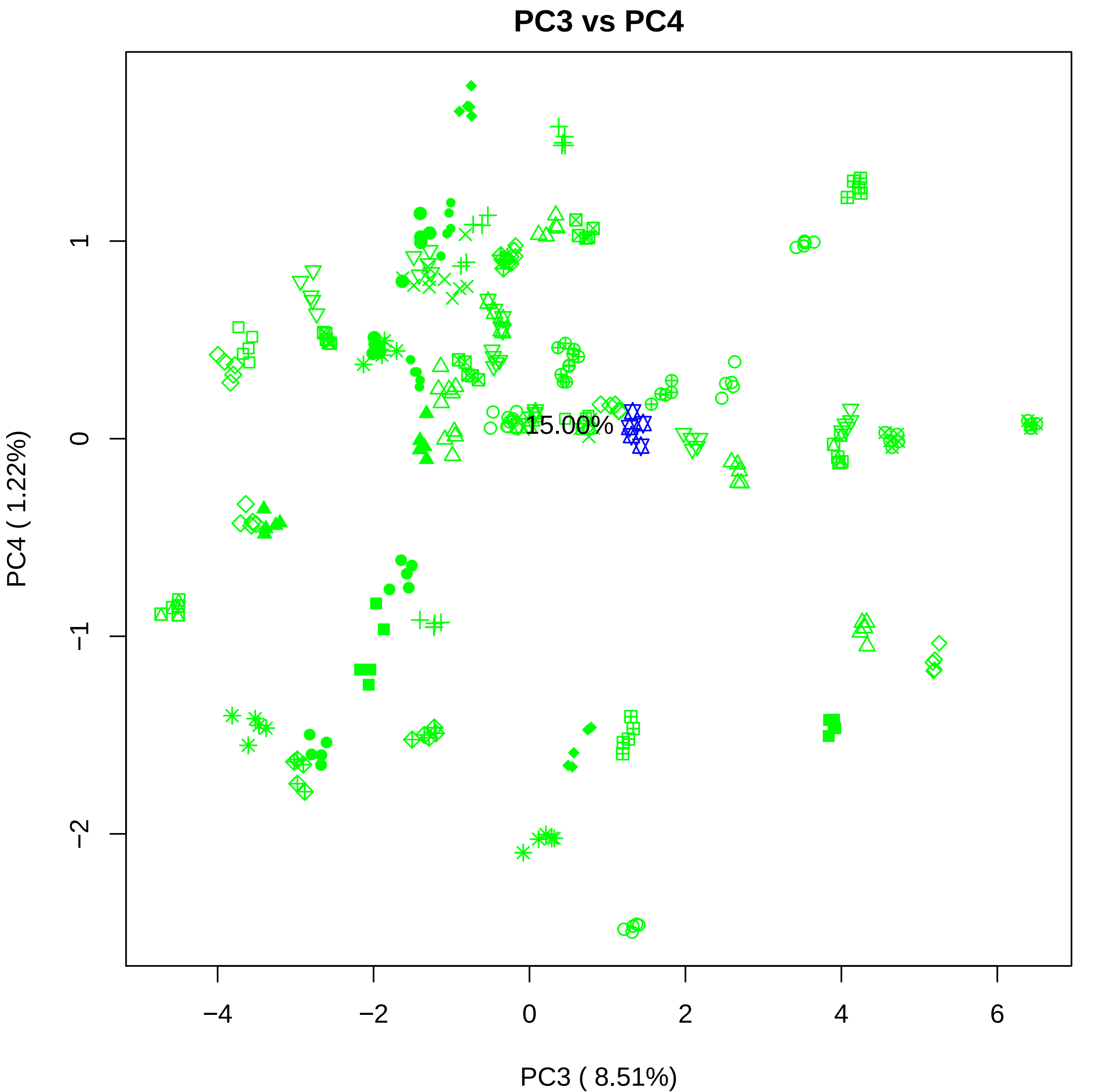

PC3 vs PC4
15.00%
PC4 ( 1.22%)
PC3 ( 8.51%)
−4
−2
0
2
4
6
1
0
−1
−2

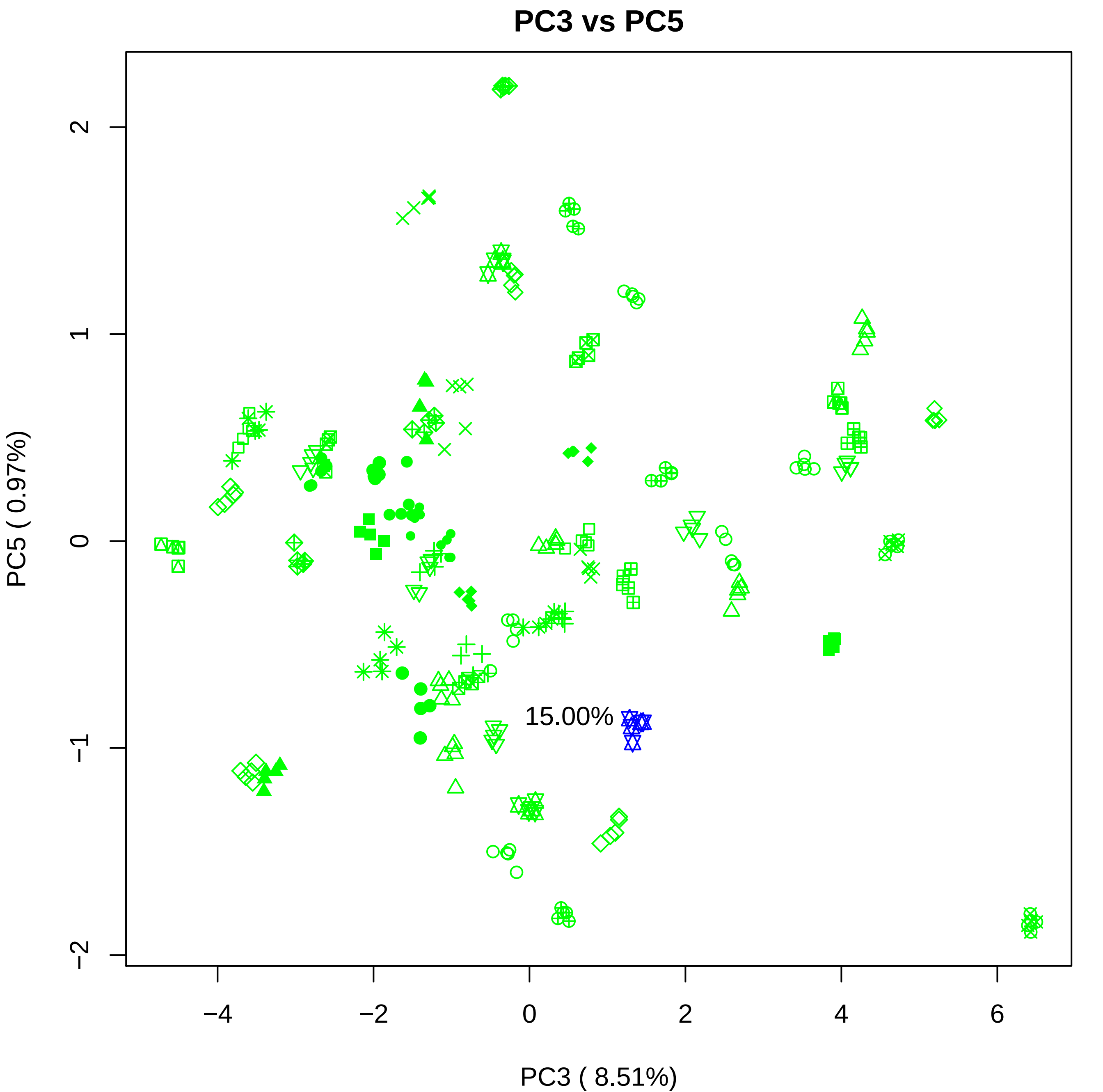
PC3 vs PC5
PC5 ( 0.97%)
PC3 ( 8.51%)
15.00%
2
1
0
−1
−2
−4
−2
0
2
4
6

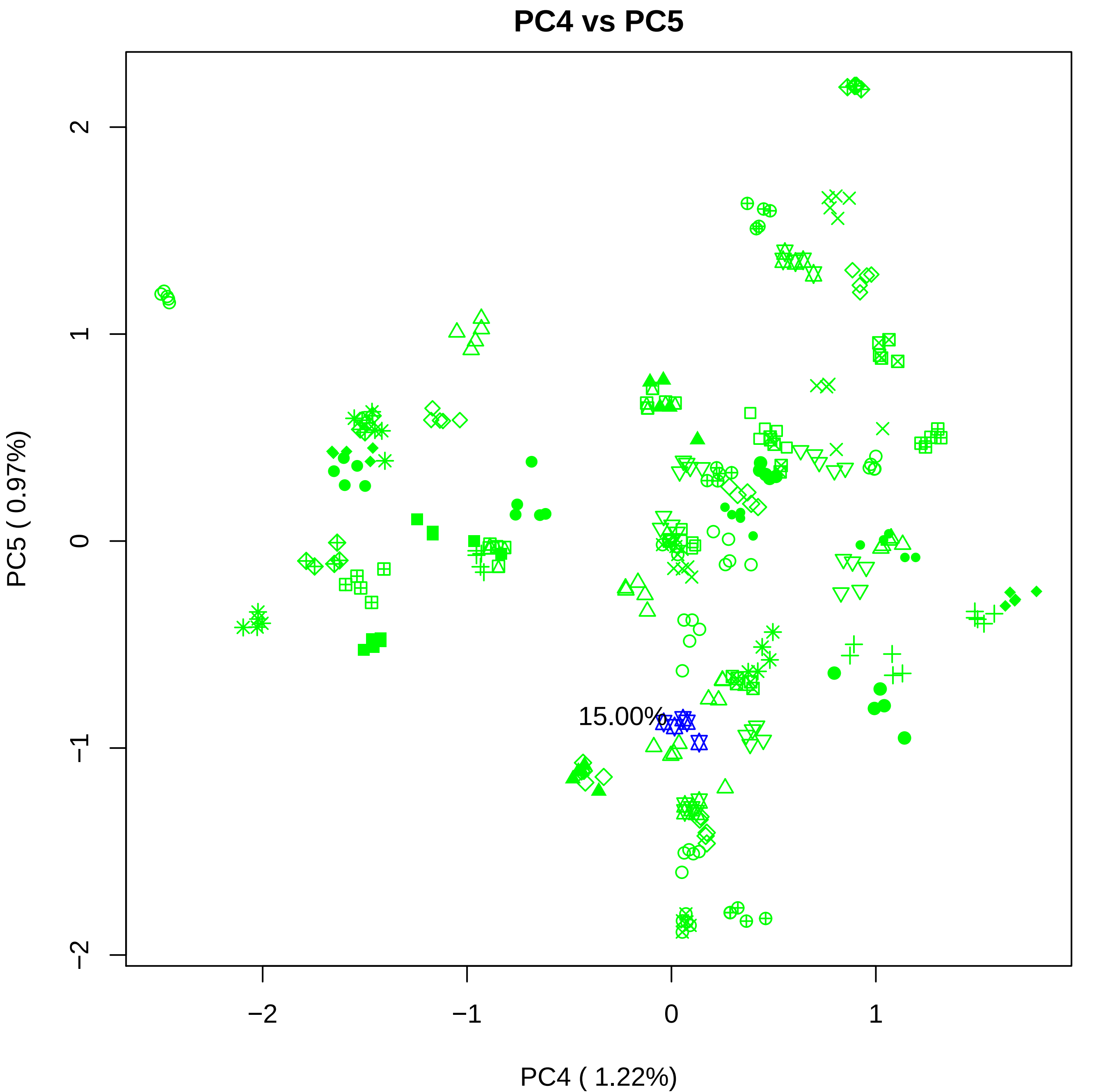
PC4 vs PC5
PC5 ( 0.97%)
PC4 ( 1.22%)
2
1
0
−1
−2
−2
−1
0
1
15.00%

### PCA of 20%  BMH(d) vs PMH(d)

Scaled Matrix PCAed

7th order polynomial baseline correction performed

Raw spectra smoothed to 21 Neighbours, 2nd order PN, Savtisky Golay

Normalisation between 0 and 100

D2 spectra smoothed to 41 Neighbours, 2nd order PN, Savtisky Golay

Regions Analysed: 3601−2501 and 2001−700cm−1

Samples Analysed: 1 bContam's 0 BLH's 0 BMH's 0 OMH's 0 pContam's 0 PLH's 60 PMH's

Keele University AD

Fri Dec 14 14:23:14 2018

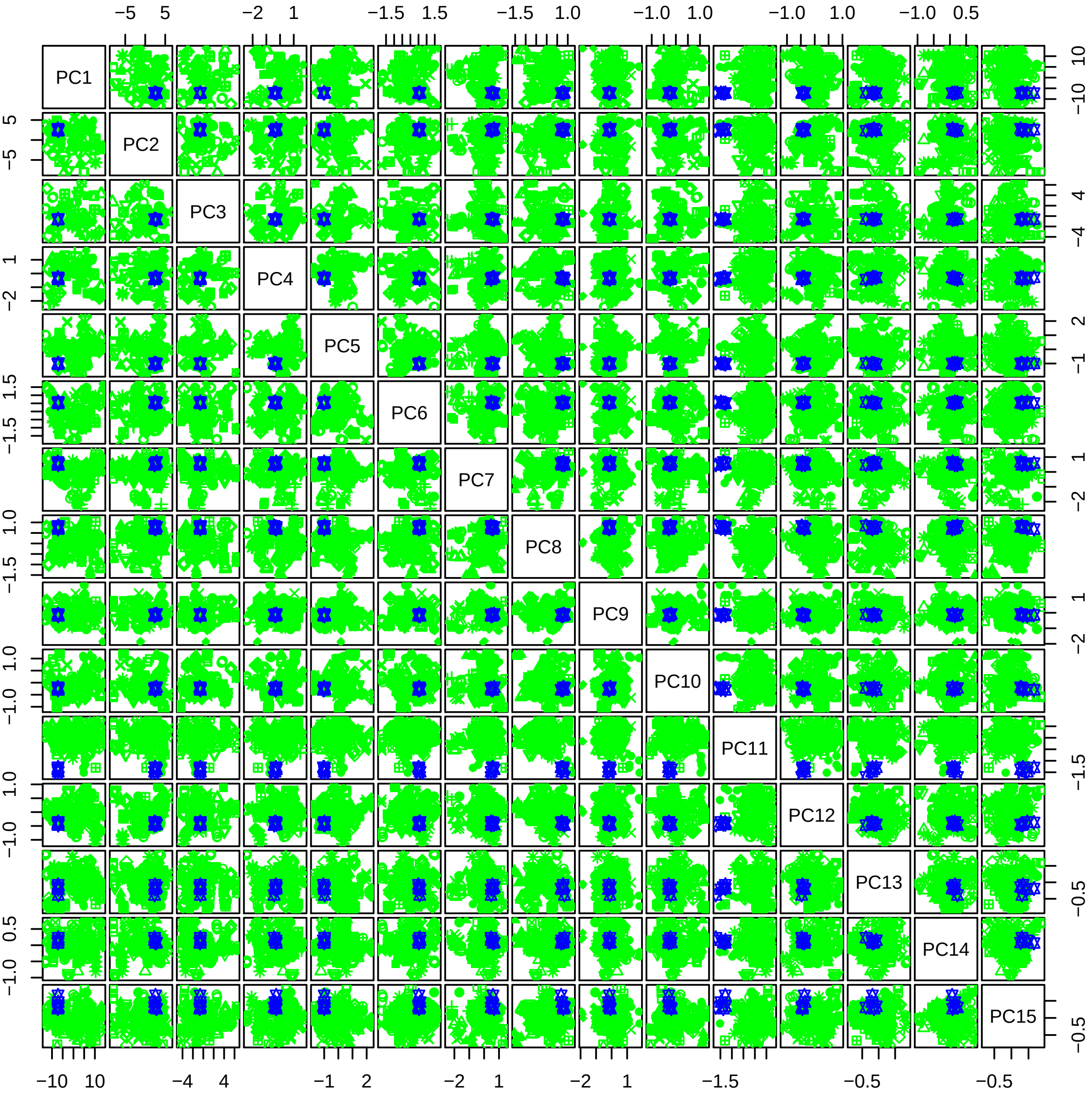
PC1
PC2
PC3
PC4
PC5
PC6
PC7
PC8
PC9
PC10
PC11
PC12
PC13
PC14
PC15

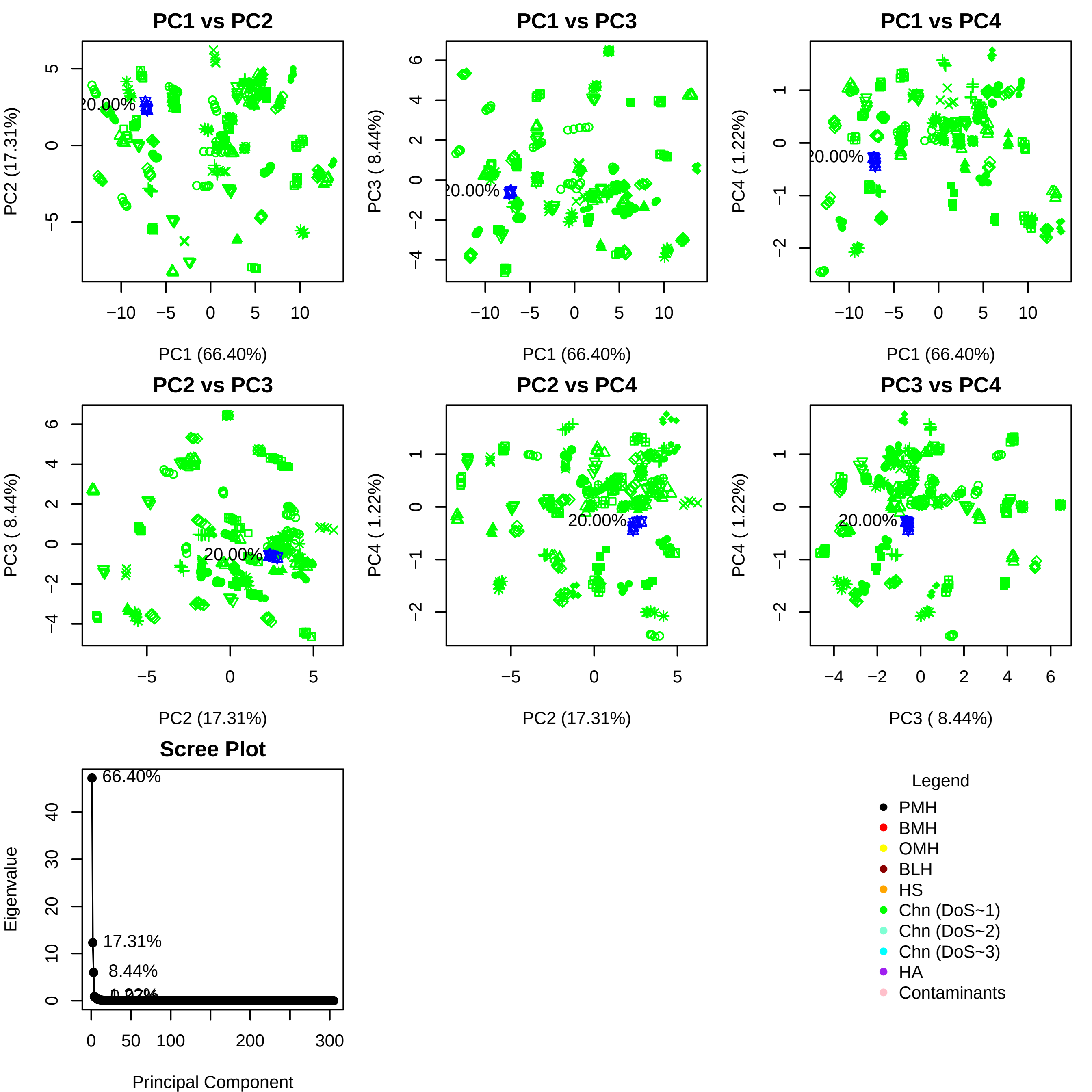

PC1 vs PC2
PC1 vs PC3
PC1 vs PC4
PC2 vs PC3
PC2 vs PC4
PC3 vs PC4
PC1 (66.40%)
PC2 (17.31%)
PC3 ( 8.44%)
PC4 ( 1.22%)
20.00%
Scree Plot
Eigenvalue
Principal Component
66.40%
17.31%
8.44%
Legend
PMH
BMH
OMH
BLH
HS
Chn (DoS~1)
Chn (DoS~2)
Chn (DoS~3)
HA
Contaminants

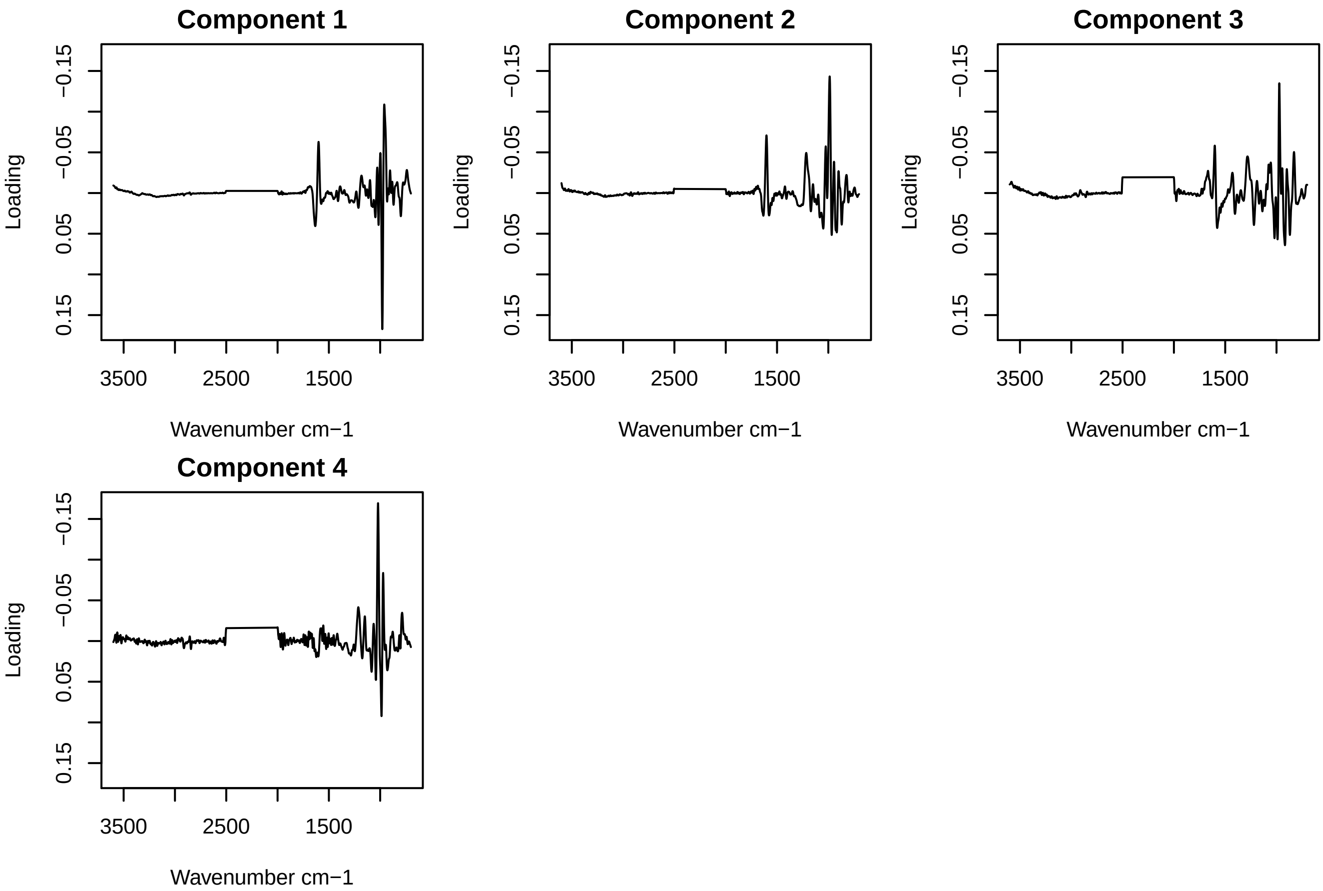
Component 1
Component 2
Component 3
Component 4
Loading
−0.15
−0.05
0.05
0.15
3500
2500
1500
Wavenumber cm−1

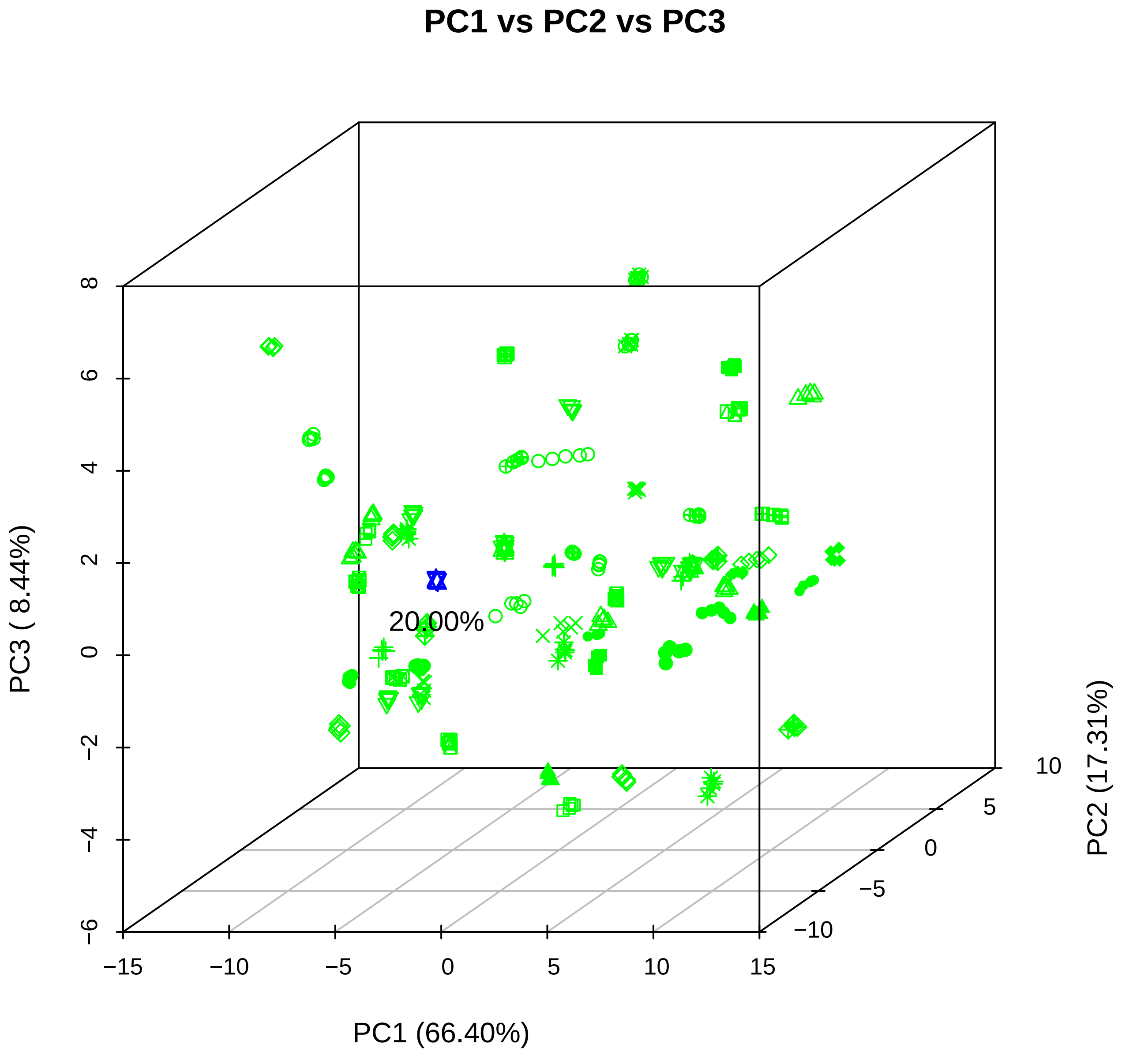

PC1 vs PC2 vs PC3
20.00%
PC1 (66.40%)
PC2 (17.31%)
PC3 ( 8.44%)
−15
−10
−5
0
5
10
15
−10
−5
0
5
10
−6
−4
−2
0
2
4
6
8

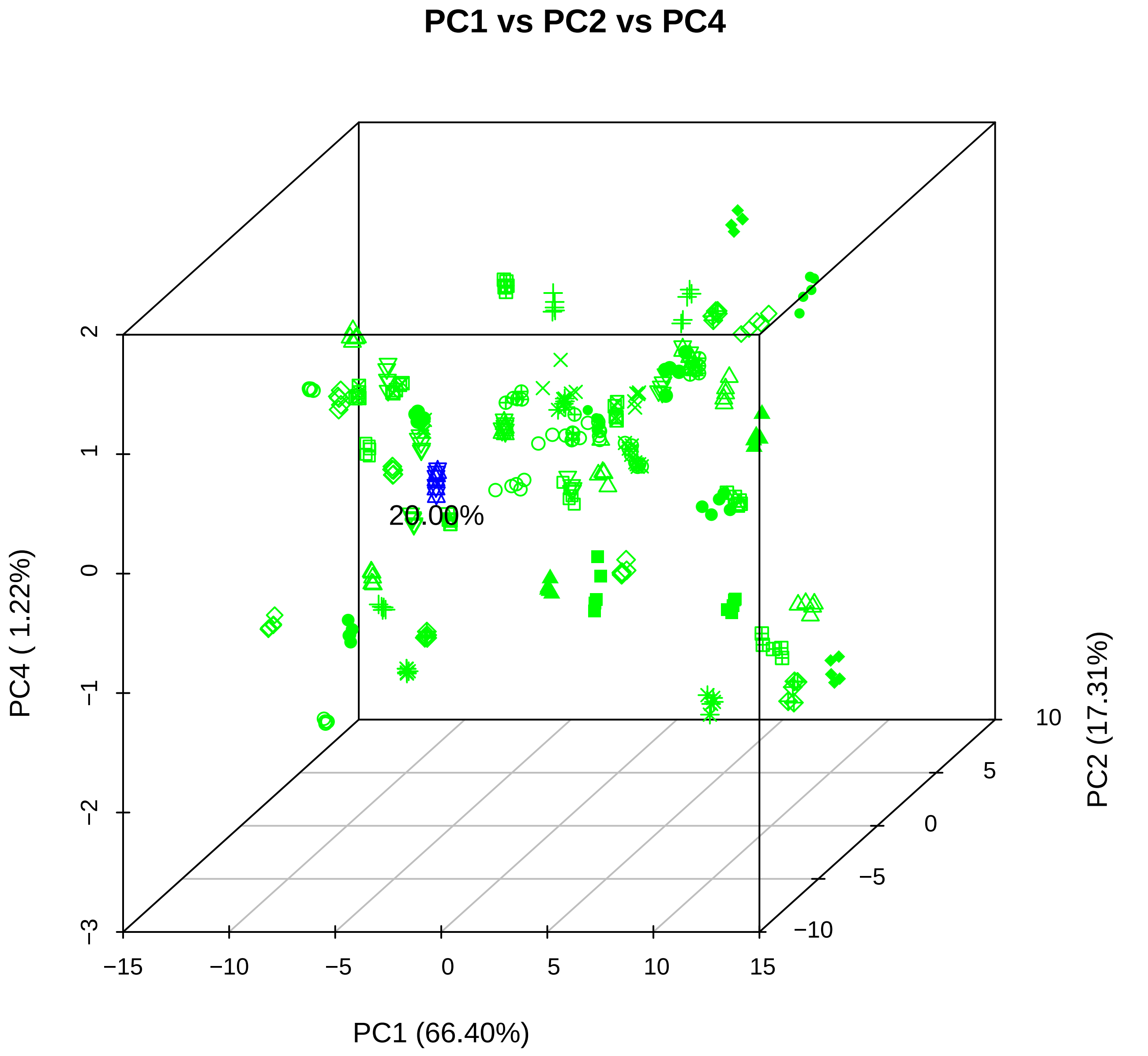
PC1 vs PC2 vs PC4
20.00%
PC1 (66.40%)
PC2 (17.31%)
PC4 ( 1.22%)
−15
−10
−5
0
5
10
15
−3
−2
−1
0
1
2
−10
−5
0
5
10

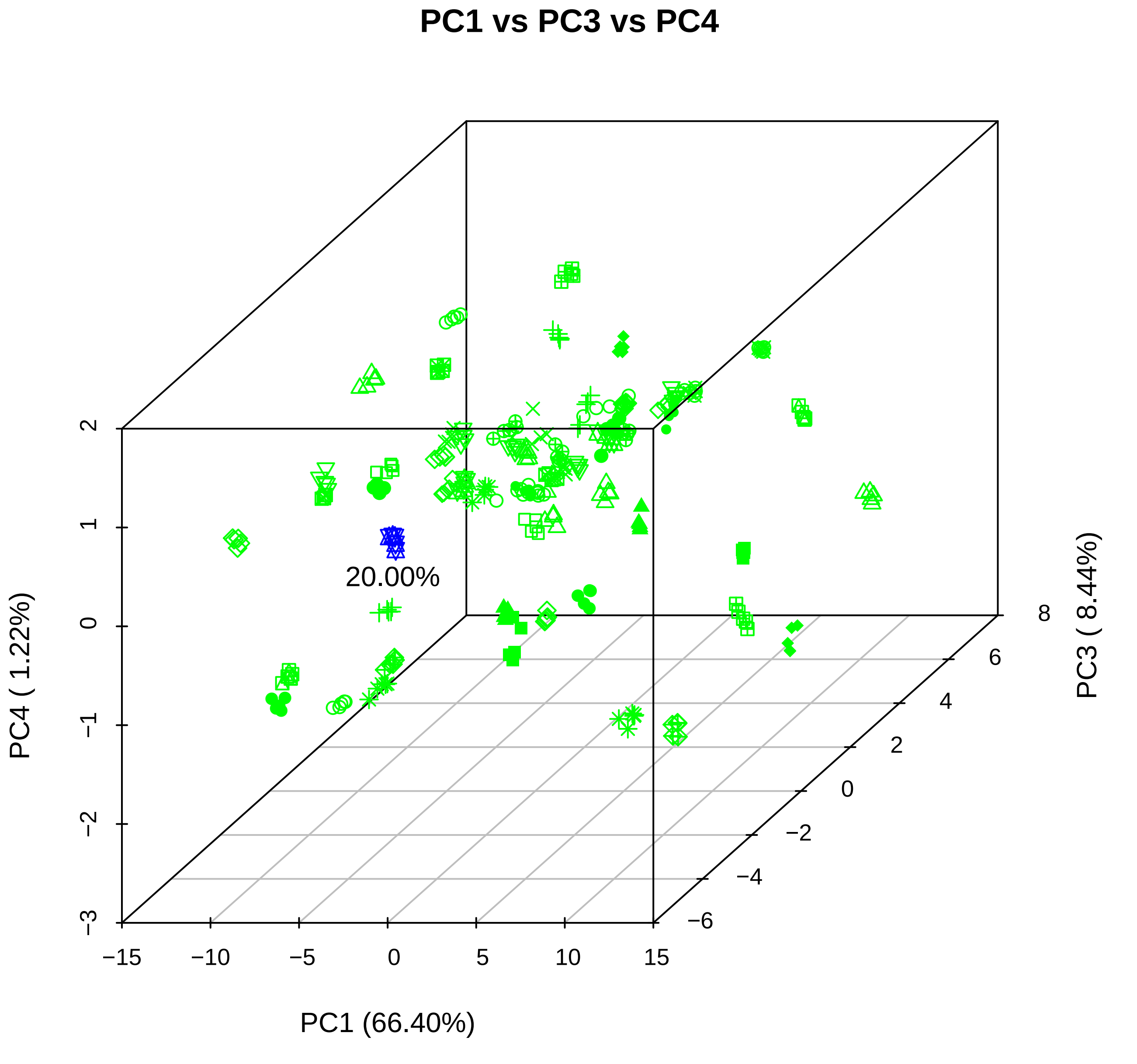
PC1 vs PC3 vs PC4
20.00%
PC1 (66.40%)
PC4 ( 1.22%)
PC3 ( 8.44%)
−15
−10
−5
0
5
10
15
−3
−2
−1
0
1
2
−6
−4
−2
0
2
4
6
8

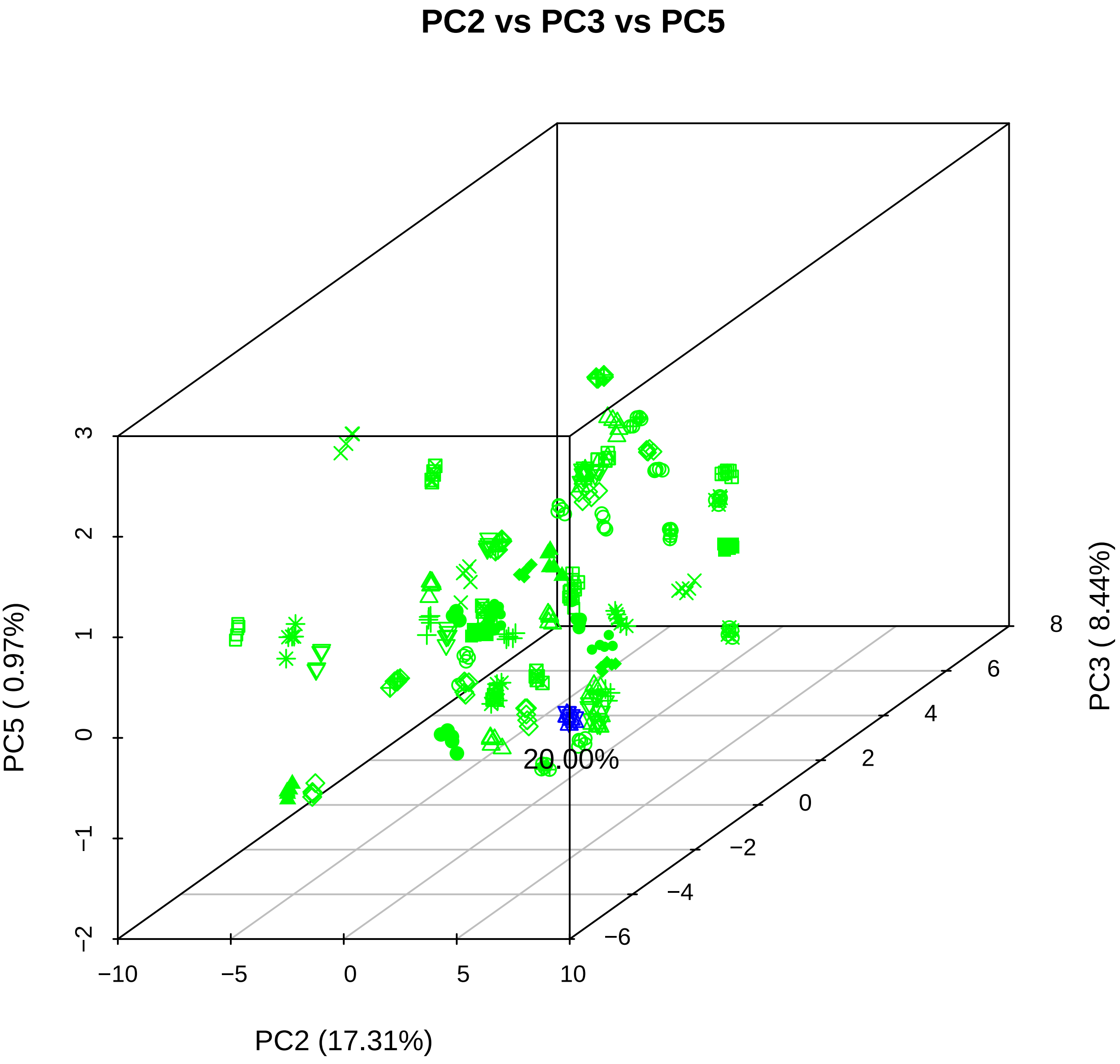
PC2 vs PC3 vs PC5
PC2 (17.31%)
PC5 ( 0.97%)
PC3 ( 8.44%)
20.00%
−10
−5
0
5
10
−2
−1
0
1
2
3
−6
−4
−2
0
2
4
6
8

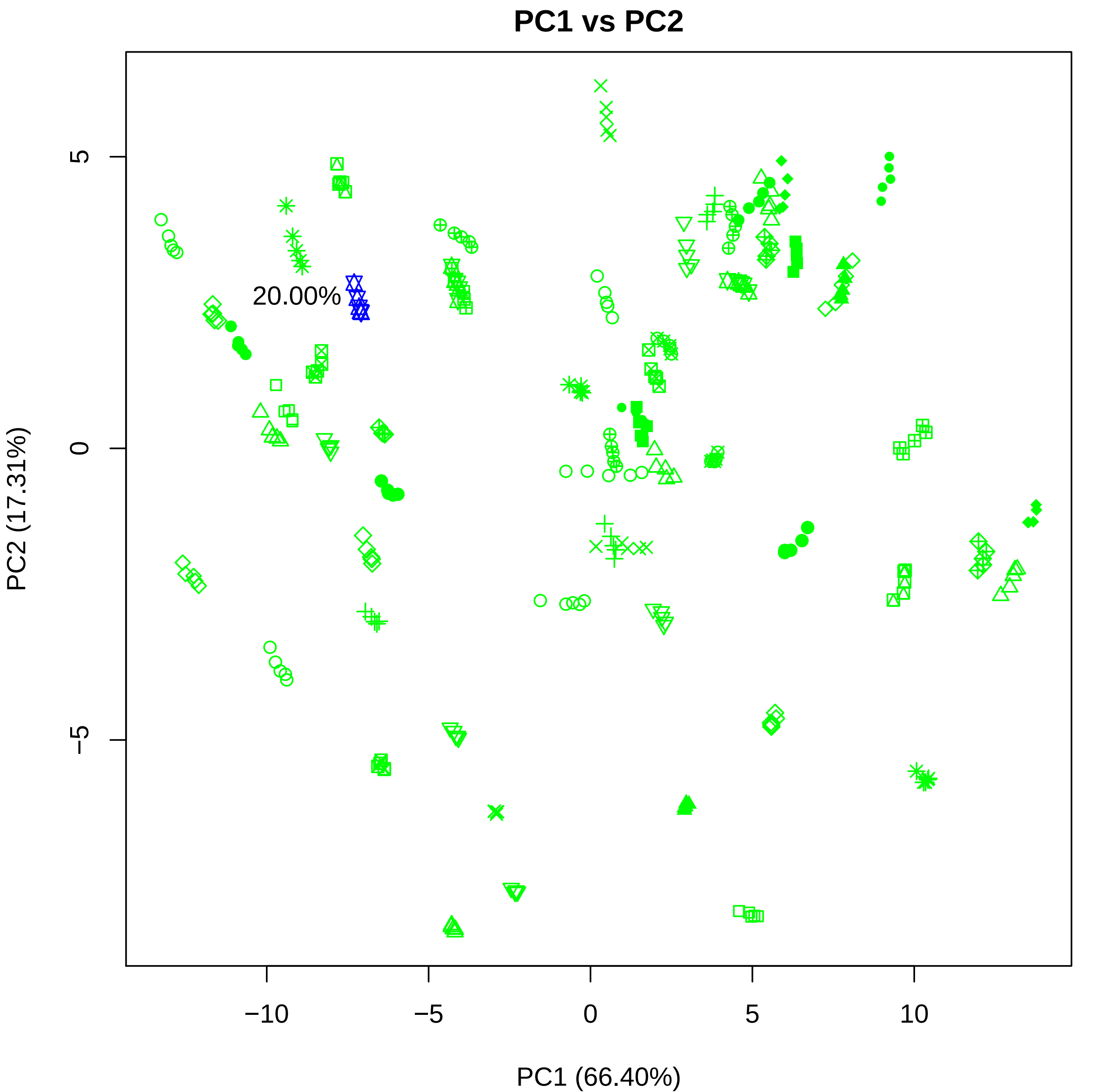
PC1 vs PC2
20.00%
PC2 (17.31%)
PC1 (66.40%)
5
0
−5
−10
−5
0
5
10

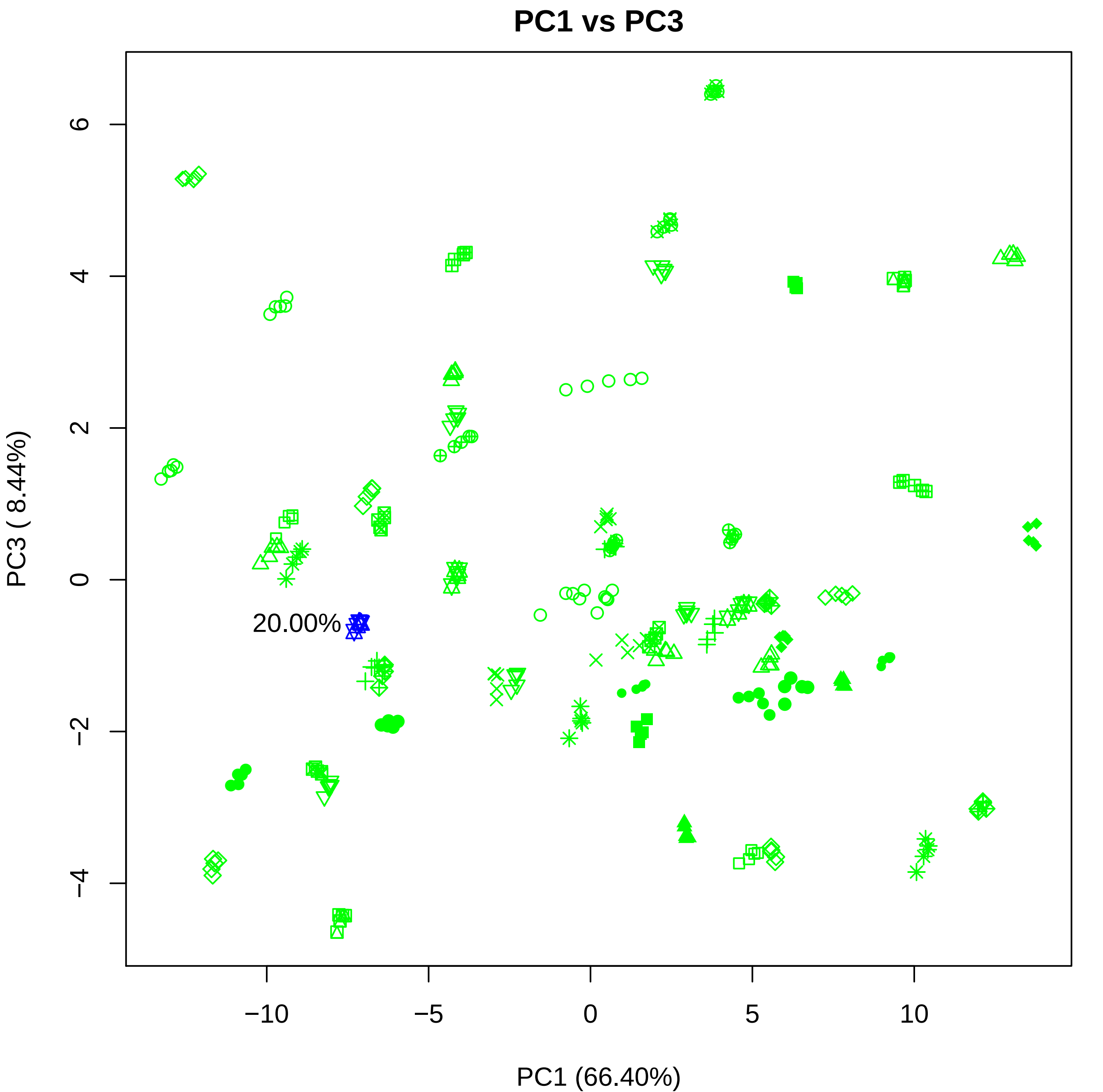
PC1 vs PC3
20.00%
PC3 ( 8.44%)
PC1 (66.40%)
6
4
2
0
−2
−4
−10
−5
0
5
10

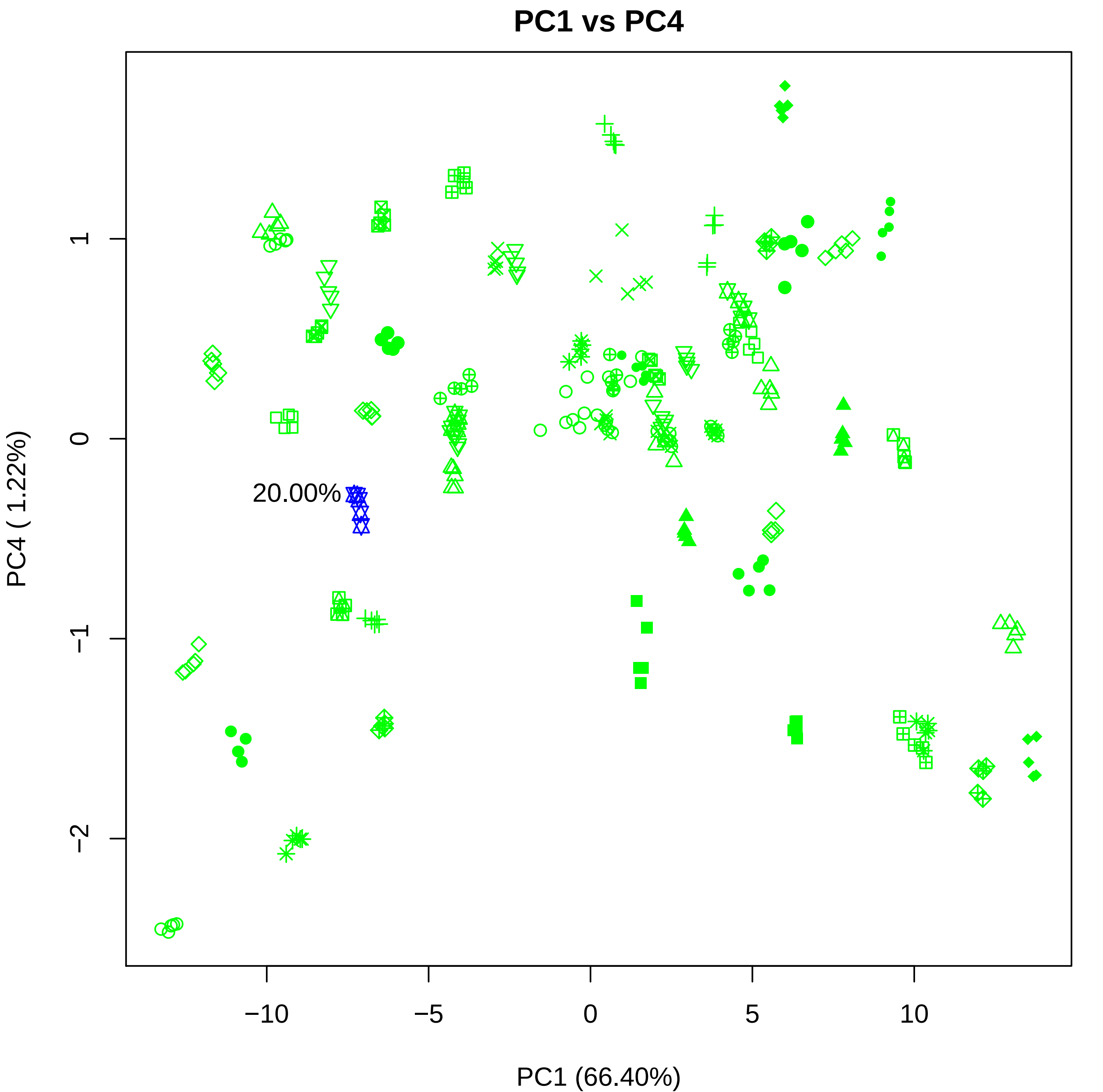
PC1 vs PC4
20.00%
PC4 ( 1.22%)
PC1 (66.40%)
−10
−5
0
5
10
1
0
−1
−2

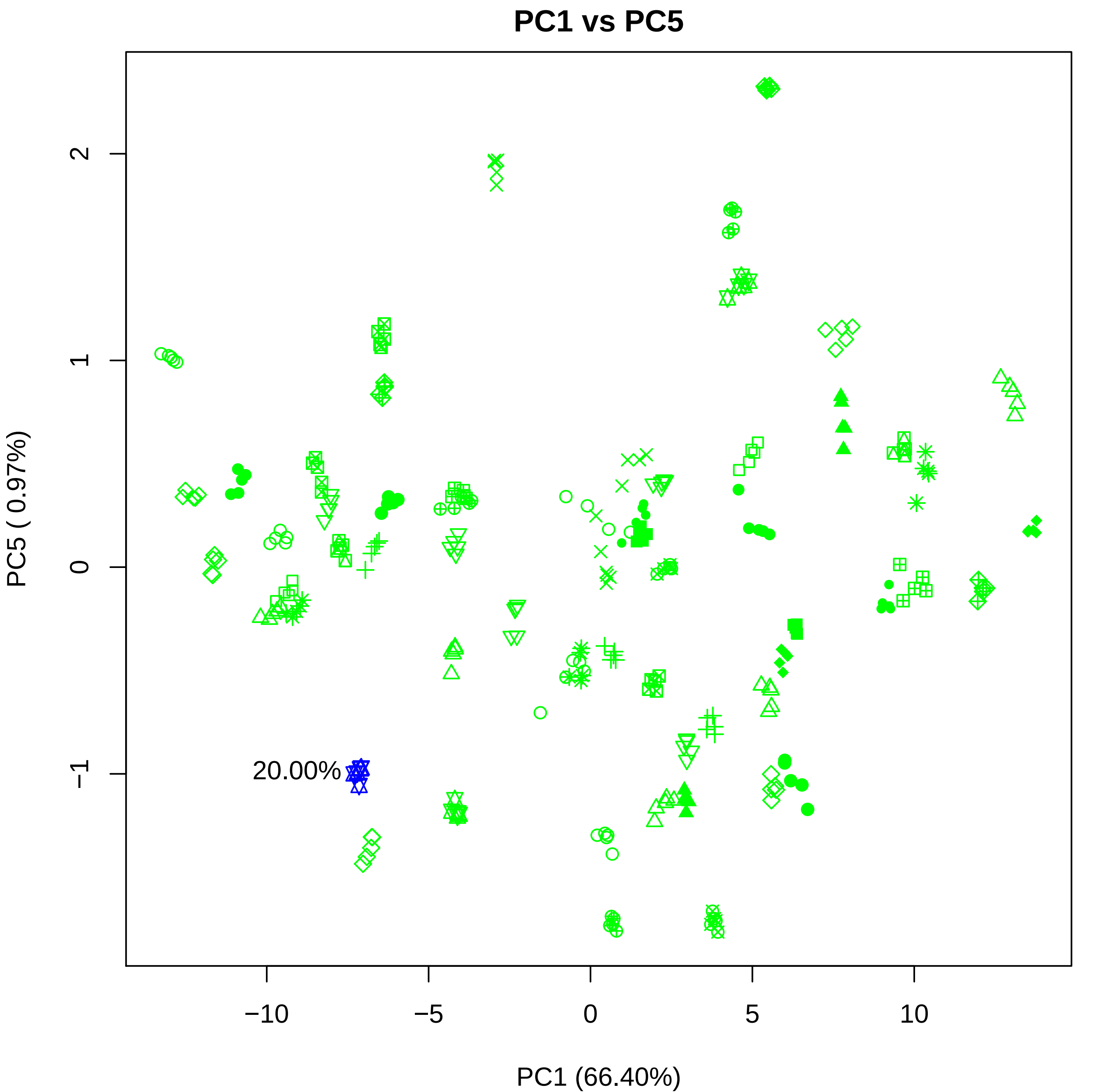

PC1 vs PC5
PC5 ( 0.97%)
PC1 (66.40%)
20.00%
2
1
0
−1
−10
−5
0
5
10

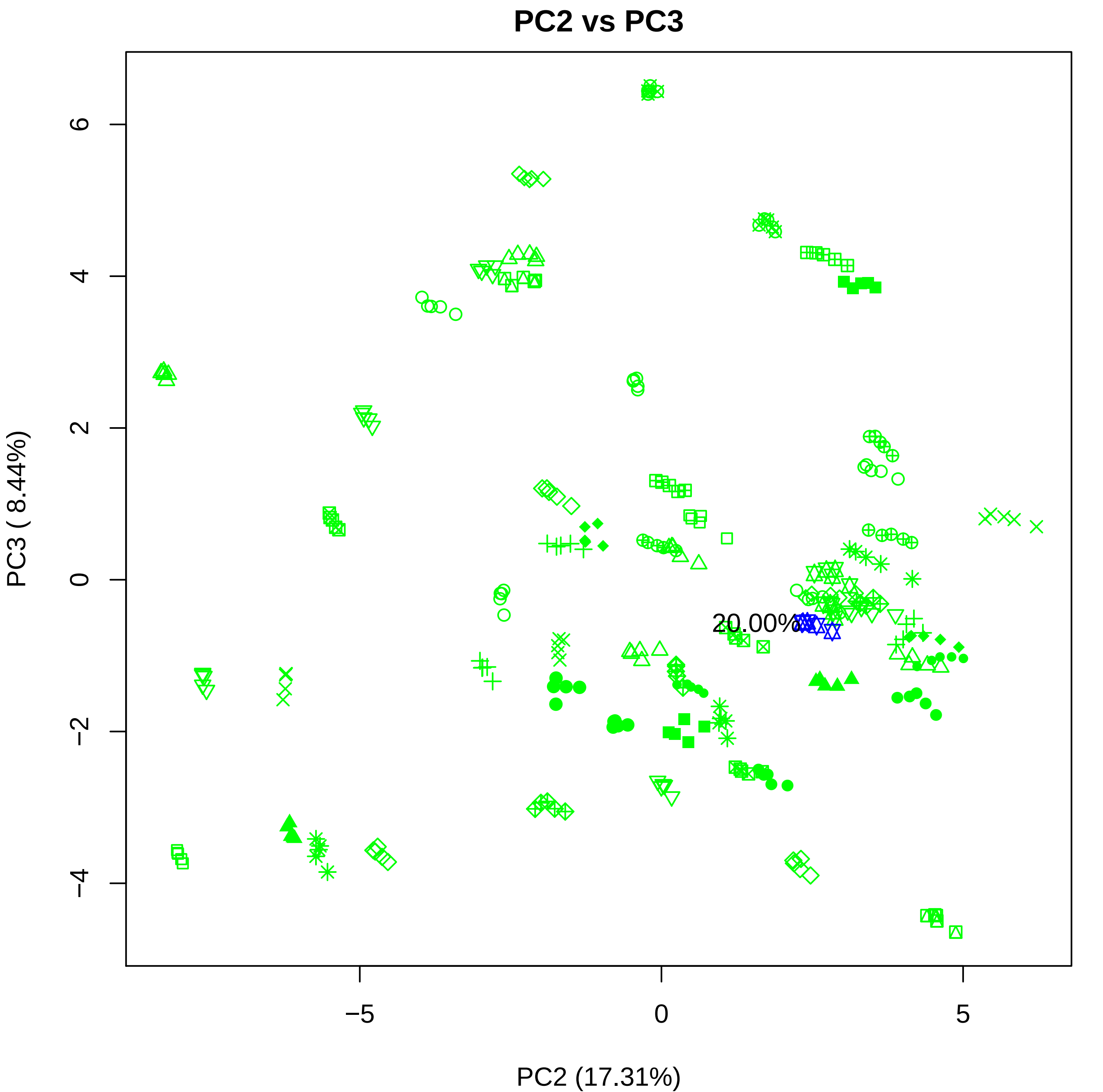

PC2 vs PC3
20.00%
PC3 ( 8.44%)
PC2 (17.31%)
6
4
2
0
−2
−4
−5
0
5

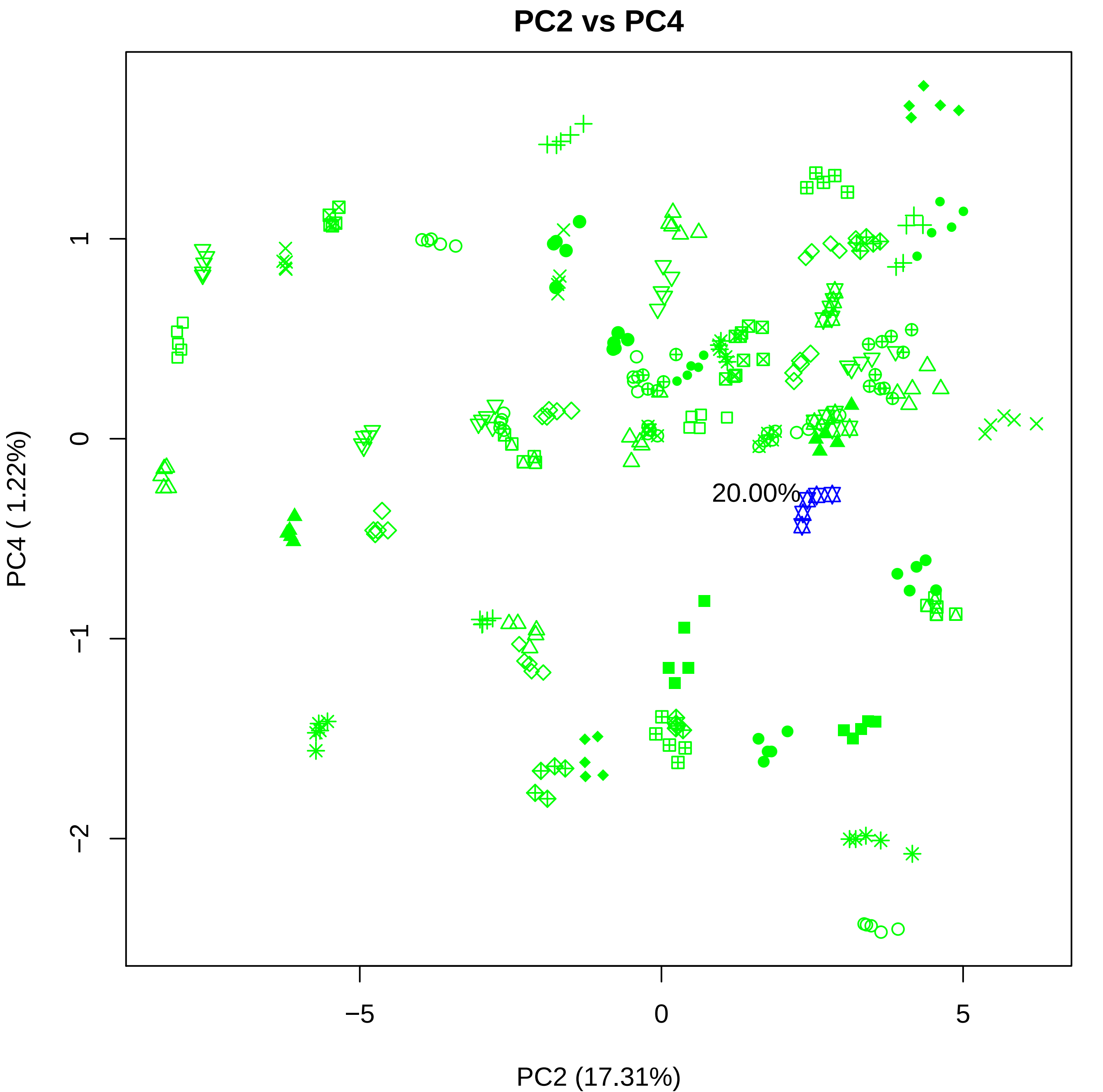

PC2 vs PC4
PC4 ( 1.22%)
PC2 (17.31%)
1
0
−1
−2
−5
0
5
20.00%

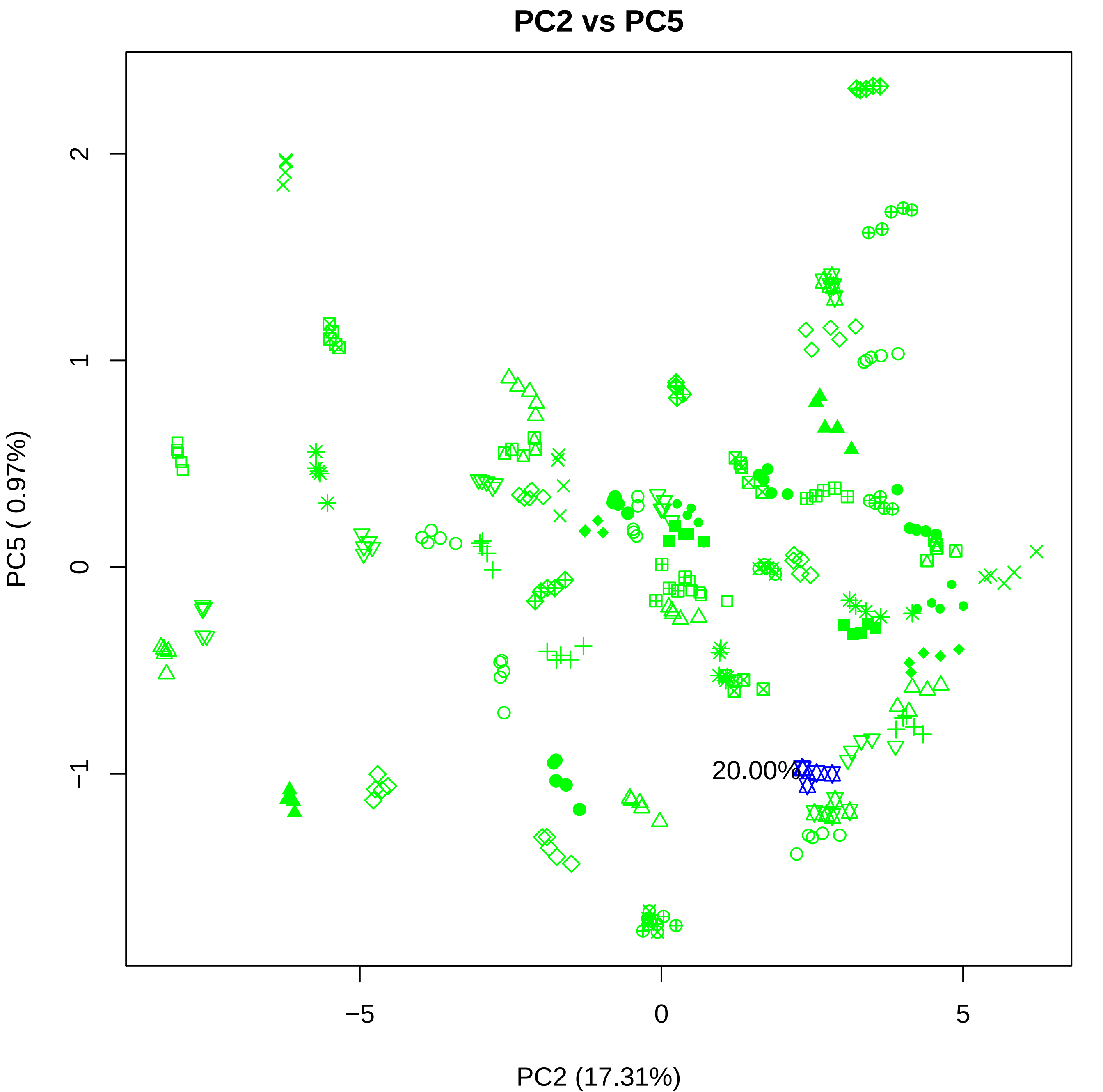
PC2 vs PC5
PC5 ( 0.97%)
PC2 (17.31%)
2
1
0
−1
−5
0
5
20.00%

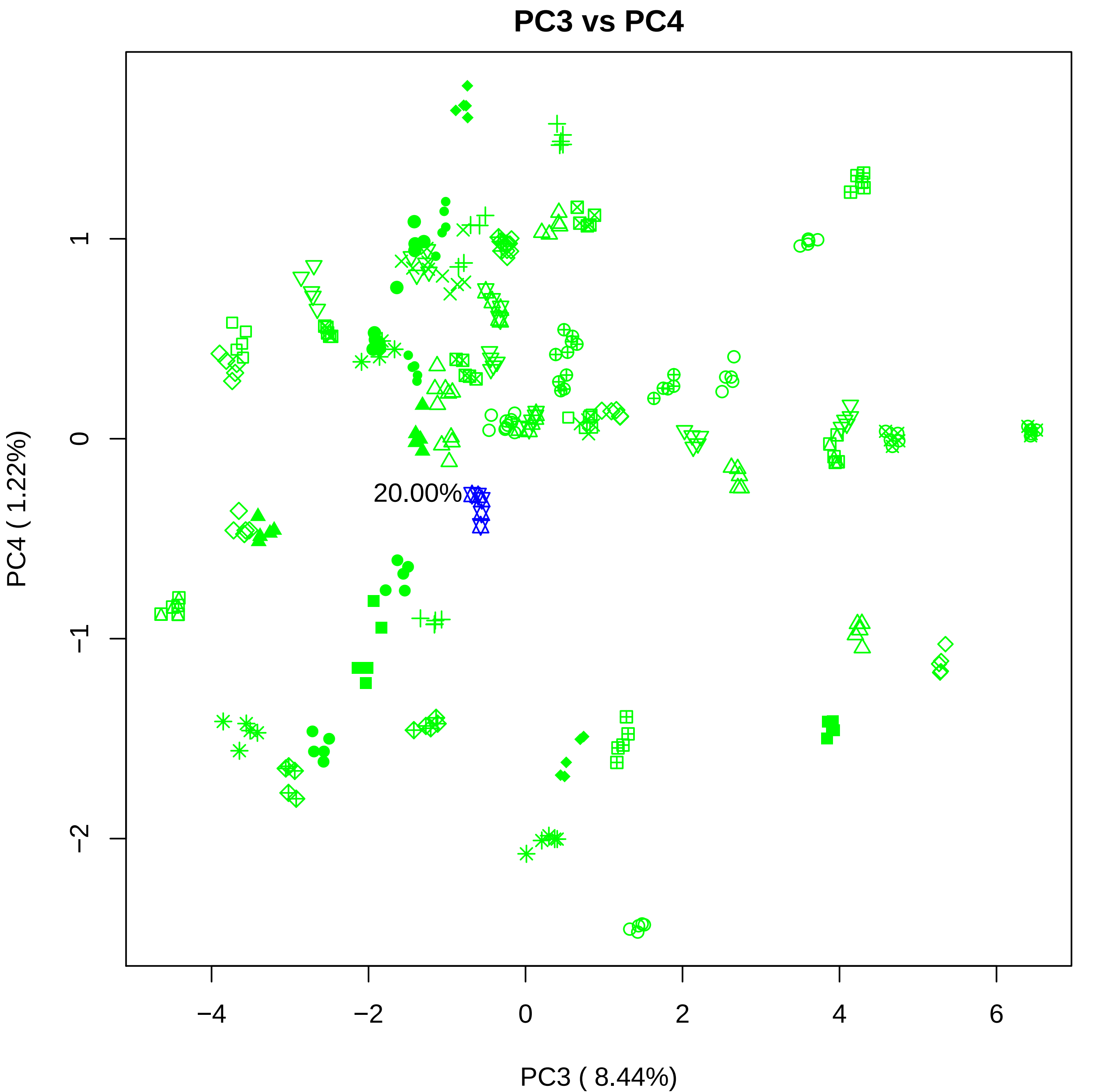
PC3 vs PC4
20.00%
PC4 ( 1.22%)
PC3 ( 8.44%)
1
0
−1
−2
−4
−2
0
2
4
6

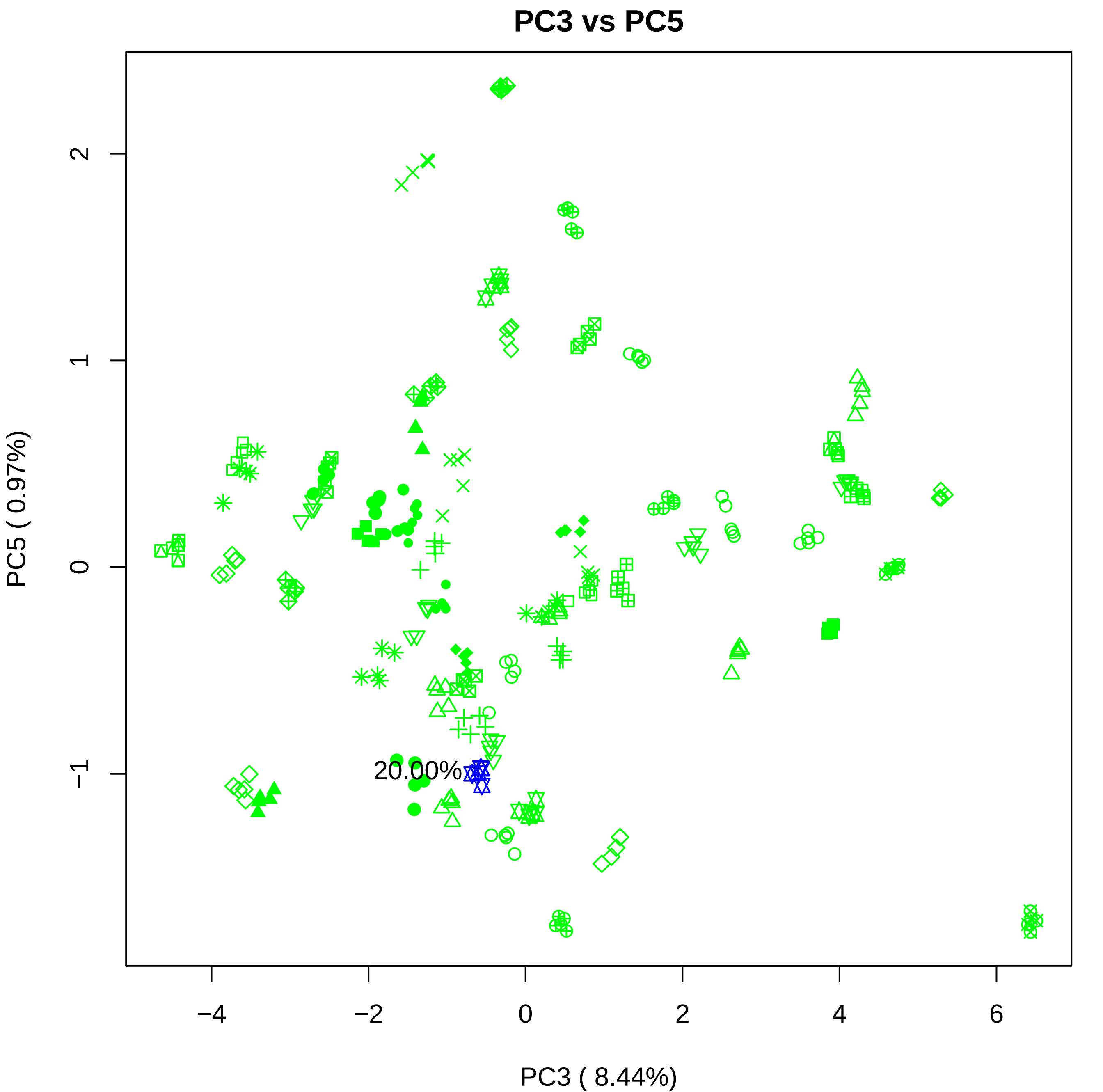

PC3 vs PC5
PC5 ( 0.97%)
PC3 ( 8.44%)
20.00%
2
1
0
−1
−4
−2
0
2
4
6

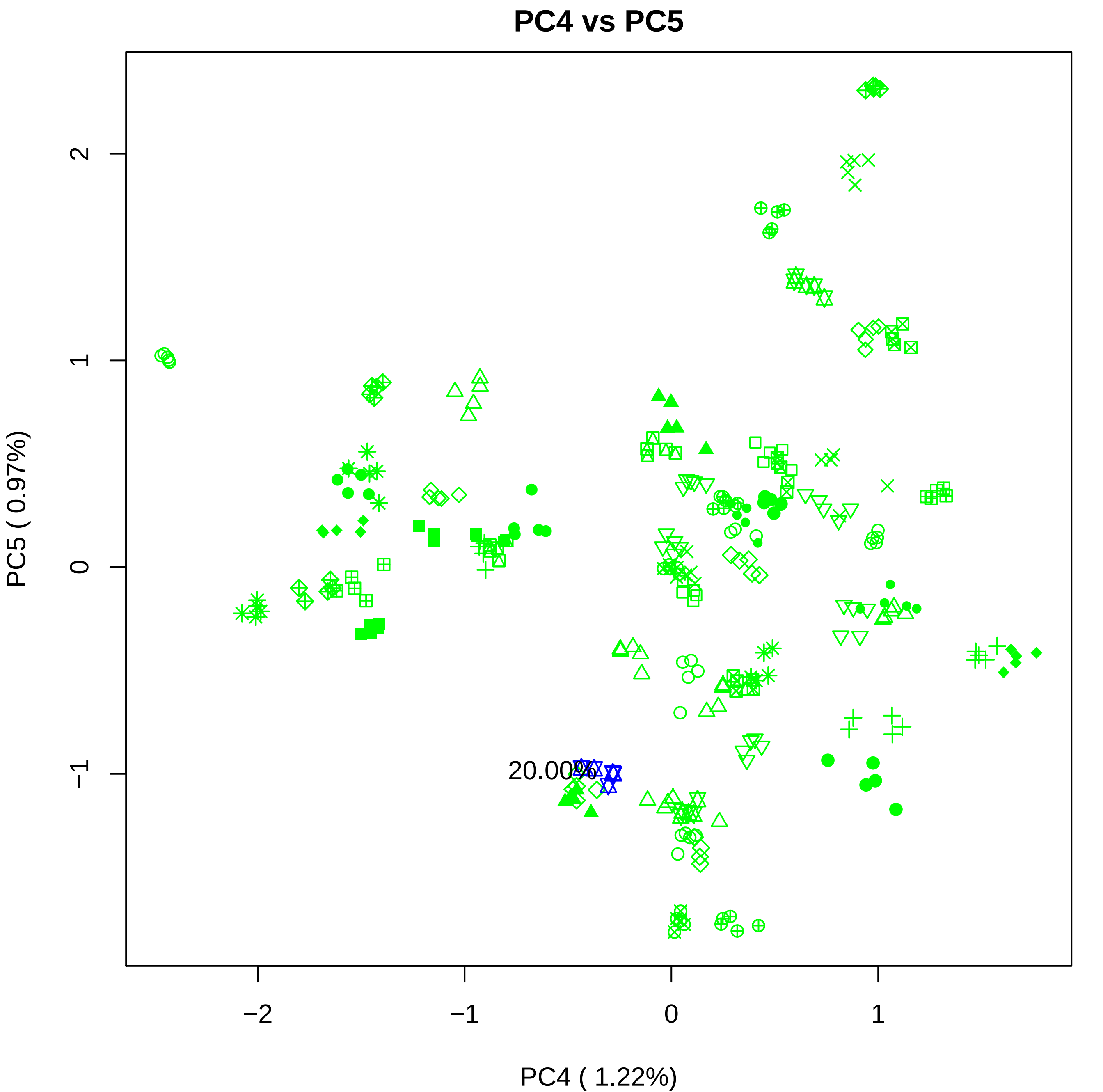

PC4 vs PC5
PC5 ( 0.97%)
PC4 ( 1.22%)
−2
−1
0
1
2
20.00%

### PCA of 40%  BMH(d) vs PMH(d)

Scaled Matrix PCAed

7th order polynomial baseline correction performed

Raw spectra smoothed to 21 Neighbours, 2nd order PN, Savtisky Golay

Normalisation between 0 and 100

D2 spectra smoothed to 41 Neighbours, 2nd order PN, Savtisky Golay

Regions Analysed: 3601−2501 and 2001−700cm−1

Samples Analysed: 1 bContam's 0 BLH's 0 BMH's 0 OMH's 0 pContam's 0 PLH's 60 PMH's

Keele University AD    Fri Dec 14 14:23:23 2018

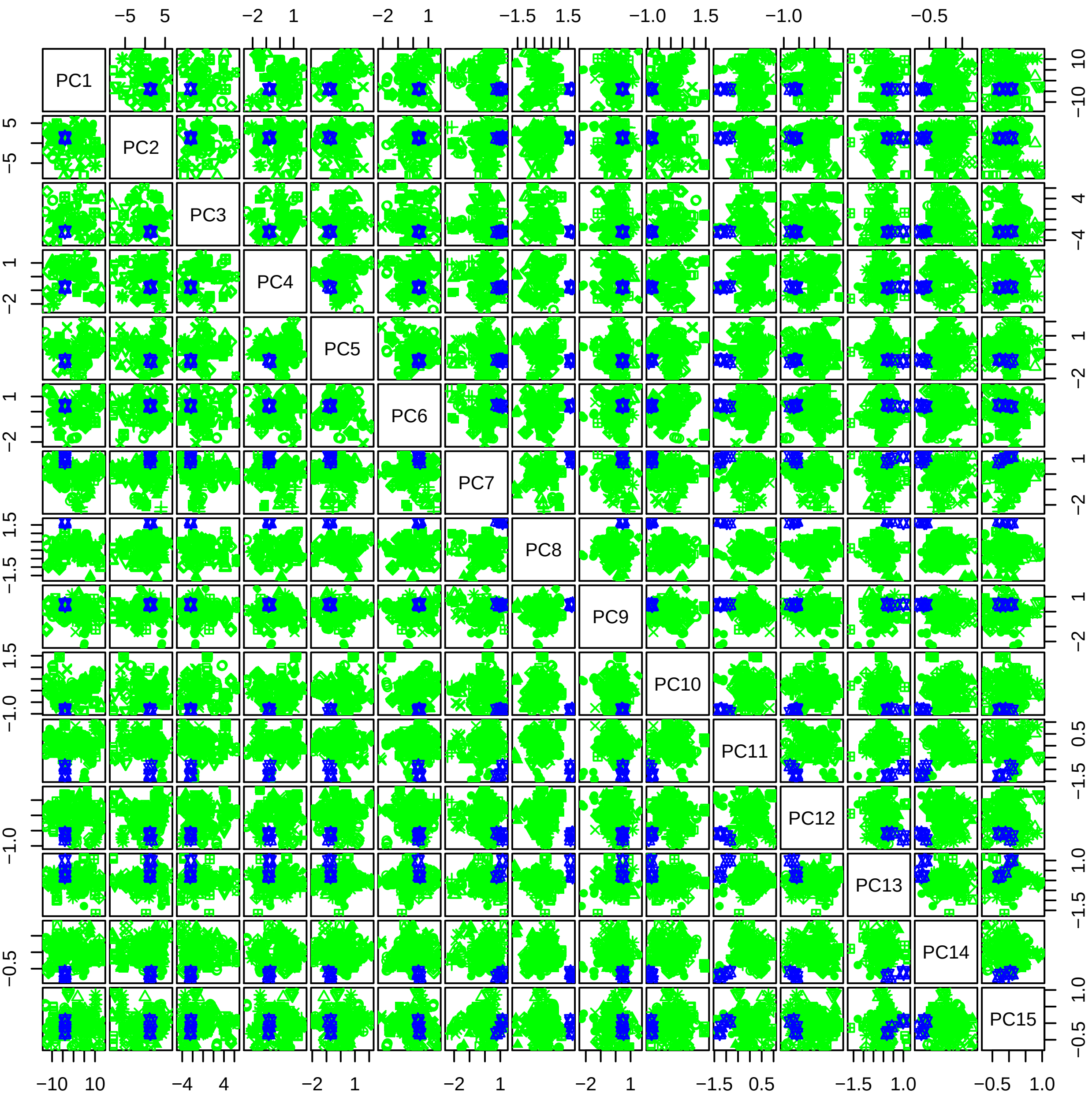

PC1
PC2
PC3
PC4
PC5
PC6
PC7
PC8
PC9
PC10
PC11
PC12
PC13
PC14
PC15

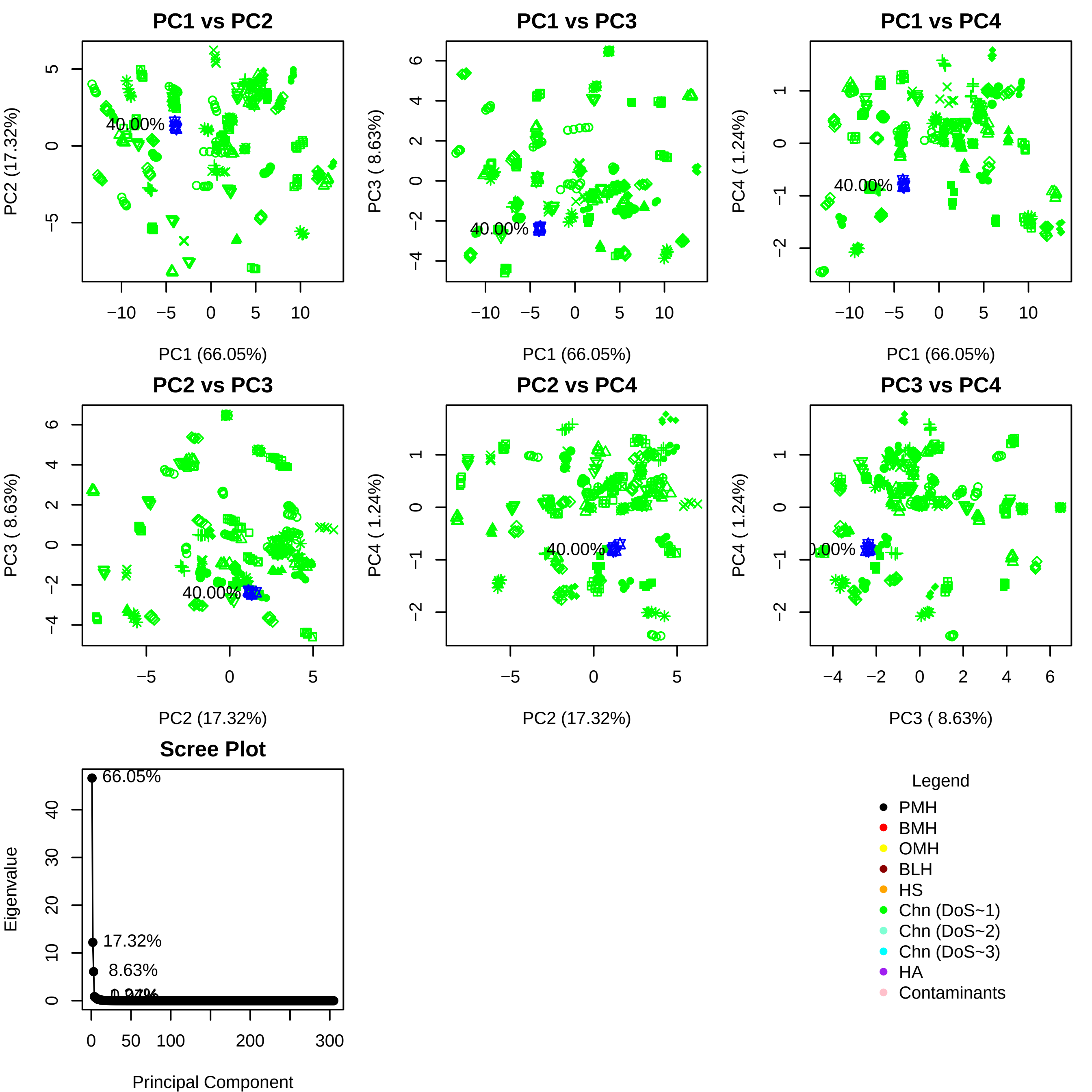

PC1 vs PC2
PC1 vs PC3
PC1 vs PC4
PC2 vs PC3
PC2 vs PC4
PC3 vs PC4
Scree Plot
PC1 (66.05%)
PC2 (17.32%)
PC3 ( 8.63%)
PC4 ( 1.24%)
40.00%
66.05%
17.32%
8.63%
Eigenvalue
Principal Component
Legend
PMH
BMH
OMH
BLH
HS
Chn (DoS~1)
Chn (DoS~2)
Chn (DoS~3)
HA
Contaminants

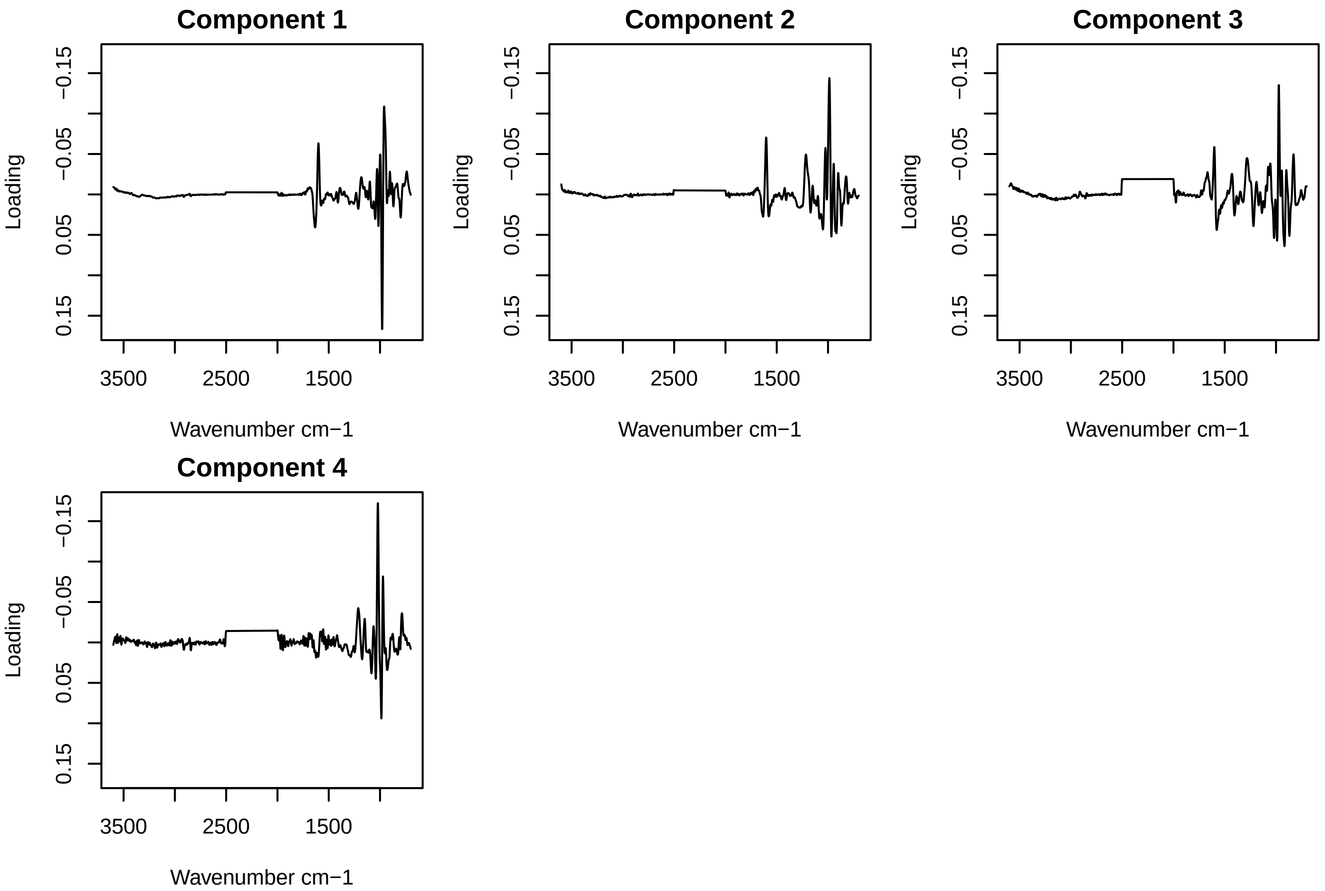
Component 1
Component 2
Component 3
Component 4
Loading
−0.15
−0.05
0.05
0.15
3500
2500
1500
Wavenumber cm−1

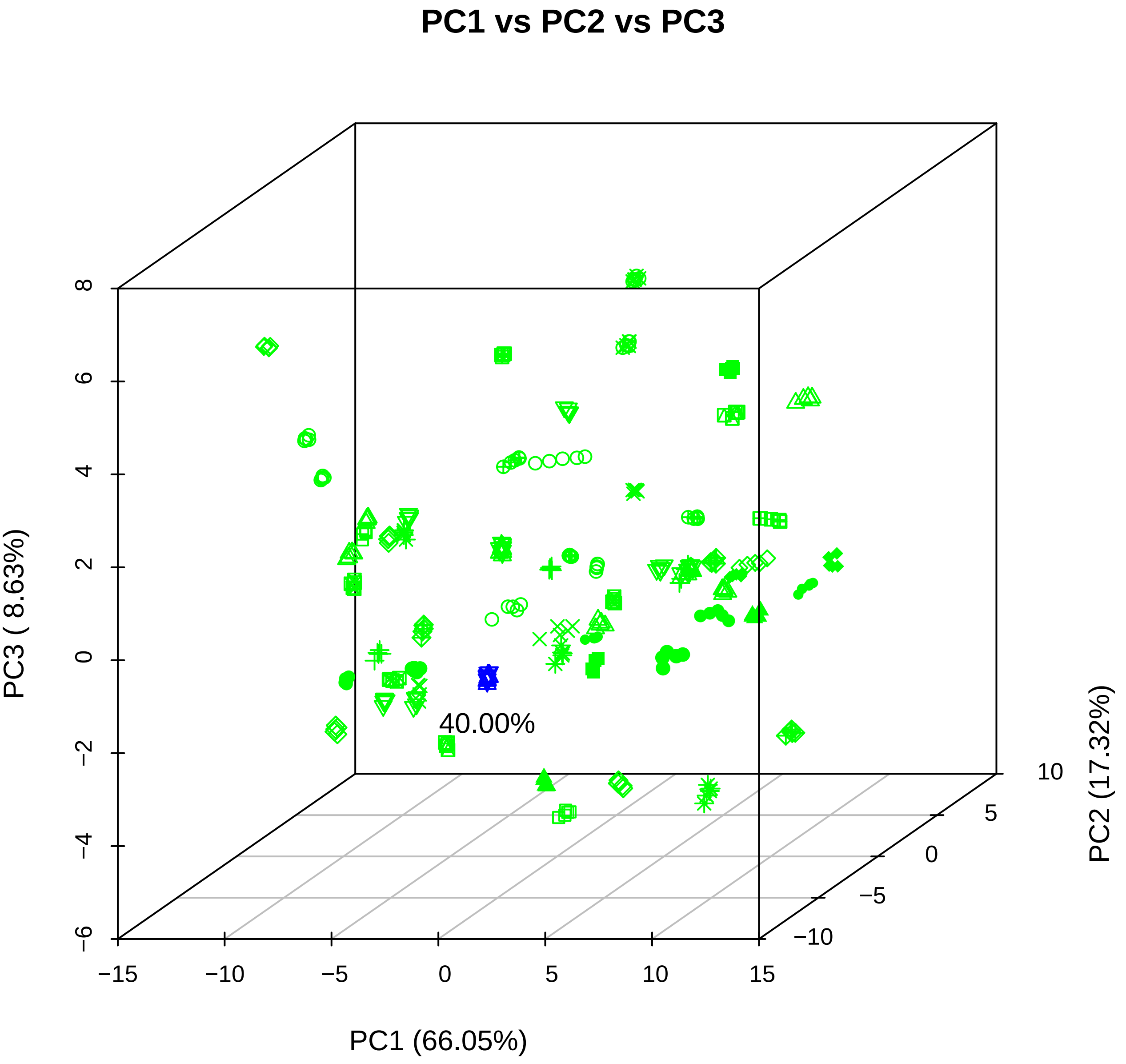
PC1 vs PC2 vs PC3
40.00%
PC1 (66.05%)
PC2 (17.32%)
PC3 ( 8.63%)
−15
−10
−5
0
5
10
15
−6
−4
−2
0
2
4
6
8
−10
−5
0
5
10

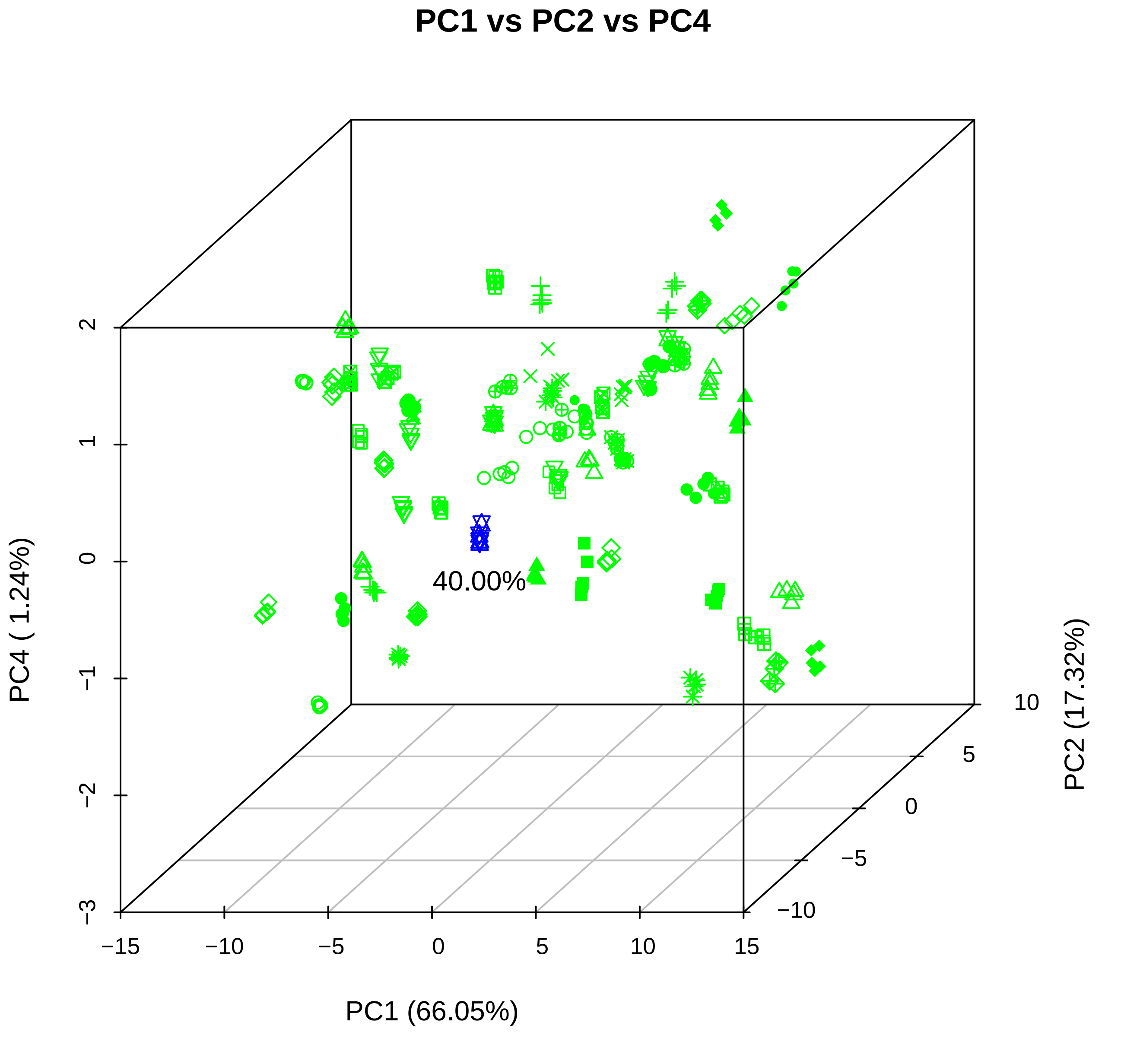

PC1 vs PC2 vs PC4
40.00%
PC1 (66.05%)
PC2 (17.32%)
PC4 ( 1.24%)
−15
−10
−5
0
5
10
15
−10
−5
0
5
10
−3
−2
−1
0
1
2

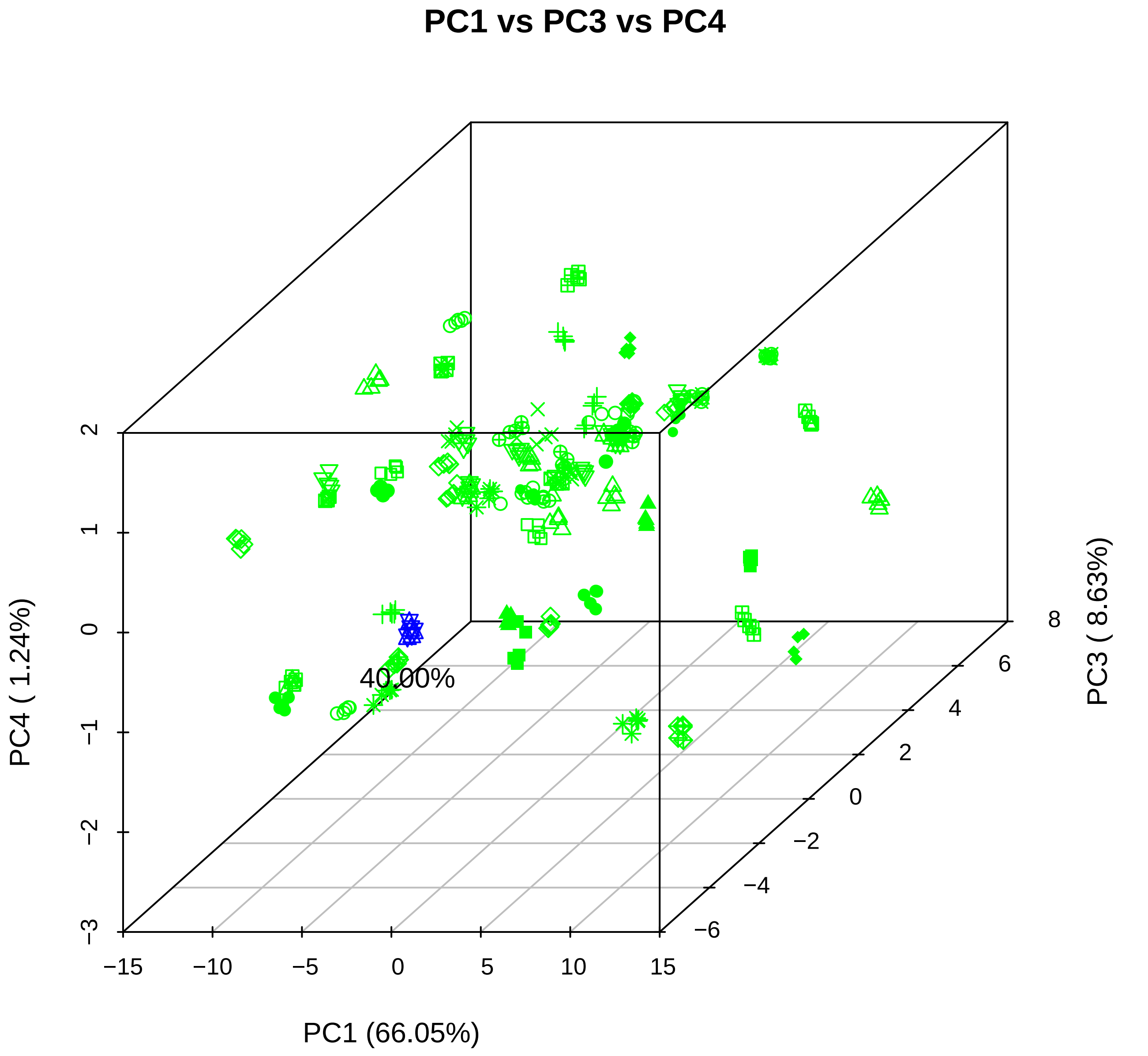

PC1 vs PC3 vs PC4
PC1 (66.05%)
PC4 ( 1.24%)
PC3 ( 8.63%)
40.00%
−15
−10
−5
0
5
10
15
−3
−2
−1
0
1
2
−6
−4
−2
0
2
4
6
8

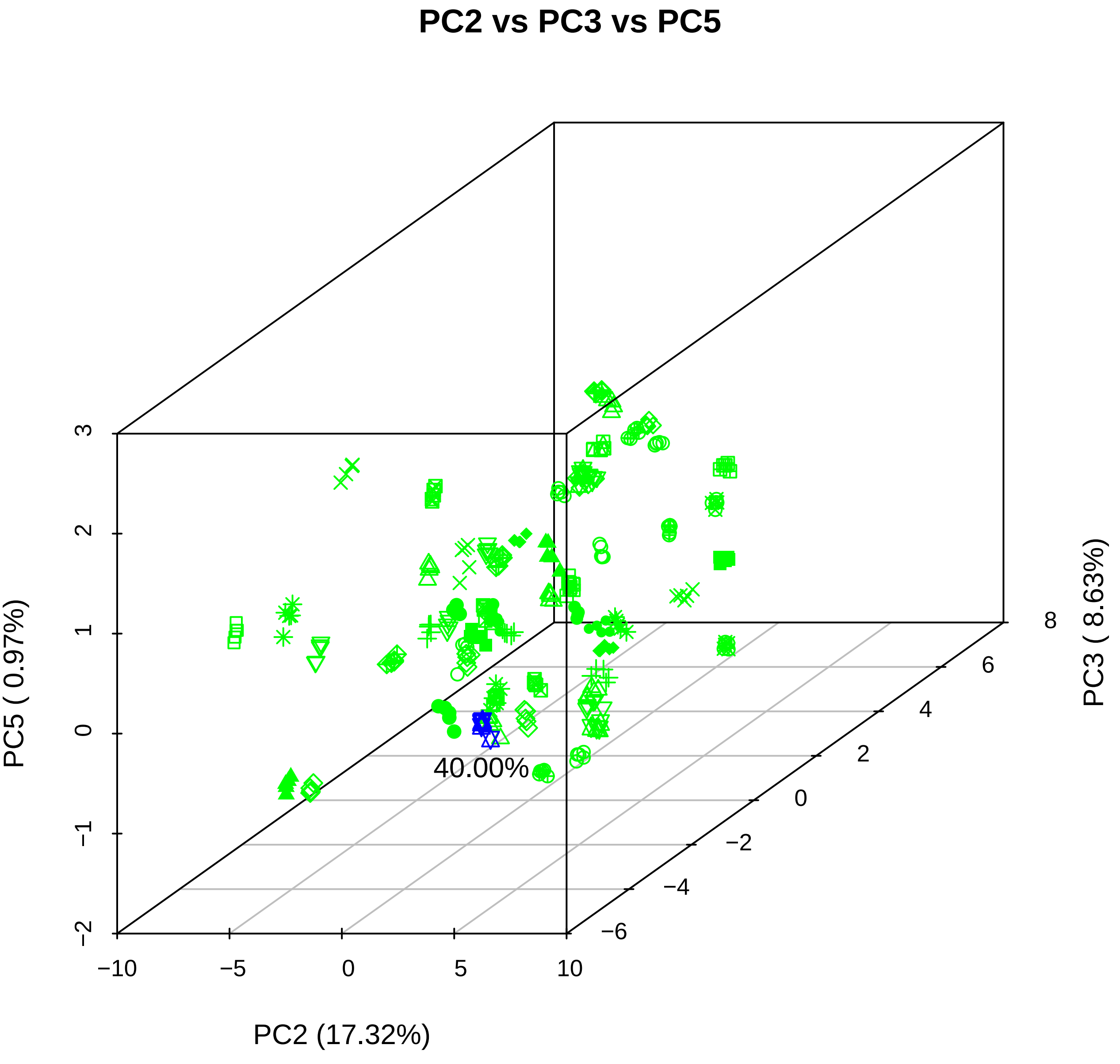

PC2 vs PC3 vs PC5
40.00%
PC2 (17.32%)
PC3 ( 8.63%)
PC5 ( 0.97%)
−10
−5
0
5
10
−6
−4
−2
0
2
4
6
8
−2
−1
0
1
2
3

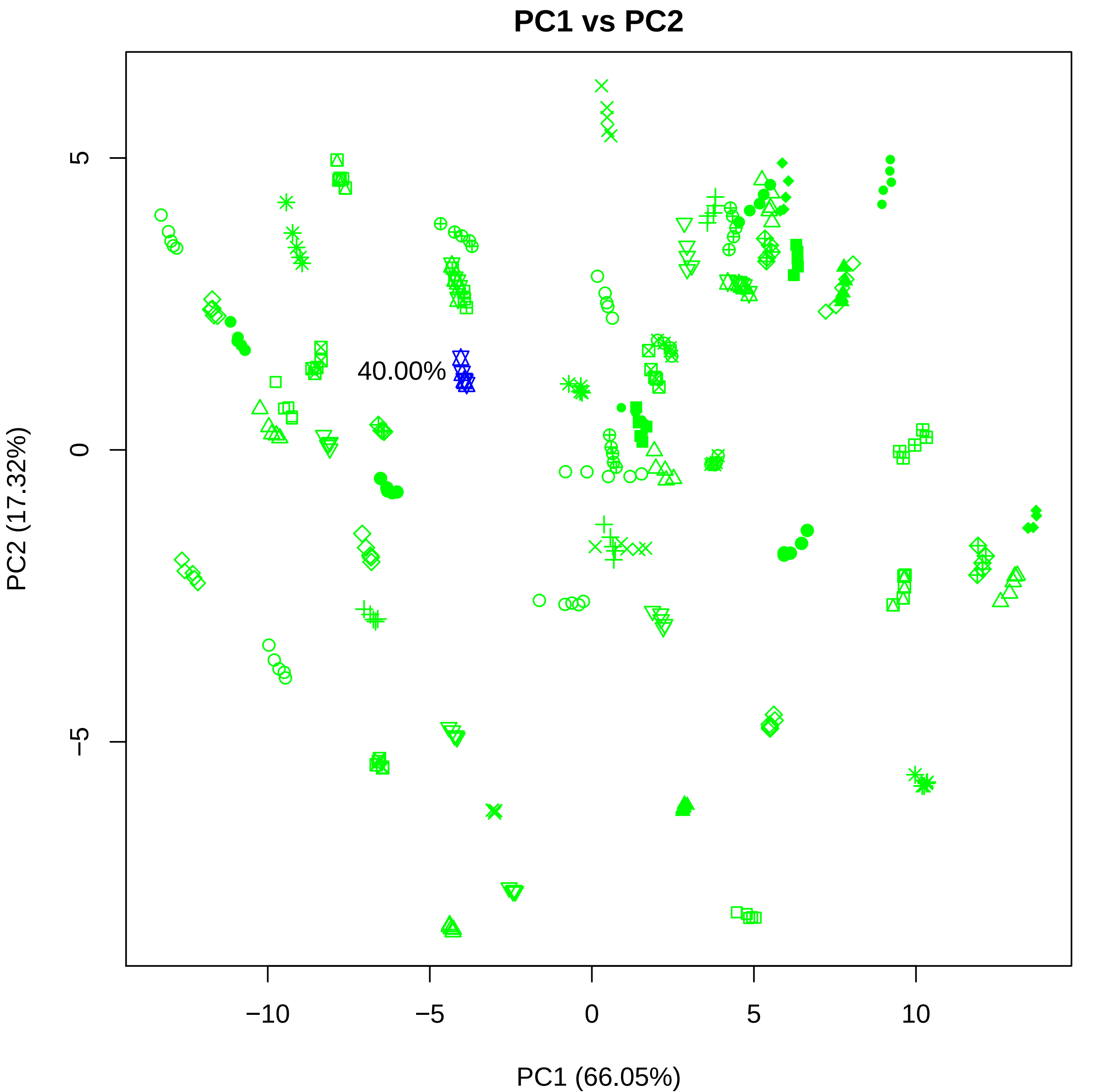
PC1 vs PC2
PC2 (17.32%)
PC1 (66.05%)
40.00%
5
0
−5
−10
−5
0
5
10

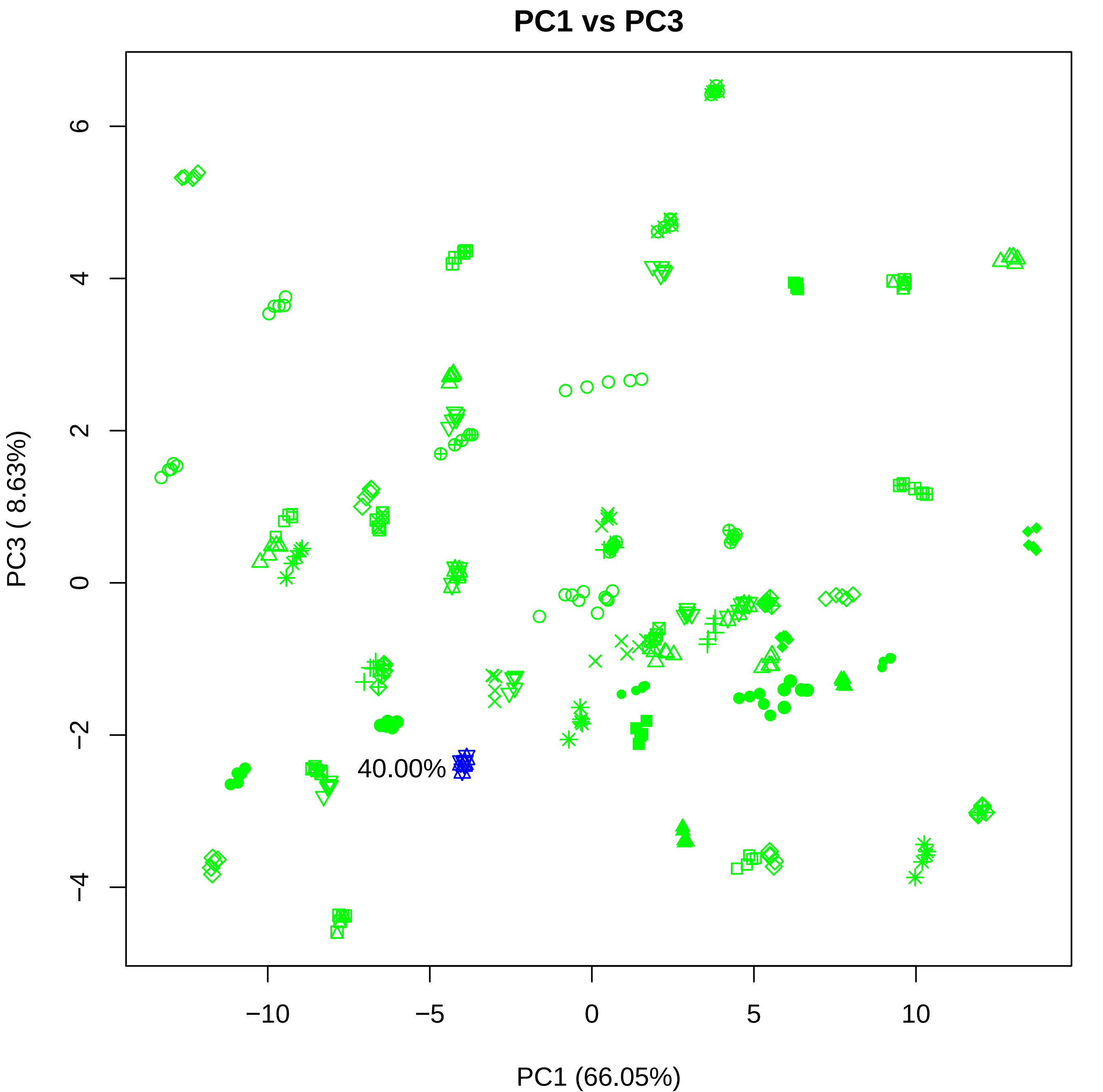

PC1 vs PC3
PC3 ( 8.63%)
PC1 (66.05%)
40.00%
6
4
2
0
−2
−4
−10
−5
0
5
10

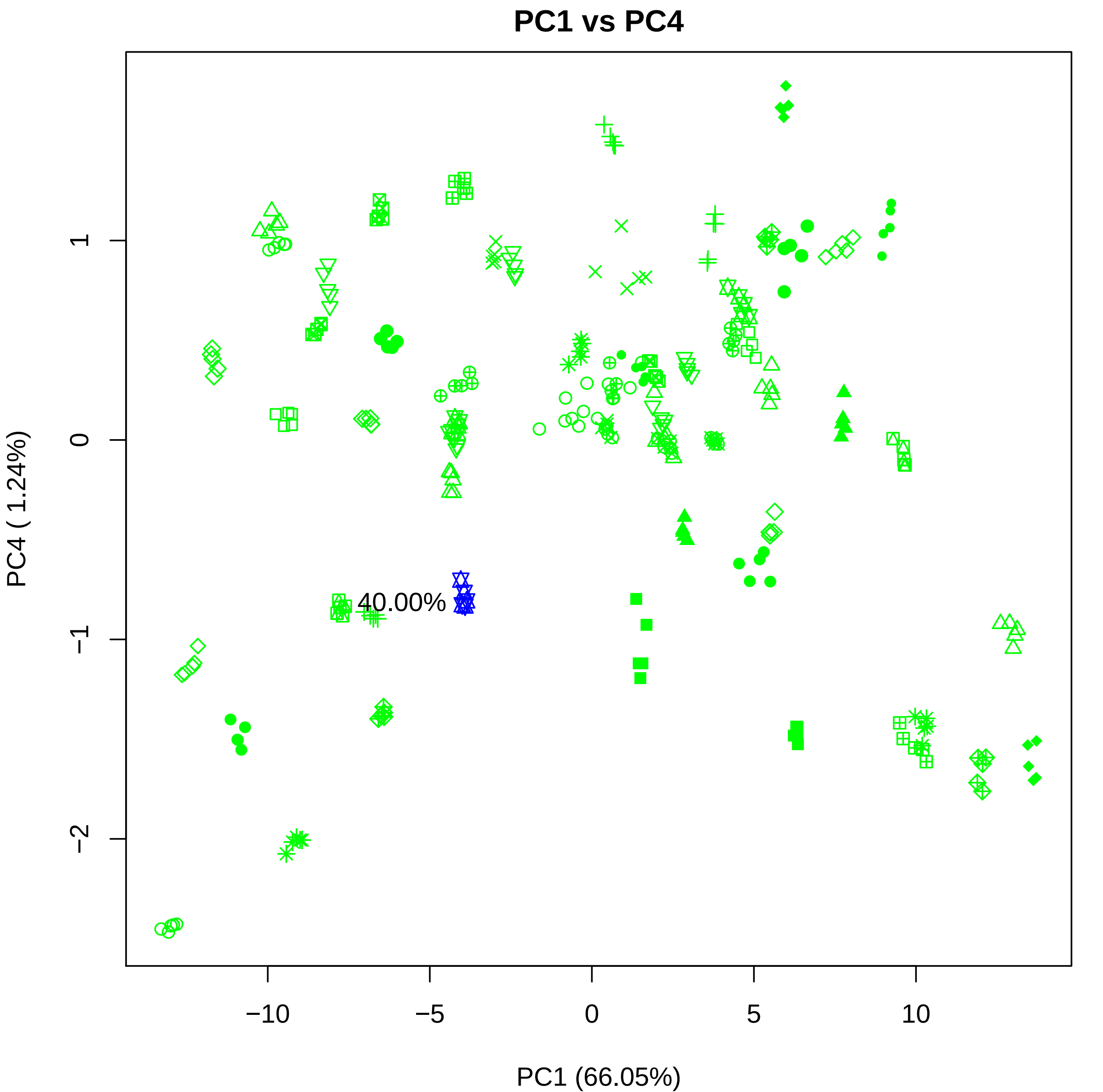
PC1 vs PC4
PC4 ( 1.24%)
PC1 (66.05%)
40.00%
1
0
−1
−2
−10
−5
0
5
10

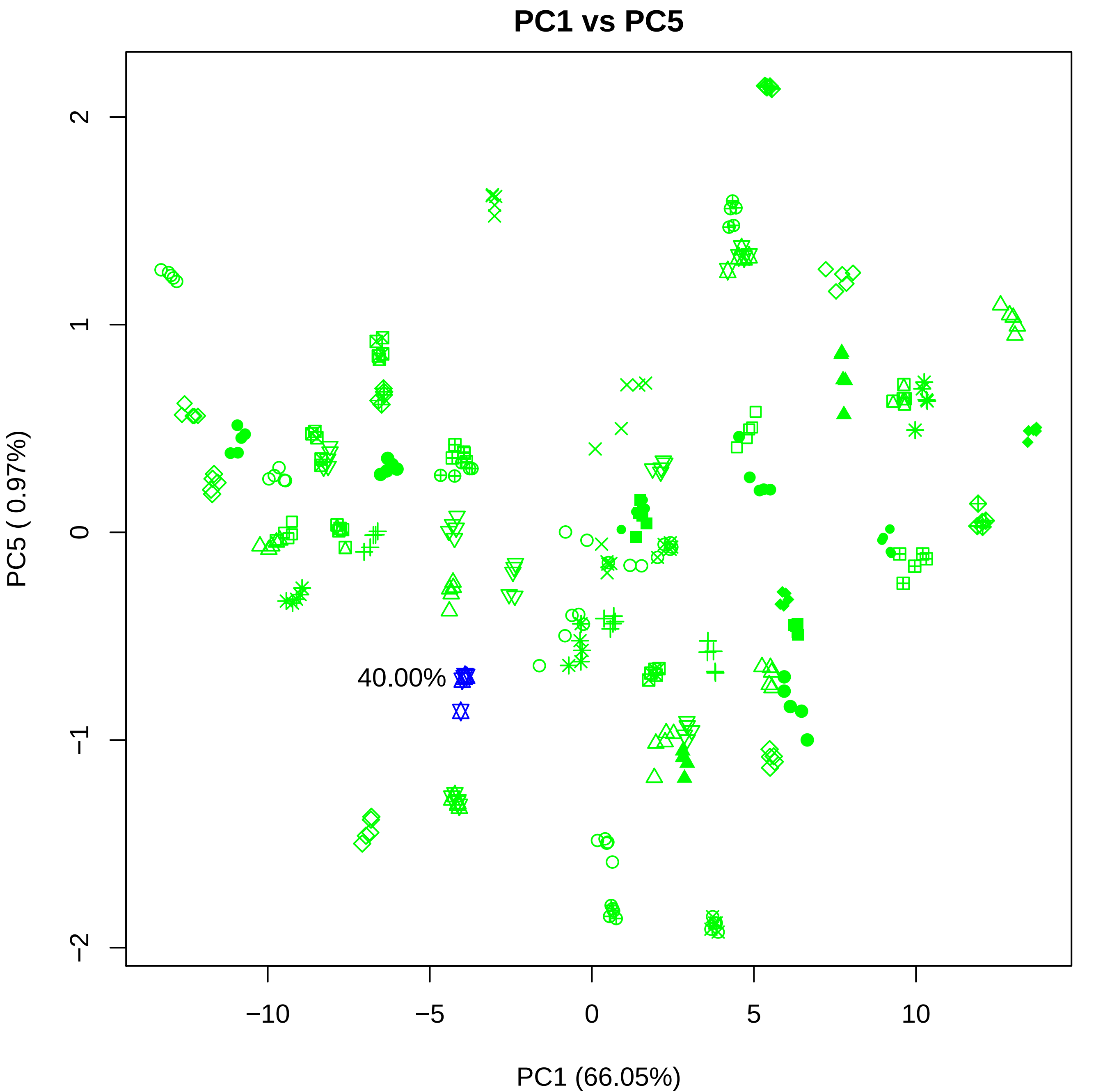
PC1 vs PC5
PC5 ( 0.97%)
PC1 (66.05%)
40.00%
2
1
0
−1
−2
−10
−5
0
5
10

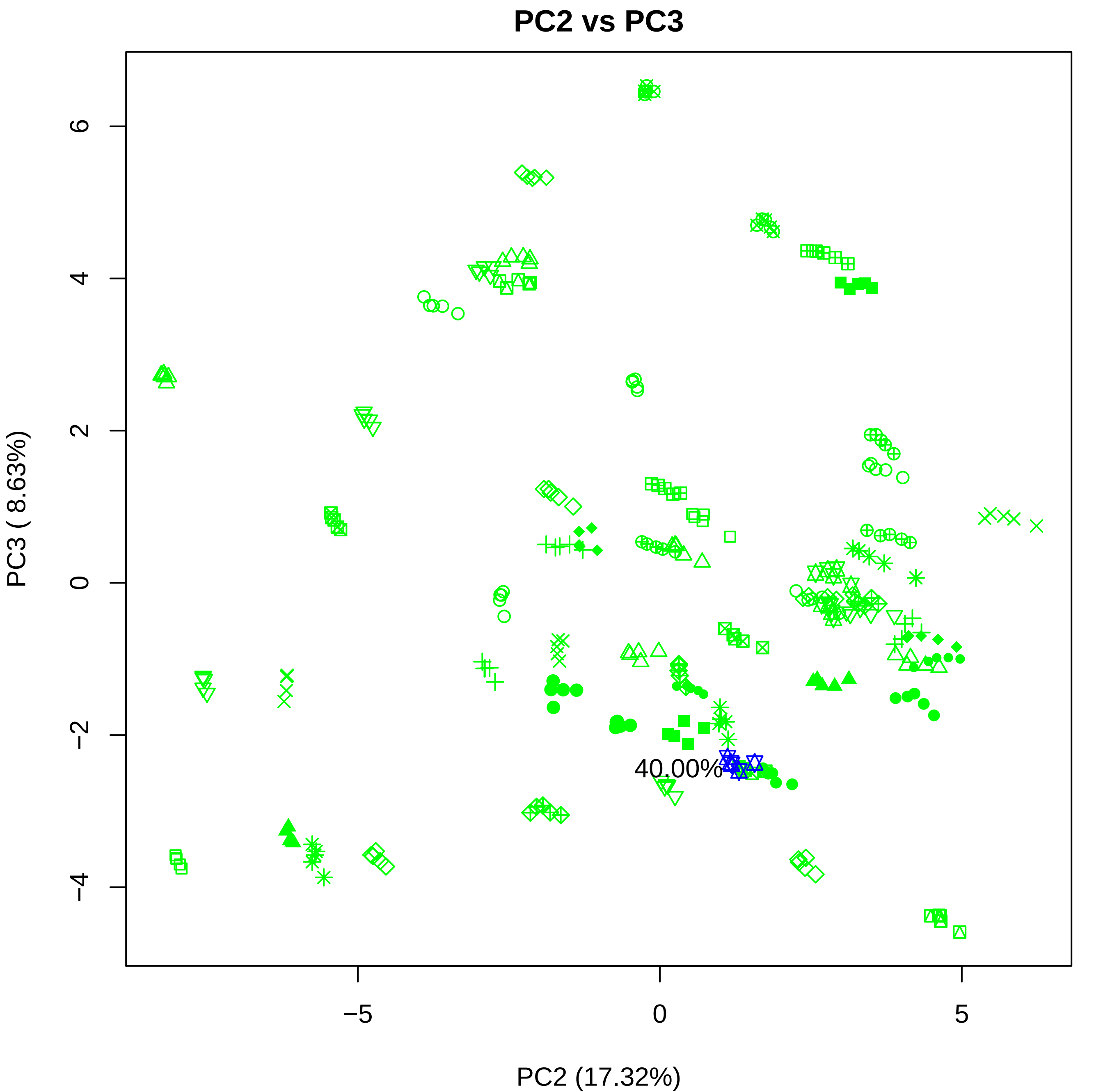
PC2 vs PC3
PC3 ( 8.63%)
PC2 (17.32%)
40.00%
6
4
2
0
−2
−4
−5
0
5

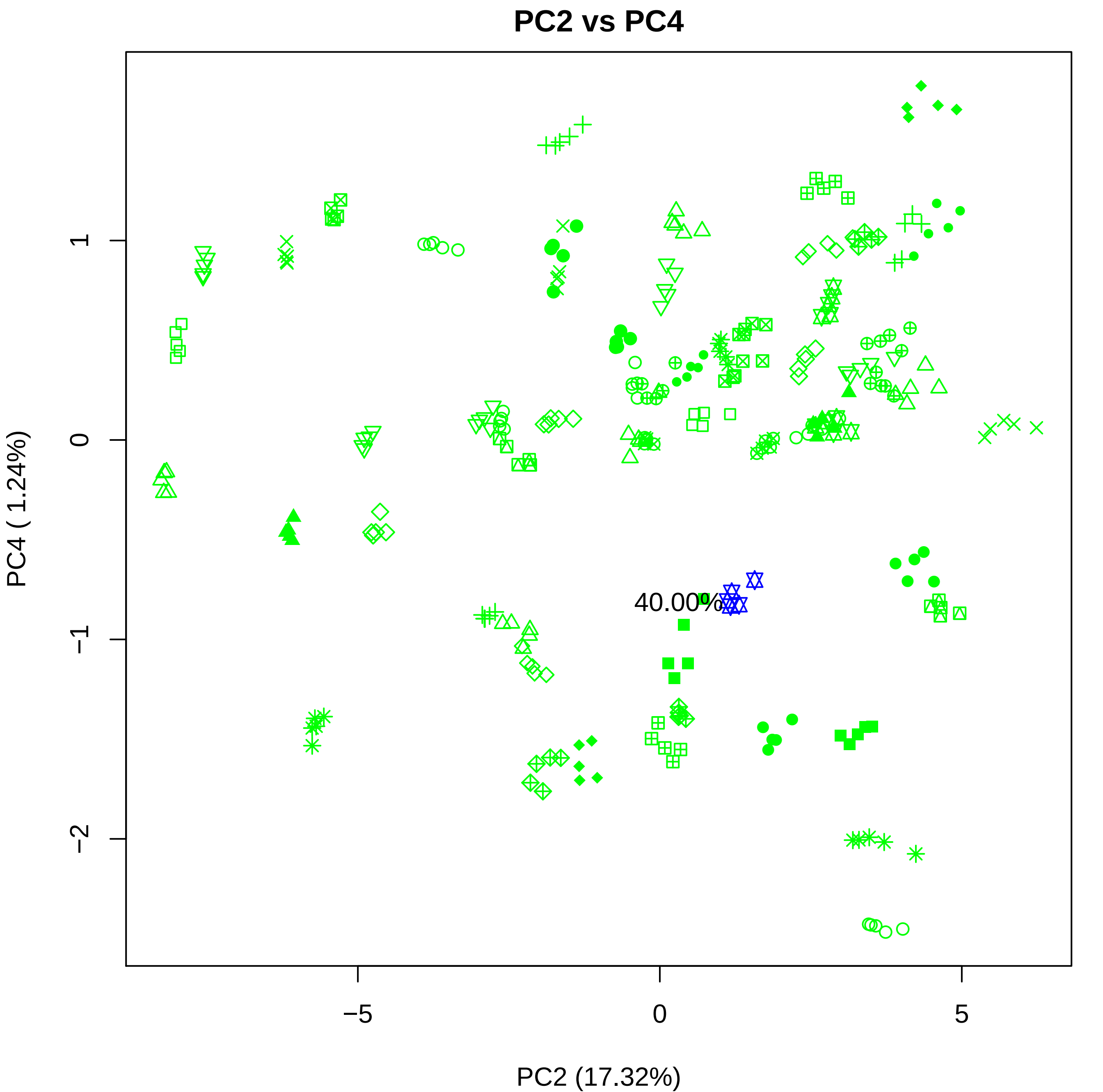
PC2 vs PC4
40.00%
PC4 ( 1.24%)
PC2 (17.32%)
−5
0
5
1
0
−1
−2

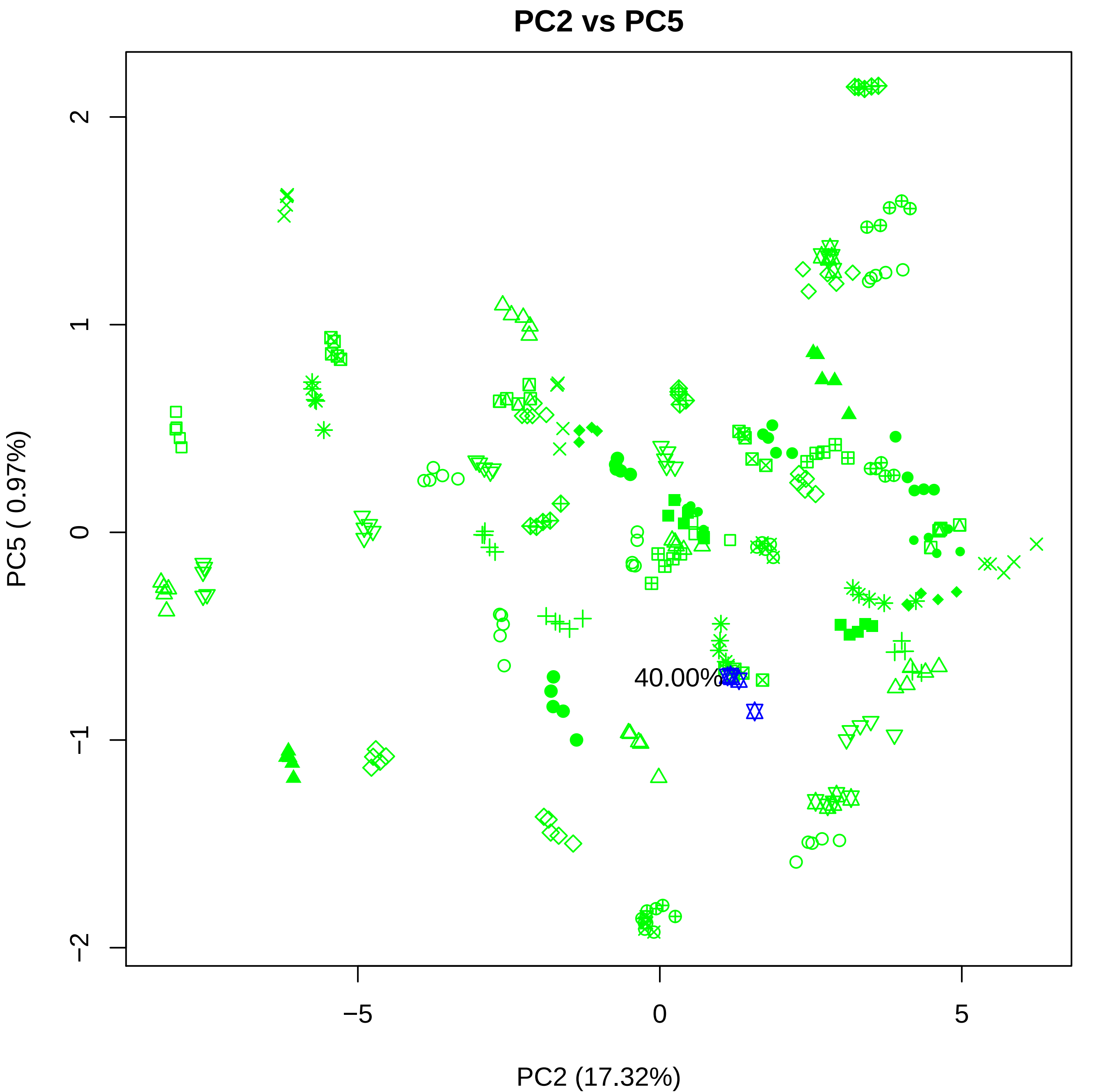
PC2 vs PC5
40.00%
2
1
0
−1
−2
−5
0
5
PC5 ( 0.97%)
PC2 (17.32%)

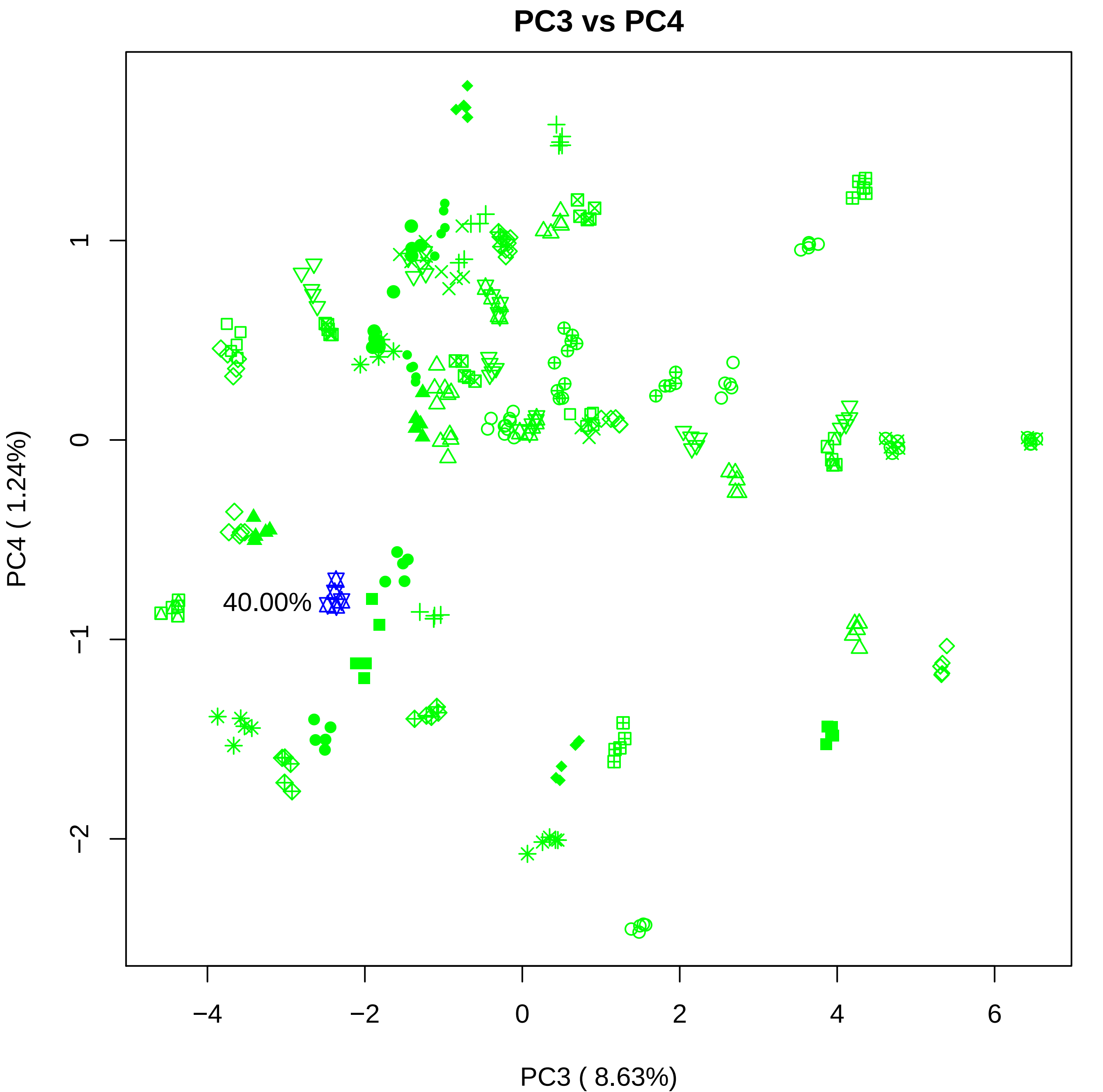
PC3 vs PC4
PC4 ( 1.24%)
PC3 ( 8.63%)
40.00%
1
0
−1
−2
−4
−2
0
2
4
6

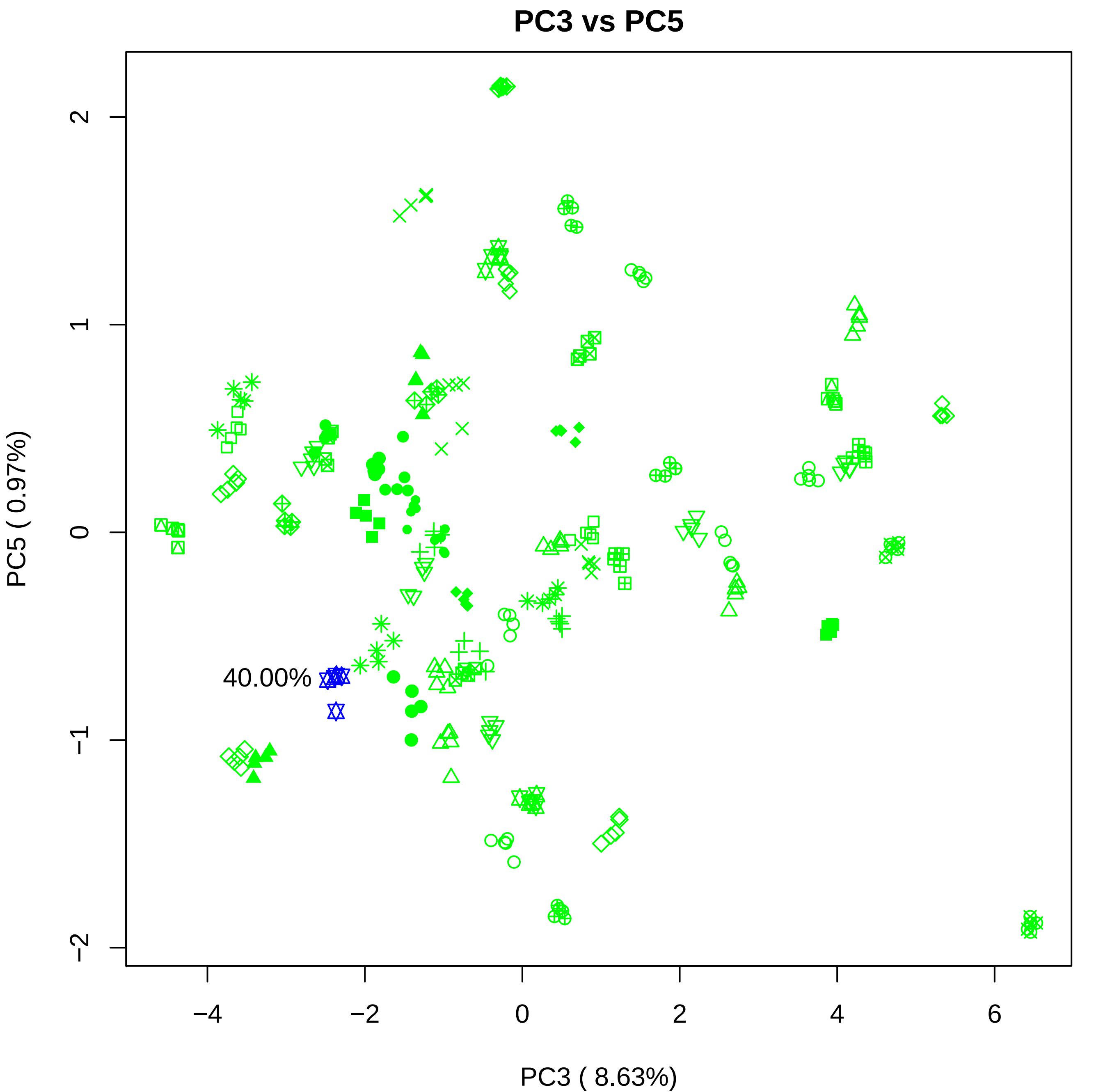
PC3 vs PC5
40.00%
PC5 ( 0.97%)
PC3 ( 8.63%)
2
1
0
−1
−2
−4
−2
0
2
4
6

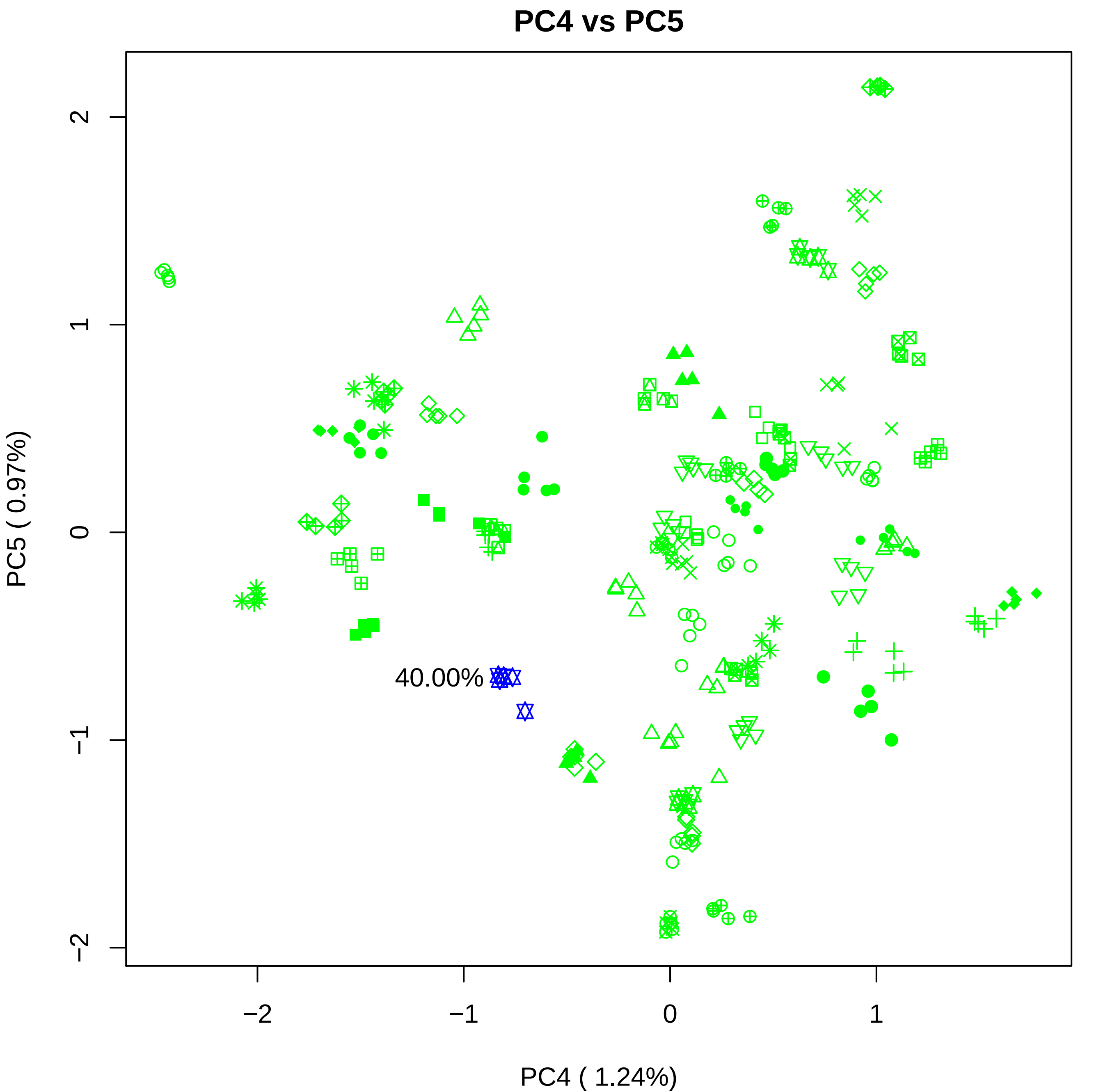

PC4 vs PC5
40.00%
2
1
0
−1
−2
PC5 ( 0.97%)
−2
−1
0
1
PC4 ( 1.24%)
